# Liver-specific Mettl14 deletion induces nuclear heterotypia and dysregulates RNA export machinery

**DOI:** 10.1101/2024.06.17.599413

**Authors:** Keith A Berggren, Saloni Sinha, Aaron E Lin, Michael P Schwoerer, Stephanie Maya, Abhishek Biswas, Thomas R Cafiero, Yongzhen Liu, Hans P Gertje, Saori Suzuki, Andrew R. Berneshawi, Sebastian Carver, Brigitte Heller, Nora Hassan, Qazi Ali, Daniel Beard, Danyang Wang, John M Cullen, Ralph E Kleiner, Nicholas A Crossland, Robert E Schwartz, Alexander Ploss

## Abstract

Modification of RNA with N^6^-methyladenosine (m^6^A) has gained attention in recent years as a general mechanism of gene regulation. In the liver, m^6^A, along with its associated machinery, has been studied as a potential biomarker of disease and cancer, with impacts on metabolism, cell cycle regulation, and pro-cancer state signaling. However these observational data have yet to be causally examined *in vivo.* For example, neither perturbation of the key m^6^A writers *Mettl3* and *Mettl14*, nor the m^6^A readers *Ythdf1* and *Ythdf2* have been thoroughly mechanistically characterized *in vivo* as they have been *in vitro*. To understand the functions of these machineries, we developed mouse models and found that deleting *Mettl14* led to progressive liver injury characterized by nuclear heterotypia, with changes in mRNA splicing, processing and export leading to increases in mRNA surveillance and recycling.

## Introduction

N^6^-methyladenosine (m^6^A) RNA modification is a critical gene regulatory mechanism, based on extensive *in vitro*, cell culture, and patient studies (*1*). RNA modification with m^6^A is implicated in cellular differentiation, metabolism, and cell-cycle regulation. Moreover, dysregulation of m^6^A or the m^6^A ‘writer’ enzymes *Mettl3* and *Mettl14* contribute to the development and malignancy of many cancers including hepatocellular carcinoma (HCC) (*2–4*). Clearly deciphering the impacts of m^6^A modification on the RNAs it is placed on is important, as this modification can lead to either RNA stability in some cases and degradation in others through various mechanisms(*5–7*). The function of m^6^A on specifically modified transcripts, and the associated machinery including *Mettl3* and *Mettl14* are important to understand not just for diagnostic and prognostic utility, but also as potential causative mechanisms which could serve as therapeutic targets. This interest, along with availability of better tools, has recently led to a flurry of studies on the impacts of disrupting m^6^A writers in liver tissue *in vivo* in mouse models (*8–11*).

RNA modification by m^6^A has been implicated in liver diseases, inflammation, and injury response, as well as viral infection (*1*). Metabolically associated steatohepatitis (MASH) has been correlated with global increases in m^6^A as well as increased levels of writer complex proteins *Mettl14* and *Mettl3* (*12*). Specific sites on interferon (IFN) beta mRNA have been shown to bear m^6^A modifications, and knockdown of *Mettl14* leads to increased IFN expression and interleukin-17RA, concurrently with increased inflammatory pathway activation and metabolic reprogramming in the liver (*13*, *14*). m^6^A modification of viral RNAs has also been reported on hepatitis B and delta virus RNAs, appearing to impact the viral replicative cycle and the switch from translation of protein and replication to packaging (*15*, *16*). Hepatitis B Virus (HBV) has also been shown to increase m^6^A levels in liver, causing a feed-forward loop of inflammation and leading towards PTEN and innate immunity changes which lead towards HCC development(*17*).

Despite this wealth of observational data, it has been difficult to pinpoint causal or mechanistic roles for m6A or its associated machinery. First, new studies have cast doubt on the accuracy and reproducibility of commonly used m^6^A sequencing techniques (*18*). Second, recent work has provided evidence for unexpected feedback loops where m^6^A-modification leads to chromatin remodeling (*19*). Third, m^6^A writers may have alternate functions, acting directly on DNA or impacting RNA splicing, nuclear export, and localization (*20–25*). Taken together, these data are difficult to integrate into a comprehensive understanding of how dysregulation of m^6^A and its associated machinery might contribute to or cause liver disease *in vivo*.

To better understand the effects of m^6^A modification machinery dysregulation in the context of liver tissue, we generated a mouse model of hepatocyte-specific deletion of m^6^A writer *Mettl14* to assess transcriptomic changes in the liver during mature steady-state tissue maintenance. We additionally developed a dual-deletion model of the ‘reader’ proteins *Ythdf1* and *Ythdf2* to disentangle the impacts of the canonical pathway of m^6^A-modified mRNA degradation from other potential outcomes of *Mettl14* deletion. Since the impacts of m^6^A modification are particularly important to temporal regulation of cellular differentiation and cell-cycle regulation during development or regeneration, we also assessed regenerative capacity and liver architecture following an extensive array of injury models: surgical, chemical, or chronic infection. Deletion of *Mettl14* or *Ythdf1*/*Ythdf2* together led to worse liver outcomes, depending on the type of challenge, highlighting the important roles and different functions of these genes in liver homeostasis and regeneration.

## Results

### m6A machinery defects impact post-natal liver maintenance, leading to injury

To study the impacts of *Mettl14* deletion on proper liver development, we developed a hepatocyte-specific deletion of *Mettl14* via a cross of mice in which exons 7-9 are flanked by loxP sites (*Mettl14^[fl/fl]^*) (*26*) with mice bearing a Cre transgene under the hepatocyte-specific albumin promoter (Alb-Cre) (*27*) (**Fig. 1A**). We further developed a hepatocyte-specific dual *Ythdf1* and *Ythdf2* deletion mouse by crossing full-body *Ythdf1* knockout (*Ythdf1^-/-^*) mice with *Ythdf2* floxed (*Ythdf2^[fl/fl]^*) mice (*26*). The subsequent mice were crossed with the Alb-Cre expressing mice to establish dual knockout in hepatocytes (**Fig. 1A**). These liver-specific deletions were necessary as whole-body knockouts of *Mettl14* or *Ythdf2* are embryonic lethal (*28*, *29*). We confirmed reduced *Mettl14* transcript and protein levels in the *Mettl14*^[fl/fl]^ Alb-Cre liver tissue by reverse-transcription qPCR and western blot, respectively (**Fig. S1E,F**).

**Fig. 1.**
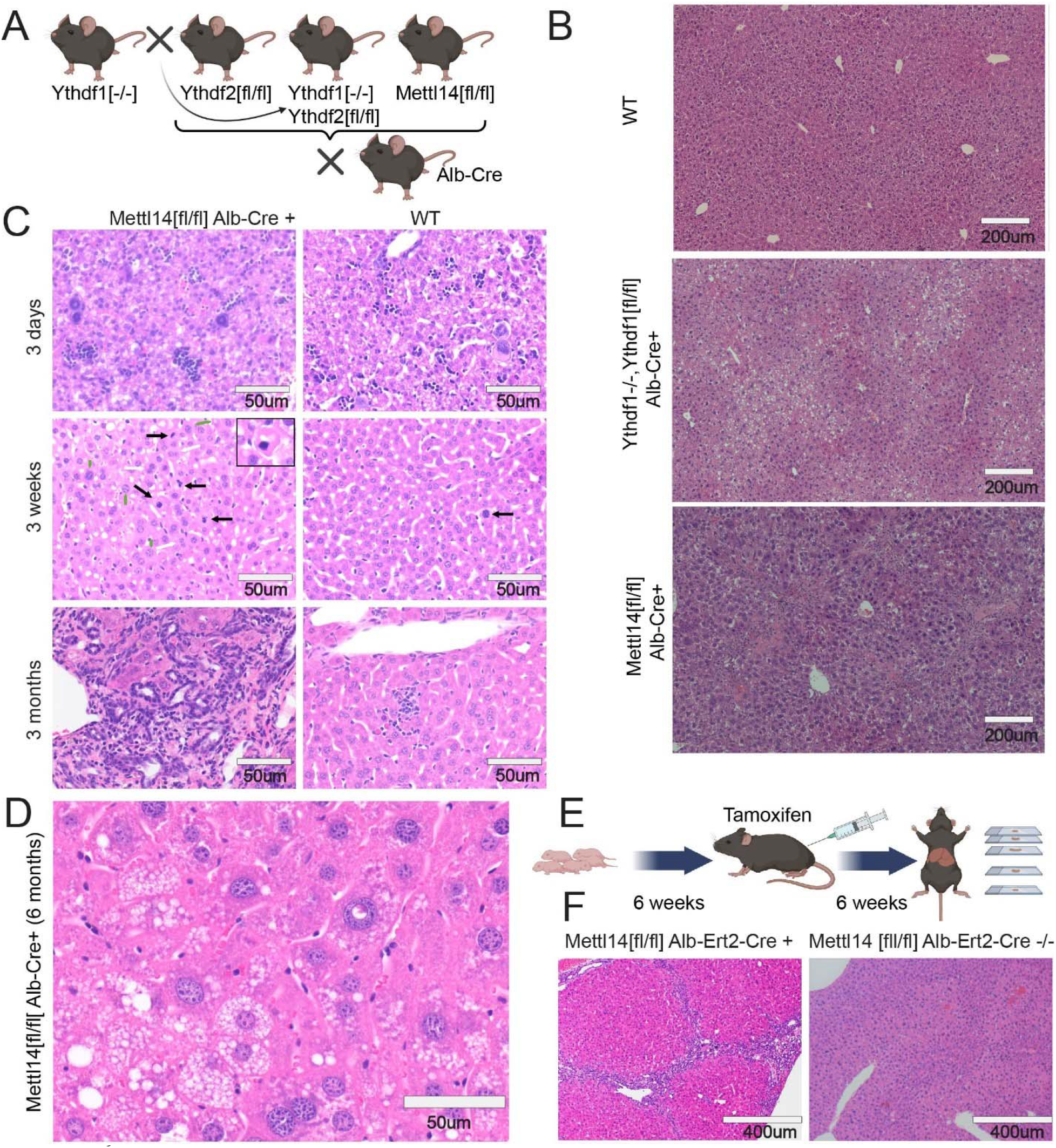
m^6^A writer and reader deficiencies lead to progressive liver damage. **(A)** Schematic showing crosses used to generate gene deletions in hepatocytes for this study (Created with BioRender.com). **(B)** Dual *Ythdf1*/*Ythdf2* deletion in liver tissue as well as *Mettl14* deletion leads to liver injury, but each with distinctly different histology. **(C)** *Mettl14*-deficient liver tissue exhibits progressive damage emerging first at 3 days and progressing in severity through 3 months of age and older. Black arrows indicate apoptotic cells with condensed nuclei, green arrows indicate enlarged nuclei, and white arrows indicate mitotic events. **(D)** Representative image of the advanced liver injury phenotype seen in 6-month-old *Mettl14* deficient liver tissue, showing steatosis and nuclear heterotypia. **(E)** Schematic of tamoxifen-induction timeline where mice were injected with tamoxifen over 6 weeks before sacrifice for experiments (Created with BioRender.com). **(F)** Liver injury is recapitulated in this inducible *Mettl14* model, showing similar levels of fibrosis and nuclear heterotypia.

The liver-specific deletion mice appeared grossly normal, although some liver-specific *Mettl14* individuals displayed lower weights and slowed growth (**Fig S1C**). We initially collected tissues from these mice at 8 weeks of age to assess impacts on mature liver development and maintenance by histology imaging of hematoxylin and eosin (H&E)-stained sections. Both the dual *Ythdf1*^-/-^/*Ythdf2*^[fl/fl]^/Alb-Cre and the *Mettl14*^[fl/fl]^/Alb-Cre mice showed liver injury with histological similarity to steatohepatitis, but *Mettl14*^[fl/fl]^/Alb-Cre mice showed additional inflammation, regions of necrosis, and signs of liver fibrosis (**Fig. 1B**). Deletion of either *Ythdf1* (in the whole animal) or *Ythdf2* (specifically in the liver) on its own did not lead to any significant histological changes (**Fig S1A**).

To assess the timeframe of onset of this damage in *Mettl14*-deleted mice, we analyzed liver tissue architecture of *Mettl14*^[fl/fl]^/Alb-Cre mice during growth and maturation (postnatal day 3, week 3, and month 3). By 3 weeks of age, individual hepatocytes could be seen undergoing apoptosis, and the rate of mitotic events was increased relative to the wild-type control (**Fig. 1C**), likely to replace the dying cells. This damage progressed over time, with *Mettl14*^[fl/fl]^/Alb-Cre livers at 3 months of age displaying pronounced immune infiltrate and signs of fibrosis. This liver injury phenotype, while variable, was worse in males than females (**Fig. S1D**): male liver injury steadily progressed to a stark phenotype of advanced steatohepatitis with immune infiltrate and pronounced nuclear heterotypia by 6 months of age (**Fig. 1D**). More broad areas of fibrosis and further necrosis and apoptosis can also be seen in mice at older ages **(Fig. S1B**).

We aimed to characterize global molecular changes underlying the liver pathology observed in *Mettl14*-deleted mice. We performed bulk-RNA sequencing of liver tissue from *Mettl14*^[fl/fl]^/Alb-Cre and *Mettl14*^[fl/fl]^ mice (**Supplemental data 1**). A volcano plot showed good distribution of upregulated and downregulated transcripts, with a relatively low percentage of genes differentially expressed at statistically significant levels (**Fig. 2A**). We compared the set of mRNA transcripts with number of known m^6^A modification sites per transcript with our list of differentially expressed genes and found a correlation between modification and upregulation of transcripts **(Fig. 2B)**. Differentially expressed genes (DEGs) represented approximately 1% of the total detected genes (**Fig. 2C**), and 84% of DEGs were known to be m^6^A modified at one or more sites, with a slight skew towards down-regulated genes (*30*) **(Fig. 2D).**

**Fig. 2.**
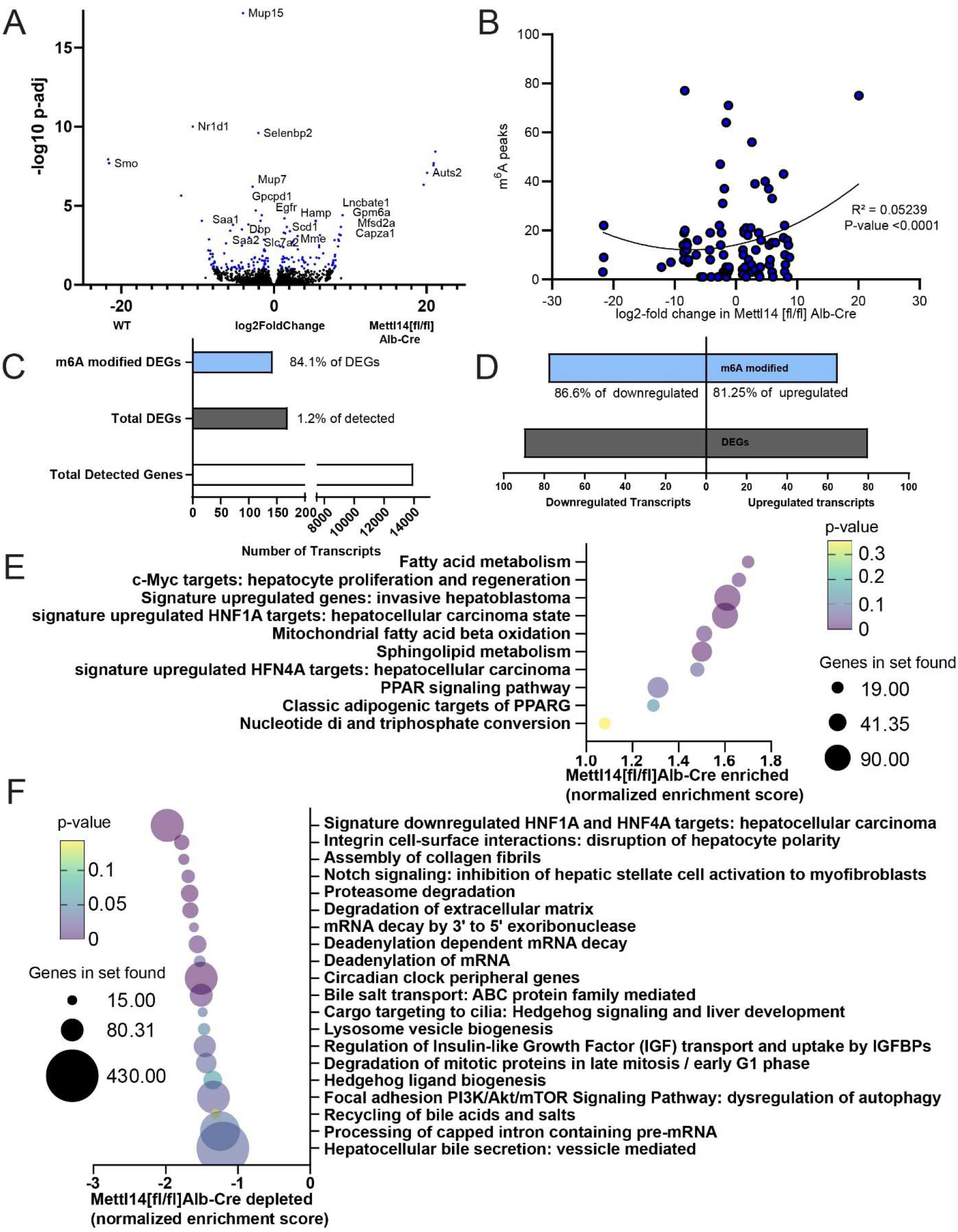
Bulk RNAseq and GSEA analysis reveal key pathways of liver damage. **(A)** Volcano plot of transcripts upregulated and downregulated between *Mettl14*^[fl/fl]^/Alb-Cre and *Mettl14*^[fl/fl]^ male mice (n = 3) Hits with a significant adjusted p-value below 0.1 are highlighted in blue. Specific genes with particularly significant p-values or high levels of up-regulation or down-regulation, as well as those with particularly interesting functions to pathways seen in the GSEA are marked with gene names next to their data points. **(B)** Comparison of known m^6^A modified sites on transcripts with significantly upregulated or downregulated transcripts in *Mettl14* deletion mice. **(C)** The ratio of enrichment of transcripts published to be m^6^A modified versus those without evidence for modification among differentially expressed genes (DEGs) seen in our data **(D)** and breakdown of m^6^A enrichment among DEGs in those upregulated and downregulated. **(E)** Gene set enrichment analysis (GSEA) of bulk transcriptomic data, revealing which pathways and functions are over-represented in the *Mettl14* deletion mice. **(F)** GSEA results of which pathways and functions are downregulated in *Mettl14* deletion mice.

Gene set enrichment analysis (GSEA) revealed dysregulation of pathways that confirmed our histological findings of liver pathology **(Supplemental Data 2)**. Livers of *Mettl14*-deleted mice had major lipid metabolism and stress response dysregulation, modulated by hepatocyte nuclear factor 1 alpha (*Hnf1a*) and *Hnf4a*. The phenotypes of these GSEA pathways, fibrosis and pro- hepatocellular carcinoma states approximated the steatohepatitis we observed in our mice (**Fig. 2E,F**) (*31–33*). Notably, *Smoothened* (*Smo)*, a member of the *Hedgehog* signaling pathway, showed remarkably decreased expression in liver tissue from *Mettl14*^[fl/fl]^/Alb-Cre as compared to *Mettl14*^[fl/fl]^ mice. *Smo* transcript expression was not detected in any knockout mouse samples (**Fig. 2A**, **Fig. S2)**, which correlated with *Hedgehog* pathway signaling changes seen in the GSEA **(Fig. 2E,F)**. *Smo* and *Hedgehog* signaling have direct impacts on hepatic insulin resistance and metabolism regulation, and decreased *Hedgehog* signaling is correlated with increased susceptibility of fatty liver disease progression to metabolically associated steatohepatitis (MASH) and fibrosis (*34–36*), similar to the phenotype observed in our mice.

To determine whether the damage seen in *Mettl14* mice is due to defects in pre-natal development versus post-natal metabolism and maintenance defects, we separately developed a liver-specific inducible *Mettl14* deletion mouse by crossing our *Mettl14*^[fl/fl]^ mice with a tamoxifen-inducible Cre, termed Alb-ERT2-Cre (*37*) expressed under the albumin promoter. At 6 weeks of age, we induced gene-specific deletion in these *Mettl14*^[fl/fl]^/Alb-ERT2-Cre mice with tamoxifen for a total of 6 weeks to allow time for any potential resulting effects to be histologically evident before sacrificing mice for histological analysis (**Fig. 1E**). Our results showed that inducible *Mettl14* deletion for 6 weeks in adulthood lead to similar fibrosis and necrosis as was seen in the constitutive deletion model at 6 weeks of age (**Fig. 1F**). Therefore, we concluded that *Mettl14* was required for proper post-natal liver growth and maintenance.

### Functional liver regeneration is impeded in *Ythdf1*/*Ythdf2* dual deletion

The liver possesses an impressive capacity for regeneration after injury, reconstituting not only tissue mass, but also the specific functional architecture of liver tissue with zonation around portal vein tracts and bile ducts (*38*, *39*). This allows rapid response to injury, toxicity, or acute infection to maintain the scope of liver function necessary to maintain metabolic processes. In wild-type C57BL/6 mice, liver regeneration occurs over a period of approximately seven days after experimentally induced injury via two-thirds partial hepatectomy (*40*). This process requires tight temporal control of cell-cycle, motility, differentiation, and coordination of developmental pathways to reconstitute tissue architecture. All of these processes have been shown in various settings to be impacted or directly regulated by m^6^A modification as a mechanism of gene regulation (*41–45*). As we initially showed that faulty m^6^A machinery affects liver maintenance at steady-state conditions, we wanted to further know whether m^6^A perturbation would also affect liver regeneration.

To better understand the impacts of m6A and it’s readers following injury, we assessed the capacity of *Mettl14*, *Ythdf1*, *Yhtdf2*, and dual *Ythdf1/Ythdf2* deficient mice to recover after two-thirds partial hepatectomy surgery. We weighed mice at 6 weeks of age, and then performed surgery to remove approximately two-thirds of total liver mass (**Fig. 3A**). Tissue was weighed after removal to confirm the appropriate amount of tissue loss. Any individual animals who experienced significant blood loss, deviated significantly from the target amount of tissue removed, or who experienced post-surgical complications were removed from the study to avoid skew of results due to any surgical technique variability. We assessed two phenotypes: (1) liver mass, as a function of the liver’s ability to regenerate, and (2) whole body mass, as reduced liver function leads to slower weight recovery. We weighed mice daily for 2 weeks post-surgery before sacrificing them for experimental sample collection and analysis. Total liver mass was assessed at time of sacrifice and compared as a ratio of liver mass to body mass to control for relative variability in overall animal size.

**Fig. 3.**
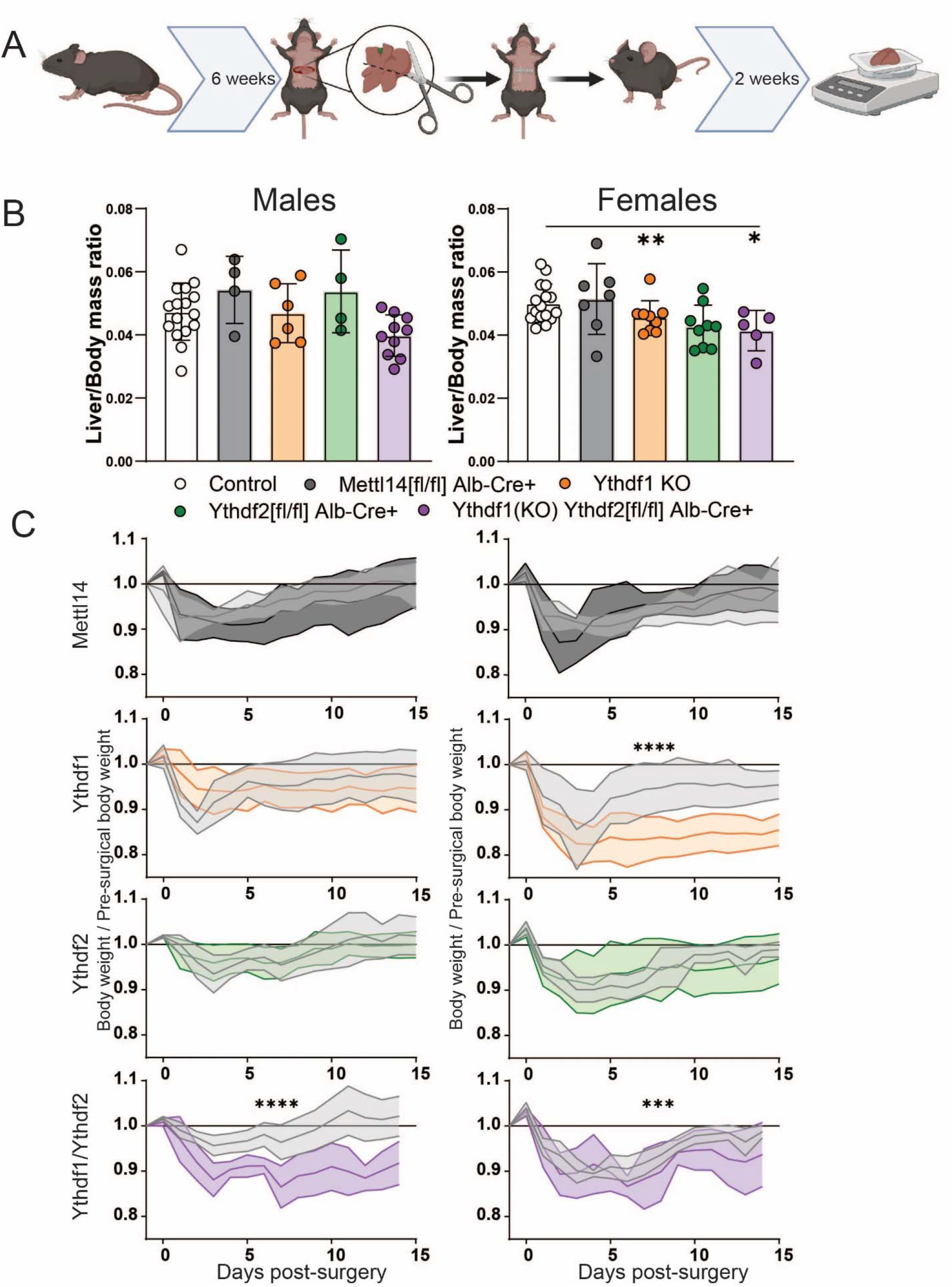
Significantly slower liver regeneration in m^6^A reader-deficient mice following partial hepatectomy. **(A)** Schematic of timeline for two-thirds hepatectomy surgery and recovery prior to sacrifice and sample collection (Created with BioRender.com). **(B)** Males (left) and females (right): comparison of liver /body mass ratios at 2 weeks post-surgery across C57BL/6 (N=16 males, 16 females) *Mettl14* (N=4 males, 7 females), *Ythdf1*(N=6 males, 9 females), *Ythdf2* (N=4 males, 9 females), and dual *Ythdf1*/*Ythdf2* deletion mice (N=10 males, 5 females). **(C)** Males (left) and females (right): weight recovery curves after surgery.

In *Ythdf1*^-/-^/*Ythdf2*^[fl/fl]^/Alb-Cre and *Ythdf1*^-/-^ mice, overall liver mass regeneration was significantly reduced compared to control animals after surgery (**Fig. 3B**). An expected sexual dimorphism in regenerative capacity was also noted, with males regenerating more liver tissue than females of the same genotype (*46*). As a proxy of liver function, the whole body weight of *Ythdf1*^-/-^/*Ythdf2*^[fl/fl]^/Alb-Cre mice recovered significantly more slowly, irrespective of sex (**Fig. 3C**). Deficiency of *Ythdf1* alone also resulted in significantly reduced weight recovery in female mice. The close agreement between which groups displayed significant reduction of regenerated liver mass and overall weight recovery was noted. These results suggest that deletion of the reader genes *Ythdf1* and *Ythdf2* slowed liver regeneration and function following surgical injury.

### Acute liver injury is exacerbated by m^6^A reader or writer deletion

Different type of liver damage can reveal different aspects of injury response and recovery. Probing these m6A reader and writer deficient mice with experimental liver injury allows us to better understand the actual nature of their defects and chronic injury phenotypes. Two separate toxicity-based injury models of liver damage response are used frequently in liver research to assess response to cholestasis-mediated liver injury as well as general hepatocyte toxicity. 1,4- dihydro 2,4,6-trimethyl 3,5-pyridinedicarboxylic acid diethyl ester (DDC) administration causes protoporphyrin plugs and stones, blocking bile ducts and impeding bile drainage from the liver parenchyma (*47*). This leads to cholestasis as bile buildup damages cholangiocytes, leading to sclerosing cholangitis and biliary fibrosis. A second model of liver injury is carbon tetrachloride (CCl_4_). CCl_4_ directly damages hepatocytes by inducing a severe state of oxidative stress by binding to triacylglycerols and phospholipids, leading to lipid peroxidation (*48*). These two distinctly different modalities of liver injury were used here as models to assess liver injury response to both post-necrotic hepatocellular fibrosis and cholestatic biliary fibrosis. In the context of our hepatocyte-specific deletion models, this allows us to probe the differences between hepatocyte response to direct damage, and response to cholangiocyte damage leading to liver injury.

We compared livers from control-chow and DDC-chow fed animals of each previously described genotype after 21 days of treatment. Histology of picrosirius red-stained liver sections displayed increased fibrosis in all m^6^A-perturbed genotypes relative to wild-type **(Fig. S3A**). Dual *Ythdf1*/*Ythdf2* deficient livers developed extensive fibrosis bridging from bile ducts to portal vein tracts (**Fig. 4A**). *Mettl14* deficient livers exhibit some fibrosis even in mock-treated conditions but developed bridging fibrosis as well as general diffuse fibrosis throughout the bulk of the tissue under DDC treatment (**Fig. 4A**, left). Quantification of the area of fibrosis staining by picrosirius red shows that all genotypes (*Ythdf1*^-/-^, *Ythdf2*^[fl/fl]^/Alb-Cre, dual *Ythdf1*^-/-^/*Ythdf2*^[fl/fl]^/Alb-Cre, and *Mettl14*^[fl/fl]^/Alb-Cre) progress to a significantly higher level of fibrosis than wild-type mice under DDC treatment, but dual *Ythdf1*/*Ythdf2*- and *Mettl14*-deficient mice both exhibited the highest-percentage areas of fibrosis (**Fig. 4B**, left). Quantification of blocked bile ducts in DDC treated liver tissue show ductular reaction and response to injury. In our model mice, *Mettl14* mice recovered similarly to wild-type mice, while *Ythdf1*/*Ythdf2* mice showed significantly reduced numbers of unblocked bile ducts per HPF **(Fig. S3B**).

**Fig. 4.**
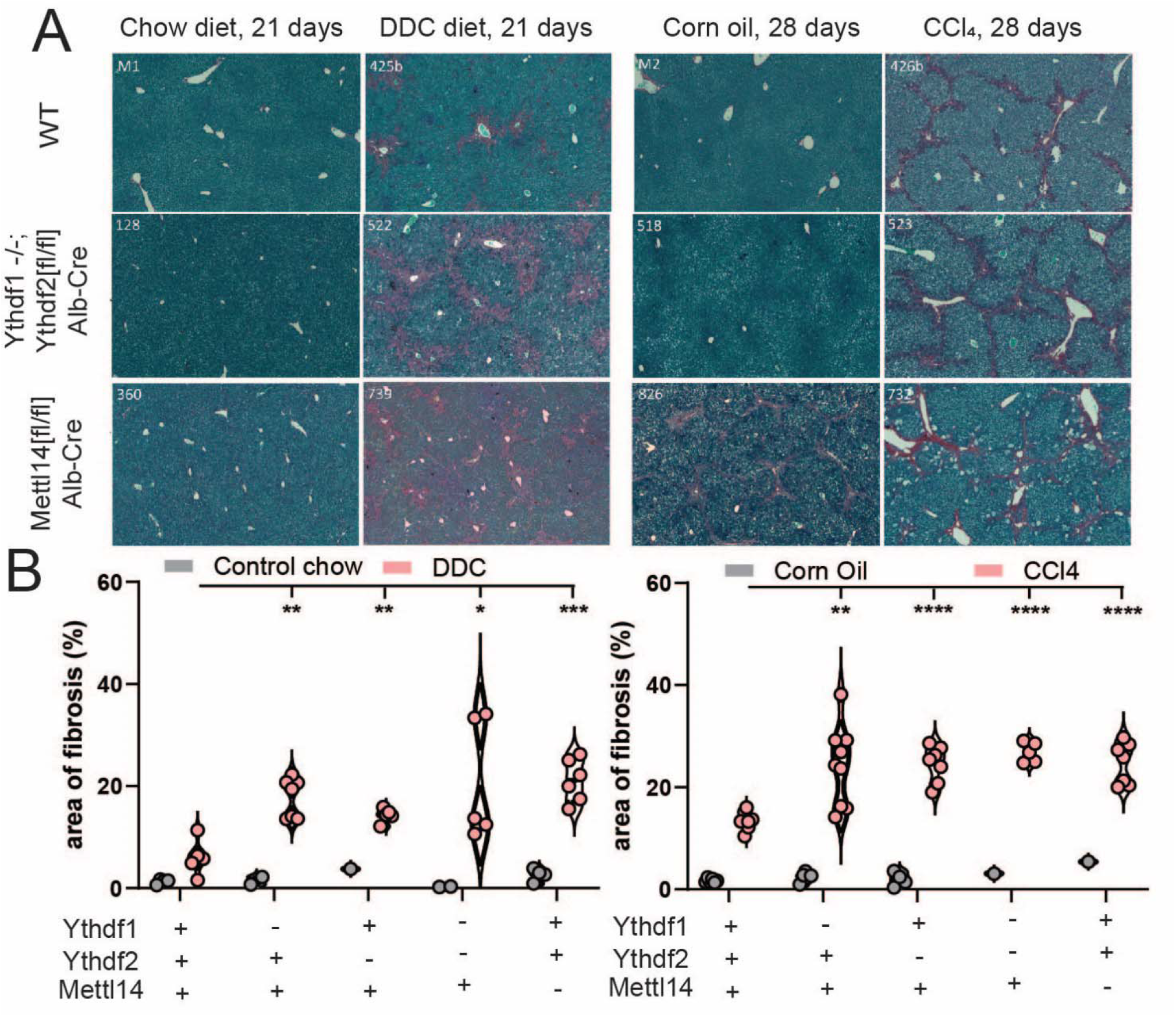
m^6^A reader and writer deficiency increases liver fibrosis response to toxicity. **(A)** Histological staining with picrosirius red of mock (chow diet) and DDC diet-treated (left) male mice reveals increased liver injury and areas of fibrosis (red). Similar staining of mock (corn oil) and CCl4-treated male mice (right) reveals the same pattern of liver injury and fibrosis advancement in m^6^A reader- and writer-perturbed mice. **(B)** Quantitative image analysis of areas of fibrosis in DDC diet-(left) and CCl_4_-(right) treated mice with mock comparison.

We similarly analyzed tissues from mice after 28 days of CCl_4_ treatment. All genotypes displayed increased fibrosis relative to control wild-type mice, with evident bridging fibrosis (**Fig. S3C**). Dual *Ythdf1*/*Ythdf2* deficient mice showed pervasive bridging fibrosis throughout the liver, and histologic analysis of *Mettl14* deficient mice revealed foci of necrosis throughout the liver in addition to the fibrosis (**Fig. 4A**, right). Automated image analysis demonstrated that while the percent area of fibrosis was increased in all perturbed genotypes relative to wild type, the dual *Ythdf1*/*Ythdf2* deficient mice had the highest average percent area of fibrosis after CCl_4_ treatment, with just over 25% of all liver tissue area staining positive for fibrosis (**Fig. 4B**, right). The number of mitotic cells counted per high-powered microscope field (HPF) was significantly elevated relative to wild type mice in *Ythdf1* and *Ythdf2* mice, but not *Mettl14* mice **(Fig. S3D**). Therefore, deletion of either the readers or writers led to worse liver injury following direct damage to hepatocytes or indirect damage via bile duct blockade.

### HBV genome induced hepatic inflammation is worsened in *Mettl14* deficient mice

Chronic HBV infection is a major cause of liver fibrosis and progression to hepatocellular carcinoma (*49*). Given the known role of both m^6^A and HBV in HCC development and progression, and the known role of m^6^A on HBV replication cycle and protein translation, we were interested in whether disruption of m^6^A modification in *Mettl14*^[fl/fl]^/Alb-Cre mice would impact outcomes of HBV exposure (*15*, *50–52*).

To assess the potential changes in HBV replication, translation, and injury, we crossed our *Mettl14*^[fl/fl]^/Alb-Cre mice with mice bearing the HBV 1.3x length genome as a transgene (1.3x HBV tg). These mice produce all gene products of HBV and produce packaged viral particles (*53*). To allow time for significant liver injury to accumulate, we waited until 3 months of age to subject tissues from these animals for analysis. Histological comparison of H&E-stained sections showed that while HBV expression leads to some minor injury and inflammation, HBV expression in *Mettl14*^[fl/fl]^/Alb-Cre mice exacerbates hepatic injury and fibrosis beyond that normally seen in *Mettl14*^[fl/fl]^/Alb-Cre mice (**Fig. 5A**). Bridging fibrosis was noted with especially broad regions of fibrosis and immune infiltrate bridging between portal vein tracts and bile ducts.

**Fig. 5.**
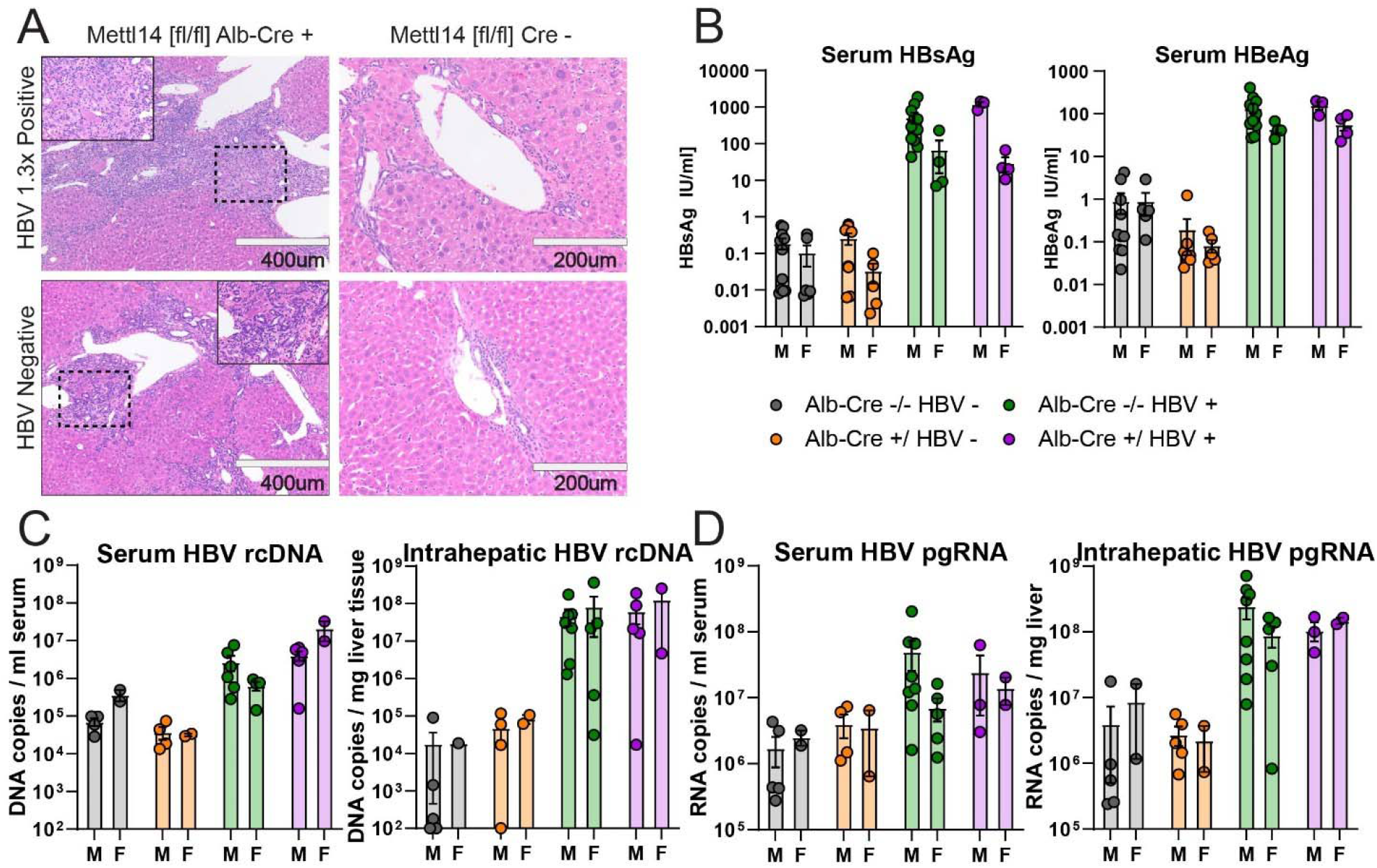
*Mettl14* deficiency impacts liver injury but not HBV translation or replication. **(A)** Histology of H&E-stained sections from male *Mettl14*^[fl/fl]^/Alb-Cre and *Mettl14*^[fl/fl]^ control mice expressing HBV 1.3x genome transgene, and comparison with control HBV negative mice. **(B)** ELISA assays showing blood serum HBsAg (left) and HBeAg (right) protein levels of HBV expressing mice in comparison to control HBV negative animals to establish baseline. **(C)** qPCR of HBV genomic rcDNA extracted from blood serum (left) and liver tissue (right). **(D)** RT-qPCR of HBV pre-genomic RNA extracted from blood serum (left) and liver tissue (right).

To assess the level of packaged and released viral particles in the blood, we collected serum from blood samples and measured HBV S antigen (HBsAg) levels by ELISA assay. This antigen is an integral part of secreted subviral and infectious HBV particles. We found no significant difference between either males or females expressing HBV with and without *Mettl14* defects (**Fig. 5B**, left). We further quantified HBV E antigen (HBeAg) levels in blood serum, a marker of ongoing virus replication, by ELISA. We again found no significant differences in HBV-expressing mice (**Fig. 5B**, right).

To thoroughly assess levels of viral replication and transcription, we also quantified levels of HBV genomic relaxed circular DNA (rcDNA) and pre-genomic RNA (pgRNA) in both serum and liver tissue by (RT)qPCR. There was no significant difference in rcDNA levels among *Mettl14*^[fl/fl]^/Alb-Cre/1.3x HBV tg and *Mettl14*^[fl/fl]^/1.3x HBV tg control mice in liver tissue or serum (**Fig. 5C**). Similarly, we saw no significant differences in pgRNA in either serum or liver tissue (**Fig. 5D**). Collectively, these data demonstrate that while HBV genome expression worsens liver disease in *Mettl14*-deleted mice, which cannot be attributed to elevated levels of HBV replication intermediates or viral proteins.

### *Mettl14* deletion leads to pronounced nuclear heterotypia and increased polyploidy in hepatocytes

One striking phenotype caused by Mettl14 deletion was nuclear heterotypia, which we observed in steady state **(Fig. 1)** and following various injuries. GSEA analysis of transcriptomic analysis of *Mettl14*-deletion mice highlighted several potential explanatory mechanisms of the nuclear heterotypia we observed in these mice **(Fig. 2E,F)**. Cell-cycle regulatory pathways of c-Myc signaling, circadian clock-related genes, and late mitosis / early G1 phase cell cycle regulation genes were all dysregulated. RNA metabolism, processing, splicing, and degradation were also evidently disrupted, with upregulated nucleotide di- and tri-phosphate conversion and downregulation of mRNA decay mechanisms including deadenylation, as well as processing and splicing of capped intron-containing pre-mRNA.

Dysregulation of cell cycle, RNA processing and export, as well as signs of oxidative stress can all lead towards nuclear heterotypia. To confirm our original histological H&E staining and perform further quantification of the nuclear heterotypia to better describe the phenotype and narrow down potential contributing mechanisms, we stained liver sections from *Mettl14*-deletion and wild-type mice with Hoechst-33342 (**Fig. 6A**). Quantitative image analysis across 3 individuals of each group showed that there was a significant increase in the mean size and size variability of nuclei in *Mettl14*-deletion liver (**Fig. 6B**, top). To assess any change in polyploidy occurring alongside nuclear size changes we isolated nuclei from liver tissue samples and performed flow cytometry, which showed a marked shift of hepatocyte ploidy towards 8n and higher ploidies (**Fig. 6B**, bottom). Representative histograms of Hoechst-33342 signal from the flow cytometry data collection show clear separation between each peak, indicating distinct populations of nuclei (**Fig. 6C**).

**Fig. 6.**
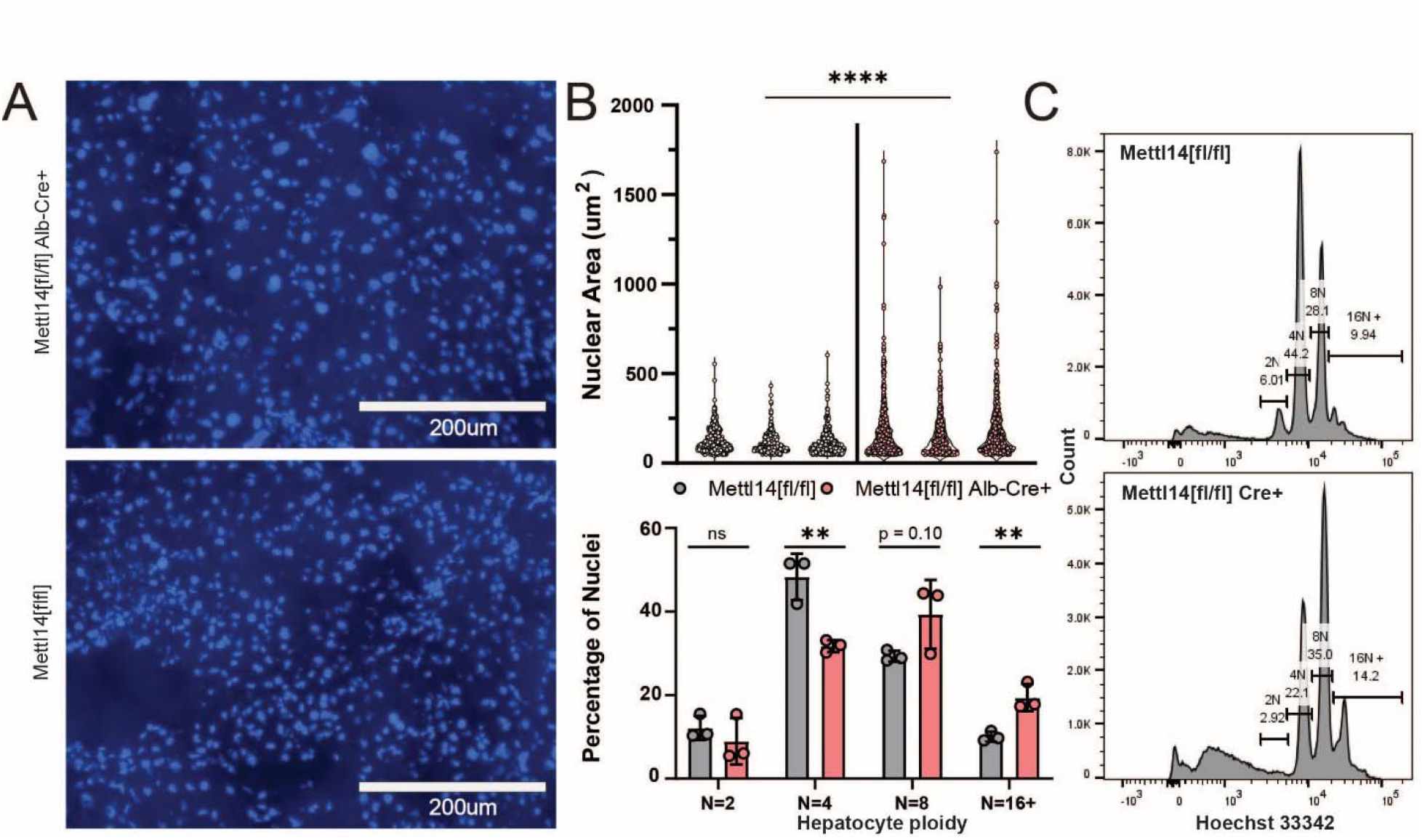
*Mettl14* deletion-related nuclear heterotypia is concurrent with increased ploidy. **(A)** Hoechst-3342-stained histological sections reveal an increase in nuclear size in *Mettl14* deletion mice (top) relative to wild type control mice (bottom). **(B)** Quantitative analysis (top) of imaging data from separate animals shows consistently increased nuclear size and greater range of size in *Mettl14*^[fl/fl]^/Alb-Cre mice vs *Mettl14*^[fl/fl]^ controls. Quantitative analysis of hepatocyte ploidy via flow cytometry (bottom) of Hoechst-33342 stained hepatocyte nuclei. **(C)** Representative histograms of Hoechst-33342 staining intensity from flow cytometry data of stained hepatocyte nuclei.

### Nuclear accumulation of the TREX complex member *Alyref* reveals impacts of *Mettl14* on RNA trafficking machinery

To gain more mechanistic insight into the nuclear heterotypia phenotype, we further explored the results of our gene set enrichment analysis of our transcriptomic data. We observed upregulation of mRNA splicing, trafficking, and activation for translation, as well as pathways involved in metabolic and oxidative stress, mitochondrial biogenesis, apoptosis, and fatty acid oxidation (**Fig. 5E,F**). RNA metabolism, processing, splicing, and degradation were also evidently disrupted, with upregulated nucleotide di- and tri-phosphate conversion and downregulation of mRNA decay mechanisms including deadenylation, as well as processing and splicing of capped intron-containing pre-mRNA. We further saw that cell-cycle regulatory pathways of circadian clock-related genes, and late mitosis / early G1 phase cell cycle regulation genes were all dysregulated.

We identified a number of differentially expressed transcripts with direct ties to the m^6^A machinery (**Fig. S2**). In Mettl14^[fl/fl]^/Alb-Cre mice, we found upregulation of *Brf2, R3hdm4, Taf1a,* and *Zfp747*, among other genes which play roles in RNA transcription initiation and regulation. We also saw upregulation of *Srpk1*, a key member involved in spliceosomal complex assembly which is responsible for mRNA processing, splicing, and export processes which all assemble within nuclear speckles. Notably, the transcription/export complex (TREX), which is the major mechanism of mRNA nuclear export in a m^6^A, assembles on m^6^A modified mRNAs via binding to the m^6^A modifying machinery within nuclear speckles (*21*, *23*). Therefore, we wondered whether *Mettl14*-deletion might dysregulate the mRNA export machinery, in turn leading to nuclear heterotopia, cell cycle defects, and liver pathology.

To test the TREX mechanism of m^6^A-related mRNA export suggested by the transcriptomic data, we wanted to quantify the nuclear concentration of the key TREX complex member Aly/REF export factor (*Alyref*). Since liver tissue produces high autofluorescence that interferes with immunofluorescence, we instead derived a mouse embryonic fibroblast line from *Mettl14*^[fl/fl]^ mice, and stably transduced them with both simian virus 40 (SV40) large T antigen. To disrupt *Mettl14*, we transduced a population of cells with Cre expressing lentivirus and compared them to cells without Cre (**Fig. S4A**). *Mettl14* was efficiently deleted in the presence of Cre, as judged by confocal microscopy **(Fig. 7A)**, reverse transcription qPCR **(Fig. 7B)**, and western blot **(Fig. S4B)**, while the other m^6^A writer *Mettl3* was unaffected. Though *Alyref* mRNA levels were unchanged **(Fig. 7B)**, *Mettl14*-deletion led to an increase in nuclear *Alyref* protein levels **(Fig. 7A,C)**. These data support a mechanism by which *Mettl14* deletion alters nuclear mRNA export machinery, causing downstream effects on nuclei, cell cycle, and liver injury.

**Fig. 7.**
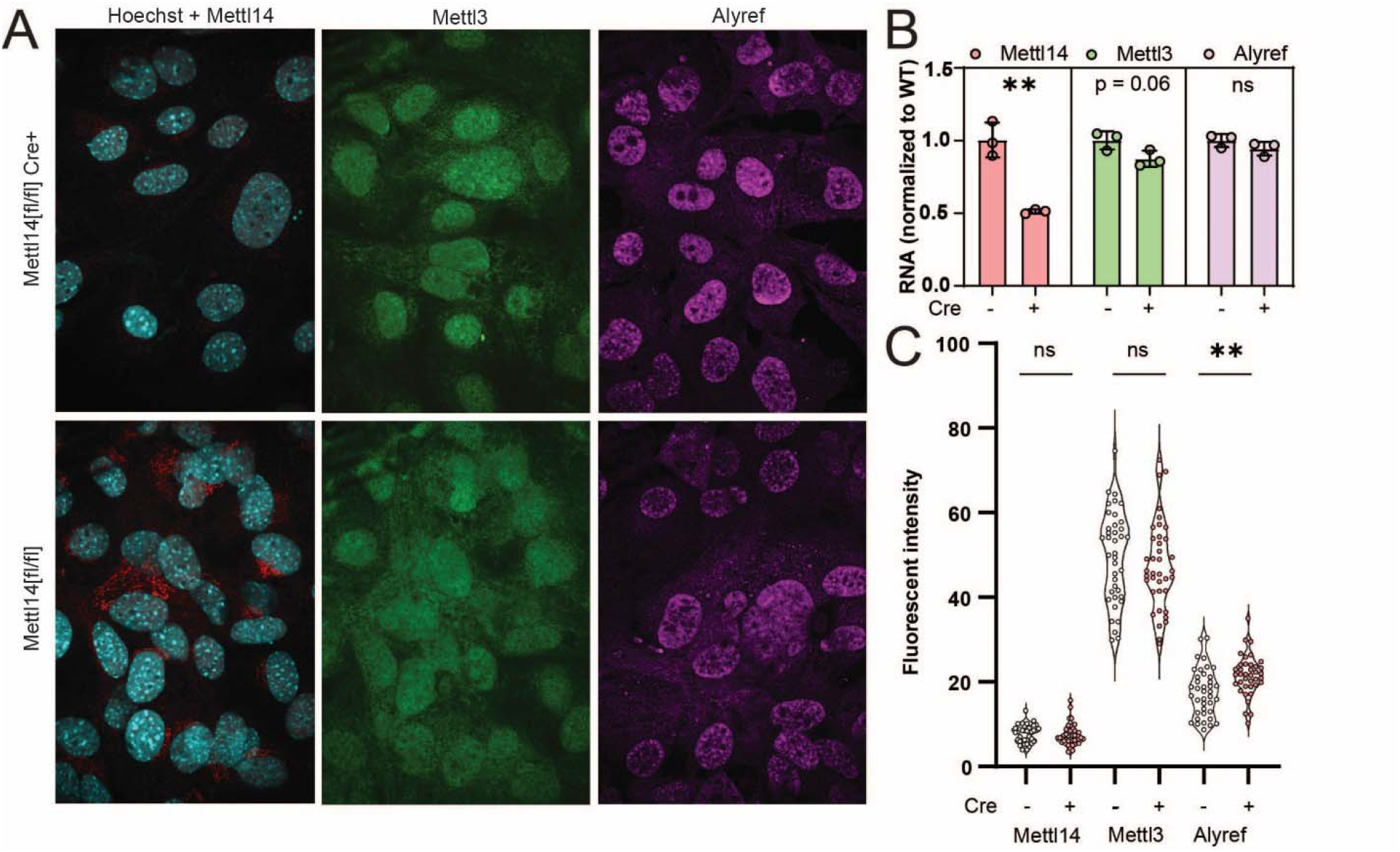
TREX complex localization changes reveal RNA trafficking machinery defects. **(A)** Representative images from *Mettl14*^[fl/fl]^/Cre (top) and *Mettl14*^[fl/fl]^/Cre MEFs (bottom) show differences in *Mettl14* signal in red (left), *Mettl3* signal in green (middle) and TREX complex marker *Alyref* in pink (right). **(B)** RT-qPCR data show gene expression of *Mettl14* is significantly reduced approximately 50%, but *Mettl3* and *Alyref* levels are unchanged. **(C)** Quantitative analysis of multiple replicate slides showing nuclear localized signal by co-localization with Hoechst-33342 signal demonstrates an increase in nuclear *Alyref* signal.

## Discussion

Many experimental studies in cell culture and observational clinical studies have shown that disruption of normal m^6^A modification impacts a layer of gene regulation leading to defects in cell-cycle regulation and important metabolic processes; however, it has been challenging to experimentally study these effects *in vivo*, as full-body knockouts of the key m6A writers and readers are embryonic lethal (*4*, *11*, *12*, *54*). Here, we generated and thoroughly characterized novel mouse mouse models with liver-specific genetic ablation of *Mettl14*, *Ythdf2*, and dual deletion of *Ythdf1* and *Ythdf2*. Though these mice are viable, both *Mettl14* and dual *Ythdf1/Ythdf2* deletion causes their livers to undergo progressive injury with steatohepatitis, and in the case of *Mettl14*, nuclear hetereotypia, which can be further exacerbated by surgical, chemical, and infectious challenges. This suggests a critical role for m^6^A in post-natal liver maintenance and regeneration.

We leveraged transcriptomic data to identify dysregulated nuclear mRNA export as a potential molecular mechanism driving nuclear heterotypia. *Mettl14*^[fl/fl]^/Alb-Cre mice showed significant changes in abundance of transcription, splicing, and nuclear export machinery mRNA. Recent literature has shown that TREX mediated mRNA export is enhanced by m^6^A modification of mRNAs, with direct TREX complex binding m^6^A modification machinery scaffold proteins vir like m^6^A methyltransferase associated (*Virma*) and WT associated protein (*Wtap*) (*21*, *23*).

Disruption of the m^6^A modification complex was shown to significantly impact the ratio nuclear to cytosolic abundances of known m^6^A modified transcripts, but not control transcripts without m^6^A sites. Assembly of the m^6^A complex occurs dynamically within phase-separated nuclear speckles, colocalizing with the TREX complex assembly and binding (*22*). This phase separation is driven by *Mettl3* when bound to *Mettl14* during assembly, but *Mettl3* homodimerization can drive phase separation and assembly in the absence of *Mettl14* (*22*). *Mettl14* is not capable of driving the same phase separation, even when homodimerization is forced by experimental fusion protein expression (*22*).

*Mettl3* self-interaction is normally capable of driving phase separation and assembly of the m^6^A modification machinery complex, which along with WTAP binding, promotes phase separation of the mRNA bound complex (*22*). This phase separation puts the m^6^A complex-bound mRNA in close contact with TREX complex, promoting binding and nuclear export via *Nxf1* recruitment and mRNA handover. Simultaneously, m^6^A modified mRNAs are recognized by *Ythdc1*, leading to export via *Srsf3* binding (*55*). TREX complex members, along with potentially other mRNA export factors, shuttle back and forth from nucleus to cytoplasm to affect mRNA export (*21*, *23*). TREX binding to non-m^6^A modified transcripts also leads to mRNA export, but through binding at alternate sites, leading to low efficiency of export of normally m^6^A modified transcripts (*23*). *Mettl14* plays a *Mettl3*-independent role in chromatin openness through direct interaction with H3K27me3 and recruiting K*dm6b* to induce H3K27me3 demethylation (*56*). This promotes transcription, with *Mettl14*, but not *Mettl3* deletion exhibiting a global decrease in transcription rate, which promotes binding to the TREX complex within phase separated nuclear speckles. These bound complexes are likely not properly bound to mRNAs due to the absence of *Mettl14*.

Taken together with our data, this suggests a possible mechanism of mRNA trafficking defects unique to *Mettl14* deletion models not seen under *Mettl3* deletion (**Fig. 8**). Perturbation of the above mechanisms in *Mettl14* deficient cells results in nuclear sequestration of the associated TREX complex via some potential mechanism of association with m^6^A machinery components in the absence of *Mettl14* (**Fig. 8, right**). Specifically, the reports of *Mettl3* self-interaction and association with *Wtap* driving phase separation of bound complex, and the known role of *Mettl3* in binding to the TREX complex (*22*, *23*), nuclear retention of RNAs in *Mettl14* deletion (*57*) and our current report of TREX nuclear sequestration in *Mettl14* deletion together could all contribute to differences in outcome of *Mettl3* deletion and *Mettl14* deletion models via *Mettl14* independent interaction of *Mettl3.* This would lead not only to dysregulation of normally m^6^A modified transcripts, but all transcripts which utilize the TREX mechanism of nuclear export.

**Fig. 8.**
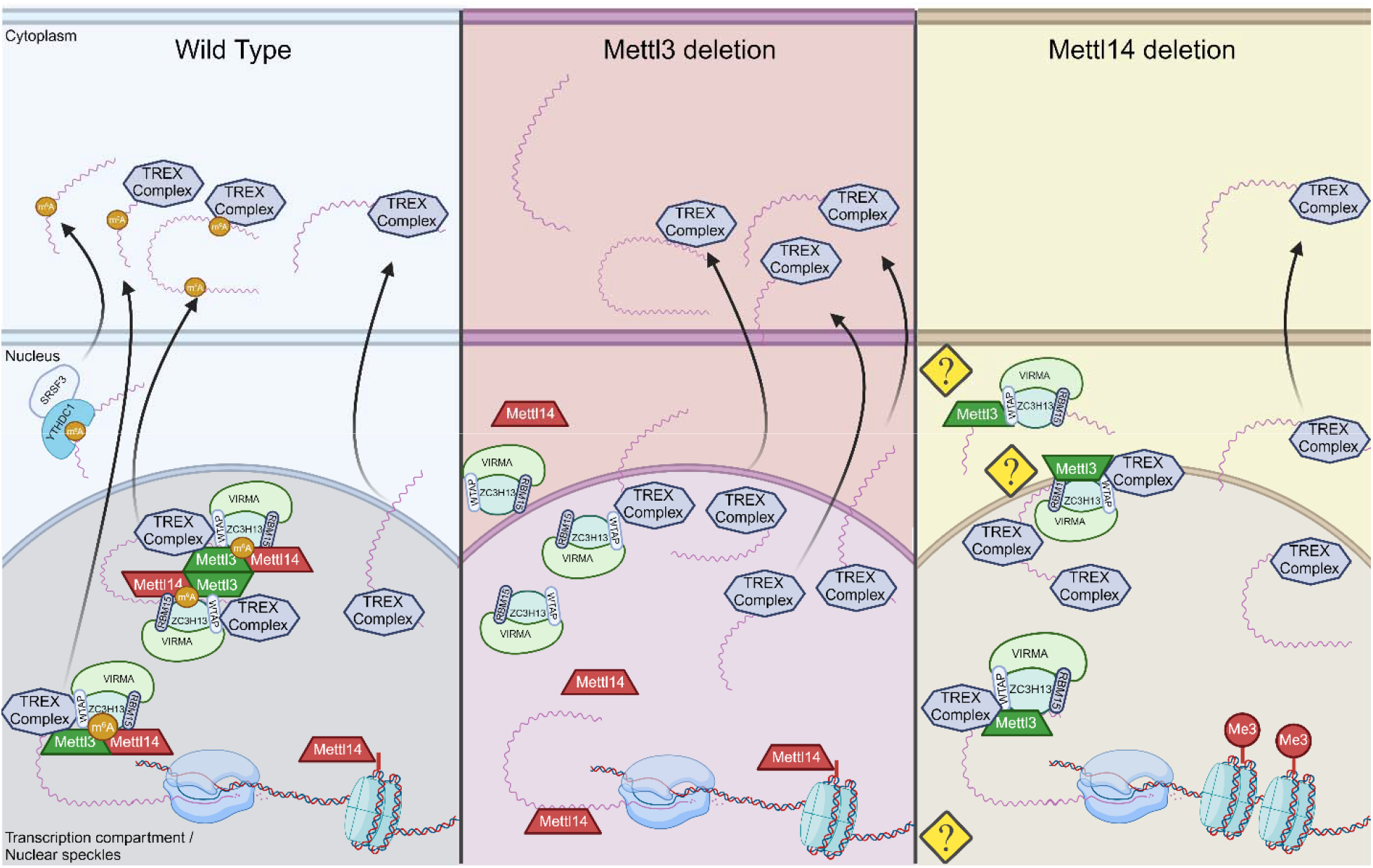
Proposed mechanisms for nuclear maintenance of TREX in *Mettl14* deletion. In wild-type cells (left), *Mettl3* and *Mettl14* function as a complex to place m^6^A modifications on mRNA transcripts. In *Mettl3*-deficient cells (middle), export of normally m^6^A-modified transcripts is slowed, but transcription rates are maintained or increased, allowing alternate pathways of TREX-mediated mRNA export to function at capacity (*56*). In *Mettl14*-deficient cells, m^6^A-mediated mechanisms of mRNA export are impaired similarly to *Mettl3* deletion, but TREX complex shuttling is also impaired through nuclear retention by mechanisms not yet understood. At the same time, global transcription rates are impacted by loss of *Mettl14* chromatin binding (Created with BioRender.com).

At the same time, pre-mRNAs are not properly m^6^A modified, reducing the processes of mRNA splicing and processing, and leading to increased nuclear mRNA surveillance recognition and degradation of these overabundant and improperly spliced pre-mRNAs, as seen represented in our data presented here. Furthermore, the co-opting of the mRNA surveillance and degradation pathways might competitively inhibit this process from regulating other pre-mRNAs which might be frequently alternatively spliced or improperly processed as a regulatory mechanism.

Overactivation of mRNA degradation and nucleotide scavenging processes due to mRNA processing and trafficking machinery defects could also lead to a cascade of apoptotic, stress, and immune responses which could contribute to nuclear heterotypia and overall liver damage.

Significant changes in *Hedgehog*, *PPAR*γ, *c-Myc*, and *PI3K/mTOR/Akt* signaling axes were detected and impacts of m^6^A on signaling deserve special consideration in follow-up work. In particular, *Smo* mRNA has been found in several studies to have multiple m^6^A modification sites, and expression is likely regulated by this mechanism (*30*). Our GSEA results confirmed the important dysregulation of *Hedgehog* signaling, with several gene sets involved not just with *Smo* itself, but also *Hedgehog* ligand synthesis and trafficking pathways significantly reduced (**Fig. 2E,F**). Noted changes to overall lipid metabolism and subsequent pre-diabetic and pro-fibrotic responses as well as cell cycle dysregulation leading towards an HCC like state are all observed possible downstream effects of *Hedgehog* signaling dysregulation (*34–36*). Our transcriptomics data showed changes in ciliary trafficking of *Hedgehog* signaling ligands, despite hepatocytes lacking cilia. Taken together with the significant changes in bile acid salt metabolism and secretion, there are likely changes in *Hedgehog* signaling cholangiocytes and potentially between hepatocytes and peri-biliary portal fibroblasts (*58*).

The results of this study merit follow-up work investigating this model mechanism. Changes in *Mettl14* and m^6^A modification abundances have been widely reported and considered as disease biomarkers in various liver diseases including MASH and HCC, and work is ongoing for potential therapeutic applications targeting *Mettl14* and overall m^6^A regulation (*1*, *2*, *59*, *60*). It is therefore urgent that we understand the outcomes of changes to *Mettl14* and *Mettl3* expression *in vivo* thoroughly, both together and in isolation, to inform drug target development. The scope of bulk-RNAseq limits this study, and follow-up studies utilizing spatial transcriptomics would strengthen our understanding of the role of *Hedgehog* signaling. This signaling occurs both within hepatocytes and between hepatocytes, peri-biliary portal fibroblasts, and cholangiocytes. Understanding this signaling more thoroughly would allow us to understand the contribution of non-hepatocyte cell types in general in this liver-injury state in *Mettl14* deletion. Follow-up work could also clarify the transcriptomic and proteomic state of *Mettl14* deletion mice at earlier timepoints during the initial onset of liver injury, and of the dual *Ythdf1*^-/-^/*Ythdf2*^[fl/fl]^/Alb-Cre mouse model in comparison to *Mettl14* deletion to further confirm and specify the gene dysregulation responsible for the differences in injury phenotypes.

## Materials and Methods

### Experimental Design

Experiments were designed to specifically assess the impacts of m^6^A within the context of an *in vivo* system. Mouse models for study were obtained as described in the relevant materials and methods section below. Study of the steady state impacts in adult mice was conducted initially via histologic analysis of H&E stained liver sections to determine any gross defects, and assess the nature of any evident changes. Due to the evident gross morphologic changes in *Mettl14* deficient liver tissue, we aimed to fully characterize the nature of the defects by determining whether this injury was due to improper liver organogenesis and development, or tissue maintenance and metabolism defects. To distinguish between these two effect types, we performed similar histology analysis of *Mettl14* model mice over a time course of early post- natal development. Simultaneously, we developed an inducible model of gene deletion for *Mettl14*, described in the relevant materials and methods section, to clearly exclude any developmental effects from impacting tissue architecture or damage. Our subsequent experimental designs were focused on two specific questions: the roles of m^6^A machinery components in injury response and regeneration, and the mechanisms leading to the unique nuclear heterotypia phenotype seen specifically in *Mettl14* deletion liver tissue.

To further probe the roles of m^6^A machinery components in injury response and regeneration, we imposed a suite of injuries and insults to the liver with various mechanisms of damage. Physical injury was induced via two-thirds partial hepatectomy, while chemical injury to hepatocytes was modeled by CCl4 treatment. DDC was utilized to model cholestatic disease via injury by blocking bile ducts. Finally, we modeled components of chronic hepatitis B infection by developing an HBV expressing *Mettl14* deficient mouse line. While the HBV expressing mouse model does not fully model chronic infection, as these animals are tolerized to HBV and express the genome themselves rather than supporting true viral infection, aspects of chronic HBV infection such as HBV protein toxicity are represented in this model.

To better understand the mechanisms underlying the nuclear heterotypia observed in *Mettl14* deletion liver tissue, we first aimed to characterize the state of these mice by transcriptomic analysis. To get better data on any changes of nuclear localization, nuclear heterotypia was further explored by flow cytometry analysis of hepatocyte nuclei ploidy. Since this can skew the data, as larger nuclei may be more fragile and less likely to be cleanly isolated from liver tissue, image analysis of confocal images of liver sections was used to quantify the nuclei size distribution *in situ*. Finally, due to the high autofluorescence background making antibody-based fluorescent imaging difficult in liver tissue, we derived a mouse embryonic fibroblast line with similar levels of *Mettl14* knockdown in order to image subcellular localization and expression levels of *Mettl14*, *Mettl3*, and *Alyref*, a component of the TREX mRNA transcription and export complex.

### Mice

C57BL/6 and B6.Cg-^Tg(Alb-cre)21Mgn/J^ (Alb-cre) were obtained from the Jackson Laboratory (Bar Harbor, ME) (*27*). *Mettl14*[fl/fl], *Ythdf1*-/- and *Ythdf1*2[fl/fl] (all on the C57BL/6 background) were kindly provided by Dr. Chuan He (University of Chicago, HHMI) (*26*) Alb^tm1(cre/ERT2)Mtz^ by Dr. Pierre Chambon (INSERM, Universite Louis Pasteur) (*37*) and 1.3x HBV transgenic mice by Dr. Frank Chisari (Scripps Research) (*61*). *Mettl14*^[fl/fl]^/Alb-Cre, *Ythdf2*^[fl/fl]/^Alb-Cre and *Ythdf2*^[fl/fl]^/Alb-Cre *Ythdf1*^-/-^, *Mettl14*^[fl/fl]^/Alb-ERT2-Cre, *Mettl14*^[fl/fl]^/Alb-Cre/1.3x HBV tg mice were generated by intercrossing mice harboring the respective alleles and typing offspring with primer combinations distinguishing wild-type and mutant alleles (typing information are available upon request).

Animal experiments were performed in accordance to a protocol (number 3063) reviewed and approved by the Institutional Animal Care and Use Committee (IACUC) at Princeton University and in accordance to IACUC protocol 2016-0047 reviewed and approved by the Weill Cornell Medical College IACUC.

### Tamoxifen induction experiments

Tamoxifen induction of Alb-ERT2-Cre expressing mice was performed using a protocol adapted from the Jackson Laboratory. Tamoxifen (Sigma-Aldrich, St. Louis, MO) was dissolved in corn oil at a concentration of 20 mg/ml by shaking overnight at 37°C. Tamoxifen solution was administered by intra-peritoneal injection at a dose of approximately 75 mg/kg body mass, a standard dose of 100 ul per mouse. 5 days of consecutive once daily administration were performed to induce recombination, followed by once weekly injection for 6 weeks of total time before animals were sacrificed for experimental analysis. All materials, animal bedding, and waste was handed appropriately to avoid exposure of personnel.

### Histology

During sample collection, mouse livers were perfused with PBS (Life Technologies, Carlsbad, CA) using a BD Vacutainer SafetyLok butterfly needle 23 gauge, 2/4” needle length, 12” tubing length (Becton, Dickinson and Company, Franklin Lakes, NJ) via the portal vein prior to removal to clear the liver tissue of blood and achieve cleaner histology sections. Samples were collected and placed in 4% [w/vol] PFA prepared from a 10% [w/vol] neutral buffered formalin solution (Sigma-Aldrich, St. Louis, MO) for fixation prior to paraffin embedding for histologic sectioning and staining.

Samples from DDC and CCl_4_ experiments were sent to Saffron Scientific Histology Services, LLC (Carbondale, IL) for paraffin-embedding and hematoxylin and eosin (H&E) or Picrosirius Red staining. Separate samples were placed in OCT in plastic cassettes and frozen at -20°C. Cryostat sectioning using a CM3050S Cryostat (Leica Biosystems, Wetzlar, Germany) was performed to obtain ∼5um thick sections and samples were mounted on glass slides and stained with Hoechst33342 (Invitrogen, Waltham, MA) at 5ug/ml [w/vol] for 30 minutes at room temperature prior to being sealed under glass coverslips with ProLong™ Gold Antifade Mountant (Invitrogen, Carlsbad, CA). These liver-section slides were imaged using a Nikon Ti-E microscope with Spinning Disc and Photomanipulation Module (Minato City, Tokyo, Japan), and nuclear area was analyzed using Fiji image analysis to set regions of interest around the nuclei (*62*).

### Partial hepatectomies

After weighing animals and recording pre-operative weight, we conducted surgeries under isoflurane induction of anesthesia. Following approved IACUC protocol (number 3063), we used a surgical technique adapted from Nevzorova et al. to remove 3 lobes from the liver, representing approximately two thirds of liver mass (*40*). Removed tissue was weighed to confirm the amount of liver mass loss, and the peritoneal wall were closed after application of analgesic medication using discontinuous 4/0 vicryl sutures (Ethicon surgical technologies, Bridgewater, NJ). Skin was closed using surgical wound clips (Stoelting, Wood Dale, IL) rather than sutures to prevent wound re-opening from animals licking or chewing on the incision site. Animals were weighed post-operatively, and daily thereafter, with recorded weights corrected for the weight of surgical staples used. Analgesic medication was administered twice daily, in accordance with the timeframes in the approved protocol. At 2 weeks post-surgery, animals were sacrificed for collection of liver samples and analysis.

### DDC and CCl_4_ toxicity experiments

Mice arriving from Princeton University were housed in the quarantine facility of Weill Cornell Medicine for 6 weeks before being used for liver injury experiments. All mice were under a 12- hour light: dark cycle with free access to regular food and water. Mice used for fibrosis or injury models were used at ages 10-12 months unless otherwise indicated.

For CCl_4_ experiments, mice received biweekly injections of 25% [w/vol] CCL4 (Sigma-Aldrich, St. Louis, MO), diluted in corn oil at a dose of 2 μl/g, for a total of 4 weeks. 0.1% [w/w] 3,5-diethoxycarbonyl-1,4-dihydrocollidine (DDC) (Sigma-Aldrich, St. Louis, MO), was mixed with 5053, Purina Picolab Rodent Diet 20 (Envigo, Indianapolis, IN) and given for 21 days. Animals were randomly assigned to groups. Blinding could not be performed given the nature of the experiments. All animal experiments were performed on at least two separate occasions and in accordance with the guidelines set by the Institutional Animal Care and Use Committee at Weill Cornell Medicine and approved in IACUC protocol 2016-0047.

Liver tissues were fixed in 4% [vol/vol] paraformaldehyde and sent to Saffron Scientific Histology Services, LLC (Carbondale, IL) for paraffin-embedding and hematoxylin and eosin (H&E), and Picrosirius Red staining. Stained liver sections were observed and imaged using the Axioscan 7 Slide Scanner (Zeiss, Jena, Germany) and analyzed for percent area fibrosis using ImageJ software (https://imagej.net/Image).

### HBV assays

HBsAg and HBeAg antigen levels were quantified as previously described (*63*) from serum samples obtained by submandibular bleeds of experimental mice. Chemiluminescence immunoassays (CLIA) for both antigens were performed using HBsAg and HBeAg CLIA kits from Autobio Diagnostics (Zhengzhou, Henan, China) according to manufacturer instructions using 50μl of serum.

HBV rcDNA and pgRNA were extracted from both mouse liver tissue and serum samples using a Quick-DNA/RNA Microprep Plus Kit (Zymo Research, Irvine, CA) following the manufacturer’s instructions. Briefly, liver samples and serum samples were resuspended in 300 μl DNA/RNA Shield. Liver samples were homogenized using a TissueLyser LT bead mill (Qiagen, Venlo, The Netherlands) for three separate 2 minute cycles followed by digestion with 15 μl Proteinase K (20 mg/ml) for 30 min. 300 μl DNA/RNA lysis buffer was then added to both liver and serum samples. Samples were loaded into Zymo-Spin IC-XM columns to collect the DNA and flow-through was saved. An equal volume of ethanol was added to the flow- through to purify RNA by using the Zymo-Spin IC column. Finally, the DNA/RNA was eluted from the columns with 30 l of nuclease-free water and concentrations were measured using a Nanodrop spectrophotometer (Thermo Fischer Scientific, Waltham, MA).

HBV rcDNA was quantified from 2 μl aliquots of HBV DNA isolated either from liver samples or blood serum was used per reaction well. We used a well-characterized HBV rcDNA qPCR system with HBV-qF (nt 1776–1797, numbered based on gt D with GenBank accession no. U95551.1): 5 -GGAGGCTGTAGGCATAAATTGG-3, HBV-qR (nt 1881-1862, numbered based on gt D with GenBank accession no. U95551.1): 5 -CACAGCTTGGAGGCTTGAAC-3 covering the conserved region of HBV(LLD 1.0E + 3 copies/mL) (*63*). Primers were kept at a final concentration of 500 nM in a 20 μl reaction volume. On a Step One Plus qPCR machine (Life Technologies), we ran the following program: denature 95 °C for 10 min, followed by 40 cycles of 95 °C for 30 s, 60 °C for 30 s, and 72 °C for 25 s.

HBV pgRNA was quantified from HBV RNA extracted from liver tissue or serum as described above. 7.5 μl of the resultant sample was treated by DNase I (Thermo Fisher Scientific, Waltham, MA, USA) followed by reverse transcription with a specific HBV primer (5 - CGAGATTGAGATCTTCTGCGAC-3, nt 2415–2436, numbered based on gt D with GenBank accession no. U95551.1) located in precure/core region (*64*) using RevertAidTM First Strand DNA Synthesis Kit (Thermo Fisher Scientific, Waltham, MA, USA). For absolute quantification, standards with 1-mer HBV target template were cloned into the TOPO-Blunt Cloning vector (Thermo Fisher, Waltham, MA, USA #450245) and copy number was calculated based on the vector molecular weight and concentration. A master mix was created containing 15 μl 2 × Taqman reaction mix (Applied Biosystems, Waltham, MA, USA), 500 nM forward and reverse primers, 200 nM probe and 3 μl synthesized cDNA in a 30 μL reaction. This master mix was then added to the samples and 10-fold serial dilution standards and the following cycling program was used to run the qPCR: 95 °C for 10 min; 45 cycles of 95 °C 15L and 58 °C for 45L

### RT-qPCRs for cellular transcripts

To assess RNA levels of *Mettl14*, *Mettl3*, and *Alyref*, as well as Cre and simian virus 40 (SV40) large T antigen (LT) transgenes, reverse transcription real-time qPCR was performed on RNA samples from mouse liver samples and mouse embryonic fibroblast cell culture samples. All qPCRs were performed using the Luna® Universal One-Step RT-qPCR Kit (New England Biolabs, Ipswich, MA) and a Step One Plus qPCR machine (Life Technologies, Carlsbad, CA). *Mettl14* was analyzed using the forward primer (5’- GACTGGCATCACTGCGAATGA-3’) and reverse primer (5’- AGGTCCAATCCTTCCCCAGAA-3’). *Mettl3* was measured using the forward primer (5’- CTGGGCACTTGGATTTAAGGAA-3’) and reverse primer (5’- TGAGAGGTGGTGTAGCAACTT-3’). *Alyref* was measured using forward primer (5’- GGCACCGTACAGTAGACCG-3’) and reverse primer (5’- AAGTCCAGGTTTGACACGAGC-3’). Cre levels were measured using forward primer (5’- GCGGTCTGGCAGTAAAAACTATC-3’) and reverse primer (5’- GTGAAACAGCATTGCTGTCACTT-3’). LT levels were assessed with forward primer (5’- CTGACTTTGGAGGCTTCTGG -3’) and reverse primer (5’- GGAAAGTCCTTGGGGTCTTC-3’). All transcript levels were normalized to housekeeping gene standard GAPDH, which was measured using the forward primer (5’- CCATGGAGAAGGCTGGGGC -3’) and reverse primer (5’- ATGACGAACATGGGGGCATCAG -3’). All primers were commercially obtained from Eton Biosciences (San Diego, CA). Standard reaction programs were run using the Step One software and Tm recommendations.

### Transcriptomics

Liver tissue was collected from animals after perfusion with PBS via the portal vein to remove blood from tissue. RNA was extracted from bulk liver tissue using the Monarch total RNA miniprep kit (New England Biolabs, Ipswich, MA) after homogenization with steel beads using a TissueLyser LT bead mill (Qiagen, Venlo, The Netherlands). After extracting total RNA, we verified high-RNA quality by Bioanalyzer RNA Nano/Pico assay (Agilent Technologies, Santa Clara, CA).

We used 50 ng total RNA per sample for gene expression profiling. We performed bulk RNA-barcoding and sequencing (BRB-Seq) (*65*) with minor modifications to the reverse transcription (RT) step. We used Template Switching RT Enzyme Mix (NEB, Ipswich, MA), along with a uniquely barcoded oligo(dT)30 primer for each sample, modified to use the Illumina TruSeq Read 1 priming site instead of Nextera Read 1 (*66*). We performed the remainder of BRB-Seq per protocol: we pooled up to 24 first-strand cDNAs into a single tube, performed Gubler-Hoffman nick translation cDNA synthesis, and tagmented cDNA with in-house-produced Tn5 (*67*). We amplified cDNAs with 17 PCR cycles using a P5-containing primer and a distinct multiplexed i7 indexing primer (Chromium i7 Multiplex Kit, 10X Genomics, Pleasanton, CA). We performed size-selection using sequential 0.55X and then 0.75X SPRIselect (Beckman Coulter, Brea, CA), and sequenced libraries on one lane of a NovaSeq SP v1.5 flowcell (Illumina, San Diego, CA) with 28 cycles Read 1, 8 cycles Read i7, and 102 cycles Read 2.

### Nuclei Isolation

Nuclei for and flow cytometry analysis were extracted from frozen liver tissue samples as previously described (*68*). Briefly, Samples were prepared by incubating freshly obtained liver tissue samples of approximately 1 gram in HypoThermosol® FRS solution (Sigma-Aldrich, St. Louis, MO) for 15 minutes on ice, followed by 30 minutes in CryoStor® CS10 cryopreservation medium (STEMCELL technologies, Vancouver, BC, Canada) on ice. Samples were then frozen overnight at -80 °C in a in a Mr. Frosty cryo-freezing container (Thermo Scientific, Waltham, MA). Tissue was then briefly washed in ice-cold DPBS (Thermo Scientific, Waltham, MA), minced using surgical scissors in a petri dish into small pieces, and homogenized using a glass tissue grinder dounce with the small sized pestle A (DWK Life Sciences, Wertheim, Germany). Nuclei were briefly fixed with 0.1% [w/vol] PFA (Electron Microscopy Sciences, Hatfield, PA) and separated by centrifugation at 500g for 5 minutes. Nuclei were then washed and prepared for downstream applications as appropriate.

### Gene set enrichment analysis

Transcriptomic data was analyzed by GSEA software (*69*, *70*). Analyses were performed using the MSigDB M2 curated mouse gene set database (https://www.gsea-msigdb.org/gsea/msigdb/). This gene set was chosen since we were looking for specific mechanisms of RNA metabolism or specific steps of metabolism important to liver function and cell cycle regulation, rather than general signaling pathways and pro-cancer gene sets which were strongly represented in other gene set databases such as the hallmark MH gene set database.

### Flow cytometry

Nuclei, isolated as described above, were prepared for flow cytometry by staining with Hoechst- 3342 (Invitrogen, Waltham, MA) at 5ug/ml [w/vol] for 30 minutes at room temperature, followed by 3 washes in DPBS (Thermo Scientific, Waltham, MA) supplemented with 5% fetal bovine serum. Flow cytometry data collection was performed using an LSRII Flow Cytometer (BD Biosciences). Data were analyzed using FlowJo software (TreeStar).

### Mouse embryonic fibroblast [MEF] generation

MEFs were generated as previously described (*71*). Briefly, In brief, the skin biopsies were scraped to remove connective tissue, cut into smaller pieces, and digested overnight at 4°C in HBSS without Ca2+ and Mg2+ (Thermo Fisher Scientific), containing 1 ml dispase (5,000 caseinolytic units/ml; Corning) for every 9 ml of HBSS containing final concentrations of 100 mg/ml streptomycin, 100 U/ml penicillin, and 250 ng/ml amphotericin B (HyClone). After digestion, the epidermis was removed and discarded, whereas the remaining dermis was cut into smaller pieces less than a few square millimeters in area. These pieces were moistened with DMEM and pressed into a six-well plate scored with a razor blade. The dermis was maintained in DMEM containing 10% FBS and 1% vol/vol penicillin/streptomycin solution at 37°C, 5% CO2. Media was changed every 4–5 d and fibroblast growth was typically observed within 1 wk of culture. Once sufficient outgrowth had occurred, the dermis was removed from the plate and the fibroblasts expanded into larger cultures.

To generate the immortalized dermal fibroblast cell line, γ-retroviral pseudoparticles containing a transfer plasmid encoding Simian virus 40 (SV40) large T antigen were produced in HEK293T cells. Cells were cultured on poly-L-lysine−coated 10 cm plates at 37 °C, 5% (vol/vol) CO2 in 10% FBS DMEM. At ∼80% confluency, Xtremegene HP DNA transfection reagent (MilliporeSigma, 6366244001) was used per manufacturer’s directions to cotransfect the cells with 4 μg of pBABE-neo-SV40 large T, a generous gift from B. Weinberg (Addgene plasmid no. 1780); 4 μg of a plasmid containing the genes for Moloney murine leukaemia virus gag-pol; and 0.57 μg a plasmid containing the gene for the G envelope protein of vesicular stomatitis virus. Supernatants were harvested 24, 48 and 72 h post-transfection, stored at 4 °C then pooled before passing through a 0.45 μm membrane filter (MilliporeSigma, HAWP02500). Polybrene (Sigma-Aldrich, TR-1003; final concentration, 4 μg ml−1) and HEPES (Gibco, 15630080; final concentration, 2 mM) were added to the ered supernatants; aliquots were prepared and at −80°C until needed. Primary dermal fibroblasts were seeded in six-well plates for transduction so that cell confluency was 30–40% at the time of transduction. The cells were ‘spinoculated’ in a centrifuge at 37 °C, 931 relative centrifugal force (r.c.f.) for 2 h with 2 ml of thawed, undiluted γ-retroviral pseudoparticles per well. The cells were subsequently kept at 37 °C, 5% (vol/vol) CO2 and the media replaced with 10% FBS DMEM 6 h post-spinoculation. The transduced cells were pooled once they achieved ∼80% confluency in the six-well plate and subsequently expanded to prepare immortalized cell stocks. Cells were verified as negative for mycoplasma by testing with the MycoAlert Mycoplasma Detection Assay kit (Lonza, LT07–318) per the manufacturer’s instructions.

To establish the *Mettl14* deficient MEF cell line, MEFs were transduced with VSV-G pseudotyped lentiviral particles expressing CRE recombinase. Lentivirus was generated as described above. The CRE expressing lentiviral backbone was obtained as CSW-CRE plasmid, a generous gift of Dr. Charles M. Rice, The Rockefeller University).

### Immunofluorescence imaging and Image analysis

MEF cells were seeded and grown overnight on glass coverslips before being fixed with 4% PFA [w/vol.] at room temperature for 30 minutes. After fixation, cells were washed with PBS and then permeabilized at −20°C for 10 min in ice-cold 90% (v/v) methanol. Cells were washed again in PBS and blocked at room temperature for 1 hour with IF buffer [PBS supplemented with 10 % (v/v) FBS and 2 mM EDTA]. Cells were incubated overnight at 4°C in primary antibody diluted in IF-T buffer (IF buffer with 0.3 % Triton X-100). The following day, cells were washed three times in IF-T buffer, incubated at room temperature for 1 hour in secondary antibody diluted 1:100 in IF-T buffer, washed three times again, and then imaged with a confocal microscope. A polyclonal antibody was used at 1:500 for *Mettl14* (Invitrogen, Waltham, MA). Monoclonal antibodies were used at 1:250 for *Mettl3* [EPR18810] (Abcam, Cambridge, UK) and *Alyref* [EPR17942] (Abcam, Cambridge, UK).

The Hoechst-3342 channel from the images was extracted, and Cellpose 2.0 (*72*) was used to generate segmentation, outlining the nuclei. These outlined nuclei images were then imported into the Tissue Analyzer (*73*) plugin of Fiji for manual inspection and correction. After corrections, the outline images were transformed into labeled images by assigning a unique label to each pixel within the nuclei boundaries using Python’s scikit-image and OpenCV2 libraries. These labeled segmentation masks were utilized to calculate the size and mean intensities for each nucleus across all other channels. This was accomplished through a Python script that iterates over each labeled region, extracting masked pixels and computing their size and mean using NumPy.

### Western Blot

Cells or liver tissue samples were lysed in ice-cold RIPA buffer [1% Nonidet P-40, 0.5% Deoxycholate, 150 mM NaCl, 50 mM Tris-HCl, pH 7.4, 1.5 mM MgCl2, 1 mM EGTA, 10% (v/v) glycerol] supplemented with 1x protease inhibitor cocktail (Sigma-Aldrich, St. Louis, MO), spun at 12,000 rpm for 10 min, and pellet discarded. Protein amounts were quantified with a Pierce BCA kit (Thermo Fisher Scientific, Somerset, NJ), mixed with 6x Laemmli buffer, heated at 95°C for 5 min, loaded into a 10% polyacrylamide gel, and ran at 170 V for 1 hour. Gels were transferred to a nitrocellulose membrane with a Genie Blotter (Idea Scientific, Minneapolis, Minnesota), blocked in Tris-buffered saline with 0.1% [v/v] Tween-20 with 5% [w/v] milk for 30 min at room temperature, and incubated with primary antibody for 1 hour at 4°C. Membranes were washed three times for 5 min in TBST and incubated for 30 min in either IRDye 680CW or 800CW secondary antibodies (Licor, Lincoln, NE) (1:20,000). Imaging was performed with the Li-Cor Odyssey Infrared Imaging System (Licor, Lincoln, NE).

### Statistical analysis

An unpaired-student’s T-test was used to compare groups to wild-type in figures 3B, 4B, 5B,C,D, 6B, 7B,C. A paired student’s T-test was used to compare weight changes over time to wild-type baseline in partial hepatectomy recoveries in figure 3C. P-values of 0.05 or less were considered significant. Transcriptomic analysis using DESeq2 determined significance by using Benjamini and Hochberg method-corrected Wald Test P values (Fig 2, Supplemental Data 1). P- adj values of 0.1 or less were considered as hits for this analysis. The regression analysis to compare known m^6^A modified sites (*30*) with DEG fold-change expression was done using internal statistics tools in Graphpad Prism software to determine non-linear regression to a second order polynomial (quadratic), to determine a best fit model to the data. A P-value of <0.0001 and R-squared of 0.05239 was recorded for the alternative hypothesis (B0 unconstrained), and a R-squared value of -0.4864 was recorded for the null hypothesis (B0 = 0). GSEA software was used for gene set enrichment analysis (*70*), and P-values are derived by permutation using the standard 100 permutation default setting. We included data from gene-sets reported with p-values over the significance cut off as well to give a more complete picture of the pathways represented by significant DEGs, even when the number of related DEGs was somewhat low for a pathway. P-values and number of DEGS found for each gene set listed are shown in the figures (Fig. 2E,F). All graphs of plotted data were plotted in GraphPad Prism 10.

## Acknowledgments

We thank Dr. Gary Laevsky and Sha Wang of the Princeton Imaging Core which is a Nikon Center of Excellence, Christina de Coste of the Flow cytometry Core and Dr. Wei Wang of the Genomics core for outstanding technical support. We are grateful to Dr. Frank Chisari (Scripps Research) for generously sharing the 1.3x HBV tg mice, Dr. Chuan He (University of Chicago/HHMI) for kindly providing the Ythdf1^-/-^, Ythdf2^[fl/fl]^, and Mettl14^[fl/fl]^ and Dr. Pierre Chambon (INSERM) for the Alb-ERT2-Cre mouse lines. We further thank Dr. Yuri Pritykin for useful discussion of the RNAseq data analysis and all members of the Ploss lab for critical discussion of the study and manuscript.

## Author contributions

Conceptualization: KAB, AP

Methodology: KAB, AP

Investigation: KAB, SS, AEL, MPS, SM, TRC, YL, HPG, SS, ARB, SC, BH, NH, QA DB, DW, JMC, AP

Visualization: KAB, SS, AEL

Supervision: AP, RES, NAC, REK, JMC

Writing—original draft: KAB, AP

Writing—review & editing: All authors

## Data and materials availability

1.3x HBV tg mice are available under MTA from Dr. Frank Chisari (Scripps Research), the Ythdf1^-/-^, Ythdf2^[fl/fl]^, and Mettl14^[fl/fl]^ under MTA from Dr. Chuan He (University of Chicago/HHMI) and the Alb-ERT2-Cre mouse lines under MTA from Dr. Pierre Chambon.

All data are available in the main text or the supplementary materials. Raw RNA-Seq reads and the processed counts matrix can be accessed at NCBI GEO accession: GSE265879.

