## Supplemental Figures for "Liver-specific Mettl14 deletion induces nuclear heterotypia and dysregulates RNA export machinery"

**SUPPLEMENTARY INFORMATION**

**FOR**

### Disruption of the $N^6$-methyladenosine RNA modification machinery in hepatocytes induces nuclear heterotypia and progressive liver disease

Keith A Berggren[1], Saloni Sinha[2], Aaron E Lin[1], Michael P Schwoerer[1], Stephanie Maya[1], Abhishek Biswas[1,3], Thomas R Cafiero[1], Yongzhen Liu[1], Hans P Gertje[4] Saori Suzuki[1]†, Andrew R. Berneshawi[1]‡, Sebastian Carver[1], Brigitte Heller[1], Nora Hassan[2], Qazi Ali[2], Daniel Beard[1], Danyang Wang[5], John M Cullen[6], Ralph E Kleiner[5], Nicholas A Crossland[4,7], Robert E Schwartz[2,8,9], Alexander Ploss[1*]

[1] Department of Molecular Biology, Princeton University, Princeton, NJ, USA.

[2] Department of Medicine, Weill Cornell Medicine, NY, USA.

[3] Research Computing, Office of Information Technology, Princeton University, Princeton, NJ, 08544, USA.

[4] National Emerging Infectious Diseases Laboratories, Boston University, Boston, MA, 02118, USA.

[5] Department of Chemistry, Princeton University, Princeton, NJ 08544, USA.

[6] Department of Population Health and Pathobiology, North Carolina State University College of Veterinary Medicine, Raleigh, NC 27607, USA.

[7] Department of Pathology and Laboratory Medicine, Boston University Chobanian & Avedisian School of Medicine, Boston, MA, 02118, USA.

[8] Department of Physiology, Biophysics, and Systems Biology, Weill Cornell Medicine, NY, USA.

[9] Department of Biomedical Engineering, Cornell University, Ithaca, NY, USA.

† Present address: Department of Microbiology and Immunology, Hokkaido University, Japan.

‡ Present address: Stanford University School of Medicine, Stanford, CA, USA.

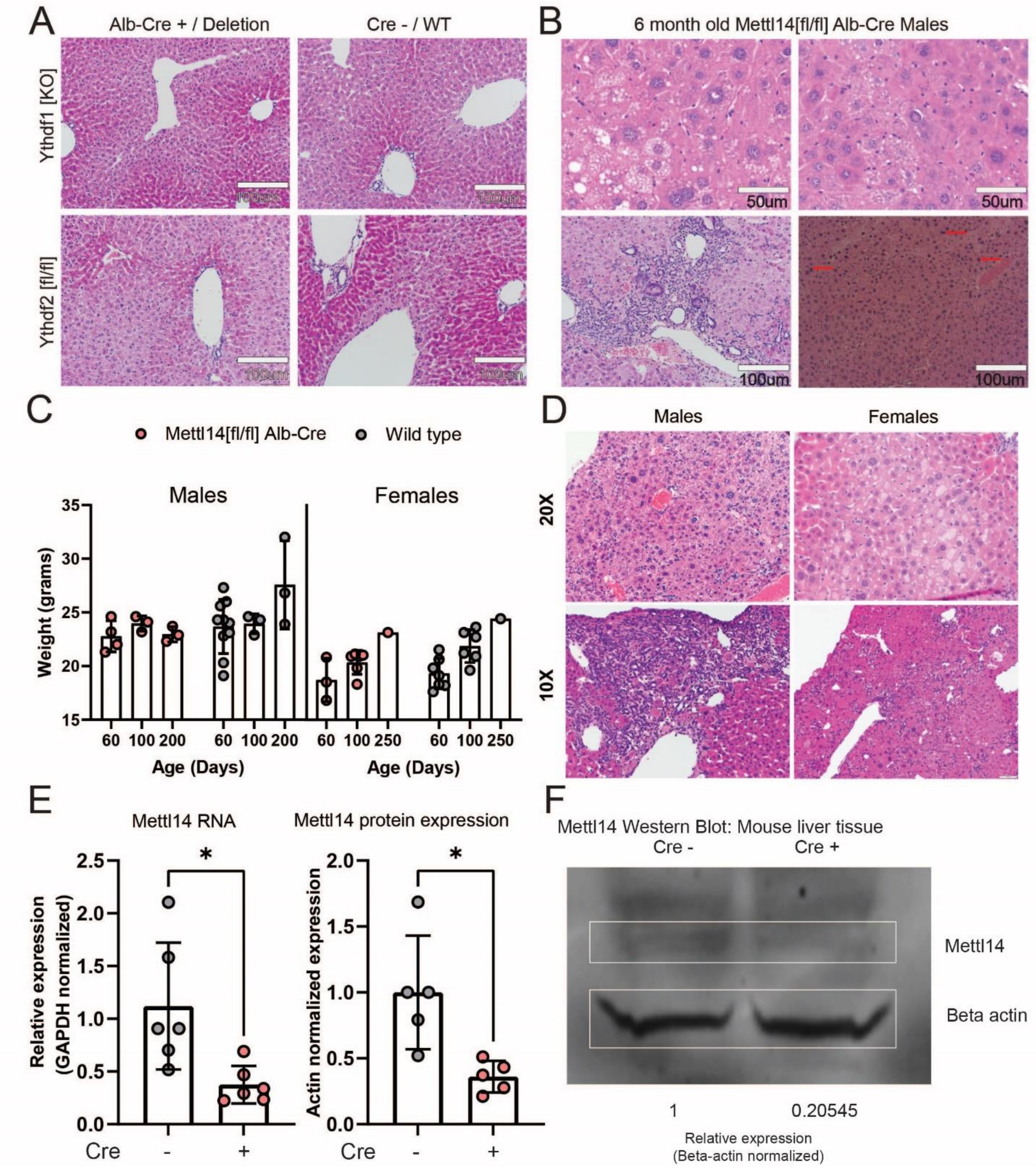


**Supplemental Fig. 1: Mettl14[fl/fl] Alb-Cre mice display a sexually dimorphic liver injury phenotype, reduced weights, and reduced Mettl14 RNA and protein levels.**

Unlike dual Ythdf1/Ythdf2, neither Ythdf1 or Ythdf2 deletion alone leads to significant histological changes in liver tissue (A). The observed injury phenotype in Mettl14 Males continues to progress with age, leading to marked steatosis (B, top), fibrosis (B, bottom left), and high levels of nuclear heterotypia (B, bottom right, red arrows). Male and female liver specific Mettl14 deletion mice show generally reduced weight throughout adulthood compared to C57BL/6 mice (C). Males show more pronounced liver damage than females, as shown by representative images (D). Reverse-transcription qPCR and western blot quantification both confirm significant reduction of Mettl14 expression by ~50% in our Mettl14[fl/fl] Alb-Cre model (E,F).

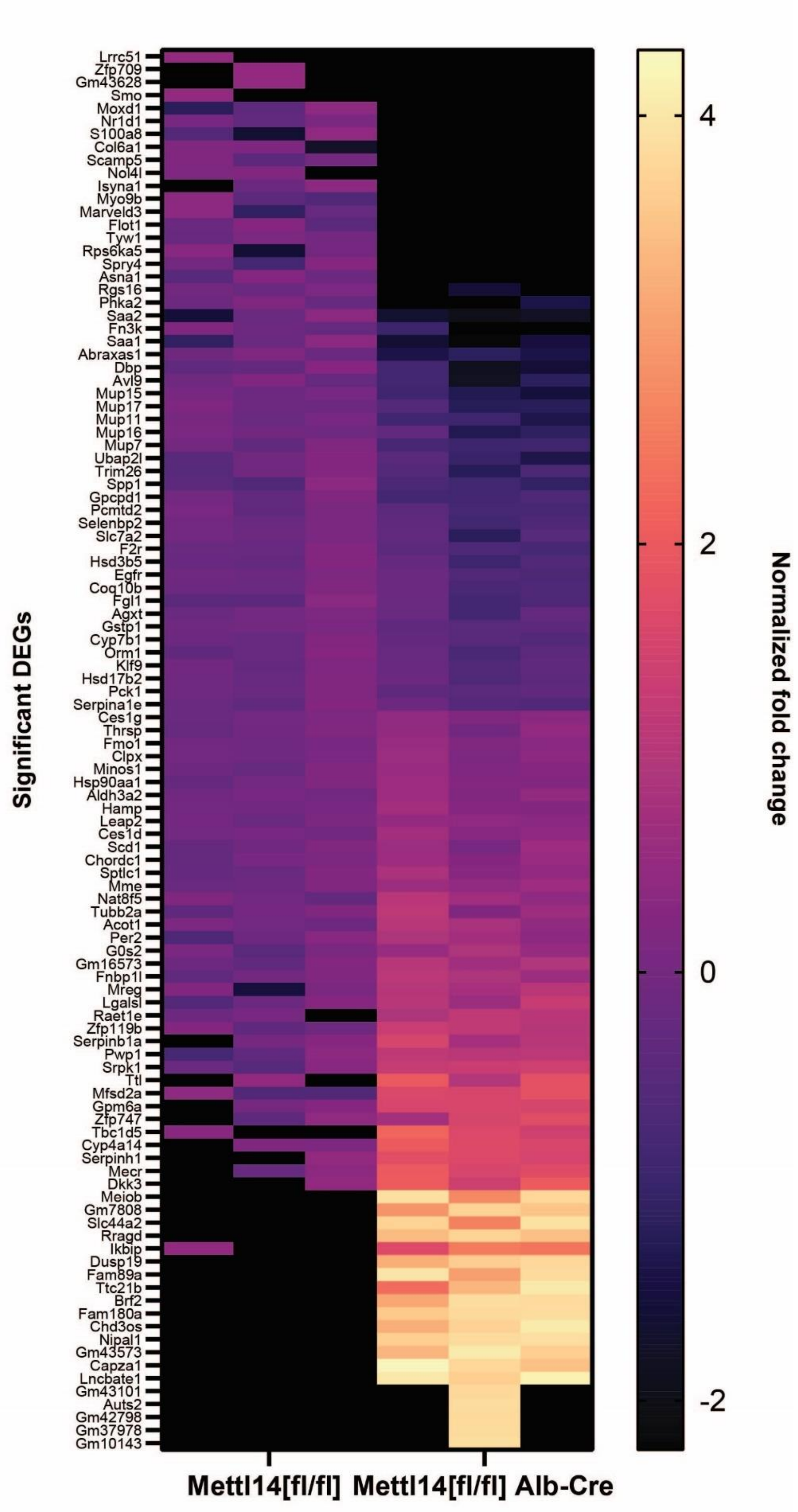
Significant DEGs
Lrrc51
Zfp709
Gm43628
Smo
Moxd1
Nr1d1
S100a8
Col6a1
Scamp5
Nol4l
Isyna1
Myo9b
Marveld3
Flot1
Tyw1
Rps6ka5
Spry4
Asna1
Rgs16
Phka2
Saa2
Fn3k
Saa1
Abraxas1
Dbp
Avl9
Mup15
Mup17
Mup11
Mup16
Mup7
Ubap2l
Trim26
Spp1
Gpcpd1
Pcmtd2
Selenbp2
Slc7a2
F2r
Hsd3b5
Egfr
Coq10b
Fgl1
Agxt
Gstp1
Cyp7b1
Orm1
Klf9
Hsd17b2
Pck1
Serpina1e
Ces1g
Thrsp
Fmo1
Clpx
Minos1
Hsp90aa1
Aldh3a2
Hamp
Leap2
Ces1d
Scd1
Chordc1
Sptlc1
Mme
Nat8f5
Tubb2a
Acot1
Per2
G0s2
Gm16573
Fnbp1l
Mreg
Lgalsl
Raet1e
Zfp119b
Serpinb1a
Pwp1
Srpk1
Ttl
Mfsd2a
Gpm6a
Zfp747
Tbc1d5
Cyp4a14
Serpinh1
Mecr
Dkk3
Meiob
Gm7808
Slc44a2
Rragd
Ikbip
Dusp19
Fam89a
Ttc21b
Brf2
Fam180a
Chd3os
Nipal1
Gm43573
Capza1
Lncbate1
Gm43101
Auts2
Gm42798
Gm37978
Gm10143
Mettl14[fl/fl]
Mettl14[fl/fl] Alb-Cre
4
2
0
-2
Normalized fold change

**Supplemental Fig. 2: Heatmap of expression changes in DEGs between wild-type and Mettl14[fl/fl] Alb-Cre expressing mice.** RNAseq count data for DEGs determined by DeSEQ2 was normalized to the mean of wild-type samples. Values were then plotted by taking the base-10 log. Where transcripts were not detected, boxes are colored black, and where values exceeded 4, values are binned into one group with the same light-yellow color at the top of the scale bar.

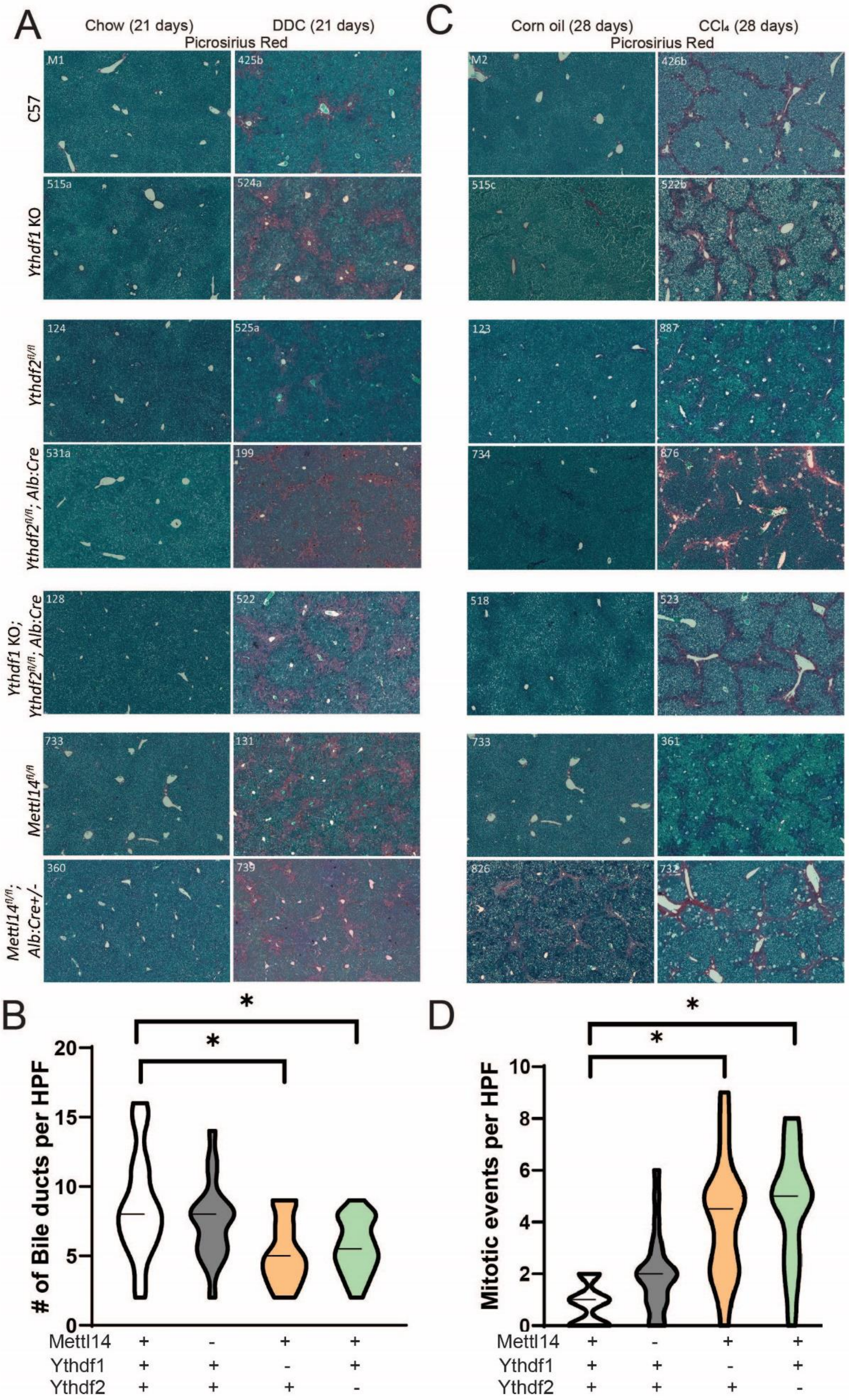
A
Chow (21 days)
DDC (21 days)
Picrosirius Red
C
Corn oil (28 days)
CCl₄ (28 days)
Picrosirius Red
C57
Ythdf1 KO
Ythdf2fl/fl
Ythdf2fl/fl; Alb:Cre
Ythdf1 KO; Ythdf2fl/fl; Alb:Cre
Mettl14fl/fl
Mettl14fl/fl; Alb:Cre+/-
M1
425b
515a
524a
124
525a
531a
199
128
522
733
131
360
739
M2
426b
515c
522b
123
887
734
876
518
523
733
361
826
732
B
### of Bile ducts per HPF
0
5
10
15
20
*
*
D
Mitotic events per HPF
0
2
4
6
8
10
*
*
Mettl14 + - + +
Ythdf1 + + - +
Ythdf2 + + + -
Mettl14 + - + +
Ythdf1 + + - +
Ythdf2 + + + -

**Supplemental Fig. 3: Liver damage is exacerbated in Ythdf1, Ythdf2 and Mettl14 deficient liver tissue**
Deletion of Ythdf1 and Ythdf2 both separately contribute to worsened injury response to DDC as seen by picrosirius red staining to show areas of fibrosis (A). Bile duct regeneration after blockage by DDC treatment is impaired significantly in Ythdf1 and Ythdf2 deletion mice (N=20 fields of view: 4 fields of view for each of 5 animals for all groups) (B). Response to $CCl_4$ is worsened in Ythdf1 and Ythdf2 mice leading to further liver fibrosis (C). This direct injury to hepatocytes leads to significantly increased rates of cell division, with higher mitotic events captured by histology imaging (N=20 fields of view: 4 fields of view for each of 5 animals for all groups) (D).

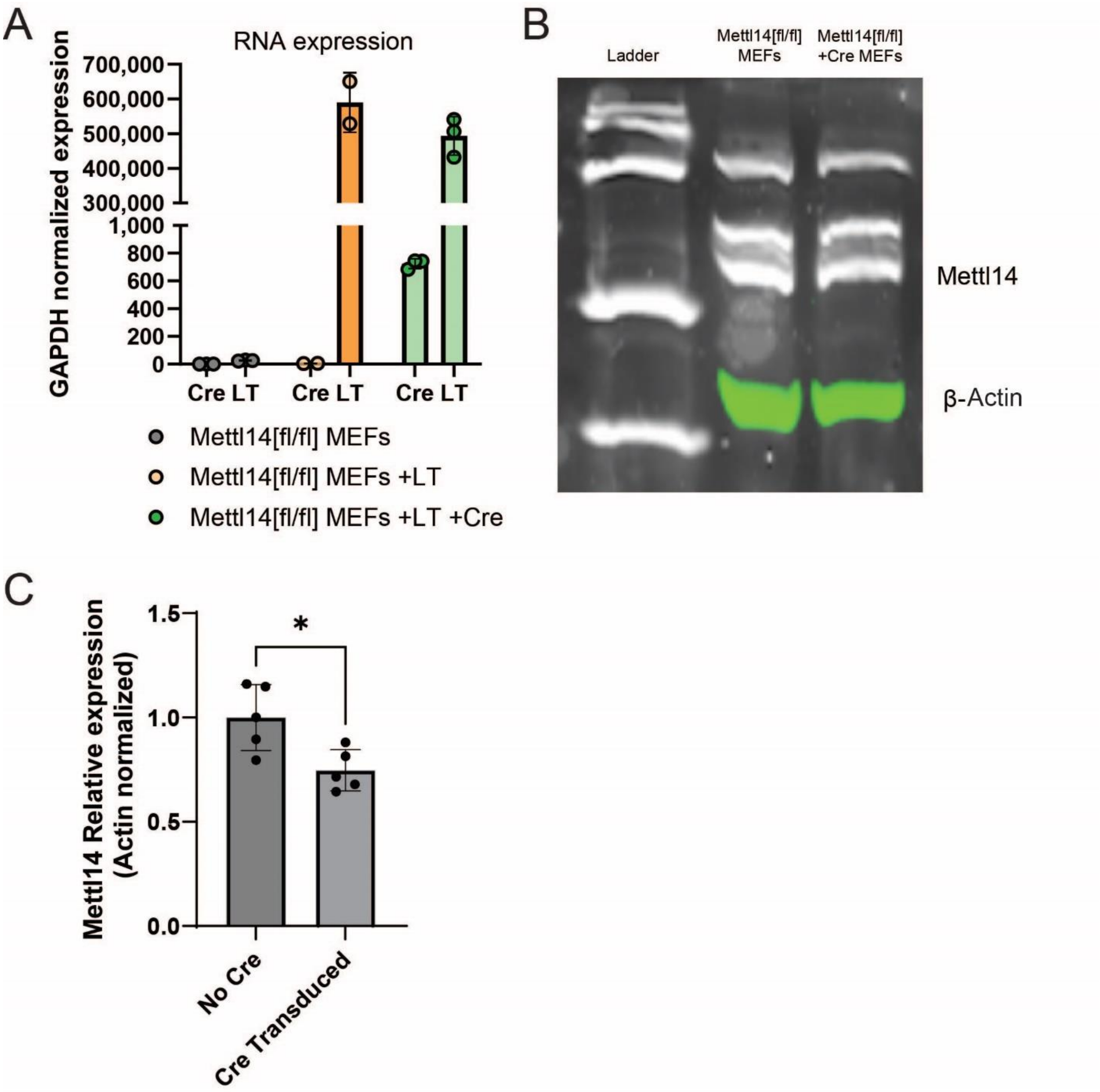


**Supplemental Fig. 4: Confirmation of Mettl14 expression changes in mouse embryonic fibroblasts**
RT-qPCR confirmed that SV40 large T antigen expression and Cre expression were present in transduced MEF cells (A). Western blot confirmed protein expression changes of Mettl14 in Cre-transduced MEF cells (B), with quantification of multiple replicates showing ~25% reduction in Mettl14 levels (C).
