## Supplemental Data 1 for "Liver-specific Mettl14 deletion induces nuclear heterotypia and dysregulates RNA export machinery"

| | baseMean | log2FoldCl | lfcSE | stat | pvalue | padj | -log10 padj |
|---|---|---|---|---|---|---|---|
| ENSMUSG00000096674.2 | 876.6743 | -4.05379 | 0.421678 | -9.61347 | 7.01E-22 | 6.65E-18 | 17.1772 |
| ENSMUSG00000020889.12 | 136.7002 | -10.6245 | 1.389649 | -7.64543 | 2.08E-14 | 9.87E-11 | 10.00566 |
| ENSMUSG00000068877.13 | 2039.568 | -2.03847 | 0.272804 | -7.47227 | 7.88E-14 | 2.49E-10 | 9.603661 |
| ENSMUSG00000064032.3 | 19.18609 | 21.14807 | 2.993193 | 7.065389 | 1.60E-12 | 3.80E-09 | 8.420679 |
| ENSMUSG00000064307.14 | 15.51242 | -21.7043 | 3.155615 | -6.87799 | 6.07E-12 | 1.15E-08 | 7.938955 |
| ENSMUSG00000001761.8 | 14.49186 | -21.5896 | 3.216747 | -6.71161 | 1.92E-11 | 2.03E-08 | 7.693032 |
| ENSMUSG00000056019.13 | 14.48839 | -21.5924 | 3.215077 | -6.71599 | 1.87E-11 | 2.03E-08 | 7.693032 |
| ENSMUSG00000103132.2 | 16.48036 | 20.93947 | 3.115788 | 6.720442 | 1.81E-11 | 2.03E-08 | 7.693032 |
| ENSMUSG00000105935.2 | 14.48839 | -21.5924 | 3.215077 | -6.71599 | 1.87E-11 | 2.03E-08 | 7.693032 |
| ENSMUSG00000106717.2 | 15.98841 | 20.89651 | 3.141656 | 6.651429 | 2.90E-11 | 2.75E-08 | 7.560404 |
| ENSMUSG00000029673.20 | 16.72634 | 20.08749 | 3.10332 | 6.472904 | 9.61E-11 | 8.29E-08 | 7.081694 |
| ENSMUSG00000106754.2 | 15.49646 | 19.61875 | 3.168853 | 6.19112 | 5.97E-10 | 4.72E-07 | 6.326121 |
| ENSMUSG00000073842.11 | 408.5183 | -2.7826 | 0.453521 | -6.13555 | 8.49E-10 | 6.19E-07 | 6.208401 |
| ENSMUSG00000020000.8 | 393.1731 | -12.1493 | 2.054272 | -5.91415 | 3.34E-09 | 2.26E-06 | 5.646094 |
| ENSMUSG00000027346.16 | 805.6587 | -2.36596 | 0.427717 | -5.53161 | 3.17E-08 | 2.01E-05 | 4.6978 |
| ENSMUSG00000020122.17 | 2417.107 | -1.59256 | 0.294967 | -5.39913 | 6.70E-08 | 3.94E-05 | 4.404697 |
| ENSMUSG00000110613.2 | 45.52601 | 8.998316 | 1.669577 | 5.38958 | 7.06E-08 | 3.94E-05 | 4.404697 |
| ENSMUSG00000050440.9 | 17410.82 | 1.385164 | 0.261695 | 5.293038 | 1.20E-07 | 6.34E-05 | 4.198196 |
| ENSMUSG00000031517.9 | 72.51598 | 5.496587 | 1.054216 | 5.213909 | 1.85E-07 | 8.86E-05 | 4.052787 |
| ENSMUSG00000056054.10 | 59.60386 | -9.42978 | 1.809248 | -5.21199 | 1.87E-07 | 8.86E-05 | 4.052787 |
| ENSMUSG00000028655.12 | 91.32863 | 5.455991 | 1.050472 | 5.193846 | 2.06E-07 | 8.88E-05 | 4.051761 |
| ENSMUSG00000038092.7 | 1276.13 | -1.75246 | 0.337407 | -5.19389 | 2.06E-07 | 8.88E-05 | 4.051761 |
| ENSMUSG00000096688.2 | 18689.05 | -3.32642 | 0.654293 | -5.084 | 3.70E-07 | 0.000152 | 3.817216 |
| ENSMUSG00000074115.6 | 2944.632 | -5.36221 | 1.058544 | -5.06564 | 4.07E-07 | 0.000161 | 3.793785 |
| ENSMUSG00000037071.4 | 21165.49 | 1.684563 | 0.337522 | 4.99097 | 6.01E-07 | 0.00022 | 3.658486 |
| ENSMUSG00000070372.12 | 38.72228 | 8.765647 | 1.756454 | 4.990538 | 6.02E-07 | 0.00022 | 3.658486 |
| ENSMUSG00000059824.13 | 96.16015 | -4.18846 | 0.852542 | -4.91291 | 8.97E-07 | 0.000315 | 3.501596 |
| ENSMUSG00000057465.6 | 2515.804 | -5.71211 | 1.173695 | -4.86677 | 1.13E-06 | 0.000384 | 3.415606 |
| ENSMUSG00000031596.16 | 1020.383 | -2.00641 | 0.413549 | -4.85169 | 1.22E-06 | 0.0004 | 3.397752 |
| ENSMUSG00000027820.13 | 379.2817 | 2.094346 | 0.43263 | 4.840962 | 1.29E-06 | 0.000408 | 3.38901 |
| ENSMUSG00000021270.14 | 3850.495 | 1.304699 | 0.272366 | 4.790235 | 1.67E-06 | 0.000509 | 3.292915 |
| ENSMUSG00000056973.7 | 4468.029 | 1.483025 | 0.310899 | 4.770109 | 1.84E-06 | 0.000545 | 3.263227 |
| ENSMUSG00000067219.10 | 34.0516 | 8.576913 | 1.814216 | 4.727613 | 2.27E-06 | 0.000653 | 3.185346 |
| ENSMUSG00000028910.13 | 45.23919 | 6.035115 | 1.286127 | 4.692472 | 2.70E-06 | 0.000731 | 3.136019 |
| ENSMUSG00000104786.2 | 38.15753 | 8.739759 | 1.861969 | 4.693827 | 2.68E-06 | 0.000731 | 3.136019 |
| ENSMUSG00000090038.8 | 120.6695 | 3.107061 | 0.666115 | 4.664449 | 3.09E-06 | 0.000815 | 3.088911 |
| ENSMUSG00000073834.11 | 40136.3 | -3.24291 | 0.70336 | -4.61059 | 4.02E-06 | 0.001029 | 2.987687 |
| ENSMUSG00000040722.8 | 31.41672 | -8.50226 | 1.869156 | -4.54872 | 5.40E-06 | 0.001347 | 2.870792 |
| ENSMUSG00000047420.9 | 31.31195 | 8.45574 | 1.862412 | 4.540209 | 5.62E-06 | 0.001366 | 2.864531 |
| ENSMUSG00000039519.7 | 3558.892 | -1.30455 | 0.288123 | -4.52774 | 5.96E-06 | 0.001413 | 2.849885 |
| ENSMUSG00000019975.13 | 43.89902 | 7.953904 | 1.761837 | 4.514552 | 6.35E-06 | 0.001467 | 2.833538 |
| ENSMUSG00000043419.12 | 32.90015 | 8.527907 | 1.90544 | 4.475558 | 7.62E-06 | 0.001686 | 2.773216 |
| ENSMUSG00000072949.7 | 397.6229 | 2.55803 | 0.571645 | 4.474859 | 7.65E-06 | 0.001686 | 2.773216 |

| | | | | | | | |
|---|---|---|---|---|---|---|---|
| ENSMUSG00000058672.8 | 298.9842 | 2.432583 | 0.550131 | 4.421822 | 9.79E-06 | 0.002109 | 2.675986 |
| ENSMUSG00000031295.14 | 51.63395 | -6.27156 | 1.4289 | -4.38908 | 1.14E-05 | 0.002398 | 2.620152 |
| ENSMUSG00000009633.4 | 640.8613 | 2.615719 | 0.605758 | 4.318093 | 1.57E-05 | 0.003199 | 2.494991 |
| ENSMUSG00000028715.3 | 691.0732 | 5.922455 | 1.372084 | 4.316395 | 1.59E-05 | 0.003199 | 2.494991 |
| ENSMUSG00000029787.11 | 74.40579 | -4.18464 | 0.973001 | -4.30076 | 1.70E-05 | 0.003362 | 2.473431 |
| ENSMUSG00000031487.6 | 30.02927 | 8.394784 | 1.963095 | 4.276301 | 1.90E-05 | 0.003676 | 2.434578 |
| ENSMUSG00000025934.16 | 17969.18 | 0.832976 | 0.195015 | 4.271347 | 1.94E-05 | 0.003684 | 2.433699 |
| ENSMUSG00000050608.12 | 1178.812 | 1.221717 | 0.288053 | 4.241299 | 2.22E-05 | 0.004131 | 2.38396 |
| ENSMUSG00000070436.13 | 45.16592 | 5.928488 | 1.400525 | 4.233047 | 2.31E-05 | 0.004203 | 2.376438 |
| ENSMUSG00000039196.3 | 6123.751 | -1.26555 | 0.304449 | -4.15686 | 3.23E-05 | 0.005664 | 2.246857 |
| ENSMUSG00000039735.17 | 104.9648 | 3.12596 | 0.751333 | 4.160551 | 3.17E-05 | 0.005664 | 2.246857 |
| ENSMUSG00000001119.8 | 33.31334 | -8.58617 | 2.086032 | -4.11603 | 3.85E-05 | 0.006644 | 2.177581 |
| ENSMUSG00000004677.18 | 28.14968 | -8.3428 | 2.036066 | -4.09751 | 4.18E-05 | 0.00707 | 2.1506 |
| ENSMUSG00000060803.7 | 1556.089 | -1.3051 | 0.318837 | -4.09333 | 4.25E-05 | 0.007072 | 2.150444 |
| ENSMUSG00000072849.11 | 55614.31 | -1.14798 | 0.281521 | -4.07776 | 4.55E-05 | 0.007432 | 2.128881 |
| ENSMUSG00000026475.8 | 35.29123 | -6.4939 | 1.625499 | -3.99502 | 6.47E-05 | 0.010279 | 1.98807 |
| ENSMUSG00000030772.7 | 34.00102 | 6.461448 | 1.617916 | 3.993686 | 6.51E-05 | 0.010279 | 1.98807 |
| ENSMUSG00000001672.15 | 27.7095 | -8.32059 | 2.093671 | -3.97416 | 7.06E-05 | 0.010976 | 1.959547 |
| ENSMUSG00000056310.15 | 22.46037 | -8.01821 | 2.024225 | -3.96113 | 7.46E-05 | 0.011406 | 1.942866 |
| ENSMUSG00000043068.7 | 22.93056 | 8.011101 | 2.036583 | 3.9336 | 8.37E-05 | 0.012592 | 1.899895 |
| ENSMUSG00000031594.10 | 4229.95 | -1.51289 | 0.389589 | -3.8833 | 0.000103 | 0.015264 | 1.816339 |
| ENSMUSG00000015357.11 | 781.3897 | 1.216308 | 0.314726 | 3.864655 | 0.000111 | 0.016225 | 1.78982 |
| ENSMUSG00000027001.11 | 22.67356 | 7.989995 | 2.071062 | 3.857922 | 0.000114 | 0.016426 | 1.78448 |
| ENSMUSG00000059714.14 | 22.74932 | -8.03576 | 2.091448 | -3.8422 | 0.000122 | 0.017253 | 1.763133 |
| ENSMUSG00000079494.3 | 357.3146 | 2.362856 | 0.61715 | 3.828657 | 0.000129 | 0.017962 | 1.745634 |
| ENSMUSG00000029304.15 | 276.9755 | -2.48515 | 0.650099 | -3.82272 | 0.000132 | 0.018134 | 1.74151 |
| ENSMUSG00000053219.16 | 65.09858 | 3.752763 | 0.982732 | 3.818706 | 0.000134 | 0.018168 | 1.740688 |
| ENSMUSG00000054381.6 | 43.58019 | 5.57893 | 1.46564 | 3.80648 | 0.000141 | 0.018821 | 1.725359 |
| ENSMUSG00000004865.17 | 36.35552 | 4.784784 | 1.258255 | 3.802715 | 0.000143 | 0.018844 | 1.724827 |
| ENSMUSG00000001774.9 | 384.8866 | 1.696875 | 0.450963 | 3.762784 | 0.000168 | 0.020695 | 1.684145 |
| ENSMUSG00000022149.19 | 4773.445 | -0.99371 | 0.264095 | -3.7627 | 0.000168 | 0.020695 | 1.684145 |
| ENSMUSG00000023923.11 | 30.10662 | 5.913734 | 1.569972 | 3.766778 | 0.000165 | 0.020695 | 1.684145 |
| ENSMUSG00000024457.17 | 151.2804 | -2.53491 | 0.672756 | -3.76794 | 0.000165 | 0.020695 | 1.684145 |
| ENSMUSG00000036216.6 | 957.9767 | 1.412891 | 0.374833 | 3.769387 | 0.000164 | 0.020695 | 1.684145 |
| ENSMUSG00000059908.10 | 19475.71 | -0.81498 | 0.21698 | -3.75604 | 0.000173 | 0.020981 | 1.678179 |
| ENSMUSG00000019139.11 | 29.97157 | -8.43721 | 2.251457 | -3.74744 | 0.000179 | 0.021438 | 1.668823 |
| ENSMUSG00000040181.15 | 847.657 | 1.166246 | 0.313878 | 3.7156 | 0.000203 | 0.023726 | 1.624776 |
| ENSMUSG00000057074.7 | 1341.577 | 1.076132 | 0.289606 | 3.715849 | 0.000203 | 0.023726 | 1.624776 |
| ENSMUSG00000042363.15 | 62.81429 | 3.715055 | 1.00336 | 3.702615 | 0.000213 | 0.024083 | 1.618284 |
| ENSMUSG00000048376.7 | 238.6158 | -1.89355 | 0.51141 | -3.70261 | 0.000213 | 0.024083 | 1.618284 |
| ENSMUSG00000061411.13 | 30.36742 | -8.45287 | 2.282202 | -3.70382 | 0.000212 | 0.024083 | 1.618284 |
| ENSMUSG00000021468.9 | 198.0033 | 1.941894 | 0.5259 | 3.692514 | 0.000222 | 0.024636 | 1.608433 |
| ENSMUSG00000078675.10 | 6419.852 | -2.89843 | 0.785298 | -3.69087 | 0.000223 | 0.024636 | 1.608433 |
| ENSMUSG00000034848.18 | 24.27744 | 8.090294 | 2.212975 | 3.655846 | 0.000256 | 0.027932 | 1.553902 |

| | | | | | | | |
|---|---|---|---|---|---|---|---|
| ENSMUSG00000025981.14 | 345.304 | -1.57628 | 0.432767 | -3.64234 | 0.00027 | 0.028459 | 1.545788 |
| ENSMUSG00000028278.15 | 19.83405 | 7.795964 | 2.14008 | 3.642838 | 0.00027 | 0.028459 | 1.545788 |
| ENSMUSG00000039395.9 | 163.5549 | 3.256876 | 0.893037 | 3.646967 | 0.000265 | 0.028459 | 1.545788 |
| ENSMUSG00000031844.9 | 784.9246 | -1.16356 | 0.320361 | -3.63202 | 0.000281 | 0.029187 | 1.534817 |
| ENSMUSG00000062101.13 | 51.62303 | 3.872071 | 1.066639 | 3.63016 | 0.000283 | 0.029187 | 1.534817 |
| ENSMUSG00000025175.13 | 39.1877 | -5.49287 | 1.518847 | -3.61647 | 0.000299 | 0.030443 | 1.516518 |
| ENSMUSG00000026272.7 | 1012.604 | -1.31479 | 0.363992 | -3.61213 | 0.000304 | 0.030627 | 1.513894 |
| ENSMUSG00000031767.14 | 4444.768 | -0.78565 | 0.218221 | -3.60024 | 0.000318 | 0.031395 | 1.503144 |
| ENSMUSG00000033863.3 | 1062.696 | -1.20146 | 0.333558 | -3.60195 | 0.000316 | 0.031395 | 1.503144 |
| ENSMUSG00000024427.8 | 19.18433 | -7.79308 | 2.176395 | -3.58073 | 0.000343 | 0.033486 | 1.475131 |
| ENSMUSG00000010025.20 | 867.1998 | 1.355303 | 0.380526 | 3.561658 | 0.000369 | 0.035649 | 1.447956 |
| ENSMUSG00000055866.10 | 161.5595 | 2.608913 | 0.734015 | 3.554303 | 0.000379 | 0.03629 | 1.440207 |
| ENSMUSG00000027589.15 | 179.7408 | -2.14145 | 0.603295 | -3.54959 | 0.000386 | 0.036577 | 1.436795 |
| ENSMUSG00000035686.9 | 5838.723 | 1.125873 | 0.317512 | 3.545929 | 0.000391 | 0.036721 | 1.43508 |
| ENSMUSG00000042520.17 | 131.2835 | -2.67636 | 0.75567 | -3.5417 | 0.000398 | 0.036949 | 1.432398 |
| ENSMUSG00000001785.14 | 43.14535 | 4.120421 | 1.165303 | 3.535923 | 0.000406 | 0.0374 | 1.427125 |
| ENSMUSG00000024155.9 | 18.95559 | 7.736749 | 2.193427 | 3.527243 | 0.00042 | 0.038276 | 1.417068 |
| ENSMUSG00000044734.17 | 58.67611 | 3.984725 | 1.13159 | 3.521351 | 0.000429 | 0.038764 | 1.411566 |
| ENSMUSG00000035234.19 | 39.35822 | -4.27883 | 1.217127 | -3.51552 | 0.000439 | 0.039157 | 1.407193 |
| ENSMUSG00000052456.10 | 18.50757 | -7.73878 | 2.202476 | -3.51367 | 0.000442 | 0.039157 | 1.407193 |
| ENSMUSG00000057193.9 | 19.37088 | 7.767026 | 2.214572 | 3.507236 | 0.000453 | 0.039745 | 1.400721 |
| ENSMUSG00000027513.12 | 1491.345 | -1.15335 | 0.331841 | -3.4756 | 0.00051 | 0.04433 | 1.353298 |
| ENSMUSG00000021180.10 | 21.23249 | -7.93704 | 2.288104 | -3.46883 | 0.000523 | 0.04505 | 1.346302 |
| ENSMUSG00000091460.3 | 19.16515 | 7.746111 | 2.239619 | 3.458674 | 0.000543 | 0.046362 | 1.333842 |
| ENSMUSG00000027394.14 | 39.20622 | 5.358508 | 1.558098 | 3.439133 | 0.000584 | 0.049396 | 1.306309 |
| ENSMUSG00000052684.5 | 153.0865 | -3.30694 | 0.963717 | -3.43144 | 0.0006 | 0.050368 | 1.297845 |
| ENSMUSG00000024378.10 | 583.9785 | -1.12757 | 0.329493 | -3.42214 | 0.000621 | 0.050596 | 1.295888 |
| ENSMUSG00000031010.18 | 28.03046 | -6.25644 | 1.825594 | -3.42707 | 0.00061 | 0.050596 | 1.295888 |
| ENSMUSG00000038224.13 | 1115.727 | -1.25765 | 0.367635 | -3.42091 | 0.000624 | 0.050596 | 1.295888 |
| ENSMUSG00000045775.16 | 25.54186 | 7.163535 | 2.094129 | 3.420771 | 0.000624 | 0.050596 | 1.295888 |
| ENSMUSG00000049764.9 | 68.22626 | 4.448087 | 1.304293 | 3.410344 | 0.000649 | 0.052125 | 1.282956 |
| ENSMUSG00000035781.15 | 19.18755 | 7.751513 | 2.275782 | 3.406088 | 0.000659 | 0.052362 | 1.280984 |
| ENSMUSG00000036106.16 | 25.62734 | -8.20813 | 2.410952 | -3.40452 | 0.000663 | 0.052362 | 1.280984 |
| ENSMUSG00000028833.14 | 25.94382 | -8.22621 | 2.421316 | -3.39741 | 0.00068 | 0.053297 | 1.2733 |
| ENSMUSG00000086437.3 | 17.63516 | 7.626077 | 2.259329 | 3.375372 | 0.000737 | 0.057281 | 1.24199 |
| ENSMUSG00000022779.17 | 28.92913 | -4.62407 | 1.373385 | -3.36692 | 0.00076 | 0.058586 | 1.232206 |
| ENSMUSG00000019143.16 | 88.02681 | -2.65533 | 0.789291 | -3.3642 | 0.000768 | 0.058689 | 1.23144 |
| ENSMUSG00000028088.15 | 5109.918 | 0.846692 | 0.252363 | 3.355059 | 0.000793 | 0.060178 | 1.220565 |
| ENSMUSG00000050587.15 | 23.40293 | 8.039541 | 2.404165 | 3.344006 | 0.000826 | 0.06213 | 1.206698 |
| ENSMUSG00000025880.12 | 109.5705 | 3.689076 | 1.112353 | 3.31646 | 0.000912 | 0.06752 | 1.17057 |
| ENSMUSG00000032224.16 | 20.41623 | -7.87932 | 2.3746 | -3.31817 | 0.000906 | 0.06752 | 1.17057 |
| ENSMUSG00000071076.9 | 59.05057 | -3.28857 | 0.996625 | -3.29971 | 0.000968 | 0.070579 | 1.151325 |
| ENSMUSG00000094793.2 | 3123.929 | -1.34567 | 0.407554 | -3.30181 | 0.000961 | 0.070579 | 1.151325 |
| ENSMUSG00000036103.10 | 205.222 | 1.985815 | 0.604039 | 3.287559 | 0.001011 | 0.07258 | 1.139186 |

| | | | | | | | |
|---|---|---|---|---|---|---|---|
| ENSMUSG00000074794.11 | 1183.248 | -1.3215 | 0.401761 | -3.28926 | 0.001004 | 0.07258 | 1.139186 |
| ENSMUSG00000017264.17 | 24.40662 | -6.13218 | 1.871067 | -3.27737 | 0.001048 | 0.073105 | 1.136051 |
| ENSMUSG00000022865.15 | 769.7953 | 1.053514 | 0.321189 | 3.280039 | 0.001038 | 0.073105 | 1.136051 |
| ENSMUSG00000034855.14 | 72.0665 | -4.12718 | 1.259723 | -3.27626 | 0.001052 | 0.073105 | 1.136051 |
| ENSMUSG00000068086.7 | 3595.025 | -0.74475 | 0.227337 | -3.27595 | 0.001053 | 0.073105 | 1.136051 |
| ENSMUSG00000072258.12 | 19.42449 | 6.759494 | 2.06524 | 3.272982 | 0.001064 | 0.073105 | 1.136051 |
| ENSMUSG00000097706.2 | 23.00955 | 8.011112 | 2.44713 | 3.273676 | 0.001062 | 0.073105 | 1.136051 |
| ENSMUSG00000055707.14 | 22.14535 | -8.0016 | 2.450564 | -3.26521 | 0.001094 | 0.074602 | 1.127251 |
| ENSMUSG00000028044.7 | 38.83675 | -5.1925 | 1.594344 | -3.25683 | 0.001127 | 0.07579 | 1.120386 |
| ENSMUSG00000039686.15 | 15.41697 | -7.47475 | 2.295211 | -3.25667 | 0.001127 | 0.07579 | 1.120386 |
| ENSMUSG00000041135.16 | 33.44993 | -4.55751 | 1.401012 | -3.25301 | 0.001142 | 0.076233 | 1.117856 |
| ENSMUSG00000003421.14 | 32.1204 | -5.04988 | 1.554803 | -3.24792 | 0.001163 | 0.077067 | 1.113131 |
| ENSMUSG00000039384.9 | 22.92329 | -5.49626 | 1.697209 | -3.23841 | 0.001202 | 0.079129 | 1.101663 |
| ENSMUSG00000048200.13 | 15.0738 | -7.44233 | 2.303214 | -3.23128 | 0.001232 | 0.080572 | 1.093818 |
| ENSMUSG00000031574.9 | 29.14542 | -6.87514 | 2.138804 | -3.21448 | 0.001307 | 0.084854 | 1.071329 |
| ENSMUSG00000022512.4 | 261.7269 | 1.578796 | 0.492085 | 3.208381 | 0.001335 | 0.086084 | 1.065078 |
| ENSMUSG00000038754.6 | 667.188 | 1.57348 | 0.49254 | 3.194625 | 0.0014 | 0.089684 | 1.047286 |
| ENSMUSG00000015837.16 | 596.5479 | -1.4332 | 0.449559 | -3.18801 | 0.001433 | 0.091145 | 1.040269 |
| ENSMUSG00000010651.5 | 4904.217 | 1.089245 | 0.342463 | 3.180624 | 0.00147 | 0.091554 | 1.038323 |
| ENSMUSG00000025372.17 | 45.22161 | 3.965414 | 1.247348 | 3.179076 | 0.001477 | 0.091554 | 1.038323 |
| ENSMUSG00000031529.6 | 51.42872 | -4.06229 | 1.277834 | -3.17905 | 0.001478 | 0.091554 | 1.038323 |
| ENSMUSG00000042831.14 | 21.00264 | 7.884017 | 2.478875 | 3.180482 | 0.00147 | 0.091554 | 1.038323 |
| ENSMUSG00000030670.17 | 271.8189 | 1.73322 | 0.546022 | 3.174265 | 0.001502 | 0.092471 | 1.033995 |
| ENSMUSG00000024743.17 | 20.17505 | -7.86649 | 2.490513 | -3.15858 | 0.001585 | 0.096664 | 1.014734 |
| ENSMUSG00000054932.7 | 24.43245 | -5.59034 | 1.771286 | -3.15609 | 0.001599 | 0.096664 | 1.014734 |
| ENSMUSG00000110206.2 | 22.24011 | -6.00118 | 1.901667 | -3.15575 | 0.001601 | 0.096664 | 1.014734 |
| ENSMUSG00000033157.18 | 42.56825 | -4.3889 | 1.392411 | -3.15202 | 0.001621 | 0.097289 | 1.011938 |
| ENSMUSG00000049800.14 | 109.6627 | -2.5482 | 0.809543 | -3.1477 | 0.001646 | 0.098116 | 1.008262 |
| ENSMUSG00000039263.17 | 37.0091 | -4.86909 | 1.549489 | -3.14239 | 0.001676 | 0.099289 | 1.003097 |
| ENSMUSG00000019261.4 | 15.67077 | -7.50268 | 2.397247 | -3.12971 | 0.00175 | 0.099364 | 1.002769 |
| ENSMUSG00000025040.14 | 95.86022 | 2.303861 | 0.734325 | 3.137387 | 0.001705 | 0.099364 | 1.002769 |
| ENSMUSG00000029048.4 | 903.3612 | 0.977747 | 0.312419 | 3.129606 | 0.00175 | 0.099364 | 1.002769 |
| ENSMUSG00000037379.8 | 21.80362 | -6.01204 | 1.919724 | -3.13172 | 0.001738 | 0.099364 | 1.002769 |
| ENSMUSG00000041930.8 | 14.08735 | 7.306314 | 2.333515 | 3.131034 | 0.001742 | 0.099364 | 1.002769 |
| ENSMUSG00000050199.14 | 197.4859 | -1.99196 | 0.634937 | -3.13726 | 0.001705 | 0.099364 | 1.002769 |
| ENSMUSG00000063235.19 | 783.546 | 0.95412 | 0.304573 | 3.132651 | 0.001732 | 0.099364 | 1.002769 |
| ENSMUSG00000021959.17 | 104.2227 | 2.530885 | 0.809786 | 3.125377 | 0.001776 | 0.099607 | 1.001709 |
| ENSMUSG00000049091.8 | 1221.544 | -1.2066 | 0.385959 | -3.12625 | 0.001771 | 0.099607 | 1.001709 |
| ENSMUSG00000066682.12 | 14.67169 | -7.40257 | 2.369845 | -3.12365 | 0.001786 | 0.099607 | 1.001709 |
| ENSMUSG00000032491.15 | 21.6594 | -7.96524 | 2.55406 | -3.11866 | 0.001817 | 0.100719 | 0.996887 |
| ENSMUSG00000038388.15 | 723.1424 | 1.000842 | 0.321582 | 3.112239 | 0.001857 | 0.102337 | 0.989969 |
| ENSMUSG00000021091.9 | 548.5052 | -1.33328 | 0.428855 | -3.10892 | 0.001878 | 0.102895 | 0.987607 |
| ENSMUSG00000053063.12 | 95.30047 | 2.713245 | 0.873256 | 3.107044 | 0.00189 | 0.102955 | 0.987352 |
| ENSMUSG00000026839.18 | 1673.691 | -1.02675 | 0.330839 | -3.10347 | 0.001913 | 0.10361 | 0.9846 |

| | | | | | | | |
|---|---|---|---|---|---|---|---|
| ENSMUSG00000000686.12 | 26.41437 | -5.58591 | 1.802263 | -3.09939 | 0.001939 | 0.104452 | 0.981082 |
| ENSMUSG00000069456.5 | 424.8009 | 1.636857 | 0.529441 | 3.091668 | 0.00199 | 0.106602 | 0.972235 |
| ENSMUSG00000013275.10 | 15.22173 | -7.46219 | 2.422937 | -3.07981 | 0.002071 | 0.106718 | 0.971761 |
| ENSMUSG00000029505.18 | 166.4457 | 2.15738 | 0.698849 | 3.087046 | 0.002022 | 0.106718 | 0.971761 |
| ENSMUSG00000036285.11 | 75.00492 | 2.482225 | 0.804987 | 3.083561 | 0.002045 | 0.106718 | 0.971761 |
| ENSMUSG00000043418.5 | 13.8349 | -7.31862 | 2.375988 | -3.08024 | 0.002068 | 0.106718 | 0.971761 |
| ENSMUSG00000073418.5 | 499.0485 | -1.52852 | 0.495087 | -3.08738 | 0.002019 | 0.106718 | 0.971761 |
| ENSMUSG00000097327.8 | 13.80916 | 7.27652 | 2.357745 | 3.08622 | 0.002027 | 0.106718 | 0.971761 |
| ENSMUSG00000118017.2 | 319.098 | 1.80806 | 0.586877 | 3.080817 | 0.002064 | 0.106718 | 0.971761 |
| ENSMUSG00000032014.7 | 495.4918 | -1.24314 | 0.404942 | -3.06993 | 0.002141 | 0.109716 | 0.959729 |
| ENSMUSG00000030060.15 | 14.03625 | 7.298439 | 2.379409 | 3.067332 | 0.00216 | 0.11008 | 0.958293 |
| ENSMUSG00000042812.6 | 13.75969 | 7.269747 | 2.374292 | 3.061859 | 0.0022 | 0.111013 | 0.954627 |
| ENSMUSG00000056476.14 | 19.03243 | -7.78018 | 2.541209 | -3.06161 | 0.002202 | 0.111013 | 0.954627 |
| ENSMUSG00000049988.6 | 19.10653 | -7.78524 | 2.545858 | -3.058 | 0.002228 | 0.111763 | 0.951703 |
| ENSMUSG00000003355.8 | 193.9098 | -1.96198 | 0.642998 | -3.05131 | 0.002278 | 0.113683 | 0.944304 |
| ENSMUSG00000028307.10 | 27877.83 | 0.569406 | 0.187259 | 3.040734 | 0.00236 | 0.115325 | 0.938077 |
| ENSMUSG00000042423.10 | 15.25425 | -7.46513 | 2.4518 | -3.04476 | 0.002329 | 0.115325 | 0.938077 |
| ENSMUSG00000046792.9 | 21.62581 | -5.59062 | 1.837053 | -3.04325 | 0.00234 | 0.115325 | 0.938077 |
| ENSMUSG00000090925.3 | 33.13927 | 6.509541 | 2.140609 | 3.040976 | 0.002358 | 0.115325 | 0.938077 |
| ENSMUSG00000018727.20 | 16.82096 | 6.548868 | 2.163051 | 3.027607 | 0.002465 | 0.119081 | 0.924156 |
| ENSMUSG00000039795.15 | 29.43909 | 4.453425 | 1.470926 | 3.027634 | 0.002465 | 0.119081 | 0.924156 |
| ENSMUSG00000046402.11 | 264.1124 | -1.64425 | 0.543296 | -3.02643 | 0.002475 | 0.119081 | 0.924156 |
| ENSMUSG00000019960.9 | 469.3074 | -1.63141 | 0.540858 | -3.01634 | 0.002558 | 0.122005 | 0.913623 |
| ENSMUSG00000069520.7 | 486.2273 | -1.17068 | 0.388154 | -3.01603 | 0.002561 | 0.122005 | 0.913623 |
| ENSMUSG00000057836.13 | 52.25235 | -3.21854 | 1.070552 | -3.00643 | 0.002643 | 0.124764 | 0.903911 |
| ENSMUSG00000071456.7 | 13.48286 | -7.28216 | 2.422377 | -3.0062 | 0.002645 | 0.124764 | 0.903911 |
| ENSMUSG00000031557.17 | 24.91357 | -6.19392 | 2.062421 | -3.00323 | 0.002671 | 0.125368 | 0.901813 |
| ENSMUSG00000030364.7 | 46.15479 | -3.45287 | 1.153031 | -2.9946 | 0.002748 | 0.128065 | 0.89257 |
| ENSMUSG00000059447.14 | 521.7785 | 1.221235 | 0.40793 | 2.993738 | 0.002756 | 0.128065 | 0.89257 |
| ENSMUSG00000031371.15 | 86.6513 | -2.793 | 0.933839 | -2.99088 | 0.002782 | 0.128641 | 0.890622 |
| ENSMUSG00000025014.14 | 17.83507 | 7.648897 | 2.559874 | 2.987998 | 0.002808 | 0.129228 | 0.888644 |
| ENSMUSG00000048796.8 | 19.52806 | -7.81554 | 2.619336 | -2.98379 | 0.002847 | 0.130387 | 0.884766 |
| ENSMUSG00000020462.15 | 145.435 | 1.919322 | 0.644012 | 2.98026 | 0.00288 | 0.130759 | 0.883529 |
| ENSMUSG00000026874.11 | 2785.039 | -0.72396 | 0.24294 | -2.97997 | 0.002883 | 0.130759 | 0.883529 |
| ENSMUSG00000034973.17 | 15.46904 | 7.446387 | 2.503663 | 2.974197 | 0.002938 | 0.13137 | 0.881503 |
| ENSMUSG00000072974.5 | 18.28467 | 7.680121 | 2.581341 | 2.975244 | 0.002928 | 0.13137 | 0.881503 |
| ENSMUSG00000078762.11 | 17.64585 | -7.67252 | 2.579717 | -2.97417 | 0.002938 | 0.13137 | 0.881503 |
| ENSMUSG00000033082.11 | 18.03431 | 7.663557 | 2.578302 | 2.972328 | 0.002956 | 0.131541 | 0.880939 |
| ENSMUSG00000052915.15 | 223.7051 | 1.740524 | 0.586268 | 2.968818 | 0.002989 | 0.131815 | 0.880035 |
| ENSMUSG00000074884.13 | 13.11645 | -7.24124 | 2.438636 | -2.96938 | 0.002984 | 0.131815 | 0.880035 |
| ENSMUSG00000024339.13 | 30.06295 | -4.49873 | 1.516916 | -2.9657 | 0.00302 | 0.13254 | 0.877653 |
| ENSMUSG00000037025.12 | 224.6285 | 1.669065 | 0.563297 | 2.963028 | 0.003046 | 0.133082 | 0.875881 |
| ENSMUSG00000052974.9 | 6751.306 | -0.67861 | 0.229344 | -2.9589 | 0.003087 | 0.134258 | 0.872061 |
| ENSMUSG00000026600.13 | 22.17702 | 6.949292 | 2.350409 | 2.956631 | 0.00311 | 0.134633 | 0.870847 |

| | | | | | | | |
|---|---|---|---|---|---|---|---|
| ENSMUSG00000003992.16 | 25.7297 | 4.435201 | 1.501868 | 2.953123 | 0.003146 | 0.134941 | 0.869858 |
| ENSMUSG00000020777.17 | 8429.554 | 0.65862 | 0.223009 | 2.953338 | 0.003144 | 0.134941 | 0.869858 |
| ENSMUSG00000022787.11 | 24.10672 | 6.016175 | 2.040244 | 2.948752 | 0.003191 | 0.135499 | 0.868064 |
| ENSMUSG00000026708.17 | 74.90305 | 2.434068 | 0.826922 | 2.943528 | 0.003245 | 0.135499 | 0.868064 |
| ENSMUSG00000030004.7 | 239.4364 | 1.925337 | 0.654386 | 2.942206 | 0.003259 | 0.135499 | 0.868064 |
| ENSMUSG00000030161.9 | 773.7287 | -1.66301 | 0.564082 | -2.94817 | 0.003197 | 0.135499 | 0.868064 |
| ENSMUSG00000030711.17 | 2879.225 | 0.660054 | 0.223944 | 2.947403 | 0.003205 | 0.135499 | 0.868064 |
| ENSMUSG00000048327.7 | 12.75152 | -7.20394 | 2.446525 | -2.94456 | 0.003234 | 0.135499 | 0.868064 |
| ENSMUSG00000062515.4 | 103.3056 | 2.572839 | 0.873277 | 2.946188 | 0.003217 | 0.135499 | 0.868064 |
| ENSMUSG00000020268.15 | 12.87649 | 7.173842 | 2.440033 | 2.940059 | 0.003281 | 0.135845 | 0.866956 |
| ENSMUSG00000040124.6 | 44.34277 | 4.886968 | 1.665152 | 2.934848 | 0.003337 | 0.137547 | 0.861549 |
| ENSMUSG00000032375.16 | 41.01434 | -3.76846 | 1.289007 | -2.92354 | 0.003461 | 0.142027 | 0.847629 |
| ENSMUSG00000016024.10 | 107.8125 | -2.56543 | 0.878908 | -2.91889 | 0.003513 | 0.142925 | 0.844891 |
| ENSMUSG00000028771.14 | 103.9797 | -2.00853 | 0.687928 | -2.91968 | 0.003504 | 0.142925 | 0.844891 |
| ENSMUSG00000033577.19 | 362.4643 | -1.32003 | 0.453044 | -2.91369 | 0.003572 | 0.143035 | 0.844559 |
| ENSMUSG00000035392.18 | 14.65276 | -7.40127 | 2.541626 | -2.91202 | 0.003591 | 0.143035 | 0.844559 |
| ENSMUSG00000074476.7 | 106.8392 | 2.229406 | 0.765578 | 2.912054 | 0.003591 | 0.143035 | 0.844559 |
| ENSMUSG00000096956.3 | 16.9054 | -7.61104 | 2.61294 | -2.91283 | 0.003582 | 0.143035 | 0.844559 |
| ENSMUSG00000101225.2 | 43.40403 | 4.130941 | 1.416695 | 2.915901 | 0.003547 | 0.143035 | 0.844559 |
| ENSMUSG00000015656.18 | 3086.396 | 0.851436 | 0.292556 | 2.910333 | 0.00361 | 0.143209 | 0.84403 |
| ENSMUSG00000047139.10 | 12.22643 | -7.14116 | 2.457562 | -2.90579 | 0.003663 | 0.144698 | 0.839536 |
| ENSMUSG00000027284.17 | 19.94085 | -4.99512 | 1.724952 | -2.8958 | 0.003782 | 0.146338 | 0.834644 |
| ENSMUSG00000030555.17 | 72.75004 | -2.60938 | 0.900296 | -2.89836 | 0.003751 | 0.146338 | 0.834644 |
| ENSMUSG00000031722.11 | 7319.776 | -1.12028 | 0.386793 | -2.89633 | 0.003776 | 0.146338 | 0.834644 |
| ENSMUSG00000034994.11 | 645.9466 | -1.46419 | 0.505218 | -2.89813 | 0.003754 | 0.146338 | 0.834644 |
| ENSMUSG00000039956.10 | 194.8083 | 1.730273 | 0.596904 | 2.898745 | 0.003747 | 0.146338 | 0.834644 |
| ENSMUSG00000098098.8 | 16.7567 | 7.558385 | 2.611432 | 2.894345 | 0.0038 | 0.14642 | 0.8344 |
| ENSMUSG00000004508.7 | 17.03723 | 7.57899 | 2.623488 | 2.888898 | 0.003866 | 0.147797 | 0.830333 |
| ENSMUSG00000024544.10 | 17.35381 | -7.64515 | 2.646426 | -2.88886 | 0.003866 | 0.147797 | 0.830333 |
| ENSMUSG00000089810.2 | 65.30347 | -2.67783 | 0.928675 | -2.8835 | 0.003933 | 0.149732 | 0.824684 |
| ENSMUSG00000013701.14 | 29.38127 | 4.287307 | 1.487654 | 2.881924 | 0.003953 | 0.149881 | 0.824255 |
| ENSMUSG00000097093.3 | 12.34105 | 7.112409 | 2.46928 | 2.880357 | 0.003972 | 0.150028 | 0.823829 |
| ENSMUSG00000070544.7 | 1240.084 | 0.989635 | 0.344102 | 2.875989 | 0.004028 | 0.150917 | 0.821262 |
| ENSMUSG00000085525.8 | 16.88451 | 7.56608 | 2.629663 | 2.877205 | 0.004012 | 0.150917 | 0.821262 |
| ENSMUSG00000033192.6 | 17.83593 | 7.646821 | 2.663579 | 2.870882 | 0.004093 | 0.15158 | 0.819359 |
| ENSMUSG00000068270.16 | 17.63225 | 7.626276 | 2.655655 | 2.871712 | 0.004083 | 0.15158 | 0.819359 |
| ENSMUSG00000115946.3 | 12.60881 | -7.18564 | 2.501563 | -2.87246 | 0.004073 | 0.15158 | 0.819359 |
| ENSMUSG00000068267.6 | 17.09454 | -6.58122 | 2.29355 | -2.86945 | 0.004112 | 0.151676 | 0.819084 |
| ENSMUSG00000023963.11 | 149.9263 | 2.226971 | 0.776468 | 2.868077 | 0.00413 | 0.151744 | 0.818888 |
| ENSMUSG00000040591.19 | 16.78314 | 7.556927 | 2.638513 | 2.864085 | 0.004182 | 0.152488 | 0.816765 |
| ENSMUSG00000057229.12 | 12.28213 | 7.106215 | 2.48108 | 2.864161 | 0.004181 | 0.152488 | 0.816765 |
| ENSMUSG00000054263.13 | 1626.996 | -0.83211 | 0.290702 | -2.86243 | 0.004204 | 0.152699 | 0.816164 |
| ENSMUSG00000011256.17 | 17.0043 | -7.61604 | 2.666352 | -2.85635 | 0.004285 | 0.154469 | 0.811159 |
| ENSMUSG00000057551.11 | 70.91454 | 3.285645 | 1.149966 | 2.857166 | 0.004274 | 0.154469 | 0.811159 |

| | | | | | | | |
|---|---|---|---|---|---|---|---|
| ENSMUSG00000020287.16 | 55.1146 | 2.854593 | 1.000893 | 2.852045 | 0.004344 | 0.154747 | 0.810378 |
| ENSMUSG00000035357.18 | 23.77376 | 5.304089 | 1.860562 | 2.8508 | 0.004361 | 0.154747 | 0.810378 |
| ENSMUSG00000037916.15 | 412.5429 | -1.44467 | 0.50691 | -2.84995 | 0.004373 | 0.154747 | 0.810378 |
| ENSMUSG00000057103.12 | 1716.565 | -0.70114 | 0.246031 | -2.8498 | 0.004375 | 0.154747 | 0.810378 |
| ENSMUSG00000107390.2 | 17.75214 | 7.635797 | 2.676638 | 2.852757 | 0.004334 | 0.154747 | 0.810378 |
| ENSMUSG00000049657.10 | 12.64849 | 7.146126 | 2.509099 | 2.848085 | 0.004398 | 0.155004 | 0.809657 |
| ENSMUSG00000047554.15 | 71.88494 | 2.49281 | 0.876413 | 2.844334 | 0.00445 | 0.15626 | 0.806152 |
| ENSMUSG00000022770.18 | 202.885 | -1.4314 | 0.50362 | -2.84223 | 0.00448 | 0.156716 | 0.804887 |
| ENSMUSG00000023079.15 | 16.03651 | 7.496298 | 2.645283 | 2.833836 | 0.004599 | 0.159968 | 0.795966 |
| ENSMUSG00000035455.13 | 17.29398 | 7.59815 | 2.68171 | 2.833323 | 0.004607 | 0.159968 | 0.795966 |
| ENSMUSG00000042419.9 | 15.95315 | -7.52689 | 2.659493 | -2.8302 | 0.004652 | 0.16095 | 0.793309 |
| ENSMUSG00000028882.14 | 24.22004 | 4.500898 | 1.591049 | 2.828888 | 0.004671 | 0.161022 | 0.793114 |
| ENSMUSG00000024747.4 | 2757.621 | 0.718399 | 0.254055 | 2.827724 | 0.004688 | 0.161023 | 0.793111 |
| ENSMUSG00000017707.10 | 2063.112 | 0.750374 | 0.265699 | 2.824146 | 0.004741 | 0.162244 | 0.789831 |
| ENSMUSG00000013787.16 | 11.72964 | -7.08269 | 2.513647 | -2.81769 | 0.004837 | 0.164411 | 0.78407 |
| ENSMUSG00000039210.17 | 49.36068 | -2.80386 | 0.995535 | -2.81643 | 0.004856 | 0.164411 | 0.78407 |
| ENSMUSG00000050103.19 | 416.0402 | 1.121425 | 0.398152 | 2.816572 | 0.004854 | 0.164411 | 0.78407 |
| ENSMUSG00000028318.15 | 12.4196 | -7.16346 | 2.545866 | -2.81376 | 0.004897 | 0.164616 | 0.783527 |
| ENSMUSG00000045064.6 | 17.03681 | 7.576512 | 2.69268 | 2.813744 | 0.004897 | 0.164616 | 0.783527 |
| ENSMUSG00000054072.13 | 1677.955 | -0.73162 | 0.260389 | -2.8097 | 0.004959 | 0.165799 | 0.780419 |
| ENSMUSG00000108414.2 | 68.37389 | 2.750156 | 0.978993 | 2.809169 | 0.004967 | 0.165799 | 0.780419 |
| ENSMUSG00000022591.6 | 12.24139 | 7.106618 | 2.531787 | 2.806957 | 0.005001 | 0.166355 | 0.778963 |
| ENSMUSG00000036775.14 | 1373.152 | 0.760108 | 0.271282 | 2.801908 | 0.00508 | 0.168064 | 0.774524 |
| ENSMUSG00000037685.16 | 35.95349 | 4.571072 | 1.631706 | 2.801407 | 0.005088 | 0.168064 | 0.774524 |
| ENSMUSG00000067928.7 | 16.40815 | -7.56426 | 2.706029 | -2.79534 | 0.005185 | 0.170659 | 0.767871 |
| ENSMUSG00000018559.17 | 41.88486 | -3.64828 | 1.307427 | -2.79043 | 0.005264 | 0.172668 | 0.762787 |
| ENSMUSG00000003955.9 | 786.2858 | 0.829933 | 0.297801 | 2.786869 | 0.005322 | 0.173377 | 0.76101 |
| ENSMUSG00000038859.8 | 67.81312 | 2.426711 | 0.870721 | 2.787013 | 0.00532 | 0.173377 | 0.76101 |
| ENSMUSG00000012187.14 | 15.45875 | 7.443268 | 2.673232 | 2.78437 | 0.005363 | 0.17412 | 0.759152 |
| ENSMUSG00000045464.6 | 12.47017 | 7.13166 | 2.565218 | 2.780138 | 0.005434 | 0.175803 | 0.754973 |
| ENSMUSG00000031858.17 | 15.52273 | -7.4887 | 2.695392 | -2.77833 | 0.005464 | 0.176181 | 0.754042 |
| ENSMUSG00000047215.15 | 3017.591 | 0.662094 | 0.238963 | 2.770699 | 0.005594 | 0.179754 | 0.745322 |
| ENSMUSG00000037849.9 | 15.28166 | -7.4641 | 2.695634 | -2.76896 | 0.005624 | 0.179833 | 0.745131 |
| ENSMUSG00000041815.15 | 33.11383 | -4.84724 | 1.750946 | -2.76836 | 0.005634 | 0.179833 | 0.745131 |
| ENSMUSG00000019590.17 | 54.02845 | -3.59549 | 1.299653 | -2.7665 | 0.005666 | 0.180252 | 0.74412 |
| ENSMUSG00000103653.2 | 2853.965 | -1.0257 | 0.371344 | -2.76214 | 0.005742 | 0.182064 | 0.739775 |
| ENSMUSG00000039801.8 | 21.16338 | -6.08815 | 2.205929 | -2.7599 | 0.005782 | 0.182706 | 0.738247 |
| ENSMUSG00000023025.17 | 288.1133 | -1.73301 | 0.628561 | -2.75711 | 0.005831 | 0.183055 | 0.737419 |
| ENSMUSG00000027678.18 | 81.92214 | 3.028657 | 1.098236 | 2.757746 | 0.00582 | 0.183055 | 0.737419 |
| ENSMUSG00000015748.14 | 14.15076 | -6.30387 | 2.291716 | -2.75072 | 0.005946 | 0.18352 | 0.736316 |
| ENSMUSG00000020238.15 | 18.21975 | -6.79162 | 2.470805 | -2.74875 | 0.005982 | 0.18352 | 0.736316 |
| ENSMUSG00000021134.18 | 106.4061 | -1.94823 | 0.707385 | -2.75413 | 0.005885 | 0.18352 | 0.736316 |
| ENSMUSG00000021775.12 | 434.153 | -1.42772 | 0.519394 | -2.74881 | 0.005981 | 0.18352 | 0.736316 |
| ENSMUSG00000024781.17 | 1337.576 | 0.934539 | 0.340115 | 2.747717 | 0.006001 | 0.18352 | 0.736316 |

| | | | | | | | |
|---|---|---|---|---|---|---|---|
| ENSMUSG00000028195.5 | 55.95731 | -3.55906 | 1.295243 | -2.74779 | 0.006 | 0.18352 | 0.736316 |
| ENSMUSG00000047022.19 | 42.09023 | -3.33423 | 1.211648 | -2.75181 | 0.005927 | 0.18352 | 0.736316 |
| ENSMUSG00000078674.3 | 4313.176 | -2.04416 | 0.743801 | -2.74826 | 0.005991 | 0.18352 | 0.736316 |
| ENSMUSG00000002763.17 | 11.28624 | 6.98892 | 2.545383 | 2.745724 | 0.006038 | 0.184045 | 0.735077 |
| ENSMUSG00000028741.14 | 98.27948 | 2.053886 | 0.749225 | 2.741348 | 0.006119 | 0.185917 | 0.730682 |
| ENSMUSG00000032582.15 | 41.18786 | -3.68438 | 1.345572 | -2.73816 | 0.006178 | 0.187131 | 0.727854 |
| ENSMUSG00000030156.6 | 15.09975 | -7.44563 | 2.724472 | -2.73287 | 0.006278 | 0.188355 | 0.725023 |
| ENSMUSG00000038773.12 | 14.88524 | 7.388047 | 2.702175 | 2.734111 | 0.006255 | 0.188355 | 0.725023 |
| ENSMUSG00000049225.15 | 15.85192 | 7.472612 | 2.733532 | 2.733683 | 0.006263 | 0.188355 | 0.725023 |
| ENSMUSG00000018160.16 | 128.5206 | -1.78546 | 0.654281 | -2.72889 | 0.006355 | 0.189658 | 0.722029 |
| ENSMUSG00000021474.10 | 996.6705 | 0.894054 | 0.32767 | 2.728519 | 0.006362 | 0.189658 | 0.722029 |
| ENSMUSG00000039148.6 | 60.11496 | -2.93391 | 1.076147 | -2.72631 | 0.006405 | 0.190335 | 0.720482 |
| ENSMUSG00000020472.15 | 55.49966 | -2.91303 | 1.069654 | -2.72334 | 0.006463 | 0.190961 | 0.719054 |
| ENSMUSG00000032192.10 | 15.75866 | 7.464052 | 2.740954 | 2.723159 | 0.006466 | 0.190961 | 0.719054 |
| ENSMUSG00000086561.2 | 13.82398 | 7.273544 | 2.67249 | 2.721636 | 0.006496 | 0.191248 | 0.718404 |
| ENSMUSG00000025511.16 | 78.75043 | 2.318251 | 0.853432 | 2.716386 | 0.0066 | 0.193706 | 0.712858 |
| ENSMUSG00000067873.12 | 546.844 | 1.093052 | 0.402715 | 2.714205 | 0.006643 | 0.194384 | 0.71134 |
| ENSMUSG00000027184.15 | 440.5382 | 1.062912 | 0.392246 | 2.709809 | 0.006732 | 0.196373 | 0.706918 |
| ENSMUSG00000051864.11 | 63.4366 | -3.59512 | 1.32814 | -2.70688 | 0.006792 | 0.197505 | 0.704423 |
| ENSMUSG00000029228.16 | 15.09877 | -7.44975 | 2.755118 | -2.70397 | 0.006852 | 0.19803 | 0.703268 |
| ENSMUSG00000034785.16 | 791.5772 | -0.84041 | 0.31077 | -2.70429 | 0.006845 | 0.19803 | 0.703268 |
| ENSMUSG00000020038.11 | 125.3527 | 1.841623 | 0.681667 | 2.701645 | 0.0069 | 0.198358 | 0.702551 |
| ENSMUSG00000035266.17 | 14.21042 | 6.371936 | 2.358755 | 2.701398 | 0.006905 | 0.198358 | 0.702551 |
| ENSMUSG00000026177.12 | 15.29642 | -7.4686 | 2.766823 | -2.69934 | 0.006948 | 0.198984 | 0.701181 |
| ENSMUSG00000026017.14 | 28.86159 | 4.170355 | 1.546834 | 2.696059 | 0.007017 | 0.200351 | 0.698208 |
| ENSMUSG00000019715.15 | 77.09087 | -2.34705 | 0.872949 | -2.68865 | 0.007174 | 0.201216 | 0.696338 |
| ENSMUSG00000024812.12 | 86.41405 | 2.114442 | 0.78502 | 2.693488 | 0.007071 | 0.201216 | 0.696338 |
| ENSMUSG00000031605.9 | 219.7817 | -1.47685 | 0.548794 | -2.69109 | 0.007122 | 0.201216 | 0.696338 |
| ENSMUSG00000038418.8 | 428.2981 | -7.73926 | 2.876627 | -2.69039 | 0.007137 | 0.201216 | 0.696338 |
| ENSMUSG00000062181.15 | 2030.548 | -0.71497 | 0.265827 | -2.68961 | 0.007154 | 0.201216 | 0.696338 |
| ENSMUSG00000063019.6 | 12.41604 | -6.23268 | 2.314756 | -2.69259 | 0.00709 | 0.201216 | 0.696338 |
| ENSMUSG00000031075.20 | 14.36428 | 7.336384 | 2.734371 | 2.683025 | 0.007296 | 0.204029 | 0.690309 |
| ENSMUSG00000054312.7 | 221.2749 | 1.519694 | 0.566903 | 2.680694 | 0.007347 | 0.204851 | 0.688562 |
| ENSMUSG00000002944.16 | 129.6798 | 2.280097 | 0.851889 | 2.676519 | 0.007439 | 0.204869 | 0.688523 |
| ENSMUSG00000023170.15 | 18.52565 | 5.624775 | 2.102111 | 2.675774 | 0.007456 | 0.204869 | 0.688523 |
| ENSMUSG00000028553.13 | 5549.677 | 0.54229 | 0.202454 | 2.678583 | 0.007393 | 0.204869 | 0.688523 |
| ENSMUSG00000028744.16 | 19.5157 | -5.49434 | 2.051694 | -2.67795 | 0.007407 | 0.204869 | 0.688523 |
| ENSMUSG00000038301.16 | 285.9187 | -1.3679 | 0.511178 | -2.67598 | 0.007451 | 0.204869 | 0.688523 |
| ENSMUSG00000002227.16 | 69.0622 | -3.24916 | 1.216814 | -2.67022 | 0.00758 | 0.206496 | 0.685089 |
| ENSMUSG00000030067.18 | 63.37457 | -2.32529 | 0.870736 | -2.67048 | 0.007574 | 0.206496 | 0.685089 |
| ENSMUSG00000116858.2 | 16.73105 | -4.52032 | 1.692309 | -2.6711 | 0.00756 | 0.206496 | 0.685089 |
| ENSMUSG00000024525.8 | 22.4035 | 4.974417 | 1.869308 | 2.661101 | 0.007789 | 0.210358 | 0.677042 |
| ENSMUSG00000030750.14 | 46.85111 | -2.92381 | 1.098473 | -2.6617 | 0.007775 | 0.210358 | 0.677042 |
| ENSMUSG00000030970.17 | 17.1579 | -5.6499 | 2.121851 | -2.66272 | 0.007751 | 0.210358 | 0.677042 |

| | | | | | | | |
|---|---|---|---|---|---|---|---|
| ENSMUSG00000026095.16 | 130.3523 | 1.862345 | 0.702526 | 2.650925 | 0.008027 | 0.215606 | 0.66634 |
| ENSMUSG00000048486.7 | 84.96082 | 2.458818 | 0.92755 | 2.650875 | 0.008028 | 0.215606 | 0.66634 |
| ENSMUSG00000042660.9 | 137.2219 | -2.02267 | 0.763819 | -2.6481 | 0.008095 | 0.216771 | 0.663998 |
| ENSMUSG00000022853.8 | 354.886 | 1.1467 | 0.433329 | 2.64626 | 0.008139 | 0.217338 | 0.662863 |
| ENSMUSG00000049922.9 | 76.58008 | 2.724085 | 1.029891 | 2.645022 | 0.008169 | 0.217522 | 0.662496 |
| ENSMUSG00000036298.11 | 13.93424 | -7.33085 | 2.773494 | -2.64318 | 0.008213 | 0.218094 | 0.661356 |
| ENSMUSG00000022678.17 | 44.90898 | 3.398633 | 1.286658 | 2.641442 | 0.008255 | 0.21836 | 0.660827 |
| ENSMUSG00000028150.15 | 142.5395 | 1.987377 | 0.752544 | 2.640879 | 0.008269 | 0.21836 | 0.660827 |
| ENSMUSG00000029022.19 | 25.61441 | -5.58728 | 2.11726 | -2.63892 | 0.008317 | 0.218409 | 0.66073 |
| ENSMUSG00000036894.4 | 16.73602 | -6.53905 | 2.477314 | -2.63957 | 0.008301 | 0.218409 | 0.66073 |
| ENSMUSG00000009733.10 | 17.49889 | 5.170606 | 1.962204 | 2.635102 | 0.008411 | 0.219791 | 0.65799 |
| ENSMUSG00000028410.14 | 1664.375 | 0.815345 | 0.309659 | 2.63304 | 0.008462 | 0.219791 | 0.65799 |
| ENSMUSG00000032598.9 | 14.24189 | -7.36067 | 2.795456 | -2.63308 | 0.008461 | 0.219791 | 0.65799 |
| ENSMUSG00000045751.17 | 14.33508 | -7.36994 | 2.798751 | -2.6333 | 0.008456 | 0.219791 | 0.65799 |
| ENSMUSG00000006522.18 | 2922.59 | -0.7884 | 0.299728 | -2.63038 | 0.008529 | 0.220914 | 0.655777 |
| ENSMUSG00000026434.13 | 826.9322 | 0.808039 | 0.307657 | 2.626432 | 0.008629 | 0.222884 | 0.651921 |
| ENSMUSG00000066406.17 | 130.5852 | -1.67134 | 0.638253 | -2.61862 | 0.008829 | 0.227434 | 0.643146 |
| ENSMUSG00000044505.6 | 13.96117 | 7.294158 | 2.786995 | 2.617212 | 0.008865 | 0.227754 | 0.642533 |
| ENSMUSG00000022386.13 | 14.44033 | -7.38063 | 2.833689 | -2.6046 | 0.009198 | 0.231588 | 0.635283 |
| ENSMUSG00000026003.6 | 1152.156 | 0.823268 | 0.315518 | 2.609253 | 0.009074 | 0.231588 | 0.635283 |
| ENSMUSG00000030104.10 | 788.1247 | 0.811919 | 0.311777 | 2.604167 | 0.00921 | 0.231588 | 0.635283 |
| ENSMUSG00000031775.6 | 32.4619 | 3.841648 | 1.474689 | 2.605056 | 0.009186 | 0.231588 | 0.635283 |
| ENSMUSG00000034171.14 | 195.6516 | -1.484 | 0.568979 | -2.60818 | 0.009102 | 0.231588 | 0.635283 |
| ENSMUSG00000036641.10 | 10.61925 | -6.93607 | 2.657 | -2.61049 | 0.009041 | 0.231588 | 0.635283 |
| ENSMUSG00000073412.7 | 16.81019 | 6.540162 | 2.509737 | 2.605915 | 0.009163 | 0.231588 | 0.635283 |
| ENSMUSG00000107091.2 | 11.74411 | 7.041032 | 2.700575 | 2.607234 | 0.009128 | 0.231588 | 0.635283 |
| ENSMUSG00000032418.16 | 646.8098 | 1.267621 | 0.487294 | 2.601349 | 0.009286 | 0.232882 | 0.632864 |
| ENSMUSG00000061353.12 | 1661.947 | 0.882746 | 0.339522 | 2.599965 | 0.009323 | 0.233206 | 0.63226 |
| ENSMUSG00000026116.12 | 46.82439 | 2.668392 | 1.028054 | 2.595576 | 0.009443 | 0.234966 | 0.628996 |
| ENSMUSG00000041431.17 | 13.78406 | -7.31286 | 2.816788 | -2.59617 | 0.009427 | 0.234966 | 0.628996 |
| ENSMUSG00000045917.18 | 10.59042 | -6.93879 | 2.680174 | -2.58893 | 0.009627 | 0.238298 | 0.62288 |
| ENSMUSG00000064225.7 | 1596.81 | 0.85468 | 0.330117 | 2.589021 | 0.009625 | 0.238298 | 0.62288 |
| ENSMUSG00000039943.17 | 26.93599 | 4.725107 | 1.82689 | 2.586422 | 0.009698 | 0.239128 | 0.621369 |
| ENSMUSG00000047123.9 | 13.5678 | 7.253069 | 2.805782 | 2.585044 | 0.009737 | 0.239128 | 0.621369 |
| ENSMUSG00000069922.13 | 6150.449 | -0.64547 | 0.249659 | -2.58542 | 0.009726 | 0.239128 | 0.621369 |
| ENSMUSG00000024213.15 | 31.7522 | -3.59149 | 1.391445 | -2.58112 | 0.009848 | 0.239383 | 0.620907 |
| ENSMUSG00000029518.16 | 34.03222 | -3.97159 | 1.53824 | -2.58191 | 0.009826 | 0.239383 | 0.620907 |
| ENSMUSG00000042650.4 | 142.7303 | -1.73237 | 0.670484 | -2.58376 | 0.009773 | 0.239383 | 0.620907 |
| ENSMUSG00000047446.19 | 253.4457 | 1.389969 | 0.538338 | 2.581964 | 0.009824 | 0.239383 | 0.620907 |
| ENSMUSG00000004558.16 | 2910.672 | -0.69668 | 0.270774 | -2.57293 | 0.010084 | 0.239861 | 0.62004 |
| ENSMUSG00000010663.6 | 1242.49 | 0.727868 | 0.282596 | 2.575649 | 0.010005 | 0.239861 | 0.62004 |
| ENSMUSG00000021257.15 | 10.17797 | -6.87558 | 2.67182 | -2.57337 | 0.010071 | 0.239861 | 0.62004 |
| ENSMUSG00000022707.17 | 1050.933 | 0.804206 | 0.312717 | 2.571677 | 0.010121 | 0.239861 | 0.62004 |
| ENSMUSG00000022868.7 | 16916.47 | -0.63569 | 0.247026 | -2.57336 | 0.010072 | 0.239861 | 0.62004 |

| | | | | | | | |
|---|---|---|---|---|---|---|---|
| ENSMUSG00000030849.19 | 93.06574 | -2.73345 | 1.061262 | -2.57566 | 0.010005 | 0.239861 | 0.62004 |
| ENSMUSG00000042429.9 | 36.89835 | 3.935805 | 1.5273 | 2.576969 | 0.009967 | 0.239861 | 0.62004 |
| ENSMUSG00000057933.11 | 526.6884 | 1.433108 | 0.556715 | 2.57422 | 0.010047 | 0.239861 | 0.62004 |
| ENSMUSG00000063047.10 | 24.11289 | 3.632323 | 1.412123 | 2.572242 | 0.010104 | 0.239861 | 0.62004 |
| ENSMUSG00000107012.2 | 22.81346 | 5.916416 | 2.294759 | 2.57823 | 0.009931 | 0.239861 | 0.62004 |
| ENSMUSG00000003099.12 | 10.42435 | 6.873443 | 2.675444 | 2.569085 | 0.010197 | 0.239864 | 0.620035 |
| ENSMUSG00000022419.17 | 1366.941 | -0.74492 | 0.289885 | -2.56971 | 0.010178 | 0.239864 | 0.620035 |
| ENSMUSG00000108738.2 | 14.94431 | -6.3762 | 2.480286 | -2.57075 | 0.010148 | 0.239864 | 0.620035 |
| ENSMUSG00000020386.6 | 1717.811 | 0.697299 | 0.271705 | 2.566386 | 0.010276 | 0.24114 | 0.617731 |
| ENSMUSG00000024371.15 | 637.0374 | -1.05668 | 0.41195 | -2.56508 | 0.010315 | 0.241453 | 0.617167 |
| ENSMUSG00000082609.2 | 12.16148 | -7.13226 | 2.783928 | -2.56194 | 0.010409 | 0.243045 | 0.614314 |
| ENSMUSG00000020022.18 | 490.2043 | 0.971036 | 0.379394 | 2.559441 | 0.010484 | 0.244199 | 0.612256 |
| ENSMUSG00000029311.17 | 1613.369 | 0.710918 | 0.278189 | 2.55552 | 0.010603 | 0.246362 | 0.608426 |
| ENSMUSG00000090659.10 | 10.99799 | 6.946541 | 2.72641 | 2.547871 | 0.010838 | 0.251214 | 0.599955 |
| ENSMUSG00000010755.18 | 22.00808 | -4.69215 | 1.846828 | -2.54065 | 0.011065 | 0.252976 | 0.596921 |
| ENSMUSG00000034528.16 | 3334.63 | 0.573527 | 0.225729 | 2.540777 | 0.011061 | 0.252976 | 0.596921 |
| ENSMUSG00000041220.11 | 973.6187 | 0.855057 | 0.336571 | 2.540495 | 0.01107 | 0.252976 | 0.596921 |
| ENSMUSG00000042323.18 | 399.6622 | -1.21762 | 0.479314 | -2.54034 | 0.011074 | 0.252976 | 0.596921 |
| ENSMUSG00000056888.12 | 13.5922 | -7.2925 | 2.869165 | -2.54168 | 0.011032 | 0.252976 | 0.596921 |
| ENSMUSG00000079215.9 | 155.9023 | 1.720215 | 0.67604 | 2.544546 | 0.010942 | 0.252976 | 0.596921 |
| ENSMUSG00000030876.8 | 528.9112 | 1.350495 | 0.531931 | 2.538855 | 0.011122 | 0.253444 | 0.596118 |
| ENSMUSG00000020773.12 | 13.42199 | 7.236906 | 2.851529 | 2.537904 | 0.011152 | 0.253525 | 0.59598 |
| ENSMUSG00000037608.17 | 220.7598 | -1.37662 | 0.543192 | -2.53431 | 0.011267 | 0.255523 | 0.592569 |
| ENSMUSG00000032042.12 | 254.2745 | -1.54764 | 0.611871 | -2.52935 | 0.011427 | 0.258547 | 0.587461 |
| ENSMUSG00000021930.15 | 287.885 | 1.272088 | 0.503586 | 2.52606 | 0.011535 | 0.260361 | 0.584424 |
| ENSMUSG00000066154.12 | 47872.45 | -0.6678 | 0.26451 | -2.52466 | 0.011581 | 0.260783 | 0.583721 |
| ENSMUSG00000033781.8 | 179.2104 | 1.385446 | 0.549607 | 2.520792 | 0.011709 | 0.262417 | 0.581008 |
| ENSMUSG00000068220.7 | 128.4244 | 2.07413 | 0.822631 | 2.521338 | 0.011691 | 0.262417 | 0.581008 |
| ENSMUSG00000001783.4 | 53.78204 | -2.40822 | 0.956537 | -2.51765 | 0.011814 | 0.262673 | 0.580585 |
| ENSMUSG00000021268.18 | 26.47815 | -4.51561 | 1.792556 | -2.51909 | 0.011766 | 0.262673 | 0.580585 |
| ENSMUSG00000059173.20 | 14.03232 | 7.300325 | 2.900252 | 2.517135 | 0.011831 | 0.262673 | 0.580585 |
| ENSMUSG00000060708.5 | 99.16776 | 2.007539 | 0.79731 | 2.51789 | 0.011806 | 0.262673 | 0.580585 |
| ENSMUSG00000051978.10 | 15.53259 | -5.51532 | 2.192003 | -2.51611 | 0.011866 | 0.262821 | 0.58034 |
| ENSMUSG00000030494.14 | 33.17436 | -3.08999 | 1.229588 | -2.51303 | 0.01197 | 0.263895 | 0.578569 |
| ENSMUSG00000039706.12 | 19.82584 | 4.256721 | 1.693738 | 2.513211 | 0.011964 | 0.263895 | 0.578569 |
| ENSMUSG00000030678.8 | 12.68569 | 7.156732 | 2.849103 | 2.511925 | 0.012007 | 0.264109 | 0.578217 |
| ENSMUSG00000003808.19 | 12.49838 | -7.17593 | 2.872121 | -2.49848 | 0.012473 | 0.270358 | 0.568061 |
| ENSMUSG00000004771.13 | 575.5064 | -1.00917 | 0.403998 | -2.49795 | 0.012491 | 0.270358 | 0.568061 |
| ENSMUSG00000007872.4 | 605.5242 | 1.175944 | 0.470424 | 2.499752 | 0.012428 | 0.270358 | 0.568061 |
| ENSMUSG00000020027.19 | 91.83632 | -2.05791 | 0.822793 | -2.50113 | 0.01238 | 0.270358 | 0.568061 |
| ENSMUSG00000031444.17 | 1451.789 | -0.68772 | 0.274829 | -2.50238 | 0.012336 | 0.270358 | 0.568061 |
| ENSMUSG00000062624.6 | 1103.261 | 0.909689 | 0.363714 | 2.501115 | 0.01238 | 0.270358 | 0.568061 |
| ENSMUSG00000115919.2 | 269.0022 | -1.70023 | 0.680271 | -2.49935 | 0.012442 | 0.270358 | 0.568061 |
| ENSMUSG00000021693.21 | 19.80698 | 4.784642 | 1.917422 | 2.495351 | 0.012583 | 0.271364 | 0.566448 |

| | | | | | | | |
|---|---|---|---|---|---|---|---|
| ENSMUSG00000037580.11 | 975.9175 | -0.91235 | 0.365851 | -2.49379 | 0.012639 | 0.271364 | 0.566448 |
| ENSMUSG00000048546.9 | 159.0106 | -1.79902 | 0.721111 | -2.49479 | 0.012603 | 0.271364 | 0.566448 |
| ENSMUSG00000057440.9 | 45.98702 | -3.49131 | 1.400664 | -2.49261 | 0.012681 | 0.271364 | 0.566448 |
| ENSMUSG00000065847.3 | 12.63313 | 7.150153 | 2.867865 | 2.493197 | 0.01266 | 0.271364 | 0.566448 |
| ENSMUSG00000005677.15 | 129.4677 | 1.8527 | 0.743958 | 2.490327 | 0.012763 | 0.272498 | 0.564637 |
| ENSMUSG00000006445.4 | 12.72993 | 7.157133 | 2.879671 | 2.485399 | 0.012941 | 0.275679 | 0.559597 |
| ENSMUSG00000026688.6 | 19.1752 | 4.747065 | 1.91227 | 2.482424 | 0.013049 | 0.276748 | 0.557916 |
| ENSMUSG00000049404.8 | 971.5989 | -0.80512 | 0.324252 | -2.483 | 0.013028 | 0.276748 | 0.557916 |
| ENSMUSG00000026526.15 | 2593.198 | 0.651715 | 0.262875 | 2.479178 | 0.013169 | 0.278099 | 0.5558 |
| ENSMUSG00000048856.10 | 449.2222 | -1.37854 | 0.556067 | -2.4791 | 0.013172 | 0.278099 | 0.5558 |
| ENSMUSG00000038541.9 | 13.44634 | -7.27699 | 2.939097 | -2.47593 | 0.013289 | 0.279957 | 0.552908 |
| ENSMUSG00000015850.12 | 12.47708 | -7.1704 | 2.900142 | -2.47243 | 0.01342 | 0.281458 | 0.550586 |
| ENSMUSG00000032515.9 | 12.63016 | -7.18769 | 2.906847 | -2.47268 | 0.013411 | 0.281458 | 0.550586 |
| ENSMUSG00000042677.8 | 10.11099 | -6.86918 | 2.779472 | -2.4714 | 0.013459 | 0.281651 | 0.550289 |
| ENSMUSG00000035811.11 | 1653.864 | -0.79425 | 0.321928 | -2.46718 | 0.013618 | 0.284364 | 0.546125 |
| ENSMUSG00000022462.8 | 968.5074 | -0.93705 | 0.380132 | -2.46507 | 0.013699 | 0.28479 | 0.545475 |
| ENSMUSG00000032310.5 | 3004.582 | -0.72909 | 0.295739 | -2.46532 | 0.013689 | 0.28479 | 0.545475 |
| ENSMUSG00000078671.12 | 73.94458 | -2.30482 | 0.936143 | -2.46204 | 0.013815 | 0.286578 | 0.542758 |
| ENSMUSG00000035585.17 | 16.33905 | -6.05659 | 2.463397 | -2.45863 | 0.013947 | 0.288679 | 0.539585 |
| ENSMUSG00000025764.15 | 317.7819 | -1.10604 | 0.450346 | -2.45597 | 0.01405 | 0.290191 | 0.537317 |
| ENSMUSG00000020803.4 | 903.2295 | 0.777159 | 0.317116 | 2.450706 | 0.014258 | 0.292916 | 0.533256 |
| ENSMUSG00000033159.15 | 98.58718 | -2.22756 | 0.908689 | -2.4514 | 0.01423 | 0.292916 | 0.533256 |
| ENSMUSG00000074227.13 | 301.7329 | -1.45433 | 0.593541 | -2.45027 | 0.014275 | 0.292916 | 0.533256 |
| ENSMUSG00000031214.14 | 18.9271 | 5.719113 | 2.335775 | 2.448487 | 0.014346 | 0.293732 | 0.532049 |
| ENSMUSG00000055538.8 | 382.9682 | -0.96285 | 0.393384 | -2.44761 | 0.014381 | 0.29381 | 0.531933 |
| ENSMUSG00000020492.12 | 12.68694 | 7.150999 | 2.924427 | 2.445265 | 0.014475 | 0.294461 | 0.530972 |
| ENSMUSG00000060419.9 | 334.9758 | 1.024831 | 0.418984 | 2.445992 | 0.014445 | 0.294461 | 0.530972 |
| ENSMUSG00000027668.14 | 82.98988 | -1.80007 | 0.736953 | -2.44258 | 0.014583 | 0.296024 | 0.528673 |
| ENSMUSG00000006021.6 | 12.11352 | -7.12752 | 2.920533 | -2.44049 | 0.014667 | 0.296217 | 0.52839 |
| ENSMUSG00000020572.9 | 539.079 | 0.957632 | 0.392647 | 2.438913 | 0.014732 | 0.296217 | 0.52839 |
| ENSMUSG00000026958.14 | 48.19109 | 3.14605 | 1.290158 | 2.4385 | 0.014748 | 0.296217 | 0.52839 |
| ENSMUSG00000030340.17 | 35.67261 | 3.196383 | 1.309486 | 2.440945 | 0.014649 | 0.296217 | 0.52839 |
| ENSMUSG00000030790.16 | 12.85835 | -5.70182 | 2.337606 | -2.43917 | 0.014721 | 0.296217 | 0.52839 |
| ENSMUSG00000022090.11 | 12.16205 | -7.13337 | 2.929955 | -2.43464 | 0.014907 | 0.298767 | 0.524668 |
| ENSMUSG00000020538.16 | 307.0925 | -1.30846 | 0.538053 | -2.43184 | 0.015022 | 0.29918 | 0.524067 |
| ENSMUSG00000025856.16 | 47.77422 | 2.486008 | 1.022213 | 2.431987 | 0.015016 | 0.29918 | 0.524067 |
| ENSMUSG00000096917.3 | 12.82569 | -7.20875 | 2.962636 | -2.43322 | 0.014965 | 0.29918 | 0.524067 |
| ENSMUSG00000063382.7 | 26.88065 | 3.338501 | 1.373259 | 2.431079 | 0.015054 | 0.299185 | 0.52406 |
| ENSMUSG00000025510.15 | 121.385 | -1.57389 | 0.648663 | -2.42637 | 0.015251 | 0.302465 | 0.519325 |
| ENSMUSG00000039069.20 | 40.17873 | -4.19274 | 1.730226 | -2.42323 | 0.015383 | 0.304451 | 0.516482 |
| ENSMUSG00000060183.7 | 12.97191 | 7.182618 | 2.966428 | 2.421302 | 0.015465 | 0.305434 | 0.515082 |
| ENSMUSG00000018169.13 | 12.90103 | 7.179406 | 2.96636 | 2.420275 | 0.015509 | 0.305662 | 0.514759 |
| ENSMUSG00000020743.16 | 133.1685 | 1.976762 | 0.817784 | 2.417218 | 0.01564 | 0.307185 | 0.5126 |
| ENSMUSG00000027340.16 | 48.89833 | -2.40918 | 0.996781 | -2.41696 | 0.015651 | 0.307185 | 0.5126 |

| | | | | | | | |
|---|---|---|---|---|---|---|---|
| ENSMUSG00000000631.21 | 57.02902 | -2.38666 | 0.988396 | -2.41468 | 0.015749 | 0.308471 | 0.510785 |
| ENSMUSG00000010307.8 | 21.81371 | -4.98108 | 2.064021 | -2.41329 | 0.015809 | 0.308964 | 0.510093 |
| ENSMUSG00000039982.9 | 19.32066 | -4.91136 | 2.035715 | -2.4126 | 0.015839 | 0.308964 | 0.510093 |
| ENSMUSG00000085704.2 | 16.16586 | -6.04079 | 2.505498 | -2.41101 | 0.015908 | 0.309038 | 0.509987 |
| ENSMUSG00000110838.2 | 12.117 | 7.086064 | 2.938264 | 2.41165 | 0.015881 | 0.309038 | 0.509987 |
| ENSMUSG00000023150.15 | 388.0233 | 1.093992 | 0.454059 | 2.40936 | 0.015981 | 0.309807 | 0.508909 |
| ENSMUSG00000024943.10 | 94.32081 | -2.26042 | 0.938954 | -2.40739 | 0.016067 | 0.310851 | 0.507448 |
| ENSMUSG00000027188.9 | 20.13698 | 5.685197 | 2.363957 | 2.40495 | 0.016175 | 0.312293 | 0.505437 |
| ENSMUSG00000114638.2 | 11.61021 | -7.06987 | 2.945136 | -2.40053 | 0.016372 | 0.315452 | 0.501067 |
| ENSMUSG00000025757.14 | 196.9052 | 1.340311 | 0.558803 | 2.398538 | 0.016461 | 0.316525 | 0.499591 |
| ENSMUSG00000021552.8 | 146.0382 | 1.585923 | 0.662605 | 2.393468 | 0.01669 | 0.319987 | 0.494868 |
| ENSMUSG00000044786.7 | 205.6636 | -1.94885 | 0.814373 | -2.39307 | 0.016708 | 0.319987 | 0.494868 |
| ENSMUSG00000078234.7 | 41.6163 | -3.38986 | 1.417122 | -2.39207 | 0.016753 | 0.320207 | 0.494569 |
| ENSMUSG00000074909.6 | 108.641 | 2.056606 | 0.860295 | 2.390582 | 0.016822 | 0.320864 | 0.493679 |
| ENSMUSG00000029469.15 | 11.662 | 7.034653 | 2.94762 | 2.386554 | 0.017007 | 0.32375 | 0.48979 |
| ENSMUSG00000024507.7 | 2626.383 | 0.634688 | 0.266166 | 2.384562 | 0.017099 | 0.324206 | 0.489179 |
| ENSMUSG00000036553.17 | 9.945501 | -6.84223 | 2.868625 | -2.3852 | 0.01707 | 0.324206 | 0.489179 |
| ENSMUSG00000026617.17 | 218.5871 | 1.193322 | 0.500678 | 2.383413 | 0.017153 | 0.32457 | 0.488691 |
| ENSMUSG00000043542.13 | 20.45169 | -5.58021 | 2.347291 | -2.3773 | 0.01744 | 0.329346 | 0.482347 |
| ENSMUSG00000020186.8 | 138.9693 | 1.939201 | 0.816566 | 2.374825 | 0.017557 | 0.3309 | 0.480303 |
| ENSMUSG00000026806.16 | 17.44872 | 5.499996 | 2.316992 | 2.373765 | 0.017608 | 0.331193 | 0.479918 |
| ENSMUSG00000032590.15 | 92.72962 | -2.32157 | 0.978758 | -2.37195 | 0.017694 | 0.331748 | 0.479192 |
| ENSMUSG00000061887.15 | 110.9016 | -1.9144 | 0.80719 | -2.37168 | 0.017707 | 0.331748 | 0.479192 |
| ENSMUSG00000019370.11 | 29.589 | 3.980089 | 1.680153 | 2.368885 | 0.017842 | 0.333394 | 0.477043 |
| ENSMUSG00000022554.8 | 11.32665 | -7.03135 | 2.969577 | -2.3678 | 0.017894 | 0.333394 | 0.477043 |
| ENSMUSG00000024866.11 | 159.634 | -1.3111 | 0.553922 | -2.36694 | 0.017936 | 0.333394 | 0.477043 |
| ENSMUSG00000070462.5 | 13.54249 | 6.297605 | 2.659942 | 2.367572 | 0.017905 | 0.333394 | 0.477043 |
| ENSMUSG00000113204.2 | 22.69878 | -3.85557 | 1.63028 | -2.36497 | 0.018031 | 0.334516 | 0.475583 |
| ENSMUSG00000027670.5 | 11.31059 | 6.993348 | 2.962164 | 2.360892 | 0.018231 | 0.336901 | 0.472498 |
| ENSMUSG00000031986.7 | 28.18607 | -3.62375 | 1.534617 | -2.36134 | 0.018209 | 0.336901 | 0.472498 |
| ENSMUSG00000050957.5 | 11.62274 | -7.06759 | 2.995805 | -2.35916 | 0.018316 | 0.337817 | 0.471319 |
| ENSMUSG00000035847.16 | 110.094 | 1.638709 | 0.694829 | 2.358436 | 0.018352 | 0.337822 | 0.471312 |
| ENSMUSG00000020232.18 | 35.03642 | -3.71337 | 1.575696 | -2.35665 | 0.01844 | 0.338789 | 0.47007 |
| ENSMUSG00000020173.18 | 39.00027 | -3.79876 | 1.61379 | -2.35394 | 0.018576 | 0.33996 | 0.468573 |
| ENSMUSG00000044258.11 | 42.25303 | 3.706537 | 1.574183 | 2.354579 | 0.018544 | 0.33996 | 0.468573 |
| ENSMUSG00000024052.18 | 663.8196 | -0.92545 | 0.393949 | -2.34916 | 0.018816 | 0.343027 | 0.464672 |
| ENSMUSG00000030980.18 | 161.1788 | 1.490703 | 0.63439 | 2.349819 | 0.018783 | 0.343027 | 0.464672 |
| ENSMUSG00000027935.15 | 44.67211 | -3.54567 | 1.511958 | -2.34508 | 0.019023 | 0.346135 | 0.460755 |
| ENSMUSG00000024177.10 | 26.20288 | -4.26601 | 1.820987 | -2.34269 | 0.019145 | 0.347029 | 0.459634 |
| ENSMUSG00000035509.18 | 12.15557 | -7.13182 | 3.043487 | -2.34331 | 0.019114 | 0.347029 | 0.459634 |
| ENSMUSG00000006717.8 | 1295.533 | 0.652052 | 0.278421 | 2.341966 | 0.019182 | 0.347041 | 0.459619 |
| ENSMUSG00000054083.9 | 11.83174 | 7.053952 | 3.016122 | 2.338749 | 0.019348 | 0.348713 | 0.457532 |
| ENSMUSG00000070934.7 | 237.569 | -1.12991 | 0.483059 | -2.33907 | 0.019332 | 0.348713 | 0.457532 |
| ENSMUSG00000004268.12 | 260.5562 | 1.074065 | 0.459818 | 2.335846 | 0.019499 | 0.35016 | 0.455733 |

| | | | | | | | |
|---|---|---|---|---|---|---|---|
| ENSMUSG00000024963.13 | 28.50998 | -4.91266 | 2.103219 | -2.33578 | 0.019503 | 0.35016 | 0.455733 |
| ENSMUSG00000020235.17 | 11.20552 | -7.01393 | 3.006187 | -2.33316 | 0.01964 | 0.351952 | 0.453517 |
| ENSMUSG00000020534.15 | 694.9573 | 0.799759 | 0.3435 | 2.328266 | 0.019898 | 0.353495 | 0.451617 |
| ENSMUSG00000026197.13 | 17.70038 | -4.49021 | 1.927163 | -2.32996 | 0.019808 | 0.353495 | 0.451617 |
| ENSMUSG00000031862.12 | 34.56556 | -3.28058 | 1.408629 | -2.32892 | 0.019864 | 0.353495 | 0.451617 |
| ENSMUSG00000048550.18 | 25.56937 | -4.40981 | 1.894151 | -2.32812 | 0.019906 | 0.353495 | 0.451617 |
| ENSMUSG00000102349.2 | 34.20575 | -3.22322 | 1.384545 | -2.328 | 0.019912 | 0.353495 | 0.451617 |
| ENSMUSG00000036295.6 | 11.04355 | 6.956393 | 2.98906 | 2.327285 | 0.01995 | 0.353509 | 0.4516 |
| ENSMUSG00000038872.11 | 97.11253 | -1.6206 | 0.696649 | -2.32628 | 0.020004 | 0.353797 | 0.451246 |
| ENSMUSG00000109764.2 | 1889.591 | -0.6881 | 0.296011 | -2.32456 | 0.020095 | 0.354755 | 0.450071 |
| ENSMUSG00000026773.20 | 28.12622 | -3.86423 | 1.664876 | -2.32103 | 0.020285 | 0.356778 | 0.447602 |
| ENSMUSG00000053128.16 | 25.552 | -3.78607 | 1.631196 | -2.32104 | 0.020285 | 0.356778 | 0.447602 |
| ENSMUSG00000011427.17 | 19.58749 | -4.01675 | 1.742244 | -2.3055 | 0.021139 | 0.363027 | 0.440061 |
| ENSMUSG00000017667.10 | 11.12623 | -7.00357 | 3.035201 | -2.30745 | 0.02103 | 0.363027 | 0.440061 |
| ENSMUSG00000017723.12 | 18.58964 | 4.563731 | 1.973512 | 2.312492 | 0.020751 | 0.363027 | 0.440061 |
| ENSMUSG00000018930.4 | 10.83669 | 6.929386 | 3.001744 | 2.308453 | 0.020974 | 0.363027 | 0.440061 |
| ENSMUSG00000019578.12 | 76.31677 | -2.7215 | 1.177424 | -2.3114 | 0.020811 | 0.363027 | 0.440061 |
| ENSMUSG00000020473.14 | 10.85553 | -6.97348 | 3.023557 | -2.30638 | 0.021089 | 0.363027 | 0.440061 |
| ENSMUSG00000025326.13 | 402.9734 | -0.96327 | 0.418047 | -2.30421 | 0.021211 | 0.363027 | 0.440061 |
| ENSMUSG00000029656.14 | 1387.007 | -0.63377 | 0.275057 | -2.30414 | 0.021215 | 0.363027 | 0.440061 |
| ENSMUSG00000030895.10 | 4161.063 | -0.8286 | 0.35876 | -2.30961 | 0.02091 | 0.363027 | 0.440061 |
| ENSMUSG00000036334.13 | 10.83423 | -6.96703 | 3.023403 | -2.30437 | 0.021202 | 0.363027 | 0.440061 |
| ENSMUSG00000038957.14 | 21.58734 | -5.09841 | 2.206386 | -2.31075 | 0.020847 | 0.363027 | 0.440061 |
| ENSMUSG00000041477.15 | 14.53964 | -6.3205 | 2.733369 | -2.31235 | 0.020758 | 0.363027 | 0.440061 |
| ENSMUSG00000045980.14 | 17.28709 | 5.594592 | 2.422345 | 2.309577 | 0.020912 | 0.363027 | 0.440061 |
| ENSMUSG00000048087.7 | 804.7125 | 0.760648 | 0.329456 | 2.308804 | 0.020954 | 0.363027 | 0.440061 |
| ENSMUSG00000062170.13 | 10.82706 | -6.96947 | 3.019882 | -2.30786 | 0.021007 | 0.363027 | 0.440061 |
| ENSMUSG00000007038.6 | 247.2857 | -2.04009 | 0.887987 | -2.29743 | 0.021594 | 0.363607 | 0.439367 |
| ENSMUSG00000015335.17 | 19.01524 | -4.3668 | 1.898684 | -2.29991 | 0.021453 | 0.363607 | 0.439367 |
| ENSMUSG00000020484.20 | 1108.108 | -0.76316 | 0.331978 | -2.29882 | 0.021515 | 0.363607 | 0.439367 |
| ENSMUSG00000021999.10 | 2924.601 | -0.53169 | 0.231412 | -2.29758 | 0.021586 | 0.363607 | 0.439367 |
| ENSMUSG00000022032.15 | 91.24506 | -1.91287 | 0.83248 | -2.2978 | 0.021573 | 0.363607 | 0.439367 |
| ENSMUSG00000024941.10 | 59.13313 | -2.52212 | 1.096631 | -2.29988 | 0.021455 | 0.363607 | 0.439367 |
| ENSMUSG00000029461.18 | 101.4014 | -1.65327 | 0.719194 | -2.29878 | 0.021517 | 0.363607 | 0.439367 |
| ENSMUSG00000046798.15 | 350.0263 | -1.02246 | 0.444303 | -2.30128 | 0.021376 | 0.363607 | 0.439367 |
| ENSMUSG00000052914.5 | 509.625 | 0.885658 | 0.3854 | 2.298023 | 0.021561 | 0.363607 | 0.439367 |
| ENSMUSG00000064294.13 | 1661.293 | -0.73267 | 0.319347 | -2.29428 | 0.021774 | 0.365436 | 0.437189 |
| ENSMUSG00000097194.3 | 14.75853 | 7.369808 | 3.212381 | 2.294188 | 0.02178 | 0.365436 | 0.437189 |
| ENSMUSG00000078349.4 | 10.65992 | 6.907422 | 3.012046 | 2.293266 | 0.021833 | 0.365679 | 0.436901 |
| ENSMUSG00000005262.14 | 155.6174 | 1.461836 | 0.638266 | 2.290324 | 0.022003 | 0.365967 | 0.436558 |
| ENSMUSG00000022414.9 | 42.81904 | -8.9499 | 3.907812 | -2.29026 | 0.022006 | 0.365967 | 0.436558 |
| ENSMUSG00000036752.5 | 381.1081 | -1.32088 | 0.576897 | -2.28963 | 0.022043 | 0.365967 | 0.436558 |
| ENSMUSG00000052392.6 | 219.5916 | 1.18474 | 0.516919 | 2.291928 | 0.02191 | 0.365967 | 0.436558 |
| ENSMUSG00000058331.16 | 20.40306 | -4.26535 | 1.861285 | -2.29161 | 0.021928 | 0.365967 | 0.436558 |

| | | | | | | | |
|---|---|---|---|---|---|---|---|
| ENSMUSG00000016319.4 | 4409.849 | 0.549204 | 0.240199 | 2.286455 | 0.022228 | 0.367527 | 0.434711 |
| ENSMUSG00000029752.13 | 38.2981 | -4.38135 | 1.915705 | -2.28707 | 0.022192 | 0.367527 | 0.434711 |
| ENSMUSG00000078656.12 | 10.55655 | -6.93065 | 3.031758 | -2.28602 | 0.022253 | 0.367527 | 0.434711 |
| ENSMUSG00000002413.16 | 54.36254 | -2.31222 | 1.012627 | -2.28339 | 0.022407 | 0.368788 | 0.433224 |
| ENSMUSG00000021097.16 | 250.743 | -1.53871 | 0.674066 | -2.28273 | 0.022446 | 0.368788 | 0.433224 |
| ENSMUSG00000054499.10 | 18.37702 | -4.62721 | 2.02655 | -2.2833 | 0.022413 | 0.368788 | 0.433224 |
| ENSMUSG00000036533.10 | 10.73645 | 6.912217 | 3.030926 | 2.280562 | 0.022574 | 0.370251 | 0.431504 |
| ENSMUSG00000008435.16 | 24.95996 | 4.94051 | 2.168465 | 2.278345 | 0.022706 | 0.371184 | 0.430411 |
| ENSMUSG00000019295.9 | 58.06929 | -2.6946 | 1.182731 | -2.27829 | 0.02271 | 0.371184 | 0.430411 |
| ENSMUSG00000016283.14 | 14.51256 | 7.345539 | 3.227685 | 2.275792 | 0.022858 | 0.372322 | 0.429082 |
| ENSMUSG00000057530.15 | 187.4971 | -1.5426 | 0.678023 | -2.27515 | 0.022897 | 0.372322 | 0.429082 |
| ENSMUSG00000066057.8 | 14.51256 | 7.345539 | 3.227685 | 2.275792 | 0.022858 | 0.372322 | 0.429082 |
| ENSMUSG00000024505.17 | 10.94063 | -6.97966 | 3.071623 | -2.2723 | 0.023068 | 0.374463 | 0.426591 |
| ENSMUSG00000037563.16 | 513.9326 | 0.960905 | 0.42368 | 2.267996 | 0.023329 | 0.377709 | 0.422843 |
| ENSMUSG00000073802.6 | 10.83915 | 6.924135 | 3.05338 | 2.267695 | 0.023348 | 0.377709 | 0.422843 |
| ENSMUSG00000020741.18 | 318.9986 | -1.2858 | 0.567842 | -2.26437 | 0.023551 | 0.379707 | 0.420552 |
| ENSMUSG00000068606.7 | 14.08364 | -7.34428 | 3.243196 | -2.26452 | 0.023542 | 0.379707 | 0.420552 |
| ENSMUSG00000034269.13 | 92.88842 | -2.00516 | 0.887089 | -2.26038 | 0.023798 | 0.383024 | 0.416775 |
| ENSMUSG00000002031.14 | 40.689 | -3.38711 | 1.501092 | -2.25643 | 0.024044 | 0.384379 | 0.41524 |
| ENSMUSG00000020877.12 | 34.91598 | -2.79803 | 1.239565 | -2.25726 | 0.023992 | 0.384379 | 0.41524 |
| ENSMUSG00000026896.15 | 134.5091 | -1.55382 | 0.689104 | -2.25484 | 0.024143 | 0.384379 | 0.41524 |
| ENSMUSG00000029490.5 | 14.26658 | 7.320855 | 3.243415 | 2.257144 | 0.023999 | 0.384379 | 0.41524 |
| ENSMUSG00000031700.12 | 1021.483 | 0.754392 | 0.334375 | 2.256129 | 0.024063 | 0.384379 | 0.41524 |
| ENSMUSG00000036613.9 | 34.70624 | -3.11921 | 1.382878 | -2.25559 | 0.024096 | 0.384379 | 0.41524 |
| ENSMUSG00000052566.9 | 59.03037 | -2.20627 | 0.978612 | -2.25449 | 0.024166 | 0.384379 | 0.41524 |
| ENSMUSG00000003721.15 | 1362.489 | 0.77439 | 0.344333 | 2.248959 | 0.024515 | 0.384999 | 0.41454 |
| ENSMUSG00000009112.5 | 86.62569 | 1.629892 | 0.72475 | 2.248904 | 0.024519 | 0.384999 | 0.41454 |
| ENSMUSG00000025077.15 | 144.6047 | 1.604219 | 0.714398 | 2.245555 | 0.024733 | 0.384999 | 0.41454 |
| ENSMUSG00000027782.11 | 177.023 | 1.879539 | 0.835053 | 2.250801 | 0.024398 | 0.384999 | 0.41454 |
| ENSMUSG00000028542.18 | 64.83574 | -2.59694 | 1.152791 | -2.25274 | 0.024275 | 0.384999 | 0.41454 |
| ENSMUSG00000040505.14 | 164.9729 | -1.55645 | 0.692909 | -2.24625 | 0.024688 | 0.384999 | 0.41454 |
| ENSMUSG00000047735.15 | 50.69044 | 3.353965 | 1.493056 | 2.246376 | 0.02468 | 0.384999 | 0.41454 |
| ENSMUSG00000050144.14 | 157.7271 | -1.32227 | 0.587629 | -2.25017 | 0.024438 | 0.384999 | 0.41454 |
| ENSMUSG00000061959.14 | 1053.815 | 0.672732 | 0.299523 | 2.246013 | 0.024703 | 0.384999 | 0.41454 |
| ENSMUSG00000063001.11 | 11.68143 | 6.079562 | 2.705446 | 2.247157 | 0.02463 | 0.384999 | 0.41454 |
| ENSMUSG00000068206.14 | 10.30381 | -6.89522 | 3.061743 | -2.25206 | 0.024319 | 0.384999 | 0.41454 |
| ENSMUSG00000074622.5 | 270.6421 | -1.47551 | 0.656491 | -2.24757 | 0.024604 | 0.384999 | 0.41454 |
| ENSMUSG00000078877.12 | 39.66479 | 8.799178 | 3.907895 | 2.251641 | 0.024345 | 0.384999 | 0.41454 |
| ENSMUSG00000058396.8 | 52.44537 | 2.800823 | 1.247861 | 2.244499 | 0.0248 | 0.385422 | 0.414064 |
| ENSMUSG00000039449.15 | 188.7257 | -1.40533 | 0.626539 | -2.24301 | 0.024896 | 0.386281 | 0.413097 |
| ENSMUSG00000044906.7 | 16.51776 | -4.33086 | 1.931759 | -2.24192 | 0.024966 | 0.386733 | 0.412589 |
| ENSMUSG00000028187.11 | 148.5594 | 1.343601 | 0.599633 | 2.240706 | 0.025045 | 0.387321 | 0.411929 |
| ENSMUSG00000052299.10 | 195.3353 | -1.16459 | 0.519891 | -2.24006 | 0.025087 | 0.387338 | 0.41191 |
| ENSMUSG00000043702.10 | 156.9411 | 1.663616 | 0.743198 | 2.238456 | 0.025191 | 0.388315 | 0.410816 |

| | | | | | | | |
|---|---|---|---|---|---|---|---|
| ENSMUSG00000100691.2 | 10.27061 | -6.8895 | 3.079268 | -2.23738 | 0.025261 | 0.388763 | 0.410315 |
| ENSMUSG00000000876.12 | 638.9192 | 0.799851 | 0.358866 | 2.228832 | 0.025825 | 0.389993 | 0.408944 |
| ENSMUSG00000018820.14 | 25.72246 | -3.53335 | 1.582018 | -2.23345 | 0.025519 | 0.389993 | 0.408944 |
| ENSMUSG00000022451.13 | 499.9671 | 0.924981 | 0.414714 | 2.230406 | 0.025721 | 0.389993 | 0.408944 |
| ENSMUSG00000024479.4 | 403.7712 | 0.955992 | 0.428654 | 2.23022 | 0.025733 | 0.389993 | 0.408944 |
| ENSMUSG00000026277.15 | 51.99339 | -2.78713 | 1.250364 | -2.22906 | 0.02581 | 0.389993 | 0.408944 |
| ENSMUSG00000026820.6 | 23.37739 | -4.81317 | 2.155099 | -2.23339 | 0.025523 | 0.389993 | 0.408944 |
| ENSMUSG00000027067.17 | 138.0987 | -1.61321 | 0.722067 | -2.23416 | 0.025473 | 0.389993 | 0.408944 |
| ENSMUSG00000028070.8 | 260.4601 | -1.15687 | 0.518935 | -2.22932 | 0.025793 | 0.389993 | 0.408944 |
| ENSMUSG00000039680.11 | 66.99494 | 2.245114 | 1.004904 | 2.234157 | 0.025473 | 0.389993 | 0.408944 |
| ENSMUSG00000062061.4 | 10.28508 | 6.854373 | 3.075525 | 2.228684 | 0.025835 | 0.389993 | 0.408944 |
| ENSMUSG00000063558.5 | 236.7249 | 1.202907 | 0.539558 | 2.229431 | 0.025785 | 0.389993 | 0.408944 |
| ENSMUSG00000066684.11 | 11.89884 | 6.114092 | 2.739819 | 2.231568 | 0.025644 | 0.389993 | 0.408944 |
| ENSMUSG00000031729.7 | 187.7251 | -1.2363 | 0.555056 | -2.22734 | 0.025924 | 0.390721 | 0.408133 |
| ENSMUSG00000005580.12 | 15.23263 | -5.51877 | 2.478629 | -2.22654 | 0.025978 | 0.390905 | 0.407929 |
| ENSMUSG00000018459.16 | 10.02613 | -6.85698 | 3.081911 | -2.22491 | 0.026087 | 0.391928 | 0.406794 |
| ENSMUSG00000005566.15 | 62.07471 | -2.40581 | 1.083075 | -2.22128 | 0.026332 | 0.393497 | 0.405059 |
| ENSMUSG00000006529.17 | 1392.344 | -0.72446 | 0.3262 | -2.2209 | 0.026358 | 0.393497 | 0.405059 |
| ENSMUSG00000027889.18 | 35.80001 | 3.387052 | 1.524934 | 2.221113 | 0.026343 | 0.393497 | 0.405059 |
| ENSMUSG00000031776.18 | 45.08636 | -3.77024 | 1.69642 | -2.22247 | 0.026252 | 0.393497 | 0.405059 |
| ENSMUSG00000006800.15 | 60.7503 | 2.347264 | 1.058332 | 2.21789 | 0.026562 | 0.393892 | 0.404623 |
| ENSMUSG00000015290.16 | 156.8395 | 1.519826 | 0.685081 | 2.218462 | 0.026523 | 0.393892 | 0.404623 |
| ENSMUSG00000022412.15 | 63.53685 | -2.6336 | 1.187993 | -2.21685 | 0.026633 | 0.393892 | 0.404623 |
| ENSMUSG00000032883.16 | 161.4978 | 1.266891 | 0.570751 | 2.21969 | 0.02644 | 0.393892 | 0.404623 |
| ENSMUSG00000048100.14 | 351.3864 | 1.106025 | 0.49867 | 2.21795 | 0.026558 | 0.393892 | 0.404623 |
| ENSMUSG00000074925.5 | 85.78093 | 2.311158 | 1.042478 | 2.216984 | 0.026624 | 0.393892 | 0.404623 |
| ENSMUSG00000112794.2 | 29.55218 | -4.51024 | 2.035189 | -2.21613 | 0.026683 | 0.394009 | 0.404494 |
| ENSMUSG00000022389.16 | 282.0455 | -1.1991 | 0.541239 | -2.21547 | 0.026728 | 0.39406 | 0.404438 |
| ENSMUSG00000032294.18 | 10.31748 | 6.854555 | 3.095876 | 2.214092 | 0.026822 | 0.394227 | 0.404253 |
| ENSMUSG00000036845.13 | 15.72434 | 4.459594 | 2.014012 | 2.214284 | 0.026809 | 0.394227 | 0.404253 |
| ENSMUSG00000022848.9 | 184.3259 | -1.32764 | 0.600341 | -2.21148 | 0.027003 | 0.395653 | 0.402685 |
| ENSMUSG00000029234.14 | 72.21448 | 2.308642 | 1.04367 | 2.212042 | 0.026964 | 0.395653 | 0.402685 |
| ENSMUSG00000035067.10 | 21.67908 | 4.163603 | 1.883362 | 2.210729 | 0.027055 | 0.395799 | 0.402526 |
| ENSMUSG00000003038.16 | 18.78155 | 3.763091 | 1.705979 | 2.205825 | 0.027396 | 0.397994 | 0.400124 |
| ENSMUSG00000019790.18 | 37.72147 | -3.0213 | 1.369852 | -2.20556 | 0.027415 | 0.397994 | 0.400124 |
| ENSMUSG00000020017.15 | 1019.542 | -0.79383 | 0.3596 | -2.20753 | 0.027277 | 0.397994 | 0.400124 |
| ENSMUSG00000029279.16 | 12.0111 | -6.04862 | 2.741153 | -2.2066 | 0.027342 | 0.397994 | 0.400124 |
| ENSMUSG00000078672.3 | 19178.12 | -0.75298 | 0.341392 | -2.20561 | 0.027411 | 0.397994 | 0.400124 |
| ENSMUSG00000054863.10 | 10.22689 | 6.841672 | 3.104771 | 2.203599 | 0.027553 | 0.399385 | 0.398608 |
| ENSMUSG00000010064.16 | 412.2399 | -0.88183 | 0.400741 | -2.20051 | 0.027771 | 0.399881 | 0.398069 |
| ENSMUSG00000020471.12 | 10.26317 | 6.849726 | 3.113474 | 2.200027 | 0.027805 | 0.399881 | 0.398069 |
| ENSMUSG00000026037.15 | 59.65902 | -2.17184 | 0.987411 | -2.19954 | 0.02784 | 0.399881 | 0.398069 |
| ENSMUSG00000049532.12 | 13.52865 | 7.244167 | 3.293363 | 2.199626 | 0.027833 | 0.399881 | 0.398069 |
| ENSMUSG00000074305.10 | 22.03617 | -4.00373 | 1.818683 | -2.20145 | 0.027704 | 0.399881 | 0.398069 |

| | | | | | | | |
|---|---|---|---|---|---|---|---|
| ENSMUSG00000113488.2 | 13.52865 | 7.244167 | 3.293363 | 2.199626 | 0.027833 | 0.399881 | 0.398069 |
| ENSMUSG00000021179.9 | 11.58565 | 5.999717 | 2.730545 | 2.19726 | 0.028002 | 0.401601 | 0.396206 |
| ENSMUSG00000021922.18 | 5886.92 | -0.68806 | 0.313433 | -2.19524 | 0.028146 | 0.403063 | 0.394627 |
| ENSMUSG00000047496.7 | 97.84531 | -1.63477 | 0.744894 | -2.19464 | 0.02819 | 0.403072 | 0.394617 |
| ENSMUSG00000041236.9 | 169.3701 | -1.20023 | 0.547128 | -2.1937 | 0.028257 | 0.403427 | 0.394235 |
| ENSMUSG00000022292.17 | 12.6579 | 5.007269 | 2.28325 | 2.193044 | 0.028304 | 0.403494 | 0.394162 |
| ENSMUSG00000028256.17 | 39.47881 | 2.535607 | 1.156759 | 2.191992 | 0.02838 | 0.403969 | 0.393652 |
| ENSMUSG00000056144.15 | 43.34792 | -2.40852 | 1.099666 | -2.19023 | 0.028507 | 0.405171 | 0.392362 |
| ENSMUSG00000040771.15 | 232.3146 | 1.118734 | 0.51116 | 2.188619 | 0.028625 | 0.406229 | 0.391229 |
| ENSMUSG00000033618.8 | 72.88512 | -2.70742 | 1.23962 | -2.18408 | 0.028957 | 0.410183 | 0.387022 |
| ENSMUSG00000060807.8 | 1289.79 | 0.646442 | 0.296041 | 2.183626 | 0.02899 | 0.410183 | 0.387022 |
| ENSMUSG00000000149.11 | 168.872 | -1.4491 | 0.664779 | -2.17982 | 0.029271 | 0.412311 | 0.384775 |
| ENSMUSG00000021306.12 | 13.28268 | 7.217672 | 3.311004 | 2.179904 | 0.029265 | 0.412311 | 0.384775 |
| ENSMUSG00000025350.16 | 34.43974 | -4.30662 | 1.974829 | -2.18076 | 0.029201 | 0.412311 | 0.384775 |
| ENSMUSG00000026305.16 | 346.6787 | 1.434013 | 0.658193 | 2.178714 | 0.029353 | 0.412857 | 0.3842 |
| ENSMUSG00000010122.15 | 352.3304 | 0.971501 | 0.44664 | 2.175131 | 0.02962 | 0.414729 | 0.382235 |
| ENSMUSG00000020261.16 | 133.2253 | -1.93047 | 0.887979 | -2.17401 | 0.029705 | 0.414729 | 0.382235 |
| ENSMUSG00000038843.19 | 9.922831 | 6.797338 | 3.126262 | 2.17427 | 0.029685 | 0.414729 | 0.382235 |
| ENSMUSG00000051232.14 | 93.77947 | -2.11719 | 0.973357 | -2.17514 | 0.029619 | 0.414729 | 0.382235 |
| ENSMUSG00000073792.12 | 50.04955 | 2.34657 | 1.079327 | 2.174105 | 0.029697 | 0.414729 | 0.382235 |
| ENSMUSG00000042182.17 | 12.92208 | -7.22045 | 3.323902 | -2.17228 | 0.029834 | 0.415317 | 0.38162 |
| ENSMUSG00000050973.12 | 10.37678 | 6.865307 | 3.160169 | 2.17245 | 0.029822 | 0.415317 | 0.38162 |
| ENSMUSG00000034707.8 | 463.8667 | -1.04098 | 0.479736 | -2.1699 | 0.030014 | 0.415378 | 0.381556 |
| ENSMUSG00000040464.16 | 49.06869 | 2.78524 | 1.283191 | 2.170558 | 0.029965 | 0.415378 | 0.381556 |
| ENSMUSG00000047866.21 | 1396.83 | -0.60519 | 0.278717 | -2.17134 | 0.029906 | 0.415378 | 0.381556 |
| ENSMUSG00000097039.9 | 23.13484 | 3.605248 | 1.661088 | 2.170414 | 0.029976 | 0.415378 | 0.381556 |
| ENSMUSG00000025223.16 | 16.94198 | 4.576754 | 2.111719 | 2.167311 | 0.030211 | 0.416887 | 0.379982 |
| ENSMUSG00000039018.16 | 46.13288 | -2.30274 | 1.062252 | -2.16779 | 0.030174 | 0.416887 | 0.379982 |
| ENSMUSG00000046688.15 | 95.50072 | -1.62833 | 0.75238 | -2.16424 | 0.030446 | 0.417693 | 0.379142 |
| ENSMUSG00000046982.12 | 180.9014 | -1.4883 | 0.687581 | -2.16454 | 0.030423 | 0.417693 | 0.379142 |
| ENSMUSG00000052291.9 | 12.85898 | -7.21298 | 3.330594 | -2.16567 | 0.030336 | 0.417693 | 0.379142 |
| ENSMUSG00000097589.10 | 154.3455 | 1.330845 | 0.6146 | 2.165382 | 0.030358 | 0.417693 | 0.379142 |
| ENSMUSG00000015757.11 | 96.54207 | -1.76592 | 0.816681 | -2.16232 | 0.030594 | 0.418511 | 0.378293 |
| ENSMUSG00000032079.13 | 4978.068 | 0.59319 | 0.274307 | 2.162501 | 0.03058 | 0.418511 | 0.378293 |
| ENSMUSG00000000320.11 | 13.0367 | 7.190683 | 3.329182 | 2.159895 | 0.030781 | 0.419859 | 0.376896 |
| ENSMUSG00000020573.18 | 13.0367 | 7.190683 | 3.329182 | 2.159895 | 0.030781 | 0.419859 | 0.376896 |
| ENSMUSG00000034993.8 | 34.56687 | -3.06338 | 1.418753 | -2.1592 | 0.030834 | 0.419986 | 0.376766 |
| ENSMUSG00000021598.10 | 76.75396 | -1.98006 | 0.917517 | -2.15806 | 0.030923 | 0.420348 | 0.376391 |
| ENSMUSG00000032977.10 | 51.12586 | -2.43148 | 1.126874 | -2.15772 | 0.03095 | 0.420348 | 0.376391 |
| ENSMUSG00000029416.18 | 38.23707 | 2.893331 | 1.343118 | 2.15419 | 0.031225 | 0.42288 | 0.373783 |
| ENSMUSG00000036636.11 | 49.28083 | -2.30584 | 1.070296 | -2.15439 | 0.031209 | 0.42288 | 0.373783 |
| ENSMUSG00000063316.14 | 1799.125 | 0.768312 | 0.357118 | 2.151424 | 0.031443 | 0.425218 | 0.371389 |
| ENSMUSG00000031200.17 | 14.71474 | -5.47055 | 2.545082 | -2.14946 | 0.031598 | 0.426708 | 0.369869 |
| ENSMUSG00000020541.13 | 106.804 | -2.18935 | 1.018938 | -2.14866 | 0.031662 | 0.426959 | 0.369614 |

| | | | | | | | |
|---|---|---|---|---|---|---|---|
| ENSMUSG00000021189.12 | 15.07858 | -3.90454 | 1.819325 | -2.14615 | 0.031861 | 0.428549 | 0.367999 |
| ENSMUSG00000030660.10 | 235.1836 | 1.070901 | 0.499013 | 2.146038 | 0.03187 | 0.428549 | 0.367999 |
| ENSMUSG00000016833.15 | 328.7822 | 1.128296 | 0.52623 | 2.144111 | 0.032024 | 0.428691 | 0.367855 |
| ENSMUSG00000028957.13 | 17.23954 | -3.95916 | 1.846888 | -2.14369 | 0.032058 | 0.428691 | 0.367855 |
| ENSMUSG00000029147.12 | 59.73559 | -2.6737 | 1.247267 | -2.14364 | 0.032061 | 0.428691 | 0.367855 |
| ENSMUSG00000073988.14 | 4474.383 | 0.56956 | 0.265651 | 2.144018 | 0.032031 | 0.428691 | 0.367855 |
| ENSMUSG00000026648.19 | 15.80919 | -4.8192 | 2.250567 | -2.14133 | 0.032248 | 0.429364 | 0.367174 |
| ENSMUSG00000034947.14 | 35.08942 | 3.777612 | 1.763951 | 2.141564 | 0.032229 | 0.429364 | 0.367174 |
| ENSMUSG00000037440.9 | 348.3266 | 1.138242 | 0.531518 | 2.141493 | 0.032234 | 0.429364 | 0.367174 |
| ENSMUSG00000025381.4 | 997.2815 | 0.667226 | 0.311914 | 2.139136 | 0.032425 | 0.431116 | 0.365406 |
| ENSMUSG00000008845.10 | 91.97077 | -2.08396 | 0.975776 | -2.13569 | 0.032705 | 0.433524 | 0.362987 |
| ENSMUSG00000020329.13 | 21.69178 | 3.578373 | 1.675013 | 2.136325 | 0.032653 | 0.433524 | 0.362987 |
| ENSMUSG00000028383.18 | 420.3568 | 0.883542 | 0.413963 | 2.134348 | 0.032814 | 0.433524 | 0.362987 |
| ENSMUSG00000035356.18 | 131.0438 | -2.14465 | 1.004449 | -2.13515 | 0.032749 | 0.433524 | 0.362987 |
| ENSMUSG00000037935.17 | 146.4718 | -1.6249 | 0.761395 | -2.1341 | 0.032834 | 0.433524 | 0.362987 |
| ENSMUSG00000015016.9 | 24.0481 | -3.8366 | 1.799822 | -2.13165 | 0.033035 | 0.435568 | 0.360944 |
| ENSMUSG00000068263.12 | 12.45076 | -7.16642 | 3.362785 | -2.1311 | 0.033081 | 0.435568 | 0.360944 |
| ENSMUSG00000020964.15 | 617.2585 | 0.866866 | 0.407148 | 2.12912 | 0.033244 | 0.43711 | 0.359409 |
| ENSMUSG00000058317.13 | 216.5593 | 1.193171 | 0.560863 | 2.127384 | 0.033388 | 0.438394 | 0.358136 |
| ENSMUSG00000027048.16 | 1382.505 | -0.72341 | 0.340193 | -2.12646 | 0.033465 | 0.438794 | 0.357739 |
| ENSMUSG00000025277.15 | 259.5019 | 1.104118 | 0.519499 | 2.12535 | 0.033557 | 0.439398 | 0.357142 |
| ENSMUSG00000024592.16 | 122.8578 | 1.481825 | 0.698293 | 2.122067 | 0.033832 | 0.439409 | 0.357131 |
| ENSMUSG00000025899.15 | 54.16534 | 2.800889 | 1.319919 | 2.122016 | 0.033836 | 0.439409 | 0.357131 |
| ENSMUSG00000031984.9 | 168.9261 | 1.512257 | 0.712095 | 2.123672 | 0.033698 | 0.439409 | 0.357131 |
| ENSMUSG00000062526.5 | 87.01707 | -2.12251 | 0.999747 | -2.12304 | 0.03375 | 0.439409 | 0.357131 |
| ENSMUSG00000084910.2 | 26.2034 | 4.359822 | 2.051961 | 2.12471 | 0.033611 | 0.439409 | 0.357131 |
| ENSMUSG00000116505.2 | 12.33471 | -7.15331 | 3.370397 | -2.12239 | 0.033805 | 0.439409 | 0.357131 |
| ENSMUSG00000024561.12 | 186.8175 | -1.6146 | 0.761217 | -2.12108 | 0.033915 | 0.439833 | 0.356712 |
| ENSMUSG00000024413.15 | 260.3278 | -1.13397 | 0.535508 | -2.11756 | 0.034212 | 0.441958 | 0.354619 |
| ENSMUSG00000027556.16 | 88.55387 | -1.8225 | 0.861343 | -2.11588 | 0.034355 | 0.441958 | 0.354619 |
| ENSMUSG00000035158.16 | 12.3857 | 5.007553 | 2.365654 | 2.116773 | 0.034279 | 0.441958 | 0.354619 |
| ENSMUSG00000054027.9 | 134.485 | -1.41558 | 0.669039 | -2.11583 | 0.034359 | 0.441958 | 0.354619 |
| ENSMUSG00000057133.15 | 73.48567 | -2.21767 | 1.047558 | -2.11699 | 0.034261 | 0.441958 | 0.354619 |
| ENSMUSG00000093930.3 | 2406.686 | -0.83406 | 0.394077 | -2.11649 | 0.034303 | 0.441958 | 0.354619 |
| ENSMUSG00000036112.16 | 510.877 | -0.75514 | 0.35705 | -2.11494 | 0.034435 | 0.442342 | 0.354242 |
| ENSMUSG00000021025.9 | 538.7206 | -1.08421 | 0.51296 | -2.11363 | 0.034547 | 0.442737 | 0.353854 |
| ENSMUSG00000059866.14 | 12.24665 | -7.14257 | 3.379528 | -2.11348 | 0.03456 | 0.442737 | 0.353854 |
| ENSMUSG00000020580.11 | 144.4832 | -1.25552 | 0.594749 | -2.11101 | 0.034771 | 0.444849 | 0.351787 |
| ENSMUSG00000025036.8 | 128.9745 | 1.724911 | 0.817515 | 2.109945 | 0.034863 | 0.44542 | 0.35123 |
| ENSMUSG00000039826.14 | 66.31219 | -2.27816 | 1.08049 | -2.10845 | 0.034992 | 0.446463 | 0.350215 |
| ENSMUSG00000006215.13 | 13.46452 | 4.965859 | 2.357429 | 2.106472 | 0.035163 | 0.44771 | 0.349003 |
| ENSMUSG00000030109.16 | 141.1843 | -1.58383 | 0.752508 | -2.10474 | 0.035314 | 0.44771 | 0.349003 |
| ENSMUSG00000053560.6 | 199.6858 | -2.3176 | 1.100378 | -2.10618 | 0.035189 | 0.44771 | 0.349003 |
| ENSMUSG00000058207.7 | 40114.15 | -0.44981 | 0.213724 | -2.10461 | 0.035326 | 0.44771 | 0.349003 |

| | | | | | | | |
|---|---|---|---|---|---|---|---|
| ENSMUSG00000073197.10 | 12.13892 | -7.13022 | 3.386699 | -2.10536 | 0.03526 | 0.44771 | 0.349003 |
| ENSMUSG00000093904.3 | 52.01873 | -2.11515 | 1.005868 | -2.10281 | 0.035483 | 0.449101 | 0.347656 |
| ENSMUSG00000020357.4 | 65.6207 | -2.39047 | 1.137498 | -2.10152 | 0.035596 | 0.449819 | 0.346962 |
| ENSMUSG00000069805.11 | 3855.284 | 0.570301 | 0.271433 | 2.101075 | 0.035634 | 0.449819 | 0.346962 |
| ENSMUSG00000027954.10 | 190.5831 | -1.34584 | 0.641114 | -2.09922 | 0.035798 | 0.451279 | 0.345555 |
| ENSMUSG00000003528.16 | 21.30196 | -4.68294 | 2.231963 | -2.09813 | 0.035894 | 0.451892 | 0.344966 |
| ENSMUSG00000032855.7 | 18.74249 | -4.89747 | 2.335614 | -2.09687 | 0.036005 | 0.452093 | 0.344773 |
| ENSMUSG00000045854.5 | 100.7346 | 1.992543 | 0.950103 | 2.097186 | 0.035977 | 0.452093 | 0.344773 |
| ENSMUSG00000039205.17 | 11.6064 | 6.072063 | 2.896829 | 2.096106 | 0.036073 | 0.452341 | 0.344534 |
| ENSMUSG00000014767.18 | 28.4691 | -2.9658 | 1.416222 | -2.09416 | 0.036246 | 0.452926 | 0.343973 |
| ENSMUSG00000015776.13 | 18.04 | 3.677399 | 1.757013 | 2.092984 | 0.036351 | 0.452926 | 0.343973 |
| ENSMUSG00000020044.14 | 127.5899 | 1.413419 | 0.674685 | 2.094932 | 0.036177 | 0.452926 | 0.343973 |
| ENSMUSG00000020640.11 | 187.009 | -1.34685 | 0.643475 | -2.09309 | 0.036341 | 0.452926 | 0.343973 |
| ENSMUSG00000032193.10 | 723.18 | 0.844139 | 0.403438 | 2.092363 | 0.036406 | 0.452926 | 0.343973 |
| ENSMUSG00000057363.13 | 36.37938 | 2.472992 | 1.181769 | 2.092619 | 0.036383 | 0.452926 | 0.343973 |
| ENSMUSG00000010607.9 | 57.13452 | -2.61761 | 1.251974 | -2.09078 | 0.036547 | 0.453389 | 0.343529 |
| ENSMUSG00000028102.16 | 83.11353 | 2.003348 | 0.958381 | 2.090345 | 0.036587 | 0.453389 | 0.343529 |
| ENSMUSG00000053898.13 | 897.3444 | 0.675806 | 0.323203 | 2.090965 | 0.036531 | 0.453389 | 0.343529 |
| ENSMUSG00000008496.20 | 11.42436 | -6.11667 | 2.930419 | -2.0873 | 0.036861 | 0.453817 | 0.343119 |
| ENSMUSG00000028051.11 | 11.94314 | -7.10675 | 3.40343 | -2.08811 | 0.036787 | 0.453817 | 0.343119 |
| ENSMUSG00000049680.18 | 136.3461 | 2.232945 | 1.069358 | 2.088118 | 0.036787 | 0.453817 | 0.343119 |
| ENSMUSG00000063415.13 | 50.24305 | 2.290446 | 1.097084 | 2.087759 | 0.03682 | 0.453817 | 0.343119 |
| ENSMUSG00000071528.5 | 1558.094 | 0.651199 | 0.311846 | 2.088206 | 0.036779 | 0.453817 | 0.343119 |
| ENSMUSG00000033396.14 | 24.0839 | -3.54092 | 1.697157 | -2.08638 | 0.036944 | 0.45425 | 0.342705 |
| ENSMUSG00000005043.14 | 14.39391 | 4.574845 | 2.194064 | 2.085101 | 0.03706 | 0.454502 | 0.342464 |
| ENSMUSG00000057605.5 | 37.62115 | -2.4995 | 1.198692 | -2.08519 | 0.037052 | 0.454502 | 0.342464 |
| ENSMUSG00000033831.6 | 27418.52 | -0.48919 | 0.234929 | -2.0823 | 0.037315 | 0.455275 | 0.341726 |
| ENSMUSG00000050088.10 | 160.1642 | 1.528096 | 0.733356 | 2.083703 | 0.037187 | 0.455275 | 0.341726 |
| ENSMUSG00000060002.15 | 2903.944 | 0.574603 | 0.275885 | 2.082762 | 0.037273 | 0.455275 | 0.341726 |
| ENSMUSG00000110353.2 | 29.79201 | -3.068 | 1.472939 | -2.08291 | 0.037259 | 0.455275 | 0.341726 |
| ENSMUSG00000031482.10 | 1394.769 | -0.56695 | 0.272496 | -2.08057 | 0.037473 | 0.456611 | 0.340454 |
| ENSMUSG00000022663.4 | 893.4436 | 0.668971 | 0.322199 | 2.076269 | 0.037869 | 0.458492 | 0.338669 |
| ENSMUSG00000028403.16 | 39.64315 | -2.44838 | 1.179082 | -2.07652 | 0.037846 | 0.458492 | 0.338669 |
| ENSMUSG00000029657.16 | 433.7154 | 1.293084 | 0.622647 | 2.076753 | 0.037824 | 0.458492 | 0.338669 |
| ENSMUSG00000037253.9 | 125.0322 | 1.35704 | 0.653551 | 2.076411 | 0.037856 | 0.458492 | 0.338669 |
| ENSMUSG00000060935.7 | 112.4362 | 1.731752 | 0.833965 | 2.076528 | 0.037845 | 0.458492 | 0.338669 |
| ENSMUSG00000001247.17 | 419.1227 | -1.10949 | 0.535302 | -2.07264 | 0.038206 | 0.460812 | 0.336476 |
| ENSMUSG00000004044.10 | 23.35614 | 3.13289 | 1.511423 | 2.072809 | 0.03819 | 0.460812 | 0.336476 |
| ENSMUSG00000024975.14 | 327.2406 | -0.98311 | 0.474381 | -2.07241 | 0.038227 | 0.460812 | 0.336476 |
| ENSMUSG00000026568.7 | 3786.179 | 0.617997 | 0.298245 | 2.07211 | 0.038255 | 0.460812 | 0.336476 |
| ENSMUSG00000036513.16 | 119.5386 | 1.586873 | 0.766026 | 2.071564 | 0.038306 | 0.46084 | 0.33645 |
| ENSMUSG00000002279.20 | 37.16749 | -3.15623 | 1.524696 | -2.07007 | 0.038446 | 0.461931 | 0.335422 |
| ENSMUSG00000059534.9 | 1688.797 | 0.589421 | 0.28502 | 2.067999 | 0.03864 | 0.463681 | 0.333781 |
| ENSMUSG00000022450.7 | 3643.957 | 0.530491 | 0.256973 | 2.064386 | 0.038981 | 0.464832 | 0.332704 |

| | | | | | | | |
|---|---|---|---|---|---|---|---|
| ENSMUSG00000024800.9 | 38.34938 | 2.336946 | 1.132023 | 2.064398 | 0.03898 | 0.464832 | 0.332704 |
| ENSMUSG00000026822.15 | 213.7673 | -4.92627 | 2.385384 | -2.06519 | 0.038905 | 0.464832 | 0.332704 |
| ENSMUSG00000030275.7 | 443.7704 | 0.882972 | 0.42762 | 2.064853 | 0.038937 | 0.464832 | 0.332704 |
| ENSMUSG00000030671.10 | 491.1022 | 0.804481 | 0.389352 | 2.066208 | 0.038809 | 0.464832 | 0.332704 |
| ENSMUSG00000023971.10 | 20.97394 | 3.617897 | 1.753438 | 2.063316 | 0.039083 | 0.465019 | 0.33253 |
| ENSMUSG00000074071.12 | 52.24548 | 3.196643 | 1.549372 | 2.063186 | 0.039095 | 0.465019 | 0.33253 |
| ENSMUSG00000028001.17 | 9977.224 | -0.49665 | 0.240885 | -2.06178 | 0.039228 | 0.466022 | 0.331594 |
| ENSMUSG00000031645.5 | 321.7781 | -1.04568 | 0.507893 | -2.05885 | 0.039508 | 0.468758 | 0.329051 |
| ENSMUSG00000049115.16 | 513.6837 | -0.82139 | 0.399249 | -2.05734 | 0.039654 | 0.469899 | 0.327995 |
| ENSMUSG00000033107.7 | 278.8014 | 1.0405 | 0.505879 | 2.056814 | 0.039704 | 0.469906 | 0.327989 |
| ENSMUSG00000004319.16 | 209.3212 | 1.088889 | 0.529586 | 2.056112 | 0.039772 | 0.469973 | 0.327927 |
| ENSMUSG00000015854.8 | 274.7794 | -1.0518 | 0.51187 | -2.05481 | 0.039897 | 0.469973 | 0.327927 |
| ENSMUSG00000021102.5 | 532.4702 | 0.734895 | 0.357512 | 2.055583 | 0.039823 | 0.469973 | 0.327927 |
| ENSMUSG00000022181.17 | 1151.834 | -0.55554 | 0.270822 | -2.05132 | 0.040236 | 0.469973 | 0.327927 |
| ENSMUSG00000022742.7 | 861.6195 | 0.707119 | 0.345287 | 2.047917 | 0.040568 | 0.469973 | 0.327927 |
| ENSMUSG00000026500.7 | 658.5255 | 1.061018 | 0.517372 | 2.050786 | 0.040288 | 0.469973 | 0.327927 |
| ENSMUSG00000029309.8 | 43.11224 | 2.608114 | 1.273282 | 2.048339 | 0.040527 | 0.469973 | 0.327927 |
| ENSMUSG00000030762.12 | 166.8864 | 1.5055 | 0.73281 | 2.054421 | 0.039935 | 0.469973 | 0.327927 |
| ENSMUSG00000032489.18 | 11.61156 | 7.027349 | 3.429862 | 2.048872 | 0.040475 | 0.469973 | 0.327927 |
| ENSMUSG00000037798.8 | 4295.65 | 0.494117 | 0.241211 | 2.048487 | 0.040512 | 0.469973 | 0.327927 |
| ENSMUSG00000039427.15 | 68.94373 | 2.064677 | 1.00544 | 2.053505 | 0.040024 | 0.469973 | 0.327927 |
| ENSMUSG00000044149.8 | 12.87322 | 4.12527 | 2.014715 | 2.04757 | 0.040602 | 0.469973 | 0.327927 |
| ENSMUSG00000047514.7 | 30.01164 | -3.3894 | 1.654677 | -2.04838 | 0.040523 | 0.469973 | 0.327927 |
| ENSMUSG00000055172.11 | 803.0716 | -0.69696 | 0.339907 | -2.05044 | 0.040321 | 0.469973 | 0.327927 |
| ENSMUSG00000062127.12 | 14.36364 | -4.26167 | 2.079046 | -2.04982 | 0.040382 | 0.469973 | 0.327927 |
| ENSMUSG00000070305.11 | 28.13438 | -3.05229 | 1.488302 | -2.05085 | 0.040281 | 0.469973 | 0.327927 |
| ENSMUSG00000073236.5 | 11.55156 | -7.05864 | 3.438257 | -2.05297 | 0.040075 | 0.469973 | 0.327927 |
| ENSMUSG00000094806.3 | 3081.923 | -0.49612 | 0.242046 | -2.0497 | 0.040394 | 0.469973 | 0.327927 |
| ENSMUSG00000001418.14 | 68.64394 | -1.80899 | 0.884304 | -2.04566 | 0.040789 | 0.471566 | 0.326458 |
| ENSMUSG00000087413.2 | 25.68309 | -3.99814 | 1.956291 | -2.04374 | 0.040979 | 0.473186 | 0.324968 |
| ENSMUSG00000027201.17 | 30.86907 | 3.124202 | 1.530916 | 2.040741 | 0.041277 | 0.473211 | 0.324945 |
| ENSMUSG00000027615.15 | 38.22974 | 2.984099 | 1.462101 | 2.040966 | 0.041254 | 0.473211 | 0.324945 |
| ENSMUSG00000028356.5 | 19313.32 | -0.41941 | 0.205487 | -2.04103 | 0.041247 | 0.473211 | 0.324945 |
| ENSMUSG00000029470.16 | 110.1528 | -1.72673 | 0.846039 | -2.04096 | 0.041255 | 0.473211 | 0.324945 |
| ENSMUSG00000040345.11 | 11.4302 | -7.04299 | 3.45127 | -2.04069 | 0.041281 | 0.473211 | 0.324945 |
| ENSMUSG00000058056.17 | 289.6844 | 1.004408 | 0.491754 | 2.042502 | 0.041102 | 0.473211 | 0.324945 |
| ENSMUSG00000021606.9 | 1419.991 | 0.590038 | 0.289373 | 2.039017 | 0.041448 | 0.473905 | 0.324308 |
| ENSMUSG00000024253.10 | 11.44328 | 7.006287 | 3.445296 | 2.03358 | 0.041994 | 0.473905 | 0.324308 |
| ENSMUSG00000026574.6 | 42.60615 | 2.884709 | 1.415638 | 2.037745 | 0.041575 | 0.473905 | 0.324308 |
| ENSMUSG00000026748.14 | 11.56085 | 7.017204 | 3.451137 | 2.033302 | 0.042022 | 0.473905 | 0.324308 |
| ENSMUSG00000028643.12 | 89.37162 | -1.94456 | 0.955507 | -2.03511 | 0.04184 | 0.473905 | 0.324308 |
| ENSMUSG00000031812.17 | 489.4472 | -0.80789 | 0.397132 | -2.03432 | 0.041919 | 0.473905 | 0.324308 |
| ENSMUSG00000035495.16 | 51.2035 | 2.752075 | 1.351702 | 2.036007 | 0.04175 | 0.473905 | 0.324308 |
| ENSMUSG00000042178.7 | 13.48004 | -4.89522 | 2.407754 | -2.03311 | 0.042042 | 0.473905 | 0.324308 |

| | | | | | | | |
|---|---|---|---|---|---|---|---|
| ENSMUSG00000042377.9 | 11.56085 | 7.017204 | 3.451137 | 2.033302 | 0.042022 | 0.473905 | 0.324308 |
| ENSMUSG00000043445.8 | 94.60617 | 1.657462 | 0.813305 | 2.037933 | 0.041557 | 0.473905 | 0.324308 |
| ENSMUSG00000046005.5 | 11.44328 | 7.006287 | 3.445296 | 2.03358 | 0.041994 | 0.473905 | 0.324308 |
| ENSMUSG00000061894.16 | 11.35577 | -7.03397 | 3.456393 | -2.03506 | 0.041845 | 0.473905 | 0.324308 |
| ENSMUSG00000066153.3 | 4714.807 | -0.78488 | 0.385111 | -2.03806 | 0.041544 | 0.473905 | 0.324308 |
| ENSMUSG00000066278.7 | 14.63569 | 5.205107 | 2.558638 | 2.034327 | 0.041919 | 0.473905 | 0.324308 |
| ENSMUSG00000022514.15 | 931.1896 | 0.632685 | 0.311625 | 2.030275 | 0.042329 | 0.476573 | 0.32187 |
| ENSMUSG00000020585.5 | 1312.105 | 0.582074 | 0.287053 | 2.027756 | 0.042585 | 0.478326 | 0.320276 |
| ENSMUSG00000020917.18 | 993.3947 | 0.783286 | 0.386245 | 2.027953 | 0.042565 | 0.478326 | 0.320276 |
| ENSMUSG00000035235.14 | 13.34331 | -5.33184 | 2.630687 | -2.02679 | 0.042684 | 0.478872 | 0.319781 |
| ENSMUSG00000003234.17 | 76.38408 | -2.16789 | 1.070041 | -2.02599 | 0.042766 | 0.479223 | 0.319463 |
| ENSMUSG00000010936.12 | 103.8188 | 1.793176 | 0.886355 | 2.023091 | 0.043064 | 0.480853 | 0.317988 |
| ENSMUSG00000024222.18 | 184.8271 | 1.573995 | 0.777908 | 2.023368 | 0.043035 | 0.480853 | 0.317988 |
| ENSMUSG00000059743.13 | 61.77893 | 2.235528 | 1.104647 | 2.023749 | 0.042996 | 0.480853 | 0.317988 |
| ENSMUSG00000021226.8 | 52.53043 | 2.663152 | 1.317668 | 2.02111 | 0.043268 | 0.481982 | 0.316969 |
| ENSMUSG00000022295.9 | 238.2305 | -1.03638 | 0.512898 | -2.02064 | 0.043317 | 0.481982 | 0.316969 |
| ENSMUSG00000024816.13 | 66.24851 | -2.63783 | 1.305272 | -2.0209 | 0.04329 | 0.481982 | 0.316969 |
| ENSMUSG00000020840.11 | 153.5309 | -1.2872 | 0.638754 | -2.01517 | 0.043886 | 0.482773 | 0.316257 |
| ENSMUSG00000023988.9 | 19.35515 | 3.022392 | 1.499434 | 2.015689 | 0.043833 | 0.482773 | 0.316257 |
| ENSMUSG00000024960.10 | 11.275 | 6.984912 | 3.4611 | 2.018119 | 0.043579 | 0.482773 | 0.316257 |
| ENSMUSG00000035236.18 | 43.98096 | -3.37213 | 1.670315 | -2.01886 | 0.043502 | 0.482773 | 0.316257 |
| ENSMUSG00000057069.8 | 404.4829 | -1.09153 | 0.541387 | -2.01617 | 0.043782 | 0.482773 | 0.316257 |
| ENSMUSG00000057228.7 | 884.579 | -0.7764 | 0.385297 | -2.01507 | 0.043898 | 0.482773 | 0.316257 |
| ENSMUSG00000073830.12 | 158.6608 | -1.2678 | 0.628981 | -2.01563 | 0.043838 | 0.482773 | 0.316257 |
| ENSMUSG00000095180.11 | 11.275 | 6.984912 | 3.4611 | 2.018119 | 0.043579 | 0.482773 | 0.316257 |
| ENSMUSG00000100075.2 | 35.76119 | -3.34859 | 1.660598 | -2.01649 | 0.043748 | 0.482773 | 0.316257 |
| ENSMUSG00000116518.2 | 9.908914 | 5.843808 | 2.896756 | 2.017363 | 0.043658 | 0.482773 | 0.316257 |
| ENSMUSG00000037095.9 | 1150.745 | -0.82924 | 0.411938 | -2.01303 | 0.044112 | 0.484563 | 0.31465 |
| ENSMUSG00000022871.14 | 568.1882 | -1.1286 | 0.561456 | -2.01013 | 0.044418 | 0.485002 | 0.314256 |
| ENSMUSG00000027353.15 | 10.86741 | 4.817181 | 2.397802 | 2.008999 | 0.044537 | 0.485002 | 0.314256 |
| ENSMUSG00000029076.15 | 308.3068 | -1.20271 | 0.598782 | -2.0086 | 0.04458 | 0.485002 | 0.314256 |
| ENSMUSG00000032322.15 | 11.31487 | 6.98615 | 3.473928 | 2.011023 | 0.044323 | 0.485002 | 0.314256 |
| ENSMUSG00000038909.17 | 32.70844 | -3.03346 | 1.509738 | -2.00926 | 0.044509 | 0.485002 | 0.314256 |
| ENSMUSG00000054422.8 | 81030.85 | 0.521185 | 0.259255 | 2.010313 | 0.044398 | 0.485002 | 0.314256 |
| ENSMUSG00000068196.6 | 11.31487 | 6.98615 | 3.473928 | 2.011023 | 0.044323 | 0.485002 | 0.314256 |
| ENSMUSG00000086332.3 | 43.55538 | -2.33496 | 1.161516 | -2.01027 | 0.044403 | 0.485002 | 0.314256 |
| ENSMUSG00000108763.2 | 29.82593 | 3.389368 | 1.687685 | 2.008295 | 0.044612 | 0.485002 | 0.314256 |
| ENSMUSG00000024131.9 | 353.8255 | -1.08214 | 0.539617 | -2.00539 | 0.044922 | 0.487251 | 0.312247 |
| ENSMUSG00000062866.17 | 38.10294 | 2.850856 | 1.421534 | 2.005478 | 0.044912 | 0.487251 | 0.312247 |
| ENSMUSG00000030287.16 | 84.61877 | -2.0808 | 1.038342 | -2.00396 | 0.045074 | 0.48835 | 0.311269 |
| ENSMUSG00000028008.11 | 12.15761 | -4.53842 | 2.26673 | -2.00219 | 0.045264 | 0.489289 | 0.310435 |
| ENSMUSG00000041654.16 | 212.3937 | -1.22382 | 0.611222 | -2.00225 | 0.045258 | 0.489289 | 0.310435 |
| ENSMUSG00000029007.9 | 36.30787 | 3.361846 | 1.680187 | 2.000876 | 0.045406 | 0.490258 | 0.309575 |
| ENSMUSG00000058230.13 | 233.6473 | -1.03161 | 0.515833 | -1.99989 | 0.045512 | 0.490503 | 0.309358 |

| | | | | | | | |
|---|---|---|---|---|---|---|---|
| ENSMUSG00000076437.12 | 112.3711 | 1.542849 | 0.771538 | 1.999707 | 0.045532 | 0.490503 | 0.309358 |
| ENSMUSG00000022051.16 | 462.6286 | -0.81836 | 0.409472 | -1.99858 | 0.045654 | 0.490698 | 0.309186 |
| ENSMUSG00000029394.12 | 161.8522 | -1.29616 | 0.648446 | -1.99887 | 0.045623 | 0.490698 | 0.309186 |
| ENSMUSG00000039804.16 | 17.21384 | 4.192019 | 2.100162 | 1.996045 | 0.045929 | 0.4931 | 0.307065 |
| ENSMUSG00000021759.15 | 129.6149 | 1.36634 | 0.685885 | 1.992083 | 0.046362 | 0.495503 | 0.304954 |
| ENSMUSG00000035472.15 | 141.1912 | -1.54954 | 0.777831 | -1.99213 | 0.046357 | 0.495503 | 0.304954 |
| ENSMUSG00000037353.10 | 116.5041 | -1.79257 | 0.899476 | -1.9929 | 0.046272 | 0.495503 | 0.304954 |
| ENSMUSG00000046805.11 | 181.0618 | -1.54586 | 0.775576 | -1.99317 | 0.046243 | 0.495503 | 0.304954 |
| ENSMUSG00000032423.13 | 282.6186 | 1.138188 | 0.571958 | 1.989985 | 0.046593 | 0.497407 | 0.303288 |
| ENSMUSG00000022641.16 | 81.72712 | -1.67351 | 0.842757 | -1.98575 | 0.047061 | 0.500211 | 0.300846 |
| ENSMUSG00000031070.16 | 10.93843 | 6.94119 | 3.493778 | 1.986729 | 0.046952 | 0.500211 | 0.300846 |
| ENSMUSG00000078676.10 | 23.42035 | 3.631405 | 1.828131 | 1.986403 | 0.046989 | 0.500211 | 0.300846 |
| ENSMUSG00000091537.3 | 243.3424 | 1.172098 | 0.590268 | 1.985703 | 0.047066 | 0.500211 | 0.300846 |
| ENSMUSG00000021747.13 | 10.81787 | -6.96352 | 3.510636 | -1.98355 | 0.047306 | 0.500514 | 0.300584 |
| ENSMUSG00000024675.20 | 22.40112 | -4.01337 | 2.022597 | -1.98427 | 0.047226 | 0.500514 | 0.300584 |
| ENSMUSG00000035004.4 | 10.58685 | -5.16867 | 2.60396 | -1.98493 | 0.047153 | 0.500514 | 0.300584 |
| ENSMUSG00000084844.4 | 10.81787 | -6.96352 | 3.510636 | -1.98355 | 0.047306 | 0.500514 | 0.300584 |
| ENSMUSG00000022177.10 | 29.89726 | 2.453598 | 1.237436 | 1.982809 | 0.047389 | 0.500832 | 0.300308 |
| ENSMUSG00000023977.16 | 114.5648 | 1.311669 | 0.662381 | 1.980232 | 0.047677 | 0.502761 | 0.298638 |
| ENSMUSG00000033487.15 | 288.0418 | 0.918041 | 0.463514 | 1.980612 | 0.047635 | 0.502761 | 0.298638 |
| ENSMUSG00000079316.11 | 216.3197 | 1.687598 | 0.85245 | 1.979704 | 0.047737 | 0.502828 | 0.298581 |
| ENSMUSG00000023089.13 | 1026.296 | 0.638478 | 0.322902 | 1.977311 | 0.048007 | 0.505107 | 0.296616 |
| ENSMUSG00000038696.15 | 129.1599 | -1.65176 | 0.835885 | -1.97606 | 0.048148 | 0.506034 | 0.29582 |
| ENSMUSG00000029250.6 | 464.2792 | 0.889687 | 0.450518 | 1.974811 | 0.04829 | 0.50696 | 0.295026 |
| ENSMUSG00000003617.17 | 3963.342 | -0.57783 | 0.293479 | -1.9689 | 0.048965 | 0.50817 | 0.293991 |
| ENSMUSG00000018381.16 | 10.77015 | 6.918822 | 3.510787 | 1.970732 | 0.048755 | 0.50817 | 0.293991 |
| ENSMUSG00000024070.16 | 1036.203 | -0.80688 | 0.409964 | -1.96817 | 0.049048 | 0.50817 | 0.293991 |
| ENSMUSG00000024986.13 | 960.9188 | 0.736082 | 0.373602 | 1.970228 | 0.048812 | 0.50817 | 0.293991 |
| ENSMUSG00000026104.15 | 161.7368 | -1.49245 | 0.756446 | -1.97298 | 0.048498 | 0.50817 | 0.293991 |
| ENSMUSG00000027805.17 | 9.922484 | 5.855036 | 2.974595 | 1.968347 | 0.049028 | 0.50817 | 0.293991 |
| ENSMUSG00000032437.11 | 651.0965 | 0.809142 | 0.41051 | 1.971064 | 0.048717 | 0.50817 | 0.293991 |
| ENSMUSG00000033960.7 | 14.78645 | -3.75471 | 1.907418 | -1.96848 | 0.049013 | 0.50817 | 0.293991 |
| ENSMUSG00000047649.10 | 34.47772 | -2.59039 | 1.313618 | -1.97195 | 0.048615 | 0.50817 | 0.293991 |
| ENSMUSG00000074412.5 | 456.1124 | 0.781027 | 0.396663 | 1.968993 | 0.048954 | 0.50817 | 0.293991 |
| ENSMUSG00000094156.3 | 47.54886 | 2.59243 | 1.316112 | 1.969764 | 0.048865 | 0.50817 | 0.293991 |
| ENSMUSG00000113774.2 | 10.77015 | 6.918822 | 3.510787 | 1.970732 | 0.048755 | 0.50817 | 0.293991 |
| ENSMUSG00000042487.7 | 188.1231 | 1.519915 | 0.77253 | 1.96745 | 0.049131 | 0.508477 | 0.293729 |
| ENSMUSG00000028436.9 | 106.6071 | -1.41933 | 0.72197 | -1.96591 | 0.049309 | 0.508993 | 0.293288 |
| ENSMUSG00000032184.6 | 10.82292 | 6.921967 | 3.521993 | 1.965355 | 0.049373 | 0.508993 | 0.293288 |
| ENSMUSG00000037962.8 | 10.82292 | 6.921967 | 3.521993 | 1.965355 | 0.049373 | 0.508993 | 0.293288 |
| ENSMUSG00000064337.1 | 14511.29 | -0.47411 | 0.241259 | -1.96516 | 0.049396 | 0.508993 | 0.293288 |
| ENSMUSG00000025153.10 | 826.3468 | 0.897624 | 0.457186 | 1.963365 | 0.049604 | 0.51058 | 0.291936 |
| ENSMUSG00000102014.2 | 12.12995 | -5.63786 | 2.87284 | -1.96247 | 0.049708 | 0.511095 | 0.291498 |
| ENSMUSG00000031171.12 | 162.3269 | 1.445231 | 0.736739 | 1.961659 | 0.049802 | 0.511155 | 0.291447 |

| | | | | | | | |
|---|---|---|---|---|---|---|---|
| ENSMUSG00000111055.2 | 10.57261 | -6.93085 | 3.533454 | -1.96149 | 0.049821 | 0.511155 | 0.291447 |
| ENSMUSG00000005583.17 | 22.18017 | 3.837649 | 1.960597 | 1.957388 | 0.050302 | 0.515525 | 0.28775 |
| ENSMUSG00000021149.9 | 329.5246 | -0.88768 | 0.45361 | -1.95693 | 0.050356 | 0.515525 | 0.28775 |
| ENSMUSG00000038069.7 | 48.79959 | -1.93038 | 0.987655 | -1.95451 | 0.050641 | 0.517879 | 0.285772 |
| ENSMUSG00000019088.15 | 44.72956 | -2.38827 | 1.223796 | -1.95152 | 0.050995 | 0.520939 | 0.283213 |
| ENSMUSG00000001909.17 | 34.57163 | -3.45666 | 1.771946 | -1.95077 | 0.051084 | 0.521291 | 0.28292 |
| ENSMUSG00000029328.16 | 373.8735 | -0.82413 | 0.42267 | -1.94982 | 0.051197 | 0.521882 | 0.282428 |
| ENSMUSG00000025780.8 | 148.8855 | -1.89505 | 0.972948 | -1.94774 | 0.051447 | 0.52386 | 0.280785 |
| ENSMUSG00000030451.17 | 89.86077 | -1.46474 | 0.752749 | -1.94585 | 0.051672 | 0.525032 | 0.279814 |
| ENSMUSG00000034330.11 | 14.06604 | -5.86018 | 3.011404 | -1.946 | 0.051655 | 0.525032 | 0.279814 |
| ENSMUSG00000011257.20 | 73.19918 | 1.569963 | 0.807144 | 1.945085 | 0.051765 | 0.525407 | 0.279504 |
| ENSMUSG00000027014.15 | 24.06807 | -3.33479 | 1.715438 | -1.94399 | 0.051897 | 0.525933 | 0.27907 |
| ENSMUSG00000030966.14 | 35.78254 | 3.520097 | 1.810997 | 1.943734 | 0.051928 | 0.525933 | 0.27907 |
| ENSMUSG00000028261.14 | 379.7436 | 0.894565 | 0.460721 | 1.941665 | 0.052178 | 0.526778 | 0.278372 |
| ENSMUSG00000045312.13 | 25.92654 | 3.465036 | 1.784198 | 1.942069 | 0.052129 | 0.526778 | 0.278372 |
| ENSMUSG00000089235.3 | 10.57695 | 6.888772 | 3.547272 | 1.941992 | 0.052138 | 0.526778 | 0.278372 |
| ENSMUSG00000002326.7 | 42.64436 | 2.391412 | 1.233964 | 1.937992 | 0.052624 | 0.529594 | 0.276057 |
| ENSMUSG00000044066.15 | 14.20362 | -4.05718 | 2.093229 | -1.93824 | 0.052594 | 0.529594 | 0.276057 |
| ENSMUSG00000073856.12 | 10.43358 | 6.873017 | 3.54546 | 1.93854 | 0.052557 | 0.529594 | 0.276057 |
| ENSMUSG00000024990.14 | 26418.59 | 0.375392 | 0.193792 | 1.937091 | 0.052734 | 0.530138 | 0.275611 |
| ENSMUSG00000001674.12 | 75.88381 | -2.33264 | 1.205234 | -1.93543 | 0.052938 | 0.530498 | 0.275316 |
| ENSMUSG00000021494.10 | 23.65571 | -3.82002 | 1.973229 | -1.93592 | 0.052877 | 0.530498 | 0.275316 |
| ENSMUSG00000086158.3 | 31.36473 | -3.19731 | 1.651686 | -1.93579 | 0.052894 | 0.530498 | 0.275316 |
| ENSMUSG00000026814.17 | 27.83679 | 2.774434 | 1.435726 | 1.932426 | 0.053307 | 0.53307 | 0.273216 |
| ENSMUSG00000028919.12 | 13.31025 | -4.37889 | 2.265643 | -1.93273 | 0.053269 | 0.53307 | 0.273216 |
| ENSMUSG00000003031.15 | 24.53163 | 3.055753 | 1.581941 | 1.931648 | 0.053403 | 0.533467 | 0.272893 |
| ENSMUSG00000028653.17 | 58.27157 | -1.86605 | 0.966893 | -1.92995 | 0.053614 | 0.533882 | 0.272554 |
| ENSMUSG00000032115.15 | 25.78452 | 3.797762 | 1.967219 | 1.930523 | 0.053542 | 0.533882 | 0.272554 |
| ENSMUSG00000041762.17 | 183.7707 | 1.144204 | 0.592847 | 1.930017 | 0.053605 | 0.533882 | 0.272554 |
| ENSMUSG00000063445.15 | 19.71062 | -4.40301 | 2.282193 | -1.92929 | 0.053695 | 0.534132 | 0.272351 |
| ENSMUSG00000021754.18 | 28.26082 | -2.80321 | 1.454402 | -1.9274 | 0.05393 | 0.53423 | 0.272272 |
| ENSMUSG00000044197.9 | 223.1155 | -1.1145 | 0.57817 | -1.92763 | 0.053901 | 0.53423 | 0.272272 |
| ENSMUSG00000048264.17 | 79.38499 | -2.23553 | 1.159489 | -1.92803 | 0.053852 | 0.53423 | 0.272272 |
| ENSMUSG00000071172.14 | 718.0929 | 0.681581 | 0.353594 | 1.927582 | 0.053907 | 0.53423 | 0.272272 |
| ENSMUSG00000026491.14 | 156.2481 | 1.157199 | 0.600993 | 1.925479 | 0.054169 | 0.53604 | 0.270802 |
| ENSMUSG00000024696.10 | 10.20554 | -6.87942 | 3.575183 | -1.92422 | 0.054328 | 0.536484 | 0.270443 |
| ENSMUSG00000060671.13 | 10.20554 | -6.87942 | 3.575183 | -1.92422 | 0.054328 | 0.536484 | 0.270443 |
| ENSMUSG00000027429.17 | 218.4012 | 1.229738 | 0.639236 | 1.923762 | 0.054384 | 0.536488 | 0.27044 |
| ENSMUSG00000028609.7 | 186.8019 | -1.32467 | 0.688768 | -1.92324 | 0.054449 | 0.53657 | 0.270374 |
| ENSMUSG00000004032.11 | 26.50719 | -2.8125 | 1.464902 | -1.91993 | 0.054867 | 0.537128 | 0.269922 |
| ENSMUSG00000020752.12 | 10.33097 | 6.854794 | 3.572869 | 1.918569 | 0.055039 | 0.537128 | 0.269922 |
| ENSMUSG00000021636.16 | 40.43604 | 2.06203 | 1.074753 | 1.918608 | 0.055034 | 0.537128 | 0.269922 |
| ENSMUSG00000026121.14 | 10.33097 | 6.854794 | 3.572869 | 1.918569 | 0.055039 | 0.537128 | 0.269922 |
| ENSMUSG00000031060.17 | 86.79863 | -1.54226 | 0.803586 | -1.91922 | 0.054957 | 0.537128 | 0.269922 |

| | | | | | | | |
|---|---|---|---|---|---|---|---|
| ENSMUSG00000036572.17 | 272.6969 | 0.94432 | 0.491947 | 1.919554 | 0.054914 | 0.537128 | 0.269922 |
| ENSMUSG00000052605.13 | 32.74508 | -3.85316 | 2.00863 | -1.9183 | 0.055073 | 0.537128 | 0.269922 |
| ENSMUSG00000058600.14 | 3820.121 | 0.540704 | 0.281735 | 1.919196 | 0.054959 | 0.537128 | 0.269922 |
| ENSMUSG00000082741.3 | 273.0093 | 0.9799 | 0.510461 | 1.919638 | 0.054904 | 0.537128 | 0.269922 |
| ENSMUSG00000089809.10 | 52.16441 | -2.02796 | 1.056766 | -1.91903 | 0.054981 | 0.537128 | 0.269922 |
| ENSMUSG00000035031.16 | 1684.809 | -0.58561 | 0.30535 | -1.91784 | 0.055131 | 0.537147 | 0.269907 |
| ENSMUSG00000051439.8 | 10.91597 | 5.394686 | 2.815219 | 1.916258 | 0.055332 | 0.538552 | 0.268772 |
| ENSMUSG00000027248.14 | 972.9613 | -0.67663 | 0.35319 | -1.91576 | 0.055396 | 0.538621 | 0.268717 |
| ENSMUSG00000025555.15 | 68.59021 | -1.75481 | 0.916429 | -1.91483 | 0.055514 | 0.538662 | 0.268684 |
| ENSMUSG00000027427.14 | 54.20943 | -1.84844 | 0.965111 | -1.91526 | 0.055459 | 0.538662 | 0.268684 |
| ENSMUSG00000068551.13 | 85.40914 | 1.629404 | 0.851149 | 1.914357 | 0.055575 | 0.538698 | 0.268655 |
| ENSMUSG00000023176.10 | 1291.137 | -0.60891 | 0.318268 | -1.91321 | 0.055722 | 0.539571 | 0.267951 |
| ENSMUSG00000014243.6 | 42.60546 | 2.135041 | 1.116737 | 1.911856 | 0.055895 | 0.539594 | 0.267933 |
| ENSMUSG00000019362.9 | 114.3132 | 1.374102 | 0.718539 | 1.912355 | 0.055831 | 0.539594 | 0.267933 |
| ENSMUSG00000024219.17 | 19.73838 | 3.06002 | 1.60039 | 1.912046 | 0.05587 | 0.539594 | 0.267933 |
| ENSMUSG00000031242.8 | 149.9745 | -1.22468 | 0.640774 | -1.91125 | 0.055972 | 0.53979 | 0.267775 |
| ENSMUSG00000066440.6 | 65.40018 | -1.62182 | 0.849142 | -1.90995 | 0.056139 | 0.540856 | 0.266918 |
| ENSMUSG00000081051.5 | 56.00072 | -1.87902 | 0.984102 | -1.90937 | 0.056214 | 0.541024 | 0.266783 |
| ENSMUSG00000028820.14 | 504.9495 | -0.74137 | 0.388591 | -1.90785 | 0.056411 | 0.541815 | 0.266149 |
| ENSMUSG00000078861.9 | 58.69428 | 2.580986 | 1.352633 | 1.90812 | 0.056376 | 0.541815 | 0.266149 |
| ENSMUSG00000041698.12 | 1265.315 | -0.65316 | 0.342654 | -1.90618 | 0.056626 | 0.543337 | 0.264931 |
| ENSMUSG00000021360.17 | 135.7922 | -1.33572 | 0.701658 | -1.90366 | 0.056955 | 0.544289 | 0.264171 |
| ENSMUSG00000026504.18 | 14.55148 | -4.80677 | 2.523979 | -1.90444 | 0.056852 | 0.544289 | 0.264171 |
| ENSMUSG00000027582.17 | 42.07651 | -3.10631 | 1.63064 | -1.90496 | 0.056785 | 0.544289 | 0.264171 |
| ENSMUSG00000112539.2 | 10.00143 | -6.85027 | 3.59839 | -1.9037 | 0.056949 | 0.544289 | 0.264171 |
| ENSMUSG00000021884.19 | 528.7711 | 0.730497 | 0.384288 | 1.900911 | 0.057314 | 0.547164 | 0.261882 |
| ENSMUSG00000026670.16 | 242.4404 | 1.018169 | 0.536469 | 1.897909 | 0.057708 | 0.550375 | 0.259341 |
| ENSMUSG00000036875.16 | 30.67101 | 3.315015 | 1.748088 | 1.896366 | 0.057912 | 0.551761 | 0.258249 |
| ENSMUSG00000040785.20 | 156.4337 | 1.116803 | 0.589323 | 1.895059 | 0.058085 | 0.552853 | 0.25739 |
| ENSMUSG00000113361.2 | 10.085 | 6.820001 | 3.600353 | 1.894259 | 0.058191 | 0.553307 | 0.257034 |
| ENSMUSG00000014907.11 | 46.15493 | -1.99318 | 1.052508 | -1.89375 | 0.058259 | 0.553346 | 0.257003 |
| ENSMUSG00000019710.14 | 424.8404 | -0.82516 | 0.436629 | -1.88985 | 0.058778 | 0.553346 | 0.257003 |
| ENSMUSG00000020526.13 | 95.90246 | 2.415303 | 1.27745 | 1.890721 | 0.058662 | 0.553346 | 0.257003 |
| ENSMUSG00000029550.12 | 83.78977 | -1.66964 | 0.8822 | -1.89259 | 0.058413 | 0.553346 | 0.257003 |
| ENSMUSG00000033688.11 | 2211.127 | -0.47228 | 0.249756 | -1.89098 | 0.058626 | 0.553346 | 0.257003 |
| ENSMUSG00000036390.9 | 87.75078 | -1.48977 | 0.787457 | -1.89188 | 0.058507 | 0.553346 | 0.257003 |
| ENSMUSG00000037262.8 | 99.11258 | -1.69921 | 0.899073 | -1.88996 | 0.058763 | 0.553346 | 0.257003 |
| ENSMUSG00000040466.17 | 626.5905 | 0.609798 | 0.322548 | 1.890565 | 0.058682 | 0.553346 | 0.257003 |
| ENSMUSG00000045252.12 | 10.13075 | -5.3776 | 2.840332 | -1.8933 | 0.058318 | 0.553346 | 0.257003 |
| ENSMUSG00000053510.13 | 187.6948 | -1.88713 | 0.997684 | -1.89151 | 0.058556 | 0.553346 | 0.257003 |
| ENSMUSG00000031502.12 | 63.4269 | -1.83265 | 0.970388 | -1.88857 | 0.058949 | 0.553784 | 0.25666 |
| ENSMUSG00000056536.15 | 20.32512 | -2.97187 | 1.574286 | -1.88776 | 0.059059 | 0.553784 | 0.25666 |
| ENSMUSG00000061619.4 | 9.928728 | 6.801462 | 3.602215 | 1.888133 | 0.059008 | 0.553784 | 0.25666 |
| ENSMUSG00000071335.6 | 120.4684 | 1.738003 | 0.920465 | 1.888179 | 0.059002 | 0.553784 | 0.25666 |

| | | | | | | | |
|---|---|---|---|---|---|---|---|
| ENSMUSG00000001289.13 | 524.4085 | 0.753349 | 0.399551 | 1.88549 | 0.059364 | 0.555546 | 0.25528 |
| ENSMUSG00000026623.17 | 936.2501 | 0.610151 | 0.323556 | 1.885767 | 0.059326 | 0.555546 | 0.25528 |
| ENSMUSG00000003161.16 | 279.233 | 1.064151 | 0.564615 | 1.884737 | 0.059465 | 0.555791 | 0.255089 |
| ENSMUSG00000038894.8 | 103.2698 | -2.0882 | 1.108389 | -1.88399 | 0.059566 | 0.555791 | 0.255089 |
| ENSMUSG00000113543.2 | 11.93406 | 4.329365 | 2.297486 | 1.884392 | 0.059512 | 0.555791 | 0.255089 |
| ENSMUSG00000025936.16 | 217.3371 | 1.139041 | 0.6048 | 1.883335 | 0.059655 | 0.556076 | 0.254866 |
| ENSMUSG00000071042.13 | 35.01918 | -3.42073 | 1.817005 | -1.88262 | 0.059752 | 0.55643 | 0.254589 |
| ENSMUSG00000046814.4 | 700.6308 | 0.587561 | 0.312288 | 1.881472 | 0.059908 | 0.557335 | 0.253883 |
| ENSMUSG00000014226.11 | 426.2745 | 0.886048 | 0.471473 | 1.87932 | 0.060201 | 0.557872 | 0.253465 |
| ENSMUSG00000029417.10 | 50.8074 | -2.58597 | 1.375435 | -1.88011 | 0.060093 | 0.557872 | 0.253465 |
| ENSMUSG00000060675.14 | 23.31226 | -3.1234 | 1.661735 | -1.8796 | 0.060162 | 0.557872 | 0.253465 |
| ENSMUSG00000091086.3 | 15.10074 | -4.58337 | 2.437778 | -1.88014 | 0.060089 | 0.557872 | 0.253465 |
| ENSMUSG00000021877.13 | 810.7931 | 0.569847 | 0.303534 | 1.877376 | 0.060467 | 0.559788 | 0.251977 |
| ENSMUSG00000038671.16 | 86.6077 | -1.66892 | 0.889187 | -1.8769 | 0.060531 | 0.559839 | 0.251937 |
| ENSMUSG00000000303.13 | 209.1743 | -1.23705 | 0.659501 | -1.87573 | 0.060692 | 0.560342 | 0.251547 |
| ENSMUSG00000009535.14 | 80.9985 | -1.41747 | 0.756027 | -1.87489 | 0.060807 | 0.560342 | 0.251547 |
| ENSMUSG00000024566.11 | 14.64847 | -4.10971 | 2.191805 | -1.87503 | 0.060788 | 0.560342 | 0.251547 |
| ENSMUSG00000116836.2 | 15.56552 | 5.476567 | 2.921167 | 1.874788 | 0.060822 | 0.560342 | 0.251547 |
| ENSMUSG00000045954.8 | 73.8182 | 1.679772 | 0.89638 | 1.873952 | 0.060937 | 0.560788 | 0.251202 |
| ENSMUSG00000063172.14 | 56.95436 | -2.0499 | 1.09411 | -1.87358 | 0.060989 | 0.560788 | 0.251202 |
| ENSMUSG00000026739.14 | 258.712 | -1.13541 | 0.606262 | -1.8728 | 0.061097 | 0.560988 | 0.251046 |
| ENSMUSG00000032235.16 | 97.51328 | 2.675593 | 1.42884 | 1.872563 | 0.061129 | 0.560988 | 0.251046 |
| ENSMUSG00000037820.16 | 205.5568 | -1.13823 | 0.60845 | -1.87071 | 0.061386 | 0.562802 | 0.249644 |
| ENSMUSG00000023572.17 | 122.6874 | -1.21968 | 0.653125 | -1.86746 | 0.061838 | 0.56585 | 0.247299 |
| ENSMUSG00000068735.15 | 20.55569 | -3.78969 | 2.029253 | -1.86753 | 0.061828 | 0.56585 | 0.247299 |
| ENSMUSG00000003849.9 | 20.30465 | -3.81234 | 2.044143 | -1.86501 | 0.06218 | 0.566958 | 0.246449 |
| ENSMUSG00000030512.14 | 209.2989 | 1.300639 | 0.697207 | 1.865498 | 0.062112 | 0.566958 | 0.246449 |
| ENSMUSG00000030614.9 | 216.6203 | 1.149818 | 0.616599 | 1.864773 | 0.062213 | 0.566958 | 0.246449 |
| ENSMUSG00000033720.13 | 24.16196 | -3.41625 | 1.832721 | -1.86403 | 0.062318 | 0.566958 | 0.246449 |
| ENSMUSG00000041567.4 | 727.5614 | -0.89463 | 0.479878 | -1.86429 | 0.062281 | 0.566958 | 0.246449 |
| ENSMUSG00000056501.4 | 634.3176 | -0.83966 | 0.450455 | -1.86403 | 0.062318 | 0.566958 | 0.246449 |
| ENSMUSG00000030545.10 | 402.7517 | 1.038977 | 0.558306 | 1.860947 | 0.062752 | 0.569814 | 0.244267 |
| ENSMUSG00000068040.11 | 125.4222 | -1.72132 | 0.924848 | -1.86119 | 0.062718 | 0.569814 | 0.244267 |
| ENSMUSG00000019854.18 | 24.9808 | 3.029316 | 1.628594 | 1.86008 | 0.062874 | 0.57038 | 0.243836 |
| ENSMUSG00000018574.15 | 508.096 | -0.65904 | 0.35455 | -1.8588 | 0.063055 | 0.571239 | 0.243182 |
| ENSMUSG00000029440.16 | 98.57465 | 2.056422 | 1.106712 | 1.858136 | 0.06315 | 0.571239 | 0.243182 |
| ENSMUSG00000040659.4 | 172.6557 | 1.057877 | 0.569279 | 1.858275 | 0.06313 | 0.571239 | 0.243182 |
| ENSMUSG00000072704.7 | 642.6663 | 0.663495 | 0.357204 | 1.857471 | 0.063244 | 0.571548 | 0.242947 |
| ENSMUSG00000005547.15 | 4370.015 | 0.473056 | 0.255565 | 1.851023 | 0.064166 | 0.572306 | 0.242371 |
| ENSMUSG00000020116.8 | 525.6163 | 0.779465 | 0.420691 | 1.85282 | 0.063908 | 0.572306 | 0.242371 |
| ENSMUSG00000026628.14 | 17.57309 | -4.40556 | 2.378746 | -1.85205 | 0.064019 | 0.572306 | 0.242371 |
| ENSMUSG00000028788.15 | 802.5492 | 0.598761 | 0.323171 | 1.852767 | 0.063916 | 0.572306 | 0.242371 |
| ENSMUSG00000028920.10 | 12.18171 | 3.884737 | 2.097555 | 1.852031 | 0.064021 | 0.572306 | 0.242371 |
| ENSMUSG00000030615.13 | 224.6948 | 1.240979 | 0.668804 | 1.855519 | 0.063522 | 0.572306 | 0.242371 |

| | | | | | | | |
|---|---|---|---|---|---|---|---|
| ENSMUSG00000033793.13 | 140.0894 | -1.3758 | 0.742277 | -1.85348 | 0.063813 | 0.572306 | 0.242371 |
| ENSMUSG00000034820.5 | 33.39836 | -3.07264 | 1.658256 | -1.85294 | 0.063891 | 0.572306 | 0.242371 |
| ENSMUSG00000036160.14 | 56.75469 | -2.15554 | 1.164592 | -1.85089 | 0.064185 | 0.572306 | 0.242371 |
| ENSMUSG00000037395.17 | 50.56481 | -2.33141 | 1.258762 | -1.85215 | 0.064005 | 0.572306 | 0.242371 |
| ENSMUSG00000037938.5 | 66.05795 | 1.903605 | 1.028667 | 1.850555 | 0.064234 | 0.572306 | 0.242371 |
| ENSMUSG00000044813.16 | 112.7351 | -1.50599 | 0.811492 | -1.85583 | 0.063477 | 0.572306 | 0.242371 |
| ENSMUSG00000047721.9 | 260.5318 | 0.901662 | 0.486975 | 1.851556 | 0.06409 | 0.572306 | 0.242371 |
| ENSMUSG00000058587.10 | 91.79809 | -1.38075 | 0.745787 | -1.8514 | 0.064112 | 0.572306 | 0.242371 |
| ENSMUSG00000112388.2 | 15.55177 | 4.479052 | 2.415609 | 1.854212 | 0.063709 | 0.572306 | 0.242371 |
| ENSMUSG00000047767.18 | 25.01951 | -2.75042 | 1.487888 | -1.84854 | 0.064525 | 0.573597 | 0.241393 |
| ENSMUSG00000047793.14 | 18.25041 | -3.65188 | 1.974949 | -1.8491 | 0.064443 | 0.573597 | 0.241393 |
| ENSMUSG00000070044.16 | 141.8646 | -1.27861 | 0.692093 | -1.84746 | 0.064681 | 0.573597 | 0.241393 |
| ENSMUSG00000074749.11 | 73.36755 | 1.715619 | 0.928361 | 1.848008 | 0.064601 | 0.573597 | 0.241393 |
| ENSMUSG00000085404.2 | 90.07227 | 1.767058 | 0.956381 | 1.847652 | 0.064653 | 0.573597 | 0.241393 |
| ENSMUSG00000003526.13 | 149.8228 | -1.86743 | 1.011772 | -1.8457 | 0.064936 | 0.575317 | 0.240093 |
| ENSMUSG00000034940.16 | 86.66231 | -1.73372 | 0.9404 | -1.84359 | 0.065242 | 0.577496 | 0.238451 |
| ENSMUSG00000026368.7 | 940.3661 | -0.55741 | 0.302433 | -1.84309 | 0.065316 | 0.577606 | 0.238368 |
| ENSMUSG00000031245.9 | 1014.16 | 0.584732 | 0.317801 | 1.839934 | 0.065778 | 0.581151 | 0.235711 |
| ENSMUSG00000022175.10 | 27.85591 | 2.944738 | 1.601682 | 1.838529 | 0.065984 | 0.58139 | 0.235533 |
| ENSMUSG00000030062.8 | 207.5759 | -1.66083 | 0.903659 | -1.83789 | 0.066078 | 0.58139 | 0.235533 |
| ENSMUSG00000031871.10 | 42.83467 | -2.23919 | 1.218446 | -1.83774 | 0.066101 | 0.58139 | 0.235533 |
| ENSMUSG00000055980.3 | 95.18175 | 1.540763 | 0.838435 | 1.837666 | 0.066112 | 0.58139 | 0.235533 |
| ENSMUSG00000113318.2 | 11.13422 | 5.9314 | 3.225538 | 1.838887 | 0.065932 | 0.58139 | 0.235533 |
| ENSMUSG00000008763.17 | 193.0701 | -0.96437 | 0.525241 | -1.83604 | 0.066351 | 0.582669 | 0.234578 |
| ENSMUSG00000024018.19 | 39.4828 | -2.06564 | 1.125422 | -1.83543 | 0.066441 | 0.582669 | 0.234578 |
| ENSMUSG00000056770.16 | 166.6007 | 1.085825 | 0.591538 | 1.835597 | 0.066417 | 0.582669 | 0.234578 |
| ENSMUSG00000018428.16 | 279.6726 | 0.80469 | 0.4387 | 1.834259 | 0.066615 | 0.583655 | 0.233844 |
| ENSMUSG00000020628.17 | 82.29574 | 2.494855 | 1.36054 | 1.833725 | 0.066695 | 0.583699 | 0.233811 |
| ENSMUSG00000026172.13 | 36.01346 | 2.421008 | 1.321241 | 1.832374 | 0.066896 | 0.583699 | 0.233811 |
| ENSMUSG00000032808.3 | 93.36074 | 1.450628 | 0.791907 | 1.831817 | 0.066979 | 0.583699 | 0.233811 |
| ENSMUSG00000042572.13 | 227.4962 | -1.30419 | 0.711997 | -1.83174 | 0.06699 | 0.583699 | 0.233811 |
| ENSMUSG00000043671.15 | 47.76622 | -2.06135 | 1.124508 | -1.83311 | 0.066786 | 0.583699 | 0.233811 |
| ENSMUSG00000055730.17 | 1907.289 | -0.60622 | 0.330738 | -1.83292 | 0.066814 | 0.583699 | 0.233811 |
| ENSMUSG00000003623.5 | 2928.216 | -0.57846 | 0.316159 | -1.82964 | 0.067304 | 0.585894 | 0.232181 |
| ENSMUSG00000022037.16 | 6612.452 | -0.4103 | 0.224331 | -1.829 | 0.0674 | 0.586196 | 0.231957 |
| ENSMUSG00000003814.9 | 961.7851 | -0.63533 | 0.34793 | -1.82604 | 0.067844 | 0.587065 | 0.231314 |
| ENSMUSG00000020549.15 | 28.61931 | -3.00341 | 1.644405 | -1.82644 | 0.067784 | 0.587065 | 0.231314 |
| ENSMUSG00000022914.18 | 91.23461 | -1.64875 | 0.902928 | -1.826 | 0.06785 | 0.587065 | 0.231314 |
| ENSMUSG00000024012.19 | 77.13909 | -1.4866 | 0.813973 | -1.82636 | 0.067797 | 0.587065 | 0.231314 |
| ENSMUSG00000038594.10 | 137.1255 | -1.14929 | 0.629455 | -1.82586 | 0.067872 | 0.587065 | 0.231314 |
| ENSMUSG00000055782.10 | 65.5763 | 2.744112 | 1.50244 | 1.826437 | 0.067784 | 0.587065 | 0.231314 |
| ENSMUSG00000009376.16 | 430.8235 | -0.70235 | 0.384769 | -1.82539 | 0.067942 | 0.587136 | 0.231261 |
| ENSMUSG00000057110.16 | 119.065 | 1.328382 | 0.728042 | 1.824596 | 0.068062 | 0.587639 | 0.23089 |
| ENSMUSG00000001665.12 | 388.6891 | 0.857042 | 0.469968 | 1.823618 | 0.06821 | 0.58804 | 0.230593 |

| | | | | | | | |
|---|---|---|---|---|---|---|---|
| ENSMUSG00000020483.15 | 525.4873 | -0.72829 | 0.3994 | -1.82347 | 0.068232 | 0.58804 | 0.230593 |
| ENSMUSG00000036256.14 | 934.4175 | 0.70639 | 0.387605 | 1.822449 | 0.068387 | 0.588294 | 0.230406 |
| ENSMUSG00000039041.16 | 196.9455 | -1.61427 | 0.885967 | -1.82205 | 0.068448 | 0.588294 | 0.230406 |
| ENSMUSG00000040661.7 | 20.92958 | -3.0986 | 1.700094 | -1.82261 | 0.068363 | 0.588294 | 0.230406 |
| ENSMUSG00000024425.5 | 2692.169 | 0.433001 | 0.238021 | 1.81917 | 0.068885 | 0.591516 | 0.228033 |
| ENSMUSG00000021492.16 | 510.8178 | -0.7543 | 0.41518 | -1.8168 | 0.069248 | 0.594092 | 0.226146 |
| ENSMUSG00000022235.16 | 1556.243 | 0.460136 | 0.253504 | 1.815107 | 0.069507 | 0.595778 | 0.224916 |
| ENSMUSG00000027808.9 | 3044.05 | 0.480554 | 0.264921 | 1.81395 | 0.069685 | 0.596226 | 0.224589 |
| ENSMUSG00000042626.14 | 91.13675 | -1.55816 | 0.858958 | -1.81401 | 0.069677 | 0.596226 | 0.224589 |
| ENSMUSG00000041560.13 | 157.2969 | -1.38057 | 0.761597 | -1.81273 | 0.069874 | 0.597301 | 0.223807 |
| ENSMUSG00000055053.19 | 275.1593 | 0.986527 | 0.544877 | 1.81055 | 0.070211 | 0.599636 | 0.222112 |
| ENSMUSG00000000127.16 | 56.68834 | -2.34314 | 1.296185 | -1.80772 | 0.07065 | 0.601818 | 0.220534 |
| ENSMUSG00000000278.11 | 135.3446 | -1.25646 | 0.695068 | -1.80768 | 0.070657 | 0.601818 | 0.220534 |
| ENSMUSG00000020108.5 | 12.16032 | 3.578416 | 1.979508 | 1.80773 | 0.070649 | 0.601818 | 0.220534 |
| ENSMUSG00000031634.14 | 295.0486 | -0.81168 | 0.449181 | -1.80703 | 0.070758 | 0.602141 | 0.220302 |
| ENSMUSG00000038766.17 | 27.96198 | 2.701593 | 1.496556 | 1.805207 | 0.071042 | 0.603167 | 0.219562 |
| ENSMUSG00000061540.4 | 881.9465 | -2.02559 | 1.122191 | -1.80503 | 0.071069 | 0.603167 | 0.219562 |
| ENSMUSG00000096215.9 | 15.45203 | -3.96697 | 2.197679 | -1.80507 | 0.071063 | 0.603167 | 0.219562 |
| ENSMUSG00000044783.17 | 62.88102 | -1.72874 | 0.959175 | -1.80232 | 0.071496 | 0.606243 | 0.217353 |
| ENSMUSG00000016194.15 | 6579.323 | 0.42818 | 0.238027 | 1.798873 | 0.072039 | 0.606385 | 0.217251 |
| ENSMUSG00000021990.17 | 92.56711 | -1.83324 | 1.019404 | -1.79835 | 0.072122 | 0.606385 | 0.217251 |
| ENSMUSG00000025759.12 | 96.23648 | -1.36437 | 0.758159 | -1.79958 | 0.071926 | 0.606385 | 0.217251 |
| ENSMUSG00000026189.14 | 1431.398 | -0.52589 | 0.292483 | -1.79802 | 0.072174 | 0.606385 | 0.217251 |
| ENSMUSG00000028107.15 | 58.59938 | 2.1064 | 1.171146 | 1.79858 | 0.072085 | 0.606385 | 0.217251 |
| ENSMUSG00000031016.10 | 34.77637 | -2.51675 | 1.399945 | -1.79775 | 0.072216 | 0.606385 | 0.217251 |
| ENSMUSG00000060548.14 | 19.04637 | -3.68771 | 2.046842 | -1.80166 | 0.071599 | 0.606385 | 0.217251 |
| ENSMUSG00000061244.15 | 169.603 | -1.14797 | 0.637752 | -1.80003 | 0.071856 | 0.606385 | 0.217251 |
| ENSMUSG00000062980.16 | 240.8334 | -0.95831 | 0.53245 | -1.79981 | 0.071891 | 0.606385 | 0.217251 |
| ENSMUSG00000068246.7 | 329.7041 | -0.79357 | 0.441029 | -1.79936 | 0.071962 | 0.606385 | 0.217251 |
| ENSMUSG00000087470.3 | 20.43419 | 2.670044 | 1.482317 | 1.801263 | 0.071661 | 0.606385 | 0.217251 |
| ENSMUSG00000031077.8 | 21.91991 | 2.917549 | 1.625721 | 1.794619 | 0.072714 | 0.60949 | 0.215034 |
| ENSMUSG00000038172.15 | 176.3794 | -1.01902 | 0.5678 | -1.79467 | 0.072706 | 0.60949 | 0.215034 |
| ENSMUSG00000029713.16 | 55.9276 | -1.86531 | 1.039662 | -1.79415 | 0.072789 | 0.609578 | 0.214971 |
| ENSMUSG00000027170.13 | 544.7086 | 0.663586 | 0.370036 | 1.793299 | 0.072925 | 0.610176 | 0.214545 |
| ENSMUSG00000029012.12 | 54.65273 | 1.904655 | 1.063712 | 1.790575 | 0.073362 | 0.613287 | 0.212336 |
| ENSMUSG00000100801.2 | 146.9696 | 1.295369 | 0.723645 | 1.790062 | 0.073444 | 0.613435 | 0.212231 |
| ENSMUSG00000038365.9 | 56.90408 | 1.776265 | 0.993055 | 1.788686 | 0.073665 | 0.614743 | 0.211307 |
| ENSMUSG00000042506.16 | 11.03064 | -4.11551 | 2.303672 | -1.7865 | 0.074019 | 0.617148 | 0.209611 |
| ENSMUSG00000034118.16 | 116.8415 | 1.268079 | 0.710131 | 1.785698 | 0.074148 | 0.61735 | 0.209468 |
| ENSMUSG00000040850.18 | 552.8865 | 0.66712 | 0.373623 | 1.785543 | 0.074173 | 0.61735 | 0.209468 |
| ENSMUSG00000006585.4 | 13.53042 | -3.91962 | 2.195825 | -1.78503 | 0.074256 | 0.617495 | 0.209367 |
| ENSMUSG00000013160.8 | 473.5629 | -0.85988 | 0.482461 | -1.78228 | 0.074704 | 0.61851 | 0.208653 |
| ENSMUSG00000019929.17 | 1445.779 | 0.574806 | 0.322385 | 1.782979 | 0.07459 | 0.61851 | 0.208653 |
| ENSMUSG00000024357.11 | 127.1769 | 1.251994 | 0.70243 | 1.782374 | 0.074688 | 0.61851 | 0.208653 |

| | | | | | | | |
|---|---|---|---|---|---|---|---|
| ENSMUSG00000045441.6 | 41.4821 | -2.09758 | 1.176909 | -1.78228 | 0.074704 | 0.61851 | 0.208653 |
| ENSMUSG00000059005.14 | 187.5234 | -1.241 | 0.695864 | -1.78339 | 0.074522 | 0.61851 | 0.208653 |
| ENSMUSG00000024966.10 | 34.04417 | 2.294607 | 1.287775 | 1.781838 | 0.074776 | 0.618563 | 0.208616 |
| ENSMUSG00000041992.10 | 24.67018 | 2.41314 | 1.355041 | 1.780861 | 0.074935 | 0.619342 | 0.208069 |
| ENSMUSG00000027198.17 | 73.6049 | -1.53914 | 0.86484 | -1.77968 | 0.075128 | 0.619972 | 0.207628 |
| ENSMUSG00000031783.11 | 102.338 | 1.333403 | 0.749441 | 1.779196 | 0.075208 | 0.619972 | 0.207628 |
| ENSMUSG00000118366.2 | 811.5475 | 0.804621 | 0.452149 | 1.779547 | 0.07515 | 0.619972 | 0.207628 |
| ENSMUSG00000025137.16 | 74.79164 | -1.384 | 0.778514 | -1.77774 | 0.075447 | 0.620324 | 0.207382 |
| ENSMUSG00000033294.12 | 29.74171 | -3.77798 | 2.124816 | -1.77803 | 0.075399 | 0.620324 | 0.207382 |
| ENSMUSG00000042742.8 | 108.9868 | 1.217796 | 0.684911 | 1.778036 | 0.075398 | 0.620324 | 0.207382 |
| ENSMUSG00000031660.14 | 174.2981 | -1.21612 | 0.684589 | -1.77642 | 0.075663 | 0.621316 | 0.206687 |
| ENSMUSG00000033060.17 | 9.981929 | -4.42046 | 2.489172 | -1.77588 | 0.075753 | 0.621316 | 0.206687 |
| ENSMUSG00000039354.17 | 26.68731 | 2.796304 | 1.574662 | 1.775812 | 0.075764 | 0.621316 | 0.206687 |
| ENSMUSG00000025076.13 | 90.59733 | 1.32087 | 0.744247 | 1.774773 | 0.075935 | 0.621646 | 0.206457 |
| ENSMUSG00000060601.14 | 14.7146 | -4.55418 | 2.565562 | -1.77512 | 0.075878 | 0.621646 | 0.206457 |
| ENSMUSG00000024515.14 | 135.6579 | -1.2687 | 0.715057 | -1.77426 | 0.07602 | 0.621801 | 0.206349 |
| ENSMUSG00000024132.7 | 1066.593 | 0.576949 | 0.325343 | 1.773357 | 0.07617 | 0.62249 | 0.205868 |
| ENSMUSG00000053436.16 | 433.382 | 0.794762 | 0.448449 | 1.772245 | 0.076354 | 0.623458 | 0.205193 |
| ENSMUSG00000020387.16 | 16.4246 | 3.658896 | 2.066301 | 1.770747 | 0.076603 | 0.624952 | 0.204153 |
| ENSMUSG00000078584.10 | 162.4614 | 1.042011 | 0.588598 | 1.770328 | 0.076673 | 0.624983 | 0.204131 |
| ENSMUSG00000031198.5 | 34.24397 | -2.2651 | 1.280656 | -1.7687 | 0.076944 | 0.626653 | 0.202973 |
| ENSMUSG00000008305.19 | 298.7276 | -0.91452 | 0.51754 | -1.76705 | 0.07722 | 0.627287 | 0.202533 |
| ENSMUSG00000027597.16 | 1644.798 | 0.546038 | 0.308965 | 1.767312 | 0.077176 | 0.627287 | 0.202533 |
| ENSMUSG00000029063.17 | 1189.833 | 0.492415 | 0.278611 | 1.767396 | 0.077162 | 0.627287 | 0.202533 |
| ENSMUSG00000004535.13 | 787.9928 | -0.69514 | 0.393706 | -1.76562 | 0.077459 | 0.628695 | 0.20156 |
| ENSMUSG00000020843.16 | 73.22384 | 1.677162 | 0.950939 | 1.76369 | 0.077784 | 0.629235 | 0.201187 |
| ENSMUSG00000024556.5 | 12.01173 | -4.83649 | 2.742177 | -1.76374 | 0.077776 | 0.629235 | 0.201187 |
| ENSMUSG00000031485.16 | 216.5392 | -1.36615 | 0.774615 | -1.76365 | 0.077791 | 0.629235 | 0.201187 |
| ENSMUSG00000066000.13 | 54.55174 | 1.79922 | 1.019665 | 1.76452 | 0.077644 | 0.629235 | 0.201187 |
| ENSMUSG00000050445.5 | 763.0511 | 0.72365 | 0.410509 | 1.762813 | 0.077932 | 0.629835 | 0.200773 |
| ENSMUSG00000004040.17 | 167.5932 | -1.5907 | 0.903245 | -1.76109 | 0.078223 | 0.631175 | 0.19985 |
| ENSMUSG00000010453.13 | 192.3762 | -1.1196 | 0.635762 | -1.76104 | 0.078231 | 0.631175 | 0.19985 |
| ENSMUSG00000010095.14 | 169.1525 | 1.459577 | 0.831446 | 1.75547 | 0.079179 | 0.631463 | 0.199652 |
| ENSMUSG00000028978.13 | 11.26314 | 4.80431 | 2.73791 | 1.754736 | 0.079304 | 0.631463 | 0.199652 |
| ENSMUSG00000029559.16 | 31.08787 | -2.87892 | 1.637636 | -1.75797 | 0.078752 | 0.631463 | 0.199652 |
| ENSMUSG00000029992.15 | 87.68284 | 1.544735 | 0.880304 | 1.754774 | 0.079298 | 0.631463 | 0.199652 |
| ENSMUSG00000031568.17 | 107.8851 | 1.597838 | 0.910564 | 1.754777 | 0.079297 | 0.631463 | 0.199652 |
| ENSMUSG00000031641.15 | 179.6286 | -1.1757 | 0.670077 | -1.75457 | 0.079333 | 0.631463 | 0.199652 |
| ENSMUSG00000042605.17 | 247.5671 | -0.90414 | 0.515004 | -1.7556 | 0.079158 | 0.631463 | 0.199652 |
| ENSMUSG00000052560.16 | 34.80494 | -2.99879 | 1.706335 | -1.75745 | 0.078842 | 0.631463 | 0.199652 |
| ENSMUSG00000055093.11 | 194.4795 | 0.985235 | 0.560632 | 1.757364 | 0.078856 | 0.631463 | 0.199652 |
| ENSMUSG00000057143.16 | 26.88514 | -2.74602 | 1.563067 | -1.75681 | 0.07895 | 0.631463 | 0.199652 |
| ENSMUSG00000062232.15 | 14.75147 | -3.47785 | 1.980033 | -1.75646 | 0.07901 | 0.631463 | 0.199652 |
| ENSMUSG00000063439.8 | 21.86891 | -4.02407 | 2.286715 | -1.75976 | 0.078448 | 0.631463 | 0.199652 |

| | | | | | | | |
|---|---|---|---|---|---|---|---|
| ENSMUSG00000068262.5 | 93.0108 | 1.457025 | 0.828748 | 1.758103 | 0.07873 | 0.631463 | 0.199652 |
| ENSMUSG00000069729.15 | 44.52467 | 2.066816 | 1.176053 | 1.757418 | 0.078847 | 0.631463 | 0.199652 |
| ENSMUSG00000075595.10 | 61.64163 | -1.61666 | 0.919098 | -1.75896 | 0.078585 | 0.631463 | 0.199652 |
| ENSMUSG00000090213.2 | 139.5234 | 1.408947 | 0.800915 | 1.759173 | 0.078548 | 0.631463 | 0.199652 |
| ENSMUSG00000000916.16 | 15.45417 | 4.248191 | 2.422998 | 1.753279 | 0.079554 | 0.632314 | 0.199067 |
| ENSMUSG00000035875.16 | 2297.176 | -0.50032 | 0.285382 | -1.75317 | 0.079573 | 0.632314 | 0.199067 |
| ENSMUSG00000017950.17 | 953.3674 | -0.58996 | 0.336619 | -1.75261 | 0.079669 | 0.632545 | 0.198909 |
| ENSMUSG00000097354.8 | 164.5351 | -1.63574 | 0.933701 | -1.75189 | 0.079793 | 0.633003 | 0.198594 |
| ENSMUSG00000015759.12 | 263.0925 | -0.87348 | 0.499467 | -1.74882 | 0.080322 | 0.634542 | 0.197539 |
| ENSMUSG00000029034.10 | 70.14513 | 1.662081 | 0.949708 | 1.750096 | 0.080102 | 0.634542 | 0.197539 |
| ENSMUSG00000032766.10 | 65.52457 | 2.303098 | 1.316635 | 1.74923 | 0.080251 | 0.634542 | 0.197539 |
| ENSMUSG00000035069.4 | 68.18728 | -1.52131 | 0.869674 | -1.74928 | 0.080242 | 0.634542 | 0.197539 |
| ENSMUSG00000037922.14 | 19.78244 | 3.762549 | 2.151296 | 1.748969 | 0.080296 | 0.634542 | 0.197539 |
| ENSMUSG00000026385.17 | 12783.04 | 0.391571 | 0.224106 | 1.747256 | 0.080593 | 0.636154 | 0.196438 |
| ENSMUSG00000020954.17 | 142.1174 | -1.16824 | 0.669707 | -1.7444 | 0.081089 | 0.639002 | 0.194498 |
| ENSMUSG00000039428.12 | 549.4366 | 0.64973 | 0.37246 | 1.744427 | 0.081085 | 0.639002 | 0.194498 |
| ENSMUSG00000037526.8 | 46.58088 | -2.27636 | 1.306102 | -1.74286 | 0.081358 | 0.640589 | 0.19342 |
| ENSMUSG00000021876.16 | 10494.62 | -0.45061 | 0.258825 | -1.741 | 0.081684 | 0.64103 | 0.193122 |
| ENSMUSG00000022160.19 | 48.67653 | -1.91917 | 1.102283 | -1.74108 | 0.081669 | 0.64103 | 0.193122 |
| ENSMUSG00000022228.15 | 119.2384 | -1.08459 | 0.622746 | -1.74163 | 0.081574 | 0.64103 | 0.193122 |
| ENSMUSG00000027639.17 | 51.8305 | -2.00316 | 1.149895 | -1.74204 | 0.081502 | 0.64103 | 0.193122 |
| ENSMUSG00000022098.11 | 162.7516 | -1.10958 | 0.637852 | -1.73956 | 0.081937 | 0.641602 | 0.192734 |
| ENSMUSG00000086429.10 | 131.075 | 1.377673 | 0.791727 | 1.740086 | 0.081844 | 0.641602 | 0.192734 |
| ENSMUSG00000087269.2 | 10.7982 | 4.939917 | 2.83997 | 1.739426 | 0.08196 | 0.641602 | 0.192734 |
| ENSMUSG00000029918.10 | 52.89706 | 1.731717 | 0.995886 | 1.73887 | 0.082058 | 0.641836 | 0.192576 |
| ENSMUSG00000019977.17 | 254.9731 | -0.96426 | 0.554883 | -1.73778 | 0.082249 | 0.642536 | 0.192102 |
| ENSMUSG00000026355.12 | 25.19852 | -2.79121 | 1.606368 | -1.73759 | 0.082283 | 0.642536 | 0.192102 |
| ENSMUSG00000022559.9 | 19.55818 | -3.41151 | 1.964585 | -1.73651 | 0.082475 | 0.642976 | 0.191805 |
| ENSMUSG00000113660.2 | 17.72544 | -3.90714 | 2.249546 | -1.73686 | 0.082412 | 0.642976 | 0.191805 |
| ENSMUSG00000001065.16 | 23.73836 | 3.495639 | 2.018261 | 1.732006 | 0.083273 | 0.642986 | 0.191799 |
| ENSMUSG00000015214.15 | 139.5922 | -1.14367 | 0.661407 | -1.72915 | 0.083783 | 0.642986 | 0.191799 |
| ENSMUSG00000016534.16 | 2163.71 | 0.424341 | 0.244696 | 1.734159 | 0.08289 | 0.642986 | 0.191799 |
| ENSMUSG00000021756.13 | 307.5663 | -0.7706 | 0.444399 | -1.73402 | 0.082915 | 0.642986 | 0.191799 |
| ENSMUSG00000023927.16 | 12.45567 | -4.56999 | 2.645105 | -1.72772 | 0.084039 | 0.642986 | 0.191799 |
| ENSMUSG00000024191.16 | 37.26971 | 2.828224 | 1.632908 | 1.732017 | 0.083271 | 0.642986 | 0.191799 |
| ENSMUSG00000024921.18 | 423.8078 | -0.83168 | 0.48089 | -1.72947 | 0.083725 | 0.642986 | 0.191799 |
| ENSMUSG00000025405.10 | 429.2315 | -0.77298 | 0.447491 | -1.72736 | 0.084104 | 0.642986 | 0.191799 |
| ENSMUSG00000027487.12 | 59.20536 | -2.14704 | 1.240826 | -1.73033 | 0.08357 | 0.642986 | 0.191799 |
| ENSMUSG00000028431.12 | 52.7365 | -2.00679 | 1.159322 | -1.731 | 0.083452 | 0.642986 | 0.191799 |
| ENSMUSG00000028563.17 | 135.4688 | 1.42894 | 0.823848 | 1.73447 | 0.082835 | 0.642986 | 0.191799 |
| ENSMUSG00000029458.15 | 302.2806 | -0.7644 | 0.442131 | -1.72891 | 0.083826 | 0.642986 | 0.191799 |
| ENSMUSG00000029591.15 | 20.80583 | 3.90092 | 2.252732 | 1.731639 | 0.083338 | 0.642986 | 0.191799 |
| ENSMUSG00000033488.12 | 41.1715 | -2.83772 | 1.642736 | -1.72744 | 0.08409 | 0.642986 | 0.191799 |
| ENSMUSG00000036391.17 | 886.7936 | 0.6774 | 0.392097 | 1.727635 | 0.084054 | 0.642986 | 0.191799 |

| | | | | | | | |
|---|---|---|---|---|---|---|---|
| ENSMUSG00000037361.9 | 546.553 | 0.769845 | 0.443466 | 1.735974 | 0.082569 | 0.642986 | 0.191799 |
| ENSMUSG00000038615.18 | 117.1823 | 1.299649 | 0.749543 | 1.733923 | 0.082932 | 0.642986 | 0.191799 |
| ENSMUSG00000038806.8 | 49.78376 | 2.325256 | 1.343574 | 1.730651 | 0.083514 | 0.642986 | 0.191799 |
| ENSMUSG00000040451.19 | 64.40441 | 1.637847 | 0.946961 | 1.729582 | 0.083705 | 0.642986 | 0.191799 |
| ENSMUSG00000040479.13 | 48.94983 | -2.19957 | 1.270297 | -1.73154 | 0.083355 | 0.642986 | 0.191799 |
| ENSMUSG00000042485.8 | 36.33635 | -3.08152 | 1.782672 | -1.7286 | 0.083881 | 0.642986 | 0.191799 |
| ENSMUSG00000055044.13 | 93.60139 | 1.40685 | 0.812002 | 1.732569 | 0.083172 | 0.642986 | 0.191799 |
| ENSMUSG00000062901.4 | 368.321 | -1.23963 | 0.717276 | -1.72825 | 0.083943 | 0.642986 | 0.191799 |
| ENSMUSG00000074882.4 | 900.0419 | 0.599432 | 0.345751 | 1.733713 | 0.082969 | 0.642986 | 0.191799 |
| ENSMUSG00000033499.15 | 1572.685 | -0.45932 | 0.266021 | -1.72663 | 0.084234 | 0.643461 | 0.191478 |
| ENSMUSG00000004341.9 | 15.29711 | 3.104432 | 1.799263 | 1.725391 | 0.084457 | 0.644524 | 0.190761 |
| ENSMUSG00000021254.10 | 44.38117 | -2.59139 | 1.502164 | -1.7251 | 0.084509 | 0.644524 | 0.190761 |
| ENSMUSG00000041438.9 | 127.1263 | 1.160972 | 0.673552 | 1.723658 | 0.08477 | 0.645994 | 0.189772 |
| ENSMUSG00000037656.10 | 131.4868 | -1.35941 | 0.788981 | -1.72299 | 0.08489 | 0.646394 | 0.189502 |
| ENSMUSG00000020638.9 | 17.56654 | -3.21921 | 1.869223 | -1.72222 | 0.08503 | 0.646935 | 0.18914 |
| ENSMUSG00000053580.17 | 36.84329 | 2.104047 | 1.222455 | 1.721165 | 0.085221 | 0.64787 | 0.188512 |
| ENSMUSG00000024002.19 | 141.9963 | -1.83328 | 1.06635 | -1.71921 | 0.085576 | 0.650047 | 0.187055 |
| ENSMUSG00000027514.15 | 23.90282 | -3.37293 | 1.964591 | -1.71686 | 0.086005 | 0.65278 | 0.185233 |
| ENSMUSG00000020902.13 | 59.00481 | 2.184321 | 1.272655 | 1.716349 | 0.086098 | 0.652968 | 0.185108 |
| ENSMUSG00000031725.9 | 3540.657 | 0.366004 | 0.213378 | 1.715287 | 0.086293 | 0.653398 | 0.184822 |
| ENSMUSG00000036323.15 | 721.9495 | -0.54318 | 0.316639 | -1.71544 | 0.086264 | 0.653398 | 0.184822 |
| ENSMUSG00000094786.2 | 175.4222 | 1.237311 | 0.722124 | 1.713433 | 0.086633 | 0.655451 | 0.18346 |
| ENSMUSG00000043587.16 | 36.72887 | -2.53025 | 1.477657 | -1.71234 | 0.086835 | 0.656453 | 0.182796 |
| ENSMUSG00000024197.11 | 87.95319 | -1.35341 | 0.790949 | -1.71112 | 0.087059 | 0.657274 | 0.182253 |
| ENSMUSG00000025217.16 | 20.95398 | 3.302696 | 1.930644 | 1.710671 | 0.087142 | 0.657274 | 0.182253 |
| ENSMUSG00000032187.17 | 96.92296 | 1.733503 | 1.013377 | 1.71062 | 0.087151 | 0.657274 | 0.182253 |
| ENSMUSG00000020460.16 | 5174.953 | 0.47641 | 0.279353 | 1.705404 | 0.088119 | 0.664025 | 0.177816 |
| ENSMUSG00000020647.11 | 107.127 | -1.18191 | 0.693185 | -1.70504 | 0.088186 | 0.664025 | 0.177816 |
| ENSMUSG00000074785.6 | 81.50707 | 1.452805 | 0.852501 | 1.704169 | 0.08835 | 0.664725 | 0.177358 |
| ENSMUSG00000027479.15 | 160.0266 | -1.01733 | 0.597602 | -1.70236 | 0.088689 | 0.664841 | 0.177282 |
| ENSMUSG00000031303.9 | 28.09732 | 3.314433 | 1.94756 | 1.701838 | 0.088786 | 0.664841 | 0.177282 |
| ENSMUSG00000038173.16 | 22.98877 | 2.814809 | 1.653093 | 1.702753 | 0.088614 | 0.664841 | 0.177282 |
| ENSMUSG00000039637.16 | 11.494 | -3.98247 | 2.340035 | -1.70189 | 0.088777 | 0.664841 | 0.177282 |
| ENSMUSG00000040706.5 | 727.4168 | -0.71927 | 0.422514 | -1.70235 | 0.08869 | 0.664841 | 0.177282 |
| ENSMUSG00000073471.4 | 14.18285 | 5.129841 | 3.012553 | 1.702822 | 0.088601 | 0.664841 | 0.177282 |
| ENSMUSG00000021947.12 | 73.8359 | -1.51395 | 0.889946 | -1.70118 | 0.08891 | 0.665246 | 0.177018 |
| ENSMUSG00000006782.17 | 17.93461 | -4.06766 | 2.394126 | -1.69902 | 0.089316 | 0.666182 | 0.176407 |
| ENSMUSG00000014232.15 | 65.02363 | 1.639696 | 0.964931 | 1.699289 | 0.089265 | 0.666182 | 0.176407 |
| ENSMUSG00000038683.17 | 338.0922 | 0.758918 | 0.446478 | 1.69979 | 0.089171 | 0.666182 | 0.176407 |
| ENSMUSG00000047126.18 | 568.495 | -0.71104 | 0.418463 | -1.69917 | 0.089287 | 0.666182 | 0.176407 |
| ENSMUSG00000022516.11 | 90.12741 | -1.33731 | 0.78733 | -1.69854 | 0.089405 | 0.666322 | 0.176316 |
| ENSMUSG00000020949.10 | 329.4766 | 0.735682 | 0.433661 | 1.696444 | 0.089802 | 0.668752 | 0.174735 |
| ENSMUSG00000038722.18 | 461.7511 | -0.80118 | 0.472736 | -1.69477 | 0.09012 | 0.670593 | 0.173541 |
| ENSMUSG00000027522.16 | 27.46374 | 2.656709 | 1.567952 | 1.694381 | 0.090193 | 0.67061 | 0.17353 |

| | | | | | | | |
|---|---|---|---|---|---|---|---|
| ENSMUSG00000002588.14 | 6843.167 | 0.393136 | 0.232157 | 1.693406 | 0.090378 | 0.671462 | 0.172979 |
| ENSMUSG00000019437.18 | 92.70798 | 1.669443 | 0.986327 | 1.692585 | 0.090534 | 0.672096 | 0.172569 |
| ENSMUSG00000041697.10 | 4409.782 | 0.418058 | 0.247128 | 1.691665 | 0.09071 | 0.672871 | 0.172068 |
| ENSMUSG00000079427.11 | 93.75784 | 1.35454 | 0.800948 | 1.691172 | 0.090804 | 0.673042 | 0.171958 |
| ENSMUSG00000001576.16 | 210.688 | -1.067 | 0.631402 | -1.68988 | 0.09105 | 0.673814 | 0.17146 |
| ENSMUSG00000046794.10 | 1658.222 | 1.40132 | 0.829085 | 1.6902 | 0.09099 | 0.673814 | 0.17146 |
| ENSMUSG00000003585.14 | 549.2324 | 0.761132 | 0.45114 | 1.68713 | 0.091578 | 0.675679 | 0.170259 |
| ENSMUSG00000020167.15 | 13.49167 | 4.272938 | 2.532302 | 1.687373 | 0.091532 | 0.675679 | 0.170259 |
| ENSMUSG00000036606.18 | 32.45788 | -2.13559 | 1.265631 | -1.68737 | 0.091533 | 0.675679 | 0.170259 |
| ENSMUSG00000048495.17 | 86.95742 | -1.29948 | 0.77042 | -1.68671 | 0.091659 | 0.675679 | 0.170259 |
| ENSMUSG00000054757.15 | 727.4845 | 0.536281 | 0.317917 | 1.686858 | 0.091631 | 0.675679 | 0.170259 |
| ENSMUSG00000001910.6 | 11.71874 | -4.48044 | 2.660687 | -1.68394 | 0.092193 | 0.677513 | 0.169083 |
| ENSMUSG00000005374.14 | 47.22989 | -2.30153 | 1.366057 | -1.6848 | 0.092028 | 0.677513 | 0.169083 |
| ENSMUSG00000026712.4 | 358.4472 | 0.739909 | 0.439356 | 1.684077 | 0.092167 | 0.677513 | 0.169083 |
| ENSMUSG00000071267.12 | 47.19577 | -1.84293 | 1.094223 | -1.68424 | 0.092135 | 0.677513 | 0.169083 |
| ENSMUSG00000037443.14 | 51.99975 | -1.63631 | 0.971995 | -1.68346 | 0.092287 | 0.677674 | 0.168979 |
| ENSMUSG00000027900.16 | 517.2282 | 0.706627 | 0.419917 | 1.682777 | 0.092418 | 0.677899 | 0.168835 |
| ENSMUSG00000039987.16 | 45.10131 | -2.30465 | 1.369836 | -1.68243 | 0.092486 | 0.677899 | 0.168835 |
| ENSMUSG00000041733.11 | 150.0286 | -1.181 | 0.70206 | -1.68219 | 0.092532 | 0.677899 | 0.168835 |
| ENSMUSG00000044452.11 | 24.29864 | 2.27874 | 1.355488 | 1.681122 | 0.092739 | 0.67837 | 0.168533 |
| ENSMUSG00000058655.10 | 205.8111 | -0.9133 | 0.54322 | -1.68127 | 0.092711 | 0.67837 | 0.168533 |
| ENSMUSG00000020961.16 | 9.946179 | -4.82154 | 2.870731 | -1.67955 | 0.093044 | 0.679158 | 0.168029 |
| ENSMUSG00000026126.16 | 18.93198 | -3.52242 | 2.097349 | -1.67946 | 0.093062 | 0.679158 | 0.168029 |
| ENSMUSG00000046876.17 | 62.35149 | -1.7886 | 1.064969 | -1.67948 | 0.093058 | 0.679158 | 0.168029 |
| ENSMUSG00000004667.19 | 92.62153 | -1.24058 | 0.739791 | -1.67693 | 0.093556 | 0.681577 | 0.166485 |
| ENSMUSG00000020109.14 | 62.592 | 1.594703 | 0.950748 | 1.677315 | 0.093481 | 0.681577 | 0.166485 |
| ENSMUSG00000097042.3 | 39.97169 | 2.076598 | 1.238533 | 1.67666 | 0.093609 | 0.681577 | 0.166485 |
| ENSMUSG00000026869.13 | 252.4286 | 1.076426 | 0.642299 | 1.675895 | 0.093759 | 0.68162 | 0.166458 |
| ENSMUSG00000031386.15 | 56.81512 | -1.68591 | 1.00588 | -1.67605 | 0.093728 | 0.68162 | 0.166458 |
| ENSMUSG00000035125.11 | 17.93889 | -3.97115 | 2.371195 | -1.67474 | 0.093984 | 0.682737 | 0.165746 |
| ENSMUSG00000009470.18 | 349.2311 | -0.90316 | 0.539639 | -1.67364 | 0.094201 | 0.68274 | 0.165744 |
| ENSMUSG00000031451.7 | 33.87951 | 2.294578 | 1.370607 | 1.674132 | 0.094105 | 0.68274 | 0.165744 |
| ENSMUSG00000038400.16 | 13.33399 | 4.253776 | 2.541625 | 1.673644 | 0.094201 | 0.68274 | 0.165744 |
| ENSMUSG00000029771.13 | 48.44595 | -2.36868 | 1.415844 | -1.67298 | 0.094331 | 0.682783 | 0.165717 |
| ENSMUSG00000111375.5 | 10.35029 | -4.87479 | 2.914008 | -1.67288 | 0.094351 | 0.682783 | 0.165717 |
| ENSMUSG00000032615.15 | 53.59482 | 1.851953 | 1.107574 | 1.672081 | 0.094508 | 0.683402 | 0.165324 |
| ENSMUSG00000002064.11 | 353.3233 | -0.74263 | 0.444261 | -1.67162 | 0.0946 | 0.683477 | 0.165276 |
| ENSMUSG00000038518.16 | 44.44999 | -2.11774 | 1.267123 | -1.6713 | 0.094663 | 0.683477 | 0.165276 |
| ENSMUSG00000003949.17 | 282.5519 | 0.892703 | 0.534387 | 1.670517 | 0.094817 | 0.684068 | 0.164901 |
| ENSMUSG00000036990.14 | 121.5755 | -1.14492 | 0.685521 | -1.67014 | 0.094891 | 0.684082 | 0.164892 |
| ENSMUSG00000032898.10 | 93.52493 | -1.87091 | 1.121128 | -1.66878 | 0.095162 | 0.684253 | 0.164783 |
| ENSMUSG00000040713.13 | 3174.527 | 0.489295 | 0.293105 | 1.669348 | 0.095048 | 0.684253 | 0.164783 |
| ENSMUSG00000048758.15 | 259.3578 | 0.90731 | 0.543767 | 1.668565 | 0.095204 | 0.684253 | 0.164783 |
| ENSMUSG00000115497.2 | 28.17635 | 2.578689 | 1.545178 | 1.668862 | 0.095145 | 0.684253 | 0.164783 |

| | | | | | | | |
|---|---|---|---|---|---|---|---|
| ENSMUSG00000030498.16 | 227.8239 | 0.834711 | 0.500804 | 1.666741 | 0.095566 | 0.685833 | 0.163781 |
| ENSMUSG00000063077.16 | 462.3815 | -0.77826 | 0.466941 | -1.66673 | 0.095568 | 0.685833 | 0.163781 |
| ENSMUSG00000003037.17 | 88.24309 | 1.4257 | 0.855643 | 1.666232 | 0.095667 | 0.686025 | 0.16366 |
| ENSMUSG00000067279.4 | 211.6796 | 1.143174 | 0.686816 | 1.664455 | 0.096021 | 0.688045 | 0.162383 |
| ENSMUSG00000009291.14 | 1035.894 | 0.530643 | 0.319043 | 1.663235 | 0.096265 | 0.688843 | 0.161879 |
| ENSMUSG00000036009.17 | 30.69469 | 2.364713 | 1.42212 | 1.662808 | 0.096351 | 0.688843 | 0.161879 |
| ENSMUSG00000045680.9 | 16.427 | 3.309156 | 1.989927 | 1.662954 | 0.096322 | 0.688843 | 0.161879 |
| ENSMUSG00000030086.17 | 13.78118 | 4.309698 | 2.593735 | 1.66158 | 0.096597 | 0.688974 | 0.161797 |
| ENSMUSG00000030834.8 | 104.2155 | -1.20212 | 0.723774 | -1.6609 | 0.096733 | 0.688974 | 0.161797 |
| ENSMUSG00000033382.16 | 70.7928 | 1.838464 | 1.106861 | 1.660971 | 0.096719 | 0.688974 | 0.161797 |
| ENSMUSG00000033430.11 | 65.66879 | 1.838834 | 1.106904 | 1.661241 | 0.096665 | 0.688974 | 0.161797 |
| ENSMUSG00000047228.10 | 511.9391 | -0.70612 | 0.424824 | -1.66214 | 0.096485 | 0.688974 | 0.161797 |
| ENSMUSG00000035834.12 | 52.82597 | 2.707723 | 1.630837 | 1.660327 | 0.096849 | 0.689283 | 0.161602 |
| ENSMUSG00000020571.14 | 719.1078 | -0.55034 | 0.331645 | -1.65941 | 0.097033 | 0.689649 | 0.161372 |
| ENSMUSG00000058756.14 | 15.0238 | -2.93796 | 1.770548 | -1.65935 | 0.097046 | 0.689649 | 0.161372 |
| ENSMUSG00000032252.15 | 175.5616 | 0.869384 | 0.524124 | 1.658736 | 0.097169 | 0.690009 | 0.161145 |
| ENSMUSG00000028948.17 | 37.98683 | -1.96576 | 1.18602 | -1.65744 | 0.09743 | 0.690946 | 0.160556 |
| ENSMUSG00000030978.11 | 57.52621 | 2.077816 | 1.253831 | 1.657174 | 0.097484 | 0.690946 | 0.160556 |
| ENSMUSG00000048731.16 | 11.03222 | 3.478167 | 2.099076 | 1.656999 | 0.09752 | 0.690946 | 0.160556 |
| ENSMUSG00000041718.17 | 43.71152 | -1.94226 | 1.172548 | -1.65645 | 0.097632 | 0.691223 | 0.160382 |
| ENSMUSG00000020372.16 | 1941.704 | 0.405337 | 0.245018 | 1.654312 | 0.098064 | 0.69364 | 0.158866 |
| ENSMUSG00000051615.15 | 166.1777 | 1.017732 | 0.615434 | 1.65368 | 0.098193 | 0.69364 | 0.158866 |
| ENSMUSG00000100594.2 | 15.50928 | -4.12457 | 2.494123 | -1.65371 | 0.098186 | 0.69364 | 0.158866 |
| ENSMUSG00000037085.9 | 25.69879 | -2.82681 | 1.710601 | -1.65253 | 0.098427 | 0.694782 | 0.158151 |
| ENSMUSG00000000711.4 | 10.59383 | 4.366352 | 2.647713 | 1.649103 | 0.099127 | 0.696104 | 0.157326 |
| ENSMUSG00000008301.12 | 123.5072 | -1.2129 | 0.73555 | -1.64897 | 0.099153 | 0.696104 | 0.157326 |
| ENSMUSG00000021385.11 | 39.49735 | 2.543337 | 1.540676 | 1.650792 | 0.098781 | 0.696104 | 0.157326 |
| ENSMUSG00000024858.15 | 37.31762 | 2.347633 | 1.425367 | 1.647038 | 0.09955 | 0.696104 | 0.157326 |
| ENSMUSG00000028138.9 | 4034.969 | 0.389388 | 0.236305 | 1.64782 | 0.09939 | 0.696104 | 0.157326 |
| ENSMUSG00000039533.9 | 13.50186 | -4.2058 | 2.55056 | -1.64897 | 0.099153 | 0.696104 | 0.157326 |
| ENSMUSG00000039958.18 | 297.944 | -1.06778 | 0.648307 | -1.64702 | 0.099553 | 0.696104 | 0.157326 |
| ENSMUSG00000040761.17 | 56.45687 | -1.61752 | 0.981391 | -1.64819 | 0.099313 | 0.696104 | 0.157326 |
| ENSMUSG00000054426.12 | 67.99999 | -1.94209 | 1.176835 | -1.65026 | 0.098889 | 0.696104 | 0.157326 |
| ENSMUSG00000056459.15 | 27.04512 | -3.56032 | 2.157513 | -1.6502 | 0.098902 | 0.696104 | 0.157326 |
| ENSMUSG00000058594.16 | 47.71354 | 2.435261 | 1.478653 | 1.646945 | 0.099569 | 0.696104 | 0.157326 |
| ENSMUSG00000060613.11 | 4425.624 | -0.43407 | 0.263532 | -1.64712 | 0.099533 | 0.696104 | 0.157326 |
| ENSMUSG00000086136.2 | 26.80547 | 3.621221 | 2.198479 | 1.647148 | 0.099528 | 0.696104 | 0.157326 |
| ENSMUSG00000107994.2 | 42.35614 | 2.376516 | 1.443505 | 1.646351 | 0.099692 | 0.696445 | 0.157113 |
| ENSMUSG00000039512.13 | 75.67514 | 1.696313 | 1.03081 | 1.645611 | 0.099844 | 0.696995 | 0.15677 |
| ENSMUSG00000032712.17 | 73.67341 | -1.85397 | 1.126903 | -1.64519 | 0.099931 | 0.697089 | 0.156712 |
| ENSMUSG00000073542.4 | 22.02322 | 3.126546 | 1.90228 | 1.643578 | 0.100263 | 0.698895 | 0.155588 |
| ENSMUSG00000019883.12 | 540.7342 | 0.794071 | 0.483285 | 1.643068 | 0.100369 | 0.699117 | 0.15545 |
| ENSMUSG00000026437.12 | 62.20436 | -2.22516 | 1.355411 | -1.64169 | 0.100655 | 0.700513 | 0.154584 |
| ENSMUSG00000027763.14 | 578.81 | -0.71712 | 0.436991 | -1.64104 | 0.100788 | 0.700513 | 0.154584 |

| | | | | | | | |
|---|---|---|---|---|---|---|---|
| ENSMUSG00000030107.11 | 65.97357 | -2.1321 | 1.299242 | -1.64103 | 0.100791 | 0.700513 | 0.154584 |
| ENSMUSG00000020024.8 | 32.27532 | -1.87888 | 1.145478 | -1.64026 | 0.100952 | 0.701022 | 0.154269 |
| ENSMUSG00000034345.15 | 345.8603 | 0.708208 | 0.431843 | 1.639966 | 0.101012 | 0.701022 | 0.154269 |
| ENSMUSG00000021639.18 | 65.99529 | -2.00583 | 1.224566 | -1.63799 | 0.101424 | 0.701932 | 0.153705 |
| ENSMUSG00000022517.18 | 79.4057 | -1.33615 | 0.815588 | -1.63826 | 0.101367 | 0.701932 | 0.153705 |
| ENSMUSG00000025499.19 | 30.21886 | -1.99809 | 1.219475 | -1.63848 | 0.101321 | 0.701932 | 0.153705 |
| ENSMUSG00000042364.12 | 68.87871 | -1.6667 | 1.017574 | -1.63791 | 0.10144 | 0.701932 | 0.153705 |
| ENSMUSG00000034459.9 | 133.0484 | -3.53057 | 2.156604 | -1.6371 | 0.10161 | 0.702597 | 0.153294 |
| ENSMUSG00000001768.17 | 85.76216 | 1.568494 | 0.959315 | 1.635015 | 0.102046 | 0.70276 | 0.153193 |
| ENSMUSG00000022136.9 | 1764.983 | -0.38788 | 0.237159 | -1.63553 | 0.101939 | 0.70276 | 0.153193 |
| ENSMUSG00000024687.12 | 165.0467 | -0.93431 | 0.571414 | -1.63508 | 0.102032 | 0.70276 | 0.153193 |
| ENSMUSG00000028608.16 | 100.9462 | 1.530814 | 0.93618 | 1.63517 | 0.102013 | 0.70276 | 0.153193 |
| ENSMUSG00000052707.9 | 157.7324 | -0.96037 | 0.586914 | -1.63631 | 0.101776 | 0.70276 | 0.153193 |
| ENSMUSG00000090877.4 | 213.5783 | 1.298691 | 0.794374 | 1.634862 | 0.102078 | 0.70276 | 0.153193 |
| ENSMUSG00000048109.11 | 102.5554 | -1.62801 | 0.996483 | -1.63375 | 0.10231 | 0.703848 | 0.152521 |
| ENSMUSG00000101970.9 | 164.9813 | 1.051832 | 0.64445 | 1.632139 | 0.10265 | 0.705673 | 0.151396 |
| ENSMUSG00000098188.2 | 17.53435 | -3.40302 | 2.08581 | -1.63151 | 0.102783 | 0.706074 | 0.15115 |
| ENSMUSG00000020034.8 | 181.3583 | -1.29424 | 0.79452 | -1.62896 | 0.103321 | 0.706168 | 0.151092 |
| ENSMUSG00000020553.9 | 571.0675 | 0.749041 | 0.460022 | 1.628273 | 0.103467 | 0.706168 | 0.151092 |
| ENSMUSG00000021067.9 | 82.78712 | -1.44158 | 0.884711 | -1.62944 | 0.10322 | 0.706168 | 0.151092 |
| ENSMUSG00000029053.17 | 59.36805 | 2.004223 | 1.229895 | 1.629589 | 0.103188 | 0.706168 | 0.151092 |
| ENSMUSG00000031792.16 | 19.99236 | -3.79564 | 2.328447 | -1.63011 | 0.103077 | 0.706168 | 0.151092 |
| ENSMUSG00000032047.6 | 1781.758 | 0.446449 | 0.274185 | 1.628279 | 0.103466 | 0.706168 | 0.151092 |
| ENSMUSG00000036452.19 | 61.23004 | 1.490303 | 0.914845 | 1.629024 | 0.103308 | 0.706168 | 0.151092 |
| ENSMUSG00000078771.11 | 16.57682 | -3.94807 | 2.424356 | -1.6285 | 0.103419 | 0.706168 | 0.151092 |
| ENSMUSG00000091264.4 | 85.98449 | -1.43984 | 0.883416 | -1.62985 | 0.103133 | 0.706168 | 0.151092 |
| ENSMUSG00000067825.12 | 69.72827 | -1.38151 | 0.848685 | -1.62783 | 0.103561 | 0.706302 | 0.151009 |
| ENSMUSG00000038538.18 | 49.47223 | -1.87355 | 1.1513 | -1.62733 | 0.103667 | 0.706512 | 0.15088 |
| ENSMUSG00000009566.13 | 144.0303 | 1.099044 | 0.675886 | 1.626079 | 0.103933 | 0.70731 | 0.15039 |
| ENSMUSG00000025982.14 | 485.2358 | 0.702853 | 0.432227 | 1.626118 | 0.103925 | 0.70731 | 0.15039 |
| ENSMUSG00000027424.12 | 26.69484 | 2.631137 | 1.618777 | 1.625386 | 0.10408 | 0.707806 | 0.150086 |
| ENSMUSG00000032553.15 | 127.1257 | 1.080753 | 0.66605 | 1.622629 | 0.104669 | 0.709357 | 0.149135 |
| ENSMUSG00000039568.7 | 18.60556 | -2.46055 | 1.515728 | -1.62334 | 0.104516 | 0.709357 | 0.149135 |
| ENSMUSG00000040250.15 | 105.3631 | 1.314562 | 0.809607 | 1.623703 | 0.104439 | 0.709357 | 0.149135 |
| ENSMUSG00000057054.12 | 35.93155 | -2.41816 | 1.490031 | -1.6229 | 0.104612 | 0.709357 | 0.149135 |
| ENSMUSG00000071369.12 | 64.67927 | -1.58261 | 0.975374 | -1.62256 | 0.104683 | 0.709357 | 0.149135 |
| ENSMUSG00000029291.12 | 236.8676 | -0.83207 | 0.513216 | -1.62128 | 0.104958 | 0.710713 | 0.148306 |
| ENSMUSG00000025558.20 | 68.97166 | 1.587512 | 0.979969 | 1.619963 | 0.10524 | 0.711652 | 0.147732 |
| ENSMUSG00000066551.13 | 464.7583 | -0.79875 | 0.493073 | -1.61993 | 0.105246 | 0.711652 | 0.147732 |
| ENSMUSG00000000594.8 | 418.1748 | -0.93558 | 0.577946 | -1.61881 | 0.105488 | 0.711763 | 0.147665 |
| ENSMUSG00000017286.16 | 166.5386 | -1.09923 | 0.678906 | -1.61912 | 0.105423 | 0.711763 | 0.147665 |
| ENSMUSG00000038213.8 | 110.2501 | -1.28961 | 0.796624 | -1.61884 | 0.105481 | 0.711763 | 0.147665 |
| ENSMUSG00000038736.11 | 41.72848 | 1.649793 | 1.020671 | 1.61638 | 0.106012 | 0.714791 | 0.145821 |
| ENSMUSG00000002661.15 | 39.9623 | 1.86132 | 1.151795 | 1.616017 | 0.106091 | 0.714811 | 0.145809 |

| | | | | | | | |
|---|---|---|---|---|---|---|---|
| ENSMUSG00000025428.16 | 5667.092 | 0.356853 | 0.220895 | 1.615487 | 0.106205 | 0.714823 | 0.145801 |
| ENSMUSG00000048696.12 | 12.66217 | 3.314652 | 2.052021 | 1.615311 | 0.106243 | 0.714823 | 0.145801 |
| ENSMUSG00000032606.8 | 130.296 | 1.195038 | 0.740148 | 1.614594 | 0.106399 | 0.714854 | 0.145783 |
| ENSMUSG00000089991.3 | 33.0586 | 2.791795 | 1.728877 | 1.614803 | 0.106353 | 0.714854 | 0.145783 |
| ENSMUSG00000019470.13 | 33.30185 | -2.33371 | 1.446425 | -1.61343 | 0.106651 | 0.716041 | 0.145062 |
| ENSMUSG00000032905.6 | 285.1363 | 0.950945 | 0.589528 | 1.613063 | 0.106731 | 0.716071 | 0.145044 |
| ENSMUSG00000017144.9 | 81.1539 | -1.52001 | 0.944069 | -1.61006 | 0.107385 | 0.716796 | 0.144605 |
| ENSMUSG00000021906.15 | 185.5294 | 1.048909 | 0.651306 | 1.610471 | 0.107295 | 0.716796 | 0.144605 |
| ENSMUSG00000027762.7 | 382.256 | 0.807902 | 0.501512 | 1.610931 | 0.107195 | 0.716796 | 0.144605 |
| ENSMUSG00000033335.18 | 33.20715 | -2.14853 | 1.334952 | -1.60944 | 0.107519 | 0.716796 | 0.144605 |
| ENSMUSG00000034867.17 | 43.30602 | -2.12949 | 1.323109 | -1.60946 | 0.107516 | 0.716796 | 0.144605 |
| ENSMUSG00000037111.10 | 20.19551 | -3.47903 | 2.158241 | -1.61198 | 0.106967 | 0.716796 | 0.144605 |
| ENSMUSG00000052926.17 | 34.15471 | -2.48441 | 1.542727 | -1.6104 | 0.10731 | 0.716796 | 0.144605 |
| ENSMUSG00000078786.10 | 27.73666 | -3.06034 | 1.90053 | -1.61025 | 0.107343 | 0.716796 | 0.144605 |
| ENSMUSG00000101397.7 | 243.3576 | 1.018261 | 0.631781 | 1.611729 | 0.107021 | 0.716796 | 0.144605 |
| ENSMUSG00000028191.12 | 81.09538 | -1.51142 | 0.939725 | -1.60837 | 0.107754 | 0.717858 | 0.143962 |
| ENSMUSG00000033917.16 | 630.5194 | 0.657748 | 0.409168 | 1.607524 | 0.10794 | 0.71831 | 0.143688 |
| ENSMUSG00000035007.6 | 32.519 | -2.58865 | 1.610488 | -1.60737 | 0.107974 | 0.71831 | 0.143688 |
| ENSMUSG00000003437.15 | 119.1715 | -1.2907 | 0.803865 | -1.60561 | 0.108359 | 0.720366 | 0.142447 |
| ENSMUSG00000037686.7 | 120.4829 | -1.05699 | 0.659234 | -1.60336 | 0.108855 | 0.721135 | 0.141984 |
| ENSMUSG00000041168.11 | 343.2004 | -0.88743 | 0.553123 | -1.60439 | 0.108628 | 0.721135 | 0.141984 |
| ENSMUSG00000041215.17 | 24.33001 | -2.59103 | 1.615979 | -1.60338 | 0.108851 | 0.721135 | 0.141984 |
| ENSMUSG00000043421.9 | 18.15112 | 3.632567 | 2.264937 | 1.603827 | 0.108752 | 0.721135 | 0.141984 |
| ENSMUSG00000071671.6 | 47.7936 | 1.659986 | 1.034721 | 1.604283 | 0.108652 | 0.721135 | 0.141984 |
| ENSMUSG00000023330.13 | 37.44664 | 2.370593 | 1.479211 | 1.602607 | 0.109022 | 0.721735 | 0.141622 |
| ENSMUSG00000036585.17 | 342.5011 | 0.845566 | 0.527886 | 1.601796 | 0.109201 | 0.721918 | 0.141512 |
| ENSMUSG00000059689.16 | 65.65851 | 1.39261 | 0.869407 | 1.601793 | 0.109201 | 0.721918 | 0.141512 |
| ENSMUSG00000019579.14 | 204.7961 | -0.90823 | 0.567273 | -1.60104 | 0.109367 | 0.722509 | 0.141157 |
| ENSMUSG00000020778.13 | 43.89246 | 1.750404 | 1.093687 | 1.600463 | 0.109496 | 0.722857 | 0.140948 |
| ENSMUSG00000040249.16 | 847.4107 | -0.60184 | 0.376165 | -1.59993 | 0.109614 | 0.72313 | 0.140784 |
| ENSMUSG00000021725.10 | 11.62991 | 5.060004 | 3.16382 | 1.599334 | 0.109746 | 0.723502 | 0.14056 |
| ENSMUSG00000006299.14 | 128.871 | -1.11912 | 0.701439 | -1.59546 | 0.110608 | 0.72505 | 0.139632 |
| ENSMUSG00000008393.10 | 288.6607 | -0.74143 | 0.464455 | -1.59634 | 0.110412 | 0.72505 | 0.139632 |
| ENSMUSG00000020375.10 | 72.29855 | -1.6636 | 1.042251 | -1.59616 | 0.110453 | 0.72505 | 0.139632 |
| ENSMUSG00000020923.18 | 27.54674 | -2.44762 | 1.533372 | -1.59623 | 0.110437 | 0.72505 | 0.139632 |
| ENSMUSG00000026678.11 | 63.6844 | -1.92204 | 1.203193 | -1.59745 | 0.110165 | 0.72505 | 0.139632 |
| ENSMUSG00000027397.15 | 153.1756 | -1.25733 | 0.788201 | -1.59519 | 0.11067 | 0.72505 | 0.139632 |
| ENSMUSG00000039983.14 | 26.1189 | -3.14158 | 1.968707 | -1.59576 | 0.110543 | 0.72505 | 0.139632 |
| ENSMUSG00000085208.4 | 136.8083 | -1.13348 | 0.709679 | -1.59718 | 0.110226 | 0.72505 | 0.139632 |
| ENSMUSG00000089917.8 | 51.08138 | -1.8522 | 1.159743 | -1.59707 | 0.110249 | 0.72505 | 0.139632 |
| ENSMUSG00000026915.17 | 230.7166 | -1.0196 | 0.639444 | -1.59451 | 0.110821 | 0.725544 | 0.139337 |
| ENSMUSG00000074063.11 | 128.7199 | -1.22695 | 0.770058 | -1.59333 | 0.111087 | 0.726279 | 0.138896 |
| ENSMUSG00000109291.2 | 12.65821 | 3.746335 | 2.351047 | 1.593475 | 0.111054 | 0.726279 | 0.138896 |
| ENSMUSG00000030814.18 | 11.52279 | -3.6698 | 2.304276 | -1.5926 | 0.111249 | 0.726339 | 0.13886 |

| | | | | | | | |
|---|---|---|---|---|---|---|---|
| ENSMUSG00000042050.9 | 62.88486 | -2.56707 | 1.61169 | -1.59278 | 0.111209 | 0.726339 | 0.13886 |
| ENSMUSG00000048498.8 | 16.12719 | -3.9051 | 2.452961 | -1.59199 | 0.111386 | 0.726733 | 0.138625 |
| ENSMUSG00000036138.17 | 889.6076 | 0.616669 | 0.387566 | 1.591133 | 0.11158 | 0.727494 | 0.138171 |
| ENSMUSG00000036006.21 | 16.12654 | 4.550152 | 2.862222 | 1.589727 | 0.111896 | 0.729057 | 0.137239 |
| ENSMUSG00000015095.13 | 124.1687 | -1.22791 | 0.773381 | -1.58772 | 0.112349 | 0.729131 | 0.137194 |
| ENSMUSG00000015846.16 | 329.5256 | -0.80417 | 0.506583 | -1.58745 | 0.112412 | 0.729131 | 0.137194 |
| ENSMUSG00000021611.10 | 28.49741 | -3.38607 | 2.131556 | -1.58854 | 0.112164 | 0.729131 | 0.137194 |
| ENSMUSG00000021809.11 | 42.11213 | 1.828283 | 1.151486 | 1.587759 | 0.112341 | 0.729131 | 0.137194 |
| ENSMUSG00000036580.16 | 89.15398 | -1.43199 | 0.901464 | -1.58852 | 0.11217 | 0.729131 | 0.137194 |
| ENSMUSG00000037149.11 | 285.5288 | -0.7757 | 0.488694 | -1.58729 | 0.112446 | 0.729131 | 0.137194 |
| ENSMUSG00000045160.15 | 644.2798 | 0.547023 | 0.344398 | 1.588345 | 0.112208 | 0.729131 | 0.137194 |
| ENSMUSG00000022636.14 | 323.8906 | -0.94279 | 0.59439 | -1.58616 | 0.112704 | 0.729153 | 0.137181 |
| ENSMUSG00000046352.8 | 1817.284 | -0.43588 | 0.274766 | -1.58637 | 0.112655 | 0.729153 | 0.137181 |
| ENSMUSG00000050965.15 | 22.95567 | 3.08641 | 1.945857 | 1.586145 | 0.112707 | 0.729153 | 0.137181 |
| ENSMUSG00000056313.6 | 74.90435 | 1.466583 | 0.924751 | 1.585921 | 0.112757 | 0.729153 | 0.137181 |
| ENSMUSG00000006932.18 | 1033.009 | 0.472582 | 0.298174 | 1.584917 | 0.112985 | 0.730129 | 0.1366 |
| ENSMUSG00000023832.15 | 419.4284 | 0.648031 | 0.409073 | 1.584146 | 0.11316 | 0.730266 | 0.136519 |
| ENSMUSG00000030374.16 | 85.80208 | 1.849061 | 1.167169 | 1.584227 | 0.113142 | 0.730266 | 0.136519 |
| ENSMUSG00000036943.10 | 12.99877 | 3.779376 | 2.386262 | 1.583806 | 0.113238 | 0.730269 | 0.136517 |
| ENSMUSG00000109901.3 | 224.3007 | -0.75752 | 0.478914 | -1.58176 | 0.113705 | 0.732784 | 0.135024 |
| ENSMUSG00000032119.6 | 85.2763 | -1.79192 | 1.134042 | -1.58012 | 0.11408 | 0.733859 | 0.134387 |
| ENSMUSG00000053175.18 | 56.71989 | -1.57005 | 0.993509 | -1.58031 | 0.114037 | 0.733859 | 0.134387 |
| ENSMUSG00000058715.12 | 59.04255 | -1.6933 | 1.071699 | -1.58001 | 0.114104 | 0.733859 | 0.134387 |
| ENSMUSG00000027131.4 | 81.52187 | 1.350718 | 0.856145 | 1.577674 | 0.11464 | 0.736476 | 0.132841 |
| ENSMUSG00000070644.14 | 651.312 | -0.56258 | 0.356612 | -1.57756 | 0.114667 | 0.736476 | 0.132841 |
| ENSMUSG00000024608.12 | 780.8186 | 0.590029 | 0.374234 | 1.576634 | 0.11488 | 0.737346 | 0.132328 |
| ENSMUSG00000037750.17 | 45.69968 | 1.876072 | 1.190298 | 1.576136 | 0.114994 | 0.737583 | 0.132189 |
| ENSMUSG00000027018.12 | 94.23956 | -1.43077 | 0.908125 | -1.57552 | 0.115137 | 0.737997 | 0.131946 |
| ENSMUSG00000051177.17 | 233.6853 | 0.791233 | 0.502924 | 1.573265 | 0.115658 | 0.740834 | 0.130279 |
| ENSMUSG00000003226.8 | 129.3374 | 0.998407 | 0.635574 | 1.570876 | 0.116211 | 0.742959 | 0.129035 |
| ENSMUSG00000025228.6 | 90.53613 | -1.88717 | 1.201533 | -1.57064 | 0.116267 | 0.742959 | 0.129035 |
| ENSMUSG00000027618.18 | 282.1641 | -0.75963 | 0.483901 | -1.56981 | 0.11646 | 0.742959 | 0.129035 |
| ENSMUSG00000029482.5 | 369.8484 | 0.688644 | 0.438495 | 1.570472 | 0.116305 | 0.742959 | 0.129035 |
| ENSMUSG00000030887.5 | 86.12747 | 1.597398 | 1.017342 | 1.570168 | 0.116376 | 0.742959 | 0.129035 |
| ENSMUSG00000072596.5 | 11.57115 | -4.20056 | 2.675595 | -1.56995 | 0.116426 | 0.742959 | 0.129035 |
| ENSMUSG00000100975.3 | 20.45536 | -2.71328 | 1.728881 | -1.56939 | 0.116558 | 0.743086 | 0.128961 |
| ENSMUSG00000020309.7 | 348.6987 | 0.706284 | 0.451327 | 1.564905 | 0.117605 | 0.74336 | 0.128801 |
| ENSMUSG00000020455.17 | 25.48668 | -2.40519 | 1.534018 | -1.5679 | 0.116905 | 0.74336 | 0.128801 |
| ENSMUSG00000027552.15 | 75.75296 | -1.69424 | 1.082053 | -1.56576 | 0.117404 | 0.74336 | 0.128801 |
| ENSMUSG00000028869.14 | 165.9517 | 0.872518 | 0.557213 | 1.565861 | 0.117381 | 0.74336 | 0.128801 |
| ENSMUSG00000030979.14 | 90.09425 | 1.432089 | 0.915166 | 1.564841 | 0.11762 | 0.74336 | 0.128801 |
| ENSMUSG00000034780.7 | 155.4458 | -1.47805 | 0.94278 | -1.56776 | 0.116938 | 0.74336 | 0.128801 |
| ENSMUSG00000037826.6 | 577.162 | -0.6273 | 0.400108 | -1.56783 | 0.11692 | 0.74336 | 0.128801 |
| ENSMUSG00000038604.10 | 14.6398 | 3.351587 | 2.137571 | 1.567942 | 0.116895 | 0.74336 | 0.128801 |

| | | | | | | | |
|---|---|---|---|---|---|---|---|
| ENSMUSG00000039615.10 | 23.29814 | -2.54917 | 1.627895 | -1.56593 | 0.117364 | 0.74336 | 0.128801 |
| ENSMUSG00000049606.17 | 114.0716 | 1.200046 | 0.766422 | 1.565776 | 0.117401 | 0.74336 | 0.128801 |
| ENSMUSG00000061292.11 | 258.6706 | -0.98861 | 0.631599 | -1.56525 | 0.117525 | 0.74336 | 0.128801 |
| ENSMUSG00000062410.15 | 1018.333 | -0.5923 | 0.378334 | -1.56554 | 0.117455 | 0.74336 | 0.128801 |
| ENSMUSG00000114797.2 | 89.29858 | -1.38862 | 0.886918 | -1.56567 | 0.117426 | 0.74336 | 0.128801 |
| ENSMUSG00000037400.18 | 127.3746 | 1.097718 | 0.70207 | 1.563545 | 0.117925 | 0.744787 | 0.127968 |
| ENSMUSG00000027466.16 | 17.31249 | -2.93556 | 1.878198 | -1.56296 | 0.118061 | 0.745151 | 0.127755 |
| ENSMUSG00000028536.13 | 38.04899 | -1.95537 | 1.251766 | -1.56209 | 0.118266 | 0.745732 | 0.127417 |
| ENSMUSG00000057778.15 | 53.77317 | 1.830604 | 1.172159 | 1.561738 | 0.11835 | 0.745732 | 0.127417 |
| ENSMUSG00000067288.14 | 958.4181 | 0.508268 | 0.325485 | 1.561572 | 0.118389 | 0.745732 | 0.127417 |
| ENSMUSG00000058952.13 | 1732.768 | -0.41479 | 0.265704 | -1.56108 | 0.118505 | 0.745968 | 0.12728 |
| ENSMUSG00000038212.17 | 143.7532 | 1.289169 | 0.826481 | 1.559829 | 0.1188 | 0.747331 | 0.126487 |
| ENSMUSG00000072082.8 | 104.6829 | -1.15691 | 0.741902 | -1.55938 | 0.118906 | 0.7475 | 0.126389 |
| ENSMUSG00000018068.14 | 33.51176 | 2.063026 | 1.323685 | 1.558547 | 0.119104 | 0.747792 | 0.126219 |
| ENSMUSG00000054404.14 | 32.93996 | -2.28324 | 1.465009 | -1.55852 | 0.11911 | 0.747792 | 0.126219 |
| ENSMUSG00000030315.16 | 17.48563 | 3.817815 | 2.451493 | 1.557342 | 0.119389 | 0.748057 | 0.126065 |
| ENSMUSG00000050213.7 | 28.65704 | -2.01575 | 1.294171 | -1.55756 | 0.119336 | 0.748057 | 0.126065 |
| ENSMUSG00000056116.19 | 168.4399 | -1.16413 | 0.747414 | -1.55754 | 0.119343 | 0.748057 | 0.126065 |
| ENSMUSG00000037669.16 | 191.5757 | 1.171346 | 0.752427 | 1.556757 | 0.119528 | 0.748432 | 0.125848 |
| ENSMUSG00000090112.9 | 95.62801 | 1.27654 | 0.820393 | 1.55601 | 0.119706 | 0.749049 | 0.125489 |
| ENSMUSG00000005078.17 | 230.5495 | 1.058613 | 0.680786 | 1.554987 | 0.119949 | 0.749643 | 0.125145 |
| ENSMUSG00000019132.12 | 858.5764 | 0.491242 | 0.316081 | 1.554163 | 0.120145 | 0.749643 | 0.125145 |
| ENSMUSG00000028086.16 | 32.83411 | -2.21378 | 1.424615 | -1.55395 | 0.120196 | 0.749643 | 0.125145 |
| ENSMUSG00000029634.16 | 288.2285 | -0.80374 | 0.516967 | -1.55472 | 0.120013 | 0.749643 | 0.125145 |
| ENSMUSG00000078853.9 | 44.02189 | 1.624189 | 1.044895 | 1.554405 | 0.120088 | 0.749643 | 0.125145 |
| ENSMUSG00000031781.15 | 51.61646 | -1.9918 | 1.282796 | -1.5527 | 0.120495 | 0.751012 | 0.124353 |
| ENSMUSG00000015597.18 | 84.66093 | -1.37005 | 0.884516 | -1.54893 | 0.121398 | 0.752192 | 0.123671 |
| ENSMUSG00000019880.11 | 74.89713 | 1.569766 | 1.012452 | 1.550459 | 0.121031 | 0.752192 | 0.123671 |
| ENSMUSG00000024713.17 | 34.30129 | 1.860308 | 1.200839 | 1.549174 | 0.12134 | 0.752192 | 0.123671 |
| ENSMUSG00000024892.18 | 356.1075 | -0.85645 | 0.552752 | -1.54944 | 0.121277 | 0.752192 | 0.123671 |
| ENSMUSG00000029500.17 | 42.04622 | -2.17107 | 1.399624 | -1.55118 | 0.120858 | 0.752192 | 0.123671 |
| ENSMUSG00000036591.17 | 73.20638 | -1.26853 | 0.818456 | -1.54991 | 0.121163 | 0.752192 | 0.123671 |
| ENSMUSG00000040913.12 | 32.47426 | -2.36138 | 1.524263 | -1.54919 | 0.121335 | 0.752192 | 0.123671 |
| ENSMUSG00000044496.7 | 41.53746 | -2.59658 | 1.675305 | -1.54992 | 0.121161 | 0.752192 | 0.123671 |
| ENSMUSG00000078673.11 | 33.28401 | -1.89329 | 1.221188 | -1.55037 | 0.121053 | 0.752192 | 0.123671 |
| ENSMUSG00000059119.10 | 104.6944 | 1.453852 | 0.939177 | 1.548007 | 0.121621 | 0.753079 | 0.123159 |
| ENSMUSG00000037260.9 | 231.4225 | 0.952471 | 0.615759 | 1.546826 | 0.121905 | 0.75407 | 0.122588 |
| ENSMUSG00000053279.9 | 15810.44 | 0.317497 | 0.205276 | 1.546682 | 0.12194 | 0.75407 | 0.122588 |
| ENSMUSG00000021039.10 | 295.4024 | -0.76267 | 0.493552 | -1.54528 | 0.12228 | 0.755186 | 0.121946 |
| ENSMUSG00000063354.8 | 98.89309 | -1.72639 | 1.117145 | -1.54536 | 0.12226 | 0.755186 | 0.121946 |
| ENSMUSG00000031996.18 | 786.0215 | 0.516205 | 0.334245 | 1.544391 | 0.122494 | 0.756015 | 0.12147 |
| ENSMUSG00000032468.12 | 157.5731 | 0.854867 | 0.553737 | 1.543815 | 0.122633 | 0.756384 | 0.121258 |
| ENSMUSG00000024732.8 | 25.49443 | 2.614944 | 1.694334 | 1.543346 | 0.122747 | 0.756469 | 0.121209 |
| ENSMUSG00000029238.12 | 126.8678 | 1.181737 | 0.765961 | 1.542818 | 0.122875 | 0.756469 | 0.121209 |

| | | | | | | | |
|---|---|---|---|---|---|---|---|
| ENSMUSG00000064325.5 | 29.59539 | 2.825744 | 1.831603 | 1.542771 | 0.122886 | 0.756469 | 0.121209 |
| ENSMUSG00000027752.16 | 106.2663 | -1.03608 | 0.671927 | -1.54196 | 0.123083 | 0.75719 | 0.120795 |
| ENSMUSG00000019122.9 | 608.8385 | -0.52066 | 0.337772 | -1.54144 | 0.12321 | 0.757475 | 0.120632 |
| ENSMUSG00000028479.19 | 108.9746 | -1.30724 | 0.848877 | -1.53996 | 0.12357 | 0.759036 | 0.119738 |
| ENSMUSG00000039176.18 | 40.65056 | -1.90829 | 1.239464 | -1.53961 | 0.123655 | 0.759036 | 0.119738 |
| ENSMUSG00000078650.3 | 951.4324 | -0.83622 | 0.543209 | -1.53941 | 0.123704 | 0.759036 | 0.119738 |
| ENSMUSG00000024789.14 | 66.74849 | -1.53166 | 0.995496 | -1.53859 | 0.123905 | 0.759783 | 0.119311 |
| ENSMUSG00000025630.9 | 562.87 | 0.610251 | 0.396933 | 1.537414 | 0.124192 | 0.761048 | 0.118588 |
| ENSMUSG00000026589.15 | 236.2158 | 0.769833 | 0.500845 | 1.537069 | 0.124276 | 0.761073 | 0.118574 |
| ENSMUSG00000022808.6 | 191.5926 | -0.80199 | 0.522186 | -1.53583 | 0.12458 | 0.762343 | 0.117849 |
| ENSMUSG00000038871.6 | 107.2834 | -1.59238 | 1.036998 | -1.53557 | 0.124645 | 0.762343 | 0.117849 |
| ENSMUSG00000040760.12 | 87.1839 | -1.76084 | 1.147253 | -1.53483 | 0.124826 | 0.762894 | 0.117536 |
| ENSMUSG00000074183.4 | 25.85362 | 3.338714 | 2.175703 | 1.534545 | 0.124896 | 0.762894 | 0.117536 |
| ENSMUSG00000027765.7 | 68.64461 | 1.381746 | 0.901042 | 1.533497 | 0.125153 | 0.76306 | 0.117441 |
| ENSMUSG00000038174.15 | 240.1152 | -0.74535 | 0.485898 | -1.53397 | 0.125036 | 0.76306 | 0.117441 |
| ENSMUSG00000038387.10 | 13.64702 | 3.869666 | 2.523499 | 1.533452 | 0.125164 | 0.76306 | 0.117441 |
| ENSMUSG00000012428.10 | 886.7987 | -0.52021 | 0.339562 | -1.532 | 0.125523 | 0.764013 | 0.1169 |
| ENSMUSG00000023829.10 | 806.2518 | -0.62971 | 0.411167 | -1.53151 | 0.125643 | 0.764013 | 0.1169 |
| ENSMUSG00000026319.14 | 57.12013 | -1.87197 | 1.222087 | -1.53178 | 0.125577 | 0.764013 | 0.1169 |
| ENSMUSG00000034501.8 | 25.11682 | -2.37007 | 1.546601 | -1.53244 | 0.125414 | 0.764013 | 0.1169 |
| ENSMUSG00000049044.17 | 887.1897 | -0.67618 | 0.441606 | -1.53118 | 0.125724 | 0.764017 | 0.116897 |
| ENSMUSG00000010609.16 | 1697.968 | 0.48387 | 0.316456 | 1.52903 | 0.126257 | 0.765557 | 0.116023 |
| ENSMUSG00000027706.9 | 2173.724 | 0.488712 | 0.319659 | 1.528854 | 0.126301 | 0.765557 | 0.116023 |
| ENSMUSG00000041124.12 | 156.4386 | 1.001812 | 0.655238 | 1.52893 | 0.126282 | 0.765557 | 0.116023 |
| ENSMUSG00000116130.2 | 56.79304 | -2.31321 | 1.513001 | -1.52889 | 0.126292 | 0.765557 | 0.116023 |
| ENSMUSG00000078813.4 | 63.79396 | -1.91182 | 1.250945 | -1.5283 | 0.126439 | 0.765903 | 0.115826 |
| ENSMUSG00000086628.8 | 197.3734 | 0.979062 | 0.641895 | 1.525269 | 0.127192 | 0.769975 | 0.113524 |
| ENSMUSG00000038539.16 | 146.9274 | -1.57514 | 1.033441 | -1.52417 | 0.127466 | 0.771143 | 0.112865 |
| ENSMUSG00000007050.18 | 66.45941 | -1.6177 | 1.061786 | -1.52356 | 0.127619 | 0.771199 | 0.112833 |
| ENSMUSG00000009378.5 | 670.4778 | 0.643667 | 0.4228 | 1.522394 | 0.12791 | 0.771199 | 0.112833 |
| ENSMUSG00000021036.11 | 75.12782 | 1.247402 | 0.819371 | 1.522391 | 0.127911 | 0.771199 | 0.112833 |
| ENSMUSG00000032342.14 | 116.5018 | -1.6817 | 1.104582 | -1.52248 | 0.127889 | 0.771199 | 0.112833 |
| ENSMUSG00000051675.14 | 46.03941 | 1.991928 | 1.308602 | 1.522181 | 0.127964 | 0.771199 | 0.112833 |
| ENSMUSG00000079941.2 | 113.7369 | 1.089046 | 0.715073 | 1.522986 | 0.127762 | 0.771199 | 0.112833 |
| ENSMUSG00000031843.3 | 36.75972 | 2.954859 | 1.941622 | 1.521851 | 0.128046 | 0.771207 | 0.112829 |
| ENSMUSG00000024697.5 | 42.3799 | -2.41091 | 1.584957 | -1.52112 | 0.128229 | 0.771328 | 0.112761 |
| ENSMUSG00000038366.16 | 592.3413 | 0.550642 | 0.361941 | 1.52136 | 0.12817 | 0.771328 | 0.112761 |
| ENSMUSG00000021686.7 | 127.3465 | -1.14176 | 0.751149 | -1.52002 | 0.128505 | 0.7725 | 0.112102 |
| ENSMUSG00000025579.15 | 119.9887 | -1.13308 | 0.746051 | -1.51878 | 0.128819 | 0.773083 | 0.111774 |
| ENSMUSG00000035047.10 | 14.11602 | 4.552888 | 2.997957 | 1.518664 | 0.128847 | 0.773083 | 0.111774 |
| ENSMUSG00000039745.9 | 146.3974 | -0.97222 | 0.639984 | -1.51913 | 0.128729 | 0.773083 | 0.111774 |
| ENSMUSG00000020189.16 | 329.4762 | 0.71148 | 0.46884 | 1.517534 | 0.129132 | 0.774115 | 0.111194 |
| ENSMUSG00000032434.10 | 366.707 | -0.76256 | 0.502596 | -1.51723 | 0.129207 | 0.774115 | 0.111194 |
| ENSMUSG00000086494.2 | 21.54753 | 2.878827 | 1.897698 | 1.51701 | 0.129264 | 0.774115 | 0.111194 |

| | | | | | | | |
|---|---|---|---|---|---|---|---|
| ENSMUSG00000022360.9 | 44.00806 | -2.05619 | 1.357139 | -1.51509 | 0.129749 | 0.77638 | 0.109926 |
| ENSMUSG00000038299.17 | 116.4651 | 0.988207 | 0.652339 | 1.514867 | 0.129806 | 0.77638 | 0.109926 |
| ENSMUSG00000030884.13 | 2053.243 | 0.434935 | 0.28723 | 1.514243 | 0.129964 | 0.776835 | 0.109671 |
| ENSMUSG00000022751.10 | 830.6616 | -0.5847 | 0.386298 | -1.5136 | 0.130127 | 0.77716 | 0.10949 |
| ENSMUSG00000029826.15 | 82.33291 | -1.20812 | 0.79829 | -1.51338 | 0.130183 | 0.77716 | 0.10949 |
| ENSMUSG00000022472.17 | 188.6283 | 1.007368 | 0.666617 | 1.511164 | 0.130747 | 0.778077 | 0.108977 |
| ENSMUSG00000025236.12 | 25.7263 | 2.995185 | 1.980436 | 1.512386 | 0.130436 | 0.778077 | 0.108977 |
| ENSMUSG00000032673.6 | 251.5644 | 0.895049 | 0.592023 | 1.511849 | 0.130572 | 0.778077 | 0.108977 |
| ENSMUSG00000040550.18 | 174.8539 | 0.982544 | 0.649928 | 1.511774 | 0.130591 | 0.778077 | 0.108977 |
| ENSMUSG00000062198.6 | 36.15097 | -2.16426 | 1.432112 | -1.51124 | 0.130728 | 0.778077 | 0.108977 |
| ENSMUSG00000022883.12 | 60.02098 | 1.570332 | 1.040131 | 1.509744 | 0.131109 | 0.779743 | 0.108048 |
| ENSMUSG00000020792.16 | 54.00763 | 1.487368 | 0.985457 | 1.509319 | 0.131217 | 0.7799 | 0.107961 |
| ENSMUSG00000073424.10 | 353.6497 | -0.78841 | 0.522709 | -1.50832 | 0.131471 | 0.780921 | 0.107393 |
| ENSMUSG00000004798.15 | 494.8716 | -0.64481 | 0.427604 | -1.50797 | 0.131562 | 0.780967 | 0.107367 |
| ENSMUSG00000062908.13 | 1748.298 | 0.491033 | 0.325895 | 1.506722 | 0.131882 | 0.782379 | 0.106583 |
| ENSMUSG00000030291.13 | 308.5292 | 0.732152 | 0.486515 | 1.50489 | 0.132352 | 0.784678 | 0.105308 |
| ENSMUSG00000056666.14 | 1007.348 | 0.608508 | 0.404516 | 1.504285 | 0.132508 | 0.78511 | 0.105069 |
| ENSMUSG00000061780.7 | 14.16787 | 3.750141 | 2.493602 | 1.503905 | 0.132606 | 0.785199 | 0.105021 |
| ENSMUSG00000031432.4 | 86.03025 | 1.247494 | 0.829823 | 1.503326 | 0.132755 | 0.785591 | 0.104803 |
| ENSMUSG00000069917.8 | 62.89487 | -2.08238 | 1.386057 | -1.50238 | 0.133 | 0.786548 | 0.104275 |
| ENSMUSG00000009569.15 | 62.39493 | 1.47591 | 0.982668 | 1.501941 | 0.133112 | 0.786724 | 0.104178 |
| ENSMUSG00000049807.17 | 21.38628 | -2.99867 | 1.998174 | -1.50071 | 0.133431 | 0.787163 | 0.103935 |
| ENSMUSG00000061906.6 | 267.0088 | -1.23393 | 0.822032 | -1.50108 | 0.133336 | 0.787163 | 0.103935 |
| ENSMUSG00000072115.12 | 1592.167 | -0.52721 | 0.35131 | -1.50069 | 0.133436 | 0.787163 | 0.103935 |
| ENSMUSG00000028792.15 | 1198.439 | -0.69355 | 0.462652 | -1.49908 | 0.133854 | 0.787853 | 0.103555 |
| ENSMUSG00000059481.6 | 625.0731 | -0.60468 | 0.403396 | -1.49897 | 0.133882 | 0.787853 | 0.103555 |
| ENSMUSG00000078817.5 | 204.5038 | 0.932382 | 0.622021 | 1.498955 | 0.133885 | 0.787853 | 0.103555 |
| ENSMUSG00000095098.3 | 49.75749 | 2.001094 | 1.334601 | 1.499395 | 0.133771 | 0.787853 | 0.103555 |
| ENSMUSG00000006850.9 | 11.03722 | 4.238246 | 2.829271 | 1.497999 | 0.134133 | 0.788016 | 0.103465 |
| ENSMUSG00000025196.5 | 342.7441 | -0.62862 | 0.419569 | -1.49825 | 0.134069 | 0.788016 | 0.103465 |
| ENSMUSG00000028577.14 | 331.7048 | -0.73693 | 0.491978 | -1.49789 | 0.134162 | 0.788016 | 0.103465 |
| ENSMUSG00000005054.8 | 304.1808 | 0.658067 | 0.439432 | 1.497543 | 0.134252 | 0.788056 | 0.103443 |
| ENSMUSG00000031848.16 | 136.1534 | -1.22622 | 0.81931 | -1.49665 | 0.134485 | 0.788934 | 0.102959 |
| ENSMUSG00000038383.17 | 10.43034 | 4.153041 | 2.776114 | 1.495991 | 0.134656 | 0.78945 | 0.102676 |
| ENSMUSG00000014313.16 | 4602.655 | 0.420355 | 0.281759 | 1.491897 | 0.135726 | 0.790656 | 0.102012 |
| ENSMUSG00000017801.16 | 91.09939 | -1.55542 | 1.042503 | -1.49201 | 0.135698 | 0.790656 | 0.102012 |
| ENSMUSG00000021115.16 | 59.323 | -2.21485 | 1.48291 | -1.49358 | 0.135285 | 0.790656 | 0.102012 |
| ENSMUSG00000023272.5 | 154.3174 | 0.973608 | 0.652578 | 1.491942 | 0.135714 | 0.790656 | 0.102012 |
| ENSMUSG00000024072.10 | 244.6333 | -0.91656 | 0.615483 | -1.48917 | 0.136442 | 0.790656 | 0.102012 |
| ENSMUSG00000024725.14 | 214.3379 | 0.800787 | 0.537311 | 1.49036 | 0.13613 | 0.790656 | 0.102012 |
| ENSMUSG00000024953.18 | 824.9249 | -0.48644 | 0.326658 | -1.48916 | 0.136447 | 0.790656 | 0.102012 |
| ENSMUSG00000025479.10 | 21577.87 | 0.300448 | 0.20169 | 1.489655 | 0.136315 | 0.790656 | 0.102012 |
| ENSMUSG00000026554.16 | 184.7672 | -1.05482 | 0.705727 | -1.49466 | 0.135004 | 0.790656 | 0.102012 |
| ENSMUSG00000027829.16 | 128.947 | -1.03644 | 0.695655 | -1.48987 | 0.136258 | 0.790656 | 0.102012 |

| | | | | | | | |
|---|---|---|---|---|---|---|---|
| ENSMUSG00000029199.12 | 259.1472 | 1.228739 | 0.824951 | 1.48947 | 0.136364 | 0.790656 | 0.102012 |
| ENSMUSG00000031627.10 | 164.8344 | 1.023394 | 0.6865 | 1.49074 | 0.13603 | 0.790656 | 0.102012 |
| ENSMUSG00000032788.16 | 125.4212 | -1.17771 | 0.789233 | -1.49222 | 0.135643 | 0.790656 | 0.102012 |
| ENSMUSG00000037236.17 | 313.1347 | 0.878158 | 0.589672 | 1.489231 | 0.136427 | 0.790656 | 0.102012 |
| ENSMUSG00000037652.16 | 86.91881 | -1.5137 | 1.015676 | -1.49034 | 0.136136 | 0.790656 | 0.102012 |
| ENSMUSG00000042293.9 | 88.09888 | -1.23632 | 0.829388 | -1.49064 | 0.136056 | 0.790656 | 0.102012 |
| ENSMUSG00000053094.16 | 182.3085 | -1.18545 | 0.793988 | -1.49303 | 0.135429 | 0.790656 | 0.102012 |
| ENSMUSG00000054509.8 | 40.69843 | -2.05738 | 1.377451 | -1.49361 | 0.135277 | 0.790656 | 0.102012 |
| ENSMUSG00000064367.1 | 10200.75 | 0.36506 | 0.244665 | 1.492084 | 0.135677 | 0.790656 | 0.102012 |
| ENSMUSG00000003166.21 | 138.2021 | -1.3654 | 0.917767 | -1.48775 | 0.136818 | 0.791846 | 0.101359 |
| ENSMUSG00000031540.15 | 49.01436 | 2.091793 | 1.406019 | 1.487742 | 0.136819 | 0.791846 | 0.101359 |
| ENSMUSG00000001552.15 | 28.70972 | 2.438152 | 1.644137 | 1.482937 | 0.138091 | 0.792476 | 0.101014 |
| ENSMUSG00000002455.15 | 165.9168 | -0.99873 | 0.673437 | -1.48304 | 0.138065 | 0.792476 | 0.101014 |
| ENSMUSG00000002475.17 | 415.4823 | 0.642892 | 0.433551 | 1.482852 | 0.138114 | 0.792476 | 0.101014 |
| ENSMUSG00000022711.17 | 25.27314 | 2.983893 | 2.012425 | 1.482735 | 0.138145 | 0.792476 | 0.101014 |
| ENSMUSG00000022856.10 | 73.86407 | 1.177468 | 0.794361 | 1.482282 | 0.138265 | 0.792476 | 0.101014 |
| ENSMUSG00000024384.5 | 138.1524 | -1.03491 | 0.696525 | -1.48582 | 0.137327 | 0.792476 | 0.101014 |
| ENSMUSG00000025940.7 | 548.4143 | 0.529647 | 0.356821 | 1.484351 | 0.137716 | 0.792476 | 0.101014 |
| ENSMUSG00000027177.15 | 321.132 | -0.69658 | 0.469187 | -1.48465 | 0.137636 | 0.792476 | 0.101014 |
| ENSMUSG00000029388.15 | 145.6612 | 1.100685 | 0.74253 | 1.482344 | 0.138249 | 0.792476 | 0.101014 |
| ENSMUSG00000036644.16 | 85.69578 | -1.36979 | 0.922352 | -1.48511 | 0.137515 | 0.792476 | 0.101014 |
| ENSMUSG00000038145.18 | 73.29737 | 1.419093 | 0.954966 | 1.486014 | 0.137275 | 0.792476 | 0.101014 |
| ENSMUSG00000040389.16 | 13.06982 | 4.503981 | 3.036127 | 1.483462 | 0.137952 | 0.792476 | 0.101014 |
| ENSMUSG00000040410.15 | 102.272 | -1.12948 | 0.760359 | -1.48545 | 0.137425 | 0.792476 | 0.101014 |
| ENSMUSG00000041881.14 | 523.6771 | 0.64645 | 0.435381 | 1.484793 | 0.137599 | 0.792476 | 0.101014 |
| ENSMUSG00000066175.3 | 24.0257 | 2.546146 | 1.716601 | 1.483248 | 0.138008 | 0.792476 | 0.101014 |
| ENSMUSG00000112793.2 | 36.20065 | -1.89689 | 1.276796 | -1.48566 | 0.137368 | 0.792476 | 0.101014 |
| ENSMUSG00000021748.10 | 741.7631 | 0.45505 | 0.307238 | 1.481101 | 0.13858 | 0.793499 | 0.100453 |
| ENSMUSG00000023092.17 | 27.33557 | 2.44925 | 1.653801 | 1.480982 | 0.138611 | 0.793499 | 0.100453 |
| ENSMUSG00000027433.6 | 198.5225 | -0.96048 | 0.648897 | -1.48017 | 0.138828 | 0.794116 | 0.100116 |
| ENSMUSG00000045374.19 | 30.54058 | -2.39209 | 1.616328 | -1.47995 | 0.138886 | 0.794116 | 0.100116 |
| ENSMUSG00000046329.15 | 92.3843 | -1.09927 | 0.743421 | -1.47866 | 0.139232 | 0.795612 | 0.099299 |
| ENSMUSG00000079469.11 | 29.89182 | -2.4541 | 1.660151 | -1.47824 | 0.139343 | 0.795768 | 0.099213 |
| ENSMUSG00000020128.16 | 233.4772 | 0.97031 | 0.657206 | 1.476417 | 0.139832 | 0.798078 | 0.097954 |
| ENSMUSG00000044167.7 | 162.9673 | -1.15809 | 0.784875 | -1.47552 | 0.140074 | 0.798409 | 0.097775 |
| ENSMUSG00000053553.12 | 31.87687 | -2.61805 | 1.774593 | -1.4753 | 0.140133 | 0.798409 | 0.097775 |
| ENSMUSG00000079057.5 | 1536.365 | -0.44966 | 0.304801 | -1.47526 | 0.140143 | 0.798409 | 0.097775 |
| ENSMUSG00000019528.19 | 25.74324 | -2.44692 | 1.660359 | -1.47373 | 0.140554 | 0.799422 | 0.097224 |
| ENSMUSG00000019797.12 | 145.1117 | 0.938221 | 0.638077 | 1.470389 | 0.141457 | 0.799422 | 0.097224 |
| ENSMUSG00000020219.8 | 139.0519 | 0.922891 | 0.626639 | 1.472763 | 0.140815 | 0.799422 | 0.097224 |
| ENSMUSG00000021215.16 | 219.9924 | 0.820519 | 0.558316 | 1.469631 | 0.141662 | 0.799422 | 0.097224 |
| ENSMUSG00000024426.18 | 32.89454 | 2.217395 | 1.504764 | 1.473583 | 0.140594 | 0.799422 | 0.097224 |
| ENSMUSG00000026153.16 | 57.8973 | 1.545931 | 1.049864 | 1.472505 | 0.140885 | 0.799422 | 0.097224 |
| ENSMUSG00000029068.17 | 289.7081 | -0.67427 | 0.458492 | -1.47062 | 0.141394 | 0.799422 | 0.097224 |

| | | | | | | | |
|---|---|---|---|---|---|---|---|
| ENSMUSG00000030059.16 | 100.6034 | -1.11182 | 0.755319 | -1.47199 | 0.141024 | 0.799422 | 0.097224 |
| ENSMUSG00000032897.18 | 58.05951 | -1.87294 | 1.273282 | -1.47095 | 0.141303 | 0.799422 | 0.097224 |
| ENSMUSG00000033429.11 | 593.3412 | 0.581526 | 0.3947 | 1.473337 | 0.14066 | 0.799422 | 0.097224 |
| ENSMUSG00000036781.14 | 3629.331 | 0.533957 | 0.363334 | 1.469602 | 0.14167 | 0.799422 | 0.097224 |
| ENSMUSG00000042228.15 | 124.723 | -0.97325 | 0.662171 | -1.46979 | 0.141618 | 0.799422 | 0.097224 |
| ENSMUSG00000042608.16 | 14.39143 | 3.779084 | 2.567453 | 1.471919 | 0.141043 | 0.799422 | 0.097224 |
| ENSMUSG00000057147.14 | 34.69912 | -2.29711 | 1.558153 | -1.47425 | 0.140413 | 0.799422 | 0.097224 |
| ENSMUSG00000073835.6 | 20.94956 | -3.15248 | 2.142911 | -1.47112 | 0.141259 | 0.799422 | 0.097224 |
| ENSMUSG00000116284.2 | 39.986 | 2.479956 | 1.68534 | 1.471487 | 0.14116 | 0.799422 | 0.097224 |
| ENSMUSG00000030681.19 | 28.91389 | -2.53801 | 1.72807 | -1.46869 | 0.141916 | 0.800333 | 0.096729 |
| ENSMUSG00000001419.18 | 13.75305 | 2.821418 | 1.929007 | 1.462627 | 0.143569 | 0.800484 | 0.096647 |
| ENSMUSG00000003948.18 | 580.204 | 0.57 | 0.390099 | 1.461169 | 0.143969 | 0.800484 | 0.096647 |
| ENSMUSG00000020634.13 | 95.44021 | -1.22313 | 0.836933 | -1.46145 | 0.143893 | 0.800484 | 0.096647 |
| ENSMUSG00000020641.17 | 32.87178 | -2.08934 | 1.427724 | -1.46341 | 0.143356 | 0.800484 | 0.096647 |
| ENSMUSG00000021619.7 | 40.97137 | 1.627983 | 1.110029 | 1.466613 | 0.142481 | 0.800484 | 0.096647 |
| ENSMUSG00000021948.18 | 30.582 | -2.32868 | 1.589178 | -1.46533 | 0.14283 | 0.800484 | 0.096647 |
| ENSMUSG00000022946.11 | 24.85485 | -3.17264 | 2.168828 | -1.46284 | 0.143512 | 0.800484 | 0.096647 |
| ENSMUSG00000025626.17 | 45.49837 | 1.946 | 1.329305 | 1.463923 | 0.143215 | 0.800484 | 0.096647 |
| ENSMUSG00000027364.15 | 166.2191 | 0.81357 | 0.556576 | 1.46174 | 0.143812 | 0.800484 | 0.096647 |
| ENSMUSG00000028420.14 | 261.9473 | 0.676849 | 0.461723 | 1.465922 | 0.14267 | 0.800484 | 0.096647 |
| ENSMUSG00000028483.14 | 45.07941 | 1.960067 | 1.33677 | 1.466271 | 0.142574 | 0.800484 | 0.096647 |
| ENSMUSG00000029556.13 | 55.1983 | 2.041509 | 1.394183 | 1.464305 | 0.143111 | 0.800484 | 0.096647 |
| ENSMUSG00000032602.7 | 555.8524 | 0.650832 | 0.443855 | 1.466315 | 0.142562 | 0.800484 | 0.096647 |
| ENSMUSG00000033585.6 | 14.34113 | -3.72364 | 2.544192 | -1.46358 | 0.143308 | 0.800484 | 0.096647 |
| ENSMUSG00000034297.15 | 103.1839 | -1.12212 | 0.765732 | -1.46542 | 0.142807 | 0.800484 | 0.096647 |
| ENSMUSG00000039114.17 | 848.0544 | 0.498971 | 0.34112 | 1.462744 | 0.143538 | 0.800484 | 0.096647 |
| ENSMUSG00000040767.11 | 22.94795 | -3.01763 | 2.064816 | -1.46145 | 0.143892 | 0.800484 | 0.096647 |
| ENSMUSG00000041444.15 | 10.18927 | 4.12754 | 2.818878 | 1.46425 | 0.143126 | 0.800484 | 0.096647 |
| ENSMUSG00000042073.16 | 508.0773 | -0.66419 | 0.453818 | -1.46355 | 0.143317 | 0.800484 | 0.096647 |
| ENSMUSG00000044005.14 | 1066.02 | -0.60168 | 0.410568 | -1.46549 | 0.142786 | 0.800484 | 0.096647 |
| ENSMUSG00000049382.11 | 289.5993 | -0.94747 | 0.647047 | -1.4643 | 0.143113 | 0.800484 | 0.096647 |
| ENSMUSG00000056629.17 | 395.108 | -0.75145 | 0.514112 | -1.46165 | 0.143836 | 0.800484 | 0.096647 |
| ENSMUSG00000079012.12 | 923.4443 | 0.638891 | 0.436384 | 1.464058 | 0.143178 | 0.800484 | 0.096647 |
| ENSMUSG00000113063.2 | 30.79094 | -2.29559 | 1.565762 | -1.46611 | 0.142617 | 0.800484 | 0.096647 |
| ENSMUSG00000021508.12 | 23.61323 | -2.48369 | 1.700513 | -1.46055 | 0.144138 | 0.800557 | 0.096608 |
| ENSMUSG00000050390.13 | 248.5627 | -0.78436 | 0.537046 | -1.46051 | 0.144151 | 0.800557 | 0.096608 |
| ENSMUSG00000020160.19 | 21.2985 | -2.79342 | 1.915743 | -1.45814 | 0.144802 | 0.801358 | 0.096174 |
| ENSMUSG00000022744.13 | 67.12032 | -1.73584 | 1.19036 | -1.45825 | 0.144772 | 0.801358 | 0.096174 |
| ENSMUSG00000025194.6 | 1115.174 | -0.41999 | 0.287911 | -1.45874 | 0.144637 | 0.801358 | 0.096174 |
| ENSMUSG00000034022.9 | 97.64656 | -1.80793 | 1.239188 | -1.45896 | 0.144575 | 0.801358 | 0.096174 |
| ENSMUSG00000046667.15 | 24.81931 | 2.292244 | 1.570785 | 1.459299 | 0.144483 | 0.801358 | 0.096174 |
| ENSMUSG00000051451.7 | 57.26305 | 1.831413 | 1.255901 | 1.458246 | 0.144773 | 0.801358 | 0.096174 |
| ENSMUSG00000021040.16 | 581.8275 | 0.586688 | 0.402543 | 1.457456 | 0.144991 | 0.801932 | 0.095863 |
| ENSMUSG00000031696.10 | 690.5411 | 0.601391 | 0.413077 | 1.455883 | 0.145425 | 0.803194 | 0.095179 |

| | | | | | | | |
|---|---|---|---|---|---|---|---|
| ENSMUSG00000034731.12 | 16.78066 | 3.168842 | 2.176837 | 1.455709 | 0.145473 | 0.803194 | 0.095179 |
| ENSMUSG00000056917.14 | 21.92073 | 3.286531 | 2.256929 | 1.456196 | 0.145339 | 0.803194 | 0.095179 |
| ENSMUSG00000019791.13 | 392.5357 | -0.68793 | 0.473621 | -1.4525 | 0.146363 | 0.804131 | 0.094673 |
| ENSMUSG00000020255.9 | 36.43293 | -2.70055 | 1.857862 | -1.45358 | 0.146062 | 0.804131 | 0.094673 |
| ENSMUSG00000021238.12 | 1859.762 | 0.397363 | 0.273286 | 1.454016 | 0.145942 | 0.804131 | 0.094673 |
| ENSMUSG00000024039.16 | 323.5352 | -0.64551 | 0.443792 | -1.45452 | 0.145801 | 0.804131 | 0.094673 |
| ENSMUSG00000027164.9 | 78.33846 | -1.3069 | 0.899217 | -1.45337 | 0.146121 | 0.804131 | 0.094673 |
| ENSMUSG00000027933.12 | 133.5444 | 1.154308 | 0.794309 | 1.453224 | 0.146162 | 0.804131 | 0.094673 |
| ENSMUSG00000029623.15 | 323.4123 | 0.668987 | 0.460626 | 1.452343 | 0.146406 | 0.804131 | 0.094673 |
| ENSMUSG00000032051.10 | 1186.84 | 0.398973 | 0.274624 | 1.452799 | 0.146279 | 0.804131 | 0.094673 |
| ENSMUSG00000087578.2 | 13.13212 | 4.44259 | 3.058681 | 1.452453 | 0.146376 | 0.804131 | 0.094673 |
| ENSMUSG00000024493.10 | 51.83557 | -1.93105 | 1.330512 | -1.45136 | 0.146679 | 0.804602 | 0.094419 |
| ENSMUSG00000032330.8 | 2751.687 | 0.458332 | 0.315867 | 1.451028 | 0.146772 | 0.804602 | 0.094419 |
| ENSMUSG00000061410.7 | 149.6024 | -1.2113 | 0.834429 | -1.45165 | 0.146599 | 0.804602 | 0.094419 |
| ENSMUSG00000078919.11 | 36.13249 | 1.84605 | 1.27269 | 1.45051 | 0.146916 | 0.804602 | 0.094419 |
| ENSMUSG00000085175.8 | 12.68131 | -3.81596 | 2.630697 | -1.45055 | 0.146905 | 0.804602 | 0.094419 |
| ENSMUSG00000019471.11 | 58.71251 | -1.34161 | 0.925353 | -1.44983 | 0.147106 | 0.805174 | 0.094111 |
| ENSMUSG00000004096.10 | 448.808 | -0.60825 | 0.419959 | -1.44835 | 0.147518 | 0.805346 | 0.094018 |
| ENSMUSG00000007817.16 | 142.277 | 0.975037 | 0.675303 | 1.443852 | 0.148781 | 0.805346 | 0.094018 |
| ENSMUSG00000015843.11 | 54.80064 | 2.38876 | 1.65004 | 1.447698 | 0.147701 | 0.805346 | 0.094018 |
| ENSMUSG00000016256.11 | 481.7326 | -0.80325 | 0.555799 | -1.44522 | 0.148396 | 0.805346 | 0.094018 |
| ENSMUSG00000024442.7 | 90.6151 | 1.186038 | 0.820058 | 1.446286 | 0.148097 | 0.805346 | 0.094018 |
| ENSMUSG00000025102.15 | 21.6836 | -2.10646 | 1.453948 | -1.44879 | 0.147397 | 0.805346 | 0.094018 |
| ENSMUSG00000025132.14 | 40.60901 | -2.01116 | 1.390354 | -1.44651 | 0.148034 | 0.805346 | 0.094018 |
| ENSMUSG00000029597.15 | 187.2029 | -0.94361 | 0.653383 | -1.44419 | 0.148685 | 0.805346 | 0.094018 |
| ENSMUSG00000031490.7 | 120.4169 | -1.46077 | 1.010201 | -1.44602 | 0.148172 | 0.805346 | 0.094018 |
| ENSMUSG00000036376.4 | 95.28794 | -1.32198 | 0.915205 | -1.44446 | 0.148609 | 0.805346 | 0.094018 |
| ENSMUSG00000039067.6 | 406.6874 | -0.68107 | 0.47177 | -1.44365 | 0.148836 | 0.805346 | 0.094018 |
| ENSMUSG00000040264.11 | 10.90976 | -3.58 | 2.479677 | -1.44374 | 0.148813 | 0.805346 | 0.094018 |
| ENSMUSG00000041594.19 | 15.02372 | -3.57424 | 2.473289 | -1.44513 | 0.14842 | 0.805346 | 0.094018 |
| ENSMUSG00000042349.14 | 22.62374 | -2.50666 | 1.733209 | -1.44625 | 0.148107 | 0.805346 | 0.094018 |
| ENSMUSG00000051234.7 | 308.5789 | -0.71969 | 0.497792 | -1.44576 | 0.148245 | 0.805346 | 0.094018 |
| ENSMUSG00000052310.10 | 27.87551 | -2.20826 | 1.526319 | -1.44679 | 0.147956 | 0.805346 | 0.094018 |
| ENSMUSG00000052712.18 | 341.626 | -0.6235 | 0.431614 | -1.44458 | 0.148577 | 0.805346 | 0.094018 |
| ENSMUSG00000062202.15 | 30.34469 | -2.29794 | 1.589322 | -1.44586 | 0.148217 | 0.805346 | 0.094018 |
| ENSMUSG00000062937.8 | 109.3945 | 1.524339 | 1.052583 | 1.44819 | 0.147564 | 0.805346 | 0.094018 |
| ENSMUSG00000097445.2 | 45.80068 | -2.43535 | 1.684355 | -1.44586 | 0.148215 | 0.805346 | 0.094018 |
| ENSMUSG00000039254.17 | 56.01444 | 2.244758 | 1.555584 | 1.443032 | 0.149011 | 0.805835 | 0.093754 |
| ENSMUSG00000026471.15 | 22.284 | -2.64338 | 1.835482 | -1.44015 | 0.149824 | 0.807688 | 0.092756 |
| ENSMUSG00000032666.17 | 22.96383 | -2.37898 | 1.651 | -1.44093 | 0.149604 | 0.807688 | 0.092756 |
| ENSMUSG00000033623.14 | 120.1166 | -1.07928 | 0.749111 | -1.44075 | 0.149655 | 0.807688 | 0.092756 |
| ENSMUSG00000034190.10 | 56.36982 | -1.34731 | 0.935407 | -1.44035 | 0.14977 | 0.807688 | 0.092756 |
| ENSMUSG00000040048.15 | 1357.605 | 0.456656 | 0.317187 | 1.439706 | 0.149951 | 0.807688 | 0.092756 |
| ENSMUSG00000078713.9 | 126.9399 | 1.118982 | 0.776871 | 1.44037 | 0.149763 | 0.807688 | 0.092756 |

| | | | | | | | |
|---|---|---|---|---|---|---|---|
| ENSMUSG00000116571.2 | 11.02191 | 4.578827 | 3.180018 | 1.439875 | 0.149903 | 0.807688 | 0.092756 |
| ENSMUSG00000021286.9 | 240.511 | 0.858022 | 0.596422 | 1.438616 | 0.150259 | 0.808432 | 0.092356 |
| ENSMUSG00000067203.9 | 14.31946 | 3.730895 | 2.593139 | 1.438756 | 0.15022 | 0.808432 | 0.092356 |
| ENSMUSG00000026102.10 | 52.72241 | -1.40741 | 0.979316 | -1.43713 | 0.15068 | 0.808684 | 0.092221 |
| ENSMUSG00000028837.9 | 556.519 | -0.6184 | 0.430243 | -1.43733 | 0.150625 | 0.808684 | 0.092221 |
| ENSMUSG00000042842.12 | 41.70365 | 2.06938 | 1.440121 | 1.436949 | 0.150733 | 0.808684 | 0.092221 |
| ENSMUSG00000055116.9 | 33.79796 | 2.573738 | 1.789783 | 1.438016 | 0.150429 | 0.808684 | 0.092221 |
| ENSMUSG00000079508.9 | 40.63394 | 2.112787 | 1.469724 | 1.43754 | 0.150565 | 0.808684 | 0.092221 |
| ENSMUSG00000021671.10 | 32.36227 | 1.74973 | 1.218437 | 1.436045 | 0.150989 | 0.808744 | 0.092189 |
| ENSMUSG00000037808.14 | 19.13838 | -2.63126 | 1.832019 | -1.43626 | 0.150928 | 0.808744 | 0.092189 |
| ENSMUSG00000061273.4 | 114.524 | -1.28815 | 0.897034 | -1.43601 | 0.151 | 0.808744 | 0.092189 |
| ENSMUSG00000018983.10 | 12.88614 | 4.413665 | 3.075038 | 1.43532 | 0.151196 | 0.809337 | 0.091871 |
| ENSMUSG00000028243.6 | 31.7663 | 2.250772 | 1.569132 | 1.434406 | 0.151456 | 0.809818 | 0.091612 |
| ENSMUSG00000041237.13 | 255.0533 | 1.017738 | 0.709446 | 1.434553 | 0.151414 | 0.809818 | 0.091612 |
| ENSMUSG00000026112.8 | 621.2109 | 0.488476 | 0.340664 | 1.433895 | 0.151602 | 0.810141 | 0.091439 |
| ENSMUSG00000074634.13 | 60.24455 | -1.38268 | 0.964489 | -1.43359 | 0.15169 | 0.810152 | 0.091433 |
| ENSMUSG00000021707.6 | 449.9531 | -0.62911 | 0.439065 | -1.43285 | 0.1519 | 0.81024 | 0.091386 |
| ENSMUSG00000024276.8 | 48.33009 | 1.511058 | 1.054741 | 1.432633 | 0.151963 | 0.81024 | 0.091386 |
| ENSMUSG00000031684.12 | 40.94112 | -2.28639 | 1.595312 | -1.43319 | 0.151803 | 0.81024 | 0.091386 |
| ENSMUSG00000002844.10 | 22.06519 | -2.55853 | 1.789453 | -1.42978 | 0.152779 | 0.811747 | 0.090579 |
| ENSMUSG00000004263.16 | 22.68738 | 2.43089 | 1.701161 | 1.428959 | 0.153016 | 0.811747 | 0.090579 |
| ENSMUSG00000016554.10 | 213.1906 | -0.86429 | 0.603903 | -1.43118 | 0.152379 | 0.811747 | 0.090579 |
| ENSMUSG00000021932.14 | 132.0758 | 1.332631 | 0.932182 | 1.429581 | 0.152837 | 0.811747 | 0.090579 |
| ENSMUSG00000022772.13 | 89.9949 | -1.25104 | 0.875263 | -1.42933 | 0.152909 | 0.811747 | 0.090579 |
| ENSMUSG00000027800.15 | 42.83641 | -1.77334 | 1.23994 | -1.43019 | 0.152664 | 0.811747 | 0.090579 |
| ENSMUSG00000036905.9 | 40.00204 | -1.89786 | 1.327857 | -1.42927 | 0.152928 | 0.811747 | 0.090579 |
| ENSMUSG00000040209.13 | 14.08419 | -3.24359 | 2.268876 | -1.4296 | 0.152832 | 0.811747 | 0.090579 |
| ENSMUSG00000043140.11 | 39.04137 | 2.652668 | 1.856031 | 1.429215 | 0.152942 | 0.811747 | 0.090579 |
| ENSMUSG00000028048.12 | 134.8423 | 0.883175 | 0.618264 | 1.428476 | 0.153155 | 0.81203 | 0.090428 |
| ENSMUSG00000021136.14 | 170.9899 | -0.82529 | 0.578019 | -1.42778 | 0.153354 | 0.81263 | 0.090107 |
| ENSMUSG00000039063.6 | 311.1389 | 0.783786 | 0.549204 | 1.427131 | 0.153542 | 0.813173 | 0.089817 |
| ENSMUSG00000003346.15 | 48.29413 | 1.585982 | 1.112251 | 1.425921 | 0.153891 | 0.813205 | 0.0898 |
| ENSMUSG00000022339.10 | 180.9868 | -0.79104 | 0.554561 | -1.42642 | 0.153746 | 0.813205 | 0.0898 |
| ENSMUSG00000022361.15 | 160.0251 | 1.457946 | 1.022077 | 1.426454 | 0.153737 | 0.813205 | 0.0898 |
| ENSMUSG00000047631.5 | 4792.921 | -0.29158 | 0.204478 | -1.42599 | 0.153871 | 0.813205 | 0.0898 |
| ENSMUSG00000019312.11 | 185.187 | 1.037582 | 0.728578 | 1.42412 | 0.154412 | 0.815327 | 0.088668 |
| ENSMUSG00000064326.14 | 26.74609 | 2.490267 | 1.748861 | 1.423936 | 0.154465 | 0.815327 | 0.088668 |
| ENSMUSG00000057880.13 | 2249.02 | -0.39009 | 0.274035 | -1.4235 | 0.154592 | 0.815546 | 0.088551 |
| ENSMUSG00000022972.11 | 75.15048 | 1.759039 | 1.238183 | 1.420662 | 0.155415 | 0.815866 | 0.088381 |
| ENSMUSG00000024712.10 | 320.5277 | 0.660729 | 0.465197 | 1.420323 | 0.155514 | 0.815866 | 0.088381 |
| ENSMUSG00000033106.7 | 20.04455 | -2.75366 | 1.93569 | -1.42257 | 0.154861 | 0.815866 | 0.088381 |
| ENSMUSG00000034211.15 | 586.1474 | 0.640716 | 0.450761 | 1.421409 | 0.155198 | 0.815866 | 0.088381 |
| ENSMUSG00000036880.11 | 6260.392 | 0.283848 | 0.199729 | 1.421166 | 0.155268 | 0.815866 | 0.088381 |
| ENSMUSG00000038582.16 | 144.2981 | -0.98241 | 0.691468 | -1.42076 | 0.155386 | 0.815866 | 0.088381 |

| | | | | | | | |
|---|---|---|---|---|---|---|---|
| ENSMUSG00000038838.16 | 18.27427 | -2.63365 | 1.851704 | -1.42228 | 0.154944 | 0.815866 | 0.088381 |
| ENSMUSG00000062169.14 | 121.482 | 0.900534 | 0.633336 | 1.42189 | 0.155058 | 0.815866 | 0.088381 |
| ENSMUSG00000081344.6 | 100.1367 | 1.024388 | 0.721102 | 1.420587 | 0.155437 | 0.815866 | 0.088381 |
| ENSMUSG00000086432.2 | 100.1287 | 1.05246 | 0.74054 | 1.421207 | 0.155257 | 0.815866 | 0.088381 |
| ENSMUSG00000001506.11 | 13.91759 | -3.58855 | 2.529432 | -1.41872 | 0.155981 | 0.816034 | 0.088292 |
| ENSMUSG00000018239.10 | 37.34003 | -2.51813 | 1.775653 | -1.41815 | 0.156148 | 0.816034 | 0.088292 |
| ENSMUSG00000029260.16 | 2986.286 | 0.380984 | 0.268573 | 1.418548 | 0.156031 | 0.816034 | 0.088292 |
| ENSMUSG00000030530.16 | 93.89384 | -1.03231 | 0.727094 | -1.41977 | 0.155674 | 0.816034 | 0.088292 |
| ENSMUSG00000039934.13 | 156.2674 | -1.35026 | 0.951578 | -1.41897 | 0.155908 | 0.816034 | 0.088292 |
| ENSMUSG00000043872.15 | 17.19772 | 3.106834 | 2.190614 | 1.418248 | 0.156118 | 0.816034 | 0.088292 |
| ENSMUSG00000056050.16 | 1118.303 | -0.43032 | 0.303297 | -1.41882 | 0.155952 | 0.816034 | 0.088292 |
| ENSMUSG00000073609.10 | 115.8677 | 0.972561 | 0.685981 | 1.417766 | 0.156259 | 0.816163 | 0.088223 |
| ENSMUSG00000047907.12 | 97.8563 | -1.23635 | 0.872935 | -1.41631 | 0.156684 | 0.817934 | 0.087282 |
| ENSMUSG00000029428.14 | 22.58004 | 2.248565 | 1.588492 | 1.415534 | 0.156912 | 0.818671 | 0.08689 |
| ENSMUSG00000001056.4 | 149.2839 | -0.98185 | 0.694096 | -1.41457 | 0.157194 | 0.81879 | 0.086827 |
| ENSMUSG00000006527.17 | 145.6696 | -0.9538 | 0.674176 | -1.41476 | 0.157138 | 0.81879 | 0.086827 |
| ENSMUSG00000007476.19 | 86.68242 | 1.077437 | 0.761548 | 1.414799 | 0.157127 | 0.81879 | 0.086827 |
| ENSMUSG00000031604.11 | 2414.587 | -0.45807 | 0.324022 | -1.4137 | 0.157449 | 0.819219 | 0.0866 |
| ENSMUSG00000051390.11 | 29.54414 | -2.63229 | 1.861886 | -1.41378 | 0.157427 | 0.819219 | 0.0866 |
| ENSMUSG00000021962.10 | 33.01755 | -2.44431 | 1.730305 | -1.41265 | 0.15776 | 0.819362 | 0.086524 |
| ENSMUSG00000032560.15 | 113.6109 | 1.265686 | 0.896009 | 1.412583 | 0.157778 | 0.819362 | 0.086524 |
| ENSMUSG00000034126.6 | 29.41867 | -2.23553 | 1.582749 | -1.41243 | 0.157822 | 0.819362 | 0.086524 |
| ENSMUSG00000066072.14 | 770.5833 | 2.690082 | 1.904322 | 1.412619 | 0.157768 | 0.819362 | 0.086524 |
| ENSMUSG00000025903.15 | 1343.115 | 0.452077 | 0.320222 | 1.411761 | 0.15802 | 0.819942 | 0.086217 |
| ENSMUSG00000021594.12 | 398.1157 | 0.705732 | 0.500054 | 1.411314 | 0.158152 | 0.820176 | 0.086093 |
| ENSMUSG00000009907.18 | 483.9459 | 0.914227 | 0.648106 | 1.410613 | 0.158359 | 0.82035 | 0.086001 |
| ENSMUSG00000073062.4 | 56.86682 | 2.359798 | 1.672837 | 1.410656 | 0.158346 | 0.82035 | 0.086001 |
| ENSMUSG00000022875.19 | 13041.82 | -0.29794 | 0.211364 | -1.4096 | 0.158657 | 0.821141 | 0.085583 |
| ENSMUSG00000028240.3 | 469.3662 | 0.851882 | 0.604382 | 1.40951 | 0.158685 | 0.821141 | 0.085583 |
| ENSMUSG00000032412.9 | 263.0581 | 0.749256 | 0.531726 | 1.409101 | 0.158805 | 0.821317 | 0.085489 |
| ENSMUSG00000022214.16 | 448.813 | -0.88555 | 0.629129 | -1.40758 | 0.159255 | 0.822455 | 0.084888 |
| ENSMUSG00000028757.5 | 276.5995 | -0.89106 | 0.632959 | -1.40776 | 0.159201 | 0.822455 | 0.084888 |
| ENSMUSG00000042213.18 | 131.6847 | 0.920234 | 0.653943 | 1.407209 | 0.159365 | 0.822455 | 0.084888 |
| ENSMUSG00000060519.12 | 19.5205 | -2.31227 | 1.643191 | -1.40719 | 0.159372 | 0.822455 | 0.084888 |
| ENSMUSG00000006169.21 | 375.505 | 0.570054 | 0.407576 | 1.398644 | 0.16192 | 0.823006 | 0.084597 |
| ENSMUSG00000018293.5 | 138.2265 | -1.34659 | 0.962604 | -1.39891 | 0.161841 | 0.823006 | 0.084597 |
| ENSMUSG00000019718.9 | 213.7513 | 0.820337 | 0.584984 | 1.402322 | 0.160819 | 0.823006 | 0.084597 |
| ENSMUSG00000019878.9 | 71.63956 | -1.62025 | 1.157757 | -1.39947 | 0.161671 | 0.823006 | 0.084597 |
| ENSMUSG00000020271.18 | 89.04298 | 1.066498 | 0.762821 | 1.398098 | 0.162084 | 0.823006 | 0.084597 |
| ENSMUSG00000020782.19 | 10.94026 | -3.87357 | 2.76679 | -1.40002 | 0.161506 | 0.823006 | 0.084597 |
| ENSMUSG00000021670.15 | 688.7572 | 0.619034 | 0.442295 | 1.399594 | 0.161635 | 0.823006 | 0.084597 |
| ENSMUSG00000022241.7 | 741.419 | 0.53291 | 0.380071 | 1.402135 | 0.160875 | 0.823006 | 0.084597 |
| ENSMUSG00000022371.17 | 118.0682 | -0.97529 | 0.695038 | -1.40322 | 0.160552 | 0.823006 | 0.084597 |
| ENSMUSG00000023026.17 | 10.47783 | -2.92607 | 2.091811 | -1.39882 | 0.161866 | 0.823006 | 0.084597 |

| | | | | | | | |
|---|---|---|---|---|---|---|---|
| ENSMUSG00000023460.14 | 102.4876 | -1.17777 | 0.838373 | -1.40483 | 0.160073 | 0.823006 | 0.084597 |
| ENSMUSG00000024084.9 | 93.93767 | 1.070113 | 0.76121 | 1.405805 | 0.159782 | 0.823006 | 0.084597 |
| ENSMUSG00000026398.15 | 278.2743 | -0.77667 | 0.554354 | -1.40103 | 0.161205 | 0.823006 | 0.084597 |
| ENSMUSG00000026427.17 | 64.35873 | -1.70685 | 1.217147 | -1.40233 | 0.160816 | 0.823006 | 0.084597 |
| ENSMUSG00000028009.8 | 10.37288 | 3.584302 | 2.562009 | 1.39902 | 0.161807 | 0.823006 | 0.084597 |
| ENSMUSG00000028089.6 | 36.66305 | -2.12933 | 1.517809 | -1.4029 | 0.160647 | 0.823006 | 0.084597 |
| ENSMUSG00000029672.17 | 247.2187 | -0.75781 | 0.541693 | -1.39896 | 0.161824 | 0.823006 | 0.084597 |
| ENSMUSG00000036646.14 | 39.53708 | -2.52137 | 1.802492 | -1.39883 | 0.161865 | 0.823006 | 0.084597 |
| ENSMUSG00000038393.15 | 240.1464 | -0.73038 | 0.521064 | -1.4017 | 0.161005 | 0.823006 | 0.084597 |
| ENSMUSG00000042599.9 | 125.902 | -0.97772 | 0.696753 | -1.40325 | 0.160543 | 0.823006 | 0.084597 |
| ENSMUSG00000046031.8 | 19.34322 | -2.55756 | 1.8258 | -1.40079 | 0.161278 | 0.823006 | 0.084597 |
| ENSMUSG00000051154.12 | 406.808 | 0.620221 | 0.442143 | 1.402761 | 0.160688 | 0.823006 | 0.084597 |
| ENSMUSG00000052302.7 | 56.81542 | -2.29847 | 1.635197 | -1.40562 | 0.159837 | 0.823006 | 0.084597 |
| ENSMUSG00000052957.9 | 145.4697 | 1.085955 | 0.77336 | 1.404205 | 0.160258 | 0.823006 | 0.084597 |
| ENSMUSG00000060636.15 | 968.7444 | 0.484715 | 0.345063 | 1.404714 | 0.160106 | 0.823006 | 0.084597 |
| ENSMUSG00000062944.5 | 11.81314 | 3.439876 | 2.459999 | 1.398325 | 0.162016 | 0.823006 | 0.084597 |
| ENSMUSG00000074170.6 | 24.46427 | 2.782381 | 1.986735 | 1.40048 | 0.16137 | 0.823006 | 0.084597 |
| ENSMUSG00000080738.2 | 76.12569 | -1.10026 | 0.783967 | -1.40346 | 0.160481 | 0.823006 | 0.084597 |
| ENSMUSG00000111361.2 | 20.7321 | -3.21848 | 2.298173 | -1.40045 | 0.161378 | 0.823006 | 0.084597 |
| ENSMUSG00000115200.2 | 10.7707 | 3.547102 | 2.531435 | 1.401222 | 0.161148 | 0.823006 | 0.084597 |
| ENSMUSG00000021009.16 | 67.09869 | 1.201797 | 0.860847 | 1.396063 | 0.162695 | 0.823669 | 0.084248 |
| ENSMUSG00000024826.16 | 60.03696 | -1.4526 | 1.041141 | -1.3952 | 0.162957 | 0.823669 | 0.084248 |
| ENSMUSG00000026439.15 | 32.03863 | 2.55522 | 1.830674 | 1.395781 | 0.16278 | 0.823669 | 0.084248 |
| ENSMUSG00000031925.18 | 34.50919 | 2.203375 | 1.579732 | 1.394778 | 0.163083 | 0.823669 | 0.084248 |
| ENSMUSG00000038910.8 | 42.03181 | -1.47921 | 1.059607 | -1.396 | 0.162714 | 0.823669 | 0.084248 |
| ENSMUSG00000039809.11 | 153.4362 | -1.04715 | 0.749638 | -1.39688 | 0.16245 | 0.823669 | 0.084248 |
| ENSMUSG00000048490.15 | 61.84535 | 1.410392 | 1.009812 | 1.396688 | 0.162507 | 0.823669 | 0.084248 |
| ENSMUSG00000048521.9 | 17.67023 | -3.29598 | 2.361636 | -1.39563 | 0.162825 | 0.823669 | 0.084248 |
| ENSMUSG00000057982.9 | 130.77 | 1.090473 | 0.780757 | 1.396687 | 0.162508 | 0.823669 | 0.084248 |
| ENSMUSG00000079659.6 | 255.2163 | 0.791412 | 0.567348 | 1.394933 | 0.163036 | 0.823669 | 0.084248 |
| ENSMUSG00000002205.17 | 36.00641 | -2.82646 | 2.027701 | -1.39393 | 0.16334 | 0.824349 | 0.083889 |
| ENSMUSG00000031216.14 | 28.57033 | 1.716387 | 1.231656 | 1.39356 | 0.16345 | 0.824349 | 0.083889 |
| ENSMUSG00000031938.16 | 1160.789 | 0.429464 | 0.308198 | 1.393467 | 0.163479 | 0.824349 | 0.083889 |
| ENSMUSG00000005575.17 | 21.59885 | 2.917135 | 2.09458 | 1.392706 | 0.163709 | 0.824478 | 0.083821 |
| ENSMUSG00000021823.11 | 269.9342 | 0.92133 | 0.661462 | 1.392869 | 0.163659 | 0.824478 | 0.083821 |
| ENSMUSG00000028494.13 | 733.2633 | 0.455125 | 0.326847 | 1.392472 | 0.163779 | 0.824478 | 0.083821 |
| ENSMUSG00000030788.17 | 290.572 | -0.71477 | 0.513448 | -1.3921 | 0.163892 | 0.824478 | 0.083821 |
| ENSMUSG00000092545.8 | 408.8744 | -0.62619 | 0.449864 | -1.39195 | 0.163939 | 0.824478 | 0.083821 |
| ENSMUSG00000043252.9 | 251.8562 | 0.720208 | 0.517634 | 1.391347 | 0.16412 | 0.824515 | 0.083801 |
| ENSMUSG00000046865.8 | 171.0056 | 0.870842 | 0.625866 | 1.39142 | 0.164098 | 0.824515 | 0.083801 |
| ENSMUSG00000020154.11 | 361.2869 | 0.909632 | 0.654249 | 1.390346 | 0.164424 | 0.824752 | 0.083677 |
| ENSMUSG00000024993.16 | 104.3201 | 1.051589 | 0.756685 | 1.389731 | 0.164611 | 0.824752 | 0.083677 |
| ENSMUSG00000029434.13 | 66.2472 | -1.41858 | 1.020952 | -1.38947 | 0.164689 | 0.824752 | 0.083677 |
| ENSMUSG00000037306.14 | 10.67895 | -3.89329 | 2.801531 | -1.3897 | 0.16462 | 0.824752 | 0.083677 |

| | | | | | | | |
|---|---|---|---|---|---|---|---|
| ENSMUSG00000039759.11 | 12.11109 | -4.29019 | 3.086658 | -1.38991 | 0.164555 | 0.824752 | 0.083677 |
| ENSMUSG00000057137.10 | 106.8965 | -1.2345 | 0.887763 | -1.39058 | 0.164353 | 0.824752 | 0.083677 |
| ENSMUSG00000018593.14 | 341.5793 | -0.59977 | 0.432339 | -1.38727 | 0.16536 | 0.825182 | 0.083451 |
| ENSMUSG00000026238.15 | 1051.058 | 0.455641 | 0.328464 | 1.387188 | 0.165384 | 0.825182 | 0.083451 |
| ENSMUSG00000028911.17 | 116.9277 | -1.09042 | 0.785752 | -1.38774 | 0.165216 | 0.825182 | 0.083451 |
| ENSMUSG00000037797.15 | 594.4968 | -0.47989 | 0.34552 | -1.38889 | 0.164865 | 0.825182 | 0.083451 |
| ENSMUSG00000049422.8 | 4183.619 | 0.340527 | 0.245365 | 1.38784 | 0.165186 | 0.825182 | 0.083451 |
| ENSMUSG00000060938.15 | 2795.794 | 0.374148 | 0.269532 | 1.388139 | 0.165095 | 0.825182 | 0.083451 |
| ENSMUSG00000112758.2 | 24.63968 | -2.6626 | 1.919376 | -1.38722 | 0.165375 | 0.825182 | 0.083451 |
| ENSMUSG00000021211.5 | 794.1051 | 0.521751 | 0.376294 | 1.386551 | 0.165579 | 0.825397 | 0.083337 |
| ENSMUSG00000022215.7 | 102.4585 | 1.767799 | 1.275479 | 1.385988 | 0.16575 | 0.825397 | 0.083337 |
| ENSMUSG00000026131.21 | 155.9705 | 0.850878 | 0.613897 | 1.386028 | 0.165738 | 0.825397 | 0.083337 |
| ENSMUSG00000039195.5 | 37.12932 | -2.02057 | 1.457946 | -1.3859 | 0.165776 | 0.825397 | 0.083337 |
| ENSMUSG00000027010.17 | 64.25577 | 1.145055 | 0.826631 | 1.385206 | 0.16599 | 0.826027 | 0.083006 |
| ENSMUSG00000019368.14 | 245.8483 | 0.9872 | 0.713204 | 1.384176 | 0.166304 | 0.826726 | 0.082639 |
| ENSMUSG00000074698.11 | 189.0143 | -1.0502 | 0.758631 | -1.38433 | 0.166257 | 0.826726 | 0.082639 |
| ENSMUSG00000032386.16 | 385.2748 | 0.636194 | 0.459949 | 1.383185 | 0.166608 | 0.827802 | 0.082074 |
| ENSMUSG00000032002.11 | 452.892 | 0.699109 | 0.505814 | 1.382145 | 0.166927 | 0.828518 | 0.081698 |
| ENSMUSG00000040274.12 | 109.6227 | -1.0368 | 0.750073 | -1.38227 | 0.16689 | 0.828518 | 0.081698 |
| ENSMUSG00000055762.17 | 434.6665 | 0.524954 | 0.379892 | 1.381851 | 0.167018 | 0.828533 | 0.08169 |
| ENSMUSG00000030737.18 | 414.3034 | 0.908792 | 0.658364 | 1.380378 | 0.16747 | 0.829657 | 0.081101 |
| ENSMUSG00000035051.16 | 14.81103 | -3.34245 | 2.421544 | -1.3803 | 0.167495 | 0.829657 | 0.081101 |
| ENSMUSG00000054484.15 | 43.01744 | -1.58775 | 1.150329 | -1.38026 | 0.167507 | 0.829657 | 0.081101 |
| ENSMUSG00000026761.13 | 135.5119 | 0.898188 | 0.651018 | 1.379667 | 0.167689 | 0.830127 | 0.080856 |
| ENSMUSG00000050312.13 | 97.56577 | -1.39829 | 1.013743 | -1.37934 | 0.167791 | 0.830198 | 0.080818 |
| ENSMUSG00000022671.14 | 27.9361 | 1.825073 | 1.324369 | 1.378071 | 0.168182 | 0.830432 | 0.080696 |
| ENSMUSG00000028207.19 | 187.2416 | 0.956668 | 0.693838 | 1.378807 | 0.167954 | 0.830432 | 0.080696 |
| ENSMUSG00000032740.18 | 12.29144 | 2.577094 | 1.869982 | 1.378138 | 0.168161 | 0.830432 | 0.080696 |
| ENSMUSG00000036550.17 | 229.5633 | -0.79453 | 0.576562 | -1.37805 | 0.168189 | 0.830432 | 0.080696 |
| ENSMUSG00000049755.15 | 51.92435 | 1.476759 | 1.07196 | 1.377624 | 0.168319 | 0.830644 | 0.080585 |
| ENSMUSG00000022607.19 | 43.12319 | -1.46016 | 1.060876 | -1.37637 | 0.168706 | 0.831687 | 0.08004 |
| ENSMUSG00000030088.16 | 2488.162 | -0.5755 | 0.418066 | -1.37659 | 0.16864 | 0.831687 | 0.08004 |
| ENSMUSG00000035954.11 | 169.3631 | -1.02215 | 0.743053 | -1.3756 | 0.168945 | 0.831997 | 0.079878 |
| ENSMUSG00000047789.6 | 13.54559 | -3.73596 | 2.715681 | -1.3757 | 0.168914 | 0.831997 | 0.079878 |
| ENSMUSG00000022601.12 | 29.64303 | -1.79017 | 1.301699 | -1.37526 | 0.169052 | 0.832096 | 0.079827 |
| ENSMUSG00000029178.15 | 177.423 | 0.859774 | 0.62581 | 1.373859 | 0.169485 | 0.833362 | 0.079166 |
| ENSMUSG00000034285.16 | 642.4242 | 0.599508 | 0.436311 | 1.37404 | 0.169429 | 0.833362 | 0.079166 |
| ENSMUSG00000112302.2 | 17.3814 | 2.581425 | 1.879441 | 1.373507 | 0.169595 | 0.833468 | 0.079111 |
| ENSMUSG00000028703.15 | 10.05882 | 3.537991 | 2.576486 | 1.373184 | 0.169695 | 0.833528 | 0.07908 |
| ENSMUSG00000021767.20 | 33.16939 | -1.9726 | 1.436874 | -1.37284 | 0.169803 | 0.833624 | 0.07903 |
| ENSMUSG00000041362.18 | 99.32279 | -1.05624 | 0.769666 | -1.37234 | 0.169959 | 0.833958 | 0.078856 |
| ENSMUSG00000091625.9 | 35.75852 | 2.136885 | 1.557847 | 1.371691 | 0.17016 | 0.834512 | 0.078567 |
| ENSMUSG00000001025.9 | 22.17109 | -2.00068 | 1.459309 | -1.37098 | 0.170381 | 0.834546 | 0.07855 |
| ENSMUSG00000018669.15 | 73.38239 | -1.40789 | 1.02689 | -1.37102 | 0.170369 | 0.834546 | 0.07855 |

| | | | | | | | |
|---|---|---|---|---|---|---|---|
| ENSMUSG00000020801.9 | 63.6799 | 1.437787 | 1.04885 | 1.370822 | 0.17043 | 0.834546 | 0.07855 |
| ENSMUSG00000028029.12 | 274.3945 | -0.77616 | 0.56641 | -1.37031 | 0.17059 | 0.834894 | 0.078369 |
| ENSMUSG00000035960.15 | 127.7797 | -0.90063 | 0.657609 | -1.36956 | 0.170825 | 0.835428 | 0.078091 |
| ENSMUSG00000040557.15 | 31.66744 | -2.52551 | 1.844248 | -1.3694 | 0.170875 | 0.835428 | 0.078091 |
| ENSMUSG00000025533.16 | 2220.433 | 0.436083 | 0.31867 | 1.368448 | 0.171172 | 0.83599 | 0.077799 |
| ENSMUSG00000026667.15 | 225.8745 | 0.763 | 0.557899 | 1.367632 | 0.171427 | 0.83599 | 0.077799 |
| ENSMUSG00000039183.7 | 57.68434 | -1.74451 | 1.275562 | -1.36764 | 0.171426 | 0.83599 | 0.077799 |
| ENSMUSG00000044857.9 | 41.80081 | -1.98565 | 1.451606 | -1.3679 | 0.171343 | 0.83599 | 0.077799 |
| ENSMUSG00000052798.14 | 15.46796 | -2.59842 | 1.899959 | -1.36762 | 0.171431 | 0.83599 | 0.077799 |
| ENSMUSG00000021365.16 | 46.75846 | -2.18248 | 1.596697 | -1.36687 | 0.171665 | 0.836049 | 0.077768 |
| ENSMUSG00000027968.12 | 143.2316 | 1.091525 | 0.798635 | 1.366738 | 0.171708 | 0.836049 | 0.077768 |
| ENSMUSG00000029864.8 | 1044.376 | 0.398417 | 0.29149 | 1.36683 | 0.171678 | 0.836049 | 0.077768 |
| ENSMUSG00000046985.12 | 255.2691 | -0.63559 | 0.465256 | -1.36612 | 0.171903 | 0.83657 | 0.077498 |
| ENSMUSG00000001999.16 | 45.29674 | -1.75308 | 1.285973 | -1.36323 | 0.17281 | 0.837331 | 0.077103 |
| ENSMUSG00000008036.12 | 36.09745 | -1.8748 | 1.379583 | -1.35896 | 0.17416 | 0.837331 | 0.077103 |
| ENSMUSG00000011148.15 | 107.4756 | 1.181815 | 0.869858 | 1.35863 | 0.174264 | 0.837331 | 0.077103 |
| ENSMUSG00000018217.13 | 50.30627 | 1.791721 | 1.317701 | 1.359733 | 0.173914 | 0.837331 | 0.077103 |
| ENSMUSG00000018322.11 | 15.22856 | -2.73526 | 2.014915 | -1.35751 | 0.174621 | 0.837331 | 0.077103 |
| ENSMUSG00000019774.9 | 66.60011 | 1.144947 | 0.843041 | 1.358116 | 0.174427 | 0.837331 | 0.077103 |
| ENSMUSG00000020441.7 | 23.87947 | -1.91793 | 1.408341 | -1.36184 | 0.17325 | 0.837331 | 0.077103 |
| ENSMUSG00000021210.17 | 13006.91 | -0.30228 | 0.222006 | -1.3616 | 0.173325 | 0.837331 | 0.077103 |
| ENSMUSG00000022940.18 | 174.1951 | 1.015223 | 0.745476 | 1.361846 | 0.173246 | 0.837331 | 0.077103 |
| ENSMUSG00000024164.16 | 13981.84 | -0.37937 | 0.278368 | -1.36285 | 0.172931 | 0.837331 | 0.077103 |
| ENSMUSG00000024661.8 | 24178.62 | 0.261461 | 0.191906 | 1.362444 | 0.173058 | 0.837331 | 0.077103 |
| ENSMUSG00000025728.17 | 31.37647 | -1.85225 | 1.363108 | -1.35884 | 0.174197 | 0.837331 | 0.077103 |
| ENSMUSG00000026614.8 | 214.704 | 0.912957 | 0.67205 | 1.358466 | 0.174316 | 0.837331 | 0.077103 |
| ENSMUSG00000026656.16 | 303.2373 | 0.920889 | 0.675304 | 1.363666 | 0.172673 | 0.837331 | 0.077103 |
| ENSMUSG00000027304.10 | 299.6999 | -0.70345 | 0.517335 | -1.35977 | 0.173904 | 0.837331 | 0.077103 |
| ENSMUSG00000028898.16 | 84.85531 | 1.010453 | 0.740364 | 1.364806 | 0.172314 | 0.837331 | 0.077103 |
| ENSMUSG00000028906.18 | 200.8787 | -0.80935 | 0.596004 | -1.35795 | 0.174479 | 0.837331 | 0.077103 |
| ENSMUSG00000029502.15 | 64.82583 | 1.338489 | 0.983471 | 1.360984 | 0.173519 | 0.837331 | 0.077103 |
| ENSMUSG00000032171.8 | 13.12305 | -3.49934 | 2.569802 | -1.36171 | 0.173288 | 0.837331 | 0.077103 |
| ENSMUSG00000034308.14 | 372.5498 | -0.58186 | 0.427335 | -1.3616 | 0.173323 | 0.837331 | 0.077103 |
| ENSMUSG00000034789.9 | 91.83389 | 1.138062 | 0.837988 | 1.358088 | 0.174436 | 0.837331 | 0.077103 |
| ENSMUSG00000037845.15 | 44.13766 | -1.86024 | 1.367589 | -1.36024 | 0.173756 | 0.837331 | 0.077103 |
| ENSMUSG00000038102.16 | 299.0852 | 0.893402 | 0.655416 | 1.363107 | 0.172849 | 0.837331 | 0.077103 |
| ENSMUSG00000043716.14 | 1838.643 | 0.424137 | 0.311744 | 1.36053 | 0.173662 | 0.837331 | 0.077103 |
| ENSMUSG00000049580.13 | 193.5566 | 0.903455 | 0.661838 | 1.36507 | 0.172231 | 0.837331 | 0.077103 |
| ENSMUSG00000067931.7 | 20.02711 | 2.596837 | 1.903787 | 1.364037 | 0.172556 | 0.837331 | 0.077103 |
| ENSMUSG00000083563.2 | 91.27836 | -1.13629 | 0.836993 | -1.35758 | 0.174596 | 0.837331 | 0.077103 |
| ENSMUSG00000098747.8 | 94.15165 | 1.300328 | 0.95393 | 1.363128 | 0.172842 | 0.837331 | 0.077103 |
| ENSMUSG00000107383.2 | 38.07603 | 1.945752 | 1.431459 | 1.35928 | 0.174058 | 0.837331 | 0.077103 |
| ENSMUSG00000025497.16 | 83.58722 | 1.16211 | 0.856548 | 1.356737 | 0.174865 | 0.838078 | 0.076715 |
| ENSMUSG00000032497.20 | 90.35031 | -1.08021 | 0.796648 | -1.35594 | 0.175117 | 0.838613 | 0.076438 |

| | | | | | | | |
|---|---|---|---|---|---|---|---|
| ENSMUSG00000036693.13 | 146.4455 | 0.980423 | 0.723277 | 1.355529 | 0.175249 | 0.838613 | 0.076438 |
| ENSMUSG00000036873.14 | 75.38304 | 1.21509 | 0.896278 | 1.355706 | 0.175193 | 0.838613 | 0.076438 |
| ENSMUSG00000038121.7 | 366.6983 | 0.689146 | 0.508492 | 1.355274 | 0.17533 | 0.838613 | 0.076438 |
| ENSMUSG00000021242.10 | 284.2214 | 0.636454 | 0.470044 | 1.35403 | 0.175727 | 0.838935 | 0.076272 |
| ENSMUSG00000032253.14 | 151.4491 | 1.029227 | 0.760165 | 1.353952 | 0.175752 | 0.838935 | 0.076272 |
| ENSMUSG00000043090.19 | 15.88624 | -3.3214 | 2.452163 | -1.35448 | 0.175584 | 0.838935 | 0.076272 |
| ENSMUSG00000044894.15 | 2735.068 | 0.396407 | 0.292659 | 1.354499 | 0.175577 | 0.838935 | 0.076272 |
| ENSMUSG00000020444.20 | 142.0862 | 0.965687 | 0.713455 | 1.353536 | 0.175885 | 0.839147 | 0.076162 |
| ENSMUSG00000028582.15 | 47.87271 | -2.09758 | 1.550191 | -1.35311 | 0.17602 | 0.839371 | 0.076046 |
| ENSMUSG00000035575.12 | 101.7875 | -1.05435 | 0.77943 | -1.35272 | 0.176146 | 0.839549 | 0.075954 |
| ENSMUSG00000022323.12 | 8969.342 | 0.3283 | 0.242858 | 1.351819 | 0.176433 | 0.840496 | 0.075464 |
| ENSMUSG00000041143.17 | 16.52383 | -3.01032 | 2.228455 | -1.35085 | 0.176742 | 0.841545 | 0.074923 |
| ENSMUSG00000021666.17 | 142.3023 | -0.90974 | 0.674323 | -1.34912 | 0.177299 | 0.841691 | 0.074848 |
| ENSMUSG00000027247.17 | 39.62246 | 2.442406 | 1.8104 | 1.349098 | 0.177305 | 0.841691 | 0.074848 |
| ENSMUSG00000032301.14 | 636.7247 | -0.4562 | 0.338071 | -1.34943 | 0.177198 | 0.841691 | 0.074848 |
| ENSMUSG00000069495.13 | 52.40309 | -2.34471 | 1.737985 | -1.3491 | 0.177305 | 0.841691 | 0.074848 |
| ENSMUSG00000097906.2 | 14.83295 | -2.60158 | 1.926952 | -1.3501 | 0.176983 | 0.841691 | 0.074848 |
| ENSMUSG00000110344.2 | 71.35983 | 1.403936 | 1.040631 | 1.34912 | 0.177298 | 0.841691 | 0.074848 |
| ENSMUSG00000058239.14 | 116.8803 | -1.11129 | 0.824276 | -1.3482 | 0.177594 | 0.84264 | 0.074358 |
| ENSMUSG00000029614.14 | 871.4459 | 0.472401 | 0.350547 | 1.347612 | 0.177783 | 0.843113 | 0.074114 |
| ENSMUSG00000041671.14 | 85.08172 | -1.2624 | 0.937142 | -1.34707 | 0.177957 | 0.843518 | 0.073906 |
| ENSMUSG00000020124.11 | 178.8485 | 0.768904 | 0.571132 | 1.346281 | 0.178212 | 0.844052 | 0.073631 |
| ENSMUSG00000078970.6 | 48.11897 | 1.307622 | 0.971366 | 1.346169 | 0.178248 | 0.844052 | 0.073631 |
| ENSMUSG00000020661.16 | 32.00428 | -2.08816 | 1.554476 | -1.34332 | 0.179169 | 0.845249 | 0.073015 |
| ENSMUSG00000022708.17 | 1301.587 | -0.41634 | 0.309887 | -1.34353 | 0.1791 | 0.845249 | 0.073015 |
| ENSMUSG00000026202.14 | 561.7092 | -0.53808 | 0.400559 | -1.34332 | 0.179168 | 0.845249 | 0.073015 |
| ENSMUSG00000036368.9 | 504.3689 | -0.56359 | 0.419045 | -1.34493 | 0.178649 | 0.845249 | 0.073015 |
| ENSMUSG00000037966.16 | 167.499 | -1.07218 | 0.797833 | -1.34386 | 0.178992 | 0.845249 | 0.073015 |
| ENSMUSG00000039886.9 | 14.84774 | 2.912491 | 2.168583 | 1.343039 | 0.179259 | 0.845249 | 0.073015 |
| ENSMUSG00000057572.16 | 341.7068 | 0.641183 | 0.47746 | 1.342903 | 0.179303 | 0.845249 | 0.073015 |
| ENSMUSG00000081400.2 | 23.74493 | 2.177462 | 1.620071 | 1.344053 | 0.178931 | 0.845249 | 0.073015 |
| ENSMUSG00000095532.2 | 28.99289 | 2.072886 | 1.541548 | 1.344678 | 0.178729 | 0.845249 | 0.073015 |
| ENSMUSG00000011306.8 | 34.57924 | -2.0067 | 1.494778 | -1.34247 | 0.179443 | 0.845487 | 0.072893 |
| ENSMUSG00000020265.17 | 391.8562 | -0.67249 | 0.5013 | -1.34149 | 0.179763 | 0.846153 | 0.072551 |
| ENSMUSG00000038759.16 | 36.76347 | 2.099203 | 1.564713 | 1.34159 | 0.179729 | 0.846153 | 0.072551 |
| ENSMUSG00000024750.12 | 412.6669 | -0.53637 | 0.400012 | -1.34089 | 0.179956 | 0.846221 | 0.072516 |
| ENSMUSG00000030630.17 | 1747.276 | -0.45067 | 0.336039 | -1.34114 | 0.179875 | 0.846221 | 0.072516 |
| ENSMUSG00000027422.16 | 1289.768 | -0.56512 | 0.421658 | -1.34022 | 0.180173 | 0.846822 | 0.072208 |
| ENSMUSG00000030082.15 | 136.5105 | -1.0909 | 0.81468 | -1.33905 | 0.180555 | 0.848195 | 0.071504 |
| ENSMUSG00000028145.10 | 128.8613 | 0.974468 | 0.728004 | 1.338548 | 0.180718 | 0.848542 | 0.071327 |
| ENSMUSG00000020412.17 | 127.6385 | -0.9825 | 0.734284 | -1.33804 | 0.180885 | 0.848905 | 0.071141 |
| ENSMUSG00000028003.7 | 70.14879 | 1.237274 | 0.927564 | 1.333896 | 0.182238 | 0.853144 | 0.068978 |
| ENSMUSG00000028514.16 | 250.8283 | 0.923902 | 0.692368 | 1.334409 | 0.18207 | 0.853144 | 0.068978 |
| ENSMUSG00000032353.10 | 50.61855 | 1.397512 | 1.046909 | 1.334893 | 0.181911 | 0.853144 | 0.068978 |

| | | | | | | | |
|---|---|---|---|---|---|---|---|
| ENSMUSG00000052253.7 | 72.82076 | -1.16531 | 0.873505 | -1.33406 | 0.182184 | 0.853144 | 0.068978 |
| ENSMUSG00000108435.2 | 38.78544 | 1.757808 | 1.317204 | 1.334499 | 0.18204 | 0.853144 | 0.068978 |
| ENSMUSG00000006315.10 | 460.5403 | 0.488678 | 0.367223 | 1.330739 | 0.183275 | 0.853304 | 0.068896 |
| ENSMUSG00000017288.16 | 48.99477 | 1.544341 | 1.160958 | 1.33023 | 0.183442 | 0.853304 | 0.068896 |
| ENSMUSG00000029246.15 | 104.9932 | -0.96018 | 0.720844 | -1.33203 | 0.182852 | 0.853304 | 0.068896 |
| ENSMUSG00000029780.15 | 130.5651 | 1.132215 | 0.850122 | 1.331827 | 0.182917 | 0.853304 | 0.068896 |
| ENSMUSG00000032040.16 | 12.2742 | -2.59851 | 1.952692 | -1.33073 | 0.183277 | 0.853304 | 0.068896 |
| ENSMUSG00000037902.19 | 53.98902 | -1.22993 | 0.923702 | -1.33153 | 0.183016 | 0.853304 | 0.068896 |
| ENSMUSG00000038068.16 | 193.7331 | -1.04481 | 0.783923 | -1.33279 | 0.1826 | 0.853304 | 0.068896 |
| ENSMUSG00000039382.13 | 194.4708 | -0.72754 | 0.546527 | -1.33121 | 0.183121 | 0.853304 | 0.068896 |
| ENSMUSG00000039976.5 | 11.50187 | -3.15221 | 2.36662 | -1.33195 | 0.182878 | 0.853304 | 0.068896 |
| ENSMUSG00000044952.6 | 29.90065 | 1.700286 | 1.276283 | 1.332216 | 0.182789 | 0.853304 | 0.068896 |
| ENSMUSG00000050628.7 | 34.01537 | -2.46685 | 1.85412 | -1.33047 | 0.183363 | 0.853304 | 0.068896 |
| ENSMUSG00000067787.11 | 15.91023 | -3.33027 | 2.500421 | -1.33189 | 0.182898 | 0.853304 | 0.068896 |
| ENSMUSG00000074247.11 | 27.24843 | -2.09076 | 1.567981 | -1.33341 | 0.182398 | 0.853304 | 0.068896 |
| ENSMUSG00000024603.9 | 83.77658 | 1.199076 | 0.90208 | 1.329235 | 0.18377 | 0.853652 | 0.068719 |
| ENSMUSG00000024949.18 | 80.10193 | -1.51216 | 1.137715 | -1.32912 | 0.183809 | 0.853652 | 0.068719 |
| ENSMUSG00000029632.8 | 6272.742 | 0.352228 | 0.264889 | 1.329717 | 0.183612 | 0.853652 | 0.068719 |
| ENSMUSG00000034435.7 | 297.1456 | -0.77857 | 0.585867 | -1.32891 | 0.183877 | 0.853652 | 0.068719 |
| ENSMUSG00000045205.17 | 47.29613 | 1.736976 | 1.307791 | 1.328176 | 0.18412 | 0.85436 | 0.068359 |
| ENSMUSG00000005674.10 | 35.03674 | 2.060832 | 1.552262 | 1.327631 | 0.1843 | 0.854459 | 0.068309 |
| ENSMUSG00000022552.17 | 30.56721 | -2.12696 | 1.602847 | -1.32699 | 0.184512 | 0.854459 | 0.068309 |
| ENSMUSG00000030094.9 | 25.20652 | 2.183468 | 1.645141 | 1.327222 | 0.184435 | 0.854459 | 0.068309 |
| ENSMUSG00000039178.10 | 12.85119 | -2.97316 | 2.240939 | -1.32675 | 0.184592 | 0.854459 | 0.068309 |
| ENSMUSG00000039653.10 | 1543.178 | 0.362655 | 0.273244 | 1.327218 | 0.184437 | 0.854459 | 0.068309 |
| ENSMUSG00000024168.9 | 33.70233 | 1.676874 | 1.265347 | 1.325229 | 0.185095 | 0.854608 | 0.068233 |
| ENSMUSG00000027007.17 | 191.497 | 0.690338 | 0.520643 | 1.325935 | 0.184861 | 0.854608 | 0.068233 |
| ENSMUSG00000028180.19 | 108.1628 | 1.293835 | 0.976069 | 1.325557 | 0.184986 | 0.854608 | 0.068233 |
| ENSMUSG00000030647.9 | 977.2722 | 0.432265 | 0.326279 | 1.324832 | 0.185227 | 0.854608 | 0.068233 |
| ENSMUSG00000032702.17 | 227.3939 | -0.77224 | 0.582936 | -1.32475 | 0.185255 | 0.854608 | 0.068233 |
| ENSMUSG00000042962.6 | 174.8366 | 0.776375 | 0.585562 | 1.325864 | 0.184885 | 0.854608 | 0.068233 |
| ENSMUSG00000089862.9 | 122.849 | 0.863013 | 0.651306 | 1.32505 | 0.185155 | 0.854608 | 0.068233 |
| ENSMUSG00000017119.21 | 628.7317 | -0.4572 | 0.345465 | -1.32344 | 0.185689 | 0.854815 | 0.068128 |
| ENSMUSG00000023057.6 | 544.0915 | 0.598063 | 0.451752 | 1.323875 | 0.185545 | 0.854815 | 0.068128 |
| ENSMUSG00000024429.10 | 33.65203 | -1.9753 | 1.491652 | -1.32424 | 0.185425 | 0.854815 | 0.068128 |
| ENSMUSG00000028540.7 | 33.53497 | -1.9429 | 1.468365 | -1.32317 | 0.185778 | 0.854815 | 0.068128 |
| ENSMUSG00000044465.20 | 34.42392 | 2.255976 | 1.705569 | 1.322712 | 0.185931 | 0.854815 | 0.068128 |
| ENSMUSG00000071054.13 | 103.8889 | 1.191448 | 0.900729 | 1.322761 | 0.185915 | 0.854815 | 0.068128 |
| ENSMUSG00000071072.13 | 603.9301 | 0.517998 | 0.391487 | 1.323154 | 0.185784 | 0.854815 | 0.068128 |
| ENSMUSG00000029135.11 | 28.3652 | -2.82643 | 2.13802 | -1.32199 | 0.186173 | 0.855158 | 0.067954 |
| ENSMUSG00000031886.15 | 1200.06 | 0.443758 | 0.336098 | 1.320322 | 0.186728 | 0.855158 | 0.067954 |
| ENSMUSG00000033557.18 | 10.01684 | -3.76448 | 2.849178 | -1.32125 | 0.186417 | 0.855158 | 0.067954 |
| ENSMUSG00000046707.10 | 69.66281 | -1.26653 | 0.958476 | -1.3214 | 0.186367 | 0.855158 | 0.067954 |
| ENSMUSG00000050919.11 | 11.76608 | -3.40067 | 2.574629 | -1.32084 | 0.186555 | 0.855158 | 0.067954 |

| | | | | | | | |
|---|---|---|---|---|---|---|---|
| ENSMUSG00000060512.8 | 251.9941 | -0.73261 | 0.554301 | -1.32169 | 0.186271 | 0.855158 | 0.067954 |
| ENSMUSG00000078566.9 | 850.9888 | 0.463809 | 0.351275 | 1.320358 | 0.186715 | 0.855158 | 0.067954 |
| ENSMUSG00000083282.4 | 36.69565 | 2.212226 | 1.674595 | 1.321051 | 0.186484 | 0.855158 | 0.067954 |
| ENSMUSG00000040964.17 | 23.45428 | -2.2715 | 1.721417 | -1.31955 | 0.186984 | 0.85592 | 0.067567 |
| ENSMUSG00000001482.16 | 27.42531 | -2.0841 | 1.579723 | -1.31928 | 0.187075 | 0.855923 | 0.067565 |
| ENSMUSG00000036438.15 | 755.6726 | -0.57818 | 0.438594 | -1.31826 | 0.187417 | 0.856658 | 0.067192 |
| ENSMUSG00000039048.16 | 98.5245 | 1.06136 | 0.804997 | 1.318466 | 0.187348 | 0.856658 | 0.067192 |
| ENSMUSG00000028478.19 | 678.6457 | -0.50326 | 0.381878 | -1.31786 | 0.187552 | 0.856864 | 0.067088 |
| ENSMUSG00000039648.15 | 19.4629 | -2.29114 | 1.739273 | -1.3173 | 0.187739 | 0.857307 | 0.066864 |
| ENSMUSG00000020321.17 | 7657.206 | 0.343413 | 0.261461 | 1.31344 | 0.189035 | 0.857356 | 0.066839 |
| ENSMUSG00000021918.11 | 59.56107 | 1.390021 | 1.05861 | 1.313063 | 0.189162 | 0.857356 | 0.066839 |
| ENSMUSG00000026014.16 | 244.9504 | -0.6693 | 0.509141 | -1.31457 | 0.188654 | 0.857356 | 0.066839 |
| ENSMUSG00000026034.18 | 238.3649 | 0.612623 | 0.465806 | 1.315189 | 0.188446 | 0.857356 | 0.066839 |
| ENSMUSG00000027199.15 | 35.30855 | 1.611763 | 1.22686 | 1.313731 | 0.188937 | 0.857356 | 0.066839 |
| ENSMUSG00000028452.8 | 883.7318 | -0.42912 | 0.326756 | -1.31327 | 0.189091 | 0.857356 | 0.066839 |
| ENSMUSG00000032207.11 | 887.9323 | -0.4164 | 0.316783 | -1.31447 | 0.188689 | 0.857356 | 0.066839 |
| ENSMUSG00000034842.17 | 15.48102 | 2.86818 | 2.181517 | 1.314764 | 0.188589 | 0.857356 | 0.066839 |
| ENSMUSG00000035476.10 | 19.31786 | 2.207017 | 1.676523 | 1.316425 | 0.188031 | 0.857356 | 0.066839 |
| ENSMUSG00000054280.15 | 37.08694 | -2.40605 | 1.832276 | -1.31315 | 0.189132 | 0.857356 | 0.066839 |
| ENSMUSG00000054920.13 | 123.4279 | -1.07745 | 0.818798 | -1.31589 | 0.18821 | 0.857356 | 0.066839 |
| ENSMUSG00000063694.6 | 337.1351 | 0.544443 | 0.414278 | 1.314197 | 0.18878 | 0.857356 | 0.066839 |
| ENSMUSG00000075590.5 | 100.2971 | -1.03995 | 0.789923 | -1.31653 | 0.187998 | 0.857356 | 0.066839 |
| ENSMUSG00000078812.11 | 150.3924 | -0.99985 | 0.76096 | -1.31393 | 0.188869 | 0.857356 | 0.066839 |
| ENSMUSG00000090081.2 | 18.43404 | -3.19004 | 2.42966 | -1.31296 | 0.189197 | 0.857356 | 0.066839 |
| ENSMUSG00000118506.2 | 16.48606 | -3.30378 | 2.512051 | -1.31517 | 0.188452 | 0.857356 | 0.066839 |
| ENSMUSG00000029247.18 | 1383.148 | -0.34122 | 0.260204 | -1.31135 | 0.18974 | 0.859406 | 0.065801 |
| ENSMUSG00000014349.8 | 160.3961 | 0.778003 | 0.593546 | 1.310771 | 0.189935 | 0.859468 | 0.06577 |
| ENSMUSG00000023367.15 | 680.2805 | -0.62989 | 0.480461 | -1.31102 | 0.189852 | 0.859468 | 0.06577 |
| ENSMUSG00000032212.11 | 586.3793 | 0.548445 | 0.418584 | 1.310238 | 0.190115 | 0.859872 | 0.065566 |
| ENSMUSG00000026848.15 | 184.8053 | 0.747181 | 0.570568 | 1.309537 | 0.190352 | 0.860445 | 0.065277 |
| ENSMUSG00000027165.17 | 201.3535 | -0.86406 | 0.659926 | -1.30933 | 0.190423 | 0.860445 | 0.065277 |
| ENSMUSG00000021428.10 | 39.52705 | -1.60734 | 1.22801 | -1.3089 | 0.190569 | 0.860695 | 0.065151 |
| ENSMUSG00000047822.9 | 63.40998 | 1.700385 | 1.299526 | 1.308466 | 0.190715 | 0.860944 | 0.065025 |
| ENSMUSG00000048661.15 | 93.29764 | -0.94319 | 0.721104 | -1.30797 | 0.190882 | 0.861286 | 0.064853 |
| ENSMUSG00000000563.18 | 5952.761 | 0.380636 | 0.292506 | 1.301293 | 0.193158 | 0.861429 | 0.064781 |
| ENSMUSG00000020246.14 | 56.39399 | -1.23397 | 0.945519 | -1.30507 | 0.191868 | 0.861429 | 0.064781 |
| ENSMUSG00000022195.8 | 23.47949 | -2.12897 | 1.633571 | -1.30326 | 0.192485 | 0.861429 | 0.064781 |
| ENSMUSG00000022704.17 | 30.13941 | -2.43048 | 1.867913 | -1.30117 | 0.193199 | 0.861429 | 0.064781 |
| ENSMUSG00000023104.10 | 87.73184 | 1.483704 | 1.135727 | 1.306392 | 0.191419 | 0.861429 | 0.064781 |
| ENSMUSG00000025138.15 | 57.75935 | -1.39171 | 1.068493 | -1.3025 | 0.192747 | 0.861429 | 0.064781 |
| ENSMUSG00000025203.7 | 11.50612 | 3.104357 | 2.375683 | 1.306722 | 0.191307 | 0.861429 | 0.064781 |
| ENSMUSG00000025240.11 | 544.4401 | -0.475 | 0.364026 | -1.30485 | 0.191945 | 0.861429 | 0.064781 |
| ENSMUSG00000025809.16 | 341.9063 | -0.52445 | 0.403211 | -1.30068 | 0.193367 | 0.861429 | 0.064781 |
| ENSMUSG00000028848.14 | 43.416 | -1.6511 | 1.264027 | -1.30622 | 0.191477 | 0.861429 | 0.064781 |

| | | | | | | | |
|---|---|---|---|---|---|---|---|
| ENSMUSG00000028955.4 | 70.32553 | -1.47948 | 1.13692 | -1.3013 | 0.193155 | 0.861429 | 0.064781 |
| ENSMUSG00000031229.17 | 465.2487 | -0.57786 | 0.442107 | -1.30707 | 0.19119 | 0.861429 | 0.064781 |
| ENSMUSG00000031665.8 | 23.51321 | -1.97589 | 1.518867 | -1.3009 | 0.193293 | 0.861429 | 0.064781 |
| ENSMUSG00000033533.15 | 630.1931 | -0.41487 | 0.317739 | -1.30569 | 0.191659 | 0.861429 | 0.064781 |
| ENSMUSG00000034327.18 | 58.94501 | -1.62201 | 1.245756 | -1.30203 | 0.192908 | 0.861429 | 0.064781 |
| ENSMUSG00000037740.9 | 96.85993 | 0.990674 | 0.761066 | 1.301691 | 0.193022 | 0.861429 | 0.064781 |
| ENSMUSG00000037818.17 | 141.8769 | -0.85366 | 0.655112 | -1.30307 | 0.192551 | 0.861429 | 0.064781 |
| ENSMUSG00000038751.6 | 12.2141 | -3.4759 | 2.666761 | -1.30342 | 0.192432 | 0.861429 | 0.064781 |
| ENSMUSG00000040640.12 | 54.20335 | -1.72305 | 1.322955 | -1.30242 | 0.192771 | 0.861429 | 0.064781 |
| ENSMUSG00000041355.14 | 44.47062 | 1.962644 | 1.503672 | 1.305235 | 0.191813 | 0.861429 | 0.064781 |
| ENSMUSG00000045316.6 | 463.1166 | 0.56863 | 0.435801 | 1.304794 | 0.191963 | 0.861429 | 0.064781 |
| ENSMUSG00000049969.4 | 250.5198 | 0.771508 | 0.591938 | 1.303359 | 0.192452 | 0.861429 | 0.064781 |
| ENSMUSG00000057425.4 | 26.57539 | -1.85964 | 1.425565 | -1.30449 | 0.192066 | 0.861429 | 0.064781 |
| ENSMUSG00000059890.17 | 37.94391 | 1.937406 | 1.486618 | 1.30323 | 0.192496 | 0.861429 | 0.064781 |
| ENSMUSG00000060441.16 | 17.57077 | -2.83814 | 2.174897 | -1.30495 | 0.191909 | 0.861429 | 0.064781 |
| ENSMUSG00000062190.13 | 64.17521 | 1.661068 | 1.272641 | 1.305213 | 0.19182 | 0.861429 | 0.064781 |
| ENSMUSG00000068011.5 | 154.1762 | 0.891391 | 0.683723 | 1.303732 | 0.192325 | 0.861429 | 0.064781 |
| ENSMUSG00000031879.15 | 163.1963 | 0.804534 | 0.618749 | 1.30026 | 0.193512 | 0.861669 | 0.06466 |
| ENSMUSG00000020440.14 | 32.98665 | -1.96112 | 1.508972 | -1.29964 | 0.193724 | 0.861803 | 0.064592 |
| ENSMUSG00000027176.9 | 125.4828 | 0.935991 | 0.720092 | 1.299822 | 0.193662 | 0.861803 | 0.064592 |
| ENSMUSG00000031845.17 | 127.8332 | -1.4547 | 1.11968 | -1.29921 | 0.193873 | 0.86206 | 0.064462 |
| ENSMUSG00000028974.14 | 31.25394 | -2.09485 | 1.613091 | -1.29865 | 0.194063 | 0.862502 | 0.06424 |
| ENSMUSG00000022797.17 | 105.8388 | 1.006831 | 0.775818 | 1.297767 | 0.194367 | 0.862642 | 0.06417 |
| ENSMUSG00000027109.17 | 142.1102 | 0.875328 | 0.674482 | 1.297778 | 0.194364 | 0.862642 | 0.06417 |
| ENSMUSG00000027381.17 | 23.24333 | -2.1813 | 1.680339 | -1.29813 | 0.194242 | 0.862642 | 0.06417 |
| ENSMUSG00000018796.14 | 4328.305 | 0.368522 | 0.284193 | 1.296732 | 0.194723 | 0.86301 | 0.063984 |
| ENSMUSG00000051098.7 | 97.72454 | 1.155188 | 0.890637 | 1.297035 | 0.194619 | 0.86301 | 0.063984 |
| ENSMUSG00000078572.4 | 178.0078 | 0.847987 | 0.653933 | 1.296749 | 0.194718 | 0.86301 | 0.063984 |
| ENSMUSG00000024338.16 | 74.34018 | -1.05442 | 0.813687 | -1.29586 | 0.195023 | 0.863129 | 0.063925 |
| ENSMUSG00000029390.14 | 261.8727 | 0.707458 | 0.545917 | 1.295907 | 0.195008 | 0.863129 | 0.063925 |
| ENSMUSG00000037295.8 | 36.94213 | -2.02269 | 1.560375 | -1.29628 | 0.194878 | 0.863129 | 0.063925 |
| ENSMUSG00000004730.16 | 47.29772 | 1.736108 | 1.343044 | 1.292666 | 0.196126 | 0.864263 | 0.063354 |
| ENSMUSG00000006494.12 | 388.6684 | 0.619228 | 0.4793 | 1.291942 | 0.196377 | 0.864263 | 0.063354 |
| ENSMUSG00000018042.19 | 926.5263 | -0.6257 | 0.483947 | -1.29291 | 0.196041 | 0.864263 | 0.063354 |
| ENSMUSG00000019232.15 | 565.5835 | 0.497344 | 0.384725 | 1.292726 | 0.196106 | 0.864263 | 0.063354 |
| ENSMUSG00000022008.5 | 84.79765 | 1.353329 | 1.047185 | 1.29235 | 0.196236 | 0.864263 | 0.063354 |
| ENSMUSG00000028470.11 | 458.0509 | -0.56947 | 0.440023 | -1.29418 | 0.195603 | 0.864263 | 0.063354 |
| ENSMUSG00000029708.10 | 33.81324 | -1.78047 | 1.378286 | -1.2918 | 0.196426 | 0.864263 | 0.063354 |
| ENSMUSG00000031352.11 | 107.1948 | 1.092785 | 0.844757 | 1.293608 | 0.195801 | 0.864263 | 0.063354 |
| ENSMUSG00000032204.14 | 1830.103 | -0.33297 | 0.257624 | -1.29248 | 0.196192 | 0.864263 | 0.063354 |
| ENSMUSG00000036820.13 | 13.42781 | 2.807314 | 2.173217 | 1.291778 | 0.196434 | 0.864263 | 0.063354 |
| ENSMUSG00000037894.14 | 208.1187 | 0.685207 | 0.530474 | 1.291689 | 0.196465 | 0.864263 | 0.063354 |
| ENSMUSG00000042207.18 | 26.69565 | -2.09263 | 1.61794 | -1.29339 | 0.195876 | 0.864263 | 0.063354 |
| ENSMUSG00000074656.13 | 1135.447 | 0.507525 | 0.392125 | 1.294295 | 0.195563 | 0.864263 | 0.063354 |

| | | | | | | | |
|---|---|---|---|---|---|---|---|
| ENSMUSG00000031917.13 | 82.24638 | 1.044835 | 0.809141 | 1.291289 | 0.196604 | 0.864472 | 0.063249 |
| ENSMUSG00000028381.9 | 158.8426 | -0.81887 | 0.63453 | -1.29051 | 0.196873 | 0.864854 | 0.063057 |
| ENSMUSG00000046324.13 | 173.0702 | 0.907246 | 0.702938 | 1.290649 | 0.196826 | 0.864854 | 0.063057 |
| ENSMUSG00000000266.12 | 21.62988 | 2.323782 | 1.801916 | 1.289618 | 0.197183 | 0.865016 | 0.062976 |
| ENSMUSG00000024098.7 | 391.1055 | 0.63384 | 0.491448 | 1.289739 | 0.197141 | 0.865016 | 0.062976 |
| ENSMUSG00000031370.13 | 85.51931 | 1.01221 | 0.784753 | 1.289845 | 0.197105 | 0.865016 | 0.062976 |
| ENSMUSG00000009614.17 | 850.6222 | -0.49315 | 0.382972 | -1.2877 | 0.19785 | 0.865537 | 0.062714 |
| ENSMUSG00000019943.11 | 339.4406 | -0.6045 | 0.469241 | -1.28826 | 0.197657 | 0.865537 | 0.062714 |
| ENSMUSG00000020677.8 | 221.0743 | 0.796307 | 0.618127 | 1.288258 | 0.197656 | 0.865537 | 0.062714 |
| ENSMUSG00000033629.7 | 820.0809 | -0.43632 | 0.33855 | -1.28878 | 0.197474 | 0.865537 | 0.062714 |
| ENSMUSG00000033799.11 | 36.22499 | -1.51117 | 1.17339 | -1.28787 | 0.197793 | 0.865537 | 0.062714 |
| ENSMUSG00000036966.16 | 18.90735 | -2.69644 | 2.093156 | -1.28822 | 0.19767 | 0.865537 | 0.062714 |
| ENSMUSG00000047213.15 | 256.2705 | -0.59896 | 0.465252 | -1.28739 | 0.197957 | 0.865607 | 0.062679 |
| ENSMUSG00000026603.14 | 94.86189 | 1.262865 | 0.981646 | 1.286477 | 0.198277 | 0.86567 | 0.062648 |
| ENSMUSG00000029368.11 | 541880.5 | 0.234128 | 0.181968 | 1.286644 | 0.198218 | 0.86567 | 0.062648 |
| ENSMUSG00000030921.18 | 59.31796 | -1.31712 | 1.023954 | -1.2863 | 0.198337 | 0.86567 | 0.062648 |
| ENSMUSG00000036435.14 | 36.11207 | -1.58342 | 1.23025 | -1.28707 | 0.198069 | 0.86567 | 0.062648 |
| ENSMUSG00000020883.19 | 34.56446 | -1.6852 | 1.311098 | -1.28534 | 0.198674 | 0.866087 | 0.062439 |
| ENSMUSG00000024335.21 | 135.1656 | -1.1543 | 0.898067 | -1.28531 | 0.198683 | 0.866087 | 0.062439 |
| ENSMUSG00000040385.8 | 55.67376 | 1.53721 | 1.196044 | 1.285245 | 0.198707 | 0.866087 | 0.062439 |
| ENSMUSG00000020319.10 | 14.04662 | 2.731265 | 2.127368 | 1.283871 | 0.199187 | 0.866227 | 0.062368 |
| ENSMUSG00000028530.15 | 303.6636 | -0.69367 | 0.540212 | -1.28407 | 0.199117 | 0.866227 | 0.062368 |
| ENSMUSG00000043019.13 | 135.6281 | 1.000442 | 0.778795 | 1.284603 | 0.198931 | 0.866227 | 0.062368 |
| ENSMUSG00000058440.15 | 16.2303 | -2.23558 | 1.741313 | -1.28385 | 0.199196 | 0.866227 | 0.062368 |
| ENSMUSG00000066366.15 | 20584.93 | -0.23929 | 0.18626 | -1.2847 | 0.198896 | 0.866227 | 0.062368 |
| ENSMUSG00000075705.14 | 959.0095 | -0.53254 | 0.415005 | -1.28321 | 0.199419 | 0.866801 | 0.062081 |
| ENSMUSG00000021143.11 | 17.84015 | 2.237146 | 1.744613 | 1.282316 | 0.199732 | 0.866967 | 0.061997 |
| ENSMUSG00000024331.12 | 166.261 | -0.74933 | 0.584305 | -1.28242 | 0.199694 | 0.866967 | 0.061997 |
| ENSMUSG00000052459.14 | 375.2709 | -0.57347 | 0.447122 | -1.28259 | 0.199637 | 0.866967 | 0.061997 |
| ENSMUSG00000028709.4 | 28.75008 | -1.51335 | 1.181176 | -1.28122 | 0.200115 | 0.867438 | 0.061762 |
| ENSMUSG00000028999.16 | 84.4682 | -1.1158 | 0.870607 | -1.28164 | 0.199969 | 0.867438 | 0.061762 |
| ENSMUSG00000038768.9 | 175.4171 | -0.82956 | 0.647389 | -1.28139 | 0.200057 | 0.867438 | 0.061762 |
| ENSMUSG00000020346.17 | 108.585 | -0.84741 | 0.661608 | -1.28084 | 0.200251 | 0.867632 | 0.061665 |
| ENSMUSG00000002733.9 | 158.7026 | -0.84046 | 0.65636 | -1.28048 | 0.200375 | 0.867772 | 0.061594 |
| ENSMUSG00000050188.9 | 43.21967 | 1.816384 | 1.419637 | 1.279471 | 0.200731 | 0.868522 | 0.061219 |
| ENSMUSG00000067144.8 | 109.3371 | -0.99672 | 0.778866 | -1.2797 | 0.200649 | 0.868522 | 0.061219 |
| ENSMUSG00000022587.15 | 765.6633 | -0.68638 | 0.536925 | -1.27835 | 0.201126 | 0.868919 | 0.061021 |
| ENSMUSG00000031320.10 | 2094.482 | 0.317749 | 0.248666 | 1.277812 | 0.201316 | 0.868919 | 0.061021 |
| ENSMUSG00000040325.17 | 81.6391 | -1.37738 | 1.07806 | -1.27765 | 0.201373 | 0.868919 | 0.061021 |
| ENSMUSG00000041997.17 | 143.734 | -0.87681 | 0.686229 | -1.27772 | 0.20135 | 0.868919 | 0.061021 |
| ENSMUSG00000070871.11 | 40.85841 | -2.27525 | 1.779794 | -1.27838 | 0.201116 | 0.868919 | 0.061021 |
| ENSMUSG00000105599.2 | 30.72594 | 1.474253 | 1.153141 | 1.278466 | 0.201085 | 0.868919 | 0.061021 |
| ENSMUSG00000024130.17 | 155.8897 | -0.91362 | 0.715263 | -1.27732 | 0.201488 | 0.869021 | 0.06097 |
| ENSMUSG00000049553.12 | 18.95247 | -2.70614 | 2.119346 | -1.27687 | 0.201647 | 0.869309 | 0.060826 |

| | | | | | | | |
|---|---|---|---|---|---|---|---|
| ENSMUSG00000028229.12 | 162.446 | 0.728549 | 0.570939 | 1.276055 | 0.201936 | 0.869766 | 0.060598 |
| ENSMUSG00000046138.16 | 157.6633 | -1.05388 | 0.825787 | -1.27622 | 0.201879 | 0.869766 | 0.060598 |
| ENSMUSG00000024487.6 | 422.901 | 0.579328 | 0.454211 | 1.275461 | 0.202146 | 0.870232 | 0.060365 |
| ENSMUSG00000028683.15 | 62.77074 | 1.6582 | 1.300315 | 1.275229 | 0.202228 | 0.870232 | 0.060365 |
| ENSMUSG00000026117.11 | 29.5773 | 2.238362 | 1.755632 | 1.274961 | 0.202323 | 0.870247 | 0.060358 |
| ENSMUSG00000023045.14 | 57.89048 | -1.49088 | 1.170064 | -1.27419 | 0.202597 | 0.870456 | 0.060253 |
| ENSMUSG00000028284.14 | 133.2636 | 0.945552 | 0.741894 | 1.274512 | 0.202482 | 0.870456 | 0.060253 |
| ENSMUSG00000039968.10 | 93.91217 | 1.280497 | 1.005064 | 1.274045 | 0.202647 | 0.870456 | 0.060253 |
| ENSMUSG00000039852.18 | 48.00751 | -1.84029 | 1.444819 | -1.27371 | 0.202764 | 0.870565 | 0.060199 |
| ENSMUSG00000037321.18 | 35.63428 | -2.12224 | 1.666596 | -1.2734 | 0.202876 | 0.870648 | 0.060157 |
| ENSMUSG00000058546.9 | 3130.184 | 0.328645 | 0.258302 | 1.272332 | 0.203255 | 0.871882 | 0.059542 |
| ENSMUSG00000032536.13 | 133.834 | -0.94898 | 0.746116 | -1.27189 | 0.203413 | 0.872163 | 0.059402 |
| ENSMUSG00000003604.15 | 22.70967 | 2.717563 | 2.137706 | 1.271251 | 0.203639 | 0.872304 | 0.059332 |
| ENSMUSG00000055491.15 | 13.1516 | -3.33293 | 2.621519 | -1.27137 | 0.203596 | 0.872304 | 0.059332 |
| ENSMUSG00000062822.9 | 240.1653 | -0.765 | 0.601878 | -1.27102 | 0.203722 | 0.872304 | 0.059332 |
| ENSMUSG00000001445.11 | 177.1941 | -0.88513 | 0.699268 | -1.26579 | 0.205588 | 0.87252 | 0.059224 |
| ENSMUSG00000005102.14 | 51.0163 | 1.302197 | 1.027308 | 1.267582 | 0.204947 | 0.87252 | 0.059224 |
| ENSMUSG00000020015.11 | 47.95218 | 1.461784 | 1.154685 | 1.26596 | 0.205528 | 0.87252 | 0.059224 |
| ENSMUSG00000020102.16 | 177.7087 | 0.727799 | 0.574981 | 1.265778 | 0.205593 | 0.87252 | 0.059224 |
| ENSMUSG00000021532.12 | 40.61543 | -1.49208 | 1.177462 | -1.2672 | 0.205084 | 0.87252 | 0.059224 |
| ENSMUSG00000021905.15 | 173.0328 | 0.853903 | 0.673818 | 1.267261 | 0.205062 | 0.87252 | 0.059224 |
| ENSMUSG00000024078.8 | 35.71492 | -1.83594 | 1.45049 | -1.26574 | 0.205606 | 0.87252 | 0.059224 |
| ENSMUSG00000025290.18 | 3266.072 | 0.365556 | 0.288672 | 1.266335 | 0.205393 | 0.87252 | 0.059224 |
| ENSMUSG00000026134.12 | 17.55698 | -2.76984 | 2.185316 | -1.26748 | 0.204984 | 0.87252 | 0.059224 |
| ENSMUSG00000028521.18 | 611.5849 | -0.4376 | 0.345628 | -1.2661 | 0.205476 | 0.87252 | 0.059224 |
| ENSMUSG00000028538.13 | 24.45243 | -2.1446 | 1.694586 | -1.26556 | 0.20567 | 0.87252 | 0.059224 |
| ENSMUSG00000029798.12 | 34.48226 | -1.60094 | 1.265024 | -1.26554 | 0.205679 | 0.87252 | 0.059224 |
| ENSMUSG00000030602.14 | 11.73678 | 3.47678 | 2.738282 | 1.269694 | 0.204194 | 0.87252 | 0.059224 |
| ENSMUSG00000032127.10 | 17.33351 | -2.63917 | 2.085955 | -1.26521 | 0.205797 | 0.87252 | 0.059224 |
| ENSMUSG00000032350.10 | 1257.05 | 0.355899 | 0.280863 | 1.26716 | 0.205098 | 0.87252 | 0.059224 |
| ENSMUSG00000036180.16 | 25.70228 | -1.95448 | 1.540689 | -1.26858 | 0.204592 | 0.87252 | 0.059224 |
| ENSMUSG00000037062.14 | 387.4079 | -0.59473 | 0.469027 | -1.26801 | 0.204794 | 0.87252 | 0.059224 |
| ENSMUSG00000040268.18 | 70.56645 | 1.110238 | 0.87738 | 1.265401 | 0.205728 | 0.87252 | 0.059224 |
| ENSMUSG00000048581.13 | 26.51989 | -2.22584 | 1.754386 | -1.26873 | 0.204539 | 0.87252 | 0.059224 |
| ENSMUSG00000049321.18 | 48.97704 | 1.20666 | 0.949962 | 1.27022 | 0.204006 | 0.87252 | 0.059224 |
| ENSMUSG00000057236.11 | 430.3011 | -0.53244 | 0.419748 | -1.26846 | 0.204632 | 0.87252 | 0.059224 |
| ENSMUSG00000117869.2 | 74.6511 | 1.85308 | 1.458893 | 1.270197 | 0.204015 | 0.87252 | 0.059224 |
| ENSMUSG00000025785.6 | 158.1899 | 1.032383 | 0.816504 | 1.264394 | 0.206089 | 0.872976 | 0.058998 |
| ENSMUSG00000060121.16 | 88.53386 | -1.06008 | 0.83836 | -1.26447 | 0.206061 | 0.872976 | 0.058998 |
| ENSMUSG00000017344.5 | 2529.438 | -0.41854 | 0.331158 | -1.26387 | 0.206278 | 0.873048 | 0.058962 |
| ENSMUSG00000027104.19 | 141.0019 | -0.91597 | 0.724893 | -1.26359 | 0.206376 | 0.873048 | 0.058962 |
| ENSMUSG00000032383.8 | 590.0033 | -0.61393 | 0.485929 | -1.26342 | 0.206437 | 0.873048 | 0.058962 |
| ENSMUSG00000059142.17 | 92.70251 | -0.98132 | 0.776779 | -1.26332 | 0.206474 | 0.873048 | 0.058962 |
| ENSMUSG00000087141.2 | 529.7717 | -0.59807 | 0.473531 | -1.263 | 0.206588 | 0.873138 | 0.058917 |

| | | | | | | | |
|---|---|---|---|---|---|---|---|
| ENSMUSG00000070565.12 | 63.16552 | -1.19101 | 0.943543 | -1.26227 | 0.20685 | 0.873746 | 0.058615 |
| ENSMUSG00000101655.2 | 40.85972 | -1.98649 | 1.573966 | -1.26209 | 0.206916 | 0.873746 | 0.058615 |
| ENSMUSG00000025898.7 | 121.6739 | -0.869 | 0.689351 | -1.2606 | 0.207452 | 0.875622 | 0.057684 |
| ENSMUSG00000066361.4 | 87.72838 | 1.312335 | 1.041383 | 1.260185 | 0.207603 | 0.875867 | 0.057562 |
| ENSMUSG00000032388.8 | 42.15075 | 1.484371 | 1.178621 | 1.259414 | 0.207881 | 0.876651 | 0.057173 |
| ENSMUSG00000005370.5 | 73.4136 | -1.13298 | 0.90104 | -1.25741 | 0.208605 | 0.876972 | 0.057014 |
| ENSMUSG00000025959.14 | 16.0814 | 2.733771 | 2.173729 | 1.257641 | 0.208521 | 0.876972 | 0.057014 |
| ENSMUSG00000027628.11 | 18.30832 | -2.82924 | 2.249628 | -1.25765 | 0.208519 | 0.876972 | 0.057014 |
| ENSMUSG00000028108.17 | 68.95763 | -1.81604 | 1.443651 | -1.25795 | 0.208411 | 0.876972 | 0.057014 |
| ENSMUSG00000040888.11 | 303.5039 | -0.98909 | 0.786577 | -1.25746 | 0.208586 | 0.876972 | 0.057014 |
| ENSMUSG00000058407.13 | 225.7457 | 0.783187 | 0.622423 | 1.258287 | 0.208288 | 0.876972 | 0.057014 |
| ENSMUSG00000079480.4 | 96.77648 | 0.980429 | 0.779277 | 1.258127 | 0.208346 | 0.876972 | 0.057014 |
| ENSMUSG00000046027.18 | 459.4262 | -0.49074 | 0.390469 | -1.2568 | 0.208826 | 0.877515 | 0.056745 |
| ENSMUSG00000024065.9 | 148.6101 | -0.84614 | 0.673554 | -1.25624 | 0.20903 | 0.877591 | 0.056708 |
| ENSMUSG00000024423.7 | 23.23736 | 1.769254 | 1.408104 | 1.25648 | 0.208942 | 0.877591 | 0.056708 |
| ENSMUSG00000020134.14 | 204.6648 | 0.626157 | 0.498933 | 1.254992 | 0.209482 | 0.878593 | 0.056212 |
| ENSMUSG00000027195.11 | 941.4818 | 0.373493 | 0.297648 | 1.254814 | 0.209546 | 0.878593 | 0.056212 |
| ENSMUSG00000027995.11 | 11.84514 | -3.34111 | 2.661569 | -1.25532 | 0.209364 | 0.878593 | 0.056212 |
| ENSMUSG00000040855.16 | 190.7766 | -0.75871 | 0.604795 | -1.25449 | 0.209665 | 0.878702 | 0.056159 |
| ENSMUSG00000024240.14 | 47.86633 | 1.186785 | 0.946266 | 1.254176 | 0.209778 | 0.878787 | 0.056117 |
| ENSMUSG00000074746.4 | 151.7352 | -0.88448 | 0.705443 | -1.2538 | 0.209915 | 0.878973 | 0.056025 |
| ENSMUSG00000001435.16 | 884.9147 | -0.37913 | 0.302765 | -1.25223 | 0.210488 | 0.879217 | 0.055904 |
| ENSMUSG00000007867.10 | 28.47249 | 2.397911 | 1.914731 | 1.252349 | 0.210443 | 0.879217 | 0.055904 |
| ENSMUSG00000018678.14 | 11.50451 | 3.42744 | 2.736647 | 1.252423 | 0.210416 | 0.879217 | 0.055904 |
| ENSMUSG00000028274.18 | 72.10517 | 1.060789 | 0.847202 | 1.25211 | 0.21053 | 0.879217 | 0.055904 |
| ENSMUSG00000054894.7 | 35.04428 | -1.51499 | 1.20939 | -1.25269 | 0.210318 | 0.879217 | 0.055904 |
| ENSMUSG00000092416.3 | 57.71083 | -1.24661 | 0.995134 | -1.25271 | 0.210312 | 0.879217 | 0.055904 |
| ENSMUSG00000073002.10 | 53.10899 | -1.60603 | 1.284014 | -1.25079 | 0.21101 | 0.880835 | 0.055105 |
| ENSMUSG00000024391.8 | 1103.168 | -0.39416 | 0.315531 | -1.24918 | 0.211598 | 0.881347 | 0.054853 |
| ENSMUSG00000027708.15 | 241.0946 | -0.69397 | 0.555232 | -1.24988 | 0.211344 | 0.881347 | 0.054853 |
| ENSMUSG00000035248.10 | 346.5051 | -0.5064 | 0.405287 | -1.24948 | 0.211491 | 0.881347 | 0.054853 |
| ENSMUSG00000036568.8 | 43.14497 | 1.657897 | 1.327031 | 1.249328 | 0.211545 | 0.881347 | 0.054853 |
| ENSMUSG00000038880.14 | 189.3504 | 0.904882 | 0.724104 | 1.249658 | 0.211425 | 0.881347 | 0.054853 |
| ENSMUSG00000014956.16 | 1008.472 | 0.447393 | 0.358288 | 1.248694 | 0.211777 | 0.881707 | 0.054676 |
| ENSMUSG00000032741.10 | 100.6672 | -1.09657 | 0.878364 | -1.24842 | 0.211877 | 0.881735 | 0.054662 |
| ENSMUSG00000033773.15 | 42.18293 | -1.6087 | 1.289237 | -1.24779 | 0.212107 | 0.882304 | 0.054382 |
| ENSMUSG00000020111.16 | 17.7428 | -2.31464 | 1.856011 | -1.24711 | 0.212358 | 0.882349 | 0.05436 |
| ENSMUSG00000020745.16 | 402.4142 | 0.71927 | 0.576719 | 1.247176 | 0.212333 | 0.882349 | 0.05436 |
| ENSMUSG00000029763.18 | 118.3131 | -0.81194 | 0.651113 | -1.247 | 0.212397 | 0.882349 | 0.05436 |
| ENSMUSG00000032803.16 | 358.9022 | 0.576568 | 0.462583 | 1.246412 | 0.212613 | 0.882862 | 0.054107 |
| ENSMUSG00000070576.5 | 45.95845 | -1.51991 | 1.219762 | -1.24607 | 0.212738 | 0.882995 | 0.054042 |
| ENSMUSG00000016181.11 | 19.43227 | -2.35731 | 1.893758 | -1.24478 | 0.213212 | 0.884575 | 0.053265 |
| ENSMUSG00000027365.15 | 154.8189 | -0.73512 | 0.590812 | -1.24426 | 0.213404 | 0.884594 | 0.053256 |
| ENSMUSG00000041229.16 | 26.08321 | -1.79652 | 1.443587 | -1.24449 | 0.213321 | 0.884594 | 0.053256 |

| | | | | | | | |
|---|---|---|---|---|---|---|---|
| ENSMUSG00000032280.17 | 50.7851 | -1.77702 | 1.42901 | -1.24353 | 0.213672 | 0.885321 | 0.052899 |
| ENSMUSG00000022680.15 | 216.962 | -0.68951 | 0.554719 | -1.24298 | 0.213874 | 0.885384 | 0.052868 |
| ENSMUSG00000074064.7 | 199.4014 | -0.76479 | 0.615254 | -1.24305 | 0.21385 | 0.885384 | 0.052868 |
| ENSMUSG00000020078.17 | 259.8761 | -0.64606 | 0.520703 | -1.24075 | 0.214698 | 0.885897 | 0.052617 |
| ENSMUSG00000020085.16 | 65.07298 | 1.812673 | 1.460408 | 1.24121 | 0.214528 | 0.885897 | 0.052617 |
| ENSMUSG00000021236.17 | 1068.579 | 0.433308 | 0.349006 | 1.241546 | 0.214404 | 0.885897 | 0.052617 |
| ENSMUSG00000022312.12 | 1214.521 | 0.36399 | 0.293236 | 1.241288 | 0.214499 | 0.885897 | 0.052617 |
| ENSMUSG00000025156.18 | 15.19786 | -2.63283 | 2.121546 | -1.241 | 0.214607 | 0.885897 | 0.052617 |
| ENSMUSG00000027709.10 | 257.8781 | -0.73826 | 0.594959 | -1.24086 | 0.214658 | 0.885897 | 0.052617 |
| ENSMUSG00000048787.14 | 38.84294 | 1.594178 | 1.284984 | 1.240621 | 0.214746 | 0.885897 | 0.052617 |
| ENSMUSG00000064350.1 | 16.64111 | -3.02755 | 2.439803 | -1.2409 | 0.214643 | 0.885897 | 0.052617 |
| ENSMUSG00000073676.5 | 2853.324 | 0.327848 | 0.264578 | 1.239134 | 0.215296 | 0.887779 | 0.051695 |
| ENSMUSG00000015890.3 | 507.0841 | -0.58531 | 0.472928 | -1.23763 | 0.215853 | 0.888446 | 0.051369 |
| ENSMUSG00000024853.11 | 241.8279 | -0.58247 | 0.470998 | -1.23668 | 0.216207 | 0.888446 | 0.051369 |
| ENSMUSG00000032397.8 | 37.81447 | -1.55394 | 1.256377 | -1.23684 | 0.216145 | 0.888446 | 0.051369 |
| ENSMUSG00000034591.6 | 78.92892 | -1.34568 | 1.088074 | -1.23675 | 0.216178 | 0.888446 | 0.051369 |
| ENSMUSG00000044122.16 | 32.36398 | 2.205051 | 1.781691 | 1.237617 | 0.215858 | 0.888446 | 0.051369 |
| ENSMUSG00000071379.3 | 39.0989 | -1.34893 | 1.089583 | -1.23803 | 0.215706 | 0.888446 | 0.051369 |
| ENSMUSG00000087107.9 | 24.32371 | 2.806969 | 2.267991 | 1.237645 | 0.215848 | 0.888446 | 0.051369 |
| ENSMUSG00000103711.2 | 277.3942 | 0.634671 | 0.512933 | 1.237337 | 0.215962 | 0.888446 | 0.051369 |
| ENSMUSG00000022965.9 | 62.7402 | -1.18376 | 0.959193 | -1.23412 | 0.217157 | 0.888801 | 0.051195 |
| ENSMUSG00000024483.20 | 152.2328 | -0.79571 | 0.644217 | -1.23515 | 0.216773 | 0.888801 | 0.051195 |
| ENSMUSG00000025078.10 | 71.48371 | 1.307475 | 1.058717 | 1.234962 | 0.216845 | 0.888801 | 0.051195 |
| ENSMUSG00000026201.13 | 277.4071 | 0.662665 | 0.536889 | 1.234268 | 0.217103 | 0.888801 | 0.051195 |
| ENSMUSG00000026248.10 | 138.6294 | -0.87385 | 0.707251 | -1.23556 | 0.216622 | 0.888801 | 0.051195 |
| ENSMUSG00000028719.9 | 639.0711 | 0.434859 | 0.352272 | 1.234438 | 0.21704 | 0.888801 | 0.051195 |
| ENSMUSG00000029635.16 | 114.5526 | -0.87368 | 0.707698 | -1.23453 | 0.217005 | 0.888801 | 0.051195 |
| ENSMUSG00000030753.16 | 103.0142 | 1.119601 | 0.90735 | 1.233924 | 0.217231 | 0.888801 | 0.051195 |
| ENSMUSG00000041483.15 | 109.8337 | -0.89204 | 0.721828 | -1.2358 | 0.216532 | 0.888801 | 0.051195 |
| ENSMUSG00000046861.10 | 91.29637 | -0.96668 | 0.783294 | -1.23412 | 0.217158 | 0.888801 | 0.051195 |
| ENSMUSG00000022048.9 | 76.63167 | 1.077333 | 0.873336 | 1.233584 | 0.217358 | 0.888936 | 0.05113 |
| ENSMUSG00000001240.14 | 19.20515 | -2.16267 | 1.754311 | -1.23278 | 0.21766 | 0.889438 | 0.050884 |
| ENSMUSG00000022887.10 | 380.3309 | -0.48129 | 0.39056 | -1.23232 | 0.21783 | 0.889438 | 0.050884 |
| ENSMUSG00000025933.16 | 52.61135 | 1.575795 | 1.27843 | 1.232602 | 0.217724 | 0.889438 | 0.050884 |
| ENSMUSG00000085793.4 | 33.04557 | -2.06236 | 1.673653 | -1.23225 | 0.217856 | 0.889438 | 0.050884 |
| ENSMUSG00000028884.15 | 27.02833 | -2.03595 | 1.652676 | -1.23191 | 0.217982 | 0.889571 | 0.05082 |
| ENSMUSG00000002043.18 | 30.718 | 2.007441 | 1.644922 | 1.220387 | 0.222318 | 0.889694 | 0.05076 |
| ENSMUSG00000007610.16 | 30.68682 | 1.623883 | 1.328874 | 1.221999 | 0.221708 | 0.889694 | 0.05076 |
| ENSMUSG00000008540.12 | 50519.86 | 0.259513 | 0.210808 | 1.231041 | 0.218307 | 0.889694 | 0.05076 |
| ENSMUSG00000008999.8 | 15.53739 | -3.00414 | 2.460973 | -1.22071 | 0.222194 | 0.889694 | 0.05076 |
| ENSMUSG00000010067.14 | 43.73035 | 1.510009 | 1.227957 | 1.229692 | 0.218812 | 0.889694 | 0.05076 |
| ENSMUSG00000015087.15 | 63.57399 | -1.72609 | 1.412113 | -1.22234 | 0.221577 | 0.889694 | 0.05076 |
| ENSMUSG00000017781.18 | 126.1641 | -0.95338 | 0.779446 | -1.22315 | 0.221273 | 0.889694 | 0.05076 |
| ENSMUSG00000020780.16 | 183.7192 | 0.878257 | 0.716559 | 1.22566 | 0.220327 | 0.889694 | 0.05076 |

| | | | | | | | |
|---|---|---|---|---|---|---|---|
| ENSMUSG00000020993.10 | 136.8938 | 0.906703 | 0.73889 | 1.227115 | 0.219779 | 0.889694 | 0.05076 |
| ENSMUSG00000021327.20 | 214.7468 | -0.65877 | 0.539508 | -1.22105 | 0.222067 | 0.889694 | 0.05076 |
| ENSMUSG00000021969.9 | 73.11015 | -1.36761 | 1.116934 | -1.22443 | 0.220789 | 0.889694 | 0.05076 |
| ENSMUSG00000022237.18 | 47.10906 | 1.489298 | 1.211409 | 1.229393 | 0.218924 | 0.889694 | 0.05076 |
| ENSMUSG00000022706.17 | 169.1014 | 0.903559 | 0.735203 | 1.228993 | 0.219074 | 0.889694 | 0.05076 |
| ENSMUSG00000023048.14 | 156.2801 | -0.83252 | 0.679722 | -1.22479 | 0.220653 | 0.889694 | 0.05076 |
| ENSMUSG00000023143.11 | 92.15631 | 0.987753 | 0.802684 | 1.230563 | 0.218486 | 0.889694 | 0.05076 |
| ENSMUSG00000025980.15 | 2838.523 | 0.296911 | 0.242219 | 1.225795 | 0.220276 | 0.889694 | 0.05076 |
| ENSMUSG00000026193.16 | 4733.906 | -0.26826 | 0.219365 | -1.22291 | 0.221363 | 0.889694 | 0.05076 |
| ENSMUSG00000026307.13 | 189.2454 | 0.731196 | 0.594005 | 1.230959 | 0.218338 | 0.889694 | 0.05076 |
| ENSMUSG00000027079.16 | 48.21267 | 1.25141 | 1.023141 | 1.223106 | 0.22129 | 0.889694 | 0.05076 |
| ENSMUSG00000028565.19 | 337.572 | 0.621556 | 0.505997 | 1.228379 | 0.219305 | 0.889694 | 0.05076 |
| ENSMUSG00000028991.16 | 121.7266 | -0.83633 | 0.683491 | -1.22362 | 0.221096 | 0.889694 | 0.05076 |
| ENSMUSG00000030714.15 | 77.533 | -0.9504 | 0.777266 | -1.22275 | 0.221424 | 0.889694 | 0.05076 |
| ENSMUSG00000030880.15 | 56.42438 | -1.26424 | 1.031014 | -1.22621 | 0.220121 | 0.889694 | 0.05076 |
| ENSMUSG00000031877.10 | 182.3655 | -0.81103 | 0.663036 | -1.2232 | 0.221253 | 0.889694 | 0.05076 |
| ENSMUSG00000032348.7 | 1140.94 | 0.538162 | 0.440099 | 1.222819 | 0.221398 | 0.889694 | 0.05076 |
| ENSMUSG00000032420.9 | 27.00339 | 2.409917 | 1.973355 | 1.221228 | 0.222 | 0.889694 | 0.05076 |
| ENSMUSG00000033111.17 | 73.06224 | -1.16082 | 0.946016 | -1.22706 | 0.2198 | 0.889694 | 0.05076 |
| ENSMUSG00000035078.7 | 171.7679 | -0.70288 | 0.576095 | -1.22008 | 0.222435 | 0.889694 | 0.05076 |
| ENSMUSG00000037999.14 | 92.12546 | -1.3266 | 1.082806 | -1.22515 | 0.22052 | 0.889694 | 0.05076 |
| ENSMUSG00000039770.7 | 262.1233 | 0.607453 | 0.494941 | 1.227325 | 0.219701 | 0.889694 | 0.05076 |
| ENSMUSG00000041319.15 | 26.58245 | -2.7174 | 2.210364 | -1.22939 | 0.218925 | 0.889694 | 0.05076 |
| ENSMUSG00000043262.17 | 16.08968 | 2.644601 | 2.149398 | 1.230392 | 0.218551 | 0.889694 | 0.05076 |
| ENSMUSG00000044715.8 | 53.62253 | -1.1305 | 0.926335 | -1.2204 | 0.222312 | 0.889694 | 0.05076 |
| ENSMUSG00000045294.12 | 279.8111 | 0.650094 | 0.530336 | 1.225816 | 0.220268 | 0.889694 | 0.05076 |
| ENSMUSG00000045776.4 | 97.03454 | 1.019127 | 0.8347 | 1.22095 | 0.222105 | 0.889694 | 0.05076 |
| ENSMUSG00000046516.11 | 234.5861 | 0.730358 | 0.59861 | 1.220089 | 0.222431 | 0.889694 | 0.05076 |
| ENSMUSG00000048755.10 | 99.96488 | 1.062905 | 0.866311 | 1.226933 | 0.219848 | 0.889694 | 0.05076 |
| ENSMUSG00000048915.13 | 40.14946 | 1.663867 | 1.362392 | 1.221284 | 0.221979 | 0.889694 | 0.05076 |
| ENSMUSG00000051671.8 | 461.7869 | 0.46288 | 0.377815 | 1.225149 | 0.220519 | 0.889694 | 0.05076 |
| ENSMUSG00000053801.19 | 10.93167 | -3.58302 | 2.921271 | -1.22653 | 0.22 | 0.889694 | 0.05076 |
| ENSMUSG00000055553.17 | 192.2575 | 0.74053 | 0.603235 | 1.227598 | 0.219598 | 0.889694 | 0.05076 |
| ENSMUSG00000063849.7 | 17.57419 | 2.931377 | 2.400753 | 1.221024 | 0.222077 | 0.889694 | 0.05076 |
| ENSMUSG00000066880.16 | 35.04064 | -1.5346 | 1.25356 | -1.22419 | 0.22088 | 0.889694 | 0.05076 |
| ENSMUSG00000076612.9 | 20.76454 | 2.603931 | 2.127721 | 1.223812 | 0.221023 | 0.889694 | 0.05076 |
| ENSMUSG00000078453.4 | 118.5177 | -0.84919 | 0.695105 | -1.22168 | 0.22183 | 0.889694 | 0.05076 |
| ENSMUSG00000096727.4 | 114.765 | -1.22101 | 1.000579 | -1.2203 | 0.222351 | 0.889694 | 0.05076 |
| ENSMUSG00000099707.7 | 161.6231 | -0.83594 | 0.684967 | -1.2204 | 0.222312 | 0.889694 | 0.05076 |
| ENSMUSG00000113769.2 | 39.08784 | -1.75198 | 1.436209 | -1.21986 | 0.222517 | 0.889694 | 0.05076 |
| ENSMUSG00000001098.16 | 26.48818 | -2.06248 | 1.692179 | -1.21883 | 0.222908 | 0.889906 | 0.050656 |
| ENSMUSG00000020315.19 | 220.1059 | 0.626625 | 0.514185 | 1.218674 | 0.222968 | 0.889906 | 0.050656 |
| ENSMUSG00000020429.8 | 404.1064 | -1.58834 | 1.303536 | -1.21849 | 0.22304 | 0.889906 | 0.050656 |
| ENSMUSG00000021501.5 | 113.1478 | 0.786015 | 0.64458 | 1.219423 | 0.222684 | 0.889906 | 0.050656 |

| | | | | | | | |
|---|---|---|---|---|---|---|---|
| ENSMUSG00000048379.10 | 137.9488 | -0.80584 | 0.661161 | -1.21882 | 0.222913 | 0.889906 | 0.050656 |
| ENSMUSG00000001670.14 | 4734.747 | 0.370078 | 0.30531 | 1.212139 | 0.225459 | 0.891095 | 0.050076 |
| ENSMUSG00000002477.6 | 264.5099 | 0.653365 | 0.537904 | 1.214649 | 0.2245 | 0.891095 | 0.050076 |
| ENSMUSG00000015501.11 | 17.14595 | -2.36204 | 1.946217 | -1.21366 | 0.224879 | 0.891095 | 0.050076 |
| ENSMUSG00000020029.7 | 1408.521 | -0.38939 | 0.320689 | -1.21422 | 0.224665 | 0.891095 | 0.050076 |
| ENSMUSG00000020611.15 | 147.6354 | 0.744757 | 0.612395 | 1.216139 | 0.223932 | 0.891095 | 0.050076 |
| ENSMUSG00000024091.10 | 394.1466 | -0.49498 | 0.407233 | -1.21548 | 0.224182 | 0.891095 | 0.050076 |
| ENSMUSG00000024856.11 | 254.0366 | 0.740555 | 0.609534 | 1.214954 | 0.224384 | 0.891095 | 0.050076 |
| ENSMUSG00000025935.11 | 1196.56 | -0.35853 | 0.295583 | -1.21295 | 0.225148 | 0.891095 | 0.050076 |
| ENSMUSG00000027133.4 | 623.9226 | 0.458512 | 0.377934 | 1.213207 | 0.225051 | 0.891095 | 0.050076 |
| ENSMUSG00000027940.19 | 191.1221 | -0.74521 | 0.613584 | -1.21452 | 0.22455 | 0.891095 | 0.050076 |
| ENSMUSG00000030105.9 | 265.2639 | -0.63878 | 0.527138 | -1.21179 | 0.225594 | 0.891095 | 0.050076 |
| ENSMUSG00000030243.17 | 30.20791 | 1.689804 | 1.391804 | 1.21411 | 0.224706 | 0.891095 | 0.050076 |
| ENSMUSG00000031513.9 | 111.0285 | -1.08339 | 0.893961 | -1.21189 | 0.225553 | 0.891095 | 0.050076 |
| ENSMUSG00000033004.17 | 186.8522 | -0.78015 | 0.643158 | -1.213 | 0.225128 | 0.891095 | 0.050076 |
| ENSMUSG00000034636.10 | 228.3101 | 0.629791 | 0.519564 | 1.212154 | 0.225454 | 0.891095 | 0.050076 |
| ENSMUSG00000035762.12 | 26.69311 | 1.621302 | 1.332728 | 1.216529 | 0.223783 | 0.891095 | 0.050076 |
| ENSMUSG00000040525.14 | 19.42289 | -2.5903 | 2.128269 | -1.21709 | 0.223569 | 0.891095 | 0.050076 |
| ENSMUSG00000045671.18 | 14.19482 | -2.18658 | 1.796587 | -1.21708 | 0.223575 | 0.891095 | 0.050076 |
| ENSMUSG00000054716.5 | 25.20371 | -2.40014 | 1.980214 | -1.21206 | 0.225489 | 0.891095 | 0.050076 |
| ENSMUSG00000059027.15 | 75.80595 | 1.142513 | 0.942598 | 1.21209 | 0.225478 | 0.891095 | 0.050076 |
| ENSMUSG00000060371.13 | 25.07478 | 2.030975 | 1.673708 | 1.213458 | 0.224955 | 0.891095 | 0.050076 |
| ENSMUSG00000060730.9 | 54.11168 | 1.16873 | 0.963445 | 1.213074 | 0.225101 | 0.891095 | 0.050076 |
| ENSMUSG00000087361.8 | 11.43999 | -3.28726 | 2.709988 | -1.21301 | 0.225124 | 0.891095 | 0.050076 |
| ENSMUSG00000117226.2 | 28.3301 | -2.3421 | 1.931233 | -1.21275 | 0.225225 | 0.891095 | 0.050076 |
| ENSMUSG00000056342.17 | 421.7194 | -0.48208 | 0.397961 | -1.21138 | 0.22575 | 0.891343 | 0.049955 |
| ENSMUSG00000006676.17 | 81.58665 | -1.04608 | 0.86475 | -1.20969 | 0.226398 | 0.891526 | 0.049866 |
| ENSMUSG00000010290.8 | 42.23038 | 1.422185 | 1.181993 | 1.203209 | 0.228895 | 0.891526 | 0.049866 |
| ENSMUSG00000018196.19 | 249.2982 | -0.82253 | 0.682369 | -1.20541 | 0.228045 | 0.891526 | 0.049866 |
| ENSMUSG00000020986.14 | 534.4017 | 0.467074 | 0.388166 | 1.203286 | 0.228866 | 0.891526 | 0.049866 |
| ENSMUSG00000021113.6 | 64.06487 | 1.00433 | 0.834587 | 1.203386 | 0.228827 | 0.891526 | 0.049866 |
| ENSMUSG00000021266.17 | 15.88875 | 2.358245 | 1.955389 | 1.206023 | 0.227808 | 0.891526 | 0.049866 |
| ENSMUSG00000021540.17 | 97.67859 | -0.91594 | 0.760596 | -1.20424 | 0.228496 | 0.891526 | 0.049866 |
| ENSMUSG00000021820.19 | 58.0912 | 1.845934 | 1.530323 | 1.206238 | 0.227726 | 0.891526 | 0.049866 |
| ENSMUSG00000021893.15 | 196.7959 | -0.67684 | 0.560026 | -1.20859 | 0.226819 | 0.891526 | 0.049866 |
| ENSMUSG00000022354.5 | 2019.435 | 0.302218 | 0.250341 | 1.207222 | 0.227347 | 0.891526 | 0.049866 |
| ENSMUSG00000023262.14 | 43.59757 | -1.61989 | 1.341077 | -1.2079 | 0.227084 | 0.891526 | 0.049866 |
| ENSMUSG00000024620.12 | 11.11475 | -3.87309 | 3.20812 | -1.20728 | 0.227326 | 0.891526 | 0.049866 |
| ENSMUSG00000025198.17 | 197.7526 | 0.636078 | 0.528657 | 1.203197 | 0.2289 | 0.891526 | 0.049866 |
| ENSMUSG00000025591.7 | 13.45757 | 2.748982 | 2.283169 | 1.20402 | 0.228582 | 0.891526 | 0.049866 |
| ENSMUSG00000027406.14 | 722.3369 | 0.372594 | 0.308455 | 1.207937 | 0.227072 | 0.891526 | 0.049866 |
| ENSMUSG00000031392.19 | 454.8627 | 0.440198 | 0.365354 | 1.204855 | 0.228259 | 0.891526 | 0.049866 |
| ENSMUSG00000031930.12 | 40.43556 | -1.68542 | 1.39372 | -1.20929 | 0.22655 | 0.891526 | 0.049866 |
| ENSMUSG00000032121.10 | 11.30233 | -3.26805 | 2.713056 | -1.20456 | 0.228372 | 0.891526 | 0.049866 |

| | | | | | | | |
|---|---|---|---|---|---|---|---|
| ENSMUSG00000038332.14 | 42.80037 | -1.36827 | 1.136335 | -1.20411 | 0.228547 | 0.891526 | 0.049866 |
| ENSMUSG00000039117.17 | 39.8428 | -1.52448 | 1.263395 | -1.20665 | 0.227566 | 0.891526 | 0.049866 |
| ENSMUSG00000040688.17 | 42.7781 | 1.615239 | 1.335329 | 1.209619 | 0.226425 | 0.891526 | 0.049866 |
| ENSMUSG00000042901.11 | 73.15573 | 1.307502 | 1.079812 | 1.210861 | 0.225949 | 0.891526 | 0.049866 |
| ENSMUSG00000047547.15 | 179.0995 | -0.77161 | 0.638661 | -1.20817 | 0.226982 | 0.891526 | 0.049866 |
| ENSMUSG00000056383.11 | 203.2853 | -0.70169 | 0.581259 | -1.20718 | 0.227361 | 0.891526 | 0.049866 |
| ENSMUSG00000056999.17 | 583.3543 | -0.40507 | 0.336489 | -1.20383 | 0.228656 | 0.891526 | 0.049866 |
| ENSMUSG00000059486.12 | 159.8781 | -0.8886 | 0.737476 | -1.20492 | 0.228234 | 0.891526 | 0.049866 |
| ENSMUSG00000061825.11 | 25.39345 | -2.70858 | 2.247171 | -1.20533 | 0.228076 | 0.891526 | 0.049866 |
| ENSMUSG00000070426.13 | 55.36408 | 1.274419 | 1.052698 | 1.210622 | 0.22604 | 0.891526 | 0.049866 |
| ENSMUSG00000071632.11 | 222.8755 | -0.67235 | 0.557111 | -1.20686 | 0.227487 | 0.891526 | 0.049866 |
| ENSMUSG00000075010.6 | 1143.608 | -0.37453 | 0.310234 | -1.20726 | 0.227332 | 0.891526 | 0.049866 |
| ENSMUSG00000085972.2 | 11.38959 | 3.257764 | 2.694653 | 1.208973 | 0.226673 | 0.891526 | 0.049866 |
| ENSMUSG00000096433.3 | 24.78478 | 1.996069 | 1.649618 | 1.210019 | 0.226272 | 0.891526 | 0.049866 |
| ENSMUSG00000109783.2 | 72.65367 | 1.183563 | 0.983573 | 1.20333 | 0.228849 | 0.891526 | 0.049866 |
| ENSMUSG00000005949.10 | 24.48715 | 2.010599 | 1.672572 | 1.2021 | 0.229325 | 0.89259 | 0.049348 |
| ENSMUSG00000034917.9 | 32.43718 | 1.89501 | 1.576541 | 1.202004 | 0.229362 | 0.89259 | 0.049348 |
| ENSMUSG00000028019.10 | 18.23835 | -2.56238 | 2.135253 | -1.20003 | 0.230126 | 0.893302 | 0.049002 |
| ENSMUSG00000030869.12 | 1003.727 | 0.377995 | 0.315048 | 1.199802 | 0.230216 | 0.893302 | 0.049002 |
| ENSMUSG00000032402.13 | 15.67895 | -2.61002 | 2.175461 | -1.19976 | 0.230234 | 0.893302 | 0.049002 |
| ENSMUSG00000034259.9 | 89.34578 | 0.997322 | 0.831386 | 1.19959 | 0.230298 | 0.893302 | 0.049002 |
| ENSMUSG00000040599.14 | 146.9858 | -0.83076 | 0.692218 | -1.20015 | 0.230083 | 0.893302 | 0.049002 |
| ENSMUSG00000047036.9 | 686.2772 | -0.53398 | 0.445019 | -1.1999 | 0.23018 | 0.893302 | 0.049002 |
| ENSMUSG00000068284.15 | 186.4008 | -0.62069 | 0.517186 | -1.20013 | 0.230089 | 0.893302 | 0.049002 |
| ENSMUSG00000071180.6 | 272.4205 | 0.612961 | 0.510352 | 1.201056 | 0.22973 | 0.893302 | 0.049002 |
| ENSMUSG00000010911.13 | 244.3097 | 0.722165 | 0.602615 | 1.198385 | 0.230767 | 0.894081 | 0.048623 |
| ENSMUSG00000020132.11 | 303.6303 | -0.59931 | 0.49999 | -1.19864 | 0.230666 | 0.894081 | 0.048623 |
| ENSMUSG00000023883.14 | 40.23292 | 1.285085 | 1.072382 | 1.198346 | 0.230782 | 0.894081 | 0.048623 |
| ENSMUSG00000022010.20 | 459.4496 | -0.43735 | 0.365224 | -1.19748 | 0.23112 | 0.894292 | 0.048521 |
| ENSMUSG00000026950.19 | 79.66024 | 1.219448 | 1.018171 | 1.197685 | 0.23104 | 0.894292 | 0.048521 |
| ENSMUSG00000076613.5 | 58.83986 | -1.82715 | 1.525521 | -1.19772 | 0.231025 | 0.894292 | 0.048521 |
| ENSMUSG00000007655.17 | 23.83195 | -2.54584 | 2.128183 | -1.19625 | 0.231598 | 0.894634 | 0.048355 |
| ENSMUSG00000019804.13 | 1074.338 | 0.320954 | 0.268218 | 1.196616 | 0.231456 | 0.894634 | 0.048355 |
| ENSMUSG00000024735.14 | 210.6895 | -0.8333 | 0.696716 | -1.19604 | 0.23168 | 0.894634 | 0.048355 |
| ENSMUSG00000036898.17 | 12.24561 | -2.99955 | 2.507697 | -1.19614 | 0.231644 | 0.894634 | 0.048355 |
| ENSMUSG00000078695.9 | 100.6498 | 0.950469 | 0.794176 | 1.196799 | 0.231385 | 0.894634 | 0.048355 |
| ENSMUSG00000001155.14 | 403.0865 | -0.57468 | 0.481023 | -1.19471 | 0.2322 | 0.895851 | 0.047764 |
| ENSMUSG00000024782.8 | 1941.945 | 0.311509 | 0.260799 | 1.194441 | 0.232305 | 0.895851 | 0.047764 |
| ENSMUSG00000025234.12 | 209.6454 | -0.6447 | 0.540253 | -1.19333 | 0.232741 | 0.895851 | 0.047764 |
| ENSMUSG00000027244.15 | 58.30855 | -1.20576 | 1.010324 | -1.19344 | 0.232696 | 0.895851 | 0.047764 |
| ENSMUSG00000029128.13 | 186.9658 | 1.010582 | 0.845927 | 1.194644 | 0.232226 | 0.895851 | 0.047764 |
| ENSMUSG00000039145.17 | 206.1618 | -0.72212 | 0.605041 | -1.1935 | 0.232673 | 0.895851 | 0.047764 |
| ENSMUSG00000042111.10 | 61.86464 | -1.1547 | 0.967442 | -1.19356 | 0.232651 | 0.895851 | 0.047764 |
| ENSMUSG00000074896.4 | 12.2676 | -2.3422 | 1.962789 | -1.1933 | 0.232751 | 0.895851 | 0.047764 |

| | | | | | | | |
|---|---|---|---|---|---|---|---|
| ENSMUSG00000062619.7 | 219.1923 | 0.65094 | 0.545836 | 1.192556 | 0.233043 | 0.896611 | 0.047396 |
| ENSMUSG00000032228.17 | 123.7122 | 0.898832 | 0.75386 | 1.192307 | 0.233141 | 0.896623 | 0.04739 |
| ENSMUSG00000030701.18 | 38.46017 | 1.658128 | 1.393569 | 1.189843 | 0.234108 | 0.89852 | 0.046472 |
| ENSMUSG00000038642.11 | 273.1195 | -0.6481 | 0.544623 | -1.19 | 0.234047 | 0.89852 | 0.046472 |
| ENSMUSG00000049550.18 | 156.0894 | -0.77133 | 0.647985 | -1.19036 | 0.233906 | 0.89852 | 0.046472 |
| ENSMUSG00000061024.9 | 151.999 | 0.824059 | 0.692134 | 1.190607 | 0.233808 | 0.89852 | 0.046472 |
| ENSMUSG00000086938.10 | 15.64197 | -2.37419 | 1.995119 | -1.19 | 0.234047 | 0.89852 | 0.046472 |
| ENSMUSG00000009035.16 | 74.01071 | 1.086896 | 0.915296 | 1.187481 | 0.235038 | 0.898814 | 0.04633 |
| ENSMUSG00000018509.9 | 51.20522 | -1.20647 | 1.015654 | -1.18787 | 0.234884 | 0.898814 | 0.04633 |
| ENSMUSG00000021244.16 | 79.68408 | -1.07966 | 0.908892 | -1.18789 | 0.234878 | 0.898814 | 0.04633 |
| ENSMUSG00000028796.18 | 20.99688 | -1.9903 | 1.674849 | -1.18835 | 0.234697 | 0.898814 | 0.04633 |
| ENSMUSG00000038056.16 | 146.079 | -0.8129 | 0.683612 | -1.18912 | 0.234392 | 0.898814 | 0.04633 |
| ENSMUSG00000051469.15 | 97.87449 | -0.81635 | 0.687403 | -1.18758 | 0.235 | 0.898814 | 0.04633 |
| ENSMUSG00000058076.13 | 644.2829 | -0.47834 | 0.402525 | -1.18836 | 0.234691 | 0.898814 | 0.04633 |
| ENSMUSG00000060149.8 | 18.78416 | 2.61641 | 2.202338 | 1.188015 | 0.234828 | 0.898814 | 0.04633 |
| ENSMUSG00000071226.12 | 54.55451 | -1.13197 | 0.952398 | -1.18855 | 0.234618 | 0.898814 | 0.04633 |
| ENSMUSG00000008892.13 | 268.1132 | 0.557963 | 0.469981 | 1.187203 | 0.235147 | 0.89887 | 0.046303 |
| ENSMUSG00000026470.15 | 22.78692 | -1.92709 | 1.624671 | -1.18614 | 0.235566 | 0.899751 | 0.045878 |
| ENSMUSG00000030131.9 | 3656.996 | -0.33226 | 0.280161 | -1.18596 | 0.235637 | 0.899751 | 0.045878 |
| ENSMUSG00000090523.3 | 62.8363 | 1.355804 | 1.143272 | 1.185898 | 0.235663 | 0.899751 | 0.045878 |
| ENSMUSG00000028684.15 | 373.559 | -0.49748 | 0.419723 | -1.18526 | 0.235914 | 0.900348 | 0.04559 |
| ENSMUSG00000004018.10 | 72.25424 | 1.112507 | 0.940386 | 1.183033 | 0.236796 | 0.900577 | 0.045479 |
| ENSMUSG00000014778.11 | 16.41588 | -2.41866 | 2.042635 | -1.18409 | 0.236378 | 0.900577 | 0.045479 |
| ENSMUSG00000020267.7 | 1252.17 | 0.326545 | 0.276027 | 1.18302 | 0.236801 | 0.900577 | 0.045479 |
| ENSMUSG00000024404.8 | 235.8771 | -0.7208 | 0.60836 | -1.18482 | 0.236089 | 0.900577 | 0.045479 |
| ENSMUSG00000025531.15 | 60.03472 | -1.16954 | 0.988683 | -1.18293 | 0.236836 | 0.900577 | 0.045479 |
| ENSMUSG00000025584.18 | 197.8682 | 0.814603 | 0.688551 | 1.183069 | 0.236782 | 0.900577 | 0.045479 |
| ENSMUSG00000029346.13 | 22.99435 | -2.72077 | 2.300313 | -1.18278 | 0.236895 | 0.900577 | 0.045479 |
| ENSMUSG00000050323.15 | 130.7597 | -1.16958 | 0.987428 | -1.18447 | 0.236227 | 0.900577 | 0.045479 |
| ENSMUSG00000054021.7 | 62.14575 | -1.6254 | 1.373924 | -1.18303 | 0.236796 | 0.900577 | 0.045479 |
| ENSMUSG00000058709.12 | 51.64855 | 1.405312 | 1.188213 | 1.182711 | 0.236924 | 0.900577 | 0.045479 |
| ENSMUSG00000026270.12 | 58.05584 | -1.48482 | 1.256414 | -1.1818 | 0.237287 | 0.901233 | 0.045163 |
| ENSMUSG00000053846.6 | 18.38946 | -2.5445 | 2.15294 | -1.18187 | 0.237256 | 0.901233 | 0.045163 |
| ENSMUSG00000055912.9 | 115.7916 | 0.861384 | 0.729041 | 1.181531 | 0.237392 | 0.901272 | 0.045144 |
| ENSMUSG00000039089.16 | 19.15232 | 2.355547 | 1.994608 | 1.180957 | 0.23762 | 0.901776 | 0.044902 |
| ENSMUSG00000025357.9 | 14.70605 | -2.29559 | 1.945235 | -1.18011 | 0.237957 | 0.902387 | 0.044607 |
| ENSMUSG00000032396.18 | 33.26697 | -2.48272 | 2.103868 | -1.18007 | 0.237971 | 0.902387 | 0.044607 |
| ENSMUSG00000007837.12 | 46.4433 | -1.36359 | 1.1558 | -1.17978 | 0.238086 | 0.902462 | 0.044571 |
| ENSMUSG00000015937.16 | 370.6735 | 0.511975 | 0.434643 | 1.177921 | 0.238828 | 0.903108 | 0.04426 |
| ENSMUSG00000021520.5 | 5437.97 | 0.350912 | 0.297823 | 1.178256 | 0.238694 | 0.903108 | 0.04426 |
| ENSMUSG00000028629.10 | 12.03505 | -2.79498 | 2.372676 | -1.17799 | 0.238802 | 0.903108 | 0.04426 |
| ENSMUSG00000028669.16 | 45.07234 | 1.646309 | 1.39668 | 1.17873 | 0.238506 | 0.903108 | 0.04426 |
| ENSMUSG00000030020.14 | 11.00665 | 2.944557 | 2.498059 | 1.178738 | 0.238503 | 0.903108 | 0.04426 |
| ENSMUSG00000059070.17 | 519.0497 | 0.563781 | 0.47848 | 1.178275 | 0.238687 | 0.903108 | 0.04426 |

| | | | | | | | |
|---|---|---|---|---|---|---|---|
| ENSMUSG00000030603.17 | 368.3946 | -0.58108 | 0.494048 | -1.17615 | 0.239534 | 0.903373 | 0.044133 |
| ENSMUSG00000032915.7 | 35.43978 | -1.91701 | 1.629679 | -1.17631 | 0.23947 | 0.903373 | 0.044133 |
| ENSMUSG00000036278.8 | 65.38317 | -0.99665 | 0.84708 | -1.17657 | 0.239367 | 0.903373 | 0.044133 |
| ENSMUSG00000042082.8 | 178.9877 | 0.659793 | 0.560945 | 1.176217 | 0.239508 | 0.903373 | 0.044133 |
| ENSMUSG00000046722.15 | 71.56436 | 1.206854 | 1.025569 | 1.176765 | 0.239289 | 0.903373 | 0.044133 |
| ENSMUSG00000058997.9 | 603.7387 | -0.44347 | 0.377079 | -1.17607 | 0.239565 | 0.903373 | 0.044133 |
| ENSMUSG00000062646.13 | 59.18217 | 1.36985 | 1.164028 | 1.176818 | 0.239268 | 0.903373 | 0.044133 |
| ENSMUSG00000022434.9 | 14.59665 | -2.99358 | 2.546446 | -1.17559 | 0.239758 | 0.903389 | 0.044125 |
| ENSMUSG00000079317.11 | 23.49508 | 2.219846 | 1.888288 | 1.175587 | 0.23976 | 0.903389 | 0.044125 |
| ENSMUSG00000029650.11 | 289.5999 | -0.62646 | 0.533126 | -1.17506 | 0.239969 | 0.903818 | 0.043919 |
| ENSMUSG00000010047.13 | 28.52547 | -1.82913 | 1.565972 | -1.16805 | 0.242788 | 0.904385 | 0.043646 |
| ENSMUSG00000014554.14 | 48.64051 | 1.224844 | 1.042972 | 1.174379 | 0.240243 | 0.904385 | 0.043646 |
| ENSMUSG00000017778.15 | 5152.333 | 0.328893 | 0.280949 | 1.170651 | 0.241739 | 0.904385 | 0.043646 |
| ENSMUSG00000018733.11 | 55.56653 | -1.17887 | 1.008643 | -1.16877 | 0.242496 | 0.904385 | 0.043646 |
| ENSMUSG00000019055.16 | 52.8156 | -1.44781 | 1.239108 | -1.16843 | 0.242633 | 0.904385 | 0.043646 |
| ENSMUSG00000020123.7 | 147.7612 | -0.68424 | 0.58391 | -1.17183 | 0.241266 | 0.904385 | 0.043646 |
| ENSMUSG00000022257.5 | 109.008 | 0.864397 | 0.739121 | 1.169493 | 0.242205 | 0.904385 | 0.043646 |
| ENSMUSG00000023050.10 | 33.79569 | 1.538848 | 1.314494 | 1.170677 | 0.241729 | 0.904385 | 0.043646 |
| ENSMUSG00000024067.15 | 237.3505 | 0.614072 | 0.525343 | 1.168897 | 0.242445 | 0.904385 | 0.043646 |
| ENSMUSG00000024589.19 | 94.57689 | 1.195554 | 1.02207 | 1.169738 | 0.242106 | 0.904385 | 0.043646 |
| ENSMUSG00000025283.16 | 212.2397 | -0.61547 | 0.526333 | -1.16936 | 0.24226 | 0.904385 | 0.043646 |
| ENSMUSG00000030314.17 | 100.6534 | -0.94584 | 0.808609 | -1.16971 | 0.242119 | 0.904385 | 0.043646 |
| ENSMUSG00000031309.16 | 95.97716 | 0.986293 | 0.844205 | 1.168309 | 0.242682 | 0.904385 | 0.043646 |
| ENSMUSG00000032349.14 | 648.0041 | 0.482699 | 0.413013 | 1.168727 | 0.242514 | 0.904385 | 0.043646 |
| ENSMUSG00000032525.16 | 119.7111 | 1.036389 | 0.884974 | 1.171096 | 0.24156 | 0.904385 | 0.043646 |
| ENSMUSG00000033965.11 | 437.8016 | -0.45853 | 0.39142 | -1.17144 | 0.241421 | 0.904385 | 0.043646 |
| ENSMUSG00000034055.17 | 16.3459 | 2.895447 | 2.473697 | 1.170494 | 0.241802 | 0.904385 | 0.043646 |
| ENSMUSG00000035569.18 | 324.2888 | -0.7077 | 0.604179 | -1.17135 | 0.24146 | 0.904385 | 0.043646 |
| ENSMUSG00000038462.4 | 2842.8 | 0.320628 | 0.274487 | 1.168098 | 0.242767 | 0.904385 | 0.043646 |
| ENSMUSG00000038845.12 | 43.56637 | -1.75206 | 1.499999 | -1.16804 | 0.242791 | 0.904385 | 0.043646 |
| ENSMUSG00000038970.8 | 13.81214 | 1.931871 | 1.653758 | 1.16817 | 0.242738 | 0.904385 | 0.043646 |
| ENSMUSG00000039536.17 | 154.3684 | -0.86601 | 0.739924 | -1.1704 | 0.24184 | 0.904385 | 0.043646 |
| ENSMUSG00000040123.18 | 290.264 | 0.547042 | 0.466412 | 1.172873 | 0.240847 | 0.904385 | 0.043646 |
| ENSMUSG00000060950.13 | 37.13753 | 2.14927 | 1.838595 | 1.168974 | 0.242414 | 0.904385 | 0.043646 |
| ENSMUSG00000063785.13 | 56.25818 | 1.150736 | 0.983737 | 1.16976 | 0.242098 | 0.904385 | 0.043646 |
| ENSMUSG00000079225.4 | 19.06764 | -2.0107 | 1.717382 | -1.17079 | 0.241682 | 0.904385 | 0.043646 |
| ENSMUSG00000091780.4 | 291.3462 | -0.60769 | 0.519239 | -1.17035 | 0.241861 | 0.904385 | 0.043646 |
| ENSMUSG00000118665.1 | 20.36809 | -1.93831 | 1.658817 | -1.16849 | 0.242609 | 0.904385 | 0.043646 |
| ENSMUSG00000049517.9 | 1929.728 | 0.273655 | 0.23438 | 1.167574 | 0.242979 | 0.904728 | 0.043482 |
| ENSMUSG00000001687.16 | 836.8493 | 0.474717 | 0.406987 | 1.166417 | 0.243446 | 0.905322 | 0.043197 |
| ENSMUSG00000032555.11 | 27.21156 | -1.58187 | 1.356306 | -1.16631 | 0.243491 | 0.905322 | 0.043197 |
| ENSMUSG00000049502.18 | 124.9723 | 0.721116 | 0.618104 | 1.166658 | 0.243348 | 0.905322 | 0.043197 |
| ENSMUSG00000070343.4 | 214.1219 | -0.59196 | 0.507593 | -1.16621 | 0.243528 | 0.905322 | 0.043197 |
| ENSMUSG00000101939.2 | 22.50603 | -1.58962 | 1.363315 | -1.166 | 0.243616 | 0.905322 | 0.043197 |

| | | | | | | | |
|---|---|---|---|---|---|---|---|
| ENSMUSG00000024277.15 | 80.29402 | 1.038865 | 0.891763 | 1.164957 | 0.244037 | 0.905821 | 0.042958 |
| ENSMUSG00000041852.16 | 27.07837 | -1.50939 | 1.295494 | -1.16511 | 0.243976 | 0.905821 | 0.042958 |
| ENSMUSG00000101517.7 | 207.5039 | 0.770477 | 0.661342 | 1.165021 | 0.244011 | 0.905821 | 0.042958 |
| ENSMUSG00000004609.12 | 12.50597 | 3.568249 | 3.075773 | 1.160115 | 0.246002 | 0.907558 | 0.042126 |
| ENSMUSG00000005936.20 | 28.47305 | -1.47014 | 1.267164 | -1.16018 | 0.245976 | 0.907558 | 0.042126 |
| ENSMUSG00000015478.14 | 201.6088 | 0.567356 | 0.48847 | 1.161495 | 0.24544 | 0.907558 | 0.042126 |
| ENSMUSG00000018548.16 | 34.18803 | -1.52131 | 1.309518 | -1.16173 | 0.245344 | 0.907558 | 0.042126 |
| ENSMUSG00000018567.15 | 190.6763 | 0.662961 | 0.571725 | 1.159582 | 0.246219 | 0.907558 | 0.042126 |
| ENSMUSG00000020288.14 | 242.1025 | 0.657034 | 0.565935 | 1.16097 | 0.245654 | 0.907558 | 0.042126 |
| ENSMUSG00000024981.7 | 1419.019 | 0.339791 | 0.292497 | 1.161691 | 0.245361 | 0.907558 | 0.042126 |
| ENSMUSG00000025059.17 | 436.9593 | 0.479556 | 0.413435 | 1.15993 | 0.246077 | 0.907558 | 0.042126 |
| ENSMUSG00000025068.9 | 664.2999 | 0.381349 | 0.329007 | 1.159091 | 0.246419 | 0.907558 | 0.042126 |
| ENSMUSG00000027566.16 | 953.8662 | -0.43788 | 0.376608 | -1.1627 | 0.244951 | 0.907558 | 0.042126 |
| ENSMUSG00000029103.17 | 173.3062 | -0.64096 | 0.552981 | -1.15909 | 0.246418 | 0.907558 | 0.042126 |
| ENSMUSG00000030236.11 | 9469.402 | -0.2459 | 0.211679 | -1.16164 | 0.245381 | 0.907558 | 0.042126 |
| ENSMUSG00000032599.13 | 21.14692 | 2.361852 | 2.030805 | 1.163013 | 0.244824 | 0.907558 | 0.042126 |
| ENSMUSG00000040620.16 | 43.15301 | -1.46763 | 1.264977 | -1.16021 | 0.245965 | 0.907558 | 0.042126 |
| ENSMUSG00000043535.14 | 83.47256 | -1.3491 | 1.163516 | -1.15951 | 0.24625 | 0.907558 | 0.042126 |
| ENSMUSG00000045962.17 | 169.9123 | -0.91408 | 0.786822 | -1.16174 | 0.245343 | 0.907558 | 0.042126 |
| ENSMUSG00000046994.10 | 19.54339 | -1.99403 | 1.717109 | -1.16127 | 0.245532 | 0.907558 | 0.042126 |
| ENSMUSG00000055435.7 | 211.1239 | 0.855064 | 0.736465 | 1.161038 | 0.245626 | 0.907558 | 0.042126 |
| ENSMUSG00000056260.16 | 72.14908 | 1.466993 | 1.262725 | 1.161768 | 0.24533 | 0.907558 | 0.042126 |
| ENSMUSG00000115708.2 | 26.64139 | 1.630246 | 1.405612 | 1.159813 | 0.246125 | 0.907558 | 0.042126 |
| ENSMUSG00000002395.15 | 463.1621 | 0.4761 | 0.412191 | 1.155046 | 0.248071 | 0.907633 | 0.04209 |
| ENSMUSG00000002458.14 | 13.4562 | -2.96248 | 2.574012 | -1.15092 | 0.249765 | 0.907633 | 0.04209 |
| ENSMUSG00000006705.14 | 45.05591 | -1.68557 | 1.459217 | -1.15512 | 0.248042 | 0.907633 | 0.04209 |
| ENSMUSG00000018906.15 | 30.93752 | -2.16519 | 1.871068 | -1.1572 | 0.247193 | 0.907633 | 0.04209 |
| ENSMUSG00000020225.11 | 722.3968 | 0.432928 | 0.37647 | 1.149967 | 0.250157 | 0.907633 | 0.04209 |
| ENSMUSG00000021228.16 | 55.2972 | 1.59956 | 1.385951 | 1.154124 | 0.248449 | 0.907633 | 0.04209 |
| ENSMUSG00000021901.11 | 25.9614 | -1.45341 | 1.262717 | -1.15102 | 0.249725 | 0.907633 | 0.04209 |
| ENSMUSG00000022664.12 | 75.78287 | 1.062876 | 0.923266 | 1.151213 | 0.249645 | 0.907633 | 0.04209 |
| ENSMUSG00000022890.14 | 5368.395 | 0.301628 | 0.261241 | 1.154594 | 0.248257 | 0.907633 | 0.04209 |
| ENSMUSG00000027509.12 | 60.78831 | -1.28914 | 1.118658 | -1.1524 | 0.249159 | 0.907633 | 0.04209 |
| ENSMUSG00000028124.16 | 1734.349 | -0.34303 | 0.296621 | -1.15644 | 0.247501 | 0.907633 | 0.04209 |
| ENSMUSG00000029014.15 | 226.4921 | -0.63584 | 0.552402 | -1.15105 | 0.249714 | 0.907633 | 0.04209 |
| ENSMUSG00000029810.16 | 995.2113 | -0.46958 | 0.40692 | -1.15397 | 0.248511 | 0.907633 | 0.04209 |
| ENSMUSG00000030218.3 | 19.01336 | 2.685738 | 2.333315 | 1.15104 | 0.249716 | 0.907633 | 0.04209 |
| ENSMUSG00000031311.18 | 99.23456 | 0.816162 | 0.706585 | 1.155079 | 0.248058 | 0.907633 | 0.04209 |
| ENSMUSG00000031864.17 | 20.2648 | 1.850713 | 1.605897 | 1.152448 | 0.249137 | 0.907633 | 0.04209 |
| ENSMUSG00000033632.17 | 15.41347 | 2.631452 | 2.288304 | 1.149958 | 0.250161 | 0.907633 | 0.04209 |
| ENSMUSG00000033972.10 | 68.89934 | -1.17922 | 1.025889 | -1.14946 | 0.250365 | 0.907633 | 0.04209 |
| ENSMUSG00000034382.15 | 11.31631 | 3.240862 | 2.812318 | 1.152381 | 0.249164 | 0.907633 | 0.04209 |
| ENSMUSG00000035105.6 | 35.9032 | 1.491098 | 1.297109 | 1.149555 | 0.250327 | 0.907633 | 0.04209 |
| ENSMUSG00000035311.17 | 40.53544 | -1.84578 | 1.596894 | -1.15586 | 0.24774 | 0.907633 | 0.04209 |

| | | | | | | | |
|---|---|---|---|---|---|---|---|
| ENSMUSG00000037370.14 | 73.09871 | 0.959227 | 0.834276 | 1.149771 | 0.250238 | 0.907633 | 0.04209 |
| ENSMUSG00000037461.11 | 36.85036 | -1.63805 | 1.416774 | -1.15619 | 0.247605 | 0.907633 | 0.04209 |
| ENSMUSG00000037851.14 | 222.4104 | 0.732578 | 0.634518 | 1.154543 | 0.248278 | 0.907633 | 0.04209 |
| ENSMUSG00000037971.11 | 299.0165 | 0.708389 | 0.615476 | 1.150962 | 0.249748 | 0.907633 | 0.04209 |
| ENSMUSG00000038250.10 | 87.99747 | 0.980858 | 0.847898 | 1.156811 | 0.24735 | 0.907633 | 0.04209 |
| ENSMUSG00000041037.9 | 33.17606 | 1.548236 | 1.342289 | 1.15343 | 0.248734 | 0.907633 | 0.04209 |
| ENSMUSG00000043991.6 | 432.9094 | 0.640129 | 0.556551 | 1.15017 | 0.250074 | 0.907633 | 0.04209 |
| ENSMUSG00000046562.6 | 67.96418 | 1.293356 | 1.125146 | 1.149501 | 0.250349 | 0.907633 | 0.04209 |
| ENSMUSG00000046826.8 | 23.41823 | 1.862196 | 1.619339 | 1.149973 | 0.250155 | 0.907633 | 0.04209 |
| ENSMUSG00000055200.8 | 22.17579 | -2.72951 | 2.358568 | -1.15727 | 0.247161 | 0.907633 | 0.04209 |
| ENSMUSG00000055401.15 | 84.74628 | -1.01512 | 0.87806 | -1.15609 | 0.247644 | 0.907633 | 0.04209 |
| ENSMUSG00000059498.14 | 24.66468 | -2.55066 | 2.203538 | -1.15753 | 0.247056 | 0.907633 | 0.04209 |
| ENSMUSG00000059708.13 | 24.47666 | 1.754915 | 1.519016 | 1.155297 | 0.247969 | 0.907633 | 0.04209 |
| ENSMUSG00000063273.12 | 251.6844 | 0.634199 | 0.550605 | 1.151822 | 0.249394 | 0.907633 | 0.04209 |
| ENSMUSG00000066900.12 | 156.937 | -0.78108 | 0.6742 | -1.15852 | 0.24665 | 0.907633 | 0.04209 |
| ENSMUSG00000072692.8 | 310.1201 | 0.522561 | 0.451138 | 1.158319 | 0.246734 | 0.907633 | 0.04209 |
| ENSMUSG00000079641.4 | 2234.29 | 0.453356 | 0.393363 | 1.152513 | 0.24911 | 0.907633 | 0.04209 |
| ENSMUSG00000086784.3 | 34.12215 | 1.599529 | 1.387273 | 1.153002 | 0.248909 | 0.907633 | 0.04209 |
| ENSMUSG00000086841.2 | 71.21882 | 1.171279 | 1.012591 | 1.156714 | 0.247389 | 0.907633 | 0.04209 |
| ENSMUSG00000112189.2 | 34.97693 | -1.43115 | 1.241509 | -1.15275 | 0.249012 | 0.907633 | 0.04209 |
| ENSMUSG00000007836.7 | 342.5671 | 0.533877 | 0.464688 | 1.148895 | 0.250599 | 0.907784 | 0.042017 |
| ENSMUSG00000019813.17 | 19.66814 | 1.742416 | 1.518331 | 1.147586 | 0.25114 | 0.907784 | 0.042017 |
| ENSMUSG00000021208.10 | 75.43318 | 1.273675 | 1.110541 | 1.146896 | 0.251425 | 0.907784 | 0.042017 |
| ENSMUSG00000022185.20 | 68.13773 | -1.04874 | 0.913025 | -1.14864 | 0.250704 | 0.907784 | 0.042017 |
| ENSMUSG00000022476.9 | 48.11456 | 1.512905 | 1.318161 | 1.147739 | 0.251076 | 0.907784 | 0.042017 |
| ENSMUSG00000025949.17 | 61.91266 | -1.23342 | 1.075506 | -1.14683 | 0.251453 | 0.907784 | 0.042017 |
| ENSMUSG00000027263.13 | 120.8455 | 0.81839 | 0.713234 | 1.147435 | 0.251202 | 0.907784 | 0.042017 |
| ENSMUSG00000028962.15 | 14.64632 | -2.72361 | 2.372415 | -1.14803 | 0.250955 | 0.907784 | 0.042017 |
| ENSMUSG00000030224.11 | 697.1699 | 0.393112 | 0.342205 | 1.148762 | 0.250654 | 0.907784 | 0.042017 |
| ENSMUSG00000034663.14 | 42.65702 | 1.432105 | 1.248772 | 1.14681 | 0.25146 | 0.907784 | 0.042017 |
| ENSMUSG00000048970.10 | 266.8632 | 0.586089 | 0.51087 | 1.147237 | 0.251284 | 0.907784 | 0.042017 |
| ENSMUSG00000097119.3 | 39.50011 | 1.581808 | 1.3798 | 1.146404 | 0.251628 | 0.908044 | 0.041893 |
| ENSMUSG00000032314.15 | 2932.331 | 0.332551 | 0.290164 | 1.146079 | 0.251763 | 0.908185 | 0.041826 |
| ENSMUSG00000005533.11 | 10.66178 | 3.154397 | 2.753546 | 1.145576 | 0.25197 | 0.908195 | 0.041821 |
| ENSMUSG00000007036.16 | 51.93524 | -1.45055 | 1.266628 | -1.14521 | 0.252124 | 0.908195 | 0.041821 |
| ENSMUSG00000026042.17 | 13.80122 | -2.63099 | 2.297516 | -1.14515 | 0.252149 | 0.908195 | 0.041821 |
| ENSMUSG00000078350.12 | 19.77417 | -2.40498 | 2.10002 | -1.14522 | 0.252119 | 0.908195 | 0.041821 |
| ENSMUSG00000000378.16 | 22.32919 | -2.11288 | 1.845464 | -1.14491 | 0.252248 | 0.90821 | 0.041814 |
| ENSMUSG00000027823.10 | 216.5704 | 0.747765 | 0.653283 | 1.144625 | 0.252364 | 0.908282 | 0.041779 |
| ENSMUSG00000033446.8 | 161.1789 | -0.83307 | 0.728238 | -1.14395 | 0.252645 | 0.908601 | 0.041627 |
| ENSMUSG00000041781.10 | 124.4743 | -0.78893 | 0.689637 | -1.14398 | 0.25263 | 0.908601 | 0.041627 |
| ENSMUSG00000024319.19 | 16.80364 | -2.29571 | 2.008149 | -1.1432 | 0.252957 | 0.909036 | 0.041419 |
| ENSMUSG00000024424.16 | 1264.622 | -0.31478 | 0.27535 | -1.14321 | 0.25295 | 0.909036 | 0.041419 |
| ENSMUSG00000029536.9 | 49.87346 | -1.49835 | 1.311654 | -1.14233 | 0.253315 | 0.909976 | 0.04097 |

| | | | | | | | |
|---|---|---|---|---|---|---|---|
| ENSMUSG00000020003.17 | 757.3066 | 0.442056 | 0.387214 | 1.141633 | 0.253606 | 0.910067 | 0.040927 |
| ENSMUSG00000028423.16 | 94.58737 | -0.93969 | 0.823148 | -1.14158 | 0.253628 | 0.910067 | 0.040927 |
| ENSMUSG00000058833.12 | 68.29688 | -1.32329 | 1.158929 | -1.14182 | 0.253528 | 0.910067 | 0.040927 |
| ENSMUSG00000021427.11 | 502.4028 | -0.49327 | 0.432427 | -1.1407 | 0.253996 | 0.910666 | 0.040641 |
| ENSMUSG00000026737.13 | 10.00173 | -3.25561 | 2.85479 | -1.1404 | 0.254119 | 0.910666 | 0.040641 |
| ENSMUSG00000031788.15 | 38.48817 | 1.343735 | 1.178611 | 1.140101 | 0.254244 | 0.910666 | 0.040641 |
| ENSMUSG00000044442.12 | 43.95835 | -1.49385 | 1.309331 | -1.14092 | 0.253902 | 0.910666 | 0.040641 |
| ENSMUSG00000085028.3 | 11.7829 | -3.1491 | 2.762309 | -1.14003 | 0.254276 | 0.910666 | 0.040641 |
| ENSMUSG00000008200.16 | 68.25976 | -1.26491 | 1.110533 | -1.13901 | 0.254697 | 0.911142 | 0.040414 |
| ENSMUSG00000011179.9 | 70.35875 | 1.332244 | 1.169216 | 1.139434 | 0.254522 | 0.911142 | 0.040414 |
| ENSMUSG00000022052.10 | 140.2997 | 0.796702 | 0.699374 | 1.139163 | 0.254635 | 0.911142 | 0.040414 |
| ENSMUSG00000030822.16 | 24.91807 | -1.99375 | 1.751292 | -1.13845 | 0.254934 | 0.911582 | 0.040204 |
| ENSMUSG00000054051.8 | 25.20202 | 1.552784 | 1.364174 | 1.138259 | 0.255012 | 0.911582 | 0.040204 |
| ENSMUSG00000019794.14 | 145.0186 | -1.02678 | 0.904009 | -1.13581 | 0.256037 | 0.911808 | 0.040097 |
| ENSMUSG00000026349.15 | 66.69663 | -1.13965 | 1.002258 | -1.13709 | 0.255502 | 0.911808 | 0.040097 |
| ENSMUSG00000026895.8 | 745.27 | 0.399165 | 0.351102 | 1.136894 | 0.255583 | 0.911808 | 0.040097 |
| ENSMUSG00000029512.12 | 98.36144 | -0.8973 | 0.78971 | -1.13624 | 0.255858 | 0.911808 | 0.040097 |
| ENSMUSG00000032018.15 | 3768.817 | -0.2937 | 0.258526 | -1.13606 | 0.25593 | 0.911808 | 0.040097 |
| ENSMUSG00000035692.8 | 30.50046 | -1.86885 | 1.644551 | -1.13639 | 0.255795 | 0.911808 | 0.040097 |
| ENSMUSG00000038641.13 | 500.5879 | 0.448455 | 0.394505 | 1.136754 | 0.255641 | 0.911808 | 0.040097 |
| ENSMUSG00000040396.13 | 112.9539 | 0.838101 | 0.736871 | 1.137379 | 0.25538 | 0.911808 | 0.040097 |
| ENSMUSG00000062328.9 | 3585.131 | 0.255936 | 0.224959 | 1.137698 | 0.255247 | 0.911808 | 0.040097 |
| ENSMUSG00000109599.2 | 16.72081 | -2.48914 | 2.19147 | -1.13583 | 0.256028 | 0.911808 | 0.040097 |
| ENSMUSG00000000902.14 | 40.36749 | 1.654597 | 1.462246 | 1.131545 | 0.257826 | 0.91214 | 0.039939 |
| ENSMUSG00000001150.8 | 66.86903 | -1.32404 | 1.17306 | -1.12871 | 0.259021 | 0.91214 | 0.039939 |
| ENSMUSG00000001211.16 | 318.389 | -0.66479 | 0.586285 | -1.1339 | 0.256837 | 0.91214 | 0.039939 |
| ENSMUSG00000001542.8 | 557.422 | -0.48569 | 0.429466 | -1.13091 | 0.258091 | 0.91214 | 0.039939 |
| ENSMUSG00000002393.15 | 131.1586 | 0.793553 | 0.708446 | 1.120133 | 0.262657 | 0.91214 | 0.039939 |
| ENSMUSG00000002486.16 | 12.16968 | -3.46512 | 3.111758 | -1.11356 | 0.265469 | 0.91214 | 0.039939 |
| ENSMUSG00000002699.14 | 16.75913 | -2.34412 | 2.104036 | -1.11411 | 0.265233 | 0.91214 | 0.039939 |
| ENSMUSG00000002778.15 | 135.5225 | -1.03409 | 0.923959 | -1.11919 | 0.263058 | 0.91214 | 0.039939 |
| ENSMUSG00000003382.19 | 89.28604 | -1.06285 | 0.945396 | -1.12424 | 0.26091 | 0.91214 | 0.039939 |
| ENSMUSG00000003518.14 | 105.1729 | -0.92787 | 0.831833 | -1.11545 | 0.264658 | 0.91214 | 0.039939 |
| ENSMUSG00000004270.14 | 229.236 | -0.72431 | 0.64929 | -1.11554 | 0.264619 | 0.91214 | 0.039939 |
| ENSMUSG00000004655.6 | 123.4751 | 0.796555 | 0.716134 | 1.112298 | 0.26601 | 0.91214 | 0.039939 |
| ENSMUSG00000004951.11 | 57.61709 | 1.547526 | 1.387604 | 1.11525 | 0.264743 | 0.91214 | 0.039939 |
| ENSMUSG00000005681.13 | 33882.88 | 0.266808 | 0.236078 | 1.130166 | 0.258406 | 0.91214 | 0.039939 |
| ENSMUSG00000009995.18 | 96.44873 | -1.47319 | 1.318686 | -1.11717 | 0.263923 | 0.91214 | 0.039939 |
| ENSMUSG00000015839.7 | 539.36 | 0.463439 | 0.416206 | 1.113485 | 0.2655 | 0.91214 | 0.039939 |
| ENSMUSG00000018287.13 | 198.2182 | -0.76732 | 0.688857 | -1.11391 | 0.265319 | 0.91214 | 0.039939 |
| ENSMUSG00000018481.10 | 103.0212 | 0.740531 | 0.662053 | 1.118536 | 0.263338 | 0.91214 | 0.039939 |
| ENSMUSG00000018761.15 | 70.67505 | 1.262174 | 1.122103 | 1.124828 | 0.260662 | 0.91214 | 0.039939 |
| ENSMUSG00000019082.19 | 106.7828 | -0.80242 | 0.716481 | -1.11994 | 0.262738 | 0.91214 | 0.039939 |
| ENSMUSG00000020311.18 | 256.8369 | 0.55414 | 0.48951 | 1.132029 | 0.257622 | 0.91214 | 0.039939 |

| | | | | | | | |
|---|---|---|---|---|---|---|---|
| ENSMUSG00000020519.6 | 151.8552 | 0.715695 | 0.6355 | 1.126193 | 0.260084 | 0.91214 | 0.039939 |
| ENSMUSG00000020794.14 | 181.9688 | 0.744786 | 0.658044 | 1.131817 | 0.257711 | 0.91214 | 0.039939 |
| ENSMUSG00000021366.9 | 36.45409 | 1.338812 | 1.184094 | 1.130663 | 0.258197 | 0.91214 | 0.039939 |
| ENSMUSG00000021716.15 | 304.2709 | 0.634819 | 0.566146 | 1.1213 | 0.26216 | 0.91214 | 0.039939 |
| ENSMUSG00000021737.5 | 619.2071 | -0.45073 | 0.405147 | -1.11251 | 0.265918 | 0.91214 | 0.039939 |
| ENSMUSG00000021958.6 | 22.77233 | -1.77702 | 1.587332 | -1.1195 | 0.262927 | 0.91214 | 0.039939 |
| ENSMUSG00000022261.7 | 1074.926 | -0.34262 | 0.30659 | -1.11752 | 0.263773 | 0.91214 | 0.039939 |
| ENSMUSG00000022353.11 | 829.759 | -0.34326 | 0.308209 | -1.11373 | 0.265397 | 0.91214 | 0.039939 |
| ENSMUSG00000023915.6 | 18.13656 | -2.11749 | 1.898498 | -1.11535 | 0.2647 | 0.91214 | 0.039939 |
| ENSMUSG00000023965.14 | 104.567 | -0.91908 | 0.814825 | -1.12795 | 0.259342 | 0.91214 | 0.039939 |
| ENSMUSG00000024254.16 | 188.3878 | -0.89328 | 0.796185 | -1.12195 | 0.261885 | 0.91214 | 0.039939 |
| ENSMUSG00000024610.16 | 114.2507 | -1.04515 | 0.924697 | -1.13026 | 0.258366 | 0.91214 | 0.039939 |
| ENSMUSG00000024694.10 | 1682.706 | 0.301033 | 0.270058 | 1.1147 | 0.264979 | 0.91214 | 0.039939 |
| ENSMUSG00000025037.7 | 54.3432 | 1.320009 | 1.17293 | 1.125394 | 0.260422 | 0.91214 | 0.039939 |
| ENSMUSG00000025862.15 | 222.8094 | 0.634993 | 0.565924 | 1.122046 | 0.261843 | 0.91214 | 0.039939 |
| ENSMUSG00000026249.11 | 128.0431 | 0.757839 | 0.681347 | 1.112267 | 0.266023 | 0.91214 | 0.039939 |
| ENSMUSG00000027531.17 | 238.4206 | -0.57091 | 0.504933 | -1.13067 | 0.258193 | 0.91214 | 0.039939 |
| ENSMUSG00000027999.16 | 124.1714 | -0.93608 | 0.825382 | -1.13411 | 0.256747 | 0.91214 | 0.039939 |
| ENSMUSG00000028266.18 | 179.9546 | 0.631465 | 0.564474 | 1.118679 | 0.263277 | 0.91214 | 0.039939 |
| ENSMUSG00000028345.16 | 36.54008 | -1.69376 | 1.502966 | -1.12694 | 0.259766 | 0.91214 | 0.039939 |
| ENSMUSG00000028790.14 | 97.73691 | 0.896236 | 0.795901 | 1.126065 | 0.260138 | 0.91214 | 0.039939 |
| ENSMUSG00000028803.19 | 26.90426 | -2.17249 | 1.915329 | -1.13427 | 0.256683 | 0.91214 | 0.039939 |
| ENSMUSG00000028932.9 | 644.741 | -0.52859 | 0.469921 | -1.12486 | 0.26065 | 0.91214 | 0.039939 |
| ENSMUSG00000029064.16 | 219.4503 | 0.765211 | 0.679165 | 1.126693 | 0.259872 | 0.91214 | 0.039939 |
| ENSMUSG00000029175.18 | 82.0063 | 1.41487 | 1.260239 | 1.1227 | 0.261565 | 0.91214 | 0.039939 |
| ENSMUSG00000029504.6 | 59.42198 | -1.4208 | 1.262225 | -1.12563 | 0.260322 | 0.91214 | 0.039939 |
| ENSMUSG00000030042.16 | 306.1474 | 0.606173 | 0.537525 | 1.127711 | 0.259442 | 0.91214 | 0.039939 |
| ENSMUSG00000030061.17 | 250.2267 | -0.54685 | 0.489811 | -1.11646 | 0.264226 | 0.91214 | 0.039939 |
| ENSMUSG00000030509.18 | 61.83011 | 1.352845 | 1.195326 | 1.131779 | 0.257727 | 0.91214 | 0.039939 |
| ENSMUSG00000030649.19 | 11.40577 | -3.02463 | 2.693951 | -1.12275 | 0.261544 | 0.91214 | 0.039939 |
| ENSMUSG00000031105.17 | 37.88861 | 1.327314 | 1.189619 | 1.115747 | 0.26453 | 0.91214 | 0.039939 |
| ENSMUSG00000031133.13 | 10.7134 | -2.82809 | 2.497675 | -1.13229 | 0.257514 | 0.91214 | 0.039939 |
| ENSMUSG00000031681.17 | 194.2727 | 0.903368 | 0.811664 | 1.112983 | 0.265716 | 0.91214 | 0.039939 |
| ENSMUSG00000032120.12 | 54.80468 | -1.5044 | 1.342376 | -1.1207 | 0.262415 | 0.91214 | 0.039939 |
| ENSMUSG00000032384.17 | 12.66826 | -2.61583 | 2.321077 | -1.12699 | 0.259746 | 0.91214 | 0.039939 |
| ENSMUSG00000032485.15 | 87.88948 | -1.23417 | 1.09989 | -1.12208 | 0.261827 | 0.91214 | 0.039939 |
| ENSMUSG00000033350.8 | 12.23579 | -3.47318 | 3.108634 | -1.11727 | 0.26388 | 0.91214 | 0.039939 |
| ENSMUSG00000034321.15 | 37.5715 | 1.360079 | 1.211066 | 1.123043 | 0.261419 | 0.91214 | 0.039939 |
| ENSMUSG00000034708.13 | 420.9568 | 0.433974 | 0.390244 | 1.11206 | 0.266113 | 0.91214 | 0.039939 |
| ENSMUSG00000036980.14 | 24.21531 | 1.612325 | 1.424685 | 1.131707 | 0.257758 | 0.91214 | 0.039939 |
| ENSMUSG00000037296.8 | 175.3025 | -0.73884 | 0.660971 | -1.11781 | 0.263649 | 0.91214 | 0.039939 |
| ENSMUSG00000037364.13 | 78.67201 | -1.40885 | 1.244268 | -1.13227 | 0.257521 | 0.91214 | 0.039939 |
| ENSMUSG00000038500.16 | 20.78018 | 1.894246 | 1.676698 | 1.129748 | 0.258583 | 0.91214 | 0.039939 |
| ENSMUSG00000038522.8 | 437.8901 | 0.499515 | 0.449181 | 1.112056 | 0.266114 | 0.91214 | 0.039939 |

| | | | | | | | |
|---|---|---|---|---|---|---|---|
| ENSMUSG00000039087.18 | 203.3666 | -0.69938 | 0.625775 | -1.11762 | 0.263729 | 0.91214 | 0.039939 |
| ENSMUSG00000039841.15 | 21.08255 | -1.77185 | 1.592009 | -1.11297 | 0.265723 | 0.91214 | 0.039939 |
| ENSMUSG00000040282.14 | 17.38603 | -2.29059 | 2.046059 | -1.11951 | 0.262922 | 0.91214 | 0.039939 |
| ENSMUSG00000040423.12 | 114.0086 | -0.9371 | 0.836509 | -1.12025 | 0.262609 | 0.91214 | 0.039939 |
| ENSMUSG00000040481.18 | 233.7388 | -0.65011 | 0.576367 | -1.12794 | 0.259345 | 0.91214 | 0.039939 |
| ENSMUSG00000040891.7 | 86.59001 | -1.26265 | 1.118279 | -1.1291 | 0.258856 | 0.91214 | 0.039939 |
| ENSMUSG00000041650.16 | 212.6056 | 0.693666 | 0.623799 | 1.112003 | 0.266137 | 0.91214 | 0.039939 |
| ENSMUSG00000041702.8 | 26.56326 | -1.88579 | 1.692512 | -1.11419 | 0.265196 | 0.91214 | 0.039939 |
| ENSMUSG00000042055.14 | 105.3738 | -0.8397 | 0.753953 | -1.11373 | 0.265393 | 0.91214 | 0.039939 |
| ENSMUSG00000042647.14 | 20.90983 | -1.6351 | 1.453831 | -1.12468 | 0.260724 | 0.91214 | 0.039939 |
| ENSMUSG00000043241.15 | 152.6999 | -0.6956 | 0.623178 | -1.11621 | 0.264333 | 0.91214 | 0.039939 |
| ENSMUSG00000045287.7 | 34.89632 | 1.466637 | 1.305947 | 1.123044 | 0.261419 | 0.91214 | 0.039939 |
| ENSMUSG00000045314.6 | 20.46318 | 1.944996 | 1.745378 | 1.114369 | 0.265121 | 0.91214 | 0.039939 |
| ENSMUSG00000046785.9 | 109.6646 | 0.964979 | 0.850136 | 1.135087 | 0.256339 | 0.91214 | 0.039939 |
| ENSMUSG00000048234.14 | 61.06887 | 1.303859 | 1.15919 | 1.124803 | 0.260673 | 0.91214 | 0.039939 |
| ENSMUSG00000051223.15 | 494.5138 | 0.574757 | 0.512987 | 1.120412 | 0.262538 | 0.91214 | 0.039939 |
| ENSMUSG00000051705.14 | 17.81546 | 2.474625 | 2.207356 | 1.121081 | 0.262253 | 0.91214 | 0.039939 |
| ENSMUSG00000051716.7 | 1951.22 | -0.31603 | 0.279518 | -1.13062 | 0.258215 | 0.91214 | 0.039939 |
| ENSMUSG00000053110.14 | 148.6545 | 0.673441 | 0.604691 | 1.113695 | 0.26541 | 0.91214 | 0.039939 |
| ENSMUSG00000057037.9 | 1302.168 | 0.448065 | 0.39837 | 1.124745 | 0.260697 | 0.91214 | 0.039939 |
| ENSMUSG00000058093.14 | 24.90225 | -1.57171 | 1.406767 | -1.11725 | 0.263887 | 0.91214 | 0.039939 |
| ENSMUSG00000059975.16 | 25.98435 | 2.266518 | 2.028146 | 1.117532 | 0.263767 | 0.91214 | 0.039939 |
| ENSMUSG00000062488.10 | 18.92134 | 2.435569 | 2.156513 | 1.129402 | 0.258728 | 0.91214 | 0.039939 |
| ENSMUSG00000062963.11 | 263.6121 | 0.589594 | 0.521365 | 1.130865 | 0.258112 | 0.91214 | 0.039939 |
| ENSMUSG00000062981.6 | 779.5444 | 0.417461 | 0.372352 | 1.121146 | 0.262226 | 0.91214 | 0.039939 |
| ENSMUSG00000063275.16 | 84.8729 | 0.872672 | 0.774956 | 1.126092 | 0.260127 | 0.91214 | 0.039939 |
| ENSMUSG00000066442.18 | 199.5685 | -0.63449 | 0.56981 | -1.11351 | 0.265491 | 0.91214 | 0.039939 |
| ENSMUSG00000067194.7 | 208.4869 | 0.611419 | 0.544273 | 1.123369 | 0.261281 | 0.91214 | 0.039939 |
| ENSMUSG00000069094.13 | 30.62856 | 1.411458 | 1.245319 | 1.13341 | 0.257042 | 0.91214 | 0.039939 |
| ENSMUSG00000069792.6 | 305.637 | 0.775335 | 0.688601 | 1.125958 | 0.260183 | 0.91214 | 0.039939 |
| ENSMUSG00000078683.10 | 41.40791 | -1.46406 | 1.30201 | -1.12446 | 0.260816 | 0.91214 | 0.039939 |
| ENSMUSG00000078685.7 | 25.57624 | 2.329277 | 2.052062 | 1.135091 | 0.256337 | 0.91214 | 0.039939 |
| ENSMUSG00000085400.2 | 11.20944 | 3.172331 | 2.851613 | 1.112469 | 0.265936 | 0.91214 | 0.039939 |
| ENSMUSG00000090150.11 | 1053.323 | 0.342852 | 0.302531 | 1.133278 | 0.257097 | 0.91214 | 0.039939 |
| ENSMUSG00000096210.3 | 548.5738 | 0.486172 | 0.430415 | 1.129541 | 0.25867 | 0.91214 | 0.039939 |
| ENSMUSG00000097290.10 | 35.63712 | 1.482471 | 1.322242 | 1.121179 | 0.262212 | 0.91214 | 0.039939 |
| ENSMUSG00000102827.2 | 13.31224 | -2.8146 | 2.5294 | -1.11275 | 0.265814 | 0.91214 | 0.039939 |
| ENSMUSG00000105879.2 | 39.27764 | 1.171515 | 1.048872 | 1.116929 | 0.264025 | 0.91214 | 0.039939 |
| ENSMUSG00000106788.2 | 20.88236 | 2.400372 | 2.11625 | 1.134257 | 0.256687 | 0.91214 | 0.039939 |
| ENSMUSG00000019814.6 | 147.0178 | 0.745523 | 0.670893 | 1.11124 | 0.266465 | 0.912401 | 0.039814 |
| ENSMUSG00000068823.11 | 758.4032 | 0.430983 | 0.38787 | 1.111154 | 0.266502 | 0.912401 | 0.039814 |
| ENSMUSG00000109118.2 | 12.32792 | 2.527562 | 2.274584 | 1.11122 | 0.266474 | 0.912401 | 0.039814 |
| ENSMUSG00000042790.17 | 34.04651 | -1.63532 | 1.472831 | -1.11032 | 0.266859 | 0.913132 | 0.039466 |
| ENSMUSG00000063049.5 | 98.35242 | 1.012013 | 0.911551 | 1.110211 | 0.266908 | 0.913132 | 0.039466 |

| | | | | | | | |
|---|---|---|---|---|---|---|---|
| ENSMUSG00000029551.14 | 71.95347 | 1.245237 | 1.122229 | 1.109611 | 0.267167 | 0.913358 | 0.039359 |
| ENSMUSG00000054455.6 | 198.6841 | 0.742546 | 0.669138 | 1.109705 | 0.267126 | 0.913358 | 0.039359 |
| ENSMUSG00000020850.15 | 131.063 | 0.889465 | 0.803502 | 1.106985 | 0.2683 | 0.913374 | 0.039351 |
| ENSMUSG00000024843.16 | 65.22201 | 1.053463 | 0.951654 | 1.106981 | 0.268302 | 0.913374 | 0.039351 |
| ENSMUSG00000026185.9 | 82.46371 | 0.939677 | 0.849053 | 1.106735 | 0.268409 | 0.913374 | 0.039351 |
| ENSMUSG00000028037.14 | 37.62394 | -1.64425 | 1.485105 | -1.10716 | 0.268224 | 0.913374 | 0.039351 |
| ENSMUSG00000028907.8 | 127.5949 | -0.93459 | 0.842912 | -1.10876 | 0.267533 | 0.913374 | 0.039351 |
| ENSMUSG00000029681.13 | 92.89212 | 0.918947 | 0.829494 | 1.10784 | 0.267931 | 0.913374 | 0.039351 |
| ENSMUSG00000039914.7 | 30.64794 | 1.759535 | 1.58934 | 1.107086 | 0.268257 | 0.913374 | 0.039351 |
| ENSMUSG00000041841.9 | 2934.884 | 0.295739 | 0.26702 | 1.107554 | 0.268054 | 0.913374 | 0.039351 |
| ENSMUSG00000042569.14 | 78.75737 | -1.68473 | 1.519938 | -1.10842 | 0.267681 | 0.913374 | 0.039351 |
| ENSMUSG00000061286.8 | 18.84787 | -2.29253 | 2.066802 | -1.10922 | 0.267336 | 0.913374 | 0.039351 |
| ENSMUSG00000063334.17 | 165.8054 | 1.039466 | 0.93718 | 1.109142 | 0.267369 | 0.913374 | 0.039351 |
| ENSMUSG00000078139.4 | 44.55152 | 1.09223 | 0.985158 | 1.108686 | 0.267566 | 0.913374 | 0.039351 |
| ENSMUSG00000078689.9 | 26.16248 | -1.44515 | 1.305824 | -1.1067 | 0.268424 | 0.913374 | 0.039351 |
| ENSMUSG00000024145.7 | 21.19667 | 1.935514 | 1.749539 | 1.106299 | 0.268597 | 0.913635 | 0.039227 |
| ENSMUSG00000022108.9 | 8558.844 | 0.24006 | 0.217151 | 1.105496 | 0.268945 | 0.91449 | 0.038821 |
| ENSMUSG00000001313.13 | 32.04748 | -2.02008 | 1.836972 | -1.09968 | 0.271473 | 0.915724 | 0.038236 |
| ENSMUSG00000001911.17 | 159.5944 | -0.60154 | 0.546967 | -1.09978 | 0.271428 | 0.915724 | 0.038236 |
| ENSMUSG00000008153.13 | 17.41148 | 2.086578 | 1.891698 | 1.103019 | 0.270019 | 0.915724 | 0.038236 |
| ENSMUSG00000018411.18 | 11.91684 | 2.964842 | 2.691109 | 1.101718 | 0.270584 | 0.915724 | 0.038236 |
| ENSMUSG00000020376.18 | 363.9908 | 0.563736 | 0.510646 | 1.103967 | 0.269608 | 0.915724 | 0.038236 |
| ENSMUSG00000021460.18 | 275.344 | -0.49598 | 0.44957 | -1.10323 | 0.269929 | 0.915724 | 0.038236 |
| ENSMUSG00000021786.14 | 209.5635 | -0.62061 | 0.564011 | -1.10036 | 0.271177 | 0.915724 | 0.038236 |
| ENSMUSG00000021892.15 | 334.2269 | 0.539075 | 0.49023 | 1.099636 | 0.271491 | 0.915724 | 0.038236 |
| ENSMUSG00000022718.13 | 19.11143 | 1.95126 | 1.771777 | 1.101301 | 0.270766 | 0.915724 | 0.038236 |
| ENSMUSG00000024962.14 | 21.54062 | 2.235623 | 2.028773 | 1.101958 | 0.27048 | 0.915724 | 0.038236 |
| ENSMUSG00000025544.14 | 500.7066 | 0.488666 | 0.444326 | 1.09979 | 0.271423 | 0.915724 | 0.038236 |
| ENSMUSG00000025962.16 | 87.32328 | -0.92452 | 0.840728 | -1.09967 | 0.271476 | 0.915724 | 0.038236 |
| ENSMUSG00000026072.13 | 53.8305 | -1.11275 | 1.00982 | -1.10193 | 0.270492 | 0.915724 | 0.038236 |
| ENSMUSG00000027602.10 | 30.46955 | 1.534273 | 1.391989 | 1.102216 | 0.270368 | 0.915724 | 0.038236 |
| ENSMUSG00000027624.21 | 13.54703 | -3.30588 | 2.997484 | -1.10289 | 0.270077 | 0.915724 | 0.038236 |
| ENSMUSG00000028224.15 | 130.1147 | 0.770787 | 0.698752 | 1.103092 | 0.269987 | 0.915724 | 0.038236 |
| ENSMUSG00000029195.11 | 62.27317 | -1.14952 | 1.04545 | -1.09955 | 0.271529 | 0.915724 | 0.038236 |
| ENSMUSG00000030830.19 | 42.81481 | -1.75687 | 1.595184 | -1.10136 | 0.27074 | 0.915724 | 0.038236 |
| ENSMUSG00000034473.15 | 106.0391 | 0.821108 | 0.746097 | 1.100538 | 0.271098 | 0.915724 | 0.038236 |
| ENSMUSG00000038217.14 | 1150.525 | 0.400291 | 0.362575 | 1.104022 | 0.269584 | 0.915724 | 0.038236 |
| ENSMUSG00000055319.9 | 134.9512 | 0.694811 | 0.631138 | 1.100886 | 0.270946 | 0.915724 | 0.038236 |
| ENSMUSG00000061544.14 | 14.55389 | 2.487593 | 2.253756 | 1.103754 | 0.2697 | 0.915724 | 0.038236 |
| ENSMUSG00000117284.2 | 48.08734 | 1.285712 | 1.166903 | 1.101816 | 0.270542 | 0.915724 | 0.038236 |
| ENSMUSG00000058603.11 | 66.16922 | 0.988227 | 0.899336 | 1.098841 | 0.271838 | 0.916437 | 0.037897 |
| ENSMUSG00000079614.5 | 179.5244 | -0.64411 | 0.586517 | -1.09819 | 0.272121 | 0.917066 | 0.0376 |
| ENSMUSG00000035699.9 | 15.94002 | -2.4418 | 2.224187 | -1.09784 | 0.272274 | 0.917257 | 0.037509 |
| ENSMUSG00000009207.16 | 18.67933 | 2.38121 | 2.174567 | 1.095027 | 0.273505 | 0.918872 | 0.036745 |

| | | | | | | | |
|---|---|---|---|---|---|---|---|
| ENSMUSG00000010914.11 | 182.1455 | -0.62568 | 0.571756 | -1.09431 | 0.273818 | 0.918872 | 0.036745 |
| ENSMUSG00000016409.5 | 114.0339 | -1.2361 | 1.128197 | -1.09564 | 0.273236 | 0.918872 | 0.036745 |
| ENSMUSG00000018449.13 | 52.89502 | -1.02991 | 0.941155 | -1.09431 | 0.27382 | 0.918872 | 0.036745 |
| ENSMUSG00000023106.16 | 156.6575 | 0.820564 | 0.748961 | 1.095603 | 0.273253 | 0.918872 | 0.036745 |
| ENSMUSG00000024143.16 | 37.78345 | 1.563984 | 1.428732 | 1.094666 | 0.273663 | 0.918872 | 0.036745 |
| ENSMUSG00000027447.7 | 933.8877 | 0.364484 | 0.333028 | 1.094452 | 0.273757 | 0.918872 | 0.036745 |
| ENSMUSG00000031556.7 | 400.3549 | 0.445737 | 0.407125 | 1.09484 | 0.273587 | 0.918872 | 0.036745 |
| ENSMUSG00000031753.17 | 95.90976 | -0.79807 | 0.728021 | -1.09622 | 0.272981 | 0.918872 | 0.036745 |
| ENSMUSG00000034602.16 | 276.8705 | 0.641287 | 0.585853 | 1.094621 | 0.273683 | 0.918872 | 0.036745 |
| ENSMUSG00000039438.8 | 564.7619 | -0.68864 | 0.628057 | -1.09645 | 0.27288 | 0.918872 | 0.036745 |
| ENSMUSG00000018442.14 | 114.1802 | -0.81566 | 0.746253 | -1.09301 | 0.274388 | 0.920126 | 0.036153 |
| ENSMUSG00000025246.14 | 127.3213 | -1.27204 | 1.16371 | -1.09309 | 0.274355 | 0.920126 | 0.036153 |
| ENSMUSG00000021338.18 | 12.04021 | 2.410021 | 2.206568 | 1.092203 | 0.274744 | 0.920756 | 0.035855 |
| ENSMUSG00000033918.17 | 291.6564 | 0.538421 | 0.492995 | 1.092143 | 0.27477 | 0.920756 | 0.035855 |
| ENSMUSG00000033307.8 | 596.7712 | 0.47129 | 0.431651 | 1.091831 | 0.274907 | 0.920891 | 0.035792 |
| ENSMUSG00000006005.19 | 558.0939 | 0.361405 | 0.331626 | 1.089796 | 0.275803 | 0.922689 | 0.034944 |
| ENSMUSG00000018143.11 | 22.27039 | -1.85536 | 1.703359 | -1.08924 | 0.276049 | 0.922689 | 0.034944 |
| ENSMUSG00000022973.19 | 64.97337 | -1.17223 | 1.076319 | -1.08911 | 0.276105 | 0.922689 | 0.034944 |
| ENSMUSG00000031232.18 | 709.5441 | 0.371559 | 0.341173 | 1.089064 | 0.276126 | 0.922689 | 0.034944 |
| ENSMUSG00000032698.16 | 30.96401 | 1.413948 | 1.298245 | 1.089123 | 0.2761 | 0.922689 | 0.034944 |
| ENSMUSG00000039159.17 | 148.0555 | -0.78001 | 0.715673 | -1.0899 | 0.275756 | 0.922689 | 0.034944 |
| ENSMUSG00000061947.11 | 998.7331 | -0.41693 | 0.382431 | -1.09022 | 0.275616 | 0.922689 | 0.034944 |
| ENSMUSG00000079045.4 | 13.19674 | -2.31492 | 2.126273 | -1.08872 | 0.276276 | 0.922868 | 0.03486 |
| ENSMUSG00000032328.13 | 1747.183 | 0.346896 | 0.318823 | 1.088052 | 0.276572 | 0.92353 | 0.034549 |
| ENSMUSG00000048285.10 | 33.22942 | 1.772567 | 1.630271 | 1.087283 | 0.276912 | 0.924339 | 0.034169 |
| ENSMUSG00000078964.3 | 305.8456 | 0.488474 | 0.449455 | 1.086815 | 0.277119 | 0.924705 | 0.033997 |
| ENSMUSG00000002580.8 | 403.5503 | 0.478943 | 0.441486 | 1.084843 | 0.277991 | 0.925902 | 0.033435 |
| ENSMUSG00000006906.11 | 36.36282 | 1.251351 | 1.155616 | 1.082843 | 0.278878 | 0.925902 | 0.033435 |
| ENSMUSG00000016503.19 | 19.62661 | 2.541054 | 2.34462 | 1.083781 | 0.278462 | 0.925902 | 0.033435 |
| ENSMUSG00000016918.16 | 21.87495 | 2.018215 | 1.862733 | 1.08347 | 0.2786 | 0.925902 | 0.033435 |
| ENSMUSG00000022445.8 | 1800.255 | 0.301406 | 0.277792 | 1.085007 | 0.277919 | 0.925902 | 0.033435 |
| ENSMUSG00000025016.12 | 791.9251 | -0.32546 | 0.300054 | -1.08467 | 0.278069 | 0.925902 | 0.033435 |
| ENSMUSG00000025491.15 | 33.45921 | 1.601429 | 1.479067 | 1.082729 | 0.278929 | 0.925902 | 0.033435 |
| ENSMUSG00000026792.17 | 26.12709 | -2.04224 | 1.884189 | -1.08388 | 0.278417 | 0.925902 | 0.033435 |
| ENSMUSG00000028402.19 | 117.6258 | 0.786765 | 0.725397 | 1.0846 | 0.278099 | 0.925902 | 0.033435 |
| ENSMUSG00000030795.19 | 137.6938 | 0.829744 | 0.766076 | 1.083108 | 0.27876 | 0.925902 | 0.033435 |
| ENSMUSG00000031731.17 | 208.9669 | -0.61283 | 0.564856 | -1.08494 | 0.27795 | 0.925902 | 0.033435 |
| ENSMUSG00000033047.6 | 510.1084 | -0.43567 | 0.402392 | -1.0827 | 0.278943 | 0.925902 | 0.033435 |
| ENSMUSG00000050335.18 | 43.34556 | 1.256981 | 1.160595 | 1.083049 | 0.278787 | 0.925902 | 0.033435 |
| ENSMUSG00000063015.13 | 646.0204 | 0.443882 | 0.409826 | 1.0831 | 0.278764 | 0.925902 | 0.033435 |
| ENSMUSG00000074918.5 | 60.88385 | -1.38633 | 1.278912 | -1.08399 | 0.278369 | 0.925902 | 0.033435 |
| ENSMUSG00000032405.11 | 48.94412 | 1.310207 | 1.210481 | 1.082386 | 0.279081 | 0.926037 | 0.033372 |
| ENSMUSG00000021514.19 | 20.69839 | 2.17437 | 2.009807 | 1.08188 | 0.279306 | 0.926225 | 0.033283 |
| ENSMUSG00000022885.17 | 186.4812 | -0.67922 | 0.62794 | -1.08167 | 0.2794 | 0.926225 | 0.033283 |

| | | | | | | | |
|---|---|---|---|---|---|---|---|
| ENSMUSG00000024663.21 | 15.30667 | 2.718621 | 2.515424 | 1.08078 | 0.279795 | 0.926225 | 0.033283 |
| ENSMUSG00000026890.20 | 26.26999 | -2.42541 | 2.243598 | -1.08104 | 0.27968 | 0.926225 | 0.033283 |
| ENSMUSG00000031023.6 | 144.6702 | 0.927454 | 0.8578 | 1.081201 | 0.279608 | 0.926225 | 0.033283 |
| ENSMUSG00000036430.9 | 18.56633 | -2.08498 | 1.929252 | -1.08072 | 0.279822 | 0.926225 | 0.033283 |
| ENSMUSG00000040213.14 | 918.0926 | -0.34509 | 0.319105 | -1.08142 | 0.279512 | 0.926225 | 0.033283 |
| ENSMUSG00000031231.5 | 5707.901 | 0.281209 | 0.260316 | 1.080258 | 0.280027 | 0.926582 | 0.033116 |
| ENSMUSG00000031950.8 | 218.81 | 0.628422 | 0.58209 | 1.079596 | 0.280322 | 0.927234 | 0.03281 |
| ENSMUSG00000022200.9 | 280.5432 | -0.54195 | 0.50236 | -1.0788 | 0.280676 | 0.927314 | 0.032773 |
| ENSMUSG00000025538.16 | 17.62183 | 1.919537 | 1.779406 | 1.078752 | 0.280698 | 0.927314 | 0.032773 |
| ENSMUSG00000029474.8 | 56.72271 | -1.16009 | 1.075489 | -1.07866 | 0.280738 | 0.927314 | 0.032773 |
| ENSMUSG00000056216.10 | 261.6449 | -0.52346 | 0.485115 | -1.07904 | 0.280571 | 0.927314 | 0.032773 |
| ENSMUSG00000048368.10 | 47.22242 | 1.432287 | 1.32888 | 1.077816 | 0.281116 | 0.927965 | 0.032468 |
| ENSMUSG00000051007.16 | 55.73337 | 1.307242 | 1.212899 | 1.077783 | 0.28113 | 0.927965 | 0.032468 |
| ENSMUSG00000024590.9 | 33.35389 | -1.81619 | 1.686145 | -1.07712 | 0.281425 | 0.92829 | 0.032316 |
| ENSMUSG00000030056.9 | 109.9184 | 0.869378 | 0.807027 | 1.077261 | 0.281364 | 0.92829 | 0.032316 |
| ENSMUSG00000030204.11 | 110.9968 | 0.861539 | 0.800058 | 1.076846 | 0.281549 | 0.928378 | 0.032275 |
| ENSMUSG00000041164.16 | 12.9715 | 2.480222 | 2.303919 | 1.076523 | 0.281693 | 0.928531 | 0.032204 |
| ENSMUSG00000021555.16 | 267.9661 | -0.56412 | 0.524137 | -1.07629 | 0.2818 | 0.928558 | 0.032191 |
| ENSMUSG00000073616.11 | 552.6042 | -0.37235 | 0.346194 | -1.07555 | 0.282127 | 0.929314 | 0.031837 |
| ENSMUSG00000004815.13 | 17.97068 | 1.883933 | 1.75441 | 1.073827 | 0.2829 | 0.930009 | 0.031513 |
| ENSMUSG00000014905.5 | 779.699 | 0.616426 | 0.573771 | 1.074342 | 0.282669 | 0.930009 | 0.031513 |
| ENSMUSG00000015247.11 | 587.3806 | 0.436898 | 0.406962 | 1.073558 | 0.283021 | 0.930009 | 0.031513 |
| ENSMUSG00000020361.14 | 753.1588 | -0.35723 | 0.332756 | -1.07355 | 0.283025 | 0.930009 | 0.031513 |
| ENSMUSG00000031885.15 | 130.8106 | 0.931683 | 0.867511 | 1.073973 | 0.282835 | 0.930009 | 0.031513 |
| ENSMUSG00000047879.18 | 278.9546 | -0.58312 | 0.542535 | -1.07481 | 0.282462 | 0.930009 | 0.031513 |
| ENSMUSG00000066319.7 | 193.2597 | 0.672762 | 0.626469 | 1.073896 | 0.282869 | 0.930009 | 0.031513 |
| ENSMUSG00000028645.12 | 22.21677 | -1.73861 | 1.620017 | -1.0732 | 0.28318 | 0.930196 | 0.031426 |
| ENSMUSG00000002289.17 | 155.2369 | -0.74847 | 0.697927 | -1.07242 | 0.28353 | 0.930271 | 0.031391 |
| ENSMUSG00000016495.13 | 34.0098 | -2.24161 | 2.090396 | -1.07234 | 0.283567 | 0.930271 | 0.031391 |
| ENSMUSG00000021963.18 | 166.4987 | 0.599358 | 0.558844 | 1.072496 | 0.283497 | 0.930271 | 0.031391 |
| ENSMUSG00000116506.2 | 27.64029 | -1.57966 | 1.473178 | -1.07228 | 0.283595 | 0.930271 | 0.031391 |
| ENSMUSG00000001151.11 | 25.96884 | 1.693752 | 1.58155 | 1.070944 | 0.284194 | 0.930414 | 0.031324 |
| ENSMUSG00000021275.17 | 12.06088 | -2.88629 | 2.702947 | -1.06783 | 0.285596 | 0.930414 | 0.031324 |
| ENSMUSG00000021417.16 | 1161.243 | 0.35297 | 0.330472 | 1.068078 | 0.285485 | 0.930414 | 0.031324 |
| ENSMUSG00000022131.4 | 204.0124 | 0.559109 | 0.524049 | 1.066903 | 0.286016 | 0.930414 | 0.031324 |
| ENSMUSG00000025413.14 | 70.23064 | -1.32202 | 1.239554 | -1.06653 | 0.286184 | 0.930414 | 0.031324 |
| ENSMUSG00000025745.13 | 436.7393 | 0.564029 | 0.528485 | 1.067256 | 0.285856 | 0.930414 | 0.031324 |
| ENSMUSG00000026273.14 | 64.34667 | -1.37278 | 1.28107 | -1.07159 | 0.283904 | 0.930414 | 0.031324 |
| ENSMUSG00000026361.10 | 127.8445 | -0.73331 | 0.687501 | -1.06664 | 0.286135 | 0.930414 | 0.031324 |
| ENSMUSG00000027746.14 | 383.4398 | 0.455002 | 0.426743 | 1.06622 | 0.286324 | 0.930414 | 0.031324 |
| ENSMUSG00000028759.14 | 384.1764 | -0.5699 | 0.534477 | -1.06628 | 0.286298 | 0.930414 | 0.031324 |
| ENSMUSG00000029406.16 | 43.0193 | -1.14408 | 1.06738 | -1.07185 | 0.283786 | 0.930414 | 0.031324 |
| ENSMUSG00000029802.14 | 713.56 | 0.374489 | 0.35104 | 1.066799 | 0.286062 | 0.930414 | 0.031324 |
| ENSMUSG00000030352.16 | 13.82095 | -2.47254 | 2.315913 | -1.06763 | 0.285688 | 0.930414 | 0.031324 |

| | | | | | | | |
|---|---|---|---|---|---|---|---|
| ENSMUSG00000030852.19 | 49.69659 | -1.78074 | 1.662856 | -1.07089 | 0.284219 | 0.930414 | 0.031324 |
| ENSMUSG00000032534.19 | 17.89752 | -2.4274 | 2.27231 | -1.06825 | 0.285407 | 0.930414 | 0.031324 |
| ENSMUSG00000032594.12 | 45.68083 | -1.99174 | 1.859583 | -1.07107 | 0.284138 | 0.930414 | 0.031324 |
| ENSMUSG00000032624.17 | 31.5824 | 1.336222 | 1.247835 | 1.070833 | 0.284245 | 0.930414 | 0.031324 |
| ENSMUSG00000033760.18 | 73.38975 | -1.18546 | 1.110305 | -1.06769 | 0.285661 | 0.930414 | 0.031324 |
| ENSMUSG00000038150.8 | 181.6801 | 0.662386 | 0.620062 | 1.068258 | 0.285404 | 0.930414 | 0.031324 |
| ENSMUSG00000038178.17 | 14.96495 | 2.417953 | 2.267029 | 1.066573 | 0.286165 | 0.930414 | 0.031324 |
| ENSMUSG00000040028.11 | 175.5582 | 0.656967 | 0.616371 | 1.065864 | 0.286485 | 0.930414 | 0.031324 |
| ENSMUSG00000044098.15 | 21.6651 | 1.714464 | 1.603017 | 1.069523 | 0.284834 | 0.930414 | 0.031324 |
| ENSMUSG00000045503.17 | 270.4523 | 0.539282 | 0.504297 | 1.069375 | 0.284901 | 0.930414 | 0.031324 |
| ENSMUSG00000047394.10 | 38.73042 | -1.59453 | 1.495889 | -1.06594 | 0.286449 | 0.930414 | 0.031324 |
| ENSMUSG00000060012.10 | 111.4774 | 0.771222 | 0.720769 | 1.069999 | 0.28462 | 0.930414 | 0.031324 |
| ENSMUSG00000068463.5 | 581.5543 | 0.399874 | 0.373901 | 1.069466 | 0.28486 | 0.930414 | 0.031324 |
| ENSMUSG00000078578.10 | 1937.659 | 0.296725 | 0.277657 | 1.068676 | 0.285216 | 0.930414 | 0.031324 |
| ENSMUSG00000097121.3 | 144.3216 | -0.86124 | 0.806596 | -1.06775 | 0.285633 | 0.930414 | 0.031324 |
| ENSMUSG00000102095.2 | 51.46339 | 1.508878 | 1.414246 | 1.066913 | 0.286011 | 0.930414 | 0.031324 |
| ENSMUSG00000025464.16 | 98.10339 | -1.08457 | 1.017904 | -1.06549 | 0.286653 | 0.930641 | 0.031218 |
| ENSMUSG00000003812.14 | 70.79432 | 0.924125 | 0.868284 | 1.064312 | 0.287188 | 0.931379 | 0.030873 |
| ENSMUSG00000024477.5 | 208.3196 | 0.582122 | 0.546712 | 1.064768 | 0.286981 | 0.931379 | 0.030873 |
| ENSMUSG00000026229.18 | 561.0731 | -0.39144 | 0.367702 | -1.06455 | 0.287079 | 0.931379 | 0.030873 |
| ENSMUSG00000059456.14 | 29.97422 | -1.90279 | 1.788133 | -1.06412 | 0.287273 | 0.931379 | 0.030873 |
| ENSMUSG00000027997.10 | 92.14249 | -1.17779 | 1.107106 | -1.06385 | 0.287397 | 0.93144 | 0.030845 |
| ENSMUSG00000063801.8 | 53.76283 | 1.578818 | 1.484408 | 1.063601 | 0.287509 | 0.93144 | 0.030845 |
| ENSMUSG00000097061.3 | 15.98077 | -2.19213 | 2.061381 | -1.06343 | 0.287587 | 0.93144 | 0.030845 |
| ENSMUSG00000003824.13 | 33.82399 | 1.779792 | 1.676226 | 1.061785 | 0.288333 | 0.931629 | 0.030757 |
| ENSMUSG00000021770.12 | 140.4427 | -0.70626 | 0.664916 | -1.06217 | 0.288157 | 0.931629 | 0.030757 |
| ENSMUSG00000024758.16 | 354.653 | -0.44534 | 0.419394 | -1.06188 | 0.288292 | 0.931629 | 0.030757 |
| ENSMUSG00000028716.16 | 11.78375 | 2.941807 | 2.769147 | 1.062352 | 0.288076 | 0.931629 | 0.030757 |
| ENSMUSG00000032179.8 | 36.98758 | 1.295727 | 1.219497 | 1.062509 | 0.288004 | 0.931629 | 0.030757 |
| ENSMUSG00000032776.10 | 68.63375 | -1.11246 | 1.047427 | -1.06209 | 0.288196 | 0.931629 | 0.030757 |
| ENSMUSG00000054428.13 | 184.6532 | -0.72605 | 0.683752 | -1.06186 | 0.2883 | 0.931629 | 0.030757 |
| ENSMUSG00000017561.17 | 45.96619 | -1.82985 | 1.725921 | -1.06021 | 0.289048 | 0.93226 | 0.030463 |
| ENSMUSG00000019809.17 | 171.8984 | -0.74852 | 0.706549 | -1.05941 | 0.289414 | 0.93226 | 0.030463 |
| ENSMUSG00000019970.16 | 118.9068 | 0.869055 | 0.8194 | 1.0606 | 0.288872 | 0.93226 | 0.030463 |
| ENSMUSG00000027349.6 | 49.56141 | -1.08699 | 1.025492 | -1.05997 | 0.28916 | 0.93226 | 0.030463 |
| ENSMUSG00000029338.14 | 66.56636 | -1.10076 | 1.038157 | -1.0603 | 0.289007 | 0.93226 | 0.030463 |
| ENSMUSG00000036330.13 | 39.6588 | -1.69287 | 1.595875 | -1.06078 | 0.288792 | 0.93226 | 0.030463 |
| ENSMUSG00000075327.11 | 19.33955 | -2.48151 | 2.341656 | -1.05972 | 0.289271 | 0.93226 | 0.030463 |
| ENSMUSG00000096145.3 | 1462.198 | 0.323562 | 0.305204 | 1.060149 | 0.289077 | 0.93226 | 0.030463 |
| ENSMUSG00000115358.2 | 83.32004 | 0.819898 | 0.773896 | 1.059442 | 0.289399 | 0.93226 | 0.030463 |
| ENSMUSG00000021710.12 | 152.5654 | -0.71802 | 0.678391 | -1.05842 | 0.289865 | 0.933255 | 0.03 |
| ENSMUSG00000027166.15 | 48.35837 | -1.49748 | 1.415182 | -1.05815 | 0.289985 | 0.933255 | 0.03 |
| ENSMUSG00000043284.13 | 117.6979 | 0.778757 | 0.736008 | 1.058082 | 0.290018 | 0.933255 | 0.03 |
| ENSMUSG00000006050.13 | 720.0926 | 0.371585 | 0.352541 | 1.054018 | 0.291875 | 0.933506 | 0.029883 |

| | | | | | | | |
|---|---|---|---|---|---|---|---|
| ENSMUSG00000019802.14 | 1114.768 | -0.33832 | 0.320911 | -1.05426 | 0.291763 | 0.933506 | 0.029883 |
| ENSMUSG00000019868.17 | 140.5653 | 0.725388 | 0.68648 | 1.056677 | 0.290659 | 0.933506 | 0.029883 |
| ENSMUSG00000021638.13 | 61.81209 | -1.1149 | 1.058965 | -1.05282 | 0.292422 | 0.933506 | 0.029883 |
| ENSMUSG00000022971.19 | 64.83797 | 0.993663 | 0.939606 | 1.057531 | 0.290269 | 0.933506 | 0.029883 |
| ENSMUSG00000026259.15 | 29.08512 | -1.71012 | 1.624777 | -1.05253 | 0.292557 | 0.933506 | 0.029883 |
| ENSMUSG00000027835.12 | 201.229 | -0.63734 | 0.605491 | -1.0526 | 0.292525 | 0.933506 | 0.029883 |
| ENSMUSG00000028822.12 | 970.5828 | -0.35885 | 0.340699 | -1.05329 | 0.292209 | 0.933506 | 0.029883 |
| ENSMUSG00000029028.15 | 34.32551 | -1.19245 | 1.130148 | -1.05513 | 0.291366 | 0.933506 | 0.029883 |
| ENSMUSG00000029762.7 | 22.83973 | 1.77889 | 1.684255 | 1.056188 | 0.290882 | 0.933506 | 0.029883 |
| ENSMUSG00000030035.15 | 13.41562 | 2.742207 | 2.598354 | 1.055363 | 0.291259 | 0.933506 | 0.029883 |
| ENSMUSG00000030447.16 | 320.0256 | -0.55609 | 0.527253 | -1.05469 | 0.291569 | 0.933506 | 0.029883 |
| ENSMUSG00000033596.6 | 27.34868 | -1.98094 | 1.878647 | -1.05445 | 0.291676 | 0.933506 | 0.029883 |
| ENSMUSG00000034216.13 | 23.7537 | -1.95279 | 1.851044 | -1.05497 | 0.29144 | 0.933506 | 0.029883 |
| ENSMUSG00000037847.15 | 187.7958 | 0.663198 | 0.629577 | 1.053403 | 0.292157 | 0.933506 | 0.029883 |
| ENSMUSG00000051510.14 | 64.64958 | -1.10617 | 1.048692 | -1.05481 | 0.291514 | 0.933506 | 0.029883 |
| ENSMUSG00000056025.13 | 12.57984 | -2.47207 | 2.33885 | -1.05696 | 0.290529 | 0.933506 | 0.029883 |
| ENSMUSG00000063904.5 | 82.56437 | 0.819866 | 0.776222 | 1.056226 | 0.290865 | 0.933506 | 0.029883 |
| ENSMUSG00000067851.12 | 438.2314 | 0.496697 | 0.470352 | 1.056011 | 0.290963 | 0.933506 | 0.029883 |
| ENSMUSG00000069895.5 | 22.81689 | -1.79215 | 1.696907 | -1.05612 | 0.290911 | 0.933506 | 0.029883 |
| ENSMUSG00000075543.5 | 47.32071 | 1.672392 | 1.587836 | 1.053252 | 0.292225 | 0.933506 | 0.029883 |
| ENSMUSG00000082065.2 | 35.4275 | -2.22796 | 2.116597 | -1.05261 | 0.292518 | 0.933506 | 0.029883 |
| ENSMUSG00000086859.5 | 10.39829 | 2.562113 | 2.430815 | 1.054014 | 0.291877 | 0.933506 | 0.029883 |
| ENSMUSG00000093661.2 | 20.63423 | -1.58245 | 1.499216 | -1.05552 | 0.291188 | 0.933506 | 0.029883 |
| ENSMUSG00000109509.2 | 207.9282 | 0.648853 | 0.616239 | 1.052924 | 0.292376 | 0.933506 | 0.029883 |
| ENSMUSG00000000561.15 | 108.2755 | -1.19363 | 1.139989 | -1.04705 | 0.295076 | 0.933791 | 0.02975 |
| ENSMUSG00000004931.12 | 21.2771 | 1.983284 | 1.892719 | 1.047849 | 0.294708 | 0.933791 | 0.02975 |
| ENSMUSG00000014850.16 | 81.4299 | 0.943285 | 0.898479 | 1.049868 | 0.293779 | 0.933791 | 0.02975 |
| ENSMUSG00000020107.11 | 154.8644 | 0.702176 | 0.670684 | 1.046956 | 0.29512 | 0.933791 | 0.02975 |
| ENSMUSG00000021340.14 | 584.5069 | -0.51165 | 0.488357 | -1.0477 | 0.294779 | 0.933791 | 0.02975 |
| ENSMUSG00000022139.18 | 266.1912 | 0.518502 | 0.495413 | 1.046605 | 0.295282 | 0.933791 | 0.02975 |
| ENSMUSG00000022338.9 | 1039.221 | 0.343636 | 0.328681 | 1.045501 | 0.295791 | 0.933791 | 0.02975 |
| ENSMUSG00000024074.9 | 74.74617 | -1.20252 | 1.146968 | -1.04843 | 0.29444 | 0.933791 | 0.02975 |
| ENSMUSG00000024217.11 | 32.64989 | -1.54301 | 1.470943 | -1.049 | 0.29418 | 0.933791 | 0.02975 |
| ENSMUSG00000024400.17 | 130.7155 | -0.68379 | 0.653413 | -1.04649 | 0.295334 | 0.933791 | 0.02975 |
| ENSMUSG00000025742.6 | 85.15159 | 0.863107 | 0.824792 | 1.046454 | 0.295351 | 0.933791 | 0.02975 |
| ENSMUSG00000025967.17 | 3160.289 | 0.248406 | 0.237163 | 1.047407 | 0.294912 | 0.933791 | 0.02975 |
| ENSMUSG00000026283.14 | 21.73815 | -2.47434 | 2.3518 | -1.05211 | 0.29275 | 0.933791 | 0.02975 |
| ENSMUSG00000026965.13 | 42.18183 | 1.179654 | 1.125019 | 1.048564 | 0.294379 | 0.933791 | 0.02975 |
| ENSMUSG00000027374.13 | 32.70165 | 1.257503 | 1.195568 | 1.051804 | 0.292889 | 0.933791 | 0.02975 |
| ENSMUSG00000027637.4 | 920.9676 | 0.319844 | 0.305162 | 1.048112 | 0.294587 | 0.933791 | 0.02975 |
| ENSMUSG00000027651.17 | 60.38763 | -1.22218 | 1.169485 | -1.04506 | 0.295996 | 0.933791 | 0.02975 |
| ENSMUSG00000031960.15 | 597.7588 | -0.40719 | 0.389152 | -1.04636 | 0.295397 | 0.933791 | 0.02975 |
| ENSMUSG00000035764.6 | 47.77201 | -1.26069 | 1.205851 | -1.04548 | 0.295801 | 0.933791 | 0.02975 |
| ENSMUSG00000038170.16 | 309.6087 | -0.51718 | 0.493772 | -1.04741 | 0.294913 | 0.933791 | 0.02975 |

| | | | | | | | |
|---|---|---|---|---|---|---|---|
| ENSMUSG00000040560.11 | 58.84774 | -1.13917 | 1.088259 | -1.04678 | 0.295201 | 0.933791 | 0.02975 |
| ENSMUSG00000042312.11 | 191.9949 | 0.558701 | 0.534438 | 1.045399 | 0.295838 | 0.933791 | 0.02975 |
| ENSMUSG00000042688.17 | 93.52583 | -0.84795 | 0.807334 | -1.05031 | 0.293574 | 0.933791 | 0.02975 |
| ENSMUSG00000044147.9 | 66.36165 | -0.88546 | 0.84261 | -1.05085 | 0.293327 | 0.933791 | 0.02975 |
| ENSMUSG00000044768.17 | 112.2516 | -0.75378 | 0.718802 | -1.04867 | 0.294331 | 0.933791 | 0.02975 |
| ENSMUSG00000044986.11 | 496.004 | 0.472525 | 0.450985 | 1.047763 | 0.294748 | 0.933791 | 0.02975 |
| ENSMUSG00000049106.8 | 136.4572 | -0.73282 | 0.700365 | -1.04634 | 0.295406 | 0.933791 | 0.02975 |
| ENSMUSG00000050705.16 | 37.69809 | -1.62837 | 1.557586 | -1.04545 | 0.295817 | 0.933791 | 0.02975 |
| ENSMUSG00000056941.18 | 111.8178 | 0.927754 | 0.887653 | 1.045176 | 0.295941 | 0.933791 | 0.02975 |
| ENSMUSG00000064202.16 | 12.5225 | 2.764278 | 2.638383 | 1.047717 | 0.294769 | 0.933791 | 0.02975 |
| ENSMUSG00000066477.14 | 100.7128 | -0.86884 | 0.829052 | -1.04799 | 0.294641 | 0.933791 | 0.02975 |
| ENSMUSG00000068015.9 | 28.95115 | -1.38381 | 1.320336 | -1.04807 | 0.294606 | 0.933791 | 0.02975 |
| ENSMUSG00000073702.12 | 2222.992 | 0.281134 | 0.268553 | 1.046849 | 0.295169 | 0.933791 | 0.02975 |
| ENSMUSG00000085438.2 | 305.7688 | -0.53221 | 0.508502 | -1.04663 | 0.295269 | 0.933791 | 0.02975 |
| ENSMUSG00000030244.8 | 1575.954 | 0.408914 | 0.39154 | 1.044374 | 0.296313 | 0.933857 | 0.029719 |
| ENSMUSG00000045410.19 | 312.6752 | 0.441946 | 0.423128 | 1.044474 | 0.296266 | 0.933857 | 0.029719 |
| ENSMUSG00000064181.7 | 145.0944 | -0.62906 | 0.602241 | -1.04453 | 0.296238 | 0.933857 | 0.029719 |
| ENSMUSG00000005823.10 | 77.26547 | -1.28821 | 1.233914 | -1.04401 | 0.296482 | 0.934081 | 0.029615 |
| ENSMUSG00000004043.15 | 14.30636 | -2.14152 | 2.054229 | -1.04249 | 0.297184 | 0.934286 | 0.02952 |
| ENSMUSG00000015013.10 | 174.5308 | 0.806756 | 0.773491 | 1.043006 | 0.296946 | 0.934286 | 0.02952 |
| ENSMUSG00000035674.14 | 933.1236 | 0.391615 | 0.375425 | 1.043123 | 0.296891 | 0.934286 | 0.02952 |
| ENSMUSG00000038279.11 | 22.14394 | 1.973482 | 1.892956 | 1.04254 | 0.297161 | 0.934286 | 0.02952 |
| ENSMUSG00000091721.5 | 47.30481 | -1.47345 | 1.413548 | -1.04238 | 0.297237 | 0.934286 | 0.02952 |
| ENSMUSG00000094483.3 | 270.2212 | -0.68312 | 0.655208 | -1.0426 | 0.297135 | 0.934286 | 0.02952 |
| ENSMUSG00000106895.2 | 61.81495 | 1.318755 | 1.264465 | 1.042936 | 0.296978 | 0.934286 | 0.02952 |
| ENSMUSG00000000826.17 | 249.6164 | -0.74301 | 0.717623 | -1.03537 | 0.300495 | 0.934602 | 0.029373 |
| ENSMUSG00000001300.17 | 41.62685 | 1.623835 | 1.571294 | 1.033438 | 0.301399 | 0.934602 | 0.029373 |
| ENSMUSG00000002028.14 | 25.03994 | 1.507158 | 1.449667 | 1.039658 | 0.298499 | 0.934602 | 0.029373 |
| ENSMUSG00000002372.10 | 32.37955 | 1.427324 | 1.382387 | 1.032506 | 0.301835 | 0.934602 | 0.029373 |
| ENSMUSG00000004071.14 | 163.6855 | 0.609151 | 0.586979 | 1.037772 | 0.299376 | 0.934602 | 0.029373 |
| ENSMUSG00000006699.18 | 995.5722 | 0.371527 | 0.357846 | 1.03823 | 0.299163 | 0.934602 | 0.029373 |
| ENSMUSG00000007589.10 | 54.66634 | -1.08556 | 1.050428 | -1.03344 | 0.301397 | 0.934602 | 0.029373 |
| ENSMUSG00000015305.7 | 93.07242 | -0.72859 | 0.705654 | -1.0325 | 0.301837 | 0.934602 | 0.029373 |
| ENSMUSG00000018765.6 | 12.20568 | -2.5216 | 2.441387 | -1.03286 | 0.301671 | 0.934602 | 0.029373 |
| ENSMUSG00000020300.15 | 220.0108 | -0.57248 | 0.550036 | -1.0408 | 0.29797 | 0.934602 | 0.029373 |
| ENSMUSG00000020696.19 | 31.1054 | 1.629665 | 1.578803 | 1.032216 | 0.301971 | 0.934602 | 0.029373 |
| ENSMUSG00000020946.14 | 106.6264 | -0.75222 | 0.725998 | -1.03611 | 0.30015 | 0.934602 | 0.029373 |
| ENSMUSG00000021712.16 | 83.54651 | 1.097074 | 1.057759 | 1.037168 | 0.299658 | 0.934602 | 0.029373 |
| ENSMUSG00000022070.8 | 12.79179 | 2.788618 | 2.687704 | 1.037547 | 0.299481 | 0.934602 | 0.029373 |
| ENSMUSG00000022129.5 | 52.17894 | -1.39664 | 1.34211 | -1.04063 | 0.298048 | 0.934602 | 0.029373 |
| ENSMUSG00000022253.18 | 366.0473 | 0.518578 | 0.501594 | 1.03386 | 0.301202 | 0.934602 | 0.029373 |
| ENSMUSG00000022951.17 | 45.60627 | -1.07609 | 1.035129 | -1.03957 | 0.298538 | 0.934602 | 0.029373 |
| ENSMUSG00000024013.16 | 11.11612 | 2.849925 | 2.746567 | 1.037632 | 0.299442 | 0.934602 | 0.029373 |
| ENSMUSG00000024309.16 | 223.7908 | 0.657128 | 0.632456 | 1.03901 | 0.2988 | 0.934602 | 0.029373 |

| | | | | | | | |
|---|---|---|---|---|---|---|---|
| ENSMUSG00000024978.12 | 1356.659 | 0.349366 | 0.337346 | 1.035632 | 0.300374 | 0.934602 | 0.029373 |
| ENSMUSG00000026558.14 | 15.31398 | -2.1507 | 2.073399 | -1.03728 | 0.299605 | 0.934602 | 0.029373 |
| ENSMUSG00000029345.14 | 36.03974 | -1.84828 | 1.783379 | -1.03639 | 0.30002 | 0.934602 | 0.029373 |
| ENSMUSG00000029599.14 | 54.96959 | 1.087993 | 1.048006 | 1.038156 | 0.299197 | 0.934602 | 0.029373 |
| ENSMUSG00000029661.17 | 55.55413 | -1.09824 | 1.062213 | -1.03391 | 0.301176 | 0.934602 | 0.029373 |
| ENSMUSG00000030064.17 | 119.5552 | -0.86807 | 0.839402 | -1.03416 | 0.301062 | 0.934602 | 0.029373 |
| ENSMUSG00000031012.18 | 117.5849 | 1.033447 | 1.00002 | 1.033426 | 0.301405 | 0.934602 | 0.029373 |
| ENSMUSG00000031367.16 | 35.20114 | -1.86572 | 1.804631 | -1.03385 | 0.301205 | 0.934602 | 0.029373 |
| ENSMUSG00000031626.19 | 150.4283 | -0.61268 | 0.59111 | -1.03649 | 0.299976 | 0.934602 | 0.029373 |
| ENSMUSG00000033478.6 | 190.7319 | -0.66048 | 0.634206 | -1.04143 | 0.297676 | 0.934602 | 0.029373 |
| ENSMUSG00000034868.10 | 57.83181 | 1.126679 | 1.091183 | 1.03253 | 0.301824 | 0.934602 | 0.029373 |
| ENSMUSG00000035451.8 | 97.98507 | 0.80783 | 0.782092 | 1.032908 | 0.301647 | 0.934602 | 0.029373 |
| ENSMUSG00000036896.6 | 77.03726 | 1.159707 | 1.114642 | 1.040431 | 0.29814 | 0.934602 | 0.029373 |
| ENSMUSG00000037254.19 | 3440.46 | -0.25218 | 0.243459 | -1.03583 | 0.300282 | 0.934602 | 0.029373 |
| ENSMUSG00000037343.9 | 44.28986 | 1.126312 | 1.090544 | 1.032799 | 0.301698 | 0.934602 | 0.029373 |
| ENSMUSG00000039157.13 | 87.41196 | 0.993176 | 0.955572 | 1.039351 | 0.298641 | 0.934602 | 0.029373 |
| ENSMUSG00000039652.17 | 67.11515 | -1.39146 | 1.342899 | -1.03616 | 0.300126 | 0.934602 | 0.029373 |
| ENSMUSG00000040253.16 | 77.43287 | -0.93178 | 0.895239 | -1.04081 | 0.297962 | 0.934602 | 0.029373 |
| ENSMUSG00000040865.16 | 33.06282 | -1.5372 | 1.477502 | -1.04041 | 0.298152 | 0.934602 | 0.029373 |
| ENSMUSG00000043067.16 | 389.7756 | 0.498714 | 0.481074 | 1.036667 | 0.299891 | 0.934602 | 0.029373 |
| ENSMUSG00000044042.20 | 43.2881 | 1.87089 | 1.800358 | 1.039176 | 0.298723 | 0.934602 | 0.029373 |
| ENSMUSG00000044475.15 | 16.40543 | -2.31032 | 2.233341 | -1.03447 | 0.300917 | 0.934602 | 0.029373 |
| ENSMUSG00000057068.16 | 195.7114 | -0.70699 | 0.684424 | -1.03297 | 0.301616 | 0.934602 | 0.029373 |
| ENSMUSG00000057841.6 | 4317.63 | 0.239905 | 0.232124 | 1.033518 | 0.301361 | 0.934602 | 0.029373 |
| ENSMUSG00000059409.10 | 40.29152 | -3.78396 | 3.64384 | -1.03846 | 0.299058 | 0.934602 | 0.029373 |
| ENSMUSG00000059772.13 | 34.72752 | 1.434767 | 1.381546 | 1.038523 | 0.299027 | 0.934602 | 0.029373 |
| ENSMUSG00000080848.4 | 355.4137 | 0.447563 | 0.433539 | 1.032347 | 0.301909 | 0.934602 | 0.029373 |
| ENSMUSG00000112909.2 | 15.03608 | 2.308942 | 2.217082 | 1.041432 | 0.297675 | 0.934602 | 0.029373 |
| ENSMUSG00000007415.16 | 257.0835 | 0.492674 | 0.477821 | 1.031084 | 0.302501 | 0.934966 | 0.029204 |
| ENSMUSG00000023031.9 | 395.0808 | -0.50782 | 0.49245 | -1.03121 | 0.302442 | 0.934966 | 0.029204 |
| ENSMUSG00000027366.13 | 736.8065 | 0.492977 | 0.47815 | 1.031008 | 0.302537 | 0.934966 | 0.029204 |
| ENSMUSG00000028412.18 | 251.0792 | -0.5988 | 0.580898 | -1.03082 | 0.302627 | 0.934966 | 0.029204 |
| ENSMUSG00000033166.12 | 26.67667 | 1.293604 | 1.255071 | 1.030703 | 0.30268 | 0.934966 | 0.029204 |
| ENSMUSG00000044950.8 | 20.67456 | -1.60516 | 1.556176 | -1.03148 | 0.302316 | 0.934966 | 0.029204 |
| ENSMUSG00000001105.16 | 901.3413 | -0.31199 | 0.302953 | -1.02982 | 0.303096 | 0.935102 | 0.029141 |
| ENSMUSG00000032557.14 | 416.7554 | -0.4734 | 0.459746 | -1.0297 | 0.303152 | 0.935102 | 0.029141 |
| ENSMUSG00000041328.17 | 100.8796 | 0.792786 | 0.770183 | 1.029348 | 0.303316 | 0.935102 | 0.029141 |
| ENSMUSG00000074211.5 | 66.09661 | 1.47835 | 1.436122 | 1.029404 | 0.30329 | 0.935102 | 0.029141 |
| ENSMUSG00000084983.8 | 228.4642 | 0.702468 | 0.682043 | 1.029947 | 0.303035 | 0.935102 | 0.029141 |
| ENSMUSG00000085148.2 | 264.3312 | -0.67872 | 0.659056 | -1.02984 | 0.303083 | 0.935102 | 0.029141 |
| ENSMUSG00000003559.9 | 486.8364 | 0.44468 | 0.433054 | 1.026846 | 0.304493 | 0.935257 | 0.029069 |
| ENSMUSG00000005397.9 | 41.92798 | 1.667598 | 1.624034 | 1.026825 | 0.304503 | 0.935257 | 0.029069 |
| ENSMUSG00000019777.17 | 163.9483 | 0.686969 | 0.668186 | 1.02811 | 0.303898 | 0.935257 | 0.029069 |
| ENSMUSG00000029273.11 | 1123.34 | 0.361718 | 0.352268 | 1.026825 | 0.304503 | 0.935257 | 0.029069 |

| | | | | | | | |
|---|---|---|---|---|---|---|---|
| ENSMUSG00000029622.17 | 250.3975 | -0.57649 | 0.560347 | -1.0288 | 0.303572 | 0.935257 | 0.029069 |
| ENSMUSG00000032425.16 | 13.78273 | 2.366332 | 2.300276 | 1.028716 | 0.303613 | 0.935257 | 0.029069 |
| ENSMUSG00000034108.7 | 334.2459 | -0.5578 | 0.543139 | -1.02699 | 0.304426 | 0.935257 | 0.029069 |
| ENSMUSG00000038412.9 | 904.719 | 0.395902 | 0.385597 | 1.026724 | 0.304551 | 0.935257 | 0.029069 |
| ENSMUSG00000042043.7 | 1030.529 | 0.332055 | 0.32316 | 1.027524 | 0.304174 | 0.935257 | 0.029069 |
| ENSMUSG00000045975.9 | 299.0096 | -0.54995 | 0.534764 | -1.0284 | 0.303762 | 0.935257 | 0.029069 |
| ENSMUSG00000046351.12 | 117.247 | -0.7539 | 0.733514 | -1.02779 | 0.304047 | 0.935257 | 0.029069 |
| ENSMUSG00000068397.5 | 138.9813 | 0.662267 | 0.644981 | 1.0268 | 0.304515 | 0.935257 | 0.029069 |
| ENSMUSG00000029821.16 | 16.51839 | 2.518729 | 2.454432 | 1.026196 | 0.304799 | 0.935717 | 0.028855 |
| ENSMUSG00000031431.14 | 1159.312 | 0.420144 | 0.409523 | 1.025935 | 0.304922 | 0.935793 | 0.02882 |
| ENSMUSG00000003053.18 | 4306.972 | -0.28449 | 0.277733 | -1.02432 | 0.305684 | 0.93614 | 0.028659 |
| ENSMUSG00000024644.18 | 98.61622 | 0.840335 | 0.819638 | 1.025252 | 0.305244 | 0.93614 | 0.028659 |
| ENSMUSG00000025366.9 | 54.92027 | -1.84668 | 1.802502 | -1.02451 | 0.305594 | 0.93614 | 0.028659 |
| ENSMUSG00000032462.15 | 93.38498 | -0.85723 | 0.836953 | -1.02423 | 0.305727 | 0.93614 | 0.028659 |
| ENSMUSG00000042426.7 | 90.10313 | 0.829895 | 0.810126 | 1.024402 | 0.305645 | 0.93614 | 0.028659 |
| ENSMUSG00000043687.17 | 19.0014 | -2.14494 | 2.093623 | -1.02451 | 0.305595 | 0.93614 | 0.028659 |
| ENSMUSG00000045795.11 | 62.15115 | -1.34915 | 1.316707 | -1.02464 | 0.305532 | 0.93614 | 0.028659 |
| ENSMUSG00000015672.5 | 341.9842 | 0.441858 | 0.431518 | 1.023961 | 0.305854 | 0.936227 | 0.028619 |
| ENSMUSG00000000538.19 | 24.80245 | -1.96962 | 1.92935 | -1.02087 | 0.307315 | 0.936437 | 0.028521 |
| ENSMUSG00000024095.16 | 80.41004 | -0.92617 | 0.906944 | -1.02119 | 0.307162 | 0.936437 | 0.028521 |
| ENSMUSG00000024360.9 | 593.3648 | -0.41828 | 0.409273 | -1.02201 | 0.306777 | 0.936437 | 0.028521 |
| ENSMUSG00000031134.17 | 28.63802 | -1.38005 | 1.350823 | -1.02164 | 0.306952 | 0.936437 | 0.028521 |
| ENSMUSG00000031523.17 | 240.119 | 0.502498 | 0.492323 | 1.020667 | 0.307412 | 0.936437 | 0.028521 |
| ENSMUSG00000031608.14 | 14.18174 | 2.326073 | 2.272814 | 1.023433 | 0.306103 | 0.936437 | 0.028521 |
| ENSMUSG00000037242.9 | 159.788 | 0.676654 | 0.661195 | 1.023381 | 0.306128 | 0.936437 | 0.028521 |
| ENSMUSG00000040651.10 | 124.9394 | -0.70644 | 0.692551 | -1.02006 | 0.307701 | 0.936437 | 0.028521 |
| ENSMUSG00000041075.9 | 54.9813 | 1.150939 | 1.128246 | 1.020114 | 0.307674 | 0.936437 | 0.028521 |
| ENSMUSG00000041268.18 | 72.60108 | 0.878511 | 0.859187 | 1.022492 | 0.306548 | 0.936437 | 0.028521 |
| ENSMUSG00000046447.4 | 389.9983 | 0.688347 | 0.674758 | 1.020139 | 0.307662 | 0.936437 | 0.028521 |
| ENSMUSG00000056211.14 | 64.99642 | -1.09543 | 1.072601 | -1.02129 | 0.307118 | 0.936437 | 0.028521 |
| ENSMUSG00000061518.12 | 2188.157 | 0.277893 | 0.271936 | 1.021906 | 0.306825 | 0.936437 | 0.028521 |
| ENSMUSG00000063511.12 | 152.5078 | 0.642497 | 0.627948 | 1.023169 | 0.306228 | 0.936437 | 0.028521 |
| ENSMUSG00000071650.7 | 289.311 | -0.60946 | 0.596411 | -1.02188 | 0.306837 | 0.936437 | 0.028521 |
| ENSMUSG00000073555.2 | 616.3068 | 0.370503 | 0.363088 | 1.020423 | 0.307528 | 0.936437 | 0.028521 |
| ENSMUSG00000084111.5 | 193.1182 | 0.570851 | 0.55868 | 1.021786 | 0.306882 | 0.936437 | 0.028521 |
| ENSMUSG00000095440.3 | 22.07305 | -2.04778 | 2.00318 | -1.02227 | 0.306655 | 0.936437 | 0.028521 |
| ENSMUSG00000027465.14 | 31.57078 | 1.38231 | 1.355489 | 1.019787 | 0.30783 | 0.936529 | 0.028479 |
| ENSMUSG00000003123.16 | 24.05453 | 2.302369 | 2.259936 | 1.018776 | 0.308309 | 0.93664 | 0.028427 |
| ENSMUSG00000009927.10 | 1544.62 | 0.287794 | 0.282449 | 1.018922 | 0.30824 | 0.93664 | 0.028427 |
| ENSMUSG00000026374.15 | 494.0456 | -0.4719 | 0.463045 | -1.01913 | 0.30814 | 0.93664 | 0.028427 |
| ENSMUSG00000034783.8 | 31.58368 | 1.377866 | 1.353167 | 1.018253 | 0.308558 | 0.93664 | 0.028427 |
| ENSMUSG00000041763.15 | 147.822 | -0.70766 | 0.694596 | -1.01881 | 0.308294 | 0.93664 | 0.028427 |
| ENSMUSG00000051065.10 | 68.74643 | 0.997414 | 0.979492 | 1.018297 | 0.308537 | 0.93664 | 0.028427 |
| ENSMUSG00000055850.13 | 624.5069 | 0.371422 | 0.364653 | 1.018564 | 0.30841 | 0.93664 | 0.028427 |

| | | | | | | | |
|---|---|---|---|---|---|---|---|
| ENSMUSG00000005510.10 | 348.3921 | 0.486399 | 0.477783 | 1.018035 | 0.308661 | 0.936655 | 0.02842 |
| ENSMUSG00000026853.16 | 59.40689 | 1.376245 | 1.352555 | 1.017515 | 0.308909 | 0.937105 | 0.028212 |
| ENSMUSG00000026768.11 | 20.41538 | -2.09493 | 2.062665 | -1.01564 | 0.309799 | 0.937333 | 0.028106 |
| ENSMUSG00000035354.10 | 162.0906 | -0.68542 | 0.675097 | -1.01529 | 0.309967 | 0.937333 | 0.028106 |
| ENSMUSG00000038274.14 | 988.2686 | 0.39762 | 0.390934 | 1.017104 | 0.309104 | 0.937333 | 0.028106 |
| ENSMUSG00000041264.17 | 64.51434 | -1.36971 | 1.348327 | -1.01586 | 0.309696 | 0.937333 | 0.028106 |
| ENSMUSG00000047221.6 | 54.82264 | 1.214983 | 1.19619 | 1.015711 | 0.309767 | 0.937333 | 0.028106 |
| ENSMUSG00000048782.16 | 93.36908 | 1.094157 | 1.077528 | 1.015433 | 0.3099 | 0.937333 | 0.028106 |
| ENSMUSG00000052906.8 | 158.7336 | 0.716309 | 0.704951 | 1.016112 | 0.309576 | 0.937333 | 0.028106 |
| ENSMUSG00000058900.6 | 13.37001 | 2.474136 | 2.434697 | 1.016199 | 0.309535 | 0.937333 | 0.028106 |
| ENSMUSG00000061808.5 | 96353.63 | 0.200873 | 0.197697 | 1.016063 | 0.3096 | 0.937333 | 0.028106 |
| ENSMUSG00000100680.2 | 46.24745 | -1.34135 | 1.321166 | -1.01528 | 0.309972 | 0.937333 | 0.028106 |
| ENSMUSG00000001280.14 | 96.1249 | -0.80617 | 0.799074 | -1.00888 | 0.313031 | 0.937551 | 0.028005 |
| ENSMUSG00000002900.18 | 27.95565 | 1.914525 | 1.897873 | 1.008774 | 0.313083 | 0.937551 | 0.028005 |
| ENSMUSG00000003198.11 | 31.31115 | 1.304547 | 1.288097 | 1.012771 | 0.31117 | 0.937551 | 0.028005 |
| ENSMUSG00000005481.19 | 65.42988 | 1.375264 | 1.355971 | 1.014228 | 0.310474 | 0.937551 | 0.028005 |
| ENSMUSG00000006818.6 | 1651.047 | 0.297002 | 0.29483 | 1.007368 | 0.313758 | 0.937551 | 0.028005 |
| ENSMUSG00000009013.6 | 212.559 | 0.650235 | 0.646361 | 1.005993 | 0.314419 | 0.937551 | 0.028005 |
| ENSMUSG00000012117.14 | 31.85464 | 1.440577 | 1.429989 | 1.007404 | 0.313741 | 0.937551 | 0.028005 |
| ENSMUSG00000014633.16 | 63.02706 | 0.924811 | 0.914234 | 1.01157 | 0.311744 | 0.937551 | 0.028005 |
| ENSMUSG00000015126.10 | 100.2056 | 0.763766 | 0.75742 | 1.008378 | 0.313273 | 0.937551 | 0.028005 |
| ENSMUSG00000017679.17 | 51.08562 | -1.27149 | 1.264934 | -1.00518 | 0.31481 | 0.937551 | 0.028005 |
| ENSMUSG00000018474.19 | 172.8615 | -0.96493 | 0.952066 | -1.01351 | 0.310815 | 0.937551 | 0.028005 |
| ENSMUSG00000020224.16 | 300.2067 | 0.469179 | 0.464938 | 1.009121 | 0.312917 | 0.937551 | 0.028005 |
| ENSMUSG00000021890.7 | 227.6429 | -0.57247 | 0.569788 | -1.0047 | 0.315041 | 0.937551 | 0.028005 |
| ENSMUSG00000022217.15 | 105.3252 | 0.841831 | 0.837325 | 1.005381 | 0.314713 | 0.937551 | 0.028005 |
| ENSMUSG00000022391.17 | 164.0989 | -0.72404 | 0.719932 | -1.00571 | 0.314556 | 0.937551 | 0.028005 |
| ENSMUSG00000022550.19 | 12.11963 | -2.20137 | 2.184844 | -1.00756 | 0.313665 | 0.937551 | 0.028005 |
| ENSMUSG00000024370.18 | 20.51072 | -1.70972 | 1.687713 | -1.01304 | 0.311041 | 0.937551 | 0.028005 |
| ENSMUSG00000025571.14 | 57.50346 | -0.89583 | 0.883048 | -1.01447 | 0.310357 | 0.937551 | 0.028005 |
| ENSMUSG00000025766.15 | 26.99505 | -1.30985 | 1.301802 | -1.00618 | 0.314328 | 0.937551 | 0.028005 |
| ENSMUSG00000026098.14 | 36.12403 | -1.30221 | 1.291393 | -1.00838 | 0.313274 | 0.937551 | 0.028005 |
| ENSMUSG00000026278.15 | 64.63467 | -1.54002 | 1.525346 | -1.00962 | 0.312677 | 0.937551 | 0.028005 |
| ENSMUSG00000026495.9 | 22.05575 | -1.69084 | 1.685204 | -1.00334 | 0.315695 | 0.937551 | 0.028005 |
| ENSMUSG00000026743.17 | 100.9869 | 0.75262 | 0.744425 | 1.011008 | 0.312012 | 0.937551 | 0.028005 |
| ENSMUSG00000026781.18 | 1643.263 | -0.30517 | 0.301475 | -1.01224 | 0.311422 | 0.937551 | 0.028005 |
| ENSMUSG00000027002.14 | 359.6425 | -0.54679 | 0.540758 | -1.01116 | 0.311939 | 0.937551 | 0.028005 |
| ENSMUSG00000027185.16 | 32.43201 | -1.59357 | 1.580362 | -1.00836 | 0.313281 | 0.937551 | 0.028005 |
| ENSMUSG00000028619.16 | 32.19266 | 1.434212 | 1.417041 | 1.012118 | 0.311482 | 0.937551 | 0.028005 |
| ENSMUSG00000029190.20 | 269.738 | 0.530276 | 0.523704 | 1.012549 | 0.311276 | 0.937551 | 0.028005 |
| ENSMUSG00000029310.14 | 155.1918 | -0.65796 | 0.652656 | -1.00812 | 0.313396 | 0.937551 | 0.028005 |
| ENSMUSG00000029513.15 | 80.69661 | 0.941382 | 0.935795 | 1.00597 | 0.31443 | 0.937551 | 0.028005 |
| ENSMUSG00000029528.20 | 34.40749 | 1.39222 | 1.378683 | 1.009819 | 0.312582 | 0.937551 | 0.028005 |
| ENSMUSG00000030046.7 | 13.80642 | -2.01518 | 1.998965 | -1.00811 | 0.313402 | 0.937551 | 0.028005 |

| | | | | | | | |
|---|---|---|---|---|---|---|---|
| ENSMUSG00000030245.11 | 11.70982 | 2.914874 | 2.900716 | 1.004881 | 0.314954 | 0.937551 | 0.028005 |
| ENSMUSG00000030282.15 | 284.8142 | 0.51743 | 0.515829 | 1.003103 | 0.315811 | 0.937551 | 0.028005 |
| ENSMUSG00000030301.18 | 115.7342 | 0.830702 | 0.823013 | 1.009343 | 0.31281 | 0.937551 | 0.028005 |
| ENSMUSG00000030659.15 | 63.23369 | -1.17357 | 1.168841 | -1.00405 | 0.315356 | 0.937551 | 0.028005 |
| ENSMUSG00000030846.16 | 159.3544 | -0.71217 | 0.71007 | -1.00296 | 0.31588 | 0.937551 | 0.028005 |
| ENSMUSG00000031563.9 | 57.7133 | -1.23944 | 1.233382 | -1.00491 | 0.314938 | 0.937551 | 0.028005 |
| ENSMUSG00000031803.9 | 10.369 | 3.281449 | 3.235886 | 1.01408 | 0.310544 | 0.937551 | 0.028005 |
| ENSMUSG00000034037.15 | 12.88616 | 2.443662 | 2.434455 | 1.003782 | 0.315484 | 0.937551 | 0.028005 |
| ENSMUSG00000034487.5 | 56.75976 | -1.33852 | 1.333639 | -1.00366 | 0.315541 | 0.937551 | 0.028005 |
| ENSMUSG00000034574.11 | 103.2493 | 0.830709 | 0.821328 | 1.011423 | 0.311814 | 0.937551 | 0.028005 |
| ENSMUSG00000035237.8 | 504.1395 | 0.364908 | 0.36125 | 1.010127 | 0.312434 | 0.937551 | 0.028005 |
| ENSMUSG00000036110.15 | 667.7953 | -0.35607 | 0.354569 | -1.00422 | 0.315271 | 0.937551 | 0.028005 |
| ENSMUSG00000038369.15 | 119.4332 | 0.755338 | 0.749779 | 1.007415 | 0.313735 | 0.937551 | 0.028005 |
| ENSMUSG00000039349.6 | 64.33735 | -0.9727 | 0.960522 | -1.01268 | 0.311214 | 0.937551 | 0.028005 |
| ENSMUSG00000040321.4 | 26.58139 | 1.695943 | 1.683413 | 1.007444 | 0.313722 | 0.937551 | 0.028005 |
| ENSMUSG00000045377.6 | 11.51355 | 2.405731 | 2.388233 | 1.007327 | 0.313778 | 0.937551 | 0.028005 |
| ENSMUSG00000046020.14 | 47.09164 | -1.14555 | 1.136784 | -1.00771 | 0.313594 | 0.937551 | 0.028005 |
| ENSMUSG00000046441.6 | 19.7553 | 1.847599 | 1.838328 | 1.005043 | 0.314876 | 0.937551 | 0.028005 |
| ENSMUSG00000046997.6 | 36.17272 | -1.23097 | 1.225493 | -1.00447 | 0.315151 | 0.937551 | 0.028005 |
| ENSMUSG00000048429.11 | 283.6841 | 0.522638 | 0.520903 | 1.00333 | 0.315702 | 0.937551 | 0.028005 |
| ENSMUSG00000050174.16 | 79.87566 | 0.957716 | 0.951273 | 1.006773 | 0.314044 | 0.937551 | 0.028005 |
| ENSMUSG00000059040.6 | 186.1788 | 0.550357 | 0.54793 | 1.004429 | 0.315172 | 0.937551 | 0.028005 |
| ENSMUSG00000063410.9 | 66.24144 | 0.88969 | 0.880531 | 1.010402 | 0.312303 | 0.937551 | 0.028005 |
| ENSMUSG00000068856.4 | 80.74688 | 0.902546 | 0.896604 | 1.006627 | 0.314114 | 0.937551 | 0.028005 |
| ENSMUSG00000070407.6 | 238.8932 | -0.49206 | 0.486621 | -1.01117 | 0.311933 | 0.937551 | 0.028005 |
| ENSMUSG00000071151.4 | 26.88238 | -1.30862 | 1.302347 | -1.00481 | 0.314986 | 0.937551 | 0.028005 |
| ENSMUSG00000090015.9 | 50.17466 | -1.72441 | 1.715299 | -1.00531 | 0.314747 | 0.937551 | 0.028005 |
| ENSMUSG00000022724.16 | 75.75782 | -0.92966 | 0.927231 | -1.00262 | 0.316043 | 0.937733 | 0.027921 |
| ENSMUSG00000042460.6 | 320.8627 | -0.44825 | 0.447168 | -1.00242 | 0.316139 | 0.937733 | 0.027921 |
| ENSMUSG00000035401.10 | 46.75468 | 1.407186 | 1.404077 | 1.002214 | 0.31624 | 0.937741 | 0.027917 |
| ENSMUSG00000001604.15 | 764.4113 | -0.31262 | 0.312603 | -1.00006 | 0.317282 | 0.937912 | 0.027838 |
| ENSMUSG00000004070.16 | 226.125 | -0.54765 | 0.547814 | -0.99971 | 0.317451 | 0.937912 | 0.027838 |
| ENSMUSG00000004451.15 | 30.21352 | -1.46295 | 1.461685 | -1.00087 | 0.316892 | 0.937912 | 0.027838 |
| ENSMUSG00000006333.15 | 1451.601 | 0.361736 | 0.36138 | 1.000985 | 0.316834 | 0.937912 | 0.027838 |
| ENSMUSG00000019947.11 | 271.4297 | 0.615696 | 0.615918 | 0.999639 | 0.317485 | 0.937912 | 0.027838 |
| ENSMUSG00000021577.15 | 3504.783 | 0.225859 | 0.225597 | 1.001165 | 0.316747 | 0.937912 | 0.027838 |
| ENSMUSG00000025075.15 | 340.61 | -0.51853 | 0.517828 | -1.00136 | 0.316653 | 0.937912 | 0.027838 |
| ENSMUSG00000036890.14 | 16.9224 | 1.960383 | 1.957195 | 1.001629 | 0.316523 | 0.937912 | 0.027838 |
| ENSMUSG00000037852.9 | 15.60743 | -2.48068 | 2.479213 | -1.00059 | 0.317025 | 0.937912 | 0.027838 |
| ENSMUSG00000038633.6 | 898.7455 | 0.409899 | 0.410028 | 0.999685 | 0.317463 | 0.937912 | 0.027838 |
| ENSMUSG00000040945.14 | 49.01568 | -1.42143 | 1.421332 | -1.00007 | 0.317278 | 0.937912 | 0.027838 |
| ENSMUSG00000118671.1 | 27.40432 | -1.73976 | 1.739202 | -1.00032 | 0.317155 | 0.937912 | 0.027838 |
| ENSMUSG00000022673.6 | 40.11954 | 1.204744 | 1.205734 | 0.999179 | 0.317708 | 0.937986 | 0.027804 |
| ENSMUSG00000035671.7 | 10.13464 | -3.25782 | 3.259991 | -0.99933 | 0.317633 | 0.937986 | 0.027804 |

| | | | | | | | |
|---|---|---|---|---|---|---|---|
| ENSMUSG00000020180.11 | 221.0183 | -0.61523 | 0.617032 | -0.99707 | 0.318729 | 0.938952 | 0.027357 |
| ENSMUSG00000029462.19 | 406.9204 | 0.410803 | 0.411974 | 0.997159 | 0.318687 | 0.938952 | 0.027357 |
| ENSMUSG00000032038.8 | 144.1461 | -0.63944 | 0.64117 | -0.9973 | 0.31862 | 0.938952 | 0.027357 |
| ENSMUSG00000033860.14 | 25372.57 | -0.26478 | 0.265408 | -0.99763 | 0.318461 | 0.938952 | 0.027357 |
| ENSMUSG00000047067.8 | 43.21335 | 1.077865 | 1.081028 | 0.997074 | 0.318729 | 0.938952 | 0.027357 |
| ENSMUSG00000062078.16 | 412.7406 | 0.432561 | 0.43371 | 0.997351 | 0.318594 | 0.938952 | 0.027357 |
| ENSMUSG00000108231.2 | 10.7745 | 2.250581 | 2.256751 | 0.997266 | 0.318635 | 0.938952 | 0.027357 |
| ENSMUSG00000027163.14 | 24.37967 | 1.581544 | 1.586735 | 0.996729 | 0.318896 | 0.939067 | 0.027303 |
| ENSMUSG00000030225.12 | 322.6412 | -0.424 | 0.425456 | -0.99659 | 0.318966 | 0.939067 | 0.027303 |
| ENSMUSG00000029833.18 | 183.1758 | -0.59175 | 0.594588 | -0.99523 | 0.319625 | 0.939618 | 0.027049 |
| ENSMUSG00000032498.10 | 87.31485 | 1.108366 | 1.113734 | 0.995181 | 0.319648 | 0.939618 | 0.027049 |
| ENSMUSG00000036983.8 | 37.33076 | 1.400747 | 1.407298 | 0.995345 | 0.319568 | 0.939618 | 0.027049 |
| ENSMUSG00000052160.8 | 20.6741 | 1.822239 | 1.830787 | 0.995331 | 0.319575 | 0.939618 | 0.027049 |
| ENSMUSG00000111080.2 | 147.1094 | 0.659756 | 0.662769 | 0.995454 | 0.319516 | 0.939618 | 0.027049 |
| ENSMUSG00000022299.10 | 155.5988 | -0.59671 | 0.599785 | -0.99487 | 0.319802 | 0.939777 | 0.026975 |
| ENSMUSG00000011831.17 | 716.2989 | 0.303585 | 0.305292 | 0.994409 | 0.320024 | 0.939819 | 0.026956 |
| ENSMUSG00000021824.14 | 460.5593 | 0.447693 | 0.450376 | 0.994043 | 0.320202 | 0.939819 | 0.026956 |
| ENSMUSG00000027282.18 | 1121.446 | 0.319123 | 0.320992 | 0.994178 | 0.320136 | 0.939819 | 0.026956 |
| ENSMUSG00000029003.12 | 30.5654 | 1.813306 | 1.826831 | 0.992597 | 0.320906 | 0.939819 | 0.026956 |
| ENSMUSG00000029231.16 | 17.20009 | -2.13954 | 2.152503 | -0.99398 | 0.320234 | 0.939819 | 0.026956 |
| ENSMUSG00000030220.14 | 42.26135 | 1.409893 | 1.419575 | 0.993179 | 0.320623 | 0.939819 | 0.026956 |
| ENSMUSG00000031631.16 | 83.26507 | -0.83125 | 0.836733 | -0.99344 | 0.320494 | 0.939819 | 0.026956 |
| ENSMUSG00000038697.16 | 36.06644 | 1.612128 | 1.622247 | 0.993762 | 0.320339 | 0.939819 | 0.026956 |
| ENSMUSG00000074797.12 | 339.7019 | 0.461778 | 0.46522 | 0.992603 | 0.320904 | 0.939819 | 0.026956 |
| ENSMUSG00000105255.2 | 56.77289 | -0.95277 | 0.959627 | -0.99286 | 0.320779 | 0.939819 | 0.026956 |
| ENSMUSG00000105837.5 | 53.15625 | 1.01741 | 1.024788 | 0.992801 | 0.320807 | 0.939819 | 0.026956 |
| ENSMUSG00000006057.16 | 772.4553 | 0.349907 | 0.352789 | 0.991832 | 0.32128 | 0.940621 | 0.026585 |
| ENSMUSG00000048924.15 | 36.40256 | 1.472852 | 1.485564 | 0.991442 | 0.32147 | 0.940887 | 0.026463 |
| ENSMUSG00000003444.9 | 28.5707 | 1.515549 | 1.532546 | 0.988909 | 0.322707 | 0.941329 | 0.026259 |
| ENSMUSG00000005534.11 | 380.1708 | -0.43287 | 0.437659 | -0.98907 | 0.322631 | 0.941329 | 0.026259 |
| ENSMUSG00000017747.14 | 86.12487 | 0.756776 | 0.765217 | 0.988969 | 0.322678 | 0.941329 | 0.026259 |
| ENSMUSG00000025220.9 | 166.969 | -0.66519 | 0.672664 | -0.98889 | 0.322715 | 0.941329 | 0.026259 |
| ENSMUSG00000025512.16 | 58.6534 | -1.24235 | 1.256026 | -0.98911 | 0.32261 | 0.941329 | 0.026259 |
| ENSMUSG00000029430.14 | 593.3685 | 0.404256 | 0.408614 | 0.989334 | 0.3225 | 0.941329 | 0.026259 |
| ENSMUSG00000034617.8 | 22.00038 | -1.84691 | 1.867957 | -0.98873 | 0.322793 | 0.941329 | 0.026259 |
| ENSMUSG00000038776.14 | 1014.941 | -0.34523 | 0.348714 | -0.99 | 0.322175 | 0.941329 | 0.026259 |
| ENSMUSG00000042476.13 | 747.2448 | -0.3833 | 0.38697 | -0.99052 | 0.321919 | 0.941329 | 0.026259 |
| ENSMUSG00000045948.9 | 154.6412 | -0.65462 | 0.661609 | -0.98943 | 0.322452 | 0.941329 | 0.026259 |
| ENSMUSG00000058291.15 | 66.76754 | 0.938608 | 0.949341 | 0.988695 | 0.322812 | 0.941329 | 0.026259 |
| ENSMUSG00000071655.13 | 125.5451 | -0.60737 | 0.614135 | -0.98898 | 0.322673 | 0.941329 | 0.026259 |
| ENSMUSG00000038780.15 | 47.18466 | -1.3032 | 1.318842 | -0.98814 | 0.323086 | 0.941837 | 0.026024 |
| ENSMUSG00000030335.7 | 458.8133 | -0.42564 | 0.430874 | -0.98786 | 0.323219 | 0.941937 | 0.025978 |
| ENSMUSG00000004500.10 | 20.07524 | -2.06798 | 2.098268 | -0.98556 | 0.324347 | 0.942294 | 0.025813 |
| ENSMUSG00000018648.16 | 20.35165 | 1.840779 | 1.868931 | 0.984937 | 0.324655 | 0.942294 | 0.025813 |

| | | | | | | | |
|---|---|---|---|---|---|---|---|
| ENSMUSG00000021411.17 | 388.8959 | 0.416883 | 0.423258 | 0.984938 | 0.324655 | 0.942294 | 0.025813 |
| ENSMUSG00000023073.3 | 298.9808 | 0.441965 | 0.448889 | 0.984574 | 0.324833 | 0.942294 | 0.025813 |
| ENSMUSG00000024083.17 | 234.6208 | 0.574297 | 0.583041 | 0.985004 | 0.324622 | 0.942294 | 0.025813 |
| ENSMUSG00000024392.18 | 39.42721 | -1.38395 | 1.40344 | -0.98611 | 0.324079 | 0.942294 | 0.025813 |
| ENSMUSG00000026851.10 | 36.06066 | -1.36359 | 1.384188 | -0.98512 | 0.324565 | 0.942294 | 0.025813 |
| ENSMUSG00000027806.10 | 127.4042 | -0.929 | 0.943536 | -0.9846 | 0.324821 | 0.942294 | 0.025813 |
| ENSMUSG00000028211.12 | 166.0797 | -0.74269 | 0.752383 | -0.98712 | 0.323586 | 0.942294 | 0.025813 |
| ENSMUSG00000031796.8 | 219.4606 | -0.50115 | 0.508788 | -0.98499 | 0.324628 | 0.942294 | 0.025813 |
| ENSMUSG00000031901.13 | 17.3766 | -2.03055 | 2.062019 | -0.98474 | 0.324753 | 0.942294 | 0.025813 |
| ENSMUSG00000035901.15 | 70.64838 | -0.92357 | 0.936083 | -0.98663 | 0.323825 | 0.942294 | 0.025813 |
| ENSMUSG00000036478.9 | 28.43685 | -1.2469 | 1.266394 | -0.9846 | 0.324819 | 0.942294 | 0.025813 |
| ENSMUSG00000044030.5 | 26.33413 | -1.53211 | 1.552514 | -0.98686 | 0.323712 | 0.942294 | 0.025813 |
| ENSMUSG00000057101.9 | 48.40319 | 0.919913 | 0.932535 | 0.986465 | 0.323905 | 0.942294 | 0.025813 |
| ENSMUSG00000019889.11 | 137.5444 | 0.586016 | 0.595599 | 0.98391 | 0.32516 | 0.942377 | 0.025776 |
| ENSMUSG00000036504.7 | 39.46737 | 1.394954 | 1.417601 | 0.984024 | 0.325104 | 0.942377 | 0.025776 |
| ENSMUSG00000049401.11 | 37.41176 | -1.44725 | 1.470877 | -0.98393 | 0.325148 | 0.942377 | 0.025776 |
| ENSMUSG00000063524.14 | 56.69797 | -1.38538 | 1.408608 | -0.98351 | 0.325356 | 0.942656 | 0.025647 |
| ENSMUSG00000014301.14 | 131.3458 | -0.77758 | 0.791117 | -0.98289 | 0.325663 | 0.94297 | 0.025502 |
| ENSMUSG00000032103.11 | 55.89016 | -1.2186 | 1.239658 | -0.98301 | 0.325603 | 0.94297 | 0.025502 |
| ENSMUSG00000024150.13 | 1794.265 | 0.259323 | 0.264074 | 0.98201 | 0.326095 | 0.943855 | 0.025095 |
| ENSMUSG00000024269.12 | 21.6801 | 1.682679 | 1.714045 | 0.981701 | 0.326247 | 0.943855 | 0.025095 |
| ENSMUSG00000024974.11 | 227.725 | -0.55849 | 0.568928 | -0.98166 | 0.326267 | 0.943855 | 0.025095 |
| ENSMUSG00000027878.12 | 134.1114 | 0.90163 | 0.919214 | 0.980872 | 0.326656 | 0.943947 | 0.025052 |
| ENSMUSG00000027963.15 | 19.65267 | 1.481567 | 1.509983 | 0.981181 | 0.326503 | 0.943947 | 0.025052 |
| ENSMUSG00000032688.10 | 27.74636 | -1.66241 | 1.694122 | -0.98128 | 0.326454 | 0.943947 | 0.025052 |
| ENSMUSG00000090306.6 | 45.81712 | -1.66481 | 1.697419 | -0.98079 | 0.326697 | 0.943947 | 0.025052 |
| ENSMUSG00000036309.15 | 561.1937 | 0.413757 | 0.421986 | 0.980499 | 0.32684 | 0.944071 | 0.024995 |
| ENSMUSG00000023110.13 | 30.05367 | -1.818 | 1.854881 | -0.98012 | 0.327028 | 0.944326 | 0.024878 |
| ENSMUSG00000000881.13 | 68.99614 | 0.934298 | 0.95909 | 0.974151 | 0.329982 | 0.944381 | 0.024853 |
| ENSMUSG00000001128.8 | 64.60789 | -1.17161 | 1.210193 | -0.96812 | 0.332987 | 0.944381 | 0.024853 |
| ENSMUSG00000002379.8 | 536.0126 | 0.43005 | 0.44106 | 0.975037 | 0.329542 | 0.944381 | 0.024853 |
| ENSMUSG00000006273.15 | 122.493 | 0.79595 | 0.821944 | 0.968376 | 0.332857 | 0.944381 | 0.024853 |
| ENSMUSG00000014075.16 | 16.80366 | -1.95547 | 2.015736 | -0.9701 | 0.331997 | 0.944381 | 0.024853 |
| ENSMUSG00000017404.13 | 726.0457 | -0.31835 | 0.327964 | -0.9707 | 0.3317 | 0.944381 | 0.024853 |
| ENSMUSG00000020427.12 | 50.04229 | -1.06924 | 1.095783 | -0.97577 | 0.329177 | 0.944381 | 0.024853 |
| ENSMUSG00000021335.14 | 58.27621 | -1.12715 | 1.151732 | -0.97866 | 0.327749 | 0.944381 | 0.024853 |
| ENSMUSG00000022100.15 | 182.6405 | 0.515405 | 0.531327 | 0.970035 | 0.332029 | 0.944381 | 0.024853 |
| ENSMUSG00000022752.10 | 936.0692 | -0.3375 | 0.348502 | -0.96843 | 0.33283 | 0.944381 | 0.024853 |
| ENSMUSG00000022811.18 | 213.5119 | 0.539278 | 0.557355 | 0.967566 | 0.333261 | 0.944381 | 0.024853 |
| ENSMUSG00000023393.16 | 74.54584 | -0.81159 | 0.832408 | -0.97499 | 0.329567 | 0.944381 | 0.024853 |
| ENSMUSG00000026426.11 | 54.12699 | 1.116282 | 1.146996 | 0.973223 | 0.330443 | 0.944381 | 0.024853 |
| ENSMUSG00000027210.21 | 18.16412 | 2.166829 | 2.224371 | 0.974131 | 0.329991 | 0.944381 | 0.024853 |
| ENSMUSG00000028049.16 | 94.80528 | 0.880673 | 0.901715 | 0.976665 | 0.328735 | 0.944381 | 0.024853 |
| ENSMUSG00000028394.10 | 188.5001 | 0.691235 | 0.708427 | 0.975733 | 0.329197 | 0.944381 | 0.024853 |

| | | | | | | | |
|---|---|---|---|---|---|---|---|
| ENSMUSG00000028443.7 | 89.28055 | 0.777911 | 0.798429 | 0.974303 | 0.329906 | 0.944381 | 0.024853 |
| ENSMUSG00000028578.9 | 87.44946 | -1.2644 | 1.291942 | -0.97868 | 0.327736 | 0.944381 | 0.024853 |
| ENSMUSG00000029227.16 | 181.9733 | -0.53886 | 0.556585 | -0.96816 | 0.332965 | 0.944381 | 0.024853 |
| ENSMUSG00000029314.15 | 12.83656 | 2.596801 | 2.674087 | 0.971098 | 0.331499 | 0.944381 | 0.024853 |
| ENSMUSG00000029447.14 | 505.0888 | -0.43804 | 0.451674 | -0.96981 | 0.332139 | 0.944381 | 0.024853 |
| ENSMUSG00000029777.11 | 186.3194 | -0.59029 | 0.608938 | -0.96938 | 0.332356 | 0.944381 | 0.024853 |
| ENSMUSG00000030231.12 | 36.88962 | 1.119973 | 1.157764 | 0.967359 | 0.333364 | 0.944381 | 0.024853 |
| ENSMUSG00000030493.14 | 13.64698 | -2.17797 | 2.24092 | -0.97191 | 0.331095 | 0.944381 | 0.024853 |
| ENSMUSG00000031393.17 | 43.81802 | 1.345281 | 1.380872 | 0.974226 | 0.329944 | 0.944381 | 0.024853 |
| ENSMUSG00000031703.9 | 413.8378 | 0.467586 | 0.480683 | 0.972754 | 0.330676 | 0.944381 | 0.024853 |
| ENSMUSG00000031770.17 | 2406.716 | -0.32259 | 0.332132 | -0.97128 | 0.331407 | 0.944381 | 0.024853 |
| ENSMUSG00000032086.13 | 54.56741 | 1.118347 | 1.149715 | 0.972717 | 0.330694 | 0.944381 | 0.024853 |
| ENSMUSG00000032116.19 | 463.471 | -0.38561 | 0.395483 | -0.97504 | 0.329543 | 0.944381 | 0.024853 |
| ENSMUSG00000033174.18 | 301.4272 | 0.546627 | 0.56211 | 0.972456 | 0.330824 | 0.944381 | 0.024853 |
| ENSMUSG00000033444.15 | 44.86552 | 1.104793 | 1.135674 | 0.972808 | 0.330649 | 0.944381 | 0.024853 |
| ENSMUSG00000034210.15 | 101.1073 | -0.69106 | 0.707604 | -0.97662 | 0.328755 | 0.944381 | 0.024853 |
| ENSMUSG00000035770.9 | 74.57422 | -0.81568 | 0.84032 | -0.97068 | 0.331708 | 0.944381 | 0.024853 |
| ENSMUSG00000035958.4 | 117.7147 | 0.82265 | 0.849278 | 0.968647 | 0.332721 | 0.944381 | 0.024853 |
| ENSMUSG00000038206.14 | 392.728 | -0.40104 | 0.410847 | -0.97613 | 0.329 | 0.944381 | 0.024853 |
| ENSMUSG00000038422.3 | 10.87318 | 2.420226 | 2.485417 | 0.973771 | 0.33017 | 0.944381 | 0.024853 |
| ENSMUSG00000039016.6 | 808.079 | 0.344197 | 0.355138 | 0.969195 | 0.332448 | 0.944381 | 0.024853 |
| ENSMUSG00000039191.13 | 29.90166 | -1.51478 | 1.565134 | -0.96783 | 0.33313 | 0.944381 | 0.024853 |
| ENSMUSG00000039230.15 | 60.75399 | -1.09579 | 1.126162 | -0.97303 | 0.330538 | 0.944381 | 0.024853 |
| ENSMUSG00000039244.11 | 15.16916 | -2.21049 | 2.264911 | -0.97597 | 0.329078 | 0.944381 | 0.024853 |
| ENSMUSG00000039910.11 | 238.8504 | 0.658413 | 0.677165 | 0.972308 | 0.330897 | 0.944381 | 0.024853 |
| ENSMUSG00000041333.7 | 36.70976 | 1.295087 | 1.323089 | 0.978836 | 0.327661 | 0.944381 | 0.024853 |
| ENSMUSG00000043323.18 | 10.75687 | -2.25889 | 2.320918 | -0.97327 | 0.330418 | 0.944381 | 0.024853 |
| ENSMUSG00000043881.10 | 65.36958 | 0.977518 | 1.009519 | 0.968301 | 0.332894 | 0.944381 | 0.024853 |
| ENSMUSG00000045639.15 | 12.74365 | 2.996669 | 3.098155 | 0.967243 | 0.333422 | 0.944381 | 0.024853 |
| ENSMUSG00000049624.15 | 206.857 | -0.57501 | 0.590747 | -0.97336 | 0.330373 | 0.944381 | 0.024853 |
| ENSMUSG00000049950.7 | 22.83562 | 1.280912 | 1.317844 | 0.971976 | 0.331063 | 0.944381 | 0.024853 |
| ENSMUSG00000052056.15 | 13.46059 | -2.21118 | 2.269906 | -0.97413 | 0.329993 | 0.944381 | 0.024853 |
| ENSMUSG00000054435.17 | 59.45565 | 1.063416 | 1.097603 | 0.968853 | 0.332618 | 0.944381 | 0.024853 |
| ENSMUSG00000058558.13 | 2448.668 | 0.287407 | 0.293953 | 0.977733 | 0.328207 | 0.944381 | 0.024853 |
| ENSMUSG00000062040.15 | 10.19468 | -3.18335 | 3.282185 | -0.96989 | 0.332103 | 0.944381 | 0.024853 |
| ENSMUSG00000063457.15 | 1121.663 | -0.28405 | 0.292277 | -0.97183 | 0.331133 | 0.944381 | 0.024853 |
| ENSMUSG00000064262.8 | 12.73573 | 2.223677 | 2.291957 | 0.970209 | 0.331942 | 0.944381 | 0.024853 |
| ENSMUSG00000067813.4 | 82.8419 | -0.83524 | 0.862388 | -0.96851 | 0.332787 | 0.944381 | 0.024853 |
| ENSMUSG00000068921.15 | 117.9399 | -0.84722 | 0.870196 | -0.9736 | 0.330256 | 0.944381 | 0.024853 |
| ENSMUSG00000070972.14 | 251.8196 | 0.56468 | 0.578785 | 0.97563 | 0.329248 | 0.944381 | 0.024853 |
| ENSMUSG00000071350.13 | 28.11573 | 1.254281 | 1.288671 | 0.973314 | 0.330397 | 0.944381 | 0.024853 |
| ENSMUSG00000071708.12 | 27.01841 | -1.84146 | 1.902601 | -0.96786 | 0.333112 | 0.944381 | 0.024853 |
| ENSMUSG00000075054.6 | 34.90785 | -1.46604 | 1.496984 | -0.97933 | 0.327416 | 0.944381 | 0.024853 |
| ENSMUSG00000078597.3 | 302.239 | -0.556 | 0.57229 | -0.97154 | 0.33128 | 0.944381 | 0.024853 |

| | | | | | | | |
|---|---|---|---|---|---|---|---|
| ENSMUSG00000079283.3 | 52.40525 | 1.321504 | 1.358934 | 0.972456 | 0.330824 | 0.944381 | 0.024853 |
| ENSMUSG00000091957.4 | 379.608 | 0.43611 | 0.450873 | 0.967257 | 0.333416 | 0.944381 | 0.024853 |
| ENSMUSG00000107552.2 | 23.99339 | -1.75014 | 1.788196 | -0.97872 | 0.327719 | 0.944381 | 0.024853 |
| ENSMUSG00000108621.2 | 114.8063 | 1.013209 | 1.043957 | 0.970547 | 0.331774 | 0.944381 | 0.024853 |
| ENSMUSG00000002280.12 | 25.56139 | -1.4882 | 1.541035 | -0.96571 | 0.334187 | 0.945124 | 0.024511 |
| ENSMUSG00000007564.16 | 66.24644 | -1.05928 | 1.096529 | -0.96603 | 0.334031 | 0.945124 | 0.024511 |
| ENSMUSG00000019948.5 | 93.29229 | 0.816184 | 0.845851 | 0.964926 | 0.334582 | 0.945124 | 0.024511 |
| ENSMUSG00000027642.16 | 886.176 | -0.32553 | 0.337347 | -0.96496 | 0.334564 | 0.945124 | 0.024511 |
| ENSMUSG00000028378.7 | 14.30218 | 1.666024 | 1.725734 | 0.9654 | 0.334344 | 0.945124 | 0.024511 |
| ENSMUSG00000028557.11 | 838.8075 | 0.421235 | 0.436307 | 0.965454 | 0.334317 | 0.945124 | 0.024511 |
| ENSMUSG00000029670.13 | 36.4319 | 1.61573 | 1.67419 | 0.965082 | 0.334504 | 0.945124 | 0.024511 |
| ENSMUSG00000052738.15 | 1457.293 | 0.272232 | 0.28195 | 0.965533 | 0.334278 | 0.945124 | 0.024511 |
| ENSMUSG00000107881.2 | 472.9208 | 0.437909 | 0.453081 | 0.966514 | 0.333787 | 0.945124 | 0.024511 |
| ENSMUSG00000000171.6 | 951.7427 | 0.313199 | 0.325262 | 0.96291 | 0.335592 | 0.945144 | 0.024502 |
| ENSMUSG00000002731.7 | 64.58738 | 0.915496 | 0.949937 | 0.963744 | 0.335174 | 0.945144 | 0.024502 |
| ENSMUSG00000003660.11 | 139.2545 | -0.74554 | 0.777922 | -0.95837 | 0.337875 | 0.945144 | 0.024502 |
| ENSMUSG00000004945.16 | 63.87376 | -1.17152 | 1.224651 | -0.95661 | 0.338763 | 0.945144 | 0.024502 |
| ENSMUSG00000007041.10 | 47.65725 | 1.037818 | 1.080674 | 0.960343 | 0.336883 | 0.945144 | 0.024502 |
| ENSMUSG00000015671.12 | 1190.383 | 0.358408 | 0.373475 | 0.959658 | 0.337228 | 0.945144 | 0.024502 |
| ENSMUSG00000017978.19 | 55.91989 | 0.939514 | 0.974485 | 0.964114 | 0.334989 | 0.945144 | 0.024502 |
| ENSMUSG00000020592.15 | 418.7054 | 0.511867 | 0.534542 | 0.957581 | 0.338274 | 0.945144 | 0.024502 |
| ENSMUSG00000020593.17 | 450.5771 | -0.47476 | 0.495959 | -0.95725 | 0.338441 | 0.945144 | 0.024502 |
| ENSMUSG00000021539.10 | 675.113 | -0.3508 | 0.365903 | -0.95874 | 0.337692 | 0.945144 | 0.024502 |
| ENSMUSG00000021765.10 | 27.81013 | -1.52513 | 1.581339 | -0.96445 | 0.334819 | 0.945144 | 0.024502 |
| ENSMUSG00000022498.17 | 104.9936 | -0.90656 | 0.947484 | -0.95681 | 0.338663 | 0.945144 | 0.024502 |
| ENSMUSG00000023886.11 | 64.07165 | -1.26794 | 1.324789 | -0.95709 | 0.338523 | 0.945144 | 0.024502 |
| ENSMUSG00000025354.6 | 48.83414 | 1.546102 | 1.613831 | 0.958032 | 0.338047 | 0.945144 | 0.024502 |
| ENSMUSG00000025509.16 | 68.31266 | -0.88669 | 0.926142 | -0.9574 | 0.338366 | 0.945144 | 0.024502 |
| ENSMUSG00000025939.20 | 109.3741 | 0.891488 | 0.927132 | 0.961555 | 0.336273 | 0.945144 | 0.024502 |
| ENSMUSG00000026353.10 | 1535.708 | 0.339732 | 0.354619 | 0.958022 | 0.338051 | 0.945144 | 0.024502 |
| ENSMUSG00000026750.7 | 733.559 | -0.35639 | 0.371961 | -0.95813 | 0.337997 | 0.945144 | 0.024502 |
| ENSMUSG00000027088.11 | 208.6577 | 0.622615 | 0.648168 | 0.960577 | 0.336765 | 0.945144 | 0.024502 |
| ENSMUSG00000028034.16 | 132.5509 | 0.66002 | 0.688426 | 0.958737 | 0.337691 | 0.945144 | 0.024502 |
| ENSMUSG00000028057.15 | 36.04602 | -1.33075 | 1.391428 | -0.95639 | 0.338876 | 0.945144 | 0.024502 |
| ENSMUSG00000031157.11 | 60.94177 | 0.919663 | 0.957129 | 0.960855 | 0.336625 | 0.945144 | 0.024502 |
| ENSMUSG00000033099.11 | 32.69969 | -1.6903 | 1.762792 | -0.95888 | 0.337621 | 0.945144 | 0.024502 |
| ENSMUSG00000033216.10 | 25.33138 | -1.65093 | 1.721597 | -0.95895 | 0.337583 | 0.945144 | 0.024502 |
| ENSMUSG00000033970.16 | 31.08561 | -1.64567 | 1.710008 | -0.96238 | 0.33586 | 0.945144 | 0.024502 |
| ENSMUSG00000036473.17 | 148.6789 | -0.74524 | 0.772842 | -0.96429 | 0.3349 | 0.945144 | 0.024502 |
| ENSMUSG00000036835.8 | 93.11439 | -0.69924 | 0.726002 | -0.96314 | 0.335475 | 0.945144 | 0.024502 |
| ENSMUSG00000036885.15 | 121.4253 | -0.81886 | 0.850879 | -0.96236 | 0.335867 | 0.945144 | 0.024502 |
| ENSMUSG00000038520.16 | 45.30192 | -1.48576 | 1.548563 | -0.95944 | 0.337335 | 0.945144 | 0.024502 |
| ENSMUSG00000039704.9 | 181.8806 | -0.57581 | 0.597761 | -0.96327 | 0.335411 | 0.945144 | 0.024502 |
| ENSMUSG00000041324.15 | 74.05484 | -1.02722 | 1.070343 | -0.95971 | 0.337201 | 0.945144 | 0.024502 |

| | | | | | | | |
|---|---|---|---|---|---|---|---|
| ENSMUSG00000042305.13 | 492.4053 | -0.4191 | 0.438176 | -0.95647 | 0.338833 | 0.945144 | 0.024502 |
| ENSMUSG00000049658.14 | 95.82828 | 0.859988 | 0.898648 | 0.956981 | 0.338577 | 0.945144 | 0.024502 |
| ENSMUSG00000055301.9 | 22.08272 | 1.512274 | 1.569403 | 0.963599 | 0.335247 | 0.945144 | 0.024502 |
| ENSMUSG00000060073.10 | 590.6093 | 0.312518 | 0.324176 | 0.964037 | 0.335027 | 0.945144 | 0.024502 |
| ENSMUSG00000061684.7 | 113.567 | -0.73934 | 0.771804 | -0.95794 | 0.338095 | 0.945144 | 0.024502 |
| ENSMUSG00000063060.7 | 31.83693 | 1.912999 | 1.987658 | 0.962439 | 0.335829 | 0.945144 | 0.024502 |
| ENSMUSG00000073563.4 | 248.3516 | 0.614091 | 0.641988 | 0.956546 | 0.338796 | 0.945144 | 0.024502 |
| ENSMUSG00000079557.12 | 137.9061 | -0.61556 | 0.639754 | -0.96217 | 0.335962 | 0.945144 | 0.024502 |
| ENSMUSG00000086370.10 | 18.36359 | 2.063105 | 2.148209 | 0.960383 | 0.336862 | 0.945144 | 0.024502 |
| ENSMUSG00000090942.2 | 66.73851 | -1.30752 | 1.359297 | -0.96191 | 0.336094 | 0.945144 | 0.024502 |
| ENSMUSG00000095041.8 | 117.422 | -0.67951 | 0.707884 | -0.95992 | 0.337095 | 0.945144 | 0.024502 |
| ENSMUSG00000117780.2 | 56.63449 | -1.23803 | 1.292834 | -0.95761 | 0.338259 | 0.945144 | 0.024502 |
| ENSMUSG00000022314.11 | 300.307 | -0.49159 | 0.514457 | -0.95555 | 0.339298 | 0.945466 | 0.024354 |
| ENSMUSG00000029823.17 | 252.5576 | -0.45788 | 0.479372 | -0.95516 | 0.339496 | 0.945466 | 0.024354 |
| ENSMUSG00000030304.12 | 405.9154 | 0.491049 | 0.513879 | 0.955573 | 0.339288 | 0.945466 | 0.024354 |
| ENSMUSG00000031666.16 | 130.663 | -0.65516 | 0.685743 | -0.9554 | 0.339376 | 0.945466 | 0.024354 |
| ENSMUSG00000052889.12 | 10.15519 | -2.03962 | 2.135344 | -0.95517 | 0.339491 | 0.945466 | 0.024354 |
| ENSMUSG00000099590.2 | 32.49155 | 1.515019 | 1.586448 | 0.954976 | 0.33959 | 0.945466 | 0.024354 |
| ENSMUSG00000112276.2 | 15.37466 | -2.42255 | 2.539302 | -0.95402 | 0.340072 | 0.946531 | 0.023865 |
| ENSMUSG00000071866.13 | 915.0543 | 0.287337 | 0.301421 | 0.953275 | 0.340451 | 0.947306 | 0.02351 |
| ENSMUSG00000000275.17 | 151.5941 | -0.71001 | 0.746273 | -0.95141 | 0.341395 | 0.947819 | 0.023275 |
| ENSMUSG00000005907.15 | 78.76248 | -0.79176 | 0.833837 | -0.94954 | 0.342347 | 0.947819 | 0.023275 |
| ENSMUSG00000007815.14 | 233.377 | 0.612247 | 0.642684 | 0.952641 | 0.340772 | 0.947819 | 0.023275 |
| ENSMUSG00000015143.16 | 72.18553 | -0.96524 | 1.017939 | -0.94823 | 0.343014 | 0.947819 | 0.023275 |
| ENSMUSG00000020392.9 | 17.94697 | 1.877157 | 1.976076 | 0.949942 | 0.342142 | 0.947819 | 0.023275 |
| ENSMUSG00000021258.11 | 83.17498 | -1.12743 | 1.191381 | -0.94632 | 0.343985 | 0.947819 | 0.023275 |
| ENSMUSG00000022667.19 | 11.17533 | 2.38142 | 2.506041 | 0.950272 | 0.341974 | 0.947819 | 0.023275 |
| ENSMUSG00000022855.10 | 67.77706 | 1.140492 | 1.201001 | 0.949618 | 0.342307 | 0.947819 | 0.023275 |
| ENSMUSG00000022898.14 | 67.05234 | 0.794638 | 0.839971 | 0.946031 | 0.344133 | 0.947819 | 0.023275 |
| ENSMUSG00000024410.16 | 49.09174 | -1.1951 | 1.263014 | -0.94623 | 0.344032 | 0.947819 | 0.023275 |
| ENSMUSG00000025034.10 | 66.32578 | -1.02248 | 1.077546 | -0.94889 | 0.342675 | 0.947819 | 0.023275 |
| ENSMUSG00000025503.6 | 494.538 | -0.45207 | 0.477858 | -0.94603 | 0.344134 | 0.947819 | 0.023275 |
| ENSMUSG00000026123.12 | 90.54634 | -0.88325 | 0.9324 | -0.94729 | 0.34349 | 0.947819 | 0.023275 |
| ENSMUSG00000026499.6 | 127.7735 | -0.73807 | 0.778431 | -0.94815 | 0.343054 | 0.947819 | 0.023275 |
| ENSMUSG00000027498.15 | 53.62 | -0.98909 | 1.04404 | -0.94737 | 0.34345 | 0.947819 | 0.023275 |
| ENSMUSG00000029563.17 | 15.97563 | -1.47171 | 1.551547 | -0.94854 | 0.342854 | 0.947819 | 0.023275 |
| ENSMUSG00000029716.14 | 709.4015 | -0.40274 | 0.424563 | -0.9486 | 0.342824 | 0.947819 | 0.023275 |
| ENSMUSG00000030341.18 | 79.36464 | -0.85115 | 0.896048 | -0.94989 | 0.342166 | 0.947819 | 0.023275 |
| ENSMUSG00000030378.16 | 7640.166 | -0.22905 | 0.240956 | -0.95059 | 0.341814 | 0.947819 | 0.023275 |
| ENSMUSG00000031375.18 | 167.2884 | -0.69801 | 0.737282 | -0.94674 | 0.343773 | 0.947819 | 0.023275 |
| ENSMUSG00000031565.19 | 53.07228 | -1.14587 | 1.203735 | -0.95193 | 0.341132 | 0.947819 | 0.023275 |
| ENSMUSG00000032010.16 | 301.8071 | -0.48194 | 0.506461 | -0.95159 | 0.341305 | 0.947819 | 0.023275 |
| ENSMUSG00000035929.12 | 15.78085 | -2.39609 | 2.528786 | -0.94752 | 0.343371 | 0.947819 | 0.023275 |
| ENSMUSG00000036023.7 | 38.74942 | -1.16478 | 1.230409 | -0.94666 | 0.343813 | 0.947819 | 0.023275 |

| | | | | | | | |
|---|---|---|---|---|---|---|---|
| ENSMUSG00000036027.14 | 12.48549 | 2.068033 | 2.174822 | 0.950897 | 0.341656 | 0.947819 | 0.023275 |
| ENSMUSG00000036617.18 | 50.00581 | 0.972625 | 1.024989 | 0.948912 | 0.342665 | 0.947819 | 0.023275 |
| ENSMUSG00000037936.16 | 138.7936 | -0.79152 | 0.833275 | -0.94989 | 0.342168 | 0.947819 | 0.023275 |
| ENSMUSG00000039844.21 | 41.426 | 1.070121 | 1.129264 | 0.947627 | 0.343319 | 0.947819 | 0.023275 |
| ENSMUSG00000042406.9 | 290.5418 | -0.59519 | 0.627998 | -0.94777 | 0.343249 | 0.947819 | 0.023275 |
| ENSMUSG00000043162.8 | 59.79825 | 1.05246 | 1.105818 | 0.951749 | 0.341225 | 0.947819 | 0.023275 |
| ENSMUSG00000045392.9 | 38.73841 | 1.391166 | 1.464313 | 0.950047 | 0.342089 | 0.947819 | 0.023275 |
| ENSMUSG00000052676.17 | 32.29148 | 1.412468 | 1.490064 | 0.947925 | 0.343168 | 0.947819 | 0.023275 |
| ENSMUSG00000062456.5 | 423.3164 | 0.44707 | 0.471283 | 0.948624 | 0.342812 | 0.947819 | 0.023275 |
| ENSMUSG00000072763.3 | 35.58147 | 1.515571 | 1.594129 | 0.95072 | 0.341746 | 0.947819 | 0.023275 |
| ENSMUSG00000086914.2 | 37.01882 | 1.468154 | 1.550815 | 0.946698 | 0.343793 | 0.947819 | 0.023275 |
| ENSMUSG00000013539.15 | 126.9244 | -0.65644 | 0.695966 | -0.94321 | 0.345573 | 0.948185 | 0.023107 |
| ENSMUSG00000021660.15 | 198.9242 | 0.596933 | 0.631999 | 0.944515 | 0.344907 | 0.948185 | 0.023107 |
| ENSMUSG00000022095.10 | 67.42391 | -0.90976 | 0.964156 | -0.94359 | 0.345382 | 0.948185 | 0.023107 |
| ENSMUSG00000022305.14 | 16.30284 | 2.057294 | 2.18133 | 0.943137 | 0.345611 | 0.948185 | 0.023107 |
| ENSMUSG00000022781.9 | 152.2298 | -0.63704 | 0.674897 | -0.94391 | 0.345217 | 0.948185 | 0.023107 |
| ENSMUSG00000026866.17 | 551.4536 | -0.36591 | 0.387702 | -0.94378 | 0.345282 | 0.948185 | 0.023107 |
| ENSMUSG00000031349.5 | 411.6895 | -0.46361 | 0.490711 | -0.94477 | 0.344774 | 0.948185 | 0.023107 |
| ENSMUSG00000031388.15 | 66.65315 | -1.03878 | 1.099085 | -0.94513 | 0.344592 | 0.948185 | 0.023107 |
| ENSMUSG00000031939.17 | 85.78542 | -0.88461 | 0.938183 | -0.94289 | 0.345735 | 0.948185 | 0.023107 |
| ENSMUSG00000034006.18 | 368.2062 | -0.39162 | 0.415222 | -0.94315 | 0.345605 | 0.948185 | 0.023107 |
| ENSMUSG00000035649.18 | 56.80882 | -0.87885 | 0.931015 | -0.94397 | 0.345185 | 0.948185 | 0.023107 |
| ENSMUSG00000039671.19 | 22.40927 | 1.657495 | 1.757998 | 0.942831 | 0.345767 | 0.948185 | 0.023107 |
| ENSMUSG00000040236.5 | 142.4912 | 0.808949 | 0.857176 | 0.943737 | 0.345304 | 0.948185 | 0.023107 |
| ENSMUSG00000046879.8 | 241.5606 | -0.48601 | 0.514829 | -0.94401 | 0.345163 | 0.948185 | 0.023107 |
| ENSMUSG00000052040.11 | 49.18496 | -0.97156 | 1.029302 | -0.9439 | 0.34522 | 0.948185 | 0.023107 |
| ENSMUSG00000000759.15 | 26.18342 | 1.285816 | 1.365455 | 0.941675 | 0.346359 | 0.94832 | 0.023045 |
| ENSMUSG00000027215.14 | 14.4656 | -2.52747 | 2.683146 | -0.94198 | 0.346202 | 0.94832 | 0.023045 |
| ENSMUSG00000028234.7 | 2003.313 | 0.30788 | 0.326699 | 0.942397 | 0.345989 | 0.94832 | 0.023045 |
| ENSMUSG00000037499.10 | 72.69909 | -0.9723 | 1.032494 | -0.9417 | 0.346347 | 0.94832 | 0.023045 |
| ENSMUSG00000045996.13 | 54.62213 | 0.898209 | 0.953956 | 0.941562 | 0.346417 | 0.94832 | 0.023045 |
| ENSMUSG00000075225.16 | 48.94216 | -1.09814 | 1.165519 | -0.94219 | 0.346096 | 0.94832 | 0.023045 |
| ENSMUSG00000018841.18 | 64.53944 | -1.31564 | 1.397923 | -0.94114 | 0.346632 | 0.948462 | 0.02298 |
| ENSMUSG00000024833.16 | 15.71414 | -2.0409 | 2.168701 | -0.94107 | 0.346669 | 0.948462 | 0.02298 |
| ENSMUSG00000002957.12 | 38.85311 | -1.21607 | 1.293323 | -0.94027 | 0.347081 | 0.948588 | 0.022922 |
| ENSMUSG00000028980.15 | 590.5064 | -0.44135 | 0.469412 | -0.94021 | 0.34711 | 0.948588 | 0.022922 |
| ENSMUSG00000036501.8 | 10.10926 | 2.033259 | 2.162582 | 0.9402 | 0.347115 | 0.948588 | 0.022922 |
| ENSMUSG00000042642.14 | 52.14479 | -1.15642 | 1.229752 | -0.94037 | 0.347028 | 0.948588 | 0.022922 |
| ENSMUSG00000064717.3 | 16.42384 | 1.599197 | 1.701354 | 0.939956 | 0.34724 | 0.948657 | 0.022891 |
| ENSMUSG00000084349.4 | 175.7047 | 0.501886 | 0.534302 | 0.93933 | 0.347561 | 0.948987 | 0.02274 |
| ENSMUSG00000113555.2 | 16.23005 | -1.9403 | 2.065548 | -0.93936 | 0.347544 | 0.948987 | 0.02274 |
| ENSMUSG00000097203.3 | 94.01391 | 0.876781 | 0.933794 | 0.938945 | 0.347759 | 0.949253 | 0.022618 |
| ENSMUSG00000090386.11 | 30.9437 | 1.786372 | 1.903035 | 0.938696 | 0.347887 | 0.949329 | 0.022583 |
| ENSMUSG00000000168.11 | 289.363 | 0.488301 | 0.522098 | 0.935267 | 0.349651 | 0.949599 | 0.02246 |

| | | | | | | | |
|---|---|---|---|---|---|---|---|
| ENSMUSG00000003546.11 | 60.69068 | -1.17708 | 1.255114 | -0.93783 | 0.348332 | 0.949599 | 0.02246 |
| ENSMUSG00000014074.18 | 62.84094 | -1.23314 | 1.319422 | -0.93461 | 0.349989 | 0.949599 | 0.02246 |
| ENSMUSG00000018433.15 | 81.15393 | 1.016574 | 1.086643 | 0.935518 | 0.349522 | 0.949599 | 0.02246 |
| ENSMUSG00000020326.8 | 259.2195 | 0.555573 | 0.592234 | 0.938096 | 0.348195 | 0.949599 | 0.02246 |
| ENSMUSG00000024887.11 | 126.1748 | -0.75872 | 0.809317 | -0.93748 | 0.348513 | 0.949599 | 0.02246 |
| ENSMUSG00000028630.10 | 379.2101 | 0.400782 | 0.428405 | 0.935522 | 0.349519 | 0.949599 | 0.02246 |
| ENSMUSG00000028975.17 | 12.37992 | -2.23371 | 2.384613 | -0.93672 | 0.348903 | 0.949599 | 0.02246 |
| ENSMUSG00000029405.17 | 514.4669 | 0.316427 | 0.338564 | 0.934616 | 0.349986 | 0.949599 | 0.02246 |
| ENSMUSG00000030733.11 | 25.30929 | -1.93613 | 2.070069 | -0.9353 | 0.349634 | 0.949599 | 0.02246 |
| ENSMUSG00000030946.14 | 295.8606 | -0.47576 | 0.508337 | -0.93591 | 0.349318 | 0.949599 | 0.02246 |
| ENSMUSG00000031903.8 | 17.13061 | 2.156591 | 2.299962 | 0.937664 | 0.348417 | 0.949599 | 0.02246 |
| ENSMUSG00000032458.7 | 639.8244 | -0.32891 | 0.351764 | -0.93503 | 0.349773 | 0.949599 | 0.02246 |
| ENSMUSG00000033910.14 | 45.14792 | -1.02233 | 1.093555 | -0.93486 | 0.349858 | 0.949599 | 0.02246 |
| ENSMUSG00000039789.8 | 34.2875 | 1.338732 | 1.429786 | 0.936316 | 0.34911 | 0.949599 | 0.02246 |
| ENSMUSG00000050503.10 | 13.72113 | -2.81183 | 3.000138 | -0.93723 | 0.348638 | 0.949599 | 0.02246 |
| ENSMUSG00000055923.17 | 12.58205 | -2.53953 | 2.711352 | -0.93663 | 0.34895 | 0.949599 | 0.02246 |
| ENSMUSG00000067365.12 | 121.3326 | 0.690006 | 0.737563 | 0.935521 | 0.34952 | 0.949599 | 0.02246 |
| ENSMUSG00000069899.4 | 51.88969 | -1.05102 | 1.121703 | -0.93699 | 0.348764 | 0.949599 | 0.02246 |
| ENSMUSG00000071041.5 | 15.54239 | 1.641154 | 1.753299 | 0.936038 | 0.349254 | 0.949599 | 0.02246 |
| ENSMUSG00000015971.6 | 161.1856 | 0.69591 | 0.745753 | 0.933164 | 0.350735 | 0.94972 | 0.022404 |
| ENSMUSG00000024955.16 | 24.10316 | -1.48014 | 1.584917 | -0.93389 | 0.35036 | 0.94972 | 0.022404 |
| ENSMUSG00000027495.5 | 259.6297 | 0.459451 | 0.492175 | 0.933512 | 0.350556 | 0.94972 | 0.022404 |
| ENSMUSG00000038079.15 | 12.53569 | 1.960861 | 2.099333 | 0.93404 | 0.350283 | 0.94972 | 0.022404 |
| ENSMUSG00000056836.9 | 12.83334 | 2.037476 | 2.1831 | 0.933295 | 0.350668 | 0.94972 | 0.022404 |
| ENSMUSG00000061132.14 | 19.21024 | -2.23748 | 2.396059 | -0.93382 | 0.350398 | 0.94972 | 0.022404 |
| ENSMUSG00000081049.2 | 29.81924 | 1.102102 | 1.180682 | 0.933445 | 0.35059 | 0.94972 | 0.022404 |
| ENSMUSG00000007739.11 | 712.68 | -0.31278 | 0.335757 | -0.93157 | 0.351561 | 0.950259 | 0.022158 |
| ENSMUSG00000020733.4 | 321.4039 | -0.53055 | 0.570249 | -0.93039 | 0.35217 | 0.950259 | 0.022158 |
| ENSMUSG00000026155.14 | 257.2846 | -0.47881 | 0.514652 | -0.93036 | 0.352186 | 0.950259 | 0.022158 |
| ENSMUSG00000026395.17 | 16.08019 | 1.804983 | 1.938241 | 0.931248 | 0.351725 | 0.950259 | 0.022158 |
| ENSMUSG00000028199.19 | 372.2004 | -0.42967 | 0.461363 | -0.93131 | 0.351691 | 0.950259 | 0.022158 |
| ENSMUSG00000028411.16 | 21.23983 | -1.8764 | 2.016664 | -0.93045 | 0.352139 | 0.950259 | 0.022158 |
| ENSMUSG00000029148.16 | 12.97585 | -2.46436 | 2.649118 | -0.93026 | 0.352237 | 0.950259 | 0.022158 |
| ENSMUSG00000030826.20 | 17.68182 | 1.648557 | 1.770889 | 0.930921 | 0.351895 | 0.950259 | 0.022158 |
| ENSMUSG00000036737.13 | 83.06233 | 0.918441 | 0.986463 | 0.931045 | 0.35183 | 0.950259 | 0.022158 |
| ENSMUSG00000040813.17 | 81.08882 | -0.79081 | 0.848434 | -0.93208 | 0.351296 | 0.950259 | 0.022158 |
| ENSMUSG00000049504.13 | 34.75751 | -1.69241 | 1.818485 | -0.93067 | 0.352025 | 0.950259 | 0.022158 |
| ENSMUSG00000062075.14 | 11.46862 | 2.515233 | 2.699809 | 0.931634 | 0.351526 | 0.950259 | 0.022158 |
| ENSMUSG00000064120.15 | 112.8512 | -0.70366 | 0.755347 | -0.93157 | 0.351561 | 0.950259 | 0.022158 |
| ENSMUSG00000027387.12 | 48.42171 | -1.25713 | 1.352527 | -0.92947 | 0.352647 | 0.950974 | 0.021831 |
| ENSMUSG00000032311.18 | 81.34625 | 1.130123 | 1.216024 | 0.929359 | 0.352703 | 0.950974 | 0.021831 |
| ENSMUSG00000002102.16 | 389.9957 | -0.49707 | 0.536679 | -0.92619 | 0.354348 | 0.951462 | 0.021609 |
| ENSMUSG00000004100.13 | 14.7132 | 2.047489 | 2.209332 | 0.926746 | 0.354059 | 0.951462 | 0.021609 |
| ENSMUSG00000015522.19 | 102.7607 | 0.760184 | 0.818825 | 0.928384 | 0.353209 | 0.951462 | 0.021609 |

| | | | | | | | |
|---|---|---|---|---|---|---|---|
| ENSMUSG00000020448.17 | 135.2135 | -0.71428 | 0.771037 | -0.92639 | 0.354245 | 0.951462 | 0.021609 |
| ENSMUSG00000024516.14 | 463.055 | -0.36568 | 0.394632 | -0.92664 | 0.354112 | 0.951462 | 0.021609 |
| ENSMUSG00000027312.15 | 142.8573 | -0.61915 | 0.668092 | -0.92675 | 0.354057 | 0.951462 | 0.021609 |
| ENSMUSG00000029141.17 | 20.74368 | -1.75715 | 1.896543 | -0.9265 | 0.354186 | 0.951462 | 0.021609 |
| ENSMUSG00000029422.16 | 447.0026 | 0.478021 | 0.516161 | 0.926109 | 0.35439 | 0.951462 | 0.021609 |
| ENSMUSG00000029714.12 | 26.79228 | -1.30864 | 1.412833 | -0.92625 | 0.354315 | 0.951462 | 0.021609 |
| ENSMUSG00000029761.17 | 2001.425 | -0.25971 | 0.280068 | -0.9273 | 0.353772 | 0.951462 | 0.021609 |
| ENSMUSG00000031090.15 | 39.87849 | -1.39661 | 1.505313 | -0.92778 | 0.353519 | 0.951462 | 0.021609 |
| ENSMUSG00000032244.10 | 55.24355 | -1.19991 | 1.29294 | -0.92805 | 0.353382 | 0.951462 | 0.021609 |
| ENSMUSG00000034761.16 | 107.5605 | -0.86403 | 0.931474 | -0.9276 | 0.353616 | 0.951462 | 0.021609 |
| ENSMUSG00000047867.12 | 47.80372 | 1.206548 | 1.301897 | 0.926762 | 0.35405 | 0.951462 | 0.021609 |
| ENSMUSG00000059474.14 | 39.4029 | -1.00966 | 1.088818 | -0.9273 | 0.353771 | 0.951462 | 0.021609 |
| ENSMUSG00000031196.14 | 59.98907 | -1.31299 | 1.418352 | -0.92571 | 0.354594 | 0.951612 | 0.02154 |
| ENSMUSG00000059461.8 | 104.5021 | 0.731875 | 0.79069 | 0.925615 | 0.354646 | 0.951612 | 0.02154 |
| ENSMUSG00000027907.5 | 33.21487 | 1.439676 | 1.55575 | 0.925391 | 0.354763 | 0.951655 | 0.02152 |
| ENSMUSG00000025317.8 | 74.65926 | 1.257714 | 1.360338 | 0.92456 | 0.355195 | 0.951767 | 0.021469 |
| ENSMUSG00000030101.16 | 114.5937 | 0.723512 | 0.782701 | 0.924378 | 0.35529 | 0.951767 | 0.021469 |
| ENSMUSG00000031691.15 | 41.42064 | -1.35809 | 1.469514 | -0.92418 | 0.355393 | 0.951767 | 0.021469 |
| ENSMUSG00000034168.8 | 88.87547 | 0.819698 | 0.886388 | 0.924762 | 0.355089 | 0.951767 | 0.021469 |
| ENSMUSG00000034449.8 | 68.49455 | -0.92292 | 0.998808 | -0.92402 | 0.355477 | 0.951767 | 0.021469 |
| ENSMUSG00000037197.12 | 158.1101 | -0.72616 | 0.78583 | -0.92406 | 0.355455 | 0.951767 | 0.021469 |
| ENSMUSG00000040429.7 | 11.67093 | -2.06392 | 2.233777 | -0.92396 | 0.355507 | 0.951767 | 0.021469 |
| ENSMUSG00000002948.20 | 27.3593 | 1.416601 | 1.538881 | 0.920539 | 0.357291 | 0.951775 | 0.021466 |
| ENSMUSG00000021715.13 | 40.86012 | 1.20838 | 1.312711 | 0.920523 | 0.3573 | 0.951775 | 0.021466 |
| ENSMUSG00000022106.15 | 290.7335 | -0.4275 | 0.465225 | -0.91891 | 0.358143 | 0.951775 | 0.021466 |
| ENSMUSG00000022332.9 | 104.0533 | -0.66263 | 0.717747 | -0.92321 | 0.355897 | 0.951775 | 0.021466 |
| ENSMUSG00000025130.12 | 2124.376 | -0.29934 | 0.324566 | -0.92227 | 0.356388 | 0.951775 | 0.021466 |
| ENSMUSG00000025613.14 | 1016.224 | 0.281527 | 0.306187 | 0.919463 | 0.357854 | 0.951775 | 0.021466 |
| ENSMUSG00000027359.17 | 2611.805 | 0.225524 | 0.244636 | 0.921877 | 0.356593 | 0.951775 | 0.021466 |
| ENSMUSG00000029472.14 | 137.3695 | 0.717596 | 0.781979 | 0.917667 | 0.358793 | 0.951775 | 0.021466 |
| ENSMUSG00000029695.14 | 2264.927 | 0.309549 | 0.335483 | 0.922694 | 0.356167 | 0.951775 | 0.021466 |
| ENSMUSG00000031221.8 | 285.6214 | -0.45655 | 0.496175 | -0.92015 | 0.357495 | 0.951775 | 0.021466 |
| ENSMUSG00000031445.6 | 1501.401 | -0.28175 | 0.306235 | -0.92006 | 0.357542 | 0.951775 | 0.021466 |
| ENSMUSG00000032786.17 | 1754.932 | 0.379486 | 0.41356 | 0.91761 | 0.358823 | 0.951775 | 0.021466 |
| ENSMUSG00000034450.9 | 702.2851 | -0.46178 | 0.502132 | -0.91964 | 0.357759 | 0.951775 | 0.021466 |
| ENSMUSG00000035164.9 | 20.28809 | 1.321082 | 1.430619 | 0.923434 | 0.355781 | 0.951775 | 0.021466 |
| ENSMUSG00000035933.6 | 53.19484 | -1.07919 | 1.173773 | -0.91942 | 0.357875 | 0.951775 | 0.021466 |
| ENSMUSG00000036552.20 | 48.33003 | -1.25466 | 1.365712 | -0.91868 | 0.358261 | 0.951775 | 0.021466 |
| ENSMUSG00000038188.18 | 19.51663 | -2.24354 | 2.443713 | -0.91809 | 0.358574 | 0.951775 | 0.021466 |
| ENSMUSG00000039405.8 | 15.84883 | 1.713326 | 1.865317 | 0.918517 | 0.358348 | 0.951775 | 0.021466 |
| ENSMUSG00000039509.9 | 54.53645 | -0.97269 | 1.059456 | -0.91811 | 0.358562 | 0.951775 | 0.021466 |
| ENSMUSG00000039988.15 | 137.4812 | -0.59268 | 0.643973 | -0.92036 | 0.357386 | 0.951775 | 0.021466 |
| ENSMUSG00000040549.17 | 57.89609 | 1.137954 | 1.233473 | 0.922561 | 0.356236 | 0.951775 | 0.021466 |
| ENSMUSG00000042632.18 | 25.07669 | 1.807413 | 1.961384 | 0.921499 | 0.35679 | 0.951775 | 0.021466 |

| | | | | | | | |
|---|---|---|---|---|---|---|---|
| ENSMUSG00000049038.4 | 32.33125 | 1.438407 | 1.565939 | 0.918558 | 0.358327 | 0.951775 | 0.021466 |
| ENSMUSG00000051236.14 | 86.5619 | -0.73855 | 0.80379 | -0.91884 | 0.358179 | 0.951775 | 0.021466 |
| ENSMUSG00000052062.15 | 14.54599 | -2.12825 | 2.317856 | -0.9182 | 0.358515 | 0.951775 | 0.021466 |
| ENSMUSG00000057278.9 | 229.4641 | 0.635755 | 0.692504 | 0.918053 | 0.358591 | 0.951775 | 0.021466 |
| ENSMUSG00000058492.6 | 1234.965 | 0.335095 | 0.363131 | 0.922795 | 0.356114 | 0.951775 | 0.021466 |
| ENSMUSG00000058886.10 | 12.57882 | -2.54737 | 2.759958 | -0.92297 | 0.356021 | 0.951775 | 0.021466 |
| ENSMUSG00000061559.16 | 417.8778 | -0.45556 | 0.496394 | -0.91773 | 0.358759 | 0.951775 | 0.021466 |
| ENSMUSG00000066233.7 | 32.29127 | 1.761297 | 1.915479 | 0.919507 | 0.35783 | 0.951775 | 0.021466 |
| ENSMUSG00000068742.12 | 67.55645 | -0.82739 | 0.899353 | -0.91998 | 0.357584 | 0.951775 | 0.021466 |
| ENSMUSG00000095567.9 | 29.99071 | 1.986655 | 2.157742 | 0.92071 | 0.357202 | 0.951775 | 0.021466 |
| ENSMUSG00000098234.8 | 278.6312 | 0.488384 | 0.528762 | 0.923638 | 0.355675 | 0.951775 | 0.021466 |
| ENSMUSG00000021595.19 | 269.44 | -0.51026 | 0.556242 | -0.91733 | 0.358971 | 0.951902 | 0.021408 |
| ENSMUSG00000028636.15 | 208.4537 | 0.571781 | 0.623519 | 0.917023 | 0.35913 | 0.952057 | 0.021337 |
| ENSMUSG00000015217.12 | 108.4485 | 0.863949 | 0.943297 | 0.915883 | 0.359728 | 0.952596 | 0.021091 |
| ENSMUSG00000020131.15 | 54.20115 | 1.413906 | 1.543115 | 0.916267 | 0.359527 | 0.952596 | 0.021091 |
| ENSMUSG00000026078.11 | 139.5096 | 0.561584 | 0.613017 | 0.916099 | 0.359615 | 0.952596 | 0.021091 |
| ENSMUSG00000026888.15 | 297.4449 | 0.414913 | 0.453063 | 0.915796 | 0.359774 | 0.952596 | 0.021091 |
| ENSMUSG00000042505.7 | 128.5622 | 0.728487 | 0.795572 | 0.915677 | 0.359836 | 0.952596 | 0.021091 |
| ENSMUSG00000066838.8 | 18.97543 | 2.57024 | 2.808371 | 0.915207 | 0.360083 | 0.952983 | 0.020915 |
| ENSMUSG00000038811.14 | 35.3745 | 1.460105 | 1.595736 | 0.915004 | 0.360189 | 0.952999 | 0.020908 |
| ENSMUSG00000006611.16 | 99.35588 | 0.696329 | 0.764717 | 0.910571 | 0.362521 | 0.95315 | 0.020839 |
| ENSMUSG00000014294.5 | 1737.764 | 0.280454 | 0.308214 | 0.909932 | 0.362858 | 0.95315 | 0.020839 |
| ENSMUSG00000019841.16 | 122.2267 | -0.85339 | 0.937717 | -0.91007 | 0.362787 | 0.95315 | 0.020839 |
| ENSMUSG00000021482.11 | 492.3962 | 0.366929 | 0.40325 | 0.909928 | 0.362861 | 0.95315 | 0.020839 |
| ENSMUSG00000021550.7 | 200.6242 | -0.52996 | 0.581649 | -0.91114 | 0.362222 | 0.95315 | 0.020839 |
| ENSMUSG00000024109.19 | 67.22412 | -0.8683 | 0.949614 | -0.91437 | 0.36052 | 0.95315 | 0.020839 |
| ENSMUSG00000024165.10 | 68.55611 | -0.86225 | 0.944261 | -0.91315 | 0.361162 | 0.95315 | 0.020839 |
| ENSMUSG00000024354.14 | 236.2713 | 0.555384 | 0.609355 | 0.911429 | 0.362069 | 0.95315 | 0.020839 |
| ENSMUSG00000026227.12 | 64.6854 | -0.92762 | 1.019177 | -0.91017 | 0.362735 | 0.95315 | 0.020839 |
| ENSMUSG00000026577.14 | 46.70155 | -1.17585 | 1.291214 | -0.91066 | 0.362477 | 0.95315 | 0.020839 |
| ENSMUSG00000028249.16 | 330.0779 | -0.40237 | 0.440491 | -0.91345 | 0.361005 | 0.95315 | 0.020839 |
| ENSMUSG00000028551.15 | 55.60583 | -0.89782 | 0.985022 | -0.91147 | 0.362049 | 0.95315 | 0.020839 |
| ENSMUSG00000030002.16 | 55.66064 | -0.99172 | 1.088261 | -0.91129 | 0.362142 | 0.95315 | 0.020839 |
| ENSMUSG00000031980.11 | 852.3992 | -0.29652 | 0.325156 | -0.91194 | 0.361803 | 0.95315 | 0.020839 |
| ENSMUSG00000041757.18 | 90.66271 | -0.79674 | 0.872019 | -0.91367 | 0.360892 | 0.95315 | 0.020839 |
| ENSMUSG00000053617.13 | 23.1913 | 1.595303 | 1.747614 | 0.912846 | 0.361323 | 0.95315 | 0.020839 |
| ENSMUSG00000054766.14 | 355.7655 | 0.439103 | 0.480642 | 0.913575 | 0.36094 | 0.95315 | 0.020839 |
| ENSMUSG00000055312.3 | 81.69858 | 0.99197 | 1.087145 | 0.912454 | 0.36153 | 0.95315 | 0.020839 |
| ENSMUSG00000059159.7 | 77.92408 | 0.756845 | 0.831167 | 0.910581 | 0.362516 | 0.95315 | 0.020839 |
| ENSMUSG00000060152.15 | 323.9205 | 0.422252 | 0.461674 | 0.914612 | 0.360395 | 0.95315 | 0.020839 |
| ENSMUSG00000060429.13 | 236.2921 | 0.472841 | 0.519007 | 0.91105 | 0.362269 | 0.95315 | 0.020839 |
| ENSMUSG00000060586.12 | 42.56659 | -1.34827 | 1.481049 | -0.91035 | 0.362638 | 0.95315 | 0.020839 |
| ENSMUSG00000064128.9 | 13.74719 | 1.803947 | 1.974383 | 0.913676 | 0.360887 | 0.95315 | 0.020839 |
| ENSMUSG00000069769.14 | 38.76186 | 1.089542 | 1.195949 | 0.911027 | 0.362281 | 0.95315 | 0.020839 |

| | | | | | | | |
|---|---|---|---|---|---|---|---|
| ENSMUSG00000079227.11 | 27.50053 | 1.419502 | 1.559834 | 0.910034 | 0.362805 | 0.95315 | 0.020839 |
| ENSMUSG00000095432.10 | 16.99665 | 1.545206 | 1.690416 | 0.914098 | 0.360665 | 0.95315 | 0.020839 |
| ENSMUSG00000000532.12 | 16.3386 | 1.952575 | 2.175233 | 0.897639 | 0.369378 | 0.953413 | 0.020719 |
| ENSMUSG00000000568.16 | 193.2966 | -0.58699 | 0.65343 | -0.89832 | 0.369016 | 0.953413 | 0.020719 |
| ENSMUSG00000002103.17 | 182.6872 | -0.48685 | 0.538784 | -0.90361 | 0.366203 | 0.953413 | 0.020719 |
| ENSMUSG00000002297.17 | 10.10752 | -2.32804 | 2.562746 | -0.90842 | 0.363659 | 0.953413 | 0.020719 |
| ENSMUSG00000005982.15 | 521.8373 | 0.419908 | 0.46188 | 0.909129 | 0.363282 | 0.953413 | 0.020719 |
| ENSMUSG00000008855.18 | 38.22381 | 1.502801 | 1.663427 | 0.903437 | 0.366294 | 0.953413 | 0.020719 |
| ENSMUSG00000017493.13 | 2541.097 | -0.2949 | 0.325022 | -0.90733 | 0.364234 | 0.953413 | 0.020719 |
| ENSMUSG00000020485.15 | 107.7677 | -0.87195 | 0.968459 | -0.90035 | 0.367936 | 0.953413 | 0.020719 |
| ENSMUSG00000020955.10 | 129.4593 | 0.684027 | 0.756145 | 0.904624 | 0.365664 | 0.953413 | 0.020719 |
| ENSMUSG00000021076.7 | 188.1668 | -0.49515 | 0.55024 | -0.89988 | 0.368184 | 0.953413 | 0.020719 |
| ENSMUSG00000021681.9 | 223.3041 | -0.44538 | 0.49392 | -0.90172 | 0.367207 | 0.953413 | 0.020719 |
| ENSMUSG00000021703.10 | 150.8562 | 0.670429 | 0.746247 | 0.898401 | 0.368972 | 0.953413 | 0.020719 |
| ENSMUSG00000021832.9 | 1101.085 | -0.33245 | 0.367816 | -0.90386 | 0.366072 | 0.953413 | 0.020719 |
| ENSMUSG00000021998.17 | 437.8484 | -0.43054 | 0.474583 | -0.90719 | 0.364308 | 0.953413 | 0.020719 |
| ENSMUSG00000022404.9 | 256.6992 | 0.518761 | 0.57794 | 0.897603 | 0.369397 | 0.953413 | 0.020719 |
| ENSMUSG00000022555.13 | 11.24029 | 2.253012 | 2.482468 | 0.907569 | 0.364106 | 0.953413 | 0.020719 |
| ENSMUSG00000022807.10 | 75.67267 | 0.790511 | 0.88015 | 0.898154 | 0.369103 | 0.953413 | 0.020719 |
| ENSMUSG00000023845.8 | 98.84886 | -0.77188 | 0.857817 | -0.89982 | 0.368214 | 0.953413 | 0.020719 |
| ENSMUSG00000023994.14 | 18.2063 | 1.888523 | 2.078263 | 0.908702 | 0.363507 | 0.953413 | 0.020719 |
| ENSMUSG00000024571.16 | 227.663 | 0.463806 | 0.510753 | 0.908082 | 0.363835 | 0.953413 | 0.020719 |
| ENSMUSG00000025157.9 | 78.70981 | 0.853215 | 0.949098 | 0.898974 | 0.368666 | 0.953413 | 0.020719 |
| ENSMUSG00000025270.14 | 120.4959 | -0.69291 | 0.769118 | -0.90091 | 0.367634 | 0.953413 | 0.020719 |
| ENSMUSG00000025810.10 | 245.4619 | -0.6169 | 0.682025 | -0.90451 | 0.365725 | 0.953413 | 0.020719 |
| ENSMUSG00000026159.14 | 131.9075 | -0.57408 | 0.635879 | -0.90281 | 0.366629 | 0.953413 | 0.020719 |
| ENSMUSG00000026669.15 | 389.3129 | -0.39168 | 0.436091 | -0.89815 | 0.369105 | 0.953413 | 0.020719 |
| ENSMUSG00000026687.15 | 1181.436 | 0.280514 | 0.310204 | 0.904288 | 0.365843 | 0.953413 | 0.020719 |
| ENSMUSG00000027384.7 | 27.86318 | 1.311662 | 1.445248 | 0.907569 | 0.364106 | 0.953413 | 0.020719 |
| ENSMUSG00000028015.4 | 242.8488 | 0.483865 | 0.533914 | 0.906262 | 0.364797 | 0.953413 | 0.020719 |
| ENSMUSG00000029263.16 | 69.52243 | -0.7954 | 0.881628 | -0.9022 | 0.366952 | 0.953413 | 0.020719 |
| ENSMUSG00000030407.3 | 44.67115 | -1.41 | 1.561008 | -0.90326 | 0.366387 | 0.953413 | 0.020719 |
| ENSMUSG00000030894.6 | 414.9965 | 0.532019 | 0.586919 | 0.906462 | 0.364691 | 0.953413 | 0.020719 |
| ENSMUSG00000031714.10 | 66.27625 | 0.768431 | 0.848158 | 0.905999 | 0.364936 | 0.953413 | 0.020719 |
| ENSMUSG00000032231.15 | 14.05607 | 2.114578 | 2.328323 | 0.908198 | 0.363774 | 0.953413 | 0.020719 |
| ENSMUSG00000036446.6 | 83.0415 | 0.985706 | 1.090986 | 0.9035 | 0.36626 | 0.953413 | 0.020719 |
| ENSMUSG00000036594.16 | 57.03594 | -1.15662 | 1.27974 | -0.90379 | 0.366106 | 0.953413 | 0.020719 |
| ENSMUSG00000037058.17 | 926.6204 | 0.261487 | 0.290002 | 0.901673 | 0.367231 | 0.953413 | 0.020719 |
| ENSMUSG00000037824.7 | 110.5588 | 0.649853 | 0.723138 | 0.898658 | 0.368835 | 0.953413 | 0.020719 |
| ENSMUSG00000038384.17 | 49.30462 | -0.86605 | 0.961285 | -0.90093 | 0.367628 | 0.953413 | 0.020719 |
| ENSMUSG00000038622.9 | 57.43209 | -0.94255 | 1.043146 | -0.90357 | 0.366224 | 0.953413 | 0.020719 |
| ENSMUSG00000039108.15 | 48.10647 | -1.17613 | 1.300286 | -0.90452 | 0.365722 | 0.953413 | 0.020719 |
| ENSMUSG00000039316.15 | 24.2759 | 1.828646 | 2.032021 | 0.899915 | 0.368165 | 0.953413 | 0.020719 |
| ENSMUSG00000041272.12 | 19.99725 | -1.53674 | 1.710426 | -0.89845 | 0.368944 | 0.953413 | 0.020719 |

| | | | | | | | |
|---|---|---|---|---|---|---|---|
| ENSMUSG00000042246.6 | 12.93156 | 2.429291 | 2.67222 | 0.909091 | 0.363302 | 0.953413 | 0.020719 |
| ENSMUSG00000043013.11 | 26.25491 | -1.52151 | 1.678749 | -0.90633 | 0.364759 | 0.953413 | 0.020719 |
| ENSMUSG00000045092.9 | 379.4207 | 0.363141 | 0.400463 | 0.906802 | 0.364511 | 0.953413 | 0.020719 |
| ENSMUSG00000049339.17 | 165.8399 | 0.510854 | 0.568447 | 0.898683 | 0.368821 | 0.953413 | 0.020719 |
| ENSMUSG00000052305.7 | 315.8117 | -0.46678 | 0.518662 | -0.89998 | 0.368133 | 0.953413 | 0.020719 |
| ENSMUSG00000053841.16 | 53.64729 | -1.19041 | 1.320645 | -0.90138 | 0.367385 | 0.953413 | 0.020719 |
| ENSMUSG00000056201.9 | 734.2791 | -0.29493 | 0.327556 | -0.9004 | 0.367908 | 0.953413 | 0.020719 |
| ENSMUSG00000058267.14 | 441.0378 | 0.350045 | 0.388367 | 0.901326 | 0.367415 | 0.953413 | 0.020719 |
| ENSMUSG00000059475.14 | 24.73726 | 1.539059 | 1.703047 | 0.903709 | 0.36615 | 0.953413 | 0.020719 |
| ENSMUSG00000061232.17 | 7745.958 | -0.17855 | 0.196492 | -0.9087 | 0.363509 | 0.953413 | 0.020719 |
| ENSMUSG00000061607.16 | 21.76794 | 1.71063 | 1.884333 | 0.907817 | 0.363975 | 0.953413 | 0.020719 |
| ENSMUSG00000061650.7 | 31.0096 | 1.472631 | 1.636309 | 0.899971 | 0.368136 | 0.953413 | 0.020719 |
| ENSMUSG00000062352.16 | 116.393 | -0.75555 | 0.841633 | -0.89771 | 0.369338 | 0.953413 | 0.020719 |
| ENSMUSG00000063145.11 | 19.87286 | -1.46187 | 1.616939 | -0.9041 | 0.365944 | 0.953413 | 0.020719 |
| ENSMUSG00000063590.4 | 140.6826 | -0.60591 | 0.671909 | -0.90177 | 0.367181 | 0.953413 | 0.020719 |
| ENSMUSG00000066152.12 | 60.95893 | 0.874145 | 0.971238 | 0.900031 | 0.368104 | 0.953413 | 0.020719 |
| ENSMUSG00000071014.11 | 2063.337 | -0.23162 | 0.256443 | -0.90319 | 0.366424 | 0.953413 | 0.020719 |
| ENSMUSG00000078185.4 | 16.53932 | 1.69123 | 1.870573 | 0.904124 | 0.36593 | 0.953413 | 0.020719 |
| ENSMUSG00000079419.5 | 13.7459 | -2.40418 | 2.676595 | -0.89822 | 0.369066 | 0.953413 | 0.020719 |
| ENSMUSG00000091337.4 | 13.67025 | -1.73213 | 1.924477 | -0.90005 | 0.368092 | 0.953413 | 0.020719 |
| ENSMUSG00000102758.4 | 43.33686 | 1.052703 | 1.167117 | 0.901969 | 0.367073 | 0.953413 | 0.020719 |
| ENSMUSG00000114019.2 | 19.71582 | 1.921118 | 2.138852 | 0.8982 | 0.369079 | 0.953413 | 0.020719 |
| ENSMUSG00000025666.17 | 60.37533 | 0.913382 | 1.017811 | 0.897398 | 0.369506 | 0.953435 | 0.020709 |
| ENSMUSG00000005621.12 | 37.35561 | -1.22735 | 1.369623 | -0.89612 | 0.370187 | 0.953556 | 0.020654 |
| ENSMUSG00000025508.14 | 1927.875 | 0.219002 | 0.24456 | 0.895496 | 0.370522 | 0.953556 | 0.020654 |
| ENSMUSG00000026819.16 | 278.5548 | -0.7602 | 0.848977 | -0.89543 | 0.370559 | 0.953556 | 0.020654 |
| ENSMUSG00000029729.13 | 157.7321 | -0.84604 | 0.943488 | -0.89671 | 0.369872 | 0.953556 | 0.020654 |
| ENSMUSG00000035473.11 | 657.4655 | -0.37153 | 0.414405 | -0.89653 | 0.36997 | 0.953556 | 0.020654 |
| ENSMUSG00000035505.15 | 78.02174 | -0.75721 | 0.845318 | -0.89576 | 0.370379 | 0.953556 | 0.020654 |
| ENSMUSG00000049354.9 | 58.69869 | -0.81539 | 0.91005 | -0.89599 | 0.370259 | 0.953556 | 0.020654 |
| ENSMUSG00000070493.4 | 4647.639 | 0.205641 | 0.229511 | 0.895995 | 0.370256 | 0.953556 | 0.020654 |
| ENSMUSG00000074639.7 | 384.0545 | -0.3757 | 0.419112 | -0.89643 | 0.370025 | 0.953556 | 0.020654 |
| ENSMUSG00000078134.2 | 24.55336 | 1.573253 | 1.753931 | 0.896987 | 0.369726 | 0.953556 | 0.020654 |
| ENSMUSG00000003477.6 | 2963.027 | 0.269611 | 0.301782 | 0.893395 | 0.371646 | 0.954058 | 0.020425 |
| ENSMUSG00000005413.9 | 39.79521 | -1.21797 | 1.363904 | -0.893 | 0.371857 | 0.954058 | 0.020425 |
| ENSMUSG00000006519.12 | 13.93271 | -2.28835 | 2.564022 | -0.89248 | 0.372134 | 0.954058 | 0.020425 |
| ENSMUSG00000022041.11 | 10.93908 | 1.938579 | 2.172351 | 0.892388 | 0.372185 | 0.954058 | 0.020425 |
| ENSMUSG00000022800.15 | 131.7589 | -0.5485 | 0.613958 | -0.89338 | 0.371655 | 0.954058 | 0.020425 |
| ENSMUSG00000025732.7 | 23.21845 | 1.502641 | 1.68266 | 0.893015 | 0.371849 | 0.954058 | 0.020425 |
| ENSMUSG00000026455.15 | 50.24514 | -1.35387 | 1.513285 | -0.89465 | 0.370972 | 0.954058 | 0.020425 |
| ENSMUSG00000028016.10 | 50.19536 | -1.18472 | 1.3278 | -0.89224 | 0.372264 | 0.954058 | 0.020425 |
| ENSMUSG00000029817.12 | 158.6988 | 0.518634 | 0.580175 | 0.893927 | 0.371361 | 0.954058 | 0.020425 |
| ENSMUSG00000030057.16 | 2177.048 | 0.26999 | 0.302156 | 0.893544 | 0.371566 | 0.954058 | 0.020425 |
| ENSMUSG00000039286.13 | 155.5427 | -0.57866 | 0.646767 | -0.89469 | 0.370953 | 0.954058 | 0.020425 |

| | | | | | | | |
|---|---|---|---|---|---|---|---|
| ENSMUSG00000040472.16 | 30.52909 | 1.387953 | 1.552325 | 0.894112 | 0.371262 | 0.954058 | 0.020425 |
| ENSMUSG00000042328.14 | 42.32793 | -1.4731 | 1.649718 | -0.89294 | 0.37189 | 0.954058 | 0.020425 |
| ENSMUSG00000051339.11 | 208.2395 | -0.65175 | 0.730441 | -0.89226 | 0.372252 | 0.954058 | 0.020425 |
| ENSMUSG00000055817.19 | 24.64793 | -1.67012 | 1.869337 | -0.89343 | 0.371628 | 0.954058 | 0.020425 |
| ENSMUSG00000020482.5 | 151.1066 | -0.61768 | 0.69334 | -0.89087 | 0.372997 | 0.954457 | 0.020243 |
| ENSMUSG00000021395.12 | 280.622 | 0.501583 | 0.563055 | 0.890825 | 0.373023 | 0.954457 | 0.020243 |
| ENSMUSG00000021957.8 | 417.9489 | 0.491507 | 0.551527 | 0.891176 | 0.372835 | 0.954457 | 0.020243 |
| ENSMUSG00000024480.10 | 80.49811 | -0.77125 | 0.865485 | -0.89112 | 0.372863 | 0.954457 | 0.020243 |
| ENSMUSG00000031158.12 | 46.2278 | 0.89535 | 1.005081 | 0.890824 | 0.373024 | 0.954457 | 0.020243 |
| ENSMUSG00000083992.2 | 155.2003 | 0.526899 | 0.590955 | 0.891606 | 0.372604 | 0.954457 | 0.020243 |
| ENSMUSG00000032475.16 | 147.0258 | 0.655999 | 0.736583 | 0.890598 | 0.373145 | 0.95451 | 0.02022 |
| ENSMUSG00000002108.11 | 181.1276 | -0.61693 | 0.694776 | -0.88796 | 0.374563 | 0.954696 | 0.020135 |
| ENSMUSG00000005268.21 | 370.676 | 0.407297 | 0.45806 | 0.889177 | 0.373908 | 0.954696 | 0.020135 |
| ENSMUSG00000005312.16 | 669.3026 | -0.36842 | 0.417377 | -0.8827 | 0.3774 | 0.954696 | 0.020135 |
| ENSMUSG00000018770.10 | 5027.535 | 0.192101 | 0.216686 | 0.886541 | 0.375326 | 0.954696 | 0.020135 |
| ENSMUSG00000020074.8 | 626.501 | 0.399894 | 0.450599 | 0.887472 | 0.374825 | 0.954696 | 0.020135 |
| ENSMUSG00000020423.7 | 118.749 | -1.14588 | 1.293566 | -0.88583 | 0.37571 | 0.954696 | 0.020135 |
| ENSMUSG00000021669.16 | 263.2107 | -0.53486 | 0.604355 | -0.885 | 0.376156 | 0.954696 | 0.020135 |
| ENSMUSG00000022272.18 | 99.61401 | -0.66561 | 0.752386 | -0.88467 | 0.376337 | 0.954696 | 0.020135 |
| ENSMUSG00000023723.18 | 73.01317 | 0.934622 | 1.05893 | 0.88261 | 0.377447 | 0.954696 | 0.020135 |
| ENSMUSG00000024474.8 | 502.7011 | -0.42815 | 0.48458 | -0.88354 | 0.376945 | 0.954696 | 0.020135 |
| ENSMUSG00000025374.14 | 72.67306 | 0.904053 | 1.022084 | 0.88452 | 0.376416 | 0.954696 | 0.020135 |
| ENSMUSG00000027357.17 | 262.2897 | -0.47555 | 0.538752 | -0.8827 | 0.3774 | 0.954696 | 0.020135 |
| ENSMUSG00000027901.13 | 53.21512 | 0.948058 | 1.067457 | 0.888146 | 0.374462 | 0.954696 | 0.020135 |
| ENSMUSG00000027937.16 | 573.6548 | -0.33744 | 0.380168 | -0.88761 | 0.374752 | 0.954696 | 0.020135 |
| ENSMUSG00000028060.15 | 86.30058 | -0.78293 | 0.886618 | -0.88305 | 0.37721 | 0.954696 | 0.020135 |
| ENSMUSG00000028082.15 | 176.5582 | -0.50582 | 0.570319 | -0.88691 | 0.375129 | 0.954696 | 0.020135 |
| ENSMUSG00000028550.16 | 80.73304 | 0.822462 | 0.929378 | 0.884959 | 0.376179 | 0.954696 | 0.020135 |
| ENSMUSG00000028990.14 | 13.08772 | 2.466554 | 2.782452 | 0.886468 | 0.375366 | 0.954696 | 0.020135 |
| ENSMUSG00000029467.16 | 470.9748 | 0.359842 | 0.4072 | 0.883697 | 0.37686 | 0.954696 | 0.020135 |
| ENSMUSG00000030844.12 | 14.84821 | -2.26291 | 2.557201 | -0.88492 | 0.376202 | 0.954696 | 0.020135 |
| ENSMUSG00000031521.6 | 222.7771 | 0.456917 | 0.516137 | 0.885262 | 0.376015 | 0.954696 | 0.020135 |
| ENSMUSG00000031546.7 | 138.475 | -0.82077 | 0.923777 | -0.8885 | 0.374274 | 0.954696 | 0.020135 |
| ENSMUSG00000032481.18 | 52.48628 | 1.238186 | 1.393026 | 0.888846 | 0.374086 | 0.954696 | 0.020135 |
| ENSMUSG00000033054.8 | 18.5309 | 1.692496 | 1.909996 | 0.886125 | 0.37555 | 0.954696 | 0.020135 |
| ENSMUSG00000034575.14 | 97.06828 | 0.75788 | 0.858619 | 0.882674 | 0.377412 | 0.954696 | 0.020135 |
| ENSMUSG00000035949.16 | 130.3626 | 0.724646 | 0.816227 | 0.887799 | 0.374649 | 0.954696 | 0.020135 |
| ENSMUSG00000036450.5 | 33.6188 | 1.333979 | 1.510892 | 0.882908 | 0.377286 | 0.954696 | 0.020135 |
| ENSMUSG00000038034.16 | 10.81564 | -2.00579 | 2.266235 | -0.88508 | 0.376115 | 0.954696 | 0.020135 |
| ENSMUSG00000038515.11 | 198.3465 | 0.681488 | 0.765636 | 0.890094 | 0.373415 | 0.954696 | 0.020135 |
| ENSMUSG00000039047.18 | 161.5787 | 0.477758 | 0.540769 | 0.883478 | 0.376978 | 0.954696 | 0.020135 |
| ENSMUSG00000044148.7 | 21.07426 | 1.52819 | 1.72347 | 0.886694 | 0.375244 | 0.954696 | 0.020135 |
| ENSMUSG00000044206.4 | 230.5211 | -0.52894 | 0.594934 | -0.88907 | 0.373965 | 0.954696 | 0.020135 |
| ENSMUSG00000050541.15 | 145.5296 | 0.600498 | 0.676963 | 0.887047 | 0.375054 | 0.954696 | 0.020135 |

| | | | | | | | |
|---|---|---|---|---|---|---|---|
| ENSMUSG00000052534.16 | 70.74611 | 0.747509 | 0.84404 | 0.885633 | 0.375815 | 0.954696 | 0.020135 |
| ENSMUSG00000053062.17 | 70.02446 | 1.169008 | 1.323056 | 0.883567 | 0.37693 | 0.954696 | 0.020135 |
| ENSMUSG00000054612.5 | 23.32793 | 1.18036 | 1.335356 | 0.883929 | 0.376734 | 0.954696 | 0.020135 |
| ENSMUSG00000055447.20 | 517.3707 | -0.37625 | 0.426028 | -0.88316 | 0.377151 | 0.954696 | 0.020135 |
| ENSMUSG00000055917.16 | 113.9006 | -0.639 | 0.719971 | -0.88754 | 0.37479 | 0.954696 | 0.020135 |
| ENSMUSG00000060961.17 | 194.0029 | -0.65624 | 0.740007 | -0.88681 | 0.375184 | 0.954696 | 0.020135 |
| ENSMUSG00000068457.15 | 20.21475 | -1.64532 | 1.862182 | -0.88354 | 0.376942 | 0.954696 | 0.020135 |
| ENSMUSG00000068663.15 | 26.53967 | -1.95009 | 2.192312 | -0.88951 | 0.373727 | 0.954696 | 0.020135 |
| ENSMUSG00000095385.4 | 31.87436 | 1.657599 | 1.871277 | 0.885811 | 0.375719 | 0.954696 | 0.020135 |
| ENSMUSG00000021477.10 | 4927.58 | -0.18484 | 0.209763 | -0.88117 | 0.378228 | 0.955108 | 0.019947 |
| ENSMUSG00000022035.7 | 564.0369 | 0.3038 | 0.344814 | 0.881054 | 0.378289 | 0.955108 | 0.019947 |
| ENSMUSG00000024810.17 | 29.69432 | 1.422708 | 1.615212 | 0.880818 | 0.378416 | 0.955108 | 0.019947 |
| ENSMUSG00000026970.17 | 291.5287 | -0.48366 | 0.548507 | -0.88177 | 0.377899 | 0.955108 | 0.019947 |
| ENSMUSG00000028868.14 | 108.2451 | -0.6773 | 0.768234 | -0.88163 | 0.377975 | 0.955108 | 0.019947 |
| ENSMUSG00000034744.14 | 40.32534 | -1.42452 | 1.616563 | -0.8812 | 0.378208 | 0.955108 | 0.019947 |
| ENSMUSG00000092241.2 | 27.79161 | -1.97053 | 2.235282 | -0.88156 | 0.378016 | 0.955108 | 0.019947 |
| ENSMUSG00000109703.2 | 13.94304 | 2.646809 | 3.004385 | 0.880982 | 0.378328 | 0.955108 | 0.019947 |
| ENSMUSG00000004846.11 | 46.6541 | -1.29735 | 1.476034 | -0.87894 | 0.379432 | 0.955564 | 0.01974 |
| ENSMUSG00000016018.5 | 93.55774 | 1.035772 | 1.178517 | 0.878877 | 0.379468 | 0.955564 | 0.01974 |
| ENSMUSG00000017999.17 | 42.06912 | -1.06328 | 1.209794 | -0.87889 | 0.379458 | 0.955564 | 0.01974 |
| ENSMUSG00000020184.16 | 107.1251 | 0.787971 | 0.897203 | 0.878253 | 0.379807 | 0.955564 | 0.01974 |
| ENSMUSG00000021209.13 | 29.42476 | -1.68391 | 1.914157 | -0.87972 | 0.379014 | 0.955564 | 0.01974 |
| ENSMUSG00000021621.16 | 73.11545 | 0.797562 | 0.907524 | 0.878834 | 0.379492 | 0.955564 | 0.01974 |
| ENSMUSG00000032182.16 | 25.2418 | 1.782354 | 2.024744 | 0.880286 | 0.378704 | 0.955564 | 0.01974 |
| ENSMUSG00000032288.10 | 364.3085 | 0.486711 | 0.553944 | 0.878627 | 0.379603 | 0.955564 | 0.01974 |
| ENSMUSG00000032724.6 | 56.53309 | 1.10566 | 1.257279 | 0.879407 | 0.379181 | 0.955564 | 0.01974 |
| ENSMUSG00000034154.16 | 13.71789 | -2.13569 | 2.429915 | -0.87892 | 0.379447 | 0.955564 | 0.01974 |
| ENSMUSG00000061787.16 | 4743.522 | 0.216951 | 0.247003 | 0.878336 | 0.379761 | 0.955564 | 0.01974 |
| ENSMUSG00000072501.6 | 231.3501 | 0.500895 | 0.570119 | 0.87858 | 0.379629 | 0.955564 | 0.01974 |
| ENSMUSG00000000184.13 | 79.61461 | -0.79175 | 0.903623 | -0.87619 | 0.380927 | 0.955592 | 0.019727 |
| ENSMUSG00000000743.10 | 35.18128 | -1.19299 | 1.359091 | -0.87779 | 0.38006 | 0.955592 | 0.019727 |
| ENSMUSG00000002105.16 | 16.7928 | 1.886652 | 2.151781 | 0.876786 | 0.380603 | 0.955592 | 0.019727 |
| ENSMUSG00000021782.15 | 22.83529 | -1.77889 | 2.029529 | -0.87651 | 0.380755 | 0.955592 | 0.019727 |
| ENSMUSG00000024387.14 | 519.8829 | -0.35084 | 0.399999 | -0.8771 | 0.380433 | 0.955592 | 0.019727 |
| ENSMUSG00000027593.16 | 14.77751 | -2.12855 | 2.427222 | -0.87695 | 0.380515 | 0.955592 | 0.019727 |
| ENSMUSG00000029538.15 | 211.1328 | 0.477989 | 0.545114 | 0.876862 | 0.380562 | 0.955592 | 0.019727 |
| ENSMUSG00000037523.14 | 274.6784 | -0.46218 | 0.527302 | -0.8765 | 0.380756 | 0.955592 | 0.019727 |
| ENSMUSG00000044763.9 | 134.6497 | 0.609408 | 0.695001 | 0.876845 | 0.380571 | 0.955592 | 0.019727 |
| ENSMUSG00000054874.13 | 24.4451 | 1.82687 | 2.083071 | 0.877008 | 0.380482 | 0.955592 | 0.019727 |
| ENSMUSG00000057421.13 | 400.8788 | 0.361804 | 0.412871 | 0.876312 | 0.38086 | 0.955592 | 0.019727 |
| ENSMUSG00000013846.11 | 77.71639 | 0.856162 | 0.978928 | 0.874591 | 0.381796 | 0.955871 | 0.019601 |
| ENSMUSG00000022003.8 | 117.2077 | 0.770925 | 0.881266 | 0.874793 | 0.381687 | 0.955871 | 0.019601 |
| ENSMUSG00000022247.13 | 300.2219 | 0.468 | 0.534817 | 0.875066 | 0.381538 | 0.955871 | 0.019601 |
| ENSMUSG00000027332.12 | 483.1161 | 0.418321 | 0.477855 | 0.875414 | 0.381349 | 0.955871 | 0.019601 |

| | | | | | | | |
|---|---|---|---|---|---|---|---|
| ENSMUSG00000030359.15 | 27715.24 | -0.20589 | 0.235317 | -0.87493 | 0.381613 | 0.955871 | 0.019601 |
| ENSMUSG00000039197.11 | 4139.739 | 0.197268 | 0.225349 | 0.875387 | 0.381363 | 0.955871 | 0.019601 |
| ENSMUSG00000051391.10 | 254.5755 | 0.47634 | 0.544462 | 0.874881 | 0.381638 | 0.955871 | 0.019601 |
| ENSMUSG00000068240.6 | 78.59369 | 0.681394 | 0.779179 | 0.874503 | 0.381844 | 0.955871 | 0.019601 |
| ENSMUSG00000006567.12 | 79.53502 | -0.77702 | 0.888988 | -0.87405 | 0.38209 | 0.955888 | 0.019593 |
| ENSMUSG00000085735.2 | 11.69762 | -2.742 | 3.137111 | -0.87405 | 0.382089 | 0.955888 | 0.019593 |
| ENSMUSG00000106568.2 | 26.72124 | -1.11536 | 1.276256 | -0.87393 | 0.382154 | 0.955888 | 0.019593 |
| ENSMUSG00000019478.18 | 136.6697 | 0.635705 | 0.727573 | 0.873733 | 0.382264 | 0.955912 | 0.019582 |
| ENSMUSG00000009654.5 | 128.98 | 0.758838 | 0.869396 | 0.872833 | 0.382754 | 0.956004 | 0.01954 |
| ENSMUSG00000025200.8 | 89.79863 | -0.70902 | 0.812931 | -0.87218 | 0.383111 | 0.956004 | 0.01954 |
| ENSMUSG00000026295.13 | 1528.433 | 0.229557 | 0.2632 | 0.872179 | 0.383111 | 0.956004 | 0.01954 |
| ENSMUSG00000034667.9 | 329.5718 | 0.472907 | 0.542101 | 0.87236 | 0.383012 | 0.956004 | 0.01954 |
| ENSMUSG00000042271.14 | 275.394 | -0.46153 | 0.529205 | -0.87212 | 0.383143 | 0.956004 | 0.01954 |
| ENSMUSG00000053347.16 | 42.87536 | 0.917322 | 1.051141 | 0.872692 | 0.382831 | 0.956004 | 0.01954 |
| ENSMUSG00000053641.10 | 88.74894 | -0.71833 | 0.823768 | -0.872 | 0.383208 | 0.956004 | 0.01954 |
| ENSMUSG00000054008.10 | 186.6404 | -0.60713 | 0.695328 | -0.87315 | 0.382579 | 0.956004 | 0.01954 |
| ENSMUSG00000058254.13 | 144.1302 | 0.635674 | 0.728841 | 0.872172 | 0.383115 | 0.956004 | 0.01954 |
| ENSMUSG00000001750.17 | 24.32325 | 1.65237 | 1.895338 | 0.871808 | 0.383313 | 0.956014 | 0.019536 |
| ENSMUSG00000001918.18 | 16.38641 | 1.919155 | 2.211044 | 0.867986 | 0.385402 | 0.956026 | 0.01953 |
| ENSMUSG00000006010.15 | 341.5723 | -0.56788 | 0.657054 | -0.86429 | 0.387429 | 0.956026 | 0.01953 |
| ENSMUSG00000018189.13 | 493.2005 | -0.30495 | 0.352637 | -0.86476 | 0.387172 | 0.956026 | 0.01953 |
| ENSMUSG00000021178.10 | 266.8598 | -0.46048 | 0.529223 | -0.8701 | 0.384245 | 0.956026 | 0.01953 |
| ENSMUSG00000022443.18 | 70.20618 | -0.83465 | 0.960953 | -0.86856 | 0.385088 | 0.956026 | 0.01953 |
| ENSMUSG00000022812.15 | 142.4542 | 0.550785 | 0.633188 | 0.86986 | 0.384377 | 0.956026 | 0.01953 |
| ENSMUSG00000024646.15 | 9792.374 | 0.198926 | 0.229158 | 0.868073 | 0.385354 | 0.956026 | 0.01953 |
| ENSMUSG00000024997.8 | 591.208 | -0.34567 | 0.399443 | -0.86537 | 0.386836 | 0.956026 | 0.01953 |
| ENSMUSG00000025195.10 | 22.24931 | -1.58626 | 1.829027 | -0.86727 | 0.385795 | 0.956026 | 0.01953 |
| ENSMUSG00000025224.17 | 78.77644 | -1.05056 | 1.207863 | -0.86977 | 0.384426 | 0.956026 | 0.01953 |
| ENSMUSG00000028066.16 | 15.84826 | 1.81572 | 2.085007 | 0.870846 | 0.383838 | 0.956026 | 0.01953 |
| ENSMUSG00000028525.17 | 58.37018 | 0.846326 | 0.979104 | 0.864389 | 0.387374 | 0.956026 | 0.01953 |
| ENSMUSG00000029207.17 | 37.34213 | -1.44235 | 1.664005 | -0.8668 | 0.386054 | 0.956026 | 0.01953 |
| ENSMUSG00000032382.15 | 33.74 | -1.37258 | 1.580232 | -0.8686 | 0.385068 | 0.956026 | 0.01953 |
| ENSMUSG00000032456.14 | 49.28305 | -1.38598 | 1.603689 | -0.86425 | 0.387453 | 0.956026 | 0.01953 |
| ENSMUSG00000033022.9 | 10491.42 | -0.20297 | 0.233167 | -0.87048 | 0.384041 | 0.956026 | 0.01953 |
| ENSMUSG00000033149.18 | 341.5028 | -0.37002 | 0.425068 | -0.87049 | 0.384035 | 0.956026 | 0.01953 |
| ENSMUSG00000034042.17 | 38.3347 | -1.20543 | 1.39248 | -0.86567 | 0.386672 | 0.956026 | 0.01953 |
| ENSMUSG00000036863.13 | 35.87211 | -1.34591 | 1.554782 | -0.86566 | 0.386679 | 0.956026 | 0.01953 |
| ENSMUSG00000038628.9 | 160.2843 | 0.676716 | 0.780713 | 0.866793 | 0.386056 | 0.956026 | 0.01953 |
| ENSMUSG00000040520.8 | 88.27873 | 0.768633 | 0.885386 | 0.868133 | 0.385321 | 0.956026 | 0.01953 |
| ENSMUSG00000040565.9 | 125.7666 | 0.643781 | 0.743258 | 0.866161 | 0.386402 | 0.956026 | 0.01953 |
| ENSMUSG00000044700.16 | 14.84438 | -2.20143 | 2.539528 | -0.86687 | 0.386015 | 0.956026 | 0.01953 |
| ENSMUSG00000045210.9 | 191.8301 | 0.619557 | 0.714943 | 0.866583 | 0.386171 | 0.956026 | 0.01953 |
| ENSMUSG00000046312.5 | 43.16903 | -1.03196 | 1.1916 | -0.86603 | 0.386474 | 0.956026 | 0.01953 |
| ENSMUSG00000049327.18 | 149.7571 | -0.52616 | 0.607266 | -0.86644 | 0.386248 | 0.956026 | 0.01953 |

| | | | | | | | |
|---|---|---|---|---|---|---|---|
| ENSMUSG00000052397.9 | 23.15082 | 1.165566 | 1.343411 | 0.867617 | 0.385604 | 0.956026 | 0.01953 |
| ENSMUSG00000052595.9 | 110.8315 | 0.682154 | 0.788689 | 0.864921 | 0.387082 | 0.956026 | 0.01953 |
| ENSMUSG00000055839.7 | 472.6245 | -0.32915 | 0.377975 | -0.87084 | 0.383844 | 0.956026 | 0.01953 |
| ENSMUSG00000056579.19 | 38.51342 | 1.106029 | 1.271225 | 0.870049 | 0.384273 | 0.956026 | 0.01953 |
| ENSMUSG00000057541.15 | 89.18868 | -0.63104 | 0.725533 | -0.86976 | 0.384432 | 0.956026 | 0.01953 |
| ENSMUSG00000060032.7 | 168.6931 | 0.525306 | 0.607538 | 0.864647 | 0.387232 | 0.956026 | 0.01953 |
| ENSMUSG00000060126.15 | 5399.718 | 0.189709 | 0.219427 | 0.864566 | 0.387277 | 0.956026 | 0.01953 |
| ENSMUSG00000061740.9 | 612.7725 | -0.33626 | 0.38804 | -0.86656 | 0.386182 | 0.956026 | 0.01953 |
| ENSMUSG00000062949.14 | 1107.583 | -0.31867 | 0.367873 | -0.86624 | 0.386357 | 0.956026 | 0.01953 |
| ENSMUSG00000063663.12 | 35.93113 | -1.06654 | 1.227238 | -0.86906 | 0.384815 | 0.956026 | 0.01953 |
| ENSMUSG00000063870.13 | 113.3821 | 0.693419 | 0.801012 | 0.865678 | 0.386666 | 0.956026 | 0.01953 |
| ENSMUSG00000065990.13 | 344.8641 | 0.41716 | 0.481943 | 0.865579 | 0.386721 | 0.956026 | 0.01953 |
| ENSMUSG00000073889.11 | 157.8319 | 0.586997 | 0.67732 | 0.866645 | 0.386136 | 0.956026 | 0.01953 |
| ENSMUSG00000074166.8 | 14.82512 | 2.114353 | 2.443812 | 0.865186 | 0.386936 | 0.956026 | 0.01953 |
| ENSMUSG00000078851.5 | 48.5109 | -1.10726 | 1.274166 | -0.86901 | 0.384844 | 0.956026 | 0.01953 |
| ENSMUSG00000074623.5 | 13.07601 | 2.033787 | 2.353905 | 0.864005 | 0.387585 | 0.956103 | 0.019495 |
| ENSMUSG00000028466.16 | 86.1825 | 0.987957 | 1.144923 | 0.862903 | 0.388191 | 0.957349 | 0.01893 |
| ENSMUSG00000021615.14 | 36.31233 | 1.182921 | 1.372379 | 0.861949 | 0.388715 | 0.95791 | 0.018675 |
| ENSMUSG00000025287.16 | 38.32011 | -1.0313 | 1.196252 | -0.86211 | 0.388628 | 0.95791 | 0.018675 |
| ENSMUSG00000030435.17 | 27.85182 | -1.0742 | 1.246266 | -0.86194 | 0.388722 | 0.95791 | 0.018675 |
| ENSMUSG00000024085.14 | 928.5587 | 0.276091 | 0.320478 | 0.861498 | 0.388964 | 0.958257 | 0.018518 |
| ENSMUSG00000038503.16 | 207.5118 | 0.538417 | 0.625329 | 0.861014 | 0.38923 | 0.958416 | 0.018446 |
| ENSMUSG00000099966.2 | 41.69422 | 1.303445 | 1.513632 | 0.861137 | 0.389162 | 0.958416 | 0.018446 |
| ENSMUSG00000003868.15 | 20.21642 | 2.061909 | 2.397845 | 0.859901 | 0.389844 | 0.958462 | 0.018425 |
| ENSMUSG00000005882.19 | 94.50019 | -0.7817 | 0.909078 | -0.85988 | 0.389855 | 0.958462 | 0.018425 |
| ENSMUSG00000018340.14 | 30.69543 | -1.12632 | 1.309403 | -0.86018 | 0.38969 | 0.958462 | 0.018425 |
| ENSMUSG00000019906.16 | 50.01225 | 0.988852 | 1.149644 | 0.860138 | 0.389713 | 0.958462 | 0.018425 |
| ENSMUSG00000032607.13 | 66.34257 | -1.06882 | 1.242352 | -0.86032 | 0.389614 | 0.958462 | 0.018425 |
| ENSMUSG00000038323.4 | 308.2963 | 0.512595 | 0.595538 | 0.860726 | 0.389389 | 0.958462 | 0.018425 |
| ENSMUSG00000019158.10 | 52.81463 | 1.174247 | 1.367144 | 0.858905 | 0.390393 | 0.959037 | 0.018165 |
| ENSMUSG00000021583.8 | 367.8177 | -0.38959 | 0.453558 | -0.85897 | 0.390356 | 0.959037 | 0.018165 |
| ENSMUSG00000067150.7 | 46.67946 | -1.30534 | 1.519416 | -0.85911 | 0.390282 | 0.959037 | 0.018165 |
| ENSMUSG00000003235.14 | 121.5542 | 0.720078 | 0.844576 | 0.852591 | 0.393886 | 0.959488 | 0.01796 |
| ENSMUSG00000003555.9 | 38.70833 | -1.56184 | 1.820795 | -0.85778 | 0.391015 | 0.959488 | 0.01796 |
| ENSMUSG00000004085.15 | 65.59258 | -0.77442 | 0.912835 | -0.84837 | 0.396233 | 0.959488 | 0.01796 |
| ENSMUSG00000005803.15 | 1339.738 | 0.249409 | 0.293423 | 0.849996 | 0.395327 | 0.959488 | 0.01796 |
| ENSMUSG00000016942.7 | 307.7537 | -0.66687 | 0.786237 | -0.84818 | 0.396336 | 0.959488 | 0.01796 |
| ENSMUSG00000017418.15 | 131.0491 | -0.56174 | 0.659719 | -0.85149 | 0.394496 | 0.959488 | 0.01796 |
| ENSMUSG00000020079.16 | 50.88244 | 0.92645 | 1.080794 | 0.857194 | 0.391337 | 0.959488 | 0.01796 |
| ENSMUSG00000020982.10 | 613.2201 | -0.31438 | 0.36748 | -0.85551 | 0.392269 | 0.959488 | 0.01796 |
| ENSMUSG00000021987.10 | 131.7773 | 0.719499 | 0.845188 | 0.851289 | 0.394609 | 0.959488 | 0.01796 |
| ENSMUSG00000022216.18 | 646.3482 | 0.288026 | 0.338715 | 0.85035 | 0.395131 | 0.959488 | 0.01796 |
| ENSMUSG00000022471.15 | 35.69135 | -1.06378 | 1.243864 | -0.85522 | 0.392429 | 0.959488 | 0.01796 |
| ENSMUSG00000024271.9 | 106.4589 | 0.628384 | 0.736079 | 0.853692 | 0.393276 | 0.959488 | 0.01796 |

| | | | | | | | |
|---|---|---|---|---|---|---|---|
| ENSMUSG00000024298.16 | 155.3023 | 0.506687 | 0.593359 | 0.853929 | 0.393144 | 0.959488 | 0.01796 |
| ENSMUSG00000024683.7 | 197.3969 | 0.531455 | 0.624432 | 0.851102 | 0.394712 | 0.959488 | 0.01796 |
| ENSMUSG00000024799.17 | 398.2806 | 0.487725 | 0.57432 | 0.849222 | 0.395758 | 0.959488 | 0.01796 |
| ENSMUSG00000025208.10 | 352.02 | 0.458968 | 0.540639 | 0.848936 | 0.395917 | 0.959488 | 0.01796 |
| ENSMUSG00000026049.12 | 82.47544 | 0.731074 | 0.861733 | 0.848377 | 0.396228 | 0.959488 | 0.01796 |
| ENSMUSG00000026917.16 | 29.26144 | -1.12411 | 1.321733 | -0.85048 | 0.395056 | 0.959488 | 0.01796 |
| ENSMUSG00000028232.14 | 49.14289 | -0.99646 | 1.168035 | -0.85311 | 0.393598 | 0.959488 | 0.01796 |
| ENSMUSG00000028329.14 | 253.0241 | 0.46779 | 0.550295 | 0.850071 | 0.395286 | 0.959488 | 0.01796 |
| ENSMUSG00000028416.14 | 663.7953 | -0.35972 | 0.420784 | -0.85488 | 0.392618 | 0.959488 | 0.01796 |
| ENSMUSG00000028461.13 | 453.2098 | 0.356001 | 0.41841 | 0.850842 | 0.394857 | 0.959488 | 0.01796 |
| ENSMUSG00000029071.17 | 91.9866 | -0.61466 | 0.719593 | -0.85418 | 0.393006 | 0.959488 | 0.01796 |
| ENSMUSG00000029125.15 | 54.42066 | -0.97846 | 1.145462 | -0.8542 | 0.392992 | 0.959488 | 0.01796 |
| ENSMUSG00000029433.17 | 99.26814 | -0.63502 | 0.746033 | -0.8512 | 0.394658 | 0.959488 | 0.01796 |
| ENSMUSG00000030522.15 | 122.3825 | -0.60404 | 0.711016 | -0.84954 | 0.395579 | 0.959488 | 0.01796 |
| ENSMUSG00000030754.10 | 330.4424 | 0.374539 | 0.441025 | 0.849246 | 0.395744 | 0.959488 | 0.01796 |
| ENSMUSG00000030870.13 | 97.36222 | 0.951927 | 1.122306 | 0.848188 | 0.396333 | 0.959488 | 0.01796 |
| ENSMUSG00000031021.14 | 231.2927 | -0.48824 | 0.573716 | -0.85101 | 0.394765 | 0.959488 | 0.01796 |
| ENSMUSG00000031246.15 | 624.0972 | -0.28754 | 0.336804 | -0.85372 | 0.393259 | 0.959488 | 0.01796 |
| ENSMUSG00000031314.19 | 74.29793 | 1.131275 | 1.327699 | 0.852057 | 0.394183 | 0.959488 | 0.01796 |
| ENSMUSG00000031853.6 | 21.41051 | 1.819768 | 2.140408 | 0.850197 | 0.395216 | 0.959488 | 0.01796 |
| ENSMUSG00000032409.16 | 89.56706 | -0.74195 | 0.872187 | -0.85067 | 0.394951 | 0.959488 | 0.01796 |
| ENSMUSG00000033634.8 | 1786.673 | -0.27978 | 0.326295 | -0.85746 | 0.391193 | 0.959488 | 0.01796 |
| ENSMUSG00000033713.13 | 380.2705 | -0.38727 | 0.453695 | -0.85358 | 0.393335 | 0.959488 | 0.01796 |
| ENSMUSG00000034173.14 | 19.78533 | 1.499009 | 1.752965 | 0.855128 | 0.39248 | 0.959488 | 0.01796 |
| ENSMUSG00000035469.18 | 294.0176 | -0.4832 | 0.565575 | -0.85435 | 0.392909 | 0.959488 | 0.01796 |
| ENSMUSG00000036073.19 | 272.0319 | -0.54258 | 0.635714 | -0.8535 | 0.39338 | 0.959488 | 0.01796 |
| ENSMUSG00000036078.17 | 382.5286 | 0.429722 | 0.506649 | 0.848166 | 0.396346 | 0.959488 | 0.01796 |
| ENSMUSG00000037366.15 | 289.9004 | -0.46925 | 0.547571 | -0.85696 | 0.391464 | 0.959488 | 0.01796 |
| ENSMUSG00000038060.16 | 63.03246 | -0.91314 | 1.06919 | -0.85405 | 0.393079 | 0.959488 | 0.01796 |
| ENSMUSG00000038587.11 | 14.89795 | 1.874306 | 2.196586 | 0.853282 | 0.393503 | 0.959488 | 0.01796 |
| ENSMUSG00000038762.16 | 456.7236 | -0.36036 | 0.420233 | -0.85753 | 0.39115 | 0.959488 | 0.01796 |
| ENSMUSG00000039967.15 | 91.35967 | 0.745652 | 0.877534 | 0.849713 | 0.395485 | 0.959488 | 0.01796 |
| ENSMUSG00000040147.15 | 790.1552 | -0.40492 | 0.475064 | -0.85235 | 0.39402 | 0.959488 | 0.01796 |
| ENSMUSG00000040234.17 | 337.2703 | 0.364877 | 0.42865 | 0.851223 | 0.394646 | 0.959488 | 0.01796 |
| ENSMUSG00000045767.10 | 321.1361 | -0.47694 | 0.556631 | -0.85683 | 0.391538 | 0.959488 | 0.01796 |
| ENSMUSG00000045930.4 | 38.77727 | 1.188219 | 1.386347 | 0.857086 | 0.391397 | 0.959488 | 0.01796 |
| ENSMUSG00000045991.20 | 498.2786 | -0.37429 | 0.437957 | -0.85462 | 0.39276 | 0.959488 | 0.01796 |
| ENSMUSG00000046949.17 | 680.8681 | 0.375348 | 0.441575 | 0.850019 | 0.395314 | 0.959488 | 0.01796 |
| ENSMUSG00000052139.19 | 60.84936 | -1.09201 | 1.28043 | -0.85285 | 0.393743 | 0.959488 | 0.01796 |
| ENSMUSG00000052609.10 | 87.67919 | -1.00717 | 1.176959 | -0.85574 | 0.39214 | 0.959488 | 0.01796 |
| ENSMUSG00000062762.18 | 326.2957 | -0.42373 | 0.496491 | -0.85345 | 0.393411 | 0.959488 | 0.01796 |
| ENSMUSG00000071657.13 | 85.38456 | -0.90176 | 1.063094 | -0.84824 | 0.396303 | 0.959488 | 0.01796 |
| ENSMUSG00000073639.7 | 1041.156 | -0.30662 | 0.361051 | -0.84923 | 0.395751 | 0.959488 | 0.01796 |
| ENSMUSG00000076617.10 | 72.6465 | -1.35216 | 1.593993 | -0.84829 | 0.396278 | 0.959488 | 0.01796 |

| | | | | | | | |
|---|---|---|---|---|---|---|---|
| ENSMUSG00000078427.4 | 78.78377 | -0.70478 | 0.828016 | -0.85117 | 0.394675 | 0.959488 | 0.01796 |
| ENSMUSG00000041828.16 | 243.6253 | -0.49963 | 0.58925 | -0.84791 | 0.396488 | 0.959587 | 0.017916 |
| ENSMUSG00000026254.18 | 204.2879 | -0.54442 | 0.642236 | -0.84769 | 0.39661 | 0.959637 | 0.017893 |
| ENSMUSG00000029397.16 | 217.0094 | 0.457668 | 0.540236 | 0.847163 | 0.396904 | 0.959733 | 0.01785 |
| ENSMUSG00000037257.9 | 66.80175 | -0.7506 | 0.886112 | -0.84708 | 0.396953 | 0.959733 | 0.01785 |
| ENSMUSG00000045140.9 | 16.07741 | -1.7955 | 2.119071 | -0.8473 | 0.396826 | 0.959733 | 0.01785 |
| ENSMUSG00000003134.11 | 36.82437 | -1.35912 | 1.607429 | -0.84552 | 0.39782 | 0.960786 | 0.017373 |
| ENSMUSG00000003923.15 | 262.4342 | 0.474933 | 0.56238 | 0.844505 | 0.398387 | 0.960786 | 0.017373 |
| ENSMUSG00000007827.11 | 57.93383 | 1.139288 | 1.349659 | 0.84413 | 0.398597 | 0.960786 | 0.017373 |
| ENSMUSG00000018401.18 | 129.2496 | 0.580025 | 0.686939 | 0.844361 | 0.398468 | 0.960786 | 0.017373 |
| ENSMUSG00000019820.12 | 128.1093 | -0.61384 | 0.727023 | -0.84432 | 0.39849 | 0.960786 | 0.017373 |
| ENSMUSG00000019945.11 | 19.30527 | -2.03923 | 2.415045 | -0.84439 | 0.398453 | 0.960786 | 0.017373 |
| ENSMUSG00000020076.8 | 234.8639 | 0.552279 | 0.653839 | 0.844671 | 0.398295 | 0.960786 | 0.017373 |
| ENSMUSG00000020680.16 | 23.64833 | -1.38791 | 1.643042 | -0.84472 | 0.398267 | 0.960786 | 0.017373 |
| ENSMUSG00000030264.15 | 121.0604 | 0.714463 | 0.846403 | 0.844116 | 0.398605 | 0.960786 | 0.017373 |
| ENSMUSG00000034300.18 | 27.01526 | -1.62751 | 1.924123 | -0.84585 | 0.397638 | 0.960786 | 0.017373 |
| ENSMUSG00000047388.11 | 37.37543 | -1.48547 | 1.755763 | -0.84605 | 0.397523 | 0.960786 | 0.017373 |
| ENSMUSG00000074466.13 | 62.90977 | -0.88525 | 1.046922 | -0.84557 | 0.397792 | 0.960786 | 0.017373 |
| ENSMUSG00000021585.11 | 294.9644 | -0.39827 | 0.472284 | -0.84329 | 0.399068 | 0.961038 | 0.01726 |
| ENSMUSG00000021709.15 | 342.3896 | 0.405266 | 0.480627 | 0.843204 | 0.399114 | 0.961038 | 0.01726 |
| ENSMUSG00000031161.16 | 22.3941 | -1.22889 | 1.4565 | -0.84373 | 0.39882 | 0.961038 | 0.01726 |
| ENSMUSG00000074238.7 | 238.9009 | -0.41083 | 0.487168 | -0.8433 | 0.399061 | 0.961038 | 0.01726 |
| ENSMUSG00000002846.10 | 88.16071 | -0.68923 | 0.818549 | -0.84202 | 0.399779 | 0.961089 | 0.017236 |
| ENSMUSG00000018848.5 | 145.4529 | 0.566002 | 0.673597 | 0.840268 | 0.400758 | 0.961089 | 0.017236 |
| ENSMUSG00000020697.17 | 79.6823 | -0.71968 | 0.855571 | -0.84117 | 0.400251 | 0.961089 | 0.017236 |
| ENSMUSG00000021094.11 | 315.6854 | 0.481474 | 0.572861 | 0.840473 | 0.400643 | 0.961089 | 0.017236 |
| ENSMUSG00000024785.8 | 605.7366 | -0.37133 | 0.44102 | -0.84198 | 0.399801 | 0.961089 | 0.017236 |
| ENSMUSG00000025907.15 | 311.4819 | -0.51899 | 0.617198 | -0.84089 | 0.400411 | 0.961089 | 0.017236 |
| ENSMUSG00000028484.17 | 315.6549 | -0.53216 | 0.631507 | -0.84268 | 0.399408 | 0.961089 | 0.017236 |
| ENSMUSG00000030521.12 | 193.884 | -0.61691 | 0.733822 | -0.84068 | 0.400527 | 0.961089 | 0.017236 |
| ENSMUSG00000034957.11 | 116.1359 | 0.613365 | 0.728385 | 0.84209 | 0.399738 | 0.961089 | 0.017236 |
| ENSMUSG00000035325.17 | 509.4924 | -0.30486 | 0.362432 | -0.84114 | 0.40027 | 0.961089 | 0.017236 |
| ENSMUSG00000036257.16 | 493.8492 | -0.33222 | 0.395319 | -0.84039 | 0.400688 | 0.961089 | 0.017236 |
| ENSMUSG00000039662.17 | 37.17513 | -1.17788 | 1.401287 | -0.84057 | 0.400589 | 0.961089 | 0.017236 |
| ENSMUSG00000040351.12 | 130.0379 | -0.53863 | 0.639377 | -0.84243 | 0.399548 | 0.961089 | 0.017236 |
| ENSMUSG00000047030.16 | 10.83792 | -1.79624 | 2.134412 | -0.84156 | 0.400032 | 0.961089 | 0.017236 |
| ENSMUSG00000056429.5 | 1037.751 | -0.24481 | 0.290605 | -0.84242 | 0.399552 | 0.961089 | 0.017236 |
| ENSMUSG00000058799.17 | 378.1178 | 0.411184 | 0.489214 | 0.840499 | 0.400628 | 0.961089 | 0.017236 |
| ENSMUSG00000026791.15 | 24.78185 | -1.64 | 1.953078 | -0.8397 | 0.401077 | 0.96161 | 0.017001 |
| ENSMUSG00000024414.14 | 586.6103 | -0.31186 | 0.371509 | -0.83945 | 0.401219 | 0.961709 | 0.016956 |
| ENSMUSG00000002504.16 | 19.06765 | 1.815258 | 2.163127 | 0.839182 | 0.401367 | 0.961714 | 0.016954 |
| ENSMUSG00000109933.2 | 79.44526 | 0.719807 | 0.857852 | 0.83908 | 0.401424 | 0.961714 | 0.016954 |
| ENSMUSG00000004069.18 | 307.467 | 0.420639 | 0.501764 | 0.838322 | 0.40185 | 0.961783 | 0.016923 |
| ENSMUSG00000022280.10 | 31.71657 | 1.259145 | 1.502011 | 0.838306 | 0.401859 | 0.961783 | 0.016923 |

| | | | | | | | |
|---|---|---|---|---|---|---|---|
| ENSMUSG00000053897.16 | 237.3599 | 0.544506 | 0.649339 | 0.838554 | 0.401719 | 0.961783 | 0.016923 |
| ENSMUSG00000104208.2 | 11.00776 | -2.01409 | 2.401496 | -0.83868 | 0.401648 | 0.961783 | 0.016923 |
| ENSMUSG00000026842.17 | 43.22834 | -1.07103 | 1.278169 | -0.83794 | 0.402065 | 0.962034 | 0.01681 |
| ENSMUSG00000039308.7 | 23.11009 | -1.70643 | 2.038188 | -0.83723 | 0.402465 | 0.962747 | 0.016488 |
| ENSMUSG00000038298.15 | 238.286 | -0.49106 | 0.586735 | -0.83694 | 0.402627 | 0.962892 | 0.016422 |
| ENSMUSG00000002897.6 | 61.24721 | -0.79627 | 0.954702 | -0.83405 | 0.404255 | 0.962899 | 0.016419 |
| ENSMUSG00000015869.17 | 248.6897 | -0.52485 | 0.627543 | -0.83635 | 0.402955 | 0.962899 | 0.016419 |
| ENSMUSG00000023075.10 | 172.2528 | -0.61625 | 0.738707 | -0.83423 | 0.40415 | 0.962899 | 0.016419 |
| ENSMUSG00000027189.17 | 97.20759 | -0.66 | 0.790361 | -0.83507 | 0.403681 | 0.962899 | 0.016419 |
| ENSMUSG00000029577.14 | 81.50726 | 0.726492 | 0.869097 | 0.835916 | 0.403202 | 0.962899 | 0.016419 |
| ENSMUSG00000032126.17 | 20.72469 | -1.70361 | 2.038736 | -0.83562 | 0.403368 | 0.962899 | 0.016419 |
| ENSMUSG00000032578.8 | 47.08072 | 1.474444 | 1.765893 | 0.834956 | 0.403742 | 0.962899 | 0.016419 |
| ENSMUSG00000032604.17 | 51.26402 | -0.99433 | 1.191943 | -0.83421 | 0.404165 | 0.962899 | 0.016419 |
| ENSMUSG00000038181.17 | 29.44481 | -1.56868 | 1.875447 | -0.83643 | 0.402912 | 0.962899 | 0.016419 |
| ENSMUSG00000052656.13 | 494.6064 | -0.3411 | 0.408787 | -0.83443 | 0.404038 | 0.962899 | 0.016419 |
| ENSMUSG00000052934.15 | 43.43693 | -0.97778 | 1.171988 | -0.83429 | 0.404118 | 0.962899 | 0.016419 |
| ENSMUSG00000058064.7 | 41.8756 | 0.904162 | 1.082958 | 0.8349 | 0.403774 | 0.962899 | 0.016419 |
| ENSMUSG00000058351.12 | 170.1701 | -0.54914 | 0.656437 | -0.83654 | 0.402849 | 0.962899 | 0.016419 |
| ENSMUSG00000073838.11 | 29.07485 | -1.80085 | 2.155681 | -0.8354 | 0.403495 | 0.962899 | 0.016419 |
| ENSMUSG00000090272.10 | 71.06495 | -0.80607 | 0.964913 | -0.83538 | 0.403502 | 0.962899 | 0.016419 |
| ENSMUSG00000090862.4 | 2488.523 | 0.230027 | 0.275333 | 0.83545 | 0.403464 | 0.962899 | 0.016419 |
| ENSMUSG00000025911.15 | 1103.487 | 0.243539 | 0.292141 | 0.833634 | 0.404487 | 0.962968 | 0.016388 |
| ENSMUSG00000035559.10 | 30.52322 | -1.26309 | 1.51493 | -0.83376 | 0.404414 | 0.962968 | 0.016388 |
| ENSMUSG00000002222.15 | 549.3627 | -0.29092 | 0.350472 | -0.83009 | 0.406489 | 0.963131 | 0.016315 |
| ENSMUSG00000013822.8 | 38.40606 | 1.48743 | 1.792783 | 0.829676 | 0.406722 | 0.963131 | 0.016315 |
| ENSMUSG00000019487.12 | 10.38279 | -2.36474 | 2.84383 | -0.83153 | 0.405671 | 0.963131 | 0.016315 |
| ENSMUSG00000019837.9 | 349.1372 | 0.42535 | 0.511634 | 0.831356 | 0.405772 | 0.963131 | 0.016315 |
| ENSMUSG00000020205.9 | 1225.41 | 0.44148 | 0.530911 | 0.831551 | 0.405662 | 0.963131 | 0.016315 |
| ENSMUSG00000022774.14 | 132.2384 | 0.647112 | 0.780218 | 0.8294 | 0.406878 | 0.963131 | 0.016315 |
| ENSMUSG00000025396.8 | 118.4501 | -0.66211 | 0.797045 | -0.8307 | 0.406142 | 0.963131 | 0.016315 |
| ENSMUSG00000026279.15 | 88.74167 | -0.68911 | 0.828851 | -0.83141 | 0.405744 | 0.963131 | 0.016315 |
| ENSMUSG00000028233.7 | 73.10221 | 0.844743 | 1.01393 | 0.833138 | 0.404767 | 0.963131 | 0.016315 |
| ENSMUSG00000028960.21 | 106.4869 | -0.74059 | 0.89158 | -0.83065 | 0.406172 | 0.963131 | 0.016315 |
| ENSMUSG00000029276.14 | 12.55314 | -1.9787 | 2.385102 | -0.82961 | 0.40676 | 0.963131 | 0.016315 |
| ENSMUSG00000029571.15 | 1245.971 | 0.29509 | 0.355737 | 0.829517 | 0.406812 | 0.963131 | 0.016315 |
| ENSMUSG00000030045.11 | 85.39312 | -0.85763 | 1.033595 | -0.82976 | 0.406677 | 0.963131 | 0.016315 |
| ENSMUSG00000031078.16 | 131.535 | -0.56376 | 0.67964 | -0.8295 | 0.406821 | 0.963131 | 0.016315 |
| ENSMUSG00000032501.10 | 239.6037 | -0.64357 | 0.77597 | -0.82937 | 0.406892 | 0.963131 | 0.016315 |
| ENSMUSG00000037270.19 | 48.36422 | 1.113387 | 1.340783 | 0.830401 | 0.406312 | 0.963131 | 0.016315 |
| ENSMUSG00000040105.4 | 40.82183 | 1.02769 | 1.238733 | 0.82963 | 0.406748 | 0.963131 | 0.016315 |
| ENSMUSG00000040564.15 | 35715.47 | 0.171689 | 0.206662 | 0.830774 | 0.406101 | 0.963131 | 0.016315 |
| ENSMUSG00000041849.8 | 20.56949 | -1.38138 | 1.664556 | -0.82988 | 0.406608 | 0.963131 | 0.016315 |
| ENSMUSG00000042185.15 | 13.2581 | -1.93896 | 2.331501 | -0.83163 | 0.405615 | 0.963131 | 0.016315 |
| ENSMUSG00000074030.4 | 56.88455 | -0.94516 | 1.137493 | -0.83092 | 0.406021 | 0.963131 | 0.016315 |

| | | | | | | | |
|---|---|---|---|---|---|---|---|
| ENSMUSG00000075266.6 | 32.96651 | 1.151439 | 1.38472 | 0.831532 | 0.405673 | 0.963131 | 0.016315 |
| ENSMUSG00000075602.11 | 82.28171 | -0.97318 | 1.173381 | -0.82938 | 0.406889 | 0.963131 | 0.016315 |
| ENSMUSG00000042167.7 | 117.3677 | -0.55526 | 0.670034 | -0.8287 | 0.407274 | 0.963794 | 0.016016 |
| ENSMUSG00000005672.13 | 11.27816 | -1.79032 | 2.162701 | -0.82782 | 0.407774 | 0.964089 | 0.015883 |
| ENSMUSG00000008958.15 | 10.45305 | -2.37222 | 2.875449 | -0.82499 | 0.409377 | 0.964089 | 0.015883 |
| ENSMUSG00000014426.10 | 12.8569 | -2.02168 | 2.4431 | -0.82751 | 0.40795 | 0.964089 | 0.015883 |
| ENSMUSG00000015542.18 | 40.07302 | -1.17848 | 1.436919 | -0.82015 | 0.412133 | 0.964089 | 0.015883 |
| ENSMUSG00000017309.12 | 18.98581 | -1.30392 | 1.586275 | -0.822 | 0.411075 | 0.964089 | 0.015883 |
| ENSMUSG00000018900.8 | 17.65981 | 1.614097 | 1.967493 | 0.820382 | 0.411998 | 0.964089 | 0.015883 |
| ENSMUSG00000019782.11 | 154.2819 | 0.661222 | 0.800774 | 0.825728 | 0.408958 | 0.964089 | 0.015883 |
| ENSMUSG00000020248.19 | 242.7251 | -0.47836 | 0.581573 | -0.82254 | 0.410772 | 0.964089 | 0.015883 |
| ENSMUSG00000021891.9 | 33.51512 | -1.26897 | 1.547733 | -0.81989 | 0.41228 | 0.964089 | 0.015883 |
| ENSMUSG00000022551.9 | 851.0638 | -0.25889 | 0.313291 | -0.82636 | 0.408602 | 0.964089 | 0.015883 |
| ENSMUSG00000023094.15 | 46.08158 | 0.823597 | 0.994777 | 0.827921 | 0.407715 | 0.964089 | 0.015883 |
| ENSMUSG00000024691.14 | 20.1425 | 1.726405 | 2.101206 | 0.821626 | 0.41129 | 0.964089 | 0.015883 |
| ENSMUSG00000024927.9 | 24.24745 | -1.62464 | 1.972454 | -0.82367 | 0.410129 | 0.964089 | 0.015883 |
| ENSMUSG00000025027.19 | 64.07953 | -0.80194 | 0.974629 | -0.82281 | 0.410613 | 0.964089 | 0.015883 |
| ENSMUSG00000025347.5 | 2057.894 | 0.264912 | 0.320687 | 0.826078 | 0.40876 | 0.964089 | 0.015883 |
| ENSMUSG00000025609.16 | 203.8515 | -0.48738 | 0.592847 | -0.82211 | 0.411016 | 0.964089 | 0.015883 |
| ENSMUSG00000026315.14 | 15.64716 | 1.851345 | 2.242366 | 0.825621 | 0.409019 | 0.964089 | 0.015883 |
| ENSMUSG00000027775.10 | 529.2859 | -0.33036 | 0.401769 | -0.82226 | 0.410929 | 0.964089 | 0.015883 |
| ENSMUSG00000028035.14 | 245.262 | 0.515094 | 0.623921 | 0.825576 | 0.409045 | 0.964089 | 0.015883 |
| ENSMUSG00000028334.11 | 39.04202 | 1.089702 | 1.32314 | 0.823573 | 0.410182 | 0.964089 | 0.015883 |
| ENSMUSG00000029815.13 | 321.0216 | 0.451709 | 0.549612 | 0.82187 | 0.411151 | 0.964089 | 0.015883 |
| ENSMUSG00000030286.7 | 148.7814 | 0.88722 | 1.075798 | 0.824708 | 0.409537 | 0.964089 | 0.015883 |
| ENSMUSG00000031176.9 | 503.6861 | 0.357754 | 0.434934 | 0.82255 | 0.410764 | 0.964089 | 0.015883 |
| ENSMUSG00000034088.16 | 727.706 | -0.30468 | 0.369261 | -0.8251 | 0.409315 | 0.964089 | 0.015883 |
| ENSMUSG00000034243.18 | 197.6821 | -0.46884 | 0.568344 | -0.82493 | 0.409414 | 0.964089 | 0.015883 |
| ENSMUSG00000035623.15 | 91.23672 | -0.81001 | 0.986522 | -0.82108 | 0.4116 | 0.964089 | 0.015883 |
| ENSMUSG00000037339.18 | 80.0463 | 0.714281 | 0.867563 | 0.823319 | 0.410327 | 0.964089 | 0.015883 |
| ENSMUSG00000039242.11 | 146.1211 | 0.604028 | 0.73442 | 0.822456 | 0.410818 | 0.964089 | 0.015883 |
| ENSMUSG00000039531.9 | 24.83314 | 1.239726 | 1.511091 | 0.820418 | 0.411978 | 0.964089 | 0.015883 |
| ENSMUSG00000041548.5 | 732.459 | -0.29036 | 0.35222 | -0.82436 | 0.409736 | 0.964089 | 0.015883 |
| ENSMUSG00000041845.12 | 39.94356 | -1.12116 | 1.3565 | -0.82651 | 0.408515 | 0.964089 | 0.015883 |
| ENSMUSG00000041926.16 | 56.75245 | 1.320999 | 1.610685 | 0.820147 | 0.412132 | 0.964089 | 0.015883 |
| ENSMUSG00000042249.12 | 23.49453 | -1.933 | 2.342292 | -0.82526 | 0.409225 | 0.964089 | 0.015883 |
| ENSMUSG00000042709.16 | 55.92073 | -1.08031 | 1.317431 | -0.82001 | 0.412209 | 0.964089 | 0.015883 |
| ENSMUSG00000048351.15 | 70.45971 | -0.75 | 0.912424 | -0.82199 | 0.411083 | 0.964089 | 0.015883 |
| ENSMUSG00000053870.8 | 60.52376 | -0.99205 | 1.207151 | -0.82181 | 0.411183 | 0.964089 | 0.015883 |
| ENSMUSG00000055436.19 | 304.7642 | 0.505374 | 0.616153 | 0.82021 | 0.412097 | 0.964089 | 0.015883 |
| ENSMUSG00000057858.9 | 46.27047 | -1.46758 | 1.784698 | -0.82231 | 0.4109 | 0.964089 | 0.015883 |
| ENSMUSG00000059273.10 | 33.68777 | -1.32426 | 1.606969 | -0.82407 | 0.409899 | 0.964089 | 0.015883 |
| ENSMUSG00000059834.13 | 11.59436 | -1.80084 | 2.180395 | -0.82592 | 0.408847 | 0.964089 | 0.015883 |
| ENSMUSG00000066571.14 | 38.6834 | 1.077234 | 1.30863 | 0.823177 | 0.410408 | 0.964089 | 0.015883 |

| | | | | | | | |
|---|---|---|---|---|---|---|---|
| ENSMUSG00000067722.5 | 172.2722 | 0.565185 | 0.688905 | 0.820411 | 0.411982 | 0.964089 | 0.015883 |
| ENSMUSG00000067995.5 | 65.70719 | -0.80095 | 0.970483 | -0.82531 | 0.409194 | 0.964089 | 0.015883 |
| ENSMUSG00000079104.5 | 27.62054 | -1.42545 | 1.731106 | -0.82343 | 0.410261 | 0.964089 | 0.015883 |
| ENSMUSG00000082895.2 | 132.2181 | -0.56026 | 0.681449 | -0.82215 | 0.410989 | 0.964089 | 0.015883 |
| ENSMUSG00000097691.2 | 68.0108 | 0.89942 | 1.086627 | 0.827717 | 0.407831 | 0.964089 | 0.015883 |
| ENSMUSG00000109511.2 | 33.3945 | -1.09662 | 1.333512 | -0.82235 | 0.410876 | 0.964089 | 0.015883 |
| ENSMUSG00000117476.2 | 90.9184 | 0.601718 | 0.73346 | 0.820383 | 0.411998 | 0.964089 | 0.015883 |
| ENSMUSG00000003746.17 | 922.9911 | 0.274802 | 0.335632 | 0.818761 | 0.412923 | 0.964214 | 0.015826 |
| ENSMUSG00000019851.9 | 1069.156 | -0.23529 | 0.287206 | -0.81924 | 0.412648 | 0.964214 | 0.015826 |
| ENSMUSG00000021248.10 | 377.9489 | -0.37476 | 0.457849 | -0.81852 | 0.41306 | 0.964214 | 0.015826 |
| ENSMUSG00000021928.10 | 855.2067 | 0.313951 | 0.383382 | 0.818898 | 0.412845 | 0.964214 | 0.015826 |
| ENSMUSG00000041415.12 | 117.9579 | -0.86739 | 1.059359 | -0.81879 | 0.412907 | 0.964214 | 0.015826 |
| ENSMUSG00000049090.4 | 379.0177 | -0.39649 | 0.48449 | -0.81837 | 0.413148 | 0.964214 | 0.015826 |
| ENSMUSG00000062580.10 | 224.4141 | -0.46139 | 0.563795 | -0.81837 | 0.413146 | 0.964214 | 0.015826 |
| ENSMUSG00000085396.8 | 35.02377 | -1.31619 | 1.607285 | -0.81889 | 0.41285 | 0.964214 | 0.015826 |
| ENSMUSG00000040883.19 | 1158.458 | -0.28447 | 0.347828 | -0.81784 | 0.413449 | 0.96468 | 0.015617 |
| ENSMUSG00000011958.18 | 170.0677 | -0.50609 | 0.619097 | -0.81746 | 0.413663 | 0.964942 | 0.015499 |
| ENSMUSG00000000555.9 | 11.08965 | -2.07646 | 2.541235 | -0.81711 | 0.413867 | 0.964949 | 0.015495 |
| ENSMUSG00000029364.16 | 209.5219 | 0.467514 | 0.572266 | 0.816952 | 0.413956 | 0.964949 | 0.015495 |
| ENSMUSG00000064284.13 | 17.561 | 1.558127 | 1.907308 | 0.816925 | 0.413971 | 0.964949 | 0.015495 |
| ENSMUSG00000006014.17 | 453.7696 | -0.424 | 0.519209 | -0.81662 | 0.414146 | 0.965119 | 0.015419 |
| ENSMUSG00000002524.18 | 98.03654 | -0.75565 | 0.952908 | -0.79299 | 0.427783 | 0.965663 | 0.015175 |
| ENSMUSG00000002781.20 | 26.81579 | 1.781126 | 2.218557 | 0.802831 | 0.422073 | 0.965663 | 0.015175 |
| ENSMUSG00000002804.5 | 13.42197 | -1.8892 | 2.332229 | -0.81004 | 0.417917 | 0.965663 | 0.015175 |
| ENSMUSG00000005107.14 | 99.03218 | -0.59914 | 0.751239 | -0.79753 | 0.425143 | 0.965663 | 0.015175 |
| ENSMUSG00000006262.16 | 303.0651 | -0.4122 | 0.517024 | -0.79725 | 0.425307 | 0.965663 | 0.015175 |
| ENSMUSG00000006281.9 | 15.68211 | 2.049487 | 2.585106 | 0.792806 | 0.427891 | 0.965663 | 0.015175 |
| ENSMUSG00000006456.11 | 22.2196 | 1.573917 | 1.99015 | 0.790853 | 0.42903 | 0.965663 | 0.015175 |
| ENSMUSG00000007029.17 | 41.12834 | -0.93334 | 1.175075 | -0.79428 | 0.427034 | 0.965663 | 0.015175 |
| ENSMUSG00000008398.17 | 19.116 | 1.312882 | 1.661163 | 0.790339 | 0.42933 | 0.965663 | 0.015175 |
| ENSMUSG00000009828.16 | 55.8059 | 0.865644 | 1.08413 | 0.798469 | 0.424599 | 0.965663 | 0.015175 |
| ENSMUSG00000010097.16 | 57.90074 | 0.764772 | 0.953833 | 0.801788 | 0.422676 | 0.965663 | 0.015175 |
| ENSMUSG00000010406.13 | 643.8013 | 0.372001 | 0.468694 | 0.793697 | 0.427372 | 0.965663 | 0.015175 |
| ENSMUSG00000018171.10 | 833.0653 | 0.256505 | 0.318475 | 0.805416 | 0.42058 | 0.965663 | 0.015175 |
| ENSMUSG00000018736.15 | 197.8872 | 0.51191 | 0.640084 | 0.799755 | 0.423853 | 0.965663 | 0.015175 |
| ENSMUSG00000019838.12 | 231.9677 | 0.391431 | 0.489588 | 0.799511 | 0.423994 | 0.965663 | 0.015175 |
| ENSMUSG00000019876.16 | 10.61051 | -2.31819 | 2.87848 | -0.80535 | 0.420617 | 0.965663 | 0.015175 |
| ENSMUSG00000020258.17 | 476.0471 | -0.32751 | 0.4015 | -0.81573 | 0.414656 | 0.965663 | 0.015175 |
| ENSMUSG00000020260.10 | 57.14254 | 0.787948 | 0.987637 | 0.797812 | 0.42498 | 0.965663 | 0.015175 |
| ENSMUSG00000020262.16 | 90.9394 | -0.73781 | 0.91502 | -0.80633 | 0.420051 | 0.965663 | 0.015175 |
| ENSMUSG00000021000.18 | 1156.892 | 0.222659 | 0.276381 | 0.805622 | 0.420461 | 0.965663 | 0.015175 |
| ENSMUSG00000021079.17 | 333.2758 | 0.366567 | 0.461566 | 0.794181 | 0.42709 | 0.965663 | 0.015175 |
| ENSMUSG00000021127.8 | 584.2211 | 0.292942 | 0.363497 | 0.8059 | 0.420301 | 0.965663 | 0.015175 |
| ENSMUSG00000021193.11 | 180.3737 | 0.449864 | 0.557384 | 0.8071 | 0.419609 | 0.965663 | 0.015175 |

| | | | | | | | |
|---|---|---|---|---|---|---|---|
| ENSMUSG00000021290.7 | 1234.447 | 0.235844 | 0.295661 | 0.797682 | 0.425055 | 0.965663 | 0.015175 |
| ENSMUSG00000021476.10 | 218.5625 | 0.468459 | 0.588659 | 0.795808 | 0.426143 | 0.965663 | 0.015175 |
| ENSMUSG00000021629.11 | 285.15 | 0.382977 | 0.482319 | 0.794033 | 0.427176 | 0.965663 | 0.015175 |
| ENSMUSG00000021668.16 | 11.9529 | -1.97209 | 2.479411 | -0.79539 | 0.426388 | 0.965663 | 0.015175 |
| ENSMUSG00000021792.16 | 671.0675 | 0.336 | 0.414895 | 0.809843 | 0.41803 | 0.965663 | 0.015175 |
| ENSMUSG00000021810.4 | 78.19692 | -0.76735 | 0.959869 | -0.79943 | 0.424041 | 0.965663 | 0.015175 |
| ENSMUSG00000022246.15 | 104.9139 | 0.78131 | 0.976001 | 0.800522 | 0.423408 | 0.965663 | 0.015175 |
| ENSMUSG00000022844.9 | 254.3879 | -0.4801 | 0.606305 | -0.79185 | 0.428449 | 0.965663 | 0.015175 |
| ENSMUSG00000023959.11 | 13.19906 | -1.85559 | 2.319657 | -0.79994 | 0.423745 | 0.965663 | 0.015175 |
| ENSMUSG00000024055.16 | 74.94557 | 0.78828 | 0.96897 | 0.813523 | 0.415918 | 0.965663 | 0.015175 |
| ENSMUSG00000024248.15 | 207.5758 | -0.48697 | 0.598594 | -0.81352 | 0.415918 | 0.965663 | 0.015175 |
| ENSMUSG00000024587.11 | 323.5247 | 0.327742 | 0.407733 | 0.803815 | 0.421504 | 0.965663 | 0.015175 |
| ENSMUSG00000024622.9 | 15.15536 | 1.667591 | 2.107461 | 0.791279 | 0.428781 | 0.965663 | 0.015175 |
| ENSMUSG00000024665.9 | 1131.435 | 0.31848 | 0.397487 | 0.801232 | 0.422997 | 0.965663 | 0.015175 |
| ENSMUSG00000024764.11 | 15.84468 | -2.0292 | 2.493933 | -0.81366 | 0.415842 | 0.965663 | 0.015175 |
| ENSMUSG00000024778.14 | 84.4172 | -0.69315 | 0.870364 | -0.7964 | 0.425802 | 0.965663 | 0.015175 |
| ENSMUSG00000025226.12 | 11.79105 | -2.3083 | 2.849522 | -0.81007 | 0.417902 | 0.965663 | 0.015175 |
| ENSMUSG00000025316.17 | 30.98396 | 1.767764 | 2.222924 | 0.795243 | 0.426472 | 0.965663 | 0.015175 |
| ENSMUSG00000025817.13 | 44.87703 | 0.944483 | 1.163095 | 0.812043 | 0.416767 | 0.965663 | 0.015175 |
| ENSMUSG00000026005.16 | 375.6936 | -0.44641 | 0.552406 | -0.80812 | 0.419022 | 0.965663 | 0.015175 |
| ENSMUSG00000026341.17 | 159.2001 | -0.55195 | 0.684162 | -0.80675 | 0.419808 | 0.965663 | 0.015175 |
| ENSMUSG00000026348.8 | 110.5603 | 0.617753 | 0.776456 | 0.795607 | 0.426261 | 0.965663 | 0.015175 |
| ENSMUSG00000026753.6 | 196.8646 | -0.4345 | 0.545124 | -0.79706 | 0.425416 | 0.965663 | 0.015175 |
| ENSMUSG00000026798.11 | 57.19745 | -0.91723 | 1.158793 | -0.79154 | 0.428628 | 0.965663 | 0.015175 |
| ENSMUSG00000026966.7 | 74.35346 | -0.81293 | 1.011822 | -0.80344 | 0.421723 | 0.965663 | 0.015175 |
| ENSMUSG00000027006.14 | 82.37055 | 0.710029 | 0.878545 | 0.808188 | 0.418982 | 0.965663 | 0.015175 |
| ENSMUSG00000027801.8 | 228.8487 | -0.45086 | 0.562508 | -0.80152 | 0.422832 | 0.965663 | 0.015175 |
| ENSMUSG00000027822.17 | 632.0941 | -0.31546 | 0.390516 | -0.80781 | 0.419197 | 0.965663 | 0.015175 |
| ENSMUSG00000027875.13 | 3851.315 | 0.208417 | 0.260604 | 0.799747 | 0.423857 | 0.965663 | 0.015175 |
| ENSMUSG00000027984.9 | 942.6431 | 0.252257 | 0.314806 | 0.801311 | 0.422952 | 0.965663 | 0.015175 |
| ENSMUSG00000028024.15 | 445.4604 | 0.387492 | 0.485569 | 0.798017 | 0.424861 | 0.965663 | 0.015175 |
| ENSMUSG00000028186.15 | 9407.698 | -0.18207 | 0.226366 | -0.8043 | 0.421221 | 0.965663 | 0.015175 |
| ENSMUSG00000028426.11 | 825.6415 | -0.23176 | 0.28997 | -0.79924 | 0.424149 | 0.965663 | 0.015175 |
| ENSMUSG00000028476.14 | 19.69083 | -1.99272 | 2.496282 | -0.79828 | 0.42471 | 0.965663 | 0.015175 |
| ENSMUSG00000028601.19 | 560.2024 | 0.330447 | 0.411569 | 0.802897 | 0.422034 | 0.965663 | 0.015175 |
| ENSMUSG00000028745.19 | 279.0088 | -0.39926 | 0.503406 | -0.79312 | 0.427706 | 0.965663 | 0.015175 |
| ENSMUSG00000028889.18 | 82.44937 | -0.72856 | 0.909818 | -0.80077 | 0.423263 | 0.965663 | 0.015175 |
| ENSMUSG00000029401.9 | 45.16783 | 0.909621 | 1.127628 | 0.806668 | 0.419858 | 0.965663 | 0.015175 |
| ENSMUSG00000029466.12 | 25.13496 | -1.25872 | 1.570773 | -0.80134 | 0.422935 | 0.965663 | 0.015175 |
| ENSMUSG00000029578.16 | 103.6853 | -0.7381 | 0.92901 | -0.7945 | 0.426906 | 0.965663 | 0.015175 |
| ENSMUSG00000029627.13 | 37.04791 | -1.45339 | 1.802104 | -0.8065 | 0.419956 | 0.965663 | 0.015175 |
| ENSMUSG00000029642.12 | 438.6814 | 0.317656 | 0.389919 | 0.814673 | 0.41526 | 0.965663 | 0.015175 |
| ENSMUSG00000029863.14 | 18.40776 | 1.729675 | 2.152114 | 0.80371 | 0.421565 | 0.965663 | 0.015175 |
| ENSMUSG00000030122.13 | 158.2109 | -0.76169 | 0.938964 | -0.8112 | 0.417251 | 0.965663 | 0.015175 |

| | | | | | | | |
|---|---|---|---|---|---|---|---|
| ENSMUSG00000030168.14 | 574.0954 | -0.32432 | 0.399181 | -0.81246 | 0.416528 | 0.965663 | 0.015175 |
| ENSMUSG00000030423.8 | 26.13269 | 1.267346 | 1.602292 | 0.790958 | 0.428969 | 0.965663 | 0.015175 |
| ENSMUSG00000030513.15 | 485.6389 | 0.341869 | 0.432247 | 0.790912 | 0.428996 | 0.965663 | 0.015175 |
| ENSMUSG00000031227.9 | 16.60828 | -1.80335 | 2.214468 | -0.81435 | 0.415445 | 0.965663 | 0.015175 |
| ENSMUSG00000031271.15 | 123.6462 | -0.56756 | 0.70307 | -0.80726 | 0.419519 | 0.965663 | 0.015175 |
| ENSMUSG00000031443.8 | 188.303 | -0.71221 | 0.893896 | -0.79675 | 0.425598 | 0.965663 | 0.015175 |
| ENSMUSG00000031570.18 | 37.55235 | -0.99916 | 1.22558 | -0.81525 | 0.414928 | 0.965663 | 0.015175 |
| ENSMUSG00000031628.10 | 134.5182 | 0.536079 | 0.667446 | 0.80318 | 0.421871 | 0.965663 | 0.015175 |
| ENSMUSG00000032246.15 | 16.85785 | -2.13969 | 2.626296 | -0.81472 | 0.415233 | 0.965663 | 0.015175 |
| ENSMUSG00000032290.8 | 34.64257 | 1.053898 | 1.308573 | 0.805379 | 0.420601 | 0.965663 | 0.015175 |
| ENSMUSG00000032320.8 | 69.57085 | -0.76075 | 0.934906 | -0.81372 | 0.415804 | 0.965663 | 0.015175 |
| ENSMUSG00000033308.17 | 2162.115 | 0.244843 | 0.308562 | 0.793495 | 0.427489 | 0.965663 | 0.015175 |
| ENSMUSG00000033671.19 | 129.7589 | -0.56764 | 0.711792 | -0.79747 | 0.425176 | 0.965663 | 0.015175 |
| ENSMUSG00000033728.15 | 11.8324 | -2.0193 | 2.483299 | -0.81315 | 0.41613 | 0.965663 | 0.015175 |
| ENSMUSG00000033845.14 | 551.9145 | 0.365036 | 0.459249 | 0.794856 | 0.426697 | 0.965663 | 0.015175 |
| ENSMUSG00000034109.16 | 428.3623 | 0.388336 | 0.483823 | 0.802641 | 0.422182 | 0.965663 | 0.015175 |
| ENSMUSG00000034563.17 | 210.6138 | -0.44073 | 0.540774 | -0.815 | 0.415071 | 0.965663 | 0.015175 |
| ENSMUSG00000034714.10 | 132.4392 | -0.5612 | 0.695617 | -0.80677 | 0.419801 | 0.965663 | 0.015175 |
| ENSMUSG00000035048.14 | 493.4426 | 0.317969 | 0.392708 | 0.809683 | 0.418123 | 0.965663 | 0.015175 |
| ENSMUSG00000035202.9 | 41.92485 | -0.90176 | 1.134428 | -0.79491 | 0.426668 | 0.965663 | 0.015175 |
| ENSMUSG00000035212.15 | 72.60397 | -0.97033 | 1.204413 | -0.80565 | 0.420446 | 0.965663 | 0.015175 |
| ENSMUSG00000035530.14 | 1342.059 | 0.246235 | 0.308465 | 0.798258 | 0.424721 | 0.965663 | 0.015175 |
| ENSMUSG00000035637.15 | 556.2857 | -0.47335 | 0.588138 | -0.80482 | 0.420921 | 0.965663 | 0.015175 |
| ENSMUSG00000036099.17 | 34.8953 | -1.15703 | 1.436269 | -0.80558 | 0.420483 | 0.965663 | 0.015175 |
| ENSMUSG00000036403.16 | 25.29743 | -1.63242 | 2.010122 | -0.8121 | 0.416735 | 0.965663 | 0.015175 |
| ENSMUSG00000036764.13 | 137.6932 | -0.49593 | 0.626367 | -0.79175 | 0.428505 | 0.965663 | 0.015175 |
| ENSMUSG00000036854.15 | 35.55298 | 0.97386 | 1.231275 | 0.790936 | 0.428981 | 0.965663 | 0.015175 |
| ENSMUSG00000036918.17 | 96.46624 | -0.66749 | 0.839647 | -0.79496 | 0.426634 | 0.965663 | 0.015175 |
| ENSMUSG00000037210.17 | 11.83142 | 1.70353 | 2.154457 | 0.7907 | 0.429119 | 0.965663 | 0.015175 |
| ENSMUSG00000037221.14 | 62.94522 | 0.886452 | 1.096582 | 0.808377 | 0.418873 | 0.965663 | 0.015175 |
| ENSMUSG00000037234.18 | 189.2582 | 0.549815 | 0.682759 | 0.805284 | 0.420656 | 0.965663 | 0.015175 |
| ENSMUSG00000037351.10 | 73.93588 | -0.66173 | 0.821947 | -0.80508 | 0.420773 | 0.965663 | 0.015175 |
| ENSMUSG00000037876.18 | 182.6459 | -0.54395 | 0.680244 | -0.79963 | 0.423923 | 0.965663 | 0.015175 |
| ENSMUSG00000038084.17 | 376.2049 | 0.305364 | 0.383892 | 0.795443 | 0.426356 | 0.965663 | 0.015175 |
| ENSMUSG00000038235.5 | 529.3071 | 0.320977 | 0.400522 | 0.801396 | 0.422902 | 0.965663 | 0.015175 |
| ENSMUSG00000038525.17 | 20.85216 | -1.30971 | 1.618966 | -0.80898 | 0.418527 | 0.965663 | 0.015175 |
| ENSMUSG00000040177.11 | 16.88011 | 2.034239 | 2.506936 | 0.811445 | 0.41711 | 0.965663 | 0.015175 |
| ENSMUSG00000040195.12 | 88.08368 | -0.82694 | 1.036089 | -0.79814 | 0.42479 | 0.965663 | 0.015175 |
| ENSMUSG00000041044.10 | 13.64675 | -1.58295 | 1.997847 | -0.79233 | 0.428168 | 0.965663 | 0.015175 |
| ENSMUSG00000041633.15 | 13.14782 | 2.455237 | 3.06877 | 0.800072 | 0.423669 | 0.965663 | 0.015175 |
| ENSMUSG00000041939.15 | 27.01802 | -1.54723 | 1.923983 | -0.80418 | 0.421292 | 0.965663 | 0.015175 |
| ENSMUSG00000041958.11 | 115.7155 | -0.74353 | 0.938863 | -0.79194 | 0.428393 | 0.965663 | 0.015175 |
| ENSMUSG00000042102.8 | 1922.901 | 0.220575 | 0.277312 | 0.795405 | 0.426378 | 0.965663 | 0.015175 |
| ENSMUSG00000042286.14 | 57.67832 | 1.066413 | 1.314762 | 0.811107 | 0.417304 | 0.965663 | 0.015175 |

| | | | | | | | |
|---|---|---|---|---|---|---|---|
| ENSMUSG00000042557.15 | 56.65885 | -0.89752 | 1.135055 | -0.79073 | 0.4291 | 0.965663 | 0.015175 |
| ENSMUSG00000042670.6 | 145.7217 | -0.56939 | 0.698827 | -0.81478 | 0.4152 | 0.965663 | 0.015175 |
| ENSMUSG00000044600.15 | 192.6464 | 0.56433 | 0.69405 | 0.813096 | 0.416163 | 0.965663 | 0.015175 |
| ENSMUSG00000044709.7 | 62.56677 | 0.948499 | 1.197291 | 0.792204 | 0.428242 | 0.965663 | 0.015175 |
| ENSMUSG00000044881.8 | 56.32976 | -0.85809 | 1.071067 | -0.80115 | 0.423043 | 0.965663 | 0.015175 |
| ENSMUSG00000047675.16 | 4578.635 | 0.19692 | 0.243538 | 0.808581 | 0.418756 | 0.965663 | 0.015175 |
| ENSMUSG00000049047.12 | 22.0967 | -1.41063 | 1.729463 | -0.81564 | 0.414703 | 0.965663 | 0.015175 |
| ENSMUSG00000051041.9 | 27.55275 | -1.42598 | 1.758409 | -0.81095 | 0.417396 | 0.965663 | 0.015175 |
| ENSMUSG00000052117.6 | 22.85686 | -1.64478 | 2.059545 | -0.79862 | 0.424513 | 0.965663 | 0.015175 |
| ENSMUSG00000053644.14 | 2182.384 | -0.22412 | 0.282678 | -0.79284 | 0.427873 | 0.965663 | 0.015175 |
| ENSMUSG00000054630.8 | 11010.67 | 0.222929 | 0.281979 | 0.790587 | 0.429185 | 0.965663 | 0.015175 |
| ENSMUSG00000054808.16 | 235.1788 | 0.493551 | 0.624512 | 0.790299 | 0.429353 | 0.965663 | 0.015175 |
| ENSMUSG00000056493.10 | 38.99518 | 0.894513 | 1.114509 | 0.802607 | 0.422202 | 0.965663 | 0.015175 |
| ENSMUSG00000057788.14 | 43.41754 | 1.368047 | 1.698461 | 0.805463 | 0.420553 | 0.965663 | 0.015175 |
| ENSMUSG00000058446.15 | 355.4848 | 0.357959 | 0.44649 | 0.801719 | 0.422715 | 0.965663 | 0.015175 |
| ENSMUSG00000060224.5 | 42.57284 | 0.926263 | 1.153436 | 0.803046 | 0.421948 | 0.965663 | 0.015175 |
| ENSMUSG00000068328.11 | 54.41463 | -1.32204 | 1.632179 | -0.80998 | 0.41795 | 0.965663 | 0.015175 |
| ENSMUSG00000068686.13 | 17.99821 | -1.7135 | 2.139605 | -0.80085 | 0.423219 | 0.965663 | 0.015175 |
| ENSMUSG00000069255.14 | 62.14132 | -0.7663 | 0.950262 | -0.80641 | 0.420009 | 0.965663 | 0.015175 |
| ENSMUSG00000069516.9 | 605.3467 | -0.42635 | 0.52892 | -0.80608 | 0.420199 | 0.965663 | 0.015175 |
| ENSMUSG00000069833.14 | 139.3273 | -0.60596 | 0.761523 | -0.79572 | 0.426193 | 0.965663 | 0.015175 |
| ENSMUSG00000070661.8 | 18.57308 | -1.69029 | 2.085624 | -0.81045 | 0.417683 | 0.965663 | 0.015175 |
| ENSMUSG00000073147.4 | 58.26571 | 1.138079 | 1.413217 | 0.805311 | 0.42064 | 0.965663 | 0.015175 |
| ENSMUSG00000073775.5 | 129.4715 | 0.568994 | 0.719353 | 0.790981 | 0.428955 | 0.965663 | 0.015175 |
| ENSMUSG00000075284.11 | 24.37428 | -1.65266 | 2.034901 | -0.81216 | 0.416702 | 0.965663 | 0.015175 |
| ENSMUSG00000080717.11 | 16.13826 | 1.488661 | 1.839043 | 0.809476 | 0.418242 | 0.965663 | 0.015175 |
| ENSMUSG00000087380.8 | 178.4419 | -0.55475 | 0.686711 | -0.80783 | 0.419187 | 0.965663 | 0.015175 |
| ENSMUSG00000089704.10 | 77.58248 | 0.814923 | 1.013664 | 0.803938 | 0.421433 | 0.965663 | 0.015175 |
| ENSMUSG00000091474.3 | 38.10488 | 1.084166 | 1.329537 | 0.815446 | 0.414817 | 0.965663 | 0.015175 |
| ENSMUSG00000091811.3 | 12.72276 | 2.226598 | 2.783068 | 0.800051 | 0.423681 | 0.965663 | 0.015175 |
| ENSMUSG00000097177.9 | 13.68741 | -2.06125 | 2.585097 | -0.79736 | 0.425244 | 0.965663 | 0.015175 |
| ENSMUSG00000097616.8 | 23.40839 | 1.301559 | 1.614898 | 0.80597 | 0.42026 | 0.965663 | 0.015175 |
| ENSMUSG00000097944.2 | 15.49459 | 1.80087 | 2.240256 | 0.803868 | 0.421473 | 0.965663 | 0.015175 |
| ENSMUSG00000104204.2 | 12.36751 | -2.22334 | 2.743352 | -0.81045 | 0.417683 | 0.965663 | 0.015175 |
| ENSMUSG00000109628.2 | 695.1156 | -0.3201 | 0.395933 | -0.80847 | 0.418821 | 0.965663 | 0.015175 |
| ENSMUSG00000116564.2 | 149.2604 | -0.63166 | 0.775775 | -0.81424 | 0.415509 | 0.965663 | 0.015175 |
| ENSMUSG00000035478.15 | 40.48049 | -0.82885 | 1.049093 | -0.79007 | 0.429489 | 0.965739 | 0.01514 |
| ENSMUSG00000033161.11 | 212.4349 | -0.44983 | 0.569818 | -0.78943 | 0.429862 | 0.966303 | 0.014887 |
| ENSMUSG00000062083.2 | 165.0652 | 0.435987 | 0.55238 | 0.789288 | 0.429944 | 0.966303 | 0.014887 |
| ENSMUSG00000020091.16 | 346.4909 | -0.36065 | 0.457142 | -0.78893 | 0.430156 | 0.96655 | 0.014776 |
| ENSMUSG00000025894.12 | 323.5667 | 0.381062 | 0.48346 | 0.788197 | 0.430582 | 0.96673 | 0.014695 |
| ENSMUSG00000027679.14 | 966.0679 | 0.240491 | 0.305156 | 0.788091 | 0.430643 | 0.96673 | 0.014695 |
| ENSMUSG00000039753.17 | 63.57862 | 0.780887 | 0.990203 | 0.788614 | 0.430338 | 0.96673 | 0.014695 |
| ENSMUSG00000040430.19 | 136.07 | 0.765599 | 0.971362 | 0.78817 | 0.430597 | 0.96673 | 0.014695 |

| | | | | | | | |
|---|---|---|---|---|---|---|---|
| ENSMUSG00000000838.18 | 119.1198 | -0.60909 | 0.777402 | -0.7835 | 0.433333 | 0.966926 | 0.014607 |
| ENSMUSG00000006307.17 | 14.7469 | 1.733825 | 2.21414 | 0.783069 | 0.433587 | 0.966926 | 0.014607 |
| ENSMUSG00000008475.14 | 746.6196 | 0.238298 | 0.302666 | 0.787331 | 0.431088 | 0.966926 | 0.014607 |
| ENSMUSG00000020810.6 | 26.08401 | 1.400086 | 1.78599 | 0.783927 | 0.433083 | 0.966926 | 0.014607 |
| ENSMUSG00000024359.11 | 2867.445 | 0.213006 | 0.271938 | 0.783288 | 0.433458 | 0.966926 | 0.014607 |
| ENSMUSG00000026544.7 | 78.37031 | 0.995987 | 1.270193 | 0.784122 | 0.432968 | 0.966926 | 0.014607 |
| ENSMUSG00000026872.19 | 76.42766 | -0.74302 | 0.943464 | -0.78754 | 0.430966 | 0.966926 | 0.014607 |
| ENSMUSG00000028637.17 | 36.30543 | -0.93507 | 1.190764 | -0.78527 | 0.432297 | 0.966926 | 0.014607 |
| ENSMUSG00000029701.16 | 93.34648 | -0.72033 | 0.918837 | -0.78396 | 0.433066 | 0.966926 | 0.014607 |
| ENSMUSG00000029994.18 | 181.5073 | -0.46783 | 0.596356 | -0.78449 | 0.432754 | 0.966926 | 0.014607 |
| ENSMUSG00000030036.9 | 163.3519 | 0.520605 | 0.663107 | 0.7851 | 0.432395 | 0.966926 | 0.014607 |
| ENSMUSG00000030682.5 | 130.0551 | -0.60031 | 0.764874 | -0.78484 | 0.432545 | 0.966926 | 0.014607 |
| ENSMUSG00000032096.17 | 157.6169 | -0.44954 | 0.573055 | -0.78446 | 0.432772 | 0.966926 | 0.014607 |
| ENSMUSG00000032743.16 | 12.51806 | 1.642139 | 2.095166 | 0.783775 | 0.433172 | 0.966926 | 0.014607 |
| ENSMUSG00000035064.18 | 40.10418 | -1.12127 | 1.431645 | -0.7832 | 0.433509 | 0.966926 | 0.014607 |
| ENSMUSG00000035104.15 | 851.157 | -0.31864 | 0.405465 | -0.78586 | 0.431947 | 0.966926 | 0.014607 |
| ENSMUSG00000035284.11 | 11.92535 | -1.95492 | 2.48956 | -0.78525 | 0.43231 | 0.966926 | 0.014607 |
| ENSMUSG00000039050.9 | 180.9085 | -0.46831 | 0.597158 | -0.78423 | 0.432907 | 0.966926 | 0.014607 |
| ENSMUSG00000041453.13 | 2141.523 | 0.181907 | 0.232274 | 0.783159 | 0.433534 | 0.966926 | 0.014607 |
| ENSMUSG00000041716.8 | 44.86058 | 0.882647 | 1.122261 | 0.78649 | 0.43158 | 0.966926 | 0.014607 |
| ENSMUSG00000041798.16 | 179.7064 | 0.589929 | 0.752911 | 0.783531 | 0.433315 | 0.966926 | 0.014607 |
| ENSMUSG00000046836.14 | 56.19323 | 0.7704 | 0.982357 | 0.784236 | 0.432902 | 0.966926 | 0.014607 |
| ENSMUSG00000054309.10 | 85.69841 | -0.67016 | 0.852077 | -0.7865 | 0.431572 | 0.966926 | 0.014607 |
| ENSMUSG00000059897.6 | 31.37659 | 1.266008 | 1.615018 | 0.783897 | 0.433101 | 0.966926 | 0.014607 |
| ENSMUSG00000063931.15 | 375.3686 | -0.33567 | 0.426151 | -0.78767 | 0.430889 | 0.966926 | 0.014607 |
| ENSMUSG00000067235.15 | 9153.902 | -0.19107 | 0.243287 | -0.78538 | 0.432233 | 0.966926 | 0.014607 |
| ENSMUSG00000078435.6 | 22.87236 | 1.429277 | 1.820914 | 0.784923 | 0.432499 | 0.966926 | 0.014607 |
| ENSMUSG00000115124.2 | 35.10142 | 1.100559 | 1.399279 | 0.786519 | 0.431564 | 0.966926 | 0.014607 |
| ENSMUSG00000035620.16 | 44.79401 | 1.158195 | 1.480059 | 0.782533 | 0.433901 | 0.967172 | 0.014496 |
| ENSMUSG00000036427.6 | 122.2228 | -0.6069 | 0.77556 | -0.78254 | 0.433899 | 0.967172 | 0.014496 |
| ENSMUSG00000021044.16 | 41.70169 | -1.06519 | 1.361616 | -0.7823 | 0.43404 | 0.967254 | 0.014459 |
| ENSMUSG00000000827.19 | 197.8067 | 0.486182 | 0.621725 | 0.781989 | 0.434221 | 0.96743 | 0.014381 |
| ENSMUSG00000030557.18 | 94.57328 | -0.63322 | 0.810868 | -0.78091 | 0.434854 | 0.96793 | 0.014156 |
| ENSMUSG00000035329.8 | 283.7563 | 0.430623 | 0.551244 | 0.781184 | 0.434694 | 0.96793 | 0.014156 |
| ENSMUSG00000044150.13 | 12.52399 | 2.121571 | 2.716266 | 0.781062 | 0.434766 | 0.96793 | 0.014156 |
| ENSMUSG00000073987.5 | 171.7971 | 0.462231 | 0.591897 | 0.780932 | 0.434843 | 0.96793 | 0.014156 |
| ENSMUSG00000026698.9 | 27.7564 | -1.1644 | 1.491573 | -0.78065 | 0.435008 | 0.967937 | 0.014153 |
| ENSMUSG00000030726.17 | 32.79179 | -1.20522 | 1.544395 | -0.78039 | 0.435163 | 0.967937 | 0.014153 |
| ENSMUSG00000035024.17 | 16.28721 | -1.2483 | 1.599316 | -0.78052 | 0.435085 | 0.967937 | 0.014153 |
| ENSMUSG00000030674.6 | 381.5985 | 0.408474 | 0.523933 | 0.77963 | 0.435609 | 0.968701 | 0.01381 |
| ENSMUSG00000021486.8 | 258.1711 | -0.44625 | 0.572679 | -0.77924 | 0.435838 | 0.968755 | 0.013786 |
| ENSMUSG00000066800.12 | 28.09942 | 1.718197 | 2.204485 | 0.77941 | 0.435738 | 0.968755 | 0.013786 |
| ENSMUSG00000001924.16 | 396.7197 | -0.41194 | 0.535104 | -0.76984 | 0.441396 | 0.968944 | 0.013701 |
| ENSMUSG00000007617.19 | 145.7556 | 0.589979 | 0.761271 | 0.774992 | 0.438345 | 0.968944 | 0.013701 |

| | | | | | | | |
|---|---|---|---|---|---|---|---|
| ENSMUSG00000010045.3 | 26.80029 | -1.72454 | 2.231359 | -0.77287 | 0.439601 | 0.968944 | 0.013701 |
| ENSMUSG00000012429.10 | 98.14255 | 0.759941 | 0.976607 | 0.778144 | 0.436484 | 0.968944 | 0.013701 |
| ENSMUSG00000014039.19 | 18.78482 | -1.86531 | 2.402124 | -0.77653 | 0.437439 | 0.968944 | 0.013701 |
| ENSMUSG00000015806.13 | 2200.107 | 0.194802 | 0.252322 | 0.772037 | 0.440093 | 0.968944 | 0.013701 |
| ENSMUSG00000020100.16 | 18.32215 | -1.65771 | 2.155599 | -0.76903 | 0.441877 | 0.968944 | 0.013701 |
| ENSMUSG00000021548.11 | 259.8815 | 0.46541 | 0.601454 | 0.773808 | 0.439044 | 0.968944 | 0.013701 |
| ENSMUSG00000021771.15 | 628.1142 | 0.282814 | 0.366218 | 0.772256 | 0.439963 | 0.968944 | 0.013701 |
| ENSMUSG00000022000.12 | 226.4308 | -0.44797 | 0.582712 | -0.76877 | 0.44203 | 0.968944 | 0.013701 |
| ENSMUSG00000022031.7 | 48.47013 | 0.780993 | 1.015639 | 0.768967 | 0.441913 | 0.968944 | 0.013701 |
| ENSMUSG00000022130.11 | 28.77453 | -1.11584 | 1.44416 | -0.77266 | 0.439726 | 0.968944 | 0.013701 |
| ENSMUSG00000022151.17 | 116.3816 | 0.572223 | 0.745545 | 0.767523 | 0.442771 | 0.968944 | 0.013701 |
| ENSMUSG00000023224.13 | 2823.455 | -0.2321 | 0.298654 | -0.77717 | 0.437059 | 0.968944 | 0.013701 |
| ENSMUSG00000024069.15 | 87.75107 | 0.621203 | 0.79878 | 0.77769 | 0.436752 | 0.968944 | 0.013701 |
| ENSMUSG00000024659.16 | 60.20705 | 0.936527 | 1.207275 | 0.775736 | 0.437905 | 0.968944 | 0.013701 |
| ENSMUSG00000025144.18 | 71.72981 | -1.02763 | 1.327898 | -0.77388 | 0.439005 | 0.968944 | 0.013701 |
| ENSMUSG00000025403.6 | 90.82967 | -0.74266 | 0.962791 | -0.77137 | 0.44049 | 0.968944 | 0.013701 |
| ENSMUSG00000025534.18 | 119.3053 | 0.640509 | 0.832136 | 0.769716 | 0.441468 | 0.968944 | 0.013701 |
| ENSMUSG00000027965.16 | 38.34169 | 0.947902 | 1.23441 | 0.767898 | 0.442548 | 0.968944 | 0.013701 |
| ENSMUSG00000028273.16 | 299.7223 | -0.46494 | 0.597327 | -0.77837 | 0.43635 | 0.968944 | 0.013701 |
| ENSMUSG00000028433.15 | 53.35299 | 0.793945 | 1.023804 | 0.775485 | 0.438053 | 0.968944 | 0.013701 |
| ENSMUSG00000028447.12 | 79.98461 | 0.593567 | 0.763124 | 0.777812 | 0.43668 | 0.968944 | 0.013701 |
| ENSMUSG00000028672.14 | 373.1352 | 0.366383 | 0.47733 | 0.767567 | 0.442744 | 0.968944 | 0.013701 |
| ENSMUSG00000028729.16 | 217.1013 | -0.38797 | 0.504206 | -0.76947 | 0.441615 | 0.968944 | 0.013701 |
| ENSMUSG00000029020.14 | 243.7339 | 0.397543 | 0.513464 | 0.774236 | 0.438791 | 0.968944 | 0.013701 |
| ENSMUSG00000029911.15 | 201.8361 | 0.453437 | 0.584859 | 0.775294 | 0.438166 | 0.968944 | 0.013701 |
| ENSMUSG00000031373.5 | 18.96736 | 1.647176 | 2.137936 | 0.770452 | 0.441032 | 0.968944 | 0.013701 |
| ENSMUSG00000031949.18 | 12.82085 | -1.93059 | 2.50994 | -0.76918 | 0.441789 | 0.968944 | 0.013701 |
| ENSMUSG00000033356.4 | 12.96036 | 2.106046 | 2.734665 | 0.770129 | 0.441223 | 0.968944 | 0.013701 |
| ENSMUSG00000033628.16 | 47.75786 | -0.86476 | 1.11365 | -0.77651 | 0.437446 | 0.968944 | 0.013701 |
| ENSMUSG00000034252.15 | 245.2912 | -0.43378 | 0.56139 | -0.7727 | 0.439703 | 0.968944 | 0.013701 |
| ENSMUSG00000034926.4 | 724.9892 | 0.323802 | 0.416178 | 0.778036 | 0.436548 | 0.968944 | 0.013701 |
| ENSMUSG00000035885.6 | 966.8345 | 0.282503 | 0.367505 | 0.768706 | 0.442068 | 0.968944 | 0.013701 |
| ENSMUSG00000036908.18 | 17.9056 | -1.41776 | 1.825912 | -0.77647 | 0.437474 | 0.968944 | 0.013701 |
| ENSMUSG00000038406.18 | 15.44783 | 1.546339 | 2.001442 | 0.772612 | 0.439752 | 0.968944 | 0.013701 |
| ENSMUSG00000039616.11 | 163.7308 | -0.44947 | 0.584104 | -0.7695 | 0.441595 | 0.968944 | 0.013701 |
| ENSMUSG00000040586.9 | 22.13531 | -1.41967 | 1.827895 | -0.77667 | 0.437354 | 0.968944 | 0.013701 |
| ENSMUSG00000040605.8 | 28.55065 | 1.175022 | 1.510084 | 0.778117 | 0.4365 | 0.968944 | 0.013701 |
| ENSMUSG00000041084.9 | 999.5719 | 0.269206 | 0.349719 | 0.76978 | 0.441431 | 0.968944 | 0.013701 |
| ENSMUSG00000041870.18 | 46.6383 | -1.01405 | 1.319243 | -0.76866 | 0.442097 | 0.968944 | 0.013701 |
| ENSMUSG00000041959.15 | 1413.125 | 0.236059 | 0.305331 | 0.773125 | 0.439449 | 0.968944 | 0.013701 |
| ENSMUSG00000042834.16 | 407.3591 | -0.34244 | 0.445828 | -0.7681 | 0.442426 | 0.968944 | 0.013701 |
| ENSMUSG00000046096.8 | 72.52236 | 0.783344 | 1.015566 | 0.771337 | 0.440507 | 0.968944 | 0.013701 |
| ENSMUSG00000046404.6 | 81.87398 | 0.651789 | 0.849007 | 0.767708 | 0.44266 | 0.968944 | 0.013701 |
| ENSMUSG00000048534.8 | 9.904431 | -1.93296 | 2.515429 | -0.76844 | 0.442225 | 0.968944 | 0.013701 |

| | | | | | | | |
|---|---|---|---|---|---|---|---|
| ENSMUSG00000048833.10 | 140.1623 | -0.53279 | 0.690855 | -0.7712 | 0.440589 | 0.968944 | 0.013701 |
| ENSMUSG00000054676.18 | 153.9687 | -0.47345 | 0.607963 | -0.77875 | 0.436125 | 0.968944 | 0.013701 |
| ENSMUSG00000055531.13 | 168.5102 | 0.536203 | 0.693148 | 0.773577 | 0.439181 | 0.968944 | 0.013701 |
| ENSMUSG00000056665.3 | 13.52015 | 2.336477 | 3.036927 | 0.769356 | 0.441682 | 0.968944 | 0.013701 |
| ENSMUSG00000056812.15 | 16.08464 | -2.01766 | 2.608232 | -0.77357 | 0.439184 | 0.968944 | 0.013701 |
| ENSMUSG00000061032.10 | 119.6992 | -0.70614 | 0.908376 | -0.77737 | 0.436942 | 0.968944 | 0.013701 |
| ENSMUSG00000062604.12 | 95.433 | 0.705183 | 0.91772 | 0.768408 | 0.442245 | 0.968944 | 0.013701 |
| ENSMUSG00000066043.14 | 95.7318 | 0.601984 | 0.780349 | 0.771429 | 0.440453 | 0.968944 | 0.013701 |
| ENSMUSG00000069208.10 | 96.17488 | -0.71733 | 0.928856 | -0.77227 | 0.439952 | 0.968944 | 0.013701 |
| ENSMUSG00000071646.10 | 37.00407 | -1.41456 | 1.830361 | -0.77283 | 0.439623 | 0.968944 | 0.013701 |
| ENSMUSG00000072640.12 | 35.83281 | -1.13324 | 1.473939 | -0.76885 | 0.44198 | 0.968944 | 0.013701 |
| ENSMUSG00000074829.11 | 78.7111 | 0.71046 | 0.925018 | 0.768051 | 0.442457 | 0.968944 | 0.013701 |
| ENSMUSG00000074922.2 | 52.68847 | 0.992015 | 1.288863 | 0.769682 | 0.441489 | 0.968944 | 0.013701 |
| ENSMUSG00000079563.11 | 353.0952 | -0.42167 | 0.547488 | -0.77019 | 0.441186 | 0.968944 | 0.013701 |
| ENSMUSG00000081434.4 | 88.78043 | 0.637777 | 0.821556 | 0.776304 | 0.437569 | 0.968944 | 0.013701 |
| ENSMUSG00000083022.2 | 333.5929 | 0.331872 | 0.430484 | 0.770929 | 0.440749 | 0.968944 | 0.013701 |
| ENSMUSG00000084235.3 | 64.26361 | 0.714438 | 0.918799 | 0.777578 | 0.436818 | 0.968944 | 0.013701 |
| ENSMUSG00000084878.2 | 11.02668 | 2.154372 | 2.779335 | 0.775139 | 0.438257 | 0.968944 | 0.013701 |
| ENSMUSG00000101249.2 | 10.51831 | 1.603995 | 2.066973 | 0.776012 | 0.437742 | 0.968944 | 0.013701 |
| ENSMUSG00000106943.3 | 13.59389 | -2.31973 | 3.018358 | -0.76854 | 0.442167 | 0.968944 | 0.013701 |
| ENSMUSG00000107283.4 | 317.4204 | -0.37007 | 0.476798 | -0.77615 | 0.437661 | 0.968944 | 0.013701 |
| ENSMUSG00000038552.15 | 26.97197 | 1.062415 | 1.384954 | 0.767112 | 0.443015 | 0.969031 | 0.013662 |
| ENSMUSG00000071654.7 | 199.869 | 0.497061 | 0.64786 | 0.767236 | 0.442941 | 0.969031 | 0.013662 |
| ENSMUSG00000030126.18 | 164.8596 | 0.439618 | 0.573231 | 0.766913 | 0.443133 | 0.969067 | 0.013646 |
| ENSMUSG00000022911.12 | 26.25174 | -1.35618 | 1.769814 | -0.76628 | 0.443508 | 0.969242 | 0.013568 |
| ENSMUSG00000031333.8 | 259.637 | -0.36723 | 0.479252 | -0.76626 | 0.44352 | 0.969242 | 0.013568 |
| ENSMUSG00000067369.13 | 142.7582 | -0.48377 | 0.631276 | -0.76633 | 0.443479 | 0.969242 | 0.013568 |
| ENSMUSG00000002769.10 | 4819.306 | 0.216098 | 0.282355 | 0.765342 | 0.444068 | 0.969557 | 0.013427 |
| ENSMUSG00000020023.19 | 123.7654 | 0.58583 | 0.766365 | 0.764427 | 0.444613 | 0.969557 | 0.013427 |
| ENSMUSG00000025871.19 | 351.6405 | 0.37664 | 0.492248 | 0.765143 | 0.444187 | 0.969557 | 0.013427 |
| ENSMUSG00000026775.10 | 370.5605 | 0.36288 | 0.474313 | 0.765064 | 0.444233 | 0.969557 | 0.013427 |
| ENSMUSG00000028042.16 | 16.42625 | -1.68808 | 2.208496 | -0.76436 | 0.444655 | 0.969557 | 0.013427 |
| ENSMUSG00000037531.9 | 90.72968 | -0.62236 | 0.814284 | -0.7643 | 0.444687 | 0.969557 | 0.013427 |
| ENSMUSG00000040818.11 | 56.08684 | -0.74258 | 0.970728 | -0.76497 | 0.444291 | 0.969557 | 0.013427 |
| ENSMUSG00000041688.17 | 59.07288 | -0.83859 | 1.096499 | -0.76479 | 0.444399 | 0.969557 | 0.013427 |
| ENSMUSG00000045519.16 | 69.05151 | -0.68389 | 0.894425 | -0.76462 | 0.444498 | 0.969557 | 0.013427 |
| ENSMUSG00000049577.16 | 33.33272 | -1.41252 | 1.845437 | -0.76541 | 0.444026 | 0.969557 | 0.013427 |
| ENSMUSG00000031422.17 | 322.9387 | 0.374841 | 0.490816 | 0.76371 | 0.44504 | 0.969709 | 0.013359 |
| ENSMUSG00000038954.15 | 14.81909 | 2.265764 | 2.966232 | 0.763852 | 0.444955 | 0.969709 | 0.013359 |
| ENSMUSG00000115987.2 | 371.9161 | -0.35887 | 0.469922 | -0.76367 | 0.445064 | 0.969709 | 0.013359 |
| ENSMUSG00000005233.17 | 35.93364 | -1.19345 | 1.563217 | -0.76346 | 0.445192 | 0.969765 | 0.013334 |
| ENSMUSG00000024146.10 | 400.5848 | -0.35822 | 0.469375 | -0.76318 | 0.445353 | 0.969894 | 0.013276 |
| ENSMUSG00000040022.15 | 45.49466 | 0.850073 | 1.114188 | 0.762953 | 0.445492 | 0.969973 | 0.013241 |
| ENSMUSG00000020397.16 | 217.705 | 0.484358 | 0.635559 | 0.762097 | 0.446002 | 0.970595 | 0.012962 |

| | | | | | | | |
|---|---|---|---|---|---|---|---|
| ENSMUSG00000026111.16 | 278.2929 | 0.436965 | 0.573476 | 0.761959 | 0.446085 | 0.970595 | 0.012962 |
| ENSMUSG00000086593.3 | 59.68958 | -0.77953 | 1.022934 | -0.76205 | 0.446029 | 0.970595 | 0.012962 |
| ENSMUSG00000037275.15 | 14.06676 | -1.71533 | 2.25249 | -0.76153 | 0.446342 | 0.97084 | 0.012852 |
| ENSMUSG00000039206.14 | 145.444 | 0.673908 | 0.885059 | 0.761427 | 0.446402 | 0.97084 | 0.012852 |
| ENSMUSG00000002949.16 | 48.78857 | -0.85243 | 1.120329 | -0.76088 | 0.446729 | 0.971016 | 0.012774 |
| ENSMUSG00000022765.11 | 177.8689 | -0.47059 | 0.618558 | -0.76079 | 0.446785 | 0.971016 | 0.012774 |
| ENSMUSG00000039656.18 | 14.84114 | 1.705821 | 2.242209 | 0.760777 | 0.44679 | 0.971016 | 0.012774 |
| ENSMUSG00000001632.17 | 27.70551 | -1.33259 | 1.753309 | -0.76004 | 0.44723 | 0.971304 | 0.012645 |
| ENSMUSG00000020190.14 | 126.6762 | -0.67564 | 0.888772 | -0.7602 | 0.447137 | 0.971304 | 0.012645 |
| ENSMUSG00000030729.18 | 22.00482 | 1.441848 | 1.896354 | 0.760326 | 0.447059 | 0.971304 | 0.012645 |
| ENSMUSG00000034377.11 | 58.80365 | 0.867082 | 1.141102 | 0.759864 | 0.447336 | 0.971311 | 0.012642 |
| ENSMUSG00000051256.11 | 251.589 | 0.382194 | 0.503173 | 0.759567 | 0.447514 | 0.971474 | 0.012569 |
| ENSMUSG00000016510.12 | 52.67587 | 1.031896 | 1.358873 | 0.759377 | 0.447627 | 0.971499 | 0.012558 |
| ENSMUSG00000032216.16 | 856.5499 | 0.304475 | 0.401099 | 0.759101 | 0.447792 | 0.971542 | 0.012538 |
| ENSMUSG00000036053.18 | 45.65543 | -0.87492 | 1.152779 | -0.75896 | 0.447875 | 0.971542 | 0.012538 |
| ENSMUSG00000037214.13 | 23.02668 | -1.2207 | 1.608677 | -0.75882 | 0.447959 | 0.971542 | 0.012538 |
| ENSMUSG00000042453.15 | 222.1346 | -0.4337 | 0.571667 | -0.75866 | 0.448057 | 0.971542 | 0.012538 |
| ENSMUSG00000026730.13 | 595.5283 | 0.301332 | 0.397553 | 0.757967 | 0.448471 | 0.971772 | 0.012436 |
| ENSMUSG00000046364.15 | 361.8198 | 0.313741 | 0.413863 | 0.758079 | 0.448403 | 0.971772 | 0.012436 |
| ENSMUSG00000090266.11 | 192.7159 | 0.431488 | 0.569176 | 0.758093 | 0.448395 | 0.971772 | 0.012436 |
| ENSMUSG00000000782.17 | 12.71695 | 1.61966 | 2.138139 | 0.757509 | 0.448745 | 0.971922 | 0.012369 |
| ENSMUSG00000050148.10 | 152.4301 | -0.49038 | 0.64723 | -0.75766 | 0.448652 | 0.971922 | 0.012369 |
| ENSMUSG00000000078.8 | 32.48408 | 1.357752 | 1.808114 | 0.750921 | 0.4527 | 0.971927 | 0.012366 |
| ENSMUSG00000000339.15 | 98.50423 | 0.531048 | 0.724386 | 0.733101 | 0.463497 | 0.971927 | 0.012366 |
| ENSMUSG00000000340.11 | 1612.801 | 0.19083 | 0.254049 | 0.751156 | 0.452559 | 0.971927 | 0.012366 |
| ENSMUSG00000000374.9 | 45.25691 | -0.92086 | 1.243164 | -0.74074 | 0.45885 | 0.971927 | 0.012366 |
| ENSMUSG00000000439.10 | 69.68126 | -0.97945 | 1.331927 | -0.73537 | 0.462117 | 0.971927 | 0.012366 |
| ENSMUSG00000000605.10 | 489.3303 | 0.310895 | 0.422272 | 0.736243 | 0.461583 | 0.971927 | 0.012366 |
| ENSMUSG00000001017.16 | 250.1807 | -0.39884 | 0.541585 | -0.73643 | 0.461469 | 0.971927 | 0.012366 |
| ENSMUSG00000001127.13 | 92.05946 | -0.57875 | 0.7879 | -0.73454 | 0.462618 | 0.971927 | 0.012366 |
| ENSMUSG00000004038.10 | 640.732 | 0.317719 | 0.425584 | 0.746549 | 0.455336 | 0.971927 | 0.012366 |
| ENSMUSG00000004748.6 | 223.6082 | 0.463499 | 0.61632 | 0.752043 | 0.452025 | 0.971927 | 0.012366 |
| ENSMUSG00000004937.7 | 15.80044 | -1.86862 | 2.507862 | -0.74511 | 0.456207 | 0.971927 | 0.012366 |
| ENSMUSG00000005034.16 | 311.2063 | -0.35719 | 0.48678 | -0.73378 | 0.463086 | 0.971927 | 0.012366 |
| ENSMUSG00000007670.11 | 112.8125 | -0.62117 | 0.826694 | -0.75139 | 0.452415 | 0.971927 | 0.012366 |
| ENSMUSG00000009905.6 | 140.6743 | 0.512932 | 0.688495 | 0.745005 | 0.456268 | 0.971927 | 0.012366 |
| ENSMUSG00000010376.16 | 211.6007 | 0.531917 | 0.721365 | 0.737375 | 0.460894 | 0.971927 | 0.012366 |
| ENSMUSG00000015083.12 | 2761.098 | -0.22393 | 0.297673 | -0.75226 | 0.451896 | 0.971927 | 0.012366 |
| ENSMUSG00000015804.15 | 188.0163 | 0.488581 | 0.654394 | 0.746616 | 0.455296 | 0.971927 | 0.012366 |
| ENSMUSG00000016253.13 | 40.14295 | 1.275691 | 1.73963 | 0.733311 | 0.463369 | 0.971927 | 0.012366 |
| ENSMUSG00000018398.19 | 12.44082 | -2.02467 | 2.679929 | -0.75549 | 0.449953 | 0.971927 | 0.012366 |
| ENSMUSG00000018501.18 | 896.1468 | -0.23678 | 0.316579 | -0.74792 | 0.454509 | 0.971927 | 0.012366 |
| ENSMUSG00000018750.15 | 23.85879 | -1.30712 | 1.75078 | -0.74659 | 0.45531 | 0.971927 | 0.012366 |
| ENSMUSG00000019659.9 | 51.63272 | 0.814018 | 1.110103 | 0.733281 | 0.463387 | 0.971927 | 0.012366 |

| | | | | | | | |
|---|---|---|---|---|---|---|---|
| ENSMUSG00000019773.8 | 12.9766 | 2.332704 | 3.096523 | 0.75333 | 0.451252 | 0.971927 | 0.012366 |
| ENSMUSG00000020092.10 | 14.00413 | -1.37768 | 1.867342 | -0.73777 | 0.460652 | 0.971927 | 0.012366 |
| ENSMUSG00000020280.13 | 162.7522 | 0.700087 | 0.951097 | 0.736084 | 0.46168 | 0.971927 | 0.012366 |
| ENSMUSG00000020358.18 | 1025.695 | 0.25528 | 0.343521 | 0.743126 | 0.457405 | 0.971927 | 0.012366 |
| ENSMUSG00000020362.14 | 277.9761 | 0.373786 | 0.498218 | 0.750245 | 0.453107 | 0.971927 | 0.012366 |
| ENSMUSG00000020390.13 | 817.7113 | -0.25962 | 0.346157 | -0.75002 | 0.453245 | 0.971927 | 0.012366 |
| ENSMUSG00000020547.6 | 51.25278 | 0.936299 | 1.24072 | 0.754642 | 0.450464 | 0.971927 | 0.012366 |
| ENSMUSG00000021116.11 | 84.58927 | -0.68721 | 0.91857 | -0.74813 | 0.45438 | 0.971927 | 0.012366 |
| ENSMUSG00000021243.15 | 224.1007 | -0.42105 | 0.570138 | -0.73851 | 0.460206 | 0.971927 | 0.012366 |
| ENSMUSG00000021252.12 | 83.51453 | 0.891406 | 1.183515 | 0.753185 | 0.451339 | 0.971927 | 0.012366 |
| ENSMUSG00000021281.16 | 32.3618 | 1.096097 | 1.467583 | 0.746872 | 0.455141 | 0.971927 | 0.012366 |
| ENSMUSG00000021371.6 | 461.712 | 0.32668 | 0.434845 | 0.751257 | 0.452498 | 0.971927 | 0.012366 |
| ENSMUSG00000021483.9 | 11.73361 | -1.81981 | 2.411328 | -0.75469 | 0.450433 | 0.971927 | 0.012366 |
| ENSMUSG00000021495.11 | 10.74747 | 1.572667 | 2.13707 | 0.735899 | 0.461792 | 0.971927 | 0.012366 |
| ENSMUSG00000021549.15 | 185.7101 | 0.525376 | 0.705296 | 0.744902 | 0.456331 | 0.971927 | 0.012366 |
| ENSMUSG00000021902.8 | 35.51879 | 1.222852 | 1.615792 | 0.756812 | 0.449162 | 0.971927 | 0.012366 |
| ENSMUSG00000021952.17 | 42.86808 | 0.962959 | 1.307843 | 0.736295 | 0.461551 | 0.971927 | 0.012366 |
| ENSMUSG00000022234.16 | 787.5978 | 0.237058 | 0.321818 | 0.736622 | 0.461352 | 0.971927 | 0.012366 |
| ENSMUSG00000022452.17 | 772.3715 | -0.27831 | 0.372695 | -0.74676 | 0.45521 | 0.971927 | 0.012366 |
| ENSMUSG00000022864.15 | 138.0481 | 0.4814 | 0.648682 | 0.742121 | 0.458014 | 0.971927 | 0.012366 |
| ENSMUSG00000022905.13 | 164.2952 | 0.417675 | 0.569386 | 0.733554 | 0.463221 | 0.971927 | 0.012366 |
| ENSMUSG00000023020.4 | 172.7768 | 0.497993 | 0.673545 | 0.739361 | 0.459688 | 0.971927 | 0.012366 |
| ENSMUSG00000024294.15 | 311.3565 | -0.46113 | 0.614338 | -0.75061 | 0.452886 | 0.971927 | 0.012366 |
| ENSMUSG00000024513.17 | 270.8701 | 0.327748 | 0.444722 | 0.736972 | 0.461139 | 0.971927 | 0.012366 |
| ENSMUSG00000024976.15 | 91.37341 | 0.854624 | 1.148992 | 0.743803 | 0.456996 | 0.971927 | 0.012366 |
| ENSMUSG00000025035.10 | 111.8822 | -0.58782 | 0.795843 | -0.73861 | 0.460146 | 0.971927 | 0.012366 |
| ENSMUSG00000025266.12 | 107.1786 | 0.538349 | 0.730025 | 0.737439 | 0.460855 | 0.971927 | 0.012366 |
| ENSMUSG00000025793.18 | 49.85492 | -0.74678 | 1.002242 | -0.74511 | 0.456203 | 0.971927 | 0.012366 |
| ENSMUSG00000026509.16 | 149.9382 | -0.7105 | 0.968772 | -0.73341 | 0.463311 | 0.971927 | 0.012366 |
| ENSMUSG00000026510.11 | 92.34223 | -0.58819 | 0.778486 | -0.75556 | 0.449914 | 0.971927 | 0.012366 |
| ENSMUSG00000026836.16 | 66.2148 | 0.770294 | 1.042642 | 0.73879 | 0.460034 | 0.971927 | 0.012366 |
| ENSMUSG00000027714.12 | 158.9124 | 0.552048 | 0.751817 | 0.734286 | 0.462775 | 0.971927 | 0.012366 |
| ENSMUSG00000027739.10 | 19.84503 | 1.428052 | 1.895731 | 0.753299 | 0.451271 | 0.971927 | 0.012366 |
| ENSMUSG00000027890.18 | 52.68543 | -0.98982 | 1.342545 | -0.73727 | 0.460956 | 0.971927 | 0.012366 |
| ENSMUSG00000027947.12 | 112.9944 | -0.56953 | 0.766667 | -0.74287 | 0.457563 | 0.971927 | 0.012366 |
| ENSMUSG00000028053.14 | 181.5808 | 0.43264 | 0.589645 | 0.733729 | 0.463114 | 0.971927 | 0.012366 |
| ENSMUSG00000028399.19 | 927.5412 | 0.25055 | 0.334687 | 0.748611 | 0.454092 | 0.971927 | 0.012366 |
| ENSMUSG00000028455.16 | 182.466 | 0.447231 | 0.593772 | 0.753203 | 0.451328 | 0.971927 | 0.012366 |
| ENSMUSG00000028467.16 | 12.0797 | -1.86298 | 2.529647 | -0.73646 | 0.461451 | 0.971927 | 0.012366 |
| ENSMUSG00000028617.11 | 96.88274 | 0.561365 | 0.760895 | 0.737769 | 0.460655 | 0.971927 | 0.012366 |
| ENSMUSG00000028894.19 | 37.22414 | 1.112666 | 1.496676 | 0.743425 | 0.457224 | 0.971927 | 0.012366 |
| ENSMUSG00000029036.19 | 10.33175 | -1.88227 | 2.533869 | -0.74284 | 0.457576 | 0.971927 | 0.012366 |
| ENSMUSG00000029455.15 | 2398.496 | 0.212608 | 0.286765 | 0.7414 | 0.458451 | 0.971927 | 0.012366 |
| ENSMUSG00000029534.18 | 31.44039 | -0.94638 | 1.268298 | -0.74618 | 0.455557 | 0.971927 | 0.012366 |

| | | | | | | | |
|---|---|---|---|---|---|---|---|
| ENSMUSG00000029920.10 | 41.33309 | 0.949445 | 1.29106 | 0.735399 | 0.462096 | 0.971927 | 0.012366 |
| ENSMUSG00000030284.12 | 41.94525 | -1.01502 | 1.373865 | -0.7388 | 0.460026 | 0.971927 | 0.012366 |
| ENSMUSG00000030347.7 | 17.75231 | 1.383078 | 1.885414 | 0.733567 | 0.463213 | 0.971927 | 0.012366 |
| ENSMUSG00000031403.15 | 141.2914 | -0.50679 | 0.674538 | -0.75132 | 0.452461 | 0.971927 | 0.012366 |
| ENSMUSG00000031438.12 | 896.1951 | -0.23025 | 0.31315 | -0.73528 | 0.462166 | 0.971927 | 0.012366 |
| ENSMUSG00000031592.11 | 41.42107 | -0.81357 | 1.100316 | -0.7394 | 0.459666 | 0.971927 | 0.012366 |
| ENSMUSG00000031618.14 | 81.35351 | -0.73023 | 0.976718 | -0.74763 | 0.454681 | 0.971927 | 0.012366 |
| ENSMUSG00000032097.11 | 239.3011 | -0.44313 | 0.602237 | -0.73581 | 0.461848 | 0.971927 | 0.012366 |
| ENSMUSG00000032422.19 | 82.26731 | -0.64494 | 0.853826 | -0.75535 | 0.450037 | 0.971927 | 0.012366 |
| ENSMUSG00000032591.16 | 274.5709 | 0.37682 | 0.506458 | 0.74403 | 0.456858 | 0.971927 | 0.012366 |
| ENSMUSG00000032870.9 | 58.81815 | -0.67845 | 0.910043 | -0.74552 | 0.455959 | 0.971927 | 0.012366 |
| ENSMUSG00000034024.9 | 707.9611 | 0.240248 | 0.324088 | 0.741304 | 0.458509 | 0.971927 | 0.012366 |
| ENSMUSG00000034066.14 | 38.51286 | -1.10748 | 1.469961 | -0.7534 | 0.451207 | 0.971927 | 0.012366 |
| ENSMUSG00000034203.9 | 64.88813 | -0.63116 | 0.859675 | -0.73418 | 0.462838 | 0.971927 | 0.012366 |
| ENSMUSG00000034320.15 | 28.44892 | -1.27169 | 1.709607 | -0.74385 | 0.456967 | 0.971927 | 0.012366 |
| ENSMUSG00000034430.17 | 13.53578 | -1.72825 | 2.315525 | -0.74638 | 0.45544 | 0.971927 | 0.012366 |
| ENSMUSG00000035041.9 | 368.6863 | -0.35397 | 0.47913 | -0.73878 | 0.460041 | 0.971927 | 0.012366 |
| ENSMUSG00000035504.17 | 2375.79 | 0.222671 | 0.297914 | 0.747436 | 0.454801 | 0.971927 | 0.012366 |
| ENSMUSG00000036442.9 | 64.35999 | 0.708101 | 0.936521 | 0.756097 | 0.449591 | 0.971927 | 0.012366 |
| ENSMUSG00000037075.8 | 357.1161 | 0.359879 | 0.476559 | 0.755161 | 0.450152 | 0.971927 | 0.012366 |
| ENSMUSG00000037493.7 | 14.00046 | 1.72592 | 2.301489 | 0.749915 | 0.453306 | 0.971927 | 0.012366 |
| ENSMUSG00000038080.18 | 95.25082 | 0.663456 | 0.89826 | 0.738601 | 0.460149 | 0.971927 | 0.012366 |
| ENSMUSG00000038127.16 | 157.3113 | -0.55476 | 0.739509 | -0.75017 | 0.453152 | 0.971927 | 0.012366 |
| ENSMUSG00000038546.10 | 219.053 | 0.454887 | 0.612431 | 0.742756 | 0.457629 | 0.971927 | 0.012366 |
| ENSMUSG00000039473.9 | 98.14025 | -0.76309 | 1.028354 | -0.74205 | 0.458056 | 0.971927 | 0.012366 |
| ENSMUSG00000039697.17 | 12.04393 | -1.75883 | 2.369564 | -0.74226 | 0.45793 | 0.971927 | 0.012366 |
| ENSMUSG00000041132.12 | 301.009 | 0.336433 | 0.452505 | 0.743491 | 0.457185 | 0.971927 | 0.012366 |
| ENSMUSG00000041679.10 | 12.47819 | -1.95905 | 2.672365 | -0.73308 | 0.463511 | 0.971927 | 0.012366 |
| ENSMUSG00000042524.15 | 75.78128 | 0.675015 | 0.894011 | 0.755041 | 0.450225 | 0.971927 | 0.012366 |
| ENSMUSG00000042541.11 | 2145.674 | 0.226959 | 0.30005 | 0.756405 | 0.449406 | 0.971927 | 0.012366 |
| ENSMUSG00000042745.10 | 18.38481 | 1.141207 | 1.533669 | 0.744103 | 0.456814 | 0.971927 | 0.012366 |
| ENSMUSG00000043467.20 | 25.32063 | 1.361491 | 1.842582 | 0.738904 | 0.459965 | 0.971927 | 0.012366 |
| ENSMUSG00000044155.12 | 45.82905 | 0.806232 | 1.073652 | 0.750925 | 0.452698 | 0.971927 | 0.012366 |
| ENSMUSG00000044252.19 | 639.3187 | 0.27934 | 0.369899 | 0.755179 | 0.450141 | 0.971927 | 0.012366 |
| ENSMUSG00000045983.17 | 246.7515 | -0.47734 | 0.643725 | -0.74153 | 0.458371 | 0.971927 | 0.012366 |
| ENSMUSG00000046179.18 | 21.78156 | 1.373497 | 1.820878 | 0.754305 | 0.450666 | 0.971927 | 0.012366 |
| ENSMUSG00000046434.17 | 40.66002 | -0.83123 | 1.128924 | -0.73631 | 0.461544 | 0.971927 | 0.012366 |
| ENSMUSG00000046567.11 | 56.78008 | 0.738026 | 0.990951 | 0.744765 | 0.456414 | 0.971927 | 0.012366 |
| ENSMUSG00000046675.8 | 63.94497 | 0.787425 | 1.065412 | 0.73908 | 0.459858 | 0.971927 | 0.012366 |
| ENSMUSG00000047492.6 | 154.7275 | -0.69103 | 0.927336 | -0.74518 | 0.456163 | 0.971927 | 0.012366 |
| ENSMUSG00000048007.5 | 43.46056 | 0.884754 | 1.203356 | 0.735239 | 0.462194 | 0.971927 | 0.012366 |
| ENSMUSG00000050846.10 | 10.24667 | 1.889254 | 2.52194 | 0.749127 | 0.453781 | 0.971927 | 0.012366 |
| ENSMUSG00000051910.14 | 122.5833 | 0.652232 | 0.886718 | 0.735558 | 0.462 | 0.971927 | 0.012366 |
| ENSMUSG00000052102.16 | 12.97033 | -1.558 | 2.080327 | -0.74892 | 0.453904 | 0.971927 | 0.012366 |

| | | | | | | | |
|---|---|---|---|---|---|---|---|
| ENSMUSG00000052155.6 | 149.706 | 0.493462 | 0.666019 | 0.740912 | 0.458747 | 0.971927 | 0.012366 |
| ENSMUSG00000053012.18 | 718.8402 | -0.24629 | 0.326888 | -0.75345 | 0.451178 | 0.971927 | 0.012366 |
| ENSMUSG00000053838.15 | 61.30047 | -1.21646 | 1.619609 | -0.75109 | 0.452601 | 0.971927 | 0.012366 |
| ENSMUSG00000054091.10 | 322.9665 | 0.350385 | 0.476214 | 0.735771 | 0.46187 | 0.971927 | 0.012366 |
| ENSMUSG00000055137.8 | 322.8271 | -0.42783 | 0.575886 | -0.74291 | 0.457534 | 0.971927 | 0.012366 |
| ENSMUSG00000057409.8 | 38.55639 | 1.223148 | 1.649446 | 0.741551 | 0.45836 | 0.971927 | 0.012366 |
| ENSMUSG00000058022.15 | 880.3385 | 0.245924 | 0.334885 | 0.734355 | 0.462732 | 0.971927 | 0.012366 |
| ENSMUSG00000058135.13 | 9299.004 | 0.169428 | 0.228307 | 0.742106 | 0.458023 | 0.971927 | 0.012366 |
| ENSMUSG00000059182.9 | 121.2354 | -0.52559 | 0.7062 | -0.74425 | 0.456723 | 0.971927 | 0.012366 |
| ENSMUSG00000060477.15 | 43.3896 | 0.829988 | 1.130467 | 0.734199 | 0.462827 | 0.971927 | 0.012366 |
| ENSMUSG00000061613.14 | 269.7339 | 0.385333 | 0.522509 | 0.737467 | 0.460838 | 0.971927 | 0.012366 |
| ENSMUSG00000062797.15 | 220.385 | 0.431873 | 0.577885 | 0.747332 | 0.454863 | 0.971927 | 0.012366 |
| ENSMUSG00000063480.9 | 110.1408 | 0.534649 | 0.710237 | 0.752775 | 0.451585 | 0.971927 | 0.012366 |
| ENSMUSG00000064373.14 | 36344.84 | 0.166402 | 0.222144 | 0.749072 | 0.453814 | 0.971927 | 0.012366 |
| ENSMUSG00000066071.7 | 772.9026 | -0.25182 | 0.341119 | -0.7382 | 0.46039 | 0.971927 | 0.012366 |
| ENSMUSG00000068798.11 | 411.9842 | 0.304798 | 0.408458 | 0.746217 | 0.455537 | 0.971927 | 0.012366 |
| ENSMUSG00000069539.12 | 118.0958 | 0.561093 | 0.744374 | 0.753778 | 0.450982 | 0.971927 | 0.012366 |
| ENSMUSG00000069743.6 | 12.5261 | 2.039786 | 2.74606 | 0.742805 | 0.4576 | 0.971927 | 0.012366 |
| ENSMUSG00000069844.13 | 13.93756 | -1.46471 | 1.977836 | -0.74056 | 0.458958 | 0.971927 | 0.012366 |
| ENSMUSG00000070639.6 | 11.53988 | -2.1029 | 2.843717 | -0.73949 | 0.459609 | 0.971927 | 0.012366 |
| ENSMUSG00000074129.15 | 161.9518 | 0.496677 | 0.670719 | 0.740514 | 0.458988 | 0.971927 | 0.012366 |
| ENSMUSG00000074207.11 | 12585.56 | 0.155846 | 0.210253 | 0.741229 | 0.458555 | 0.971927 | 0.012366 |
| ENSMUSG00000074582.11 | 99.33218 | -0.56948 | 0.775404 | -0.73443 | 0.462686 | 0.971927 | 0.012366 |
| ENSMUSG00000074649.7 | 17.31184 | -1.61816 | 2.19955 | -0.73568 | 0.461927 | 0.971927 | 0.012366 |
| ENSMUSG00000074826.11 | 21.10209 | 1.26651 | 1.727531 | 0.733133 | 0.463478 | 0.971927 | 0.012366 |
| ENSMUSG00000078154.4 | 53.45732 | 0.716538 | 0.956491 | 0.749132 | 0.453778 | 0.971927 | 0.012366 |
| ENSMUSG00000081406.5 | 231.953 | 0.358176 | 0.485002 | 0.738504 | 0.460208 | 0.971927 | 0.012366 |
| ENSMUSG00000089911.5 | 216.0918 | 0.445973 | 0.603517 | 0.738958 | 0.459933 | 0.971927 | 0.012366 |
| ENSMUSG00000092558.4 | 36.54249 | -1.14167 | 1.540885 | -0.74092 | 0.458743 | 0.971927 | 0.012366 |
| ENSMUSG00000096199.3 | 77.71878 | -0.61357 | 0.831513 | -0.73789 | 0.46058 | 0.971927 | 0.012366 |
| ENSMUSG00000109644.2 | 539.4976 | -0.32673 | 0.435468 | -0.75029 | 0.453082 | 0.971927 | 0.012366 |
| ENSMUSG00000110755.2 | 64.00718 | 0.999792 | 1.324305 | 0.754956 | 0.450276 | 0.971927 | 0.012366 |
| ENSMUSG00000113831.2 | 10.07745 | -1.69334 | 2.252776 | -0.75167 | 0.452251 | 0.971927 | 0.012366 |
| ENSMUSG00000117962.2 | 50.04594 | 0.704777 | 0.959879 | 0.734235 | 0.462805 | 0.971927 | 0.012366 |
| ENSMUSG00000028158.15 | 1932.257 | 0.200943 | 0.2742 | 0.732834 | 0.46366 | 0.972025 | 0.012323 |
| ENSMUSG00000004633.18 | 112.6407 | 0.536315 | 0.737348 | 0.727357 | 0.467008 | 0.972638 | 0.012049 |
| ENSMUSG00000006356.11 | 203.5753 | 0.561563 | 0.769956 | 0.729345 | 0.465791 | 0.972638 | 0.012049 |
| ENSMUSG00000011305.12 | 125.8339 | 0.798129 | 1.095653 | 0.728451 | 0.466338 | 0.972638 | 0.012049 |
| ENSMUSG00000014077.14 | 2327.254 | 0.188486 | 0.258641 | 0.728756 | 0.466151 | 0.972638 | 0.012049 |
| ENSMUSG00000015202.14 | 15.36244 | 1.710319 | 2.341127 | 0.730554 | 0.465052 | 0.972638 | 0.012049 |
| ENSMUSG00000018425.19 | 103.0843 | -0.53924 | 0.741647 | -0.72708 | 0.467176 | 0.972638 | 0.012049 |
| ENSMUSG00000018774.14 | 20.69538 | 1.074551 | 1.477201 | 0.727424 | 0.466967 | 0.972638 | 0.012049 |
| ENSMUSG00000020171.9 | 74.3454 | -0.87996 | 1.204651 | -0.73047 | 0.465105 | 0.972638 | 0.012049 |
| ENSMUSG00000020755.10 | 27.64812 | 1.377241 | 1.889314 | 0.728963 | 0.466024 | 0.972638 | 0.012049 |

| | | | | | | | |
|---|---|---|---|---|---|---|---|
| ENSMUSG00000020829.10 | 146.7356 | -0.53 | 0.727224 | -0.7288 | 0.466122 | 0.972638 | 0.012049 |
| ENSMUSG00000024026.14 | 1044.776 | -0.24663 | 0.337987 | -0.72971 | 0.465567 | 0.972638 | 0.012049 |
| ENSMUSG00000025261.19 | 151.5544 | 0.456353 | 0.626184 | 0.728784 | 0.466134 | 0.972638 | 0.012049 |
| ENSMUSG00000025337.11 | 195.8849 | -0.41347 | 0.565482 | -0.73119 | 0.464663 | 0.972638 | 0.012049 |
| ENSMUSG00000026333.15 | 86.00202 | 0.623436 | 0.857566 | 0.726984 | 0.467236 | 0.972638 | 0.012049 |
| ENSMUSG00000026766.17 | 487.97 | 0.293552 | 0.401329 | 0.731451 | 0.464504 | 0.972638 | 0.012049 |
| ENSMUSG00000027809.15 | 1209.954 | -0.21769 | 0.298777 | -0.72861 | 0.466239 | 0.972638 | 0.012049 |
| ENSMUSG00000029059.10 | 119.9242 | -0.67862 | 0.931158 | -0.7288 | 0.466126 | 0.972638 | 0.012049 |
| ENSMUSG00000029484.13 | 36.40907 | 0.828119 | 1.136537 | 0.728634 | 0.466226 | 0.972638 | 0.012049 |
| ENSMUSG00000031838.11 | 185.6877 | -0.57138 | 0.785575 | -0.72734 | 0.467015 | 0.972638 | 0.012049 |
| ENSMUSG00000032469.13 | 63.68166 | -0.69836 | 0.957399 | -0.72944 | 0.465734 | 0.972638 | 0.012049 |
| ENSMUSG00000035798.15 | 21.26166 | -1.34033 | 1.833816 | -0.7309 | 0.464841 | 0.972638 | 0.012049 |
| ENSMUSG00000037355.15 | 36.08614 | -1.13376 | 1.549702 | -0.7316 | 0.464412 | 0.972638 | 0.012049 |
| ENSMUSG00000042729.12 | 99.08116 | 0.743087 | 1.021226 | 0.727642 | 0.466833 | 0.972638 | 0.012049 |
| ENSMUSG00000057522.16 | 504.216 | -0.27099 | 0.371598 | -0.72925 | 0.465851 | 0.972638 | 0.012049 |
| ENSMUSG00000058806.16 | 37.23313 | -1.09764 | 1.509702 | -0.72706 | 0.467189 | 0.972638 | 0.012049 |
| ENSMUSG00000070733.14 | 22.54891 | -1.24921 | 1.715491 | -0.72819 | 0.466495 | 0.972638 | 0.012049 |
| ENSMUSG00000073664.12 | 189.4199 | 0.498356 | 0.68143 | 0.731338 | 0.464572 | 0.972638 | 0.012049 |
| ENSMUSG00000079418.6 | 14.25266 | 1.689467 | 2.322005 | 0.72759 | 0.466865 | 0.972638 | 0.012049 |
| ENSMUSG00000089824.11 | 21.47841 | -1.25381 | 1.72137 | -0.72838 | 0.466382 | 0.972638 | 0.012049 |
| ENSMUSG00000091764.4 | 13.29557 | -2.21129 | 3.039086 | -0.72762 | 0.466849 | 0.972638 | 0.012049 |
| ENSMUSG00000100153.3 | 44.74495 | -0.76505 | 1.047532 | -0.73033 | 0.465186 | 0.972638 | 0.012049 |
| ENSMUSG00000108366.2 | 253.9176 | -0.35915 | 0.492338 | -0.72948 | 0.465709 | 0.972638 | 0.012049 |
| ENSMUSG00000003199.17 | 73.79915 | -0.79831 | 1.099261 | -0.72623 | 0.467699 | 0.972707 | 0.012018 |
| ENSMUSG00000005687.15 | 268.6659 | 0.368865 | 0.509673 | 0.723729 | 0.469232 | 0.972707 | 0.012018 |
| ENSMUSG00000009073.17 | 24.95336 | 1.18521 | 1.638489 | 0.723355 | 0.469462 | 0.972707 | 0.012018 |
| ENSMUSG00000020059.10 | 19.5291 | 1.556035 | 2.141582 | 0.726582 | 0.467482 | 0.972707 | 0.012018 |
| ENSMUSG00000020413.12 | 30.7759 | 0.900132 | 1.242889 | 0.724225 | 0.468927 | 0.972707 | 0.012018 |
| ENSMUSG00000024096.18 | 163.0625 | 0.545316 | 0.751115 | 0.726008 | 0.467834 | 0.972707 | 0.012018 |
| ENSMUSG00000024870.7 | 89.34439 | -0.79706 | 1.100783 | -0.72409 | 0.469013 | 0.972707 | 0.012018 |
| ENSMUSG00000026675.13 | 289.8947 | -0.40477 | 0.558698 | -0.72448 | 0.468771 | 0.972707 | 0.012018 |
| ENSMUSG00000027508.16 | 28.58819 | 0.983239 | 1.355024 | 0.725625 | 0.468069 | 0.972707 | 0.012018 |
| ENSMUSG00000028945.10 | 286.6085 | 0.387319 | 0.533924 | 0.725421 | 0.468194 | 0.972707 | 0.012018 |
| ENSMUSG00000031378.15 | 16.48693 | -1.63441 | 2.252038 | -0.72575 | 0.467994 | 0.972707 | 0.012018 |
| ENSMUSG00000032064.18 | 30.19625 | -1.20735 | 1.668143 | -0.72377 | 0.469208 | 0.972707 | 0.012018 |
| ENSMUSG00000032403.9 | 150.6997 | 0.503635 | 0.695089 | 0.724561 | 0.468721 | 0.972707 | 0.012018 |
| ENSMUSG00000035614.12 | 86.19246 | 0.76967 | 1.062927 | 0.724104 | 0.469002 | 0.972707 | 0.012018 |
| ENSMUSG00000036620.15 | 15.38877 | -1.6071 | 2.218734 | -0.72433 | 0.468862 | 0.972707 | 0.012018 |
| ENSMUSG00000041912.13 | 17.33233 | -1.32882 | 1.836575 | -0.72353 | 0.469353 | 0.972707 | 0.012018 |
| ENSMUSG00000047109.14 | 27.66709 | 1.398741 | 1.930454 | 0.724566 | 0.468718 | 0.972707 | 0.012018 |
| ENSMUSG00000053613.7 | 10.38576 | 2.049049 | 2.833767 | 0.723083 | 0.469629 | 0.972707 | 0.012018 |
| ENSMUSG00000056153.16 | 123.5801 | 0.532844 | 0.736366 | 0.723613 | 0.469303 | 0.972707 | 0.012018 |
| ENSMUSG00000056692.14 | 166.7591 | -0.46831 | 0.647049 | -0.72377 | 0.469208 | 0.972707 | 0.012018 |
| ENSMUSG00000058793.15 | 147.4893 | -0.52067 | 0.716431 | -0.72676 | 0.467372 | 0.972707 | 0.012018 |

| | | | | | | | |
|---|---|---|---|---|---|---|---|
| ENSMUSG00000087370.4 | 117.9294 | -0.7154 | 0.989364 | -0.72309 | 0.469624 | 0.972707 | 0.012018 |
| ENSMUSG00000097328.8 | 16.52017 | -1.53119 | 2.117173 | -0.72322 | 0.469542 | 0.972707 | 0.012018 |
| ENSMUSG00000021733.12 | 58.79143 | 0.682636 | 0.944711 | 0.722588 | 0.469933 | 0.973125 | 0.011831 |
| ENSMUSG00000027650.13 | 56.24456 | 0.717886 | 0.994501 | 0.721856 | 0.470383 | 0.973631 | 0.011605 |
| ENSMUSG00000062070.13 | 134.9179 | 0.574996 | 0.796438 | 0.721959 | 0.470319 | 0.973631 | 0.011605 |
| ENSMUSG00000054277.9 | 46.34522 | 1.312176 | 1.818378 | 0.721619 | 0.470529 | 0.973721 | 0.011566 |
| ENSMUSG00000018347.17 | 27.18749 | -1.29317 | 1.793043 | -0.72121 | 0.470778 | 0.973876 | 0.011496 |
| ENSMUSG00000020903.14 | 97.34618 | -0.54503 | 0.756109 | -0.72084 | 0.47101 | 0.973876 | 0.011496 |
| ENSMUSG00000024395.10 | 51.05609 | -0.97637 | 1.354449 | -0.72086 | 0.470994 | 0.973876 | 0.011496 |
| ENSMUSG00000062270.15 | 103.0885 | -0.80802 | 1.12096 | -0.72083 | 0.471015 | 0.973876 | 0.011496 |
| ENSMUSG00000020604.14 | 48.03846 | -0.777 | 1.079268 | -0.71993 | 0.471565 | 0.973919 | 0.011477 |
| ENSMUSG00000028556.16 | 39.85022 | 0.963559 | 1.33866 | 0.719793 | 0.471652 | 0.973919 | 0.011477 |
| ENSMUSG00000037470.15 | 183.7932 | 0.440704 | 0.61156 | 0.720623 | 0.471142 | 0.973919 | 0.011477 |
| ENSMUSG00000039763.11 | 51.89214 | -0.81802 | 1.136449 | -0.7198 | 0.471647 | 0.973919 | 0.011477 |
| ENSMUSG00000057130.13 | 221.4243 | 0.41858 | 0.581443 | 0.719898 | 0.471588 | 0.973919 | 0.011477 |
| ENSMUSG00000091586.3 | 11.37085 | 1.539726 | 2.137188 | 0.720445 | 0.471251 | 0.973919 | 0.011477 |
| ENSMUSG00000051124.7 | 149.1675 | 0.549785 | 0.764196 | 0.71943 | 0.471876 | 0.97417 | 0.011365 |
| ENSMUSG00000000420.16 | 129.632 | 0.451451 | 0.631137 | 0.715298 | 0.474425 | 0.974715 | 0.011122 |
| ENSMUSG00000001249.15 | 365.75 | -0.28355 | 0.395143 | -0.71758 | 0.473018 | 0.974715 | 0.011122 |
| ENSMUSG00000003662.11 | 143.8231 | 0.53754 | 0.74934 | 0.717352 | 0.473157 | 0.974715 | 0.011122 |
| ENSMUSG00000004996.10 | 41.22679 | 1.265378 | 1.769195 | 0.715228 | 0.474468 | 0.974715 | 0.011122 |
| ENSMUSG00000005779.13 | 1117.258 | -0.22926 | 0.319013 | -0.71866 | 0.472353 | 0.974715 | 0.011122 |
| ENSMUSG00000006392.17 | 168.8085 | 0.437031 | 0.609266 | 0.717308 | 0.473184 | 0.974715 | 0.011122 |
| ENSMUSG00000020097.15 | 253.9968 | -0.33733 | 0.471296 | -0.71576 | 0.474141 | 0.974715 | 0.011122 |
| ENSMUSG00000021302.11 | 20.73585 | -1.25741 | 1.751021 | -0.7181 | 0.472696 | 0.974715 | 0.011122 |
| ENSMUSG00000021661.17 | 43.51643 | -1.04654 | 1.456171 | -0.71869 | 0.47233 | 0.974715 | 0.011122 |
| ENSMUSG00000024797.15 | 10.19604 | -1.83834 | 2.569489 | -0.71545 | 0.474332 | 0.974715 | 0.011122 |
| ENSMUSG00000026317.8 | 35.59065 | -0.94882 | 1.3233 | -0.71701 | 0.473367 | 0.974715 | 0.011122 |
| ENSMUSG00000028121.9 | 360.9989 | 0.369799 | 0.514881 | 0.718222 | 0.47262 | 0.974715 | 0.011122 |
| ENSMUSG00000029587.17 | 36.016 | -1.1024 | 1.54069 | -0.71552 | 0.474287 | 0.974715 | 0.011122 |
| ENSMUSG00000029999.15 | 96.98 | -0.62644 | 0.874486 | -0.71635 | 0.473776 | 0.974715 | 0.011122 |
| ENSMUSG00000031667.17 | 33.94026 | 0.9417 | 1.31637 | 0.715376 | 0.474376 | 0.974715 | 0.011122 |
| ENSMUSG00000033715.16 | 2825.501 | 0.25919 | 0.36145 | 0.717085 | 0.473321 | 0.974715 | 0.011122 |
| ENSMUSG00000035824.8 | 59.32679 | -0.69844 | 0.9757 | -0.71583 | 0.474094 | 0.974715 | 0.011122 |
| ENSMUSG00000038718.16 | 35.07695 | -1.35726 | 1.891542 | -0.71754 | 0.473041 | 0.974715 | 0.011122 |
| ENSMUSG00000042202.16 | 85.7407 | -0.55629 | 0.775351 | -0.71746 | 0.473087 | 0.974715 | 0.011122 |
| ENSMUSG00000042675.16 | 75.97018 | -0.61798 | 0.862569 | -0.71644 | 0.47372 | 0.974715 | 0.011122 |
| ENSMUSG00000048799.9 | 41.60653 | -0.79066 | 1.105819 | -0.715 | 0.474608 | 0.974715 | 0.011122 |
| ENSMUSG00000050530.15 | 18.60029 | 1.720587 | 2.405865 | 0.715164 | 0.474508 | 0.974715 | 0.011122 |
| ENSMUSG00000078622.9 | 491.2867 | -0.25904 | 0.362271 | -0.71505 | 0.474576 | 0.974715 | 0.011122 |
| ENSMUSG00000079017.4 | 64.30642 | -0.75869 | 1.060186 | -0.71562 | 0.474227 | 0.974715 | 0.011122 |
| ENSMUSG00000000290.14 | 13.57452 | -1.44123 | 2.017399 | -0.7144 | 0.47498 | 0.974755 | 0.011105 |
| ENSMUSG00000000326.14 | 4538.931 | -0.187 | 0.264123 | -0.70799 | 0.478951 | 0.974755 | 0.011105 |
| ENSMUSG00000002625.11 | 20.73475 | -1.6978 | 2.405101 | -0.70592 | 0.480241 | 0.974755 | 0.011105 |

| | | | | | | | |
|---|---|---|---|---|---|---|---|
| ENSMUSG00000003039.10 | 449.1043 | 0.33434 | 0.472304 | 0.707892 | 0.479013 | 0.974755 | 0.011105 |
| ENSMUSG00000007458.15 | 452.3571 | 0.304651 | 0.428821 | 0.710438 | 0.477433 | 0.974755 | 0.011105 |
| ENSMUSG00000012422.15 | 705.0125 | 0.269314 | 0.38132 | 0.706266 | 0.480022 | 0.974755 | 0.011105 |
| ENSMUSG00000013593.13 | 879.841 | 0.2248 | 0.316637 | 0.709961 | 0.477728 | 0.974755 | 0.011105 |
| ENSMUSG00000013997.11 | 296.4456 | -0.33643 | 0.473671 | -0.71026 | 0.477542 | 0.974755 | 0.011105 |
| ENSMUSG00000017132.18 | 94.85028 | -0.58245 | 0.823624 | -0.70718 | 0.479454 | 0.974755 | 0.011105 |
| ENSMUSG00000017929.14 | 205.9289 | 0.483736 | 0.680437 | 0.710919 | 0.477134 | 0.974755 | 0.011105 |
| ENSMUSG00000019917.11 | 66.36728 | 0.724341 | 1.021852 | 0.708851 | 0.478417 | 0.974755 | 0.011105 |
| ENSMUSG00000020385.17 | 166.3094 | -0.47033 | 0.659547 | -0.71311 | 0.475777 | 0.974755 | 0.011105 |
| ENSMUSG00000020918.15 | 21.79191 | 1.714961 | 2.412659 | 0.710818 | 0.477197 | 0.974755 | 0.011105 |
| ENSMUSG00000020948.10 | 28.69445 | -0.90768 | 1.277579 | -0.71047 | 0.477413 | 0.974755 | 0.011105 |
| ENSMUSG00000021368.16 | 55.70091 | 0.681688 | 0.959616 | 0.710375 | 0.477471 | 0.974755 | 0.011105 |
| ENSMUSG00000021807.7 | 771.1169 | -0.25628 | 0.361352 | -0.70923 | 0.478183 | 0.974755 | 0.011105 |
| ENSMUSG00000022383.14 | 67.52026 | -0.82279 | 1.159469 | -0.70963 | 0.477934 | 0.974755 | 0.011105 |
| ENSMUSG00000022564.8 | 105.1651 | 0.622201 | 0.882521 | 0.705027 | 0.480794 | 0.974755 | 0.011105 |
| ENSMUSG00000022749.9 | 27.34749 | 1.080745 | 1.515094 | 0.713319 | 0.475648 | 0.974755 | 0.011105 |
| ENSMUSG00000022799.5 | 76.02662 | 0.645263 | 0.912494 | 0.707142 | 0.479478 | 0.974755 | 0.011105 |
| ENSMUSG00000023070.7 | 4540.771 | -0.15404 | 0.217173 | -0.70931 | 0.478129 | 0.974755 | 0.011105 |
| ENSMUSG00000023939.8 | 349.3035 | -0.31779 | 0.449799 | -0.70651 | 0.479868 | 0.974755 | 0.011105 |
| ENSMUSG00000024180.15 | 23.00264 | -1.16482 | 1.64425 | -0.70842 | 0.478683 | 0.974755 | 0.011105 |
| ENSMUSG00000024231.16 | 268.9412 | -0.33294 | 0.470916 | -0.707 | 0.479564 | 0.974755 | 0.011105 |
| ENSMUSG00000024863.7 | 1834.63 | 0.209798 | 0.297295 | 0.705689 | 0.480382 | 0.974755 | 0.011105 |
| ENSMUSG00000026154.5 | 225.526 | 0.419059 | 0.59478 | 0.704561 | 0.481083 | 0.974755 | 0.011105 |
| ENSMUSG00000027944.14 | 85.55504 | -0.70988 | 0.993582 | -0.71446 | 0.474942 | 0.974755 | 0.011105 |
| ENSMUSG00000028322.12 | 55.48885 | 0.820624 | 1.150193 | 0.713466 | 0.475558 | 0.974755 | 0.011105 |
| ENSMUSG00000029106.15 | 36.13919 | 0.897445 | 1.267788 | 0.707883 | 0.479018 | 0.974755 | 0.011105 |
| ENSMUSG00000029152.14 | 394.9156 | 0.378709 | 0.533205 | 0.710251 | 0.477548 | 0.974755 | 0.011105 |
| ENSMUSG00000029177.10 | 19.67597 | -1.20165 | 1.698765 | -0.70737 | 0.479339 | 0.974755 | 0.011105 |
| ENSMUSG00000029192.18 | 35.8994 | -1.18307 | 1.673623 | -0.70689 | 0.479636 | 0.974755 | 0.011105 |
| ENSMUSG00000029547.11 | 11.42115 | -1.75613 | 2.460934 | -0.7136 | 0.475472 | 0.974755 | 0.011105 |
| ENSMUSG00000031138.5 | 1154.1 | -0.25396 | 0.356974 | -0.71143 | 0.476818 | 0.974755 | 0.011105 |
| ENSMUSG00000031534.16 | 107.4026 | 0.506622 | 0.715509 | 0.708058 | 0.478909 | 0.974755 | 0.011105 |
| ENSMUSG00000031616.8 | 20.76711 | 1.504171 | 2.125754 | 0.707594 | 0.479197 | 0.974755 | 0.011105 |
| ENSMUSG00000032067.7 | 206.1718 | 0.453907 | 0.644005 | 0.704819 | 0.480923 | 0.974755 | 0.011105 |
| ENSMUSG00000032580.13 | 195.173 | 0.473565 | 0.671717 | 0.705006 | 0.480806 | 0.974755 | 0.011105 |
| ENSMUSG00000034245.11 | 84.09195 | -0.56604 | 0.801071 | -0.7066 | 0.479813 | 0.974755 | 0.011105 |
| ENSMUSG00000034854.9 | 13.67323 | 1.689127 | 2.39645 | 0.704845 | 0.480907 | 0.974755 | 0.011105 |
| ENSMUSG00000035572.17 | 75.50461 | 0.605892 | 0.859687 | 0.704782 | 0.480946 | 0.974755 | 0.011105 |
| ENSMUSG00000035936.7 | 202.4075 | 0.366599 | 0.516824 | 0.709329 | 0.47812 | 0.974755 | 0.011105 |
| ENSMUSG00000036992.11 | 91.58232 | -0.5945 | 0.838286 | -0.70918 | 0.478212 | 0.974755 | 0.011105 |
| ENSMUSG00000038619.13 | 64.66306 | -0.64877 | 0.910772 | -0.71233 | 0.47626 | 0.974755 | 0.011105 |
| ENSMUSG00000039294.15 | 26.27917 | 1.387765 | 1.953573 | 0.710373 | 0.477473 | 0.974755 | 0.011105 |
| ENSMUSG00000041235.13 | 28.84799 | -1.1947 | 1.694415 | -0.70508 | 0.480761 | 0.974755 | 0.011105 |
| ENSMUSG00000041777.13 | 143.0458 | -0.48661 | 0.686491 | -0.70884 | 0.478423 | 0.974755 | 0.011105 |

| | | | | | | | |
|---|---|---|---|---|---|---|---|
| ENSMUSG00000042978.12 | 43.46878 | -1.15146 | 1.617288 | -0.71197 | 0.476483 | 0.974755 | 0.011105 |
| ENSMUSG00000044469.9 | 130.0266 | 0.506861 | 0.713493 | 0.710394 | 0.47746 | 0.974755 | 0.011105 |
| ENSMUSG00000047045.18 | 18.3243 | -1.56734 | 2.224663 | -0.70453 | 0.481103 | 0.974755 | 0.011105 |
| ENSMUSG00000047090.14 | 14.27886 | -1.89068 | 2.657883 | -0.71135 | 0.476868 | 0.974755 | 0.011105 |
| ENSMUSG00000053754.15 | 42.76577 | -0.71815 | 1.013785 | -0.70838 | 0.478707 | 0.974755 | 0.011105 |
| ENSMUSG00000056228.11 | 23.28345 | 1.235298 | 1.751205 | 0.705399 | 0.480562 | 0.974755 | 0.011105 |
| ENSMUSG00000058070.15 | 16.41919 | -1.78305 | 2.516628 | -0.70851 | 0.478629 | 0.974755 | 0.011105 |
| ENSMUSG00000058569.9 | 455.637 | 0.32575 | 0.457597 | 0.71187 | 0.476545 | 0.974755 | 0.011105 |
| ENSMUSG00000059316.3 | 14.32724 | 1.84234 | 2.614372 | 0.704697 | 0.480999 | 0.974755 | 0.011105 |
| ENSMUSG00000061536.14 | 68.33922 | -0.59636 | 0.835372 | -0.71389 | 0.475296 | 0.974755 | 0.011105 |
| ENSMUSG00000061731.10 | 224.3224 | 0.530848 | 0.753483 | 0.704526 | 0.481105 | 0.974755 | 0.011105 |
| ENSMUSG00000061904.13 | 2905.671 | 0.181961 | 0.256337 | 0.709849 | 0.477798 | 0.974755 | 0.011105 |
| ENSMUSG00000068184.8 | 218.5779 | 0.380302 | 0.538318 | 0.706464 | 0.4799 | 0.974755 | 0.011105 |
| ENSMUSG00000070394.11 | 609.6059 | 0.274536 | 0.385171 | 0.712764 | 0.475992 | 0.974755 | 0.011105 |
| ENSMUSG00000100094.2 | 387.6496 | 0.371147 | 0.521531 | 0.711648 | 0.476683 | 0.974755 | 0.011105 |
| ENSMUSG00000110444.3 | 104.11 | -0.61225 | 0.865104 | -0.70772 | 0.479116 | 0.974755 | 0.011105 |
| ENSMUSG00000020454.18 | 73.28581 | 0.676864 | 0.962907 | 0.702939 | 0.482094 | 0.975091 | 0.010955 |
| ENSMUSG00000025736.16 | 71.3941 | -0.88805 | 1.261707 | -0.70385 | 0.481529 | 0.975091 | 0.010955 |
| ENSMUSG00000027580.18 | 111.592 | 0.500899 | 0.712528 | 0.702989 | 0.482062 | 0.975091 | 0.010955 |
| ENSMUSG00000028568.17 | 62.69353 | 0.673743 | 0.957985 | 0.703292 | 0.481874 | 0.975091 | 0.010955 |
| ENSMUSG00000045284.10 | 61.83062 | -0.78377 | 1.11436 | -0.70334 | 0.481844 | 0.975091 | 0.010955 |
| ENSMUSG00000049999.5 | 10.945 | -2.00286 | 2.848085 | -0.70323 | 0.481912 | 0.975091 | 0.010955 |
| ENSMUSG00000058240.15 | 27.4848 | 0.854176 | 1.214833 | 0.703122 | 0.48198 | 0.975091 | 0.010955 |
| ENSMUSG00000092274.4 | 321.6193 | 0.31707 | 0.450423 | 0.703938 | 0.481471 | 0.975091 | 0.010955 |
| ENSMUSG00000001123.16 | 1256.505 | -0.20743 | 0.296432 | -0.69975 | 0.484082 | 0.975608 | 0.010725 |
| ENSMUSG00000006024.14 | 64.60392 | -0.62756 | 0.894421 | -0.70164 | 0.482905 | 0.975608 | 0.010725 |
| ENSMUSG00000018001.19 | 33.05222 | -1.18662 | 1.693435 | -0.70072 | 0.483477 | 0.975608 | 0.010725 |
| ENSMUSG00000019080.9 | 24.00691 | -1.60999 | 2.298846 | -0.70035 | 0.483711 | 0.975608 | 0.010725 |
| ENSMUSG00000020063.17 | 63.96442 | 0.697892 | 0.994923 | 0.701453 | 0.48302 | 0.975608 | 0.010725 |
| ENSMUSG00000021484.9 | 691.6018 | 0.312184 | 0.445091 | 0.701394 | 0.483057 | 0.975608 | 0.010725 |
| ENSMUSG00000022982.11 | 8778.236 | 0.185529 | 0.26492 | 0.700322 | 0.483726 | 0.975608 | 0.010725 |
| ENSMUSG00000028030.13 | 65.79184 | -0.66555 | 0.950756 | -0.70003 | 0.483911 | 0.975608 | 0.010725 |
| ENSMUSG00000039007.11 | 549.1092 | -0.26126 | 0.373223 | -0.70001 | 0.483923 | 0.975608 | 0.010725 |
| ENSMUSG00000039046.16 | 30.43949 | -1.2904 | 1.843018 | -0.70016 | 0.483828 | 0.975608 | 0.010725 |
| ENSMUSG00000041445.10 | 52.54652 | -0.75542 | 1.080353 | -0.69923 | 0.484408 | 0.975608 | 0.010725 |
| ENSMUSG00000046201.10 | 95.66565 | 0.537118 | 0.766305 | 0.700918 | 0.483354 | 0.975608 | 0.010725 |
| ENSMUSG00000058173.13 | 56.6856 | 1.013418 | 1.448622 | 0.699574 | 0.484193 | 0.975608 | 0.010725 |
| ENSMUSG00000058216.14 | 49.89217 | -0.77405 | 1.10517 | -0.70039 | 0.483682 | 0.975608 | 0.010725 |
| ENSMUSG00000059586.11 | 62.29062 | -0.86611 | 1.233641 | -0.70208 | 0.482632 | 0.975608 | 0.010725 |
| ENSMUSG00000073436.12 | 32.87047 | 1.561947 | 2.23356 | 0.699308 | 0.484359 | 0.975608 | 0.010725 |
| ENSMUSG00000073940.4 | 47.73738 | -0.86814 | 1.240208 | -0.69999 | 0.483931 | 0.975608 | 0.010725 |
| ENSMUSG00000078920.4 | 179.5453 | -0.42592 | 0.607066 | -0.7016 | 0.482931 | 0.975608 | 0.010725 |
| ENSMUSG00000093565.2 | 25.07591 | 1.218251 | 1.741841 | 0.699404 | 0.4843 | 0.975608 | 0.010725 |
| ENSMUSG00000096935.6 | 23.48714 | -1.40933 | 2.012488 | -0.70029 | 0.483743 | 0.975608 | 0.010725 |

| | | | | | | | |
|---|---|---|---|---|---|---|---|
| ENSMUSG00000000049.12 | 16008.19 | 0.144246 | 0.206504 | 0.698516 | 0.484854 | 0.97616 | 0.010479 |
| ENSMUSG00000002249.22 | 11.23721 | 1.690255 | 2.422877 | 0.697623 | 0.485413 | 0.97616 | 0.010479 |
| ENSMUSG00000030541.17 | 283.0574 | 0.348517 | 0.499373 | 0.697908 | 0.485234 | 0.97616 | 0.010479 |
| ENSMUSG00000030972.7 | 383.2756 | 0.270222 | 0.387257 | 0.697784 | 0.485312 | 0.97616 | 0.010479 |
| ENSMUSG00000032527.14 | 619.6159 | -0.26126 | 0.37458 | -0.69747 | 0.485506 | 0.97616 | 0.010479 |
| ENSMUSG00000033114.12 | 265.7296 | 0.394575 | 0.565541 | 0.697694 | 0.485369 | 0.97616 | 0.010479 |
| ENSMUSG00000033610.17 | 1307.43 | -0.18571 | 0.266178 | -0.69768 | 0.48538 | 0.97616 | 0.010479 |
| ENSMUSG00000049792.7 | 59.30532 | -0.71344 | 1.022788 | -0.69755 | 0.48546 | 0.97616 | 0.010479 |
| ENSMUSG00000025134.3 | 14.38452 | 1.665982 | 2.39064 | 0.696877 | 0.48588 | 0.976462 | 0.010345 |
| ENSMUSG00000026484.14 | 53.0104 | -0.62737 | 0.900382 | -0.69678 | 0.485941 | 0.976462 | 0.010345 |
| ENSMUSG00000046314.17 | 127.7221 | -0.51103 | 0.733462 | -0.69674 | 0.485965 | 0.976462 | 0.010345 |
| ENSMUSG00000066568.13 | 115.4084 | 0.503737 | 0.723509 | 0.696242 | 0.486277 | 0.976883 | 0.010158 |
| ENSMUSG00000078348.5 | 196.6245 | 0.371167 | 0.533321 | 0.695954 | 0.486458 | 0.977038 | 0.010089 |
| ENSMUSG00000028271.10 | 304.2685 | -0.3212 | 0.461809 | -0.69552 | 0.486729 | 0.97708 | 0.01007 |
| ENSMUSG00000030556.14 | 121.5234 | -0.56348 | 0.810575 | -0.69516 | 0.486954 | 0.97708 | 0.01007 |
| ENSMUSG00000030987.6 | 21.84834 | 1.291978 | 1.858701 | 0.695098 | 0.486994 | 0.97708 | 0.01007 |
| ENSMUSG00000032849.15 | 17.30243 | 1.352824 | 1.945841 | 0.695239 | 0.486906 | 0.97708 | 0.01007 |
| ENSMUSG00000038612.17 | 307.4568 | -0.40121 | 0.576797 | -0.69558 | 0.486692 | 0.97708 | 0.01007 |
| ENSMUSG00000003581.15 | 88.98653 | -0.67404 | 0.97008 | -0.69482 | 0.487165 | 0.977124 | 0.01005 |
| ENSMUSG00000021027.18 | 36.09251 | -0.93497 | 1.346119 | -0.69457 | 0.487325 | 0.977124 | 0.01005 |
| ENSMUSG00000039202.13 | 86.33198 | 0.827873 | 1.191912 | 0.694576 | 0.487321 | 0.977124 | 0.01005 |
| ENSMUSG00000006941.6 | 243.3604 | 0.374868 | 0.539851 | 0.694392 | 0.487436 | 0.97714 | 0.010043 |
| ENSMUSG00000026728.10 | 204.7339 | 0.461806 | 0.66654 | 0.69284 | 0.48841 | 0.977644 | 0.009819 |
| ENSMUSG00000029136.10 | 81.78594 | -0.6368 | 0.919028 | -0.69291 | 0.488367 | 0.977644 | 0.009819 |
| ENSMUSG00000030339.8 | 141.2566 | -0.68596 | 0.989602 | -0.69317 | 0.488203 | 0.977644 | 0.009819 |
| ENSMUSG00000033880.12 | 80.90799 | 0.589998 | 0.851433 | 0.692946 | 0.488343 | 0.977644 | 0.009819 |
| ENSMUSG00000034422.15 | 31.79119 | -0.92655 | 1.337175 | -0.69292 | 0.48836 | 0.977644 | 0.009819 |
| ENSMUSG00000038095.16 | 249.7478 | -0.34832 | 0.502262 | -0.6935 | 0.487995 | 0.977644 | 0.009819 |
| ENSMUSG00000038240.14 | 123.3864 | 0.618965 | 0.893165 | 0.693001 | 0.488309 | 0.977644 | 0.009819 |
| ENSMUSG00000032028.16 | 58.82709 | -0.71431 | 1.032234 | -0.69201 | 0.488934 | 0.978487 | 0.009445 |
| ENSMUSG00000023944.15 | 4122.595 | 0.273624 | 0.395559 | 0.69174 | 0.489101 | 0.978542 | 0.00942 |
| ENSMUSG00000026921.21 | 35.87489 | 1.122967 | 1.623662 | 0.691626 | 0.489172 | 0.978542 | 0.00942 |
| ENSMUSG00000038884.15 | 92.67561 | 0.557426 | 0.806148 | 0.691469 | 0.489271 | 0.978542 | 0.00942 |
| ENSMUSG00000003299.11 | 19.01029 | 1.358317 | 1.966776 | 0.690631 | 0.489797 | 0.979388 | 0.009045 |
| ENSMUSG00000033658.17 | 28.52228 | 0.868588 | 1.258218 | 0.690332 | 0.489986 | 0.979558 | 0.00897 |
| ENSMUSG00000114828.2 | 174.3145 | -0.48652 | 0.70533 | -0.68978 | 0.490334 | 0.980047 | 0.008753 |
| ENSMUSG00000001525.11 | 122.4627 | -0.72182 | 1.055577 | -0.68382 | 0.494091 | 0.980357 | 0.008616 |
| ENSMUSG00000004394.13 | 404.9099 | 0.265807 | 0.386119 | 0.688407 | 0.491197 | 0.980357 | 0.008616 |
| ENSMUSG00000010054.5 | 131.3124 | -0.51594 | 0.757299 | -0.68129 | 0.495685 | 0.980357 | 0.008616 |
| ENSMUSG00000015749.13 | 322.6531 | 0.398484 | 0.582783 | 0.68376 | 0.494127 | 0.980357 | 0.008616 |
| ENSMUSG00000020289.16 | 28.77695 | -0.8064 | 1.176067 | -0.68568 | 0.492916 | 0.980357 | 0.008616 |
| ENSMUSG00000020644.10 | 1813.537 | 0.24342 | 0.355919 | 0.683921 | 0.494025 | 0.980357 | 0.008616 |
| ENSMUSG00000020935.9 | 11.70138 | 1.283923 | 1.885291 | 0.681021 | 0.495858 | 0.980357 | 0.008616 |
| ENSMUSG00000021054.7 | 700.9337 | -0.26063 | 0.383574 | -0.67948 | 0.496831 | 0.980357 | 0.008616 |

| | | | | | | | |
|---|---|---|---|---|---|---|---|
| ENSMUSG00000021589.14 | 23.99602 | -1.14942 | 1.674244 | -0.68653 | 0.492377 | 0.980357 | 0.008616 |
| ENSMUSG00000021840.18 | 174.691 | -0.47999 | 0.704662 | -0.68117 | 0.495764 | 0.980357 | 0.008616 |
| ENSMUSG00000022191.17 | 14.52628 | 1.550559 | 2.282845 | 0.679222 | 0.496997 | 0.980357 | 0.008616 |
| ENSMUSG00000022538.21 | 54.18064 | -0.63018 | 0.923697 | -0.68224 | 0.495087 | 0.980357 | 0.008616 |
| ENSMUSG00000022635.10 | 735.0502 | 0.294987 | 0.431889 | 0.683016 | 0.494597 | 0.980357 | 0.008616 |
| ENSMUSG00000022656.16 | 259.6618 | 0.313505 | 0.456578 | 0.686641 | 0.492309 | 0.980357 | 0.008616 |
| ENSMUSG00000023186.15 | 108.8238 | 0.461073 | 0.678982 | 0.679066 | 0.497096 | 0.980357 | 0.008616 |
| ENSMUSG00000023973.14 | 17.70763 | -1.27198 | 1.851707 | -0.68692 | 0.492131 | 0.980357 | 0.008616 |
| ENSMUSG00000024006.18 | 221.8643 | -0.39045 | 0.566587 | -0.68913 | 0.490741 | 0.980357 | 0.008616 |
| ENSMUSG00000024127.16 | 31.27193 | -1.08287 | 1.586656 | -0.68248 | 0.494933 | 0.980357 | 0.008616 |
| ENSMUSG00000024181.10 | 117.8336 | 0.479893 | 0.706092 | 0.679646 | 0.496728 | 0.980357 | 0.008616 |
| ENSMUSG00000024228.13 | 386.9475 | 0.364352 | 0.529196 | 0.688501 | 0.491137 | 0.980357 | 0.008616 |
| ENSMUSG00000024369.17 | 88.03884 | -0.64668 | 0.946344 | -0.68334 | 0.494391 | 0.980357 | 0.008616 |
| ENSMUSG00000024527.11 | 125.1168 | -0.43045 | 0.633585 | -0.67939 | 0.496888 | 0.980357 | 0.008616 |
| ENSMUSG00000024535.17 | 92.9549 | 0.488412 | 0.719433 | 0.678885 | 0.497211 | 0.980357 | 0.008616 |
| ENSMUSG00000026179.15 | 131.4765 | 0.624807 | 0.910488 | 0.686233 | 0.492566 | 0.980357 | 0.008616 |
| ENSMUSG00000026718.18 | 92.56787 | 0.597099 | 0.875365 | 0.682115 | 0.495166 | 0.980357 | 0.008616 |
| ENSMUSG00000027130.16 | 10.70852 | 1.477408 | 2.173132 | 0.679852 | 0.496598 | 0.980357 | 0.008616 |
| ENSMUSG00000027193.13 | 169.6736 | 0.391585 | 0.574546 | 0.681556 | 0.495519 | 0.980357 | 0.008616 |
| ENSMUSG00000028136.16 | 100.3946 | -0.62037 | 0.912124 | -0.68014 | 0.496415 | 0.980357 | 0.008616 |
| ENSMUSG00000028344.13 | 45.93088 | 1.102088 | 1.602587 | 0.687693 | 0.491646 | 0.980357 | 0.008616 |
| ENSMUSG00000028367.6 | 2249.984 | 0.188782 | 0.276441 | 0.682901 | 0.494669 | 0.980357 | 0.008616 |
| ENSMUSG00000028419.6 | 393.018 | 0.33605 | 0.494408 | 0.6797 | 0.496694 | 0.980357 | 0.008616 |
| ENSMUSG00000029427.12 | 48.42403 | 0.985244 | 1.436862 | 0.685692 | 0.492908 | 0.980357 | 0.008616 |
| ENSMUSG00000029840.9 | 1078.642 | 0.256869 | 0.375583 | 0.683922 | 0.494025 | 0.980357 | 0.008616 |
| ENSMUSG00000030103.12 | 591.1696 | 0.292049 | 0.426312 | 0.68506 | 0.493306 | 0.980357 | 0.008616 |
| ENSMUSG00000031483.9 | 131.1955 | 0.63173 | 0.919582 | 0.686975 | 0.492098 | 0.980357 | 0.008616 |
| ENSMUSG00000031824.16 | 55.09424 | 0.953379 | 1.391106 | 0.685339 | 0.49313 | 0.980357 | 0.008616 |
| ENSMUSG00000031967.16 | 152.9707 | -0.70571 | 1.02947 | -0.68551 | 0.493025 | 0.980357 | 0.008616 |
| ENSMUSG00000032059.14 | 71.31898 | 0.796561 | 1.161799 | 0.685627 | 0.492948 | 0.980357 | 0.008616 |
| ENSMUSG00000032410.14 | 50.91007 | -0.81962 | 1.205048 | -0.68016 | 0.496405 | 0.980357 | 0.008616 |
| ENSMUSG00000032554.16 | 14720.31 | -0.20434 | 0.296813 | -0.68846 | 0.49116 | 0.980357 | 0.008616 |
| ENSMUSG00000033364.14 | 11.20865 | -1.56232 | 2.285669 | -0.68353 | 0.494273 | 0.980357 | 0.008616 |
| ENSMUSG00000033813.16 | 17.55429 | 1.237691 | 1.822565 | 0.679093 | 0.497079 | 0.980357 | 0.008616 |
| ENSMUSG00000034620.18 | 148.7622 | -0.42459 | 0.622275 | -0.68232 | 0.495039 | 0.980357 | 0.008616 |
| ENSMUSG00000035168.17 | 48.21264 | -1.18329 | 1.732872 | -0.68285 | 0.494701 | 0.980357 | 0.008616 |
| ENSMUSG00000036678.9 | 20.39001 | 1.66736 | 2.44328 | 0.682427 | 0.494969 | 0.980357 | 0.008616 |
| ENSMUSG00000037348.16 | 44.52459 | 0.942566 | 1.385754 | 0.680183 | 0.496389 | 0.980357 | 0.008616 |
| ENSMUSG00000040296.16 | 119.7443 | -0.74434 | 1.096393 | -0.6789 | 0.497202 | 0.980357 | 0.008616 |
| ENSMUSG00000041765.7 | 60.62531 | -0.63652 | 0.931861 | -0.68307 | 0.494566 | 0.980357 | 0.008616 |
| ENSMUSG00000041769.14 | 119.8484 | 0.528893 | 0.77661 | 0.681027 | 0.495854 | 0.980357 | 0.008616 |
| ENSMUSG00000041846.16 | 193.0363 | 0.358626 | 0.521406 | 0.687805 | 0.491576 | 0.980357 | 0.008616 |
| ENSMUSG00000044876.16 | 22.00569 | -1.25065 | 1.826579 | -0.6847 | 0.493536 | 0.980357 | 0.008616 |
| ENSMUSG00000045103.20 | 95.23327 | 0.60161 | 0.875301 | 0.687319 | 0.491882 | 0.980357 | 0.008616 |

| | | | | | | | |
|---|---|---|---|---|---|---|---|
| ENSMUSG00000048371.9 | 226.6967 | -0.46092 | 0.676121 | -0.68171 | 0.495424 | 0.980357 | 0.008616 |
| ENSMUSG00000050270.13 | 200.0411 | 0.413842 | 0.607708 | 0.680989 | 0.495879 | 0.980357 | 0.008616 |
| ENSMUSG00000050912.16 | 268.828 | 0.334757 | 0.489691 | 0.683608 | 0.494223 | 0.980357 | 0.008616 |
| ENSMUSG00000056167.9 | 56.10705 | -0.85039 | 1.243245 | -0.68401 | 0.493969 | 0.980357 | 0.008616 |
| ENSMUSG00000059669.9 | 42.19349 | -0.87514 | 1.286619 | -0.68019 | 0.496384 | 0.980357 | 0.008616 |
| ENSMUSG00000062110.15 | 41.45301 | 0.910883 | 1.336666 | 0.681459 | 0.495581 | 0.980357 | 0.008616 |
| ENSMUSG00000072941.6 | 49.7935 | 0.705809 | 1.032244 | 0.683762 | 0.494126 | 0.980357 | 0.008616 |
| ENSMUSG00000073633.10 | 40.03456 | 0.861707 | 1.258733 | 0.684583 | 0.493607 | 0.980357 | 0.008616 |
| ENSMUSG00000077450.14 | 113.9737 | -0.52679 | 0.771468 | -0.68285 | 0.494703 | 0.980357 | 0.008616 |
| ENSMUSG00000082884.2 | 31.13759 | 0.841741 | 1.226944 | 0.686047 | 0.492684 | 0.980357 | 0.008616 |
| ENSMUSG00000084786.10 | 572.3723 | 0.23601 | 0.34278 | 0.688518 | 0.491127 | 0.980357 | 0.008616 |
| ENSMUSG00000087213.3 | 37.83431 | 0.72949 | 1.062678 | 0.686464 | 0.492421 | 0.980357 | 0.008616 |
| ENSMUSG00000090110.9 | 43.17917 | 0.959414 | 1.405489 | 0.68262 | 0.494847 | 0.980357 | 0.008616 |
| ENSMUSG00000019428.17 | 70.0941 | 0.712408 | 1.050344 | 0.678262 | 0.497606 | 0.980738 | 0.008447 |
| ENSMUSG00000024383.10 | 82.44803 | -0.59814 | 0.882452 | -0.67781 | 0.497891 | 0.980738 | 0.008447 |
| ENSMUSG00000025486.18 | 235.1359 | -0.41198 | 0.60759 | -0.67806 | 0.497734 | 0.980738 | 0.008447 |
| ENSMUSG00000062115.16 | 52.49926 | 0.788364 | 1.163182 | 0.677765 | 0.497921 | 0.980738 | 0.008447 |
| ENSMUSG00000063268.13 | 16.02825 | -1.48878 | 2.19645 | -0.67781 | 0.497891 | 0.980738 | 0.008447 |
| ENSMUSG00000034621.15 | 68.63072 | -0.65819 | 0.971731 | -0.67734 | 0.498191 | 0.980788 | 0.008425 |
| ENSMUSG00000036282.16 | 215.5849 | 0.448921 | 0.662732 | 0.677378 | 0.498166 | 0.980788 | 0.008425 |
| ENSMUSG00000050890.16 | 61.98657 | -0.68693 | 1.014322 | -0.67723 | 0.498257 | 0.980788 | 0.008425 |
| ENSMUSG00000000355.14 | 537.1507 | 0.25417 | 0.378449 | 0.671609 | 0.501833 | 0.981123 | 0.008276 |
| ENSMUSG00000015970.20 | 371.2508 | 0.341089 | 0.507804 | 0.671694 | 0.501778 | 0.981123 | 0.008276 |
| ENSMUSG00000020098.8 | 2107.331 | 0.18615 | 0.27546 | 0.675779 | 0.499181 | 0.981123 | 0.008276 |
| ENSMUSG00000020669.16 | 40.25355 | 0.801786 | 1.193381 | 0.671861 | 0.501672 | 0.981123 | 0.008276 |
| ENSMUSG00000020739.12 | 41.46323 | 1.004857 | 1.495694 | 0.671834 | 0.50169 | 0.981123 | 0.008276 |
| ENSMUSG00000021497.11 | 827.2688 | 0.214292 | 0.31845 | 0.672923 | 0.500996 | 0.981123 | 0.008276 |
| ENSMUSG00000021690.11 | 53.32493 | -0.65951 | 0.98058 | -0.67257 | 0.50122 | 0.981123 | 0.008276 |
| ENSMUSG00000022828.11 | 121.4805 | 0.545385 | 0.806284 | 0.676419 | 0.498775 | 0.981123 | 0.008276 |
| ENSMUSG00000024238.16 | 97.12828 | 0.554869 | 0.822053 | 0.67498 | 0.499688 | 0.981123 | 0.008276 |
| ENSMUSG00000025437.16 | 255.7127 | -0.30647 | 0.453493 | -0.6758 | 0.499165 | 0.981123 | 0.008276 |
| ENSMUSG00000027883.16 | 116.5177 | 0.454791 | 0.67418 | 0.674584 | 0.49994 | 0.981123 | 0.008276 |
| ENSMUSG00000031668.15 | 20.00972 | -1.0172 | 1.513361 | -0.67215 | 0.501489 | 0.981123 | 0.008276 |
| ENSMUSG00000031889.10 | 79.06026 | -0.82657 | 1.222936 | -0.67589 | 0.499112 | 0.981123 | 0.008276 |
| ENSMUSG00000032340.9 | 10.14989 | -1.50674 | 2.236171 | -0.67381 | 0.500435 | 0.981123 | 0.008276 |
| ENSMUSG00000032667.18 | 378.5297 | 0.371264 | 0.551601 | 0.673066 | 0.500905 | 0.981123 | 0.008276 |
| ENSMUSG00000034932.5 | 103.8064 | -0.4805 | 0.713027 | -0.67389 | 0.500383 | 0.981123 | 0.008276 |
| ENSMUSG00000035152.15 | 22.78527 | 0.999705 | 1.488556 | 0.671594 | 0.501842 | 0.981123 | 0.008276 |
| ENSMUSG00000036087.19 | 268.6256 | -0.37985 | 0.564925 | -0.67239 | 0.501339 | 0.981123 | 0.008276 |
| ENSMUSG00000036819.15 | 27.2545 | 1.206607 | 1.79045 | 0.673912 | 0.500367 | 0.981123 | 0.008276 |
| ENSMUSG00000037640.16 | 20.11549 | -1.35651 | 2.016766 | -0.67262 | 0.501191 | 0.981123 | 0.008276 |
| ENSMUSG00000038116.17 | 19.07562 | 1.040416 | 1.54902 | 0.671661 | 0.5018 | 0.981123 | 0.008276 |
| ENSMUSG00000040025.18 | 189.5491 | -0.41835 | 0.620556 | -0.67416 | 0.500209 | 0.981123 | 0.008276 |
| ENSMUSG00000041720.15 | 87.022 | 0.723868 | 1.07141 | 0.675622 | 0.499281 | 0.981123 | 0.008276 |

| | | | | | | | |
|---|---|---|---|---|---|---|---|
| ENSMUSG00000042508.17 | 46.04002 | 0.763647 | 1.130443 | 0.675529 | 0.49934 | 0.981123 | 0.008276 |
| ENSMUSG00000042682.11 | 1056.671 | -0.21654 | 0.320799 | -0.67501 | 0.49967 | 0.981123 | 0.008276 |
| ENSMUSG00000043091.10 | 489.7008 | 0.353703 | 0.524973 | 0.673755 | 0.500467 | 0.981123 | 0.008276 |
| ENSMUSG00000044712.16 | 22.52175 | -1.09446 | 1.626195 | -0.67302 | 0.500935 | 0.981123 | 0.008276 |
| ENSMUSG00000047407.18 | 125.3535 | 0.492376 | 0.729165 | 0.675259 | 0.499511 | 0.981123 | 0.008276 |
| ENSMUSG00000047989.12 | 114.3562 | -0.50098 | 0.743866 | -0.67348 | 0.500642 | 0.981123 | 0.008276 |
| ENSMUSG00000050697.10 | 78.0771 | -0.61611 | 0.912674 | -0.67506 | 0.499638 | 0.981123 | 0.008276 |
| ENSMUSG00000051730.16 | 58.69209 | -0.84138 | 1.24788 | -0.67425 | 0.500152 | 0.981123 | 0.008276 |
| ENSMUSG00000052435.8 | 31.02364 | 1.198516 | 1.779088 | 0.673669 | 0.500522 | 0.981123 | 0.008276 |
| ENSMUSG00000055320.19 | 69.31169 | 0.690397 | 1.024221 | 0.67407 | 0.500267 | 0.981123 | 0.008276 |
| ENSMUSG00000045193.14 | 28.70915 | -1.06491 | 1.586932 | -0.67105 | 0.502191 | 0.981602 | 0.008064 |
| ENSMUSG00000060181.7 | 48.20418 | -0.75648 | 1.127847 | -0.67073 | 0.50239 | 0.98179 | 0.007982 |
| ENSMUSG00000006740.15 | 659.4213 | -0.32435 | 0.484722 | -0.66915 | 0.5034 | 0.982173 | 0.007812 |
| ENSMUSG00000020458.17 | 700.058 | -0.20775 | 0.310388 | -0.66932 | 0.503289 | 0.982173 | 0.007812 |
| ENSMUSG00000021944.16 | 46.72223 | 1.056358 | 1.578715 | 0.669125 | 0.503415 | 0.982173 | 0.007812 |
| ENSMUSG00000028847.9 | 25.25417 | -1.40478 | 2.097822 | -0.66964 | 0.50309 | 0.982173 | 0.007812 |
| ENSMUSG00000033676.14 | 13.63154 | -1.45025 | 2.166961 | -0.66925 | 0.503333 | 0.982173 | 0.007812 |
| ENSMUSG00000034807.10 | 211.9939 | -0.36308 | 0.542484 | -0.66929 | 0.503313 | 0.982173 | 0.007812 |
| ENSMUSG00000038544.15 | 12.95197 | 1.639515 | 2.448257 | 0.669666 | 0.503071 | 0.982173 | 0.007812 |
| ENSMUSG00000099083.8 | 12.97575 | 1.562278 | 2.332133 | 0.669892 | 0.502926 | 0.982173 | 0.007812 |
| ENSMUSG00000022913.17 | 311.5946 | 0.365852 | 0.547517 | 0.668202 | 0.504005 | 0.982337 | 0.007739 |
| ENSMUSG00000025487.16 | 92.24047 | -0.5728 | 0.856944 | -0.66842 | 0.503864 | 0.982337 | 0.007739 |
| ENSMUSG00000030180.16 | 219.0369 | 0.397973 | 0.595606 | 0.668181 | 0.504018 | 0.982337 | 0.007739 |
| ENSMUSG00000031327.11 | 18.10701 | -1.19151 | 1.782745 | -0.66836 | 0.503906 | 0.982337 | 0.007739 |
| ENSMUSG00000061950.18 | 91.33102 | 0.592802 | 0.886592 | 0.668631 | 0.503731 | 0.982337 | 0.007739 |
| ENSMUSG00000015341.18 | 80.03359 | 0.56479 | 0.845895 | 0.667683 | 0.504336 | 0.982385 | 0.007718 |
| ENSMUSG00000021264.13 | 333.5282 | -0.32506 | 0.487655 | -0.66657 | 0.505045 | 0.982385 | 0.007718 |
| ENSMUSG00000021830.15 | 17.48465 | -1.51306 | 2.271948 | -0.66597 | 0.505428 | 0.982385 | 0.007718 |
| ENSMUSG00000026234.13 | 1647.032 | 0.29046 | 0.435132 | 0.667523 | 0.504438 | 0.982385 | 0.007718 |
| ENSMUSG00000028106.14 | 78.042 | 0.569535 | 0.854167 | 0.666773 | 0.504917 | 0.982385 | 0.007718 |
| ENSMUSG00000028832.12 | 14.59746 | 1.55797 | 2.33501 | 0.667222 | 0.50463 | 0.982385 | 0.007718 |
| ENSMUSG00000031807.11 | 23.91637 | 1.08952 | 1.632215 | 0.66751 | 0.504446 | 0.982385 | 0.007718 |
| ENSMUSG00000035790.16 | 60.99858 | 0.620265 | 0.930909 | 0.666301 | 0.505219 | 0.982385 | 0.007718 |
| ENSMUSG00000040043.18 | 12.10389 | 1.834011 | 2.755998 | 0.665462 | 0.505755 | 0.982385 | 0.007718 |
| ENSMUSG00000042248.5 | 1048.385 | 0.264828 | 0.397729 | 0.665851 | 0.505507 | 0.982385 | 0.007718 |
| ENSMUSG00000044676.11 | 39.08199 | -1.15743 | 1.739064 | -0.66555 | 0.5057 | 0.982385 | 0.007718 |
| ENSMUSG00000046962.17 | 20.31143 | 1.078126 | 1.617529 | 0.666527 | 0.505075 | 0.982385 | 0.007718 |
| ENSMUSG00000054237.7 | 99.52321 | -0.53122 | 0.797579 | -0.66604 | 0.505385 | 0.982385 | 0.007718 |
| ENSMUSG00000085826.2 | 62.45584 | 0.597281 | 0.897011 | 0.665857 | 0.505503 | 0.982385 | 0.007718 |
| ENSMUSG00000089847.9 | 106.2714 | -0.61617 | 0.92603 | -0.66539 | 0.505804 | 0.982385 | 0.007718 |
| ENSMUSG00000097167.2 | 26.73061 | 1.351129 | 2.027574 | 0.666377 | 0.50517 | 0.982385 | 0.007718 |
| ENSMUSG00000099881.2 | 38.58795 | -0.94164 | 1.412371 | -0.66671 | 0.504957 | 0.982385 | 0.007718 |
| ENSMUSG00000020687.14 | 53.73793 | 0.687631 | 1.03398 | 0.665033 | 0.506029 | 0.982424 | 0.007701 |
| ENSMUSG00000040648.16 | 241.5809 | 0.37064 | 0.557328 | 0.66503 | 0.506031 | 0.982424 | 0.007701 |

| | | | | | | | |
|---|---|---|---|---|---|---|---|
| ENSMUSG00000062300.15 | 67.83099 | -0.62362 | 0.938509 | -0.66448 | 0.506385 | 0.9828 | 0.007535 |
| ENSMUSG00000062627.10 | 137.0301 | 0.473482 | 0.712642 | 0.664404 | 0.506432 | 0.9828 | 0.007535 |
| ENSMUSG00000020737.13 | 35.52541 | -1.11533 | 1.684559 | -0.66209 | 0.507913 | 0.983192 | 0.007362 |
| ENSMUSG00000031613.10 | 503.6684 | 0.266893 | 0.403079 | 0.662137 | 0.507883 | 0.983192 | 0.007362 |
| ENSMUSG00000032046.16 | 27.21087 | -1.2424 | 1.875775 | -0.66234 | 0.507754 | 0.983192 | 0.007362 |
| ENSMUSG00000032199.14 | 382.0149 | -0.35657 | 0.538002 | -0.66277 | 0.507477 | 0.983192 | 0.007362 |
| ENSMUSG00000034610.15 | 90.57128 | -0.63553 | 0.960038 | -0.66198 | 0.507983 | 0.983192 | 0.007362 |
| ENSMUSG00000035878.14 | 288.1447 | -0.37252 | 0.561301 | -0.66367 | 0.506903 | 0.983192 | 0.007362 |
| ENSMUSG00000041258.19 | 52.40942 | -0.68608 | 1.034642 | -0.66311 | 0.507261 | 0.983192 | 0.007362 |
| ENSMUSG00000042625.17 | 142.0489 | 0.466025 | 0.702755 | 0.66314 | 0.507241 | 0.983192 | 0.007362 |
| ENSMUSG00000047150.9 | 10.96269 | 1.728784 | 2.609105 | 0.662597 | 0.507589 | 0.983192 | 0.007362 |
| ENSMUSG00000060036.15 | 2465.654 | 0.171759 | 0.259311 | 0.662368 | 0.507735 | 0.983192 | 0.007362 |
| ENSMUSG00000071633.12 | 366.8577 | 0.338223 | 0.510184 | 0.662943 | 0.507367 | 0.983192 | 0.007362 |
| ENSMUSG00000083380.2 | 27.41779 | 0.863438 | 1.303959 | 0.662167 | 0.507864 | 0.983192 | 0.007362 |
| ENSMUSG00000116835.2 | 20.13987 | -0.95017 | 1.43225 | -0.66341 | 0.507068 | 0.983192 | 0.007362 |
| ENSMUSG00000000159.17 | 202.7711 | -0.43394 | 0.664684 | -0.65284 | 0.513857 | 0.983423 | 0.00726 |
| ENSMUSG00000000567.6 | 51.16534 | -0.8025 | 1.215291 | -0.66033 | 0.509041 | 0.983423 | 0.00726 |
| ENSMUSG00000000751.14 | 55.2207 | -0.91655 | 1.633039 | -0.56126 | 0.574623 | 0.983423 | 0.00726 |
| ENSMUSG00000000811.15 | 12.6928 | -1.55186 | 2.397057 | -0.6474 | 0.517371 | 0.983423 | 0.00726 |
| ENSMUSG00000000823.17 | 23.95256 | -1.009 | 1.746843 | -0.57761 | 0.563526 | 0.983423 | 0.00726 |
| ENSMUSG00000001054.10 | 79.31176 | -0.54068 | 0.837804 | -0.64535 | 0.518701 | 0.983423 | 0.00726 |
| ENSMUSG00000001134.10 | 205.8043 | -0.48881 | 0.86566 | -0.56466 | 0.572304 | 0.983423 | 0.00726 |
| ENSMUSG00000001157.14 | 93.34965 | -0.78989 | 1.200593 | -0.65792 | 0.510591 | 0.983423 | 0.00726 |
| ENSMUSG00000001229.10 | 55.81081 | 0.553751 | 0.972504 | 0.569407 | 0.56908 | 0.983423 | 0.00726 |
| ENSMUSG00000001376.18 | 44.69465 | -0.72843 | 1.174611 | -0.62015 | 0.535161 | 0.983423 | 0.00726 |
| ENSMUSG00000001518.13 | 13.99833 | -1.40502 | 2.262459 | -0.62101 | 0.53459 | 0.983423 | 0.00726 |
| ENSMUSG00000001663.11 | 843.2339 | 0.261609 | 0.433538 | 0.603427 | 0.546225 | 0.983423 | 0.00726 |
| ENSMUSG00000001700.11 | 264.8256 | 0.386896 | 0.59659 | 0.648513 | 0.516653 | 0.983423 | 0.00726 |
| ENSMUSG00000001763.15 | 31.12982 | -0.911 | 1.63905 | -0.55581 | 0.578341 | 0.983423 | 0.00726 |
| ENSMUSG00000001767.6 | 86.21396 | 0.537185 | 0.883055 | 0.608325 | 0.542972 | 0.983423 | 0.00726 |
| ENSMUSG00000001855.16 | 16.50674 | -1.40998 | 2.165114 | -0.65123 | 0.5149 | 0.983423 | 0.00726 |
| ENSMUSG00000001983.8 | 36.5583 | 0.77475 | 1.291187 | 0.600029 | 0.548487 | 0.983423 | 0.00726 |
| ENSMUSG00000002017.11 | 93.12541 | -0.52958 | 0.819438 | -0.64627 | 0.518102 | 0.983423 | 0.00726 |
| ENSMUSG00000002083.14 | 12.29904 | 1.553989 | 2.491852 | 0.623628 | 0.532872 | 0.983423 | 0.00726 |
| ENSMUSG00000002107.19 | 138.6843 | 0.406911 | 0.61592 | 0.660655 | 0.508833 | 0.983423 | 0.00726 |
| ENSMUSG00000002109.15 | 21.4886 | 1.043583 | 1.81202 | 0.575923 | 0.564668 | 0.983423 | 0.00726 |
| ENSMUSG00000002325.16 | 141.5853 | -0.44334 | 0.798178 | -0.55544 | 0.578591 | 0.983423 | 0.00726 |
| ENSMUSG00000002679.15 | 16.88194 | 1.150526 | 1.931482 | 0.59567 | 0.551396 | 0.983423 | 0.00726 |
| ENSMUSG00000002732.15 | 27.7436 | 0.690183 | 1.234574 | 0.559046 | 0.57613 | 0.983423 | 0.00726 |
| ENSMUSG00000002803.15 | 10.86684 | -1.27686 | 2.096881 | -0.60893 | 0.542569 | 0.983423 | 0.00726 |
| ENSMUSG00000002981.11 | 33.43527 | -0.80184 | 1.243275 | -0.64494 | 0.518964 | 0.983423 | 0.00726 |
| ENSMUSG00000002996.18 | 336.6006 | -0.25761 | 0.456668 | -0.56411 | 0.572677 | 0.983423 | 0.00726 |
| ENSMUSG00000003062.14 | 61.31134 | 0.775631 | 1.195446 | 0.648822 | 0.516454 | 0.983423 | 0.00726 |
| ENSMUSG00000003344.15 | 14.41573 | -1.35283 | 2.08952 | -0.64744 | 0.51735 | 0.983423 | 0.00726 |

| | | | | | | | |
|---|---|---|---|---|---|---|---|
| ENSMUSG00000003345.17 | 47.13746 | -0.91474 | 1.400761 | -0.65303 | 0.513734 | 0.983423 | 0.00726 |
| ENSMUSG00000003438.18 | 34.90013 | -0.98406 | 1.608207 | -0.6119 | 0.540604 | 0.983423 | 0.00726 |
| ENSMUSG00000003778.15 | 37.7768 | -0.69212 | 1.148781 | -0.60248 | 0.546853 | 0.983423 | 0.00726 |
| ENSMUSG00000004285.13 | 308.8317 | -0.41348 | 0.644338 | -0.64172 | 0.521056 | 0.983423 | 0.00726 |
| ENSMUSG00000004393.11 | 43.51345 | -0.81441 | 1.422286 | -0.57261 | 0.56691 | 0.983423 | 0.00726 |
| ENSMUSG00000004768.15 | 64.59562 | 0.658249 | 1.021541 | 0.644369 | 0.519336 | 0.983423 | 0.00726 |
| ENSMUSG00000004936.9 | 207.7766 | -0.38863 | 0.652016 | -0.59604 | 0.55115 | 0.983423 | 0.00726 |
| ENSMUSG00000004980.17 | 3019.833 | 0.175329 | 0.293053 | 0.598285 | 0.54965 | 0.983423 | 0.00726 |
| ENSMUSG00000005142.11 | 63.14901 | -0.73614 | 1.165848 | -0.63142 | 0.527764 | 0.983423 | 0.00726 |
| ENSMUSG00000005373.14 | 259.1837 | 0.324431 | 0.544654 | 0.595665 | 0.551399 | 0.983423 | 0.00726 |
| ENSMUSG00000005609.17 | 117.2204 | 0.61157 | 0.936828 | 0.652809 | 0.513879 | 0.983423 | 0.00726 |
| ENSMUSG00000005625.16 | 425.8011 | -0.25025 | 0.445089 | -0.56226 | 0.57394 | 0.983423 | 0.00726 |
| ENSMUSG00000005683.9 | 407.4333 | -0.26161 | 0.401208 | -0.65206 | 0.514361 | 0.983423 | 0.00726 |
| ENSMUSG00000005698.16 | 92.36248 | 0.708438 | 1.264032 | 0.560459 | 0.575166 | 0.983423 | 0.00726 |
| ENSMUSG00000005897.15 | 45.30532 | 0.622182 | 1.069069 | 0.581985 | 0.560576 | 0.983423 | 0.00726 |
| ENSMUSG00000006191.18 | 33.83487 | -0.85548 | 1.353145 | -0.63222 | 0.527246 | 0.983423 | 0.00726 |
| ENSMUSG00000006526.14 | 32.56768 | -0.91723 | 1.626451 | -0.56395 | 0.572791 | 0.983423 | 0.00726 |
| ENSMUSG00000007570.17 | 17.93064 | 1.269634 | 2.068186 | 0.613888 | 0.53929 | 0.983423 | 0.00726 |
| ENSMUSG00000007613.16 | 102.8184 | -0.55459 | 0.864451 | -0.64155 | 0.521163 | 0.983423 | 0.00726 |
| ENSMUSG00000007656.14 | 928.8342 | 0.213077 | 0.362359 | 0.588028 | 0.556513 | 0.983423 | 0.00726 |
| ENSMUSG00000007721.7 | 46.22561 | -0.59283 | 1.055088 | -0.56188 | 0.574197 | 0.983423 | 0.00726 |
| ENSMUSG00000007833.16 | 109.7277 | 0.616235 | 0.991786 | 0.621339 | 0.534377 | 0.983423 | 0.00726 |
| ENSMUSG00000007880.17 | 105.3471 | -0.58447 | 0.938665 | -0.62266 | 0.533511 | 0.983423 | 0.00726 |
| ENSMUSG00000009630.11 | 33.69274 | -0.69233 | 1.137215 | -0.60879 | 0.542661 | 0.983423 | 0.00726 |
| ENSMUSG00000009640.12 | 18.06126 | -1.37274 | 2.202584 | -0.62324 | 0.533126 | 0.983423 | 0.00726 |
| ENSMUSG00000010392.9 | 30.49618 | 0.82796 | 1.337148 | 0.619198 | 0.535786 | 0.983423 | 0.00726 |
| ENSMUSG00000011382.9 | 779.4818 | 0.1962 | 0.314199 | 0.624446 | 0.532335 | 0.983423 | 0.00726 |
| ENSMUSG00000013662.7 | 134.9942 | 0.347057 | 0.609875 | 0.569062 | 0.569314 | 0.983423 | 0.00726 |
| ENSMUSG00000013663.8 | 442.9304 | 0.231577 | 0.357452 | 0.647855 | 0.517079 | 0.983423 | 0.00726 |
| ENSMUSG00000013698.13 | 72.93446 | -0.46819 | 0.813946 | -0.57521 | 0.565152 | 0.983423 | 0.00726 |
| ENSMUSG00000014470.5 | 30.59096 | 0.759823 | 1.178893 | 0.644523 | 0.519237 | 0.983423 | 0.00726 |
| ENSMUSG00000014542.4 | 88.74183 | -0.52512 | 0.828844 | -0.63355 | 0.526372 | 0.983423 | 0.00726 |
| ENSMUSG00000014856.13 | 254.1808 | 0.359222 | 0.633182 | 0.567329 | 0.570491 | 0.983423 | 0.00726 |
| ENSMUSG00000015002.19 | 313.2727 | -0.26384 | 0.473086 | -0.5577 | 0.577051 | 0.983423 | 0.00726 |
| ENSMUSG00000015189.13 | 81.07898 | -0.73316 | 1.130884 | -0.64831 | 0.516787 | 0.983423 | 0.00726 |
| ENSMUSG00000015354.9 | 11.0579 | 1.537763 | 2.484341 | 0.618982 | 0.535928 | 0.983423 | 0.00726 |
| ENSMUSG00000015733.14 | 492.0534 | -0.21209 | 0.353601 | -0.59979 | 0.548647 | 0.983423 | 0.00726 |
| ENSMUSG00000015882.19 | 107.6936 | 0.627759 | 0.975187 | 0.643732 | 0.519749 | 0.983423 | 0.00726 |
| ENSMUSG00000015932.9 | 58.73204 | -0.61375 | 0.949194 | -0.6466 | 0.51789 | 0.983423 | 0.00726 |
| ENSMUSG00000016239.12 | 12.12209 | 1.561907 | 2.686627 | 0.581364 | 0.560995 | 0.983423 | 0.00726 |
| ENSMUSG00000016481.16 | 321.4566 | 0.330975 | 0.510798 | 0.647956 | 0.517014 | 0.983423 | 0.00726 |
| ENSMUSG00000016520.8 | 39.39646 | 0.838045 | 1.412597 | 0.593265 | 0.553004 | 0.983423 | 0.00726 |
| ENSMUSG00000017421.19 | 201.3235 | -0.466 | 0.707102 | -0.65902 | 0.509882 | 0.983423 | 0.00726 |
| ENSMUSG00000018166.9 | 240.0394 | -0.3417 | 0.569359 | -0.60015 | 0.548404 | 0.983423 | 0.00726 |

| | | | | | | | |
|---|---|---|---|---|---|---|---|
| ENSMUSG00000018541.11 | 74.41705 | -0.68929 | 1.127357 | -0.61142 | 0.540923 | 0.983423 | 0.00726 |
| ENSMUSG00000018565.18 | 59.51753 | -0.63168 | 1.051796 | -0.60057 | 0.548127 | 0.983423 | 0.00726 |
| ENSMUSG00000018604.19 | 45.6357 | -0.83075 | 1.381313 | -0.60142 | 0.547559 | 0.983423 | 0.00726 |
| ENSMUSG00000018654.18 | 22.807 | -1.46783 | 2.319558 | -0.63281 | 0.52686 | 0.983423 | 0.00726 |
| ENSMUSG00000018659.13 | 192.5099 | 0.381457 | 0.675971 | 0.564309 | 0.572544 | 0.983423 | 0.00726 |
| ENSMUSG00000018661.18 | 28.095 | 1.037261 | 1.746329 | 0.593967 | 0.552534 | 0.983423 | 0.00726 |
| ENSMUSG00000018861.9 | 26.60835 | -0.95766 | 1.580368 | -0.60597 | 0.544532 | 0.983423 | 0.00726 |
| ENSMUSG00000018882.13 | 244.5719 | 0.455559 | 0.785648 | 0.579851 | 0.562015 | 0.983423 | 0.00726 |
| ENSMUSG00000018920.12 | 35.49168 | 0.950279 | 1.630291 | 0.582889 | 0.559968 | 0.983423 | 0.00726 |
| ENSMUSG00000018932.10 | 299.4714 | -0.32069 | 0.562176 | -0.57044 | 0.568382 | 0.983423 | 0.00726 |
| ENSMUSG00000019505.8 | 1136.564 | 0.163337 | 0.280355 | 0.582607 | 0.560158 | 0.983423 | 0.00726 |
| ENSMUSG00000019577.7 | 42.04833 | 0.843684 | 1.51657 | 0.556311 | 0.577998 | 0.983423 | 0.00726 |
| ENSMUSG00000019779.16 | 45.1476 | 0.958188 | 1.473188 | 0.650418 | 0.515422 | 0.983423 | 0.00726 |
| ENSMUSG00000019874.12 | 27.18952 | -1.06519 | 1.799707 | -0.59187 | 0.553937 | 0.983423 | 0.00726 |
| ENSMUSG00000019920.19 | 89.3124 | 0.693497 | 1.209634 | 0.573312 | 0.566434 | 0.983423 | 0.00726 |
| ENSMUSG00000019951.11 | 387.7893 | -0.25781 | 0.458861 | -0.56184 | 0.574225 | 0.983423 | 0.00726 |
| ENSMUSG00000019952.16 | 14.23978 | -1.41886 | 2.439457 | -0.58163 | 0.560817 | 0.983423 | 0.00726 |
| ENSMUSG00000019961.18 | 250.234 | 0.327901 | 0.508123 | 0.645318 | 0.518721 | 0.983423 | 0.00726 |
| ENSMUSG00000019969.15 | 98.15245 | -0.43041 | 0.768233 | -0.56026 | 0.575302 | 0.983423 | 0.00726 |
| ENSMUSG00000019987.10 | 10771.69 | 0.158344 | 0.249128 | 0.635591 | 0.525043 | 0.983423 | 0.00726 |
| ENSMUSG00000019998.7 | 85.33758 | 0.537143 | 0.900462 | 0.596519 | 0.550829 | 0.983423 | 0.00726 |
| ENSMUSG00000020089.9 | 1810.501 | 0.181906 | 0.319315 | 0.569674 | 0.568899 | 0.983423 | 0.00726 |
| ENSMUSG00000020105.10 | 19.26401 | -1.05941 | 1.736832 | -0.60997 | 0.541884 | 0.983423 | 0.00726 |
| ENSMUSG00000020114.13 | 193.1349 | -0.37468 | 0.668072 | -0.56083 | 0.574913 | 0.983423 | 0.00726 |
| ENSMUSG00000020149.18 | 888.4153 | 0.218136 | 0.370098 | 0.5894 | 0.555593 | 0.983423 | 0.00726 |
| ENSMUSG00000020153.15 | 261.1442 | 0.349916 | 0.538286 | 0.650056 | 0.515656 | 0.983423 | 0.00726 |
| ENSMUSG00000020263.15 | 51.36942 | -0.63712 | 1.065293 | -0.59807 | 0.549796 | 0.983423 | 0.00726 |
| ENSMUSG00000020328.11 | 340.3605 | 0.287349 | 0.505469 | 0.568479 | 0.56971 | 0.983423 | 0.00726 |
| ENSMUSG00000020359.14 | 140.1708 | -0.4748 | 0.729204 | -0.65112 | 0.514966 | 0.983423 | 0.00726 |
| ENSMUSG00000020402.12 | 777.6708 | -0.23784 | 0.382134 | -0.6224 | 0.533678 | 0.983423 | 0.00726 |
| ENSMUSG00000020409.11 | 207.2973 | -0.39212 | 0.641366 | -0.61139 | 0.540943 | 0.983423 | 0.00726 |
| ENSMUSG00000020463.16 | 283.3415 | -0.35354 | 0.584286 | -0.60509 | 0.545122 | 0.983423 | 0.00726 |
| ENSMUSG00000020467.16 | 109.4087 | 0.451811 | 0.78952 | 0.57226 | 0.567146 | 0.983423 | 0.00726 |
| ENSMUSG00000020477.11 | 644.1655 | -0.22623 | 0.379652 | -0.59589 | 0.551248 | 0.983423 | 0.00726 |
| ENSMUSG00000020496.11 | 244.1471 | 0.333719 | 0.538655 | 0.61954 | 0.535561 | 0.983423 | 0.00726 |
| ENSMUSG00000020522.14 | 29.39559 | 0.855417 | 1.35923 | 0.629339 | 0.529127 | 0.983423 | 0.00726 |
| ENSMUSG00000020672.15 | 26.85523 | 0.76119 | 1.370029 | 0.555602 | 0.578483 | 0.983423 | 0.00726 |
| ENSMUSG00000020709.16 | 16.73318 | 1.243294 | 2.005552 | 0.619926 | 0.535307 | 0.983423 | 0.00726 |
| ENSMUSG00000020717.20 | 27.29237 | 1.166479 | 1.897434 | 0.614767 | 0.538709 | 0.983423 | 0.00726 |
| ENSMUSG00000020774.10 | 122.2156 | -0.45971 | 0.705948 | -0.6512 | 0.514919 | 0.983423 | 0.00726 |
| ENSMUSG00000020775.14 | 115.5414 | 0.389974 | 0.696633 | 0.559798 | 0.575617 | 0.983423 | 0.00726 |
| ENSMUSG00000020817.17 | 214.3374 | 0.456668 | 0.745567 | 0.612511 | 0.540199 | 0.983423 | 0.00726 |
| ENSMUSG00000020821.18 | 240.3026 | 0.309302 | 0.504966 | 0.61252 | 0.540193 | 0.983423 | 0.00726 |
| ENSMUSG00000020859.17 | 251.6572 | -0.35153 | 0.568837 | -0.61797 | 0.536594 | 0.983423 | 0.00726 |

| | | | | | | | |
|---|---|---|---|---|---|---|---|
| ENSMUSG00000020869.9 | 265.1119 | 0.30283 | 0.524723 | 0.577125 | 0.563855 | 0.983423 | 0.00726 |
| ENSMUSG00000021018.9 | 21.15827 | -1.40002 | 2.348582 | -0.59611 | 0.551101 | 0.983423 | 0.00726 |
| ENSMUSG00000021037.8 | 53.27156 | 0.860707 | 1.492075 | 0.576852 | 0.564039 | 0.983423 | 0.00726 |
| ENSMUSG00000021111.16 | 430.9174 | 0.243192 | 0.418785 | 0.58071 | 0.561436 | 0.983423 | 0.00726 |
| ENSMUSG00000021135.9 | 6300.886 | 0.16171 | 0.268092 | 0.60319 | 0.546382 | 0.983423 | 0.00726 |
| ENSMUSG00000021186.10 | 26.0968 | -1.02684 | 1.553738 | -0.66088 | 0.508686 | 0.983423 | 0.00726 |
| ENSMUSG00000021262.16 | 11.78952 | -1.72367 | 2.757912 | -0.62499 | 0.531976 | 0.983423 | 0.00726 |
| ENSMUSG00000021277.17 | 50.64715 | 1.125492 | 1.790111 | 0.628728 | 0.529527 | 0.983423 | 0.00726 |
| ENSMUSG00000021288.20 | 14.84733 | -1.47394 | 2.265514 | -0.6506 | 0.515307 | 0.983423 | 0.00726 |
| ENSMUSG00000021361.8 | 591.1479 | 0.305441 | 0.482786 | 0.632664 | 0.526953 | 0.983423 | 0.00726 |
| ENSMUSG00000021500.18 | 235.0958 | -0.37608 | 0.659736 | -0.57004 | 0.56865 | 0.983423 | 0.00726 |
| ENSMUSG00000021518.5 | 110.5505 | -0.48492 | 0.770406 | -0.62944 | 0.529061 | 0.983423 | 0.00726 |
| ENSMUSG00000021537.10 | 370.6815 | 0.318028 | 0.489735 | 0.649389 | 0.516087 | 0.983423 | 0.00726 |
| ENSMUSG00000021557.16 | 72.30987 | 0.538061 | 0.915315 | 0.587842 | 0.556638 | 0.983423 | 0.00726 |
| ENSMUSG00000021676.11 | 1278.033 | 0.169223 | 0.285623 | 0.592469 | 0.553536 | 0.983423 | 0.00726 |
| ENSMUSG00000021694.19 | 13.63029 | -1.35036 | 2.351871 | -0.57416 | 0.565857 | 0.983423 | 0.00726 |
| ENSMUSG00000021711.18 | 149.1757 | -0.47289 | 0.742781 | -0.63664 | 0.524356 | 0.983423 | 0.00726 |
| ENSMUSG00000021751.14 | 967.6459 | -0.20583 | 0.357862 | -0.57516 | 0.565182 | 0.983423 | 0.00726 |
| ENSMUSG00000021794.17 | 3728.809 | -0.1454 | 0.221361 | -0.65685 | 0.511276 | 0.983423 | 0.00726 |
| ENSMUSG00000021895.11 | 149.0611 | -0.37234 | 0.655464 | -0.56805 | 0.569999 | 0.983423 | 0.00726 |
| ENSMUSG00000021972.15 | 50.83897 | -0.63668 | 1.145208 | -0.55596 | 0.578241 | 0.983423 | 0.00726 |
| ENSMUSG00000021973.10 | 241.2658 | -0.48888 | 0.781839 | -0.6253 | 0.531774 | 0.983423 | 0.00726 |
| ENSMUSG00000021981.11 | 278.1452 | 0.331704 | 0.582821 | 0.569135 | 0.569264 | 0.983423 | 0.00726 |
| ENSMUSG00000021982.15 | 167.5769 | -0.3807 | 0.641125 | -0.59381 | 0.552641 | 0.983423 | 0.00726 |
| ENSMUSG00000021993.11 | 94.1612 | -0.51384 | 0.778032 | -0.66044 | 0.508974 | 0.983423 | 0.00726 |
| ENSMUSG00000022009.2 | 27.48918 | -0.86954 | 1.397944 | -0.62201 | 0.533932 | 0.983423 | 0.00726 |
| ENSMUSG00000022013.5 | 405.1415 | 0.26066 | 0.431355 | 0.604281 | 0.545657 | 0.983423 | 0.00726 |
| ENSMUSG00000022019.17 | 64.34618 | 0.591109 | 0.980796 | 0.602683 | 0.54672 | 0.983423 | 0.00726 |
| ENSMUSG00000022020.16 | 161.4425 | 0.530753 | 0.84397 | 0.628876 | 0.52943 | 0.983423 | 0.00726 |
| ENSMUSG00000022022.4 | 92.78519 | 0.591063 | 1.045915 | 0.565116 | 0.571995 | 0.983423 | 0.00726 |
| ENSMUSG00000022204.10 | 171.1275 | 0.331392 | 0.580252 | 0.571118 | 0.56792 | 0.983423 | 0.00726 |
| ENSMUSG00000022210.8 | 894.7945 | 0.175844 | 0.309129 | 0.568837 | 0.569467 | 0.983423 | 0.00726 |
| ENSMUSG00000022244.8 | 730.5219 | -0.23596 | 0.397997 | -0.59286 | 0.553274 | 0.983423 | 0.00726 |
| ENSMUSG00000022255.17 | 694.566 | 0.259026 | 0.408567 | 0.633986 | 0.52609 | 0.983423 | 0.00726 |
| ENSMUSG00000022263.11 | 55.76958 | 0.697527 | 1.175865 | 0.593204 | 0.553045 | 0.983423 | 0.00726 |
| ENSMUSG00000022336.4 | 850.2722 | -0.19076 | 0.334096 | -0.57097 | 0.568023 | 0.983423 | 0.00726 |
| ENSMUSG00000022370.14 | 576.7513 | -0.28866 | 0.452461 | -0.63798 | 0.523485 | 0.983423 | 0.00726 |
| ENSMUSG00000022377.17 | 59.2416 | -0.7668 | 1.217285 | -0.62993 | 0.528742 | 0.983423 | 0.00726 |
| ENSMUSG00000022390.12 | 36.33273 | -1.00572 | 1.550675 | -0.64857 | 0.516616 | 0.983423 | 0.00726 |
| ENSMUSG00000022401.13 | 73.54984 | 0.712009 | 1.101973 | 0.646122 | 0.518201 | 0.983423 | 0.00726 |
| ENSMUSG00000022427.5 | 315.9152 | -0.32526 | 0.515581 | -0.63087 | 0.528126 | 0.983423 | 0.00726 |
| ENSMUSG00000022561.14 | 19.28417 | -1.29231 | 2.120841 | -0.60934 | 0.542299 | 0.983423 | 0.00726 |
| ENSMUSG00000022568.18 | 11.27404 | 1.465096 | 2.407456 | 0.608566 | 0.542812 | 0.983423 | 0.00726 |
| ENSMUSG00000022668.11 | 53.11391 | 0.616577 | 1.022551 | 0.602979 | 0.546522 | 0.983423 | 0.00726 |

| | | | | | | | |
|---|---|---|---|---|---|---|---|
| ENSMUSG00000022682.9 | 36.92424 | -0.68168 | 1.143339 | -0.59622 | 0.551031 | 0.983423 | 0.00726 |
| ENSMUSG00000022710.18 | 85.29402 | -0.45092 | 0.80287 | -0.56164 | 0.574363 | 0.983423 | 0.00726 |
| ENSMUSG00000022723.17 | 39.09512 | -0.79263 | 1.381867 | -0.57359 | 0.566243 | 0.983423 | 0.00726 |
| ENSMUSG00000022892.12 | 117.0779 | 0.411625 | 0.707144 | 0.582095 | 0.560503 | 0.983423 | 0.00726 |
| ENSMUSG00000022893.15 | 55.55986 | 0.849603 | 1.296542 | 0.655284 | 0.512285 | 0.983423 | 0.00726 |
| ENSMUSG00000022897.16 | 89.58511 | -0.50133 | 0.856107 | -0.58559 | 0.558152 | 0.983423 | 0.00726 |
| ENSMUSG00000022956.12 | 943.3414 | 0.191439 | 0.332202 | 0.576272 | 0.564431 | 0.983423 | 0.00726 |
| ENSMUSG00000022967.14 | 60.53168 | -0.56602 | 0.929427 | -0.609 | 0.542524 | 0.983423 | 0.00726 |
| ENSMUSG00000023004.9 | 83.44529 | -0.47489 | 0.803029 | -0.59137 | 0.554272 | 0.983423 | 0.00726 |
| ENSMUSG00000023147.18 | 32.69854 | -1.09359 | 1.877161 | -0.58258 | 0.560179 | 0.983423 | 0.00726 |
| ENSMUSG00000023175.16 | 488.3073 | 0.25207 | 0.395373 | 0.63755 | 0.523767 | 0.983423 | 0.00726 |
| ENSMUSG00000023348.12 | 19.87687 | -0.91321 | 1.441944 | -0.63332 | 0.526527 | 0.983423 | 0.00726 |
| ENSMUSG00000023932.10 | 133.6506 | -0.498 | 0.784164 | -0.63507 | 0.525381 | 0.983423 | 0.00726 |
| ENSMUSG00000023961.17 | 72.93682 | -0.63339 | 1.041046 | -0.60842 | 0.542911 | 0.983423 | 0.00726 |
| ENSMUSG00000023980.8 | 25.10309 | 1.057077 | 1.685593 | 0.627124 | 0.530578 | 0.983423 | 0.00726 |
| ENSMUSG00000024066.10 | 355.2413 | -0.32223 | 0.542235 | -0.59427 | 0.552331 | 0.983423 | 0.00726 |
| ENSMUSG00000024140.11 | 776.3897 | 0.227736 | 0.408565 | 0.557406 | 0.57725 | 0.983423 | 0.00726 |
| ENSMUSG00000024242.16 | 162.6284 | 0.390784 | 0.660751 | 0.591424 | 0.554236 | 0.983423 | 0.00726 |
| ENSMUSG00000024258.5 | 45.19578 | -0.90388 | 1.47252 | -0.61383 | 0.539328 | 0.983423 | 0.00726 |
| ENSMUSG00000024287.9 | 147.7089 | 0.454027 | 0.740918 | 0.61279 | 0.540015 | 0.983423 | 0.00726 |
| ENSMUSG00000024381.17 | 22.72656 | -1.01112 | 1.810034 | -0.55862 | 0.576423 | 0.983423 | 0.00726 |
| ENSMUSG00000024397.15 | 30.71014 | -1.16483 | 1.903003 | -0.6121 | 0.540472 | 0.983423 | 0.00726 |
| ENSMUSG00000024411.12 | 22.02038 | 1.22468 | 1.963387 | 0.623759 | 0.532786 | 0.983423 | 0.00726 |
| ENSMUSG00000024446.14 | 84.11607 | 0.683412 | 1.193884 | 0.572427 | 0.567032 | 0.983423 | 0.00726 |
| ENSMUSG00000024456.17 | 60.2717 | 0.922882 | 1.418884 | 0.650428 | 0.515416 | 0.983423 | 0.00726 |
| ENSMUSG00000024560.8 | 43.08468 | 0.728458 | 1.254921 | 0.580481 | 0.56159 | 0.983423 | 0.00726 |
| ENSMUSG00000024580.9 | 105.146 | 0.825776 | 1.354616 | 0.609601 | 0.542126 | 0.983423 | 0.00726 |
| ENSMUSG00000024581.14 | 84.69694 | -0.54643 | 0.846236 | -0.64572 | 0.518462 | 0.983423 | 0.00726 |
| ENSMUSG00000024604.8 | 64.25768 | 0.639482 | 1.073143 | 0.595897 | 0.551244 | 0.983423 | 0.00726 |
| ENSMUSG00000024642.18 | 64.98481 | -0.56959 | 0.910333 | -0.62569 | 0.531518 | 0.983423 | 0.00726 |
| ENSMUSG00000024654.9 | 49.73838 | 1.007495 | 1.747953 | 0.576386 | 0.564355 | 0.983423 | 0.00726 |
| ENSMUSG00000024772.10 | 22.84073 | -0.85915 | 1.503001 | -0.57162 | 0.567578 | 0.983423 | 0.00726 |
| ENSMUSG00000024790.9 | 47.28293 | 0.870894 | 1.406103 | 0.619367 | 0.535675 | 0.983423 | 0.00726 |
| ENSMUSG00000024811.12 | 212.6438 | 0.378991 | 0.657097 | 0.576766 | 0.564098 | 0.983423 | 0.00726 |
| ENSMUSG00000024818.16 | 42.19293 | -0.77169 | 1.286527 | -0.59982 | 0.548624 | 0.983423 | 0.00726 |
| ENSMUSG00000024829.13 | 202.7896 | -0.37976 | 0.63725 | -0.59593 | 0.55122 | 0.983423 | 0.00726 |
| ENSMUSG00000024841.9 | 214.4128 | -0.36556 | 0.553325 | -0.66066 | 0.508833 | 0.983423 | 0.00726 |
| ENSMUSG00000024875.6 | 112.7334 | -0.70753 | 1.070913 | -0.66068 | 0.508818 | 0.983423 | 0.00726 |
| ENSMUSG00000024998.18 | 26.7782 | -1.12363 | 1.710475 | -0.65691 | 0.511237 | 0.983423 | 0.00726 |
| ENSMUSG00000025047.10 | 83.9061 | -0.50124 | 0.829055 | -0.6046 | 0.545447 | 0.983423 | 0.00726 |
| ENSMUSG00000025151.17 | 305.2017 | -0.3077 | 0.483178 | -0.63682 | 0.524244 | 0.983423 | 0.00726 |
| ENSMUSG00000025207.17 | 97.75849 | -0.41539 | 0.742068 | -0.55978 | 0.57563 | 0.983423 | 0.00726 |
| ENSMUSG00000025264.13 | 27.48632 | -1.07818 | 1.662279 | -0.64862 | 0.516586 | 0.983423 | 0.00726 |
| ENSMUSG00000025271.14 | 82.41088 | 0.632432 | 1.04529 | 0.60503 | 0.545159 | 0.983423 | 0.00726 |

| | | | | | | | |
|---|---|---|---|---|---|---|---|
| ENSMUSG00000025314.17 | 59.19892 | -0.57446 | 0.938632 | -0.61202 | 0.540523 | 0.983423 | 0.00726 |
| ENSMUSG00000025358.17 | 19.54748 | -0.91768 | 1.62705 | -0.56401 | 0.572746 | 0.983423 | 0.00726 |
| ENSMUSG00000025395.15 | 38.63595 | -1.1231 | 1.878221 | -0.59796 | 0.549866 | 0.983423 | 0.00726 |
| ENSMUSG00000025498.16 | 87.66025 | -0.60964 | 1.03538 | -0.5888 | 0.555992 | 0.983423 | 0.00726 |
| ENSMUSG00000025575.15 | 49.01232 | 0.586469 | 0.93233 | 0.629036 | 0.529326 | 0.983423 | 0.00726 |
| ENSMUSG00000025647.17 | 22.80704 | 1.035193 | 1.780587 | 0.581377 | 0.560986 | 0.983423 | 0.00726 |
| ENSMUSG00000025786.18 | 50.32953 | -0.95218 | 1.539859 | -0.61835 | 0.536343 | 0.983423 | 0.00726 |
| ENSMUSG00000025872.6 | 105.2832 | -0.47038 | 0.754929 | -0.62307 | 0.533236 | 0.983423 | 0.00726 |
| ENSMUSG00000025887.11 | 22.38493 | -1.17679 | 1.890131 | -0.6226 | 0.53355 | 0.983423 | 0.00726 |
| ENSMUSG00000025956.11 | 27.73395 | 0.992815 | 1.71938 | 0.577426 | 0.563651 | 0.983423 | 0.00726 |
| ENSMUSG00000025995.17 | 85.50133 | 0.515025 | 0.890331 | 0.578464 | 0.562951 | 0.983423 | 0.00726 |
| ENSMUSG00000026000.17 | 65.20892 | -0.52447 | 0.893225 | -0.58716 | 0.557093 | 0.983423 | 0.00726 |
| ENSMUSG00000026031.16 | 164.3959 | -0.33485 | 0.543475 | -0.61612 | 0.537812 | 0.983423 | 0.00726 |
| ENSMUSG00000026043.19 | 79.71924 | -0.56887 | 0.895291 | -0.6354 | 0.525166 | 0.983423 | 0.00726 |
| ENSMUSG00000026097.10 | 299.6442 | -0.30346 | 0.530993 | -0.5715 | 0.567661 | 0.983423 | 0.00726 |
| ENSMUSG00000026150.15 | 207.2059 | 0.34125 | 0.529659 | 0.644283 | 0.519392 | 0.983423 | 0.00726 |
| ENSMUSG00000026181.14 | 29.58529 | -0.88484 | 1.496627 | -0.59122 | 0.55437 | 0.983423 | 0.00726 |
| ENSMUSG00000026240.13 | 33.05464 | -1.01321 | 1.732631 | -0.58478 | 0.558694 | 0.983423 | 0.00726 |
| ENSMUSG00000026269.15 | 162.0957 | -0.35761 | 0.597868 | -0.59815 | 0.54974 | 0.983423 | 0.00726 |
| ENSMUSG00000026360.10 | 66.06044 | 0.742984 | 1.261716 | 0.588868 | 0.55595 | 0.983423 | 0.00726 |
| ENSMUSG00000026390.8 | 52.06609 | -1.07839 | 1.737219 | -0.62076 | 0.534759 | 0.983423 | 0.00726 |
| ENSMUSG00000026456.19 | 36.86149 | -0.84514 | 1.340035 | -0.63069 | 0.528246 | 0.983423 | 0.00726 |
| ENSMUSG00000026496.12 | 96.93407 | 0.762715 | 1.290271 | 0.591127 | 0.554435 | 0.983423 | 0.00726 |
| ENSMUSG00000026579.9 | 3497.758 | -0.15392 | 0.269228 | -0.57172 | 0.56751 | 0.983423 | 0.00726 |
| ENSMUSG00000026621.14 | 2004.898 | -0.15269 | 0.233671 | -0.65346 | 0.513461 | 0.983423 | 0.00726 |
| ENSMUSG00000026740.13 | 271.3852 | -0.33353 | 0.576651 | -0.5784 | 0.562996 | 0.983423 | 0.00726 |
| ENSMUSG00000026812.17 | 36.54361 | 0.626718 | 1.11245 | 0.563367 | 0.573185 | 0.983423 | 0.00726 |
| ENSMUSG00000026867.20 | 230.1324 | -0.28823 | 0.510689 | -0.5644 | 0.57248 | 0.983423 | 0.00726 |
| ENSMUSG00000026870.20 | 145.2761 | 0.435517 | 0.734048 | 0.593308 | 0.552975 | 0.983423 | 0.00726 |
| ENSMUSG00000026887.10 | 87.97068 | 0.509841 | 0.849023 | 0.600503 | 0.548171 | 0.983423 | 0.00726 |
| ENSMUSG00000026932.15 | 57.65216 | -0.63622 | 1.100046 | -0.57836 | 0.563022 | 0.983423 | 0.00726 |
| ENSMUSG00000026933.18 | 21.76616 | -1.158 | 2.036656 | -0.56858 | 0.569643 | 0.983423 | 0.00726 |
| ENSMUSG00000027011.15 | 286.7591 | 0.289929 | 0.51467 | 0.56333 | 0.57321 | 0.983423 | 0.00726 |
| ENSMUSG00000027035.11 | 42.33872 | 0.67102 | 1.199257 | 0.55953 | 0.5758 | 0.983423 | 0.00726 |
| ENSMUSG00000027086.17 | 48.36853 | -0.72181 | 1.114715 | -0.64753 | 0.517288 | 0.983423 | 0.00726 |
| ENSMUSG00000027175.11 | 58.51316 | -0.51979 | 0.926632 | -0.56094 | 0.574838 | 0.983423 | 0.00726 |
| ENSMUSG00000027180.13 | 646.7351 | 0.246337 | 0.378281 | 0.6512 | 0.514917 | 0.983423 | 0.00726 |
| ENSMUSG00000027194.17 | 80.14506 | -0.57612 | 0.977878 | -0.58915 | 0.55576 | 0.983423 | 0.00726 |
| ENSMUSG00000027200.18 | 45.43443 | 0.821608 | 1.254988 | 0.654674 | 0.512678 | 0.983423 | 0.00726 |
| ENSMUSG00000027227.8 | 4757.785 | 0.157276 | 0.241489 | 0.651276 | 0.514868 | 0.983423 | 0.00726 |
| ENSMUSG00000027243.9 | 62.70891 | 0.761348 | 1.360548 | 0.559589 | 0.57576 | 0.983423 | 0.00726 |
| ENSMUSG00000027274.17 | 43.70194 | -0.67243 | 1.144144 | -0.58772 | 0.556722 | 0.983423 | 0.00726 |
| ENSMUSG00000027367.17 | 788.2154 | 0.233392 | 0.407743 | 0.572401 | 0.56705 | 0.983423 | 0.00726 |
| ENSMUSG00000027371.11 | 19.81832 | 1.30562 | 2.043144 | 0.639025 | 0.522807 | 0.983423 | 0.00726 |

| | | | | | | | |
|---|---|---|---|---|---|---|---|
| ENSMUSG00000027398.14 | 27.19281 | -1.25075 | 1.970319 | -0.6348 | 0.525562 | 0.983423 | 0.00726 |
| ENSMUSG00000027455.17 | 95.73112 | -0.55356 | 0.879105 | -0.62968 | 0.528904 | 0.983423 | 0.00726 |
| ENSMUSG00000027523.21 | 3834.439 | 0.142735 | 0.228937 | 0.623469 | 0.532977 | 0.983423 | 0.00726 |
| ENSMUSG00000027536.7 | 9.942076 | 1.791942 | 2.862624 | 0.625979 | 0.531329 | 0.983423 | 0.00726 |
| ENSMUSG00000027559.6 | 28116.05 | 0.143716 | 0.250674 | 0.57332 | 0.566428 | 0.983423 | 0.00726 |
| ENSMUSG00000027575.20 | 28.52352 | -1.2709 | 1.993141 | -0.63764 | 0.52371 | 0.983423 | 0.00726 |
| ENSMUSG00000027598.17 | 372.4115 | -0.26352 | 0.453844 | -0.58063 | 0.561487 | 0.983423 | 0.00726 |
| ENSMUSG00000027690.14 | 1749.81 | -0.17727 | 0.288491 | -0.61446 | 0.538913 | 0.983423 | 0.00726 |
| ENSMUSG00000027710.15 | 37.15049 | 0.981756 | 1.492636 | 0.657733 | 0.51071 | 0.983423 | 0.00726 |
| ENSMUSG00000027750.17 | 31.554 | -0.8154 | 1.357815 | -0.60053 | 0.548155 | 0.983423 | 0.00726 |
| ENSMUSG00000027751.14 | 31.81756 | 0.897847 | 1.546373 | 0.580615 | 0.5615 | 0.983423 | 0.00726 |
| ENSMUSG00000027761.5 | 6127.422 | 0.154907 | 0.235033 | 0.659083 | 0.509842 | 0.983423 | 0.00726 |
| ENSMUSG00000027770.6 | 208.6527 | -0.35466 | 0.569069 | -0.62323 | 0.533131 | 0.983423 | 0.00726 |
| ENSMUSG00000027784.11 | 49.66541 | -0.68206 | 1.223263 | -0.55757 | 0.577135 | 0.983423 | 0.00726 |
| ENSMUSG00000027803.15 | 52.41915 | 0.730305 | 1.163715 | 0.627564 | 0.53029 | 0.983423 | 0.00726 |
| ENSMUSG00000027881.15 | 340.4682 | -0.27973 | 0.45187 | -0.61905 | 0.535881 | 0.983423 | 0.00726 |
| ENSMUSG00000027957.14 | 389.1471 | -0.31559 | 0.505922 | -0.6238 | 0.532762 | 0.983423 | 0.00726 |
| ENSMUSG00000027983.14 | 141.3243 | -0.43956 | 0.669417 | -0.65663 | 0.511419 | 0.983423 | 0.00726 |
| ENSMUSG00000027993.17 | 246.6797 | 0.510641 | 0.77458 | 0.659249 | 0.509736 | 0.983423 | 0.00726 |
| ENSMUSG00000028011.17 | 12215.38 | -0.16396 | 0.294347 | -0.55702 | 0.577517 | 0.983423 | 0.00726 |
| ENSMUSG00000028059.17 | 35.43735 | -1.19342 | 1.909351 | -0.62504 | 0.531944 | 0.983423 | 0.00726 |
| ENSMUSG00000028081.7 | 2923.22 | 0.161988 | 0.251418 | 0.644297 | 0.519383 | 0.983423 | 0.00726 |
| ENSMUSG00000028096.14 | 96.16846 | 0.430506 | 0.762031 | 0.564946 | 0.57211 | 0.983423 | 0.00726 |
| ENSMUSG00000028127.11 | 3176.566 | 0.158149 | 0.271509 | 0.582479 | 0.560244 | 0.983423 | 0.00726 |
| ENSMUSG00000028184.13 | 146.8032 | -0.42866 | 0.730891 | -0.58649 | 0.557549 | 0.983423 | 0.00726 |
| ENSMUSG00000028221.4 | 129.8487 | 0.534013 | 0.862143 | 0.619402 | 0.535651 | 0.983423 | 0.00726 |
| ENSMUSG00000028252.21 | 109.6115 | 0.411621 | 0.726058 | 0.566926 | 0.570765 | 0.983423 | 0.00726 |
| ENSMUSG00000028268.15 | 39.92462 | -0.88877 | 1.411778 | -0.62954 | 0.528994 | 0.983423 | 0.00726 |
| ENSMUSG00000028270.13 | 32.60521 | 1.082612 | 1.695862 | 0.638385 | 0.523223 | 0.983423 | 0.00726 |
| ENSMUSG00000028277.14 | 157.0161 | -0.34652 | 0.585806 | -0.59152 | 0.554169 | 0.983423 | 0.00726 |
| ENSMUSG00000028295.15 | 171.3449 | 0.41609 | 0.699202 | 0.595092 | 0.551782 | 0.983423 | 0.00726 |
| ENSMUSG00000028560.12 | 86.22817 | 0.528749 | 0.905368 | 0.584016 | 0.55921 | 0.983423 | 0.00726 |
| ENSMUSG00000028646.17 | 77.2476 | 0.525455 | 0.808223 | 0.650136 | 0.515604 | 0.983423 | 0.00726 |
| ENSMUSG00000028690.5 | 59.35844 | 0.613742 | 0.945209 | 0.649319 | 0.516132 | 0.983423 | 0.00726 |
| ENSMUSG00000028706.15 | 75.2343 | -0.65797 | 1.042913 | -0.6309 | 0.528105 | 0.983423 | 0.00726 |
| ENSMUSG00000028776.15 | 54.34699 | -0.7414 | 1.308806 | -0.56647 | 0.571074 | 0.983423 | 0.00726 |
| ENSMUSG00000028779.17 | 36.87579 | 0.700703 | 1.150464 | 0.609061 | 0.542484 | 0.983423 | 0.00726 |
| ENSMUSG00000028793.16 | 48.76255 | -0.61263 | 1.033625 | -0.5927 | 0.55338 | 0.983423 | 0.00726 |
| ENSMUSG00000028797.21 | 277.0159 | 0.309343 | 0.542796 | 0.569907 | 0.568741 | 0.983423 | 0.00726 |
| ENSMUSG00000028842.16 | 23.05671 | -1.06403 | 1.694301 | -0.628 | 0.530001 | 0.983423 | 0.00726 |
| ENSMUSG00000028878.12 | 67.4751 | 0.571173 | 0.91908 | 0.621462 | 0.534295 | 0.983423 | 0.00726 |
| ENSMUSG00000028914.14 | 11.72874 | 1.776074 | 2.752361 | 0.645291 | 0.518739 | 0.983423 | 0.00726 |
| ENSMUSG00000028936.16 | 2200.745 | 0.17142 | 0.273949 | 0.625737 | 0.531487 | 0.983423 | 0.00726 |
| ENSMUSG00000028952.13 | 13.75665 | 1.387637 | 2.399379 | 0.578332 | 0.56304 | 0.983423 | 0.00726 |

| | | | | | | | |
|---|---|---|---|---|---|---|---|
| ENSMUSG00000028973.19 | 29.75743 | 1.036827 | 1.735676 | 0.597362 | 0.550266 | 0.983423 | 0.00726 |
| ENSMUSG00000029098.18 | 43.94563 | -0.89073 | 1.47892 | -0.60228 | 0.546987 | 0.983423 | 0.00726 |
| ENSMUSG00000029163.10 | 13.8281 | -1.34424 | 2.321232 | -0.57911 | 0.562517 | 0.983423 | 0.00726 |
| ENSMUSG00000029176.14 | 83.5555 | 0.745208 | 1.242298 | 0.599863 | 0.548598 | 0.983423 | 0.00726 |
| ENSMUSG00000029186.13 | 141.0405 | 0.409379 | 0.686096 | 0.596679 | 0.550722 | 0.983423 | 0.00726 |
| ENSMUSG00000029270.11 | 118.4993 | 0.482813 | 0.811407 | 0.595032 | 0.551822 | 0.983423 | 0.00726 |
| ENSMUSG00000029344.16 | 169.6526 | 0.432109 | 0.70206 | 0.615488 | 0.538233 | 0.983423 | 0.00726 |
| ENSMUSG00000029439.15 | 60.43072 | 0.618616 | 1.105961 | 0.559347 | 0.575925 | 0.983423 | 0.00726 |
| ENSMUSG00000029569.10 | 92.19705 | 0.497078 | 0.781765 | 0.635841 | 0.52488 | 0.983423 | 0.00726 |
| ENSMUSG00000029613.16 | 98.44391 | 0.486812 | 0.765037 | 0.636325 | 0.524565 | 0.983423 | 0.00726 |
| ENSMUSG00000029616.10 | 210.9276 | -0.37433 | 0.61019 | -0.61346 | 0.539571 | 0.983423 | 0.00726 |
| ENSMUSG00000029726.17 | 41.63567 | 0.997301 | 1.718517 | 0.580327 | 0.561694 | 0.983423 | 0.00726 |
| ENSMUSG00000029776.12 | 1525.437 | 0.16517 | 0.264485 | 0.624498 | 0.532301 | 0.983423 | 0.00726 |
| ENSMUSG00000030029.15 | 47.87245 | -0.65404 | 1.107935 | -0.59032 | 0.554976 | 0.983423 | 0.00726 |
| ENSMUSG00000030058.18 | 219.1692 | 0.328703 | 0.535329 | 0.61402 | 0.539202 | 0.983423 | 0.00726 |
| ENSMUSG00000030079.16 | 72.30663 | 0.643903 | 1.016977 | 0.633154 | 0.526633 | 0.983423 | 0.00726 |
| ENSMUSG00000030087.12 | 1290.152 | -0.18549 | 0.332244 | -0.55829 | 0.576646 | 0.983423 | 0.00726 |
| ENSMUSG00000030172.16 | 79.13016 | 0.636451 | 1.045531 | 0.608735 | 0.5427 | 0.983423 | 0.00726 |
| ENSMUSG00000030189.16 | 102.1604 | -0.51674 | 0.886421 | -0.58295 | 0.559926 | 0.983423 | 0.00726 |
| ENSMUSG00000030214.8 | 224.5272 | -0.49752 | 0.762351 | -0.65261 | 0.51401 | 0.983423 | 0.00726 |
| ENSMUSG00000030222.14 | 14.0163 | 1.095348 | 1.760426 | 0.622206 | 0.533806 | 0.983423 | 0.00726 |
| ENSMUSG00000030465.20 | 191.6488 | -0.37603 | 0.59057 | -0.63673 | 0.524299 | 0.983423 | 0.00726 |
| ENSMUSG00000030470.16 | 89.17343 | -0.73396 | 1.166248 | -0.62934 | 0.52913 | 0.983423 | 0.00726 |
| ENSMUSG00000030515.10 | 81.31061 | 0.493918 | 0.80307 | 0.615037 | 0.53853 | 0.983423 | 0.00726 |
| ENSMUSG00000030612.7 | 141.9324 | 0.382872 | 0.654201 | 0.585251 | 0.558379 | 0.983423 | 0.00726 |
| ENSMUSG00000030652.12 | 401.0357 | 0.268992 | 0.461367 | 0.583033 | 0.559871 | 0.983423 | 0.00726 |
| ENSMUSG00000030689.16 | 11.1989 | 1.711329 | 2.843122 | 0.601919 | 0.547228 | 0.983423 | 0.00726 |
| ENSMUSG00000030748.10 | 27.90012 | -1.00295 | 1.745146 | -0.57471 | 0.56549 | 0.983423 | 0.00726 |
| ENSMUSG00000030770.16 | 245.3442 | 0.427297 | 0.699036 | 0.611266 | 0.541024 | 0.983423 | 0.00726 |
| ENSMUSG00000030787.5 | 22.85738 | 1.048575 | 1.80622 | 0.580536 | 0.561553 | 0.983423 | 0.00726 |
| ENSMUSG00000030801.11 | 31.88178 | -0.78842 | 1.303164 | -0.605 | 0.545176 | 0.983423 | 0.00726 |
| ENSMUSG00000030839.13 | 29.48905 | 1.196078 | 1.854333 | 0.645018 | 0.518916 | 0.983423 | 0.00726 |
| ENSMUSG00000030847.9 | 117.683 | 0.471746 | 0.725409 | 0.650318 | 0.515487 | 0.983423 | 0.00726 |
| ENSMUSG00000030850.18 | 165.0138 | 0.394668 | 0.637555 | 0.619035 | 0.535894 | 0.983423 | 0.00726 |
| ENSMUSG00000030868.7 | 42.49243 | 0.738185 | 1.194053 | 0.618218 | 0.536431 | 0.983423 | 0.00726 |
| ENSMUSG00000030909.7 | 393.1654 | -0.35309 | 0.556943 | -0.63397 | 0.526098 | 0.983423 | 0.00726 |
| ENSMUSG00000030942.9 | 87.23465 | 0.587678 | 0.90682 | 0.648064 | 0.516944 | 0.983423 | 0.00726 |
| ENSMUSG00000030965.20 | 67.24342 | -0.58754 | 0.954373 | -0.61563 | 0.538137 | 0.983423 | 0.00726 |
| ENSMUSG00000031119.5 | 115.8496 | -0.50985 | 0.804747 | -0.63355 | 0.526373 | 0.983423 | 0.00726 |
| ENSMUSG00000031174.18 | 15.79733 | 1.333666 | 2.243248 | 0.594524 | 0.552161 | 0.983423 | 0.00726 |
| ENSMUSG00000031310.17 | 10.75657 | -1.28294 | 2.251776 | -0.56975 | 0.56885 | 0.983423 | 0.00726 |
| ENSMUSG00000031357.4 | 136.8735 | 0.375268 | 0.650975 | 0.576471 | 0.564297 | 0.983423 | 0.00726 |
| ENSMUSG00000031402.13 | 106.923 | 0.450608 | 0.752274 | 0.598994 | 0.549177 | 0.983423 | 0.00726 |
| ENSMUSG00000031590.9 | 58.98649 | 0.763537 | 1.283288 | 0.594985 | 0.551853 | 0.983423 | 0.00726 |

| | | | | | | | |
|---|---|---|---|---|---|---|---|
| ENSMUSG00000031644.20 | 43.39792 | -0.88787 | 1.354851 | -0.65533 | 0.512258 | 0.983423 | 0.00726 |
| ENSMUSG00000031661.13 | 14.94794 | -1.24506 | 1.963559 | -0.63408 | 0.526027 | 0.983423 | 0.00726 |
| ENSMUSG00000031708.18 | 280.0272 | -0.30156 | 0.470776 | -0.64056 | 0.521805 | 0.983423 | 0.00726 |
| ENSMUSG00000031730.19 | 26.23742 | 1.027845 | 1.752185 | 0.586607 | 0.557467 | 0.983423 | 0.00726 |
| ENSMUSG00000031818.13 | 1952.187 | -0.19714 | 0.340083 | -0.57969 | 0.562126 | 0.983423 | 0.00726 |
| ENSMUSG00000031924.5 | 617.2034 | -0.2292 | 0.3929 | -0.58336 | 0.559648 | 0.983423 | 0.00726 |
| ENSMUSG00000031948.15 | 156.5873 | -0.41537 | 0.712101 | -0.58331 | 0.559685 | 0.983423 | 0.00726 |
| ENSMUSG00000031954.7 | 455.809 | -0.2302 | 0.403094 | -0.57107 | 0.567952 | 0.983423 | 0.00726 |
| ENSMUSG00000031955.11 | 35.93107 | 0.771997 | 1.255537 | 0.614874 | 0.538638 | 0.983423 | 0.00726 |
| ENSMUSG00000031958.17 | 470.2938 | -0.26033 | 0.447398 | -0.58187 | 0.560652 | 0.983423 | 0.00726 |
| ENSMUSG00000031987.7 | 136.1385 | 0.4517 | 0.76379 | 0.591393 | 0.554257 | 0.983423 | 0.00726 |
| ENSMUSG00000031993.8 | 138.8813 | -0.41834 | 0.676181 | -0.61868 | 0.536125 | 0.983423 | 0.00726 |
| ENSMUSG00000032012.10 | 74.14237 | 0.852552 | 1.504755 | 0.566572 | 0.571005 | 0.983423 | 0.00726 |
| ENSMUSG00000032035.17 | 56.8956 | 0.847702 | 1.318284 | 0.643034 | 0.520202 | 0.983423 | 0.00726 |
| ENSMUSG00000032076.21 | 144.1663 | -0.42116 | 0.748781 | -0.56246 | 0.573804 | 0.983423 | 0.00726 |
| ENSMUSG00000032123.6 | 94.84787 | 0.52008 | 0.926632 | 0.561259 | 0.574621 | 0.983423 | 0.00726 |
| ENSMUSG00000032178.15 | 35.93185 | -0.92893 | 1.423071 | -0.65277 | 0.513908 | 0.983423 | 0.00726 |
| ENSMUSG00000032220.11 | 75.95716 | -0.72135 | 1.192716 | -0.60479 | 0.545316 | 0.983423 | 0.00726 |
| ENSMUSG00000032261.17 | 83.98035 | -0.52749 | 0.898956 | -0.58678 | 0.557355 | 0.983423 | 0.00726 |
| ENSMUSG00000032263.15 | 397.7581 | 0.258032 | 0.398709 | 0.647167 | 0.517524 | 0.983423 | 0.00726 |
| ENSMUSG00000032285.16 | 34.91367 | -1.06828 | 1.817564 | -0.58776 | 0.556696 | 0.983423 | 0.00726 |
| ENSMUSG00000032377.9 | 12.64183 | 1.134328 | 1.979538 | 0.573027 | 0.566626 | 0.983423 | 0.00726 |
| ENSMUSG00000032436.18 | 10.51146 | -1.79081 | 2.848619 | -0.62866 | 0.529573 | 0.983423 | 0.00726 |
| ENSMUSG00000032480.18 | 56.38711 | -0.66502 | 1.080468 | -0.61549 | 0.538228 | 0.983423 | 0.00726 |
| ENSMUSG00000032526.12 | 516.1923 | -0.26613 | 0.429886 | -0.61908 | 0.535863 | 0.983423 | 0.00726 |
| ENSMUSG00000032733.7 | 17.41304 | 1.339479 | 2.122374 | 0.631123 | 0.52796 | 0.983423 | 0.00726 |
| ENSMUSG00000032757.7 | 564.5949 | 0.270989 | 0.447757 | 0.605215 | 0.545036 | 0.983423 | 0.00726 |
| ENSMUSG00000032842.14 | 10.78215 | 1.539656 | 2.665743 | 0.577571 | 0.563554 | 0.983423 | 0.00726 |
| ENSMUSG00000033170.15 | 20.92904 | -1.21513 | 2.00162 | -0.60707 | 0.543804 | 0.983423 | 0.00726 |
| ENSMUSG00000033184.15 | 503.8984 | 0.254362 | 0.445048 | 0.571539 | 0.567634 | 0.983423 | 0.00726 |
| ENSMUSG00000033207.8 | 24.76278 | 1.504546 | 2.292935 | 0.656166 | 0.511717 | 0.983423 | 0.00726 |
| ENSMUSG00000033355.7 | 168.3732 | 0.399826 | 0.682969 | 0.585423 | 0.558263 | 0.983423 | 0.00726 |
| ENSMUSG00000033386.11 | 213.9809 | -0.35131 | 0.596451 | -0.58901 | 0.555856 | 0.983423 | 0.00726 |
| ENSMUSG00000033416.16 | 461.4146 | 0.241843 | 0.435575 | 0.555227 | 0.57874 | 0.983423 | 0.00726 |
| ENSMUSG00000033669.9 | 13.18852 | 1.857269 | 3.096058 | 0.599882 | 0.548585 | 0.983423 | 0.00726 |
| ENSMUSG00000033701.14 | 117.3576 | 0.409772 | 0.735833 | 0.556882 | 0.577608 | 0.983423 | 0.00726 |
| ENSMUSG00000033732.11 | 97.20476 | -0.42498 | 0.730657 | -0.58164 | 0.560808 | 0.983423 | 0.00726 |
| ENSMUSG00000033933.14 | 68.85268 | 0.593465 | 1.012656 | 0.586048 | 0.557843 | 0.983423 | 0.00726 |
| ENSMUSG00000033938.6 | 416.3168 | -0.26962 | 0.423465 | -0.63669 | 0.524326 | 0.983423 | 0.00726 |
| ENSMUSG00000034163.11 | 69.41133 | -0.59129 | 1.000314 | -0.59111 | 0.554448 | 0.983423 | 0.00726 |
| ENSMUSG00000034187.19 | 77.50734 | -0.52553 | 0.899243 | -0.58441 | 0.558945 | 0.983423 | 0.00726 |
| ENSMUSG00000034312.16 | 34.40608 | 1.029931 | 1.563375 | 0.658787 | 0.510032 | 0.983423 | 0.00726 |
| ENSMUSG00000034445.9 | 46.3598 | 0.872285 | 1.328042 | 0.65682 | 0.511297 | 0.983423 | 0.00726 |
| ENSMUSG00000034560.7 | 84.88243 | 0.460642 | 0.735016 | 0.62671 | 0.530849 | 0.983423 | 0.00726 |

| | | | | | | | |
|---|---|---|---|---|---|---|---|
| ENSMUSG00000034738.9 | 84.02163 | -0.5619 | 1.010806 | -0.55589 | 0.578287 | 0.983423 | 0.00726 |
| ENSMUSG00000034751.18 | 11.86239 | -1.37892 | 2.443908 | -0.56423 | 0.572599 | 0.983423 | 0.00726 |
| ENSMUSG00000034774.7 | 12.85237 | -1.34532 | 2.35012 | -0.57245 | 0.56702 | 0.983423 | 0.00726 |
| ENSMUSG00000034781.6 | 83.51229 | -0.60688 | 1.043157 | -0.58177 | 0.560723 | 0.983423 | 0.00726 |
| ENSMUSG00000034880.8 | 174.5202 | 0.474645 | 0.727108 | 0.652784 | 0.513895 | 0.983423 | 0.00726 |
| ENSMUSG00000034892.9 | 2340.518 | 0.161486 | 0.273779 | 0.589842 | 0.555297 | 0.983423 | 0.00726 |
| ENSMUSG00000034893.10 | 58.67231 | 0.572711 | 0.902817 | 0.634359 | 0.525846 | 0.983423 | 0.00726 |
| ENSMUSG00000034928.17 | 310.1619 | 0.313557 | 0.531543 | 0.589899 | 0.555258 | 0.983423 | 0.00726 |
| ENSMUSG00000034936.3 | 57.22702 | -1.071 | 1.628579 | -0.65763 | 0.510776 | 0.983423 | 0.00726 |
| ENSMUSG00000035093.17 | 210.1873 | 0.338973 | 0.525612 | 0.64491 | 0.518985 | 0.983423 | 0.00726 |
| ENSMUSG00000035133.10 | 463.4355 | 0.230102 | 0.381111 | 0.603766 | 0.545999 | 0.983423 | 0.00726 |
| ENSMUSG00000035181.7 | 67.12737 | -0.52823 | 0.923824 | -0.57178 | 0.567468 | 0.983423 | 0.00726 |
| ENSMUSG00000035268.15 | 33.8697 | -0.76874 | 1.384568 | -0.55522 | 0.578746 | 0.983423 | 0.00726 |
| ENSMUSG00000035696.16 | 10.47691 | -1.54049 | 2.497608 | -0.61679 | 0.537376 | 0.983423 | 0.00726 |
| ENSMUSG00000035754.9 | 138.0478 | -0.48563 | 0.793362 | -0.61212 | 0.540458 | 0.983423 | 0.00726 |
| ENSMUSG00000035757.18 | 169.4973 | -0.36628 | 0.618913 | -0.59182 | 0.553972 | 0.983423 | 0.00726 |
| ENSMUSG00000035765.11 | 74.03337 | -0.65755 | 1.034672 | -0.63551 | 0.525095 | 0.983423 | 0.00726 |
| ENSMUSG00000035780.3 | 2778.281 | 0.184912 | 0.287745 | 0.642624 | 0.520468 | 0.983423 | 0.00726 |
| ENSMUSG00000035836.3 | 3000.358 | -0.18662 | 0.315335 | -0.59183 | 0.553966 | 0.983423 | 0.00726 |
| ENSMUSG00000035868.10 | 25.66741 | -0.80138 | 1.421718 | -0.56367 | 0.572979 | 0.983423 | 0.00726 |
| ENSMUSG00000035890.10 | 20.51268 | 1.225398 | 2.07774 | 0.589775 | 0.555342 | 0.983423 | 0.00726 |
| ENSMUSG00000035944.18 | 340.653 | -0.25786 | 0.427336 | -0.60341 | 0.546234 | 0.983423 | 0.00726 |
| ENSMUSG00000035992.16 | 219.5918 | -0.40642 | 0.686675 | -0.59187 | 0.55394 | 0.983423 | 0.00726 |
| ENSMUSG00000036002.13 | 107.2348 | -0.53636 | 0.896811 | -0.59807 | 0.549792 | 0.983423 | 0.00726 |
| ENSMUSG00000036019.9 | 36.64402 | 1.057589 | 1.800367 | 0.58743 | 0.556915 | 0.983423 | 0.00726 |
| ENSMUSG00000036188.5 | 57.76384 | -0.53098 | 0.931901 | -0.56979 | 0.568823 | 0.983423 | 0.00726 |
| ENSMUSG00000036199.10 | 1169.077 | -0.18286 | 0.327156 | -0.55893 | 0.576208 | 0.983423 | 0.00726 |
| ENSMUSG00000036270.17 | 15.10393 | 1.353535 | 2.353564 | 0.5751 | 0.565224 | 0.983423 | 0.00726 |
| ENSMUSG00000036315.14 | 178.4156 | 0.478211 | 0.743107 | 0.643529 | 0.519881 | 0.983423 | 0.00726 |
| ENSMUSG00000036362.3 | 14.30971 | 1.774958 | 2.730965 | 0.649938 | 0.515732 | 0.983423 | 0.00726 |
| ENSMUSG00000036655.9 | 116.4897 | 0.437705 | 0.724106 | 0.604477 | 0.545527 | 0.983423 | 0.00726 |
| ENSMUSG00000036698.12 | 123.3737 | -0.46106 | 0.826133 | -0.55809 | 0.57678 | 0.983423 | 0.00726 |
| ENSMUSG00000036769.15 | 27.51063 | 0.774637 | 1.36715 | 0.566607 | 0.570981 | 0.983423 | 0.00726 |
| ENSMUSG00000036779.13 | 85.37861 | 0.445821 | 0.799449 | 0.55766 | 0.577076 | 0.983423 | 0.00726 |
| ENSMUSG00000036782.14 | 10.57973 | -1.58921 | 2.804628 | -0.56664 | 0.570961 | 0.983423 | 0.00726 |
| ENSMUSG00000036792.13 | 36.51961 | 0.972605 | 1.540367 | 0.631411 | 0.527772 | 0.983423 | 0.00726 |
| ENSMUSG00000036840.6 | 39.78241 | -0.77243 | 1.364418 | -0.56612 | 0.57131 | 0.983423 | 0.00726 |
| ENSMUSG00000036850.5 | 152.7194 | 0.399329 | 0.626518 | 0.637379 | 0.523878 | 0.983423 | 0.00726 |
| ENSMUSG00000036860.15 | 128.4457 | 0.382644 | 0.634991 | 0.602597 | 0.546777 | 0.983423 | 0.00726 |
| ENSMUSG00000036940.16 | 32.92866 | -0.78234 | 1.241782 | -0.63001 | 0.528687 | 0.983423 | 0.00726 |
| ENSMUSG00000037022.4 | 115.8923 | -0.43517 | 0.70241 | -0.61953 | 0.535564 | 0.983423 | 0.00726 |
| ENSMUSG00000037072.17 | 1344.364 | -0.16298 | 0.288368 | -0.5652 | 0.571941 | 0.983423 | 0.00726 |
| ENSMUSG00000037089.12 | 89.96442 | -0.54356 | 0.914891 | -0.59412 | 0.552429 | 0.983423 | 0.00726 |
| ENSMUSG00000037172.15 | 19.33789 | 1.116664 | 1.881549 | 0.593481 | 0.552859 | 0.983423 | 0.00726 |

| | | | | | | | |
|---|---|---|---|---|---|---|---|
| ENSMUSG00000037369.18 | 59.44942 | 0.703813 | 1.211664 | 0.580865 | 0.561331 | 0.983423 | 0.00726 |
| ENSMUSG00000037406.8 | 13.01396 | -1.51594 | 2.511818 | -0.60352 | 0.546161 | 0.983423 | 0.00726 |
| ENSMUSG00000037410.14 | 9.940674 | -1.26435 | 1.953759 | -0.64713 | 0.517545 | 0.983423 | 0.00726 |
| ENSMUSG00000037415.8 | 145.0583 | 0.421085 | 0.65718 | 0.640746 | 0.521688 | 0.983423 | 0.00726 |
| ENSMUSG00000037416.14 | 108.2044 | -0.53903 | 0.819244 | -0.65796 | 0.510565 | 0.983423 | 0.00726 |
| ENSMUSG00000037455.17 | 31.33911 | 1.063564 | 1.888561 | 0.563161 | 0.573325 | 0.983423 | 0.00726 |
| ENSMUSG00000037583.3 | 143.6119 | -0.69457 | 1.067657 | -0.65056 | 0.515331 | 0.983423 | 0.00726 |
| ENSMUSG00000037773.15 | 20.62729 | -0.96199 | 1.594927 | -0.60316 | 0.546405 | 0.983423 | 0.00726 |
| ENSMUSG00000037805.16 | 342.3645 | 0.280481 | 0.486032 | 0.577085 | 0.563882 | 0.983423 | 0.00726 |
| ENSMUSG00000037815.8 | 506.5538 | -0.29911 | 0.492461 | -0.60738 | 0.543597 | 0.983423 | 0.00726 |
| ENSMUSG00000037926.16 | 54.08433 | 0.793395 | 1.369942 | 0.579145 | 0.562491 | 0.983423 | 0.00726 |
| ENSMUSG00000038005.16 | 67.34766 | 0.591264 | 0.904902 | 0.653401 | 0.513498 | 0.983423 | 0.00726 |
| ENSMUSG00000038195.7 | 30.00326 | -0.99574 | 1.761967 | -0.56513 | 0.571986 | 0.983423 | 0.00726 |
| ENSMUSG00000038225.16 | 29.97459 | 0.743323 | 1.310354 | 0.567269 | 0.570531 | 0.983423 | 0.00726 |
| ENSMUSG00000038312.11 | 58.01793 | -0.58139 | 0.994744 | -0.58446 | 0.558911 | 0.983423 | 0.00726 |
| ENSMUSG00000038344.15 | 76.76604 | -0.60322 | 1.070448 | -0.56352 | 0.573082 | 0.983423 | 0.00726 |
| ENSMUSG00000038374.11 | 244.061 | 0.367124 | 0.566567 | 0.64798 | 0.516998 | 0.983423 | 0.00726 |
| ENSMUSG00000038403.11 | 308.2592 | 0.319595 | 0.491067 | 0.650818 | 0.515164 | 0.983423 | 0.00726 |
| ENSMUSG00000038417.5 | 67.58833 | -0.58059 | 0.978134 | -0.59357 | 0.552803 | 0.983423 | 0.00726 |
| ENSMUSG00000038526.15 | 173.3268 | -0.39286 | 0.673883 | -0.58297 | 0.559911 | 0.983423 | 0.00726 |
| ENSMUSG00000038591.4 | 165.1086 | -0.46519 | 0.790898 | -0.58818 | 0.556414 | 0.983423 | 0.00726 |
| ENSMUSG00000038648.6 | 76.82564 | 0.511809 | 0.850523 | 0.601757 | 0.547336 | 0.983423 | 0.00726 |
| ENSMUSG00000038650.16 | 151.6928 | -0.60136 | 0.943528 | -0.63735 | 0.523897 | 0.983423 | 0.00726 |
| ENSMUSG00000038745.16 | 123.2588 | -0.44959 | 0.704987 | -0.63773 | 0.523651 | 0.983423 | 0.00726 |
| ENSMUSG00000038812.17 | 77.20609 | 0.503276 | 0.89775 | 0.560598 | 0.575072 | 0.983423 | 0.00726 |
| ENSMUSG00000038886.11 | 76.38349 | -0.77199 | 1.282251 | -0.60206 | 0.547135 | 0.983423 | 0.00726 |
| ENSMUSG00000039103.13 | 37.81282 | 0.813713 | 1.437694 | 0.565985 | 0.571404 | 0.983423 | 0.00726 |
| ENSMUSG00000039105.7 | 304.5919 | 0.308068 | 0.469902 | 0.655601 | 0.512081 | 0.983423 | 0.00726 |
| ENSMUSG00000039163.11 | 286.7315 | 0.312364 | 0.534188 | 0.584746 | 0.558719 | 0.983423 | 0.00726 |
| ENSMUSG00000039218.18 | 189.1949 | -0.43736 | 0.740786 | -0.5904 | 0.554923 | 0.983423 | 0.00726 |
| ENSMUSG00000039219.19 | 288.1213 | 0.343543 | 0.540464 | 0.635645 | 0.525008 | 0.983423 | 0.00726 |
| ENSMUSG00000039233.13 | 114.5396 | -0.44762 | 0.742085 | -0.6032 | 0.546378 | 0.983423 | 0.00726 |
| ENSMUSG00000039246.9 | 90.67466 | -0.54635 | 0.853385 | -0.64022 | 0.522031 | 0.983423 | 0.00726 |
| ENSMUSG00000039262.17 | 86.53212 | -0.47785 | 0.851381 | -0.56126 | 0.574619 | 0.983423 | 0.00726 |
| ENSMUSG00000039356.16 | 36.69275 | 0.874755 | 1.492468 | 0.586113 | 0.557799 | 0.983423 | 0.00726 |
| ENSMUSG00000039782.16 | 105.6092 | -0.53446 | 0.851137 | -0.62794 | 0.530045 | 0.983423 | 0.00726 |
| ENSMUSG00000039849.7 | 67.19663 | -0.69032 | 1.19571 | -0.57733 | 0.563718 | 0.983423 | 0.00726 |
| ENSMUSG00000039865.9 | 19.90142 | 1.437663 | 2.192661 | 0.65567 | 0.512036 | 0.983423 | 0.00726 |
| ENSMUSG00000039997.17 | 18.1457 | -1.38418 | 2.16081 | -0.64058 | 0.521793 | 0.983423 | 0.00726 |
| ENSMUSG00000040297.13 | 27.60063 | -0.82113 | 1.29324 | -0.63494 | 0.525468 | 0.983423 | 0.00726 |
| ENSMUSG00000040462.14 | 130.2543 | -0.54118 | 0.925244 | -0.5849 | 0.558613 | 0.983423 | 0.00726 |
| ENSMUSG00000040488.19 | 38.81787 | -0.7582 | 1.267005 | -0.59842 | 0.549563 | 0.983423 | 0.00726 |
| ENSMUSG00000040511.15 | 24.6711 | -1.07028 | 1.758529 | -0.60862 | 0.542775 | 0.983423 | 0.00726 |
| ENSMUSG00000040552.9 | 32.89407 | 1.115723 | 1.938189 | 0.575652 | 0.56485 | 0.983423 | 0.00726 |

| | | | | | | | |
|---|---|---|---|---|---|---|---|
| ENSMUSG00000040584.9 | 20.17492 | -1.13031 | 1.71369 | -0.65958 | 0.509526 | 0.983423 | 0.00726 |
| ENSMUSG00000040624.19 | 13.70878 | -1.88473 | 3.024167 | -0.62322 | 0.533137 | 0.983423 | 0.00726 |
| ENSMUSG00000040697.11 | 61.21093 | -0.67435 | 1.1493 | -0.58675 | 0.557372 | 0.983423 | 0.00726 |
| ENSMUSG00000040724.6 | 23.22298 | -1.42778 | 2.303089 | -0.61994 | 0.535297 | 0.983423 | 0.00726 |
| ENSMUSG00000040746.16 | 53.06253 | -0.98008 | 1.493439 | -0.65625 | 0.51166 | 0.983423 | 0.00726 |
| ENSMUSG00000040774.16 | 341.9339 | 0.338829 | 0.59764 | 0.566945 | 0.570752 | 0.983423 | 0.00726 |
| ENSMUSG00000040848.12 | 103.8395 | 0.50702 | 0.905877 | 0.5597 | 0.575684 | 0.983423 | 0.00726 |
| ENSMUSG00000041025.16 | 20.14113 | -1.13006 | 1.900631 | -0.59457 | 0.552132 | 0.983423 | 0.00726 |
| ENSMUSG00000041035.10 | 19.15468 | 1.052816 | 1.860237 | 0.565958 | 0.571422 | 0.983423 | 0.00726 |
| ENSMUSG00000041115.17 | 16.7701 | -1.0005 | 1.683621 | -0.59425 | 0.552342 | 0.983423 | 0.00726 |
| ENSMUSG00000041119.13 | 51.67661 | 0.641666 | 1.1205 | 0.572661 | 0.566875 | 0.983423 | 0.00726 |
| ENSMUSG00000041293.6 | 21.03657 | -1.24363 | 2.15521 | -0.57704 | 0.563916 | 0.983423 | 0.00726 |
| ENSMUSG00000041360.9 | 124.7151 | 0.505764 | 0.84152 | 0.601012 | 0.547832 | 0.983423 | 0.00726 |
| ENSMUSG00000041408.19 | 441.0413 | 0.264903 | 0.473627 | 0.559306 | 0.575953 | 0.983423 | 0.00726 |
| ENSMUSG00000041530.18 | 42.12095 | 0.716493 | 1.257443 | 0.569802 | 0.568812 | 0.983423 | 0.00726 |
| ENSMUSG00000041977.19 | 11.98341 | -1.14265 | 1.890334 | -0.60447 | 0.545533 | 0.983423 | 0.00726 |
| ENSMUSG00000042340.7 | 22.80643 | -1.02262 | 1.807042 | -0.56591 | 0.571457 | 0.983423 | 0.00726 |
| ENSMUSG00000042354.8 | 281.1733 | -0.41133 | 0.671455 | -0.6126 | 0.54014 | 0.983423 | 0.00726 |
| ENSMUSG00000042380.9 | 53.13769 | -1.03263 | 1.652382 | -0.62493 | 0.532014 | 0.983423 | 0.00726 |
| ENSMUSG00000042410.17 | 52.11986 | -0.66511 | 1.142148 | -0.58233 | 0.560343 | 0.983423 | 0.00726 |
| ENSMUSG00000042446.17 | 20.72362 | -0.82011 | 1.471137 | -0.55747 | 0.577209 | 0.983423 | 0.00726 |
| ENSMUSG00000042595.8 | 91.65488 | -0.4743 | 0.846285 | -0.56045 | 0.575172 | 0.983423 | 0.00726 |
| ENSMUSG00000042628.9 | 38.67689 | -0.79672 | 1.367362 | -0.58267 | 0.560116 | 0.983423 | 0.00726 |
| ENSMUSG00000042659.16 | 21.04722 | 1.131961 | 1.827607 | 0.619368 | 0.535674 | 0.983423 | 0.00726 |
| ENSMUSG00000042719.18 | 153.046 | 0.595349 | 0.959691 | 0.620355 | 0.535024 | 0.983423 | 0.00726 |
| ENSMUSG00000042744.17 | 44.29637 | 0.630522 | 1.090335 | 0.578283 | 0.563073 | 0.983423 | 0.00726 |
| ENSMUSG00000042747.13 | 775.601 | -0.23919 | 0.426449 | -0.56088 | 0.574876 | 0.983423 | 0.00726 |
| ENSMUSG00000042797.10 | 202.4026 | 0.360777 | 0.576263 | 0.626063 | 0.531274 | 0.983423 | 0.00726 |
| ENSMUSG00000043154.16 | 14.26406 | -1.20743 | 2.044763 | -0.5905 | 0.554855 | 0.983423 | 0.00726 |
| ENSMUSG00000043207.11 | 244.9299 | -0.33797 | 0.53057 | -0.63699 | 0.524128 | 0.983423 | 0.00726 |
| ENSMUSG00000043510.13 | 12.8951 | -1.40984 | 2.410935 | -0.58477 | 0.558704 | 0.983423 | 0.00726 |
| ENSMUSG00000044345.10 | 22.15406 | 1.011911 | 1.730393 | 0.584787 | 0.558691 | 0.983423 | 0.00726 |
| ENSMUSG00000044477.13 | 186.9425 | -0.41825 | 0.676771 | -0.61801 | 0.536568 | 0.983423 | 0.00726 |
| ENSMUSG00000044749.14 | 396.5609 | -0.32777 | 0.562505 | -0.5827 | 0.560095 | 0.983423 | 0.00726 |
| ENSMUSG00000044982.8 | 21.76488 | 1.237863 | 1.939807 | 0.638137 | 0.523384 | 0.983423 | 0.00726 |
| ENSMUSG00000045038.15 | 50.98667 | -0.97612 | 1.490283 | -0.65499 | 0.512472 | 0.983423 | 0.00726 |
| ENSMUSG00000045094.9 | 14.98302 | -1.25021 | 2.246383 | -0.55654 | 0.57784 | 0.983423 | 0.00726 |
| ENSMUSG00000045211.6 | 26.16283 | 0.900692 | 1.423724 | 0.632631 | 0.526975 | 0.983423 | 0.00726 |
| ENSMUSG00000045435.10 | 75.5132 | -0.46571 | 0.789105 | -0.59018 | 0.55507 | 0.983423 | 0.00726 |
| ENSMUSG00000045482.17 | 19.05352 | 1.039431 | 1.604438 | 0.647848 | 0.517084 | 0.983423 | 0.00726 |
| ENSMUSG00000045624.16 | 312.1738 | -0.30116 | 0.487689 | -0.61752 | 0.536895 | 0.983423 | 0.00726 |
| ENSMUSG00000045658.17 | 869.2572 | -0.18813 | 0.301746 | -0.62348 | 0.532972 | 0.983423 | 0.00726 |
| ENSMUSG00000045896.15 | 13.10643 | 1.353973 | 2.062278 | 0.656542 | 0.511475 | 0.983423 | 0.00726 |
| ENSMUSG00000046079.17 | 110.5851 | -0.46416 | 0.835844 | -0.55532 | 0.578674 | 0.983423 | 0.00726 |

| | | | | | | | |
|---|---|---|---|---|---|---|---|
| ENSMUSG00000046139.9 | 60.18502 | 0.600701 | 0.920082 | 0.652878 | 0.513835 | 0.983423 | 0.00726 |
| ENSMUSG00000046519.16 | 27.45757 | -0.86534 | 1.389955 | -0.62257 | 0.533568 | 0.983423 | 0.00726 |
| ENSMUSG00000046791.9 | 37.78948 | -1.09934 | 1.837237 | -0.59837 | 0.549595 | 0.983423 | 0.00726 |
| ENSMUSG00000047728.14 | 320.968 | -0.29334 | 0.47503 | -0.61753 | 0.536888 | 0.983423 | 0.00726 |
| ENSMUSG00000047843.17 | 17.97272 | 1.173393 | 1.963892 | 0.597484 | 0.550185 | 0.983423 | 0.00726 |
| ENSMUSG00000047888.11 | 97.06537 | -0.41596 | 0.702909 | -0.59178 | 0.554 | 0.983423 | 0.00726 |
| ENSMUSG00000048047.4 | 71.10862 | -0.63792 | 1.036148 | -0.61566 | 0.538117 | 0.983423 | 0.00726 |
| ENSMUSG00000048126.17 | 31.19905 | -0.97031 | 1.572175 | -0.61718 | 0.537119 | 0.983423 | 0.00726 |
| ENSMUSG00000048489.13 | 253.0879 | -0.43354 | 0.738718 | -0.58688 | 0.557286 | 0.983423 | 0.00726 |
| ENSMUSG00000048772.16 | 50.57486 | 0.748348 | 1.184524 | 0.631771 | 0.527537 | 0.983423 | 0.00726 |
| ENSMUSG00000049421.14 | 84.51997 | -0.65602 | 1.16452 | -0.56334 | 0.573205 | 0.983423 | 0.00726 |
| ENSMUSG00000049482.17 | 42.25536 | 0.802523 | 1.385939 | 0.579046 | 0.562558 | 0.983423 | 0.00726 |
| ENSMUSG00000049489.6 | 37.62411 | 0.834602 | 1.374766 | 0.607086 | 0.543794 | 0.983423 | 0.00726 |
| ENSMUSG00000050299.9 | 235.5766 | -0.30773 | 0.496575 | -0.61971 | 0.53545 | 0.983423 | 0.00726 |
| ENSMUSG00000050627.14 | 43.83475 | 0.702049 | 1.117878 | 0.628019 | 0.529991 | 0.983423 | 0.00726 |
| ENSMUSG00000050730.18 | 232.4964 | -0.37606 | 0.577994 | -0.65063 | 0.515288 | 0.983423 | 0.00726 |
| ENSMUSG00000050953.11 | 38.69782 | 0.885952 | 1.505662 | 0.588414 | 0.556255 | 0.983423 | 0.00726 |
| ENSMUSG00000051147.11 | 102.9484 | 0.407436 | 0.708641 | 0.574954 | 0.565322 | 0.983423 | 0.00726 |
| ENSMUSG00000051285.18 | 287.9635 | 0.356736 | 0.573041 | 0.622531 | 0.533593 | 0.983423 | 0.00726 |
| ENSMUSG00000051316.9 | 18.38832 | 1.336874 | 2.237103 | 0.597591 | 0.550113 | 0.983423 | 0.00726 |
| ENSMUSG00000051495.9 | 110.3323 | -0.52926 | 0.860042 | -0.61539 | 0.538297 | 0.983423 | 0.00726 |
| ENSMUSG00000051586.18 | 19.76217 | 1.13421 | 1.932452 | 0.586928 | 0.557252 | 0.983423 | 0.00726 |
| ENSMUSG00000051721.7 | 50.96539 | 0.643238 | 1.125706 | 0.571409 | 0.567723 | 0.983423 | 0.00726 |
| ENSMUSG00000052369.9 | 21.00957 | 1.096495 | 1.722091 | 0.636723 | 0.524305 | 0.983423 | 0.00726 |
| ENSMUSG00000052794.14 | 27.28522 | -1.08008 | 1.925826 | -0.56084 | 0.574906 | 0.983423 | 0.00726 |
| ENSMUSG00000053134.14 | 46.83881 | -0.72094 | 1.152219 | -0.6257 | 0.531511 | 0.983423 | 0.00726 |
| ENSMUSG00000053565.11 | 113.472 | -0.41301 | 0.672546 | -0.6141 | 0.539151 | 0.983423 | 0.00726 |
| ENSMUSG00000053768.14 | 555.7921 | 0.208023 | 0.359094 | 0.5793 | 0.562387 | 0.983423 | 0.00726 |
| ENSMUSG00000054203.8 | 25.42326 | -1.38392 | 2.284864 | -0.60569 | 0.544722 | 0.983423 | 0.00726 |
| ENSMUSG00000054619.8 | 2371.584 | -0.16969 | 0.263855 | -0.64312 | 0.520146 | 0.983423 | 0.00726 |
| ENSMUSG00000054843.10 | 111.557 | 0.44248 | 0.689219 | 0.642002 | 0.520872 | 0.983423 | 0.00726 |
| ENSMUSG00000055013.17 | 63.48259 | -0.55266 | 0.901061 | -0.61335 | 0.539648 | 0.983423 | 0.00726 |
| ENSMUSG00000055184.5 | 13.13168 | 1.647985 | 2.660289 | 0.619476 | 0.535603 | 0.983423 | 0.00726 |
| ENSMUSG00000055720.11 | 72.27755 | -0.60936 | 1.093917 | -0.55704 | 0.5775 | 0.983423 | 0.00726 |
| ENSMUSG00000055900.15 | 31.57353 | 0.989569 | 1.527861 | 0.647682 | 0.51719 | 0.983423 | 0.00726 |
| ENSMUSG00000056131.14 | 114.6715 | 0.532408 | 0.853496 | 0.623797 | 0.532761 | 0.983423 | 0.00726 |
| ENSMUSG00000056608.15 | 78.52165 | -0.60846 | 0.941449 | -0.6463 | 0.518086 | 0.983423 | 0.00726 |
| ENSMUSG00000056851.15 | 1185.787 | 0.165621 | 0.289794 | 0.571512 | 0.567653 | 0.983423 | 0.00726 |
| ENSMUSG00000056899.11 | 171.5302 | 0.382967 | 0.630114 | 0.607775 | 0.543337 | 0.983423 | 0.00726 |
| ENSMUSG00000056938.17 | 42.51266 | 0.94396 | 1.439494 | 0.655758 | 0.51198 | 0.983423 | 0.00726 |
| ENSMUSG00000057113.14 | 601.6277 | 0.223817 | 0.350562 | 0.638451 | 0.52318 | 0.983423 | 0.00726 |
| ENSMUSG00000057342.16 | 80.56514 | 0.484155 | 0.799056 | 0.605909 | 0.544575 | 0.983423 | 0.00726 |
| ENSMUSG00000057469.9 | 88.56932 | 0.636713 | 1.021999 | 0.623008 | 0.533279 | 0.983423 | 0.00726 |
| ENSMUSG00000057561.11 | 599.2971 | 0.270595 | 0.421613 | 0.641808 | 0.520998 | 0.983423 | 0.00726 |

| | | | | | | | |
|---|---|---|---|---|---|---|---|
| ENSMUSG00000057691.9 | 39.53614 | -0.77274 | 1.199315 | -0.64431 | 0.519372 | 0.983423 | 0.00726 |
| ENSMUSG00000057835.8 | 26.44024 | 0.954814 | 1.618729 | 0.589854 | 0.555289 | 0.983423 | 0.00726 |
| ENSMUSG00000058835.15 | 215.7449 | 0.32755 | 0.562199 | 0.582623 | 0.560147 | 0.983423 | 0.00726 |
| ENSMUSG00000058921.4 | 93.17681 | 0.683942 | 1.188608 | 0.575415 | 0.565011 | 0.983423 | 0.00726 |
| ENSMUSG00000058934.9 | 9.944938 | 1.624925 | 2.9217 | 0.556157 | 0.578103 | 0.983423 | 0.00726 |
| ENSMUSG00000059263.10 | 355.6611 | 0.257325 | 0.428857 | 0.600026 | 0.548489 | 0.983423 | 0.00726 |
| ENSMUSG00000059278.10 | 359.8268 | 0.373385 | 0.616789 | 0.605369 | 0.544934 | 0.983423 | 0.00726 |
| ENSMUSG00000059325.15 | 1222.044 | -0.20244 | 0.359956 | -0.56241 | 0.573837 | 0.983423 | 0.00726 |
| ENSMUSG00000059482.16 | 32.11146 | 0.978249 | 1.586781 | 0.616499 | 0.537565 | 0.983423 | 0.00726 |
| ENSMUSG00000059552.14 | 43.03228 | -1.13066 | 1.746003 | -0.64757 | 0.517261 | 0.983423 | 0.00726 |
| ENSMUSG00000059734.8 | 379.8227 | -0.22312 | 0.392234 | -0.56885 | 0.569458 | 0.983423 | 0.00726 |
| ENSMUSG00000059811.14 | 725.0358 | 0.180685 | 0.311066 | 0.580856 | 0.561337 | 0.983423 | 0.00726 |
| ENSMUSG00000060407.7 | 1925.113 | 0.14451 | 0.253242 | 0.570639 | 0.568244 | 0.983423 | 0.00726 |
| ENSMUSG00000060591.10 | 366.0611 | -0.24954 | 0.410016 | -0.6086 | 0.542787 | 0.983423 | 0.00726 |
| ENSMUSG00000060639.6 | 11.89807 | 1.611022 | 2.441686 | 0.659799 | 0.509383 | 0.983423 | 0.00726 |
| ENSMUSG00000060904.15 | 516.8148 | 0.251492 | 0.38542 | 0.652515 | 0.514069 | 0.983423 | 0.00726 |
| ENSMUSG00000061119.8 | 16.2472 | -1.41672 | 2.263568 | -0.62588 | 0.531395 | 0.983423 | 0.00726 |
| ENSMUSG00000061313.11 | 83.95212 | -0.49812 | 0.837776 | -0.59457 | 0.552128 | 0.983423 | 0.00726 |
| ENSMUSG00000061315.15 | 1485.668 | 0.18707 | 0.294994 | 0.634149 | 0.525984 | 0.983423 | 0.00726 |
| ENSMUSG00000061455.14 | 92.30501 | -0.57038 | 0.966257 | -0.5903 | 0.554991 | 0.983423 | 0.00726 |
| ENSMUSG00000061665.8 | 226.4146 | 0.430495 | 0.68619 | 0.627369 | 0.530417 | 0.983423 | 0.00726 |
| ENSMUSG00000061689.16 | 27.86424 | -1.06848 | 1.889221 | -0.56557 | 0.571689 | 0.983423 | 0.00726 |
| ENSMUSG00000061778.11 | 97.29317 | 0.56483 | 0.974084 | 0.579858 | 0.56201 | 0.983423 | 0.00726 |
| ENSMUSG00000061838.8 | 646.0893 | -0.26376 | 0.405545 | -0.65039 | 0.515443 | 0.983423 | 0.00726 |
| ENSMUSG00000062210.14 | 18.51694 | -1.30059 | 2.198739 | -0.59152 | 0.554174 | 0.983423 | 0.00726 |
| ENSMUSG00000062585.12 | 12.19312 | 1.400225 | 2.41414 | 0.58001 | 0.561908 | 0.983423 | 0.00726 |
| ENSMUSG00000062611.5 | 31.39363 | 0.93565 | 1.425031 | 0.656582 | 0.51145 | 0.983423 | 0.00726 |
| ENSMUSG00000062691.11 | 95.7288 | 0.40004 | 0.704374 | 0.567938 | 0.570077 | 0.983423 | 0.00726 |
| ENSMUSG00000062997.7 | 277.3253 | 0.413189 | 0.705893 | 0.585343 | 0.558317 | 0.983423 | 0.00726 |
| ENSMUSG00000063229.16 | 2656.563 | 0.14306 | 0.245974 | 0.581606 | 0.560832 | 0.983423 | 0.00726 |
| ENSMUSG00000063488.7 | 27.27736 | 0.801558 | 1.319568 | 0.60744 | 0.543559 | 0.983423 | 0.00726 |
| ENSMUSG00000063787.5 | 542.7932 | 0.249942 | 0.406678 | 0.614594 | 0.538823 | 0.983423 | 0.00726 |
| ENSMUSG00000063808.15 | 70.99303 | 0.634673 | 1.059274 | 0.599159 | 0.549067 | 0.983423 | 0.00726 |
| ENSMUSG00000063888.8 | 278.6018 | -0.36802 | 0.574057 | -0.64109 | 0.521466 | 0.983423 | 0.00726 |
| ENSMUSG00000063953.4 | 19.91712 | -0.86492 | 1.405713 | -0.61529 | 0.538363 | 0.983423 | 0.00726 |
| ENSMUSG00000064068.16 | 53.48871 | -0.99671 | 1.519854 | -0.65579 | 0.511957 | 0.983423 | 0.00726 |
| ENSMUSG00000064343.1 | 12.74771 | 1.551569 | 2.690402 | 0.576705 | 0.564139 | 0.983423 | 0.00726 |
| ENSMUSG00000064368.1 | 1734.186 | 0.193548 | 0.303563 | 0.637587 | 0.523743 | 0.983423 | 0.00726 |
| ENSMUSG00000066441.15 | 326.9305 | 0.319769 | 0.485592 | 0.658515 | 0.510208 | 0.983423 | 0.00726 |
| ENSMUSG00000067148.16 | 130.2058 | -0.41601 | 0.731206 | -0.56894 | 0.569396 | 0.983423 | 0.00726 |
| ENSMUSG00000067225.3 | 1553.172 | -0.16896 | 0.284636 | -0.59361 | 0.552772 | 0.983423 | 0.00726 |
| ENSMUSG00000067567.13 | 19.28163 | 1.408424 | 2.486741 | 0.566373 | 0.57114 | 0.983423 | 0.00726 |
| ENSMUSG00000068036.17 | 284.7155 | -0.28688 | 0.495019 | -0.57954 | 0.562226 | 0.983423 | 0.00726 |
| ENSMUSG00000068391.9 | 70.66311 | 0.525876 | 0.946405 | 0.555657 | 0.578446 | 0.983423 | 0.00726 |

| | | | | | | | |
|---|---|---|---|---|---|---|---|
| ENSMUSG00000068732.6 | 52.7999 | -0.81734 | 1.281156 | -0.63797 | 0.523494 | 0.983423 | 0.00726 |
| ENSMUSG00000068762.12 | 274.5877 | -0.42949 | 0.664231 | -0.6466 | 0.517893 | 0.983423 | 0.00726 |
| ENSMUSG00000068854.8 | 42.92987 | -2.20234 | 3.732439 | -0.59005 | 0.555154 | 0.983423 | 0.00726 |
| ENSMUSG00000068882.14 | 768.2126 | 0.196323 | 0.353443 | 0.555458 | 0.578581 | 0.983423 | 0.00726 |
| ENSMUSG00000068923.15 | 10.82624 | -1.41237 | 2.240111 | -0.63049 | 0.528374 | 0.983423 | 0.00726 |
| ENSMUSG00000069135.12 | 121.771 | 0.476406 | 0.731535 | 0.651242 | 0.51489 | 0.983423 | 0.00726 |
| ENSMUSG00000069633.13 | 112.7076 | -0.51777 | 0.783701 | -0.66068 | 0.508819 | 0.983423 | 0.00726 |
| ENSMUSG00000070315.13 | 36.3264 | -0.91215 | 1.499657 | -0.60824 | 0.543028 | 0.983423 | 0.00726 |
| ENSMUSG00000070427.5 | 36.28981 | 0.75334 | 1.345334 | 0.559965 | 0.575503 | 0.983423 | 0.00726 |
| ENSMUSG00000070594.11 | 706.8241 | -1.12028 | 1.895975 | -0.59087 | 0.554605 | 0.983423 | 0.00726 |
| ENSMUSG00000070730.12 | 242.8796 | -0.3675 | 0.594249 | -0.61843 | 0.536291 | 0.983423 | 0.00726 |
| ENSMUSG00000070953.14 | 88.61782 | -0.5465 | 0.967386 | -0.56492 | 0.572129 | 0.983423 | 0.00726 |
| ENSMUSG00000071415.7 | 4782.786 | 0.155021 | 0.261488 | 0.592843 | 0.553286 | 0.983423 | 0.00726 |
| ENSMUSG00000071454.14 | 24.6184 | -1.1127 | 1.93626 | -0.57466 | 0.565518 | 0.983423 | 0.00726 |
| ENSMUSG00000071497.5 | 16.89645 | -0.91127 | 1.549453 | -0.58812 | 0.55645 | 0.983423 | 0.00726 |
| ENSMUSG00000071662.5 | 93.33627 | 0.470215 | 0.722786 | 0.650559 | 0.515331 | 0.983423 | 0.00726 |
| ENSMUSG00000072889.10 | 32.15563 | 0.97433 | 1.6414 | 0.593597 | 0.552782 | 0.983423 | 0.00726 |
| ENSMUSG00000072964.15 | 13.16464 | 1.302054 | 2.331061 | 0.558567 | 0.576457 | 0.983423 | 0.00726 |
| ENSMUSG00000073481.10 | 3666.447 | -0.13419 | 0.219093 | -0.6125 | 0.540206 | 0.983423 | 0.00726 |
| ENSMUSG00000073640.4 | 479.4494 | 0.251036 | 0.386915 | 0.648815 | 0.516458 | 0.983423 | 0.00726 |
| ENSMUSG00000074088.7 | 21.24947 | -1.16481 | 1.785136 | -0.6525 | 0.514076 | 0.983423 | 0.00726 |
| ENSMUSG00000074151.14 | 12.26681 | -1.48427 | 2.448178 | -0.60628 | 0.544331 | 0.983423 | 0.00726 |
| ENSMUSG00000074284.7 | 28.90966 | 0.967715 | 1.706262 | 0.567155 | 0.570609 | 0.983423 | 0.00726 |
| ENSMUSG00000074781.6 | 44.41012 | 0.688903 | 1.108704 | 0.621359 | 0.534364 | 0.983423 | 0.00726 |
| ENSMUSG00000074825.5 | 23.08261 | -1.24514 | 2.105265 | -0.59144 | 0.554224 | 0.983423 | 0.00726 |
| ENSMUSG00000075227.7 | 38.19061 | 0.720246 | 1.15303 | 0.624655 | 0.532197 | 0.983423 | 0.00726 |
| ENSMUSG00000075229.12 | 391.4784 | 0.291724 | 0.479674 | 0.608171 | 0.543074 | 0.983423 | 0.00726 |
| ENSMUSG00000075376.11 | 184.6912 | 0.326875 | 0.518413 | 0.630529 | 0.528349 | 0.983423 | 0.00726 |
| ENSMUSG00000075415.14 | 48.29953 | -0.66334 | 1.11703 | -0.59384 | 0.552618 | 0.983423 | 0.00726 |
| ENSMUSG00000075467.5 | 41.18453 | -0.81223 | 1.253473 | -0.64798 | 0.516996 | 0.983423 | 0.00726 |
| ENSMUSG00000075517.6 | 84.14588 | 0.564411 | 0.867339 | 0.650739 | 0.515215 | 0.983423 | 0.00726 |
| ENSMUSG00000075704.18 | 47.33836 | 0.69486 | 1.100502 | 0.631403 | 0.527777 | 0.983423 | 0.00726 |
| ENSMUSG00000076441.10 | 2223.885 | -0.19729 | 0.327826 | -0.60182 | 0.547293 | 0.983423 | 0.00726 |
| ENSMUSG00000078153.4 | 119.7022 | 0.490634 | 0.846261 | 0.579767 | 0.562072 | 0.983423 | 0.00726 |
| ENSMUSG00000078517.13 | 149.1935 | -0.70136 | 1.13832 | -0.61613 | 0.537806 | 0.983423 | 0.00726 |
| ENSMUSG00000078681.11 | 111.8294 | 0.547059 | 0.896411 | 0.610277 | 0.541679 | 0.983423 | 0.00726 |
| ENSMUSG00000078784.9 | 233.5335 | 0.321526 | 0.542108 | 0.593103 | 0.553112 | 0.983423 | 0.00726 |
| ENSMUSG00000079184.11 | 284.2312 | 0.312321 | 0.559268 | 0.558446 | 0.57654 | 0.983423 | 0.00726 |
| ENSMUSG00000079357.10 | 10.32041 | -1.6044 | 2.546003 | -0.63016 | 0.528588 | 0.983423 | 0.00726 |
| ENSMUSG00000080727.3 | 19.76582 | 1.287062 | 2.314226 | 0.556152 | 0.578107 | 0.983423 | 0.00726 |
| ENSMUSG00000080904.2 | 442.422 | 0.216115 | 0.38822 | 0.556682 | 0.577744 | 0.983423 | 0.00726 |
| ENSMUSG00000081207.5 | 521.5651 | -0.25743 | 0.408544 | -0.63011 | 0.528625 | 0.983423 | 0.00726 |
| ENSMUSG00000082363.2 | 17.73339 | -0.8799 | 1.487846 | -0.59139 | 0.554257 | 0.983423 | 0.00726 |
| ENSMUSG00000084309.2 | 13.1034 | -1.50884 | 2.690106 | -0.56089 | 0.574876 | 0.983423 | 0.00726 |

| | | | | | | | |
|---|---|---|---|---|---|---|---|
| ENSMUSG00000084771.9 | 11.36227 | -1.93271 | 3.204319 | -0.60316 | 0.546404 | 0.983423 | 0.00726 |
| ENSMUSG00000085037.2 | 22.7819 | -1.20301 | 2.018228 | -0.59607 | 0.551128 | 0.983423 | 0.00726 |
| ENSMUSG00000086054.3 | 36.4218 | 0.926699 | 1.648183 | 0.562255 | 0.573942 | 0.983423 | 0.00726 |
| ENSMUSG00000086290.10 | 68.57453 | -0.5855 | 1.012076 | -0.57852 | 0.562915 | 0.983423 | 0.00726 |
| ENSMUSG00000087260.6 | 279.8714 | -0.32228 | 0.520769 | -0.61885 | 0.536016 | 0.983423 | 0.00726 |
| ENSMUSG00000087412.5 | 53.93931 | -0.64053 | 1.081508 | -0.59226 | 0.553677 | 0.983423 | 0.00726 |
| ENSMUSG00000089774.3 | 15.74779 | 1.293839 | 2.197626 | 0.588744 | 0.556033 | 0.983423 | 0.00726 |
| ENSMUSG00000089902.2 | 12.83901 | 1.563017 | 2.626818 | 0.595023 | 0.551828 | 0.983423 | 0.00726 |
| ENSMUSG00000090035.4 | 74.19611 | -0.4665 | 0.797917 | -0.58465 | 0.558781 | 0.983423 | 0.00726 |
| ENSMUSG00000090555.2 | 79.09057 | -0.60411 | 0.996842 | -0.60603 | 0.544497 | 0.983423 | 0.00726 |
| ENSMUSG00000091387.3 | 23.44792 | -0.96601 | 1.719304 | -0.56186 | 0.574211 | 0.983423 | 0.00726 |
| ENSMUSG00000091509.8 | 232.8338 | 0.387237 | 0.654735 | 0.591441 | 0.554225 | 0.983423 | 0.00726 |
| ENSMUSG00000092203.8 | 84.71174 | 0.548684 | 0.847043 | 0.647765 | 0.517137 | 0.983423 | 0.00726 |
| ENSMUSG00000097048.8 | 39.62641 | 0.729267 | 1.13332 | 0.643479 | 0.519914 | 0.983423 | 0.00726 |
| ENSMUSG00000097378.9 | 24.48591 | -1.2433 | 2.076408 | -0.59878 | 0.549322 | 0.983423 | 0.00726 |
| ENSMUSG00000097503.3 | 33.04241 | 1.270011 | 2.137248 | 0.594228 | 0.55236 | 0.983423 | 0.00726 |
| ENSMUSG00000097730.4 | 25.50244 | 1.015632 | 1.569192 | 0.647233 | 0.517481 | 0.983423 | 0.00726 |
| ENSMUSG00000098557.2 | 62.43281 | 0.611677 | 0.973061 | 0.628611 | 0.529603 | 0.983423 | 0.00726 |
| ENSMUSG00000099471.2 | 60.33786 | 0.655889 | 1.047163 | 0.626349 | 0.531086 | 0.983423 | 0.00726 |
| ENSMUSG00000100826.7 | 15.58004 | -1.44793 | 2.343457 | -0.61786 | 0.536668 | 0.983423 | 0.00726 |
| ENSMUSG00000102644.6 | 42.78212 | 0.73837 | 1.222956 | 0.603759 | 0.546004 | 0.983423 | 0.00726 |
| ENSMUSG00000104459.2 | 10.80228 | 1.725145 | 2.862835 | 0.6026 | 0.546775 | 0.983423 | 0.00726 |
| ENSMUSG00000106864.4 | 162.5395 | -0.62477 | 0.949438 | -0.65804 | 0.51051 | 0.983423 | 0.00726 |
| ENSMUSG00000106990.2 | 15.90402 | -1.21183 | 2.067463 | -0.58615 | 0.557778 | 0.983423 | 0.00726 |
| ENSMUSG00000108659.2 | 27.21408 | 1.335395 | 2.28377 | 0.584733 | 0.558728 | 0.983423 | 0.00726 |
| ENSMUSG00000109482.3 | 147.8672 | 0.403002 | 0.632828 | 0.636827 | 0.524238 | 0.983423 | 0.00726 |
| ENSMUSG00000109881.2 | 70.31774 | -0.48343 | 0.835565 | -0.57857 | 0.562882 | 0.983423 | 0.00726 |
| ENSMUSG00000110165.2 | 15.48315 | 1.180277 | 1.96112 | 0.601838 | 0.547282 | 0.983423 | 0.00726 |
| ENSMUSG00000111013.2 | 35.40569 | 1.031506 | 1.833269 | 0.56266 | 0.573667 | 0.983423 | 0.00726 |
| ENSMUSG00000111074.2 | 10.22705 | 1.669393 | 2.961057 | 0.563783 | 0.572902 | 0.983423 | 0.00726 |
| ENSMUSG00000112006.2 | 218.1871 | -0.31894 | 0.545815 | -0.58434 | 0.55899 | 0.983423 | 0.00726 |
| ENSMUSG00000112796.2 | 47.37747 | -0.71705 | 1.18878 | -0.60318 | 0.546387 | 0.983423 | 0.00726 |
| ENSMUSG00000113061.2 | 169.7353 | 0.428221 | 0.670411 | 0.638744 | 0.522989 | 0.983423 | 0.00726 |
| ENSMUSG00000113450.2 | 11.01725 | 1.818396 | 2.783657 | 0.65324 | 0.513602 | 0.983423 | 0.00726 |
| ENSMUSG00000114154.2 | 10.91024 | -1.57881 | 2.807347 | -0.56239 | 0.573853 | 0.983423 | 0.00726 |
| ENSMUSG00000115938.2 | 19.61789 | 1.331889 | 2.027396 | 0.656946 | 0.511216 | 0.983423 | 0.00726 |
| ENSMUSG00000026986.15 | 300.055 | 0.293956 | 0.529976 | 0.554659 | 0.579128 | 0.983718 | 0.007129 |
| ENSMUSG00000037674.16 | 56.136 | 0.681346 | 1.228394 | 0.554664 | 0.579124 | 0.983718 | 0.007129 |
| ENSMUSG00000026280.18 | 62.17077 | -0.54161 | 0.977686 | -0.55397 | 0.579602 | 0.984171 | 0.006929 |
| ENSMUSG00000116097.2 | 12.81941 | -1.17427 | 2.119186 | -0.55411 | 0.579502 | 0.984171 | 0.006929 |
| ENSMUSG00000059355.14 | 176.6337 | 0.346088 | 0.625146 | 0.553611 | 0.579845 | 0.984407 | 0.006825 |
| ENSMUSG00000019188.17 | 148.1359 | -0.41163 | 0.744409 | -0.55297 | 0.580286 | 0.984472 | 0.006797 |
| ENSMUSG00000024187.16 | 104.0241 | 0.464516 | 0.839563 | 0.553283 | 0.58007 | 0.984472 | 0.006797 |
| ENSMUSG00000029998.15 | 623.98 | 0.186817 | 0.337855 | 0.552949 | 0.580298 | 0.984472 | 0.006797 |

| | | | | | | | |
|---|---|---|---|---|---|---|---|
| ENSMUSG00000079477.10 | 631.4109 | -0.2135 | 0.386047 | -0.55304 | 0.580234 | 0.984472 | 0.006797 |
| ENSMUSG00000032601.14 | 483.4011 | 0.202257 | 0.366266 | 0.552213 | 0.580802 | 0.985151 | 0.006497 |
| ENSMUSG00000022205.16 | 975.1249 | 0.221256 | 0.400803 | 0.552032 | 0.580926 | 0.985185 | 0.006482 |
| ENSMUSG00000029464.11 | 33.8309 | -0.64838 | 1.174883 | -0.55187 | 0.581039 | 0.985199 | 0.006476 |
| ENSMUSG00000045973.19 | 166.9083 | -0.34183 | 0.619591 | -0.5517 | 0.581153 | 0.985216 | 0.006468 |
| ENSMUSG00000027951.17 | 26.0446 | 1.059139 | 1.920485 | 0.551495 | 0.581294 | 0.98528 | 0.006441 |
| ENSMUSG00000001175.17 | 3735.299 | 0.129436 | 0.236298 | 0.547766 | 0.583853 | 0.985311 | 0.006427 |
| ENSMUSG00000003233.13 | 36.48564 | 0.975427 | 1.783648 | 0.546872 | 0.584467 | 0.985311 | 0.006427 |
| ENSMUSG00000004105.9 | 20.53596 | 0.99717 | 1.827052 | 0.545781 | 0.585217 | 0.985311 | 0.006427 |
| ENSMUSG00000006732.15 | 51.73092 | -0.69586 | 1.268782 | -0.54845 | 0.583385 | 0.985311 | 0.006427 |
| ENSMUSG00000019373.12 | 411.7288 | 0.23538 | 0.427138 | 0.551064 | 0.58159 | 0.985311 | 0.006427 |
| ENSMUSG00000019762.5 | 851.9032 | 0.177945 | 0.32335 | 0.550319 | 0.582101 | 0.985311 | 0.006427 |
| ENSMUSG00000019843.15 | 27.26441 | 0.923217 | 1.68986 | 0.546328 | 0.584841 | 0.985311 | 0.006427 |
| ENSMUSG00000020415.17 | 103.4676 | -0.4448 | 0.81149 | -0.54813 | 0.583601 | 0.985311 | 0.006427 |
| ENSMUSG00000020516.16 | 222.5207 | -0.32561 | 0.591013 | -0.55094 | 0.581672 | 0.985311 | 0.006427 |
| ENSMUSG00000021023.15 | 25.64551 | 1.061066 | 1.944362 | 0.545714 | 0.585262 | 0.985311 | 0.006427 |
| ENSMUSG00000023206.17 | 168.3202 | 0.403286 | 0.738784 | 0.545878 | 0.58515 | 0.985311 | 0.006427 |
| ENSMUSG00000025495.16 | 11.28666 | -1.44265 | 2.63565 | -0.54736 | 0.584131 | 0.985311 | 0.006427 |
| ENSMUSG00000027291.16 | 12.43455 | 1.317629 | 2.401637 | 0.548638 | 0.583254 | 0.985311 | 0.006427 |
| ENSMUSG00000029110.14 | 367.1939 | -0.26237 | 0.479391 | -0.5473 | 0.584171 | 0.985311 | 0.006427 |
| ENSMUSG00000029202.13 | 127.1279 | 0.389461 | 0.711137 | 0.54766 | 0.583926 | 0.985311 | 0.006427 |
| ENSMUSG00000029592.12 | 27.02562 | 0.953192 | 1.746488 | 0.545777 | 0.58522 | 0.985311 | 0.006427 |
| ENSMUSG00000029686.16 | 282.4894 | -0.27782 | 0.505584 | -0.54951 | 0.582654 | 0.985311 | 0.006427 |
| ENSMUSG00000031549.17 | 372.5489 | 0.287249 | 0.526175 | 0.545919 | 0.585121 | 0.985311 | 0.006427 |
| ENSMUSG00000032381.6 | 107.0827 | -0.36022 | 0.658591 | -0.54696 | 0.584409 | 0.985311 | 0.006427 |
| ENSMUSG00000033377.15 | 88.9167 | -0.40204 | 0.732934 | -0.54853 | 0.583328 | 0.985311 | 0.006427 |
| ENSMUSG00000034729.17 | 202.1891 | -0.32428 | 0.590116 | -0.54952 | 0.582647 | 0.985311 | 0.006427 |
| ENSMUSG00000036611.6 | 78.53319 | -0.56554 | 1.03464 | -0.54661 | 0.584649 | 0.985311 | 0.006427 |
| ENSMUSG00000037780.4 | 1134.363 | -0.17091 | 0.311047 | -0.54946 | 0.582693 | 0.985311 | 0.006427 |
| ENSMUSG00000040225.16 | 150.9109 | -0.43136 | 0.785904 | -0.54887 | 0.583093 | 0.985311 | 0.006427 |
| ENSMUSG00000041417.16 | 262.0545 | -0.3087 | 0.563375 | -0.54795 | 0.583727 | 0.985311 | 0.006427 |
| ENSMUSG00000041645.15 | 261.9051 | -0.26478 | 0.484751 | -0.54622 | 0.584917 | 0.985311 | 0.006427 |
| ENSMUSG00000044037.16 | 43.60209 | -0.63544 | 1.155071 | -0.55013 | 0.582232 | 0.985311 | 0.006427 |
| ENSMUSG00000045594.11 | 25.58169 | -0.76025 | 1.384698 | -0.54903 | 0.582982 | 0.985311 | 0.006427 |
| ENSMUSG00000054640.16 | 10.68393 | -1.54177 | 2.799666 | -0.5507 | 0.581842 | 0.985311 | 0.006427 |
| ENSMUSG00000054693.15 | 211.4159 | 0.302034 | 0.548423 | 0.550731 | 0.581818 | 0.985311 | 0.006427 |
| ENSMUSG00000055334.5 | 21.88695 | 1.294099 | 2.363043 | 0.547641 | 0.583939 | 0.985311 | 0.006427 |
| ENSMUSG00000056091.13 | 523.1998 | 0.253533 | 0.463722 | 0.546734 | 0.584561 | 0.985311 | 0.006427 |
| ENSMUSG00000058126.8 | 34.52457 | -0.8458 | 1.545831 | -0.54715 | 0.584277 | 0.985311 | 0.006427 |
| ENSMUSG00000063065.14 | 10.33587 | -1.54566 | 2.825594 | -0.54702 | 0.584363 | 0.985311 | 0.006427 |
| ENSMUSG00000066148.4 | 15.78158 | 0.875981 | 1.598817 | 0.547893 | 0.583765 | 0.985311 | 0.006427 |
| ENSMUSG00000075470.3 | 131.6205 | -0.39999 | 0.725581 | -0.55127 | 0.581447 | 0.985311 | 0.006427 |
| ENSMUSG00000091896.9 | 164.7395 | 0.325165 | 0.592002 | 0.549264 | 0.582825 | 0.985311 | 0.006427 |
| ENSMUSG00000092075.2 | 141.7787 | 0.634793 | 1.160822 | 0.546848 | 0.584483 | 0.985311 | 0.006427 |

| | | | | | | | |
|---|---|---|---|---|---|---|---|
| ENSMUSG00000029446.15 | 48.18438 | -0.79241 | 1.453201 | -0.54528 | 0.585558 | 0.985634 | 0.006284 |
| ENSMUSG00000035898.14 | 35.23786 | -0.68646 | 1.25959 | -0.54499 | 0.58576 | 0.9858 | 0.006211 |
| ENSMUSG00000005968.15 | 34.05113 | -0.82101 | 1.507073 | -0.54477 | 0.585909 | 0.985827 | 0.006199 |
| ENSMUSG00000024563.17 | 109.9214 | 0.46886 | 0.860825 | 0.544664 | 0.585985 | 0.985827 | 0.006199 |
| ENSMUSG00000021024.15 | 954.9545 | -0.15763 | 0.289608 | -0.54427 | 0.586254 | 0.986023 | 0.006113 |
| ENSMUSG00000055546.7 | 32.4965 | -1.05597 | 1.94068 | -0.54412 | 0.586356 | 0.986023 | 0.006113 |
| ENSMUSG00000063412.5 | 46.96692 | -0.55347 | 1.017331 | -0.54404 | 0.586413 | 0.986023 | 0.006113 |
| ENSMUSG00000020790.15 | 118.6465 | 0.600029 | 1.103382 | 0.543809 | 0.586573 | 0.986117 | 0.006072 |
| ENSMUSG00000020720.14 | 768.0535 | 0.170787 | 0.314265 | 0.54345 | 0.58682 | 0.986182 | 0.006043 |
| ENSMUSG00000039990.17 | 38.98883 | -0.73095 | 1.344809 | -0.54354 | 0.586761 | 0.986182 | 0.006043 |
| ENSMUSG00000000673.10 | 814.4226 | -0.18869 | 0.358269 | -0.52666 | 0.598431 | 0.986277 | 0.006001 |
| ENSMUSG00000001052.16 | 101.4632 | 0.549563 | 1.044474 | 0.526163 | 0.598775 | 0.986277 | 0.006001 |
| ENSMUSG00000001741.13 | 30.03898 | -1.04334 | 1.995419 | -0.52287 | 0.601066 | 0.986277 | 0.006001 |
| ENSMUSG00000002396.9 | 42.63114 | -0.51858 | 0.990754 | -0.52342 | 0.600679 | 0.986277 | 0.006001 |
| ENSMUSG00000002847.8 | 226.7435 | -0.28011 | 0.528255 | -0.53025 | 0.595939 | 0.986277 | 0.006001 |
| ENSMUSG00000002985.17 | 32703.29 | -0.14528 | 0.277212 | -0.52409 | 0.600218 | 0.986277 | 0.006001 |
| ENSMUSG00000003423.16 | 95.24177 | 0.458092 | 0.855075 | 0.535733 | 0.592143 | 0.986277 | 0.006001 |
| ENSMUSG00000003644.18 | 31.19078 | -0.95854 | 1.829631 | -0.5239 | 0.60035 | 0.986277 | 0.006001 |
| ENSMUSG00000004207.15 | 185.808 | -0.95058 | 1.791564 | -0.53059 | 0.595706 | 0.986277 | 0.006001 |
| ENSMUSG00000008822.16 | 73.48345 | 0.549748 | 1.058831 | 0.519203 | 0.603619 | 0.986277 | 0.006001 |
| ENSMUSG00000011114.19 | 33.17378 | -0.7012 | 1.300229 | -0.53929 | 0.589687 | 0.986277 | 0.006001 |
| ENSMUSG00000013465.20 | 40.05714 | -0.78574 | 1.488662 | -0.52782 | 0.597626 | 0.986277 | 0.006001 |
| ENSMUSG00000013833.16 | 15.51557 | -1.24915 | 2.347116 | -0.5322 | 0.594584 | 0.986277 | 0.006001 |
| ENSMUSG00000013973.13 | 43.38202 | 0.76359 | 1.452916 | 0.525557 | 0.599196 | 0.986277 | 0.006001 |
| ENSMUSG00000014164.16 | 28.01952 | -1.04688 | 1.961295 | -0.53377 | 0.593502 | 0.986277 | 0.006001 |
| ENSMUSG00000014353.16 | 86.21621 | -0.37936 | 0.723614 | -0.52426 | 0.600098 | 0.986277 | 0.006001 |
| ENSMUSG00000014402.16 | 124.9392 | 0.380477 | 0.712176 | 0.534245 | 0.593172 | 0.986277 | 0.006001 |
| ENSMUSG00000014601.14 | 17.67619 | -0.97818 | 1.851085 | -0.52844 | 0.597197 | 0.986277 | 0.006001 |
| ENSMUSG00000014932.16 | 118.5667 | 0.399698 | 0.75156 | 0.531825 | 0.594847 | 0.986277 | 0.006001 |
| ENSMUSG00000015745.10 | 15.9004 | -1.15731 | 2.188334 | -0.52886 | 0.596905 | 0.986277 | 0.006001 |
| ENSMUSG00000016252.8 | 1779.774 | 0.144725 | 0.267986 | 0.540048 | 0.589164 | 0.986277 | 0.006001 |
| ENSMUSG00000016637.8 | 28.87104 | -0.66554 | 1.262609 | -0.52712 | 0.598113 | 0.986277 | 0.006001 |
| ENSMUSG00000017291.14 | 296.0846 | 0.239146 | 0.457331 | 0.522916 | 0.601032 | 0.986277 | 0.006001 |
| ENSMUSG00000017390.16 | 54.56391 | -0.7018 | 1.293867 | -0.5424 | 0.58754 | 0.986277 | 0.006001 |
| ENSMUSG00000018583.6 | 313.158 | -0.2469 | 0.469368 | -0.52603 | 0.598865 | 0.986277 | 0.006001 |
| ENSMUSG00000018697.15 | 63.56432 | 0.498475 | 0.933698 | 0.533872 | 0.59343 | 0.986277 | 0.006001 |
| ENSMUSG00000018921.12 | 38.46088 | -0.90103 | 1.687256 | -0.53402 | 0.593326 | 0.986277 | 0.006001 |
| ENSMUSG00000019087.14 | 91.61228 | 0.483701 | 0.905875 | 0.53396 | 0.593369 | 0.986277 | 0.006001 |
| ENSMUSG00000019189.14 | 96.897 | 0.440553 | 0.828738 | 0.531595 | 0.595007 | 0.986277 | 0.006001 |
| ENSMUSG00000019857.8 | 131.1874 | -0.42107 | 0.784613 | -0.53666 | 0.591502 | 0.986277 | 0.006001 |
| ENSMUSG00000020290.15 | 183.2151 | 0.285781 | 0.532006 | 0.537177 | 0.591145 | 0.986277 | 0.006001 |
| ENSMUSG00000020864.14 | 113.3647 | 0.393138 | 0.733527 | 0.535955 | 0.591989 | 0.986277 | 0.006001 |
| ENSMUSG00000021470.20 | 33.75254 | -0.59461 | 1.124885 | -0.52859 | 0.597088 | 0.986277 | 0.006001 |
| ENSMUSG00000022092.12 | 14.34589 | -1.307 | 2.434678 | -0.53683 | 0.591386 | 0.986277 | 0.006001 |

| | | | | | | | |
|---|---|---|---|---|---|---|---|
| ENSMUSG00000022125.8 | 149.1077 | 0.385211 | 0.71445 | 0.539171 | 0.589769 | 0.986277 | 0.006001 |
| ENSMUSG00000022174.9 | 255.4748 | 0.256439 | 0.482447 | 0.531539 | 0.595045 | 0.986277 | 0.006001 |
| ENSMUSG00000022621.17 | 13.52793 | 1.205729 | 2.315091 | 0.520813 | 0.602497 | 0.986277 | 0.006001 |
| ENSMUSG00000023019.13 | 287.832 | -0.33292 | 0.627 | -0.53098 | 0.595433 | 0.986277 | 0.006001 |
| ENSMUSG00000023495.14 | 17.53533 | -1.29729 | 2.448964 | -0.52973 | 0.596298 | 0.986277 | 0.006001 |
| ENSMUSG00000023861.18 | 185.2802 | -0.36547 | 0.679849 | -0.53757 | 0.590874 | 0.986277 | 0.006001 |
| ENSMUSG00000024097.12 | 199.7549 | 0.308624 | 0.594242 | 0.519357 | 0.603512 | 0.986277 | 0.006001 |
| ENSMUSG00000024188.17 | 102.1979 | 0.361407 | 0.695165 | 0.519886 | 0.603143 | 0.986277 | 0.006001 |
| ENSMUSG00000024273.9 | 88.79544 | -0.51067 | 0.974429 | -0.52407 | 0.600232 | 0.986277 | 0.006001 |
| ENSMUSG00000024740.11 | 119.3521 | 0.4226 | 0.80213 | 0.526848 | 0.598299 | 0.986277 | 0.006001 |
| ENSMUSG00000025019.18 | 91.63393 | -0.44136 | 0.849014 | -0.51985 | 0.603168 | 0.986277 | 0.006001 |
| ENSMUSG00000025255.19 | 79.57021 | -0.50693 | 0.96068 | -0.52768 | 0.597724 | 0.986277 | 0.006001 |
| ENSMUSG00000025451.16 | 59.55416 | -0.46655 | 0.888814 | -0.52491 | 0.599644 | 0.986277 | 0.006001 |
| ENSMUSG00000025492.7 | 2249.612 | -0.15439 | 0.295824 | -0.52191 | 0.601735 | 0.986277 | 0.006001 |
| ENSMUSG00000026176.14 | 151.5203 | -0.36036 | 0.671576 | -0.53659 | 0.591551 | 0.986277 | 0.006001 |
| ENSMUSG00000026578.7 | 26.19158 | 0.796308 | 1.504048 | 0.529443 | 0.596498 | 0.986277 | 0.006001 |
| ENSMUSG00000026594.15 | 31.81884 | 0.770881 | 1.424417 | 0.541191 | 0.588376 | 0.986277 | 0.006001 |
| ENSMUSG00000026924.18 | 237.2894 | 0.297109 | 0.550657 | 0.539554 | 0.589504 | 0.986277 | 0.006001 |
| ENSMUSG00000027259.16 | 69.57008 | 0.568963 | 1.066088 | 0.533692 | 0.593555 | 0.986277 | 0.006001 |
| ENSMUSG00000027698.15 | 121.2234 | 0.41601 | 0.768133 | 0.541586 | 0.588104 | 0.986277 | 0.006001 |
| ENSMUSG00000027787.15 | 306.9297 | 0.326667 | 0.618702 | 0.527989 | 0.597507 | 0.986277 | 0.006001 |
| ENSMUSG00000027882.19 | 193.5157 | -0.32705 | 0.605106 | -0.54049 | 0.58886 | 0.986277 | 0.006001 |
| ENSMUSG00000028076.13 | 1328.416 | 0.167034 | 0.312537 | 0.534446 | 0.593033 | 0.986277 | 0.006001 |
| ENSMUSG00000028414.18 | 42.92889 | -0.59174 | 1.134269 | -0.5217 | 0.601882 | 0.986277 | 0.006001 |
| ENSMUSG00000028552.14 | 248.0886 | 0.337717 | 0.636127 | 0.530896 | 0.595491 | 0.986277 | 0.006001 |
| ENSMUSG00000028580.16 | 432.5588 | 0.222046 | 0.414835 | 0.535263 | 0.592468 | 0.986277 | 0.006001 |
| ENSMUSG00000028647.14 | 128.6882 | -0.3749 | 0.706202 | -0.53087 | 0.59551 | 0.986277 | 0.006001 |
| ENSMUSG00000028700.15 | 29.84857 | -0.8969 | 1.668634 | -0.53751 | 0.590918 | 0.986277 | 0.006001 |
| ENSMUSG00000028737.16 | 645.8135 | 0.217507 | 0.404076 | 0.538283 | 0.590382 | 0.986277 | 0.006001 |
| ENSMUSG00000028811.13 | 84.22719 | -0.46643 | 0.875646 | -0.53267 | 0.594261 | 0.986277 | 0.006001 |
| ENSMUSG00000028899.12 | 142.2011 | 0.381465 | 0.709852 | 0.537386 | 0.591001 | 0.986277 | 0.006001 |
| ENSMUSG00000029017.14 | 642.551 | 0.190536 | 0.36114 | 0.527594 | 0.597781 | 0.986277 | 0.006001 |
| ENSMUSG00000029299.15 | 14.38166 | 0.927802 | 1.773594 | 0.52312 | 0.600891 | 0.986277 | 0.006001 |
| ENSMUSG00000029392.13 | 25.78662 | -1.08646 | 2.034502 | -0.53402 | 0.593329 | 0.986277 | 0.006001 |
| ENSMUSG00000029403.15 | 29.71437 | -0.90306 | 1.739407 | -0.51918 | 0.603637 | 0.986277 | 0.006001 |
| ENSMUSG00000029445.14 | 3799.537 | -0.14149 | 0.266024 | -0.53189 | 0.594804 | 0.986277 | 0.006001 |
| ENSMUSG00000029617.11 | 235.1477 | 0.309347 | 0.577915 | 0.535281 | 0.592455 | 0.986277 | 0.006001 |
| ENSMUSG00000029649.11 | 1677.456 | 0.130056 | 0.250202 | 0.519804 | 0.6032 | 0.986277 | 0.006001 |
| ENSMUSG00000029705.18 | 107.9716 | -0.43855 | 0.817866 | -0.53622 | 0.591808 | 0.986277 | 0.006001 |
| ENSMUSG00000030096.9 | 268.4398 | 0.338854 | 0.646592 | 0.524062 | 0.600236 | 0.986277 | 0.006001 |
| ENSMUSG00000030201.16 | 119.5331 | 0.336503 | 0.625023 | 0.538385 | 0.590312 | 0.986277 | 0.006001 |
| ENSMUSG00000030313.16 | 357.4734 | -0.23557 | 0.442172 | -0.53276 | 0.594197 | 0.986277 | 0.006001 |
| ENSMUSG00000030533.17 | 15.43496 | 1.116678 | 2.11372 | 0.5283 | 0.597291 | 0.986277 | 0.006001 |
| ENSMUSG00000030727.13 | 25.07733 | -1.05256 | 2.006475 | -0.52458 | 0.599875 | 0.986277 | 0.006001 |

| | | | | | | | |
|---|---|---|---|---|---|---|---|
| ENSMUSG00000030747.6 | 880.131 | 0.163841 | 0.314227 | 0.521409 | 0.602082 | 0.986277 | 0.006001 |
| ENSMUSG00000030929.18 | 53.53606 | 0.652461 | 1.207443 | 0.540366 | 0.588945 | 0.986277 | 0.006001 |
| ENSMUSG00000031520.7 | 18.59522 | -0.99613 | 1.898908 | -0.52458 | 0.599876 | 0.986277 | 0.006001 |
| ENSMUSG00000031527.8 | 59.60217 | 0.629261 | 1.17801 | 0.534173 | 0.593222 | 0.986277 | 0.006001 |
| ENSMUSG00000031601.17 | 162.5768 | -0.30194 | 0.570619 | -0.52915 | 0.596702 | 0.986277 | 0.006001 |
| ENSMUSG00000031609.4 | 35.56199 | 0.801469 | 1.496139 | 0.535691 | 0.592172 | 0.986277 | 0.006001 |
| ENSMUSG00000031931.5 | 80.85946 | 0.515966 | 0.956726 | 0.539304 | 0.589677 | 0.986277 | 0.006001 |
| ENSMUSG00000032376.13 | 79.34252 | 0.55072 | 1.050663 | 0.524165 | 0.600164 | 0.986277 | 0.006001 |
| ENSMUSG00000032387.16 | 235.8765 | 0.272045 | 0.507461 | 0.53609 | 0.591896 | 0.986277 | 0.006001 |
| ENSMUSG00000032548.15 | 178.2009 | -0.29659 | 0.563285 | -0.52654 | 0.598516 | 0.986277 | 0.006001 |
| ENSMUSG00000033809.16 | 23.21208 | -0.94721 | 1.825446 | -0.51889 | 0.603835 | 0.986277 | 0.006001 |
| ENSMUSG00000034064.15 | 55.08172 | 0.5495 | 1.045516 | 0.525578 | 0.599182 | 0.986277 | 0.006001 |
| ENSMUSG00000034349.15 | 107.9064 | -0.42374 | 0.807567 | -0.52471 | 0.599783 | 0.986277 | 0.006001 |
| ENSMUSG00000034525.8 | 30.70471 | 0.719254 | 1.384183 | 0.519623 | 0.603326 | 0.986277 | 0.006001 |
| ENSMUSG00000034826.15 | 32.0849 | 0.664216 | 1.273377 | 0.521617 | 0.601937 | 0.986277 | 0.006001 |
| ENSMUSG00000034998.19 | 49.85249 | 0.797444 | 1.490638 | 0.534968 | 0.592672 | 0.986277 | 0.006001 |
| ENSMUSG00000035517.18 | 82.93637 | 0.455374 | 0.841318 | 0.541263 | 0.588326 | 0.986277 | 0.006001 |
| ENSMUSG00000036503.14 | 935.1099 | 0.160381 | 0.309002 | 0.519029 | 0.603741 | 0.986277 | 0.006001 |
| ENSMUSG00000037541.23 | 24.92504 | 0.985129 | 1.888188 | 0.521732 | 0.601857 | 0.986277 | 0.006001 |
| ENSMUSG00000038023.13 | 40.34879 | -0.57168 | 1.085881 | -0.52647 | 0.598563 | 0.986277 | 0.006001 |
| ENSMUSG00000038175.11 | 19.69072 | -0.98952 | 1.843935 | -0.53663 | 0.591521 | 0.986277 | 0.006001 |
| ENSMUSG00000038563.15 | 37.10033 | 0.865215 | 1.608809 | 0.537798 | 0.590716 | 0.986277 | 0.006001 |
| ENSMUSG00000039001.13 | 4221.796 | 0.140049 | 0.264187 | 0.530112 | 0.596034 | 0.986277 | 0.006001 |
| ENSMUSG00000039182.7 | 148.921 | -0.4097 | 0.761081 | -0.53831 | 0.590362 | 0.986277 | 0.006001 |
| ENSMUSG00000039715.9 | 14.18998 | 1.23926 | 2.360579 | 0.524981 | 0.599596 | 0.986277 | 0.006001 |
| ENSMUSG00000040269.6 | 266.87 | -0.29991 | 0.556687 | -0.53875 | 0.59006 | 0.986277 | 0.006001 |
| ENSMUSG00000040354.15 | 173.5172 | -0.40556 | 0.779196 | -0.52049 | 0.602725 | 0.986277 | 0.006001 |
| ENSMUSG00000040370.14 | 514.9957 | 0.181448 | 0.340044 | 0.533602 | 0.593617 | 0.986277 | 0.006001 |
| ENSMUSG00000040447.16 | 37.97639 | -0.79422 | 1.514541 | -0.5244 | 0.600001 | 0.986277 | 0.006001 |
| ENSMUSG00000040521.12 | 55.56277 | -0.65092 | 1.224506 | -0.53158 | 0.595016 | 0.986277 | 0.006001 |
| ENSMUSG00000040721.10 | 28.11482 | 1.057713 | 1.955432 | 0.54091 | 0.588569 | 0.986277 | 0.006001 |
| ENSMUSG00000041079.13 | 19.75075 | 1.127412 | 2.147891 | 0.524893 | 0.599658 | 0.986277 | 0.006001 |
| ENSMUSG00000041426.13 | 556.2362 | -0.20392 | 0.375812 | -0.5426 | 0.587404 | 0.986277 | 0.006001 |
| ENSMUSG00000041570.15 | 38.27435 | 0.759556 | 1.428785 | 0.53161 | 0.594996 | 0.986277 | 0.006001 |
| ENSMUSG00000041879.14 | 92.7678 | 0.450207 | 0.853274 | 0.527623 | 0.597761 | 0.986277 | 0.006001 |
| ENSMUSG00000042079.14 | 335.2611 | 0.223109 | 0.418265 | 0.533414 | 0.593747 | 0.986277 | 0.006001 |
| ENSMUSG00000042284.11 | 260.758 | 0.323138 | 0.596529 | 0.541697 | 0.588027 | 0.986277 | 0.006001 |
| ENSMUSG00000042348.11 | 14.61076 | -0.91763 | 1.760676 | -0.52118 | 0.602241 | 0.986277 | 0.006001 |
| ENSMUSG00000042447.14 | 76.30924 | 0.464 | 0.864822 | 0.536527 | 0.591594 | 0.986277 | 0.006001 |
| ENSMUSG00000043183.19 | 50.37801 | 0.506783 | 0.949459 | 0.53376 | 0.593508 | 0.986277 | 0.006001 |
| ENSMUSG00000044018.4 | 325.9043 | -0.24052 | 0.457343 | -0.5259 | 0.598955 | 0.986277 | 0.006001 |
| ENSMUSG00000045665.6 | 40.60391 | -0.66484 | 1.247157 | -0.53309 | 0.593973 | 0.986277 | 0.006001 |
| ENSMUSG00000046111.19 | 17.29523 | -1.11653 | 2.119479 | -0.52679 | 0.598337 | 0.986277 | 0.006001 |
| ENSMUSG00000046753.11 | 12.83548 | -0.9204 | 1.771649 | -0.51951 | 0.603402 | 0.986277 | 0.006001 |

| | | | | | | | |
|---|---|---|---|---|---|---|---|
| ENSMUSG00000047635.7 | 36.36293 | -0.6539 | 1.255932 | -0.52065 | 0.60261 | 0.986277 | 0.006001 |
| ENSMUSG00000051518.9 | 99.57039 | 0.438489 | 0.830016 | 0.52829 | 0.597298 | 0.986277 | 0.006001 |
| ENSMUSG00000052562.16 | 1514.698 | 0.14793 | 0.284055 | 0.52078 | 0.60252 | 0.986277 | 0.006001 |
| ENSMUSG00000053730.16 | 17.51903 | 1.210759 | 2.254846 | 0.536959 | 0.591296 | 0.986277 | 0.006001 |
| ENSMUSG00000053819.17 | 23.09042 | 1.049937 | 2.013709 | 0.521395 | 0.602092 | 0.986277 | 0.006001 |
| ENSMUSG00000057696.5 | 10.19956 | 1.074824 | 1.997344 | 0.538126 | 0.59049 | 0.986277 | 0.006001 |
| ENSMUSG00000057802.9 | 13.66821 | 1.430241 | 2.64264 | 0.541217 | 0.588358 | 0.986277 | 0.006001 |
| ENSMUSG00000058013.12 | 24.16395 | 0.856794 | 1.603469 | 0.534338 | 0.593108 | 0.986277 | 0.006001 |
| ENSMUSG00000058325.7 | 55.81579 | -0.72072 | 1.384935 | -0.5204 | 0.602783 | 0.986277 | 0.006001 |
| ENSMUSG00000058454.16 | 247.3485 | -0.33151 | 0.638077 | -0.51954 | 0.603381 | 0.986277 | 0.006001 |
| ENSMUSG00000058706.6 | 206.0239 | -0.33113 | 0.632728 | -0.52334 | 0.600738 | 0.986277 | 0.006001 |
| ENSMUSG00000059796.17 | 404.6007 | -0.22514 | 0.417341 | -0.53947 | 0.589561 | 0.986277 | 0.006001 |
| ENSMUSG00000060227.16 | 21.35654 | -0.85728 | 1.578325 | -0.54316 | 0.587019 | 0.986277 | 0.006001 |
| ENSMUSG00000060261.17 | 201.2285 | 0.308362 | 0.581609 | 0.530188 | 0.595982 | 0.986277 | 0.006001 |
| ENSMUSG00000060538.15 | 98.52201 | -0.55394 | 1.061659 | -0.52177 | 0.601834 | 0.986277 | 0.006001 |
| ENSMUSG00000060923.6 | 40.04387 | 0.767256 | 1.416278 | 0.541741 | 0.587997 | 0.986277 | 0.006001 |
| ENSMUSG00000061306.17 | 44.92849 | -0.82181 | 1.548464 | -0.53073 | 0.595609 | 0.986277 | 0.006001 |
| ENSMUSG00000061758.14 | 13.87746 | 1.265954 | 2.377381 | 0.5325 | 0.59438 | 0.986277 | 0.006001 |
| ENSMUSG00000062098.12 | 18.17439 | 0.973706 | 1.835266 | 0.530553 | 0.595729 | 0.986277 | 0.006001 |
| ENSMUSG00000062960.11 | 385.8425 | 0.251935 | 0.483944 | 0.520588 | 0.602654 | 0.986277 | 0.006001 |
| ENSMUSG00000066026.15 | 243.0301 | -0.29872 | 0.574757 | -0.51973 | 0.603249 | 0.986277 | 0.006001 |
| ENSMUSG00000068290.12 | 147.1305 | 0.426009 | 0.788339 | 0.540388 | 0.58893 | 0.986277 | 0.006001 |
| ENSMUSG00000069972.7 | 181.5216 | 0.304527 | 0.563277 | 0.540635 | 0.588759 | 0.986277 | 0.006001 |
| ENSMUSG00000071074.10 | 58.66398 | -0.77132 | 1.452075 | -0.53119 | 0.595289 | 0.986277 | 0.006001 |
| ENSMUSG00000075254.13 | 28.79901 | 0.84726 | 1.590777 | 0.532608 | 0.594305 | 0.986277 | 0.006001 |
| ENSMUSG00000078612.10 | 36.35095 | 0.850832 | 1.608757 | 0.528875 | 0.596892 | 0.986277 | 0.006001 |
| ENSMUSG00000079179.10 | 28.31023 | 0.838747 | 1.59601 | 0.525528 | 0.599216 | 0.986277 | 0.006001 |
| ENSMUSG00000079435.10 | 1826.268 | 0.13343 | 0.254086 | 0.525136 | 0.599489 | 0.986277 | 0.006001 |
| ENSMUSG00000084289.2 | 19.80259 | 0.763354 | 1.467294 | 0.520246 | 0.602892 | 0.986277 | 0.006001 |
| ENSMUSG00000084416.4 | 221.1336 | -0.27975 | 0.5292 | -0.52864 | 0.597058 | 0.986277 | 0.006001 |
| ENSMUSG00000085023.2 | 94.26758 | 0.48831 | 0.934033 | 0.522798 | 0.601115 | 0.986277 | 0.006001 |
| ENSMUSG00000087174.8 | 13.34753 | 1.269243 | 2.389127 | 0.531258 | 0.59524 | 0.986277 | 0.006001 |
| ENSMUSG00000090733.7 | 1850.016 | 0.136198 | 0.261564 | 0.520708 | 0.602571 | 0.986277 | 0.006001 |
| ENSMUSG00000092072.4 | 68.37898 | 0.497448 | 0.922115 | 0.539464 | 0.589567 | 0.986277 | 0.006001 |
| ENSMUSG00000101892.2 | 78.18415 | -0.63541 | 1.224417 | -0.51895 | 0.603795 | 0.986277 | 0.006001 |
| ENSMUSG00000105388.2 | 91.1225 | 0.389316 | 0.744985 | 0.522583 | 0.601265 | 0.986277 | 0.006001 |
| ENSMUSG00000107176.2 | 698.3826 | 0.200005 | 0.374723 | 0.533742 | 0.59352 | 0.986277 | 0.006001 |
| ENSMUSG00000107198.2 | 47.13282 | -1.96806 | 3.731129 | -0.52747 | 0.597867 | 0.986277 | 0.006001 |
| ENSMUSG00000112099.2 | 426.2129 | 0.210487 | 0.40072 | 0.525273 | 0.599393 | 0.986277 | 0.006001 |
| ENSMUSG00000114358.2 | 17.67842 | -0.99552 | 1.893723 | -0.5257 | 0.5991 | 0.986277 | 0.006001 |
| ENSMUSG00000115338.3 | 463.6307 | 0.249132 | 0.470008 | 0.530058 | 0.596072 | 0.986277 | 0.006001 |
| ENSMUSG00000000088.8 | 3151.683 | 0.137996 | 0.266784 | 0.517257 | 0.604977 | 0.986396 | 0.005949 |
| ENSMUSG00000001472.18 | 227.8618 | -0.29664 | 0.574732 | -0.51613 | 0.605762 | 0.986396 | 0.005949 |
| ENSMUSG00000015790.17 | 229.5126 | -0.35697 | 0.690916 | -0.51666 | 0.605395 | 0.986396 | 0.005949 |

| | | | | | | | |
|---|---|---|---|---|---|---|---|
| ENSMUSG00000017210.15 | 13.89563 | 1.013467 | 1.967506 | 0.515103 | 0.606481 | 0.986396 | 0.005949 |
| ENSMUSG00000020037.16 | 16.61539 | 1.134065 | 2.187561 | 0.518415 | 0.604169 | 0.986396 | 0.005949 |
| ENSMUSG00000020674.18 | 13.63853 | -1.06474 | 2.061364 | -0.51652 | 0.605488 | 0.986396 | 0.005949 |
| ENSMUSG00000022313.11 | 47.26034 | 0.632229 | 1.226177 | 0.51561 | 0.606127 | 0.986396 | 0.005949 |
| ENSMUSG00000023967.10 | 39.99496 | 0.748748 | 1.4537 | 0.515064 | 0.606509 | 0.986396 | 0.005949 |
| ENSMUSG00000025085.17 | 92.72055 | 0.469102 | 0.90963 | 0.515706 | 0.60606 | 0.986396 | 0.005949 |
| ENSMUSG00000025921.8 | 235.909 | 0.27893 | 0.540981 | 0.515601 | 0.606133 | 0.986396 | 0.005949 |
| ENSMUSG00000026088.16 | 96.46576 | 0.473634 | 0.914435 | 0.517953 | 0.604491 | 0.986396 | 0.005949 |
| ENSMUSG00000028359.5 | 102.7522 | -1.39859 | 2.699293 | -0.51813 | 0.604365 | 0.986396 | 0.005949 |
| ENSMUSG00000028756.13 | 318.3127 | 0.260603 | 0.504019 | 0.51705 | 0.605121 | 0.986396 | 0.005949 |
| ENSMUSG00000029073.10 | 46.10021 | 0.846326 | 1.632529 | 0.518414 | 0.60417 | 0.986396 | 0.005949 |
| ENSMUSG00000031029.7 | 630.5856 | -0.2095 | 0.405518 | -0.51662 | 0.605424 | 0.986396 | 0.005949 |
| ENSMUSG00000031754.11 | 97.41522 | 0.398884 | 0.770306 | 0.517826 | 0.60458 | 0.986396 | 0.005949 |
| ENSMUSG00000031918.17 | 46.43961 | 0.666982 | 1.292884 | 0.515887 | 0.605934 | 0.986396 | 0.005949 |
| ENSMUSG00000032479.16 | 67.23416 | -0.54885 | 1.062071 | -0.51677 | 0.605313 | 0.986396 | 0.005949 |
| ENSMUSG00000038902.15 | 17.15849 | -0.95293 | 1.850081 | -0.51507 | 0.606502 | 0.986396 | 0.005949 |
| ENSMUSG00000039270.10 | 126.8566 | -0.32378 | 0.625121 | -0.51796 | 0.604489 | 0.986396 | 0.005949 |
| ENSMUSG00000042042.15 | 75.89425 | 0.483851 | 0.937059 | 0.516351 | 0.60561 | 0.986396 | 0.005949 |
| ENSMUSG00000055128.16 | 216.2619 | 0.291267 | 0.563952 | 0.516475 | 0.605523 | 0.986396 | 0.005949 |
| ENSMUSG00000055760.13 | 26.09881 | 0.841245 | 1.62839 | 0.516612 | 0.605427 | 0.986396 | 0.005949 |
| ENSMUSG00000063253.12 | 86.15349 | 0.462527 | 0.89488 | 0.516859 | 0.605254 | 0.986396 | 0.005949 |
| ENSMUSG00000110686.2 | 36.1374 | 0.583006 | 1.131281 | 0.51535 | 0.606308 | 0.986396 | 0.005949 |
| ENSMUSG00000029322.13 | 59.78697 | 0.615649 | 1.197075 | 0.514295 | 0.607046 | 0.98671 | 0.00581 |
| ENSMUSG00000035184.16 | 46.93435 | -0.71475 | 1.390052 | -0.51419 | 0.607118 | 0.98671 | 0.00581 |
| ENSMUSG00000038764.15 | 232.3099 | -0.30619 | 0.595436 | -0.51424 | 0.607087 | 0.98671 | 0.00581 |
| ENSMUSG00000115867.2 | 25.86602 | -0.83825 | 1.629238 | -0.5145 | 0.606901 | 0.98671 | 0.00581 |
| ENSMUSG00000079015.3 | 46663.99 | -0.10128 | 0.19714 | -0.51372 | 0.607444 | 0.987071 | 0.005652 |
| ENSMUSG00000016257.16 | 243.141 | -0.27903 | 0.543403 | -0.51348 | 0.607615 | 0.987179 | 0.005604 |
| ENSMUSG00000021038.18 | 18.00964 | -1.14231 | 2.229485 | -0.51237 | 0.608394 | 0.987261 | 0.005568 |
| ENSMUSG00000024151.14 | 27.49191 | -1.04736 | 2.04408 | -0.51239 | 0.60838 | 0.987261 | 0.005568 |
| ENSMUSG00000024422.15 | 82.6125 | -0.4846 | 0.945258 | -0.51267 | 0.608185 | 0.987261 | 0.005568 |
| ENSMUSG00000029389.18 | 25.48473 | -1.08623 | 2.119613 | -0.51247 | 0.608324 | 0.987261 | 0.005568 |
| ENSMUSG00000036461.17 | 73.72672 | -0.467 | 0.910967 | -0.51265 | 0.608198 | 0.987261 | 0.005568 |
| ENSMUSG00000063358.18 | 437.5278 | 0.2759 | 0.537999 | 0.512827 | 0.608072 | 0.987261 | 0.005568 |
| ENSMUSG00000111403.2 | 47.40448 | 0.637114 | 1.242808 | 0.512641 | 0.608203 | 0.987261 | 0.005568 |
| ENSMUSG00000024792.10 | 18.80286 | -1.08157 | 2.111576 | -0.51221 | 0.608503 | 0.987269 | 0.005564 |
| ENSMUSG00000029471.14 | 35.63197 | 0.95974 | 1.874391 | 0.512028 | 0.608632 | 0.987308 | 0.005547 |
| ENSMUSG00000033964.13 | 100.7385 | 0.402049 | 0.785757 | 0.51167 | 0.608882 | 0.987545 | 0.005443 |
| ENSMUSG00000034640.10 | 23.92794 | 0.987093 | 1.930243 | 0.511383 | 0.609083 | 0.987702 | 0.005374 |
| ENSMUSG00000020464.16 | 90.83959 | 0.440577 | 0.862278 | 0.510945 | 0.60939 | 0.987906 | 0.005285 |
| ENSMUSG00000032727.15 | 65.00852 | -0.5321 | 1.041486 | -0.51091 | 0.609417 | 0.987906 | 0.005285 |
| ENSMUSG00000032596.15 | 27.66404 | 1.004678 | 1.968269 | 0.510437 | 0.609745 | 0.988269 | 0.005125 |
| ENSMUSG00000001016.13 | 20.72737 | 0.895936 | 1.756159 | 0.510168 | 0.609934 | 0.988295 | 0.005114 |
| ENSMUSG00000003429.12 | 1548.747 | -0.13114 | 0.257748 | -0.50878 | 0.610908 | 0.988295 | 0.005114 |

| | | | | | | | |
|---|---|---|---|---|---|---|---|
| ENSMUSG00000021806.5 | 21.64317 | -0.84861 | 1.664415 | -0.50985 | 0.610155 | 0.988295 | 0.005114 |
| ENSMUSG00000023156.5 | 82.84502 | -0.41474 | 0.814846 | -0.50898 | 0.610763 | 0.988295 | 0.005114 |
| ENSMUSG00000023800.16 | 64.62676 | 0.513909 | 1.009992 | 0.508825 | 0.610875 | 0.988295 | 0.005114 |
| ENSMUSG00000026880.12 | 44.28771 | -0.94584 | 1.855602 | -0.50972 | 0.610249 | 0.988295 | 0.005114 |
| ENSMUSG00000028407.5 | 217.3966 | 0.304861 | 0.598746 | 0.509166 | 0.610636 | 0.988295 | 0.005114 |
| ENSMUSG00000029767.17 | 208.9478 | -0.28544 | 0.560673 | -0.5091 | 0.610684 | 0.988295 | 0.005114 |
| ENSMUSG00000037622.15 | 27.29914 | -0.89973 | 1.768075 | -0.50887 | 0.61084 | 0.988295 | 0.005114 |
| ENSMUSG00000044641.8 | 53.34327 | 0.562091 | 1.103511 | 0.509366 | 0.610496 | 0.988295 | 0.005114 |
| ENSMUSG00000050234.8 | 10.32055 | 1.6868 | 3.314676 | 0.508888 | 0.610831 | 0.988295 | 0.005114 |
| ENSMUSG00000048076.15 | 482.4767 | -0.19521 | 0.383859 | -0.50854 | 0.611076 | 0.988397 | 0.005068 |
| ENSMUSG00000032372.15 | 165.5508 | 0.336542 | 0.662055 | 0.50833 | 0.611222 | 0.988466 | 0.005038 |
| ENSMUSG00000024528.9 | 27.49683 | -0.77249 | 1.520392 | -0.50808 | 0.611394 | 0.988575 | 0.00499 |
| ENSMUSG00000028668.6 | 142.7125 | -0.34015 | 0.669824 | -0.50783 | 0.611575 | 0.988699 | 0.004936 |
| ENSMUSG00000016756.18 | 456.5595 | 0.229731 | 0.452944 | 0.507195 | 0.612018 | 0.988874 | 0.004859 |
| ENSMUSG00000022814.7 | 137.5035 | 0.449821 | 0.887086 | 0.507077 | 0.6121 | 0.988874 | 0.004859 |
| ENSMUSG00000024082.6 | 86.09744 | -0.45431 | 0.895589 | -0.50727 | 0.611965 | 0.988874 | 0.004859 |
| ENSMUSG00000035772.12 | 98.54699 | -0.36675 | 0.723213 | -0.50711 | 0.612076 | 0.988874 | 0.004859 |
| ENSMUSG00000000001.5 | 608.5477 | 0.139176 | 0.401558 | 0.34659 | 0.7289 | 0.98928 | 0.004681 |
| ENSMUSG00000000131.16 | 66.09521 | -0.41133 | 0.94521 | -0.43517 | 0.663436 | 0.98928 | 0.004681 |
| ENSMUSG00000000247.12 | 36.73536 | 0.794264 | 1.960997 | 0.405031 | 0.685455 | 0.98928 | 0.004681 |
| ENSMUSG00000000346.10 | 827.1522 | 0.195013 | 0.417046 | 0.467605 | 0.640067 | 0.98928 | 0.004681 |
| ENSMUSG00000000384.16 | 39.66119 | 0.32925 | 1.021414 | 0.322347 | 0.74719 | 0.98928 | 0.004681 |
| ENSMUSG00000000581.9 | 742.0302 | 0.183893 | 0.372906 | 0.493134 | 0.621918 | 0.98928 | 0.004681 |
| ENSMUSG00000000776.14 | 53.15021 | 0.557317 | 1.298006 | 0.429364 | 0.667658 | 0.98928 | 0.004681 |
| ENSMUSG00000000787.13 | 731.0091 | -0.13869 | 0.345099 | -0.40189 | 0.687767 | 0.98928 | 0.004681 |
| ENSMUSG00000000934.10 | 36.94431 | 0.908006 | 1.812152 | 0.501065 | 0.616325 | 0.98928 | 0.004681 |
| ENSMUSG00000001138.14 | 29.27473 | -0.49511 | 1.22389 | -0.40454 | 0.685816 | 0.98928 | 0.004681 |
| ENSMUSG00000001285.13 | 35.92662 | 0.760384 | 1.541045 | 0.493421 | 0.621715 | 0.98928 | 0.004681 |
| ENSMUSG00000001323.18 | 260.1635 | 0.21154 | 0.449837 | 0.470259 | 0.63817 | 0.98928 | 0.004681 |
| ENSMUSG00000001348.16 | 357.8552 | 0.165068 | 0.444729 | 0.371166 | 0.710514 | 0.98928 | 0.004681 |
| ENSMUSG00000001366.15 | 695.1757 | -0.14058 | 0.425831 | -0.33013 | 0.7413 | 0.98928 | 0.004681 |
| ENSMUSG00000001416.14 | 270.977 | 0.140156 | 0.437784 | 0.320149 | 0.748855 | 0.98928 | 0.004681 |
| ENSMUSG00000001424.15 | 196.5192 | -0.36363 | 0.775159 | -0.46911 | 0.638993 | 0.98928 | 0.004681 |
| ENSMUSG00000001440.6 | 317.6772 | 0.239058 | 0.498377 | 0.479672 | 0.631461 | 0.98928 | 0.004681 |
| ENSMUSG00000001467.6 | 1108.919 | -0.16999 | 0.3496 | -0.48625 | 0.626791 | 0.98928 | 0.004681 |
| ENSMUSG00000001729.15 | 16.41709 | 0.752142 | 1.867842 | 0.402679 | 0.687184 | 0.98928 | 0.004681 |
| ENSMUSG00000001751.10 | 18.90346 | 0.948015 | 2.072841 | 0.457351 | 0.647419 | 0.98928 | 0.004681 |
| ENSMUSG00000001755.13 | 389.9369 | -0.17183 | 0.42693 | -0.40248 | 0.687332 | 0.98928 | 0.004681 |
| ENSMUSG00000001833.18 | 389.3629 | 0.184885 | 0.405421 | 0.456031 | 0.648367 | 0.98928 | 0.004681 |
| ENSMUSG00000001844.11 | 30.87275 | 0.517373 | 1.462133 | 0.353848 | 0.723453 | 0.98928 | 0.004681 |
| ENSMUSG00000001847.15 | 720.0203 | -0.11547 | 0.342287 | -0.33736 | 0.735846 | 0.98928 | 0.004681 |
| ENSMUSG00000001942.9 | 197.2177 | -0.30376 | 0.666322 | -0.45587 | 0.648483 | 0.98928 | 0.004681 |
| ENSMUSG00000001946.15 | 32.58352 | -0.76436 | 1.710102 | -0.44697 | 0.654897 | 0.98928 | 0.004681 |
| ENSMUSG00000001962.9 | 189.5932 | -0.26183 | 0.645712 | -0.4055 | 0.685111 | 0.98928 | 0.004681 |

| | | | | | | | |
|---|---|---|---|---|---|---|---|
| ENSMUSG00000001964.15 | 32.72786 | -0.77524 | 2.027593 | -0.38235 | 0.702205 | 0.98928 | 0.004681 |
| ENSMUSG00000002032.18 | 30.47231 | 0.656217 | 1.337993 | 0.490449 | 0.623816 | 0.98928 | 0.004681 |
| ENSMUSG00000002329.8 | 50.29408 | 0.570179 | 1.370163 | 0.41614 | 0.677308 | 0.98928 | 0.004681 |
| ENSMUSG00000002496.11 | 48.60985 | -0.47176 | 1.298477 | -0.36332 | 0.716369 | 0.98928 | 0.004681 |
| ENSMUSG00000002550.17 | 46.01214 | 0.663749 | 1.378202 | 0.481605 | 0.630086 | 0.98928 | 0.004681 |
| ENSMUSG00000002602.17 | 33.19746 | -0.45118 | 1.415212 | -0.31881 | 0.749872 | 0.98928 | 0.004681 |
| ENSMUSG00000002608.9 | 21.15368 | -1.09727 | 2.434864 | -0.45065 | 0.652241 | 0.98928 | 0.004681 |
| ENSMUSG00000002660.9 | 137.2561 | 0.317847 | 0.66481 | 0.478103 | 0.632577 | 0.98928 | 0.004681 |
| ENSMUSG00000002718.15 | 269.4564 | 0.193158 | 0.445224 | 0.433845 | 0.664401 | 0.98928 | 0.004681 |
| ENSMUSG00000002748.8 | 136.5223 | -0.27636 | 0.639837 | -0.43192 | 0.665799 | 0.98928 | 0.004681 |
| ENSMUSG00000002825.9 | 35.43916 | 0.817176 | 1.979476 | 0.412825 | 0.679735 | 0.98928 | 0.004681 |
| ENSMUSG00000002845.15 | 58.47133 | -0.37451 | 0.9233 | -0.40562 | 0.685023 | 0.98928 | 0.004681 |
| ENSMUSG00000002881.15 | 767.5484 | 0.140435 | 0.360647 | 0.389398 | 0.696981 | 0.98928 | 0.004681 |
| ENSMUSG00000002963.18 | 24.86147 | 0.510856 | 1.536254 | 0.332534 | 0.739486 | 0.98928 | 0.004681 |
| ENSMUSG00000003072.16 | 228.5377 | 0.257564 | 0.619334 | 0.415873 | 0.677503 | 0.98928 | 0.004681 |
| ENSMUSG00000003119.15 | 49.99723 | 0.366015 | 1.048466 | 0.349096 | 0.727017 | 0.98928 | 0.004681 |
| ENSMUSG00000003131.8 | 126.2955 | 0.286506 | 0.744396 | 0.384884 | 0.700323 | 0.98928 | 0.004681 |
| ENSMUSG00000003135.16 | 28.3411 | 0.531451 | 1.349729 | 0.393746 | 0.693768 | 0.98928 | 0.004681 |
| ENSMUSG00000003184.16 | 72.57499 | 0.577977 | 1.266002 | 0.456537 | 0.648004 | 0.98928 | 0.004681 |
| ENSMUSG00000003190.13 | 15.59752 | -0.85196 | 1.912467 | -0.44548 | 0.655973 | 0.98928 | 0.004681 |
| ENSMUSG00000003308.16 | 102.2943 | -0.50098 | 1.018791 | -0.49174 | 0.622902 | 0.98928 | 0.004681 |
| ENSMUSG00000003316.15 | 101.9336 | 0.486715 | 1.090391 | 0.446367 | 0.655332 | 0.98928 | 0.004681 |
| ENSMUSG00000003435.10 | 92.87275 | -0.27641 | 0.749187 | -0.36895 | 0.712166 | 0.98928 | 0.004681 |
| ENSMUSG00000003527.10 | 18.63342 | 0.825086 | 2.196242 | 0.375681 | 0.707154 | 0.98928 | 0.004681 |
| ENSMUSG00000003549.10 | 16.98559 | -0.8635 | 2.247009 | -0.38429 | 0.700765 | 0.98928 | 0.004681 |
| ENSMUSG00000003731.14 | 94.45855 | 0.40166 | 0.826089 | 0.486219 | 0.626812 | 0.98928 | 0.004681 |
| ENSMUSG00000003813.16 | 135.2326 | -0.42602 | 0.865999 | -0.49194 | 0.622759 | 0.98928 | 0.004681 |
| ENSMUSG00000003873.12 | 31.10907 | -0.77743 | 1.86756 | -0.41628 | 0.677205 | 0.98928 | 0.004681 |
| ENSMUSG00000004056.16 | 47.96429 | 0.447515 | 1.243433 | 0.359903 | 0.71892 | 0.98928 | 0.004681 |
| ENSMUSG00000004266.16 | 41.37566 | 0.379984 | 1.197507 | 0.317312 | 0.751007 | 0.98928 | 0.004681 |
| ENSMUSG00000004356.9 | 93.07846 | -0.36461 | 0.756352 | -0.48206 | 0.629763 | 0.98928 | 0.004681 |
| ENSMUSG00000004446.13 | 96.36546 | 0.444356 | 1.259649 | 0.352762 | 0.724267 | 0.98928 | 0.004681 |
| ENSMUSG00000004460.18 | 616.8345 | 0.185262 | 0.383149 | 0.483524 | 0.628723 | 0.98928 | 0.004681 |
| ENSMUSG00000004462.8 | 86.63248 | -0.2749 | 0.769214 | -0.35738 | 0.720805 | 0.98928 | 0.004681 |
| ENSMUSG00000004473.11 | 38.4252 | -0.75167 | 1.999396 | -0.37595 | 0.706957 | 0.98928 | 0.004681 |
| ENSMUSG00000004567.17 | 119.104 | -0.33467 | 0.964662 | -0.34693 | 0.728642 | 0.98928 | 0.004681 |
| ENSMUSG00000004637.16 | 25.47191 | -0.70933 | 1.685987 | -0.42072 | 0.673958 | 0.98928 | 0.004681 |
| ENSMUSG00000004665.11 | 10.32029 | -1.30065 | 2.956249 | -0.43997 | 0.659962 | 0.98928 | 0.004681 |
| ENSMUSG00000004897.17 | 389.9028 | -0.13506 | 0.407296 | -0.33159 | 0.740198 | 0.98928 | 0.004681 |
| ENSMUSG00000005103.13 | 175.0376 | 0.177563 | 0.561314 | 0.316334 | 0.751749 | 0.98928 | 0.004681 |
| ENSMUSG00000005198.16 | 18.64365 | -0.84004 | 1.933172 | -0.43454 | 0.663897 | 0.98928 | 0.004681 |
| ENSMUSG00000005204.13 | 14.92 | -0.78021 | 1.679545 | -0.46454 | 0.642261 | 0.98928 | 0.004681 |
| ENSMUSG00000005225.16 | 92.63135 | 0.454091 | 0.990264 | 0.458556 | 0.646553 | 0.98928 | 0.004681 |
| ENSMUSG00000005299.7 | 554.1085 | -0.16682 | 0.369323 | -0.4517 | 0.651487 | 0.98928 | 0.004681 |

| | | | | | | | |
|---|---|---|---|---|---|---|---|
| ENSMUSG00000005371.16 | 175.8409 | -0.24575 | 0.613335 | -0.40068 | 0.688658 | 0.98928 | 0.004681 |
| ENSMUSG00000005469.11 | 54.18919 | 0.327804 | 0.999845 | 0.327855 | 0.743021 | 0.98928 | 0.004681 |
| ENSMUSG00000005514.15 | 223.6818 | -0.28021 | 0.556096 | -0.50389 | 0.614337 | 0.98928 | 0.004681 |
| ENSMUSG00000005615.16 | 283.6503 | 0.314424 | 0.677289 | 0.464239 | 0.642476 | 0.98928 | 0.004681 |
| ENSMUSG00000005656.10 | 135.3116 | 0.327429 | 1.008118 | 0.324792 | 0.745338 | 0.98928 | 0.004681 |
| ENSMUSG00000005732.15 | 262.8467 | -0.27242 | 0.550864 | -0.49453 | 0.62093 | 0.98928 | 0.004681 |
| ENSMUSG00000005774.13 | 21.52843 | -0.75157 | 2.128416 | -0.35311 | 0.724003 | 0.98928 | 0.004681 |
| ENSMUSG00000005813.13 | 188.3824 | -0.24335 | 0.723203 | -0.33649 | 0.7365 | 0.98928 | 0.004681 |
| ENSMUSG00000005871.16 | 104.4165 | 0.316657 | 0.911606 | 0.347362 | 0.728319 | 0.98928 | 0.004681 |
| ENSMUSG00000006058.11 | 138.1109 | -0.36966 | 0.77599 | -0.47637 | 0.633812 | 0.98928 | 0.004681 |
| ENSMUSG00000006289.15 | 43.49654 | -0.36675 | 1.018664 | -0.36003 | 0.718821 | 0.98928 | 0.004681 |
| ENSMUSG00000006373.11 | 4814.325 | -0.0748 | 0.218951 | -0.34165 | 0.732613 | 0.98928 | 0.004681 |
| ENSMUSG00000006395.17 | 190.3394 | 0.33009 | 0.681352 | 0.484463 | 0.628058 | 0.98928 | 0.004681 |
| ENSMUSG00000006442.11 | 75.34301 | 0.335638 | 0.9154 | 0.366657 | 0.713875 | 0.98928 | 0.004681 |
| ENSMUSG00000006463.14 | 16.96709 | 1.124646 | 2.243413 | 0.50131 | 0.616153 | 0.98928 | 0.004681 |
| ENSMUSG00000006715.12 | 85.26176 | -0.53078 | 1.103099 | -0.48117 | 0.630397 | 0.98928 | 0.004681 |
| ENSMUSG00000006731.11 | 11.76284 | 1.187171 | 2.453381 | 0.483892 | 0.628463 | 0.98928 | 0.004681 |
| ENSMUSG00000006763.15 | 17.77046 | -0.77959 | 1.985536 | -0.39264 | 0.694589 | 0.98928 | 0.004681 |
| ENSMUSG00000007812.17 | 214.3044 | 0.273332 | 0.67656 | 0.404003 | 0.686211 | 0.98928 | 0.004681 |
| ENSMUSG00000007850.17 | 268.9267 | 0.287959 | 0.627029 | 0.459243 | 0.646059 | 0.98928 | 0.004681 |
| ENSMUSG00000008035.13 | 840.0913 | 0.175259 | 0.520069 | 0.336991 | 0.736124 | 0.98928 | 0.004681 |
| ENSMUSG00000008167.15 | 55.89883 | -0.48494 | 0.985493 | -0.49208 | 0.622664 | 0.98928 | 0.004681 |
| ENSMUSG00000008226.15 | 404.356 | 0.177033 | 0.426391 | 0.415189 | 0.678004 | 0.98928 | 0.004681 |
| ENSMUSG00000008429.9 | 250.1499 | 0.280065 | 0.631042 | 0.443814 | 0.657177 | 0.98928 | 0.004681 |
| ENSMUSG00000008668.16 | 3233.314 | -0.08324 | 0.245836 | -0.33858 | 0.734926 | 0.98928 | 0.004681 |
| ENSMUSG00000008682.14 | 1002.879 | -0.16454 | 0.424404 | -0.38769 | 0.698245 | 0.98928 | 0.004681 |
| ENSMUSG00000008690.16 | 46.69479 | -0.43634 | 1.379327 | -0.31634 | 0.751744 | 0.98928 | 0.004681 |
| ENSMUSG00000009079.17 | 289.2222 | 0.222786 | 0.550741 | 0.40452 | 0.68583 | 0.98928 | 0.004681 |
| ENSMUSG00000009293.18 | 137.1009 | 0.365925 | 0.88374 | 0.414064 | 0.678827 | 0.98928 | 0.004681 |
| ENSMUSG00000009563.17 | 104.4885 | 0.226929 | 0.664247 | 0.341633 | 0.732627 | 0.98928 | 0.004681 |
| ENSMUSG00000009772.16 | 24.73019 | 0.565914 | 1.506703 | 0.375597 | 0.707216 | 0.98928 | 0.004681 |
| ENSMUSG00000009863.15 | 996.5028 | -0.11987 | 0.318314 | -0.37657 | 0.706492 | 0.98928 | 0.004681 |
| ENSMUSG00000009894.16 | 164.9323 | -0.2622 | 0.565773 | -0.46343 | 0.643058 | 0.98928 | 0.004681 |
| ENSMUSG00000010051.16 | 23.26715 | 0.755955 | 1.521404 | 0.49688 | 0.619274 | 0.98928 | 0.004681 |
| ENSMUSG00000010057.9 | 40.60116 | -0.52581 | 1.132512 | -0.46429 | 0.642441 | 0.98928 | 0.004681 |
| ENSMUSG00000010110.18 | 49.15278 | -0.47176 | 1.12548 | -0.41916 | 0.675097 | 0.98928 | 0.004681 |
| ENSMUSG00000010175.14 | 234.1361 | 0.218984 | 0.655679 | 0.333981 | 0.738394 | 0.98928 | 0.004681 |
| ENSMUSG00000010517.8 | 165.0254 | 0.330942 | 0.864473 | 0.382825 | 0.70185 | 0.98928 | 0.004681 |
| ENSMUSG00000010554.15 | 66.89126 | -0.44853 | 1.070121 | -0.41914 | 0.675113 | 0.98928 | 0.004681 |
| ENSMUSG00000010601.15 | 634.5099 | 0.136395 | 0.344983 | 0.395367 | 0.692572 | 0.98928 | 0.004681 |
| ENSMUSG00000011096.18 | 36.38553 | -0.80887 | 1.794837 | -0.45067 | 0.65223 | 0.98928 | 0.004681 |
| ENSMUSG00000011158.9 | 37.18356 | -0.54202 | 1.618019 | -0.33499 | 0.737635 | 0.98928 | 0.004681 |
| ENSMUSG00000011960.13 | 96.21121 | 0.388613 | 0.902189 | 0.430744 | 0.666654 | 0.98928 | 0.004681 |
| ENSMUSG00000012076.9 | 12.91418 | 0.913291 | 2.316826 | 0.394199 | 0.693434 | 0.98928 | 0.004681 |

| | | | | | | | |
|---|---|---|---|---|---|---|---|
| ENSMUSG00000012405.17 | 167.8599 | 0.274319 | 0.560509 | 0.48941 | 0.624551 | 0.98928 | 0.004681 |
| ENSMUSG00000012483.5 | 34.95402 | 0.464379 | 1.336988 | 0.347332 | 0.728342 | 0.98928 | 0.004681 |
| ENSMUSG00000013622.16 | 413.3093 | -0.20916 | 0.449968 | -0.46483 | 0.642057 | 0.98928 | 0.004681 |
| ENSMUSG00000013736.17 | 40.52535 | -0.4262 | 1.143431 | -0.37274 | 0.709342 | 0.98928 | 0.004681 |
| ENSMUSG00000014496.9 | 125.2372 | -0.24222 | 0.626302 | -0.38674 | 0.698946 | 0.98928 | 0.004681 |
| ENSMUSG00000014504.17 | 299.9159 | 0.156022 | 0.48273 | 0.323207 | 0.746539 | 0.98928 | 0.004681 |
| ENSMUSG00000014547.11 | 29.22685 | 0.514699 | 1.54855 | 0.332375 | 0.739606 | 0.98928 | 0.004681 |
| ENSMUSG00000014606.15 | 429.9574 | -0.16703 | 0.377933 | -0.44195 | 0.658524 | 0.98928 | 0.004681 |
| ENSMUSG00000014748.14 | 170.5343 | 0.270496 | 0.654156 | 0.413504 | 0.679237 | 0.98928 | 0.004681 |
| ENSMUSG00000014771.14 | 72.67726 | 0.528116 | 1.127916 | 0.468223 | 0.639625 | 0.98928 | 0.004681 |
| ENSMUSG00000014791.11 | 23.25581 | 0.541049 | 1.507747 | 0.358846 | 0.71971 | 0.98928 | 0.004681 |
| ENSMUSG00000014980.15 | 72.39589 | 0.421324 | 0.958231 | 0.439689 | 0.660162 | 0.98928 | 0.004681 |
| ENSMUSG00000015023.8 | 133.5007 | -0.31139 | 0.841205 | -0.37017 | 0.711256 | 0.98928 | 0.004681 |
| ENSMUSG00000015243.5 | 1219.113 | -0.18814 | 0.387244 | -0.48583 | 0.627086 | 0.98928 | 0.004681 |
| ENSMUSG00000015337.6 | 190.816 | -0.23522 | 0.632923 | -0.37165 | 0.710157 | 0.98928 | 0.004681 |
| ENSMUSG00000015342.7 | 18.72161 | 0.68902 | 1.834995 | 0.375489 | 0.707297 | 0.98928 | 0.004681 |
| ENSMUSG00000015451.17 | 54.65939 | 0.530859 | 1.528087 | 0.347401 | 0.72829 | 0.98928 | 0.004681 |
| ENSMUSG00000015474.17 | 49.92592 | -0.48393 | 1.374557 | -0.35206 | 0.72479 | 0.98928 | 0.004681 |
| ENSMUSG00000015536.15 | 1122.24 | 0.143353 | 0.321602 | 0.445746 | 0.65578 | 0.98928 | 0.004681 |
| ENSMUSG00000015653.14 | 51.44676 | -0.61743 | 1.507029 | -0.4097 | 0.682024 | 0.98928 | 0.004681 |
| ENSMUSG00000015714.12 | 1740.116 | -0.15842 | 0.362558 | -0.43695 | 0.662149 | 0.98928 | 0.004681 |
| ENSMUSG00000015755.17 | 251.9807 | 0.208389 | 0.483551 | 0.430955 | 0.666501 | 0.98928 | 0.004681 |
| ENSMUSG00000015942.10 | 30.58451 | 0.747221 | 2.234141 | 0.334455 | 0.738036 | 0.98928 | 0.004681 |
| ENSMUSG00000015961.9 | 191.6892 | -0.38573 | 0.779399 | -0.49491 | 0.620662 | 0.98928 | 0.004681 |
| ENSMUSG00000015966.18 | 19.30504 | 0.64943 | 1.944964 | 0.333903 | 0.738452 | 0.98928 | 0.004681 |
| ENSMUSG00000015994.11 | 184.3071 | 0.219714 | 0.579992 | 0.378822 | 0.70482 | 0.98928 | 0.004681 |
| ENSMUSG00000016128.15 | 46.96176 | 0.421969 | 1.039756 | 0.405835 | 0.684864 | 0.98928 | 0.004681 |
| ENSMUSG00000016308.13 | 319.2778 | -0.14403 | 0.438802 | -0.32823 | 0.742739 | 0.98928 | 0.004681 |
| ENSMUSG00000016487.16 | 105.3878 | -0.39862 | 0.959807 | -0.41531 | 0.677914 | 0.98928 | 0.004681 |
| ENSMUSG00000016528.11 | 37.83191 | -0.6533 | 1.710362 | -0.38196 | 0.702487 | 0.98928 | 0.004681 |
| ENSMUSG00000016664.17 | 185.0011 | -0.23449 | 0.643959 | -0.36414 | 0.715752 | 0.98928 | 0.004681 |
| ENSMUSG00000016940.19 | 21.04447 | -0.65123 | 1.495484 | -0.43547 | 0.663223 | 0.98928 | 0.004681 |
| ENSMUSG00000016984.8 | 33.85653 | -0.49 | 1.37371 | -0.3567 | 0.721319 | 0.98928 | 0.004681 |
| ENSMUSG00000017485.12 | 328.0171 | 0.149938 | 0.441026 | 0.339975 | 0.733875 | 0.98928 | 0.004681 |
| ENSMUSG00000017802.15 | 37.32646 | 0.507552 | 1.399957 | 0.362549 | 0.716942 | 0.98928 | 0.004681 |
| ENSMUSG00000017831.9 | 12.43698 | -0.77456 | 2.257954 | -0.34303 | 0.731572 | 0.98928 | 0.004681 |
| ENSMUSG00000017858.13 | 93.63942 | 0.401766 | 0.833763 | 0.48187 | 0.629898 | 0.98928 | 0.004681 |
| ENSMUSG00000017868.17 | 58.38658 | -0.44138 | 0.903271 | -0.48864 | 0.625094 | 0.98928 | 0.004681 |
| ENSMUSG00000018286.7 | 656.3722 | -0.1663 | 0.361858 | -0.45957 | 0.645827 | 0.98928 | 0.004681 |
| ENSMUSG00000018326.10 | 648.7717 | -0.12646 | 0.319426 | -0.3959 | 0.692179 | 0.98928 | 0.004681 |
| ENSMUSG00000018362.15 | 44.61005 | 0.497861 | 1.132143 | 0.439751 | 0.660117 | 0.98928 | 0.004681 |
| ENSMUSG00000018363.17 | 178.2339 | 0.261287 | 0.628767 | 0.415555 | 0.677736 | 0.98928 | 0.004681 |
| ENSMUSG00000018387.13 | 14.63452 | 1.087392 | 2.238194 | 0.485834 | 0.627085 | 0.98928 | 0.004681 |
| ENSMUSG00000018427.9 | 78.33091 | 0.481926 | 0.99677 | 0.483488 | 0.628749 | 0.98928 | 0.004681 |

| | | | | | | | |
|---|---|---|---|---|---|---|---|
| ENSMUSG00000018585.10 | 1053.331 | -0.1269 | 0.31194 | -0.4068 | 0.684157 | 0.98928 | 0.004681 |
| ENSMUSG00000018599.6 | 15.42544 | -0.88208 | 2.20972 | -0.39918 | 0.68976 | 0.98928 | 0.004681 |
| ENSMUSG00000018846.9 | 224.9427 | -0.21941 | 0.553167 | -0.39664 | 0.691634 | 0.98928 | 0.004681 |
| ENSMUSG00000018849.7 | 190.9232 | 0.258261 | 0.559607 | 0.461503 | 0.644438 | 0.98928 | 0.004681 |
| ENSMUSG00000018927.4 | 39.46771 | 0.914384 | 1.855897 | 0.492691 | 0.622231 | 0.98928 | 0.004681 |
| ENSMUSG00000018965.12 | 131.7434 | 0.352595 | 0.823347 | 0.428246 | 0.668472 | 0.98928 | 0.004681 |
| ENSMUSG00000018974.8 | 37.76866 | 0.531983 | 1.329967 | 0.399997 | 0.689159 | 0.98928 | 0.004681 |
| ENSMUSG00000018995.13 | 78.62612 | -0.3857 | 0.876394 | -0.4401 | 0.659868 | 0.98928 | 0.004681 |
| ENSMUSG00000019054.14 | 76.09547 | 0.325964 | 0.921968 | 0.353553 | 0.723674 | 0.98928 | 0.004681 |
| ENSMUSG00000019297.10 | 66.20451 | 0.354902 | 0.891686 | 0.398012 | 0.690621 | 0.98928 | 0.004681 |
| ENSMUSG00000019302.17 | 11.49709 | -1.04027 | 2.502313 | -0.41572 | 0.677611 | 0.98928 | 0.004681 |
| ENSMUSG00000019432.16 | 157.2925 | -0.30954 | 0.689909 | -0.44867 | 0.653667 | 0.98928 | 0.004681 |
| ENSMUSG00000019518.12 | 39.18197 | 0.926433 | 1.992384 | 0.464987 | 0.641941 | 0.98928 | 0.004681 |
| ENSMUSG00000019763.12 | 154.7482 | -0.26119 | 0.726333 | -0.3596 | 0.719146 | 0.98928 | 0.004681 |
| ENSMUSG00000019795.18 | 225.0633 | 0.211083 | 0.555262 | 0.38015 | 0.703834 | 0.98928 | 0.004681 |
| ENSMUSG00000019810.16 | 33.39045 | 0.730702 | 1.721691 | 0.424409 | 0.671267 | 0.98928 | 0.004681 |
| ENSMUSG00000019818.16 | 1547.98 | -0.14984 | 0.327137 | -0.45804 | 0.646921 | 0.98928 | 0.004681 |
| ENSMUSG00000019849.12 | 107.6901 | 0.362467 | 0.771586 | 0.469769 | 0.63852 | 0.98928 | 0.004681 |
| ENSMUSG00000019863.8 | 19.39809 | -0.87474 | 2.09859 | -0.41682 | 0.676808 | 0.98928 | 0.004681 |
| ENSMUSG00000019866.16 | 43.93515 | -0.66486 | 1.440529 | -0.46154 | 0.644413 | 0.98928 | 0.004681 |
| ENSMUSG00000019872.14 | 368.9259 | 0.277895 | 0.553322 | 0.50223 | 0.615506 | 0.98928 | 0.004681 |
| ENSMUSG00000019877.11 | 1291.703 | -0.15347 | 0.329081 | -0.46635 | 0.640967 | 0.98928 | 0.004681 |
| ENSMUSG00000019907.11 | 121.2661 | 0.414972 | 0.847931 | 0.489393 | 0.624563 | 0.98928 | 0.004681 |
| ENSMUSG00000019927.7 | 77.96522 | -0.39354 | 0.817851 | -0.48119 | 0.630383 | 0.98928 | 0.004681 |
| ENSMUSG00000019935.17 | 63.27779 | 0.642431 | 1.302792 | 0.493119 | 0.621929 | 0.98928 | 0.004681 |
| ENSMUSG00000019944.15 | 182.2963 | -0.29445 | 0.807452 | -0.36466 | 0.715363 | 0.98928 | 0.004681 |
| ENSMUSG00000019966.19 | 28.44753 | 0.671423 | 1.683702 | 0.398778 | 0.690057 | 0.98928 | 0.004681 |
| ENSMUSG00000019984.17 | 22.87654 | 0.693316 | 1.502405 | 0.461471 | 0.644461 | 0.98928 | 0.004681 |
| ENSMUSG00000019988.8 | 61.44598 | -0.33341 | 0.95919 | -0.3476 | 0.728143 | 0.98928 | 0.004681 |
| ENSMUSG00000019989.9 | 194.4045 | -0.23249 | 0.553437 | -0.42008 | 0.674425 | 0.98928 | 0.004681 |
| ENSMUSG00000020010.8 | 101.022 | 0.313589 | 0.689762 | 0.454634 | 0.649372 | 0.98928 | 0.004681 |
| ENSMUSG00000020019.5 | 39.22951 | -0.70414 | 1.538239 | -0.45776 | 0.647125 | 0.98928 | 0.004681 |
| ENSMUSG00000020048.14 | 3524.657 | -0.1023 | 0.255793 | -0.39992 | 0.689218 | 0.98928 | 0.004681 |
| ENSMUSG00000020051.18 | 7981.102 | 0.10653 | 0.211149 | 0.504524 | 0.613893 | 0.98928 | 0.004681 |
| ENSMUSG00000020069.17 | 46.09292 | 0.471755 | 1.390018 | 0.339388 | 0.734317 | 0.98928 | 0.004681 |
| ENSMUSG00000020072.16 | 764.3222 | -0.13299 | 0.342192 | -0.38865 | 0.697534 | 0.98928 | 0.004681 |
| ENSMUSG00000020077.15 | 40.2 | 0.558595 | 1.516232 | 0.36841 | 0.712568 | 0.98928 | 0.004681 |
| ENSMUSG00000020115.4 | 106.4765 | -0.28545 | 0.789615 | -0.36151 | 0.717717 | 0.98928 | 0.004681 |
| ENSMUSG00000020196.11 | 23.11185 | -0.82502 | 2.144399 | -0.38473 | 0.700434 | 0.98928 | 0.004681 |
| ENSMUSG00000020198.9 | 104.4384 | -0.40019 | 0.814141 | -0.49155 | 0.623039 | 0.98928 | 0.004681 |
| ENSMUSG00000020366.19 | 170.4402 | -0.19888 | 0.534989 | -0.37175 | 0.710077 | 0.98928 | 0.004681 |
| ENSMUSG00000020368.16 | 1455.083 | 0.092789 | 0.27042 | 0.343131 | 0.7315 | 0.98928 | 0.004681 |
| ENSMUSG00000020380.17 | 95.29985 | -0.41959 | 0.902004 | -0.46518 | 0.641806 | 0.98928 | 0.004681 |
| ENSMUSG00000020381.11 | 33.53968 | 0.703162 | 2.038665 | 0.344913 | 0.73016 | 0.98928 | 0.004681 |

| | | | | | | | |
|---|---|---|---|---|---|---|---|
| ENSMUSG00000020456.18 | 358.4486 | -0.17585 | 0.48039 | -0.36605 | 0.714325 | 0.98928 | 0.004681 |
| ENSMUSG00000020515.14 | 189.2375 | -0.33797 | 0.738338 | -0.45774 | 0.647138 | 0.98928 | 0.004681 |
| ENSMUSG00000020521.8 | 159.9464 | 0.243968 | 0.769148 | 0.317193 | 0.751098 | 0.98928 | 0.004681 |
| ENSMUSG00000020530.15 | 892.8333 | 0.142624 | 0.30807 | 0.462958 | 0.643394 | 0.98928 | 0.004681 |
| ENSMUSG00000020532.19 | 145.997 | 0.484416 | 1.070296 | 0.452601 | 0.650836 | 0.98928 | 0.004681 |
| ENSMUSG00000020537.10 | 52.42557 | -0.3804 | 1.192078 | -0.31911 | 0.749646 | 0.98928 | 0.004681 |
| ENSMUSG00000020544.15 | 74.86987 | -0.35369 | 0.77907 | -0.45399 | 0.649839 | 0.98928 | 0.004681 |
| ENSMUSG00000020561.9 | 171.9627 | -0.26765 | 0.680548 | -0.39329 | 0.694105 | 0.98928 | 0.004681 |
| ENSMUSG00000020570.16 | 734.7624 | -0.17293 | 0.372568 | -0.46416 | 0.642535 | 0.98928 | 0.004681 |
| ENSMUSG00000020590.17 | 128.658 | -0.30082 | 0.666361 | -0.45144 | 0.651675 | 0.98928 | 0.004681 |
| ENSMUSG00000020608.8 | 242.8775 | -0.18729 | 0.570623 | -0.32821 | 0.742752 | 0.98928 | 0.004681 |
| ENSMUSG00000020610.18 | 45.28002 | -0.60048 | 1.196509 | -0.50186 | 0.615765 | 0.98928 | 0.004681 |
| ENSMUSG00000020620.15 | 191.0145 | -0.21392 | 0.579764 | -0.36898 | 0.71214 | 0.98928 | 0.004681 |
| ENSMUSG00000020621.7 | 366.7485 | 0.244597 | 0.487817 | 0.501412 | 0.616081 | 0.98928 | 0.004681 |
| ENSMUSG00000020664.11 | 686.1214 | 0.114512 | 0.358329 | 0.319572 | 0.749293 | 0.98928 | 0.004681 |
| ENSMUSG00000020694.18 | 53.65193 | -0.31753 | 0.981455 | -0.32353 | 0.74629 | 0.98928 | 0.004681 |
| ENSMUSG00000020705.7 | 107.4665 | -0.33612 | 0.70673 | -0.4756 | 0.634358 | 0.98928 | 0.004681 |
| ENSMUSG00000020707.7 | 53.48873 | 0.440349 | 1.217155 | 0.361785 | 0.717512 | 0.98928 | 0.004681 |
| ENSMUSG00000020719.15 | 2034.504 | -0.10593 | 0.318699 | -0.33238 | 0.739602 | 0.98928 | 0.004681 |
| ENSMUSG00000020736.13 | 105.0093 | -0.31267 | 0.942751 | -0.33166 | 0.740144 | 0.98928 | 0.004681 |
| ENSMUSG00000020783.15 | 29.12917 | -0.62243 | 1.878341 | -0.33137 | 0.740362 | 0.98928 | 0.004681 |
| ENSMUSG00000020818.17 | 11.2647 | -1.1001 | 2.868073 | -0.38357 | 0.701298 | 0.98928 | 0.004681 |
| ENSMUSG00000020823.17 | 30.40373 | -1.01106 | 2.239969 | -0.45137 | 0.651722 | 0.98928 | 0.004681 |
| ENSMUSG00000020830.4 | 37.67744 | -0.64058 | 1.801799 | -0.35552 | 0.722197 | 0.98928 | 0.004681 |
| ENSMUSG00000020863.16 | 1149.739 | 0.096371 | 0.301234 | 0.319921 | 0.749028 | 0.98928 | 0.004681 |
| ENSMUSG00000020865.17 | 357.8073 | -0.24992 | 0.539519 | -0.46323 | 0.643197 | 0.98928 | 0.004681 |
| ENSMUSG00000020922.12 | 76.9369 | -0.43443 | 0.906215 | -0.47939 | 0.631663 | 0.98928 | 0.004681 |
| ENSMUSG00000020925.17 | 60.53961 | -0.40423 | 0.86503 | -0.4673 | 0.640286 | 0.98928 | 0.004681 |
| ENSMUSG00000020929.15 | 33.881 | -0.72501 | 1.487072 | -0.48754 | 0.625876 | 0.98928 | 0.004681 |
| ENSMUSG00000020952.11 | 256.9507 | -0.19795 | 0.604024 | -0.32772 | 0.743122 | 0.98928 | 0.004681 |
| ENSMUSG00000020962.15 | 84.19987 | 0.396437 | 0.789947 | 0.501853 | 0.615771 | 0.98928 | 0.004681 |
| ENSMUSG00000020994.6 | 282.0188 | -0.19953 | 0.456775 | -0.43682 | 0.662241 | 0.98928 | 0.004681 |
| ENSMUSG00000021012.17 | 427.8604 | 0.154961 | 0.482262 | 0.321322 | 0.747967 | 0.98928 | 0.004681 |
| ENSMUSG00000021028.9 | 17.51696 | 0.690385 | 2.026546 | 0.340671 | 0.733351 | 0.98928 | 0.004681 |
| ENSMUSG00000021069.18 | 2325.033 | -0.10614 | 0.280944 | -0.37779 | 0.705587 | 0.98928 | 0.004681 |
| ENSMUSG00000021087.19 | 10.72773 | -0.96538 | 2.154695 | -0.44803 | 0.654129 | 0.98928 | 0.004681 |
| ENSMUSG00000021103.13 | 119.6246 | 0.250034 | 0.68972 | 0.362516 | 0.716967 | 0.98928 | 0.004681 |
| ENSMUSG00000021109.14 | 178.3205 | -0.29751 | 0.786038 | -0.37849 | 0.705065 | 0.98928 | 0.004681 |
| ENSMUSG00000021124.14 | 323.4205 | 0.238267 | 0.542538 | 0.439171 | 0.660538 | 0.98928 | 0.004681 |
| ENSMUSG00000021131.15 | 26.46294 | -0.67632 | 2.111347 | -0.32033 | 0.74872 | 0.98928 | 0.004681 |
| ENSMUSG00000021133.10 | 43.27968 | -0.43959 | 1.205276 | -0.36472 | 0.71532 | 0.98928 | 0.004681 |
| ENSMUSG00000021187.15 | 16.91841 | -0.62884 | 1.918972 | -0.3277 | 0.74314 | 0.98928 | 0.004681 |
| ENSMUSG00000021218.11 | 2147.638 | -0.11886 | 0.260009 | -0.45712 | 0.647583 | 0.98928 | 0.004681 |
| ENSMUSG00000021241.8 | 164.4622 | 0.281628 | 0.596475 | 0.472153 | 0.636817 | 0.98928 | 0.004681 |

| | | | | | | | |
|---|---|---|---|---|---|---|---|
| ENSMUSG00000021279.6 | 26.49839 | 0.969577 | 2.279405 | 0.425364 | 0.670571 | 0.98928 | 0.004681 |
| ENSMUSG00000021282.18 | 1944.521 | 0.12504 | 0.273939 | 0.456453 | 0.648064 | 0.98928 | 0.004681 |
| ENSMUSG00000021285.16 | 17.03152 | 0.610826 | 1.91425 | 0.319094 | 0.749655 | 0.98928 | 0.004681 |
| ENSMUSG00000021311.10 | 19.93951 | 0.995373 | 2.813347 | 0.353804 | 0.723486 | 0.98928 | 0.004681 |
| ENSMUSG00000021339.5 | 77.31257 | 0.441115 | 0.892829 | 0.494064 | 0.621261 | 0.98928 | 0.004681 |
| ENSMUSG00000021357.15 | 65.08405 | -0.66853 | 1.351986 | -0.49448 | 0.620967 | 0.98928 | 0.004681 |
| ENSMUSG00000021364.17 | 1890.801 | 0.11184 | 0.259883 | 0.430348 | 0.666943 | 0.98928 | 0.004681 |
| ENSMUSG00000021466.13 | 128.8043 | 0.304467 | 0.815144 | 0.373513 | 0.708767 | 0.98928 | 0.004681 |
| ENSMUSG00000021488.9 | 57.1153 | 0.393754 | 1.082187 | 0.363851 | 0.71597 | 0.98928 | 0.004681 |
| ENSMUSG00000021504.15 | 16.66742 | 0.755424 | 2.308322 | 0.327261 | 0.74347 | 0.98928 | 0.004681 |
| ENSMUSG00000021510.12 | 15.73348 | -0.72664 | 1.724178 | -0.42144 | 0.673434 | 0.98928 | 0.004681 |
| ENSMUSG00000021559.15 | 224.534 | 0.246362 | 0.626707 | 0.393106 | 0.694241 | 0.98928 | 0.004681 |
| ENSMUSG00000021576.7 | 734.0209 | 0.113524 | 0.325772 | 0.348477 | 0.727482 | 0.98928 | 0.004681 |
| ENSMUSG00000021592.19 | 32.33015 | -0.53786 | 1.299018 | -0.41405 | 0.678838 | 0.98928 | 0.004681 |
| ENSMUSG00000021597.17 | 20.55554 | 0.548636 | 1.728616 | 0.317385 | 0.750952 | 0.98928 | 0.004681 |
| ENSMUSG00000021607.10 | 199.7656 | -0.2349 | 0.649628 | -0.36159 | 0.717655 | 0.98928 | 0.004681 |
| ENSMUSG00000021610.9 | 309.4882 | 0.179437 | 0.505814 | 0.35475 | 0.722777 | 0.98928 | 0.004681 |
| ENSMUSG00000021620.5 | 429.3425 | -0.15877 | 0.500169 | -0.31743 | 0.750916 | 0.98928 | 0.004681 |
| ENSMUSG00000021645.17 | 42.63887 | 0.459911 | 1.271586 | 0.361683 | 0.717589 | 0.98928 | 0.004681 |
| ENSMUSG00000021665.10 | 142.3979 | 0.374405 | 1.010168 | 0.370637 | 0.710908 | 0.98928 | 0.004681 |
| ENSMUSG00000021728.9 | 22.94515 | 0.670793 | 1.874415 | 0.357868 | 0.720442 | 0.98928 | 0.004681 |
| ENSMUSG00000021752.15 | 86.53355 | -0.31506 | 0.786904 | -0.40038 | 0.688874 | 0.98928 | 0.004681 |
| ENSMUSG00000021814.18 | 471.4137 | -0.11553 | 0.35202 | -0.3282 | 0.742763 | 0.98928 | 0.004681 |
| ENSMUSG00000021816.12 | 268.3934 | 0.187638 | 0.479052 | 0.391686 | 0.69529 | 0.98928 | 0.004681 |
| ENSMUSG00000021866.16 | 67.64387 | -0.40698 | 0.962578 | -0.4228 | 0.67244 | 0.98928 | 0.004681 |
| ENSMUSG00000021870.18 | 329.9355 | 0.140556 | 0.435904 | 0.322447 | 0.747114 | 0.98928 | 0.004681 |
| ENSMUSG00000021900.11 | 66.12248 | -0.486 | 1.193543 | -0.40719 | 0.683868 | 0.98928 | 0.004681 |
| ENSMUSG00000021916.17 | 39.87328 | -0.75583 | 1.768264 | -0.42744 | 0.669057 | 0.98928 | 0.004681 |
| ENSMUSG00000021917.17 | 787.6936 | -0.12232 | 0.370632 | -0.33002 | 0.741385 | 0.98928 | 0.004681 |
| ENSMUSG00000021975.10 | 16.91617 | -1.16379 | 2.527369 | -0.46047 | 0.645176 | 0.98928 | 0.004681 |
| ENSMUSG00000021996.17 | 2756.713 | 0.077747 | 0.238389 | 0.326134 | 0.744323 | 0.98928 | 0.004681 |
| ENSMUSG00000022016.18 | 96.76424 | 0.344126 | 0.929374 | 0.370277 | 0.711176 | 0.98928 | 0.004681 |
| ENSMUSG00000022023.7 | 152.8862 | -0.34386 | 0.820441 | -0.41912 | 0.675129 | 0.98928 | 0.004681 |
| ENSMUSG00000022024.11 | 276.7132 | -0.19765 | 0.450378 | -0.43886 | 0.660766 | 0.98928 | 0.004681 |
| ENSMUSG00000022025.14 | 97.4331 | 0.34272 | 1.07188 | 0.319738 | 0.749167 | 0.98928 | 0.004681 |
| ENSMUSG00000022057.9 | 18.48393 | -0.80812 | 2.498346 | -0.32346 | 0.746346 | 0.98928 | 0.004681 |
| ENSMUSG00000022109.7 | 25.75389 | -0.55345 | 1.626174 | -0.34034 | 0.733602 | 0.98928 | 0.004681 |
| ENSMUSG00000022110.14 | 1207.371 | 0.167285 | 0.374484 | 0.446709 | 0.655086 | 0.98928 | 0.004681 |
| ENSMUSG00000022141.8 | 337.2924 | -0.20359 | 0.570484 | -0.35688 | 0.721182 | 0.98928 | 0.004681 |
| ENSMUSG00000022178.12 | 21.16888 | 0.711507 | 2.165663 | 0.32854 | 0.742503 | 0.98928 | 0.004681 |
| ENSMUSG00000022193.8 | 103.9688 | 0.437481 | 0.980426 | 0.446216 | 0.655441 | 0.98928 | 0.004681 |
| ENSMUSG00000022197.17 | 10.2262 | 1.520413 | 3.311403 | 0.459145 | 0.64613 | 0.98928 | 0.004681 |
| ENSMUSG00000022219.11 | 1598.4 | -0.10075 | 0.264149 | -0.38142 | 0.702895 | 0.98928 | 0.004681 |
| ENSMUSG00000022236.11 | 165.0441 | 0.242487 | 0.74889 | 0.323795 | 0.746093 | 0.98928 | 0.004681 |

| | | | | | | | |
|---|---|---|---|---|---|---|---|
| ENSMUSG00000022265.8 | 82.27077 | -0.35342 | 0.939245 | -0.37628 | 0.706709 | 0.98928 | 0.004681 |
| ENSMUSG00000022283.16 | 1734.062 | -0.14553 | 0.303716 | -0.47918 | 0.631814 | 0.98928 | 0.004681 |
| ENSMUSG00000022300.11 | 152.5965 | -0.21766 | 0.61739 | -0.35255 | 0.724426 | 0.98928 | 0.004681 |
| ENSMUSG00000022337.8 | 319.926 | 0.230799 | 0.468296 | 0.492849 | 0.622119 | 0.98928 | 0.004681 |
| ENSMUSG00000022351.15 | 819.9124 | -0.17736 | 0.40221 | -0.44096 | 0.659243 | 0.98928 | 0.004681 |
| ENSMUSG00000022359.11 | 54.47459 | -0.39461 | 1.001426 | -0.39405 | 0.693544 | 0.98928 | 0.004681 |
| ENSMUSG00000022364.15 | 20.83589 | -0.47533 | 1.429617 | -0.33249 | 0.739523 | 0.98928 | 0.004681 |
| ENSMUSG00000022425.17 | 671.8701 | 0.147976 | 0.399546 | 0.37036 | 0.711114 | 0.98928 | 0.004681 |
| ENSMUSG00000022453.9 | 16.71634 | 0.987912 | 2.120155 | 0.465962 | 0.641243 | 0.98928 | 0.004681 |
| ENSMUSG00000022477.14 | 555.1441 | 0.173769 | 0.479219 | 0.362608 | 0.716898 | 0.98928 | 0.004681 |
| ENSMUSG00000022503.11 | 111.4379 | 0.349312 | 0.70833 | 0.49315 | 0.621907 | 0.98928 | 0.004681 |
| ENSMUSG00000022507.7 | 136.7648 | -0.24094 | 0.694241 | -0.34705 | 0.728552 | 0.98928 | 0.004681 |
| ENSMUSG00000022528.9 | 26.48335 | -0.77912 | 2.193371 | -0.35522 | 0.722427 | 0.98928 | 0.004681 |
| ENSMUSG00000022533.15 | 483.7153 | 0.134425 | 0.393975 | 0.341201 | 0.732953 | 0.98928 | 0.004681 |
| ENSMUSG00000022544.15 | 50.67395 | 0.455546 | 1.121962 | 0.406026 | 0.684723 | 0.98928 | 0.004681 |
| ENSMUSG00000022556.12 | 23.55484 | 0.684922 | 1.895551 | 0.361331 | 0.717852 | 0.98928 | 0.004681 |
| ENSMUSG00000022557.12 | 37.41023 | 0.575715 | 1.481435 | 0.38862 | 0.697557 | 0.98928 | 0.004681 |
| ENSMUSG00000022672.9 | 14.44778 | 0.660469 | 1.752872 | 0.376792 | 0.706328 | 0.98928 | 0.004681 |
| ENSMUSG00000022677.16 | 161.462 | 0.283442 | 0.789448 | 0.359038 | 0.719567 | 0.98928 | 0.004681 |
| ENSMUSG00000022679.14 | 190.5292 | -0.25744 | 0.64249 | -0.40069 | 0.688647 | 0.98928 | 0.004681 |
| ENSMUSG00000022684.16 | 819.3476 | -0.11195 | 0.336699 | -0.33249 | 0.739522 | 0.98928 | 0.004681 |
| ENSMUSG00000022685.10 | 21.36092 | -0.82428 | 1.691823 | -0.48722 | 0.626105 | 0.98928 | 0.004681 |
| ENSMUSG00000022698.18 | 696.3937 | 0.163808 | 0.335518 | 0.488225 | 0.62539 | 0.98928 | 0.004681 |
| ENSMUSG00000022701.19 | 39.57523 | 0.58987 | 1.492371 | 0.395257 | 0.692653 | 0.98928 | 0.004681 |
| ENSMUSG00000022747.18 | 11.25524 | -1.17826 | 2.51382 | -0.46871 | 0.639274 | 0.98928 | 0.004681 |
| ENSMUSG00000022748.9 | 24.34864 | -0.75734 | 2.155713 | -0.35132 | 0.725349 | 0.98928 | 0.004681 |
| ENSMUSG00000022761.13 | 64.32836 | -0.54724 | 1.409835 | -0.38816 | 0.697899 | 0.98928 | 0.004681 |
| ENSMUSG00000022816.12 | 11.26247 | 1.211295 | 2.509637 | 0.482657 | 0.629339 | 0.98928 | 0.004681 |
| ENSMUSG00000022820.5 | 240.8117 | 0.200615 | 0.544426 | 0.368488 | 0.712509 | 0.98928 | 0.004681 |
| ENSMUSG00000022877.10 | 9218.425 | 0.091774 | 0.211054 | 0.434837 | 0.663681 | 0.98928 | 0.004681 |
| ENSMUSG00000022881.16 | 51.78058 | 0.543236 | 1.405045 | 0.386632 | 0.699028 | 0.98928 | 0.004681 |
| ENSMUSG00000022957.21 | 32.95644 | -0.58252 | 1.611102 | -0.36156 | 0.717678 | 0.98928 | 0.004681 |
| ENSMUSG00000022962.16 | 87.68359 | 0.283331 | 0.848224 | 0.334028 | 0.738358 | 0.98928 | 0.004681 |
| ENSMUSG00000022974.13 | 69.49732 | -0.45243 | 0.994147 | -0.4551 | 0.649039 | 0.98928 | 0.004681 |
| ENSMUSG00000022978.12 | 12.66287 | 1.031229 | 2.356791 | 0.437557 | 0.661708 | 0.98928 | 0.004681 |
| ENSMUSG00000022983.17 | 28.2172 | 0.598405 | 1.66099 | 0.36027 | 0.718645 | 0.98928 | 0.004681 |
| ENSMUSG00000022995.17 | 10.0298 | 1.482904 | 3.303631 | 0.448871 | 0.653525 | 0.98928 | 0.004681 |
| ENSMUSG00000023043.8 | 611.0777 | -0.20542 | 0.439733 | -0.46714 | 0.640402 | 0.98928 | 0.004681 |
| ENSMUSG00000023122.7 | 22.76401 | 0.586292 | 1.838513 | 0.318894 | 0.749807 | 0.98928 | 0.004681 |
| ENSMUSG00000023235.15 | 92.65415 | 0.353779 | 0.870423 | 0.406445 | 0.684415 | 0.98928 | 0.004681 |
| ENSMUSG00000023249.16 | 129.3801 | 0.269233 | 0.819979 | 0.328342 | 0.742653 | 0.98928 | 0.004681 |
| ENSMUSG00000023286.17 | 91.99412 | 0.355916 | 0.780244 | 0.45616 | 0.648275 | 0.98928 | 0.004681 |
| ENSMUSG00000023456.17 | 312.8047 | -0.17992 | 0.449248 | -0.40048 | 0.688801 | 0.98928 | 0.004681 |
| ENSMUSG00000023882.16 | 16.45282 | -0.75797 | 1.754949 | -0.4319 | 0.665813 | 0.98928 | 0.004681 |

| | | | | | | | |
|---|---|---|---|---|---|---|---|
| ENSMUSG00000023892.9 | 11.29557 | 0.840884 | 1.858293 | 0.452504 | 0.650906 | 0.98928 | 0.004681 |
| ENSMUSG00000023938.9 | 16.11103 | 0.995495 | 2.168039 | 0.459169 | 0.646113 | 0.98928 | 0.004681 |
| ENSMUSG00000023947.9 | 17.78492 | 0.856144 | 2.383334 | 0.359221 | 0.71943 | 0.98928 | 0.004681 |
| ENSMUSG00000024007.16 | 13.93494 | 0.949789 | 2.345961 | 0.404861 | 0.68558 | 0.98928 | 0.004681 |
| ENSMUSG00000024037.14 | 10.0758 | -1.07006 | 2.892477 | -0.36995 | 0.711423 | 0.98928 | 0.004681 |
| ENSMUSG00000024038.18 | 1195.14 | -0.13798 | 0.331992 | -0.4156 | 0.677704 | 0.98928 | 0.004681 |
| ENSMUSG00000024045.7 | 126.6423 | 0.296852 | 0.69777 | 0.425429 | 0.670524 | 0.98928 | 0.004681 |
| ENSMUSG00000024077.16 | 24.62151 | 0.640063 | 1.527932 | 0.418908 | 0.675283 | 0.98928 | 0.004681 |
| ENSMUSG00000024079.5 | 79.99331 | -0.39802 | 0.832882 | -0.47788 | 0.632737 | 0.98928 | 0.004681 |
| ENSMUSG00000024081.10 | 281.4387 | -0.19434 | 0.494444 | -0.39305 | 0.694286 | 0.98928 | 0.004681 |
| ENSMUSG00000024104.12 | 110.188 | -0.46771 | 1.187302 | -0.39393 | 0.693635 | 0.98928 | 0.004681 |
| ENSMUSG00000024122.17 | 86.1597 | -0.34142 | 0.734098 | -0.46509 | 0.64187 | 0.98928 | 0.004681 |
| ENSMUSG00000024194.17 | 108.5204 | 0.310127 | 0.804833 | 0.38533 | 0.699993 | 0.98928 | 0.004681 |
| ENSMUSG00000024205.8 | 181.9589 | 0.210232 | 0.569721 | 0.36901 | 0.712121 | 0.98928 | 0.004681 |
| ENSMUSG00000024208.17 | 60.16467 | 0.558592 | 1.71546 | 0.325622 | 0.74471 | 0.98928 | 0.004681 |
| ENSMUSG00000024218.14 | 52.57941 | -0.50262 | 1.097261 | -0.45806 | 0.646906 | 0.98928 | 0.004681 |
| ENSMUSG00000024260.15 | 13.86231 | -0.80783 | 2.163928 | -0.37332 | 0.708914 | 0.98928 | 0.004681 |
| ENSMUSG00000024290.10 | 570.6455 | -0.198 | 0.440479 | -0.44952 | 0.653055 | 0.98928 | 0.004681 |
| ENSMUSG00000024292.16 | 1144.545 | 0.11427 | 0.295182 | 0.387118 | 0.698669 | 0.98928 | 0.004681 |
| ENSMUSG00000024537.6 | 83.9004 | 0.361246 | 0.840445 | 0.429827 | 0.667322 | 0.98928 | 0.004681 |
| ENSMUSG00000024583.4 | 1232.131 | 0.139028 | 0.297308 | 0.467622 | 0.640055 | 0.98928 | 0.004681 |
| ENSMUSG00000024601.10 | 687.5565 | 0.127643 | 0.334928 | 0.381105 | 0.703125 | 0.98928 | 0.004681 |
| ENSMUSG00000024614.8 | 150.572 | 0.420607 | 0.858443 | 0.489965 | 0.624158 | 0.98928 | 0.004681 |
| ENSMUSG00000024621.17 | 92.18817 | -0.30645 | 0.798027 | -0.38402 | 0.700967 | 0.98928 | 0.004681 |
| ENSMUSG00000024645.6 | 154.6388 | 0.334202 | 0.670019 | 0.498795 | 0.617924 | 0.98928 | 0.004681 |
| ENSMUSG00000024666.7 | 14.08676 | -1.31088 | 2.643752 | -0.49584 | 0.620006 | 0.98928 | 0.004681 |
| ENSMUSG00000024668.9 | 200.9199 | -0.29163 | 0.774851 | -0.37636 | 0.706646 | 0.98928 | 0.004681 |
| ENSMUSG00000024678.7 | 21.34789 | 0.71156 | 1.839583 | 0.386805 | 0.6989 | 0.98928 | 0.004681 |
| ENSMUSG00000024754.14 | 15.45868 | -0.90757 | 2.208731 | -0.4109 | 0.681145 | 0.98928 | 0.004681 |
| ENSMUSG00000024759.15 | 393.7239 | -0.22831 | 0.493035 | -0.46307 | 0.643316 | 0.98928 | 0.004681 |
| ENSMUSG00000024780.8 | 168.9089 | -0.35052 | 0.751786 | -0.46624 | 0.641042 | 0.98928 | 0.004681 |
| ENSMUSG00000024805.17 | 71.16838 | 0.382026 | 0.999141 | 0.382355 | 0.702198 | 0.98928 | 0.004681 |
| ENSMUSG00000024830.18 | 13.26254 | 0.88807 | 2.481516 | 0.357874 | 0.720438 | 0.98928 | 0.004681 |
| ENSMUSG00000024845.20 | 99.9797 | 0.299289 | 0.891539 | 0.335699 | 0.737098 | 0.98928 | 0.004681 |
| ENSMUSG00000024896.10 | 41.29537 | -0.38072 | 1.075013 | -0.35415 | 0.723227 | 0.98928 | 0.004681 |
| ENSMUSG00000024914.17 | 58.46545 | 0.318806 | 0.930298 | 0.342692 | 0.73183 | 0.98928 | 0.004681 |
| ENSMUSG00000024947.18 | 33.99833 | -0.58164 | 1.571244 | -0.37018 | 0.71125 | 0.98928 | 0.004681 |
| ENSMUSG00000024982.11 | 263.4463 | -0.25962 | 0.535336 | -0.48496 | 0.627703 | 0.98928 | 0.004681 |
| ENSMUSG00000025017.11 | 117.7648 | 0.342204 | 0.760145 | 0.450183 | 0.652579 | 0.98928 | 0.004681 |
| ENSMUSG00000025041.18 | 53.27783 | 0.386413 | 0.902224 | 0.42829 | 0.66844 | 0.98928 | 0.004681 |
| ENSMUSG00000025044.17 | 43.88196 | 0.807451 | 1.618688 | 0.498831 | 0.617899 | 0.98928 | 0.004681 |
| ENSMUSG00000025050.9 | 46.65688 | -0.50015 | 1.308656 | -0.38218 | 0.702326 | 0.98928 | 0.004681 |
| ENSMUSG00000025066.11 | 156.6335 | 0.308237 | 0.635938 | 0.484696 | 0.627892 | 0.98928 | 0.004681 |
| ENSMUSG00000025162.19 | 136.6157 | -0.3611 | 0.714247 | -0.50557 | 0.613162 | 0.98928 | 0.004681 |

| | | | | | | | |
|---|---|---|---|---|---|---|---|
| ENSMUSG00000025176.15 | 611.6715 | -0.15586 | 0.385445 | -0.40438 | 0.685937 | 0.98928 | 0.004681 |
| ENSMUSG00000025204.11 | 420.0801 | 0.199477 | 0.46606 | 0.428006 | 0.668647 | 0.98928 | 0.004681 |
| ENSMUSG00000025239.4 | 181.1653 | 0.262957 | 0.586092 | 0.448662 | 0.653675 | 0.98928 | 0.004681 |
| ENSMUSG00000025245.15 | 40.93296 | 0.39541 | 1.215422 | 0.325327 | 0.744933 | 0.98928 | 0.004681 |
| ENSMUSG00000025289.16 | 1134.679 | 0.105448 | 0.308791 | 0.341488 | 0.732736 | 0.98928 | 0.004681 |
| ENSMUSG00000025323.11 | 37.0181 | 0.48012 | 1.233434 | 0.389255 | 0.697088 | 0.98928 | 0.004681 |
| ENSMUSG00000025351.15 | 11.37463 | -0.8838 | 2.475182 | -0.35706 | 0.721044 | 0.98928 | 0.004681 |
| ENSMUSG00000025353.6 | 146.6863 | 0.327067 | 0.810694 | 0.403441 | 0.686624 | 0.98928 | 0.004681 |
| ENSMUSG00000025355.8 | 188.6378 | 0.262615 | 0.53946 | 0.48681 | 0.626393 | 0.98928 | 0.004681 |
| ENSMUSG00000025364.14 | 122.0766 | 0.359071 | 0.748213 | 0.479905 | 0.631295 | 0.98928 | 0.004681 |
| ENSMUSG00000025371.13 | 18.15303 | -0.73363 | 2.028579 | -0.36165 | 0.717615 | 0.98928 | 0.004681 |
| ENSMUSG00000025393.13 | 3819.116 | 0.082014 | 0.215169 | 0.381161 | 0.703084 | 0.98928 | 0.004681 |
| ENSMUSG00000025409.15 | 25.09517 | 0.796208 | 1.911483 | 0.416539 | 0.677016 | 0.98928 | 0.004681 |
| ENSMUSG00000025421.16 | 14.02129 | 1.079398 | 2.405003 | 0.448814 | 0.653566 | 0.98928 | 0.004681 |
| ENSMUSG00000025470.12 | 12.77939 | 0.877342 | 2.461845 | 0.356376 | 0.721559 | 0.98928 | 0.004681 |
| ENSMUSG00000025504.15 | 53.2421 | 0.5626 | 1.234146 | 0.455862 | 0.648489 | 0.98928 | 0.004681 |
| ENSMUSG00000025525.13 | 305.1269 | -0.2088 | 0.570041 | -0.36629 | 0.714146 | 0.98928 | 0.004681 |
| ENSMUSG00000025545.11 | 75.8667 | 0.581139 | 1.218268 | 0.477021 | 0.633347 | 0.98928 | 0.004681 |
| ENSMUSG00000025588.5 | 26.9975 | -0.73863 | 1.806101 | -0.40896 | 0.682567 | 0.98928 | 0.004681 |
| ENSMUSG00000025612.6 | 116.7941 | -0.26253 | 0.640008 | -0.4102 | 0.681662 | 0.98928 | 0.004681 |
| ENSMUSG00000025702.16 | 410.8259 | -0.15341 | 0.476709 | -0.32182 | 0.747589 | 0.98928 | 0.004681 |
| ENSMUSG00000025747.13 | 34.99213 | 0.497492 | 1.265352 | 0.393165 | 0.694198 | 0.98928 | 0.004681 |
| ENSMUSG00000025758.11 | 14.26671 | -0.8975 | 2.321265 | -0.38664 | 0.699021 | 0.98928 | 0.004681 |
| ENSMUSG00000025792.10 | 31.675 | 0.601704 | 1.578285 | 0.381239 | 0.703026 | 0.98928 | 0.004681 |
| ENSMUSG00000025794.10 | 711.3565 | 0.114061 | 0.341573 | 0.333929 | 0.738433 | 0.98928 | 0.004681 |
| ENSMUSG00000025795.9 | 354.5401 | -0.16811 | 0.42613 | -0.39449 | 0.693217 | 0.98928 | 0.004681 |
| ENSMUSG00000025812.19 | 293.0046 | -0.24088 | 0.504826 | -0.47716 | 0.63325 | 0.98928 | 0.004681 |
| ENSMUSG00000025815.14 | 191.7988 | -0.19577 | 0.606852 | -0.3226 | 0.746997 | 0.98928 | 0.004681 |
| ENSMUSG00000025816.16 | 42.3705 | 0.724882 | 1.459583 | 0.496637 | 0.619445 | 0.98928 | 0.004681 |
| ENSMUSG00000025878.17 | 53.4598 | -0.47496 | 1.276796 | -0.372 | 0.709896 | 0.98928 | 0.004681 |
| ENSMUSG00000025917.10 | 375.32 | -0.1894 | 0.394806 | -0.47973 | 0.631421 | 0.98928 | 0.004681 |
| ENSMUSG00000025920.20 | 19.49212 | -0.86599 | 1.751468 | -0.49444 | 0.620997 | 0.98928 | 0.004681 |
| ENSMUSG00000025937.7 | 1186.385 | 0.125616 | 0.320806 | 0.391564 | 0.69538 | 0.98928 | 0.004681 |
| ENSMUSG00000025969.16 | 95.80023 | 0.353864 | 1.063737 | 0.332661 | 0.73939 | 0.98928 | 0.004681 |
| ENSMUSG00000025971.7 | 85.32919 | 0.400841 | 0.924857 | 0.433408 | 0.664718 | 0.98928 | 0.004681 |
| ENSMUSG00000025993.11 | 682.9355 | -0.16432 | 0.344299 | -0.47727 | 0.633168 | 0.98928 | 0.004681 |
| ENSMUSG00000026028.13 | 46.16555 | -0.42579 | 0.97812 | -0.43531 | 0.663335 | 0.98928 | 0.004681 |
| ENSMUSG00000026035.16 | 34.77762 | -0.76988 | 1.803948 | -0.42678 | 0.669541 | 0.98928 | 0.004681 |
| ENSMUSG00000026047.14 | 31.26097 | -0.66147 | 1.71138 | -0.38651 | 0.699116 | 0.98928 | 0.004681 |
| ENSMUSG00000026096.15 | 103.2995 | -0.32358 | 0.730494 | -0.44297 | 0.65779 | 0.98928 | 0.004681 |
| ENSMUSG00000026219.15 | 332.3199 | -0.28796 | 0.575774 | -0.50013 | 0.616983 | 0.98928 | 0.004681 |
| ENSMUSG00000026245.17 | 172.8787 | -0.23101 | 0.603506 | -0.38278 | 0.701881 | 0.98928 | 0.004681 |
| ENSMUSG00000026260.13 | 793.0847 | 0.144729 | 0.334577 | 0.432574 | 0.665324 | 0.98928 | 0.004681 |
| ENSMUSG00000026377.13 | 178.2237 | -0.27357 | 0.636471 | -0.42982 | 0.667327 | 0.98928 | 0.004681 |

| | | | | | | | |
|---|---|---|---|---|---|---|---|
| ENSMUSG00000026384.14 | 36.06797 | 0.433173 | 1.187282 | 0.364844 | 0.715228 | 0.98928 | 0.004681 |
| ENSMUSG00000026411.18 | 83.19108 | -0.35982 | 0.937463 | -0.38383 | 0.701107 | 0.98928 | 0.004681 |
| ENSMUSG00000026417.14 | 10974.28 | 0.100454 | 0.21635 | 0.464311 | 0.642425 | 0.98928 | 0.004681 |
| ENSMUSG00000026433.14 | 58.11046 | -0.37139 | 0.983902 | -0.37746 | 0.705829 | 0.98928 | 0.004681 |
| ENSMUSG00000026457.15 | 179.3478 | 0.207321 | 0.592489 | 0.349916 | 0.726402 | 0.98928 | 0.004681 |
| ENSMUSG00000026466.17 | 142.9683 | -0.29346 | 0.720683 | -0.4072 | 0.683865 | 0.98928 | 0.004681 |
| ENSMUSG00000026490.19 | 88.63946 | -0.36957 | 0.898466 | -0.41133 | 0.680828 | 0.98928 | 0.004681 |
| ENSMUSG00000026502.14 | 108.6269 | 0.322241 | 0.726581 | 0.443503 | 0.657402 | 0.98928 | 0.004681 |
| ENSMUSG00000026516.9 | 47.45151 | -0.33452 | 0.97491 | -0.34313 | 0.731497 | 0.98928 | 0.004681 |
| ENSMUSG00000026542.7 | 854.066 | -0.21817 | 0.524985 | -0.41558 | 0.677718 | 0.98928 | 0.004681 |
| ENSMUSG00000026571.13 | 50.11247 | -0.50674 | 1.082657 | -0.46805 | 0.63975 | 0.98928 | 0.004681 |
| ENSMUSG00000026576.13 | 280.1831 | 0.177453 | 0.534929 | 0.331732 | 0.740092 | 0.98928 | 0.004681 |
| ENSMUSG00000026585.14 | 40.2979 | -0.60655 | 1.251036 | -0.48484 | 0.627793 | 0.98928 | 0.004681 |
| ENSMUSG00000026615.15 | 720.6052 | 0.126445 | 0.313488 | 0.403348 | 0.686692 | 0.98928 | 0.004681 |
| ENSMUSG00000026618.12 | 62.61503 | -0.32802 | 0.949781 | -0.34536 | 0.729822 | 0.98928 | 0.004681 |
| ENSMUSG00000026662.14 | 148.4661 | -0.35166 | 0.845906 | -0.41572 | 0.677612 | 0.98928 | 0.004681 |
| ENSMUSG00000026664.8 | 17650.42 | 0.065309 | 0.199138 | 0.327957 | 0.742944 | 0.98928 | 0.004681 |
| ENSMUSG00000026672.12 | 247.1084 | -0.2227 | 0.629189 | -0.35395 | 0.723379 | 0.98928 | 0.004681 |
| ENSMUSG00000026694.20 | 16.21406 | -1.11363 | 2.565376 | -0.4341 | 0.664217 | 0.98928 | 0.004681 |
| ENSMUSG00000026696.16 | 102.0314 | -0.44439 | 0.964082 | -0.46094 | 0.64484 | 0.98928 | 0.004681 |
| ENSMUSG00000026701.16 | 3132.119 | 0.088135 | 0.26877 | 0.327921 | 0.742971 | 0.98928 | 0.004681 |
| ENSMUSG00000026705.17 | 17.96088 | 0.826122 | 1.79376 | 0.460553 | 0.645119 | 0.98928 | 0.004681 |
| ENSMUSG00000026709.11 | 89.10935 | 0.429765 | 1.145647 | 0.375129 | 0.707565 | 0.98928 | 0.004681 |
| ENSMUSG00000026754.17 | 96.35132 | -0.44884 | 1.326313 | -0.33841 | 0.735055 | 0.98928 | 0.004681 |
| ENSMUSG00000026755.15 | 442.3135 | -0.17437 | 0.413717 | -0.42148 | 0.673406 | 0.98928 | 0.004681 |
| ENSMUSG00000026827.13 | 250.6649 | 0.230826 | 0.508512 | 0.453923 | 0.649884 | 0.98928 | 0.004681 |
| ENSMUSG00000026843.16 | 63.25146 | 0.360023 | 1.021552 | 0.352427 | 0.724518 | 0.98928 | 0.004681 |
| ENSMUSG00000026864.14 | 3989.674 | 0.11537 | 0.249722 | 0.461993 | 0.644086 | 0.98928 | 0.004681 |
| ENSMUSG00000026914.16 | 633.5433 | 0.144356 | 0.351963 | 0.410147 | 0.681698 | 0.98928 | 0.004681 |
| ENSMUSG00000026922.14 | 539.9155 | 0.139369 | 0.438999 | 0.31747 | 0.750887 | 0.98928 | 0.004681 |
| ENSMUSG00000026926.15 | 258.1981 | -0.25251 | 0.514127 | -0.49114 | 0.623327 | 0.98928 | 0.004681 |
| ENSMUSG00000026938.11 | 62.87699 | -0.40842 | 1.2085 | -0.33796 | 0.735394 | 0.98928 | 0.004681 |
| ENSMUSG00000026974.12 | 12.71239 | -1.02871 | 2.181184 | -0.47163 | 0.637191 | 0.98928 | 0.004681 |
| ENSMUSG00000026987.17 | 131.6738 | -0.21091 | 0.6605 | -0.31932 | 0.749487 | 0.98928 | 0.004681 |
| ENSMUSG00000026991.21 | 172.9159 | -0.26357 | 0.650402 | -0.40523 | 0.685305 | 0.98928 | 0.004681 |
| ENSMUSG00000026999.15 | 56.80209 | 0.624459 | 1.370729 | 0.455567 | 0.648701 | 0.98928 | 0.004681 |
| ENSMUSG00000027004.4 | 14.0523 | 1.060766 | 2.252089 | 0.471014 | 0.637631 | 0.98928 | 0.004681 |
| ENSMUSG00000027012.16 | 184.9078 | 0.238987 | 0.550677 | 0.433987 | 0.664298 | 0.98928 | 0.004681 |
| ENSMUSG00000027091.15 | 678.6743 | 0.140545 | 0.380683 | 0.369192 | 0.711985 | 0.98928 | 0.004681 |
| ENSMUSG00000027099.10 | 381.4961 | -0.18094 | 0.420379 | -0.43042 | 0.666888 | 0.98928 | 0.004681 |
| ENSMUSG00000027122.16 | 54.12838 | -0.58396 | 1.176777 | -0.49624 | 0.619725 | 0.98928 | 0.004681 |
| ENSMUSG00000027132.4 | 41.73198 | 0.505266 | 1.053476 | 0.479618 | 0.631499 | 0.98928 | 0.004681 |
| ENSMUSG00000027160.17 | 14.92951 | 1.001483 | 2.031305 | 0.493024 | 0.621995 | 0.98928 | 0.004681 |
| ENSMUSG00000027207.16 | 57.09586 | -0.5488 | 1.53949 | -0.35648 | 0.721482 | 0.98928 | 0.004681 |

| | | | | | | | |
|---|---|---|---|---|---|---|---|
| ENSMUSG00000027236.9 | 295.4613 | -0.18772 | 0.484551 | -0.38741 | 0.698453 | 0.98928 | 0.004681 |
| ENSMUSG00000027285.14 | 70.37358 | -0.40242 | 1.205272 | -0.33389 | 0.738466 | 0.98928 | 0.004681 |
| ENSMUSG00000027286.17 | 37.50625 | 0.741009 | 1.517821 | 0.488206 | 0.625404 | 0.98928 | 0.004681 |
| ENSMUSG00000027305.10 | 183.5974 | 0.232416 | 0.592889 | 0.392007 | 0.695053 | 0.98928 | 0.004681 |
| ENSMUSG00000027309.19 | 154.214 | 0.228142 | 0.711165 | 0.3208 | 0.748362 | 0.98928 | 0.004681 |
| ENSMUSG00000027341.11 | 127.7178 | 0.27173 | 0.691034 | 0.393222 | 0.694155 | 0.98928 | 0.004681 |
| ENSMUSG00000027342.15 | 123.918 | 0.309848 | 0.674429 | 0.459423 | 0.64593 | 0.98928 | 0.004681 |
| ENSMUSG00000027347.19 | 20.42311 | 0.770033 | 2.191824 | 0.351321 | 0.725348 | 0.98928 | 0.004681 |
| ENSMUSG00000027411.18 | 35.99098 | -0.51705 | 1.602249 | -0.3227 | 0.746922 | 0.98928 | 0.004681 |
| ENSMUSG00000027423.16 | 328.3593 | 0.22469 | 0.620861 | 0.3619 | 0.717427 | 0.98928 | 0.004681 |
| ENSMUSG00000027435.9 | 28.02006 | 0.70966 | 2.213371 | 0.320624 | 0.748495 | 0.98928 | 0.004681 |
| ENSMUSG00000027439.10 | 233.315 | 0.190627 | 0.531878 | 0.358403 | 0.720042 | 0.98928 | 0.004681 |
| ENSMUSG00000027546.16 | 12.66103 | 1.278044 | 2.666376 | 0.479319 | 0.631712 | 0.98928 | 0.004681 |
| ENSMUSG00000027569.16 | 24.17796 | -0.73052 | 1.699047 | -0.42996 | 0.667227 | 0.98928 | 0.004681 |
| ENSMUSG00000027599.10 | 331.3463 | 0.215617 | 0.493245 | 0.43714 | 0.66201 | 0.98928 | 0.004681 |
| ENSMUSG00000027601.14 | 342.7216 | -0.22157 | 0.46494 | -0.47656 | 0.633679 | 0.98928 | 0.004681 |
| ENSMUSG00000027610.18 | 80.74222 | -0.37901 | 0.958381 | -0.39547 | 0.692498 | 0.98928 | 0.004681 |
| ENSMUSG00000027613.16 | 119.0251 | 0.379363 | 0.806045 | 0.470648 | 0.637892 | 0.98928 | 0.004681 |
| ENSMUSG00000027620.17 | 596.1711 | -0.15895 | 0.343545 | -0.46269 | 0.643589 | 0.98928 | 0.004681 |
| ENSMUSG00000027630.14 | 151.0847 | -0.28638 | 0.772339 | -0.37079 | 0.710792 | 0.98928 | 0.004681 |
| ENSMUSG00000027636.12 | 135.7572 | 0.335897 | 0.840108 | 0.399827 | 0.689284 | 0.98928 | 0.004681 |
| ENSMUSG00000027660.17 | 52.02604 | -0.46221 | 1.17027 | -0.39496 | 0.692874 | 0.98928 | 0.004681 |
| ENSMUSG00000027665.14 | 205.9457 | 0.18274 | 0.499294 | 0.365997 | 0.714367 | 0.98928 | 0.004681 |
| ENSMUSG00000027671.15 | 53.94106 | 0.392961 | 1.06891 | 0.367628 | 0.713151 | 0.98928 | 0.004681 |
| ENSMUSG00000027673.13 | 1612.324 | 0.166039 | 0.398891 | 0.416251 | 0.677226 | 0.98928 | 0.004681 |
| ENSMUSG00000027677.18 | 199.0175 | 0.313122 | 0.674032 | 0.46455 | 0.642254 | 0.98928 | 0.004681 |
| ENSMUSG00000027742.14 | 197.524 | 0.200422 | 0.616507 | 0.325094 | 0.74511 | 0.98928 | 0.004681 |
| ENSMUSG00000027792.12 | 272.7783 | 0.226769 | 0.582955 | 0.388999 | 0.697277 | 0.98928 | 0.004681 |
| ENSMUSG00000027796.3 | 69.06524 | -0.32424 | 0.804534 | -0.40302 | 0.686935 | 0.98928 | 0.004681 |
| ENSMUSG00000027810.15 | 270.1374 | -0.27499 | 0.615932 | -0.44647 | 0.655261 | 0.98928 | 0.004681 |
| ENSMUSG00000027852.15 | 122.1679 | -0.39841 | 0.801695 | -0.49696 | 0.619217 | 0.98928 | 0.004681 |
| ENSMUSG00000027860.16 | 29.8714 | 0.551539 | 1.615383 | 0.341429 | 0.732781 | 0.98928 | 0.004681 |
| ENSMUSG00000027884.17 | 232.9276 | 0.214659 | 0.642064 | 0.334326 | 0.738133 | 0.98928 | 0.004681 |
| ENSMUSG00000027905.16 | 78.68058 | 0.486583 | 1.309078 | 0.371699 | 0.710117 | 0.98928 | 0.004681 |
| ENSMUSG00000027936.19 | 12.09727 | -0.75135 | 2.375393 | -0.31631 | 0.75177 | 0.98928 | 0.004681 |
| ENSMUSG00000027998.12 | 81.27739 | -0.29216 | 0.883449 | -0.3307 | 0.74087 | 0.98928 | 0.004681 |
| ENSMUSG00000028007.14 | 136.1166 | 0.300196 | 0.743691 | 0.403657 | 0.686465 | 0.98928 | 0.004681 |
| ENSMUSG00000028013.17 | 395.8651 | 0.161366 | 0.508964 | 0.317049 | 0.751207 | 0.98928 | 0.004681 |
| ENSMUSG00000028080.17 | 120.3857 | 0.407221 | 0.995313 | 0.409138 | 0.682438 | 0.98928 | 0.004681 |
| ENSMUSG00000028085.13 | 120.3763 | -0.44192 | 1.12977 | -0.39116 | 0.695678 | 0.98928 | 0.004681 |
| ENSMUSG00000028093.16 | 140.3568 | -0.21468 | 0.654604 | -0.32795 | 0.742946 | 0.98928 | 0.004681 |
| ENSMUSG00000028099.9 | 39.70237 | -0.59581 | 1.545147 | -0.3856 | 0.699791 | 0.98928 | 0.004681 |
| ENSMUSG00000028104.15 | 21.46036 | -0.90551 | 1.915643 | -0.47269 | 0.636432 | 0.98928 | 0.004681 |
| ENSMUSG00000028134.12 | 64.03747 | -0.51398 | 1.148217 | -0.44763 | 0.654418 | 0.98928 | 0.004681 |

| | | | | | | | |
|---|---|---|---|---|---|---|---|
| ENSMUSG00000028149.13 | 113.4674 | 0.236412 | 0.661937 | 0.357151 | 0.720979 | 0.98928 | 0.004681 |
| ENSMUSG00000028161.18 | 318.0173 | -0.17536 | 0.432015 | -0.40592 | 0.684801 | 0.98928 | 0.004681 |
| ENSMUSG00000028165.10 | 295.799 | 0.237298 | 0.61335 | 0.386889 | 0.698838 | 0.98928 | 0.004681 |
| ENSMUSG00000028185.13 | 137.0533 | 0.334147 | 0.722809 | 0.46229 | 0.643874 | 0.98928 | 0.004681 |
| ENSMUSG00000028212.17 | 21.1728 | 0.853816 | 2.121911 | 0.402381 | 0.687404 | 0.98928 | 0.004681 |
| ENSMUSG00000028228.6 | 351.5626 | -0.21416 | 0.445001 | -0.48125 | 0.630338 | 0.98928 | 0.004681 |
| ENSMUSG00000028245.16 | 70.66559 | -0.38142 | 1.094741 | -0.34841 | 0.727533 | 0.98928 | 0.004681 |
| ENSMUSG00000028309.15 | 186.6022 | 0.365858 | 0.741353 | 0.4935 | 0.621659 | 0.98928 | 0.004681 |
| ENSMUSG00000028312.20 | 22.72174 | 0.849796 | 1.715789 | 0.49528 | 0.620402 | 0.98928 | 0.004681 |
| ENSMUSG00000028327.15 | 327.7137 | -0.24178 | 0.536635 | -0.45054 | 0.65232 | 0.98928 | 0.004681 |
| ENSMUSG00000028333.11 | 472.7173 | 0.188132 | 0.38301 | 0.491193 | 0.62329 | 0.98928 | 0.004681 |
| ENSMUSG00000028343.11 | 815.6116 | -0.15408 | 0.361732 | -0.42595 | 0.670143 | 0.98928 | 0.004681 |
| ENSMUSG00000028397.14 | 20.6678 | -0.74259 | 2.063966 | -0.35979 | 0.719004 | 0.98928 | 0.004681 |
| ENSMUSG00000028398.9 | 272.8037 | -0.20111 | 0.581705 | -0.34572 | 0.729549 | 0.98928 | 0.004681 |
| ENSMUSG00000028405.10 | 1288.499 | 0.106604 | 0.323239 | 0.3298 | 0.741551 | 0.98928 | 0.004681 |
| ENSMUSG00000028409.12 | 239.1538 | -0.18244 | 0.527894 | -0.34559 | 0.729648 | 0.98928 | 0.004681 |
| ENSMUSG00000028437.15 | 171.7792 | 0.35196 | 0.840156 | 0.418922 | 0.675273 | 0.98928 | 0.004681 |
| ENSMUSG00000028465.17 | 14.66288 | -0.89385 | 2.435256 | -0.36705 | 0.713585 | 0.98928 | 0.004681 |
| ENSMUSG00000028468.16 | 32.39759 | -0.59019 | 1.639886 | -0.3599 | 0.718923 | 0.98928 | 0.004681 |
| ENSMUSG00000028517.9 | 690.9055 | 0.203001 | 0.462418 | 0.438998 | 0.660663 | 0.98928 | 0.004681 |
| ENSMUSG00000028518.9 | 288.4825 | -0.19213 | 0.461459 | -0.41636 | 0.677144 | 0.98928 | 0.004681 |
| ENSMUSG00000028522.17 | 158.9459 | -0.37519 | 0.778419 | -0.48199 | 0.629815 | 0.98928 | 0.004681 |
| ENSMUSG00000028549.18 | 37.44964 | -0.65978 | 1.421647 | -0.46409 | 0.64258 | 0.98928 | 0.004681 |
| ENSMUSG00000028607.17 | 653.0393 | -0.14118 | 0.328358 | -0.42997 | 0.667218 | 0.98928 | 0.004681 |
| ENSMUSG00000028618.12 | 1526.118 | -0.10415 | 0.282783 | -0.3683 | 0.712647 | 0.98928 | 0.004681 |
| ENSMUSG00000028639.15 | 1006.353 | 0.165758 | 0.384843 | 0.430717 | 0.666674 | 0.98928 | 0.004681 |
| ENSMUSG00000028648.14 | 172.5144 | -0.288 | 0.796265 | -0.36169 | 0.717586 | 0.98928 | 0.004681 |
| ENSMUSG00000028692.15 | 1979.146 | -0.12435 | 0.285673 | -0.43529 | 0.663352 | 0.98928 | 0.004681 |
| ENSMUSG00000028696.13 | 66.06835 | 0.493606 | 1.060472 | 0.465459 | 0.641603 | 0.98928 | 0.004681 |
| ENSMUSG00000028708.17 | 110.9311 | -0.29716 | 0.91537 | -0.32463 | 0.745461 | 0.98928 | 0.004681 |
| ENSMUSG00000028713.18 | 43.07242 | 0.577463 | 1.159921 | 0.497847 | 0.618592 | 0.98928 | 0.004681 |
| ENSMUSG00000028766.11 | 22.8709 | 0.731883 | 2.028635 | 0.360776 | 0.718267 | 0.98928 | 0.004681 |
| ENSMUSG00000028772.20 | 149.4087 | -0.36239 | 0.791861 | -0.45764 | 0.647211 | 0.98928 | 0.004681 |
| ENSMUSG00000028798.17 | 795.8494 | 0.205389 | 0.44579 | 0.460731 | 0.644992 | 0.98928 | 0.004681 |
| ENSMUSG00000028809.17 | 145.7916 | 0.295124 | 0.601249 | 0.490852 | 0.623531 | 0.98928 | 0.004681 |
| ENSMUSG00000028821.9 | 107.1567 | 0.341296 | 0.6854 | 0.497951 | 0.618518 | 0.98928 | 0.004681 |
| ENSMUSG00000028830.15 | 67.20553 | 0.621367 | 1.348875 | 0.460656 | 0.645045 | 0.98928 | 0.004681 |
| ENSMUSG00000028840.11 | 28.51836 | 0.586422 | 1.408552 | 0.41633 | 0.677169 | 0.98928 | 0.004681 |
| ENSMUSG00000028851.7 | 101.7183 | 0.427074 | 1.158517 | 0.368639 | 0.712397 | 0.98928 | 0.004681 |
| ENSMUSG00000028886.16 | 12.33409 | 0.894907 | 2.248538 | 0.397995 | 0.690634 | 0.98928 | 0.004681 |
| ENSMUSG00000028902.5 | 105.8013 | 0.275985 | 0.815374 | 0.338476 | 0.735005 | 0.98928 | 0.004681 |
| ENSMUSG00000028961.16 | 56.09698 | 0.350944 | 0.983721 | 0.356751 | 0.721278 | 0.98928 | 0.004681 |
| ENSMUSG00000028967.11 | 3682.015 | -0.19011 | 0.376483 | -0.50495 | 0.613594 | 0.98928 | 0.004681 |
| ENSMUSG00000028998.7 | 666.3529 | 0.149703 | 0.385699 | 0.388133 | 0.697917 | 0.98928 | 0.004681 |

| | | | | | | | |
|---|---|---|---|---|---|---|---|
| ENSMUSG00000029030.15 | 306.9058 | -0.27187 | 0.550869 | -0.49353 | 0.621637 | 0.98928 | 0.004681 |
| ENSMUSG00000029050.16 | 48.27466 | 0.45313 | 1.022967 | 0.442957 | 0.657797 | 0.98928 | 0.004681 |
| ENSMUSG00000029070.10 | 10.91579 | -1.0435 | 2.507737 | -0.41611 | 0.677328 | 0.98928 | 0.004681 |
| ENSMUSG00000029090.13 | 182.0279 | 0.251988 | 0.639793 | 0.393858 | 0.693686 | 0.98928 | 0.004681 |
| ENSMUSG00000029102.10 | 330.635 | -0.31069 | 0.780156 | -0.39824 | 0.69045 | 0.98928 | 0.004681 |
| ENSMUSG00000029145.13 | 145.5786 | -0.29458 | 0.725592 | -0.40599 | 0.684752 | 0.98928 | 0.004681 |
| ENSMUSG00000029153.12 | 638.6486 | 0.17668 | 0.35916 | 0.491926 | 0.622772 | 0.98928 | 0.004681 |
| ENSMUSG00000029169.12 | 279.0167 | 0.256229 | 0.619081 | 0.413886 | 0.678958 | 0.98928 | 0.004681 |
| ENSMUSG00000029179.15 | 15.7166 | 0.940439 | 1.974437 | 0.476307 | 0.633855 | 0.98928 | 0.004681 |
| ENSMUSG00000029185.15 | 102.6816 | 0.358565 | 0.873085 | 0.410688 | 0.681301 | 0.98928 | 0.004681 |
| ENSMUSG00000029198.4 | 509.9526 | -0.16125 | 0.390281 | -0.41316 | 0.679488 | 0.98928 | 0.004681 |
| ENSMUSG00000029201.15 | 866.2062 | 0.131725 | 0.369226 | 0.356759 | 0.721272 | 0.98928 | 0.004681 |
| ENSMUSG00000029203.17 | 594.767 | -0.20794 | 0.479013 | -0.43411 | 0.664211 | 0.98928 | 0.004681 |
| ENSMUSG00000029208.17 | 136.7262 | -0.28382 | 0.729447 | -0.38909 | 0.697207 | 0.98928 | 0.004681 |
| ENSMUSG00000029213.12 | 179.9478 | 0.229979 | 0.60164 | 0.382254 | 0.702273 | 0.98928 | 0.004681 |
| ENSMUSG00000029229.9 | 202.4798 | -0.23368 | 0.57369 | -0.40733 | 0.683764 | 0.98928 | 0.004681 |
| ENSMUSG00000029233.16 | 62.17877 | 0.540071 | 1.171637 | 0.460954 | 0.644832 | 0.98928 | 0.004681 |
| ENSMUSG00000029249.16 | 67.80786 | -0.51184 | 1.108724 | -0.46165 | 0.644332 | 0.98928 | 0.004681 |
| ENSMUSG00000029265.5 | 147.3896 | 0.23396 | 0.65354 | 0.357988 | 0.720352 | 0.98928 | 0.004681 |
| ENSMUSG00000029313.19 | 127.6985 | -0.41665 | 0.875815 | -0.47572 | 0.634271 | 0.98928 | 0.004681 |
| ENSMUSG00000029369.18 | 2146.619 | 0.135559 | 0.314085 | 0.431599 | 0.666033 | 0.98928 | 0.004681 |
| ENSMUSG00000029381.17 | 21.4715 | 0.74933 | 2.098701 | 0.357045 | 0.721058 | 0.98928 | 0.004681 |
| ENSMUSG00000029387.11 | 56.27356 | 0.375235 | 1.081309 | 0.347019 | 0.728577 | 0.98928 | 0.004681 |
| ENSMUSG00000029404.16 | 127.8815 | -0.25186 | 0.722334 | -0.34868 | 0.72733 | 0.98928 | 0.004681 |
| ENSMUSG00000029415.5 | 25.95241 | -0.4275 | 1.32784 | -0.32195 | 0.747489 | 0.98928 | 0.004681 |
| ENSMUSG00000029426.9 | 342.6542 | -0.24347 | 0.570295 | -0.42692 | 0.669436 | 0.98928 | 0.004681 |
| ENSMUSG00000029432.13 | 26.67099 | 0.64424 | 1.369395 | 0.470456 | 0.638029 | 0.98928 | 0.004681 |
| ENSMUSG00000029456.13 | 91.84556 | -0.42481 | 1.169009 | -0.36339 | 0.716312 | 0.98928 | 0.004681 |
| ENSMUSG00000029463.6 | 45.03335 | -0.66087 | 1.733346 | -0.38127 | 0.703004 | 0.98928 | 0.004681 |
| ENSMUSG00000029486.14 | 185.254 | -0.22549 | 0.622773 | -0.36208 | 0.717291 | 0.98928 | 0.004681 |
| ENSMUSG00000029499.12 | 1323.33 | 0.162485 | 0.326246 | 0.498045 | 0.618453 | 0.98928 | 0.004681 |
| ENSMUSG00000029598.13 | 75.52787 | -0.45418 | 0.975018 | -0.46581 | 0.641349 | 0.98928 | 0.004681 |
| ENSMUSG00000029621.15 | 426.7742 | 0.224147 | 0.443572 | 0.505322 | 0.613333 | 0.98928 | 0.004681 |
| ENSMUSG00000029625.17 | 39.66229 | 0.641224 | 1.42882 | 0.448778 | 0.653591 | 0.98928 | 0.004681 |
| ENSMUSG00000029629.18 | 259.0345 | 0.158442 | 0.487662 | 0.324901 | 0.745256 | 0.98928 | 0.004681 |
| ENSMUSG00000029648.14 | 85.82027 | -0.25689 | 0.811791 | -0.31645 | 0.751665 | 0.98928 | 0.004681 |
| ENSMUSG00000029655.18 | 218.2253 | 0.218696 | 0.519856 | 0.420686 | 0.673984 | 0.98928 | 0.004681 |
| ENSMUSG00000029664.11 | 443.7701 | -0.16531 | 0.434834 | -0.38017 | 0.703821 | 0.98928 | 0.004681 |
| ENSMUSG00000029669.15 | 953.795 | -0.17209 | 0.362119 | -0.47524 | 0.634615 | 0.98928 | 0.004681 |
| ENSMUSG00000029684.15 | 384.7295 | 0.191456 | 0.429095 | 0.446185 | 0.655464 | 0.98928 | 0.004681 |
| ENSMUSG00000029687.17 | 19.63429 | 0.79316 | 2.469683 | 0.321159 | 0.74809 | 0.98928 | 0.004681 |
| ENSMUSG00000029703.11 | 28.87169 | -0.90722 | 1.978119 | -0.45863 | 0.646502 | 0.98928 | 0.004681 |
| ENSMUSG00000029735.18 | 138.125 | 0.211393 | 0.652771 | 0.323839 | 0.74606 | 0.98928 | 0.004681 |
| ENSMUSG00000029781.8 | 96.95973 | -0.27605 | 0.715599 | -0.38576 | 0.699672 | 0.98928 | 0.004681 |

| | | | | | | | |
|---|---|---|---|---|---|---|---|
| ENSMUSG00000029782.21 | 33.2593 | -0.51657 | 1.524108 | -0.33893 | 0.73466 | 0.98928 | 0.004681 |
| ENSMUSG00000029822.16 | 29.01241 | 0.586755 | 1.569875 | 0.373759 | 0.708583 | 0.98928 | 0.004681 |
| ENSMUSG00000029859.7 | 25.09893 | -0.94521 | 1.99171 | -0.47457 | 0.635092 | 0.98928 | 0.004681 |
| ENSMUSG00000029860.17 | 27.93948 | -0.50012 | 1.398849 | -0.35752 | 0.720701 | 0.98928 | 0.004681 |
| ENSMUSG00000030016.15 | 130.2484 | -0.3242 | 0.749043 | -0.43282 | 0.665144 | 0.98928 | 0.004681 |
| ENSMUSG00000030083.12 | 15.3692 | 1.071222 | 2.949404 | 0.363199 | 0.716456 | 0.98928 | 0.004681 |
| ENSMUSG00000030095.11 | 29.31953 | 0.678153 | 1.371799 | 0.494353 | 0.621057 | 0.98928 | 0.004681 |
| ENSMUSG00000030157.6 | 404.9472 | 0.18654 | 0.549305 | 0.339592 | 0.734163 | 0.98928 | 0.004681 |
| ENSMUSG00000030159.16 | 49.71066 | -0.45157 | 1.173027 | -0.38497 | 0.700263 | 0.98928 | 0.004681 |
| ENSMUSG00000030177.15 | 44.60356 | -0.83209 | 1.929241 | -0.4313 | 0.666248 | 0.98928 | 0.004681 |
| ENSMUSG00000030216.13 | 39.10471 | 0.368507 | 1.085384 | 0.339517 | 0.73422 | 0.98928 | 0.004681 |
| ENSMUSG00000030228.20 | 39.2372 | 0.595499 | 1.638178 | 0.363513 | 0.716221 | 0.98928 | 0.004681 |
| ENSMUSG00000030259.13 | 103.9921 | 0.349204 | 0.873332 | 0.399852 | 0.689266 | 0.98928 | 0.004681 |
| ENSMUSG00000030269.15 | 30.32808 | 0.675031 | 1.570916 | 0.429705 | 0.66741 | 0.98928 | 0.004681 |
| ENSMUSG00000030298.11 | 281.9558 | -0.22023 | 0.524257 | -0.42007 | 0.674433 | 0.98928 | 0.004681 |
| ENSMUSG00000030327.9 | 151.7599 | -0.3157 | 0.69918 | -0.45152 | 0.651611 | 0.98928 | 0.004681 |
| ENSMUSG00000030342.9 | 40.45063 | 0.423241 | 1.18465 | 0.357271 | 0.720889 | 0.98928 | 0.004681 |
| ENSMUSG00000030357.11 | 206.7501 | -0.22071 | 0.595922 | -0.37036 | 0.711113 | 0.98928 | 0.004681 |
| ENSMUSG00000030403.10 | 13.16793 | 1.148742 | 2.668632 | 0.430461 | 0.66686 | 0.98928 | 0.004681 |
| ENSMUSG00000030431.9 | 58.63661 | -0.47342 | 0.986007 | -0.48014 | 0.63113 | 0.98928 | 0.004681 |
| ENSMUSG00000030446.18 | 13.72473 | -1.50698 | 3.029981 | -0.49736 | 0.618939 | 0.98928 | 0.004681 |
| ENSMUSG00000030471.18 | 20.21455 | -0.83029 | 1.732308 | -0.4793 | 0.631728 | 0.98928 | 0.004681 |
| ENSMUSG00000030516.15 | 208.1529 | 0.270506 | 0.649952 | 0.416195 | 0.677267 | 0.98928 | 0.004681 |
| ENSMUSG00000030534.14 | 24.60599 | -0.69011 | 1.745748 | -0.39531 | 0.692613 | 0.98928 | 0.004681 |
| ENSMUSG00000030536.11 | 24.06863 | 0.757014 | 1.992614 | 0.37991 | 0.704012 | 0.98928 | 0.004681 |
| ENSMUSG00000030560.18 | 546.5759 | -0.18496 | 0.466 | -0.39692 | 0.691428 | 0.98928 | 0.004681 |
| ENSMUSG00000030587.6 | 213.3593 | 0.264266 | 0.636646 | 0.415091 | 0.678075 | 0.98928 | 0.004681 |
| ENSMUSG00000030588.13 | 18.42177 | -0.77483 | 2.105585 | -0.36799 | 0.712881 | 0.98928 | 0.004681 |
| ENSMUSG00000030591.18 | 319.0522 | -0.32992 | 0.673808 | -0.48963 | 0.624394 | 0.98928 | 0.004681 |
| ENSMUSG00000030611.11 | 32.80092 | 0.45058 | 1.399435 | 0.321973 | 0.747473 | 0.98928 | 0.004681 |
| ENSMUSG00000030613.14 | 51.81667 | 0.399349 | 1.006372 | 0.39682 | 0.6915 | 0.98928 | 0.004681 |
| ENSMUSG00000030619.11 | 95.4267 | -0.42441 | 0.855841 | -0.4959 | 0.619964 | 0.98928 | 0.004681 |
| ENSMUSG00000030655.16 | 38.4282 | 0.652973 | 1.509938 | 0.43245 | 0.665414 | 0.98928 | 0.004681 |
| ENSMUSG00000030662.11 | 143.7163 | 0.196141 | 0.60144 | 0.326119 | 0.744334 | 0.98928 | 0.004681 |
| ENSMUSG00000030688.16 | 212.4808 | -0.20006 | 0.517007 | -0.38695 | 0.69879 | 0.98928 | 0.004681 |
| ENSMUSG00000030691.16 | 49.66214 | -0.37634 | 0.960679 | -0.39175 | 0.695246 | 0.98928 | 0.004681 |
| ENSMUSG00000030697.8 | 27.39006 | 0.506978 | 1.583149 | 0.320234 | 0.748791 | 0.98928 | 0.004681 |
| ENSMUSG00000030718.10 | 14.2439 | 1.0965 | 2.316964 | 0.473249 | 0.636036 | 0.98928 | 0.004681 |
| ENSMUSG00000030744.14 | 1917.74 | 0.089356 | 0.267012 | 0.334652 | 0.737888 | 0.98928 | 0.004681 |
| ENSMUSG00000030751.19 | 1198.628 | 0.127249 | 0.335345 | 0.379459 | 0.704347 | 0.98928 | 0.004681 |
| ENSMUSG00000030759.17 | 12.2113 | -0.98592 | 2.401177 | -0.4106 | 0.681368 | 0.98928 | 0.004681 |
| ENSMUSG00000030763.8 | 88.4461 | -0.33854 | 0.853138 | -0.39682 | 0.691498 | 0.98928 | 0.004681 |
| ENSMUSG00000030805.15 | 60.30278 | -0.52303 | 1.451479 | -0.36034 | 0.718593 | 0.98928 | 0.004681 |
| ENSMUSG00000030824.18 | 127.646 | -0.2291 | 0.656216 | -0.34912 | 0.726999 | 0.98928 | 0.004681 |

| | | | | | | | |
|---|---|---|---|---|---|---|---|
| ENSMUSG00000030835.7 | 23.47079 | -0.72269 | 2.05441 | -0.35177 | 0.725008 | 0.98928 | 0.004681 |
| ENSMUSG00000030888.15 | 65.35128 | 0.511705 | 1.257757 | 0.40684 | 0.684126 | 0.98928 | 0.004681 |
| ENSMUSG00000030930.15 | 27.99844 | -0.91816 | 1.920197 | -0.47816 | 0.632535 | 0.98928 | 0.004681 |
| ENSMUSG00000030935.16 | 557.7692 | -0.18381 | 0.422292 | -0.43527 | 0.663365 | 0.98928 | 0.004681 |
| ENSMUSG00000030982.20 | 59.979 | 0.46019 | 1.418585 | 0.324401 | 0.745635 | 0.98928 | 0.004681 |
| ENSMUSG00000031015.9 | 18.45944 | -0.54386 | 1.656069 | -0.32841 | 0.742604 | 0.98928 | 0.004681 |
| ENSMUSG00000031059.10 | 826.4346 | -0.21472 | 0.469945 | -0.45691 | 0.647736 | 0.98928 | 0.004681 |
| ENSMUSG00000031072.15 | 23.25912 | 0.754072 | 1.778118 | 0.424084 | 0.671504 | 0.98928 | 0.004681 |
| ENSMUSG00000031166.14 | 52.10713 | 0.535538 | 1.156923 | 0.462898 | 0.643437 | 0.98928 | 0.004681 |
| ENSMUSG00000031168.15 | 900.5931 | -0.1216 | 0.313955 | -0.38733 | 0.698511 | 0.98928 | 0.004681 |
| ENSMUSG00000031207.17 | 48.42573 | 0.597545 | 1.349971 | 0.442635 | 0.65803 | 0.98928 | 0.004681 |
| ENSMUSG00000031278.13 | 561.802 | 0.194027 | 0.399054 | 0.486217 | 0.626814 | 0.98928 | 0.004681 |
| ENSMUSG00000031358.18 | 131.7411 | 0.249639 | 0.642011 | 0.388838 | 0.697396 | 0.98928 | 0.004681 |
| ENSMUSG00000031379.14 | 287.5875 | 0.225928 | 0.480758 | 0.469941 | 0.638397 | 0.98928 | 0.004681 |
| ENSMUSG00000031399.16 | 28.93372 | -0.60287 | 1.685316 | -0.35772 | 0.720555 | 0.98928 | 0.004681 |
| ENSMUSG00000031516.12 | 67.03402 | 0.415165 | 1.086758 | 0.382022 | 0.702445 | 0.98928 | 0.004681 |
| ENSMUSG00000031545.7 | 364.8632 | -0.15639 | 0.448165 | -0.34897 | 0.727116 | 0.98928 | 0.004681 |
| ENSMUSG00000031561.17 | 13.25942 | 0.826781 | 2.147277 | 0.385037 | 0.70021 | 0.98928 | 0.004681 |
| ENSMUSG00000031577.11 | 34.76815 | -0.64751 | 1.480441 | -0.43738 | 0.661839 | 0.98928 | 0.004681 |
| ENSMUSG00000031578.6 | 100.5848 | -0.32289 | 0.999602 | -0.32302 | 0.746678 | 0.98928 | 0.004681 |
| ENSMUSG00000031583.14 | 57.13914 | -0.48234 | 0.996751 | -0.48391 | 0.628449 | 0.98928 | 0.004681 |
| ENSMUSG00000031584.17 | 563.5518 | -0.1304 | 0.366892 | -0.35543 | 0.72227 | 0.98928 | 0.004681 |
| ENSMUSG00000031585.14 | 46.78687 | -0.52009 | 1.075672 | -0.4835 | 0.628737 | 0.98928 | 0.004681 |
| ENSMUSG00000031600.12 | 296.1038 | -0.16726 | 0.485296 | -0.34465 | 0.730355 | 0.98928 | 0.004681 |
| ENSMUSG00000031633.5 | 42.63196 | 0.678493 | 1.440364 | 0.471056 | 0.637601 | 0.98928 | 0.004681 |
| ENSMUSG00000031671.12 | 145.3418 | 0.265363 | 0.630487 | 0.420886 | 0.673839 | 0.98928 | 0.004681 |
| ENSMUSG00000031701.7 | 871.1197 | -0.14148 | 0.323678 | -0.43709 | 0.662043 | 0.98928 | 0.004681 |
| ENSMUSG00000031711.8 | 26.46713 | -0.7304 | 1.494605 | -0.48869 | 0.625059 | 0.98928 | 0.004681 |
| ENSMUSG00000031782.16 | 296.8864 | -0.20289 | 0.44124 | -0.45982 | 0.645647 | 0.98928 | 0.004681 |
| ENSMUSG00000031799.11 | 130.7829 | -0.30118 | 0.721642 | -0.41736 | 0.676417 | 0.98928 | 0.004681 |
| ENSMUSG00000031813.9 | 91.81859 | -0.35986 | 0.949411 | -0.37903 | 0.704664 | 0.98928 | 0.004681 |
| ENSMUSG00000031816.16 | 21.54364 | 0.972269 | 2.356868 | 0.412526 | 0.679954 | 0.98928 | 0.004681 |
| ENSMUSG00000031822.21 | 25.53575 | 0.5283 | 1.441406 | 0.366517 | 0.713979 | 0.98928 | 0.004681 |
| ENSMUSG00000031826.21 | 138.0548 | -0.33701 | 0.850971 | -0.39603 | 0.692086 | 0.98928 | 0.004681 |
| ENSMUSG00000031839.8 | 563.2339 | 0.139601 | 0.341577 | 0.408695 | 0.682764 | 0.98928 | 0.004681 |
| ENSMUSG00000031875.8 | 19.04571 | -0.77353 | 1.614518 | -0.47911 | 0.63186 | 0.98928 | 0.004681 |
| ENSMUSG00000031878.19 | 241.6392 | -0.20501 | 0.596231 | -0.34384 | 0.730964 | 0.98928 | 0.004681 |
| ENSMUSG00000031902.11 | 98.88887 | -0.42088 | 0.883465 | -0.4764 | 0.633789 | 0.98928 | 0.004681 |
| ENSMUSG00000031906.10 | 16.45663 | 0.732379 | 2.274825 | 0.32195 | 0.747491 | 0.98928 | 0.004681 |
| ENSMUSG00000031913.10 | 17.52836 | 1.135042 | 2.265068 | 0.501107 | 0.616296 | 0.98928 | 0.004681 |
| ENSMUSG00000031916.18 | 78.86024 | -0.41333 | 0.935191 | -0.44197 | 0.658509 | 0.98928 | 0.004681 |
| ENSMUSG00000031922.13 | 47.11145 | 0.591832 | 1.212608 | 0.488066 | 0.625503 | 0.98928 | 0.004681 |
| ENSMUSG00000031979.18 | 28.72081 | -0.54776 | 1.431795 | -0.38257 | 0.702039 | 0.98928 | 0.004681 |
| ENSMUSG00000031988.11 | 114.4042 | -0.48412 | 0.985414 | -0.49129 | 0.623222 | 0.98928 | 0.004681 |

| | | | | | | | |
|---|---|---|---|---|---|---|---|
| ENSMUSG00000032009.9 | 117.9069 | -0.30673 | 0.933516 | -0.32858 | 0.742476 | 0.98928 | 0.004681 |
| ENSMUSG00000032041.16 | 64.70313 | 0.51841 | 1.033776 | 0.501472 | 0.616039 | 0.98928 | 0.004681 |
| ENSMUSG00000032044.5 | 41.74552 | 0.415709 | 1.252303 | 0.331956 | 0.739923 | 0.98928 | 0.004681 |
| ENSMUSG00000032050.19 | 3253.948 | -0.08144 | 0.246845 | -0.32994 | 0.741449 | 0.98928 | 0.004681 |
| ENSMUSG00000032080.7 | 287.1251 | 0.373539 | 0.765773 | 0.487793 | 0.625697 | 0.98928 | 0.004681 |
| ENSMUSG00000032081.11 | 12866.42 | -0.10224 | 0.234783 | -0.43546 | 0.663228 | 0.98928 | 0.004681 |
| ENSMUSG00000032083.9 | 48424.44 | -0.09996 | 0.214451 | -0.46613 | 0.641119 | 0.98928 | 0.004681 |
| ENSMUSG00000032092.6 | 284.918 | -0.14972 | 0.46513 | -0.32188 | 0.747543 | 0.98928 | 0.004681 |
| ENSMUSG00000032112.11 | 104.0433 | 0.322926 | 0.815756 | 0.395861 | 0.692208 | 0.98928 | 0.004681 |
| ENSMUSG00000032114.10 | 845.9128 | 0.163456 | 0.357774 | 0.45687 | 0.647764 | 0.98928 | 0.004681 |
| ENSMUSG00000032122.16 | 12.12465 | 1.363549 | 2.747713 | 0.496249 | 0.619719 | 0.98928 | 0.004681 |
| ENSMUSG00000032198.10 | 60.38032 | 0.681909 | 1.498148 | 0.455168 | 0.648988 | 0.98928 | 0.004681 |
| ENSMUSG00000032217.14 | 68.25642 | 0.328328 | 0.867485 | 0.378483 | 0.705072 | 0.98928 | 0.004681 |
| ENSMUSG00000032238.18 | 427.0298 | -0.20477 | 0.430346 | -0.47582 | 0.634206 | 0.98928 | 0.004681 |
| ENSMUSG00000032249.15 | 353.588 | 0.224887 | 0.499068 | 0.450615 | 0.652267 | 0.98928 | 0.004681 |
| ENSMUSG00000032271.14 | 770.5303 | 0.147124 | 0.387574 | 0.379602 | 0.704241 | 0.98928 | 0.004681 |
| ENSMUSG00000032299.11 | 69.57108 | 0.485164 | 1.239926 | 0.391285 | 0.695587 | 0.98928 | 0.004681 |
| ENSMUSG00000032307.17 | 61.16884 | -0.63371 | 1.410546 | -0.44927 | 0.653238 | 0.98928 | 0.004681 |
| ENSMUSG00000032309.16 | 158.0193 | 0.310066 | 0.62277 | 0.497881 | 0.618568 | 0.98928 | 0.004681 |
| ENSMUSG00000032312.8 | 108.2232 | -0.3268 | 0.85478 | -0.38232 | 0.702223 | 0.98928 | 0.004681 |
| ENSMUSG00000032329.5 | 46.65473 | -0.59251 | 1.394754 | -0.42481 | 0.670973 | 0.98928 | 0.004681 |
| ENSMUSG00000032332.18 | 12.30424 | -1.05942 | 2.440904 | -0.43403 | 0.664268 | 0.98928 | 0.004681 |
| ENSMUSG00000032336.18 | 440.7891 | 0.188864 | 0.410116 | 0.460515 | 0.645147 | 0.98928 | 0.004681 |
| ENSMUSG00000032359.15 | 792.3454 | -0.17933 | 0.389904 | -0.45993 | 0.645563 | 0.98928 | 0.004681 |
| ENSMUSG00000032370.9 | 568.5348 | -0.15656 | 0.354878 | -0.44115 | 0.659103 | 0.98928 | 0.004681 |
| ENSMUSG00000032440.14 | 80.91972 | 0.243655 | 0.768585 | 0.317018 | 0.75123 | 0.98928 | 0.004681 |
| ENSMUSG00000032463.11 | 106.3177 | 0.316567 | 0.88361 | 0.358266 | 0.720144 | 0.98928 | 0.004681 |
| ENSMUSG00000032500.11 | 25.94678 | 0.689931 | 2.033567 | 0.339271 | 0.734405 | 0.98928 | 0.004681 |
| ENSMUSG00000032504.16 | 367.3248 | 0.154782 | 0.434885 | 0.355915 | 0.721904 | 0.98928 | 0.004681 |
| ENSMUSG00000032512.8 | 15.7121 | -0.91662 | 2.223567 | -0.41223 | 0.680172 | 0.98928 | 0.004681 |
| ENSMUSG00000032518.7 | 1025.663 | 0.126821 | 0.340603 | 0.372342 | 0.709639 | 0.98928 | 0.004681 |
| ENSMUSG00000032540.17 | 128.2719 | -0.32158 | 0.734677 | -0.43772 | 0.66159 | 0.98928 | 0.004681 |
| ENSMUSG00000032652.14 | 185.0514 | 0.24364 | 0.571065 | 0.426641 | 0.66964 | 0.98928 | 0.004681 |
| ENSMUSG00000032679.13 | 554.3609 | 0.127607 | 0.363091 | 0.351446 | 0.725254 | 0.98928 | 0.004681 |
| ENSMUSG00000032705.10 | 94.29869 | -0.37202 | 0.979086 | -0.37997 | 0.70397 | 0.98928 | 0.004681 |
| ENSMUSG00000032725.11 | 51.03971 | -0.6192 | 1.400748 | -0.44205 | 0.658456 | 0.98928 | 0.004681 |
| ENSMUSG00000032745.19 | 307.6467 | 0.267457 | 0.583093 | 0.458687 | 0.646459 | 0.98928 | 0.004681 |
| ENSMUSG00000032763.12 | 85.0945 | -0.28656 | 0.896493 | -0.31965 | 0.749235 | 0.98928 | 0.004681 |
| ENSMUSG00000032902.2 | 230.4251 | 0.214354 | 0.637834 | 0.336065 | 0.736822 | 0.98928 | 0.004681 |
| ENSMUSG00000033014.13 | 123.5223 | 0.378413 | 1.040142 | 0.363809 | 0.716001 | 0.98928 | 0.004681 |
| ENSMUSG00000033083.17 | 16.82031 | -1.03452 | 2.166447 | -0.47752 | 0.632994 | 0.98928 | 0.004681 |
| ENSMUSG00000033105.13 | 129.6178 | 0.258898 | 0.818483 | 0.316315 | 0.751764 | 0.98928 | 0.004681 |
| ENSMUSG00000033147.17 | 43.87627 | 0.436461 | 1.294078 | 0.337276 | 0.735909 | 0.98928 | 0.004681 |
| ENSMUSG00000033237.20 | 56.98926 | 0.452914 | 1.001646 | 0.45217 | 0.651147 | 0.98928 | 0.004681 |

| | | | | | | | |
|---|---|---|---|---|---|---|---|
| ENSMUSG00000033257.15 | 28.72737 | -0.55168 | 1.248278 | -0.44196 | 0.65852 | 0.98928 | 0.004681 |
| ENSMUSG00000033295.15 | 510.6376 | -0.19972 | 0.507465 | -0.39357 | 0.6939 | 0.98928 | 0.004681 |
| ENSMUSG00000033352.12 | 177.4977 | -0.26903 | 0.559278 | -0.48104 | 0.630491 | 0.98928 | 0.004681 |
| ENSMUSG00000033565.19 | 35.31863 | 0.732071 | 1.86918 | 0.391654 | 0.695314 | 0.98928 | 0.004681 |
| ENSMUSG00000033684.15 | 55.94263 | -0.4234 | 1.123448 | -0.37688 | 0.706264 | 0.98928 | 0.004681 |
| ENSMUSG00000033685.14 | 23.20211 | -0.77322 | 1.56981 | -0.49256 | 0.622325 | 0.98928 | 0.004681 |
| ENSMUSG00000033735.10 | 324.9808 | 0.184424 | 0.469836 | 0.392529 | 0.694668 | 0.98928 | 0.004681 |
| ENSMUSG00000033751.6 | 120.1455 | 0.338959 | 0.710549 | 0.477038 | 0.633335 | 0.98928 | 0.004681 |
| ENSMUSG00000033767.15 | 34.24356 | 0.476999 | 1.107015 | 0.430887 | 0.66655 | 0.98928 | 0.004681 |
| ENSMUSG00000033790.12 | 22.92839 | -0.78633 | 1.830882 | -0.42948 | 0.667575 | 0.98928 | 0.004681 |
| ENSMUSG00000033931.10 | 113.305 | -0.33581 | 0.727879 | -0.46136 | 0.644543 | 0.98928 | 0.004681 |
| ENSMUSG00000033953.11 | 45.31676 | -0.34454 | 1.016241 | -0.33904 | 0.734581 | 0.98928 | 0.004681 |
| ENSMUSG00000033983.15 | 21.1609 | 0.881525 | 1.949752 | 0.452122 | 0.651181 | 0.98928 | 0.004681 |
| ENSMUSG00000033991.10 | 57.31183 | -0.50007 | 1.386502 | -0.36067 | 0.718345 | 0.98928 | 0.004681 |
| ENSMUSG00000034135.16 | 172.0895 | 0.29043 | 0.748324 | 0.388107 | 0.697937 | 0.98928 | 0.004681 |
| ENSMUSG00000034158.10 | 184.5941 | 0.219714 | 0.639874 | 0.343371 | 0.731319 | 0.98928 | 0.004681 |
| ENSMUSG00000034247.10 | 39.50912 | -1.58085 | 3.731305 | -0.42367 | 0.671804 | 0.98928 | 0.004681 |
| ENSMUSG00000034248.8 | 301.743 | -0.27949 | 0.614529 | -0.4548 | 0.64925 | 0.98928 | 0.004681 |
| ENSMUSG00000034263.14 | 50.76011 | -0.4396 | 1.342734 | -0.32739 | 0.743374 | 0.98928 | 0.004681 |
| ENSMUSG00000034274.12 | 46.59594 | -0.5107 | 1.405693 | -0.36331 | 0.716374 | 0.98928 | 0.004681 |
| ENSMUSG00000034278.4 | 11.27188 | -0.90489 | 2.798113 | -0.32339 | 0.746398 | 0.98928 | 0.004681 |
| ENSMUSG00000034334.9 | 64.34439 | -0.31189 | 0.947864 | -0.32904 | 0.742124 | 0.98928 | 0.004681 |
| ENSMUSG00000034342.10 | 21.76563 | -0.73816 | 1.713792 | -0.43072 | 0.666672 | 0.98928 | 0.004681 |
| ENSMUSG00000034354.18 | 77.27806 | -0.52172 | 1.319734 | -0.39532 | 0.692604 | 0.98928 | 0.004681 |
| ENSMUSG00000034371.10 | 547.0828 | 0.219009 | 0.490347 | 0.44664 | 0.655135 | 0.98928 | 0.004681 |
| ENSMUSG00000034379.10 | 14.80374 | 0.944103 | 2.045151 | 0.46163 | 0.644347 | 0.98928 | 0.004681 |
| ENSMUSG00000034432.12 | 88.14031 | 0.524105 | 1.164347 | 0.450128 | 0.652618 | 0.98928 | 0.004681 |
| ENSMUSG00000034453.9 | 48.6972 | -0.47088 | 1.349848 | -0.34884 | 0.727211 | 0.98928 | 0.004681 |
| ENSMUSG00000034456.16 | 762.3409 | 0.104393 | 0.309248 | 0.33757 | 0.735687 | 0.98928 | 0.004681 |
| ENSMUSG00000034462.10 | 38.78469 | 0.455543 | 1.146927 | 0.397186 | 0.691231 | 0.98928 | 0.004681 |
| ENSMUSG00000034484.9 | 286.1575 | 0.218435 | 0.475173 | 0.459694 | 0.645736 | 0.98928 | 0.004681 |
| ENSMUSG00000034601.18 | 14.04466 | -0.81523 | 2.270456 | -0.35906 | 0.719552 | 0.98928 | 0.004681 |
| ENSMUSG00000034653.12 | 45.44295 | -0.74171 | 1.757159 | -0.42211 | 0.672947 | 0.98928 | 0.004681 |
| ENSMUSG00000034673.15 | 27.75733 | -0.53521 | 1.618157 | -0.33075 | 0.74083 | 0.98928 | 0.004681 |
| ENSMUSG00000034709.9 | 45.53126 | 0.604887 | 1.419626 | 0.426089 | 0.670043 | 0.98928 | 0.004681 |
| ENSMUSG00000034723.12 | 160.1933 | -0.23257 | 0.57961 | -0.40125 | 0.688238 | 0.98928 | 0.004681 |
| ENSMUSG00000034724.19 | 155.5695 | -0.25289 | 0.605397 | -0.41773 | 0.676143 | 0.98928 | 0.004681 |
| ENSMUSG00000034801.14 | 101.4977 | -0.32095 | 0.818421 | -0.39216 | 0.694941 | 0.98928 | 0.004681 |
| ENSMUSG00000034858.17 | 232.6095 | 0.239655 | 0.549084 | 0.436463 | 0.662501 | 0.98928 | 0.004681 |
| ENSMUSG00000034908.17 | 79.88427 | 0.45379 | 0.963481 | 0.47099 | 0.637648 | 0.98928 | 0.004681 |
| ENSMUSG00000034931.16 | 32.07234 | 0.759546 | 1.575386 | 0.482133 | 0.629711 | 0.98928 | 0.004681 |
| ENSMUSG00000035000.9 | 563.8797 | 0.170455 | 0.377533 | 0.451497 | 0.651632 | 0.98928 | 0.004681 |
| ENSMUSG00000035021.14 | 147.6598 | -0.24014 | 0.664732 | -0.36126 | 0.717905 | 0.98928 | 0.004681 |
| ENSMUSG00000035049.5 | 23.01618 | 0.465984 | 1.461256 | 0.318893 | 0.749808 | 0.98928 | 0.004681 |

| | | | | | | | |
|---|---|---|---|---|---|---|---|
| ENSMUSG00000035139.15 | 77.59338 | 0.674888 | 1.392072 | 0.484808 | 0.627812 | 0.98928 | 0.004681 |
| ENSMUSG00000035150.16 | 63.96384 | 0.483474 | 1.141099 | 0.423691 | 0.671791 | 0.98928 | 0.004681 |
| ENSMUSG00000035161.8 | 111.1256 | 0.383633 | 0.907215 | 0.422869 | 0.672391 | 0.98928 | 0.004681 |
| ENSMUSG00000035186.7 | 10.15198 | 0.844842 | 2.237056 | 0.377658 | 0.705685 | 0.98928 | 0.004681 |
| ENSMUSG00000035215.10 | 33.72044 | -0.60931 | 1.650948 | -0.36907 | 0.712077 | 0.98928 | 0.004681 |
| ENSMUSG00000035227.8 | 883.4151 | 0.142861 | 0.286996 | 0.497781 | 0.618638 | 0.98928 | 0.004681 |
| ENSMUSG00000035242.16 | 814.5631 | 0.150503 | 0.384611 | 0.391311 | 0.695567 | 0.98928 | 0.004681 |
| ENSMUSG00000035273.15 | 33.17932 | -0.59255 | 1.437762 | -0.41213 | 0.680244 | 0.98928 | 0.004681 |
| ENSMUSG00000035367.10 | 40.79553 | 0.68242 | 1.396519 | 0.488657 | 0.625084 | 0.98928 | 0.004681 |
| ENSMUSG00000035372.3 | 49.68109 | -0.3685 | 1.071789 | -0.34382 | 0.730985 | 0.98928 | 0.004681 |
| ENSMUSG00000035413.9 | 32.0128 | -0.99089 | 2.24183 | -0.442 | 0.658489 | 0.98928 | 0.004681 |
| ENSMUSG00000035443.11 | 134.4916 | -0.294 | 0.65621 | -0.44803 | 0.654134 | 0.98928 | 0.004681 |
| ENSMUSG00000035498.11 | 25.52099 | -0.97182 | 2.115352 | -0.45941 | 0.645936 | 0.98928 | 0.004681 |
| ENSMUSG00000035596.15 | 86.33891 | -0.36892 | 1.025144 | -0.35987 | 0.718945 | 0.98928 | 0.004681 |
| ENSMUSG00000035597.20 | 103.5491 | -0.45435 | 0.948306 | -0.47912 | 0.631855 | 0.98928 | 0.004681 |
| ENSMUSG00000035629.20 | 37.59534 | -0.55029 | 1.461812 | -0.37644 | 0.706587 | 0.98928 | 0.004681 |
| ENSMUSG00000035864.15 | 36.64969 | 0.671017 | 1.687687 | 0.397596 | 0.690928 | 0.98928 | 0.004681 |
| ENSMUSG00000036083.16 | 462.606 | -0.16275 | 0.48477 | -0.33573 | 0.737077 | 0.98928 | 0.004681 |
| ENSMUSG00000036093.8 | 259.8323 | -0.26928 | 0.636093 | -0.42333 | 0.672053 | 0.98928 | 0.004681 |
| ENSMUSG00000036098.16 | 17.49422 | -0.54816 | 1.559294 | -0.35154 | 0.72518 | 0.98928 | 0.004681 |
| ENSMUSG00000036104.13 | 55.26834 | -0.42222 | 0.911547 | -0.46319 | 0.64323 | 0.98928 | 0.004681 |
| ENSMUSG00000036114.4 | 50.72886 | -0.59203 | 1.424103 | -0.41572 | 0.677612 | 0.98928 | 0.004681 |
| ENSMUSG00000036167.17 | 18.52691 | 1.210831 | 2.407324 | 0.502978 | 0.61498 | 0.98928 | 0.004681 |
| ENSMUSG00000036197.17 | 114.8536 | -0.28595 | 0.788843 | -0.36249 | 0.716987 | 0.98928 | 0.004681 |
| ENSMUSG00000036214.15 | 210.0313 | -0.24312 | 0.597464 | -0.40692 | 0.684067 | 0.98928 | 0.004681 |
| ENSMUSG00000036327.19 | 20.84611 | -0.7441 | 2.142519 | -0.3473 | 0.728365 | 0.98928 | 0.004681 |
| ENSMUSG00000036339.19 | 137.5296 | -0.2054 | 0.645273 | -0.31831 | 0.750248 | 0.98928 | 0.004681 |
| ENSMUSG00000036398.14 | 23.37322 | 0.690495 | 1.971233 | 0.350286 | 0.726124 | 0.98928 | 0.004681 |
| ENSMUSG00000036402.14 | 238.214 | -0.25612 | 0.677259 | -0.37818 | 0.705299 | 0.98928 | 0.004681 |
| ENSMUSG00000036528.17 | 136.6128 | -0.44038 | 0.909993 | -0.48394 | 0.628427 | 0.98928 | 0.004681 |
| ENSMUSG00000036676.15 | 21.03264 | -0.5829 | 1.775822 | -0.32824 | 0.742729 | 0.98928 | 0.004681 |
| ENSMUSG00000036748.17 | 31.8376 | -0.85595 | 1.705553 | -0.50186 | 0.615765 | 0.98928 | 0.004681 |
| ENSMUSG00000036751.8 | 1216.096 | -0.13095 | 0.31119 | -0.42079 | 0.673908 | 0.98928 | 0.004681 |
| ENSMUSG00000036810.17 | 76.91139 | 0.328541 | 0.827212 | 0.397167 | 0.691244 | 0.98928 | 0.004681 |
| ENSMUSG00000036813.12 | 190.5956 | 0.19951 | 0.581172 | 0.343288 | 0.731382 | 0.98928 | 0.004681 |
| ENSMUSG00000036817.15 | 31.95477 | 0.659411 | 1.840463 | 0.358285 | 0.72013 | 0.98928 | 0.004681 |
| ENSMUSG00000036822.7 | 194.2781 | 0.183484 | 0.56654 | 0.323867 | 0.746039 | 0.98928 | 0.004681 |
| ENSMUSG00000036867.8 | 16.90495 | -0.55858 | 1.690708 | -0.33038 | 0.741111 | 0.98928 | 0.004681 |
| ENSMUSG00000036893.18 | 64.57406 | -0.37107 | 0.92716 | -0.40022 | 0.688994 | 0.98928 | 0.004681 |
| ENSMUSG00000036904.7 | 120.6867 | -0.29404 | 0.662511 | -0.44382 | 0.657171 | 0.98928 | 0.004681 |
| ENSMUSG00000036934.10 | 105.9507 | 0.418583 | 0.832312 | 0.502916 | 0.615023 | 0.98928 | 0.004681 |
| ENSMUSG00000037053.7 | 9032.636 | -0.07711 | 0.194632 | -0.3962 | 0.691956 | 0.98928 | 0.004681 |
| ENSMUSG00000037119.5 | 375.8121 | 0.168781 | 0.45167 | 0.373681 | 0.708642 | 0.98928 | 0.004681 |
| ENSMUSG00000037190.13 | 52.49363 | -0.32848 | 1.005985 | -0.32652 | 0.74403 | 0.98928 | 0.004681 |

| | | | | | | | |
|---|---|---|---|---|---|---|---|
| ENSMUSG00000037211.13 | 15.36401 | 0.593876 | 1.564319 | 0.379639 | 0.704214 | 0.98928 | 0.004681 |
| ENSMUSG00000037266.19 | 1138.944 | 0.161335 | 0.373256 | 0.432236 | 0.66557 | 0.98928 | 0.004681 |
| ENSMUSG00000037278.10 | 172.2062 | -0.53925 | 1.077688 | -0.50038 | 0.616809 | 0.98928 | 0.004681 |
| ENSMUSG00000037286.16 | 111.2168 | 0.284303 | 0.838605 | 0.33902 | 0.734595 | 0.98928 | 0.004681 |
| ENSMUSG00000037326.11 | 32.93299 | -0.94593 | 1.894674 | -0.49926 | 0.617599 | 0.98928 | 0.004681 |
| ENSMUSG00000037331.16 | 352.948 | -0.16808 | 0.474975 | -0.35388 | 0.72343 | 0.98928 | 0.004681 |
| ENSMUSG00000037405.9 | 13.08495 | -0.86375 | 2.710229 | -0.3187 | 0.749953 | 0.98928 | 0.004681 |
| ENSMUSG00000037475.16 | 336.291 | 0.266405 | 0.573862 | 0.464232 | 0.642482 | 0.98928 | 0.004681 |
| ENSMUSG00000037486.19 | 108.7014 | -0.31583 | 0.799891 | -0.39484 | 0.692964 | 0.98928 | 0.004681 |
| ENSMUSG00000037514.19 | 127.4839 | -0.27866 | 0.615199 | -0.45296 | 0.650574 | 0.98928 | 0.004681 |
| ENSMUSG00000037542.14 | 2153.811 | -0.08936 | 0.249201 | -0.35859 | 0.719899 | 0.98928 | 0.004681 |
| ENSMUSG00000037573.6 | 796.1487 | 0.180554 | 0.403235 | 0.447763 | 0.654324 | 0.98928 | 0.004681 |
| ENSMUSG00000037638.6 | 14.64353 | 1.005971 | 2.632507 | 0.382134 | 0.702362 | 0.98928 | 0.004681 |
| ENSMUSG00000037646.11 | 230.4497 | 0.169674 | 0.484365 | 0.350302 | 0.726112 | 0.98928 | 0.004681 |
| ENSMUSG00000037697.20 | 108.3444 | 0.321189 | 0.671065 | 0.478625 | 0.632205 | 0.98928 | 0.004681 |
| ENSMUSG00000037706.18 | 505.2955 | -0.1622 | 0.409385 | -0.39622 | 0.691946 | 0.98928 | 0.004681 |
| ENSMUSG00000037712.18 | 411.7166 | -0.20395 | 0.465213 | -0.43841 | 0.66109 | 0.98928 | 0.004681 |
| ENSMUSG00000037722.9 | 504.589 | 0.172184 | 0.376978 | 0.456747 | 0.647853 | 0.98928 | 0.004681 |
| ENSMUSG00000037772.14 | 18.91574 | 0.682647 | 1.986452 | 0.343651 | 0.731109 | 0.98928 | 0.004681 |
| ENSMUSG00000037787.15 | 149.551 | -0.32403 | 0.695511 | -0.46589 | 0.641296 | 0.98928 | 0.004681 |
| ENSMUSG00000037857.17 | 104.9589 | -0.34044 | 0.857382 | -0.39706 | 0.691321 | 0.98928 | 0.004681 |
| ENSMUSG00000037896.18 | 125.8788 | -0.29191 | 0.657014 | -0.4443 | 0.656827 | 0.98928 | 0.004681 |
| ENSMUSG00000037921.16 | 9.959858 | 0.901703 | 2.657259 | 0.339336 | 0.734357 | 0.98928 | 0.004681 |
| ENSMUSG00000037965.16 | 125.3194 | -0.24686 | 0.725069 | -0.34046 | 0.733509 | 0.98928 | 0.004681 |
| ENSMUSG00000038002.10 | 25.64716 | -0.83922 | 1.749191 | -0.47978 | 0.631385 | 0.98928 | 0.004681 |
| ENSMUSG00000038009.8 | 183.1625 | -0.27811 | 0.687857 | -0.40431 | 0.685982 | 0.98928 | 0.004681 |
| ENSMUSG00000038013.15 | 19.47535 | -0.66988 | 1.733805 | -0.38636 | 0.699227 | 0.98928 | 0.004681 |
| ENSMUSG00000038014.8 | 618.9505 | -0.12741 | 0.39331 | -0.32395 | 0.745977 | 0.98928 | 0.004681 |
| ENSMUSG00000038055.13 | 80.32432 | -0.35822 | 0.790584 | -0.45311 | 0.650473 | 0.98928 | 0.004681 |
| ENSMUSG00000038122.10 | 19.37497 | -0.82617 | 2.103382 | -0.39278 | 0.694481 | 0.98928 | 0.004681 |
| ENSMUSG00000038160.8 | 262.4079 | -0.2121 | 0.531469 | -0.39909 | 0.689828 | 0.98928 | 0.004681 |
| ENSMUSG00000038187.15 | 52.26339 | -0.63755 | 1.436417 | -0.44385 | 0.657151 | 0.98928 | 0.004681 |
| ENSMUSG00000038214.16 | 21.08053 | -0.64566 | 1.457599 | -0.44296 | 0.657791 | 0.98928 | 0.004681 |
| ENSMUSG00000038267.16 | 220.7058 | -0.28385 | 0.570859 | -0.49723 | 0.619024 | 0.98928 | 0.004681 |
| ENSMUSG00000038290.16 | 51.33643 | -0.86739 | 1.839235 | -0.4716 | 0.637212 | 0.98928 | 0.004681 |
| ENSMUSG00000038335.14 | 146.9242 | 0.307402 | 0.700198 | 0.439022 | 0.660645 | 0.98928 | 0.004681 |
| ENSMUSG00000038459.11 | 192.9725 | -0.30166 | 0.600207 | -0.50259 | 0.61525 | 0.98928 | 0.004681 |
| ENSMUSG00000038467.16 | 167.1678 | -0.27256 | 0.576519 | -0.47276 | 0.636381 | 0.98928 | 0.004681 |
| ENSMUSG00000038495.15 | 78.05242 | 0.408997 | 0.829928 | 0.49281 | 0.622147 | 0.98928 | 0.004681 |
| ENSMUSG00000038550.11 | 65.55534 | -0.5947 | 1.358559 | -0.43774 | 0.661572 | 0.98928 | 0.004681 |
| ENSMUSG00000038564.12 | 10.89514 | -1.05885 | 2.917425 | -0.36294 | 0.716649 | 0.98928 | 0.004681 |
| ENSMUSG00000038611.18 | 11.25159 | 0.984785 | 2.81687 | 0.349603 | 0.726637 | 0.98928 | 0.004681 |
| ENSMUSG00000038658.16 | 32.80235 | -0.49075 | 1.262929 | -0.38858 | 0.697586 | 0.98928 | 0.004681 |
| ENSMUSG00000038685.19 | 21.97185 | -0.8774 | 2.021361 | -0.43406 | 0.664242 | 0.98928 | 0.004681 |

| | | | | | | | |
|---|---|---|---|---|---|---|---|
| ENSMUSG00000038705.14 | 16.41675 | -0.73892 | 2.229019 | -0.3315 | 0.740266 | 0.98928 | 0.004681 |
| ENSMUSG00000038708.11 | 346.2634 | -0.25471 | 0.512761 | -0.49674 | 0.619374 | 0.98928 | 0.004681 |
| ENSMUSG00000038784.14 | 62.11994 | -0.3097 | 0.905649 | -0.34196 | 0.732378 | 0.98928 | 0.004681 |
| ENSMUSG00000038866.16 | 143.1147 | -0.30068 | 0.62719 | -0.47941 | 0.631648 | 0.98928 | 0.004681 |
| ENSMUSG00000038876.13 | 154.2724 | 0.284271 | 0.624005 | 0.455559 | 0.648707 | 0.98928 | 0.004681 |
| ENSMUSG00000038900.18 | 1910.598 | 0.119454 | 0.294056 | 0.40623 | 0.684574 | 0.98928 | 0.004681 |
| ENSMUSG00000038914.16 | 78.31285 | 0.420575 | 1.124615 | 0.373972 | 0.708425 | 0.98928 | 0.004681 |
| ENSMUSG00000038967.14 | 57.96035 | 0.397949 | 1.197191 | 0.332403 | 0.739585 | 0.98928 | 0.004681 |
| ENSMUSG00000039000.9 | 125.2755 | 0.298305 | 0.705072 | 0.423085 | 0.672233 | 0.98928 | 0.004681 |
| ENSMUSG00000039031.17 | 66.66496 | -0.4512 | 1.107254 | -0.4075 | 0.683643 | 0.98928 | 0.004681 |
| ENSMUSG00000039065.12 | 77.72859 | 0.367966 | 1.012937 | 0.363267 | 0.716406 | 0.98928 | 0.004681 |
| ENSMUSG00000039128.14 | 205.005 | -0.16484 | 0.519869 | -0.31709 | 0.751177 | 0.98928 | 0.004681 |
| ENSMUSG00000039166.17 | 40.82992 | 0.584368 | 1.564364 | 0.37355 | 0.708739 | 0.98928 | 0.004681 |
| ENSMUSG00000039167.13 | 36.35896 | -0.55472 | 1.179737 | -0.47021 | 0.638207 | 0.98928 | 0.004681 |
| ENSMUSG00000039318.13 | 42.83127 | -0.51095 | 1.051537 | -0.48591 | 0.627032 | 0.98928 | 0.004681 |
| ENSMUSG00000039345.17 | 28.40902 | 0.838868 | 2.34589 | 0.35759 | 0.72065 | 0.98928 | 0.004681 |
| ENSMUSG00000039361.12 | 569.7193 | 0.192063 | 0.381215 | 0.503818 | 0.614389 | 0.98928 | 0.004681 |
| ENSMUSG00000039456.11 | 58.98 | 0.456588 | 0.95836 | 0.476427 | 0.63377 | 0.98928 | 0.004681 |
| ENSMUSG00000039457.5 | 49.30691 | 0.56071 | 1.435778 | 0.390527 | 0.696147 | 0.98928 | 0.004681 |
| ENSMUSG00000039501.15 | 69.80204 | -0.62111 | 1.253794 | -0.49538 | 0.62033 | 0.98928 | 0.004681 |
| ENSMUSG00000039630.11 | 1008.109 | 0.098599 | 0.306424 | 0.321774 | 0.747624 | 0.98928 | 0.004681 |
| ENSMUSG00000039682.13 | 1949.166 | 0.099666 | 0.279558 | 0.356511 | 0.721458 | 0.98928 | 0.004681 |
| ENSMUSG00000039701.12 | 43.90122 | -0.53952 | 1.337246 | -0.40346 | 0.686613 | 0.98928 | 0.004681 |
| ENSMUSG00000039703.16 | 48.3248 | -0.43361 | 1.131957 | -0.38306 | 0.701676 | 0.98928 | 0.004681 |
| ENSMUSG00000039737.12 | 24.0107 | 0.777719 | 2.285048 | 0.340351 | 0.733592 | 0.98928 | 0.004681 |
| ENSMUSG00000039740.7 | 58.50773 | -0.40346 | 0.912253 | -0.44227 | 0.658296 | 0.98928 | 0.004681 |
| ENSMUSG00000039768.17 | 86.74901 | 0.3919 | 0.933044 | 0.420023 | 0.674469 | 0.98928 | 0.004681 |
| ENSMUSG00000039840.9 | 83.95688 | -0.28633 | 0.790238 | -0.36233 | 0.717102 | 0.98928 | 0.004681 |
| ENSMUSG00000039887.12 | 71.31784 | -0.29064 | 0.870539 | -0.33386 | 0.738484 | 0.98928 | 0.004681 |
| ENSMUSG00000040018.10 | 173.3266 | 0.374241 | 0.896294 | 0.417543 | 0.676281 | 0.98928 | 0.004681 |
| ENSMUSG00000040026.9 | 27.92984 | -0.68744 | 1.566907 | -0.43872 | 0.660862 | 0.98928 | 0.004681 |
| ENSMUSG00000040034.16 | 41.43124 | -0.67053 | 1.432861 | -0.46797 | 0.639809 | 0.98928 | 0.004681 |
| ENSMUSG00000040054.18 | 68.30232 | 0.433076 | 1.357742 | 0.318967 | 0.749751 | 0.98928 | 0.004681 |
| ENSMUSG00000040139.15 | 44.21432 | -0.52949 | 1.125049 | -0.47064 | 0.6379 | 0.98928 | 0.004681 |
| ENSMUSG00000040146.10 | 34.36526 | 0.740382 | 1.728869 | 0.428246 | 0.668472 | 0.98928 | 0.004681 |
| ENSMUSG00000040152.9 | 19.30347 | -0.96116 | 2.492743 | -0.38558 | 0.699806 | 0.98928 | 0.004681 |
| ENSMUSG00000040167.17 | 64.24278 | 0.361066 | 1.037023 | 0.348176 | 0.727708 | 0.98928 | 0.004681 |
| ENSMUSG00000040174.15 | 102.624 | -0.35675 | 1.06867 | -0.33382 | 0.738513 | 0.98928 | 0.004681 |
| ENSMUSG00000040263.17 | 20.78016 | -0.65405 | 2.036745 | -0.32112 | 0.748116 | 0.98928 | 0.004681 |
| ENSMUSG00000040359.15 | 207.8151 | -0.18956 | 0.487266 | -0.38903 | 0.697257 | 0.98928 | 0.004681 |
| ENSMUSG00000040363.16 | 17.80149 | 0.485015 | 1.494662 | 0.324498 | 0.745561 | 0.98928 | 0.004681 |
| ENSMUSG00000040374.14 | 363.327 | 0.216574 | 0.475065 | 0.455882 | 0.648475 | 0.98928 | 0.004681 |
| ENSMUSG00000040433.17 | 170.219 | -0.25213 | 0.595424 | -0.42344 | 0.671972 | 0.98928 | 0.004681 |
| ENSMUSG00000040463.17 | 121.3175 | 0.480842 | 1.073539 | 0.447903 | 0.654223 | 0.98928 | 0.004681 |

| | | | | | | | |
|---|---|---|---|---|---|---|---|
| ENSMUSG00000040548.17 | 352.169 | -0.17426 | 0.493054 | -0.35344 | 0.723761 | 0.98928 | 0.004681 |
| ENSMUSG00000040596.16 | 13.61952 | -1.05972 | 2.309674 | -0.45882 | 0.646365 | 0.98928 | 0.004681 |
| ENSMUSG00000040681.17 | 664.4809 | -0.14857 | 0.338336 | -0.43912 | 0.660577 | 0.98928 | 0.004681 |
| ENSMUSG00000040723.15 | 13.84451 | 1.346906 | 3.050221 | 0.441576 | 0.658796 | 0.98928 | 0.004681 |
| ENSMUSG00000040732.21 | 23.59141 | -0.67152 | 1.586363 | -0.42331 | 0.672069 | 0.98928 | 0.004681 |
| ENSMUSG00000040782.13 | 190.1069 | 0.350302 | 0.704067 | 0.49754 | 0.618808 | 0.98928 | 0.004681 |
| ENSMUSG00000040820.17 | 72.91555 | 0.379762 | 1.059855 | 0.358315 | 0.720107 | 0.98928 | 0.004681 |
| ENSMUSG00000040824.4 | 159.4958 | 0.366957 | 0.762812 | 0.481059 | 0.630475 | 0.98928 | 0.004681 |
| ENSMUSG00000040918.13 | 447.2453 | 0.189588 | 0.445528 | 0.425535 | 0.670447 | 0.98928 | 0.004681 |
| ENSMUSG00000040928.16 | 53.39358 | 0.809136 | 1.681239 | 0.481274 | 0.630322 | 0.98928 | 0.004681 |
| ENSMUSG00000040936.16 | 75.86481 | 0.436183 | 0.9202 | 0.474009 | 0.635493 | 0.98928 | 0.004681 |
| ENSMUSG00000040952.17 | 390.1533 | 0.20503 | 0.410709 | 0.49921 | 0.617631 | 0.98928 | 0.004681 |
| ENSMUSG00000040997.15 | 18.98973 | -0.63215 | 1.892783 | -0.33398 | 0.738395 | 0.98928 | 0.004681 |
| ENSMUSG00000041057.11 | 86.49534 | -0.30445 | 0.961014 | -0.31681 | 0.751391 | 0.98928 | 0.004681 |
| ENSMUSG00000041058.16 | 573.1416 | 0.130067 | 0.342916 | 0.379297 | 0.704468 | 0.98928 | 0.004681 |
| ENSMUSG00000041126.17 | 318.8765 | 0.150553 | 0.439566 | 0.342505 | 0.731971 | 0.98928 | 0.004681 |
| ENSMUSG00000041238.17 | 14.31181 | 0.826709 | 2.362817 | 0.349883 | 0.726427 | 0.98928 | 0.004681 |
| ENSMUSG00000041263.15 | 14.43685 | 1.328099 | 2.688772 | 0.493943 | 0.621347 | 0.98928 | 0.004681 |
| ENSMUSG00000041303.8 | 105.4861 | 0.40645 | 1.036776 | 0.392032 | 0.695034 | 0.98928 | 0.004681 |
| ENSMUSG00000041341.11 | 65.55213 | -0.45473 | 1.155155 | -0.39366 | 0.693835 | 0.98928 | 0.004681 |
| ENSMUSG00000041481.18 | 15.95892 | -0.72397 | 2.246184 | -0.32231 | 0.747217 | 0.98928 | 0.004681 |
| ENSMUSG00000041540.17 | 66.46907 | -0.30149 | 0.908969 | -0.33168 | 0.740131 | 0.98928 | 0.004681 |
| ENSMUSG00000041629.8 | 108.1237 | 0.227724 | 0.685364 | 0.332267 | 0.739687 | 0.98928 | 0.004681 |
| ENSMUSG00000041684.12 | 48.80055 | -0.43171 | 1.093702 | -0.39472 | 0.693046 | 0.98928 | 0.004681 |
| ENSMUSG00000041685.17 | 72.80468 | 0.381729 | 1.134843 | 0.336371 | 0.736591 | 0.98928 | 0.004681 |
| ENSMUSG00000041740.17 | 30.65978 | -0.69047 | 1.491898 | -0.46281 | 0.643497 | 0.98928 | 0.004681 |
| ENSMUSG00000041891.17 | 879.8742 | -0.15553 | 0.308247 | -0.50456 | 0.613867 | 0.98928 | 0.004681 |
| ENSMUSG00000041895.16 | 40.12954 | 0.660746 | 1.51717 | 0.435512 | 0.663191 | 0.98928 | 0.004681 |
| ENSMUSG00000041915.10 | 82.89758 | -0.50025 | 1.258876 | -0.39738 | 0.691086 | 0.98928 | 0.004681 |
| ENSMUSG00000041920.15 | 19.38766 | 0.93683 | 2.416788 | 0.387634 | 0.698287 | 0.98928 | 0.004681 |
| ENSMUSG00000041936.19 | 39.20923 | -0.53047 | 1.326103 | -0.40002 | 0.68914 | 0.98928 | 0.004681 |
| ENSMUSG00000041961.15 | 28.41395 | 0.512247 | 1.214153 | 0.421897 | 0.6731 | 0.98928 | 0.004681 |
| ENSMUSG00000041966.19 | 119.0536 | -0.22789 | 0.688442 | -0.33102 | 0.740633 | 0.98928 | 0.004681 |
| ENSMUSG00000041974.11 | 17.80339 | 1.063905 | 2.54881 | 0.417412 | 0.676377 | 0.98928 | 0.004681 |
| ENSMUSG00000042010.17 | 45.09213 | -0.55643 | 1.432805 | -0.38835 | 0.697758 | 0.98928 | 0.004681 |
| ENSMUSG00000042015.18 | 13.0395 | 0.823957 | 1.890764 | 0.43578 | 0.662996 | 0.98928 | 0.004681 |
| ENSMUSG00000042041.8 | 13.83471 | 0.898625 | 1.997947 | 0.449774 | 0.652873 | 0.98928 | 0.004681 |
| ENSMUSG00000042165.9 | 40.80256 | -0.36163 | 1.02014 | -0.35449 | 0.722969 | 0.98928 | 0.004681 |
| ENSMUSG00000042197.14 | 15.01346 | 0.936049 | 1.976458 | 0.4736 | 0.635786 | 0.98928 | 0.004681 |
| ENSMUSG00000042225.4 | 15.33897 | -0.79207 | 2.196225 | -0.36065 | 0.718361 | 0.98928 | 0.004681 |
| ENSMUSG00000042229.10 | 38.4902 | -0.38774 | 1.214752 | -0.31919 | 0.749583 | 0.98928 | 0.004681 |
| ENSMUSG00000042350.14 | 80.31904 | 0.269687 | 0.801047 | 0.336669 | 0.736367 | 0.98928 | 0.004681 |
| ENSMUSG00000042389.14 | 24.28557 | 0.678102 | 1.965502 | 0.345002 | 0.730093 | 0.98928 | 0.004681 |
| ENSMUSG00000042472.12 | 34.01294 | -0.74947 | 1.973931 | -0.37968 | 0.704182 | 0.98928 | 0.004681 |

| | | | | | | | |
|---|---|---|---|---|---|---|---|
| ENSMUSG00000042680.9 | 29.53354 | -0.58616 | 1.548421 | -0.37855 | 0.705021 | 0.98928 | 0.004681 |
| ENSMUSG00000042712.11 | 79.5486 | -0.34576 | 0.912178 | -0.37905 | 0.704653 | 0.98928 | 0.004681 |
| ENSMUSG00000042770.9 | 1531.159 | 0.093408 | 0.266416 | 0.35061 | 0.725881 | 0.98928 | 0.004681 |
| ENSMUSG00000042810.15 | 21.89148 | 0.78187 | 1.798333 | 0.434775 | 0.663726 | 0.98928 | 0.004681 |
| ENSMUSG00000042988.11 | 76.55871 | -0.49603 | 1.14293 | -0.434 | 0.664289 | 0.98928 | 0.004681 |
| ENSMUSG00000043131.15 | 125.0441 | 0.343422 | 0.76813 | 0.447088 | 0.654812 | 0.98928 | 0.004681 |
| ENSMUSG00000043336.15 | 24.78513 | -0.80445 | 1.672313 | -0.48104 | 0.630487 | 0.98928 | 0.004681 |
| ENSMUSG00000043681.6 | 517.5877 | -0.25761 | 0.515601 | -0.49964 | 0.61733 | 0.98928 | 0.004681 |
| ENSMUSG00000043733.15 | 175.7108 | -0.24809 | 0.750323 | -0.33064 | 0.740917 | 0.98928 | 0.004681 |
| ENSMUSG00000043789.9 | 142.8527 | -0.27012 | 0.685033 | -0.39431 | 0.693352 | 0.98928 | 0.004681 |
| ENSMUSG00000043885.15 | 31.1911 | 0.527616 | 1.555386 | 0.339219 | 0.734445 | 0.98928 | 0.004681 |
| ENSMUSG00000043909.17 | 25.97874 | 0.916996 | 1.989907 | 0.460824 | 0.644925 | 0.98928 | 0.004681 |
| ENSMUSG00000043940.16 | 147.2684 | 0.218151 | 0.672368 | 0.324452 | 0.745596 | 0.98928 | 0.004681 |
| ENSMUSG00000044033.17 | 17.72432 | -1.04494 | 2.298981 | -0.45453 | 0.649451 | 0.98928 | 0.004681 |
| ENSMUSG00000044068.8 | 116.8034 | -0.23938 | 0.751458 | -0.31856 | 0.750063 | 0.98928 | 0.004681 |
| ENSMUSG00000044221.15 | 454.2958 | -0.17503 | 0.42552 | -0.41134 | 0.680827 | 0.98928 | 0.004681 |
| ENSMUSG00000044224.17 | 314.6417 | -0.19809 | 0.550324 | -0.35995 | 0.718881 | 0.98928 | 0.004681 |
| ENSMUSG00000044254.7 | 76.88868 | 0.32276 | 0.786948 | 0.410141 | 0.681703 | 0.98928 | 0.004681 |
| ENSMUSG00000044339.13 | 22.71591 | -0.49593 | 1.423846 | -0.34831 | 0.727611 | 0.98928 | 0.004681 |
| ENSMUSG00000044533.16 | 2498.024 | -0.10045 | 0.295379 | -0.34007 | 0.733803 | 0.98928 | 0.004681 |
| ENSMUSG00000044548.12 | 12.59006 | 0.588128 | 1.719351 | 0.342064 | 0.732303 | 0.98928 | 0.004681 |
| ENSMUSG00000044617.9 | 15.91322 | 0.698971 | 2.101632 | 0.332585 | 0.739448 | 0.98928 | 0.004681 |
| ENSMUSG00000044791.17 | 84.36516 | 0.430546 | 0.972063 | 0.44292 | 0.657824 | 0.98928 | 0.004681 |
| ENSMUSG00000044827.11 | 14.06235 | -1.00136 | 2.620897 | -0.38207 | 0.702411 | 0.98928 | 0.004681 |
| ENSMUSG00000044934.14 | 87.12164 | 0.389535 | 0.896195 | 0.434654 | 0.663814 | 0.98928 | 0.004681 |
| ENSMUSG00000045100.12 | 19.73383 | 0.789189 | 2.064317 | 0.3823 | 0.702239 | 0.98928 | 0.004681 |
| ENSMUSG00000045106.13 | 12.03799 | -0.88554 | 2.423088 | -0.36546 | 0.714767 | 0.98928 | 0.004681 |
| ENSMUSG00000045180.14 | 25.3291 | -0.89398 | 1.941124 | -0.46055 | 0.645123 | 0.98928 | 0.004681 |
| ENSMUSG00000045237.6 | 40.5897 | 0.533035 | 1.204065 | 0.442696 | 0.657985 | 0.98928 | 0.004681 |
| ENSMUSG00000045538.7 | 67.7195 | 0.476811 | 1.382041 | 0.345005 | 0.73009 | 0.98928 | 0.004681 |
| ENSMUSG00000045551.8 | 12.66467 | 1.07749 | 3.129066 | 0.344349 | 0.730584 | 0.98928 | 0.004681 |
| ENSMUSG00000045636.17 | 337.4996 | 0.172609 | 0.427735 | 0.403542 | 0.686549 | 0.98928 | 0.004681 |
| ENSMUSG00000045672.16 | 75.48338 | 0.367581 | 0.901949 | 0.407541 | 0.683611 | 0.98928 | 0.004681 |
| ENSMUSG00000045694.5 | 34.36124 | 0.534723 | 1.167382 | 0.458053 | 0.646914 | 0.98928 | 0.004681 |
| ENSMUSG00000045752.14 | 11.19998 | 0.795184 | 2.420413 | 0.328532 | 0.742509 | 0.98928 | 0.004681 |
| ENSMUSG00000045827.11 | 38.90243 | 0.601303 | 1.515078 | 0.396879 | 0.691456 | 0.98928 | 0.004681 |
| ENSMUSG00000045867.11 | 36.28755 | 0.72353 | 1.530005 | 0.472894 | 0.636289 | 0.98928 | 0.004681 |
| ENSMUSG00000046410.11 | 25.75493 | 0.746568 | 1.926015 | 0.387623 | 0.698295 | 0.98928 | 0.004681 |
| ENSMUSG00000046546.4 | 18.46466 | -0.9523 | 1.915463 | -0.49716 | 0.619073 | 0.98928 | 0.004681 |
| ENSMUSG00000046598.16 | 622.327 | 0.179508 | 0.374541 | 0.479274 | 0.631744 | 0.98928 | 0.004681 |
| ENSMUSG00000046718.9 | 877.7456 | 0.156374 | 0.338492 | 0.461973 | 0.644101 | 0.98928 | 0.004681 |
| ENSMUSG00000046727.14 | 20.31715 | -0.86784 | 1.858291 | -0.46701 | 0.640491 | 0.98928 | 0.004681 |
| ENSMUSG00000046756.11 | 407.6405 | 0.17178 | 0.488387 | 0.35173 | 0.725041 | 0.98928 | 0.004681 |
| ENSMUSG00000046761.14 | 27.85043 | -1.03155 | 2.261676 | -0.4561 | 0.648318 | 0.98928 | 0.004681 |

| | | | | | | | |
|---|---|---|---|---|---|---|---|
| ENSMUSG00000046897.18 | 24.94223 | 0.940153 | 2.033082 | 0.462428 | 0.643775 | 0.98928 | 0.004681 |
| ENSMUSG00000047084.9 | 174.7072 | 0.357149 | 0.720994 | 0.495356 | 0.620349 | 0.98928 | 0.004681 |
| ENSMUSG00000047153.5 | 73.01087 | -0.99954 | 2.720024 | -0.36747 | 0.713265 | 0.98928 | 0.004681 |
| ENSMUSG00000047180.9 | 17.32213 | -0.9632 | 2.134563 | -0.45124 | 0.651817 | 0.98928 | 0.004681 |
| ENSMUSG00000047187.10 | 615.6711 | -0.13498 | 0.340253 | -0.39672 | 0.691575 | 0.98928 | 0.004681 |
| ENSMUSG00000047230.7 | 173.394 | 0.244923 | 0.572021 | 0.428171 | 0.668527 | 0.98928 | 0.004681 |
| ENSMUSG00000047250.14 | 19.71496 | -0.78649 | 1.770708 | -0.44417 | 0.656921 | 0.98928 | 0.004681 |
| ENSMUSG00000047260.5 | 526.2511 | 0.177086 | 0.394754 | 0.448597 | 0.653722 | 0.98928 | 0.004681 |
| ENSMUSG00000047371.8 | 17.56239 | 0.826961 | 2.159836 | 0.382882 | 0.701808 | 0.98928 | 0.004681 |
| ENSMUSG00000047412.17 | 305.1585 | 0.206817 | 0.448544 | 0.461086 | 0.644737 | 0.98928 | 0.004681 |
| ENSMUSG00000047423.11 | 14.28696 | -0.70836 | 1.848587 | -0.38319 | 0.701578 | 0.98928 | 0.004681 |
| ENSMUSG00000047454.13 | 226.3274 | -0.26876 | 0.57885 | -0.46429 | 0.642437 | 0.98928 | 0.004681 |
| ENSMUSG00000047459.15 | 312.127 | -0.34793 | 0.841419 | -0.4135 | 0.679241 | 0.98928 | 0.004681 |
| ENSMUSG00000047603.10 | 24.39694 | 0.633568 | 1.760485 | 0.359883 | 0.718935 | 0.98928 | 0.004681 |
| ENSMUSG00000047613.11 | 91.65905 | -0.25815 | 0.791789 | -0.32604 | 0.744395 | 0.98928 | 0.004681 |
| ENSMUSG00000047648.14 | 107.8206 | -0.34986 | 0.824964 | -0.4241 | 0.671495 | 0.98928 | 0.004681 |
| ENSMUSG00000047731.18 | 293.4385 | -0.18465 | 0.540783 | -0.34145 | 0.732767 | 0.98928 | 0.004681 |
| ENSMUSG00000047881.15 | 105.8071 | 0.357262 | 0.744839 | 0.479649 | 0.631477 | 0.98928 | 0.004681 |
| ENSMUSG00000047963.8 | 389.0785 | -0.19776 | 0.391525 | -0.5051 | 0.61349 | 0.98928 | 0.004681 |
| ENSMUSG00000048175.14 | 97.36916 | 0.350958 | 0.790905 | 0.443742 | 0.657229 | 0.98928 | 0.004681 |
| ENSMUSG00000048222.4 | 56.50246 | -0.40731 | 1.056938 | -0.38537 | 0.699963 | 0.98928 | 0.004681 |
| ENSMUSG00000048277.16 | 54.07011 | -0.61106 | 1.36686 | -0.44705 | 0.654836 | 0.98928 | 0.004681 |
| ENSMUSG00000048307.9 | 89.40307 | 0.325795 | 0.891229 | 0.365557 | 0.714695 | 0.98928 | 0.004681 |
| ENSMUSG00000048388.4 | 13.83977 | -0.87977 | 2.401204 | -0.36639 | 0.714077 | 0.98928 | 0.004681 |
| ENSMUSG00000048920.9 | 30.80797 | 0.444582 | 1.394485 | 0.318815 | 0.749867 | 0.98928 | 0.004681 |
| ENSMUSG00000049086.9 | 28.85337 | -0.65188 | 1.651346 | -0.39476 | 0.693021 | 0.98928 | 0.004681 |
| ENSMUSG00000049323.9 | 23.3031 | -0.80902 | 1.699523 | -0.47603 | 0.634053 | 0.98928 | 0.004681 |
| ENSMUSG00000049470.14 | 238.5437 | -0.2504 | 0.566128 | -0.4423 | 0.658269 | 0.98928 | 0.004681 |
| ENSMUSG00000049516.10 | 150.6158 | -0.25639 | 0.679297 | -0.37743 | 0.705855 | 0.98928 | 0.004681 |
| ENSMUSG00000049686.15 | 45.01487 | -0.59213 | 1.312846 | -0.45103 | 0.651971 | 0.98928 | 0.004681 |
| ENSMUSG00000049739.11 | 15.31698 | 0.945636 | 2.661098 | 0.355355 | 0.722323 | 0.98928 | 0.004681 |
| ENSMUSG00000049791.5 | 172.9846 | -0.22648 | 0.609342 | -0.37169 | 0.710126 | 0.98928 | 0.004681 |
| ENSMUSG00000049940.8 | 331.4981 | 0.205421 | 0.499871 | 0.410947 | 0.681111 | 0.98928 | 0.004681 |
| ENSMUSG00000049957.15 | 30.06172 | -0.5329 | 1.586072 | -0.33599 | 0.736881 | 0.98928 | 0.004681 |
| ENSMUSG00000049960.5 | 275.6725 | -0.21747 | 0.520236 | -0.41803 | 0.675927 | 0.98928 | 0.004681 |
| ENSMUSG00000049971.18 | 51.2612 | -0.51746 | 1.373098 | -0.37686 | 0.706279 | 0.98928 | 0.004681 |
| ENSMUSG00000050017.12 | 287.0204 | 0.175438 | 0.553914 | 0.316725 | 0.751452 | 0.98928 | 0.004681 |
| ENSMUSG00000050069.4 | 38.9702 | -0.70729 | 1.658602 | -0.42644 | 0.669787 | 0.98928 | 0.004681 |
| ENSMUSG00000050229.4 | 26.4853 | 0.651857 | 1.852595 | 0.351861 | 0.724942 | 0.98928 | 0.004681 |
| ENSMUSG00000050310.10 | 33.61425 | -0.44407 | 1.259925 | -0.35246 | 0.724496 | 0.98928 | 0.004681 |
| ENSMUSG00000050552.9 | 89.07665 | -0.45825 | 0.953647 | -0.48053 | 0.630851 | 0.98928 | 0.004681 |
| ENSMUSG00000050663.9 | 12.98954 | 1.020347 | 2.719619 | 0.37518 | 0.707527 | 0.98928 | 0.004681 |
| ENSMUSG00000050668.11 | 68.86699 | 0.588668 | 1.260083 | 0.467166 | 0.640381 | 0.98928 | 0.004681 |
| ENSMUSG00000050796.5 | 34.79896 | -0.65999 | 1.653427 | -0.39917 | 0.68977 | 0.98928 | 0.004681 |

| | | | | | | | |
|---|---|---|---|---|---|---|---|
| ENSMUSG00000050891.11 | 139.3712 | 0.334758 | 0.787357 | 0.425166 | 0.670716 | 0.98928 | 0.004681 |
| ENSMUSG00000050931.8 | 276.2582 | 0.201543 | 0.466162 | 0.432345 | 0.665491 | 0.98928 | 0.004681 |
| ENSMUSG00000051169.15 | 34.80593 | -0.90173 | 1.800075 | -0.50094 | 0.616414 | 0.98928 | 0.004681 |
| ENSMUSG00000051185.10 | 152.1033 | 0.308993 | 0.625542 | 0.49396 | 0.621334 | 0.98928 | 0.004681 |
| ENSMUSG00000051355.19 | 183.2632 | 0.229844 | 0.663106 | 0.346617 | 0.728879 | 0.98928 | 0.004681 |
| ENSMUSG00000051413.18 | 10.2384 | 1.032013 | 2.198104 | 0.469502 | 0.638711 | 0.98928 | 0.004681 |
| ENSMUSG00000051579.11 | 19.51214 | -0.72155 | 2.068528 | -0.34882 | 0.727221 | 0.98928 | 0.004681 |
| ENSMUSG00000052085.8 | 117.0524 | -0.38789 | 0.774049 | -0.50112 | 0.616288 | 0.98928 | 0.004681 |
| ENSMUSG00000052144.9 | 226.1301 | 0.245172 | 0.528599 | 0.463815 | 0.642781 | 0.98928 | 0.004681 |
| ENSMUSG00000052151.13 | 19.27447 | -0.59871 | 1.746384 | -0.34283 | 0.731728 | 0.98928 | 0.004681 |
| ENSMUSG00000052298.13 | 37.62129 | 0.705061 | 1.513721 | 0.46578 | 0.641373 | 0.98928 | 0.004681 |
| ENSMUSG00000052336.8 | 13.61875 | -1.02685 | 2.654221 | -0.38687 | 0.69885 | 0.98928 | 0.004681 |
| ENSMUSG00000052395.18 | 54.56592 | 0.439724 | 0.919721 | 0.478106 | 0.632575 | 0.98928 | 0.004681 |
| ENSMUSG00000052419.8 | 68.28335 | 0.383883 | 0.850369 | 0.451431 | 0.651679 | 0.98928 | 0.004681 |
| ENSMUSG00000052488.8 | 37.702 | 0.586259 | 1.593822 | 0.367832 | 0.712998 | 0.98928 | 0.004681 |
| ENSMUSG00000052512.18 | 46.89354 | 0.50017 | 1.090125 | 0.458819 | 0.646364 | 0.98928 | 0.004681 |
| ENSMUSG00000052751.17 | 61.29895 | 0.549746 | 1.325632 | 0.414705 | 0.678358 | 0.98928 | 0.004681 |
| ENSMUSG00000052812.6 | 70.59559 | -0.35739 | 0.93908 | -0.38057 | 0.703519 | 0.98928 | 0.004681 |
| ENSMUSG00000052928.10 | 11.9048 | -1.11494 | 2.501574 | -0.44569 | 0.655818 | 0.98928 | 0.004681 |
| ENSMUSG00000052962.4 | 367.2298 | -0.19826 | 0.415611 | -0.47703 | 0.633343 | 0.98928 | 0.004681 |
| ENSMUSG00000053253.17 | 239.4935 | -0.24078 | 0.628785 | -0.38293 | 0.701775 | 0.98928 | 0.004681 |
| ENSMUSG00000053286.12 | 103.0864 | 0.313975 | 0.905566 | 0.346716 | 0.728804 | 0.98928 | 0.004681 |
| ENSMUSG00000053289.9 | 101.1921 | -0.29206 | 0.910213 | -0.32087 | 0.748307 | 0.98928 | 0.004681 |
| ENSMUSG00000053333.16 | 14.48068 | 0.722799 | 2.077761 | 0.347874 | 0.727935 | 0.98928 | 0.004681 |
| ENSMUSG00000053536.8 | 98.70773 | -0.35149 | 0.817102 | -0.43017 | 0.667072 | 0.98928 | 0.004681 |
| ENSMUSG00000053646.14 | 47.98722 | -0.43631 | 1.338583 | -0.32595 | 0.744465 | 0.98928 | 0.004681 |
| ENSMUSG00000053769.6 | 24.8007 | 0.697543 | 1.796937 | 0.388185 | 0.697879 | 0.98928 | 0.004681 |
| ENSMUSG00000053950.11 | 40.28566 | 0.56536 | 1.536073 | 0.368055 | 0.712832 | 0.98928 | 0.004681 |
| ENSMUSG00000054252.19 | 125.2339 | -0.28045 | 0.705697 | -0.3974 | 0.691069 | 0.98928 | 0.004681 |
| ENSMUSG00000054302.16 | 167.4677 | 0.322399 | 0.761388 | 0.423436 | 0.671977 | 0.98928 | 0.004681 |
| ENSMUSG00000054321.8 | 13.0632 | -1.06722 | 2.370211 | -0.45026 | 0.65252 | 0.98928 | 0.004681 |
| ENSMUSG00000054387.14 | 68.94402 | -0.42022 | 0.927163 | -0.45323 | 0.650381 | 0.98928 | 0.004681 |
| ENSMUSG00000054408.10 | 238.7309 | -0.23575 | 0.560812 | -0.42037 | 0.674216 | 0.98928 | 0.004681 |
| ENSMUSG00000054452.11 | 312.5939 | 0.26205 | 0.609537 | 0.429917 | 0.667256 | 0.98928 | 0.004681 |
| ENSMUSG00000054469.15 | 116.4049 | -0.21721 | 0.67999 | -0.31943 | 0.749403 | 0.98928 | 0.004681 |
| ENSMUSG00000054474.16 | 92.50919 | -0.38648 | 1.028874 | -0.37564 | 0.707186 | 0.98928 | 0.004681 |
| ENSMUSG00000054477.17 | 59.43898 | 0.445376 | 0.901936 | 0.4938 | 0.621448 | 0.98928 | 0.004681 |
| ENSMUSG00000054604.8 | 286.2149 | -0.24761 | 0.490168 | -0.50516 | 0.613447 | 0.98928 | 0.004681 |
| ENSMUSG00000054611.18 | 72.4485 | -0.42159 | 0.900619 | -0.46811 | 0.639705 | 0.98928 | 0.004681 |
| ENSMUSG00000054715.12 | 13.18728 | 0.957104 | 2.348115 | 0.407605 | 0.683564 | 0.98928 | 0.004681 |
| ENSMUSG00000054723.7 | 67.60265 | -0.77093 | 1.547046 | -0.49833 | 0.618255 | 0.98928 | 0.004681 |
| ENSMUSG00000054814.15 | 75.96143 | -0.42588 | 0.999832 | -0.42595 | 0.670145 | 0.98928 | 0.004681 |
| ENSMUSG00000054827.13 | 4954.199 | -0.10213 | 0.232319 | -0.43961 | 0.660216 | 0.98928 | 0.004681 |
| ENSMUSG00000054836.13 | 18.13604 | -0.931 | 1.961138 | -0.47472 | 0.634985 | 0.98928 | 0.004681 |

| | | | | | | | |
|---|---|---|---|---|---|---|---|
| ENSMUSG00000055065.8 | 168.3449 | -0.27622 | 0.586803 | -0.47072 | 0.637843 | 0.98928 | 0.004681 |
| ENSMUSG00000055148.8 | 57.35392 | 0.737316 | 1.633883 | 0.451266 | 0.651798 | 0.98928 | 0.004681 |
| ENSMUSG00000055240.18 | 15.8102 | 0.867357 | 2.543488 | 0.341011 | 0.733095 | 0.98928 | 0.004681 |
| ENSMUSG00000055296.15 | 62.1283 | 0.358225 | 1.058885 | 0.338304 | 0.735134 | 0.98928 | 0.004681 |
| ENSMUSG00000055302.6 | 1150.695 | 0.101613 | 0.300188 | 0.338497 | 0.734988 | 0.98928 | 0.004681 |
| ENSMUSG00000055322.17 | 100.5833 | -0.32121 | 0.776919 | -0.41344 | 0.679283 | 0.98928 | 0.004681 |
| ENSMUSG00000055541.19 | 36.06165 | -0.42243 | 1.271828 | -0.33214 | 0.739781 | 0.98928 | 0.004681 |
| ENSMUSG00000055652.15 | 42.50369 | -0.63928 | 1.567149 | -0.40793 | 0.683328 | 0.98928 | 0.004681 |
| ENSMUSG00000055660.10 | 50.80963 | -0.48302 | 1.007938 | -0.47922 | 0.631784 | 0.98928 | 0.004681 |
| ENSMUSG00000055681.15 | 530.5156 | -0.12442 | 0.38976 | -0.31922 | 0.74956 | 0.98928 | 0.004681 |
| ENSMUSG00000055737.13 | 3275.393 | -0.10113 | 0.253289 | -0.39925 | 0.689707 | 0.98928 | 0.004681 |
| ENSMUSG00000056035.9 | 28857.48 | 0.313853 | 0.877192 | 0.357793 | 0.720498 | 0.98928 | 0.004681 |
| ENSMUSG00000056076.14 | 43.09383 | -0.67258 | 1.536662 | -0.43769 | 0.66161 | 0.98928 | 0.004681 |
| ENSMUSG00000056204.9 | 129.9616 | 0.351049 | 0.842822 | 0.416516 | 0.677032 | 0.98928 | 0.004681 |
| ENSMUSG00000056515.10 | 32.17814 | 0.662578 | 1.954746 | 0.338959 | 0.734641 | 0.98928 | 0.004681 |
| ENSMUSG00000056602.14 | 14.28552 | 1.248685 | 3.017639 | 0.413795 | 0.679024 | 0.98928 | 0.004681 |
| ENSMUSG00000056643.6 | 42.84304 | -0.46589 | 0.998176 | -0.46674 | 0.640683 | 0.98928 | 0.004681 |
| ENSMUSG00000056820.8 | 155.4506 | -0.36103 | 0.844417 | -0.42754 | 0.668983 | 0.98928 | 0.004681 |
| ENSMUSG00000056919.10 | 12.96733 | 0.896331 | 2.468358 | 0.363128 | 0.716509 | 0.98928 | 0.004681 |
| ENSMUSG00000057036.8 | 298.605 | 0.1627 | 0.419918 | 0.387458 | 0.698417 | 0.98928 | 0.004681 |
| ENSMUSG00000057134.7 | 41.31445 | -0.50615 | 1.332199 | -0.37994 | 0.703993 | 0.98928 | 0.004681 |
| ENSMUSG00000057156.11 | 13.03401 | 0.925034 | 2.746425 | 0.336814 | 0.736257 | 0.98928 | 0.004681 |
| ENSMUSG00000057177.13 | 44.64649 | 0.62131 | 1.396629 | 0.444864 | 0.656418 | 0.98928 | 0.004681 |
| ENSMUSG00000057234.10 | 28.1274 | -1.03719 | 2.083292 | -0.49786 | 0.618582 | 0.98928 | 0.004681 |
| ENSMUSG00000057497.9 | 258.7245 | 0.188305 | 0.476546 | 0.395144 | 0.692736 | 0.98928 | 0.004681 |
| ENSMUSG00000057506.6 | 16.01594 | 0.947097 | 2.258442 | 0.419359 | 0.674954 | 0.98928 | 0.004681 |
| ENSMUSG00000057531.15 | 133.8455 | -0.32216 | 0.782804 | -0.41154 | 0.680675 | 0.98928 | 0.004681 |
| ENSMUSG00000057657.5 | 175.4537 | 0.210013 | 0.548886 | 0.382616 | 0.702004 | 0.98928 | 0.004681 |
| ENSMUSG00000057863.7 | 2032.297 | -0.1239 | 0.286529 | -0.43243 | 0.665427 | 0.98928 | 0.004681 |
| ENSMUSG00000058586.13 | 24.75348 | 0.903432 | 2.06032 | 0.438491 | 0.66103 | 0.98928 | 0.004681 |
| ENSMUSG00000058704.10 | 192.9885 | 0.232631 | 0.707467 | 0.328822 | 0.74229 | 0.98928 | 0.004681 |
| ENSMUSG00000058748.10 | 21.70494 | 0.701749 | 1.459912 | 0.480679 | 0.630744 | 0.98928 | 0.004681 |
| ENSMUSG00000058905.6 | 28.915 | -0.54875 | 1.203004 | -0.45615 | 0.648284 | 0.98928 | 0.004681 |
| ENSMUSG00000059089.5 | 70.7664 | 0.387877 | 1.223279 | 0.31708 | 0.751183 | 0.98928 | 0.004681 |
| ENSMUSG00000059183.13 | 80.42202 | -0.42615 | 0.88335 | -0.48243 | 0.6295 | 0.98928 | 0.004681 |
| ENSMUSG00000059208.16 | 315.4407 | -0.3167 | 0.6739 | -0.46994 | 0.638395 | 0.98928 | 0.004681 |
| ENSMUSG00000059291.16 | 2408.642 | 0.110404 | 0.293151 | 0.376611 | 0.706463 | 0.98928 | 0.004681 |
| ENSMUSG00000059434.10 | 136.7553 | -0.29081 | 0.80333 | -0.36201 | 0.717346 | 0.98928 | 0.004681 |
| ENSMUSG00000059495.15 | 374.0484 | -0.18597 | 0.39847 | -0.46672 | 0.640701 | 0.98928 | 0.004681 |
| ENSMUSG00000059518.15 | 98.03187 | -0.51499 | 1.159825 | -0.44402 | 0.657026 | 0.98928 | 0.004681 |
| ENSMUSG00000059820.15 | 129.3174 | -0.3797 | 0.750554 | -0.5059 | 0.612929 | 0.98928 | 0.004681 |
| ENSMUSG00000059839.10 | 29.62993 | -0.6397 | 1.606748 | -0.39813 | 0.690531 | 0.98928 | 0.004681 |
| ENSMUSG00000059920.10 | 225.2103 | 0.263477 | 0.583767 | 0.451339 | 0.651745 | 0.98928 | 0.004681 |
| ENSMUSG00000059923.14 | 133.9624 | -0.29231 | 0.677358 | -0.43155 | 0.666068 | 0.98928 | 0.004681 |

| | | | | | | | |
|---|---|---|---|---|---|---|---|
| ENSMUSG00000059970.8 | 24.35855 | 0.666502 | 1.594558 | 0.417985 | 0.675958 | 0.98928 | 0.004681 |
| ENSMUSG00000060038.15 | 38.54278 | -0.75803 | 1.848686 | -0.41004 | 0.681779 | 0.98928 | 0.004681 |
| ENSMUSG00000060143.8 | 91.90591 | -0.28031 | 0.848875 | -0.33021 | 0.741243 | 0.98928 | 0.004681 |
| ENSMUSG00000060216.16 | 11.60498 | 0.958871 | 2.236748 | 0.42869 | 0.668149 | 0.98928 | 0.004681 |
| ENSMUSG00000060373.16 | 372.1659 | -0.19102 | 0.420082 | -0.45473 | 0.649306 | 0.98928 | 0.004681 |
| ENSMUSG00000060376.8 | 111.337 | 0.41186 | 1.035468 | 0.397753 | 0.690812 | 0.98928 | 0.004681 |
| ENSMUSG00000060377.5 | 35.02304 | 0.49129 | 1.398605 | 0.351271 | 0.725385 | 0.98928 | 0.004681 |
| ENSMUSG00000060397.7 | 10.5096 | 1.371833 | 3.286646 | 0.417396 | 0.676389 | 0.98928 | 0.004681 |
| ENSMUSG00000060450.15 | 169.8688 | -0.31481 | 0.874471 | -0.36 | 0.718845 | 0.98928 | 0.004681 |
| ENSMUSG00000060579.14 | 45.65539 | 0.418812 | 1.223747 | 0.342238 | 0.732172 | 0.98928 | 0.004681 |
| ENSMUSG00000060678.6 | 10.67526 | 0.992176 | 2.943361 | 0.337089 | 0.736049 | 0.98928 | 0.004681 |
| ENSMUSG00000060703.13 | 2480.432 | 0.129475 | 0.282055 | 0.459041 | 0.646205 | 0.98928 | 0.004681 |
| ENSMUSG00000060733.14 | 216.1659 | 0.272823 | 0.593186 | 0.459929 | 0.645567 | 0.98928 | 0.004681 |
| ENSMUSG00000060743.13 | 164.1788 | -0.18047 | 0.563721 | -0.32015 | 0.748855 | 0.98928 | 0.004681 |
| ENSMUSG00000060802.9 | 33468 | 0.128845 | 0.259339 | 0.496822 | 0.619314 | 0.98928 | 0.004681 |
| ENSMUSG00000061079.15 | 22.31022 | -0.59752 | 1.825985 | -0.32723 | 0.743492 | 0.98928 | 0.004681 |
| ENSMUSG00000061136.17 | 517.8813 | -0.17085 | 0.380411 | -0.44912 | 0.653342 | 0.98928 | 0.004681 |
| ENSMUSG00000061175.13 | 46.73379 | -0.4544 | 1.209005 | -0.37584 | 0.707032 | 0.98928 | 0.004681 |
| ENSMUSG00000061374.15 | 13.91012 | 0.766402 | 2.37928 | 0.322115 | 0.747366 | 0.98928 | 0.004681 |
| ENSMUSG00000061479.16 | 90.67746 | 0.57426 | 1.314958 | 0.436714 | 0.662319 | 0.98928 | 0.004681 |
| ENSMUSG00000061983.8 | 5099.916 | 0.091266 | 0.221859 | 0.411368 | 0.680802 | 0.98928 | 0.004681 |
| ENSMUSG00000061992.3 | 13.28967 | -0.91293 | 2.007427 | -0.45477 | 0.649272 | 0.98928 | 0.004681 |
| ENSMUSG00000062203.16 | 152.7086 | 0.283514 | 0.610063 | 0.464729 | 0.642125 | 0.98928 | 0.004681 |
| ENSMUSG00000062234.15 | 45.64453 | -0.56068 | 1.308036 | -0.42864 | 0.668186 | 0.98928 | 0.004681 |
| ENSMUSG00000062248.6 | 72.39597 | -0.50822 | 1.080249 | -0.47046 | 0.638024 | 0.98928 | 0.004681 |
| ENSMUSG00000062376.8 | 61.74751 | 0.469147 | 1.232808 | 0.380552 | 0.703536 | 0.98928 | 0.004681 |
| ENSMUSG00000062647.17 | 2161.268 | 0.106591 | 0.239301 | 0.445426 | 0.656012 | 0.98928 | 0.004681 |
| ENSMUSG00000062825.16 | 35.97277 | 0.458531 | 1.143476 | 0.400997 | 0.688422 | 0.98928 | 0.004681 |
| ENSMUSG00000063108.16 | 24.71738 | 0.469421 | 1.382444 | 0.339559 | 0.734189 | 0.98928 | 0.004681 |
| ENSMUSG00000063236.7 | 14.31107 | -0.81001 | 2.061813 | -0.39286 | 0.69442 | 0.98928 | 0.004681 |
| ENSMUSG00000063406.12 | 941.7578 | 0.144877 | 0.332075 | 0.436278 | 0.662635 | 0.98928 | 0.004681 |
| ENSMUSG00000063576.13 | 38.0657 | -0.57664 | 1.427505 | -0.40395 | 0.686248 | 0.98928 | 0.004681 |
| ENSMUSG00000063659.12 | 78.47356 | 0.412692 | 0.927428 | 0.444986 | 0.65633 | 0.98928 | 0.004681 |
| ENSMUSG00000063683.13 | 2475 | 0.073001 | 0.228505 | 0.319473 | 0.749368 | 0.98928 | 0.004681 |
| ENSMUSG00000063856.9 | 10680.76 | -0.10411 | 0.287527 | -0.36209 | 0.717283 | 0.98928 | 0.004681 |
| ENSMUSG00000063882.13 | 3952.835 | -0.07889 | 0.216188 | -0.3649 | 0.715183 | 0.98928 | 0.004681 |
| ENSMUSG00000063889.17 | 123.1059 | 0.232606 | 0.613691 | 0.379027 | 0.704668 | 0.98928 | 0.004681 |
| ENSMUSG00000063894.16 | 10.54964 | -0.96446 | 2.525246 | -0.38193 | 0.702515 | 0.98928 | 0.004681 |
| ENSMUSG00000064037.14 | 63.11019 | 0.457564 | 1.074118 | 0.42599 | 0.670115 | 0.98928 | 0.004681 |
| ENSMUSG00000064127.14 | 84.84114 | 0.424973 | 0.979004 | 0.434087 | 0.664225 | 0.98928 | 0.004681 |
| ENSMUSG00000064138.15 | 41.60244 | 0.609981 | 1.617614 | 0.377087 | 0.706109 | 0.98928 | 0.004681 |
| ENSMUSG00000064302.14 | 31.7159 | 0.557362 | 1.721842 | 0.323701 | 0.746164 | 0.98928 | 0.004681 |
| ENSMUSG00000064339.1 | 92963.96 | -0.07858 | 0.240624 | -0.32657 | 0.743996 | 0.98928 | 0.004681 |
| ENSMUSG00000064342.1 | 463.2562 | -0.16178 | 0.408389 | -0.39613 | 0.692006 | 0.98928 | 0.004681 |

| | | | | | | | |
|---|---|---|---|---|---|---|---|
| ENSMUSG00000064351.1 | 7410.065 | -0.09913 | 0.198071 | -0.50048 | 0.616736 | 0.98928 | 0.004681 |
| ENSMUSG00000065952.14 | 19.55421 | 0.90048 | 2.053746 | 0.438458 | 0.661055 | 0.98928 | 0.004681 |
| ENSMUSG00000065979.13 | 185.6472 | -0.2217 | 0.658974 | -0.33644 | 0.73654 | 0.98928 | 0.004681 |
| ENSMUSG00000066036.15 | 112.6859 | -0.2944 | 0.782052 | -0.37644 | 0.706588 | 0.98928 | 0.004681 |
| ENSMUSG00000066149.12 | 198.3396 | 0.283046 | 0.68295 | 0.414446 | 0.678548 | 0.98928 | 0.004681 |
| ENSMUSG00000066151.13 | 57.2394 | 0.524694 | 1.194966 | 0.439087 | 0.660599 | 0.98928 | 0.004681 |
| ENSMUSG00000066232.5 | 352.1322 | 0.222625 | 0.490923 | 0.453481 | 0.650202 | 0.98928 | 0.004681 |
| ENSMUSG00000066443.6 | 14.35692 | 0.631819 | 1.743104 | 0.362468 | 0.717003 | 0.98928 | 0.004681 |
| ENSMUSG00000066595.10 | 26.00356 | 0.548666 | 1.636219 | 0.335325 | 0.73738 | 0.98928 | 0.004681 |
| ENSMUSG00000066798.4 | 57.41297 | -0.40794 | 1.15892 | -0.352 | 0.724838 | 0.98928 | 0.004681 |
| ENSMUSG00000066839.13 | 91.95485 | -0.43908 | 1.03195 | -0.42548 | 0.670485 | 0.98928 | 0.004681 |
| ENSMUSG00000066979.7 | 173.7432 | 0.264373 | 0.590785 | 0.447494 | 0.654518 | 0.98928 | 0.004681 |
| ENSMUSG00000067071.9 | 113.5804 | -0.43198 | 0.879134 | -0.49137 | 0.623166 | 0.98928 | 0.004681 |
| ENSMUSG00000067212.9 | 340.3065 | -0.24918 | 0.559001 | -0.44576 | 0.655769 | 0.98928 | 0.004681 |
| ENSMUSG00000067274.11 | 1292.283 | -0.11555 | 0.309028 | -0.37393 | 0.708456 | 0.98928 | 0.004681 |
| ENSMUSG00000067870.6 | 390.2448 | 0.186552 | 0.484548 | 0.385001 | 0.700236 | 0.98928 | 0.004681 |
| ENSMUSG00000067889.8 | 9.960154 | -1.26526 | 2.752241 | -0.45972 | 0.645717 | 0.98928 | 0.004681 |
| ENSMUSG00000068749.10 | 570.6262 | 0.163944 | 0.346393 | 0.47329 | 0.636006 | 0.98928 | 0.004681 |
| ENSMUSG00000068874.14 | 528.207 | 0.176401 | 0.441994 | 0.399104 | 0.689817 | 0.98928 | 0.004681 |
| ENSMUSG00000069020.10 | 18.49916 | -0.79451 | 2.164013 | -0.36715 | 0.71351 | 0.98928 | 0.004681 |
| ENSMUSG00000069045.12 | 800.2621 | 0.136017 | 0.409439 | 0.332203 | 0.739736 | 0.98928 | 0.004681 |
| ENSMUSG00000069049.12 | 191.4858 | -0.20849 | 0.526188 | -0.39622 | 0.691941 | 0.98928 | 0.004681 |
| ENSMUSG00000069170.15 | 32.72805 | 1.035692 | 2.159113 | 0.479684 | 0.631452 | 0.98928 | 0.004681 |
| ENSMUSG00000069184.5 | 16.73454 | 0.616444 | 1.843438 | 0.334399 | 0.738079 | 0.98928 | 0.004681 |
| ENSMUSG00000069324.7 | 3101.261 | -0.13589 | 0.351922 | -0.38614 | 0.69939 | 0.98928 | 0.004681 |
| ENSMUSG00000069565.13 | 12.04024 | -0.98668 | 2.170605 | -0.45457 | 0.649422 | 0.98928 | 0.004681 |
| ENSMUSG00000069835.11 | 28.15469 | -0.5871 | 1.646948 | -0.35648 | 0.721481 | 0.98928 | 0.004681 |
| ENSMUSG00000069862.7 | 585.0876 | 0.175888 | 0.379146 | 0.463905 | 0.642716 | 0.98928 | 0.004681 |
| ENSMUSG00000070056.7 | 82.13528 | 0.381231 | 0.963917 | 0.395502 | 0.692473 | 0.98928 | 0.004681 |
| ENSMUSG00000070319.7 | 326.6493 | -0.23865 | 0.500838 | -0.4765 | 0.63372 | 0.98928 | 0.004681 |
| ENSMUSG00000070692.7 | 15.91515 | -0.88483 | 1.830138 | -0.48348 | 0.628756 | 0.98928 | 0.004681 |
| ENSMUSG00000070713.6 | 20.15491 | 0.618359 | 1.378262 | 0.448651 | 0.653683 | 0.98928 | 0.004681 |
| ENSMUSG00000070738.11 | 15.18843 | -0.96849 | 2.063915 | -0.46925 | 0.638893 | 0.98928 | 0.004681 |
| ENSMUSG00000070780.12 | 95.11599 | -0.27475 | 0.727276 | -0.37778 | 0.705596 | 0.98928 | 0.004681 |
| ENSMUSG00000070923.5 | 181.2713 | 0.245599 | 0.59361 | 0.413739 | 0.679066 | 0.98928 | 0.004681 |
| ENSMUSG00000071281.13 | 27.99809 | -0.74648 | 1.931194 | -0.38654 | 0.699099 | 0.98928 | 0.004681 |
| ENSMUSG00000071451.11 | 153.1347 | 0.25665 | 0.561008 | 0.457481 | 0.647325 | 0.98928 | 0.004681 |
| ENSMUSG00000071533.12 | 318.6986 | 0.190374 | 0.523495 | 0.36366 | 0.716112 | 0.98928 | 0.004681 |
| ENSMUSG00000071551.14 | 862.0411 | 0.159576 | 0.402338 | 0.396621 | 0.691647 | 0.98928 | 0.004681 |
| ENSMUSG00000071637.6 | 70.59478 | -0.5036 | 1.434557 | -0.35105 | 0.72555 | 0.98928 | 0.004681 |
| ENSMUSG00000071757.11 | 15.34187 | -1.00012 | 2.199516 | -0.4547 | 0.649325 | 0.98928 | 0.004681 |
| ENSMUSG00000072940.6 | 20.0511 | 0.633099 | 1.473069 | 0.429783 | 0.667354 | 0.98928 | 0.004681 |
| ENSMUSG00000073096.12 | 20.2461 | 0.883158 | 1.799608 | 0.49075 | 0.623603 | 0.98928 | 0.004681 |
| ENSMUSG00000073411.13 | 1939.06 | 0.11604 | 0.271641 | 0.427181 | 0.669247 | 0.98928 | 0.004681 |

| | | | | | | | |
|---|---|---|---|---|---|---|---|
| ENSMUSG00000073421.7 | 35.34213 | -0.62639 | 1.387634 | -0.45141 | 0.651698 | 0.98928 | 0.004681 |
| ENSMUSG00000073478.7 | 26.33938 | -0.792 | 2.24183 | -0.35328 | 0.723878 | 0.98928 | 0.004681 |
| ENSMUSG00000073565.6 | 267.694 | 0.222869 | 0.494948 | 0.450288 | 0.652503 | 0.98928 | 0.004681 |
| ENSMUSG00000073647.5 | 11.29923 | 1.05322 | 3.242609 | 0.324806 | 0.745328 | 0.98928 | 0.004681 |
| ENSMUSG00000073678.5 | 170.5271 | -0.26161 | 0.745366 | -0.35099 | 0.725597 | 0.98928 | 0.004681 |
| ENSMUSG00000073684.14 | 32.1333 | 0.597694 | 1.549172 | 0.385815 | 0.699634 | 0.98928 | 0.004681 |
| ENSMUSG00000074182.9 | 107.1333 | -0.36836 | 1.144716 | -0.32179 | 0.747612 | 0.98928 | 0.004681 |
| ENSMUSG00000074264.13 | 1538.511 | 0.115795 | 0.304819 | 0.379883 | 0.704032 | 0.98928 | 0.004681 |
| ENSMUSG00000074336.6 | 1742.793 | 0.104413 | 0.251872 | 0.414549 | 0.678472 | 0.98928 | 0.004681 |
| ENSMUSG00000074513.10 | 58.83299 | 0.350136 | 1.040413 | 0.336536 | 0.736467 | 0.98928 | 0.004681 |
| ENSMUSG00000074521.10 | 22.97513 | 0.604042 | 1.851793 | 0.326193 | 0.744278 | 0.98928 | 0.004681 |
| ENSMUSG00000074591.9 | 13.72864 | -1.01281 | 3.06442 | -0.33051 | 0.741017 | 0.98928 | 0.004681 |
| ENSMUSG00000074733.15 | 163.7245 | 0.35455 | 0.81473 | 0.435174 | 0.663436 | 0.98928 | 0.004681 |
| ENSMUSG00000075279.6 | 47.07814 | 0.312791 | 0.940198 | 0.332686 | 0.739371 | 0.98928 | 0.004681 |
| ENSMUSG00000075324.14 | 64.80753 | 0.547957 | 1.153698 | 0.474957 | 0.634817 | 0.98928 | 0.004681 |
| ENSMUSG00000075486.11 | 443.9865 | 0.195312 | 0.446526 | 0.437402 | 0.661819 | 0.98928 | 0.004681 |
| ENSMUSG00000075700.11 | 1694.313 | 0.100076 | 0.291451 | 0.343371 | 0.731319 | 0.98928 | 0.004681 |
| ENSMUSG00000076609.3 | 381.4551 | 0.234941 | 0.685415 | 0.342771 | 0.73177 | 0.98928 | 0.004681 |
| ENSMUSG00000078240.3 | 63.41927 | -0.41653 | 0.933257 | -0.44632 | 0.655365 | 0.98928 | 0.004681 |
| ENSMUSG00000078570.11 | 66.58073 | -0.57096 | 1.238602 | -0.46097 | 0.644818 | 0.98928 | 0.004681 |
| ENSMUSG00000078636.5 | 27.42141 | -0.46164 | 1.304944 | -0.35376 | 0.723519 | 0.98928 | 0.004681 |
| ENSMUSG00000078652.10 | 247.38 | -0.18846 | 0.478782 | -0.39363 | 0.693855 | 0.98928 | 0.004681 |
| ENSMUSG00000078686.12 | 3287.164 | 0.70797 | 1.588076 | 0.445804 | 0.655739 | 0.98928 | 0.004681 |
| ENSMUSG00000078941.10 | 120.109 | 0.333299 | 0.787512 | 0.423231 | 0.672127 | 0.98928 | 0.004681 |
| ENSMUSG00000079036.11 | 26.70373 | -0.54682 | 1.710937 | -0.3196 | 0.749271 | 0.98928 | 0.004681 |
| ENSMUSG00000079111.4 | 282.7557 | 0.290754 | 0.578303 | 0.502771 | 0.615125 | 0.98928 | 0.004681 |
| ENSMUSG00000079144.8 | 17.91648 | -0.97526 | 1.944154 | -0.50164 | 0.615921 | 0.98928 | 0.004681 |
| ENSMUSG00000079164.11 | 35.41617 | -1.03429 | 2.137284 | -0.48393 | 0.628437 | 0.98928 | 0.004681 |
| ENSMUSG00000079197.11 | 556.2193 | -0.15145 | 0.42795 | -0.35391 | 0.723408 | 0.98928 | 0.004681 |
| ENSMUSG00000079293.12 | 17.41997 | 0.766553 | 2.221567 | 0.345051 | 0.730056 | 0.98928 | 0.004681 |
| ENSMUSG00000079429.10 | 15.96862 | -0.78029 | 1.569173 | -0.49726 | 0.619004 | 0.98928 | 0.004681 |
| ENSMUSG00000079555.3 | 39.45826 | 0.406783 | 1.252292 | 0.324831 | 0.745309 | 0.98928 | 0.004681 |
| ENSMUSG00000079562.8 | 179.9505 | -0.24039 | 0.650833 | -0.36935 | 0.711863 | 0.98928 | 0.004681 |
| ENSMUSG00000079658.10 | 680.5332 | -0.14411 | 0.392317 | -0.36734 | 0.713363 | 0.98928 | 0.004681 |
| ENSMUSG00000079942.6 | 10.7121 | -0.79742 | 1.883493 | -0.42337 | 0.672025 | 0.98928 | 0.004681 |
| ENSMUSG00000081214.4 | 284.5793 | 0.204927 | 0.481545 | 0.425561 | 0.670428 | 0.98928 | 0.004681 |
| ENSMUSG00000081534.4 | 60.31985 | -0.4404 | 0.916105 | -0.48073 | 0.630708 | 0.98928 | 0.004681 |
| ENSMUSG00000081684.5 | 49.95226 | 0.443648 | 1.080994 | 0.410407 | 0.681507 | 0.98928 | 0.004681 |
| ENSMUSG00000081824.6 | 70.91647 | -0.64327 | 1.281677 | -0.50189 | 0.615742 | 0.98928 | 0.004681 |
| ENSMUSG00000081926.2 | 107.3775 | -0.30965 | 0.651166 | -0.47553 | 0.634412 | 0.98928 | 0.004681 |
| ENSMUSG00000082193.2 | 199.5921 | 0.224764 | 0.605525 | 0.371188 | 0.710497 | 0.98928 | 0.004681 |
| ENSMUSG00000082465.2 | 60.41629 | -0.41112 | 0.850947 | -0.48313 | 0.629004 | 0.98928 | 0.004681 |
| ENSMUSG00000083621.2 | 441.6648 | 0.186655 | 0.406727 | 0.458919 | 0.646292 | 0.98928 | 0.004681 |
| ENSMUSG00000084106.5 | 35.69572 | 0.455051 | 1.263782 | 0.360071 | 0.718794 | 0.98928 | 0.004681 |

| | | | | | | | |
|---|---|---|---|---|---|---|---|
| ENSMUSG00000084128.11 | 135.4944 | -0.2912 | 0.868508 | -0.33529 | 0.737404 | 0.98928 | 0.004681 |
| ENSMUSG00000084384.4 | 14.13378 | -0.89879 | 1.79264 | -0.50138 | 0.616104 | 0.98928 | 0.004681 |
| ENSMUSG00000085287.9 | 15.40216 | -0.93954 | 1.959577 | -0.47946 | 0.631611 | 0.98928 | 0.004681 |
| ENSMUSG00000085385.8 | 35.32545 | 0.679711 | 1.538696 | 0.441745 | 0.658674 | 0.98928 | 0.004681 |
| ENSMUSG00000085442.2 | 254.4767 | 0.164781 | 0.492297 | 0.33472 | 0.737837 | 0.98928 | 0.004681 |
| ENSMUSG00000085492.8 | 27.32856 | -0.63945 | 1.328754 | -0.48124 | 0.630344 | 0.98928 | 0.004681 |
| ENSMUSG00000085655.6 | 15.14478 | -0.67098 | 1.683101 | -0.39866 | 0.690146 | 0.98928 | 0.004681 |
| ENSMUSG00000086447.3 | 48.25505 | -0.47259 | 1.482583 | -0.31876 | 0.749907 | 0.98928 | 0.004681 |
| ENSMUSG00000086583.4 | 195.1293 | -0.17241 | 0.533176 | -0.32337 | 0.746415 | 0.98928 | 0.004681 |
| ENSMUSG00000086968.9 | 17.52318 | -1.2208 | 2.513455 | -0.48571 | 0.627176 | 0.98928 | 0.004681 |
| ENSMUSG00000087385.9 | 10.13078 | -1.01031 | 2.44364 | -0.41344 | 0.679282 | 0.98928 | 0.004681 |
| ENSMUSG00000087579.8 | 483.8005 | -0.1907 | 0.515616 | -0.36985 | 0.711495 | 0.98928 | 0.004681 |
| ENSMUSG00000087687.4 | 33.38661 | -0.86659 | 1.75202 | -0.49462 | 0.620866 | 0.98928 | 0.004681 |
| ENSMUSG00000089665.3 | 43.61139 | -0.68808 | 1.732003 | -0.39727 | 0.691166 | 0.98928 | 0.004681 |
| ENSMUSG00000090083.12 | 46.76116 | 0.377763 | 1.133401 | 0.333301 | 0.738907 | 0.98928 | 0.004681 |
| ENSMUSG00000090394.9 | 47.51502 | 0.477028 | 1.209589 | 0.394372 | 0.693307 | 0.98928 | 0.004681 |
| ENSMUSG00000090553.9 | 323.8406 | -0.15499 | 0.489755 | -0.31647 | 0.751647 | 0.98928 | 0.004681 |
| ENSMUSG00000092607.10 | 61.54201 | -0.44581 | 1.121855 | -0.39739 | 0.691081 | 0.98928 | 0.004681 |
| ENSMUSG00000094185.3 | 74.7361 | -0.30296 | 0.81812 | -0.37032 | 0.711148 | 0.98928 | 0.004681 |
| ENSMUSG00000094344.2 | 161.416 | -0.19748 | 0.613135 | -0.32209 | 0.747384 | 0.98928 | 0.004681 |
| ENSMUSG00000094870.9 | 146.8158 | -0.30693 | 0.639613 | -0.47986 | 0.631324 | 0.98928 | 0.004681 |
| ENSMUSG00000094936.9 | 16.46835 | -1.05716 | 2.493954 | -0.42389 | 0.671647 | 0.98928 | 0.004681 |
| ENSMUSG00000095253.5 | 37.30982 | -0.38436 | 1.059442 | -0.3628 | 0.716757 | 0.98928 | 0.004681 |
| ENSMUSG00000095597.3 | 63.73442 | -0.28769 | 0.897163 | -0.32066 | 0.748467 | 0.98928 | 0.004681 |
| ENSMUSG00000095930.2 | 11.09328 | -1.13503 | 2.797073 | -0.40579 | 0.684895 | 0.98928 | 0.004681 |
| ENSMUSG00000096188.2 | 94.2214 | 0.415313 | 0.826401 | 0.502556 | 0.615276 | 0.98928 | 0.004681 |
| ENSMUSG00000096910.3 | 64.72895 | -0.44045 | 1.381496 | -0.31882 | 0.749862 | 0.98928 | 0.004681 |
| ENSMUSG00000096929.8 | 17.10914 | 0.920127 | 2.493517 | 0.369008 | 0.712122 | 0.98928 | 0.004681 |
| ENSMUSG00000097148.3 | 12.65405 | -0.68804 | 2.115491 | -0.32524 | 0.745001 | 0.98928 | 0.004681 |
| ENSMUSG00000097187.2 | 26.48212 | 0.697172 | 2.013702 | 0.346214 | 0.729182 | 0.98928 | 0.004681 |
| ENSMUSG00000097277.8 | 18.81063 | 0.66281 | 1.876969 | 0.353128 | 0.723992 | 0.98928 | 0.004681 |
| ENSMUSG00000097482.3 | 135.3499 | 0.322612 | 0.674245 | 0.478478 | 0.63231 | 0.98928 | 0.004681 |
| ENSMUSG00000097750.3 | 10.29815 | 0.892331 | 2.485621 | 0.358997 | 0.719597 | 0.98928 | 0.004681 |
| ENSMUSG00000098104.2 | 22.2229 | -0.59049 | 1.477602 | -0.39962 | 0.689433 | 0.98928 | 0.004681 |
| ENSMUSG00000098923.2 | 10.60568 | -1.02589 | 2.548452 | -0.40255 | 0.687276 | 0.98928 | 0.004681 |
| ENSMUSG00000099764.2 | 22.93548 | -0.67517 | 1.354944 | -0.4983 | 0.618271 | 0.98928 | 0.004681 |
| ENSMUSG00000100235.9 | 15.00598 | 1.044445 | 2.710416 | 0.385345 | 0.699982 | 0.98928 | 0.004681 |
| ENSMUSG00000101111.2 | 66.03488 | 0.298508 | 0.936836 | 0.318634 | 0.750004 | 0.98928 | 0.004681 |
| ENSMUSG00000101514.2 | 55.80992 | -0.53537 | 1.087901 | -0.49211 | 0.622641 | 0.98928 | 0.004681 |
| ENSMUSG00000102059.2 | 12.72619 | -1.35497 | 2.705886 | -0.50075 | 0.616548 | 0.98928 | 0.004681 |
| ENSMUSG00000102070.2 | 30.57193 | 0.623524 | 1.24451 | 0.50102 | 0.616357 | 0.98928 | 0.004681 |
| ENSMUSG00000102562.2 | 15.97912 | -1.39379 | 2.956257 | -0.47147 | 0.637305 | 0.98928 | 0.004681 |
| ENSMUSG00000103502.2 | 18.93807 | -1.0693 | 2.136791 | -0.50042 | 0.616779 | 0.98928 | 0.004681 |
| ENSMUSG00000104394.3 | 82.04886 | 0.47998 | 1.274976 | 0.376462 | 0.706574 | 0.98928 | 0.004681 |

| | | | | | | | |
|---|---|---|---|---|---|---|---|
| ENSMUSG00000104445.2 | 19.19427 | -0.97902 | 2.232427 | -0.43854 | 0.660992 | 0.98928 | 0.004681 |
| ENSMUSG00000104548.2 | 12.90202 | 0.988308 | 2.090937 | 0.472663 | 0.636454 | 0.98928 | 0.004681 |
| ENSMUSG00000104950.2 | 10.82015 | 1.094411 | 2.590652 | 0.422446 | 0.672699 | 0.98928 | 0.004681 |
| ENSMUSG00000105509.2 | 12.54772 | 0.775619 | 2.4513 | 0.316411 | 0.75169 | 0.98928 | 0.004681 |
| ENSMUSG00000105810.2 | 10.63467 | 1.163949 | 3.269129 | 0.356043 | 0.721809 | 0.98928 | 0.004681 |
| ENSMUSG00000106087.2 | 12.62954 | 0.961347 | 2.731713 | 0.351921 | 0.724898 | 0.98928 | 0.004681 |
| ENSMUSG00000106539.2 | 17.28122 | -0.73634 | 1.639359 | -0.44916 | 0.653315 | 0.98928 | 0.004681 |
| ENSMUSG00000106926.2 | 248.3343 | 0.194338 | 0.477126 | 0.407311 | 0.68378 | 0.98928 | 0.004681 |
| ENSMUSG00000107160.2 | 61.71294 | -0.43965 | 0.924998 | -0.4753 | 0.634572 | 0.98928 | 0.004681 |
| ENSMUSG00000107451.2 | 32.74388 | 0.597027 | 1.210933 | 0.493031 | 0.621991 | 0.98928 | 0.004681 |
| ENSMUSG00000107470.3 | 125.6899 | -0.23027 | 0.707965 | -0.32526 | 0.744984 | 0.98928 | 0.004681 |
| ENSMUSG00000109841.3 | 47.85067 | -0.48374 | 1.432681 | -0.33765 | 0.735629 | 0.98928 | 0.004681 |
| ENSMUSG00000110035.2 | 13.95797 | -1.12024 | 2.769463 | -0.4045 | 0.685846 | 0.98928 | 0.004681 |
| ENSMUSG00000110397.3 | 15.68423 | 0.9824 | 2.935975 | 0.334608 | 0.737921 | 0.98928 | 0.004681 |
| ENSMUSG00000110834.2 | 38.58226 | 0.794766 | 1.972563 | 0.40291 | 0.687014 | 0.98928 | 0.004681 |
| ENSMUSG00000110841.2 | 55.81206 | 0.37112 | 1.051353 | 0.352993 | 0.724094 | 0.98928 | 0.004681 |
| ENSMUSG00000112249.2 | 43.41472 | 0.438648 | 1.270759 | 0.345186 | 0.729955 | 0.98928 | 0.004681 |
| ENSMUSG00000112294.2 | 17.94589 | -1.06154 | 2.247218 | -0.47238 | 0.636656 | 0.98928 | 0.004681 |
| ENSMUSG00000113198.2 | 169.9449 | 0.215186 | 0.616812 | 0.348868 | 0.727188 | 0.98928 | 0.004681 |
| ENSMUSG00000113625.2 | 121.7617 | 0.224827 | 0.632415 | 0.355506 | 0.722211 | 0.98928 | 0.004681 |
| ENSMUSG00000113948.2 | 134.0862 | 0.271188 | 0.58756 | 0.46155 | 0.644404 | 0.98928 | 0.004681 |
| ENSMUSG00000114255.2 | 55.34652 | 0.682402 | 1.373193 | 0.496945 | 0.619228 | 0.98928 | 0.004681 |
| ENSMUSG00000115852.3 | 49.44958 | -0.62212 | 1.546278 | -0.40234 | 0.687437 | 0.98928 | 0.004681 |
| ENSMUSG00000116508.2 | 13.61799 | 1.164212 | 2.443625 | 0.476428 | 0.633769 | 0.98928 | 0.004681 |
| ENSMUSG00000117105.2 | 55.24321 | 0.65481 | 1.718736 | 0.380984 | 0.703215 | 0.98928 | 0.004681 |
| ENSMUSG00000117403.2 | 18.34552 | 0.504636 | 1.508631 | 0.334499 | 0.738003 | 0.98928 | 0.004681 |
| ENSMUSG00000117621.2 | 47.95698 | 0.474234 | 1.058497 | 0.448025 | 0.654135 | 0.98928 | 0.004681 |
| ENSMUSG00000117662.2 | 51.76608 | -0.73215 | 1.448997 | -0.50528 | 0.613363 | 0.98928 | 0.004681 |
| ENSMUSG00000117879.2 | 39.18703 | 0.621075 | 1.461488 | 0.424961 | 0.670865 | 0.98928 | 0.004681 |
| ENSMUSG00000118052.2 | 35.03018 | 0.400147 | 1.174609 | 0.340664 | 0.733356 | 0.98928 | 0.004681 |
| ENSMUSG00000118537.2 | 23.91246 | -0.72136 | 1.773491 | -0.40674 | 0.684195 | 0.98928 | 0.004681 |
| ENSMUSG00000118664.1 | 134.4792 | 0.243254 | 0.686316 | 0.354434 | 0.723013 | 0.98928 | 0.004681 |
| __ambiguous | 80222.88 | 0.066986 | 0.177394 | 0.377612 | 0.705719 | 0.98928 | 0.004681 |
| ENSMUSG00000001707.9 | 42.23292 | 0.340605 | 1.079938 | 0.315393 | 0.752463 | 0.989797 | 0.004454 |
| ENSMUSG00000022721.20 | 59.14774 | 0.302843 | 0.959571 | 0.315603 | 0.752304 | 0.989797 | 0.004454 |
| ENSMUSG00000031197.12 | 149.656 | -0.35841 | 1.136465 | -0.31537 | 0.75248 | 0.989797 | 0.004454 |
| ENSMUSG00000039660.17 | 13.63815 | 0.656165 | 2.081483 | 0.315239 | 0.75258 | 0.989797 | 0.004454 |
| ENSMUSG00000034641.3 | 39.47767 | -0.52231 | 1.657872 | -0.31505 | 0.752727 | 0.989853 | 0.004429 |
| ENSMUSG00000006134.5 | 183.3363 | -0.2034 | 0.646644 | -0.31455 | 0.753106 | 0.990047 | 0.004344 |
| ENSMUSG00000021774.14 | 326.8649 | 0.159638 | 0.508409 | 0.313995 | 0.753525 | 0.990047 | 0.004344 |
| ENSMUSG00000022186.15 | 25.67305 | -0.51152 | 1.626111 | -0.31457 | 0.753092 | 0.990047 | 0.004344 |
| ENSMUSG00000022681.16 | 15.76507 | 0.689941 | 2.206741 | 0.312652 | 0.754545 | 0.990047 | 0.004344 |
| ENSMUSG00000023018.6 | 11.11166 | 0.785651 | 2.509396 | 0.313084 | 0.754217 | 0.990047 | 0.004344 |
| ENSMUSG00000024667.13 | 40.13441 | 0.351088 | 1.118046 | 0.314019 | 0.753506 | 0.990047 | 0.004344 |

| | | | | | | | |
|---|---|---|---|---|---|---|---|
| ENSMUSG00000032366.16 | 49.03889 | 0.473803 | 1.513498 | 0.313052 | 0.754241 | 0.990047 | 0.004344 |
| ENSMUSG00000033318.8 | 285.3994 | 0.196409 | 0.627843 | 0.312831 | 0.754409 | 0.990047 | 0.004344 |
| ENSMUSG00000035941.16 | 268.0086 | -0.14467 | 0.461886 | -0.31321 | 0.754121 | 0.990047 | 0.004344 |
| ENSMUSG00000037791.16 | 43.48446 | 0.508711 | 1.619409 | 0.314134 | 0.753419 | 0.990047 | 0.004344 |
| ENSMUSG00000039771.14 | 90.12795 | -0.37484 | 1.198837 | -0.31267 | 0.754533 | 0.990047 | 0.004344 |
| ENSMUSG00000039908.15 | 28.80412 | -0.39604 | 1.265093 | -0.31305 | 0.754243 | 0.990047 | 0.004344 |
| ENSMUSG00000057649.7 | 38.40987 | 0.333545 | 1.063906 | 0.31351 | 0.753893 | 0.990047 | 0.004344 |
| ENSMUSG00000058258.15 | 38.18486 | 0.389043 | 1.2418 | 0.313289 | 0.754061 | 0.990047 | 0.004344 |
| ENSMUSG00000079037.10 | 334.9601 | -0.15791 | 0.503394 | -0.3137 | 0.753751 | 0.990047 | 0.004344 |
| ENSMUSG00000095362.8 | 13.15747 | -0.76215 | 2.431713 | -0.31342 | 0.753962 | 0.990047 | 0.004344 |
| ENSMUSG00000024014.9 | 22.04039 | -0.67666 | 2.167185 | -0.31223 | 0.754866 | 0.99033 | 0.00422 |
| ENSMUSG00000022604.19 | 21.09922 | 0.699804 | 2.259042 | 0.309779 | 0.756729 | 0.990732 | 0.004044 |
| ENSMUSG00000022676.8 | 38.05471 | -0.36789 | 1.183114 | -0.31095 | 0.755835 | 0.990732 | 0.004044 |
| ENSMUSG00000028992.14 | 81.74451 | -0.27615 | 0.89172 | -0.30969 | 0.7568 | 0.990732 | 0.004044 |
| ENSMUSG00000030579.11 | 80.32691 | -0.33423 | 1.073992 | -0.3112 | 0.755648 | 0.990732 | 0.004044 |
| ENSMUSG00000032621.9 | 270.7286 | -0.18645 | 0.601168 | -0.31014 | 0.756455 | 0.990732 | 0.004044 |
| ENSMUSG00000032959.13 | 554.7291 | -0.16632 | 0.533795 | -0.31157 | 0.755364 | 0.990732 | 0.004044 |
| ENSMUSG00000033411.17 | 12.47824 | -0.55181 | 1.781088 | -0.30982 | 0.7567 | 0.990732 | 0.004044 |
| ENSMUSG00000035199.7 | 36.93859 | -0.34726 | 1.115587 | -0.31128 | 0.755586 | 0.990732 | 0.004044 |
| ENSMUSG00000037949.9 | 16.2712 | -0.91474 | 2.952925 | -0.30977 | 0.756733 | 0.990732 | 0.004044 |
| ENSMUSG00000041660.9 | 1480.957 | 0.105577 | 0.340057 | 0.31047 | 0.756204 | 0.990732 | 0.004044 |
| ENSMUSG00000048878.7 | 137.4282 | 0.251449 | 0.809375 | 0.31067 | 0.756051 | 0.990732 | 0.004044 |
| ENSMUSG00000050777.8 | 147.1737 | 0.187142 | 0.602 | 0.310867 | 0.755902 | 0.990732 | 0.004044 |
| ENSMUSG00000055884.9 | 22.472 | 0.739299 | 2.382369 | 0.310321 | 0.756317 | 0.990732 | 0.004044 |
| ENSMUSG00000074272.11 | 237.856 | 0.156747 | 0.506197 | 0.309655 | 0.756823 | 0.990732 | 0.004044 |
| ENSMUSG00000102101.2 | 56.3481 | -0.39612 | 1.273616 | -0.31102 | 0.755783 | 0.990732 | 0.004044 |
| ENSMUSG00000108884.2 | 16.74445 | -0.69242 | 2.236283 | -0.30963 | 0.756844 | 0.990732 | 0.004044 |
| ENSMUSG00000028251.6 | 261.2167 | -0.17037 | 0.550945 | -0.30923 | 0.757144 | 0.990787 | 0.00402 |
| ENSMUSG00000050471.18 | 43.23748 | 0.550672 | 1.781188 | 0.30916 | 0.7572 | 0.990787 | 0.00402 |
| ENSMUSG00000071659.5 | 295.308 | -0.1402 | 0.453476 | -0.30918 | 0.757187 | 0.990787 | 0.00402 |
| ENSMUSG00000021738.6 | 66.79634 | 0.302644 | 0.979426 | 0.309001 | 0.757321 | 0.990809 | 0.00401 |
| ENSMUSG00000000154.17 | 361.0994 | -0.15427 | 0.500417 | -0.30829 | 0.757865 | 0.990817 | 0.004007 |
| ENSMUSG00000001441.14 | 107.536 | -0.24848 | 0.80685 | -0.30796 | 0.758111 | 0.990817 | 0.004007 |
| ENSMUSG00000015165.17 | 48.91621 | -0.40449 | 1.316151 | -0.30732 | 0.758596 | 0.990817 | 0.004007 |
| ENSMUSG00000015711.9 | 22.02608 | 0.589735 | 1.91537 | 0.307896 | 0.758161 | 0.990817 | 0.004007 |
| ENSMUSG00000021171.9 | 184.4201 | 0.208666 | 0.675821 | 0.308758 | 0.757505 | 0.990817 | 0.004007 |
| ENSMUSG00000021835.16 | 17.98392 | -0.66926 | 2.175456 | -0.30764 | 0.758357 | 0.990817 | 0.004007 |
| ENSMUSG00000027261.3 | 1902.492 | 0.098211 | 0.318164 | 0.308681 | 0.757564 | 0.990817 | 0.004007 |
| ENSMUSG00000028456.19 | 21.67771 | 0.657928 | 2.142889 | 0.307029 | 0.758822 | 0.990817 | 0.004007 |
| ENSMUSG00000028559.18 | 425.1822 | -0.14328 | 0.465763 | -0.30762 | 0.758373 | 0.990817 | 0.004007 |
| ENSMUSG00000030199.17 | 31.5183 | 0.440984 | 1.434224 | 0.307472 | 0.758484 | 0.990817 | 0.004007 |
| ENSMUSG00000037795.15 | 131.5654 | 0.259838 | 0.84694 | 0.306796 | 0.758999 | 0.990817 | 0.004007 |
| ENSMUSG00000041203.10 | 155.8938 | 0.236406 | 0.770226 | 0.30693 | 0.758896 | 0.990817 | 0.004007 |
| ENSMUSG00000041625.16 | 84.6393 | -0.31401 | 1.023306 | -0.30686 | 0.758951 | 0.990817 | 0.004007 |

| | | | | | | | |
|---|---|---|---|---|---|---|---|
| ENSMUSG00000042700.17 | 25.63856 | 0.514612 | 1.670299 | 0.308096 | 0.758009 | 0.990817 | 0.004007 |
| ENSMUSG00000064345.1 | 41669.24 | 0.093575 | 0.304884 | 0.306918 | 0.758905 | 0.990817 | 0.004007 |
| ENSMUSG00000066258.14 | 38.71084 | 0.549259 | 1.787213 | 0.307327 | 0.758594 | 0.990817 | 0.004007 |
| ENSMUSG00000000959.8 | 77.82513 | -0.2517 | 0.821701 | -0.30632 | 0.759364 | 0.990883 | 0.003978 |
| ENSMUSG00000018379.18 | 507.3597 | -0.10518 | 0.343328 | -0.30635 | 0.759338 | 0.990883 | 0.003978 |
| ENSMUSG00000035342.15 | 12.99654 | -0.8428 | 2.751381 | -0.30632 | 0.759361 | 0.990883 | 0.003978 |
| ENSMUSG00000031897.10 | 55.11442 | -0.30413 | 0.993814 | -0.30603 | 0.759585 | 0.991035 | 0.003911 |
| ENSMUSG00000032407.15 | 284.9882 | 0.163141 | 0.53337 | 0.305868 | 0.759705 | 0.991056 | 0.003902 |
| ENSMUSG00000000740.13 | 1254.85 | 0.086807 | 0.288341 | 0.301056 | 0.763372 | 0.99126 | 0.003813 |
| ENSMUSG00000006711.16 | 16.02413 | 0.678936 | 2.262633 | 0.300065 | 0.764128 | 0.99126 | 0.003813 |
| ENSMUSG00000014177.11 | 151.9049 | -0.18024 | 0.591678 | -0.30462 | 0.760653 | 0.99126 | 0.003813 |
| ENSMUSG00000020130.10 | 255.1384 | -0.15924 | 0.527619 | -0.30181 | 0.762795 | 0.99126 | 0.003813 |
| ENSMUSG00000020349.5 | 660.7588 | 0.101886 | 0.33974 | 0.299895 | 0.764257 | 0.99126 | 0.003813 |
| ENSMUSG00000020630.10 | 11.91697 | 0.842037 | 2.772783 | 0.303679 | 0.761372 | 0.99126 | 0.003813 |
| ENSMUSG00000021713.10 | 10.58676 | -0.86 | 2.867859 | -0.29987 | 0.764273 | 0.99126 | 0.003813 |
| ENSMUSG00000021910.16 | 84.77213 | 0.382101 | 1.27176 | 0.300451 | 0.763833 | 0.99126 | 0.003813 |
| ENSMUSG00000021929.10 | 159.6813 | 0.180451 | 0.600287 | 0.300608 | 0.763713 | 0.99126 | 0.003813 |
| ENSMUSG00000021967.5 | 180.4774 | -0.20955 | 0.688587 | -0.30432 | 0.760883 | 0.99126 | 0.003813 |
| ENSMUSG00000022463.9 | 19.56739 | 0.647069 | 2.149581 | 0.301021 | 0.763398 | 0.99126 | 0.003813 |
| ENSMUSG00000022750.19 | 23.16293 | -0.50785 | 1.682057 | -0.30192 | 0.76271 | 0.99126 | 0.003813 |
| ENSMUSG00000024135.11 | 50.28899 | -0.30371 | 1.013734 | -0.2996 | 0.764483 | 0.99126 | 0.003813 |
| ENSMUSG00000025190.14 | 1274.742 | -0.11106 | 0.369603 | -0.30048 | 0.763813 | 0.99126 | 0.003813 |
| ENSMUSG00000025885.19 | 45.70331 | -0.41694 | 1.37945 | -0.30225 | 0.762458 | 0.99126 | 0.003813 |
| ENSMUSG00000025925.15 | 51.64682 | -0.3751 | 1.243438 | -0.30166 | 0.76291 | 0.99126 | 0.003813 |
| ENSMUSG00000027472.15 | 51.90917 | 0.388007 | 1.285452 | 0.301845 | 0.76277 | 0.99126 | 0.003813 |
| ENSMUSG00000028863.14 | 18.98117 | 0.663757 | 2.193286 | 0.302632 | 0.762171 | 0.99126 | 0.003813 |
| ENSMUSG00000028890.14 | 42.80001 | 0.545688 | 1.802537 | 0.302733 | 0.762093 | 0.99126 | 0.003813 |
| ENSMUSG00000030562.18 | 222.0102 | -0.19365 | 0.63831 | -0.30338 | 0.761604 | 0.99126 | 0.003813 |
| ENSMUSG00000030629.16 | 696.8549 | -0.12726 | 0.424932 | -0.29949 | 0.764566 | 0.99126 | 0.003813 |
| ENSMUSG00000030934.10 | 4032.842 | -0.09067 | 0.299654 | -0.30258 | 0.762208 | 0.99126 | 0.003813 |
| ENSMUSG00000031505.12 | 124.4763 | 0.201926 | 0.663456 | 0.304355 | 0.760857 | 0.99126 | 0.003813 |
| ENSMUSG00000031652.12 | 28.82969 | 0.482075 | 1.588218 | 0.303532 | 0.761485 | 0.99126 | 0.003813 |
| ENSMUSG00000032030.17 | 146.7692 | -0.17274 | 0.575332 | -0.30024 | 0.763993 | 0.99126 | 0.003813 |
| ENSMUSG00000032846.13 | 21.48893 | 0.623721 | 2.048833 | 0.304427 | 0.760802 | 0.99126 | 0.003813 |
| ENSMUSG00000034101.15 | 57.73409 | 0.283351 | 0.942146 | 0.300751 | 0.763605 | 0.99126 | 0.003813 |
| ENSMUSG00000037710.9 | 2319.859 | 0.079338 | 0.264669 | 0.299763 | 0.764358 | 0.99126 | 0.003813 |
| ENSMUSG00000038481.14 | 14.86204 | -0.62968 | 2.066954 | -0.30464 | 0.76064 | 0.99126 | 0.003813 |
| ENSMUSG00000039357.3 | 24.75352 | 0.692009 | 2.284305 | 0.302941 | 0.761935 | 0.99126 | 0.003813 |
| ENSMUSG00000039901.11 | 29.98377 | -0.66411 | 2.184009 | -0.30408 | 0.761067 | 0.99126 | 0.003813 |
| ENSMUSG00000040536.16 | 376.2141 | -0.17108 | 0.565019 | -0.30278 | 0.762056 | 0.99126 | 0.003813 |
| ENSMUSG00000041012.5 | 268.8271 | -0.14014 | 0.463169 | -0.30257 | 0.762214 | 0.99126 | 0.003813 |
| ENSMUSG00000046721.12 | 129.589 | 0.190827 | 0.625626 | 0.305018 | 0.760353 | 0.99126 | 0.003813 |
| ENSMUSG00000052748.15 | 79.24302 | 0.242483 | 0.801069 | 0.302699 | 0.762119 | 0.99126 | 0.003813 |
| ENSMUSG00000055371.18 | 70.68277 | -0.34076 | 1.12656 | -0.30248 | 0.762287 | 0.99126 | 0.003813 |

| | | | | | | | |
|---|---|---|---|---|---|---|---|
| ENSMUSG00000055862.8 | 27.15266 | 0.694602 | 2.306285 | 0.301178 | 0.763279 | 0.99126 | 0.003813 |
| ENSMUSG00000056492.7 | 121.5359 | 0.20625 | 0.681037 | 0.302846 | 0.762007 | 0.99126 | 0.003813 |
| ENSMUSG00000058443.6 | 577.5806 | 0.103328 | 0.344288 | 0.300121 | 0.764085 | 0.99126 | 0.003813 |
| ENSMUSG00000059974.11 | 15.00624 | -0.90608 | 2.995407 | -0.30249 | 0.76228 | 0.99126 | 0.003813 |
| ENSMUSG00000060438.6 | 70.06286 | 0.254423 | 0.837394 | 0.303828 | 0.761259 | 0.99126 | 0.003813 |
| ENSMUSG00000064341.1 | 41167.93 | 0.064286 | 0.214027 | 0.300364 | 0.7639 | 0.99126 | 0.003813 |
| ENSMUSG00000090841.3 | 109.1936 | -0.34625 | 1.137039 | -0.30452 | 0.760732 | 0.99126 | 0.003813 |
| ENSMUSG00000104649.2 | 579.8446 | 0.103813 | 0.346195 | 0.29987 | 0.764276 | 0.99126 | 0.003813 |
| ENSMUSG00000118346.2 | 73.85245 | -0.37241 | 1.223549 | -0.30437 | 0.760845 | 0.99126 | 0.003813 |
| ENSMUSG00000000738.19 | 102.6247 | 0.227626 | 0.777141 | 0.292902 | 0.769597 | 0.991295 | 0.003797 |
| ENSMUSG00000002768.9 | 64.80949 | -0.43698 | 1.47941 | -0.29537 | 0.767708 | 0.991295 | 0.003797 |
| ENSMUSG00000004264.18 | 428.0071 | 0.123071 | 0.411599 | 0.299006 | 0.764936 | 0.991295 | 0.003797 |
| ENSMUSG00000005506.17 | 556.9649 | 0.108178 | 0.365133 | 0.296269 | 0.767024 | 0.991295 | 0.003797 |
| ENSMUSG00000005610.18 | 2196.902 | 0.073435 | 0.251418 | 0.292084 | 0.770222 | 0.991295 | 0.003797 |
| ENSMUSG00000005804.15 | 162.8718 | 0.204868 | 0.697245 | 0.293825 | 0.768892 | 0.991295 | 0.003797 |
| ENSMUSG00000006095.13 | 171.9491 | 0.20607 | 0.690629 | 0.298381 | 0.765413 | 0.991295 | 0.003797 |
| ENSMUSG00000006423.16 | 12.14156 | 0.530561 | 1.824604 | 0.290781 | 0.771218 | 0.991295 | 0.003797 |
| ENSMUSG00000008683.17 | 2732.703 | 0.083525 | 0.289649 | 0.288368 | 0.773065 | 0.991295 | 0.003797 |
| ENSMUSG00000014592.22 | 49.37453 | -0.46366 | 1.57377 | -0.29461 | 0.768289 | 0.991295 | 0.003797 |
| ENSMUSG00000014959.13 | 437.2269 | 0.117732 | 0.404936 | 0.290743 | 0.771248 | 0.991295 | 0.003797 |
| ENSMUSG00000015149.15 | 77.45874 | -0.27327 | 0.958812 | -0.28501 | 0.775636 | 0.991295 | 0.003797 |
| ENSMUSG00000015889.9 | 275.6217 | 0.140658 | 0.476049 | 0.295469 | 0.767636 | 0.991295 | 0.003797 |
| ENSMUSG00000017002.15 | 14.35203 | -0.75806 | 2.630148 | -0.28822 | 0.773179 | 0.991295 | 0.003797 |
| ENSMUSG00000017670.17 | 18.48544 | 0.540371 | 1.863276 | 0.290011 | 0.771808 | 0.991295 | 0.003797 |
| ENSMUSG00000018102.6 | 2081.003 | 0.095174 | 0.329682 | 0.288684 | 0.772823 | 0.991295 | 0.003797 |
| ENSMUSG00000018672.16 | 46.70938 | 0.425084 | 1.478412 | 0.287527 | 0.773708 | 0.991295 | 0.003797 |
| ENSMUSG00000019494.15 | 206.6387 | 0.161969 | 0.549815 | 0.294589 | 0.768308 | 0.991295 | 0.003797 |
| ENSMUSG00000020021.5 | 38.03036 | -0.35235 | 1.201183 | -0.29334 | 0.769265 | 0.991295 | 0.003797 |
| ENSMUSG00000020716.17 | 34.85223 | 0.435273 | 1.500074 | 0.290168 | 0.771688 | 0.991295 | 0.003797 |
| ENSMUSG00000020884.16 | 2255.077 | -0.06986 | 0.24426 | -0.28599 | 0.774887 | 0.991295 | 0.003797 |
| ENSMUSG00000021147.18 | 61.95594 | -0.30445 | 1.052303 | -0.28932 | 0.772335 | 0.991295 | 0.003797 |
| ENSMUSG00000021190.15 | 179.9408 | 0.192337 | 0.674575 | 0.285123 | 0.77555 | 0.991295 | 0.003797 |
| ENSMUSG00000021831.10 | 66.75722 | 0.312147 | 1.04668 | 0.298225 | 0.765531 | 0.991295 | 0.003797 |
| ENSMUSG00000022637.12 | 18.66016 | 0.492732 | 1.648245 | 0.298943 | 0.764983 | 0.991295 | 0.003797 |
| ENSMUSG00000022757.20 | 410.1648 | -0.12929 | 0.444928 | -0.29059 | 0.771363 | 0.991295 | 0.003797 |
| ENSMUSG00000022792.17 | 95.38272 | -0.28427 | 0.9976 | -0.28496 | 0.775678 | 0.991295 | 0.003797 |
| ENSMUSG00000022809.5 | 65.32353 | 0.383732 | 1.32621 | 0.289345 | 0.772317 | 0.991295 | 0.003797 |
| ENSMUSG00000022889.8 | 290.2238 | 0.157913 | 0.532552 | 0.296521 | 0.766832 | 0.991295 | 0.003797 |
| ENSMUSG00000023010.16 | 4328.743 | -0.06523 | 0.222096 | -0.2937 | 0.768985 | 0.991295 | 0.003797 |
| ENSMUSG00000023452.20 | 142.7004 | 0.176268 | 0.617322 | 0.285537 | 0.775233 | 0.991295 | 0.003797 |
| ENSMUSG00000024431.16 | 537.3436 | 0.105349 | 0.363763 | 0.289608 | 0.772116 | 0.991295 | 0.003797 |
| ENSMUSG00000024844.16 | 165.023 | -0.21865 | 0.762896 | -0.2866 | 0.774419 | 0.991295 | 0.003797 |
| ENSMUSG00000024878.10 | 32.34695 | -0.42175 | 1.46246 | -0.28838 | 0.773052 | 0.991295 | 0.003797 |
| ENSMUSG00000026019.16 | 173.313 | -0.23865 | 0.817037 | -0.2921 | 0.770213 | 0.991295 | 0.003797 |

| | | | | | | | |
|---|---|---|---|---|---|---|---|
| ENSMUSG00000026918.17 | 68.56877 | -0.35397 | 1.214016 | -0.29157 | 0.770613 | 0.991295 | 0.003797 |
| ENSMUSG00000026939.13 | 18.57109 | 0.686629 | 2.301139 | 0.298387 | 0.765408 | 0.991295 | 0.003797 |
| ENSMUSG00000027506.16 | 154.0014 | -0.19337 | 0.66099 | -0.29255 | 0.769865 | 0.991295 | 0.003797 |
| ENSMUSG00000027952.17 | 92.76402 | 0.26309 | 0.891266 | 0.295187 | 0.767851 | 0.991295 | 0.003797 |
| ENSMUSG00000028010.17 | 34.86813 | 0.392991 | 1.366748 | 0.287537 | 0.773701 | 0.991295 | 0.003797 |
| ENSMUSG00000028126.17 | 25.32054 | -0.48223 | 1.675159 | -0.28787 | 0.773446 | 0.991295 | 0.003797 |
| ENSMUSG00000028140.14 | 126.8587 | -0.22519 | 0.764863 | -0.29442 | 0.768434 | 0.991295 | 0.003797 |
| ENSMUSG00000028282.13 | 39.06444 | -0.45686 | 1.595616 | -0.28632 | 0.774634 | 0.991295 | 0.003797 |
| ENSMUSG00000028641.17 | 25.93171 | 0.660048 | 2.287073 | 0.2886 | 0.772888 | 0.991295 | 0.003797 |
| ENSMUSG00000028671.17 | 21.13493 | 0.658761 | 2.227789 | 0.295702 | 0.767458 | 0.991295 | 0.003797 |
| ENSMUSG00000028673.11 | 220.5116 | -0.18835 | 0.656347 | -0.28697 | 0.774139 | 0.991295 | 0.003797 |
| ENSMUSG00000028879.5 | 177.1567 | -0.16602 | 0.557264 | -0.29792 | 0.765765 | 0.991295 | 0.003797 |
| ENSMUSG00000028944.15 | 13.77794 | 0.689169 | 2.327834 | 0.296056 | 0.767187 | 0.991295 | 0.003797 |
| ENSMUSG00000029156.12 | 12.58844 | 0.532516 | 1.811302 | 0.293996 | 0.768761 | 0.991295 | 0.003797 |
| ENSMUSG00000029545.14 | 41.77992 | 0.41139 | 1.415257 | 0.290682 | 0.771295 | 0.991295 | 0.003797 |
| ENSMUSG00000029723.17 | 29.35399 | -0.48409 | 1.681116 | -0.28796 | 0.77338 | 0.991295 | 0.003797 |
| ENSMUSG00000030138.9 | 107.6817 | -0.37083 | 1.268883 | -0.29225 | 0.770098 | 0.991295 | 0.003797 |
| ENSMUSG00000030213.13 | 36.0957 | 0.358119 | 1.208458 | 0.296343 | 0.766968 | 0.991295 | 0.003797 |
| ENSMUSG00000030643.15 | 39.74844 | 0.509487 | 1.771814 | 0.287551 | 0.773691 | 0.991295 | 0.003797 |
| ENSMUSG00000031226.14 | 139.2979 | 0.203541 | 0.695798 | 0.292528 | 0.769883 | 0.991295 | 0.003797 |
| ENSMUSG00000032293.9 | 387.3873 | -0.11722 | 0.395857 | -0.29612 | 0.767139 | 0.991295 | 0.003797 |
| ENSMUSG00000032570.18 | 142.2579 | 0.178076 | 0.618011 | 0.288144 | 0.773236 | 0.991295 | 0.003797 |
| ENSMUSG00000032575.17 | 461.0028 | 0.132551 | 0.455465 | 0.291023 | 0.771034 | 0.991295 | 0.003797 |
| ENSMUSG00000032737.14 | 17.52755 | 0.714582 | 2.466308 | 0.289738 | 0.772017 | 0.991295 | 0.003797 |
| ENSMUSG00000032946.17 | 95.19893 | 0.242615 | 0.810902 | 0.299192 | 0.764793 | 0.991295 | 0.003797 |
| ENSMUSG00000033021.17 | 12.3662 | -0.60732 | 2.096923 | -0.28962 | 0.772105 | 0.991295 | 0.003797 |
| ENSMUSG00000033228.9 | 228.2555 | 0.165666 | 0.578007 | 0.286616 | 0.774406 | 0.991295 | 0.003797 |
| ENSMUSG00000034152.7 | 97.4482 | 0.253041 | 0.873933 | 0.289543 | 0.772166 | 0.991295 | 0.003797 |
| ENSMUSG00000034424.6 | 639.0623 | 0.115006 | 0.39417 | 0.291768 | 0.770464 | 0.991295 | 0.003797 |
| ENSMUSG00000034480.19 | 52.70439 | -0.28668 | 1.003032 | -0.28581 | 0.775021 | 0.991295 | 0.003797 |
| ENSMUSG00000034518.15 | 13.52318 | -0.67658 | 2.310478 | -0.29283 | 0.769652 | 0.991295 | 0.003797 |
| ENSMUSG00000036046.15 | 40.43267 | 0.437785 | 1.517542 | 0.288483 | 0.772977 | 0.991295 | 0.003797 |
| ENSMUSG00000036299.6 | 408.4591 | 0.195218 | 0.668804 | 0.291891 | 0.77037 | 0.991295 | 0.003797 |
| ENSMUSG00000037134.18 | 61.33379 | 0.40065 | 1.39708 | 0.286777 | 0.774283 | 0.991295 | 0.003797 |
| ENSMUSG00000037679.10 | 32.32871 | 0.47709 | 1.671447 | 0.285436 | 0.77531 | 0.991295 | 0.003797 |
| ENSMUSG00000037742.15 | 2299.267 | -0.09122 | 0.316905 | -0.28786 | 0.773457 | 0.991295 | 0.003797 |
| ENSMUSG00000038039.14 | 242.0727 | 0.16765 | 0.566661 | 0.295856 | 0.76734 | 0.991295 | 0.003797 |
| ENSMUSG00000038058.15 | 26.19486 | -0.40145 | 1.349044 | -0.29758 | 0.766022 | 0.991295 | 0.003797 |
| ENSMUSG00000038949.9 | 11.66514 | -0.63039 | 2.179812 | -0.2892 | 0.772431 | 0.991295 | 0.003797 |
| ENSMUSG00000039115.14 | 41.86346 | 0.335822 | 1.135068 | 0.29586 | 0.767337 | 0.991295 | 0.003797 |
| ENSMUSG00000039221.11 | 869.5577 | -0.10252 | 0.352429 | -0.29091 | 0.771123 | 0.991295 | 0.003797 |
| ENSMUSG00000039304.12 | 11.98181 | -0.5766 | 1.988164 | -0.29001 | 0.771806 | 0.991295 | 0.003797 |
| ENSMUSG00000039497.9 | 19.28751 | 0.606432 | 2.069327 | 0.293058 | 0.769478 | 0.991295 | 0.003797 |
| ENSMUSG00000040612.14 | 125.3819 | -0.25046 | 0.850067 | -0.29463 | 0.768275 | 0.991295 | 0.003797 |

| | | | | | | | |
|---|---|---|---|---|---|---|---|
| ENSMUSG00000040738.11 | 57.85291 | 0.28879 | 0.973783 | 0.296565 | 0.766799 | 0.991295 | 0.003797 |
| ENSMUSG00000041390.19 | 58.53283 | 0.254552 | 0.881333 | 0.288827 | 0.772714 | 0.991295 | 0.003797 |
| ENSMUSG00000041577.6 | 213.6257 | 0.164204 | 0.575199 | 0.285472 | 0.775282 | 0.991295 | 0.003797 |
| ENSMUSG00000041827.16 | 93.3399 | 0.235214 | 0.789398 | 0.297967 | 0.765728 | 0.991295 | 0.003797 |
| ENSMUSG00000042997.14 | 39.73741 | -0.37196 | 1.284827 | -0.2895 | 0.772196 | 0.991295 | 0.003797 |
| ENSMUSG00000044501.18 | 12.84934 | 0.703224 | 2.375098 | 0.296082 | 0.767167 | 0.991295 | 0.003797 |
| ENSMUSG00000044573.17 | 440.2003 | 0.14158 | 0.482583 | 0.293379 | 0.769232 | 0.991295 | 0.003797 |
| ENSMUSG00000046185.11 | 27.04878 | -0.48044 | 1.676019 | -0.28665 | 0.774377 | 0.991295 | 0.003797 |
| ENSMUSG00000047638.16 | 460.7529 | 0.134414 | 0.46084 | 0.291672 | 0.770538 | 0.991295 | 0.003797 |
| ENSMUSG00000049164.8 | 34.66718 | -0.35614 | 1.245878 | -0.28585 | 0.77499 | 0.991295 | 0.003797 |
| ENSMUSG00000050708.17 | 179.1309 | 0.194432 | 0.678269 | 0.286659 | 0.774374 | 0.991295 | 0.003797 |
| ENSMUSG00000053886.5 | 18.47729 | 0.757325 | 2.580075 | 0.293528 | 0.769118 | 0.991295 | 0.003797 |
| ENSMUSG00000056305.5 | 75.32376 | -0.31994 | 1.120597 | -0.28551 | 0.775253 | 0.991295 | 0.003797 |
| ENSMUSG00000058761.6 | 133.0529 | 0.199796 | 0.687939 | 0.290427 | 0.77149 | 0.991295 | 0.003797 |
| ENSMUSG00000060679.15 | 24.91444 | -0.45375 | 1.569302 | -0.28914 | 0.772472 | 0.991295 | 0.003797 |
| ENSMUSG00000062931.16 | 67.74859 | 0.288281 | 0.984849 | 0.292716 | 0.769739 | 0.991295 | 0.003797 |
| ENSMUSG00000067713.8 | 25.61978 | 0.431532 | 1.470876 | 0.293384 | 0.769228 | 0.991295 | 0.003797 |
| ENSMUSG00000072625.2 | 35.91961 | -0.5052 | 1.712003 | -0.2951 | 0.767921 | 0.991295 | 0.003797 |
| ENSMUSG00000075014.2 | 180.0965 | 0.166827 | 0.582465 | 0.286415 | 0.77456 | 0.991295 | 0.003797 |
| ENSMUSG00000075232.7 | 23.89369 | 0.658485 | 2.309247 | 0.285152 | 0.775528 | 0.991295 | 0.003797 |
| ENSMUSG00000075273.5 | 15.12096 | 0.544264 | 1.890809 | 0.287847 | 0.773464 | 0.991295 | 0.003797 |
| ENSMUSG00000078317.7 | 33.63503 | -0.63662 | 2.162336 | -0.29441 | 0.768444 | 0.991295 | 0.003797 |
| ENSMUSG00000084319.4 | 69.0176 | 0.240436 | 0.835901 | 0.287637 | 0.773624 | 0.991295 | 0.003797 |
| ENSMUSG00000087497.8 | 34.95313 | -0.64419 | 2.179248 | -0.2956 | 0.767534 | 0.991295 | 0.003797 |
| ENSMUSG00000091931.6 | 69.76799 | 0.334755 | 1.120675 | 0.298708 | 0.765163 | 0.991295 | 0.003797 |
| ENSMUSG00000110275.2 | 187.9826 | -0.16616 | 0.567758 | -0.29267 | 0.769776 | 0.991295 | 0.003797 |
| ENSMUSG00000111966.2 | 136.104 | 0.194789 | 0.656831 | 0.296559 | 0.766803 | 0.991295 | 0.003797 |
| ENSMUSG00000030265.15 | 182.4201 | 0.167657 | 0.589464 | 0.284423 | 0.776086 | 0.991548 | 0.003686 |
| ENSMUSG00000051224.14 | 24.1986 | -0.487 | 1.71304 | -0.28429 | 0.77619 | 0.991548 | 0.003686 |
| ENSMUSG00000052539.16 | 72.76861 | -0.29123 | 1.024161 | -0.28436 | 0.776134 | 0.991548 | 0.003686 |
| ENSMUSG00000000296.9 | 101.5547 | -0.24822 | 0.918195 | -0.27034 | 0.786901 | 0.991777 | 0.003586 |
| ENSMUSG00000001666.9 | 854.1673 | -0.12009 | 0.438257 | -0.27403 | 0.784063 | 0.991777 | 0.003586 |
| ENSMUSG00000002416.14 | 517.4175 | 0.103684 | 0.366319 | 0.283044 | 0.777143 | 0.991777 | 0.003586 |
| ENSMUSG00000004127.13 | 66.89571 | -0.26877 | 0.97211 | -0.27648 | 0.782181 | 0.991777 | 0.003586 |
| ENSMUSG00000005501.15 | 100.777 | 0.215848 | 0.807462 | 0.267317 | 0.789225 | 0.991777 | 0.003586 |
| ENSMUSG00000006301.18 | 94.18318 | -0.3367 | 1.241006 | -0.27131 | 0.78615 | 0.991777 | 0.003586 |
| ENSMUSG00000006736.10 | 1265.209 | -0.0861 | 0.309191 | -0.27847 | 0.780649 | 0.991777 | 0.003586 |
| ENSMUSG00000008575.18 | 845.7392 | -0.10167 | 0.381315 | -0.26662 | 0.78976 | 0.991777 | 0.003586 |
| ENSMUSG00000012848.16 | 2298.434 | 0.067218 | 0.244267 | 0.275181 | 0.783177 | 0.991777 | 0.003586 |
| ENSMUSG00000014418.17 | 31.26383 | 0.374494 | 1.346302 | 0.278165 | 0.780886 | 0.991777 | 0.003586 |
| ENSMUSG00000014763.9 | 76.22194 | -0.23899 | 0.89239 | -0.26781 | 0.788847 | 0.991777 | 0.003586 |
| ENSMUSG00000015092.10 | 241.0992 | -0.1527 | 0.556951 | -0.27417 | 0.783958 | 0.991777 | 0.003586 |
| ENSMUSG00000015127.15 | 29.30583 | -0.392 | 1.434946 | -0.27318 | 0.784713 | 0.991777 | 0.003586 |
| ENSMUSG00000016541.11 | 230.1594 | 0.148525 | 0.539888 | 0.275103 | 0.783237 | 0.991777 | 0.003586 |

| | | | | | | | |
|---|---|---|---|---|---|---|---|
| ENSMUSG00000016933.18 | 31.83096 | -0.43981 | 1.622553 | -0.27106 | 0.786344 | 0.991777 | 0.003586 |
| ENSMUSG00000017548.16 | 95.81532 | 0.221481 | 0.787245 | 0.281337 | 0.778452 | 0.991777 | 0.003586 |
| ENSMUSG00000017760.17 | 203.771 | -0.19787 | 0.701608 | -0.28203 | 0.777923 | 0.991777 | 0.003586 |
| ENSMUSG00000018412.17 | 73.69838 | 0.249441 | 0.898487 | 0.277623 | 0.781301 | 0.991777 | 0.003586 |
| ENSMUSG00000019726.13 | 25.27544 | 0.434608 | 1.621248 | 0.26807 | 0.788646 | 0.991777 | 0.003586 |
| ENSMUSG00000020018.7 | 216.0132 | 0.184272 | 0.684365 | 0.26926 | 0.78773 | 0.991777 | 0.003586 |
| ENSMUSG00000020064.10 | 196.9803 | 0.147763 | 0.536288 | 0.275529 | 0.78291 | 0.991777 | 0.003586 |
| ENSMUSG00000020137.6 | 48.58062 | 0.272798 | 1.009149 | 0.270325 | 0.78691 | 0.991777 | 0.003586 |
| ENSMUSG00000020220.17 | 84.14608 | 0.263315 | 0.956769 | 0.275213 | 0.783153 | 0.991777 | 0.003586 |
| ENSMUSG00000020275.10 | 24.72832 | -0.39309 | 1.443412 | -0.27233 | 0.785368 | 0.991777 | 0.003586 |
| ENSMUSG00000020307.16 | 66.12939 | -0.30567 | 1.117949 | -0.27342 | 0.784533 | 0.991777 | 0.003586 |
| ENSMUSG00000020849.14 | 1214.708 | 0.074513 | 0.271389 | 0.274562 | 0.783653 | 0.991777 | 0.003586 |
| ENSMUSG00000020910.17 | 122.0484 | -0.20926 | 0.769578 | -0.27192 | 0.785686 | 0.991777 | 0.003586 |
| ENSMUSG00000020978.11 | 96.49016 | 0.211486 | 0.780512 | 0.270959 | 0.786423 | 0.991777 | 0.003586 |
| ENSMUSG00000021496.13 | 227.8946 | 0.19517 | 0.709587 | 0.275048 | 0.78328 | 0.991777 | 0.003586 |
| ENSMUSG00000021796.16 | 135.7216 | -0.19853 | 0.732758 | -0.27094 | 0.786439 | 0.991777 | 0.003586 |
| ENSMUSG00000022387.18 | 48.82476 | 0.330269 | 1.203812 | 0.274353 | 0.783813 | 0.991777 | 0.003586 |
| ENSMUSG00000022529.12 | 35.56495 | 0.516715 | 1.860044 | 0.277797 | 0.781168 | 0.991777 | 0.003586 |
| ENSMUSG00000022847.16 | 39.47856 | 0.399405 | 1.454087 | 0.274678 | 0.783564 | 0.991777 | 0.003586 |
| ENSMUSG00000022858.17 | 185.3324 | 0.192471 | 0.691179 | 0.278467 | 0.780654 | 0.991777 | 0.003586 |
| ENSMUSG00000022895.17 | 127.0561 | -0.23512 | 0.838722 | -0.28034 | 0.77922 | 0.991777 | 0.003586 |
| ENSMUSG00000023307.15 | 548.5004 | 0.098667 | 0.366979 | 0.268863 | 0.788035 | 0.991777 | 0.003586 |
| ENSMUSG00000025199.17 | 764.9485 | 0.090883 | 0.329038 | 0.276208 | 0.782388 | 0.991777 | 0.003586 |
| ENSMUSG00000025362.7 | 3555.473 | 0.071663 | 0.254157 | 0.281964 | 0.777971 | 0.991777 | 0.003586 |
| ENSMUSG00000025466.20 | 136.996 | 0.204116 | 0.728038 | 0.280364 | 0.779198 | 0.991777 | 0.003586 |
| ENSMUSG00000025583.16 | 77.44872 | -0.31764 | 1.135549 | -0.27972 | 0.77969 | 0.991777 | 0.003586 |
| ENSMUSG00000025823.10 | 168.6353 | 0.16502 | 0.593758 | 0.277925 | 0.78107 | 0.991777 | 0.003586 |
| ENSMUSG00000025857.11 | 17.14052 | -0.53224 | 1.965635 | -0.27077 | 0.786566 | 0.991777 | 0.003586 |
| ENSMUSG00000025873.17 | 208.2398 | -0.17472 | 0.63341 | -0.27584 | 0.78267 | 0.991777 | 0.003586 |
| ENSMUSG00000026276.21 | 14.43591 | 0.5748 | 2.093559 | 0.274556 | 0.783657 | 0.991777 | 0.003586 |
| ENSMUSG00000026281.16 | 179.6565 | -0.1562 | 0.565444 | -0.27624 | 0.782361 | 0.991777 | 0.003586 |
| ENSMUSG00000026383.15 | 283.7346 | 0.142713 | 0.534342 | 0.267081 | 0.789407 | 0.991777 | 0.003586 |
| ENSMUSG00000026473.17 | 2269.506 | -0.08649 | 0.311886 | -0.27731 | 0.781543 | 0.991777 | 0.003586 |
| ENSMUSG00000026575.16 | 29.76234 | -0.44539 | 1.627895 | -0.2736 | 0.784391 | 0.991777 | 0.003586 |
| ENSMUSG00000026584.15 | 93.65551 | 0.222742 | 0.829698 | 0.268462 | 0.788344 | 0.991777 | 0.003586 |
| ENSMUSG00000026767.13 | 159.4156 | 0.155525 | 0.581759 | 0.267336 | 0.78921 | 0.991777 | 0.003586 |
| ENSMUSG00000026849.19 | 256.1597 | -0.1635 | 0.607781 | -0.26902 | 0.787917 | 0.991777 | 0.003586 |
| ENSMUSG00000026858.19 | 54.30116 | 0.273991 | 0.998529 | 0.274395 | 0.783781 | 0.991777 | 0.003586 |
| ENSMUSG00000026977.18 | 167.391 | -0.15457 | 0.574797 | -0.26891 | 0.788 | 0.991777 | 0.003586 |
| ENSMUSG00000027245.12 | 55.32841 | 0.325323 | 1.160245 | 0.280392 | 0.779177 | 0.991777 | 0.003586 |
| ENSMUSG00000027428.10 | 138.7455 | -0.1806 | 0.662564 | -0.27257 | 0.785183 | 0.991777 | 0.003586 |
| ENSMUSG00000028238.7 | 24.48063 | -0.46647 | 1.746503 | -0.26709 | 0.789401 | 0.991777 | 0.003586 |
| ENSMUSG00000028393.11 | 448.9317 | -0.14134 | 0.506308 | -0.27915 | 0.780128 | 0.991777 | 0.003586 |
| ENSMUSG00000028675.13 | 202.5507 | 0.211548 | 0.74565 | 0.28371 | 0.776633 | 0.991777 | 0.003586 |

| | | | | | | | |
|---|---|---|---|---|---|---|---|
| ENSMUSG00000029082.18 | 39.36987 | 1.03354 | 3.730381 | 0.27706 | 0.781734 | 0.991777 | 0.003586 |
| ENSMUSG00000029189.11 | 37.99305 | 0.420081 | 1.516851 | 0.276943 | 0.781824 | 0.991777 | 0.003586 |
| ENSMUSG00000029535.5 | 73.91153 | -0.24486 | 0.893311 | -0.2741 | 0.784006 | 0.991777 | 0.003586 |
| ENSMUSG00000029722.16 | 19.30864 | -0.42364 | 1.556357 | -0.2722 | 0.785471 | 0.991777 | 0.003586 |
| ENSMUSG00000030663.13 | 306.3895 | -0.1404 | 0.500964 | -0.28027 | 0.779271 | 0.991777 | 0.003586 |
| ENSMUSG00000031109.17 | 48.85357 | -0.28114 | 1.017358 | -0.27634 | 0.782284 | 0.991777 | 0.003586 |
| ENSMUSG00000031441.16 | 27.56446 | -0.40263 | 1.46125 | -0.27553 | 0.782905 | 0.991777 | 0.003586 |
| ENSMUSG00000031672.9 | 433.8646 | -0.12284 | 0.443513 | -0.27697 | 0.781802 | 0.991777 | 0.003586 |
| ENSMUSG00000031715.15 | 181.6693 | -0.18035 | 0.672216 | -0.2683 | 0.788472 | 0.991777 | 0.003586 |
| ENSMUSG00000032000.18 | 67.71331 | 0.252842 | 0.908709 | 0.278243 | 0.780826 | 0.991777 | 0.003586 |
| ENSMUSG00000032528.6 | 69.03585 | -0.22994 | 0.862373 | -0.26664 | 0.789746 | 0.991777 | 0.003586 |
| ENSMUSG00000032966.15 | 159.0413 | 0.166912 | 0.595645 | 0.280221 | 0.779308 | 0.991777 | 0.003586 |
| ENSMUSG00000034021.16 | 134.3332 | 0.225425 | 0.81613 | 0.276213 | 0.782385 | 0.991777 | 0.003586 |
| ENSMUSG00000034189.6 | 52.11646 | -0.34372 | 1.267189 | -0.27125 | 0.7862 | 0.991777 | 0.003586 |
| ENSMUSG00000035305.6 | 10.05907 | 0.842711 | 3.026046 | 0.278486 | 0.780639 | 0.991777 | 0.003586 |
| ENSMUSG00000035566.8 | 27.88251 | -0.46737 | 1.684426 | -0.27746 | 0.781425 | 0.991777 | 0.003586 |
| ENSMUSG00000036707.17 | 255.1036 | -0.13991 | 0.500377 | -0.27962 | 0.779772 | 0.991777 | 0.003586 |
| ENSMUSG00000036712.16 | 136.7894 | -0.17802 | 0.653974 | -0.27222 | 0.785454 | 0.991777 | 0.003586 |
| ENSMUSG00000038690.16 | 2410.343 | 0.071354 | 0.264499 | 0.269771 | 0.787337 | 0.991777 | 0.003586 |
| ENSMUSG00000038827.12 | 12.39171 | 0.741938 | 2.698154 | 0.27498 | 0.783332 | 0.991777 | 0.003586 |
| ENSMUSG00000038828.14 | 21.19494 | -0.55572 | 2.079896 | -0.26719 | 0.789326 | 0.991777 | 0.003586 |
| ENSMUSG00000038990.15 | 169.5043 | -0.20645 | 0.764425 | -0.27007 | 0.787108 | 0.991777 | 0.003586 |
| ENSMUSG00000039033.12 | 92.55362 | -0.25534 | 0.95423 | -0.26759 | 0.789018 | 0.991777 | 0.003586 |
| ENSMUSG00000039217.14 | 59.18468 | -0.25695 | 0.93852 | -0.27378 | 0.78425 | 0.991777 | 0.003586 |
| ENSMUSG00000039646.6 | 46.79553 | 0.410049 | 1.533321 | 0.267425 | 0.789142 | 0.991777 | 0.003586 |
| ENSMUSG00000039879.10 | 151.4215 | -0.19915 | 0.737717 | -0.26995 | 0.787201 | 0.991777 | 0.003586 |
| ENSMUSG00000040667.12 | 193.8807 | -0.17421 | 0.636976 | -0.2735 | 0.784471 | 0.991777 | 0.003586 |
| ENSMUSG00000040842.18 | 70.36268 | -0.31181 | 1.142951 | -0.27281 | 0.784997 | 0.991777 | 0.003586 |
| ENSMUSG00000040859.15 | 59.09304 | -0.33663 | 1.189751 | -0.28294 | 0.77722 | 0.991777 | 0.003586 |
| ENSMUSG00000042302.15 | 12.69404 | -0.76807 | 2.719015 | -0.28248 | 0.777575 | 0.991777 | 0.003586 |
| ENSMUSG00000042396.11 | 307.2262 | 0.169389 | 0.602639 | 0.281078 | 0.778651 | 0.991777 | 0.003586 |
| ENSMUSG00000043683.5 | 61.57659 | -0.26554 | 0.99454 | -0.267 | 0.78947 | 0.991777 | 0.003586 |
| ENSMUSG00000043923.9 | 35.12251 | 0.455593 | 1.639913 | 0.277815 | 0.781154 | 0.991777 | 0.003586 |
| ENSMUSG00000043929.17 | 24.18393 | -0.43255 | 1.58207 | -0.2734 | 0.784542 | 0.991777 | 0.003586 |
| ENSMUSG00000045282.8 | 410.7503 | -0.12266 | 0.442969 | -0.2769 | 0.781859 | 0.991777 | 0.003586 |
| ENSMUSG00000045414.8 | 296.0168 | -0.14923 | 0.534026 | -0.27944 | 0.779905 | 0.991777 | 0.003586 |
| ENSMUSG00000045886.8 | 17.12867 | 0.492085 | 1.803645 | 0.272828 | 0.784985 | 0.991777 | 0.003586 |
| ENSMUSG00000046010.8 | 27.22885 | -0.61155 | 2.268505 | -0.26958 | 0.787482 | 0.991777 | 0.003586 |
| ENSMUSG00000046811.4 | 84.98116 | 0.275092 | 1.013749 | 0.271361 | 0.786113 | 0.991777 | 0.003586 |
| ENSMUSG00000049555.11 | 33.5751 | -0.33491 | 1.195977 | -0.28003 | 0.779453 | 0.991777 | 0.003586 |
| ENSMUSG00000050029.8 | 235.3449 | 0.143496 | 0.532558 | 0.269447 | 0.787586 | 0.991777 | 0.003586 |
| ENSMUSG00000051695.7 | 502.236 | -0.12828 | 0.473061 | -0.27117 | 0.786258 | 0.991777 | 0.003586 |
| ENSMUSG00000052296.8 | 19.14199 | -0.48582 | 1.814422 | -0.26775 | 0.788889 | 0.991777 | 0.003586 |
| ENSMUSG00000054364.6 | 37.70546 | 0.313744 | 1.114971 | 0.281392 | 0.77841 | 0.991777 | 0.003586 |

| | | | | | | | |
|---|---|---|---|---|---|---|---|
| ENSMUSG00000054770.17 | 18.725 | -0.58024 | 2.130607 | -0.27234 | 0.785364 | 0.991777 | 0.003586 |
| ENSMUSG00000056209.8 | 58.5233 | 0.37091 | 1.362647 | 0.272198 | 0.785469 | 0.991777 | 0.003586 |
| ENSMUSG00000056537.12 | 107.7685 | -0.19699 | 0.724635 | -0.27184 | 0.785742 | 0.991777 | 0.003586 |
| ENSMUSG00000057388.10 | 683.4182 | 0.095935 | 0.352326 | 0.272289 | 0.7854 | 0.991777 | 0.003586 |
| ENSMUSG00000058402.15 | 14.54194 | 0.673787 | 2.469019 | 0.272897 | 0.784933 | 0.991777 | 0.003586 |
| ENSMUSG00000062929.9 | 332.8819 | 0.12732 | 0.47217 | 0.26965 | 0.78743 | 0.991777 | 0.003586 |
| ENSMUSG00000063232.17 | 38.07771 | -0.31601 | 1.155669 | -0.27344 | 0.784514 | 0.991777 | 0.003586 |
| ENSMUSG00000064344.1 | 19.02548 | 0.403166 | 1.432654 | 0.281412 | 0.778394 | 0.991777 | 0.003586 |
| ENSMUSG00000069755.6 | 35.49818 | -0.47937 | 1.713764 | -0.27972 | 0.779696 | 0.991777 | 0.003586 |
| ENSMUSG00000070520.5 | 110.9203 | 0.201499 | 0.733664 | 0.274647 | 0.783587 | 0.991777 | 0.003586 |
| ENSMUSG00000070737.11 | 27.29739 | 0.493662 | 1.775448 | 0.278049 | 0.780975 | 0.991777 | 0.003586 |
| ENSMUSG00000071252.7 | 10.11452 | 0.819779 | 2.972629 | 0.275776 | 0.78272 | 0.991777 | 0.003586 |
| ENSMUSG00000071713.7 | 9.987561 | -0.71872 | 2.68107 | -0.26807 | 0.788643 | 0.991777 | 0.003586 |
| ENSMUSG00000072676.13 | 1180.451 | 0.090707 | 0.334811 | 0.270921 | 0.786452 | 0.991777 | 0.003586 |
| ENSMUSG00000075511.10 | 20.1667 | -0.45629 | 1.703579 | -0.26784 | 0.78882 | 0.991777 | 0.003586 |
| ENSMUSG00000078515.5 | 500.4426 | -0.10099 | 0.378658 | -0.2667 | 0.789701 | 0.991777 | 0.003586 |
| ENSMUSG00000079509.11 | 84.02955 | -0.25023 | 0.91318 | -0.27402 | 0.784068 | 0.991777 | 0.003586 |
| ENSMUSG00000080877.2 | 10.78924 | -0.51537 | 1.867134 | -0.27602 | 0.782532 | 0.991777 | 0.003586 |
| ENSMUSG00000083337.2 | 64.33813 | -0.234 | 0.863494 | -0.27099 | 0.786396 | 0.991777 | 0.003586 |
| ENSMUSG00000083863.2 | 481.3817 | -0.11603 | 0.413692 | -0.28048 | 0.77911 | 0.991777 | 0.003586 |
| ENSMUSG00000086448.2 | 38.98543 | -0.53106 | 1.965384 | -0.27021 | 0.787002 | 0.991777 | 0.003586 |
| ENSMUSG00000097375.4 | 44.06093 | -0.32699 | 1.226401 | -0.26663 | 0.789757 | 0.991777 | 0.003586 |
| ENSMUSG00000097392.5 | 223.1222 | -0.15928 | 0.572332 | -0.2783 | 0.780782 | 0.991777 | 0.003586 |
| ENSMUSG00000097574.3 | 10.18201 | -0.80651 | 2.90063 | -0.27805 | 0.780977 | 0.991777 | 0.003586 |
| ENSMUSG00000105315.2 | 105.6224 | -0.23203 | 0.860104 | -0.26977 | 0.787338 | 0.991777 | 0.003586 |
| ENSMUSG00000112129.2 | 544.2956 | -0.10188 | 0.376019 | -0.27095 | 0.786428 | 0.991777 | 0.003586 |
| ENSMUSG00000026749.12 | 194.2671 | 0.183978 | 0.690707 | 0.266361 | 0.789961 | 0.991898 | 0.003533 |
| ENSMUSG00000001089.15 | 34.46893 | -0.40928 | 1.550685 | -0.26394 | 0.79183 | 0.991981 | 0.003496 |
| ENSMUSG00000001383.10 | 196.3875 | -0.16989 | 0.642852 | -0.26428 | 0.791563 | 0.991981 | 0.003496 |
| ENSMUSG00000002741.10 | 125.0014 | -0.16597 | 0.637435 | -0.26038 | 0.794573 | 0.991981 | 0.003496 |
| ENSMUSG00000002812.5 | 73.69517 | -0.22167 | 0.847821 | -0.26145 | 0.793743 | 0.991981 | 0.003496 |
| ENSMUSG00000003402.15 | 136.066 | 0.193874 | 0.744802 | 0.260303 | 0.79463 | 0.991981 | 0.003496 |
| ENSMUSG00000003847.17 | 80.809 | 0.215571 | 0.814985 | 0.264509 | 0.791388 | 0.991981 | 0.003496 |
| ENSMUSG00000004561.15 | 42.7755 | -0.28128 | 1.084618 | -0.25934 | 0.795377 | 0.991981 | 0.003496 |
| ENSMUSG00000007411.17 | 152.9546 | -0.1555 | 0.610078 | -0.25488 | 0.798817 | 0.991981 | 0.003496 |
| ENSMUSG00000007891.17 | 255.2213 | 0.156973 | 0.597854 | 0.262561 | 0.792889 | 0.991981 | 0.003496 |
| ENSMUSG00000015568.17 | 225.1856 | -0.21136 | 0.816313 | -0.25892 | 0.795699 | 0.991981 | 0.003496 |
| ENSMUSG00000015697.15 | 61.84193 | 0.236308 | 0.917266 | 0.257623 | 0.796698 | 0.991981 | 0.003496 |
| ENSMUSG00000016619.11 | 168.1659 | 0.211804 | 0.801106 | 0.26439 | 0.79148 | 0.991981 | 0.003496 |
| ENSMUSG00000018417.15 | 763.1822 | -0.09522 | 0.359793 | -0.26466 | 0.791269 | 0.991981 | 0.003496 |
| ENSMUSG00000020176.18 | 38.4709 | -0.41951 | 1.620679 | -0.25885 | 0.795753 | 0.991981 | 0.003496 |
| ENSMUSG00000020629.13 | 288.03 | 0.169147 | 0.658906 | 0.256708 | 0.797404 | 0.991981 | 0.003496 |
| ENSMUSG00000021144.15 | 15.42844 | 0.433339 | 1.673047 | 0.259012 | 0.795626 | 0.991981 | 0.003496 |
| ENSMUSG00000021418.16 | 84.88633 | 0.233013 | 0.887609 | 0.262518 | 0.792922 | 0.991981 | 0.003496 |

| | | | | | | | |
|---|---|---|---|---|---|---|---|
| ENSMUSG00000022346.18 | 40.88096 | 0.417064 | 1.62484 | 0.25668 | 0.797426 | 0.991981 | 0.003496 |
| ENSMUSG00000024491.17 | 87.74723 | 0.219923 | 0.838515 | 0.262276 | 0.793108 | 0.991981 | 0.003496 |
| ENSMUSG00000024539.18 | 65.61847 | 0.272066 | 1.027379 | 0.264816 | 0.791151 | 0.991981 | 0.003496 |
| ENSMUSG00000025791.19 | 1283.001 | 0.093278 | 0.357053 | 0.261245 | 0.793904 | 0.991981 | 0.003496 |
| ENSMUSG00000026435.16 | 63.41566 | 0.257074 | 0.978467 | 0.262732 | 0.792757 | 0.991981 | 0.003496 |
| ENSMUSG00000026860.17 | 34.15942 | 0.407265 | 1.568085 | 0.259721 | 0.795079 | 0.991981 | 0.003496 |
| ENSMUSG00000027016.18 | 134.1944 | -0.16993 | 0.6427 | -0.26441 | 0.791466 | 0.991981 | 0.003496 |
| ENSMUSG00000027108.16 | 329.5906 | 0.136735 | 0.535213 | 0.255477 | 0.798355 | 0.991981 | 0.003496 |
| ENSMUSG00000027206.14 | 387.2574 | 0.149222 | 0.57485 | 0.259585 | 0.795184 | 0.991981 | 0.003496 |
| ENSMUSG00000027404.17 | 133.1037 | -0.20744 | 0.78763 | -0.26337 | 0.792268 | 0.991981 | 0.003496 |
| ENSMUSG00000027573.12 | 143.7697 | -0.18891 | 0.722753 | -0.26137 | 0.793805 | 0.991981 | 0.003496 |
| ENSMUSG00000028133.18 | 43.36633 | -0.37337 | 1.436228 | -0.25997 | 0.794889 | 0.991981 | 0.003496 |
| ENSMUSG00000028189.13 | 29.8526 | -0.43774 | 1.708063 | -0.25628 | 0.797734 | 0.991981 | 0.003496 |
| ENSMUSG00000028247.3 | 328.6011 | -0.15281 | 0.58734 | -0.26018 | 0.794728 | 0.991981 | 0.003496 |
| ENSMUSG00000028293.15 | 126.4384 | 0.220413 | 0.854512 | 0.25794 | 0.796453 | 0.991981 | 0.003496 |
| ENSMUSG00000028976.11 | 21.52563 | -0.53821 | 2.098596 | -0.25646 | 0.797595 | 0.991981 | 0.003496 |
| ENSMUSG00000028986.13 | 101.1656 | 0.286142 | 1.089998 | 0.262516 | 0.792923 | 0.991981 | 0.003496 |
| ENSMUSG00000031147.9 | 14.81659 | -0.68311 | 2.652599 | -0.25753 | 0.796773 | 0.991981 | 0.003496 |
| ENSMUSG00000031488.15 | 30.50883 | 0.449948 | 1.761454 | 0.255441 | 0.798383 | 0.991981 | 0.003496 |
| ENSMUSG00000031657.17 | 139.0243 | 0.161365 | 0.628669 | 0.256678 | 0.797428 | 0.991981 | 0.003496 |
| ENSMUSG00000031808.8 | 14.46956 | 0.596526 | 2.317491 | 0.257402 | 0.796869 | 0.991981 | 0.003496 |
| ENSMUSG00000032215.16 | 119.9578 | -0.19364 | 0.744288 | -0.26016 | 0.794738 | 0.991981 | 0.003496 |
| ENSMUSG00000032251.13 | 28.95488 | 0.518773 | 2.005581 | 0.258665 | 0.795894 | 0.991981 | 0.003496 |
| ENSMUSG00000032579.15 | 21.82473 | 0.494139 | 1.89644 | 0.260561 | 0.794431 | 0.991981 | 0.003496 |
| ENSMUSG00000034832.16 | 17.33817 | -0.52313 | 1.998051 | -0.26182 | 0.793462 | 0.991981 | 0.003496 |
| ENSMUSG00000034850.16 | 81.13 | -0.22329 | 0.862062 | -0.25902 | 0.795621 | 0.991981 | 0.003496 |
| ENSMUSG00000035086.14 | 184.7951 | -0.15905 | 0.620868 | -0.25617 | 0.79782 | 0.991981 | 0.003496 |
| ENSMUSG00000035578.17 | 13.78024 | 0.591104 | 2.318957 | 0.254901 | 0.7988 | 0.991981 | 0.003496 |
| ENSMUSG00000035948.14 | 78.44233 | -0.27706 | 1.067451 | -0.25955 | 0.795209 | 0.991981 | 0.003496 |
| ENSMUSG00000036615.8 | 185.5791 | -0.17806 | 0.693018 | -0.25693 | 0.797234 | 0.991981 | 0.003496 |
| ENSMUSG00000037110.21 | 64.91334 | -0.21998 | 0.860332 | -0.25569 | 0.798186 | 0.991981 | 0.003496 |
| ENSMUSG00000037300.20 | 38.41997 | 0.37955 | 1.429744 | 0.265467 | 0.79065 | 0.991981 | 0.003496 |
| ENSMUSG00000038774.10 | 86.87224 | -0.32025 | 1.241253 | -0.258 | 0.796404 | 0.991981 | 0.003496 |
| ENSMUSG00000040102.11 | 20.07209 | -0.495 | 1.908387 | -0.25938 | 0.79534 | 0.991981 | 0.003496 |
| ENSMUSG00000040532.15 | 34.39772 | -0.38385 | 1.458725 | -0.26314 | 0.79244 | 0.991981 | 0.003496 |
| ENSMUSG00000041040.13 | 13.89801 | 0.615986 | 2.382314 | 0.258566 | 0.79597 | 0.991981 | 0.003496 |
| ENSMUSG00000041741.11 | 14.33797 | 0.783472 | 3.006122 | 0.260626 | 0.794381 | 0.991981 | 0.003496 |
| ENSMUSG00000041957.16 | 266.8091 | -0.12724 | 0.480811 | -0.26463 | 0.791295 | 0.991981 | 0.003496 |
| ENSMUSG00000044350.15 | 27.14525 | 0.448835 | 1.707231 | 0.262902 | 0.792626 | 0.991981 | 0.003496 |
| ENSMUSG00000045098.20 | 47.4224 | 0.365147 | 1.426774 | 0.255925 | 0.798009 | 0.991981 | 0.003496 |
| ENSMUSG00000047617.12 | 40.27288 | -0.3779 | 1.480444 | -0.25526 | 0.798519 | 0.991981 | 0.003496 |
| ENSMUSG00000048106.7 | 12.23114 | -0.68697 | 2.586589 | -0.26559 | 0.790557 | 0.991981 | 0.003496 |
| ENSMUSG00000048865.17 | 13.84274 | 0.626547 | 2.403958 | 0.260631 | 0.794377 | 0.991981 | 0.003496 |
| ENSMUSG00000049287.6 | 11.258 | -0.63605 | 2.461432 | -0.25841 | 0.796093 | 0.991981 | 0.003496 |

| | | | | | | | |
|---|---|---|---|---|---|---|---|
| ENSMUSG00000052917.16 | 23.68776 | -0.39265 | 1.531624 | -0.25636 | 0.797671 | 0.991981 | 0.003496 |
| ENSMUSG00000053332.14 | 1505.867 | -0.0964 | 0.376162 | -0.25627 | 0.797741 | 0.991981 | 0.003496 |
| ENSMUSG00000054733.10 | 805.673 | 0.088641 | 0.343409 | 0.25812 | 0.796314 | 0.991981 | 0.003496 |
| ENSMUSG00000054889.11 | 92.8159 | 0.22443 | 0.851272 | 0.263641 | 0.792057 | 0.991981 | 0.003496 |
| ENSMUSG00000056268.16 | 146.6073 | 0.152791 | 0.598873 | 0.255132 | 0.798621 | 0.991981 | 0.003496 |
| ENSMUSG00000056673.15 | 30.77406 | -0.40849 | 1.542188 | -0.26487 | 0.791106 | 0.991981 | 0.003496 |
| ENSMUSG00000057346.13 | 114.6516 | -0.19893 | 0.773982 | -0.25702 | 0.79716 | 0.991981 | 0.003496 |
| ENSMUSG00000059033.8 | 11.79438 | 0.546815 | 2.092978 | 0.261262 | 0.793891 | 0.991981 | 0.003496 |
| ENSMUSG00000060739.9 | 168.159 | -0.16789 | 0.654283 | -0.2566 | 0.797484 | 0.991981 | 0.003496 |
| ENSMUSG00000061130.14 | 506.9073 | 0.128583 | 0.492649 | 0.261002 | 0.794091 | 0.991981 | 0.003496 |
| ENSMUSG00000062006.13 | 213.2272 | -0.14751 | 0.573399 | -0.25725 | 0.796988 | 0.991981 | 0.003496 |
| ENSMUSG00000066621.13 | 30.79102 | 0.488388 | 1.839827 | 0.265453 | 0.790661 | 0.991981 | 0.003496 |
| ENSMUSG00000066724.4 | 22.54745 | 0.33984 | 1.322677 | 0.256934 | 0.79723 | 0.991981 | 0.003496 |
| ENSMUSG00000068876.17 | 36.10543 | 0.472969 | 1.778742 | 0.265901 | 0.790316 | 0.991981 | 0.003496 |
| ENSMUSG00000074024.6 | 14.99583 | -0.53164 | 2.032643 | -0.26155 | 0.793667 | 0.991981 | 0.003496 |
| ENSMUSG00000074578.15 | 137.1207 | -0.15115 | 0.58435 | -0.25867 | 0.795892 | 0.991981 | 0.003496 |
| ENSMUSG00000079426.14 | 24.52967 | 0.499264 | 1.95114 | 0.255883 | 0.798041 | 0.991981 | 0.003496 |
| ENSMUSG00000079470.9 | 54.63389 | 0.264194 | 1.013232 | 0.260743 | 0.79429 | 0.991981 | 0.003496 |
| ENSMUSG00000082419.4 | 281.4468 | -0.12332 | 0.483626 | -0.25499 | 0.798728 | 0.991981 | 0.003496 |
| ENSMUSG00000085241.8 | 55.52616 | -0.3748 | 1.41205 | -0.26543 | 0.790677 | 0.991981 | 0.003496 |
| ENSMUSG00000091512.6 | 12.49434 | 0.469737 | 1.787327 | 0.262815 | 0.792693 | 0.991981 | 0.003496 |
| ENSMUSG00000117924.2 | 55.20224 | 0.348842 | 1.346331 | 0.259106 | 0.795553 | 0.991981 | 0.003496 |
| ENSMUSG00000118155.2 | 24.5428 | -0.46455 | 1.763822 | -0.26338 | 0.792258 | 0.991981 | 0.003496 |
| ENSMUSG00000006673.15 | 37.79437 | -0.31291 | 1.240885 | -0.25216 | 0.800914 | 0.992063 | 0.003461 |
| ENSMUSG00000015488.15 | 19.69704 | -0.55755 | 2.214643 | -0.25176 | 0.801228 | 0.992063 | 0.003461 |
| ENSMUSG00000016028.12 | 34.19278 | -0.40911 | 1.625465 | -0.25169 | 0.801283 | 0.992063 | 0.003461 |
| ENSMUSG00000020612.17 | 458.2775 | 0.128738 | 0.509548 | 0.252651 | 0.800538 | 0.992063 | 0.003461 |
| ENSMUSG00000020721.17 | 29.56016 | -0.38175 | 1.511712 | -0.25253 | 0.80063 | 0.992063 | 0.003461 |
| ENSMUSG00000021951.8 | 132.923 | -0.19109 | 0.753198 | -0.25371 | 0.79972 | 0.992063 | 0.003461 |
| ENSMUSG00000024099.16 | 572.2559 | 0.11027 | 0.437973 | 0.251773 | 0.801216 | 0.992063 | 0.003461 |
| ENSMUSG00000025860.16 | 529.2616 | -0.10727 | 0.423392 | -0.25336 | 0.79999 | 0.992063 | 0.003461 |
| ENSMUSG00000027257.14 | 116.9312 | -0.184 | 0.724591 | -0.25393 | 0.799548 | 0.992063 | 0.003461 |
| ENSMUSG00000029191.17 | 67.9922 | 0.237795 | 0.944247 | 0.251836 | 0.801168 | 0.992063 | 0.003461 |
| ENSMUSG00000029924.13 | 75.57148 | -0.37206 | 1.477204 | -0.25187 | 0.801142 | 0.992063 | 0.003461 |
| ENSMUSG00000030706.18 | 183.8361 | -0.26514 | 1.044538 | -0.25384 | 0.799623 | 0.992063 | 0.003461 |
| ENSMUSG00000030986.14 | 159.0564 | 0.153952 | 0.605372 | 0.25431 | 0.799256 | 0.992063 | 0.003461 |
| ENSMUSG00000031479.10 | 260.1681 | -0.16009 | 0.633927 | -0.25254 | 0.800625 | 0.992063 | 0.003461 |
| ENSMUSG00000033400.15 | 203.4793 | 0.133662 | 0.530997 | 0.251718 | 0.801259 | 0.992063 | 0.003461 |
| ENSMUSG00000033530.9 | 67.23594 | -0.23168 | 0.911366 | -0.25421 | 0.799335 | 0.992063 | 0.003461 |
| ENSMUSG00000038215.16 | 12.54356 | 0.53625 | 2.127457 | 0.252062 | 0.800994 | 0.992063 | 0.003461 |
| ENSMUSG00000041375.20 | 11.93174 | -0.53844 | 2.129092 | -0.25289 | 0.800349 | 0.992063 | 0.003461 |
| ENSMUSG00000042046.16 | 12.98922 | 0.512768 | 2.035595 | 0.251901 | 0.801118 | 0.992063 | 0.003461 |
| ENSMUSG00000046062.6 | 254.3233 | 0.133168 | 0.528868 | 0.251798 | 0.801197 | 0.992063 | 0.003461 |
| ENSMUSG00000060288.14 | 19.13528 | 0.371035 | 1.474246 | 0.251678 | 0.80129 | 0.992063 | 0.003461 |

| | | | | | | | |
|---|---|---|---|---|---|---|---|
| ENSMUSG00000105135.2 | 65.5636 | 0.286725 | 1.134337 | 0.252769 | 0.800447 | 0.992063 | 0.003461 |
| ENSMUSG00000108350.2 | 9.998993 | 0.737952 | 2.917593 | 0.252932 | 0.800321 | 0.992063 | 0.003461 |
| ENSMUSG00000019647.17 | 24.63696 | 0.447613 | 1.78036 | 0.251417 | 0.801491 | 0.992183 | 0.003408 |
| ENSMUSG00000015575.16 | 253.6746 | 0.128211 | 0.512349 | 0.250242 | 0.8024 | 0.992214 | 0.003395 |
| ENSMUSG00000020150.14 | 1077.321 | -0.08669 | 0.345914 | -0.2506 | 0.802122 | 0.992214 | 0.003395 |
| ENSMUSG00000022142.9 | 59.71293 | -0.29979 | 1.199004 | -0.25003 | 0.802563 | 0.992214 | 0.003395 |
| ENSMUSG00000025474.10 | 20.55309 | -0.44552 | 1.780352 | -0.25024 | 0.802398 | 0.992214 | 0.003395 |
| ENSMUSG00000027351.15 | 56.50747 | -0.23106 | 0.920405 | -0.25104 | 0.801782 | 0.992214 | 0.003395 |
| ENSMUSG00000031575.19 | 46.02611 | -0.30197 | 1.203625 | -0.25089 | 0.801902 | 0.992214 | 0.003395 |
| ENSMUSG00000033285.16 | 164.9539 | -0.17208 | 0.688032 | -0.2501 | 0.802511 | 0.992214 | 0.003395 |
| ENSMUSG00000036372.15 | 288.3894 | -0.13445 | 0.537212 | -0.25027 | 0.802382 | 0.992214 | 0.003395 |
| ENSMUSG00000050052.13 | 66.50787 | 0.234528 | 0.937224 | 0.250237 | 0.802404 | 0.992214 | 0.003395 |
| ENSMUSG00000052428.11 | 1099.091 | -0.07562 | 0.301017 | -0.25121 | 0.801653 | 0.992214 | 0.003395 |
| ENSMUSG00000016477.19 | 20.63329 | 0.527118 | 2.110446 | 0.249766 | 0.802768 | 0.992338 | 0.00334 |
| ENSMUSG00000019210.11 | 529.876 | -0.0895 | 0.359828 | -0.24872 | 0.803579 | 0.992646 | 0.003206 |
| ENSMUSG00000019738.16 | 254.8126 | 0.159785 | 0.642303 | 0.248769 | 0.80354 | 0.992646 | 0.003206 |
| ENSMUSG00000025147.8 | 22.71427 | 0.510266 | 2.049249 | 0.249001 | 0.80336 | 0.992646 | 0.003206 |
| ENSMUSG00000036632.10 | 379.7413 | -0.1151 | 0.462069 | -0.24909 | 0.80329 | 0.992646 | 0.003206 |
| ENSMUSG00000039599.17 | 127.5146 | 0.161289 | 0.648705 | 0.248633 | 0.803645 | 0.992646 | 0.003206 |
| ENSMUSG00000070704.11 | 5805.622 | -0.07107 | 0.285824 | -0.24864 | 0.803639 | 0.992646 | 0.003206 |
| ENSMUSG00000020614.14 | 47.50284 | -0.24559 | 1.001531 | -0.24522 | 0.806289 | 0.992718 | 0.003174 |
| ENSMUSG00000020671.10 | 361.8092 | -0.12507 | 0.505829 | -0.24726 | 0.80471 | 0.992718 | 0.003174 |
| ENSMUSG00000023852.15 | 64.93321 | 0.238148 | 0.969498 | 0.24564 | 0.805961 | 0.992718 | 0.003174 |
| ENSMUSG00000025173.14 | 264.0258 | -0.12089 | 0.492301 | -0.24557 | 0.806018 | 0.992718 | 0.003174 |
| ENSMUSG00000025958.15 | 105.5183 | -0.29838 | 1.217852 | -0.245 | 0.806456 | 0.992718 | 0.003174 |
| ENSMUSG00000026663.7 | 215.6031 | -0.13716 | 0.552931 | -0.24806 | 0.80409 | 0.992718 | 0.003174 |
| ENSMUSG00000027695.17 | 38.72159 | -0.31905 | 1.285402 | -0.24821 | 0.80397 | 0.992718 | 0.003174 |
| ENSMUSG00000032125.22 | 12.15379 | 0.611691 | 2.483879 | 0.246265 | 0.805477 | 0.992718 | 0.003174 |
| ENSMUSG00000032470.18 | 17.03436 | 0.488848 | 1.987439 | 0.245969 | 0.805706 | 0.992718 | 0.003174 |
| ENSMUSG00000033427.15 | 1199.166 | 0.092485 | 0.375175 | 0.246511 | 0.805287 | 0.992718 | 0.003174 |
| ENSMUSG00000033885.16 | 123.3378 | 0.179276 | 0.72985 | 0.245634 | 0.805966 | 0.992718 | 0.003174 |
| ENSMUSG00000037709.14 | 73.40821 | 0.205753 | 0.834257 | 0.246631 | 0.805194 | 0.992718 | 0.003174 |
| ENSMUSG00000038244.15 | 37.96658 | -0.35598 | 1.436631 | -0.24779 | 0.804296 | 0.992718 | 0.003174 |
| ENSMUSG00000038398.9 | 125.5547 | -0.19172 | 0.777734 | -0.24652 | 0.805283 | 0.992718 | 0.003174 |
| ENSMUSG00000038712.17 | 51.48214 | 0.264773 | 1.067811 | 0.247958 | 0.804167 | 0.992718 | 0.003174 |
| ENSMUSG00000040613.15 | 34.51764 | 0.321842 | 1.311452 | 0.245409 | 0.80614 | 0.992718 | 0.003174 |
| ENSMUSG00000040747.10 | 19.77917 | 0.527807 | 2.149728 | 0.245523 | 0.806052 | 0.992718 | 0.003174 |
| ENSMUSG00000049037.9 | 14.42894 | -0.6461 | 2.616371 | -0.24695 | 0.80495 | 0.992718 | 0.003174 |
| ENSMUSG00000052833.10 | 68.02929 | -0.25687 | 1.044352 | -0.24596 | 0.805714 | 0.992718 | 0.003174 |
| ENSMUSG00000063428.9 | 107.3801 | 0.212404 | 0.858445 | 0.247429 | 0.804576 | 0.992718 | 0.003174 |
| ENSMUSG00000064254.7 | 204.1302 | 0.153166 | 0.625013 | 0.245061 | 0.806409 | 0.992718 | 0.003174 |
| ENSMUSG00000064289.16 | 121.0836 | 0.172544 | 0.704539 | 0.244904 | 0.806531 | 0.992718 | 0.003174 |
| ENSMUSG00000066037.15 | 156.6055 | -0.15019 | 0.609188 | -0.24654 | 0.805267 | 0.992718 | 0.003174 |
| ENSMUSG00000067336.7 | 234.3967 | 0.162263 | 0.660317 | 0.245735 | 0.805887 | 0.992718 | 0.003174 |

| | | | | | | | |
|---|---|---|---|---|---|---|---|
| ENSMUSG00000073131.13 | 251.5457 | -0.13986 | 0.564897 | -0.24758 | 0.804457 | 0.992718 | 0.003174 |
| ENSMUSG00000075701.11 | 725.6003 | 0.087494 | 0.354224 | 0.247001 | 0.804907 | 0.992718 | 0.003174 |
| ENSMUSG00000097162.3 | 142.509 | 0.164196 | 0.662547 | 0.247825 | 0.80427 | 0.992718 | 0.003174 |
| ENSMUSG00000021273.12 | 295.8145 | 0.140415 | 0.574105 | 0.244581 | 0.806781 | 0.992768 | 0.003152 |
| ENSMUSG00000081382.2 | 13.97524 | 0.40601 | 1.659649 | 0.244636 | 0.806738 | 0.992768 | 0.003152 |
| ENSMUSG00000005802.13 | 35.46609 | 0.319494 | 1.309708 | 0.243943 | 0.807275 | 0.99305 | 0.003029 |
| ENSMUSG00000025574.14 | 22.63114 | -0.43589 | 1.786337 | -0.24402 | 0.807219 | 0.99305 | 0.003029 |
| ENSMUSG00000061288.14 | 130.9936 | 0.229515 | 0.941101 | 0.243879 | 0.807324 | 0.99305 | 0.003029 |
| ENSMUSG00000019996.18 | 66.38688 | 0.214653 | 0.882116 | 0.243339 | 0.807743 | 0.993178 | 0.002973 |
| ENSMUSG00000054226.14 | 698.776 | -0.11052 | 0.453862 | -0.24351 | 0.807612 | 0.993178 | 0.002973 |
| ENSMUSG00000070697.5 | 464.6978 | -0.09793 | 0.40239 | -0.24336 | 0.807728 | 0.993178 | 0.002973 |
| ENSMUSG00000000957.12 | 107.9296 | -0.16655 | 0.783572 | -0.21255 | 0.831679 | 0.993188 | 0.002969 |
| ENSMUSG00000000976.14 | 26.98639 | -0.29337 | 1.296918 | -0.22621 | 0.821041 | 0.993188 | 0.002969 |
| ENSMUSG00000001158.14 | 227.9288 | 0.139494 | 0.598001 | 0.233268 | 0.815554 | 0.993188 | 0.002969 |
| ENSMUSG00000001569.11 | 54.18484 | -0.22283 | 1.041577 | -0.21394 | 0.830597 | 0.993188 | 0.002969 |
| ENSMUSG00000002010.18 | 202.1228 | -0.19501 | 0.85919 | -0.22697 | 0.820448 | 0.993188 | 0.002969 |
| ENSMUSG00000002015.6 | 423.8937 | 0.096414 | 0.398718 | 0.241811 | 0.808927 | 0.993188 | 0.002969 |
| ENSMUSG00000002059.19 | 21.07439 | -0.54977 | 2.479698 | -0.22171 | 0.824541 | 0.993188 | 0.002969 |
| ENSMUSG00000002129.12 | 87.00584 | -0.19296 | 0.891245 | -0.2165 | 0.828596 | 0.993188 | 0.002969 |
| ENSMUSG00000003283.16 | 18.60174 | -0.39702 | 1.849717 | -0.21464 | 0.83005 | 0.993188 | 0.002969 |
| ENSMUSG00000003420.9 | 158.3207 | 0.137759 | 0.616472 | 0.223463 | 0.823175 | 0.993188 | 0.002969 |
| ENSMUSG00000003970.10 | 1158.275 | -0.07172 | 0.316207 | -0.22681 | 0.820568 | 0.993188 | 0.002969 |
| ENSMUSG00000005846.13 | 105.4344 | 0.166722 | 0.774118 | 0.215371 | 0.829478 | 0.993188 | 0.002969 |
| ENSMUSG00000005881.14 | 85.00014 | 0.21227 | 0.993778 | 0.213599 | 0.83086 | 0.993188 | 0.002969 |
| ENSMUSG00000005899.15 | 45.87397 | 0.353501 | 1.465883 | 0.241152 | 0.809437 | 0.993188 | 0.002969 |
| ENSMUSG00000006390.16 | 96.49971 | -0.18976 | 0.898201 | -0.21127 | 0.832676 | 0.993188 | 0.002969 |
| ENSMUSG00000008090.15 | 35.20737 | -0.3432 | 1.512288 | -0.22694 | 0.820472 | 0.993188 | 0.002969 |
| ENSMUSG00000008730.18 | 216.3334 | 0.105886 | 0.484193 | 0.218686 | 0.826895 | 0.993188 | 0.002969 |
| ENSMUSG00000009281.7 | 3997.562 | 0.054489 | 0.253051 | 0.215329 | 0.829511 | 0.993188 | 0.002969 |
| ENSMUSG00000012114.18 | 29.23708 | -0.36588 | 1.566812 | -0.23352 | 0.815359 | 0.993188 | 0.002969 |
| ENSMUSG00000014859.10 | 23.94713 | -0.35853 | 1.641332 | -0.21844 | 0.82709 | 0.993188 | 0.002969 |
| ENSMUSG00000016427.8 | 1357.709 | -0.05729 | 0.254625 | -0.22499 | 0.821986 | 0.993188 | 0.002969 |
| ENSMUSG00000017428.17 | 466.6859 | 0.100086 | 0.443425 | 0.22571 | 0.821427 | 0.993188 | 0.002969 |
| ENSMUSG00000018377.11 | 121.7345 | 0.170724 | 0.712919 | 0.239472 | 0.810739 | 0.993188 | 0.002969 |
| ENSMUSG00000018415.15 | 50.76851 | 0.251141 | 1.146082 | 0.21913 | 0.826549 | 0.993188 | 0.002969 |
| ENSMUSG00000018899.18 | 39.94844 | 0.24904 | 1.079531 | 0.230693 | 0.817554 | 0.993188 | 0.002969 |
| ENSMUSG00000018999.15 | 119.9188 | -0.16232 | 0.765092 | -0.21216 | 0.831986 | 0.993188 | 0.002969 |
| ENSMUSG00000019254.17 | 38.24865 | -0.34029 | 1.59147 | -0.21382 | 0.830688 | 0.993188 | 0.002969 |
| ENSMUSG00000019832.6 | 239.049 | -0.14942 | 0.660776 | -0.22612 | 0.821105 | 0.993188 | 0.002969 |
| ENSMUSG00000019971.11 | 16.54909 | -0.52651 | 2.274134 | -0.23152 | 0.816909 | 0.993188 | 0.002969 |
| ENSMUSG00000019978.17 | 65.00316 | -0.22619 | 1.076256 | -0.21016 | 0.833543 | 0.993188 | 0.002969 |
| ENSMUSG00000020053.19 | 6871.158 | 0.048166 | 0.208227 | 0.231317 | 0.817069 | 0.993188 | 0.002969 |
| ENSMUSG00000020393.17 | 20.24278 | -0.39007 | 1.725481 | -0.22606 | 0.821152 | 0.993188 | 0.002969 |
| ENSMUSG00000020766.5 | 57.36425 | 0.28656 | 1.304311 | 0.219702 | 0.826103 | 0.993188 | 0.002969 |

| | | | | | | | |
|---|---|---|---|---|---|---|---|
| ENSMUSG00000020873.16 | 405.2774 | 0.090118 | 0.412334 | 0.218555 | 0.826997 | 0.993188 | 0.002969 |
| ENSMUSG00000021114.10 | 331.0133 | 0.140061 | 0.65529 | 0.213739 | 0.83075 | 0.993188 | 0.002969 |
| ENSMUSG00000021156.18 | 205.2295 | 0.124527 | 0.586191 | 0.212434 | 0.831768 | 0.993188 | 0.002969 |
| ENSMUSG00000021509.6 | 60.22737 | -0.28127 | 1.33993 | -0.20992 | 0.833733 | 0.993188 | 0.002969 |
| ENSMUSG00000021811.8 | 21.32475 | 0.523376 | 2.405489 | 0.217576 | 0.82776 | 0.993188 | 0.002969 |
| ENSMUSG00000021846.10 | 111.854 | 0.153819 | 0.724407 | 0.212338 | 0.831844 | 0.993188 | 0.002969 |
| ENSMUSG00000021868.8 | 119.8525 | 0.187588 | 0.826381 | 0.226999 | 0.820425 | 0.993188 | 0.002969 |
| ENSMUSG00000021911.17 | 10.72447 | 0.493388 | 2.16687 | 0.227696 | 0.819882 | 0.993188 | 0.002969 |
| ENSMUSG00000022089.10 | 43.26787 | -0.25251 | 1.049855 | -0.24052 | 0.809925 | 0.993188 | 0.002969 |
| ENSMUSG00000022122.16 | 49.84736 | 0.303058 | 1.38844 | 0.218272 | 0.827217 | 0.993188 | 0.002969 |
| ENSMUSG00000022248.15 | 24.7732 | 0.363088 | 1.711872 | 0.2121 | 0.832029 | 0.993188 | 0.002969 |
| ENSMUSG00000022270.17 | 370.7959 | -0.13922 | 0.612261 | -0.22739 | 0.820122 | 0.993188 | 0.002969 |
| ENSMUSG00000022350.8 | 57.7254 | 0.291394 | 1.375452 | 0.211853 | 0.832222 | 0.993188 | 0.002969 |
| ENSMUSG00000022372.15 | 13.06919 | 0.563816 | 2.678658 | 0.210484 | 0.83329 | 0.993188 | 0.002969 |
| ENSMUSG00000022378.15 | 35.2162 | -0.28226 | 1.199703 | -0.23527 | 0.813997 | 0.993188 | 0.002969 |
| ENSMUSG00000022403.16 | 1004.68 | -0.06883 | 0.29644 | -0.23219 | 0.816392 | 0.993188 | 0.002969 |
| ENSMUSG00000022570.8 | 27.83965 | 0.375227 | 1.558371 | 0.240782 | 0.809724 | 0.993188 | 0.002969 |
| ENSMUSG00000022766.15 | 1524.4 | -0.06846 | 0.292301 | -0.23421 | 0.814819 | 0.993188 | 0.002969 |
| ENSMUSG00000022841.16 | 167.5988 | 0.151259 | 0.654675 | 0.231044 | 0.817281 | 0.993188 | 0.002969 |
| ENSMUSG00000023336.7 | 20.68414 | -0.42571 | 1.967859 | -0.21633 | 0.828729 | 0.993188 | 0.002969 |
| ENSMUSG00000023951.19 | 290.7482 | -0.14227 | 0.655966 | -0.21689 | 0.828294 | 0.993188 | 0.002969 |
| ENSMUSG00000024030.8 | 13.41943 | -0.46727 | 2.216252 | -0.21084 | 0.833013 | 0.993188 | 0.002969 |
| ENSMUSG00000024120.13 | 1298.905 | 0.066246 | 0.311352 | 0.21277 | 0.831506 | 0.993188 | 0.002969 |
| ENSMUSG00000024158.18 | 1512.376 | 0.070963 | 0.323204 | 0.219561 | 0.826213 | 0.993188 | 0.002969 |
| ENSMUSG00000024241.8 | 243.7597 | 0.118804 | 0.543528 | 0.218579 | 0.826978 | 0.993188 | 0.002969 |
| ENSMUSG00000024259.10 | 358.3052 | -0.11593 | 0.53709 | -0.21584 | 0.82911 | 0.993188 | 0.002969 |
| ENSMUSG00000024317.16 | 105.9147 | -0.26602 | 1.164883 | -0.22836 | 0.819365 | 0.993188 | 0.002969 |
| ENSMUSG00000024454.17 | 22.07725 | 0.456367 | 2.097449 | 0.217582 | 0.827755 | 0.993188 | 0.002969 |
| ENSMUSG00000024594.10 | 170.5634 | 0.141853 | 0.64413 | 0.220224 | 0.825697 | 0.993188 | 0.002969 |
| ENSMUSG00000024899.8 | 1219.701 | -0.09151 | 0.403129 | -0.22699 | 0.82043 | 0.993188 | 0.002969 |
| ENSMUSG00000025002.6 | 92.35474 | -0.21302 | 0.881085 | -0.24177 | 0.808956 | 0.993188 | 0.002969 |
| ENSMUSG00000025135.13 | 85.62317 | 0.249275 | 1.105338 | 0.225519 | 0.821575 | 0.993188 | 0.002969 |
| ENSMUSG00000025209.6 | 65.15849 | 0.205964 | 0.942242 | 0.218589 | 0.82697 | 0.993188 | 0.002969 |
| ENSMUSG00000025241.17 | 140.6195 | -0.15798 | 0.690379 | -0.22883 | 0.819002 | 0.993188 | 0.002969 |
| ENSMUSG00000025494.9 | 14.2453 | -0.4766 | 2.113822 | -0.22547 | 0.821616 | 0.993188 | 0.002969 |
| ENSMUSG00000025722.17 | 67.96299 | -0.22565 | 1.022225 | -0.22074 | 0.825295 | 0.993188 | 0.002969 |
| ENSMUSG00000025733.18 | 107.1386 | -0.21517 | 0.898709 | -0.23942 | 0.81078 | 0.993188 | 0.002969 |
| ENSMUSG00000025868.7 | 372.8909 | 0.120686 | 0.49989 | 0.241424 | 0.809226 | 0.993188 | 0.002969 |
| ENSMUSG00000026127.14 | 72.03044 | 0.227241 | 1.075092 | 0.211369 | 0.8326 | 0.993188 | 0.002969 |
| ENSMUSG00000026170.7 | 1017.367 | -0.07553 | 0.31114 | -0.24276 | 0.80819 | 0.993188 | 0.002969 |
| ENSMUSG00000026289.16 | 57.60892 | 0.38409 | 1.634648 | 0.234968 | 0.814234 | 0.993188 | 0.002969 |
| ENSMUSG00000026626.12 | 358.8559 | -0.11702 | 0.502567 | -0.23284 | 0.815884 | 0.993188 | 0.002969 |
| ENSMUSG00000026788.14 | 44.75976 | -0.27927 | 1.268852 | -0.2201 | 0.825796 | 0.993188 | 0.002969 |
| ENSMUSG00000026889.13 | 32.03504 | -0.35334 | 1.538755 | -0.22963 | 0.818379 | 0.993188 | 0.002969 |

| | | | | | | | |
|---|---|---|---|---|---|---|---|
| ENSMUSG00000026960.7 | 45.26221 | 0.283518 | 1.222567 | 0.231904 | 0.816613 | 0.993188 | 0.002969 |
| ENSMUSG00000027087.12 | 136.7246 | -0.1963 | 0.83031 | -0.23641 | 0.813113 | 0.993188 | 0.002969 |
| ENSMUSG00000027249.16 | 6591.762 | -0.04647 | 0.221253 | -0.21002 | 0.833655 | 0.993188 | 0.002969 |
| ENSMUSG00000027712.14 | 372.1137 | 0.101927 | 0.423959 | 0.240417 | 0.810007 | 0.993188 | 0.002969 |
| ENSMUSG00000027778.16 | 17.81663 | -0.49771 | 2.316645 | -0.21484 | 0.82989 | 0.993188 | 0.002969 |
| ENSMUSG00000028062.16 | 738.4257 | 0.095422 | 0.417284 | 0.228673 | 0.819123 | 0.993188 | 0.002969 |
| ENSMUSG00000028064.18 | 140.7275 | -0.14858 | 0.639043 | -0.2325 | 0.816148 | 0.993188 | 0.002969 |
| ENSMUSG00000028464.17 | 14.57929 | 0.495901 | 2.277668 | 0.217723 | 0.827645 | 0.993188 | 0.002969 |
| ENSMUSG00000028603.16 | 3701.166 | -0.05667 | 0.267072 | -0.21217 | 0.831971 | 0.993188 | 0.002969 |
| ENSMUSG00000028691.13 | 10902.31 | 0.045631 | 0.214276 | 0.212955 | 0.831362 | 0.993188 | 0.002969 |
| ENSMUSG00000028760.17 | 280.6551 | 0.110761 | 0.488818 | 0.226588 | 0.820744 | 0.993188 | 0.002969 |
| ENSMUSG00000028800.16 | 34.71967 | -0.42706 | 1.782271 | -0.23961 | 0.810629 | 0.993188 | 0.002969 |
| ENSMUSG00000028901.14 | 18.04875 | -0.47147 | 2.122583 | -0.22212 | 0.824219 | 0.993188 | 0.002969 |
| ENSMUSG00000028954.12 | 73.09615 | 0.260635 | 1.229524 | 0.21198 | 0.832123 | 0.993188 | 0.002969 |
| ENSMUSG00000029004.16 | 289.6282 | -0.10046 | 0.461223 | -0.21781 | 0.827578 | 0.993188 | 0.002969 |
| ENSMUSG00000029066.13 | 468.7587 | -0.09178 | 0.421904 | -0.21753 | 0.827796 | 0.993188 | 0.002969 |
| ENSMUSG00000029167.14 | 24.11458 | -0.29967 | 1.343258 | -0.22309 | 0.823465 | 0.993188 | 0.002969 |
| ENSMUSG00000029269.12 | 305.2503 | 0.11575 | 0.550157 | 0.210394 | 0.83336 | 0.993188 | 0.002969 |
| ENSMUSG00000029385.15 | 109.1459 | 0.176618 | 0.792112 | 0.222972 | 0.823558 | 0.993188 | 0.002969 |
| ENSMUSG00000029402.9 | 25.40762 | 0.40474 | 1.719931 | 0.235324 | 0.813958 | 0.993188 | 0.002969 |
| ENSMUSG00000029580.15 | 445.3139 | -0.1402 | 0.639154 | -0.21935 | 0.826374 | 0.993188 | 0.002969 |
| ENSMUSG00000029610.14 | 65.88496 | 0.229843 | 0.974656 | 0.23582 | 0.813573 | 0.993188 | 0.002969 |
| ENSMUSG00000029759.10 | 357.0338 | -0.09515 | 0.451514 | -0.21073 | 0.833098 | 0.993188 | 0.002969 |
| ENSMUSG00000030505.18 | 58.33575 | 0.274171 | 1.14469 | 0.239515 | 0.810706 | 0.993188 | 0.002969 |
| ENSMUSG00000030538.16 | 161.7906 | -0.16113 | 0.70215 | -0.22948 | 0.818493 | 0.993188 | 0.002969 |
| ENSMUSG00000030695.17 | 94.88981 | -0.19349 | 0.838825 | -0.23067 | 0.81757 | 0.993188 | 0.002969 |
| ENSMUSG00000030738.12 | 710.4041 | -0.09499 | 0.440292 | -0.21575 | 0.82918 | 0.993188 | 0.002969 |
| ENSMUSG00000030760.15 | 63.6559 | 0.20434 | 0.889465 | 0.229734 | 0.818299 | 0.993188 | 0.002969 |
| ENSMUSG00000030871.12 | 17.28122 | -0.45972 | 2.007679 | -0.22898 | 0.818885 | 0.993188 | 0.002969 |
| ENSMUSG00000030890.17 | 82.20994 | 0.601391 | 2.745219 | 0.219069 | 0.826597 | 0.993188 | 0.002969 |
| ENSMUSG00000030968.4 | 28.40908 | 0.403846 | 1.888071 | 0.213893 | 0.83063 | 0.993188 | 0.002969 |
| ENSMUSG00000031292.15 | 16.83203 | 0.433939 | 1.941503 | 0.223507 | 0.823141 | 0.993188 | 0.002969 |
| ENSMUSG00000031453.17 | 18.2078 | -0.52533 | 2.222754 | -0.23634 | 0.813166 | 0.993188 | 0.002969 |
| ENSMUSG00000031508.15 | 46.00281 | 0.251527 | 1.066816 | 0.235773 | 0.813609 | 0.993188 | 0.002969 |
| ENSMUSG00000031642.10 | 41.43557 | 0.385789 | 1.692993 | 0.227874 | 0.819744 | 0.993188 | 0.002969 |
| ENSMUSG00000031827.14 | 28.88669 | -0.42564 | 1.891177 | -0.22507 | 0.821929 | 0.993188 | 0.002969 |
| ENSMUSG00000031828.10 | 13.4038 | 0.716681 | 3.075906 | 0.232999 | 0.815763 | 0.993188 | 0.002969 |
| ENSMUSG00000031935.10 | 82.34841 | 0.220972 | 1.023197 | 0.215962 | 0.829017 | 0.993188 | 0.002969 |
| ENSMUSG00000031974.9 | 293.7097 | -0.13393 | 0.574706 | -0.23304 | 0.815727 | 0.993188 | 0.002969 |
| ENSMUSG00000031976.15 | 11.80018 | -0.5278 | 2.426258 | -0.21754 | 0.827791 | 0.993188 | 0.002969 |
| ENSMUSG00000032026.8 | 287.4441 | 0.109439 | 0.460944 | 0.237424 | 0.812328 | 0.993188 | 0.002969 |
| ENSMUSG00000032279.12 | 132.8345 | 0.148048 | 0.702866 | 0.210635 | 0.833172 | 0.993188 | 0.002969 |
| ENSMUSG00000032306.15 | 64.02206 | -0.31353 | 1.348013 | -0.23259 | 0.816081 | 0.993188 | 0.002969 |
| ENSMUSG00000032392.12 | 86.76319 | -0.19196 | 0.872881 | -0.21992 | 0.825934 | 0.993188 | 0.002969 |

| | | | | | | | |
|---|---|---|---|---|---|---|---|
| ENSMUSG00000032411.16 | 43.80501 | 0.232053 | 1.104374 | 0.210122 | 0.833572 | 0.993188 | 0.002969 |
| ENSMUSG00000032431.7 | 13.06592 | -0.57444 | 2.362639 | -0.24313 | 0.807901 | 0.993188 | 0.002969 |
| ENSMUSG00000032435.10 | 121.9512 | -0.1596 | 0.693162 | -0.23025 | 0.817894 | 0.993188 | 0.002969 |
| ENSMUSG00000032478.15 | 51.10406 | -0.2862 | 1.349588 | -0.21206 | 0.832057 | 0.993188 | 0.002969 |
| ENSMUSG00000032637.16 | 40.44878 | 0.287767 | 1.257782 | 0.228789 | 0.819033 | 0.993188 | 0.002969 |
| ENSMUSG00000032952.12 | 18.15593 | -0.45872 | 1.982704 | -0.23136 | 0.817036 | 0.993188 | 0.002969 |
| ENSMUSG00000032998.17 | 38.52394 | -0.29703 | 1.3182 | -0.22533 | 0.82172 | 0.993188 | 0.002969 |
| ENSMUSG00000033306.16 | 272.265 | -0.11331 | 0.51485 | -0.22008 | 0.825808 | 0.993188 | 0.002969 |
| ENSMUSG00000033454.7 | 38.56115 | -0.26111 | 1.184457 | -0.22045 | 0.825524 | 0.993188 | 0.002969 |
| ENSMUSG00000033542.14 | 29.19259 | -0.28527 | 1.356973 | -0.21022 | 0.833493 | 0.993188 | 0.002969 |
| ENSMUSG00000033900.14 | 19.22315 | 0.559831 | 2.509677 | 0.223069 | 0.823482 | 0.993188 | 0.002969 |
| ENSMUSG00000033943.16 | 35.6614 | -0.39061 | 1.621233 | -0.24094 | 0.809604 | 0.993188 | 0.002969 |
| ENSMUSG00000034075.11 | 120.0486 | -0.17423 | 0.769574 | -0.22639 | 0.820895 | 0.993188 | 0.002969 |
| ENSMUSG00000034343.15 | 352.8922 | 0.104057 | 0.460735 | 0.225851 | 0.821317 | 0.993188 | 0.002969 |
| ENSMUSG00000034557.15 | 40.63813 | 0.286516 | 1.275132 | 0.224695 | 0.822216 | 0.993188 | 0.002969 |
| ENSMUSG00000034903.19 | 92.6625 | -0.19028 | 0.841796 | -0.22604 | 0.821171 | 0.993188 | 0.002969 |
| ENSMUSG00000035151.13 | 58.55965 | -0.24011 | 1.074701 | -0.22342 | 0.823212 | 0.993188 | 0.002969 |
| ENSMUSG00000035382.10 | 46.20543 | -0.32723 | 1.547628 | -0.21144 | 0.832544 | 0.993188 | 0.002969 |
| ENSMUSG00000035521.14 | 126.1373 | -0.16048 | 0.678537 | -0.23651 | 0.813041 | 0.993188 | 0.002969 |
| ENSMUSG00000035561.6 | 178.0957 | -0.12756 | 0.532345 | -0.23963 | 0.81062 | 0.993188 | 0.002969 |
| ENSMUSG00000035840.8 | 199.0788 | -0.11942 | 0.563072 | -0.21208 | 0.832046 | 0.993188 | 0.002969 |
| ENSMUSG00000036833.17 | 774.9285 | 0.079226 | 0.354785 | 0.223307 | 0.823297 | 0.993188 | 0.002969 |
| ENSMUSG00000036975.6 | 23.89714 | -0.2973 | 1.291975 | -0.23011 | 0.818002 | 0.993188 | 0.002969 |
| ENSMUSG00000037013.18 | 186.0411 | -0.1734 | 0.719941 | -0.24085 | 0.80967 | 0.993188 | 0.002969 |
| ENSMUSG00000037070.11 | 99.75023 | -0.20754 | 0.887071 | -0.23396 | 0.815016 | 0.993188 | 0.002969 |
| ENSMUSG00000037376.15 | 63.92476 | 0.25638 | 1.221334 | 0.209918 | 0.833732 | 0.993188 | 0.002969 |
| ENSMUSG00000037465.11 | 57.71274 | -0.36511 | 1.722806 | -0.21193 | 0.832161 | 0.993188 | 0.002969 |
| ENSMUSG00000037730.14 | 20.81708 | 0.333655 | 1.441586 | 0.23145 | 0.816965 | 0.993188 | 0.002969 |
| ENSMUSG00000038028.10 | 29.12484 | -0.42612 | 1.865018 | -0.22848 | 0.819271 | 0.993188 | 0.002969 |
| ENSMUSG00000038425.19 | 37.08293 | -0.30887 | 1.455249 | -0.21225 | 0.831915 | 0.993188 | 0.002969 |
| ENSMUSG00000038489.9 | 164.6426 | 0.1218 | 0.568713 | 0.214168 | 0.830416 | 0.993188 | 0.002969 |
| ENSMUSG00000038510.13 | 130.2269 | -0.19958 | 0.862593 | -0.23138 | 0.817022 | 0.993188 | 0.002969 |
| ENSMUSG00000038733.15 | 581.2864 | 0.107102 | 0.459873 | 0.232895 | 0.815843 | 0.993188 | 0.002969 |
| ENSMUSG00000038831.17 | 17.08313 | 0.501355 | 2.146963 | 0.233518 | 0.815359 | 0.993188 | 0.002969 |
| ENSMUSG00000038888.9 | 12.89405 | -0.62207 | 2.723576 | -0.2284 | 0.819333 | 0.993188 | 0.002969 |
| ENSMUSG00000038991.18 | 222.3979 | 0.126709 | 0.571235 | 0.221816 | 0.824457 | 0.993188 | 0.002969 |
| ENSMUSG00000039062.16 | 27.26887 | -0.31249 | 1.418451 | -0.2203 | 0.825636 | 0.993188 | 0.002969 |
| ENSMUSG00000039234.12 | 206.0673 | 0.127264 | 0.591213 | 0.21526 | 0.829565 | 0.993188 | 0.002969 |
| ENSMUSG00000039323.19 | 1387.227 | 0.088978 | 0.379492 | 0.234465 | 0.814624 | 0.993188 | 0.002969 |
| ENSMUSG00000039480.15 | 153.0732 | -0.1769 | 0.840658 | -0.21043 | 0.83333 | 0.993188 | 0.002969 |
| ENSMUSG00000039523.17 | 15.79454 | -0.42508 | 1.766687 | -0.24061 | 0.809858 | 0.993188 | 0.002969 |
| ENSMUSG00000039529.10 | 134.5988 | 0.182452 | 0.860378 | 0.212061 | 0.83206 | 0.993188 | 0.002969 |
| ENSMUSG00000039640.8 | 215.5293 | -0.14347 | 0.627181 | -0.22876 | 0.819057 | 0.993188 | 0.002969 |
| ENSMUSG00000040112.10 | 464.9037 | -0.11564 | 0.528962 | -0.21862 | 0.826947 | 0.993188 | 0.002969 |

| | | | | | | | |
|---|---|---|---|---|---|---|---|
| ENSMUSG00000040270.17 | 26.93088 | -0.35983 | 1.647999 | -0.21834 | 0.827164 | 0.993188 | 0.002969 |
| ENSMUSG00000040356.17 | 133.4773 | 0.192222 | 0.813899 | 0.236174 | 0.813298 | 0.993188 | 0.002969 |
| ENSMUSG00000040471.18 | 83.46295 | 0.203638 | 0.899858 | 0.2263 | 0.820968 | 0.993188 | 0.002969 |
| ENSMUSG00000041736.8 | 188.0705 | -0.14474 | 0.604905 | -0.23927 | 0.810894 | 0.993188 | 0.002969 |
| ENSMUSG00000041840.9 | 11.06459 | 0.537886 | 2.394467 | 0.224637 | 0.822261 | 0.993188 | 0.002969 |
| ENSMUSG00000041975.18 | 44.87946 | 0.338369 | 1.568345 | 0.215749 | 0.829183 | 0.993188 | 0.002969 |
| ENSMUSG00000042121.17 | 15.20409 | -0.59002 | 2.575093 | -0.22913 | 0.81877 | 0.993188 | 0.002969 |
| ENSMUSG00000042138.10 | 53.92102 | 0.247949 | 1.100647 | 0.225276 | 0.821765 | 0.993188 | 0.002969 |
| ENSMUSG00000042548.15 | 40.77337 | 0.349616 | 1.646896 | 0.212288 | 0.831882 | 0.993188 | 0.002969 |
| ENSMUSG00000042807.16 | 31.62991 | -0.38865 | 1.625739 | -0.23906 | 0.811057 | 0.993188 | 0.002969 |
| ENSMUSG00000043190.15 | 146.2692 | 0.138343 | 0.637263 | 0.217089 | 0.828139 | 0.993188 | 0.002969 |
| ENSMUSG00000043391.11 | 19.44538 | -0.33489 | 1.463412 | -0.22884 | 0.818994 | 0.993188 | 0.002969 |
| ENSMUSG00000043411.16 | 162.8207 | 0.148302 | 0.64341 | 0.230494 | 0.817708 | 0.993188 | 0.002969 |
| ENSMUSG00000043415.6 | 15.76832 | -0.49077 | 2.18361 | -0.22475 | 0.822171 | 0.993188 | 0.002969 |
| ENSMUSG00000043770.11 | 24.76407 | -0.3304 | 1.522768 | -0.21697 | 0.82823 | 0.993188 | 0.002969 |
| ENSMUSG00000044456.17 | 15.1753 | 0.519409 | 2.346301 | 0.221373 | 0.824802 | 0.993188 | 0.002969 |
| ENSMUSG00000044627.12 | 723.8001 | 0.100891 | 0.442811 | 0.227842 | 0.819769 | 0.993188 | 0.002969 |
| ENSMUSG00000044788.11 | 157.3596 | 0.163201 | 0.700543 | 0.232964 | 0.815789 | 0.993188 | 0.002969 |
| ENSMUSG00000045409.17 | 28.87328 | 0.436456 | 1.864087 | 0.234139 | 0.814877 | 0.993188 | 0.002969 |
| ENSMUSG00000045969.9 | 152.9519 | -0.13074 | 0.602756 | -0.2169 | 0.828289 | 0.993188 | 0.002969 |
| ENSMUSG00000046873.19 | 37.66469 | 0.270877 | 1.169293 | 0.231659 | 0.816803 | 0.993188 | 0.002969 |
| ENSMUSG00000047044.8 | 69.73119 | 0.260013 | 1.222424 | 0.212703 | 0.831559 | 0.993188 | 0.002969 |
| ENSMUSG00000047539.10 | 27.6542 | -0.37805 | 1.645816 | -0.2297 | 0.818324 | 0.993188 | 0.002969 |
| ENSMUSG00000047719.3 | 28.41274 | -0.33634 | 1.597541 | -0.21053 | 0.833251 | 0.993188 | 0.002969 |
| ENSMUSG00000048000.16 | 143.8124 | -0.13895 | 0.625056 | -0.2223 | 0.824079 | 0.993188 | 0.002969 |
| ENSMUSG00000048578.12 | 88.07247 | -0.20845 | 0.913102 | -0.22829 | 0.819421 | 0.993188 | 0.002969 |
| ENSMUSG00000048701.14 | 72.72571 | 0.170318 | 0.790692 | 0.215403 | 0.829453 | 0.993188 | 0.002969 |
| ENSMUSG00000049305.7 | 16.29139 | -0.58121 | 2.678932 | -0.21696 | 0.828242 | 0.993188 | 0.002969 |
| ENSMUSG00000049659.16 | 199.49 | -0.12033 | 0.569906 | -0.21113 | 0.832785 | 0.993188 | 0.002969 |
| ENSMUSG00000049717.10 | 77.00587 | 0.210648 | 0.929758 | 0.226563 | 0.820764 | 0.993188 | 0.002969 |
| ENSMUSG00000049878.14 | 39.45022 | 0.258644 | 1.182805 | 0.21867 | 0.826907 | 0.993188 | 0.002969 |
| ENSMUSG00000050079.11 | 79.10874 | -0.19153 | 0.857911 | -0.22326 | 0.823336 | 0.993188 | 0.002969 |
| ENSMUSG00000050493.9 | 15.89608 | -0.58283 | 2.535395 | -0.22988 | 0.818188 | 0.993188 | 0.002969 |
| ENSMUSG00000050854.10 | 24.01992 | -0.40067 | 1.660437 | -0.2413 | 0.809321 | 0.993188 | 0.002969 |
| ENSMUSG00000051483.10 | 152.476 | 0.14059 | 0.626411 | 0.224437 | 0.822417 | 0.993188 | 0.002969 |
| ENSMUSG00000051989.12 | 226.2064 | 0.186885 | 0.782455 | 0.238845 | 0.811226 | 0.993188 | 0.002969 |
| ENSMUSG00000052997.16 | 165.9659 | -0.13967 | 0.604659 | -0.23098 | 0.817329 | 0.993188 | 0.002969 |
| ENSMUSG00000053317.12 | 1463.809 | -0.08139 | 0.365693 | -0.22257 | 0.823873 | 0.993188 | 0.002969 |
| ENSMUSG00000053334.7 | 25.74702 | 0.312847 | 1.440596 | 0.217165 | 0.82808 | 0.993188 | 0.002969 |
| ENSMUSG00000053931.12 | 475.5662 | 0.08139 | 0.355452 | 0.228975 | 0.818889 | 0.993188 | 0.002969 |
| ENSMUSG00000054079.13 | 52.69166 | 0.214355 | 0.982731 | 0.218122 | 0.827334 | 0.993188 | 0.002969 |
| ENSMUSG00000054115.13 | 21.29643 | -0.5616 | 2.371309 | -0.23683 | 0.812788 | 0.993188 | 0.002969 |
| ENSMUSG00000056234.16 | 23.44629 | -0.4373 | 1.966535 | -0.22237 | 0.824027 | 0.993188 | 0.002969 |
| ENSMUSG00000057322.13 | 939.9886 | 0.071003 | 0.300105 | 0.236593 | 0.812973 | 0.993188 | 0.002969 |

| | | | | | | | |
|---|---|---|---|---|---|---|---|
| ENSMUSG00000057400.15 | 8628.974 | 0.044729 | 0.207513 | 0.215548 | 0.82934 | 0.993188 | 0.002969 |
| ENSMUSG00000058318.16 | 50.36675 | -0.26958 | 1.117434 | -0.24125 | 0.809363 | 0.993188 | 0.002969 |
| ENSMUSG00000060771.16 | 14.61109 | 0.511306 | 2.260149 | 0.226227 | 0.821025 | 0.993188 | 0.002969 |
| ENSMUSG00000061111.9 | 45.59221 | 0.427767 | 1.945919 | 0.219828 | 0.826005 | 0.993188 | 0.002969 |
| ENSMUSG00000061118.9 | 101.09 | -0.22307 | 0.943622 | -0.23639 | 0.813127 | 0.993188 | 0.002969 |
| ENSMUSG00000063800.6 | 39.26206 | -0.28251 | 1.187175 | -0.23796 | 0.811909 | 0.993188 | 0.002969 |
| ENSMUSG00000066613.15 | 29.22993 | -0.34736 | 1.619573 | -0.21448 | 0.830174 | 0.993188 | 0.002969 |
| ENSMUSG00000066637.3 | 195.7122 | 0.118569 | 0.538876 | 0.22003 | 0.825848 | 0.993188 | 0.002969 |
| ENSMUSG00000066735.9 | 174.8235 | 0.181913 | 0.755218 | 0.240875 | 0.809652 | 0.993188 | 0.002969 |
| ENSMUSG00000066878.6 | 20.47541 | -0.32286 | 1.413628 | -0.22839 | 0.81934 | 0.993188 | 0.002969 |
| ENSMUSG00000070284.11 | 28.50688 | 0.390163 | 1.778883 | 0.21933 | 0.826393 | 0.993188 | 0.002969 |
| ENSMUSG00000071177.5 | 33581.19 | 0.045123 | 0.207568 | 0.217391 | 0.827904 | 0.993188 | 0.002969 |
| ENSMUSG00000071178.12 | 40887.35 | -0.03902 | 0.18547 | -0.2104 | 0.833352 | 0.993188 | 0.002969 |
| ENSMUSG00000071253.9 | 324.3794 | 0.13931 | 0.608393 | 0.22898 | 0.818885 | 0.993188 | 0.002969 |
| ENSMUSG00000071644.11 | 455.0206 | -0.12483 | 0.550629 | -0.22671 | 0.820653 | 0.993188 | 0.002969 |
| ENSMUSG00000071711.13 | 50.71342 | -0.32316 | 1.506504 | -0.21451 | 0.830147 | 0.993188 | 0.002969 |
| ENSMUSG00000072582.10 | 110.6782 | 0.205047 | 0.868952 | 0.23597 | 0.813456 | 0.993188 | 0.002969 |
| ENSMUSG00000073489.7 | 50.10152 | -0.32762 | 1.515835 | -0.21613 | 0.828887 | 0.993188 | 0.002969 |
| ENSMUSG00000074102.5 | 17.20244 | 0.529728 | 2.241293 | 0.236349 | 0.813162 | 0.993188 | 0.002969 |
| ENSMUSG00000074748.4 | 359.8549 | -0.1161 | 0.508577 | -0.22828 | 0.81943 | 0.993188 | 0.002969 |
| ENSMUSG00000081058.6 | 16.24535 | -0.39774 | 1.870558 | -0.21263 | 0.831615 | 0.993188 | 0.002969 |
| ENSMUSG00000083087.2 | 27.47334 | 0.288025 | 1.366732 | 0.21074 | 0.83309 | 0.993188 | 0.002969 |
| ENSMUSG00000084936.2 | 14.62256 | 0.517525 | 2.419778 | 0.213873 | 0.830646 | 0.993188 | 0.002969 |
| ENSMUSG00000086382.2 | 34.95598 | -0.28114 | 1.259122 | -0.22329 | 0.823313 | 0.993188 | 0.002969 |
| ENSMUSG00000087165.8 | 13.48265 | -0.64615 | 2.683381 | -0.2408 | 0.809712 | 0.993188 | 0.002969 |
| ENSMUSG00000090330.3 | 17.63904 | 0.542354 | 2.234251 | 0.242745 | 0.808203 | 0.993188 | 0.002969 |
| ENSMUSG00000091119.3 | 3173.412 | 0.072111 | 0.340996 | 0.211471 | 0.83252 | 0.993188 | 0.002969 |
| ENSMUSG00000095304.10 | 20.0667 | -0.40082 | 1.843443 | -0.21743 | 0.827873 | 0.993188 | 0.002969 |
| ENSMUSG00000096546.3 | 492.6456 | -0.09245 | 0.432557 | -0.21373 | 0.830754 | 0.993188 | 0.002969 |
| ENSMUSG00000096768.9 | 31.74657 | -0.4419 | 1.902387 | -0.23229 | 0.816316 | 0.993188 | 0.002969 |
| ENSMUSG00000097321.6 | 33.95805 | 0.293587 | 1.224322 | 0.239796 | 0.810489 | 0.993188 | 0.002969 |
| ENSMUSG00000100510.8 | 28.13222 | 0.429968 | 1.94435 | 0.221137 | 0.824986 | 0.993188 | 0.002969 |
| ENSMUSG00000100865.3 | 58.1357 | -0.20663 | 0.866368 | -0.2385 | 0.811493 | 0.993188 | 0.002969 |
| ENSMUSG00000101795.3 | 52.9221 | 0.234015 | 0.992014 | 0.235899 | 0.813511 | 0.993188 | 0.002969 |
| ENSMUSG00000109093.2 | 142.0772 | 0.1513 | 0.700006 | 0.21614 | 0.828878 | 0.993188 | 0.002969 |
| ENSMUSG00000110899.2 | 15.38373 | 0.450265 | 2.011086 | 0.223892 | 0.822842 | 0.993188 | 0.002969 |
| ENSMUSG00000111709.2 | 11.20698 | 0.683453 | 3.237693 | 0.211093 | 0.832815 | 0.993188 | 0.002969 |
| ENSMUSG00000116903.2 | 39.83214 | 0.359658 | 1.64719 | 0.218346 | 0.827159 | 0.993188 | 0.002969 |
| ENSMUSG00000119584.1 | 440.3654 | -0.12387 | 0.577361 | -0.21454 | 0.830127 | 0.993188 | 0.002969 |
| ENSMUSG00000030249.16 | 97.82171 | 0.157808 | 0.752987 | 0.209576 | 0.833998 | 0.993379 | 0.002885 |
| ENSMUSG00000036026.16 | 12.58579 | 0.51249 | 2.449336 | 0.209236 | 0.834264 | 0.99357 | 0.002801 |
| ENSMUSG00000043059.17 | 10.86943 | -0.55271 | 2.645145 | -0.20895 | 0.834486 | 0.99371 | 0.00274 |
| ENSMUSG00000007892.9 | 4907.312 | -0.05085 | 0.246946 | -0.20591 | 0.836858 | 0.993713 | 0.002739 |
| ENSMUSG00000009549.15 | 154.9962 | -0.14644 | 0.703646 | -0.20811 | 0.835141 | 0.993713 | 0.002739 |

| | | | | | | | |
|---|---|---|---|---|---|---|---|
| ENSMUSG00000012640.17 | 35.21288 | 0.250564 | 1.211146 | 0.206882 | 0.836102 | 0.993713 | 0.002739 |
| ENSMUSG00000020609.15 | 9402.352 | -0.05984 | 0.290769 | -0.20579 | 0.836957 | 0.993713 | 0.002739 |
| ENSMUSG00000024236.19 | 101.4594 | -0.15184 | 0.729588 | -0.20812 | 0.835139 | 0.993713 | 0.002739 |
| ENSMUSG00000024902.6 | 211.7099 | 0.116519 | 0.566377 | 0.205727 | 0.837004 | 0.993713 | 0.002739 |
| ENSMUSG00000025025.15 | 59.42774 | -0.2557 | 1.225649 | -0.20863 | 0.834741 | 0.993713 | 0.002739 |
| ENSMUSG00000025888.7 | 13.5192 | 0.559257 | 2.710481 | 0.206331 | 0.836532 | 0.993713 | 0.002739 |
| ENSMUSG00000026608.14 | 26.17237 | 0.343826 | 1.649261 | 0.208473 | 0.83486 | 0.993713 | 0.002739 |
| ENSMUSG00000028163.18 | 72.00113 | -0.25222 | 1.221882 | -0.20642 | 0.836464 | 0.993713 | 0.002739 |
| ENSMUSG00000028649.22 | 34.67266 | -0.29509 | 1.421866 | -0.20753 | 0.835593 | 0.993713 | 0.002739 |
| ENSMUSG00000031537.16 | 162.949 | 0.128437 | 0.622808 | 0.206222 | 0.836618 | 0.993713 | 0.002739 |
| ENSMUSG00000032551.8 | 125.6843 | -0.15602 | 0.755623 | -0.20648 | 0.836413 | 0.993713 | 0.002739 |
| ENSMUSG00000033706.14 | 43.72545 | -0.34413 | 1.657695 | -0.20759 | 0.835546 | 0.993713 | 0.002739 |
| ENSMUSG00000035828.12 | 15.26138 | 0.549745 | 2.648641 | 0.207557 | 0.835575 | 0.993713 | 0.002739 |
| ENSMUSG00000035845.18 | 27.10116 | 0.267428 | 1.294083 | 0.206655 | 0.83628 | 0.993713 | 0.002739 |
| ENSMUSG00000043061.11 | 73.08245 | 0.177478 | 0.857819 | 0.206895 | 0.836092 | 0.993713 | 0.002739 |
| ENSMUSG00000046167.6 | 13.94326 | 0.552388 | 2.679902 | 0.206122 | 0.836695 | 0.993713 | 0.002739 |
| ENSMUSG00000047909.12 | 24.70724 | 0.284343 | 1.371557 | 0.207314 | 0.835765 | 0.993713 | 0.002739 |
| ENSMUSG00000053483.16 | 11.30706 | 0.454998 | 2.208264 | 0.206043 | 0.836757 | 0.993713 | 0.002739 |
| ENSMUSG00000053799.9 | 82.19178 | -0.18806 | 0.905079 | -0.20779 | 0.835396 | 0.993713 | 0.002739 |
| ENSMUSG00000055493.5 | 24.58581 | 0.311031 | 1.506538 | 0.206454 | 0.836436 | 0.993713 | 0.002739 |
| ENSMUSG00000062014.14 | 161.251 | -0.1278 | 0.620333 | -0.20601 | 0.83678 | 0.993713 | 0.002739 |
| ENSMUSG00000070939.10 | 71.86588 | 0.229246 | 1.101887 | 0.208048 | 0.835191 | 0.993713 | 0.002739 |
| ENSMUSG00000022961.19 | 844.0379 | -0.06 | 0.292454 | -0.20515 | 0.837457 | 0.993737 | 0.002729 |
| ENSMUSG00000027804.14 | 333.526 | -0.08679 | 0.422956 | -0.20521 | 0.837411 | 0.993737 | 0.002729 |
| ENSMUSG00000028330.9 | 84.94511 | 0.22176 | 1.081792 | 0.204993 | 0.837578 | 0.993737 | 0.002729 |
| ENSMUSG00000038903.14 | 14.06349 | -0.43886 | 2.141868 | -0.2049 | 0.837653 | 0.993737 | 0.002729 |
| ENSMUSG00000062480.13 | 984.9896 | -0.07279 | 0.354345 | -0.20543 | 0.83724 | 0.993737 | 0.002729 |
| ENSMUSG00000066829.8 | 34.51205 | -0.33232 | 1.621506 | -0.20495 | 0.837613 | 0.993737 | 0.002729 |
| ENSMUSG00000006304.15 | 475.7025 | -0.08235 | 0.402402 | -0.20465 | 0.837849 | 0.993845 | 0.002681 |
| ENSMUSG00000002346.17 | 81.38001 | 0.19865 | 0.976026 | 0.203529 | 0.838721 | 0.993927 | 0.002645 |
| ENSMUSG00000009418.17 | 17.82085 | 0.512619 | 2.53187 | 0.202467 | 0.839552 | 0.993927 | 0.002645 |
| ENSMUSG00000017677.12 | 91.44312 | -0.18995 | 0.938149 | -0.20247 | 0.839547 | 0.993927 | 0.002645 |
| ENSMUSG00000020956.15 | 50.86957 | 0.240988 | 1.191493 | 0.202257 | 0.839716 | 0.993927 | 0.002645 |
| ENSMUSG00000021591.9 | 141.9273 | -0.15036 | 0.743468 | -0.20224 | 0.839732 | 0.993927 | 0.002645 |
| ENSMUSG00000022634.10 | 186.9246 | 0.158403 | 0.781234 | 0.20276 | 0.839322 | 0.993927 | 0.002645 |
| ENSMUSG00000024472.10 | 64.50172 | 0.19251 | 0.957963 | 0.200958 | 0.840731 | 0.993927 | 0.002645 |
| ENSMUSG00000024807.19 | 61.6772 | 0.261348 | 1.295754 | 0.201696 | 0.840155 | 0.993927 | 0.002645 |
| ENSMUSG00000025139.15 | 124.5165 | -0.16741 | 0.833176 | -0.20094 | 0.840749 | 0.993927 | 0.002645 |
| ENSMUSG00000026655.16 | 265.0718 | -0.09981 | 0.492899 | -0.2025 | 0.839523 | 0.993927 | 0.002645 |
| ENSMUSG00000026676.8 | 11.17044 | -0.56771 | 2.797211 | -0.20296 | 0.839169 | 0.993927 | 0.002645 |
| ENSMUSG00000026980.16 | 11.84055 | 0.596257 | 2.930137 | 0.203491 | 0.838751 | 0.993927 | 0.002645 |
| ENSMUSG00000028063.16 | 15.98731 | 0.464719 | 2.309723 | 0.201201 | 0.840541 | 0.993927 | 0.002645 |
| ENSMUSG00000029272.9 | 18.92251 | 0.44431 | 2.195198 | 0.202401 | 0.839603 | 0.993927 | 0.002645 |
| ENSMUSG00000030432.13 | 893.8641 | -0.0824 | 0.408778 | -0.20157 | 0.840255 | 0.993927 | 0.002645 |

| | | | | | | | |
|---|---|---|---|---|---|---|---|
| ENSMUSG00000031467.11 | 47.10813 | 0.219609 | 1.080527 | 0.203242 | 0.838946 | 0.993927 | 0.002645 |
| ENSMUSG00000033554.18 | 82.91495 | 0.226328 | 1.108002 | 0.204267 | 0.838145 | 0.993927 | 0.002645 |
| ENSMUSG00000035704.18 | 24.07805 | 0.409305 | 2.033241 | 0.201306 | 0.840459 | 0.993927 | 0.002645 |
| ENSMUSG00000035726.9 | 77.49234 | 0.203474 | 1.001254 | 0.203219 | 0.838964 | 0.993927 | 0.002645 |
| ENSMUSG00000036721.15 | 66.50599 | 0.187015 | 0.928108 | 0.201501 | 0.840307 | 0.993927 | 0.002645 |
| ENSMUSG00000038072.15 | 36.33285 | 0.23749 | 1.18088 | 0.201113 | 0.84061 | 0.993927 | 0.002645 |
| ENSMUSG00000039450.12 | 114.798 | 0.136928 | 0.673837 | 0.203206 | 0.838974 | 0.993927 | 0.002645 |
| ENSMUSG00000048874.16 | 178.3035 | 0.144973 | 0.711359 | 0.203797 | 0.838512 | 0.993927 | 0.002645 |
| ENSMUSG00000054405.15 | 160.3692 | 0.198401 | 0.973911 | 0.203716 | 0.838575 | 0.993927 | 0.002645 |
| ENSMUSG00000057181.16 | 110.4347 | 0.181786 | 0.902147 | 0.201504 | 0.840304 | 0.993927 | 0.002645 |
| ENSMUSG00000067847.14 | 399.2915 | -0.09609 | 0.47563 | -0.20203 | 0.839894 | 0.993927 | 0.002645 |
| ENSMUSG00000085156.9 | 36.21045 | -0.30795 | 1.512861 | -0.20356 | 0.838699 | 0.993927 | 0.002645 |
| ENSMUSG00000002728.15 | 23.69911 | -0.29252 | 1.518079 | -0.19269 | 0.847203 | 0.994112 | 0.002565 |
| ENSMUSG00000002767.14 | 182.2145 | -0.11647 | 0.603718 | -0.19291 | 0.847027 | 0.994112 | 0.002565 |
| ENSMUSG00000006412.11 | 369.4723 | 0.08499 | 0.44103 | 0.192709 | 0.847187 | 0.994112 | 0.002565 |
| ENSMUSG00000007777.10 | 286.7027 | -0.12583 | 0.640922 | -0.19633 | 0.844356 | 0.994112 | 0.002565 |
| ENSMUSG00000008333.13 | 194.1382 | -0.12692 | 0.642194 | -0.19763 | 0.843332 | 0.994112 | 0.002565 |
| ENSMUSG00000008348.10 | 3071.191 | 0.060862 | 0.311774 | 0.195213 | 0.845226 | 0.994112 | 0.002565 |
| ENSMUSG00000019173.12 | 152.8757 | 0.130481 | 0.651273 | 0.200347 | 0.841209 | 0.994112 | 0.002565 |
| ENSMUSG00000019826.12 | 54.70597 | -0.17995 | 0.922889 | -0.19499 | 0.845402 | 0.994112 | 0.002565 |
| ENSMUSG00000020257.14 | 116.8943 | -0.13106 | 0.662592 | -0.19781 | 0.843196 | 0.994112 | 0.002565 |
| ENSMUSG00000020514.9 | 108.2484 | 0.173058 | 0.886954 | 0.195115 | 0.845303 | 0.994112 | 0.002565 |
| ENSMUSG00000021374.11 | 29.83068 | -0.35892 | 1.853402 | -0.19365 | 0.846448 | 0.994112 | 0.002565 |
| ENSMUSG00000021687.15 | 283.6591 | 0.108354 | 0.543691 | 0.199294 | 0.842033 | 0.994112 | 0.002565 |
| ENSMUSG00000021938.13 | 12.63004 | -0.41289 | 2.131995 | -0.19367 | 0.846438 | 0.994112 | 0.002565 |
| ENSMUSG00000022119.16 | 39.48199 | 0.229949 | 1.181585 | 0.194611 | 0.845698 | 0.994112 | 0.002565 |
| ENSMUSG00000022437.8 | 284.8234 | -0.09455 | 0.480049 | -0.19696 | 0.843862 | 0.994112 | 0.002565 |
| ENSMUSG00000022571.10 | 41.28709 | 0.297428 | 1.544578 | 0.192563 | 0.847301 | 0.994112 | 0.002565 |
| ENSMUSG00000024576.15 | 492.6985 | -0.07436 | 0.384728 | -0.19327 | 0.846746 | 0.994112 | 0.002565 |
| ENSMUSG00000024985.21 | 141.8292 | -0.14474 | 0.751234 | -0.19267 | 0.84722 | 0.994112 | 0.002565 |
| ENSMUSG00000024987.7 | 222.0562 | 0.168138 | 0.864244 | 0.194549 | 0.845746 | 0.994112 | 0.002565 |
| ENSMUSG00000025158.8 | 18.65102 | 0.447566 | 2.267414 | 0.197391 | 0.843522 | 0.994112 | 0.002565 |
| ENSMUSG00000025169.7 | 16.54514 | 0.444808 | 2.247101 | 0.197948 | 0.843086 | 0.994112 | 0.002565 |
| ENSMUSG00000025232.9 | 76.82311 | 0.200162 | 1.01365 | 0.197466 | 0.843463 | 0.994112 | 0.002565 |
| ENSMUSG00000025408.16 | 28.99628 | -0.33582 | 1.723621 | -0.19483 | 0.845524 | 0.994112 | 0.002565 |
| ENSMUSG00000026192.14 | 32.04798 | -0.27816 | 1.393654 | -0.19959 | 0.841801 | 0.994112 | 0.002565 |
| ENSMUSG00000026482.14 | 10.78651 | 0.523133 | 2.669145 | 0.195993 | 0.844616 | 0.994112 | 0.002565 |
| ENSMUSG00000026492.12 | 35.63706 | -0.33089 | 1.656108 | -0.1998 | 0.841636 | 0.994112 | 0.002565 |
| ENSMUSG00000027475.10 | 33.29211 | -0.30179 | 1.55947 | -0.19352 | 0.846552 | 0.994112 | 0.002565 |
| ENSMUSG00000028382.16 | 416.7315 | 0.074676 | 0.386729 | 0.193097 | 0.846883 | 0.994112 | 0.002565 |
| ENSMUSG00000028964.15 | 460.0168 | 0.080786 | 0.413542 | 0.19535 | 0.845119 | 0.994112 | 0.002565 |
| ENSMUSG00000029910.15 | 17.63197 | -0.41077 | 2.126569 | -0.19316 | 0.846834 | 0.994112 | 0.002565 |
| ENSMUSG00000029993.17 | 191.183 | -0.11126 | 0.558367 | -0.19926 | 0.84206 | 0.994112 | 0.002565 |
| ENSMUSG00000031007.3 | 192.3903 | -0.12463 | 0.629381 | -0.19802 | 0.843031 | 0.994112 | 0.002565 |

| | | | | | | | |
|---|---|---|---|---|---|---|---|
| ENSMUSG00000031173.14 | 4087.887 | -0.05833 | 0.293111 | -0.19901 | 0.842257 | 0.994112 | 0.002565 |
| ENSMUSG00000031446.15 | 459.637 | -0.09855 | 0.5015 | -0.19651 | 0.844211 | 0.994112 | 0.002565 |
| ENSMUSG00000031532.8 | 1141.35 | -0.05267 | 0.263066 | -0.20021 | 0.841316 | 0.994112 | 0.002565 |
| ENSMUSG00000032571.15 | 102.6881 | -0.15875 | 0.801592 | -0.19805 | 0.843007 | 0.994112 | 0.002565 |
| ENSMUSG00000033898.18 | 5447.236 | 0.057298 | 0.295546 | 0.193873 | 0.846275 | 0.994112 | 0.002565 |
| ENSMUSG00000034218.17 | 28.59656 | 0.402249 | 2.028046 | 0.198343 | 0.842777 | 0.994112 | 0.002565 |
| ENSMUSG00000034875.6 | 344.472 | -0.09363 | 0.485091 | -0.19302 | 0.846942 | 0.994112 | 0.002565 |
| ENSMUSG00000036879.16 | 133.1765 | -0.17092 | 0.860605 | -0.19861 | 0.842572 | 0.994112 | 0.002565 |
| ENSMUSG00000038375.16 | 610.8964 | -0.07886 | 0.40362 | -0.19539 | 0.845088 | 0.994112 | 0.002565 |
| ENSMUSG00000038485.7 | 11.02584 | 0.481011 | 2.491232 | 0.193082 | 0.846895 | 0.994112 | 0.002565 |
| ENSMUSG00000038704.13 | 53.17448 | -0.2043 | 1.034549 | -0.19747 | 0.843458 | 0.994112 | 0.002565 |
| ENSMUSG00000038965.18 | 102.3924 | 0.15136 | 0.771672 | 0.196145 | 0.844497 | 0.994112 | 0.002565 |
| ENSMUSG00000039068.17 | 90.70777 | -0.15004 | 0.761591 | -0.19701 | 0.843821 | 0.994112 | 0.002565 |
| ENSMUSG00000039285.13 | 242.1975 | -0.10229 | 0.517938 | -0.1975 | 0.843438 | 0.994112 | 0.002565 |
| ENSMUSG00000040007.9 | 23.9283 | -0.33313 | 1.720737 | -0.1936 | 0.84649 | 0.994112 | 0.002565 |
| ENSMUSG00000041225.17 | 125.9304 | 0.154035 | 0.796981 | 0.193273 | 0.846746 | 0.994112 | 0.002565 |
| ENSMUSG00000042198.13 | 250.1991 | 0.112731 | 0.562305 | 0.20048 | 0.841106 | 0.994112 | 0.002565 |
| ENSMUSG00000044502.17 | 145.6465 | -0.14837 | 0.770346 | -0.1926 | 0.847276 | 0.994112 | 0.002565 |
| ENSMUSG00000046668.10 | 66.19173 | -0.22192 | 1.15325 | -0.19243 | 0.847406 | 0.994112 | 0.002565 |
| ENSMUSG00000048170.15 | 137.7421 | 0.132143 | 0.664726 | 0.198793 | 0.842424 | 0.994112 | 0.002565 |
| ENSMUSG00000051748.3 | 13813.76 | -0.04698 | 0.244142 | -0.19243 | 0.847407 | 0.994112 | 0.002565 |
| ENSMUSG00000053460.9 | 104.5081 | -0.14088 | 0.71944 | -0.19581 | 0.844757 | 0.994112 | 0.002565 |
| ENSMUSG00000057230.15 | 21.3479 | -0.35258 | 1.815755 | -0.19418 | 0.846036 | 0.994112 | 0.002565 |
| ENSMUSG00000060098.12 | 63.61759 | -0.18064 | 0.919385 | -0.19648 | 0.844233 | 0.994112 | 0.002565 |
| ENSMUSG00000063179.14 | 105.266 | 0.214007 | 1.084023 | 0.197419 | 0.843499 | 0.994112 | 0.002565 |
| ENSMUSG00000072825.12 | 25.12628 | -0.33003 | 1.655502 | -0.19935 | 0.841987 | 0.994112 | 0.002565 |
| ENSMUSG00000072946.13 | 184.4116 | 0.133132 | 0.687117 | 0.193755 | 0.846368 | 0.994112 | 0.002565 |
| ENSMUSG00000073910.11 | 173.4381 | -0.12085 | 0.616944 | -0.19588 | 0.844703 | 0.994112 | 0.002565 |
| ENSMUSG00000092417.4 | 20.14044 | 0.404083 | 2.096542 | 0.192738 | 0.847164 | 0.994112 | 0.002565 |
| ENSMUSG00000102091.3 | 18.77025 | 0.297849 | 1.497088 | 0.198953 | 0.8423 | 0.994112 | 0.002565 |
| ENSMUSG00000000441.18 | 145.2177 | -0.13045 | 0.723196 | -0.18037 | 0.856858 | 0.99419 | 0.002531 |
| ENSMUSG00000002014.13 | 1418.499 | -0.04756 | 0.25582 | -0.1859 | 0.852525 | 0.99419 | 0.002531 |
| ENSMUSG00000002342.18 | 27.93496 | -0.30106 | 1.6109 | -0.18689 | 0.851745 | 0.99419 | 0.002531 |
| ENSMUSG00000003464.14 | 337.0426 | 0.099836 | 0.531023 | 0.188007 | 0.850871 | 0.99419 | 0.002531 |
| ENSMUSG00000004035.13 | 238.5174 | 0.145279 | 0.784844 | 0.185106 | 0.853146 | 0.99419 | 0.002531 |
| ENSMUSG00000004789.11 | 483.8655 | 0.074957 | 0.41217 | 0.18186 | 0.855693 | 0.99419 | 0.002531 |
| ENSMUSG00000007594.11 | 17.77506 | 0.351844 | 1.955737 | 0.179903 | 0.857228 | 0.99419 | 0.002531 |
| ENSMUSG00000009621.19 | 50.00334 | -0.28328 | 1.532469 | -0.18485 | 0.853346 | 0.99419 | 0.002531 |
| ENSMUSG00000010048.11 | 41.8915 | 0.260464 | 1.424765 | 0.182812 | 0.854945 | 0.99419 | 0.002531 |
| ENSMUSG00000016344.15 | 70.97842 | -0.17513 | 0.980741 | -0.17857 | 0.858275 | 0.99419 | 0.002531 |
| ENSMUSG00000017057.10 | 283.5908 | 0.089665 | 0.490215 | 0.182909 | 0.854869 | 0.99419 | 0.002531 |
| ENSMUSG00000017453.5 | 1070.312 | 0.050451 | 0.27064 | 0.186414 | 0.85212 | 0.99419 | 0.002531 |
| ENSMUSG00000017466.10 | 40.74827 | -0.28553 | 1.567402 | -0.18217 | 0.855449 | 0.99419 | 0.002531 |
| ENSMUSG00000017756.10 | 29.48748 | 0.310176 | 1.638127 | 0.189348 | 0.84982 | 0.99419 | 0.002531 |

| | | | | | | | |
|---|---|---|---|---|---|---|---|
| ENSMUSG00000020163.13 | 1140.857 | 0.062631 | 0.335558 | 0.186646 | 0.851938 | 0.99419 | 0.002531 |
| ENSMUSG00000020228.12 | 25.67576 | 0.417116 | 2.310347 | 0.180543 | 0.856727 | 0.99419 | 0.002531 |
| ENSMUSG00000020283.6 | 346.7944 | 0.089169 | 0.501611 | 0.177766 | 0.858907 | 0.99419 | 0.002531 |
| ENSMUSG00000020900.16 | 51.84858 | 0.327037 | 1.716597 | 0.190515 | 0.848906 | 0.99419 | 0.002531 |
| ENSMUSG00000021224.16 | 109.3258 | -0.12815 | 0.70096 | -0.18282 | 0.854936 | 0.99419 | 0.002531 |
| ENSMUSG00000021326.14 | 116.8234 | -0.14174 | 0.775106 | -0.18287 | 0.854903 | 0.99419 | 0.002531 |
| ENSMUSG00000021779.20 | 215.0426 | 0.107574 | 0.575101 | 0.187052 | 0.85162 | 0.99419 | 0.002531 |
| ENSMUSG00000022111.10 | 223.6763 | -0.09073 | 0.505584 | -0.17945 | 0.857581 | 0.99419 | 0.002531 |
| ENSMUSG00000022285.18 | 93.8968 | -0.14739 | 0.767365 | -0.19207 | 0.847688 | 0.99419 | 0.002531 |
| ENSMUSG00000022442.17 | 12.17914 | -0.46042 | 2.449431 | -0.18797 | 0.850899 | 0.99419 | 0.002531 |
| ENSMUSG00000022906.16 | 126.9488 | 0.136628 | 0.747543 | 0.18277 | 0.854979 | 0.99419 | 0.002531 |
| ENSMUSG00000022912.9 | 452.2933 | 0.079456 | 0.413453 | 0.192177 | 0.847603 | 0.99419 | 0.002531 |
| ENSMUSG00000024048.16 | 419.9765 | 0.078227 | 0.410929 | 0.190365 | 0.849023 | 0.99419 | 0.002531 |
| ENSMUSG00000024190.8 | 312.3776 | 0.152283 | 0.84049 | 0.181184 | 0.856223 | 0.99419 | 0.002531 |
| ENSMUSG00000024386.10 | 1245.82 | -0.06357 | 0.340014 | -0.18696 | 0.851688 | 0.99419 | 0.002531 |
| ENSMUSG00000024570.8 | 41.60006 | 0.299075 | 1.685832 | 0.177405 | 0.85919 | 0.99419 | 0.002531 |
| ENSMUSG00000025332.15 | 13.86084 | -0.39407 | 2.189346 | -0.17999 | 0.857158 | 0.99419 | 0.002531 |
| ENSMUSG00000025645.9 | 27.60774 | -0.29976 | 1.645377 | -0.18219 | 0.855437 | 0.99419 | 0.002531 |
| ENSMUSG00000026029.15 | 33.8929 | 0.285891 | 1.581868 | 0.18073 | 0.856579 | 0.99419 | 0.002531 |
| ENSMUSG00000026032.9 | 999.5654 | 0.05903 | 0.330928 | 0.178377 | 0.858427 | 0.99419 | 0.002531 |
| ENSMUSG00000026107.12 | 93.23892 | -0.16197 | 0.903127 | -0.17935 | 0.857665 | 0.99419 | 0.002531 |
| ENSMUSG00000026275.14 | 123.7976 | -0.15813 | 0.84752 | -0.18658 | 0.851988 | 0.99419 | 0.002531 |
| ENSMUSG00000026469.15 | 156.7111 | 0.118433 | 0.62021 | 0.190956 | 0.84856 | 0.99419 | 0.002531 |
| ENSMUSG00000026565.19 | 48.20885 | 0.236781 | 1.25055 | 0.189341 | 0.849825 | 0.99419 | 0.002531 |
| ENSMUSG00000026659.14 | 103.1772 | 0.14248 | 0.794222 | 0.179396 | 0.857627 | 0.99419 | 0.002531 |
| ENSMUSG00000026782.16 | 21.96652 | 0.356207 | 1.918899 | 0.185631 | 0.852734 | 0.99419 | 0.002531 |
| ENSMUSG00000026893.5 | 22.6416 | -0.2497 | 1.394989 | -0.179 | 0.85794 | 0.99419 | 0.002531 |
| ENSMUSG00000027076.11 | 135.5368 | -0.14539 | 0.792991 | -0.18334 | 0.854533 | 0.99419 | 0.002531 |
| ENSMUSG00000027303.19 | 115.1125 | 0.151745 | 0.814204 | 0.186372 | 0.852153 | 0.99419 | 0.002531 |
| ENSMUSG00000028018.16 | 24.33695 | -0.2886 | 1.523688 | -0.18941 | 0.84977 | 0.99419 | 0.002531 |
| ENSMUSG00000028132.16 | 460.2026 | -0.07507 | 0.40603 | -0.18489 | 0.853312 | 0.99419 | 0.002531 |
| ENSMUSG00000028194.16 | 576.3899 | -0.10101 | 0.554895 | -0.18204 | 0.855551 | 0.99419 | 0.002531 |
| ENSMUSG00000028223.9 | 1090.702 | 0.060893 | 0.322631 | 0.18874 | 0.850297 | 0.99419 | 0.002531 |
| ENSMUSG00000028469.16 | 86.13803 | -0.2047 | 1.144373 | -0.17887 | 0.858038 | 0.99419 | 0.002531 |
| ENSMUSG00000028572.14 | 394.9421 | 0.075841 | 0.426503 | 0.17782 | 0.858864 | 0.99419 | 0.002531 |
| ENSMUSG00000028576.13 | 40.77727 | 0.301873 | 1.611187 | 0.18736 | 0.851378 | 0.99419 | 0.002531 |
| ENSMUSG00000029290.13 | 101.3047 | 0.143515 | 0.809133 | 0.177369 | 0.859219 | 0.99419 | 0.002531 |
| ENSMUSG00000029319.9 | 287.141 | -0.09487 | 0.502287 | -0.18888 | 0.850188 | 0.99419 | 0.002531 |
| ENSMUSG00000030203.18 | 129.3324 | 0.145017 | 0.799609 | 0.18136 | 0.856085 | 0.99419 | 0.002531 |
| ENSMUSG00000030551.15 | 271.0817 | -0.10642 | 0.578691 | -0.18389 | 0.8541 | 0.99419 | 0.002531 |
| ENSMUSG00000031148.15 | 46.42828 | -0.23471 | 1.226659 | -0.19134 | 0.848257 | 0.99419 | 0.002531 |
| ENSMUSG00000031533.5 | 135.3411 | -0.11168 | 0.617402 | -0.18089 | 0.856456 | 0.99419 | 0.002531 |
| ENSMUSG00000031959.16 | 11.68372 | 0.402066 | 2.135287 | 0.188296 | 0.850645 | 0.99419 | 0.002531 |
| ENSMUSG00000032062.4 | 54.5275 | 0.238181 | 1.289698 | 0.18468 | 0.85348 | 0.99419 | 0.002531 |

| | | | | | | | |
|---|---|---|---|---|---|---|---|
| ENSMUSG00000033883.12 | 22.52783 | 0.479801 | 2.540126 | 0.188889 | 0.85018 | 0.99419 | 0.002531 |
| ENSMUSG00000033940.9 | 70.14612 | -0.18943 | 1.046169 | -0.18107 | 0.856315 | 0.99419 | 0.002531 |
| ENSMUSG00000035011.16 | 133.4254 | 0.158031 | 0.832555 | 0.189815 | 0.849454 | 0.99419 | 0.002531 |
| ENSMUSG00000035203.17 | 55.83104 | 0.17413 | 0.922127 | 0.188835 | 0.850222 | 0.99419 | 0.002531 |
| ENSMUSG00000036291.7 | 61.35185 | 0.170705 | 0.956292 | 0.178508 | 0.858324 | 0.99419 | 0.002531 |
| ENSMUSG00000037235.14 | 28.6351 | 0.290433 | 1.634519 | 0.177687 | 0.858969 | 0.99419 | 0.002531 |
| ENSMUSG00000038506.14 | 88.9289 | 0.145108 | 0.770043 | 0.188441 | 0.850531 | 0.99419 | 0.002531 |
| ENSMUSG00000038535.18 | 41.30386 | 0.320851 | 1.75516 | 0.182805 | 0.854951 | 0.99419 | 0.002531 |
| ENSMUSG00000038664.17 | 100.0512 | -0.12794 | 0.711247 | -0.17989 | 0.857243 | 0.99419 | 0.002531 |
| ENSMUSG00000038803.8 | 123.7788 | 0.124918 | 0.670044 | 0.186432 | 0.852106 | 0.99419 | 0.002531 |
| ENSMUSG00000039156.20 | 74.87352 | 0.241363 | 1.291991 | 0.186815 | 0.851806 | 0.99419 | 0.002531 |
| ENSMUSG00000039960.6 | 199.7054 | -0.1288 | 0.710634 | -0.18124 | 0.856178 | 0.99419 | 0.002531 |
| ENSMUSG00000039989.8 | 155.9328 | 0.123802 | 0.683392 | 0.181158 | 0.856244 | 0.99419 | 0.002531 |
| ENSMUSG00000040021.14 | 91.19136 | -0.15275 | 0.851883 | -0.1793 | 0.857699 | 0.99419 | 0.002531 |
| ENSMUSG00000040029.16 | 111.7975 | 0.143754 | 0.761674 | 0.188735 | 0.850301 | 0.99419 | 0.002531 |
| ENSMUSG00000040033.17 | 62.46272 | -0.18364 | 0.996555 | -0.18427 | 0.853799 | 0.99419 | 0.002531 |
| ENSMUSG00000040061.18 | 15.88492 | 0.54755 | 2.95753 | 0.185138 | 0.853121 | 0.99419 | 0.002531 |
| ENSMUSG00000040407.19 | 350.2932 | -0.1124 | 0.620353 | -0.18118 | 0.856224 | 0.99419 | 0.002531 |
| ENSMUSG00000040963.16 | 1006.811 | -0.06992 | 0.367373 | -0.19031 | 0.849065 | 0.99419 | 0.002531 |
| ENSMUSG00000041153.10 | 16.12459 | 0.552829 | 2.963477 | 0.186548 | 0.852015 | 0.99419 | 0.002531 |
| ENSMUSG00000044340.8 | 58.71564 | 0.164564 | 0.914279 | 0.179993 | 0.857158 | 0.99419 | 0.002531 |
| ENSMUSG00000045087.9 | 48.05529 | -0.18831 | 1.05684 | -0.17818 | 0.858581 | 0.99419 | 0.002531 |
| ENSMUSG00000046959.17 | 306.4313 | -0.0877 | 0.475019 | -0.18462 | 0.85353 | 0.99419 | 0.002531 |
| ENSMUSG00000048271.15 | 139.9921 | -0.16001 | 0.857022 | -0.18671 | 0.851891 | 0.99419 | 0.002531 |
| ENSMUSG00000049152.12 | 853.6553 | 0.060567 | 0.323771 | 0.187067 | 0.851608 | 0.99419 | 0.002531 |
| ENSMUSG00000049232.4 | 68.54405 | -0.20598 | 1.144045 | -0.18005 | 0.857117 | 0.99419 | 0.002531 |
| ENSMUSG00000050619.15 | 21.82273 | -0.32575 | 1.736308 | -0.18761 | 0.851181 | 0.99419 | 0.002531 |
| ENSMUSG00000050812.19 | 269.8578 | -0.11493 | 0.603537 | -0.19043 | 0.84897 | 0.99419 | 0.002531 |
| ENSMUSG00000053453.10 | 684.3591 | -0.06771 | 0.353691 | -0.19144 | 0.848182 | 0.99419 | 0.002531 |
| ENSMUSG00000053600.9 | 28.143 | 0.31236 | 1.631575 | 0.191447 | 0.848176 | 0.99419 | 0.002531 |
| ENSMUSG00000054517.10 | 10.35216 | 0.513535 | 2.88847 | 0.177788 | 0.85889 | 0.99419 | 0.002531 |
| ENSMUSG00000057406.17 | 44.15258 | -0.20467 | 1.134085 | -0.18047 | 0.856784 | 0.99419 | 0.002531 |
| ENSMUSG00000057894.11 | 24.03086 | 0.309673 | 1.670618 | 0.185365 | 0.852943 | 0.99419 | 0.002531 |
| ENSMUSG00000057963.10 | 11.13336 | 0.458814 | 2.49052 | 0.184224 | 0.853838 | 0.99419 | 0.002531 |
| ENSMUSG00000058355.9 | 452.7698 | 0.084149 | 0.456287 | 0.184421 | 0.853683 | 0.99419 | 0.002531 |
| ENSMUSG00000059363.11 | 102.9108 | 0.15478 | 0.83871 | 0.184545 | 0.853586 | 0.99419 | 0.002531 |
| ENSMUSG00000063362.14 | 164.768 | 0.117348 | 0.618761 | 0.18965 | 0.849583 | 0.99419 | 0.002531 |
| ENSMUSG00000063816.4 | 16.75392 | 0.278752 | 1.535787 | 0.181504 | 0.855972 | 0.99419 | 0.002531 |
| ENSMUSG00000064363.1 | 87122.05 | 0.04854 | 0.265143 | 0.183071 | 0.854742 | 0.99419 | 0.002531 |
| ENSMUSG00000066150.13 | 1560.906 | -0.04763 | 0.26585 | -0.17916 | 0.857813 | 0.99419 | 0.002531 |
| ENSMUSG00000067149.7 | 37.12664 | -0.28889 | 1.583534 | -0.18243 | 0.855244 | 0.99419 | 0.002531 |
| ENSMUSG00000068396.10 | 97.91517 | 0.135248 | 0.745372 | 0.18145 | 0.856015 | 0.99419 | 0.002531 |
| ENSMUSG00000073468.13 | 62.69543 | -0.18798 | 0.982004 | -0.19143 | 0.848191 | 0.99419 | 0.002531 |
| ENSMUSG00000074212.8 | 184.9749 | 0.116439 | 0.641356 | 0.181551 | 0.855935 | 0.99419 | 0.002531 |

| | | | | | | | |
|---|---|---|---|---|---|---|---|
| ENSMUSG00000074221.13 | 14.356 | 0.313934 | 1.661473 | 0.188949 | 0.850133 | 0.99419 | 0.002531 |
| ENSMUSG00000075703.17 | 168.3345 | 0.111078 | 0.605708 | 0.183386 | 0.854495 | 0.99419 | 0.002531 |
| ENSMUSG00000082543.2 | 27.35569 | -0.30124 | 1.654442 | -0.18208 | 0.855521 | 0.99419 | 0.002531 |
| ENSMUSG00000084819.2 | 31.64873 | 0.372889 | 1.945258 | 0.191691 | 0.847984 | 0.99419 | 0.002531 |
| ENSMUSG00000085945.4 | 13.66649 | -0.55768 | 3.049532 | -0.18287 | 0.854897 | 0.99419 | 0.002531 |
| ENSMUSG00000090935.11 | 849.3774 | -0.05479 | 0.302314 | -0.18123 | 0.856185 | 0.99419 | 0.002531 |
| ENSMUSG00000093674.8 | 4294.263 | 0.037364 | 0.209405 | 0.17843 | 0.858385 | 0.99419 | 0.002531 |
| ENSMUSG00000095526.3 | 69.73194 | 0.155891 | 0.878002 | 0.177552 | 0.859075 | 0.99419 | 0.002531 |
| ENSMUSG00000096971.3 | 19.9598 | -0.32429 | 1.771562 | -0.18306 | 0.854754 | 0.99419 | 0.002531 |
| ENSMUSG00000110185.3 | 115.7077 | -0.11195 | 0.629791 | -0.17775 | 0.858919 | 0.99419 | 0.002531 |
| ENSMUSG00000047694.13 | 185.5135 | 0.098246 | 0.554731 | 0.177106 | 0.859425 | 0.994307 | 0.00248 |
| ENSMUSG00000002885.15 | 60.72326 | 0.179919 | 1.01853 | 0.176645 | 0.859787 | 0.994532 | 0.002381 |
| ENSMUSG00000028567.9 | 325.4506 | 0.102116 | 0.5787 | 0.176458 | 0.859934 | 0.994532 | 0.002381 |
| ENSMUSG00000030779.17 | 369.899 | -0.08175 | 0.463221 | -0.17648 | 0.859917 | 0.994532 | 0.002381 |
| ENSMUSG00000028693.16 | 131.7589 | -0.13497 | 0.765491 | -0.17632 | 0.860046 | 0.99454 | 0.002378 |
| ENSMUSG00000006517.6 | 10.50631 | -0.38077 | 2.182112 | -0.1745 | 0.861475 | 0.994705 | 0.002306 |
| ENSMUSG00000020576.11 | 20.44831 | -0.36227 | 2.077353 | -0.17439 | 0.86156 | 0.994705 | 0.002306 |
| ENSMUSG00000021033.12 | 2449.67 | -0.04464 | 0.256538 | -0.17402 | 0.861847 | 0.994705 | 0.002306 |
| ENSMUSG00000022500.16 | 475.1331 | 0.071193 | 0.408122 | 0.174441 | 0.861519 | 0.994705 | 0.002306 |
| ENSMUSG00000022789.16 | 227.8243 | -0.1043 | 0.595977 | -0.17501 | 0.861071 | 0.994705 | 0.002306 |
| ENSMUSG00000024827.11 | 207.8895 | -0.10587 | 0.607679 | -0.17421 | 0.861698 | 0.994705 | 0.002306 |
| ENSMUSG00000025280.9 | 34.80979 | 0.357334 | 2.031199 | 0.175923 | 0.860354 | 0.994705 | 0.002306 |
| ENSMUSG00000028032.14 | 43.74247 | -0.18105 | 1.035622 | -0.17482 | 0.861221 | 0.994705 | 0.002306 |
| ENSMUSG00000028849.18 | 13.18127 | 0.306258 | 1.752055 | 0.174799 | 0.861238 | 0.994705 | 0.002306 |
| ENSMUSG00000030780.16 | 10.30152 | -0.43701 | 2.488035 | -0.17565 | 0.860572 | 0.994705 | 0.002306 |
| ENSMUSG00000039367.12 | 132.2942 | -0.13095 | 0.746481 | -0.17542 | 0.860749 | 0.994705 | 0.002306 |
| ENSMUSG00000040017.9 | 2074.556 | -0.06148 | 0.353272 | -0.17404 | 0.861837 | 0.994705 | 0.002306 |
| ENSMUSG00000049235.9 | 11.88453 | 0.305884 | 1.75798 | 0.173997 | 0.861868 | 0.994705 | 0.002306 |
| ENSMUSG00000060475.14 | 31.57185 | 0.264194 | 1.512329 | 0.174694 | 0.86132 | 0.994705 | 0.002306 |
| ENSMUSG00000063320.12 | 113.0192 | 0.154111 | 0.883764 | 0.17438 | 0.861567 | 0.994705 | 0.002306 |
| ENSMUSG00000070327.15 | 76.28838 | -0.17034 | 0.975853 | -0.17456 | 0.861426 | 0.994705 | 0.002306 |
| ENSMUSG00000000440.13 | 56.57729 | -0.15792 | 1.000225 | -0.15789 | 0.874547 | 0.994712 | 0.002303 |
| ENSMUSG00000001100.11 | 256.1027 | -0.10182 | 0.65541 | -0.15536 | 0.876538 | 0.994712 | 0.002303 |
| ENSMUSG00000001627.13 | 156.2608 | 0.159012 | 0.970805 | 0.163794 | 0.869894 | 0.994712 | 0.002303 |
| ENSMUSG00000001630.14 | 101.6728 | 0.132132 | 0.850322 | 0.15539 | 0.876514 | 0.994712 | 0.002303 |
| ENSMUSG00000001794.14 | 332.8709 | -0.06543 | 0.411932 | -0.15884 | 0.873796 | 0.994712 | 0.002303 |
| ENSMUSG00000002307.17 | 20.17015 | 0.318843 | 1.963279 | 0.162403 | 0.870988 | 0.994712 | 0.002303 |
| ENSMUSG00000002365.11 | 134.6539 | -0.11603 | 0.745498 | -0.15564 | 0.876321 | 0.994712 | 0.002303 |
| ENSMUSG00000005087.18 | 53.90985 | 0.172239 | 1.035052 | 0.166406 | 0.867838 | 0.994712 | 0.002303 |
| ENSMUSG00000005682.10 | 55.55999 | -0.22297 | 1.361647 | -0.16375 | 0.869927 | 0.994712 | 0.002303 |
| ENSMUSG00000006641.13 | 33.26222 | -0.32139 | 2.072373 | -0.15508 | 0.876758 | 0.994712 | 0.002303 |
| ENSMUSG00000008859.9 | 292.1332 | 0.08798 | 0.507924 | 0.173214 | 0.862483 | 0.994712 | 0.002303 |
| ENSMUSG00000009647.14 | 23.03756 | -0.31205 | 1.822784 | -0.1712 | 0.86407 | 0.994712 | 0.002303 |
| ENSMUSG00000014361.6 | 35.78943 | -0.1791 | 1.107572 | -0.16171 | 0.871538 | 0.994712 | 0.002303 |

| | | | | | | | |
|---|---|---|---|---|---|---|---|
| ENSMUSG00000015340.11 | 61.73829 | 0.196947 | 1.27835 | 0.154064 | 0.87756 | 0.994712 | 0.002303 |
| ENSMUSG00000016559.15 | 1233.238 | -0.04537 | 0.288561 | -0.15721 | 0.875075 | 0.994712 | 0.002303 |
| ENSMUSG00000017376.16 | 47.68536 | 0.216417 | 1.355755 | 0.159628 | 0.873174 | 0.994712 | 0.002303 |
| ENSMUSG00000018923.14 | 95.36452 | -0.15468 | 0.964295 | -0.16041 | 0.872561 | 0.994712 | 0.002303 |
| ENSMUSG00000020142.13 | 15.41557 | -0.40062 | 2.555535 | -0.15676 | 0.87543 | 0.994712 | 0.002303 |
| ENSMUSG00000020253.16 | 33.76632 | -0.21676 | 1.363898 | -0.15892 | 0.873728 | 0.994712 | 0.002303 |
| ENSMUSG00000020459.15 | 135.9793 | -0.14293 | 0.872178 | -0.16388 | 0.869829 | 0.994712 | 0.002303 |
| ENSMUSG00000020605.10 | 37.28605 | -0.3216 | 1.903148 | -0.16898 | 0.86581 | 0.994712 | 0.002303 |
| ENSMUSG00000020689.5 | 10.46038 | 0.514237 | 3.252649 | 0.158098 | 0.87438 | 0.994712 | 0.002303 |
| ENSMUSG00000020718.13 | 27.72129 | 0.227882 | 1.319259 | 0.172735 | 0.86286 | 0.994712 | 0.002303 |
| ENSMUSG00000020806.16 | 11.12804 | -0.45832 | 2.836662 | -0.16157 | 0.871645 | 0.994712 | 0.002303 |
| ENSMUSG00000020988.10 | 340.7412 | 0.070531 | 0.453176 | 0.155638 | 0.876318 | 0.994712 | 0.002303 |
| ENSMUSG00000021048.8 | 855.3449 | 0.068178 | 0.403615 | 0.168919 | 0.865861 | 0.994712 | 0.002303 |
| ENSMUSG00000021051.11 | 268.6239 | 0.086492 | 0.526593 | 0.164248 | 0.869536 | 0.994712 | 0.002303 |
| ENSMUSG00000021140.10 | 23.10099 | 0.270256 | 1.671756 | 0.16166 | 0.871574 | 0.994712 | 0.002303 |
| ENSMUSG00000021213.13 | 746.0827 | 0.05278 | 0.338691 | 0.155835 | 0.876163 | 0.994712 | 0.002303 |
| ENSMUSG00000021458.18 | 94.67325 | -0.14721 | 0.88256 | -0.1668 | 0.867525 | 0.994712 | 0.002303 |
| ENSMUSG00000021643.16 | 13.08692 | 0.376244 | 2.341082 | 0.160714 | 0.872319 | 0.994712 | 0.002303 |
| ENSMUSG00000021731.11 | 154.0247 | 0.115703 | 0.740295 | 0.156293 | 0.875802 | 0.994712 | 0.002303 |
| ENSMUSG00000021936.15 | 118.5234 | 0.139841 | 0.87862 | 0.15916 | 0.873543 | 0.994712 | 0.002303 |
| ENSMUSG00000022201.17 | 401.6056 | 0.075346 | 0.464533 | 0.162197 | 0.871151 | 0.994712 | 0.002303 |
| ENSMUSG00000022307.17 | 489.4167 | 0.07766 | 0.487008 | 0.159464 | 0.873303 | 0.994712 | 0.002303 |
| ENSMUSG00000022400.10 | 681.9695 | -0.0562 | 0.35391 | -0.1588 | 0.873829 | 0.994712 | 0.002303 |
| ENSMUSG00000022579.11 | 15.92796 | -0.27591 | 1.777862 | -0.15519 | 0.876671 | 0.994712 | 0.002303 |
| ENSMUSG00000022702.17 | 31.79774 | 0.351538 | 2.206889 | 0.159291 | 0.87344 | 0.994712 | 0.002303 |
| ENSMUSG00000022760.18 | 33.70005 | -0.27186 | 1.686439 | -0.1612 | 0.871933 | 0.994712 | 0.002303 |
| ENSMUSG00000024312.11 | 89.53745 | 0.160605 | 0.946466 | 0.16969 | 0.865254 | 0.994712 | 0.002303 |
| ENSMUSG00000024346.7 | 60.19335 | -0.15567 | 0.939938 | -0.16562 | 0.868455 | 0.994712 | 0.002303 |
| ENSMUSG00000024679.11 | 14.64428 | 0.439726 | 2.617087 | 0.168021 | 0.866567 | 0.994712 | 0.002303 |
| ENSMUSG00000024726.11 | 57.33767 | 0.210043 | 1.222663 | 0.171791 | 0.863602 | 0.994712 | 0.002303 |
| ENSMUSG00000025006.19 | 64.72171 | 0.258279 | 1.612914 | 0.160132 | 0.872777 | 0.994712 | 0.002303 |
| ENSMUSG00000025155.16 | 140.1731 | -0.10098 | 0.592517 | -0.17042 | 0.864679 | 0.994712 | 0.002303 |
| ENSMUSG00000025197.10 | 2317.102 | 0.045882 | 0.264814 | 0.173262 | 0.862445 | 0.994712 | 0.002303 |
| ENSMUSG00000025481.13 | 4483.253 | 0.036604 | 0.2125 | 0.172254 | 0.863238 | 0.994712 | 0.002303 |
| ENSMUSG00000025532.14 | 38.6582 | 0.231161 | 1.477346 | 0.15647 | 0.875662 | 0.994712 | 0.002303 |
| ENSMUSG00000025616.15 | 268.2006 | -0.10301 | 0.597689 | -0.17235 | 0.863165 | 0.994712 | 0.002303 |
| ENSMUSG00000025724.13 | 508.2498 | -0.06138 | 0.375799 | -0.16333 | 0.870256 | 0.994712 | 0.002303 |
| ENSMUSG00000025743.15 | 94.27518 | -0.13351 | 0.796548 | -0.16761 | 0.866888 | 0.994712 | 0.002303 |
| ENSMUSG00000026036.18 | 100.284 | -0.15239 | 0.91839 | -0.16593 | 0.86821 | 0.994712 | 0.002303 |
| ENSMUSG00000026142.16 | 106.4526 | -0.11956 | 0.692179 | -0.17273 | 0.862863 | 0.994712 | 0.002303 |
| ENSMUSG00000026203.17 | 59.66185 | -0.18833 | 1.198163 | -0.15719 | 0.875098 | 0.994712 | 0.002303 |
| ENSMUSG00000026393.11 | 207.5618 | -0.1126 | 0.655111 | -0.17189 | 0.863528 | 0.994712 | 0.002303 |
| ENSMUSG00000026489.14 | 150.8653 | 0.143347 | 0.899158 | 0.159424 | 0.873335 | 0.994712 | 0.002303 |
| ENSMUSG00000026638.16 | 131.398 | -0.13109 | 0.788563 | -0.16624 | 0.867967 | 0.994712 | 0.002303 |

| | | | | | | | |
|---|---|---|---|---|---|---|---|
| ENSMUSG00000026715.13 | 6921.551 | -0.03421 | 0.206568 | -0.1656 | 0.868472 | 0.994712 | 0.002303 |
| ENSMUSG00000026923.16 | 55.46428 | 0.169288 | 0.987971 | 0.171349 | 0.863949 | 0.994712 | 0.002303 |
| ENSMUSG00000027078.15 | 20.51741 | -0.35529 | 2.126014 | -0.16711 | 0.86728 | 0.994712 | 0.002303 |
| ENSMUSG00000027173.8 | 321.9772 | -0.07503 | 0.438762 | -0.171 | 0.864227 | 0.994712 | 0.002303 |
| ENSMUSG00000027540.14 | 115.7626 | 0.122316 | 0.710483 | 0.17216 | 0.863312 | 0.994712 | 0.002303 |
| ENSMUSG00000027605.19 | 940.6992 | 0.07335 | 0.439685 | 0.166823 | 0.867509 | 0.994712 | 0.002303 |
| ENSMUSG00000027649.16 | 12.04065 | -0.34983 | 2.077531 | -0.16839 | 0.866278 | 0.994712 | 0.002303 |
| ENSMUSG00000028098.14 | 149.7808 | -0.09659 | 0.611977 | -0.15783 | 0.874593 | 0.994712 | 0.002303 |
| ENSMUSG00000028622.12 | 37.95238 | -0.27639 | 1.776381 | -0.15559 | 0.876354 | 0.994712 | 0.002303 |
| ENSMUSG00000028763.19 | 25.94504 | 0.391788 | 2.329231 | 0.168205 | 0.866422 | 0.994712 | 0.002303 |
| ENSMUSG00000029162.16 | 427.6506 | 0.088632 | 0.521312 | 0.170017 | 0.864997 | 0.994712 | 0.002303 |
| ENSMUSG00000029366.11 | 22.67609 | 0.256639 | 1.555818 | 0.164954 | 0.86898 | 0.994712 | 0.002303 |
| ENSMUSG00000030108.15 | 198.0239 | 0.106626 | 0.670535 | 0.159016 | 0.873656 | 0.994712 | 0.002303 |
| ENSMUSG00000030246.13 | 26.30817 | -0.37811 | 2.289889 | -0.16512 | 0.868847 | 0.994712 | 0.002303 |
| ENSMUSG00000030382.16 | 4130.041 | -0.03745 | 0.241485 | -0.15507 | 0.876763 | 0.994712 | 0.002303 |
| ENSMUSG00000030386.20 | 33.68133 | 0.200766 | 1.195241 | 0.167971 | 0.866606 | 0.994712 | 0.002303 |
| ENSMUSG00000030417.16 | 229.63 | 0.091034 | 0.589184 | 0.154509 | 0.877208 | 0.994712 | 0.002303 |
| ENSMUSG00000031095.16 | 112.2913 | 0.119224 | 0.771248 | 0.154586 | 0.877148 | 0.994712 | 0.002303 |
| ENSMUSG00000031365.14 | 10.25564 | 0.444936 | 2.636827 | 0.168739 | 0.866002 | 0.994712 | 0.002303 |
| ENSMUSG00000031928.16 | 12.04432 | 0.30088 | 1.903648 | 0.158055 | 0.874414 | 0.994712 | 0.002303 |
| ENSMUSG00000032324.12 | 26.12154 | 0.267869 | 1.542854 | 0.173619 | 0.862165 | 0.994712 | 0.002303 |
| ENSMUSG00000032633.13 | 45.0819 | 0.175664 | 1.031346 | 0.170325 | 0.864754 | 0.994712 | 0.002303 |
| ENSMUSG00000033319.6 | 78.72879 | -0.12827 | 0.83274 | -0.15404 | 0.877582 | 0.994712 | 0.002303 |
| ENSMUSG00000033543.18 | 358.9895 | 0.085274 | 0.526741 | 0.161889 | 0.871393 | 0.994712 | 0.002303 |
| ENSMUSG00000034120.19 | 513.4513 | -0.06097 | 0.359742 | -0.16948 | 0.865418 | 0.994712 | 0.002303 |
| ENSMUSG00000034290.10 | 207.8824 | -0.08443 | 0.531752 | -0.15877 | 0.873848 | 0.994712 | 0.002303 |
| ENSMUSG00000034543.16 | 21.83839 | 0.332253 | 2.017105 | 0.164718 | 0.869166 | 0.994712 | 0.002303 |
| ENSMUSG00000035142.19 | 233.4223 | 0.101284 | 0.622045 | 0.162823 | 0.870657 | 0.994712 | 0.002303 |
| ENSMUSG00000035278.10 | 27.96279 | 0.328497 | 1.905133 | 0.172428 | 0.863101 | 0.994712 | 0.002303 |
| ENSMUSG00000036202.17 | 136.1607 | 0.166745 | 0.962314 | 0.173275 | 0.862435 | 0.994712 | 0.002303 |
| ENSMUSG00000036333.12 | 131.3564 | -0.12167 | 0.70993 | -0.17139 | 0.863917 | 0.994712 | 0.002303 |
| ENSMUSG00000036932.15 | 302.4765 | -0.08246 | 0.489216 | -0.16855 | 0.866149 | 0.994712 | 0.002303 |
| ENSMUSG00000037204.8 | 26.58712 | 0.23697 | 1.482203 | 0.159877 | 0.872978 | 0.994712 | 0.002303 |
| ENSMUSG00000037487.8 | 348.9781 | -0.08628 | 0.521259 | -0.16552 | 0.868533 | 0.994712 | 0.002303 |
| ENSMUSG00000037822.13 | 651.33 | 0.06555 | 0.416204 | 0.157494 | 0.874855 | 0.994712 | 0.002303 |
| ENSMUSG00000038576.16 | 205.3014 | -0.11606 | 0.707201 | -0.16411 | 0.869647 | 0.994712 | 0.002303 |
| ENSMUSG00000039917.11 | 15.98434 | 0.354373 | 2.077521 | 0.170575 | 0.864558 | 0.994712 | 0.002303 |
| ENSMUSG00000040151.10 | 113.3159 | 0.13194 | 0.820065 | 0.160889 | 0.872181 | 0.994712 | 0.002303 |
| ENSMUSG00000040658.8 | 55.5341 | 0.222242 | 1.43098 | 0.155308 | 0.876579 | 0.994712 | 0.002303 |
| ENSMUSG00000041935.11 | 146.3394 | -0.10075 | 0.610673 | -0.16498 | 0.868958 | 0.994712 | 0.002303 |
| ENSMUSG00000042105.19 | 71.7602 | -0.15967 | 1.021576 | -0.1563 | 0.875796 | 0.994712 | 0.002303 |
| ENSMUSG00000042133.17 | 565.8613 | -0.06061 | 0.375554 | -0.16139 | 0.871789 | 0.994712 | 0.002303 |
| ENSMUSG00000043398.6 | 16.48876 | -0.37598 | 2.24472 | -0.16749 | 0.866982 | 0.994712 | 0.002303 |
| ENSMUSG00000043998.11 | 489.9257 | 0.072839 | 0.431327 | 0.168872 | 0.865897 | 0.994712 | 0.002303 |

| | | | | | | | |
|---|---|---|---|---|---|---|---|
| ENSMUSG00000044026.4 | 146.039 | -0.10158 | 0.63909 | -0.15894 | 0.873712 | 0.994712 | 0.002303 |
| ENSMUSG00000044378.13 | 34.98671 | 0.241082 | 1.514242 | 0.15921 | 0.873504 | 0.994712 | 0.002303 |
| ENSMUSG00000044408.8 | 453.7986 | -0.07253 | 0.462784 | -0.15673 | 0.875461 | 0.994712 | 0.002303 |
| ENSMUSG00000044792.8 | 71.23046 | 0.13801 | 0.868757 | 0.158859 | 0.87378 | 0.994712 | 0.002303 |
| ENSMUSG00000045071.15 | 21.05297 | -0.25547 | 1.606013 | -0.15907 | 0.873611 | 0.994712 | 0.002303 |
| ENSMUSG00000045659.19 | 18.94032 | -0.42695 | 2.466735 | -0.17308 | 0.862586 | 0.994712 | 0.002303 |
| ENSMUSG00000046070.10 | 356.9067 | 0.093462 | 0.588254 | 0.15888 | 0.873763 | 0.994712 | 0.002303 |
| ENSMUSG00000046230.12 | 56.22004 | 0.198696 | 1.176332 | 0.168912 | 0.865866 | 0.994712 | 0.002303 |
| ENSMUSG00000046318.17 | 37.2883 | -0.27442 | 1.655587 | -0.16575 | 0.868352 | 0.994712 | 0.002303 |
| ENSMUSG00000046679.17 | 36.00403 | -0.19059 | 1.160496 | -0.16423 | 0.869547 | 0.994712 | 0.002303 |
| ENSMUSG00000047141.6 | 57.68359 | 0.138929 | 0.901307 | 0.154141 | 0.877498 | 0.994712 | 0.002303 |
| ENSMUSG00000047714.15 | 164.5656 | -0.10719 | 0.64332 | -0.16662 | 0.867672 | 0.994712 | 0.002303 |
| ENSMUSG00000048410.18 | 30.07668 | -0.23232 | 1.368782 | -0.16973 | 0.865226 | 0.994712 | 0.002303 |
| ENSMUSG00000049858.9 | 302.7889 | -0.09077 | 0.548641 | -0.16545 | 0.868587 | 0.994712 | 0.002303 |
| ENSMUSG00000051306.8 | 11.77116 | -0.30356 | 1.877408 | -0.16169 | 0.871549 | 0.994712 | 0.002303 |
| ENSMUSG00000051674.16 | 109.2156 | -0.16864 | 1.069975 | -0.15762 | 0.87476 | 0.994712 | 0.002303 |
| ENSMUSG00000052446.18 | 71.76008 | 0.161759 | 1.022401 | 0.158215 | 0.874287 | 0.994712 | 0.002303 |
| ENSMUSG00000052520.8 | 4308.832 | -0.03794 | 0.246374 | -0.154 | 0.877613 | 0.994712 | 0.002303 |
| ENSMUSG00000053168.7 | 43.17649 | 0.266656 | 1.677957 | 0.158917 | 0.873734 | 0.994712 | 0.002303 |
| ENSMUSG00000054717.8 | 23.19882 | -0.26908 | 1.584703 | -0.1698 | 0.865167 | 0.994712 | 0.002303 |
| ENSMUSG00000055943.6 | 619.5094 | -0.06311 | 0.376196 | -0.16776 | 0.866768 | 0.994712 | 0.002303 |
| ENSMUSG00000056121.17 | 228.4268 | 0.116973 | 0.673443 | 0.173694 | 0.862106 | 0.994712 | 0.002303 |
| ENSMUSG00000056763.17 | 52.94598 | 0.181146 | 1.139722 | 0.158939 | 0.873717 | 0.994712 | 0.002303 |
| ENSMUSG00000057596.14 | 23.53759 | -0.23546 | 1.510762 | -0.15586 | 0.876145 | 0.994712 | 0.002303 |
| ENSMUSG00000057738.15 | 29.93399 | -0.21758 | 1.337661 | -0.16266 | 0.870787 | 0.994712 | 0.002303 |
| ENSMUSG00000059248.14 | 17.54564 | -0.36996 | 2.174262 | -0.17015 | 0.864889 | 0.994712 | 0.002303 |
| ENSMUSG00000059883.17 | 19.73898 | 0.299777 | 1.884716 | 0.159057 | 0.873624 | 0.994712 | 0.002303 |
| ENSMUSG00000060681.16 | 70.25676 | -0.14425 | 0.932417 | -0.1547 | 0.877054 | 0.994712 | 0.002303 |
| ENSMUSG00000060992.10 | 404.5256 | 0.072858 | 0.435186 | 0.167417 | 0.867042 | 0.994712 | 0.002303 |
| ENSMUSG00000061272.9 | 18.26523 | 0.261316 | 1.51029 | 0.173024 | 0.862633 | 0.994712 | 0.002303 |
| ENSMUSG00000061360.9 | 246.3419 | 0.097089 | 0.577324 | 0.168171 | 0.866449 | 0.994712 | 0.002303 |
| ENSMUSG00000063450.15 | 48.96452 | 0.17228 | 1.087039 | 0.158486 | 0.874074 | 0.994712 | 0.002303 |
| ENSMUSG00000068114.8 | 12.30357 | 0.416588 | 2.421551 | 0.172034 | 0.863411 | 0.994712 | 0.002303 |
| ENSMUSG00000068394.5 | 11.16874 | -0.44918 | 2.8332 | -0.15854 | 0.874029 | 0.994712 | 0.002303 |
| ENSMUSG00000070348.6 | 205.1716 | 0.112887 | 0.667517 | 0.169115 | 0.865706 | 0.994712 | 0.002303 |
| ENSMUSG00000073139.10 | 100.0045 | 0.13924 | 0.85287 | 0.163261 | 0.870313 | 0.994712 | 0.002303 |
| ENSMUSG00000073155.13 | 679.1761 | -0.05887 | 0.379921 | -0.15494 | 0.876867 | 0.994712 | 0.002303 |
| ENSMUSG00000076435.4 | 11.03136 | 0.400288 | 2.52011 | 0.158838 | 0.873797 | 0.994712 | 0.002303 |
| ENSMUSG00000079043.4 | 52.81122 | 0.207628 | 1.242244 | 0.16714 | 0.86726 | 0.994712 | 0.002303 |
| ENSMUSG00000085894.2 | 20.96091 | 0.340085 | 2.034761 | 0.167138 | 0.867262 | 0.994712 | 0.002303 |
| ENSMUSG00000087590.3 | 120.1317 | -0.10955 | 0.68239 | -0.16054 | 0.872453 | 0.994712 | 0.002303 |
| ENSMUSG00000095325.4 | 12.66951 | 0.364871 | 2.350459 | 0.155234 | 0.876637 | 0.994712 | 0.002303 |
| ENSMUSG00000098332.3 | 199.8605 | 0.103716 | 0.673042 | 0.154101 | 0.87753 | 0.994712 | 0.002303 |
| ENSMUSG00000104623.2 | 192.2983 | 0.089263 | 0.532668 | 0.167576 | 0.866917 | 0.994712 | 0.002303 |

| | | | | | | | |
|---|---|---|---|---|---|---|---|
| ENSMUSG00000106258.2 | 234.6965 | 0.078814 | 0.465321 | 0.169375 | 0.865502 | 0.994712 | 0.002303 |
| ENSMUSG00000106918.4 | 468.756 | 0.070319 | 0.448765 | 0.156695 | 0.875485 | 0.994712 | 0.002303 |
| ENSMUSG00000109336.2 | 20.34966 | -0.30117 | 1.812981 | -0.16612 | 0.868063 | 0.994712 | 0.002303 |
| ENSMUSG00000110277.3 | 18.15696 | 0.336461 | 2.150487 | 0.156458 | 0.875672 | 0.994712 | 0.002303 |
| ENSMUSG00000116993.2 | 116.3755 | -0.13757 | 0.846853 | -0.16245 | 0.870951 | 0.994712 | 0.002303 |
| ENSMUSG00000085172.2 | 16.97128 | -0.29815 | 1.943449 | -0.15341 | 0.878072 | 0.994994 | 0.00218 |
| ENSMUSG00000086587.8 | 63.95591 | -0.20304 | 1.323133 | -0.15345 | 0.87804 | 0.994994 | 0.00218 |
| ENSMUSG00000000804.15 | 187.9524 | 0.121881 | 0.810361 | 0.150403 | 0.880447 | 0.995045 | 0.002157 |
| ENSMUSG00000002332.17 | 656.7478 | 0.055758 | 0.372813 | 0.14956 | 0.881111 | 0.995045 | 0.002157 |
| ENSMUSG00000002658.10 | 58.21963 | 0.200351 | 1.3297 | 0.150674 | 0.880233 | 0.995045 | 0.002157 |
| ENSMUSG00000003068.18 | 25.38069 | -0.25936 | 1.710613 | -0.15162 | 0.879486 | 0.995045 | 0.002157 |
| ENSMUSG00000005836.11 | 32.25905 | -0.25458 | 1.712652 | -0.14865 | 0.881834 | 0.995045 | 0.002157 |
| ENSMUSG00000019923.15 | 222.4951 | 0.088547 | 0.591028 | 0.149819 | 0.880908 | 0.995045 | 0.002157 |
| ENSMUSG00000020152.8 | 557.6322 | -0.06238 | 0.407517 | -0.15307 | 0.878345 | 0.995045 | 0.002157 |
| ENSMUSG00000020272.9 | 14.89188 | -0.38395 | 2.58433 | -0.14857 | 0.881893 | 0.995045 | 0.002157 |
| ENSMUSG00000020527.15 | 25.65874 | 0.324438 | 2.140096 | 0.1516 | 0.879503 | 0.995045 | 0.002157 |
| ENSMUSG00000024073.17 | 125.5286 | 0.101172 | 0.677773 | 0.149272 | 0.881339 | 0.995045 | 0.002157 |
| ENSMUSG00000026927.18 | 103.0523 | -0.14367 | 0.952633 | -0.15081 | 0.880125 | 0.995045 | 0.002157 |
| ENSMUSG00000027438.15 | 14.9516 | -0.33574 | 2.251156 | -0.14914 | 0.881442 | 0.995045 | 0.002157 |
| ENSMUSG00000027981.15 | 17.82782 | -0.28219 | 1.860984 | -0.15163 | 0.879475 | 0.995045 | 0.002157 |
| ENSMUSG00000030452.17 | 147.2631 | -0.12901 | 0.848935 | -0.15197 | 0.879214 | 0.995045 | 0.002157 |
| ENSMUSG00000030766.16 | 22.35992 | -0.26533 | 1.756686 | -0.15104 | 0.879946 | 0.995045 | 0.002157 |
| ENSMUSG00000031591.15 | 185.9412 | 0.098135 | 0.649071 | 0.151194 | 0.879823 | 0.995045 | 0.002157 |
| ENSMUSG00000032058.15 | 38.20961 | -0.18521 | 1.221076 | -0.15168 | 0.879439 | 0.995045 | 0.002157 |
| ENSMUSG00000032860.6 | 54.17724 | -0.17143 | 1.144588 | -0.14978 | 0.880941 | 0.995045 | 0.002157 |
| ENSMUSG00000033009.16 | 33.47335 | 0.219349 | 1.433025 | 0.153067 | 0.878346 | 0.995045 | 0.002157 |
| ENSMUSG00000034390.17 | 57.6876 | 0.182751 | 1.195948 | 0.152809 | 0.878549 | 0.995045 | 0.002157 |
| ENSMUSG00000036352.17 | 35.56833 | 0.240159 | 1.601708 | 0.149939 | 0.880813 | 0.995045 | 0.002157 |
| ENSMUSG00000036639.13 | 93.75929 | -0.11852 | 0.796873 | -0.14873 | 0.88177 | 0.995045 | 0.002157 |
| ENSMUSG00000037001.11 | 19.72127 | 0.27083 | 1.793141 | 0.151037 | 0.879947 | 0.995045 | 0.002157 |
| ENSMUSG00000037519.18 | 48.073 | -0.23952 | 1.593633 | -0.1503 | 0.880532 | 0.995045 | 0.002157 |
| ENSMUSG00000037720.17 | 616.5457 | 0.065972 | 0.438172 | 0.150562 | 0.880321 | 0.995045 | 0.002157 |
| ENSMUSG00000038346.19 | 11.26945 | 0.381962 | 2.502228 | 0.152649 | 0.878675 | 0.995045 | 0.002157 |
| ENSMUSG00000039756.13 | 140.6983 | -0.10576 | 0.701817 | -0.15069 | 0.880218 | 0.995045 | 0.002157 |
| ENSMUSG00000040687.17 | 24.92244 | -0.31328 | 2.068437 | -0.15146 | 0.879615 | 0.995045 | 0.002157 |
| ENSMUSG00000054720.13 | 13.96462 | 0.319622 | 2.108632 | 0.151578 | 0.87952 | 0.995045 | 0.002157 |
| ENSMUSG00000057375.14 | 172.7634 | 0.114634 | 0.762937 | 0.150254 | 0.880564 | 0.995045 | 0.002157 |
| ENSMUSG00000061859.18 | 29.98859 | -0.23912 | 1.605661 | -0.14892 | 0.881613 | 0.995045 | 0.002157 |
| ENSMUSG00000063052.10 | 124.5595 | -0.17366 | 1.162726 | -0.14936 | 0.881273 | 0.995045 | 0.002157 |
| ENSMUSG00000066513.5 | 47.53665 | 0.226463 | 1.517044 | 0.149279 | 0.881333 | 0.995045 | 0.002157 |
| ENSMUSG00000068566.14 | 44.62711 | 0.176574 | 1.18852 | 0.148567 | 0.881896 | 0.995045 | 0.002157 |
| ENSMUSG00000090625.2 | 188.9154 | 0.095863 | 0.636434 | 0.150626 | 0.880271 | 0.995045 | 0.002157 |
| ENSMUSG00000101013.2 | 58.48757 | -0.13975 | 0.92208 | -0.15156 | 0.879534 | 0.995045 | 0.002157 |
| ENSMUSG00000024983.11 | 27.7884 | -0.27823 | 1.87574 | -0.14833 | 0.882084 | 0.995089 | 0.002138 |

| | | | | | | | |
|---|---|---|---|---|---|---|---|
| ENSMUSG00000028218.20 | 92.37149 | -0.11465 | 0.773334 | -0.14825 | 0.882144 | 0.995089 | 0.002138 |
| ENSMUSG00000000301.17 | 1071.517 | -0.05595 | 0.392936 | -0.14239 | 0.886769 | 0.995175 | 0.0021 |
| ENSMUSG00000001380.8 | 201.9745 | 0.091311 | 0.665182 | 0.137272 | 0.890816 | 0.995175 | 0.0021 |
| ENSMUSG00000004843.8 | 230.4437 | -0.08354 | 0.600371 | -0.13915 | 0.88933 | 0.995175 | 0.0021 |
| ENSMUSG00000006599.14 | 195.4595 | -0.07301 | 0.516184 | -0.14145 | 0.887518 | 0.995175 | 0.0021 |
| ENSMUSG00000008140.19 | 154.979 | 0.097838 | 0.667558 | 0.146561 | 0.883479 | 0.995175 | 0.0021 |
| ENSMUSG00000009646.14 | 234.8895 | 0.079203 | 0.547152 | 0.144755 | 0.884904 | 0.995175 | 0.0021 |
| ENSMUSG00000014769.11 | 1768.665 | -0.03647 | 0.256664 | -0.1421 | 0.886997 | 0.995175 | 0.0021 |
| ENSMUSG00000015355.14 | 23.78027 | 0.310817 | 2.183824 | 0.142327 | 0.886822 | 0.995175 | 0.0021 |
| ENSMUSG00000017686.18 | 179.5897 | 0.092512 | 0.656246 | 0.140971 | 0.887892 | 0.995175 | 0.0021 |
| ENSMUSG00000017754.14 | 94.49419 | -0.1391 | 0.993534 | -0.14 | 0.88866 | 0.995175 | 0.0021 |
| ENSMUSG00000019861.16 | 159.7811 | 0.087002 | 0.597883 | 0.145516 | 0.884303 | 0.995175 | 0.0021 |
| ENSMUSG00000019873.9 | 453.5334 | 0.062724 | 0.446585 | 0.140453 | 0.888302 | 0.995175 | 0.0021 |
| ENSMUSG00000019997.12 | 27.12054 | -0.28391 | 2.005618 | -0.14156 | 0.887431 | 0.995175 | 0.0021 |
| ENSMUSG00000020182.17 | 238.8293 | -0.08074 | 0.559749 | -0.14424 | 0.885312 | 0.995175 | 0.0021 |
| ENSMUSG00000020430.13 | 26.8787 | -0.24302 | 1.707492 | -0.14232 | 0.886825 | 0.995175 | 0.0021 |
| ENSMUSG00000021376.12 | 459.2342 | -0.07383 | 0.517268 | -0.14273 | 0.886505 | 0.995175 | 0.0021 |
| ENSMUSG00000021392.9 | 122.6349 | -0.09856 | 0.709381 | -0.13894 | 0.889496 | 0.995175 | 0.0021 |
| ENSMUSG00000021431.16 | 104.3899 | -0.12353 | 0.872329 | -0.14161 | 0.887389 | 0.995175 | 0.0021 |
| ENSMUSG00000021519.12 | 95.24374 | 0.126476 | 0.900901 | 0.140388 | 0.888354 | 0.995175 | 0.0021 |
| ENSMUSG00000023044.3 | 594.0438 | 0.054254 | 0.394006 | 0.137698 | 0.890479 | 0.995175 | 0.0021 |
| ENSMUSG00000023068.17 | 371.6128 | -0.05708 | 0.408573 | -0.1397 | 0.888899 | 0.995175 | 0.0021 |
| ENSMUSG00000023074.12 | 37.96129 | 0.206053 | 1.411265 | 0.146006 | 0.883917 | 0.995175 | 0.0021 |
| ENSMUSG00000023942.16 | 294.6153 | 0.108178 | 0.738902 | 0.146404 | 0.883603 | 0.995175 | 0.0021 |
| ENSMUSG00000024382.14 | 62.67939 | -0.22044 | 1.597168 | -0.13802 | 0.890223 | 0.995175 | 0.0021 |
| ENSMUSG00000025505.17 | 24.04933 | -0.29407 | 2.062594 | -0.14257 | 0.886627 | 0.995175 | 0.0021 |
| ENSMUSG00000026004.16 | 28.91183 | 0.225906 | 1.596631 | 0.141489 | 0.887484 | 0.995175 | 0.0021 |
| ENSMUSG00000026222.17 | 21.4826 | -0.29298 | 1.983478 | -0.14771 | 0.882571 | 0.995175 | 0.0021 |
| ENSMUSG00000028114.17 | 44.6247 | 0.188518 | 1.288981 | 0.146253 | 0.883721 | 0.995175 | 0.0021 |
| ENSMUSG00000028248.16 | 300.2492 | -0.07541 | 0.528259 | -0.14275 | 0.886485 | 0.995175 | 0.0021 |
| ENSMUSG00000028527.19 | 390.1468 | 0.064153 | 0.449622 | 0.142683 | 0.886541 | 0.995175 | 0.0021 |
| ENSMUSG00000028657.15 | 133.1266 | 0.124287 | 0.85152 | 0.145959 | 0.883954 | 0.995175 | 0.0021 |
| ENSMUSG00000028743.8 | 226.7117 | -0.10742 | 0.732293 | -0.14669 | 0.883376 | 0.995175 | 0.0021 |
| ENSMUSG00000029062.15 | 164.3119 | 0.125603 | 0.901931 | 0.13926 | 0.889245 | 0.995175 | 0.0021 |
| ENSMUSG00000029465.15 | 235.5879 | 0.07173 | 0.509728 | 0.140723 | 0.888089 | 0.995175 | 0.0021 |
| ENSMUSG00000029727.8 | 857.7572 | -0.0456 | 0.317661 | -0.14355 | 0.885858 | 0.995175 | 0.0021 |
| ENSMUSG00000029922.16 | 81.04181 | 0.15474 | 1.076465 | 0.143749 | 0.885699 | 0.995175 | 0.0021 |
| ENSMUSG00000030007.9 | 169.9753 | 0.090292 | 0.628133 | 0.143746 | 0.885701 | 0.995175 | 0.0021 |
| ENSMUSG00000030654.10 | 1743.918 | -0.03943 | 0.282628 | -0.1395 | 0.889056 | 0.995175 | 0.0021 |
| ENSMUSG00000031536.7 | 45.33092 | 0.195637 | 1.389665 | 0.14078 | 0.888044 | 0.995175 | 0.0021 |
| ENSMUSG00000031697.13 | 14.25552 | 0.312341 | 2.264223 | 0.137946 | 0.890283 | 0.995175 | 0.0021 |
| ENSMUSG00000031953.3 | 26.54521 | 0.218979 | 1.575114 | 0.139024 | 0.889431 | 0.995175 | 0.0021 |
| ENSMUSG00000032175.12 | 27.10062 | -0.23871 | 1.673816 | -0.14262 | 0.886593 | 0.995175 | 0.0021 |
| ENSMUSG00000032735.16 | 46.10148 | -0.1578 | 1.108781 | -0.14232 | 0.886831 | 0.995175 | 0.0021 |

| | | | | | | | |
|---|---|---|---|---|---|---|---|
| ENSMUSG00000032872.17 | 67.59046 | 0.145841 | 1.030585 | 0.141513 | 0.887465 | 0.995175 | 0.0021 |
| ENSMUSG00000032932.16 | 195.3346 | -0.0866 | 0.622939 | -0.13902 | 0.889438 | 0.995175 | 0.0021 |
| ENSMUSG00000033909.18 | 19.35575 | 0.283136 | 1.965466 | 0.144056 | 0.885457 | 0.995175 | 0.0021 |
| ENSMUSG00000033985.18 | 87.11143 | -0.15141 | 1.043447 | -0.14511 | 0.884628 | 0.995175 | 0.0021 |
| ENSMUSG00000034083.17 | 16.66503 | 0.269117 | 1.875651 | 0.143479 | 0.885912 | 0.995175 | 0.0021 |
| ENSMUSG00000035595.7 | 10.47652 | 0.417341 | 2.897132 | 0.144053 | 0.885459 | 0.995175 | 0.0021 |
| ENSMUSG00000038291.17 | 115.5992 | -0.12506 | 0.907888 | -0.13775 | 0.890439 | 0.995175 | 0.0021 |
| ENSMUSG00000038618.13 | 36.30707 | 0.222969 | 1.582776 | 0.140872 | 0.887971 | 0.995175 | 0.0021 |
| ENSMUSG00000039004.7 | 31.65166 | -0.23244 | 1.617739 | -0.14368 | 0.885754 | 0.995175 | 0.0021 |
| ENSMUSG00000039458.16 | 91.45143 | 0.156159 | 1.093674 | 0.142784 | 0.886461 | 0.995175 | 0.0021 |
| ENSMUSG00000040652.17 | 47.67238 | -0.16406 | 1.186821 | -0.13823 | 0.890057 | 0.995175 | 0.0021 |
| ENSMUSG00000040720.16 | 84.18076 | 0.147635 | 1.038532 | 0.142158 | 0.886955 | 0.995175 | 0.0021 |
| ENSMUSG00000040731.14 | 752.6386 | -0.04555 | 0.321616 | -0.14164 | 0.887367 | 0.995175 | 0.0021 |
| ENSMUSG00000041180.14 | 18.85096 | 0.26725 | 1.91194 | 0.13978 | 0.888834 | 0.995175 | 0.0021 |
| ENSMUSG00000041632.8 | 37.1876 | -0.21388 | 1.463782 | -0.14611 | 0.883832 | 0.995175 | 0.0021 |
| ENSMUSG00000042118.8 | 759.4118 | 0.054007 | 0.370862 | 0.145627 | 0.884216 | 0.995175 | 0.0021 |
| ENSMUSG00000042298.19 | 108.2341 | -0.10129 | 0.722846 | -0.14012 | 0.888565 | 0.995175 | 0.0021 |
| ENSMUSG00000045078.13 | 37.53624 | -0.18676 | 1.347548 | -0.1386 | 0.88977 | 0.995175 | 0.0021 |
| ENSMUSG00000045257.11 | 23.18995 | 0.233761 | 1.701702 | 0.137369 | 0.890739 | 0.995175 | 0.0021 |
| ENSMUSG00000047466.11 | 40.32125 | 0.211568 | 1.467717 | 0.144147 | 0.885384 | 0.995175 | 0.0021 |
| ENSMUSG00000048249.15 | 85.59534 | -0.16329 | 1.179959 | -0.13838 | 0.889938 | 0.995175 | 0.0021 |
| ENSMUSG00000049076.13 | 70.92205 | 0.147386 | 1.022637 | 0.144123 | 0.885403 | 0.995175 | 0.0021 |
| ENSMUSG00000050565.17 | 418.244 | -0.058 | 0.404632 | -0.14334 | 0.886021 | 0.995175 | 0.0021 |
| ENSMUSG00000051403.10 | 24.92965 | 0.232631 | 1.626648 | 0.143013 | 0.88628 | 0.995175 | 0.0021 |
| ENSMUSG00000052516.20 | 12.2423 | 0.341417 | 2.360699 | 0.144625 | 0.885007 | 0.995175 | 0.0021 |
| ENSMUSG00000053470.14 | 78.60038 | -0.12454 | 0.899094 | -0.13852 | 0.88983 | 0.995175 | 0.0021 |
| ENSMUSG00000054823.18 | 210.4781 | -0.07612 | 0.554622 | -0.13726 | 0.890829 | 0.995175 | 0.0021 |
| ENSMUSG00000057315.15 | 15.95065 | -0.28398 | 1.975695 | -0.14374 | 0.885709 | 0.995175 | 0.0021 |
| ENSMUSG00000063730.14 | 63.57741 | 0.141398 | 1.001718 | 0.141155 | 0.887747 | 0.995175 | 0.0021 |
| ENSMUSG00000066363.13 | 11.68154 | 0.382012 | 2.780525 | 0.137388 | 0.890724 | 0.995175 | 0.0021 |
| ENSMUSG00000068039.14 | 870.964 | -0.0534 | 0.363201 | -0.14702 | 0.88312 | 0.995175 | 0.0021 |
| ENSMUSG00000069744.8 | 380.6907 | -0.08293 | 0.575452 | -0.14411 | 0.885417 | 0.995175 | 0.0021 |
| ENSMUSG00000071660.11 | 57.94334 | -0.17315 | 1.220402 | -0.14188 | 0.887178 | 0.995175 | 0.0021 |
| ENSMUSG00000072294.6 | 50.05676 | -0.17463 | 1.266844 | -0.13785 | 0.89036 | 0.995175 | 0.0021 |
| ENSMUSG00000074165.11 | 10.27569 | 0.481095 | 3.295101 | 0.146003 | 0.883919 | 0.995175 | 0.0021 |
| ENSMUSG00000074491.10 | 258.5302 | -0.08229 | 0.559713 | -0.14702 | 0.88312 | 0.995175 | 0.0021 |
| ENSMUSG00000079083.4 | 10.91325 | 0.415044 | 2.952226 | 0.140587 | 0.888196 | 0.995175 | 0.0021 |
| ENSMUSG00000089678.9 | 776.6831 | 0.05397 | 0.387566 | 0.139253 | 0.88925 | 0.995175 | 0.0021 |
| ENSMUSG00000090136.5 | 70.64951 | -0.12262 | 0.840835 | -0.14584 | 0.884051 | 0.995175 | 0.0021 |
| ENSMUSG00000038742.8 | 32.10528 | 0.20566 | 1.501256 | 0.136992 | 0.891037 | 0.995291 | 0.00205 |
| ENSMUSG00000028712.13 | 26.20351 | -0.22719 | 1.664051 | -0.13653 | 0.891403 | 0.995465 | 0.001974 |
| ENSMUSG00000037958.14 | 241.4553 | 0.08069 | 0.590543 | 0.136637 | 0.891318 | 0.995465 | 0.001974 |
| ENSMUSG00000024286.9 | 209.6191 | 0.074327 | 0.544976 | 0.136386 | 0.891516 | 0.995474 | 0.00197 |
| ENSMUSG00000015750.16 | 19.92704 | -0.23197 | 1.710479 | -0.13562 | 0.892124 | 0.995917 | 0.001777 |

| | | | | | | | |
|---|---|---|---|---|---|---|---|
| ENSMUSG00000045095.17 | 52.1309 | -0.13578 | 1.00064 | -0.13569 | 0.892064 | 0.995917 | 0.001777 |
| ENSMUSG00000051346.10 | 124.7489 | -0.10279 | 0.758973 | -0.13543 | 0.892274 | 0.995968 | 0.001754 |
| ENSMUSG00000053929.18 | 70.80331 | -0.1671 | 1.235744 | -0.13522 | 0.892438 | 0.996034 | 0.001726 |
| ENSMUSG00000001018.16 | 101.1243 | -0.09567 | 0.735115 | -0.13014 | 0.896457 | 0.996165 | 0.001669 |
| ENSMUSG00000003154.16 | 51.36058 | -0.14052 | 1.064747 | -0.13197 | 0.895007 | 0.996165 | 0.001669 |
| ENSMUSG00000014668.16 | 17.30078 | -0.28032 | 2.186614 | -0.1282 | 0.897993 | 0.996165 | 0.001669 |
| ENSMUSG00000014867.10 | 912.7288 | 0.036501 | 0.280813 | 0.129984 | 0.896579 | 0.996165 | 0.001669 |
| ENSMUSG00000015027.12 | 34.73849 | 0.290247 | 2.161442 | 0.134284 | 0.893178 | 0.996165 | 0.001669 |
| ENSMUSG00000015668.14 | 48.04357 | 0.134115 | 1.046102 | 0.128204 | 0.897987 | 0.996165 | 0.001669 |
| ENSMUSG00000019039.15 | 94.82645 | -0.10401 | 0.803156 | -0.1295 | 0.896962 | 0.996165 | 0.001669 |
| ENSMUSG00000019792.9 | 49.59392 | 0.132532 | 1.010646 | 0.131135 | 0.895668 | 0.996165 | 0.001669 |
| ENSMUSG00000020250.12 | 1190.419 | 0.042128 | 0.328719 | 0.128157 | 0.898025 | 0.996165 | 0.001669 |
| ENSMUSG00000020656.17 | 13.75524 | 0.313627 | 2.419486 | 0.129626 | 0.896863 | 0.996165 | 0.001669 |
| ENSMUSG00000021245.16 | 13.04322 | 0.304457 | 2.327863 | 0.130788 | 0.895943 | 0.996165 | 0.001669 |
| ENSMUSG00000022788.17 | 56.81422 | -0.13879 | 1.037794 | -0.13374 | 0.893612 | 0.996165 | 0.001669 |
| ENSMUSG00000024817.14 | 114.3001 | 0.086759 | 0.67195 | 0.129115 | 0.897267 | 0.996165 | 0.001669 |
| ENSMUSG00000025278.10 | 144.7089 | 0.095626 | 0.740676 | 0.129106 | 0.897274 | 0.996165 | 0.001669 |
| ENSMUSG00000025369.16 | 59.34989 | 0.125046 | 0.933803 | 0.13391 | 0.893474 | 0.996165 | 0.001669 |
| ENSMUSG00000025731.17 | 608.781 | -0.0553 | 0.416498 | -0.13277 | 0.894376 | 0.996165 | 0.001669 |
| ENSMUSG00000025893.9 | 43.10302 | 0.220683 | 1.640734 | 0.134503 | 0.893005 | 0.996165 | 0.001669 |
| ENSMUSG00000026027.14 | 43.47811 | -0.17038 | 1.282396 | -0.13286 | 0.894301 | 0.996165 | 0.001669 |
| ENSMUSG00000026048.17 | 36.04475 | -0.16932 | 1.265727 | -0.13377 | 0.893582 | 0.996165 | 0.001669 |
| ENSMUSG00000026339.19 | 38.67082 | -0.2154 | 1.612038 | -0.13362 | 0.893704 | 0.996165 | 0.001669 |
| ENSMUSG00000028179.13 | 2791.91 | 0.038375 | 0.300667 | 0.127633 | 0.89844 | 0.996165 | 0.001669 |
| ENSMUSG00000028495.15 | 1219.8 | -0.03856 | 0.289508 | -0.13318 | 0.894047 | 0.996165 | 0.001669 |
| ENSMUSG00000031337.17 | 57.86669 | -0.17706 | 1.349832 | -0.13117 | 0.895641 | 0.996165 | 0.001669 |
| ENSMUSG00000032020.16 | 10.08736 | 0.424502 | 3.287607 | 0.129122 | 0.897261 | 0.996165 | 0.001669 |
| ENSMUSG00000032459.10 | 202.4305 | 0.074084 | 0.578434 | 0.128077 | 0.898088 | 0.996165 | 0.001669 |
| ENSMUSG00000032563.17 | 119.4334 | 0.092892 | 0.721838 | 0.128688 | 0.897604 | 0.996165 | 0.001669 |
| ENSMUSG00000034341.18 | 226.2574 | 0.078657 | 0.594706 | 0.132262 | 0.894777 | 0.996165 | 0.001669 |
| ENSMUSG00000034566.11 | 2861.894 | -0.02962 | 0.223768 | -0.13236 | 0.8947 | 0.996165 | 0.001669 |
| ENSMUSG00000035776.15 | 56.79654 | 0.187943 | 1.403141 | 0.133944 | 0.893447 | 0.996165 | 0.001669 |
| ENSMUSG00000035953.15 | 15.42853 | 0.211171 | 1.649386 | 0.12803 | 0.898125 | 0.996165 | 0.001669 |
| ENSMUSG00000037942.6 | 1810.601 | -0.03623 | 0.272525 | -0.13295 | 0.89423 | 0.996165 | 0.001669 |
| ENSMUSG00000039810.12 | 13.37987 | -0.28893 | 2.220438 | -0.13012 | 0.896469 | 0.996165 | 0.001669 |
| ENSMUSG00000040459.12 | 348.9103 | -0.0625 | 0.46476 | -0.13447 | 0.89303 | 0.996165 | 0.001669 |
| ENSMUSG00000041278.11 | 74.93605 | 0.17121 | 1.285847 | 0.133149 | 0.894075 | 0.996165 | 0.001669 |
| ENSMUSG00000043279.10 | 42.62793 | 0.161122 | 1.251045 | 0.12879 | 0.897524 | 0.996165 | 0.001669 |
| ENSMUSG00000045216.8 | 237.8553 | 0.068425 | 0.516358 | 0.132514 | 0.894578 | 0.996165 | 0.001669 |
| ENSMUSG00000049439.14 | 174.9193 | 0.076154 | 0.595395 | 0.127904 | 0.898225 | 0.996165 | 0.001669 |
| ENSMUSG00000050114.8 | 33.62442 | 0.153621 | 1.169031 | 0.131409 | 0.895452 | 0.996165 | 0.001669 |
| ENSMUSG00000050549.9 | 17.75243 | -0.23906 | 1.835351 | -0.13025 | 0.896367 | 0.996165 | 0.001669 |
| ENSMUSG00000051319.7 | 208.7076 | -0.07669 | 0.592146 | -0.12952 | 0.896948 | 0.996165 | 0.001669 |
| ENSMUSG00000051652.5 | 70.03793 | 0.111726 | 0.870004 | 0.128421 | 0.897816 | 0.996165 | 0.001669 |

| | | | | | | | |
|---|---|---|---|---|---|---|---|
| ENSMUSG00000053825.16 | 14.16397 | 0.342502 | 2.637748 | 0.129846 | 0.896688 | 0.996165 | 0.001669 |
| ENSMUSG00000054199.19 | 33.45599 | -0.22604 | 1.712698 | -0.13198 | 0.895001 | 0.996165 | 0.001669 |
| ENSMUSG00000055239.11 | 223.392 | 0.066789 | 0.518685 | 0.128766 | 0.897543 | 0.996165 | 0.001669 |
| ENSMUSG00000060147.16 | 35.87229 | 0.210418 | 1.615582 | 0.130243 | 0.896374 | 0.996165 | 0.001669 |
| ENSMUSG00000069206.15 | 106.825 | -0.11349 | 0.845095 | -0.13429 | 0.893173 | 0.996165 | 0.001669 |
| ENSMUSG00000069601.15 | 76.6031 | 0.104911 | 0.814364 | 0.128825 | 0.897496 | 0.996165 | 0.001669 |
| ENSMUSG00000073725.9 | 411.054 | 0.050386 | 0.394415 | 0.127748 | 0.898348 | 0.996165 | 0.001669 |
| ENSMUSG00000074415.15 | 17.18508 | -0.33355 | 2.582903 | -0.12914 | 0.897248 | 0.996165 | 0.001669 |
| ENSMUSG00000074547.5 | 40.98044 | 0.153504 | 1.194505 | 0.128509 | 0.897747 | 0.996165 | 0.001669 |
| ENSMUSG00000074754.5 | 280.383 | -0.08281 | 0.648147 | -0.12777 | 0.898335 | 0.996165 | 0.001669 |
| ENSMUSG00000078184.2 | 15.39451 | -0.23269 | 1.772088 | -0.13131 | 0.895531 | 0.996165 | 0.001669 |
| ENSMUSG00000085995.3 | 172.0911 | -0.13711 | 1.061989 | -0.12911 | 0.897273 | 0.996165 | 0.001669 |
| ENSMUSG00000097195.11 | 39.07797 | -0.15804 | 1.228098 | -0.12869 | 0.897604 | 0.996165 | 0.001669 |
| ENSMUSG00000097908.8 | 14.81371 | -0.39448 | 3.010254 | -0.13105 | 0.895739 | 0.996165 | 0.001669 |
| ENSMUSG00000112023.2 | 18.11708 | -0.29631 | 2.307248 | -0.12843 | 0.897811 | 0.996165 | 0.001669 |
| ENSMUSG00000000600.16 | 65.56236 | 0.123123 | 1.081368 | 0.113859 | 0.90935 | 0.996254 | 0.00163 |
| ENSMUSG00000000708.15 | 316.6162 | -0.05827 | 0.518946 | -0.11228 | 0.9106 | 0.996254 | 0.00163 |
| ENSMUSG00000000753.16 | 1070.051 | 0.038188 | 0.354428 | 0.107746 | 0.914197 | 0.996254 | 0.00163 |
| ENSMUSG00000001829.18 | 143.0593 | 0.088063 | 0.731212 | 0.120435 | 0.904139 | 0.996254 | 0.00163 |
| ENSMUSG00000001870.17 | 21.29552 | -0.21259 | 1.857875 | -0.11442 | 0.908901 | 0.996254 | 0.00163 |
| ENSMUSG00000002635.12 | 36.18499 | 0.120547 | 1.240481 | 0.097178 | 0.922585 | 0.996254 | 0.00163 |
| ENSMUSG00000002797.10 | 90.05857 | -0.05942 | 0.879414 | -0.06757 | 0.946127 | 0.996254 | 0.00163 |
| ENSMUSG00000003033.16 | 33.78545 | 0.206074 | 1.901973 | 0.108347 | 0.91372 | 0.996254 | 0.00163 |
| ENSMUSG00000003458.13 | 78.59133 | -0.0598 | 0.843784 | -0.07087 | 0.9435 | 0.996254 | 0.00163 |
| ENSMUSG00000003809.15 | 1710.691 | -0.02923 | 0.300182 | -0.09739 | 0.922417 | 0.996254 | 0.00163 |
| ENSMUSG00000004221.17 | 257.6951 | 0.061572 | 0.590756 | 0.104225 | 0.916991 | 0.996254 | 0.00163 |
| ENSMUSG00000004364.15 | 428.9841 | 0.051156 | 0.438694 | 0.116609 | 0.90717 | 0.996254 | 0.00163 |
| ENSMUSG00000004591.17 | 294.3504 | -0.05083 | 0.476917 | -0.10657 | 0.91513 | 0.996254 | 0.00163 |
| ENSMUSG00000005069.13 | 205.0703 | -0.06437 | 0.556079 | -0.11575 | 0.907851 | 0.996254 | 0.00163 |
| ENSMUSG00000005320.10 | 13.38182 | 0.202187 | 2.061663 | 0.09807 | 0.921877 | 0.996254 | 0.00163 |
| ENSMUSG00000005417.18 | 112.7291 | -0.08108 | 0.780213 | -0.10392 | 0.917236 | 0.996254 | 0.00163 |
| ENSMUSG00000005505.13 | 85.18825 | -0.09085 | 0.871447 | -0.10426 | 0.916967 | 0.996254 | 0.00163 |
| ENSMUSG00000005873.7 | 58.22302 | -0.12123 | 1.219393 | -0.09942 | 0.920803 | 0.996254 | 0.00163 |
| ENSMUSG00000006378.16 | 86.79584 | -0.10612 | 1.110262 | -0.09558 | 0.923856 | 0.996254 | 0.00163 |
| ENSMUSG00000006386.16 | 45.37653 | 0.109646 | 1.076444 | 0.101859 | 0.918869 | 0.996254 | 0.00163 |
| ENSMUSG00000006418.18 | 164.8037 | 0.083418 | 0.86014 | 0.096982 | 0.922741 | 0.996254 | 0.00163 |
| ENSMUSG00000006498.17 | 39.15358 | 0.138078 | 1.175863 | 0.117427 | 0.906522 | 0.996254 | 0.00163 |
| ENSMUSG00000006589.9 | 161.6285 | 0.090036 | 0.880683 | 0.102234 | 0.918571 | 0.996254 | 0.00163 |
| ENSMUSG00000006728.13 | 68.66699 | -0.12429 | 1.037236 | -0.11983 | 0.90462 | 0.996254 | 0.00163 |
| ENSMUSG00000008373.17 | 33.63971 | 0.159541 | 1.560028 | 0.102268 | 0.918544 | 0.996254 | 0.00163 |
| ENSMUSG00000008976.17 | 121.2276 | 0.089325 | 0.757296 | 0.117953 | 0.906105 | 0.996254 | 0.00163 |
| ENSMUSG00000009030.15 | 94.77823 | 0.052877 | 0.756565 | 0.069891 | 0.94428 | 0.996254 | 0.00163 |
| ENSMUSG00000009090.18 | 63.39881 | 0.108227 | 1.012658 | 0.106874 | 0.914889 | 0.996254 | 0.00163 |
| ENSMUSG00000009575.15 | 68.42288 | 0.075066 | 1.008869 | 0.074406 | 0.940687 | 0.996254 | 0.00163 |

| | | | | | | | |
|---|---|---|---|---|---|---|---|
| ENSMUSG00000009741.16 | 246.4539 | -0.03567 | 0.521362 | -0.06842 | 0.945452 | 0.996254 | 0.00163 |
| ENSMUSG00000010277.8 | 120.01 | -0.0627 | 0.87086 | -0.072 | 0.942604 | 0.996254 | 0.00163 |
| ENSMUSG00000012535.15 | 177.6554 | -0.05803 | 0.614684 | -0.09441 | 0.924781 | 0.996254 | 0.00163 |
| ENSMUSG00000014195.17 | 169.6926 | 0.065661 | 0.667916 | 0.098308 | 0.921688 | 0.996254 | 0.00163 |
| ENSMUSG00000014550.15 | 44.49853 | -0.11447 | 1.214584 | -0.09425 | 0.924911 | 0.996254 | 0.00163 |
| ENSMUSG00000014873.16 | 65.11804 | 0.157821 | 1.386143 | 0.113856 | 0.909352 | 0.996254 | 0.00163 |
| ENSMUSG00000015943.12 | 159.5107 | 0.105796 | 0.835585 | 0.126613 | 0.899247 | 0.996254 | 0.00163 |
| ENSMUSG00000016386.16 | 16.77714 | -0.35408 | 2.935058 | -0.12064 | 0.903979 | 0.996254 | 0.00163 |
| ENSMUSG00000017188.4 | 150.5438 | -0.06921 | 0.94306 | -0.07338 | 0.9415 | 0.996254 | 0.00163 |
| ENSMUSG00000017386.11 | 23.54545 | -0.17702 | 1.489201 | -0.11887 | 0.905376 | 0.996254 | 0.00163 |
| ENSMUSG00000017615.13 | 149.5437 | -0.06111 | 0.690267 | -0.08853 | 0.929454 | 0.996254 | 0.00163 |
| ENSMUSG00000017776.16 | 214.8705 | 0.041641 | 0.520446 | 0.08001 | 0.936229 | 0.996254 | 0.00163 |
| ENSMUSG00000018395.20 | 16.91422 | 0.233361 | 2.517448 | 0.092697 | 0.926144 | 0.996254 | 0.00163 |
| ENSMUSG00000018446.9 | 508.7108 | 0.02894 | 0.402193 | 0.071955 | 0.942638 | 0.996254 | 0.00163 |
| ENSMUSG00000018666.14 | 169.2913 | 0.048776 | 0.677124 | 0.072034 | 0.942575 | 0.996254 | 0.00163 |
| ENSMUSG00000018858.16 | 60.27055 | -0.14188 | 1.271019 | -0.11163 | 0.911119 | 0.996254 | 0.00163 |
| ENSMUSG00000019179.11 | 817.6724 | 0.030876 | 0.317404 | 0.097276 | 0.922507 | 0.996254 | 0.00163 |
| ENSMUSG00000019731.14 | 71.60467 | -0.07271 | 1.00456 | -0.07238 | 0.9423 | 0.996254 | 0.00163 |
| ENSMUSG00000019897.9 | 139.3125 | -0.06214 | 0.680325 | -0.09134 | 0.927223 | 0.996254 | 0.00163 |
| ENSMUSG00000020009.14 | 13.63173 | -0.21482 | 1.902799 | -0.1129 | 0.910112 | 0.996254 | 0.00163 |
| ENSMUSG00000020056.17 | 180.2516 | 0.057097 | 0.618744 | 0.092279 | 0.926476 | 0.996254 | 0.00163 |
| ENSMUSG00000020088.11 | 509.5182 | 0.040673 | 0.40045 | 0.101568 | 0.9191 | 0.996254 | 0.00163 |
| ENSMUSG00000020166.16 | 152.2727 | -0.05736 | 0.622918 | -0.09208 | 0.926638 | 0.996254 | 0.00163 |
| ENSMUSG00000020170.5 | 78.47507 | -0.10069 | 0.9529 | -0.10567 | 0.915846 | 0.996254 | 0.00163 |
| ENSMUSG00000020305.14 | 22.70204 | -0.27483 | 2.386122 | -0.11518 | 0.908305 | 0.996254 | 0.00163 |
| ENSMUSG00000020308.8 | 39.3275 | -0.13812 | 1.570762 | -0.08793 | 0.929929 | 0.996254 | 0.00163 |
| ENSMUSG00000020364.15 | 19.27742 | 0.188697 | 1.602063 | 0.117784 | 0.906239 | 0.996254 | 0.00163 |
| ENSMUSG00000020422.15 | 160.7471 | -0.0732 | 0.634806 | -0.11531 | 0.908198 | 0.996254 | 0.00163 |
| ENSMUSG00000020432.13 | 35.48052 | 0.121851 | 1.263562 | 0.096434 | 0.923176 | 0.996254 | 0.00163 |
| ENSMUSG00000020457.14 | 136.0231 | -0.06725 | 0.942269 | -0.07137 | 0.943101 | 0.996254 | 0.00163 |
| ENSMUSG00000020476.15 | 92.99048 | -0.09287 | 1.080342 | -0.08596 | 0.931497 | 0.996254 | 0.00163 |
| ENSMUSG00000020523.15 | 94.11663 | -0.11354 | 0.953383 | -0.1191 | 0.905199 | 0.996254 | 0.00163 |
| ENSMUSG00000020650.16 | 136.7437 | -0.08011 | 0.832704 | -0.09621 | 0.923357 | 0.996254 | 0.00163 |
| ENSMUSG00000020691.14 | 10.03599 | 0.27718 | 2.503592 | 0.110713 | 0.911844 | 0.996254 | 0.00163 |
| ENSMUSG00000020715.10 | 184.1262 | 0.065421 | 0.726141 | 0.090093 | 0.928213 | 0.996254 | 0.00163 |
| ENSMUSG00000020919.12 | 80.99558 | -0.11246 | 0.903415 | -0.12449 | 0.900929 | 0.996254 | 0.00163 |
| ENSMUSG00000020936.9 | 95.75084 | 0.068746 | 0.808367 | 0.085043 | 0.932227 | 0.996254 | 0.00163 |
| ENSMUSG00000021072.13 | 230.3427 | 0.059838 | 0.556304 | 0.107563 | 0.914343 | 0.996254 | 0.00163 |
| ENSMUSG00000021096.12 | 573.8649 | -0.05097 | 0.401431 | -0.12697 | 0.898962 | 0.996254 | 0.00163 |
| ENSMUSG00000021185.17 | 75.93518 | 0.080108 | 0.892249 | 0.089782 | 0.92846 | 0.996254 | 0.00163 |
| ENSMUSG00000021271.6 | 38.36903 | -0.13409 | 1.246006 | -0.10762 | 0.914301 | 0.996254 | 0.00163 |
| ENSMUSG00000021336.18 | 110.6754 | 0.081524 | 0.829502 | 0.09828 | 0.92171 | 0.996254 | 0.00163 |
| ENSMUSG00000021375.12 | 44.70387 | -0.12245 | 1.532193 | -0.07992 | 0.936304 | 0.996254 | 0.00163 |
| ENSMUSG00000021377.16 | 646.7572 | -0.03349 | 0.465047 | -0.07201 | 0.942596 | 0.996254 | 0.00163 |

| | | | | | | | |
|---|---|---|---|---|---|---|---|
| ENSMUSG00000021420.14 | 144.751 | -0.06464 | 0.677127 | -0.09547 | 0.923943 | 0.996254 | 0.00163 |
| ENSMUSG00000021423.7 | 16.46331 | -0.198 | 1.86328 | -0.10626 | 0.915375 | 0.996254 | 0.00163 |
| ENSMUSG00000021546.19 | 1244.415 | -0.02102 | 0.283492 | -0.07414 | 0.940902 | 0.996254 | 0.00163 |
| ENSMUSG00000021646.10 | 290.2951 | -0.04791 | 0.477609 | -0.10031 | 0.920095 | 0.996254 | 0.00163 |
| ENSMUSG00000021706.15 | 46.62485 | -0.11596 | 0.987844 | -0.11739 | 0.90655 | 0.996254 | 0.00163 |
| ENSMUSG00000021772.16 | 146.7115 | -0.07275 | 0.651604 | -0.11165 | 0.911099 | 0.996254 | 0.00163 |
| ENSMUSG00000021785.9 | 228.5281 | 0.073402 | 0.642345 | 0.114273 | 0.909022 | 0.996254 | 0.00163 |
| ENSMUSG00000021943.8 | 42.06382 | 0.146098 | 1.403821 | 0.104072 | 0.917112 | 0.996254 | 0.00163 |
| ENSMUSG00000021945.10 | 163.8879 | 0.072881 | 0.659863 | 0.110449 | 0.912054 | 0.996254 | 0.00163 |
| ENSMUSG00000022040.9 | 3593.431 | -0.01759 | 0.254639 | -0.06908 | 0.944929 | 0.996254 | 0.00163 |
| ENSMUSG00000022094.16 | 114.3674 | 0.082229 | 0.726719 | 0.113151 | 0.909911 | 0.996254 | 0.00163 |
| ENSMUSG00000022124.16 | 164.4996 | -0.05448 | 0.629959 | -0.08648 | 0.931083 | 0.996254 | 0.00163 |
| ENSMUSG00000022148.17 | 30.76313 | 0.107067 | 1.413605 | 0.07574 | 0.939626 | 0.996254 | 0.00163 |
| ENSMUSG00000022304.14 | 712.5638 | 0.041951 | 0.397035 | 0.105661 | 0.915852 | 0.996254 | 0.00163 |
| ENSMUSG00000022329.16 | 46.41336 | -0.13409 | 1.219441 | -0.10996 | 0.91244 | 0.996254 | 0.00163 |
| ENSMUSG00000022365.7 | 302.1708 | 0.060949 | 0.551773 | 0.11046 | 0.912045 | 0.996254 | 0.00163 |
| ENSMUSG00000022466.7 | 29.18032 | -0.13911 | 1.301661 | -0.10687 | 0.914892 | 0.996254 | 0.00163 |
| ENSMUSG00000022521.11 | 148.0967 | 0.060595 | 0.700911 | 0.086452 | 0.931107 | 0.996254 | 0.00163 |
| ENSMUSG00000022574.8 | 39.23418 | -0.10677 | 1.512141 | -0.07061 | 0.943711 | 0.996254 | 0.00163 |
| ENSMUSG00000022620.15 | 65.98396 | -0.14364 | 1.167395 | -0.12305 | 0.902071 | 0.996254 | 0.00163 |
| ENSMUSG00000022629.18 | 443.5534 | 0.041926 | 0.481337 | 0.087104 | 0.930589 | 0.996254 | 0.00163 |
| ENSMUSG00000022674.16 | 91.05233 | -0.08498 | 0.852197 | -0.09971 | 0.920572 | 0.996254 | 0.00163 |
| ENSMUSG00000022769.10 | 87.43325 | -0.10541 | 0.960884 | -0.1097 | 0.91265 | 0.996254 | 0.00163 |
| ENSMUSG00000022867.12 | 96.7091 | 0.090631 | 0.805237 | 0.112552 | 0.910386 | 0.996254 | 0.00163 |
| ENSMUSG00000022884.17 | 410.522 | -0.04542 | 0.413154 | -0.10992 | 0.912469 | 0.996254 | 0.00163 |
| ENSMUSG00000022969.14 | 42.70962 | -0.18289 | 1.579238 | -0.11581 | 0.907804 | 0.996254 | 0.00163 |
| ENSMUSG00000023027.14 | 150.9372 | -0.06341 | 0.636546 | -0.09962 | 0.920649 | 0.996254 | 0.00163 |
| ENSMUSG00000023030.17 | 112.1373 | -0.04809 | 0.655343 | -0.07338 | 0.941506 | 0.996254 | 0.00163 |
| ENSMUSG00000023087.16 | 246.485 | 0.073579 | 0.744006 | 0.098896 | 0.921221 | 0.996254 | 0.00163 |
| ENSMUSG00000023904.11 | 294.2688 | 0.066831 | 0.641702 | 0.104147 | 0.917053 | 0.996254 | 0.00163 |
| ENSMUSG00000023921.10 | 1156.042 | 0.032189 | 0.340303 | 0.094591 | 0.92464 | 0.996254 | 0.00163 |
| ENSMUSG00000023960.14 | 61.68426 | 0.069135 | 1.021527 | 0.067678 | 0.946042 | 0.996254 | 0.00163 |
| ENSMUSG00000024101.16 | 25.77121 | -0.16519 | 2.049641 | -0.0806 | 0.935764 | 0.996254 | 0.00163 |
| ENSMUSG00000024118.15 | 138.1122 | -0.05649 | 0.59783 | -0.0945 | 0.924713 | 0.996254 | 0.00163 |
| ENSMUSG00000024182.18 | 21.7474 | -0.20424 | 2.077274 | -0.09832 | 0.921678 | 0.996254 | 0.00163 |
| ENSMUSG00000024247.15 | 30.33233 | 0.185361 | 2.112668 | 0.087738 | 0.930085 | 0.996254 | 0.00163 |
| ENSMUSG00000024293.17 | 15.86102 | -0.19462 | 2.235942 | -0.08704 | 0.930637 | 0.996254 | 0.00163 |
| ENSMUSG00000024308.16 | 82.42088 | -0.10994 | 1.301657 | -0.08446 | 0.932692 | 0.996254 | 0.00163 |
| ENSMUSG00000024430.15 | 25.88989 | -0.18431 | 2.094157 | -0.08801 | 0.929869 | 0.996254 | 0.00163 |
| ENSMUSG00000024588.11 | 2564.895 | 0.035866 | 0.307804 | 0.116523 | 0.907238 | 0.996254 | 0.00163 |
| ENSMUSG00000024769.8 | 25.01124 | -0.22254 | 2.039546 | -0.10911 | 0.913114 | 0.996254 | 0.00163 |
| ENSMUSG00000024847.16 | 25.28683 | 0.099722 | 1.30401 | 0.076473 | 0.939043 | 0.996254 | 0.00163 |
| ENSMUSG00000024911.8 | 56.14197 | -0.1198 | 1.158408 | -0.10341 | 0.917634 | 0.996254 | 0.00163 |
| ENSMUSG00000024925.12 | 150.3449 | 0.046798 | 0.61081 | 0.076616 | 0.938929 | 0.996254 | 0.00163 |

| | | | | | | | |
|---|---|---|---|---|---|---|---|
| ENSMUSG00000024999.8 | 14.15576 | -0.18722 | 1.998929 | -0.09366 | 0.925378 | 0.996254 | 0.00163 |
| ENSMUSG00000025004.17 | 25.40744 | -0.18437 | 1.786193 | -0.10322 | 0.917787 | 0.996254 | 0.00163 |
| ENSMUSG00000025024.8 | 88.60952 | 0.063437 | 0.850511 | 0.074587 | 0.940543 | 0.996254 | 0.00163 |
| ENSMUSG00000025026.16 | 59.84576 | 0.14831 | 1.659595 | 0.089365 | 0.928792 | 0.996254 | 0.00163 |
| ENSMUSG00000025103.9 | 244.3439 | -0.03587 | 0.50891 | -0.07049 | 0.943807 | 0.996254 | 0.00163 |
| ENSMUSG00000025133.10 | 56.18813 | 0.100275 | 1.224777 | 0.081872 | 0.934749 | 0.996254 | 0.00163 |
| ENSMUSG00000025192.14 | 20.83853 | 0.164297 | 2.095774 | 0.078395 | 0.937514 | 0.996254 | 0.00163 |
| ENSMUSG00000025193.15 | 162.7758 | -0.05938 | 0.741839 | -0.08005 | 0.936199 | 0.996254 | 0.00163 |
| ENSMUSG00000025268.16 | 11.5128 | 0.239268 | 3.216866 | 0.074379 | 0.940709 | 0.996254 | 0.00163 |
| ENSMUSG00000025279.8 | 187.3152 | 0.058927 | 0.665757 | 0.088511 | 0.92947 | 0.996254 | 0.00163 |
| ENSMUSG00000025410.11 | 153.4949 | 0.079978 | 0.672529 | 0.118922 | 0.905337 | 0.996254 | 0.00163 |
| ENSMUSG00000025439.6 | 250.8138 | 0.074427 | 0.882291 | 0.084357 | 0.932773 | 0.996254 | 0.00163 |
| ENSMUSG00000025521.16 | 34.56933 | 0.228885 | 1.800498 | 0.127123 | 0.898843 | 0.996254 | 0.00163 |
| ENSMUSG00000025762.17 | 633.6025 | 0.042143 | 0.384308 | 0.109658 | 0.91268 | 0.996254 | 0.00163 |
| ENSMUSG00000025782.13 | 56.96031 | -0.09323 | 1.108295 | -0.08412 | 0.932959 | 0.996254 | 0.00163 |
| ENSMUSG00000025813.15 | 338.2153 | 0.045982 | 0.541345 | 0.08494 | 0.932309 | 0.996254 | 0.00163 |
| ENSMUSG00000025854.16 | 142.0321 | -0.07706 | 0.607625 | -0.12682 | 0.899084 | 0.996254 | 0.00163 |
| ENSMUSG00000025869.12 | 105.1593 | 0.076463 | 0.836445 | 0.091414 | 0.927163 | 0.996254 | 0.00163 |
| ENSMUSG00000025915.15 | 65.88364 | -0.07532 | 0.915535 | -0.08227 | 0.934435 | 0.996254 | 0.00163 |
| ENSMUSG00000025968.17 | 892.608 | -0.03067 | 0.3667 | -0.08363 | 0.933347 | 0.996254 | 0.00163 |
| ENSMUSG00000025979.14 | 426.4352 | 0.036902 | 0.468174 | 0.078821 | 0.937175 | 0.996254 | 0.00163 |
| ENSMUSG00000025991.9 | 7500.927 | 0.022844 | 0.222849 | 0.10251 | 0.918352 | 0.996254 | 0.00163 |
| ENSMUSG00000026021.16 | 259.4054 | 0.071186 | 0.663632 | 0.107268 | 0.914577 | 0.996254 | 0.00163 |
| ENSMUSG00000026087.12 | 603.1807 | 0.028484 | 0.404651 | 0.070391 | 0.943883 | 0.996254 | 0.00163 |
| ENSMUSG00000026174.10 | 29.08273 | 0.12363 | 1.822638 | 0.06783 | 0.945921 | 0.996254 | 0.00163 |
| ENSMUSG00000026187.10 | 98.46517 | -0.07342 | 0.836357 | -0.08779 | 0.930042 | 0.996254 | 0.00163 |
| ENSMUSG00000026304.14 | 55.74686 | -0.17978 | 1.485106 | -0.12106 | 0.903646 | 0.996254 | 0.00163 |
| ENSMUSG00000026309.15 | 70.77701 | -0.07967 | 0.929893 | -0.08568 | 0.931723 | 0.996254 | 0.00163 |
| ENSMUSG00000026356.16 | 521.1609 | -0.03924 | 0.499121 | -0.07862 | 0.937332 | 0.996254 | 0.00163 |
| ENSMUSG00000026365.16 | 4474.304 | -0.03421 | 0.276352 | -0.12378 | 0.901491 | 0.996254 | 0.00163 |
| ENSMUSG00000026399.13 | 109.0827 | -0.08991 | 1.239761 | -0.07253 | 0.942184 | 0.996254 | 0.00163 |
| ENSMUSG00000026405.15 | 5349.358 | -0.02578 | 0.279417 | -0.09226 | 0.926491 | 0.996254 | 0.00163 |
| ENSMUSG00000026430.17 | 107.1932 | 0.111957 | 0.985514 | 0.113602 | 0.909553 | 0.996254 | 0.00163 |
| ENSMUSG00000026480.13 | 13.46975 | 0.283268 | 2.41963 | 0.117071 | 0.906804 | 0.996254 | 0.00163 |
| ENSMUSG00000026553.18 | 276.5302 | -0.05995 | 0.506255 | -0.11841 | 0.90574 | 0.996254 | 0.00163 |
| ENSMUSG00000026563.14 | 58.25066 | 0.092105 | 0.956753 | 0.096269 | 0.923307 | 0.996254 | 0.00163 |
| ENSMUSG00000026640.13 | 35.0745 | 0.117132 | 1.384665 | 0.084592 | 0.932586 | 0.996254 | 0.00163 |
| ENSMUSG00000026643.17 | 67.96534 | -0.09678 | 1.051434 | -0.09205 | 0.92666 | 0.996254 | 0.00163 |
| ENSMUSG00000026674.10 | 34.66153 | 0.142048 | 1.952893 | 0.072737 | 0.942015 | 0.996254 | 0.00163 |
| ENSMUSG00000026721.17 | 159.0783 | 0.049788 | 0.705718 | 0.07055 | 0.943756 | 0.996254 | 0.00163 |
| ENSMUSG00000026879.15 | 79.07614 | -0.0933 | 1.318914 | -0.07074 | 0.943603 | 0.996254 | 0.00163 |
| ENSMUSG00000026946.10 | 250.2065 | -0.03574 | 0.482419 | -0.07408 | 0.940948 | 0.996254 | 0.00163 |
| ENSMUSG00000027082.16 | 40.84578 | -0.13874 | 1.107953 | -0.12522 | 0.900348 | 0.996254 | 0.00163 |
| ENSMUSG00000027187.11 | 16154.61 | -0.02971 | 0.285897 | -0.10393 | 0.917223 | 0.996254 | 0.00163 |

| | | | | | | | |
|---|---|---|---|---|---|---|---|
| ENSMUSG00000027272.6 | 181.5805 | 0.044159 | 0.63103 | 0.069979 | 0.94421 | 0.996254 | 0.00163 |
| ENSMUSG00000027276.8 | 32.2955 | 0.155099 | 1.409541 | 0.110035 | 0.912382 | 0.996254 | 0.00163 |
| ENSMUSG00000027288.17 | 296.8254 | 0.073951 | 0.756942 | 0.097698 | 0.922172 | 0.996254 | 0.00163 |
| ENSMUSG00000027425.19 | 15.51603 | 0.238658 | 2.096142 | 0.113856 | 0.909352 | 0.996254 | 0.00163 |
| ENSMUSG00000027430.13 | 52.51202 | 0.119957 | 1.282249 | 0.093552 | 0.925465 | 0.996254 | 0.00163 |
| ENSMUSG00000027502.12 | 97.78368 | 0.107147 | 0.959888 | 0.111624 | 0.911121 | 0.996254 | 0.00163 |
| ENSMUSG00000027533.11 | 53.86033 | 0.175393 | 1.591848 | 0.110182 | 0.912265 | 0.996254 | 0.00163 |
| ENSMUSG00000027534.17 | 18.57232 | -0.14161 | 1.609028 | -0.08801 | 0.929871 | 0.996254 | 0.00163 |
| ENSMUSG00000027652.16 | 46.22397 | 0.169742 | 1.418977 | 0.119623 | 0.904782 | 0.996254 | 0.00163 |
| ENSMUSG00000027680.16 | 251.6592 | 0.062723 | 0.641152 | 0.097828 | 0.922069 | 0.996254 | 0.00163 |
| ENSMUSG00000027774.17 | 375.4326 | -0.0539 | 0.425676 | -0.12661 | 0.899245 | 0.996254 | 0.00163 |
| ENSMUSG00000027854.13 | 84.28324 | -0.06002 | 0.880323 | -0.06818 | 0.945642 | 0.996254 | 0.00163 |
| ENSMUSG00000027879.10 | 395.1952 | -0.04556 | 0.41558 | -0.10963 | 0.9127 | 0.996254 | 0.00163 |
| ENSMUSG00000027893.15 | 329.5514 | 0.04906 | 0.502776 | 0.097578 | 0.922268 | 0.996254 | 0.00163 |
| ENSMUSG00000027942.18 | 216.08 | -0.06393 | 0.548448 | -0.11657 | 0.907204 | 0.996254 | 0.00163 |
| ENSMUSG00000027962.15 | 76.63178 | -0.0773 | 0.964103 | -0.08018 | 0.936096 | 0.996254 | 0.00163 |
| ENSMUSG00000028164.16 | 21.27361 | -0.1655 | 1.375737 | -0.1203 | 0.904248 | 0.996254 | 0.00163 |
| ENSMUSG00000028291.8 | 109.311 | -0.08164 | 0.709539 | -0.11506 | 0.9084 | 0.996254 | 0.00163 |
| ENSMUSG00000028300.15 | 57.73508 | -0.14491 | 1.39875 | -0.1036 | 0.917488 | 0.996254 | 0.00163 |
| ENSMUSG00000028434.13 | 426.0719 | -0.04355 | 0.415198 | -0.10489 | 0.91646 | 0.996254 | 0.00163 |
| ENSMUSG00000028454.17 | 10.05787 | -0.19895 | 2.590812 | -0.07679 | 0.938791 | 0.996254 | 0.00163 |
| ENSMUSG00000028573.19 | 470.3218 | -0.0482 | 0.438111 | -0.11003 | 0.912388 | 0.996254 | 0.00163 |
| ENSMUSG00000028599.11 | 33.13197 | -0.22162 | 1.831968 | -0.12098 | 0.90371 | 0.996254 | 0.00163 |
| ENSMUSG00000028621.18 | 22.15135 | -0.17052 | 2.394619 | -0.07121 | 0.943231 | 0.996254 | 0.00163 |
| ENSMUSG00000028670.15 | 12.2961 | -0.23334 | 2.128887 | -0.10961 | 0.912721 | 0.996254 | 0.00163 |
| ENSMUSG00000028710.19 | 10.882 | 0.224308 | 2.330361 | 0.096255 | 0.923318 | 0.996254 | 0.00163 |
| ENSMUSG00000028826.15 | 191.7395 | 0.100242 | 0.80223 | 0.124954 | 0.90056 | 0.996254 | 0.00163 |
| ENSMUSG00000028861.14 | 151.9871 | -0.06219 | 0.834977 | -0.07448 | 0.940631 | 0.996254 | 0.00163 |
| ENSMUSG00000028937.15 | 80.76797 | -0.14084 | 1.336898 | -0.10535 | 0.916097 | 0.996254 | 0.00163 |
| ENSMUSG00000028979.18 | 926.4513 | -0.04837 | 0.390905 | -0.12373 | 0.901531 | 0.996254 | 0.00163 |
| ENSMUSG00000028988.14 | 68.58411 | -0.12437 | 1.019231 | -0.12202 | 0.902882 | 0.996254 | 0.00163 |
| ENSMUSG00000029056.14 | 22.60536 | -0.17914 | 2.080685 | -0.0861 | 0.931391 | 0.996254 | 0.00163 |
| ENSMUSG00000029084.6 | 98.65244 | -0.11435 | 1.004546 | -0.11384 | 0.909368 | 0.996254 | 0.00163 |
| ENSMUSG00000029131.15 | 330.3863 | -0.03964 | 0.528507 | -0.075 | 0.940218 | 0.996254 | 0.00163 |
| ENSMUSG00000029166.15 | 12.9958 | 0.224755 | 2.386993 | 0.094158 | 0.924983 | 0.996254 | 0.00163 |
| ENSMUSG00000029221.16 | 218.7355 | -0.06164 | 0.584601 | -0.10543 | 0.916034 | 0.996254 | 0.00163 |
| ENSMUSG00000029253.13 | 29.04933 | 0.150307 | 1.219675 | 0.123235 | 0.901921 | 0.996254 | 0.00163 |
| ENSMUSG00000029287.15 | 30.53503 | 0.145645 | 1.730327 | 0.084172 | 0.93292 | 0.996254 | 0.00163 |
| ENSMUSG00000029407.11 | 810.8155 | -0.03481 | 0.289827 | -0.12011 | 0.904396 | 0.996254 | 0.00163 |
| ENSMUSG00000029501.15 | 109.3575 | 0.10055 | 0.871264 | 0.115407 | 0.908122 | 0.996254 | 0.00163 |
| ENSMUSG00000029507.17 | 27.72829 | -0.15312 | 1.745107 | -0.08774 | 0.930084 | 0.996254 | 0.00163 |
| ENSMUSG00000029554.16 | 18.24892 | -0.24332 | 2.159704 | -0.11266 | 0.910297 | 0.996254 | 0.00163 |
| ENSMUSG00000029715.6 | 20.98673 | -0.10985 | 1.604237 | -0.06847 | 0.945408 | 0.996254 | 0.00163 |
| ENSMUSG00000029772.18 | 137.7917 | 0.060796 | 0.818143 | 0.07431 | 0.940764 | 0.996254 | 0.00163 |

| | | | | | | | |
|---|---|---|---|---|---|---|---|
| ENSMUSG00000030120.15 | 448.4929 | 0.033815 | 0.4351 | 0.077718 | 0.938052 | 0.996254 | 0.00163 |
| ENSMUSG00000030127.16 | 159.8047 | -0.05885 | 0.566523 | -0.10388 | 0.917266 | 0.996254 | 0.00163 |
| ENSMUSG00000030166.15 | 30.54074 | -0.13443 | 1.713186 | -0.07847 | 0.937455 | 0.996254 | 0.00163 |
| ENSMUSG00000030232.19 | 70.39264 | -0.08974 | 0.941872 | -0.09528 | 0.924093 | 0.996254 | 0.00163 |
| ENSMUSG00000030704.15 | 460.5114 | -0.03934 | 0.358841 | -0.10964 | 0.912696 | 0.996254 | 0.00163 |
| ENSMUSG00000030802.15 | 14.72957 | -0.1641 | 2.296546 | -0.07146 | 0.943034 | 0.996254 | 0.00163 |
| ENSMUSG00000030815.12 | 23.17758 | 0.206309 | 2.061315 | 0.100086 | 0.920276 | 0.996254 | 0.00163 |
| ENSMUSG00000030823.7 | 13.67751 | 0.3102 | 2.647414 | 0.117171 | 0.906725 | 0.996254 | 0.00163 |
| ENSMUSG00000030879.11 | 279.8488 | -0.07063 | 0.584746 | -0.12078 | 0.903866 | 0.996254 | 0.00163 |
| ENSMUSG00000030922.13 | 18.71861 | -0.20058 | 1.642559 | -0.12211 | 0.902809 | 0.996254 | 0.00163 |
| ENSMUSG00000030960.17 | 22.87588 | 0.226509 | 2.00397 | 0.11303 | 0.910007 | 0.996254 | 0.00163 |
| ENSMUSG00000030990.19 | 93.15924 | -0.06355 | 0.869355 | -0.0731 | 0.94173 | 0.996254 | 0.00163 |
| ENSMUSG00000031068.9 | 531.7641 | -0.0427 | 0.359206 | -0.11886 | 0.905384 | 0.996254 | 0.00163 |
| ENSMUSG00000031156.19 | 94.22001 | 0.081338 | 0.940229 | 0.086508 | 0.931062 | 0.996254 | 0.00163 |
| ENSMUSG00000031299.11 | 734.601 | -0.03563 | 0.398965 | -0.0893 | 0.928844 | 0.996254 | 0.00163 |
| ENSMUSG00000031353.14 | 347.1972 | -0.05427 | 0.490137 | -0.11073 | 0.911834 | 0.996254 | 0.00163 |
| ENSMUSG00000031447.8 | 2114.888 | -0.0178 | 0.259426 | -0.06861 | 0.945303 | 0.996254 | 0.00163 |
| ENSMUSG00000031458.9 | 17.55279 | 0.154983 | 2.155006 | 0.071918 | 0.942667 | 0.996254 | 0.00163 |
| ENSMUSG00000031503.14 | 33.63507 | 0.156345 | 1.48627 | 0.105193 | 0.916223 | 0.996254 | 0.00163 |
| ENSMUSG00000031555.16 | 213.5361 | 0.064665 | 0.57054 | 0.11334 | 0.909761 | 0.996254 | 0.00163 |
| ENSMUSG00000031683.17 | 439.1007 | 0.04311 | 0.442044 | 0.097525 | 0.92231 | 0.996254 | 0.00163 |
| ENSMUSG00000031790.9 | 73.63103 | 0.07385 | 0.880084 | 0.083912 | 0.933126 | 0.996254 | 0.00163 |
| ENSMUSG00000031983.8 | 24.50115 | -0.17919 | 1.671703 | -0.10719 | 0.914636 | 0.996254 | 0.00163 |
| ENSMUSG00000031985.10 | 151.2303 | -0.07624 | 0.750908 | -0.10153 | 0.919126 | 0.996254 | 0.00163 |
| ENSMUSG00000032078.18 | 75.76739 | -0.08358 | 1.004255 | -0.08322 | 0.933675 | 0.996254 | 0.00163 |
| ENSMUSG00000032239.11 | 534.6292 | 0.041596 | 0.479418 | 0.086764 | 0.930859 | 0.996254 | 0.00163 |
| ENSMUSG00000032380.10 | 65.43131 | -0.08079 | 0.98763 | -0.0818 | 0.934805 | 0.996254 | 0.00163 |
| ENSMUSG00000032393.16 | 74.57051 | -0.09776 | 1.033971 | -0.09455 | 0.924672 | 0.996254 | 0.00163 |
| ENSMUSG00000032398.8 | 243.7849 | 0.046871 | 0.583329 | 0.080351 | 0.935958 | 0.996254 | 0.00163 |
| ENSMUSG00000032415.15 | 13.12098 | -0.2999 | 3.056282 | -0.09812 | 0.921834 | 0.996254 | 0.00163 |
| ENSMUSG00000032492.15 | 21.05943 | 0.173665 | 1.960994 | 0.088559 | 0.929432 | 0.996254 | 0.00163 |
| ENSMUSG00000032519.12 | 29.91254 | 0.155788 | 1.359635 | 0.114581 | 0.908777 | 0.996254 | 0.00163 |
| ENSMUSG00000032531.16 | 91.25388 | 0.068762 | 0.830403 | 0.082806 | 0.934006 | 0.996254 | 0.00163 |
| ENSMUSG00000032547.13 | 218.0539 | -0.05204 | 0.521729 | -0.09975 | 0.920545 | 0.996254 | 0.00163 |
| ENSMUSG00000032612.15 | 216.1086 | -0.05117 | 0.567392 | -0.09019 | 0.928139 | 0.996254 | 0.00163 |
| ENSMUSG00000032869.16 | 107.9768 | 0.081484 | 0.774092 | 0.105264 | 0.916167 | 0.996254 | 0.00163 |
| ENSMUSG00000033102.16 | 159.4668 | 0.068865 | 0.614767 | 0.112018 | 0.910809 | 0.996254 | 0.00163 |
| ENSMUSG00000033417.17 | 203.3478 | 0.083505 | 0.666354 | 0.125316 | 0.900273 | 0.996254 | 0.00163 |
| ENSMUSG00000033545.15 | 66.08433 | -0.15831 | 1.347796 | -0.11746 | 0.906496 | 0.996254 | 0.00163 |
| ENSMUSG00000033752.8 | 33.32855 | -0.12904 | 1.483427 | -0.08699 | 0.930683 | 0.996254 | 0.00163 |
| ENSMUSG00000034329.17 | 14.70973 | 0.185814 | 2.225176 | 0.083505 | 0.93345 | 0.996254 | 0.00163 |
| ENSMUSG00000034403.17 | 241.8231 | -0.07325 | 0.582417 | -0.12576 | 0.899919 | 0.996254 | 0.00163 |
| ENSMUSG00000034429.16 | 29.38851 | -0.15271 | 2.019298 | -0.07563 | 0.939716 | 0.996254 | 0.00163 |
| ENSMUSG00000034647.16 | 140.5691 | -0.04741 | 0.669143 | -0.07086 | 0.943511 | 0.996254 | 0.00163 |

| | | | | | | | |
|---|---|---|---|---|---|---|---|
| ENSMUSG00000034659.15 | 69.05402 | -0.0799 | 1.082842 | -0.07379 | 0.94118 | 0.996254 | 0.00163 |
| ENSMUSG00000034845.7 | 13.95949 | 0.297287 | 2.392194 | 0.124274 | 0.901098 | 0.996254 | 0.00163 |
| ENSMUSG00000034951.11 | 120.1617 | 0.078318 | 0.807253 | 0.097018 | 0.922712 | 0.996254 | 0.00163 |
| ENSMUSG00000035173.15 | 30.9668 | -0.18568 | 2.18377 | -0.08503 | 0.932241 | 0.996254 | 0.00163 |
| ENSMUSG00000035351.17 | 31.47151 | -0.10211 | 1.195176 | -0.08543 | 0.931916 | 0.996254 | 0.00163 |
| ENSMUSG00000035376.11 | 384.2061 | 0.036913 | 0.390176 | 0.094607 | 0.924627 | 0.996254 | 0.00163 |
| ENSMUSG00000035378.18 | 20.74792 | -0.1272 | 1.460479 | -0.08709 | 0.930598 | 0.996254 | 0.00163 |
| ENSMUSG00000035437.16 | 59.08693 | -0.09671 | 0.94497 | -0.10234 | 0.918489 | 0.996254 | 0.00163 |
| ENSMUSG00000035441.15 | 63.02885 | -0.12244 | 1.043392 | -0.11735 | 0.906583 | 0.996254 | 0.00163 |
| ENSMUSG00000035493.11 | 312.5349 | -0.06039 | 0.606851 | -0.09951 | 0.920732 | 0.996254 | 0.00163 |
| ENSMUSG00000035529.11 | 30.30601 | 0.121905 | 1.577527 | 0.077276 | 0.938404 | 0.996254 | 0.00163 |
| ENSMUSG00000035545.14 | 25.76423 | 0.201554 | 1.830049 | 0.110136 | 0.912302 | 0.996254 | 0.00163 |
| ENSMUSG00000035769.10 | 363.5841 | 0.064308 | 0.546028 | 0.117774 | 0.906246 | 0.996254 | 0.00163 |
| ENSMUSG00000035877.18 | 59.79868 | 0.152746 | 1.3181 | 0.115883 | 0.907745 | 0.996254 | 0.00163 |
| ENSMUSG00000036151.16 | 53.39738 | 0.133045 | 1.436835 | 0.092596 | 0.926225 | 0.996254 | 0.00163 |
| ENSMUSG00000036181.3 | 175.4085 | 0.072261 | 0.677539 | 0.106651 | 0.915065 | 0.996254 | 0.00163 |
| ENSMUSG00000036241.4 | 1169.174 | -0.02646 | 0.290681 | -0.09101 | 0.927481 | 0.996254 | 0.00163 |
| ENSMUSG00000036570.15 | 449.5135 | 0.035524 | 0.495988 | 0.071623 | 0.942902 | 0.996254 | 0.00163 |
| ENSMUSG00000036661.15 | 11.05287 | 0.235075 | 2.94472 | 0.079829 | 0.936373 | 0.996254 | 0.00163 |
| ENSMUSG00000036892.13 | 271.3829 | 0.039725 | 0.578422 | 0.068678 | 0.945246 | 0.996254 | 0.00163 |
| ENSMUSG00000036955.17 | 95.95858 | -0.0802 | 0.899829 | -0.08913 | 0.928979 | 0.996254 | 0.00163 |
| ENSMUSG00000036986.17 | 13.84853 | 0.246742 | 2.01898 | 0.122211 | 0.902732 | 0.996254 | 0.00163 |
| ENSMUSG00000037017.16 | 104.815 | -0.06766 | 0.920008 | -0.07354 | 0.941376 | 0.996254 | 0.00163 |
| ENSMUSG00000037152.12 | 772.2748 | 0.049443 | 0.419482 | 0.117866 | 0.906174 | 0.996254 | 0.00163 |
| ENSMUSG00000037316.10 | 203.6417 | 0.082086 | 0.647777 | 0.12672 | 0.899162 | 0.996254 | 0.00163 |
| ENSMUSG00000037434.8 | 227.5984 | 0.061316 | 0.587623 | 0.104347 | 0.916894 | 0.996254 | 0.00163 |
| ENSMUSG00000037458.16 | 490.3642 | 0.045349 | 0.406573 | 0.111539 | 0.911189 | 0.996254 | 0.00163 |
| ENSMUSG00000037503.13 | 91.16118 | -0.11927 | 0.964064 | -0.12371 | 0.901544 | 0.996254 | 0.00163 |
| ENSMUSG00000037533.21 | 80.67459 | -0.0741 | 1.025596 | -0.07225 | 0.9424 | 0.996254 | 0.00163 |
| ENSMUSG00000037536.15 | 114.0888 | 0.05357 | 0.743522 | 0.072049 | 0.942563 | 0.996254 | 0.00163 |
| ENSMUSG00000037905.14 | 137.7013 | -0.06115 | 0.664387 | -0.09204 | 0.92667 | 0.996254 | 0.00163 |
| ENSMUSG00000037957.15 | 194.274 | 0.069502 | 0.562506 | 0.123558 | 0.901666 | 0.996254 | 0.00163 |
| ENSMUSG00000037993.9 | 20.60016 | 0.239564 | 2.124844 | 0.112744 | 0.910233 | 0.996254 | 0.00163 |
| ENSMUSG00000038025.9 | 16.25578 | -0.20885 | 2.539888 | -0.08223 | 0.934465 | 0.996254 | 0.00163 |
| ENSMUSG00000038046.5 | 11.64476 | 0.171574 | 2.19476 | 0.078174 | 0.937689 | 0.996254 | 0.00163 |
| ENSMUSG00000038280.13 | 85.43443 | 0.081519 | 0.843796 | 0.09661 | 0.923036 | 0.996254 | 0.00163 |
| ENSMUSG00000038286.12 | 778.4799 | -0.03141 | 0.322731 | -0.09733 | 0.922461 | 0.996254 | 0.00163 |
| ENSMUSG00000038324.14 | 57.58374 | -0.1254 | 1.01301 | -0.12379 | 0.901478 | 0.996254 | 0.00163 |
| ENSMUSG00000038368.17 | 35.2592 | 0.140147 | 1.554569 | 0.090152 | 0.928166 | 0.996254 | 0.00163 |
| ENSMUSG00000038371.16 | 49.67197 | -0.11393 | 1.116058 | -0.10209 | 0.918689 | 0.996254 | 0.00163 |
| ENSMUSG00000038415.11 | 204.4688 | -0.05569 | 0.777865 | -0.07159 | 0.942929 | 0.996254 | 0.00163 |
| ENSMUSG00000038446.9 | 85.34087 | 0.061536 | 0.902741 | 0.068166 | 0.945653 | 0.996254 | 0.00163 |
| ENSMUSG00000038482.12 | 49.74112 | 0.099571 | 1.097429 | 0.090731 | 0.927707 | 0.996254 | 0.00163 |
| ENSMUSG00000038527.10 | 246.1918 | -0.06927 | 0.590166 | -0.11738 | 0.906558 | 0.996254 | 0.00163 |

| | | | | | | | |
|---|---|---|---|---|---|---|---|
| ENSMUSG00000038533.16 | 17.27937 | 0.131257 | 1.799639 | 0.072935 | 0.941858 | 0.996254 | 0.00163 |
| ENSMUSG00000038578.16 | 20.78606 | 0.151843 | 1.542664 | 0.098429 | 0.921592 | 0.996254 | 0.00163 |
| ENSMUSG00000038848.15 | 147.6586 | 0.064919 | 0.794586 | 0.081702 | 0.934884 | 0.996254 | 0.00163 |
| ENSMUSG00000038982.11 | 187.4469 | 0.064667 | 0.700927 | 0.092259 | 0.926492 | 0.996254 | 0.00163 |
| ENSMUSG00000039164.3 | 19.41237 | 0.269446 | 2.822636 | 0.095459 | 0.92395 | 0.996254 | 0.00163 |
| ENSMUSG00000039461.13 | 182.032 | -0.04394 | 0.648213 | -0.06779 | 0.945952 | 0.996254 | 0.00163 |
| ENSMUSG00000039633.8 | 61.54753 | -0.14006 | 1.168636 | -0.11985 | 0.904605 | 0.996254 | 0.00163 |
| ENSMUSG00000039831.17 | 197.0107 | 0.04741 | 0.519333 | 0.091291 | 0.927262 | 0.996254 | 0.00163 |
| ENSMUSG00000039952.16 | 284.423 | -0.03384 | 0.465861 | -0.07265 | 0.942085 | 0.996254 | 0.00163 |
| ENSMUSG00000040044.12 | 75.1058 | 0.05821 | 0.859581 | 0.067719 | 0.946009 | 0.996254 | 0.00163 |
| ENSMUSG00000040128.10 | 584.5388 | 0.046065 | 0.439335 | 0.104851 | 0.916494 | 0.996254 | 0.00163 |
| ENSMUSG00000040134.9 | 2109.132 | 0.033127 | 0.266484 | 0.124311 | 0.901069 | 0.996254 | 0.00163 |
| ENSMUSG00000040170.14 | 103.733 | -0.10107 | 1.371941 | -0.07367 | 0.941275 | 0.996254 | 0.00163 |
| ENSMUSG00000040365.15 | 101.1681 | -0.05999 | 0.832932 | -0.07202 | 0.942583 | 0.996254 | 0.00163 |
| ENSMUSG00000040414.9 | 60.63282 | -0.13054 | 1.254169 | -0.10409 | 0.9171 | 0.996254 | 0.00163 |
| ENSMUSG00000040725.14 | 127.6217 | 0.082476 | 0.737232 | 0.111872 | 0.910925 | 0.996254 | 0.00163 |
| ENSMUSG00000040843.11 | 148.5416 | -0.06712 | 0.570494 | -0.11765 | 0.906345 | 0.996254 | 0.00163 |
| ENSMUSG00000041130.11 | 217.8486 | 0.08718 | 0.778867 | 0.111932 | 0.910877 | 0.996254 | 0.00163 |
| ENSMUSG00000041133.12 | 556.6102 | 0.03921 | 0.424461 | 0.092377 | 0.926399 | 0.996254 | 0.00163 |
| ENSMUSG00000041439.16 | 28.76521 | 0.08578 | 1.245988 | 0.068845 | 0.945113 | 0.996254 | 0.00163 |
| ENSMUSG00000041459.16 | 392.4971 | -0.03605 | 0.379287 | -0.09506 | 0.92427 | 0.996254 | 0.00163 |
| ENSMUSG00000041471.10 | 157.7335 | -0.04682 | 0.610497 | -0.0767 | 0.938864 | 0.996254 | 0.00163 |
| ENSMUSG00000041598.8 | 108.7119 | -0.06565 | 0.895289 | -0.07332 | 0.941549 | 0.996254 | 0.00163 |
| ENSMUSG00000041712.8 | 36.56112 | -0.11371 | 1.556276 | -0.07306 | 0.941756 | 0.996254 | 0.00163 |
| ENSMUSG00000041747.4 | 31.10252 | -0.21745 | 1.893088 | -0.11487 | 0.908552 | 0.996254 | 0.00163 |
| ENSMUSG00000041775.8 | 33.05363 | 0.104538 | 1.428088 | 0.073201 | 0.941646 | 0.996254 | 0.00163 |
| ENSMUSG00000041837.5 | 37.75858 | 0.155717 | 1.294975 | 0.120247 | 0.904288 | 0.996254 | 0.00163 |
| ENSMUSG00000042032.14 | 223.2871 | 0.079688 | 0.654926 | 0.121676 | 0.903156 | 0.996254 | 0.00163 |
| ENSMUSG00000042148.9 | 57.50835 | 0.112941 | 1.129864 | 0.099959 | 0.920376 | 0.996254 | 0.00163 |
| ENSMUSG00000042208.16 | 156.4675 | 0.082052 | 0.701274 | 0.117004 | 0.906857 | 0.996254 | 0.00163 |
| ENSMUSG00000042462.5 | 107.8456 | -0.06198 | 0.905095 | -0.06848 | 0.945406 | 0.996254 | 0.00163 |
| ENSMUSG00000042613.10 | 22.01236 | 0.20791 | 2.761819 | 0.07528 | 0.939992 | 0.996254 | 0.00163 |
| ENSMUSG00000042699.12 | 248.1851 | -0.06695 | 0.54109 | -0.12374 | 0.901523 | 0.996254 | 0.00163 |
| ENSMUSG00000042737.10 | 269.8944 | 0.045987 | 0.592156 | 0.077659 | 0.938099 | 0.996254 | 0.00163 |
| ENSMUSG00000043964.15 | 38.08715 | 0.169224 | 1.694298 | 0.099879 | 0.920441 | 0.996254 | 0.00163 |
| ENSMUSG00000044080.10 | 443.9756 | -0.04549 | 0.478065 | -0.09516 | 0.924184 | 0.996254 | 0.00163 |
| ENSMUSG00000044279.16 | 31.13125 | 0.14562 | 1.492278 | 0.097582 | 0.922264 | 0.996254 | 0.00163 |
| ENSMUSG00000044308.18 | 623.6324 | -0.03432 | 0.41271 | -0.08316 | 0.933726 | 0.996254 | 0.00163 |
| ENSMUSG00000045005.10 | 30.26331 | -0.16188 | 1.305443 | -0.124 | 0.901313 | 0.996254 | 0.00163 |
| ENSMUSG00000045128.10 | 935.6828 | -0.02716 | 0.400704 | -0.06778 | 0.945958 | 0.996254 | 0.00163 |
| ENSMUSG00000045302.14 | 174.9294 | 0.080436 | 0.780089 | 0.103112 | 0.917874 | 0.996254 | 0.00163 |
| ENSMUSG00000045438.13 | 193.7297 | -0.06681 | 0.580898 | -0.11502 | 0.908432 | 0.996254 | 0.00163 |
| ENSMUSG00000045576.17 | 177.6467 | 0.087613 | 0.818214 | 0.107078 | 0.914727 | 0.996254 | 0.00163 |
| ENSMUSG00000046034.9 | 210.5772 | -0.06389 | 0.664092 | -0.09621 | 0.923358 | 0.996254 | 0.00163 |

| | | | | | | | |
|---|---|---|---|---|---|---|---|
| ENSMUSG00000046330.11 | 1690.934 | -0.02555 | 0.342673 | -0.07455 | 0.940573 | 0.996254 | 0.00163 |
| ENSMUSG00000046862.15 | 28.42076 | -0.18874 | 1.59094 | -0.11864 | 0.905563 | 0.996254 | 0.00163 |
| ENSMUSG00000046916.10 | 38.08807 | 0.147245 | 1.414503 | 0.104097 | 0.917093 | 0.996254 | 0.00163 |
| ENSMUSG00000047583.10 | 13.9435 | -0.18787 | 2.682019 | -0.07005 | 0.944154 | 0.996254 | 0.00163 |
| ENSMUSG00000047804.16 | 18.29142 | -0.20347 | 1.807668 | -0.11256 | 0.910382 | 0.996254 | 0.00163 |
| ENSMUSG00000048217.12 | 161.65 | 0.051722 | 0.60637 | 0.085297 | 0.932025 | 0.996254 | 0.00163 |
| ENSMUSG00000048482.15 | 29.31192 | 0.120444 | 1.372208 | 0.087774 | 0.930057 | 0.996254 | 0.00163 |
| ENSMUSG00000048720.9 | 78.65418 | -0.08442 | 0.875817 | -0.0964 | 0.923206 | 0.996254 | 0.00163 |
| ENSMUSG00000048756.12 | 143.3099 | 0.052027 | 0.744221 | 0.069908 | 0.944267 | 0.996254 | 0.00163 |
| ENSMUSG00000048960.14 | 40.21298 | 0.111955 | 1.165355 | 0.096069 | 0.923466 | 0.996254 | 0.00163 |
| ENSMUSG00000049300.12 | 40.47317 | -0.14582 | 1.831245 | -0.07963 | 0.936533 | 0.996254 | 0.00163 |
| ENSMUSG00000049751.7 | 82.84236 | -0.07472 | 0.830364 | -0.08998 | 0.928303 | 0.996254 | 0.00163 |
| ENSMUSG00000049775.17 | 1182.806 | -0.0233 | 0.303739 | -0.07672 | 0.93885 | 0.996254 | 0.00163 |
| ENSMUSG00000049881.14 | 17.72763 | -0.15987 | 2.23647 | -0.07149 | 0.943012 | 0.996254 | 0.00163 |
| ENSMUSG00000050567.17 | 51.74199 | 0.092416 | 1.018352 | 0.090751 | 0.927691 | 0.996254 | 0.00163 |
| ENSMUSG00000050621.8 | 124.2823 | -0.04993 | 0.61748 | -0.08086 | 0.93555 | 0.996254 | 0.00163 |
| ENSMUSG00000050732.10 | 304.489 | 0.049189 | 0.666675 | 0.073783 | 0.941183 | 0.996254 | 0.00163 |
| ENSMUSG00000050786.9 | 25.90665 | -0.14635 | 1.823783 | -0.08024 | 0.936043 | 0.996254 | 0.00163 |
| ENSMUSG00000051238.7 | 21.05723 | 0.222187 | 2.0427 | 0.108771 | 0.913384 | 0.996254 | 0.00163 |
| ENSMUSG00000051723.8 | 21.43931 | -0.13267 | 1.361504 | -0.09745 | 0.922372 | 0.996254 | 0.00163 |
| ENSMUSG00000052214.10 | 75.0826 | 0.078734 | 1.167445 | 0.067441 | 0.946231 | 0.996254 | 0.00163 |
| ENSMUSG00000052337.16 | 374.1933 | -0.05389 | 0.452571 | -0.11907 | 0.905217 | 0.996254 | 0.00163 |
| ENSMUSG00000052632.17 | 30.45351 | 0.198727 | 1.577113 | 0.126007 | 0.899727 | 0.996254 | 0.00163 |
| ENSMUSG00000053119.12 | 115.093 | -0.07765 | 0.766356 | -0.10133 | 0.919291 | 0.996254 | 0.00163 |
| ENSMUSG00000053205.10 | 33.44733 | -0.08511 | 1.25495 | -0.06782 | 0.945932 | 0.996254 | 0.00163 |
| ENSMUSG00000053477.19 | 127.1536 | -0.08489 | 0.746459 | -0.11373 | 0.909455 | 0.996254 | 0.00163 |
| ENSMUSG00000053604.6 | 29.27581 | 0.111353 | 1.650725 | 0.067457 | 0.946218 | 0.996254 | 0.00163 |
| ENSMUSG00000053907.11 | 217.5767 | 0.052495 | 0.530569 | 0.098941 | 0.921185 | 0.996254 | 0.00163 |
| ENSMUSG00000054414.5 | 156.461 | 0.069471 | 0.657489 | 0.105661 | 0.915851 | 0.996254 | 0.00163 |
| ENSMUSG00000055024.13 | 33.38524 | 0.084498 | 1.204911 | 0.070128 | 0.944092 | 0.996254 | 0.00163 |
| ENSMUSG00000055027.18 | 11.70082 | -0.22893 | 2.470158 | -0.09268 | 0.92616 | 0.996254 | 0.00163 |
| ENSMUSG00000055204.16 | 346.3325 | 0.037989 | 0.490064 | 0.077519 | 0.93821 | 0.996254 | 0.00163 |
| ENSMUSG00000056978.9 | 142.1267 | -0.092 | 0.877293 | -0.10486 | 0.916483 | 0.996254 | 0.00163 |
| ENSMUSG00000057329.8 | 24.82665 | -0.18356 | 1.738578 | -0.10558 | 0.915917 | 0.996254 | 0.00163 |
| ENSMUSG00000057335.12 | 25.83313 | -0.11015 | 1.331491 | -0.08273 | 0.934069 | 0.996254 | 0.00163 |
| ENSMUSG00000057554.14 | 402.374 | -0.0503 | 0.403012 | -0.12482 | 0.900665 | 0.996254 | 0.00163 |
| ENSMUSG00000057637.14 | 28.48779 | -0.17373 | 1.574473 | -0.11034 | 0.912139 | 0.996254 | 0.00163 |
| ENSMUSG00000059423.14 | 11.11367 | 0.242485 | 2.591287 | 0.093577 | 0.925445 | 0.996254 | 0.00163 |
| ENSMUSG00000059436.14 | 48.07179 | -0.12678 | 1.36377 | -0.09296 | 0.925934 | 0.996254 | 0.00163 |
| ENSMUSG00000059439.16 | 24.65551 | 0.145228 | 1.451726 | 0.100038 | 0.920314 | 0.996254 | 0.00163 |
| ENSMUSG00000059588.14 | 19.8771 | -0.18552 | 1.551759 | -0.11956 | 0.904835 | 0.996254 | 0.00163 |
| ENSMUSG00000060019.7 | 558.1672 | -0.04395 | 0.348571 | -0.12608 | 0.899667 | 0.996254 | 0.00163 |
| ENSMUSG00000060301.7 | 32.00266 | 0.123064 | 1.313363 | 0.093701 | 0.925347 | 0.996254 | 0.00163 |
| ENSMUSG00000060459.15 | 1510.741 | -0.02803 | 0.29782 | -0.09412 | 0.925015 | 0.996254 | 0.00163 |

| | | | | | | | |
|---|---|---|---|---|---|---|---|
| ENSMUSG00000060510.15 | 70.2088 | -0.06936 | 0.840074 | -0.08257 | 0.934195 | 0.996254 | 0.00163 |
| ENSMUSG00000061461.12 | 75.622 | -0.07649 | 0.831099 | -0.09204 | 0.926668 | 0.996254 | 0.00163 |
| ENSMUSG00000061751.17 | 35.9684 | -0.14373 | 1.476046 | -0.09737 | 0.922429 | 0.996254 | 0.00163 |
| ENSMUSG00000061755.11 | 234.1764 | 0.054237 | 0.662867 | 0.081822 | 0.934788 | 0.996254 | 0.00163 |
| ENSMUSG00000061759.16 | 124.9351 | -0.07928 | 0.730573 | -0.10851 | 0.91359 | 0.996254 | 0.00163 |
| ENSMUSG00000061981.15 | 79.40488 | 0.132162 | 1.208674 | 0.109345 | 0.912929 | 0.996254 | 0.00163 |
| ENSMUSG00000062397.9 | 942.0319 | -0.02569 | 0.314128 | -0.08178 | 0.934823 | 0.996254 | 0.00163 |
| ENSMUSG00000062563.16 | 10.80353 | 0.184609 | 1.88923 | 0.097717 | 0.922157 | 0.996254 | 0.00163 |
| ENSMUSG00000062867.14 | 168.3917 | -0.07489 | 0.732045 | -0.10231 | 0.918512 | 0.996254 | 0.00163 |
| ENSMUSG00000063171.5 | 38.75592 | -0.10547 | 1.558619 | -0.06767 | 0.946049 | 0.996254 | 0.00163 |
| ENSMUSG00000063952.17 | 44.40295 | -0.16183 | 1.293691 | -0.12509 | 0.90045 | 0.996254 | 0.00163 |
| ENSMUSG00000064145.13 | 135.5908 | -0.05729 | 0.763102 | -0.07507 | 0.94016 | 0.996254 | 0.00163 |
| ENSMUSG00000064215.14 | 524.3087 | -0.03527 | 0.473276 | -0.07453 | 0.94059 | 0.996254 | 0.00163 |
| ENSMUSG00000064346.1 | 10.03141 | 0.250564 | 2.583122 | 0.097 | 0.922726 | 0.996254 | 0.00163 |
| ENSMUSG00000065126.3 | 68.25774 | 0.118743 | 1.167885 | 0.101674 | 0.919016 | 0.996254 | 0.00163 |
| ENSMUSG00000066235.8 | 12.95182 | -0.26133 | 3.131704 | -0.08345 | 0.933495 | 0.996254 | 0.00163 |
| ENSMUSG00000066357.7 | 42.44948 | -0.11682 | 1.180448 | -0.09896 | 0.921168 | 0.996254 | 0.00163 |
| ENSMUSG00000068083.3 | 248.5578 | 0.054665 | 0.530687 | 0.103009 | 0.917956 | 0.996254 | 0.00163 |
| ENSMUSG00000068250.14 | 98.56759 | -0.09977 | 1.010118 | -0.09877 | 0.921323 | 0.996254 | 0.00163 |
| ENSMUSG00000068523.13 | 261.4045 | 0.05304 | 0.476419 | 0.11133 | 0.911355 | 0.996254 | 0.00163 |
| ENSMUSG00000068587.11 | 719.4346 | -0.02872 | 0.344106 | -0.08345 | 0.93349 | 0.996254 | 0.00163 |
| ENSMUSG00000069089.9 | 114.1971 | 0.080196 | 0.96858 | 0.082797 | 0.934013 | 0.996254 | 0.00163 |
| ENSMUSG00000069237.9 | 469.5442 | -0.04172 | 0.376356 | -0.11084 | 0.911744 | 0.996254 | 0.00163 |
| ENSMUSG00000069678.11 | 13.0108 | -0.20403 | 2.381979 | -0.08566 | 0.931739 | 0.996254 | 0.00163 |
| ENSMUSG00000069733.12 | 27.16153 | -0.13011 | 1.511776 | -0.08606 | 0.931418 | 0.996254 | 0.00163 |
| ENSMUSG00000069919.8 | 112.5947 | 0.06327 | 0.909183 | 0.06959 | 0.94452 | 0.996254 | 0.00163 |
| ENSMUSG00000070047.15 | 11.52441 | -0.27862 | 2.790758 | -0.09984 | 0.920473 | 0.996254 | 0.00163 |
| ENSMUSG00000071337.12 | 55.37992 | 0.138785 | 1.327373 | 0.104556 | 0.916728 | 0.996254 | 0.00163 |
| ENSMUSG00000071856.11 | 100.7763 | -0.06652 | 0.691422 | -0.09621 | 0.923353 | 0.996254 | 0.00163 |
| ENSMUSG00000073295.5 | 12.76099 | 0.225737 | 3.144494 | 0.071788 | 0.942771 | 0.996254 | 0.00163 |
| ENSMUSG00000073700.4 | 84.03934 | 0.083489 | 0.864996 | 0.09652 | 0.923108 | 0.996254 | 0.00163 |
| ENSMUSG00000073888.14 | 54.83166 | -0.1335 | 1.258258 | -0.1061 | 0.915503 | 0.996254 | 0.00163 |
| ENSMUSG00000074093.5 | 118.8877 | -0.08408 | 0.933473 | -0.09008 | 0.928227 | 0.996254 | 0.00163 |
| ENSMUSG00000074800.5 | 103.9396 | 0.060321 | 0.710679 | 0.084877 | 0.932359 | 0.996254 | 0.00163 |
| ENSMUSG00000074802.12 | 12.77675 | 0.273534 | 2.45897 | 0.111239 | 0.911427 | 0.996254 | 0.00163 |
| ENSMUSG00000074994.7 | 87.05331 | -0.06369 | 0.841599 | -0.07567 | 0.939679 | 0.996254 | 0.00163 |
| ENSMUSG00000078193.3 | 315.222 | -0.04979 | 0.444547 | -0.112 | 0.910822 | 0.996254 | 0.00163 |
| ENSMUSG00000078619.11 | 137.3691 | -0.0739 | 0.710015 | -0.10409 | 0.9171 | 0.996254 | 0.00163 |
| ENSMUSG00000079020.10 | 10.80025 | -0.27236 | 3.263912 | -0.08345 | 0.933497 | 0.996254 | 0.00163 |
| ENSMUSG00000080006.5 | 11.66326 | -0.22277 | 2.48455 | -0.08966 | 0.928554 | 0.996254 | 0.00163 |
| ENSMUSG00000083844.9 | 73.39961 | 0.089035 | 1.313805 | 0.067769 | 0.94597 | 0.996254 | 0.00163 |
| ENSMUSG00000084265.2 | 12.21344 | 0.299259 | 3.165692 | 0.094532 | 0.924687 | 0.996254 | 0.00163 |
| ENSMUSG00000084350.2 | 11.25625 | -0.2523 | 3.226855 | -0.07819 | 0.937679 | 0.996254 | 0.00163 |
| ENSMUSG00000084790.2 | 16.92927 | 0.236604 | 2.281454 | 0.103708 | 0.917401 | 0.996254 | 0.00163 |

| | | | | | | | |
|---|---|---|---|---|---|---|---|
| ENSMUSG00000085795.9 | 10.34222 | 0.300875 | 3.288789 | 0.091485 | 0.927107 | 0.996254 | 0.00163 |
| ENSMUSG00000086316.9 | 17.00608 | -0.15264 | 1.878383 | -0.08126 | 0.935234 | 0.996254 | 0.00163 |
| ENSMUSG00000086390.8 | 37.4268 | -0.15768 | 1.566346 | -0.10067 | 0.919813 | 0.996254 | 0.00163 |
| ENSMUSG00000086567.2 | 10.98858 | 0.310887 | 2.952236 | 0.105306 | 0.916133 | 0.996254 | 0.00163 |
| ENSMUSG00000087163.2 | 11.33791 | 0.313921 | 2.788732 | 0.112568 | 0.910373 | 0.996254 | 0.00163 |
| ENSMUSG00000089682.11 | 53.37658 | -0.08858 | 0.974174 | -0.09093 | 0.927546 | 0.996254 | 0.00163 |
| ENSMUSG00000090000.3 | 412.5221 | -0.04358 | 0.531691 | -0.08196 | 0.934677 | 0.996254 | 0.00163 |
| ENSMUSG00000090290.3 | 11.86977 | 0.174919 | 2.475999 | 0.070646 | 0.94368 | 0.996254 | 0.00163 |
| ENSMUSG00000092220.2 | 44.54183 | 0.126369 | 1.280296 | 0.098703 | 0.921374 | 0.996254 | 0.00163 |
| ENSMUSG00000092626.2 | 20.30499 | -0.3074 | 2.450179 | -0.12546 | 0.900159 | 0.996254 | 0.00163 |
| ENSMUSG00000094410.9 | 11.79971 | -0.27526 | 2.762434 | -0.09964 | 0.920628 | 0.996254 | 0.00163 |
| ENSMUSG00000094557.2 | 10.78645 | -0.23168 | 1.868603 | -0.12399 | 0.901326 | 0.996254 | 0.00163 |
| ENSMUSG00000097080.9 | 14.75817 | 0.292478 | 2.616047 | 0.111802 | 0.910981 | 0.996254 | 0.00163 |
| ENSMUSG00000097333.8 | 19.94807 | -0.13655 | 1.6277 | -0.08389 | 0.933145 | 0.996254 | 0.00163 |
| ENSMUSG00000097571.10 | 24.51098 | -0.122 | 1.389411 | -0.08781 | 0.930029 | 0.996254 | 0.00163 |
| ENSMUSG00000098134.2 | 61.28043 | -0.08008 | 1.167887 | -0.06856 | 0.945336 | 0.996254 | 0.00163 |
| ENSMUSG00000100164.2 | 18.25813 | 0.227928 | 2.446979 | 0.093147 | 0.925787 | 0.996254 | 0.00163 |
| ENSMUSG00000102869.3 | 14.59325 | -0.17949 | 2.339862 | -0.07671 | 0.938854 | 0.996254 | 0.00163 |
| ENSMUSG00000106352.5 | 176.6462 | -0.06712 | 0.675304 | -0.09939 | 0.920832 | 0.996254 | 0.00163 |
| ENSMUSG00000107002.2 | 491.8327 | 0.059294 | 0.478333 | 0.123959 | 0.901348 | 0.996254 | 0.00163 |
| ENSMUSG00000109089.2 | 47.91767 | 0.099873 | 1.087009 | 0.091878 | 0.926795 | 0.996254 | 0.00163 |
| ENSMUSG00000112567.2 | 41.00327 | 0.087153 | 1.05215 | 0.082833 | 0.933984 | 0.996254 | 0.00163 |
| ENSMUSG00000113249.2 | 27.765 | 0.157597 | 1.326265 | 0.118828 | 0.905412 | 0.996254 | 0.00163 |
| ENSMUSG00000113902.2 | 2364.671 | -0.0186 | 0.269255 | -0.06909 | 0.94492 | 0.996254 | 0.00163 |
| ENSMUSG00000114103.2 | 17.42256 | -0.36615 | 2.910678 | -0.1258 | 0.899893 | 0.996254 | 0.00163 |
| ENSMUSG00000115100.2 | 11.54817 | -0.33497 | 2.844167 | -0.11778 | 0.906245 | 0.996254 | 0.00163 |
| ENSMUSG00000117327.2 | 27.22004 | -0.10937 | 1.355206 | -0.0807 | 0.93568 | 0.996254 | 0.00163 |
| ENSMUSG00000118516.2 | 40.76099 | -0.08915 | 1.271032 | -0.07014 | 0.94408 | 0.996254 | 0.00163 |
| ENSMUSG00000118669.2 | 88.64531 | 0.113812 | 1.429554 | 0.079613 | 0.936545 | 0.996254 | 0.00163 |
| ENSMUSG00000002984.18 | 19.59865 | -0.16453 | 2.469262 | -0.06663 | 0.946875 | 0.996547 | 0.001502 |
| ENSMUSG00000014329.15 | 32.93427 | 0.098861 | 1.495051 | 0.066125 | 0.947278 | 0.996547 | 0.001502 |
| ENSMUSG00000025404.16 | 85.87987 | -0.06354 | 0.98182 | -0.06471 | 0.948401 | 0.996547 | 0.001502 |
| ENSMUSG00000031429.15 | 173.0077 | -0.05085 | 0.779066 | -0.06526 | 0.947964 | 0.996547 | 0.001502 |
| ENSMUSG00000034007.11 | 55.25119 | -0.09395 | 1.425973 | -0.06588 | 0.947471 | 0.996547 | 0.001502 |
| ENSMUSG00000034127.16 | 12.9081 | -0.20639 | 3.097607 | -0.06663 | 0.946878 | 0.996547 | 0.001502 |
| ENSMUSG00000034292.14 | 34.17232 | -0.099 | 1.504213 | -0.06582 | 0.947525 | 0.996547 | 0.001502 |
| ENSMUSG00000034757.16 | 24.30739 | -0.1104 | 1.703659 | -0.0648 | 0.94833 | 0.996547 | 0.001502 |
| ENSMUSG00000035258.16 | 13.10291 | -0.13394 | 2.06632 | -0.06482 | 0.948317 | 0.996547 | 0.001502 |
| ENSMUSG00000038623.10 | 13.45889 | -0.16478 | 2.51956 | -0.0654 | 0.947854 | 0.996547 | 0.001502 |
| ENSMUSG00000039670.3 | 31.87438 | 0.111039 | 1.708008 | 0.065011 | 0.948165 | 0.996547 | 0.001502 |
| ENSMUSG00000040242.15 | 349.7939 | 0.037937 | 0.5791 | 0.06551 | 0.947768 | 0.996547 | 0.001502 |
| ENSMUSG00000044285.9 | 45.59124 | 0.068864 | 1.033046 | 0.066661 | 0.946851 | 0.996547 | 0.001502 |
| ENSMUSG00000062353.7 | 86.49442 | -0.04835 | 0.739679 | -0.06537 | 0.947883 | 0.996547 | 0.001502 |
| ENSMUSG00000063972.14 | 40.67395 | 0.072932 | 1.124887 | 0.064835 | 0.948306 | 0.996547 | 0.001502 |

| | | | | | | | |
|---|---|---|---|---|---|---|---|
| ENSMUSG00000066415.5 | 87.25944 | 0.060157 | 0.915848 | 0.065684 | 0.947629 | 0.996547 | 0.001502 |
| ENSMUSG00000070002.8 | 64.03087 | -0.07047 | 1.076532 | -0.06546 | 0.94781 | 0.996547 | 0.001502 |
| ENSMUSG00002075795.1 | 29.11494 | 0.112477 | 1.736715 | 0.064764 | 0.948362 | 0.996547 | 0.001502 |
| ENSMUSG00000005161.16 | 649.7984 | 0.023321 | 0.362021 | 0.064418 | 0.948638 | 0.996574 | 0.00149 |
| ENSMUSG00000021432.16 | 43.4087 | 0.079934 | 1.238715 | 0.06453 | 0.948548 | 0.996574 | 0.00149 |
| ENSMUSG00000032399.9 | 4179.758 | -0.01396 | 0.218171 | -0.064 | 0.948968 | 0.996723 | 0.001425 |
| ENSMUSG00000064370.1 | 62661.4 | 0.012653 | 0.197784 | 0.063976 | 0.94899 | 0.996723 | 0.001425 |
| ENSMUSG00000090946.4 | 19.07871 | -0.11088 | 1.740125 | -0.06372 | 0.949192 | 0.996825 | 0.001381 |
| ENSMUSG00000034111.7 | 28.61874 | 0.127243 | 2.005345 | 0.063452 | 0.949406 | 0.99694 | 0.001331 |
| ENSMUSG00000000056.8 | 75.67506 | 0.050142 | 0.906011 | 0.055344 | 0.955864 | 0.997337 | 0.001158 |
| ENSMUSG00000001036.18 | 12.76722 | -0.15364 | 2.79974 | -0.05488 | 0.956237 | 0.997337 | 0.001158 |
| ENSMUSG00000001891.17 | 1582.927 | 0.01675 | 0.289075 | 0.057943 | 0.953794 | 0.997337 | 0.001158 |
| ENSMUSG00000002345.19 | 17.98794 | 0.106448 | 2.358815 | 0.045128 | 0.964006 | 0.997337 | 0.001158 |
| ENSMUSG00000003380.12 | 35.16695 | 0.061007 | 1.175198 | 0.051912 | 0.958599 | 0.997337 | 0.001158 |
| ENSMUSG00000003541.7 | 10.33266 | 0.188175 | 3.265906 | 0.057618 | 0.954053 | 0.997337 | 0.001158 |
| ENSMUSG00000004455.17 | 11.27443 | -0.12816 | 2.836517 | -0.04518 | 0.963963 | 0.997337 | 0.001158 |
| ENSMUSG00000004788.12 | 24.58116 | -0.07535 | 1.755566 | -0.04292 | 0.965765 | 0.997337 | 0.001158 |
| ENSMUSG00000005089.16 | 666.751 | 0.014504 | 0.374514 | 0.038726 | 0.969109 | 0.997337 | 0.001158 |
| ENSMUSG00000005354.17 | 640.5427 | 0.012253 | 0.335048 | 0.036571 | 0.970827 | 0.997337 | 0.001158 |
| ENSMUSG00000005362.15 | 176.4027 | -0.0274 | 0.721217 | -0.03799 | 0.969694 | 0.997337 | 0.001158 |
| ENSMUSG00000005378.18 | 28.11551 | -0.06758 | 1.649237 | -0.04098 | 0.967313 | 0.997337 | 0.001158 |
| ENSMUSG00000005886.15 | 205.7999 | -0.03261 | 0.704719 | -0.04628 | 0.963089 | 0.997337 | 0.001158 |
| ENSMUSG00000005973.7 | 14.72268 | 0.126808 | 2.075994 | 0.061083 | 0.951293 | 0.997337 | 0.001158 |
| ENSMUSG00000006998.17 | 272.6946 | 0.025589 | 0.498532 | 0.051328 | 0.959064 | 0.997337 | 0.001158 |
| ENSMUSG00000014355.11 | 153.4264 | 0.032606 | 0.772604 | 0.042203 | 0.966337 | 0.997337 | 0.001158 |
| ENSMUSG00000014551.9 | 116.239 | -0.05155 | 0.843206 | -0.06114 | 0.951251 | 0.997337 | 0.001158 |
| ENSMUSG00000015112.16 | 725.0787 | -0.02703 | 0.494133 | -0.05469 | 0.956384 | 0.997337 | 0.001158 |
| ENSMUSG00000015176.12 | 104.942 | -0.03484 | 0.826055 | -0.04218 | 0.966354 | 0.997337 | 0.001158 |
| ENSMUSG00000016382.16 | 295.3863 | -0.0264 | 0.449307 | -0.05876 | 0.953147 | 0.997337 | 0.001158 |
| ENSMUSG00000017221.15 | 26.31969 | -0.07729 | 1.721261 | -0.0449 | 0.964186 | 0.997337 | 0.001158 |
| ENSMUSG00000018572.11 | 31.09431 | 0.06343 | 1.380137 | 0.045959 | 0.963343 | 0.997337 | 0.001158 |
| ENSMUSG00000018677.10 | 374.419 | 0.026683 | 0.528603 | 0.050478 | 0.959741 | 0.997337 | 0.001158 |
| ENSMUSG00000018707.14 | 122.8268 | -0.04371 | 0.943613 | -0.04633 | 0.96305 | 0.997337 | 0.001158 |
| ENSMUSG00000019256.18 | 218.3217 | 0.032123 | 0.579354 | 0.055447 | 0.955782 | 0.997337 | 0.001158 |
| ENSMUSG00000019761.11 | 63.27874 | 0.058274 | 1.075136 | 0.054202 | 0.956775 | 0.997337 | 0.001158 |
| ENSMUSG00000019916.15 | 22.42679 | 0.06268 | 1.751714 | 0.035782 | 0.971456 | 0.997337 | 0.001158 |
| ENSMUSG00000020486.20 | 12.98477 | -0.09907 | 2.021476 | -0.04901 | 0.960913 | 0.997337 | 0.001158 |
| ENSMUSG00000020659.17 | 31.05757 | -0.10035 | 2.022972 | -0.04961 | 0.960435 | 0.997337 | 0.001158 |
| ENSMUSG00000021022.10 | 15.35441 | -0.08366 | 2.169849 | -0.03856 | 0.969245 | 0.997337 | 0.001158 |
| ENSMUSG00000021065.17 | 18.56461 | -0.08194 | 1.737205 | -0.04717 | 0.962381 | 0.997337 | 0.001158 |
| ENSMUSG00000021192.17 | 43.26831 | 0.049235 | 1.128653 | 0.043623 | 0.965205 | 0.997337 | 0.001158 |
| ENSMUSG00000021400.9 | 228.2251 | 0.025895 | 0.518351 | 0.049957 | 0.960156 | 0.997337 | 0.001158 |
| ENSMUSG00000021408.16 | 69.03799 | -0.0444 | 0.961299 | -0.04618 | 0.963164 | 0.997337 | 0.001158 |
| ENSMUSG00000021635.15 | 100.856 | 0.043947 | 0.734916 | 0.059799 | 0.952316 | 0.997337 | 0.001158 |

| | | | | | | | |
|---|---|---|---|---|---|---|---|
| ENSMUSG00000021843.18 | 156.1441 | -0.02536 | 0.642829 | -0.03945 | 0.968529 | 0.997337 | 0.001158 |
| ENSMUSG00000022150.17 | 78.3023 | -0.05193 | 1.374721 | -0.03777 | 0.969869 | 0.997337 | 0.001158 |
| ENSMUSG00000022407.11 | 86.84466 | -0.06096 | 1.101468 | -0.05535 | 0.955863 | 0.997337 | 0.001158 |
| ENSMUSG00000022464.15 | 2637.654 | -0.01329 | 0.269778 | -0.04925 | 0.960723 | 0.997337 | 0.001158 |
| ENSMUSG00000022546.6 | 652.0476 | 0.015349 | 0.345123 | 0.044475 | 0.964526 | 0.997337 | 0.001158 |
| ENSMUSG00000022575.6 | 43.18618 | 0.069639 | 1.26091 | 0.055229 | 0.955956 | 0.997337 | 0.001158 |
| ENSMUSG00000022791.20 | 21.96862 | 0.074791 | 2.08653 | 0.035845 | 0.971406 | 0.997337 | 0.001158 |
| ENSMUSG00000022801.14 | 13.65167 | 0.108684 | 1.819242 | 0.059741 | 0.952362 | 0.997337 | 0.001158 |
| ENSMUSG00000022821.14 | 3474.286 | 0.008907 | 0.232424 | 0.038323 | 0.96943 | 0.997337 | 0.001158 |
| ENSMUSG00000023022.15 | 184.0959 | -0.03314 | 0.653682 | -0.0507 | 0.959567 | 0.997337 | 0.001158 |
| ENSMUSG00000023828.4 | 46.4249 | -0.04597 | 1.078099 | -0.04264 | 0.965985 | 0.997337 | 0.001158 |
| ENSMUSG00000023919.10 | 14.04437 | 0.118277 | 2.547174 | 0.046435 | 0.962964 | 0.997337 | 0.001158 |
| ENSMUSG00000024068.9 | 217.124 | 0.028365 | 0.64126 | 0.044233 | 0.964719 | 0.997337 | 0.001158 |
| ENSMUSG00000024137.10 | 55.63414 | -0.05621 | 1.30841 | -0.04296 | 0.965734 | 0.997337 | 0.001158 |
| ENSMUSG00000024234.8 | 41.53758 | -0.07781 | 1.642983 | -0.04736 | 0.962228 | 0.997337 | 0.001158 |
| ENSMUSG00000024283.16 | 100.5262 | -0.03248 | 0.775526 | -0.04188 | 0.966593 | 0.997337 | 0.001158 |
| ENSMUSG00000024498.17 | 105.0222 | -0.05161 | 1.02257 | -0.05047 | 0.959744 | 0.997337 | 0.001158 |
| ENSMUSG00000024639.6 | 114.8981 | 0.03281 | 0.741439 | 0.044251 | 0.964704 | 0.997337 | 0.001158 |
| ENSMUSG00000024695.17 | 203.9429 | 0.031322 | 0.545623 | 0.057405 | 0.954223 | 0.997337 | 0.001158 |
| ENSMUSG00000024787.15 | 74.14839 | 0.037131 | 0.95477 | 0.03889 | 0.968978 | 0.997337 | 0.001158 |
| ENSMUSG00000024900.6 | 1180.984 | 0.014292 | 0.264862 | 0.05396 | 0.956967 | 0.997337 | 0.001158 |
| ENSMUSG00000025049.7 | 36.01485 | 0.067848 | 1.566799 | 0.043304 | 0.965459 | 0.997337 | 0.001158 |
| ENSMUSG00000025086.14 | 113.6286 | -0.02721 | 0.746646 | -0.03645 | 0.970925 | 0.997337 | 0.001158 |
| ENSMUSG00000025089.16 | 890.404 | 0.017732 | 0.290178 | 0.061107 | 0.951274 | 0.997337 | 0.001158 |
| ENSMUSG00000025127.16 | 360.6577 | 0.028869 | 0.561611 | 0.051404 | 0.959004 | 0.997337 | 0.001158 |
| ENSMUSG00000025159.9 | 31.50558 | 0.086865 | 1.667447 | 0.052095 | 0.958453 | 0.997337 | 0.001158 |
| ENSMUSG00000025178.10 | 90.11053 | -0.02824 | 0.764703 | -0.03693 | 0.970538 | 0.997337 | 0.001158 |
| ENSMUSG00000025243.18 | 35.30359 | 0.064531 | 1.70098 | 0.037938 | 0.969737 | 0.997337 | 0.001158 |
| ENSMUSG00000025485.14 | 80.88676 | 0.0788 | 1.384101 | 0.056932 | 0.954599 | 0.997337 | 0.001158 |
| ENSMUSG00000025580.16 | 82.00659 | -0.04399 | 1.22356 | -0.03596 | 0.971318 | 0.997337 | 0.001158 |
| ENSMUSG00000025607.16 | 36.45824 | -0.11111 | 1.825291 | -0.06087 | 0.951463 | 0.997337 | 0.001158 |
| ENSMUSG00000025781.15 | 3661.575 | 0.011272 | 0.244267 | 0.046146 | 0.963194 | 0.997337 | 0.001158 |
| ENSMUSG00000025825.13 | 165.6859 | -0.03778 | 0.680531 | -0.05552 | 0.955725 | 0.997337 | 0.001158 |
| ENSMUSG00000025950.17 | 2211.03 | 0.014095 | 0.253833 | 0.055527 | 0.955719 | 0.997337 | 0.001158 |
| ENSMUSG00000026020.10 | 400.5993 | 0.023642 | 0.51209 | 0.046167 | 0.963177 | 0.997337 | 0.001158 |
| ENSMUSG00000026082.12 | 53.32759 | 0.086647 | 1.464743 | 0.059155 | 0.952829 | 0.997337 | 0.001158 |
| ENSMUSG00000026223.16 | 93.85597 | -0.06601 | 1.203712 | -0.05484 | 0.956267 | 0.997337 | 0.001158 |
| ENSMUSG00000026380.11 | 10.76298 | 0.168074 | 2.877114 | 0.058418 | 0.953416 | 0.997337 | 0.001158 |
| ENSMUSG00000026634.17 | 98.44856 | 0.036844 | 0.924067 | 0.039871 | 0.968196 | 0.997337 | 0.001158 |
| ENSMUSG00000026727.11 | 91.21971 | -0.0455 | 0.751926 | -0.06051 | 0.951746 | 0.997337 | 0.001158 |
| ENSMUSG00000026810.13 | 11.47596 | 0.079089 | 2.201374 | 0.035927 | 0.97134 | 0.997337 | 0.001158 |
| ENSMUSG00000027030.16 | 14.99619 | -0.11442 | 2.597181 | -0.04406 | 0.96486 | 0.997337 | 0.001158 |
| ENSMUSG00000027080.7 | 30.25187 | 0.087729 | 1.674214 | 0.0524 | 0.95821 | 0.997337 | 0.001158 |
| ENSMUSG00000027361.16 | 32.61233 | 0.077398 | 1.554743 | 0.049782 | 0.960296 | 0.997337 | 0.001158 |

| | | | | | | | |
|---|---|---|---|---|---|---|---|
| ENSMUSG00000027363.16 | 184.834 | -0.03022 | 0.691621 | -0.0437 | 0.965147 | 0.997337 | 0.001158 |
| ENSMUSG00000027519.11 | 108.4347 | -0.04496 | 0.724368 | -0.06207 | 0.950506 | 0.997337 | 0.001158 |
| ENSMUSG00000027562.13 | 34.25797 | 0.049778 | 1.352657 | 0.0368 | 0.970644 | 0.997337 | 0.001158 |
| ENSMUSG00000027663.13 | 43.13834 | -0.06101 | 1.541894 | -0.03957 | 0.968436 | 0.997337 | 0.001158 |
| ENSMUSG00000027667.14 | 120.1241 | 0.049797 | 0.851647 | 0.058471 | 0.953373 | 0.997337 | 0.001158 |
| ENSMUSG00000028069.17 | 135.7811 | -0.04409 | 0.903077 | -0.04883 | 0.961058 | 0.997337 | 0.001158 |
| ENSMUSG00000028156.13 | 256.3235 | 0.032509 | 0.671499 | 0.048413 | 0.961387 | 0.997337 | 0.001158 |
| ENSMUSG00000028167.16 | 114.6084 | 0.031681 | 0.848028 | 0.037359 | 0.970199 | 0.997337 | 0.001158 |
| ENSMUSG00000028458.13 | 43.3521 | -0.07478 | 1.31456 | -0.05688 | 0.954638 | 0.997337 | 0.001158 |
| ENSMUSG00000028614.15 | 69.80987 | 0.048824 | 1.067974 | 0.045717 | 0.963536 | 0.997337 | 0.001158 |
| ENSMUSG00000028676.18 | 124.2773 | -0.0362 | 0.746759 | -0.04848 | 0.961336 | 0.997337 | 0.001158 |
| ENSMUSG00000028688.14 | 60.31084 | 0.049713 | 1.014103 | 0.049022 | 0.960902 | 0.997337 | 0.001158 |
| ENSMUSG00000028982.10 | 84.96333 | -0.02764 | 0.760676 | -0.03634 | 0.971015 | 0.997337 | 0.001158 |
| ENSMUSG00000029038.10 | 420.3785 | 0.024275 | 0.397034 | 0.06114 | 0.951248 | 0.997337 | 0.001158 |
| ENSMUSG00000029173.13 | 334.8966 | -0.02559 | 0.459099 | -0.05575 | 0.955544 | 0.997337 | 0.001158 |
| ENSMUSG00000029267.18 | 42.12546 | 0.062266 | 1.214559 | 0.051266 | 0.959113 | 0.997337 | 0.001158 |
| ENSMUSG00000029370.11 | 20.15598 | -0.08613 | 1.456464 | -0.05914 | 0.952844 | 0.997337 | 0.001158 |
| ENSMUSG00000029775.15 | 110.4537 | 0.031116 | 0.822108 | 0.037849 | 0.969808 | 0.997337 | 0.001158 |
| ENSMUSG00000030217.3 | 43.96559 | -0.07116 | 1.180727 | -0.06027 | 0.951942 | 0.997337 | 0.001158 |
| ENSMUSG00000031154.16 | 50.05014 | 0.092568 | 1.580768 | 0.058559 | 0.953303 | 0.997337 | 0.001158 |
| ENSMUSG00000031586.18 | 174.5843 | -0.03509 | 0.706812 | -0.04964 | 0.960407 | 0.997337 | 0.001158 |
| ENSMUSG00000031617.17 | 87.36925 | -0.03887 | 0.985185 | -0.03945 | 0.968529 | 0.997337 | 0.001158 |
| ENSMUSG00000031622.17 | 96.30984 | -0.05351 | 0.949964 | -0.05633 | 0.955077 | 0.997337 | 0.001158 |
| ENSMUSG00000031639.13 | 19.3901 | -0.11021 | 2.067184 | -0.05332 | 0.95748 | 0.997337 | 0.001158 |
| ENSMUSG00000031887.14 | 33.89734 | 0.0716 | 1.51219 | 0.047349 | 0.962235 | 0.997337 | 0.001158 |
| ENSMUSG00000031969.17 | 313.5011 | 0.018848 | 0.468706 | 0.040212 | 0.967924 | 0.997337 | 0.001158 |
| ENSMUSG00000032369.14 | 22.85657 | 0.092748 | 1.736814 | 0.053401 | 0.957412 | 0.997337 | 0.001158 |
| ENSMUSG00000032508.9 | 17.80587 | -0.09435 | 2.502434 | -0.0377 | 0.969923 | 0.997337 | 0.001158 |
| ENSMUSG00000032562.14 | 129.7603 | 0.054738 | 0.911853 | 0.060029 | 0.952132 | 0.997337 | 0.001158 |
| ENSMUSG00000032754.15 | 12.18922 | -0.09241 | 2.548511 | -0.03626 | 0.971073 | 0.997337 | 0.001158 |
| ENSMUSG00000033096.8 | 60.07307 | 0.058027 | 1.33883 | 0.043341 | 0.965429 | 0.997337 | 0.001158 |
| ENSMUSG00000033186.10 | 158.9549 | 0.042589 | 0.719459 | 0.059195 | 0.952797 | 0.997337 | 0.001158 |
| ENSMUSG00000033220.8 | 10.40705 | -0.10533 | 2.557898 | -0.04118 | 0.967152 | 0.997337 | 0.001158 |
| ENSMUSG00000033379.14 | 61.66603 | -0.05656 | 1.180246 | -0.04792 | 0.961779 | 0.997337 | 0.001158 |
| ENSMUSG00000033589.5 | 12.26068 | -0.08478 | 2.085604 | -0.04065 | 0.967576 | 0.997337 | 0.001158 |
| ENSMUSG00000033916.6 | 899.1151 | 0.017976 | 0.397456 | 0.045228 | 0.963926 | 0.997337 | 0.001158 |
| ENSMUSG00000034157.13 | 176.5083 | -0.02732 | 0.528577 | -0.05169 | 0.958777 | 0.997337 | 0.001158 |
| ENSMUSG00000034201.14 | 46.82387 | -0.0496 | 0.947649 | -0.05234 | 0.958256 | 0.997337 | 0.001158 |
| ENSMUSG00000034795.15 | 141.0869 | 0.03422 | 0.942674 | 0.036301 | 0.971043 | 0.997337 | 0.001158 |
| ENSMUSG00000035027.19 | 299.4885 | -0.02219 | 0.549377 | -0.04039 | 0.967781 | 0.997337 | 0.001158 |
| ENSMUSG00000035198.10 | 64.78251 | 0.07006 | 1.194152 | 0.05867 | 0.953215 | 0.997337 | 0.001158 |
| ENSMUSG00000035247.17 | 476.8687 | 0.01752 | 0.367023 | 0.047737 | 0.961926 | 0.997337 | 0.001158 |
| ENSMUSG00000035540.13 | 48420.42 | -0.00817 | 0.186167 | -0.04388 | 0.965002 | 0.997337 | 0.001158 |
| ENSMUSG00000035642.17 | 179.8696 | 0.104084 | 1.840566 | 0.05655 | 0.954904 | 0.997337 | 0.001158 |

| | | | | | | | |
|---|---|---|---|---|---|---|---|
| ENSMUSG00000036155.14 | 18.80142 | -0.06958 | 1.80377 | -0.03857 | 0.969231 | 0.997337 | 0.001158 |
| ENSMUSG00000036371.7 | 1731.802 | -0.01363 | 0.260915 | -0.05225 | 0.958331 | 0.997337 | 0.001158 |
| ENSMUSG00000036825.13 | 15.38381 | -0.11293 | 2.324363 | -0.04858 | 0.961251 | 0.997337 | 0.001158 |
| ENSMUSG00000036968.16 | 15.44046 | 0.14894 | 2.614876 | 0.056959 | 0.954578 | 0.997337 | 0.001158 |
| ENSMUSG00000036985.13 | 90.14707 | 0.050851 | 0.827443 | 0.061455 | 0.950997 | 0.997337 | 0.001158 |
| ENSMUSG00000037007.18 | 29.74926 | 0.088211 | 1.947707 | 0.04529 | 0.963876 | 0.997337 | 0.001158 |
| ENSMUSG00000037174.19 | 55.12257 | -0.04779 | 1.326191 | -0.03604 | 0.971254 | 0.997337 | 0.001158 |
| ENSMUSG00000037885.19 | 10.4897 | -0.10853 | 2.509096 | -0.04325 | 0.9655 | 0.997337 | 0.001158 |
| ENSMUSG00000038074.17 | 45.45172 | -0.06266 | 1.115392 | -0.05618 | 0.955198 | 0.997337 | 0.001158 |
| ENSMUSG00000038451.14 | 16.2527 | 0.082566 | 2.209853 | 0.037363 | 0.970196 | 0.997337 | 0.001158 |
| ENSMUSG00000038646.14 | 212.0461 | 0.028583 | 0.492074 | 0.058087 | 0.95368 | 0.997337 | 0.001158 |
| ENSMUSG00000039100.11 | 550.641 | -0.01817 | 0.409685 | -0.04434 | 0.964631 | 0.997337 | 0.001158 |
| ENSMUSG00000039168.16 | 1421.468 | -0.01624 | 0.270095 | -0.06012 | 0.95206 | 0.997337 | 0.001158 |
| ENSMUSG00000039231.19 | 21.10486 | -0.09082 | 1.899616 | -0.04781 | 0.96187 | 0.997337 | 0.001158 |
| ENSMUSG00000039238.7 | 23.14829 | 0.112478 | 2.096747 | 0.053644 | 0.957219 | 0.997337 | 0.001158 |
| ENSMUSG00000040006.14 | 392.2926 | -0.02244 | 0.498512 | -0.04502 | 0.964093 | 0.997337 | 0.001158 |
| ENSMUSG00000040302.18 | 47.19236 | -0.09621 | 1.781893 | -0.05399 | 0.95694 | 0.997337 | 0.001158 |
| ENSMUSG00000040383.17 | 21.179 | -0.08601 | 1.76131 | -0.04884 | 0.96105 | 0.997337 | 0.001158 |
| ENSMUSG00000040413.16 | 783.2784 | 0.021318 | 0.366585 | 0.058152 | 0.953627 | 0.997337 | 0.001158 |
| ENSMUSG00000040455.18 | 73.6963 | -0.0488 | 1.194903 | -0.04084 | 0.967426 | 0.997337 | 0.001158 |
| ENSMUSG00000041261.10 | 704.6816 | 0.018336 | 0.333711 | 0.054946 | 0.956182 | 0.997337 | 0.001158 |
| ENSMUSG00000041297.9 | 82.98479 | 0.032393 | 0.732617 | 0.044215 | 0.964733 | 0.997337 | 0.001158 |
| ENSMUSG00000041396.14 | 18.33904 | 0.104261 | 1.906653 | 0.054683 | 0.956391 | 0.997337 | 0.001158 |
| ENSMUSG00000041429.4 | 29.42802 | -0.07467 | 1.899215 | -0.03932 | 0.968638 | 0.997337 | 0.001158 |
| ENSMUSG00000041528.16 | 33.10691 | 0.063546 | 1.643903 | 0.038656 | 0.969165 | 0.997337 | 0.001158 |
| ENSMUSG00000042063.12 | 34.22657 | 0.106657 | 1.814497 | 0.05878 | 0.953127 | 0.997337 | 0.001158 |
| ENSMUSG00000042251.13 | 313.5303 | 0.025727 | 0.555557 | 0.046308 | 0.963065 | 0.997337 | 0.001158 |
| ENSMUSG00000042390.10 | 115.9287 | 0.042238 | 0.804134 | 0.052526 | 0.95811 | 0.997337 | 0.001158 |
| ENSMUSG00000042590.18 | 92.82418 | 0.042643 | 0.830473 | 0.051348 | 0.959048 | 0.997337 | 0.001158 |
| ENSMUSG00000042814.9 | 57.99243 | 0.074508 | 1.399233 | 0.053249 | 0.957534 | 0.997337 | 0.001158 |
| ENSMUSG00000043866.12 | 250.8735 | 0.029113 | 0.559417 | 0.052042 | 0.958495 | 0.997337 | 0.001158 |
| ENSMUSG00000043962.17 | 34.73088 | 0.061246 | 1.441879 | 0.042477 | 0.966119 | 0.997337 | 0.001158 |
| ENSMUSG00000044393.16 | 489.804 | -0.01846 | 0.364178 | -0.05069 | 0.959574 | 0.997337 | 0.001158 |
| ENSMUSG00000044468.15 | 39.80519 | -0.0509 | 1.133982 | -0.04489 | 0.964197 | 0.997337 | 0.001158 |
| ENSMUSG00000046532.9 | 74.97134 | 0.053631 | 1.271186 | 0.04219 | 0.966348 | 0.997337 | 0.001158 |
| ENSMUSG00000046603.17 | 70.70404 | -0.04268 | 1.158661 | -0.03684 | 0.970614 | 0.997337 | 0.001158 |
| ENSMUSG00000046909.10 | 150.4153 | -0.04132 | 0.775918 | -0.05325 | 0.957534 | 0.997337 | 0.001158 |
| ENSMUSG00000047368.5 | 210.6655 | -0.03416 | 0.718303 | -0.04756 | 0.96207 | 0.997337 | 0.001158 |
| ENSMUSG00000048332.14 | 60.71109 | -0.04112 | 0.952247 | -0.04318 | 0.965558 | 0.997337 | 0.001158 |
| ENSMUSG00000050002.15 | 26.2131 | 0.072653 | 1.712927 | 0.042414 | 0.966168 | 0.997337 | 0.001158 |
| ENSMUSG00000051043.17 | 70.88171 | -0.06743 | 1.497673 | -0.04502 | 0.96409 | 0.997337 | 0.001158 |
| ENSMUSG00000051435.12 | 57.81606 | -0.08257 | 1.462423 | -0.05646 | 0.954976 | 0.997337 | 0.001158 |
| ENSMUSG00000052033.14 | 124.7202 | -0.0307 | 0.709423 | -0.04328 | 0.96548 | 0.997337 | 0.001158 |
| ENSMUSG00000052593.17 | 49.14984 | -0.05606 | 1.20063 | -0.0467 | 0.962756 | 0.997337 | 0.001158 |

| | | | | | | | |
|---|---|---|---|---|---|---|---|
| ENSMUSG00000053684.9 | 11.7502 | 0.079301 | 2.140465 | 0.037048 | 0.970446 | 0.997337 | 0.001158 |
| ENSMUSG00000055723.7 | 76.98866 | 0.036749 | 0.992113 | 0.037042 | 0.970452 | 0.997337 | 0.001158 |
| ENSMUSG00000055771.3 | 13.77554 | 0.09983 | 1.956018 | 0.051037 | 0.959296 | 0.997337 | 0.001158 |
| ENSMUSG00000056698.12 | 28.63489 | -0.08419 | 1.482075 | -0.0568 | 0.954701 | 0.997337 | 0.001158 |
| ENSMUSG00000056749.8 | 60.39653 | -0.08186 | 1.37083 | -0.05972 | 0.952381 | 0.997337 | 0.001158 |
| ENSMUSG00000058152.9 | 13.02746 | -0.1468 | 2.460424 | -0.05967 | 0.952422 | 0.997337 | 0.001158 |
| ENSMUSG00000058388.15 | 27.17889 | -0.10929 | 2.078647 | -0.05258 | 0.958069 | 0.997337 | 0.001158 |
| ENSMUSG00000058444.9 | 63.49459 | -0.05593 | 1.39469 | -0.0401 | 0.968013 | 0.997337 | 0.001158 |
| ENSMUSG00000058638.15 | 103.2466 | 0.068531 | 1.12735 | 0.060789 | 0.951527 | 0.997337 | 0.001158 |
| ENSMUSG00000060791.16 | 27.77209 | -0.12211 | 2.282844 | -0.05349 | 0.95734 | 0.997337 | 0.001158 |
| ENSMUSG00000061207.12 | 15.24226 | 0.109642 | 2.20064 | 0.049823 | 0.960264 | 0.997337 | 0.001158 |
| ENSMUSG00000062382.10 | 170.6848 | -0.03628 | 0.58887 | -0.0616 | 0.95088 | 0.997337 | 0.001158 |
| ENSMUSG00000063281.10 | 47.68567 | 0.07209 | 1.321476 | 0.054552 | 0.956495 | 0.997337 | 0.001158 |
| ENSMUSG00000064372.1 | 17.65473 | 0.139931 | 2.483198 | 0.056351 | 0.955062 | 0.997337 | 0.001158 |
| ENSMUSG00000066324.3 | 500.0203 | 0.020564 | 0.442446 | 0.046479 | 0.962929 | 0.997337 | 0.001158 |
| ENSMUSG00000068134.14 | 94.52515 | 0.058009 | 0.932805 | 0.062188 | 0.950413 | 0.997337 | 0.001158 |
| ENSMUSG00000068479.6 | 54.11725 | 0.067661 | 1.14369 | 0.05916 | 0.952824 | 0.997337 | 0.001158 |
| ENSMUSG00000069874.8 | 88.88147 | -0.05106 | 0.864932 | -0.05904 | 0.952922 | 0.997337 | 0.001158 |
| ENSMUSG00000070283.5 | 76.77596 | -0.06048 | 1.238234 | -0.04884 | 0.961046 | 0.997337 | 0.001158 |
| ENSMUSG00000070690.8 | 45.69883 | 0.061122 | 1.191946 | 0.051279 | 0.959103 | 0.997337 | 0.001158 |
| ENSMUSG00000071204.11 | 56.04167 | -0.07904 | 1.39405 | -0.0567 | 0.954785 | 0.997337 | 0.001158 |
| ENSMUSG00000071302.11 | 11.53369 | 0.091527 | 2.0794 | 0.044016 | 0.964891 | 0.997337 | 0.001158 |
| ENSMUSG00000073643.12 | 45.85413 | 0.055944 | 1.167411 | 0.047921 | 0.961779 | 0.997337 | 0.001158 |
| ENSMUSG00000074768.7 | 21473.2 | -0.01254 | 0.344806 | -0.03636 | 0.970993 | 0.997337 | 0.001158 |
| ENSMUSG00000078201.6 | 71.54225 | 0.065991 | 1.185641 | 0.055658 | 0.955614 | 0.997337 | 0.001158 |
| ENSMUSG00000078429.10 | 49.24757 | -0.0451 | 1.135588 | -0.03971 | 0.968321 | 0.997337 | 0.001158 |
| ENSMUSG00000079334.9 | 10.93783 | 0.175007 | 2.825784 | 0.061932 | 0.950617 | 0.997337 | 0.001158 |
| ENSMUSG00000086219.8 | 13.62588 | -0.11005 | 2.659353 | -0.04138 | 0.966992 | 0.997337 | 0.001158 |
| ENSMUSG00000087132.9 | 14.19703 | -0.07566 | 2.015557 | -0.03754 | 0.970057 | 0.997337 | 0.001158 |
| ENSMUSG00000090258.8 | 219.4698 | -0.02814 | 0.547269 | -0.05142 | 0.958987 | 0.997337 | 0.001158 |
| ENSMUSG00000092021.9 | 14.03797 | -0.08156 | 1.946375 | -0.0419 | 0.966576 | 0.997337 | 0.001158 |
| ENSMUSG00000092341.4 | 2983.273 | -0.02683 | 0.434577 | -0.06174 | 0.950772 | 0.997337 | 0.001158 |
| ENSMUSG00000097412.3 | 43.94224 | -0.07974 | 1.276782 | -0.06245 | 0.950202 | 0.997337 | 0.001158 |
| ENSMUSG00000099146.8 | 119.8849 | -0.04923 | 0.886873 | -0.05551 | 0.955734 | 0.997337 | 0.001158 |
| ENSMUSG00000101609.3 | 50.46403 | -0.0644 | 1.680173 | -0.03833 | 0.969424 | 0.997337 | 0.001158 |
| ENSMUSG00000104960.2 | 319.9899 | 0.016252 | 0.44049 | 0.036895 | 0.970568 | 0.997337 | 0.001158 |
| ENSMUSG00000105814.3 | 671.5902 | 0.021335 | 0.41041 | 0.051984 | 0.958542 | 0.997337 | 0.001158 |
| ENSMUSG00000109865.2 | 115.584 | 0.045858 | 0.757634 | 0.060528 | 0.951735 | 0.997337 | 0.001158 |
| ENSMUSG00000110234.3 | 16.10958 | -0.10941 | 2.951353 | -0.03707 | 0.970429 | 0.997337 | 0.001158 |
| ENSMUSG00000026436.16 | 211.1374 | -0.01969 | 0.553256 | -0.0356 | 0.971604 | 0.99738 | 0.001139 |
| ENSMUSG00000000399.11 | 505.5725 | -0.01608 | 0.46375 | -0.03468 | 0.972333 | 0.997498 | 0.001088 |
| ENSMUSG00000005951.14 | 137.4248 | -0.02672 | 0.773735 | -0.03453 | 0.972455 | 0.997498 | 0.001088 |
| ENSMUSG00000022843.18 | 60.1215 | 0.044551 | 1.273035 | 0.034996 | 0.972083 | 0.997498 | 0.001088 |
| ENSMUSG00000037373.13 | 51.68291 | 0.062697 | 1.812595 | 0.03459 | 0.972407 | 0.997498 | 0.001088 |

| | | | | | | | |
|---|---|---|---|---|---|---|---|
| ENSMUSG00000039783.16 | 483.1028 | -0.01667 | 0.480185 | -0.03473 | 0.972298 | 0.997498 | 0.001088 |
| ENSMUSG00000040127.14 | 205.2716 | -0.02397 | 0.681943 | -0.03515 | 0.971956 | 0.997498 | 0.001088 |
| ENSMUSG00000061848.6 | 66.7544 | 0.037467 | 1.064148 | 0.035208 | 0.971914 | 0.997498 | 0.001088 |
| ENSMUSG00000010608.16 | 1531.226 | 0.008583 | 0.253525 | 0.033854 | 0.972994 | 0.997632 | 0.00103 |
| ENSMUSG00000021764.9 | 512.7478 | -0.01414 | 0.42052 | -0.03362 | 0.973176 | 0.997632 | 0.00103 |
| ENSMUSG00000022536.15 | 159.0736 | -0.0207 | 0.614429 | -0.03369 | 0.973127 | 0.997632 | 0.00103 |
| ENSMUSG00000028386.2 | 18.41094 | -0.05466 | 1.618245 | -0.03378 | 0.973055 | 0.997632 | 0.00103 |
| ENSMUSG00000031266.8 | 18.59945 | 0.073873 | 2.177797 | 0.033921 | 0.97294 | 0.997632 | 0.00103 |
| ENSMUSG00000032802.9 | 62.67185 | -0.03647 | 1.086385 | -0.03357 | 0.973217 | 0.997632 | 0.00103 |
| ENSMUSG00000001305.6 | 25.6061 | 0.048502 | 1.506255 | 0.032201 | 0.974312 | 0.998073 | 0.000838 |
| ENSMUSG00000025060.16 | 132.3906 | -0.02198 | 0.681943 | -0.03223 | 0.974286 | 0.998073 | 0.000838 |
| ENSMUSG00000035666.15 | 65.79072 | -0.02687 | 0.823645 | -0.03262 | 0.97398 | 0.998073 | 0.000838 |
| ENSMUSG00000036275.14 | 273.049 | -0.01752 | 0.533007 | -0.03287 | 0.973781 | 0.998073 | 0.000838 |
| ENSMUSG00000041241.13 | 107.6475 | -0.03905 | 1.216256 | -0.03211 | 0.974385 | 0.998073 | 0.000838 |
| ENSMUSG00000071266.14 | 33.21651 | 0.046517 | 1.439923 | 0.032305 | 0.974229 | 0.998073 | 0.000838 |
| ENSMUSG00000074240.6 | 43.98634 | 0.056832 | 1.763618 | 0.032225 | 0.974293 | 0.998073 | 0.000838 |
| ENSMUSG00000036432.9 | 80.56656 | 0.030031 | 0.94412 | 0.031809 | 0.974625 | 0.998151 | 0.000804 |
| ENSMUSG00000047905.11 | 14.69862 | 0.049738 | 1.566564 | 0.03175 | 0.974672 | 0.998151 | 0.000804 |
| ENSMUSG00000065954.12 | 113.3758 | 0.029281 | 0.927545 | 0.031569 | 0.974816 | 0.998191 | 0.000786 |
| ENSMUSG00000005150.17 | 62.06304 | 0.031145 | 1.00133 | 0.031103 | 0.975187 | 0.998192 | 0.000786 |
| ENSMUSG00000031774.9 | 95.628 | -0.02421 | 0.780045 | -0.03104 | 0.975238 | 0.998192 | 0.000786 |
| ENSMUSG00000039835.17 | 37.74868 | -0.04358 | 1.403165 | -0.03106 | 0.975222 | 0.998192 | 0.000786 |
| ENSMUSG00000074890.5 | 50.61269 | 0.116297 | 3.728688 | 0.03119 | 0.975118 | 0.998192 | 0.000786 |
| ENSMUSG00000030872.15 | 60.60421 | -0.03827 | 1.242024 | -0.03081 | 0.975422 | 0.998194 | 0.000785 |
| ENSMUSG00000091144.3 | 26.13014 | -0.06406 | 2.081598 | -0.03077 | 0.97545 | 0.998194 | 0.000785 |
| ENSMUSG00000019689.5 | 431.3686 | -0.01444 | 0.488752 | -0.02955 | 0.976424 | 0.998343 | 0.00072 |
| ENSMUSG00000020914.18 | 13.49138 | 0.085008 | 2.813786 | 0.030211 | 0.975898 | 0.998343 | 0.00072 |
| ENSMUSG00000021578.18 | 125.5725 | -0.02577 | 0.848907 | -0.03036 | 0.975784 | 0.998343 | 0.00072 |
| ENSMUSG00000021701.9 | 26.07103 | 0.050756 | 1.718111 | 0.029542 | 0.976432 | 0.998343 | 0.00072 |
| ENSMUSG00000027358.7 | 102.1831 | 0.021091 | 0.708762 | 0.029758 | 0.97626 | 0.998343 | 0.00072 |
| ENSMUSG00000030102.12 | 244.6461 | -0.01731 | 0.580624 | -0.02981 | 0.976221 | 0.998343 | 0.00072 |
| ENSMUSG00000030483.15 | 65.27534 | -0.09168 | 3.05751 | -0.02998 | 0.976079 | 0.998343 | 0.00072 |
| ENSMUSG00000033282.15 | 12.56385 | 0.083084 | 2.813143 | 0.029534 | 0.976438 | 0.998343 | 0.00072 |
| ENSMUSG00000005125.14 | 20.06547 | 0.054634 | 1.913576 | 0.028551 | 0.977223 | 0.998379 | 0.000705 |
| ENSMUSG00000020594.15 | 231.3814 | -0.01667 | 0.572425 | -0.02912 | 0.976767 | 0.998379 | 0.000705 |
| ENSMUSG00000030800.11 | 11.03065 | -0.09277 | 3.232937 | -0.0287 | 0.977107 | 0.998379 | 0.000705 |
| ENSMUSG00000041353.13 | 57.12047 | -0.03562 | 1.216581 | -0.02928 | 0.976643 | 0.998379 | 0.000705 |
| ENSMUSG00000044674.7 | 27.53801 | -0.03709 | 1.286157 | -0.02884 | 0.976995 | 0.998379 | 0.000705 |
| ENSMUSG00000057594.13 | 12.38052 | 0.088986 | 3.12954 | 0.028434 | 0.977316 | 0.998379 | 0.000705 |
| ENSMUSG00000062054.8 | 735.1348 | 0.009839 | 0.342504 | 0.028725 | 0.977084 | 0.998379 | 0.000705 |
| ENSMUSG00000063317.8 | 15.19533 | -0.06152 | 2.126635 | -0.02893 | 0.97692 | 0.998379 | 0.000705 |
| ENSMUSG00000020841.6 | 131.4893 | -0.0224 | 0.793083 | -0.02824 | 0.977469 | 0.998427 | 0.000684 |
| ENSMUSG00000002910.12 | 22.63125 | -0.05273 | 2.019053 | -0.02612 | 0.979164 | 0.998452 | 0.000673 |
| ENSMUSG00000015289.5 | 96.8121 | 0.018534 | 0.696598 | 0.026607 | 0.978773 | 0.998452 | 0.000673 |

| | | | | | | | |
|---|---|---|---|---|---|---|---|
| ENSMUSG00000018821.4 | 219.6173 | -0.01374 | 0.496784 | -0.02767 | 0.977928 | 0.998452 | 0.000673 |
| ENSMUSG00000019768.17 | 306.431 | 0.016934 | 0.637462 | 0.026565 | 0.978807 | 0.998452 | 0.000673 |
| ENSMUSG00000020706.14 | 114.9731 | -0.01975 | 0.769935 | -0.02565 | 0.979539 | 0.998452 | 0.000673 |
| ENSMUSG00000020708.13 | 232.4072 | 0.016119 | 0.633033 | 0.025463 | 0.979686 | 0.998452 | 0.000673 |
| ENSMUSG00000022014.16 | 22.6898 | 0.06438 | 2.392077 | 0.026914 | 0.978529 | 0.998452 | 0.000673 |
| ENSMUSG00000022836.12 | 157.2466 | -0.01963 | 0.775551 | -0.02531 | 0.97981 | 0.998452 | 0.000673 |
| ENSMUSG00000023243.10 | 53.13058 | 0.035226 | 1.289411 | 0.027319 | 0.978205 | 0.998452 | 0.000673 |
| ENSMUSG00000023830.15 | 115.5512 | -0.02039 | 0.767855 | -0.02655 | 0.978815 | 0.998452 | 0.000673 |
| ENSMUSG00000024991.9 | 2373.64 | 0.008235 | 0.303085 | 0.02717 | 0.978324 | 0.998452 | 0.000673 |
| ENSMUSG00000025465.14 | 1019.102 | -0.00841 | 0.326756 | -0.02575 | 0.979458 | 0.998452 | 0.000673 |
| ENSMUSG00000027478.16 | 16.71256 | -0.06552 | 2.53907 | -0.02581 | 0.979413 | 0.998452 | 0.000673 |
| ENSMUSG00000027828.13 | 1584.336 | 0.008328 | 0.307138 | 0.027116 | 0.978367 | 0.998452 | 0.000673 |
| ENSMUSG00000029089.9 | 15.33436 | -0.06481 | 2.379824 | -0.02723 | 0.978275 | 0.998452 | 0.000673 |
| ENSMUSG00000030237.15 | 230.1183 | 0.017062 | 0.674106 | 0.025311 | 0.979807 | 0.998452 | 0.000673 |
| ENSMUSG00000030595.16 | 32.47781 | 0.039562 | 1.509295 | 0.026212 | 0.979088 | 0.998452 | 0.000673 |
| ENSMUSG00000032513.9 | 31.22608 | 0.049278 | 1.756367 | 0.028057 | 0.977617 | 0.998452 | 0.000673 |
| ENSMUSG00000035297.15 | 355.9988 | 0.01312 | 0.471889 | 0.027804 | 0.977819 | 0.998452 | 0.000673 |
| ENSMUSG00000037287.16 | 150.0778 | -0.02163 | 0.782396 | -0.02765 | 0.977941 | 0.998452 | 0.000673 |
| ENSMUSG00000037325.11 | 14.72725 | -0.06716 | 2.631063 | -0.02553 | 0.979634 | 0.998452 | 0.000673 |
| ENSMUSG00000038521.18 | 1515.92 | -0.00783 | 0.303934 | -0.02576 | 0.979451 | 0.998452 | 0.000673 |
| ENSMUSG00000035851.15 | 155.0868 | 0.018545 | 0.737362 | 0.025151 | 0.979935 | 0.998472 | 0.000664 |
| ENSMUSG00000037933.17 | 48.127 | -0.03484 | 1.405903 | -0.02478 | 0.980227 | 0.998662 | 0.000581 |
| ENSMUSG00000061477.5 | 1143.053 | 0.007307 | 0.297469 | 0.024564 | 0.980402 | 0.998734 | 0.00055 |
| ENSMUSG00000002833.12 | 46.9834 | 0.028252 | 1.177536 | 0.023992 | 0.980859 | 0.998768 | 0.000535 |
| ENSMUSG00000015363.14 | 45.414 | 0.027837 | 1.153419 | 0.024134 | 0.980745 | 0.998768 | 0.000535 |
| ENSMUSG00000021704.9 | 52.65612 | -0.03288 | 1.351685 | -0.02432 | 0.980594 | 0.998768 | 0.000535 |
| ENSMUSG00000022790.15 | 104.0188 | -0.01744 | 0.727332 | -0.02397 | 0.980873 | 0.998768 | 0.000535 |
| ENSMUSG00000023791.19 | 133.4016 | 0.018874 | 0.792593 | 0.023813 | 0.981002 | 0.998768 | 0.000535 |
| ENSMUSG00000026158.12 | 157.2942 | -0.0146 | 0.622572 | -0.02345 | 0.981294 | 0.998768 | 0.000535 |
| ENSMUSG00000041571.11 | 167.0743 | -0.01422 | 0.606779 | -0.02344 | 0.981302 | 0.998768 | 0.000535 |
| ENSMUSG00000068706.4 | 275.981 | 0.011762 | 0.504081 | 0.023333 | 0.981384 | 0.998768 | 0.000535 |
| ENSMUSG00000076432.14 | 323.3439 | 0.010137 | 0.432586 | 0.023433 | 0.981305 | 0.998768 | 0.000535 |
| ENSMUSG00000005483.11 | 105.4501 | -0.01628 | 0.765761 | -0.02126 | 0.983041 | 0.999126 | 0.00038 |
| ENSMUSG00000016921.15 | 342.1071 | -0.00925 | 0.42821 | -0.02161 | 0.98276 | 0.999126 | 0.00038 |
| ENSMUSG00000024063.14 | 36.73003 | 0.041139 | 1.880513 | 0.021877 | 0.982546 | 0.999126 | 0.00038 |
| ENSMUSG00000024742.16 | 22.50947 | -0.0303 | 1.412699 | -0.02145 | 0.982888 | 0.999126 | 0.00038 |
| ENSMUSG00000025436.13 | 15.74917 | -0.0471 | 2.101951 | -0.02241 | 0.982123 | 0.999126 | 0.00038 |
| ENSMUSG00000026342.11 | 271.5502 | 0.012055 | 0.566655 | 0.021275 | 0.983027 | 0.999126 | 0.00038 |
| ENSMUSG00000026511.8 | 544.8584 | 0.008389 | 0.374045 | 0.022428 | 0.982106 | 0.999126 | 0.00038 |
| ENSMUSG00000039220.17 | 16.70736 | -0.04457 | 2.023175 | -0.02203 | 0.982423 | 0.999126 | 0.00038 |
| ENSMUSG00000042594.17 | 69.24086 | 0.022565 | 1.066778 | 0.021152 | 0.983124 | 0.999126 | 0.00038 |
| ENSMUSG00000048039.9 | 49.01793 | 0.025997 | 1.229744 | 0.02114 | 0.983134 | 0.999126 | 0.00038 |
| ENSMUSG00000051853.11 | 43.46256 | 0.022678 | 1.071544 | 0.021164 | 0.983115 | 0.999126 | 0.00038 |
| ENSMUSG00000058690.15 | 288.4701 | -0.00927 | 0.440457 | -0.02104 | 0.983211 | 0.999126 | 0.00038 |

| | | | | | | | |
|---|---|---|---|---|---|---|---|
| ENSMUSG00000092187.2 | 13.00879 | -0.06089 | 2.711579 | -0.02245 | 0.982086 | 0.999126 | 0.00038 |
| ENSMUSG00000097772.8 | 55.18572 | -0.01925 | 0.901522 | -0.02135 | 0.982963 | 0.999126 | 0.00038 |
| ENSMUSG00000002428.13 | 52.34931 | 0.024116 | 1.193409 | 0.020208 | 0.983878 | 0.99916 | 0.000365 |
| ENSMUSG00000042705.10 | 13.77818 | -0.04617 | 2.277796 | -0.02027 | 0.983827 | 0.99916 | 0.000365 |
| ENSMUSG00000048706.4 | 91.46287 | -0.01918 | 0.93418 | -0.02053 | 0.983623 | 0.99916 | 0.000365 |
| ENSMUSG00000055932.16 | 112.2976 | 0.01304 | 0.63382 | 0.020574 | 0.983585 | 0.99916 | 0.000365 |
| ENSMUSG00000063200.5 | 525.1469 | -0.0079 | 0.386431 | -0.02045 | 0.983684 | 0.99916 | 0.000365 |
| ENSMUSG00000071645.7 | 19.18131 | 0.040829 | 1.976672 | 0.020655 | 0.983521 | 0.99916 | 0.000365 |
| ENSMUSG00000070473.5 | 81.42212 | 0.024984 | 1.267103 | 0.019717 | 0.984269 | 0.999451 | 0.000239 |
| ENSMUSG00000017009.4 | 356.7817 | -0.01157 | 0.597374 | -0.01936 | 0.984554 | 0.999473 | 0.000229 |
| ENSMUSG00000025651.15 | 263.0327 | -0.01294 | 0.666546 | -0.01942 | 0.984509 | 0.999473 | 0.000229 |
| ENSMUSG00000055897.14 | 25.83591 | -0.03449 | 1.787882 | -0.01929 | 0.984608 | 0.999473 | 0.000229 |
| ENSMUSG00000020087.6 | 96.1759 | 0.014118 | 0.751613 | 0.018784 | 0.985013 | 0.999475 | 0.000228 |
| ENSMUSG00000021939.9 | 2401.779 | 0.005889 | 0.309137 | 0.019049 | 0.984802 | 0.999475 | 0.000228 |
| ENSMUSG00000027162.8 | 317.9524 | 0.008221 | 0.437837 | 0.018777 | 0.985019 | 0.999475 | 0.000228 |
| ENSMUSG00000029196.10 | 157.7077 | -0.01525 | 0.812589 | -0.01876 | 0.985031 | 0.999475 | 0.000228 |
| ENSMUSG00000041231.16 | 38.72557 | -0.02222 | 1.214987 | -0.01829 | 0.98541 | 0.999539 | 0.0002 |
| ENSMUSG00000050856.17 | 2375.584 | -0.00505 | 0.274409 | -0.01839 | 0.985325 | 0.999539 | 0.0002 |
| ENSMUSG00000081855.5 | 72.09135 | -0.01642 | 0.896279 | -0.01832 | 0.985383 | 0.999539 | 0.0002 |
| ENSMUSG00000032194.10 | 102.5194 | -0.01745 | 0.963861 | -0.01811 | 0.985555 | 0.999578 | 0.000183 |
| ENSMUSG00000000295.14 | 12.85947 | 0.029512 | 2.384592 | 0.012376 | 0.990126 | 0.999655 | 0.00015 |
| ENSMUSG00000003531.16 | 65.1607 | 0.014015 | 0.985975 | 0.014215 | 0.988659 | 0.999655 | 0.00015 |
| ENSMUSG00000004610.5 | 1355.374 | 0.003403 | 0.268749 | 0.012662 | 0.989898 | 0.999655 | 0.00015 |
| ENSMUSG00000004880.12 | 127.1733 | -0.00476 | 0.803744 | -0.00592 | 0.995275 | 0.999655 | 0.00015 |
| ENSMUSG00000005981.12 | 165.1524 | -0.0037 | 0.9284 | -0.00399 | 0.99682 | 0.999655 | 0.00015 |
| ENSMUSG00000005983.16 | 18.87667 | -0.00796 | 2.07905 | -0.00383 | 0.996944 | 0.999655 | 0.00015 |
| ENSMUSG00000007338.11 | 119.9146 | 0.006799 | 0.674929 | 0.010074 | 0.991962 | 0.999655 | 0.00015 |
| ENSMUSG00000008384.9 | 38.61761 | -0.00652 | 1.342999 | -0.00485 | 0.996128 | 0.999655 | 0.00015 |
| ENSMUSG00000015291.11 | 144.4981 | 0.011791 | 0.71479 | 0.016496 | 0.986839 | 0.999655 | 0.00015 |
| ENSMUSG00000016831.13 | 120.2602 | 0.009673 | 1.111808 | 0.008701 | 0.993058 | 0.999655 | 0.00015 |
| ENSMUSG00000017299.14 | 26.18483 | 0.009796 | 1.580379 | 0.006199 | 0.995054 | 0.999655 | 0.00015 |
| ENSMUSG00000017478.16 | 26.25514 | -0.02191 | 1.445732 | -0.01516 | 0.987907 | 0.999655 | 0.00015 |
| ENSMUSG00000017843.15 | 227.9429 | -0.0067 | 0.593502 | -0.01129 | 0.990992 | 0.999655 | 0.00015 |
| ENSMUSG00000019806.14 | 96.12539 | -0.01272 | 0.890301 | -0.01429 | 0.988598 | 0.999655 | 0.00015 |
| ENSMUSG00000020075.9 | 104.6235 | -0.00754 | 0.882625 | -0.00854 | 0.993182 | 0.999655 | 0.00015 |
| ENSMUSG00000020528.15 | 11.78651 | 0.009349 | 2.50144 | 0.003737 | 0.997018 | 0.999655 | 0.00015 |
| ENSMUSG00000020589.18 | 21.44349 | 0.008699 | 1.484756 | 0.005859 | 0.995325 | 0.999655 | 0.00015 |
| ENSMUSG00000021112.10 | 87.65957 | 0.003256 | 0.883885 | 0.003684 | 0.997061 | 0.999655 | 0.00015 |
| ENSMUSG00000021413.10 | 133.9386 | -0.00595 | 0.776902 | -0.00766 | 0.993888 | 0.999655 | 0.00015 |
| ENSMUSG00000021650.8 | 78.86388 | -0.01454 | 0.893216 | -0.01627 | 0.987016 | 0.999655 | 0.00015 |
| ENSMUSG00000022184.9 | 69.09548 | 0.009577 | 0.92006 | 0.010409 | 0.991695 | 0.999655 | 0.00015 |
| ENSMUSG00000022827.15 | 36.09829 | -0.01068 | 1.95561 | -0.00546 | 0.995644 | 0.999655 | 0.00015 |
| ENSMUSG00000023707.14 | 68.76562 | -0.02208 | 1.457877 | -0.01514 | 0.987918 | 0.999655 | 0.00015 |
| ENSMUSG00000023764.19 | 18.10082 | 0.009288 | 2.306622 | 0.004027 | 0.996787 | 0.999655 | 0.00015 |

| | | | | | | | |
|---|---|---|---|---|---|---|---|
| ENSMUSG00000024597.12 | 48.67718 | -0.0181 | 1.43977 | -0.01257 | 0.989969 | 0.999655 | 0.00015 |
| ENSMUSG00000025260.11 | 664.4066 | -0.0078 | 0.483485 | -0.01614 | 0.987125 | 0.999655 | 0.00015 |
| ENSMUSG00000025423.17 | 107.9887 | 0.008397 | 0.708691 | 0.011849 | 0.990546 | 0.999655 | 0.00015 |
| ENSMUSG00000026209.16 | 68.41995 | -0.02723 | 1.608547 | -0.01693 | 0.986494 | 0.999655 | 0.00015 |
| ENSMUSG00000026803.18 | 40.9382 | 0.018494 | 1.181896 | 0.015648 | 0.987515 | 0.999655 | 0.00015 |
| ENSMUSG00000026878.18 | 1049.454 | 0.003673 | 0.347055 | 0.010583 | 0.991556 | 0.999655 | 0.00015 |
| ENSMUSG00000027203.16 | 103.1193 | 0.005323 | 0.889249 | 0.005986 | 0.995224 | 0.999655 | 0.00015 |
| ENSMUSG00000027287.15 | 211.4648 | -0.00747 | 0.615691 | -0.01214 | 0.990315 | 0.999655 | 0.00015 |
| ENSMUSG00000027488.13 | 19.34578 | -0.03722 | 2.465232 | -0.0151 | 0.987954 | 0.999655 | 0.00015 |
| ENSMUSG00000027865.11 | 186.59 | 0.002275 | 0.582963 | 0.003903 | 0.996886 | 0.999655 | 0.00015 |
| ENSMUSG00000028292.15 | 48.79093 | 0.024588 | 1.413878 | 0.017391 | 0.986125 | 0.999655 | 0.00015 |
| ENSMUSG00000028413.14 | 109.2953 | -0.0044 | 0.854976 | -0.00514 | 0.995896 | 0.999655 | 0.00015 |
| ENSMUSG00000029111.8 | 10.16131 | -0.03453 | 2.256365 | -0.0153 | 0.987789 | 0.999655 | 0.00015 |
| ENSMUSG00000029376.9 | 15.38473 | 0.026604 | 2.390022 | 0.011131 | 0.991119 | 0.999655 | 0.00015 |
| ENSMUSG00000029630.16 | 4757.233 | -0.00423 | 0.367295 | -0.01153 | 0.990803 | 0.999655 | 0.00015 |
| ENSMUSG00000029647.16 | 40.23825 | 0.027474 | 1.8696 | 0.014695 | 0.988275 | 0.999655 | 0.00015 |
| ENSMUSG00000030279.16 | 13.13558 | -0.02154 | 2.735648 | -0.00788 | 0.993716 | 0.999655 | 0.00015 |
| ENSMUSG00000030421.10 | 170.0072 | 0.014507 | 0.879963 | 0.016485 | 0.986847 | 0.999655 | 0.00015 |
| ENSMUSG00000030469.11 | 21.03929 | -0.03111 | 2.071855 | -0.01502 | 0.988018 | 0.999655 | 0.00015 |
| ENSMUSG00000030861.16 | 399.069 | 0.002735 | 0.391723 | 0.006982 | 0.994429 | 0.999655 | 0.00015 |
| ENSMUSG00000030967.16 | 435.4499 | -0.00425 | 0.454688 | -0.00934 | 0.992549 | 0.999655 | 0.00015 |
| ENSMUSG00000030983.5 | 534.0183 | 0.004688 | 0.38579 | 0.012151 | 0.990305 | 0.999655 | 0.00015 |
| ENSMUSG00000031201.18 | 253.408 | 0.003016 | 0.463809 | 0.006502 | 0.994812 | 0.999655 | 0.00015 |
| ENSMUSG00000031256.12 | 32.2515 | -0.00725 | 1.216936 | -0.00596 | 0.995245 | 0.999655 | 0.00015 |
| ENSMUSG00000031347.13 | 104.3926 | 0.011375 | 0.853544 | 0.013327 | 0.989367 | 0.999655 | 0.00015 |
| ENSMUSG00000031751.15 | 1569.267 | 0.000954 | 0.254278 | 0.00375 | 0.997008 | 0.999655 | 0.00015 |
| ENSMUSG00000031820.10 | 204.3198 | -0.00852 | 0.513882 | -0.01658 | 0.986769 | 0.999655 | 0.00015 |
| ENSMUSG00000031835.17 | 166.7557 | 0.003001 | 0.631736 | 0.00475 | 0.99621 | 0.999655 | 0.00015 |
| ENSMUSG00000032316.9 | 26.75116 | -0.0316 | 1.879071 | -0.01681 | 0.986585 | 0.999655 | 0.00015 |
| ENSMUSG00000032561.16 | 23.06308 | -0.02083 | 2.157478 | -0.00965 | 0.992297 | 0.999655 | 0.00015 |
| ENSMUSG00000032565.10 | 52.20264 | -0.01248 | 1.089009 | -0.01146 | 0.990858 | 0.999655 | 0.00015 |
| ENSMUSG00000033016.17 | 18.59944 | 0.018577 | 2.020536 | 0.009194 | 0.992664 | 0.999655 | 0.00015 |
| ENSMUSG00000033763.15 | 11.42007 | 0.011039 | 2.793432 | 0.003952 | 0.996847 | 0.999655 | 0.00015 |
| ENSMUSG00000034160.14 | 195.4411 | 0.008563 | 0.661515 | 0.012944 | 0.989672 | 0.999655 | 0.00015 |
| ENSMUSG00000034544.18 | 230.9499 | -0.00582 | 0.712848 | -0.00817 | 0.993484 | 0.999655 | 0.00015 |
| ENSMUSG00000036097.9 | 19.70996 | 0.008736 | 1.538941 | 0.005677 | 0.995471 | 0.999655 | 0.00015 |
| ENSMUSG00000036499.10 | 278.5704 | 0.005466 | 0.568317 | 0.009618 | 0.992326 | 0.999655 | 0.00015 |
| ENSMUSG00000037601.12 | 711.3599 | -0.00434 | 0.380961 | -0.01139 | 0.990912 | 0.999655 | 0.00015 |
| ENSMUSG00000037643.15 | 65.14399 | 0.009339 | 1.116414 | 0.008365 | 0.993326 | 0.999655 | 0.00015 |
| ENSMUSG00000038507.7 | 94.78893 | -0.01299 | 1.187843 | -0.01094 | 0.991271 | 0.999655 | 0.00015 |
| ENSMUSG00000038717.10 | 34.35922 | -0.00762 | 1.387661 | -0.00549 | 0.995619 | 0.999655 | 0.00015 |
| ENSMUSG00000038975.15 | 322.9948 | -0.00628 | 0.518664 | -0.0121 | 0.990344 | 0.999655 | 0.00015 |
| ENSMUSG00000039130.19 | 30.87259 | -0.0111 | 1.929336 | -0.00575 | 0.995408 | 0.999655 | 0.00015 |
| ENSMUSG00000040188.11 | 30.41261 | 0.020863 | 1.650388 | 0.012641 | 0.989914 | 0.999655 | 0.00015 |

| | | | | | | | |
|---|---|---|---|---|---|---|---|
| ENSMUSG00000040331.17 | 291.9352 | 0.005135 | 0.496284 | 0.010347 | 0.991744 | 0.999655 | 0.00015 |
| ENSMUSG00000040482.17 | 10.91583 | 0.014812 | 2.543293 | 0.005824 | 0.995353 | 0.999655 | 0.00015 |
| ENSMUSG00000040562.13 | 92.12646 | -0.00459 | 1.039133 | -0.00442 | 0.996473 | 0.999655 | 0.00015 |
| ENSMUSG00000040822.8 | 111.5664 | -0.00902 | 0.751823 | -0.01199 | 0.99043 | 0.999655 | 0.00015 |
| ENSMUSG00000042211.8 | 61.96935 | 0.018781 | 1.314908 | 0.014283 | 0.988604 | 0.999655 | 0.00015 |
| ENSMUSG00000042444.11 | 163.8351 | -0.00842 | 0.786816 | -0.0107 | 0.991461 | 0.999655 | 0.00015 |
| ENSMUSG00000045107.6 | 53.37442 | -0.00956 | 1.430204 | -0.00668 | 0.994668 | 0.999655 | 0.00015 |
| ENSMUSG00000046311.14 | 80.6486 | -0.00306 | 0.794801 | -0.00385 | 0.996931 | 0.999655 | 0.00015 |
| ENSMUSG00000046671.19 | 203.2533 | -0.00365 | 0.540959 | -0.00674 | 0.994621 | 0.999655 | 0.00015 |
| ENSMUSG00000047797.15 | 110.0841 | -0.00443 | 0.826541 | -0.00536 | 0.995727 | 0.999655 | 0.00015 |
| ENSMUSG00000049299.15 | 181.9829 | -0.00524 | 0.631683 | -0.0083 | 0.993376 | 0.999655 | 0.00015 |
| ENSMUSG00000049760.7 | 172.5571 | 0.006413 | 0.916612 | 0.006996 | 0.994418 | 0.999655 | 0.00015 |
| ENSMUSG00000050043.17 | 157.2735 | 0.007143 | 0.782588 | 0.009127 | 0.992718 | 0.999655 | 0.00015 |
| ENSMUSG00000051359.16 | 12.55161 | -0.0226 | 2.43132 | -0.00929 | 0.992585 | 0.999655 | 0.00015 |
| ENSMUSG00000052406.15 | 71.38936 | -0.01373 | 1.33521 | -0.01028 | 0.991797 | 0.999655 | 0.00015 |
| ENSMUSG00000052681.9 | 423.6992 | -0.00536 | 0.50619 | -0.01058 | 0.991557 | 0.999655 | 0.00015 |
| ENSMUSG00000053329.8 | 231.242 | -0.01138 | 0.707967 | -0.01608 | 0.987174 | 0.999655 | 0.00015 |
| ENSMUSG00000053581.14 | 20.07461 | -0.01884 | 1.808382 | -0.01042 | 0.991689 | 0.999655 | 0.00015 |
| ENSMUSG00000054648.9 | 64.68699 | -0.01905 | 1.097839 | -0.01735 | 0.986156 | 0.999655 | 0.00015 |
| ENSMUSG00000055991.16 | 52.07443 | 0.008252 | 1.367353 | 0.006035 | 0.995185 | 0.999655 | 0.00015 |
| ENSMUSG00000056962.12 | 17.4877 | -0.03425 | 2.011962 | -0.01702 | 0.986419 | 0.999655 | 0.00015 |
| ENSMUSG00000057842.15 | 28.26741 | -0.02884 | 1.898475 | -0.01519 | 0.98788 | 0.999655 | 0.00015 |
| ENSMUSG00000058624.14 | 230.3494 | 0.002711 | 0.586067 | 0.004625 | 0.99631 | 0.999655 | 0.00015 |
| ENSMUSG00000060090.17 | 41.00252 | 0.017739 | 1.067496 | 0.016618 | 0.986742 | 0.999655 | 0.00015 |
| ENSMUSG00000061104.6 | 44.20108 | -0.01676 | 1.275296 | -0.01314 | 0.989516 | 0.999655 | 0.00015 |
| ENSMUSG00000061898.11 | 15.05037 | 0.007121 | 1.975851 | 0.003604 | 0.997124 | 0.999655 | 0.00015 |
| ENSMUSG00000062519.14 | 11.67344 | 0.0279 | 2.383332 | 0.011706 | 0.99066 | 0.999655 | 0.00015 |
| ENSMUSG00000063550.9 | 31.68703 | 0.006421 | 1.427716 | 0.004497 | 0.996412 | 0.999655 | 0.00015 |
| ENSMUSG00000063727.4 | 18.08814 | 0.038121 | 2.209025 | 0.017257 | 0.986231 | 0.999655 | 0.00015 |
| ENSMUSG00000063760.10 | 212.359 | -0.00667 | 0.526909 | -0.01267 | 0.989894 | 0.999655 | 0.00015 |
| ENSMUSG00000067367.10 | 136.9997 | -0.00578 | 0.603617 | -0.00958 | 0.992359 | 0.999655 | 0.00015 |
| ENSMUSG00000069662.6 | 193.5363 | -0.00358 | 0.687754 | -0.0052 | 0.995849 | 0.999655 | 0.00015 |
| ENSMUSG00000069682.7 | 28.89858 | -0.02374 | 1.530793 | -0.01551 | 0.987628 | 0.999655 | 0.00015 |
| ENSMUSG00000071359.14 | 25.07333 | -0.01221 | 1.48204 | -0.00824 | 0.993426 | 0.999655 | 0.00015 |
| ENSMUSG00000084957.4 | 151.2916 | -0.01342 | 0.77677 | -0.01727 | 0.986217 | 0.999655 | 0.00015 |
| ENSMUSG00000087679.3 | 79.63805 | -0.01225 | 0.933605 | -0.01312 | 0.989532 | 0.999655 | 0.00015 |
| ENSMUSG00000093622.2 | 18.33642 | -0.02424 | 2.042358 | -0.01187 | 0.990532 | 0.999655 | 0.00015 |
| ENSMUSG00000107838.2 | 115.1718 | -0.00848 | 0.858718 | -0.00987 | 0.992125 | 0.999655 | 0.00015 |
| ENSMUSG00000110057.2 | 166.995 | 0.002735 | 0.563725 | 0.004852 | 0.996128 | 0.999655 | 0.00015 |
| ENSMUSG00000112090.2 | 19.45395 | -0.01144 | 2.512509 | -0.00455 | 0.996368 | 0.999655 | 0.00015 |
| __no_feature | 384638.3 | -0.00143 | 0.187624 | -0.00763 | 0.993915 | 0.999655 | 0.00015 |
| ENSMUSG00000020649.12 | 47.77587 | -0.00262 | 0.970771 | -0.0027 | 0.997848 | 0.99967 | 0.000143 |
| ENSMUSG00000021188.15 | 224.5905 | -0.00159 | 0.530105 | -0.003 | 0.997608 | 0.99967 | 0.000143 |
| ENSMUSG00000022665.16 | 31.23016 | -0.00532 | 1.683968 | -0.00316 | 0.997477 | 0.99967 | 0.000143 |

| | | | | | | | |
|---|---|---|---|---|---|---|---|
| ENSMUSG00000024304.15 | 176.876 | -0.00172 | 0.5202 | -0.0033 | 0.997364 | 0.99967 | 0.000143 |
| ENSMUSG00000025184.11 | 25.02547 | -0.00345 | 1.849148 | -0.00187 | 0.99851 | 0.99967 | 0.000143 |
| ENSMUSG00000026771.15 | 65.33316 | 0.002597 | 1.035958 | 0.002507 | 0.998 | 0.99967 | 0.000143 |
| ENSMUSG00000027074.15 | 103.8212 | 0.002467 | 0.79545 | 0.003101 | 0.997525 | 0.99967 | 0.000143 |
| ENSMUSG00000041028.17 | 1934.591 | 0.000653 | 0.283044 | 0.002308 | 0.998158 | 0.99967 | 0.000143 |
| ENSMUSG00000046032.17 | 14.534 | 0.004422 | 2.275345 | 0.001944 | 0.998449 | 0.99967 | 0.000143 |
| ENSMUSG00000046573.8 | 46.78764 | 0.003454 | 1.579579 | 0.002187 | 0.998255 | 0.99967 | 0.000143 |
| ENSMUSG00000055670.15 | 87.21432 | 0.001651 | 0.782681 | 0.002109 | 0.998317 | 0.99967 | 0.000143 |
| ENSMUSG00000062031.14 | 38.03453 | 0.00494 | 1.756583 | 0.002812 | 0.997756 | 0.99967 | 0.000143 |
| ENSMUSG00000066306.14 | 10.49233 | 0.005692 | 2.532814 | 0.002247 | 0.998207 | 0.99967 | 0.000143 |
| ENSMUSG00000019822.13 | 60.99801 | -0.00162 | 1.200574 | -0.00135 | 0.998925 | 0.999701 | 0.00013 |
| ENSMUSG00000036292.15 | 109.3139 | -0.0012 | 0.812138 | -0.00147 | 0.998826 | 0.999701 | 0.00013 |
| ENSMUSG00000043987.19 | 39.99116 | 0.002225 | 1.511234 | 0.001472 | 0.998825 | 0.999701 | 0.00013 |
| ENSMUSG00000057367.13 | 55.80612 | 0.001426 | 1.097448 | 0.0013 | 0.998963 | 0.999701 | 0.00013 |
| ENSMUSG00000026425.16 | 46.1268 | 0.001221 | 1.095558 | 0.001114 | 0.999111 | 0.999744 | 0.000111 |
| ENSMUSG00000048118.18 | 314.9672 | -0.00054 | 0.556404 | -0.00097 | 0.999226 | 0.999753 | 0.000107 |
| ENSMUSG00000002221.12 | 30.4773 | 0.000604 | 1.181537 | 0.000511 | 0.999592 | 0.999876 | 5.37E-05 |
| ENSMUSG00000026083.13 | 1604.673 | -9.29E-05 | 0.258678 | -0.00036 | 0.999713 | 0.999876 | 5.37E-05 |
| ENSMUSG00000030842.13 | 60.50911 | 0.000535 | 0.971922 | 0.000551 | 0.99956 | 0.999876 | 5.37E-05 |
| ENSMUSG00000031504.7 | 42.14899 | 0.000216 | 1.397845 | 0.000155 | 0.999876 | 0.999876 | 5.37E-05 |
| ENSMUSG00000038007.15 | 68.93657 | 0.000359 | 1.362012 | 0.000264 | 0.99979 | 0.999876 | 5.37E-05 |
| ENSMUSG00000000003.16 | 0 | NA | NA | NA | NA | NA | #VALUE! |
| ENSMUSG00000000028.16 | 0 | NA | NA | NA | NA | NA | #VALUE! |
| ENSMUSG00000000031.17 | 0 | NA | NA | NA | NA | NA | #VALUE! |
| ENSMUSG00000000037.18 | 0 | NA | NA | NA | NA | NA | #VALUE! |
| ENSMUSG00000000058.7 | 9.90362 | 3.299945 | 2.691822 | 1.225915 | 0.220231 | NA | #VALUE! |
| ENSMUSG00000000085.17 | 12.79015 | 0.896927 | 3.099281 | 0.289398 | NA | NA | #VALUE! |
| ENSMUSG00000000093.7 | 7.572759 | 6.410662 | 3.9127 | 1.638424 | 0.101333 | NA | #VALUE! |
| ENSMUSG00000000094.13 | 0 | NA | NA | NA | NA | NA | #VALUE! |
| ENSMUSG00000000103.13 | 0 | NA | NA | NA | NA | NA | #VALUE! |
| ENSMUSG00000000120.7 | 0.093255 | 0.868093 | 4.072464 | 0.213162 | 0.831201 | NA | #VALUE! |
| ENSMUSG00000000125.6 | 0 | NA | NA | NA | NA | NA | #VALUE! |
| ENSMUSG00000000126.12 | 4.129939 | 5.537758 | 3.917624 | 1.41355 | 0.157494 | NA | #VALUE! |
| ENSMUSG00000000134.18 | 8.470884 | 0.087862 | 3.488522 | 0.025186 | 0.979907 | NA | #VALUE! |
| ENSMUSG00000000142.16 | 3.125261 | 0.561961 | 3.786092 | 0.148428 | 0.882005 | NA | #VALUE! |
| ENSMUSG00000000148.18 | 9.084167 | -1.65882 | 2.983165 | -0.55606 | 0.57817 | NA | #VALUE! |
| ENSMUSG00000000157.17 | 0 | NA | NA | NA | NA | NA | #VALUE! |
| ENSMUSG00000000167.15 | 2.520315 | -4.8742 | 3.924358 | -1.24204 | 0.214223 | NA | #VALUE! |
| ENSMUSG00000000182.10 | 0 | NA | NA | NA | NA | NA | #VALUE! |
| ENSMUSG00000000183.7 | 0 | NA | NA | NA | NA | NA | #VALUE! |
| ENSMUSG00000000194.14 | 24.13529 | 0.839049 | 1.951797 | 0.429885 | NA | NA | #VALUE! |
| ENSMUSG00000000197.9 | 1.865104 | 4.40601 | 3.930481 | 1.120985 | 0.262294 | NA | #VALUE! |
| ENSMUSG00000000202.10 | 0 | NA | NA | NA | NA | NA | #VALUE! |
| ENSMUSG00000000204.17 | 0 | NA | NA | NA | NA | NA | #VALUE! |

| | | | | | | | |
|---|---|---|---|---|---|---|---|
| ENSMUSG00000000214.12 | 0 | NA | NA | NA | NA | NA | #VALUE! |
| ENSMUSG00000000215.12 | 0 | NA | NA | NA | NA | NA | #VALUE! |
| ENSMUSG00000000216.9 | 0 | NA | NA | NA | NA | NA | #VALUE! |
| ENSMUSG00000000223.14 | 0 | NA | NA | NA | NA | NA | #VALUE! |
| ENSMUSG00000000244.18 | 0 | NA | NA | NA | NA | NA | #VALUE! |
| ENSMUSG00000000248.17 | 0 | NA | NA | NA | NA | NA | #VALUE! |
| ENSMUSG00000000253.14 | 0 | NA | NA | NA | NA | NA | #VALUE! |
| ENSMUSG00000000263.16 | 0 | NA | NA | NA | NA | NA | #VALUE! |
| ENSMUSG00000000276.12 | 0.408222 | -2.20205 | 4.016842 | -0.54821 | 0.583551 | NA | #VALUE! |
| ENSMUSG00000000282.13 | 1.923902 | 4.423325 | 3.9302 | 1.125471 | 0.26039 | NA | #VALUE! |
| ENSMUSG00000000305.13 | 0 | NA | NA | NA | NA | NA | #VALUE! |
| ENSMUSG00000000308.15 | 0 | NA | NA | NA | NA | NA | #VALUE! |
| ENSMUSG00000000317.12 | 0 | NA | NA | NA | NA | NA | #VALUE! |
| ENSMUSG00000000318.17 | 0 | NA | NA | NA | NA | NA | #VALUE! |
| ENSMUSG00000000359.4 | 0.373021 | 2.140748 | 4.018182 | 0.532765 | 0.594196 | NA | #VALUE! |
| ENSMUSG00000000365.10 | 0 | NA | NA | NA | NA | NA | #VALUE! |
| ENSMUSG00000000381.11 | 0 | NA | NA | NA | NA | NA | #VALUE! |
| ENSMUSG00000000385.9 | 3.338328 | 0.513707 | 3.783137 | 0.135789 | 0.891988 | NA | #VALUE! |
| ENSMUSG00000000386.17 | 3.340873 | -4.12431 | 3.906995 | -1.05562 | 0.291141 | NA | #VALUE! |
| ENSMUSG00000000392.18 | 0 | NA | NA | NA | NA | NA | #VALUE! |
| ENSMUSG00000000394.16 | 0 | NA | NA | NA | NA | NA | #VALUE! |
| ENSMUSG00000000402.3 | 0 | NA | NA | NA | NA | NA | #VALUE! |
| ENSMUSG00000000409.15 | 0 | NA | NA | NA | NA | NA | #VALUE! |
| ENSMUSG00000000411.8 | 0 | NA | NA | NA | NA | NA | #VALUE! |
| ENSMUSG00000000416.17 | 0 | NA | NA | NA | NA | NA | #VALUE! |
| ENSMUSG00000000435.2 | 0 | NA | NA | NA | NA | NA | #VALUE! |
| ENSMUSG00000000486.14 | 0.828518 | 3.195613 | 3.961197 | 0.806729 | 0.419823 | NA | #VALUE! |
| ENSMUSG00000000489.8 | 0 | NA | NA | NA | NA | NA | #VALUE! |
| ENSMUSG00000000530.17 | 1.967804 | 4.449735 | 3.929778 | 1.132312 | 0.257503 | NA | #VALUE! |
| ENSMUSG00000000531.6 | 8.750743 | 6.619253 | 3.754899 | 1.762831 | 0.077929 | NA | #VALUE! |
| ENSMUSG00000000544.15 | 0 | NA | NA | NA | NA | NA | #VALUE! |
| ENSMUSG00000000552.11 | 0 | NA | NA | NA | NA | NA | #VALUE! |
| ENSMUSG00000000560.10 | 0 | NA | NA | NA | NA | NA | #VALUE! |
| ENSMUSG00000000562.6 | 0 | NA | NA | NA | NA | NA | #VALUE! |
| ENSMUSG00000000579.15 | 0 | NA | NA | NA | NA | NA | #VALUE! |
| ENSMUSG00000000606.19 | 0 | NA | NA | NA | NA | NA | #VALUE! |
| ENSMUSG00000000617.15 | 0 | NA | NA | NA | NA | NA | #VALUE! |
| ENSMUSG00000000627.16 | 0 | NA | NA | NA | NA | NA | #VALUE! |
| ENSMUSG00000000628.11 | 9.184984 | -6.72736 | 3.700099 | -1.81816 | 0.06904 | NA | #VALUE! |
| ENSMUSG00000000632.14 | 0 | NA | NA | NA | NA | NA | #VALUE! |
| ENSMUSG00000000682.8 | 23.7316 | -0.54624 | 2.028918 | -0.26923 | NA | NA | #VALUE! |
| ENSMUSG00000000690.6 | 0 | NA | NA | NA | NA | NA | #VALUE! |
| ENSMUSG00000000693.11 | 4.480072 | -5.69697 | 3.916727 | -1.45452 | 0.145801 | NA | #VALUE! |
| ENSMUSG00000000701.9 | 0 | NA | NA | NA | NA | NA | #VALUE! |

| | | | | | | | |
|---|---|---|---|---|---|---|---|
| ENSMUSG00000000706.17 | 0 | NA | NA | NA | NA | NA | #VALUE! |
| ENSMUSG00000000724.11 | 0 | NA | NA | NA | NA | NA | #VALUE! |
| ENSMUSG00000000730.14 | 0 | NA | NA | NA | NA | NA | #VALUE! |
| ENSMUSG00000000731.16 | 0 | NA | NA | NA | NA | NA | #VALUE! |
| ENSMUSG00000000732.10 | 0 | NA | NA | NA | NA | NA | #VALUE! |
| ENSMUSG00000000739.15 | 15.97617 | -1.75382 | 2.922889 | -0.60003 | NA | NA | #VALUE! |
| ENSMUSG00000000766.19 | 0 | NA | NA | NA | NA | NA | #VALUE! |
| ENSMUSG00000000791.10 | 0.18651 | 1.401769 | 4.056354 | 0.345574 | 0.729663 | NA | #VALUE! |
| ENSMUSG00000000792.3 | 0 | NA | NA | NA | NA | NA | #VALUE! |
| ENSMUSG00000000794.10 | 0 | NA | NA | NA | NA | NA | #VALUE! |
| ENSMUSG00000000805.19 | 0 | NA | NA | NA | NA | NA | #VALUE! |
| ENSMUSG00000000817.11 | 1.767477 | -4.3586 | 3.931877 | -1.10853 | 0.267633 | NA | #VALUE! |
| ENSMUSG00000000861.16 | 3.232286 | -5.22438 | 3.920577 | -1.33255 | 0.182678 | NA | #VALUE! |
| ENSMUSG00000000869.13 | 0 | NA | NA | NA | NA | NA | #VALUE! |
| ENSMUSG00000000884.18 | 0 | NA | NA | NA | NA | NA | #VALUE! |
| ENSMUSG00000000889.9 | 0 | NA | NA | NA | NA | NA | #VALUE! |
| ENSMUSG00000000901.18 | 0 | NA | NA | NA | NA | NA | #VALUE! |
| ENSMUSG00000000903.4 | 4.68165 | 5.712018 | 3.916391 | 1.45849 | 0.144705 | NA | #VALUE! |
| ENSMUSG00000000915.16 | 0 | NA | NA | NA | NA | NA | #VALUE! |
| ENSMUSG00000000938.12 | 0 | NA | NA | NA | NA | NA | #VALUE! |
| ENSMUSG00000000942.11 | 0 | NA | NA | NA | NA | NA | #VALUE! |
| ENSMUSG00000000948.17 | 0.336567 | 1.944925 | 4.029936 | 0.482619 | 0.629366 | NA | #VALUE! |
| ENSMUSG00000000958.11 | 6.72084 | -6.28266 | 3.913407 | -1.60542 | 0.108402 | NA | #VALUE! |
| ENSMUSG00000000982.6 | 0 | NA | NA | NA | NA | NA | #VALUE! |
| ENSMUSG00000000983.14 | 0 | NA | NA | NA | NA | NA | #VALUE! |
| ENSMUSG00000001014.7 | 0 | NA | NA | NA | NA | NA | #VALUE! |
| ENSMUSG00000001020.9 | 5.978686 | -6.10788 | 3.847895 | -1.58733 | 0.112438 | NA | #VALUE! |
| ENSMUSG00000001021.14 | 0 | NA | NA | NA | NA | NA | #VALUE! |
| ENSMUSG00000001023.11 | 0 | NA | NA | NA | NA | NA | #VALUE! |
| ENSMUSG00000001027.8 | 0 | NA | NA | NA | NA | NA | #VALUE! |
| ENSMUSG00000001029.16 | 3.555455 | 4.352892 | 3.915336 | 1.111754 | 0.266244 | NA | #VALUE! |
| ENSMUSG00000001034.18 | 3.030092 | -4.17196 | 3.91369 | -1.06599 | 0.286427 | NA | #VALUE! |
| ENSMUSG00000001039.13 | 0 | NA | NA | NA | NA | NA | #VALUE! |
| ENSMUSG00000001053.16 | 0 | NA | NA | NA | NA | NA | #VALUE! |
| ENSMUSG00000001062.19 | 8.861548 | 0.577264 | 2.720544 | 0.212187 | 0.831961 | NA | #VALUE! |
| ENSMUSG00000001076.8 | 0.408222 | -2.20205 | 4.016842 | -0.54821 | 0.583551 | NA | #VALUE! |
| ENSMUSG00000001082.13 | 0 | NA | NA | NA | NA | NA | #VALUE! |
| ENSMUSG00000001095.6 | 7.337452 | -5.47573 | 3.65159 | -1.49955 | 0.133732 | NA | #VALUE! |
| ENSMUSG00000001103.8 | 2.520315 | -4.8742 | 3.924358 | -1.24204 | 0.214223 | NA | #VALUE! |
| ENSMUSG00000001120.16 | 0 | NA | NA | NA | NA | NA | #VALUE! |
| ENSMUSG00000001131.12 | 0 | NA | NA | NA | NA | NA | #VALUE! |
| ENSMUSG00000001143.14 | 0.932552 | 3.421684 | 3.953399 | 0.865504 | 0.386762 | NA | #VALUE! |
| ENSMUSG00000001156.10 | 8.296037 | -6.5859 | 3.798445 | -1.73384 | NA | NA | #VALUE! |
| ENSMUSG00000001166.18 | 0 | NA | NA | NA | NA | NA | #VALUE! |

| | | | | | | | |
|---|---|---|---|---|---|---|---|
| ENSMUSG00000001168.17 | 0 | NA | NA | NA | NA | NA | #VALUE! |
| ENSMUSG00000001173.16 | 2.502152 | -3.30121 | 3.900949 | -0.84626 | 0.397408 | NA | #VALUE! |
| ENSMUSG00000001225.13 | 0 | NA | NA | NA | NA | NA | #VALUE! |
| ENSMUSG00000001227.13 | 0 | NA | NA | NA | NA | NA | #VALUE! |
| ENSMUSG00000001228.15 | 0 | NA | NA | NA | NA | NA | #VALUE! |
| ENSMUSG00000001248.16 | 0 | NA | NA | NA | NA | NA | #VALUE! |
| ENSMUSG00000001260.11 | 0 | NA | NA | NA | NA | NA | #VALUE! |
| ENSMUSG00000001270.10 | 7.43506 | 1.495862 | 2.777555 | 0.538554 | 0.590195 | NA | #VALUE! |
| ENSMUSG00000001281.11 | 0 | NA | NA | NA | NA | NA | #VALUE! |
| ENSMUSG00000001288.16 | 0.612332 | -2.79827 | 3.980109 | -0.70306 | 0.482016 | NA | #VALUE! |
| ENSMUSG00000001333.10 | 0 | NA | NA | NA | NA | NA | #VALUE! |
| ENSMUSG00000001334.10 | 0 | NA | NA | NA | NA | NA | #VALUE! |
| ENSMUSG00000001349.6 | 0 | NA | NA | NA | NA | NA | #VALUE! |
| ENSMUSG00000001403.14 | 0 | NA | NA | NA | NA | NA | #VALUE! |
| ENSMUSG00000001415.11 | 1.128341 | 2.40607 | 3.945995 | 0.60975 | 0.542028 | NA | #VALUE! |
| ENSMUSG00000001420.14 | 0 | NA | NA | NA | NA | NA | #VALUE! |
| ENSMUSG00000001436.16 | 7.668942 | -6.46689 | 3.457681 | -1.8703 | 0.061443 | NA | #VALUE! |
| ENSMUSG00000001444.3 | 0 | NA | NA | NA | NA | NA | #VALUE! |
| ENSMUSG00000001473.8 | 8.241622 | -1.0232 | 3.117344 | -0.32823 | 0.74274 | NA | #VALUE! |
| ENSMUSG00000001493.10 | 0 | NA | NA | NA | NA | NA | #VALUE! |
| ENSMUSG00000001494.7 | 0 | NA | NA | NA | NA | NA | #VALUE! |
| ENSMUSG00000001496.16 | 0 | NA | NA | NA | NA | NA | #VALUE! |
| ENSMUSG00000001497.19 | 0 | NA | NA | NA | NA | NA | #VALUE! |
| ENSMUSG00000001504.11 | 0 | NA | NA | NA | NA | NA | #VALUE! |
| ENSMUSG00000001507.17 | 0.315039 | -1.93549 | 4.038621 | -0.47925 | 0.631763 | NA | #VALUE! |
| ENSMUSG00000001508.16 | 0 | NA | NA | NA | NA | NA | #VALUE! |
| ENSMUSG00000001510.9 | 0 | NA | NA | NA | NA | NA | #VALUE! |
| ENSMUSG00000001517.15 | 0 | NA | NA | NA | NA | NA | #VALUE! |
| ENSMUSG00000001520.13 | 0 | NA | NA | NA | NA | NA | #VALUE! |
| ENSMUSG00000001521.13 | 2.425886 | -2.77737 | 3.882111 | -0.71543 | 0.474346 | NA | #VALUE! |
| ENSMUSG00000001524.15 | 7.955712 | -0.3695 | 3.146721 | -0.11742 | 0.906524 | NA | #VALUE! |
| ENSMUSG00000001555.11 | 0 | NA | NA | NA | NA | NA | #VALUE! |
| ENSMUSG00000001558.6 | 0 | NA | NA | NA | NA | NA | #VALUE! |
| ENSMUSG00000001566.10 | 0 | NA | NA | NA | NA | NA | #VALUE! |
| ENSMUSG00000001583.14 | 0 | NA | NA | NA | NA | NA | #VALUE! |
| ENSMUSG00000001588.13 | 0 | NA | NA | NA | NA | NA | #VALUE! |
| ENSMUSG00000001622.16 | 0 | NA | NA | NA | NA | NA | #VALUE! |
| ENSMUSG00000001642.19 | 5.51099 | -5.9895 | 3.914902 | -1.52992 | 0.126035 | NA | #VALUE! |
| ENSMUSG00000001655.7 | 0 | NA | NA | NA | NA | NA | #VALUE! |
| ENSMUSG00000001656.4 | 0 | NA | NA | NA | NA | NA | #VALUE! |
| ENSMUSG00000001657.8 | 0 | NA | NA | NA | NA | NA | #VALUE! |
| ENSMUSG00000001661.6 | 0 | NA | NA | NA | NA | NA | #VALUE! |
| ENSMUSG00000001739.15 | 0.491951 | 2.402185 | 4.000147 | 0.600524 | 0.548157 | NA | #VALUE! |
| ENSMUSG00000001773.15 | 0 | NA | NA | NA | NA | NA | #VALUE! |

| | | | | | | | |
|---|---|---|---|---|---|---|---|
| ENSMUSG00000001786.15 | 8.736877 | -4.02945 | 2.984334 | -1.3502 | 0.176952 | NA | #VALUE! |
| ENSMUSG00000001802.17 | 4.432747 | -5.67814 | 3.916857 | -1.44967 | 0.147151 | NA | #VALUE! |
| ENSMUSG00000001804.10 | 0 | NA | NA | NA | NA | NA | #VALUE! |
| ENSMUSG00000001815.16 | 0 | NA | NA | NA | NA | NA | #VALUE! |
| ENSMUSG00000001819.5 | 0 | NA | NA | NA | NA | NA | #VALUE! |
| ENSMUSG00000001823.6 | 0 | NA | NA | NA | NA | NA | #VALUE! |
| ENSMUSG00000001827.13 | 0 | NA | NA | NA | NA | NA | #VALUE! |
| ENSMUSG00000001864.14 | 0 | NA | NA | NA | NA | NA | #VALUE! |
| ENSMUSG00000001865.3 | 0 | NA | NA | NA | NA | NA | #VALUE! |
| ENSMUSG00000001901.16 | 0 | NA | NA | NA | NA | NA | #VALUE! |
| ENSMUSG00000001930.18 | 1.370524 | -3.97623 | 3.939448 | -1.00934 | 0.312813 | NA | #VALUE! |
| ENSMUSG00000001943.9 | 0 | NA | NA | NA | NA | NA | #VALUE! |
| ENSMUSG00000001948.11 | 4.18556 | 3.547143 | 3.874759 | 0.915449 | 0.359956 | NA | #VALUE! |
| ENSMUSG00000001985.10 | 0 | NA | NA | NA | NA | NA | #VALUE! |
| ENSMUSG00000001986.17 | 0.525066 | -2.64687 | 3.988095 | -0.66369 | 0.506887 | NA | #VALUE! |
| ENSMUSG00000001988.10 | 0 | NA | NA | NA | NA | NA | #VALUE! |
| ENSMUSG00000001995.10 | 4.837167 | 2.699997 | 3.807497 | 0.709126 | 0.478246 | NA | #VALUE! |
| ENSMUSG00000001998.16 | 19.90601 | -2.33069 | 2.743294 | -0.8496 | NA | NA | #VALUE! |
| ENSMUSG00000002006.13 | 0 | NA | NA | NA | NA | NA | #VALUE! |
| ENSMUSG00000002007.6 | 0 | NA | NA | NA | NA | NA | #VALUE! |
| ENSMUSG00000002012.14 | 0 | NA | NA | NA | NA | NA | #VALUE! |
| ENSMUSG00000002020.16 | 0 | NA | NA | NA | NA | NA | #VALUE! |
| ENSMUSG00000002033.15 | 6.761843 | -6.28552 | 3.691268 | -1.70281 | 0.088604 | NA | #VALUE! |
| ENSMUSG00000002052.7 | 5.569989 | 4.100633 | 3.217921 | 1.274311 | 0.202553 | NA | #VALUE! |
| ENSMUSG00000002055.10 | 1.597059 | -2.96633 | 3.924337 | -0.75588 | 0.449721 | NA | #VALUE! |
| ENSMUSG00000002057.5 | 0 | NA | NA | NA | NA | NA | #VALUE! |
| ENSMUSG00000002058.14 | 5.468767 | 4.388352 | 3.639778 | 1.205665 | 0.227947 | NA | #VALUE! |
| ENSMUSG00000002068.17 | 0.364073 | 0.012025 | 4.059931 | 0.002962 | 0.997637 | NA | #VALUE! |
| ENSMUSG00000002076.13 | 1.053643 | -0.25287 | 3.897552 | -0.06488 | 0.94827 | NA | #VALUE! |
| ENSMUSG00000002100.16 | 0 | NA | NA | NA | NA | NA | #VALUE! |
| ENSMUSG00000002104.12 | 0 | NA | NA | NA | NA | NA | #VALUE! |
| ENSMUSG00000002111.9 | 0.420053 | -2.33493 | 4.007343 | -0.58266 | 0.56012 | NA | #VALUE! |
| ENSMUSG00000002147.19 | 31.47485 | -0.9124 | 1.671007 | -0.54602 | NA | NA | #VALUE! |
| ENSMUSG00000002190.14 | 0 | NA | NA | NA | NA | NA | #VALUE! |
| ENSMUSG00000002204.10 | 0 | NA | NA | NA | NA | NA | #VALUE! |
| ENSMUSG00000002210.12 | 6.218482 | -3.05137 | 3.793267 | -0.80442 | 0.421156 | NA | #VALUE! |
| ENSMUSG00000002228.8 | 0 | NA | NA | NA | NA | NA | #VALUE! |
| ENSMUSG00000002233.14 | 7.253781 | 1.182222 | 3.651007 | 0.323807 | 0.746084 | NA | #VALUE! |
| ENSMUSG00000002240.13 | 0 | NA | NA | NA | NA | NA | #VALUE! |
| ENSMUSG00000002250.17 | 0.771041 | -1.57901 | 3.974438 | -0.39729 | 0.691152 | NA | #VALUE! |
| ENSMUSG00000002257.9 | 0 | NA | NA | NA | NA | NA | #VALUE! |
| ENSMUSG00000002265.18 | 6.539701 | -5.12233 | 3.371746 | -1.51919 | 0.128714 | NA | #VALUE! |
| ENSMUSG00000002266.18 | 0 | NA | NA | NA | NA | NA | #VALUE! |
| ENSMUSG00000002274.14 | 4.757613 | -1.55322 | 3.735173 | -0.41584 | 0.67753 | NA | #VALUE! |

| | | | | | | | |
|---|---|---|---|---|---|---|---|
| ENSMUSG00000002308.17 | 1.579384 | 0.791289 | 3.850326 | 0.205512 | 0.837172 | NA | #VALUE! |
| ENSMUSG00000002319.17 | 41.41393 | -1.18227 | 1.746533 | -0.67692 | NA | NA | #VALUE! |
| ENSMUSG00000002320.16 | 1.174735 | -3.75246 | 3.944889 | -0.95122 | 0.341493 | NA | #VALUE! |
| ENSMUSG00000002324.9 | 2.349469 | -4.75734 | 3.925837 | -1.2118 | 0.225588 | NA | #VALUE! |
| ENSMUSG00000002341.9 | 0 | NA | NA | NA | NA | NA | #VALUE! |
| ENSMUSG00000002343.12 | 1.444189 | -3.07104 | 3.931081 | -0.78122 | 0.434673 | NA | #VALUE! |
| ENSMUSG00000002384.3 | 0 | NA | NA | NA | NA | NA | #VALUE! |
| ENSMUSG00000002409.19 | 0 | NA | NA | NA | NA | NA | #VALUE! |
| ENSMUSG00000002459.18 | 0.587367 | -2.74297 | 3.982932 | -0.68868 | 0.491024 | NA | #VALUE! |
| ENSMUSG00000002489.17 | 0 | NA | NA | NA | NA | NA | #VALUE! |
| ENSMUSG00000002500.16 | 0 | NA | NA | NA | NA | NA | #VALUE! |
| ENSMUSG00000002546.18 | 4.931192 | -1.73861 | 3.402395 | -0.511 | 0.609354 | NA | #VALUE! |
| ENSMUSG00000002565.17 | 0 | NA | NA | NA | NA | NA | #VALUE! |
| ENSMUSG00000002578.19 | 0 | NA | NA | NA | NA | NA | #VALUE! |
| ENSMUSG00000002603.16 | 0 | NA | NA | NA | NA | NA | #VALUE! |
| ENSMUSG00000002617.15 | 1.626034 | 4.194606 | 3.934197 | 1.066191 | 0.286337 | NA | #VALUE! |
| ENSMUSG00000002633.5 | 0 | NA | NA | NA | NA | NA | #VALUE! |
| ENSMUSG00000002664.3 | 0 | NA | NA | NA | NA | NA | #VALUE! |
| ENSMUSG00000002668.9 | 3.265772 | -5.23273 | 3.920498 | -1.33471 | 0.181971 | NA | #VALUE! |
| ENSMUSG00000002688.9 | 0 | NA | NA | NA | NA | NA | #VALUE! |
| ENSMUSG00000002771.13 | 0 | NA | NA | NA | NA | NA | #VALUE! |
| ENSMUSG00000002791.18 | 0 | NA | NA | NA | NA | NA | #VALUE! |
| ENSMUSG00000002799.7 | 0 | NA | NA | NA | NA | NA | #VALUE! |
| ENSMUSG00000002808.8 | 0 | NA | NA | NA | NA | NA | #VALUE! |
| ENSMUSG00000002814.16 | 8.275181 | -1.32603 | 3.528742 | -0.37578 | 0.707081 | NA | #VALUE! |
| ENSMUSG00000002820.7 | 7.868604 | 4.067313 | 3.03609 | 1.339655 | 0.180358 | NA | #VALUE! |
| ENSMUSG00000002831.15 | 12.12318 | 7.093299 | 3.358357 | 2.112134 | NA | NA | #VALUE! |
| ENSMUSG00000002835.10 | 1.957891 | -4.49332 | 3.929649 | -1.14344 | 0.252855 | NA | #VALUE! |
| ENSMUSG00000002870.9 | 8.801548 | -0.8825 | 2.991706 | -0.29498 | 0.768008 | NA | #VALUE! |
| ENSMUSG00000002871.15 | 5.052611 | 0.817057 | 3.693315 | 0.221226 | 0.824917 | NA | #VALUE! |
| ENSMUSG00000002908.18 | 4.382994 | 5.629097 | 3.916959 | 1.437109 | 0.150687 | NA | #VALUE! |
| ENSMUSG00000002930.7 | 0 | NA | NA | NA | NA | NA | #VALUE! |
| ENSMUSG00000002968.10 | 13.63097 | 7.258679 | 3.26914 | 2.220364 | NA | NA | #VALUE! |
| ENSMUSG00000002980.15 | 0 | NA | NA | NA | NA | NA | #VALUE! |
| ENSMUSG00000002983.18 | 1.050131 | -3.6251 | 3.948379 | -0.91812 | 0.358554 | NA | #VALUE! |
| ENSMUSG00000002992.18 | 0 | NA | NA | NA | NA | NA | #VALUE! |
| ENSMUSG00000002997.16 | 0.279766 | 1.808031 | 4.036658 | 0.447903 | 0.654223 | NA | #VALUE! |
| ENSMUSG00000003032.9 | 9.199413 | -5.64931 | 3.377277 | -1.67274 | NA | NA | #VALUE! |
| ENSMUSG00000003051.14 | 0 | NA | NA | NA | NA | NA | #VALUE! |
| ENSMUSG00000003070.7 | 0 | NA | NA | NA | NA | NA | #VALUE! |
| ENSMUSG00000003153.11 | 0 | NA | NA | NA | NA | NA | #VALUE! |
| ENSMUSG00000003200.11 | 1.724765 | -4.31574 | 3.932631 | -1.09742 | 0.272459 | NA | #VALUE! |
| ENSMUSG00000003206.8 | 0 | NA | NA | NA | NA | NA | #VALUE! |
| ENSMUSG00000003208.10 | 6.194543 | -5.03944 | 3.404521 | -1.48022 | 0.138815 | NA | #VALUE! |

| | | | | | | | |
|---|---|---|---|---|---|---|---|
| ENSMUSG00000003227.5 | 0 | NA | NA | NA | NA | NA | #VALUE! |
| ENSMUSG00000003228.10 | 1.350934 | -3.97933 | 3.939378 | -1.01014 | 0.312427 | NA | #VALUE! |
| ENSMUSG00000003269.18 | 17.05362 | -0.35275 | 2.886074 | -0.12222 | NA | NA | #VALUE! |
| ENSMUSG00000003271.18 | 0 | NA | NA | NA | NA | NA | #VALUE! |
| ENSMUSG00000003273.15 | 0 | NA | NA | NA | NA | NA | #VALUE! |
| ENSMUSG00000003279.18 | 8.431504 | -0.37506 | 3.090953 | -0.12134 | 0.90342 | NA | #VALUE! |
| ENSMUSG00000003282.10 | 0 | NA | NA | NA | NA | NA | #VALUE! |
| ENSMUSG00000003309.15 | 0 | NA | NA | NA | NA | NA | #VALUE! |
| ENSMUSG00000003341.10 | 0 | NA | NA | NA | NA | NA | #VALUE! |
| ENSMUSG00000003348.11 | 0.210026 | -1.52494 | 4.058692 | -0.37572 | 0.707123 | NA | #VALUE! |
| ENSMUSG00000003352.15 | 0 | NA | NA | NA | NA | NA | #VALUE! |
| ENSMUSG00000003354.7 | 0 | NA | NA | NA | NA | NA | #VALUE! |
| ENSMUSG00000003360.16 | 6.062396 | 2.612052 | 3.788012 | 0.689557 | 0.490473 | NA | #VALUE! |
| ENSMUSG00000003363.16 | 5.242805 | -3.62231 | 3.5204 | -1.02895 | 0.303503 | NA | #VALUE! |
| ENSMUSG00000003378.10 | 0 | NA | NA | NA | NA | NA | #VALUE! |
| ENSMUSG00000003379.8 | 0 | NA | NA | NA | NA | NA | #VALUE! |
| ENSMUSG00000003410.9 | 0 | NA | NA | NA | NA | NA | #VALUE! |
| ENSMUSG00000003411.11 | 0.735092 | -3.12016 | 3.965612 | -0.7868 | 0.431397 | NA | #VALUE! |
| ENSMUSG00000003418.12 | 0 | NA | NA | NA | NA | NA | #VALUE! |
| ENSMUSG00000003436.12 | 0 | NA | NA | NA | NA | NA | #VALUE! |
| ENSMUSG00000003452.16 | 0.315039 | -1.93549 | 4.038621 | -0.47925 | 0.631763 | NA | #VALUE! |
| ENSMUSG00000003469.10 | 0 | NA | NA | NA | NA | NA | #VALUE! |
| ENSMUSG00000003476.17 | 0 | NA | NA | NA | NA | NA | #VALUE! |
| ENSMUSG00000003484.5 | 0 | NA | NA | NA | NA | NA | #VALUE! |
| ENSMUSG00000003500.14 | 0 | NA | NA | NA | NA | NA | #VALUE! |
| ENSMUSG00000003505.16 | 0 | NA | NA | NA | NA | NA | #VALUE! |
| ENSMUSG00000003534.18 | 0 | NA | NA | NA | NA | NA | #VALUE! |
| ENSMUSG00000003545.4 | 3.36042 | -5.28663 | 3.919996 | -1.34863 | 0.177455 | NA | #VALUE! |
| ENSMUSG00000003573.16 | 1.771849 | 4.332864 | 3.931707 | 1.102031 | 0.270448 | NA | #VALUE! |
| ENSMUSG00000003575.14 | 0 | NA | NA | NA | NA | NA | #VALUE! |
| ENSMUSG00000003657.10 | 0 | NA | NA | NA | NA | NA | #VALUE! |
| ENSMUSG00000003665.8 | 0 | NA | NA | NA | NA | NA | #VALUE! |
| ENSMUSG00000003680.17 | 0 | NA | NA | NA | NA | NA | #VALUE! |
| ENSMUSG00000003752.10 | 0.630079 | -2.90291 | 3.975047 | -0.73028 | 0.465217 | NA | #VALUE! |
| ENSMUSG00000003762.14 | 3.962644 | -1.48363 | 3.785349 | -0.39194 | 0.695103 | NA | #VALUE! |
| ENSMUSG00000003779.17 | 3.447843 | 0.775256 | 3.782727 | 0.204946 | 0.837614 | NA | #VALUE! |
| ENSMUSG00000003810.14 | 8.423389 | 1.579197 | 3.125548 | 0.505254 | 0.61338 | NA | #VALUE! |
| ENSMUSG00000003848.15 | 23.39002 | 0.339853 | 2.071824 | 0.164036 | NA | NA | #VALUE! |
| ENSMUSG00000003863.19 | 0 | NA | NA | NA | NA | NA | #VALUE! |
| ENSMUSG00000003865.17 | 1.025807 | 3.556426 | 3.949287 | 0.900523 | 0.367842 | NA | #VALUE! |
| ENSMUSG00000003872.10 | 0 | NA | NA | NA | NA | NA | #VALUE! |
| ENSMUSG00000003882.6 | 0 | NA | NA | NA | NA | NA | #VALUE! |
| ENSMUSG00000003929.12 | 7.01722 | -1.85769 | 3.350187 | -0.5545 | 0.579233 | NA | #VALUE! |
| ENSMUSG00000003934.6 | 0.336567 | 1.944925 | 4.029936 | 0.482619 | 0.629366 | NA | #VALUE! |

| | | | | | | | |
|---|---|---|---|---|---|---|---|
| ENSMUSG00000003974.7 | 0 | NA | NA | NA | NA | NA | #VALUE! |
| ENSMUSG00000004031.14 | 0 | NA | NA | NA | NA | NA | #VALUE! |
| ENSMUSG00000004054.10 | 4.808529 | 4.683455 | 3.687365 | 1.270136 | 0.204036 | NA | #VALUE! |
| ENSMUSG00000004098.8 | 4.935617 | -5.8387 | 3.915796 | -1.49106 | 0.135945 | NA | #VALUE! |
| ENSMUSG00000004099.17 | 5.916003 | -1.87464 | 3.013813 | -0.62201 | 0.533932 | NA | #VALUE! |
| ENSMUSG00000004110.18 | 0 | NA | NA | NA | NA | NA | #VALUE! |
| ENSMUSG00000004113.19 | 0 | NA | NA | NA | NA | NA | #VALUE! |
| ENSMUSG00000004151.18 | 1.048192 | 2.22288 | 3.952233 | 0.562436 | 0.573819 | NA | #VALUE! |
| ENSMUSG00000004187.12 | 2.81658 | -1.95702 | 3.825053 | -0.51163 | 0.608909 | NA | #VALUE! |
| ENSMUSG00000004231.16 | 0 | NA | NA | NA | NA | NA | #VALUE! |
| ENSMUSG00000004233.15 | 4.083735 | -2.59981 | 3.817756 | -0.68098 | 0.495885 | NA | #VALUE! |
| ENSMUSG00000004267.17 | 0 | NA | NA | NA | NA | NA | #VALUE! |
| ENSMUSG00000004296.15 | 0 | NA | NA | NA | NA | NA | #VALUE! |
| ENSMUSG00000004317.15 | 5.493385 | -5.98712 | 3.398778 | -1.76155 | 0.078145 | NA | #VALUE! |
| ENSMUSG00000004328.16 | 0 | NA | NA | NA | NA | NA | #VALUE! |
| ENSMUSG00000004344.13 | 0 | NA | NA | NA | NA | NA | #VALUE! |
| ENSMUSG00000004347.18 | 1.425667 | 2.74624 | 3.936277 | 0.697675 | 0.485381 | NA | #VALUE! |
| ENSMUSG00000004359.17 | 0.673134 | 2.918386 | 3.972531 | 0.734641 | 0.462558 | NA | #VALUE! |
| ENSMUSG00000004360.10 | 0 | NA | NA | NA | NA | NA | #VALUE! |
| ENSMUSG00000004366.5 | 0 | NA | NA | NA | NA | NA | #VALUE! |
| ENSMUSG00000004371.16 | 0 | NA | NA | NA | NA | NA | #VALUE! |
| ENSMUSG00000004383.19 | 8.77681 | -0.91062 | 2.693968 | -0.33802 | 0.735345 | NA | #VALUE! |
| ENSMUSG00000004415.16 | 0 | NA | NA | NA | NA | NA | #VALUE! |
| ENSMUSG00000004530.13 | 3.213238 | 0.516597 | 3.784533 | 0.136502 | 0.891424 | NA | #VALUE! |
| ENSMUSG00000004540.8 | 0 | NA | NA | NA | NA | NA | #VALUE! |
| ENSMUSG00000004542.16 | 0 | NA | NA | NA | NA | NA | #VALUE! |
| ENSMUSG00000004552.17 | 2.797656 | 4.98547 | 3.922669 | 1.270938 | 0.203751 | NA | #VALUE! |
| ENSMUSG00000004562.17 | 1.48846 | -1.88314 | 3.898866 | -0.483 | 0.629099 | NA | #VALUE! |
| ENSMUSG00000004565.16 | 4.824759 | 4.692676 | 3.766214 | 1.245993 | 0.212767 | NA | #VALUE! |
| ENSMUSG00000004568.14 | 5.250656 | -5.92764 | 3.915257 | -1.51398 | 0.13003 | NA | #VALUE! |
| ENSMUSG00000004612.10 | 3.254185 | -0.21235 | 3.782348 | -0.05614 | 0.955229 | NA | #VALUE! |
| ENSMUSG00000004626.15 | 5.410418 | 1.143578 | 3.764939 | 0.303744 | 0.761323 | NA | #VALUE! |
| ENSMUSG00000004630.19 | 0.983902 | 3.433476 | 3.953024 | 0.868569 | 0.385083 | NA | #VALUE! |
| ENSMUSG00000004631.16 | 6.946192 | -3.01418 | 3.012113 | -1.00069 | 0.316979 | NA | #VALUE! |
| ENSMUSG00000004642.14 | 1.222896 | 0.900279 | 3.885843 | 0.231682 | 0.816785 | NA | #VALUE! |
| ENSMUSG00000004651.7 | 0 | NA | NA | NA | NA | NA | #VALUE! |
| ENSMUSG00000004654.7 | 0 | NA | NA | NA | NA | NA | #VALUE! |
| ENSMUSG00000004661.16 | 4.712294 | -5.76437 | 3.916273 | -1.4719 | 0.141048 | NA | #VALUE! |
| ENSMUSG00000004668.15 | 0 | NA | NA | NA | NA | NA | #VALUE! |
| ENSMUSG00000004698.12 | 1.305573 | 3.898287 | 3.940396 | 0.989314 | 0.32251 | NA | #VALUE! |
| ENSMUSG00000004707.15 | 4.193144 | -5.59513 | 3.917454 | -1.42826 | 0.153218 | NA | #VALUE! |
| ENSMUSG00000004709.15 | 0.210026 | -1.52494 | 4.058692 | -0.37572 | 0.707123 | NA | #VALUE! |
| ENSMUSG00000004791.8 | 0 | NA | NA | NA | NA | NA | #VALUE! |
| ENSMUSG00000004814.11 | 1.549202 | -3.17824 | 3.928755 | -0.80897 | 0.418533 | NA | #VALUE! |

| | | | | | | | |
|---|---|---|---|---|---|---|---|
| ENSMUSG00000004821.2 | 0 | NA | NA | NA | NA | NA | #VALUE! |
| ENSMUSG00000004837.3 | 0 | NA | NA | NA | NA | NA | #VALUE! |
| ENSMUSG00000004842.19 | 0 | NA | NA | NA | NA | NA | #VALUE! |
| ENSMUSG00000004849.15 | 1.817209 | -1.48924 | 3.853033 | -0.38651 | 0.699118 | NA | #VALUE! |
| ENSMUSG00000004864.14 | 0 | NA | NA | NA | NA | NA | #VALUE! |
| ENSMUSG00000004872.16 | 0 | NA | NA | NA | NA | NA | #VALUE! |
| ENSMUSG00000004885.6 | 0 | NA | NA | NA | NA | NA | #VALUE! |
| ENSMUSG00000004891.17 | 9.760444 | 6.776799 | 3.622161 | 1.870927 | 0.061355 | NA | #VALUE! |
| ENSMUSG00000004892.14 | 0 | NA | NA | NA | NA | NA | #VALUE! |
| ENSMUSG00000004894.11 | 0 | NA | NA | NA | NA | NA | #VALUE! |
| ENSMUSG00000004895.10 | 0 | NA | NA | NA | NA | NA | #VALUE! |
| ENSMUSG00000004896.17 | 6.079004 | -5.20772 | 3.859413 | -1.34936 | 0.177223 | NA | #VALUE! |
| ENSMUSG00000004902.8 | 0 | NA | NA | NA | NA | NA | #VALUE! |
| ENSMUSG00000004929.13 | 1.025807 | 3.556426 | 3.949287 | 0.900523 | 0.367842 | NA | #VALUE! |
| ENSMUSG00000004933.18 | 0 | NA | NA | NA | NA | NA | #VALUE! |
| ENSMUSG00000004934.15 | 0.408222 | -2.20205 | 4.016842 | -0.54821 | 0.583551 | NA | #VALUE! |
| ENSMUSG00000004939.8 | 0 | NA | NA | NA | NA | NA | #VALUE! |
| ENSMUSG00000004947.16 | 0 | NA | NA | NA | NA | NA | #VALUE! |
| ENSMUSG00000004948.13 | 0 | NA | NA | NA | NA | NA | #VALUE! |
| ENSMUSG00000004952.14 | 2.936837 | -5.08026 | 3.922024 | -1.29532 | 0.195212 | NA | #VALUE! |
| ENSMUSG00000004961.8 | 0 | NA | NA | NA | NA | NA | #VALUE! |
| ENSMUSG00000004988.10 | 0 | NA | NA | NA | NA | NA | #VALUE! |
| ENSMUSG00000004994.13 | 5.889284 | 0.149349 | 3.027035 | 0.049338 | 0.96065 | NA | #VALUE! |
| ENSMUSG00000005045.17 | 0 | NA | NA | NA | NA | NA | #VALUE! |
| ENSMUSG00000005057.14 | 1.865104 | 4.40601 | 3.930481 | 1.120985 | 0.262294 | NA | #VALUE! |
| ENSMUSG00000005124.11 | 0.210026 | -1.52494 | 4.058692 | -0.37572 | 0.707123 | NA | #VALUE! |
| ENSMUSG00000005131.5 | 0 | NA | NA | NA | NA | NA | #VALUE! |
| ENSMUSG00000005148.9 | 0.525066 | -2.64687 | 3.988095 | -0.66369 | 0.506887 | NA | #VALUE! |
| ENSMUSG00000005202.3 | 0 | NA | NA | NA | NA | NA | #VALUE! |
| ENSMUSG00000005220.11 | 0 | NA | NA | NA | NA | NA | #VALUE! |
| ENSMUSG00000005232.12 | 0 | NA | NA | NA | NA | NA | #VALUE! |
| ENSMUSG00000005237.15 | 0 | NA | NA | NA | NA | NA | #VALUE! |
| ENSMUSG00000005251.16 | 42.68246 | 1.322905 | 1.506393 | 0.878193 | NA | NA | #VALUE! |
| ENSMUSG00000005267.14 | 1.323133 | 3.872684 | 3.940993 | 0.982667 | 0.325771 | NA | #VALUE! |
| ENSMUSG00000005338.15 | 0.093255 | 0.868093 | 4.072464 | 0.213162 | 0.831201 | NA | #VALUE! |
| ENSMUSG00000005339.11 | 0 | NA | NA | NA | NA | NA | #VALUE! |
| ENSMUSG00000005355.10 | 0 | NA | NA | NA | NA | NA | #VALUE! |
| ENSMUSG00000005357.10 | 0 | NA | NA | NA | NA | NA | #VALUE! |
| ENSMUSG00000005360.15 | 0 | NA | NA | NA | NA | NA | #VALUE! |
| ENSMUSG00000005364.12 | 0 | NA | NA | NA | NA | NA | #VALUE! |
| ENSMUSG00000005410.7 | 2.524253 | 4.825754 | 3.924522 | 1.229641 | 0.218831 | NA | #VALUE! |
| ENSMUSG00000005442.14 | 7.574173 | 3.069576 | 3.173792 | 0.967164 | 0.333462 | NA | #VALUE! |
| ENSMUSG00000005447.13 | 0 | NA | NA | NA | NA | NA | #VALUE! |
| ENSMUSG00000005465.10 | 0 | NA | NA | NA | NA | NA | #VALUE! |

| | | | | | | | |
|---|---|---|---|---|---|---|---|
| ENSMUSG00000005470.9 | 0.339231 | 1.823695 | 4.035859 | 0.451873 | 0.651361 | NA | #VALUE! |
| ENSMUSG00000005474.10 | 0 | NA | NA | NA | NA | NA | #VALUE! |
| ENSMUSG00000005493.16 | 0.420053 | -2.33493 | 4.007343 | -0.58266 | 0.56012 | NA | #VALUE! |
| ENSMUSG00000005503.9 | 0 | NA | NA | NA | NA | NA | #VALUE! |
| ENSMUSG00000005540.11 | 0 | NA | NA | NA | NA | NA | #VALUE! |
| ENSMUSG00000005553.15 | 3.332975 | 0.491719 | 3.782626 | 0.129994 | 0.896571 | NA | #VALUE! |
| ENSMUSG00000005611.16 | 2.287083 | -3.48404 | 3.914877 | -0.88995 | 0.373494 | NA | #VALUE! |
| ENSMUSG00000005628.13 | 0 | NA | NA | NA | NA | NA | #VALUE! |
| ENSMUSG00000005640.12 | 0 | NA | NA | NA | NA | NA | #VALUE! |
| ENSMUSG00000005649.18 | 0 | NA | NA | NA | NA | NA | #VALUE! |
| ENSMUSG00000005667.9 | 3.523415 | -3.01313 | 3.856001 | -0.78141 | 0.43456 | NA | #VALUE! |
| ENSMUSG00000005686.18 | 3.045381 | -5.14543 | 3.921352 | -1.31216 | 0.189467 | NA | #VALUE! |
| ENSMUSG00000005696.12 | 0 | NA | NA | NA | NA | NA | #VALUE! |
| ENSMUSG00000005699.17 | 1.836997 | -4.39907 | 3.931186 | -1.11902 | 0.263133 | NA | #VALUE! |
| ENSMUSG00000005705.11 | 0 | NA | NA | NA | NA | NA | #VALUE! |
| ENSMUSG00000005716.17 | 0 | NA | NA | NA | NA | NA | #VALUE! |
| ENSMUSG00000005718.9 | 3.451764 | 0.285874 | 3.779456 | 0.075639 | 0.939706 | NA | #VALUE! |
| ENSMUSG00000005763.16 | 0 | NA | NA | NA | NA | NA | #VALUE! |
| ENSMUSG00000005800.4 | 0.816443 | -3.21914 | 3.961753 | -0.81255 | 0.416474 | NA | #VALUE! |
| ENSMUSG00000005824.8 | 0 | NA | NA | NA | NA | NA | #VALUE! |
| ENSMUSG00000005864.8 | 0 | NA | NA | NA | NA | NA | #VALUE! |
| ENSMUSG00000005883.16 | 0.093255 | 0.868093 | 4.072464 | 0.213162 | 0.831201 | NA | #VALUE! |
| ENSMUSG00000005891.8 | 0 | NA | NA | NA | NA | NA | #VALUE! |
| ENSMUSG00000005892.5 | 0 | NA | NA | NA | NA | NA | #VALUE! |
| ENSMUSG00000005893.15 | 28.65983 | -0.95598 | 1.92584 | -0.4964 | NA | NA | #VALUE! |
| ENSMUSG00000005917.16 | 0 | NA | NA | NA | NA | NA | #VALUE! |
| ENSMUSG00000005947.12 | 2.219898 | 4.653556 | 3.926761 | 1.185088 | 0.235983 | NA | #VALUE! |
| ENSMUSG00000005950.15 | 0 | NA | NA | NA | NA | NA | #VALUE! |
| ENSMUSG00000005952.16 | 0 | NA | NA | NA | NA | NA | #VALUE! |
| ENSMUSG00000005958.16 | 3.328415 | -5.26129 | 3.920229 | -1.34209 | 0.179567 | NA | #VALUE! |
| ENSMUSG00000005980.16 | 6.836354 | 5.307382 | 3.738714 | 1.419574 | 0.155732 | NA | #VALUE! |
| ENSMUSG00000005986.17 | 0 | NA | NA | NA | NA | NA | #VALUE! |
| ENSMUSG00000005994.15 | 0 | NA | NA | NA | NA | NA | #VALUE! |
| ENSMUSG00000006007.15 | 0 | NA | NA | NA | NA | NA | #VALUE! |
| ENSMUSG00000006019.16 | 13.90426 | 6.340954 | 3.03245 | 2.091033 | NA | NA | #VALUE! |
| ENSMUSG00000006056.14 | 0 | NA | NA | NA | NA | NA | #VALUE! |
| ENSMUSG00000006127.10 | 0 | NA | NA | NA | NA | NA | #VALUE! |
| ENSMUSG00000006143.13 | 0 | NA | NA | NA | NA | NA | #VALUE! |
| ENSMUSG00000006154.14 | 0.491951 | 2.402185 | 4.000147 | 0.600524 | 0.548157 | NA | #VALUE! |
| ENSMUSG00000006179.10 | 0 | NA | NA | NA | NA | NA | #VALUE! |
| ENSMUSG00000006200.4 | 0 | NA | NA | NA | NA | NA | #VALUE! |
| ENSMUSG00000006204.8 | 0 | NA | NA | NA | NA | NA | #VALUE! |
| ENSMUSG00000006205.14 | 4.698938 | -5.75981 | 3.916303 | -1.47073 | 0.141365 | NA | #VALUE! |
| ENSMUSG00000006216.11 | 0 | NA | NA | NA | NA | NA | #VALUE! |

| | | | | | | | | |
|---|---|---|---|---|---|---|---|---|
| ENSMUSG00000006218.5 | 0 | NA | NA | NA | NA | NA | | #VALUE! |
| ENSMUSG00000006219.13 | 9.005834 | 3.935232 | 2.921013 | 1.347215 | 0.177911 | NA | | #VALUE! |
| ENSMUSG00000006221.8 | 0 | NA | NA | NA | NA | NA | | #VALUE! |
| ENSMUSG00000006235.7 | 4.694547 | -5.7577 | 3.916317 | -1.47018 | 0.141512 | NA | | #VALUE! |
| ENSMUSG00000006241.17 | 0 | NA | NA | NA | NA | NA | | #VALUE! |
| ENSMUSG00000006269.8 | 0 | NA | NA | NA | NA | NA | | #VALUE! |
| ENSMUSG00000006270.8 | 0 | NA | NA | NA | NA | NA | | #VALUE! |
| ENSMUSG00000006276.11 | 4.076285 | 0.020851 | 3.542121 | 0.005887 | 0.995303 | NA | | #VALUE! |
| ENSMUSG00000006288.9 | 36.85383 | -1.47516 | 2.438318 | -0.60499 | NA | NA | | #VALUE! |
| ENSMUSG00000006310.11 | 0.525066 | -2.64687 | 3.988095 | -0.66369 | 0.506887 | NA | | #VALUE! |
| ENSMUSG00000006311.9 | 0 | NA | NA | NA | NA | NA | | #VALUE! |
| ENSMUSG00000006313.6 | 0 | NA | NA | NA | NA | NA | | #VALUE! |
| ENSMUSG00000006335.17 | 9.643852 | -5.70098 | 3.370977 | -1.6912 | 0.090799 | NA | | #VALUE! |
| ENSMUSG00000006342.16 | 0 | NA | NA | NA | NA | NA | | #VALUE! |
| ENSMUSG00000006344.18 | 2.259413 | 2.554725 | 3.886298 | 0.657367 | 0.510945 | NA | | #VALUE! |
| ENSMUSG00000006345.11 | 0 | NA | NA | NA | NA | NA | | #VALUE! |
| ENSMUSG00000006360.12 | 9.347568 | -2.79113 | 2.672854 | -1.04425 | 0.29637 | NA | | #VALUE! |
| ENSMUSG00000006362.17 | 16.15541 | -1.09779 | 2.887693 | -0.38016 | NA | NA | | #VALUE! |
| ENSMUSG00000006369.15 | 9.451182 | -6.77386 | 3.637007 | -1.86248 | NA | NA | | #VALUE! |
| ENSMUSG00000006389.14 | 0 | NA | NA | NA | NA | NA | | #VALUE! |
| ENSMUSG00000006398.16 | 3.061661 | -5.13931 | 3.921414 | -1.31058 | 0.190001 | NA | | #VALUE! |
| ENSMUSG00000006403.14 | 0 | NA | NA | NA | NA | NA | | #VALUE! |
| ENSMUSG00000006411.13 | 0.391578 | -2.1489 | 4.020883 | -0.53444 | 0.59304 | NA | | #VALUE! |
| ENSMUSG00000006435.17 | 0 | NA | NA | NA | NA | NA | | #VALUE! |
| ENSMUSG00000006457.5 | 0 | NA | NA | NA | NA | NA | | #VALUE! |
| ENSMUSG00000006462.7 | 4.06631 | 4.426787 | 3.910893 | 1.131912 | 0.257671 | NA | | #VALUE! |
| ENSMUSG00000006464.10 | 0 | NA | NA | NA | NA | NA | | #VALUE! |
| ENSMUSG00000006469.14 | 0 | NA | NA | NA | NA | NA | | #VALUE! |
| ENSMUSG00000006471.19 | 5.573571 | 3.219076 | 3.819298 | 0.842845 | 0.399315 | NA | | #VALUE! |
| ENSMUSG00000006476.20 | 7.351906 | -2.23064 | 2.160204 | -1.03261 | 0.301789 | NA | | #VALUE! |
| ENSMUSG00000006488.18 | 0 | NA | NA | NA | NA | NA | | #VALUE! |
| ENSMUSG00000006490.4 | 0 | NA | NA | NA | NA | NA | | #VALUE! |
| ENSMUSG00000006538.13 | 0 | NA | NA | NA | NA | NA | | #VALUE! |
| ENSMUSG00000006542.14 | 0 | NA | NA | NA | NA | NA | | #VALUE! |
| ENSMUSG00000006546.4 | 0 | NA | NA | NA | NA | NA | | #VALUE! |
| ENSMUSG00000006570.4 | 0 | NA | NA | NA | NA | NA | | #VALUE! |
| ENSMUSG00000006574.16 | 0 | NA | NA | NA | NA | NA | | #VALUE! |
| ENSMUSG00000006575.15 | 0 | NA | NA | NA | NA | NA | | #VALUE! |
| ENSMUSG00000006576.17 | 0 | NA | NA | NA | NA | NA | | #VALUE! |
| ENSMUSG00000006586.16 | 0 | NA | NA | NA | NA | NA | | #VALUE! |
| ENSMUSG00000006587.7 | 0 | NA | NA | NA | NA | NA | | #VALUE! |
| ENSMUSG00000006638.16 | 2.21378 | 4.621487 | 3.927208 | 1.176787 | 0.239281 | NA | | #VALUE! |
| ENSMUSG00000006642.4 | 0 | NA | NA | NA | NA | NA | | #VALUE! |
| ENSMUSG00000006649.18 | 0 | NA | NA | NA | NA | NA | | #VALUE! |

| | | | | | | | |
|---|---|---|---|---|---|---|---|
| ENSMUSG00000006651.9 | 0 | NA | NA | NA | NA | NA | #VALUE! |
| ENSMUSG00000006675.11 | 0 | NA | NA | NA | NA | NA | #VALUE! |
| ENSMUSG00000006678.7 | 31.29887 | 0.457869 | 1.889801 | 0.242284 | NA | NA | #VALUE! |
| ENSMUSG00000006720.17 | 0 | NA | NA | NA | NA | NA | #VALUE! |
| ENSMUSG00000006724.13 | 0 | NA | NA | NA | NA | NA | #VALUE! |
| ENSMUSG00000006764.9 | 0 | NA | NA | NA | NA | NA | #VALUE! |
| ENSMUSG00000006777.8 | 2.888772 | -5.05981 | 3.922241 | -1.29003 | 0.19704 | NA | #VALUE! |
| ENSMUSG00000006784.15 | 9.241155 | -6.74146 | 3.664007 | -1.83991 | NA | NA | #VALUE! |
| ENSMUSG00000006920.15 | 2.489267 | 1.576424 | 3.822543 | 0.412402 | 0.680045 | NA | #VALUE! |
| ENSMUSG00000006930.16 | 0 | NA | NA | NA | NA | NA | #VALUE! |
| ENSMUSG00000006931.16 | 7.19954 | 6.337151 | 3.473764 | 1.82429 | 0.068108 | NA | #VALUE! |
| ENSMUSG00000006948.8 | 0 | NA | NA | NA | NA | NA | #VALUE! |
| ENSMUSG00000006958.15 | 8.677337 | 0.512691 | 2.747202 | 0.186623 | 0.851956 | NA | #VALUE! |
| ENSMUSG00000007021.6 | 0 | NA | NA | NA | NA | NA | #VALUE! |
| ENSMUSG00000007030.9 | 0 | NA | NA | NA | NA | NA | #VALUE! |
| ENSMUSG00000007033.5 | 0 | NA | NA | NA | NA | NA | #VALUE! |
| ENSMUSG00000007034.16 | 0 | NA | NA | NA | NA | NA | #VALUE! |
| ENSMUSG00000007035.16 | 0 | NA | NA | NA | NA | NA | #VALUE! |
| ENSMUSG00000007039.13 | 6.934289 | -3.43731 | 3.188091 | -1.07817 | 0.280957 | NA | #VALUE! |
| ENSMUSG00000007080.15 | 1.566313 | -4.16993 | 3.935366 | -1.0596 | 0.289325 | NA | #VALUE! |
| ENSMUSG00000007097.15 | 0 | NA | NA | NA | NA | NA | #VALUE! |
| ENSMUSG00000007107.7 | 0 | NA | NA | NA | NA | NA | #VALUE! |
| ENSMUSG00000007122.12 | 0 | NA | NA | NA | NA | NA | #VALUE! |
| ENSMUSG00000007207.11 | 0 | NA | NA | NA | NA | NA | #VALUE! |
| ENSMUSG00000007209.5 | 0 | NA | NA | NA | NA | NA | #VALUE! |
| ENSMUSG00000007216.7 | 0.336567 | 1.944925 | 4.029936 | 0.482619 | 0.629366 | NA | #VALUE! |
| ENSMUSG00000007279.15 | 0 | NA | NA | NA | NA | NA | #VALUE! |
| ENSMUSG00000007379.16 | 0 | NA | NA | NA | NA | NA | #VALUE! |
| ENSMUSG00000007440.11 | 0 | NA | NA | NA | NA | NA | #VALUE! |
| ENSMUSG00000007457.4 | 0 | NA | NA | NA | NA | NA | #VALUE! |
| ENSMUSG00000007480.6 | 0 | NA | NA | NA | NA | NA | #VALUE! |
| ENSMUSG00000007591.16 | 0 | NA | NA | NA | NA | NA | #VALUE! |
| ENSMUSG00000007603.10 | 6.912441 | -1.30976 | 2.900129 | -0.45162 | 0.651542 | NA | #VALUE! |
| ENSMUSG00000007646.14 | 0.737927 | 3.007825 | 3.968641 | 0.757898 | 0.448512 | NA | #VALUE! |
| ENSMUSG00000007653.13 | 0 | NA | NA | NA | NA | NA | #VALUE! |
| ENSMUSG00000007659.19 | 1.043407 | 2.257396 | 3.950998 | 0.571348 | 0.567763 | NA | #VALUE! |
| ENSMUSG00000007682.8 | 0 | NA | NA | NA | NA | NA | #VALUE! |
| ENSMUSG00000007783.11 | 0 | NA | NA | NA | NA | NA | #VALUE! |
| ENSMUSG00000007805.5 | 0 | NA | NA | NA | NA | NA | #VALUE! |
| ENSMUSG00000007877.3 | 0 | NA | NA | NA | NA | NA | #VALUE! |
| ENSMUSG00000007888.16 | 0 | NA | NA | NA | NA | NA | #VALUE! |
| ENSMUSG00000007907.5 | 0 | NA | NA | NA | NA | NA | #VALUE! |
| ENSMUSG00000007908.15 | 0.18651 | 1.401769 | 4.056354 | 0.345574 | 0.729663 | NA | #VALUE! |
| ENSMUSG00000007944.9 | 0 | NA | NA | NA | NA | NA | #VALUE! |

| | | | | | | | |
|---|---|---|---|---|---|---|---|
| ENSMUSG00000007946.12 | 0 | NA | NA | NA | NA | NA | #VALUE! |
| ENSMUSG00000007950.10 | 1.893071 | -0.71247 | 3.828881 | -0.18608 | 0.852384 | NA | #VALUE! |
| ENSMUSG00000007987.13 | 3.550231 | -2.60238 | 3.831279 | -0.67925 | 0.496982 | NA | #VALUE! |
| ENSMUSG00000007989.8 | 0.168284 | 1.027628 | 4.067126 | 0.252667 | 0.800526 | NA | #VALUE! |
| ENSMUSG00000008028.16 | 0 | NA | NA | NA | NA | NA | #VALUE! |
| ENSMUSG00000008129.15 | 0 | NA | NA | NA | NA | NA | #VALUE! |
| ENSMUSG00000008136.15 | 0.279766 | 1.808031 | 4.036658 | 0.447903 | 0.654223 | NA | #VALUE! |
| ENSMUSG00000008193.14 | 0 | NA | NA | NA | NA | NA | #VALUE! |
| ENSMUSG00000008206.17 | 5.902263 | -4.02629 | 3.862072 | -1.04252 | 0.29717 | NA | #VALUE! |
| ENSMUSG00000008307.12 | 0 | NA | NA | NA | NA | NA | #VALUE! |
| ENSMUSG00000008318.11 | 1.957891 | -4.49332 | 3.929649 | -1.14344 | 0.252855 | NA | #VALUE! |
| ENSMUSG00000008438.4 | 0 | NA | NA | NA | NA | NA | #VALUE! |
| ENSMUSG00000008450.9 | 0 | NA | NA | NA | NA | NA | #VALUE! |
| ENSMUSG00000008461.7 | 0 | NA | NA | NA | NA | NA | #VALUE! |
| ENSMUSG00000008482.10 | 0 | NA | NA | NA | NA | NA | #VALUE! |
| ENSMUSG00000008489.19 | 0 | NA | NA | NA | NA | NA | #VALUE! |
| ENSMUSG00000008590.5 | 0 | NA | NA | NA | NA | NA | #VALUE! |
| ENSMUSG00000008601.13 | 0 | NA | NA | NA | NA | NA | #VALUE! |
| ENSMUSG00000008604.12 | 3.870356 | 0.170608 | 2.803464 | 0.060856 | 0.951474 | NA | #VALUE! |
| ENSMUSG00000008658.17 | 0 | NA | NA | NA | NA | NA | #VALUE! |
| ENSMUSG00000008734.10 | 9.51203 | 6.743611 | 3.627857 | 1.858842 | NA | NA | #VALUE! |
| ENSMUSG00000008789.11 | 0 | NA | NA | NA | NA | NA | #VALUE! |
| ENSMUSG00000008813.14 | 0 | NA | NA | NA | NA | NA | #VALUE! |
| ENSMUSG00000008843.5 | 0 | NA | NA | NA | NA | NA | #VALUE! |
| ENSMUSG00000008874.4 | 0 | NA | NA | NA | NA | NA | #VALUE! |
| ENSMUSG00000008932.11 | 0 | NA | NA | NA | NA | NA | #VALUE! |
| ENSMUSG00000009047.5 | 0 | NA | NA | NA | NA | NA | #VALUE! |
| ENSMUSG00000009070.15 | 0 | NA | NA | NA | NA | NA | #VALUE! |
| ENSMUSG00000009075.3 | 0 | NA | NA | NA | NA | NA | #VALUE! |
| ENSMUSG00000009076.11 | 0.466276 | 2.449889 | 3.997179 | 0.612904 | 0.53994 | NA | #VALUE! |
| ENSMUSG00000009092.6 | 0 | NA | NA | NA | NA | NA | #VALUE! |
| ENSMUSG00000009093.8 | 0 | NA | NA | NA | NA | NA | #VALUE! |
| ENSMUSG00000009097.11 | 0 | NA | NA | NA | NA | NA | #VALUE! |
| ENSMUSG00000009108.16 | 0.18651 | 1.401769 | 4.056354 | 0.345574 | 0.729663 | NA | #VALUE! |
| ENSMUSG00000009114.18 | 0 | NA | NA | NA | NA | NA | #VALUE! |
| ENSMUSG00000009115.6 | 0 | NA | NA | NA | NA | NA | #VALUE! |
| ENSMUSG00000009145.7 | 6.195775 | -6.16564 | 3.913968 | -1.57529 | 0.115189 | NA | #VALUE! |
| ENSMUSG00000009185.3 | 0 | NA | NA | NA | NA | NA | #VALUE! |
| ENSMUSG00000009210.11 | 0 | NA | NA | NA | NA | NA | #VALUE! |
| ENSMUSG00000009214.10 | 0 | NA | NA | NA | NA | NA | #VALUE! |
| ENSMUSG00000009216.8 | 0 | NA | NA | NA | NA | NA | #VALUE! |
| ENSMUSG00000009246.15 | 0 | NA | NA | NA | NA | NA | #VALUE! |
| ENSMUSG00000009248.7 | 0 | NA | NA | NA | NA | NA | #VALUE! |
| ENSMUSG00000009292.19 | 2.210091 | -1.83783 | 3.845912 | -0.47787 | 0.632745 | NA | #VALUE! |

| | | | | | | | |
|---|---|---|---|---|---|---|---|
| ENSMUSG00000009350.14 | 0 | NA | NA | NA | NA | NA | #VALUE! |
| ENSMUSG00000009356.13 | 0 | NA | NA | NA | NA | NA | #VALUE! |
| ENSMUSG00000009394.14 | 0 | NA | NA | NA | NA | NA | #VALUE! |
| ENSMUSG00000009406.14 | 0 | NA | NA | NA | NA | NA | #VALUE! |
| ENSMUSG00000009471.5 | 0 | NA | NA | NA | NA | NA | #VALUE! |
| ENSMUSG00000009487.11 | 0 | NA | NA | NA | NA | NA | #VALUE! |
| ENSMUSG00000009545.15 | 2.355969 | 4.72622 | 3.925784 | 1.203892 | 0.228631 | NA | #VALUE! |
| ENSMUSG00000009551.3 | 0 | NA | NA | NA | NA | NA | #VALUE! |
| ENSMUSG00000009555.17 | 17.13044 | 3.355578 | 2.236586 | 1.500313 | NA | NA | #VALUE! |
| ENSMUSG00000009580.16 | 0 | NA | NA | NA | NA | NA | #VALUE! |
| ENSMUSG00000009585.19 | 8.409554 | -1.5069 | 3.497313 | -0.43087 | 0.666561 | NA | #VALUE! |
| ENSMUSG00000009588.10 | 0 | NA | NA | NA | NA | NA | #VALUE! |
| ENSMUSG00000009596.10 | 0 | NA | NA | NA | NA | NA | #VALUE! |
| ENSMUSG00000009628.15 | 0 | NA | NA | NA | NA | NA | #VALUE! |
| ENSMUSG00000009670.12 | 0 | NA | NA | NA | NA | NA | #VALUE! |
| ENSMUSG00000009681.11 | 2.664509 | -4.94119 | 3.923563 | -1.25936 | 0.207899 | NA | #VALUE! |
| ENSMUSG00000009687.15 | 2.770284 | -4.99742 | 3.922923 | -1.2739 | 0.202698 | NA | #VALUE! |
| ENSMUSG00000009731.5 | 0 | NA | NA | NA | NA | NA | #VALUE! |
| ENSMUSG00000009734.19 | 0 | NA | NA | NA | NA | NA | #VALUE! |
| ENSMUSG00000009739.18 | 2.529648 | 3.847701 | 3.920736 | 0.981372 | 0.326409 | NA | #VALUE! |
| ENSMUSG00000009876.14 | 0 | NA | NA | NA | NA | NA | #VALUE! |
| ENSMUSG00000009900.8 | 0 | NA | NA | NA | NA | NA | #VALUE! |
| ENSMUSG00000009941.11 | 0 | NA | NA | NA | NA | NA | #VALUE! |
| ENSMUSG00000010021.14 | 0 | NA | NA | NA | NA | NA | #VALUE! |
| ENSMUSG00000010044.13 | 0 | NA | NA | NA | NA | NA | #VALUE! |
| ENSMUSG00000010066.16 | 0 | NA | NA | NA | NA | NA | #VALUE! |
| ENSMUSG00000010080.16 | 0 | NA | NA | NA | NA | NA | #VALUE! |
| ENSMUSG00000010086.18 | 0 | NA | NA | NA | NA | NA | #VALUE! |
| ENSMUSG00000010136.15 | 0 | NA | NA | NA | NA | NA | #VALUE! |
| ENSMUSG00000010142.13 | 0 | NA | NA | NA | NA | NA | #VALUE! |
| ENSMUSG00000010154.8 | 0 | NA | NA | NA | NA | NA | #VALUE! |
| ENSMUSG00000010205.12 | 0 | NA | NA | NA | NA | NA | #VALUE! |
| ENSMUSG00000010311.16 | 0 | NA | NA | NA | NA | NA | #VALUE! |
| ENSMUSG00000010342.17 | 0 | NA | NA | NA | NA | NA | #VALUE! |
| ENSMUSG00000010358.14 | 1.685563 | -4.2806 | 3.933265 | -1.08831 | 0.276459 | NA | #VALUE! |
| ENSMUSG00000010362.10 | 3.552976 | 4.231018 | 3.912577 | 1.081389 | 0.279524 | NA | #VALUE! |
| ENSMUSG00000010435.8 | 0 | NA | NA | NA | NA | NA | #VALUE! |
| ENSMUSG00000010461.17 | 0 | NA | NA | NA | NA | NA | #VALUE! |
| ENSMUSG00000010476.15 | 0 | NA | NA | NA | NA | NA | #VALUE! |
| ENSMUSG00000010492.11 | 0 | NA | NA | NA | NA | NA | #VALUE! |
| ENSMUSG00000010505.17 | 0 | NA | NA | NA | NA | NA | #VALUE! |
| ENSMUSG00000010529.7 | 0 | NA | NA | NA | NA | NA | #VALUE! |
| ENSMUSG00000010538.15 | 6.15226 | -1.27818 | 2.936619 | -0.43526 | 0.663377 | NA | #VALUE! |
| ENSMUSG00000010592.10 | 0 | NA | NA | NA | NA | NA | #VALUE! |

| | | | | | | | |
|---|---|---|---|---|---|---|---|
| ENSMUSG00000010660.4 | 0 | NA | NA | NA | NA | NA | #VALUE! |
| ENSMUSG00000010721.16 | 73.75845 | 1.65103 | 2.886147 | 0.572053 | NA | NA | #VALUE! |
| ENSMUSG00000010751.16 | 1.082609 | 3.624054 | 3.947361 | 0.918096 | 0.358569 | NA | #VALUE! |
| ENSMUSG00000010760.6 | 1.050131 | -3.6251 | 3.948379 | -0.91812 | 0.358554 | NA | #VALUE! |
| ENSMUSG00000010796.12 | 0 | NA | NA | NA | NA | NA | #VALUE! |
| ENSMUSG00000010797.7 | 4.6198 | 1.178027 | 3.753179 | 0.313874 | 0.753616 | NA | #VALUE! |
| ENSMUSG00000010803.14 | 0 | NA | NA | NA | NA | NA | #VALUE! |
| ENSMUSG00000010825.18 | 0 | NA | NA | NA | NA | NA | #VALUE! |
| ENSMUSG00000010830.9 | 0 | NA | NA | NA | NA | NA | #VALUE! |
| ENSMUSG00000010841.14 | 0 | NA | NA | NA | NA | NA | #VALUE! |
| ENSMUSG00000011008.14 | 0 | NA | NA | NA | NA | NA | #VALUE! |
| ENSMUSG00000011034.7 | 0 | NA | NA | NA | NA | NA | #VALUE! |
| ENSMUSG00000011118.5 | 0 | NA | NA | NA | NA | NA | #VALUE! |
| ENSMUSG00000011154.18 | 0 | NA | NA | NA | NA | NA | #VALUE! |
| ENSMUSG00000011171.12 | 0 | NA | NA | NA | NA | NA | #VALUE! |
| ENSMUSG00000011254.17 | 0 | NA | NA | NA | NA | NA | #VALUE! |
| ENSMUSG00000011263.17 | 4.791237 | -3.84984 | 3.771008 | -1.0209 | 0.3073 | NA | #VALUE! |
| ENSMUSG00000011267.9 | 6.700625 | -3.62299 | 3.723234 | -0.97308 | 0.330516 | NA | #VALUE! |
| ENSMUSG00000011349.6 | 0 | NA | NA | NA | NA | NA | #VALUE! |
| ENSMUSG00000011350.16 | 0 | NA | NA | NA | NA | NA | #VALUE! |
| ENSMUSG00000011463.6 | 0 | NA | NA | NA | NA | NA | #VALUE! |
| ENSMUSG00000011486.15 | 0 | NA | NA | NA | NA | NA | #VALUE! |
| ENSMUSG00000011589.10 | 0 | NA | NA | NA | NA | NA | #VALUE! |
| ENSMUSG00000011632.8 | 0 | NA | NA | NA | NA | NA | #VALUE! |
| ENSMUSG00000011658.17 | 2.545958 | -2.00884 | 3.838843 | -0.52329 | 0.60077 | NA | #VALUE! |
| ENSMUSG00000011751.17 | 0 | NA | NA | NA | NA | NA | #VALUE! |
| ENSMUSG00000011752.8 | 2.979548 | -5.10391 | 3.921776 | -1.30143 | 0.193112 | NA | #VALUE! |
| ENSMUSG00000011832.17 | 0.204111 | -1.05581 | 4.073302 | -0.2592 | 0.795479 | NA | #VALUE! |
| ENSMUSG00000011837.5 | 9.694545 | 1.091627 | 2.553855 | 0.427443 | 0.669057 | NA | #VALUE! |
| ENSMUSG00000011877.14 | 5.354121 | 5.915943 | 3.915124 | 1.511049 | 0.130776 | NA | #VALUE! |
| ENSMUSG00000011884.14 | 12.18086 | 1.59918 | 2.852542 | 0.560616 | NA | NA | #VALUE! |
| ENSMUSG00000012017.5 | 13.61526 | 20.68194 | 3.243083 | 6.377245 | NA | NA | #VALUE! |
| ENSMUSG00000012042.10 | 1.196212 | 3.752416 | 3.943939 | 0.951439 | 0.341382 | NA | #VALUE! |
| ENSMUSG00000012123.18 | 2.424635 | 4.780725 | 3.925082 | 1.217993 | 0.223226 | NA | #VALUE! |
| ENSMUSG00000012126.17 | 0 | NA | NA | NA | NA | NA | #VALUE! |
| ENSMUSG00000012211.14 | 0 | NA | NA | NA | NA | NA | #VALUE! |
| ENSMUSG00000012282.3 | 0 | NA | NA | NA | NA | NA | #VALUE! |
| ENSMUSG00000012296.17 | 7.244197 | -6.38517 | 3.912952 | -1.6318 | 0.102721 | NA | #VALUE! |
| ENSMUSG00000012350.16 | 0 | NA | NA | NA | NA | NA | #VALUE! |
| ENSMUSG00000012396.13 | 0 | NA | NA | NA | NA | NA | #VALUE! |
| ENSMUSG00000012443.5 | 1.388554 | 2.257526 | 3.92166 | 0.575656 | 0.564848 | NA | #VALUE! |
| ENSMUSG00000012519.15 | 0 | NA | NA | NA | NA | NA | #VALUE! |
| ENSMUSG00000012520.11 | 0 | NA | NA | NA | NA | NA | #VALUE! |
| ENSMUSG00000012609.20 | 0.735092 | -3.12016 | 3.965612 | -0.7868 | 0.431397 | NA | #VALUE! |

| | | | | | | | |
|---|---|---|---|---|---|---|---|
| ENSMUSG00000012705.17 | 0 | NA | NA | NA | NA | NA | #VALUE! |
| ENSMUSG00000012777.15 | 0 | NA | NA | NA | NA | NA | #VALUE! |
| ENSMUSG00000012819.17 | 0 | NA | NA | NA | NA | NA | #VALUE! |
| ENSMUSG00000012889.9 | 0 | NA | NA | NA | NA | NA | #VALUE! |
| ENSMUSG00000013033.17 | 5.553356 | 5.96322 | 3.914855 | 1.523229 | 0.127701 | NA | #VALUE! |
| ENSMUSG00000013076.18 | 4.012742 | 0.231694 | 3.464757 | 0.066872 | 0.946684 | NA | #VALUE! |
| ENSMUSG00000013083.9 | 0 | NA | NA | NA | NA | NA | #VALUE! |
| ENSMUSG00000013089.16 | 3.674651 | -1.72975 | 3.795259 | -0.45577 | 0.648557 | NA | #VALUE! |
| ENSMUSG00000013091.6 | 0 | NA | NA | NA | NA | NA | #VALUE! |
| ENSMUSG00000013150.16 | 0.877871 | 3.316761 | 3.95687 | 0.838228 | 0.401902 | NA | #VALUE! |
| ENSMUSG00000013155.11 | 0 | NA | NA | NA | NA | NA | #VALUE! |
| ENSMUSG00000013158.7 | 0 | NA | NA | NA | NA | NA | #VALUE! |
| ENSMUSG00000013236.18 | 0.204111 | -1.05581 | 4.073302 | -0.2592 | 0.795479 | NA | #VALUE! |
| ENSMUSG00000013338.17 | 0 | NA | NA | NA | NA | NA | #VALUE! |
| ENSMUSG00000013353.11 | 0 | NA | NA | NA | NA | NA | #VALUE! |
| ENSMUSG00000013367.6 | 0 | NA | NA | NA | NA | NA | #VALUE! |
| ENSMUSG00000013415.10 | 0.373021 | 2.140748 | 4.018182 | 0.532765 | 0.594196 | NA | #VALUE! |
| ENSMUSG00000013418.9 | 0 | NA | NA | NA | NA | NA | #VALUE! |
| ENSMUSG00000013419.8 | 0 | NA | NA | NA | NA | NA | #VALUE! |
| ENSMUSG00000013483.15 | 0 | NA | NA | NA | NA | NA | #VALUE! |
| ENSMUSG00000013495.16 | 71.57086 | 0.434639 | 1.33021 | 0.326745 | NA | NA | #VALUE! |
| ENSMUSG00000013523.14 | 0 | NA | NA | NA | NA | NA | #VALUE! |
| ENSMUSG00000013562.13 | 0 | NA | NA | NA | NA | NA | #VALUE! |
| ENSMUSG00000013584.6 | 0 | NA | NA | NA | NA | NA | #VALUE! |
| ENSMUSG00000013611.15 | 0 | NA | NA | NA | NA | NA | #VALUE! |
| ENSMUSG00000013629.17 | 8.631267 | 2.119685 | 3.011604 | 0.703839 | 0.481533 | NA | #VALUE! |
| ENSMUSG00000013643.14 | 0 | NA | NA | NA | NA | NA | #VALUE! |
| ENSMUSG00000013646.18 | 28.10317 | -4.22126 | 2.529169 | -1.66903 | NA | NA | #VALUE! |
| ENSMUSG00000013653.3 | 0 | NA | NA | NA | NA | NA | #VALUE! |
| ENSMUSG00000013668.10 | 0 | NA | NA | NA | NA | NA | #VALUE! |
| ENSMUSG00000013707.4 | 2.355969 | 4.72622 | 3.925784 | 1.203892 | 0.228631 | NA | #VALUE! |
| ENSMUSG00000013766.12 | 0 | NA | NA | NA | NA | NA | #VALUE! |
| ENSMUSG00000013858.15 | 14.46906 | 0.632816 | 2.334317 | 0.271093 | NA | NA | #VALUE! |
| ENSMUSG00000013878.19 | 21.42181 | -1.37814 | 1.931106 | -0.71365 | NA | NA | #VALUE! |
| ENSMUSG00000013921.16 | 0 | NA | NA | NA | NA | NA | #VALUE! |
| ENSMUSG00000013928.14 | 0 | NA | NA | NA | NA | NA | #VALUE! |
| ENSMUSG00000013936.13 | 0 | NA | NA | NA | NA | NA | #VALUE! |
| ENSMUSG00000013974.4 | 0.315039 | -1.93549 | 4.038621 | -0.47925 | 0.631763 | NA | #VALUE! |
| ENSMUSG00000014030.16 | 0 | NA | NA | NA | NA | NA | #VALUE! |
| ENSMUSG00000014104.13 | 0 | NA | NA | NA | NA | NA | #VALUE! |
| ENSMUSG00000014158.13 | 0 | NA | NA | NA | NA | NA | #VALUE! |
| ENSMUSG00000014198.16 | 0 | NA | NA | NA | NA | NA | #VALUE! |
| ENSMUSG00000014245.9 | 1.967804 | 4.449735 | 3.929778 | 1.132312 | 0.257503 | NA | #VALUE! |
| ENSMUSG00000014303.14 | 3.36567 | 5.240782 | 3.920098 | 1.336901 | 0.181255 | NA | #VALUE! |

| | | | | | | | |
|---|---|---|---|---|---|---|---|
| ENSMUSG00000014351.13 | 0 | NA | NA | NA | NA | NA | #VALUE! |
| ENSMUSG00000014444.18 | 0 | NA | NA | NA | NA | NA | #VALUE! |
| ENSMUSG00000014453.5 | 0 | NA | NA | NA | NA | NA | #VALUE! |
| ENSMUSG00000014498.10 | 2.741047 | -4.98044 | 3.923113 | -1.26951 | 0.204259 | NA | #VALUE! |
| ENSMUSG00000014503.16 | 0 | NA | NA | NA | NA | NA | #VALUE! |
| ENSMUSG00000014529.10 | 0 | NA | NA | NA | NA | NA | #VALUE! |
| ENSMUSG00000014543.15 | 0 | NA | NA | NA | NA | NA | #VALUE! |
| ENSMUSG00000014599.11 | 8.147751 | -1.29748 | 3.087735 | -0.42021 | 0.674336 | NA | #VALUE! |
| ENSMUSG00000014602.16 | 0 | NA | NA | NA | NA | NA | #VALUE! |
| ENSMUSG00000014603.4 | 0 | NA | NA | NA | NA | NA | #VALUE! |
| ENSMUSG00000014609.17 | 0 | NA | NA | NA | NA | NA | #VALUE! |
| ENSMUSG00000014686.9 | 0 | NA | NA | NA | NA | NA | #VALUE! |
| ENSMUSG00000014704.11 | 0 | NA | NA | NA | NA | NA | #VALUE! |
| ENSMUSG00000014725.17 | 0 | NA | NA | NA | NA | NA | #VALUE! |
| ENSMUSG00000014747.7 | 0 | NA | NA | NA | NA | NA | #VALUE! |
| ENSMUSG00000014773.14 | 1.282998 | 2.793116 | 3.940291 | 0.70886 | 0.478411 | NA | #VALUE! |
| ENSMUSG00000014776.8 | 1.428775 | -4.03418 | 3.938169 | -1.02438 | 0.305656 | NA | #VALUE! |
| ENSMUSG00000014782.17 | 0 | NA | NA | NA | NA | NA | #VALUE! |
| ENSMUSG00000014786.9 | 0 | NA | NA | NA | NA | NA | #VALUE! |
| ENSMUSG00000014813.10 | 0 | NA | NA | NA | NA | NA | #VALUE! |
| ENSMUSG00000014837.16 | 14.97723 | 20.81131 | 3.181896 | 6.540536 | NA | NA | #VALUE! |
| ENSMUSG00000014846.13 | 0 | NA | NA | NA | NA | NA | #VALUE! |
| ENSMUSG00000014852.17 | 0 | NA | NA | NA | NA | NA | #VALUE! |
| ENSMUSG00000015001.18 | 0 | NA | NA | NA | NA | NA | #VALUE! |
| ENSMUSG00000015053.15 | 1.924702 | -1.25408 | 3.837748 | -0.32677 | 0.743838 | NA | #VALUE! |
| ENSMUSG00000015085.9 | 0 | NA | NA | NA | NA | NA | #VALUE! |
| ENSMUSG00000015090.14 | 2.58797 | -4.90094 | 3.924036 | -1.24895 | 0.211682 | NA | #VALUE! |
| ENSMUSG00000015093.16 | 0 | NA | NA | NA | NA | NA | #VALUE! |
| ENSMUSG00000015094.17 | 3.727774 | -5.42754 | 3.911086 | -1.38773 | 0.165219 | NA | #VALUE! |
| ENSMUSG00000015120.16 | 4.270245 | -2.19657 | 3.798684 | -0.57825 | 0.563098 | NA | #VALUE! |
| ENSMUSG00000015133.18 | 8.910734 | -1.13656 | 2.646179 | -0.42951 | 0.667552 | NA | #VALUE! |
| ENSMUSG00000015134.16 | 5.30688 | -5.93495 | 3.915215 | -1.51587 | 0.129553 | NA | #VALUE! |
| ENSMUSG00000015222.19 | 0 | NA | NA | NA | NA | NA | #VALUE! |
| ENSMUSG00000015224.11 | 8.266599 | -2.74309 | 2.482058 | -1.10517 | 0.269087 | NA | #VALUE! |
| ENSMUSG00000015242.15 | 0 | NA | NA | NA | NA | NA | #VALUE! |
| ENSMUSG00000015312.10 | 0.093255 | 0.868093 | 4.072464 | 0.213162 | 0.831201 | NA | #VALUE! |
| ENSMUSG00000015314.11 | 0.504851 | 2.502843 | 3.993993 | 0.626652 | 0.530887 | NA | #VALUE! |
| ENSMUSG00000015316.12 | 0 | NA | NA | NA | NA | NA | #VALUE! |
| ENSMUSG00000015365.16 | 0 | NA | NA | NA | NA | NA | #VALUE! |
| ENSMUSG00000015377.11 | 6.875472 | 1.276857 | 3.714626 | 0.343738 | 0.731044 | NA | #VALUE! |
| ENSMUSG00000015396.5 | 12.62267 | -2.12579 | 3.057413 | -0.69529 | NA | NA | #VALUE! |
| ENSMUSG00000015401.13 | 0 | NA | NA | NA | NA | NA | #VALUE! |
| ENSMUSG00000015405.16 | 0 | NA | NA | NA | NA | NA | #VALUE! |
| ENSMUSG00000015437.6 | 0 | NA | NA | NA | NA | NA | #VALUE! |

| | | | | | | | |
|---|---|---|---|---|---|---|---|
| ENSMUSG00000015441.5 | 0 | NA | NA | NA | NA | NA | #VALUE! |
| ENSMUSG00000015443.10 | 0 | NA | NA | NA | NA | NA | #VALUE! |
| ENSMUSG00000015452.15 | 0 | NA | NA | NA | NA | NA | #VALUE! |
| ENSMUSG00000015461.16 | 3.466746 | -1.65193 | 3.797748 | -0.43498 | 0.663579 | NA | #VALUE! |
| ENSMUSG00000015467.15 | 0 | NA | NA | NA | NA | NA | #VALUE! |
| ENSMUSG00000015468.15 | 4.040622 | 4.524985 | 3.913885 | 1.156136 | 0.247625 | NA | #VALUE! |
| ENSMUSG00000015476.7 | 0 | NA | NA | NA | NA | NA | #VALUE! |
| ENSMUSG00000015484.4 | 0 | NA | NA | NA | NA | NA | #VALUE! |
| ENSMUSG00000015519.13 | 0 | NA | NA | NA | NA | NA | #VALUE! |
| ENSMUSG00000015533.10 | 0 | NA | NA | NA | NA | NA | #VALUE! |
| ENSMUSG00000015579.6 | 0 | NA | NA | NA | NA | NA | #VALUE! |
| ENSMUSG00000015599.10 | 0 | NA | NA | NA | NA | NA | #VALUE! |
| ENSMUSG00000015605.7 | 2.389531 | 0.299422 | 3.803025 | 0.078733 | 0.937245 | NA | #VALUE! |
| ENSMUSG00000015619.11 | 0 | NA | NA | NA | NA | NA | #VALUE! |
| ENSMUSG00000015627.6 | 0 | NA | NA | NA | NA | NA | #VALUE! |
| ENSMUSG00000015647.10 | 0.315039 | -1.93549 | 4.038621 | -0.47925 | 0.631763 | NA | #VALUE! |
| ENSMUSG00000015652.10 | 0.093255 | 0.868093 | 4.072464 | 0.213162 | 0.831201 | NA | #VALUE! |
| ENSMUSG00000015659.13 | 3.689633 | 5.364344 | 3.919007 | 1.368802 | 0.171061 | NA | #VALUE! |
| ENSMUSG00000015665.9 | 0 | NA | NA | NA | NA | NA | #VALUE! |
| ENSMUSG00000015702.14 | 0 | NA | NA | NA | NA | NA | #VALUE! |
| ENSMUSG00000015709.10 | 0 | NA | NA | NA | NA | NA | #VALUE! |
| ENSMUSG00000015721.19 | 0 | NA | NA | NA | NA | NA | #VALUE! |
| ENSMUSG00000015747.6 | 1.981875 | 2.848137 | 3.909095 | 0.728592 | 0.466251 | NA | #VALUE! |
| ENSMUSG00000015766.15 | 7.205854 | -3.47318 | 2.715743 | -1.27891 | 0.20093 | NA | #VALUE! |
| ENSMUSG00000015787.16 | 0 | NA | NA | NA | NA | NA | #VALUE! |
| ENSMUSG00000015812.9 | 4.095512 | -5.57062 | 3.917637 | -1.42193 | 0.155045 | NA | #VALUE! |
| ENSMUSG00000015829.14 | 0 | NA | NA | NA | NA | NA | #VALUE! |
| ENSMUSG00000015852.14 | 0 | NA | NA | NA | NA | NA | #VALUE! |
| ENSMUSG00000015879.9 | 0 | NA | NA | NA | NA | NA | #VALUE! |
| ENSMUSG00000015880.14 | 0 | NA | NA | NA | NA | NA | #VALUE! |
| ENSMUSG00000015944.10 | 0.408295 | -1.02718 | 4.03053 | -0.25485 | 0.79884 | NA | #VALUE! |
| ENSMUSG00000015947.11 | 19.74247 | -22.0018 | 2.927309 | -7.51607 | NA | NA | #VALUE! |
| ENSMUSG00000015950.14 | 6.679098 | -5.17173 | 3.740362 | -1.38268 | 0.166763 | NA | #VALUE! |
| ENSMUSG00000015957.13 | 0 | NA | NA | NA | NA | NA | #VALUE! |
| ENSMUSG00000015962.6 | 0 | NA | NA | NA | NA | NA | #VALUE! |
| ENSMUSG00000015968.19 | 0.673134 | 2.918386 | 3.972531 | 0.734641 | 0.462558 | NA | #VALUE! |
| ENSMUSG00000015980.15 | 0 | NA | NA | NA | NA | NA | #VALUE! |
| ENSMUSG00000015981.13 | 0 | NA | NA | NA | NA | NA | #VALUE! |
| ENSMUSG00000016087.14 | 1.346268 | 3.918821 | 3.939925 | 0.994644 | 0.31991 | NA | #VALUE! |
| ENSMUSG00000016150.16 | 0 | NA | NA | NA | NA | NA | #VALUE! |
| ENSMUSG00000016179.12 | 0 | NA | NA | NA | NA | NA | #VALUE! |
| ENSMUSG00000016200.15 | 0 | NA | NA | NA | NA | NA | #VALUE! |
| ENSMUSG00000016206.7 | 15.99745 | 2.875251 | 2.568157 | 1.119578 | NA | NA | #VALUE! |
| ENSMUSG00000016255.6 | 0 | NA | NA | NA | NA | NA | #VALUE! |

| | | | | | | | |
|---|---|---|---|---|---|---|---|
| ENSMUSG00000016262.15 | 7.824139 | 2.168223 | 3.591012 | 0.603792 | 0.545982 | NA | #VALUE! |
| ENSMUSG00000016327.10 | 0 | NA | NA | NA | NA | NA | #VALUE! |
| ENSMUSG00000016346.18 | 0 | NA | NA | NA | NA | NA | #VALUE! |
| ENSMUSG00000016349.11 | 0 | NA | NA | NA | NA | NA | #VALUE! |
| ENSMUSG00000016356.19 | 0 | NA | NA | NA | NA | NA | #VALUE! |
| ENSMUSG00000016458.16 | 0 | NA | NA | NA | NA | NA | #VALUE! |
| ENSMUSG00000016493.14 | 8.550431 | -2.22038 | 3.451102 | -0.64338 | 0.519975 | NA | #VALUE! |
| ENSMUSG00000016494.10 | 9.592161 | 6.751707 | 3.642661 | 1.85351 | 0.063809 | NA | #VALUE! |
| ENSMUSG00000016496.8 | 1.117245 | -3.68056 | 3.946822 | -0.93254 | 0.351058 | NA | #VALUE! |
| ENSMUSG00000016498.11 | 0 | NA | NA | NA | NA | NA | #VALUE! |
| ENSMUSG00000016524.13 | 0 | NA | NA | NA | NA | NA | #VALUE! |
| ENSMUSG00000016526.9 | 2.824742 | 2.606062 | 3.859052 | 0.675312 | 0.499478 | NA | #VALUE! |
| ENSMUSG00000016529.6 | 0 | NA | NA | NA | NA | NA | #VALUE! |
| ENSMUSG00000016552.14 | 0.735092 | -3.12016 | 3.965612 | -0.7868 | 0.431397 | NA | #VALUE! |
| ENSMUSG00000016624.16 | 4.305538 | -5.64245 | 3.91711 | -1.44046 | 0.149737 | NA | #VALUE! |
| ENSMUSG00000016626.11 | 0 | NA | NA | NA | NA | NA | #VALUE! |
| ENSMUSG00000016757.12 | 0.559531 | 2.703958 | 3.982863 | 0.678898 | 0.497202 | NA | #VALUE! |
| ENSMUSG00000016758.5 | 1.0383 | -3.59394 | 3.949279 | -0.91003 | 0.362809 | NA | #VALUE! |
| ENSMUSG00000016763.16 | 0 | NA | NA | NA | NA | NA | #VALUE! |
| ENSMUSG00000016946.7 | 2.311601 | -0.82308 | 3.811215 | -0.21596 | 0.829016 | NA | #VALUE! |
| ENSMUSG00000016982.7 | 0 | NA | NA | NA | NA | NA | #VALUE! |
| ENSMUSG00000016995.18 | 0 | NA | NA | NA | NA | NA | #VALUE! |
| ENSMUSG00000016998.3 | 0 | NA | NA | NA | NA | NA | #VALUE! |
| ENSMUSG00000017000.3 | 0 | NA | NA | NA | NA | NA | #VALUE! |
| ENSMUSG00000017003.8 | 0 | NA | NA | NA | NA | NA | #VALUE! |
| ENSMUSG00000017004.8 | 0 | NA | NA | NA | NA | NA | #VALUE! |
| ENSMUSG00000017007.10 | 0 | NA | NA | NA | NA | NA | #VALUE! |
| ENSMUSG00000017049.5 | 0 | NA | NA | NA | NA | NA | #VALUE! |
| ENSMUSG00000017064.5 | 0 | NA | NA | NA | NA | NA | #VALUE! |
| ENSMUSG00000017146.13 | 0 | NA | NA | NA | NA | NA | #VALUE! |
| ENSMUSG00000017165.2 | 0 | NA | NA | NA | NA | NA | #VALUE! |
| ENSMUSG00000017167.7 | 0.525066 | -2.64687 | 3.988095 | -0.66369 | 0.506887 | NA | #VALUE! |
| ENSMUSG00000017176.18 | 8.303425 | 1.439824 | 3.523305 | 0.408657 | 0.682791 | NA | #VALUE! |
| ENSMUSG00000017195.16 | 0 | NA | NA | NA | NA | NA | #VALUE! |
| ENSMUSG00000017204.5 | 0 | NA | NA | NA | NA | NA | #VALUE! |
| ENSMUSG00000017211.15 | 0 | NA | NA | NA | NA | NA | #VALUE! |
| ENSMUSG00000017300.10 | 0 | NA | NA | NA | NA | NA | #VALUE! |
| ENSMUSG00000017307.17 | 0 | NA | NA | NA | NA | NA | #VALUE! |
| ENSMUSG00000017310.8 | 0 | NA | NA | NA | NA | NA | #VALUE! |
| ENSMUSG00000017311.16 | 0 | NA | NA | NA | NA | NA | #VALUE! |
| ENSMUSG00000017314.13 | 0 | NA | NA | NA | NA | NA | #VALUE! |
| ENSMUSG00000017316.14 | 0 | NA | NA | NA | NA | NA | #VALUE! |
| ENSMUSG00000017400.11 | 0 | NA | NA | NA | NA | NA | #VALUE! |
| ENSMUSG00000017405.15 | 0.466276 | 2.449889 | 3.997179 | 0.612904 | 0.53994 | NA | #VALUE! |

| | | | | | | | |
|---|---|---|---|---|---|---|---|
| ENSMUSG00000017412.16 | 2.415302 | -4.81324 | 3.925115 | -1.22627 | 0.220098 | NA | #VALUE! |
| ENSMUSG00000017417.15 | 1.475853 | 4.029233 | 3.937502 | 1.023297 | 0.306167 | NA | #VALUE! |
| ENSMUSG00000017446.15 | 0 | NA | NA | NA | NA | NA | #VALUE! |
| ENSMUSG00000017491.10 | 8.338287 | 6.554739 | 3.357032 | 1.95254 | 0.050874 | NA | #VALUE! |
| ENSMUSG00000017499.16 | 0 | NA | NA | NA | NA | NA | #VALUE! |
| ENSMUSG00000017550.15 | 8.660759 | 1.879826 | 2.723153 | 0.690312 | 0.489998 | NA | #VALUE! |
| ENSMUSG00000017588.3 | 0 | NA | NA | NA | NA | NA | #VALUE! |
| ENSMUSG00000017607.10 | 0 | NA | NA | NA | NA | NA | #VALUE! |
| ENSMUSG00000017631.20 | 3.36042 | -5.28663 | 3.919996 | -1.34863 | 0.177455 | NA | #VALUE! |
| ENSMUSG00000017639.14 | 0 | NA | NA | NA | NA | NA | #VALUE! |
| ENSMUSG00000017652.17 | 7.926691 | 6.480757 | 3.856882 | 1.68031 | NA | NA | #VALUE! |
| ENSMUSG00000017664.17 | 0 | NA | NA | NA | NA | NA | #VALUE! |
| ENSMUSG00000017688.15 | 0 | NA | NA | NA | NA | NA | #VALUE! |
| ENSMUSG00000017692.9 | 0 | NA | NA | NA | NA | NA | #VALUE! |
| ENSMUSG00000017697.4 | 23.5237 | -2.91936 | 2.625345 | -1.11199 | NA | NA | #VALUE! |
| ENSMUSG00000017713.10 | 3.719993 | -5.42213 | 3.918812 | -1.38361 | 0.166476 | NA | #VALUE! |
| ENSMUSG00000017715.12 | 6.772178 | -3.16574 | 3.25703 | -0.97197 | 0.331065 | NA | #VALUE! |
| ENSMUSG00000017716.16 | 1.930385 | -3.27282 | 3.918337 | -0.83526 | 0.403573 | NA | #VALUE! |
| ENSMUSG00000017718.16 | 22.66721 | 1.400193 | 2.383049 | 0.587564 | NA | NA | #VALUE! |
| ENSMUSG00000017720.3 | 0 | NA | NA | NA | NA | NA | #VALUE! |
| ENSMUSG00000017721.16 | 0 | NA | NA | NA | NA | NA | #VALUE! |
| ENSMUSG00000017724.15 | 0 | NA | NA | NA | NA | NA | #VALUE! |
| ENSMUSG00000017733.5 | 0 | NA | NA | NA | NA | NA | #VALUE! |
| ENSMUSG00000017734.16 | 1.365171 | -3.99822 | 3.938957 | -1.01505 | 0.310084 | NA | #VALUE! |
| ENSMUSG00000017737.3 | 0 | NA | NA | NA | NA | NA | #VALUE! |
| ENSMUSG00000017740.18 | 3.082339 | 0.324871 | 3.785537 | 0.085819 | 0.93161 | NA | #VALUE! |
| ENSMUSG00000017764.3 | 2.473283 | -2.4323 | 3.86308 | -0.62963 | 0.528939 | NA | #VALUE! |
| ENSMUSG00000017765.18 | 8.192782 | -0.29444 | 3.562829 | -0.08264 | 0.934137 | NA | #VALUE! |
| ENSMUSG00000017767.4 | 0 | NA | NA | NA | NA | NA | #VALUE! |
| ENSMUSG00000017774.20 | 6.174475 | -6.15438 | 3.80551 | -1.61723 | 0.105829 | NA | #VALUE! |
| ENSMUSG00000017817.12 | 0 | NA | NA | NA | NA | NA | #VALUE! |
| ENSMUSG00000017830.16 | 7.942436 | -2.71684 | 3.159555 | -0.85988 | 0.389855 | NA | #VALUE! |
| ENSMUSG00000017832.3 | 0 | NA | NA | NA | NA | NA | #VALUE! |
| ENSMUSG00000017837.13 | 1.819051 | -4.39655 | 3.931229 | -1.11837 | 0.263411 | NA | #VALUE! |
| ENSMUSG00000017861.12 | 0 | NA | NA | NA | NA | NA | #VALUE! |
| ENSMUSG00000017897.19 | 0 | NA | NA | NA | NA | NA | #VALUE! |
| ENSMUSG00000017922.15 | 0 | NA | NA | NA | NA | NA | #VALUE! |
| ENSMUSG00000017943.16 | 0 | NA | NA | NA | NA | NA | #VALUE! |
| ENSMUSG00000017969.14 | 6.327433 | -6.18917 | 3.913851 | -1.58135 | 0.113798 | NA | #VALUE! |
| ENSMUSG00000018008.9 | 1.521083 | -0.11467 | 3.848703 | -0.02979 | 0.976231 | NA | #VALUE! |
| ENSMUSG00000018012.12 | 0 | NA | NA | NA | NA | NA | #VALUE! |
| ENSMUSG00000018040.10 | 3.524204 | -5.34395 | 3.919482 | -1.36343 | 0.172746 | NA | #VALUE! |
| ENSMUSG00000018076.13 | 5.904632 | 1.350245 | 2.764015 | 0.488509 | 0.62519 | NA | #VALUE! |
| ENSMUSG00000018126.16 | 0 | NA | NA | NA | NA | NA | #VALUE! |

| | | | | | | | |
|---|---|---|---|---|---|---|---|
| ENSMUSG00000018167.11 | 1.260158 | -3.88425 | 3.941584 | -0.98545 | 0.324401 | NA | #VALUE! |
| ENSMUSG00000018168.9 | 0 | NA | NA | NA | NA | NA | #VALUE! |
| ENSMUSG00000018199.10 | 20.0178 | -0.17897 | 1.953593 | -0.09161 | NA | NA | #VALUE! |
| ENSMUSG00000018209.16 | 52.9862 | 2.159489 | 1.339765 | 1.611842 | NA | NA | #VALUE! |
| ENSMUSG00000018211.14 | 1.632886 | -4.22813 | 3.934241 | -1.0747 | 0.282509 | NA | #VALUE! |
| ENSMUSG00000018238.7 | 0 | NA | NA | NA | NA | NA | #VALUE! |
| ENSMUSG00000018259.14 | 0 | NA | NA | NA | NA | NA | #VALUE! |
| ENSMUSG00000018263.12 | 0 | NA | NA | NA | NA | NA | #VALUE! |
| ENSMUSG00000018334.19 | 0 | NA | NA | NA | NA | NA | #VALUE! |
| ENSMUSG00000018339.13 | 6.011214 | -0.14752 | 3.756128 | -0.03927 | 0.968672 | NA | #VALUE! |
| ENSMUSG00000018341.13 | 0 | NA | NA | NA | NA | NA | #VALUE! |
| ENSMUSG00000018372.14 | 4.828908 | 1.704465 | 3.728873 | 0.457099 | 0.6476 | NA | #VALUE! |
| ENSMUSG00000018378.14 | 0 | NA | NA | NA | NA | NA | #VALUE! |
| ENSMUSG00000018405.8 | 1.155144 | -3.7605 | 3.944679 | -0.95331 | 0.340433 | NA | #VALUE! |
| ENSMUSG00000018451.8 | 0 | NA | NA | NA | NA | NA | #VALUE! |
| ENSMUSG00000018470.9 | 0 | NA | NA | NA | NA | NA | #VALUE! |
| ENSMUSG00000018476.8 | 3.962235 | -0.26551 | 3.548751 | -0.07482 | 0.94036 | NA | #VALUE! |
| ENSMUSG00000018479.13 | 0 | NA | NA | NA | NA | NA | #VALUE! |
| ENSMUSG00000018486.3 | 0 | NA | NA | NA | NA | NA | #VALUE! |
| ENSMUSG00000018500.3 | 0 | NA | NA | NA | NA | NA | #VALUE! |
| ENSMUSG00000018507.17 | 0.279766 | 1.808031 | 4.036658 | 0.447903 | 0.654223 | NA | #VALUE! |
| ENSMUSG00000018537.17 | 0 | NA | NA | NA | NA | NA | #VALUE! |
| ENSMUSG00000018543.9 | 0 | NA | NA | NA | NA | NA | #VALUE! |
| ENSMUSG00000018547.9 | 31.80824 | -2.14506 | 1.535587 | -1.3969 | NA | NA | #VALUE! |
| ENSMUSG00000018554.14 | 0 | NA | NA | NA | NA | NA | #VALUE! |
| ENSMUSG00000018566.15 | 0 | NA | NA | NA | NA | NA | #VALUE! |
| ENSMUSG00000018569.13 | 0 | NA | NA | NA | NA | NA | #VALUE! |
| ENSMUSG00000018570.18 | 0 | NA | NA | NA | NA | NA | #VALUE! |
| ENSMUSG00000018581.16 | 0 | NA | NA | NA | NA | NA | #VALUE! |
| ENSMUSG00000018589.9 | 0 | NA | NA | NA | NA | NA | #VALUE! |
| ENSMUSG00000018595.7 | 0 | NA | NA | NA | NA | NA | #VALUE! |
| ENSMUSG00000018620.4 | 0 | NA | NA | NA | NA | NA | #VALUE! |
| ENSMUSG00000018623.10 | 0 | NA | NA | NA | NA | NA | #VALUE! |
| ENSMUSG00000018634.11 | 0 | NA | NA | NA | NA | NA | #VALUE! |
| ENSMUSG00000018651.15 | 3.485686 | 5.292617 | 3.919629 | 1.350285 | 0.176925 | NA | #VALUE! |
| ENSMUSG00000018656.6 | 0 | NA | NA | NA | NA | NA | #VALUE! |
| ENSMUSG00000018698.16 | 0 | NA | NA | NA | NA | NA | #VALUE! |
| ENSMUSG00000018740.15 | 0 | NA | NA | NA | NA | NA | #VALUE! |
| ENSMUSG00000018752.20 | 0 | NA | NA | NA | NA | NA | #VALUE! |
| ENSMUSG00000018776.11 | 0 | NA | NA | NA | NA | NA | #VALUE! |
| ENSMUSG00000018800.15 | 4.854718 | 5.763343 | 3.916055 | 1.471722 | 0.141096 | NA | #VALUE! |
| ENSMUSG00000018809.3 | 3.029103 | 5.088783 | 3.921574 | 1.297638 | 0.194412 | NA | #VALUE! |
| ENSMUSG00000018819.11 | 0.105013 | -0.97324 | 4.076293 | -0.23876 | 0.811295 | NA | #VALUE! |
| ENSMUSG00000018822.8 | 0 | NA | NA | NA | NA | NA | #VALUE! |

| | | | | | | | |
|---|---|---|---|---|---|---|---|
| ENSMUSG00000018830.11 | 7.964337 | 4.307005 | 3.551535 | 1.212717 | 0.225238 | NA | #VALUE! |
| ENSMUSG00000018844.7 | 0 | NA | NA | NA | NA | NA | #VALUE! |
| ENSMUSG00000018845.15 | 9.488463 | -0.34982 | 3.412902 | -0.1025 | 0.918361 | NA | #VALUE! |
| ENSMUSG00000018862.12 | 0 | NA | NA | NA | NA | NA | #VALUE! |
| ENSMUSG00000018865.10 | 0 | NA | NA | NA | NA | NA | #VALUE! |
| ENSMUSG00000018868.5 | 0 | NA | NA | NA | NA | NA | #VALUE! |
| ENSMUSG00000018893.16 | 0 | NA | NA | NA | NA | NA | #VALUE! |
| ENSMUSG00000018907.11 | 0 | NA | NA | NA | NA | NA | #VALUE! |
| ENSMUSG00000018909.16 | 0.336567 | 1.944925 | 4.029936 | 0.482619 | 0.629366 | NA | #VALUE! |
| ENSMUSG00000018914.6 | 0 | NA | NA | NA | NA | NA | #VALUE! |
| ENSMUSG00000018916.6 | 0 | NA | NA | NA | NA | NA | #VALUE! |
| ENSMUSG00000018919.11 | 0 | NA | NA | NA | NA | NA | #VALUE! |
| ENSMUSG00000018924.7 | 0 | NA | NA | NA | NA | NA | #VALUE! |
| ENSMUSG00000018925.4 | 0 | NA | NA | NA | NA | NA | #VALUE! |
| ENSMUSG00000018931.4 | 0 | NA | NA | NA | NA | NA | #VALUE! |
| ENSMUSG00000018973.3 | 0 | NA | NA | NA | NA | NA | #VALUE! |
| ENSMUSG00000018986.11 | 0 | NA | NA | NA | NA | NA | #VALUE! |
| ENSMUSG00000019027.9 | 0 | NA | NA | NA | NA | NA | #VALUE! |
| ENSMUSG00000019066.14 | 8.456928 | -3.73763 | 2.968433 | -1.25912 | 0.207985 | NA | #VALUE! |
| ENSMUSG00000019102.11 | 0 | NA | NA | NA | NA | NA | #VALUE! |
| ENSMUSG00000019124.11 | 0 | NA | NA | NA | NA | NA | #VALUE! |
| ENSMUSG00000019146.3 | 0 | NA | NA | NA | NA | NA | #VALUE! |
| ENSMUSG00000019178.18 | 0 | NA | NA | NA | NA | NA | #VALUE! |
| ENSMUSG00000019194.16 | 1.470184 | -4.10383 | 3.936698 | -1.04245 | 0.297201 | NA | #VALUE! |
| ENSMUSG00000019214.15 | 0 | NA | NA | NA | NA | NA | #VALUE! |
| ENSMUSG00000019230.15 | 0 | NA | NA | NA | NA | NA | #VALUE! |
| ENSMUSG00000019235.10 | 0 | NA | NA | NA | NA | NA | #VALUE! |
| ENSMUSG00000019278.6 | 1.212318 | 3.793111 | 3.942915 | 0.962007 | 0.336046 | NA | #VALUE! |
| ENSMUSG00000019301.6 | 0 | NA | NA | NA | NA | NA | #VALUE! |
| ENSMUSG00000019303.15 | 0.612332 | -2.79827 | 3.980109 | -0.70306 | 0.482016 | NA | #VALUE! |
| ENSMUSG00000019320.15 | 0 | NA | NA | NA | NA | NA | #VALUE! |
| ENSMUSG00000019326.15 | 0 | NA | NA | NA | NA | NA | #VALUE! |
| ENSMUSG00000019338.14 | 3.135876 | -0.72698 | 3.788384 | -0.1919 | 0.847824 | NA | #VALUE! |
| ENSMUSG00000019359.15 | 0 | NA | NA | NA | NA | NA | #VALUE! |
| ENSMUSG00000019429.8 | 0 | NA | NA | NA | NA | NA | #VALUE! |
| ENSMUSG00000019433.10 | 0 | NA | NA | NA | NA | NA | #VALUE! |
| ENSMUSG00000019461.13 | 1.707091 | -1.99111 | 3.887076 | -0.51224 | 0.608484 | NA | #VALUE! |
| ENSMUSG00000019464.7 | 0 | NA | NA | NA | NA | NA | #VALUE! |
| ENSMUSG00000019467.16 | 3.935608 | 5.458007 | 3.918239 | 1.392975 | 0.163627 | NA | #VALUE! |
| ENSMUSG00000019489.8 | 0 | NA | NA | NA | NA | NA | #VALUE! |
| ENSMUSG00000019539.12 | 3.719827 | -0.55405 | 3.776773 | -0.1467 | 0.883369 | NA | #VALUE! |
| ENSMUSG00000019558.15 | 0 | NA | NA | NA | NA | NA | #VALUE! |
| ENSMUSG00000019564.13 | 0.840105 | -3.30885 | 3.958472 | -0.83589 | 0.403216 | NA | #VALUE! |
| ENSMUSG00000019699.17 | 1.890236 | -4.46257 | 3.93014 | -1.13547 | 0.256177 | NA | #VALUE! |

| | | | | | | | |
|---|---|---|---|---|---|---|---|
| ENSMUSG00000019732.15 | 1.310688 | 1.250168 | 3.893942 | 0.321055 | 0.748169 | NA | #VALUE! |
| ENSMUSG00000019734.17 | 6.251567 | 2.651073 | 3.787776 | 0.699902 | 0.483988 | NA | #VALUE! |
| ENSMUSG00000019737.15 | 1.441709 | -4.06896 | 3.937426 | -1.03341 | 0.301414 | NA | #VALUE! |
| ENSMUSG00000019756.11 | 0 | NA | NA | NA | NA | NA | #VALUE! |
| ENSMUSG00000019767.13 | 0 | NA | NA | NA | NA | NA | #VALUE! |
| ENSMUSG00000019772.6 | 0 | NA | NA | NA | NA | NA | #VALUE! |
| ENSMUSG00000019775.18 | 0 | NA | NA | NA | NA | NA | #VALUE! |
| ENSMUSG00000019785.15 | 0 | NA | NA | NA | NA | NA | #VALUE! |
| ENSMUSG00000019787.10 | 0 | NA | NA | NA | NA | NA | #VALUE! |
| ENSMUSG00000019789.10 | 0 | NA | NA | NA | NA | NA | #VALUE! |
| ENSMUSG00000019796.13 | 1.305573 | 3.898287 | 3.940396 | 0.989314 | 0.32251 | NA | #VALUE! |
| ENSMUSG00000019803.12 | 0 | NA | NA | NA | NA | NA | #VALUE! |
| ENSMUSG00000019808.9 | 9.212943 | -0.35627 | 2.694633 | -0.13222 | 0.894814 | NA | #VALUE! |
| ENSMUSG00000019815.8 | 0 | NA | NA | NA | NA | NA | #VALUE! |
| ENSMUSG00000019817.19 | 0.525066 | -2.64687 | 3.988095 | -0.66369 | 0.506887 | NA | #VALUE! |
| ENSMUSG00000019823.18 | 0 | NA | NA | NA | NA | NA | #VALUE! |
| ENSMUSG00000019828.14 | 0 | NA | NA | NA | NA | NA | #VALUE! |
| ENSMUSG00000019831.15 | 2.180952 | -0.16387 | 3.808817 | -0.04302 | 0.965683 | NA | #VALUE! |
| ENSMUSG00000019834.16 | 0 | NA | NA | NA | NA | NA | #VALUE! |
| ENSMUSG00000019836.6 | 0 | NA | NA | NA | NA | NA | #VALUE! |
| ENSMUSG00000019842.7 | 20.16628 | -22.0541 | 2.940979 | -7.4989 | NA | NA | #VALUE! |
| ENSMUSG00000019845.9 | 1.957891 | -4.49332 | 3.929649 | -1.14344 | 0.252855 | NA | #VALUE! |
| ENSMUSG00000019846.12 | 0 | NA | NA | NA | NA | NA | #VALUE! |
| ENSMUSG00000019848.15 | 0 | NA | NA | NA | NA | NA | #VALUE! |
| ENSMUSG00000019850.12 | 56.84927 | -4.34436 | 3.743446 | -1.16052 | NA | NA | #VALUE! |
| ENSMUSG00000019852.8 | 0 | NA | NA | NA | NA | NA | #VALUE! |
| ENSMUSG00000019853.7 | 0.195789 | -1.05581 | 4.073302 | -0.2592 | 0.795479 | NA | #VALUE! |
| ENSMUSG00000019856.15 | 0 | NA | NA | NA | NA | NA | #VALUE! |
| ENSMUSG00000019864.15 | 8.983511 | 0.814689 | 2.630887 | 0.309663 | 0.756817 | NA | #VALUE! |
| ENSMUSG00000019865.10 | 0 | NA | NA | NA | NA | NA | #VALUE! |
| ENSMUSG00000019867.5 | 0 | NA | NA | NA | NA | NA | #VALUE! |
| ENSMUSG00000019888.17 | 0 | NA | NA | NA | NA | NA | #VALUE! |
| ENSMUSG00000019890.5 | 0 | NA | NA | NA | NA | NA | #VALUE! |
| ENSMUSG00000019891.16 | 16.91087 | 4.831536 | 2.83933 | 1.701646 | NA | NA | #VALUE! |
| ENSMUSG00000019892.14 | 0 | NA | NA | NA | NA | NA | #VALUE! |
| ENSMUSG00000019893.14 | 0 | NA | NA | NA | NA | NA | #VALUE! |
| ENSMUSG00000019894.15 | 0 | NA | NA | NA | NA | NA | #VALUE! |
| ENSMUSG00000019899.18 | 2.940368 | -5.09511 | 3.921868 | -1.29915 | 0.193891 | NA | #VALUE! |
| ENSMUSG00000019900.20 | 0 | NA | NA | NA | NA | NA | #VALUE! |
| ENSMUSG00000019905.9 | 0 | NA | NA | NA | NA | NA | #VALUE! |
| ENSMUSG00000019909.5 | 0 | NA | NA | NA | NA | NA | #VALUE! |
| ENSMUSG00000019913.5 | 0 | NA | NA | NA | NA | NA | #VALUE! |
| ENSMUSG00000019932.9 | 0 | NA | NA | NA | NA | NA | #VALUE! |
| ENSMUSG00000019933.8 | 0 | NA | NA | NA | NA | NA | #VALUE! |

| | | | | | | | |
|---|---|---|---|---|---|---|---|
| ENSMUSG00000019936.11 | 0 | NA | NA | NA | NA | NA | #VALUE! |
| ENSMUSG00000019942.14 | 6.892082 | 4.285797 | 3.200426 | 1.339133 | 0.180527 | NA | #VALUE! |
| ENSMUSG00000019979.13 | 0.652786 | 2.919659 | 3.972474 | 0.734972 | 0.462356 | NA | #VALUE! |
| ENSMUSG00000019982.16 | 0 | NA | NA | NA | NA | NA | #VALUE! |
| ENSMUSG00000019986.18 | 0 | NA | NA | NA | NA | NA | #VALUE! |
| ENSMUSG00000019990.16 | 3.507052 | -0.08596 | 3.778297 | -0.02275 | 0.981849 | NA | #VALUE! |
| ENSMUSG00000019992.9 | 0 | NA | NA | NA | NA | NA | #VALUE! |
| ENSMUSG00000020007.5 | 0 | NA | NA | NA | NA | NA | #VALUE! |
| ENSMUSG00000020014.19 | 0 | NA | NA | NA | NA | NA | #VALUE! |
| ENSMUSG00000020020.9 | 0 | NA | NA | NA | NA | NA | #VALUE! |
| ENSMUSG00000020032.15 | 3.25745 | -5.2291 | 3.920532 | -1.33377 | 0.182278 | NA | #VALUE! |
| ENSMUSG00000020033.3 | 0 | NA | NA | NA | NA | NA | #VALUE! |
| ENSMUSG00000020042.16 | 0.525066 | -2.64687 | 3.988095 | -0.66369 | 0.506887 | NA | #VALUE! |
| ENSMUSG00000020052.10 | 0 | NA | NA | NA | NA | NA | #VALUE! |
| ENSMUSG00000020057.3 | 0 | NA | NA | NA | NA | NA | #VALUE! |
| ENSMUSG00000020061.19 | 0 | NA | NA | NA | NA | NA | #VALUE! |
| ENSMUSG00000020062.8 | 0 | NA | NA | NA | NA | NA | #VALUE! |
| ENSMUSG00000020067.9 | 0 | NA | NA | NA | NA | NA | #VALUE! |
| ENSMUSG00000020070.19 | 6.718128 | 0.842761 | 2.289547 | 0.368091 | 0.712806 | NA | #VALUE! |
| ENSMUSG00000020080.9 | 0 | NA | NA | NA | NA | NA | #VALUE! |
| ENSMUSG00000020081.6 | 0 | NA | NA | NA | NA | NA | #VALUE! |
| ENSMUSG00000020083.14 | 3.804437 | 0.536594 | 3.775567 | 0.142123 | 0.886983 | NA | #VALUE! |
| ENSMUSG00000020086.7 | 0 | NA | NA | NA | NA | NA | #VALUE! |
| ENSMUSG00000020090.8 | 0 | NA | NA | NA | NA | NA | #VALUE! |
| ENSMUSG00000020096.22 | 0 | NA | NA | NA | NA | NA | #VALUE! |
| ENSMUSG00000020099.9 | 6.609479 | -2.25788 | 3.379932 | -0.66803 | 0.504117 | NA | #VALUE! |
| ENSMUSG00000020101.15 | 7.86197 | -0.10427 | 2.473484 | -0.04216 | 0.966374 | NA | #VALUE! |
| ENSMUSG00000020120.16 | 20.71259 | -4.40451 | 2.149441 | -2.04914 | NA | NA | #VALUE! |
| ENSMUSG00000020121.16 | 1.119062 | 3.679627 | 3.945843 | 0.932533 | 0.351061 | NA | #VALUE! |
| ENSMUSG00000020125.8 | 0.168284 | 1.027628 | 4.067126 | 0.252667 | 0.800526 | NA | #VALUE! |
| ENSMUSG00000020133.9 | 5.756648 | 4.463979 | 3.461432 | 1.289633 | 0.197178 | NA | #VALUE! |
| ENSMUSG00000020135.14 | 0 | NA | NA | NA | NA | NA | #VALUE! |
| ENSMUSG00000020140.16 | 19.86508 | 2.585196 | 2.245998 | 1.151023 | NA | NA | #VALUE! |
| ENSMUSG00000020143.16 | 2.29759 | 4.700848 | 3.92612 | 1.197327 | 0.231179 | NA | #VALUE! |
| ENSMUSG00000020151.17 | 0 | NA | NA | NA | NA | NA | #VALUE! |
| ENSMUSG00000020155.3 | 0 | NA | NA | NA | NA | NA | #VALUE! |
| ENSMUSG00000020156.17 | 8.868096 | -3.80859 | 2.947423 | -1.29218 | 0.196296 | NA | #VALUE! |
| ENSMUSG00000020159.9 | 0 | NA | NA | NA | NA | NA | #VALUE! |
| ENSMUSG00000020164.8 | 0 | NA | NA | NA | NA | NA | #VALUE! |
| ENSMUSG00000020168.6 | 0 | NA | NA | NA | NA | NA | #VALUE! |
| ENSMUSG00000020169.5 | 0 | NA | NA | NA | NA | NA | #VALUE! |
| ENSMUSG00000020175.16 | 3.780473 | -5.45568 | 3.918536 | -1.39228 | 0.163839 | NA | #VALUE! |
| ENSMUSG00000020177.13 | 0 | NA | NA | NA | NA | NA | #VALUE! |
| ENSMUSG00000020178.7 | 0 | NA | NA | NA | NA | NA | #VALUE! |

| | | | | | | | |
|---|---|---|---|---|---|---|---|
| ENSMUSG00000020181.19 | 0 | NA | NA | NA | NA | NA | #VALUE! |
| ENSMUSG00000020183.12 | 0.989353 | 3.494937 | 3.951117 | 0.884544 | 0.376403 | NA | #VALUE! |
| ENSMUSG00000020185.17 | 0 | NA | NA | NA | NA | NA | #VALUE! |
| ENSMUSG00000020191.12 | 0 | NA | NA | NA | NA | NA | #VALUE! |
| ENSMUSG00000020193.4 | 0 | NA | NA | NA | NA | NA | #VALUE! |
| ENSMUSG00000020211.16 | 0.210026 | -1.52494 | 4.058692 | -0.37572 | 0.707123 | NA | #VALUE! |
| ENSMUSG00000020212.15 | 0 | NA | NA | NA | NA | NA | #VALUE! |
| ENSMUSG00000020213.7 | 0 | NA | NA | NA | NA | NA | #VALUE! |
| ENSMUSG00000020214.11 | 0 | NA | NA | NA | NA | NA | #VALUE! |
| ENSMUSG00000020216.14 | 0 | NA | NA | NA | NA | NA | #VALUE! |
| ENSMUSG00000020218.12 | 0 | NA | NA | NA | NA | NA | #VALUE! |
| ENSMUSG00000020226.10 | 0 | NA | NA | NA | NA | NA | #VALUE! |
| ENSMUSG00000020227.11 | 0.525066 | -2.64687 | 3.988095 | -0.66369 | 0.506887 | NA | #VALUE! |
| ENSMUSG00000020229.5 | 0 | NA | NA | NA | NA | NA | #VALUE! |
| ENSMUSG00000020230.16 | 0.816443 | -3.21914 | 3.961753 | -0.81255 | 0.416474 | NA | #VALUE! |
| ENSMUSG00000020231.16 | 3.61434 | 0.629536 | 3.779107 | 0.166583 | 0.867698 | NA | #VALUE! |
| ENSMUSG00000020234.11 | 0 | NA | NA | NA | NA | NA | #VALUE! |
| ENSMUSG00000020241.14 | 0 | NA | NA | NA | NA | NA | #VALUE! |
| ENSMUSG00000020251.15 | 0 | NA | NA | NA | NA | NA | #VALUE! |
| ENSMUSG00000020256.15 | 0 | NA | NA | NA | NA | NA | #VALUE! |
| ENSMUSG00000020264.6 | 0 | NA | NA | NA | NA | NA | #VALUE! |
| ENSMUSG00000020270.10 | 0 | NA | NA | NA | NA | NA | #VALUE! |
| ENSMUSG00000020273.14 | 6.766336 | -6.28729 | 3.151877 | -1.99478 | 0.046067 | NA | #VALUE! |
| ENSMUSG00000020277.11 | 8.105115 | 5.460563 | 3.544659 | 1.540504 | 0.123437 | NA | #VALUE! |
| ENSMUSG00000020279.11 | 0 | NA | NA | NA | NA | NA | #VALUE! |
| ENSMUSG00000020282.19 | 7.347987 | -6.40523 | 3.912867 | -1.63697 | 0.101638 | NA | #VALUE! |
| ENSMUSG00000020284.17 | 6.709009 | -6.2804 | 3.63513 | -1.7277 | 0.084043 | NA | #VALUE! |
| ENSMUSG00000020286.14 | 0 | NA | NA | NA | NA | NA | #VALUE! |
| ENSMUSG00000020295.2 | 0 | NA | NA | NA | NA | NA | #VALUE! |
| ENSMUSG00000020297.11 | 0 | NA | NA | NA | NA | NA | #VALUE! |
| ENSMUSG00000020299.7 | 0 | NA | NA | NA | NA | NA | #VALUE! |
| ENSMUSG00000020303.3 | 0 | NA | NA | NA | NA | NA | #VALUE! |
| ENSMUSG00000020310.10 | 0 | NA | NA | NA | NA | NA | #VALUE! |
| ENSMUSG00000020312.13 | 0 | NA | NA | NA | NA | NA | #VALUE! |
| ENSMUSG00000020317.15 | 0 | NA | NA | NA | NA | NA | #VALUE! |
| ENSMUSG00000020323.15 | 0 | NA | NA | NA | NA | NA | #VALUE! |
| ENSMUSG00000020325.11 | 0 | NA | NA | NA | NA | NA | #VALUE! |
| ENSMUSG00000020327.5 | 0 | NA | NA | NA | NA | NA | #VALUE! |
| ENSMUSG00000020330.17 | 0 | NA | NA | NA | NA | NA | #VALUE! |
| ENSMUSG00000020331.10 | 0 | NA | NA | NA | NA | NA | #VALUE! |
| ENSMUSG00000020332.8 | 0 | NA | NA | NA | NA | NA | #VALUE! |
| ENSMUSG00000020333.18 | 2.325214 | 0.471535 | 3.807234 | 0.123852 | 0.901432 | NA | #VALUE! |
| ENSMUSG00000020334.7 | 2.206407 | -0.89933 | 3.817824 | -0.23556 | 0.813772 | NA | #VALUE! |
| ENSMUSG00000020335.14 | 9.388576 | -0.0248 | 2.656962 | -0.00933 | 0.992553 | NA | #VALUE! |

| | | | | | | | |
|---|---|---|---|---|---|---|---|
| ENSMUSG00000020340.17 | 1.177985 | 3.72614 | 3.944615 | 0.944614 | 0.344856 | NA | #VALUE! |
| ENSMUSG00000020354.16 | 0 | NA | NA | NA | NA | NA | #VALUE! |
| ENSMUSG00000020363.7 | 0 | NA | NA | NA | NA | NA | #VALUE! |
| ENSMUSG00000020374.17 | 0 | NA | NA | NA | NA | NA | #VALUE! |
| ENSMUSG00000020377.16 | 0 | NA | NA | NA | NA | NA | #VALUE! |
| ENSMUSG00000020383.2 | 0 | NA | NA | NA | NA | NA | #VALUE! |
| ENSMUSG00000020388.13 | 4.606341 | -5.73912 | 3.916441 | -1.46539 | 0.142814 | NA | #VALUE! |
| ENSMUSG00000020389.20 | 0 | NA | NA | NA | NA | NA | #VALUE! |
| ENSMUSG00000020395.14 | 0.204111 | -1.05581 | 4.073302 | -0.2592 | 0.795479 | NA | #VALUE! |
| ENSMUSG00000020396.9 | 0 | NA | NA | NA | NA | NA | #VALUE! |
| ENSMUSG00000020399.15 | 0 | NA | NA | NA | NA | NA | #VALUE! |
| ENSMUSG00000020400.18 | 19.14001 | 1.91519 | 2.324907 | 0.823771 | NA | NA | #VALUE! |
| ENSMUSG00000020401.7 | 0 | NA | NA | NA | NA | NA | #VALUE! |
| ENSMUSG00000020405.5 | 0 | NA | NA | NA | NA | NA | #VALUE! |
| ENSMUSG00000020407.14 | 1.154336 | 1.377753 | 3.915234 | 0.351895 | 0.724917 | NA | #VALUE! |
| ENSMUSG00000020411.3 | 0 | NA | NA | NA | NA | NA | #VALUE! |
| ENSMUSG00000020419.12 | 3.175859 | 3.111058 | 3.878883 | 0.80205 | 0.422524 | NA | #VALUE! |
| ENSMUSG00000020420.9 | 7.871217 | 6.462163 | 3.906699 | 1.654124 | 0.098102 | NA | #VALUE! |
| ENSMUSG00000020424.4 | 9.234924 | 6.695938 | 3.181822 | 2.104435 | 0.03534 | NA | #VALUE! |
| ENSMUSG00000020428.14 | 0 | NA | NA | NA | NA | NA | #VALUE! |
| ENSMUSG00000020431.6 | 0 | NA | NA | NA | NA | NA | #VALUE! |
| ENSMUSG00000020434.5 | 0 | NA | NA | NA | NA | NA | #VALUE! |
| ENSMUSG00000020435.18 | 0 | NA | NA | NA | NA | NA | #VALUE! |
| ENSMUSG00000020436.18 | 0 | NA | NA | NA | NA | NA | #VALUE! |
| ENSMUSG00000020437.13 | 0 | NA | NA | NA | NA | NA | #VALUE! |
| ENSMUSG00000020439.18 | 0.673134 | 2.918386 | 3.972531 | 0.734641 | 0.462558 | NA | #VALUE! |
| ENSMUSG00000020447.6 | 0 | NA | NA | NA | NA | NA | #VALUE! |
| ENSMUSG00000020451.18 | 2.347825 | -4.75738 | 3.925836 | -1.21181 | 0.225584 | NA | #VALUE! |
| ENSMUSG00000020453.18 | 7.467271 | -0.63155 | 3.639435 | -0.17353 | 0.862236 | NA | #VALUE! |
| ENSMUSG00000020461.11 | 0 | NA | NA | NA | NA | NA | #VALUE! |
| ENSMUSG00000020469.9 | 1.851119 | 4.378291 | 3.930938 | 1.113803 | 0.265364 | NA | #VALUE! |
| ENSMUSG00000020474.12 | 9.832167 | -6.82601 | 3.179551 | -2.14685 | 0.031805 | NA | #VALUE! |
| ENSMUSG00000020475.4 | 0 | NA | NA | NA | NA | NA | #VALUE! |
| ENSMUSG00000020481.16 | 0 | NA | NA | NA | NA | NA | #VALUE! |
| ENSMUSG00000020490.17 | 0 | NA | NA | NA | NA | NA | #VALUE! |
| ENSMUSG00000020491.16 | 6.418526 | 1.503392 | 3.08172 | 0.487842 | 0.625662 | NA | #VALUE! |
| ENSMUSG00000020493.9 | 0 | NA | NA | NA | NA | NA | #VALUE! |
| ENSMUSG00000020495.14 | 25.03455 | -1.5681 | 1.767543 | -0.88716 | NA | NA | #VALUE! |
| ENSMUSG00000020513.16 | 7.32199 | -1.16084 | 3.225209 | -0.35993 | 0.718901 | NA | #VALUE! |
| ENSMUSG00000020520.15 | 3.855789 | -0.63923 | 3.775604 | -0.16931 | 0.865556 | NA | #VALUE! |
| ENSMUSG00000020524.17 | 0 | NA | NA | NA | NA | NA | #VALUE! |
| ENSMUSG00000020525.18 | 8.270028 | -3.27195 | 2.992659 | -1.09332 | 0.274251 | NA | #VALUE! |
| ENSMUSG00000020536.13 | 20.62764 | -0.83421 | 2.032526 | -0.41043 | NA | NA | #VALUE! |
| ENSMUSG00000020542.19 | 0 | NA | NA | NA | NA | NA | #VALUE! |

| | | | | | | | |
|---|---|---|---|---|---|---|---|
| ENSMUSG00000020545.15 | 0 | NA | NA | NA | NA | NA | #VALUE! |
| ENSMUSG00000020546.15 | 7.129831 | -0.96777 | 2.881642 | -0.33584 | 0.736992 | NA | #VALUE! |
| ENSMUSG00000020548.11 | 0 | NA | NA | NA | NA | NA | #VALUE! |
| ENSMUSG00000020562.8 | 0 | NA | NA | NA | NA | NA | #VALUE! |
| ENSMUSG00000020564.18 | 4.278884 | 0.856593 | 3.443844 | 0.248732 | 0.803568 | NA | #VALUE! |
| ENSMUSG00000020566.20 | 8.85694 | -0.57154 | 3.042093 | -0.18788 | 0.850974 | NA | #VALUE! |
| ENSMUSG00000020577.18 | 3.263932 | 5.206263 | 3.920419 | 1.327986 | 0.184183 | NA | #VALUE! |
| ENSMUSG00000020581.12 | 0 | NA | NA | NA | NA | NA | #VALUE! |
| ENSMUSG00000020583.5 | 0 | NA | NA | NA | NA | NA | #VALUE! |
| ENSMUSG00000020591.12 | 0 | NA | NA | NA | NA | NA | #VALUE! |
| ENSMUSG00000020598.17 | 0 | NA | NA | NA | NA | NA | #VALUE! |
| ENSMUSG00000020599.14 | 0 | NA | NA | NA | NA | NA | #VALUE! |
| ENSMUSG00000020600.14 | 0 | NA | NA | NA | NA | NA | #VALUE! |
| ENSMUSG00000020601.9 | 2.460982 | 3.793196 | 3.921438 | 0.967297 | 0.333396 | NA | #VALUE! |
| ENSMUSG00000020607.8 | 0 | NA | NA | NA | NA | NA | #VALUE! |
| ENSMUSG00000020617.14 | 0 | NA | NA | NA | NA | NA | #VALUE! |
| ENSMUSG00000020622.18 | 0 | NA | NA | NA | NA | NA | #VALUE! |
| ENSMUSG00000020623.12 | 25.88632 | -2.03782 | 2.181019 | -0.93434 | NA | NA | #VALUE! |
| ENSMUSG00000020624.4 | 0 | NA | NA | NA | NA | NA | #VALUE! |
| ENSMUSG00000020627.11 | 0 | NA | NA | NA | NA | NA | #VALUE! |
| ENSMUSG00000020633.15 | 0 | NA | NA | NA | NA | NA | #VALUE! |
| ENSMUSG00000020635.9 | 0 | NA | NA | NA | NA | NA | #VALUE! |
| ENSMUSG00000020636.15 | 0 | NA | NA | NA | NA | NA | #VALUE! |
| ENSMUSG00000020639.15 | 6.294991 | 3.937065 | 3.851261 | 1.022279 | 0.306649 | NA | #VALUE! |
| ENSMUSG00000020642.14 | 5.222291 | 5.88069 | 3.915331 | 1.501965 | 0.133106 | NA | #VALUE! |
| ENSMUSG00000020646.18 | 0 | NA | NA | NA | NA | NA | #VALUE! |
| ENSMUSG00000020648.4 | 6.583835 | -0.52751 | 3.754557 | -0.1405 | 0.888265 | NA | #VALUE! |
| ENSMUSG00000020651.9 | 5.408801 | 5.931107 | 3.915037 | 1.514955 | 0.129784 | NA | #VALUE! |
| ENSMUSG00000020652.18 | 4.47837 | 5.658661 | 3.916753 | 1.444733 | 0.148533 | NA | #VALUE! |
| ENSMUSG00000020653.13 | 7.479888 | -3.78891 | 2.808171 | -1.34924 | 0.177259 | NA | #VALUE! |
| ENSMUSG00000020654.16 | 1.721829 | 4.254739 | 3.933085 | 1.081782 | 0.27935 | NA | #VALUE! |
| ENSMUSG00000020657.17 | 0 | NA | NA | NA | NA | NA | #VALUE! |
| ENSMUSG00000020658.11 | 4.390036 | -5.66275 | 3.687207 | -1.53578 | 0.124592 | NA | #VALUE! |
| ENSMUSG00000020660.7 | 0 | NA | NA | NA | NA | NA | #VALUE! |
| ENSMUSG00000020668.11 | 2.21378 | 4.621487 | 3.927208 | 1.176787 | 0.239281 | NA | #VALUE! |
| ENSMUSG00000020673.15 | 0 | NA | NA | NA | NA | NA | #VALUE! |
| ENSMUSG00000020676.3 | 0 | NA | NA | NA | NA | NA | #VALUE! |
| ENSMUSG00000020679.12 | 9.894962 | -0.03499 | 2.620761 | -0.01335 | 0.989347 | NA | #VALUE! |
| ENSMUSG00000020681.15 | 6.123323 | -6.14178 | 3.914088 | -1.56915 | 0.116614 | NA | #VALUE! |
| ENSMUSG00000020682.18 | 0 | NA | NA | NA | NA | NA | #VALUE! |
| ENSMUSG00000020684.15 | 0 | NA | NA | NA | NA | NA | #VALUE! |
| ENSMUSG00000020686.13 | 0 | NA | NA | NA | NA | NA | #VALUE! |
| ENSMUSG00000020690.13 | 0 | NA | NA | NA | NA | NA | #VALUE! |
| ENSMUSG00000020692.15 | 0.559531 | 2.703958 | 3.982863 | 0.678898 | 0.497202 | NA | #VALUE! |

| | | | | | | | |
|---|---|---|---|---|---|---|---|
| ENSMUSG00000020695.15 | 0 | NA | NA | NA | NA | NA | #VALUE! |
| ENSMUSG00000020698.12 | 0.279766 | 1.808031 | 4.036658 | 0.447903 | 0.654223 | NA | #VALUE! |
| ENSMUSG00000020700.12 | 0 | NA | NA | NA | NA | NA | #VALUE! |
| ENSMUSG00000020701.13 | 5.690884 | 2.278906 | 3.782777 | 0.602443 | 0.54688 | NA | #VALUE! |
| ENSMUSG00000020702.14 | 0 | NA | NA | NA | NA | NA | #VALUE! |
| ENSMUSG00000020703.4 | 0 | NA | NA | NA | NA | NA | #VALUE! |
| ENSMUSG00000020704.14 | 0 | NA | NA | NA | NA | NA | #VALUE! |
| ENSMUSG00000020712.12 | 0 | NA | NA | NA | NA | NA | #VALUE! |
| ENSMUSG00000020713.7 | 0 | NA | NA | NA | NA | NA | #VALUE! |
| ENSMUSG00000020722.6 | 0 | NA | NA | NA | NA | NA | #VALUE! |
| ENSMUSG00000020723.4 | 0 | NA | NA | NA | NA | NA | #VALUE! |
| ENSMUSG00000020728.18 | 0 | NA | NA | NA | NA | NA | #VALUE! |
| ENSMUSG00000020732.14 | 0 | NA | NA | NA | NA | NA | #VALUE! |
| ENSMUSG00000020734.14 | 0 | NA | NA | NA | NA | NA | #VALUE! |
| ENSMUSG00000020738.17 | 3.935608 | 5.458007 | 3.918239 | 1.392975 | 0.163627 | NA | #VALUE! |
| ENSMUSG00000020740.14 | 7.403004 | 3.19716 | 2.843867 | 1.12423 | 0.260916 | NA | #VALUE! |
| ENSMUSG00000020744.14 | 5.242097 | -0.41133 | 3.564366 | -0.1154 | 0.908127 | NA | #VALUE! |
| ENSMUSG00000020747.19 | 3.432215 | 5.27787 | 3.91976 | 1.346478 | 0.178149 | NA | #VALUE! |
| ENSMUSG00000020758.16 | 0 | NA | NA | NA | NA | NA | #VALUE! |
| ENSMUSG00000020770.14 | 4.830604 | -5.80779 | 3.915992 | -1.4831 | 0.138049 | NA | #VALUE! |
| ENSMUSG00000020776.19 | 0 | NA | NA | NA | NA | NA | #VALUE! |
| ENSMUSG00000020781.15 | 3.150394 | -5.19405 | 3.92087 | -1.32472 | 0.185265 | NA | #VALUE! |
| ENSMUSG00000020785.18 | 0 | NA | NA | NA | NA | NA | #VALUE! |
| ENSMUSG00000020787.15 | 0 | NA | NA | NA | NA | NA | #VALUE! |
| ENSMUSG00000020788.16 | 0 | NA | NA | NA | NA | NA | #VALUE! |
| ENSMUSG00000020793.6 | 0 | NA | NA | NA | NA | NA | #VALUE! |
| ENSMUSG00000020798.15 | 0 | NA | NA | NA | NA | NA | #VALUE! |
| ENSMUSG00000020799.17 | 0 | NA | NA | NA | NA | NA | #VALUE! |
| ENSMUSG00000020802.9 | 0.504851 | 2.502843 | 3.993993 | 0.626652 | 0.530887 | NA | #VALUE! |
| ENSMUSG00000020804.13 | 0 | NA | NA | NA | NA | NA | #VALUE! |
| ENSMUSG00000020805.15 | 0.315039 | -1.93549 | 4.038621 | -0.47925 | 0.631763 | NA | #VALUE! |
| ENSMUSG00000020807.16 | 0.587367 | -2.74297 | 3.982932 | -0.68868 | 0.491024 | NA | #VALUE! |
| ENSMUSG00000020808.4 | 0 | NA | NA | NA | NA | NA | #VALUE! |
| ENSMUSG00000020811.17 | 0 | NA | NA | NA | NA | NA | #VALUE! |
| ENSMUSG00000020812.19 | 0.336567 | 1.944925 | 4.029936 | 0.482619 | 0.629366 | NA | #VALUE! |
| ENSMUSG00000020814.14 | 3.357187 | 5.246639 | 3.920044 | 1.338413 | 0.180762 | NA | #VALUE! |
| ENSMUSG00000020826.10 | 0 | NA | NA | NA | NA | NA | #VALUE! |
| ENSMUSG00000020827.19 | 0 | NA | NA | NA | NA | NA | #VALUE! |
| ENSMUSG00000020828.14 | 0 | NA | NA | NA | NA | NA | #VALUE! |
| ENSMUSG00000020831.19 | 0 | NA | NA | NA | NA | NA | #VALUE! |
| ENSMUSG00000020832.15 | 8.385382 | 1.745387 | 2.642221 | 0.660576 | 0.508884 | NA | #VALUE! |
| ENSMUSG00000020834.18 | 1.723265 | -0.66651 | 3.83672 | -0.17372 | 0.862087 | NA | #VALUE! |
| ENSMUSG00000020836.16 | 0 | NA | NA | NA | NA | NA | #VALUE! |
| ENSMUSG00000020838.13 | 0 | NA | NA | NA | NA | NA | #VALUE! |

| | | | | | | | |
|---|---|---|---|---|---|---|---|
| ENSMUSG00000020839.17 | 0 | NA | NA | NA | NA | NA | #VALUE! |
| ENSMUSG00000020844.7 | 3.077421 | 5.121959 | 3.921238 | 1.30621 | 0.191481 | NA | #VALUE! |
| ENSMUSG00000020846.7 | 0 | NA | NA | NA | NA | NA | #VALUE! |
| ENSMUSG00000020847.16 | 2.797656 | 4.98547 | 3.922669 | 1.270938 | 0.203751 | NA | #VALUE! |
| ENSMUSG00000020848.8 | 0 | NA | NA | NA | NA | NA | #VALUE! |
| ENSMUSG00000020857.12 | 0 | NA | NA | NA | NA | NA | #VALUE! |
| ENSMUSG00000020866.18 | 0 | NA | NA | NA | NA | NA | #VALUE! |
| ENSMUSG00000020867.14 | 0 | NA | NA | NA | NA | NA | #VALUE! |
| ENSMUSG00000020868.16 | 2.321964 | -3.56351 | 3.9137 | -0.91052 | 0.362547 | NA | #VALUE! |
| ENSMUSG00000020870.8 | 0 | NA | NA | NA | NA | NA | #VALUE! |
| ENSMUSG00000020871.9 | 0 | NA | NA | NA | NA | NA | #VALUE! |
| ENSMUSG00000020872.5 | 0 | NA | NA | NA | NA | NA | #VALUE! |
| ENSMUSG00000020875.10 | 0 | NA | NA | NA | NA | NA | #VALUE! |
| ENSMUSG00000020876.15 | 0 | NA | NA | NA | NA | NA | #VALUE! |
| ENSMUSG00000020878.7 | 3.496698 | -4.19344 | 3.906333 | -1.0735 | 0.283048 | NA | #VALUE! |
| ENSMUSG00000020882.18 | 0 | NA | NA | NA | NA | NA | #VALUE! |
| ENSMUSG00000020886.18 | 0 | NA | NA | NA | NA | NA | #VALUE! |
| ENSMUSG00000020887.4 | 0 | NA | NA | NA | NA | NA | #VALUE! |
| ENSMUSG00000020888.17 | 6.792164 | -0.46019 | 3.288091 | -0.13996 | 0.888695 | NA | #VALUE! |
| ENSMUSG00000020890.12 | 0 | NA | NA | NA | NA | NA | #VALUE! |
| ENSMUSG00000020891.12 | 0 | NA | NA | NA | NA | NA | #VALUE! |
| ENSMUSG00000020892.11 | 0 | NA | NA | NA | NA | NA | #VALUE! |
| ENSMUSG00000020893.18 | 9.353565 | -6.75721 | 2.729608 | -2.47552 | 0.013304 | NA | #VALUE! |
| ENSMUSG00000020894.17 | 0.300802 | -1.80634 | 4.050033 | -0.44601 | 0.655592 | NA | #VALUE! |
| ENSMUSG00000020895.15 | 4.626967 | 2.317509 | 3.736264 | 0.620274 | 0.535077 | NA | #VALUE! |
| ENSMUSG00000020897.13 | 1.762102 | -4.34067 | 3.93219 | -1.10388 | 0.269645 | NA | #VALUE! |
| ENSMUSG00000020898.19 | 1.236642 | -2.18313 | 3.923447 | -0.55643 | 0.577915 | NA | #VALUE! |
| ENSMUSG00000020899.17 | 6.521142 | -1.87231 | 3.045522 | -0.61477 | 0.538704 | NA | #VALUE! |
| ENSMUSG00000020901.14 | 0 | NA | NA | NA | NA | NA | #VALUE! |
| ENSMUSG00000020904.12 | 0 | NA | NA | NA | NA | NA | #VALUE! |
| ENSMUSG00000020905.12 | 0 | NA | NA | NA | NA | NA | #VALUE! |
| ENSMUSG00000020907.2 | 0 | NA | NA | NA | NA | NA | #VALUE! |
| ENSMUSG00000020908.15 | 0 | NA | NA | NA | NA | NA | #VALUE! |
| ENSMUSG00000020911.15 | 0 | NA | NA | NA | NA | NA | #VALUE! |
| ENSMUSG00000020912.5 | 0 | NA | NA | NA | NA | NA | #VALUE! |
| ENSMUSG00000020913.4 | 0 | NA | NA | NA | NA | NA | #VALUE! |
| ENSMUSG00000020916.9 | 0 | NA | NA | NA | NA | NA | #VALUE! |
| ENSMUSG00000020921.7 | 4.09888 | 4.56066 | 3.898478 | 1.169857 | 0.242059 | NA | #VALUE! |
| ENSMUSG00000020926.17 | 0 | NA | NA | NA | NA | NA | #VALUE! |
| ENSMUSG00000020928.15 | 0 | NA | NA | NA | NA | NA | #VALUE! |
| ENSMUSG00000020930.10 | 0 | NA | NA | NA | NA | NA | #VALUE! |
| ENSMUSG00000020932.15 | 0 | NA | NA | NA | NA | NA | #VALUE! |
| ENSMUSG00000020937.15 | 0 | NA | NA | NA | NA | NA | #VALUE! |
| ENSMUSG00000020940.14 | 0 | NA | NA | NA | NA | NA | #VALUE! |

| | | | | | | | |
|---|---|---|---|---|---|---|---|
| ENSMUSG00000020941.8 | 5.222291 | 5.88069 | 3.915331 | 1.501965 | 0.133106 | NA | #VALUE! |
| ENSMUSG00000020945.14 | 0 | NA | NA | NA | NA | NA | #VALUE! |
| ENSMUSG00000020950.11 | 0 | NA | NA | NA | NA | NA | #VALUE! |
| ENSMUSG00000020953.18 | 0 | NA | NA | NA | NA | NA | #VALUE! |
| ENSMUSG00000020963.16 | 0 | NA | NA | NA | NA | NA | #VALUE! |
| ENSMUSG00000020973.8 | 5.502056 | 5.955668 | 3.914897 | 1.521283 | 0.128189 | NA | #VALUE! |
| ENSMUSG00000020974.7 | 0 | NA | NA | NA | NA | NA | #VALUE! |
| ENSMUSG00000020990.6 | 0.735092 | -3.12016 | 3.965612 | -0.7868 | 0.431397 | NA | #VALUE! |
| ENSMUSG00000020992.6 | 0 | NA | NA | NA | NA | NA | #VALUE! |
| ENSMUSG00000021003.11 | 9.594705 | 5.71846 | 2.545674 | 2.246345 | 0.024682 | NA | #VALUE! |
| ENSMUSG00000021007.15 | 9.885104 | 0.492457 | 2.980522 | 0.165225 | 0.868767 | NA | #VALUE! |
| ENSMUSG00000021010.10 | 5.411462 | 5.919712 | 3.915102 | 1.51202 | 0.130529 | NA | #VALUE! |
| ENSMUSG00000021013.17 | 2.430165 | 0.973684 | 3.810369 | 0.255535 | 0.79831 | NA | #VALUE! |
| ENSMUSG00000021032.14 | 0 | NA | NA | NA | NA | NA | #VALUE! |
| ENSMUSG00000021047.8 | 0 | NA | NA | NA | NA | NA | #VALUE! |
| ENSMUSG00000021055.15 | 0 | NA | NA | NA | NA | NA | #VALUE! |
| ENSMUSG00000021056.9 | 0 | NA | NA | NA | NA | NA | #VALUE! |
| ENSMUSG00000021057.16 | 0 | NA | NA | NA | NA | NA | #VALUE! |
| ENSMUSG00000021061.16 | 0 | NA | NA | NA | NA | NA | #VALUE! |
| ENSMUSG00000021062.16 | 0 | NA | NA | NA | NA | NA | #VALUE! |
| ENSMUSG00000021066.10 | 3.423691 | -4.17323 | 3.906523 | -1.06827 | 0.285398 | NA | #VALUE! |
| ENSMUSG00000021068.18 | 0.804831 | -0.79862 | 3.943253 | -0.20253 | 0.839504 | NA | #VALUE! |
| ENSMUSG00000021070.7 | 0 | NA | NA | NA | NA | NA | #VALUE! |
| ENSMUSG00000021071.18 | 0 | NA | NA | NA | NA | NA | #VALUE! |
| ENSMUSG00000021078.6 | 1.75848 | -0.49659 | 3.83513 | -0.12948 | 0.896974 | NA | #VALUE! |
| ENSMUSG00000021081.13 | 0 | NA | NA | NA | NA | NA | #VALUE! |
| ENSMUSG00000021086.6 | 1.212318 | 3.793111 | 3.942915 | 0.962007 | 0.336046 | NA | #VALUE! |
| ENSMUSG00000021090.17 | 3.885486 | -5.49502 | 3.91822 | -1.40243 | 0.160787 | NA | #VALUE! |
| ENSMUSG00000021095.6 | 0 | NA | NA | NA | NA | NA | #VALUE! |
| ENSMUSG00000021098.15 | 0 | NA | NA | NA | NA | NA | #VALUE! |
| ENSMUSG00000021099.7 | 0 | NA | NA | NA | NA | NA | #VALUE! |
| ENSMUSG00000021101.3 | 0 | NA | NA | NA | NA | NA | #VALUE! |
| ENSMUSG00000021108.19 | 7.148609 | 0.429164 | 3.652244 | 0.117507 | 0.906458 | NA | #VALUE! |
| ENSMUSG00000021118.8 | 0 | NA | NA | NA | NA | NA | #VALUE! |
| ENSMUSG00000021120.12 | 6.394774 | 6.166745 | 3.913792 | 1.575645 | 0.115108 | NA | #VALUE! |
| ENSMUSG00000021123.13 | 0 | NA | NA | NA | NA | NA | #VALUE! |
| ENSMUSG00000021125.7 | 0.746042 | 3.107177 | 3.964586 | 0.783733 | 0.433197 | NA | #VALUE! |
| ENSMUSG00000021130.9 | 0 | NA | NA | NA | NA | NA | #VALUE! |
| ENSMUSG00000021139.18 | 0 | NA | NA | NA | NA | NA | #VALUE! |
| ENSMUSG00000021148.12 | 0 | NA | NA | NA | NA | NA | #VALUE! |
| ENSMUSG00000021175.16 | 0 | NA | NA | NA | NA | NA | #VALUE! |
| ENSMUSG00000021176.7 | 3.061661 | -5.13931 | 3.921414 | -1.31058 | 0.190001 | NA | #VALUE! |
| ENSMUSG00000021177.17 | 9.883798 | 4.602229 | 2.958214 | 1.555746 | NA | NA | #VALUE! |
| ENSMUSG00000021182.17 | 2.940368 | -5.09511 | 3.921868 | -1.29915 | 0.193891 | NA | #VALUE! |

| | | | | | | | |
|---|---|---|---|---|---|---|---|
| ENSMUSG00000021194.7 | 0 | NA | NA | NA | NA | NA | #VALUE! |
| ENSMUSG00000021196.15 | 6.253129 | -5.0797 | 3.452679 | -1.47123 | 0.141228 | NA | #VALUE! |
| ENSMUSG00000021198.17 | 0 | NA | NA | NA | NA | NA | #VALUE! |
| ENSMUSG00000021200.15 | 3.990499 | -5.53332 | 3.917921 | -1.41231 | 0.157859 | NA | #VALUE! |
| ENSMUSG00000021203.16 | 1.989334 | -4.5335 | 3.929023 | -1.15385 | 0.248562 | NA | #VALUE! |
| ENSMUSG00000021207.10 | 0 | NA | NA | NA | NA | NA | #VALUE! |
| ENSMUSG00000021214.15 | 0 | NA | NA | NA | NA | NA | #VALUE! |
| ENSMUSG00000021216.8 | 0 | NA | NA | NA | NA | NA | #VALUE! |
| ENSMUSG00000021217.8 | 4.410551 | -5.67707 | 3.916865 | -1.44939 | 0.147228 | NA | #VALUE! |
| ENSMUSG00000021219.17 | 2.940368 | -5.09511 | 3.921868 | -1.29915 | 0.193891 | NA | #VALUE! |
| ENSMUSG00000021221.16 | 0 | NA | NA | NA | NA | NA | #VALUE! |
| ENSMUSG00000021222.10 | 5.697487 | 0.878627 | 3.760835 | 0.233625 | 0.815276 | NA | #VALUE! |
| ENSMUSG00000021223.14 | 0 | NA | NA | NA | NA | NA | #VALUE! |
| ENSMUSG00000021234.10 | 1.836997 | -4.39907 | 3.931186 | -1.11902 | 0.263133 | NA | #VALUE! |
| ENSMUSG00000021235.14 | 7.222697 | 0.988527 | 2.967803 | 0.333084 | 0.739071 | NA | #VALUE! |
| ENSMUSG00000021239.13 | 0 | NA | NA | NA | NA | NA | #VALUE! |
| ENSMUSG00000021240.7 | 9.720351 | -0.95665 | 2.587708 | -0.36969 | 0.711613 | NA | #VALUE! |
| ENSMUSG00000021250.14 | 368.2245 | -5.44792 | 3.73081 | -1.46025 | NA | NA | #VALUE! |
| ENSMUSG00000021253.8 | 0 | NA | NA | NA | NA | NA | #VALUE! |
| ENSMUSG00000021255.18 | 2.763323 | 4.962925 | 3.922918 | 1.265111 | 0.205832 | NA | #VALUE! |
| ENSMUSG00000021256.6 | 0 | NA | NA | NA | NA | NA | #VALUE! |
| ENSMUSG00000021259.6 | 0 | NA | NA | NA | NA | NA | #VALUE! |
| ENSMUSG00000021260.9 | 3.870521 | 5.442409 | 3.918364 | 1.388949 | 0.164848 | NA | #VALUE! |
| ENSMUSG00000021263.12 | 0 | NA | NA | NA | NA | NA | #VALUE! |
| ENSMUSG00000021265.4 | 0 | NA | NA | NA | NA | NA | #VALUE! |
| ENSMUSG00000021276.11 | 45.61904 | -0.28656 | 1.376117 | -0.20824 | NA | NA | #VALUE! |
| ENSMUSG00000021278.8 | 0 | NA | NA | NA | NA | NA | #VALUE! |
| ENSMUSG00000021280.10 | 0.612332 | -2.79827 | 3.980109 | -0.70306 | 0.482016 | NA | #VALUE! |
| ENSMUSG00000021287.13 | 1.212318 | 3.793111 | 3.942915 | 0.962007 | 0.336046 | NA | #VALUE! |
| ENSMUSG00000021294.8 | 0 | NA | NA | NA | NA | NA | #VALUE! |
| ENSMUSG00000021298.9 | 0 | NA | NA | NA | NA | NA | #VALUE! |
| ENSMUSG00000021301.10 | 0 | NA | NA | NA | NA | NA | #VALUE! |
| ENSMUSG00000021303.15 | 0 | NA | NA | NA | NA | NA | #VALUE! |
| ENSMUSG00000021313.17 | 0 | NA | NA | NA | NA | NA | #VALUE! |
| ENSMUSG00000021314.14 | 0 | NA | NA | NA | NA | NA | #VALUE! |
| ENSMUSG00000021318.16 | 1.365171 | -3.99822 | 3.938957 | -1.01505 | 0.310084 | NA | #VALUE! |
| ENSMUSG00000021319.8 | 0 | NA | NA | NA | NA | NA | #VALUE! |
| ENSMUSG00000021322.9 | 9.728416 | -1.39053 | 3.334959 | -0.41696 | 0.67671 | NA | #VALUE! |
| ENSMUSG00000021337.9 | 0 | NA | NA | NA | NA | NA | #VALUE! |
| ENSMUSG00000021342.15 | 0 | NA | NA | NA | NA | NA | #VALUE! |
| ENSMUSG00000021345.17 | 0 | NA | NA | NA | NA | NA | #VALUE! |
| ENSMUSG00000021346.15 | 0 | NA | NA | NA | NA | NA | #VALUE! |
| ENSMUSG00000021347.4 | 0 | NA | NA | NA | NA | NA | #VALUE! |
| ENSMUSG00000021348.4 | 0 | NA | NA | NA | NA | NA | #VALUE! |

| | | | | | | | |
|---|---|---|---|---|---|---|---|
| ENSMUSG00000021356.11 | 0 | NA | NA | NA | NA | NA | #VALUE! |
| ENSMUSG00000021359.17 | 0 | NA | NA | NA | NA | NA | #VALUE! |
| ENSMUSG00000021362.8 | 0 | NA | NA | NA | NA | NA | #VALUE! |
| ENSMUSG00000021363.15 | 0 | NA | NA | NA | NA | NA | #VALUE! |
| ENSMUSG00000021367.9 | 0 | NA | NA | NA | NA | NA | #VALUE! |
| ENSMUSG00000021373.17 | 0 | NA | NA | NA | NA | NA | #VALUE! |
| ENSMUSG00000021379.3 | 2.99874 | 1.344426 | 3.524602 | 0.381441 | 0.702876 | NA | #VALUE! |
| ENSMUSG00000021381.5 | 0 | NA | NA | NA | NA | NA | #VALUE! |
| ENSMUSG00000021384.15 | 0 | NA | NA | NA | NA | NA | #VALUE! |
| ENSMUSG00000021388.15 | 0 | NA | NA | NA | NA | NA | #VALUE! |
| ENSMUSG00000021390.7 | 5.373305 | 1.499085 | 3.153266 | 0.475407 | 0.634497 | NA | #VALUE! |
| ENSMUSG00000021391.11 | 0 | NA | NA | NA | NA | NA | #VALUE! |
| ENSMUSG00000021396.6 | 0 | NA | NA | NA | NA | NA | #VALUE! |
| ENSMUSG00000021403.4 | 0 | NA | NA | NA | NA | NA | #VALUE! |
| ENSMUSG00000021404.13 | 0 | NA | NA | NA | NA | NA | #VALUE! |
| ENSMUSG00000021414.9 | 2.21378 | 4.621487 | 3.927208 | 1.176787 | 0.239281 | NA | #VALUE! |
| ENSMUSG00000021415.14 | 0 | NA | NA | NA | NA | NA | #VALUE! |
| ENSMUSG00000021416.12 | 0.18651 | 1.401769 | 4.056354 | 0.345574 | 0.729663 | NA | #VALUE! |
| ENSMUSG00000021439.9 | 0 | NA | NA | NA | NA | NA | #VALUE! |
| ENSMUSG00000021440.8 | 0 | NA | NA | NA | NA | NA | #VALUE! |
| ENSMUSG00000021441.5 | 0 | NA | NA | NA | NA | NA | #VALUE! |
| ENSMUSG00000021448.9 | 0 | NA | NA | NA | NA | NA | #VALUE! |
| ENSMUSG00000021451.17 | 0 | NA | NA | NA | NA | NA | #VALUE! |
| ENSMUSG00000021453.3 | 76.31943 | -2.13538 | 1.675077 | -1.27479 | NA | NA | #VALUE! |
| ENSMUSG00000021456.9 | 0 | NA | NA | NA | NA | NA | #VALUE! |
| ENSMUSG00000021457.15 | 0.420053 | -2.33493 | 4.007343 | -0.58266 | 0.56012 | NA | #VALUE! |
| ENSMUSG00000021461.18 | 0.978946 | -3.48749 | 3.952505 | -0.88235 | 0.377588 | NA | #VALUE! |
| ENSMUSG00000021464.16 | 0 | NA | NA | NA | NA | NA | #VALUE! |
| ENSMUSG00000021469.10 | 0 | NA | NA | NA | NA | NA | #VALUE! |
| ENSMUSG00000021478.7 | 0 | NA | NA | NA | NA | NA | #VALUE! |
| ENSMUSG00000021481.12 | 6.969343 | -6.32942 | 3.540516 | -1.78771 | 0.073823 | NA | #VALUE! |
| ENSMUSG00000021485.15 | 0 | NA | NA | NA | NA | NA | #VALUE! |
| ENSMUSG00000021490.12 | 0 | NA | NA | NA | NA | NA | #VALUE! |
| ENSMUSG00000021493.16 | 0 | NA | NA | NA | NA | NA | #VALUE! |
| ENSMUSG00000021499.13 | 0.336567 | 1.944925 | 4.029936 | 0.482619 | 0.629366 | NA | #VALUE! |
| ENSMUSG00000021506.8 | 0 | NA | NA | NA | NA | NA | #VALUE! |
| ENSMUSG00000021534.9 | 0 | NA | NA | NA | NA | NA | #VALUE! |
| ENSMUSG00000021536.8 | 0 | NA | NA | NA | NA | NA | #VALUE! |
| ENSMUSG00000021538.4 | 0 | NA | NA | NA | NA | NA | #VALUE! |
| ENSMUSG00000021541.15 | 0 | NA | NA | NA | NA | NA | #VALUE! |
| ENSMUSG00000021545.6 | 0 | NA | NA | NA | NA | NA | #VALUE! |
| ENSMUSG00000021553.15 | 0 | NA | NA | NA | NA | NA | #VALUE! |
| ENSMUSG00000021556.13 | 0.510828 | -2.58279 | 3.991728 | -0.64704 | 0.517609 | NA | #VALUE! |
| ENSMUSG00000021565.17 | 0 | NA | NA | NA | NA | NA | #VALUE! |

| | | | | | | | |
|---|---|---|---|---|---|---|---|
| ENSMUSG00000021566.8 | 0 | NA | NA | NA | NA | NA | #VALUE! |
| ENSMUSG00000021567.16 | 0 | NA | NA | NA | NA | NA | #VALUE! |
| ENSMUSG00000021569.11 | 5.647193 | -4.54206 | 3.889786 | -1.16769 | 0.242932 | NA | #VALUE! |
| ENSMUSG00000021572.10 | 2.352719 | 0.013891 | 3.804191 | 0.003652 | 0.997086 | NA | #VALUE! |
| ENSMUSG00000021573.16 | 37.34293 | -0.60566 | 1.813619 | -0.33395 | NA | NA | #VALUE! |
| ENSMUSG00000021575.17 | 0 | NA | NA | NA | NA | NA | #VALUE! |
| ENSMUSG00000021579.5 | 0 | NA | NA | NA | NA | NA | #VALUE! |
| ENSMUSG00000021587.6 | 0 | NA | NA | NA | NA | NA | #VALUE! |
| ENSMUSG00000021590.8 | 0 | NA | NA | NA | NA | NA | #VALUE! |
| ENSMUSG00000021596.17 | 0.841418 | 3.240539 | 3.959551 | 0.818411 | 0.413123 | NA | #VALUE! |
| ENSMUSG00000021600.6 | 0 | NA | NA | NA | NA | NA | #VALUE! |
| ENSMUSG00000021604.16 | 0 | NA | NA | NA | NA | NA | #VALUE! |
| ENSMUSG00000021608.16 | 8.097824 | 3.795844 | 3.494521 | 1.086227 | 0.277378 | NA | #VALUE! |
| ENSMUSG00000021609.7 | 0 | NA | NA | NA | NA | NA | #VALUE! |
| ENSMUSG00000021612.16 | 0 | NA | NA | NA | NA | NA | #VALUE! |
| ENSMUSG00000021613.10 | 0 | NA | NA | NA | NA | NA | #VALUE! |
| ENSMUSG00000021614.17 | 0 | NA | NA | NA | NA | NA | #VALUE! |
| ENSMUSG00000021622.4 | 0 | NA | NA | NA | NA | NA | #VALUE! |
| ENSMUSG00000021624.10 | 0 | NA | NA | NA | NA | NA | #VALUE! |
| ENSMUSG00000021631.9 | 0 | NA | NA | NA | NA | NA | #VALUE! |
| ENSMUSG00000021640.7 | 5.715101 | -6.04207 | 3.914611 | -1.54347 | 0.122718 | NA | #VALUE! |
| ENSMUSG00000021647.8 | 0 | NA | NA | NA | NA | NA | #VALUE! |
| ENSMUSG00000021662.12 | 2.520315 | -4.8742 | 3.924358 | -1.24204 | 0.214223 | NA | #VALUE! |
| ENSMUSG00000021675.5 | 0 | NA | NA | NA | NA | NA | #VALUE! |
| ENSMUSG00000021678.10 | 0 | NA | NA | NA | NA | NA | #VALUE! |
| ENSMUSG00000021679.5 | 0 | NA | NA | NA | NA | NA | #VALUE! |
| ENSMUSG00000021680.9 | 0 | NA | NA | NA | NA | NA | #VALUE! |
| ENSMUSG00000021684.18 | 1.110947 | 3.632223 | 3.947134 | 0.920218 | 0.357459 | NA | #VALUE! |
| ENSMUSG00000021685.13 | 0 | NA | NA | NA | NA | NA | #VALUE! |
| ENSMUSG00000021692.10 | 4.127485 | -1.14189 | 3.511754 | -0.32516 | 0.745058 | NA | #VALUE! |
| ENSMUSG00000021696.10 | 0 | NA | NA | NA | NA | NA | #VALUE! |
| ENSMUSG00000021697.13 | 0 | NA | NA | NA | NA | NA | #VALUE! |
| ENSMUSG00000021699.18 | 0 | NA | NA | NA | NA | NA | #VALUE! |
| ENSMUSG00000021700.11 | 0 | NA | NA | NA | NA | NA | #VALUE! |
| ENSMUSG00000021702.8 | 0 | NA | NA | NA | NA | NA | #VALUE! |
| ENSMUSG00000021708.19 | 0.84294 | 2.074799 | 3.964343 | 0.523365 | 0.60072 | NA | #VALUE! |
| ENSMUSG00000021714.16 | 7.143308 | 0.150293 | 3.338446 | 0.045019 | 0.964092 | NA | #VALUE! |
| ENSMUSG00000021718.6 | 0 | NA | NA | NA | NA | NA | #VALUE! |
| ENSMUSG00000021719.10 | 1.585338 | 4.174397 | 3.934581 | 1.060951 | 0.288712 | NA | #VALUE! |
| ENSMUSG00000021720.12 | 0.587367 | -2.74297 | 3.982932 | -0.68868 | 0.491024 | NA | #VALUE! |
| ENSMUSG00000021721.6 | 0 | NA | NA | NA | NA | NA | #VALUE! |
| ENSMUSG00000021730.9 | 0 | NA | NA | NA | NA | NA | #VALUE! |
| ENSMUSG00000021732.15 | 0 | NA | NA | NA | NA | NA | #VALUE! |
| ENSMUSG00000021741.5 | 0.840707 | 0.109948 | 3.943962 | 0.027877 | 0.97776 | NA | #VALUE! |

| | | | | | | | |
|---|---|---|---|---|---|---|---|
| ENSMUSG00000021743.7 | 0 | NA | NA | NA | NA | NA | #VALUE! |
| ENSMUSG00000021749.7 | 0 | NA | NA | NA | NA | NA | #VALUE! |
| ENSMUSG00000021750.17 | 0 | NA | NA | NA | NA | NA | #VALUE! |
| ENSMUSG00000021758.15 | 0 | NA | NA | NA | NA | NA | #VALUE! |
| ENSMUSG00000021760.6 | 1.271783 | 3.851779 | 3.941488 | 0.97724 | 0.32845 | NA | #VALUE! |
| ENSMUSG00000021763.18 | 0 | NA | NA | NA | NA | NA | #VALUE! |
| ENSMUSG00000021768.16 | 0 | NA | NA | NA | NA | NA | #VALUE! |
| ENSMUSG00000021773.12 | 2.524253 | 4.825754 | 3.924522 | 1.229641 | 0.218831 | NA | #VALUE! |
| ENSMUSG00000021789.15 | 0.816443 | -3.21914 | 3.961753 | -0.81255 | 0.416474 | NA | #VALUE! |
| ENSMUSG00000021790.13 | 0 | NA | NA | NA | NA | NA | #VALUE! |
| ENSMUSG00000021791.8 | 0 | NA | NA | NA | NA | NA | #VALUE! |
| ENSMUSG00000021795.11 | 0 | NA | NA | NA | NA | NA | #VALUE! |
| ENSMUSG00000021797.3 | 0 | NA | NA | NA | NA | NA | #VALUE! |
| ENSMUSG00000021798.15 | 0 | NA | NA | NA | NA | NA | #VALUE! |
| ENSMUSG00000021799.10 | 0 | NA | NA | NA | NA | NA | #VALUE! |
| ENSMUSG00000021803.11 | 0 | NA | NA | NA | NA | NA | #VALUE! |
| ENSMUSG00000021804.7 | 0 | NA | NA | NA | NA | NA | #VALUE! |
| ENSMUSG00000021815.5 | 0 | NA | NA | NA | NA | NA | #VALUE! |
| ENSMUSG00000021819.12 | 0.420053 | -2.33493 | 4.007343 | -0.58266 | 0.56012 | NA | #VALUE! |
| ENSMUSG00000021822.4 | 0 | NA | NA | NA | NA | NA | #VALUE! |
| ENSMUSG00000021838.18 | 4.91951 | 5.781601 | 3.915938 | 1.476428 | 0.139829 | NA | #VALUE! |
| ENSMUSG00000021848.17 | 0 | NA | NA | NA | NA | NA | #VALUE! |
| ENSMUSG00000021850.15 | 0 | NA | NA | NA | NA | NA | #VALUE! |
| ENSMUSG00000021852.15 | 0 | NA | NA | NA | NA | NA | #VALUE! |
| ENSMUSG00000021871.20 | 0 | NA | NA | NA | NA | NA | #VALUE! |
| ENSMUSG00000021872.9 | 4.66276 | 5.717927 | 3.916352 | 1.460014 | 0.144286 | NA | #VALUE! |
| ENSMUSG00000021874.13 | 0 | NA | NA | NA | NA | NA | #VALUE! |
| ENSMUSG00000021879.14 | 0 | NA | NA | NA | NA | NA | #VALUE! |
| ENSMUSG00000021880.8 | 0.420053 | -2.33493 | 4.007343 | -0.58266 | 0.56012 | NA | #VALUE! |
| ENSMUSG00000021886.8 | 8.689418 | -4.06413 | 3.419321 | -1.18858 | 0.234606 | NA | #VALUE! |
| ENSMUSG00000021898.17 | 0 | NA | NA | NA | NA | NA | #VALUE! |
| ENSMUSG00000021903.12 | 3.443657 | 5.264174 | 3.919884 | 1.342941 | 0.179291 | NA | #VALUE! |
| ENSMUSG00000021904.7 | 0 | NA | NA | NA | NA | NA | #VALUE! |
| ENSMUSG00000021907.14 | 0 | NA | NA | NA | NA | NA | #VALUE! |
| ENSMUSG00000021908.11 | 0 | NA | NA | NA | NA | NA | #VALUE! |
| ENSMUSG00000021913.9 | 0 | NA | NA | NA | NA | NA | #VALUE! |
| ENSMUSG00000021919.9 | 0 | NA | NA | NA | NA | NA | #VALUE! |
| ENSMUSG00000021927.8 | 0 | NA | NA | NA | NA | NA | #VALUE! |
| ENSMUSG00000021933.12 | 0 | NA | NA | NA | NA | NA | #VALUE! |
| ENSMUSG00000021940.11 | 0 | NA | NA | NA | NA | NA | #VALUE! |
| ENSMUSG00000021950.16 | 0 | NA | NA | NA | NA | NA | #VALUE! |
| ENSMUSG00000021953.15 | 0 | NA | NA | NA | NA | NA | #VALUE! |
| ENSMUSG00000021961.7 | 0 | NA | NA | NA | NA | NA | #VALUE! |
| ENSMUSG00000021965.4 | 50.82358 | -23.3141 | 3.907647 | -5.96627 | NA | NA | #VALUE! |

| | | | | | | | |
|---|---|---|---|---|---|---|---|
| ENSMUSG00000021966.8 | 0 | NA | NA | NA | NA | NA | #VALUE! |
| ENSMUSG00000021974.16 | 0 | NA | NA | NA | NA | NA | #VALUE! |
| ENSMUSG00000021977.12 | 0 | NA | NA | NA | NA | NA | #VALUE! |
| ENSMUSG00000021978.10 | 6.510992 | -6.23349 | 3.116906 | -1.9999 | 0.045511 | NA | #VALUE! |
| ENSMUSG00000021983.17 | 0 | NA | NA | NA | NA | NA | #VALUE! |
| ENSMUSG00000021986.10 | 0.391578 | -2.1489 | 4.020883 | -0.53444 | 0.59304 | NA | #VALUE! |
| ENSMUSG00000021991.10 | 0 | NA | NA | NA | NA | NA | #VALUE! |
| ENSMUSG00000021994.16 | 5.944689 | -1.67164 | 3.415987 | -0.48936 | 0.624588 | NA | #VALUE! |
| ENSMUSG00000021997.5 | 0 | NA | NA | NA | NA | NA | #VALUE! |
| ENSMUSG00000022002.3 | 0 | NA | NA | NA | NA | NA | #VALUE! |
| ENSMUSG00000022012.10 | 0 | NA | NA | NA | NA | NA | #VALUE! |
| ENSMUSG00000022015.9 | 0 | NA | NA | NA | NA | NA | #VALUE! |
| ENSMUSG00000022018.8 | 0 | NA | NA | NA | NA | NA | #VALUE! |
| ENSMUSG00000022021.15 | 1.050131 | -3.6251 | 3.948379 | -0.91812 | 0.358554 | NA | #VALUE! |
| ENSMUSG00000022026.8 | 0 | NA | NA | NA | NA | NA | #VALUE! |
| ENSMUSG00000022033.10 | 0 | NA | NA | NA | NA | NA | #VALUE! |
| ENSMUSG00000022034.11 | 0.783156 | -3.1627 | 3.963922 | -0.79787 | 0.424945 | NA | #VALUE! |
| ENSMUSG00000022039.7 | 0 | NA | NA | NA | NA | NA | #VALUE! |
| ENSMUSG00000022043.20 | 9.752739 | 1.490537 | 2.941372 | 0.506749 | 0.612331 | NA | #VALUE! |
| ENSMUSG00000022044.15 | 0 | NA | NA | NA | NA | NA | #VALUE! |
| ENSMUSG00000022053.14 | 0 | NA | NA | NA | NA | NA | #VALUE! |
| ENSMUSG00000022054.12 | 0 | NA | NA | NA | NA | NA | #VALUE! |
| ENSMUSG00000022055.8 | 0 | NA | NA | NA | NA | NA | #VALUE! |
| ENSMUSG00000022056.10 | 0 | NA | NA | NA | NA | NA | #VALUE! |
| ENSMUSG00000022061.9 | 0 | NA | NA | NA | NA | NA | #VALUE! |
| ENSMUSG00000022064.11 | 2.061063 | -1.71234 | 3.848282 | -0.44496 | 0.656347 | NA | #VALUE! |
| ENSMUSG00000022066.17 | 0 | NA | NA | NA | NA | NA | #VALUE! |
| ENSMUSG00000022074.7 | 3.878104 | -5.48139 | 3.918329 | -1.39891 | 0.16184 | NA | #VALUE! |
| ENSMUSG00000022075.8 | 6.306529 | 2.5603 | 3.456945 | 0.740625 | 0.458921 | NA | #VALUE! |
| ENSMUSG00000022076.11 | 0 | NA | NA | NA | NA | NA | #VALUE! |
| ENSMUSG00000022085.5 | 0 | NA | NA | NA | NA | NA | #VALUE! |
| ENSMUSG00000022091.7 | 0 | NA | NA | NA | NA | NA | #VALUE! |
| ENSMUSG00000022096.15 | 0 | NA | NA | NA | NA | NA | #VALUE! |
| ENSMUSG00000022097.5 | 0 | NA | NA | NA | NA | NA | #VALUE! |
| ENSMUSG00000022099.18 | 0 | NA | NA | NA | NA | NA | #VALUE! |
| ENSMUSG00000022101.7 | 0 | NA | NA | NA | NA | NA | #VALUE! |
| ENSMUSG00000022102.14 | 1.365171 | -3.99822 | 3.938957 | -1.01505 | 0.310084 | NA | #VALUE! |
| ENSMUSG00000022103.11 | 0 | NA | NA | NA | NA | NA | #VALUE! |
| ENSMUSG00000022105.7 | 41.63182 | -0.30906 | 1.712335 | -0.18049 | NA | NA | #VALUE! |
| ENSMUSG00000022112.15 | 0 | NA | NA | NA | NA | NA | #VALUE! |
| ENSMUSG00000022113.7 | 0 | NA | NA | NA | NA | NA | #VALUE! |
| ENSMUSG00000022114.6 | 2.59073 | -1.14796 | 3.807663 | -0.30149 | 0.763042 | NA | #VALUE! |
| ENSMUSG00000022116.8 | 0 | NA | NA | NA | NA | NA | #VALUE! |
| ENSMUSG00000022120.5 | 0 | NA | NA | NA | NA | NA | #VALUE! |

| | | | | | | | |
|---|---|---|---|---|---|---|---|
| ENSMUSG00000022123.10 | 0 | NA | NA | NA | NA | NA | #VALUE! |
| ENSMUSG00000022126.7 | 3.524204 | -5.34395 | 3.919482 | -1.36343 | 0.172746 | NA | #VALUE! |
| ENSMUSG00000022132.16 | 0 | NA | NA | NA | NA | NA | #VALUE! |
| ENSMUSG00000022144.5 | 0 | NA | NA | NA | NA | NA | #VALUE! |
| ENSMUSG00000022146.13 | 12.06762 | 1.694648 | 3.105917 | 0.545619 | NA | NA | #VALUE! |
| ENSMUSG00000022155.10 | 0 | NA | NA | NA | NA | NA | #VALUE! |
| ENSMUSG00000022156.9 | 0 | NA | NA | NA | NA | NA | #VALUE! |
| ENSMUSG00000022157.9 | 0 | NA | NA | NA | NA | NA | #VALUE! |
| ENSMUSG00000022159.17 | 3.357187 | 5.246639 | 3.920044 | 1.338413 | 0.180762 | NA | #VALUE! |
| ENSMUSG00000022176.12 | 0 | NA | NA | NA | NA | NA | #VALUE! |
| ENSMUSG00000022179.6 | 6.957686 | 6.293813 | 3.58814 | 1.75406 | 0.07942 | NA | #VALUE! |
| ENSMUSG00000022180.8 | 1.900378 | 2.279131 | 3.891159 | 0.58572 | 0.558063 | NA | #VALUE! |
| ENSMUSG00000022187.3 | 0 | NA | NA | NA | NA | NA | #VALUE! |
| ENSMUSG00000022194.16 | 0 | NA | NA | NA | NA | NA | #VALUE! |
| ENSMUSG00000022199.13 | 0 | NA | NA | NA | NA | NA | #VALUE! |
| ENSMUSG00000022203.8 | 0 | NA | NA | NA | NA | NA | #VALUE! |
| ENSMUSG00000022206.8 | 1.356923 | 3.919485 | 3.93991 | 0.994816 | 0.319826 | NA | #VALUE! |
| ENSMUSG00000022208.13 | 0 | NA | NA | NA | NA | NA | #VALUE! |
| ENSMUSG00000022209.13 | 0 | NA | NA | NA | NA | NA | #VALUE! |
| ENSMUSG00000022211.10 | 0 | NA | NA | NA | NA | NA | #VALUE! |
| ENSMUSG00000022212.16 | 0 | NA | NA | NA | NA | NA | #VALUE! |
| ENSMUSG00000022218.17 | 8.077609 | 6.503772 | 3.856709 | 1.686353 | 0.091728 | NA | #VALUE! |
| ENSMUSG00000022220.15 | 1.305573 | 3.898287 | 3.940396 | 0.989314 | 0.32251 | NA | #VALUE! |
| ENSMUSG00000022221.15 | 6.057585 | -0.1136 | 3.498134 | -0.03247 | 0.974093 | NA | #VALUE! |
| ENSMUSG00000022223.10 | 15.91018 | 7.486057 | 2.797199 | 2.676269 | NA | NA | #VALUE! |
| ENSMUSG00000022225.8 | 0 | NA | NA | NA | NA | NA | #VALUE! |
| ENSMUSG00000022226.7 | 0 | NA | NA | NA | NA | NA | #VALUE! |
| ENSMUSG00000022227.6 | 0 | NA | NA | NA | NA | NA | #VALUE! |
| ENSMUSG00000022229.4 | 0 | NA | NA | NA | NA | NA | #VALUE! |
| ENSMUSG00000022231.11 | 0 | NA | NA | NA | NA | NA | #VALUE! |
| ENSMUSG00000022240.11 | 0.168284 | 1.027628 | 4.067126 | 0.252667 | 0.800526 | NA | #VALUE! |
| ENSMUSG00000022243.11 | 0.673134 | 2.918386 | 3.972531 | 0.734641 | 0.462558 | NA | #VALUE! |
| ENSMUSG00000022245.16 | 0 | NA | NA | NA | NA | NA | #VALUE! |
| ENSMUSG00000022249.15 | 0 | NA | NA | NA | NA | NA | #VALUE! |
| ENSMUSG00000022262.9 | 1.346268 | 3.918821 | 3.939925 | 0.994644 | 0.31991 | NA | #VALUE! |
| ENSMUSG00000022269.14 | 0 | NA | NA | NA | NA | NA | #VALUE! |
| ENSMUSG00000022286.17 | 0 | NA | NA | NA | NA | NA | #VALUE! |
| ENSMUSG00000022288.2 | 0 | NA | NA | NA | NA | NA | #VALUE! |
| ENSMUSG00000022296.11 | 0.408222 | -2.20205 | 4.016842 | -0.54821 | 0.583551 | NA | #VALUE! |
| ENSMUSG00000022297.16 | 3.344143 | 3.193518 | 3.878005 | 0.823495 | 0.410227 | NA | #VALUE! |
| ENSMUSG00000022303.10 | 0 | NA | NA | NA | NA | NA | #VALUE! |
| ENSMUSG00000022306.10 | 1.738059 | 4.300351 | 3.932271 | 1.093605 | 0.274128 | NA | #VALUE! |
| ENSMUSG00000022309.10 | 0.746042 | 3.107177 | 3.964586 | 0.783733 | 0.433197 | NA | #VALUE! |
| ENSMUSG00000022311.16 | 0 | NA | NA | NA | NA | NA | #VALUE! |

| | | | | | | | |
|---|---|---|---|---|---|---|---|
| ENSMUSG00000022315.4 | 0 | NA | NA | NA | NA | NA | #VALUE! |
| ENSMUSG00000022321.16 | 0 | NA | NA | NA | NA | NA | #VALUE! |
| ENSMUSG00000022322.9 | 0 | NA | NA | NA | NA | NA | #VALUE! |
| ENSMUSG00000022324.16 | 2.60352 | -1.05992 | 3.806558 | -0.27845 | 0.78067 | NA | #VALUE! |
| ENSMUSG00000022325.15 | 0.504851 | 2.502843 | 3.993993 | 0.626652 | 0.530887 | NA | #VALUE! |
| ENSMUSG00000022330.6 | 0 | NA | NA | NA | NA | NA | #VALUE! |
| ENSMUSG00000022335.18 | 0.541304 | 2.641495 | 3.986162 | 0.662666 | 0.507544 | NA | #VALUE! |
| ENSMUSG00000022340.16 | 0.18651 | 1.401769 | 4.056354 | 0.345574 | 0.729663 | NA | #VALUE! |
| ENSMUSG00000022342.7 | 0 | NA | NA | NA | NA | NA | #VALUE! |
| ENSMUSG00000022347.9 | 0 | NA | NA | NA | NA | NA | #VALUE! |
| ENSMUSG00000022357.3 | 0 | NA | NA | NA | NA | NA | #VALUE! |
| ENSMUSG00000022358.8 | 0.652786 | 2.919659 | 3.972474 | 0.734972 | 0.462356 | NA | #VALUE! |
| ENSMUSG00000022362.14 | 0 | NA | NA | NA | NA | NA | #VALUE! |
| ENSMUSG00000022366.13 | 0 | NA | NA | NA | NA | NA | #VALUE! |
| ENSMUSG00000022367.8 | 0 | NA | NA | NA | NA | NA | #VALUE! |
| ENSMUSG00000022369.14 | 0 | NA | NA | NA | NA | NA | #VALUE! |
| ENSMUSG00000022375.7 | 0 | NA | NA | NA | NA | NA | #VALUE! |
| ENSMUSG00000022376.9 | 0 | NA | NA | NA | NA | NA | #VALUE! |
| ENSMUSG00000022382.16 | 0 | NA | NA | NA | NA | NA | #VALUE! |
| ENSMUSG00000022385.12 | 0 | NA | NA | NA | NA | NA | #VALUE! |
| ENSMUSG00000022388.15 | 0 | NA | NA | NA | NA | NA | #VALUE! |
| ENSMUSG00000022394.15 | 9.752977 | 0.421622 | 2.598606 | 0.162249 | 0.87111 | NA | #VALUE! |
| ENSMUSG00000022408.8 | 3.552856 | 0.661292 | 3.779989 | 0.174946 | 0.861122 | NA | #VALUE! |
| ENSMUSG00000022415.13 | 0 | NA | NA | NA | NA | NA | #VALUE! |
| ENSMUSG00000022416.16 | 0 | NA | NA | NA | NA | NA | #VALUE! |
| ENSMUSG00000022420.17 | 7.831564 | -6.49776 | 3.90065 | -1.66581 | 0.095751 | NA | #VALUE! |
| ENSMUSG00000022421.21 | 0 | NA | NA | NA | NA | NA | #VALUE! |
| ENSMUSG00000022422.14 | 0 | NA | NA | NA | NA | NA | #VALUE! |
| ENSMUSG00000022426.7 | 49.8359 | -1.91754 | 1.655051 | -1.1586 | NA | NA | #VALUE! |
| ENSMUSG00000022428.9 | 0.841418 | 3.240539 | 3.959551 | 0.818411 | 0.413123 | NA | #VALUE! |
| ENSMUSG00000022429.12 | 0 | NA | NA | NA | NA | NA | #VALUE! |
| ENSMUSG00000022431.4 | 0 | NA | NA | NA | NA | NA | #VALUE! |
| ENSMUSG00000022432.8 | 0.373021 | 2.140748 | 4.018182 | 0.532765 | 0.594196 | NA | #VALUE! |
| ENSMUSG00000022433.22 | 7.89703 | 1.554574 | 2.821296 | 0.551014 | 0.581624 | NA | #VALUE! |
| ENSMUSG00000022435.7 | 0 | NA | NA | NA | NA | NA | #VALUE! |
| ENSMUSG00000022436.17 | 0 | NA | NA | NA | NA | NA | #VALUE! |
| ENSMUSG00000022438.7 | 7.07718 | -0.13485 | 3.689854 | -0.03655 | 0.970846 | NA | #VALUE! |
| ENSMUSG00000022439.10 | 0 | NA | NA | NA | NA | NA | #VALUE! |
| ENSMUSG00000022440.10 | 2.692536 | 4.918862 | 3.923417 | 1.253719 | 0.209944 | NA | #VALUE! |
| ENSMUSG00000022441.18 | 0 | NA | NA | NA | NA | NA | #VALUE! |
| ENSMUSG00000022449.15 | 0 | NA | NA | NA | NA | NA | #VALUE! |
| ENSMUSG00000022454.18 | 0 | NA | NA | NA | NA | NA | #VALUE! |
| ENSMUSG00000022455.5 | 0 | NA | NA | NA | NA | NA | #VALUE! |
| ENSMUSG00000022456.19 | 0 | NA | NA | NA | NA | NA | #VALUE! |

| | | | | | | | |
|---|---|---|---|---|---|---|---|
| ENSMUSG00000022468.13 | 0 | NA | NA | NA | NA | NA | #VALUE! |
| ENSMUSG00000022469.18 | 5.315546 | 5.90612 | 3.915181 | 1.508518 | 0.131422 | NA | #VALUE! |
| ENSMUSG00000022474.16 | 0 | NA | NA | NA | NA | NA | #VALUE! |
| ENSMUSG00000022475.20 | 1.846078 | -2.32169 | 3.889915 | -0.59685 | 0.550608 | NA | #VALUE! |
| ENSMUSG00000022479.16 | 0 | NA | NA | NA | NA | NA | #VALUE! |
| ENSMUSG00000022483.18 | 0.105013 | -0.97324 | 4.076293 | -0.23876 | 0.811295 | NA | #VALUE! |
| ENSMUSG00000022484.8 | 0 | NA | NA | NA | NA | NA | #VALUE! |
| ENSMUSG00000022485.4 | 0 | NA | NA | NA | NA | NA | #VALUE! |
| ENSMUSG00000022487.15 | 0 | NA | NA | NA | NA | NA | #VALUE! |
| ENSMUSG00000022488.10 | 1.865104 | 4.40601 | 3.930481 | 1.120985 | 0.262294 | NA | #VALUE! |
| ENSMUSG00000022489.7 | 1.492083 | 4.088114 | 3.936282 | 1.038572 | 0.299004 | NA | #VALUE! |
| ENSMUSG00000022490.8 | 0 | NA | NA | NA | NA | NA | #VALUE! |
| ENSMUSG00000022491.6 | 0 | NA | NA | NA | NA | NA | #VALUE! |
| ENSMUSG00000022494.16 | 0 | NA | NA | NA | NA | NA | #VALUE! |
| ENSMUSG00000022496.6 | 0 | NA | NA | NA | NA | NA | #VALUE! |
| ENSMUSG00000022501.7 | 0 | NA | NA | NA | NA | NA | #VALUE! |
| ENSMUSG00000022504.11 | 0 | NA | NA | NA | NA | NA | #VALUE! |
| ENSMUSG00000022505.10 | 9.609403 | -0.48975 | 1.993399 | -0.24569 | 0.805925 | NA | #VALUE! |
| ENSMUSG00000022508.6 | 273.6576 | -1.08599 | 0.938974 | -1.15658 | NA | NA | #VALUE! |
| ENSMUSG00000022510.15 | 0 | NA | NA | NA | NA | NA | #VALUE! |
| ENSMUSG00000022515.7 | 8.219557 | -1.87354 | 2.704548 | -0.69274 | 0.488475 | NA | #VALUE! |
| ENSMUSG00000022518.17 | 0 | NA | NA | NA | NA | NA | #VALUE! |
| ENSMUSG00000022519.15 | 0 | NA | NA | NA | NA | NA | #VALUE! |
| ENSMUSG00000022523.11 | 0.932552 | 3.421684 | 3.953399 | 0.865504 | 0.386762 | NA | #VALUE! |
| ENSMUSG00000022525.14 | 0 | NA | NA | NA | NA | NA | #VALUE! |
| ENSMUSG00000022526.9 | 1.958359 | 4.475621 | 3.929371 | 1.139017 | 0.254696 | NA | #VALUE! |
| ENSMUSG00000022534.15 | 0 | NA | NA | NA | NA | NA | #VALUE! |
| ENSMUSG00000022537.19 | 1.550873 | 0.949778 | 3.853765 | 0.246455 | 0.80533 | NA | #VALUE! |
| ENSMUSG00000022540.17 | 8.761596 | 2.941003 | 3.440077 | 0.854924 | 0.392593 | NA | #VALUE! |
| ENSMUSG00000022542.8 | 0 | NA | NA | NA | NA | NA | #VALUE! |
| ENSMUSG00000022543.10 | 0 | NA | NA | NA | NA | NA | #VALUE! |
| ENSMUSG00000022545.14 | 3.839813 | 5.426526 | 3.9039 | 1.390027 | 0.164521 | NA | #VALUE! |
| ENSMUSG00000022548.15 | 0 | NA | NA | NA | NA | NA | #VALUE! |
| ENSMUSG00000022553.17 | 4.03474 | -0.46636 | 3.772321 | -0.12363 | 0.90161 | NA | #VALUE! |
| ENSMUSG00000022558.17 | 4.289739 | 5.598228 | 3.917179 | 1.429148 | 0.152962 | NA | #VALUE! |
| ENSMUSG00000022560.10 | 17.62102 | -21.6187 | 3.045977 | -7.09745 | NA | NA | #VALUE! |
| ENSMUSG00000022562.16 | 7.360199 | 3.09362 | 3.143341 | 0.984182 | 0.325026 | NA | #VALUE! |
| ENSMUSG00000022565.16 | 4.173909 | 4.587149 | 3.867894 | 1.185955 | 0.23564 | NA | #VALUE! |
| ENSMUSG00000022577.18 | 0 | NA | NA | NA | NA | NA | #VALUE! |
| ENSMUSG00000022580.14 | 0 | NA | NA | NA | NA | NA | #VALUE! |
| ENSMUSG00000022582.5 | 0 | NA | NA | NA | NA | NA | #VALUE! |
| ENSMUSG00000022583.13 | 0 | NA | NA | NA | NA | NA | #VALUE! |
| ENSMUSG00000022584.15 | 0 | NA | NA | NA | NA | NA | #VALUE! |
| ENSMUSG00000022586.12 | 0 | NA | NA | NA | NA | NA | #VALUE! |

| | | | | | | | |
|---|---|---|---|---|---|---|---|
| ENSMUSG00000022589.9 | 0 | NA | NA | NA | NA | NA | #VALUE! |
| ENSMUSG00000022594.15 | 0 | NA | NA | NA | NA | NA | #VALUE! |
| ENSMUSG00000022595.8 | 0 | NA | NA | NA | NA | NA | #VALUE! |
| ENSMUSG00000022596.4 | 0 | NA | NA | NA | NA | NA | #VALUE! |
| ENSMUSG00000022598.7 | 0 | NA | NA | NA | NA | NA | #VALUE! |
| ENSMUSG00000022602.15 | 0 | NA | NA | NA | NA | NA | #VALUE! |
| ENSMUSG00000022603.10 | 0 | NA | NA | NA | NA | NA | #VALUE! |
| ENSMUSG00000022610.11 | 4.057005 | 3.066667 | 3.842827 | 0.798024 | 0.424857 | NA | #VALUE! |
| ENSMUSG00000022613.9 | 0.210026 | -1.52494 | 4.058692 | -0.37572 | 0.707123 | NA | #VALUE! |
| ENSMUSG00000022614.9 | 9.094654 | -1.28381 | 3.013962 | -0.42596 | 0.67014 | NA | #VALUE! |
| ENSMUSG00000022615.8 | 0 | NA | NA | NA | NA | NA | #VALUE! |
| ENSMUSG00000022617.16 | 4.267911 | 0.136123 | 3.482608 | 0.039086 | 0.968822 | NA | #VALUE! |
| ENSMUSG00000022619.6 | 0 | NA | NA | NA | NA | NA | #VALUE! |
| ENSMUSG00000022622.6 | 0 | NA | NA | NA | NA | NA | #VALUE! |
| ENSMUSG00000022623.18 | 0.105013 | -0.97324 | 4.076293 | -0.23876 | 0.811295 | NA | #VALUE! |
| ENSMUSG00000022639.15 | 5.927728 | 0.218112 | 3.756654 | 0.05806 | 0.953701 | NA | #VALUE! |
| ENSMUSG00000022650.8 | 0 | NA | NA | NA | NA | NA | #VALUE! |
| ENSMUSG00000022651.7 | 11.13139 | -7.00979 | 3.449286 | -2.03224 | NA | NA | #VALUE! |
| ENSMUSG00000022652.8 | 0 | NA | NA | NA | NA | NA | #VALUE! |
| ENSMUSG00000022657.10 | 0 | NA | NA | NA | NA | NA | #VALUE! |
| ENSMUSG00000022658.11 | 0 | NA | NA | NA | NA | NA | #VALUE! |
| ENSMUSG00000022659.14 | 0 | NA | NA | NA | NA | NA | #VALUE! |
| ENSMUSG00000022661.15 | 7.301574 | 6.352735 | 3.612118 | 1.758729 | 0.078624 | NA | #VALUE! |
| ENSMUSG00000022683.14 | 0 | NA | NA | NA | NA | NA | #VALUE! |
| ENSMUSG00000022686.15 | 0 | NA | NA | NA | NA | NA | #VALUE! |
| ENSMUSG00000022687.13 | 4.28478 | 5.588653 | 3.917248 | 1.426678 | 0.153673 | NA | #VALUE! |
| ENSMUSG00000022696.18 | 0 | NA | NA | NA | NA | NA | #VALUE! |
| ENSMUSG00000022705.6 | 0 | NA | NA | NA | NA | NA | #VALUE! |
| ENSMUSG00000022715.4 | 0 | NA | NA | NA | NA | NA | #VALUE! |
| ENSMUSG00000022722.19 | 9.318463 | 0.916165 | 2.557311 | 0.358253 | 0.720154 | NA | #VALUE! |
| ENSMUSG00000022738.8 | 0 | NA | NA | NA | NA | NA | #VALUE! |
| ENSMUSG00000022753.16 | 0 | NA | NA | NA | NA | NA | #VALUE! |
| ENSMUSG00000022754.7 | 0 | NA | NA | NA | NA | NA | #VALUE! |
| ENSMUSG00000022755.5 | 0 | NA | NA | NA | NA | NA | #VALUE! |
| ENSMUSG00000022756.18 | 0 | NA | NA | NA | NA | NA | #VALUE! |
| ENSMUSG00000022758.15 | 0 | NA | NA | NA | NA | NA | #VALUE! |
| ENSMUSG00000022759.16 | 0 | NA | NA | NA | NA | NA | #VALUE! |
| ENSMUSG00000022762.19 | 7.924234 | 1.449048 | 2.686701 | 0.539341 | 0.589651 | NA | #VALUE! |
| ENSMUSG00000022763.19 | 0.978946 | -3.48749 | 3.952505 | -0.88235 | 0.377588 | NA | #VALUE! |
| ENSMUSG00000022768.18 | 0 | NA | NA | NA | NA | NA | #VALUE! |
| ENSMUSG00000022771.18 | 33.35571 | -0.98457 | 1.561675 | -0.63046 | NA | NA | #VALUE! |
| ENSMUSG00000022773.20 | 0 | NA | NA | NA | NA | NA | #VALUE! |
| ENSMUSG00000022780.6 | 0 | NA | NA | NA | NA | NA | #VALUE! |
| ENSMUSG00000022783.12 | 0 | NA | NA | NA | NA | NA | #VALUE! |

| | | | | | | | |
|---|---|---|---|---|---|---|---|
| ENSMUSG00000022793.18 | 0.279766 | 1.808031 | 4.036658 | 0.447903 | 0.654223 | NA | #VALUE! |
| ENSMUSG00000022798.9 | 0 | NA | NA | NA | NA | NA | #VALUE! |
| ENSMUSG00000022802.3 | 0 | NA | NA | NA | NA | NA | #VALUE! |
| ENSMUSG00000022803.13 | 0 | NA | NA | NA | NA | NA | #VALUE! |
| ENSMUSG00000022805.18 | 0 | NA | NA | NA | NA | NA | #VALUE! |
| ENSMUSG00000022817.16 | 48.81241 | 1.001594 | 1.365239 | 0.73364 | NA | NA | #VALUE! |
| ENSMUSG00000022818.14 | 0 | NA | NA | NA | NA | NA | #VALUE! |
| ENSMUSG00000022822.17 | 5.346785 | -5.95166 | 3.88155 | -1.53332 | 0.125197 | NA | #VALUE! |
| ENSMUSG00000022824.13 | 0 | NA | NA | NA | NA | NA | #VALUE! |
| ENSMUSG00000022829.15 | 0 | NA | NA | NA | NA | NA | #VALUE! |
| ENSMUSG00000022831.15 | 6.512103 | 0.186813 | 3.753821 | 0.049766 | 0.960309 | NA | #VALUE! |
| ENSMUSG00000022832.12 | 0 | NA | NA | NA | NA | NA | #VALUE! |
| ENSMUSG00000022833.8 | 0 | NA | NA | NA | NA | NA | #VALUE! |
| ENSMUSG00000022837.15 | 14.38471 | -0.22118 | 2.608478 | -0.08479 | NA | NA | #VALUE! |
| ENSMUSG00000022838.15 | 8.645873 | -5.56491 | 3.089949 | -1.80097 | 0.071707 | NA | #VALUE! |
| ENSMUSG00000022840.10 | 0.18651 | 1.401769 | 4.056354 | 0.345574 | 0.729663 | NA | #VALUE! |
| ENSMUSG00000022842.19 | 0 | NA | NA | NA | NA | NA | #VALUE! |
| ENSMUSG00000022849.6 | 7.723358 | 6.439591 | 3.085745 | 2.086884 | 0.036899 | NA | #VALUE! |
| ENSMUSG00000022857.14 | 0 | NA | NA | NA | NA | NA | #VALUE! |
| ENSMUSG00000022860.16 | 0 | NA | NA | NA | NA | NA | #VALUE! |
| ENSMUSG00000022861.18 | 0 | NA | NA | NA | NA | NA | #VALUE! |
| ENSMUSG00000022863.16 | 2.103097 | -0.95019 | 3.822986 | -0.24855 | 0.803712 | NA | #VALUE! |
| ENSMUSG00000022876.19 | 8.329762 | 0.636785 | 3.092261 | 0.205928 | 0.836847 | NA | #VALUE! |
| ENSMUSG00000022878.6 | 0 | NA | NA | NA | NA | NA | #VALUE! |
| ENSMUSG00000022886.6 | 0 | NA | NA | NA | NA | NA | #VALUE! |
| ENSMUSG00000022894.7 | 18.33747 | 0.902158 | 2.205056 | 0.409132 | NA | NA | #VALUE! |
| ENSMUSG00000022899.11 | 7.268645 | 3.474674 | 3.297945 | 1.053588 | 0.292072 | NA | #VALUE! |
| ENSMUSG00000022900.15 | 0 | NA | NA | NA | NA | NA | #VALUE! |
| ENSMUSG00000022901.14 | 2.167917 | -4.64264 | 3.927408 | -1.18211 | 0.237161 | NA | #VALUE! |
| ENSMUSG00000022902.9 | 0 | NA | NA | NA | NA | NA | #VALUE! |
| ENSMUSG00000022931.8 | 0 | NA | NA | NA | NA | NA | #VALUE! |
| ENSMUSG00000022935.17 | 0 | NA | NA | NA | NA | NA | #VALUE! |
| ENSMUSG00000022938.9 | 0 | NA | NA | NA | NA | NA | #VALUE! |
| ENSMUSG00000022941.9 | 7.478917 | 2.436911 | 3.275738 | 0.743927 | 0.456921 | NA | #VALUE! |
| ENSMUSG00000022945.13 | 3.089749 | -0.07325 | 3.784982 | -0.01935 | 0.984559 | NA | #VALUE! |
| ENSMUSG00000022947.9 | 0.18651 | 1.401769 | 4.056354 | 0.345574 | 0.729663 | NA | #VALUE! |
| ENSMUSG00000022948.17 | 7.655509 | -2.64135 | 2.774444 | -0.95203 | 0.341082 | NA | #VALUE! |
| ENSMUSG00000022949.10 | 0 | NA | NA | NA | NA | NA | #VALUE! |
| ENSMUSG00000022952.18 | 0 | NA | NA | NA | NA | NA | #VALUE! |
| ENSMUSG00000022960.14 | 8.062049 | 2.557344 | 3.549618 | 0.720456 | 0.471244 | NA | #VALUE! |
| ENSMUSG00000022964.15 | 6.646412 | 5.169685 | 2.866147 | 1.803706 | 0.071277 | NA | #VALUE! |
| ENSMUSG00000022986.6 | 0 | NA | NA | NA | NA | NA | #VALUE! |
| ENSMUSG00000022987.13 | 0 | NA | NA | NA | NA | NA | #VALUE! |
| ENSMUSG00000022991.6 | 0 | NA | NA | NA | NA | NA | #VALUE! |

| | | | | | | | |
|---|---|---|---|---|---|---|---|
| ENSMUSG00000022992.18 | 5.914563 | 6.055898 | 3.252763 | 1.86177 | 0.062635 | NA | #VALUE! |
| ENSMUSG00000022993.8 | 2.951706 | 5.040208 | 3.922079 | 1.285086 | 0.198762 | NA | #VALUE! |
| ENSMUSG00000022994.10 | 8.864364 | 1.900697 | 2.676554 | 0.710128 | 0.477625 | NA | #VALUE! |
| ENSMUSG00000022996.16 | 0 | NA | NA | NA | NA | NA | #VALUE! |
| ENSMUSG00000022997.10 | 0 | NA | NA | NA | NA | NA | #VALUE! |
| ENSMUSG00000022999.16 | 0.783156 | -3.1627 | 3.963922 | -0.79787 | 0.424945 | NA | #VALUE! |
| ENSMUSG00000023000.6 | 0 | NA | NA | NA | NA | NA | #VALUE! |
| ENSMUSG00000023007.16 | 0 | NA | NA | NA | NA | NA | #VALUE! |
| ENSMUSG00000023008.19 | 0 | NA | NA | NA | NA | NA | #VALUE! |
| ENSMUSG00000023009.15 | 0 | NA | NA | NA | NA | NA | #VALUE! |
| ENSMUSG00000023011.9 | 0 | NA | NA | NA | NA | NA | #VALUE! |
| ENSMUSG00000023013.6 | 0 | NA | NA | NA | NA | NA | #VALUE! |
| ENSMUSG00000023015.15 | 0 | NA | NA | NA | NA | NA | #VALUE! |
| ENSMUSG00000023017.11 | 0 | NA | NA | NA | NA | NA | #VALUE! |
| ENSMUSG00000023021.16 | 0.903375 | -2.05231 | 3.951877 | -0.51933 | 0.603534 | NA | #VALUE! |
| ENSMUSG00000023032.13 | 3.499948 | -0.67234 | 3.781341 | -0.1778 | 0.858877 | NA | #VALUE! |
| ENSMUSG00000023033.15 | 0.168284 | 1.027628 | 4.067126 | 0.252667 | 0.800526 | NA | #VALUE! |
| ENSMUSG00000023034.8 | 4.620578 | -5.74391 | 3.916409 | -1.46663 | 0.142478 | NA | #VALUE! |
| ENSMUSG00000023036.15 | 0 | NA | NA | NA | NA | NA | #VALUE! |
| ENSMUSG00000023039.18 | 0 | NA | NA | NA | NA | NA | #VALUE! |
| ENSMUSG00000023041.8 | 0 | NA | NA | NA | NA | NA | #VALUE! |
| ENSMUSG00000023046.7 | 0.778147 | 1.98536 | 3.968237 | 0.500313 | 0.616855 | NA | #VALUE! |
| ENSMUSG00000023047.12 | 0 | NA | NA | NA | NA | NA | #VALUE! |
| ENSMUSG00000023051.12 | 0.168284 | 1.027628 | 4.067126 | 0.252667 | 0.800526 | NA | #VALUE! |
| ENSMUSG00000023052.9 | 0 | NA | NA | NA | NA | NA | #VALUE! |
| ENSMUSG00000023055.11 | 2.741291 | -1.39248 | 3.808428 | -0.36563 | 0.71464 | NA | #VALUE! |
| ENSMUSG00000023064.5 | 0 | NA | NA | NA | NA | NA | #VALUE! |
| ENSMUSG00000023066.11 | 1.962601 | 4.464983 | 3.929537 | 1.136262 | 0.255847 | NA | #VALUE! |
| ENSMUSG00000023067.15 | 2.34307 | 4.715369 | 3.925927 | 1.201084 | 0.229719 | NA | #VALUE! |
| ENSMUSG00000023072.15 | 4.698938 | -5.75981 | 3.916303 | -1.47073 | 0.141365 | NA | #VALUE! |
| ENSMUSG00000023078.7 | 3.113993 | 2.052276 | 3.820255 | 0.537209 | 0.591123 | NA | #VALUE! |
| ENSMUSG00000023083.8 | 0 | NA | NA | NA | NA | NA | #VALUE! |
| ENSMUSG00000023084.12 | 0 | NA | NA | NA | NA | NA | #VALUE! |
| ENSMUSG00000023088.18 | 0 | NA | NA | NA | NA | NA | #VALUE! |
| ENSMUSG00000023093.8 | 0 | NA | NA | NA | NA | NA | #VALUE! |
| ENSMUSG00000023118.16 | 3.947787 | -5.51641 | 3.918052 | -1.40795 | 0.159147 | NA | #VALUE! |
| ENSMUSG00000023120.10 | 0 | NA | NA | NA | NA | NA | #VALUE! |
| ENSMUSG00000023132.9 | 0.630079 | -2.90291 | 3.975047 | -0.73028 | 0.465217 | NA | #VALUE! |
| ENSMUSG00000023140.5 | 0 | NA | NA | NA | NA | NA | #VALUE! |
| ENSMUSG00000023151.10 | 0 | NA | NA | NA | NA | NA | #VALUE! |
| ENSMUSG00000023153.10 | 0 | NA | NA | NA | NA | NA | #VALUE! |
| ENSMUSG00000023159.9 | 0 | NA | NA | NA | NA | NA | #VALUE! |
| ENSMUSG00000023165.14 | 0 | NA | NA | NA | NA | NA | #VALUE! |
| ENSMUSG00000023169.16 | 4.811391 | -5.79422 | 3.916079 | -1.4796 | 0.138981 | NA | #VALUE! |

| | | | | | | | |
|---|---|---|---|---|---|---|---|
| ENSMUSG00000023185.5 | 0 | NA | NA | NA | NA | NA | #VALUE! |
| ENSMUSG00000023191.10 | 0 | NA | NA | NA | NA | NA | #VALUE! |
| ENSMUSG00000023192.13 | 0 | NA | NA | NA | NA | NA | #VALUE! |
| ENSMUSG00000023210.3 | 0 | NA | NA | NA | NA | NA | #VALUE! |
| ENSMUSG00000023216.14 | 0 | NA | NA | NA | NA | NA | #VALUE! |
| ENSMUSG00000023232.18 | 8.634221 | 2.182754 | 3.466761 | 0.629623 | 0.528941 | NA | #VALUE! |
| ENSMUSG00000023236.8 | 0 | NA | NA | NA | NA | NA | #VALUE! |
| ENSMUSG00000023247.3 | 0 | NA | NA | NA | NA | NA | #VALUE! |
| ENSMUSG00000023257.4 | 0 | NA | NA | NA | NA | NA | #VALUE! |
| ENSMUSG00000023259.18 | 0 | NA | NA | NA | NA | NA | #VALUE! |
| ENSMUSG00000023263.4 | 0 | NA | NA | NA | NA | NA | #VALUE! |
| ENSMUSG00000023266.12 | 0.680622 | -1.83467 | 3.974726 | -0.46158 | 0.644379 | NA | #VALUE! |
| ENSMUSG00000023267.11 | 0 | NA | NA | NA | NA | NA | #VALUE! |
| ENSMUSG00000023274.15 | 0 | NA | NA | NA | NA | NA | #VALUE! |
| ENSMUSG00000023277.14 | 5.071565 | -2.95312 | 3.563005 | -0.82883 | 0.407202 | NA | #VALUE! |
| ENSMUSG00000023279.8 | 0 | NA | NA | NA | NA | NA | #VALUE! |
| ENSMUSG00000023284.16 | 0 | NA | NA | NA | NA | NA | #VALUE! |
| ENSMUSG00000023289.12 | 0 | NA | NA | NA | NA | NA | #VALUE! |
| ENSMUSG00000023328.15 | 0 | NA | NA | NA | NA | NA | #VALUE! |
| ENSMUSG00000023333.9 | 0 | NA | NA | NA | NA | NA | #VALUE! |
| ENSMUSG00000023341.16 | 8.466138 | -3.9782 | 2.602616 | -1.52854 | 0.126379 | NA | #VALUE! |
| ENSMUSG00000023345.18 | 3.395294 | -0.42562 | 3.781015 | -0.11257 | 0.910373 | NA | #VALUE! |
| ENSMUSG00000023349.15 | 0 | NA | NA | NA | NA | NA | #VALUE! |
| ENSMUSG00000023350.7 | 0 | NA | NA | NA | NA | NA | #VALUE! |
| ENSMUSG00000023353.15 | 0.105013 | -0.97324 | 4.076293 | -0.23876 | 0.811295 | NA | #VALUE! |
| ENSMUSG00000023387.9 | 0 | NA | NA | NA | NA | NA | #VALUE! |
| ENSMUSG00000023391.9 | 0 | NA | NA | NA | NA | NA | #VALUE! |
| ENSMUSG00000023403.15 | 0 | NA | NA | NA | NA | NA | #VALUE! |
| ENSMUSG00000023411.13 | 0 | NA | NA | NA | NA | NA | #VALUE! |
| ENSMUSG00000023433.9 | 0 | NA | NA | NA | NA | NA | #VALUE! |
| ENSMUSG00000023439.12 | 0 | NA | NA | NA | NA | NA | #VALUE! |
| ENSMUSG00000023443.14 | 0 | NA | NA | NA | NA | NA | #VALUE! |
| ENSMUSG00000023467.19 | 0 | NA | NA | NA | NA | NA | #VALUE! |
| ENSMUSG00000023473.14 | 0 | NA | NA | NA | NA | NA | #VALUE! |
| ENSMUSG00000023484.15 | 0 | NA | NA | NA | NA | NA | #VALUE! |
| ENSMUSG00000023505.14 | 8.353343 | -0.9664 | 2.736412 | -0.35316 | 0.723966 | NA | #VALUE! |
| ENSMUSG00000023571.5 | 3.788977 | 0.45947 | 3.775558 | 0.121696 | 0.90314 | NA | #VALUE! |
| ENSMUSG00000023737.15 | 0 | NA | NA | NA | NA | NA | #VALUE! |
| ENSMUSG00000023755.11 | 5.102769 | -5.87825 | 3.915553 | -1.50126 | 0.133289 | NA | #VALUE! |
| ENSMUSG00000023781.3 | 0 | NA | NA | NA | NA | NA | #VALUE! |
| ENSMUSG00000023795.17 | 0 | NA | NA | NA | NA | NA | #VALUE! |
| ENSMUSG00000023802.5 | 0 | NA | NA | NA | NA | NA | #VALUE! |
| ENSMUSG00000023805.18 | 6.935092 | -1.73546 | 3.311843 | -0.52402 | 0.600266 | NA | #VALUE! |
| ENSMUSG00000023806.11 | 3.357187 | 5.246639 | 3.920044 | 1.338413 | 0.180762 | NA | #VALUE! |

| | | | | | | | |
|---|---|---|---|---|---|---|---|
| ENSMUSG00000023809.11 | 0 | NA | NA | NA | NA | NA | #VALUE! |
| ENSMUSG00000023826.17 | 3.140947 | -5.17738 | 3.921033 | -1.32041 | 0.186698 | NA | #VALUE! |
| ENSMUSG00000023827.9 | 0 | NA | NA | NA | NA | NA | #VALUE! |
| ENSMUSG00000023868.18 | 2.671552 | -2.19294 | 3.840214 | -0.57105 | 0.567968 | NA | #VALUE! |
| ENSMUSG00000023873.14 | 0.210026 | -1.52494 | 4.058692 | -0.37572 | 0.707123 | NA | #VALUE! |
| ENSMUSG00000023885.10 | 2.277174 | -1.93851 | 3.848056 | -0.50376 | 0.614428 | NA | #VALUE! |
| ENSMUSG00000023902.19 | 0 | NA | NA | NA | NA | NA | #VALUE! |
| ENSMUSG00000023903.9 | 0 | NA | NA | NA | NA | NA | #VALUE! |
| ENSMUSG00000023905.16 | 9.803931 | -0.99337 | 2.514775 | -0.39502 | 0.692832 | NA | #VALUE! |
| ENSMUSG00000023906.4 | 0 | NA | NA | NA | NA | NA | #VALUE! |
| ENSMUSG00000023908.9 | 0.420053 | -2.33493 | 4.007343 | -0.58266 | 0.56012 | NA | #VALUE! |
| ENSMUSG00000023909.5 | 0 | NA | NA | NA | NA | NA | #VALUE! |
| ENSMUSG00000023911.13 | 2.898625 | -3.92103 | 3.909132 | -1.00304 | 0.31584 | NA | #VALUE! |
| ENSMUSG00000023912.11 | 0 | NA | NA | NA | NA | NA | #VALUE! |
| ENSMUSG00000023913.18 | 9.105927 | -0.64443 | 2.433532 | -0.26481 | 0.791154 | NA | #VALUE! |
| ENSMUSG00000023914.17 | 0 | NA | NA | NA | NA | NA | #VALUE! |
| ENSMUSG00000023918.13 | 0 | NA | NA | NA | NA | NA | #VALUE! |
| ENSMUSG00000023926.9 | 0 | NA | NA | NA | NA | NA | #VALUE! |
| ENSMUSG00000023930.15 | 0 | NA | NA | NA | NA | NA | #VALUE! |
| ENSMUSG00000023931.6 | 0 | NA | NA | NA | NA | NA | #VALUE! |
| ENSMUSG00000023935.11 | 0 | NA | NA | NA | NA | NA | #VALUE! |
| ENSMUSG00000023940.15 | 2.936837 | -5.08026 | 3.922024 | -1.29532 | 0.195212 | NA | #VALUE! |
| ENSMUSG00000023943.8 | 0 | NA | NA | NA | NA | NA | #VALUE! |
| ENSMUSG00000023945.8 | 0 | NA | NA | NA | NA | NA | #VALUE! |
| ENSMUSG00000023949.8 | 0 | NA | NA | NA | NA | NA | #VALUE! |
| ENSMUSG00000023952.14 | 6.667044 | 0.357651 | 3.753455 | 0.095286 | 0.924088 | NA | #VALUE! |
| ENSMUSG00000023953.10 | 3.384555 | -1.55998 | 3.797272 | -0.41082 | 0.681207 | NA | #VALUE! |
| ENSMUSG00000023964.16 | 0 | NA | NA | NA | NA | NA | #VALUE! |
| ENSMUSG00000023966.9 | 0 | NA | NA | NA | NA | NA | #VALUE! |
| ENSMUSG00000023968.16 | 0 | NA | NA | NA | NA | NA | #VALUE! |
| ENSMUSG00000023972.11 | 0 | NA | NA | NA | NA | NA | #VALUE! |
| ENSMUSG00000023978.6 | 0 | NA | NA | NA | NA | NA | #VALUE! |
| ENSMUSG00000023979.15 | 0 | NA | NA | NA | NA | NA | #VALUE! |
| ENSMUSG00000023982.9 | 0 | NA | NA | NA | NA | NA | #VALUE! |
| ENSMUSG00000023984.18 | 0 | NA | NA | NA | NA | NA | #VALUE! |
| ENSMUSG00000023987.15 | 0 | NA | NA | NA | NA | NA | #VALUE! |
| ENSMUSG00000023990.19 | 0.093255 | 0.868093 | 4.072464 | 0.213162 | 0.831201 | NA | #VALUE! |
| ENSMUSG00000023991.17 | 5.250656 | -5.92764 | 3.915257 | -1.51398 | 0.13003 | NA | #VALUE! |
| ENSMUSG00000023992.15 | 7.209824 | -2.41927 | 3.614442 | -0.66933 | 0.503282 | NA | #VALUE! |
| ENSMUSG00000023993.8 | 0 | NA | NA | NA | NA | NA | #VALUE! |
| ENSMUSG00000023995.13 | 0 | NA | NA | NA | NA | NA | #VALUE! |
| ENSMUSG00000023999.15 | 0 | NA | NA | NA | NA | NA | #VALUE! |
| ENSMUSG00000024008.17 | 0 | NA | NA | NA | NA | NA | #VALUE! |
| ENSMUSG00000024011.18 | 0 | NA | NA | NA | NA | NA | #VALUE! |

| | | | | | | | |
|---|---|---|---|---|---|---|---|
| ENSMUSG00000024019.19 | 8.988793 | -4.76834 | 2.928763 | -1.62811 | 0.103502 | NA | #VALUE! |
| ENSMUSG00000024027.9 | 0 | NA | NA | NA | NA | NA | #VALUE! |
| ENSMUSG00000024028.8 | 0 | NA | NA | NA | NA | NA | #VALUE! |
| ENSMUSG00000024029.5 | 154.2895 | -2.59771 | 2.793331 | -0.92997 | NA | NA | #VALUE! |
| ENSMUSG00000024032.8 | 0 | NA | NA | NA | NA | NA | #VALUE! |
| ENSMUSG00000024033.11 | 0 | NA | NA | NA | NA | NA | #VALUE! |
| ENSMUSG00000024034.14 | 0 | NA | NA | NA | NA | NA | #VALUE! |
| ENSMUSG00000024036.17 | 1.866574 | -4.43493 | 3.93059 | -1.12831 | 0.259189 | NA | #VALUE! |
| ENSMUSG00000024041.11 | 0.525066 | -2.64687 | 3.988095 | -0.66369 | 0.506887 | NA | #VALUE! |
| ENSMUSG00000024042.8 | 0.612332 | -2.79827 | 3.980109 | -0.70306 | 0.482016 | NA | #VALUE! |
| ENSMUSG00000024043.14 | 0 | NA | NA | NA | NA | NA | #VALUE! |
| ENSMUSG00000024044.20 | 6.900884 | 6.28129 | 3.913256 | 1.605131 | NA | NA | #VALUE! |
| ENSMUSG00000024049.16 | 9.296947 | 0.462166 | 3.406331 | 0.135679 | 0.892075 | NA | #VALUE! |
| ENSMUSG00000024050.19 | 1.632886 | -4.22813 | 3.934241 | -1.0747 | 0.282509 | NA | #VALUE! |
| ENSMUSG00000024053.12 | 2.835354 | -5.04297 | 3.922422 | -1.28568 | 0.198556 | NA | #VALUE! |
| ENSMUSG00000024054.15 | 9.591857 | 3.315824 | 2.665241 | 1.244099 | 0.213463 | NA | #VALUE! |
| ENSMUSG00000024056.11 | 0 | NA | NA | NA | NA | NA | #VALUE! |
| ENSMUSG00000024059.11 | 4.082215 | -5.55558 | 3.91775 | -1.41805 | 0.156175 | NA | #VALUE! |
| ENSMUSG00000024064.15 | 0 | NA | NA | NA | NA | NA | #VALUE! |
| ENSMUSG00000024076.11 | 0 | NA | NA | NA | NA | NA | #VALUE! |
| ENSMUSG00000024087.5 | 4.448138 | -3.58169 | 3.864998 | -0.9267 | 0.354083 | NA | #VALUE! |
| ENSMUSG00000024088.5 | 0 | NA | NA | NA | NA | NA | #VALUE! |
| ENSMUSG00000024105.4 | 0 | NA | NA | NA | NA | NA | #VALUE! |
| ENSMUSG00000024107.8 | 0 | NA | NA | NA | NA | NA | #VALUE! |
| ENSMUSG00000024112.17 | 0 | NA | NA | NA | NA | NA | #VALUE! |
| ENSMUSG00000024114.16 | 0 | NA | NA | NA | NA | NA | #VALUE! |
| ENSMUSG00000024116.7 | 0 | NA | NA | NA | NA | NA | #VALUE! |
| ENSMUSG00000024121.14 | 0 | NA | NA | NA | NA | NA | #VALUE! |
| ENSMUSG00000024124.11 | 1.967804 | 4.449735 | 3.929778 | 1.132312 | 0.257503 | NA | #VALUE! |
| ENSMUSG00000024125.3 | 0 | NA | NA | NA | NA | NA | #VALUE! |
| ENSMUSG00000024128.14 | 0 | NA | NA | NA | NA | NA | #VALUE! |
| ENSMUSG00000024134.12 | 0 | NA | NA | NA | NA | NA | #VALUE! |
| ENSMUSG00000024136.15 | 6.747486 | -6.28211 | 3.621855 | -1.7345 | 0.08283 | NA | #VALUE! |
| ENSMUSG00000024142.16 | 1.119062 | 3.679627 | 3.945843 | 0.932533 | 0.351061 | NA | #VALUE! |
| ENSMUSG00000024154.12 | 0 | NA | NA | NA | NA | NA | #VALUE! |
| ENSMUSG00000024160.17 | 5.781822 | 6.026936 | 3.914506 | 1.539642 | 0.123648 | NA | #VALUE! |
| ENSMUSG00000024163.18 | 3.427735 | 2.122768 | 3.812879 | 0.556736 | 0.577708 | NA | #VALUE! |
| ENSMUSG00000024169.17 | 0.978946 | -3.48749 | 3.952505 | -0.88235 | 0.377588 | NA | #VALUE! |
| ENSMUSG00000024170.16 | 0.391578 | -2.1489 | 4.020883 | -0.53444 | 0.59304 | NA | #VALUE! |
| ENSMUSG00000024171.6 | 0 | NA | NA | NA | NA | NA | #VALUE! |
| ENSMUSG00000024172.11 | 0 | NA | NA | NA | NA | NA | #VALUE! |
| ENSMUSG00000024173.12 | 0 | NA | NA | NA | NA | NA | #VALUE! |
| ENSMUSG00000024174.10 | 1.9661 | 1.181914 | 3.835285 | 0.308169 | 0.757954 | NA | #VALUE! |
| ENSMUSG00000024175.3 | 0 | NA | NA | NA | NA | NA | #VALUE! |

| | | | | | | | |
|---|---|---|---|---|---|---|---|
| ENSMUSG00000024176.11 | 0 | NA | NA | NA | NA | NA | #VALUE! |
| ENSMUSG00000024184.16 | 0 | NA | NA | NA | NA | NA | #VALUE! |
| ENSMUSG00000024186.16 | 0 | NA | NA | NA | NA | NA | #VALUE! |
| ENSMUSG00000024193.9 | 0.210026 | -1.52494 | 4.058692 | -0.37572 | 0.707123 | NA | #VALUE! |
| ENSMUSG00000024201.13 | 8.003735 | -6.53264 | 2.890166 | -2.2603 | 0.023803 | NA | #VALUE! |
| ENSMUSG00000024206.16 | 4.30736 | -5.63406 | 3.91717 | -1.4383 | 0.15035 | NA | #VALUE! |
| ENSMUSG00000024207.9 | 0 | NA | NA | NA | NA | NA | #VALUE! |
| ENSMUSG00000024209.10 | 0 | NA | NA | NA | NA | NA | #VALUE! |
| ENSMUSG00000024210.4 | 0 | NA | NA | NA | NA | NA | #VALUE! |
| ENSMUSG00000024211.16 | 2.168234 | 1.077989 | 3.82397 | 0.281903 | 0.778018 | NA | #VALUE! |
| ENSMUSG00000024212.10 | 38.8391 | -6.30146 | 3.820876 | -1.64922 | NA | NA | #VALUE! |
| ENSMUSG00000024215.15 | 0 | NA | NA | NA | NA | NA | #VALUE! |
| ENSMUSG00000024220.15 | 2.12297 | -1.14396 | 3.824639 | -0.2991 | 0.764863 | NA | #VALUE! |
| ENSMUSG00000024223.4 | 0 | NA | NA | NA | NA | NA | #VALUE! |
| ENSMUSG00000024224.6 | 0 | NA | NA | NA | NA | NA | #VALUE! |
| ENSMUSG00000024225.6 | 0 | NA | NA | NA | NA | NA | #VALUE! |
| ENSMUSG00000024227.15 | 0 | NA | NA | NA | NA | NA | #VALUE! |
| ENSMUSG00000024232.3 | 6.982398 | 2.634193 | 3.218196 | 0.818531 | 0.413054 | NA | #VALUE! |
| ENSMUSG00000024233.11 | 0 | NA | NA | NA | NA | NA | #VALUE! |
| ENSMUSG00000024235.12 | 27.17479 | 0.69652 | 2.075007 | 0.335671 | NA | NA | #VALUE! |
| ENSMUSG00000024245.5 | 7.26845 | 6.347413 | 3.493571 | 1.816884 | 0.069235 | NA | #VALUE! |
| ENSMUSG00000024246.10 | 5.496332 | -5.98658 | 3.914918 | -1.52917 | 0.126222 | NA | #VALUE! |
| ENSMUSG00000024251.11 | 13.89733 | 0.1501 | 2.413434 | 0.062193 | NA | NA | #VALUE! |
| ENSMUSG00000024256.8 | 0 | NA | NA | NA | NA | NA | #VALUE! |
| ENSMUSG00000024261.7 | 0 | NA | NA | NA | NA | NA | #VALUE! |
| ENSMUSG00000024266.9 | 0 | NA | NA | NA | NA | NA | #VALUE! |
| ENSMUSG00000024268.17 | 0 | NA | NA | NA | NA | NA | #VALUE! |
| ENSMUSG00000024270.12 | 3.517207 | -5.34584 | 3.919465 | -1.36392 | 0.172593 | NA | #VALUE! |
| ENSMUSG00000024274.11 | 0 | NA | NA | NA | NA | NA | #VALUE! |
| ENSMUSG00000024295.10 | 2.310289 | -4.7496 | 3.925939 | -1.2098 | 0.226355 | NA | #VALUE! |
| ENSMUSG00000024299.18 | 0 | NA | NA | NA | NA | NA | #VALUE! |
| ENSMUSG00000024300.18 | 7.710428 | -0.13214 | 3.289794 | -0.04017 | 0.967961 | NA | #VALUE! |
| ENSMUSG00000024301.11 | 0 | NA | NA | NA | NA | NA | #VALUE! |
| ENSMUSG00000024302.17 | 0 | NA | NA | NA | NA | NA | #VALUE! |
| ENSMUSG00000024306.14 | 0 | NA | NA | NA | NA | NA | #VALUE! |
| ENSMUSG00000024313.9 | 0 | NA | NA | NA | NA | NA | #VALUE! |
| ENSMUSG00000024325.9 | 2.695818 | -1.44529 | 3.813395 | -0.379 | 0.704685 | NA | #VALUE! |
| ENSMUSG00000024327.17 | 0 | NA | NA | NA | NA | NA | #VALUE! |
| ENSMUSG00000024330.18 | 0 | NA | NA | NA | NA | NA | #VALUE! |
| ENSMUSG00000024334.10 | 0 | NA | NA | NA | NA | NA | #VALUE! |
| ENSMUSG00000024340.16 | 0 | NA | NA | NA | NA | NA | #VALUE! |
| ENSMUSG00000024347.17 | 0 | NA | NA | NA | NA | NA | #VALUE! |
| ENSMUSG00000024349.11 | 3.36567 | 5.240782 | 3.920098 | 1.336901 | 0.181255 | NA | #VALUE! |
| ENSMUSG00000024350.7 | 3.885486 | -5.49502 | 3.91822 | -1.40243 | 0.160787 | NA | #VALUE! |

| | | | | | | | |
|---|---|---|---|---|---|---|---|
| ENSMUSG00000024352.13 | 1.787848 | 3.30776 | 3.929002 | 0.841883 | 0.399853 | NA | #VALUE! |
| ENSMUSG00000024353.6 | 0 | NA | NA | NA | NA | NA | #VALUE! |
| ENSMUSG00000024365.10 | 2.100263 | -4.61322 | 3.927832 | -1.17449 | 0.240197 | NA | #VALUE! |
| ENSMUSG00000024366.9 | 0 | NA | NA | NA | NA | NA | #VALUE! |
| ENSMUSG00000024376.8 | 0 | NA | NA | NA | NA | NA | #VALUE! |
| ENSMUSG00000024379.5 | 8.855119 | 6.632223 | 3.756168 | 1.765689 | 0.077448 | NA | #VALUE! |
| ENSMUSG00000024388.12 | 0 | NA | NA | NA | NA | NA | #VALUE! |
| ENSMUSG00000024393.15 | 1.514552 | 4.088766 | 3.936269 | 1.038742 | 0.298925 | NA | #VALUE! |
| ENSMUSG00000024399.6 | 0 | NA | NA | NA | NA | NA | #VALUE! |
| ENSMUSG00000024401.15 | 0 | NA | NA | NA | NA | NA | #VALUE! |
| ENSMUSG00000024402.6 | 0 | NA | NA | NA | NA | NA | #VALUE! |
| ENSMUSG00000024403.17 | 0 | NA | NA | NA | NA | NA | #VALUE! |
| ENSMUSG00000024406.17 | 0 | NA | NA | NA | NA | NA | #VALUE! |
| ENSMUSG00000024409.9 | 0 | NA | NA | NA | NA | NA | #VALUE! |
| ENSMUSG00000024420.10 | 0 | NA | NA | NA | NA | NA | #VALUE! |
| ENSMUSG00000024421.17 | 8.03727 | -1.39435 | 2.516319 | -0.55412 | 0.579496 | NA | #VALUE! |
| ENSMUSG00000024436.17 | 3.263932 | 5.206263 | 3.920419 | 1.327986 | 0.184183 | NA | #VALUE! |
| ENSMUSG00000024437.7 | 0 | NA | NA | NA | NA | NA | #VALUE! |
| ENSMUSG00000024440.14 | 0.658075 | 0.886597 | 3.979906 | 0.222768 | 0.823716 | NA | #VALUE! |
| ENSMUSG00000024448.15 | 0 | NA | NA | NA | NA | NA | #VALUE! |
| ENSMUSG00000024451.10 | 2.710126 | -2.13997 | 3.836897 | -0.55774 | 0.577025 | NA | #VALUE! |
| ENSMUSG00000024459.19 | 0 | NA | NA | NA | NA | NA | #VALUE! |
| ENSMUSG00000024462.18 | 2.898625 | -3.92103 | 3.909132 | -1.00304 | 0.31584 | NA | #VALUE! |
| ENSMUSG00000024471.13 | 0 | NA | NA | NA | NA | NA | #VALUE! |
| ENSMUSG00000024481.6 | 0 | NA | NA | NA | NA | NA | #VALUE! |
| ENSMUSG00000024485.17 | 0 | NA | NA | NA | NA | NA | #VALUE! |
| ENSMUSG00000024486.7 | 2.424635 | 4.780725 | 3.925082 | 1.217993 | 0.223226 | NA | #VALUE! |
| ENSMUSG00000024497.5 | 0 | NA | NA | NA | NA | NA | #VALUE! |
| ENSMUSG00000024500.21 | 0 | NA | NA | NA | NA | NA | #VALUE! |
| ENSMUSG00000024501.21 | 2.041473 | -1.71214 | 3.848287 | -0.44491 | 0.656385 | NA | #VALUE! |
| ENSMUSG00000024502.11 | 0 | NA | NA | NA | NA | NA | #VALUE! |
| ENSMUSG00000024503.4 | 0 | NA | NA | NA | NA | NA | #VALUE! |
| ENSMUSG00000024510.7 | 0 | NA | NA | NA | NA | NA | #VALUE! |
| ENSMUSG00000024511.16 | 7.11476 | -0.635 | 3.271486 | -0.1941 | 0.846097 | NA | #VALUE! |
| ENSMUSG00000024512.5 | 0 | NA | NA | NA | NA | NA | #VALUE! |
| ENSMUSG00000024517.18 | 0 | NA | NA | NA | NA | NA | #VALUE! |
| ENSMUSG00000024518.5 | 0 | NA | NA | NA | NA | NA | #VALUE! |
| ENSMUSG00000024519.7 | 0 | NA | NA | NA | NA | NA | #VALUE! |
| ENSMUSG00000024521.9 | 9.833352 | 2.480198 | 3.318027 | 0.747492 | 0.454767 | NA | #VALUE! |
| ENSMUSG00000024524.18 | 2.645118 | -4.92765 | 3.923721 | -1.25586 | 0.209166 | NA | #VALUE! |
| ENSMUSG00000024526.10 | 12.58945 | 20.57345 | 3.319966 | 6.196885 | NA | NA | #VALUE! |
| ENSMUSG00000024529.15 | 0 | NA | NA | NA | NA | NA | #VALUE! |
| ENSMUSG00000024530.9 | 0 | NA | NA | NA | NA | NA | #VALUE! |
| ENSMUSG00000024532.14 | 0 | NA | NA | NA | NA | NA | #VALUE! |

| | | | | | | | |
|---|---|---|---|---|---|---|---|
| ENSMUSG00000024533.18 | 0.18651 | 1.401769 | 4.056354 | 0.345574 | 0.729663 | NA | #VALUE! |
| ENSMUSG00000024534.16 | 0.336567 | 1.944925 | 4.029936 | 0.482619 | 0.629366 | NA | #VALUE! |
| ENSMUSG00000024538.9 | 5.119513 | -3.19934 | 3.671029 | -0.87151 | 0.383476 | NA | #VALUE! |
| ENSMUSG00000024542.10 | 3.689633 | 5.364344 | 3.919007 | 1.368802 | 0.171061 | NA | #VALUE! |
| ENSMUSG00000024546.7 | 0 | NA | NA | NA | NA | NA | #VALUE! |
| ENSMUSG00000024548.12 | 2.908511 | 3.921326 | 3.915749 | 1.001424 | 0.316622 | NA | #VALUE! |
| ENSMUSG00000024552.17 | 0 | NA | NA | NA | NA | NA | #VALUE! |
| ENSMUSG00000024553.9 | 0 | NA | NA | NA | NA | NA | #VALUE! |
| ENSMUSG00000024558.13 | 4.219063 | -4.664 | 3.90928 | -1.19306 | 0.232846 | NA | #VALUE! |
| ENSMUSG00000024565.11 | 0 | NA | NA | NA | NA | NA | #VALUE! |
| ENSMUSG00000024575.17 | 0 | NA | NA | NA | NA | NA | #VALUE! |
| ENSMUSG00000024578.4 | 0 | NA | NA | NA | NA | NA | #VALUE! |
| ENSMUSG00000024579.8 | 0.195789 | -1.05581 | 4.073302 | -0.2592 | 0.795479 | NA | #VALUE! |
| ENSMUSG00000024593.16 | 0 | NA | NA | NA | NA | NA | #VALUE! |
| ENSMUSG00000024598.10 | 0 | NA | NA | NA | NA | NA | #VALUE! |
| ENSMUSG00000024600.9 | 0 | NA | NA | NA | NA | NA | #VALUE! |
| ENSMUSG00000024613.17 | 2.431196 | 0.71356 | 3.805862 | 0.18749 | 0.851277 | NA | #VALUE! |
| ENSMUSG00000024617.17 | 0 | NA | NA | NA | NA | NA | #VALUE! |
| ENSMUSG00000024619.9 | 0 | NA | NA | NA | NA | NA | #VALUE! |
| ENSMUSG00000024640.10 | 0.168284 | 1.027628 | 4.067126 | 0.252667 | 0.800526 | NA | #VALUE! |
| ENSMUSG00000024647.15 | 0 | NA | NA | NA | NA | NA | #VALUE! |
| ENSMUSG00000024650.6 | 0 | NA | NA | NA | NA | NA | #VALUE! |
| ENSMUSG00000024653.3 | 0 | NA | NA | NA | NA | NA | #VALUE! |
| ENSMUSG00000024658.8 | 0 | NA | NA | NA | NA | NA | #VALUE! |
| ENSMUSG00000024660.10 | 0 | NA | NA | NA | NA | NA | #VALUE! |
| ENSMUSG00000024664.6 | 0 | NA | NA | NA | NA | NA | #VALUE! |
| ENSMUSG00000024669.9 | 0 | NA | NA | NA | NA | NA | #VALUE! |
| ENSMUSG00000024670.18 | 0 | NA | NA | NA | NA | NA | #VALUE! |
| ENSMUSG00000024672.12 | 0 | NA | NA | NA | NA | NA | #VALUE! |
| ENSMUSG00000024673.10 | 0.816443 | -3.21914 | 3.961753 | -0.81255 | 0.416474 | NA | #VALUE! |
| ENSMUSG00000024677.14 | 3.663702 | -4.46534 | 3.910883 | -1.14177 | 0.253548 | NA | #VALUE! |
| ENSMUSG00000024680.13 | 0 | NA | NA | NA | NA | NA | #VALUE! |
| ENSMUSG00000024681.12 | 0 | NA | NA | NA | NA | NA | #VALUE! |
| ENSMUSG00000024682.4 | 0 | NA | NA | NA | NA | NA | #VALUE! |
| ENSMUSG00000024727.10 | 0 | NA | NA | NA | NA | NA | #VALUE! |
| ENSMUSG00000024728.14 | 0 | NA | NA | NA | NA | NA | #VALUE! |
| ENSMUSG00000024729.8 | 0 | NA | NA | NA | NA | NA | #VALUE! |
| ENSMUSG00000024730.8 | 3.543697 | 5.324148 | 3.919351 | 1.358426 | 0.174329 | NA | #VALUE! |
| ENSMUSG00000024731.16 | 0 | NA | NA | NA | NA | NA | #VALUE! |
| ENSMUSG00000024734.9 | 0 | NA | NA | NA | NA | NA | #VALUE! |
| ENSMUSG00000024736.16 | 0 | NA | NA | NA | NA | NA | #VALUE! |
| ENSMUSG00000024737.3 | 6.017077 | -3.82536 | 3.375586 | -1.13324 | 0.257112 | NA | #VALUE! |
| ENSMUSG00000024738.7 | 9.255594 | 6.700175 | 3.686258 | 1.817609 | 0.069124 | NA | #VALUE! |
| ENSMUSG00000024749.10 | 0 | NA | NA | NA | NA | NA | #VALUE! |

| | | | | | | | |
|---|---|---|---|---|---|---|---|
| ENSMUSG00000024757.8 | 0 | NA | NA | NA | NA | NA | #VALUE! |
| ENSMUSG00000024761.4 | 0 | NA | NA | NA | NA | NA | #VALUE! |
| ENSMUSG00000024766.16 | 4.313159 | 2.689139 | 3.41832 | 0.786684 | 0.431467 | NA | #VALUE! |
| ENSMUSG00000024767.12 | 1.339103 | -3.95748 | 3.939873 | -1.00447 | 0.315153 | NA | #VALUE! |
| ENSMUSG00000024768.6 | 0 | NA | NA | NA | NA | NA | #VALUE! |
| ENSMUSG00000024770.15 | 0 | NA | NA | NA | NA | NA | #VALUE! |
| ENSMUSG00000024771.8 | 0 | NA | NA | NA | NA | NA | #VALUE! |
| ENSMUSG00000024773.16 | 2.814712 | -4.06782 | 3.91483 | -1.03908 | 0.298768 | NA | #VALUE! |
| ENSMUSG00000024774.17 | 0 | NA | NA | NA | NA | NA | #VALUE! |
| ENSMUSG00000024776.19 | 0 | NA | NA | NA | NA | NA | #VALUE! |
| ENSMUSG00000024777.11 | 0 | NA | NA | NA | NA | NA | #VALUE! |
| ENSMUSG00000024784.14 | 0 | NA | NA | NA | NA | NA | #VALUE! |
| ENSMUSG00000024786.10 | 0 | NA | NA | NA | NA | NA | #VALUE! |
| ENSMUSG00000024791.11 | 8.368541 | -6.59301 | 3.815764 | -1.72783 | 0.084018 | NA | #VALUE! |
| ENSMUSG00000024793.15 | 0 | NA | NA | NA | NA | NA | #VALUE! |
| ENSMUSG00000024795.12 | 0 | NA | NA | NA | NA | NA | #VALUE! |
| ENSMUSG00000024798.16 | 0 | NA | NA | NA | NA | NA | #VALUE! |
| ENSMUSG00000024803.10 | 0.841418 | 3.240539 | 3.959551 | 0.818411 | 0.413123 | NA | #VALUE! |
| ENSMUSG00000024806.5 | 0 | NA | NA | NA | NA | NA | #VALUE! |
| ENSMUSG00000024815.6 | 0 | NA | NA | NA | NA | NA | #VALUE! |
| ENSMUSG00000024824.8 | 0 | NA | NA | NA | NA | NA | #VALUE! |
| ENSMUSG00000024831.15 | 2.509934 | -0.24668 | 3.799553 | -0.06492 | 0.948236 | NA | #VALUE! |
| ENSMUSG00000024835.16 | 0 | NA | NA | NA | NA | NA | #VALUE! |
| ENSMUSG00000024837.16 | 0 | NA | NA | NA | NA | NA | #VALUE! |
| ENSMUSG00000024842.9 | 0.746042 | 3.107177 | 3.964586 | 0.783733 | 0.433197 | NA | #VALUE! |
| ENSMUSG00000024846.7 | 0 | NA | NA | NA | NA | NA | #VALUE! |
| ENSMUSG00000024851.15 | 0 | NA | NA | NA | NA | NA | #VALUE! |
| ENSMUSG00000024854.5 | 23.74543 | 1.143747 | 1.942485 | 0.588806 | NA | NA | #VALUE! |
| ENSMUSG00000024855.11 | 4.503149 | -5.6983 | 3.916718 | -1.45487 | 0.145706 | NA | #VALUE! |
| ENSMUSG00000024857.17 | 5.090517 | -5.87548 | 3.91557 | -1.50054 | 0.133474 | NA | #VALUE! |
| ENSMUSG00000024862.17 | 0 | NA | NA | NA | NA | NA | #VALUE! |
| ENSMUSG00000024867.15 | 0 | NA | NA | NA | NA | NA | #VALUE! |
| ENSMUSG00000024868.9 | 0 | NA | NA | NA | NA | NA | #VALUE! |
| ENSMUSG00000024869.11 | 0 | NA | NA | NA | NA | NA | #VALUE! |
| ENSMUSG00000024871.12 | 0 | NA | NA | NA | NA | NA | #VALUE! |
| ENSMUSG00000024873.8 | 0 | NA | NA | NA | NA | NA | #VALUE! |
| ENSMUSG00000024883.9 | 0 | NA | NA | NA | NA | NA | #VALUE! |
| ENSMUSG00000024885.10 | 0 | NA | NA | NA | NA | NA | #VALUE! |
| ENSMUSG00000024889.6 | 2.355969 | 4.72622 | 3.925784 | 1.203892 | 0.228631 | NA | #VALUE! |
| ENSMUSG00000024891.7 | 0 | NA | NA | NA | NA | NA | #VALUE! |
| ENSMUSG00000024897.10 | 0 | NA | NA | NA | NA | NA | #VALUE! |
| ENSMUSG00000024901.16 | 5.165486 | 5.852308 | 3.9155 | 1.494651 | 0.135005 | NA | #VALUE! |
| ENSMUSG00000024903.11 | 0 | NA | NA | NA | NA | NA | #VALUE! |
| ENSMUSG00000024905.16 | 0 | NA | NA | NA | NA | NA | #VALUE! |

| | | | | | | | |
|---|---|---|---|---|---|---|---|
| ENSMUSG00000024906.11 | 1.105487 | -2.62506 | 3.942814 | -0.66578 | 0.505549 | NA | #VALUE! |
| ENSMUSG00000024907.8 | 0 | NA | NA | NA | NA | NA | #VALUE! |
| ENSMUSG00000024908.16 | 113.9974 | 0.152924 | 1.090868 | 0.140185 | NA | NA | #VALUE! |
| ENSMUSG00000024909.16 | 0.210026 | -1.52494 | 4.058692 | -0.37572 | 0.707123 | NA | #VALUE! |
| ENSMUSG00000024910.6 | 0.168284 | 1.027628 | 4.067126 | 0.252667 | 0.800526 | NA | #VALUE! |
| ENSMUSG00000024912.7 | 0 | NA | NA | NA | NA | NA | #VALUE! |
| ENSMUSG00000024913.17 | 5.062091 | -2.11087 | 3.574539 | -0.59053 | 0.554836 | NA | #VALUE! |
| ENSMUSG00000024922.3 | 0 | NA | NA | NA | NA | NA | #VALUE! |
| ENSMUSG00000024924.15 | 0.587367 | -2.74297 | 3.982932 | -0.68868 | 0.491024 | NA | #VALUE! |
| ENSMUSG00000024926.11 | 15.27403 | -6.53786 | 2.727622 | -2.39691 | NA | NA | #VALUE! |
| ENSMUSG00000024935.12 | 0 | NA | NA | NA | NA | NA | #VALUE! |
| ENSMUSG00000024936.9 | 0 | NA | NA | NA | NA | NA | #VALUE! |
| ENSMUSG00000024937.16 | 3.00106 | 5.067676 | 3.921791 | 1.292184 | 0.196293 | NA | #VALUE! |
| ENSMUSG00000024939.7 | 0 | NA | NA | NA | NA | NA | #VALUE! |
| ENSMUSG00000024940.12 | 5.050868 | 1.068155 | 3.667482 | 0.29125 | 0.77086 | NA | #VALUE! |
| ENSMUSG00000024942.20 | 0 | NA | NA | NA | NA | NA | #VALUE! |
| ENSMUSG00000024944.7 | 2.871886 | -0.23143 | 3.789927 | -0.06107 | 0.951307 | NA | #VALUE! |
| ENSMUSG00000024948.16 | 8.098419 | -0.6368 | 2.81402 | -0.22629 | 0.820973 | NA | #VALUE! |
| ENSMUSG00000024957.10 | 0 | NA | NA | NA | NA | NA | #VALUE! |
| ENSMUSG00000024958.14 | 0 | NA | NA | NA | NA | NA | #VALUE! |
| ENSMUSG00000024959.15 | 9.659553 | -1.41514 | 3.342751 | -0.42335 | 0.672043 | NA | #VALUE! |
| ENSMUSG00000024965.10 | 9.440469 | -6.76665 | 2.752567 | -2.45831 | 0.013959 | NA | #VALUE! |
| ENSMUSG00000024968.15 | 1.865104 | 4.40601 | 3.930481 | 1.120985 | 0.262294 | NA | #VALUE! |
| ENSMUSG00000024969.16 | 0 | NA | NA | NA | NA | NA | #VALUE! |
| ENSMUSG00000024970.6 | 0 | NA | NA | NA | NA | NA | #VALUE! |
| ENSMUSG00000024972.17 | 0 | NA | NA | NA | NA | NA | #VALUE! |
| ENSMUSG00000024973.18 | 0 | NA | NA | NA | NA | NA | #VALUE! |
| ENSMUSG00000024979.14 | 0 | NA | NA | NA | NA | NA | #VALUE! |
| ENSMUSG00000024989.16 | 0 | NA | NA | NA | NA | NA | #VALUE! |
| ENSMUSG00000024992.14 | 9.389095 | -6.75908 | 3.672758 | -1.84033 | 0.06572 | NA | #VALUE! |
| ENSMUSG00000025001.12 | 9.282274 | -5.8205 | 2.684572 | -2.16813 | 0.030149 | NA | #VALUE! |
| ENSMUSG00000025003.7 | 0.737927 | 3.007825 | 3.968641 | 0.757898 | 0.448512 | NA | #VALUE! |
| ENSMUSG00000025007.14 | 3.915782 | -5.49628 | 3.91821 | -1.40275 | 0.16069 | NA | #VALUE! |
| ENSMUSG00000025008.16 | 1.632886 | -4.22813 | 3.934241 | -1.0747 | 0.282509 | NA | #VALUE! |
| ENSMUSG00000025010.15 | 0 | NA | NA | NA | NA | NA | #VALUE! |
| ENSMUSG00000025013.16 | 0 | NA | NA | NA | NA | NA | #VALUE! |
| ENSMUSG00000025020.12 | 0 | NA | NA | NA | NA | NA | #VALUE! |
| ENSMUSG00000025038.8 | 0 | NA | NA | NA | NA | NA | #VALUE! |
| ENSMUSG00000025043.4 | 0 | NA | NA | NA | NA | NA | #VALUE! |
| ENSMUSG00000025051.2 | 0 | NA | NA | NA | NA | NA | #VALUE! |
| ENSMUSG00000025056.5 | 0 | NA | NA | NA | NA | NA | #VALUE! |
| ENSMUSG00000025058.5 | 0.168284 | 1.027628 | 4.067126 | 0.252667 | 0.800526 | NA | #VALUE! |
| ENSMUSG00000025064.16 | 0 | NA | NA | NA | NA | NA | #VALUE! |
| ENSMUSG00000025069.16 | 0 | NA | NA | NA | NA | NA | #VALUE! |

| | | | | | | | |
|---|---|---|---|---|---|---|---|
| ENSMUSG00000025081.14 | 0 | NA | NA | NA | NA | NA | #VALUE! |
| ENSMUSG00000025082.9 | 0 | NA | NA | NA | NA | NA | #VALUE! |
| ENSMUSG00000025083.19 | 0 | NA | NA | NA | NA | NA | #VALUE! |
| ENSMUSG00000025090.10 | 0 | NA | NA | NA | NA | NA | #VALUE! |
| ENSMUSG00000025091.5 | 0 | NA | NA | NA | NA | NA | #VALUE! |
| ENSMUSG00000025092.8 | 7.236192 | 6.345076 | 3.912976 | 1.621547 | 0.1049 | NA | #VALUE! |
| ENSMUSG00000025094.9 | 3.925428 | -5.50357 | 3.918153 | -1.40463 | 0.16013 | NA | #VALUE! |
| ENSMUSG00000025104.14 | 1.489577 | -0.26054 | 3.850482 | -0.06767 | 0.946052 | NA | #VALUE! |
| ENSMUSG00000025105.11 | 5.411462 | 5.919712 | 3.915102 | 1.51202 | 0.130529 | NA | #VALUE! |
| ENSMUSG00000025128.8 | 0 | NA | NA | NA | NA | NA | #VALUE! |
| ENSMUSG00000025129.3 | 0 | NA | NA | NA | NA | NA | #VALUE! |
| ENSMUSG00000025140.16 | 0 | NA | NA | NA | NA | NA | #VALUE! |
| ENSMUSG00000025141.4 | 0 | NA | NA | NA | NA | NA | #VALUE! |
| ENSMUSG00000025142.19 | 9.097205 | -2.15635 | 2.645514 | -0.8151 | 0.415016 | NA | #VALUE! |
| ENSMUSG00000025145.14 | 8.381193 | -3.40063 | 2.775121 | -1.2254 | 0.220425 | NA | #VALUE! |
| ENSMUSG00000025150.11 | 0 | NA | NA | NA | NA | NA | #VALUE! |
| ENSMUSG00000025154.16 | 0.384779 | 0.875002 | 4.032432 | 0.216991 | 0.828215 | NA | #VALUE! |
| ENSMUSG00000025161.17 | 0 | NA | NA | NA | NA | NA | #VALUE! |
| ENSMUSG00000025163.7 | 0 | NA | NA | NA | NA | NA | #VALUE! |
| ENSMUSG00000025165.13 | 0.210026 | -1.52494 | 4.058692 | -0.37572 | 0.707123 | NA | #VALUE! |
| ENSMUSG00000025170.14 | 0 | NA | NA | NA | NA | NA | #VALUE! |
| ENSMUSG00000025171.3 | 1.174735 | -3.75246 | 3.944889 | -0.95122 | 0.341493 | NA | #VALUE! |
| ENSMUSG00000025172.4 | 2.187686 | 4.619304 | 3.927239 | 1.176222 | 0.239506 | NA | #VALUE! |
| ENSMUSG00000025185.15 | 0 | NA | NA | NA | NA | NA | #VALUE! |
| ENSMUSG00000025188.16 | 9.866025 | 2.420484 | 3.301721 | 0.733098 | NA | NA | #VALUE! |
| ENSMUSG00000025189.10 | 0 | NA | NA | NA | NA | NA | #VALUE! |
| ENSMUSG00000025201.8 | 0 | NA | NA | NA | NA | NA | #VALUE! |
| ENSMUSG00000025202.9 | 0 | NA | NA | NA | NA | NA | #VALUE! |
| ENSMUSG00000025212.18 | 5.480157 | 1.437827 | 3.767925 | 0.381596 | 0.702761 | NA | #VALUE! |
| ENSMUSG00000025213.12 | 3.625764 | 0.713248 | 3.77924 | 0.188728 | 0.850306 | NA | #VALUE! |
| ENSMUSG00000025215.11 | 0 | NA | NA | NA | NA | NA | #VALUE! |
| ENSMUSG00000025216.10 | 0 | NA | NA | NA | NA | NA | #VALUE! |
| ENSMUSG00000025218.7 | 6.555206 | -6.24085 | 3.626871 | -1.72072 | 0.085301 | NA | #VALUE! |
| ENSMUSG00000025219.14 | 0 | NA | NA | NA | NA | NA | #VALUE! |
| ENSMUSG00000025221.17 | 0 | NA | NA | NA | NA | NA | #VALUE! |
| ENSMUSG00000025225.16 | 0 | NA | NA | NA | NA | NA | #VALUE! |
| ENSMUSG00000025227.15 | 0 | NA | NA | NA | NA | NA | #VALUE! |
| ENSMUSG00000025229.16 | 0 | NA | NA | NA | NA | NA | #VALUE! |
| ENSMUSG00000025231.18 | 8.400653 | -1.09381 | 2.834567 | -0.38588 | 0.699584 | NA | #VALUE! |
| ENSMUSG00000025235.9 | 2.918463 | 1.33982 | 3.802756 | 0.352329 | 0.724592 | NA | #VALUE! |
| ENSMUSG00000025237.17 | 5.664188 | 1.387613 | 3.540662 | 0.391908 | 0.695126 | NA | #VALUE! |
| ENSMUSG00000025257.6 | 3.974733 | -2.77742 | 3.829577 | -0.72525 | 0.468296 | NA | #VALUE! |
| ENSMUSG00000025262.9 | 2.415302 | -4.81324 | 3.925115 | -1.22627 | 0.220098 | NA | #VALUE! |
| ENSMUSG00000025265.15 | 0 | NA | NA | NA | NA | NA | #VALUE! |

| | | | | | | | |
|---|---|---|---|---|---|---|---|
| ENSMUSG00000025269.17 | 6.089115 | -1.05609 | 3.440071 | -0.307 | 0.758845 | NA | #VALUE! |
| ENSMUSG00000025272.17 | 0 | NA | NA | NA | NA | NA | #VALUE! |
| ENSMUSG00000025288.2 | 0 | NA | NA | NA | NA | NA | #VALUE! |
| ENSMUSG00000025318.14 | 0 | NA | NA | NA | NA | NA | #VALUE! |
| ENSMUSG00000025321.15 | 7.789638 | 0.719076 | 3.552532 | 0.202412 | 0.839594 | NA | #VALUE! |
| ENSMUSG00000025324.9 | 0 | NA | NA | NA | NA | NA | #VALUE! |
| ENSMUSG00000025328.10 | 0 | NA | NA | NA | NA | NA | #VALUE! |
| ENSMUSG00000025329.4 | 0 | NA | NA | NA | NA | NA | #VALUE! |
| ENSMUSG00000025330.7 | 0 | NA | NA | NA | NA | NA | #VALUE! |
| ENSMUSG00000025333.11 | 0 | NA | NA | NA | NA | NA | #VALUE! |
| ENSMUSG00000025340.15 | 1.50649 | -3.12458 | 3.929898 | -0.79508 | 0.426568 | NA | #VALUE! |
| ENSMUSG00000025348.10 | 0 | NA | NA | NA | NA | NA | #VALUE! |
| ENSMUSG00000025352.7 | 0 | NA | NA | NA | NA | NA | #VALUE! |
| ENSMUSG00000025359.11 | 0 | NA | NA | NA | NA | NA | #VALUE! |
| ENSMUSG00000025370.8 | 0 | NA | NA | NA | NA | NA | #VALUE! |
| ENSMUSG00000025373.15 | 3.378989 | -0.37282 | 3.781043 | -0.0986 | 0.921454 | NA | #VALUE! |
| ENSMUSG00000025375.16 | 18.1245 | 7.665798 | 2.795954 | 2.741746 | NA | NA | #VALUE! |
| ENSMUSG00000025377.15 | 3.003638 | -3.97512 | 3.908534 | -1.01704 | 0.309137 | NA | #VALUE! |
| ENSMUSG00000025380.3 | 0 | NA | NA | NA | NA | NA | #VALUE! |
| ENSMUSG00000025383.3 | 0 | NA | NA | NA | NA | NA | #VALUE! |
| ENSMUSG00000025384.16 | 4.468627 | 5.649855 | 3.916814 | 1.442462 | 0.149172 | NA | #VALUE! |
| ENSMUSG00000025386.3 | 0 | NA | NA | NA | NA | NA | #VALUE! |
| ENSMUSG00000025389.8 | 0 | NA | NA | NA | NA | NA | #VALUE! |
| ENSMUSG00000025400.12 | 0 | NA | NA | NA | NA | NA | #VALUE! |
| ENSMUSG00000025401.10 | 0 | NA | NA | NA | NA | NA | #VALUE! |
| ENSMUSG00000025402.13 | 6.358949 | -1.96222 | 3.00254 | -0.65352 | 0.513421 | NA | #VALUE! |
| ENSMUSG00000025407.8 | 0 | NA | NA | NA | NA | NA | #VALUE! |
| ENSMUSG00000025417.9 | 1.282865 | 0.056953 | 3.8641 | 0.014739 | 0.98824 | NA | #VALUE! |
| ENSMUSG00000025418.8 | 0 | NA | NA | NA | NA | NA | #VALUE! |
| ENSMUSG00000025420.14 | 0 | NA | NA | NA | NA | NA | #VALUE! |
| ENSMUSG00000025422.11 | 0 | NA | NA | NA | NA | NA | #VALUE! |
| ENSMUSG00000025425.19 | 0 | NA | NA | NA | NA | NA | #VALUE! |
| ENSMUSG00000025427.16 | 0 | NA | NA | NA | NA | NA | #VALUE! |
| ENSMUSG00000025429.10 | 1.177985 | 3.72614 | 3.944615 | 0.944614 | 0.344856 | NA | #VALUE! |
| ENSMUSG00000025431.5 | 0 | NA | NA | NA | NA | NA | #VALUE! |
| ENSMUSG00000025432.12 | 0 | NA | NA | NA | NA | NA | #VALUE! |
| ENSMUSG00000025433.8 | 0 | NA | NA | NA | NA | NA | #VALUE! |
| ENSMUSG00000025450.5 | 0 | NA | NA | NA | NA | NA | #VALUE! |
| ENSMUSG00000025453.19 | 5.2602 | -1.78924 | 3.776041 | -0.47384 | 0.635615 | NA | #VALUE! |
| ENSMUSG00000025461.12 | 0 | NA | NA | NA | NA | NA | #VALUE! |
| ENSMUSG00000025467.9 | 0 | NA | NA | NA | NA | NA | #VALUE! |
| ENSMUSG00000025468.16 | 0 | NA | NA | NA | NA | NA | #VALUE! |
| ENSMUSG00000025469.11 | 0 | NA | NA | NA | NA | NA | #VALUE! |
| ENSMUSG00000025473.17 | 0 | NA | NA | NA | NA | NA | #VALUE! |

| | | | | | | | |
|---|---|---|---|---|---|---|---|
| ENSMUSG00000025475.18 | 0 | NA | NA | NA | NA | NA | #VALUE! |
| ENSMUSG00000025477.15 | 28.87343 | 0.215181 | 2.29987 | 0.093562 | NA | NA | #VALUE! |
| ENSMUSG00000025478.16 | 0 | NA | NA | NA | NA | NA | #VALUE! |
| ENSMUSG00000025480.6 | 0 | NA | NA | NA | NA | NA | #VALUE! |
| ENSMUSG00000025482.12 | 0 | NA | NA | NA | NA | NA | #VALUE! |
| ENSMUSG00000025484.10 | 19.27369 | 2.100897 | 2.256681 | 0.930967 | NA | NA | #VALUE! |
| ENSMUSG00000025488.10 | 2.051614 | 4.542018 | 3.92836 | 1.156212 | 0.247594 | NA | #VALUE! |
| ENSMUSG00000025489.6 | 0 | NA | NA | NA | NA | NA | #VALUE! |
| ENSMUSG00000025496.6 | 0 | NA | NA | NA | NA | NA | #VALUE! |
| ENSMUSG00000025500.13 | 0.630079 | -2.90291 | 3.975047 | -0.73028 | 0.465217 | NA | #VALUE! |
| ENSMUSG00000025507.14 | 5.51099 | -5.9895 | 3.914902 | -1.52992 | 0.126035 | NA | #VALUE! |
| ENSMUSG00000025515.17 | 0 | NA | NA | NA | NA | NA | #VALUE! |
| ENSMUSG00000025519.6 | 0 | NA | NA | NA | NA | NA | #VALUE! |
| ENSMUSG00000025527.10 | 0 | NA | NA | NA | NA | NA | #VALUE! |
| ENSMUSG00000025528.11 | 0 | NA | NA | NA | NA | NA | #VALUE! |
| ENSMUSG00000025529.15 | 0 | NA | NA | NA | NA | NA | #VALUE! |
| ENSMUSG00000025537.13 | 0 | NA | NA | NA | NA | NA | #VALUE! |
| ENSMUSG00000025551.14 | 0 | NA | NA | NA | NA | NA | #VALUE! |
| ENSMUSG00000025557.11 | 0 | NA | NA | NA | NA | NA | #VALUE! |
| ENSMUSG00000025572.19 | 9.625181 | 3.336483 | 2.867238 | 1.163658 | 0.244563 | NA | #VALUE! |
| ENSMUSG00000025573.5 | 0 | NA | NA | NA | NA | NA | #VALUE! |
| ENSMUSG00000025576.18 | 0 | NA | NA | NA | NA | NA | #VALUE! |
| ENSMUSG00000025577.8 | 1.365171 | -3.99822 | 3.938957 | -1.01505 | 0.310084 | NA | #VALUE! |
| ENSMUSG00000025578.10 | 0 | NA | NA | NA | NA | NA | #VALUE! |
| ENSMUSG00000025582.5 | 0 | NA | NA | NA | NA | NA | #VALUE! |
| ENSMUSG00000025586.18 | 2.769352 | 0.060301 | 3.79189 | 0.015903 | 0.987312 | NA | #VALUE! |
| ENSMUSG00000025592.18 | 0 | NA | NA | NA | NA | NA | #VALUE! |
| ENSMUSG00000025597.14 | 1.358374 | 0.349112 | 3.861208 | 0.090415 | 0.927957 | NA | #VALUE! |
| ENSMUSG00000025602.14 | 0 | NA | NA | NA | NA | NA | #VALUE! |
| ENSMUSG00000025608.10 | 7.281241 | 0.488505 | 2.840526 | 0.171977 | 0.863456 | NA | #VALUE! |
| ENSMUSG00000025610.8 | 0 | NA | NA | NA | NA | NA | #VALUE! |
| ENSMUSG00000025644.11 | 0 | NA | NA | NA | NA | NA | #VALUE! |
| ENSMUSG00000025646.10 | 1.050131 | -3.6251 | 3.948379 | -0.91812 | 0.358554 | NA | #VALUE! |
| ENSMUSG00000025648.19 | 0 | NA | NA | NA | NA | NA | #VALUE! |
| ENSMUSG00000025650.14 | 0 | NA | NA | NA | NA | NA | #VALUE! |
| ENSMUSG00000025652.7 | 0 | NA | NA | NA | NA | NA | #VALUE! |
| ENSMUSG00000025656.18 | 1.875374 | -1.43319 | 3.849798 | -0.37228 | 0.709688 | NA | #VALUE! |
| ENSMUSG00000025658.17 | 0 | NA | NA | NA | NA | NA | #VALUE! |
| ENSMUSG00000025665.17 | 0 | NA | NA | NA | NA | NA | #VALUE! |
| ENSMUSG00000025701.13 | 0 | NA | NA | NA | NA | NA | #VALUE! |
| ENSMUSG00000025716.15 | 0 | NA | NA | NA | NA | NA | #VALUE! |
| ENSMUSG00000025723.13 | 0 | NA | NA | NA | NA | NA | #VALUE! |
| ENSMUSG00000025726.12 | 0 | NA | NA | NA | NA | NA | #VALUE! |
| ENSMUSG00000025727.9 | 0 | NA | NA | NA | NA | NA | #VALUE! |

| | | | | | | | |
|---|---|---|---|---|---|---|---|
| ENSMUSG00000025730.15 | 36.12112 | 0.912132 | 2.10784 | 0.432733 | NA | NA | #VALUE! |
| ENSMUSG00000025735.14 | 0 | NA | NA | NA | NA | NA | #VALUE! |
| ENSMUSG00000025737.6 | 4.916405 | -5.82558 | 3.915879 | -1.48768 | 0.136835 | NA | #VALUE! |
| ENSMUSG00000025738.9 | 0 | NA | NA | NA | NA | NA | #VALUE! |
| ENSMUSG00000025739.14 | 0 | NA | NA | NA | NA | NA | #VALUE! |
| ENSMUSG00000025746.12 | 0 | NA | NA | NA | NA | NA | #VALUE! |
| ENSMUSG00000025754.12 | 0 | NA | NA | NA | NA | NA | #VALUE! |
| ENSMUSG00000025774.15 | 0 | NA | NA | NA | NA | NA | #VALUE! |
| ENSMUSG00000025776.14 | 0 | NA | NA | NA | NA | NA | #VALUE! |
| ENSMUSG00000025777.9 | 0 | NA | NA | NA | NA | NA | #VALUE! |
| ENSMUSG00000025779.11 | 8.632827 | -2.32593 | 2.701271 | -0.86105 | 0.389211 | NA | #VALUE! |
| ENSMUSG00000025783.3 | 0 | NA | NA | NA | NA | NA | #VALUE! |
| ENSMUSG00000025784.6 | 0 | NA | NA | NA | NA | NA | #VALUE! |
| ENSMUSG00000025787.7 | 0 | NA | NA | NA | NA | NA | #VALUE! |
| ENSMUSG00000025789.10 | 0 | NA | NA | NA | NA | NA | #VALUE! |
| ENSMUSG00000025790.15 | 2.858699 | 5.008545 | 3.922417 | 1.276903 | 0.201637 | NA | #VALUE! |
| ENSMUSG00000025804.6 | 0 | NA | NA | NA | NA | NA | #VALUE! |
| ENSMUSG00000025808.18 | 0 | NA | NA | NA | NA | NA | #VALUE! |
| ENSMUSG00000025821.10 | 9.264157 | 4.711497 | 2.889423 | 1.630602 | 0.102974 | NA | #VALUE! |
| ENSMUSG00000025838.13 | 0 | NA | NA | NA | NA | NA | #VALUE! |
| ENSMUSG00000025839.4 | 0 | NA | NA | NA | NA | NA | #VALUE! |
| ENSMUSG00000025855.14 | 0.168284 | 1.027628 | 4.067126 | 0.252667 | 0.800526 | NA | #VALUE! |
| ENSMUSG00000025858.13 | 0 | NA | NA | NA | NA | NA | #VALUE! |
| ENSMUSG00000025867.9 | 0 | NA | NA | NA | NA | NA | #VALUE! |
| ENSMUSG00000025870.11 | 1.728275 | -4.32474 | 3.932471 | -1.09975 | 0.27144 | NA | #VALUE! |
| ENSMUSG00000025875.15 | 1.728275 | -4.32474 | 3.932471 | -1.09975 | 0.27144 | NA | #VALUE! |
| ENSMUSG00000025876.16 | 0.315039 | -1.93549 | 4.038621 | -0.47925 | 0.631763 | NA | #VALUE! |
| ENSMUSG00000025877.15 | 1.137398 | -3.71943 | 3.945765 | -0.94264 | 0.345865 | NA | #VALUE! |
| ENSMUSG00000025889.14 | 0.903375 | -2.05231 | 3.951877 | -0.51933 | 0.603534 | NA | #VALUE! |
| ENSMUSG00000025892.17 | 0 | NA | NA | NA | NA | NA | #VALUE! |
| ENSMUSG00000025900.14 | 0 | NA | NA | NA | NA | NA | #VALUE! |
| ENSMUSG00000025902.14 | 8.312954 | 4.156194 | 3.117277 | 1.333277 | NA | NA | #VALUE! |
| ENSMUSG00000025905.15 | 0 | NA | NA | NA | NA | NA | #VALUE! |
| ENSMUSG00000025909.17 | 0 | NA | NA | NA | NA | NA | #VALUE! |
| ENSMUSG00000025912.17 | 0 | NA | NA | NA | NA | NA | #VALUE! |
| ENSMUSG00000025916.11 | 0 | NA | NA | NA | NA | NA | #VALUE! |
| ENSMUSG00000025922.10 | 0.245976 | 1.027628 | 4.067126 | 0.252667 | 0.800526 | NA | #VALUE! |
| ENSMUSG00000025927.14 | 0 | NA | NA | NA | NA | NA | #VALUE! |
| ENSMUSG00000025929.5 | 0 | NA | NA | NA | NA | NA | #VALUE! |
| ENSMUSG00000025930.7 | 0 | NA | NA | NA | NA | NA | #VALUE! |
| ENSMUSG00000025931.16 | 0.245976 | 1.027628 | 4.067126 | 0.252667 | 0.800526 | NA | #VALUE! |
| ENSMUSG00000025932.15 | 0 | NA | NA | NA | NA | NA | #VALUE! |
| ENSMUSG00000025938.17 | 0 | NA | NA | NA | NA | NA | #VALUE! |
| ENSMUSG00000025945.12 | 0 | NA | NA | NA | NA | NA | #VALUE! |

| | | | | | | | |
|---|---|---|---|---|---|---|---|
| ENSMUSG00000025946.14 | 0 | NA | NA | NA | NA | NA | #VALUE! |
| ENSMUSG00000025952.16 | 0 | NA | NA | NA | NA | NA | #VALUE! |
| ENSMUSG00000025955.14 | 0 | NA | NA | NA | NA | NA | #VALUE! |
| ENSMUSG00000025961.2 | 0 | NA | NA | NA | NA | NA | #VALUE! |
| ENSMUSG00000025963.6 | 0 | NA | NA | NA | NA | NA | #VALUE! |
| ENSMUSG00000025964.16 | 9.075653 | 3.521846 | 2.619374 | 1.344537 | 0.178775 | NA | #VALUE! |
| ENSMUSG00000025977.16 | 0 | NA | NA | NA | NA | NA | #VALUE! |
| ENSMUSG00000025978.15 | 0 | NA | NA | NA | NA | NA | #VALUE! |
| ENSMUSG00000025983.12 | 0 | NA | NA | NA | NA | NA | #VALUE! |
| ENSMUSG00000025986.7 | 7.420539 | -0.06257 | 2.441157 | -0.02563 | 0.979553 | NA | #VALUE! |
| ENSMUSG00000025997.14 | 0.195789 | -1.05581 | 4.073302 | -0.2592 | 0.795479 | NA | #VALUE! |
| ENSMUSG00000026009.15 | 0.408222 | -2.20205 | 4.016842 | -0.54821 | 0.583551 | NA | #VALUE! |
| ENSMUSG00000026011.14 | 0 | NA | NA | NA | NA | NA | #VALUE! |
| ENSMUSG00000026012.3 | 13.96753 | 20.71215 | 3.246055 | 6.380715 | NA | NA | #VALUE! |
| ENSMUSG00000026018.13 | 0 | NA | NA | NA | NA | NA | #VALUE! |
| ENSMUSG00000026023.17 | 2.940368 | -5.09511 | 3.921868 | -1.29915 | 0.193891 | NA | #VALUE! |
| ENSMUSG00000026024.15 | 9.826079 | -0.97073 | 2.640057 | -0.36769 | 0.713102 | NA | #VALUE! |
| ENSMUSG00000026039.10 | 0 | NA | NA | NA | NA | NA | #VALUE! |
| ENSMUSG00000026051.9 | 0 | NA | NA | NA | NA | NA | #VALUE! |
| ENSMUSG00000026058.12 | 0 | NA | NA | NA | NA | NA | #VALUE! |
| ENSMUSG00000026062.13 | 0 | NA | NA | NA | NA | NA | #VALUE! |
| ENSMUSG00000026063.5 | 0 | NA | NA | NA | NA | NA | #VALUE! |
| ENSMUSG00000026064.17 | 0 | NA | NA | NA | NA | NA | #VALUE! |
| ENSMUSG00000026065.9 | 0 | NA | NA | NA | NA | NA | #VALUE! |
| ENSMUSG00000026068.12 | 0 | NA | NA | NA | NA | NA | #VALUE! |
| ENSMUSG00000026069.16 | 0 | NA | NA | NA | NA | NA | #VALUE! |
| ENSMUSG00000026070.16 | 8.368541 | -6.59301 | 3.815764 | -1.72783 | 0.084018 | NA | #VALUE! |
| ENSMUSG00000026073.14 | 0 | NA | NA | NA | NA | NA | #VALUE! |
| ENSMUSG00000026074.16 | 3.296726 | 2.825219 | 3.85244 | 0.733358 | 0.46334 | NA | #VALUE! |
| ENSMUSG00000026077.16 | 0 | NA | NA | NA | NA | NA | #VALUE! |
| ENSMUSG00000026080.15 | 0 | NA | NA | NA | NA | NA | #VALUE! |
| ENSMUSG00000026085.12 | 0 | NA | NA | NA | NA | NA | #VALUE! |
| ENSMUSG00000026090.17 | 0 | NA | NA | NA | NA | NA | #VALUE! |
| ENSMUSG00000026094.15 | 36.15203 | 0.39416 | 1.873164 | 0.210425 | NA | NA | #VALUE! |
| ENSMUSG00000026100.7 | 0 | NA | NA | NA | NA | NA | #VALUE! |
| ENSMUSG00000026103.15 | 4.395677 | -0.84928 | 3.068557 | -0.27677 | 0.781957 | NA | #VALUE! |
| ENSMUSG00000026109.15 | 4.579147 | -5.72326 | 3.630757 | -1.57633 | 0.11495 | NA | #VALUE! |
| ENSMUSG00000026110.16 | 0 | NA | NA | NA | NA | NA | #VALUE! |
| ENSMUSG00000026113.18 | 2.590054 | -2.90137 | 3.880378 | -0.7477 | 0.454639 | NA | #VALUE! |
| ENSMUSG00000026114.15 | 0 | NA | NA | NA | NA | NA | #VALUE! |
| ENSMUSG00000026124.5 | 0 | NA | NA | NA | NA | NA | #VALUE! |
| ENSMUSG00000026125.10 | 0 | NA | NA | NA | NA | NA | #VALUE! |
| ENSMUSG00000026135.18 | 4.42189 | 0.976587 | 3.77227 | 0.258886 | 0.795723 | NA | #VALUE! |
| ENSMUSG00000026141.14 | 0 | NA | NA | NA | NA | NA | #VALUE! |

| | | | | | | | |
|---|---|---|---|---|---|---|---|
| ENSMUSG00000026147.17 | 0 | NA | NA | NA | NA | NA | #VALUE! |
| ENSMUSG00000026149.3 | 0.932552 | 3.421684 | 3.953399 | 0.865504 | 0.386762 | NA | #VALUE! |
| ENSMUSG00000026156.9 | 8.272073 | 4.380919 | 3.019927 | 1.450671 | 0.146872 | NA | #VALUE! |
| ENSMUSG00000026162.9 | 3.87292 | -1.63322 | 3.789477 | -0.43099 | 0.666477 | NA | #VALUE! |
| ENSMUSG00000026163.18 | 0 | NA | NA | NA | NA | NA | #VALUE! |
| ENSMUSG00000026166.15 | 0 | NA | NA | NA | NA | NA | #VALUE! |
| ENSMUSG00000026167.15 | 0 | NA | NA | NA | NA | NA | #VALUE! |
| ENSMUSG00000026171.13 | 1.470184 | -4.10383 | 3.936698 | -1.04245 | 0.297201 | NA | #VALUE! |
| ENSMUSG00000026173.16 | 0 | NA | NA | NA | NA | NA | #VALUE! |
| ENSMUSG00000026175.13 | 0 | NA | NA | NA | NA | NA | #VALUE! |
| ENSMUSG00000026180.9 | 1.865104 | 4.40601 | 3.930481 | 1.120985 | 0.262294 | NA | #VALUE! |
| ENSMUSG00000026182.7 | 0 | NA | NA | NA | NA | NA | #VALUE! |
| ENSMUSG00000026188.12 | 0 | NA | NA | NA | NA | NA | #VALUE! |
| ENSMUSG00000026196.8 | 3.570446 | -5.37363 | 3.919223 | -1.3711 | 0.170345 | NA | #VALUE! |
| ENSMUSG00000026198.16 | 5.677884 | -6.03326 | 3.914659 | -1.5412 | 0.123269 | NA | #VALUE! |
| ENSMUSG00000026199.18 | 4.889375 | -5.81945 | 3.915918 | -1.4861 | 0.137252 | NA | #VALUE! |
| ENSMUSG00000026200.14 | 4.382994 | 5.629097 | 3.916959 | 1.437109 | 0.150687 | NA | #VALUE! |
| ENSMUSG00000026204.16 | 0 | NA | NA | NA | NA | NA | #VALUE! |
| ENSMUSG00000026205.9 | 0 | NA | NA | NA | NA | NA | #VALUE! |
| ENSMUSG00000026207.17 | 0 | NA | NA | NA | NA | NA | #VALUE! |
| ENSMUSG00000026208.10 | 0 | NA | NA | NA | NA | NA | #VALUE! |
| ENSMUSG00000026211.19 | 0.391578 | -2.1489 | 4.020883 | -0.53444 | 0.59304 | NA | #VALUE! |
| ENSMUSG00000026213.16 | 1.514552 | 4.088766 | 3.936269 | 1.038742 | 0.298925 | NA | #VALUE! |
| ENSMUSG00000026220.7 | 0 | NA | NA | NA | NA | NA | #VALUE! |
| ENSMUSG00000026224.12 | 0 | NA | NA | NA | NA | NA | #VALUE! |
| ENSMUSG00000026226.17 | 0 | NA | NA | NA | NA | NA | #VALUE! |
| ENSMUSG00000026228.7 | 0 | NA | NA | NA | NA | NA | #VALUE! |
| ENSMUSG00000026235.15 | 3.265772 | -5.23273 | 3.920498 | -1.33471 | 0.181971 | NA | #VALUE! |
| ENSMUSG00000026237.7 | 0 | NA | NA | NA | NA | NA | #VALUE! |
| ENSMUSG00000026239.15 | 7.155977 | 0.33823 | 2.842813 | 0.118977 | 0.905293 | NA | #VALUE! |
| ENSMUSG00000026241.6 | 0 | NA | NA | NA | NA | NA | #VALUE! |
| ENSMUSG00000026246.13 | 0 | NA | NA | NA | NA | NA | #VALUE! |
| ENSMUSG00000026247.14 | 0 | NA | NA | NA | NA | NA | #VALUE! |
| ENSMUSG00000026251.14 | 0 | NA | NA | NA | NA | NA | #VALUE! |
| ENSMUSG00000026253.15 | 0 | NA | NA | NA | NA | NA | #VALUE! |
| ENSMUSG00000026255.16 | 0 | NA | NA | NA | NA | NA | #VALUE! |
| ENSMUSG00000026258.6 | 0 | NA | NA | NA | NA | NA | #VALUE! |
| ENSMUSG00000026271.16 | 0 | NA | NA | NA | NA | NA | #VALUE! |
| ENSMUSG00000026274.12 | 0 | NA | NA | NA | NA | NA | #VALUE! |
| ENSMUSG00000026285.9 | 0 | NA | NA | NA | NA | NA | #VALUE! |
| ENSMUSG00000026288.15 | 5.863564 | -0.76685 | 3.471569 | -0.22089 | 0.825175 | NA | #VALUE! |
| ENSMUSG00000026301.17 | 0 | NA | NA | NA | NA | NA | #VALUE! |
| ENSMUSG00000026303.7 | 5.060913 | 0.742363 | 3.764111 | 0.197221 | 0.843654 | NA | #VALUE! |
| ENSMUSG00000026308.9 | 0 | NA | NA | NA | NA | NA | #VALUE! |

| | | | | | | | |
|---|---|---|---|---|---|---|---|
| ENSMUSG00000026311.17 | 5.318633 | -0.98677 | 3.551843 | -0.27782 | 0.781151 | NA | #VALUE! |
| ENSMUSG00000026312.18 | 0 | NA | NA | NA | NA | NA | #VALUE! |
| ENSMUSG00000026313.17 | 2.846977 | 0.816203 | 3.795756 | 0.21503 | 0.829744 | NA | #VALUE! |
| ENSMUSG00000026321.9 | 0.673134 | 2.918386 | 3.972531 | 0.734641 | 0.462558 | NA | #VALUE! |
| ENSMUSG00000026322.10 | 0 | NA | NA | NA | NA | NA | #VALUE! |
| ENSMUSG00000026327.9 | 0 | NA | NA | NA | NA | NA | #VALUE! |
| ENSMUSG00000026331.14 | 0 | NA | NA | NA | NA | NA | #VALUE! |
| ENSMUSG00000026335.17 | 5.128272 | 0.032122 | 3.292887 | 0.009755 | 0.992217 | NA | #VALUE! |
| ENSMUSG00000026336.14 | 0 | NA | NA | NA | NA | NA | #VALUE! |
| ENSMUSG00000026343.7 | 9.471594 | 0.330116 | 2.606697 | 0.126642 | 0.899224 | NA | #VALUE! |
| ENSMUSG00000026344.10 | 0 | NA | NA | NA | NA | NA | #VALUE! |
| ENSMUSG00000026347.14 | 0 | NA | NA | NA | NA | NA | #VALUE! |
| ENSMUSG00000026354.9 | 0 | NA | NA | NA | NA | NA | #VALUE! |
| ENSMUSG00000026357.4 | 0 | NA | NA | NA | NA | NA | #VALUE! |
| ENSMUSG00000026358.14 | 2.310289 | -4.7496 | 3.925939 | -1.2098 | 0.226355 | NA | #VALUE! |
| ENSMUSG00000026387.16 | 0 | NA | NA | NA | NA | NA | #VALUE! |
| ENSMUSG00000026388.16 | 4.775076 | 3.314085 | 3.66705 | 0.903747 | 0.36613 | NA | #VALUE! |
| ENSMUSG00000026389.17 | 9.836306 | 1.447696 | 2.615546 | 0.553497 | 0.579923 | NA | #VALUE! |
| ENSMUSG00000026394.6 | 0 | NA | NA | NA | NA | NA | #VALUE! |
| ENSMUSG00000026401.16 | 0 | NA | NA | NA | NA | NA | #VALUE! |
| ENSMUSG00000026404.13 | 23.85907 | -2.34142 | 2.221139 | -1.05416 | NA | NA | #VALUE! |
| ENSMUSG00000026407.18 | 0 | NA | NA | NA | NA | NA | #VALUE! |
| ENSMUSG00000026409.15 | 7.887287 | 1.825605 | 3.25312 | 0.561186 | 0.574671 | NA | #VALUE! |
| ENSMUSG00000026413.13 | 0 | NA | NA | NA | NA | NA | #VALUE! |
| ENSMUSG00000026414.14 | 0 | NA | NA | NA | NA | NA | #VALUE! |
| ENSMUSG00000026415.12 | 0 | NA | NA | NA | NA | NA | #VALUE! |
| ENSMUSG00000026416.13 | 0 | NA | NA | NA | NA | NA | #VALUE! |
| ENSMUSG00000026418.17 | 0 | NA | NA | NA | NA | NA | #VALUE! |
| ENSMUSG00000026420.17 | 0 | NA | NA | NA | NA | NA | #VALUE! |
| ENSMUSG00000026421.15 | 6.72529 | -6.28056 | 3.553441 | -1.76746 | 0.077151 | NA | #VALUE! |
| ENSMUSG00000026424.9 | 0 | NA | NA | NA | NA | NA | #VALUE! |
| ENSMUSG00000026429.10 | 0.105013 | -0.97324 | 4.076293 | -0.23876 | 0.811295 | NA | #VALUE! |
| ENSMUSG00000026432.8 | 0 | NA | NA | NA | NA | NA | #VALUE! |
| ENSMUSG00000026442.15 | 0 | NA | NA | NA | NA | NA | #VALUE! |
| ENSMUSG00000026443.4 | 0 | NA | NA | NA | NA | NA | #VALUE! |
| ENSMUSG00000026447.18 | 3.029103 | 5.088783 | 3.921574 | 1.297638 | 0.194412 | NA | #VALUE! |
| ENSMUSG00000026450.15 | 0 | NA | NA | NA | NA | NA | #VALUE! |
| ENSMUSG00000026452.16 | 0 | NA | NA | NA | NA | NA | #VALUE! |
| ENSMUSG00000026458.14 | 4.338617 | 0.725749 | 3.77116 | 0.192447 | 0.847392 | NA | #VALUE! |
| ENSMUSG00000026459.6 | 0 | NA | NA | NA | NA | NA | #VALUE! |
| ENSMUSG00000026463.18 | 5.818102 | -0.89813 | 3.500968 | -0.25654 | 0.797535 | NA | #VALUE! |
| ENSMUSG00000026468.15 | 0 | NA | NA | NA | NA | NA | #VALUE! |
| ENSMUSG00000026478.15 | 9.755104 | 0.756212 | 2.659326 | 0.284362 | 0.776133 | NA | #VALUE! |
| ENSMUSG00000026479.14 | 0 | NA | NA | NA | NA | NA | #VALUE! |

| | | | | | | | |
|---|---|---|---|---|---|---|---|
| ENSMUSG00000026483.14 | 8.56213 | -0.71741 | 2.711463 | -0.26459 | 0.791328 | NA | #VALUE! |
| ENSMUSG00000026494.13 | 1.230875 | 1.164322 | 3.896933 | 0.298779 | 0.765109 | NA | #VALUE! |
| ENSMUSG00000026497.8 | 0 | NA | NA | NA | NA | NA | #VALUE! |
| ENSMUSG00000026514.15 | 0 | NA | NA | NA | NA | NA | #VALUE! |
| ENSMUSG00000026519.17 | 2.195288 | -4.65832 | 3.927186 | -1.18617 | 0.235554 | NA | #VALUE! |
| ENSMUSG00000026520.9 | 35.49908 | 0.266695 | 2.487749 | 0.107203 | NA | NA | #VALUE! |
| ENSMUSG00000026523.15 | 0 | NA | NA | NA | NA | NA | #VALUE! |
| ENSMUSG00000026525.10 | 0.18651 | 1.401769 | 4.056354 | 0.345574 | 0.729663 | NA | #VALUE! |
| ENSMUSG00000026527.14 | 0 | NA | NA | NA | NA | NA | #VALUE! |
| ENSMUSG00000026531.5 | 0 | NA | NA | NA | NA | NA | #VALUE! |
| ENSMUSG00000026532.8 | 0.105013 | -0.97324 | 4.076293 | -0.23876 | 0.811295 | NA | #VALUE! |
| ENSMUSG00000026535.10 | 0 | NA | NA | NA | NA | NA | #VALUE! |
| ENSMUSG00000026536.10 | 6.957951 | -3.4433 | 3.173717 | -1.08494 | 0.277948 | NA | #VALUE! |
| ENSMUSG00000026546.17 | 0 | NA | NA | NA | NA | NA | #VALUE! |
| ENSMUSG00000026547.16 | 3.381948 | -3.1603 | 3.868735 | -0.81688 | 0.413995 | NA | #VALUE! |
| ENSMUSG00000026548.5 | 1.131482 | -3.70524 | 3.946148 | -0.93895 | 0.347756 | NA | #VALUE! |
| ENSMUSG00000026556.16 | 1.762102 | -4.34067 | 3.93219 | -1.10388 | 0.269645 | NA | #VALUE! |
| ENSMUSG00000026560.14 | 0 | NA | NA | NA | NA | NA | #VALUE! |
| ENSMUSG00000026564.10 | 0 | NA | NA | NA | NA | NA | #VALUE! |
| ENSMUSG00000026566.16 | 0.291524 | 0.468739 | 4.052149 | 0.115677 | 0.907909 | NA | #VALUE! |
| ENSMUSG00000026567.17 | 0 | NA | NA | NA | NA | NA | #VALUE! |
| ENSMUSG00000026572.12 | 0 | NA | NA | NA | NA | NA | #VALUE! |
| ENSMUSG00000026573.8 | 0 | NA | NA | NA | NA | NA | #VALUE! |
| ENSMUSG00000026580.17 | 0.245976 | 1.027628 | 4.067126 | 0.252667 | 0.800526 | NA | #VALUE! |
| ENSMUSG00000026581.15 | 0.420053 | -2.33493 | 4.007343 | -0.58266 | 0.56012 | NA | #VALUE! |
| ENSMUSG00000026582.7 | 0 | NA | NA | NA | NA | NA | #VALUE! |
| ENSMUSG00000026586.17 | 1.492083 | 4.088114 | 3.936282 | 1.038572 | 0.299004 | NA | #VALUE! |
| ENSMUSG00000026587.16 | 0.727815 | 3.063145 | 3.96635 | 0.772283 | 0.439947 | NA | #VALUE! |
| ENSMUSG00000026592.14 | 0 | NA | NA | NA | NA | NA | #VALUE! |
| ENSMUSG00000026596.14 | 31.35146 | -3.09876 | 1.503818 | -2.0606 | NA | NA | #VALUE! |
| ENSMUSG00000026601.16 | 0 | NA | NA | NA | NA | NA | #VALUE! |
| ENSMUSG00000026602.10 | 0 | NA | NA | NA | NA | NA | #VALUE! |
| ENSMUSG00000026604.18 | 0 | NA | NA | NA | NA | NA | #VALUE! |
| ENSMUSG00000026605.15 | 0.273297 | 0.094602 | 4.062933 | 0.023284 | 0.981424 | NA | #VALUE! |
| ENSMUSG00000026609.16 | 0 | NA | NA | NA | NA | NA | #VALUE! |
| ENSMUSG00000026610.14 | 1.970287 | -1.62992 | 3.849953 | -0.42336 | 0.672033 | NA | #VALUE! |
| ENSMUSG00000026611.18 | 0 | NA | NA | NA | NA | NA | #VALUE! |
| ENSMUSG00000026616.17 | 0 | NA | NA | NA | NA | NA | #VALUE! |
| ENSMUSG00000026620.12 | 2.220253 | -4.67428 | 3.926962 | -1.1903 | 0.233927 | NA | #VALUE! |
| ENSMUSG00000026622.16 | 6.064057 | -1.08476 | 3.016492 | -0.35961 | 0.719138 | NA | #VALUE! |
| ENSMUSG00000026627.8 | 3.277306 | -0.25259 | 3.781625 | -0.06679 | 0.946746 | NA | #VALUE! |
| ENSMUSG00000026630.10 | 1.967804 | 4.449735 | 3.929778 | 1.132312 | 0.257503 | NA | #VALUE! |
| ENSMUSG00000026632.18 | 3.276654 | -0.21169 | 3.782335 | -0.05597 | 0.955367 | NA | #VALUE! |
| ENSMUSG00000026637.15 | 1.721829 | 4.254739 | 3.933085 | 1.081782 | 0.27935 | NA | #VALUE! |

| | | | | | | | |
|---|---|---|---|---|---|---|---|
| ENSMUSG00000026639.19 | 0.168284 | 1.027628 | 4.067126 | 0.252667 | 0.800526 | NA | #VALUE! |
| ENSMUSG00000026641.14 | 0 | NA | NA | NA | NA | NA | #VALUE! |
| ENSMUSG00000026644.8 | 0 | NA | NA | NA | NA | NA | #VALUE! |
| ENSMUSG00000026645.12 | 0 | NA | NA | NA | NA | NA | #VALUE! |
| ENSMUSG00000026646.17 | 6.574331 | 1.853099 | 2.945687 | 0.629089 | 0.529291 | NA | #VALUE! |
| ENSMUSG00000026649.15 | 1.81242 | 4.333085 | 3.931703 | 1.102089 | 0.270423 | NA | #VALUE! |
| ENSMUSG00000026650.16 | 0.598106 | 2.755755 | 3.98023 | 0.692361 | 0.488711 | NA | #VALUE! |
| ENSMUSG00000026657.18 | 2.209391 | -3.48404 | 3.914877 | -0.88995 | 0.373494 | NA | #VALUE! |
| ENSMUSG00000026668.11 | 0 | NA | NA | NA | NA | NA | #VALUE! |
| ENSMUSG00000026679.4 | 0 | NA | NA | NA | NA | NA | #VALUE! |
| ENSMUSG00000026683.15 | 0 | NA | NA | NA | NA | NA | #VALUE! |
| ENSMUSG00000026686.15 | 0 | NA | NA | NA | NA | NA | #VALUE! |
| ENSMUSG00000026691.11 | 1.823199 | 4.354872 | 3.931331 | 1.107735 | 0.267976 | NA | #VALUE! |
| ENSMUSG00000026692.13 | 6.614879 | -0.22552 | 2.453891 | -0.0919 | 0.926775 | NA | #VALUE! |
| ENSMUSG00000026697.11 | 0 | NA | NA | NA | NA | NA | #VALUE! |
| ENSMUSG00000026700.5 | 0 | NA | NA | NA | NA | NA | #VALUE! |
| ENSMUSG00000026707.16 | 7.658684 | 5.478477 | 3.578267 | 1.531042 | NA | NA | #VALUE! |
| ENSMUSG00000026723.11 | 8.314173 | 3.2254 | 3.165428 | 1.018946 | 0.308229 | NA | #VALUE! |
| ENSMUSG00000026725.18 | 0 | NA | NA | NA | NA | NA | #VALUE! |
| ENSMUSG00000026726.11 | 0 | NA | NA | NA | NA | NA | #VALUE! |
| ENSMUSG00000026729.10 | 0 | NA | NA | NA | NA | NA | #VALUE! |
| ENSMUSG00000026734.10 | 0 | NA | NA | NA | NA | NA | #VALUE! |
| ENSMUSG00000026735.3 | 0 | NA | NA | NA | NA | NA | #VALUE! |
| ENSMUSG00000026736.3 | 0 | NA | NA | NA | NA | NA | #VALUE! |
| ENSMUSG00000026751.15 | 0 | NA | NA | NA | NA | NA | #VALUE! |
| ENSMUSG00000026764.16 | 0 | NA | NA | NA | NA | NA | #VALUE! |
| ENSMUSG00000026765.13 | 0 | NA | NA | NA | NA | NA | #VALUE! |
| ENSMUSG00000026770.6 | 0 | NA | NA | NA | NA | NA | #VALUE! |
| ENSMUSG00000026774.12 | 0 | NA | NA | NA | NA | NA | #VALUE! |
| ENSMUSG00000026778.14 | 0.245976 | 1.027628 | 4.067126 | 0.252667 | 0.800526 | NA | #VALUE! |
| ENSMUSG00000026779.7 | 0.504851 | 2.502843 | 3.993993 | 0.626652 | 0.530887 | NA | #VALUE! |
| ENSMUSG00000026784.15 | 6.253168 | -0.36523 | 2.620369 | -0.13938 | 0.889148 | NA | #VALUE! |
| ENSMUSG00000026785.15 | 0 | NA | NA | NA | NA | NA | #VALUE! |
| ENSMUSG00000026786.15 | 14.40304 | 0.067267 | 2.982889 | 0.022551 | NA | NA | #VALUE! |
| ENSMUSG00000026787.4 | 0 | NA | NA | NA | NA | NA | #VALUE! |
| ENSMUSG00000026790.20 | 3.382319 | -0.33426 | 3.780217 | -0.08842 | 0.929541 | NA | #VALUE! |
| ENSMUSG00000026796.17 | 0 | NA | NA | NA | NA | NA | #VALUE! |
| ENSMUSG00000026797.17 | 0 | NA | NA | NA | NA | NA | #VALUE! |
| ENSMUSG00000026799.16 | 9.560056 | -1.38148 | 2.268885 | -0.60888 | 0.542604 | NA | #VALUE! |
| ENSMUSG00000026805.15 | 0 | NA | NA | NA | NA | NA | #VALUE! |
| ENSMUSG00000026807.9 | 0 | NA | NA | NA | NA | NA | #VALUE! |
| ENSMUSG00000026809.16 | 0 | NA | NA | NA | NA | NA | #VALUE! |
| ENSMUSG00000026811.19 | 0.841418 | 3.240539 | 3.959551 | 0.818411 | 0.413123 | NA | #VALUE! |
| ENSMUSG00000026815.15 | 0 | NA | NA | NA | NA | NA | #VALUE! |

| | | | | | | | |
|---|---|---|---|---|---|---|---|
| ENSMUSG00000026816.15 | 2.53199 | -3.6997 | 3.911826 | -0.94577 | 0.344264 | NA | #VALUE! |
| ENSMUSG00000026817.15 | 0 | NA | NA | NA | NA | NA | #VALUE! |
| ENSMUSG00000026818.6 | 0 | NA | NA | NA | NA | NA | #VALUE! |
| ENSMUSG00000026821.17 | 3.132626 | -5.17361 | 3.921071 | -1.31944 | 0.187023 | NA | #VALUE! |
| ENSMUSG00000026824.12 | 0 | NA | NA | NA | NA | NA | #VALUE! |
| ENSMUSG00000026825.18 | 0 | NA | NA | NA | NA | NA | #VALUE! |
| ENSMUSG00000026826.14 | 5.775722 | -6.06468 | 3.914489 | -1.54929 | 0.121312 | NA | #VALUE! |
| ENSMUSG00000026828.12 | 0 | NA | NA | NA | NA | NA | #VALUE! |
| ENSMUSG00000026829.10 | 0 | NA | NA | NA | NA | NA | #VALUE! |
| ENSMUSG00000026830.10 | 0 | NA | NA | NA | NA | NA | #VALUE! |
| ENSMUSG00000026831.17 | 0 | NA | NA | NA | NA | NA | #VALUE! |
| ENSMUSG00000026832.13 | 9.102477 | -2.52394 | 3.406913 | -0.74083 | 0.458797 | NA | #VALUE! |
| ENSMUSG00000026833.19 | 17.79569 | 3.346033 | 2.535835 | 1.319499 | NA | NA | #VALUE! |
| ENSMUSG00000026834.14 | 0 | NA | NA | NA | NA | NA | #VALUE! |
| ENSMUSG00000026835.16 | 0 | NA | NA | NA | NA | NA | #VALUE! |
| ENSMUSG00000026837.17 | 16.43979 | -7.56721 | 2.786113 | -2.71605 | NA | NA | #VALUE! |
| ENSMUSG00000026840.15 | 0 | NA | NA | NA | NA | NA | #VALUE! |
| ENSMUSG00000026841.8 | 0 | NA | NA | NA | NA | NA | #VALUE! |
| ENSMUSG00000026854.17 | 1.174735 | -3.75246 | 3.944889 | -0.95122 | 0.341493 | NA | #VALUE! |
| ENSMUSG00000026856.15 | 7.913052 | 0.679062 | 3.147162 | 0.21577 | 0.829167 | NA | #VALUE! |
| ENSMUSG00000026857.10 | 7.010646 | -0.68454 | 2.874486 | -0.23814 | 0.811769 | NA | #VALUE! |
| ENSMUSG00000026873.10 | 0.245976 | 1.027628 | 4.067126 | 0.252667 | 0.800526 | NA | #VALUE! |
| ENSMUSG00000026875.15 | 0 | NA | NA | NA | NA | NA | #VALUE! |
| ENSMUSG00000026882.2 | 0 | NA | NA | NA | NA | NA | #VALUE! |
| ENSMUSG00000026883.18 | 4.5321 | 2.614725 | 3.010974 | 0.868398 | 0.385176 | NA | #VALUE! |
| ENSMUSG00000026885.14 | 0 | NA | NA | NA | NA | NA | #VALUE! |
| ENSMUSG00000026894.5 | 0 | NA | NA | NA | NA | NA | #VALUE! |
| ENSMUSG00000026904.18 | 0 | NA | NA | NA | NA | NA | #VALUE! |
| ENSMUSG00000026919.2 | 0 | NA | NA | NA | NA | NA | #VALUE! |
| ENSMUSG00000026925.17 | 8.564121 | -2.9799 | 2.759127 | -1.08002 | 0.280135 | NA | #VALUE! |
| ENSMUSG00000026928.15 | 0 | NA | NA | NA | NA | NA | #VALUE! |
| ENSMUSG00000026930.16 | 0 | NA | NA | NA | NA | NA | #VALUE! |
| ENSMUSG00000026931.11 | 0 | NA | NA | NA | NA | NA | #VALUE! |
| ENSMUSG00000026934.16 | 0 | NA | NA | NA | NA | NA | #VALUE! |
| ENSMUSG00000026936.5 | 0 | NA | NA | NA | NA | NA | #VALUE! |
| ENSMUSG00000026937.13 | 0 | NA | NA | NA | NA | NA | #VALUE! |
| ENSMUSG00000026940.4 | 0 | NA | NA | NA | NA | NA | #VALUE! |
| ENSMUSG00000026941.17 | 0 | NA | NA | NA | NA | NA | #VALUE! |
| ENSMUSG00000026942.14 | 6.807629 | 6.261727 | 3.913345 | 1.600096 | NA | NA | #VALUE! |
| ENSMUSG00000026943.13 | 0 | NA | NA | NA | NA | NA | #VALUE! |
| ENSMUSG00000026944.19 | 3.524204 | -5.34395 | 3.919482 | -1.36343 | 0.172746 | NA | #VALUE! |
| ENSMUSG00000026955.14 | 0 | NA | NA | NA | NA | NA | #VALUE! |
| ENSMUSG00000026956.16 | 5.048506 | 5.825722 | 3.915663 | 1.4878 | 0.136804 | NA | #VALUE! |
| ENSMUSG00000026959.14 | 0 | NA | NA | NA | NA | NA | #VALUE! |

| | | | | | | | |
|---|---|---|---|---|---|---|---|
| ENSMUSG00000026961.7 | 0 | NA | NA | NA | NA | NA | #VALUE! |
| ENSMUSG00000026963.5 | 0 | NA | NA | NA | NA | NA | #VALUE! |
| ENSMUSG00000026969.4 | 0 | NA | NA | NA | NA | NA | #VALUE! |
| ENSMUSG00000026971.16 | 0 | NA | NA | NA | NA | NA | #VALUE! |
| ENSMUSG00000026972.16 | 0.315039 | -1.93549 | 4.038621 | -0.47925 | 0.631763 | NA | #VALUE! |
| ENSMUSG00000026975.11 | 7.362172 | -1.42834 | 3.253703 | -0.43899 | 0.66067 | NA | #VALUE! |
| ENSMUSG00000026976.16 | 0 | NA | NA | NA | NA | NA | #VALUE! |
| ENSMUSG00000026979.17 | 0.714877 | 1.018111 | 3.971736 | 0.256339 | 0.797689 | NA | #VALUE! |
| ENSMUSG00000026981.16 | 7.679178 | 0.093091 | 2.868965 | 0.032448 | 0.974115 | NA | #VALUE! |
| ENSMUSG00000026983.11 | 0 | NA | NA | NA | NA | NA | #VALUE! |
| ENSMUSG00000026984.5 | 0 | NA | NA | NA | NA | NA | #VALUE! |
| ENSMUSG00000026985.2 | 0 | NA | NA | NA | NA | NA | #VALUE! |
| ENSMUSG00000026988.16 | 9.33082 | 3.181499 | 3.016478 | 1.054706 | 0.29156 | NA | #VALUE! |
| ENSMUSG00000026989.6 | 0 | NA | NA | NA | NA | NA | #VALUE! |
| ENSMUSG00000026994.10 | 2.625328 | -4.93268 | 3.923662 | -1.25716 | 0.208695 | NA | #VALUE! |
| ENSMUSG00000027009.19 | 1.585338 | 4.174397 | 3.934581 | 1.060951 | 0.288712 | NA | #VALUE! |
| ENSMUSG00000027015.5 | 0 | NA | NA | NA | NA | NA | #VALUE! |
| ENSMUSG00000027022.15 | 0 | NA | NA | NA | NA | NA | #VALUE! |
| ENSMUSG00000027068.7 | 8.383607 | -2.64977 | 3.102208 | -0.85416 | 0.393018 | NA | #VALUE! |
| ENSMUSG00000027070.15 | 4.865945 | 1.197489 | 3.668852 | 0.326393 | 0.744127 | NA | #VALUE! |
| ENSMUSG00000027071.15 | 0 | NA | NA | NA | NA | NA | #VALUE! |
| ENSMUSG00000027072.14 | 0 | NA | NA | NA | NA | NA | #VALUE! |
| ENSMUSG00000027073.6 | 0 | NA | NA | NA | NA | NA | #VALUE! |
| ENSMUSG00000027075.17 | 0.932552 | 3.421684 | 3.953399 | 0.865504 | 0.386762 | NA | #VALUE! |
| ENSMUSG00000027077.8 | 0 | NA | NA | NA | NA | NA | #VALUE! |
| ENSMUSG00000027102.5 | 0 | NA | NA | NA | NA | NA | #VALUE! |
| ENSMUSG00000027107.4 | 0 | NA | NA | NA | NA | NA | #VALUE! |
| ENSMUSG00000027111.17 | 0.735092 | -3.12016 | 3.965612 | -0.7868 | 0.431397 | NA | #VALUE! |
| ENSMUSG00000027115.15 | 2.415302 | -4.81324 | 3.925115 | -1.22627 | 0.220098 | NA | #VALUE! |
| ENSMUSG00000027120.7 | 0 | NA | NA | NA | NA | NA | #VALUE! |
| ENSMUSG00000027134.5 | 0 | NA | NA | NA | NA | NA | #VALUE! |
| ENSMUSG00000027157.11 | 0 | NA | NA | NA | NA | NA | #VALUE! |
| ENSMUSG00000027167.12 | 7.345074 | 0.765086 | 2.963611 | 0.25816 | 0.796283 | NA | #VALUE! |
| ENSMUSG00000027168.22 | 0 | NA | NA | NA | NA | NA | #VALUE! |
| ENSMUSG00000027171.11 | 0 | NA | NA | NA | NA | NA | #VALUE! |
| ENSMUSG00000027186.16 | 0 | NA | NA | NA | NA | NA | #VALUE! |
| ENSMUSG00000027196.12 | 0 | NA | NA | NA | NA | NA | #VALUE! |
| ENSMUSG00000027202.13 | 0 | NA | NA | NA | NA | NA | #VALUE! |
| ENSMUSG00000027204.14 | 8.269385 | -6.57869 | 3.296086 | -1.99591 | 0.045944 | NA | #VALUE! |
| ENSMUSG00000027208.15 | 0 | NA | NA | NA | NA | NA | #VALUE! |
| ENSMUSG00000027209.18 | 0 | NA | NA | NA | NA | NA | #VALUE! |
| ENSMUSG00000027217.14 | 0 | NA | NA | NA | NA | NA | #VALUE! |
| ENSMUSG00000027219.14 | 0 | NA | NA | NA | NA | NA | #VALUE! |
| ENSMUSG00000027220.3 | 0 | NA | NA | NA | NA | NA | #VALUE! |

| | | | | | | | |
|---|---|---|---|---|---|---|---|
| ENSMUSG00000027221.6 | 0 | NA | NA | NA | NA | NA | #VALUE! |
| ENSMUSG00000027222.15 | 7.769262 | -6.48621 | 3.440761 | -1.88511 | 0.059415 | NA | #VALUE! |
| ENSMUSG00000027223.16 | 0 | NA | NA | NA | NA | NA | #VALUE! |
| ENSMUSG00000027224.15 | 0 | NA | NA | NA | NA | NA | #VALUE! |
| ENSMUSG00000027225.8 | 0 | NA | NA | NA | NA | NA | #VALUE! |
| ENSMUSG00000027229.6 | 0 | NA | NA | NA | NA | NA | #VALUE! |
| ENSMUSG00000027230.10 | 0 | NA | NA | NA | NA | NA | #VALUE! |
| ENSMUSG00000027233.5 | 0 | NA | NA | NA | NA | NA | #VALUE! |
| ENSMUSG00000027238.18 | 0 | NA | NA | NA | NA | NA | #VALUE! |
| ENSMUSG00000027239.15 | 0 | NA | NA | NA | NA | NA | #VALUE! |
| ENSMUSG00000027242.15 | 6.188375 | -0.85363 | 3.039401 | -0.28085 | 0.778822 | NA | #VALUE! |
| ENSMUSG00000027246.13 | 0 | NA | NA | NA | NA | NA | #VALUE! |
| ENSMUSG00000027253.16 | 32.21219 | 0.014191 | 1.961348 | 0.007235 | NA | NA | #VALUE! |
| ENSMUSG00000027254.14 | 0 | NA | NA | NA | NA | NA | #VALUE! |
| ENSMUSG00000027255.15 | 9.67957 | -1.12229 | 2.98688 | -0.37574 | 0.707111 | NA | #VALUE! |
| ENSMUSG00000027270.15 | 0 | NA | NA | NA | NA | NA | #VALUE! |
| ENSMUSG00000027273.14 | 0 | NA | NA | NA | NA | NA | #VALUE! |
| ENSMUSG00000027281.17 | 11.01746 | -3.67009 | 2.909391 | -1.26146 | NA | NA | #VALUE! |
| ENSMUSG00000027293.14 | 2.857411 | -1.70827 | 3.595033 | -0.47517 | 0.634663 | NA | #VALUE! |
| ENSMUSG00000027296.8 | 0 | NA | NA | NA | NA | NA | #VALUE! |
| ENSMUSG00000027297.16 | 0 | NA | NA | NA | NA | NA | #VALUE! |
| ENSMUSG00000027298.18 | 0 | NA | NA | NA | NA | NA | #VALUE! |
| ENSMUSG00000027300.11 | 9.425241 | -1.30357 | 3.343423 | -0.38989 | 0.696616 | NA | #VALUE! |
| ENSMUSG00000027301.8 | 0 | NA | NA | NA | NA | NA | #VALUE! |
| ENSMUSG00000027306.16 | 1.025807 | 3.556426 | 3.949287 | 0.900523 | 0.367842 | NA | #VALUE! |
| ENSMUSG00000027313.4 | 10.31374 | 2.977891 | 3.247425 | 0.917001 | NA | NA | #VALUE! |
| ENSMUSG00000027314.7 | 1.074494 | 3.572911 | 3.94881 | 0.904807 | 0.365568 | NA | #VALUE! |
| ENSMUSG00000027315.14 | 0 | NA | NA | NA | NA | NA | #VALUE! |
| ENSMUSG00000027316.16 | 0 | NA | NA | NA | NA | NA | #VALUE! |
| ENSMUSG00000027317.9 | 0 | NA | NA | NA | NA | NA | #VALUE! |
| ENSMUSG00000027318.18 | 0 | NA | NA | NA | NA | NA | #VALUE! |
| ENSMUSG00000027322.12 | 6.090761 | -6.14106 | 3.914091 | -1.56896 | 0.116657 | NA | #VALUE! |
| ENSMUSG00000027323.11 | 4.006779 | -5.53111 | 3.917938 | -1.41174 | 0.158026 | NA | #VALUE! |
| ENSMUSG00000027324.2 | 0 | NA | NA | NA | NA | NA | #VALUE! |
| ENSMUSG00000027326.14 | 2.730341 | -4.98888 | 3.923018 | -1.27169 | 0.203482 | NA | #VALUE! |
| ENSMUSG00000027327.17 | 4.529511 | 1.901048 | 3.721024 | 0.510894 | 0.609425 | NA | #VALUE! |
| ENSMUSG00000027329.10 | 0.315039 | -1.93549 | 4.038621 | -0.47925 | 0.631763 | NA | #VALUE! |
| ENSMUSG00000027330.17 | 0.735092 | -3.12016 | 3.965612 | -0.7868 | 0.431397 | NA | #VALUE! |
| ENSMUSG00000027331.16 | 0.491951 | 2.402185 | 4.000147 | 0.600524 | 0.548157 | NA | #VALUE! |
| ENSMUSG00000027333.19 | 9.547272 | -3.16408 | 3.320977 | -0.95275 | 0.340714 | NA | #VALUE! |
| ENSMUSG00000027335.10 | 0 | NA | NA | NA | NA | NA | #VALUE! |
| ENSMUSG00000027338.17 | 0 | NA | NA | NA | NA | NA | #VALUE! |
| ENSMUSG00000027339.16 | 1.077157 | 3.572282 | 3.948828 | 0.904644 | 0.365654 | NA | #VALUE! |
| ENSMUSG00000027344.15 | 0 | NA | NA | NA | NA | NA | #VALUE! |

| | | | | | | | |
|---|---|---|---|---|---|---|---|
| ENSMUSG00000027345.7 | 0 | NA | NA | NA | NA | NA | #VALUE! |
| ENSMUSG00000027350.9 | 0 | NA | NA | NA | NA | NA | #VALUE! |
| ENSMUSG00000027355.16 | 0 | NA | NA | NA | NA | NA | #VALUE! |
| ENSMUSG00000027356.9 | 0 | NA | NA | NA | NA | NA | #VALUE! |
| ENSMUSG00000027360.6 | 11.43996 | 0.505558 | 3.17119 | 0.159422 | NA | NA | #VALUE! |
| ENSMUSG00000027368.7 | 1.632886 | -4.22813 | 3.934241 | -1.0747 | 0.282509 | NA | #VALUE! |
| ENSMUSG00000027375.15 | 1.995249 | -4.53986 | 3.928926 | -1.1555 | 0.247887 | NA | #VALUE! |
| ENSMUSG00000027376.16 | 0 | NA | NA | NA | NA | NA | #VALUE! |
| ENSMUSG00000027377.3 | 0 | NA | NA | NA | NA | NA | #VALUE! |
| ENSMUSG00000027378.17 | 9.119571 | -4.09275 | 3.392826 | -1.20629 | 0.227704 | NA | #VALUE! |
| ENSMUSG00000027379.15 | 0 | NA | NA | NA | NA | NA | #VALUE! |
| ENSMUSG00000027380.11 | 0 | NA | NA | NA | NA | NA | #VALUE! |
| ENSMUSG00000027386.10 | 0 | NA | NA | NA | NA | NA | #VALUE! |
| ENSMUSG00000027395.16 | 5.735158 | 0.689139 | 3.148264 | 0.218895 | 0.826732 | NA | #VALUE! |
| ENSMUSG00000027399.2 | 6.735655 | -6.27951 | 3.913422 | -1.60461 | 0.10858 | NA | #VALUE! |
| ENSMUSG00000027400.12 | 0 | NA | NA | NA | NA | NA | #VALUE! |
| ENSMUSG00000027401.10 | 0 | NA | NA | NA | NA | NA | #VALUE! |
| ENSMUSG00000027403.13 | 0 | NA | NA | NA | NA | NA | #VALUE! |
| ENSMUSG00000027405.17 | 7.435413 | 0.949789 | 2.837776 | 0.334695 | 0.737855 | NA | #VALUE! |
| ENSMUSG00000027408.8 | 0.578089 | -0.70693 | 3.996403 | -0.17689 | 0.859593 | NA | #VALUE! |
| ENSMUSG00000027409.6 | 0 | NA | NA | NA | NA | NA | #VALUE! |
| ENSMUSG00000027412.13 | 0.204111 | -1.05581 | 4.073302 | -0.2592 | 0.795479 | NA | #VALUE! |
| ENSMUSG00000027416.3 | 0 | NA | NA | NA | NA | NA | #VALUE! |
| ENSMUSG00000027419.10 | 0 | NA | NA | NA | NA | NA | #VALUE! |
| ENSMUSG00000027420.14 | 0 | NA | NA | NA | NA | NA | #VALUE! |
| ENSMUSG00000027431.6 | 0 | NA | NA | NA | NA | NA | #VALUE! |
| ENSMUSG00000027434.12 | 0 | NA | NA | NA | NA | NA | #VALUE! |
| ENSMUSG00000027442.11 | 0 | NA | NA | NA | NA | NA | #VALUE! |
| ENSMUSG00000027443.11 | 0 | NA | NA | NA | NA | NA | #VALUE! |
| ENSMUSG00000027444.3 | 0 | NA | NA | NA | NA | NA | #VALUE! |
| ENSMUSG00000027445.4 | 0 | NA | NA | NA | NA | NA | #VALUE! |
| ENSMUSG00000027446.3 | 0 | NA | NA | NA | NA | NA | #VALUE! |
| ENSMUSG00000027452.12 | 0 | NA | NA | NA | NA | NA | #VALUE! |
| ENSMUSG00000027454.11 | 9.555795 | 0.168149 | 2.953494 | 0.056932 | 0.954599 | NA | #VALUE! |
| ENSMUSG00000027456.9 | 0 | NA | NA | NA | NA | NA | #VALUE! |
| ENSMUSG00000027457.16 | 0 | NA | NA | NA | NA | NA | #VALUE! |
| ENSMUSG00000027459.17 | 0 | NA | NA | NA | NA | NA | #VALUE! |
| ENSMUSG00000027460.8 | 0 | NA | NA | NA | NA | NA | #VALUE! |
| ENSMUSG00000027463.15 | 0 | NA | NA | NA | NA | NA | #VALUE! |
| ENSMUSG00000027468.3 | 0 | NA | NA | NA | NA | NA | #VALUE! |
| ENSMUSG00000027469.17 | 9.09526 | 1.703152 | 2.980777 | 0.571379 | 0.567743 | NA | #VALUE! |
| ENSMUSG00000027470.10 | 0 | NA | NA | NA | NA | NA | #VALUE! |
| ENSMUSG00000027474.14 | 1.855357 | -3.43237 | 3.923877 | -0.87474 | 0.381716 | NA | #VALUE! |
| ENSMUSG00000027480.13 | 0 | NA | NA | NA | NA | NA | #VALUE! |

| | | | | | | | |
|---|---|---|---|---|---|---|---|
| ENSMUSG00000027481.3 | 0 | NA | NA | NA | NA | NA | #VALUE! |
| ENSMUSG00000027482.13 | 0 | NA | NA | NA | NA | NA | #VALUE! |
| ENSMUSG00000027483.8 | 0 | NA | NA | NA | NA | NA | #VALUE! |
| ENSMUSG00000027484.3 | 0 | NA | NA | NA | NA | NA | #VALUE! |
| ENSMUSG00000027485.16 | 0 | NA | NA | NA | NA | NA | #VALUE! |
| ENSMUSG00000027489.16 | 0.195789 | -1.05581 | 4.073302 | -0.2592 | 0.795479 | NA | #VALUE! |
| ENSMUSG00000027490.18 | 0 | NA | NA | NA | NA | NA | #VALUE! |
| ENSMUSG00000027496.16 | 2.704401 | 4.936946 | 3.92321 | 1.258394 | 0.208249 | NA | #VALUE! |
| ENSMUSG00000027499.13 | 2.529648 | 3.847701 | 3.920736 | 0.981372 | 0.326409 | NA | #VALUE! |
| ENSMUSG00000027500.11 | 0 | NA | NA | NA | NA | NA | #VALUE! |
| ENSMUSG00000027505.3 | 0 | NA | NA | NA | NA | NA | #VALUE! |
| ENSMUSG00000027510.18 | 0 | NA | NA | NA | NA | NA | #VALUE! |
| ENSMUSG00000027517.15 | 0 | NA | NA | NA | NA | NA | #VALUE! |
| ENSMUSG00000027518.4 | 0 | NA | NA | NA | NA | NA | #VALUE! |
| ENSMUSG00000027520.17 | 0 | NA | NA | NA | NA | NA | #VALUE! |
| ENSMUSG00000027524.10 | 0 | NA | NA | NA | NA | NA | #VALUE! |
| ENSMUSG00000027525.19 | 0 | NA | NA | NA | NA | NA | #VALUE! |
| ENSMUSG00000027528.13 | 0 | NA | NA | NA | NA | NA | #VALUE! |
| ENSMUSG00000027530.16 | 0 | NA | NA | NA | NA | NA | #VALUE! |
| ENSMUSG00000027544.17 | 0 | NA | NA | NA | NA | NA | #VALUE! |
| ENSMUSG00000027547.18 | 0 | NA | NA | NA | NA | NA | #VALUE! |
| ENSMUSG00000027550.15 | 4.537155 | -0.09783 | 3.766315 | -0.02598 | 0.979277 | NA | #VALUE! |
| ENSMUSG00000027551.16 | 9.850512 | 3.908422 | 2.966542 | 1.317501 | NA | NA | #VALUE! |
| ENSMUSG00000027555.9 | 0 | NA | NA | NA | NA | NA | #VALUE! |
| ENSMUSG00000027560.5 | 0 | NA | NA | NA | NA | NA | #VALUE! |
| ENSMUSG00000027564.5 | 0 | NA | NA | NA | NA | NA | #VALUE! |
| ENSMUSG00000027568.10 | 0 | NA | NA | NA | NA | NA | #VALUE! |
| ENSMUSG00000027570.16 | 0 | NA | NA | NA | NA | NA | #VALUE! |
| ENSMUSG00000027574.16 | 4.769959 | -1.11211 | 3.770096 | -0.29498 | 0.768007 | NA | #VALUE! |
| ENSMUSG00000027577.15 | 5.119044 | 5.841908 | 3.457752 | 1.68951 | 0.091122 | NA | #VALUE! |
| ENSMUSG00000027579.7 | 0.848575 | 1.904055 | 3.965106 | 0.480203 | 0.631083 | NA | #VALUE! |
| ENSMUSG00000027581.13 | 0 | NA | NA | NA | NA | NA | #VALUE! |
| ENSMUSG00000027583.14 | 5.954265 | -0.11924 | 3.453611 | -0.03453 | 0.972458 | NA | #VALUE! |
| ENSMUSG00000027584.18 | 0 | NA | NA | NA | NA | NA | #VALUE! |
| ENSMUSG00000027596.11 | 0 | NA | NA | NA | NA | NA | #VALUE! |
| ENSMUSG00000027603.16 | 0 | NA | NA | NA | NA | NA | #VALUE! |
| ENSMUSG00000027606.13 | 0 | NA | NA | NA | NA | NA | #VALUE! |
| ENSMUSG00000027611.13 | 0.309124 | -1.83395 | 4.047982 | -0.45305 | 0.650511 | NA | #VALUE! |
| ENSMUSG00000027612.8 | 0 | NA | NA | NA | NA | NA | #VALUE! |
| ENSMUSG00000027634.15 | 23.53864 | -1.14882 | 2.433299 | -0.47212 | NA | NA | #VALUE! |
| ENSMUSG00000027635.16 | 2.821117 | -5.03482 | 3.922511 | -1.28357 | 0.199292 | NA | #VALUE! |
| ENSMUSG00000027641.14 | 15.56897 | -5.56129 | 2.577675 | -2.15748 | NA | NA | #VALUE! |
| ENSMUSG00000027643.10 | 0 | NA | NA | NA | NA | NA | #VALUE! |
| ENSMUSG00000027646.16 | 0 | NA | NA | NA | NA | NA | #VALUE! |

| | | | | | | | |
|---|---|---|---|---|---|---|---|
| ENSMUSG00000027654.3 | 2.205276 | -4.68302 | 3.926841 | -1.19257 | 0.233039 | NA | #VALUE! |
| ENSMUSG00000027655.15 | 0.876054 | -1.83507 | 3.961344 | -0.46324 | 0.64319 | NA | #VALUE! |
| ENSMUSG00000027656.8 | 0 | NA | NA | NA | NA | NA | #VALUE! |
| ENSMUSG00000027661.5 | 0 | NA | NA | NA | NA | NA | #VALUE! |
| ENSMUSG00000027669.15 | 4.398796 | -2.71833 | 3.354007 | -0.81047 | 0.417668 | NA | #VALUE! |
| ENSMUSG00000027674.17 | 0 | NA | NA | NA | NA | NA | #VALUE! |
| ENSMUSG00000027676.12 | 0 | NA | NA | NA | NA | NA | #VALUE! |
| ENSMUSG00000027684.17 | 2.355969 | 4.72622 | 3.925784 | 1.203892 | 0.228631 | NA | #VALUE! |
| ENSMUSG00000027692.17 | 4.718209 | -5.76658 | 3.916259 | -1.47247 | 0.140893 | NA | #VALUE! |
| ENSMUSG00000027694.10 | 0 | NA | NA | NA | NA | NA | #VALUE! |
| ENSMUSG00000027699.20 | 8.54454 | -6.62462 | 3.267831 | -2.02722 | 0.04264 | NA | #VALUE! |
| ENSMUSG00000027702.8 | 0 | NA | NA | NA | NA | NA | #VALUE! |
| ENSMUSG00000027703.17 | 0 | NA | NA | NA | NA | NA | #VALUE! |
| ENSMUSG00000027713.7 | 0 | NA | NA | NA | NA | NA | #VALUE! |
| ENSMUSG00000027715.10 | 2.402649 | -0.04371 | 3.802655 | -0.01149 | 0.990829 | NA | #VALUE! |
| ENSMUSG00000027716.14 | 0.373021 | 2.140748 | 4.018182 | 0.532765 | 0.594196 | NA | #VALUE! |
| ENSMUSG00000027718.9 | 0 | NA | NA | NA | NA | NA | #VALUE! |
| ENSMUSG00000027719.16 | 0 | NA | NA | NA | NA | NA | #VALUE! |
| ENSMUSG00000027720.8 | 0 | NA | NA | NA | NA | NA | #VALUE! |
| ENSMUSG00000027722.15 | 5.361314 | -0.39828 | 3.227643 | -0.1234 | 0.901792 | NA | #VALUE! |
| ENSMUSG00000027737.11 | 0 | NA | NA | NA | NA | NA | #VALUE! |
| ENSMUSG00000027744.5 | 0 | NA | NA | NA | NA | NA | #VALUE! |
| ENSMUSG00000027748.12 | 0 | NA | NA | NA | NA | NA | #VALUE! |
| ENSMUSG00000027776.13 | 0 | NA | NA | NA | NA | NA | #VALUE! |
| ENSMUSG00000027777.16 | 0 | NA | NA | NA | NA | NA | #VALUE! |
| ENSMUSG00000027788.7 | 0 | NA | NA | NA | NA | NA | #VALUE! |
| ENSMUSG00000027790.15 | 0 | NA | NA | NA | NA | NA | #VALUE! |
| ENSMUSG00000027793.7 | 0 | NA | NA | NA | NA | NA | #VALUE! |
| ENSMUSG00000027794.5 | 0 | NA | NA | NA | NA | NA | #VALUE! |
| ENSMUSG00000027797.16 | 0 | NA | NA | NA | NA | NA | #VALUE! |
| ENSMUSG00000027799.13 | 0 | NA | NA | NA | NA | NA | #VALUE! |
| ENSMUSG00000027811.10 | 7.589113 | -2.4556 | 2.892037 | -0.84909 | 0.395831 | NA | #VALUE! |
| ENSMUSG00000027824.6 | 0 | NA | NA | NA | NA | NA | #VALUE! |
| ENSMUSG00000027827.18 | 0 | NA | NA | NA | NA | NA | #VALUE! |
| ENSMUSG00000027831.10 | 0 | NA | NA | NA | NA | NA | #VALUE! |
| ENSMUSG00000027832.6 | 0 | NA | NA | NA | NA | NA | #VALUE! |
| ENSMUSG00000027833.17 | 0 | NA | NA | NA | NA | NA | #VALUE! |
| ENSMUSG00000027834.16 | 4.894728 | -5.8188 | 3.915922 | -1.48593 | 0.137296 | NA | #VALUE! |
| ENSMUSG00000027840.6 | 0 | NA | NA | NA | NA | NA | #VALUE! |
| ENSMUSG00000027843.14 | 4.378546 | -5.65989 | 3.916986 | -1.44496 | 0.148469 | NA | #VALUE! |
| ENSMUSG00000027845.16 | 7.155652 | -2.70774 | 2.843879 | -0.95213 | 0.341031 | NA | #VALUE! |
| ENSMUSG00000027848.13 | 9.689906 | 3.114176 | 2.869039 | 1.085442 | 0.277726 | NA | #VALUE! |
| ENSMUSG00000027849.20 | 0 | NA | NA | NA | NA | NA | #VALUE! |
| ENSMUSG00000027855.14 | 0 | NA | NA | NA | NA | NA | #VALUE! |

| | | | | | | | |
|---|---|---|---|---|---|---|---|
| ENSMUSG00000027857.11 | 0 | NA | NA | NA | NA | NA | #VALUE! |
| ENSMUSG00000027858.14 | 2.529648 | 3.847701 | 3.920736 | 0.981372 | 0.326409 | NA | #VALUE! |
| ENSMUSG00000027859.11 | 0 | NA | NA | NA | NA | NA | #VALUE! |
| ENSMUSG00000027861.14 | 1.009701 | 3.503687 | 3.950852 | 0.886818 | 0.375177 | NA | #VALUE! |
| ENSMUSG00000027863.9 | 0 | NA | NA | NA | NA | NA | #VALUE! |
| ENSMUSG00000027864.10 | 2.943337 | 2.023459 | 3.824297 | 0.529106 | 0.596732 | NA | #VALUE! |
| ENSMUSG00000027867.11 | 0 | NA | NA | NA | NA | NA | #VALUE! |
| ENSMUSG00000027868.12 | 0 | NA | NA | NA | NA | NA | #VALUE! |
| ENSMUSG00000027869.15 | 0 | NA | NA | NA | NA | NA | #VALUE! |
| ENSMUSG00000027870.9 | 0 | NA | NA | NA | NA | NA | #VALUE! |
| ENSMUSG00000027871.17 | 0 | NA | NA | NA | NA | NA | #VALUE! |
| ENSMUSG00000027876.5 | 0 | NA | NA | NA | NA | NA | #VALUE! |
| ENSMUSG00000027880.7 | 0 | NA | NA | NA | NA | NA | #VALUE! |
| ENSMUSG00000027886.8 | 0 | NA | NA | NA | NA | NA | #VALUE! |
| ENSMUSG00000027887.12 | 0 | NA | NA | NA | NA | NA | #VALUE! |
| ENSMUSG00000027894.15 | 0 | NA | NA | NA | NA | NA | #VALUE! |
| ENSMUSG00000027895.11 | 0 | NA | NA | NA | NA | NA | #VALUE! |
| ENSMUSG00000027896.16 | 0 | NA | NA | NA | NA | NA | #VALUE! |
| ENSMUSG00000027902.9 | 0 | NA | NA | NA | NA | NA | #VALUE! |
| ENSMUSG00000027908.3 | 0 | NA | NA | NA | NA | NA | #VALUE! |
| ENSMUSG00000027912.9 | 0 | NA | NA | NA | NA | NA | #VALUE! |
| ENSMUSG00000027913.6 | 0 | NA | NA | NA | NA | NA | #VALUE! |
| ENSMUSG00000027919.6 | 0 | NA | NA | NA | NA | NA | #VALUE! |
| ENSMUSG00000027923.4 | 0 | NA | NA | NA | NA | NA | #VALUE! |
| ENSMUSG00000027925.3 | 0 | NA | NA | NA | NA | NA | #VALUE! |
| ENSMUSG00000027927.6 | 0 | NA | NA | NA | NA | NA | #VALUE! |
| ENSMUSG00000027931.13 | 7.498461 | 1.236271 | 3.223277 | 0.383545 | 0.701316 | NA | #VALUE! |
| ENSMUSG00000027932.15 | 0 | NA | NA | NA | NA | NA | #VALUE! |
| ENSMUSG00000027938.12 | 0.315039 | -1.93549 | 4.038621 | -0.47925 | 0.631763 | NA | #VALUE! |
| ENSMUSG00000027939.12 | 0 | NA | NA | NA | NA | NA | #VALUE! |
| ENSMUSG00000027950.8 | 0 | NA | NA | NA | NA | NA | #VALUE! |
| ENSMUSG00000027953.14 | 4.373289 | -2.13842 | 3.794904 | -0.5635 | 0.573095 | NA | #VALUE! |
| ENSMUSG00000027955.17 | 0 | NA | NA | NA | NA | NA | #VALUE! |
| ENSMUSG00000027956.12 | 17.50593 | 2.239574 | 2.566014 | 0.872783 | NA | NA | #VALUE! |
| ENSMUSG00000027959.15 | 2.019402 | 4.503826 | 3.928936 | 1.146322 | 0.251662 | NA | #VALUE! |
| ENSMUSG00000027961.8 | 0.373021 | 2.140748 | 4.018182 | 0.532765 | 0.594196 | NA | #VALUE! |
| ENSMUSG00000027966.21 | 0 | NA | NA | NA | NA | NA | #VALUE! |
| ENSMUSG00000027967.9 | 0.195789 | -1.05581 | 4.073302 | -0.2592 | 0.795479 | NA | #VALUE! |
| ENSMUSG00000027971.17 | 0 | NA | NA | NA | NA | NA | #VALUE! |
| ENSMUSG00000027973.12 | 0 | NA | NA | NA | NA | NA | #VALUE! |
| ENSMUSG00000027977.16 | 0 | NA | NA | NA | NA | NA | #VALUE! |
| ENSMUSG00000027978.10 | 0 | NA | NA | NA | NA | NA | #VALUE! |
| ENSMUSG00000027985.15 | 0 | NA | NA | NA | NA | NA | #VALUE! |
| ENSMUSG00000027994.15 | 0 | NA | NA | NA | NA | NA | #VALUE! |

| | | | | | | | |
|---|---|---|---|---|---|---|---|
| ENSMUSG00000027996.14 | 0 | NA | NA | NA | NA | NA | #VALUE! |
| ENSMUSG00000028004.13 | 0 | NA | NA | NA | NA | NA | #VALUE! |
| ENSMUSG00000028005.14 | 6.994725 | -3.01223 | 3.033706 | -0.99292 | 0.320748 | NA | #VALUE! |
| ENSMUSG00000028012.16 | 0 | NA | NA | NA | NA | NA | #VALUE! |
| ENSMUSG00000028017.8 | 0 | NA | NA | NA | NA | NA | #VALUE! |
| ENSMUSG00000028020.17 | 0.652786 | 2.919659 | 3.972474 | 0.734972 | 0.462356 | NA | #VALUE! |
| ENSMUSG00000028023.17 | 0 | NA | NA | NA | NA | NA | #VALUE! |
| ENSMUSG00000028028.12 | 4.048188 | -5.54974 | 3.917795 | -1.41655 | 0.156616 | NA | #VALUE! |
| ENSMUSG00000028031.7 | 0 | NA | NA | NA | NA | NA | #VALUE! |
| ENSMUSG00000028033.17 | 0 | NA | NA | NA | NA | NA | #VALUE! |
| ENSMUSG00000028036.7 | 0 | NA | NA | NA | NA | NA | #VALUE! |
| ENSMUSG00000028039.12 | 0 | NA | NA | NA | NA | NA | #VALUE! |
| ENSMUSG00000028040.8 | 0 | NA | NA | NA | NA | NA | #VALUE! |
| ENSMUSG00000028041.18 | 0 | NA | NA | NA | NA | NA | #VALUE! |
| ENSMUSG00000028047.12 | 0 | NA | NA | NA | NA | NA | #VALUE! |
| ENSMUSG00000028068.15 | 0 | NA | NA | NA | NA | NA | #VALUE! |
| ENSMUSG00000028071.13 | 0 | NA | NA | NA | NA | NA | #VALUE! |
| ENSMUSG00000028072.7 | 0 | NA | NA | NA | NA | NA | #VALUE! |
| ENSMUSG00000028073.16 | 9.978306 | 6.812606 | 3.570725 | 1.907906 | NA | NA | #VALUE! |
| ENSMUSG00000028078.15 | 0 | NA | NA | NA | NA | NA | #VALUE! |
| ENSMUSG00000028100.12 | 0 | NA | NA | NA | NA | NA | #VALUE! |
| ENSMUSG00000028101.19 | 7.239806 | -6.38344 | 3.494309 | -1.82681 | 0.067728 | NA | #VALUE! |
| ENSMUSG00000028109.15 | 0 | NA | NA | NA | NA | NA | #VALUE! |
| ENSMUSG00000028111.5 | 5.60448 | -2.27023 | 3.783435 | -0.60004 | 0.548476 | NA | #VALUE! |
| ENSMUSG00000028115.17 | 0 | NA | NA | NA | NA | NA | #VALUE! |
| ENSMUSG00000028116.14 | 0 | NA | NA | NA | NA | NA | #VALUE! |
| ENSMUSG00000028125.15 | 0 | NA | NA | NA | NA | NA | #VALUE! |
| ENSMUSG00000028128.14 | 6.026592 | -2.2672 | 2.744295 | -0.82615 | 0.40872 | NA | #VALUE! |
| ENSMUSG00000028137.10 | 0 | NA | NA | NA | NA | NA | #VALUE! |
| ENSMUSG00000028139.6 | 0 | NA | NA | NA | NA | NA | #VALUE! |
| ENSMUSG00000028141.13 | 0 | NA | NA | NA | NA | NA | #VALUE! |
| ENSMUSG00000028148.12 | 0 | NA | NA | NA | NA | NA | #VALUE! |
| ENSMUSG00000028152.11 | 3.935608 | 5.458007 | 3.918239 | 1.392975 | 0.163627 | NA | #VALUE! |
| ENSMUSG00000028159.15 | 0 | NA | NA | NA | NA | NA | #VALUE! |
| ENSMUSG00000028172.6 | 0 | NA | NA | NA | NA | NA | #VALUE! |
| ENSMUSG00000028173.11 | 6.501643 | 2.182485 | 3.313611 | 0.658642 | 0.510125 | NA | #VALUE! |
| ENSMUSG00000028174.13 | 0 | NA | NA | NA | NA | NA | #VALUE! |
| ENSMUSG00000028175.16 | 0 | NA | NA | NA | NA | NA | #VALUE! |
| ENSMUSG00000028176.14 | 0 | NA | NA | NA | NA | NA | #VALUE! |
| ENSMUSG00000028177.14 | 4.984434 | -1.19149 | 3.769338 | -0.3161 | 0.751925 | NA | #VALUE! |
| ENSMUSG00000028182.15 | 7.09395 | 0.676555 | 3.233171 | 0.209254 | 0.83425 | NA | #VALUE! |
| ENSMUSG00000028188.14 | 2.390978 | -0.40159 | 3.802891 | -0.1056 | 0.915899 | NA | #VALUE! |
| ENSMUSG00000028197.5 | 0 | NA | NA | NA | NA | NA | #VALUE! |
| ENSMUSG00000028214.14 | 1.09228 | -3.64955 | 3.947685 | -0.92448 | 0.355237 | NA | #VALUE! |

| | | | | | | | |
|---|---|---|---|---|---|---|---|
| ENSMUSG00000028217.12 | 0 | NA | NA | NA | NA | NA | #VALUE! |
| ENSMUSG00000028222.3 | 0 | NA | NA | NA | NA | NA | #VALUE! |
| ENSMUSG00000028226.16 | 0 | NA | NA | NA | NA | NA | #VALUE! |
| ENSMUSG00000028236.4 | 0 | NA | NA | NA | NA | NA | #VALUE! |
| ENSMUSG00000028244.7 | 0 | NA | NA | NA | NA | NA | #VALUE! |
| ENSMUSG00000028246.14 | 0 | NA | NA | NA | NA | NA | #VALUE! |
| ENSMUSG00000028255.7 | 0 | NA | NA | NA | NA | NA | #VALUE! |
| ENSMUSG00000028259.14 | 0 | NA | NA | NA | NA | NA | #VALUE! |
| ENSMUSG00000028262.14 | 0 | NA | NA | NA | NA | NA | #VALUE! |
| ENSMUSG00000028264.17 | 0 | NA | NA | NA | NA | NA | #VALUE! |
| ENSMUSG00000028280.10 | 0 | NA | NA | NA | NA | NA | #VALUE! |
| ENSMUSG00000028287.5 | 0 | NA | NA | NA | NA | NA | #VALUE! |
| ENSMUSG00000028289.13 | 1.938301 | -4.49059 | 3.929692 | -1.14273 | 0.253149 | NA | #VALUE! |
| ENSMUSG00000028294.16 | 0 | NA | NA | NA | NA | NA | #VALUE! |
| ENSMUSG00000028298.11 | 0 | NA | NA | NA | NA | NA | #VALUE! |
| ENSMUSG00000028310.3 | 0 | NA | NA | NA | NA | NA | #VALUE! |
| ENSMUSG00000028314.7 | 0 | NA | NA | NA | NA | NA | #VALUE! |
| ENSMUSG00000028328.14 | 0 | NA | NA | NA | NA | NA | #VALUE! |
| ENSMUSG00000028331.13 | 1.177985 | 3.72614 | 3.944615 | 0.944614 | 0.344856 | NA | #VALUE! |
| ENSMUSG00000028332.14 | 0 | NA | NA | NA | NA | NA | #VALUE! |
| ENSMUSG00000028337.15 | 2.58392 | -4.90098 | 3.924036 | -1.24896 | 0.211678 | NA | #VALUE! |
| ENSMUSG00000028339.18 | 7.850364 | -0.73948 | 3.608091 | -0.20495 | 0.83761 | NA | #VALUE! |
| ENSMUSG00000028341.10 | 4.095512 | -5.57062 | 3.917637 | -1.42193 | 0.155045 | NA | #VALUE! |
| ENSMUSG00000028347.15 | 0 | NA | NA | NA | NA | NA | #VALUE! |
| ENSMUSG00000028348.8 | 0 | NA | NA | NA | NA | NA | #VALUE! |
| ENSMUSG00000028351.6 | 0 | NA | NA | NA | NA | NA | #VALUE! |
| ENSMUSG00000028354.14 | 5.449071 | -1.16935 | 3.609377 | -0.32398 | 0.745956 | NA | #VALUE! |
| ENSMUSG00000028357.14 | 0 | NA | NA | NA | NA | NA | #VALUE! |
| ENSMUSG00000028358.16 | 0 | NA | NA | NA | NA | NA | #VALUE! |
| ENSMUSG00000028360.11 | 0 | NA | NA | NA | NA | NA | #VALUE! |
| ENSMUSG00000028362.3 | 0 | NA | NA | NA | NA | NA | #VALUE! |
| ENSMUSG00000028364.16 | 0 | NA | NA | NA | NA | NA | #VALUE! |
| ENSMUSG00000028369.16 | 0 | NA | NA | NA | NA | NA | #VALUE! |
| ENSMUSG00000028370.8 | 0 | NA | NA | NA | NA | NA | #VALUE! |
| ENSMUSG00000028373.17 | 2.21378 | 4.621487 | 3.927208 | 1.176787 | 0.239281 | NA | #VALUE! |
| ENSMUSG00000028385.15 | 6.908645 | 5.322371 | 2.853306 | 1.865335 | 0.062135 | NA | #VALUE! |
| ENSMUSG00000028389.13 | 0 | NA | NA | NA | NA | NA | #VALUE! |
| ENSMUSG00000028391.17 | 0 | NA | NA | NA | NA | NA | #VALUE! |
| ENSMUSG00000028392.16 | 0 | NA | NA | NA | NA | NA | #VALUE! |
| ENSMUSG00000028396.6 | 0 | NA | NA | NA | NA | NA | #VALUE! |
| ENSMUSG00000028415.5 | 0 | NA | NA | NA | NA | NA | #VALUE! |
| ENSMUSG00000028417.4 | 0 | NA | NA | NA | NA | NA | #VALUE! |
| ENSMUSG00000028427.14 | 0 | NA | NA | NA | NA | NA | #VALUE! |
| ENSMUSG00000028430.15 | 7.090504 | -0.91152 | 2.815166 | -0.32379 | 0.746099 | NA | #VALUE! |

| | | | | | | | |
|---|---|---|---|---|---|---|---|
| ENSMUSG00000028435.9 | 0 | NA | NA | NA | NA | NA | #VALUE! |
| ENSMUSG00000028438.17 | 0 | NA | NA | NA | NA | NA | #VALUE! |
| ENSMUSG00000028439.15 | 1.050131 | -3.6251 | 3.948379 | -0.91812 | 0.358554 | NA | #VALUE! |
| ENSMUSG00000028441.13 | 0 | NA | NA | NA | NA | NA | #VALUE! |
| ENSMUSG00000028444.18 | 0 | NA | NA | NA | NA | NA | #VALUE! |
| ENSMUSG00000028445.8 | 9.171792 | -6.72804 | 3.178841 | -2.11651 | 0.034302 | NA | #VALUE! |
| ENSMUSG00000028451.13 | 0 | NA | NA | NA | NA | NA | #VALUE! |
| ENSMUSG00000028453.11 | 3.533954 | 5.311169 | 3.919465 | 1.355075 | 0.175394 | NA | #VALUE! |
| ENSMUSG00000028457.19 | 9.095712 | -3.00593 | 3.371971 | -0.89144 | NA | NA | #VALUE! |
| ENSMUSG00000028459.12 | 9.760099 | -6.81468 | 3.14792 | -2.16482 | 0.030401 | NA | #VALUE! |
| ENSMUSG00000028460.7 | 0 | NA | NA | NA | NA | NA | #VALUE! |
| ENSMUSG00000028463.15 | 0.315039 | -1.93549 | 4.038621 | -0.47925 | 0.631763 | NA | #VALUE! |
| ENSMUSG00000028475.10 | 0 | NA | NA | NA | NA | NA | #VALUE! |
| ENSMUSG00000028480.15 | 0 | NA | NA | NA | NA | NA | #VALUE! |
| ENSMUSG00000028487.19 | 0 | NA | NA | NA | NA | NA | #VALUE! |
| ENSMUSG00000028488.16 | 0 | NA | NA | NA | NA | NA | #VALUE! |
| ENSMUSG00000028492.14 | 0 | NA | NA | NA | NA | NA | #VALUE! |
| ENSMUSG00000028496.18 | 9.529962 | -1.60822 | 2.915405 | -0.55163 | 0.581203 | NA | #VALUE! |
| ENSMUSG00000028497.13 | 2.144869 | 4.605508 | 3.927435 | 1.17265 | 0.240936 | NA | #VALUE! |
| ENSMUSG00000028519.17 | 0 | NA | NA | NA | NA | NA | #VALUE! |
| ENSMUSG00000028520.13 | 0 | NA | NA | NA | NA | NA | #VALUE! |
| ENSMUSG00000028523.15 | 0 | NA | NA | NA | NA | NA | #VALUE! |
| ENSMUSG00000028524.22 | 0.373021 | 2.140748 | 4.018182 | 0.532765 | 0.594196 | NA | #VALUE! |
| ENSMUSG00000028528.18 | 0 | NA | NA | NA | NA | NA | #VALUE! |
| ENSMUSG00000028532.15 | 0 | NA | NA | NA | NA | NA | #VALUE! |
| ENSMUSG00000028533.11 | 0 | NA | NA | NA | NA | NA | #VALUE! |
| ENSMUSG00000028539.15 | 0 | NA | NA | NA | NA | NA | #VALUE! |
| ENSMUSG00000028541.15 | 0 | NA | NA | NA | NA | NA | #VALUE! |
| ENSMUSG00000028544.15 | 0 | NA | NA | NA | NA | NA | #VALUE! |
| ENSMUSG00000028545.14 | 0.587367 | -2.74297 | 3.982932 | -0.68868 | 0.491024 | NA | #VALUE! |
| ENSMUSG00000028546.18 | 0 | NA | NA | NA | NA | NA | #VALUE! |
| ENSMUSG00000028555.16 | 3.146614 | 0.383512 | 3.78538 | 0.101314 | 0.919301 | NA | #VALUE! |
| ENSMUSG00000028558.15 | 0 | NA | NA | NA | NA | NA | #VALUE! |
| ENSMUSG00000028571.16 | 0 | NA | NA | NA | NA | NA | #VALUE! |
| ENSMUSG00000028575.12 | 0.840105 | -3.30885 | 3.958472 | -0.83589 | 0.403216 | NA | #VALUE! |
| ENSMUSG00000028581.18 | 5.95689 | -6.10215 | 3.752695 | -1.62607 | 0.103935 | NA | #VALUE! |
| ENSMUSG00000028583.15 | 1.229878 | 3.761857 | 3.943699 | 0.95389 | 0.340139 | NA | #VALUE! |
| ENSMUSG00000028584.4 | 0 | NA | NA | NA | NA | NA | #VALUE! |
| ENSMUSG00000028587.19 | 0.105013 | -0.97324 | 4.076293 | -0.23876 | 0.811295 | NA | #VALUE! |
| ENSMUSG00000028589.3 | 0 | NA | NA | NA | NA | NA | #VALUE! |
| ENSMUSG00000028590.7 | 0 | NA | NA | NA | NA | NA | #VALUE! |
| ENSMUSG00000028591.12 | 0 | NA | NA | NA | NA | NA | #VALUE! |
| ENSMUSG00000028593.6 | 0 | NA | NA | NA | NA | NA | #VALUE! |
| ENSMUSG00000028597.12 | 8.365007 | 2.03067 | 3.079595 | 0.659395 | NA | NA | #VALUE! |

| | | | | | | | |
|---|---|---|---|---|---|---|---|
| ENSMUSG00000028600.16 | 1.00278 | -0.14045 | 3.909579 | -0.03592 | 0.971343 | NA | #VALUE! |
| ENSMUSG00000028602.13 | 0 | NA | NA | NA | NA | NA | #VALUE! |
| ENSMUSG00000028610.17 | 0 | NA | NA | NA | NA | NA | #VALUE! |
| ENSMUSG00000028612.11 | 0 | NA | NA | NA | NA | NA | #VALUE! |
| ENSMUSG00000028613.16 | 0 | NA | NA | NA | NA | NA | #VALUE! |
| ENSMUSG00000028626.6 | 0 | NA | NA | NA | NA | NA | #VALUE! |
| ENSMUSG00000028631.8 | 0 | NA | NA | NA | NA | NA | #VALUE! |
| ENSMUSG00000028633.8 | 7.826996 | -2.06224 | 2.274142 | -0.90682 | 0.364502 | NA | #VALUE! |
| ENSMUSG00000028634.18 | 0.766389 | 3.110667 | 3.964448 | 0.784641 | 0.432664 | NA | #VALUE! |
| ENSMUSG00000028635.8 | 0 | NA | NA | NA | NA | NA | #VALUE! |
| ENSMUSG00000028640.12 | 0 | NA | NA | NA | NA | NA | #VALUE! |
| ENSMUSG00000028642.8 | 0 | NA | NA | NA | NA | NA | #VALUE! |
| ENSMUSG00000028644.18 | 0 | NA | NA | NA | NA | NA | #VALUE! |
| ENSMUSG00000028651.13 | 9.825716 | -1.73592 | 2.894808 | -0.59967 | 0.548728 | NA | #VALUE! |
| ENSMUSG00000028654.14 | 7.317396 | 2.034648 | 3.238606 | 0.628248 | 0.529842 | NA | #VALUE! |
| ENSMUSG00000028656.15 | 0 | NA | NA | NA | NA | NA | #VALUE! |
| ENSMUSG00000028661.9 | 0 | NA | NA | NA | NA | NA | #VALUE! |
| ENSMUSG00000028664.15 | 0 | NA | NA | NA | NA | NA | #VALUE! |
| ENSMUSG00000028677.20 | 2.671009 | 2.830954 | 3.882263 | 0.729202 | 0.465878 | NA | #VALUE! |
| ENSMUSG00000028678.14 | 0 | NA | NA | NA | NA | NA | #VALUE! |
| ENSMUSG00000028680.15 | 2.464018 | 0.934075 | 3.807352 | 0.245335 | 0.806197 | NA | #VALUE! |
| ENSMUSG00000028681.12 | 6.735655 | -6.27951 | 3.913422 | -1.60461 | 0.10858 | NA | #VALUE! |
| ENSMUSG00000028687.18 | 0 | NA | NA | NA | NA | NA | #VALUE! |
| ENSMUSG00000028689.15 | 7.698519 | -6.47341 | 3.404538 | -1.90141 | 0.057249 | NA | #VALUE! |
| ENSMUSG00000028698.14 | 9.241582 | 4.300842 | 2.897614 | 1.48427 | 0.137737 | NA | #VALUE! |
| ENSMUSG00000028699.10 | 0 | NA | NA | NA | NA | NA | #VALUE! |
| ENSMUSG00000028701.5 | 0 | NA | NA | NA | NA | NA | #VALUE! |
| ENSMUSG00000028702.16 | 4.286326 | -5.62615 | 3.917227 | -1.43626 | 0.150929 | NA | #VALUE! |
| ENSMUSG00000028707.16 | 0 | NA | NA | NA | NA | NA | #VALUE! |
| ENSMUSG00000028717.13 | 0 | NA | NA | NA | NA | NA | #VALUE! |
| ENSMUSG00000028718.17 | 0 | NA | NA | NA | NA | NA | #VALUE! |
| ENSMUSG00000028730.15 | 0 | NA | NA | NA | NA | NA | #VALUE! |
| ENSMUSG00000028736.14 | 0 | NA | NA | NA | NA | NA | #VALUE! |
| ENSMUSG00000028738.15 | 0 | NA | NA | NA | NA | NA | #VALUE! |
| ENSMUSG00000028747.11 | 0 | NA | NA | NA | NA | NA | #VALUE! |
| ENSMUSG00000028749.8 | 0 | NA | NA | NA | NA | NA | #VALUE! |
| ENSMUSG00000028750.13 | 0 | NA | NA | NA | NA | NA | #VALUE! |
| ENSMUSG00000028751.14 | 0 | NA | NA | NA | NA | NA | #VALUE! |
| ENSMUSG00000028753.13 | 0 | NA | NA | NA | NA | NA | #VALUE! |
| ENSMUSG00000028755.4 | 3.047873 | 5.100691 | 3.921452 | 1.300715 | 0.193356 | NA | #VALUE! |
| ENSMUSG00000028758.13 | 0 | NA | NA | NA | NA | NA | #VALUE! |
| ENSMUSG00000028773.9 | 0 | NA | NA | NA | NA | NA | #VALUE! |
| ENSMUSG00000028777.9 | 0 | NA | NA | NA | NA | NA | #VALUE! |
| ENSMUSG00000028778.16 | 0 | NA | NA | NA | NA | NA | #VALUE! |

| | | | | | | | |
|---|---|---|---|---|---|---|---|
| ENSMUSG00000028780.14 | 0.630079 | -2.90291 | 3.975047 | -0.73028 | 0.465217 | NA | #VALUE! |
| ENSMUSG00000028782.15 | 0 | NA | NA | NA | NA | NA | #VALUE! |
| ENSMUSG00000028784.15 | 0 | NA | NA | NA | NA | NA | #VALUE! |
| ENSMUSG00000028785.14 | 0 | NA | NA | NA | NA | NA | #VALUE! |
| ENSMUSG00000028786.16 | 0 | NA | NA | NA | NA | NA | #VALUE! |
| ENSMUSG00000028789.17 | 0 | NA | NA | NA | NA | NA | #VALUE! |
| ENSMUSG00000028794.14 | 0 | NA | NA | NA | NA | NA | #VALUE! |
| ENSMUSG00000028795.16 | 3.055303 | -0.39721 | 3.78672 | -0.1049 | 0.916459 | NA | #VALUE! |
| ENSMUSG00000028799.16 | 2.741047 | -4.98044 | 3.923113 | -1.26951 | 0.204259 | NA | #VALUE! |
| ENSMUSG00000028801.15 | 0 | NA | NA | NA | NA | NA | #VALUE! |
| ENSMUSG00000028804.21 | 0 | NA | NA | NA | NA | NA | #VALUE! |
| ENSMUSG00000028807.3 | 3.224875 | -1.93185 | 3.811719 | -0.50682 | 0.612281 | NA | #VALUE! |
| ENSMUSG00000028813.3 | 0 | NA | NA | NA | NA | NA | #VALUE! |
| ENSMUSG00000028825.8 | 0 | NA | NA | NA | NA | NA | #VALUE! |
| ENSMUSG00000028834.14 | 0 | NA | NA | NA | NA | NA | #VALUE! |
| ENSMUSG00000028836.15 | 0 | NA | NA | NA | NA | NA | #VALUE! |
| ENSMUSG00000028838.12 | 2.167917 | -4.64264 | 3.927408 | -1.18211 | 0.237161 | NA | #VALUE! |
| ENSMUSG00000028841.15 | 0 | NA | NA | NA | NA | NA | #VALUE! |
| ENSMUSG00000028843.9 | 6.521619 | 3.29889 | 2.917591 | 1.13069 | 0.258186 | NA | #VALUE! |
| ENSMUSG00000028845.16 | 0 | NA | NA | NA | NA | NA | #VALUE! |
| ENSMUSG00000028850.3 | 0 | NA | NA | NA | NA | NA | #VALUE! |
| ENSMUSG00000028854.10 | 3.823463 | 5.43312 | 3.918438 | 1.386552 | 0.165578 | NA | #VALUE! |
| ENSMUSG00000028857.17 | 4.086379 | -0.38986 | 3.771319 | -0.10337 | 0.917666 | NA | #VALUE! |
| ENSMUSG00000028859.15 | 1.890236 | -4.46257 | 3.93014 | -1.13547 | 0.256177 | NA | #VALUE! |
| ENSMUSG00000028860.14 | 0 | NA | NA | NA | NA | NA | #VALUE! |
| ENSMUSG00000028862.7 | 0 | NA | NA | NA | NA | NA | #VALUE! |
| ENSMUSG00000028864.8 | 5.710652 | -6.04563 | 3.780477 | -1.59917 | 0.109783 | NA | #VALUE! |
| ENSMUSG00000028865.15 | 0 | NA | NA | NA | NA | NA | #VALUE! |
| ENSMUSG00000028871.8 | 0 | NA | NA | NA | NA | NA | #VALUE! |
| ENSMUSG00000028873.17 | 0 | NA | NA | NA | NA | NA | #VALUE! |
| ENSMUSG00000028874.15 | 0 | NA | NA | NA | NA | NA | #VALUE! |
| ENSMUSG00000028876.8 | 0.279766 | 1.808031 | 4.036658 | 0.447903 | 0.654223 | NA | #VALUE! |
| ENSMUSG00000028883.20 | 0 | NA | NA | NA | NA | NA | #VALUE! |
| ENSMUSG00000028885.9 | 0 | NA | NA | NA | NA | NA | #VALUE! |
| ENSMUSG00000028893.9 | 2.545258 | -4.8732 | 3.92437 | -1.24178 | 0.214318 | NA | #VALUE! |
| ENSMUSG00000028896.14 | 0 | NA | NA | NA | NA | NA | #VALUE! |
| ENSMUSG00000028909.18 | 0 | NA | NA | NA | NA | NA | #VALUE! |
| ENSMUSG00000028917.15 | 5.042143 | 5.82709 | 3.915654 | 1.488152 | 0.136711 | NA | #VALUE! |
| ENSMUSG00000028923.15 | 3.450833 | -5.31342 | 3.919753 | -1.35555 | 0.175242 | NA | #VALUE! |
| ENSMUSG00000028926.16 | 3.100758 | 2.757021 | 3.857193 | 0.714774 | 0.474749 | NA | #VALUE! |
| ENSMUSG00000028927.7 | 0 | NA | NA | NA | NA | NA | #VALUE! |
| ENSMUSG00000028931.13 | 3.150394 | -5.19405 | 3.92087 | -1.32472 | 0.185265 | NA | #VALUE! |
| ENSMUSG00000028933.12 | 16.12475 | -21.7525 | 3.121981 | -6.96755 | NA | NA | #VALUE! |
| ENSMUSG00000028938.10 | 0 | NA | NA | NA | NA | NA | #VALUE! |

| | | | | | | | |
|---|---|---|---|---|---|---|---|
| ENSMUSG00000028940.3 | 0 | NA | NA | NA | NA | NA | #VALUE! |
| ENSMUSG00000028943.19 | 0 | NA | NA | NA | NA | NA | #VALUE! |
| ENSMUSG00000028946.8 | 0 | NA | NA | NA | NA | NA | #VALUE! |
| ENSMUSG00000028949.14 | 0 | NA | NA | NA | NA | NA | #VALUE! |
| ENSMUSG00000028950.4 | 0 | NA | NA | NA | NA | NA | #VALUE! |
| ENSMUSG00000028953.11 | 9.34707 | 6.710288 | 3.689587 | 1.81871 | 0.068956 | NA | #VALUE! |
| ENSMUSG00000028958.16 | 0.195789 | -1.05581 | 4.073302 | -0.2592 | 0.795479 | NA | #VALUE! |
| ENSMUSG00000028959.15 | 1.721829 | 4.254739 | 3.933085 | 1.081782 | 0.27935 | NA | #VALUE! |
| ENSMUSG00000028963.2 | 0 | NA | NA | NA | NA | NA | #VALUE! |
| ENSMUSG00000028965.14 | 0 | NA | NA | NA | NA | NA | #VALUE! |
| ENSMUSG00000028969.11 | 8.669542 | -0.10442 | 3.051633 | -0.03422 | 0.972703 | NA | #VALUE! |
| ENSMUSG00000028970.10 | 1.865104 | 4.40601 | 3.930481 | 1.120985 | 0.262294 | NA | #VALUE! |
| ENSMUSG00000028971.5 | 0 | NA | NA | NA | NA | NA | #VALUE! |
| ENSMUSG00000028972.12 | 0 | NA | NA | NA | NA | NA | #VALUE! |
| ENSMUSG00000028977.18 | 0 | NA | NA | NA | NA | NA | #VALUE! |
| ENSMUSG00000028989.4 | 0 | NA | NA | NA | NA | NA | #VALUE! |
| ENSMUSG00000028995.15 | 32.44958 | -1.6179 | 1.535811 | -1.05345 | NA | NA | #VALUE! |
| ENSMUSG00000028996.10 | 0 | NA | NA | NA | NA | NA | #VALUE! |
| ENSMUSG00000029001.16 | 1.771849 | 4.332864 | 3.931707 | 1.102031 | 0.270448 | NA | #VALUE! |
| ENSMUSG00000029005.5 | 0 | NA | NA | NA | NA | NA | #VALUE! |
| ENSMUSG00000029009.18 | 6.771175 | 6.254637 | 3.621597 | 1.727038 | 0.084161 | NA | #VALUE! |
| ENSMUSG00000029015.10 | 0 | NA | NA | NA | NA | NA | #VALUE! |
| ENSMUSG00000029016.14 | 6.153439 | -4.23341 | 3.846836 | -1.10049 | 0.271118 | NA | #VALUE! |
| ENSMUSG00000029019.9 | 0 | NA | NA | NA | NA | NA | #VALUE! |
| ENSMUSG00000029026.18 | 0 | NA | NA | NA | NA | NA | #VALUE! |
| ENSMUSG00000029027.10 | 4.243542 | 5.575812 | 3.917342 | 1.423366 | 0.15463 | NA | #VALUE! |
| ENSMUSG00000029029.15 | 5.83343 | -0.91853 | 3.066887 | -0.2995 | 0.76456 | NA | #VALUE! |
| ENSMUSG00000029032.13 | 0 | NA | NA | NA | NA | NA | #VALUE! |
| ENSMUSG00000029033.17 | 0 | NA | NA | NA | NA | NA | #VALUE! |
| ENSMUSG00000029044.10 | 0 | NA | NA | NA | NA | NA | #VALUE! |
| ENSMUSG00000029047.14 | 0 | NA | NA | NA | NA | NA | #VALUE! |
| ENSMUSG00000029049.15 | 4.111571 | -5.56681 | 3.917665 | -1.42095 | 0.155331 | NA | #VALUE! |
| ENSMUSG00000029054.9 | 0 | NA | NA | NA | NA | NA | #VALUE! |
| ENSMUSG00000029055.18 | 0 | NA | NA | NA | NA | NA | #VALUE! |
| ENSMUSG00000029060.18 | 0 | NA | NA | NA | NA | NA | #VALUE! |
| ENSMUSG00000029061.3 | 1.602939 | 2.980712 | 3.930773 | 0.758302 | 0.44827 | NA | #VALUE! |
| ENSMUSG00000029072.4 | 2.611145 | 4.886731 | 3.92379 | 1.245411 | 0.212981 | NA | #VALUE! |
| ENSMUSG00000029074.15 | 0 | NA | NA | NA | NA | NA | #VALUE! |
| ENSMUSG00000029075.7 | 0 | NA | NA | NA | NA | NA | #VALUE! |
| ENSMUSG00000029086.16 | 0 | NA | NA | NA | NA | NA | #VALUE! |
| ENSMUSG00000029088.17 | 0 | NA | NA | NA | NA | NA | #VALUE! |
| ENSMUSG00000029092.10 | 0 | NA | NA | NA | NA | NA | #VALUE! |
| ENSMUSG00000029093.15 | 0 | NA | NA | NA | NA | NA | #VALUE! |
| ENSMUSG00000029094.13 | 0.210026 | -1.52494 | 4.058692 | -0.37572 | 0.707123 | NA | #VALUE! |

| | | | | | | | |
|---|---|---|---|---|---|---|---|
| ENSMUSG00000029095.18 | 6.248098 | 6.138416 | 3.913931 | 1.56835 | 0.116799 | NA | #VALUE! |
| ENSMUSG00000029096.16 | 2.21378 | 4.621487 | 3.927208 | 1.176787 | 0.239281 | NA | #VALUE! |
| ENSMUSG00000029097.12 | 0.279766 | 1.808031 | 4.036658 | 0.447903 | 0.654223 | NA | #VALUE! |
| ENSMUSG00000029101.17 | 1.940132 | 4.460891 | 3.929601 | 1.135202 | 0.256291 | NA | #VALUE! |
| ENSMUSG00000029104.16 | 3.737971 | 0.371018 | 3.616199 | 0.102599 | 0.918281 | NA | #VALUE! |
| ENSMUSG00000029108.15 | 7.571633 | -2.80206 | 2.767227 | -1.01259 | 0.311258 | NA | #VALUE! |
| ENSMUSG00000029112.6 | 0 | NA | NA | NA | NA | NA | #VALUE! |
| ENSMUSG00000029119.10 | 3.798088 | -0.46849 | 3.523274 | -0.13297 | 0.894217 | NA | #VALUE! |
| ENSMUSG00000029120.11 | 0 | NA | NA | NA | NA | NA | #VALUE! |
| ENSMUSG00000029121.17 | 0 | NA | NA | NA | NA | NA | #VALUE! |
| ENSMUSG00000029122.12 | 7.143876 | -6.36453 | 3.913041 | -1.62649 | 0.103845 | NA | #VALUE! |
| ENSMUSG00000029123.9 | 0 | NA | NA | NA | NA | NA | #VALUE! |
| ENSMUSG00000029126.8 | 0 | NA | NA | NA | NA | NA | #VALUE! |
| ENSMUSG00000029127.16 | 4.703895 | -1.94845 | 3.785923 | -0.51466 | 0.606794 | NA | #VALUE! |
| ENSMUSG00000029130.13 | 0 | NA | NA | NA | NA | NA | #VALUE! |
| ENSMUSG00000029134.15 | 0 | NA | NA | NA | NA | NA | #VALUE! |
| ENSMUSG00000029138.5 | 0 | NA | NA | NA | NA | NA | #VALUE! |
| ENSMUSG00000029146.15 | 3.339608 | -0.29512 | 3.780882 | -0.07806 | 0.937783 | NA | #VALUE! |
| ENSMUSG00000029149.15 | 0 | NA | NA | NA | NA | NA | #VALUE! |
| ENSMUSG00000029151.15 | 0 | NA | NA | NA | NA | NA | #VALUE! |
| ENSMUSG00000029154.14 | 0 | NA | NA | NA | NA | NA | #VALUE! |
| ENSMUSG00000029155.17 | 0 | NA | NA | NA | NA | NA | #VALUE! |
| ENSMUSG00000029158.10 | 0 | NA | NA | NA | NA | NA | #VALUE! |
| ENSMUSG00000029161.9 | 0 | NA | NA | NA | NA | NA | #VALUE! |
| ENSMUSG00000029165.18 | 6.269769 | 3.226041 | 3.810009 | 0.846728 | 0.397147 | NA | #VALUE! |
| ENSMUSG00000029168.15 | 0 | NA | NA | NA | NA | NA | #VALUE! |
| ENSMUSG00000029171.13 | 7.708218 | -0.71848 | 3.153386 | -0.22784 | 0.819768 | NA | #VALUE! |
| ENSMUSG00000029174.19 | 8.566593 | -5.54611 | 2.611871 | -2.12343 | 0.033718 | NA | #VALUE! |
| ENSMUSG00000029182.11 | 0 | NA | NA | NA | NA | NA | #VALUE! |
| ENSMUSG00000029184.6 | 0 | NA | NA | NA | NA | NA | #VALUE! |
| ENSMUSG00000029188.15 | 7.706391 | 6.440911 | 3.507221 | 1.836471 | 0.066288 | NA | #VALUE! |
| ENSMUSG00000029193.8 | 0 | NA | NA | NA | NA | NA | #VALUE! |
| ENSMUSG00000029204.8 | 2.688599 | -3.80635 | 3.910477 | -0.97337 | 0.330369 | NA | #VALUE! |
| ENSMUSG00000029205.13 | 0 | NA | NA | NA | NA | NA | #VALUE! |
| ENSMUSG00000029206.14 | 0 | NA | NA | NA | NA | NA | #VALUE! |
| ENSMUSG00000029209.16 | 6.812178 | 3.663034 | 2.388521 | 1.533599 | 0.125128 | NA | #VALUE! |
| ENSMUSG00000029211.12 | 0 | NA | NA | NA | NA | NA | #VALUE! |
| ENSMUSG00000029212.12 | 0 | NA | NA | NA | NA | NA | #VALUE! |
| ENSMUSG00000029217.17 | 7.34784 | 0.411228 | 2.844696 | 0.14456 | 0.885059 | NA | #VALUE! |
| ENSMUSG00000029219.11 | 0 | NA | NA | NA | NA | NA | #VALUE! |
| ENSMUSG00000029223.13 | 0 | NA | NA | NA | NA | NA | #VALUE! |
| ENSMUSG00000029235.15 | 0 | NA | NA | NA | NA | NA | #VALUE! |
| ENSMUSG00000029236.5 | 0 | NA | NA | NA | NA | NA | #VALUE! |
| ENSMUSG00000029245.17 | 0 | NA | NA | NA | NA | NA | #VALUE! |

| | | | | | | | |
|---|---|---|---|---|---|---|---|
| ENSMUSG00000029248.15 | 0 | NA | NA | NA | NA | NA | #VALUE! |
| ENSMUSG00000029254.17 | 1.052966 | 1.112538 | 3.925595 | 0.283406 | 0.776866 | NA | #VALUE! |
| ENSMUSG00000029255.9 | 0 | NA | NA | NA | NA | NA | #VALUE! |
| ENSMUSG00000029268.15 | 0 | NA | NA | NA | NA | NA | #VALUE! |
| ENSMUSG00000029275.19 | 0 | NA | NA | NA | NA | NA | #VALUE! |
| ENSMUSG00000029280.14 | 0 | NA | NA | NA | NA | NA | #VALUE! |
| ENSMUSG00000029281.14 | 0 | NA | NA | NA | NA | NA | #VALUE! |
| ENSMUSG00000029282.4 | 0 | NA | NA | NA | NA | NA | #VALUE! |
| ENSMUSG00000029283.18 | 0 | NA | NA | NA | NA | NA | #VALUE! |
| ENSMUSG00000029286.13 | 0 | NA | NA | NA | NA | NA | #VALUE! |
| ENSMUSG00000029288.12 | 0 | NA | NA | NA | NA | NA | #VALUE! |
| ENSMUSG00000029298.16 | 3.914993 | -3.18002 | 3.854438 | -0.82503 | 0.409355 | NA | #VALUE! |
| ENSMUSG00000029306.9 | 0 | NA | NA | NA | NA | NA | #VALUE! |
| ENSMUSG00000029307.8 | 0 | NA | NA | NA | NA | NA | #VALUE! |
| ENSMUSG00000029312.13 | 9.275316 | 1.183007 | 2.973083 | 0.397906 | 0.6907 | NA | #VALUE! |
| ENSMUSG00000029320.12 | 0 | NA | NA | NA | NA | NA | #VALUE! |
| ENSMUSG00000029321.2 | 0 | NA | NA | NA | NA | NA | #VALUE! |
| ENSMUSG00000029326.14 | 1.831447 | -1.50814 | 3.852603 | -0.39146 | 0.695458 | NA | #VALUE! |
| ENSMUSG00000029330.9 | 0.840105 | -3.30885 | 3.958472 | -0.83589 | 0.403216 | NA | #VALUE! |
| ENSMUSG00000029331.4 | 0 | NA | NA | NA | NA | NA | #VALUE! |
| ENSMUSG00000029334.15 | 1.958359 | 4.475621 | 3.929371 | 1.139017 | 0.254696 | NA | #VALUE! |
| ENSMUSG00000029335.6 | 0 | NA | NA | NA | NA | NA | #VALUE! |
| ENSMUSG00000029337.3 | 0 | NA | NA | NA | NA | NA | #VALUE! |
| ENSMUSG00000029343.18 | 0 | NA | NA | NA | NA | NA | #VALUE! |
| ENSMUSG00000029348.12 | 0 | NA | NA | NA | NA | NA | #VALUE! |
| ENSMUSG00000029352.15 | 0.105013 | -0.97324 | 4.076293 | -0.23876 | 0.811295 | NA | #VALUE! |
| ENSMUSG00000029359.14 | 0.559531 | 2.703958 | 3.982863 | 0.678898 | 0.497202 | NA | #VALUE! |
| ENSMUSG00000029360.4 | 6.731341 | 6.240743 | 3.913441 | 1.594695 | 0.110781 | NA | #VALUE! |
| ENSMUSG00000029361.19 | 0 | NA | NA | NA | NA | NA | #VALUE! |
| ENSMUSG00000029363.15 | 22.34389 | -2.02782 | 2.198132 | -0.92252 | NA | NA | #VALUE! |
| ENSMUSG00000029371.8 | 0 | NA | NA | NA | NA | NA | #VALUE! |
| ENSMUSG00000029372.8 | 0.81989 | 0.607554 | 3.951222 | 0.153764 | 0.877796 | NA | #VALUE! |
| ENSMUSG00000029373.8 | 1.114714 | 2.570661 | 3.946534 | 0.651372 | 0.514806 | NA | #VALUE! |
| ENSMUSG00000029375.7 | 0 | NA | NA | NA | NA | NA | #VALUE! |
| ENSMUSG00000029377.6 | 0 | NA | NA | NA | NA | NA | #VALUE! |
| ENSMUSG00000029378.6 | 0 | NA | NA | NA | NA | NA | #VALUE! |
| ENSMUSG00000029379.11 | 0 | NA | NA | NA | NA | NA | #VALUE! |
| ENSMUSG00000029380.12 | 323.3605 | -3.86113 | 1.790029 | -2.15702 | NA | NA | #VALUE! |
| ENSMUSG00000029384.6 | 0.168284 | 1.027628 | 4.067126 | 0.252667 | 0.800526 | NA | #VALUE! |
| ENSMUSG00000029408.14 | 0.420053 | -2.33493 | 4.007343 | -0.58266 | 0.56012 | NA | #VALUE! |
| ENSMUSG00000029409.8 | 0 | NA | NA | NA | NA | NA | #VALUE! |
| ENSMUSG00000029410.12 | 1.865104 | 4.40601 | 3.930481 | 1.120985 | 0.262294 | NA | #VALUE! |
| ENSMUSG00000029413.15 | 9.5643 | 0.941605 | 2.922196 | 0.322225 | 0.747282 | NA | #VALUE! |
| ENSMUSG00000029414.12 | 0 | NA | NA | NA | NA | NA | #VALUE! |

| | | | | | | | |
|---|---|---|---|---|---|---|---|
| ENSMUSG00000029419.9 | 0 | NA | NA | NA | NA | NA | #VALUE! |
| ENSMUSG00000029420.15 | 0 | NA | NA | NA | NA | NA | #VALUE! |
| ENSMUSG00000029423.11 | 0 | NA | NA | NA | NA | NA | #VALUE! |
| ENSMUSG00000029431.6 | 0 | NA | NA | NA | NA | NA | #VALUE! |
| ENSMUSG00000029436.10 | 0 | NA | NA | NA | NA | NA | #VALUE! |
| ENSMUSG00000029437.13 | 0 | NA | NA | NA | NA | NA | #VALUE! |
| ENSMUSG00000029438.10 | 4.877439 | 4.697347 | 3.692952 | 1.271976 | 0.203382 | NA | #VALUE! |
| ENSMUSG00000029442.19 | 0 | NA | NA | NA | NA | NA | #VALUE! |
| ENSMUSG00000029449.12 | 0 | NA | NA | NA | NA | NA | #VALUE! |
| ENSMUSG00000029451.15 | 0 | NA | NA | NA | NA | NA | #VALUE! |
| ENSMUSG00000029452.19 | 0 | NA | NA | NA | NA | NA | #VALUE! |
| ENSMUSG00000029454.16 | 0 | NA | NA | NA | NA | NA | #VALUE! |
| ENSMUSG00000029468.18 | 1.738059 | 4.300351 | 3.932271 | 1.093605 | 0.274128 | NA | #VALUE! |
| ENSMUSG00000029475.18 | 10.94246 | -1.87061 | 3.189273 | -0.58653 | NA | NA | #VALUE! |
| ENSMUSG00000029477.15 | 0 | NA | NA | NA | NA | NA | #VALUE! |
| ENSMUSG00000029478.17 | 0.373021 | 2.140748 | 4.018182 | 0.532765 | 0.594196 | NA | #VALUE! |
| ENSMUSG00000029480.14 | 3.300909 | -4.10576 | 3.907177 | -1.05083 | 0.293339 | NA | #VALUE! |
| ENSMUSG00000029491.8 | 0 | NA | NA | NA | NA | NA | #VALUE! |
| ENSMUSG00000029503.17 | 0 | NA | NA | NA | NA | NA | #VALUE! |
| ENSMUSG00000029510.16 | 0 | NA | NA | NA | NA | NA | #VALUE! |
| ENSMUSG00000029516.20 | 12.32629 | 0.479731 | 3.106961 | 0.154405 | NA | NA | #VALUE! |
| ENSMUSG00000029517.14 | 0 | NA | NA | NA | NA | NA | #VALUE! |
| ENSMUSG00000029521.8 | 0.50155 | -0.4935 | 4.014251 | -0.12294 | 0.902157 | NA | #VALUE! |
| ENSMUSG00000029522.13 | 0 | NA | NA | NA | NA | NA | #VALUE! |
| ENSMUSG00000029524.17 | 5.678054 | -2.04688 | 2.724156 | -0.75138 | 0.452424 | NA | #VALUE! |
| ENSMUSG00000029526.6 | 0 | NA | NA | NA | NA | NA | #VALUE! |
| ENSMUSG00000029530.17 | 0 | NA | NA | NA | NA | NA | #VALUE! |
| ENSMUSG00000029541.13 | 0 | NA | NA | NA | NA | NA | #VALUE! |
| ENSMUSG00000029544.17 | 0 | NA | NA | NA | NA | NA | #VALUE! |
| ENSMUSG00000029546.13 | 0 | NA | NA | NA | NA | NA | #VALUE! |
| ENSMUSG00000029552.20 | 5.883423 | 0.594035 | 3.758208 | 0.158063 | 0.874407 | NA | #VALUE! |
| ENSMUSG00000029553.11 | 3.132626 | -5.17361 | 3.921071 | -1.31944 | 0.187023 | NA | #VALUE! |
| ENSMUSG00000029557.8 | 29.30214 | 2.462654 | 2.288447 | 1.076125 | NA | NA | #VALUE! |
| ENSMUSG00000029560.13 | 5.78817 | -0.44613 | 3.758294 | -0.11871 | 0.905509 | NA | #VALUE! |
| ENSMUSG00000029561.18 | 6.902725 | 0.169094 | 3.722033 | 0.04543 | 0.963764 | NA | #VALUE! |
| ENSMUSG00000029564.13 | 0 | NA | NA | NA | NA | NA | #VALUE! |
| ENSMUSG00000029570.6 | 0.51915 | -2.6028 | 3.990576 | -0.65224 | 0.514248 | NA | #VALUE! |
| ENSMUSG00000029575.18 | 45.45713 | -0.42676 | 1.458621 | -0.29258 | NA | NA | #VALUE! |
| ENSMUSG00000029576.18 | 0 | NA | NA | NA | NA | NA | #VALUE! |
| ENSMUSG00000029581.15 | 0 | NA | NA | NA | NA | NA | #VALUE! |
| ENSMUSG00000029586.10 | 0 | NA | NA | NA | NA | NA | #VALUE! |
| ENSMUSG00000029594.13 | 7.361706 | -1.04353 | 2.891763 | -0.36086 | 0.718201 | NA | #VALUE! |
| ENSMUSG00000029595.10 | 0 | NA | NA | NA | NA | NA | #VALUE! |
| ENSMUSG00000029596.14 | 17.49029 | -2.17624 | 2.874041 | -0.75721 | NA | NA | #VALUE! |

| | | | | | | | |
|---|---|---|---|---|---|---|---|
| ENSMUSG00000029600.11 | 1.762102 | -4.34067 | 3.93219 | -1.10388 | 0.269645 | NA | #VALUE! |
| ENSMUSG00000029601.14 | 0 | NA | NA | NA | NA | NA | #VALUE! |
| ENSMUSG00000029602.12 | 0 | NA | NA | NA | NA | NA | #VALUE! |
| ENSMUSG00000029603.16 | 0 | NA | NA | NA | NA | NA | #VALUE! |
| ENSMUSG00000029605.17 | 0 | NA | NA | NA | NA | NA | #VALUE! |
| ENSMUSG00000029607.15 | 0 | NA | NA | NA | NA | NA | #VALUE! |
| ENSMUSG00000029608.11 | 0 | NA | NA | NA | NA | NA | #VALUE! |
| ENSMUSG00000029618.14 | 0 | NA | NA | NA | NA | NA | #VALUE! |
| ENSMUSG00000029620.5 | 0 | NA | NA | NA | NA | NA | #VALUE! |
| ENSMUSG00000029624.11 | 11.73113 | 2.71354 | 3.133323 | 0.866026 | NA | NA | #VALUE! |
| ENSMUSG00000029633.10 | 0 | NA | NA | NA | NA | NA | #VALUE! |
| ENSMUSG00000029636.12 | 0 | NA | NA | NA | NA | NA | #VALUE! |
| ENSMUSG00000029638.18 | 0 | NA | NA | NA | NA | NA | #VALUE! |
| ENSMUSG00000029640.18 | 5.968332 | 6.072569 | 3.914265 | 1.551394 | 0.120807 | NA | #VALUE! |
| ENSMUSG00000029641.9 | 0 | NA | NA | NA | NA | NA | #VALUE! |
| ENSMUSG00000029644.8 | 0 | NA | NA | NA | NA | NA | #VALUE! |
| ENSMUSG00000029646.4 | 0 | NA | NA | NA | NA | NA | #VALUE! |
| ENSMUSG00000029651.17 | 0.18651 | 1.401769 | 4.056354 | 0.345574 | 0.729663 | NA | #VALUE! |
| ENSMUSG00000029658.18 | 0 | NA | NA | NA | NA | NA | #VALUE! |
| ENSMUSG00000029659.17 | 0 | NA | NA | NA | NA | NA | #VALUE! |
| ENSMUSG00000029660.11 | 0 | NA | NA | NA | NA | NA | #VALUE! |
| ENSMUSG00000029663.11 | 4.116497 | 5.530815 | 3.917676 | 1.411759 | 0.158021 | NA | #VALUE! |
| ENSMUSG00000029671.16 | 0 | NA | NA | NA | NA | NA | #VALUE! |
| ENSMUSG00000029674.14 | 0 | NA | NA | NA | NA | NA | #VALUE! |
| ENSMUSG00000029675.13 | 2.144869 | 4.605508 | 3.927435 | 1.17265 | 0.240936 | NA | #VALUE! |
| ENSMUSG00000029676.16 | 6.881733 | 5.222324 | 2.853593 | 1.830087 | 0.067237 | NA | #VALUE! |
| ENSMUSG00000029678.9 | 0 | NA | NA | NA | NA | NA | #VALUE! |
| ENSMUSG00000029679.8 | 0 | NA | NA | NA | NA | NA | #VALUE! |
| ENSMUSG00000029680.3 | 0 | NA | NA | NA | NA | NA | #VALUE! |
| ENSMUSG00000029682.6 | 0 | NA | NA | NA | NA | NA | #VALUE! |
| ENSMUSG00000029683.8 | 0.673134 | 2.918386 | 3.972531 | 0.734641 | 0.462558 | NA | #VALUE! |
| ENSMUSG00000029685.16 | 0 | NA | NA | NA | NA | NA | #VALUE! |
| ENSMUSG00000029697.7 | 0 | NA | NA | NA | NA | NA | #VALUE! |
| ENSMUSG00000029699.14 | 1.957891 | -4.49332 | 3.929649 | -1.14344 | 0.252855 | NA | #VALUE! |
| ENSMUSG00000029700.12 | 0 | NA | NA | NA | NA | NA | #VALUE! |
| ENSMUSG00000029706.16 | 0 | NA | NA | NA | NA | NA | #VALUE! |
| ENSMUSG00000029707.7 | 0 | NA | NA | NA | NA | NA | #VALUE! |
| ENSMUSG00000029710.16 | 0.093255 | 0.868093 | 4.072464 | 0.213162 | 0.831201 | NA | #VALUE! |
| ENSMUSG00000029711.14 | 0 | NA | NA | NA | NA | NA | #VALUE! |
| ENSMUSG00000029712.15 | 0 | NA | NA | NA | NA | NA | #VALUE! |
| ENSMUSG00000029718.15 | 15.21719 | -0.14666 | 2.965573 | -0.04945 | NA | NA | #VALUE! |
| ENSMUSG00000029720.10 | 0 | NA | NA | NA | NA | NA | #VALUE! |
| ENSMUSG00000029725.11 | 3.532525 | -5.3473 | 3.919452 | -1.3643 | 0.172474 | NA | #VALUE! |
| ENSMUSG00000029730.17 | 7.30558 | 0.904584 | 2.780965 | 0.325277 | 0.744971 | NA | #VALUE! |

| | | | | | | | |
|---|---|---|---|---|---|---|---|
| ENSMUSG00000029736.16 | 0 | NA | NA | NA | NA | NA | #VALUE! |
| ENSMUSG00000029754.14 | 0 | NA | NA | NA | NA | NA | #VALUE! |
| ENSMUSG00000029755.11 | 0 | NA | NA | NA | NA | NA | #VALUE! |
| ENSMUSG00000029757.17 | 0 | NA | NA | NA | NA | NA | #VALUE! |
| ENSMUSG00000029765.13 | 3.294828 | 4.227288 | 3.785689 | 1.116649 | 0.264144 | NA | #VALUE! |
| ENSMUSG00000029766.8 | 0 | NA | NA | NA | NA | NA | #VALUE! |
| ENSMUSG00000029769.18 | 0 | NA | NA | NA | NA | NA | #VALUE! |
| ENSMUSG00000029778.13 | 5.677884 | -6.03326 | 3.914659 | -1.5412 | 0.123269 | NA | #VALUE! |
| ENSMUSG00000029784.14 | 0 | NA | NA | NA | NA | NA | #VALUE! |
| ENSMUSG00000029788.14 | 0 | NA | NA | NA | NA | NA | #VALUE! |
| ENSMUSG00000029790.17 | 7.687371 | -6.47239 | 3.392544 | -1.90783 | 0.056414 | NA | #VALUE! |
| ENSMUSG00000029797.14 | 0 | NA | NA | NA | NA | NA | #VALUE! |
| ENSMUSG00000029804.18 | 2.499316 | -3.64018 | 3.912624 | -0.93037 | 0.35218 | NA | #VALUE! |
| ENSMUSG00000029811.15 | 0 | NA | NA | NA | NA | NA | #VALUE! |
| ENSMUSG00000029813.9 | 0 | NA | NA | NA | NA | NA | #VALUE! |
| ENSMUSG00000029814.11 | 0 | NA | NA | NA | NA | NA | #VALUE! |
| ENSMUSG00000029816.11 | 0 | NA | NA | NA | NA | NA | #VALUE! |
| ENSMUSG00000029819.7 | 0 | NA | NA | NA | NA | NA | #VALUE! |
| ENSMUSG00000029828.5 | 0.746042 | 3.107177 | 3.964586 | 0.783733 | 0.433197 | NA | #VALUE! |
| ENSMUSG00000029829.5 | 0 | NA | NA | NA | NA | NA | #VALUE! |
| ENSMUSG00000029830.16 | 0 | NA | NA | NA | NA | NA | #VALUE! |
| ENSMUSG00000029831.8 | 0 | NA | NA | NA | NA | NA | #VALUE! |
| ENSMUSG00000029832.17 | 9.346168 | -6.75775 | 3.650385 | -1.85124 | NA | NA | #VALUE! |
| ENSMUSG00000029836.16 | 0 | NA | NA | NA | NA | NA | #VALUE! |
| ENSMUSG00000029837.3 | 0 | NA | NA | NA | NA | NA | #VALUE! |
| ENSMUSG00000029838.12 | 0 | NA | NA | NA | NA | NA | #VALUE! |
| ENSMUSG00000029843.5 | 3.197387 | 5.166784 | 3.920797 | 1.317789 | 0.187574 | NA | #VALUE! |
| ENSMUSG00000029844.10 | 0 | NA | NA | NA | NA | NA | #VALUE! |
| ENSMUSG00000029847.14 | 0 | NA | NA | NA | NA | NA | #VALUE! |
| ENSMUSG00000029848.12 | 0 | NA | NA | NA | NA | NA | #VALUE! |
| ENSMUSG00000029851.6 | 1.878624 | 3.225164 | 3.925897 | 0.82151 | 0.411356 | NA | #VALUE! |
| ENSMUSG00000029861.18 | 0 | NA | NA | NA | NA | NA | #VALUE! |
| ENSMUSG00000029862.17 | 0 | NA | NA | NA | NA | NA | #VALUE! |
| ENSMUSG00000029865.5 | 0 | NA | NA | NA | NA | NA | #VALUE! |
| ENSMUSG00000029866.14 | 0 | NA | NA | NA | NA | NA | #VALUE! |
| ENSMUSG00000029867.6 | 0 | NA | NA | NA | NA | NA | #VALUE! |
| ENSMUSG00000029868.15 | 0 | NA | NA | NA | NA | NA | #VALUE! |
| ENSMUSG00000029869.8 | 0 | NA | NA | NA | NA | NA | #VALUE! |
| ENSMUSG00000029875.6 | 0 | NA | NA | NA | NA | NA | #VALUE! |
| ENSMUSG00000029878.7 | 0 | NA | NA | NA | NA | NA | #VALUE! |
| ENSMUSG00000029882.6 | 0 | NA | NA | NA | NA | NA | #VALUE! |
| ENSMUSG00000029883.13 | 0 | NA | NA | NA | NA | NA | #VALUE! |
| ENSMUSG00000029885.4 | 0 | NA | NA | NA | NA | NA | #VALUE! |
| ENSMUSG00000029909.7 | 0 | NA | NA | NA | NA | NA | #VALUE! |

| | | | | | | | |
|---|---|---|---|---|---|---|---|
| ENSMUSG00000029913.15 | 0 | NA | NA | NA | NA | NA | #VALUE! |
| ENSMUSG00000029915.15 | 0 | NA | NA | NA | NA | NA | #VALUE! |
| ENSMUSG00000029916.12 | 9.083549 | -1.21887 | 2.962081 | -0.41149 | 0.680712 | NA | #VALUE! |
| ENSMUSG00000029917.16 | 0 | NA | NA | NA | NA | NA | #VALUE! |
| ENSMUSG00000029919.6 | 0.245976 | 1.027628 | 4.067126 | 0.252667 | 0.800526 | NA | #VALUE! |
| ENSMUSG00000029923.5 | 0.373021 | 2.140748 | 4.018182 | 0.532765 | 0.594196 | NA | #VALUE! |
| ENSMUSG00000029925.14 | 4.830604 | -5.80779 | 3.915992 | -1.4831 | 0.138049 | NA | #VALUE! |
| ENSMUSG00000030000.11 | 0 | NA | NA | NA | NA | NA | #VALUE! |
| ENSMUSG00000030001.5 | 0 | NA | NA | NA | NA | NA | #VALUE! |
| ENSMUSG00000030008.10 | 3.136211 | 2.280084 | 3.827631 | 0.595691 | 0.551382 | NA | #VALUE! |
| ENSMUSG00000030017.3 | 0 | NA | NA | NA | NA | NA | #VALUE! |
| ENSMUSG00000030019.7 | 2.998401 | -1.78345 | 3.813727 | -0.46764 | 0.640042 | NA | #VALUE! |
| ENSMUSG00000030022.15 | 8.277603 | -2.85374 | 2.726903 | -1.04651 | 0.295325 | NA | #VALUE! |
| ENSMUSG00000030030.6 | 0 | NA | NA | NA | NA | NA | #VALUE! |
| ENSMUSG00000030031.15 | 9.292502 | -3.41282 | 2.572686 | -1.32656 | 0.184654 | NA | #VALUE! |
| ENSMUSG00000030032.15 | 0 | NA | NA | NA | NA | NA | #VALUE! |
| ENSMUSG00000030034.13 | 2.210018 | -2.21814 | 3.866491 | -0.57368 | 0.566182 | NA | #VALUE! |
| ENSMUSG00000030037.10 | 0 | NA | NA | NA | NA | NA | #VALUE! |
| ENSMUSG00000030041.10 | 0 | NA | NA | NA | NA | NA | #VALUE! |
| ENSMUSG00000030043.12 | 0 | NA | NA | NA | NA | NA | #VALUE! |
| ENSMUSG00000030047.15 | 2.763866 | 4.966032 | 3.922883 | 1.265914 | 0.205544 | NA | #VALUE! |
| ENSMUSG00000030048.6 | 0 | NA | NA | NA | NA | NA | #VALUE! |
| ENSMUSG00000030049.6 | 0 | NA | NA | NA | NA | NA | #VALUE! |
| ENSMUSG00000030050.5 | 0 | NA | NA | NA | NA | NA | #VALUE! |
| ENSMUSG00000030051.11 | 9.226933 | 6.694717 | 2.761375 | 2.424415 | 0.015333 | NA | #VALUE! |
| ENSMUSG00000030054.5 | 0 | NA | NA | NA | NA | NA | #VALUE! |
| ENSMUSG00000030055.17 | 5.315546 | 5.90612 | 3.915181 | 1.508518 | 0.131422 | NA | #VALUE! |
| ENSMUSG00000030068.5 | 0 | NA | NA | NA | NA | NA | #VALUE! |
| ENSMUSG00000030069.16 | 0 | NA | NA | NA | NA | NA | #VALUE! |
| ENSMUSG00000030074.10 | 0 | NA | NA | NA | NA | NA | #VALUE! |
| ENSMUSG00000030075.11 | 0 | NA | NA | NA | NA | NA | #VALUE! |
| ENSMUSG00000030077.12 | 0 | NA | NA | NA | NA | NA | #VALUE! |
| ENSMUSG00000030084.12 | 0 | NA | NA | NA | NA | NA | #VALUE! |
| ENSMUSG00000030089.16 | 0 | NA | NA | NA | NA | NA | #VALUE! |
| ENSMUSG00000030091.18 | 3.763981 | -1.21709 | 3.783402 | -0.32169 | 0.747686 | NA | #VALUE! |
| ENSMUSG00000030092.15 | 0 | NA | NA | NA | NA | NA | #VALUE! |
| ENSMUSG00000030093.8 | 0 | NA | NA | NA | NA | NA | #VALUE! |
| ENSMUSG00000030098.16 | 0 | NA | NA | NA | NA | NA | #VALUE! |
| ENSMUSG00000030110.14 | 0 | NA | NA | NA | NA | NA | #VALUE! |
| ENSMUSG00000030111.10 | 0.105013 | -0.97324 | 4.076293 | -0.23876 | 0.811295 | NA | #VALUE! |
| ENSMUSG00000030114.9 | 0 | NA | NA | NA | NA | NA | #VALUE! |
| ENSMUSG00000030116.15 | 0 | NA | NA | NA | NA | NA | #VALUE! |
| ENSMUSG00000030117.6 | 0 | NA | NA | NA | NA | NA | #VALUE! |
| ENSMUSG00000030123.16 | 2.986766 | -5.10416 | 3.921774 | -1.30149 | 0.19309 | NA | #VALUE! |

| | | | | | | | |
|---|---|---|---|---|---|---|---|
| ENSMUSG00000030124.3 | 0.932552 | 3.421684 | 3.953399 | 0.865504 | 0.386762 | NA | #VALUE! |
| ENSMUSG00000030125.12 | 0 | NA | NA | NA | NA | NA | #VALUE! |
| ENSMUSG00000030134.12 | 0 | NA | NA | NA | NA | NA | #VALUE! |
| ENSMUSG00000030137.9 | 0 | NA | NA | NA | NA | NA | #VALUE! |
| ENSMUSG00000030142.11 | 0 | NA | NA | NA | NA | NA | #VALUE! |
| ENSMUSG00000030143.8 | 0 | NA | NA | NA | NA | NA | #VALUE! |
| ENSMUSG00000030144.5 | 0 | NA | NA | NA | NA | NA | #VALUE! |
| ENSMUSG00000030145.15 | 0 | NA | NA | NA | NA | NA | #VALUE! |
| ENSMUSG00000030147.13 | 4.324641 | 0.949326 | 3.772573 | 0.251639 | 0.80132 | NA | #VALUE! |
| ENSMUSG00000030148.16 | 8.750743 | 6.619253 | 3.754899 | 1.762831 | 0.077929 | NA | #VALUE! |
| ENSMUSG00000030149.16 | 2.663947 | 1.783953 | 3.824003 | 0.466515 | 0.640847 | NA | #VALUE! |
| ENSMUSG00000030154.11 | 0 | NA | NA | NA | NA | NA | #VALUE! |
| ENSMUSG00000030155.8 | 0.559531 | 2.703958 | 3.982863 | 0.678898 | 0.497202 | NA | #VALUE! |
| ENSMUSG00000030158.13 | 0 | NA | NA | NA | NA | NA | #VALUE! |
| ENSMUSG00000030160.8 | 0 | NA | NA | NA | NA | NA | #VALUE! |
| ENSMUSG00000030162.15 | 0 | NA | NA | NA | NA | NA | #VALUE! |
| ENSMUSG00000030165.17 | 1.100835 | 3.652188 | 3.946585 | 0.925405 | 0.354755 | NA | #VALUE! |
| ENSMUSG00000030167.16 | 0 | NA | NA | NA | NA | NA | #VALUE! |
| ENSMUSG00000030170.15 | 5.210668 | 2.937019 | 3.587182 | 0.818754 | 0.412927 | NA | #VALUE! |
| ENSMUSG00000030173.16 | 0 | NA | NA | NA | NA | NA | #VALUE! |
| ENSMUSG00000030178.16 | 0 | NA | NA | NA | NA | NA | #VALUE! |
| ENSMUSG00000030187.16 | 2.520315 | -4.8742 | 3.924358 | -1.24204 | 0.214223 | NA | #VALUE! |
| ENSMUSG00000030188.13 | 9.467099 | -6.77293 | 2.797222 | -2.42131 | 0.015465 | NA | #VALUE! |
| ENSMUSG00000030194.3 | 0 | NA | NA | NA | NA | NA | #VALUE! |
| ENSMUSG00000030196.4 | 0 | NA | NA | NA | NA | NA | #VALUE! |
| ENSMUSG00000030200.14 | 0 | NA | NA | NA | NA | NA | #VALUE! |
| ENSMUSG00000030205.16 | 0 | NA | NA | NA | NA | NA | #VALUE! |
| ENSMUSG00000030206.14 | 0 | NA | NA | NA | NA | NA | #VALUE! |
| ENSMUSG00000030207.16 | 29.5307 | -0.46896 | 1.691406 | -0.27726 | NA | NA | #VALUE! |
| ENSMUSG00000030208.16 | 0.983902 | 3.433476 | 3.953024 | 0.868569 | 0.385083 | NA | #VALUE! |
| ENSMUSG00000030209.15 | 0 | NA | NA | NA | NA | NA | #VALUE! |
| ENSMUSG00000030219.14 | 0 | NA | NA | NA | NA | NA | #VALUE! |
| ENSMUSG00000030223.15 | 0 | NA | NA | NA | NA | NA | #VALUE! |
| ENSMUSG00000030226.13 | 0 | NA | NA | NA | NA | NA | #VALUE! |
| ENSMUSG00000030230.16 | 0 | NA | NA | NA | NA | NA | #VALUE! |
| ENSMUSG00000030235.18 | 0 | NA | NA | NA | NA | NA | #VALUE! |
| ENSMUSG00000030247.10 | 0.981067 | -2.05231 | 3.951877 | -0.51933 | 0.603534 | NA | #VALUE! |
| ENSMUSG00000030254.17 | 2.990405 | 5.064792 | 3.921821 | 1.291439 | 0.196552 | NA | #VALUE! |
| ENSMUSG00000030255.14 | 0.604011 | -2.78003 | 3.981028 | -0.69832 | 0.484977 | NA | #VALUE! |
| ENSMUSG00000030256.12 | 6.642138 | -5.16559 | 3.285484 | -1.57225 | 0.115893 | NA | #VALUE! |
| ENSMUSG00000030257.17 | 9.014396 | 0.64959 | 2.993881 | 0.216973 | 0.82823 | NA | #VALUE! |
| ENSMUSG00000030263.14 | 1.346268 | 3.918821 | 3.939925 | 0.994644 | 0.31991 | NA | #VALUE! |
| ENSMUSG00000030268.18 | 0 | NA | NA | NA | NA | NA | #VALUE! |
| ENSMUSG00000030270.12 | 0 | NA | NA | NA | NA | NA | #VALUE! |

| | | | | | | | |
|---|---|---|---|---|---|---|---|
| ENSMUSG00000030271.15 | 8.659141 | -6.64822 | 3.303531 | -2.01246 | 0.044172 | NA | #VALUE! |
| ENSMUSG00000030272.16 | 6.557243 | -1.01967 | 3.404492 | -0.29951 | 0.764552 | NA | #VALUE! |
| ENSMUSG00000030276.20 | 0.504851 | 2.502843 | 3.993993 | 0.626652 | 0.530887 | NA | #VALUE! |
| ENSMUSG00000030278.12 | 0.934673 | 3.395605 | 3.954239 | 0.858725 | 0.390492 | NA | #VALUE! |
| ENSMUSG00000030281.17 | 0.195789 | -1.05581 | 4.073302 | -0.2592 | 0.795479 | NA | #VALUE! |
| ENSMUSG00000030283.8 | 0 | NA | NA | NA | NA | NA | #VALUE! |
| ENSMUSG00000030292.11 | 0 | NA | NA | NA | NA | NA | #VALUE! |
| ENSMUSG00000030302.18 | 0.525066 | -2.64687 | 3.988095 | -0.66369 | 0.506887 | NA | #VALUE! |
| ENSMUSG00000030303.16 | 1.236642 | -2.18313 | 3.923447 | -0.55643 | 0.577915 | NA | #VALUE! |
| ENSMUSG00000030306.15 | 1.836997 | -4.39907 | 3.931186 | -1.11902 | 0.263133 | NA | #VALUE! |
| ENSMUSG00000030307.9 | 0 | NA | NA | NA | NA | NA | #VALUE! |
| ENSMUSG00000030309.17 | 7.027373 | -6.34048 | 3.529732 | -1.79631 | 0.072446 | NA | #VALUE! |
| ENSMUSG00000030310.11 | 0 | NA | NA | NA | NA | NA | #VALUE! |
| ENSMUSG00000030316.14 | 6.876106 | 2.089377 | 2.551548 | 0.818867 | 0.412863 | NA | #VALUE! |
| ENSMUSG00000030317.9 | 0 | NA | NA | NA | NA | NA | #VALUE! |
| ENSMUSG00000030319.9 | 0 | NA | NA | NA | NA | NA | #VALUE! |
| ENSMUSG00000030321.16 | 0 | NA | NA | NA | NA | NA | #VALUE! |
| ENSMUSG00000030322.13 | 3.617088 | 0.980605 | 3.78279 | 0.259228 | 0.795459 | NA | #VALUE! |
| ENSMUSG00000030323.14 | 9.063905 | -1.67256 | 3.0204 | -0.55375 | 0.579747 | NA | #VALUE! |
| ENSMUSG00000030324.9 | 0 | NA | NA | NA | NA | NA | #VALUE! |
| ENSMUSG00000030325.17 | 0 | NA | NA | NA | NA | NA | #VALUE! |
| ENSMUSG00000030329.12 | 4.698938 | -5.75981 | 3.916303 | -1.47073 | 0.141365 | NA | #VALUE! |
| ENSMUSG00000030330.16 | 2.625328 | -4.93268 | 3.923662 | -1.25716 | 0.208695 | NA | #VALUE! |
| ENSMUSG00000030336.15 | 8.438432 | 2.376533 | 3.487689 | 0.681406 | 0.495615 | NA | #VALUE! |
| ENSMUSG00000030337.17 | 0 | NA | NA | NA | NA | NA | #VALUE! |
| ENSMUSG00000030344.12 | 0 | NA | NA | NA | NA | NA | #VALUE! |
| ENSMUSG00000030345.17 | 0 | NA | NA | NA | NA | NA | #VALUE! |
| ENSMUSG00000030346.17 | 0 | NA | NA | NA | NA | NA | #VALUE! |
| ENSMUSG00000030350.9 | 0 | NA | NA | NA | NA | NA | #VALUE! |
| ENSMUSG00000030351.6 | 0 | NA | NA | NA | NA | NA | #VALUE! |
| ENSMUSG00000030353.16 | 0 | NA | NA | NA | NA | NA | #VALUE! |
| ENSMUSG00000030361.17 | 0 | NA | NA | NA | NA | NA | #VALUE! |
| ENSMUSG00000030363.9 | 0 | NA | NA | NA | NA | NA | #VALUE! |
| ENSMUSG00000030365.12 | 0.420053 | -2.33493 | 4.007343 | -0.58266 | 0.56012 | NA | #VALUE! |
| ENSMUSG00000030366.11 | 0 | NA | NA | NA | NA | NA | #VALUE! |
| ENSMUSG00000030368.11 | 0 | NA | NA | NA | NA | NA | #VALUE! |
| ENSMUSG00000030373.7 | 0 | NA | NA | NA | NA | NA | #VALUE! |
| ENSMUSG00000030376.9 | 0 | NA | NA | NA | NA | NA | #VALUE! |
| ENSMUSG00000030380.18 | 0 | NA | NA | NA | NA | NA | #VALUE! |
| ENSMUSG00000030385.5 | 0 | NA | NA | NA | NA | NA | #VALUE! |
| ENSMUSG00000030393.9 | 0 | NA | NA | NA | NA | NA | #VALUE! |
| ENSMUSG00000030397.12 | 2.997175 | -1.54826 | 3.806097 | -0.40678 | 0.684167 | NA | #VALUE! |
| ENSMUSG00000030399.3 | 0 | NA | NA | NA | NA | NA | #VALUE! |
| ENSMUSG00000030400.18 | 8.223142 | -6.56817 | 3.836124 | -1.71219 | 0.086862 | NA | #VALUE! |

| | | | | | | | |
|---|---|---|---|---|---|---|---|
| ENSMUSG00000030401.17 | 0 | NA | NA | NA | NA | NA | #VALUE! |
| ENSMUSG00000030402.6 | 0 | NA | NA | NA | NA | NA | #VALUE! |
| ENSMUSG00000030406.8 | 0 | NA | NA | NA | NA | NA | #VALUE! |
| ENSMUSG00000030409.17 | 2.273078 | 3.536985 | 3.920751 | 0.902119 | 0.366994 | NA | #VALUE! |
| ENSMUSG00000030410.17 | 0.523077 | 2.574714 | 3.989843 | 0.645317 | 0.518722 | NA | #VALUE! |
| ENSMUSG00000030411.17 | 0 | NA | NA | NA | NA | NA | #VALUE! |
| ENSMUSG00000030413.8 | 0 | NA | NA | NA | NA | NA | #VALUE! |
| ENSMUSG00000030424.16 | 0.652786 | 2.919659 | 3.972474 | 0.734972 | 0.462356 | NA | #VALUE! |
| ENSMUSG00000030427.18 | 0 | NA | NA | NA | NA | NA | #VALUE! |
| ENSMUSG00000030428.17 | 2.86082 | 5.006323 | 3.922441 | 1.276328 | 0.201839 | NA | #VALUE! |
| ENSMUSG00000030433.16 | 0 | NA | NA | NA | NA | NA | #VALUE! |
| ENSMUSG00000030443.17 | 0 | NA | NA | NA | NA | NA | #VALUE! |
| ENSMUSG00000030450.12 | 0 | NA | NA | NA | NA | NA | #VALUE! |
| ENSMUSG00000030463.6 | 0 | NA | NA | NA | NA | NA | #VALUE! |
| ENSMUSG00000030468.13 | 0 | NA | NA | NA | NA | NA | #VALUE! |
| ENSMUSG00000030472.10 | 0 | NA | NA | NA | NA | NA | #VALUE! |
| ENSMUSG00000030474.10 | 0 | NA | NA | NA | NA | NA | #VALUE! |
| ENSMUSG00000030484.6 | 0 | NA | NA | NA | NA | NA | #VALUE! |
| ENSMUSG00000030486.10 | 4.66276 | 5.717927 | 3.916352 | 1.460014 | 0.144286 | NA | #VALUE! |
| ENSMUSG00000030491.17 | 0 | NA | NA | NA | NA | NA | #VALUE! |
| ENSMUSG00000030492.17 | 0 | NA | NA | NA | NA | NA | #VALUE! |
| ENSMUSG00000030495.13 | 0 | NA | NA | NA | NA | NA | #VALUE! |
| ENSMUSG00000030499.10 | 4.725591 | -5.7762 | 3.916196 | -1.47495 | 0.140225 | NA | #VALUE! |
| ENSMUSG00000030500.7 | 0 | NA | NA | NA | NA | NA | #VALUE! |
| ENSMUSG00000030507.7 | 0 | NA | NA | NA | NA | NA | #VALUE! |
| ENSMUSG00000030510.12 | 0 | NA | NA | NA | NA | NA | #VALUE! |
| ENSMUSG00000030518.18 | 0 | NA | NA | NA | NA | NA | #VALUE! |
| ENSMUSG00000030519.15 | 0 | NA | NA | NA | NA | NA | #VALUE! |
| ENSMUSG00000030523.19 | 0 | NA | NA | NA | NA | NA | #VALUE! |
| ENSMUSG00000030525.9 | 0 | NA | NA | NA | NA | NA | #VALUE! |
| ENSMUSG00000030527.16 | 1.719289 | 4.275716 | 3.932707 | 1.087219 | 0.27694 | NA | #VALUE! |
| ENSMUSG00000030528.13 | 3.976013 | 4.38472 | 3.911236 | 1.121057 | 0.262263 | NA | #VALUE! |
| ENSMUSG00000030532.7 | 7.249386 | 6.345536 | 3.497336 | 1.814391 | 0.069617 | NA | #VALUE! |
| ENSMUSG00000030539.14 | 1.594787 | -4.20243 | 3.934732 | -1.06804 | 0.285505 | NA | #VALUE! |
| ENSMUSG00000030543.11 | 0 | NA | NA | NA | NA | NA | #VALUE! |
| ENSMUSG00000030544.6 | 0 | NA | NA | NA | NA | NA | #VALUE! |
| ENSMUSG00000030546.15 | 0 | NA | NA | NA | NA | NA | #VALUE! |
| ENSMUSG00000030549.7 | 0 | NA | NA | NA | NA | NA | #VALUE! |
| ENSMUSG00000030553.4 | 0 | NA | NA | NA | NA | NA | #VALUE! |
| ENSMUSG00000030554.17 | 3.616127 | 0.882172 | 3.781216 | 0.233304 | 0.815526 | NA | #VALUE! |
| ENSMUSG00000030559.9 | 6.142329 | -3.09319 | 3.509093 | -0.88148 | 0.378059 | NA | #VALUE! |
| ENSMUSG00000030577.15 | 0 | NA | NA | NA | NA | NA | #VALUE! |
| ENSMUSG00000030583.17 | 6.840668 | -6.30166 | 3.678891 | -1.71292 | 0.086727 | NA | #VALUE! |
| ENSMUSG00000030584.15 | 0 | NA | NA | NA | NA | NA | #VALUE! |

| | | | | | | | |
|---|---|---|---|---|---|---|---|
| ENSMUSG00000030589.16 | 0 | NA | NA | NA | NA | NA | #VALUE! |
| ENSMUSG00000030590.16 | 4.095512 | -5.57062 | 3.917637 | -1.42193 | 0.155045 | NA | #VALUE! |
| ENSMUSG00000030592.19 | 0.093255 | 0.868093 | 4.072464 | 0.213162 | 0.831201 | NA | #VALUE! |
| ENSMUSG00000030598.16 | 0.983902 | 3.433476 | 3.953024 | 0.868569 | 0.385083 | NA | #VALUE! |
| ENSMUSG00000030600.16 | 0 | NA | NA | NA | NA | NA | #VALUE! |
| ENSMUSG00000030604.10 | 7.98334 | -0.23841 | 3.196725 | -0.07458 | 0.940548 | NA | #VALUE! |
| ENSMUSG00000030605.16 | 5.020615 | 3.822481 | 3.578002 | 1.068328 | 0.285372 | NA | #VALUE! |
| ENSMUSG00000030606.9 | 0 | NA | NA | NA | NA | NA | #VALUE! |
| ENSMUSG00000030607.8 | 0 | NA | NA | NA | NA | NA | #VALUE! |
| ENSMUSG00000030609.19 | 9.306026 | 0.725183 | 3.389393 | 0.213957 | 0.830581 | NA | #VALUE! |
| ENSMUSG00000030610.14 | 9.197797 | 3.910245 | 2.901139 | 1.347831 | 0.177713 | NA | #VALUE! |
| ENSMUSG00000030616.17 | 1.177985 | 3.72614 | 3.944615 | 0.944614 | 0.344856 | NA | #VALUE! |
| ENSMUSG00000030617.14 | 0 | NA | NA | NA | NA | NA | #VALUE! |
| ENSMUSG00000030621.18 | 0.245976 | 1.027628 | 4.067126 | 0.252667 | 0.800526 | NA | #VALUE! |
| ENSMUSG00000030623.5 | 0 | NA | NA | NA | NA | NA | #VALUE! |
| ENSMUSG00000030636.16 | 0 | NA | NA | NA | NA | NA | #VALUE! |
| ENSMUSG00000030638.14 | 0 | NA | NA | NA | NA | NA | #VALUE! |
| ENSMUSG00000030641.11 | 0 | NA | NA | NA | NA | NA | #VALUE! |
| ENSMUSG00000030650.19 | 0 | NA | NA | NA | NA | NA | #VALUE! |
| ENSMUSG00000030651.6 | 0 | NA | NA | NA | NA | NA | #VALUE! |
| ENSMUSG00000030653.18 | 0 | NA | NA | NA | NA | NA | #VALUE! |
| ENSMUSG00000030657.12 | 0 | NA | NA | NA | NA | NA | #VALUE! |
| ENSMUSG00000030664.4 | 5.385073 | 5.918828 | 3.915107 | 1.511792 | 0.130587 | NA | #VALUE! |
| ENSMUSG00000030666.12 | 0 | NA | NA | NA | NA | NA | #VALUE! |
| ENSMUSG00000030669.14 | 0 | NA | NA | NA | NA | NA | #VALUE! |
| ENSMUSG00000030672.13 | 1.575197 | -4.20222 | 3.934736 | -1.06798 | 0.285529 | NA | #VALUE! |
| ENSMUSG00000030677.9 | 0 | NA | NA | NA | NA | NA | #VALUE! |
| ENSMUSG00000030680.7 | 0 | NA | NA | NA | NA | NA | #VALUE! |
| ENSMUSG00000030683.12 | 0 | NA | NA | NA | NA | NA | #VALUE! |
| ENSMUSG00000030685.6 | 1.566313 | -4.16993 | 3.935366 | -1.0596 | 0.289325 | NA | #VALUE! |
| ENSMUSG00000030693.11 | 0 | NA | NA | NA | NA | NA | #VALUE! |
| ENSMUSG00000030699.17 | 0 | NA | NA | NA | NA | NA | #VALUE! |
| ENSMUSG00000030703.9 | 0 | NA | NA | NA | NA | NA | #VALUE! |
| ENSMUSG00000030707.16 | 2.15368 | -4.63136 | 3.92757 | -1.17919 | 0.238321 | NA | #VALUE! |
| ENSMUSG00000030708.15 | 0 | NA | NA | NA | NA | NA | #VALUE! |
| ENSMUSG00000030713.7 | 0 | NA | NA | NA | NA | NA | #VALUE! |
| ENSMUSG00000030717.10 | 0.204111 | -1.05581 | 4.073302 | -0.2592 | 0.795479 | NA | #VALUE! |
| ENSMUSG00000030720.17 | 7.131947 | 1.836067 | 2.832433 | 0.64823 | 0.516836 | NA | #VALUE! |
| ENSMUSG00000030722.8 | 4.843657 | -1.63359 | 3.774341 | -0.43281 | 0.66515 | NA | #VALUE! |
| ENSMUSG00000030724.8 | 5.408801 | 5.931107 | 3.915037 | 1.514955 | 0.129784 | NA | #VALUE! |
| ENSMUSG00000030725.5 | 9.545896 | -1.75621 | 2.524483 | -0.69567 | 0.486634 | NA | #VALUE! |
| ENSMUSG00000030730.13 | 0 | NA | NA | NA | NA | NA | #VALUE! |
| ENSMUSG00000030731.14 | 10.63109 | 6.903964 | 3.498501 | 1.973406 | NA | NA | #VALUE! |
| ENSMUSG00000030732.14 | 2.705731 | 4.913666 | 3.923477 | 1.252375 | 0.210433 | NA | #VALUE! |

| | | | | | | | |
|---|---|---|---|---|---|---|---|
| ENSMUSG00000030735.7 | 0 | NA | NA | NA | NA | NA | #VALUE! |
| ENSMUSG00000030739.19 | 0 | NA | NA | NA | NA | NA | #VALUE! |
| ENSMUSG00000030741.16 | 6.056086 | 6.089083 | 3.71983 | 1.636925 | 0.101646 | NA | #VALUE! |
| ENSMUSG00000030742.8 | 0 | NA | NA | NA | NA | NA | #VALUE! |
| ENSMUSG00000030745.10 | 0 | NA | NA | NA | NA | NA | #VALUE! |
| ENSMUSG00000030752.9 | 2.369303 | -1.08658 | 3.813797 | -0.28491 | 0.775716 | NA | #VALUE! |
| ENSMUSG00000030757.14 | 0 | NA | NA | NA | NA | NA | #VALUE! |
| ENSMUSG00000030761.17 | 35.10982 | 0.869545 | 2.471565 | 0.35182 | NA | NA | #VALUE! |
| ENSMUSG00000030768.13 | 3.719993 | -5.42213 | 3.918812 | -1.38361 | 0.166476 | NA | #VALUE! |
| ENSMUSG00000030769.16 | 0 | NA | NA | NA | NA | NA | #VALUE! |
| ENSMUSG00000030774.14 | 1.177852 | 0.312998 | 3.877522 | 0.080721 | 0.935664 | NA | #VALUE! |
| ENSMUSG00000030775.12 | 3.045381 | -5.14543 | 3.921352 | -1.31216 | 0.189467 | NA | #VALUE! |
| ENSMUSG00000030781.14 | 0 | NA | NA | NA | NA | NA | #VALUE! |
| ENSMUSG00000030782.18 | 0 | NA | NA | NA | NA | NA | #VALUE! |
| ENSMUSG00000030785.9 | 0 | NA | NA | NA | NA | NA | #VALUE! |
| ENSMUSG00000030786.19 | 3.36042 | -5.28663 | 3.919996 | -1.34863 | 0.177455 | NA | #VALUE! |
| ENSMUSG00000030789.10 | 0 | NA | NA | NA | NA | NA | #VALUE! |
| ENSMUSG00000030792.9 | 0 | NA | NA | NA | NA | NA | #VALUE! |
| ENSMUSG00000030793.5 | 5.376347 | 0.331736 | 3.554223 | 0.093336 | 0.925637 | NA | #VALUE! |
| ENSMUSG00000030796.18 | 0 | NA | NA | NA | NA | NA | #VALUE! |
| ENSMUSG00000030798.16 | 0.466276 | 2.449889 | 3.997179 | 0.612904 | 0.53994 | NA | #VALUE! |
| ENSMUSG00000030804.9 | 0 | NA | NA | NA | NA | NA | #VALUE! |
| ENSMUSG00000030806.7 | 8.581318 | 0.715369 | 3.474354 | 0.2059 | 0.836869 | NA | #VALUE! |
| ENSMUSG00000030811.15 | 0 | NA | NA | NA | NA | NA | #VALUE! |
| ENSMUSG00000030816.9 | 4.475134 | -1.7869 | 3.784469 | -0.47217 | 0.636808 | NA | #VALUE! |
| ENSMUSG00000030825.20 | 0 | NA | NA | NA | NA | NA | #VALUE! |
| ENSMUSG00000030827.6 | 3.061661 | -5.13931 | 3.921414 | -1.31058 | 0.190001 | NA | #VALUE! |
| ENSMUSG00000030838.21 | 0 | NA | NA | NA | NA | NA | #VALUE! |
| ENSMUSG00000030851.16 | 0 | NA | NA | NA | NA | NA | #VALUE! |
| ENSMUSG00000030854.18 | 0 | NA | NA | NA | NA | NA | #VALUE! |
| ENSMUSG00000030858.11 | 0 | NA | NA | NA | NA | NA | #VALUE! |
| ENSMUSG00000030859.9 | 0 | NA | NA | NA | NA | NA | #VALUE! |
| ENSMUSG00000030862.14 | 0 | NA | NA | NA | NA | NA | #VALUE! |
| ENSMUSG00000030865.5 | 0 | NA | NA | NA | NA | NA | #VALUE! |
| ENSMUSG00000030866.9 | 0 | NA | NA | NA | NA | NA | #VALUE! |
| ENSMUSG00000030867.8 | 1.398828 | 3.99633 | 3.938205 | 1.014759 | 0.310221 | NA | #VALUE! |
| ENSMUSG00000030873.10 | 0 | NA | NA | NA | NA | NA | #VALUE! |
| ENSMUSG00000030877.12 | 0 | NA | NA | NA | NA | NA | #VALUE! |
| ENSMUSG00000030878.12 | 0 | NA | NA | NA | NA | NA | #VALUE! |
| ENSMUSG00000030881.16 | 1.113402 | -2.40779 | 3.939098 | -0.61126 | 0.541031 | NA | #VALUE! |
| ENSMUSG00000030882.20 | 0 | NA | NA | NA | NA | NA | #VALUE! |
| ENSMUSG00000030889.15 | 0 | NA | NA | NA | NA | NA | #VALUE! |
| ENSMUSG00000030897.6 | 0.195789 | -1.05581 | 4.073302 | -0.2592 | 0.795479 | NA | #VALUE! |
| ENSMUSG00000030898.16 | 0 | NA | NA | NA | NA | NA | #VALUE! |

| | | | | | | | | |
|---|---|---|---|---|---|---|---|---|
| ENSMUSG00000030905.6 | 5.51099 | -5.9895 | 3.914902 | -1.52992 | 0.126035 | NA | | #VALUE! |
| ENSMUSG00000030911.8 | 0 | NA | NA | NA | NA | NA | | #VALUE! |
| ENSMUSG00000030917.14 | 9.856269 | -0.63462 | 3.332443 | -0.19044 | 0.848967 | NA | | #VALUE! |
| ENSMUSG00000030924.17 | 9.333431 | -0.43074 | 2.952814 | -0.14587 | 0.884021 | NA | | #VALUE! |
| ENSMUSG00000030945.18 | 4.999741 | -5.84964 | 3.915728 | -1.49388 | 0.135206 | NA | | #VALUE! |
| ENSMUSG00000030954.12 | 0 | NA | NA | NA | NA | NA | | #VALUE! |
| ENSMUSG00000030956.16 | 8.094176 | 2.073503 | 3.514595 | 0.589969 | NA | NA | | #VALUE! |
| ENSMUSG00000030963.7 | 0 | NA | NA | NA | NA | NA | | #VALUE! |
| ENSMUSG00000030976.4 | 0 | NA | NA | NA | NA | NA | | #VALUE! |
| ENSMUSG00000030981.10 | 0 | NA | NA | NA | NA | NA | | #VALUE! |
| ENSMUSG00000030994.17 | 0 | NA | NA | NA | NA | NA | | #VALUE! |
| ENSMUSG00000030996.9 | 0 | NA | NA | NA | NA | NA | | #VALUE! |
| ENSMUSG00000031004.9 | 0 | NA | NA | NA | NA | NA | | #VALUE! |
| ENSMUSG00000031022.16 | 0 | NA | NA | NA | NA | NA | | #VALUE! |
| ENSMUSG00000031024.14 | 4.611068 | 3.40522 | 3.720522 | 0.915253 | 0.360059 | NA | | #VALUE! |
| ENSMUSG00000031026.16 | 0 | NA | NA | NA | NA | NA | | #VALUE! |
| ENSMUSG00000031027.16 | 0 | NA | NA | NA | NA | NA | | #VALUE! |
| ENSMUSG00000031028.15 | 0 | NA | NA | NA | NA | NA | | #VALUE! |
| ENSMUSG00000031065.15 | 0.816443 | -3.21914 | 3.961753 | -0.81255 | 0.416474 | NA | | #VALUE! |
| ENSMUSG00000031066.11 | 2.545258 | -4.8732 | 3.92437 | -1.24178 | 0.214318 | NA | | #VALUE! |
| ENSMUSG00000031073.7 | 0 | NA | NA | NA | NA | NA | | #VALUE! |
| ENSMUSG00000031074.15 | 0 | NA | NA | NA | NA | NA | | #VALUE! |
| ENSMUSG00000031079.13 | 0 | NA | NA | NA | NA | NA | | #VALUE! |
| ENSMUSG00000031085.17 | 0 | NA | NA | NA | NA | NA | | #VALUE! |
| ENSMUSG00000031089.8 | 0 | NA | NA | NA | NA | NA | | #VALUE! |
| ENSMUSG00000031093.15 | 4.1981 | 0.58058 | 3.770855 | 0.153965 | 0.877637 | NA | | #VALUE! |
| ENSMUSG00000031097.16 | 0 | NA | NA | NA | NA | NA | | #VALUE! |
| ENSMUSG00000031098.19 | 0 | NA | NA | NA | NA | NA | | #VALUE! |
| ENSMUSG00000031099.17 | 0.195789 | -1.05581 | 4.073302 | -0.2592 | 0.795479 | NA | | #VALUE! |
| ENSMUSG00000031101.7 | 0.466276 | 2.449889 | 3.997179 | 0.612904 | 0.53994 | NA | | #VALUE! |
| ENSMUSG00000031103.13 | 6.615827 | -6.26 | 3.913512 | -1.59959 | 0.10969 | NA | | #VALUE! |
| ENSMUSG00000031104.3 | 0 | NA | NA | NA | NA | NA | | #VALUE! |
| ENSMUSG00000031107.7 | 2.100263 | -4.61322 | 3.927832 | -1.17449 | 0.240197 | NA | | #VALUE! |
| ENSMUSG00000031111.17 | 0 | NA | NA | NA | NA | NA | | #VALUE! |
| ENSMUSG00000031112.11 | 1.370524 | -3.97623 | 3.939448 | -1.00934 | 0.312813 | NA | | #VALUE! |
| ENSMUSG00000031118.9 | 0 | NA | NA | NA | NA | NA | | #VALUE! |
| ENSMUSG00000031125.3 | 0 | NA | NA | NA | NA | NA | | #VALUE! |
| ENSMUSG00000031129.10 | 6.43337 | -0.23329 | 3.341229 | -0.06982 | 0.944335 | NA | | #VALUE! |
| ENSMUSG00000031130.4 | 0 | NA | NA | NA | NA | NA | | #VALUE! |
| ENSMUSG00000031131.12 | 0 | NA | NA | NA | NA | NA | | #VALUE! |
| ENSMUSG00000031132.2 | 0 | NA | NA | NA | NA | NA | | #VALUE! |
| ENSMUSG00000031137.18 | 0 | NA | NA | NA | NA | NA | | #VALUE! |
| ENSMUSG00000031139.17 | 0 | NA | NA | NA | NA | NA | | #VALUE! |
| ENSMUSG00000031142.18 | 0 | NA | NA | NA | NA | NA | | #VALUE! |

| | | | | | | | |
|---|---|---|---|---|---|---|---|
| ENSMUSG00000031143.5 | 0.630079 | -2.90291 | 3.975047 | -0.73028 | 0.465217 | NA | #VALUE! |
| ENSMUSG00000031144.16 | 0 | NA | NA | NA | NA | NA | #VALUE! |
| ENSMUSG00000031145.16 | 0 | NA | NA | NA | NA | NA | #VALUE! |
| ENSMUSG00000031146.7 | 4.009973 | 5.501447 | 3.9179 | 1.404183 | 0.160264 | NA | #VALUE! |
| ENSMUSG00000031149.8 | 1.428775 | -4.03418 | 3.938169 | -1.02438 | 0.305656 | NA | #VALUE! |
| ENSMUSG00000031150.14 | 0 | NA | NA | NA | NA | NA | #VALUE! |
| ENSMUSG00000031153.17 | 5.278487 | -2.1293 | 3.608499 | -0.59008 | 0.555138 | NA | #VALUE! |
| ENSMUSG00000031155.18 | 3.823463 | 5.43312 | 3.918438 | 1.386552 | 0.165578 | NA | #VALUE! |
| ENSMUSG00000031160.5 | 0 | NA | NA | NA | NA | NA | #VALUE! |
| ENSMUSG00000031162.15 | 0 | NA | NA | NA | NA | NA | #VALUE! |
| ENSMUSG00000031163.3 | 0 | NA | NA | NA | NA | NA | #VALUE! |
| ENSMUSG00000031165.7 | 0 | NA | NA | NA | NA | NA | #VALUE! |
| ENSMUSG00000031167.17 | 9.729062 | 0.596738 | 2.540834 | 0.234859 | 0.814318 | NA | #VALUE! |
| ENSMUSG00000031169.14 | 5.553356 | 5.96322 | 3.914855 | 1.523229 | 0.127701 | NA | #VALUE! |
| ENSMUSG00000031170.15 | 0 | NA | NA | NA | NA | NA | #VALUE! |
| ENSMUSG00000031179.15 | 0 | NA | NA | NA | NA | NA | #VALUE! |
| ENSMUSG00000031181.3 | 0 | NA | NA | NA | NA | NA | #VALUE! |
| ENSMUSG00000031182.3 | 0 | NA | NA | NA | NA | NA | #VALUE! |
| ENSMUSG00000031189.13 | 0 | NA | NA | NA | NA | NA | #VALUE! |
| ENSMUSG00000031194.2 | 0 | NA | NA | NA | NA | NA | #VALUE! |
| ENSMUSG00000031202.6 | 0 | NA | NA | NA | NA | NA | #VALUE! |
| ENSMUSG00000031204.4 | 0 | NA | NA | NA | NA | NA | #VALUE! |
| ENSMUSG00000031209.15 | 1.771849 | 4.332864 | 3.931707 | 1.102031 | 0.270448 | NA | #VALUE! |
| ENSMUSG00000031210.11 | 0 | NA | NA | NA | NA | NA | #VALUE! |
| ENSMUSG00000031212.4 | 0 | NA | NA | NA | NA | NA | #VALUE! |
| ENSMUSG00000031217.9 | 6.9244 | 4.754328 | 3.690283 | 1.288337 | NA | NA | #VALUE! |
| ENSMUSG00000031220.15 | 0 | NA | NA | NA | NA | NA | #VALUE! |
| ENSMUSG00000031224.7 | 0 | NA | NA | NA | NA | NA | #VALUE! |
| ENSMUSG00000031230.4 | 0 | NA | NA | NA | NA | NA | #VALUE! |
| ENSMUSG00000031233.7 | 0 | NA | NA | NA | NA | NA | #VALUE! |
| ENSMUSG00000031239.6 | 0 | NA | NA | NA | NA | NA | #VALUE! |
| ENSMUSG00000031241.18 | 0 | NA | NA | NA | NA | NA | #VALUE! |
| ENSMUSG00000031250.9 | 0 | NA | NA | NA | NA | NA | #VALUE! |
| ENSMUSG00000031253.16 | 1.174735 | -3.75246 | 3.944889 | -0.95122 | 0.341493 | NA | #VALUE! |
| ENSMUSG00000031255.15 | 3.255407 | -5.24108 | 3.920419 | -1.33687 | 0.181266 | NA | #VALUE! |
| ENSMUSG00000031257.15 | 0 | NA | NA | NA | NA | NA | #VALUE! |
| ENSMUSG00000031258.5 | 0 | NA | NA | NA | NA | NA | #VALUE! |
| ENSMUSG00000031262.13 | 0.525066 | -2.64687 | 3.988095 | -0.66369 | 0.506887 | NA | #VALUE! |
| ENSMUSG00000031264.14 | 0 | NA | NA | NA | NA | NA | #VALUE! |
| ENSMUSG00000031270.2 | 0.364073 | 0.012025 | 4.059931 | 0.002962 | 0.997637 | NA | #VALUE! |
| ENSMUSG00000031273.17 | 0 | NA | NA | NA | NA | NA | #VALUE! |
| ENSMUSG00000031274.17 | 2.377944 | -4.77767 | 3.925571 | -1.21706 | 0.22358 | NA | #VALUE! |
| ENSMUSG00000031283.17 | 0 | NA | NA | NA | NA | NA | #VALUE! |
| ENSMUSG00000031284.17 | 0 | NA | NA | NA | NA | NA | #VALUE! |

| | | | | | | | |
|---|---|---|---|---|---|---|---|
| ENSMUSG00000031285.15 | 0 | NA | NA | NA | NA | NA | #VALUE! |
| ENSMUSG00000031286.7 | 6.811564 | 2.097476 | 2.910988 | 0.720538 | 0.471194 | NA | #VALUE! |
| ENSMUSG00000031289.11 | 1.876862 | 3.39984 | 3.927368 | 0.865679 | 0.386666 | NA | #VALUE! |
| ENSMUSG00000031290.15 | 1.967804 | 4.449735 | 3.929778 | 1.132312 | 0.257503 | NA | #VALUE! |
| ENSMUSG00000031293.14 | 0 | NA | NA | NA | NA | NA | #VALUE! |
| ENSMUSG00000031294.10 | 0 | NA | NA | NA | NA | NA | #VALUE! |
| ENSMUSG00000031297.15 | 0 | NA | NA | NA | NA | NA | #VALUE! |
| ENSMUSG00000031298.16 | 2.704401 | 4.936946 | 3.92321 | 1.258394 | 0.208249 | NA | #VALUE! |
| ENSMUSG00000031302.17 | 0 | NA | NA | NA | NA | NA | #VALUE! |
| ENSMUSG00000031304.19 | 0 | NA | NA | NA | NA | NA | #VALUE! |
| ENSMUSG00000031312.6 | 3.702238 | 5.378281 | 3.918889 | 1.372399 | 0.169939 | NA | #VALUE! |
| ENSMUSG00000031323.9 | 0 | NA | NA | NA | NA | NA | #VALUE! |
| ENSMUSG00000031325.16 | 0 | NA | NA | NA | NA | NA | #VALUE! |
| ENSMUSG00000031326.3 | 0 | NA | NA | NA | NA | NA | #VALUE! |
| ENSMUSG00000031328.17 | 14.97022 | 1.049939 | 2.972506 | 0.353217 | NA | NA | #VALUE! |
| ENSMUSG00000031329.5 | 0 | NA | NA | NA | NA | NA | #VALUE! |
| ENSMUSG00000031330.9 | 0 | NA | NA | NA | NA | NA | #VALUE! |
| ENSMUSG00000031340.9 | 0 | NA | NA | NA | NA | NA | #VALUE! |
| ENSMUSG00000031342.18 | 0.336567 | 1.944925 | 4.029936 | 0.482619 | 0.629366 | NA | #VALUE! |
| ENSMUSG00000031343.14 | 1.666686 | -1.31006 | 3.861741 | -0.33924 | 0.734428 | NA | #VALUE! |
| ENSMUSG00000031344.12 | 0.466276 | 2.449889 | 3.997179 | 0.612904 | 0.53994 | NA | #VALUE! |
| ENSMUSG00000031351.16 | 0 | NA | NA | NA | NA | NA | #VALUE! |
| ENSMUSG00000031354.17 | 0 | NA | NA | NA | NA | NA | #VALUE! |
| ENSMUSG00000031355.17 | 7.995905 | -2.01591 | 2.345883 | -0.85934 | 0.390153 | NA | #VALUE! |
| ENSMUSG00000031360.15 | 21.46836 | -2.43832 | 2.43212 | -1.00255 | NA | NA | #VALUE! |
| ENSMUSG00000031362.18 | 4.711939 | 5.72619 | 3.916297 | 1.462144 | 0.143702 | NA | #VALUE! |
| ENSMUSG00000031364.3 | 0 | NA | NA | NA | NA | NA | #VALUE! |
| ENSMUSG00000031372.8 | 0 | NA | NA | NA | NA | NA | #VALUE! |
| ENSMUSG00000031374.14 | 0 | NA | NA | NA | NA | NA | #VALUE! |
| ENSMUSG00000031376.16 | 0 | NA | NA | NA | NA | NA | #VALUE! |
| ENSMUSG00000031377.12 | 0 | NA | NA | NA | NA | NA | #VALUE! |
| ENSMUSG00000031380.11 | 0 | NA | NA | NA | NA | NA | #VALUE! |
| ENSMUSG00000031381.17 | 7.803122 | -0.45868 | 2.094616 | -0.21898 | 0.826665 | NA | #VALUE! |
| ENSMUSG00000031382.15 | 0 | NA | NA | NA | NA | NA | #VALUE! |
| ENSMUSG00000031383.9 | 0 | NA | NA | NA | NA | NA | #VALUE! |
| ENSMUSG00000031384.3 | 0 | NA | NA | NA | NA | NA | #VALUE! |
| ENSMUSG00000031385.15 | 0 | NA | NA | NA | NA | NA | #VALUE! |
| ENSMUSG00000031387.15 | 6.358506 | -5.26683 | 3.827333 | -1.37611 | 0.168787 | NA | #VALUE! |
| ENSMUSG00000031389.18 | 0 | NA | NA | NA | NA | NA | #VALUE! |
| ENSMUSG00000031390.9 | 0 | NA | NA | NA | NA | NA | #VALUE! |
| ENSMUSG00000031391.19 | 3.780473 | -5.45568 | 3.918536 | -1.39228 | 0.163839 | NA | #VALUE! |
| ENSMUSG00000031394.12 | 0 | NA | NA | NA | NA | NA | #VALUE! |
| ENSMUSG00000031397.12 | 0 | NA | NA | NA | NA | NA | #VALUE! |
| ENSMUSG00000031398.14 | 0 | NA | NA | NA | NA | NA | #VALUE! |

| | | | | | | | |
|---|---|---|---|---|---|---|---|
| ENSMUSG00000031400.11 | 3.636953 | 5.361394 | 3.919032 | 1.36804 | 0.171299 | NA | #VALUE! |
| ENSMUSG00000031409.2 | 0 | NA | NA | NA | NA | NA | #VALUE! |
| ENSMUSG00000031410.15 | 0 | NA | NA | NA | NA | NA | #VALUE! |
| ENSMUSG00000031411.12 | 0 | NA | NA | NA | NA | NA | #VALUE! |
| ENSMUSG00000031424.17 | 0 | NA | NA | NA | NA | NA | #VALUE! |
| ENSMUSG00000031425.16 | 0.484833 | -1.24061 | 4.012754 | -0.30917 | 0.757196 | NA | #VALUE! |
| ENSMUSG00000031428.12 | 0 | NA | NA | NA | NA | NA | #VALUE! |
| ENSMUSG00000031430.9 | 0 | NA | NA | NA | NA | NA | #VALUE! |
| ENSMUSG00000031433.16 | 0 | NA | NA | NA | NA | NA | #VALUE! |
| ENSMUSG00000031434.16 | 0.315039 | -1.93549 | 4.038621 | -0.47925 | 0.631763 | NA | #VALUE! |
| ENSMUSG00000031442.22 | 0 | NA | NA | NA | NA | NA | #VALUE! |
| ENSMUSG00000031448.13 | 0 | NA | NA | NA | NA | NA | #VALUE! |
| ENSMUSG00000031449.11 | 0 | NA | NA | NA | NA | NA | #VALUE! |
| ENSMUSG00000031450.9 | 0 | NA | NA | NA | NA | NA | #VALUE! |
| ENSMUSG00000031452.16 | 0 | NA | NA | NA | NA | NA | #VALUE! |
| ENSMUSG00000031461.5 | 0 | NA | NA | NA | NA | NA | #VALUE! |
| ENSMUSG00000031465.7 | 1.501652 | 4.069382 | 3.936665 | 1.033713 | 0.30127 | NA | #VALUE! |
| ENSMUSG00000031471.4 | 0 | NA | NA | NA | NA | NA | #VALUE! |
| ENSMUSG00000031478.17 | 0 | NA | NA | NA | NA | NA | #VALUE! |
| ENSMUSG00000031480.16 | 0 | NA | NA | NA | NA | NA | #VALUE! |
| ENSMUSG00000031481.9 | 0 | NA | NA | NA | NA | NA | #VALUE! |
| ENSMUSG00000031486.16 | 6.055582 | 1.365157 | 3.76317 | 0.362768 | 0.716778 | NA | #VALUE! |
| ENSMUSG00000031489.16 | 4.903049 | -5.82123 | 3.915906 | -1.48656 | 0.137131 | NA | #VALUE! |
| ENSMUSG00000031491.11 | 0 | NA | NA | NA | NA | NA | #VALUE! |
| ENSMUSG00000031492.14 | 0 | NA | NA | NA | NA | NA | #VALUE! |
| ENSMUSG00000031493.11 | 0 | NA | NA | NA | NA | NA | #VALUE! |
| ENSMUSG00000031494.8 | 0 | NA | NA | NA | NA | NA | #VALUE! |
| ENSMUSG00000031495.9 | 0 | NA | NA | NA | NA | NA | #VALUE! |
| ENSMUSG00000031497.11 | 3.689633 | 5.364344 | 3.919007 | 1.368802 | 0.171061 | NA | #VALUE! |
| ENSMUSG00000031506.12 | 0 | NA | NA | NA | NA | NA | #VALUE! |
| ENSMUSG00000031509.11 | 0 | NA | NA | NA | NA | NA | #VALUE! |
| ENSMUSG00000031510.6 | 0 | NA | NA | NA | NA | NA | #VALUE! |
| ENSMUSG00000031511.17 | 14.58165 | -6.47277 | 2.992849 | -2.16274 | NA | NA | #VALUE! |
| ENSMUSG00000031512.16 | 0 | NA | NA | NA | NA | NA | #VALUE! |
| ENSMUSG00000031518.7 | 0 | NA | NA | NA | NA | NA | #VALUE! |
| ENSMUSG00000031519.5 | 0 | NA | NA | NA | NA | NA | #VALUE! |
| ENSMUSG00000031530.7 | 0.373021 | 2.140748 | 4.018182 | 0.532765 | 0.594196 | NA | #VALUE! |
| ENSMUSG00000031535.8 | 0 | NA | NA | NA | NA | NA | #VALUE! |
| ENSMUSG00000031538.7 | 0 | NA | NA | NA | NA | NA | #VALUE! |
| ENSMUSG00000031539.7 | 0 | NA | NA | NA | NA | NA | #VALUE! |
| ENSMUSG00000031543.19 | 0 | NA | NA | NA | NA | NA | #VALUE! |
| ENSMUSG00000031548.8 | 0 | NA | NA | NA | NA | NA | #VALUE! |
| ENSMUSG00000031551.13 | 0 | NA | NA | NA | NA | NA | #VALUE! |
| ENSMUSG00000031552.14 | 0 | NA | NA | NA | NA | NA | #VALUE! |

| | | | | | | | |
|---|---|---|---|---|---|---|---|
| ENSMUSG00000031553.17 | 0 | NA | NA | NA | NA | NA | #VALUE! |
| ENSMUSG00000031554.19 | 0 | NA | NA | NA | NA | NA | #VALUE! |
| ENSMUSG00000031558.16 | 0 | NA | NA | NA | NA | NA | #VALUE! |
| ENSMUSG00000031559.16 | 0 | NA | NA | NA | NA | NA | #VALUE! |
| ENSMUSG00000031562.17 | 0 | NA | NA | NA | NA | NA | #VALUE! |
| ENSMUSG00000031576.17 | 0 | NA | NA | NA | NA | NA | #VALUE! |
| ENSMUSG00000031595.10 | 0 | NA | NA | NA | NA | NA | #VALUE! |
| ENSMUSG00000031603.15 | 0 | NA | NA | NA | NA | NA | #VALUE! |
| ENSMUSG00000031610.4 | 0 | NA | NA | NA | NA | NA | #VALUE! |
| ENSMUSG00000031620.17 | 0 | NA | NA | NA | NA | NA | #VALUE! |
| ENSMUSG00000031621.10 | 0 | NA | NA | NA | NA | NA | #VALUE! |
| ENSMUSG00000031629.15 | 0 | NA | NA | NA | NA | NA | #VALUE! |
| ENSMUSG00000031635.15 | 0 | NA | NA | NA | NA | NA | #VALUE! |
| ENSMUSG00000031636.8 | 0 | NA | NA | NA | NA | NA | #VALUE! |
| ENSMUSG00000031637.14 | 6.751137 | -6.28598 | 3.547512 | -1.77194 | 0.076405 | NA | #VALUE! |
| ENSMUSG00000031640.12 | 0 | NA | NA | NA | NA | NA | #VALUE! |
| ENSMUSG00000031647.11 | 0 | NA | NA | NA | NA | NA | #VALUE! |
| ENSMUSG00000031651.6 | 0 | NA | NA | NA | NA | NA | #VALUE! |
| ENSMUSG00000031654.17 | 0 | NA | NA | NA | NA | NA | #VALUE! |
| ENSMUSG00000031659.14 | 4.289739 | 5.598228 | 3.917179 | 1.429148 | 0.152962 | NA | #VALUE! |
| ENSMUSG00000031662.9 | 2.205276 | -4.68302 | 3.926841 | -1.19257 | 0.233039 | NA | #VALUE! |
| ENSMUSG00000031669.11 | 2.415302 | -4.81324 | 3.925115 | -1.22627 | 0.220098 | NA | #VALUE! |
| ENSMUSG00000031673.7 | 0.839297 | 3.273019 | 3.958392 | 0.826856 | 0.408319 | NA | #VALUE! |
| ENSMUSG00000031682.6 | 0 | NA | NA | NA | NA | NA | #VALUE! |
| ENSMUSG00000031688.5 | 0 | NA | NA | NA | NA | NA | #VALUE! |
| ENSMUSG00000031698.15 | 0 | NA | NA | NA | NA | NA | #VALUE! |
| ENSMUSG00000031706.8 | 5.691105 | -4.93055 | 3.514403 | -1.40295 | 0.16063 | NA | #VALUE! |
| ENSMUSG00000031709.16 | 5.513814 | 4.998085 | 3.910679 | 1.278061 | 0.201228 | NA | #VALUE! |
| ENSMUSG00000031710.5 | 0 | NA | NA | NA | NA | NA | #VALUE! |
| ENSMUSG00000031712.11 | 5.095077 | -0.68796 | 3.634954 | -0.18926 | 0.849887 | NA | #VALUE! |
| ENSMUSG00000031723.9 | 0.279766 | 1.808031 | 4.036658 | 0.447903 | 0.654223 | NA | #VALUE! |
| ENSMUSG00000031727.9 | 0 | NA | NA | NA | NA | NA | #VALUE! |
| ENSMUSG00000031728.11 | 0 | NA | NA | NA | NA | NA | #VALUE! |
| ENSMUSG00000031732.13 | 6.241313 | 6.131012 | 3.660809 | 1.674769 | 0.093979 | NA | #VALUE! |
| ENSMUSG00000031734.14 | 0 | NA | NA | NA | NA | NA | #VALUE! |
| ENSMUSG00000031736.12 | 0 | NA | NA | NA | NA | NA | #VALUE! |
| ENSMUSG00000031737.12 | 0 | NA | NA | NA | NA | NA | #VALUE! |
| ENSMUSG00000031738.15 | 0 | NA | NA | NA | NA | NA | #VALUE! |
| ENSMUSG00000031740.9 | 0 | NA | NA | NA | NA | NA | #VALUE! |
| ENSMUSG00000031748.17 | 0 | NA | NA | NA | NA | NA | #VALUE! |
| ENSMUSG00000031749.13 | 0 | NA | NA | NA | NA | NA | #VALUE! |
| ENSMUSG00000031750.16 | 0.336567 | 1.944925 | 4.029936 | 0.482619 | 0.629366 | NA | #VALUE! |
| ENSMUSG00000031755.7 | 4.277 | 2.661584 | 3.817339 | 0.697235 | 0.485656 | NA | #VALUE! |
| ENSMUSG00000031756.6 | 0 | NA | NA | NA | NA | NA | #VALUE! |

| | | | | | | | |
|---|---|---|---|---|---|---|---|
| ENSMUSG00000031757.8 | 0 | NA | NA | NA | NA | NA | #VALUE! |
| ENSMUSG00000031758.11 | 0.18651 | 1.401769 | 4.056354 | 0.345574 | 0.729663 | NA | #VALUE! |
| ENSMUSG00000031760.10 | 0 | NA | NA | NA | NA | NA | #VALUE! |
| ENSMUSG00000031762.8 | 180.2476 | -3.62465 | 3.308638 | -1.09551 | NA | NA | #VALUE! |
| ENSMUSG00000031765.9 | 762.5367 | -2.90572 | 1.110606 | -2.61634 | NA | NA | #VALUE! |
| ENSMUSG00000031766.5 | 0 | NA | NA | NA | NA | NA | #VALUE! |
| ENSMUSG00000031772.18 | 0 | NA | NA | NA | NA | NA | #VALUE! |
| ENSMUSG00000031778.14 | 0 | NA | NA | NA | NA | NA | #VALUE! |
| ENSMUSG00000031779.4 | 0.168284 | 1.027628 | 4.067126 | 0.252667 | 0.800526 | NA | #VALUE! |
| ENSMUSG00000031780.3 | 0 | NA | NA | NA | NA | NA | #VALUE! |
| ENSMUSG00000031785.17 | 4.075121 | 0.307452 | 3.771211 | 0.081526 | 0.935024 | NA | #VALUE! |
| ENSMUSG00000031786.8 | 0 | NA | NA | NA | NA | NA | #VALUE! |
| ENSMUSG00000031787.9 | 0.18651 | 1.401769 | 4.056354 | 0.345574 | 0.729663 | NA | #VALUE! |
| ENSMUSG00000031789.18 | 0 | NA | NA | NA | NA | NA | #VALUE! |
| ENSMUSG00000031791.9 | 7.995514 | -0.12189 | 3.153849 | -0.03865 | 0.969171 | NA | #VALUE! |
| ENSMUSG00000031802.10 | 1.585338 | 4.174397 | 3.934581 | 1.060951 | 0.288712 | NA | #VALUE! |
| ENSMUSG00000031805.18 | 0 | NA | NA | NA | NA | NA | #VALUE! |
| ENSMUSG00000031809.11 | 0 | NA | NA | NA | NA | NA | #VALUE! |
| ENSMUSG00000031819.14 | 104.2497 | 0.839732 | 1.007781 | 0.833249 | NA | NA | #VALUE! |
| ENSMUSG00000031821.12 | 0 | NA | NA | NA | NA | NA | #VALUE! |
| ENSMUSG00000031823.15 | 2.224866 | -4.68666 | 3.926791 | -1.19351 | 0.23267 | NA | #VALUE! |
| ENSMUSG00000031825.17 | 8.937297 | 2.030913 | 3.015699 | 0.673447 | 0.500663 | NA | #VALUE! |
| ENSMUSG00000031831.7 | 0 | NA | NA | NA | NA | NA | #VALUE! |
| ENSMUSG00000031832.16 | 0 | NA | NA | NA | NA | NA | #VALUE! |
| ENSMUSG00000031833.11 | 6.836737 | -6.30055 | 3.215662 | -1.95933 | 0.050074 | NA | #VALUE! |
| ENSMUSG00000031834.16 | 1.704401 | -2.1343 | 3.892719 | -0.54828 | 0.583499 | NA | #VALUE! |
| ENSMUSG00000031837.15 | 0 | NA | NA | NA | NA | NA | #VALUE! |
| ENSMUSG00000031840.15 | 1.050131 | -3.6251 | 3.948379 | -0.91812 | 0.358554 | NA | #VALUE! |
| ENSMUSG00000031841.20 | 9.731163 | -0.63911 | 2.274356 | -0.28101 | 0.778704 | NA | #VALUE! |
| ENSMUSG00000031842.16 | 3.353942 | -5.27766 | 3.920078 | -1.34632 | 0.178201 | NA | #VALUE! |
| ENSMUSG00000031847.9 | 0 | NA | NA | NA | NA | NA | #VALUE! |
| ENSMUSG00000031849.10 | 0 | NA | NA | NA | NA | NA | #VALUE! |
| ENSMUSG00000031851.15 | 40.03844 | 0.591585 | 1.493442 | 0.396122 | NA | NA | #VALUE! |
| ENSMUSG00000031860.18 | 0 | NA | NA | NA | NA | NA | #VALUE! |
| ENSMUSG00000031861.17 | 0 | NA | NA | NA | NA | NA | #VALUE! |
| ENSMUSG00000031865.17 | 0 | NA | NA | NA | NA | NA | #VALUE! |
| ENSMUSG00000031870.17 | 0 | NA | NA | NA | NA | NA | #VALUE! |
| ENSMUSG00000031872.15 | 0 | NA | NA | NA | NA | NA | #VALUE! |
| ENSMUSG00000031876.19 | 0 | NA | NA | NA | NA | NA | #VALUE! |
| ENSMUSG00000031880.8 | 0 | NA | NA | NA | NA | NA | #VALUE! |
| ENSMUSG00000031881.7 | 0 | NA | NA | NA | NA | NA | #VALUE! |
| ENSMUSG00000031883.14 | 0 | NA | NA | NA | NA | NA | #VALUE! |
| ENSMUSG00000031884.11 | 0 | NA | NA | NA | NA | NA | #VALUE! |
| ENSMUSG00000031891.7 | 0 | NA | NA | NA | NA | NA | #VALUE! |

| | | | | | | | |
|---|---|---|---|---|---|---|---|
| ENSMUSG00000031893.8 | 0 | NA | NA | NA | NA | NA | #VALUE! |
| ENSMUSG00000031896.8 | 0 | NA | NA | NA | NA | NA | #VALUE! |
| ENSMUSG00000031898.10 | 0 | NA | NA | NA | NA | NA | #VALUE! |
| ENSMUSG00000031904.6 | 0.391578 | -2.1489 | 4.020883 | -0.53444 | 0.59304 | NA | #VALUE! |
| ENSMUSG00000031907.10 | 3.060811 | -0.27912 | 3.78569 | -0.07373 | 0.941224 | NA | #VALUE! |
| ENSMUSG00000031910.15 | 0 | NA | NA | NA | NA | NA | #VALUE! |
| ENSMUSG00000031919.7 | 0 | NA | NA | NA | NA | NA | #VALUE! |
| ENSMUSG00000031921.18 | 14.81133 | 0.022406 | 2.961553 | 0.007566 | NA | NA | #VALUE! |
| ENSMUSG00000031927.10 | 0 | NA | NA | NA | NA | NA | #VALUE! |
| ENSMUSG00000031932.15 | 0 | NA | NA | NA | NA | NA | #VALUE! |
| ENSMUSG00000031933.18 | 0 | NA | NA | NA | NA | NA | #VALUE! |
| ENSMUSG00000031934.15 | 1.794317 | 4.336098 | 3.931651 | 1.102869 | 0.270084 | NA | #VALUE! |
| ENSMUSG00000031936.10 | 0 | NA | NA | NA | NA | NA | #VALUE! |
| ENSMUSG00000031937.8 | 0 | NA | NA | NA | NA | NA | #VALUE! |
| ENSMUSG00000031951.9 | 2.041108 | -4.55188 | 3.928743 | -1.15861 | 0.246615 | NA | #VALUE! |
| ENSMUSG00000031952.6 | 0 | NA | NA | NA | NA | NA | #VALUE! |
| ENSMUSG00000031957.7 | 0 | NA | NA | NA | NA | NA | #VALUE! |
| ENSMUSG00000031962.7 | 0 | NA | NA | NA | NA | NA | #VALUE! |
| ENSMUSG00000031963.9 | 0 | NA | NA | NA | NA | NA | #VALUE! |
| ENSMUSG00000031965.16 | 8.955468 | -2.08919 | 2.277597 | -0.91728 | 0.358998 | NA | #VALUE! |
| ENSMUSG00000031966.8 | 0 | NA | NA | NA | NA | NA | #VALUE! |
| ENSMUSG00000031970.18 | 0 | NA | NA | NA | NA | NA | #VALUE! |
| ENSMUSG00000031971.16 | 1.020554 | -3.54462 | 3.950745 | -0.8972 | 0.36961 | NA | #VALUE! |
| ENSMUSG00000031972.6 | 0 | NA | NA | NA | NA | NA | #VALUE! |
| ENSMUSG00000031981.8 | 0 | NA | NA | NA | NA | NA | #VALUE! |
| ENSMUSG00000031982.6 | 22.4341 | -6.9711 | 2.414873 | -2.88674 | NA | NA | #VALUE! |
| ENSMUSG00000031990.16 | 0 | NA | NA | NA | NA | NA | #VALUE! |
| ENSMUSG00000031991.11 | 0 | NA | NA | NA | NA | NA | #VALUE! |
| ENSMUSG00000031994.14 | 0 | NA | NA | NA | NA | NA | #VALUE! |
| ENSMUSG00000031995.10 | 4.490437 | -5.69342 | 3.916751 | -1.45361 | 0.146055 | NA | #VALUE! |
| ENSMUSG00000031997.10 | 0 | NA | NA | NA | NA | NA | #VALUE! |
| ENSMUSG00000031998.6 | 0 | NA | NA | NA | NA | NA | #VALUE! |
| ENSMUSG00000032006.16 | 7.415349 | -1.58751 | 3.172079 | -0.50046 | 0.616749 | NA | #VALUE! |
| ENSMUSG00000032011.6 | 0.504851 | 2.502843 | 3.993993 | 0.626652 | 0.530887 | NA | #VALUE! |
| ENSMUSG00000032013.7 | 0 | NA | NA | NA | NA | NA | #VALUE! |
| ENSMUSG00000032015.17 | 0 | NA | NA | NA | NA | NA | #VALUE! |
| ENSMUSG00000032017.15 | 0 | NA | NA | NA | NA | NA | #VALUE! |
| ENSMUSG00000032021.15 | 0 | NA | NA | NA | NA | NA | #VALUE! |
| ENSMUSG00000032023.8 | 0 | NA | NA | NA | NA | NA | #VALUE! |
| ENSMUSG00000032024.11 | 0.652786 | 2.919659 | 3.972474 | 0.734972 | 0.462356 | NA | #VALUE! |
| ENSMUSG00000032033.12 | 0 | NA | NA | NA | NA | NA | #VALUE! |
| ENSMUSG00000032034.7 | 0 | NA | NA | NA | NA | NA | #VALUE! |
| ENSMUSG00000032036.16 | 0 | NA | NA | NA | NA | NA | #VALUE! |
| ENSMUSG00000032048.4 | 0 | NA | NA | NA | NA | NA | #VALUE! |

| | | | | | | | |
|---|---|---|---|---|---|---|---|
| ENSMUSG00000032053.9 | 0.093255 | 0.868093 | 4.072464 | 0.213162 | 0.831201 | NA | #VALUE! |
| ENSMUSG00000032056.11 | 0 | NA | NA | NA | NA | NA | #VALUE! |
| ENSMUSG00000032057.10 | 0 | NA | NA | NA | NA | NA | #VALUE! |
| ENSMUSG00000032060.11 | 3.544601 | 0.229379 | 3.666651 | 0.062558 | 0.950118 | NA | #VALUE! |
| ENSMUSG00000032065.7 | 9.067035 | -1.51132 | 3.406189 | -0.4437 | 0.657261 | NA | #VALUE! |
| ENSMUSG00000032066.10 | 45.37494 | -0.2095 | 1.525895 | -0.13729 | NA | NA | #VALUE! |
| ENSMUSG00000032068.15 | 18.64143 | -0.24652 | 2.114194 | -0.1166 | NA | NA | #VALUE! |
| ENSMUSG00000032077.6 | 5.116212 | -4.39156 | 3.679326 | -1.19358 | 0.232643 | NA | #VALUE! |
| ENSMUSG00000032085.6 | 1.619565 | 3.155742 | 3.931935 | 0.802593 | 0.42221 | NA | #VALUE! |
| ENSMUSG00000032087.12 | 0 | NA | NA | NA | NA | NA | #VALUE! |
| ENSMUSG00000032089.17 | 0 | NA | NA | NA | NA | NA | #VALUE! |
| ENSMUSG00000032091.16 | 0 | NA | NA | NA | NA | NA | #VALUE! |
| ENSMUSG00000032093.8 | 0 | NA | NA | NA | NA | NA | #VALUE! |
| ENSMUSG00000032094.9 | 3.748467 | -5.43432 | 3.918711 | -1.38676 | 0.165514 | NA | #VALUE! |
| ENSMUSG00000032098.14 | 0 | NA | NA | NA | NA | NA | #VALUE! |
| ENSMUSG00000032099.4 | 0 | NA | NA | NA | NA | NA | #VALUE! |
| ENSMUSG00000032101.7 | 0 | NA | NA | NA | NA | NA | #VALUE! |
| ENSMUSG00000032105.6 | 0 | NA | NA | NA | NA | NA | #VALUE! |
| ENSMUSG00000032108.4 | 0 | NA | NA | NA | NA | NA | #VALUE! |
| ENSMUSG00000032109.16 | 0.737927 | 3.007825 | 3.968641 | 0.757898 | 0.448512 | NA | #VALUE! |
| ENSMUSG00000032110.5 | 0 | NA | NA | NA | NA | NA | #VALUE! |
| ENSMUSG00000032113.17 | 4.039291 | 4.535958 | 3.913798 | 1.158966 | 0.24647 | NA | #VALUE! |
| ENSMUSG00000032118.17 | 0 | NA | NA | NA | NA | NA | #VALUE! |
| ENSMUSG00000032128.16 | 0 | NA | NA | NA | NA | NA | #VALUE! |
| ENSMUSG00000032131.17 | 0 | NA | NA | NA | NA | NA | #VALUE! |
| ENSMUSG00000032135.16 | 3.893626 | -0.94669 | 3.58274 | -0.26424 | 0.791597 | NA | #VALUE! |
| ENSMUSG00000032172.9 | 0 | NA | NA | NA | NA | NA | #VALUE! |
| ENSMUSG00000032174.6 | 0 | NA | NA | NA | NA | NA | #VALUE! |
| ENSMUSG00000032177.18 | 0 | NA | NA | NA | NA | NA | #VALUE! |
| ENSMUSG00000032180.12 | 4.32494 | 3.305846 | 3.851848 | 0.858249 | 0.390755 | NA | #VALUE! |
| ENSMUSG00000032181.8 | 0 | NA | NA | NA | NA | NA | #VALUE! |
| ENSMUSG00000032185.16 | 28.14443 | -1.06599 | 2.147554 | -0.49638 | NA | NA | #VALUE! |
| ENSMUSG00000032186.16 | 0 | NA | NA | NA | NA | NA | #VALUE! |
| ENSMUSG00000032191.7 | 0 | NA | NA | NA | NA | NA | #VALUE! |
| ENSMUSG00000032202.12 | 1.575872 | -1.01852 | 3.85513 | -0.2642 | 0.791627 | NA | #VALUE! |
| ENSMUSG00000032218.8 | 8.590398 | -6.63067 | 3.322252 | -1.99584 | 0.045952 | NA | #VALUE! |
| ENSMUSG00000032221.15 | 0 | NA | NA | NA | NA | NA | #VALUE! |
| ENSMUSG00000032226.6 | 0 | NA | NA | NA | NA | NA | #VALUE! |
| ENSMUSG00000032232.15 | 99.43204 | -0.55657 | 1.017228 | -0.54714 | NA | NA | #VALUE! |
| ENSMUSG00000032243.9 | 0 | NA | NA | NA | NA | NA | #VALUE! |
| ENSMUSG00000032245.13 | 8.447406 | -6.60687 | 3.357826 | -1.9676 | 0.049114 | NA | #VALUE! |
| ENSMUSG00000032254.11 | 2.245218 | -4.69006 | 3.926744 | -1.19439 | 0.232326 | NA | #VALUE! |
| ENSMUSG00000032257.7 | 0 | NA | NA | NA | NA | NA | #VALUE! |
| ENSMUSG00000032258.16 | 0 | NA | NA | NA | NA | NA | #VALUE! |

| | | | | | | | |
|---|---|---|---|---|---|---|---|
| ENSMUSG00000032259.9 | 0 | NA | NA | NA | NA | NA | #VALUE! |
| ENSMUSG00000032262.14 | 0 | NA | NA | NA | NA | NA | #VALUE! |
| ENSMUSG00000032264.10 | 7.478371 | 0.636341 | 3.197666 | 0.199002 | 0.842261 | NA | #VALUE! |
| ENSMUSG00000032265.15 | 15.48199 | -2.15111 | 2.402156 | -0.89549 | NA | NA | #VALUE! |
| ENSMUSG00000032267.9 | 14.57834 | 0.915628 | 2.677091 | 0.342024 | NA | NA | #VALUE! |
| ENSMUSG00000032268.14 | 0 | NA | NA | NA | NA | NA | #VALUE! |
| ENSMUSG00000032269.9 | 0 | NA | NA | NA | NA | NA | #VALUE! |
| ENSMUSG00000032274.10 | 0 | NA | NA | NA | NA | NA | #VALUE! |
| ENSMUSG00000032278.12 | 0 | NA | NA | NA | NA | NA | #VALUE! |
| ENSMUSG00000032281.12 | 0 | NA | NA | NA | NA | NA | #VALUE! |
| ENSMUSG00000032289.16 | 0 | NA | NA | NA | NA | NA | #VALUE! |
| ENSMUSG00000032291.9 | 0 | NA | NA | NA | NA | NA | #VALUE! |
| ENSMUSG00000032292.9 | 0 | NA | NA | NA | NA | NA | #VALUE! |
| ENSMUSG00000032295.16 | 9.658021 | 2.109378 | 2.484346 | 0.849068 | 0.395844 | NA | #VALUE! |
| ENSMUSG00000032297.12 | 0 | NA | NA | NA | NA | NA | #VALUE! |
| ENSMUSG00000032298.14 | 9.000941 | -1.35457 | 2.58781 | -0.52344 | 0.600667 | NA | #VALUE! |
| ENSMUSG00000032300.8 | 1.796981 | 4.336699 | 3.931641 | 1.103025 | 0.270016 | NA | #VALUE! |
| ENSMUSG00000032303.9 | 0 | NA | NA | NA | NA | NA | #VALUE! |
| ENSMUSG00000032305.16 | 6.114268 | 6.100707 | 3.690589 | 1.653044 | 0.098322 | NA | #VALUE! |
| ENSMUSG00000032308.10 | 9.045754 | 6.671149 | 3.688539 | 1.808616 | NA | NA | #VALUE! |
| ENSMUSG00000032313.12 | 0 | NA | NA | NA | NA | NA | #VALUE! |
| ENSMUSG00000032315.7 | 1.229878 | 3.761857 | 3.943699 | 0.95389 | 0.340139 | NA | #VALUE! |
| ENSMUSG00000032318.13 | 0 | NA | NA | NA | NA | NA | #VALUE! |
| ENSMUSG00000032323.14 | 0 | NA | NA | NA | NA | NA | #VALUE! |
| ENSMUSG00000032327.15 | 0 | NA | NA | NA | NA | NA | #VALUE! |
| ENSMUSG00000032333.7 | 0 | NA | NA | NA | NA | NA | #VALUE! |
| ENSMUSG00000032334.11 | 0 | NA | NA | NA | NA | NA | #VALUE! |
| ENSMUSG00000032338.10 | 0 | NA | NA | NA | NA | NA | #VALUE! |
| ENSMUSG00000032343.16 | 0 | NA | NA | NA | NA | NA | #VALUE! |
| ENSMUSG00000032344.18 | 8.443465 | -1.22646 | 3.508703 | -0.34955 | 0.726678 | NA | #VALUE! |
| ENSMUSG00000032346.8 | 0 | NA | NA | NA | NA | NA | #VALUE! |
| ENSMUSG00000032352.17 | 0 | NA | NA | NA | NA | NA | #VALUE! |
| ENSMUSG00000032355.17 | 0 | NA | NA | NA | NA | NA | #VALUE! |
| ENSMUSG00000032356.13 | 0 | NA | NA | NA | NA | NA | #VALUE! |
| ENSMUSG00000032357.13 | 0 | NA | NA | NA | NA | NA | #VALUE! |
| ENSMUSG00000032358.13 | 0 | NA | NA | NA | NA | NA | #VALUE! |
| ENSMUSG00000032360.17 | 2.251499 | -1.89082 | 3.844972 | -0.49176 | 0.622886 | NA | #VALUE! |
| ENSMUSG00000032363.16 | 0 | NA | NA | NA | NA | NA | #VALUE! |
| ENSMUSG00000032368.15 | 0 | NA | NA | NA | NA | NA | #VALUE! |
| ENSMUSG00000032373.16 | 0 | NA | NA | NA | NA | NA | #VALUE! |
| ENSMUSG00000032374.15 | 0 | NA | NA | NA | NA | NA | #VALUE! |
| ENSMUSG00000032394.7 | 0 | NA | NA | NA | NA | NA | #VALUE! |
| ENSMUSG00000032400.19 | 0.559531 | 2.703958 | 3.982863 | 0.678898 | 0.497202 | NA | #VALUE! |
| ENSMUSG00000032401.16 | 0 | NA | NA | NA | NA | NA | #VALUE! |

| | | | | | | | |
|---|---|---|---|---|---|---|---|
| ENSMUSG00000032413.13 | 6.148963 | 2.029741 | 3.463711 | 0.586002 | 0.557874 | NA | #VALUE! |
| ENSMUSG00000032417.11 | 0 | NA | NA | NA | NA | NA | #VALUE! |
| ENSMUSG00000032419.9 | 0 | NA | NA | NA | NA | NA | #VALUE! |
| ENSMUSG00000032443.18 | 1.785223 | -4.3809 | 3.931494 | -1.11431 | 0.265146 | NA | #VALUE! |
| ENSMUSG00000032446.15 | 0 | NA | NA | NA | NA | NA | #VALUE! |
| ENSMUSG00000032449.14 | 9.585987 | -0.42586 | 3.09639 | -0.13753 | 0.890609 | NA | #VALUE! |
| ENSMUSG00000032451.7 | 0 | NA | NA | NA | NA | NA | #VALUE! |
| ENSMUSG00000032452.13 | 0.420053 | -2.33493 | 4.007343 | -0.58266 | 0.56012 | NA | #VALUE! |
| ENSMUSG00000032454.10 | 0 | NA | NA | NA | NA | NA | #VALUE! |
| ENSMUSG00000032473.14 | 2.019402 | 4.503826 | 3.928936 | 1.146322 | 0.251662 | NA | #VALUE! |
| ENSMUSG00000032477.15 | 3.357187 | 5.246639 | 3.920044 | 1.338413 | 0.180762 | NA | #VALUE! |
| ENSMUSG00000032482.10 | 9.285609 | -1.49366 | 3.394752 | -0.43999 | 0.659943 | NA | #VALUE! |
| ENSMUSG00000032484.9 | 8.164275 | -0.3791 | 3.503361 | -0.10821 | 0.913828 | NA | #VALUE! |
| ENSMUSG00000032487.9 | 0 | NA | NA | NA | NA | NA | #VALUE! |
| ENSMUSG00000032493.9 | 0 | NA | NA | NA | NA | NA | #VALUE! |
| ENSMUSG00000032494.13 | 0 | NA | NA | NA | NA | NA | #VALUE! |
| ENSMUSG00000032495.9 | 0 | NA | NA | NA | NA | NA | #VALUE! |
| ENSMUSG00000032496.8 | 0 | NA | NA | NA | NA | NA | #VALUE! |
| ENSMUSG00000032502.9 | 0 | NA | NA | NA | NA | NA | #VALUE! |
| ENSMUSG00000032503.19 | 0 | NA | NA | NA | NA | NA | #VALUE! |
| ENSMUSG00000032507.16 | 2.51789 | 4.834703 | 3.924413 | 1.231956 | 0.217966 | NA | #VALUE! |
| ENSMUSG00000032511.18 | 0 | NA | NA | NA | NA | NA | #VALUE! |
| ENSMUSG00000032514.12 | 0 | NA | NA | NA | NA | NA | #VALUE! |
| ENSMUSG00000032517.17 | 0 | NA | NA | NA | NA | NA | #VALUE! |
| ENSMUSG00000032523.12 | 0 | NA | NA | NA | NA | NA | #VALUE! |
| ENSMUSG00000032530.15 | 0 | NA | NA | NA | NA | NA | #VALUE! |
| ENSMUSG00000032532.8 | 0 | NA | NA | NA | NA | NA | #VALUE! |
| ENSMUSG00000032537.16 | 0 | NA | NA | NA | NA | NA | #VALUE! |
| ENSMUSG00000032549.8 | 2.611145 | 4.886731 | 3.92379 | 1.245411 | 0.212981 | NA | #VALUE! |
| ENSMUSG00000032556.12 | 0 | NA | NA | NA | NA | NA | #VALUE! |
| ENSMUSG00000032558.20 | 7.417292 | 1.043463 | 3.255015 | 0.320571 | 0.748536 | NA | #VALUE! |
| ENSMUSG00000032564.17 | 0 | NA | NA | NA | NA | NA | #VALUE! |
| ENSMUSG00000032566.5 | 0 | NA | NA | NA | NA | NA | #VALUE! |
| ENSMUSG00000032567.17 | 6.858789 | 4.414104 | 2.602342 | 1.696205 | 0.089847 | NA | #VALUE! |
| ENSMUSG00000032572.10 | 0 | NA | NA | NA | NA | NA | #VALUE! |
| ENSMUSG00000032577.17 | 1.428775 | -4.03418 | 3.938169 | -1.02438 | 0.305656 | NA | #VALUE! |
| ENSMUSG00000032583.8 | 6.756911 | -6.28439 | 3.217564 | -1.95315 | 0.050801 | NA | #VALUE! |
| ENSMUSG00000032584.13 | 0 | NA | NA | NA | NA | NA | #VALUE! |
| ENSMUSG00000032586.10 | 0 | NA | NA | NA | NA | NA | #VALUE! |
| ENSMUSG00000032589.15 | 0 | NA | NA | NA | NA | NA | #VALUE! |
| ENSMUSG00000032593.6 | 5.874609 | 1.014572 | 3.760833 | 0.269773 | 0.787335 | NA | #VALUE! |
| ENSMUSG00000032595.11 | 0 | NA | NA | NA | NA | NA | #VALUE! |
| ENSMUSG00000032609.13 | 2.238125 | 4.666312 | 3.926586 | 1.188389 | 0.23468 | NA | #VALUE! |
| ENSMUSG00000032611.10 | 0 | NA | NA | NA | NA | NA | #VALUE! |

| | | | | | | | |
|---|---|---|---|---|---|---|---|
| ENSMUSG00000032623.10 | 0 | NA | NA | NA | NA | NA | #VALUE! |
| ENSMUSG00000032625.15 | 0 | NA | NA | NA | NA | NA | #VALUE! |
| ENSMUSG00000032640.11 | 8.868391 | -0.84428 | 3.450943 | -0.24465 | 0.806726 | NA | #VALUE! |
| ENSMUSG00000032641.19 | 5.286306 | -5.93001 | 3.915244 | -1.5146 | 0.129875 | NA | #VALUE! |
| ENSMUSG00000032643.13 | 0.735092 | -3.12016 | 3.965612 | -0.7868 | 0.431397 | NA | #VALUE! |
| ENSMUSG00000032648.15 | 2.524253 | 4.825754 | 3.924522 | 1.229641 | 0.218831 | NA | #VALUE! |
| ENSMUSG00000032649.15 | 2.951706 | 5.040208 | 3.922079 | 1.285086 | 0.198762 | NA | #VALUE! |
| ENSMUSG00000032656.15 | 0 | NA | NA | NA | NA | NA | #VALUE! |
| ENSMUSG00000032657.16 | 0.945118 | -3.47564 | 3.952879 | -0.87927 | 0.379256 | NA | #VALUE! |
| ENSMUSG00000032661.10 | 3.935608 | 5.458007 | 3.918239 | 1.392975 | 0.163627 | NA | #VALUE! |
| ENSMUSG00000032671.6 | 0 | NA | NA | NA | NA | NA | #VALUE! |
| ENSMUSG00000032680.12 | 0 | NA | NA | NA | NA | NA | #VALUE! |
| ENSMUSG00000032690.17 | 4.935617 | -5.8387 | 3.915796 | -1.49106 | 0.135945 | NA | #VALUE! |
| ENSMUSG00000032691.15 | 0.195789 | -1.05581 | 4.073302 | -0.2592 | 0.795479 | NA | #VALUE! |
| ENSMUSG00000032714.7 | 1.723265 | -0.66651 | 3.83672 | -0.17372 | 0.862087 | NA | #VALUE! |
| ENSMUSG00000032715.10 | 12.64376 | -0.7702 | 3.082801 | -0.24984 | NA | NA | #VALUE! |
| ENSMUSG00000032717.15 | 0 | NA | NA | NA | NA | NA | #VALUE! |
| ENSMUSG00000032718.5 | 1.020554 | -3.54462 | 3.950745 | -0.8972 | 0.36961 | NA | #VALUE! |
| ENSMUSG00000032719.5 | 0 | NA | NA | NA | NA | NA | #VALUE! |
| ENSMUSG00000032726.13 | 0 | NA | NA | NA | NA | NA | #VALUE! |
| ENSMUSG00000032739.17 | 0 | NA | NA | NA | NA | NA | #VALUE! |
| ENSMUSG00000032744.5 | 5.28944 | -2.16877 | 3.784161 | -0.57312 | 0.566565 | NA | #VALUE! |
| ENSMUSG00000032750.14 | 0 | NA | NA | NA | NA | NA | #VALUE! |
| ENSMUSG00000032758.5 | 0 | NA | NA | NA | NA | NA | #VALUE! |
| ENSMUSG00000032769.6 | 0 | NA | NA | NA | NA | NA | #VALUE! |
| ENSMUSG00000032773.10 | 0 | NA | NA | NA | NA | NA | #VALUE! |
| ENSMUSG00000032777.10 | 9.875899 | 2.782103 | 2.543274 | 1.093906 | 0.273996 | NA | #VALUE! |
| ENSMUSG00000032782.20 | 1.824007 | 0.164829 | 3.82696 | 0.043071 | 0.965645 | NA | #VALUE! |
| ENSMUSG00000032783.10 | 0 | NA | NA | NA | NA | NA | #VALUE! |
| ENSMUSG00000032796.9 | 3.00047 | 1.588138 | 3.807058 | 0.417156 | 0.676564 | NA | #VALUE! |
| ENSMUSG00000032806.14 | 5.226332 | -2.01035 | 3.780523 | -0.53177 | 0.594888 | NA | #VALUE! |
| ENSMUSG00000032807.6 | 0 | NA | NA | NA | NA | NA | #VALUE! |
| ENSMUSG00000032812.19 | 4.664805 | -0.63193 | 3.044735 | -0.20755 | 0.835581 | NA | #VALUE! |
| ENSMUSG00000032815.17 | 0 | NA | NA | NA | NA | NA | #VALUE! |
| ENSMUSG00000032816.16 | 0 | NA | NA | NA | NA | NA | #VALUE! |
| ENSMUSG00000032818.17 | 0 | NA | NA | NA | NA | NA | #VALUE! |
| ENSMUSG00000032826.20 | 0 | NA | NA | NA | NA | NA | #VALUE! |
| ENSMUSG00000032827.16 | 7.636379 | -0.0338 | 2.60887 | -0.01296 | 0.989663 | NA | #VALUE! |
| ENSMUSG00000032834.12 | 7.749712 | 4.907553 | 3.102615 | 1.581747 | 0.113707 | NA | #VALUE! |
| ENSMUSG00000032839.14 | 0.093255 | 0.868093 | 4.072464 | 0.213162 | 0.831201 | NA | #VALUE! |
| ENSMUSG00000032840.8 | 8.538182 | -6.62223 | 3.360884 | -1.97038 | 0.048795 | NA | #VALUE! |
| ENSMUSG00000032841.16 | 1.260158 | -3.88425 | 3.941584 | -0.98545 | 0.324401 | NA | #VALUE! |
| ENSMUSG00000032845.17 | 0 | NA | NA | NA | NA | NA | #VALUE! |
| ENSMUSG00000032850.17 | 0 | NA | NA | NA | NA | NA | #VALUE! |

| | | | | | | | |
|---|---|---|---|---|---|---|---|
| ENSMUSG00000032852.4 | 0 | NA | NA | NA | NA | NA | #VALUE! |
| ENSMUSG00000032854.13 | 0.491951 | 2.402185 | 4.000147 | 0.600524 | 0.548157 | NA | #VALUE! |
| ENSMUSG00000032864.5 | 0 | NA | NA | NA | NA | NA | #VALUE! |
| ENSMUSG00000032867.9 | 31.0125 | -4.20511 | 2.304623 | -1.82464 | NA | NA | #VALUE! |
| ENSMUSG00000032875.9 | 3.699366 | 4.275669 | 3.912172 | 1.092914 | 0.274431 | NA | #VALUE! |
| ENSMUSG00000032878.17 | 0 | NA | NA | NA | NA | NA | #VALUE! |
| ENSMUSG00000032889.8 | 0 | NA | NA | NA | NA | NA | #VALUE! |
| ENSMUSG00000032890.18 | 0 | NA | NA | NA | NA | NA | #VALUE! |
| ENSMUSG00000032892.15 | 0 | NA | NA | NA | NA | NA | #VALUE! |
| ENSMUSG00000032894.14 | 0 | NA | NA | NA | NA | NA | #VALUE! |
| ENSMUSG00000032899.15 | 0 | NA | NA | NA | NA | NA | #VALUE! |
| ENSMUSG00000032900.16 | 0 | NA | NA | NA | NA | NA | #VALUE! |
| ENSMUSG00000032908.10 | 0 | NA | NA | NA | NA | NA | #VALUE! |
| ENSMUSG00000032911.7 | 0 | NA | NA | NA | NA | NA | #VALUE! |
| ENSMUSG00000032913.14 | 2.938512 | 5.044044 | 3.922038 | 1.286077 | 0.198416 | NA | #VALUE! |
| ENSMUSG00000032921.14 | 0 | NA | NA | NA | NA | NA | #VALUE! |
| ENSMUSG00000032925.17 | 1.995249 | -4.53986 | 3.928926 | -1.1555 | 0.247887 | NA | #VALUE! |
| ENSMUSG00000032936.14 | 0 | NA | NA | NA | NA | NA | #VALUE! |
| ENSMUSG00000032937.6 | 0 | NA | NA | NA | NA | NA | #VALUE! |
| ENSMUSG00000032939.15 | 6.971357 | 0.190402 | 2.928009 | 0.065028 | 0.948152 | NA | #VALUE! |
| ENSMUSG00000032940.17 | 0 | NA | NA | NA | NA | NA | #VALUE! |
| ENSMUSG00000032942.15 | 0 | NA | NA | NA | NA | NA | #VALUE! |
| ENSMUSG00000032948.11 | 0 | NA | NA | NA | NA | NA | #VALUE! |
| ENSMUSG00000032965.12 | 8.46066 | 1.311151 | 3.041236 | 0.431124 | 0.666378 | NA | #VALUE! |
| ENSMUSG00000032968.5 | 0 | NA | NA | NA | NA | NA | #VALUE! |
| ENSMUSG00000032978.15 | 0 | NA | NA | NA | NA | NA | #VALUE! |
| ENSMUSG00000032985.16 | 0 | NA | NA | NA | NA | NA | #VALUE! |
| ENSMUSG00000032987.10 | 0 | NA | NA | NA | NA | NA | #VALUE! |
| ENSMUSG00000032988.5 | 0 | NA | NA | NA | NA | NA | #VALUE! |
| ENSMUSG00000032997.17 | 0.673134 | 2.918386 | 3.972531 | 0.734641 | 0.462558 | NA | #VALUE! |
| ENSMUSG00000032999.11 | 0 | NA | NA | NA | NA | NA | #VALUE! |
| ENSMUSG00000033006.11 | 0 | NA | NA | NA | NA | NA | #VALUE! |
| ENSMUSG00000033007.16 | 0 | NA | NA | NA | NA | NA | #VALUE! |
| ENSMUSG00000033015.17 | 0 | NA | NA | NA | NA | NA | #VALUE! |
| ENSMUSG00000033020.8 | 7.562949 | 5.362913 | 3.596849 | 1.491003 | NA | NA | #VALUE! |
| ENSMUSG00000033024.17 | 0 | NA | NA | NA | NA | NA | #VALUE! |
| ENSMUSG00000033027.16 | 0 | NA | NA | NA | NA | NA | #VALUE! |
| ENSMUSG00000033029.13 | 0 | NA | NA | NA | NA | NA | #VALUE! |
| ENSMUSG00000033031.14 | 0.391578 | -2.1489 | 4.020883 | -0.53444 | 0.59304 | NA | #VALUE! |
| ENSMUSG00000033032.16 | 0.336567 | 1.944925 | 4.029936 | 0.482619 | 0.629366 | NA | #VALUE! |
| ENSMUSG00000033033.11 | 20.27656 | -2.88014 | 2.756487 | -1.04486 | NA | NA | #VALUE! |
| ENSMUSG00000033036.10 | 0 | NA | NA | NA | NA | NA | #VALUE! |
| ENSMUSG00000033039.11 | 9.803232 | -6.82101 | 3.24275 | -2.10347 | 0.035425 | NA | #VALUE! |
| ENSMUSG00000033044.13 | 0 | NA | NA | NA | NA | NA | #VALUE! |

| | | | | | | | |
|---|---|---|---|---|---|---|---|
| ENSMUSG00000033053.5 | 0 | NA | NA | NA | NA | NA | #VALUE! |
| ENSMUSG00000033055.13 | 10.11349 | 2.801462 | 2.948595 | 0.9501 | NA | NA | #VALUE! |
| ENSMUSG00000033059.8 | 0 | NA | NA | NA | NA | NA | #VALUE! |
| ENSMUSG00000033061.16 | 0 | NA | NA | NA | NA | NA | #VALUE! |
| ENSMUSG00000033063.9 | 0 | NA | NA | NA | NA | NA | #VALUE! |
| ENSMUSG00000033065.15 | 0.484833 | -1.24061 | 4.012754 | -0.30917 | 0.757196 | NA | #VALUE! |
| ENSMUSG00000033066.18 | 7.807914 | 1.406943 | 3.132362 | 0.449163 | 0.653314 | NA | #VALUE! |
| ENSMUSG00000033068.17 | 4.207088 | 5.562698 | 3.917438 | 1.419984 | 0.155612 | NA | #VALUE! |
| ENSMUSG00000033075.18 | 5.093587 | 3.412527 | 3.840166 | 0.888641 | 0.374196 | NA | #VALUE! |
| ENSMUSG00000033080.10 | 0 | NA | NA | NA | NA | NA | #VALUE! |
| ENSMUSG00000033088.20 | 0 | NA | NA | NA | NA | NA | #VALUE! |
| ENSMUSG00000033122.15 | 0 | NA | NA | NA | NA | NA | #VALUE! |
| ENSMUSG00000033124.17 | 0 | NA | NA | NA | NA | NA | #VALUE! |
| ENSMUSG00000033126.6 | 18.1004 | -0.05959 | 2.818721 | -0.02114 | NA | NA | #VALUE! |
| ENSMUSG00000033128.10 | 6.463782 | -3.95687 | 3.749485 | -1.05531 | 0.291284 | NA | #VALUE! |
| ENSMUSG00000033152.14 | 0 | NA | NA | NA | NA | NA | #VALUE! |
| ENSMUSG00000033156.10 | 0 | NA | NA | NA | NA | NA | #VALUE! |
| ENSMUSG00000033177.14 | 0 | NA | NA | NA | NA | NA | #VALUE! |
| ENSMUSG00000033182.13 | 4.52893 | 5.664391 | 3.916713 | 1.44621 | 0.148118 | NA | #VALUE! |
| ENSMUSG00000033187.9 | 0 | NA | NA | NA | NA | NA | #VALUE! |
| ENSMUSG00000033191.15 | 2.039617 | 1.482119 | 3.840422 | 0.385926 | 0.699551 | NA | #VALUE! |
| ENSMUSG00000033196.18 | 0 | NA | NA | NA | NA | NA | #VALUE! |
| ENSMUSG00000033200.17 | 0 | NA | NA | NA | NA | NA | #VALUE! |
| ENSMUSG00000033208.8 | 0 | NA | NA | NA | NA | NA | #VALUE! |
| ENSMUSG00000033209.18 | 1.224665 | -3.81006 | 3.943408 | -0.96618 | 0.333952 | NA | #VALUE! |
| ENSMUSG00000033210.17 | 0 | NA | NA | NA | NA | NA | #VALUE! |
| ENSMUSG00000033213.17 | 0 | NA | NA | NA | NA | NA | #VALUE! |
| ENSMUSG00000033214.11 | 0 | NA | NA | NA | NA | NA | #VALUE! |
| ENSMUSG00000033219.9 | 0 | NA | NA | NA | NA | NA | #VALUE! |
| ENSMUSG00000033222.17 | 8.606014 | 6.595146 | 2.976276 | 2.215905 | 0.026698 | NA | #VALUE! |
| ENSMUSG00000033227.8 | 0 | NA | NA | NA | NA | NA | #VALUE! |
| ENSMUSG00000033233.18 | 0 | NA | NA | NA | NA | NA | #VALUE! |
| ENSMUSG00000033249.11 | 0.210026 | -1.52494 | 4.058692 | -0.37572 | 0.707123 | NA | #VALUE! |
| ENSMUSG00000033253.19 | 0 | NA | NA | NA | NA | NA | #VALUE! |
| ENSMUSG00000033255.9 | 0 | NA | NA | NA | NA | NA | #VALUE! |
| ENSMUSG00000033256.15 | 0.414259 | 2.174227 | 4.015696 | 0.541432 | 0.58821 | NA | #VALUE! |
| ENSMUSG00000033268.9 | 0 | NA | NA | NA | NA | NA | #VALUE! |
| ENSMUSG00000033272.13 | 5.665133 | 5.987759 | 3.363569 | 1.78018 | 0.075046 | NA | #VALUE! |
| ENSMUSG00000033276.19 | 0 | NA | NA | NA | NA | NA | #VALUE! |
| ENSMUSG00000033278.11 | 41.1098 | 2.180136 | 1.754002 | 1.242949 | NA | NA | #VALUE! |
| ENSMUSG00000033287.18 | 3.197387 | 5.166784 | 3.920797 | 1.317789 | 0.187574 | NA | #VALUE! |
| ENSMUSG00000033313.12 | 5.715935 | 0.331737 | 3.758087 | 0.088273 | 0.92966 | NA | #VALUE! |
| ENSMUSG00000033316.15 | 0 | NA | NA | NA | NA | NA | #VALUE! |
| ENSMUSG00000033323.15 | 2.731654 | -1.18183 | 3.80444 | -0.31064 | 0.756071 | NA | #VALUE! |

| | | | | | | | |
|---|---|---|---|---|---|---|---|
| ENSMUSG00000033326.16 | 7.971644 | 1.384711 | 2.39706 | 0.577671 | 0.563487 | NA | #VALUE! |
| ENSMUSG00000033327.19 | 3.249947 | 5.194551 | 3.92053 | 1.324961 | 0.185184 | NA | #VALUE! |
| ENSMUSG00000033342.16 | 2.238125 | 4.666312 | 3.926586 | 1.188389 | 0.23468 | NA | #VALUE! |
| ENSMUSG00000033343.2 | 0 | NA | NA | NA | NA | NA | #VALUE! |
| ENSMUSG00000033361.14 | 0 | NA | NA | NA | NA | NA | #VALUE! |
| ENSMUSG00000033365.15 | 4.783158 | -1.84354 | 3.651553 | -0.50487 | 0.613653 | NA | #VALUE! |
| ENSMUSG00000033368.9 | 0 | NA | NA | NA | NA | NA | #VALUE! |
| ENSMUSG00000033373.17 | 0 | NA | NA | NA | NA | NA | #VALUE! |
| ENSMUSG00000033383.7 | 0 | NA | NA | NA | NA | NA | #VALUE! |
| ENSMUSG00000033389.17 | 0.315039 | -1.93549 | 4.038621 | -0.47925 | 0.631763 | NA | #VALUE! |
| ENSMUSG00000033392.17 | 4.103229 | 5.534434 | 3.917649 | 1.412693 | 0.157746 | NA | #VALUE! |
| ENSMUSG00000033405.5 | 16.21016 | 0.269249 | 2.89453 | 0.09302 | NA | NA | #VALUE! |
| ENSMUSG00000033409.15 | 0 | NA | NA | NA | NA | NA | #VALUE! |
| ENSMUSG00000033419.17 | 0 | NA | NA | NA | NA | NA | #VALUE! |
| ENSMUSG00000033420.13 | 4.597673 | 4.606838 | 3.788644 | 1.21596 | 0.224 | NA | #VALUE! |
| ENSMUSG00000033423.17 | 4.51137 | 5.662692 | 3.916725 | 1.445772 | 0.148241 | NA | #VALUE! |
| ENSMUSG00000033434.16 | 38.82799 | 0.445326 | 3.729925 | 0.119393 | NA | NA | #VALUE! |
| ENSMUSG00000033436.14 | 1.343018 | -2.68461 | 3.931081 | -0.68292 | 0.494658 | NA | #VALUE! |
| ENSMUSG00000033439.13 | 4.029295 | 0.242503 | 3.55345 | 0.068244 | 0.945591 | NA | #VALUE! |
| ENSMUSG00000033450.9 | 0 | NA | NA | NA | NA | NA | #VALUE! |
| ENSMUSG00000033453.9 | 0 | NA | NA | NA | NA | NA | #VALUE! |
| ENSMUSG00000033458.18 | 7.048408 | -6.3456 | 3.913124 | -1.62162 | 0.104885 | NA | #VALUE! |
| ENSMUSG00000033460.15 | 19.83513 | -2.05812 | 2.249068 | -0.9151 | NA | NA | #VALUE! |
| ENSMUSG00000033467.13 | 6.974864 | 5.239524 | 2.863022 | 1.830068 | 0.06724 | NA | #VALUE! |
| ENSMUSG00000033470.13 | 0 | NA | NA | NA | NA | NA | #VALUE! |
| ENSMUSG00000033475.16 | 0.783156 | -3.1627 | 3.963922 | -0.79787 | 0.424945 | NA | #VALUE! |
| ENSMUSG00000033486.15 | 5.438519 | 1.789014 | 3.081609 | 0.580545 | 0.561547 | NA | #VALUE! |
| ENSMUSG00000033491.14 | 0 | NA | NA | NA | NA | NA | #VALUE! |
| ENSMUSG00000033498.15 | 0 | NA | NA | NA | NA | NA | #VALUE! |
| ENSMUSG00000033501.6 | 0 | NA | NA | NA | NA | NA | #VALUE! |
| ENSMUSG00000033502.16 | 5.064889 | 1.91004 | 3.238863 | 0.589726 | 0.555375 | NA | #VALUE! |
| ENSMUSG00000033508.9 | 0 | NA | NA | NA | NA | NA | #VALUE! |
| ENSMUSG00000033510.15 | 0 | NA | NA | NA | NA | NA | #VALUE! |
| ENSMUSG00000033520.3 | 0 | NA | NA | NA | NA | NA | #VALUE! |
| ENSMUSG00000033526.17 | 9.224213 | 1.164068 | 3.080076 | 0.377935 | 0.705479 | NA | #VALUE! |
| ENSMUSG00000033538.14 | 3.439269 | -0.55881 | 3.78176 | -0.14776 | 0.882529 | NA | #VALUE! |
| ENSMUSG00000033540.19 | 6.217674 | -1.64788 | 3.765718 | -0.4376 | 0.661675 | NA | #VALUE! |
| ENSMUSG00000033544.7 | 0 | NA | NA | NA | NA | NA | #VALUE! |
| ENSMUSG00000033569.18 | 0 | NA | NA | NA | NA | NA | #VALUE! |
| ENSMUSG00000033576.13 | 0 | NA | NA | NA | NA | NA | #VALUE! |
| ENSMUSG00000033578.8 | 0 | NA | NA | NA | NA | NA | #VALUE! |
| ENSMUSG00000033579.17 | 0 | NA | NA | NA | NA | NA | #VALUE! |
| ENSMUSG00000033581.18 | 3.570446 | -5.37363 | 3.919223 | -1.3711 | 0.170345 | NA | #VALUE! |
| ENSMUSG00000033590.9 | 0 | NA | NA | NA | NA | NA | #VALUE! |

| | | | | | | | |
|---|---|---|---|---|---|---|---|
| ENSMUSG00000033594.12 | 1.224665 | -3.81006 | 3.943408 | -0.96618 | 0.333952 | NA | #VALUE! |
| ENSMUSG00000033595.8 | 0 | NA | NA | NA | NA | NA | #VALUE! |
| ENSMUSG00000033597.10 | 0 | NA | NA | NA | NA | NA | #VALUE! |
| ENSMUSG00000033615.10 | 0 | NA | NA | NA | NA | NA | #VALUE! |
| ENSMUSG00000033624.11 | 0.773878 | -1.301 | 3.958218 | -0.32868 | 0.742396 | NA | #VALUE! |
| ENSMUSG00000033633.16 | 0 | NA | NA | NA | NA | NA | #VALUE! |
| ENSMUSG00000033644.6 | 1.260158 | -3.88425 | 3.941584 | -0.98545 | 0.324401 | NA | #VALUE! |
| ENSMUSG00000033653.19 | 4.757577 | -3.14479 | 3.73954 | -0.84096 | 0.400373 | NA | #VALUE! |
| ENSMUSG00000033697.17 | 1.119062 | 3.679627 | 3.945843 | 0.932533 | 0.351061 | NA | #VALUE! |
| ENSMUSG00000033703.10 | 5.984525 | 0.544673 | 2.648042 | 0.205689 | 0.837034 | NA | #VALUE! |
| ENSMUSG00000033705.18 | 7.672992 | -6.46726 | 3.432471 | -1.88414 | 0.059546 | NA | #VALUE! |
| ENSMUSG00000033707.10 | 0 | NA | NA | NA | NA | NA | #VALUE! |
| ENSMUSG00000033712.7 | 1.020554 | -3.54462 | 3.950745 | -0.8972 | 0.36961 | NA | #VALUE! |
| ENSMUSG00000033717.7 | 0 | NA | NA | NA | NA | NA | #VALUE! |
| ENSMUSG00000033721.17 | 6.073346 | 5.139602 | 3.863135 | 1.330423 | 0.183379 | NA | #VALUE! |
| ENSMUSG00000033722.10 | 0 | NA | NA | NA | NA | NA | #VALUE! |
| ENSMUSG00000033726.9 | 0 | NA | NA | NA | NA | NA | #VALUE! |
| ENSMUSG00000033730.5 | 3.271688 | -5.23618 | 3.920465 | -1.3356 | 0.181679 | NA | #VALUE! |
| ENSMUSG00000033731.10 | 0 | NA | NA | NA | NA | NA | #VALUE! |
| ENSMUSG00000033737.15 | 0 | NA | NA | NA | NA | NA | #VALUE! |
| ENSMUSG00000033739.9 | 0.532356 | 0.929314 | 4.022674 | 0.231019 | 0.8173 | NA | #VALUE! |
| ENSMUSG00000033740.18 | 0 | NA | NA | NA | NA | NA | #VALUE! |
| ENSMUSG00000033762.9 | 2.981986 | 0.410692 | 3.788795 | 0.108396 | 0.913681 | NA | #VALUE! |
| ENSMUSG00000033765.6 | 0 | NA | NA | NA | NA | NA | #VALUE! |
| ENSMUSG00000033768.18 | 7.18065 | 6.338444 | 3.913004 | 1.619841 | NA | NA | #VALUE! |
| ENSMUSG00000033769.17 | 4.430052 | 5.638116 | 3.916896 | 1.439435 | 0.150027 | NA | #VALUE! |
| ENSMUSG00000033770.14 | 0 | NA | NA | NA | NA | NA | #VALUE! |
| ENSMUSG00000033774.5 | 0 | NA | NA | NA | NA | NA | #VALUE! |
| ENSMUSG00000033777.5 | 0 | NA | NA | NA | NA | NA | #VALUE! |
| ENSMUSG00000033788.16 | 0 | NA | NA | NA | NA | NA | #VALUE! |
| ENSMUSG00000033792.13 | 5.269388 | -1.0673 | 3.573042 | -0.29871 | 0.765162 | NA | #VALUE! |
| ENSMUSG00000033794.7 | 0 | NA | NA | NA | NA | NA | #VALUE! |
| ENSMUSG00000033805.13 | 6.46104 | -6.21992 | 3.913702 | -1.58927 | 0.112 | NA | #VALUE! |
| ENSMUSG00000033808.17 | 1.212318 | 3.793111 | 3.942915 | 0.962007 | 0.336046 | NA | #VALUE! |
| ENSMUSG00000033819.17 | 0.945118 | -3.47564 | 3.952879 | -0.87927 | 0.379256 | NA | #VALUE! |
| ENSMUSG00000033825.11 | 0 | NA | NA | NA | NA | NA | #VALUE! |
| ENSMUSG00000033826.11 | 0 | NA | NA | NA | NA | NA | #VALUE! |
| ENSMUSG00000033834.8 | 0 | NA | NA | NA | NA | NA | #VALUE! |
| ENSMUSG00000033837.6 | 0 | NA | NA | NA | NA | NA | #VALUE! |
| ENSMUSG00000033847.16 | 0 | NA | NA | NA | NA | NA | #VALUE! |
| ENSMUSG00000033849.4 | 0 | NA | NA | NA | NA | NA | #VALUE! |
| ENSMUSG00000033850.6 | 0 | NA | NA | NA | NA | NA | #VALUE! |
| ENSMUSG00000033852.17 | 0 | NA | NA | NA | NA | NA | #VALUE! |
| ENSMUSG00000033854.11 | 5.506082 | 1.839813 | 3.597631 | 0.511396 | 0.609074 | NA | #VALUE! |

| | | | | | | | |
|---|---|---|---|---|---|---|---|
| ENSMUSG00000033855.16 | 0 | NA | NA | NA | NA | NA | #VALUE! |
| ENSMUSG00000033857.13 | 4.562852 | 1.198518 | 3.773399 | 0.317623 | 0.750771 | NA | #VALUE! |
| ENSMUSG00000033862.8 | 9.004712 | -3.26127 | 2.629488 | -1.24027 | 0.214876 | NA | #VALUE! |
| ENSMUSG00000033871.15 | 2.33138 | 4.724655 | 3.925805 | 1.203487 | 0.228788 | NA | #VALUE! |
| ENSMUSG00000033872.7 | 0 | NA | NA | NA | NA | NA | #VALUE! |
| ENSMUSG00000033882.16 | 0 | NA | NA | NA | NA | NA | #VALUE! |
| ENSMUSG00000033902.13 | 0 | NA | NA | NA | NA | NA | #VALUE! |
| ENSMUSG00000033904.17 | 9.145246 | 4.357554 | 2.309617 | 1.886699 | 0.059201 | NA | #VALUE! |
| ENSMUSG00000033906.6 | 7.625835 | 1.196173 | 3.570719 | 0.334995 | 0.737629 | NA | #VALUE! |
| ENSMUSG00000033948.4 | 10.4681 | 5.355119 | 3.241546 | 1.652026 | NA | NA | #VALUE! |
| ENSMUSG00000033949.13 | 0 | NA | NA | NA | NA | NA | #VALUE! |
| ENSMUSG00000033952.15 | 0 | NA | NA | NA | NA | NA | #VALUE! |
| ENSMUSG00000033955.14 | 3.36567 | 5.240782 | 3.920098 | 1.336901 | 0.181255 | NA | #VALUE! |
| ENSMUSG00000033961.17 | 5.002123 | 1.403382 | 3.771461 | 0.372106 | 0.709814 | NA | #VALUE! |
| ENSMUSG00000033963.17 | 0 | NA | NA | NA | NA | NA | #VALUE! |
| ENSMUSG00000033966.9 | 0 | NA | NA | NA | NA | NA | #VALUE! |
| ENSMUSG00000033967.5 | 0 | NA | NA | NA | NA | NA | #VALUE! |
| ENSMUSG00000033981.15 | 0 | NA | NA | NA | NA | NA | #VALUE! |
| ENSMUSG00000033987.17 | 0 | NA | NA | NA | NA | NA | #VALUE! |
| ENSMUSG00000033998.10 | 0 | NA | NA | NA | NA | NA | #VALUE! |
| ENSMUSG00000034000.16 | 0 | NA | NA | NA | NA | NA | #VALUE! |
| ENSMUSG00000034009.15 | 0 | NA | NA | NA | NA | NA | #VALUE! |
| ENSMUSG00000034023.17 | 0 | NA | NA | NA | NA | NA | #VALUE! |
| ENSMUSG00000034028.17 | 2.56008 | 3.7106 | 3.918328 | 0.946986 | 0.343646 | NA | #VALUE! |
| ENSMUSG00000034031.6 | 0 | NA | NA | NA | NA | NA | #VALUE! |
| ENSMUSG00000034032.16 | 9.130484 | 0.848456 | 3.38194 | 0.250878 | 0.801908 | NA | #VALUE! |
| ENSMUSG00000034035.19 | 1.995249 | -4.53986 | 3.928926 | -1.1555 | 0.247887 | NA | #VALUE! |
| ENSMUSG00000034039.9 | 0 | NA | NA | NA | NA | NA | #VALUE! |
| ENSMUSG00000034040.18 | 0 | NA | NA | NA | NA | NA | #VALUE! |
| ENSMUSG00000034041.6 | 0.559531 | 2.703958 | 3.982863 | 0.678898 | 0.497202 | NA | #VALUE! |
| ENSMUSG00000034057.9 | 0 | NA | NA | NA | NA | NA | #VALUE! |
| ENSMUSG00000034059.15 | 0 | NA | NA | NA | NA | NA | #VALUE! |
| ENSMUSG00000034063.9 | 0 | NA | NA | NA | NA | NA | #VALUE! |
| ENSMUSG00000034071.16 | 0.336567 | 1.944925 | 4.029936 | 0.482619 | 0.629366 | NA | #VALUE! |
| ENSMUSG00000034087.15 | 0 | NA | NA | NA | NA | NA | #VALUE! |
| ENSMUSG00000034098.15 | 0 | NA | NA | NA | NA | NA | #VALUE! |
| ENSMUSG00000034105.10 | 6.959156 | -6.32598 | 3.121809 | -2.02638 | 0.042726 | NA | #VALUE! |
| ENSMUSG00000034107.11 | 0 | NA | NA | NA | NA | NA | #VALUE! |
| ENSMUSG00000034110.9 | 1.009701 | 3.503687 | 3.950852 | 0.886818 | 0.375177 | NA | #VALUE! |
| ENSMUSG00000034112.10 | 0 | NA | NA | NA | NA | NA | #VALUE! |
| ENSMUSG00000034115.11 | 0 | NA | NA | NA | NA | NA | #VALUE! |
| ENSMUSG00000034116.18 | 6.154843 | 6.1168 | 3.914039 | 1.562784 | 0.118103 | NA | #VALUE! |
| ENSMUSG00000034117.4 | 1.224665 | -3.81006 | 3.943408 | -0.96618 | 0.333952 | NA | #VALUE! |
| ENSMUSG00000034121.13 | 0 | NA | NA | NA | NA | NA | #VALUE! |

| | | | | | | | |
|---|---|---|---|---|---|---|---|
| ENSMUSG00000034122.18 | 0 | NA | NA | NA | NA | NA | #VALUE! |
| ENSMUSG00000034139.5 | 0 | NA | NA | NA | NA | NA | #VALUE! |
| ENSMUSG00000034145.15 | 0 | NA | NA | NA | NA | NA | #VALUE! |
| ENSMUSG00000034151.14 | 0 | NA | NA | NA | NA | NA | #VALUE! |
| ENSMUSG00000034156.17 | 0 | NA | NA | NA | NA | NA | #VALUE! |
| ENSMUSG00000034159.13 | 0 | NA | NA | NA | NA | NA | #VALUE! |
| ENSMUSG00000034161.9 | 0 | NA | NA | NA | NA | NA | #VALUE! |
| ENSMUSG00000034164.18 | 0 | NA | NA | NA | NA | NA | #VALUE! |
| ENSMUSG00000034165.17 | 3.536618 | 5.311848 | 3.919459 | 1.35525 | 0.175338 | NA | #VALUE! |
| ENSMUSG00000034175.14 | 8.729216 | 4.650448 | 3.43032 | 1.355689 | 0.175198 | NA | #VALUE! |
| ENSMUSG00000034177.16 | 9.133785 | -1.94854 | 2.70513 | -0.72031 | 0.471333 | NA | #VALUE! |
| ENSMUSG00000034185.10 | 0 | NA | NA | NA | NA | NA | #VALUE! |
| ENSMUSG00000034192.6 | 5.191678 | -0.27742 | 3.620238 | -0.07663 | 0.938918 | NA | #VALUE! |
| ENSMUSG00000034194.19 | 5.34709 | -0.87909 | 3.211671 | -0.27372 | 0.784302 | NA | #VALUE! |
| ENSMUSG00000034205.17 | 0.916644 | -3.41541 | 3.954826 | -0.8636 | 0.387805 | NA | #VALUE! |
| ENSMUSG00000034206.16 | 0 | NA | NA | NA | NA | NA | #VALUE! |
| ENSMUSG00000034209.5 | 0 | NA | NA | NA | NA | NA | #VALUE! |
| ENSMUSG00000034212.15 | 0 | NA | NA | NA | NA | NA | #VALUE! |
| ENSMUSG00000034219.16 | 0 | NA | NA | NA | NA | NA | #VALUE! |
| ENSMUSG00000034220.8 | 7.297421 | 4.831037 | 3.626092 | 1.332299 | NA | NA | #VALUE! |
| ENSMUSG00000034224.15 | 0 | NA | NA | NA | NA | NA | #VALUE! |
| ENSMUSG00000034226.8 | 0 | NA | NA | NA | NA | NA | #VALUE! |
| ENSMUSG00000034227.8 | 0 | NA | NA | NA | NA | NA | #VALUE! |
| ENSMUSG00000034235.18 | 6.563057 | 6.204219 | 3.913612 | 1.585292 | 0.1129 | NA | #VALUE! |
| ENSMUSG00000034239.15 | 0 | NA | NA | NA | NA | NA | #VALUE! |
| ENSMUSG00000034254.18 | 0 | NA | NA | NA | NA | NA | #VALUE! |
| ENSMUSG00000034255.19 | 0 | NA | NA | NA | NA | NA | #VALUE! |
| ENSMUSG00000034258.5 | 0.624163 | -2.86979 | 3.97661 | -0.72167 | 0.470499 | NA | #VALUE! |
| ENSMUSG00000034265.9 | 0.210026 | -1.52494 | 4.058692 | -0.37572 | 0.707123 | NA | #VALUE! |
| ENSMUSG00000034266.7 | 7.14619 | 6.321751 | 3.579523 | 1.766087 | 0.077381 | NA | #VALUE! |
| ENSMUSG00000034271.17 | 7.188181 | -0.33721 | 3.684332 | -0.09153 | 0.927075 | NA | #VALUE! |
| ENSMUSG00000034275.19 | 0 | NA | NA | NA | NA | NA | #VALUE! |
| ENSMUSG00000034282.4 | 0 | NA | NA | NA | NA | NA | #VALUE! |
| ENSMUSG00000034295.11 | 4.543655 | 5.673725 | 3.916649 | 1.448617 | 0.147445 | NA | #VALUE! |
| ENSMUSG00000034303.9 | 0 | NA | NA | NA | NA | NA | #VALUE! |
| ENSMUSG00000034310.9 | 0 | NA | NA | NA | NA | NA | #VALUE! |
| ENSMUSG00000034311.4 | 0 | NA | NA | NA | NA | NA | #VALUE! |
| ENSMUSG00000034317.15 | 0 | NA | NA | NA | NA | NA | #VALUE! |
| ENSMUSG00000034324.17 | 0 | NA | NA | NA | NA | NA | #VALUE! |
| ENSMUSG00000034333.16 | 2.663947 | 1.783953 | 3.824003 | 0.466515 | 0.640847 | NA | #VALUE! |
| ENSMUSG00000034336.4 | 0 | NA | NA | NA | NA | NA | #VALUE! |
| ENSMUSG00000034339.9 | 0 | NA | NA | NA | NA | NA | #VALUE! |
| ENSMUSG00000034353.15 | 5.790219 | -0.9618 | 3.046684 | -0.31569 | 0.75224 | NA | #VALUE! |
| ENSMUSG00000034359.17 | 0 | NA | NA | NA | NA | NA | #VALUE! |

| | | | | | | | |
|---|---|---|---|---|---|---|---|
| ENSMUSG00000034361.11 | 8.360745 | 0.189627 | 2.810674 | 0.067467 | 0.94621 | NA | #VALUE! |
| ENSMUSG00000034362.8 | 0 | NA | NA | NA | NA | NA | #VALUE! |
| ENSMUSG00000034384.12 | 0 | NA | NA | NA | NA | NA | #VALUE! |
| ENSMUSG00000034387.14 | 0 | NA | NA | NA | NA | NA | #VALUE! |
| ENSMUSG00000034391.13 | 0 | NA | NA | NA | NA | NA | #VALUE! |
| ENSMUSG00000034394.15 | 0 | NA | NA | NA | NA | NA | #VALUE! |
| ENSMUSG00000034401.17 | 0 | NA | NA | NA | NA | NA | #VALUE! |
| ENSMUSG00000034402.4 | 0 | NA | NA | NA | NA | NA | #VALUE! |
| ENSMUSG00000034412.14 | 1.632886 | -4.22813 | 3.934241 | -1.0747 | 0.282509 | NA | #VALUE! |
| ENSMUSG00000034413.15 | 3.673994 | -5.40317 | 3.918971 | -1.37872 | 0.167981 | NA | #VALUE! |
| ENSMUSG00000034416.19 | 0 | NA | NA | NA | NA | NA | #VALUE! |
| ENSMUSG00000034427.18 | 0 | NA | NA | NA | NA | NA | #VALUE! |
| ENSMUSG00000034437.9 | 0 | NA | NA | NA | NA | NA | #VALUE! |
| ENSMUSG00000034438.17 | 1.540956 | -0.39117 | 3.84553 | -0.10172 | 0.918978 | NA | #VALUE! |
| ENSMUSG00000034442.12 | 25.73661 | 1.095274 | 1.731732 | 0.632473 | NA | NA | #VALUE! |
| ENSMUSG00000034452.8 | 0 | NA | NA | NA | NA | NA | #VALUE! |
| ENSMUSG00000034457.11 | 0 | NA | NA | NA | NA | NA | #VALUE! |
| ENSMUSG00000034460.10 | 0 | NA | NA | NA | NA | NA | #VALUE! |
| ENSMUSG00000034463.5 | 0.245976 | 1.027628 | 4.067126 | 0.252667 | 0.800526 | NA | #VALUE! |
| ENSMUSG00000034467.8 | 0 | NA | NA | NA | NA | NA | #VALUE! |
| ENSMUSG00000034471.13 | 2.041108 | -4.55188 | 3.928743 | -1.15861 | 0.246615 | NA | #VALUE! |
| ENSMUSG00000034472.14 | 0 | NA | NA | NA | NA | NA | #VALUE! |
| ENSMUSG00000034482.3 | 0 | NA | NA | NA | NA | NA | #VALUE! |
| ENSMUSG00000034485.11 | 9.178469 | -6.7296 | 2.73759 | -2.45822 | 0.013963 | NA | #VALUE! |
| ENSMUSG00000034486.9 | 0 | NA | NA | NA | NA | NA | #VALUE! |
| ENSMUSG00000034488.15 | 0 | NA | NA | NA | NA | NA | #VALUE! |
| ENSMUSG00000034493.2 | 0 | NA | NA | NA | NA | NA | #VALUE! |
| ENSMUSG00000034509.10 | 80.83121 | -1.28973 | 1.086486 | -1.18706 | NA | NA | #VALUE! |
| ENSMUSG00000034520.15 | 1.020554 | -3.54462 | 3.950745 | -0.8972 | 0.36961 | NA | #VALUE! |
| ENSMUSG00000034522.11 | 5.372118 | 1.93088 | 2.812463 | 0.686544 | 0.49237 | NA | #VALUE! |
| ENSMUSG00000034532.10 | 0 | NA | NA | NA | NA | NA | #VALUE! |
| ENSMUSG00000034533.11 | 0 | NA | NA | NA | NA | NA | #VALUE! |
| ENSMUSG00000034538.9 | 1.085272 | 3.623578 | 3.947374 | 0.917972 | 0.358634 | NA | #VALUE! |
| ENSMUSG00000034551.13 | 0 | NA | NA | NA | NA | NA | #VALUE! |
| ENSMUSG00000034552.9 | 0 | NA | NA | NA | NA | NA | #VALUE! |
| ENSMUSG00000034555.15 | 0 | NA | NA | NA | NA | NA | #VALUE! |
| ENSMUSG00000034570.13 | 0 | NA | NA | NA | NA | NA | #VALUE! |
| ENSMUSG00000034573.15 | 0.652786 | 2.919659 | 3.972474 | 0.734972 | 0.462356 | NA | #VALUE! |
| ENSMUSG00000034579.18 | 0 | NA | NA | NA | NA | NA | #VALUE! |
| ENSMUSG00000034583.10 | 0 | NA | NA | NA | NA | NA | #VALUE! |
| ENSMUSG00000034584.4 | 3.850862 | 3.554228 | 3.605543 | 0.985768 | 0.324247 | NA | #VALUE! |
| ENSMUSG00000034586.15 | 0.630079 | -2.90291 | 3.975047 | -0.73028 | 0.465217 | NA | #VALUE! |
| ENSMUSG00000034587.9 | 1.632886 | -4.22813 | 3.934241 | -1.0747 | 0.282509 | NA | #VALUE! |
| ENSMUSG00000034593.17 | 0.587367 | -2.74297 | 3.982932 | -0.68868 | 0.491024 | NA | #VALUE! |

| | | | | | | | |
|---|---|---|---|---|---|---|---|
| ENSMUSG00000034595.18 | 7.601511 | 2.313843 | 3.555168 | 0.650839 | 0.51515 | NA | #VALUE! |
| ENSMUSG00000034607.7 | 0 | NA | NA | NA | NA | NA | #VALUE! |
| ENSMUSG00000034612.8 | 0 | NA | NA | NA | NA | NA | #VALUE! |
| ENSMUSG00000034613.13 | 0.504851 | 2.502843 | 3.993993 | 0.626652 | 0.530887 | NA | #VALUE! |
| ENSMUSG00000034614.15 | 0 | NA | NA | NA | NA | NA | #VALUE! |
| ENSMUSG00000034616.18 | 9.495335 | -1.74153 | 2.951112 | -0.59013 | 0.555107 | NA | #VALUE! |
| ENSMUSG00000034623.15 | 0 | NA | NA | NA | NA | NA | #VALUE! |
| ENSMUSG00000034634.8 | 1.119062 | 3.679627 | 3.945843 | 0.932533 | 0.351061 | NA | #VALUE! |
| ENSMUSG00000034639.8 | 0.587367 | -2.74297 | 3.982932 | -0.68868 | 0.491024 | NA | #VALUE! |
| ENSMUSG00000034645.15 | 6.394774 | 6.166745 | 3.913792 | 1.575645 | 0.115108 | NA | #VALUE! |
| ENSMUSG00000034648.10 | 0 | NA | NA | NA | NA | NA | #VALUE! |
| ENSMUSG00000034652.13 | 8.238339 | -1.31841 | 3.108558 | -0.42412 | 0.671476 | NA | #VALUE! |
| ENSMUSG00000034656.19 | 4.162826 | 1.027458 | 3.77524 | 0.272157 | 0.785501 | NA | #VALUE! |
| ENSMUSG00000034660.9 | 0 | NA | NA | NA | NA | NA | #VALUE! |
| ENSMUSG00000034664.14 | 2.491841 | -4.85553 | 3.924586 | -1.23721 | 0.21601 | NA | #VALUE! |
| ENSMUSG00000034674.19 | 0 | NA | NA | NA | NA | NA | #VALUE! |
| ENSMUSG00000034675.18 | 0 | NA | NA | NA | NA | NA | #VALUE! |
| ENSMUSG00000034677.10 | 0 | NA | NA | NA | NA | NA | #VALUE! |
| ENSMUSG00000034681.18 | 15.41558 | -3.1969 | 2.922955 | -1.09372 | NA | NA | #VALUE! |
| ENSMUSG00000034683.13 | 0 | NA | NA | NA | NA | NA | #VALUE! |
| ENSMUSG00000034684.13 | 0.279766 | 1.808031 | 4.036658 | 0.447903 | 0.654223 | NA | #VALUE! |
| ENSMUSG00000034685.5 | 0 | NA | NA | NA | NA | NA | #VALUE! |
| ENSMUSG00000034686.9 | 0 | NA | NA | NA | NA | NA | #VALUE! |
| ENSMUSG00000034687.9 | 1.26147 | 0.945811 | 3.884142 | 0.243506 | 0.807614 | NA | #VALUE! |
| ENSMUSG00000034689.3 | 0 | NA | NA | NA | NA | NA | #VALUE! |
| ENSMUSG00000034690.14 | 0 | NA | NA | NA | NA | NA | #VALUE! |
| ENSMUSG00000034701.10 | 0 | NA | NA | NA | NA | NA | #VALUE! |
| ENSMUSG00000034706.17 | 0 | NA | NA | NA | NA | NA | #VALUE! |
| ENSMUSG00000034730.18 | 0 | NA | NA | NA | NA | NA | #VALUE! |
| ENSMUSG00000034732.5 | 0 | NA | NA | NA | NA | NA | #VALUE! |
| ENSMUSG00000034739.18 | 0 | NA | NA | NA | NA | NA | #VALUE! |
| ENSMUSG00000034748.17 | 0 | NA | NA | NA | NA | NA | #VALUE! |
| ENSMUSG00000034755.19 | 0 | NA | NA | NA | NA | NA | #VALUE! |
| ENSMUSG00000034758.13 | 4.490437 | -5.69342 | 3.916751 | -1.45361 | 0.146055 | NA | #VALUE! |
| ENSMUSG00000034762.10 | 0 | NA | NA | NA | NA | NA | #VALUE! |
| ENSMUSG00000034764.16 | 0 | NA | NA | NA | NA | NA | #VALUE! |
| ENSMUSG00000034765.7 | 0.315039 | -1.93549 | 4.038621 | -0.47925 | 0.631763 | NA | #VALUE! |
| ENSMUSG00000034768.5 | 0 | NA | NA | NA | NA | NA | #VALUE! |
| ENSMUSG00000034771.16 | 3.343284 | 2.931512 | 3.86004 | 0.759451 | 0.447583 | NA | #VALUE! |
| ENSMUSG00000034773.17 | 0 | NA | NA | NA | NA | NA | #VALUE! |
| ENSMUSG00000034777.3 | 0 | NA | NA | NA | NA | NA | #VALUE! |
| ENSMUSG00000034786.18 | 0 | NA | NA | NA | NA | NA | #VALUE! |
| ENSMUSG00000034792.9 | 0 | NA | NA | NA | NA | NA | #VALUE! |
| ENSMUSG00000034793.16 | 0.598106 | 2.755755 | 3.98023 | 0.692361 | 0.488711 | NA | #VALUE! |

| | | | | | | | |
|---|---|---|---|---|---|---|---|
| ENSMUSG00000034796.15 | 0 | NA | NA | NA | NA | NA | #VALUE! |
| ENSMUSG00000034799.17 | 0 | NA | NA | NA | NA | NA | #VALUE! |
| ENSMUSG00000034800.16 | 3.636953 | 5.361394 | 3.919032 | 1.36804 | 0.171299 | NA | #VALUE! |
| ENSMUSG00000034810.8 | 0 | NA | NA | NA | NA | NA | #VALUE! |
| ENSMUSG00000034813.20 | 0 | NA | NA | NA | NA | NA | #VALUE! |
| ENSMUSG00000034818.18 | 0 | NA | NA | NA | NA | NA | #VALUE! |
| ENSMUSG00000034825.15 | 0 | NA | NA | NA | NA | NA | #VALUE! |
| ENSMUSG00000034829.9 | 0 | NA | NA | NA | NA | NA | #VALUE! |
| ENSMUSG00000034833.11 | 0.841418 | 3.240539 | 3.959551 | 0.818411 | 0.413123 | NA | #VALUE! |
| ENSMUSG00000034837.13 | 14.26356 | -0.12946 | 2.990733 | -0.04329 | NA | NA | #VALUE! |
| ENSMUSG00000034839.6 | 0 | NA | NA | NA | NA | NA | #VALUE! |
| ENSMUSG00000034853.17 | 4.749621 | 1.460351 | 3.320089 | 0.439853 | 0.660044 | NA | #VALUE! |
| ENSMUSG00000034863.10 | 0 | NA | NA | NA | NA | NA | #VALUE! |
| ENSMUSG00000034871.3 | 0 | NA | NA | NA | NA | NA | #VALUE! |
| ENSMUSG00000034872.7 | 0 | NA | NA | NA | NA | NA | #VALUE! |
| ENSMUSG00000034881.9 | 0 | NA | NA | NA | NA | NA | #VALUE! |
| ENSMUSG00000034883.10 | 0.783156 | -3.1627 | 3.963922 | -0.79787 | 0.424945 | NA | #VALUE! |
| ENSMUSG00000034889.9 | 21.57419 | -2.53074 | 2.231383 | -1.13416 | NA | NA | #VALUE! |
| ENSMUSG00000034891.14 | 0 | NA | NA | NA | NA | NA | #VALUE! |
| ENSMUSG00000034898.18 | 4.756015 | 5.746359 | 3.916165 | 1.467344 | 0.142283 | NA | #VALUE! |
| ENSMUSG00000034902.18 | 4.148367 | -5.58054 | 3.85697 | -1.44687 | 0.147933 | NA | #VALUE! |
| ENSMUSG00000034906.16 | 0 | NA | NA | NA | NA | NA | #VALUE! |
| ENSMUSG00000034910.6 | 0.093255 | 0.868093 | 4.072464 | 0.213162 | 0.831201 | NA | #VALUE! |
| ENSMUSG00000034911.9 | 0 | NA | NA | NA | NA | NA | #VALUE! |
| ENSMUSG00000034912.19 | 0.204111 | -1.05581 | 4.073302 | -0.2592 | 0.795479 | NA | #VALUE! |
| ENSMUSG00000034913.11 | 0 | NA | NA | NA | NA | NA | #VALUE! |
| ENSMUSG00000034918.9 | 0 | NA | NA | NA | NA | NA | #VALUE! |
| ENSMUSG00000034919.3 | 0 | NA | NA | NA | NA | NA | #VALUE! |
| ENSMUSG00000034923.10 | 0 | NA | NA | NA | NA | NA | #VALUE! |
| ENSMUSG00000034930.17 | 5.759445 | 0.140111 | 3.757465 | 0.037289 | 0.970255 | NA | #VALUE! |
| ENSMUSG00000034949.19 | 0 | NA | NA | NA | NA | NA | #VALUE! |
| ENSMUSG00000034958.12 | 0 | NA | NA | NA | NA | NA | #VALUE! |
| ENSMUSG00000034959.9 | 0 | NA | NA | NA | NA | NA | #VALUE! |
| ENSMUSG00000034968.4 | 0 | NA | NA | NA | NA | NA | #VALUE! |
| ENSMUSG00000034974.14 | 0.093255 | 0.868093 | 4.072464 | 0.213162 | 0.831201 | NA | #VALUE! |
| ENSMUSG00000034981.10 | 0 | NA | NA | NA | NA | NA | #VALUE! |
| ENSMUSG00000034987.5 | 0 | NA | NA | NA | NA | NA | #VALUE! |
| ENSMUSG00000034990.16 | 0 | NA | NA | NA | NA | NA | #VALUE! |
| ENSMUSG00000034997.6 | 0 | NA | NA | NA | NA | NA | #VALUE! |
| ENSMUSG00000035020.12 | 0 | NA | NA | NA | NA | NA | #VALUE! |
| ENSMUSG00000035032.13 | 0 | NA | NA | NA | NA | NA | #VALUE! |
| ENSMUSG00000035033.16 | 0 | NA | NA | NA | NA | NA | #VALUE! |
| ENSMUSG00000035042.3 | 1.177985 | 3.72614 | 3.944615 | 0.944614 | 0.344856 | NA | #VALUE! |
| ENSMUSG00000035045.18 | 0 | NA | NA | NA | NA | NA | #VALUE! |

| | | | | | | | |
|---|---|---|---|---|---|---|---|
| ENSMUSG00000035057.8 | 0 | NA | NA | NA | NA | NA | #VALUE! |
| ENSMUSG00000035062.14 | 0 | NA | NA | NA | NA | NA | #VALUE! |
| ENSMUSG00000035085.6 | 0 | NA | NA | NA | NA | NA | #VALUE! |
| ENSMUSG00000035095.12 | 1.229878 | 3.761857 | 3.943699 | 0.95389 | 0.340139 | NA | #VALUE! |
| ENSMUSG00000035107.14 | 6.216394 | -1.13635 | 3.410824 | -0.33316 | 0.739013 | NA | #VALUE! |
| ENSMUSG00000035109.15 | 0 | NA | NA | NA | NA | NA | #VALUE! |
| ENSMUSG00000035112.18 | 0 | NA | NA | NA | NA | NA | #VALUE! |
| ENSMUSG00000035121.7 | 0 | NA | NA | NA | NA | NA | #VALUE! |
| ENSMUSG00000035126.20 | 0 | NA | NA | NA | NA | NA | #VALUE! |
| ENSMUSG00000035129.7 | 0 | NA | NA | NA | NA | NA | #VALUE! |
| ENSMUSG00000035131.15 | 0 | NA | NA | NA | NA | NA | #VALUE! |
| ENSMUSG00000035148.5 | 0 | NA | NA | NA | NA | NA | #VALUE! |
| ENSMUSG00000035165.15 | 9.557138 | 1.089061 | 2.93362 | 0.371235 | 0.710463 | NA | #VALUE! |
| ENSMUSG00000035171.10 | 1.492083 | 4.088114 | 3.936282 | 1.038572 | 0.299004 | NA | #VALUE! |
| ENSMUSG00000035172.16 | 7.258434 | -6.38785 | 3.566455 | -1.79109 | 0.073279 | NA | #VALUE! |
| ENSMUSG00000035177.9 | 0 | NA | NA | NA | NA | NA | #VALUE! |
| ENSMUSG00000035179.5 | 0 | NA | NA | NA | NA | NA | #VALUE! |
| ENSMUSG00000035183.15 | 5.63667 | 5.979364 | 3.402102 | 1.75755 | 0.078824 | NA | #VALUE! |
| ENSMUSG00000035187.11 | 0 | NA | NA | NA | NA | NA | #VALUE! |
| ENSMUSG00000035189.18 | 0 | NA | NA | NA | NA | NA | #VALUE! |
| ENSMUSG00000035191.16 | 0 | NA | NA | NA | NA | NA | #VALUE! |
| ENSMUSG00000035200.8 | 0 | NA | NA | NA | NA | NA | #VALUE! |
| ENSMUSG00000035201.16 | 0 | NA | NA | NA | NA | NA | #VALUE! |
| ENSMUSG00000035206.11 | 4.625944 | -1.726 | 3.273062 | -0.52734 | 0.597961 | NA | #VALUE! |
| ENSMUSG00000035208.17 | 1.749949 | -2.2556 | 3.891263 | -0.57966 | 0.562146 | NA | #VALUE! |
| ENSMUSG00000035211.10 | 0 | NA | NA | NA | NA | NA | #VALUE! |
| ENSMUSG00000035226.6 | 0 | NA | NA | NA | NA | NA | #VALUE! |
| ENSMUSG00000035228.16 | 0 | NA | NA | NA | NA | NA | #VALUE! |
| ENSMUSG00000035232.9 | 9.389095 | -6.75908 | 3.672758 | -1.84033 | 0.06572 | NA | #VALUE! |
| ENSMUSG00000035238.5 | 0 | NA | NA | NA | NA | NA | #VALUE! |
| ENSMUSG00000035239.7 | 7.511715 | 6.400543 | 3.414427 | 1.874559 | 0.060853 | NA | #VALUE! |
| ENSMUSG00000035245.13 | 5.412423 | 1.852544 | 3.508238 | 0.528055 | 0.597461 | NA | #VALUE! |
| ENSMUSG00000035246.17 | 0 | NA | NA | NA | NA | NA | #VALUE! |
| ENSMUSG00000035262.6 | 0 | NA | NA | NA | NA | NA | #VALUE! |
| ENSMUSG00000035270.16 | 4.034298 | 1.492991 | 3.78405 | 0.394548 | 0.693176 | NA | #VALUE! |
| ENSMUSG00000035274.14 | 0 | NA | NA | NA | NA | NA | #VALUE! |
| ENSMUSG00000035275.15 | 0.746042 | 3.107177 | 3.964586 | 0.783733 | 0.433197 | NA | #VALUE! |
| ENSMUSG00000035277.16 | 0 | NA | NA | NA | NA | NA | #VALUE! |
| ENSMUSG00000035279.9 | 1.682835 | 4.240781 | 3.933339 | 1.078163 | 0.280961 | NA | #VALUE! |
| ENSMUSG00000035283.6 | 1.155144 | -3.7605 | 3.944679 | -0.95331 | 0.340433 | NA | #VALUE! |
| ENSMUSG00000035285.7 | 3.916718 | 5.467688 | 3.918163 | 1.395472 | 0.162873 | NA | #VALUE! |
| ENSMUSG00000035293.14 | 5.740952 | 0.497877 | 2.738128 | 0.181831 | 0.855715 | NA | #VALUE! |
| ENSMUSG00000035295.11 | 0 | NA | NA | NA | NA | NA | #VALUE! |
| ENSMUSG00000035296.15 | 0 | NA | NA | NA | NA | NA | #VALUE! |

| | | | | | | | |
|---|---|---|---|---|---|---|---|
| ENSMUSG00000035298.9 | 0 | NA | NA | NA | NA | NA | #VALUE! |
| ENSMUSG00000035299.17 | 0 | NA | NA | NA | NA | NA | #VALUE! |
| ENSMUSG00000035314.11 | 0.839297 | 3.273019 | 3.958392 | 0.826856 | 0.408319 | NA | #VALUE! |
| ENSMUSG00000035337.4 | 2.3255 | -0.11767 | 3.804698 | -0.03093 | 0.975327 | NA | #VALUE! |
| ENSMUSG00000035352.4 | 0 | NA | NA | NA | NA | NA | #VALUE! |
| ENSMUSG00000035355.16 | 0 | NA | NA | NA | NA | NA | #VALUE! |
| ENSMUSG00000035365.16 | 0 | NA | NA | NA | NA | NA | #VALUE! |
| ENSMUSG00000035370.10 | 0 | NA | NA | NA | NA | NA | #VALUE! |
| ENSMUSG00000035371.9 | 0 | NA | NA | NA | NA | NA | #VALUE! |
| ENSMUSG00000035373.3 | 0 | NA | NA | NA | NA | NA | #VALUE! |
| ENSMUSG00000035383.8 | 0 | NA | NA | NA | NA | NA | #VALUE! |
| ENSMUSG00000035385.6 | 10.93413 | -1.22778 | 2.823815 | -0.4348 | NA | NA | #VALUE! |
| ENSMUSG00000035387.6 | 0 | NA | NA | NA | NA | NA | #VALUE! |
| ENSMUSG00000035390.17 | 2.524253 | 4.825754 | 3.924522 | 1.229641 | 0.218831 | NA | #VALUE! |
| ENSMUSG00000035394.12 | 0 | NA | NA | NA | NA | NA | #VALUE! |
| ENSMUSG00000035395.12 | 0 | NA | NA | NA | NA | NA | #VALUE! |
| ENSMUSG00000035397.9 | 1.68021 | -4.29432 | 3.933015 | -1.09187 | 0.274892 | NA | #VALUE! |
| ENSMUSG00000035399.13 | 8.959697 | 0.189767 | 2.688096 | 0.070595 | 0.94372 | NA | #VALUE! |
| ENSMUSG00000035403.11 | 0 | NA | NA | NA | NA | NA | #VALUE! |
| ENSMUSG00000035407.9 | 0 | NA | NA | NA | NA | NA | #VALUE! |
| ENSMUSG00000035420.8 | 0 | NA | NA | NA | NA | NA | #VALUE! |
| ENSMUSG00000035427.19 | 0 | NA | NA | NA | NA | NA | #VALUE! |
| ENSMUSG00000035429.14 | 0 | NA | NA | NA | NA | NA | #VALUE! |
| ENSMUSG00000035431.5 | 0 | NA | NA | NA | NA | NA | #VALUE! |
| ENSMUSG00000035435.18 | 0 | NA | NA | NA | NA | NA | #VALUE! |
| ENSMUSG00000035439.14 | 0.279766 | 1.808031 | 4.036658 | 0.447903 | 0.654223 | NA | #VALUE! |
| ENSMUSG00000035448.10 | 0 | NA | NA | NA | NA | NA | #VALUE! |
| ENSMUSG00000035454.2 | 0 | NA | NA | NA | NA | NA | #VALUE! |
| ENSMUSG00000035456.10 | 0 | NA | NA | NA | NA | NA | #VALUE! |
| ENSMUSG00000035458.16 | 0.093255 | 0.868093 | 4.072464 | 0.213162 | 0.831201 | NA | #VALUE! |
| ENSMUSG00000035459.19 | 79.96161 | 0.280241 | 1.073376 | 0.261084 | NA | NA | #VALUE! |
| ENSMUSG00000035486.15 | 7.439986 | -6.42369 | 3.91279 | -1.64172 | 0.100649 | NA | #VALUE! |
| ENSMUSG00000035491.6 | 0 | NA | NA | NA | NA | NA | #VALUE! |
| ENSMUSG00000035506.17 | 0 | NA | NA | NA | NA | NA | #VALUE! |
| ENSMUSG00000035513.20 | 0 | NA | NA | NA | NA | NA | #VALUE! |
| ENSMUSG00000035522.4 | 0 | NA | NA | NA | NA | NA | #VALUE! |
| ENSMUSG00000035523.9 | 0 | NA | NA | NA | NA | NA | #VALUE! |
| ENSMUSG00000035528.5 | 0 | NA | NA | NA | NA | NA | #VALUE! |
| ENSMUSG00000035539.15 | 0 | NA | NA | NA | NA | NA | #VALUE! |
| ENSMUSG00000035547.15 | 0 | NA | NA | NA | NA | NA | #VALUE! |
| ENSMUSG00000035551.7 | 0 | NA | NA | NA | NA | NA | #VALUE! |
| ENSMUSG00000035557.10 | 0 | NA | NA | NA | NA | NA | #VALUE! |
| ENSMUSG00000035560.6 | 0 | NA | NA | NA | NA | NA | #VALUE! |
| ENSMUSG00000035576.14 | 0 | NA | NA | NA | NA | NA | #VALUE! |

| | | | | | | | |
|---|---|---|---|---|---|---|---|
| ENSMUSG00000035580.11 | 0 | NA | NA | NA | NA | NA | #VALUE! |
| ENSMUSG00000035582.14 | 0 | NA | NA | NA | NA | NA | #VALUE! |
| ENSMUSG00000035592.3 | 0 | NA | NA | NA | NA | NA | #VALUE! |
| ENSMUSG00000035594.11 | 0 | NA | NA | NA | NA | NA | #VALUE! |
| ENSMUSG00000035601.10 | 0.245976 | 1.027628 | 4.067126 | 0.252667 | 0.800526 | NA | #VALUE! |
| ENSMUSG00000035606.9 | 0 | NA | NA | NA | NA | NA | #VALUE! |
| ENSMUSG00000035615.13 | 0 | NA | NA | NA | NA | NA | #VALUE! |
| ENSMUSG00000035621.14 | 0.391578 | -2.1489 | 4.020883 | -0.53444 | 0.59304 | NA | #VALUE! |
| ENSMUSG00000035626.5 | 0 | NA | NA | NA | NA | NA | #VALUE! |
| ENSMUSG00000035632.17 | 0 | NA | NA | NA | NA | NA | #VALUE! |
| ENSMUSG00000035638.15 | 0 | NA | NA | NA | NA | NA | #VALUE! |
| ENSMUSG00000035640.19 | 4.158016 | -2.92202 | 3.832089 | -0.76251 | 0.445754 | NA | #VALUE! |
| ENSMUSG00000035651.8 | 0 | NA | NA | NA | NA | NA | #VALUE! |
| ENSMUSG00000035653.17 | 0 | NA | NA | NA | NA | NA | #VALUE! |
| ENSMUSG00000035673.11 | 0 | NA | NA | NA | NA | NA | #VALUE! |
| ENSMUSG00000035678.9 | 0 | NA | NA | NA | NA | NA | #VALUE! |
| ENSMUSG00000035681.8 | 0 | NA | NA | NA | NA | NA | #VALUE! |
| ENSMUSG00000035683.14 | 0 | NA | NA | NA | NA | NA | #VALUE! |
| ENSMUSG00000035694.16 | 0 | NA | NA | NA | NA | NA | #VALUE! |
| ENSMUSG00000035697.15 | 2.881802 | 0.497636 | 3.790981 | 0.131268 | 0.895563 | NA | #VALUE! |
| ENSMUSG00000035711.6 | 0 | NA | NA | NA | NA | NA | #VALUE! |
| ENSMUSG00000035713.16 | 0 | NA | NA | NA | NA | NA | #VALUE! |
| ENSMUSG00000035722.16 | 0 | NA | NA | NA | NA | NA | #VALUE! |
| ENSMUSG00000035725.14 | 2.07098 | -0.52901 | 3.816115 | -0.13862 | 0.889747 | NA | #VALUE! |
| ENSMUSG00000035735.11 | 0 | NA | NA | NA | NA | NA | #VALUE! |
| ENSMUSG00000035745.10 | 0 | NA | NA | NA | NA | NA | #VALUE! |
| ENSMUSG00000035759.5 | 4.395341 | 1.394877 | 3.777962 | 0.369214 | 0.711968 | NA | #VALUE! |
| ENSMUSG00000035773.7 | 0 | NA | NA | NA | NA | NA | #VALUE! |
| ENSMUSG00000035775.3 | 0 | NA | NA | NA | NA | NA | #VALUE! |
| ENSMUSG00000035778.18 | 3.721071 | -0.62062 | 3.777957 | -0.16427 | 0.869515 | NA | #VALUE! |
| ENSMUSG00000035783.10 | 4.368871 | -0.21331 | 3.768084 | -0.05661 | 0.954856 | NA | #VALUE! |
| ENSMUSG00000035785.7 | 0 | NA | NA | NA | NA | NA | #VALUE! |
| ENSMUSG00000035799.7 | 0 | NA | NA | NA | NA | NA | #VALUE! |
| ENSMUSG00000035804.6 | 0 | NA | NA | NA | NA | NA | #VALUE! |
| ENSMUSG00000035805.14 | 0 | NA | NA | NA | NA | NA | #VALUE! |
| ENSMUSG00000035818.15 | 0 | NA | NA | NA | NA | NA | #VALUE! |
| ENSMUSG00000035829.14 | 0 | NA | NA | NA | NA | NA | #VALUE! |
| ENSMUSG00000035831.3 | 0 | NA | NA | NA | NA | NA | #VALUE! |
| ENSMUSG00000035835.15 | 0 | NA | NA | NA | NA | NA | #VALUE! |
| ENSMUSG00000035842.11 | 4.413069 | 0.496483 | 3.052032 | 0.162673 | 0.870776 | NA | #VALUE! |
| ENSMUSG00000035849.15 | 0 | NA | NA | NA | NA | NA | #VALUE! |
| ENSMUSG00000035852.12 | 0 | NA | NA | NA | NA | NA | #VALUE! |
| ENSMUSG00000035860.10 | 0 | NA | NA | NA | NA | NA | #VALUE! |
| ENSMUSG00000035861.7 | 0 | NA | NA | NA | NA | NA | #VALUE! |

| | | | | | | | |
|---|---|---|---|---|---|---|---|
| ENSMUSG00000035863.15 | 0.840105 | -3.30885 | 3.958472 | -0.83589 | 0.403216 | NA | #VALUE! |
| ENSMUSG00000035873.9 | 0.93336 | -2.40055 | 3.950215 | -0.6077 | 0.543385 | NA | #VALUE! |
| ENSMUSG00000035891.17 | 0 | NA | NA | NA | NA | NA | #VALUE! |
| ENSMUSG00000035896.6 | 0 | NA | NA | NA | NA | NA | #VALUE! |
| ENSMUSG00000035900.19 | 8.592549 | 0.863265 | 3.046355 | 0.283376 | 0.776888 | NA | #VALUE! |
| ENSMUSG00000035910.16 | 8.611889 | -1.6122 | 2.720437 | -0.59262 | 0.553433 | NA | #VALUE! |
| ENSMUSG00000035914.12 | 7.74015 | -2.02641 | 2.76884 | -0.73186 | 0.464253 | NA | #VALUE! |
| ENSMUSG00000035916.9 | 0 | NA | NA | NA | NA | NA | #VALUE! |
| ENSMUSG00000035919.17 | 9.757002 | 3.171 | 2.863655 | 1.107326 | 0.268153 | NA | #VALUE! |
| ENSMUSG00000035923.5 | 0 | NA | NA | NA | NA | NA | #VALUE! |
| ENSMUSG00000035930.10 | 0 | NA | NA | NA | NA | NA | #VALUE! |
| ENSMUSG00000035932.7 | 0 | NA | NA | NA | NA | NA | #VALUE! |
| ENSMUSG00000035934.17 | 0 | NA | NA | NA | NA | NA | #VALUE! |
| ENSMUSG00000035946.8 | 0 | NA | NA | NA | NA | NA | #VALUE! |
| ENSMUSG00000035951.3 | 0 | NA | NA | NA | NA | NA | #VALUE! |
| ENSMUSG00000035963.9 | 0 | NA | NA | NA | NA | NA | #VALUE! |
| ENSMUSG00000035964.9 | 0 | NA | NA | NA | NA | NA | #VALUE! |
| ENSMUSG00000035967.16 | 1.68021 | -4.29432 | 3.933015 | -1.09187 | 0.274892 | NA | #VALUE! |
| ENSMUSG00000035969.16 | 2.019402 | 4.503826 | 3.928936 | 1.146322 | 0.251662 | NA | #VALUE! |
| ENSMUSG00000035983.5 | 0 | NA | NA | NA | NA | NA | #VALUE! |
| ENSMUSG00000035984.15 | 0 | NA | NA | NA | NA | NA | #VALUE! |
| ENSMUSG00000036013.13 | 0 | NA | NA | NA | NA | NA | #VALUE! |
| ENSMUSG00000036022.16 | 4.742728 | -0.40164 | 3.241199 | -0.12392 | 0.901381 | NA | #VALUE! |
| ENSMUSG00000036030.10 | 0 | NA | NA | NA | NA | NA | #VALUE! |
| ENSMUSG00000036036.16 | 0 | NA | NA | NA | NA | NA | #VALUE! |
| ENSMUSG00000036040.15 | 2.465992 | 1.118287 | 3.81229 | 0.293337 | 0.769264 | NA | #VALUE! |
| ENSMUSG00000036045.7 | 0 | NA | NA | NA | NA | NA | #VALUE! |
| ENSMUSG00000036052.15 | 0.279766 | 1.808031 | 4.036658 | 0.447903 | 0.654223 | NA | #VALUE! |
| ENSMUSG00000036054.16 | 1.958359 | 4.475621 | 3.929371 | 1.139017 | 0.254696 | NA | #VALUE! |
| ENSMUSG00000036057.12 | 7.115096 | -4.31408 | 3.206667 | -1.34535 | 0.178513 | NA | #VALUE! |
| ENSMUSG00000036061.11 | 0.559531 | 2.703958 | 3.982863 | 0.678898 | 0.497202 | NA | #VALUE! |
| ENSMUSG00000036062.14 | 0 | NA | NA | NA | NA | NA | #VALUE! |
| ENSMUSG00000036067.13 | 0 | NA | NA | NA | NA | NA | #VALUE! |
| ENSMUSG00000036086.17 | 9.825568 | 1.442307 | 2.92407 | 0.493253 | NA | NA | #VALUE! |
| ENSMUSG00000036091.9 | 1.260158 | -3.88425 | 3.941584 | -0.98545 | 0.324401 | NA | #VALUE! |
| ENSMUSG00000036095.12 | 0 | NA | NA | NA | NA | NA | #VALUE! |
| ENSMUSG00000036109.18 | 5.048506 | 5.825722 | 3.915663 | 1.4878 | 0.136804 | NA | #VALUE! |
| ENSMUSG00000036111.9 | 0 | NA | NA | NA | NA | NA | #VALUE! |
| ENSMUSG00000036117.3 | 0 | NA | NA | NA | NA | NA | #VALUE! |
| ENSMUSG00000036120.11 | 4.2033 | 1.047458 | 3.775775 | 0.277415 | 0.781461 | NA | #VALUE! |
| ENSMUSG00000036123.9 | 0 | NA | NA | NA | NA | NA | #VALUE! |
| ENSMUSG00000036131.13 | 0 | NA | NA | NA | NA | NA | #VALUE! |
| ENSMUSG00000036136.10 | 0.18651 | 1.401769 | 4.056354 | 0.345574 | 0.729663 | NA | #VALUE! |
| ENSMUSG00000036139.7 | 0 | NA | NA | NA | NA | NA | #VALUE! |

| | | | | | | | |
|---|---|---|---|---|---|---|---|
| ENSMUSG00000036144.7 | 0 | NA | NA | NA | NA | NA | #VALUE! |
| ENSMUSG00000036158.13 | 2.547797 | 0.10283 | 3.798897 | 0.027068 | 0.978405 | NA | #VALUE! |
| ENSMUSG00000036162.4 | 0 | NA | NA | NA | NA | NA | #VALUE! |
| ENSMUSG00000036168.16 | 0.848575 | 1.904055 | 3.965106 | 0.480203 | 0.631083 | NA | #VALUE! |
| ENSMUSG00000036169.7 | 0 | NA | NA | NA | NA | NA | #VALUE! |
| ENSMUSG00000036172.18 | 0 | NA | NA | NA | NA | NA | #VALUE! |
| ENSMUSG00000036185.10 | 0 | NA | NA | NA | NA | NA | #VALUE! |
| ENSMUSG00000036186.6 | 0 | NA | NA | NA | NA | NA | #VALUE! |
| ENSMUSG00000036192.16 | 0 | NA | NA | NA | NA | NA | #VALUE! |
| ENSMUSG00000036196.17 | 0 | NA | NA | NA | NA | NA | #VALUE! |
| ENSMUSG00000036198.13 | 0 | NA | NA | NA | NA | NA | #VALUE! |
| ENSMUSG00000036206.13 | 0 | NA | NA | NA | NA | NA | #VALUE! |
| ENSMUSG00000036208.14 | 9.529681 | 0.788253 | 2.975084 | 0.264952 | 0.791047 | NA | #VALUE! |
| ENSMUSG00000036211.5 | 0 | NA | NA | NA | NA | NA | #VALUE! |
| ENSMUSG00000036218.17 | 0 | NA | NA | NA | NA | NA | #VALUE! |
| ENSMUSG00000036223.17 | 0 | NA | NA | NA | NA | NA | #VALUE! |
| ENSMUSG00000036225.16 | 0 | NA | NA | NA | NA | NA | #VALUE! |
| ENSMUSG00000036231.6 | 0 | NA | NA | NA | NA | NA | #VALUE! |
| ENSMUSG00000036242.16 | 0 | NA | NA | NA | NA | NA | #VALUE! |
| ENSMUSG00000036244.6 | 0 | NA | NA | NA | NA | NA | #VALUE! |
| ENSMUSG00000036246.15 | 0 | NA | NA | NA | NA | NA | #VALUE! |
| ENSMUSG00000036249.17 | 4.496671 | 1.789514 | 3.784377 | 0.472869 | 0.636307 | NA | #VALUE! |
| ENSMUSG00000036251.17 | 0 | NA | NA | NA | NA | NA | #VALUE! |
| ENSMUSG00000036264.10 | 0 | NA | NA | NA | NA | NA | #VALUE! |
| ENSMUSG00000036273.16 | 16.12475 | -21.7525 | 3.121981 | -6.96755 | NA | NA | #VALUE! |
| ENSMUSG00000036281.14 | 3.437988 | 0.386109 | 3.780274 | 0.102138 | 0.918647 | NA | #VALUE! |
| ENSMUSG00000036304.15 | 1.212318 | 3.793111 | 3.942915 | 0.962007 | 0.336046 | NA | #VALUE! |
| ENSMUSG00000036305.8 | 0.795232 | 1.195798 | 3.961772 | 0.301834 | 0.762779 | NA | #VALUE! |
| ENSMUSG00000036306.14 | 0 | NA | NA | NA | NA | NA | #VALUE! |
| ENSMUSG00000036322.12 | 0 | NA | NA | NA | NA | NA | #VALUE! |
| ENSMUSG00000036353.14 | 9.505006 | 3.72914 | 2.450122 | 1.522022 | 0.128003 | NA | #VALUE! |
| ENSMUSG00000036356.16 | 0.612332 | -2.79827 | 3.980109 | -0.70306 | 0.482016 | NA | #VALUE! |
| ENSMUSG00000036357.6 | 0 | NA | NA | NA | NA | NA | #VALUE! |
| ENSMUSG00000036377.20 | 0 | NA | NA | NA | NA | NA | #VALUE! |
| ENSMUSG00000036381.14 | 0.315039 | -1.93549 | 4.038621 | -0.47925 | 0.631763 | NA | #VALUE! |
| ENSMUSG00000036395.16 | 0 | NA | NA | NA | NA | NA | #VALUE! |
| ENSMUSG00000036401.6 | 0 | NA | NA | NA | NA | NA | #VALUE! |
| ENSMUSG00000036411.11 | 0 | NA | NA | NA | NA | NA | #VALUE! |
| ENSMUSG00000036412.6 | 0 | NA | NA | NA | NA | NA | #VALUE! |
| ENSMUSG00000036422.11 | 0 | NA | NA | NA | NA | NA | #VALUE! |
| ENSMUSG00000036437.7 | 0 | NA | NA | NA | NA | NA | #VALUE! |
| ENSMUSG00000036449.7 | 0 | NA | NA | NA | NA | NA | #VALUE! |
| ENSMUSG00000036459.16 | 3.499223 | 0.278415 | 3.778384 | 0.073686 | 0.94126 | NA | #VALUE! |
| ENSMUSG00000036463.9 | 0 | NA | NA | NA | NA | NA | #VALUE! |

| | | | | | | | |
|---|---|---|---|---|---|---|---|
| ENSMUSG00000036466.18 | 0 | NA | NA | NA | NA | NA | #VALUE! |
| ENSMUSG00000036469.17 | 4.371519 | 0.800994 | 3.771094 | 0.212403 | 0.831792 | NA | #VALUE! |
| ENSMUSG00000036480.10 | 0 | NA | NA | NA | NA | NA | #VALUE! |
| ENSMUSG00000036492.13 | 0 | NA | NA | NA | NA | NA | #VALUE! |
| ENSMUSG00000036500.5 | 0 | NA | NA | NA | NA | NA | #VALUE! |
| ENSMUSG00000036502.15 | 7.234242 | 6.340002 | 3.069586 | 2.065426 | 0.038883 | NA | #VALUE! |
| ENSMUSG00000036510.18 | 0 | NA | NA | NA | NA | NA | #VALUE! |
| ENSMUSG00000036521.6 | 0 | NA | NA | NA | NA | NA | #VALUE! |
| ENSMUSG00000036523.17 | 0 | NA | NA | NA | NA | NA | #VALUE! |
| ENSMUSG00000036526.9 | 0 | NA | NA | NA | NA | NA | #VALUE! |
| ENSMUSG00000036529.18 | 6.978891 | 1.572727 | 3.717031 | 0.423114 | 0.672212 | NA | #VALUE! |
| ENSMUSG00000036534.7 | 2.995194 | 0.273676 | 3.787301 | 0.072261 | 0.942394 | NA | #VALUE! |
| ENSMUSG00000036537.7 | 0.783156 | -3.1627 | 3.963922 | -0.79787 | 0.424945 | NA | #VALUE! |
| ENSMUSG00000036545.10 | 0 | NA | NA | NA | NA | NA | #VALUE! |
| ENSMUSG00000036551.7 | 0 | NA | NA | NA | NA | NA | #VALUE! |
| ENSMUSG00000036555.15 | 1.785223 | -4.3809 | 3.931494 | -1.11431 | 0.265146 | NA | #VALUE! |
| ENSMUSG00000036557.9 | 0 | NA | NA | NA | NA | NA | #VALUE! |
| ENSMUSG00000036560.15 | 0 | NA | NA | NA | NA | NA | #VALUE! |
| ENSMUSG00000036561.10 | 2.205276 | -4.68302 | 3.926841 | -1.19257 | 0.233039 | NA | #VALUE! |
| ENSMUSG00000036564.18 | 0 | NA | NA | NA | NA | NA | #VALUE! |
| ENSMUSG00000036565.8 | 3.150394 | -5.19405 | 3.92087 | -1.32472 | 0.185265 | NA | #VALUE! |
| ENSMUSG00000036574.6 | 0 | NA | NA | NA | NA | NA | #VALUE! |
| ENSMUSG00000036578.8 | 0 | NA | NA | NA | NA | NA | #VALUE! |
| ENSMUSG00000036586.7 | 0 | NA | NA | NA | NA | NA | #VALUE! |
| ENSMUSG00000036587.14 | 0 | NA | NA | NA | NA | NA | #VALUE! |
| ENSMUSG00000036596.7 | 0 | NA | NA | NA | NA | NA | #VALUE! |
| ENSMUSG00000036598.5 | 0 | NA | NA | NA | NA | NA | #VALUE! |
| ENSMUSG00000036599.11 | 0 | NA | NA | NA | NA | NA | #VALUE! |
| ENSMUSG00000036602.15 | 0 | NA | NA | NA | NA | NA | #VALUE! |
| ENSMUSG00000036622.16 | 8.167508 | -5.08141 | 3.496494 | -1.45329 | NA | NA | #VALUE! |
| ENSMUSG00000036634.16 | 0 | NA | NA | NA | NA | NA | #VALUE! |
| ENSMUSG00000036647.6 | 0 | NA | NA | NA | NA | NA | #VALUE! |
| ENSMUSG00000036658.5 | 0 | NA | NA | NA | NA | NA | #VALUE! |
| ENSMUSG00000036667.15 | 0.606563 | -0.89296 | 3.98278 | -0.22421 | 0.822597 | NA | #VALUE! |
| ENSMUSG00000036672.6 | 6.123446 | 0.346781 | 3.03381 | 0.114305 | 0.908996 | NA | #VALUE! |
| ENSMUSG00000036686.17 | 0 | NA | NA | NA | NA | NA | #VALUE! |
| ENSMUSG00000036687.14 | 6.059714 | -1.15638 | 3.463877 | -0.33384 | 0.738501 | NA | #VALUE! |
| ENSMUSG00000036699.13 | 0 | NA | NA | NA | NA | NA | #VALUE! |
| ENSMUSG00000036718.18 | 0 | NA | NA | NA | NA | NA | #VALUE! |
| ENSMUSG00000036731.8 | 0 | NA | NA | NA | NA | NA | #VALUE! |
| ENSMUSG00000036733.17 | 0 | NA | NA | NA | NA | NA | #VALUE! |
| ENSMUSG00000036743.5 | 0 | NA | NA | NA | NA | NA | #VALUE! |
| ENSMUSG00000036744.14 | 1.398828 | 3.99633 | 3.938205 | 1.014759 | 0.310221 | NA | #VALUE! |
| ENSMUSG00000036745.16 | 2.2543 | -2.65698 | 3.883942 | -0.68409 | 0.493915 | NA | #VALUE! |

| | | | | | | | |
|---|---|---|---|---|---|---|---|
| ENSMUSG00000036749.16 | 0 | NA | NA | NA | NA | NA | #VALUE! |
| ENSMUSG00000036760.8 | 0 | NA | NA | NA | NA | NA | #VALUE! |
| ENSMUSG00000036766.13 | 0 | NA | NA | NA | NA | NA | #VALUE! |
| ENSMUSG00000036768.7 | 1.771849 | 4.332864 | 3.931707 | 1.102031 | 0.270448 | NA | #VALUE! |
| ENSMUSG00000036770.12 | 0 | NA | NA | NA | NA | NA | #VALUE! |
| ENSMUSG00000036777.9 | 0 | NA | NA | NA | NA | NA | #VALUE! |
| ENSMUSG00000036790.6 | 0 | NA | NA | NA | NA | NA | #VALUE! |
| ENSMUSG00000036800.9 | 0 | NA | NA | NA | NA | NA | #VALUE! |
| ENSMUSG00000036805.14 | 0 | NA | NA | NA | NA | NA | #VALUE! |
| ENSMUSG00000036814.14 | 0 | NA | NA | NA | NA | NA | #VALUE! |
| ENSMUSG00000036815.18 | 0.168284 | 1.027628 | 4.067126 | 0.252667 | 0.800526 | NA | #VALUE! |
| ENSMUSG00000036816.5 | 0 | NA | NA | NA | NA | NA | #VALUE! |
| ENSMUSG00000036826.15 | 0 | NA | NA | NA | NA | NA | #VALUE! |
| ENSMUSG00000036832.6 | 0 | NA | NA | NA | NA | NA | #VALUE! |
| ENSMUSG00000036834.17 | 0 | NA | NA | NA | NA | NA | #VALUE! |
| ENSMUSG00000036853.6 | 0 | NA | NA | NA | NA | NA | #VALUE! |
| ENSMUSG00000036855.4 | 0 | NA | NA | NA | NA | NA | #VALUE! |
| ENSMUSG00000036856.5 | 0 | NA | NA | NA | NA | NA | #VALUE! |
| ENSMUSG00000036858.9 | 0 | NA | NA | NA | NA | NA | #VALUE! |
| ENSMUSG00000036862.16 | 0 | NA | NA | NA | NA | NA | #VALUE! |
| ENSMUSG00000036864.17 | 0 | NA | NA | NA | NA | NA | #VALUE! |
| ENSMUSG00000036872.17 | 0 | NA | NA | NA | NA | NA | #VALUE! |
| ENSMUSG00000036882.8 | 1.398828 | 3.99633 | 3.938205 | 1.014759 | 0.310221 | NA | #VALUE! |
| ENSMUSG00000036887.6 | 0.840105 | -3.30885 | 3.958472 | -0.83589 | 0.403216 | NA | #VALUE! |
| ENSMUSG00000036899.11 | 0 | NA | NA | NA | NA | NA | #VALUE! |
| ENSMUSG00000036902.12 | 3.543697 | 5.324148 | 3.919351 | 1.358426 | 0.174329 | NA | #VALUE! |
| ENSMUSG00000036907.12 | 0 | NA | NA | NA | NA | NA | #VALUE! |
| ENSMUSG00000036912.19 | 0 | NA | NA | NA | NA | NA | #VALUE! |
| ENSMUSG00000036913.16 | 0 | NA | NA | NA | NA | NA | #VALUE! |
| ENSMUSG00000036915.18 | 0 | NA | NA | NA | NA | NA | #VALUE! |
| ENSMUSG00000036916.14 | 6.908328 | -0.20407 | 2.610495 | -0.07817 | 0.93769 | NA | #VALUE! |
| ENSMUSG00000036921.3 | 0 | NA | NA | NA | NA | NA | #VALUE! |
| ENSMUSG00000036923.13 | 0 | NA | NA | NA | NA | NA | #VALUE! |
| ENSMUSG00000036924.4 | 0 | NA | NA | NA | NA | NA | #VALUE! |
| ENSMUSG00000036925.9 | 0 | NA | NA | NA | NA | NA | #VALUE! |
| ENSMUSG00000036928.15 | 0 | NA | NA | NA | NA | NA | #VALUE! |
| ENSMUSG00000036931.16 | 0 | NA | NA | NA | NA | NA | #VALUE! |
| ENSMUSG00000036938.18 | 0 | NA | NA | NA | NA | NA | #VALUE! |
| ENSMUSG00000036941.3 | 3.289873 | -2.24474 | 3.821387 | -0.58741 | 0.556925 | NA | #VALUE! |
| ENSMUSG00000036944.7 | 6.173125 | -2.20673 | 3.43023 | -0.64332 | 0.520018 | NA | #VALUE! |
| ENSMUSG00000036948.18 | 0 | NA | NA | NA | NA | NA | #VALUE! |
| ENSMUSG00000036949.17 | 0 | NA | NA | NA | NA | NA | #VALUE! |
| ENSMUSG00000036951.3 | 0 | NA | NA | NA | NA | NA | #VALUE! |
| ENSMUSG00000036957.9 | 9.850172 | 6.787803 | 2.668762 | 2.543428 | 0.010977 | NA | #VALUE! |

| | | | | | | | |
|---|---|---|---|---|---|---|---|
| ENSMUSG00000036958.6 | 0 | NA | NA | NA | NA | NA | #VALUE! |
| ENSMUSG00000036959.16 | 7.802523 | -2.02642 | 2.776654 | -0.72981 | 0.465508 | NA | #VALUE! |
| ENSMUSG00000036960.11 | 0 | NA | NA | NA | NA | NA | #VALUE! |
| ENSMUSG00000036961.6 | 0.195789 | -1.05581 | 4.073302 | -0.2592 | 0.795479 | NA | #VALUE! |
| ENSMUSG00000036962.13 | 0 | NA | NA | NA | NA | NA | #VALUE! |
| ENSMUSG00000036964.15 | 0 | NA | NA | NA | NA | NA | #VALUE! |
| ENSMUSG00000036972.15 | 0 | NA | NA | NA | NA | NA | #VALUE! |
| ENSMUSG00000036977.9 | 47.69235 | 1.903826 | 1.617587 | 1.176954 | NA | NA | #VALUE! |
| ENSMUSG00000036989.17 | 0 | NA | NA | NA | NA | NA | #VALUE! |
| ENSMUSG00000036995.8 | 0.630079 | -2.90291 | 3.975047 | -0.73028 | 0.465217 | NA | #VALUE! |
| ENSMUSG00000037003.17 | 7.951672 | 1.225243 | 2.747101 | 0.446013 | 0.655588 | NA | #VALUE! |
| ENSMUSG00000037005.17 | 0 | NA | NA | NA | NA | NA | #VALUE! |
| ENSMUSG00000037010.8 | 0 | NA | NA | NA | NA | NA | #VALUE! |
| ENSMUSG00000037012.19 | 0 | NA | NA | NA | NA | NA | #VALUE! |
| ENSMUSG00000037014.5 | 0 | NA | NA | NA | NA | NA | #VALUE! |
| ENSMUSG00000037016.12 | 0 | NA | NA | NA | NA | NA | #VALUE! |
| ENSMUSG00000037020.17 | 0.391578 | -2.1489 | 4.020883 | -0.53444 | 0.59304 | NA | #VALUE! |
| ENSMUSG00000037028.8 | 0 | NA | NA | NA | NA | NA | #VALUE! |
| ENSMUSG00000037029.9 | 0.315039 | -1.93549 | 4.038621 | -0.47925 | 0.631763 | NA | #VALUE! |
| ENSMUSG00000037031.11 | 8.953679 | 2.59857 | 2.685468 | 0.967641 | 0.333223 | NA | #VALUE! |
| ENSMUSG00000037032.17 | 0 | NA | NA | NA | NA | NA | #VALUE! |
| ENSMUSG00000037033.10 | 0 | NA | NA | NA | NA | NA | #VALUE! |
| ENSMUSG00000037034.17 | 0 | NA | NA | NA | NA | NA | #VALUE! |
| ENSMUSG00000037035.6 | 0 | NA | NA | NA | NA | NA | #VALUE! |
| ENSMUSG00000037049.10 | 6.637239 | -6.25967 | 3.596208 | -1.74063 | 0.081748 | NA | #VALUE! |
| ENSMUSG00000037060.4 | 0.18651 | 1.401769 | 4.056354 | 0.345574 | 0.729663 | NA | #VALUE! |
| ENSMUSG00000037064.7 | 0 | NA | NA | NA | NA | NA | #VALUE! |
| ENSMUSG00000037086.4 | 0 | NA | NA | NA | NA | NA | #VALUE! |
| ENSMUSG00000037096.7 | 3.662079 | 3.060124 | 3.725588 | 0.82138 | 0.41143 | NA | #VALUE! |
| ENSMUSG00000037098.19 | 0.523077 | 2.574714 | 3.989843 | 0.645317 | 0.518722 | NA | #VALUE! |
| ENSMUSG00000037101.17 | 0 | NA | NA | NA | NA | NA | #VALUE! |
| ENSMUSG00000037103.9 | 0 | NA | NA | NA | NA | NA | #VALUE! |
| ENSMUSG00000037104.10 | 9.132536 | 0.750111 | 2.341655 | 0.320334 | 0.748715 | NA | #VALUE! |
| ENSMUSG00000037106.9 | 0 | NA | NA | NA | NA | NA | #VALUE! |
| ENSMUSG00000037108.14 | 0.840105 | -3.30885 | 3.958472 | -0.83589 | 0.403216 | NA | #VALUE! |
| ENSMUSG00000037112.17 | 3.407413 | 3.682059 | 3.898122 | 0.944572 | 0.344877 | NA | #VALUE! |
| ENSMUSG00000037124.7 | 0 | NA | NA | NA | NA | NA | #VALUE! |
| ENSMUSG00000037126.17 | 0 | NA | NA | NA | NA | NA | #VALUE! |
| ENSMUSG00000037129.8 | 0 | NA | NA | NA | NA | NA | #VALUE! |
| ENSMUSG00000037130.5 | 0 | NA | NA | NA | NA | NA | #VALUE! |
| ENSMUSG00000037138.19 | 0 | NA | NA | NA | NA | NA | #VALUE! |
| ENSMUSG00000037139.16 | 0.637369 | 0.178787 | 3.999109 | 0.044707 | 0.964341 | NA | #VALUE! |
| ENSMUSG00000037140.7 | 0 | NA | NA | NA | NA | NA | #VALUE! |
| ENSMUSG00000037143.18 | 0 | NA | NA | NA | NA | NA | #VALUE! |

| | | | | | | | |
|---|---|---|---|---|---|---|---|
| ENSMUSG00000037145.15 | 0 | NA | NA | NA | NA | NA | #VALUE! |
| ENSMUSG00000037148.9 | 9.53466 | -3.65757 | 2.578691 | -1.41838 | 0.156079 | NA | #VALUE! |
| ENSMUSG00000037151.9 | 17.10172 | -0.16904 | 2.171343 | -0.07785 | NA | NA | #VALUE! |
| ENSMUSG00000037157.9 | 0 | NA | NA | NA | NA | NA | #VALUE! |
| ENSMUSG00000037159.12 | 0 | NA | NA | NA | NA | NA | #VALUE! |
| ENSMUSG00000037161.15 | 0 | NA | NA | NA | NA | NA | #VALUE! |
| ENSMUSG00000037166.6 | 3.24117 | -5.23408 | 3.920485 | -1.33506 | 0.181857 | NA | #VALUE! |
| ENSMUSG00000037167.6 | 0 | NA | NA | NA | NA | NA | #VALUE! |
| ENSMUSG00000037169.16 | 5.418326 | -0.02402 | 3.589532 | -0.00669 | 0.994661 | NA | #VALUE! |
| ENSMUSG00000037171.7 | 0 | NA | NA | NA | NA | NA | #VALUE! |
| ENSMUSG00000037185.10 | 0 | NA | NA | NA | NA | NA | #VALUE! |
| ENSMUSG00000037188.8 | 0.245976 | 1.027628 | 4.067126 | 0.252667 | 0.800526 | NA | #VALUE! |
| ENSMUSG00000037196.7 | 0 | NA | NA | NA | NA | NA | #VALUE! |
| ENSMUSG00000037202.7 | 0 | NA | NA | NA | NA | NA | #VALUE! |
| ENSMUSG00000037206.16 | 9.446011 | -0.60741 | 2.638799 | -0.23019 | 0.817947 | NA | #VALUE! |
| ENSMUSG00000037216.6 | 18.74459 | 1.047888 | 2.838796 | 0.369131 | NA | NA | #VALUE! |
| ENSMUSG00000037217.16 | 0 | NA | NA | NA | NA | NA | #VALUE! |
| ENSMUSG00000037224.16 | 0 | NA | NA | NA | NA | NA | #VALUE! |
| ENSMUSG00000037225.14 | 0.587367 | -2.74297 | 3.982932 | -0.68868 | 0.491024 | NA | #VALUE! |
| ENSMUSG00000037239.9 | 0 | NA | NA | NA | NA | NA | #VALUE! |
| ENSMUSG00000037243.18 | 9.897554 | 2.487127 | 2.881274 | 0.863204 | 0.388026 | NA | #VALUE! |
| ENSMUSG00000037246.7 | 0 | NA | NA | NA | NA | NA | #VALUE! |
| ENSMUSG00000037247.4 | 0 | NA | NA | NA | NA | NA | #VALUE! |
| ENSMUSG00000037251.6 | 6.657173 | -1.43348 | 2.84758 | -0.5034 | 0.61468 | NA | #VALUE! |
| ENSMUSG00000037259.16 | 6.22649 | 6.128273 | 3.913982 | 1.565739 | 0.11741 | NA | #VALUE! |
| ENSMUSG00000037263.14 | 0 | NA | NA | NA | NA | NA | #VALUE! |
| ENSMUSG00000037279.14 | 0 | NA | NA | NA | NA | NA | #VALUE! |
| ENSMUSG00000037280.13 | 0 | NA | NA | NA | NA | NA | #VALUE! |
| ENSMUSG00000037307.12 | 0 | NA | NA | NA | NA | NA | #VALUE! |
| ENSMUSG00000037313.17 | 0 | NA | NA | NA | NA | NA | #VALUE! |
| ENSMUSG00000037315.16 | 8.596387 | -5.70374 | 3.039795 | -1.87636 | 0.060606 | NA | #VALUE! |
| ENSMUSG00000037318.11 | 0.652786 | 2.919659 | 3.972474 | 0.734972 | 0.462356 | NA | #VALUE! |
| ENSMUSG00000037334.3 | 0 | NA | NA | NA | NA | NA | #VALUE! |
| ENSMUSG00000037335.14 | 0 | NA | NA | NA | NA | NA | #VALUE! |
| ENSMUSG00000037336.15 | 0.408222 | -2.20205 | 4.016842 | -0.54821 | 0.583551 | NA | #VALUE! |
| ENSMUSG00000037337.12 | 0 | NA | NA | NA | NA | NA | #VALUE! |
| ENSMUSG00000037341.15 | 0 | NA | NA | NA | NA | NA | #VALUE! |
| ENSMUSG00000037344.15 | 0.902407 | -3.38379 | 3.95588 | -0.85538 | 0.39234 | NA | #VALUE! |
| ENSMUSG00000037346.6 | 0 | NA | NA | NA | NA | NA | #VALUE! |
| ENSMUSG00000037347.8 | 5.287242 | 1.53166 | 3.771071 | 0.406161 | 0.684625 | NA | #VALUE! |
| ENSMUSG00000037349.10 | 6.071856 | 0.938096 | 2.841197 | 0.330176 | 0.741267 | NA | #VALUE! |
| ENSMUSG00000037358.7 | 1.678593 | 4.255795 | 3.933066 | 1.082055 | 0.279228 | NA | #VALUE! |
| ENSMUSG00000037362.9 | 0.840105 | -3.30885 | 3.958472 | -0.83589 | 0.403216 | NA | #VALUE! |
| ENSMUSG00000037363.10 | 3.689633 | 5.364344 | 3.919007 | 1.368802 | 0.171061 | NA | #VALUE! |

| | | | | | | | |
|---|---|---|---|---|---|---|---|
| ENSMUSG00000037375.17 | 0.093255 | 0.868093 | 4.072464 | 0.213162 | 0.831201 | NA | #VALUE! |
| ENSMUSG00000037386.17 | 0 | NA | NA | NA | NA | NA | #VALUE! |
| ENSMUSG00000037390.14 | 0 | NA | NA | NA | NA | NA | #VALUE! |
| ENSMUSG00000037393.3 | 0 | NA | NA | NA | NA | NA | #VALUE! |
| ENSMUSG00000037408.11 | 8.427374 | 0.217117 | 2.657834 | 0.08169 | 0.934894 | NA | #VALUE! |
| ENSMUSG00000037411.11 | 20.47756 | -22.0806 | 2.899942 | -7.61414 | NA | NA | #VALUE! |
| ENSMUSG00000037418.7 | 0 | NA | NA | NA | NA | NA | #VALUE! |
| ENSMUSG00000037419.10 | 8.522066 | 1.898949 | 3.000343 | 0.632911 | 0.526792 | NA | #VALUE! |
| ENSMUSG00000037424.7 | 0 | NA | NA | NA | NA | NA | #VALUE! |
| ENSMUSG00000037426.19 | 0.587367 | -2.74297 | 3.982932 | -0.68868 | 0.491024 | NA | #VALUE! |
| ENSMUSG00000037428.15 | 0 | NA | NA | NA | NA | NA | #VALUE! |
| ENSMUSG00000037432.16 | 0 | NA | NA | NA | NA | NA | #VALUE! |
| ENSMUSG00000037437.19 | 0 | NA | NA | NA | NA | NA | #VALUE! |
| ENSMUSG00000037438.7 | 1.099645 | -1.94663 | 3.930833 | -0.49522 | 0.620444 | NA | #VALUE! |
| ENSMUSG00000037446.15 | 0 | NA | NA | NA | NA | NA | #VALUE! |
| ENSMUSG00000037447.17 | 0 | NA | NA | NA | NA | NA | #VALUE! |
| ENSMUSG00000037451.9 | 0 | NA | NA | NA | NA | NA | #VALUE! |
| ENSMUSG00000037463.15 | 0 | NA | NA | NA | NA | NA | #VALUE! |
| ENSMUSG00000037466.14 | 0 | NA | NA | NA | NA | NA | #VALUE! |
| ENSMUSG00000037469.12 | 0 | NA | NA | NA | NA | NA | #VALUE! |
| ENSMUSG00000037474.14 | 0 | NA | NA | NA | NA | NA | #VALUE! |
| ENSMUSG00000037477.9 | 0 | NA | NA | NA | NA | NA | #VALUE! |
| ENSMUSG00000037482.3 | 0 | NA | NA | NA | NA | NA | #VALUE! |
| ENSMUSG00000037490.6 | 0 | NA | NA | NA | NA | NA | #VALUE! |
| ENSMUSG00000037492.17 | 0 | NA | NA | NA | NA | NA | #VALUE! |
| ENSMUSG00000037509.22 | 0.559531 | 2.703958 | 3.982863 | 0.678898 | 0.497202 | NA | #VALUE! |
| ENSMUSG00000037525.6 | 6.629313 | 0.532323 | 3.323719 | 0.160159 | 0.872756 | NA | #VALUE! |
| ENSMUSG00000037529.14 | 0 | NA | NA | NA | NA | NA | #VALUE! |
| ENSMUSG00000037535.6 | 0 | NA | NA | NA | NA | NA | #VALUE! |
| ENSMUSG00000037537.7 | 0 | NA | NA | NA | NA | NA | #VALUE! |
| ENSMUSG00000037544.15 | 0 | NA | NA | NA | NA | NA | #VALUE! |
| ENSMUSG00000037548.16 | 0 | NA | NA | NA | NA | NA | #VALUE! |
| ENSMUSG00000037552.18 | 6.085912 | 2.24356 | 3.77779 | 0.593882 | 0.552591 | NA | #VALUE! |
| ENSMUSG00000037553.15 | 0.841418 | 3.240539 | 3.959551 | 0.818411 | 0.413123 | NA | #VALUE! |
| ENSMUSG00000037568.13 | 0 | NA | NA | NA | NA | NA | #VALUE! |
| ENSMUSG00000037570.17 | 0 | NA | NA | NA | NA | NA | #VALUE! |
| ENSMUSG00000037572.18 | 1.177985 | 3.72614 | 3.944615 | 0.944614 | 0.344856 | NA | #VALUE! |
| ENSMUSG00000037577.14 | 0 | NA | NA | NA | NA | NA | #VALUE! |
| ENSMUSG00000037578.5 | 0 | NA | NA | NA | NA | NA | #VALUE! |
| ENSMUSG00000037579.8 | 0 | NA | NA | NA | NA | NA | #VALUE! |
| ENSMUSG00000037593.13 | 0 | NA | NA | NA | NA | NA | #VALUE! |
| ENSMUSG00000037594.12 | 0 | NA | NA | NA | NA | NA | #VALUE! |
| ENSMUSG00000037600.17 | 3.476117 | 5.293931 | 3.919617 | 1.350625 | 0.176816 | NA | #VALUE! |
| ENSMUSG00000037603.8 | 0 | NA | NA | NA | NA | NA | #VALUE! |

| | | | | | | | |
|---|---|---|---|---|---|---|---|
| ENSMUSG00000037605.17 | 0 | NA | NA | NA | NA | NA | #VALUE! |
| ENSMUSG00000037606.19 | 0 | NA | NA | NA | NA | NA | #VALUE! |
| ENSMUSG00000037610.16 | 0 | NA | NA | NA | NA | NA | #VALUE! |
| ENSMUSG00000037613.17 | 0 | NA | NA | NA | NA | NA | #VALUE! |
| ENSMUSG00000037617.13 | 1.958359 | 4.475621 | 3.929371 | 1.139017 | 0.254696 | NA | #VALUE! |
| ENSMUSG00000037621.9 | 5.135659 | 1.390775 | 2.847839 | 0.488362 | 0.625294 | NA | #VALUE! |
| ENSMUSG00000037624.16 | 0.576505 | -1.13421 | 4.003282 | -0.28332 | 0.776932 | NA | #VALUE! |
| ENSMUSG00000037625.8 | 0 | NA | NA | NA | NA | NA | #VALUE! |
| ENSMUSG00000037627.17 | 0 | NA | NA | NA | NA | NA | #VALUE! |
| ENSMUSG00000037628.11 | 0 | NA | NA | NA | NA | NA | #VALUE! |
| ENSMUSG00000037636.7 | 0 | NA | NA | NA | NA | NA | #VALUE! |
| ENSMUSG00000037649.11 | 0 | NA | NA | NA | NA | NA | #VALUE! |
| ENSMUSG00000037653.8 | 0 | NA | NA | NA | NA | NA | #VALUE! |
| ENSMUSG00000037660.6 | 0 | NA | NA | NA | NA | NA | #VALUE! |
| ENSMUSG00000037661.15 | 0.978946 | -3.48749 | 3.952505 | -0.88235 | 0.377588 | NA | #VALUE! |
| ENSMUSG00000037664.14 | 0.947953 | 1.523095 | 3.946242 | 0.385961 | 0.699526 | NA | #VALUE! |
| ENSMUSG00000037681.14 | 0 | NA | NA | NA | NA | NA | #VALUE! |
| ENSMUSG00000037683.15 | 0 | NA | NA | NA | NA | NA | #VALUE! |
| ENSMUSG00000037689.7 | 0 | NA | NA | NA | NA | NA | #VALUE! |
| ENSMUSG00000037692.15 | 1.392001 | 2.736801 | 3.936518 | 0.695234 | 0.486909 | NA | #VALUE! |
| ENSMUSG00000037703.15 | 1.56096 | -4.18629 | 3.935045 | -1.06385 | 0.287398 | NA | #VALUE! |
| ENSMUSG00000037705.14 | 0 | NA | NA | NA | NA | NA | #VALUE! |
| ENSMUSG00000037708.17 | 0 | NA | NA | NA | NA | NA | #VALUE! |
| ENSMUSG00000037716.16 | 0 | NA | NA | NA | NA | NA | #VALUE! |
| ENSMUSG00000037725.9 | 4.648689 | -5.74744 | 3.916385 | -1.46754 | 0.14223 | NA | #VALUE! |
| ENSMUSG00000037727.7 | 0 | NA | NA | NA | NA | NA | #VALUE! |
| ENSMUSG00000037731.6 | 7.331707 | -6.40458 | 3.444297 | -1.85947 | 0.06296 | NA | #VALUE! |
| ENSMUSG00000037735.8 | 0 | NA | NA | NA | NA | NA | #VALUE! |
| ENSMUSG00000037736.21 | 1.192284 | 0.355789 | 3.880132 | 0.091695 | 0.92694 | NA | #VALUE! |
| ENSMUSG00000037737.7 | 0 | NA | NA | NA | NA | NA | #VALUE! |
| ENSMUSG00000037738.12 | 0 | NA | NA | NA | NA | NA | #VALUE! |
| ENSMUSG00000037747.10 | 0 | NA | NA | NA | NA | NA | #VALUE! |
| ENSMUSG00000037752.5 | 9.021463 | 4.816571 | 2.761303 | 1.744311 | 0.081105 | NA | #VALUE! |
| ENSMUSG00000037754.14 | 2.869382 | -5.04758 | 3.922372 | -1.28687 | 0.19814 | NA | #VALUE! |
| ENSMUSG00000037759.7 | 4.02703 | -1.03762 | 3.777254 | -0.2747 | 0.783544 | NA | #VALUE! |
| ENSMUSG00000037761.17 | 3.109878 | 5.12106 | 3.921247 | 1.305977 | 0.19156 | NA | #VALUE! |
| ENSMUSG00000037762.7 | 1.32413 | 1.312987 | 3.891861 | 0.337367 | 0.73584 | NA | #VALUE! |
| ENSMUSG00000037771.12 | 0 | NA | NA | NA | NA | NA | #VALUE! |
| ENSMUSG00000037784.15 | 0 | NA | NA | NA | NA | NA | #VALUE! |
| ENSMUSG00000037788.15 | 7.779832 | 0.150661 | 2.866722 | 0.052555 | 0.958086 | NA | #VALUE! |
| ENSMUSG00000037790.4 | 0 | NA | NA | NA | NA | NA | #VALUE! |
| ENSMUSG00000037801.14 | 0 | NA | NA | NA | NA | NA | #VALUE! |
| ENSMUSG00000037813.14 | 0 | NA | NA | NA | NA | NA | #VALUE! |
| ENSMUSG00000037816.11 | 4.314233 | -0.38074 | 3.768763 | -0.10102 | 0.919531 | NA | #VALUE! |

| | | | | | | | |
|---|---|---|---|---|---|---|---|
| ENSMUSG00000037827.6 | 0 | NA | NA | NA | NA | NA | #VALUE! |
| ENSMUSG00000037833.15 | 0 | NA | NA | NA | NA | NA | #VALUE! |
| ENSMUSG00000037843.7 | 0 | NA | NA | NA | NA | NA | #VALUE! |
| ENSMUSG00000037846.19 | 0 | NA | NA | NA | NA | NA | #VALUE! |
| ENSMUSG00000037855.16 | 0 | NA | NA | NA | NA | NA | #VALUE! |
| ENSMUSG00000037860.16 | 5.995824 | -5.17817 | 3.49018 | -1.48364 | 0.137904 | NA | #VALUE! |
| ENSMUSG00000037868.16 | 9.451182 | -6.77386 | 3.637007 | -1.86248 | NA | NA | #VALUE! |
| ENSMUSG00000037872.6 | 1.851119 | 4.378291 | 3.930938 | 1.113803 | 0.265364 | NA | #VALUE! |
| ENSMUSG00000037884.7 | 0 | NA | NA | NA | NA | NA | #VALUE! |
| ENSMUSG00000037887.12 | 2.424635 | 4.780725 | 3.925082 | 1.217993 | 0.223226 | NA | #VALUE! |
| ENSMUSG00000037890.14 | 9.186763 | -0.13356 | 3.039916 | -0.04394 | 0.964955 | NA | #VALUE! |
| ENSMUSG00000037892.14 | 0 | NA | NA | NA | NA | NA | #VALUE! |
| ENSMUSG00000037904.15 | 0 | NA | NA | NA | NA | NA | #VALUE! |
| ENSMUSG00000037907.17 | 0 | NA | NA | NA | NA | NA | #VALUE! |
| ENSMUSG00000037910.3 | 0 | NA | NA | NA | NA | NA | #VALUE! |
| ENSMUSG00000037913.13 | 0 | NA | NA | NA | NA | NA | #VALUE! |
| ENSMUSG00000037924.7 | 0 | NA | NA | NA | NA | NA | #VALUE! |
| ENSMUSG00000037940.18 | 0 | NA | NA | NA | NA | NA | #VALUE! |
| ENSMUSG00000037944.9 | 0 | NA | NA | NA | NA | NA | #VALUE! |
| ENSMUSG00000037946.12 | 0 | NA | NA | NA | NA | NA | #VALUE! |
| ENSMUSG00000037953.8 | 0 | NA | NA | NA | NA | NA | #VALUE! |
| ENSMUSG00000037960.12 | 9.245178 | -0.75283 | 3.019701 | -0.2493 | 0.803125 | NA | #VALUE! |
| ENSMUSG00000037972.8 | 0.168284 | 1.027628 | 4.067126 | 0.252667 | 0.800526 | NA | #VALUE! |
| ENSMUSG00000037973.7 | 0 | NA | NA | NA | NA | NA | #VALUE! |
| ENSMUSG00000037974.17 | 0 | NA | NA | NA | NA | NA | #VALUE! |
| ENSMUSG00000037977.7 | 5.502056 | 5.955668 | 3.914897 | 1.521283 | 0.128189 | NA | #VALUE! |
| ENSMUSG00000037979.14 | 0 | NA | NA | NA | NA | NA | #VALUE! |
| ENSMUSG00000037982.17 | 0 | NA | NA | NA | NA | NA | #VALUE! |
| ENSMUSG00000037984.10 | 0 | NA | NA | NA | NA | NA | #VALUE! |
| ENSMUSG00000037989.16 | 0 | NA | NA | NA | NA | NA | #VALUE! |
| ENSMUSG00000037990.19 | 0.420053 | -2.33493 | 4.007343 | -0.58266 | 0.56012 | NA | #VALUE! |
| ENSMUSG00000037991.10 | 2.611145 | 4.886731 | 3.92379 | 1.245411 | 0.212981 | NA | #VALUE! |
| ENSMUSG00000037992.17 | 9.231274 | 1.011044 | 2.70326 | 0.374009 | 0.708397 | NA | #VALUE! |
| ENSMUSG00000037994.15 | 0 | NA | NA | NA | NA | NA | #VALUE! |
| ENSMUSG00000037995.16 | 0 | NA | NA | NA | NA | NA | #VALUE! |
| ENSMUSG00000037996.18 | 0 | NA | NA | NA | NA | NA | #VALUE! |
| ENSMUSG00000037997.13 | 4.744129 | -4.83717 | 3.71327 | -1.30267 | 0.192687 | NA | #VALUE! |
| ENSMUSG00000038000.11 | 3.850831 | -0.93469 | 3.778509 | -0.24737 | 0.804621 | NA | #VALUE! |
| ENSMUSG00000038010.10 | 2.649481 | 1.641124 | 3.819326 | 0.429689 | 0.667422 | NA | #VALUE! |
| ENSMUSG00000038011.16 | 0 | NA | NA | NA | NA | NA | #VALUE! |
| ENSMUSG00000038015.7 | 0 | NA | NA | NA | NA | NA | #VALUE! |
| ENSMUSG00000038020.6 | 0 | NA | NA | NA | NA | NA | #VALUE! |
| ENSMUSG00000038022.12 | 0.945118 | -3.47564 | 3.952879 | -0.87927 | 0.379256 | NA | #VALUE! |
| ENSMUSG00000038024.18 | 4.844062 | -4.29098 | 3.199819 | -1.34101 | 0.179918 | NA | #VALUE! |

| | | | | | | | | |
|---|---|---|---|---|---|---|---|---|
| ENSMUSG00000038026.13 | 0 | NA | NA | NA | NA | NA | | #VALUE! |
| ENSMUSG00000038037.6 | 0 | NA | NA | NA | NA | NA | | #VALUE! |
| ENSMUSG00000038042.12 | 1.177985 | 3.72614 | 3.944615 | 0.944614 | 0.344856 | NA | | #VALUE! |
| ENSMUSG00000038044.9 | 0 | NA | NA | NA | NA | NA | | #VALUE! |
| ENSMUSG00000038045.15 | 0 | NA | NA | NA | NA | NA | | #VALUE! |
| ENSMUSG00000038047.19 | 7.274943 | -5.29978 | 3.225142 | -1.64327 | 0.100327 | NA | | #VALUE! |
| ENSMUSG00000038048.9 | 0 | NA | NA | NA | NA | NA | | #VALUE! |
| ENSMUSG00000038057.5 | 0 | NA | NA | NA | NA | NA | | #VALUE! |
| ENSMUSG00000038059.8 | 8.467996 | 6.576988 | 3.401774 | 1.933399 | NA | NA | | #VALUE! |
| ENSMUSG00000038064.8 | 0 | NA | NA | NA | NA | NA | | #VALUE! |
| ENSMUSG00000038065.14 | 3.255407 | -5.24108 | 3.920419 | -1.33687 | 0.181266 | NA | | #VALUE! |
| ENSMUSG00000038067.3 | 0 | NA | NA | NA | NA | NA | | #VALUE! |
| ENSMUSG00000038070.16 | 4.81548 | 5.763521 | 3.916054 | 1.471768 | 0.141084 | NA | | #VALUE! |
| ENSMUSG00000038071.4 | 0 | NA | NA | NA | NA | NA | | #VALUE! |
| ENSMUSG00000038077.8 | 0 | NA | NA | NA | NA | NA | | #VALUE! |
| ENSMUSG00000038085.14 | 0 | NA | NA | NA | NA | NA | | #VALUE! |
| ENSMUSG00000038086.5 | 0 | NA | NA | NA | NA | NA | | #VALUE! |
| ENSMUSG00000038094.16 | 0 | NA | NA | NA | NA | NA | | #VALUE! |
| ENSMUSG00000038112.16 | 0 | NA | NA | NA | NA | NA | | #VALUE! |
| ENSMUSG00000038115.17 | 0.945118 | -3.47564 | 3.952879 | -0.87927 | 0.379256 | NA | | #VALUE! |
| ENSMUSG00000038119.16 | 8.855119 | 6.632223 | 3.756168 | 1.765689 | 0.077448 | NA | | #VALUE! |
| ENSMUSG00000038126.18 | 3.84518 | -5.47788 | 3.918357 | -1.398 | 0.162112 | NA | | #VALUE! |
| ENSMUSG00000038128.8 | 0 | NA | NA | NA | NA | NA | | #VALUE! |
| ENSMUSG00000038132.7 | 0 | NA | NA | NA | NA | NA | | #VALUE! |
| ENSMUSG00000038135.5 | 7.259005 | 6.348671 | 3.464089 | 1.83271 | 0.066846 | NA | | #VALUE! |
| ENSMUSG00000038141.12 | 0.315039 | -1.93549 | 4.038621 | -0.47925 | 0.631763 | NA | | #VALUE! |
| ENSMUSG00000038143.17 | 0.168284 | 1.027628 | 4.067126 | 0.252667 | 0.800526 | NA | | #VALUE! |
| ENSMUSG00000038146.9 | 0 | NA | NA | NA | NA | NA | | #VALUE! |
| ENSMUSG00000038147.15 | 9.031467 | -2.94354 | 2.358322 | -1.24815 | 0.211976 | NA | | #VALUE! |
| ENSMUSG00000038148.14 | 0 | NA | NA | NA | NA | NA | | #VALUE! |
| ENSMUSG00000038151.14 | 0.204111 | -1.05581 | 4.073302 | -0.2592 | 0.795479 | NA | | #VALUE! |
| ENSMUSG00000038152.5 | 6.015903 | -6.11562 | 3.464384 | -1.76528 | 0.077516 | NA | | #VALUE! |
| ENSMUSG00000038155.12 | 7.734808 | -0.44079 | 2.275994 | -0.19367 | 0.846436 | NA | | #VALUE! |
| ENSMUSG00000038156.17 | 1.550873 | 0.949778 | 3.853765 | 0.246455 | 0.80533 | NA | | #VALUE! |
| ENSMUSG00000038165.9 | 0 | NA | NA | NA | NA | NA | | #VALUE! |
| ENSMUSG00000038167.7 | 4.574306 | 1.688094 | 3.030578 | 0.557021 | 0.577513 | NA | | #VALUE! |
| ENSMUSG00000038168.6 | 0 | NA | NA | NA | NA | NA | | #VALUE! |
| ENSMUSG00000038179.14 | 1.119062 | 3.679627 | 3.945843 | 0.932533 | 0.351061 | NA | | #VALUE! |
| ENSMUSG00000038180.12 | 0 | NA | NA | NA | NA | NA | | #VALUE! |
| ENSMUSG00000038192.6 | 0 | NA | NA | NA | NA | NA | | #VALUE! |
| ENSMUSG00000038193.5 | 4.456556 | -4.18404 | 3.77706 | -1.10775 | 0.26797 | NA | | #VALUE! |
| ENSMUSG00000038199.13 | 0 | NA | NA | NA | NA | NA | | #VALUE! |
| ENSMUSG00000038201.11 | 0 | NA | NA | NA | NA | NA | | #VALUE! |
| ENSMUSG00000038203.22 | 0 | NA | NA | NA | NA | NA | | #VALUE! |

| | | | | | | | |
|---|---|---|---|---|---|---|---|
| ENSMUSG00000038204.14 | 0 | NA | NA | NA | NA | NA | #VALUE! |
| ENSMUSG00000038205.13 | 4.727797 | 5.726132 | 3.664137 | 1.562751 | 0.118111 | NA | #VALUE! |
| ENSMUSG00000038208.14 | 0 | NA | NA | NA | NA | NA | #VALUE! |
| ENSMUSG00000038209.13 | 0 | NA | NA | NA | NA | NA | #VALUE! |
| ENSMUSG00000038210.11 | 0 | NA | NA | NA | NA | NA | #VALUE! |
| ENSMUSG00000038216.8 | 0 | NA | NA | NA | NA | NA | #VALUE! |
| ENSMUSG00000038218.4 | 0 | NA | NA | NA | NA | NA | #VALUE! |
| ENSMUSG00000038227.16 | 0 | NA | NA | NA | NA | NA | #VALUE! |
| ENSMUSG00000038233.11 | 8.571798 | 1.16881 | 2.376293 | 0.491863 | 0.622816 | NA | #VALUE! |
| ENSMUSG00000038236.9 | 0 | NA | NA | NA | NA | NA | #VALUE! |
| ENSMUSG00000038239.12 | 0 | NA | NA | NA | NA | NA | #VALUE! |
| ENSMUSG00000038241.17 | 7.733989 | -4.55454 | 3.191728 | -1.42698 | 0.153585 | NA | #VALUE! |
| ENSMUSG00000038242.13 | 0 | NA | NA | NA | NA | NA | #VALUE! |
| ENSMUSG00000038246.8 | 0 | NA | NA | NA | NA | NA | #VALUE! |
| ENSMUSG00000038248.9 | 0.204111 | -1.05581 | 4.073302 | -0.2592 | 0.795479 | NA | #VALUE! |
| ENSMUSG00000038252.14 | 0 | NA | NA | NA | NA | NA | #VALUE! |
| ENSMUSG00000038253.7 | 7.751939 | 5.496127 | 3.564411 | 1.541945 | NA | NA | #VALUE! |
| ENSMUSG00000038255.7 | 0 | NA | NA | NA | NA | NA | #VALUE! |
| ENSMUSG00000038256.16 | 9.397877 | -6.76087 | 3.669872 | -1.84226 | 0.065437 | NA | #VALUE! |
| ENSMUSG00000038257.11 | 0 | NA | NA | NA | NA | NA | #VALUE! |
| ENSMUSG00000038259.5 | 0 | NA | NA | NA | NA | NA | #VALUE! |
| ENSMUSG00000038260.11 | 0 | NA | NA | NA | NA | NA | #VALUE! |
| ENSMUSG00000038264.9 | 0 | NA | NA | NA | NA | NA | #VALUE! |
| ENSMUSG00000038268.5 | 7.670598 | -4.27797 | 2.512669 | -1.70256 | 0.088651 | NA | #VALUE! |
| ENSMUSG00000038271.18 | 4.286326 | -5.62615 | 3.917227 | -1.43626 | 0.150929 | NA | #VALUE! |
| ENSMUSG00000038276.13 | 0 | NA | NA | NA | NA | NA | #VALUE! |
| ENSMUSG00000038292.15 | 0 | NA | NA | NA | NA | NA | #VALUE! |
| ENSMUSG00000038295.15 | 0 | NA | NA | NA | NA | NA | #VALUE! |
| ENSMUSG00000038296.15 | 0 | NA | NA | NA | NA | NA | #VALUE! |
| ENSMUSG00000038300.7 | 0 | NA | NA | NA | NA | NA | #VALUE! |
| ENSMUSG00000038302.15 | 5.417213 | -3.52101 | 3.165325 | -1.11237 | 0.26598 | NA | #VALUE! |
| ENSMUSG00000038304.15 | 0 | NA | NA | NA | NA | NA | #VALUE! |
| ENSMUSG00000038305.16 | 2.881578 | -2.32316 | 3.838449 | -0.60523 | 0.545023 | NA | #VALUE! |
| ENSMUSG00000038311.14 | 0 | NA | NA | NA | NA | NA | #VALUE! |
| ENSMUSG00000038319.15 | 0 | NA | NA | NA | NA | NA | #VALUE! |
| ENSMUSG00000038327.9 | 0 | NA | NA | NA | NA | NA | #VALUE! |
| ENSMUSG00000038330.11 | 0 | NA | NA | NA | NA | NA | #VALUE! |
| ENSMUSG00000038331.16 | 0 | NA | NA | NA | NA | NA | #VALUE! |
| ENSMUSG00000038342.16 | 9.195362 | -1.45992 | 2.945772 | -0.4956 | 0.620178 | NA | #VALUE! |
| ENSMUSG00000038347.16 | 0 | NA | NA | NA | NA | NA | #VALUE! |
| ENSMUSG00000038349.11 | 6.243063 | 0.051309 | 3.755018 | 0.013664 | 0.989098 | NA | #VALUE! |
| ENSMUSG00000038351.15 | 2.984166 | 5.077883 | 3.921685 | 1.294822 | 0.195382 | NA | #VALUE! |
| ENSMUSG00000038352.16 | 0 | NA | NA | NA | NA | NA | #VALUE! |
| ENSMUSG00000038354.14 | 0 | NA | NA | NA | NA | NA | #VALUE! |

| | | | | | | | |
|---|---|---|---|---|---|---|---|
| ENSMUSG00000038357.11 | 7.665958 | -6.47201 | 3.903408 | -1.65804 | NA | NA | #VALUE! |
| ENSMUSG00000038370.7 | 7.591947 | 1.565531 | 2.726183 | 0.574257 | 0.565794 | NA | #VALUE! |
| ENSMUSG00000038372.15 | 0 | NA | NA | NA | NA | NA | #VALUE! |
| ENSMUSG00000038379.16 | 1.428775 | -4.03418 | 3.938169 | -1.02438 | 0.305656 | NA | #VALUE! |
| ENSMUSG00000038390.12 | 0 | NA | NA | NA | NA | NA | #VALUE! |
| ENSMUSG00000038402.3 | 0 | NA | NA | NA | NA | NA | #VALUE! |
| ENSMUSG00000038408.15 | 0 | NA | NA | NA | NA | NA | #VALUE! |
| ENSMUSG00000038416.16 | 71.59737 | -1.99408 | 1.344346 | -1.48331 | NA | NA | #VALUE! |
| ENSMUSG00000038421.14 | 1.365171 | -3.99822 | 3.938957 | -1.01505 | 0.310084 | NA | #VALUE! |
| ENSMUSG00000038429.11 | 8.314438 | 3.321107 | 3.070132 | 1.081747 | 0.279365 | NA | #VALUE! |
| ENSMUSG00000038437.12 | 6.054301 | -0.47375 | 2.983793 | -0.15878 | 0.873846 | NA | #VALUE! |
| ENSMUSG00000038453.18 | 0 | NA | NA | NA | NA | NA | #VALUE! |
| ENSMUSG00000038456.10 | 7.692131 | -0.64444 | 2.894054 | -0.22268 | 0.823787 | NA | #VALUE! |
| ENSMUSG00000038457.18 | 0 | NA | NA | NA | NA | NA | #VALUE! |
| ENSMUSG00000038463.9 | 0 | NA | NA | NA | NA | NA | #VALUE! |
| ENSMUSG00000038473.15 | 0 | NA | NA | NA | NA | NA | #VALUE! |
| ENSMUSG00000038486.10 | 0 | NA | NA | NA | NA | NA | #VALUE! |
| ENSMUSG00000038491.10 | 0 | NA | NA | NA | NA | NA | #VALUE! |
| ENSMUSG00000038496.17 | 0 | NA | NA | NA | NA | NA | #VALUE! |
| ENSMUSG00000038497.8 | 9.737928 | -1.59566 | 2.310199 | -0.6907 | 0.489753 | NA | #VALUE! |
| ENSMUSG00000038498.5 | 0 | NA | NA | NA | NA | NA | #VALUE! |
| ENSMUSG00000038502.18 | 31.26195 | 0.49662 | 1.966614 | 0.252525 | NA | NA | #VALUE! |
| ENSMUSG00000038508.8 | 1.154336 | 1.377753 | 3.915234 | 0.351895 | 0.724917 | NA | #VALUE! |
| ENSMUSG00000038517.16 | 1.585338 | 4.174397 | 3.934581 | 1.060951 | 0.288712 | NA | #VALUE! |
| ENSMUSG00000038523.11 | 0 | NA | NA | NA | NA | NA | #VALUE! |
| ENSMUSG00000038524.14 | 0 | NA | NA | NA | NA | NA | #VALUE! |
| ENSMUSG00000038528.16 | 0 | NA | NA | NA | NA | NA | #VALUE! |
| ENSMUSG00000038530.12 | 3.773892 | 3.07894 | 3.853816 | 0.798933 | 0.424329 | NA | #VALUE! |
| ENSMUSG00000038534.20 | 0.105013 | -0.97324 | 4.076293 | -0.23876 | 0.811295 | NA | #VALUE! |
| ENSMUSG00000038537.2 | 0 | NA | NA | NA | NA | NA | #VALUE! |
| ENSMUSG00000038540.15 | 0 | NA | NA | NA | NA | NA | #VALUE! |
| ENSMUSG00000038542.15 | 2.364341 | 3.099744 | 3.905103 | 0.793768 | 0.427331 | NA | #VALUE! |
| ENSMUSG00000038543.14 | 2.41652 | 4.767904 | 3.925245 | 1.214677 | 0.224489 | NA | #VALUE! |
| ENSMUSG00000038545.14 | 8.988255 | -6.69859 | 3.20319 | -2.09122 | 0.036508 | NA | #VALUE! |
| ENSMUSG00000038555.8 | 0 | NA | NA | NA | NA | NA | #VALUE! |
| ENSMUSG00000038560.8 | 0 | NA | NA | NA | NA | NA | #VALUE! |
| ENSMUSG00000038567.6 | 0 | NA | NA | NA | NA | NA | #VALUE! |
| ENSMUSG00000038569.14 | 0.414137 | -2.27192 | 4.011742 | -0.56632 | 0.571177 | NA | #VALUE! |
| ENSMUSG00000038570.16 | 2.915605 | -2.32606 | 3.838411 | -0.606 | 0.544517 | NA | #VALUE! |
| ENSMUSG00000038572.8 | 0 | NA | NA | NA | NA | NA | #VALUE! |
| ENSMUSG00000038580.14 | 0 | NA | NA | NA | NA | NA | #VALUE! |
| ENSMUSG00000038583.14 | 0 | NA | NA | NA | NA | NA | #VALUE! |
| ENSMUSG00000038593.19 | 0 | NA | NA | NA | NA | NA | #VALUE! |
| ENSMUSG00000038598.10 | 0 | NA | NA | NA | NA | NA | #VALUE! |

| | | | | | | | |
|---|---|---|---|---|---|---|---|
| ENSMUSG00000038599.15 | 37.68153 | -1.60694 | 1.720586 | -0.93395 | NA | NA | #VALUE! |
| ENSMUSG00000038600.13 | 0 | NA | NA | NA | NA | NA | #VALUE! |
| ENSMUSG00000038602.8 | 0 | NA | NA | NA | NA | NA | #VALUE! |
| ENSMUSG00000038605.12 | 0 | NA | NA | NA | NA | NA | #VALUE! |
| ENSMUSG00000038607.13 | 0 | NA | NA | NA | NA | NA | #VALUE! |
| ENSMUSG00000038608.16 | 4.543655 | 5.673725 | 3.916649 | 1.448617 | 0.147445 | NA | #VALUE! |
| ENSMUSG00000038624.14 | 0 | NA | NA | NA | NA | NA | #VALUE! |
| ENSMUSG00000038630.9 | 0 | NA | NA | NA | NA | NA | #VALUE! |
| ENSMUSG00000038637.16 | 0 | NA | NA | NA | NA | NA | #VALUE! |
| ENSMUSG00000038644.15 | 3.197387 | 5.166784 | 3.920797 | 1.317789 | 0.187574 | NA | #VALUE! |
| ENSMUSG00000038651.16 | 0 | NA | NA | NA | NA | NA | #VALUE! |
| ENSMUSG00000038656.6 | 0 | NA | NA | NA | NA | NA | #VALUE! |
| ENSMUSG00000038663.8 | 0 | NA | NA | NA | NA | NA | #VALUE! |
| ENSMUSG00000038665.16 | 0 | NA | NA | NA | NA | NA | #VALUE! |
| ENSMUSG00000038668.15 | 5.35567 | -5.95611 | 3.915092 | -1.52132 | 0.12818 | NA | #VALUE! |
| ENSMUSG00000038670.12 | 0 | NA | NA | NA | NA | NA | #VALUE! |
| ENSMUSG00000038676.7 | 0 | NA | NA | NA | NA | NA | #VALUE! |
| ENSMUSG00000038677.15 | 0 | NA | NA | NA | NA | NA | #VALUE! |
| ENSMUSG00000038679.17 | 0 | NA | NA | NA | NA | NA | #VALUE! |
| ENSMUSG00000038691.6 | 0 | NA | NA | NA | NA | NA | #VALUE! |
| ENSMUSG00000038692.9 | 0.466276 | 2.449889 | 3.997179 | 0.612904 | 0.53994 | NA | #VALUE! |
| ENSMUSG00000038695.14 | 4.153201 | -5.58673 | 3.917516 | -1.42609 | 0.153843 | NA | #VALUE! |
| ENSMUSG00000038700.5 | 22.96898 | 8.007932 | 2.649631 | 3.022282 | NA | NA | #VALUE! |
| ENSMUSG00000038702.7 | 0.945118 | -3.47564 | 3.952879 | -0.87927 | 0.379256 | NA | #VALUE! |
| ENSMUSG00000038709.16 | 0 | NA | NA | NA | NA | NA | #VALUE! |
| ENSMUSG00000038721.9 | 0 | NA | NA | NA | NA | NA | #VALUE! |
| ENSMUSG00000038725.14 | 0.559531 | 2.703958 | 3.982863 | 0.678898 | 0.497202 | NA | #VALUE! |
| ENSMUSG00000038729.25 | 0 | NA | NA | NA | NA | NA | #VALUE! |
| ENSMUSG00000038732.16 | 4.382994 | 5.629097 | 3.916959 | 1.437109 | 0.150687 | NA | #VALUE! |
| ENSMUSG00000038738.16 | 0 | NA | NA | NA | NA | NA | #VALUE! |
| ENSMUSG00000038740.10 | 0 | NA | NA | NA | NA | NA | #VALUE! |
| ENSMUSG00000038750.13 | 0 | NA | NA | NA | NA | NA | #VALUE! |
| ENSMUSG00000038756.14 | 0 | NA | NA | NA | NA | NA | #VALUE! |
| ENSMUSG00000038760.15 | 0 | NA | NA | NA | NA | NA | #VALUE! |
| ENSMUSG00000038763.13 | 0 | NA | NA | NA | NA | NA | #VALUE! |
| ENSMUSG00000038765.15 | 0 | NA | NA | NA | NA | NA | #VALUE! |
| ENSMUSG00000038770.18 | 0 | NA | NA | NA | NA | NA | #VALUE! |
| ENSMUSG00000038775.15 | 0 | NA | NA | NA | NA | NA | #VALUE! |
| ENSMUSG00000038777.20 | 0 | NA | NA | NA | NA | NA | #VALUE! |
| ENSMUSG00000038781.8 | 32.40546 | -0.16607 | 1.40108 | -0.11853 | NA | NA | #VALUE! |
| ENSMUSG00000038782.8 | 0 | NA | NA | NA | NA | NA | #VALUE! |
| ENSMUSG00000038791.15 | 0 | NA | NA | NA | NA | NA | #VALUE! |
| ENSMUSG00000038793.9 | 1.68021 | -4.29432 | 3.933015 | -1.09187 | 0.274892 | NA | #VALUE! |
| ENSMUSG00000038797.14 | 0.373021 | 2.140748 | 4.018182 | 0.532765 | 0.594196 | NA | #VALUE! |

| | | | | | | | |
|---|---|---|---|---|---|---|---|
| ENSMUSG00000038801.7 | 4.410551 | -5.67707 | 3.916865 | -1.44939 | 0.147228 | NA | #VALUE! |
| ENSMUSG00000038805.11 | 0 | NA | NA | NA | NA | NA | #VALUE! |
| ENSMUSG00000038807.20 | 0 | NA | NA | NA | NA | NA | #VALUE! |
| ENSMUSG00000038816.15 | 14.45895 | -1.37354 | 2.684424 | -0.51167 | NA | NA | #VALUE! |
| ENSMUSG00000038822.16 | 0.717345 | -3.03856 | 3.968994 | -0.76558 | 0.443929 | NA | #VALUE! |
| ENSMUSG00000038836.16 | 8.802258 | 2.658716 | 2.590921 | 1.026167 | 0.304813 | NA | #VALUE! |
| ENSMUSG00000038840.8 | 0 | NA | NA | NA | NA | NA | #VALUE! |
| ENSMUSG00000038844.11 | 73.5019 | -1.26407 | 1.384917 | -0.91274 | NA | NA | #VALUE! |
| ENSMUSG00000038855.11 | 5.479042 | 5.938864 | 3.891674 | 1.526043 | 0.126999 | NA | #VALUE! |
| ENSMUSG00000038860.16 | 1.470184 | -4.10383 | 3.936698 | -1.04245 | 0.297201 | NA | #VALUE! |
| ENSMUSG00000038861.15 | 0 | NA | NA | NA | NA | NA | #VALUE! |
| ENSMUSG00000038879.9 | 0 | NA | NA | NA | NA | NA | #VALUE! |
| ENSMUSG00000038883.4 | 0 | NA | NA | NA | NA | NA | #VALUE! |
| ENSMUSG00000038891.4 | 0 | NA | NA | NA | NA | NA | #VALUE! |
| ENSMUSG00000038893.13 | 2.993193 | -0.78877 | 3.791476 | -0.20804 | 0.835199 | NA | #VALUE! |
| ENSMUSG00000038895.17 | 0 | NA | NA | NA | NA | NA | #VALUE! |
| ENSMUSG00000038916.8 | 0 | NA | NA | NA | NA | NA | #VALUE! |
| ENSMUSG00000038917.5 | 0 | NA | NA | NA | NA | NA | #VALUE! |
| ENSMUSG00000038925.16 | 0 | NA | NA | NA | NA | NA | #VALUE! |
| ENSMUSG00000038930.12 | 0 | NA | NA | NA | NA | NA | #VALUE! |
| ENSMUSG00000038932.14 | 0 | NA | NA | NA | NA | NA | #VALUE! |
| ENSMUSG00000038936.14 | 7.163089 | -6.37245 | 3.474566 | -1.83403 | 0.06665 | NA | #VALUE! |
| ENSMUSG00000038943.17 | 0 | NA | NA | NA | NA | NA | #VALUE! |
| ENSMUSG00000038963.16 | 0 | NA | NA | NA | NA | NA | #VALUE! |
| ENSMUSG00000038968.4 | 0 | NA | NA | NA | NA | NA | #VALUE! |
| ENSMUSG00000038973.3 | 0 | NA | NA | NA | NA | NA | #VALUE! |
| ENSMUSG00000038976.13 | 4.585304 | -3.79422 | 3.871444 | -0.98005 | 0.32706 | NA | #VALUE! |
| ENSMUSG00000038980.2 | 0 | NA | NA | NA | NA | NA | #VALUE! |
| ENSMUSG00000038984.5 | 0.210026 | -1.52494 | 4.058692 | -0.37572 | 0.707123 | NA | #VALUE! |
| ENSMUSG00000038987.9 | 0 | NA | NA | NA | NA | NA | #VALUE! |
| ENSMUSG00000038994.7 | 0 | NA | NA | NA | NA | NA | #VALUE! |
| ENSMUSG00000038997.5 | 0 | NA | NA | NA | NA | NA | #VALUE! |
| ENSMUSG00000039005.14 | 0 | NA | NA | NA | NA | NA | #VALUE! |
| ENSMUSG00000039013.17 | 0 | NA | NA | NA | NA | NA | #VALUE! |
| ENSMUSG00000039021.16 | 0 | NA | NA | NA | NA | NA | #VALUE! |
| ENSMUSG00000039032.3 | 0.491951 | 2.402185 | 4.000147 | 0.600524 | 0.548157 | NA | #VALUE! |
| ENSMUSG00000039037.6 | 0 | NA | NA | NA | NA | NA | #VALUE! |
| ENSMUSG00000039043.6 | 1.398828 | 3.99633 | 3.938205 | 1.014759 | 0.310221 | NA | #VALUE! |
| ENSMUSG00000039055.5 | 12.33188 | -3.15493 | 2.73832 | -1.15214 | NA | NA | #VALUE! |
| ENSMUSG00000039057.12 | 0 | NA | NA | NA | NA | NA | #VALUE! |
| ENSMUSG00000039058.12 | 0 | NA | NA | NA | NA | NA | #VALUE! |
| ENSMUSG00000039059.12 | 0 | NA | NA | NA | NA | NA | #VALUE! |
| ENSMUSG00000039070.6 | 0 | NA | NA | NA | NA | NA | #VALUE! |
| ENSMUSG00000039079.2 | 0 | NA | NA | NA | NA | NA | #VALUE! |

| | | | | | | | |
|---|---|---|---|---|---|---|---|
| ENSMUSG00000039081.11 | 4.159363 | 5.548619 | 3.764171 | 1.474061 | 0.140465 | NA | #VALUE! |
| ENSMUSG00000039084.9 | 0 | NA | NA | NA | NA | NA | #VALUE! |
| ENSMUSG00000039086.10 | 2.625328 | -4.93268 | 3.923662 | -1.25716 | 0.208695 | NA | #VALUE! |
| ENSMUSG00000039092.10 | 0 | NA | NA | NA | NA | NA | #VALUE! |
| ENSMUSG00000039095.9 | 0 | NA | NA | NA | NA | NA | #VALUE! |
| ENSMUSG00000039096.11 | 2.927558 | -3.53847 | 3.89802 | -0.90776 | 0.364004 | NA | #VALUE! |
| ENSMUSG00000039097.6 | 0 | NA | NA | NA | NA | NA | #VALUE! |
| ENSMUSG00000039098.4 | 0 | NA | NA | NA | NA | NA | #VALUE! |
| ENSMUSG00000039099.8 | 0 | NA | NA | NA | NA | NA | #VALUE! |
| ENSMUSG00000039106.7 | 0 | NA | NA | NA | NA | NA | #VALUE! |
| ENSMUSG00000039109.18 | 0 | NA | NA | NA | NA | NA | #VALUE! |
| ENSMUSG00000039110.16 | 0 | NA | NA | NA | NA | NA | #VALUE! |
| ENSMUSG00000039116.8 | 0 | NA | NA | NA | NA | NA | #VALUE! |
| ENSMUSG00000039126.11 | 2.187686 | 4.619304 | 3.927239 | 1.176222 | 0.239506 | NA | #VALUE! |
| ENSMUSG00000039131.16 | 0 | NA | NA | NA | NA | NA | #VALUE! |
| ENSMUSG00000039133.9 | 0 | NA | NA | NA | NA | NA | #VALUE! |
| ENSMUSG00000039137.19 | 0.841418 | 3.240539 | 3.959551 | 0.818411 | 0.413123 | NA | #VALUE! |
| ENSMUSG00000039138.2 | 0 | NA | NA | NA | NA | NA | #VALUE! |
| ENSMUSG00000039146.6 | 0 | NA | NA | NA | NA | NA | #VALUE! |
| ENSMUSG00000039153.18 | 0.093255 | 0.868093 | 4.072464 | 0.213162 | 0.831201 | NA | #VALUE! |
| ENSMUSG00000039154.9 | 0 | NA | NA | NA | NA | NA | #VALUE! |
| ENSMUSG00000039155.16 | 0 | NA | NA | NA | NA | NA | #VALUE! |
| ENSMUSG00000039158.12 | 6.149388 | 6.104864 | 3.9141 | 1.559711 | 0.118828 | NA | #VALUE! |
| ENSMUSG00000039174.15 | 0 | NA | NA | NA | NA | NA | #VALUE! |
| ENSMUSG00000039179.14 | 0 | NA | NA | NA | NA | NA | #VALUE! |
| ENSMUSG00000039187.17 | 0 | NA | NA | NA | NA | NA | #VALUE! |
| ENSMUSG00000039193.10 | 0 | NA | NA | NA | NA | NA | #VALUE! |
| ENSMUSG00000039194.17 | 0 | NA | NA | NA | NA | NA | #VALUE! |
| ENSMUSG00000039198.9 | 0 | NA | NA | NA | NA | NA | #VALUE! |
| ENSMUSG00000039199.7 | 0 | NA | NA | NA | NA | NA | #VALUE! |
| ENSMUSG00000039200.17 | 0 | NA | NA | NA | NA | NA | #VALUE! |
| ENSMUSG00000039201.11 | 8.662195 | -0.71999 | 2.701542 | -0.26651 | 0.789845 | NA | #VALUE! |
| ENSMUSG00000039208.16 | 11.11074 | 2.36936 | 3.163659 | 0.74893 | NA | NA | #VALUE! |
| ENSMUSG00000039209.13 | 0 | NA | NA | NA | NA | NA | #VALUE! |
| ENSMUSG00000039215.6 | 0 | NA | NA | NA | NA | NA | #VALUE! |
| ENSMUSG00000039224.2 | 0 | NA | NA | NA | NA | NA | #VALUE! |
| ENSMUSG00000039232.13 | 0 | NA | NA | NA | NA | NA | #VALUE! |
| ENSMUSG00000039236.19 | 1.995249 | -4.53986 | 3.928926 | -1.1555 | 0.247887 | NA | #VALUE! |
| ENSMUSG00000039239.15 | 0 | NA | NA | NA | NA | NA | #VALUE! |
| ENSMUSG00000039252.12 | 0 | NA | NA | NA | NA | NA | #VALUE! |
| ENSMUSG00000039253.8 | 10.07156 | 6.826017 | 3.559952 | 1.917446 | NA | NA | #VALUE! |
| ENSMUSG00000039257.12 | 0 | NA | NA | NA | NA | NA | #VALUE! |
| ENSMUSG00000039264.10 | 4.181584 | 5.545962 | 3.917562 | 1.415667 | 0.156873 | NA | #VALUE! |
| ENSMUSG00000039269.6 | 0 | NA | NA | NA | NA | NA | #VALUE! |

| | | | | | | | |
|---|---|---|---|---|---|---|---|
| ENSMUSG00000039275.10 | 146.5439 | -0.14996 | 3.292171 | -0.04555 | NA | NA | #VALUE! |
| ENSMUSG00000039278.11 | 0 | NA | NA | NA | NA | NA | #VALUE! |
| ENSMUSG00000039296.16 | 0 | NA | NA | NA | NA | NA | #VALUE! |
| ENSMUSG00000039298.17 | 2.187686 | 4.619304 | 3.927239 | 1.176222 | 0.239506 | NA | #VALUE! |
| ENSMUSG00000039307.17 | 6.837032 | 2.561221 | 2.973988 | 0.861207 | 0.389124 | NA | #VALUE! |
| ENSMUSG00000039313.15 | 0 | NA | NA | NA | NA | NA | #VALUE! |
| ENSMUSG00000039315.16 | 0 | NA | NA | NA | NA | NA | #VALUE! |
| ENSMUSG00000039321.2 | 0 | NA | NA | NA | NA | NA | #VALUE! |
| ENSMUSG00000039328.12 | 0 | NA | NA | NA | NA | NA | #VALUE! |
| ENSMUSG00000039329.9 | 0 | NA | NA | NA | NA | NA | #VALUE! |
| ENSMUSG00000039330.6 | 0 | NA | NA | NA | NA | NA | #VALUE! |
| ENSMUSG00000039335.12 | 0 | NA | NA | NA | NA | NA | #VALUE! |
| ENSMUSG00000039337.4 | 0 | NA | NA | NA | NA | NA | #VALUE! |
| ENSMUSG00000039339.5 | 0 | NA | NA | NA | NA | NA | #VALUE! |
| ENSMUSG00000039342.6 | 0 | NA | NA | NA | NA | NA | #VALUE! |
| ENSMUSG00000039347.8 | 0 | NA | NA | NA | NA | NA | #VALUE! |
| ENSMUSG00000039358.4 | 0 | NA | NA | NA | NA | NA | #VALUE! |
| ENSMUSG00000039364.14 | 0 | NA | NA | NA | NA | NA | #VALUE! |
| ENSMUSG00000039372.6 | 0 | NA | NA | NA | NA | NA | #VALUE! |
| ENSMUSG00000039375.17 | 0 | NA | NA | NA | NA | NA | #VALUE! |
| ENSMUSG00000039376.14 | 0 | NA | NA | NA | NA | NA | #VALUE! |
| ENSMUSG00000039377.8 | 3.588965 | 3.058087 | 3.858738 | 0.79251 | 0.428064 | NA | #VALUE! |
| ENSMUSG00000039385.6 | 0.420053 | -2.33493 | 4.007343 | -0.58266 | 0.56012 | NA | #VALUE! |
| ENSMUSG00000039391.12 | 0 | NA | NA | NA | NA | NA | #VALUE! |
| ENSMUSG00000039396.12 | 0 | NA | NA | NA | NA | NA | #VALUE! |
| ENSMUSG00000039410.17 | 0 | NA | NA | NA | NA | NA | #VALUE! |
| ENSMUSG00000039414.11 | 8.575921 | -0.99513 | 3.495379 | -0.2847 | 0.775874 | NA | #VALUE! |
| ENSMUSG00000039419.18 | 0 | NA | NA | NA | NA | NA | #VALUE! |
| ENSMUSG00000039431.17 | 4.711939 | 5.72619 | 3.916297 | 1.462144 | 0.143702 | NA | #VALUE! |
| ENSMUSG00000039452.7 | 1.567111 | 4.155831 | 3.934939 | 1.056136 | 0.290906 | NA | #VALUE! |
| ENSMUSG00000039462.5 | 0 | NA | NA | NA | NA | NA | #VALUE! |
| ENSMUSG00000039463.15 | 6.326359 | -0.17749 | 3.424624 | -0.05183 | 0.958665 | NA | #VALUE! |
| ENSMUSG00000039470.16 | 6.123323 | -6.14178 | 3.914088 | -1.56915 | 0.116614 | NA | #VALUE! |
| ENSMUSG00000039474.14 | 12.35304 | -1.25261 | 3.095057 | -0.40471 | NA | NA | #VALUE! |
| ENSMUSG00000039476.14 | 0 | NA | NA | NA | NA | NA | #VALUE! |
| ENSMUSG00000039477.17 | 28.78205 | -0.12509 | 1.648907 | -0.07586 | NA | NA | #VALUE! |
| ENSMUSG00000039478.16 | 11.54417 | 0.917214 | 3.15875 | 0.290372 | NA | NA | #VALUE! |
| ENSMUSG00000039481.6 | 0 | NA | NA | NA | NA | NA | #VALUE! |
| ENSMUSG00000039483.11 | 9.457945 | 0.877974 | 2.689354 | 0.326463 | 0.744074 | NA | #VALUE! |
| ENSMUSG00000039485.9 | 9.187179 | 2.359652 | 2.615641 | 0.902132 | 0.366987 | NA | #VALUE! |
| ENSMUSG00000039488.16 | 0 | NA | NA | NA | NA | NA | #VALUE! |
| ENSMUSG00000039492.8 | 0 | NA | NA | NA | NA | NA | #VALUE! |
| ENSMUSG00000039496.9 | 3.328415 | -5.26129 | 3.920229 | -1.34209 | 0.179567 | NA | #VALUE! |
| ENSMUSG00000039508.7 | 0 | NA | NA | NA | NA | NA | #VALUE! |

| | | | | | | | |
|---|---|---|---|---|---|---|---|
| ENSMUSG00000039515.12 | 7.087395 | 6.319644 | 3.913086 | 1.615003 | NA | NA | #VALUE! |
| ENSMUSG00000039518.8 | 0 | NA | NA | NA | NA | NA | #VALUE! |
| ENSMUSG00000039521.14 | 0 | NA | NA | NA | NA | NA | #VALUE! |
| ENSMUSG00000039539.14 | 0 | NA | NA | NA | NA | NA | #VALUE! |
| ENSMUSG00000039540.10 | 0 | NA | NA | NA | NA | NA | #VALUE! |
| ENSMUSG00000039542.17 | 0 | NA | NA | NA | NA | NA | #VALUE! |
| ENSMUSG00000039543.17 | 0 | NA | NA | NA | NA | NA | #VALUE! |
| ENSMUSG00000039545.17 | 0 | NA | NA | NA | NA | NA | #VALUE! |
| ENSMUSG00000039546.10 | 0 | NA | NA | NA | NA | NA | #VALUE! |
| ENSMUSG00000039552.15 | 0.420053 | -2.33493 | 4.007343 | -0.58266 | 0.56012 | NA | #VALUE! |
| ENSMUSG00000039555.16 | 0 | NA | NA | NA | NA | NA | #VALUE! |
| ENSMUSG00000039556.17 | 0 | NA | NA | NA | NA | NA | #VALUE! |
| ENSMUSG00000039563.7 | 1.009701 | 3.503687 | 3.950852 | 0.886818 | 0.375177 | NA | #VALUE! |
| ENSMUSG00000039577.18 | 0 | NA | NA | NA | NA | NA | #VALUE! |
| ENSMUSG00000039578.18 | 0 | NA | NA | NA | NA | NA | #VALUE! |
| ENSMUSG00000039579.16 | 0 | NA | NA | NA | NA | NA | #VALUE! |
| ENSMUSG00000039585.16 | 4.552227 | 0.753792 | 3.327016 | 0.226567 | 0.82076 | NA | #VALUE! |
| ENSMUSG00000039601.17 | 35.69476 | -1.34207 | 2.430906 | -0.55208 | NA | NA | #VALUE! |
| ENSMUSG00000039607.18 | 6.278386 | 2.458052 | 3.34563 | 0.734705 | 0.462519 | NA | #VALUE! |
| ENSMUSG00000039608.7 | 0 | NA | NA | NA | NA | NA | #VALUE! |
| ENSMUSG00000039611.3 | 0 | NA | NA | NA | NA | NA | #VALUE! |
| ENSMUSG00000039617.9 | 0 | NA | NA | NA | NA | NA | #VALUE! |
| ENSMUSG00000039620.18 | 13.42667 | -2.22578 | 2.748032 | -0.80995 | NA | NA | #VALUE! |
| ENSMUSG00000039621.14 | 0.637369 | 0.178787 | 3.999109 | 0.044707 | 0.964341 | NA | #VALUE! |
| ENSMUSG00000039623.14 | 8.512972 | 0.46801 | 3.458024 | 0.13534 | 0.892343 | NA | #VALUE! |
| ENSMUSG00000039628.10 | 0.737927 | 3.007825 | 3.968641 | 0.757898 | 0.448512 | NA | #VALUE! |
| ENSMUSG00000039629.15 | 0 | NA | NA | NA | NA | NA | #VALUE! |
| ENSMUSG00000039632.14 | 2.984166 | 5.077883 | 3.921685 | 1.294822 | 0.195382 | NA | #VALUE! |
| ENSMUSG00000039634.13 | 19.7056 | 0.576567 | 2.176177 | 0.264945 | NA | NA | #VALUE! |
| ENSMUSG00000039639.8 | 0 | NA | NA | NA | NA | NA | #VALUE! |
| ENSMUSG00000039661.15 | 0 | NA | NA | NA | NA | NA | #VALUE! |
| ENSMUSG00000039672.13 | 0 | NA | NA | NA | NA | NA | #VALUE! |
| ENSMUSG00000039676.5 | 0 | NA | NA | NA | NA | NA | #VALUE! |
| ENSMUSG00000039678.13 | 7.589763 | -3.36743 | 3.224463 | -1.04434 | 0.296329 | NA | #VALUE! |
| ENSMUSG00000039683.17 | 0 | NA | NA | NA | NA | NA | #VALUE! |
| ENSMUSG00000039684.10 | 0 | NA | NA | NA | NA | NA | #VALUE! |
| ENSMUSG00000039691.11 | 0 | NA | NA | NA | NA | NA | #VALUE! |
| ENSMUSG00000039693.12 | 0 | NA | NA | NA | NA | NA | #VALUE! |
| ENSMUSG00000039699.15 | 0 | NA | NA | NA | NA | NA | #VALUE! |
| ENSMUSG00000039710.15 | 0 | NA | NA | NA | NA | NA | #VALUE! |
| ENSMUSG00000039713.18 | 1.119062 | 3.679627 | 3.945843 | 0.932533 | 0.351061 | NA | #VALUE! |
| ENSMUSG00000039714.10 | 0 | NA | NA | NA | NA | NA | #VALUE! |
| ENSMUSG00000039716.13 | 0.737927 | 3.007825 | 3.968641 | 0.757898 | 0.448512 | NA | #VALUE! |
| ENSMUSG00000039717.17 | 0 | NA | NA | NA | NA | NA | #VALUE! |

| | | | | | | | |
|---|---|---|---|---|---|---|---|
| ENSMUSG00000039720.8 | 0 | NA | NA | NA | NA | NA | #VALUE! |
| ENSMUSG00000039725.7 | 8.966372 | -0.07515 | 3.037319 | -0.02474 | 0.98026 | NA | #VALUE! |
| ENSMUSG00000039728.16 | 0 | NA | NA | NA | NA | NA | #VALUE! |
| ENSMUSG00000039738.17 | 2.100263 | -4.61322 | 3.927832 | -1.17449 | 0.240197 | NA | #VALUE! |
| ENSMUSG00000039741.16 | 1.616499 | -2.90839 | 3.92562 | -0.74087 | 0.45877 | NA | #VALUE! |
| ENSMUSG00000039742.16 | 0 | NA | NA | NA | NA | NA | #VALUE! |
| ENSMUSG00000039747.12 | 0 | NA | NA | NA | NA | NA | #VALUE! |
| ENSMUSG00000039748.12 | 4.942525 | 5.801604 | 3.915812 | 1.481584 | 0.138451 | NA | #VALUE! |
| ENSMUSG00000039754.11 | 7.215172 | 1.586967 | 3.294843 | 0.481652 | 0.630053 | NA | #VALUE! |
| ENSMUSG00000039760.9 | 0 | NA | NA | NA | NA | NA | #VALUE! |
| ENSMUSG00000039765.16 | 7.89157 | -0.34751 | 2.859901 | -0.12151 | 0.903285 | NA | #VALUE! |
| ENSMUSG00000039774.13 | 0 | NA | NA | NA | NA | NA | #VALUE! |
| ENSMUSG00000039775.8 | 0 | NA | NA | NA | NA | NA | #VALUE! |
| ENSMUSG00000039781.15 | 1.613981 | 0.570769 | 3.844101 | 0.148479 | 0.881965 | NA | #VALUE! |
| ENSMUSG00000039785.6 | 0 | NA | NA | NA | NA | NA | #VALUE! |
| ENSMUSG00000039787.14 | 0 | NA | NA | NA | NA | NA | #VALUE! |
| ENSMUSG00000039798.3 | 0 | NA | NA | NA | NA | NA | #VALUE! |
| ENSMUSG00000039813.15 | 0 | NA | NA | NA | NA | NA | #VALUE! |
| ENSMUSG00000039814.16 | 0 | NA | NA | NA | NA | NA | #VALUE! |
| ENSMUSG00000039824.5 | 0 | NA | NA | NA | NA | NA | #VALUE! |
| ENSMUSG00000039828.8 | 24.0014 | 0.796307 | 1.985478 | 0.401066 | NA | NA | #VALUE! |
| ENSMUSG00000039830.10 | 0 | NA | NA | NA | NA | NA | #VALUE! |
| ENSMUSG00000039834.18 | 0 | NA | NA | NA | NA | NA | #VALUE! |
| ENSMUSG00000039838.12 | 0 | NA | NA | NA | NA | NA | #VALUE! |
| ENSMUSG00000039842.16 | 8.107723 | -5.45377 | 3.202653 | -1.70289 | 0.088589 | NA | #VALUE! |
| ENSMUSG00000039850.17 | 7.759071 | 6.445604 | 3.381039 | 1.906397 | 0.056599 | NA | #VALUE! |
| ENSMUSG00000039851.12 | 0 | NA | NA | NA | NA | NA | #VALUE! |
| ENSMUSG00000039853.19 | 0 | NA | NA | NA | NA | NA | #VALUE! |
| ENSMUSG00000039860.20 | 0 | NA | NA | NA | NA | NA | #VALUE! |
| ENSMUSG00000039873.5 | 0.612332 | -2.79827 | 3.980109 | -0.70306 | 0.482016 | NA | #VALUE! |
| ENSMUSG00000039878.14 | 0 | NA | NA | NA | NA | NA | #VALUE! |
| ENSMUSG00000039883.6 | 0 | NA | NA | NA | NA | NA | #VALUE! |
| ENSMUSG00000039891.7 | 0 | NA | NA | NA | NA | NA | #VALUE! |
| ENSMUSG00000039899.6 | 8.614231 | -1.56897 | 3.132739 | -0.50083 | 0.61649 | NA | #VALUE! |
| ENSMUSG00000039903.19 | 5.30688 | -5.93495 | 3.915215 | -1.51587 | 0.129553 | NA | #VALUE! |
| ENSMUSG00000039904.10 | 0 | NA | NA | NA | NA | NA | #VALUE! |
| ENSMUSG00000039911.14 | 0 | NA | NA | NA | NA | NA | #VALUE! |
| ENSMUSG00000039913.13 | 0 | NA | NA | NA | NA | NA | #VALUE! |
| ENSMUSG00000039929.15 | 5.70874 | 6.002521 | 3.871497 | 1.550439 | 0.121036 | NA | #VALUE! |
| ENSMUSG00000039936.19 | 0.483323 | -0.86763 | 4.025136 | -0.21555 | 0.829336 | NA | #VALUE! |
| ENSMUSG00000039942.16 | 0 | NA | NA | NA | NA | NA | #VALUE! |
| ENSMUSG00000039953.14 | 9.0214 | -3.40702 | 3.00113 | -1.13525 | 0.256272 | NA | #VALUE! |
| ENSMUSG00000039954.10 | 0 | NA | NA | NA | NA | NA | #VALUE! |
| ENSMUSG00000039959.14 | 8.772455 | -1.6981 | 3.409689 | -0.49802 | 0.618469 | NA | #VALUE! |

| | | | | | | | |
|---|---|---|---|---|---|---|---|
| ENSMUSG00000039962.4 | 0 | NA | NA | NA | NA | NA | #VALUE! |
| ENSMUSG00000039963.21 | 0 | NA | NA | NA | NA | NA | #VALUE! |
| ENSMUSG00000039977.17 | 0.559531 | 2.703958 | 3.982863 | 0.678898 | 0.497202 | NA | #VALUE! |
| ENSMUSG00000039981.7 | 0 | NA | NA | NA | NA | NA | #VALUE! |
| ENSMUSG00000039985.15 | 0 | NA | NA | NA | NA | NA | #VALUE! |
| ENSMUSG00000039994.16 | 9.371272 | -3.4249 | 2.653161 | -1.29088 | 0.196747 | NA | #VALUE! |
| ENSMUSG00000040003.19 | 0 | NA | NA | NA | NA | NA | #VALUE! |
| ENSMUSG00000040009.7 | 0 | NA | NA | NA | NA | NA | #VALUE! |
| ENSMUSG00000040010.11 | 0.420053 | -2.33493 | 4.007343 | -0.58266 | 0.56012 | NA | #VALUE! |
| ENSMUSG00000040013.12 | 0 | NA | NA | NA | NA | NA | #VALUE! |
| ENSMUSG00000040016.17 | 0.504851 | 2.502843 | 3.993993 | 0.626652 | 0.530887 | NA | #VALUE! |
| ENSMUSG00000040035.15 | 3.565687 | -0.77726 | 3.781598 | -0.20554 | 0.837153 | NA | #VALUE! |
| ENSMUSG00000040037.14 | 0 | NA | NA | NA | NA | NA | #VALUE! |
| ENSMUSG00000040040.17 | 9.119247 | -2.61636 | 2.321132 | -1.12719 | 0.259661 | NA | #VALUE! |
| ENSMUSG00000040046.15 | 0 | NA | NA | NA | NA | NA | #VALUE! |
| ENSMUSG00000040055.10 | 0 | NA | NA | NA | NA | NA | #VALUE! |
| ENSMUSG00000040065.9 | 0 | NA | NA | NA | NA | NA | #VALUE! |
| ENSMUSG00000040078.10 | 0 | NA | NA | NA | NA | NA | #VALUE! |
| ENSMUSG00000040084.10 | 0 | NA | NA | NA | NA | NA | #VALUE! |
| ENSMUSG00000040086.14 | 0 | NA | NA | NA | NA | NA | #VALUE! |
| ENSMUSG00000040093.17 | 3.226933 | -5.22704 | 3.920552 | -1.33324 | 0.182453 | NA | #VALUE! |
| ENSMUSG00000040097.17 | 7.179299 | 0.764962 | 2.494113 | 0.306707 | 0.759066 | NA | #VALUE! |
| ENSMUSG00000040111.18 | 0.210026 | -1.52494 | 4.058692 | -0.37572 | 0.707123 | NA | #VALUE! |
| ENSMUSG00000040113.15 | 0 | NA | NA | NA | NA | NA | #VALUE! |
| ENSMUSG00000040118.16 | 0 | NA | NA | NA | NA | NA | #VALUE! |
| ENSMUSG00000040121.6 | 0 | NA | NA | NA | NA | NA | #VALUE! |
| ENSMUSG00000040125.5 | 0 | NA | NA | NA | NA | NA | #VALUE! |
| ENSMUSG00000040132.7 | 0 | NA | NA | NA | NA | NA | #VALUE! |
| ENSMUSG00000040133.3 | 0 | NA | NA | NA | NA | NA | #VALUE! |
| ENSMUSG00000040136.11 | 0 | NA | NA | NA | NA | NA | #VALUE! |
| ENSMUSG00000040138.8 | 0 | NA | NA | NA | NA | NA | #VALUE! |
| ENSMUSG00000040140.16 | 0 | NA | NA | NA | NA | NA | #VALUE! |
| ENSMUSG00000040148.6 | 0 | NA | NA | NA | NA | NA | #VALUE! |
| ENSMUSG00000040154.12 | 0 | NA | NA | NA | NA | NA | #VALUE! |
| ENSMUSG00000040158.13 | 1.346268 | 3.918821 | 3.939925 | 0.994644 | 0.31991 | NA | #VALUE! |
| ENSMUSG00000040163.15 | 0 | NA | NA | NA | NA | NA | #VALUE! |
| ENSMUSG00000040164.4 | 0 | NA | NA | NA | NA | NA | #VALUE! |
| ENSMUSG00000040165.15 | 0 | NA | NA | NA | NA | NA | #VALUE! |
| ENSMUSG00000040183.14 | 0 | NA | NA | NA | NA | NA | #VALUE! |
| ENSMUSG00000040187.16 | 0 | NA | NA | NA | NA | NA | #VALUE! |
| ENSMUSG00000040189.19 | 0 | NA | NA | NA | NA | NA | #VALUE! |
| ENSMUSG00000040197.5 | 0 | NA | NA | NA | NA | NA | #VALUE! |
| ENSMUSG00000040204.7 | 0.525066 | -2.64687 | 3.988095 | -0.66369 | 0.506887 | NA | #VALUE! |
| ENSMUSG00000040205.9 | 0 | NA | NA | NA | NA | NA | #VALUE! |

| | | | | | | | |
|---|---|---|---|---|---|---|---|
| ENSMUSG00000040212.13 | 0.273297 | 0.094602 | 4.062933 | 0.023284 | 0.981424 | NA | #VALUE! |
| ENSMUSG00000040217.8 | 0 | NA | NA | NA | NA | NA | #VALUE! |
| ENSMUSG00000040219.5 | 0 | NA | NA | NA | NA | NA | #VALUE! |
| ENSMUSG00000040220.11 | 5.385073 | 5.918828 | 3.915107 | 1.511792 | 0.130587 | NA | #VALUE! |
| ENSMUSG00000040229.12 | 0 | NA | NA | NA | NA | NA | #VALUE! |
| ENSMUSG00000040231.17 | 0 | NA | NA | NA | NA | NA | #VALUE! |
| ENSMUSG00000040247.7 | 3.443657 | 5.264174 | 3.919884 | 1.342941 | 0.179291 | NA | #VALUE! |
| ENSMUSG00000040254.12 | 2.592451 | -1.58514 | 3.819833 | -0.41498 | 0.678159 | NA | #VALUE! |
| ENSMUSG00000040258.7 | 0 | NA | NA | NA | NA | NA | #VALUE! |
| ENSMUSG00000040260.9 | 0.315039 | -1.93549 | 4.038621 | -0.47925 | 0.631763 | NA | #VALUE! |
| ENSMUSG00000040265.17 | 0 | NA | NA | NA | NA | NA | #VALUE! |
| ENSMUSG00000040272.15 | 5.967152 | 4.083545 | 3.437961 | 1.187781 | 0.23492 | NA | #VALUE! |
| ENSMUSG00000040276.16 | 0.630079 | -2.90291 | 3.975047 | -0.73028 | 0.465217 | NA | #VALUE! |
| ENSMUSG00000040280.11 | 3.197387 | 5.166784 | 3.920797 | 1.317789 | 0.187574 | NA | #VALUE! |
| ENSMUSG00000040283.15 | 1.595425 | -0.85495 | 3.848186 | -0.22217 | 0.824182 | NA | #VALUE! |
| ENSMUSG00000040284.5 | 0 | NA | NA | NA | NA | NA | #VALUE! |
| ENSMUSG00000040287.10 | 0 | NA | NA | NA | NA | NA | #VALUE! |
| ENSMUSG00000040289.9 | 0.195789 | -1.05581 | 4.073302 | -0.2592 | 0.795479 | NA | #VALUE! |
| ENSMUSG00000040298.7 | 0 | NA | NA | NA | NA | NA | #VALUE! |
| ENSMUSG00000040310.13 | 0 | NA | NA | NA | NA | NA | #VALUE! |
| ENSMUSG00000040312.15 | 0 | NA | NA | NA | NA | NA | #VALUE! |
| ENSMUSG00000040314.3 | 0 | NA | NA | NA | NA | NA | #VALUE! |
| ENSMUSG00000040322.11 | 4.760753 | 1.022203 | 3.748845 | 0.272671 | 0.785106 | NA | #VALUE! |
| ENSMUSG00000040323.14 | 0 | NA | NA | NA | NA | NA | #VALUE! |
| ENSMUSG00000040327.17 | 7.236192 | 6.345076 | 3.912976 | 1.621547 | 0.1049 | NA | #VALUE! |
| ENSMUSG00000040328.17 | 0 | NA | NA | NA | NA | NA | #VALUE! |
| ENSMUSG00000040329.15 | 4.503149 | -5.6983 | 3.916718 | -1.45487 | 0.145706 | NA | #VALUE! |
| ENSMUSG00000040339.11 | 8.026022 | -3.54967 | 2.710824 | -1.30944 | 0.190384 | NA | #VALUE! |
| ENSMUSG00000040340.14 | 0 | NA | NA | NA | NA | NA | #VALUE! |
| ENSMUSG00000040350.17 | 4.974797 | -5.84639 | 3.915749 | -1.49304 | 0.135425 | NA | #VALUE! |
| ENSMUSG00000040364.9 | 0 | NA | NA | NA | NA | NA | #VALUE! |
| ENSMUSG00000040367.3 | 3.170677 | 5.164728 | 3.920817 | 1.317258 | 0.187752 | NA | #VALUE! |
| ENSMUSG00000040372.3 | 0 | NA | NA | NA | NA | NA | #VALUE! |
| ENSMUSG00000040373.13 | 0 | NA | NA | NA | NA | NA | #VALUE! |
| ENSMUSG00000040380.15 | 0 | NA | NA | NA | NA | NA | #VALUE! |
| ENSMUSG00000040387.17 | 0.18651 | 1.401769 | 4.056354 | 0.345574 | 0.729663 | NA | #VALUE! |
| ENSMUSG00000040390.14 | 3.276603 | 1.394819 | 3.795951 | 0.367449 | 0.713284 | NA | #VALUE! |
| ENSMUSG00000040405.14 | 0 | NA | NA | NA | NA | NA | #VALUE! |
| ENSMUSG00000040412.18 | 0 | NA | NA | NA | NA | NA | #VALUE! |
| ENSMUSG00000040415.16 | 1.177985 | 3.72614 | 3.944615 | 0.944614 | 0.344856 | NA | #VALUE! |
| ENSMUSG00000040420.16 | 0 | NA | NA | NA | NA | NA | #VALUE! |
| ENSMUSG00000040424.16 | 0 | NA | NA | NA | NA | NA | #VALUE! |
| ENSMUSG00000040428.20 | 0 | NA | NA | NA | NA | NA | #VALUE! |
| ENSMUSG00000040432.6 | 2.797656 | 4.98547 | 3.922669 | 1.270938 | 0.203751 | NA | #VALUE! |

| | | | | | | | |
|---|---|---|---|---|---|---|---|
| ENSMUSG00000040434.17 | 0 | NA | NA | NA | NA | NA | #VALUE! |
| ENSMUSG00000040435.13 | 4.830604 | -5.80779 | 3.915992 | -1.4831 | 0.138049 | NA | #VALUE! |
| ENSMUSG00000040441.13 | 6.99414 | 6.300596 | 3.91317 | 1.6101 | NA | NA | #VALUE! |
| ENSMUSG00000040446.5 | 6.438594 | -1.30798 | 3.018669 | -0.4333 | 0.664799 | NA | #VALUE! |
| ENSMUSG00000040452.10 | 0 | NA | NA | NA | NA | NA | #VALUE! |
| ENSMUSG00000040456.3 | 0 | NA | NA | NA | NA | NA | #VALUE! |
| ENSMUSG00000040467.2 | 0 | NA | NA | NA | NA | NA | #VALUE! |
| ENSMUSG00000040473.16 | 0 | NA | NA | NA | NA | NA | #VALUE! |
| ENSMUSG00000040478.15 | 0 | NA | NA | NA | NA | NA | #VALUE! |
| ENSMUSG00000040483.16 | 3.475798 | -5.32367 | 3.919661 | -1.3582 | 0.174401 | NA | #VALUE! |
| ENSMUSG00000040485.6 | 0 | NA | NA | NA | NA | NA | #VALUE! |
| ENSMUSG00000040489.6 | 0 | NA | NA | NA | NA | NA | #VALUE! |
| ENSMUSG00000040490.5 | 0 | NA | NA | NA | NA | NA | #VALUE! |
| ENSMUSG00000040495.5 | 0 | NA | NA | NA | NA | NA | #VALUE! |
| ENSMUSG00000040498.8 | 1.333409 | 0.404416 | 3.864119 | 0.104659 | 0.916646 | NA | #VALUE! |
| ENSMUSG00000040502.6 | 0 | NA | NA | NA | NA | NA | #VALUE! |
| ENSMUSG00000040506.17 | 6.005373 | -0.63831 | 3.75766 | -0.16987 | 0.865114 | NA | #VALUE! |
| ENSMUSG00000040514.7 | 0 | NA | NA | NA | NA | NA | #VALUE! |
| ENSMUSG00000040522.6 | 0 | NA | NA | NA | NA | NA | #VALUE! |
| ENSMUSG00000040524.10 | 6.93622 | -6.32767 | 3.535274 | -1.78987 | 0.073475 | NA | #VALUE! |
| ENSMUSG00000040528.16 | 0 | NA | NA | NA | NA | NA | #VALUE! |
| ENSMUSG00000040533.8 | 0 | NA | NA | NA | NA | NA | #VALUE! |
| ENSMUSG00000040537.18 | 0 | NA | NA | NA | NA | NA | #VALUE! |
| ENSMUSG00000040540.7 | 0 | NA | NA | NA | NA | NA | #VALUE! |
| ENSMUSG00000040541.9 | 0 | NA | NA | NA | NA | NA | #VALUE! |
| ENSMUSG00000040543.17 | 0 | NA | NA | NA | NA | NA | #VALUE! |
| ENSMUSG00000040554.11 | 0 | NA | NA | NA | NA | NA | #VALUE! |
| ENSMUSG00000040563.14 | 0 | NA | NA | NA | NA | NA | #VALUE! |
| ENSMUSG00000040569.14 | 0 | NA | NA | NA | NA | NA | #VALUE! |
| ENSMUSG00000040570.15 | 0 | NA | NA | NA | NA | NA | #VALUE! |
| ENSMUSG00000040576.9 | 0 | NA | NA | NA | NA | NA | #VALUE! |
| ENSMUSG00000040583.9 | 0 | NA | NA | NA | NA | NA | #VALUE! |
| ENSMUSG00000040592.12 | 0.612332 | -2.79827 | 3.980109 | -0.70306 | 0.482016 | NA | #VALUE! |
| ENSMUSG00000040594.20 | 0 | NA | NA | NA | NA | NA | #VALUE! |
| ENSMUSG00000040600.10 | 0 | NA | NA | NA | NA | NA | #VALUE! |
| ENSMUSG00000040601.14 | 0 | NA | NA | NA | NA | NA | #VALUE! |
| ENSMUSG00000040606.15 | 0 | NA | NA | NA | NA | NA | #VALUE! |
| ENSMUSG00000040610.8 | 0 | NA | NA | NA | NA | NA | #VALUE! |
| ENSMUSG00000040614.15 | 0 | NA | NA | NA | NA | NA | #VALUE! |
| ENSMUSG00000040616.4 | 4.189699 | 5.551587 | 3.91752 | 1.417118 | 0.156449 | NA | #VALUE! |
| ENSMUSG00000040618.8 | 1.232996 | 1.131842 | 3.89811 | 0.290357 | 0.771543 | NA | #VALUE! |
| ENSMUSG00000040621.15 | 0 | NA | NA | NA | NA | NA | #VALUE! |
| ENSMUSG00000040627.15 | 0 | NA | NA | NA | NA | NA | #VALUE! |
| ENSMUSG00000040629.9 | 0 | NA | NA | NA | NA | NA | #VALUE! |

| | | | | | | | |
|---|---|---|---|---|---|---|---|
| ENSMUSG00000040631.9 | 2.100263 | -4.61322 | 3.927832 | -1.17449 | 0.240197 | NA | #VALUE! |
| ENSMUSG00000040632.17 | 0 | NA | NA | NA | NA | NA | #VALUE! |
| ENSMUSG00000040649.16 | 0 | NA | NA | NA | NA | NA | #VALUE! |
| ENSMUSG00000040650.9 | 0 | NA | NA | NA | NA | NA | #VALUE! |
| ENSMUSG00000040653.7 | 0 | NA | NA | NA | NA | NA | #VALUE! |
| ENSMUSG00000040657.4 | 0 | NA | NA | NA | NA | NA | #VALUE! |
| ENSMUSG00000040660.7 | 6.724978 | 6.241644 | 3.544936 | 1.760721 | 0.078286 | NA | #VALUE! |
| ENSMUSG00000040663.10 | 3.533954 | 5.311169 | 3.919465 | 1.355075 | 0.175394 | NA | #VALUE! |
| ENSMUSG00000040666.20 | 0 | NA | NA | NA | NA | NA | #VALUE! |
| ENSMUSG00000040669.15 | 0 | NA | NA | NA | NA | NA | #VALUE! |
| ENSMUSG00000040675.18 | 0.630079 | -2.90291 | 3.975047 | -0.73028 | 0.465217 | NA | #VALUE! |
| ENSMUSG00000040680.10 | 0 | NA | NA | NA | NA | NA | #VALUE! |
| ENSMUSG00000040690.16 | 7.590545 | -3.2613 | 3.611337 | -0.90307 | 0.366487 | NA | #VALUE! |
| ENSMUSG00000040693.8 | 0 | NA | NA | NA | NA | NA | #VALUE! |
| ENSMUSG00000040694.5 | 0 | NA | NA | NA | NA | NA | #VALUE! |
| ENSMUSG00000040699.14 | 6.435904 | 0.874836 | 2.656153 | 0.329362 | 0.741882 | NA | #VALUE! |
| ENSMUSG00000040701.17 | 2.20361 | -4.66366 | 3.927111 | -1.18755 | 0.235009 | NA | #VALUE! |
| ENSMUSG00000040703.12 | 0 | NA | NA | NA | NA | NA | #VALUE! |
| ENSMUSG00000040705.3 | 0 | NA | NA | NA | NA | NA | #VALUE! |
| ENSMUSG00000040710.11 | 0.168284 | 1.027628 | 4.067126 | 0.252667 | 0.800526 | NA | #VALUE! |
| ENSMUSG00000040711.9 | 0 | NA | NA | NA | NA | NA | #VALUE! |
| ENSMUSG00000040712.17 | 0.466276 | 2.449889 | 3.997179 | 0.612904 | 0.53994 | NA | #VALUE! |
| ENSMUSG00000040714.15 | 0 | NA | NA | NA | NA | NA | #VALUE! |
| ENSMUSG00000040715.13 | 0 | NA | NA | NA | NA | NA | #VALUE! |
| ENSMUSG00000040717.7 | 2.33138 | 4.724655 | 3.925805 | 1.203487 | 0.228788 | NA | #VALUE! |
| ENSMUSG00000040726.11 | 0 | NA | NA | NA | NA | NA | #VALUE! |
| ENSMUSG00000040728.16 | 0 | NA | NA | NA | NA | NA | #VALUE! |
| ENSMUSG00000040729.9 | 0 | NA | NA | NA | NA | NA | #VALUE! |
| ENSMUSG00000040734.15 | 16.74709 | -7.59441 | 2.834672 | -2.67912 | NA | NA | #VALUE! |
| ENSMUSG00000040740.8 | 0 | NA | NA | NA | NA | NA | #VALUE! |
| ENSMUSG00000040749.12 | 1.212318 | 3.793111 | 3.942915 | 0.962007 | 0.336046 | NA | #VALUE! |
| ENSMUSG00000040751.13 | 3.357187 | 5.246639 | 3.920044 | 1.338413 | 0.180762 | NA | #VALUE! |
| ENSMUSG00000040752.17 | 0 | NA | NA | NA | NA | NA | #VALUE! |
| ENSMUSG00000040759.10 | 0 | NA | NA | NA | NA | NA | #VALUE! |
| ENSMUSG00000040770.6 | 0 | NA | NA | NA | NA | NA | #VALUE! |
| ENSMUSG00000040775.7 | 0 | NA | NA | NA | NA | NA | #VALUE! |
| ENSMUSG00000040794.6 | 0 | NA | NA | NA | NA | NA | #VALUE! |
| ENSMUSG00000040795.13 | 4.498776 | 1.697447 | 2.952244 | 0.574969 | 0.565313 | NA | #VALUE! |
| ENSMUSG00000040797.17 | 0 | NA | NA | NA | NA | NA | #VALUE! |
| ENSMUSG00000040808.5 | 0 | NA | NA | NA | NA | NA | #VALUE! |
| ENSMUSG00000040809.11 | 0 | NA | NA | NA | NA | NA | #VALUE! |
| ENSMUSG00000040811.16 | 0.978946 | -3.48749 | 3.952505 | -0.88235 | 0.377588 | NA | #VALUE! |
| ENSMUSG00000040812.15 | 0 | NA | NA | NA | NA | NA | #VALUE! |
| ENSMUSG00000040828.11 | 2.705731 | 4.913666 | 3.923477 | 1.252375 | 0.210433 | NA | #VALUE! |

| | | | | | | | |
|---|---|---|---|---|---|---|---|
| ENSMUSG00000040829.15 | 0 | NA | NA | NA | NA | NA | #VALUE! |
| ENSMUSG00000040836.16 | 0 | NA | NA | NA | NA | NA | #VALUE! |
| ENSMUSG00000040838.11 | 0 | NA | NA | NA | NA | NA | #VALUE! |
| ENSMUSG00000040840.8 | 0 | NA | NA | NA | NA | NA | #VALUE! |
| ENSMUSG00000040841.6 | 0.673134 | 2.918386 | 3.972531 | 0.734641 | 0.462558 | NA | #VALUE! |
| ENSMUSG00000040852.7 | 2.86082 | 5.006323 | 3.922441 | 1.276328 | 0.201839 | NA | #VALUE! |
| ENSMUSG00000040856.18 | 5.482095 | -5.98256 | 3.914941 | -1.52813 | 0.126479 | NA | #VALUE! |
| ENSMUSG00000040857.16 | 1.072839 | 0.624931 | 3.897316 | 0.160349 | 0.872606 | NA | #VALUE! |
| ENSMUSG00000040860.17 | 0 | NA | NA | NA | NA | NA | #VALUE! |
| ENSMUSG00000040866.14 | 0 | NA | NA | NA | NA | NA | #VALUE! |
| ENSMUSG00000040867.14 | 0 | NA | NA | NA | NA | NA | #VALUE! |
| ENSMUSG00000040875.13 | 0 | NA | NA | NA | NA | NA | #VALUE! |
| ENSMUSG00000040877.17 | 5.364194 | 0.633252 | 3.761287 | 0.16836 | 0.8663 | NA | #VALUE! |
| ENSMUSG00000040896.17 | 0 | NA | NA | NA | NA | NA | #VALUE! |
| ENSMUSG00000040899.15 | 0 | NA | NA | NA | NA | NA | #VALUE! |
| ENSMUSG00000040901.9 | 0 | NA | NA | NA | NA | NA | #VALUE! |
| ENSMUSG00000040904.11 | 0 | NA | NA | NA | NA | NA | #VALUE! |
| ENSMUSG00000040907.16 | 0 | NA | NA | NA | NA | NA | #VALUE! |
| ENSMUSG00000040919.14 | 0 | NA | NA | NA | NA | NA | #VALUE! |
| ENSMUSG00000040929.18 | 6.874117 | 2.006783 | 2.936181 | 0.683467 | 0.494312 | NA | #VALUE! |
| ENSMUSG00000040935.13 | 0 | NA | NA | NA | NA | NA | #VALUE! |
| ENSMUSG00000040938.17 | 1.762102 | -4.34067 | 3.93219 | -1.10388 | 0.269645 | NA | #VALUE! |
| ENSMUSG00000040940.19 | 2.816667 | 1.883324 | 3.822534 | 0.49269 | 0.622232 | NA | #VALUE! |
| ENSMUSG00000040943.13 | 2.581785 | -2.70493 | 3.8742 | -0.69819 | 0.485057 | NA | #VALUE! |
| ENSMUSG00000040950.14 | 0 | NA | NA | NA | NA | NA | #VALUE! |
| ENSMUSG00000040957.16 | 2.911246 | -3.87321 | 3.90968 | -0.99067 | 0.321846 | NA | #VALUE! |
| ENSMUSG00000040966.11 | 0 | NA | NA | NA | NA | NA | #VALUE! |
| ENSMUSG00000040969.15 | 0 | NA | NA | NA | NA | NA | #VALUE! |
| ENSMUSG00000040972.9 | 0 | NA | NA | NA | NA | NA | #VALUE! |
| ENSMUSG00000040978.5 | 0.491951 | 2.402185 | 4.000147 | 0.600524 | 0.548157 | NA | #VALUE! |
| ENSMUSG00000040985.14 | 0 | NA | NA | NA | NA | NA | #VALUE! |
| ENSMUSG00000040987.16 | 6.852619 | -6.30491 | 3.913306 | -1.61115 | 0.107148 | NA | #VALUE! |
| ENSMUSG00000040990.18 | 5.595312 | 5.979818 | 3.914763 | 1.527505 | 0.126636 | NA | #VALUE! |
| ENSMUSG00000040998.19 | 0 | NA | NA | NA | NA | NA | #VALUE! |
| ENSMUSG00000041000.9 | 0.279766 | 1.808031 | 4.036658 | 0.447903 | 0.654223 | NA | #VALUE! |
| ENSMUSG00000041009.9 | 0 | NA | NA | NA | NA | NA | #VALUE! |
| ENSMUSG00000041014.18 | 0 | NA | NA | NA | NA | NA | #VALUE! |
| ENSMUSG00000041020.15 | 0 | NA | NA | NA | NA | NA | #VALUE! |
| ENSMUSG00000041046.8 | 0 | NA | NA | NA | NA | NA | #VALUE! |
| ENSMUSG00000041052.9 | 0 | NA | NA | NA | NA | NA | #VALUE! |
| ENSMUSG00000041062.8 | 0 | NA | NA | NA | NA | NA | #VALUE! |
| ENSMUSG00000041064.15 | 0 | NA | NA | NA | NA | NA | #VALUE! |
| ENSMUSG00000041068.7 | 0 | NA | NA | NA | NA | NA | #VALUE! |
| ENSMUSG00000041073.11 | 0 | NA | NA | NA | NA | NA | #VALUE! |

| | | | | | | | |
|---|---|---|---|---|---|---|---|
| ENSMUSG00000041078.8 | 6.22649 | 6.128273 | 3.913982 | 1.565739 | 0.11741 | NA | #VALUE! |
| ENSMUSG00000041096.14 | 6.386588 | -6.20479 | 3.628121 | -1.71019 | 0.08723 | NA | #VALUE! |
| ENSMUSG00000041112.18 | 5.831945 | -6.07186 | 3.801483 | -1.59723 | 0.110214 | NA | #VALUE! |
| ENSMUSG00000041117.9 | 0 | NA | NA | NA | NA | NA | #VALUE! |
| ENSMUSG00000041120.7 | 0 | NA | NA | NA | NA | NA | #VALUE! |
| ENSMUSG00000041134.16 | 0.735092 | -3.12016 | 3.965612 | -0.7868 | 0.431397 | NA | #VALUE! |
| ENSMUSG00000041138.17 | 0 | NA | NA | NA | NA | NA | #VALUE! |
| ENSMUSG00000041141.8 | 0 | NA | NA | NA | NA | NA | #VALUE! |
| ENSMUSG00000041144.13 | 0 | NA | NA | NA | NA | NA | #VALUE! |
| ENSMUSG00000041147.11 | 3.66691 | 2.417382 | 3.819649 | 0.632881 | 0.526812 | NA | #VALUE! |
| ENSMUSG00000041161.9 | 1.224665 | -3.81006 | 3.943408 | -0.96618 | 0.333952 | NA | #VALUE! |
| ENSMUSG00000041165.7 | 0 | NA | NA | NA | NA | NA | #VALUE! |
| ENSMUSG00000041187.17 | 1.029132 | -2.09486 | 3.95018 | -0.53032 | 0.595891 | NA | #VALUE! |
| ENSMUSG00000041189.10 | 0 | NA | NA | NA | NA | NA | #VALUE! |
| ENSMUSG00000041193.16 | 0 | NA | NA | NA | NA | NA | #VALUE! |
| ENSMUSG00000041199.4 | 0 | NA | NA | NA | NA | NA | #VALUE! |
| ENSMUSG00000041202.13 | 0 | NA | NA | NA | NA | NA | #VALUE! |
| ENSMUSG00000041205.9 | 0 | NA | NA | NA | NA | NA | #VALUE! |
| ENSMUSG00000041216.16 | 0 | NA | NA | NA | NA | NA | #VALUE! |
| ENSMUSG00000041219.15 | 4.305538 | -5.64245 | 3.91711 | -1.44046 | 0.149737 | NA | #VALUE! |
| ENSMUSG00000041245.14 | 0 | NA | NA | NA | NA | NA | #VALUE! |
| ENSMUSG00000041247.9 | 0 | NA | NA | NA | NA | NA | #VALUE! |
| ENSMUSG00000041248.10 | 0 | NA | NA | NA | NA | NA | #VALUE! |
| ENSMUSG00000041255.3 | 0 | NA | NA | NA | NA | NA | #VALUE! |
| ENSMUSG00000041287.6 | 0 | NA | NA | NA | NA | NA | #VALUE! |
| ENSMUSG00000041298.16 | 7.640079 | -0.79619 | 2.430434 | -0.32759 | 0.74322 | NA | #VALUE! |
| ENSMUSG00000041301.16 | 0 | NA | NA | NA | NA | NA | #VALUE! |
| ENSMUSG00000041308.7 | 9.682164 | -0.53707 | 2.412433 | -0.22262 | 0.823828 | NA | #VALUE! |
| ENSMUSG00000041309.18 | 0 | NA | NA | NA | NA | NA | #VALUE! |
| ENSMUSG00000041313.15 | 0 | NA | NA | NA | NA | NA | #VALUE! |
| ENSMUSG00000041323.7 | 4.694547 | -5.7577 | 3.916317 | -1.47018 | 0.141512 | NA | #VALUE! |
| ENSMUSG00000041329.14 | 0 | NA | NA | NA | NA | NA | #VALUE! |
| ENSMUSG00000041343.20 | 0 | NA | NA | NA | NA | NA | #VALUE! |
| ENSMUSG00000041346.12 | 1.032385 | -3.57787 | 3.949752 | -0.90585 | 0.365018 | NA | #VALUE! |
| ENSMUSG00000041347.6 | 0 | NA | NA | NA | NA | NA | #VALUE! |
| ENSMUSG00000041351.17 | 0.168284 | 1.027628 | 4.067126 | 0.252667 | 0.800526 | NA | #VALUE! |
| ENSMUSG00000041354.17 | 10.92989 | -1.22328 | 3.20321 | -0.38189 | NA | NA | #VALUE! |
| ENSMUSG00000041358.2 | 0 | NA | NA | NA | NA | NA | #VALUE! |
| ENSMUSG00000041359.15 | 0 | NA | NA | NA | NA | NA | #VALUE! |
| ENSMUSG00000041361.14 | 0 | NA | NA | NA | NA | NA | #VALUE! |
| ENSMUSG00000041372.11 | 0 | NA | NA | NA | NA | NA | #VALUE! |
| ENSMUSG00000041377.13 | 0 | NA | NA | NA | NA | NA | #VALUE! |
| ENSMUSG00000041378.3 | 0 | NA | NA | NA | NA | NA | #VALUE! |
| ENSMUSG00000041380.14 | 0 | NA | NA | NA | NA | NA | #VALUE! |

| | | | | | | | |
|---|---|---|---|---|---|---|---|
| ENSMUSG00000041399.4 | 0 | NA | NA | NA | NA | NA | #VALUE! |
| ENSMUSG00000041406.15 | 3.36567 | 5.240782 | 3.920098 | 1.336901 | 0.181255 | NA | #VALUE! |
| ENSMUSG00000041420.19 | 0 | NA | NA | NA | NA | NA | #VALUE! |
| ENSMUSG00000041423.17 | 0 | NA | NA | NA | NA | NA | #VALUE! |
| ENSMUSG00000041440.19 | 0 | NA | NA | NA | NA | NA | #VALUE! |
| ENSMUSG00000041449.17 | 0 | NA | NA | NA | NA | NA | #VALUE! |
| ENSMUSG00000041460.17 | 0.587367 | -2.74297 | 3.982932 | -0.68868 | 0.491024 | NA | #VALUE! |
| ENSMUSG00000041468.8 | 0 | NA | NA | NA | NA | NA | #VALUE! |
| ENSMUSG00000041476.13 | 0 | NA | NA | NA | NA | NA | #VALUE! |
| ENSMUSG00000041479.15 | 0 | NA | NA | NA | NA | NA | #VALUE! |
| ENSMUSG00000041482.18 | 4.694547 | -5.7577 | 3.916317 | -1.47018 | 0.141512 | NA | #VALUE! |
| ENSMUSG00000041488.17 | 6.290288 | 5.089151 | 3.334523 | 1.526201 | 0.12696 | NA | #VALUE! |
| ENSMUSG00000041491.9 | 7.71206 | -5.38471 | 3.267249 | -1.64809 | 0.099335 | NA | #VALUE! |
| ENSMUSG00000041498.14 | 0 | NA | NA | NA | NA | NA | #VALUE! |
| ENSMUSG00000041505.16 | 0 | NA | NA | NA | NA | NA | #VALUE! |
| ENSMUSG00000041506.16 | 6.416382 | 6.177056 | 3.728672 | 1.656637 | 0.097593 | NA | #VALUE! |
| ENSMUSG00000041512.4 | 0 | NA | NA | NA | NA | NA | #VALUE! |
| ENSMUSG00000041515.11 | 3.948756 | -4.38783 | 3.904634 | -1.12375 | 0.26112 | NA | #VALUE! |
| ENSMUSG00000041523.4 | 0 | NA | NA | NA | NA | NA | #VALUE! |
| ENSMUSG00000041534.10 | 0 | NA | NA | NA | NA | NA | #VALUE! |
| ENSMUSG00000041536.14 | 0 | NA | NA | NA | NA | NA | #VALUE! |
| ENSMUSG00000041538.17 | 0 | NA | NA | NA | NA | NA | #VALUE! |
| ENSMUSG00000041544.12 | 0 | NA | NA | NA | NA | NA | #VALUE! |
| ENSMUSG00000041550.5 | 0.279766 | 1.808031 | 4.036658 | 0.447903 | 0.654223 | NA | #VALUE! |
| ENSMUSG00000041552.8 | 0 | NA | NA | NA | NA | NA | #VALUE! |
| ENSMUSG00000041556.9 | 0 | NA | NA | NA | NA | NA | #VALUE! |
| ENSMUSG00000041559.8 | 0 | NA | NA | NA | NA | NA | #VALUE! |
| ENSMUSG00000041565.18 | 0.504851 | 2.502843 | 3.993993 | 0.626652 | 0.530887 | NA | #VALUE! |
| ENSMUSG00000041566.4 | 1.785223 | -4.3809 | 3.931494 | -1.11431 | 0.265146 | NA | #VALUE! |
| ENSMUSG00000041578.16 | 0 | NA | NA | NA | NA | NA | #VALUE! |
| ENSMUSG00000041583.9 | 0 | NA | NA | NA | NA | NA | #VALUE! |
| ENSMUSG00000041592.17 | 0 | NA | NA | NA | NA | NA | #VALUE! |
| ENSMUSG00000041596.13 | 0 | NA | NA | NA | NA | NA | #VALUE! |
| ENSMUSG00000041605.17 | 0 | NA | NA | NA | NA | NA | #VALUE! |
| ENSMUSG00000041607.18 | 0 | NA | NA | NA | NA | NA | #VALUE! |
| ENSMUSG00000041608.9 | 0 | NA | NA | NA | NA | NA | #VALUE! |
| ENSMUSG00000041609.17 | 4.111571 | -5.56681 | 3.917665 | -1.42095 | 0.155331 | NA | #VALUE! |
| ENSMUSG00000041616.10 | 0 | NA | NA | NA | NA | NA | #VALUE! |
| ENSMUSG00000041617.11 | 0 | NA | NA | NA | NA | NA | #VALUE! |
| ENSMUSG00000041620.9 | 0 | NA | NA | NA | NA | NA | #VALUE! |
| ENSMUSG00000041623.18 | 8.354017 | -1.65426 | 3.077644 | -0.53751 | 0.590917 | NA | #VALUE! |
| ENSMUSG00000041624.11 | 3.543697 | 5.324148 | 3.919351 | 1.358426 | 0.174329 | NA | #VALUE! |
| ENSMUSG00000041638.19 | 0.525066 | -2.64687 | 3.988095 | -0.66369 | 0.506887 | NA | #VALUE! |
| ENSMUSG00000041642.19 | 7.553671 | 6.411285 | 3.912697 | 1.638584 | NA | NA | #VALUE! |

| | | | | | | | |
|---|---|---|---|---|---|---|---|
| ENSMUSG00000041644.15 | 0 | NA | NA | NA | NA | NA | #VALUE! |
| ENSMUSG00000041649.14 | 0 | NA | NA | NA | NA | NA | #VALUE! |
| ENSMUSG00000041653.6 | 0.737927 | 3.007825 | 3.968641 | 0.757898 | 0.448512 | NA | #VALUE! |
| ENSMUSG00000041658.13 | 0 | NA | NA | NA | NA | NA | #VALUE! |
| ENSMUSG00000041669.16 | 0 | NA | NA | NA | NA | NA | #VALUE! |
| ENSMUSG00000041670.20 | 0 | NA | NA | NA | NA | NA | #VALUE! |
| ENSMUSG00000041673.13 | 14.91742 | 1.321047 | 2.970481 | 0.444725 | NA | NA | #VALUE! |
| ENSMUSG00000041674.17 | 0 | NA | NA | NA | NA | NA | #VALUE! |
| ENSMUSG00000041681.3 | 0 | NA | NA | NA | NA | NA | #VALUE! |
| ENSMUSG00000041695.3 | 0 | NA | NA | NA | NA | NA | #VALUE! |
| ENSMUSG00000041696.15 | 0 | NA | NA | NA | NA | NA | #VALUE! |
| ENSMUSG00000041700.10 | 0 | NA | NA | NA | NA | NA | #VALUE! |
| ENSMUSG00000041703.8 | 0 | NA | NA | NA | NA | NA | #VALUE! |
| ENSMUSG00000041707.9 | 0 | NA | NA | NA | NA | NA | #VALUE! |
| ENSMUSG00000041708.13 | 0 | NA | NA | NA | NA | NA | #VALUE! |
| ENSMUSG00000041710.5 | 0 | NA | NA | NA | NA | NA | #VALUE! |
| ENSMUSG00000041722.8 | 0 | NA | NA | NA | NA | NA | #VALUE! |
| ENSMUSG00000041729.16 | 0 | NA | NA | NA | NA | NA | #VALUE! |
| ENSMUSG00000041730.15 | 0 | NA | NA | NA | NA | NA | #VALUE! |
| ENSMUSG00000041731.14 | 2.779429 | 4.97543 | 3.922779 | 1.268343 | 0.204675 | NA | #VALUE! |
| ENSMUSG00000041734.16 | 0 | NA | NA | NA | NA | NA | #VALUE! |
| ENSMUSG00000041735.9 | 0 | NA | NA | NA | NA | NA | #VALUE! |
| ENSMUSG00000041737.9 | 0.652786 | 2.919659 | 3.972474 | 0.734972 | 0.462356 | NA | #VALUE! |
| ENSMUSG00000041750.14 | 0 | NA | NA | NA | NA | NA | #VALUE! |
| ENSMUSG00000041754.6 | 0 | NA | NA | NA | NA | NA | #VALUE! |
| ENSMUSG00000041771.15 | 0 | NA | NA | NA | NA | NA | #VALUE! |
| ENSMUSG00000041773.9 | 8.452565 | 2.358857 | 3.436075 | 0.686497 | NA | NA | #VALUE! |
| ENSMUSG00000041774.17 | 0 | NA | NA | NA | NA | NA | #VALUE! |
| ENSMUSG00000041779.6 | 36.45534 | -0.72704 | 1.549356 | -0.46926 | NA | NA | #VALUE! |
| ENSMUSG00000041782.15 | 3.77488 | 1.359927 | 3.785538 | 0.359243 | 0.719413 | NA | #VALUE! |
| ENSMUSG00000041789.8 | 0 | NA | NA | NA | NA | NA | #VALUE! |
| ENSMUSG00000041791.7 | 0 | NA | NA | NA | NA | NA | #VALUE! |
| ENSMUSG00000041794.14 | 0 | NA | NA | NA | NA | NA | #VALUE! |
| ENSMUSG00000041797.8 | 3.815814 | 2.966902 | 3.84372 | 0.771883 | 0.440184 | NA | #VALUE! |
| ENSMUSG00000041801.6 | 0 | NA | NA | NA | NA | NA | #VALUE! |
| ENSMUSG00000041805.4 | 0 | NA | NA | NA | NA | NA | #VALUE! |
| ENSMUSG00000041809.6 | 0 | NA | NA | NA | NA | NA | #VALUE! |
| ENSMUSG00000041817.15 | 0.373021 | 2.140748 | 4.018182 | 0.532765 | 0.594196 | NA | #VALUE! |
| ENSMUSG00000041831.18 | 0 | NA | NA | NA | NA | NA | #VALUE! |
| ENSMUSG00000041836.11 | 5.191867 | -3.28612 | 3.829328 | -0.85815 | 0.390812 | NA | #VALUE! |
| ENSMUSG00000041842.16 | 0.840105 | -3.30885 | 3.958472 | -0.83589 | 0.403216 | NA | #VALUE! |
| ENSMUSG00000041857.11 | 0 | NA | NA | NA | NA | NA | #VALUE! |
| ENSMUSG00000041859.11 | 4.455795 | 4.682139 | 3.765274 | 1.243506 | 0.213682 | NA | #VALUE! |
| ENSMUSG00000041872.10 | 0 | NA | NA | NA | NA | NA | #VALUE! |

| | | | | | | | |
|---|---|---|---|---|---|---|---|
| ENSMUSG00000041878.4 | 0 | NA | NA | NA | NA | NA | #VALUE! |
| ENSMUSG00000041885.6 | 0 | NA | NA | NA | NA | NA | #VALUE! |
| ENSMUSG00000041886.8 | 0 | NA | NA | NA | NA | NA | #VALUE! |
| ENSMUSG00000041889.8 | 0 | NA | NA | NA | NA | NA | #VALUE! |
| ENSMUSG00000041890.18 | 6.825491 | 0.68557 | 2.561876 | 0.267605 | 0.789004 | NA | #VALUE! |
| ENSMUSG00000041907.10 | 0 | NA | NA | NA | NA | NA | #VALUE! |
| ENSMUSG00000041911.4 | 0 | NA | NA | NA | NA | NA | #VALUE! |
| ENSMUSG00000041921.17 | 8.117192 | 6.506594 | 3.865977 | 1.68304 | 0.092367 | NA | #VALUE! |
| ENSMUSG00000041923.16 | 0 | NA | NA | NA | NA | NA | #VALUE! |
| ENSMUSG00000041945.13 | 1.836997 | -4.39907 | 3.931186 | -1.11902 | 0.263133 | NA | #VALUE! |
| ENSMUSG00000041949.7 | 0 | NA | NA | NA | NA | NA | #VALUE! |
| ENSMUSG00000041954.19 | 0 | NA | NA | NA | NA | NA | #VALUE! |
| ENSMUSG00000041984.3 | 0 | NA | NA | NA | NA | NA | #VALUE! |
| ENSMUSG00000041986.17 | 0 | NA | NA | NA | NA | NA | #VALUE! |
| ENSMUSG00000041991.19 | 0 | NA | NA | NA | NA | NA | #VALUE! |
| ENSMUSG00000041995.9 | 22.88656 | 21.40094 | 2.847188 | 7.516517 | NA | NA | #VALUE! |
| ENSMUSG00000042002.12 | 0 | NA | NA | NA | NA | NA | #VALUE! |
| ENSMUSG00000042029.8 | 0.336567 | 1.944925 | 4.029936 | 0.482619 | 0.629366 | NA | #VALUE! |
| ENSMUSG00000042031.5 | 0 | NA | NA | NA | NA | NA | #VALUE! |
| ENSMUSG00000042035.12 | 0.420053 | -2.33493 | 4.007343 | -0.58266 | 0.56012 | NA | #VALUE! |
| ENSMUSG00000042045.7 | 0 | NA | NA | NA | NA | NA | #VALUE! |
| ENSMUSG00000042064.14 | 0 | NA | NA | NA | NA | NA | #VALUE! |
| ENSMUSG00000042066.17 | 0.630079 | -2.90291 | 3.975047 | -0.73028 | 0.465217 | NA | #VALUE! |
| ENSMUSG00000042078.15 | 0 | NA | NA | NA | NA | NA | #VALUE! |
| ENSMUSG00000042087.6 | 1.682835 | 4.240781 | 3.933339 | 1.078163 | 0.280961 | NA | #VALUE! |
| ENSMUSG00000042092.6 | 0 | NA | NA | NA | NA | NA | #VALUE! |
| ENSMUSG00000042096.16 | 0 | NA | NA | NA | NA | NA | #VALUE! |
| ENSMUSG00000042097.18 | 2.705731 | 4.913666 | 3.923477 | 1.252375 | 0.210433 | NA | #VALUE! |
| ENSMUSG00000042099.17 | 6.675463 | 0.851391 | 3.302058 | 0.257837 | 0.796533 | NA | #VALUE! |
| ENSMUSG00000042104.19 | 0 | NA | NA | NA | NA | NA | #VALUE! |
| ENSMUSG00000042106.6 | 3.255407 | -5.24108 | 3.920419 | -1.33687 | 0.181266 | NA | #VALUE! |
| ENSMUSG00000042109.9 | 0 | NA | NA | NA | NA | NA | #VALUE! |
| ENSMUSG00000042115.5 | 2.611145 | 4.886731 | 3.92379 | 1.245411 | 0.212981 | NA | #VALUE! |
| ENSMUSG00000042116.4 | 0.105013 | -0.97324 | 4.076293 | -0.23876 | 0.811295 | NA | #VALUE! |
| ENSMUSG00000042124.6 | 0 | NA | NA | NA | NA | NA | #VALUE! |
| ENSMUSG00000042129.9 | 0.491951 | 2.402185 | 4.000147 | 0.600524 | 0.548157 | NA | #VALUE! |
| ENSMUSG00000042155.4 | 0 | NA | NA | NA | NA | NA | #VALUE! |
| ENSMUSG00000042156.17 | 0.18651 | 1.401769 | 4.056354 | 0.345574 | 0.729663 | NA | #VALUE! |
| ENSMUSG00000042157.3 | 0 | NA | NA | NA | NA | NA | #VALUE! |
| ENSMUSG00000042179.7 | 0 | NA | NA | NA | NA | NA | #VALUE! |
| ENSMUSG00000042184.12 | 0 | NA | NA | NA | NA | NA | #VALUE! |
| ENSMUSG00000042189.6 | 0 | NA | NA | NA | NA | NA | #VALUE! |
| ENSMUSG00000042190.13 | 0.315039 | -1.93549 | 4.038621 | -0.47925 | 0.631763 | NA | #VALUE! |
| ENSMUSG00000042195.10 | 0 | NA | NA | NA | NA | NA | #VALUE! |

| | | | | | | | |
|---|---|---|---|---|---|---|---|
| ENSMUSG00000042200.4 | 0 | NA | NA | NA | NA | NA | #VALUE! |
| ENSMUSG00000042203.9 | 5.270247 | -5.93136 | 3.911985 | -1.5162 | 0.129468 | NA | #VALUE! |
| ENSMUSG00000042210.18 | 39.19938 | -0.97675 | 2.043321 | -0.47802 | NA | NA | #VALUE! |
| ENSMUSG00000042212.4 | 0 | NA | NA | NA | NA | NA | #VALUE! |
| ENSMUSG00000042215.9 | 9.026598 | 1.666364 | 2.362058 | 0.705471 | 0.480517 | NA | #VALUE! |
| ENSMUSG00000042216.14 | 0 | NA | NA | NA | NA | NA | #VALUE! |
| ENSMUSG00000042219.7 | 0 | NA | NA | NA | NA | NA | #VALUE! |
| ENSMUSG00000042240.15 | 0 | NA | NA | NA | NA | NA | #VALUE! |
| ENSMUSG00000042243.11 | 0 | NA | NA | NA | NA | NA | #VALUE! |
| ENSMUSG00000042244.5 | 0 | NA | NA | NA | NA | NA | #VALUE! |
| ENSMUSG00000042250.14 | 0 | NA | NA | NA | NA | NA | #VALUE! |
| ENSMUSG00000042254.15 | 0 | NA | NA | NA | NA | NA | #VALUE! |
| ENSMUSG00000042256.6 | 0 | NA | NA | NA | NA | NA | #VALUE! |
| ENSMUSG00000042258.14 | 0 | NA | NA | NA | NA | NA | #VALUE! |
| ENSMUSG00000042262.6 | 0 | NA | NA | NA | NA | NA | #VALUE! |
| ENSMUSG00000042265.14 | 0 | NA | NA | NA | NA | NA | #VALUE! |
| ENSMUSG00000042268.11 | 0 | NA | NA | NA | NA | NA | #VALUE! |
| ENSMUSG00000042269.16 | 0 | NA | NA | NA | NA | NA | #VALUE! |
| ENSMUSG00000042272.18 | 1.842912 | -4.40636 | 3.931064 | -1.12091 | 0.262327 | NA | #VALUE! |
| ENSMUSG00000042275.10 | 0 | NA | NA | NA | NA | NA | #VALUE! |
| ENSMUSG00000042279.14 | 0 | NA | NA | NA | NA | NA | #VALUE! |
| ENSMUSG00000042282.5 | 0 | NA | NA | NA | NA | NA | #VALUE! |
| ENSMUSG00000042289.12 | 0 | NA | NA | NA | NA | NA | #VALUE! |
| ENSMUSG00000042292.19 | 2.122016 | 0.719358 | 3.81732 | 0.188446 | 0.850527 | NA | #VALUE! |
| ENSMUSG00000042303.20 | 3.524204 | -5.34395 | 3.919482 | -1.36343 | 0.172746 | NA | #VALUE! |
| ENSMUSG00000042306.12 | 0 | NA | NA | NA | NA | NA | #VALUE! |
| ENSMUSG00000042308.15 | 0 | NA | NA | NA | NA | NA | #VALUE! |
| ENSMUSG00000042320.18 | 0 | NA | NA | NA | NA | NA | #VALUE! |
| ENSMUSG00000042331.14 | 9.541577 | -1.75484 | 2.951644 | -0.59453 | 0.552157 | NA | #VALUE! |
| ENSMUSG00000042333.17 | 0 | NA | NA | NA | NA | NA | #VALUE! |
| ENSMUSG00000042345.16 | 0 | NA | NA | NA | NA | NA | #VALUE! |
| ENSMUSG00000042351.10 | 0 | NA | NA | NA | NA | NA | #VALUE! |
| ENSMUSG00000042353.10 | 6.069462 | -6.1296 | 3.91415 | -1.56601 | 0.117346 | NA | #VALUE! |
| ENSMUSG00000042357.3 | 0 | NA | NA | NA | NA | NA | #VALUE! |
| ENSMUSG00000042359.19 | 3.730208 | 5.397701 | 3.918728 | 1.377412 | 0.168385 | NA | #VALUE! |
| ENSMUSG00000042360.13 | 0 | NA | NA | NA | NA | NA | #VALUE! |
| ENSMUSG00000042367.13 | 0 | NA | NA | NA | NA | NA | #VALUE! |
| ENSMUSG00000042369.9 | 0 | NA | NA | NA | NA | NA | #VALUE! |
| ENSMUSG00000042371.13 | 0 | NA | NA | NA | NA | NA | #VALUE! |
| ENSMUSG00000042372.6 | 0 | NA | NA | NA | NA | NA | #VALUE! |
| ENSMUSG00000042379.9 | 4.012094 | 5.500544 | 3.917907 | 1.40395 | 0.160334 | NA | #VALUE! |
| ENSMUSG00000042385.15 | 0 | NA | NA | NA | NA | NA | #VALUE! |
| ENSMUSG00000042386.8 | 0 | NA | NA | NA | NA | NA | #VALUE! |
| ENSMUSG00000042388.15 | 0 | NA | NA | NA | NA | NA | #VALUE! |

| | | | | | | | |
|---|---|---|---|---|---|---|---|
| ENSMUSG00000042401.9 | 0 | NA | NA | NA | NA | NA | #VALUE! |
| ENSMUSG00000042404.17 | 5.52905 | -3.43428 | 3.496207 | -0.98229 | 0.325958 | NA | #VALUE! |
| ENSMUSG00000042408.14 | 0.195789 | -1.05581 | 4.073302 | -0.2592 | 0.795479 | NA | #VALUE! |
| ENSMUSG00000042414.8 | 0 | NA | NA | NA | NA | NA | #VALUE! |
| ENSMUSG00000042417.6 | 0 | NA | NA | NA | NA | NA | #VALUE! |
| ENSMUSG00000042425.19 | 0 | NA | NA | NA | NA | NA | #VALUE! |
| ENSMUSG00000042428.7 | 4.490437 | -5.69342 | 3.916751 | -1.45361 | 0.146055 | NA | #VALUE! |
| ENSMUSG00000042433.3 | 0 | NA | NA | NA | NA | NA | #VALUE! |
| ENSMUSG00000042436.13 | 0 | NA | NA | NA | NA | NA | #VALUE! |
| ENSMUSG00000042439.14 | 6.941816 | -3.01569 | 3.652881 | -0.82556 | 0.409052 | NA | #VALUE! |
| ENSMUSG00000042448.6 | 0 | NA | NA | NA | NA | NA | #VALUE! |
| ENSMUSG00000042451.13 | 0 | NA | NA | NA | NA | NA | #VALUE! |
| ENSMUSG00000042459.9 | 0 | NA | NA | NA | NA | NA | #VALUE! |
| ENSMUSG00000042473.9 | 9.254055 | 0.552009 | 3.021893 | 0.18267 | 0.855057 | NA | #VALUE! |
| ENSMUSG00000042474.7 | 0 | NA | NA | NA | NA | NA | #VALUE! |
| ENSMUSG00000042477.8 | 0 | NA | NA | NA | NA | NA | #VALUE! |
| ENSMUSG00000042489.16 | 0 | NA | NA | NA | NA | NA | #VALUE! |
| ENSMUSG00000042492.13 | 3.591255 | 1.055655 | 3.784102 | 0.278971 | 0.780267 | NA | #VALUE! |
| ENSMUSG00000042496.19 | 3.232853 | -1.26493 | 3.794005 | -0.3334 | 0.738831 | NA | #VALUE! |
| ENSMUSG00000042498.17 | 0 | NA | NA | NA | NA | NA | #VALUE! |
| ENSMUSG00000042499.14 | 0 | NA | NA | NA | NA | NA | #VALUE! |
| ENSMUSG00000042500.12 | 2.704401 | 4.936946 | 3.92321 | 1.258394 | 0.208249 | NA | #VALUE! |
| ENSMUSG00000042501.13 | 0 | NA | NA | NA | NA | NA | #VALUE! |
| ENSMUSG00000042502.11 | 48.22377 | 1.366617 | 1.48179 | 0.922274 | NA | NA | #VALUE! |
| ENSMUSG00000042507.16 | 5.800084 | -2.16716 | 3.518232 | -0.61598 | 0.537909 | NA | #VALUE! |
| ENSMUSG00000042510.8 | 0 | NA | NA | NA | NA | NA | #VALUE! |
| ENSMUSG00000042514.12 | 0 | NA | NA | NA | NA | NA | #VALUE! |
| ENSMUSG00000042515.14 | 0.245976 | 1.027628 | 4.067126 | 0.252667 | 0.800526 | NA | #VALUE! |
| ENSMUSG00000042523.13 | 5.903413 | 6.04575 | 3.914406 | 1.544487 | 0.12247 | NA | #VALUE! |
| ENSMUSG00000042525.6 | 0 | NA | NA | NA | NA | NA | #VALUE! |
| ENSMUSG00000042529.15 | 0 | NA | NA | NA | NA | NA | #VALUE! |
| ENSMUSG00000042532.15 | 0 | NA | NA | NA | NA | NA | #VALUE! |
| ENSMUSG00000042535.10 | 2.763866 | 4.966032 | 3.922883 | 1.265914 | 0.205544 | NA | #VALUE! |
| ENSMUSG00000042540.13 | 0 | NA | NA | NA | NA | NA | #VALUE! |
| ENSMUSG00000042549.11 | 0 | NA | NA | NA | NA | NA | #VALUE! |
| ENSMUSG00000042554.11 | 0 | NA | NA | NA | NA | NA | #VALUE! |
| ENSMUSG00000042558.17 | 0 | NA | NA | NA | NA | NA | #VALUE! |
| ENSMUSG00000042564.14 | 0 | NA | NA | NA | NA | NA | #VALUE! |
| ENSMUSG00000042567.21 | 0 | NA | NA | NA | NA | NA | #VALUE! |
| ENSMUSG00000042570.15 | 0.466276 | 2.449889 | 3.997179 | 0.612904 | 0.53994 | NA | #VALUE! |
| ENSMUSG00000042579.3 | 5.408295 | 4.850981 | 3.569284 | 1.359091 | 0.174118 | NA | #VALUE! |
| ENSMUSG00000042581.15 | 0 | NA | NA | NA | NA | NA | #VALUE! |
| ENSMUSG00000042589.19 | 0 | NA | NA | NA | NA | NA | #VALUE! |
| ENSMUSG00000042596.8 | 0 | NA | NA | NA | NA | NA | #VALUE! |

| | | | | | | | |
|---|---|---|---|---|---|---|---|
| ENSMUSG00000042604.6 | 0 | NA | NA | NA | NA | NA | #VALUE! |
| ENSMUSG00000042606.10 | 7.387084 | -0.79846 | 3.660193 | -0.21815 | 0.827314 | NA | #VALUE! |
| ENSMUSG00000042607.16 | 0 | NA | NA | NA | NA | NA | #VALUE! |
| ENSMUSG00000042616.9 | 0 | NA | NA | NA | NA | NA | #VALUE! |
| ENSMUSG00000042622.15 | 7.897397 | -6.51141 | 3.358124 | -1.939 | 0.052501 | NA | #VALUE! |
| ENSMUSG00000042631.8 | 0 | NA | NA | NA | NA | NA | #VALUE! |
| ENSMUSG00000042638.15 | 0 | NA | NA | NA | NA | NA | #VALUE! |
| ENSMUSG00000042641.20 | 0 | NA | NA | NA | NA | NA | #VALUE! |
| ENSMUSG00000042644.10 | 0 | NA | NA | NA | NA | NA | #VALUE! |
| ENSMUSG00000042655.6 | 0 | NA | NA | NA | NA | NA | #VALUE! |
| ENSMUSG00000042662.17 | 0 | NA | NA | NA | NA | NA | #VALUE! |
| ENSMUSG00000042671.13 | 0 | NA | NA | NA | NA | NA | #VALUE! |
| ENSMUSG00000042672.16 | 0 | NA | NA | NA | NA | NA | #VALUE! |
| ENSMUSG00000042678.18 | 0 | NA | NA | NA | NA | NA | #VALUE! |
| ENSMUSG00000042684.9 | 0 | NA | NA | NA | NA | NA | #VALUE! |
| ENSMUSG00000042686.6 | 0 | NA | NA | NA | NA | NA | #VALUE! |
| ENSMUSG00000042694.18 | 1.683045 | -0.4276 | 3.837329 | -0.11143 | 0.911273 | NA | #VALUE! |
| ENSMUSG00000042707.7 | 0 | NA | NA | NA | NA | NA | #VALUE! |
| ENSMUSG00000042708.13 | 0 | NA | NA | NA | NA | NA | #VALUE! |
| ENSMUSG00000042717.6 | 0 | NA | NA | NA | NA | NA | #VALUE! |
| ENSMUSG00000042724.9 | 0 | NA | NA | NA | NA | NA | #VALUE! |
| ENSMUSG00000042726.15 | 8.868771 | -3.25256 | 3.421695 | -0.95057 | 0.341822 | NA | #VALUE! |
| ENSMUSG00000042734.7 | 0 | NA | NA | NA | NA | NA | #VALUE! |
| ENSMUSG00000042743.14 | 0 | NA | NA | NA | NA | NA | #VALUE! |
| ENSMUSG00000042750.8 | 0 | NA | NA | NA | NA | NA | #VALUE! |
| ENSMUSG00000042751.15 | 0 | NA | NA | NA | NA | NA | #VALUE! |
| ENSMUSG00000042757.17 | 4.725591 | -5.7762 | 3.916196 | -1.47495 | 0.140225 | NA | #VALUE! |
| ENSMUSG00000042759.13 | 0 | NA | NA | NA | NA | NA | #VALUE! |
| ENSMUSG00000042761.15 | 0 | NA | NA | NA | NA | NA | #VALUE! |
| ENSMUSG00000042763.10 | 0 | NA | NA | NA | NA | NA | #VALUE! |
| ENSMUSG00000042766.18 | 0 | NA | NA | NA | NA | NA | #VALUE! |
| ENSMUSG00000042772.16 | 9.209182 | -3.8421 | 3.049188 | -1.26004 | NA | NA | #VALUE! |
| ENSMUSG00000042774.9 | 0 | NA | NA | NA | NA | NA | #VALUE! |
| ENSMUSG00000042784.10 | 0 | NA | NA | NA | NA | NA | #VALUE! |
| ENSMUSG00000042787.16 | 0.420053 | -2.33493 | 4.007343 | -0.58266 | 0.56012 | NA | #VALUE! |
| ENSMUSG00000042788.13 | 0 | NA | NA | NA | NA | NA | #VALUE! |
| ENSMUSG00000042793.14 | 0 | NA | NA | NA | NA | NA | #VALUE! |
| ENSMUSG00000042796.3 | 0 | NA | NA | NA | NA | NA | #VALUE! |
| ENSMUSG00000042800.6 | 0 | NA | NA | NA | NA | NA | #VALUE! |
| ENSMUSG00000042801.6 | 0 | NA | NA | NA | NA | NA | #VALUE! |
| ENSMUSG00000042804.14 | 0 | NA | NA | NA | NA | NA | #VALUE! |
| ENSMUSG00000042808.11 | 0 | NA | NA | NA | NA | NA | #VALUE! |
| ENSMUSG00000042816.5 | 0 | NA | NA | NA | NA | NA | #VALUE! |
| ENSMUSG00000042817.16 | 0 | NA | NA | NA | NA | NA | #VALUE! |

| | | | | | | | |
|---|---|---|---|---|---|---|---|
| ENSMUSG00000042821.8 | 1.260158 | -3.88425 | 3.941584 | -0.98545 | 0.324401 | NA | #VALUE! |
| ENSMUSG00000042826.14 | 0 | NA | NA | NA | NA | NA | #VALUE! |
| ENSMUSG00000042828.13 | 0 | NA | NA | NA | NA | NA | #VALUE! |
| ENSMUSG00000042845.8 | 6.589599 | -3.64919 | 3.746823 | -0.97394 | 0.330085 | NA | #VALUE! |
| ENSMUSG00000042846.9 | 0 | NA | NA | NA | NA | NA | #VALUE! |
| ENSMUSG00000042848.5 | 0 | NA | NA | NA | NA | NA | #VALUE! |
| ENSMUSG00000042849.12 | 0 | NA | NA | NA | NA | NA | #VALUE! |
| ENSMUSG00000042851.18 | 5.616003 | -6.01693 | 3.914749 | -1.53699 | 0.124296 | NA | #VALUE! |
| ENSMUSG00000042854.10 | 0.501477 | -1.29375 | 4.008705 | -0.32274 | 0.746896 | NA | #VALUE! |
| ENSMUSG00000042857.4 | 4.515565 | -5.71088 | 3.916632 | -1.45811 | 0.14481 | NA | #VALUE! |
| ENSMUSG00000042861.8 | 0 | NA | NA | NA | NA | NA | #VALUE! |
| ENSMUSG00000042863.6 | 0 | NA | NA | NA | NA | NA | #VALUE! |
| ENSMUSG00000042869.3 | 0 | NA | NA | NA | NA | NA | #VALUE! |
| ENSMUSG00000042870.16 | 0 | NA | NA | NA | NA | NA | #VALUE! |
| ENSMUSG00000042873.12 | 0 | NA | NA | NA | NA | NA | #VALUE! |
| ENSMUSG00000042874.3 | 0 | NA | NA | NA | NA | NA | #VALUE! |
| ENSMUSG00000042888.11 | 0 | NA | NA | NA | NA | NA | #VALUE! |
| ENSMUSG00000042894.12 | 0 | NA | NA | NA | NA | NA | #VALUE! |
| ENSMUSG00000042895.7 | 0.297366 | -0.14751 | 4.065279 | -0.03629 | 0.971054 | NA | #VALUE! |
| ENSMUSG00000042903.9 | 3.259345 | 1.745807 | 3.804954 | 0.458825 | 0.64636 | NA | #VALUE! |
| ENSMUSG00000042909.5 | 0 | NA | NA | NA | NA | NA | #VALUE! |
| ENSMUSG00000042918.18 | 0 | NA | NA | NA | NA | NA | #VALUE! |
| ENSMUSG00000042938.9 | 0 | NA | NA | NA | NA | NA | #VALUE! |
| ENSMUSG00000042940.9 | 0 | NA | NA | NA | NA | NA | #VALUE! |
| ENSMUSG00000042942.19 | 0 | NA | NA | NA | NA | NA | #VALUE! |
| ENSMUSG00000042961.14 | 0 | NA | NA | NA | NA | NA | #VALUE! |
| ENSMUSG00000042976.7 | 0 | NA | NA | NA | NA | NA | #VALUE! |
| ENSMUSG00000042985.8 | 2.730341 | -4.98888 | 3.923018 | -1.27169 | 0.203482 | NA | #VALUE! |
| ENSMUSG00000042992.16 | 4.91951 | 5.781601 | 3.915938 | 1.476428 | 0.139829 | NA | #VALUE! |
| ENSMUSG00000042993.7 | 0 | NA | NA | NA | NA | NA | #VALUE! |
| ENSMUSG00000043003.15 | 0 | NA | NA | NA | NA | NA | #VALUE! |
| ENSMUSG00000043004.14 | 2.625328 | -4.93268 | 3.923662 | -1.25716 | 0.208695 | NA | #VALUE! |
| ENSMUSG00000043008.10 | 4.476249 | 5.659319 | 3.916748 | 1.444902 | 0.148485 | NA | #VALUE! |
| ENSMUSG00000043015.16 | 2.857551 | -5.03942 | 3.922461 | -1.28476 | 0.198876 | NA | #VALUE! |
| ENSMUSG00000043017.10 | 0 | NA | NA | NA | NA | NA | #VALUE! |
| ENSMUSG00000043020.14 | 0 | NA | NA | NA | NA | NA | #VALUE! |
| ENSMUSG00000043024.7 | 0 | NA | NA | NA | NA | NA | #VALUE! |
| ENSMUSG00000043029.3 | 0 | NA | NA | NA | NA | NA | #VALUE! |
| ENSMUSG00000043036.15 | 0 | NA | NA | NA | NA | NA | #VALUE! |
| ENSMUSG00000043050.9 | 0 | NA | NA | NA | NA | NA | #VALUE! |
| ENSMUSG00000043051.18 | 0 | NA | NA | NA | NA | NA | #VALUE! |
| ENSMUSG00000043060.9 | 0 | NA | NA | NA | NA | NA | #VALUE! |
| ENSMUSG00000043065.13 | 2.781426 | 4.964897 | 3.922896 | 1.26562 | 0.205649 | NA | #VALUE! |
| ENSMUSG00000043066.6 | 0 | NA | NA | NA | NA | NA | #VALUE! |

| | | | | | | | |
|---|---|---|---|---|---|---|---|
| ENSMUSG00000043073.11 | 0 | NA | NA | NA | NA | NA | #VALUE! |
| ENSMUSG00000043079.18 | 4.385419 | -0.42798 | 3.768156 | -0.11358 | 0.909572 | NA | #VALUE! |
| ENSMUSG00000043085.15 | 3.150223 | 4.166957 | 3.917109 | 1.063784 | 0.287427 | NA | #VALUE! |
| ENSMUSG00000043087.6 | 0 | NA | NA | NA | NA | NA | #VALUE! |
| ENSMUSG00000043088.17 | 0 | NA | NA | NA | NA | NA | #VALUE! |
| ENSMUSG00000043089.8 | 0 | NA | NA | NA | NA | NA | #VALUE! |
| ENSMUSG00000043099.6 | 0.105013 | -0.97324 | 4.076293 | -0.23876 | 0.811295 | NA | #VALUE! |
| ENSMUSG00000043102.3 | 0 | NA | NA | NA | NA | NA | #VALUE! |
| ENSMUSG00000043110.3 | 0 | NA | NA | NA | NA | NA | #VALUE! |
| ENSMUSG00000043119.6 | 0 | NA | NA | NA | NA | NA | #VALUE! |
| ENSMUSG00000043122.8 | 0 | NA | NA | NA | NA | NA | #VALUE! |
| ENSMUSG00000043126.6 | 0 | NA | NA | NA | NA | NA | #VALUE! |
| ENSMUSG00000043144.7 | 0 | NA | NA | NA | NA | NA | #VALUE! |
| ENSMUSG00000043145.8 | 0 | NA | NA | NA | NA | NA | #VALUE! |
| ENSMUSG00000043153.7 | 31.04615 | -0.90868 | 2.294463 | -0.39603 | NA | NA | #VALUE! |
| ENSMUSG00000043155.5 | 1.475853 | 4.029233 | 3.937502 | 1.023297 | 0.306167 | NA | #VALUE! |
| ENSMUSG00000043157.9 | 1.514552 | 4.088766 | 3.936269 | 1.038742 | 0.298925 | NA | #VALUE! |
| ENSMUSG00000043164.4 | 0 | NA | NA | NA | NA | NA | #VALUE! |
| ENSMUSG00000043165.8 | 2.019402 | 4.503826 | 3.928936 | 1.146322 | 0.251662 | NA | #VALUE! |
| ENSMUSG00000043168.13 | 0 | NA | NA | NA | NA | NA | #VALUE! |
| ENSMUSG00000043192.5 | 0 | NA | NA | NA | NA | NA | #VALUE! |
| ENSMUSG00000043219.10 | 0 | NA | NA | NA | NA | NA | #VALUE! |
| ENSMUSG00000043223.8 | 0.289044 | -0.14751 | 4.065279 | -0.03629 | 0.971054 | NA | #VALUE! |
| ENSMUSG00000043226.6 | 0 | NA | NA | NA | NA | NA | #VALUE! |
| ENSMUSG00000043230.3 | 0 | NA | NA | NA | NA | NA | #VALUE! |
| ENSMUSG00000043243.16 | 0 | NA | NA | NA | NA | NA | #VALUE! |
| ENSMUSG00000043251.7 | 0 | NA | NA | NA | NA | NA | #VALUE! |
| ENSMUSG00000043257.16 | 3.860158 | 2.438082 | 3.815944 | 0.63892 | 0.522875 | NA | #VALUE! |
| ENSMUSG00000043259.16 | 0 | NA | NA | NA | NA | NA | #VALUE! |
| ENSMUSG00000043263.14 | 0.105013 | -0.97324 | 4.076293 | -0.23876 | 0.811295 | NA | #VALUE! |
| ENSMUSG00000043267.4 | 0 | NA | NA | NA | NA | NA | #VALUE! |
| ENSMUSG00000043274.5 | 0 | NA | NA | NA | NA | NA | #VALUE! |
| ENSMUSG00000043282.4 | 0 | NA | NA | NA | NA | NA | #VALUE! |
| ENSMUSG00000043286.13 | 0 | NA | NA | NA | NA | NA | #VALUE! |
| ENSMUSG00000043289.13 | 0 | NA | NA | NA | NA | NA | #VALUE! |
| ENSMUSG00000043290.7 | 0 | NA | NA | NA | NA | NA | #VALUE! |
| ENSMUSG00000043295.7 | 0.44158 | 1.011896 | 4.025703 | 0.251359 | 0.801537 | NA | #VALUE! |
| ENSMUSG00000043298.9 | 0 | NA | NA | NA | NA | NA | #VALUE! |
| ENSMUSG00000043300.3 | 0 | NA | NA | NA | NA | NA | #VALUE! |
| ENSMUSG00000043301.17 | 0 | NA | NA | NA | NA | NA | #VALUE! |
| ENSMUSG00000043308.3 | 0 | NA | NA | NA | NA | NA | #VALUE! |
| ENSMUSG00000043310.5 | 0 | NA | NA | NA | NA | NA | #VALUE! |
| ENSMUSG00000043311.8 | 24.81327 | -1.27134 | 2.19402 | -0.57946 | NA | NA | #VALUE! |
| ENSMUSG00000043312.7 | 0 | NA | NA | NA | NA | NA | #VALUE! |

| | | | | | | | |
|---|---|---|---|---|---|---|---|
| ENSMUSG00000043313.7 | 0 | NA | NA | NA | NA | NA | #VALUE! |
| ENSMUSG00000043314.7 | 0 | NA | NA | NA | NA | NA | #VALUE! |
| ENSMUSG00000043319.6 | 0 | NA | NA | NA | NA | NA | #VALUE! |
| ENSMUSG00000043329.5 | 0 | NA | NA | NA | NA | NA | #VALUE! |
| ENSMUSG00000043331.7 | 0 | NA | NA | NA | NA | NA | #VALUE! |
| ENSMUSG00000043333.13 | 0 | NA | NA | NA | NA | NA | #VALUE! |
| ENSMUSG00000043340.4 | 0 | NA | NA | NA | NA | NA | #VALUE! |
| ENSMUSG00000043342.10 | 0 | NA | NA | NA | NA | NA | #VALUE! |
| ENSMUSG00000043346.10 | 0 | NA | NA | NA | NA | NA | #VALUE! |
| ENSMUSG00000043354.3 | 0 | NA | NA | NA | NA | NA | #VALUE! |
| ENSMUSG00000043357.6 | 0 | NA | NA | NA | NA | NA | #VALUE! |
| ENSMUSG00000043366.17 | 0 | NA | NA | NA | NA | NA | #VALUE! |
| ENSMUSG00000043372.13 | 0.279766 | 1.808031 | 4.036658 | 0.447903 | 0.654223 | NA | #VALUE! |
| ENSMUSG00000043383.6 | 0 | NA | NA | NA | NA | NA | #VALUE! |
| ENSMUSG00000043384.16 | 9.485928 | -1.82745 | 2.617522 | -0.69816 | 0.485078 | NA | #VALUE! |
| ENSMUSG00000043385.5 | 0 | NA | NA | NA | NA | NA | #VALUE! |
| ENSMUSG00000043388.9 | 0 | NA | NA | NA | NA | NA | #VALUE! |
| ENSMUSG00000043410.17 | 0 | NA | NA | NA | NA | NA | #VALUE! |
| ENSMUSG00000043424.11 | 1.119871 | -1.46061 | 3.91329 | -0.37324 | 0.708966 | NA | #VALUE! |
| ENSMUSG00000043429.8 | 0 | NA | NA | NA | NA | NA | #VALUE! |
| ENSMUSG00000043430.6 | 0 | NA | NA | NA | NA | NA | #VALUE! |
| ENSMUSG00000043432.8 | 7.181725 | -2.01621 | 3.296782 | -0.61157 | 0.540823 | NA | #VALUE! |
| ENSMUSG00000043439.6 | 28.54494 | -4.00255 | 2.525843 | -1.58464 | NA | NA | #VALUE! |
| ENSMUSG00000043441.6 | 0 | NA | NA | NA | NA | NA | #VALUE! |
| ENSMUSG00000043448.14 | 0 | NA | NA | NA | NA | NA | #VALUE! |
| ENSMUSG00000043453.6 | 0 | NA | NA | NA | NA | NA | #VALUE! |
| ENSMUSG00000043456.18 | 0 | NA | NA | NA | NA | NA | #VALUE! |
| ENSMUSG00000043458.6 | 1.492083 | 4.088114 | 3.936282 | 1.038572 | 0.299004 | NA | #VALUE! |
| ENSMUSG00000043460.8 | 0 | NA | NA | NA | NA | NA | #VALUE! |
| ENSMUSG00000043461.14 | 0 | NA | NA | NA | NA | NA | #VALUE! |
| ENSMUSG00000043463.7 | 0 | NA | NA | NA | NA | NA | #VALUE! |
| ENSMUSG00000043468.5 | 0 | NA | NA | NA | NA | NA | #VALUE! |
| ENSMUSG00000043472.11 | 0 | NA | NA | NA | NA | NA | #VALUE! |
| ENSMUSG00000043483.5 | 8.263566 | 0.541002 | 2.024111 | 0.267279 | 0.789254 | NA | #VALUE! |
| ENSMUSG00000043484.4 | 0.105013 | -0.97324 | 4.076293 | -0.23876 | 0.811295 | NA | #VALUE! |
| ENSMUSG00000043485.3 | 0 | NA | NA | NA | NA | NA | #VALUE! |
| ENSMUSG00000043487.5 | 1.851119 | 4.378291 | 3.930938 | 1.113803 | 0.265364 | NA | #VALUE! |
| ENSMUSG00000043488.6 | 22.45218 | -22.2053 | 2.865536 | -7.74908 | NA | NA | #VALUE! |
| ENSMUSG00000043496.8 | 0 | NA | NA | NA | NA | NA | #VALUE! |
| ENSMUSG00000043501.6 | 0 | NA | NA | NA | NA | NA | #VALUE! |
| ENSMUSG00000043505.9 | 7.869188 | -3.20203 | 3.102992 | -1.03192 | 0.30211 | NA | #VALUE! |
| ENSMUSG00000043518.8 | 0.245976 | 1.027628 | 4.067126 | 0.252667 | 0.800526 | NA | #VALUE! |
| ENSMUSG00000043522.9 | 0 | NA | NA | NA | NA | NA | #VALUE! |
| ENSMUSG00000043529.5 | 0 | NA | NA | NA | NA | NA | #VALUE! |

| | | | | | | | |
|---|---|---|---|---|---|---|---|
| ENSMUSG00000043531.17 | 0 | NA | NA | NA | NA | NA | #VALUE! |
| ENSMUSG00000043537.3 | 0 | NA | NA | NA | NA | NA | #VALUE! |
| ENSMUSG00000043541.17 | 0 | NA | NA | NA | NA | NA | #VALUE! |
| ENSMUSG00000043549.14 | 0 | NA | NA | NA | NA | NA | #VALUE! |
| ENSMUSG00000043556.11 | 6.327433 | -6.18917 | 3.913851 | -1.58135 | 0.113798 | NA | #VALUE! |
| ENSMUSG00000043557.17 | 0 | NA | NA | NA | NA | NA | #VALUE! |
| ENSMUSG00000043569.8 | 0 | NA | NA | NA | NA | NA | #VALUE! |
| ENSMUSG00000043572.8 | 0 | NA | NA | NA | NA | NA | #VALUE! |
| ENSMUSG00000043583.2 | 0 | NA | NA | NA | NA | NA | #VALUE! |
| ENSMUSG00000043592.16 | 0 | NA | NA | NA | NA | NA | #VALUE! |
| ENSMUSG00000043602.6 | 0.783156 | -3.1627 | 3.963922 | -0.79787 | 0.424945 | NA | #VALUE! |
| ENSMUSG00000043605.6 | 0 | NA | NA | NA | NA | NA | #VALUE! |
| ENSMUSG00000043613.9 | 0 | NA | NA | NA | NA | NA | #VALUE! |
| ENSMUSG00000043614.14 | 0 | NA | NA | NA | NA | NA | #VALUE! |
| ENSMUSG00000043618.10 | 0 | NA | NA | NA | NA | NA | #VALUE! |
| ENSMUSG00000043621.14 | 0 | NA | NA | NA | NA | NA | #VALUE! |
| ENSMUSG00000043623.8 | 0 | NA | NA | NA | NA | NA | #VALUE! |
| ENSMUSG00000043629.13 | 0 | NA | NA | NA | NA | NA | #VALUE! |
| ENSMUSG00000043631.9 | 14.97723 | 20.81131 | 3.181896 | 6.540536 | NA | NA | #VALUE! |
| ENSMUSG00000043633.7 | 0 | NA | NA | NA | NA | NA | #VALUE! |
| ENSMUSG00000043635.13 | 0 | NA | NA | NA | NA | NA | #VALUE! |
| ENSMUSG00000043639.16 | 0.204111 | -1.05581 | 4.073302 | -0.2592 | 0.795479 | NA | #VALUE! |
| ENSMUSG00000043644.5 | 7.434969 | 1.611203 | 3.274413 | 0.492059 | 0.622678 | NA | #VALUE! |
| ENSMUSG00000043648.8 | 1.68021 | -4.29432 | 3.933015 | -1.09187 | 0.274892 | NA | #VALUE! |
| ENSMUSG00000043659.12 | 0 | NA | NA | NA | NA | NA | #VALUE! |
| ENSMUSG00000043661.4 | 0 | NA | NA | NA | NA | NA | #VALUE! |
| ENSMUSG00000043664.8 | 0.204111 | -1.05581 | 4.073302 | -0.2592 | 0.795479 | NA | #VALUE! |
| ENSMUSG00000043668.14 | 0 | NA | NA | NA | NA | NA | #VALUE! |
| ENSMUSG00000043670.5 | 0 | NA | NA | NA | NA | NA | #VALUE! |
| ENSMUSG00000043673.12 | 2.310289 | -4.7496 | 3.925939 | -1.2098 | 0.226355 | NA | #VALUE! |
| ENSMUSG00000043692.7 | 0 | NA | NA | NA | NA | NA | #VALUE! |
| ENSMUSG00000043698.10 | 0 | NA | NA | NA | NA | NA | #VALUE! |
| ENSMUSG00000043705.8 | 0 | NA | NA | NA | NA | NA | #VALUE! |
| ENSMUSG00000043715.13 | 0 | NA | NA | NA | NA | NA | #VALUE! |
| ENSMUSG00000043719.15 | 0 | NA | NA | NA | NA | NA | #VALUE! |
| ENSMUSG00000043727.6 | 0 | NA | NA | NA | NA | NA | #VALUE! |
| ENSMUSG00000043740.16 | 3.043764 | 0.297782 | 3.786042 | 0.078652 | 0.937309 | NA | #VALUE! |
| ENSMUSG00000043747.13 | 0 | NA | NA | NA | NA | NA | #VALUE! |
| ENSMUSG00000043753.3 | 0.195789 | -1.05581 | 4.073302 | -0.2592 | 0.795479 | NA | #VALUE! |
| ENSMUSG00000043760.17 | 1.009701 | 3.503687 | 3.950852 | 0.886818 | 0.375177 | NA | #VALUE! |
| ENSMUSG00000043773.6 | 8.414176 | 6.562668 | 3.80431 | 1.725061 | 0.084516 | NA | #VALUE! |
| ENSMUSG00000043782.10 | 0 | NA | NA | NA | NA | NA | #VALUE! |
| ENSMUSG00000043787.9 | 0 | NA | NA | NA | NA | NA | #VALUE! |
| ENSMUSG00000043794.9 | 0.168284 | 1.027628 | 4.067126 | 0.252667 | 0.800526 | NA | #VALUE! |

| | | | | | | | |
|---|---|---|---|---|---|---|---|
| ENSMUSG00000043795.11 | 0 | NA | NA | NA | NA | NA | #VALUE! |
| ENSMUSG00000043800.12 | 0 | NA | NA | NA | NA | NA | #VALUE! |
| ENSMUSG00000043801.8 | 0.577132 | 1.125107 | 4.010901 | 0.280512 | 0.779084 | NA | #VALUE! |
| ENSMUSG00000043807.7 | 0.315039 | -1.93549 | 4.038621 | -0.47925 | 0.631763 | NA | #VALUE! |
| ENSMUSG00000043811.6 | 0 | NA | NA | NA | NA | NA | #VALUE! |
| ENSMUSG00000043822.19 | 0 | NA | NA | NA | NA | NA | #VALUE! |
| ENSMUSG00000043827.10 | 0 | NA | NA | NA | NA | NA | #VALUE! |
| ENSMUSG00000043831.13 | 3.684554 | -0.08627 | 3.775395 | -0.02285 | 0.98177 | NA | #VALUE! |
| ENSMUSG00000043832.14 | 4.91951 | 5.781601 | 3.915938 | 1.476428 | 0.139829 | NA | #VALUE! |
| ENSMUSG00000043833.7 | 0.587367 | -2.74297 | 3.982932 | -0.68868 | 0.491024 | NA | #VALUE! |
| ENSMUSG00000043843.18 | 0 | NA | NA | NA | NA | NA | #VALUE! |
| ENSMUSG00000043850.16 | 0 | NA | NA | NA | NA | NA | #VALUE! |
| ENSMUSG00000043855.8 | 0 | NA | NA | NA | NA | NA | #VALUE! |
| ENSMUSG00000043857.17 | 0 | NA | NA | NA | NA | NA | #VALUE! |
| ENSMUSG00000043859.6 | 0 | NA | NA | NA | NA | NA | #VALUE! |
| ENSMUSG00000043865.4 | 0 | NA | NA | NA | NA | NA | #VALUE! |
| ENSMUSG00000043870.10 | 0 | NA | NA | NA | NA | NA | #VALUE! |
| ENSMUSG00000043873.12 | 0 | NA | NA | NA | NA | NA | #VALUE! |
| ENSMUSG00000043880.6 | 0 | NA | NA | NA | NA | NA | #VALUE! |
| ENSMUSG00000043889.7 | 0 | NA | NA | NA | NA | NA | #VALUE! |
| ENSMUSG00000043892.4 | 0 | NA | NA | NA | NA | NA | #VALUE! |
| ENSMUSG00000043895.7 | 4.181584 | 5.545962 | 3.917562 | 1.415667 | 0.156873 | NA | #VALUE! |
| ENSMUSG00000043897.14 | 0 | NA | NA | NA | NA | NA | #VALUE! |
| ENSMUSG00000043903.5 | 0 | NA | NA | NA | NA | NA | #VALUE! |
| ENSMUSG00000043911.6 | 0 | NA | NA | NA | NA | NA | #VALUE! |
| ENSMUSG00000043913.15 | 0 | NA | NA | NA | NA | NA | #VALUE! |
| ENSMUSG00000043924.17 | 0 | NA | NA | NA | NA | NA | #VALUE! |
| ENSMUSG00000043925.8 | 0 | NA | NA | NA | NA | NA | #VALUE! |
| ENSMUSG00000043931.11 | 0.18651 | 1.401769 | 4.056354 | 0.345574 | 0.729663 | NA | #VALUE! |
| ENSMUSG00000043932.6 | 0 | NA | NA | NA | NA | NA | #VALUE! |
| ENSMUSG00000043939.17 | 0 | NA | NA | NA | NA | NA | #VALUE! |
| ENSMUSG00000043943.15 | 8.027353 | -6.53339 | 3.867678 | -1.68923 | 0.091176 | NA | #VALUE! |
| ENSMUSG00000043945.5 | 0 | NA | NA | NA | NA | NA | #VALUE! |
| ENSMUSG00000043948.3 | 0 | NA | NA | NA | NA | NA | #VALUE! |
| ENSMUSG00000043953.13 | 0.866176 | 0.314441 | 3.93423 | 0.079924 | 0.936297 | NA | #VALUE! |
| ENSMUSG00000043969.5 | 0 | NA | NA | NA | NA | NA | #VALUE! |
| ENSMUSG00000043972.9 | 0 | NA | NA | NA | NA | NA | #VALUE! |
| ENSMUSG00000043982.3 | 0 | NA | NA | NA | NA | NA | #VALUE! |
| ENSMUSG00000043986.6 | 0 | NA | NA | NA | NA | NA | #VALUE! |
| ENSMUSG00000043993.7 | 0.559531 | 2.703958 | 3.982863 | 0.678898 | 0.497202 | NA | #VALUE! |
| ENSMUSG00000043999.7 | 0 | NA | NA | NA | NA | NA | #VALUE! |
| ENSMUSG00000044006.9 | 0 | NA | NA | NA | NA | NA | #VALUE! |
| ENSMUSG00000044014.10 | 0 | NA | NA | NA | NA | NA | #VALUE! |
| ENSMUSG00000044017.17 | 0 | NA | NA | NA | NA | NA | #VALUE! |

| | | | | | | | |
|---|---|---|---|---|---|---|---|
| ENSMUSG00000044022.6 | 0 | NA | NA | NA | NA | NA | #VALUE! |
| ENSMUSG00000044024.17 | 0 | NA | NA | NA | NA | NA | #VALUE! |
| ENSMUSG00000044025.7 | 0 | NA | NA | NA | NA | NA | #VALUE! |
| ENSMUSG00000044029.7 | 0 | NA | NA | NA | NA | NA | #VALUE! |
| ENSMUSG00000044034.12 | 0.210026 | -1.52494 | 4.058692 | -0.37572 | 0.707123 | NA | #VALUE! |
| ENSMUSG00000044039.5 | 0 | NA | NA | NA | NA | NA | #VALUE! |
| ENSMUSG00000044040.4 | 0 | NA | NA | NA | NA | NA | #VALUE! |
| ENSMUSG00000044041.5 | 0 | NA | NA | NA | NA | NA | #VALUE! |
| ENSMUSG00000044043.5 | 0.652786 | 2.919659 | 3.972474 | 0.734972 | 0.462356 | NA | #VALUE! |
| ENSMUSG00000044052.4 | 0 | NA | NA | NA | NA | NA | #VALUE! |
| ENSMUSG00000044055.6 | 0 | NA | NA | NA | NA | NA | #VALUE! |
| ENSMUSG00000044056.4 | 0 | NA | NA | NA | NA | NA | #VALUE! |
| ENSMUSG00000044058.9 | 0 | NA | NA | NA | NA | NA | #VALUE! |
| ENSMUSG00000044061.8 | 0 | NA | NA | NA | NA | NA | #VALUE! |
| ENSMUSG00000044062.8 | 0 | NA | NA | NA | NA | NA | #VALUE! |
| ENSMUSG00000044065.4 | 0 | NA | NA | NA | NA | NA | #VALUE! |
| ENSMUSG00000044067.8 | 0 | NA | NA | NA | NA | NA | #VALUE! |
| ENSMUSG00000044071.10 | 5.03578 | 5.828452 | 3.915646 | 1.488503 | 0.136618 | NA | #VALUE! |
| ENSMUSG00000044072.15 | 2.984166 | 5.077883 | 3.921685 | 1.294822 | 0.195382 | NA | #VALUE! |
| ENSMUSG00000044081.8 | 2.520315 | -4.8742 | 3.924358 | -1.24204 | 0.214223 | NA | #VALUE! |
| ENSMUSG00000044083.13 | 0 | NA | NA | NA | NA | NA | #VALUE! |
| ENSMUSG00000044084.3 | 0 | NA | NA | NA | NA | NA | #VALUE! |
| ENSMUSG00000044086.9 | 3.823463 | 5.43312 | 3.918438 | 1.386552 | 0.165578 | NA | #VALUE! |
| ENSMUSG00000044092.9 | 3.885486 | -5.49502 | 3.91822 | -1.40243 | 0.160787 | NA | #VALUE! |
| ENSMUSG00000044103.5 | 0 | NA | NA | NA | NA | NA | #VALUE! |
| ENSMUSG00000044106.6 | 0 | NA | NA | NA | NA | NA | #VALUE! |
| ENSMUSG00000044117.13 | 0 | NA | NA | NA | NA | NA | #VALUE! |
| ENSMUSG00000044120.6 | 0 | NA | NA | NA | NA | NA | #VALUE! |
| ENSMUSG00000044121.4 | 0 | NA | NA | NA | NA | NA | #VALUE! |
| ENSMUSG00000044125.7 | 9.892144 | 1.782216 | 3.314229 | 0.537747 | NA | NA | #VALUE! |
| ENSMUSG00000044134.10 | 0 | NA | NA | NA | NA | NA | #VALUE! |
| ENSMUSG00000044139.13 | 0 | NA | NA | NA | NA | NA | #VALUE! |
| ENSMUSG00000044141.9 | 0 | NA | NA | NA | NA | NA | #VALUE! |
| ENSMUSG00000044145.3 | 50.18934 | 0.099302 | 1.446963 | 0.068628 | NA | NA | #VALUE! |
| ENSMUSG00000044156.15 | 0 | NA | NA | NA | NA | NA | #VALUE! |
| ENSMUSG00000044162.13 | 0 | NA | NA | NA | NA | NA | #VALUE! |
| ENSMUSG00000044164.3 | 0.612332 | -2.79827 | 3.980109 | -0.70306 | 0.482016 | NA | #VALUE! |
| ENSMUSG00000044165.13 | 0 | NA | NA | NA | NA | NA | #VALUE! |
| ENSMUSG00000044170.6 | 0 | NA | NA | NA | NA | NA | #VALUE! |
| ENSMUSG00000044172.8 | 0 | NA | NA | NA | NA | NA | #VALUE! |
| ENSMUSG00000044176.13 | 0 | NA | NA | NA | NA | NA | #VALUE! |
| ENSMUSG00000044177.5 | 0 | NA | NA | NA | NA | NA | #VALUE! |
| ENSMUSG00000044186.11 | 0 | NA | NA | NA | NA | NA | #VALUE! |
| ENSMUSG00000044199.7 | 2.362062 | -4.76758 | 3.925702 | -1.21445 | 0.224575 | NA | #VALUE! |

| | | | | | | | |
|---|---|---|---|---|---|---|---|
| ENSMUSG00000044201.11 | 0 | NA | NA | NA | NA | NA | #VALUE! |
| ENSMUSG00000044205.8 | 0 | NA | NA | NA | NA | NA | #VALUE! |
| ENSMUSG00000044211.10 | 0 | NA | NA | NA | NA | NA | #VALUE! |
| ENSMUSG00000044213.5 | 0 | NA | NA | NA | NA | NA | #VALUE! |
| ENSMUSG00000044216.8 | 0 | NA | NA | NA | NA | NA | #VALUE! |
| ENSMUSG00000044217.18 | 0 | NA | NA | NA | NA | NA | #VALUE! |
| ENSMUSG00000044220.14 | 0 | NA | NA | NA | NA | NA | #VALUE! |
| ENSMUSG00000044222.4 | 0 | NA | NA | NA | NA | NA | #VALUE! |
| ENSMUSG00000044227.6 | 0 | NA | NA | NA | NA | NA | #VALUE! |
| ENSMUSG00000044229.10 | 0 | NA | NA | NA | NA | NA | #VALUE! |
| ENSMUSG00000044231.4 | 1.470184 | -4.10383 | 3.936698 | -1.04245 | 0.297201 | NA | #VALUE! |
| ENSMUSG00000044243.4 | 0 | NA | NA | NA | NA | NA | #VALUE! |
| ENSMUSG00000044244.19 | 0 | NA | NA | NA | NA | NA | #VALUE! |
| ENSMUSG00000044248.5 | 0 | NA | NA | NA | NA | NA | #VALUE! |
| ENSMUSG00000044249.4 | 0 | NA | NA | NA | NA | NA | #VALUE! |
| ENSMUSG00000044250.9 | 0.746042 | 3.107177 | 3.964586 | 0.783733 | 0.433197 | NA | #VALUE! |
| ENSMUSG00000044265.17 | 0 | NA | NA | NA | NA | NA | #VALUE! |
| ENSMUSG00000044268.5 | 0 | NA | NA | NA | NA | NA | #VALUE! |
| ENSMUSG00000044276.3 | 0.093255 | 0.868093 | 4.072464 | 0.213162 | 0.831201 | NA | #VALUE! |
| ENSMUSG00000044286.7 | 0 | NA | NA | NA | NA | NA | #VALUE! |
| ENSMUSG00000044287.7 | 0 | NA | NA | NA | NA | NA | #VALUE! |
| ENSMUSG00000044288.7 | 0 | NA | NA | NA | NA | NA | #VALUE! |
| ENSMUSG00000044292.5 | 0 | NA | NA | NA | NA | NA | #VALUE! |
| ENSMUSG00000044293.6 | 0 | NA | NA | NA | NA | NA | #VALUE! |
| ENSMUSG00000044294.9 | 0 | NA | NA | NA | NA | NA | #VALUE! |
| ENSMUSG00000044296.11 | 6.716495 | 6.242664 | 3.586842 | 1.740434 | 0.081783 | NA | #VALUE! |
| ENSMUSG00000044303.7 | 0 | NA | NA | NA | NA | NA | #VALUE! |
| ENSMUSG00000044306.5 | 0 | NA | NA | NA | NA | NA | #VALUE! |
| ENSMUSG00000044309.9 | 0 | NA | NA | NA | NA | NA | #VALUE! |
| ENSMUSG00000044312.5 | 0 | NA | NA | NA | NA | NA | #VALUE! |
| ENSMUSG00000044313.14 | 0 | NA | NA | NA | NA | NA | #VALUE! |
| ENSMUSG00000044317.8 | 0.408222 | -2.20205 | 4.016842 | -0.54821 | 0.583551 | NA | #VALUE! |
| ENSMUSG00000044320.15 | 0 | NA | NA | NA | NA | NA | #VALUE! |
| ENSMUSG00000044322.8 | 0 | NA | NA | NA | NA | NA | #VALUE! |
| ENSMUSG00000044328.14 | 0 | NA | NA | NA | NA | NA | #VALUE! |
| ENSMUSG00000044330.9 | 6.048988 | -0.04838 | 2.678639 | -0.01806 | 0.985589 | NA | #VALUE! |
| ENSMUSG00000044337.6 | 0 | NA | NA | NA | NA | NA | #VALUE! |
| ENSMUSG00000044338.10 | 0 | NA | NA | NA | NA | NA | #VALUE! |
| ENSMUSG00000044348.15 | 0.735092 | -3.12016 | 3.965612 | -0.7868 | 0.431397 | NA | #VALUE! |
| ENSMUSG00000044349.17 | 8.027353 | -6.53339 | 3.867678 | -1.68923 | 0.091176 | NA | #VALUE! |
| ENSMUSG00000044352.7 | 3.702238 | 5.378281 | 3.918889 | 1.372399 | 0.169939 | NA | #VALUE! |
| ENSMUSG00000044359.2 | 0 | NA | NA | NA | NA | NA | #VALUE! |
| ENSMUSG00000044361.17 | 7.635775 | -6.4612 | 3.912635 | -1.65137 | 0.098663 | NA | #VALUE! |
| ENSMUSG00000044362.9 | 0 | NA | NA | NA | NA | NA | #VALUE! |

| | | | | | | | |
|---|---|---|---|---|---|---|---|
| ENSMUSG00000044364.6 | 0 | NA | NA | NA | NA | NA | #VALUE! |
| ENSMUSG00000044365.16 | 0.204111 | -1.05581 | 4.073302 | -0.2592 | 0.795479 | NA | #VALUE! |
| ENSMUSG00000044367.8 | 0 | NA | NA | NA | NA | NA | #VALUE! |
| ENSMUSG00000044375.9 | 0 | NA | NA | NA | NA | NA | #VALUE! |
| ENSMUSG00000044377.4 | 0 | NA | NA | NA | NA | NA | #VALUE! |
| ENSMUSG00000044387.6 | 0 | NA | NA | NA | NA | NA | #VALUE! |
| ENSMUSG00000044390.6 | 0 | NA | NA | NA | NA | NA | #VALUE! |
| ENSMUSG00000044400.2 | 0 | NA | NA | NA | NA | NA | #VALUE! |
| ENSMUSG00000044405.5 | 0 | NA | NA | NA | NA | NA | #VALUE! |
| ENSMUSG00000044424.5 | 1.051444 | 1.755809 | 3.9371 | 0.445965 | 0.655622 | NA | #VALUE! |
| ENSMUSG00000044429.9 | 0 | NA | NA | NA | NA | NA | #VALUE! |
| ENSMUSG00000044430.12 | 0 | NA | NA | NA | NA | NA | #VALUE! |
| ENSMUSG00000044433.17 | 9.743416 | -2.5261 | 2.983362 | -0.84673 | 0.397146 | NA | #VALUE! |
| ENSMUSG00000044434.6 | 0 | NA | NA | NA | NA | NA | #VALUE! |
| ENSMUSG00000044441.6 | 0.816443 | -3.21914 | 3.961753 | -0.81255 | 0.416474 | NA | #VALUE! |
| ENSMUSG00000044444.7 | 0 | NA | NA | NA | NA | NA | #VALUE! |
| ENSMUSG00000044447.6 | 31.07329 | -0.64575 | 1.884313 | -0.3427 | NA | NA | #VALUE! |
| ENSMUSG00000044453.6 | 0 | NA | NA | NA | NA | NA | #VALUE! |
| ENSMUSG00000044454.6 | 0 | NA | NA | NA | NA | NA | #VALUE! |
| ENSMUSG00000044457.9 | 0 | NA | NA | NA | NA | NA | #VALUE! |
| ENSMUSG00000044461.7 | 0 | NA | NA | NA | NA | NA | #VALUE! |
| ENSMUSG00000044471.13 | 0.18651 | 1.401769 | 4.056354 | 0.345574 | 0.729663 | NA | #VALUE! |
| ENSMUSG00000044485.5 | 0 | NA | NA | NA | NA | NA | #VALUE! |
| ENSMUSG00000044487.3 | 0 | NA | NA | NA | NA | NA | #VALUE! |
| ENSMUSG00000044499.12 | 0 | NA | NA | NA | NA | NA | #VALUE! |
| ENSMUSG00000044515.6 | 0 | NA | NA | NA | NA | NA | #VALUE! |
| ENSMUSG00000044518.4 | 0 | NA | NA | NA | NA | NA | #VALUE! |
| ENSMUSG00000044519.9 | 0 | NA | NA | NA | NA | NA | #VALUE! |
| ENSMUSG00000044522.9 | 0 | NA | NA | NA | NA | NA | #VALUE! |
| ENSMUSG00000044526.4 | 0 | NA | NA | NA | NA | NA | #VALUE! |
| ENSMUSG00000044528.6 | 0 | NA | NA | NA | NA | NA | #VALUE! |
| ENSMUSG00000044534.9 | 2.631025 | -1.53473 | 3.817443 | -0.40203 | 0.687661 | NA | #VALUE! |
| ENSMUSG00000044537.9 | 0 | NA | NA | NA | NA | NA | #VALUE! |
| ENSMUSG00000044538.11 | 0 | NA | NA | NA | NA | NA | #VALUE! |
| ENSMUSG00000044542.4 | 0 | NA | NA | NA | NA | NA | #VALUE! |
| ENSMUSG00000044544.5 | 0 | NA | NA | NA | NA | NA | #VALUE! |
| ENSMUSG00000044550.12 | 0 | NA | NA | NA | NA | NA | #VALUE! |
| ENSMUSG00000044551.14 | 4.865069 | -2.89831 | 3.815321 | -0.75965 | 0.447464 | NA | #VALUE! |
| ENSMUSG00000044556.4 | 0 | NA | NA | NA | NA | NA | #VALUE! |
| ENSMUSG00000044560.4 | 0 | NA | NA | NA | NA | NA | #VALUE! |
| ENSMUSG00000044562.13 | 5.983179 | 6.070861 | 3.844042 | 1.579291 | 0.114269 | NA | #VALUE! |
| ENSMUSG00000044566.16 | 4.741871 | -5.77539 | 3.916201 | -1.47474 | 0.140282 | NA | #VALUE! |
| ENSMUSG00000044574.6 | 0 | NA | NA | NA | NA | NA | #VALUE! |
| ENSMUSG00000044576.7 | 0 | NA | NA | NA | NA | NA | #VALUE! |

| | | | | | | | |
|---|---|---|---|---|---|---|---|
| ENSMUSG00000044581.9 | 0 | NA | NA | NA | NA | NA | #VALUE! |
| ENSMUSG00000044583.14 | 0.783156 | -3.1627 | 3.963922 | -0.79787 | 0.424945 | NA | #VALUE! |
| ENSMUSG00000044594.15 | 0 | NA | NA | NA | NA | NA | #VALUE! |
| ENSMUSG00000044595.11 | 2.126175 | -3.42547 | 3.915786 | -0.87479 | 0.381691 | NA | #VALUE! |
| ENSMUSG00000044597.5 | 0 | NA | NA | NA | NA | NA | #VALUE! |
| ENSMUSG00000044609.7 | 0.540678 | 0.929314 | 4.022674 | 0.231019 | 0.8173 | NA | #VALUE! |
| ENSMUSG00000044624.8 | 0 | NA | NA | NA | NA | NA | #VALUE! |
| ENSMUSG00000044626.14 | 3.780473 | -5.45568 | 3.918536 | -1.39228 | 0.163839 | NA | #VALUE! |
| ENSMUSG00000044628.6 | 0 | NA | NA | NA | NA | NA | #VALUE! |
| ENSMUSG00000044629.6 | 3.25672 | -1.5325 | 3.799249 | -0.40337 | 0.686677 | NA | #VALUE! |
| ENSMUSG00000044633.5 | 0 | NA | NA | NA | NA | NA | #VALUE! |
| ENSMUSG00000044636.7 | 24.82268 | 2.026949 | 2.662249 | 0.761367 | NA | NA | #VALUE! |
| ENSMUSG00000044639.4 | 0 | NA | NA | NA | NA | NA | #VALUE! |
| ENSMUSG00000044645.9 | 0 | NA | NA | NA | NA | NA | #VALUE! |
| ENSMUSG00000044646.16 | 4.448828 | -5.68014 | 3.916844 | -1.45018 | 0.147008 | NA | #VALUE! |
| ENSMUSG00000044647.17 | 0 | NA | NA | NA | NA | NA | #VALUE! |
| ENSMUSG00000044649.5 | 0 | NA | NA | NA | NA | NA | #VALUE! |
| ENSMUSG00000044664.8 | 0 | NA | NA | NA | NA | NA | #VALUE! |
| ENSMUSG00000044667.13 | 0 | NA | NA | NA | NA | NA | #VALUE! |
| ENSMUSG00000044678.12 | 0 | NA | NA | NA | NA | NA | #VALUE! |
| ENSMUSG00000044681.20 | 0 | NA | NA | NA | NA | NA | #VALUE! |
| ENSMUSG00000044689.6 | 0 | NA | NA | NA | NA | NA | #VALUE! |
| ENSMUSG00000044694.3 | 0 | NA | NA | NA | NA | NA | #VALUE! |
| ENSMUSG00000044697.6 | 0 | NA | NA | NA | NA | NA | #VALUE! |
| ENSMUSG00000044701.6 | 0 | NA | NA | NA | NA | NA | #VALUE! |
| ENSMUSG00000044702.14 | 0 | NA | NA | NA | NA | NA | #VALUE! |
| ENSMUSG00000044703.6 | 0 | NA | NA | NA | NA | NA | #VALUE! |
| ENSMUSG00000044705.4 | 0 | NA | NA | NA | NA | NA | #VALUE! |
| ENSMUSG00000044707.8 | 0 | NA | NA | NA | NA | NA | #VALUE! |
| ENSMUSG00000044708.6 | 0.901387 | 1.484278 | 3.947911 | 0.375966 | 0.706943 | NA | #VALUE! |
| ENSMUSG00000044716.13 | 0 | NA | NA | NA | NA | NA | #VALUE! |
| ENSMUSG00000044719.6 | 0 | NA | NA | NA | NA | NA | #VALUE! |
| ENSMUSG00000044724.10 | 0 | NA | NA | NA | NA | NA | #VALUE! |
| ENSMUSG00000044726.9 | 0 | NA | NA | NA | NA | NA | #VALUE! |
| ENSMUSG00000044730.18 | 0.408222 | -2.20205 | 4.016842 | -0.54821 | 0.583551 | NA | #VALUE! |
| ENSMUSG00000044737.7 | 0 | NA | NA | NA | NA | NA | #VALUE! |
| ENSMUSG00000044743.4 | 0 | NA | NA | NA | NA | NA | #VALUE! |
| ENSMUSG00000044748.9 | 1.387167 | -3.99301 | 3.939073 | -1.01369 | 0.310729 | NA | #VALUE! |
| ENSMUSG00000044751.6 | 2.219624 | 1.744072 | 3.842496 | 0.45389 | 0.649908 | NA | #VALUE! |
| ENSMUSG00000044757.8 | 0 | NA | NA | NA | NA | NA | #VALUE! |
| ENSMUSG00000044770.15 | 0 | NA | NA | NA | NA | NA | #VALUE! |
| ENSMUSG00000044772.3 | 0 | NA | NA | NA | NA | NA | #VALUE! |
| ENSMUSG00000044787.8 | 0 | NA | NA | NA | NA | NA | #VALUE! |
| ENSMUSG00000044795.13 | 3.823463 | 5.43312 | 3.918438 | 1.386552 | 0.165578 | NA | #VALUE! |

| | | | | | | | |
|---|---|---|---|---|---|---|---|
| ENSMUSG00000044798.8 | 0 | NA | NA | NA | NA | NA | #VALUE! |
| ENSMUSG00000044801.2 | 0 | NA | NA | NA | NA | NA | #VALUE! |
| ENSMUSG00000044807.14 | 8.960047 | 2.869106 | 2.979713 | 0.96288 | 0.335608 | NA | #VALUE! |
| ENSMUSG00000044811.14 | 8.311966 | 0.315529 | 3.083585 | 0.102326 | 0.918498 | NA | #VALUE! |
| ENSMUSG00000044814.7 | 0 | NA | NA | NA | NA | NA | #VALUE! |
| ENSMUSG00000044816.11 | 0 | NA | NA | NA | NA | NA | #VALUE! |
| ENSMUSG00000044819.6 | 0 | NA | NA | NA | NA | NA | #VALUE! |
| ENSMUSG00000044820.7 | 1.668452 | -3.29393 | 3.926429 | -0.83891 | 0.401518 | NA | #VALUE! |
| ENSMUSG00000044824.10 | 0 | NA | NA | NA | NA | NA | #VALUE! |
| ENSMUSG00000044835.16 | 0 | NA | NA | NA | NA | NA | #VALUE! |
| ENSMUSG00000044847.14 | 8.510547 | 2.804393 | 3.420579 | 0.819859 | NA | NA | #VALUE! |
| ENSMUSG00000044854.8 | 1.458426 | -3.0899 | 3.930659 | -0.7861 | 0.431807 | NA | #VALUE! |
| ENSMUSG00000044860.10 | 0 | NA | NA | NA | NA | NA | #VALUE! |
| ENSMUSG00000044863.8 | 0 | NA | NA | NA | NA | NA | #VALUE! |
| ENSMUSG00000044864.17 | 17.19968 | -0.53475 | 2.878842 | -0.18575 | NA | NA | #VALUE! |
| ENSMUSG00000044867.6 | 0 | NA | NA | NA | NA | NA | #VALUE! |
| ENSMUSG00000044897.4 | 0 | NA | NA | NA | NA | NA | #VALUE! |
| ENSMUSG00000044899.4 | 0 | NA | NA | NA | NA | NA | #VALUE! |
| ENSMUSG00000044903.16 | 0 | NA | NA | NA | NA | NA | #VALUE! |
| ENSMUSG00000044912.12 | 0 | NA | NA | NA | NA | NA | #VALUE! |
| ENSMUSG00000044916.11 | 2.936837 | -5.08026 | 3.922024 | -1.29532 | 0.195212 | NA | #VALUE! |
| ENSMUSG00000044921.6 | 0 | NA | NA | NA | NA | NA | #VALUE! |
| ENSMUSG00000044923.5 | 0 | NA | NA | NA | NA | NA | #VALUE! |
| ENSMUSG00000044927.7 | 0 | NA | NA | NA | NA | NA | #VALUE! |
| ENSMUSG00000044933.4 | 0 | NA | NA | NA | NA | NA | #VALUE! |
| ENSMUSG00000044937.16 | 0 | NA | NA | NA | NA | NA | #VALUE! |
| ENSMUSG00000044938.9 | 2.705731 | 4.913666 | 3.923477 | 1.252375 | 0.210433 | NA | #VALUE! |
| ENSMUSG00000044948.18 | 0 | NA | NA | NA | NA | NA | #VALUE! |
| ENSMUSG00000044949.5 | 0 | NA | NA | NA | NA | NA | #VALUE! |
| ENSMUSG00000044951.18 | 0 | NA | NA | NA | NA | NA | #VALUE! |
| ENSMUSG00000044957.4 | 0 | NA | NA | NA | NA | NA | #VALUE! |
| ENSMUSG00000044966.5 | 0.093255 | 0.868093 | 4.072464 | 0.213162 | 0.831201 | NA | #VALUE! |
| ENSMUSG00000044968.17 | 7.190789 | -1.42364 | 3.281962 | -0.43378 | 0.664449 | NA | #VALUE! |
| ENSMUSG00000044976.18 | 0 | NA | NA | NA | NA | NA | #VALUE! |
| ENSMUSG00000044985.7 | 0 | NA | NA | NA | NA | NA | #VALUE! |
| ENSMUSG00000044988.8 | 0 | NA | NA | NA | NA | NA | #VALUE! |
| ENSMUSG00000044991.11 | 5.761485 | -6.06101 | 3.871928 | -1.56537 | 0.117496 | NA | #VALUE! |
| ENSMUSG00000044994.11 | 0 | NA | NA | NA | NA | NA | #VALUE! |
| ENSMUSG00000044997.4 | 0 | NA | NA | NA | NA | NA | #VALUE! |
| ENSMUSG00000045004.4 | 0 | NA | NA | NA | NA | NA | #VALUE! |
| ENSMUSG00000045007.10 | 4.410551 | -5.67707 | 3.916865 | -1.44939 | 0.147228 | NA | #VALUE! |
| ENSMUSG00000045008.3 | 0 | NA | NA | NA | NA | NA | #VALUE! |
| ENSMUSG00000045009.10 | 0 | NA | NA | NA | NA | NA | #VALUE! |
| ENSMUSG00000045010.15 | 0 | NA | NA | NA | NA | NA | #VALUE! |

| | | | | | | | |
|---|---|---|---|---|---|---|---|
| ENSMUSG00000045013.8 | 0 | NA | NA | NA | NA | NA | #VALUE! |
| ENSMUSG00000045019.5 | 0 | NA | NA | NA | NA | NA | #VALUE! |
| ENSMUSG00000045022.7 | 0 | NA | NA | NA | NA | NA | #VALUE! |
| ENSMUSG00000045027.8 | 0 | NA | NA | NA | NA | NA | #VALUE! |
| ENSMUSG00000045030.9 | 2.741047 | -4.98044 | 3.923113 | -1.26951 | 0.204259 | NA | #VALUE! |
| ENSMUSG00000045031.19 | 0 | NA | NA | NA | NA | NA | #VALUE! |
| ENSMUSG00000045034.13 | 0 | NA | NA | NA | NA | NA | #VALUE! |
| ENSMUSG00000045036.16 | 0 | NA | NA | NA | NA | NA | #VALUE! |
| ENSMUSG00000045039.10 | 5.747935 | -1.71268 | 3.451547 | -0.49621 | 0.619748 | NA | #VALUE! |
| ENSMUSG00000045045.8 | 0 | NA | NA | NA | NA | NA | #VALUE! |
| ENSMUSG00000045048.8 | 0 | NA | NA | NA | NA | NA | #VALUE! |
| ENSMUSG00000045052.4 | 0 | NA | NA | NA | NA | NA | #VALUE! |
| ENSMUSG00000045053.3 | 0 | NA | NA | NA | NA | NA | #VALUE! |
| ENSMUSG00000045055.6 | 0.198268 | -0.06494 | 4.068276 | -0.01596 | 0.987265 | NA | #VALUE! |
| ENSMUSG00000045062.5 | 0 | NA | NA | NA | NA | NA | #VALUE! |
| ENSMUSG00000045065.8 | 0 | NA | NA | NA | NA | NA | #VALUE! |
| ENSMUSG00000045075.4 | 0 | NA | NA | NA | NA | NA | #VALUE! |
| ENSMUSG00000045083.15 | 0 | NA | NA | NA | NA | NA | #VALUE! |
| ENSMUSG00000045091.5 | 0 | NA | NA | NA | NA | NA | #VALUE! |
| ENSMUSG00000045102.13 | 0 | NA | NA | NA | NA | NA | #VALUE! |
| ENSMUSG00000045104.10 | 0 | NA | NA | NA | NA | NA | #VALUE! |
| ENSMUSG00000045109.7 | 0 | NA | NA | NA | NA | NA | #VALUE! |
| ENSMUSG00000045111.6 | 0 | NA | NA | NA | NA | NA | #VALUE! |
| ENSMUSG00000045114.11 | 0 | NA | NA | NA | NA | NA | #VALUE! |
| ENSMUSG00000045126.3 | 0 | NA | NA | NA | NA | NA | #VALUE! |
| ENSMUSG00000045132.4 | 0 | NA | NA | NA | NA | NA | #VALUE! |
| ENSMUSG00000045136.7 | 0 | NA | NA | NA | NA | NA | #VALUE! |
| ENSMUSG00000045148.6 | 0 | NA | NA | NA | NA | NA | #VALUE! |
| ENSMUSG00000045150.6 | 0 | NA | NA | NA | NA | NA | #VALUE! |
| ENSMUSG00000045165.7 | 0 | NA | NA | NA | NA | NA | #VALUE! |
| ENSMUSG00000045174.11 | 0 | NA | NA | NA | NA | NA | #VALUE! |
| ENSMUSG00000045176.4 | 6.668798 | -0.57006 | 3.391826 | -0.16807 | 0.866528 | NA | #VALUE! |
| ENSMUSG00000045178.11 | 0 | NA | NA | NA | NA | NA | #VALUE! |
| ENSMUSG00000045179.10 | 0 | NA | NA | NA | NA | NA | #VALUE! |
| ENSMUSG00000045201.7 | 0 | NA | NA | NA | NA | NA | #VALUE! |
| ENSMUSG00000045202.7 | 0 | NA | NA | NA | NA | NA | #VALUE! |
| ENSMUSG00000045204.5 | 0 | NA | NA | NA | NA | NA | #VALUE! |
| ENSMUSG00000045215.18 | 0 | NA | NA | NA | NA | NA | #VALUE! |
| ENSMUSG00000045217.6 | 0 | NA | NA | NA | NA | NA | #VALUE! |
| ENSMUSG00000045225.5 | 0 | NA | NA | NA | NA | NA | #VALUE! |
| ENSMUSG00000045231.6 | 0 | NA | NA | NA | NA | NA | #VALUE! |
| ENSMUSG00000045232.5 | 0 | NA | NA | NA | NA | NA | #VALUE! |
| ENSMUSG00000045236.6 | 0 | NA | NA | NA | NA | NA | #VALUE! |
| ENSMUSG00000045238.10 | 0 | NA | NA | NA | NA | NA | #VALUE! |

| | | | | | | | |
|---|---|---|---|---|---|---|---|
| ENSMUSG00000045246.12 | 0 | NA | NA | NA | NA | NA | #VALUE! |
| ENSMUSG00000045248.7 | 0.598106 | 2.755755 | 3.98023 | 0.692361 | 0.488711 | NA | #VALUE! |
| ENSMUSG00000045251.14 | 3.550791 | -0.75155 | 3.780973 | -0.19877 | 0.842441 | NA | #VALUE! |
| ENSMUSG00000045259.5 | 0 | NA | NA | NA | NA | NA | #VALUE! |
| ENSMUSG00000045267.4 | 0 | NA | NA | NA | NA | NA | #VALUE! |
| ENSMUSG00000045268.14 | 0 | NA | NA | NA | NA | NA | #VALUE! |
| ENSMUSG00000045273.9 | 0 | NA | NA | NA | NA | NA | #VALUE! |
| ENSMUSG00000045275.18 | 0 | NA | NA | NA | NA | NA | #VALUE! |
| ENSMUSG00000045281.6 | 0 | NA | NA | NA | NA | NA | #VALUE! |
| ENSMUSG00000045288.11 | 0 | NA | NA | NA | NA | NA | #VALUE! |
| ENSMUSG00000045291.9 | 0 | NA | NA | NA | NA | NA | #VALUE! |
| ENSMUSG00000045306.4 | 0 | NA | NA | NA | NA | NA | #VALUE! |
| ENSMUSG00000045318.7 | 0 | NA | NA | NA | NA | NA | #VALUE! |
| ENSMUSG00000045319.14 | 9.64462 | 0.094312 | 1.979381 | 0.047647 | 0.961997 | NA | #VALUE! |
| ENSMUSG00000045322.11 | 0 | NA | NA | NA | NA | NA | #VALUE! |
| ENSMUSG00000045326.14 | 0 | NA | NA | NA | NA | NA | #VALUE! |
| ENSMUSG00000045327.8 | 3.870521 | 5.442409 | 3.918364 | 1.388949 | 0.164848 | NA | #VALUE! |
| ENSMUSG00000045328.12 | 0.210026 | -1.52494 | 4.058692 | -0.37572 | 0.707123 | NA | #VALUE! |
| ENSMUSG00000045330.9 | 0 | NA | NA | NA | NA | NA | #VALUE! |
| ENSMUSG00000045331.4 | 0 | NA | NA | NA | NA | NA | #VALUE! |
| ENSMUSG00000045333.16 | 0 | NA | NA | NA | NA | NA | #VALUE! |
| ENSMUSG00000045336.6 | 0 | NA | NA | NA | NA | NA | #VALUE! |
| ENSMUSG00000045337.5 | 0 | NA | NA | NA | NA | NA | #VALUE! |
| ENSMUSG00000045339.8 | 0 | NA | NA | NA | NA | NA | #VALUE! |
| ENSMUSG00000045340.5 | 0 | NA | NA | NA | NA | NA | #VALUE! |
| ENSMUSG00000045341.3 | 0 | NA | NA | NA | NA | NA | #VALUE! |
| ENSMUSG00000045345.8 | 0 | NA | NA | NA | NA | NA | #VALUE! |
| ENSMUSG00000045348.17 | 0 | NA | NA | NA | NA | NA | #VALUE! |
| ENSMUSG00000045349.16 | 0 | NA | NA | NA | NA | NA | #VALUE! |
| ENSMUSG00000045350.16 | 0 | NA | NA | NA | NA | NA | #VALUE! |
| ENSMUSG00000045362.9 | 1.398828 | 3.99633 | 3.938205 | 1.014759 | 0.310221 | NA | #VALUE! |
| ENSMUSG00000045364.3 | 0 | NA | NA | NA | NA | NA | #VALUE! |
| ENSMUSG00000045378.6 | 0 | NA | NA | NA | NA | NA | #VALUE! |
| ENSMUSG00000045381.7 | 0 | NA | NA | NA | NA | NA | #VALUE! |
| ENSMUSG00000045382.7 | 0 | NA | NA | NA | NA | NA | #VALUE! |
| ENSMUSG00000045391.5 | 0 | NA | NA | NA | NA | NA | #VALUE! |
| ENSMUSG00000045394.10 | 0 | NA | NA | NA | NA | NA | #VALUE! |
| ENSMUSG00000045395.5 | 0 | NA | NA | NA | NA | NA | #VALUE! |
| ENSMUSG00000045404.17 | 2.404612 | 0.049605 | 3.803241 | 0.013043 | 0.989594 | NA | #VALUE! |
| ENSMUSG00000045411.17 | 0 | NA | NA | NA | NA | NA | #VALUE! |
| ENSMUSG00000045417.3 | 0 | NA | NA | NA | NA | NA | #VALUE! |
| ENSMUSG00000045421.3 | 0 | NA | NA | NA | NA | NA | #VALUE! |
| ENSMUSG00000045427.14 | 7.196886 | -2.79101 | 3.237198 | -0.86217 | 0.388594 | NA | #VALUE! |
| ENSMUSG00000045440.5 | 0 | NA | NA | NA | NA | NA | #VALUE! |

| | | | | | | | |
|---|---|---|---|---|---|---|---|
| ENSMUSG00000045455.9 | 1.475853 | 4.029233 | 3.937502 | 1.023297 | 0.306167 | NA | #VALUE! |
| ENSMUSG00000045463.16 | 0 | NA | NA | NA | NA | NA | #VALUE! |
| ENSMUSG00000045466.19 | 2.692536 | 4.918862 | 3.923417 | 1.253719 | 0.209944 | NA | #VALUE! |
| ENSMUSG00000045467.14 | 0 | NA | NA | NA | NA | NA | #VALUE! |
| ENSMUSG00000045471.5 | 0 | NA | NA | NA | NA | NA | #VALUE! |
| ENSMUSG00000045474.8 | 0 | NA | NA | NA | NA | NA | #VALUE! |
| ENSMUSG00000045475.6 | 0 | NA | NA | NA | NA | NA | #VALUE! |
| ENSMUSG00000045479.8 | 0 | NA | NA | NA | NA | NA | #VALUE! |
| ENSMUSG00000045493.5 | 0 | NA | NA | NA | NA | NA | #VALUE! |
| ENSMUSG00000045498.2 | 0.279766 | 1.808031 | 4.036658 | 0.447903 | 0.654223 | NA | #VALUE! |
| ENSMUSG00000045502.7 | 0 | NA | NA | NA | NA | NA | #VALUE! |
| ENSMUSG00000045506.4 | 0 | NA | NA | NA | NA | NA | #VALUE! |
| ENSMUSG00000045508.5 | 0 | NA | NA | NA | NA | NA | #VALUE! |
| ENSMUSG00000045509.8 | 0 | NA | NA | NA | NA | NA | #VALUE! |
| ENSMUSG00000045514.6 | 0 | NA | NA | NA | NA | NA | #VALUE! |
| ENSMUSG00000045515.5 | 0 | NA | NA | NA | NA | NA | #VALUE! |
| ENSMUSG00000045518.9 | 0 | NA | NA | NA | NA | NA | #VALUE! |
| ENSMUSG00000045521.8 | 0 | NA | NA | NA | NA | NA | #VALUE! |
| ENSMUSG00000045528.2 | 0 | NA | NA | NA | NA | NA | #VALUE! |
| ENSMUSG00000045532.6 | 0 | NA | NA | NA | NA | NA | #VALUE! |
| ENSMUSG00000045534.5 | 0 | NA | NA | NA | NA | NA | #VALUE! |
| ENSMUSG00000045539.4 | 0 | NA | NA | NA | NA | NA | #VALUE! |
| ENSMUSG00000045540.6 | 0 | NA | NA | NA | NA | NA | #VALUE! |
| ENSMUSG00000045545.9 | 0 | NA | NA | NA | NA | NA | #VALUE! |
| ENSMUSG00000045555.4 | 0 | NA | NA | NA | NA | NA | #VALUE! |
| ENSMUSG00000045559.7 | 0 | NA | NA | NA | NA | NA | #VALUE! |
| ENSMUSG00000045566.2 | 0 | NA | NA | NA | NA | NA | #VALUE! |
| ENSMUSG00000045569.8 | 0 | NA | NA | NA | NA | NA | #VALUE! |
| ENSMUSG00000045573.10 | 0 | NA | NA | NA | NA | NA | #VALUE! |
| ENSMUSG00000045575.3 | 0 | NA | NA | NA | NA | NA | #VALUE! |
| ENSMUSG00000045581.8 | 0 | NA | NA | NA | NA | NA | #VALUE! |
| ENSMUSG00000045584.5 | 0 | NA | NA | NA | NA | NA | #VALUE! |
| ENSMUSG00000045587.10 | 0 | NA | NA | NA | NA | NA | #VALUE! |
| ENSMUSG00000045589.8 | 0 | NA | NA | NA | NA | NA | #VALUE! |
| ENSMUSG00000045591.7 | 0 | NA | NA | NA | NA | NA | #VALUE! |
| ENSMUSG00000045598.11 | 2.485079 | -3.62954 | 3.91277 | -0.92762 | 0.353607 | NA | #VALUE! |
| ENSMUSG00000045608.8 | 0 | NA | NA | NA | NA | NA | #VALUE! |
| ENSMUSG00000045609.7 | 0 | NA | NA | NA | NA | NA | #VALUE! |
| ENSMUSG00000045613.10 | 4.382994 | 5.629097 | 3.916959 | 1.437109 | 0.150687 | NA | #VALUE! |
| ENSMUSG00000045620.8 | 0 | NA | NA | NA | NA | NA | #VALUE! |
| ENSMUSG00000045625.6 | 0 | NA | NA | NA | NA | NA | #VALUE! |
| ENSMUSG00000045629.9 | 0.588336 | -1.26709 | 3.993751 | -0.31727 | 0.75104 | NA | #VALUE! |
| ENSMUSG00000045631.11 | 0 | NA | NA | NA | NA | NA | #VALUE! |
| ENSMUSG00000045648.16 | 0 | NA | NA | NA | NA | NA | #VALUE! |

| | | | | | | | |
|---|---|---|---|---|---|---|---|
| ENSMUSG00000045655.10 | 0 | NA | NA | NA | NA | NA | #VALUE! |
| ENSMUSG00000045657.4 | 0 | NA | NA | NA | NA | NA | #VALUE! |
| ENSMUSG00000045662.18 | 0 | NA | NA | NA | NA | NA | #VALUE! |
| ENSMUSG00000045664.6 | 2.353222 | 3.581246 | 3.920106 | 0.913558 | 0.360949 | NA | #VALUE! |
| ENSMUSG00000045667.15 | 0.44158 | 1.011896 | 4.025703 | 0.251359 | 0.801537 | NA | #VALUE! |
| ENSMUSG00000045678.9 | 0 | NA | NA | NA | NA | NA | #VALUE! |
| ENSMUSG00000045679.15 | 0 | NA | NA | NA | NA | NA | #VALUE! |
| ENSMUSG00000045684.11 | 0 | NA | NA | NA | NA | NA | #VALUE! |
| ENSMUSG00000045689.4 | 0 | NA | NA | NA | NA | NA | #VALUE! |
| ENSMUSG00000045690.9 | 6.599616 | 0.76932 | 3.371243 | 0.228201 | 0.81949 | NA | #VALUE! |
| ENSMUSG00000045691.15 | 0 | NA | NA | NA | NA | NA | #VALUE! |
| ENSMUSG00000045693.9 | 0 | NA | NA | NA | NA | NA | #VALUE! |
| ENSMUSG00000045699.5 | 0 | NA | NA | NA | NA | NA | #VALUE! |
| ENSMUSG00000045708.5 | 0 | NA | NA | NA | NA | NA | #VALUE! |
| ENSMUSG00000045709.12 | 0 | NA | NA | NA | NA | NA | #VALUE! |
| ENSMUSG00000045725.6 | 0.279766 | 1.808031 | 4.036658 | 0.447903 | 0.654223 | NA | #VALUE! |
| ENSMUSG00000045730.5 | 0 | NA | NA | NA | NA | NA | #VALUE! |
| ENSMUSG00000045731.7 | 0 | NA | NA | NA | NA | NA | #VALUE! |
| ENSMUSG00000045733.8 | 2.984166 | 5.077883 | 3.921685 | 1.294822 | 0.195382 | NA | #VALUE! |
| ENSMUSG00000045744.12 | 0 | NA | NA | NA | NA | NA | #VALUE! |
| ENSMUSG00000045746.4 | 0 | NA | NA | NA | NA | NA | #VALUE! |
| ENSMUSG00000045757.7 | 4.280187 | -0.7485 | 3.771342 | -0.19847 | 0.842678 | NA | #VALUE! |
| ENSMUSG00000045761.17 | 0 | NA | NA | NA | NA | NA | #VALUE! |
| ENSMUSG00000045763.9 | 5.377114 | -2.9911 | 3.157638 | -0.94726 | 0.343507 | NA | #VALUE! |
| ENSMUSG00000045777.15 | 0 | NA | NA | NA | NA | NA | #VALUE! |
| ENSMUSG00000045780.4 | 0 | NA | NA | NA | NA | NA | #VALUE! |
| ENSMUSG00000045790.11 | 2.041108 | -4.55188 | 3.928743 | -1.15861 | 0.246615 | NA | #VALUE! |
| ENSMUSG00000045792.3 | 0 | NA | NA | NA | NA | NA | #VALUE! |
| ENSMUSG00000045794.4 | 0 | NA | NA | NA | NA | NA | #VALUE! |
| ENSMUSG00000045797.2 | 0 | NA | NA | NA | NA | NA | #VALUE! |
| ENSMUSG00000045799.7 | 0.303281 | -0.61664 | 4.050639 | -0.15223 | 0.879003 | NA | #VALUE! |
| ENSMUSG00000045802.14 | 0 | NA | NA | NA | NA | NA | #VALUE! |
| ENSMUSG00000045812.6 | 0 | NA | NA | NA | NA | NA | #VALUE! |
| ENSMUSG00000045813.8 | 0 | NA | NA | NA | NA | NA | #VALUE! |
| ENSMUSG00000045815.5 | 0 | NA | NA | NA | NA | NA | #VALUE! |
| ENSMUSG00000045817.9 | 5.502891 | 3.531047 | 3.462718 | 1.019733 | 0.307855 | NA | #VALUE! |
| ENSMUSG00000045822.5 | 1.771849 | 4.332864 | 3.931707 | 1.102031 | 0.270448 | NA | #VALUE! |
| ENSMUSG00000045824.7 | 0 | NA | NA | NA | NA | NA | #VALUE! |
| ENSMUSG00000045826.10 | 0 | NA | NA | NA | NA | NA | #VALUE! |
| ENSMUSG00000045835.6 | 0 | NA | NA | NA | NA | NA | #VALUE! |
| ENSMUSG00000045838.10 | 0 | NA | NA | NA | NA | NA | #VALUE! |
| ENSMUSG00000045842.10 | 0 | NA | NA | NA | NA | NA | #VALUE! |
| ENSMUSG00000045868.13 | 0 | NA | NA | NA | NA | NA | #VALUE! |
| ENSMUSG00000045871.6 | 0 | NA | NA | NA | NA | NA | #VALUE! |

| | | | | | | | |
|---|---|---|---|---|---|---|---|
| ENSMUSG00000045875.14 | 0 | NA | NA | NA | NA | NA | #VALUE! |
| ENSMUSG00000045876.5 | 0.983902 | 3.433476 | 3.953024 | 0.868569 | 0.385083 | NA | #VALUE! |
| ENSMUSG00000045877.3 | 0.391578 | -2.1489 | 4.020883 | -0.53444 | 0.59304 | NA | #VALUE! |
| ENSMUSG00000045883.6 | 0 | NA | NA | NA | NA | NA | #VALUE! |
| ENSMUSG00000045903.10 | 0 | NA | NA | NA | NA | NA | #VALUE! |
| ENSMUSG00000045912.10 | 0 | NA | NA | NA | NA | NA | #VALUE! |
| ENSMUSG00000045915.16 | 0 | NA | NA | NA | NA | NA | #VALUE! |
| ENSMUSG00000045928.3 | 0 | NA | NA | NA | NA | NA | #VALUE! |
| ENSMUSG00000045929.9 | 0 | NA | NA | NA | NA | NA | #VALUE! |
| ENSMUSG00000045932.13 | 7.187698 | -0.13373 | 2.861417 | -0.04673 | 0.962725 | NA | #VALUE! |
| ENSMUSG00000045934.16 | 0 | NA | NA | NA | NA | NA | #VALUE! |
| ENSMUSG00000045942.14 | 1.890236 | -4.46257 | 3.93014 | -1.13547 | 0.256177 | NA | #VALUE! |
| ENSMUSG00000045952.4 | 0 | NA | NA | NA | NA | NA | #VALUE! |
| ENSMUSG00000045967.12 | 0 | NA | NA | NA | NA | NA | #VALUE! |
| ENSMUSG00000045968.8 | 2.355969 | 4.72622 | 3.925784 | 1.203892 | 0.228631 | NA | #VALUE! |
| ENSMUSG00000045989.5 | 0 | NA | NA | NA | NA | NA | #VALUE! |
| ENSMUSG00000045994.18 | 0 | NA | NA | NA | NA | NA | #VALUE! |
| ENSMUSG00000046000.6 | 0 | NA | NA | NA | NA | NA | #VALUE! |
| ENSMUSG00000046006.7 | 0 | NA | NA | NA | NA | NA | #VALUE! |
| ENSMUSG00000046008.9 | 0 | NA | NA | NA | NA | NA | #VALUE! |
| ENSMUSG00000046016.4 | 0 | NA | NA | NA | NA | NA | #VALUE! |
| ENSMUSG00000046041.5 | 0 | NA | NA | NA | NA | NA | #VALUE! |
| ENSMUSG00000046049.8 | 0 | NA | NA | NA | NA | NA | #VALUE! |
| ENSMUSG00000046056.16 | 0 | NA | NA | NA | NA | NA | #VALUE! |
| ENSMUSG00000046057.5 | 0 | NA | NA | NA | NA | NA | #VALUE! |
| ENSMUSG00000046058.8 | 0 | NA | NA | NA | NA | NA | #VALUE! |
| ENSMUSG00000046080.18 | 5.914762 | -6.09611 | 3.730177 | -1.63427 | 0.102202 | NA | #VALUE! |
| ENSMUSG00000046082.5 | 0 | NA | NA | NA | NA | NA | #VALUE! |
| ENSMUSG00000046085.8 | 0 | NA | NA | NA | NA | NA | #VALUE! |
| ENSMUSG00000046093.10 | 0 | NA | NA | NA | NA | NA | #VALUE! |
| ENSMUSG00000046095.6 | 0 | NA | NA | NA | NA | NA | #VALUE! |
| ENSMUSG00000046101.17 | 0 | NA | NA | NA | NA | NA | #VALUE! |
| ENSMUSG00000046108.4 | 0 | NA | NA | NA | NA | NA | #VALUE! |
| ENSMUSG00000046109.7 | 0 | NA | NA | NA | NA | NA | #VALUE! |
| ENSMUSG00000046110.16 | 0 | NA | NA | NA | NA | NA | #VALUE! |
| ENSMUSG00000046119.3 | 0 | NA | NA | NA | NA | NA | #VALUE! |
| ENSMUSG00000046130.8 | 0 | NA | NA | NA | NA | NA | #VALUE! |
| ENSMUSG00000046133.4 | 0 | NA | NA | NA | NA | NA | #VALUE! |
| ENSMUSG00000046145.8 | 0 | NA | NA | NA | NA | NA | #VALUE! |
| ENSMUSG00000046150.4 | 0 | NA | NA | NA | NA | NA | #VALUE! |
| ENSMUSG00000046152.17 | 0 | NA | NA | NA | NA | NA | #VALUE! |
| ENSMUSG00000046157.14 | 0.245976 | 1.027628 | 4.067126 | 0.252667 | 0.800526 | NA | #VALUE! |
| ENSMUSG00000046159.17 | 0.630079 | -2.90291 | 3.975047 | -0.73028 | 0.465217 | NA | #VALUE! |
| ENSMUSG00000046160.7 | 7.578719 | 5.373194 | 2.93385 | 1.831448 | 0.067034 | NA | #VALUE! |

| | | | | | | | |
|---|---|---|---|---|---|---|---|
| ENSMUSG00000046168.10 | 0 | NA | NA | NA | NA | NA | #VALUE! |
| ENSMUSG00000046169.11 | 6.069462 | -6.1296 | 3.91415 | -1.56601 | 0.117346 | NA | #VALUE! |
| ENSMUSG00000046173.4 | 0 | NA | NA | NA | NA | NA | #VALUE! |
| ENSMUSG00000046178.4 | 0 | NA | NA | NA | NA | NA | #VALUE! |
| ENSMUSG00000046180.12 | 0 | NA | NA | NA | NA | NA | #VALUE! |
| ENSMUSG00000046182.9 | 0.466276 | 2.449889 | 3.997179 | 0.612904 | 0.53994 | NA | #VALUE! |
| ENSMUSG00000046186.9 | 0 | NA | NA | NA | NA | NA | #VALUE! |
| ENSMUSG00000046191.8 | 0 | NA | NA | NA | NA | NA | #VALUE! |
| ENSMUSG00000046192.5 | 0 | NA | NA | NA | NA | NA | #VALUE! |
| ENSMUSG00000046196.5 | 0 | NA | NA | NA | NA | NA | #VALUE! |
| ENSMUSG00000046203.12 | 0 | NA | NA | NA | NA | NA | #VALUE! |
| ENSMUSG00000046204.15 | 0 | NA | NA | NA | NA | NA | #VALUE! |
| ENSMUSG00000046207.15 | 0 | NA | NA | NA | NA | NA | #VALUE! |
| ENSMUSG00000046210.8 | 0 | NA | NA | NA | NA | NA | #VALUE! |
| ENSMUSG00000046213.10 | 0 | NA | NA | NA | NA | NA | #VALUE! |
| ENSMUSG00000046215.4 | 0 | NA | NA | NA | NA | NA | #VALUE! |
| ENSMUSG00000046219.10 | 0 | NA | NA | NA | NA | NA | #VALUE! |
| ENSMUSG00000046223.11 | 0 | NA | NA | NA | NA | NA | #VALUE! |
| ENSMUSG00000046229.11 | 7.740181 | 6.446428 | 3.889072 | 1.657575 | NA | NA | #VALUE! |
| ENSMUSG00000046240.9 | 0 | NA | NA | NA | NA | NA | #VALUE! |
| ENSMUSG00000046242.9 | 0 | NA | NA | NA | NA | NA | #VALUE! |
| ENSMUSG00000046245.14 | 5.909641 | -0.49543 | 3.035763 | -0.1632 | 0.870362 | NA | #VALUE! |
| ENSMUSG00000046248.10 | 0 | NA | NA | NA | NA | NA | #VALUE! |
| ENSMUSG00000046258.5 | 0 | NA | NA | NA | NA | NA | #VALUE! |
| ENSMUSG00000046259.3 | 0 | NA | NA | NA | NA | NA | #VALUE! |
| ENSMUSG00000046262.15 | 0 | NA | NA | NA | NA | NA | #VALUE! |
| ENSMUSG00000046269.18 | 4.898449 | 5.782444 | 3.915933 | 1.476645 | 0.139771 | NA | #VALUE! |
| ENSMUSG00000046272.5 | 0 | NA | NA | NA | NA | NA | #VALUE! |
| ENSMUSG00000046275.3 | 0 | NA | NA | NA | NA | NA | #VALUE! |
| ENSMUSG00000046280.10 | 0.559531 | 2.703958 | 3.982863 | 0.678898 | 0.497202 | NA | #VALUE! |
| ENSMUSG00000046282.8 | 0 | NA | NA | NA | NA | NA | #VALUE! |
| ENSMUSG00000046287.8 | 0 | NA | NA | NA | NA | NA | #VALUE! |
| ENSMUSG00000046295.14 | 0 | NA | NA | NA | NA | NA | #VALUE! |
| ENSMUSG00000046300.5 | 0 | NA | NA | NA | NA | NA | #VALUE! |
| ENSMUSG00000046317.15 | 0 | NA | NA | NA | NA | NA | #VALUE! |
| ENSMUSG00000046321.9 | 0 | NA | NA | NA | NA | NA | #VALUE! |
| ENSMUSG00000046323.9 | 0.093255 | 0.868093 | 4.072464 | 0.213162 | 0.831201 | NA | #VALUE! |
| ENSMUSG00000046334.5 | 0 | NA | NA | NA | NA | NA | #VALUE! |
| ENSMUSG00000046337.18 | 0 | NA | NA | NA | NA | NA | #VALUE! |
| ENSMUSG00000046338.5 | 0.195789 | -1.05581 | 4.073302 | -0.2592 | 0.795479 | NA | #VALUE! |
| ENSMUSG00000046341.6 | 0 | NA | NA | NA | NA | NA | #VALUE! |
| ENSMUSG00000046345.6 | 0 | NA | NA | NA | NA | NA | #VALUE! |
| ENSMUSG00000046354.4 | 0 | NA | NA | NA | NA | NA | #VALUE! |
| ENSMUSG00000046367.11 | 0 | NA | NA | NA | NA | NA | #VALUE! |

| | | | | | | | |
|---|---|---|---|---|---|---|---|
| ENSMUSG00000046372.5 | 0 | NA | NA | NA | NA | NA | #VALUE! |
| ENSMUSG00000046378.12 | 0 | NA | NA | NA | NA | NA | #VALUE! |
| ENSMUSG00000046380.4 | 0 | NA | NA | NA | NA | NA | #VALUE! |
| ENSMUSG00000046387.6 | 9.101094 | 6.671784 | 3.72269 | 1.792194 | 0.073102 | NA | #VALUE! |
| ENSMUSG00000046388.5 | 0 | NA | NA | NA | NA | NA | #VALUE! |
| ENSMUSG00000046396.8 | 0 | NA | NA | NA | NA | NA | #VALUE! |
| ENSMUSG00000046408.16 | 0 | NA | NA | NA | NA | NA | #VALUE! |
| ENSMUSG00000046411.3 | 0 | NA | NA | NA | NA | NA | #VALUE! |
| ENSMUSG00000046413.15 | 0 | NA | NA | NA | NA | NA | #VALUE! |
| ENSMUSG00000046415.12 | 0 | NA | NA | NA | NA | NA | #VALUE! |
| ENSMUSG00000046417.15 | 0.420053 | -2.33493 | 4.007343 | -0.58266 | 0.56012 | NA | #VALUE! |
| ENSMUSG00000046431.3 | 0 | NA | NA | NA | NA | NA | #VALUE! |
| ENSMUSG00000046432.13 | 0 | NA | NA | NA | NA | NA | #VALUE! |
| ENSMUSG00000046435.8 | 0 | NA | NA | NA | NA | NA | #VALUE! |
| ENSMUSG00000046438.6 | 0 | NA | NA | NA | NA | NA | #VALUE! |
| ENSMUSG00000046440.10 | 0 | NA | NA | NA | NA | NA | #VALUE! |
| ENSMUSG00000046442.5 | 0 | NA | NA | NA | NA | NA | #VALUE! |
| ENSMUSG00000046449.16 | 0 | NA | NA | NA | NA | NA | #VALUE! |
| ENSMUSG00000046450.4 | 0 | NA | NA | NA | NA | NA | #VALUE! |
| ENSMUSG00000046456.15 | 0 | NA | NA | NA | NA | NA | #VALUE! |
| ENSMUSG00000046460.16 | 0 | NA | NA | NA | NA | NA | #VALUE! |
| ENSMUSG00000046463.6 | 0 | NA | NA | NA | NA | NA | #VALUE! |
| ENSMUSG00000046470.6 | 2.051614 | 4.542018 | 3.92836 | 1.156212 | 0.247594 | NA | #VALUE! |
| ENSMUSG00000046474.10 | 0 | NA | NA | NA | NA | NA | #VALUE! |
| ENSMUSG00000046480.7 | 0 | NA | NA | NA | NA | NA | #VALUE! |
| ENSMUSG00000046486.5 | 0 | NA | NA | NA | NA | NA | #VALUE! |
| ENSMUSG00000046487.7 | 0 | NA | NA | NA | NA | NA | #VALUE! |
| ENSMUSG00000046490.8 | 0 | NA | NA | NA | NA | NA | #VALUE! |
| ENSMUSG00000046491.11 | 5.528805 | -0.85781 | 3.761648 | -0.22804 | 0.819615 | NA | #VALUE! |
| ENSMUSG00000046493.6 | 0 | NA | NA | NA | NA | NA | #VALUE! |
| ENSMUSG00000046500.9 | 0 | NA | NA | NA | NA | NA | #VALUE! |
| ENSMUSG00000046518.4 | 0 | NA | NA | NA | NA | NA | #VALUE! |
| ENSMUSG00000046523.6 | 0 | NA | NA | NA | NA | NA | #VALUE! |
| ENSMUSG00000046541.10 | 0 | NA | NA | NA | NA | NA | #VALUE! |
| ENSMUSG00000046550.9 | 0 | NA | NA | NA | NA | NA | #VALUE! |
| ENSMUSG00000046556.8 | 5.462391 | 2.603297 | 3.077135 | 0.846013 | 0.397545 | NA | #VALUE! |
| ENSMUSG00000046561.10 | 0 | NA | NA | NA | NA | NA | #VALUE! |
| ENSMUSG00000046572.11 | 1.365171 | -3.99822 | 3.938957 | -1.01505 | 0.310084 | NA | #VALUE! |
| ENSMUSG00000046574.9 | 6.395995 | -0.60648 | 3.755535 | -0.16149 | 0.871708 | NA | #VALUE! |
| ENSMUSG00000046580.11 | 0 | NA | NA | NA | NA | NA | #VALUE! |
| ENSMUSG00000046585.10 | 0 | NA | NA | NA | NA | NA | #VALUE! |
| ENSMUSG00000046589.8 | 0 | NA | NA | NA | NA | NA | #VALUE! |
| ENSMUSG00000046590.4 | 0 | NA | NA | NA | NA | NA | #VALUE! |
| ENSMUSG00000046591.11 | 0 | NA | NA | NA | NA | NA | #VALUE! |

| | | | | | | | |
|---|---|---|---|---|---|---|---|
| ENSMUSG00000046593.4 | 0 | NA | NA | NA | NA | NA | #VALUE! |
| ENSMUSG00000046605.15 | 0.525066 | -2.64687 | 3.988095 | -0.66369 | 0.506887 | NA | #VALUE! |
| ENSMUSG00000046607.7 | 0 | NA | NA | NA | NA | NA | #VALUE! |
| ENSMUSG00000046610.16 | 0 | NA | NA | NA | NA | NA | #VALUE! |
| ENSMUSG00000046613.20 | 0 | NA | NA | NA | NA | NA | #VALUE! |
| ENSMUSG00000046615.3 | 0 | NA | NA | NA | NA | NA | #VALUE! |
| ENSMUSG00000046618.8 | 0 | NA | NA | NA | NA | NA | #VALUE! |
| ENSMUSG00000046620.5 | 0.105013 | -0.97324 | 4.076293 | -0.23876 | 0.811295 | NA | #VALUE! |
| ENSMUSG00000046623.9 | 0 | NA | NA | NA | NA | NA | #VALUE! |
| ENSMUSG00000046634.13 | 0 | NA | NA | NA | NA | NA | #VALUE! |
| ENSMUSG00000046636.6 | 0 | NA | NA | NA | NA | NA | #VALUE! |
| ENSMUSG00000046637.4 | 0 | NA | NA | NA | NA | NA | #VALUE! |
| ENSMUSG00000046643.3 | 0 | NA | NA | NA | NA | NA | #VALUE! |
| ENSMUSG00000046650.5 | 0 | NA | NA | NA | NA | NA | #VALUE! |
| ENSMUSG00000046652.4 | 0 | NA | NA | NA | NA | NA | #VALUE! |
| ENSMUSG00000046658.17 | 1.616243 | -4.21378 | 3.934514 | -1.07098 | 0.284179 | NA | #VALUE! |
| ENSMUSG00000046676.5 | 0 | NA | NA | NA | NA | NA | #VALUE! |
| ENSMUSG00000046678.7 | 0 | NA | NA | NA | NA | NA | #VALUE! |
| ENSMUSG00000046683.17 | 0 | NA | NA | NA | NA | NA | #VALUE! |
| ENSMUSG00000046687.7 | 0.988358 | 0.480316 | 3.915094 | 0.122683 | 0.902358 | NA | #VALUE! |
| ENSMUSG00000046691.15 | 0 | NA | NA | NA | NA | NA | #VALUE! |
| ENSMUSG00000046694.7 | 0 | NA | NA | NA | NA | NA | #VALUE! |
| ENSMUSG00000046697.18 | 0 | NA | NA | NA | NA | NA | #VALUE! |
| ENSMUSG00000046699.15 | 0 | NA | NA | NA | NA | NA | #VALUE! |
| ENSMUSG00000046709.19 | 0 | NA | NA | NA | NA | NA | #VALUE! |
| ENSMUSG00000046711.17 | 0 | NA | NA | NA | NA | NA | #VALUE! |
| ENSMUSG00000046714.8 | 0 | NA | NA | NA | NA | NA | #VALUE! |
| ENSMUSG00000046716.8 | 0 | NA | NA | NA | NA | NA | #VALUE! |
| ENSMUSG00000046717.6 | 0 | NA | NA | NA | NA | NA | #VALUE! |
| ENSMUSG00000046719.8 | 0 | NA | NA | NA | NA | NA | #VALUE! |
| ENSMUSG00000046723.5 | 0 | NA | NA | NA | NA | NA | #VALUE! |
| ENSMUSG00000046731.6 | 1.020554 | -3.54462 | 3.950745 | -0.8972 | 0.36961 | NA | #VALUE! |
| ENSMUSG00000046733.8 | 0 | NA | NA | NA | NA | NA | #VALUE! |
| ENSMUSG00000046743.7 | 5.015502 | -5.85421 | 3.9157 | -1.49506 | 0.134899 | NA | #VALUE! |
| ENSMUSG00000046747.9 | 0 | NA | NA | NA | NA | NA | #VALUE! |
| ENSMUSG00000046748.9 | 0 | NA | NA | NA | NA | NA | #VALUE! |
| ENSMUSG00000046750.19 | 0 | NA | NA | NA | NA | NA | #VALUE! |
| ENSMUSG00000046755.6 | 0 | NA | NA | NA | NA | NA | #VALUE! |
| ENSMUSG00000046764.9 | 0 | NA | NA | NA | NA | NA | #VALUE! |
| ENSMUSG00000046768.14 | 3.851299 | 2.503502 | 3.81879 | 0.655575 | 0.512098 | NA | #VALUE! |
| ENSMUSG00000046774.18 | 0 | NA | NA | NA | NA | NA | #VALUE! |
| ENSMUSG00000046782.15 | 1.575197 | -4.20222 | 3.934736 | -1.06798 | 0.285529 | NA | #VALUE! |
| ENSMUSG00000046790.8 | 0 | NA | NA | NA | NA | NA | #VALUE! |
| ENSMUSG00000046793.9 | 0 | NA | NA | NA | NA | NA | #VALUE! |

| | | | | | | | |
|---|---|---|---|---|---|---|---|
| ENSMUSG00000046804.14 | 0 | NA | NA | NA | NA | NA | #VALUE! |
| ENSMUSG00000046806.14 | 8.894486 | -4.77263 | 3.412252 | -1.39867 | 0.161911 | NA | #VALUE! |
| ENSMUSG00000046807.11 | 0 | NA | NA | NA | NA | NA | #VALUE! |
| ENSMUSG00000046808.18 | 6.749568 | 6.244819 | 3.651237 | 1.710329 | 0.087205 | NA | #VALUE! |
| ENSMUSG00000046818.8 | 0 | NA | NA | NA | NA | NA | #VALUE! |
| ENSMUSG00000046822.15 | 3.380703 | 3.680003 | 3.898143 | 0.94404 | 0.345149 | NA | #VALUE! |
| ENSMUSG00000046828.4 | 0 | NA | NA | NA | NA | NA | #VALUE! |
| ENSMUSG00000046834.8 | 0 | NA | NA | NA | NA | NA | #VALUE! |
| ENSMUSG00000046840.17 | 5.129036 | 5.854808 | 3.915485 | 1.495296 | 0.134837 | NA | #VALUE! |
| ENSMUSG00000046841.6 | 0.924965 | -3.42737 | 3.954433 | -0.86672 | 0.386098 | NA | #VALUE! |
| ENSMUSG00000046844.7 | 0 | NA | NA | NA | NA | NA | #VALUE! |
| ENSMUSG00000046845.2 | 0 | NA | NA | NA | NA | NA | #VALUE! |
| ENSMUSG00000046846.5 | 0 | NA | NA | NA | NA | NA | #VALUE! |
| ENSMUSG00000046854.17 | 0 | NA | NA | NA | NA | NA | #VALUE! |
| ENSMUSG00000046856.14 | 0 | NA | NA | NA | NA | NA | #VALUE! |
| ENSMUSG00000046881.8 | 0 | NA | NA | NA | NA | NA | #VALUE! |
| ENSMUSG00000046886.7 | 0 | NA | NA | NA | NA | NA | #VALUE! |
| ENSMUSG00000046899.8 | 0 | NA | NA | NA | NA | NA | #VALUE! |
| ENSMUSG00000046908.6 | 0 | NA | NA | NA | NA | NA | #VALUE! |
| ENSMUSG00000046913.4 | 0 | NA | NA | NA | NA | NA | #VALUE! |
| ENSMUSG00000046922.8 | 0 | NA | NA | NA | NA | NA | #VALUE! |
| ENSMUSG00000046924.5 | 0 | NA | NA | NA | NA | NA | #VALUE! |
| ENSMUSG00000046932.3 | 0 | NA | NA | NA | NA | NA | #VALUE! |
| ENSMUSG00000046934.6 | 0 | NA | NA | NA | NA | NA | #VALUE! |
| ENSMUSG00000046942.18 | 0 | NA | NA | NA | NA | NA | #VALUE! |
| ENSMUSG00000046947.13 | 1.789449 | 3.15878 | 3.927142 | 0.804346 | 0.421197 | NA | #VALUE! |
| ENSMUSG00000046952.10 | 0 | NA | NA | NA | NA | NA | #VALUE! |
| ENSMUSG00000046957.6 | 0 | NA | NA | NA | NA | NA | #VALUE! |
| ENSMUSG00000046958.11 | 0 | NA | NA | NA | NA | NA | #VALUE! |
| ENSMUSG00000046961.9 | 0 | NA | NA | NA | NA | NA | #VALUE! |
| ENSMUSG00000046971.8 | 0 | NA | NA | NA | NA | NA | #VALUE! |
| ENSMUSG00000046974.7 | 0 | NA | NA | NA | NA | NA | #VALUE! |
| ENSMUSG00000046975.8 | 0 | NA | NA | NA | NA | NA | #VALUE! |
| ENSMUSG00000046991.13 | 3.183978 | -0.51037 | 3.784731 | -0.13485 | 0.892731 | NA | #VALUE! |
| ENSMUSG00000046993.7 | 0 | NA | NA | NA | NA | NA | #VALUE! |
| ENSMUSG00000046995.4 | 0 | NA | NA | NA | NA | NA | #VALUE! |
| ENSMUSG00000046999.3 | 0 | NA | NA | NA | NA | NA | #VALUE! |
| ENSMUSG00000047002.3 | 0 | NA | NA | NA | NA | NA | #VALUE! |
| ENSMUSG00000047003.16 | 0.84294 | 2.074799 | 3.964343 | 0.523365 | 0.60072 | NA | #VALUE! |
| ENSMUSG00000047013.16 | 0 | NA | NA | NA | NA | NA | #VALUE! |
| ENSMUSG00000047014.8 | 0 | NA | NA | NA | NA | NA | #VALUE! |
| ENSMUSG00000047021.15 | 0 | NA | NA | NA | NA | NA | #VALUE! |
| ENSMUSG00000047025.6 | 0 | NA | NA | NA | NA | NA | #VALUE! |
| ENSMUSG00000047026.10 | 0 | NA | NA | NA | NA | NA | #VALUE! |

| | | | | | | | |
|---|---|---|---|---|---|---|---|
| ENSMUSG00000047031.6 | 0 | NA | NA | NA | NA | NA | #VALUE! |
| ENSMUSG00000047033.3 | 0 | NA | NA | NA | NA | NA | #VALUE! |
| ENSMUSG00000047034.11 | 0 | NA | NA | NA | NA | NA | #VALUE! |
| ENSMUSG00000047037.6 | 0.195789 | -1.05581 | 4.073302 | -0.2592 | 0.795479 | NA | #VALUE! |
| ENSMUSG00000047039.5 | 0 | NA | NA | NA | NA | NA | #VALUE! |
| ENSMUSG00000047040.10 | 0 | NA | NA | NA | NA | NA | #VALUE! |
| ENSMUSG00000047046.9 | 0 | NA | NA | NA | NA | NA | #VALUE! |
| ENSMUSG00000047048.5 | 0 | NA | NA | NA | NA | NA | #VALUE! |
| ENSMUSG00000047050.3 | 0 | NA | NA | NA | NA | NA | #VALUE! |
| ENSMUSG00000047053.6 | 0 | NA | NA | NA | NA | NA | #VALUE! |
| ENSMUSG00000047061.7 | 0 | NA | NA | NA | NA | NA | #VALUE! |
| ENSMUSG00000047079.2 | 0 | NA | NA | NA | NA | NA | #VALUE! |
| ENSMUSG00000047085.15 | 0 | NA | NA | NA | NA | NA | #VALUE! |
| ENSMUSG00000047094.10 | 0 | NA | NA | NA | NA | NA | #VALUE! |
| ENSMUSG00000047098.19 | 7.269162 | -6.38971 | 3.545633 | -1.80214 | 0.071524 | NA | #VALUE! |
| ENSMUSG00000047102.5 | 0 | NA | NA | NA | NA | NA | #VALUE! |
| ENSMUSG00000047104.6 | 0 | NA | NA | NA | NA | NA | #VALUE! |
| ENSMUSG00000047108.5 | 0 | NA | NA | NA | NA | NA | #VALUE! |
| ENSMUSG00000047115.16 | 0 | NA | NA | NA | NA | NA | #VALUE! |
| ENSMUSG00000047117.14 | 0 | NA | NA | NA | NA | NA | #VALUE! |
| ENSMUSG00000047129.12 | 0.195789 | -1.05581 | 4.073302 | -0.2592 | 0.795479 | NA | #VALUE! |
| ENSMUSG00000047143.4 | 0 | NA | NA | NA | NA | NA | #VALUE! |
| ENSMUSG00000047146.18 | 0 | NA | NA | NA | NA | NA | #VALUE! |
| ENSMUSG00000047149.6 | 0 | NA | NA | NA | NA | NA | #VALUE! |
| ENSMUSG00000047155.14 | 0 | NA | NA | NA | NA | NA | #VALUE! |
| ENSMUSG00000047161.13 | 0 | NA | NA | NA | NA | NA | #VALUE! |
| ENSMUSG00000047168.7 | 0 | NA | NA | NA | NA | NA | #VALUE! |
| ENSMUSG00000047171.9 | 0 | NA | NA | NA | NA | NA | #VALUE! |
| ENSMUSG00000047181.13 | 3.450442 | 5.285914 | 3.919688 | 1.348555 | 0.17748 | NA | #VALUE! |
| ENSMUSG00000047182.7 | 0 | NA | NA | NA | NA | NA | #VALUE! |
| ENSMUSG00000047189.4 | 0 | NA | NA | NA | NA | NA | #VALUE! |
| ENSMUSG00000047193.17 | 8.822299 | 3.709054 | 3.069999 | 1.208162 | NA | NA | #VALUE! |
| ENSMUSG00000047197.2 | 0 | NA | NA | NA | NA | NA | #VALUE! |
| ENSMUSG00000047203.4 | 0 | NA | NA | NA | NA | NA | #VALUE! |
| ENSMUSG00000047205.13 | 0 | NA | NA | NA | NA | NA | #VALUE! |
| ENSMUSG00000047207.4 | 0 | NA | NA | NA | NA | NA | #VALUE! |
| ENSMUSG00000047216.9 | 0.168284 | 1.027628 | 4.067126 | 0.252667 | 0.800526 | NA | #VALUE! |
| ENSMUSG00000047220.6 | 0 | NA | NA | NA | NA | NA | #VALUE! |
| ENSMUSG00000047222.8 | 2.310289 | -4.7496 | 3.925939 | -1.2098 | 0.226355 | NA | #VALUE! |
| ENSMUSG00000047225.9 | 0 | NA | NA | NA | NA | NA | #VALUE! |
| ENSMUSG00000047227.6 | 0.525066 | -2.64687 | 3.988095 | -0.66369 | 0.506887 | NA | #VALUE! |
| ENSMUSG00000047237.14 | 0 | NA | NA | NA | NA | NA | #VALUE! |
| ENSMUSG00000047238.5 | 8.398953 | 3.30138 | 3.024997 | 1.091367 | 0.275112 | NA | #VALUE! |
| ENSMUSG00000047242.15 | 1.836997 | -4.39907 | 3.931186 | -1.11902 | 0.263133 | NA | #VALUE! |

| | | | | | | | |
|---|---|---|---|---|---|---|---|
| ENSMUSG00000047246.8 | 2.144869 | 4.605508 | 3.927435 | 1.17265 | 0.240936 | NA | #VALUE! |
| ENSMUSG00000047248.21 | 8.639423 | 2.520022 | 3.449718 | 0.730501 | 0.465084 | NA | #VALUE! |
| ENSMUSG00000047253.5 | 0 | NA | NA | NA | NA | NA | #VALUE! |
| ENSMUSG00000047257.14 | 0 | NA | NA | NA | NA | NA | #VALUE! |
| ENSMUSG00000047259.4 | 0 | NA | NA | NA | NA | NA | #VALUE! |
| ENSMUSG00000047261.10 | 0 | NA | NA | NA | NA | NA | #VALUE! |
| ENSMUSG00000047264.9 | 0 | NA | NA | NA | NA | NA | #VALUE! |
| ENSMUSG00000047281.4 | 0 | NA | NA | NA | NA | NA | #VALUE! |
| ENSMUSG00000047284.15 | 3.121217 | 4.147825 | 3.917305 | 1.058847 | 0.28967 | NA | #VALUE! |
| ENSMUSG00000047286.8 | 0 | NA | NA | NA | NA | NA | #VALUE! |
| ENSMUSG00000047293.8 | 0 | NA | NA | NA | NA | NA | #VALUE! |
| ENSMUSG00000047295.17 | 0 | NA | NA | NA | NA | NA | #VALUE! |
| ENSMUSG00000047298.4 | 0 | NA | NA | NA | NA | NA | #VALUE! |
| ENSMUSG00000047307.3 | 0 | NA | NA | NA | NA | NA | #VALUE! |
| ENSMUSG00000047324.6 | 0 | NA | NA | NA | NA | NA | #VALUE! |
| ENSMUSG00000047330.9 | 1.995249 | -4.53986 | 3.928926 | -1.1555 | 0.247887 | NA | #VALUE! |
| ENSMUSG00000047342.18 | 0 | NA | NA | NA | NA | NA | #VALUE! |
| ENSMUSG00000047343.5 | 0 | NA | NA | NA | NA | NA | #VALUE! |
| ENSMUSG00000047344.10 | 0 | NA | NA | NA | NA | NA | #VALUE! |
| ENSMUSG00000047347.8 | 0 | NA | NA | NA | NA | NA | #VALUE! |
| ENSMUSG00000047352.7 | 0 | NA | NA | NA | NA | NA | #VALUE! |
| ENSMUSG00000047356.3 | 0 | NA | NA | NA | NA | NA | #VALUE! |
| ENSMUSG00000047361.17 | 0 | NA | NA | NA | NA | NA | #VALUE! |
| ENSMUSG00000047363.5 | 0 | NA | NA | NA | NA | NA | #VALUE! |
| ENSMUSG00000047369.18 | 5.090517 | -5.87548 | 3.91557 | -1.50054 | 0.133474 | NA | #VALUE! |
| ENSMUSG00000047370.6 | 0 | NA | NA | NA | NA | NA | #VALUE! |
| ENSMUSG00000047379.8 | 0 | NA | NA | NA | NA | NA | #VALUE! |
| ENSMUSG00000047383.9 | 0 | NA | NA | NA | NA | NA | #VALUE! |
| ENSMUSG00000047384.4 | 0 | NA | NA | NA | NA | NA | #VALUE! |
| ENSMUSG00000047390.7 | 0 | NA | NA | NA | NA | NA | #VALUE! |
| ENSMUSG00000047409.14 | 30.21187 | -0.42891 | 2.297303 | -0.1867 | NA | NA | #VALUE! |
| ENSMUSG00000047414.7 | 0.168284 | 1.027628 | 4.067126 | 0.252667 | 0.800526 | NA | #VALUE! |
| ENSMUSG00000047415.13 | 0 | NA | NA | NA | NA | NA | #VALUE! |
| ENSMUSG00000047417.19 | 7.26072 | -6.38797 | 3.52188 | -1.8138 | 0.069709 | NA | #VALUE! |
| ENSMUSG00000047419.6 | 0 | NA | NA | NA | NA | NA | #VALUE! |
| ENSMUSG00000047428.15 | 0 | NA | NA | NA | NA | NA | #VALUE! |
| ENSMUSG00000047434.15 | 4.401589 | 5.627369 | 3.703574 | 1.519443 | 0.128651 | NA | #VALUE! |
| ENSMUSG00000047441.5 | 0 | NA | NA | NA | NA | NA | #VALUE! |
| ENSMUSG00000047443.15 | 0 | NA | NA | NA | NA | NA | #VALUE! |
| ENSMUSG00000047444.4 | 0 | NA | NA | NA | NA | NA | #VALUE! |
| ENSMUSG00000047462.10 | 0 | NA | NA | NA | NA | NA | #VALUE! |
| ENSMUSG00000047473.15 | 7.784981 | 1.220946 | 3.121957 | 0.391084 | 0.695735 | NA | #VALUE! |
| ENSMUSG00000047485.7 | 0 | NA | NA | NA | NA | NA | #VALUE! |
| ENSMUSG00000047495.16 | 0 | NA | NA | NA | NA | NA | #VALUE! |

| | | | | | | | |
|---|---|---|---|---|---|---|---|
| ENSMUSG00000047497.11 | 0 | NA | NA | NA | NA | NA | #VALUE! |
| ENSMUSG00000047501.4 | 0 | NA | NA | NA | NA | NA | #VALUE! |
| ENSMUSG00000047502.15 | 0 | NA | NA | NA | NA | NA | #VALUE! |
| ENSMUSG00000047507.13 | 0 | NA | NA | NA | NA | NA | #VALUE! |
| ENSMUSG00000047508.10 | 0 | NA | NA | NA | NA | NA | #VALUE! |
| ENSMUSG00000047509.2 | 0 | NA | NA | NA | NA | NA | #VALUE! |
| ENSMUSG00000047511.7 | 1.229878 | 3.761857 | 3.943699 | 0.95389 | 0.340139 | NA | #VALUE! |
| ENSMUSG00000047515.9 | 0 | NA | NA | NA | NA | NA | #VALUE! |
| ENSMUSG00000047517.14 | 0 | NA | NA | NA | NA | NA | #VALUE! |
| ENSMUSG00000047518.4 | 0 | NA | NA | NA | NA | NA | #VALUE! |
| ENSMUSG00000047528.14 | 0 | NA | NA | NA | NA | NA | #VALUE! |
| ENSMUSG00000047531.10 | 0 | NA | NA | NA | NA | NA | #VALUE! |
| ENSMUSG00000047534.18 | 4.529533 | 0.560003 | 3.767911 | 0.148624 | 0.88185 | NA | #VALUE! |
| ENSMUSG00000047535.12 | 0 | NA | NA | NA | NA | NA | #VALUE! |
| ENSMUSG00000047544.8 | 0 | NA | NA | NA | NA | NA | #VALUE! |
| ENSMUSG00000047545.6 | 0 | NA | NA | NA | NA | NA | #VALUE! |
| ENSMUSG00000047557.3 | 1.678593 | 4.255795 | 3.933066 | 1.082055 | 0.279228 | NA | #VALUE! |
| ENSMUSG00000047562.4 | 0 | NA | NA | NA | NA | NA | #VALUE! |
| ENSMUSG00000047564.5 | 0 | NA | NA | NA | NA | NA | #VALUE! |
| ENSMUSG00000047565.6 | 0 | NA | NA | NA | NA | NA | #VALUE! |
| ENSMUSG00000047586.5 | 0 | NA | NA | NA | NA | NA | #VALUE! |
| ENSMUSG00000047591.6 | 4.181584 | 5.545962 | 3.917562 | 1.415667 | 0.156873 | NA | #VALUE! |
| ENSMUSG00000047592.18 | 0 | NA | NA | NA | NA | NA | #VALUE! |
| ENSMUSG00000047594.5 | 0 | NA | NA | NA | NA | NA | #VALUE! |
| ENSMUSG00000047604.4 | 0 | NA | NA | NA | NA | NA | #VALUE! |
| ENSMUSG00000047606.5 | 0 | NA | NA | NA | NA | NA | #VALUE! |
| ENSMUSG00000047619.6 | 0 | NA | NA | NA | NA | NA | #VALUE! |
| ENSMUSG00000047626.5 | 0 | NA | NA | NA | NA | NA | #VALUE! |
| ENSMUSG00000047632.12 | 1.492083 | 4.088114 | 3.936282 | 1.038572 | 0.299004 | NA | #VALUE! |
| ENSMUSG00000047636.4 | 0 | NA | NA | NA | NA | NA | #VALUE! |
| ENSMUSG00000047641.9 | 0 | NA | NA | NA | NA | NA | #VALUE! |
| ENSMUSG00000047642.16 | 0 | NA | NA | NA | NA | NA | #VALUE! |
| ENSMUSG00000047643.6 | 0 | NA | NA | NA | NA | NA | #VALUE! |
| ENSMUSG00000047652.5 | 0 | NA | NA | NA | NA | NA | #VALUE! |
| ENSMUSG00000047654.7 | 0 | NA | NA | NA | NA | NA | #VALUE! |
| ENSMUSG00000047655.3 | 0 | NA | NA | NA | NA | NA | #VALUE! |
| ENSMUSG00000047656.10 | 5.380447 | 1.230005 | 3.239882 | 0.379645 | 0.704209 | NA | #VALUE! |
| ENSMUSG00000047657.7 | 0.093255 | 0.868093 | 4.072464 | 0.213162 | 0.831201 | NA | #VALUE! |
| ENSMUSG00000047658.7 | 0 | NA | NA | NA | NA | NA | #VALUE! |
| ENSMUSG00000047661.10 | 0 | NA | NA | NA | NA | NA | #VALUE! |
| ENSMUSG00000047667.6 | 0 | NA | NA | NA | NA | NA | #VALUE! |
| ENSMUSG00000047669.9 | 5.72164 | 6.006287 | 3.914617 | 1.534323 | 0.12495 | NA | #VALUE! |
| ENSMUSG00000047671.9 | 0 | NA | NA | NA | NA | NA | #VALUE! |
| ENSMUSG00000047674.3 | 0 | NA | NA | NA | NA | NA | #VALUE! |

| | | | | | | | |
|---|---|---|---|---|---|---|---|
| ENSMUSG00000047676.7 | 1.710884 | 3.083092 | 3.928631 | 0.784775 | 0.432585 | NA | #VALUE! |
| ENSMUSG00000047678.2 | 0 | NA | NA | NA | NA | NA | #VALUE! |
| ENSMUSG00000047686.10 | 0 | NA | NA | NA | NA | NA | #VALUE! |
| ENSMUSG00000047692.7 | 0 | NA | NA | NA | NA | NA | #VALUE! |
| ENSMUSG00000047696.17 | 0 | NA | NA | NA | NA | NA | #VALUE! |
| ENSMUSG00000047702.3 | 0 | NA | NA | NA | NA | NA | #VALUE! |
| ENSMUSG00000047710.9 | 37.70002 | 1.004691 | 1.597771 | 0.628808 | NA | NA | #VALUE! |
| ENSMUSG00000047712.9 | 4.492049 | -4.20049 | 3.891977 | -1.07927 | 0.280468 | NA | #VALUE! |
| ENSMUSG00000047716.5 | 0 | NA | NA | NA | NA | NA | #VALUE! |
| ENSMUSG00000047720.8 | 0 | NA | NA | NA | NA | NA | #VALUE! |
| ENSMUSG00000047730.18 | 0 | NA | NA | NA | NA | NA | #VALUE! |
| ENSMUSG00000047733.11 | 0 | NA | NA | NA | NA | NA | #VALUE! |
| ENSMUSG00000047746.15 | 0 | NA | NA | NA | NA | NA | #VALUE! |
| ENSMUSG00000047747.11 | 0 | NA | NA | NA | NA | NA | #VALUE! |
| ENSMUSG00000047749.3 | 0 | NA | NA | NA | NA | NA | #VALUE! |
| ENSMUSG00000047751.10 | 0 | NA | NA | NA | NA | NA | #VALUE! |
| ENSMUSG00000047757.18 | 4.402877 | 4.547076 | 3.909966 | 1.162945 | 0.244852 | NA | #VALUE! |
| ENSMUSG00000047759.7 | 0.336567 | 1.944925 | 4.029936 | 0.482619 | 0.629366 | NA | #VALUE! |
| ENSMUSG00000047766.16 | 0 | NA | NA | NA | NA | NA | #VALUE! |
| ENSMUSG00000047773.16 | 0 | NA | NA | NA | NA | NA | #VALUE! |
| ENSMUSG00000047777.10 | 0.504851 | 2.502843 | 3.993993 | 0.626652 | 0.530887 | NA | #VALUE! |
| ENSMUSG00000047786.13 | 0 | NA | NA | NA | NA | NA | #VALUE! |
| ENSMUSG00000047787.9 | 2.051614 | 4.542018 | 3.92836 | 1.156212 | 0.247594 | NA | #VALUE! |
| ENSMUSG00000047794.7 | 0 | NA | NA | NA | NA | NA | #VALUE! |
| ENSMUSG00000047798.16 | 6.489038 | 0.158954 | 3.753961 | 0.042343 | 0.966225 | NA | #VALUE! |
| ENSMUSG00000047799.16 | 0 | NA | NA | NA | NA | NA | #VALUE! |
| ENSMUSG00000047807.11 | 0 | NA | NA | NA | NA | NA | #VALUE! |
| ENSMUSG00000047810.10 | 0 | NA | NA | NA | NA | NA | #VALUE! |
| ENSMUSG00000047819.4 | 0 | NA | NA | NA | NA | NA | #VALUE! |
| ENSMUSG00000047821.17 | 0 | NA | NA | NA | NA | NA | #VALUE! |
| ENSMUSG00000047824.14 | 1.970117 | 3.472986 | 3.926141 | 0.88458 | 0.376383 | NA | #VALUE! |
| ENSMUSG00000047828.5 | 0 | NA | NA | NA | NA | NA | #VALUE! |
| ENSMUSG00000047832.10 | 2.386828 | -4.79366 | 3.925364 | -1.2212 | 0.22201 | NA | #VALUE! |
| ENSMUSG00000047841.9 | 0 | NA | NA | NA | NA | NA | #VALUE! |
| ENSMUSG00000047842.8 | 0 | NA | NA | NA | NA | NA | #VALUE! |
| ENSMUSG00000047844.4 | 0 | NA | NA | NA | NA | NA | #VALUE! |
| ENSMUSG00000047854.6 | 0 | NA | NA | NA | NA | NA | #VALUE! |
| ENSMUSG00000047861.3 | 0 | NA | NA | NA | NA | NA | #VALUE! |
| ENSMUSG00000047868.7 | 0 | NA | NA | NA | NA | NA | #VALUE! |
| ENSMUSG00000047875.7 | 0 | NA | NA | NA | NA | NA | #VALUE! |
| ENSMUSG00000047878.14 | 3.061661 | -5.13931 | 3.921414 | -1.31058 | 0.190001 | NA | #VALUE! |
| ENSMUSG00000047880.16 | 0 | NA | NA | NA | NA | NA | #VALUE! |
| ENSMUSG00000047884.7 | 0 | NA | NA | NA | NA | NA | #VALUE! |
| ENSMUSG00000047889.10 | 0 | NA | NA | NA | NA | NA | #VALUE! |

| | | | | | | | |
|---|---|---|---|---|---|---|---|
| ENSMUSG00000047894.3 | 0 | NA | NA | NA | NA | NA | #VALUE! |
| ENSMUSG00000047897.11 | 0 | NA | NA | NA | NA | NA | #VALUE! |
| ENSMUSG00000047898.8 | 0 | NA | NA | NA | NA | NA | #VALUE! |
| ENSMUSG00000047904.7 | 5.745699 | 0.781798 | 3.495064 | 0.223686 | 0.823001 | NA | #VALUE! |
| ENSMUSG00000047910.7 | 0 | NA | NA | NA | NA | NA | #VALUE! |
| ENSMUSG00000047911.7 | 0 | NA | NA | NA | NA | NA | #VALUE! |
| ENSMUSG00000047921.18 | 1.305573 | 3.898287 | 3.940396 | 0.989314 | 0.32251 | NA | #VALUE! |
| ENSMUSG00000047935.18 | 0 | NA | NA | NA | NA | NA | #VALUE! |
| ENSMUSG00000047938.8 | 0 | NA | NA | NA | NA | NA | #VALUE! |
| ENSMUSG00000047940.14 | 0 | NA | NA | NA | NA | NA | #VALUE! |
| ENSMUSG00000047945.7 | 35.4798 | -1.38242 | 1.596867 | -0.86571 | NA | NA | #VALUE! |
| ENSMUSG00000047953.8 | 0 | NA | NA | NA | NA | NA | #VALUE! |
| ENSMUSG00000047959.5 | 0 | NA | NA | NA | NA | NA | #VALUE! |
| ENSMUSG00000047960.5 | 0 | NA | NA | NA | NA | NA | #VALUE! |
| ENSMUSG00000047965.10 | 0.245976 | 1.027628 | 4.067126 | 0.252667 | 0.800526 | NA | #VALUE! |
| ENSMUSG00000047969.6 | 0 | NA | NA | NA | NA | NA | #VALUE! |
| ENSMUSG00000047976.5 | 0 | NA | NA | NA | NA | NA | #VALUE! |
| ENSMUSG00000047977.4 | 0 | NA | NA | NA | NA | NA | #VALUE! |
| ENSMUSG00000047980.7 | 0 | NA | NA | NA | NA | NA | #VALUE! |
| ENSMUSG00000047986.13 | 0 | NA | NA | NA | NA | NA | #VALUE! |
| ENSMUSG00000047988.2 | 0 | NA | NA | NA | NA | NA | #VALUE! |
| ENSMUSG00000047990.7 | 0 | NA | NA | NA | NA | NA | #VALUE! |
| ENSMUSG00000047992.12 | 0 | NA | NA | NA | NA | NA | #VALUE! |
| ENSMUSG00000047995.8 | 0 | NA | NA | NA | NA | NA | #VALUE! |
| ENSMUSG00000047996.17 | 3.539928 | 0.738575 | 3.709194 | 0.19912 | 0.842169 | NA | #VALUE! |
| ENSMUSG00000048001.8 | 0 | NA | NA | NA | NA | NA | #VALUE! |
| ENSMUSG00000048003.13 | 0 | NA | NA | NA | NA | NA | #VALUE! |
| ENSMUSG00000048004.16 | 0 | NA | NA | NA | NA | NA | #VALUE! |
| ENSMUSG00000048012.20 | 0 | NA | NA | NA | NA | NA | #VALUE! |
| ENSMUSG00000048013.10 | 0 | NA | NA | NA | NA | NA | #VALUE! |
| ENSMUSG00000048015.5 | 0 | NA | NA | NA | NA | NA | #VALUE! |
| ENSMUSG00000048022.12 | 5.436834 | 1.312691 | 3.767015 | 0.34847 | 0.727487 | NA | #VALUE! |
| ENSMUSG00000048027.10 | 8.91355 | -3.061 | 3.367086 | -0.9091 | NA | NA | #VALUE! |
| ENSMUSG00000048029.11 | 0 | NA | NA | NA | NA | NA | #VALUE! |
| ENSMUSG00000048031.16 | 0 | NA | NA | NA | NA | NA | #VALUE! |
| ENSMUSG00000048038.9 | 0 | NA | NA | NA | NA | NA | #VALUE! |
| ENSMUSG00000048040.9 | 0 | NA | NA | NA | NA | NA | #VALUE! |
| ENSMUSG00000048058.18 | 2.891094 | -3.8598 | 3.909837 | -0.9872 | 0.323544 | NA | #VALUE! |
| ENSMUSG00000048062.5 | 0 | NA | NA | NA | NA | NA | #VALUE! |
| ENSMUSG00000048065.9 | 0 | NA | NA | NA | NA | NA | #VALUE! |
| ENSMUSG00000048067.9 | 0 | NA | NA | NA | NA | NA | #VALUE! |
| ENSMUSG00000048070.5 | 0 | NA | NA | NA | NA | NA | #VALUE! |
| ENSMUSG00000048077.7 | 0 | NA | NA | NA | NA | NA | #VALUE! |
| ENSMUSG00000048078.17 | 0 | NA | NA | NA | NA | NA | #VALUE! |

| | | | | | | | |
|---|---|---|---|---|---|---|---|
| ENSMUSG00000048080.3 | 0 | NA | NA | NA | NA | NA | #VALUE! |
| ENSMUSG00000048096.8 | 0 | NA | NA | NA | NA | NA | #VALUE! |
| ENSMUSG00000048101.7 | 0 | NA | NA | NA | NA | NA | #VALUE! |
| ENSMUSG00000048108.14 | 0 | NA | NA | NA | NA | NA | #VALUE! |
| ENSMUSG00000048120.17 | 7.658731 | -1.90634 | 3.151627 | -0.60488 | 0.545262 | NA | #VALUE! |
| ENSMUSG00000048138.11 | 0 | NA | NA | NA | NA | NA | #VALUE! |
| ENSMUSG00000048142.10 | 0 | NA | NA | NA | NA | NA | #VALUE! |
| ENSMUSG00000048148.19 | 0 | NA | NA | NA | NA | NA | #VALUE! |
| ENSMUSG00000048153.7 | 0 | NA | NA | NA | NA | NA | #VALUE! |
| ENSMUSG00000048154.18 | 8.399978 | 1.817415 | 2.656068 | 0.68425 | 0.493817 | NA | #VALUE! |
| ENSMUSG00000048155.5 | 0 | NA | NA | NA | NA | NA | #VALUE! |
| ENSMUSG00000048163.14 | 0.466276 | 2.449889 | 3.997179 | 0.612904 | 0.53994 | NA | #VALUE! |
| ENSMUSG00000048164.10 | 2.170051 | -1.40355 | 3.831165 | -0.36635 | 0.714103 | NA | #VALUE! |
| ENSMUSG00000048173.6 | 0 | NA | NA | NA | NA | NA | #VALUE! |
| ENSMUSG00000048174.3 | 1.271783 | 3.851779 | 3.941488 | 0.97724 | 0.32845 | NA | #VALUE! |
| ENSMUSG00000048185.6 | 0 | NA | NA | NA | NA | NA | #VALUE! |
| ENSMUSG00000048186.15 | 3.306665 | -0.77396 | 3.784818 | -0.20449 | 0.837971 | NA | #VALUE! |
| ENSMUSG00000048188.7 | 0 | NA | NA | NA | NA | NA | #VALUE! |
| ENSMUSG00000048197.5 | 0 | NA | NA | NA | NA | NA | #VALUE! |
| ENSMUSG00000048206.7 | 0 | NA | NA | NA | NA | NA | #VALUE! |
| ENSMUSG00000048215.15 | 0 | NA | NA | NA | NA | NA | #VALUE! |
| ENSMUSG00000048216.14 | 0 | NA | NA | NA | NA | NA | #VALUE! |
| ENSMUSG00000048218.7 | 2.961801 | -5.09225 | 3.921898 | -1.29842 | 0.194145 | NA | #VALUE! |
| ENSMUSG00000048226.6 | 0 | NA | NA | NA | NA | NA | #VALUE! |
| ENSMUSG00000048230.7 | 0 | NA | NA | NA | NA | NA | #VALUE! |
| ENSMUSG00000048231.15 | 0 | NA | NA | NA | NA | NA | #VALUE! |
| ENSMUSG00000048232.13 | 0.105013 | -0.97324 | 4.076293 | -0.23876 | 0.811295 | NA | #VALUE! |
| ENSMUSG00000048236.9 | 0 | NA | NA | NA | NA | NA | #VALUE! |
| ENSMUSG00000048240.15 | 0 | NA | NA | NA | NA | NA | #VALUE! |
| ENSMUSG00000048251.16 | 0 | NA | NA | NA | NA | NA | #VALUE! |
| ENSMUSG00000048261.7 | 0 | NA | NA | NA | NA | NA | #VALUE! |
| ENSMUSG00000048279.20 | 2.187686 | 4.619304 | 3.927239 | 1.176222 | 0.239506 | NA | #VALUE! |
| ENSMUSG00000048280.19 | 7.532957 | -1.08805 | 2.831181 | -0.38431 | 0.700749 | NA | #VALUE! |
| ENSMUSG00000048281.10 | 0 | NA | NA | NA | NA | NA | #VALUE! |
| ENSMUSG00000048284.5 | 0 | NA | NA | NA | NA | NA | #VALUE! |
| ENSMUSG00000048292.4 | 0 | NA | NA | NA | NA | NA | #VALUE! |
| ENSMUSG00000048294.5 | 0 | NA | NA | NA | NA | NA | #VALUE! |
| ENSMUSG00000048299.9 | 0 | NA | NA | NA | NA | NA | #VALUE! |
| ENSMUSG00000048304.9 | 0 | NA | NA | NA | NA | NA | #VALUE! |
| ENSMUSG00000048310.9 | 34.71398 | 1.318355 | 1.702523 | 0.774354 | NA | NA | #VALUE! |
| ENSMUSG00000048312.8 | 0 | NA | NA | NA | NA | NA | #VALUE! |
| ENSMUSG00000048316.4 | 0 | NA | NA | NA | NA | NA | #VALUE! |
| ENSMUSG00000048329.8 | 0 | NA | NA | NA | NA | NA | #VALUE! |
| ENSMUSG00000048330.15 | 5.090517 | -5.87548 | 3.91557 | -1.50054 | 0.133474 | NA | #VALUE! |

| | | | | | | | |
|---|---|---|---|---|---|---|---|
| ENSMUSG00000048334.9 | 1.170879 | 0.767818 | 3.891098 | 0.197327 | 0.843572 | NA | #VALUE! |
| ENSMUSG00000048337.4 | 0 | NA | NA | NA | NA | NA | #VALUE! |
| ENSMUSG00000048347.6 | 0.336567 | 1.944925 | 4.029936 | 0.482619 | 0.629366 | NA | #VALUE! |
| ENSMUSG00000048349.10 | 0 | NA | NA | NA | NA | NA | #VALUE! |
| ENSMUSG00000048355.3 | 0 | NA | NA | NA | NA | NA | #VALUE! |
| ENSMUSG00000048356.7 | 0 | NA | NA | NA | NA | NA | #VALUE! |
| ENSMUSG00000048373.9 | 0 | NA | NA | NA | NA | NA | #VALUE! |
| ENSMUSG00000048377.8 | 0 | NA | NA | NA | NA | NA | #VALUE! |
| ENSMUSG00000048378.3 | 0 | NA | NA | NA | NA | NA | #VALUE! |
| ENSMUSG00000048385.10 | 0 | NA | NA | NA | NA | NA | #VALUE! |
| ENSMUSG00000048387.9 | 0 | NA | NA | NA | NA | NA | #VALUE! |
| ENSMUSG00000048391.4 | 0 | NA | NA | NA | NA | NA | #VALUE! |
| ENSMUSG00000048398.8 | 0 | NA | NA | NA | NA | NA | #VALUE! |
| ENSMUSG00000048399.5 | 0 | NA | NA | NA | NA | NA | #VALUE! |
| ENSMUSG00000048400.14 | 0 | NA | NA | NA | NA | NA | #VALUE! |
| ENSMUSG00000048402.15 | 0 | NA | NA | NA | NA | NA | #VALUE! |
| ENSMUSG00000048406.6 | 3.873185 | 5.44306 | 3.918358 | 1.389118 | 0.164797 | NA | #VALUE! |
| ENSMUSG00000048411.5 | 0 | NA | NA | NA | NA | NA | #VALUE! |
| ENSMUSG00000048416.16 | 0 | NA | NA | NA | NA | NA | #VALUE! |
| ENSMUSG00000048424.19 | 2.545258 | -4.8732 | 3.92437 | -1.24178 | 0.214318 | NA | #VALUE! |
| ENSMUSG00000048425.4 | 0 | NA | NA | NA | NA | NA | #VALUE! |
| ENSMUSG00000048439.16 | 2.255685 | 4.662059 | 3.926645 | 1.187288 | 0.235114 | NA | #VALUE! |
| ENSMUSG00000048440.17 | 3.730208 | 5.397701 | 3.918728 | 1.377412 | 0.168385 | NA | #VALUE! |
| ENSMUSG00000048442.16 | 0 | NA | NA | NA | NA | NA | #VALUE! |
| ENSMUSG00000048445.7 | 0 | NA | NA | NA | NA | NA | #VALUE! |
| ENSMUSG00000048450.11 | 0 | NA | NA | NA | NA | NA | #VALUE! |
| ENSMUSG00000048455.10 | 0 | NA | NA | NA | NA | NA | #VALUE! |
| ENSMUSG00000048456.4 | 0 | NA | NA | NA | NA | NA | #VALUE! |
| ENSMUSG00000048458.9 | 0 | NA | NA | NA | NA | NA | #VALUE! |
| ENSMUSG00000048469.7 | 0 | NA | NA | NA | NA | NA | #VALUE! |
| ENSMUSG00000048473.16 | 0 | NA | NA | NA | NA | NA | #VALUE! |
| ENSMUSG00000048478.8 | 0 | NA | NA | NA | NA | NA | #VALUE! |
| ENSMUSG00000048480.6 | 0 | NA | NA | NA | NA | NA | #VALUE! |
| ENSMUSG00000048481.16 | 0 | NA | NA | NA | NA | NA | #VALUE! |
| ENSMUSG00000048483.7 | 0 | NA | NA | NA | NA | NA | #VALUE! |
| ENSMUSG00000048484.5 | 0 | NA | NA | NA | NA | NA | #VALUE! |
| ENSMUSG00000048485.13 | 0 | NA | NA | NA | NA | NA | #VALUE! |
| ENSMUSG00000048497.12 | 0 | NA | NA | NA | NA | NA | #VALUE! |
| ENSMUSG00000048500.3 | 0 | NA | NA | NA | NA | NA | #VALUE! |
| ENSMUSG00000048501.4 | 0 | NA | NA | NA | NA | NA | #VALUE! |
| ENSMUSG00000048502.15 | 0 | NA | NA | NA | NA | NA | #VALUE! |
| ENSMUSG00000048503.8 | 7.749331 | -6.48223 | 2.930486 | -2.212 | 0.026967 | NA | #VALUE! |
| ENSMUSG00000048516.3 | 0 | NA | NA | NA | NA | NA | #VALUE! |
| ENSMUSG00000048520.17 | 0.245976 | 1.027628 | 4.067126 | 0.252667 | 0.800526 | NA | #VALUE! |

| | | | | | | | |
|---|---|---|---|---|---|---|---|
| ENSMUSG00000048528.8 | 0 | NA | NA | NA | NA | NA | #VALUE! |
| ENSMUSG00000048537.17 | 1.155144 | -3.7605 | 3.944679 | -0.95331 | 0.340433 | NA | #VALUE! |
| ENSMUSG00000048538.8 | 0 | NA | NA | NA | NA | NA | #VALUE! |
| ENSMUSG00000048540.9 | 0 | NA | NA | NA | NA | NA | #VALUE! |
| ENSMUSG00000048559.6 | 0 | NA | NA | NA | NA | NA | #VALUE! |
| ENSMUSG00000048562.8 | 0 | NA | NA | NA | NA | NA | #VALUE! |
| ENSMUSG00000048572.6 | 9.839021 | 6.784348 | 3.628918 | 1.869524 | 0.06155 | NA | #VALUE! |
| ENSMUSG00000048573.10 | 0 | NA | NA | NA | NA | NA | #VALUE! |
| ENSMUSG00000048574.10 | 0 | NA | NA | NA | NA | NA | #VALUE! |
| ENSMUSG00000048582.8 | 0 | NA | NA | NA | NA | NA | #VALUE! |
| ENSMUSG00000048583.17 | 0 | NA | NA | NA | NA | NA | #VALUE! |
| ENSMUSG00000048592.10 | 0 | NA | NA | NA | NA | NA | #VALUE! |
| ENSMUSG00000048600.8 | 0 | NA | NA | NA | NA | NA | #VALUE! |
| ENSMUSG00000048602.10 | 0 | NA | NA | NA | NA | NA | #VALUE! |
| ENSMUSG00000048603.9 | 0.831182 | 3.193261 | 3.961284 | 0.806118 | 0.420175 | NA | #VALUE! |
| ENSMUSG00000048612.17 | 9.282423 | 0.467796 | 2.941965 | 0.159008 | 0.873662 | NA | #VALUE! |
| ENSMUSG00000048616.5 | 0 | NA | NA | NA | NA | NA | #VALUE! |
| ENSMUSG00000048617.17 | 0 | NA | NA | NA | NA | NA | #VALUE! |
| ENSMUSG00000048620.7 | 0 | NA | NA | NA | NA | NA | #VALUE! |
| ENSMUSG00000048621.4 | 0.664544 | 1.770909 | 3.978581 | 0.445111 | 0.65624 | NA | #VALUE! |
| ENSMUSG00000048626.6 | 0 | NA | NA | NA | NA | NA | #VALUE! |
| ENSMUSG00000048628.6 | 0 | NA | NA | NA | NA | NA | #VALUE! |
| ENSMUSG00000048636.8 | 0 | NA | NA | NA | NA | NA | #VALUE! |
| ENSMUSG00000048643.6 | 0 | NA | NA | NA | NA | NA | #VALUE! |
| ENSMUSG00000048644.9 | 0 | NA | NA | NA | NA | NA | #VALUE! |
| ENSMUSG00000048647.10 | 7.796484 | 2.492782 | 3.562799 | 0.69967 | 0.484134 | NA | #VALUE! |
| ENSMUSG00000048652.13 | 0 | NA | NA | NA | NA | NA | #VALUE! |
| ENSMUSG00000048655.18 | 0 | NA | NA | NA | NA | NA | #VALUE! |
| ENSMUSG00000048668.17 | 0.877871 | 3.316761 | 3.95687 | 0.838228 | 0.401902 | NA | #VALUE! |
| ENSMUSG00000048677.10 | 0 | NA | NA | NA | NA | NA | #VALUE! |
| ENSMUSG00000048686.5 | 0 | NA | NA | NA | NA | NA | #VALUE! |
| ENSMUSG00000048693.6 | 0 | NA | NA | NA | NA | NA | #VALUE! |
| ENSMUSG00000048697.5 | 0 | NA | NA | NA | NA | NA | #VALUE! |
| ENSMUSG00000048699.5 | 0 | NA | NA | NA | NA | NA | #VALUE! |
| ENSMUSG00000048703.10 | 0 | NA | NA | NA | NA | NA | #VALUE! |
| ENSMUSG00000048707.10 | 0.75532 | 1.688323 | 3.975516 | 0.42468 | 0.67107 | NA | #VALUE! |
| ENSMUSG00000048709.7 | 0 | NA | NA | NA | NA | NA | #VALUE! |
| ENSMUSG00000048721.3 | 0 | NA | NA | NA | NA | NA | #VALUE! |
| ENSMUSG00000048728.16 | 0 | NA | NA | NA | NA | NA | #VALUE! |
| ENSMUSG00000048732.6 | 0 | NA | NA | NA | NA | NA | #VALUE! |
| ENSMUSG00000048745.4 | 0 | NA | NA | NA | NA | NA | #VALUE! |
| ENSMUSG00000048747.15 | 0.204111 | -1.05581 | 4.073302 | -0.2592 | 0.795479 | NA | #VALUE! |
| ENSMUSG00000048752.8 | 0 | NA | NA | NA | NA | NA | #VALUE! |
| ENSMUSG00000048763.12 | 0.983902 | 3.433476 | 3.953024 | 0.868569 | 0.385083 | NA | #VALUE! |

| | | | | | | | |
|---|---|---|---|---|---|---|---|
| ENSMUSG00000048764.17 | 0 | NA | NA | NA | NA | NA | #VALUE! |
| ENSMUSG00000048766.7 | 0 | NA | NA | NA | NA | NA | #VALUE! |
| ENSMUSG00000048775.11 | 0 | NA | NA | NA | NA | NA | #VALUE! |
| ENSMUSG00000048776.9 | 0 | NA | NA | NA | NA | NA | #VALUE! |
| ENSMUSG00000048779.6 | 2.809414 | 4.003923 | 3.918862 | 1.021706 | 0.30692 | NA | #VALUE! |
| ENSMUSG00000048794.15 | 0 | NA | NA | NA | NA | NA | #VALUE! |
| ENSMUSG00000048806.5 | 0 | NA | NA | NA | NA | NA | #VALUE! |
| ENSMUSG00000048807.3 | 0.168284 | 1.027628 | 4.067126 | 0.252667 | 0.800526 | NA | #VALUE! |
| ENSMUSG00000048810.3 | 0 | NA | NA | NA | NA | NA | #VALUE! |
| ENSMUSG00000048814.12 | 0 | NA | NA | NA | NA | NA | #VALUE! |
| ENSMUSG00000048824.8 | 0 | NA | NA | NA | NA | NA | #VALUE! |
| ENSMUSG00000048826.8 | 6.265251 | -6.17547 | 3.913919 | -1.57782 | 0.114607 | NA | #VALUE! |
| ENSMUSG00000048827.13 | 0 | NA | NA | NA | NA | NA | #VALUE! |
| ENSMUSG00000048830.4 | 0 | NA | NA | NA | NA | NA | #VALUE! |
| ENSMUSG00000048832.10 | 5.430178 | -0.39504 | 3.525943 | -0.11204 | 0.910792 | NA | #VALUE! |
| ENSMUSG00000048834.17 | 0 | NA | NA | NA | NA | NA | #VALUE! |
| ENSMUSG00000048852.14 | 0 | NA | NA | NA | NA | NA | #VALUE! |
| ENSMUSG00000048875.4 | 0 | NA | NA | NA | NA | NA | #VALUE! |
| ENSMUSG00000048895.14 | 0 | NA | NA | NA | NA | NA | #VALUE! |
| ENSMUSG00000048897.16 | 7.881957 | -2.6969 | 3.046578 | -0.88522 | 0.376036 | NA | #VALUE! |
| ENSMUSG00000048899.9 | 0 | NA | NA | NA | NA | NA | #VALUE! |
| ENSMUSG00000048904.6 | 0 | NA | NA | NA | NA | NA | #VALUE! |
| ENSMUSG00000048905.6 | 0 | NA | NA | NA | NA | NA | #VALUE! |
| ENSMUSG00000048911.16 | 9.022738 | -5.6005 | 3.440442 | -1.62784 | 0.103558 | NA | #VALUE! |
| ENSMUSG00000048916.8 | 0 | NA | NA | NA | NA | NA | #VALUE! |
| ENSMUSG00000048919.6 | 0 | NA | NA | NA | NA | NA | #VALUE! |
| ENSMUSG00000048921.12 | 2.017149 | 1.481466 | 3.840436 | 0.385755 | 0.699678 | NA | #VALUE! |
| ENSMUSG00000048922.19 | 0 | NA | NA | NA | NA | NA | #VALUE! |
| ENSMUSG00000048930.13 | 6.977879 | -5.40376 | 3.212725 | -1.68199 | 0.092571 | NA | #VALUE! |
| ENSMUSG00000048933.9 | 0 | NA | NA | NA | NA | NA | #VALUE! |
| ENSMUSG00000048938.13 | 5.066495 | 0.835217 | 2.506944 | 0.333161 | 0.739013 | NA | #VALUE! |
| ENSMUSG00000048939.14 | 0 | NA | NA | NA | NA | NA | #VALUE! |
| ENSMUSG00000048949.9 | 0 | NA | NA | NA | NA | NA | #VALUE! |
| ENSMUSG00000048965.10 | 2.984166 | 5.077883 | 3.921685 | 1.294822 | 0.195382 | NA | #VALUE! |
| ENSMUSG00000048967.17 | 0 | NA | NA | NA | NA | NA | #VALUE! |
| ENSMUSG00000048978.15 | 0 | NA | NA | NA | NA | NA | #VALUE! |
| ENSMUSG00000048981.2 | 0 | NA | NA | NA | NA | NA | #VALUE! |
| ENSMUSG00000048982.4 | 0 | NA | NA | NA | NA | NA | #VALUE! |
| ENSMUSG00000048988.9 | 0 | NA | NA | NA | NA | NA | #VALUE! |
| ENSMUSG00000048992.10 | 0 | NA | NA | NA | NA | NA | #VALUE! |
| ENSMUSG00000048994.3 | 0 | NA | NA | NA | NA | NA | #VALUE! |
| ENSMUSG00000048996.7 | 0 | NA | NA | NA | NA | NA | #VALUE! |
| ENSMUSG00000048997.18 | 0 | NA | NA | NA | NA | NA | #VALUE! |
| ENSMUSG00000049001.6 | 0 | NA | NA | NA | NA | NA | #VALUE! |

| | | | | | | | |
|---|---|---|---|---|---|---|---|
| ENSMUSG00000049008.5 | 0 | NA | NA | NA | NA | NA | #VALUE! |
| ENSMUSG00000049010.7 | 0 | NA | NA | NA | NA | NA | #VALUE! |
| ENSMUSG00000049011.3 | 0 | NA | NA | NA | NA | NA | #VALUE! |
| ENSMUSG00000049013.4 | 0 | NA | NA | NA | NA | NA | #VALUE! |
| ENSMUSG00000049015.4 | 0 | NA | NA | NA | NA | NA | #VALUE! |
| ENSMUSG00000049018.9 | 0 | NA | NA | NA | NA | NA | #VALUE! |
| ENSMUSG00000049028.9 | 0 | NA | NA | NA | NA | NA | #VALUE! |
| ENSMUSG00000049036.8 | 0 | NA | NA | NA | NA | NA | #VALUE! |
| ENSMUSG00000049041.7 | 0 | NA | NA | NA | NA | NA | #VALUE! |
| ENSMUSG00000049052.5 | 0 | NA | NA | NA | NA | NA | #VALUE! |
| ENSMUSG00000049057.5 | 0 | NA | NA | NA | NA | NA | #VALUE! |
| ENSMUSG00000049062.4 | 0 | NA | NA | NA | NA | NA | #VALUE! |
| ENSMUSG00000049073.8 | 0 | NA | NA | NA | NA | NA | #VALUE! |
| ENSMUSG00000049092.12 | 6.46754 | -0.80039 | 3.756276 | -0.21308 | 0.831264 | NA | #VALUE! |
| ENSMUSG00000049093.10 | 0 | NA | NA | NA | NA | NA | #VALUE! |
| ENSMUSG00000049094.8 | 0 | NA | NA | NA | NA | NA | #VALUE! |
| ENSMUSG00000049097.10 | 0 | NA | NA | NA | NA | NA | #VALUE! |
| ENSMUSG00000049098.5 | 0 | NA | NA | NA | NA | NA | #VALUE! |
| ENSMUSG00000049100.16 | 0 | NA | NA | NA | NA | NA | #VALUE! |
| ENSMUSG00000049103.15 | 1.936314 | -2.41366 | 3.888138 | -0.62077 | 0.534748 | NA | #VALUE! |
| ENSMUSG00000049107.14 | 2.538238 | 4.842935 | 3.924313 | 1.234085 | 0.217171 | NA | #VALUE! |
| ENSMUSG00000049109.16 | 6.882657 | 6.278181 | 3.641137 | 1.724236 | 0.084665 | NA | #VALUE! |
| ENSMUSG00000049112.10 | 0 | NA | NA | NA | NA | NA | #VALUE! |
| ENSMUSG00000049119.15 | 0 | NA | NA | NA | NA | NA | #VALUE! |
| ENSMUSG00000049122.18 | 0 | NA | NA | NA | NA | NA | #VALUE! |
| ENSMUSG00000049123.11 | 0 | NA | NA | NA | NA | NA | #VALUE! |
| ENSMUSG00000049124.5 | 1.602939 | 2.980712 | 3.930773 | 0.758302 | 0.44827 | NA | #VALUE! |
| ENSMUSG00000049128.8 | 0 | NA | NA | NA | NA | NA | #VALUE! |
| ENSMUSG00000049130.7 | 0 | NA | NA | NA | NA | NA | #VALUE! |
| ENSMUSG00000049133.8 | 0 | NA | NA | NA | NA | NA | #VALUE! |
| ENSMUSG00000049134.16 | 0 | NA | NA | NA | NA | NA | #VALUE! |
| ENSMUSG00000049148.9 | 0 | NA | NA | NA | NA | NA | #VALUE! |
| ENSMUSG00000049149.10 | 0 | NA | NA | NA | NA | NA | #VALUE! |
| ENSMUSG00000049154.13 | 0 | NA | NA | NA | NA | NA | #VALUE! |
| ENSMUSG00000049160.16 | 0 | NA | NA | NA | NA | NA | #VALUE! |
| ENSMUSG00000049168.10 | 0 | NA | NA | NA | NA | NA | #VALUE! |
| ENSMUSG00000049173.9 | 0 | NA | NA | NA | NA | NA | #VALUE! |
| ENSMUSG00000049176.17 | 0 | NA | NA | NA | NA | NA | #VALUE! |
| ENSMUSG00000049184.7 | 0.204111 | -1.05581 | 4.073302 | -0.2592 | 0.795479 | NA | #VALUE! |
| ENSMUSG00000049191.13 | 0 | NA | NA | NA | NA | NA | #VALUE! |
| ENSMUSG00000049202.6 | 0 | NA | NA | NA | NA | NA | #VALUE! |
| ENSMUSG00000049214.14 | 0 | NA | NA | NA | NA | NA | #VALUE! |
| ENSMUSG00000049217.5 | 0 | NA | NA | NA | NA | NA | #VALUE! |
| ENSMUSG00000049229.6 | 0 | NA | NA | NA | NA | NA | #VALUE! |

| | | | | | | | |
|---|---|---|---|---|---|---|---|
| ENSMUSG00000049230.9 | 0 | NA | NA | NA | NA | NA | #VALUE! |
| ENSMUSG00000049231.7 | 0.872091 | 0.242697 | 3.92879 | 0.061774 | 0.950743 | NA | #VALUE! |
| ENSMUSG00000049233.8 | 0.168284 | 1.027628 | 4.067126 | 0.252667 | 0.800526 | NA | #VALUE! |
| ENSMUSG00000049241.7 | 0 | NA | NA | NA | NA | NA | #VALUE! |
| ENSMUSG00000049246.6 | 0 | NA | NA | NA | NA | NA | #VALUE! |
| ENSMUSG00000049249.3 | 0 | NA | NA | NA | NA | NA | #VALUE! |
| ENSMUSG00000049252.18 | 0 | NA | NA | NA | NA | NA | #VALUE! |
| ENSMUSG00000049265.9 | 0 | NA | NA | NA | NA | NA | #VALUE! |
| ENSMUSG00000049280.5 | 0 | NA | NA | NA | NA | NA | #VALUE! |
| ENSMUSG00000049281.17 | 0 | NA | NA | NA | NA | NA | #VALUE! |
| ENSMUSG00000049285.5 | 12.45298 | 7.12828 | 3.357786 | 2.122911 | NA | NA | #VALUE! |
| ENSMUSG00000049288.5 | 2.173564 | -2.41394 | 3.878702 | -0.62236 | 0.533707 | NA | #VALUE! |
| ENSMUSG00000049291.7 | 0 | NA | NA | NA | NA | NA | #VALUE! |
| ENSMUSG00000049295.18 | 0 | NA | NA | NA | NA | NA | #VALUE! |
| ENSMUSG00000049303.11 | 0 | NA | NA | NA | NA | NA | #VALUE! |
| ENSMUSG00000049307.7 | 0 | NA | NA | NA | NA | NA | #VALUE! |
| ENSMUSG00000049313.9 | 3.823463 | 5.43312 | 3.918438 | 1.386552 | 0.165578 | NA | #VALUE! |
| ENSMUSG00000049314.12 | 0 | NA | NA | NA | NA | NA | #VALUE! |
| ENSMUSG00000049315.5 | 0 | NA | NA | NA | NA | NA | #VALUE! |
| ENSMUSG00000049334.4 | 0 | NA | NA | NA | NA | NA | #VALUE! |
| ENSMUSG00000049336.17 | 0.315039 | -1.93549 | 4.038621 | -0.47925 | 0.631763 | NA | #VALUE! |
| ENSMUSG00000049349.4 | 0 | NA | NA | NA | NA | NA | #VALUE! |
| ENSMUSG00000049350.7 | 0 | NA | NA | NA | NA | NA | #VALUE! |
| ENSMUSG00000049353.18 | 1.995249 | -4.53986 | 3.928926 | -1.1555 | 0.247887 | NA | #VALUE! |
| ENSMUSG00000049357.12 | 0 | NA | NA | NA | NA | NA | #VALUE! |
| ENSMUSG00000049362.7 | 0 | NA | NA | NA | NA | NA | #VALUE! |
| ENSMUSG00000049372.12 | 0 | NA | NA | NA | NA | NA | #VALUE! |
| ENSMUSG00000049387.11 | 0 | NA | NA | NA | NA | NA | #VALUE! |
| ENSMUSG00000049396.7 | 0 | NA | NA | NA | NA | NA | #VALUE! |
| ENSMUSG00000049409.10 | 0 | NA | NA | NA | NA | NA | #VALUE! |
| ENSMUSG00000049410.9 | 0 | NA | NA | NA | NA | NA | #VALUE! |
| ENSMUSG00000049411.16 | 1.428454 | 2.787556 | 3.935242 | 0.708357 | 0.478724 | NA | #VALUE! |
| ENSMUSG00000049414.8 | 0 | NA | NA | NA | NA | NA | #VALUE! |
| ENSMUSG00000049420.10 | 0 | NA | NA | NA | NA | NA | #VALUE! |
| ENSMUSG00000049427.3 | 0 | NA | NA | NA | NA | NA | #VALUE! |
| ENSMUSG00000049436.6 | 6.784111 | -5.19507 | 3.721131 | -1.3961 | 0.162685 | NA | #VALUE! |
| ENSMUSG00000049456.4 | 0 | NA | NA | NA | NA | NA | #VALUE! |
| ENSMUSG00000049460.8 | 0 | NA | NA | NA | NA | NA | #VALUE! |
| ENSMUSG00000049476.13 | 0 | NA | NA | NA | NA | NA | #VALUE! |
| ENSMUSG00000049477.6 | 0 | NA | NA | NA | NA | NA | #VALUE! |
| ENSMUSG00000049488.15 | 5.487051 | -2.22462 | 3.783755 | -0.58794 | 0.556572 | NA | #VALUE! |
| ENSMUSG00000049491.16 | 0 | NA | NA | NA | NA | NA | #VALUE! |
| ENSMUSG00000049493.14 | 1.575197 | -4.20222 | 3.934736 | -1.06798 | 0.285529 | NA | #VALUE! |
| ENSMUSG00000049498.6 | 0 | NA | NA | NA | NA | NA | #VALUE! |

| | | | | | | | |
|---|---|---|---|---|---|---|---|
| ENSMUSG00000049506.8 | 0 | NA | NA | NA | NA | NA | #VALUE! |
| ENSMUSG00000049511.6 | 0 | NA | NA | NA | NA | NA | #VALUE! |
| ENSMUSG00000049515.13 | 0 | NA | NA | NA | NA | NA | #VALUE! |
| ENSMUSG00000049521.3 | 2.051614 | 4.542018 | 3.92836 | 1.156212 | 0.247594 | NA | #VALUE! |
| ENSMUSG00000049526.9 | 0 | NA | NA | NA | NA | NA | #VALUE! |
| ENSMUSG00000049528.8 | 0 | NA | NA | NA | NA | NA | #VALUE! |
| ENSMUSG00000049530.7 | 0 | NA | NA | NA | NA | NA | #VALUE! |
| ENSMUSG00000049536.6 | 1.851119 | 4.378291 | 3.930938 | 1.113803 | 0.265364 | NA | #VALUE! |
| ENSMUSG00000049537.11 | 0 | NA | NA | NA | NA | NA | #VALUE! |
| ENSMUSG00000049538.15 | 0 | NA | NA | NA | NA | NA | #VALUE! |
| ENSMUSG00000049539.4 | 0 | NA | NA | NA | NA | NA | #VALUE! |
| ENSMUSG00000049548.10 | 0 | NA | NA | NA | NA | NA | #VALUE! |
| ENSMUSG00000049551.3 | 0 | NA | NA | NA | NA | NA | #VALUE! |
| ENSMUSG00000049556.6 | 0 | NA | NA | NA | NA | NA | #VALUE! |
| ENSMUSG00000049560.5 | 0 | NA | NA | NA | NA | NA | #VALUE! |
| ENSMUSG00000049561.10 | 0 | NA | NA | NA | NA | NA | #VALUE! |
| ENSMUSG00000049562.7 | 0 | NA | NA | NA | NA | NA | #VALUE! |
| ENSMUSG00000049565.17 | 0 | NA | NA | NA | NA | NA | #VALUE! |
| ENSMUSG00000049571.19 | 0 | NA | NA | NA | NA | NA | #VALUE! |
| ENSMUSG00000049573.4 | 0 | NA | NA | NA | NA | NA | #VALUE! |
| ENSMUSG00000049576.7 | 0 | NA | NA | NA | NA | NA | #VALUE! |
| ENSMUSG00000049583.16 | 0 | NA | NA | NA | NA | NA | #VALUE! |
| ENSMUSG00000049588.14 | 6.641339 | 6.216291 | 3.913555 | 1.5884 | 0.112196 | NA | #VALUE! |
| ENSMUSG00000049593.9 | 0 | NA | NA | NA | NA | NA | #VALUE! |
| ENSMUSG00000049598.10 | 0 | NA | NA | NA | NA | NA | #VALUE! |
| ENSMUSG00000049600.9 | 0.525066 | -2.64687 | 3.988095 | -0.66369 | 0.506887 | NA | #VALUE! |
| ENSMUSG00000049604.4 | 0 | NA | NA | NA | NA | NA | #VALUE! |
| ENSMUSG00000049605.4 | 0 | NA | NA | NA | NA | NA | #VALUE! |
| ENSMUSG00000049608.10 | 0 | NA | NA | NA | NA | NA | #VALUE! |
| ENSMUSG00000049612.11 | 0 | NA | NA | NA | NA | NA | #VALUE! |
| ENSMUSG00000049618.7 | 0 | NA | NA | NA | NA | NA | #VALUE! |
| ENSMUSG00000049620.10 | 0 | NA | NA | NA | NA | NA | #VALUE! |
| ENSMUSG00000049625.7 | 8.116727 | 1.703982 | 3.516859 | 0.484518 | 0.628018 | NA | #VALUE! |
| ENSMUSG00000049630.7 | 0 | NA | NA | NA | NA | NA | #VALUE! |
| ENSMUSG00000049641.16 | 0 | NA | NA | NA | NA | NA | #VALUE! |
| ENSMUSG00000049643.16 | 5.95944 | -2.97142 | 3.802612 | -0.78142 | 0.434558 | NA | #VALUE! |
| ENSMUSG00000049648.5 | 0 | NA | NA | NA | NA | NA | #VALUE! |
| ENSMUSG00000049649.9 | 0 | NA | NA | NA | NA | NA | #VALUE! |
| ENSMUSG00000049653.6 | 0 | NA | NA | NA | NA | NA | #VALUE! |
| ENSMUSG00000049670.10 | 0 | NA | NA | NA | NA | NA | #VALUE! |
| ENSMUSG00000049672.16 | 9.143838 | 0.218641 | 2.615644 | 0.08359 | 0.933383 | NA | #VALUE! |
| ENSMUSG00000049674.8 | 0 | NA | NA | NA | NA | NA | #VALUE! |
| ENSMUSG00000049676.15 | 0 | NA | NA | NA | NA | NA | #VALUE! |
| ENSMUSG00000049685.9 | 12.95793 | -1.7849 | 2.776176 | -0.64293 | NA | NA | #VALUE! |

| | | | | | | | |
|---|---|---|---|---|---|---|---|
| ENSMUSG00000049687.6 | 0 | NA | NA | NA | NA | NA | #VALUE! |
| ENSMUSG00000049690.16 | 0 | NA | NA | NA | NA | NA | #VALUE! |
| ENSMUSG00000049691.9 | 0 | NA | NA | NA | NA | NA | #VALUE! |
| ENSMUSG00000049692.8 | 0 | NA | NA | NA | NA | NA | #VALUE! |
| ENSMUSG00000049694.14 | 0 | NA | NA | NA | NA | NA | #VALUE! |
| ENSMUSG00000049699.5 | 0 | NA | NA | NA | NA | NA | #VALUE! |
| ENSMUSG00000049708.10 | 0 | NA | NA | NA | NA | NA | #VALUE! |
| ENSMUSG00000049709.5 | 0.491951 | 2.402185 | 4.000147 | 0.600524 | 0.548157 | NA | #VALUE! |
| ENSMUSG00000049719.15 | 0 | NA | NA | NA | NA | NA | #VALUE! |
| ENSMUSG00000049721.15 | 7.65104 | 6.420573 | 3.467297 | 1.851752 | 0.064062 | NA | #VALUE! |
| ENSMUSG00000049723.15 | 0 | NA | NA | NA | NA | NA | #VALUE! |
| ENSMUSG00000049728.15 | 3.357187 | 5.246639 | 3.920044 | 1.338413 | 0.180762 | NA | #VALUE! |
| ENSMUSG00000049734.10 | 0 | NA | NA | NA | NA | NA | #VALUE! |
| ENSMUSG00000049737.3 | 0 | NA | NA | NA | NA | NA | #VALUE! |
| ENSMUSG00000049740.3 | 0 | NA | NA | NA | NA | NA | #VALUE! |
| ENSMUSG00000049741.5 | 0 | NA | NA | NA | NA | NA | #VALUE! |
| ENSMUSG00000049744.16 | 7.871217 | 6.462163 | 3.906699 | 1.654124 | 0.098102 | NA | #VALUE! |
| ENSMUSG00000049758.8 | 0 | NA | NA | NA | NA | NA | #VALUE! |
| ENSMUSG00000049761.7 | 0 | NA | NA | NA | NA | NA | #VALUE! |
| ENSMUSG00000049796.5 | 0 | NA | NA | NA | NA | NA | #VALUE! |
| ENSMUSG00000049797.5 | 0 | NA | NA | NA | NA | NA | #VALUE! |
| ENSMUSG00000049799.14 | 0 | NA | NA | NA | NA | NA | #VALUE! |
| ENSMUSG00000049804.10 | 6.590695 | -1.32988 | 3.74146 | -0.35544 | 0.722256 | NA | #VALUE! |
| ENSMUSG00000049806.5 | 0 | NA | NA | NA | NA | NA | #VALUE! |
| ENSMUSG00000049809.3 | 0 | NA | NA | NA | NA | NA | #VALUE! |
| ENSMUSG00000049811.18 | 0 | NA | NA | NA | NA | NA | #VALUE! |
| ENSMUSG00000049815.4 | 0 | NA | NA | NA | NA | NA | #VALUE! |
| ENSMUSG00000049823.10 | 0 | NA | NA | NA | NA | NA | #VALUE! |
| ENSMUSG00000049832.7 | 1.014906 | 0.518024 | 3.910042 | 0.132486 | 0.8946 | NA | #VALUE! |
| ENSMUSG00000049843.7 | 0 | NA | NA | NA | NA | NA | #VALUE! |
| ENSMUSG00000049848.8 | 0 | NA | NA | NA | NA | NA | #VALUE! |
| ENSMUSG00000049864.3 | 0 | NA | NA | NA | NA | NA | #VALUE! |
| ENSMUSG00000049866.13 | 0.195789 | -1.05581 | 4.073302 | -0.2592 | 0.795479 | NA | #VALUE! |
| ENSMUSG00000049871.15 | 0 | NA | NA | NA | NA | NA | #VALUE! |
| ENSMUSG00000049872.9 | 1.925939 | 3.239121 | 3.925643 | 0.825119 | 0.409304 | NA | #VALUE! |
| ENSMUSG00000049882.11 | 2.07513 | 2.921282 | 3.907863 | 0.74754 | 0.454738 | NA | #VALUE! |
| ENSMUSG00000049891.5 | 0 | NA | NA | NA | NA | NA | #VALUE! |
| ENSMUSG00000049892.8 | 1.470184 | -4.10383 | 3.936698 | -1.04245 | 0.297201 | NA | #VALUE! |
| ENSMUSG00000049894.4 | 0 | NA | NA | NA | NA | NA | #VALUE! |
| ENSMUSG00000049897.15 | 0 | NA | NA | NA | NA | NA | #VALUE! |
| ENSMUSG00000049902.7 | 0 | NA | NA | NA | NA | NA | #VALUE! |
| ENSMUSG00000049904.8 | 0.466276 | 2.449889 | 3.997179 | 0.612904 | 0.53994 | NA | #VALUE! |
| ENSMUSG00000049907.9 | 2.487076 | 3.796194 | 3.921399 | 0.968071 | 0.333009 | NA | #VALUE! |
| ENSMUSG00000049908.7 | 0 | NA | NA | NA | NA | NA | #VALUE! |

| | | | | | | | |
|---|---|---|---|---|---|---|---|
| ENSMUSG00000049916.13 | 0 | NA | NA | NA | NA | NA | #VALUE! |
| ENSMUSG00000049926.6 | 0 | NA | NA | NA | NA | NA | #VALUE! |
| ENSMUSG00000049928.16 | 2.21378 | 4.621487 | 3.927208 | 1.176787 | 0.239281 | NA | #VALUE! |
| ENSMUSG00000049929.8 | 0.504851 | 2.502843 | 3.993993 | 0.626652 | 0.530887 | NA | #VALUE! |
| ENSMUSG00000049932.4 | 16.54414 | 1.108296 | 2.24382 | 0.493933 | NA | NA | #VALUE! |
| ENSMUSG00000049939.7 | 0 | NA | NA | NA | NA | NA | #VALUE! |
| ENSMUSG00000049946.8 | 0 | NA | NA | NA | NA | NA | #VALUE! |
| ENSMUSG00000049972.5 | 0 | NA | NA | NA | NA | NA | #VALUE! |
| ENSMUSG00000049985.16 | 0 | NA | NA | NA | NA | NA | #VALUE! |
| ENSMUSG00000050010.9 | 1.916454 | 4.427787 | 3.930128 | 1.126627 | 0.2599 | NA | #VALUE! |
| ENSMUSG00000050014.9 | 0 | NA | NA | NA | NA | NA | #VALUE! |
| ENSMUSG00000050015.4 | 0 | NA | NA | NA | NA | NA | #VALUE! |
| ENSMUSG00000050022.17 | 0 | NA | NA | NA | NA | NA | #VALUE! |
| ENSMUSG00000050023.5 | 0 | NA | NA | NA | NA | NA | #VALUE! |
| ENSMUSG00000050028.6 | 0 | NA | NA | NA | NA | NA | #VALUE! |
| ENSMUSG00000050030.6 | 0 | NA | NA | NA | NA | NA | #VALUE! |
| ENSMUSG00000050035.8 | 0 | NA | NA | NA | NA | NA | #VALUE! |
| ENSMUSG00000050050.18 | 6.061588 | 6.094854 | 3.914151 | 1.557133 | 0.119439 | NA | #VALUE! |
| ENSMUSG00000050058.7 | 0 | NA | NA | NA | NA | NA | #VALUE! |
| ENSMUSG00000050063.19 | 0 | NA | NA | NA | NA | NA | #VALUE! |
| ENSMUSG00000050064.15 | 2.590054 | -2.90137 | 3.880378 | -0.7477 | 0.454639 | NA | #VALUE! |
| ENSMUSG00000050066.7 | 0 | NA | NA | NA | NA | NA | #VALUE! |
| ENSMUSG00000050071.9 | 5.04063 | -5.86896 | 3.915609 | -1.49886 | 0.133909 | NA | #VALUE! |
| ENSMUSG00000050074.13 | 0 | NA | NA | NA | NA | NA | #VALUE! |
| ENSMUSG00000050075.9 | 0.504851 | 2.502843 | 3.993993 | 0.626652 | 0.530887 | NA | #VALUE! |
| ENSMUSG00000050085.6 | 0 | NA | NA | NA | NA | NA | #VALUE! |
| ENSMUSG00000050087.4 | 0 | NA | NA | NA | NA | NA | #VALUE! |
| ENSMUSG00000050089.14 | 0 | NA | NA | NA | NA | NA | #VALUE! |
| ENSMUSG00000050092.4 | 0 | NA | NA | NA | NA | NA | #VALUE! |
| ENSMUSG00000050097.7 | 7.149792 | -6.3655 | 3.586241 | -1.77498 | 0.075902 | NA | #VALUE! |
| ENSMUSG00000050100.15 | 0 | NA | NA | NA | NA | NA | #VALUE! |
| ENSMUSG00000050102.7 | 0 | NA | NA | NA | NA | NA | #VALUE! |
| ENSMUSG00000050105.6 | 0 | NA | NA | NA | NA | NA | #VALUE! |
| ENSMUSG00000050106.19 | 0.90621 | 3.318856 | 3.956799 | 0.838773 | 0.401597 | NA | #VALUE! |
| ENSMUSG00000050107.3 | 0 | NA | NA | NA | NA | NA | #VALUE! |
| ENSMUSG00000050108.13 | 0 | NA | NA | NA | NA | NA | #VALUE! |
| ENSMUSG00000050121.10 | 0 | NA | NA | NA | NA | NA | #VALUE! |
| ENSMUSG00000050122.20 | 0 | NA | NA | NA | NA | NA | #VALUE! |
| ENSMUSG00000050128.6 | 0 | NA | NA | NA | NA | NA | #VALUE! |
| ENSMUSG00000050132.14 | 0 | NA | NA | NA | NA | NA | #VALUE! |
| ENSMUSG00000050134.8 | 0 | NA | NA | NA | NA | NA | #VALUE! |
| ENSMUSG00000050138.9 | 0 | NA | NA | NA | NA | NA | #VALUE! |
| ENSMUSG00000050141.14 | 0 | NA | NA | NA | NA | NA | #VALUE! |
| ENSMUSG00000050147.10 | 0 | NA | NA | NA | NA | NA | #VALUE! |

| | | | | | | | |
|---|---|---|---|---|---|---|---|
| ENSMUSG00000050150.17 | 0 | NA | NA | NA | NA | NA | #VALUE! |
| ENSMUSG00000050157.7 | 0 | NA | NA | NA | NA | NA | #VALUE! |
| ENSMUSG00000050158.8 | 0 | NA | NA | NA | NA | NA | #VALUE! |
| ENSMUSG00000050164.12 | 0 | NA | NA | NA | NA | NA | #VALUE! |
| ENSMUSG00000050179.4 | 0 | NA | NA | NA | NA | NA | #VALUE! |
| ENSMUSG00000050189.5 | 0 | NA | NA | NA | NA | NA | #VALUE! |
| ENSMUSG00000050190.5 | 0 | NA | NA | NA | NA | NA | #VALUE! |
| ENSMUSG00000050192.9 | 0 | NA | NA | NA | NA | NA | #VALUE! |
| ENSMUSG00000050195.9 | 0 | NA | NA | NA | NA | NA | #VALUE! |
| ENSMUSG00000050197.6 | 0 | NA | NA | NA | NA | NA | #VALUE! |
| ENSMUSG00000050198.5 | 0 | NA | NA | NA | NA | NA | #VALUE! |
| ENSMUSG00000050201.10 | 0 | NA | NA | NA | NA | NA | #VALUE! |
| ENSMUSG00000050211.15 | 0 | NA | NA | NA | NA | NA | #VALUE! |
| ENSMUSG00000050212.5 | 5.102769 | -5.87825 | 3.915553 | -1.50126 | 0.133289 | NA | #VALUE! |
| ENSMUSG00000050215.7 | 0 | NA | NA | NA | NA | NA | #VALUE! |
| ENSMUSG00000050217.14 | 0 | NA | NA | NA | NA | NA | #VALUE! |
| ENSMUSG00000050222.12 | 0 | NA | NA | NA | NA | NA | #VALUE! |
| ENSMUSG00000050224.4 | 0 | NA | NA | NA | NA | NA | #VALUE! |
| ENSMUSG00000050232.7 | 0 | NA | NA | NA | NA | NA | #VALUE! |
| ENSMUSG00000050239.4 | 0 | NA | NA | NA | NA | NA | #VALUE! |
| ENSMUSG00000050240.17 | 3.175859 | 3.111058 | 3.878883 | 0.80205 | 0.422524 | NA | #VALUE! |
| ENSMUSG00000050241.9 | 0 | NA | NA | NA | NA | NA | #VALUE! |
| ENSMUSG00000050243.7 | 0 | NA | NA | NA | NA | NA | #VALUE! |
| ENSMUSG00000050244.10 | 20.38011 | -0.31811 | 2.385199 | -0.13337 | NA | NA | #VALUE! |
| ENSMUSG00000050248.12 | 0 | NA | NA | NA | NA | NA | #VALUE! |
| ENSMUSG00000050251.6 | 0 | NA | NA | NA | NA | NA | #VALUE! |
| ENSMUSG00000050266.7 | 0 | NA | NA | NA | NA | NA | #VALUE! |
| ENSMUSG00000050271.13 | 0 | NA | NA | NA | NA | NA | #VALUE! |
| ENSMUSG00000050272.11 | 0 | NA | NA | NA | NA | NA | #VALUE! |
| ENSMUSG00000050276.8 | 0 | NA | NA | NA | NA | NA | #VALUE! |
| ENSMUSG00000050281.5 | 0 | NA | NA | NA | NA | NA | #VALUE! |
| ENSMUSG00000050288.7 | 0 | NA | NA | NA | NA | NA | #VALUE! |
| ENSMUSG00000050295.5 | 0 | NA | NA | NA | NA | NA | #VALUE! |
| ENSMUSG00000050296.15 | 0 | NA | NA | NA | NA | NA | #VALUE! |
| ENSMUSG00000050304.5 | 0 | NA | NA | NA | NA | NA | #VALUE! |
| ENSMUSG00000050315.15 | 0 | NA | NA | NA | NA | NA | #VALUE! |
| ENSMUSG00000050321.4 | 0 | NA | NA | NA | NA | NA | #VALUE! |
| ENSMUSG00000050328.3 | 0 | NA | NA | NA | NA | NA | #VALUE! |
| ENSMUSG00000050332.8 | 9.093539 | -0.46715 | 3.026588 | -0.15435 | 0.877334 | NA | #VALUE! |
| ENSMUSG00000050334.13 | 0 | NA | NA | NA | NA | NA | #VALUE! |
| ENSMUSG00000050343.3 | 0 | NA | NA | NA | NA | NA | #VALUE! |
| ENSMUSG00000050345.10 | 0 | NA | NA | NA | NA | NA | #VALUE! |
| ENSMUSG00000050350.9 | 8.855119 | 6.632223 | 3.756168 | 1.765689 | 0.077448 | NA | #VALUE! |
| ENSMUSG00000050357.10 | 0 | NA | NA | NA | NA | NA | #VALUE! |

| | | | | | | | |
|---|---|---|---|---|---|---|---|
| ENSMUSG00000050359.8 | 0 | NA | NA | NA | NA | NA | #VALUE! |
| ENSMUSG00000050366.7 | 0 | NA | NA | NA | NA | NA | #VALUE! |
| ENSMUSG00000050368.5 | 0 | NA | NA | NA | NA | NA | #VALUE! |
| ENSMUSG00000050370.6 | 0 | NA | NA | NA | NA | NA | #VALUE! |
| ENSMUSG00000050373.14 | 2.520315 | -4.8742 | 3.924358 | -1.24204 | 0.214223 | NA | #VALUE! |
| ENSMUSG00000050377.9 | 0 | NA | NA | NA | NA | NA | #VALUE! |
| ENSMUSG00000050379.16 | 0 | NA | NA | NA | NA | NA | #VALUE! |
| ENSMUSG00000050382.15 | 0 | NA | NA | NA | NA | NA | #VALUE! |
| ENSMUSG00000050383.6 | 0 | NA | NA | NA | NA | NA | #VALUE! |
| ENSMUSG00000050394.15 | 0 | NA | NA | NA | NA | NA | #VALUE! |
| ENSMUSG00000050395.10 | 1.678593 | 4.255795 | 3.933066 | 1.082055 | 0.279228 | NA | #VALUE! |
| ENSMUSG00000050397.5 | 0 | NA | NA | NA | NA | NA | #VALUE! |
| ENSMUSG00000050410.16 | 1.147229 | -2.41965 | 3.938723 | -0.61432 | 0.539002 | NA | #VALUE! |
| ENSMUSG00000050423.12 | 6.405801 | -6.21359 | 3.913733 | -1.58764 | 0.112368 | NA | #VALUE! |
| ENSMUSG00000050424.10 | 0 | NA | NA | NA | NA | NA | #VALUE! |
| ENSMUSG00000050425.3 | 0 | NA | NA | NA | NA | NA | #VALUE! |
| ENSMUSG00000050428.8 | 0.491951 | 2.402185 | 4.000147 | 0.600524 | 0.548157 | NA | #VALUE! |
| ENSMUSG00000050435.3 | 0 | NA | NA | NA | NA | NA | #VALUE! |
| ENSMUSG00000050439.8 | 0 | NA | NA | NA | NA | NA | #VALUE! |
| ENSMUSG00000050447.16 | 5.731802 | 6.01122 | 3.788854 | 1.586553 | 0.112614 | NA | #VALUE! |
| ENSMUSG00000050463.10 | 0 | NA | NA | NA | NA | NA | #VALUE! |
| ENSMUSG00000050468.13 | 0 | NA | NA | NA | NA | NA | #VALUE! |
| ENSMUSG00000050473.6 | 0 | NA | NA | NA | NA | NA | #VALUE! |
| ENSMUSG00000050478.4 | 0 | NA | NA | NA | NA | NA | #VALUE! |
| ENSMUSG00000050490.8 | 3.802386 | -3.62124 | 3.689797 | -0.98142 | 0.326386 | NA | #VALUE! |
| ENSMUSG00000050504.4 | 0 | NA | NA | NA | NA | NA | #VALUE! |
| ENSMUSG00000050505.8 | 0 | NA | NA | NA | NA | NA | #VALUE! |
| ENSMUSG00000050511.2 | 0 | NA | NA | NA | NA | NA | #VALUE! |
| ENSMUSG00000050520.5 | 0 | NA | NA | NA | NA | NA | #VALUE! |
| ENSMUSG00000050526.6 | 0 | NA | NA | NA | NA | NA | #VALUE! |
| ENSMUSG00000050533.4 | 0 | NA | NA | NA | NA | NA | #VALUE! |
| ENSMUSG00000050534.9 | 0 | NA | NA | NA | NA | NA | #VALUE! |
| ENSMUSG00000050538.9 | 5.098821 | 5.840499 | 3.435766 | 1.699912 | 0.089148 | NA | #VALUE! |
| ENSMUSG00000050545.11 | 9.476647 | -1.21448 | 3.363724 | -0.36105 | 0.718061 | NA | #VALUE! |
| ENSMUSG00000050550.8 | 0 | NA | NA | NA | NA | NA | #VALUE! |
| ENSMUSG00000050553.4 | 0 | NA | NA | NA | NA | NA | #VALUE! |
| ENSMUSG00000050555.10 | 0 | NA | NA | NA | NA | NA | #VALUE! |
| ENSMUSG00000050556.10 | 8.992598 | -1.54512 | 2.630565 | -0.58737 | 0.556955 | NA | #VALUE! |
| ENSMUSG00000050558.14 | 0 | NA | NA | NA | NA | NA | #VALUE! |
| ENSMUSG00000050577.9 | 0 | NA | NA | NA | NA | NA | #VALUE! |
| ENSMUSG00000050578.11 | 0 | NA | NA | NA | NA | NA | #VALUE! |
| ENSMUSG00000050592.9 | 1.846078 | -2.32169 | 3.889915 | -0.59685 | 0.550608 | NA | #VALUE! |
| ENSMUSG00000050600.6 | 0 | NA | NA | NA | NA | NA | #VALUE! |
| ENSMUSG00000050603.2 | 0 | NA | NA | NA | NA | NA | #VALUE! |

| | | | | | | | |
|---|---|---|---|---|---|---|---|
| ENSMUSG00000050605.18 | 0 | NA | NA | NA | NA | NA | #VALUE! |
| ENSMUSG00000050612.7 | 0 | NA | NA | NA | NA | NA | #VALUE! |
| ENSMUSG00000050613.6 | 0 | NA | NA | NA | NA | NA | #VALUE! |
| ENSMUSG00000050623.6 | 0 | NA | NA | NA | NA | NA | #VALUE! |
| ENSMUSG00000050625.5 | 0 | NA | NA | NA | NA | NA | #VALUE! |
| ENSMUSG00000050635.2 | 0 | NA | NA | NA | NA | NA | #VALUE! |
| ENSMUSG00000050640.17 | 4.66276 | 5.717927 | 3.916352 | 1.460014 | 0.144286 | NA | #VALUE! |
| ENSMUSG00000050641.8 | 0 | NA | NA | NA | NA | NA | #VALUE! |
| ENSMUSG00000050645.4 | 0 | NA | NA | NA | NA | NA | #VALUE! |
| ENSMUSG00000050650.8 | 0 | NA | NA | NA | NA | NA | #VALUE! |
| ENSMUSG00000050654.6 | 0 | NA | NA | NA | NA | NA | #VALUE! |
| ENSMUSG00000050666.15 | 3.543697 | 5.324148 | 3.919351 | 1.358426 | 0.174329 | NA | #VALUE! |
| ENSMUSG00000050671.13 | 0 | NA | NA | NA | NA | NA | #VALUE! |
| ENSMUSG00000050675.8 | 0 | NA | NA | NA | NA | NA | #VALUE! |
| ENSMUSG00000050677.3 | 7.831564 | -6.49776 | 3.90065 | -1.66581 | 0.095751 | NA | #VALUE! |
| ENSMUSG00000050685.6 | 0 | NA | NA | NA | NA | NA | #VALUE! |
| ENSMUSG00000050700.12 | 0 | NA | NA | NA | NA | NA | #VALUE! |
| ENSMUSG00000050702.7 | 0 | NA | NA | NA | NA | NA | #VALUE! |
| ENSMUSG00000050704.4 | 0 | NA | NA | NA | NA | NA | #VALUE! |
| ENSMUSG00000050711.8 | 0 | NA | NA | NA | NA | NA | #VALUE! |
| ENSMUSG00000050714.10 | 4.353603 | -5.65681 | 3.917008 | -1.44417 | 0.148692 | NA | #VALUE! |
| ENSMUSG00000050721.10 | 0 | NA | NA | NA | NA | NA | #VALUE! |
| ENSMUSG00000050737.14 | 0 | NA | NA | NA | NA | NA | #VALUE! |
| ENSMUSG00000050742.6 | 0 | NA | NA | NA | NA | NA | #VALUE! |
| ENSMUSG00000050747.15 | 0 | NA | NA | NA | NA | NA | #VALUE! |
| ENSMUSG00000050751.15 | 0 | NA | NA | NA | NA | NA | #VALUE! |
| ENSMUSG00000050756.4 | 0 | NA | NA | NA | NA | NA | #VALUE! |
| ENSMUSG00000050761.5 | 0 | NA | NA | NA | NA | NA | #VALUE! |
| ENSMUSG00000050762.6 | 0 | NA | NA | NA | NA | NA | #VALUE! |
| ENSMUSG00000050763.5 | 0 | NA | NA | NA | NA | NA | #VALUE! |
| ENSMUSG00000050765.4 | 0 | NA | NA | NA | NA | NA | #VALUE! |
| ENSMUSG00000050766.3 | 0 | NA | NA | NA | NA | NA | #VALUE! |
| ENSMUSG00000050772.3 | 0 | NA | NA | NA | NA | NA | #VALUE! |
| ENSMUSG00000050776.4 | 0 | NA | NA | NA | NA | NA | #VALUE! |
| ENSMUSG00000050783.6 | 0 | NA | NA | NA | NA | NA | #VALUE! |
| ENSMUSG00000050788.6 | 0 | NA | NA | NA | NA | NA | #VALUE! |
| ENSMUSG00000050799.4 | 0 | NA | NA | NA | NA | NA | #VALUE! |
| ENSMUSG00000050803.5 | 0 | NA | NA | NA | NA | NA | #VALUE! |
| ENSMUSG00000050808.14 | 0 | NA | NA | NA | NA | NA | #VALUE! |
| ENSMUSG00000050810.9 | 0 | NA | NA | NA | NA | NA | #VALUE! |
| ENSMUSG00000050813.8 | 0 | NA | NA | NA | NA | NA | #VALUE! |
| ENSMUSG00000050815.7 | 0 | NA | NA | NA | NA | NA | #VALUE! |
| ENSMUSG00000050818.8 | 0 | NA | NA | NA | NA | NA | #VALUE! |
| ENSMUSG00000050821.15 | 0 | NA | NA | NA | NA | NA | #VALUE! |

| | | | | | | | |
|---|---|---|---|---|---|---|---|
| ENSMUSG00000050822.12 | 0 | NA | NA | NA | NA | NA | #VALUE! |
| ENSMUSG00000050824.13 | 0 | NA | NA | NA | NA | NA | #VALUE! |
| ENSMUSG00000050830.18 | 0 | NA | NA | NA | NA | NA | #VALUE! |
| ENSMUSG00000050840.9 | 0 | NA | NA | NA | NA | NA | #VALUE! |
| ENSMUSG00000050844.6 | 0 | NA | NA | NA | NA | NA | #VALUE! |
| ENSMUSG00000050853.4 | 0 | NA | NA | NA | NA | NA | #VALUE! |
| ENSMUSG00000050855.17 | 0 | NA | NA | NA | NA | NA | #VALUE! |
| ENSMUSG00000050860.7 | 0 | NA | NA | NA | NA | NA | #VALUE! |
| ENSMUSG00000050865.6 | 0 | NA | NA | NA | NA | NA | #VALUE! |
| ENSMUSG00000050866.8 | 0 | NA | NA | NA | NA | NA | #VALUE! |
| ENSMUSG00000050870.5 | 0 | NA | NA | NA | NA | NA | #VALUE! |
| ENSMUSG00000050875.12 | 0 | NA | NA | NA | NA | NA | #VALUE! |
| ENSMUSG00000050876.10 | 0 | NA | NA | NA | NA | NA | #VALUE! |
| ENSMUSG00000050896.13 | 0 | NA | NA | NA | NA | NA | #VALUE! |
| ENSMUSG00000050900.6 | 0 | NA | NA | NA | NA | NA | #VALUE! |
| ENSMUSG00000050901.7 | 0 | NA | NA | NA | NA | NA | #VALUE! |
| ENSMUSG00000050908.13 | 0 | NA | NA | NA | NA | NA | #VALUE! |
| ENSMUSG00000050910.6 | 6.825853 | -6.30496 | 3.913306 | -1.61116 | 0.107145 | NA | #VALUE! |
| ENSMUSG00000050914.17 | 8.183749 | 3.541848 | 3.456456 | 1.024705 | NA | NA | #VALUE! |
| ENSMUSG00000050917.6 | 0 | NA | NA | NA | NA | NA | #VALUE! |
| ENSMUSG00000050921.13 | 0 | NA | NA | NA | NA | NA | #VALUE! |
| ENSMUSG00000050926.6 | 0 | NA | NA | NA | NA | NA | #VALUE! |
| ENSMUSG00000050930.6 | 0 | NA | NA | NA | NA | NA | #VALUE! |
| ENSMUSG00000050933.5 | 0 | NA | NA | NA | NA | NA | #VALUE! |
| ENSMUSG00000050936.6 | 0 | NA | NA | NA | NA | NA | #VALUE! |
| ENSMUSG00000050944.15 | 0 | NA | NA | NA | NA | NA | #VALUE! |
| ENSMUSG00000050945.9 | 5.184189 | -0.70685 | 3.579963 | -0.19745 | 0.843478 | NA | #VALUE! |
| ENSMUSG00000050947.10 | 0 | NA | NA | NA | NA | NA | #VALUE! |
| ENSMUSG00000050954.16 | 0 | NA | NA | NA | NA | NA | #VALUE! |
| ENSMUSG00000050961.5 | 0 | NA | NA | NA | NA | NA | #VALUE! |
| ENSMUSG00000050963.8 | 0 | NA | NA | NA | NA | NA | #VALUE! |
| ENSMUSG00000050966.10 | 0 | NA | NA | NA | NA | NA | #VALUE! |
| ENSMUSG00000050967.6 | 1.177985 | 3.72614 | 3.944615 | 0.944614 | 0.344856 | NA | #VALUE! |
| ENSMUSG00000050974.5 | 0 | NA | NA | NA | NA | NA | #VALUE! |
| ENSMUSG00000050978.4 | 0 | NA | NA | NA | NA | NA | #VALUE! |
| ENSMUSG00000050982.14 | 0 | NA | NA | NA | NA | NA | #VALUE! |
| ENSMUSG00000050989.11 | 0 | NA | NA | NA | NA | NA | #VALUE! |
| ENSMUSG00000050994.21 | 0 | NA | NA | NA | NA | NA | #VALUE! |
| ENSMUSG00000050996.8 | 0 | NA | NA | NA | NA | NA | #VALUE! |
| ENSMUSG00000051000.18 | 5.87863 | -2.34512 | 3.782931 | -0.61992 | 0.535309 | NA | #VALUE! |
| ENSMUSG00000051002.8 | 0 | NA | NA | NA | NA | NA | #VALUE! |
| ENSMUSG00000051003.3 | 0 | NA | NA | NA | NA | NA | #VALUE! |
| ENSMUSG00000051008.6 | 0 | NA | NA | NA | NA | NA | #VALUE! |
| ENSMUSG00000051022.8 | 0 | NA | NA | NA | NA | NA | #VALUE! |

| | | | | | | | |
|---|---|---|---|---|---|---|---|
| ENSMUSG00000051029.9 | 0 | NA | NA | NA | NA | NA | #VALUE! |
| ENSMUSG00000051034.8 | 0 | NA | NA | NA | NA | NA | #VALUE! |
| ENSMUSG00000051036.14 | 0 | NA | NA | NA | NA | NA | #VALUE! |
| ENSMUSG00000051037.16 | 3.543697 | 5.324148 | 3.919351 | 1.358426 | 0.174329 | NA | #VALUE! |
| ENSMUSG00000051038.11 | 0 | NA | NA | NA | NA | NA | #VALUE! |
| ENSMUSG00000051046.8 | 0 | NA | NA | NA | NA | NA | #VALUE! |
| ENSMUSG00000051048.18 | 0 | NA | NA | NA | NA | NA | #VALUE! |
| ENSMUSG00000051050.11 | 0 | NA | NA | NA | NA | NA | #VALUE! |
| ENSMUSG00000051051.8 | 0 | NA | NA | NA | NA | NA | #VALUE! |
| ENSMUSG00000051054.5 | 0 | NA | NA | NA | NA | NA | #VALUE! |
| ENSMUSG00000051056.6 | 0 | NA | NA | NA | NA | NA | #VALUE! |
| ENSMUSG00000051062.7 | 0 | NA | NA | NA | NA | NA | #VALUE! |
| ENSMUSG00000051067.9 | 0 | NA | NA | NA | NA | NA | #VALUE! |
| ENSMUSG00000051074.5 | 0.839297 | 3.273019 | 3.958392 | 0.826856 | 0.408319 | NA | #VALUE! |
| ENSMUSG00000051076.9 | 0 | NA | NA | NA | NA | NA | #VALUE! |
| ENSMUSG00000051079.9 | 0 | NA | NA | NA | NA | NA | #VALUE! |
| ENSMUSG00000051081.8 | 0 | NA | NA | NA | NA | NA | #VALUE! |
| ENSMUSG00000051095.8 | 0 | NA | NA | NA | NA | NA | #VALUE! |
| ENSMUSG00000051111.17 | 0 | NA | NA | NA | NA | NA | #VALUE! |
| ENSMUSG00000051113.10 | 0 | NA | NA | NA | NA | NA | #VALUE! |
| ENSMUSG00000051116.7 | 0.37831 | -0.4571 | 4.045272 | -0.113 | 0.910033 | NA | #VALUE! |
| ENSMUSG00000051118.10 | 0 | NA | NA | NA | NA | NA | #VALUE! |
| ENSMUSG00000051133.4 | 0 | NA | NA | NA | NA | NA | #VALUE! |
| ENSMUSG00000051136.2 | 0 | NA | NA | NA | NA | NA | #VALUE! |
| ENSMUSG00000051146.4 | 1.912652 | -2.37989 | 3.888777 | -0.61199 | 0.540544 | NA | #VALUE! |
| ENSMUSG00000051149.16 | 0 | NA | NA | NA | NA | NA | #VALUE! |
| ENSMUSG00000051153.6 | 0 | NA | NA | NA | NA | NA | #VALUE! |
| ENSMUSG00000051156.3 | 0 | NA | NA | NA | NA | NA | #VALUE! |
| ENSMUSG00000051159.17 | 0 | NA | NA | NA | NA | NA | #VALUE! |
| ENSMUSG00000051160.5 | 0 | NA | NA | NA | NA | NA | #VALUE! |
| ENSMUSG00000051166.11 | 5.731668 | 4.942596 | 3.516244 | 1.405647 | 0.159829 | NA | #VALUE! |
| ENSMUSG00000051172.8 | 0 | NA | NA | NA | NA | NA | #VALUE! |
| ENSMUSG00000051176.7 | 0 | NA | NA | NA | NA | NA | #VALUE! |
| ENSMUSG00000051180.8 | 0 | NA | NA | NA | NA | NA | #VALUE! |
| ENSMUSG00000051182.6 | 0 | NA | NA | NA | NA | NA | #VALUE! |
| ENSMUSG00000051184.8 | 6.790738 | -6.29041 | 3.23983 | -1.94158 | 0.052187 | NA | #VALUE! |
| ENSMUSG00000051190.9 | 0 | NA | NA | NA | NA | NA | #VALUE! |
| ENSMUSG00000051198.7 | 0 | NA | NA | NA | NA | NA | #VALUE! |
| ENSMUSG00000051200.4 | 0 | NA | NA | NA | NA | NA | #VALUE! |
| ENSMUSG00000051207.8 | 0 | NA | NA | NA | NA | NA | #VALUE! |
| ENSMUSG00000051209.4 | 0 | NA | NA | NA | NA | NA | #VALUE! |
| ENSMUSG00000051212.9 | 0 | NA | NA | NA | NA | NA | #VALUE! |
| ENSMUSG00000051220.3 | 1.346268 | 3.918821 | 3.939925 | 0.994644 | 0.31991 | NA | #VALUE! |
| ENSMUSG00000051225.10 | 0.373021 | 2.140748 | 4.018182 | 0.532765 | 0.594196 | NA | #VALUE! |

| | | | | | | | |
|---|---|---|---|---|---|---|---|
| ENSMUSG00000051228.6 | 0 | NA | NA | NA | NA | NA | #VALUE! |
| ENSMUSG00000051235.11 | 3.622967 | 5.351386 | 3.919117 | 1.365457 | 0.172109 | NA | #VALUE! |
| ENSMUSG00000051237.8 | 0 | NA | NA | NA | NA | NA | #VALUE! |
| ENSMUSG00000051242.2 | 0 | NA | NA | NA | NA | NA | #VALUE! |
| ENSMUSG00000051243.15 | 0 | NA | NA | NA | NA | NA | #VALUE! |
| ENSMUSG00000051246.4 | 0 | NA | NA | NA | NA | NA | #VALUE! |
| ENSMUSG00000051251.4 | 0 | NA | NA | NA | NA | NA | #VALUE! |
| ENSMUSG00000051255.7 | 4.746404 | 4.168649 | 3.40789 | 1.223235 | 0.221241 | NA | #VALUE! |
| ENSMUSG00000051257.4 | 0 | NA | NA | NA | NA | NA | #VALUE! |
| ENSMUSG00000051258.6 | 0 | NA | NA | NA | NA | NA | #VALUE! |
| ENSMUSG00000051262.10 | 0 | NA | NA | NA | NA | NA | #VALUE! |
| ENSMUSG00000051276.5 | 0 | NA | NA | NA | NA | NA | #VALUE! |
| ENSMUSG00000051278.13 | 0.681591 | -0.6064 | 3.969202 | -0.15278 | 0.878575 | NA | #VALUE! |
| ENSMUSG00000051279.3 | 0.105013 | -0.97324 | 4.076293 | -0.23876 | 0.811295 | NA | #VALUE! |
| ENSMUSG00000051295.9 | 0 | NA | NA | NA | NA | NA | #VALUE! |
| ENSMUSG00000051297.9 | 0 | NA | NA | NA | NA | NA | #VALUE! |
| ENSMUSG00000051313.13 | 0 | NA | NA | NA | NA | NA | #VALUE! |
| ENSMUSG00000051314.12 | 0 | NA | NA | NA | NA | NA | #VALUE! |
| ENSMUSG00000051323.17 | 0 | NA | NA | NA | NA | NA | #VALUE! |
| ENSMUSG00000051329.14 | 5.255137 | -2.83303 | 3.622741 | -0.78201 | 0.434207 | NA | #VALUE! |
| ENSMUSG00000051331.17 | 0 | NA | NA | NA | NA | NA | #VALUE! |
| ENSMUSG00000051335.7 | 2.238125 | 4.666312 | 3.926586 | 1.188389 | 0.23468 | NA | #VALUE! |
| ENSMUSG00000051340.7 | 0 | NA | NA | NA | NA | NA | #VALUE! |
| ENSMUSG00000051341.7 | 7.213192 | 1.27326 | 3.192092 | 0.39888 | 0.689982 | NA | #VALUE! |
| ENSMUSG00000051343.12 | 2.100263 | -4.61322 | 3.927832 | -1.17449 | 0.240197 | NA | #VALUE! |
| ENSMUSG00000051344.14 | 1.174735 | -3.75246 | 3.944889 | -0.95122 | 0.341493 | NA | #VALUE! |
| ENSMUSG00000051351.15 | 5.270499 | 4.919287 | 3.911143 | 1.257762 | 0.208478 | NA | #VALUE! |
| ENSMUSG00000051354.15 | 0 | NA | NA | NA | NA | NA | #VALUE! |
| ENSMUSG00000051361.8 | 0.735092 | -3.12016 | 3.965612 | -0.7868 | 0.431397 | NA | #VALUE! |
| ENSMUSG00000051362.3 | 0 | NA | NA | NA | NA | NA | #VALUE! |
| ENSMUSG00000051367.9 | 0 | NA | NA | NA | NA | NA | #VALUE! |
| ENSMUSG00000051373.6 | 0 | NA | NA | NA | NA | NA | #VALUE! |
| ENSMUSG00000051375.16 | 7.233334 | -5.29022 | 3.289081 | -1.60842 | 0.107743 | NA | #VALUE! |
| ENSMUSG00000051378.11 | 0 | NA | NA | NA | NA | NA | #VALUE! |
| ENSMUSG00000051379.13 | 4.694547 | -5.7577 | 3.916317 | -1.47018 | 0.141512 | NA | #VALUE! |
| ENSMUSG00000051392.6 | 0 | NA | NA | NA | NA | NA | #VALUE! |
| ENSMUSG00000051396.16 | 1.155144 | -3.7605 | 3.944679 | -0.95331 | 0.340433 | NA | #VALUE! |
| ENSMUSG00000051397.6 | 0 | NA | NA | NA | NA | NA | #VALUE! |
| ENSMUSG00000051401.8 | 0 | NA | NA | NA | NA | NA | #VALUE! |
| ENSMUSG00000051414.6 | 0 | NA | NA | NA | NA | NA | #VALUE! |
| ENSMUSG00000051424.3 | 0 | NA | NA | NA | NA | NA | #VALUE! |
| ENSMUSG00000051427.15 | 8.241332 | -6.57267 | 3.307572 | -1.98716 | 0.046905 | NA | #VALUE! |
| ENSMUSG00000051431.8 | 0 | NA | NA | NA | NA | NA | #VALUE! |
| ENSMUSG00000051437.5 | 0 | NA | NA | NA | NA | NA | #VALUE! |

| | | | | | | | |
|---|---|---|---|---|---|---|---|
| ENSMUSG00000051444.10 | 0.204111 | -1.05581 | 4.073302 | -0.2592 | 0.795479 | NA | #VALUE! |
| ENSMUSG00000051452.9 | 5.515837 | 1.279888 | 2.87163 | 0.445701 | 0.655813 | NA | #VALUE! |
| ENSMUSG00000051455.14 | 0 | NA | NA | NA | NA | NA | #VALUE! |
| ENSMUSG00000051456.5 | 0 | NA | NA | NA | NA | NA | #VALUE! |
| ENSMUSG00000051457.8 | 0 | NA | NA | NA | NA | NA | #VALUE! |
| ENSMUSG00000051481.6 | 0 | NA | NA | NA | NA | NA | #VALUE! |
| ENSMUSG00000051486.6 | 0 | NA | NA | NA | NA | NA | #VALUE! |
| ENSMUSG00000051490.4 | 0 | NA | NA | NA | NA | NA | #VALUE! |
| ENSMUSG00000051493.8 | 0 | NA | NA | NA | NA | NA | #VALUE! |
| ENSMUSG00000051497.16 | 0.630079 | -2.90291 | 3.975047 | -0.73028 | 0.465217 | NA | #VALUE! |
| ENSMUSG00000051498.8 | 0 | NA | NA | NA | NA | NA | #VALUE! |
| ENSMUSG00000051499.8 | 0 | NA | NA | NA | NA | NA | #VALUE! |
| ENSMUSG00000051502.8 | 2.071788 | -4.59026 | 3.928168 | -1.16855 | 0.242586 | NA | #VALUE! |
| ENSMUSG00000051503.3 | 0 | NA | NA | NA | NA | NA | #VALUE! |
| ENSMUSG00000051504.19 | 3.150394 | -5.19405 | 3.92087 | -1.32472 | 0.185265 | NA | #VALUE! |
| ENSMUSG00000051506.18 | 6.621119 | 6.221784 | 3.913529 | 1.589814 | NA | NA | #VALUE! |
| ENSMUSG00000051509.7 | 0 | NA | NA | NA | NA | NA | #VALUE! |
| ENSMUSG00000051515.12 | 0 | NA | NA | NA | NA | NA | #VALUE! |
| ENSMUSG00000051517.15 | 0 | NA | NA | NA | NA | NA | #VALUE! |
| ENSMUSG00000051527.12 | 0 | NA | NA | NA | NA | NA | #VALUE! |
| ENSMUSG00000051528.5 | 0 | NA | NA | NA | NA | NA | #VALUE! |
| ENSMUSG00000051537.10 | 0 | NA | NA | NA | NA | NA | #VALUE! |
| ENSMUSG00000051548.5 | 0 | NA | NA | NA | NA | NA | #VALUE! |
| ENSMUSG00000051550.11 | 0.932552 | 3.421684 | 3.953399 | 0.865504 | 0.386762 | NA | #VALUE! |
| ENSMUSG00000051557.16 | 3.971493 | -4.38576 | 3.904651 | -1.12321 | 0.261347 | NA | #VALUE! |
| ENSMUSG00000051582.3 | 0 | NA | NA | NA | NA | NA | #VALUE! |
| ENSMUSG00000051590.11 | 0 | NA | NA | NA | NA | NA | #VALUE! |
| ENSMUSG00000051591.5 | 0 | NA | NA | NA | NA | NA | #VALUE! |
| ENSMUSG00000051592.15 | 0 | NA | NA | NA | NA | NA | #VALUE! |
| ENSMUSG00000051593.5 | 0 | NA | NA | NA | NA | NA | #VALUE! |
| ENSMUSG00000051596.16 | 0 | NA | NA | NA | NA | NA | #VALUE! |
| ENSMUSG00000051599.5 | 0 | NA | NA | NA | NA | NA | #VALUE! |
| ENSMUSG00000051606.6 | 0 | NA | NA | NA | NA | NA | #VALUE! |
| ENSMUSG00000051611.10 | 0 | NA | NA | NA | NA | NA | #VALUE! |
| ENSMUSG00000051616.15 | 0 | NA | NA | NA | NA | NA | #VALUE! |
| ENSMUSG00000051617.4 | 0 | NA | NA | NA | NA | NA | #VALUE! |
| ENSMUSG00000051618.6 | 0 | NA | NA | NA | NA | NA | #VALUE! |
| ENSMUSG00000051627.4 | 0 | NA | NA | NA | NA | NA | #VALUE! |
| ENSMUSG00000051638.3 | 0 | NA | NA | NA | NA | NA | #VALUE! |
| ENSMUSG00000051639.4 | 8.402176 | -3.44022 | 3.096425 | -1.11103 | 0.266556 | NA | #VALUE! |
| ENSMUSG00000051648.14 | 0 | NA | NA | NA | NA | NA | #VALUE! |
| ENSMUSG00000051650.12 | 0 | NA | NA | NA | NA | NA | #VALUE! |
| ENSMUSG00000051663.5 | 0 | NA | NA | NA | NA | NA | #VALUE! |
| ENSMUSG00000051669.7 | 8.942755 | 6.654041 | 3.198403 | 2.080426 | 0.037486 | NA | #VALUE! |

| | | | | | | | |
|---|---|---|---|---|---|---|---|
| ENSMUSG00000051678.5 | 0 | NA | NA | NA | NA | NA | #VALUE! |
| ENSMUSG00000051680.4 | 0 | NA | NA | NA | NA | NA | #VALUE! |
| ENSMUSG00000051682.16 | 3.213664 | -4.07757 | 3.90746 | -1.04354 | 0.2967 | NA | #VALUE! |
| ENSMUSG00000051687.3 | 0 | NA | NA | NA | NA | NA | #VALUE! |
| ENSMUSG00000051703.15 | 0 | NA | NA | NA | NA | NA | #VALUE! |
| ENSMUSG00000051706.4 | 0 | NA | NA | NA | NA | NA | #VALUE! |
| ENSMUSG00000051726.7 | 0 | NA | NA | NA | NA | NA | #VALUE! |
| ENSMUSG00000051727.15 | 0 | NA | NA | NA | NA | NA | #VALUE! |
| ENSMUSG00000051728.8 | 0 | NA | NA | NA | NA | NA | #VALUE! |
| ENSMUSG00000051729.13 | 0 | NA | NA | NA | NA | NA | #VALUE! |
| ENSMUSG00000051732.3 | 0 | NA | NA | NA | NA | NA | #VALUE! |
| ENSMUSG00000051735.8 | 0.279766 | 1.808031 | 4.036658 | 0.447903 | 0.654223 | NA | #VALUE! |
| ENSMUSG00000051736.9 | 0 | NA | NA | NA | NA | NA | #VALUE! |
| ENSMUSG00000051747.16 | 0 | NA | NA | NA | NA | NA | #VALUE! |
| ENSMUSG00000051758.7 | 0 | NA | NA | NA | NA | NA | #VALUE! |
| ENSMUSG00000051768.10 | 9.193904 | -5.80763 | 2.97901 | -1.94952 | 0.051234 | NA | #VALUE! |
| ENSMUSG00000051769.3 | 0 | NA | NA | NA | NA | NA | #VALUE! |
| ENSMUSG00000051777.7 | 0 | NA | NA | NA | NA | NA | #VALUE! |
| ENSMUSG00000051786.16 | 3.533951 | 0.032405 | 3.777587 | 0.008578 | 0.993156 | NA | #VALUE! |
| ENSMUSG00000051788.5 | 0 | NA | NA | NA | NA | NA | #VALUE! |
| ENSMUSG00000051790.16 | 0 | NA | NA | NA | NA | NA | #VALUE! |
| ENSMUSG00000051793.5 | 0 | NA | NA | NA | NA | NA | #VALUE! |
| ENSMUSG00000051802.4 | 0 | NA | NA | NA | NA | NA | #VALUE! |
| ENSMUSG00000051804.4 | 0 | NA | NA | NA | NA | NA | #VALUE! |
| ENSMUSG00000051805.2 | 0 | NA | NA | NA | NA | NA | #VALUE! |
| ENSMUSG00000051811.15 | 2.467451 | 4.794393 | 3.92491 | 1.221529 | 0.221886 | NA | #VALUE! |
| ENSMUSG00000051817.9 | 0.525066 | -2.64687 | 3.988095 | -0.66369 | 0.506887 | NA | #VALUE! |
| ENSMUSG00000051827.12 | 0 | NA | NA | NA | NA | NA | #VALUE! |
| ENSMUSG00000051832.8 | 0 | NA | NA | NA | NA | NA | #VALUE! |
| ENSMUSG00000051839.8 | 0 | NA | NA | NA | NA | NA | #VALUE! |
| ENSMUSG00000051844.14 | 0 | NA | NA | NA | NA | NA | #VALUE! |
| ENSMUSG00000051851.6 | 0 | NA | NA | NA | NA | NA | #VALUE! |
| ENSMUSG00000051855.16 | 4.429426 | 4.55397 | 3.865339 | 1.178156 | 0.238735 | NA | #VALUE! |
| ENSMUSG00000051860.14 | 0 | NA | NA | NA | NA | NA | #VALUE! |
| ENSMUSG00000051877.9 | 0 | NA | NA | NA | NA | NA | #VALUE! |
| ENSMUSG00000051879.5 | 0 | NA | NA | NA | NA | NA | #VALUE! |
| ENSMUSG00000051885.7 | 0 | NA | NA | NA | NA | NA | #VALUE! |
| ENSMUSG00000051890.14 | 0 | NA | NA | NA | NA | NA | #VALUE! |
| ENSMUSG00000051896.5 | 0 | NA | NA | NA | NA | NA | #VALUE! |
| ENSMUSG00000051900.13 | 0 | NA | NA | NA | NA | NA | #VALUE! |
| ENSMUSG00000051906.16 | 0 | NA | NA | NA | NA | NA | #VALUE! |
| ENSMUSG00000051917.4 | 0 | NA | NA | NA | NA | NA | #VALUE! |
| ENSMUSG00000051920.8 | 0 | NA | NA | NA | NA | NA | #VALUE! |
| ENSMUSG00000051934.6 | 8.702571 | -6.6524 | 2.817801 | -2.36085 | 0.018233 | NA | #VALUE! |

| | | | | | | | |
|---|---|---|---|---|---|---|---|
| ENSMUSG00000051936.5 | 0 | NA | NA | NA | NA | NA | #VALUE! |
| ENSMUSG00000051940.3 | 0 | NA | NA | NA | NA | NA | #VALUE! |
| ENSMUSG00000051950.11 | 2.617413 | -3.7591 | 3.911063 | -0.96115 | 0.336479 | NA | #VALUE! |
| ENSMUSG00000051951.6 | 0 | NA | NA | NA | NA | NA | #VALUE! |
| ENSMUSG00000051952.6 | 0 | NA | NA | NA | NA | NA | #VALUE! |
| ENSMUSG00000051956.5 | 0 | NA | NA | NA | NA | NA | #VALUE! |
| ENSMUSG00000051965.9 | 0 | NA | NA | NA | NA | NA | #VALUE! |
| ENSMUSG00000051969.8 | 0 | NA | NA | NA | NA | NA | #VALUE! |
| ENSMUSG00000051977.17 | 5.372351 | -1.79421 | 3.165103 | -0.56687 | 0.5708 | NA | #VALUE! |
| ENSMUSG00000051980.14 | 0 | NA | NA | NA | NA | NA | #VALUE! |
| ENSMUSG00000051984.14 | 4.91951 | 5.781601 | 3.915938 | 1.476428 | 0.139829 | NA | #VALUE! |
| ENSMUSG00000051985.13 | 0 | NA | NA | NA | NA | NA | #VALUE! |
| ENSMUSG00000051986.7 | 0 | NA | NA | NA | NA | NA | #VALUE! |
| ENSMUSG00000051998.15 | 0 | NA | NA | NA | NA | NA | #VALUE! |
| ENSMUSG00000052005.9 | 0 | NA | NA | NA | NA | NA | #VALUE! |
| ENSMUSG00000052012.7 | 0 | NA | NA | NA | NA | NA | #VALUE! |
| ENSMUSG00000052013.16 | 5.146956 | -2.36171 | 3.790978 | -0.62298 | 0.533297 | NA | #VALUE! |
| ENSMUSG00000052014.7 | 0 | NA | NA | NA | NA | NA | #VALUE! |
| ENSMUSG00000052025.10 | 0 | NA | NA | NA | NA | NA | #VALUE! |
| ENSMUSG00000052026.9 | 0 | NA | NA | NA | NA | NA | #VALUE! |
| ENSMUSG00000052031.9 | 8.943043 | -1.80232 | 2.676238 | -0.67345 | 0.50066 | NA | #VALUE! |
| ENSMUSG00000052058.4 | 0 | NA | NA | NA | NA | NA | #VALUE! |
| ENSMUSG00000052075.8 | 0 | NA | NA | NA | NA | NA | #VALUE! |
| ENSMUSG00000052087.15 | 0 | NA | NA | NA | NA | NA | #VALUE! |
| ENSMUSG00000052099.15 | 0 | NA | NA | NA | NA | NA | #VALUE! |
| ENSMUSG00000052105.18 | 0 | NA | NA | NA | NA | NA | #VALUE! |
| ENSMUSG00000052125.17 | 0 | NA | NA | NA | NA | NA | #VALUE! |
| ENSMUSG00000052131.8 | 1.398828 | 3.99633 | 3.938205 | 1.014759 | 0.310221 | NA | #VALUE! |
| ENSMUSG00000052133.17 | 2.60352 | -1.05992 | 3.806558 | -0.27845 | 0.78067 | NA | #VALUE! |
| ENSMUSG00000052135.9 | 0 | NA | NA | NA | NA | NA | #VALUE! |
| ENSMUSG00000052137.12 | 9.201905 | -2.55769 | 2.982213 | -0.85765 | NA | NA | #VALUE! |
| ENSMUSG00000052142.16 | 0 | NA | NA | NA | NA | NA | #VALUE! |
| ENSMUSG00000052143.9 | 0 | NA | NA | NA | NA | NA | #VALUE! |
| ENSMUSG00000052146.17 | 0 | NA | NA | NA | NA | NA | #VALUE! |
| ENSMUSG00000052154.5 | 0 | NA | NA | NA | NA | NA | #VALUE! |
| ENSMUSG00000052157.5 | 0 | NA | NA | NA | NA | NA | #VALUE! |
| ENSMUSG00000052180.16 | 0 | NA | NA | NA | NA | NA | #VALUE! |
| ENSMUSG00000052182.5 | 0 | NA | NA | NA | NA | NA | #VALUE! |
| ENSMUSG00000052187.5 | 0 | NA | NA | NA | NA | NA | #VALUE! |
| ENSMUSG00000052188.7 | 7.020572 | -3.47168 | 3.663172 | -0.94773 | 0.343269 | NA | #VALUE! |
| ENSMUSG00000052192.6 | 5.057569 | 0.679355 | 2.476853 | 0.274282 | 0.783868 | NA | #VALUE! |
| ENSMUSG00000052212.7 | 0 | NA | NA | NA | NA | NA | #VALUE! |
| ENSMUSG00000052217.7 | 0 | NA | NA | NA | NA | NA | #VALUE! |
| ENSMUSG00000052221.9 | 0 | NA | NA | NA | NA | NA | #VALUE! |

| | | | | | | | |
|---|---|---|---|---|---|---|---|
| ENSMUSG00000052229.6 | 0 | NA | NA | NA | NA | NA | #VALUE! |
| ENSMUSG00000052234.3 | 0 | NA | NA | NA | NA | NA | #VALUE! |
| ENSMUSG00000052236.8 | 0 | NA | NA | NA | NA | NA | #VALUE! |
| ENSMUSG00000052241.4 | 0 | NA | NA | NA | NA | NA | #VALUE! |
| ENSMUSG00000052248.18 | 0 | NA | NA | NA | NA | NA | #VALUE! |
| ENSMUSG00000052270.8 | 3.15829 | -2.38905 | 3.83365 | -0.62318 | 0.533167 | NA | #VALUE! |
| ENSMUSG00000052271.8 | 1.575197 | -4.20222 | 3.934736 | -1.06798 | 0.285529 | NA | #VALUE! |
| ENSMUSG00000052273.5 | 0 | NA | NA | NA | NA | NA | #VALUE! |
| ENSMUSG00000052276.9 | 0 | NA | NA | NA | NA | NA | #VALUE! |
| ENSMUSG00000052281.15 | 0 | NA | NA | NA | NA | NA | #VALUE! |
| ENSMUSG00000052293.15 | 0 | NA | NA | NA | NA | NA | #VALUE! |
| ENSMUSG00000052295.5 | 0 | NA | NA | NA | NA | NA | #VALUE! |
| ENSMUSG00000052301.15 | 0 | NA | NA | NA | NA | NA | #VALUE! |
| ENSMUSG00000052303.7 | 0 | NA | NA | NA | NA | NA | #VALUE! |
| ENSMUSG00000052316.10 | 0 | NA | NA | NA | NA | NA | #VALUE! |
| ENSMUSG00000052331.15 | 3.580588 | -0.32526 | 3.777183 | -0.08611 | 0.931378 | NA | #VALUE! |
| ENSMUSG00000052334.12 | 0 | NA | NA | NA | NA | NA | #VALUE! |
| ENSMUSG00000052353.14 | 0 | NA | NA | NA | NA | NA | #VALUE! |
| ENSMUSG00000052363.9 | 0 | NA | NA | NA | NA | NA | #VALUE! |
| ENSMUSG00000052364.5 | 2.355969 | 4.72622 | 3.925784 | 1.203892 | 0.228631 | NA | #VALUE! |
| ENSMUSG00000052368.3 | 0 | NA | NA | NA | NA | NA | #VALUE! |
| ENSMUSG00000052371.6 | 0 | NA | NA | NA | NA | NA | #VALUE! |
| ENSMUSG00000052372.11 | 0 | NA | NA | NA | NA | NA | #VALUE! |
| ENSMUSG00000052373.15 | 0 | NA | NA | NA | NA | NA | #VALUE! |
| ENSMUSG00000052374.17 | 0 | NA | NA | NA | NA | NA | #VALUE! |
| ENSMUSG00000052376.3 | 0 | NA | NA | NA | NA | NA | #VALUE! |
| ENSMUSG00000052382.6 | 0 | NA | NA | NA | NA | NA | #VALUE! |
| ENSMUSG00000052384.15 | 6.788894 | -6.29254 | 3.551696 | -1.7717 | 0.076444 | NA | #VALUE! |
| ENSMUSG00000052387.17 | 0 | NA | NA | NA | NA | NA | #VALUE! |
| ENSMUSG00000052396.8 | 0 | NA | NA | NA | NA | NA | #VALUE! |
| ENSMUSG00000052403.12 | 0 | NA | NA | NA | NA | NA | #VALUE! |
| ENSMUSG00000052407.18 | 0.816443 | -3.21914 | 3.961753 | -0.81255 | 0.416474 | NA | #VALUE! |
| ENSMUSG00000052414.10 | 0 | NA | NA | NA | NA | NA | #VALUE! |
| ENSMUSG00000052415.6 | 3.404219 | 2.783156 | 3.845534 | 0.723737 | 0.469227 | NA | #VALUE! |
| ENSMUSG00000052417.7 | 0 | NA | NA | NA | NA | NA | #VALUE! |
| ENSMUSG00000052423.16 | 1.851119 | 4.378291 | 3.930938 | 1.113803 | 0.265364 | NA | #VALUE! |
| ENSMUSG00000052426.5 | 0 | NA | NA | NA | NA | NA | #VALUE! |
| ENSMUSG00000052430.16 | 0 | NA | NA | NA | NA | NA | #VALUE! |
| ENSMUSG00000052468.8 | 0 | NA | NA | NA | NA | NA | #VALUE! |
| ENSMUSG00000052469.10 | 0 | NA | NA | NA | NA | NA | #VALUE! |
| ENSMUSG00000052477.16 | 0.315039 | -1.93549 | 4.038621 | -0.47925 | 0.631763 | NA | #VALUE! |
| ENSMUSG00000052479.6 | 0 | NA | NA | NA | NA | NA | #VALUE! |
| ENSMUSG00000052482.7 | 0 | NA | NA | NA | NA | NA | #VALUE! |
| ENSMUSG00000052485.8 | 0 | NA | NA | NA | NA | NA | #VALUE! |

| | | | | | | | |
|---|---|---|---|---|---|---|---|
| ENSMUSG00000052496.6 | 0 | NA | NA | NA | NA | NA | #VALUE! |
| ENSMUSG00000052504.8 | 5.581976 | -6.0115 | 3.811213 | -1.57732 | 0.114722 | NA | #VALUE! |
| ENSMUSG00000052508.11 | 0 | NA | NA | NA | NA | NA | #VALUE! |
| ENSMUSG00000052525.15 | 0 | NA | NA | NA | NA | NA | #VALUE! |
| ENSMUSG00000052533.15 | 9.794281 | 2.208659 | 2.603118 | 0.848467 | 0.396178 | NA | #VALUE! |
| ENSMUSG00000052537.5 | 0 | NA | NA | NA | NA | NA | #VALUE! |
| ENSMUSG00000052544.10 | 2.963075 | -1.78973 | 3.813564 | -0.46931 | 0.63885 | NA | #VALUE! |
| ENSMUSG00000052549.5 | 0 | NA | NA | NA | NA | NA | #VALUE! |
| ENSMUSG00000052551.17 | 0 | NA | NA | NA | NA | NA | #VALUE! |
| ENSMUSG00000052554.4 | 0 | NA | NA | NA | NA | NA | #VALUE! |
| ENSMUSG00000052557.12 | 8.248142 | 0.915638 | 2.659047 | 0.344348 | 0.730584 | NA | #VALUE! |
| ENSMUSG00000052563.14 | 0.746042 | 3.107177 | 3.964586 | 0.783733 | 0.433197 | NA | #VALUE! |
| ENSMUSG00000052565.8 | 4.974401 | -2.05964 | 3.785 | -0.54416 | 0.586333 | NA | #VALUE! |
| ENSMUSG00000052572.20 | 0 | NA | NA | NA | NA | NA | #VALUE! |
| ENSMUSG00000052581.14 | 0 | NA | NA | NA | NA | NA | #VALUE! |
| ENSMUSG00000052584.11 | 0 | NA | NA | NA | NA | NA | #VALUE! |
| ENSMUSG00000052613.17 | 0 | NA | NA | NA | NA | NA | #VALUE! |
| ENSMUSG00000052616.11 | 0 | NA | NA | NA | NA | NA | #VALUE! |
| ENSMUSG00000052625.9 | 0 | NA | NA | NA | NA | NA | #VALUE! |
| ENSMUSG00000052629.6 | 0 | NA | NA | NA | NA | NA | #VALUE! |
| ENSMUSG00000052631.16 | 0 | NA | NA | NA | NA | NA | #VALUE! |
| ENSMUSG00000052642.4 | 0 | NA | NA | NA | NA | NA | #VALUE! |
| ENSMUSG00000052658.6 | 9.232264 | 6.700573 | 3.663674 | 1.828922 | NA | NA | #VALUE! |
| ENSMUSG00000052673.3 | 0 | NA | NA | NA | NA | NA | #VALUE! |
| ENSMUSG00000052675.12 | 2.51789 | 4.834703 | 3.924413 | 1.231956 | 0.217966 | NA | #VALUE! |
| ENSMUSG00000052688.13 | 6.622838 | 0.579372 | 3.384561 | 0.171181 | 0.864082 | NA | #VALUE! |
| ENSMUSG00000052698.16 | 2.170545 | 4.615911 | 3.927287 | 1.175343 | 0.239857 | NA | #VALUE! |
| ENSMUSG00000052713.10 | 5.158528 | 2.496786 | 3.795459 | 0.657835 | 0.510644 | NA | #VALUE! |
| ENSMUSG00000052724.8 | 0 | NA | NA | NA | NA | NA | #VALUE! |
| ENSMUSG00000052726.17 | 7.376431 | 3.13265 | 3.61897 | 0.865619 | 0.386699 | NA | #VALUE! |
| ENSMUSG00000052727.7 | 0.168284 | 1.027628 | 4.067126 | 0.252667 | 0.800526 | NA | #VALUE! |
| ENSMUSG00000052730.8 | 0 | NA | NA | NA | NA | NA | #VALUE! |
| ENSMUSG00000052736.16 | 0 | NA | NA | NA | NA | NA | #VALUE! |
| ENSMUSG00000052749.10 | 0 | NA | NA | NA | NA | NA | #VALUE! |
| ENSMUSG00000052752.16 | 0 | NA | NA | NA | NA | NA | #VALUE! |
| ENSMUSG00000052759.8 | 0 | NA | NA | NA | NA | NA | #VALUE! |
| ENSMUSG00000052760.17 | 0.093255 | 0.868093 | 4.072464 | 0.213162 | 0.831201 | NA | #VALUE! |
| ENSMUSG00000052763.8 | 7.840648 | -6.49824 | 3.065932 | -2.1195 | 0.034048 | NA | #VALUE! |
| ENSMUSG00000052767.6 | 0 | NA | NA | NA | NA | NA | #VALUE! |
| ENSMUSG00000052769.9 | 0 | NA | NA | NA | NA | NA | #VALUE! |
| ENSMUSG00000052776.11 | 0 | NA | NA | NA | NA | NA | #VALUE! |
| ENSMUSG00000052779.5 | 0 | NA | NA | NA | NA | NA | #VALUE! |
| ENSMUSG00000052783.17 | 0.864972 | 3.275682 | 3.958298 | 0.827548 | 0.407926 | NA | #VALUE! |
| ENSMUSG00000052785.7 | 0 | NA | NA | NA | NA | NA | #VALUE! |

| | | | | | | | |
|---|---|---|---|---|---|---|---|
| ENSMUSG00000052790.7 | 0 | NA | NA | NA | NA | NA | #VALUE! |
| ENSMUSG00000052818.6 | 0 | NA | NA | NA | NA | NA | #VALUE! |
| ENSMUSG00000052819.7 | 0 | NA | NA | NA | NA | NA | #VALUE! |
| ENSMUSG00000052821.4 | 4.30736 | -5.63406 | 3.91717 | -1.4383 | 0.15035 | NA | #VALUE! |
| ENSMUSG00000052825.7 | 1.963632 | 0.748915 | 3.825786 | 0.195754 | 0.844802 | NA | #VALUE! |
| ENSMUSG00000052837.8 | 12.43529 | -4.89101 | 2.780461 | -1.75906 | NA | NA | #VALUE! |
| ENSMUSG00000052848.8 | 0 | NA | NA | NA | NA | NA | #VALUE! |
| ENSMUSG00000052850.6 | 0 | NA | NA | NA | NA | NA | #VALUE! |
| ENSMUSG00000052852.9 | 5.51099 | -5.9895 | 3.914902 | -1.52992 | 0.126035 | NA | #VALUE! |
| ENSMUSG00000052854.16 | 0 | NA | NA | NA | NA | NA | #VALUE! |
| ENSMUSG00000052861.14 | 0 | NA | NA | NA | NA | NA | #VALUE! |
| ENSMUSG00000052865.6 | 0 | NA | NA | NA | NA | NA | #VALUE! |
| ENSMUSG00000052909.9 | 0 | NA | NA | NA | NA | NA | #VALUE! |
| ENSMUSG00000052911.10 | 0.874497 | 0.288047 | 3.932198 | 0.073253 | 0.941604 | NA | #VALUE! |
| ENSMUSG00000052912.8 | 0 | NA | NA | NA | NA | NA | #VALUE! |
| ENSMUSG00000052920.17 | 0 | NA | NA | NA | NA | NA | #VALUE! |
| ENSMUSG00000052921.14 | 18.06194 | -3.86098 | 2.539612 | -1.5203 | NA | NA | #VALUE! |
| ENSMUSG00000052922.10 | 0 | NA | NA | NA | NA | NA | #VALUE! |
| ENSMUSG00000052942.14 | 7.404475 | 6.378241 | 3.912835 | 1.630082 | 0.103084 | NA | #VALUE! |
| ENSMUSG00000052949.15 | 0 | NA | NA | NA | NA | NA | #VALUE! |
| ENSMUSG00000052951.6 | 0 | NA | NA | NA | NA | NA | #VALUE! |
| ENSMUSG00000052955.6 | 0 | NA | NA | NA | NA | NA | #VALUE! |
| ENSMUSG00000052981.7 | 0 | NA | NA | NA | NA | NA | #VALUE! |
| ENSMUSG00000053004.10 | 0 | NA | NA | NA | NA | NA | #VALUE! |
| ENSMUSG00000053007.10 | 0 | NA | NA | NA | NA | NA | #VALUE! |
| ENSMUSG00000053024.15 | 0 | NA | NA | NA | NA | NA | #VALUE! |
| ENSMUSG00000053025.14 | 0 | NA | NA | NA | NA | NA | #VALUE! |
| ENSMUSG00000053030.12 | 0 | NA | NA | NA | NA | NA | #VALUE! |
| ENSMUSG00000053038.9 | 0 | NA | NA | NA | NA | NA | #VALUE! |
| ENSMUSG00000053040.14 | 0.373021 | 2.140748 | 4.018182 | 0.532765 | 0.594196 | NA | #VALUE! |
| ENSMUSG00000053044.9 | 0 | NA | NA | NA | NA | NA | #VALUE! |
| ENSMUSG00000053046.17 | 0 | NA | NA | NA | NA | NA | #VALUE! |
| ENSMUSG00000053049.4 | 0 | NA | NA | NA | NA | NA | #VALUE! |
| ENSMUSG00000053054.14 | 0 | NA | NA | NA | NA | NA | #VALUE! |
| ENSMUSG00000053070.6 | 0 | NA | NA | NA | NA | NA | #VALUE! |
| ENSMUSG00000053080.11 | 5.03578 | 5.828452 | 3.915646 | 1.488503 | 0.136618 | NA | #VALUE! |
| ENSMUSG00000053081.4 | 0 | NA | NA | NA | NA | NA | #VALUE! |
| ENSMUSG00000053091.17 | 8.963938 | 2.709455 | 2.675919 | 1.012533 | 0.311283 | NA | #VALUE! |
| ENSMUSG00000053093.17 | 0 | NA | NA | NA | NA | NA | #VALUE! |
| ENSMUSG00000053101.4 | 0 | NA | NA | NA | NA | NA | #VALUE! |
| ENSMUSG00000053111.14 | 0 | NA | NA | NA | NA | NA | #VALUE! |
| ENSMUSG00000053113.4 | 7.057407 | -4.29902 | 3.669336 | -1.17161 | 0.241355 | NA | #VALUE! |
| ENSMUSG00000053117.12 | 0 | NA | NA | NA | NA | NA | #VALUE! |
| ENSMUSG00000053121.5 | 0 | NA | NA | NA | NA | NA | #VALUE! |

| | | | | | | | |
|---|---|---|---|---|---|---|---|
| ENSMUSG00000053129.6 | 0 | NA | NA | NA | NA | NA | #VALUE! |
| ENSMUSG00000053137.8 | 0 | NA | NA | NA | NA | NA | #VALUE! |
| ENSMUSG00000053141.17 | 0 | NA | NA | NA | NA | NA | #VALUE! |
| ENSMUSG00000053146.3 | 0 | NA | NA | NA | NA | NA | #VALUE! |
| ENSMUSG00000053153.15 | 0 | NA | NA | NA | NA | NA | #VALUE! |
| ENSMUSG00000053158.11 | 1.492083 | 4.088114 | 3.936282 | 1.038572 | 0.299004 | NA | #VALUE! |
| ENSMUSG00000053161.17 | 0 | NA | NA | NA | NA | NA | #VALUE! |
| ENSMUSG00000053164.7 | 0 | NA | NA | NA | NA | NA | #VALUE! |
| ENSMUSG00000053166.15 | 0 | NA | NA | NA | NA | NA | #VALUE! |
| ENSMUSG00000053173.13 | 1.544258 | -0.11577 | 3.844104 | -0.03012 | 0.975974 | NA | #VALUE! |
| ENSMUSG00000053178.6 | 0 | NA | NA | NA | NA | NA | #VALUE! |
| ENSMUSG00000053181.2 | 0 | NA | NA | NA | NA | NA | #VALUE! |
| ENSMUSG00000053182.12 | 0 | NA | NA | NA | NA | NA | #VALUE! |
| ENSMUSG00000053184.16 | 0 | NA | NA | NA | NA | NA | #VALUE! |
| ENSMUSG00000053185.3 | 0 | NA | NA | NA | NA | NA | #VALUE! |
| ENSMUSG00000053192.10 | 0.303281 | -0.61664 | 4.050639 | -0.15223 | 0.879003 | NA | #VALUE! |
| ENSMUSG00000053194.3 | 0 | NA | NA | NA | NA | NA | #VALUE! |
| ENSMUSG00000053198.15 | 0 | NA | NA | NA | NA | NA | #VALUE! |
| ENSMUSG00000053199.14 | 0 | NA | NA | NA | NA | NA | #VALUE! |
| ENSMUSG00000053208.11 | 0 | NA | NA | NA | NA | NA | #VALUE! |
| ENSMUSG00000053211.11 | 0 | NA | NA | NA | NA | NA | #VALUE! |
| ENSMUSG00000053214.8 | 0 | NA | NA | NA | NA | NA | #VALUE! |
| ENSMUSG00000053216.16 | 0 | NA | NA | NA | NA | NA | #VALUE! |
| ENSMUSG00000053217.4 | 0 | NA | NA | NA | NA | NA | #VALUE! |
| ENSMUSG00000053226.8 | 0.210026 | -1.52494 | 4.058692 | -0.37572 | 0.707123 | NA | #VALUE! |
| ENSMUSG00000053228.13 | 0 | NA | NA | NA | NA | NA | #VALUE! |
| ENSMUSG00000053251.7 | 0 | NA | NA | NA | NA | NA | #VALUE! |
| ENSMUSG00000053263.4 | 0 | NA | NA | NA | NA | NA | #VALUE! |
| ENSMUSG00000053268.8 | 0 | NA | NA | NA | NA | NA | #VALUE! |
| ENSMUSG00000053287.5 | 0 | NA | NA | NA | NA | NA | #VALUE! |
| ENSMUSG00000053291.16 | 0 | NA | NA | NA | NA | NA | #VALUE! |
| ENSMUSG00000053293.10 | 30.44466 | -1.31783 | 1.729437 | -0.762 | NA | NA | #VALUE! |
| ENSMUSG00000053297.15 | 0 | NA | NA | NA | NA | NA | #VALUE! |
| ENSMUSG00000053303.15 | 9.418775 | 6.729407 | 3.639605 | 1.848939 | NA | NA | #VALUE! |
| ENSMUSG00000053310.12 | 1.575197 | -4.20222 | 3.934736 | -1.06798 | 0.285529 | NA | #VALUE! |
| ENSMUSG00000053318.8 | 0 | NA | NA | NA | NA | NA | #VALUE! |
| ENSMUSG00000053338.10 | 0 | NA | NA | NA | NA | NA | #VALUE! |
| ENSMUSG00000053353.6 | 0 | NA | NA | NA | NA | NA | #VALUE! |
| ENSMUSG00000053358.3 | 1.470184 | -4.10383 | 3.936698 | -1.04245 | 0.297201 | NA | #VALUE! |
| ENSMUSG00000053360.5 | 0 | NA | NA | NA | NA | NA | #VALUE! |
| ENSMUSG00000053367.16 | 0 | NA | NA | NA | NA | NA | #VALUE! |
| ENSMUSG00000053368.14 | 0 | NA | NA | NA | NA | NA | #VALUE! |
| ENSMUSG00000053375.9 | 0 | NA | NA | NA | NA | NA | #VALUE! |
| ENSMUSG00000053388.11 | 0 | NA | NA | NA | NA | NA | #VALUE! |

| | | | | | | | |
|---|---|---|---|---|---|---|---|
| ENSMUSG00000053389.5 | 0 | NA | NA | NA | NA | NA | #VALUE! |
| ENSMUSG00000053390.17 | 4.898658 | -5.81923 | 3.915919 | -1.48605 | 0.137267 | NA | #VALUE! |
| ENSMUSG00000053391.6 | 0 | NA | NA | NA | NA | NA | #VALUE! |
| ENSMUSG00000053395.16 | 0 | NA | NA | NA | NA | NA | #VALUE! |
| ENSMUSG00000053398.12 | 0 | NA | NA | NA | NA | NA | #VALUE! |
| ENSMUSG00000053399.10 | 0 | NA | NA | NA | NA | NA | #VALUE! |
| ENSMUSG00000053411.17 | 7.533319 | -1.0539 | 3.229291 | -0.32636 | 0.744155 | NA | #VALUE! |
| ENSMUSG00000053414.9 | 6.976282 | -2.09869 | 3.659452 | -0.5735 | 0.566306 | NA | #VALUE! |
| ENSMUSG00000053420.7 | 0 | NA | NA | NA | NA | NA | #VALUE! |
| ENSMUSG00000053441.6 | 0 | NA | NA | NA | NA | NA | #VALUE! |
| ENSMUSG00000053442.5 | 0 | NA | NA | NA | NA | NA | #VALUE! |
| ENSMUSG00000053461.10 | 0 | NA | NA | NA | NA | NA | #VALUE! |
| ENSMUSG00000053465.7 | 0 | NA | NA | NA | NA | NA | #VALUE! |
| ENSMUSG00000053469.15 | 0 | NA | NA | NA | NA | NA | #VALUE! |
| ENSMUSG00000053472.4 | 0 | NA | NA | NA | NA | NA | #VALUE! |
| ENSMUSG00000053475.6 | 0 | NA | NA | NA | NA | NA | #VALUE! |
| ENSMUSG00000053490.9 | 0 | NA | NA | NA | NA | NA | #VALUE! |
| ENSMUSG00000053499.5 | 0 | NA | NA | NA | NA | NA | #VALUE! |
| ENSMUSG00000053508.9 | 0 | NA | NA | NA | NA | NA | #VALUE! |
| ENSMUSG00000053519.18 | 0 | NA | NA | NA | NA | NA | #VALUE! |
| ENSMUSG00000053522.12 | 0 | NA | NA | NA | NA | NA | #VALUE! |
| ENSMUSG00000053528.7 | 0 | NA | NA | NA | NA | NA | #VALUE! |
| ENSMUSG00000053541.4 | 0 | NA | NA | NA | NA | NA | #VALUE! |
| ENSMUSG00000053545.7 | 0 | NA | NA | NA | NA | NA | #VALUE! |
| ENSMUSG00000053550.14 | 0 | NA | NA | NA | NA | NA | #VALUE! |
| ENSMUSG00000053552.15 | 0 | NA | NA | NA | NA | NA | #VALUE! |
| ENSMUSG00000053559.14 | 7.30862 | -5.28996 | 3.291752 | -1.60704 | 0.108047 | NA | #VALUE! |
| ENSMUSG00000053574.6 | 0.195789 | -1.05581 | 4.073302 | -0.2592 | 0.795479 | NA | #VALUE! |
| ENSMUSG00000053588.5 | 0.168284 | 1.027628 | 4.067126 | 0.252667 | 0.800526 | NA | #VALUE! |
| ENSMUSG00000053593.11 | 0 | NA | NA | NA | NA | NA | #VALUE! |
| ENSMUSG00000053603.5 | 0 | NA | NA | NA | NA | NA | #VALUE! |
| ENSMUSG00000053615.2 | 0 | NA | NA | NA | NA | NA | #VALUE! |
| ENSMUSG00000053624.5 | 0 | NA | NA | NA | NA | NA | #VALUE! |
| ENSMUSG00000053626.6 | 0 | NA | NA | NA | NA | NA | #VALUE! |
| ENSMUSG00000053640.3 | 0 | NA | NA | NA | NA | NA | #VALUE! |
| ENSMUSG00000053647.5 | 2.245218 | -4.69006 | 3.926744 | -1.19439 | 0.232326 | NA | #VALUE! |
| ENSMUSG00000053654.8 | 0 | NA | NA | NA | NA | NA | #VALUE! |
| ENSMUSG00000053675.9 | 0 | NA | NA | NA | NA | NA | #VALUE! |
| ENSMUSG00000053678.8 | 0 | NA | NA | NA | NA | NA | #VALUE! |
| ENSMUSG00000053687.16 | 0 | NA | NA | NA | NA | NA | #VALUE! |
| ENSMUSG00000053693.17 | 0 | NA | NA | NA | NA | NA | #VALUE! |
| ENSMUSG00000053695.4 | 0 | NA | NA | NA | NA | NA | #VALUE! |
| ENSMUSG00000053702.17 | 6.102283 | 6.103895 | 3.692297 | 1.653143 | 0.098302 | NA | #VALUE! |
| ENSMUSG00000053706.3 | 0 | NA | NA | NA | NA | NA | #VALUE! |

| | | | | | | | |
|---|---|---|---|---|---|---|---|
| ENSMUSG00000053714.9 | 0 | NA | NA | NA | NA | NA | #VALUE! |
| ENSMUSG00000053716.10 | 17.15167 | -1.59896 | 2.545831 | -0.62807 | NA | NA | #VALUE! |
| ENSMUSG00000053719.11 | 0 | NA | NA | NA | NA | NA | #VALUE! |
| ENSMUSG00000053720.5 | 0 | NA | NA | NA | NA | NA | #VALUE! |
| ENSMUSG00000053729.5 | 0 | NA | NA | NA | NA | NA | #VALUE! |
| ENSMUSG00000053740.9 | 0 | NA | NA | NA | NA | NA | #VALUE! |
| ENSMUSG00000053742.9 | 0 | NA | NA | NA | NA | NA | #VALUE! |
| ENSMUSG00000053746.9 | 5.857614 | -6.08321 | 3.748598 | -1.62279 | 0.104633 | NA | #VALUE! |
| ENSMUSG00000053747.10 | 0 | NA | NA | NA | NA | NA | #VALUE! |
| ENSMUSG00000053749.13 | 5.706127 | 6.00576 | 3.806272 | 1.577859 | 0.114598 | NA | #VALUE! |
| ENSMUSG00000053765.5 | 0 | NA | NA | NA | NA | NA | #VALUE! |
| ENSMUSG00000053773.6 | 0 | NA | NA | NA | NA | NA | #VALUE! |
| ENSMUSG00000053774.10 | 9.878542 | -2.32265 | 2.548951 | -0.91122 | 0.362181 | NA | #VALUE! |
| ENSMUSG00000053783.6 | 0 | NA | NA | NA | NA | NA | #VALUE! |
| ENSMUSG00000053790.4 | 0 | NA | NA | NA | NA | NA | #VALUE! |
| ENSMUSG00000053797.11 | 0 | NA | NA | NA | NA | NA | #VALUE! |
| ENSMUSG00000053815.6 | 0 | NA | NA | NA | NA | NA | #VALUE! |
| ENSMUSG00000053820.5 | 0 | NA | NA | NA | NA | NA | #VALUE! |
| ENSMUSG00000053821.5 | 0 | NA | NA | NA | NA | NA | #VALUE! |
| ENSMUSG00000053830.5 | 0 | NA | NA | NA | NA | NA | #VALUE! |
| ENSMUSG00000053835.18 | 0 | NA | NA | NA | NA | NA | #VALUE! |
| ENSMUSG00000053852.12 | 0 | NA | NA | NA | NA | NA | #VALUE! |
| ENSMUSG00000053856.8 | 0 | NA | NA | NA | NA | NA | #VALUE! |
| ENSMUSG00000053861.2 | 0 | NA | NA | NA | NA | NA | #VALUE! |
| ENSMUSG00000053862.7 | 0.735092 | -3.12016 | 3.965612 | -0.7868 | 0.431397 | NA | #VALUE! |
| ENSMUSG00000053863.4 | 0 | NA | NA | NA | NA | NA | #VALUE! |
| ENSMUSG00000053868.5 | 0 | NA | NA | NA | NA | NA | #VALUE! |
| ENSMUSG00000053873.2 | 0 | NA | NA | NA | NA | NA | #VALUE! |
| ENSMUSG00000053877.13 | 0.195789 | -1.05581 | 4.073302 | -0.2592 | 0.795479 | NA | #VALUE! |
| ENSMUSG00000053889.6 | 0 | NA | NA | NA | NA | NA | #VALUE! |
| ENSMUSG00000053896.15 | 0 | NA | NA | NA | NA | NA | #VALUE! |
| ENSMUSG00000053909.9 | 0 | NA | NA | NA | NA | NA | #VALUE! |
| ENSMUSG00000053914.10 | 0 | NA | NA | NA | NA | NA | #VALUE! |
| ENSMUSG00000053916.5 | 0 | NA | NA | NA | NA | NA | #VALUE! |
| ENSMUSG00000053925.5 | 0 | NA | NA | NA | NA | NA | #VALUE! |
| ENSMUSG00000053930.14 | 0 | NA | NA | NA | NA | NA | #VALUE! |
| ENSMUSG00000053957.6 | 0 | NA | NA | NA | NA | NA | #VALUE! |
| ENSMUSG00000053961.8 | 0 | NA | NA | NA | NA | NA | #VALUE! |
| ENSMUSG00000053963.8 | 0 | NA | NA | NA | NA | NA | #VALUE! |
| ENSMUSG00000053964.18 | 6.875983 | -0.93833 | 3.278462 | -0.28621 | 0.774718 | NA | #VALUE! |
| ENSMUSG00000053965.11 | 0 | NA | NA | NA | NA | NA | #VALUE! |
| ENSMUSG00000053977.14 | 0 | NA | NA | NA | NA | NA | #VALUE! |
| ENSMUSG00000053979.5 | 0 | NA | NA | NA | NA | NA | #VALUE! |
| ENSMUSG00000053980.5 | 0 | NA | NA | NA | NA | NA | #VALUE! |

| | | | | | | | |
|---|---|---|---|---|---|---|---|
| ENSMUSG00000053985.11 | 7.347667 | 0.130061 | 3.645626 | 0.035676 | 0.971541 | NA | #VALUE! |
| ENSMUSG00000054000.6 | 1.364362 | 0.666372 | 3.863094 | 0.172497 | 0.863047 | NA | #VALUE! |
| ENSMUSG00000054003.14 | 0 | NA | NA | NA | NA | NA | #VALUE! |
| ENSMUSG00000054005.5 | 0 | NA | NA | NA | NA | NA | #VALUE! |
| ENSMUSG00000054006.15 | 0 | NA | NA | NA | NA | NA | #VALUE! |
| ENSMUSG00000054013.7 | 0 | NA | NA | NA | NA | NA | #VALUE! |
| ENSMUSG00000054031.14 | 0 | NA | NA | NA | NA | NA | #VALUE! |
| ENSMUSG00000054033.7 | 0 | NA | NA | NA | NA | NA | #VALUE! |
| ENSMUSG00000054034.11 | 0 | NA | NA | NA | NA | NA | #VALUE! |
| ENSMUSG00000054036.7 | 0 | NA | NA | NA | NA | NA | #VALUE! |
| ENSMUSG00000054046.8 | 0 | NA | NA | NA | NA | NA | #VALUE! |
| ENSMUSG00000054052.6 | 0 | NA | NA | NA | NA | NA | #VALUE! |
| ENSMUSG00000054054.4 | 0 | NA | NA | NA | NA | NA | #VALUE! |
| ENSMUSG00000054057.3 | 7.474597 | 2.75255 | 3.131029 | 0.87912 | 0.379336 | NA | #VALUE! |
| ENSMUSG00000054061.10 | 0 | NA | NA | NA | NA | NA | #VALUE! |
| ENSMUSG00000054065.13 | 0 | NA | NA | NA | NA | NA | #VALUE! |
| ENSMUSG00000054069.9 | 0 | NA | NA | NA | NA | NA | #VALUE! |
| ENSMUSG00000054074.10 | 2.215933 | -2.22544 | 3.866366 | -0.57559 | 0.564893 | NA | #VALUE! |
| ENSMUSG00000054087.12 | 0 | NA | NA | NA | NA | NA | #VALUE! |
| ENSMUSG00000054099.12 | 7.340482 | 2.723543 | 3.627817 | 0.750739 | 0.45281 | NA | #VALUE! |
| ENSMUSG00000054102.16 | 0 | NA | NA | NA | NA | NA | #VALUE! |
| ENSMUSG00000054106.8 | 0 | NA | NA | NA | NA | NA | #VALUE! |
| ENSMUSG00000054116.2 | 0 | NA | NA | NA | NA | NA | #VALUE! |
| ENSMUSG00000054117.9 | 0 | NA | NA | NA | NA | NA | #VALUE! |
| ENSMUSG00000054128.17 | 0 | NA | NA | NA | NA | NA | #VALUE! |
| ENSMUSG00000054134.15 | 0 | NA | NA | NA | NA | NA | #VALUE! |
| ENSMUSG00000054135.3 | 0 | NA | NA | NA | NA | NA | #VALUE! |
| ENSMUSG00000054136.7 | 0 | NA | NA | NA | NA | NA | #VALUE! |
| ENSMUSG00000054141.6 | 0 | NA | NA | NA | NA | NA | #VALUE! |
| ENSMUSG00000054142.6 | 0 | NA | NA | NA | NA | NA | #VALUE! |
| ENSMUSG00000054146.9 | 0 | NA | NA | NA | NA | NA | #VALUE! |
| ENSMUSG00000054150.13 | 0 | NA | NA | NA | NA | NA | #VALUE! |
| ENSMUSG00000054156.9 | 0 | NA | NA | NA | NA | NA | #VALUE! |
| ENSMUSG00000054160.3 | 0 | NA | NA | NA | NA | NA | #VALUE! |
| ENSMUSG00000054161.13 | 0 | NA | NA | NA | NA | NA | #VALUE! |
| ENSMUSG00000054162.16 | 0 | NA | NA | NA | NA | NA | #VALUE! |
| ENSMUSG00000054169.8 | 0 | NA | NA | NA | NA | NA | #VALUE! |
| ENSMUSG00000054181.12 | 0 | NA | NA | NA | NA | NA | #VALUE! |
| ENSMUSG00000054191.10 | 4.024183 | 2.844486 | 3.515425 | 0.809144 | 0.418432 | NA | #VALUE! |
| ENSMUSG00000054196.8 | 0 | NA | NA | NA | NA | NA | #VALUE! |
| ENSMUSG00000054200.8 | 0 | NA | NA | NA | NA | NA | #VALUE! |
| ENSMUSG00000054204.7 | 0 | NA | NA | NA | NA | NA | #VALUE! |
| ENSMUSG00000054206.3 | 0 | NA | NA | NA | NA | NA | #VALUE! |
| ENSMUSG00000054215.3 | 0 | NA | NA | NA | NA | NA | #VALUE! |

| | | | | | | | |
|---|---|---|---|---|---|---|---|
| ENSMUSG00000054236.7 | 0 | NA | NA | NA | NA | NA | #VALUE! |
| ENSMUSG00000054247.7 | 0 | NA | NA | NA | NA | NA | #VALUE! |
| ENSMUSG00000054256.12 | 0 | NA | NA | NA | NA | NA | #VALUE! |
| ENSMUSG00000054258.10 | 0 | NA | NA | NA | NA | NA | #VALUE! |
| ENSMUSG00000054266.3 | 0 | NA | NA | NA | NA | NA | #VALUE! |
| ENSMUSG00000054272.7 | 0 | NA | NA | NA | NA | NA | #VALUE! |
| ENSMUSG00000054293.11 | 0.204111 | -1.05581 | 4.073302 | -0.2592 | 0.795479 | NA | #VALUE! |
| ENSMUSG00000054304.4 | 0 | NA | NA | NA | NA | NA | #VALUE! |
| ENSMUSG00000054310.17 | 0 | NA | NA | NA | NA | NA | #VALUE! |
| ENSMUSG00000054320.17 | 0 | NA | NA | NA | NA | NA | #VALUE! |
| ENSMUSG00000054325.6 | 0 | NA | NA | NA | NA | NA | #VALUE! |
| ENSMUSG00000054342.9 | 0 | NA | NA | NA | NA | NA | #VALUE! |
| ENSMUSG00000054351.3 | 0 | NA | NA | NA | NA | NA | #VALUE! |
| ENSMUSG00000054360.5 | 0 | NA | NA | NA | NA | NA | #VALUE! |
| ENSMUSG00000054362.10 | 0 | NA | NA | NA | NA | NA | #VALUE! |
| ENSMUSG00000054379.8 | 0 | NA | NA | NA | NA | NA | #VALUE! |
| ENSMUSG00000054383.3 | 0 | NA | NA | NA | NA | NA | #VALUE! |
| ENSMUSG00000054385.15 | 4.375372 | 5.61928 | 3.917028 | 1.434577 | 0.151408 | NA | #VALUE! |
| ENSMUSG00000054391.13 | 0 | NA | NA | NA | NA | NA | #VALUE! |
| ENSMUSG00000054400.17 | 34.15199 | 0.876065 | 2.105005 | 0.416182 | NA | NA | #VALUE! |
| ENSMUSG00000054406.10 | 0 | NA | NA | NA | NA | NA | #VALUE! |
| ENSMUSG00000054409.6 | 0 | NA | NA | NA | NA | NA | #VALUE! |
| ENSMUSG00000054412.6 | 0 | NA | NA | NA | NA | NA | #VALUE! |
| ENSMUSG00000054417.6 | 0.630079 | -2.90291 | 3.975047 | -0.73028 | 0.465217 | NA | #VALUE! |
| ENSMUSG00000054418.7 | 0 | NA | NA | NA | NA | NA | #VALUE! |
| ENSMUSG00000054423.15 | 0 | NA | NA | NA | NA | NA | #VALUE! |
| ENSMUSG00000054431.5 | 0 | NA | NA | NA | NA | NA | #VALUE! |
| ENSMUSG00000054434.15 | 4.038805 | 5.503807 | 3.917882 | 1.404792 | 0.160083 | NA | #VALUE! |
| ENSMUSG00000054446.9 | 0 | NA | NA | NA | NA | NA | #VALUE! |
| ENSMUSG00000054450.2 | 0 | NA | NA | NA | NA | NA | #VALUE! |
| ENSMUSG00000054453.12 | 0 | NA | NA | NA | NA | NA | #VALUE! |
| ENSMUSG00000054457.6 | 0 | NA | NA | NA | NA | NA | #VALUE! |
| ENSMUSG00000054459.9 | 0 | NA | NA | NA | NA | NA | #VALUE! |
| ENSMUSG00000054488.15 | 0 | NA | NA | NA | NA | NA | #VALUE! |
| ENSMUSG00000054493.3 | 2.704401 | 4.936946 | 3.92321 | 1.258394 | 0.208249 | NA | #VALUE! |
| ENSMUSG00000054497.6 | 0 | NA | NA | NA | NA | NA | #VALUE! |
| ENSMUSG00000054498.7 | 0 | NA | NA | NA | NA | NA | #VALUE! |
| ENSMUSG00000054510.6 | 0 | NA | NA | NA | NA | NA | #VALUE! |
| ENSMUSG00000054514.4 | 1.119062 | 3.679627 | 3.945843 | 0.932533 | 0.351061 | NA | #VALUE! |
| ENSMUSG00000054519.9 | 26.61762 | -2.61911 | 2.333627 | -1.12233 | NA | NA | #VALUE! |
| ENSMUSG00000054520.16 | 13.94491 | -2.41208 | 2.659493 | -0.90697 | NA | NA | #VALUE! |
| ENSMUSG00000054523.14 | 0 | NA | NA | NA | NA | NA | #VALUE! |
| ENSMUSG00000054526.6 | 0 | NA | NA | NA | NA | NA | #VALUE! |
| ENSMUSG00000054537.9 | 0 | NA | NA | NA | NA | NA | #VALUE! |

| | | | | | | | |
|---|---|---|---|---|---|---|---|
| ENSMUSG00000054545.18 | 1.566313 | -4.16993 | 3.935366 | -1.0596 | 0.289325 | NA | #VALUE! |
| ENSMUSG00000054555.12 | 1.765352 | 1.023379 | 3.843626 | 0.266254 | 0.790044 | NA | #VALUE! |
| ENSMUSG00000054556.7 | 0 | NA | NA | NA | NA | NA | #VALUE! |
| ENSMUSG00000054568.8 | 0 | NA | NA | NA | NA | NA | #VALUE! |
| ENSMUSG00000054580.15 | 8.896893 | -1.07615 | 2.659086 | -0.40471 | 0.685694 | NA | #VALUE! |
| ENSMUSG00000054582.5 | 0 | NA | NA | NA | NA | NA | #VALUE! |
| ENSMUSG00000054589.2 | 0 | NA | NA | NA | NA | NA | #VALUE! |
| ENSMUSG00000054594.14 | 0 | NA | NA | NA | NA | NA | #VALUE! |
| ENSMUSG00000054598.14 | 0 | NA | NA | NA | NA | NA | #VALUE! |
| ENSMUSG00000054618.9 | 0 | NA | NA | NA | NA | NA | #VALUE! |
| ENSMUSG00000054622.5 | 0 | NA | NA | NA | NA | NA | #VALUE! |
| ENSMUSG00000054626.12 | 0.210026 | -1.52494 | 4.058692 | -0.37572 | 0.707123 | NA | #VALUE! |
| ENSMUSG00000054641.11 | 0.559531 | 2.703958 | 3.982863 | 0.678898 | 0.497202 | NA | #VALUE! |
| ENSMUSG00000054659.14 | 0 | NA | NA | NA | NA | NA | #VALUE! |
| ENSMUSG00000054662.9 | 0 | NA | NA | NA | NA | NA | #VALUE! |
| ENSMUSG00000054666.8 | 0 | NA | NA | NA | NA | NA | #VALUE! |
| ENSMUSG00000054667.8 | 0 | NA | NA | NA | NA | NA | #VALUE! |
| ENSMUSG00000054672.14 | 0 | NA | NA | NA | NA | NA | #VALUE! |
| ENSMUSG00000054675.6 | 0 | NA | NA | NA | NA | NA | #VALUE! |
| ENSMUSG00000054679.9 | 0 | NA | NA | NA | NA | NA | #VALUE! |
| ENSMUSG00000054690.18 | 6.79987 | 1.452003 | 3.295816 | 0.440559 | 0.659532 | NA | #VALUE! |
| ENSMUSG00000054702.15 | 3.702238 | 5.378281 | 3.918889 | 1.372399 | 0.169939 | NA | #VALUE! |
| ENSMUSG00000054708.18 | 0 | NA | NA | NA | NA | NA | #VALUE! |
| ENSMUSG00000054727.3 | 0 | NA | NA | NA | NA | NA | #VALUE! |
| ENSMUSG00000054728.18 | 0 | NA | NA | NA | NA | NA | #VALUE! |
| ENSMUSG00000054737.12 | 0 | NA | NA | NA | NA | NA | #VALUE! |
| ENSMUSG00000054745.2 | 0 | NA | NA | NA | NA | NA | #VALUE! |
| ENSMUSG00000054746.19 | 0 | NA | NA | NA | NA | NA | #VALUE! |
| ENSMUSG00000054752.17 | 0 | NA | NA | NA | NA | NA | #VALUE! |
| ENSMUSG00000054753.10 | 0 | NA | NA | NA | NA | NA | #VALUE! |
| ENSMUSG00000054759.7 | 0 | NA | NA | NA | NA | NA | #VALUE! |
| ENSMUSG00000054763.10 | 0 | NA | NA | NA | NA | NA | #VALUE! |
| ENSMUSG00000054764.10 | 1.229878 | 3.761857 | 3.943699 | 0.95389 | 0.340139 | NA | #VALUE! |
| ENSMUSG00000054779.6 | 0 | NA | NA | NA | NA | NA | #VALUE! |
| ENSMUSG00000054792.16 | 6.825853 | -6.30496 | 3.913306 | -1.61116 | 0.107145 | NA | #VALUE! |
| ENSMUSG00000054793.9 | 0 | NA | NA | NA | NA | NA | #VALUE! |
| ENSMUSG00000054822.6 | 0 | NA | NA | NA | NA | NA | #VALUE! |
| ENSMUSG00000054850.5 | 0 | NA | NA | NA | NA | NA | #VALUE! |
| ENSMUSG00000054855.14 | 1.344956 | -0.7658 | 3.869263 | -0.19792 | 0.843108 | NA | #VALUE! |
| ENSMUSG00000054871.6 | 0 | NA | NA | NA | NA | NA | #VALUE! |
| ENSMUSG00000054885.12 | 0 | NA | NA | NA | NA | NA | #VALUE! |
| ENSMUSG00000054890.3 | 0 | NA | NA | NA | NA | NA | #VALUE! |
| ENSMUSG00000054892.15 | 3.730208 | 5.397701 | 3.918728 | 1.377412 | 0.168385 | NA | #VALUE! |
| ENSMUSG00000054893.16 | 5.270541 | 0.062698 | 3.593726 | 0.017447 | 0.98608 | NA | #VALUE! |

| | | | | | | | |
|---|---|---|---|---|---|---|---|
| ENSMUSG00000054901.8 | 0 | NA | NA | NA | NA | NA | #VALUE! |
| ENSMUSG00000054905.3 | 0 | NA | NA | NA | NA | NA | #VALUE! |
| ENSMUSG00000054909.12 | 0 | NA | NA | NA | NA | NA | #VALUE! |
| ENSMUSG00000054910.6 | 0 | NA | NA | NA | NA | NA | #VALUE! |
| ENSMUSG00000054931.8 | 1.995474 | 0.670616 | 3.822169 | 0.175454 | 0.860723 | NA | #VALUE! |
| ENSMUSG00000054934.11 | 0 | NA | NA | NA | NA | NA | #VALUE! |
| ENSMUSG00000054938.8 | 0 | NA | NA | NA | NA | NA | #VALUE! |
| ENSMUSG00000054939.7 | 2.344116 | -4.76364 | 3.925754 | -1.21343 | 0.224964 | NA | #VALUE! |
| ENSMUSG00000054940.7 | 0 | NA | NA | NA | NA | NA | #VALUE! |
| ENSMUSG00000054942.14 | 4.305538 | -5.64245 | 3.91711 | -1.44046 | 0.149737 | NA | #VALUE! |
| ENSMUSG00000054944.8 | 0 | NA | NA | NA | NA | NA | #VALUE! |
| ENSMUSG00000054945.5 | 0.839297 | 3.273019 | 3.958392 | 0.826856 | 0.408319 | NA | #VALUE! |
| ENSMUSG00000054951.4 | 0 | NA | NA | NA | NA | NA | #VALUE! |
| ENSMUSG00000054957.6 | 0 | NA | NA | NA | NA | NA | #VALUE! |
| ENSMUSG00000054958.7 | 0 | NA | NA | NA | NA | NA | #VALUE! |
| ENSMUSG00000054966.14 | 0 | NA | NA | NA | NA | NA | #VALUE! |
| ENSMUSG00000054967.8 | 2.705731 | 4.913666 | 3.923477 | 1.252375 | 0.210433 | NA | #VALUE! |
| ENSMUSG00000054976.15 | 0 | NA | NA | NA | NA | NA | #VALUE! |
| ENSMUSG00000054978.4 | 0 | NA | NA | NA | NA | NA | #VALUE! |
| ENSMUSG00000054986.14 | 0 | NA | NA | NA | NA | NA | #VALUE! |
| ENSMUSG00000054988.10 | 0 | NA | NA | NA | NA | NA | #VALUE! |
| ENSMUSG00000054994.3 | 0 | NA | NA | NA | NA | NA | #VALUE! |
| ENSMUSG00000054999.3 | 0 | NA | NA | NA | NA | NA | #VALUE! |
| ENSMUSG00000055003.15 | 0 | NA | NA | NA | NA | NA | #VALUE! |
| ENSMUSG00000055010.2 | 0 | NA | NA | NA | NA | NA | #VALUE! |
| ENSMUSG00000055015.6 | 0 | NA | NA | NA | NA | NA | #VALUE! |
| ENSMUSG00000055022.15 | 0 | NA | NA | NA | NA | NA | #VALUE! |
| ENSMUSG00000055026.14 | 0 | NA | NA | NA | NA | NA | #VALUE! |
| ENSMUSG00000055030.2 | 0 | NA | NA | NA | NA | NA | #VALUE! |
| ENSMUSG00000055033.5 | 0 | NA | NA | NA | NA | NA | #VALUE! |
| ENSMUSG00000055041.9 | 0 | NA | NA | NA | NA | NA | #VALUE! |
| ENSMUSG00000055045.6 | 0 | NA | NA | NA | NA | NA | #VALUE! |
| ENSMUSG00000055048.8 | 0 | NA | NA | NA | NA | NA | #VALUE! |
| ENSMUSG00000055061.6 | 0 | NA | NA | NA | NA | NA | #VALUE! |
| ENSMUSG00000055067.16 | 7.663484 | -0.28749 | 2.620194 | -0.10972 | 0.912631 | NA | #VALUE! |
| ENSMUSG00000055069.4 | 0 | NA | NA | NA | NA | NA | #VALUE! |
| ENSMUSG00000055072.3 | 0 | NA | NA | NA | NA | NA | #VALUE! |
| ENSMUSG00000055078.8 | 0 | NA | NA | NA | NA | NA | #VALUE! |
| ENSMUSG00000055088.6 | 0 | NA | NA | NA | NA | NA | #VALUE! |
| ENSMUSG00000055095.4 | 0 | NA | NA | NA | NA | NA | #VALUE! |
| ENSMUSG00000055102.16 | 0 | NA | NA | NA | NA | NA | #VALUE! |
| ENSMUSG00000055108.7 | 0 | NA | NA | NA | NA | NA | #VALUE! |
| ENSMUSG00000055109.6 | 0 | NA | NA | NA | NA | NA | #VALUE! |
| ENSMUSG00000055110.9 | 0 | NA | NA | NA | NA | NA | #VALUE! |

| | | | | | | | |
|---|---|---|---|---|---|---|---|
| ENSMUSG00000055114.11 | 0.245976 | 1.027628 | 4.067126 | 0.252667 | 0.800526 | NA | #VALUE! |
| ENSMUSG00000055125.8 | 0 | NA | NA | NA | NA | NA | #VALUE! |
| ENSMUSG00000055134.6 | 0 | NA | NA | NA | NA | NA | #VALUE! |
| ENSMUSG00000055138.11 | 0 | NA | NA | NA | NA | NA | #VALUE! |
| ENSMUSG00000055150.16 | 3.093896 | 5.116778 | 3.92129 | 1.304871 | 0.191937 | NA | #VALUE! |
| ENSMUSG00000055159.8 | 0 | NA | NA | NA | NA | NA | #VALUE! |
| ENSMUSG00000055170.5 | 0 | NA | NA | NA | NA | NA | #VALUE! |
| ENSMUSG00000055177.16 | 0 | NA | NA | NA | NA | NA | #VALUE! |
| ENSMUSG00000055188.13 | 0 | NA | NA | NA | NA | NA | #VALUE! |
| ENSMUSG00000055193.5 | 0 | NA | NA | NA | NA | NA | #VALUE! |
| ENSMUSG00000055194.4 | 0 | NA | NA | NA | NA | NA | #VALUE! |
| ENSMUSG00000055197.5 | 0 | NA | NA | NA | NA | NA | #VALUE! |
| ENSMUSG00000055198.4 | 0 | NA | NA | NA | NA | NA | #VALUE! |
| ENSMUSG00000055202.12 | 0.652786 | 2.919659 | 3.972474 | 0.734972 | 0.462356 | NA | #VALUE! |
| ENSMUSG00000055210.5 | 0 | NA | NA | NA | NA | NA | #VALUE! |
| ENSMUSG00000055214.16 | 0 | NA | NA | NA | NA | NA | #VALUE! |
| ENSMUSG00000055216.4 | 0 | NA | NA | NA | NA | NA | #VALUE! |
| ENSMUSG00000055228.9 | 0 | NA | NA | NA | NA | NA | #VALUE! |
| ENSMUSG00000055235.12 | 0 | NA | NA | NA | NA | NA | #VALUE! |
| ENSMUSG00000055254.16 | 49.78258 | 3.722678 | 3.739351 | 0.995541 | NA | NA | #VALUE! |
| ENSMUSG00000055271.6 | 0 | NA | NA | NA | NA | NA | #VALUE! |
| ENSMUSG00000055298.7 | 0 | NA | NA | NA | NA | NA | #VALUE! |
| ENSMUSG00000055305.16 | 3.470864 | 1.286181 | 3.789908 | 0.33937 | 0.734331 | NA | #VALUE! |
| ENSMUSG00000055313.16 | 5.165486 | 5.852308 | 3.9155 | 1.494651 | 0.135005 | NA | #VALUE! |
| ENSMUSG00000055323.6 | 0 | NA | NA | NA | NA | NA | #VALUE! |
| ENSMUSG00000055333.15 | 0 | NA | NA | NA | NA | NA | #VALUE! |
| ENSMUSG00000055341.11 | 0 | NA | NA | NA | NA | NA | #VALUE! |
| ENSMUSG00000055357.9 | 0 | NA | NA | NA | NA | NA | #VALUE! |
| ENSMUSG00000055360.14 | 0 | NA | NA | NA | NA | NA | #VALUE! |
| ENSMUSG00000055368.15 | 0 | NA | NA | NA | NA | NA | #VALUE! |
| ENSMUSG00000055370.4 | 0 | NA | NA | NA | NA | NA | #VALUE! |
| ENSMUSG00000055373.9 | 0 | NA | NA | NA | NA | NA | #VALUE! |
| ENSMUSG00000055385.13 | 0 | NA | NA | NA | NA | NA | #VALUE! |
| ENSMUSG00000055403.14 | 0 | NA | NA | NA | NA | NA | #VALUE! |
| ENSMUSG00000055407.15 | 0 | NA | NA | NA | NA | NA | #VALUE! |
| ENSMUSG00000055408.8 | 0 | NA | NA | NA | NA | NA | #VALUE! |
| ENSMUSG00000055409.15 | 0 | NA | NA | NA | NA | NA | #VALUE! |
| ENSMUSG00000055413.13 | 0 | NA | NA | NA | NA | NA | #VALUE! |
| ENSMUSG00000055415.9 | 0 | NA | NA | NA | NA | NA | #VALUE! |
| ENSMUSG00000055416.7 | 0 | NA | NA | NA | NA | NA | #VALUE! |
| ENSMUSG00000055421.9 | 0 | NA | NA | NA | NA | NA | #VALUE! |
| ENSMUSG00000055424.5 | 0 | NA | NA | NA | NA | NA | #VALUE! |
| ENSMUSG00000055430.5 | 0 | NA | NA | NA | NA | NA | #VALUE! |
| ENSMUSG00000055446.6 | 0 | NA | NA | NA | NA | NA | #VALUE! |

| | | | | | | | |
|---|---|---|---|---|---|---|---|
| ENSMUSG00000055452.7 | 0 | NA | NA | NA | NA | NA | #VALUE! |
| ENSMUSG00000055456.2 | 0 | NA | NA | NA | NA | NA | #VALUE! |
| ENSMUSG00000055471.8 | 0 | NA | NA | NA | NA | NA | #VALUE! |
| ENSMUSG00000055480.8 | 2.017149 | 1.481466 | 3.840436 | 0.385755 | 0.699678 | NA | #VALUE! |
| ENSMUSG00000055485.7 | 1.342688 | -0.41854 | 3.863366 | -0.10833 | 0.91373 | NA | #VALUE! |
| ENSMUSG00000055489.9 | 0 | NA | NA | NA | NA | NA | #VALUE! |
| ENSMUSG00000055494.6 | 0 | NA | NA | NA | NA | NA | #VALUE! |
| ENSMUSG00000055497.6 | 0 | NA | NA | NA | NA | NA | #VALUE! |
| ENSMUSG00000055515.9 | 0 | NA | NA | NA | NA | NA | #VALUE! |
| ENSMUSG00000055523.8 | 0 | NA | NA | NA | NA | NA | #VALUE! |
| ENSMUSG00000055540.16 | 0 | NA | NA | NA | NA | NA | #VALUE! |
| ENSMUSG00000055547.5 | 0 | NA | NA | NA | NA | NA | #VALUE! |
| ENSMUSG00000055555.2 | 0 | NA | NA | NA | NA | NA | #VALUE! |
| ENSMUSG00000055560.9 | 0 | NA | NA | NA | NA | NA | #VALUE! |
| ENSMUSG00000055561.10 | 0.291524 | 0.468739 | 4.052149 | 0.115677 | 0.907909 | NA | #VALUE! |
| ENSMUSG00000055567.19 | 1.771849 | 4.332864 | 3.931707 | 1.102031 | 0.270448 | NA | #VALUE! |
| ENSMUSG00000055571.5 | 0 | NA | NA | NA | NA | NA | #VALUE! |
| ENSMUSG00000055594.7 | 0 | NA | NA | NA | NA | NA | #VALUE! |
| ENSMUSG00000055602.18 | 0 | NA | NA | NA | NA | NA | #VALUE! |
| ENSMUSG00000055609.9 | 0 | NA | NA | NA | NA | NA | #VALUE! |
| ENSMUSG00000055610.6 | 0 | NA | NA | NA | NA | NA | #VALUE! |
| ENSMUSG00000055612.16 | 0.204111 | -1.05581 | 4.073302 | -0.2592 | 0.795479 | NA | #VALUE! |
| ENSMUSG00000055621.9 | 0 | NA | NA | NA | NA | NA | #VALUE! |
| ENSMUSG00000055629.6 | 0 | NA | NA | NA | NA | NA | #VALUE! |
| ENSMUSG00000055632.19 | 0 | NA | NA | NA | NA | NA | #VALUE! |
| ENSMUSG00000055633.7 | 0 | NA | NA | NA | NA | NA | #VALUE! |
| ENSMUSG00000055639.17 | 0 | NA | NA | NA | NA | NA | #VALUE! |
| ENSMUSG00000055643.5 | 0 | NA | NA | NA | NA | NA | #VALUE! |
| ENSMUSG00000055653.14 | 0 | NA | NA | NA | NA | NA | #VALUE! |
| ENSMUSG00000055675.7 | 0.315039 | -1.93549 | 4.038621 | -0.47925 | 0.631763 | NA | #VALUE! |
| ENSMUSG00000055676.5 | 0 | NA | NA | NA | NA | NA | #VALUE! |
| ENSMUSG00000055679.6 | 0 | NA | NA | NA | NA | NA | #VALUE! |
| ENSMUSG00000055691.6 | 0 | NA | NA | NA | NA | NA | #VALUE! |
| ENSMUSG00000055692.22 | 4.207088 | 5.562698 | 3.917438 | 1.419984 | 0.155612 | NA | #VALUE! |
| ENSMUSG00000055700.8 | 0 | NA | NA | NA | NA | NA | #VALUE! |
| ENSMUSG00000055704.5 | 0 | NA | NA | NA | NA | NA | #VALUE! |
| ENSMUSG00000055717.14 | 2.310289 | -4.7496 | 3.925939 | -1.2098 | 0.226355 | NA | #VALUE! |
| ENSMUSG00000055725.12 | 0.839297 | 3.273019 | 3.958392 | 0.826856 | 0.408319 | NA | #VALUE! |
| ENSMUSG00000055732.2 | 0 | NA | NA | NA | NA | NA | #VALUE! |
| ENSMUSG00000055733.7 | 0.204111 | -1.05581 | 4.073302 | -0.2592 | 0.795479 | NA | #VALUE! |
| ENSMUSG00000055745.5 | 0 | NA | NA | NA | NA | NA | #VALUE! |
| ENSMUSG00000055746.10 | 0 | NA | NA | NA | NA | NA | #VALUE! |
| ENSMUSG00000055748.13 | 0 | NA | NA | NA | NA | NA | #VALUE! |
| ENSMUSG00000055761.15 | 0 | NA | NA | NA | NA | NA | #VALUE! |

| | | | | | | | |
|---|---|---|---|---|---|---|---|
| ENSMUSG00000055763.10 | 0 | NA | NA | NA | NA | NA | #VALUE! |
| ENSMUSG00000055775.17 | 0 | NA | NA | NA | NA | NA | #VALUE! |
| ENSMUSG00000055780.11 | 0 | NA | NA | NA | NA | NA | #VALUE! |
| ENSMUSG00000055789.8 | 0 | NA | NA | NA | NA | NA | #VALUE! |
| ENSMUSG00000055795.7 | 0.195789 | -1.05581 | 4.073302 | -0.2592 | 0.795479 | NA | #VALUE! |
| ENSMUSG00000055799.14 | 2.403544 | -3.84131 | 3.917613 | -0.98052 | 0.326828 | NA | #VALUE! |
| ENSMUSG00000055805.16 | 3.465433 | -5.33079 | 3.919598 | -1.36004 | 0.173819 | NA | #VALUE! |
| ENSMUSG00000055809.8 | 0 | NA | NA | NA | NA | NA | #VALUE! |
| ENSMUSG00000055811.3 | 0 | NA | NA | NA | NA | NA | #VALUE! |
| ENSMUSG00000055818.3 | 0 | NA | NA | NA | NA | NA | #VALUE! |
| ENSMUSG00000055820.3 | 0 | NA | NA | NA | NA | NA | #VALUE! |
| ENSMUSG00000055826.6 | 0 | NA | NA | NA | NA | NA | #VALUE! |
| ENSMUSG00000055827.14 | 0 | NA | NA | NA | NA | NA | #VALUE! |
| ENSMUSG00000055833.15 | 0 | NA | NA | NA | NA | NA | #VALUE! |
| ENSMUSG00000055835.11 | 9.127952 | -3.24595 | 2.889868 | -1.12322 | 0.261346 | NA | #VALUE! |
| ENSMUSG00000055838.5 | 0 | NA | NA | NA | NA | NA | #VALUE! |
| ENSMUSG00000055865.9 | 0 | NA | NA | NA | NA | NA | #VALUE! |
| ENSMUSG00000055872.2 | 0 | NA | NA | NA | NA | NA | #VALUE! |
| ENSMUSG00000055874.7 | 0 | NA | NA | NA | NA | NA | #VALUE! |
| ENSMUSG00000055882.9 | 0 | NA | NA | NA | NA | NA | #VALUE! |
| ENSMUSG00000055891.8 | 0 | NA | NA | NA | NA | NA | #VALUE! |
| ENSMUSG00000055895.7 | 0 | NA | NA | NA | NA | NA | #VALUE! |
| ENSMUSG00000055926.4 | 0 | NA | NA | NA | NA | NA | #VALUE! |
| ENSMUSG00000055933.14 | 0 | NA | NA | NA | NA | NA | #VALUE! |
| ENSMUSG00000055937.2 | 0 | NA | NA | NA | NA | NA | #VALUE! |
| ENSMUSG00000055942.14 | 0 | NA | NA | NA | NA | NA | #VALUE! |
| ENSMUSG00000055944.4 | 0 | NA | NA | NA | NA | NA | #VALUE! |
| ENSMUSG00000055945.9 | 0 | NA | NA | NA | NA | NA | #VALUE! |
| ENSMUSG00000055958.4 | 0 | NA | NA | NA | NA | NA | #VALUE! |
| ENSMUSG00000055960.13 | 0 | NA | NA | NA | NA | NA | #VALUE! |
| ENSMUSG00000055961.8 | 0 | NA | NA | NA | NA | NA | #VALUE! |
| ENSMUSG00000055963.13 | 0 | NA | NA | NA | NA | NA | #VALUE! |
| ENSMUSG00000055968.2 | 0 | NA | NA | NA | NA | NA | #VALUE! |
| ENSMUSG00000055971.12 | 0 | NA | NA | NA | NA | NA | #VALUE! |
| ENSMUSG00000055972.6 | 0 | NA | NA | NA | NA | NA | #VALUE! |
| ENSMUSG00000055976.7 | 0 | NA | NA | NA | NA | NA | #VALUE! |
| ENSMUSG00000055978.6 | 0 | NA | NA | NA | NA | NA | #VALUE! |
| ENSMUSG00000055994.16 | 0 | NA | NA | NA | NA | NA | #VALUE! |
| ENSMUSG00000056004.17 | 7.314473 | 6.35544 | 3.538195 | 1.796238 | 0.072457 | NA | #VALUE! |
| ENSMUSG00000056008.2 | 0 | NA | NA | NA | NA | NA | #VALUE! |
| ENSMUSG00000056014.16 | 3.702238 | 5.378281 | 3.918889 | 1.372399 | 0.169939 | NA | #VALUE! |
| ENSMUSG00000056018.15 | 0 | NA | NA | NA | NA | NA | #VALUE! |
| ENSMUSG00000056023.4 | 0 | NA | NA | NA | NA | NA | #VALUE! |
| ENSMUSG00000056031.14 | 0 | NA | NA | NA | NA | NA | #VALUE! |

| | | | | | | | |
|---|---|---|---|---|---|---|---|
| ENSMUSG00000056032.13 | 0 | NA | NA | NA | NA | NA | #VALUE! |
| ENSMUSG00000056043.6 | 0 | NA | NA | NA | NA | NA | #VALUE! |
| ENSMUSG00000056055.14 | 0 | NA | NA | NA | NA | NA | #VALUE! |
| ENSMUSG00000056061.4 | 0 | NA | NA | NA | NA | NA | #VALUE! |
| ENSMUSG00000056069.11 | 5.822476 | 1.456973 | 3.023478 | 0.481886 | 0.629887 | NA | #VALUE! |
| ENSMUSG00000056071.13 | 57.82371 | -2.82219 | 1.780487 | -1.58507 | NA | NA | #VALUE! |
| ENSMUSG00000056073.17 | 0 | NA | NA | NA | NA | NA | #VALUE! |
| ENSMUSG00000056078.7 | 0 | NA | NA | NA | NA | NA | #VALUE! |
| ENSMUSG00000056089.7 | 0 | NA | NA | NA | NA | NA | #VALUE! |
| ENSMUSG00000056115.3 | 0 | NA | NA | NA | NA | NA | #VALUE! |
| ENSMUSG00000056124.6 | 6.734767 | 1.694428 | 3.387143 | 0.500253 | 0.616897 | NA | #VALUE! |
| ENSMUSG00000056128.4 | 0 | NA | NA | NA | NA | NA | #VALUE! |
| ENSMUSG00000056130.11 | 0 | NA | NA | NA | NA | NA | #VALUE! |
| ENSMUSG00000056133.6 | 8.08767 | -1.53167 | 3.109934 | -0.49251 | 0.62236 | NA | #VALUE! |
| ENSMUSG00000056145.6 | 0 | NA | NA | NA | NA | NA | #VALUE! |
| ENSMUSG00000056148.9 | 2.58392 | -4.90098 | 3.924036 | -1.24896 | 0.211678 | NA | #VALUE! |
| ENSMUSG00000056155.7 | 0 | NA | NA | NA | NA | NA | #VALUE! |
| ENSMUSG00000056158.15 | 0 | NA | NA | NA | NA | NA | #VALUE! |
| ENSMUSG00000056162.15 | 0 | NA | NA | NA | NA | NA | #VALUE! |
| ENSMUSG00000056174.9 | 0 | NA | NA | NA | NA | NA | #VALUE! |
| ENSMUSG00000056184.6 | 0 | NA | NA | NA | NA | NA | #VALUE! |
| ENSMUSG00000056185.6 | 0 | NA | NA | NA | NA | NA | #VALUE! |
| ENSMUSG00000056197.7 | 0 | NA | NA | NA | NA | NA | #VALUE! |
| ENSMUSG00000056203.5 | 0 | NA | NA | NA | NA | NA | #VALUE! |
| ENSMUSG00000056214.10 | 0.652786 | 2.919659 | 3.972474 | 0.734972 | 0.462356 | NA | #VALUE! |
| ENSMUSG00000056215.14 | 0 | NA | NA | NA | NA | NA | #VALUE! |
| ENSMUSG00000056219.8 | 0 | NA | NA | NA | NA | NA | #VALUE! |
| ENSMUSG00000056220.15 | 7.325348 | -2.1736 | 3.138629 | -0.69253 | 0.488603 | NA | #VALUE! |
| ENSMUSG00000056222.16 | 0 | NA | NA | NA | NA | NA | #VALUE! |
| ENSMUSG00000056223.9 | 0 | NA | NA | NA | NA | NA | #VALUE! |
| ENSMUSG00000056257.4 | 0 | NA | NA | NA | NA | NA | #VALUE! |
| ENSMUSG00000056258.10 | 0 | NA | NA | NA | NA | NA | #VALUE! |
| ENSMUSG00000056267.15 | 5.879119 | -1.31097 | 3.763575 | -0.34833 | 0.727591 | NA | #VALUE! |
| ENSMUSG00000056270.4 | 0 | NA | NA | NA | NA | NA | #VALUE! |
| ENSMUSG00000056271.14 | 0 | NA | NA | NA | NA | NA | #VALUE! |
| ENSMUSG00000056281.5 | 0 | NA | NA | NA | NA | NA | #VALUE! |
| ENSMUSG00000056288.12 | 0 | NA | NA | NA | NA | NA | #VALUE! |
| ENSMUSG00000056290.17 | 10.48241 | -2.76672 | 3.234597 | -0.85535 | NA | NA | #VALUE! |
| ENSMUSG00000056293.13 | 0 | NA | NA | NA | NA | NA | #VALUE! |
| ENSMUSG00000056296.18 | 0 | NA | NA | NA | NA | NA | #VALUE! |
| ENSMUSG00000056300.15 | 0 | NA | NA | NA | NA | NA | #VALUE! |
| ENSMUSG00000056306.7 | 0 | NA | NA | NA | NA | NA | #VALUE! |
| ENSMUSG00000056316.3 | 0 | NA | NA | NA | NA | NA | #VALUE! |
| ENSMUSG00000056328.15 | 0 | NA | NA | NA | NA | NA | #VALUE! |

| | | | | | | | |
|---|---|---|---|---|---|---|---|
| ENSMUSG00000056329.4 | 0 | NA | NA | NA | NA | NA | #VALUE! |
| ENSMUSG00000056332.5 | 0 | NA | NA | NA | NA | NA | #VALUE! |
| ENSMUSG00000056350.8 | 0 | NA | NA | NA | NA | NA | #VALUE! |
| ENSMUSG00000056359.7 | 0 | NA | NA | NA | NA | NA | #VALUE! |
| ENSMUSG00000056366.6 | 0 | NA | NA | NA | NA | NA | #VALUE! |
| ENSMUSG00000056367.15 | 0 | NA | NA | NA | NA | NA | #VALUE! |
| ENSMUSG00000056370.18 | 0 | NA | NA | NA | NA | NA | #VALUE! |
| ENSMUSG00000056379.4 | 0 | NA | NA | NA | NA | NA | #VALUE! |
| ENSMUSG00000056380.14 | 0 | NA | NA | NA | NA | NA | #VALUE! |
| ENSMUSG00000056394.21 | 7.159636 | -0.39284 | 3.676837 | -0.10684 | 0.914915 | NA | #VALUE! |
| ENSMUSG00000056399.3 | 0 | NA | NA | NA | NA | NA | #VALUE! |
| ENSMUSG00000056411.3 | 0.70064 | 1.487231 | 3.986665 | 0.373051 | 0.70911 | NA | #VALUE! |
| ENSMUSG00000056412.7 | 0.391578 | -2.1489 | 4.020883 | -0.53444 | 0.59304 | NA | #VALUE! |
| ENSMUSG00000056413.17 | 0 | NA | NA | NA | NA | NA | #VALUE! |
| ENSMUSG00000056418.4 | 0 | NA | NA | NA | NA | NA | #VALUE! |
| ENSMUSG00000056423.8 | 0 | NA | NA | NA | NA | NA | #VALUE! |
| ENSMUSG00000056426.4 | 0 | NA | NA | NA | NA | NA | #VALUE! |
| ENSMUSG00000056427.11 | 0 | NA | NA | NA | NA | NA | #VALUE! |
| ENSMUSG00000056436.5 | 0 | NA | NA | NA | NA | NA | #VALUE! |
| ENSMUSG00000056445.9 | 0 | NA | NA | NA | NA | NA | #VALUE! |
| ENSMUSG00000056457.7 | 0 | NA | NA | NA | NA | NA | #VALUE! |
| ENSMUSG00000056458.17 | 0 | NA | NA | NA | NA | NA | #VALUE! |
| ENSMUSG00000056468.4 | 0 | NA | NA | NA | NA | NA | #VALUE! |
| ENSMUSG00000056481.8 | 0 | NA | NA | NA | NA | NA | #VALUE! |
| ENSMUSG00000056486.19 | 0 | NA | NA | NA | NA | NA | #VALUE! |
| ENSMUSG00000056487.9 | 1.087334 | -2.3404 | 3.94129 | -0.59382 | 0.552635 | NA | #VALUE! |
| ENSMUSG00000056494.8 | 0 | NA | NA | NA | NA | NA | #VALUE! |
| ENSMUSG00000056498.14 | 0 | NA | NA | NA | NA | NA | #VALUE! |
| ENSMUSG00000056508.7 | 0 | NA | NA | NA | NA | NA | #VALUE! |
| ENSMUSG00000056509.11 | 0 | NA | NA | NA | NA | NA | #VALUE! |
| ENSMUSG00000056529.8 | 0.391578 | -2.1489 | 4.020883 | -0.53444 | 0.59304 | NA | #VALUE! |
| ENSMUSG00000056531.12 | 0.168284 | 1.027628 | 4.067126 | 0.252667 | 0.800526 | NA | #VALUE! |
| ENSMUSG00000056544.3 | 0 | NA | NA | NA | NA | NA | #VALUE! |
| ENSMUSG00000056553.15 | 0 | NA | NA | NA | NA | NA | #VALUE! |
| ENSMUSG00000056564.8 | 0 | NA | NA | NA | NA | NA | #VALUE! |
| ENSMUSG00000056569.12 | 0 | NA | NA | NA | NA | NA | #VALUE! |
| ENSMUSG00000056586.9 | 0 | NA | NA | NA | NA | NA | #VALUE! |
| ENSMUSG00000056592.15 | 2.520315 | -4.8742 | 3.924358 | -1.24204 | 0.214223 | NA | #VALUE! |
| ENSMUSG00000056596.9 | 0 | NA | NA | NA | NA | NA | #VALUE! |
| ENSMUSG00000056598.17 | 0 | NA | NA | NA | NA | NA | #VALUE! |
| ENSMUSG00000056600.5 | 0 | NA | NA | NA | NA | NA | #VALUE! |
| ENSMUSG00000056605.8 | 0 | NA | NA | NA | NA | NA | #VALUE! |
| ENSMUSG00000056612.8 | 7.960319 | -6.52083 | 3.880481 | -1.68042 | 0.092876 | NA | #VALUE! |
| ENSMUSG00000056617.13 | 0 | NA | NA | NA | NA | NA | #VALUE! |

| | | | | | | | |
|---|---|---|---|---|---|---|---|
| ENSMUSG00000056632.9 | 0.093255 | 0.868093 | 4.072464 | 0.213162 | 0.831201 | NA | #VALUE! |
| ENSMUSG00000056648.6 | 0 | NA | NA | NA | NA | NA | #VALUE! |
| ENSMUSG00000056656.7 | 0 | NA | NA | NA | NA | NA | #VALUE! |
| ENSMUSG00000056671.8 | 3.702238 | 5.378281 | 3.918889 | 1.372399 | 0.169939 | NA | #VALUE! |
| ENSMUSG00000056679.14 | 0 | NA | NA | NA | NA | NA | #VALUE! |
| ENSMUSG00000056682.9 | 0 | NA | NA | NA | NA | NA | #VALUE! |
| ENSMUSG00000056687.10 | 0 | NA | NA | NA | NA | NA | #VALUE! |
| ENSMUSG00000056696.6 | 0 | NA | NA | NA | NA | NA | #VALUE! |
| ENSMUSG00000056699.8 | 0 | NA | NA | NA | NA | NA | #VALUE! |
| ENSMUSG00000056706.4 | 0 | NA | NA | NA | NA | NA | #VALUE! |
| ENSMUSG00000056708.6 | 7.975098 | -1.54855 | 2.847449 | -0.54384 | 0.586554 | NA | #VALUE! |
| ENSMUSG00000056716.11 | 0 | NA | NA | NA | NA | NA | #VALUE! |
| ENSMUSG00000056718.3 | 8.299712 | 6.547057 | 3.796415 | 1.724536 | NA | NA | #VALUE! |
| ENSMUSG00000056724.16 | 8.847664 | 5.601634 | 2.620465 | 2.137649 | 0.032545 | NA | #VALUE! |
| ENSMUSG00000056728.7 | 0 | NA | NA | NA | NA | NA | #VALUE! |
| ENSMUSG00000056735.6 | 0 | NA | NA | NA | NA | NA | #VALUE! |
| ENSMUSG00000056737.15 | 5.408801 | 5.931107 | 3.915037 | 1.514955 | 0.129784 | NA | #VALUE! |
| ENSMUSG00000056738.12 | 3.823463 | 5.43312 | 3.918438 | 1.386552 | 0.165578 | NA | #VALUE! |
| ENSMUSG00000056752.17 | 0 | NA | NA | NA | NA | NA | #VALUE! |
| ENSMUSG00000056753.4 | 0 | NA | NA | NA | NA | NA | #VALUE! |
| ENSMUSG00000056755.14 | 0 | NA | NA | NA | NA | NA | #VALUE! |
| ENSMUSG00000056758.15 | 0 | NA | NA | NA | NA | NA | #VALUE! |
| ENSMUSG00000056771.8 | 0 | NA | NA | NA | NA | NA | #VALUE! |
| ENSMUSG00000056772.4 | 0 | NA | NA | NA | NA | NA | #VALUE! |
| ENSMUSG00000056777.6 | 0 | NA | NA | NA | NA | NA | #VALUE! |
| ENSMUSG00000056782.6 | 0 | NA | NA | NA | NA | NA | #VALUE! |
| ENSMUSG00000056797.7 | 0 | NA | NA | NA | NA | NA | #VALUE! |
| ENSMUSG00000056815.2 | 0 | NA | NA | NA | NA | NA | #VALUE! |
| ENSMUSG00000056821.14 | 0 | NA | NA | NA | NA | NA | #VALUE! |
| ENSMUSG00000056822.5 | 0 | NA | NA | NA | NA | NA | #VALUE! |
| ENSMUSG00000056824.16 | 0 | NA | NA | NA | NA | NA | #VALUE! |
| ENSMUSG00000056829.5 | 0 | NA | NA | NA | NA | NA | #VALUE! |
| ENSMUSG00000056832.15 | 0.660235 | 2.855092 | 3.975428 | 0.718185 | 0.472643 | NA | #VALUE! |
| ENSMUSG00000056849.7 | 0 | NA | NA | NA | NA | NA | #VALUE! |
| ENSMUSG00000056853.4 | 0 | NA | NA | NA | NA | NA | #VALUE! |
| ENSMUSG00000056854.5 | 0 | NA | NA | NA | NA | NA | #VALUE! |
| ENSMUSG00000056856.15 | 0 | NA | NA | NA | NA | NA | #VALUE! |
| ENSMUSG00000056858.2 | 0 | NA | NA | NA | NA | NA | #VALUE! |
| ENSMUSG00000056863.6 | 0 | NA | NA | NA | NA | NA | #VALUE! |
| ENSMUSG00000056870.10 | 6.265251 | -6.17547 | 3.913919 | -1.57782 | 0.114607 | NA | #VALUE! |
| ENSMUSG00000056877.3 | 0.093255 | 0.868093 | 4.072464 | 0.213162 | 0.831201 | NA | #VALUE! |
| ENSMUSG00000056880.14 | 0 | NA | NA | NA | NA | NA | #VALUE! |
| ENSMUSG00000056883.6 | 0 | NA | NA | NA | NA | NA | #VALUE! |
| ENSMUSG00000056885.6 | 0 | NA | NA | NA | NA | NA | #VALUE! |

| | | | | | | | |
|---|---|---|---|---|---|---|---|
| ENSMUSG00000056895.5 | 0 | NA | NA | NA | NA | NA | #VALUE! |
| ENSMUSG00000056900.14 | 0 | NA | NA | NA | NA | NA | #VALUE! |
| ENSMUSG00000056901.2 | 0 | NA | NA | NA | NA | NA | #VALUE! |
| ENSMUSG00000056904.4 | 0.261539 | 1.68833 | 4.043024 | 0.417591 | 0.676246 | NA | #VALUE! |
| ENSMUSG00000056910.6 | 0 | NA | NA | NA | NA | NA | #VALUE! |
| ENSMUSG00000056912.13 | 0 | NA | NA | NA | NA | NA | #VALUE! |
| ENSMUSG00000056921.3 | 0 | NA | NA | NA | NA | NA | #VALUE! |
| ENSMUSG00000056926.3 | 0 | NA | NA | NA | NA | NA | #VALUE! |
| ENSMUSG00000056943.3 | 0 | NA | NA | NA | NA | NA | #VALUE! |
| ENSMUSG00000056946.5 | 0 | NA | NA | NA | NA | NA | #VALUE! |
| ENSMUSG00000056947.6 | 0 | NA | NA | NA | NA | NA | #VALUE! |
| ENSMUSG00000056952.14 | 9.267895 | 1.239089 | 2.974124 | 0.416623 | 0.676954 | NA | #VALUE! |
| ENSMUSG00000056959.7 | 0 | NA | NA | NA | NA | NA | #VALUE! |
| ENSMUSG00000056961.3 | 0 | NA | NA | NA | NA | NA | #VALUE! |
| ENSMUSG00000056966.8 | 0 | NA | NA | NA | NA | NA | #VALUE! |
| ENSMUSG00000056972.7 | 0 | NA | NA | NA | NA | NA | #VALUE! |
| ENSMUSG00000056987.9 | 0 | NA | NA | NA | NA | NA | #VALUE! |
| ENSMUSG00000056995.4 | 0 | NA | NA | NA | NA | NA | #VALUE! |
| ENSMUSG00000057000.13 | 0 | NA | NA | NA | NA | NA | #VALUE! |
| ENSMUSG00000057003.13 | 0 | NA | NA | NA | NA | NA | #VALUE! |
| ENSMUSG00000057021.14 | 0 | NA | NA | NA | NA | NA | #VALUE! |
| ENSMUSG00000057047.6 | 0 | NA | NA | NA | NA | NA | #VALUE! |
| ENSMUSG00000057050.8 | 0 | NA | NA | NA | NA | NA | #VALUE! |
| ENSMUSG00000057058.17 | 0 | NA | NA | NA | NA | NA | #VALUE! |
| ENSMUSG00000057060.9 | 0 | NA | NA | NA | NA | NA | #VALUE! |
| ENSMUSG00000057067.5 | 0 | NA | NA | NA | NA | NA | #VALUE! |
| ENSMUSG00000057072.12 | 0 | NA | NA | NA | NA | NA | #VALUE! |
| ENSMUSG00000057092.13 | 0 | NA | NA | NA | NA | NA | #VALUE! |
| ENSMUSG00000057093.15 | 3.274827 | -0.41315 | 3.782436 | -0.10923 | 0.913022 | NA | #VALUE! |
| ENSMUSG00000057098.15 | 0 | NA | NA | NA | NA | NA | #VALUE! |
| ENSMUSG00000057116.6 | 0 | NA | NA | NA | NA | NA | #VALUE! |
| ENSMUSG00000057123.15 | 0 | NA | NA | NA | NA | NA | #VALUE! |
| ENSMUSG00000057132.17 | 4.475781 | 0.381587 | 3.767344 | 0.101288 | 0.919322 | NA | #VALUE! |
| ENSMUSG00000057135.13 | 1.050131 | -3.6251 | 3.948379 | -0.91812 | 0.358554 | NA | #VALUE! |
| ENSMUSG00000057149.5 | 0 | NA | NA | NA | NA | NA | #VALUE! |
| ENSMUSG00000057157.4 | 0 | NA | NA | NA | NA | NA | #VALUE! |
| ENSMUSG00000057160.3 | 0 | NA | NA | NA | NA | NA | #VALUE! |
| ENSMUSG00000057163.4 | 0 | NA | NA | NA | NA | NA | #VALUE! |
| ENSMUSG00000057170.7 | 0 | NA | NA | NA | NA | NA | #VALUE! |
| ENSMUSG00000057173.9 | 0 | NA | NA | NA | NA | NA | #VALUE! |
| ENSMUSG00000057174.6 | 0 | NA | NA | NA | NA | NA | #VALUE! |
| ENSMUSG00000057176.5 | 2.755285 | -4.98901 | 3.923017 | -1.27173 | 0.20347 | NA | #VALUE! |
| ENSMUSG00000057179.7 | 0 | NA | NA | NA | NA | NA | #VALUE! |
| ENSMUSG00000057182.16 | 0 | NA | NA | NA | NA | NA | #VALUE! |

| | | | | | | | |
|---|---|---|---|---|---|---|---|
| ENSMUSG00000057191.15 | 9.37927 | -6.75848 | 2.729912 | -2.47571 | 0.013297 | NA | #VALUE! |
| ENSMUSG00000057195.14 | 0 | NA | NA | NA | NA | NA | #VALUE! |
| ENSMUSG00000057203.4 | 0 | NA | NA | NA | NA | NA | #VALUE! |
| ENSMUSG00000057207.3 | 0 | NA | NA | NA | NA | NA | #VALUE! |
| ENSMUSG00000057215.7 | 0 | NA | NA | NA | NA | NA | #VALUE! |
| ENSMUSG00000057216.7 | 0 | NA | NA | NA | NA | NA | #VALUE! |
| ENSMUSG00000057219.12 | 0.373021 | 2.140748 | 4.018182 | 0.532765 | 0.594196 | NA | #VALUE! |
| ENSMUSG00000057223.7 | 4.200525 | -5.60698 | 3.917367 | -1.43131 | 0.15234 | NA | #VALUE! |
| ENSMUSG00000057240.15 | 0 | NA | NA | NA | NA | NA | #VALUE! |
| ENSMUSG00000057244.7 | 0.195789 | -1.05581 | 4.073302 | -0.2592 | 0.795479 | NA | #VALUE! |
| ENSMUSG00000057246.17 | 0 | NA | NA | NA | NA | NA | #VALUE! |
| ENSMUSG00000057262.3 | 0 | NA | NA | NA | NA | NA | #VALUE! |
| ENSMUSG00000057265.14 | 1.229878 | 3.761857 | 3.943699 | 0.95389 | 0.340139 | NA | #VALUE! |
| ENSMUSG00000057270.4 | 0 | NA | NA | NA | NA | NA | #VALUE! |
| ENSMUSG00000057280.16 | 0 | NA | NA | NA | NA | NA | #VALUE! |
| ENSMUSG00000057286.7 | 2.310289 | -4.7496 | 3.925939 | -1.2098 | 0.226355 | NA | #VALUE! |
| ENSMUSG00000057290.6 | 0 | NA | NA | NA | NA | NA | #VALUE! |
| ENSMUSG00000057321.3 | 0 | NA | NA | NA | NA | NA | #VALUE! |
| ENSMUSG00000057337.14 | 0 | NA | NA | NA | NA | NA | #VALUE! |
| ENSMUSG00000057349.3 | 0 | NA | NA | NA | NA | NA | #VALUE! |
| ENSMUSG00000057359.8 | 2.424364 | -4.80166 | 3.925262 | -1.22327 | 0.221227 | NA | #VALUE! |
| ENSMUSG00000057370.6 | 0 | NA | NA | NA | NA | NA | #VALUE! |
| ENSMUSG00000057378.16 | 0 | NA | NA | NA | NA | NA | #VALUE! |
| ENSMUSG00000057381.2 | 0 | NA | NA | NA | NA | NA | #VALUE! |
| ENSMUSG00000057396.8 | 3.615742 | -5.38042 | 3.919165 | -1.37285 | 0.1698 | NA | #VALUE! |
| ENSMUSG00000057402.6 | 0 | NA | NA | NA | NA | NA | #VALUE! |
| ENSMUSG00000057411.10 | 0 | NA | NA | NA | NA | NA | #VALUE! |
| ENSMUSG00000057417.8 | 0 | NA | NA | NA | NA | NA | #VALUE! |
| ENSMUSG00000057424.4 | 0 | NA | NA | NA | NA | NA | #VALUE! |
| ENSMUSG00000057439.13 | 0 | NA | NA | NA | NA | NA | #VALUE! |
| ENSMUSG00000057443.14 | 0 | NA | NA | NA | NA | NA | #VALUE! |
| ENSMUSG00000057444.4 | 0 | NA | NA | NA | NA | NA | #VALUE! |
| ENSMUSG00000057446.4 | 0 | NA | NA | NA | NA | NA | #VALUE! |
| ENSMUSG00000057447.3 | 0 | NA | NA | NA | NA | NA | #VALUE! |
| ENSMUSG00000057454.6 | 0 | NA | NA | NA | NA | NA | #VALUE! |
| ENSMUSG00000057455.13 | 0 | NA | NA | NA | NA | NA | #VALUE! |
| ENSMUSG00000057457.12 | 0 | NA | NA | NA | NA | NA | #VALUE! |
| ENSMUSG00000057461.4 | 0 | NA | NA | NA | NA | NA | #VALUE! |
| ENSMUSG00000057464.5 | 0 | NA | NA | NA | NA | NA | #VALUE! |
| ENSMUSG00000057499.4 | 0 | NA | NA | NA | NA | NA | #VALUE! |
| ENSMUSG00000057503.7 | 0 | NA | NA | NA | NA | NA | #VALUE! |
| ENSMUSG00000057513.4 | 0 | NA | NA | NA | NA | NA | #VALUE! |
| ENSMUSG00000057534.14 | 0 | NA | NA | NA | NA | NA | #VALUE! |
| ENSMUSG00000057540.4 | 0 | NA | NA | NA | NA | NA | #VALUE! |

| | | | | | | | |
|---|---|---|---|---|---|---|---|
| ENSMUSG00000057564.7 | 0 | NA | NA | NA | NA | NA | #VALUE! |
| ENSMUSG00000057577.3 | 0 | NA | NA | NA | NA | NA | #VALUE! |
| ENSMUSG00000057580.5 | 5.377866 | -5.9579 | 3.866125 | -1.54105 | 0.123304 | NA | #VALUE! |
| ENSMUSG00000057592.2 | 0 | NA | NA | NA | NA | NA | #VALUE! |
| ENSMUSG00000057604.10 | 0.408222 | -2.20205 | 4.016842 | -0.54821 | 0.583551 | NA | #VALUE! |
| ENSMUSG00000057606.15 | 0 | NA | NA | NA | NA | NA | #VALUE! |
| ENSMUSG00000057609.7 | 0 | NA | NA | NA | NA | NA | #VALUE! |
| ENSMUSG00000057614.7 | 8.643185 | 6.606806 | 2.941408 | 2.246137 | 0.024695 | NA | #VALUE! |
| ENSMUSG00000057615.6 | 0 | NA | NA | NA | NA | NA | #VALUE! |
| ENSMUSG00000057650.4 | 0 | NA | NA | NA | NA | NA | #VALUE! |
| ENSMUSG00000057654.7 | 0 | NA | NA | NA | NA | NA | #VALUE! |
| ENSMUSG00000057666.19 | 102.3399 | -1.67528 | 1.328812 | -1.26074 | NA | NA | #VALUE! |
| ENSMUSG00000057667.8 | 5.742679 | -4.94953 | 3.161547 | -1.56554 | 0.117457 | NA | #VALUE! |
| ENSMUSG00000057672.17 | 0 | NA | NA | NA | NA | NA | #VALUE! |
| ENSMUSG00000057674.4 | 0 | NA | NA | NA | NA | NA | #VALUE! |
| ENSMUSG00000057685.9 | 0 | NA | NA | NA | NA | NA | #VALUE! |
| ENSMUSG00000057697.3 | 1.009701 | 3.503687 | 3.950852 | 0.886818 | 0.375177 | NA | #VALUE! |
| ENSMUSG00000057699.6 | 0 | NA | NA | NA | NA | NA | #VALUE! |
| ENSMUSG00000057706.8 | 0.936195 | 2.260233 | 3.956978 | 0.571202 | 0.567863 | NA | #VALUE! |
| ENSMUSG00000057710.8 | 0 | NA | NA | NA | NA | NA | #VALUE! |
| ENSMUSG00000057715.14 | 0 | NA | NA | NA | NA | NA | #VALUE! |
| ENSMUSG00000057716.8 | 0 | NA | NA | NA | NA | NA | #VALUE! |
| ENSMUSG00000057719.12 | 0.491951 | 2.402185 | 4.000147 | 0.600524 | 0.548157 | NA | #VALUE! |
| ENSMUSG00000057722.18 | 2.161045 | -3.06325 | 3.904409 | -0.78456 | 0.432711 | NA | #VALUE! |
| ENSMUSG00000057723.4 | 0 | NA | NA | NA | NA | NA | #VALUE! |
| ENSMUSG00000057726.4 | 0 | NA | NA | NA | NA | NA | #VALUE! |
| ENSMUSG00000057729.13 | 6.930867 | -6.32693 | 3.913207 | -1.61681 | 0.105918 | NA | #VALUE! |
| ENSMUSG00000057735.7 | 0 | NA | NA | NA | NA | NA | #VALUE! |
| ENSMUSG00000057751.15 | 1.009701 | 3.503687 | 3.950852 | 0.886818 | 0.375177 | NA | #VALUE! |
| ENSMUSG00000057754.3 | 0 | NA | NA | NA | NA | NA | #VALUE! |
| ENSMUSG00000057761.2 | 0 | NA | NA | NA | NA | NA | #VALUE! |
| ENSMUSG00000057762.7 | 0.420053 | -2.33493 | 4.007343 | -0.58266 | 0.56012 | NA | #VALUE! |
| ENSMUSG00000057766.15 | 0 | NA | NA | NA | NA | NA | #VALUE! |
| ENSMUSG00000057770.6 | 0 | NA | NA | NA | NA | NA | #VALUE! |
| ENSMUSG00000057777.5 | 4.84927 | 5.774248 | 3.915985 | 1.474533 | 0.140338 | NA | #VALUE! |
| ENSMUSG00000057789.15 | 0 | NA | NA | NA | NA | NA | #VALUE! |
| ENSMUSG00000057801.4 | 0 | NA | NA | NA | NA | NA | #VALUE! |
| ENSMUSG00000057805.5 | 0 | NA | NA | NA | NA | NA | #VALUE! |
| ENSMUSG00000057816.8 | 0 | NA | NA | NA | NA | NA | #VALUE! |
| ENSMUSG00000057817.7 | 0 | NA | NA | NA | NA | NA | #VALUE! |
| ENSMUSG00000057818.9 | 0 | NA | NA | NA | NA | NA | #VALUE! |
| ENSMUSG00000057829.6 | 0 | NA | NA | NA | NA | NA | #VALUE! |
| ENSMUSG00000057855.3 | 0 | NA | NA | NA | NA | NA | #VALUE! |
| ENSMUSG00000057886.6 | 0 | NA | NA | NA | NA | NA | #VALUE! |

| | | | | | | | |
|---|---|---|---|---|---|---|---|
| ENSMUSG00000057890.6 | 0 | NA | NA | NA | NA | NA | #VALUE! |
| ENSMUSG00000057895.12 | 0 | NA | NA | NA | NA | NA | #VALUE! |
| ENSMUSG00000057897.15 | 14.74608 | 6.891385 | 2.964984 | 2.324257 | NA | NA | #VALUE! |
| ENSMUSG00000057899.8 | 0 | NA | NA | NA | NA | NA | #VALUE! |
| ENSMUSG00000057903.5 | 0 | NA | NA | NA | NA | NA | #VALUE! |
| ENSMUSG00000057913.4 | 2.187686 | 4.619304 | 3.927239 | 1.176222 | 0.239506 | NA | #VALUE! |
| ENSMUSG00000057914.16 | 0 | NA | NA | NA | NA | NA | #VALUE! |
| ENSMUSG00000057945.4 | 0 | NA | NA | NA | NA | NA | #VALUE! |
| ENSMUSG00000057948.13 | 0 | NA | NA | NA | NA | NA | #VALUE! |
| ENSMUSG00000057967.13 | 0 | NA | NA | NA | NA | NA | #VALUE! |
| ENSMUSG00000057969.16 | 0.523077 | 2.574714 | 3.989843 | 0.645317 | 0.518722 | NA | #VALUE! |
| ENSMUSG00000057977.11 | 0 | NA | NA | NA | NA | NA | #VALUE! |
| ENSMUSG00000057981.6 | 0 | NA | NA | NA | NA | NA | #VALUE! |
| ENSMUSG00000057990.5 | 0 | NA | NA | NA | NA | NA | #VALUE! |
| ENSMUSG00000057997.7 | 0 | NA | NA | NA | NA | NA | #VALUE! |
| ENSMUSG00000058005.4 | 0 | NA | NA | NA | NA | NA | #VALUE! |
| ENSMUSG00000058006.13 | 45.62838 | 1.331659 | 1.389562 | 0.95833 | NA | NA | #VALUE! |
| ENSMUSG00000058014.5 | 0 | NA | NA | NA | NA | NA | #VALUE! |
| ENSMUSG00000058017.7 | 0 | NA | NA | NA | NA | NA | #VALUE! |
| ENSMUSG00000058019.14 | 0 | NA | NA | NA | NA | NA | #VALUE! |
| ENSMUSG00000058028.15 | 0 | NA | NA | NA | NA | NA | #VALUE! |
| ENSMUSG00000058030.6 | 0 | NA | NA | NA | NA | NA | #VALUE! |
| ENSMUSG00000058046.2 | 0.652786 | 2.919659 | 3.972474 | 0.734972 | 0.462356 | NA | #VALUE! |
| ENSMUSG00000058050.6 | 0 | NA | NA | NA | NA | NA | #VALUE! |
| ENSMUSG00000058052.3 | 0 | NA | NA | NA | NA | NA | #VALUE! |
| ENSMUSG00000058057.6 | 0 | NA | NA | NA | NA | NA | #VALUE! |
| ENSMUSG00000058063.5 | 0 | NA | NA | NA | NA | NA | #VALUE! |
| ENSMUSG00000058071.5 | 0 | NA | NA | NA | NA | NA | #VALUE! |
| ENSMUSG00000058073.5 | 0 | NA | NA | NA | NA | NA | #VALUE! |
| ENSMUSG00000058078.9 | 0 | NA | NA | NA | NA | NA | #VALUE! |
| ENSMUSG00000058084.6 | 0 | NA | NA | NA | NA | NA | #VALUE! |
| ENSMUSG00000058099.17 | 6.169834 | -0.35364 | 3.755603 | -0.09416 | 0.92498 | NA | #VALUE! |
| ENSMUSG00000058101.4 | 0 | NA | NA | NA | NA | NA | #VALUE! |
| ENSMUSG00000058114.4 | 0 | NA | NA | NA | NA | NA | #VALUE! |
| ENSMUSG00000058119.7 | 0 | NA | NA | NA | NA | NA | #VALUE! |
| ENSMUSG00000058124.5 | 0 | NA | NA | NA | NA | NA | #VALUE! |
| ENSMUSG00000058132.6 | 0 | NA | NA | NA | NA | NA | #VALUE! |
| ENSMUSG00000058145.17 | 0 | NA | NA | NA | NA | NA | #VALUE! |
| ENSMUSG00000058147.2 | 0 | NA | NA | NA | NA | NA | #VALUE! |
| ENSMUSG00000058153.16 | 0 | NA | NA | NA | NA | NA | #VALUE! |
| ENSMUSG00000058159.15 | 0 | NA | NA | NA | NA | NA | #VALUE! |
| ENSMUSG00000058163.15 | 7.350919 | -6.4116 | 3.91284 | -1.6386 | NA | NA | #VALUE! |
| ENSMUSG00000058172.2 | 0 | NA | NA | NA | NA | NA | #VALUE! |
| ENSMUSG00000058174.8 | 0.105013 | -0.97324 | 4.076293 | -0.23876 | 0.811295 | NA | #VALUE! |

| | | | | | | | |
|---|---|---|---|---|---|---|---|
| ENSMUSG00000058176.4 | 0 | NA | NA | NA | NA | NA | #VALUE! |
| ENSMUSG00000058183.15 | 0 | NA | NA | NA | NA | NA | #VALUE! |
| ENSMUSG00000058186.14 | 0 | NA | NA | NA | NA | NA | #VALUE! |
| ENSMUSG00000058188.4 | 0 | NA | NA | NA | NA | NA | #VALUE! |
| ENSMUSG00000058192.17 | 6.861847 | -3.19945 | 3.29746 | -0.97028 | 0.331909 | NA | #VALUE! |
| ENSMUSG00000058194.4 | 0 | NA | NA | NA | NA | NA | #VALUE! |
| ENSMUSG00000058200.3 | 0 | NA | NA | NA | NA | NA | #VALUE! |
| ENSMUSG00000058244.5 | 0 | NA | NA | NA | NA | NA | #VALUE! |
| ENSMUSG00000058246.9 | 1.851119 | 4.378291 | 3.930938 | 1.113803 | 0.265364 | NA | #VALUE! |
| ENSMUSG00000058248.13 | 0 | NA | NA | NA | NA | NA | #VALUE! |
| ENSMUSG00000058250.3 | 0 | NA | NA | NA | NA | NA | #VALUE! |
| ENSMUSG00000058251.4 | 0 | NA | NA | NA | NA | NA | #VALUE! |
| ENSMUSG00000058252.7 | 0 | NA | NA | NA | NA | NA | #VALUE! |
| ENSMUSG00000058260.3 | 4.750079 | 1.802583 | 3.29661 | 0.546799 | 0.584517 | NA | #VALUE! |
| ENSMUSG00000058270.4 | 0 | NA | NA | NA | NA | NA | #VALUE! |
| ENSMUSG00000058275.4 | 0 | NA | NA | NA | NA | NA | #VALUE! |
| ENSMUSG00000058281.2 | 0 | NA | NA | NA | NA | NA | #VALUE! |
| ENSMUSG00000058287.6 | 0 | NA | NA | NA | NA | NA | #VALUE! |
| ENSMUSG00000058290.5 | 0 | NA | NA | NA | NA | NA | #VALUE! |
| ENSMUSG00000058295.7 | 0 | NA | NA | NA | NA | NA | #VALUE! |
| ENSMUSG00000058297.17 | 1.174735 | -3.75246 | 3.944889 | -0.95122 | 0.341493 | NA | #VALUE! |
| ENSMUSG00000058298.11 | 1.682835 | 4.240781 | 3.933339 | 1.078163 | 0.280961 | NA | #VALUE! |
| ENSMUSG00000058301.9 | 6.240473 | 1.100585 | 2.68608 | 0.409736 | 0.681999 | NA | #VALUE! |
| ENSMUSG00000058328.14 | 0 | NA | NA | NA | NA | NA | #VALUE! |
| ENSMUSG00000058349.3 | 0 | NA | NA | NA | NA | NA | #VALUE! |
| ENSMUSG00000058354.8 | 0 | NA | NA | NA | NA | NA | #VALUE! |
| ENSMUSG00000058357.3 | 0 | NA | NA | NA | NA | NA | #VALUE! |
| ENSMUSG00000058360.6 | 0 | NA | NA | NA | NA | NA | #VALUE! |
| ENSMUSG00000058368.4 | 0 | NA | NA | NA | NA | NA | #VALUE! |
| ENSMUSG00000058385.9 | 0 | NA | NA | NA | NA | NA | #VALUE! |
| ENSMUSG00000058392.14 | 3.72495 | -1.50676 | 3.790375 | -0.39752 | 0.690982 | NA | #VALUE! |
| ENSMUSG00000058398.7 | 0 | NA | NA | NA | NA | NA | #VALUE! |
| ENSMUSG00000058400.14 | 0 | NA | NA | NA | NA | NA | #VALUE! |
| ENSMUSG00000058420.9 | 0 | NA | NA | NA | NA | NA | #VALUE! |
| ENSMUSG00000058427.11 | 10.29129 | -6.89665 | 3.5367 | -1.95002 | NA | NA | #VALUE! |
| ENSMUSG00000058430.2 | 0.093255 | 0.868093 | 4.072464 | 0.213162 | 0.831201 | NA | #VALUE! |
| ENSMUSG00000058435.7 | 0 | NA | NA | NA | NA | NA | #VALUE! |
| ENSMUSG00000058441.8 | 0 | NA | NA | NA | NA | NA | #VALUE! |
| ENSMUSG00000058447.9 | 0 | NA | NA | NA | NA | NA | #VALUE! |
| ENSMUSG00000058463.4 | 0 | NA | NA | NA | NA | NA | #VALUE! |
| ENSMUSG00000058470.11 | 2.238125 | 4.666312 | 3.926586 | 1.188389 | 0.23468 | NA | #VALUE! |
| ENSMUSG00000058486.12 | 6.794054 | 2.539132 | 3.357598 | 0.756235 | 0.449509 | NA | #VALUE! |
| ENSMUSG00000058488.8 | 0 | NA | NA | NA | NA | NA | #VALUE! |
| ENSMUSG00000058491.9 | 0 | NA | NA | NA | NA | NA | #VALUE! |

| | | | | | | | |
|---|---|---|---|---|---|---|---|
| ENSMUSG00000058498.13 | 0 | NA | NA | NA | NA | NA | #VALUE! |
| ENSMUSG00000058499.10 | 0 | NA | NA | NA | NA | NA | #VALUE! |
| ENSMUSG00000058503.12 | 102.8446 | -0.51511 | 1.121969 | -0.45911 | NA | NA | #VALUE! |
| ENSMUSG00000058508.2 | 0 | NA | NA | NA | NA | NA | #VALUE! |
| ENSMUSG00000058513.8 | 0 | NA | NA | NA | NA | NA | #VALUE! |
| ENSMUSG00000058515.5 | 0 | NA | NA | NA | NA | NA | #VALUE! |
| ENSMUSG00000058523.3 | 9.378692 | -0.35139 | 1.956861 | -0.17957 | 0.857492 | NA | #VALUE! |
| ENSMUSG00000058537.5 | 0 | NA | NA | NA | NA | NA | #VALUE! |
| ENSMUSG00000058542.4 | 0.405815 | -2.2474 | 4.013505 | -0.55996 | 0.575507 | NA | #VALUE! |
| ENSMUSG00000058550.16 | 0 | NA | NA | NA | NA | NA | #VALUE! |
| ENSMUSG00000058567.7 | 0 | NA | NA | NA | NA | NA | #VALUE! |
| ENSMUSG00000058568.7 | 0 | NA | NA | NA | NA | NA | #VALUE! |
| ENSMUSG00000058571.12 | 0 | NA | NA | NA | NA | NA | #VALUE! |
| ENSMUSG00000058579.6 | 0 | NA | NA | NA | NA | NA | #VALUE! |
| ENSMUSG00000058581.8 | 0 | NA | NA | NA | NA | NA | #VALUE! |
| ENSMUSG00000058589.15 | 0 | NA | NA | NA | NA | NA | #VALUE! |
| ENSMUSG00000058590.3 | 0 | NA | NA | NA | NA | NA | #VALUE! |
| ENSMUSG00000058607.5 | 0 | NA | NA | NA | NA | NA | #VALUE! |
| ENSMUSG00000058613.7 | 2.310289 | -4.7496 | 3.925939 | -1.2098 | 0.226355 | NA | #VALUE! |
| ENSMUSG00000058618.7 | 0 | NA | NA | NA | NA | NA | #VALUE! |
| ENSMUSG00000058620.6 | 3.627209 | 5.348973 | 3.919137 | 1.364834 | 0.172305 | NA | #VALUE! |
| ENSMUSG00000058622.2 | 0 | NA | NA | NA | NA | NA | #VALUE! |
| ENSMUSG00000058625.7 | 0 | NA | NA | NA | NA | NA | #VALUE! |
| ENSMUSG00000058626.17 | 0 | NA | NA | NA | NA | NA | #VALUE! |
| ENSMUSG00000058628.5 | 0 | NA | NA | NA | NA | NA | #VALUE! |
| ENSMUSG00000058631.7 | 0 | NA | NA | NA | NA | NA | #VALUE! |
| ENSMUSG00000058643.4 | 0 | NA | NA | NA | NA | NA | #VALUE! |
| ENSMUSG00000058656.14 | 0 | NA | NA | NA | NA | NA | #VALUE! |
| ENSMUSG00000058659.7 | 0 | NA | NA | NA | NA | NA | #VALUE! |
| ENSMUSG00000058662.5 | 0 | NA | NA | NA | NA | NA | #VALUE! |
| ENSMUSG00000058665.9 | 0 | NA | NA | NA | NA | NA | #VALUE! |
| ENSMUSG00000058669.8 | 0 | NA | NA | NA | NA | NA | #VALUE! |
| ENSMUSG00000058670.6 | 0 | NA | NA | NA | NA | NA | #VALUE! |
| ENSMUSG00000058685.8 | 0 | NA | NA | NA | NA | NA | #VALUE! |
| ENSMUSG00000058692.5 | 0 | NA | NA | NA | NA | NA | #VALUE! |
| ENSMUSG00000058700.3 | 0.093255 | 0.868093 | 4.072464 | 0.213162 | 0.831201 | NA | #VALUE! |
| ENSMUSG00000058717.9 | 0 | NA | NA | NA | NA | NA | #VALUE! |
| ENSMUSG00000058725.2 | 0 | NA | NA | NA | NA | NA | #VALUE! |
| ENSMUSG00000058728.16 | 0 | NA | NA | NA | NA | NA | #VALUE! |
| ENSMUSG00000058729.14 | 8.576792 | -2.38954 | 2.151018 | -1.11089 | 0.266616 | NA | #VALUE! |
| ENSMUSG00000058740.15 | 0 | NA | NA | NA | NA | NA | #VALUE! |
| ENSMUSG00000058741.5 | 0 | NA | NA | NA | NA | NA | #VALUE! |
| ENSMUSG00000058743.7 | 0 | NA | NA | NA | NA | NA | #VALUE! |
| ENSMUSG00000058745.5 | 0 | NA | NA | NA | NA | NA | #VALUE! |

| | | | | | | | |
|---|---|---|---|---|---|---|---|
| ENSMUSG00000058755.4 | 0 | NA | NA | NA | NA | NA | #VALUE! |
| ENSMUSG00000058773.3 | 0 | NA | NA | NA | NA | NA | #VALUE! |
| ENSMUSG00000058794.13 | 0 | NA | NA | NA | NA | NA | #VALUE! |
| ENSMUSG00000058802.6 | 0 | NA | NA | NA | NA | NA | #VALUE! |
| ENSMUSG00000058807.5 | 0 | NA | NA | NA | NA | NA | #VALUE! |
| ENSMUSG00000058809.9 | 1.515952 | 0.906781 | 3.859007 | 0.234978 | 0.814226 | NA | #VALUE! |
| ENSMUSG00000058812.3 | 0 | NA | NA | NA | NA | NA | #VALUE! |
| ENSMUSG00000058816.6 | 0 | NA | NA | NA | NA | NA | #VALUE! |
| ENSMUSG00000058818.14 | 1.566313 | -4.16993 | 3.935366 | -1.0596 | 0.289325 | NA | #VALUE! |
| ENSMUSG00000058820.4 | 0 | NA | NA | NA | NA | NA | #VALUE! |
| ENSMUSG00000058831.14 | 0 | NA | NA | NA | NA | NA | #VALUE! |
| ENSMUSG00000058838.10 | 4.47273 | 1.248106 | 2.746865 | 0.454375 | 0.649559 | NA | #VALUE! |
| ENSMUSG00000058856.4 | 0 | NA | NA | NA | NA | NA | #VALUE! |
| ENSMUSG00000058869.10 | 0.105013 | -0.97324 | 4.076293 | -0.23876 | 0.811295 | NA | #VALUE! |
| ENSMUSG00000058873.2 | 0 | NA | NA | NA | NA | NA | #VALUE! |
| ENSMUSG00000058881.14 | 0.168284 | 1.027628 | 4.067126 | 0.252667 | 0.800526 | NA | #VALUE! |
| ENSMUSG00000058883.17 | 0 | NA | NA | NA | NA | NA | #VALUE! |
| ENSMUSG00000058884.7 | 0 | NA | NA | NA | NA | NA | #VALUE! |
| ENSMUSG00000058897.19 | 0 | NA | NA | NA | NA | NA | #VALUE! |
| ENSMUSG00000058904.4 | 0 | NA | NA | NA | NA | NA | #VALUE! |
| ENSMUSG00000058908.16 | 0 | NA | NA | NA | NA | NA | #VALUE! |
| ENSMUSG00000058914.13 | 0 | NA | NA | NA | NA | NA | #VALUE! |
| ENSMUSG00000058922.8 | 0 | NA | NA | NA | NA | NA | #VALUE! |
| ENSMUSG00000058925.12 | 0 | NA | NA | NA | NA | NA | #VALUE! |
| ENSMUSG00000058927.7 | 1.396653 | -0.12506 | 3.856374 | -0.03243 | 0.97413 | NA | #VALUE! |
| ENSMUSG00000058932.7 | 0 | NA | NA | NA | NA | NA | #VALUE! |
| ENSMUSG00000058933.6 | 0 | NA | NA | NA | NA | NA | #VALUE! |
| ENSMUSG00000058935.15 | 0 | NA | NA | NA | NA | NA | #VALUE! |
| ENSMUSG00000058953.4 | 0 | NA | NA | NA | NA | NA | #VALUE! |
| ENSMUSG00000058966.14 | 0 | NA | NA | NA | NA | NA | #VALUE! |
| ENSMUSG00000058975.8 | 0 | NA | NA | NA | NA | NA | #VALUE! |
| ENSMUSG00000058976.5 | 0 | NA | NA | NA | NA | NA | #VALUE! |
| ENSMUSG00000058979.8 | 9.327266 | -2.57192 | 3.35358 | -0.76692 | NA | NA | #VALUE! |
| ENSMUSG00000058981.5 | 0 | NA | NA | NA | NA | NA | #VALUE! |
| ENSMUSG00000059003.13 | 0 | NA | NA | NA | NA | NA | #VALUE! |
| ENSMUSG00000059013.13 | 9.477675 | 2.418727 | 2.655425 | 0.910862 | 0.362368 | NA | #VALUE! |
| ENSMUSG00000059014.9 | 0 | NA | NA | NA | NA | NA | #VALUE! |
| ENSMUSG00000059022.16 | 1.025807 | 3.556426 | 3.949287 | 0.900523 | 0.367842 | NA | #VALUE! |
| ENSMUSG00000059023.3 | 0 | NA | NA | NA | NA | NA | #VALUE! |
| ENSMUSG00000059030.6 | 0 | NA | NA | NA | NA | NA | #VALUE! |
| ENSMUSG00000059031.4 | 0.204111 | -1.05581 | 4.073302 | -0.2592 | 0.795479 | NA | #VALUE! |
| ENSMUSG00000059042.4 | 0 | NA | NA | NA | NA | NA | #VALUE! |
| ENSMUSG00000059043.3 | 0 | NA | NA | NA | NA | NA | #VALUE! |
| ENSMUSG00000059049.15 | 0 | NA | NA | NA | NA | NA | #VALUE! |

| | | | | | | | |
|---|---|---|---|---|---|---|---|
| ENSMUSG00000059058.8 | 0.245976 | 1.027628 | 4.067126 | 0.252667 | 0.800526 | NA | #VALUE! |
| ENSMUSG00000059060.16 | 0 | NA | NA | NA | NA | NA | #VALUE! |
| ENSMUSG00000059064.9 | 0 | NA | NA | NA | NA | NA | #VALUE! |
| ENSMUSG00000059069.6 | 0 | NA | NA | NA | NA | NA | #VALUE! |
| ENSMUSG00000059077.7 | 0 | NA | NA | NA | NA | NA | #VALUE! |
| ENSMUSG00000059087.5 | 0 | NA | NA | NA | NA | NA | #VALUE! |
| ENSMUSG00000059105.5 | 0 | NA | NA | NA | NA | NA | #VALUE! |
| ENSMUSG00000059106.6 | 0 | NA | NA | NA | NA | NA | #VALUE! |
| ENSMUSG00000059108.5 | 5.382448 | 1.124158 | 3.765107 | 0.298573 | 0.765266 | NA | #VALUE! |
| ENSMUSG00000059112.7 | 0 | NA | NA | NA | NA | NA | #VALUE! |
| ENSMUSG00000059113.3 | 0 | NA | NA | NA | NA | NA | #VALUE! |
| ENSMUSG00000059114.9 | 0 | NA | NA | NA | NA | NA | #VALUE! |
| ENSMUSG00000059128.4 | 0 | NA | NA | NA | NA | NA | #VALUE! |
| ENSMUSG00000059134.3 | 0 | NA | NA | NA | NA | NA | #VALUE! |
| ENSMUSG00000059136.5 | 0 | NA | NA | NA | NA | NA | #VALUE! |
| ENSMUSG00000059146.13 | 0 | NA | NA | NA | NA | NA | #VALUE! |
| ENSMUSG00000059149.18 | 0 | NA | NA | NA | NA | NA | #VALUE! |
| ENSMUSG00000059150.4 | 0 | NA | NA | NA | NA | NA | #VALUE! |
| ENSMUSG00000059169.10 | 0 | NA | NA | NA | NA | NA | #VALUE! |
| ENSMUSG00000059179.7 | 0 | NA | NA | NA | NA | NA | #VALUE! |
| ENSMUSG00000059187.13 | 0 | NA | NA | NA | NA | NA | #VALUE! |
| ENSMUSG00000059189.4 | 0 | NA | NA | NA | NA | NA | #VALUE! |
| ENSMUSG00000059195.11 | 0 | NA | NA | NA | NA | NA | #VALUE! |
| ENSMUSG00000059201.13 | 0 | NA | NA | NA | NA | NA | #VALUE! |
| ENSMUSG00000059203.11 | 0 | NA | NA | NA | NA | NA | #VALUE! |
| ENSMUSG00000059205.6 | 0 | NA | NA | NA | NA | NA | #VALUE! |
| ENSMUSG00000059206.5 | 0 | NA | NA | NA | NA | NA | #VALUE! |
| ENSMUSG00000059213.8 | 0 | NA | NA | NA | NA | NA | #VALUE! |
| ENSMUSG00000059218.15 | 0 | NA | NA | NA | NA | NA | #VALUE! |
| ENSMUSG00000059229.8 | 0 | NA | NA | NA | NA | NA | #VALUE! |
| ENSMUSG00000059230.3 | 0 | NA | NA | NA | NA | NA | #VALUE! |
| ENSMUSG00000059237.7 | 0 | NA | NA | NA | NA | NA | #VALUE! |
| ENSMUSG00000059244.7 | 0 | NA | NA | NA | NA | NA | #VALUE! |
| ENSMUSG00000059246.5 | 0 | NA | NA | NA | NA | NA | #VALUE! |
| ENSMUSG00000059251.2 | 0 | NA | NA | NA | NA | NA | #VALUE! |
| ENSMUSG00000059252.6 | 0.093255 | 0.868093 | 4.072464 | 0.213162 | 0.831201 | NA | #VALUE! |
| ENSMUSG00000059256.7 | 0 | NA | NA | NA | NA | NA | #VALUE! |
| ENSMUSG00000059277.13 | 0 | NA | NA | NA | NA | NA | #VALUE! |
| ENSMUSG00000059279.12 | 0 | NA | NA | NA | NA | NA | #VALUE! |
| ENSMUSG00000059280.5 | 0 | NA | NA | NA | NA | NA | #VALUE! |
| ENSMUSG00000059288.15 | 9.644778 | 1.159288 | 2.258543 | 0.51329 | 0.607748 | NA | #VALUE! |
| ENSMUSG00000059301.8 | 0 | NA | NA | NA | NA | NA | #VALUE! |
| ENSMUSG00000059303.7 | 0 | NA | NA | NA | NA | NA | #VALUE! |
| ENSMUSG00000059305.13 | 0 | NA | NA | NA | NA | NA | #VALUE! |

| | | | | | | | |
|---|---|---|---|---|---|---|---|
| ENSMUSG00000059319.4 | 0 | NA | NA | NA | NA | NA | #VALUE! |
| ENSMUSG00000059323.15 | 0 | NA | NA | NA | NA | NA | #VALUE! |
| ENSMUSG00000059326.8 | 19.50405 | -3.76816 | 2.309875 | -1.63133 | NA | NA | #VALUE! |
| ENSMUSG00000059327.10 | 0 | NA | NA | NA | NA | NA | #VALUE! |
| ENSMUSG00000059334.7 | 0 | NA | NA | NA | NA | NA | #VALUE! |
| ENSMUSG00000059336.15 | 0 | NA | NA | NA | NA | NA | #VALUE! |
| ENSMUSG00000059343.5 | 0.204111 | -1.05581 | 4.073302 | -0.2592 | 0.795479 | NA | #VALUE! |
| ENSMUSG00000059352.6 | 0.245976 | 1.027628 | 4.067126 | 0.252667 | 0.800526 | NA | #VALUE! |
| ENSMUSG00000059361.7 | 0 | NA | NA | NA | NA | NA | #VALUE! |
| ENSMUSG00000059366.4 | 0 | NA | NA | NA | NA | NA | #VALUE! |
| ENSMUSG00000059371.5 | 0 | NA | NA | NA | NA | NA | #VALUE! |
| ENSMUSG00000059375.6 | 0 | NA | NA | NA | NA | NA | #VALUE! |
| ENSMUSG00000059379.4 | 0 | NA | NA | NA | NA | NA | #VALUE! |
| ENSMUSG00000059382.3 | 0 | NA | NA | NA | NA | NA | #VALUE! |
| ENSMUSG00000059383.15 | 0 | NA | NA | NA | NA | NA | #VALUE! |
| ENSMUSG00000059395.5 | 0 | NA | NA | NA | NA | NA | #VALUE! |
| ENSMUSG00000059397.3 | 0 | NA | NA | NA | NA | NA | #VALUE! |
| ENSMUSG00000059401.15 | 1.68021 | -4.29432 | 3.933015 | -1.09187 | 0.274892 | NA | #VALUE! |
| ENSMUSG00000059406.9 | 0 | NA | NA | NA | NA | NA | #VALUE! |
| ENSMUSG00000059408.4 | 0 | NA | NA | NA | NA | NA | #VALUE! |
| ENSMUSG00000059410.6 | 0 | NA | NA | NA | NA | NA | #VALUE! |
| ENSMUSG00000059411.7 | 0 | NA | NA | NA | NA | NA | #VALUE! |
| ENSMUSG00000059412.8 | 0 | NA | NA | NA | NA | NA | #VALUE! |
| ENSMUSG00000059422.6 | 0.279766 | 1.808031 | 4.036658 | 0.447903 | 0.654223 | NA | #VALUE! |
| ENSMUSG00000059429.4 | 0 | NA | NA | NA | NA | NA | #VALUE! |
| ENSMUSG00000059430.15 | 0 | NA | NA | NA | NA | NA | #VALUE! |
| ENSMUSG00000059455.4 | 0 | NA | NA | NA | NA | NA | #VALUE! |
| ENSMUSG00000059460.8 | 0 | NA | NA | NA | NA | NA | #VALUE! |
| ENSMUSG00000059463.8 | 0 | NA | NA | NA | NA | NA | #VALUE! |
| ENSMUSG00000059473.5 | 0 | NA | NA | NA | NA | NA | #VALUE! |
| ENSMUSG00000059479.8 | 0.737927 | 3.007825 | 3.968641 | 0.757898 | 0.448512 | NA | #VALUE! |
| ENSMUSG00000059483.6 | 0 | NA | NA | NA | NA | NA | #VALUE! |
| ENSMUSG00000059488.4 | 0 | NA | NA | NA | NA | NA | #VALUE! |
| ENSMUSG00000059493.14 | 0 | NA | NA | NA | NA | NA | #VALUE! |
| ENSMUSG00000059503.3 | 0 | NA | NA | NA | NA | NA | #VALUE! |
| ENSMUSG00000059504.8 | 0 | NA | NA | NA | NA | NA | #VALUE! |
| ENSMUSG00000059540.16 | 1.865104 | 4.40601 | 3.930481 | 1.120985 | 0.262294 | NA | #VALUE! |
| ENSMUSG00000059547.4 | 0 | NA | NA | NA | NA | NA | #VALUE! |
| ENSMUSG00000059554.13 | 6.615827 | -6.26 | 3.913512 | -1.59959 | 0.10969 | NA | #VALUE! |
| ENSMUSG00000059555.7 | 1.682835 | 4.240781 | 3.933339 | 1.078163 | 0.280961 | NA | #VALUE! |
| ENSMUSG00000059562.15 | 0 | NA | NA | NA | NA | NA | #VALUE! |
| ENSMUSG00000059565.5 | 0 | NA | NA | NA | NA | NA | #VALUE! |
| ENSMUSG00000059571.8 | 0 | NA | NA | NA | NA | NA | #VALUE! |
| ENSMUSG00000059585.5 | 0 | NA | NA | NA | NA | NA | #VALUE! |

| | | | | | | | |
|---|---|---|---|---|---|---|---|
| ENSMUSG00000059595.3 | 0 | NA | NA | NA | NA | NA | #VALUE! |
| ENSMUSG00000059602.15 | 6.887315 | 6.268935 | 3.913312 | 1.601951 | 0.109166 | NA | #VALUE! |
| ENSMUSG00000059606.4 | 0 | NA | NA | NA | NA | NA | #VALUE! |
| ENSMUSG00000059610.6 | 0 | NA | NA | NA | NA | NA | #VALUE! |
| ENSMUSG00000059623.7 | 0 | NA | NA | NA | NA | NA | #VALUE! |
| ENSMUSG00000059625.7 | 0 | NA | NA | NA | NA | NA | #VALUE! |
| ENSMUSG00000059626.14 | 0 | NA | NA | NA | NA | NA | #VALUE! |
| ENSMUSG00000059631.7 | 0 | NA | NA | NA | NA | NA | #VALUE! |
| ENSMUSG00000059632.5 | 0 | NA | NA | NA | NA | NA | #VALUE! |
| ENSMUSG00000059639.3 | 0 | NA | NA | NA | NA | NA | #VALUE! |
| ENSMUSG00000059645.7 | 0 | NA | NA | NA | NA | NA | #VALUE! |
| ENSMUSG00000059647.6 | 0 | NA | NA | NA | NA | NA | #VALUE! |
| ENSMUSG00000059648.4 | 0 | NA | NA | NA | NA | NA | #VALUE! |
| ENSMUSG00000059654.8 | 0 | NA | NA | NA | NA | NA | #VALUE! |
| ENSMUSG00000059657.6 | 0 | NA | NA | NA | NA | NA | #VALUE! |
| ENSMUSG00000059658.6 | 0.204111 | -1.05581 | 4.073302 | -0.2592 | 0.795479 | NA | #VALUE! |
| ENSMUSG00000059659.8 | 0 | NA | NA | NA | NA | NA | #VALUE! |
| ENSMUSG00000059663.2 | 0 | NA | NA | NA | NA | NA | #VALUE! |
| ENSMUSG00000059668.7 | 0 | NA | NA | NA | NA | NA | #VALUE! |
| ENSMUSG00000059674.8 | 0 | NA | NA | NA | NA | NA | #VALUE! |
| ENSMUSG00000059687.5 | 0 | NA | NA | NA | NA | NA | #VALUE! |
| ENSMUSG00000059690.2 | 0 | NA | NA | NA | NA | NA | #VALUE! |
| ENSMUSG00000059695.5 | 0 | NA | NA | NA | NA | NA | #VALUE! |
| ENSMUSG00000059706.5 | 0 | NA | NA | NA | NA | NA | #VALUE! |
| ENSMUSG00000059713.13 | 7.112571 | -0.08721 | 3.260963 | -0.02674 | 0.978664 | NA | #VALUE! |
| ENSMUSG00000059729.3 | 0 | NA | NA | NA | NA | NA | #VALUE! |
| ENSMUSG00000059741.14 | 0 | NA | NA | NA | NA | NA | #VALUE! |
| ENSMUSG00000059742.11 | 0 | NA | NA | NA | NA | NA | #VALUE! |
| ENSMUSG00000059751.8 | 0.988862 | -0.67189 | 3.913491 | -0.17169 | 0.863684 | NA | #VALUE! |
| ENSMUSG00000059762.5 | 0 | NA | NA | NA | NA | NA | #VALUE! |
| ENSMUSG00000059763.5 | 0 | NA | NA | NA | NA | NA | #VALUE! |
| ENSMUSG00000059768.8 | 0 | NA | NA | NA | NA | NA | #VALUE! |
| ENSMUSG00000059775.6 | 5.221596 | -0.27435 | 2.43642 | -0.1126 | 0.910345 | NA | #VALUE! |
| ENSMUSG00000059776.8 | 0.315039 | -1.93549 | 4.038621 | -0.47925 | 0.631763 | NA | #VALUE! |
| ENSMUSG00000059791.15 | 2.459755 | 4.774963 | 3.925155 | 1.216503 | 0.223793 | NA | #VALUE! |
| ENSMUSG00000059810.19 | 0.783156 | -3.1627 | 3.963922 | -0.79787 | 0.424945 | NA | #VALUE! |
| ENSMUSG00000059816.6 | 0 | NA | NA | NA | NA | NA | #VALUE! |
| ENSMUSG00000059821.7 | 0 | NA | NA | NA | NA | NA | #VALUE! |
| ENSMUSG00000059827.6 | 0 | NA | NA | NA | NA | NA | #VALUE! |
| ENSMUSG00000059830.4 | 0 | NA | NA | NA | NA | NA | #VALUE! |
| ENSMUSG00000059832.5 | 0 | NA | NA | NA | NA | NA | #VALUE! |
| ENSMUSG00000059835.2 | 0 | NA | NA | NA | NA | NA | #VALUE! |
| ENSMUSG00000059842.13 | 1.213812 | 2.488071 | 3.943444 | 0.630939 | 0.528081 | NA | #VALUE! |
| ENSMUSG00000059845.6 | 0 | NA | NA | NA | NA | NA | #VALUE! |

| | | | | | | | |
|---|---|---|---|---|---|---|---|
| ENSMUSG00000059851.16 | 0 | NA | NA | NA | NA | NA | #VALUE! |
| ENSMUSG00000059852.8 | 0 | NA | NA | NA | NA | NA | #VALUE! |
| ENSMUSG00000059854.9 | 0 | NA | NA | NA | NA | NA | #VALUE! |
| ENSMUSG00000059857.16 | 0 | NA | NA | NA | NA | NA | #VALUE! |
| ENSMUSG00000059862.5 | 0 | NA | NA | NA | NA | NA | #VALUE! |
| ENSMUSG00000059864.3 | 0 | NA | NA | NA | NA | NA | #VALUE! |
| ENSMUSG00000059867.4 | 0 | NA | NA | NA | NA | NA | #VALUE! |
| ENSMUSG00000059873.5 | 0 | NA | NA | NA | NA | NA | #VALUE! |
| ENSMUSG00000059874.3 | 0 | NA | NA | NA | NA | NA | #VALUE! |
| ENSMUSG00000059877.4 | 0 | NA | NA | NA | NA | NA | #VALUE! |
| ENSMUSG00000059878.13 | 81.70849 | 2.423399 | 1.421205 | 1.705172 | NA | NA | #VALUE! |
| ENSMUSG00000059887.8 | 0 | NA | NA | NA | NA | NA | #VALUE! |
| ENSMUSG00000059891.15 | 0 | NA | NA | NA | NA | NA | #VALUE! |
| ENSMUSG00000059895.14 | 0.105013 | -0.97324 | 4.076293 | -0.23876 | 0.811295 | NA | #VALUE! |
| ENSMUSG00000059898.10 | 0 | NA | NA | NA | NA | NA | #VALUE! |
| ENSMUSG00000059900.15 | 0 | NA | NA | NA | NA | NA | #VALUE! |
| ENSMUSG00000059901.13 | 0 | NA | NA | NA | NA | NA | #VALUE! |
| ENSMUSG00000059910.5 | 0 | NA | NA | NA | NA | NA | #VALUE! |
| ENSMUSG00000059912.5 | 2.275337 | -4.71144 | 3.926451 | -1.19992 | 0.230169 | NA | #VALUE! |
| ENSMUSG00000059921.16 | 0.168284 | 1.027628 | 4.067126 | 0.252667 | 0.800526 | NA | #VALUE! |
| ENSMUSG00000059934.13 | 0 | NA | NA | NA | NA | NA | #VALUE! |
| ENSMUSG00000059939.14 | 0.816443 | -3.21914 | 3.961753 | -0.81255 | 0.416474 | NA | #VALUE! |
| ENSMUSG00000059956.15 | 0 | NA | NA | NA | NA | NA | #VALUE! |
| ENSMUSG00000059964.10 | 0 | NA | NA | NA | NA | NA | #VALUE! |
| ENSMUSG00000059972.8 | 0 | NA | NA | NA | NA | NA | #VALUE! |
| ENSMUSG00000059981.13 | 5.745991 | -2.54031 | 3.520462 | -0.72158 | 0.47055 | NA | #VALUE! |
| ENSMUSG00000059991.8 | 0 | NA | NA | NA | NA | NA | #VALUE! |
| ENSMUSG00000059994.11 | 0 | NA | NA | NA | NA | NA | #VALUE! |
| ENSMUSG00000059995.13 | 3.666892 | -0.60278 | 3.778215 | -0.15954 | 0.873244 | NA | #VALUE! |
| ENSMUSG00000060017.9 | 0 | NA | NA | NA | NA | NA | #VALUE! |
| ENSMUSG00000060024.3 | 0 | NA | NA | NA | NA | NA | #VALUE! |
| ENSMUSG00000060029.14 | 1.682835 | 4.240781 | 3.933339 | 1.078163 | 0.280961 | NA | #VALUE! |
| ENSMUSG00000060030.6 | 0 | NA | NA | NA | NA | NA | #VALUE! |
| ENSMUSG00000060034.6 | 0 | NA | NA | NA | NA | NA | #VALUE! |
| ENSMUSG00000060044.9 | 5.936516 | 1.165021 | 3.435384 | 0.339124 | 0.734516 | NA | #VALUE! |
| ENSMUSG00000060049.6 | 0 | NA | NA | NA | NA | NA | #VALUE! |
| ENSMUSG00000060057.5 | 0 | NA | NA | NA | NA | NA | #VALUE! |
| ENSMUSG00000060063.10 | 1.995249 | -4.53986 | 3.928926 | -1.1555 | 0.247887 | NA | #VALUE! |
| ENSMUSG00000060068.8 | 0.952962 | -0.12305 | 3.914486 | -0.03144 | 0.974922 | NA | #VALUE! |
| ENSMUSG00000060070.6 | 0 | NA | NA | NA | NA | NA | #VALUE! |
| ENSMUSG00000060081.2 | 0 | NA | NA | NA | NA | NA | #VALUE! |
| ENSMUSG00000060084.5 | 0 | NA | NA | NA | NA | NA | #VALUE! |
| ENSMUSG00000060093.7 | 8.919027 | 6.646734 | 3.731324 | 1.781334 | 0.074858 | NA | #VALUE! |
| ENSMUSG00000060096.6 | 0 | NA | NA | NA | NA | NA | #VALUE! |

| | | | | | | | |
|---|---|---|---|---|---|---|---|
| ENSMUSG00000060105.3 | 0 | NA | NA | NA | NA | NA | #VALUE! |
| ENSMUSG00000060112.5 | 0 | NA | NA | NA | NA | NA | #VALUE! |
| ENSMUSG00000060114.5 | 0 | NA | NA | NA | NA | NA | #VALUE! |
| ENSMUSG00000060128.6 | 0.289044 | -0.14751 | 4.065279 | -0.03629 | 0.971054 | NA | #VALUE! |
| ENSMUSG00000060131.12 | 3.767249 | -4.49487 | 3.910631 | -1.1494 | 0.250392 | NA | #VALUE! |
| ENSMUSG00000060161.10 | 0 | NA | NA | NA | NA | NA | #VALUE! |
| ENSMUSG00000060166.7 | 0 | NA | NA | NA | NA | NA | #VALUE! |
| ENSMUSG00000060170.4 | 0 | NA | NA | NA | NA | NA | #VALUE! |
| ENSMUSG00000060176.5 | 0 | NA | NA | NA | NA | NA | #VALUE! |
| ENSMUSG00000060177.7 | 0 | NA | NA | NA | NA | NA | #VALUE! |
| ENSMUSG00000060180.13 | 0 | NA | NA | NA | NA | NA | #VALUE! |
| ENSMUSG00000060187.5 | 0 | NA | NA | NA | NA | NA | #VALUE! |
| ENSMUSG00000060188.7 | 0 | NA | NA | NA | NA | NA | #VALUE! |
| ENSMUSG00000060198.8 | 1.268308 | 0.631835 | 3.877516 | 0.162948 | 0.870559 | NA | #VALUE! |
| ENSMUSG00000060201.6 | 0 | NA | NA | NA | NA | NA | #VALUE! |
| ENSMUSG00000060204.7 | 0 | NA | NA | NA | NA | NA | #VALUE! |
| ENSMUSG00000060205.4 | 0 | NA | NA | NA | NA | NA | #VALUE! |
| ENSMUSG00000060206.12 | 0 | NA | NA | NA | NA | NA | #VALUE! |
| ENSMUSG00000060208.6 | 0 | NA | NA | NA | NA | NA | #VALUE! |
| ENSMUSG00000060212.14 | 0 | NA | NA | NA | NA | NA | #VALUE! |
| ENSMUSG00000060240.13 | 0 | NA | NA | NA | NA | NA | #VALUE! |
| ENSMUSG00000060244.2 | 2.940114 | 0.399322 | 3.78923 | 0.105383 | 0.916072 | NA | #VALUE! |
| ENSMUSG00000060245.6 | 0 | NA | NA | NA | NA | NA | #VALUE! |
| ENSMUSG00000060254.8 | 0 | NA | NA | NA | NA | NA | #VALUE! |
| ENSMUSG00000060256.2 | 0 | NA | NA | NA | NA | NA | #VALUE! |
| ENSMUSG00000060257.3 | 0 | NA | NA | NA | NA | NA | #VALUE! |
| ENSMUSG00000060260.14 | 2.730341 | -4.98888 | 3.923018 | -1.27169 | 0.203482 | NA | #VALUE! |
| ENSMUSG00000060268.13 | 0 | NA | NA | NA | NA | NA | #VALUE! |
| ENSMUSG00000060275.15 | 0 | NA | NA | NA | NA | NA | #VALUE! |
| ENSMUSG00000060279.15 | 0 | NA | NA | NA | NA | NA | #VALUE! |
| ENSMUSG00000060284.8 | 0 | NA | NA | NA | NA | NA | #VALUE! |
| ENSMUSG00000060303.5 | 0 | NA | NA | NA | NA | NA | #VALUE! |
| ENSMUSG00000060311.12 | 0 | NA | NA | NA | NA | NA | #VALUE! |
| ENSMUSG00000060314.14 | 4.830604 | -5.80779 | 3.915992 | -1.4831 | 0.138049 | NA | #VALUE! |
| ENSMUSG00000060317.10 | 23.75797 | 1.247177 | 2.064801 | 0.604018 | NA | NA | #VALUE! |
| ENSMUSG00000060332.10 | 0 | NA | NA | NA | NA | NA | #VALUE! |
| ENSMUSG00000060335.3 | 0 | NA | NA | NA | NA | NA | #VALUE! |
| ENSMUSG00000060336.7 | 0.962867 | -0.02325 | 3.910747 | -0.00594 | 0.995257 | NA | #VALUE! |
| ENSMUSG00000060375.3 | 0 | NA | NA | NA | NA | NA | #VALUE! |
| ENSMUSG00000060380.4 | 3.029103 | 5.088783 | 3.921574 | 1.297638 | 0.194412 | NA | #VALUE! |
| ENSMUSG00000060402.9 | 0 | NA | NA | NA | NA | NA | #VALUE! |
| ENSMUSG00000060404.4 | 0 | NA | NA | NA | NA | NA | #VALUE! |
| ENSMUSG00000060411.4 | 0 | NA | NA | NA | NA | NA | #VALUE! |
| ENSMUSG00000060412.3 | 0 | NA | NA | NA | NA | NA | #VALUE! |

| | | | | | | | |
|---|---|---|---|---|---|---|---|
| ENSMUSG00000060416.3 | 0 | NA | NA | NA | NA | NA | #VALUE! |
| ENSMUSG00000060417.7 | 0 | NA | NA | NA | NA | NA | #VALUE! |
| ENSMUSG00000060424.15 | 0 | NA | NA | NA | NA | NA | #VALUE! |
| ENSMUSG00000060427.16 | 5.852502 | 5.084622 | 3.432846 | 1.481168 | 0.138562 | NA | #VALUE! |
| ENSMUSG00000060445.12 | 0 | NA | NA | NA | NA | NA | #VALUE! |
| ENSMUSG00000060461.6 | 0 | NA | NA | NA | NA | NA | #VALUE! |
| ENSMUSG00000060467.5 | 0.198268 | -0.06494 | 4.068276 | -0.01596 | 0.987265 | NA | #VALUE! |
| ENSMUSG00000060469.4 | 0 | NA | NA | NA | NA | NA | #VALUE! |
| ENSMUSG00000060470.8 | 8.616538 | 1.10075 | 2.735498 | 0.402395 | 0.687393 | NA | #VALUE! |
| ENSMUSG00000060480.6 | 0 | NA | NA | NA | NA | NA | #VALUE! |
| ENSMUSG00000060487.8 | 6.147935 | -2.00539 | 3.772236 | -0.53162 | 0.594991 | NA | #VALUE! |
| ENSMUSG00000060491.15 | 0 | NA | NA | NA | NA | NA | #VALUE! |
| ENSMUSG00000060499.8 | 0 | NA | NA | NA | NA | NA | #VALUE! |
| ENSMUSG00000060503.7 | 0 | NA | NA | NA | NA | NA | #VALUE! |
| ENSMUSG00000060508.17 | 0 | NA | NA | NA | NA | NA | #VALUE! |
| ENSMUSG00000060509.6 | 0 | NA | NA | NA | NA | NA | #VALUE! |
| ENSMUSG00000060523.6 | 0 | NA | NA | NA | NA | NA | #VALUE! |
| ENSMUSG00000060530.3 | 0 | NA | NA | NA | NA | NA | #VALUE! |
| ENSMUSG00000060534.17 | 0 | NA | NA | NA | NA | NA | #VALUE! |
| ENSMUSG00000060550.17 | 2.925612 | 5.035797 | 3.922125 | 1.283946 | 0.199161 | NA | #VALUE! |
| ENSMUSG00000060556.5 | 0 | NA | NA | NA | NA | NA | #VALUE! |
| ENSMUSG00000060560.9 | 0.105013 | -0.97324 | 4.076293 | -0.23876 | 0.811295 | NA | #VALUE! |
| ENSMUSG00000060565.6 | 0 | NA | NA | NA | NA | NA | #VALUE! |
| ENSMUSG00000060566.5 | 0.612332 | -2.79827 | 3.980109 | -0.70306 | 0.482016 | NA | #VALUE! |
| ENSMUSG00000060568.15 | 0 | NA | NA | NA | NA | NA | #VALUE! |
| ENSMUSG00000060572.18 | 0 | NA | NA | NA | NA | NA | #VALUE! |
| ENSMUSG00000060583.5 | 0 | NA | NA | NA | NA | NA | #VALUE! |
| ENSMUSG00000060594.7 | 0 | NA | NA | NA | NA | NA | #VALUE! |
| ENSMUSG00000060600.16 | 2.809222 | 4.971827 | 3.922819 | 1.267412 | 0.205008 | NA | #VALUE! |
| ENSMUSG00000060615.6 | 0 | NA | NA | NA | NA | NA | #VALUE! |
| ENSMUSG00000060621.7 | 0 | NA | NA | NA | NA | NA | #VALUE! |
| ENSMUSG00000060630.8 | 0 | NA | NA | NA | NA | NA | #VALUE! |
| ENSMUSG00000060647.3 | 0 | NA | NA | NA | NA | NA | #VALUE! |
| ENSMUSG00000060657.9 | 2.25334 | -4.70803 | 3.926497 | -1.19904 | 0.230512 | NA | #VALUE! |
| ENSMUSG00000060663.7 | 0 | NA | NA | NA | NA | NA | #VALUE! |
| ENSMUSG00000060673.3 | 0 | NA | NA | NA | NA | NA | #VALUE! |
| ENSMUSG00000060680.3 | 0 | NA | NA | NA | NA | NA | #VALUE! |
| ENSMUSG00000060685.5 | 0 | NA | NA | NA | NA | NA | #VALUE! |
| ENSMUSG00000060688.5 | 0 | NA | NA | NA | NA | NA | #VALUE! |
| ENSMUSG00000060691.5 | 0 | NA | NA | NA | NA | NA | #VALUE! |
| ENSMUSG00000060699.3 | 0 | NA | NA | NA | NA | NA | #VALUE! |
| ENSMUSG00000060701.2 | 0 | NA | NA | NA | NA | NA | #VALUE! |
| ENSMUSG00000060715.10 | 0 | NA | NA | NA | NA | NA | #VALUE! |
| ENSMUSG00000060716.8 | 0 | NA | NA | NA | NA | NA | #VALUE! |

| | | | | | | | |
|---|---|---|---|---|---|---|---|
| ENSMUSG00000060724.3 | 0 | NA | NA | NA | NA | NA | #VALUE! |
| ENSMUSG00000060726.11 | 0 | NA | NA | NA | NA | NA | #VALUE! |
| ENSMUSG00000060727.5 | 0 | NA | NA | NA | NA | NA | #VALUE! |
| ENSMUSG00000060735.5 | 0 | NA | NA | NA | NA | NA | #VALUE! |
| ENSMUSG00000060738.5 | 0 | NA | NA | NA | NA | NA | #VALUE! |
| ENSMUSG00000060742.2 | 0 | NA | NA | NA | NA | NA | #VALUE! |
| ENSMUSG00000060747.4 | 0 | NA | NA | NA | NA | NA | #VALUE! |
| ENSMUSG00000060756.6 | 0 | NA | NA | NA | NA | NA | #VALUE! |
| ENSMUSG00000060759.5 | 0 | NA | NA | NA | NA | NA | #VALUE! |
| ENSMUSG00000060765.9 | 0 | NA | NA | NA | NA | NA | #VALUE! |
| ENSMUSG00000060777.5 | 0 | NA | NA | NA | NA | NA | #VALUE! |
| ENSMUSG00000060780.3 | 0 | NA | NA | NA | NA | NA | #VALUE! |
| ENSMUSG00000060787.10 | 0 | NA | NA | NA | NA | NA | #VALUE! |
| ENSMUSG00000060794.9 | 0 | NA | NA | NA | NA | NA | #VALUE! |
| ENSMUSG00000060795.6 | 0 | NA | NA | NA | NA | NA | #VALUE! |
| ENSMUSG00000060798.8 | 5.053376 | 1.650668 | 2.901806 | 0.568842 | 0.569464 | NA | #VALUE! |
| ENSMUSG00000060808.5 | 0 | NA | NA | NA | NA | NA | #VALUE! |
| ENSMUSG00000060811.5 | 0 | NA | NA | NA | NA | NA | #VALUE! |
| ENSMUSG00000060816.4 | 0 | NA | NA | NA | NA | NA | #VALUE! |
| ENSMUSG00000060827.3 | 0 | NA | NA | NA | NA | NA | #VALUE! |
| ENSMUSG00000060843.12 | 0 | NA | NA | NA | NA | NA | #VALUE! |
| ENSMUSG00000060860.9 | 0 | NA | NA | NA | NA | NA | #VALUE! |
| ENSMUSG00000060862.11 | 3.700117 | 5.379425 | 3.91888 | 1.372695 | 0.169847 | NA | #VALUE! |
| ENSMUSG00000060863.13 | 0 | NA | NA | NA | NA | NA | #VALUE! |
| ENSMUSG00000060878.5 | 0 | NA | NA | NA | NA | NA | #VALUE! |
| ENSMUSG00000060879.4 | 0 | NA | NA | NA | NA | NA | #VALUE! |
| ENSMUSG00000060882.6 | 0 | NA | NA | NA | NA | NA | #VALUE! |
| ENSMUSG00000060888.5 | 0 | NA | NA | NA | NA | NA | #VALUE! |
| ENSMUSG00000060890.12 | 0 | NA | NA | NA | NA | NA | #VALUE! |
| ENSMUSG00000060913.7 | 0 | NA | NA | NA | NA | NA | #VALUE! |
| ENSMUSG00000060918.3 | 0 | NA | NA | NA | NA | NA | #VALUE! |
| ENSMUSG00000060924.16 | 0 | NA | NA | NA | NA | NA | #VALUE! |
| ENSMUSG00000060956.5 | 0 | NA | NA | NA | NA | NA | #VALUE! |
| ENSMUSG00000060962.13 | 0 | NA | NA | NA | NA | NA | #VALUE! |
| ENSMUSG00000060967.7 | 0 | NA | NA | NA | NA | NA | #VALUE! |
| ENSMUSG00000060969.10 | 1.370524 | -3.97623 | 3.939448 | -1.00934 | 0.312813 | NA | #VALUE! |
| ENSMUSG00000060974.4 | 0 | NA | NA | NA | NA | NA | #VALUE! |
| ENSMUSG00000060981.8 | 2.144869 | 4.605508 | 3.927435 | 1.17265 | 0.240936 | NA | #VALUE! |
| ENSMUSG00000060985.16 | 0 | NA | NA | NA | NA | NA | #VALUE! |
| ENSMUSG00000060988.13 | 0 | NA | NA | NA | NA | NA | #VALUE! |
| ENSMUSG00000060989.9 | 0 | NA | NA | NA | NA | NA | #VALUE! |
| ENSMUSG00000061000.2 | 0 | NA | NA | NA | NA | NA | #VALUE! |
| ENSMUSG00000061013.7 | 0.336567 | 1.944925 | 4.029936 | 0.482619 | 0.629366 | NA | #VALUE! |
| ENSMUSG00000061022.6 | 0 | NA | NA | NA | NA | NA | #VALUE! |

| | | | | | | | |
|---|---|---|---|---|---|---|---|
| ENSMUSG00000061028.8 | 0.245976 | 1.027628 | 4.067126 | 0.252667 | 0.800526 | NA | #VALUE! |
| ENSMUSG00000061039.6 | 1.890236 | -4.46257 | 3.93014 | -1.13547 | 0.256177 | NA | #VALUE! |
| ENSMUSG00000061046.10 | 1.27124 | 3.837908 | 3.94182 | 0.973639 | 0.330236 | NA | #VALUE! |
| ENSMUSG00000061048.9 | 0 | NA | NA | NA | NA | NA | #VALUE! |
| ENSMUSG00000061062.6 | 0 | NA | NA | NA | NA | NA | #VALUE! |
| ENSMUSG00000061065.4 | 0 | NA | NA | NA | NA | NA | #VALUE! |
| ENSMUSG00000061068.8 | 0 | NA | NA | NA | NA | NA | #VALUE! |
| ENSMUSG00000061080.12 | 0 | NA | NA | NA | NA | NA | #VALUE! |
| ENSMUSG00000061082.12 | 0 | NA | NA | NA | NA | NA | #VALUE! |
| ENSMUSG00000061086.13 | 6.160595 | 1.278583 | 3.76158 | 0.339906 | 0.733927 | NA | #VALUE! |
| ENSMUSG00000061099.13 | 0.840105 | -3.30885 | 3.958472 | -0.83589 | 0.403216 | NA | #VALUE! |
| ENSMUSG00000061100.6 | 1.155144 | -3.7605 | 3.944679 | -0.95331 | 0.340433 | NA | #VALUE! |
| ENSMUSG00000061115.5 | 0 | NA | NA | NA | NA | NA | #VALUE! |
| ENSMUSG00000061126.6 | 0 | NA | NA | NA | NA | NA | #VALUE! |
| ENSMUSG00000061143.16 | 3.19204 | -1.3127 | 3.79635 | -0.34578 | 0.729509 | NA | #VALUE! |
| ENSMUSG00000061144.7 | 0 | NA | NA | NA | NA | NA | #VALUE! |
| ENSMUSG00000061165.10 | 0 | NA | NA | NA | NA | NA | #VALUE! |
| ENSMUSG00000061167.10 | 0 | NA | NA | NA | NA | NA | #VALUE! |
| ENSMUSG00000061171.16 | 0 | NA | NA | NA | NA | NA | #VALUE! |
| ENSMUSG00000061184.9 | 0 | NA | NA | NA | NA | NA | #VALUE! |
| ENSMUSG00000061186.16 | 2.890911 | 5.03241 | 3.922161 | 1.283071 | 0.199467 | NA | #VALUE! |
| ENSMUSG00000061195.9 | 0 | NA | NA | NA | NA | NA | #VALUE! |
| ENSMUSG00000061208.7 | 0 | NA | NA | NA | NA | NA | #VALUE! |
| ENSMUSG00000061210.5 | 0 | NA | NA | NA | NA | NA | #VALUE! |
| ENSMUSG00000061242.7 | 0.91911 | 3.358864 | 3.955448 | 0.849174 | 0.395784 | NA | #VALUE! |
| ENSMUSG00000061246.7 | 0 | NA | NA | NA | NA | NA | #VALUE! |
| ENSMUSG00000061259.12 | 0 | NA | NA | NA | NA | NA | #VALUE! |
| ENSMUSG00000061295.6 | 0 | NA | NA | NA | NA | NA | #VALUE! |
| ENSMUSG00000061296.3 | 0 | NA | NA | NA | NA | NA | #VALUE! |
| ENSMUSG00000061298.15 | 0 | NA | NA | NA | NA | NA | #VALUE! |
| ENSMUSG00000061311.7 | 0 | NA | NA | NA | NA | NA | #VALUE! |
| ENSMUSG00000061322.16 | 2.415302 | -4.81324 | 3.925115 | -1.22627 | 0.220098 | NA | #VALUE! |
| ENSMUSG00000061330.9 | 0 | NA | NA | NA | NA | NA | #VALUE! |
| ENSMUSG00000061331.6 | 0 | NA | NA | NA | NA | NA | #VALUE! |
| ENSMUSG00000061336.4 | 0 | NA | NA | NA | NA | NA | #VALUE! |
| ENSMUSG00000061342.6 | 0 | NA | NA | NA | NA | NA | #VALUE! |
| ENSMUSG00000061356.14 | 0 | NA | NA | NA | NA | NA | #VALUE! |
| ENSMUSG00000061361.3 | 0 | NA | NA | NA | NA | NA | #VALUE! |
| ENSMUSG00000061367.4 | 0 | NA | NA | NA | NA | NA | #VALUE! |
| ENSMUSG00000061371.8 | 0.778147 | 1.98536 | 3.968237 | 0.500313 | 0.616855 | NA | #VALUE! |
| ENSMUSG00000061376.3 | 0 | NA | NA | NA | NA | NA | #VALUE! |
| ENSMUSG00000061387.4 | 0 | NA | NA | NA | NA | NA | #VALUE! |
| ENSMUSG00000061388.12 | 0 | NA | NA | NA | NA | NA | #VALUE! |
| ENSMUSG00000061390.7 | 1.119062 | 3.679627 | 3.945843 | 0.932533 | 0.351061 | NA | #VALUE! |

| | | | | | | | |
|---|---|---|---|---|---|---|---|
| ENSMUSG00000061392.3 | 0 | NA | NA | NA | NA | NA | #VALUE! |
| ENSMUSG00000061393.15 | 2.94128 | -2.37375 | 3.841501 | -0.61792 | 0.536626 | NA | #VALUE! |
| ENSMUSG00000061397.9 | 0 | NA | NA | NA | NA | NA | #VALUE! |
| ENSMUSG00000061414.9 | 0 | NA | NA | NA | NA | NA | #VALUE! |
| ENSMUSG00000061436.16 | 7.236322 | 2.383784 | 3.636235 | 0.655564 | 0.512105 | NA | #VALUE! |
| ENSMUSG00000061451.14 | 0.504851 | 2.502843 | 3.993993 | 0.626652 | 0.530887 | NA | #VALUE! |
| ENSMUSG00000061457.4 | 0 | NA | NA | NA | NA | NA | #VALUE! |
| ENSMUSG00000061458.10 | 3.215614 | 5.175564 | 3.920712 | 1.320057 | 0.186816 | NA | #VALUE! |
| ENSMUSG00000061462.19 | 0 | NA | NA | NA | NA | NA | #VALUE! |
| ENSMUSG00000061469.4 | 0 | NA | NA | NA | NA | NA | #VALUE! |
| ENSMUSG00000061474.12 | 8.3679 | 2.470984 | 3.039206 | 0.813036 | 0.416197 | NA | #VALUE! |
| ENSMUSG00000061482.8 | 0 | NA | NA | NA | NA | NA | #VALUE! |
| ENSMUSG00000061486.8 | 0 | NA | NA | NA | NA | NA | #VALUE! |
| ENSMUSG00000061488.6 | 0 | NA | NA | NA | NA | NA | #VALUE! |
| ENSMUSG00000061489.4 | 0 | NA | NA | NA | NA | NA | #VALUE! |
| ENSMUSG00000061490.7 | 0 | NA | NA | NA | NA | NA | #VALUE! |
| ENSMUSG00000061510.4 | 1.224665 | -3.81006 | 3.943408 | -0.96618 | 0.333952 | NA | #VALUE! |
| ENSMUSG00000061517.9 | 0 | NA | NA | NA | NA | NA | #VALUE! |
| ENSMUSG00000061520.5 | 0 | NA | NA | NA | NA | NA | #VALUE! |
| ENSMUSG00000061524.9 | 0 | NA | NA | NA | NA | NA | #VALUE! |
| ENSMUSG00000061525.3 | 0 | NA | NA | NA | NA | NA | #VALUE! |
| ENSMUSG00000061527.8 | 0 | NA | NA | NA | NA | NA | #VALUE! |
| ENSMUSG00000061531.9 | 0 | NA | NA | NA | NA | NA | #VALUE! |
| ENSMUSG00000061533.17 | 0 | NA | NA | NA | NA | NA | #VALUE! |
| ENSMUSG00000061535.12 | 2.310289 | -4.7496 | 3.925939 | -1.2098 | 0.226355 | NA | #VALUE! |
| ENSMUSG00000061549.7 | 0 | NA | NA | NA | NA | NA | #VALUE! |
| ENSMUSG00000061561.6 | 0 | NA | NA | NA | NA | NA | #VALUE! |
| ENSMUSG00000061576.16 | 0 | NA | NA | NA | NA | NA | #VALUE! |
| ENSMUSG00000061577.13 | 0 | NA | NA | NA | NA | NA | #VALUE! |
| ENSMUSG00000061578.9 | 0 | NA | NA | NA | NA | NA | #VALUE! |
| ENSMUSG00000061584.7 | 0 | NA | NA | NA | NA | NA | #VALUE! |
| ENSMUSG00000061589.15 | 2.647228 | 0.367823 | 3.795331 | 0.096915 | 0.922794 | NA | #VALUE! |
| ENSMUSG00000061601.16 | 0 | NA | NA | NA | NA | NA | #VALUE! |
| ENSMUSG00000061602.4 | 0 | NA | NA | NA | NA | NA | #VALUE! |
| ENSMUSG00000061603.9 | 0 | NA | NA | NA | NA | NA | #VALUE! |
| ENSMUSG00000061614.6 | 0 | NA | NA | NA | NA | NA | #VALUE! |
| ENSMUSG00000061615.3 | 0 | NA | NA | NA | NA | NA | #VALUE! |
| ENSMUSG00000061616.6 | 0 | NA | NA | NA | NA | NA | #VALUE! |
| ENSMUSG00000061626.7 | 0 | NA | NA | NA | NA | NA | #VALUE! |
| ENSMUSG00000061633.4 | 0 | NA | NA | NA | NA | NA | #VALUE! |
| ENSMUSG00000061637.6 | 0 | NA | NA | NA | NA | NA | #VALUE! |
| ENSMUSG00000061653.5 | 0 | NA | NA | NA | NA | NA | #VALUE! |
| ENSMUSG00000061666.7 | 1.514552 | 4.088766 | 3.936269 | 1.038742 | 0.298925 | NA | #VALUE! |
| ENSMUSG00000061669.7 | 0.168284 | 1.027628 | 4.067126 | 0.252667 | 0.800526 | NA | #VALUE! |

| | | | | | | | |
|---|---|---|---|---|---|---|---|
| ENSMUSG00000061701.12 | 0 | NA | NA | NA | NA | NA | #VALUE! |
| ENSMUSG00000061702.11 | 0 | NA | NA | NA | NA | NA | #VALUE! |
| ENSMUSG00000061707.6 | 0 | NA | NA | NA | NA | NA | #VALUE! |
| ENSMUSG00000061718.13 | 3.328415 | -5.26129 | 3.920229 | -1.34209 | 0.179567 | NA | #VALUE! |
| ENSMUSG00000061723.19 | 0 | NA | NA | NA | NA | NA | #VALUE! |
| ENSMUSG00000061724.6 | 0 | NA | NA | NA | NA | NA | #VALUE! |
| ENSMUSG00000061728.6 | 0 | NA | NA | NA | NA | NA | #VALUE! |
| ENSMUSG00000061742.13 | 0 | NA | NA | NA | NA | NA | #VALUE! |
| ENSMUSG00000061762.13 | 0 | NA | NA | NA | NA | NA | #VALUE! |
| ENSMUSG00000061769.3 | 0 | NA | NA | NA | NA | NA | #VALUE! |
| ENSMUSG00000061798.3 | 0 | NA | NA | NA | NA | NA | #VALUE! |
| ENSMUSG00000061802.7 | 0 | NA | NA | NA | NA | NA | #VALUE! |
| ENSMUSG00000061809.9 | 0 | NA | NA | NA | NA | NA | #VALUE! |
| ENSMUSG00000061815.12 | 0 | NA | NA | NA | NA | NA | #VALUE! |
| ENSMUSG00000061816.16 | 0 | NA | NA | NA | NA | NA | #VALUE! |
| ENSMUSG00000061829.5 | 0 | NA | NA | NA | NA | NA | #VALUE! |
| ENSMUSG00000061833.7 | 0 | NA | NA | NA | NA | NA | #VALUE! |
| ENSMUSG00000061843.6 | 0 | NA | NA | NA | NA | NA | #VALUE! |
| ENSMUSG00000061845.4 | 0 | NA | NA | NA | NA | NA | #VALUE! |
| ENSMUSG00000061847.3 | 0 | NA | NA | NA | NA | NA | #VALUE! |
| ENSMUSG00000061852.10 | 0 | NA | NA | NA | NA | NA | #VALUE! |
| ENSMUSG00000061863.7 | 0 | NA | NA | NA | NA | NA | #VALUE! |
| ENSMUSG00000061875.3 | 0 | NA | NA | NA | NA | NA | #VALUE! |
| ENSMUSG00000061877.14 | 0 | NA | NA | NA | NA | NA | #VALUE! |
| ENSMUSG00000061878.17 | 0 | NA | NA | NA | NA | NA | #VALUE! |
| ENSMUSG00000061882.13 | 5.394122 | -1.36385 | 3.619114 | -0.37685 | 0.706287 | NA | #VALUE! |
| ENSMUSG00000061897.6 | 0 | NA | NA | NA | NA | NA | #VALUE! |
| ENSMUSG00000061911.16 | 0 | NA | NA | NA | NA | NA | #VALUE! |
| ENSMUSG00000061923.5 | 0 | NA | NA | NA | NA | NA | #VALUE! |
| ENSMUSG00000061928.7 | 0 | NA | NA | NA | NA | NA | #VALUE! |
| ENSMUSG00000061937.8 | 0 | NA | NA | NA | NA | NA | #VALUE! |
| ENSMUSG00000061952.7 | 0 | NA | NA | NA | NA | NA | #VALUE! |
| ENSMUSG00000061958.6 | 0 | NA | NA | NA | NA | NA | #VALUE! |
| ENSMUSG00000061961.4 | 0 | NA | NA | NA | NA | NA | #VALUE! |
| ENSMUSG00000061969.9 | 0 | NA | NA | NA | NA | NA | #VALUE! |
| ENSMUSG00000061972.4 | 0 | NA | NA | NA | NA | NA | #VALUE! |
| ENSMUSG00000061974.3 | 0 | NA | NA | NA | NA | NA | #VALUE! |
| ENSMUSG00000061977.3 | 0 | NA | NA | NA | NA | NA | #VALUE! |
| ENSMUSG00000061979.9 | 7.365071 | 2.7074 | 2.88242 | 0.93928 | 0.347587 | NA | #VALUE! |
| ENSMUSG00000061984.7 | 0 | NA | NA | NA | NA | NA | #VALUE! |
| ENSMUSG00000061988.4 | 0.261539 | 1.68833 | 4.043024 | 0.417591 | 0.676246 | NA | #VALUE! |
| ENSMUSG00000061991.3 | 0 | NA | NA | NA | NA | NA | #VALUE! |
| ENSMUSG00000062007.14 | 0 | NA | NA | NA | NA | NA | #VALUE! |
| ENSMUSG00000062012.15 | 0 | NA | NA | NA | NA | NA | #VALUE! |

| | | | | | | | |
|---|---|---|---|---|---|---|---|
| ENSMUSG00000062017.16 | 0 | NA | NA | NA | NA | NA | #VALUE! |
| ENSMUSG00000062028.9 | 0 | NA | NA | NA | NA | NA | #VALUE! |
| ENSMUSG00000062036.10 | 0 | NA | NA | NA | NA | NA | #VALUE! |
| ENSMUSG00000062037.4 | 0 | NA | NA | NA | NA | NA | #VALUE! |
| ENSMUSG00000062038.6 | 0 | NA | NA | NA | NA | NA | #VALUE! |
| ENSMUSG00000062042.2 | 0 | NA | NA | NA | NA | NA | #VALUE! |
| ENSMUSG00000062044.17 | 0 | NA | NA | NA | NA | NA | #VALUE! |
| ENSMUSG00000062046.2 | 0 | NA | NA | NA | NA | NA | #VALUE! |
| ENSMUSG00000062064.7 | 0 | NA | NA | NA | NA | NA | #VALUE! |
| ENSMUSG00000062073.7 | 0 | NA | NA | NA | NA | NA | #VALUE! |
| ENSMUSG00000062074.8 | 1.714037 | -4.31023 | 3.932729 | -1.09599 | 0.273083 | NA | #VALUE! |
| ENSMUSG00000062077.15 | 0 | NA | NA | NA | NA | NA | #VALUE! |
| ENSMUSG00000062081.7 | 0 | NA | NA | NA | NA | NA | #VALUE! |
| ENSMUSG00000062082.17 | 1.009701 | 3.503687 | 3.950852 | 0.886818 | 0.375177 | NA | #VALUE! |
| ENSMUSG00000062093.6 | 6.993895 | 0.471602 | 2.386961 | 0.197574 | 0.843378 | NA | #VALUE! |
| ENSMUSG00000062103.6 | 0 | NA | NA | NA | NA | NA | #VALUE! |
| ENSMUSG00000062105.6 | 0 | NA | NA | NA | NA | NA | #VALUE! |
| ENSMUSG00000062116.11 | 1.263668 | 3.813247 | 3.942419 | 0.967236 | 0.333426 | NA | #VALUE! |
| ENSMUSG00000062121.4 | 0 | NA | NA | NA | NA | NA | #VALUE! |
| ENSMUSG00000062124.8 | 0 | NA | NA | NA | NA | NA | #VALUE! |
| ENSMUSG00000062128.9 | 0 | NA | NA | NA | NA | NA | #VALUE! |
| ENSMUSG00000062132.11 | 0 | NA | NA | NA | NA | NA | #VALUE! |
| ENSMUSG00000062142.6 | 0 | NA | NA | NA | NA | NA | #VALUE! |
| ENSMUSG00000062148.7 | 0 | NA | NA | NA | NA | NA | #VALUE! |
| ENSMUSG00000062151.14 | 0 | NA | NA | NA | NA | NA | #VALUE! |
| ENSMUSG00000062152.5 | 0 | NA | NA | NA | NA | NA | #VALUE! |
| ENSMUSG00000062154.15 | 0 | NA | NA | NA | NA | NA | #VALUE! |
| ENSMUSG00000062157.7 | 0 | NA | NA | NA | NA | NA | #VALUE! |
| ENSMUSG00000062162.3 | 0 | NA | NA | NA | NA | NA | #VALUE! |
| ENSMUSG00000062165.4 | 0 | NA | NA | NA | NA | NA | #VALUE! |
| ENSMUSG00000062168.13 | 0 | NA | NA | NA | NA | NA | #VALUE! |
| ENSMUSG00000062175.14 | 0 | NA | NA | NA | NA | NA | #VALUE! |
| ENSMUSG00000062184.12 | 0 | NA | NA | NA | NA | NA | #VALUE! |
| ENSMUSG00000062186.3 | 0 | NA | NA | NA | NA | NA | #VALUE! |
| ENSMUSG00000062199.3 | 0 | NA | NA | NA | NA | NA | #VALUE! |
| ENSMUSG00000062200.15 | 0 | NA | NA | NA | NA | NA | #VALUE! |
| ENSMUSG00000062201.5 | 0 | NA | NA | NA | NA | NA | #VALUE! |
| ENSMUSG00000062204.2 | 0 | NA | NA | NA | NA | NA | #VALUE! |
| ENSMUSG00000062209.16 | 3.044572 | -1.36981 | 3.799825 | -0.36049 | 0.718479 | NA | #VALUE! |
| ENSMUSG00000062224.7 | 0 | NA | NA | NA | NA | NA | #VALUE! |
| ENSMUSG00000062226.6 | 0 | NA | NA | NA | NA | NA | #VALUE! |
| ENSMUSG00000062235.7 | 0 | NA | NA | NA | NA | NA | #VALUE! |
| ENSMUSG00000062245.5 | 0 | NA | NA | NA | NA | NA | #VALUE! |
| ENSMUSG00000062252.10 | 0 | NA | NA | NA | NA | NA | #VALUE! |

| | | | | | | | |
|---|---|---|---|---|---|---|---|
| ENSMUSG00000062257.14 | 0 | NA | NA | NA | NA | NA | #VALUE! |
| ENSMUSG00000062272.5 | 0 | NA | NA | NA | NA | NA | #VALUE! |
| ENSMUSG00000062275.8 | 0 | NA | NA | NA | NA | NA | #VALUE! |
| ENSMUSG00000062278.3 | 0 | NA | NA | NA | NA | NA | #VALUE! |
| ENSMUSG00000062279.7 | 0 | NA | NA | NA | NA | NA | #VALUE! |
| ENSMUSG00000062282.5 | 0 | NA | NA | NA | NA | NA | #VALUE! |
| ENSMUSG00000062284.6 | 0 | NA | NA | NA | NA | NA | #VALUE! |
| ENSMUSG00000062296.9 | 0 | NA | NA | NA | NA | NA | #VALUE! |
| ENSMUSG00000062309.9 | 0 | NA | NA | NA | NA | NA | #VALUE! |
| ENSMUSG00000062310.8 | 0 | NA | NA | NA | NA | NA | #VALUE! |
| ENSMUSG00000062312.6 | 0 | NA | NA | NA | NA | NA | #VALUE! |
| ENSMUSG00000062314.4 | 0 | NA | NA | NA | NA | NA | #VALUE! |
| ENSMUSG00000062319.3 | 0 | NA | NA | NA | NA | NA | #VALUE! |
| ENSMUSG00000062327.11 | 0 | NA | NA | NA | NA | NA | #VALUE! |
| ENSMUSG00000062329.5 | 0 | NA | NA | NA | NA | NA | #VALUE! |
| ENSMUSG00000062342.4 | 0 | NA | NA | NA | NA | NA | #VALUE! |
| ENSMUSG00000062345.11 | 0 | NA | NA | NA | NA | NA | #VALUE! |
| ENSMUSG00000062365.5 | 0 | NA | NA | NA | NA | NA | #VALUE! |
| ENSMUSG00000062369.3 | 0 | NA | NA | NA | NA | NA | #VALUE! |
| ENSMUSG00000062372.14 | 0 | NA | NA | NA | NA | NA | #VALUE! |
| ENSMUSG00000062373.9 | 6.70171 | -0.36835 | 2.186469 | -0.16847 | 0.866216 | NA | #VALUE! |
| ENSMUSG00000062380.5 | 0 | NA | NA | NA | NA | NA | #VALUE! |
| ENSMUSG00000062391.4 | 0 | NA | NA | NA | NA | NA | #VALUE! |
| ENSMUSG00000062393.15 | 0 | NA | NA | NA | NA | NA | #VALUE! |
| ENSMUSG00000062400.4 | 0 | NA | NA | NA | NA | NA | #VALUE! |
| ENSMUSG00000062421.14 | 9.189274 | 0.515377 | 3.378633 | 0.15254 | 0.878761 | NA | #VALUE! |
| ENSMUSG00000062426.3 | 0 | NA | NA | NA | NA | NA | #VALUE! |
| ENSMUSG00000062432.5 | 0 | NA | NA | NA | NA | NA | #VALUE! |
| ENSMUSG00000062433.4 | 0 | NA | NA | NA | NA | NA | #VALUE! |
| ENSMUSG00000062434.3 | 0 | NA | NA | NA | NA | NA | #VALUE! |
| ENSMUSG00000062438.5 | 0 | NA | NA | NA | NA | NA | #VALUE! |
| ENSMUSG00000062444.16 | 0 | NA | NA | NA | NA | NA | #VALUE! |
| ENSMUSG00000062458.6 | 0 | NA | NA | NA | NA | NA | #VALUE! |
| ENSMUSG00000062461.8 | 0.475555 | 0.792421 | 4.029408 | 0.196659 | 0.844094 | NA | #VALUE! |
| ENSMUSG00000062464.7 | 0 | NA | NA | NA | NA | NA | #VALUE! |
| ENSMUSG00000062470.11 | 0 | NA | NA | NA | NA | NA | #VALUE! |
| ENSMUSG00000062472.7 | 0 | NA | NA | NA | NA | NA | #VALUE! |
| ENSMUSG00000062477.6 | 0 | NA | NA | NA | NA | NA | #VALUE! |
| ENSMUSG00000062478.18 | 0 | NA | NA | NA | NA | NA | #VALUE! |
| ENSMUSG00000062483.5 | 0 | NA | NA | NA | NA | NA | #VALUE! |
| ENSMUSG00000062496.10 | 0 | NA | NA | NA | NA | NA | #VALUE! |
| ENSMUSG00000062497.4 | 0 | NA | NA | NA | NA | NA | #VALUE! |
| ENSMUSG00000062510.13 | 0.168284 | 1.027628 | 4.067126 | 0.252667 | 0.800526 | NA | #VALUE! |
| ENSMUSG00000062511.7 | 0 | NA | NA | NA | NA | NA | #VALUE! |

| | | | | | | | |
|---|---|---|---|---|---|---|---|
| ENSMUSG00000062518.8 | 0 | NA | NA | NA | NA | NA | #VALUE! |
| ENSMUSG00000062524.5 | 0 | NA | NA | NA | NA | NA | #VALUE! |
| ENSMUSG00000062527.7 | 0 | NA | NA | NA | NA | NA | #VALUE! |
| ENSMUSG00000062528.3 | 0 | NA | NA | NA | NA | NA | #VALUE! |
| ENSMUSG00000062542.12 | 3.197387 | 5.166784 | 3.920797 | 1.317789 | 0.187574 | NA | #VALUE! |
| ENSMUSG00000062543.9 | 0 | NA | NA | NA | NA | NA | #VALUE! |
| ENSMUSG00000062545.5 | 0 | NA | NA | NA | NA | NA | #VALUE! |
| ENSMUSG00000062546.6 | 0 | NA | NA | NA | NA | NA | #VALUE! |
| ENSMUSG00000062551.14 | 0 | NA | NA | NA | NA | NA | #VALUE! |
| ENSMUSG00000062553.4 | 0 | NA | NA | NA | NA | NA | #VALUE! |
| ENSMUSG00000062554.8 | 0 | NA | NA | NA | NA | NA | #VALUE! |
| ENSMUSG00000062556.4 | 0 | NA | NA | NA | NA | NA | #VALUE! |
| ENSMUSG00000062561.4 | 0 | NA | NA | NA | NA | NA | #VALUE! |
| ENSMUSG00000062564.13 | 0 | NA | NA | NA | NA | NA | #VALUE! |
| ENSMUSG00000062582.7 | 4.17547 | -4.64729 | 3.909407 | -1.18875 | 0.23454 | NA | #VALUE! |
| ENSMUSG00000062588.5 | 0.168284 | 1.027628 | 4.067126 | 0.252667 | 0.800526 | NA | #VALUE! |
| ENSMUSG00000062590.14 | 1.224665 | -3.81006 | 3.943408 | -0.96618 | 0.333952 | NA | #VALUE! |
| ENSMUSG00000062591.6 | 0 | NA | NA | NA | NA | NA | #VALUE! |
| ENSMUSG00000062593.18 | 0 | NA | NA | NA | NA | NA | #VALUE! |
| ENSMUSG00000062598.4 | 0 | NA | NA | NA | NA | NA | #VALUE! |
| ENSMUSG00000062608.6 | 0 | NA | NA | NA | NA | NA | #VALUE! |
| ENSMUSG00000062609.15 | 0 | NA | NA | NA | NA | NA | #VALUE! |
| ENSMUSG00000062621.4 | 0 | NA | NA | NA | NA | NA | #VALUE! |
| ENSMUSG00000062629.5 | 0 | NA | NA | NA | NA | NA | #VALUE! |
| ENSMUSG00000062638.12 | 0 | NA | NA | NA | NA | NA | #VALUE! |
| ENSMUSG00000062649.7 | 0 | NA | NA | NA | NA | NA | #VALUE! |
| ENSMUSG00000062651.5 | 0 | NA | NA | NA | NA | NA | #VALUE! |
| ENSMUSG00000062661.7 | 0 | NA | NA | NA | NA | NA | #VALUE! |
| ENSMUSG00000062683.12 | 2.993445 | 3.96985 | 3.915206 | 1.013957 | 0.310603 | NA | #VALUE! |
| ENSMUSG00000062687.7 | 0 | NA | NA | NA | NA | NA | #VALUE! |
| ENSMUSG00000062694.8 | 0 | NA | NA | NA | NA | NA | #VALUE! |
| ENSMUSG00000062704.4 | 7.522541 | 6.398071 | 3.43319 | 1.863594 | 0.062379 | NA | #VALUE! |
| ENSMUSG00000062705.8 | 0 | NA | NA | NA | NA | NA | #VALUE! |
| ENSMUSG00000062712.4 | 0 | NA | NA | NA | NA | NA | #VALUE! |
| ENSMUSG00000062713.9 | 0 | NA | NA | NA | NA | NA | #VALUE! |
| ENSMUSG00000062727.5 | 0 | NA | NA | NA | NA | NA | #VALUE! |
| ENSMUSG00000062729.13 | 3.76629 | 1.166441 | 3.782374 | 0.308389 | 0.757787 | NA | #VALUE! |
| ENSMUSG00000062732.11 | 0 | NA | NA | NA | NA | NA | #VALUE! |
| ENSMUSG00000062737.14 | 0 | NA | NA | NA | NA | NA | #VALUE! |
| ENSMUSG00000062742.5 | 0 | NA | NA | NA | NA | NA | #VALUE! |
| ENSMUSG00000062743.10 | 0.339231 | 1.823695 | 4.035859 | 0.451873 | 0.651361 | NA | #VALUE! |
| ENSMUSG00000062751.6 | 0 | NA | NA | NA | NA | NA | #VALUE! |
| ENSMUSG00000062753.15 | 4.559363 | 0.915866 | 3.769813 | 0.242947 | 0.808046 | NA | #VALUE! |
| ENSMUSG00000062757.3 | 0 | NA | NA | NA | NA | NA | #VALUE! |

| | | | | | | | |
|---|---|---|---|---|---|---|---|
| ENSMUSG00000062758.7 | 0 | NA | NA | NA | NA | NA | #VALUE! |
| ENSMUSG00000062760.12 | 0 | NA | NA | NA | NA | NA | #VALUE! |
| ENSMUSG00000062761.8 | 3.918558 | -2.27285 | 3.807962 | -0.59687 | 0.550595 | NA | #VALUE! |
| ENSMUSG00000062773.7 | 0 | NA | NA | NA | NA | NA | #VALUE! |
| ENSMUSG00000062778.13 | 0 | NA | NA | NA | NA | NA | #VALUE! |
| ENSMUSG00000062782.5 | 0 | NA | NA | NA | NA | NA | #VALUE! |
| ENSMUSG00000062783.3 | 0 | NA | NA | NA | NA | NA | #VALUE! |
| ENSMUSG00000062785.15 | 4.894728 | -5.8188 | 3.915922 | -1.48593 | 0.137296 | NA | #VALUE! |
| ENSMUSG00000062791.5 | 0 | NA | NA | NA | NA | NA | #VALUE! |
| ENSMUSG00000062793.7 | 0 | NA | NA | NA | NA | NA | #VALUE! |
| ENSMUSG00000062794.9 | 1.470184 | -4.10383 | 3.936698 | -1.04245 | 0.297201 | NA | #VALUE! |
| ENSMUSG00000062814.10 | 0 | NA | NA | NA | NA | NA | #VALUE! |
| ENSMUSG00000062818.8 | 3.197387 | 5.166784 | 3.920797 | 1.317789 | 0.187574 | NA | #VALUE! |
| ENSMUSG00000062826.8 | 0 | NA | NA | NA | NA | NA | #VALUE! |
| ENSMUSG00000062833.3 | 0 | NA | NA | NA | NA | NA | #VALUE! |
| ENSMUSG00000062844.7 | 0 | NA | NA | NA | NA | NA | #VALUE! |
| ENSMUSG00000062846.8 | 0 | NA | NA | NA | NA | NA | #VALUE! |
| ENSMUSG00000062859.16 | 0 | NA | NA | NA | NA | NA | #VALUE! |
| ENSMUSG00000062861.9 | 0 | NA | NA | NA | NA | NA | #VALUE! |
| ENSMUSG00000062868.5 | 0 | NA | NA | NA | NA | NA | #VALUE! |
| ENSMUSG00000062873.8 | 0 | NA | NA | NA | NA | NA | #VALUE! |
| ENSMUSG00000062874.7 | 0 | NA | NA | NA | NA | NA | #VALUE! |
| ENSMUSG00000062878.4 | 0 | NA | NA | NA | NA | NA | #VALUE! |
| ENSMUSG00000062892.5 | 0 | NA | NA | NA | NA | NA | #VALUE! |
| ENSMUSG00000062896.4 | 0 | NA | NA | NA | NA | NA | #VALUE! |
| ENSMUSG00000062905.3 | 0 | NA | NA | NA | NA | NA | #VALUE! |
| ENSMUSG00000062906.13 | 3.876264 | 2.448198 | 3.815845 | 0.641587 | 0.521141 | NA | #VALUE! |
| ENSMUSG00000062914.4 | 0 | NA | NA | NA | NA | NA | #VALUE! |
| ENSMUSG00000062933.8 | 1.410586 | 2.965274 | 3.936066 | 0.75336 | 0.451234 | NA | #VALUE! |
| ENSMUSG00000062939.12 | 0 | NA | NA | NA | NA | NA | #VALUE! |
| ENSMUSG00000062952.2 | 0 | NA | NA | NA | NA | NA | #VALUE! |
| ENSMUSG00000062961.8 | 0 | NA | NA | NA | NA | NA | #VALUE! |
| ENSMUSG00000062962.6 | 0 | NA | NA | NA | NA | NA | #VALUE! |
| ENSMUSG00000062987.7 | 0 | NA | NA | NA | NA | NA | #VALUE! |
| ENSMUSG00000062991.10 | 0.105013 | -0.97324 | 4.076293 | -0.23876 | 0.811295 | NA | #VALUE! |
| ENSMUSG00000062995.13 | 0 | NA | NA | NA | NA | NA | #VALUE! |
| ENSMUSG00000063011.8 | 4.921965 | 4.241468 | 3.893596 | 1.089345 | 0.276002 | NA | #VALUE! |
| ENSMUSG00000063018.7 | 0 | NA | NA | NA | NA | NA | #VALUE! |
| ENSMUSG00000063020.3 | 0 | NA | NA | NA | NA | NA | #VALUE! |
| ENSMUSG00000063021.4 | 0 | NA | NA | NA | NA | NA | #VALUE! |
| ENSMUSG00000063058.3 | 0 | NA | NA | NA | NA | NA | #VALUE! |
| ENSMUSG00000063063.13 | 0 | NA | NA | NA | NA | NA | #VALUE! |
| ENSMUSG00000063087.16 | 0.612332 | -2.79827 | 3.980109 | -0.70306 | 0.482016 | NA | #VALUE! |
| ENSMUSG00000063089.5 | 0 | NA | NA | NA | NA | NA | #VALUE! |

| | | | | | | | |
|---|---|---|---|---|---|---|---|
| ENSMUSG00000063106.7 | 0 | NA | NA | NA | NA | NA | #VALUE! |
| ENSMUSG00000063109.14 | 0 | NA | NA | NA | NA | NA | #VALUE! |
| ENSMUSG00000063116.5 | 0 | NA | NA | NA | NA | NA | #VALUE! |
| ENSMUSG00000063120.6 | 0 | NA | NA | NA | NA | NA | #VALUE! |
| ENSMUSG00000063129.2 | 0 | NA | NA | NA | NA | NA | #VALUE! |
| ENSMUSG00000063130.5 | 0 | NA | NA | NA | NA | NA | #VALUE! |
| ENSMUSG00000063133.3 | 0 | NA | NA | NA | NA | NA | #VALUE! |
| ENSMUSG00000063137.6 | 0 | NA | NA | NA | NA | NA | #VALUE! |
| ENSMUSG00000063142.16 | 0 | NA | NA | NA | NA | NA | #VALUE! |
| ENSMUSG00000063146.12 | 0 | NA | NA | NA | NA | NA | #VALUE! |
| ENSMUSG00000063157.10 | 0 | NA | NA | NA | NA | NA | #VALUE! |
| ENSMUSG00000063160.13 | 0 | NA | NA | NA | NA | NA | #VALUE! |
| ENSMUSG00000063163.14 | 0 | NA | NA | NA | NA | NA | #VALUE! |
| ENSMUSG00000063166.5 | 0 | NA | NA | NA | NA | NA | #VALUE! |
| ENSMUSG00000063173.3 | 0 | NA | NA | NA | NA | NA | #VALUE! |
| ENSMUSG00000063176.3 | 0 | NA | NA | NA | NA | NA | #VALUE! |
| ENSMUSG00000063177.7 | 0 | NA | NA | NA | NA | NA | #VALUE! |
| ENSMUSG00000063188.5 | 0 | NA | NA | NA | NA | NA | #VALUE! |
| ENSMUSG00000063193.10 | 1.177985 | 3.72614 | 3.944615 | 0.944614 | 0.344856 | NA | #VALUE! |
| ENSMUSG00000063206.5 | 0 | NA | NA | NA | NA | NA | #VALUE! |
| ENSMUSG00000063216.4 | 0 | NA | NA | NA | NA | NA | #VALUE! |
| ENSMUSG00000063221.7 | 0 | NA | NA | NA | NA | NA | #VALUE! |
| ENSMUSG00000063225.2 | 0 | NA | NA | NA | NA | NA | #VALUE! |
| ENSMUSG00000063230.6 | 0 | NA | NA | NA | NA | NA | #VALUE! |
| ENSMUSG00000063234.6 | 0 | NA | NA | NA | NA | NA | #VALUE! |
| ENSMUSG00000063239.18 | 0 | NA | NA | NA | NA | NA | #VALUE! |
| ENSMUSG00000063240.7 | 0 | NA | NA | NA | NA | NA | #VALUE! |
| ENSMUSG00000063245.12 | 0 | NA | NA | NA | NA | NA | #VALUE! |
| ENSMUSG00000063251.7 | 0 | NA | NA | NA | NA | NA | #VALUE! |
| ENSMUSG00000063252.8 | 0 | NA | NA | NA | NA | NA | #VALUE! |
| ENSMUSG00000063260.3 | 0 | NA | NA | NA | NA | NA | #VALUE! |
| ENSMUSG00000063277.8 | 0 | NA | NA | NA | NA | NA | #VALUE! |
| ENSMUSG00000063286.8 | 5.843947 | 6.031264 | 3.853432 | 1.565167 | 0.117544 | NA | #VALUE! |
| ENSMUSG00000063296.6 | 0 | NA | NA | NA | NA | NA | #VALUE! |
| ENSMUSG00000063297.8 | 0 | NA | NA | NA | NA | NA | #VALUE! |
| ENSMUSG00000063305.11 | 0 | NA | NA | NA | NA | NA | #VALUE! |
| ENSMUSG00000063314.6 | 0 | NA | NA | NA | NA | NA | #VALUE! |
| ENSMUSG00000063328.7 | 4.728922 | 0.385579 | 3.25113 | 0.118599 | 0.905593 | NA | #VALUE! |
| ENSMUSG00000063350.7 | 0 | NA | NA | NA | NA | NA | #VALUE! |
| ENSMUSG00000063364.11 | 0 | NA | NA | NA | NA | NA | #VALUE! |
| ENSMUSG00000063374.5 | 0 | NA | NA | NA | NA | NA | #VALUE! |
| ENSMUSG00000063376.7 | 0 | NA | NA | NA | NA | NA | #VALUE! |
| ENSMUSG00000063380.4 | 0 | NA | NA | NA | NA | NA | #VALUE! |
| ENSMUSG00000063383.6 | 0.245976 | 1.027628 | 4.067126 | 0.252667 | 0.800526 | NA | #VALUE! |

| | | | | | | | |
|---|---|---|---|---|---|---|---|
| ENSMUSG00000063386.3 | 0 | NA | NA | NA | NA | NA | #VALUE! |
| ENSMUSG00000063388.6 | 5.51099 | -5.9895 | 3.914902 | -1.52992 | 0.126035 | NA | #VALUE! |
| ENSMUSG00000063394.4 | 0 | NA | NA | NA | NA | NA | #VALUE! |
| ENSMUSG00000063409.13 | 0 | NA | NA | NA | NA | NA | #VALUE! |
| ENSMUSG00000063430.10 | 0 | NA | NA | NA | NA | NA | #VALUE! |
| ENSMUSG00000063434.7 | 0 | NA | NA | NA | NA | NA | #VALUE! |
| ENSMUSG00000063442.6 | 1.836997 | -4.39907 | 3.931186 | -1.11902 | 0.263133 | NA | #VALUE! |
| ENSMUSG00000063446.5 | 0 | NA | NA | NA | NA | NA | #VALUE! |
| ENSMUSG00000063447.8 | 0 | NA | NA | NA | NA | NA | #VALUE! |
| ENSMUSG00000063455.17 | 7.134205 | 6.32387 | 3.008995 | 2.101656 | 0.035583 | NA | #VALUE! |
| ENSMUSG00000063458.14 | 0 | NA | NA | NA | NA | NA | #VALUE! |
| ENSMUSG00000063478.3 | 0 | NA | NA | NA | NA | NA | #VALUE! |
| ENSMUSG00000063506.15 | 0 | NA | NA | NA | NA | NA | #VALUE! |
| ENSMUSG00000063507.6 | 0 | NA | NA | NA | NA | NA | #VALUE! |
| ENSMUSG00000063514.4 | 0 | NA | NA | NA | NA | NA | #VALUE! |
| ENSMUSG00000063522.6 | 0 | NA | NA | NA | NA | NA | #VALUE! |
| ENSMUSG00000063529.4 | 6.069462 | -6.1296 | 3.91415 | -1.56601 | 0.117346 | NA | #VALUE! |
| ENSMUSG00000063531.8 | 0 | NA | NA | NA | NA | NA | #VALUE! |
| ENSMUSG00000063535.8 | 0 | NA | NA | NA | NA | NA | #VALUE! |
| ENSMUSG00000063543.6 | 0 | NA | NA | NA | NA | NA | #VALUE! |
| ENSMUSG00000063548.4 | 0.245976 | 1.027628 | 4.067126 | 0.252667 | 0.800526 | NA | #VALUE! |
| ENSMUSG00000063549.5 | 0 | NA | NA | NA | NA | NA | #VALUE! |
| ENSMUSG00000063556.6 | 0.728145 | -0.16377 | 3.969585 | -0.04126 | 0.967091 | NA | #VALUE! |
| ENSMUSG00000063564.14 | 0 | NA | NA | NA | NA | NA | #VALUE! |
| ENSMUSG00000063568.12 | 0 | NA | NA | NA | NA | NA | #VALUE! |
| ENSMUSG00000063582.3 | 0 | NA | NA | NA | NA | NA | #VALUE! |
| ENSMUSG00000063583.7 | 0 | NA | NA | NA | NA | NA | #VALUE! |
| ENSMUSG00000063586.2 | 0 | NA | NA | NA | NA | NA | #VALUE! |
| ENSMUSG00000063594.6 | 0 | NA | NA | NA | NA | NA | #VALUE! |
| ENSMUSG00000063600.15 | 0.336567 | 1.944925 | 4.029936 | 0.482619 | 0.629366 | NA | #VALUE! |
| ENSMUSG00000063605.6 | 0 | NA | NA | NA | NA | NA | #VALUE! |
| ENSMUSG00000063611.6 | 1.260158 | -3.88425 | 3.941584 | -0.98545 | 0.324401 | NA | #VALUE! |
| ENSMUSG00000063615.7 | 0 | NA | NA | NA | NA | NA | #VALUE! |
| ENSMUSG00000063623.5 | 3.434336 | 5.276494 | 3.919773 | 1.346123 | 0.178263 | NA | #VALUE! |
| ENSMUSG00000063626.7 | 0 | NA | NA | NA | NA | NA | #VALUE! |
| ENSMUSG00000063628.3 | 0 | NA | NA | NA | NA | NA | #VALUE! |
| ENSMUSG00000063632.7 | 0 | NA | NA | NA | NA | NA | #VALUE! |
| ENSMUSG00000063646.19 | 0 | NA | NA | NA | NA | NA | #VALUE! |
| ENSMUSG00000063651.12 | 0 | NA | NA | NA | NA | NA | #VALUE! |
| ENSMUSG00000063652.11 | 0 | NA | NA | NA | NA | NA | #VALUE! |
| ENSMUSG00000063656.7 | 0 | NA | NA | NA | NA | NA | #VALUE! |
| ENSMUSG00000063660.5 | 0 | NA | NA | NA | NA | NA | #VALUE! |
| ENSMUSG00000063661.7 | 0 | NA | NA | NA | NA | NA | #VALUE! |
| ENSMUSG00000063672.8 | 0 | NA | NA | NA | NA | NA | #VALUE! |

| | | | | | | | |
|---|---|---|---|---|---|---|---|
| ENSMUSG00000063681.15 | 0 | NA | NA | NA | NA | NA | #VALUE! |
| ENSMUSG00000063684.10 | 0 | NA | NA | NA | NA | NA | #VALUE! |
| ENSMUSG00000063687.4 | 0.195789 | -1.05581 | 4.073302 | -0.2592 | 0.795479 | NA | #VALUE! |
| ENSMUSG00000063689.5 | 0 | NA | NA | NA | NA | NA | #VALUE! |
| ENSMUSG00000063696.8 | 0.568183 | -1.10781 | 4.005278 | -0.27659 | 0.782096 | NA | #VALUE! |
| ENSMUSG00000063698.10 | 2.355969 | 4.72622 | 3.925784 | 1.203892 | 0.228631 | NA | #VALUE! |
| ENSMUSG00000063704.14 | 0 | NA | NA | NA | NA | NA | #VALUE! |
| ENSMUSG00000063713.8 | 0 | NA | NA | NA | NA | NA | #VALUE! |
| ENSMUSG00000063714.16 | 9.3163 | 0.990127 | 2.959917 | 0.334512 | 0.737993 | NA | #VALUE! |
| ENSMUSG00000063715.3 | 0 | NA | NA | NA | NA | NA | #VALUE! |
| ENSMUSG00000063728.11 | 0 | NA | NA | NA | NA | NA | #VALUE! |
| ENSMUSG00000063739.3 | 0 | NA | NA | NA | NA | NA | #VALUE! |
| ENSMUSG00000063751.7 | 0 | NA | NA | NA | NA | NA | #VALUE! |
| ENSMUSG00000063754.8 | 2.553969 | 1.691997 | 3.674788 | 0.460434 | 0.645205 | NA | #VALUE! |
| ENSMUSG00000063762.6 | 0 | NA | NA | NA | NA | NA | #VALUE! |
| ENSMUSG00000063764.3 | 0 | NA | NA | NA | NA | NA | #VALUE! |
| ENSMUSG00000063765.13 | 3.197682 | 5.156529 | 3.920897 | 1.31514 | 0.188463 | NA | #VALUE! |
| ENSMUSG00000063767.5 | 0 | NA | NA | NA | NA | NA | #VALUE! |
| ENSMUSG00000063777.4 | 0 | NA | NA | NA | NA | NA | #VALUE! |
| ENSMUSG00000063779.7 | 0 | NA | NA | NA | NA | NA | #VALUE! |
| ENSMUSG00000063780.8 | 0 | NA | NA | NA | NA | NA | #VALUE! |
| ENSMUSG00000063796.10 | 0 | NA | NA | NA | NA | NA | #VALUE! |
| ENSMUSG00000063802.6 | 1.957891 | -4.49332 | 3.929649 | -1.14344 | 0.252855 | NA | #VALUE! |
| ENSMUSG00000063804.9 | 0 | NA | NA | NA | NA | NA | #VALUE! |
| ENSMUSG00000063810.8 | 0 | NA | NA | NA | NA | NA | #VALUE! |
| ENSMUSG00000063820.12 | 0 | NA | NA | NA | NA | NA | #VALUE! |
| ENSMUSG00000063821.7 | 0 | NA | NA | NA | NA | NA | #VALUE! |
| ENSMUSG00000063823.3 | 0 | NA | NA | NA | NA | NA | #VALUE! |
| ENSMUSG00000063827.3 | 0 | NA | NA | NA | NA | NA | #VALUE! |
| ENSMUSG00000063830.6 | 0 | NA | NA | NA | NA | NA | #VALUE! |
| ENSMUSG00000063838.7 | 2.374196 | 4.738398 | 3.925625 | 1.207043 | 0.227416 | NA | #VALUE! |
| ENSMUSG00000063842.6 | 0 | NA | NA | NA | NA | NA | #VALUE! |
| ENSMUSG00000063844.6 | 0 | NA | NA | NA | NA | NA | #VALUE! |
| ENSMUSG00000063846.9 | 0 | NA | NA | NA | NA | NA | #VALUE! |
| ENSMUSG00000063851.13 | 0 | NA | NA | NA | NA | NA | #VALUE! |
| ENSMUSG00000063867.3 | 0 | NA | NA | NA | NA | NA | #VALUE! |
| ENSMUSG00000063873.11 | 0 | NA | NA | NA | NA | NA | #VALUE! |
| ENSMUSG00000063875.7 | 5.82603 | -6.07014 | 3.826235 | -1.58645 | 0.112637 | NA | #VALUE! |
| ENSMUSG00000063881.12 | 0 | NA | NA | NA | NA | NA | #VALUE! |
| ENSMUSG00000063884.7 | 8.240755 | 1.157412 | 2.693682 | 0.429677 | 0.667431 | NA | #VALUE! |
| ENSMUSG00000063885.6 | 0 | NA | NA | NA | NA | NA | #VALUE! |
| ENSMUSG00000063887.14 | 0 | NA | NA | NA | NA | NA | #VALUE! |
| ENSMUSG00000063895.8 | 0 | NA | NA | NA | NA | NA | #VALUE! |
| ENSMUSG00000063900.6 | 0 | NA | NA | NA | NA | NA | #VALUE! |

| | | | | | | | |
|---|---|---|---|---|---|---|---|
| ENSMUSG00000063902.14 | 0 | NA | NA | NA | NA | NA | #VALUE! |
| ENSMUSG00000063903.6 | 0 | NA | NA | NA | NA | NA | #VALUE! |
| ENSMUSG00000063905.3 | 0 | NA | NA | NA | NA | NA | #VALUE! |
| ENSMUSG00000063911.5 | 0 | NA | NA | NA | NA | NA | #VALUE! |
| ENSMUSG00000063919.12 | 14.07176 | -21.5772 | 3.213417 | -6.71473 | NA | NA | #VALUE! |
| ENSMUSG00000063929.7 | 0 | NA | NA | NA | NA | NA | #VALUE! |
| ENSMUSG00000063932.6 | 0 | NA | NA | NA | NA | NA | #VALUE! |
| ENSMUSG00000063935.4 | 0 | NA | NA | NA | NA | NA | #VALUE! |
| ENSMUSG00000063954.8 | 0 | NA | NA | NA | NA | NA | #VALUE! |
| ENSMUSG00000063971.8 | 0 | NA | NA | NA | NA | NA | #VALUE! |
| ENSMUSG00000063975.14 | 0 | NA | NA | NA | NA | NA | #VALUE! |
| ENSMUSG00000063994.6 | 0 | NA | NA | NA | NA | NA | #VALUE! |
| ENSMUSG00000063998.5 | 0 | NA | NA | NA | NA | NA | #VALUE! |
| ENSMUSG00000064006.7 | 0 | NA | NA | NA | NA | NA | #VALUE! |
| ENSMUSG00000064010.13 | 0 | NA | NA | NA | NA | NA | #VALUE! |
| ENSMUSG00000064016.5 | 0 | NA | NA | NA | NA | NA | #VALUE! |
| ENSMUSG00000064023.5 | 0.673134 | 2.918386 | 3.972531 | 0.734641 | 0.462558 | NA | #VALUE! |
| ENSMUSG00000064030.16 | 0 | NA | NA | NA | NA | NA | #VALUE! |
| ENSMUSG00000064036.16 | 0 | NA | NA | NA | NA | NA | #VALUE! |
| ENSMUSG00000064039.5 | 0 | NA | NA | NA | NA | NA | #VALUE! |
| ENSMUSG00000064043.16 | 0 | NA | NA | NA | NA | NA | #VALUE! |
| ENSMUSG00000064044.3 | 0 | NA | NA | NA | NA | NA | #VALUE! |
| ENSMUSG00000064052.4 | 0 | NA | NA | NA | NA | NA | #VALUE! |
| ENSMUSG00000064057.13 | 0 | NA | NA | NA | NA | NA | #VALUE! |
| ENSMUSG00000064061.14 | 2.256848 | 3.503118 | 3.921259 | 0.893366 | 0.371661 | NA | #VALUE! |
| ENSMUSG00000064063.7 | 0 | NA | NA | NA | NA | NA | #VALUE! |
| ENSMUSG00000064065.16 | 0 | NA | NA | NA | NA | NA | #VALUE! |
| ENSMUSG00000064080.13 | 0.408222 | -2.20205 | 4.016842 | -0.54821 | 0.583551 | NA | #VALUE! |
| ENSMUSG00000064081.5 | 0 | NA | NA | NA | NA | NA | #VALUE! |
| ENSMUSG00000064084.3 | 0.245976 | 1.027628 | 4.067126 | 0.252667 | 0.800526 | NA | #VALUE! |
| ENSMUSG00000064090.15 | 7.922832 | -0.27609 | 2.858644 | -0.09658 | 0.923059 | NA | #VALUE! |
| ENSMUSG00000064105.14 | 2.705871 | -1.1217 | 3.803699 | -0.2949 | 0.768072 | NA | #VALUE! |
| ENSMUSG00000064109.9 | 0 | NA | NA | NA | NA | NA | #VALUE! |
| ENSMUSG00000064110.3 | 0 | NA | NA | NA | NA | NA | #VALUE! |
| ENSMUSG00000064115.14 | 0 | NA | NA | NA | NA | NA | #VALUE! |
| ENSMUSG00000064121.5 | 0 | NA | NA | NA | NA | NA | #VALUE! |
| ENSMUSG00000064125.15 | 0 | NA | NA | NA | NA | NA | #VALUE! |
| ENSMUSG00000064129.5 | 0 | NA | NA | NA | NA | NA | #VALUE! |
| ENSMUSG00000064137.7 | 0 | NA | NA | NA | NA | NA | #VALUE! |
| ENSMUSG00000064140.4 | 0 | NA | NA | NA | NA | NA | #VALUE! |
| ENSMUSG00000064141.15 | 0 | NA | NA | NA | NA | NA | #VALUE! |
| ENSMUSG00000064147.8 | 0.420053 | -2.33493 | 4.007343 | -0.58266 | 0.56012 | NA | #VALUE! |
| ENSMUSG00000064156.9 | 0 | NA | NA | NA | NA | NA | #VALUE! |
| ENSMUSG00000064158.7 | 0 | NA | NA | NA | NA | NA | #VALUE! |

| | | | | | | | |
|---|---|---|---|---|---|---|---|
| ENSMUSG00000064165.9 | 0 | NA | NA | NA | NA | NA | #VALUE! |
| ENSMUSG00000064177.9 | 0 | NA | NA | NA | NA | NA | #VALUE! |
| ENSMUSG00000064179.14 | 2.920362 | -0.04549 | 3.788548 | -0.01201 | 0.990419 | NA | #VALUE! |
| ENSMUSG00000064193.9 | 0 | NA | NA | NA | NA | NA | #VALUE! |
| ENSMUSG00000064194.11 | 0 | NA | NA | NA | NA | NA | #VALUE! |
| ENSMUSG00000064201.9 | 0 | NA | NA | NA | NA | NA | #VALUE! |
| ENSMUSG00000064208.7 | 0 | NA | NA | NA | NA | NA | #VALUE! |
| ENSMUSG00000064210.8 | 0.945118 | -3.47564 | 3.952879 | -0.87927 | 0.379256 | NA | #VALUE! |
| ENSMUSG00000064213.6 | 0 | NA | NA | NA | NA | NA | #VALUE! |
| ENSMUSG00000064220.7 | 0 | NA | NA | NA | NA | NA | #VALUE! |
| ENSMUSG00000064223.5 | 0 | NA | NA | NA | NA | NA | #VALUE! |
| ENSMUSG00000064224.11 | 0 | NA | NA | NA | NA | NA | #VALUE! |
| ENSMUSG00000064231.2 | 0.75813 | -0.78481 | 3.957593 | -0.1983 | 0.842807 | NA | #VALUE! |
| ENSMUSG00000064232.3 | 0 | NA | NA | NA | NA | NA | #VALUE! |
| ENSMUSG00000064246.11 | 0 | NA | NA | NA | NA | NA | #VALUE! |
| ENSMUSG00000064247.15 | 5.72164 | 6.006287 | 3.914617 | 1.534323 | 0.12495 | NA | #VALUE! |
| ENSMUSG00000064252.10 | 0 | NA | NA | NA | NA | NA | #VALUE! |
| ENSMUSG00000064259.3 | 0 | NA | NA | NA | NA | NA | #VALUE! |
| ENSMUSG00000064263.7 | 0 | NA | NA | NA | NA | NA | #VALUE! |
| ENSMUSG00000064264.15 | 0 | NA | NA | NA | NA | NA | #VALUE! |
| ENSMUSG00000064267.14 | 1.009701 | 3.503687 | 3.950852 | 0.886818 | 0.375177 | NA | #VALUE! |
| ENSMUSG00000064272.4 | 0 | NA | NA | NA | NA | NA | #VALUE! |
| ENSMUSG00000064280.15 | 1.575197 | -4.20222 | 3.934736 | -1.06798 | 0.285529 | NA | #VALUE! |
| ENSMUSG00000064281.8 | 0 | NA | NA | NA | NA | NA | #VALUE! |
| ENSMUSG00000064288.5 | 0 | NA | NA | NA | NA | NA | #VALUE! |
| ENSMUSG00000064293.15 | 0 | NA | NA | NA | NA | NA | #VALUE! |
| ENSMUSG00000064299.7 | 0 | NA | NA | NA | NA | NA | #VALUE! |
| ENSMUSG00000064308.11 | 0 | NA | NA | NA | NA | NA | #VALUE! |
| ENSMUSG00000064310.11 | 0 | NA | NA | NA | NA | NA | #VALUE! |
| ENSMUSG00000064317.6 | 0.093255 | 0.868093 | 4.072464 | 0.213162 | 0.831201 | NA | #VALUE! |
| ENSMUSG00000064329.14 | 0 | NA | NA | NA | NA | NA | #VALUE! |
| ENSMUSG00000064330.10 | 0 | NA | NA | NA | NA | NA | #VALUE! |
| ENSMUSG00000064333.3 | 0 | NA | NA | NA | NA | NA | #VALUE! |
| ENSMUSG00000064336.1 | 0 | NA | NA | NA | NA | NA | #VALUE! |
| ENSMUSG00000064338.1 | 0 | NA | NA | NA | NA | NA | #VALUE! |
| ENSMUSG00000064340.1 | 0 | NA | NA | NA | NA | NA | #VALUE! |
| ENSMUSG00000064347.1 | 1.995249 | -4.53986 | 3.928926 | -1.1555 | 0.247887 | NA | #VALUE! |
| ENSMUSG00000064348.1 | 0 | NA | NA | NA | NA | NA | #VALUE! |
| ENSMUSG00000064349.1 | 0 | NA | NA | NA | NA | NA | #VALUE! |
| ENSMUSG00000064352.1 | 2.857551 | -5.03942 | 3.922461 | -1.28476 | 0.198876 | NA | #VALUE! |
| ENSMUSG00000064353.1 | 0 | NA | NA | NA | NA | NA | #VALUE! |
| ENSMUSG00000064354.1 | 0 | NA | NA | NA | NA | NA | #VALUE! |
| ENSMUSG00000064355.1 | 0 | NA | NA | NA | NA | NA | #VALUE! |
| ENSMUSG00000064356.1 | 0 | NA | NA | NA | NA | NA | #VALUE! |

| | | | | | | | |
|---|---|---|---|---|---|---|---|
| ENSMUSG00000064357.1 | 0 | NA | NA | NA | NA | NA | #VALUE! |
| ENSMUSG00000064358.1 | 0 | NA | NA | NA | NA | NA | #VALUE! |
| ENSMUSG00000064359.1 | 0 | NA | NA | NA | NA | NA | #VALUE! |
| ENSMUSG00000064360.1 | 2.289475 | 4.686787 | 3.926308 | 1.193688 | 0.2326 | NA | #VALUE! |
| ENSMUSG00000064361.1 | 0 | NA | NA | NA | NA | NA | #VALUE! |
| ENSMUSG00000064364.1 | 0 | NA | NA | NA | NA | NA | #VALUE! |
| ENSMUSG00000064365.1 | 0 | NA | NA | NA | NA | NA | #VALUE! |
| ENSMUSG00000064366.1 | 0 | NA | NA | NA | NA | NA | #VALUE! |
| ENSMUSG00000064369.1 | 8.121028 | 0.561585 | 2.699808 | 0.208009 | 0.835222 | NA | #VALUE! |
| ENSMUSG00000064371.1 | 0 | NA | NA | NA | NA | NA | #VALUE! |
| ENSMUSG00000064377.3 | 0 | NA | NA | NA | NA | NA | #VALUE! |
| ENSMUSG00000064380.3 | 0 | NA | NA | NA | NA | NA | #VALUE! |
| ENSMUSG00000064382.3 | 0 | NA | NA | NA | NA | NA | #VALUE! |
| ENSMUSG00000064386.4 | 0 | NA | NA | NA | NA | NA | #VALUE! |
| ENSMUSG00000064387.3 | 0 | NA | NA | NA | NA | NA | #VALUE! |
| ENSMUSG00000064389.3 | 0 | NA | NA | NA | NA | NA | #VALUE! |
| ENSMUSG00000064390.3 | 0 | NA | NA | NA | NA | NA | #VALUE! |
| ENSMUSG00000064391.4 | 0 | NA | NA | NA | NA | NA | #VALUE! |
| ENSMUSG00000064392.3 | 0 | NA | NA | NA | NA | NA | #VALUE! |
| ENSMUSG00000064394.3 | 0 | NA | NA | NA | NA | NA | #VALUE! |
| ENSMUSG00000064395.3 | 0 | NA | NA | NA | NA | NA | #VALUE! |
| ENSMUSG00000064398.3 | 0 | NA | NA | NA | NA | NA | #VALUE! |
| ENSMUSG00000064399.3 | 0 | NA | NA | NA | NA | NA | #VALUE! |
| ENSMUSG00000064400.4 | 0 | NA | NA | NA | NA | NA | #VALUE! |
| ENSMUSG00000064403.4 | 0 | NA | NA | NA | NA | NA | #VALUE! |
| ENSMUSG00000064405.3 | 0 | NA | NA | NA | NA | NA | #VALUE! |
| ENSMUSG00000064407.4 | 0 | NA | NA | NA | NA | NA | #VALUE! |
| ENSMUSG00000064408.3 | 0 | NA | NA | NA | NA | NA | #VALUE! |
| ENSMUSG00000064410.2 | 0 | NA | NA | NA | NA | NA | #VALUE! |
| ENSMUSG00000064413.4 | 0 | NA | NA | NA | NA | NA | #VALUE! |
| ENSMUSG00000064416.3 | 0 | NA | NA | NA | NA | NA | #VALUE! |
| ENSMUSG00000064417.3 | 0 | NA | NA | NA | NA | NA | #VALUE! |
| ENSMUSG00000064419.3 | 0 | NA | NA | NA | NA | NA | #VALUE! |
| ENSMUSG00000064427.3 | 0 | NA | NA | NA | NA | NA | #VALUE! |
| ENSMUSG00000064432.3 | 0 | NA | NA | NA | NA | NA | #VALUE! |
| ENSMUSG00000064435.3 | 0 | NA | NA | NA | NA | NA | #VALUE! |
| ENSMUSG00000064436.3 | 0 | NA | NA | NA | NA | NA | #VALUE! |
| ENSMUSG00000064437.4 | 0 | NA | NA | NA | NA | NA | #VALUE! |
| ENSMUSG00000064440.3 | 0 | NA | NA | NA | NA | NA | #VALUE! |
| ENSMUSG00000064441.3 | 0 | NA | NA | NA | NA | NA | #VALUE! |
| ENSMUSG00000064442.3 | 0 | NA | NA | NA | NA | NA | #VALUE! |
| ENSMUSG00000064443.3 | 0 | NA | NA | NA | NA | NA | #VALUE! |
| ENSMUSG00000064444.3 | 0 | NA | NA | NA | NA | NA | #VALUE! |
| ENSMUSG00000064446.3 | 0 | NA | NA | NA | NA | NA | #VALUE! |

| | | | | | | | |
|---|---|---|---|---|---|---|---|
| ENSMUSG00000064450.3 | 0 | NA | NA | NA | NA | NA | #VALUE! |
| ENSMUSG00000064451.3 | 0 | NA | NA | NA | NA | NA | #VALUE! |
| ENSMUSG00000064452.3 | 0 | NA | NA | NA | NA | NA | #VALUE! |
| ENSMUSG00000064453.3 | 0 | NA | NA | NA | NA | NA | #VALUE! |
| ENSMUSG00000064460.3 | 0 | NA | NA | NA | NA | NA | #VALUE! |
| ENSMUSG00000064463.3 | 0 | NA | NA | NA | NA | NA | #VALUE! |
| ENSMUSG00000064468.4 | 0 | NA | NA | NA | NA | NA | #VALUE! |
| ENSMUSG00000064471.3 | 0 | NA | NA | NA | NA | NA | #VALUE! |
| ENSMUSG00000064472.3 | 0 | NA | NA | NA | NA | NA | #VALUE! |
| ENSMUSG00000064476.4 | 0 | NA | NA | NA | NA | NA | #VALUE! |
| ENSMUSG00000064477.3 | 0 | NA | NA | NA | NA | NA | #VALUE! |
| ENSMUSG00000064481.3 | 0 | NA | NA | NA | NA | NA | #VALUE! |
| ENSMUSG00000064485.3 | 0 | NA | NA | NA | NA | NA | #VALUE! |
| ENSMUSG00000064486.4 | 0 | NA | NA | NA | NA | NA | #VALUE! |
| ENSMUSG00000064487.3 | 0 | NA | NA | NA | NA | NA | #VALUE! |
| ENSMUSG00000064490.3 | 0 | NA | NA | NA | NA | NA | #VALUE! |
| ENSMUSG00000064492.3 | 0 | NA | NA | NA | NA | NA | #VALUE! |
| ENSMUSG00000064493.3 | 0 | NA | NA | NA | NA | NA | #VALUE! |
| ENSMUSG00000064495.3 | 0 | NA | NA | NA | NA | NA | #VALUE! |
| ENSMUSG00000064496.3 | 0 | NA | NA | NA | NA | NA | #VALUE! |
| ENSMUSG00000064500.4 | 0 | NA | NA | NA | NA | NA | #VALUE! |
| ENSMUSG00000064511.3 | 0 | NA | NA | NA | NA | NA | #VALUE! |
| ENSMUSG00000064512.4 | 0 | NA | NA | NA | NA | NA | #VALUE! |
| ENSMUSG00000064513.3 | 0 | NA | NA | NA | NA | NA | #VALUE! |
| ENSMUSG00000064514.3 | 0 | NA | NA | NA | NA | NA | #VALUE! |
| ENSMUSG00000064515.3 | 0 | NA | NA | NA | NA | NA | #VALUE! |
| ENSMUSG00000064517.3 | 0 | NA | NA | NA | NA | NA | #VALUE! |
| ENSMUSG00000064522.3 | 0 | NA | NA | NA | NA | NA | #VALUE! |
| ENSMUSG00000064524.3 | 0 | NA | NA | NA | NA | NA | #VALUE! |
| ENSMUSG00000064525.3 | 0 | NA | NA | NA | NA | NA | #VALUE! |
| ENSMUSG00000064526.3 | 0 | NA | NA | NA | NA | NA | #VALUE! |
| ENSMUSG00000064527.4 | 0 | NA | NA | NA | NA | NA | #VALUE! |
| ENSMUSG00000064529.3 | 0 | NA | NA | NA | NA | NA | #VALUE! |
| ENSMUSG00000064530.4 | 0 | NA | NA | NA | NA | NA | #VALUE! |
| ENSMUSG00000064532.3 | 0 | NA | NA | NA | NA | NA | #VALUE! |
| ENSMUSG00000064536.3 | 0 | NA | NA | NA | NA | NA | #VALUE! |
| ENSMUSG00000064540.3 | 0 | NA | NA | NA | NA | NA | #VALUE! |
| ENSMUSG00000064541.3 | 0 | NA | NA | NA | NA | NA | #VALUE! |
| ENSMUSG00000064542.3 | 0 | NA | NA | NA | NA | NA | #VALUE! |
| ENSMUSG00000064543.3 | 0 | NA | NA | NA | NA | NA | #VALUE! |
| ENSMUSG00000064545.4 | 0 | NA | NA | NA | NA | NA | #VALUE! |
| ENSMUSG00000064548.3 | 0 | NA | NA | NA | NA | NA | #VALUE! |
| ENSMUSG00000064549.3 | 0 | NA | NA | NA | NA | NA | #VALUE! |
| ENSMUSG00000064550.3 | 0 | NA | NA | NA | NA | NA | #VALUE! |

| | | | | | | | |
|---|---|---|---|---|---|---|---|
| ENSMUSG00000064551.3 | 0 | NA | NA | NA | NA | NA | #VALUE! |
| ENSMUSG00000064553.4 | 0 | NA | NA | NA | NA | NA | #VALUE! |
| ENSMUSG00000064554.3 | 0 | NA | NA | NA | NA | NA | #VALUE! |
| ENSMUSG00000064555.3 | 0 | NA | NA | NA | NA | NA | #VALUE! |
| ENSMUSG00000064556.3 | 0 | NA | NA | NA | NA | NA | #VALUE! |
| ENSMUSG00000064558.3 | 0 | NA | NA | NA | NA | NA | #VALUE! |
| ENSMUSG00000064563.3 | 0 | NA | NA | NA | NA | NA | #VALUE! |
| ENSMUSG00000064565.3 | 0 | NA | NA | NA | NA | NA | #VALUE! |
| ENSMUSG00000064566.3 | 0 | NA | NA | NA | NA | NA | #VALUE! |
| ENSMUSG00000064567.3 | 0 | NA | NA | NA | NA | NA | #VALUE! |
| ENSMUSG00000064569.3 | 0 | NA | NA | NA | NA | NA | #VALUE! |
| ENSMUSG00000064570.3 | 0 | NA | NA | NA | NA | NA | #VALUE! |
| ENSMUSG00000064572.3 | 0 | NA | NA | NA | NA | NA | #VALUE! |
| ENSMUSG00000064573.3 | 0 | NA | NA | NA | NA | NA | #VALUE! |
| ENSMUSG00000064579.4 | 0 | NA | NA | NA | NA | NA | #VALUE! |
| ENSMUSG00000064580.3 | 0 | NA | NA | NA | NA | NA | #VALUE! |
| ENSMUSG00000064581.3 | 0 | NA | NA | NA | NA | NA | #VALUE! |
| ENSMUSG00000064582.3 | 0 | NA | NA | NA | NA | NA | #VALUE! |
| ENSMUSG00000064584.3 | 0 | NA | NA | NA | NA | NA | #VALUE! |
| ENSMUSG00000064585.3 | 0 | NA | NA | NA | NA | NA | #VALUE! |
| ENSMUSG00000064586.3 | 0 | NA | NA | NA | NA | NA | #VALUE! |
| ENSMUSG00000064587.3 | 0 | NA | NA | NA | NA | NA | #VALUE! |
| ENSMUSG00000064590.3 | 0 | NA | NA | NA | NA | NA | #VALUE! |
| ENSMUSG00000064591.3 | 0 | NA | NA | NA | NA | NA | #VALUE! |
| ENSMUSG00000064595.2 | 0 | NA | NA | NA | NA | NA | #VALUE! |
| ENSMUSG00000064596.3 | 0 | NA | NA | NA | NA | NA | #VALUE! |
| ENSMUSG00000064597.3 | 0 | NA | NA | NA | NA | NA | #VALUE! |
| ENSMUSG00000064600.3 | 0 | NA | NA | NA | NA | NA | #VALUE! |
| ENSMUSG00000064601.3 | 0 | NA | NA | NA | NA | NA | #VALUE! |
| ENSMUSG00000064602.3 | 0 | NA | NA | NA | NA | NA | #VALUE! |
| ENSMUSG00000064604.3 | 0 | NA | NA | NA | NA | NA | #VALUE! |
| ENSMUSG00000064605.5 | 0 | NA | NA | NA | NA | NA | #VALUE! |
| ENSMUSG00000064606.3 | 0 | NA | NA | NA | NA | NA | #VALUE! |
| ENSMUSG00000064609.3 | 0 | NA | NA | NA | NA | NA | #VALUE! |
| ENSMUSG00000064612.4 | 0 | NA | NA | NA | NA | NA | #VALUE! |
| ENSMUSG00000064613.3 | 0 | NA | NA | NA | NA | NA | #VALUE! |
| ENSMUSG00000064619.2 | 0 | NA | NA | NA | NA | NA | #VALUE! |
| ENSMUSG00000064620.3 | 0 | NA | NA | NA | NA | NA | #VALUE! |
| ENSMUSG00000064621.3 | 0 | NA | NA | NA | NA | NA | #VALUE! |
| ENSMUSG00000064622.3 | 0 | NA | NA | NA | NA | NA | #VALUE! |
| ENSMUSG00000064624.3 | 0 | NA | NA | NA | NA | NA | #VALUE! |
| ENSMUSG00000064630.4 | 0 | NA | NA | NA | NA | NA | #VALUE! |
| ENSMUSG00000064632.3 | 0 | NA | NA | NA | NA | NA | #VALUE! |
| ENSMUSG00000064634.3 | 0 | NA | NA | NA | NA | NA | #VALUE! |

| | | | | | | | |
|---|---|---|---|---|---|---|---|
| ENSMUSG00000064637.3 | 0 | NA | NA | NA | NA | NA | #VALUE! |
| ENSMUSG00000064644.3 | 0 | NA | NA | NA | NA | NA | #VALUE! |
| ENSMUSG00000064646.3 | 0 | NA | NA | NA | NA | NA | #VALUE! |
| ENSMUSG00000064647.3 | 0 | NA | NA | NA | NA | NA | #VALUE! |
| ENSMUSG00000064648.5 | 0 | NA | NA | NA | NA | NA | #VALUE! |
| ENSMUSG00000064652.3 | 0 | NA | NA | NA | NA | NA | #VALUE! |
| ENSMUSG00000064653.3 | 0 | NA | NA | NA | NA | NA | #VALUE! |
| ENSMUSG00000064655.3 | 0 | NA | NA | NA | NA | NA | #VALUE! |
| ENSMUSG00000064658.3 | 0 | NA | NA | NA | NA | NA | #VALUE! |
| ENSMUSG00000064659.4 | 0 | NA | NA | NA | NA | NA | #VALUE! |
| ENSMUSG00000064664.4 | 0 | NA | NA | NA | NA | NA | #VALUE! |
| ENSMUSG00000064665.3 | 0 | NA | NA | NA | NA | NA | #VALUE! |
| ENSMUSG00000064666.3 | 0 | NA | NA | NA | NA | NA | #VALUE! |
| ENSMUSG00000064669.3 | 0 | NA | NA | NA | NA | NA | #VALUE! |
| ENSMUSG00000064670.3 | 0 | NA | NA | NA | NA | NA | #VALUE! |
| ENSMUSG00000064671.3 | 0 | NA | NA | NA | NA | NA | #VALUE! |
| ENSMUSG00000064672.3 | 0 | NA | NA | NA | NA | NA | #VALUE! |
| ENSMUSG00000064673.3 | 0 | NA | NA | NA | NA | NA | #VALUE! |
| ENSMUSG00000064675.4 | 0 | NA | NA | NA | NA | NA | #VALUE! |
| ENSMUSG00000064677.3 | 0 | NA | NA | NA | NA | NA | #VALUE! |
| ENSMUSG00000064679.3 | 0 | NA | NA | NA | NA | NA | #VALUE! |
| ENSMUSG00000064682.3 | 0 | NA | NA | NA | NA | NA | #VALUE! |
| ENSMUSG00000064685.3 | 0 | NA | NA | NA | NA | NA | #VALUE! |
| ENSMUSG00000064686.3 | 0 | NA | NA | NA | NA | NA | #VALUE! |
| ENSMUSG00000064687.3 | 0 | NA | NA | NA | NA | NA | #VALUE! |
| ENSMUSG00000064688.3 | 0 | NA | NA | NA | NA | NA | #VALUE! |
| ENSMUSG00000064689.3 | 0 | NA | NA | NA | NA | NA | #VALUE! |
| ENSMUSG00000064692.3 | 0 | NA | NA | NA | NA | NA | #VALUE! |
| ENSMUSG00000064693.3 | 0 | NA | NA | NA | NA | NA | #VALUE! |
| ENSMUSG00000064697.3 | 0 | NA | NA | NA | NA | NA | #VALUE! |
| ENSMUSG00000064698.2 | 0 | NA | NA | NA | NA | NA | #VALUE! |
| ENSMUSG00000064707.4 | 0 | NA | NA | NA | NA | NA | #VALUE! |
| ENSMUSG00000064715.3 | 0 | NA | NA | NA | NA | NA | #VALUE! |
| ENSMUSG00000064719.3 | 0 | NA | NA | NA | NA | NA | #VALUE! |
| ENSMUSG00000064720.3 | 0 | NA | NA | NA | NA | NA | #VALUE! |
| ENSMUSG00000064721.6 | 0 | NA | NA | NA | NA | NA | #VALUE! |
| ENSMUSG00000064722.3 | 0 | NA | NA | NA | NA | NA | #VALUE! |
| ENSMUSG00000064724.3 | 0 | NA | NA | NA | NA | NA | #VALUE! |
| ENSMUSG00000064725.3 | 0 | NA | NA | NA | NA | NA | #VALUE! |
| ENSMUSG00000064726.3 | 0 | NA | NA | NA | NA | NA | #VALUE! |
| ENSMUSG00000064727.3 | 0 | NA | NA | NA | NA | NA | #VALUE! |
| ENSMUSG00000064731.3 | 0 | NA | NA | NA | NA | NA | #VALUE! |
| ENSMUSG00000064732.3 | 0 | NA | NA | NA | NA | NA | #VALUE! |
| ENSMUSG00000064734.3 | 1.762102 | -4.34067 | 3.93219 | -1.10388 | 0.269645 | NA | #VALUE! |

| | | | | | | | |
|---|---|---|---|---|---|---|---|
| ENSMUSG00000064735.3 | 0 | NA | NA | NA | NA | NA | #VALUE! |
| ENSMUSG00000064737.3 | 0 | NA | NA | NA | NA | NA | #VALUE! |
| ENSMUSG00000064738.3 | 0 | NA | NA | NA | NA | NA | #VALUE! |
| ENSMUSG00000064739.3 | 0 | NA | NA | NA | NA | NA | #VALUE! |
| ENSMUSG00000064741.3 | 0 | NA | NA | NA | NA | NA | #VALUE! |
| ENSMUSG00000064742.3 | 0 | NA | NA | NA | NA | NA | #VALUE! |
| ENSMUSG00000064745.3 | 0 | NA | NA | NA | NA | NA | #VALUE! |
| ENSMUSG00000064748.3 | 0 | NA | NA | NA | NA | NA | #VALUE! |
| ENSMUSG00000064751.3 | 0 | NA | NA | NA | NA | NA | #VALUE! |
| ENSMUSG00000064755.3 | 0 | NA | NA | NA | NA | NA | #VALUE! |
| ENSMUSG00000064759.4 | 0 | NA | NA | NA | NA | NA | #VALUE! |
| ENSMUSG00000064761.3 | 0 | NA | NA | NA | NA | NA | #VALUE! |
| ENSMUSG00000064762.3 | 0 | NA | NA | NA | NA | NA | #VALUE! |
| ENSMUSG00000064763.3 | 0 | NA | NA | NA | NA | NA | #VALUE! |
| ENSMUSG00000064764.3 | 0 | NA | NA | NA | NA | NA | #VALUE! |
| ENSMUSG00000064765.4 | 0 | NA | NA | NA | NA | NA | #VALUE! |
| ENSMUSG00000064766.3 | 0 | NA | NA | NA | NA | NA | #VALUE! |
| ENSMUSG00000064767.3 | 0 | NA | NA | NA | NA | NA | #VALUE! |
| ENSMUSG00000064768.3 | 0 | NA | NA | NA | NA | NA | #VALUE! |
| ENSMUSG00000064769.3 | 0 | NA | NA | NA | NA | NA | #VALUE! |
| ENSMUSG00000064772.3 | 0 | NA | NA | NA | NA | NA | #VALUE! |
| ENSMUSG00000064775.3 | 0 | NA | NA | NA | NA | NA | #VALUE! |
| ENSMUSG00000064776.4 | 0 | NA | NA | NA | NA | NA | #VALUE! |
| ENSMUSG00000064777.3 | 0 | NA | NA | NA | NA | NA | #VALUE! |
| ENSMUSG00000064778.4 | 0 | NA | NA | NA | NA | NA | #VALUE! |
| ENSMUSG00000064780.3 | 0 | NA | NA | NA | NA | NA | #VALUE! |
| ENSMUSG00000064781.4 | 0 | NA | NA | NA | NA | NA | #VALUE! |
| ENSMUSG00000064782.2 | 0 | NA | NA | NA | NA | NA | #VALUE! |
| ENSMUSG00000064783.3 | 0 | NA | NA | NA | NA | NA | #VALUE! |
| ENSMUSG00000064788.3 | 0 | NA | NA | NA | NA | NA | #VALUE! |
| ENSMUSG00000064792.3 | 0 | NA | NA | NA | NA | NA | #VALUE! |
| ENSMUSG00000064793.3 | 0 | NA | NA | NA | NA | NA | #VALUE! |
| ENSMUSG00000064795.3 | 0 | NA | NA | NA | NA | NA | #VALUE! |
| ENSMUSG00000064797.3 | 0 | NA | NA | NA | NA | NA | #VALUE! |
| ENSMUSG00000064800.3 | 0 | NA | NA | NA | NA | NA | #VALUE! |
| ENSMUSG00000064802.3 | 0 | NA | NA | NA | NA | NA | #VALUE! |
| ENSMUSG00000064805.3 | 0 | NA | NA | NA | NA | NA | #VALUE! |
| ENSMUSG00000064806.4 | 0 | NA | NA | NA | NA | NA | #VALUE! |
| ENSMUSG00000064811.3 | 0 | NA | NA | NA | NA | NA | #VALUE! |
| ENSMUSG00000064814.3 | 0 | NA | NA | NA | NA | NA | #VALUE! |
| ENSMUSG00000064816.3 | 0 | NA | NA | NA | NA | NA | #VALUE! |
| ENSMUSG00000064818.3 | 0 | NA | NA | NA | NA | NA | #VALUE! |
| ENSMUSG00000064822.2 | 0 | NA | NA | NA | NA | NA | #VALUE! |
| ENSMUSG00000064823.3 | 0 | NA | NA | NA | NA | NA | #VALUE! |

| | | | | | | | |
|---|---|---|---|---|---|---|---|
| ENSMUSG00000064829.5 | 0 | NA | NA | NA | NA | NA | #VALUE! |
| ENSMUSG00000064830.3 | 0 | NA | NA | NA | NA | NA | #VALUE! |
| ENSMUSG00000064833.3 | 0 | NA | NA | NA | NA | NA | #VALUE! |
| ENSMUSG00000064836.3 | 0 | NA | NA | NA | NA | NA | #VALUE! |
| ENSMUSG00000064837.3 | 0 | NA | NA | NA | NA | NA | #VALUE! |
| ENSMUSG00000064841.3 | 0 | NA | NA | NA | NA | NA | #VALUE! |
| ENSMUSG00000064842.3 | 0 | NA | NA | NA | NA | NA | #VALUE! |
| ENSMUSG00000064843.3 | 0 | NA | NA | NA | NA | NA | #VALUE! |
| ENSMUSG00000064844.3 | 0 | NA | NA | NA | NA | NA | #VALUE! |
| ENSMUSG00000064845.4 | 0 | NA | NA | NA | NA | NA | #VALUE! |
| ENSMUSG00000064850.3 | 0 | NA | NA | NA | NA | NA | #VALUE! |
| ENSMUSG00000064851.3 | 0 | NA | NA | NA | NA | NA | #VALUE! |
| ENSMUSG00000064853.3 | 0 | NA | NA | NA | NA | NA | #VALUE! |
| ENSMUSG00000064854.3 | 0 | NA | NA | NA | NA | NA | #VALUE! |
| ENSMUSG00000064855.3 | 0 | NA | NA | NA | NA | NA | #VALUE! |
| ENSMUSG00000064857.3 | 0 | NA | NA | NA | NA | NA | #VALUE! |
| ENSMUSG00000064858.4 | 0 | NA | NA | NA | NA | NA | #VALUE! |
| ENSMUSG00000064859.3 | 0 | NA | NA | NA | NA | NA | #VALUE! |
| ENSMUSG00000064862.3 | 0 | NA | NA | NA | NA | NA | #VALUE! |
| ENSMUSG00000064867.3 | 0 | NA | NA | NA | NA | NA | #VALUE! |
| ENSMUSG00000064871.3 | 0 | NA | NA | NA | NA | NA | #VALUE! |
| ENSMUSG00000064872.3 | 0 | NA | NA | NA | NA | NA | #VALUE! |
| ENSMUSG00000064876.3 | 0 | NA | NA | NA | NA | NA | #VALUE! |
| ENSMUSG00000064877.5 | 0 | NA | NA | NA | NA | NA | #VALUE! |
| ENSMUSG00000064880.3 | 0 | NA | NA | NA | NA | NA | #VALUE! |
| ENSMUSG00000064882.4 | 0 | NA | NA | NA | NA | NA | #VALUE! |
| ENSMUSG00000064883.3 | 0 | NA | NA | NA | NA | NA | #VALUE! |
| ENSMUSG00000064884.3 | 0 | NA | NA | NA | NA | NA | #VALUE! |
| ENSMUSG00000064889.3 | 0 | NA | NA | NA | NA | NA | #VALUE! |
| ENSMUSG00000064890.3 | 0 | NA | NA | NA | NA | NA | #VALUE! |
| ENSMUSG00000064897.3 | 0 | NA | NA | NA | NA | NA | #VALUE! |
| ENSMUSG00000064899.3 | 6.25428 | -0.35533 | 3.431044 | -0.10356 | 0.917516 | NA | #VALUE! |
| ENSMUSG00000064900.3 | 0 | NA | NA | NA | NA | NA | #VALUE! |
| ENSMUSG00000064901.3 | 0 | NA | NA | NA | NA | NA | #VALUE! |
| ENSMUSG00000064906.3 | 0 | NA | NA | NA | NA | NA | #VALUE! |
| ENSMUSG00000064909.3 | 0 | NA | NA | NA | NA | NA | #VALUE! |
| ENSMUSG00000064915.3 | 0 | NA | NA | NA | NA | NA | #VALUE! |
| ENSMUSG00000064916.3 | 0 | NA | NA | NA | NA | NA | #VALUE! |
| ENSMUSG00000064917.2 | 0.210026 | -1.52494 | 4.058692 | -0.37572 | 0.707123 | NA | #VALUE! |
| ENSMUSG00000064918.5 | 0 | NA | NA | NA | NA | NA | #VALUE! |
| ENSMUSG00000064919.3 | 0 | NA | NA | NA | NA | NA | #VALUE! |
| ENSMUSG00000064921.3 | 0 | NA | NA | NA | NA | NA | #VALUE! |
| ENSMUSG00000064923.2 | 0 | NA | NA | NA | NA | NA | #VALUE! |
| ENSMUSG00000064925.3 | 0 | NA | NA | NA | NA | NA | #VALUE! |

| | | | | | | | |
|---|---|---|---|---|---|---|---|
| ENSMUSG00000064927.3 | 0 | NA | NA | NA | NA | NA | #VALUE! |
| ENSMUSG00000064928.3 | 0 | NA | NA | NA | NA | NA | #VALUE! |
| ENSMUSG00000064930.3 | 0 | NA | NA | NA | NA | NA | #VALUE! |
| ENSMUSG00000064936.3 | 0 | NA | NA | NA | NA | NA | #VALUE! |
| ENSMUSG00000064940.4 | 0 | NA | NA | NA | NA | NA | #VALUE! |
| ENSMUSG00000064941.3 | 0 | NA | NA | NA | NA | NA | #VALUE! |
| ENSMUSG00000064943.3 | 0 | NA | NA | NA | NA | NA | #VALUE! |
| ENSMUSG00000064945.3 | 0 | NA | NA | NA | NA | NA | #VALUE! |
| ENSMUSG00000064948.4 | 0 | NA | NA | NA | NA | NA | #VALUE! |
| ENSMUSG00000064949.3 | 0 | NA | NA | NA | NA | NA | #VALUE! |
| ENSMUSG00000064952.3 | 0 | NA | NA | NA | NA | NA | #VALUE! |
| ENSMUSG00000064954.3 | 0 | NA | NA | NA | NA | NA | #VALUE! |
| ENSMUSG00000064956.2 | 0 | NA | NA | NA | NA | NA | #VALUE! |
| ENSMUSG00000064959.3 | 0 | NA | NA | NA | NA | NA | #VALUE! |
| ENSMUSG00000064961.3 | 0 | NA | NA | NA | NA | NA | #VALUE! |
| ENSMUSG00000064964.3 | 0 | NA | NA | NA | NA | NA | #VALUE! |
| ENSMUSG00000064966.3 | 0 | NA | NA | NA | NA | NA | #VALUE! |
| ENSMUSG00000064967.3 | 0 | NA | NA | NA | NA | NA | #VALUE! |
| ENSMUSG00000064968.3 | 0 | NA | NA | NA | NA | NA | #VALUE! |
| ENSMUSG00000064970.3 | 0 | NA | NA | NA | NA | NA | #VALUE! |
| ENSMUSG00000064977.3 | 0 | NA | NA | NA | NA | NA | #VALUE! |
| ENSMUSG00000064982.3 | 0 | NA | NA | NA | NA | NA | #VALUE! |
| ENSMUSG00000064984.3 | 0 | NA | NA | NA | NA | NA | #VALUE! |
| ENSMUSG00000064987.4 | 0 | NA | NA | NA | NA | NA | #VALUE! |
| ENSMUSG00000064994.3 | 0 | NA | NA | NA | NA | NA | #VALUE! |
| ENSMUSG00000064995.3 | 0 | NA | NA | NA | NA | NA | #VALUE! |
| ENSMUSG00000065004.3 | 0 | NA | NA | NA | NA | NA | #VALUE! |
| ENSMUSG00000065006.3 | 0 | NA | NA | NA | NA | NA | #VALUE! |
| ENSMUSG00000065012.3 | 0 | NA | NA | NA | NA | NA | #VALUE! |
| ENSMUSG00000065013.3 | 0 | NA | NA | NA | NA | NA | #VALUE! |
| ENSMUSG00000065014.3 | 0 | NA | NA | NA | NA | NA | #VALUE! |
| ENSMUSG00000065016.3 | 0 | NA | NA | NA | NA | NA | #VALUE! |
| ENSMUSG00000065022.3 | 0 | NA | NA | NA | NA | NA | #VALUE! |
| ENSMUSG00000065027.3 | 0 | NA | NA | NA | NA | NA | #VALUE! |
| ENSMUSG00000065028.3 | 0 | NA | NA | NA | NA | NA | #VALUE! |
| ENSMUSG00000065029.3 | 0.245976 | 1.027628 | 4.067126 | 0.252667 | 0.800526 | NA | #VALUE! |
| ENSMUSG00000065031.3 | 0 | NA | NA | NA | NA | NA | #VALUE! |
| ENSMUSG00000065034.3 | 0 | NA | NA | NA | NA | NA | #VALUE! |
| ENSMUSG00000065036.3 | 0 | NA | NA | NA | NA | NA | #VALUE! |
| ENSMUSG00000065040.2 | 0 | NA | NA | NA | NA | NA | #VALUE! |
| ENSMUSG00000065041.3 | 0 | NA | NA | NA | NA | NA | #VALUE! |
| ENSMUSG00000065042.3 | 0 | NA | NA | NA | NA | NA | #VALUE! |
| ENSMUSG00000065045.3 | 0 | NA | NA | NA | NA | NA | #VALUE! |
| ENSMUSG00000065047.3 | 0 | NA | NA | NA | NA | NA | #VALUE! |

| | | | | | | | |
|---|---|---|---|---|---|---|---|
| ENSMUSG00000065048.4 | 0 | NA | NA | NA | NA | NA | #VALUE! |
| ENSMUSG00000065049.4 | 0 | NA | NA | NA | NA | NA | #VALUE! |
| ENSMUSG00000065050.4 | 0 | NA | NA | NA | NA | NA | #VALUE! |
| ENSMUSG00000065053.2 | 0 | NA | NA | NA | NA | NA | #VALUE! |
| ENSMUSG00000065057.3 | 0 | NA | NA | NA | NA | NA | #VALUE! |
| ENSMUSG00000065061.3 | 0 | NA | NA | NA | NA | NA | #VALUE! |
| ENSMUSG00000065072.3 | 0 | NA | NA | NA | NA | NA | #VALUE! |
| ENSMUSG00000065074.3 | 0 | NA | NA | NA | NA | NA | #VALUE! |
| ENSMUSG00000065078.3 | 0 | NA | NA | NA | NA | NA | #VALUE! |
| ENSMUSG00000065079.3 | 0 | NA | NA | NA | NA | NA | #VALUE! |
| ENSMUSG00000065080.3 | 0 | NA | NA | NA | NA | NA | #VALUE! |
| ENSMUSG00000065086.3 | 0 | NA | NA | NA | NA | NA | #VALUE! |
| ENSMUSG00000065087.3 | 0 | NA | NA | NA | NA | NA | #VALUE! |
| ENSMUSG00000065089.3 | 0 | NA | NA | NA | NA | NA | #VALUE! |
| ENSMUSG00000065090.3 | 0 | NA | NA | NA | NA | NA | #VALUE! |
| ENSMUSG00000065094.3 | 0 | NA | NA | NA | NA | NA | #VALUE! |
| ENSMUSG00000065096.3 | 0 | NA | NA | NA | NA | NA | #VALUE! |
| ENSMUSG00000065097.3 | 0 | NA | NA | NA | NA | NA | #VALUE! |
| ENSMUSG00000065100.3 | 0 | NA | NA | NA | NA | NA | #VALUE! |
| ENSMUSG00000065104.3 | 0 | NA | NA | NA | NA | NA | #VALUE! |
| ENSMUSG00000065105.3 | 0 | NA | NA | NA | NA | NA | #VALUE! |
| ENSMUSG00000065107.4 | 0 | NA | NA | NA | NA | NA | #VALUE! |
| ENSMUSG00000065110.6 | 0 | NA | NA | NA | NA | NA | #VALUE! |
| ENSMUSG00000065111.3 | 0 | NA | NA | NA | NA | NA | #VALUE! |
| ENSMUSG00000065118.3 | 0 | NA | NA | NA | NA | NA | #VALUE! |
| ENSMUSG00000065122.4 | 0 | NA | NA | NA | NA | NA | #VALUE! |
| ENSMUSG00000065124.3 | 0 | NA | NA | NA | NA | NA | #VALUE! |
| ENSMUSG00000065130.3 | 0 | NA | NA | NA | NA | NA | #VALUE! |
| ENSMUSG00000065141.3 | 0 | NA | NA | NA | NA | NA | #VALUE! |
| ENSMUSG00000065143.3 | 0 | NA | NA | NA | NA | NA | #VALUE! |
| ENSMUSG00000065147.3 | 0 | NA | NA | NA | NA | NA | #VALUE! |
| ENSMUSG00000065151.3 | 0 | NA | NA | NA | NA | NA | #VALUE! |
| ENSMUSG00000065155.3 | 0 | NA | NA | NA | NA | NA | #VALUE! |
| ENSMUSG00000065159.3 | 0 | NA | NA | NA | NA | NA | #VALUE! |
| ENSMUSG00000065160.3 | 0 | NA | NA | NA | NA | NA | #VALUE! |
| ENSMUSG00000065162.3 | 0 | NA | NA | NA | NA | NA | #VALUE! |
| ENSMUSG00000065164.3 | 0 | NA | NA | NA | NA | NA | #VALUE! |
| ENSMUSG00000065173.3 | 0 | NA | NA | NA | NA | NA | #VALUE! |
| ENSMUSG00000065176.3 | 0 | NA | NA | NA | NA | NA | #VALUE! |
| ENSMUSG00000065178.3 | 0 | NA | NA | NA | NA | NA | #VALUE! |
| ENSMUSG00000065181.3 | 0 | NA | NA | NA | NA | NA | #VALUE! |
| ENSMUSG00000065182.3 | 0 | NA | NA | NA | NA | NA | #VALUE! |
| ENSMUSG00000065185.3 | 0 | NA | NA | NA | NA | NA | #VALUE! |
| ENSMUSG00000065187.3 | 0 | NA | NA | NA | NA | NA | #VALUE! |

| | | | | | | | |
|---|---|---|---|---|---|---|---|
| ENSMUSG00000065188.3 | 0 | NA | NA | NA | NA | NA | #VALUE! |
| ENSMUSG00000065191.4 | 0 | NA | NA | NA | NA | NA | #VALUE! |
| ENSMUSG00000065192.3 | 0 | NA | NA | NA | NA | NA | #VALUE! |
| ENSMUSG00000065193.4 | 0 | NA | NA | NA | NA | NA | #VALUE! |
| ENSMUSG00000065196.3 | 0 | NA | NA | NA | NA | NA | #VALUE! |
| ENSMUSG00000065198.3 | 0 | NA | NA | NA | NA | NA | #VALUE! |
| ENSMUSG00000065200.3 | 0 | NA | NA | NA | NA | NA | #VALUE! |
| ENSMUSG00000065202.3 | 0 | NA | NA | NA | NA | NA | #VALUE! |
| ENSMUSG00000065203.3 | 0 | NA | NA | NA | NA | NA | #VALUE! |
| ENSMUSG00000065204.3 | 0 | NA | NA | NA | NA | NA | #VALUE! |
| ENSMUSG00000065208.3 | 0 | NA | NA | NA | NA | NA | #VALUE! |
| ENSMUSG00000065211.5 | 0 | NA | NA | NA | NA | NA | #VALUE! |
| ENSMUSG00000065212.3 | 0 | NA | NA | NA | NA | NA | #VALUE! |
| ENSMUSG00000065215.3 | 0 | NA | NA | NA | NA | NA | #VALUE! |
| ENSMUSG00000065219.3 | 0 | NA | NA | NA | NA | NA | #VALUE! |
| ENSMUSG00000065220.3 | 0 | NA | NA | NA | NA | NA | #VALUE! |
| ENSMUSG00000065223.4 | 0 | NA | NA | NA | NA | NA | #VALUE! |
| ENSMUSG00000065226.3 | 0 | NA | NA | NA | NA | NA | #VALUE! |
| ENSMUSG00000065228.3 | 0 | NA | NA | NA | NA | NA | #VALUE! |
| ENSMUSG00000065231.3 | 0 | NA | NA | NA | NA | NA | #VALUE! |
| ENSMUSG00000065232.3 | 0.18651 | 1.401769 | 4.056354 | 0.345574 | 0.729663 | NA | #VALUE! |
| ENSMUSG00000065235.3 | 0 | NA | NA | NA | NA | NA | #VALUE! |
| ENSMUSG00000065238.3 | 0 | NA | NA | NA | NA | NA | #VALUE! |
| ENSMUSG00000065239.3 | 0 | NA | NA | NA | NA | NA | #VALUE! |
| ENSMUSG00000065246.3 | 0 | NA | NA | NA | NA | NA | #VALUE! |
| ENSMUSG00000065248.3 | 0 | NA | NA | NA | NA | NA | #VALUE! |
| ENSMUSG00000065255.3 | 0 | NA | NA | NA | NA | NA | #VALUE! |
| ENSMUSG00000065256.3 | 0 | NA | NA | NA | NA | NA | #VALUE! |
| ENSMUSG00000065258.3 | 0 | NA | NA | NA | NA | NA | #VALUE! |
| ENSMUSG00000065259.3 | 0 | NA | NA | NA | NA | NA | #VALUE! |
| ENSMUSG00000065260.3 | 0 | NA | NA | NA | NA | NA | #VALUE! |
| ENSMUSG00000065262.3 | 0 | NA | NA | NA | NA | NA | #VALUE! |
| ENSMUSG00000065265.3 | 0 | NA | NA | NA | NA | NA | #VALUE! |
| ENSMUSG00000065266.3 | 0 | NA | NA | NA | NA | NA | #VALUE! |
| ENSMUSG00000065272.3 | 0 | NA | NA | NA | NA | NA | #VALUE! |
| ENSMUSG00000065273.3 | 0 | NA | NA | NA | NA | NA | #VALUE! |
| ENSMUSG00000065274.3 | 0 | NA | NA | NA | NA | NA | #VALUE! |
| ENSMUSG00000065277.3 | 0 | NA | NA | NA | NA | NA | #VALUE! |
| ENSMUSG00000065279.3 | 0 | NA | NA | NA | NA | NA | #VALUE! |
| ENSMUSG00000065280.3 | 0 | NA | NA | NA | NA | NA | #VALUE! |
| ENSMUSG00000065281.4 | 0 | NA | NA | NA | NA | NA | #VALUE! |
| ENSMUSG00000065282.3 | 0 | NA | NA | NA | NA | NA | #VALUE! |
| ENSMUSG00000065283.3 | 0 | NA | NA | NA | NA | NA | #VALUE! |
| ENSMUSG00000065287.3 | 0 | NA | NA | NA | NA | NA | #VALUE! |

| | | | | | | | |
|---|---|---|---|---|---|---|---|
| ENSMUSG00000065289.3 | 0 | NA | NA | NA | NA | NA | #VALUE! |
| ENSMUSG00000065291.3 | 0 | NA | NA | NA | NA | NA | #VALUE! |
| ENSMUSG00000065293.3 | 0 | NA | NA | NA | NA | NA | #VALUE! |
| ENSMUSG00000065295.3 | 0 | NA | NA | NA | NA | NA | #VALUE! |
| ENSMUSG00000065296.4 | 0 | NA | NA | NA | NA | NA | #VALUE! |
| ENSMUSG00000065297.3 | 0 | NA | NA | NA | NA | NA | #VALUE! |
| ENSMUSG00000065298.3 | 0 | NA | NA | NA | NA | NA | #VALUE! |
| ENSMUSG00000065299.3 | 0 | NA | NA | NA | NA | NA | #VALUE! |
| ENSMUSG00000065301.3 | 0 | NA | NA | NA | NA | NA | #VALUE! |
| ENSMUSG00000065303.4 | 0 | NA | NA | NA | NA | NA | #VALUE! |
| ENSMUSG00000065304.3 | 0 | NA | NA | NA | NA | NA | #VALUE! |
| ENSMUSG00000065305.3 | 0 | NA | NA | NA | NA | NA | #VALUE! |
| ENSMUSG00000065309.4 | 0 | NA | NA | NA | NA | NA | #VALUE! |
| ENSMUSG00000065312.3 | 0 | NA | NA | NA | NA | NA | #VALUE! |
| ENSMUSG00000065315.3 | 0 | NA | NA | NA | NA | NA | #VALUE! |
| ENSMUSG00000065320.4 | 0 | NA | NA | NA | NA | NA | #VALUE! |
| ENSMUSG00000065322.2 | 0 | NA | NA | NA | NA | NA | #VALUE! |
| ENSMUSG00000065328.4 | 0 | NA | NA | NA | NA | NA | #VALUE! |
| ENSMUSG00000065329.4 | 0 | NA | NA | NA | NA | NA | #VALUE! |
| ENSMUSG00000065331.3 | 0 | NA | NA | NA | NA | NA | #VALUE! |
| ENSMUSG00000065333.3 | 0 | NA | NA | NA | NA | NA | #VALUE! |
| ENSMUSG00000065337.3 | 0 | NA | NA | NA | NA | NA | #VALUE! |
| ENSMUSG00000065340.3 | 0 | NA | NA | NA | NA | NA | #VALUE! |
| ENSMUSG00000065341.3 | 0 | NA | NA | NA | NA | NA | #VALUE! |
| ENSMUSG00000065344.2 | 0 | NA | NA | NA | NA | NA | #VALUE! |
| ENSMUSG00000065346.4 | 0 | NA | NA | NA | NA | NA | #VALUE! |
| ENSMUSG00000065353.3 | 0.391578 | -2.1489 | 4.020883 | -0.53444 | 0.59304 | NA | #VALUE! |
| ENSMUSG00000065360.3 | 0 | NA | NA | NA | NA | NA | #VALUE! |
| ENSMUSG00000065362.3 | 0 | NA | NA | NA | NA | NA | #VALUE! |
| ENSMUSG00000065371.3 | 0 | NA | NA | NA | NA | NA | #VALUE! |
| ENSMUSG00000065372.3 | 0 | NA | NA | NA | NA | NA | #VALUE! |
| ENSMUSG00000065373.3 | 0 | NA | NA | NA | NA | NA | #VALUE! |
| ENSMUSG00000065374.3 | 0 | NA | NA | NA | NA | NA | #VALUE! |
| ENSMUSG00000065378.3 | 0 | NA | NA | NA | NA | NA | #VALUE! |
| ENSMUSG00000065379.3 | 0 | NA | NA | NA | NA | NA | #VALUE! |
| ENSMUSG00000065380.3 | 0 | NA | NA | NA | NA | NA | #VALUE! |
| ENSMUSG00000065382.2 | 0 | NA | NA | NA | NA | NA | #VALUE! |
| ENSMUSG00000065384.4 | 0 | NA | NA | NA | NA | NA | #VALUE! |
| ENSMUSG00000065387.4 | 0 | NA | NA | NA | NA | NA | #VALUE! |
| ENSMUSG00000065388.3 | 0 | NA | NA | NA | NA | NA | #VALUE! |
| ENSMUSG00000065389.3 | 0 | NA | NA | NA | NA | NA | #VALUE! |
| ENSMUSG00000065392.3 | 0 | NA | NA | NA | NA | NA | #VALUE! |
| ENSMUSG00000065394.3 | 0 | NA | NA | NA | NA | NA | #VALUE! |
| ENSMUSG00000065395.3 | 0 | NA | NA | NA | NA | NA | #VALUE! |

| | | | | | | | |
|---|---|---|---|---|---|---|---|
| ENSMUSG00000065396.3 | 0 | NA | NA | NA | NA | NA | #VALUE! |
| ENSMUSG00000065397.3 | 0 | NA | NA | NA | NA | NA | #VALUE! |
| ENSMUSG00000065398.3 | 0 | NA | NA | NA | NA | NA | #VALUE! |
| ENSMUSG00000065399.3 | 0 | NA | NA | NA | NA | NA | #VALUE! |
| ENSMUSG00000065400.3 | 0 | NA | NA | NA | NA | NA | #VALUE! |
| ENSMUSG00000065401.3 | 0 | NA | NA | NA | NA | NA | #VALUE! |
| ENSMUSG00000065402.3 | 0 | NA | NA | NA | NA | NA | #VALUE! |
| ENSMUSG00000065403.3 | 0 | NA | NA | NA | NA | NA | #VALUE! |
| ENSMUSG00000065404.3 | 0 | NA | NA | NA | NA | NA | #VALUE! |
| ENSMUSG00000065405.5 | 0 | NA | NA | NA | NA | NA | #VALUE! |
| ENSMUSG00000065406.5 | 0 | NA | NA | NA | NA | NA | #VALUE! |
| ENSMUSG00000065407.3 | 0 | NA | NA | NA | NA | NA | #VALUE! |
| ENSMUSG00000065408.3 | 0 | NA | NA | NA | NA | NA | #VALUE! |
| ENSMUSG00000065410.3 | 0 | NA | NA | NA | NA | NA | #VALUE! |
| ENSMUSG00000065411.4 | 0 | NA | NA | NA | NA | NA | #VALUE! |
| ENSMUSG00000065412.5 | 0 | NA | NA | NA | NA | NA | #VALUE! |
| ENSMUSG00000065413.4 | 0 | NA | NA | NA | NA | NA | #VALUE! |
| ENSMUSG00000065414.3 | 0 | NA | NA | NA | NA | NA | #VALUE! |
| ENSMUSG00000065415.3 | 0 | NA | NA | NA | NA | NA | #VALUE! |
| ENSMUSG00000065416.3 | 0 | NA | NA | NA | NA | NA | #VALUE! |
| ENSMUSG00000065417.3 | 0 | NA | NA | NA | NA | NA | #VALUE! |
| ENSMUSG00000065418.3 | 0 | NA | NA | NA | NA | NA | #VALUE! |
| ENSMUSG00000065420.4 | 0 | NA | NA | NA | NA | NA | #VALUE! |
| ENSMUSG00000065421.5 | 0 | NA | NA | NA | NA | NA | #VALUE! |
| ENSMUSG00000065422.3 | 0 | NA | NA | NA | NA | NA | #VALUE! |
| ENSMUSG00000065423.3 | 0 | NA | NA | NA | NA | NA | #VALUE! |
| ENSMUSG00000065424.4 | 0 | NA | NA | NA | NA | NA | #VALUE! |
| ENSMUSG00000065425.3 | 0 | NA | NA | NA | NA | NA | #VALUE! |
| ENSMUSG00000065426.5 | 0 | NA | NA | NA | NA | NA | #VALUE! |
| ENSMUSG00000065427.3 | 0 | NA | NA | NA | NA | NA | #VALUE! |
| ENSMUSG00000065428.6 | 0 | NA | NA | NA | NA | NA | #VALUE! |
| ENSMUSG00000065429.3 | 0 | NA | NA | NA | NA | NA | #VALUE! |
| ENSMUSG00000065430.3 | 0 | NA | NA | NA | NA | NA | #VALUE! |
| ENSMUSG00000065431.3 | 0 | NA | NA | NA | NA | NA | #VALUE! |
| ENSMUSG00000065432.3 | 0 | NA | NA | NA | NA | NA | #VALUE! |
| ENSMUSG00000065433.3 | 0 | NA | NA | NA | NA | NA | #VALUE! |
| ENSMUSG00000065434.3 | 0 | NA | NA | NA | NA | NA | #VALUE! |
| ENSMUSG00000065435.3 | 0 | NA | NA | NA | NA | NA | #VALUE! |
| ENSMUSG00000065436.3 | 0 | NA | NA | NA | NA | NA | #VALUE! |
| ENSMUSG00000065437.3 | 0 | NA | NA | NA | NA | NA | #VALUE! |
| ENSMUSG00000065438.4 | 0 | NA | NA | NA | NA | NA | #VALUE! |
| ENSMUSG00000065439.3 | 0 | NA | NA | NA | NA | NA | #VALUE! |
| ENSMUSG00000065440.3 | 0 | NA | NA | NA | NA | NA | #VALUE! |
| ENSMUSG00000065441.4 | 0 | NA | NA | NA | NA | NA | #VALUE! |

| | | | | | | | |
|---|---|---|---|---|---|---|---|
| ENSMUSG00000065442.3 | 0 | NA | NA | NA | NA | NA | #VALUE! |
| ENSMUSG00000065443.3 | 0 | NA | NA | NA | NA | NA | #VALUE! |
| ENSMUSG00000065444.3 | 0 | NA | NA | NA | NA | NA | #VALUE! |
| ENSMUSG00000065445.5 | 0 | NA | NA | NA | NA | NA | #VALUE! |
| ENSMUSG00000065446.3 | 0 | NA | NA | NA | NA | NA | #VALUE! |
| ENSMUSG00000065447.3 | 0 | NA | NA | NA | NA | NA | #VALUE! |
| ENSMUSG00000065448.4 | 0 | NA | NA | NA | NA | NA | #VALUE! |
| ENSMUSG00000065449.3 | 0 | NA | NA | NA | NA | NA | #VALUE! |
| ENSMUSG00000065450.3 | 0 | NA | NA | NA | NA | NA | #VALUE! |
| ENSMUSG00000065451.3 | 0 | NA | NA | NA | NA | NA | #VALUE! |
| ENSMUSG00000065452.2 | 0 | NA | NA | NA | NA | NA | #VALUE! |
| ENSMUSG00000065453.3 | 0 | NA | NA | NA | NA | NA | #VALUE! |
| ENSMUSG00000065454.4 | 0 | NA | NA | NA | NA | NA | #VALUE! |
| ENSMUSG00000065455.3 | 0 | NA | NA | NA | NA | NA | #VALUE! |
| ENSMUSG00000065456.3 | 0 | NA | NA | NA | NA | NA | #VALUE! |
| ENSMUSG00000065457.5 | 0 | NA | NA | NA | NA | NA | #VALUE! |
| ENSMUSG00000065458.3 | 0 | NA | NA | NA | NA | NA | #VALUE! |
| ENSMUSG00000065460.3 | 0 | NA | NA | NA | NA | NA | #VALUE! |
| ENSMUSG00000065462.5 | 0 | NA | NA | NA | NA | NA | #VALUE! |
| ENSMUSG00000065463.3 | 0 | NA | NA | NA | NA | NA | #VALUE! |
| ENSMUSG00000065464.4 | 0 | NA | NA | NA | NA | NA | #VALUE! |
| ENSMUSG00000065465.3 | 0 | NA | NA | NA | NA | NA | #VALUE! |
| ENSMUSG00000065468.3 | 0 | NA | NA | NA | NA | NA | #VALUE! |
| ENSMUSG00000065469.3 | 0 | NA | NA | NA | NA | NA | #VALUE! |
| ENSMUSG00000065470.3 | 0 | NA | NA | NA | NA | NA | #VALUE! |
| ENSMUSG00000065471.3 | 0 | NA | NA | NA | NA | NA | #VALUE! |
| ENSMUSG00000065472.3 | 0 | NA | NA | NA | NA | NA | #VALUE! |
| ENSMUSG00000065473.3 | 0 | NA | NA | NA | NA | NA | #VALUE! |
| ENSMUSG00000065474.3 | 0 | NA | NA | NA | NA | NA | #VALUE! |
| ENSMUSG00000065475.4 | 0 | NA | NA | NA | NA | NA | #VALUE! |
| ENSMUSG00000065476.4 | 0 | NA | NA | NA | NA | NA | #VALUE! |
| ENSMUSG00000065477.5 | 0 | NA | NA | NA | NA | NA | #VALUE! |
| ENSMUSG00000065479.4 | 0 | NA | NA | NA | NA | NA | #VALUE! |
| ENSMUSG00000065480.3 | 0 | NA | NA | NA | NA | NA | #VALUE! |
| ENSMUSG00000065481.3 | 0 | NA | NA | NA | NA | NA | #VALUE! |
| ENSMUSG00000065482.3 | 0 | NA | NA | NA | NA | NA | #VALUE! |
| ENSMUSG00000065483.4 | 0 | NA | NA | NA | NA | NA | #VALUE! |
| ENSMUSG00000065484.3 | 0 | NA | NA | NA | NA | NA | #VALUE! |
| ENSMUSG00000065485.3 | 0 | NA | NA | NA | NA | NA | #VALUE! |
| ENSMUSG00000065486.3 | 0 | NA | NA | NA | NA | NA | #VALUE! |
| ENSMUSG00000065488.4 | 0 | NA | NA | NA | NA | NA | #VALUE! |
| ENSMUSG00000065489.3 | 0 | NA | NA | NA | NA | NA | #VALUE! |
| ENSMUSG00000065490.3 | 0 | NA | NA | NA | NA | NA | #VALUE! |
| ENSMUSG00000065491.3 | 0 | NA | NA | NA | NA | NA | #VALUE! |

| | | | | | | | |
|---|---|---|---|---|---|---|---|
| ENSMUSG00000065492.3 | 0 | NA | NA | NA | NA | NA | #VALUE! |
| ENSMUSG00000065493.3 | 0 | NA | NA | NA | NA | NA | #VALUE! |
| ENSMUSG00000065494.3 | 0 | NA | NA | NA | NA | NA | #VALUE! |
| ENSMUSG00000065495.3 | 0 | NA | NA | NA | NA | NA | #VALUE! |
| ENSMUSG00000065497.3 | 0 | NA | NA | NA | NA | NA | #VALUE! |
| ENSMUSG00000065498.6 | 0 | NA | NA | NA | NA | NA | #VALUE! |
| ENSMUSG00000065499.3 | 0 | NA | NA | NA | NA | NA | #VALUE! |
| ENSMUSG00000065500.5 | 0 | NA | NA | NA | NA | NA | #VALUE! |
| ENSMUSG00000065501.3 | 0 | NA | NA | NA | NA | NA | #VALUE! |
| ENSMUSG00000065502.3 | 0 | NA | NA | NA | NA | NA | #VALUE! |
| ENSMUSG00000065503.3 | 0 | NA | NA | NA | NA | NA | #VALUE! |
| ENSMUSG00000065505.3 | 0 | NA | NA | NA | NA | NA | #VALUE! |
| ENSMUSG00000065507.5 | 0 | NA | NA | NA | NA | NA | #VALUE! |
| ENSMUSG00000065508.3 | 0 | NA | NA | NA | NA | NA | #VALUE! |
| ENSMUSG00000065509.5 | 0 | NA | NA | NA | NA | NA | #VALUE! |
| ENSMUSG00000065510.3 | 0 | NA | NA | NA | NA | NA | #VALUE! |
| ENSMUSG00000065511.3 | 0 | NA | NA | NA | NA | NA | #VALUE! |
| ENSMUSG00000065512.5 | 0 | NA | NA | NA | NA | NA | #VALUE! |
| ENSMUSG00000065513.3 | 0 | NA | NA | NA | NA | NA | #VALUE! |
| ENSMUSG00000065514.3 | 0 | NA | NA | NA | NA | NA | #VALUE! |
| ENSMUSG00000065515.3 | 0 | NA | NA | NA | NA | NA | #VALUE! |
| ENSMUSG00000065516.3 | 0 | NA | NA | NA | NA | NA | #VALUE! |
| ENSMUSG00000065518.4 | 0 | NA | NA | NA | NA | NA | #VALUE! |
| ENSMUSG00000065519.3 | 0 | NA | NA | NA | NA | NA | #VALUE! |
| ENSMUSG00000065520.4 | 0 | NA | NA | NA | NA | NA | #VALUE! |
| ENSMUSG00000065521.3 | 0 | NA | NA | NA | NA | NA | #VALUE! |
| ENSMUSG00000065523.3 | 0 | NA | NA | NA | NA | NA | #VALUE! |
| ENSMUSG00000065524.3 | 0 | NA | NA | NA | NA | NA | #VALUE! |
| ENSMUSG00000065526.6 | 0 | NA | NA | NA | NA | NA | #VALUE! |
| ENSMUSG00000065527.3 | 0 | NA | NA | NA | NA | NA | #VALUE! |
| ENSMUSG00000065528.3 | 0 | NA | NA | NA | NA | NA | #VALUE! |
| ENSMUSG00000065529.3 | 0 | NA | NA | NA | NA | NA | #VALUE! |
| ENSMUSG00000065530.3 | 0 | NA | NA | NA | NA | NA | #VALUE! |
| ENSMUSG00000065532.3 | 0 | NA | NA | NA | NA | NA | #VALUE! |
| ENSMUSG00000065533.3 | 0 | NA | NA | NA | NA | NA | #VALUE! |
| ENSMUSG00000065534.3 | 0 | NA | NA | NA | NA | NA | #VALUE! |
| ENSMUSG00000065536.4 | 0 | NA | NA | NA | NA | NA | #VALUE! |
| ENSMUSG00000065537.4 | 0 | NA | NA | NA | NA | NA | #VALUE! |
| ENSMUSG00000065538.3 | 0 | NA | NA | NA | NA | NA | #VALUE! |
| ENSMUSG00000065539.3 | 0 | NA | NA | NA | NA | NA | #VALUE! |
| ENSMUSG00000065540.3 | 0 | NA | NA | NA | NA | NA | #VALUE! |
| ENSMUSG00000065541.3 | 0 | NA | NA | NA | NA | NA | #VALUE! |
| ENSMUSG00000065542.3 | 0 | NA | NA | NA | NA | NA | #VALUE! |
| ENSMUSG00000065543.3 | 0 | NA | NA | NA | NA | NA | #VALUE! |

| | | | | | | | |
|---|---|---|---|---|---|---|---|
| ENSMUSG00000065544.2 | 0 | NA | NA | NA | NA | NA | #VALUE! |
| ENSMUSG00000065546.3 | 0 | NA | NA | NA | NA | NA | #VALUE! |
| ENSMUSG00000065547.3 | 0 | NA | NA | NA | NA | NA | #VALUE! |
| ENSMUSG00000065548.3 | 0 | NA | NA | NA | NA | NA | #VALUE! |
| ENSMUSG00000065549.5 | 0 | NA | NA | NA | NA | NA | #VALUE! |
| ENSMUSG00000065551.3 | 0 | NA | NA | NA | NA | NA | #VALUE! |
| ENSMUSG00000065552.3 | 0 | NA | NA | NA | NA | NA | #VALUE! |
| ENSMUSG00000065553.3 | 0 | NA | NA | NA | NA | NA | #VALUE! |
| ENSMUSG00000065555.3 | 0 | NA | NA | NA | NA | NA | #VALUE! |
| ENSMUSG00000065556.3 | 0 | NA | NA | NA | NA | NA | #VALUE! |
| ENSMUSG00000065557.5 | 0 | NA | NA | NA | NA | NA | #VALUE! |
| ENSMUSG00000065559.3 | 0 | NA | NA | NA | NA | NA | #VALUE! |
| ENSMUSG00000065560.3 | 0 | NA | NA | NA | NA | NA | #VALUE! |
| ENSMUSG00000065561.4 | 0 | NA | NA | NA | NA | NA | #VALUE! |
| ENSMUSG00000065562.3 | 0 | NA | NA | NA | NA | NA | #VALUE! |
| ENSMUSG00000065563.3 | 0 | NA | NA | NA | NA | NA | #VALUE! |
| ENSMUSG00000065564.3 | 0 | NA | NA | NA | NA | NA | #VALUE! |
| ENSMUSG00000065565.3 | 0 | NA | NA | NA | NA | NA | #VALUE! |
| ENSMUSG00000065567.3 | 0 | NA | NA | NA | NA | NA | #VALUE! |
| ENSMUSG00000065568.3 | 0 | NA | NA | NA | NA | NA | #VALUE! |
| ENSMUSG00000065569.4 | 0 | NA | NA | NA | NA | NA | #VALUE! |
| ENSMUSG00000065570.3 | 0 | NA | NA | NA | NA | NA | #VALUE! |
| ENSMUSG00000065571.3 | 0 | NA | NA | NA | NA | NA | #VALUE! |
| ENSMUSG00000065572.3 | 0 | NA | NA | NA | NA | NA | #VALUE! |
| ENSMUSG00000065573.3 | 0 | NA | NA | NA | NA | NA | #VALUE! |
| ENSMUSG00000065574.3 | 0 | NA | NA | NA | NA | NA | #VALUE! |
| ENSMUSG00000065575.3 | 0 | NA | NA | NA | NA | NA | #VALUE! |
| ENSMUSG00000065577.3 | 0 | NA | NA | NA | NA | NA | #VALUE! |
| ENSMUSG00000065578.3 | 0 | NA | NA | NA | NA | NA | #VALUE! |
| ENSMUSG00000065579.3 | 0 | NA | NA | NA | NA | NA | #VALUE! |
| ENSMUSG00000065580.4 | 0 | NA | NA | NA | NA | NA | #VALUE! |
| ENSMUSG00000065581.3 | 0 | NA | NA | NA | NA | NA | #VALUE! |
| ENSMUSG00000065582.3 | 0 | NA | NA | NA | NA | NA | #VALUE! |
| ENSMUSG00000065583.3 | 0 | NA | NA | NA | NA | NA | #VALUE! |
| ENSMUSG00000065585.3 | 0 | NA | NA | NA | NA | NA | #VALUE! |
| ENSMUSG00000065586.3 | 0 | NA | NA | NA | NA | NA | #VALUE! |
| ENSMUSG00000065587.3 | 0 | NA | NA | NA | NA | NA | #VALUE! |
| ENSMUSG00000065589.3 | 0 | NA | NA | NA | NA | NA | #VALUE! |
| ENSMUSG00000065590.3 | 0 | NA | NA | NA | NA | NA | #VALUE! |
| ENSMUSG00000065591.3 | 0 | NA | NA | NA | NA | NA | #VALUE! |
| ENSMUSG00000065592.4 | 0 | NA | NA | NA | NA | NA | #VALUE! |
| ENSMUSG00000065593.3 | 0 | NA | NA | NA | NA | NA | #VALUE! |
| ENSMUSG00000065594.3 | 0 | NA | NA | NA | NA | NA | #VALUE! |
| ENSMUSG00000065596.3 | 0 | NA | NA | NA | NA | NA | #VALUE! |

| | | | | | | | |
|---|---|---|---|---|---|---|---|
| ENSMUSG00000065597.3 | 0 | NA | NA | NA | NA | NA | #VALUE! |
| ENSMUSG00000065599.4 | 0 | NA | NA | NA | NA | NA | #VALUE! |
| ENSMUSG00000065600.3 | 0 | NA | NA | NA | NA | NA | #VALUE! |
| ENSMUSG00000065601.3 | 0 | NA | NA | NA | NA | NA | #VALUE! |
| ENSMUSG00000065602.3 | 0 | NA | NA | NA | NA | NA | #VALUE! |
| ENSMUSG00000065603.3 | 0 | NA | NA | NA | NA | NA | #VALUE! |
| ENSMUSG00000065604.4 | 0 | NA | NA | NA | NA | NA | #VALUE! |
| ENSMUSG00000065606.3 | 0 | NA | NA | NA | NA | NA | #VALUE! |
| ENSMUSG00000065607.3 | 0 | NA | NA | NA | NA | NA | #VALUE! |
| ENSMUSG00000065608.3 | 0 | NA | NA | NA | NA | NA | #VALUE! |
| ENSMUSG00000065609.3 | 0 | NA | NA | NA | NA | NA | #VALUE! |
| ENSMUSG00000065610.3 | 0 | NA | NA | NA | NA | NA | #VALUE! |
| ENSMUSG00000065611.3 | 0 | NA | NA | NA | NA | NA | #VALUE! |
| ENSMUSG00000065612.3 | 0 | NA | NA | NA | NA | NA | #VALUE! |
| ENSMUSG00000065613.3 | 0 | NA | NA | NA | NA | NA | #VALUE! |
| ENSMUSG00000065616.5 | 0 | NA | NA | NA | NA | NA | #VALUE! |
| ENSMUSG00000065617.3 | 0 | NA | NA | NA | NA | NA | #VALUE! |
| ENSMUSG00000065619.4 | 0 | NA | NA | NA | NA | NA | #VALUE! |
| ENSMUSG00000065620.3 | 0 | NA | NA | NA | NA | NA | #VALUE! |
| ENSMUSG00000065622.3 | 0 | NA | NA | NA | NA | NA | #VALUE! |
| ENSMUSG00000065623.3 | 0 | NA | NA | NA | NA | NA | #VALUE! |
| ENSMUSG00000065625.3 | 0 | NA | NA | NA | NA | NA | #VALUE! |
| ENSMUSG00000065629.4 | 0 | NA | NA | NA | NA | NA | #VALUE! |
| ENSMUSG00000065630.4 | 0 | NA | NA | NA | NA | NA | #VALUE! |
| ENSMUSG00000065634.3 | 0 | NA | NA | NA | NA | NA | #VALUE! |
| ENSMUSG00000065635.3 | 0 | NA | NA | NA | NA | NA | #VALUE! |
| ENSMUSG00000065636.3 | 0 | NA | NA | NA | NA | NA | #VALUE! |
| ENSMUSG00000065637.3 | 0 | NA | NA | NA | NA | NA | #VALUE! |
| ENSMUSG00000065640.3 | 0 | NA | NA | NA | NA | NA | #VALUE! |
| ENSMUSG00000065642.3 | 0 | NA | NA | NA | NA | NA | #VALUE! |
| ENSMUSG00000065643.3 | 0 | NA | NA | NA | NA | NA | #VALUE! |
| ENSMUSG00000065647.3 | 0 | NA | NA | NA | NA | NA | #VALUE! |
| ENSMUSG00000065648.3 | 0 | NA | NA | NA | NA | NA | #VALUE! |
| ENSMUSG00000065649.3 | 0 | NA | NA | NA | NA | NA | #VALUE! |
| ENSMUSG00000065651.4 | 0 | NA | NA | NA | NA | NA | #VALUE! |
| ENSMUSG00000065653.3 | 0 | NA | NA | NA | NA | NA | #VALUE! |
| ENSMUSG00000065656.4 | 0 | NA | NA | NA | NA | NA | #VALUE! |
| ENSMUSG00000065657.3 | 0 | NA | NA | NA | NA | NA | #VALUE! |
| ENSMUSG00000065658.3 | 0 | NA | NA | NA | NA | NA | #VALUE! |
| ENSMUSG00000065663.3 | 0 | NA | NA | NA | NA | NA | #VALUE! |
| ENSMUSG00000065664.3 | 0 | NA | NA | NA | NA | NA | #VALUE! |
| ENSMUSG00000065665.3 | 0 | NA | NA | NA | NA | NA | #VALUE! |
| ENSMUSG00000065667.3 | 0 | NA | NA | NA | NA | NA | #VALUE! |
| ENSMUSG00000065669.3 | 0 | NA | NA | NA | NA | NA | #VALUE! |

| | | | | | | | |
|---|---|---|---|---|---|---|---|
| ENSMUSG00000065673.3 | 0 | NA | NA | NA | NA | NA | #VALUE! |
| ENSMUSG00000065674.3 | 0 | NA | NA | NA | NA | NA | #VALUE! |
| ENSMUSG00000065676.3 | 0 | NA | NA | NA | NA | NA | #VALUE! |
| ENSMUSG00000065678.3 | 0 | NA | NA | NA | NA | NA | #VALUE! |
| ENSMUSG00000065680.4 | 0 | NA | NA | NA | NA | NA | #VALUE! |
| ENSMUSG00000065684.3 | 0 | NA | NA | NA | NA | NA | #VALUE! |
| ENSMUSG00000065686.3 | 0 | NA | NA | NA | NA | NA | #VALUE! |
| ENSMUSG00000065687.3 | 0 | NA | NA | NA | NA | NA | #VALUE! |
| ENSMUSG00000065689.3 | 0 | NA | NA | NA | NA | NA | #VALUE! |
| ENSMUSG00000065692.3 | 0 | NA | NA | NA | NA | NA | #VALUE! |
| ENSMUSG00000065694.3 | 0 | NA | NA | NA | NA | NA | #VALUE! |
| ENSMUSG00000065698.3 | 0 | NA | NA | NA | NA | NA | #VALUE! |
| ENSMUSG00000065700.3 | 0 | NA | NA | NA | NA | NA | #VALUE! |
| ENSMUSG00000065701.3 | 0 | NA | NA | NA | NA | NA | #VALUE! |
| ENSMUSG00000065702.3 | 0 | NA | NA | NA | NA | NA | #VALUE! |
| ENSMUSG00000065705.3 | 0 | NA | NA | NA | NA | NA | #VALUE! |
| ENSMUSG00000065706.3 | 0 | NA | NA | NA | NA | NA | #VALUE! |
| ENSMUSG00000065708.3 | 0 | NA | NA | NA | NA | NA | #VALUE! |
| ENSMUSG00000065709.3 | 0 | NA | NA | NA | NA | NA | #VALUE! |
| ENSMUSG00000065714.3 | 0 | NA | NA | NA | NA | NA | #VALUE! |
| ENSMUSG00000065715.3 | 0 | NA | NA | NA | NA | NA | #VALUE! |
| ENSMUSG00000065716.4 | 0 | NA | NA | NA | NA | NA | #VALUE! |
| ENSMUSG00000065718.3 | 0 | NA | NA | NA | NA | NA | #VALUE! |
| ENSMUSG00000065721.3 | 0 | NA | NA | NA | NA | NA | #VALUE! |
| ENSMUSG00000065722.3 | 0 | NA | NA | NA | NA | NA | #VALUE! |
| ENSMUSG00000065725.3 | 0 | NA | NA | NA | NA | NA | #VALUE! |
| ENSMUSG00000065726.3 | 0 | NA | NA | NA | NA | NA | #VALUE! |
| ENSMUSG00000065727.3 | 0 | NA | NA | NA | NA | NA | #VALUE! |
| ENSMUSG00000065728.3 | 0 | NA | NA | NA | NA | NA | #VALUE! |
| ENSMUSG00000065729.3 | 0 | NA | NA | NA | NA | NA | #VALUE! |
| ENSMUSG00000065732.3 | 1.475853 | 4.029233 | 3.937502 | 1.023297 | 0.306167 | NA | #VALUE! |
| ENSMUSG00000065733.3 | 0 | NA | NA | NA | NA | NA | #VALUE! |
| ENSMUSG00000065734.3 | 0 | NA | NA | NA | NA | NA | #VALUE! |
| ENSMUSG00000065735.3 | 0 | NA | NA | NA | NA | NA | #VALUE! |
| ENSMUSG00000065737.3 | 0 | NA | NA | NA | NA | NA | #VALUE! |
| ENSMUSG00000065738.3 | 0 | NA | NA | NA | NA | NA | #VALUE! |
| ENSMUSG00000065742.4 | 0 | NA | NA | NA | NA | NA | #VALUE! |
| ENSMUSG00000065743.4 | 0 | NA | NA | NA | NA | NA | #VALUE! |
| ENSMUSG00000065745.4 | 0 | NA | NA | NA | NA | NA | #VALUE! |
| ENSMUSG00000065746.3 | 0 | NA | NA | NA | NA | NA | #VALUE! |
| ENSMUSG00000065747.3 | 0 | NA | NA | NA | NA | NA | #VALUE! |
| ENSMUSG00000065749.3 | 0 | NA | NA | NA | NA | NA | #VALUE! |
| ENSMUSG00000065750.3 | 5.236419 | -5.92346 | 3.915282 | -1.51291 | 0.130303 | NA | #VALUE! |
| ENSMUSG00000065751.3 | 0 | NA | NA | NA | NA | NA | #VALUE! |

| | | | | | | | |
|---|---|---|---|---|---|---|---|
| ENSMUSG00000065752.3 | 0 | NA | NA | NA | NA | NA | #VALUE! |
| ENSMUSG00000065757.3 | 0 | NA | NA | NA | NA | NA | #VALUE! |
| ENSMUSG00000065760.3 | 0 | NA | NA | NA | NA | NA | #VALUE! |
| ENSMUSG00000065763.3 | 0 | NA | NA | NA | NA | NA | #VALUE! |
| ENSMUSG00000065764.3 | 0 | NA | NA | NA | NA | NA | #VALUE! |
| ENSMUSG00000065768.3 | 0 | NA | NA | NA | NA | NA | #VALUE! |
| ENSMUSG00000065778.3 | 0 | NA | NA | NA | NA | NA | #VALUE! |
| ENSMUSG00000065782.3 | 0 | NA | NA | NA | NA | NA | #VALUE! |
| ENSMUSG00000065787.3 | 0 | NA | NA | NA | NA | NA | #VALUE! |
| ENSMUSG00000065792.3 | 0 | NA | NA | NA | NA | NA | #VALUE! |
| ENSMUSG00000065794.4 | 0 | NA | NA | NA | NA | NA | #VALUE! |
| ENSMUSG00000065795.3 | 0 | NA | NA | NA | NA | NA | #VALUE! |
| ENSMUSG00000065797.3 | 0 | NA | NA | NA | NA | NA | #VALUE! |
| ENSMUSG00000065799.3 | 0 | NA | NA | NA | NA | NA | #VALUE! |
| ENSMUSG00000065800.3 | 0 | NA | NA | NA | NA | NA | #VALUE! |
| ENSMUSG00000065802.3 | 0 | NA | NA | NA | NA | NA | #VALUE! |
| ENSMUSG00000065803.4 | 0 | NA | NA | NA | NA | NA | #VALUE! |
| ENSMUSG00000065804.3 | 0 | NA | NA | NA | NA | NA | #VALUE! |
| ENSMUSG00000065807.3 | 0 | NA | NA | NA | NA | NA | #VALUE! |
| ENSMUSG00000065811.4 | 0 | NA | NA | NA | NA | NA | #VALUE! |
| ENSMUSG00000065812.3 | 0 | NA | NA | NA | NA | NA | #VALUE! |
| ENSMUSG00000065815.3 | 0 | NA | NA | NA | NA | NA | #VALUE! |
| ENSMUSG00000065817.3 | 0 | NA | NA | NA | NA | NA | #VALUE! |
| ENSMUSG00000065818.3 | 0 | NA | NA | NA | NA | NA | #VALUE! |
| ENSMUSG00000065822.3 | 0 | NA | NA | NA | NA | NA | #VALUE! |
| ENSMUSG00000065825.3 | 0 | NA | NA | NA | NA | NA | #VALUE! |
| ENSMUSG00000065836.4 | 0 | NA | NA | NA | NA | NA | #VALUE! |
| ENSMUSG00000065840.3 | 0 | NA | NA | NA | NA | NA | #VALUE! |
| ENSMUSG00000065843.3 | 0 | NA | NA | NA | NA | NA | #VALUE! |
| ENSMUSG00000065845.3 | 0 | NA | NA | NA | NA | NA | #VALUE! |
| ENSMUSG00000065852.3 | 0 | NA | NA | NA | NA | NA | #VALUE! |
| ENSMUSG00000065853.3 | 0 | NA | NA | NA | NA | NA | #VALUE! |
| ENSMUSG00000065858.3 | 0 | NA | NA | NA | NA | NA | #VALUE! |
| ENSMUSG00000065859.4 | 0 | NA | NA | NA | NA | NA | #VALUE! |
| ENSMUSG00000065862.3 | 0 | NA | NA | NA | NA | NA | #VALUE! |
| ENSMUSG00000065864.3 | 0 | NA | NA | NA | NA | NA | #VALUE! |
| ENSMUSG00000065865.3 | 0 | NA | NA | NA | NA | NA | #VALUE! |
| ENSMUSG00000065866.3 | 0 | NA | NA | NA | NA | NA | #VALUE! |
| ENSMUSG00000065869.3 | 0 | NA | NA | NA | NA | NA | #VALUE! |
| ENSMUSG00000065872.3 | 0 | NA | NA | NA | NA | NA | #VALUE! |
| ENSMUSG00000065874.3 | 0 | NA | NA | NA | NA | NA | #VALUE! |
| ENSMUSG00000065876.3 | 0 | NA | NA | NA | NA | NA | #VALUE! |
| ENSMUSG00000065878.3 | 0 | NA | NA | NA | NA | NA | #VALUE! |
| ENSMUSG00000065881.3 | 0 | NA | NA | NA | NA | NA | #VALUE! |

| | | | | | | | |
|---|---|---|---|---|---|---|---|
| ENSMUSG00000065883.4 | 0 | NA | NA | NA | NA | NA | #VALUE! |
| ENSMUSG00000065887.2 | 0 | NA | NA | NA | NA | NA | #VALUE! |
| ENSMUSG00000065892.3 | 0 | NA | NA | NA | NA | NA | #VALUE! |
| ENSMUSG00000065898.3 | 0 | NA | NA | NA | NA | NA | #VALUE! |
| ENSMUSG00000065899.3 | 0 | NA | NA | NA | NA | NA | #VALUE! |
| ENSMUSG00000065903.3 | 0 | NA | NA | NA | NA | NA | #VALUE! |
| ENSMUSG00000065904.3 | 0 | NA | NA | NA | NA | NA | #VALUE! |
| ENSMUSG00000065905.3 | 0 | NA | NA | NA | NA | NA | #VALUE! |
| ENSMUSG00000065907.3 | 0 | NA | NA | NA | NA | NA | #VALUE! |
| ENSMUSG00000065916.3 | 0 | NA | NA | NA | NA | NA | #VALUE! |
| ENSMUSG00000065919.3 | 0 | NA | NA | NA | NA | NA | #VALUE! |
| ENSMUSG00000065924.3 | 0 | NA | NA | NA | NA | NA | #VALUE! |
| ENSMUSG00000065934.3 | 0 | NA | NA | NA | NA | NA | #VALUE! |
| ENSMUSG00000065936.3 | 0 | NA | NA | NA | NA | NA | #VALUE! |
| ENSMUSG00000065937.4 | 0 | NA | NA | NA | NA | NA | #VALUE! |
| ENSMUSG00000065939.3 | 0 | NA | NA | NA | NA | NA | #VALUE! |
| ENSMUSG00000065940.3 | 0.336567 | 1.944925 | 4.029936 | 0.482619 | 0.629366 | NA | #VALUE! |
| ENSMUSG00000065945.4 | 0 | NA | NA | NA | NA | NA | #VALUE! |
| ENSMUSG00000065947.1 | 0 | NA | NA | NA | NA | NA | #VALUE! |
| ENSMUSG00000065956.3 | 0 | NA | NA | NA | NA | NA | #VALUE! |
| ENSMUSG00000065968.9 | 0 | NA | NA | NA | NA | NA | #VALUE! |
| ENSMUSG00000065987.13 | 0 | NA | NA | NA | NA | NA | #VALUE! |
| ENSMUSG00000065999.14 | 0 | NA | NA | NA | NA | NA | #VALUE! |
| ENSMUSG00000066007.16 | 0 | NA | NA | NA | NA | NA | #VALUE! |
| ENSMUSG00000066009.9 | 0 | NA | NA | NA | NA | NA | #VALUE! |
| ENSMUSG00000066027.8 | 0 | NA | NA | NA | NA | NA | #VALUE! |
| ENSMUSG00000066030.7 | 0 | NA | NA | NA | NA | NA | #VALUE! |
| ENSMUSG00000066031.14 | 0 | NA | NA | NA | NA | NA | #VALUE! |
| ENSMUSG00000066042.5 | 0 | NA | NA | NA | NA | NA | #VALUE! |
| ENSMUSG00000066058.12 | 0 | NA | NA | NA | NA | NA | #VALUE! |
| ENSMUSG00000066060.6 | 0 | NA | NA | NA | NA | NA | #VALUE! |
| ENSMUSG00000066061.7 | 0 | NA | NA | NA | NA | NA | #VALUE! |
| ENSMUSG00000066068.6 | 2.10267 | -0.94636 | 3.822451 | -0.24758 | 0.80446 | NA | #VALUE! |
| ENSMUSG00000066090.6 | 0 | NA | NA | NA | NA | NA | #VALUE! |
| ENSMUSG00000066097.7 | 0 | NA | NA | NA | NA | NA | #VALUE! |
| ENSMUSG00000066100.8 | 0 | NA | NA | NA | NA | NA | #VALUE! |
| ENSMUSG00000066101.6 | 0 | NA | NA | NA | NA | NA | #VALUE! |
| ENSMUSG00000066107.7 | 0 | NA | NA | NA | NA | NA | #VALUE! |
| ENSMUSG00000066108.8 | 0 | NA | NA | NA | NA | NA | #VALUE! |
| ENSMUSG00000066113.17 | 4.456775 | -3.04323 | 3.831092 | -0.79435 | 0.426991 | NA | #VALUE! |
| ENSMUSG00000066116.4 | 1.544453 | 2.009383 | 3.909027 | 0.514037 | 0.607226 | NA | #VALUE! |
| ENSMUSG00000066122.7 | 0 | NA | NA | NA | NA | NA | #VALUE! |
| ENSMUSG00000066129.18 | 0 | NA | NA | NA | NA | NA | #VALUE! |
| ENSMUSG00000066137.13 | 0 | NA | NA | NA | NA | NA | #VALUE! |

| | | | | | | | |
|---|---|---|---|---|---|---|---|
| ENSMUSG00000066141.3 | 0 | NA | NA | NA | NA | NA | #VALUE! |
| ENSMUSG00000066158.8 | 0 | NA | NA | NA | NA | NA | #VALUE! |
| ENSMUSG00000066170.8 | 0 | NA | NA | NA | NA | NA | #VALUE! |
| ENSMUSG00000066176.4 | 0 | NA | NA | NA | NA | NA | #VALUE! |
| ENSMUSG00000066178.5 | 0 | NA | NA | NA | NA | NA | #VALUE! |
| ENSMUSG00000066180.6 | 1.819051 | -4.39655 | 3.931229 | -1.11837 | 0.263411 | NA | #VALUE! |
| ENSMUSG00000066189.10 | 0 | NA | NA | NA | NA | NA | #VALUE! |
| ENSMUSG00000066191.13 | 0 | NA | NA | NA | NA | NA | #VALUE! |
| ENSMUSG00000066196.12 | 0 | NA | NA | NA | NA | NA | #VALUE! |
| ENSMUSG00000066197.6 | 0 | NA | NA | NA | NA | NA | #VALUE! |
| ENSMUSG00000066224.15 | 0 | NA | NA | NA | NA | NA | #VALUE! |
| ENSMUSG00000066239.3 | 0 | NA | NA | NA | NA | NA | #VALUE! |
| ENSMUSG00000066240.4 | 0 | NA | NA | NA | NA | NA | #VALUE! |
| ENSMUSG00000066241.3 | 0 | NA | NA | NA | NA | NA | #VALUE! |
| ENSMUSG00000066242.4 | 0 | NA | NA | NA | NA | NA | #VALUE! |
| ENSMUSG00000066245.6 | 0 | NA | NA | NA | NA | NA | #VALUE! |
| ENSMUSG00000066257.5 | 0 | NA | NA | NA | NA | NA | #VALUE! |
| ENSMUSG00000066262.5 | 0 | NA | NA | NA | NA | NA | #VALUE! |
| ENSMUSG00000066263.11 | 0 | NA | NA | NA | NA | NA | #VALUE! |
| ENSMUSG00000066268.7 | 0 | NA | NA | NA | NA | NA | #VALUE! |
| ENSMUSG00000066269.8 | 0 | NA | NA | NA | NA | NA | #VALUE! |
| ENSMUSG00000066270.4 | 0 | NA | NA | NA | NA | NA | #VALUE! |
| ENSMUSG00000066272.6 | 0 | NA | NA | NA | NA | NA | #VALUE! |
| ENSMUSG00000066273.7 | 0 | NA | NA | NA | NA | NA | #VALUE! |
| ENSMUSG00000066279.4 | 0 | NA | NA | NA | NA | NA | #VALUE! |
| ENSMUSG00000066315.10 | 0.210026 | -1.52494 | 4.058692 | -0.37572 | 0.707123 | NA | #VALUE! |
| ENSMUSG00000066359.7 | 0 | NA | NA | NA | NA | NA | #VALUE! |
| ENSMUSG00000066362.6 | 0 | NA | NA | NA | NA | NA | #VALUE! |
| ENSMUSG00000066364.3 | 0 | NA | NA | NA | NA | NA | #VALUE! |
| ENSMUSG00000066368.2 | 0 | NA | NA | NA | NA | NA | #VALUE! |
| ENSMUSG00000066372.7 | 0 | NA | NA | NA | NA | NA | #VALUE! |
| ENSMUSG00000066378.4 | 0 | NA | NA | NA | NA | NA | #VALUE! |
| ENSMUSG00000066382.5 | 0 | NA | NA | NA | NA | NA | #VALUE! |
| ENSMUSG00000066383.8 | 0 | NA | NA | NA | NA | NA | #VALUE! |
| ENSMUSG00000066389.2 | 0 | NA | NA | NA | NA | NA | #VALUE! |
| ENSMUSG00000066392.13 | 0 | NA | NA | NA | NA | NA | #VALUE! |
| ENSMUSG00000066407.4 | 0.3999 | -2.17566 | 4.018831 | -0.54137 | 0.588256 | NA | #VALUE! |
| ENSMUSG00000066438.7 | 0 | NA | NA | NA | NA | NA | #VALUE! |
| ENSMUSG00000066456.15 | 6.704043 | -0.36169 | 2.987588 | -0.12106 | 0.90364 | NA | #VALUE! |
| ENSMUSG00000066457.7 | 0 | NA | NA | NA | NA | NA | #VALUE! |
| ENSMUSG00000066463.14 | 0 | NA | NA | NA | NA | NA | #VALUE! |
| ENSMUSG00000066475.7 | 0.372394 | 0.012025 | 4.059931 | 0.002962 | 0.997637 | NA | #VALUE! |
| ENSMUSG00000066478.4 | 0 | NA | NA | NA | NA | NA | #VALUE! |
| ENSMUSG00000066487.4 | 0.093255 | 0.868093 | 4.072464 | 0.213162 | 0.831201 | NA | #VALUE! |

| | | | | | | | |
|---|---|---|---|---|---|---|---|
| ENSMUSG00000066489.3 | 0 | NA | NA | NA | NA | NA | #VALUE! |
| ENSMUSG00000066491.5 | 5.063414 | -0.62802 | 2.490453 | -0.25217 | 0.800908 | NA | #VALUE! |
| ENSMUSG00000066500.6 | 0 | NA | NA | NA | NA | NA | #VALUE! |
| ENSMUSG00000066510.6 | 0 | NA | NA | NA | NA | NA | #VALUE! |
| ENSMUSG00000066512.4 | 0 | NA | NA | NA | NA | NA | #VALUE! |
| ENSMUSG00000066515.4 | 0 | NA | NA | NA | NA | NA | #VALUE! |
| ENSMUSG00000066516.6 | 0 | NA | NA | NA | NA | NA | #VALUE! |
| ENSMUSG00000066537.6 | 0 | NA | NA | NA | NA | NA | #VALUE! |
| ENSMUSG00000066538.6 | 0 | NA | NA | NA | NA | NA | #VALUE! |
| ENSMUSG00000066543.9 | 0.364073 | 0.012025 | 4.059931 | 0.002962 | 0.997637 | NA | #VALUE! |
| ENSMUSG00000066553.6 | 8.723965 | 0.449418 | 3.456284 | 0.130029 | 0.896543 | NA | #VALUE! |
| ENSMUSG00000066554.7 | 0.195789 | -1.05581 | 4.073302 | -0.2592 | 0.795479 | NA | #VALUE! |
| ENSMUSG00000066583.5 | 0 | NA | NA | NA | NA | NA | #VALUE! |
| ENSMUSG00000066584.11 | 0 | NA | NA | NA | NA | NA | #VALUE! |
| ENSMUSG00000066586.4 | 0 | NA | NA | NA | NA | NA | #VALUE! |
| ENSMUSG00000066592.4 | 0 | NA | NA | NA | NA | NA | #VALUE! |
| ENSMUSG00000066607.6 | 0 | NA | NA | NA | NA | NA | #VALUE! |
| ENSMUSG00000066620.3 | 0 | NA | NA | NA | NA | NA | #VALUE! |
| ENSMUSG00000066626.6 | 2.497672 | 1.050808 | 3.738884 | 0.281049 | 0.778673 | NA | #VALUE! |
| ENSMUSG00000066629.3 | 0.261539 | 1.68833 | 4.043024 | 0.417591 | 0.676246 | NA | #VALUE! |
| ENSMUSG00000066632.4 | 0 | NA | NA | NA | NA | NA | #VALUE! |
| ENSMUSG00000066640.12 | 2.407276 | 2.332576 | 3.862953 | 0.603832 | 0.545955 | NA | #VALUE! |
| ENSMUSG00000066643.13 | 8.133918 | 6.51637 | 2.905074 | 2.243099 | 0.02489 | NA | #VALUE! |
| ENSMUSG00000066647.9 | 9.516795 | 1.038731 | 2.940392 | 0.353263 | 0.723891 | NA | #VALUE! |
| ENSMUSG00000066652.6 | 0 | NA | NA | NA | NA | NA | #VALUE! |
| ENSMUSG00000066671.5 | 0 | NA | NA | NA | NA | NA | #VALUE! |
| ENSMUSG00000066672.3 | 0 | NA | NA | NA | NA | NA | #VALUE! |
| ENSMUSG00000066677.13 | 0 | NA | NA | NA | NA | NA | #VALUE! |
| ENSMUSG00000066680.5 | 0 | NA | NA | NA | NA | NA | #VALUE! |
| ENSMUSG00000066687.6 | 1.858263 | -1.08829 | 3.837713 | -0.28358 | 0.776733 | NA | #VALUE! |
| ENSMUSG00000066688.5 | 0 | NA | NA | NA | NA | NA | #VALUE! |
| ENSMUSG00000066693.3 | 0 | NA | NA | NA | NA | NA | #VALUE! |
| ENSMUSG00000066697.4 | 0 | NA | NA | NA | NA | NA | #VALUE! |
| ENSMUSG00000066704.7 | 0 | NA | NA | NA | NA | NA | #VALUE! |
| ENSMUSG00000066705.8 | 0 | NA | NA | NA | NA | NA | #VALUE! |
| ENSMUSG00000066714.7 | 0.093255 | 0.868093 | 4.072464 | 0.213162 | 0.831201 | NA | #VALUE! |
| ENSMUSG00000066720.8 | 0 | NA | NA | NA | NA | NA | #VALUE! |
| ENSMUSG00000066721.4 | 0 | NA | NA | NA | NA | NA | #VALUE! |
| ENSMUSG00000066747.7 | 0 | NA | NA | NA | NA | NA | #VALUE! |
| ENSMUSG00000066748.5 | 0 | NA | NA | NA | NA | NA | #VALUE! |
| ENSMUSG00000066749.4 | 0 | NA | NA | NA | NA | NA | #VALUE! |
| ENSMUSG00000066750.4 | 0 | NA | NA | NA | NA | NA | #VALUE! |
| ENSMUSG00000066752.5 | 0 | NA | NA | NA | NA | NA | #VALUE! |
| ENSMUSG00000066755.3 | 0 | NA | NA | NA | NA | NA | #VALUE! |

| | | | | | | | |
|---|---|---|---|---|---|---|---|
| ENSMUSG00000066756.6 | 0 | NA | NA | NA | NA | NA | #VALUE! |
| ENSMUSG00000066760.13 | 0 | NA | NA | NA | NA | NA | #VALUE! |
| ENSMUSG00000066767.7 | 0 | NA | NA | NA | NA | NA | #VALUE! |
| ENSMUSG00000066771.3 | 0 | NA | NA | NA | NA | NA | #VALUE! |
| ENSMUSG00000066772.13 | 0 | NA | NA | NA | NA | NA | #VALUE! |
| ENSMUSG00000066797.7 | 0 | NA | NA | NA | NA | NA | #VALUE! |
| ENSMUSG00000066803.3 | 0 | NA | NA | NA | NA | NA | #VALUE! |
| ENSMUSG00000066804.4 | 0 | NA | NA | NA | NA | NA | #VALUE! |
| ENSMUSG00000066807.3 | 0 | NA | NA | NA | NA | NA | #VALUE! |
| ENSMUSG00000066809.2 | 1.42396 | 3.995882 | 3.938214 | 1.014643 | 0.310276 | NA | #VALUE! |
| ENSMUSG00000066810.3 | 0 | NA | NA | NA | NA | NA | #VALUE! |
| ENSMUSG00000066820.8 | 0 | NA | NA | NA | NA | NA | #VALUE! |
| ENSMUSG00000066828.6 | 0 | NA | NA | NA | NA | NA | #VALUE! |
| ENSMUSG00000066842.19 | 0.18651 | 1.401769 | 4.056354 | 0.345574 | 0.729663 | NA | #VALUE! |
| ENSMUSG00000066850.3 | 0.105013 | -0.97324 | 4.076293 | -0.23876 | 0.811295 | NA | #VALUE! |
| ENSMUSG00000066861.15 | 0 | NA | NA | NA | NA | NA | #VALUE! |
| ENSMUSG00000066867.9 | 0 | NA | NA | NA | NA | NA | #VALUE! |
| ENSMUSG00000066877.12 | 0 | NA | NA | NA | NA | NA | #VALUE! |
| ENSMUSG00000066892.15 | 3.170677 | 5.164728 | 3.920817 | 1.317258 | 0.187752 | NA | #VALUE! |
| ENSMUSG00000066894.15 | 0 | NA | NA | NA | NA | NA | #VALUE! |
| ENSMUSG00000066896.7 | 0 | NA | NA | NA | NA | NA | #VALUE! |
| ENSMUSG00000066897.9 | 0 | NA | NA | NA | NA | NA | #VALUE! |
| ENSMUSG00000066899.7 | 0 | NA | NA | NA | NA | NA | #VALUE! |
| ENSMUSG00000066902.8 | 0.093255 | 0.868093 | 4.072464 | 0.213162 | 0.831201 | NA | #VALUE! |
| ENSMUSG00000066905.5 | 0 | NA | NA | NA | NA | NA | #VALUE! |
| ENSMUSG00000066912.4 | 0 | NA | NA | NA | NA | NA | #VALUE! |
| ENSMUSG00000066914.5 | 0 | NA | NA | NA | NA | NA | #VALUE! |
| ENSMUSG00000066936.5 | 0 | NA | NA | NA | NA | NA | #VALUE! |
| ENSMUSG00000066938.2 | 0 | NA | NA | NA | NA | NA | #VALUE! |
| ENSMUSG00000066952.12 | 7.586744 | 6.414791 | 3.404211 | 1.88437 | 0.059515 | NA | #VALUE! |
| ENSMUSG00000066958.2 | 0 | NA | NA | NA | NA | NA | #VALUE! |
| ENSMUSG00000066964.3 | 0 | NA | NA | NA | NA | NA | #VALUE! |
| ENSMUSG00000066975.13 | 0 | NA | NA | NA | NA | NA | #VALUE! |
| ENSMUSG00000066983.7 | 0.204111 | -1.05581 | 4.073302 | -0.2592 | 0.795479 | NA | #VALUE! |
| ENSMUSG00000067001.12 | 0 | NA | NA | NA | NA | NA | #VALUE! |
| ENSMUSG00000067006.13 | 0 | NA | NA | NA | NA | NA | #VALUE! |
| ENSMUSG00000067010.5 | 0 | NA | NA | NA | NA | NA | #VALUE! |
| ENSMUSG00000067017.6 | 2.24757 | 4.647237 | 3.926849 | 1.183452 | 0.23663 | NA | #VALUE! |
| ENSMUSG00000067028.12 | 0 | NA | NA | NA | NA | NA | #VALUE! |
| ENSMUSG00000067038.7 | 0.195789 | -1.05581 | 4.073302 | -0.2592 | 0.795479 | NA | #VALUE! |
| ENSMUSG00000067047.5 | 0 | NA | NA | NA | NA | NA | #VALUE! |
| ENSMUSG00000067049.9 | 0 | NA | NA | NA | NA | NA | #VALUE! |
| ENSMUSG00000067058.8 | 1.088082 | 1.385035 | 3.931522 | 0.35229 | 0.724621 | NA | #VALUE! |
| ENSMUSG00000067063.2 | 0 | NA | NA | NA | NA | NA | #VALUE! |

| | | | | | | | |
|---|---|---|---|---|---|---|---|
| ENSMUSG00000067064.5 | 0 | NA | NA | NA | NA | NA | #VALUE! |
| ENSMUSG00000067081.14 | 0 | NA | NA | NA | NA | NA | #VALUE! |
| ENSMUSG00000067085.2 | 0 | NA | NA | NA | NA | NA | #VALUE! |
| ENSMUSG00000067101.6 | 0 | NA | NA | NA | NA | NA | #VALUE! |
| ENSMUSG00000067103.7 | 0 | NA | NA | NA | NA | NA | #VALUE! |
| ENSMUSG00000067106.7 | 0 | NA | NA | NA | NA | NA | #VALUE! |
| ENSMUSG00000067121.3 | 0.780616 | -1.73043 | 3.966423 | -0.43627 | 0.662642 | NA | #VALUE! |
| ENSMUSG00000067122.3 | 0 | NA | NA | NA | NA | NA | #VALUE! |
| ENSMUSG00000067147.8 | 0.630079 | -2.90291 | 3.975047 | -0.73028 | 0.465217 | NA | #VALUE! |
| ENSMUSG00000067156.6 | 0 | NA | NA | NA | NA | NA | #VALUE! |
| ENSMUSG00000067158.10 | 6.463156 | -5.12925 | 3.416821 | -1.50117 | 0.13331 | NA | #VALUE! |
| ENSMUSG00000067161.7 | 0.315039 | -1.93549 | 4.038621 | -0.47925 | 0.631763 | NA | #VALUE! |
| ENSMUSG00000067173.4 | 0 | NA | NA | NA | NA | NA | #VALUE! |
| ENSMUSG00000067186.3 | 0 | NA | NA | NA | NA | NA | #VALUE! |
| ENSMUSG00000067189.6 | 0 | NA | NA | NA | NA | NA | #VALUE! |
| ENSMUSG00000067199.5 | 2.692536 | 4.918862 | 3.923417 | 1.253719 | 0.209944 | NA | #VALUE! |
| ENSMUSG00000067201.12 | 0 | NA | NA | NA | NA | NA | #VALUE! |
| ENSMUSG00000067206.4 | 0 | NA | NA | NA | NA | NA | #VALUE! |
| ENSMUSG00000067215.6 | 0 | NA | NA | NA | NA | NA | #VALUE! |
| ENSMUSG00000067220.13 | 0 | NA | NA | NA | NA | NA | #VALUE! |
| ENSMUSG00000067224.4 | 0 | NA | NA | NA | NA | NA | #VALUE! |
| ENSMUSG00000067229.4 | 0 | NA | NA | NA | NA | NA | #VALUE! |
| ENSMUSG00000067231.5 | 0 | NA | NA | NA | NA | NA | #VALUE! |
| ENSMUSG00000067242.12 | 0 | NA | NA | NA | NA | NA | #VALUE! |
| ENSMUSG00000067261.5 | 0 | NA | NA | NA | NA | NA | #VALUE! |
| ENSMUSG00000067262.10 | 0 | NA | NA | NA | NA | NA | #VALUE! |
| ENSMUSG00000067276.6 | 6.731341 | 6.240743 | 3.913441 | 1.594695 | 0.110781 | NA | #VALUE! |
| ENSMUSG00000067285.5 | 0 | NA | NA | NA | NA | NA | #VALUE! |
| ENSMUSG00000067292.4 | 0 | NA | NA | NA | NA | NA | #VALUE! |
| ENSMUSG00000067297.6 | 0 | NA | NA | NA | NA | NA | #VALUE! |
| ENSMUSG00000067299.12 | 0 | NA | NA | NA | NA | NA | #VALUE! |
| ENSMUSG00000067321.6 | 0 | NA | NA | NA | NA | NA | #VALUE! |
| ENSMUSG00000067331.6 | 0 | NA | NA | NA | NA | NA | #VALUE! |
| ENSMUSG00000067338.7 | 0 | NA | NA | NA | NA | NA | #VALUE! |
| ENSMUSG00000067341.7 | 0 | NA | NA | NA | NA | NA | #VALUE! |
| ENSMUSG00000067344.8 | 6.424451 | -0.19992 | 2.242732 | -0.08914 | 0.928968 | NA | #VALUE! |
| ENSMUSG00000067351.2 | 0.105013 | -0.97324 | 4.076293 | -0.23876 | 0.811295 | NA | #VALUE! |
| ENSMUSG00000067352.7 | 0 | NA | NA | NA | NA | NA | #VALUE! |
| ENSMUSG00000067356.6 | 0 | NA | NA | NA | NA | NA | #VALUE! |
| ENSMUSG00000067360.13 | 0 | NA | NA | NA | NA | NA | #VALUE! |
| ENSMUSG00000067370.5 | 0 | NA | NA | NA | NA | NA | #VALUE! |
| ENSMUSG00000067377.13 | 3.263932 | 5.206263 | 3.920419 | 1.327986 | 0.184183 | NA | #VALUE! |
| ENSMUSG00000067389.4 | 0 | NA | NA | NA | NA | NA | #VALUE! |
| ENSMUSG00000067399.11 | 0 | NA | NA | NA | NA | NA | #VALUE! |

| | | | | | | | |
|---|---|---|---|---|---|---|---|
| ENSMUSG00000067424.14 | 3.980712 | 1.263476 | 3.781164 | 0.33415 | 0.738266 | NA | #VALUE! |
| ENSMUSG00000067430.8 | 0 | NA | NA | NA | NA | NA | #VALUE! |
| ENSMUSG00000067438.6 | 0 | NA | NA | NA | NA | NA | #VALUE! |
| ENSMUSG00000067441.6 | 0 | NA | NA | NA | NA | NA | #VALUE! |
| ENSMUSG00000067455.6 | 0 | NA | NA | NA | NA | NA | #VALUE! |
| ENSMUSG00000067482.3 | 0 | NA | NA | NA | NA | NA | #VALUE! |
| ENSMUSG00000067513.2 | 0 | NA | NA | NA | NA | NA | #VALUE! |
| ENSMUSG00000067519.2 | 0 | NA | NA | NA | NA | NA | #VALUE! |
| ENSMUSG00000067522.4 | 0 | NA | NA | NA | NA | NA | #VALUE! |
| ENSMUSG00000067524.5 | 0 | NA | NA | NA | NA | NA | #VALUE! |
| ENSMUSG00000067525.4 | 0 | NA | NA | NA | NA | NA | #VALUE! |
| ENSMUSG00000067526.4 | 0 | NA | NA | NA | NA | NA | #VALUE! |
| ENSMUSG00000067528.4 | 0 | NA | NA | NA | NA | NA | #VALUE! |
| ENSMUSG00000067529.3 | 0 | NA | NA | NA | NA | NA | #VALUE! |
| ENSMUSG00000067541.8 | 0 | NA | NA | NA | NA | NA | #VALUE! |
| ENSMUSG00000067543.9 | 0 | NA | NA | NA | NA | NA | #VALUE! |
| ENSMUSG00000067545.4 | 0 | NA | NA | NA | NA | NA | #VALUE! |
| ENSMUSG00000067547.7 | 0 | NA | NA | NA | NA | NA | #VALUE! |
| ENSMUSG00000067555.3 | 0 | NA | NA | NA | NA | NA | #VALUE! |
| ENSMUSG00000067561.13 | 0 | NA | NA | NA | NA | NA | #VALUE! |
| ENSMUSG00000067562.12 | 0 | NA | NA | NA | NA | NA | #VALUE! |
| ENSMUSG00000067571.5 | 0 | NA | NA | NA | NA | NA | #VALUE! |
| ENSMUSG00000067575.6 | 1.303829 | -1.71873 | 3.904795 | -0.44016 | 0.659822 | NA | #VALUE! |
| ENSMUSG00000067577.9 | 0 | NA | NA | NA | NA | NA | #VALUE! |
| ENSMUSG00000067578.4 | 0 | NA | NA | NA | NA | NA | #VALUE! |
| ENSMUSG00000067586.5 | 4.289261 | 1.244736 | 3.776844 | 0.32957 | 0.741725 | NA | #VALUE! |
| ENSMUSG00000067591.13 | 0.105013 | -0.97324 | 4.076293 | -0.23876 | 0.811295 | NA | #VALUE! |
| ENSMUSG00000067594.7 | 0 | NA | NA | NA | NA | NA | #VALUE! |
| ENSMUSG00000067596.5 | 0 | NA | NA | NA | NA | NA | #VALUE! |
| ENSMUSG00000067597.6 | 0 | NA | NA | NA | NA | NA | #VALUE! |
| ENSMUSG00000067599.14 | 0 | NA | NA | NA | NA | NA | #VALUE! |
| ENSMUSG00000067604.9 | 0 | NA | NA | NA | NA | NA | #VALUE! |
| ENSMUSG00000067608.6 | 0.587367 | -2.74297 | 3.982932 | -0.68868 | 0.491024 | NA | #VALUE! |
| ENSMUSG00000067610.2 | 0 | NA | NA | NA | NA | NA | #VALUE! |
| ENSMUSG00000067613.7 | 0 | NA | NA | NA | NA | NA | #VALUE! |
| ENSMUSG00000067614.6 | 0 | NA | NA | NA | NA | NA | #VALUE! |
| ENSMUSG00000067615.5 | 0 | NA | NA | NA | NA | NA | #VALUE! |
| ENSMUSG00000067616.12 | 0 | NA | NA | NA | NA | NA | #VALUE! |
| ENSMUSG00000067627.3 | 0 | NA | NA | NA | NA | NA | #VALUE! |
| ENSMUSG00000067629.14 | 0 | NA | NA | NA | NA | NA | #VALUE! |
| ENSMUSG00000067642.12 | 0 | NA | NA | NA | NA | NA | #VALUE! |
| ENSMUSG00000067649.4 | 0 | NA | NA | NA | NA | NA | #VALUE! |
| ENSMUSG00000067653.13 | 0 | NA | NA | NA | NA | NA | #VALUE! |
| ENSMUSG00000067656.14 | 1.678593 | 4.255795 | 3.933066 | 1.082055 | 0.279228 | NA | #VALUE! |

| | | | | | | | |
|---|---|---|---|---|---|---|---|
| ENSMUSG00000067679.4 | 0 | NA | NA | NA | NA | NA | #VALUE! |
| ENSMUSG00000067684.2 | 0 | NA | NA | NA | NA | NA | #VALUE! |
| ENSMUSG00000067698.5 | 0 | NA | NA | NA | NA | NA | #VALUE! |
| ENSMUSG00000067700.6 | 0 | NA | NA | NA | NA | NA | #VALUE! |
| ENSMUSG00000067702.6 | 0 | NA | NA | NA | NA | NA | #VALUE! |
| ENSMUSG00000067704.2 | 0 | NA | NA | NA | NA | NA | #VALUE! |
| ENSMUSG00000067714.14 | 0 | NA | NA | NA | NA | NA | #VALUE! |
| ENSMUSG00000067716.4 | 0 | NA | NA | NA | NA | NA | #VALUE! |
| ENSMUSG00000067719.6 | 2.779437 | 0.337625 | 3.206476 | 0.105295 | 0.916142 | NA | #VALUE! |
| ENSMUSG00000067724.6 | 0 | NA | NA | NA | NA | NA | #VALUE! |
| ENSMUSG00000067736.3 | 0 | NA | NA | NA | NA | NA | #VALUE! |
| ENSMUSG00000067745.5 | 0 | NA | NA | NA | NA | NA | #VALUE! |
| ENSMUSG00000067750.4 | 0 | NA | NA | NA | NA | NA | #VALUE! |
| ENSMUSG00000067764.12 | 0 | NA | NA | NA | NA | NA | #VALUE! |
| ENSMUSG00000067767.6 | 0 | NA | NA | NA | NA | NA | #VALUE! |
| ENSMUSG00000067768.13 | 4.103229 | 5.534434 | 3.917649 | 1.412693 | 0.157746 | NA | #VALUE! |
| ENSMUSG00000067771.10 | 0 | NA | NA | NA | NA | NA | #VALUE! |
| ENSMUSG00000067773.7 | 0 | NA | NA | NA | NA | NA | #VALUE! |
| ENSMUSG00000067780.4 | 0 | NA | NA | NA | NA | NA | #VALUE! |
| ENSMUSG00000067786.17 | 0 | NA | NA | NA | NA | NA | #VALUE! |
| ENSMUSG00000067795.14 | 0 | NA | NA | NA | NA | NA | #VALUE! |
| ENSMUSG00000067818.7 | 0 | NA | NA | NA | NA | NA | #VALUE! |
| ENSMUSG00000067835.5 | 0 | NA | NA | NA | NA | NA | #VALUE! |
| ENSMUSG00000067848.13 | 0 | NA | NA | NA | NA | NA | #VALUE! |
| ENSMUSG00000067855.8 | 0 | NA | NA | NA | NA | NA | #VALUE! |
| ENSMUSG00000067860.12 | 0 | NA | NA | NA | NA | NA | #VALUE! |
| ENSMUSG00000067869.5 | 0.195789 | -1.05581 | 4.073302 | -0.2592 | 0.795479 | NA | #VALUE! |
| ENSMUSG00000067872.3 | 0 | NA | NA | NA | NA | NA | #VALUE! |
| ENSMUSG00000067878.14 | 2.019402 | 4.503826 | 3.928936 | 1.146322 | 0.251662 | NA | #VALUE! |
| ENSMUSG00000067879.4 | 0 | NA | NA | NA | NA | NA | #VALUE! |
| ENSMUSG00000067882.15 | 0 | NA | NA | NA | NA | NA | #VALUE! |
| ENSMUSG00000067909.5 | 0 | NA | NA | NA | NA | NA | #VALUE! |
| ENSMUSG00000067916.10 | 0.932552 | 3.421684 | 3.953399 | 0.865504 | 0.386762 | NA | #VALUE! |
| ENSMUSG00000067919.9 | 0 | NA | NA | NA | NA | NA | #VALUE! |
| ENSMUSG00000067924.5 | 6.85113 | 0.585141 | 3.267959 | 0.179054 | 0.857895 | NA | #VALUE! |
| ENSMUSG00000067925.5 | 0 | NA | NA | NA | NA | NA | #VALUE! |
| ENSMUSG00000067929.4 | 0 | NA | NA | NA | NA | NA | #VALUE! |
| ENSMUSG00000067941.6 | 0 | NA | NA | NA | NA | NA | #VALUE! |
| ENSMUSG00000067942.7 | 9.458824 | -1.111 | 1.983557 | -0.5601 | 0.575409 | NA | #VALUE! |
| ENSMUSG00000067949.6 | 0 | NA | NA | NA | NA | NA | #VALUE! |
| ENSMUSG00000067951.6 | 0 | NA | NA | NA | NA | NA | #VALUE! |
| ENSMUSG00000067971.6 | 0 | NA | NA | NA | NA | NA | #VALUE! |
| ENSMUSG00000067973.12 | 0 | NA | NA | NA | NA | NA | #VALUE! |
| ENSMUSG00000067996.6 | 0 | NA | NA | NA | NA | NA | #VALUE! |

| | | | | | | | |
|---|---|---|---|---|---|---|---|
| ENSMUSG00000067998.12 | 0 | NA | NA | NA | NA | NA | #VALUE! |
| ENSMUSG00000068008.9 | 0 | NA | NA | NA | NA | NA | #VALUE! |
| ENSMUSG00000068009.12 | 0 | NA | NA | NA | NA | NA | #VALUE! |
| ENSMUSG00000068014.6 | 0 | NA | NA | NA | NA | NA | #VALUE! |
| ENSMUSG00000068037.12 | 0 | NA | NA | NA | NA | NA | #VALUE! |
| ENSMUSG00000068048.9 | 0 | NA | NA | NA | NA | NA | #VALUE! |
| ENSMUSG00000068062.4 | 0 | NA | NA | NA | NA | NA | #VALUE! |
| ENSMUSG00000068067.3 | 0 | NA | NA | NA | NA | NA | #VALUE! |
| ENSMUSG00000068068.2 | 0 | NA | NA | NA | NA | NA | #VALUE! |
| ENSMUSG00000068071.3 | 0 | NA | NA | NA | NA | NA | #VALUE! |
| ENSMUSG00000068073.3 | 0 | NA | NA | NA | NA | NA | #VALUE! |
| ENSMUSG00000068074.4 | 0 | NA | NA | NA | NA | NA | #VALUE! |
| ENSMUSG00000068075.4 | 0 | NA | NA | NA | NA | NA | #VALUE! |
| ENSMUSG00000068078.5 | 0 | NA | NA | NA | NA | NA | #VALUE! |
| ENSMUSG00000068079.6 | 0 | NA | NA | NA | NA | NA | #VALUE! |
| ENSMUSG00000068082.14 | 0 | NA | NA | NA | NA | NA | #VALUE! |
| ENSMUSG00000068085.7 | 0.793895 | 1.740142 | 3.972877 | 0.438006 | 0.661382 | NA | #VALUE! |
| ENSMUSG00000068099.6 | 0 | NA | NA | NA | NA | NA | #VALUE! |
| ENSMUSG00000068101.12 | 0.420053 | -2.33493 | 4.007343 | -0.58266 | 0.56012 | NA | #VALUE! |
| ENSMUSG00000068105.12 | 0 | NA | NA | NA | NA | NA | #VALUE! |
| ENSMUSG00000068113.7 | 0 | NA | NA | NA | NA | NA | #VALUE! |
| ENSMUSG00000068115.14 | 1.75449 | 1.75698 | 3.873983 | 0.453533 | 0.650165 | NA | #VALUE! |
| ENSMUSG00000068117.11 | 0 | NA | NA | NA | NA | NA | #VALUE! |
| ENSMUSG00000068120.5 | 0 | NA | NA | NA | NA | NA | #VALUE! |
| ENSMUSG00000068122.11 | 0 | NA | NA | NA | NA | NA | #VALUE! |
| ENSMUSG00000068129.6 | 0.105013 | -0.97324 | 4.076293 | -0.23876 | 0.811295 | NA | #VALUE! |
| ENSMUSG00000068130.12 | 0 | NA | NA | NA | NA | NA | #VALUE! |
| ENSMUSG00000068141.6 | 0 | NA | NA | NA | NA | NA | #VALUE! |
| ENSMUSG00000068149.6 | 0 | NA | NA | NA | NA | NA | #VALUE! |
| ENSMUSG00000068151.8 | 0 | NA | NA | NA | NA | NA | #VALUE! |
| ENSMUSG00000068154.6 | 0 | NA | NA | NA | NA | NA | #VALUE! |
| ENSMUSG00000068165.4 | 0 | NA | NA | NA | NA | NA | #VALUE! |
| ENSMUSG00000068167.7 | 0 | NA | NA | NA | NA | NA | #VALUE! |
| ENSMUSG00000068173.13 | 0 | NA | NA | NA | NA | NA | #VALUE! |
| ENSMUSG00000068180.14 | 0 | NA | NA | NA | NA | NA | #VALUE! |
| ENSMUSG00000068181.2 | 0 | NA | NA | NA | NA | NA | #VALUE! |
| ENSMUSG00000068182.4 | 0 | NA | NA | NA | NA | NA | #VALUE! |
| ENSMUSG00000068200.7 | 0 | NA | NA | NA | NA | NA | #VALUE! |
| ENSMUSG00000068205.15 | 105.5497 | -1.79205 | 1.054368 | -1.69965 | NA | NA | #VALUE! |
| ENSMUSG00000068218.7 | 0 | NA | NA | NA | NA | NA | #VALUE! |
| ENSMUSG00000068219.9 | 0 | NA | NA | NA | NA | NA | #VALUE! |
| ENSMUSG00000068226.4 | 0 | NA | NA | NA | NA | NA | #VALUE! |
| ENSMUSG00000068227.10 | 0.210026 | -1.52494 | 4.058692 | -0.37572 | 0.707123 | NA | #VALUE! |
| ENSMUSG00000068231.6 | 0 | NA | NA | NA | NA | NA | #VALUE! |

| | | | | | | | |
|---|---|---|---|---|---|---|---|
| ENSMUSG00000068232.4 | 0 | NA | NA | NA | NA | NA | #VALUE! |
| ENSMUSG00000068234.8 | 0 | NA | NA | NA | NA | NA | #VALUE! |
| ENSMUSG00000068243.7 | 0.204111 | -1.05581 | 4.073302 | -0.2592 | 0.795479 | NA | #VALUE! |
| ENSMUSG00000068245.15 | 4.698938 | -5.75981 | 3.916303 | -1.47073 | 0.141365 | NA | #VALUE! |
| ENSMUSG00000068252.7 | 0 | NA | NA | NA | NA | NA | #VALUE! |
| ENSMUSG00000068259.5 | 0 | NA | NA | NA | NA | NA | #VALUE! |
| ENSMUSG00000068264.12 | 12.30836 | 1.486349 | 2.556981 | 0.581291 | NA | NA | #VALUE! |
| ENSMUSG00000068289.7 | 0 | NA | NA | NA | NA | NA | #VALUE! |
| ENSMUSG00000068299.12 | 1.459101 | -0.46049 | 3.8524 | -0.11953 | 0.904852 | NA | #VALUE! |
| ENSMUSG00000068302.9 | 0 | NA | NA | NA | NA | NA | #VALUE! |
| ENSMUSG00000068303.7 | 0 | NA | NA | NA | NA | NA | #VALUE! |
| ENSMUSG00000068314.6 | 0 | NA | NA | NA | NA | NA | #VALUE! |
| ENSMUSG00000068323.13 | 0 | NA | NA | NA | NA | NA | #VALUE! |
| ENSMUSG00000068324.6 | 0 | NA | NA | NA | NA | NA | #VALUE! |
| ENSMUSG00000068327.6 | 0 | NA | NA | NA | NA | NA | #VALUE! |
| ENSMUSG00000068329.13 | 1.305573 | 3.898287 | 3.940396 | 0.989314 | 0.32251 | NA | #VALUE! |
| ENSMUSG00000068335.7 | 2.019402 | 4.503826 | 3.928936 | 1.146322 | 0.251662 | NA | #VALUE! |
| ENSMUSG00000068341.9 | 0 | NA | NA | NA | NA | NA | #VALUE! |
| ENSMUSG00000068348.5 | 0 | NA | NA | NA | NA | NA | #VALUE! |
| ENSMUSG00000068349.13 | 0 | NA | NA | NA | NA | NA | #VALUE! |
| ENSMUSG00000068373.16 | 0 | NA | NA | NA | NA | NA | #VALUE! |
| ENSMUSG00000068392.4 | 0 | NA | NA | NA | NA | NA | #VALUE! |
| ENSMUSG00000068399.6 | 0 | NA | NA | NA | NA | NA | #VALUE! |
| ENSMUSG00000068407.4 | 0 | NA | NA | NA | NA | NA | #VALUE! |
| ENSMUSG00000068417.9 | 0 | NA | NA | NA | NA | NA | #VALUE! |
| ENSMUSG00000068428.8 | 0 | NA | NA | NA | NA | NA | #VALUE! |
| ENSMUSG00000068431.4 | 0 | NA | NA | NA | NA | NA | #VALUE! |
| ENSMUSG00000068437.5 | 0 | NA | NA | NA | NA | NA | #VALUE! |
| ENSMUSG00000068452.9 | 0 | NA | NA | NA | NA | NA | #VALUE! |
| ENSMUSG00000068466.7 | 0 | NA | NA | NA | NA | NA | #VALUE! |
| ENSMUSG00000068480.7 | 0 | NA | NA | NA | NA | NA | #VALUE! |
| ENSMUSG00000068487.7 | 0.195789 | -1.05581 | 4.073302 | -0.2592 | 0.795479 | NA | #VALUE! |
| ENSMUSG00000068506.4 | 0 | NA | NA | NA | NA | NA | #VALUE! |
| ENSMUSG00000068522.5 | 0 | NA | NA | NA | NA | NA | #VALUE! |
| ENSMUSG00000068535.3 | 0 | NA | NA | NA | NA | NA | #VALUE! |
| ENSMUSG00000068536.12 | 0 | NA | NA | NA | NA | NA | #VALUE! |
| ENSMUSG00000068537.3 | 0 | NA | NA | NA | NA | NA | #VALUE! |
| ENSMUSG00000068547.8 | 0 | NA | NA | NA | NA | NA | #VALUE! |
| ENSMUSG00000068574.3 | 0 | NA | NA | NA | NA | NA | #VALUE! |
| ENSMUSG00000068579.6 | 0.309124 | -1.83395 | 4.047982 | -0.45305 | 0.650511 | NA | #VALUE! |
| ENSMUSG00000068580.12 | 20.3466 | -2.34862 | 2.236218 | -1.05027 | NA | NA | #VALUE! |
| ENSMUSG00000068600.11 | 0 | NA | NA | NA | NA | NA | #VALUE! |
| ENSMUSG00000068601.3 | 6.656829 | -6.26304 | 3.913498 | -1.60037 | 0.109517 | NA | #VALUE! |
| ENSMUSG00000068604.5 | 0 | NA | NA | NA | NA | NA | #VALUE! |

| | | | | | | | |
|---|---|---|---|---|---|---|---|
| ENSMUSG00000068614.8 | 0 | NA | NA | NA | NA | NA | #VALUE! |
| ENSMUSG00000068615.5 | 0 | NA | NA | NA | NA | NA | #VALUE! |
| ENSMUSG00000068617.6 | 0 | NA | NA | NA | NA | NA | #VALUE! |
| ENSMUSG00000068629.5 | 0 | NA | NA | NA | NA | NA | #VALUE! |
| ENSMUSG00000068631.9 | 0 | NA | NA | NA | NA | NA | #VALUE! |
| ENSMUSG00000068647.3 | 0 | NA | NA | NA | NA | NA | #VALUE! |
| ENSMUSG00000068674.4 | 0.587367 | -2.74297 | 3.982932 | -0.68868 | 0.491024 | NA | #VALUE! |
| ENSMUSG00000068680.2 | 0 | NA | NA | NA | NA | NA | #VALUE! |
| ENSMUSG00000068689.9 | 0 | NA | NA | NA | NA | NA | #VALUE! |
| ENSMUSG00000068696.7 | 0 | NA | NA | NA | NA | NA | #VALUE! |
| ENSMUSG00000068697.8 | 0 | NA | NA | NA | NA | NA | #VALUE! |
| ENSMUSG00000068699.13 | 0 | NA | NA | NA | NA | NA | #VALUE! |
| ENSMUSG00000068739.14 | 5.296072 | -1.59481 | 3.77181 | -0.42282 | 0.672423 | NA | #VALUE! |
| ENSMUSG00000068740.15 | 0 | NA | NA | NA | NA | NA | #VALUE! |
| ENSMUSG00000068744.13 | 0 | NA | NA | NA | NA | NA | #VALUE! |
| ENSMUSG00000068745.7 | 0 | NA | NA | NA | NA | NA | #VALUE! |
| ENSMUSG00000068747.15 | 74.10251 | -0.79488 | 2.713766 | -0.29291 | NA | NA | #VALUE! |
| ENSMUSG00000068748.8 | 0 | NA | NA | NA | NA | NA | #VALUE! |
| ENSMUSG00000068758.9 | 0.420053 | -2.33493 | 4.007343 | -0.58266 | 0.56012 | NA | #VALUE! |
| ENSMUSG00000068776.8 | 0 | NA | NA | NA | NA | NA | #VALUE! |
| ENSMUSG00000068794.8 | 0 | NA | NA | NA | NA | NA | #VALUE! |
| ENSMUSG00000068806.6 | 0 | NA | NA | NA | NA | NA | #VALUE! |
| ENSMUSG00000068808.3 | 0 | NA | NA | NA | NA | NA | #VALUE! |
| ENSMUSG00000068809.5 | 0 | NA | NA | NA | NA | NA | #VALUE! |
| ENSMUSG00000068810.6 | 0 | NA | NA | NA | NA | NA | #VALUE! |
| ENSMUSG00000068814.3 | 0 | NA | NA | NA | NA | NA | #VALUE! |
| ENSMUSG00000068815.4 | 0 | NA | NA | NA | NA | NA | #VALUE! |
| ENSMUSG00000068816.4 | 0 | NA | NA | NA | NA | NA | #VALUE! |
| ENSMUSG00000068817.6 | 0 | NA | NA | NA | NA | NA | #VALUE! |
| ENSMUSG00000068818.6 | 0 | NA | NA | NA | NA | NA | #VALUE! |
| ENSMUSG00000068819.4 | 0 | NA | NA | NA | NA | NA | #VALUE! |
| ENSMUSG00000068824.5 | 0 | NA | NA | NA | NA | NA | #VALUE! |
| ENSMUSG00000068855.4 | 0 | NA | NA | NA | NA | NA | #VALUE! |
| ENSMUSG00000068859.6 | 0 | NA | NA | NA | NA | NA | #VALUE! |
| ENSMUSG00000068860.6 | 0 | NA | NA | NA | NA | NA | #VALUE! |
| ENSMUSG00000068879.5 | 0 | NA | NA | NA | NA | NA | #VALUE! |
| ENSMUSG00000068885.5 | 0 | NA | NA | NA | NA | NA | #VALUE! |
| ENSMUSG00000068887.5 | 0 | NA | NA | NA | NA | NA | #VALUE! |
| ENSMUSG00000068888.4 | 0 | NA | NA | NA | NA | NA | #VALUE! |
| ENSMUSG00000068889.2 | 0 | NA | NA | NA | NA | NA | #VALUE! |
| ENSMUSG00000068890.9 | 0 | NA | NA | NA | NA | NA | #VALUE! |
| ENSMUSG00000068893.6 | 0 | NA | NA | NA | NA | NA | #VALUE! |
| ENSMUSG00000068917.13 | 2.238125 | 4.666312 | 3.926586 | 1.188389 | 0.23468 | NA | #VALUE! |
| ENSMUSG00000068922.15 | 2.595048 | -0.19616 | 3.797087 | -0.05166 | 0.9588 | NA | #VALUE! |

| | | | | | | | |
|---|---|---|---|---|---|---|---|
| ENSMUSG00000068940.8 | 0 | NA | NA | NA | NA | NA | #VALUE! |
| ENSMUSG00000068947.3 | 0 | NA | NA | NA | NA | NA | #VALUE! |
| ENSMUSG00000068948.5 | 0 | NA | NA | NA | NA | NA | #VALUE! |
| ENSMUSG00000068950.4 | 0 | NA | NA | NA | NA | NA | #VALUE! |
| ENSMUSG00000068957.2 | 0 | NA | NA | NA | NA | NA | #VALUE! |
| ENSMUSG00000068959.7 | 0.88817 | -3.35104 | 3.956997 | -0.84686 | 0.397071 | NA | #VALUE! |
| ENSMUSG00000068962.8 | 0 | NA | NA | NA | NA | NA | #VALUE! |
| ENSMUSG00000068964.6 | 0 | NA | NA | NA | NA | NA | #VALUE! |
| ENSMUSG00000068966.11 | 7.029333 | 6.30364 | 3.555098 | 1.773127 | 0.076208 | NA | #VALUE! |
| ENSMUSG00000068969.4 | 2.245218 | -4.69006 | 3.926744 | -1.19439 | 0.232326 | NA | #VALUE! |
| ENSMUSG00000068999.14 | 0 | NA | NA | NA | NA | NA | #VALUE! |
| ENSMUSG00000069008.4 | 0 | NA | NA | NA | NA | NA | #VALUE! |
| ENSMUSG00000069011.6 | 0 | NA | NA | NA | NA | NA | #VALUE! |
| ENSMUSG00000069014.5 | 0.18651 | 1.401769 | 4.056354 | 0.345574 | 0.729663 | NA | #VALUE! |
| ENSMUSG00000069036.4 | 0 | NA | NA | NA | NA | NA | #VALUE! |
| ENSMUSG00000069038.3 | 0 | NA | NA | NA | NA | NA | #VALUE! |
| ENSMUSG00000069041.9 | 0 | NA | NA | NA | NA | NA | #VALUE! |
| ENSMUSG00000069044.7 | 3.255407 | -5.24108 | 3.920419 | -1.33687 | 0.181266 | NA | #VALUE! |
| ENSMUSG00000069053.12 | 0 | NA | NA | NA | NA | NA | #VALUE! |
| ENSMUSG00000069072.10 | 0 | NA | NA | NA | NA | NA | #VALUE! |
| ENSMUSG00000069074.3 | 0 | NA | NA | NA | NA | NA | #VALUE! |
| ENSMUSG00000069080.6 | 0 | NA | NA | NA | NA | NA | #VALUE! |
| ENSMUSG00000069083.5 | 0 | NA | NA | NA | NA | NA | #VALUE! |
| ENSMUSG00000069085.6 | 0 | NA | NA | NA | NA | NA | #VALUE! |
| ENSMUSG00000069114.9 | 6.267301 | 5.172612 | 3.43725 | 1.504869 | 0.132358 | NA | #VALUE! |
| ENSMUSG00000069117.5 | 0.53502 | 0.80809 | 4.028608 | 0.200588 | 0.841021 | NA | #VALUE! |
| ENSMUSG00000069118.4 | 0 | NA | NA | NA | NA | NA | #VALUE! |
| ENSMUSG00000069125.3 | 0.093255 | 0.868093 | 4.072464 | 0.213162 | 0.831201 | NA | #VALUE! |
| ENSMUSG00000069132.4 | 0 | NA | NA | NA | NA | NA | #VALUE! |
| ENSMUSG00000069136.5 | 0 | NA | NA | NA | NA | NA | #VALUE! |
| ENSMUSG00000069171.15 | 2.998348 | 1.05654 | 3.79586 | 0.27834 | 0.780751 | NA | #VALUE! |
| ENSMUSG00000069188.6 | 0.279766 | 1.808031 | 4.036658 | 0.447903 | 0.654223 | NA | #VALUE! |
| ENSMUSG00000069189.6 | 0 | NA | NA | NA | NA | NA | #VALUE! |
| ENSMUSG00000069196.7 | 0 | NA | NA | NA | NA | NA | #VALUE! |
| ENSMUSG00000069223.11 | 0 | NA | NA | NA | NA | NA | #VALUE! |
| ENSMUSG00000069227.5 | 0 | NA | NA | NA | NA | NA | #VALUE! |
| ENSMUSG00000069236.6 | 0 | NA | NA | NA | NA | NA | #VALUE! |
| ENSMUSG00000069248.12 | 0 | NA | NA | NA | NA | NA | #VALUE! |
| ENSMUSG00000069258.11 | 0 | NA | NA | NA | NA | NA | #VALUE! |
| ENSMUSG00000069259.11 | 0 | NA | NA | NA | NA | NA | #VALUE! |
| ENSMUSG00000069265.3 | 0 | NA | NA | NA | NA | NA | #VALUE! |
| ENSMUSG00000069266.6 | 0 | NA | NA | NA | NA | NA | #VALUE! |
| ENSMUSG00000069267.3 | 0 | NA | NA | NA | NA | NA | #VALUE! |
| ENSMUSG00000069268.5 | 0 | NA | NA | NA | NA | NA | #VALUE! |

| | | | | | | | |
|---|---|---|---|---|---|---|---|
| ENSMUSG00000069270.7 | 0.783156 | -3.1627 | 3.963922 | -0.79787 | 0.424945 | NA | #VALUE! |
| ENSMUSG00000069272.7 | 0 | NA | NA | NA | NA | NA | #VALUE! |
| ENSMUSG00000069273.3 | 0 | NA | NA | NA | NA | NA | #VALUE! |
| ENSMUSG00000069274.4 | 0 | NA | NA | NA | NA | NA | #VALUE! |
| ENSMUSG00000069280.3 | 0 | NA | NA | NA | NA | NA | #VALUE! |
| ENSMUSG00000069289.3 | 0 | NA | NA | NA | NA | NA | #VALUE! |
| ENSMUSG00000069292.5 | 0 | NA | NA | NA | NA | NA | #VALUE! |
| ENSMUSG00000069294.3 | 0 | NA | NA | NA | NA | NA | #VALUE! |
| ENSMUSG00000069295.3 | 0 | NA | NA | NA | NA | NA | #VALUE! |
| ENSMUSG00000069296.5 | 0 | NA | NA | NA | NA | NA | #VALUE! |
| ENSMUSG00000069297.2 | 0 | NA | NA | NA | NA | NA | #VALUE! |
| ENSMUSG00000069299.3 | 0 | NA | NA | NA | NA | NA | #VALUE! |
| ENSMUSG00000069300.4 | 0 | NA | NA | NA | NA | NA | #VALUE! |
| ENSMUSG00000069301.6 | 0 | NA | NA | NA | NA | NA | #VALUE! |
| ENSMUSG00000069302.6 | 0 | NA | NA | NA | NA | NA | #VALUE! |
| ENSMUSG00000069303.8 | 0 | NA | NA | NA | NA | NA | #VALUE! |
| ENSMUSG00000069305.4 | 0 | NA | NA | NA | NA | NA | #VALUE! |
| ENSMUSG00000069306.6 | 0 | NA | NA | NA | NA | NA | #VALUE! |
| ENSMUSG00000069307.8 | 0 | NA | NA | NA | NA | NA | #VALUE! |
| ENSMUSG00000069308.8 | 0 | NA | NA | NA | NA | NA | #VALUE! |
| ENSMUSG00000069309.3 | 0 | NA | NA | NA | NA | NA | #VALUE! |
| ENSMUSG00000069310.5 | 0 | NA | NA | NA | NA | NA | #VALUE! |
| ENSMUSG00000069315.3 | 0 | NA | NA | NA | NA | NA | #VALUE! |
| ENSMUSG00000069355.13 | 0 | NA | NA | NA | NA | NA | #VALUE! |
| ENSMUSG00000069372.4 | 0 | NA | NA | NA | NA | NA | #VALUE! |
| ENSMUSG00000069376.3 | 0 | NA | NA | NA | NA | NA | #VALUE! |
| ENSMUSG00000069378.15 | 0 | NA | NA | NA | NA | NA | #VALUE! |
| ENSMUSG00000069379.4 | 0.893055 | -2.28286 | 3.95457 | -0.57727 | 0.563756 | NA | #VALUE! |
| ENSMUSG00000069385.11 | 0 | NA | NA | NA | NA | NA | #VALUE! |
| ENSMUSG00000069390.6 | 0 | NA | NA | NA | NA | NA | #VALUE! |
| ENSMUSG00000069421.5 | 0 | NA | NA | NA | NA | NA | #VALUE! |
| ENSMUSG00000069430.4 | 0 | NA | NA | NA | NA | NA | #VALUE! |
| ENSMUSG00000069439.4 | 0 | NA | NA | NA | NA | NA | #VALUE! |
| ENSMUSG00000069441.4 | 0 | NA | NA | NA | NA | NA | #VALUE! |
| ENSMUSG00000069475.2 | 0 | NA | NA | NA | NA | NA | #VALUE! |
| ENSMUSG00000069476.14 | 0 | NA | NA | NA | NA | NA | #VALUE! |
| ENSMUSG00000069515.7 | 0 | NA | NA | NA | NA | NA | #VALUE! |
| ENSMUSG00000069518.6 | 0 | NA | NA | NA | NA | NA | #VALUE! |
| ENSMUSG00000069540.5 | 0 | NA | NA | NA | NA | NA | #VALUE! |
| ENSMUSG00000069554.5 | 0 | NA | NA | NA | NA | NA | #VALUE! |
| ENSMUSG00000069581.13 | 0 | NA | NA | NA | NA | NA | #VALUE! |
| ENSMUSG00000069582.4 | 0 | NA | NA | NA | NA | NA | #VALUE! |
| ENSMUSG00000069583.3 | 0 | NA | NA | NA | NA | NA | #VALUE! |
| ENSMUSG00000069584.5 | 0 | NA | NA | NA | NA | NA | #VALUE! |

| | | | | | | | |
|---|---|---|---|---|---|---|---|
| ENSMUSG00000069588.4 | 0 | NA | NA | NA | NA | NA | #VALUE! |
| ENSMUSG00000069607.5 | 0 | NA | NA | NA | NA | NA | #VALUE! |
| ENSMUSG00000069609.8 | 0 | NA | NA | NA | NA | NA | #VALUE! |
| ENSMUSG00000069620.7 | 0 | NA | NA | NA | NA | NA | #VALUE! |
| ENSMUSG00000069622.6 | 0 | NA | NA | NA | NA | NA | #VALUE! |
| ENSMUSG00000069631.15 | 0 | NA | NA | NA | NA | NA | #VALUE! |
| ENSMUSG00000069668.5 | 0 | NA | NA | NA | NA | NA | #VALUE! |
| ENSMUSG00000069670.9 | 1.119062 | 3.679627 | 3.945843 | 0.932533 | 0.351061 | NA | #VALUE! |
| ENSMUSG00000069706.4 | 0 | NA | NA | NA | NA | NA | #VALUE! |
| ENSMUSG00000069707.2 | 0 | NA | NA | NA | NA | NA | #VALUE! |
| ENSMUSG00000069708.4 | 0 | NA | NA | NA | NA | NA | #VALUE! |
| ENSMUSG00000069712.6 | 0 | NA | NA | NA | NA | NA | #VALUE! |
| ENSMUSG00000069713.3 | 0 | NA | NA | NA | NA | NA | #VALUE! |
| ENSMUSG00000069717.8 | 0 | NA | NA | NA | NA | NA | #VALUE! |
| ENSMUSG00000069718.5 | 0 | NA | NA | NA | NA | NA | #VALUE! |
| ENSMUSG00000069720.5 | 0 | NA | NA | NA | NA | NA | #VALUE! |
| ENSMUSG00000069721.3 | 0 | NA | NA | NA | NA | NA | #VALUE! |
| ENSMUSG00000069722.5 | 0 | NA | NA | NA | NA | NA | #VALUE! |
| ENSMUSG00000069727.6 | 0 | NA | NA | NA | NA | NA | #VALUE! |
| ENSMUSG00000069763.4 | 5.480525 | 3.957043 | 3.871863 | 1.022 | 0.306781 | NA | #VALUE! |
| ENSMUSG00000069785.5 | 0 | NA | NA | NA | NA | NA | #VALUE! |
| ENSMUSG00000069793.13 | 4.018784 | -0.18515 | 3.7715 | -0.04909 | 0.960847 | NA | #VALUE! |
| ENSMUSG00000069796.4 | 0 | NA | NA | NA | NA | NA | #VALUE! |
| ENSMUSG00000069804.3 | 0 | NA | NA | NA | NA | NA | #VALUE! |
| ENSMUSG00000069806.6 | 0 | NA | NA | NA | NA | NA | #VALUE! |
| ENSMUSG00000069808.14 | 8.230967 | -6.57382 | 3.315613 | -1.98268 | 0.047403 | NA | #VALUE! |
| ENSMUSG00000069814.7 | 0 | NA | NA | NA | NA | NA | #VALUE! |
| ENSMUSG00000069816.6 | 0 | NA | NA | NA | NA | NA | #VALUE! |
| ENSMUSG00000069818.7 | 0 | NA | NA | NA | NA | NA | #VALUE! |
| ENSMUSG00000069823.12 | 0 | NA | NA | NA | NA | NA | #VALUE! |
| ENSMUSG00000069824.6 | 0 | NA | NA | NA | NA | NA | #VALUE! |
| ENSMUSG00000069825.13 | 0 | NA | NA | NA | NA | NA | #VALUE! |
| ENSMUSG00000069830.11 | 0 | NA | NA | NA | NA | NA | #VALUE! |
| ENSMUSG00000069855.10 | 0 | NA | NA | NA | NA | NA | #VALUE! |
| ENSMUSG00000069867.4 | 0 | NA | NA | NA | NA | NA | #VALUE! |
| ENSMUSG00000069873.5 | 0 | NA | NA | NA | NA | NA | #VALUE! |
| ENSMUSG00000069890.4 | 0 | NA | NA | NA | NA | NA | #VALUE! |
| ENSMUSG00000069892.10 | 30.11452 | 2.341589 | 2.482242 | 0.943336 | NA | NA | #VALUE! |
| ENSMUSG00000069893.11 | 0 | NA | NA | NA | NA | NA | #VALUE! |
| ENSMUSG00000069910.3 | 2.299168 | 4.692706 | 3.926229 | 1.19522 | 0.232001 | NA | #VALUE! |
| ENSMUSG00000069911.12 | 0 | NA | NA | NA | NA | NA | #VALUE! |
| ENSMUSG00000069913.4 | 0 | NA | NA | NA | NA | NA | #VALUE! |
| ENSMUSG00000069920.11 | 11.5325 | -0.82971 | 3.20082 | -0.25922 | NA | NA | #VALUE! |
| ENSMUSG00000069925.7 | 0 | NA | NA | NA | NA | NA | #VALUE! |

| | | | | | | | |
|---|---|---|---|---|---|---|---|
| ENSMUSG00000069939.9 | 0 | NA | NA | NA | NA | NA | #VALUE! |
| ENSMUSG00000069962.4 | 0 | NA | NA | NA | NA | NA | #VALUE! |
| ENSMUSG00000069971.6 | 0 | NA | NA | NA | NA | NA | #VALUE! |
| ENSMUSG00000069986.5 | 0 | NA | NA | NA | NA | NA | #VALUE! |
| ENSMUSG00000069996.4 | 0 | NA | NA | NA | NA | NA | #VALUE! |
| ENSMUSG00000069998.6 | 0 | NA | NA | NA | NA | NA | #VALUE! |
| ENSMUSG00000070000.14 | 0 | NA | NA | NA | NA | NA | #VALUE! |
| ENSMUSG00000070003.8 | 0 | NA | NA | NA | NA | NA | #VALUE! |
| ENSMUSG00000070023.6 | 0 | NA | NA | NA | NA | NA | #VALUE! |
| ENSMUSG00000070025.9 | 0 | NA | NA | NA | NA | NA | #VALUE! |
| ENSMUSG00000070031.12 | 5.502056 | 5.955668 | 3.914897 | 1.521283 | 0.128189 | NA | #VALUE! |
| ENSMUSG00000070034.14 | 7.61377 | -2.56407 | 3.23206 | -0.79332 | 0.42759 | NA | #VALUE! |
| ENSMUSG00000070063.3 | 0 | NA | NA | NA | NA | NA | #VALUE! |
| ENSMUSG00000070071.3 | 0 | NA | NA | NA | NA | NA | #VALUE! |
| ENSMUSG00000070072.3 | 0 | NA | NA | NA | NA | NA | #VALUE! |
| ENSMUSG00000070073.3 | 0 | NA | NA | NA | NA | NA | #VALUE! |
| ENSMUSG00000070074.3 | 0 | NA | NA | NA | NA | NA | #VALUE! |
| ENSMUSG00000070075.3 | 0 | NA | NA | NA | NA | NA | #VALUE! |
| ENSMUSG00000070076.4 | 0 | NA | NA | NA | NA | NA | #VALUE! |
| ENSMUSG00000070077.4 | 0 | NA | NA | NA | NA | NA | #VALUE! |
| ENSMUSG00000070078.3 | 0 | NA | NA | NA | NA | NA | #VALUE! |
| ENSMUSG00000070080.3 | 0 | NA | NA | NA | NA | NA | #VALUE! |
| ENSMUSG00000070081.4 | 0 | NA | NA | NA | NA | NA | #VALUE! |
| ENSMUSG00000070084.5 | 0 | NA | NA | NA | NA | NA | #VALUE! |
| ENSMUSG00000070099.3 | 0 | NA | NA | NA | NA | NA | #VALUE! |
| ENSMUSG00000070100.3 | 0 | NA | NA | NA | NA | NA | #VALUE! |
| ENSMUSG00000070101.3 | 0 | NA | NA | NA | NA | NA | #VALUE! |
| ENSMUSG00000070102.3 | 0 | NA | NA | NA | NA | NA | #VALUE! |
| ENSMUSG00000070103.4 | 0 | NA | NA | NA | NA | NA | #VALUE! |
| ENSMUSG00000070104.3 | 0 | NA | NA | NA | NA | NA | #VALUE! |
| ENSMUSG00000070105.5 | 0 | NA | NA | NA | NA | NA | #VALUE! |
| ENSMUSG00000070106.3 | 0 | NA | NA | NA | NA | NA | #VALUE! |
| ENSMUSG00000070107.2 | 0 | NA | NA | NA | NA | NA | #VALUE! |
| ENSMUSG00000070108.4 | 0 | NA | NA | NA | NA | NA | #VALUE! |
| ENSMUSG00000070109.4 | 0 | NA | NA | NA | NA | NA | #VALUE! |
| ENSMUSG00000070110.4 | 0 | NA | NA | NA | NA | NA | #VALUE! |
| ENSMUSG00000070111.5 | 0 | NA | NA | NA | NA | NA | #VALUE! |
| ENSMUSG00000070126.3 | 0 | NA | NA | NA | NA | NA | #VALUE! |
| ENSMUSG00000070127.4 | 0 | NA | NA | NA | NA | NA | #VALUE! |
| ENSMUSG00000070128.5 | 0 | NA | NA | NA | NA | NA | #VALUE! |
| ENSMUSG00000070129.4 | 0 | NA | NA | NA | NA | NA | #VALUE! |
| ENSMUSG00000070130.3 | 0 | NA | NA | NA | NA | NA | #VALUE! |
| ENSMUSG00000070133.3 | 0 | NA | NA | NA | NA | NA | #VALUE! |
| ENSMUSG00000070136.3 | 0 | NA | NA | NA | NA | NA | #VALUE! |

| | | | | | | | |
|---|---|---|---|---|---|---|---|
| ENSMUSG00000070138.3 | 0 | NA | NA | NA | NA | NA | #VALUE! |
| ENSMUSG00000070139.3 | 0 | NA | NA | NA | NA | NA | #VALUE! |
| ENSMUSG00000070140.3 | 0 | NA | NA | NA | NA | NA | #VALUE! |
| ENSMUSG00000070141.4 | 0 | NA | NA | NA | NA | NA | #VALUE! |
| ENSMUSG00000070144.3 | 0 | NA | NA | NA | NA | NA | #VALUE! |
| ENSMUSG00000070166.3 | 0 | NA | NA | NA | NA | NA | #VALUE! |
| ENSMUSG00000070167.3 | 0 | NA | NA | NA | NA | NA | #VALUE! |
| ENSMUSG00000070170.3 | 0 | NA | NA | NA | NA | NA | #VALUE! |
| ENSMUSG00000070173.3 | 0 | NA | NA | NA | NA | NA | #VALUE! |
| ENSMUSG00000070175.3 | 0 | NA | NA | NA | NA | NA | #VALUE! |
| ENSMUSG00000070182.3 | 0 | NA | NA | NA | NA | NA | #VALUE! |
| ENSMUSG00000070203.3 | 0 | NA | NA | NA | NA | NA | #VALUE! |
| ENSMUSG00000070214.3 | 0 | NA | NA | NA | NA | NA | #VALUE! |
| ENSMUSG00000070225.3 | 0 | NA | NA | NA | NA | NA | #VALUE! |
| ENSMUSG00000070227.3 | 0 | NA | NA | NA | NA | NA | #VALUE! |
| ENSMUSG00000070235.3 | 0 | NA | NA | NA | NA | NA | #VALUE! |
| ENSMUSG00000070236.4 | 0 | NA | NA | NA | NA | NA | #VALUE! |
| ENSMUSG00000070263.3 | 0 | NA | NA | NA | NA | NA | #VALUE! |
| ENSMUSG00000070271.8 | 1.554565 | 1.99418 | 3.909407 | 0.510098 | 0.609983 | NA | #VALUE! |
| ENSMUSG00000070280.14 | 0 | NA | NA | NA | NA | NA | #VALUE! |
| ENSMUSG00000070282.7 | 0 | NA | NA | NA | NA | NA | #VALUE! |
| ENSMUSG00000070287.4 | 1.177985 | 3.72614 | 3.944615 | 0.944614 | 0.344856 | NA | #VALUE! |
| ENSMUSG00000070291.5 | 0 | NA | NA | NA | NA | NA | #VALUE! |
| ENSMUSG00000070298.7 | 0 | NA | NA | NA | NA | NA | #VALUE! |
| ENSMUSG00000070304.13 | 0 | NA | NA | NA | NA | NA | #VALUE! |
| ENSMUSG00000070306.4 | 0 | NA | NA | NA | NA | NA | #VALUE! |
| ENSMUSG00000070311.5 | 0 | NA | NA | NA | NA | NA | #VALUE! |
| ENSMUSG00000070313.9 | 0 | NA | NA | NA | NA | NA | #VALUE! |
| ENSMUSG00000070323.12 | 0 | NA | NA | NA | NA | NA | #VALUE! |
| ENSMUSG00000070324.7 | 0 | NA | NA | NA | NA | NA | #VALUE! |
| ENSMUSG00000070330.3 | 0 | NA | NA | NA | NA | NA | #VALUE! |
| ENSMUSG00000070331.15 | 0 | NA | NA | NA | NA | NA | #VALUE! |
| ENSMUSG00000070332.5 | 0 | NA | NA | NA | NA | NA | #VALUE! |
| ENSMUSG00000070334.5 | 0 | NA | NA | NA | NA | NA | #VALUE! |
| ENSMUSG00000070335.5 | 0 | NA | NA | NA | NA | NA | #VALUE! |
| ENSMUSG00000070336.4 | 0 | NA | NA | NA | NA | NA | #VALUE! |
| ENSMUSG00000070337.5 | 0 | NA | NA | NA | NA | NA | #VALUE! |
| ENSMUSG00000070342.4 | 0 | NA | NA | NA | NA | NA | #VALUE! |
| ENSMUSG00000070345.4 | 4.289739 | 5.598228 | 3.917179 | 1.429148 | 0.152962 | NA | #VALUE! |
| ENSMUSG00000070354.14 | 0 | NA | NA | NA | NA | NA | #VALUE! |
| ENSMUSG00000070360.6 | 0 | NA | NA | NA | NA | NA | #VALUE! |
| ENSMUSG00000070366.6 | 0 | NA | NA | NA | NA | NA | #VALUE! |
| ENSMUSG00000070368.10 | 0.914325 | 3.387843 | 3.954492 | 0.856708 | 0.391606 | NA | #VALUE! |
| ENSMUSG00000070369.14 | 0 | NA | NA | NA | NA | NA | #VALUE! |

| | | | | | | | |
|---|---|---|---|---|---|---|---|
| ENSMUSG00000070371.12 | 2.245218 | -4.69006 | 3.926744 | -1.19439 | 0.232326 | NA | #VALUE! |
| ENSMUSG00000070374.8 | 0 | NA | NA | NA | NA | NA | #VALUE! |
| ENSMUSG00000070375.8 | 0 | NA | NA | NA | NA | NA | #VALUE! |
| ENSMUSG00000070377.7 | 0 | NA | NA | NA | NA | NA | #VALUE! |
| ENSMUSG00000070378.4 | 0 | NA | NA | NA | NA | NA | #VALUE! |
| ENSMUSG00000070379.6 | 0 | NA | NA | NA | NA | NA | #VALUE! |
| ENSMUSG00000070380.3 | 0 | NA | NA | NA | NA | NA | #VALUE! |
| ENSMUSG00000070382.9 | 0 | NA | NA | NA | NA | NA | #VALUE! |
| ENSMUSG00000070383.6 | 0 | NA | NA | NA | NA | NA | #VALUE! |
| ENSMUSG00000070385.13 | 0 | NA | NA | NA | NA | NA | #VALUE! |
| ENSMUSG00000070388.4 | 0 | NA | NA | NA | NA | NA | #VALUE! |
| ENSMUSG00000070390.13 | 0 | NA | NA | NA | NA | NA | #VALUE! |
| ENSMUSG00000070392.6 | 0 | NA | NA | NA | NA | NA | #VALUE! |
| ENSMUSG00000070417.7 | 0 | NA | NA | NA | NA | NA | #VALUE! |
| ENSMUSG00000070419.12 | 0 | NA | NA | NA | NA | NA | #VALUE! |
| ENSMUSG00000070420.5 | 0 | NA | NA | NA | NA | NA | #VALUE! |
| ENSMUSG00000070421.5 | 0 | NA | NA | NA | NA | NA | #VALUE! |
| ENSMUSG00000070423.7 | 0 | NA | NA | NA | NA | NA | #VALUE! |
| ENSMUSG00000070424.13 | 0 | NA | NA | NA | NA | NA | #VALUE! |
| ENSMUSG00000070425.14 | 0 | NA | NA | NA | NA | NA | #VALUE! |
| ENSMUSG00000070438.6 | 0 | NA | NA | NA | NA | NA | #VALUE! |
| ENSMUSG00000070443.7 | 0 | NA | NA | NA | NA | NA | #VALUE! |
| ENSMUSG00000070448.14 | 0 | NA | NA | NA | NA | NA | #VALUE! |
| ENSMUSG00000070458.6 | 0 | NA | NA | NA | NA | NA | #VALUE! |
| ENSMUSG00000070459.7 | 0 | NA | NA | NA | NA | NA | #VALUE! |
| ENSMUSG00000070460.8 | 0 | NA | NA | NA | NA | NA | #VALUE! |
| ENSMUSG00000070461.4 | 0 | NA | NA | NA | NA | NA | #VALUE! |
| ENSMUSG00000070464.2 | 0 | NA | NA | NA | NA | NA | #VALUE! |
| ENSMUSG00000070465.10 | 0 | NA | NA | NA | NA | NA | #VALUE! |
| ENSMUSG00000070469.13 | 0 | NA | NA | NA | NA | NA | #VALUE! |
| ENSMUSG00000070471.4 | 0 | NA | NA | NA | NA | NA | #VALUE! |
| ENSMUSG00000070476.11 | 0.587367 | -2.74297 | 3.982932 | -0.68868 | 0.491024 | NA | #VALUE! |
| ENSMUSG00000070489.7 | 0 | NA | NA | NA | NA | NA | #VALUE! |
| ENSMUSG00000070490.5 | 0 | NA | NA | NA | NA | NA | #VALUE! |
| ENSMUSG00000070495.12 | 0 | NA | NA | NA | NA | NA | #VALUE! |
| ENSMUSG00000070498.4 | 0 | NA | NA | NA | NA | NA | #VALUE! |
| ENSMUSG00000070501.15 | 0 | NA | NA | NA | NA | NA | #VALUE! |
| ENSMUSG00000070504.10 | 0 | NA | NA | NA | NA | NA | #VALUE! |
| ENSMUSG00000070509.16 | 0 | NA | NA | NA | NA | NA | #VALUE! |
| ENSMUSG00000070522.5 | 0 | NA | NA | NA | NA | NA | #VALUE! |
| ENSMUSG00000070524.2 | 0 | NA | NA | NA | NA | NA | #VALUE! |
| ENSMUSG00000070526.3 | 0 | NA | NA | NA | NA | NA | #VALUE! |
| ENSMUSG00000070527.4 | 0 | NA | NA | NA | NA | NA | #VALUE! |
| ENSMUSG00000070529.8 | 0 | NA | NA | NA | NA | NA | #VALUE! |

| | | | | | | | |
|---|---|---|---|---|---|---|---|
| ENSMUSG00000070530.6 | 0 | NA | NA | NA | NA | NA | #VALUE! |
| ENSMUSG00000070531.5 | 0 | NA | NA | NA | NA | NA | #VALUE! |
| ENSMUSG00000070532.6 | 0 | NA | NA | NA | NA | NA | #VALUE! |
| ENSMUSG00000070533.12 | 0 | NA | NA | NA | NA | NA | #VALUE! |
| ENSMUSG00000070546.3 | 0 | NA | NA | NA | NA | NA | #VALUE! |
| ENSMUSG00000070547.5 | 0 | NA | NA | NA | NA | NA | #VALUE! |
| ENSMUSG00000070550.3 | 0 | NA | NA | NA | NA | NA | #VALUE! |
| ENSMUSG00000070551.4 | 0 | NA | NA | NA | NA | NA | #VALUE! |
| ENSMUSG00000070552.3 | 0 | NA | NA | NA | NA | NA | #VALUE! |
| ENSMUSG00000070563.7 | 0 | NA | NA | NA | NA | NA | #VALUE! |
| ENSMUSG00000070564.16 | 0 | NA | NA | NA | NA | NA | #VALUE! |
| ENSMUSG00000070568.13 | 0 | NA | NA | NA | NA | NA | #VALUE! |
| ENSMUSG00000070570.6 | 0 | NA | NA | NA | NA | NA | #VALUE! |
| ENSMUSG00000070572.8 | 0 | NA | NA | NA | NA | NA | #VALUE! |
| ENSMUSG00000070574.3 | 0 | NA | NA | NA | NA | NA | #VALUE! |
| ENSMUSG00000070577.6 | 0 | NA | NA | NA | NA | NA | #VALUE! |
| ENSMUSG00000070583.2 | 0 | NA | NA | NA | NA | NA | #VALUE! |
| ENSMUSG00000070601.6 | 0.093255 | 0.868093 | 4.072464 | 0.213162 | 0.831201 | NA | #VALUE! |
| ENSMUSG00000070604.6 | 1.370524 | -3.97623 | 3.939448 | -1.00934 | 0.312813 | NA | #VALUE! |
| ENSMUSG00000070605.5 | 33.73386 | 0.09771 | 1.895539 | 0.051547 | NA | NA | #VALUE! |
| ENSMUSG00000070606.5 | 0 | NA | NA | NA | NA | NA | #VALUE! |
| ENSMUSG00000070609.4 | 0 | NA | NA | NA | NA | NA | #VALUE! |
| ENSMUSG00000070610.5 | 0 | NA | NA | NA | NA | NA | #VALUE! |
| ENSMUSG00000070616.16 | 0 | NA | NA | NA | NA | NA | #VALUE! |
| ENSMUSG00000070617.3 | 0 | NA | NA | NA | NA | NA | #VALUE! |
| ENSMUSG00000070618.11 | 0 | NA | NA | NA | NA | NA | #VALUE! |
| ENSMUSG00000070619.6 | 0 | NA | NA | NA | NA | NA | #VALUE! |
| ENSMUSG00000070621.4 | 0 | NA | NA | NA | NA | NA | #VALUE! |
| ENSMUSG00000070637.3 | 0 | NA | NA | NA | NA | NA | #VALUE! |
| ENSMUSG00000070641.4 | 0 | NA | NA | NA | NA | NA | #VALUE! |
| ENSMUSG00000070643.12 | 8.111942 | 6.510777 | 3.343377 | 1.947365 | 0.051491 | NA | #VALUE! |
| ENSMUSG00000070645.6 | 0 | NA | NA | NA | NA | NA | #VALUE! |
| ENSMUSG00000070667.8 | 0.291524 | 0.468739 | 4.052149 | 0.115677 | 0.907909 | NA | #VALUE! |
| ENSMUSG00000070677.9 | 0 | NA | NA | NA | NA | NA | #VALUE! |
| ENSMUSG00000070683.8 | 0 | NA | NA | NA | NA | NA | #VALUE! |
| ENSMUSG00000070686.11 | 0 | NA | NA | NA | NA | NA | #VALUE! |
| ENSMUSG00000070687.12 | 0 | NA | NA | NA | NA | NA | #VALUE! |
| ENSMUSG00000070691.11 | 0.841418 | 3.240539 | 3.959551 | 0.818411 | 0.413123 | NA | #VALUE! |
| ENSMUSG00000070695.5 | 0 | NA | NA | NA | NA | NA | #VALUE! |
| ENSMUSG00000070699.6 | 6.615827 | -6.26 | 3.913512 | -1.59959 | 0.10969 | NA | #VALUE! |
| ENSMUSG00000070702.10 | 0 | NA | NA | NA | NA | NA | #VALUE! |
| ENSMUSG00000070705.4 | 24.35421 | 3.379248 | 2.378949 | 1.42048 | NA | NA | #VALUE! |
| ENSMUSG00000070708.6 | 0 | NA | NA | NA | NA | NA | #VALUE! |
| ENSMUSG00000070709.13 | 6.753859 | -2.77363 | 3.240933 | -0.85581 | 0.392102 | NA | #VALUE! |

| | | | | | | | |
|---|---|---|---|---|---|---|---|
| ENSMUSG00000070717.2 | 0 | NA | NA | NA | NA | NA | #VALUE! |
| ENSMUSG00000070719.5 | 0 | NA | NA | NA | NA | NA | #VALUE! |
| ENSMUSG00000070720.5 | 0 | NA | NA | NA | NA | NA | #VALUE! |
| ENSMUSG00000070729.6 | 0 | NA | NA | NA | NA | NA | #VALUE! |
| ENSMUSG00000070732.3 | 0 | NA | NA | NA | NA | NA | #VALUE! |
| ENSMUSG00000070777.6 | 0 | NA | NA | NA | NA | NA | #VALUE! |
| ENSMUSG00000070778.6 | 0 | NA | NA | NA | NA | NA | #VALUE! |
| ENSMUSG00000070794.12 | 0 | NA | NA | NA | NA | NA | #VALUE! |
| ENSMUSG00000070796.12 | 0 | NA | NA | NA | NA | NA | #VALUE! |
| ENSMUSG00000070797.5 | 0 | NA | NA | NA | NA | NA | #VALUE! |
| ENSMUSG00000070798.5 | 0 | NA | NA | NA | NA | NA | #VALUE! |
| ENSMUSG00000070799.5 | 0 | NA | NA | NA | NA | NA | #VALUE! |
| ENSMUSG00000070802.6 | 0 | NA | NA | NA | NA | NA | #VALUE! |
| ENSMUSG00000070803.7 | 0 | NA | NA | NA | NA | NA | #VALUE! |
| ENSMUSG00000070806.6 | 0 | NA | NA | NA | NA | NA | #VALUE! |
| ENSMUSG00000070808.5 | 0.18651 | 1.401769 | 4.056354 | 0.345574 | 0.729663 | NA | #VALUE! |
| ENSMUSG00000070810.11 | 0 | NA | NA | NA | NA | NA | #VALUE! |
| ENSMUSG00000070811.7 | 0 | NA | NA | NA | NA | NA | #VALUE! |
| ENSMUSG00000070814.18 | 1.177985 | 3.72614 | 3.944615 | 0.944614 | 0.344856 | NA | #VALUE! |
| ENSMUSG00000070815.4 | 0 | NA | NA | NA | NA | NA | #VALUE! |
| ENSMUSG00000070816.4 | 0 | NA | NA | NA | NA | NA | #VALUE! |
| ENSMUSG00000070817.4 | 0 | NA | NA | NA | NA | NA | #VALUE! |
| ENSMUSG00000070820.4 | 0 | NA | NA | NA | NA | NA | #VALUE! |
| ENSMUSG00000070821.7 | 0 | NA | NA | NA | NA | NA | #VALUE! |
| ENSMUSG00000070822.6 | 0 | NA | NA | NA | NA | NA | #VALUE! |
| ENSMUSG00000070828.11 | 0 | NA | NA | NA | NA | NA | #VALUE! |
| ENSMUSG00000070834.6 | 0 | NA | NA | NA | NA | NA | #VALUE! |
| ENSMUSG00000070837.7 | 0 | NA | NA | NA | NA | NA | #VALUE! |
| ENSMUSG00000070841.6 | 0 | NA | NA | NA | NA | NA | #VALUE! |
| ENSMUSG00000070844.12 | 0 | NA | NA | NA | NA | NA | #VALUE! |
| ENSMUSG00000070847.14 | 0 | NA | NA | NA | NA | NA | #VALUE! |
| ENSMUSG00000070852.3 | 0 | NA | NA | NA | NA | NA | #VALUE! |
| ENSMUSG00000070853.6 | 0 | NA | NA | NA | NA | NA | #VALUE! |
| ENSMUSG00000070855.9 | 0 | NA | NA | NA | NA | NA | #VALUE! |
| ENSMUSG00000070856.6 | 0 | NA | NA | NA | NA | NA | #VALUE! |
| ENSMUSG00000070857.5 | 0 | NA | NA | NA | NA | NA | #VALUE! |
| ENSMUSG00000070858.13 | 0 | NA | NA | NA | NA | NA | #VALUE! |
| ENSMUSG00000070866.5 | 0 | NA | NA | NA | NA | NA | #VALUE! |
| ENSMUSG00000070867.5 | 0 | NA | NA | NA | NA | NA | #VALUE! |
| ENSMUSG00000070868.12 | 0.336567 | 1.944925 | 4.029936 | 0.482619 | 0.629366 | NA | #VALUE! |
| ENSMUSG00000070870.7 | 0 | NA | NA | NA | NA | NA | #VALUE! |
| ENSMUSG00000070873.6 | 0.18651 | 1.401769 | 4.056354 | 0.345574 | 0.729663 | NA | #VALUE! |
| ENSMUSG00000070875.6 | 0 | NA | NA | NA | NA | NA | #VALUE! |
| ENSMUSG00000070877.5 | 0 | NA | NA | NA | NA | NA | #VALUE! |

| | | | | | | | |
|---|---|---|---|---|---|---|---|
| ENSMUSG00000070880.11 | 0 | NA | NA | NA | NA | NA | #VALUE! |
| ENSMUSG00000070883.4 | 5.156447 | -2.98453 | 3.566168 | -0.8369 | 0.402649 | NA | #VALUE! |
| ENSMUSG00000070890.4 | 0 | NA | NA | NA | NA | NA | #VALUE! |
| ENSMUSG00000070891.3 | 0 | NA | NA | NA | NA | NA | #VALUE! |
| ENSMUSG00000070896.3 | 0 | NA | NA | NA | NA | NA | #VALUE! |
| ENSMUSG00000070902.6 | 0 | NA | NA | NA | NA | NA | #VALUE! |
| ENSMUSG00000070904.5 | 0 | NA | NA | NA | NA | NA | #VALUE! |
| ENSMUSG00000070908.2 | 0 | NA | NA | NA | NA | NA | #VALUE! |
| ENSMUSG00000070933.11 | 0 | NA | NA | NA | NA | NA | #VALUE! |
| ENSMUSG00000070942.9 | 0 | NA | NA | NA | NA | NA | #VALUE! |
| ENSMUSG00000070943.4 | 0 | NA | NA | NA | NA | NA | #VALUE! |
| ENSMUSG00000070960.5 | 0 | NA | NA | NA | NA | NA | #VALUE! |
| ENSMUSG00000070977.5 | 0 | NA | NA | NA | NA | NA | #VALUE! |
| ENSMUSG00000070979.6 | 0 | NA | NA | NA | NA | NA | #VALUE! |
| ENSMUSG00000070980.5 | 0 | NA | NA | NA | NA | NA | #VALUE! |
| ENSMUSG00000070983.4 | 0 | NA | NA | NA | NA | NA | #VALUE! |
| ENSMUSG00000070985.4 | 0 | NA | NA | NA | NA | NA | #VALUE! |
| ENSMUSG00000070990.3 | 0 | NA | NA | NA | NA | NA | #VALUE! |
| ENSMUSG00000070995.11 | 0 | NA | NA | NA | NA | NA | #VALUE! |
| ENSMUSG00000070997.5 | 6.14722 | 6.111548 | 3.672676 | 1.664059 | 0.096101 | NA | #VALUE! |
| ENSMUSG00000070999.3 | 0 | NA | NA | NA | NA | NA | #VALUE! |
| ENSMUSG00000071000.4 | 0 | NA | NA | NA | NA | NA | #VALUE! |
| ENSMUSG00000071001.5 | 0 | NA | NA | NA | NA | NA | #VALUE! |
| ENSMUSG00000071005.8 | 0 | NA | NA | NA | NA | NA | #VALUE! |
| ENSMUSG00000071015.3 | 0 | NA | NA | NA | NA | NA | #VALUE! |
| ENSMUSG00000071019.12 | 0 | NA | NA | NA | NA | NA | #VALUE! |
| ENSMUSG00000071033.2 | 0 | NA | NA | NA | NA | NA | #VALUE! |
| ENSMUSG00000071035.6 | 0 | NA | NA | NA | NA | NA | #VALUE! |
| ENSMUSG00000071036.3 | 0 | NA | NA | NA | NA | NA | #VALUE! |
| ENSMUSG00000071037.7 | 2.076747 | -3.02061 | 3.905091 | -0.7735 | 0.439224 | NA | #VALUE! |
| ENSMUSG00000071040.3 | 0 | NA | NA | NA | NA | NA | #VALUE! |
| ENSMUSG00000071047.6 | 0 | NA | NA | NA | NA | NA | #VALUE! |
| ENSMUSG00000071052.5 | 0 | NA | NA | NA | NA | NA | #VALUE! |
| ENSMUSG00000071064.16 | 0 | NA | NA | NA | NA | NA | #VALUE! |
| ENSMUSG00000071065.7 | 0 | NA | NA | NA | NA | NA | #VALUE! |
| ENSMUSG00000071068.8 | 0 | NA | NA | NA | NA | NA | #VALUE! |
| ENSMUSG00000071073.6 | 0 | NA | NA | NA | NA | NA | #VALUE! |
| ENSMUSG00000071078.7 | 7.532692 | -0.76582 | 3.186321 | -0.24035 | 0.810062 | NA | #VALUE! |
| ENSMUSG00000071089.4 | 0 | NA | NA | NA | NA | NA | #VALUE! |
| ENSMUSG00000071102.2 | 0 | NA | NA | NA | NA | NA | #VALUE! |
| ENSMUSG00000071103.11 | 2.424635 | 4.780725 | 3.925082 | 1.217993 | 0.223226 | NA | #VALUE! |
| ENSMUSG00000071104.11 | 0 | NA | NA | NA | NA | NA | #VALUE! |
| ENSMUSG00000071112.13 | 0 | NA | NA | NA | NA | NA | #VALUE! |
| ENSMUSG00000071113.5 | 0 | NA | NA | NA | NA | NA | #VALUE! |

| | | | | | | | |
|---|---|---|---|---|---|---|---|
| ENSMUSG00000071138.5 | 0 | NA | NA | NA | NA | NA | #VALUE! |
| ENSMUSG00000071141.8 | 0.967199 | -0.19126 | 3.910416 | -0.04891 | 0.96099 | NA | #VALUE! |
| ENSMUSG00000071147.3 | 0 | NA | NA | NA | NA | NA | #VALUE! |
| ENSMUSG00000071149.4 | 0 | NA | NA | NA | NA | NA | #VALUE! |
| ENSMUSG00000071150.2 | 0 | NA | NA | NA | NA | NA | #VALUE! |
| ENSMUSG00000071158.10 | 0 | NA | NA | NA | NA | NA | #VALUE! |
| ENSMUSG00000071165.4 | 0 | NA | NA | NA | NA | NA | #VALUE! |
| ENSMUSG00000071169.2 | 0 | NA | NA | NA | NA | NA | #VALUE! |
| ENSMUSG00000071176.11 | 5.556981 | 0.997788 | 3.542446 | 0.281666 | 0.778199 | NA | #VALUE! |
| ENSMUSG00000071179.9 | 0.105013 | -0.97324 | 4.076293 | -0.23876 | 0.811295 | NA | #VALUE! |
| ENSMUSG00000071181.3 | 0 | NA | NA | NA | NA | NA | #VALUE! |
| ENSMUSG00000071185.6 | 0 | NA | NA | NA | NA | NA | #VALUE! |
| ENSMUSG00000071192.7 | 0 | NA | NA | NA | NA | NA | #VALUE! |
| ENSMUSG00000071195.5 | 0 | NA | NA | NA | NA | NA | #VALUE! |
| ENSMUSG00000071202.6 | 0 | NA | NA | NA | NA | NA | #VALUE! |
| ENSMUSG00000071203.7 | 5.882171 | 0.711543 | 3.758703 | 0.189305 | 0.849854 | NA | #VALUE! |
| ENSMUSG00000071207.5 | 0 | NA | NA | NA | NA | NA | #VALUE! |
| ENSMUSG00000071229.4 | 0 | NA | NA | NA | NA | NA | #VALUE! |
| ENSMUSG00000071230.8 | 0 | NA | NA | NA | NA | NA | #VALUE! |
| ENSMUSG00000071234.4 | 0 | NA | NA | NA | NA | NA | #VALUE! |
| ENSMUSG00000071235.13 | 0 | NA | NA | NA | NA | NA | #VALUE! |
| ENSMUSG00000071237.5 | 0 | NA | NA | NA | NA | NA | #VALUE! |
| ENSMUSG00000071256.5 | 3.029103 | 5.088783 | 3.921574 | 1.297638 | 0.194412 | NA | #VALUE! |
| ENSMUSG00000071262.11 | 0 | NA | NA | NA | NA | NA | #VALUE! |
| ENSMUSG00000071265.11 | 0 | NA | NA | NA | NA | NA | #VALUE! |
| ENSMUSG00000071273.5 | 0 | NA | NA | NA | NA | NA | #VALUE! |
| ENSMUSG00000071275.2 | 0 | NA | NA | NA | NA | NA | #VALUE! |
| ENSMUSG00000071276.2 | 0 | NA | NA | NA | NA | NA | #VALUE! |
| ENSMUSG00000071280.3 | 0 | NA | NA | NA | NA | NA | #VALUE! |
| ENSMUSG00000071291.11 | 7.385824 | 1.42404 | 3.252796 | 0.43779 | 0.661539 | NA | #VALUE! |
| ENSMUSG00000071294.3 | 0 | NA | NA | NA | NA | NA | #VALUE! |
| ENSMUSG00000071303.5 | 5.414413 | -1.86459 | 2.564951 | -0.72695 | 0.467258 | NA | #VALUE! |
| ENSMUSG00000071311.3 | 0 | NA | NA | NA | NA | NA | #VALUE! |
| ENSMUSG00000071317.5 | 0 | NA | NA | NA | NA | NA | #VALUE! |
| ENSMUSG00000071322.13 | 0 | NA | NA | NA | NA | NA | #VALUE! |
| ENSMUSG00000071324.12 | 0 | NA | NA | NA | NA | NA | #VALUE! |
| ENSMUSG00000071332.13 | 0 | NA | NA | NA | NA | NA | #VALUE! |
| ENSMUSG00000071340.3 | 0 | NA | NA | NA | NA | NA | #VALUE! |
| ENSMUSG00000071341.5 | 0 | NA | NA | NA | NA | NA | #VALUE! |
| ENSMUSG00000071342.6 | 0 | NA | NA | NA | NA | NA | #VALUE! |
| ENSMUSG00000071343.3 | 0 | NA | NA | NA | NA | NA | #VALUE! |
| ENSMUSG00000071347.4 | 0 | NA | NA | NA | NA | NA | #VALUE! |
| ENSMUSG00000071356.8 | 0 | NA | NA | NA | NA | NA | #VALUE! |
| ENSMUSG00000071361.2 | 0 | NA | NA | NA | NA | NA | #VALUE! |

| | | | | | | | |
|---|---|---|---|---|---|---|---|
| ENSMUSG00000071362.4 | 0 | NA | NA | NA | NA | NA | #VALUE! |
| ENSMUSG00000071392.7 | 0 | NA | NA | NA | NA | NA | #VALUE! |
| ENSMUSG00000071398.9 | 0 | NA | NA | NA | NA | NA | #VALUE! |
| ENSMUSG00000071414.6 | 0 | NA | NA | NA | NA | NA | #VALUE! |
| ENSMUSG00000071419.8 | 0.640033 | 0.057569 | 4.005078 | 0.014374 | 0.988532 | NA | #VALUE! |
| ENSMUSG00000071424.11 | 0 | NA | NA | NA | NA | NA | #VALUE! |
| ENSMUSG00000071428.3 | 0 | NA | NA | NA | NA | NA | #VALUE! |
| ENSMUSG00000071434.3 | 0 | NA | NA | NA | NA | NA | #VALUE! |
| ENSMUSG00000071452.5 | 0 | NA | NA | NA | NA | NA | #VALUE! |
| ENSMUSG00000071470.5 | 0 | NA | NA | NA | NA | NA | #VALUE! |
| ENSMUSG00000071471.3 | 0 | NA | NA | NA | NA | NA | #VALUE! |
| ENSMUSG00000071475.4 | 0 | NA | NA | NA | NA | NA | #VALUE! |
| ENSMUSG00000071477.11 | 0 | NA | NA | NA | NA | NA | #VALUE! |
| ENSMUSG00000071478.7 | 0 | NA | NA | NA | NA | NA | #VALUE! |
| ENSMUSG00000071481.5 | 0 | NA | NA | NA | NA | NA | #VALUE! |
| ENSMUSG00000071489.2 | 0 | NA | NA | NA | NA | NA | #VALUE! |
| ENSMUSG00000071490.6 | 0 | NA | NA | NA | NA | NA | #VALUE! |
| ENSMUSG00000071491.3 | 0 | NA | NA | NA | NA | NA | #VALUE! |
| ENSMUSG00000071493.6 | 0 | NA | NA | NA | NA | NA | #VALUE! |
| ENSMUSG00000071494.6 | 0 | NA | NA | NA | NA | NA | #VALUE! |
| ENSMUSG00000071506.5 | 0 | NA | NA | NA | NA | NA | #VALUE! |
| ENSMUSG00000071510.4 | 0 | NA | NA | NA | NA | NA | #VALUE! |
| ENSMUSG00000071516.3 | 0 | NA | NA | NA | NA | NA | #VALUE! |
| ENSMUSG00000071517.7 | 0 | NA | NA | NA | NA | NA | #VALUE! |
| ENSMUSG00000071519.7 | 0 | NA | NA | NA | NA | NA | #VALUE! |
| ENSMUSG00000071521.5 | 0 | NA | NA | NA | NA | NA | #VALUE! |
| ENSMUSG00000071522.4 | 0 | NA | NA | NA | NA | NA | #VALUE! |
| ENSMUSG00000071531.4 | 0 | NA | NA | NA | NA | NA | #VALUE! |
| ENSMUSG00000071532.7 | 0 | NA | NA | NA | NA | NA | #VALUE! |
| ENSMUSG00000071537.7 | 0 | NA | NA | NA | NA | NA | #VALUE! |
| ENSMUSG00000071540.5 | 0 | NA | NA | NA | NA | NA | #VALUE! |
| ENSMUSG00000071547.5 | 9.13385 | -0.144 | 2.65208 | -0.0543 | 0.956698 | NA | #VALUE! |
| ENSMUSG00000071550.15 | 0 | NA | NA | NA | NA | NA | #VALUE! |
| ENSMUSG00000071552.6 | 0 | NA | NA | NA | NA | NA | #VALUE! |
| ENSMUSG00000071553.11 | 0 | NA | NA | NA | NA | NA | #VALUE! |
| ENSMUSG00000071561.3 | 0 | NA | NA | NA | NA | NA | #VALUE! |
| ENSMUSG00000071562.7 | 0 | NA | NA | NA | NA | NA | #VALUE! |
| ENSMUSG00000071568.6 | 0 | NA | NA | NA | NA | NA | #VALUE! |
| ENSMUSG00000071573.15 | 0 | NA | NA | NA | NA | NA | #VALUE! |
| ENSMUSG00000071586.5 | 0 | NA | NA | NA | NA | NA | #VALUE! |
| ENSMUSG00000071591.4 | 0 | NA | NA | NA | NA | NA | #VALUE! |
| ENSMUSG00000071604.6 | 0.420053 | -2.33493 | 4.007343 | -0.58266 | 0.56012 | NA | #VALUE! |
| ENSMUSG00000071613.12 | 0 | NA | NA | NA | NA | NA | #VALUE! |
| ENSMUSG00000071629.4 | 0 | NA | NA | NA | NA | NA | #VALUE! |

| | | | | | | | |
|---|---|---|---|---|---|---|---|
| ENSMUSG00000071636.9 | 0 | NA | NA | NA | NA | NA | #VALUE! |
| ENSMUSG00000071640.4 | 0 | NA | NA | NA | NA | NA | #VALUE! |
| ENSMUSG00000071647.6 | 12.86285 | 1.678099 | 2.746108 | 0.611083 | NA | NA | #VALUE! |
| ENSMUSG00000071648.5 | 1.804813 | -4.38285 | 3.931461 | -1.11481 | 0.26493 | NA | #VALUE! |
| ENSMUSG00000071649.7 | 9.117324 | 6.674675 | 3.226447 | 2.068738 | 0.038571 | NA | #VALUE! |
| ENSMUSG00000071652.9 | 2.717775 | -0.91901 | 3.799438 | -0.24188 | 0.808872 | NA | #VALUE! |
| ENSMUSG00000071653.10 | 0 | NA | NA | NA | NA | NA | #VALUE! |
| ENSMUSG00000071656.8 | 0 | NA | NA | NA | NA | NA | #VALUE! |
| ENSMUSG00000071658.6 | 0 | NA | NA | NA | NA | NA | #VALUE! |
| ENSMUSG00000071661.8 | 0 | NA | NA | NA | NA | NA | #VALUE! |
| ENSMUSG00000071665.11 | 0 | NA | NA | NA | NA | NA | #VALUE! |
| ENSMUSG00000071669.15 | 0 | NA | NA | NA | NA | NA | #VALUE! |
| ENSMUSG00000071679.5 | 0 | NA | NA | NA | NA | NA | #VALUE! |
| ENSMUSG00000071680.7 | 0.195789 | -1.05581 | 4.073302 | -0.2592 | 0.795479 | NA | #VALUE! |
| ENSMUSG00000071686.3 | 0 | NA | NA | NA | NA | NA | #VALUE! |
| ENSMUSG00000071691.12 | 0 | NA | NA | NA | NA | NA | #VALUE! |
| ENSMUSG00000071714.8 | 2.236897 | -4.68482 | 3.926816 | -1.19303 | 0.232857 | NA | #VALUE! |
| ENSMUSG00000071715.13 | 6.380673 | -6.2032 | 3.632131 | -1.70787 | 0.087661 | NA | #VALUE! |
| ENSMUSG00000071716.6 | 0 | NA | NA | NA | NA | NA | #VALUE! |
| ENSMUSG00000071719.3 | 0.315039 | -1.93549 | 4.038621 | -0.47925 | 0.631763 | NA | #VALUE! |
| ENSMUSG00000071721.6 | 0 | NA | NA | NA | NA | NA | #VALUE! |
| ENSMUSG00000071722.5 | 0.466276 | 2.449889 | 3.997179 | 0.612904 | 0.53994 | NA | #VALUE! |
| ENSMUSG00000071723.8 | 0.408222 | -2.20205 | 4.016842 | -0.54821 | 0.583551 | NA | #VALUE! |
| ENSMUSG00000071724.6 | 0.18651 | 1.401769 | 4.056354 | 0.345574 | 0.729663 | NA | #VALUE! |
| ENSMUSG00000071726.3 | 0 | NA | NA | NA | NA | NA | #VALUE! |
| ENSMUSG00000071738.5 | 0 | NA | NA | NA | NA | NA | #VALUE! |
| ENSMUSG00000071745.2 | 0 | NA | NA | NA | NA | NA | #VALUE! |
| ENSMUSG00000071748.3 | 0.204111 | -1.05581 | 4.073302 | -0.2592 | 0.795479 | NA | #VALUE! |
| ENSMUSG00000071749.4 | 0 | NA | NA | NA | NA | NA | #VALUE! |
| ENSMUSG00000071750.4 | 0 | NA | NA | NA | NA | NA | #VALUE! |
| ENSMUSG00000071753.14 | 0 | NA | NA | NA | NA | NA | #VALUE! |
| ENSMUSG00000071763.10 | 0 | NA | NA | NA | NA | NA | #VALUE! |
| ENSMUSG00000071764.3 | 0 | NA | NA | NA | NA | NA | #VALUE! |
| ENSMUSG00000071766.13 | 0 | NA | NA | NA | NA | NA | #VALUE! |
| ENSMUSG00000071767.5 | 0 | NA | NA | NA | NA | NA | #VALUE! |
| ENSMUSG00000071769.12 | 0 | NA | NA | NA | NA | NA | #VALUE! |
| ENSMUSG00000071770.6 | 0 | NA | NA | NA | NA | NA | #VALUE! |
| ENSMUSG00000071771.5 | 0 | NA | NA | NA | NA | NA | #VALUE! |
| ENSMUSG00000071772.14 | 0 | NA | NA | NA | NA | NA | #VALUE! |
| ENSMUSG00000071773.5 | 0 | NA | NA | NA | NA | NA | #VALUE! |
| ENSMUSG00000071788.6 | 0 | NA | NA | NA | NA | NA | #VALUE! |
| ENSMUSG00000071793.13 | 0 | NA | NA | NA | NA | NA | #VALUE! |
| ENSMUSG00000071796.14 | 5.179471 | -5.90661 | 3.915382 | -1.50856 | 0.13141 | NA | #VALUE! |
| ENSMUSG00000071815.7 | 0 | NA | NA | NA | NA | NA | #VALUE! |

| | | | | | | | |
|---|---|---|---|---|---|---|---|
| ENSMUSG00000071816.6 | 0 | NA | NA | NA | NA | NA | #VALUE! |
| ENSMUSG00000071818.10 | 0 | NA | NA | NA | NA | NA | #VALUE! |
| ENSMUSG00000071847.14 | 0 | NA | NA | NA | NA | NA | #VALUE! |
| ENSMUSG00000071855.6 | 9.804362 | 3.197858 | 3.266712 | 0.978923 | NA | NA | #VALUE! |
| ENSMUSG00000071858.5 | 0 | NA | NA | NA | NA | NA | #VALUE! |
| ENSMUSG00000071862.4 | 0 | NA | NA | NA | NA | NA | #VALUE! |
| ENSMUSG00000071867.6 | 0 | NA | NA | NA | NA | NA | #VALUE! |
| ENSMUSG00000071890.5 | 0 | NA | NA | NA | NA | NA | #VALUE! |
| ENSMUSG00000071893.8 | 0.093255 | 0.868093 | 4.072464 | 0.213162 | 0.831201 | NA | #VALUE! |
| ENSMUSG00000071909.3 | 0 | NA | NA | NA | NA | NA | #VALUE! |
| ENSMUSG00000071937.5 | 0 | NA | NA | NA | NA | NA | #VALUE! |
| ENSMUSG00000071952.4 | 0 | NA | NA | NA | NA | NA | #VALUE! |
| ENSMUSG00000071984.12 | 0 | NA | NA | NA | NA | NA | #VALUE! |
| ENSMUSG00000072049.7 | 0 | NA | NA | NA | NA | NA | #VALUE! |
| ENSMUSG00000072066.8 | 0 | NA | NA | NA | NA | NA | #VALUE! |
| ENSMUSG00000072068.5 | 0 | NA | NA | NA | NA | NA | #VALUE! |
| ENSMUSG00000072100.5 | 0 | NA | NA | NA | NA | NA | #VALUE! |
| ENSMUSG00000072109.12 | 0 | NA | NA | NA | NA | NA | #VALUE! |
| ENSMUSG00000072145.4 | 0 | NA | NA | NA | NA | NA | #VALUE! |
| ENSMUSG00000072188.5 | 0 | NA | NA | NA | NA | NA | #VALUE! |
| ENSMUSG00000072214.8 | 0 | NA | NA | NA | NA | NA | #VALUE! |
| ENSMUSG00000072235.7 | 7.147281 | -2.52676 | 2.803861 | -0.90117 | 0.367497 | NA | #VALUE! |
| ENSMUSG00000072244.12 | 0.336567 | 1.944925 | 4.029936 | 0.482619 | 0.629366 | NA | #VALUE! |
| ENSMUSG00000072249.4 | 0 | NA | NA | NA | NA | NA | #VALUE! |
| ENSMUSG00000072255.4 | 0.198268 | -0.06494 | 4.068276 | -0.01596 | 0.987265 | NA | #VALUE! |
| ENSMUSG00000072259.4 | 0 | NA | NA | NA | NA | NA | #VALUE! |
| ENSMUSG00000072295.8 | 0 | NA | NA | NA | NA | NA | #VALUE! |
| ENSMUSG00000072324.5 | 0 | NA | NA | NA | NA | NA | #VALUE! |
| ENSMUSG00000072381.4 | 0 | NA | NA | NA | NA | NA | #VALUE! |
| ENSMUSG00000072387.4 | 0 | NA | NA | NA | NA | NA | #VALUE! |
| ENSMUSG00000072407.5 | 0.093255 | 0.868093 | 4.072464 | 0.213162 | 0.831201 | NA | #VALUE! |
| ENSMUSG00000072419.6 | 0 | NA | NA | NA | NA | NA | #VALUE! |
| ENSMUSG00000072421.5 | 0 | NA | NA | NA | NA | NA | #VALUE! |
| ENSMUSG00000072423.6 | 0 | NA | NA | NA | NA | NA | #VALUE! |
| ENSMUSG00000072437.5 | 0 | NA | NA | NA | NA | NA | #VALUE! |
| ENSMUSG00000072460.6 | 0 | NA | NA | NA | NA | NA | #VALUE! |
| ENSMUSG00000072473.5 | 0 | NA | NA | NA | NA | NA | #VALUE! |
| ENSMUSG00000072476.7 | 0.204111 | -1.05581 | 4.073302 | -0.2592 | 0.795479 | NA | #VALUE! |
| ENSMUSG00000072477.6 | 0 | NA | NA | NA | NA | NA | #VALUE! |
| ENSMUSG00000072479.11 | 0 | NA | NA | NA | NA | NA | #VALUE! |
| ENSMUSG00000072487.12 | 0 | NA | NA | NA | NA | NA | #VALUE! |
| ENSMUSG00000072492.6 | 0 | NA | NA | NA | NA | NA | #VALUE! |
| ENSMUSG00000072494.8 | 5.833172 | 6.038059 | 3.784953 | 1.59528 | 0.11065 | NA | #VALUE! |
| ENSMUSG00000072511.4 | 0 | NA | NA | NA | NA | NA | #VALUE! |

| | | | | | | | |
|---|---|---|---|---|---|---|---|
| ENSMUSG00000072541.3 | 0 | NA | NA | NA | NA | NA | #VALUE! |
| ENSMUSG00000072553.11 | 0 | NA | NA | NA | NA | NA | #VALUE! |
| ENSMUSG00000072568.5 | 30.27278 | 0.582744 | 2.379864 | 0.244864 | NA | NA | #VALUE! |
| ENSMUSG00000072571.4 | 0 | NA | NA | NA | NA | NA | #VALUE! |
| ENSMUSG00000072572.10 | 0 | NA | NA | NA | NA | NA | #VALUE! |
| ENSMUSG00000072573.3 | 0 | NA | NA | NA | NA | NA | #VALUE! |
| ENSMUSG00000072574.5 | 0 | NA | NA | NA | NA | NA | #VALUE! |
| ENSMUSG00000072575.5 | 0 | NA | NA | NA | NA | NA | #VALUE! |
| ENSMUSG00000072576.6 | 0 | NA | NA | NA | NA | NA | #VALUE! |
| ENSMUSG00000072584.3 | 0 | NA | NA | NA | NA | NA | #VALUE! |
| ENSMUSG00000072589.5 | 0 | NA | NA | NA | NA | NA | #VALUE! |
| ENSMUSG00000072591.11 | 0 | NA | NA | NA | NA | NA | #VALUE! |
| ENSMUSG00000072592.3 | 0 | NA | NA | NA | NA | NA | #VALUE! |
| ENSMUSG00000072594.4 | 0 | NA | NA | NA | NA | NA | #VALUE! |
| ENSMUSG00000072595.4 | 0 | NA | NA | NA | NA | NA | #VALUE! |
| ENSMUSG00000072598.4 | 0 | NA | NA | NA | NA | NA | #VALUE! |
| ENSMUSG00000072599.4 | 0 | NA | NA | NA | NA | NA | #VALUE! |
| ENSMUSG00000072600.5 | 0 | NA | NA | NA | NA | NA | #VALUE! |
| ENSMUSG00000072601.5 | 0 | NA | NA | NA | NA | NA | #VALUE! |
| ENSMUSG00000072605.11 | 0 | NA | NA | NA | NA | NA | #VALUE! |
| ENSMUSG00000072612.4 | 0 | NA | NA | NA | NA | NA | #VALUE! |
| ENSMUSG00000072616.6 | 0 | NA | NA | NA | NA | NA | #VALUE! |
| ENSMUSG00000072618.3 | 0 | NA | NA | NA | NA | NA | #VALUE! |
| ENSMUSG00000072620.4 | 0 | NA | NA | NA | NA | NA | #VALUE! |
| ENSMUSG00000072621.14 | 0 | NA | NA | NA | NA | NA | #VALUE! |
| ENSMUSG00000072623.7 | 9.540261 | -0.66318 | 3.000085 | -0.22105 | 0.825051 | NA | #VALUE! |
| ENSMUSG00000072624.4 | 0 | NA | NA | NA | NA | NA | #VALUE! |
| ENSMUSG00000072647.7 | 0 | NA | NA | NA | NA | NA | #VALUE! |
| ENSMUSG00000072653.15 | 0 | NA | NA | NA | NA | NA | #VALUE! |
| ENSMUSG00000072660.6 | 0 | NA | NA | NA | NA | NA | #VALUE! |
| ENSMUSG00000072662.3 | 0 | NA | NA | NA | NA | NA | #VALUE! |
| ENSMUSG00000072663.13 | 0 | NA | NA | NA | NA | NA | #VALUE! |
| ENSMUSG00000072664.12 | 0 | NA | NA | NA | NA | NA | #VALUE! |
| ENSMUSG00000072673.3 | 0 | NA | NA | NA | NA | NA | #VALUE! |
| ENSMUSG00000072677.5 | 0 | NA | NA | NA | NA | NA | #VALUE! |
| ENSMUSG00000072679.11 | 0 | NA | NA | NA | NA | NA | #VALUE! |
| ENSMUSG00000072683.4 | 1.428775 | -4.03418 | 3.938169 | -1.02438 | 0.305656 | NA | #VALUE! |
| ENSMUSG00000072686.3 | 0 | NA | NA | NA | NA | NA | #VALUE! |
| ENSMUSG00000072688.3 | 0 | NA | NA | NA | NA | NA | #VALUE! |
| ENSMUSG00000072693.3 | 0.816589 | -1.46094 | 3.950283 | -0.36983 | 0.711508 | NA | #VALUE! |
| ENSMUSG00000072694.9 | 3.139126 | 1.603726 | 3.804495 | 0.421534 | 0.673365 | NA | #VALUE! |
| ENSMUSG00000072707.4 | 0 | NA | NA | NA | NA | NA | #VALUE! |
| ENSMUSG00000072708.4 | 0 | NA | NA | NA | NA | NA | #VALUE! |
| ENSMUSG00000072709.4 | 0 | NA | NA | NA | NA | NA | #VALUE! |

| | | | | | | | |
|---|---|---|---|---|---|---|---|
| ENSMUSG00000072714.3 | 0 | NA | NA | NA | NA | NA | #VALUE! |
| ENSMUSG00000072717.9 | 0 | NA | NA | NA | NA | NA | #VALUE! |
| ENSMUSG00000072718.7 | 0 | NA | NA | NA | NA | NA | #VALUE! |
| ENSMUSG00000072720.10 | 0 | NA | NA | NA | NA | NA | #VALUE! |
| ENSMUSG00000072721.9 | 0 | NA | NA | NA | NA | NA | #VALUE! |
| ENSMUSG00000072722.3 | 0 | NA | NA | NA | NA | NA | #VALUE! |
| ENSMUSG00000072723.13 | 1.995249 | -4.53986 | 3.928926 | -1.1555 | 0.247887 | NA | #VALUE! |
| ENSMUSG00000072726.4 | 0 | NA | NA | NA | NA | NA | #VALUE! |
| ENSMUSG00000072731.7 | 0 | NA | NA | NA | NA | NA | #VALUE! |
| ENSMUSG00000072738.4 | 0 | NA | NA | NA | NA | NA | #VALUE! |
| ENSMUSG00000072739.5 | 0 | NA | NA | NA | NA | NA | #VALUE! |
| ENSMUSG00000072753.13 | 0 | NA | NA | NA | NA | NA | #VALUE! |
| ENSMUSG00000072754.10 | 0 | NA | NA | NA | NA | NA | #VALUE! |
| ENSMUSG00000072761.11 | 1.566313 | -4.16993 | 3.935366 | -1.0596 | 0.289325 | NA | #VALUE! |
| ENSMUSG00000072762.10 | 7.982981 | 1.901901 | 2.774642 | 0.685458 | 0.493055 | NA | #VALUE! |
| ENSMUSG00000072768.2 | 0 | NA | NA | NA | NA | NA | #VALUE! |
| ENSMUSG00000072769.7 | 0 | NA | NA | NA | NA | NA | #VALUE! |
| ENSMUSG00000072770.10 | 6.740908 | -2.44453 | 3.72366 | -0.65649 | 0.511512 | NA | #VALUE! |
| ENSMUSG00000072772.4 | 0 | NA | NA | NA | NA | NA | #VALUE! |
| ENSMUSG00000072774.11 | 1.575197 | -4.20222 | 3.934736 | -1.06798 | 0.285529 | NA | #VALUE! |
| ENSMUSG00000072778.4 | 0 | NA | NA | NA | NA | NA | #VALUE! |
| ENSMUSG00000072780.4 | 0 | NA | NA | NA | NA | NA | #VALUE! |
| ENSMUSG00000072789.5 | 0.624285 | 0.689511 | 3.993559 | 0.172656 | 0.862922 | NA | #VALUE! |
| ENSMUSG00000072791.12 | 0 | NA | NA | NA | NA | NA | #VALUE! |
| ENSMUSG00000072809.3 | 7.740181 | 6.446428 | 3.889072 | 1.657575 | NA | NA | #VALUE! |
| ENSMUSG00000072813.6 | 0 | NA | NA | NA | NA | NA | #VALUE! |
| ENSMUSG00000072814.11 | 0 | NA | NA | NA | NA | NA | #VALUE! |
| ENSMUSG00000072821.5 | 0 | NA | NA | NA | NA | NA | #VALUE! |
| ENSMUSG00000072822.3 | 0 | NA | NA | NA | NA | NA | #VALUE! |
| ENSMUSG00000072834.5 | 0 | NA | NA | NA | NA | NA | #VALUE! |
| ENSMUSG00000072837.3 | 0 | NA | NA | NA | NA | NA | #VALUE! |
| ENSMUSG00000072844.7 | 1.155144 | -3.7605 | 3.944679 | -0.95331 | 0.340433 | NA | #VALUE! |
| ENSMUSG00000072845.4 | 0 | NA | NA | NA | NA | NA | #VALUE! |
| ENSMUSG00000072847.5 | 5.136342 | -0.37618 | 3.217866 | -0.1169 | 0.906938 | NA | #VALUE! |
| ENSMUSG00000072852.5 | 0 | NA | NA | NA | NA | NA | #VALUE! |
| ENSMUSG00000072872.4 | 0 | NA | NA | NA | NA | NA | #VALUE! |
| ENSMUSG00000072874.5 | 0 | NA | NA | NA | NA | NA | #VALUE! |
| ENSMUSG00000072875.3 | 0 | NA | NA | NA | NA | NA | #VALUE! |
| ENSMUSG00000072878.5 | 0 | NA | NA | NA | NA | NA | #VALUE! |
| ENSMUSG00000072884.11 | 0 | NA | NA | NA | NA | NA | #VALUE! |
| ENSMUSG00000072888.3 | 0 | NA | NA | NA | NA | NA | #VALUE! |
| ENSMUSG00000072893.13 | 0 | NA | NA | NA | NA | NA | #VALUE! |
| ENSMUSG00000072898.4 | 0 | NA | NA | NA | NA | NA | #VALUE! |
| ENSMUSG00000072900.4 | 0 | NA | NA | NA | NA | NA | #VALUE! |

| | | | | | | | |
|---|---|---|---|---|---|---|---|
| ENSMUSG00000072902.5 | 0 | NA | NA | NA | NA | NA | #VALUE! |
| ENSMUSG00000072905.6 | 0 | NA | NA | NA | NA | NA | #VALUE! |
| ENSMUSG00000072915.4 | 8.176106 | 3.790601 | 3.078107 | 1.231471 | 0.218147 | NA | #VALUE! |
| ENSMUSG00000072919.5 | 0 | NA | NA | NA | NA | NA | #VALUE! |
| ENSMUSG00000072923.7 | 0 | NA | NA | NA | NA | NA | #VALUE! |
| ENSMUSG00000072929.7 | 0 | NA | NA | NA | NA | NA | #VALUE! |
| ENSMUSG00000072930.4 | 0 | NA | NA | NA | NA | NA | #VALUE! |
| ENSMUSG00000072931.5 | 0 | NA | NA | NA | NA | NA | #VALUE! |
| ENSMUSG00000072934.11 | 0 | NA | NA | NA | NA | NA | #VALUE! |
| ENSMUSG00000072944.12 | 0 | NA | NA | NA | NA | NA | #VALUE! |
| ENSMUSG00000072945.4 | 3.197387 | 5.166784 | 3.920797 | 1.317789 | 0.187574 | NA | #VALUE! |
| ENSMUSG00000072950.6 | 0 | NA | NA | NA | NA | NA | #VALUE! |
| ENSMUSG00000072952.8 | 0 | NA | NA | NA | NA | NA | #VALUE! |
| ENSMUSG00000072955.15 | 0 | NA | NA | NA | NA | NA | #VALUE! |
| ENSMUSG00000072962.4 | 0 | NA | NA | NA | NA | NA | #VALUE! |
| ENSMUSG00000072963.3 | 0 | NA | NA | NA | NA | NA | #VALUE! |
| ENSMUSG00000072966.12 | 0 | NA | NA | NA | NA | NA | #VALUE! |
| ENSMUSG00000072968.5 | 0 | NA | NA | NA | NA | NA | #VALUE! |
| ENSMUSG00000072969.3 | 4.764193 | -0.45807 | 3.365493 | -0.13611 | 0.891736 | NA | #VALUE! |
| ENSMUSG00000072972.4 | 1.119871 | -1.46061 | 3.91329 | -0.37324 | 0.708966 | NA | #VALUE! |
| ENSMUSG00000072978.4 | 0 | NA | NA | NA | NA | NA | #VALUE! |
| ENSMUSG00000072980.4 | 0.95144 | -2.09486 | 3.95018 | -0.53032 | 0.595891 | NA | #VALUE! |
| ENSMUSG00000072982.2 | 0 | NA | NA | NA | NA | NA | #VALUE! |
| ENSMUSG00000072983.5 | 0 | NA | NA | NA | NA | NA | #VALUE! |
| ENSMUSG00000072995.2 | 0 | NA | NA | NA | NA | NA | #VALUE! |
| ENSMUSG00000072999.10 | 0 | NA | NA | NA | NA | NA | #VALUE! |
| ENSMUSG00000073000.4 | 7.01631 | 6.298932 | 3.546075 | 1.776311 | 0.075682 | NA | #VALUE! |
| ENSMUSG00000073001.5 | 0 | NA | NA | NA | NA | NA | #VALUE! |
| ENSMUSG00000073006.3 | 0 | NA | NA | NA | NA | NA | #VALUE! |
| ENSMUSG00000073007.11 | 0 | NA | NA | NA | NA | NA | #VALUE! |
| ENSMUSG00000073008.12 | 0 | NA | NA | NA | NA | NA | #VALUE! |
| ENSMUSG00000073010.10 | 0 | NA | NA | NA | NA | NA | #VALUE! |
| ENSMUSG00000073012.3 | 0 | NA | NA | NA | NA | NA | #VALUE! |
| ENSMUSG00000073016.4 | 0 | NA | NA | NA | NA | NA | #VALUE! |
| ENSMUSG00000073018.7 | 0 | NA | NA | NA | NA | NA | #VALUE! |
| ENSMUSG00000073027.11 | 0 | NA | NA | NA | NA | NA | #VALUE! |
| ENSMUSG00000073028.5 | 0 | NA | NA | NA | NA | NA | #VALUE! |
| ENSMUSG00000073043.6 | 0 | NA | NA | NA | NA | NA | #VALUE! |
| ENSMUSG00000073045.10 | 0 | NA | NA | NA | NA | NA | #VALUE! |
| ENSMUSG00000073051.5 | 0 | NA | NA | NA | NA | NA | #VALUE! |
| ENSMUSG00000073052.5 | 0 | NA | NA | NA | NA | NA | #VALUE! |
| ENSMUSG00000073060.5 | 0 | NA | NA | NA | NA | NA | #VALUE! |
| ENSMUSG00000073063.4 | 0 | NA | NA | NA | NA | NA | #VALUE! |
| ENSMUSG00000073067.11 | 0 | NA | NA | NA | NA | NA | #VALUE! |

| | | | | | | | |
|---|---|---|---|---|---|---|---|
| ENSMUSG00000073069.3 | 0 | NA | NA | NA | NA | NA | #VALUE! |
| ENSMUSG00000073073.5 | 0 | NA | NA | NA | NA | NA | #VALUE! |
| ENSMUSG00000073077.8 | 0 | NA | NA | NA | NA | NA | #VALUE! |
| ENSMUSG00000073079.7 | 1.260158 | -3.88425 | 3.941584 | -0.98545 | 0.324401 | NA | #VALUE! |
| ENSMUSG00000073085.3 | 0 | NA | NA | NA | NA | NA | #VALUE! |
| ENSMUSG00000073094.10 | 0 | NA | NA | NA | NA | NA | #VALUE! |
| ENSMUSG00000073102.8 | 5.408801 | 5.931107 | 3.915037 | 1.514955 | 0.129784 | NA | #VALUE! |
| ENSMUSG00000073103.3 | 0 | NA | NA | NA | NA | NA | #VALUE! |
| ENSMUSG00000073110.4 | 0 | NA | NA | NA | NA | NA | #VALUE! |
| ENSMUSG00000073111.4 | 0 | NA | NA | NA | NA | NA | #VALUE! |
| ENSMUSG00000073116.5 | 0 | NA | NA | NA | NA | NA | #VALUE! |
| ENSMUSG00000073117.3 | 0 | NA | NA | NA | NA | NA | #VALUE! |
| ENSMUSG00000073119.4 | 0 | NA | NA | NA | NA | NA | #VALUE! |
| ENSMUSG00000073125.11 | 0 | NA | NA | NA | NA | NA | #VALUE! |
| ENSMUSG00000073130.11 | 0 | NA | NA | NA | NA | NA | #VALUE! |
| ENSMUSG00000073141.9 | 0 | NA | NA | NA | NA | NA | #VALUE! |
| ENSMUSG00000073144.5 | 0.105013 | -0.97324 | 4.076293 | -0.23876 | 0.811295 | NA | #VALUE! |
| ENSMUSG00000073145.7 | 0 | NA | NA | NA | NA | NA | #VALUE! |
| ENSMUSG00000073152.5 | 0 | NA | NA | NA | NA | NA | #VALUE! |
| ENSMUSG00000073154.11 | 0 | NA | NA | NA | NA | NA | #VALUE! |
| ENSMUSG00000073158.5 | 0 | NA | NA | NA | NA | NA | #VALUE! |
| ENSMUSG00000073164.12 | 0 | NA | NA | NA | NA | NA | #VALUE! |
| ENSMUSG00000073173.3 | 0 | NA | NA | NA | NA | NA | #VALUE! |
| ENSMUSG00000073174.7 | 0 | NA | NA | NA | NA | NA | #VALUE! |
| ENSMUSG00000073176.5 | 1.370524 | -3.97623 | 3.939448 | -1.00934 | 0.312813 | NA | #VALUE! |
| ENSMUSG00000073177.9 | 0 | NA | NA | NA | NA | NA | #VALUE! |
| ENSMUSG00000073179.4 | 0.195789 | -1.05581 | 4.073302 | -0.2592 | 0.795479 | NA | #VALUE! |
| ENSMUSG00000073184.5 | 0 | NA | NA | NA | NA | NA | #VALUE! |
| ENSMUSG00000073198.6 | 0.093255 | 0.868093 | 4.072464 | 0.213162 | 0.831201 | NA | #VALUE! |
| ENSMUSG00000073207.2 | 0 | NA | NA | NA | NA | NA | #VALUE! |
| ENSMUSG00000073208.13 | 0 | NA | NA | NA | NA | NA | #VALUE! |
| ENSMUSG00000073209.5 | 0 | NA | NA | NA | NA | NA | #VALUE! |
| ENSMUSG00000073226.10 | 0 | NA | NA | NA | NA | NA | #VALUE! |
| ENSMUSG00000073233.5 | 0 | NA | NA | NA | NA | NA | #VALUE! |
| ENSMUSG00000073234.5 | 0 | NA | NA | NA | NA | NA | #VALUE! |
| ENSMUSG00000073242.5 | 0 | NA | NA | NA | NA | NA | #VALUE! |
| ENSMUSG00000073243.3 | 0 | NA | NA | NA | NA | NA | #VALUE! |
| ENSMUSG00000073245.11 | 0 | NA | NA | NA | NA | NA | #VALUE! |
| ENSMUSG00000073247.4 | 0 | NA | NA | NA | NA | NA | #VALUE! |
| ENSMUSG00000073255.11 | 0 | NA | NA | NA | NA | NA | #VALUE! |
| ENSMUSG00000073257.4 | 0 | NA | NA | NA | NA | NA | #VALUE! |
| ENSMUSG00000073267.6 | 0 | NA | NA | NA | NA | NA | #VALUE! |
| ENSMUSG00000073274.5 | 0 | NA | NA | NA | NA | NA | #VALUE! |
| ENSMUSG00000073290.2 | 0 | NA | NA | NA | NA | NA | #VALUE! |

| | | | | | | | |
|---|---|---|---|---|---|---|---|
| ENSMUSG00000073291.4 | 0 | NA | NA | NA | NA | NA | #VALUE! |
| ENSMUSG00000073293.5 | 0.391578 | -2.1489 | 4.020883 | -0.53444 | 0.59304 | NA | #VALUE! |
| ENSMUSG00000073294.5 | 0 | NA | NA | NA | NA | NA | #VALUE! |
| ENSMUSG00000073367.6 | 0 | NA | NA | NA | NA | NA | #VALUE! |
| ENSMUSG00000073371.5 | 0 | NA | NA | NA | NA | NA | #VALUE! |
| ENSMUSG00000073374.4 | 3.256595 | 2.325332 | 3.827093 | 0.607597 | 0.543455 | NA | #VALUE! |
| ENSMUSG00000073375.4 | 0 | NA | NA | NA | NA | NA | #VALUE! |
| ENSMUSG00000073377.10 | 0 | NA | NA | NA | NA | NA | #VALUE! |
| ENSMUSG00000073380.3 | 0 | NA | NA | NA | NA | NA | #VALUE! |
| ENSMUSG00000073386.6 | 0 | NA | NA | NA | NA | NA | #VALUE! |
| ENSMUSG00000073394.5 | 0 | NA | NA | NA | NA | NA | #VALUE! |
| ENSMUSG00000073396.6 | 0 | NA | NA | NA | NA | NA | #VALUE! |
| ENSMUSG00000073399.10 | 0 | NA | NA | NA | NA | NA | #VALUE! |
| ENSMUSG00000073400.7 | 0 | NA | NA | NA | NA | NA | #VALUE! |
| ENSMUSG00000073402.12 | 0 | NA | NA | NA | NA | NA | #VALUE! |
| ENSMUSG00000073403.12 | 0 | NA | NA | NA | NA | NA | #VALUE! |
| ENSMUSG00000073405.7 | 0 | NA | NA | NA | NA | NA | #VALUE! |
| ENSMUSG00000073406.11 | 0 | NA | NA | NA | NA | NA | #VALUE! |
| ENSMUSG00000073407.2 | 0 | NA | NA | NA | NA | NA | #VALUE! |
| ENSMUSG00000073408.3 | 0 | NA | NA | NA | NA | NA | #VALUE! |
| ENSMUSG00000073409.13 | 0 | NA | NA | NA | NA | NA | #VALUE! |
| ENSMUSG00000073413.10 | 0 | NA | NA | NA | NA | NA | #VALUE! |
| ENSMUSG00000073414.9 | 0 | NA | NA | NA | NA | NA | #VALUE! |
| ENSMUSG00000073415.5 | 0 | NA | NA | NA | NA | NA | #VALUE! |
| ENSMUSG00000073420.11 | 0 | NA | NA | NA | NA | NA | #VALUE! |
| ENSMUSG00000073422.12 | 0 | NA | NA | NA | NA | NA | #VALUE! |
| ENSMUSG00000073423.11 | 0.315039 | -1.93549 | 4.038621 | -0.47925 | 0.631763 | NA | #VALUE! |
| ENSMUSG00000073427.5 | 1.428775 | -4.03418 | 3.938169 | -1.02438 | 0.305656 | NA | #VALUE! |
| ENSMUSG00000073430.6 | 0 | NA | NA | NA | NA | NA | #VALUE! |
| ENSMUSG00000073433.12 | 0 | NA | NA | NA | NA | NA | #VALUE! |
| ENSMUSG00000073434.13 | 0 | NA | NA | NA | NA | NA | #VALUE! |
| ENSMUSG00000073435.11 | 0 | NA | NA | NA | NA | NA | #VALUE! |
| ENSMUSG00000073437.11 | 0 | NA | NA | NA | NA | NA | #VALUE! |
| ENSMUSG00000073448.4 | 0 | NA | NA | NA | NA | NA | #VALUE! |
| ENSMUSG00000073457.5 | 0 | NA | NA | NA | NA | NA | #VALUE! |
| ENSMUSG00000073458.3 | 0 | NA | NA | NA | NA | NA | #VALUE! |
| ENSMUSG00000073460.6 | 7.390353 | 1.604874 | 3.649119 | 0.439798 | 0.660084 | NA | #VALUE! |
| ENSMUSG00000073482.3 | 0 | NA | NA | NA | NA | NA | #VALUE! |
| ENSMUSG00000073485.4 | 0 | NA | NA | NA | NA | NA | #VALUE! |
| ENSMUSG00000073486.3 | 0 | NA | NA | NA | NA | NA | #VALUE! |
| ENSMUSG00000073490.12 | 5.905284 | -4.59983 | 3.436847 | -1.33839 | 0.18077 | NA | #VALUE! |
| ENSMUSG00000073491.12 | 19.18733 | -21.9894 | 2.978667 | -7.3823 | NA | NA | #VALUE! |
| ENSMUSG00000073492.3 | 0 | NA | NA | NA | NA | NA | #VALUE! |
| ENSMUSG00000073494.6 | 0 | NA | NA | NA | NA | NA | #VALUE! |

| | | | | | | | |
|---|---|---|---|---|---|---|---|
| ENSMUSG00000073497.7 | 0 | NA | NA | NA | NA | NA | #VALUE! |
| ENSMUSG00000073510.3 | 0 | NA | NA | NA | NA | NA | #VALUE! |
| ENSMUSG00000073514.5 | 0 | NA | NA | NA | NA | NA | #VALUE! |
| ENSMUSG00000073528.8 | 0 | NA | NA | NA | NA | NA | #VALUE! |
| ENSMUSG00000073529.10 | 0 | NA | NA | NA | NA | NA | #VALUE! |
| ENSMUSG00000073530.12 | 0 | NA | NA | NA | NA | NA | #VALUE! |
| ENSMUSG00000073532.4 | 0 | NA | NA | NA | NA | NA | #VALUE! |
| ENSMUSG00000073535.6 | 0 | NA | NA | NA | NA | NA | #VALUE! |
| ENSMUSG00000073538.11 | 0 | NA | NA | NA | NA | NA | #VALUE! |
| ENSMUSG00000073551.6 | 0 | NA | NA | NA | NA | NA | #VALUE! |
| ENSMUSG00000073554.4 | 0 | NA | NA | NA | NA | NA | #VALUE! |
| ENSMUSG00000073557.13 | 0.816443 | -3.21914 | 3.961753 | -0.81255 | 0.416474 | NA | #VALUE! |
| ENSMUSG00000073559.3 | 0 | NA | NA | NA | NA | NA | #VALUE! |
| ENSMUSG00000073568.3 | 0 | NA | NA | NA | NA | NA | #VALUE! |
| ENSMUSG00000073569.3 | 0 | NA | NA | NA | NA | NA | #VALUE! |
| ENSMUSG00000073572.3 | 0 | NA | NA | NA | NA | NA | #VALUE! |
| ENSMUSG00000073573.7 | 0 | NA | NA | NA | NA | NA | #VALUE! |
| ENSMUSG00000073574.5 | 0 | NA | NA | NA | NA | NA | #VALUE! |
| ENSMUSG00000073590.4 | 0 | NA | NA | NA | NA | NA | #VALUE! |
| ENSMUSG00000073591.5 | 0 | NA | NA | NA | NA | NA | #VALUE! |
| ENSMUSG00000073594.4 | 0 | NA | NA | NA | NA | NA | #VALUE! |
| ENSMUSG00000073598.4 | 0 | NA | NA | NA | NA | NA | #VALUE! |
| ENSMUSG00000073599.11 | 10.48875 | -3.31649 | 3.198564 | -1.03687 | NA | NA | #VALUE! |
| ENSMUSG00000073600.6 | 1.680714 | 4.248361 | 3.933201 | 1.080128 | 0.280085 | NA | #VALUE! |
| ENSMUSG00000073601.3 | 0 | NA | NA | NA | NA | NA | #VALUE! |
| ENSMUSG00000073602.11 | 0 | NA | NA | NA | NA | NA | #VALUE! |
| ENSMUSG00000073607.3 | 0.840105 | -3.30885 | 3.958472 | -0.83589 | 0.403216 | NA | #VALUE! |
| ENSMUSG00000073608.5 | 0 | NA | NA | NA | NA | NA | #VALUE! |
| ENSMUSG00000073627.8 | 0 | NA | NA | NA | NA | NA | #VALUE! |
| ENSMUSG00000073628.6 | 0 | NA | NA | NA | NA | NA | #VALUE! |
| ENSMUSG00000073631.3 | 0 | NA | NA | NA | NA | NA | #VALUE! |
| ENSMUSG00000073650.9 | 0 | NA | NA | NA | NA | NA | #VALUE! |
| ENSMUSG00000073652.11 | 0 | NA | NA | NA | NA | NA | #VALUE! |
| ENSMUSG00000073656.5 | 0 | NA | NA | NA | NA | NA | #VALUE! |
| ENSMUSG00000073658.5 | 0 | NA | NA | NA | NA | NA | #VALUE! |
| ENSMUSG00000073679.3 | 0 | NA | NA | NA | NA | NA | #VALUE! |
| ENSMUSG00000073680.3 | 2.679637 | 4.909687 | 3.923523 | 1.251347 | 0.210808 | NA | #VALUE! |
| ENSMUSG00000073682.4 | 0 | NA | NA | NA | NA | NA | #VALUE! |
| ENSMUSG00000073686.3 | 0 | NA | NA | NA | NA | NA | #VALUE! |
| ENSMUSG00000073699.4 | 0 | NA | NA | NA | NA | NA | #VALUE! |
| ENSMUSG00000073705.11 | 0.504851 | 2.502843 | 3.993993 | 0.626652 | 0.530887 | NA | #VALUE! |
| ENSMUSG00000073716.7 | 0 | NA | NA | NA | NA | NA | #VALUE! |
| ENSMUSG00000073721.6 | 0 | NA | NA | NA | NA | NA | #VALUE! |
| ENSMUSG00000073722.4 | 0 | NA | NA | NA | NA | NA | #VALUE! |

| | | | | | | | |
|---|---|---|---|---|---|---|---|
| ENSMUSG00000073723.11 | 0 | NA | NA | NA | NA | NA | #VALUE! |
| ENSMUSG00000073724.13 | 0 | NA | NA | NA | NA | NA | #VALUE! |
| ENSMUSG00000073728.11 | 0 | NA | NA | NA | NA | NA | #VALUE! |
| ENSMUSG00000073730.3 | 0 | NA | NA | NA | NA | NA | #VALUE! |
| ENSMUSG00000073731.13 | 0 | NA | NA | NA | NA | NA | #VALUE! |
| ENSMUSG00000073733.5 | 0 | NA | NA | NA | NA | NA | #VALUE! |
| ENSMUSG00000073735.2 | 0 | NA | NA | NA | NA | NA | #VALUE! |
| ENSMUSG00000073737.4 | 0 | NA | NA | NA | NA | NA | #VALUE! |
| ENSMUSG00000073739.4 | 0 | NA | NA | NA | NA | NA | #VALUE! |
| ENSMUSG00000073747.3 | 0 | NA | NA | NA | NA | NA | #VALUE! |
| ENSMUSG00000073752.3 | 0 | NA | NA | NA | NA | NA | #VALUE! |
| ENSMUSG00000073753.5 | 0 | NA | NA | NA | NA | NA | #VALUE! |
| ENSMUSG00000073755.5 | 3.704894 | -3.45608 | 3.604553 | -0.95881 | 0.337655 | NA | #VALUE! |
| ENSMUSG00000073758.11 | 0 | NA | NA | NA | NA | NA | #VALUE! |
| ENSMUSG00000073759.7 | 0 | NA | NA | NA | NA | NA | #VALUE! |
| ENSMUSG00000073761.3 | 0 | NA | NA | NA | NA | NA | #VALUE! |
| ENSMUSG00000073764.3 | 0 | NA | NA | NA | NA | NA | #VALUE! |
| ENSMUSG00000073765.7 | 0 | NA | NA | NA | NA | NA | #VALUE! |
| ENSMUSG00000073768.4 | 0 | NA | NA | NA | NA | NA | #VALUE! |
| ENSMUSG00000073769.5 | 0 | NA | NA | NA | NA | NA | #VALUE! |
| ENSMUSG00000073771.12 | 3.150394 | -5.19405 | 3.92087 | -1.32472 | 0.185265 | NA | #VALUE! |
| ENSMUSG00000073774.3 | 0 | NA | NA | NA | NA | NA | #VALUE! |
| ENSMUSG00000073778.3 | 0 | NA | NA | NA | NA | NA | #VALUE! |
| ENSMUSG00000073779.8 | 0 | NA | NA | NA | NA | NA | #VALUE! |
| ENSMUSG00000073781.5 | 0 | NA | NA | NA | NA | NA | #VALUE! |
| ENSMUSG00000073785.3 | 0 | NA | NA | NA | NA | NA | #VALUE! |
| ENSMUSG00000073786.2 | 0 | NA | NA | NA | NA | NA | #VALUE! |
| ENSMUSG00000073787.5 | 0 | NA | NA | NA | NA | NA | #VALUE! |
| ENSMUSG00000073789.3 | 0 | NA | NA | NA | NA | NA | #VALUE! |
| ENSMUSG00000073791.14 | 6.149388 | 6.104864 | 3.9141 | 1.559711 | 0.118828 | NA | #VALUE! |
| ENSMUSG00000073794.4 | 0 | NA | NA | NA | NA | NA | #VALUE! |
| ENSMUSG00000073795.12 | 0 | NA | NA | NA | NA | NA | #VALUE! |
| ENSMUSG00000073797.6 | 0 | NA | NA | NA | NA | NA | #VALUE! |
| ENSMUSG00000073803.4 | 0 | NA | NA | NA | NA | NA | #VALUE! |
| ENSMUSG00000073804.6 | 0 | NA | NA | NA | NA | NA | #VALUE! |
| ENSMUSG00000073805.6 | 0 | NA | NA | NA | NA | NA | #VALUE! |
| ENSMUSG00000073807.2 | 0 | NA | NA | NA | NA | NA | #VALUE! |
| ENSMUSG00000073809.2 | 0 | NA | NA | NA | NA | NA | #VALUE! |
| ENSMUSG00000073811.5 | 0 | NA | NA | NA | NA | NA | #VALUE! |
| ENSMUSG00000073821.12 | 0 | NA | NA | NA | NA | NA | #VALUE! |
| ENSMUSG00000073840.7 | 0 | NA | NA | NA | NA | NA | #VALUE! |
| ENSMUSG00000073843.4 | 0 | NA | NA | NA | NA | NA | #VALUE! |
| ENSMUSG00000073844.6 | 0 | NA | NA | NA | NA | NA | #VALUE! |
| ENSMUSG00000073846.5 | 0 | NA | NA | NA | NA | NA | #VALUE! |

| | | | | | | | |
|---|---|---|---|---|---|---|---|
| ENSMUSG00000073860.13 | 0 | NA | NA | NA | NA | NA | #VALUE! |
| ENSMUSG00000073862.5 | 0 | NA | NA | NA | NA | NA | #VALUE! |
| ENSMUSG00000073867.3 | 0 | NA | NA | NA | NA | NA | #VALUE! |
| ENSMUSG00000073876.4 | 0 | NA | NA | NA | NA | NA | #VALUE! |
| ENSMUSG00000073877.11 | 0 | NA | NA | NA | NA | NA | #VALUE! |
| ENSMUSG00000073878.4 | 0 | NA | NA | NA | NA | NA | #VALUE! |
| ENSMUSG00000073879.5 | 0 | NA | NA | NA | NA | NA | #VALUE! |
| ENSMUSG00000073893.7 | 0 | NA | NA | NA | NA | NA | #VALUE! |
| ENSMUSG00000073894.2 | 0 | NA | NA | NA | NA | NA | #VALUE! |
| ENSMUSG00000073896.6 | 0 | NA | NA | NA | NA | NA | #VALUE! |
| ENSMUSG00000073897.6 | 0 | NA | NA | NA | NA | NA | #VALUE! |
| ENSMUSG00000073898.4 | 0 | NA | NA | NA | NA | NA | #VALUE! |
| ENSMUSG00000073899.4 | 0 | NA | NA | NA | NA | NA | #VALUE! |
| ENSMUSG00000073900.6 | 0 | NA | NA | NA | NA | NA | #VALUE! |
| ENSMUSG00000073901.5 | 0 | NA | NA | NA | NA | NA | #VALUE! |
| ENSMUSG00000073902.6 | 0 | NA | NA | NA | NA | NA | #VALUE! |
| ENSMUSG00000073906.4 | 0 | NA | NA | NA | NA | NA | #VALUE! |
| ENSMUSG00000073907.4 | 0 | NA | NA | NA | NA | NA | #VALUE! |
| ENSMUSG00000073908.6 | 0 | NA | NA | NA | NA | NA | #VALUE! |
| ENSMUSG00000073909.3 | 0 | NA | NA | NA | NA | NA | #VALUE! |
| ENSMUSG00000073913.3 | 0 | NA | NA | NA | NA | NA | #VALUE! |
| ENSMUSG00000073914.3 | 0 | NA | NA | NA | NA | NA | #VALUE! |
| ENSMUSG00000073915.6 | 0 | NA | NA | NA | NA | NA | #VALUE! |
| ENSMUSG00000073916.4 | 0 | NA | NA | NA | NA | NA | #VALUE! |
| ENSMUSG00000073917.3 | 0 | NA | NA | NA | NA | NA | #VALUE! |
| ENSMUSG00000073920.5 | 0.105013 | -0.97324 | 4.076293 | -0.23876 | 0.811295 | NA | #VALUE! |
| ENSMUSG00000073922.3 | 0 | NA | NA | NA | NA | NA | #VALUE! |
| ENSMUSG00000073923.5 | 0 | NA | NA | NA | NA | NA | #VALUE! |
| ENSMUSG00000073924.6 | 0 | NA | NA | NA | NA | NA | #VALUE! |
| ENSMUSG00000073925.6 | 0 | NA | NA | NA | NA | NA | #VALUE! |
| ENSMUSG00000073926.5 | 0 | NA | NA | NA | NA | NA | #VALUE! |
| ENSMUSG00000073927.7 | 0 | NA | NA | NA | NA | NA | #VALUE! |
| ENSMUSG00000073928.6 | 0 | NA | NA | NA | NA | NA | #VALUE! |
| ENSMUSG00000073929.13 | 0 | NA | NA | NA | NA | NA | #VALUE! |
| ENSMUSG00000073931.4 | 0 | NA | NA | NA | NA | NA | #VALUE! |
| ENSMUSG00000073932.3 | 0 | NA | NA | NA | NA | NA | #VALUE! |
| ENSMUSG00000073937.3 | 0 | NA | NA | NA | NA | NA | #VALUE! |
| ENSMUSG00000073938.4 | 0 | NA | NA | NA | NA | NA | #VALUE! |
| ENSMUSG00000073943.5 | 0 | NA | NA | NA | NA | NA | #VALUE! |
| ENSMUSG00000073944.7 | 0 | NA | NA | NA | NA | NA | #VALUE! |
| ENSMUSG00000073945.3 | 0 | NA | NA | NA | NA | NA | #VALUE! |
| ENSMUSG00000073946.6 | 0 | NA | NA | NA | NA | NA | #VALUE! |
| ENSMUSG00000073947.5 | 0 | NA | NA | NA | NA | NA | #VALUE! |
| ENSMUSG00000073948.4 | 0 | NA | NA | NA | NA | NA | #VALUE! |

| | | | | | | | |
|---|---|---|---|---|---|---|---|
| ENSMUSG00000073949.5 | 0 | NA | NA | NA | NA | NA | #VALUE! |
| ENSMUSG00000073950.7 | 0 | NA | NA | NA | NA | NA | #VALUE! |
| ENSMUSG00000073951.5 | 0 | NA | NA | NA | NA | NA | #VALUE! |
| ENSMUSG00000073952.6 | 0 | NA | NA | NA | NA | NA | #VALUE! |
| ENSMUSG00000073953.5 | 0 | NA | NA | NA | NA | NA | #VALUE! |
| ENSMUSG00000073954.6 | 0 | NA | NA | NA | NA | NA | #VALUE! |
| ENSMUSG00000073955.3 | 0 | NA | NA | NA | NA | NA | #VALUE! |
| ENSMUSG00000073956.4 | 0 | NA | NA | NA | NA | NA | #VALUE! |
| ENSMUSG00000073959.6 | 0 | NA | NA | NA | NA | NA | #VALUE! |
| ENSMUSG00000073960.2 | 0 | NA | NA | NA | NA | NA | #VALUE! |
| ENSMUSG00000073961.5 | 0 | NA | NA | NA | NA | NA | #VALUE! |
| ENSMUSG00000073962.10 | 0 | NA | NA | NA | NA | NA | #VALUE! |
| ENSMUSG00000073963.4 | 0 | NA | NA | NA | NA | NA | #VALUE! |
| ENSMUSG00000073964.4 | 0 | NA | NA | NA | NA | NA | #VALUE! |
| ENSMUSG00000073965.2 | 0 | NA | NA | NA | NA | NA | #VALUE! |
| ENSMUSG00000073966.4 | 0 | NA | NA | NA | NA | NA | #VALUE! |
| ENSMUSG00000073967.4 | 0 | NA | NA | NA | NA | NA | #VALUE! |
| ENSMUSG00000073968.5 | 0 | NA | NA | NA | NA | NA | #VALUE! |
| ENSMUSG00000073969.4 | 0 | NA | NA | NA | NA | NA | #VALUE! |
| ENSMUSG00000073970.2 | 0 | NA | NA | NA | NA | NA | #VALUE! |
| ENSMUSG00000073971.4 | 0 | NA | NA | NA | NA | NA | #VALUE! |
| ENSMUSG00000073972.3 | 0 | NA | NA | NA | NA | NA | #VALUE! |
| ENSMUSG00000073973.5 | 0 | NA | NA | NA | NA | NA | #VALUE! |
| ENSMUSG00000073974.5 | 0 | NA | NA | NA | NA | NA | #VALUE! |
| ENSMUSG00000073975.4 | 0 | NA | NA | NA | NA | NA | #VALUE! |
| ENSMUSG00000073976.11 | 0 | NA | NA | NA | NA | NA | #VALUE! |
| ENSMUSG00000073977.3 | 0 | NA | NA | NA | NA | NA | #VALUE! |
| ENSMUSG00000073978.9 | 0 | NA | NA | NA | NA | NA | #VALUE! |
| ENSMUSG00000073979.3 | 0 | NA | NA | NA | NA | NA | #VALUE! |
| ENSMUSG00000073980.6 | 0 | NA | NA | NA | NA | NA | #VALUE! |
| ENSMUSG00000073982.12 | 4.894728 | -5.8188 | 3.915922 | -1.48593 | 0.137296 | NA | #VALUE! |
| ENSMUSG00000073985.3 | 3.029103 | 5.088783 | 3.921574 | 1.297638 | 0.194412 | NA | #VALUE! |
| ENSMUSG00000073991.5 | 0 | NA | NA | NA | NA | NA | #VALUE! |
| ENSMUSG00000073997.4 | 0 | NA | NA | NA | NA | NA | #VALUE! |
| ENSMUSG00000073998.3 | 0 | NA | NA | NA | NA | NA | #VALUE! |
| ENSMUSG00000074001.4 | 0 | NA | NA | NA | NA | NA | #VALUE! |
| ENSMUSG00000074003.5 | 0 | NA | NA | NA | NA | NA | #VALUE! |
| ENSMUSG00000074004.4 | 0 | NA | NA | NA | NA | NA | #VALUE! |
| ENSMUSG00000074006.4 | 0 | NA | NA | NA | NA | NA | #VALUE! |
| ENSMUSG00000074011.5 | 0 | NA | NA | NA | NA | NA | #VALUE! |
| ENSMUSG00000074026.4 | 0 | NA | NA | NA | NA | NA | #VALUE! |
| ENSMUSG00000074028.5 | 0 | NA | NA | NA | NA | NA | #VALUE! |
| ENSMUSG00000074029.3 | 0.093255 | 0.868093 | 4.072464 | 0.213162 | 0.831201 | NA | #VALUE! |
| ENSMUSG00000074034.2 | 1.008991 | 0.587892 | 3.915273 | 0.150153 | 0.880644 | NA | #VALUE! |

| | | | | | | | |
|---|---|---|---|---|---|---|---|
| ENSMUSG00000074037.4 | 0 | NA | NA | NA | NA | NA | #VALUE! |
| ENSMUSG00000074039.11 | 0 | NA | NA | NA | NA | NA | #VALUE! |
| ENSMUSG00000074052.8 | 0 | NA | NA | NA | NA | NA | #VALUE! |
| ENSMUSG00000074059.4 | 0 | NA | NA | NA | NA | NA | #VALUE! |
| ENSMUSG00000074060.11 | 0 | NA | NA | NA | NA | NA | #VALUE! |
| ENSMUSG00000074061.5 | 0 | NA | NA | NA | NA | NA | #VALUE! |
| ENSMUSG00000074062.6 | 0 | NA | NA | NA | NA | NA | #VALUE! |
| ENSMUSG00000074065.5 | 0 | NA | NA | NA | NA | NA | #VALUE! |
| ENSMUSG00000074067.10 | 0 | NA | NA | NA | NA | NA | #VALUE! |
| ENSMUSG00000074072.3 | 0 | NA | NA | NA | NA | NA | #VALUE! |
| ENSMUSG00000074092.6 | 0 | NA | NA | NA | NA | NA | #VALUE! |
| ENSMUSG00000074108.2 | 0.195789 | -1.05581 | 4.073302 | -0.2592 | 0.795479 | NA | #VALUE! |
| ENSMUSG00000074109.5 | 0 | NA | NA | NA | NA | NA | #VALUE! |
| ENSMUSG00000074111.5 | 0 | NA | NA | NA | NA | NA | #VALUE! |
| ENSMUSG00000074113.5 | 0 | NA | NA | NA | NA | NA | #VALUE! |
| ENSMUSG00000074121.4 | 0 | NA | NA | NA | NA | NA | #VALUE! |
| ENSMUSG00000074123.2 | 0 | NA | NA | NA | NA | NA | #VALUE! |
| ENSMUSG00000074127.7 | 0 | NA | NA | NA | NA | NA | #VALUE! |
| ENSMUSG00000074128.5 | 0 | NA | NA | NA | NA | NA | #VALUE! |
| ENSMUSG00000074136.10 | 7.552098 | -6.44481 | 3.912702 | -1.64715 | 0.099527 | NA | #VALUE! |
| ENSMUSG00000074139.9 | 0 | NA | NA | NA | NA | NA | #VALUE! |
| ENSMUSG00000074141.14 | 0 | NA | NA | NA | NA | NA | #VALUE! |
| ENSMUSG00000074146.10 | 0 | NA | NA | NA | NA | NA | #VALUE! |
| ENSMUSG00000074149.10 | 0 | NA | NA | NA | NA | NA | #VALUE! |
| ENSMUSG00000074155.3 | 0 | NA | NA | NA | NA | NA | #VALUE! |
| ENSMUSG00000074156.7 | 0 | NA | NA | NA | NA | NA | #VALUE! |
| ENSMUSG00000074158.10 | 5.763964 | -5.13 | 3.906281 | -1.31327 | 0.189092 | NA | #VALUE! |
| ENSMUSG00000074164.13 | 0 | NA | NA | NA | NA | NA | #VALUE! |
| ENSMUSG00000074171.3 | 2.84792 | 4.997812 | 3.922534 | 1.274128 | 0.202618 | NA | #VALUE! |
| ENSMUSG00000074178.4 | 0 | NA | NA | NA | NA | NA | #VALUE! |
| ENSMUSG00000074179.12 | 1.470184 | -4.10383 | 3.936698 | -1.04245 | 0.297201 | NA | #VALUE! |
| ENSMUSG00000074184.6 | 0 | NA | NA | NA | NA | NA | #VALUE! |
| ENSMUSG00000074194.5 | 4.308296 | 3.358991 | 3.856061 | 0.871094 | 0.383703 | NA | #VALUE! |
| ENSMUSG00000074195.4 | 0 | NA | NA | NA | NA | NA | #VALUE! |
| ENSMUSG00000074196.11 | 0 | NA | NA | NA | NA | NA | #VALUE! |
| ENSMUSG00000074199.8 | 0 | NA | NA | NA | NA | NA | #VALUE! |
| ENSMUSG00000074203.5 | 0 | NA | NA | NA | NA | NA | #VALUE! |
| ENSMUSG00000074206.8 | 0 | NA | NA | NA | NA | NA | #VALUE! |
| ENSMUSG00000074210.4 | 0 | NA | NA | NA | NA | NA | #VALUE! |
| ENSMUSG00000074213.4 | 0 | NA | NA | NA | NA | NA | #VALUE! |
| ENSMUSG00000074215.3 | 0 | NA | NA | NA | NA | NA | #VALUE! |
| ENSMUSG00000074217.9 | 2.238125 | 4.666312 | 3.926586 | 1.188389 | 0.23468 | NA | #VALUE! |
| ENSMUSG00000074218.4 | 7.404475 | 6.378241 | 3.912835 | 1.630082 | 0.103084 | NA | #VALUE! |
| ENSMUSG00000074219.4 | 2.984166 | 5.077883 | 3.921685 | 1.294822 | 0.195382 | NA | #VALUE! |

| | | | | | | | |
|---|---|---|---|---|---|---|---|
| ENSMUSG00000074220.11 | 0 | NA | NA | NA | NA | NA | #VALUE! |
| ENSMUSG00000074224.4 | 0 | NA | NA | NA | NA | NA | #VALUE! |
| ENSMUSG00000074228.4 | 0 | NA | NA | NA | NA | NA | #VALUE! |
| ENSMUSG00000074232.4 | 0 | NA | NA | NA | NA | NA | #VALUE! |
| ENSMUSG00000074235.5 | 0 | NA | NA | NA | NA | NA | #VALUE! |
| ENSMUSG00000074236.4 | 0 | NA | NA | NA | NA | NA | #VALUE! |
| ENSMUSG00000074237.4 | 0 | NA | NA | NA | NA | NA | #VALUE! |
| ENSMUSG00000074243.5 | 0 | NA | NA | NA | NA | NA | #VALUE! |
| ENSMUSG00000074248.8 | 0 | NA | NA | NA | NA | NA | #VALUE! |
| ENSMUSG00000074250.12 | 0 | NA | NA | NA | NA | NA | #VALUE! |
| ENSMUSG00000074252.3 | 0 | NA | NA | NA | NA | NA | #VALUE! |
| ENSMUSG00000074254.5 | 5.60204 | 1.958917 | 2.522873 | 0.776463 | 0.437476 | NA | #VALUE! |
| ENSMUSG00000074256.4 | 0 | NA | NA | NA | NA | NA | #VALUE! |
| ENSMUSG00000074259.11 | 0 | NA | NA | NA | NA | NA | #VALUE! |
| ENSMUSG00000074261.4 | 3.36567 | 5.240782 | 3.920098 | 1.336901 | 0.181255 | NA | #VALUE! |
| ENSMUSG00000074268.5 | 0 | NA | NA | NA | NA | NA | #VALUE! |
| ENSMUSG00000074269.11 | 34.91654 | 1.575797 | 2.238723 | 0.703882 | NA | NA | #VALUE! |
| ENSMUSG00000074274.8 | 0 | NA | NA | NA | NA | NA | #VALUE! |
| ENSMUSG00000074277.5 | 5.048506 | 5.825722 | 3.915663 | 1.4878 | 0.136804 | NA | #VALUE! |
| ENSMUSG00000074280.7 | 0 | NA | NA | NA | NA | NA | #VALUE! |
| ENSMUSG00000074281.4 | 0 | NA | NA | NA | NA | NA | #VALUE! |
| ENSMUSG00000074282.11 | 3.532525 | -5.3473 | 3.919452 | -1.3643 | 0.172474 | NA | #VALUE! |
| ENSMUSG00000074283.6 | 0 | NA | NA | NA | NA | NA | #VALUE! |
| ENSMUSG00000074291.4 | 0 | NA | NA | NA | NA | NA | #VALUE! |
| ENSMUSG00000074292.5 | 0 | NA | NA | NA | NA | NA | #VALUE! |
| ENSMUSG00000074300.5 | 0 | NA | NA | NA | NA | NA | #VALUE! |
| ENSMUSG00000074302.2 | 0 | NA | NA | NA | NA | NA | #VALUE! |
| ENSMUSG00000074311.4 | 0 | NA | NA | NA | NA | NA | #VALUE! |
| ENSMUSG00000074322.6 | 0 | NA | NA | NA | NA | NA | #VALUE! |
| ENSMUSG00000074340.10 | 0 | NA | NA | NA | NA | NA | #VALUE! |
| ENSMUSG00000074342.5 | 0 | NA | NA | NA | NA | NA | #VALUE! |
| ENSMUSG00000074343.4 | 0 | NA | NA | NA | NA | NA | #VALUE! |
| ENSMUSG00000074344.8 | 0 | NA | NA | NA | NA | NA | #VALUE! |
| ENSMUSG00000074345.5 | 0.559531 | 2.703958 | 3.982863 | 0.678898 | 0.497202 | NA | #VALUE! |
| ENSMUSG00000074346.7 | 0 | NA | NA | NA | NA | NA | #VALUE! |
| ENSMUSG00000074354.12 | 0 | NA | NA | NA | NA | NA | #VALUE! |
| ENSMUSG00000074355.3 | 0 | NA | NA | NA | NA | NA | #VALUE! |
| ENSMUSG00000074357.4 | 0 | NA | NA | NA | NA | NA | #VALUE! |
| ENSMUSG00000074358.10 | 2.33138 | 4.724655 | 3.925805 | 1.203487 | 0.228788 | NA | #VALUE! |
| ENSMUSG00000074359.3 | 0 | NA | NA | NA | NA | NA | #VALUE! |
| ENSMUSG00000074361.12 | 0 | NA | NA | NA | NA | NA | #VALUE! |
| ENSMUSG00000074364.7 | 1.785223 | -4.3809 | 3.931494 | -1.11431 | 0.265146 | NA | #VALUE! |
| ENSMUSG00000074365.14 | 0 | NA | NA | NA | NA | NA | #VALUE! |
| ENSMUSG00000074366.10 | 0 | NA | NA | NA | NA | NA | #VALUE! |

| | | | | | | | |
|---|---|---|---|---|---|---|---|
| ENSMUSG00000074369.13 | 0 | NA | NA | NA | NA | NA | #VALUE! |
| ENSMUSG00000074373.7 | 3.524204 | -5.34395 | 3.919482 | -1.36343 | 0.172746 | NA | #VALUE! |
| ENSMUSG00000074375.4 | 0 | NA | NA | NA | NA | NA | #VALUE! |
| ENSMUSG00000074377.13 | 0 | NA | NA | NA | NA | NA | #VALUE! |
| ENSMUSG00000074378.6 | 0 | NA | NA | NA | NA | NA | #VALUE! |
| ENSMUSG00000074384.5 | 2.21378 | 4.621487 | 3.927208 | 1.176787 | 0.239281 | NA | #VALUE! |
| ENSMUSG00000074385.4 | 0 | NA | NA | NA | NA | NA | #VALUE! |
| ENSMUSG00000074388.4 | 3.36567 | 5.240782 | 3.920098 | 1.336901 | 0.181255 | NA | #VALUE! |
| ENSMUSG00000074397.6 | 0 | NA | NA | NA | NA | NA | #VALUE! |
| ENSMUSG00000074398.6 | 0 | NA | NA | NA | NA | NA | #VALUE! |
| ENSMUSG00000074403.3 | 0 | NA | NA | NA | NA | NA | #VALUE! |
| ENSMUSG00000074404.7 | 0 | NA | NA | NA | NA | NA | #VALUE! |
| ENSMUSG00000074405.9 | 1.750771 | -1.32041 | 3.852498 | -0.34274 | 0.731793 | NA | #VALUE! |
| ENSMUSG00000074406.6 | 1.188972 | -3.77568 | 3.944285 | -0.95725 | 0.338439 | NA | #VALUE! |
| ENSMUSG00000074417.10 | 0 | NA | NA | NA | NA | NA | #VALUE! |
| ENSMUSG00000074418.5 | 0 | NA | NA | NA | NA | NA | #VALUE! |
| ENSMUSG00000074419.12 | 0 | NA | NA | NA | NA | NA | #VALUE! |
| ENSMUSG00000074420.6 | 0 | NA | NA | NA | NA | NA | #VALUE! |
| ENSMUSG00000074424.4 | 0 | NA | NA | NA | NA | NA | #VALUE! |
| ENSMUSG00000074433.5 | 0 | NA | NA | NA | NA | NA | #VALUE! |
| ENSMUSG00000074434.5 | 0 | NA | NA | NA | NA | NA | #VALUE! |
| ENSMUSG00000074435.11 | 0 | NA | NA | NA | NA | NA | #VALUE! |
| ENSMUSG00000074436.3 | 0 | NA | NA | NA | NA | NA | #VALUE! |
| ENSMUSG00000074437.6 | 0 | NA | NA | NA | NA | NA | #VALUE! |
| ENSMUSG00000074439.5 | 0 | NA | NA | NA | NA | NA | #VALUE! |
| ENSMUSG00000074440.4 | 0 | NA | NA | NA | NA | NA | #VALUE! |
| ENSMUSG00000074441.4 | 0 | NA | NA | NA | NA | NA | #VALUE! |
| ENSMUSG00000074442.5 | 0 | NA | NA | NA | NA | NA | #VALUE! |
| ENSMUSG00000074443.2 | 0 | NA | NA | NA | NA | NA | #VALUE! |
| ENSMUSG00000074444.5 | 0 | NA | NA | NA | NA | NA | #VALUE! |
| ENSMUSG00000074445.5 | 0 | NA | NA | NA | NA | NA | #VALUE! |
| ENSMUSG00000074446.4 | 0 | NA | NA | NA | NA | NA | #VALUE! |
| ENSMUSG00000074447.3 | 0 | NA | NA | NA | NA | NA | #VALUE! |
| ENSMUSG00000074448.3 | 0 | NA | NA | NA | NA | NA | #VALUE! |
| ENSMUSG00000074449.4 | 0 | NA | NA | NA | NA | NA | #VALUE! |
| ENSMUSG00000074452.14 | 0 | NA | NA | NA | NA | NA | #VALUE! |
| ENSMUSG00000074454.4 | 0 | NA | NA | NA | NA | NA | #VALUE! |
| ENSMUSG00000074457.11 | 3.846979 | 3.87667 | 3.896347 | 0.99495 | 0.319761 | NA | #VALUE! |
| ENSMUSG00000074461.3 | 0 | NA | NA | NA | NA | NA | #VALUE! |
| ENSMUSG00000074465.3 | 0 | NA | NA | NA | NA | NA | #VALUE! |
| ENSMUSG00000074467.5 | 0 | NA | NA | NA | NA | NA | #VALUE! |
| ENSMUSG00000074469.7 | 0 | NA | NA | NA | NA | NA | #VALUE! |
| ENSMUSG00000074472.11 | 0.336567 | 1.944925 | 4.029936 | 0.482619 | 0.629366 | NA | #VALUE! |
| ENSMUSG00000074473.2 | 0 | NA | NA | NA | NA | NA | #VALUE! |

| | | | | | | | |
|---|---|---|---|---|---|---|---|
| ENSMUSG00000074479.4 | 3.906503 | -3.98016 | 3.893691 | -1.02221 | 0.306682 | NA | #VALUE! |
| ENSMUSG00000074480.5 | 0 | NA | NA | NA | NA | NA | #VALUE! |
| ENSMUSG00000074482.6 | 0 | NA | NA | NA | NA | NA | #VALUE! |
| ENSMUSG00000074483.3 | 0 | NA | NA | NA | NA | NA | #VALUE! |
| ENSMUSG00000074485.2 | 0 | NA | NA | NA | NA | NA | #VALUE! |
| ENSMUSG00000074486.3 | 0 | NA | NA | NA | NA | NA | #VALUE! |
| ENSMUSG00000074489.10 | 0 | NA | NA | NA | NA | NA | #VALUE! |
| ENSMUSG00000074500.12 | 0 | NA | NA | NA | NA | NA | #VALUE! |
| ENSMUSG00000074501.7 | 0 | NA | NA | NA | NA | NA | #VALUE! |
| ENSMUSG00000074502.5 | 0 | NA | NA | NA | NA | NA | #VALUE! |
| ENSMUSG00000074505.7 | 0 | NA | NA | NA | NA | NA | #VALUE! |
| ENSMUSG00000074506.5 | 0 | NA | NA | NA | NA | NA | #VALUE! |
| ENSMUSG00000074507.7 | 0 | NA | NA | NA | NA | NA | #VALUE! |
| ENSMUSG00000074508.5 | 0 | NA | NA | NA | NA | NA | #VALUE! |
| ENSMUSG00000074516.7 | 0 | NA | NA | NA | NA | NA | #VALUE! |
| ENSMUSG00000074517.3 | 0 | NA | NA | NA | NA | NA | #VALUE! |
| ENSMUSG00000074519.12 | 2.93356 | 2.671348 | 3.858224 | 0.692378 | 0.4887 | NA | #VALUE! |
| ENSMUSG00000074527.12 | 0 | NA | NA | NA | NA | NA | #VALUE! |
| ENSMUSG00000074529.11 | 0 | NA | NA | NA | NA | NA | #VALUE! |
| ENSMUSG00000074553.5 | 0.588889 | 0.014287 | 4.003809 | 0.003568 | 0.997153 | NA | #VALUE! |
| ENSMUSG00000074555.6 | 0 | NA | NA | NA | NA | NA | #VALUE! |
| ENSMUSG00000074564.4 | 0.245976 | 1.027628 | 4.067126 | 0.252667 | 0.800526 | NA | #VALUE! |
| ENSMUSG00000074569.5 | 0 | NA | NA | NA | NA | NA | #VALUE! |
| ENSMUSG00000074570.15 | 0 | NA | NA | NA | NA | NA | #VALUE! |
| ENSMUSG00000074575.5 | 0 | NA | NA | NA | NA | NA | #VALUE! |
| ENSMUSG00000074576.5 | 9.752572 | 1.275629 | 2.24928 | 0.567128 | 0.570627 | NA | #VALUE! |
| ENSMUSG00000074577.10 | 5.165486 | 5.852308 | 3.9155 | 1.494651 | 0.135005 | NA | #VALUE! |
| ENSMUSG00000074579.15 | 0 | NA | NA | NA | NA | NA | #VALUE! |
| ENSMUSG00000074580.12 | 2.40029 | 4.741481 | 3.925585 | 1.207841 | 0.227109 | NA | #VALUE! |
| ENSMUSG00000074589.5 | 0 | NA | NA | NA | NA | NA | #VALUE! |
| ENSMUSG00000074593.12 | 0 | NA | NA | NA | NA | NA | #VALUE! |
| ENSMUSG00000074594.10 | 0 | NA | NA | NA | NA | NA | #VALUE! |
| ENSMUSG00000074595.10 | 0 | NA | NA | NA | NA | NA | #VALUE! |
| ENSMUSG00000074596.4 | 0 | NA | NA | NA | NA | NA | #VALUE! |
| ENSMUSG00000074603.4 | 0 | NA | NA | NA | NA | NA | #VALUE! |
| ENSMUSG00000074604.10 | 0 | NA | NA | NA | NA | NA | #VALUE! |
| ENSMUSG00000074607.12 | 7.245906 | -6.39089 | 3.912928 | -1.63328 | 0.102411 | NA | #VALUE! |
| ENSMUSG00000074619.6 | 0 | NA | NA | NA | NA | NA | #VALUE! |
| ENSMUSG00000074620.7 | 0 | NA | NA | NA | NA | NA | #VALUE! |
| ENSMUSG00000074625.12 | 0 | NA | NA | NA | NA | NA | #VALUE! |
| ENSMUSG00000074627.12 | 0 | NA | NA | NA | NA | NA | #VALUE! |
| ENSMUSG00000074628.10 | 0 | NA | NA | NA | NA | NA | #VALUE! |
| ENSMUSG00000074629.4 | 0 | NA | NA | NA | NA | NA | #VALUE! |
| ENSMUSG00000074635.4 | 0 | NA | NA | NA | NA | NA | #VALUE! |

| | | | | | | | |
|---|---|---|---|---|---|---|---|
| ENSMUSG00000074637.8 | 0 | NA | NA | NA | NA | NA | #VALUE! |
| ENSMUSG00000074643.13 | 0 | NA | NA | NA | NA | NA | #VALUE! |
| ENSMUSG00000074646.11 | 3.780473 | -5.45568 | 3.918536 | -1.39228 | 0.163839 | NA | #VALUE! |
| ENSMUSG00000074647.5 | 0 | NA | NA | NA | NA | NA | #VALUE! |
| ENSMUSG00000074650.3 | 0 | NA | NA | NA | NA | NA | #VALUE! |
| ENSMUSG00000074651.4 | 0 | NA | NA | NA | NA | NA | #VALUE! |
| ENSMUSG00000074652.4 | 0 | NA | NA | NA | NA | NA | #VALUE! |
| ENSMUSG00000074653.11 | 0 | NA | NA | NA | NA | NA | #VALUE! |
| ENSMUSG00000074655.2 | 0 | NA | NA | NA | NA | NA | #VALUE! |
| ENSMUSG00000074657.6 | 0 | NA | NA | NA | NA | NA | #VALUE! |
| ENSMUSG00000074664.4 | 0 | NA | NA | NA | NA | NA | #VALUE! |
| ENSMUSG00000074665.12 | 0 | NA | NA | NA | NA | NA | #VALUE! |
| ENSMUSG00000074671.7 | 4.991704 | 5.809761 | 3.915761 | 1.483686 | 0.137892 | NA | #VALUE! |
| ENSMUSG00000074673.16 | 0 | NA | NA | NA | NA | NA | #VALUE! |
| ENSMUSG00000074676.3 | 0 | NA | NA | NA | NA | NA | #VALUE! |
| ENSMUSG00000074677.12 | 0 | NA | NA | NA | NA | NA | #VALUE! |
| ENSMUSG00000074678.2 | 0 | NA | NA | NA | NA | NA | #VALUE! |
| ENSMUSG00000074679.4 | 0 | NA | NA | NA | NA | NA | #VALUE! |
| ENSMUSG00000074680.2 | 0 | NA | NA | NA | NA | NA | #VALUE! |
| ENSMUSG00000074681.3 | 0 | NA | NA | NA | NA | NA | #VALUE! |
| ENSMUSG00000074682.5 | 0 | NA | NA | NA | NA | NA | #VALUE! |
| ENSMUSG00000074695.4 | 0 | NA | NA | NA | NA | NA | #VALUE! |
| ENSMUSG00000074704.10 | 0 | NA | NA | NA | NA | NA | #VALUE! |
| ENSMUSG00000074715.4 | 0.746042 | 3.107177 | 3.964586 | 0.783733 | 0.433197 | NA | #VALUE! |
| ENSMUSG00000074720.2 | 0 | NA | NA | NA | NA | NA | #VALUE! |
| ENSMUSG00000074731.4 | 0 | NA | NA | NA | NA | NA | #VALUE! |
| ENSMUSG00000074734.4 | 0 | NA | NA | NA | NA | NA | #VALUE! |
| ENSMUSG00000074735.3 | 0 | NA | NA | NA | NA | NA | #VALUE! |
| ENSMUSG00000074736.11 | 0 | NA | NA | NA | NA | NA | #VALUE! |
| ENSMUSG00000074737.3 | 0 | NA | NA | NA | NA | NA | #VALUE! |
| ENSMUSG00000074738.3 | 1.174735 | -3.75246 | 3.944889 | -0.95122 | 0.341493 | NA | #VALUE! |
| ENSMUSG00000074739.3 | 0 | NA | NA | NA | NA | NA | #VALUE! |
| ENSMUSG00000074743.5 | 26.06788 | 1.267889 | 2.320607 | 0.546361 | NA | NA | #VALUE! |
| ENSMUSG00000074753.6 | 0 | NA | NA | NA | NA | NA | #VALUE! |
| ENSMUSG00000074758.11 | 0 | NA | NA | NA | NA | NA | #VALUE! |
| ENSMUSG00000074763.6 | 0 | NA | NA | NA | NA | NA | #VALUE! |
| ENSMUSG00000074764.12 | 0 | NA | NA | NA | NA | NA | #VALUE! |
| ENSMUSG00000074766.11 | 0 | NA | NA | NA | NA | NA | #VALUE! |
| ENSMUSG00000074771.11 | 0 | NA | NA | NA | NA | NA | #VALUE! |
| ENSMUSG00000074776.5 | 0 | NA | NA | NA | NA | NA | #VALUE! |
| ENSMUSG00000074780.3 | 0 | NA | NA | NA | NA | NA | #VALUE! |
| ENSMUSG00000074782.4 | 0.841418 | 3.240539 | 3.959551 | 0.818411 | 0.413123 | NA | #VALUE! |
| ENSMUSG00000074783.11 | 0 | NA | NA | NA | NA | NA | #VALUE! |
| ENSMUSG00000074793.11 | 0.336567 | 1.944925 | 4.029936 | 0.482619 | 0.629366 | NA | #VALUE! |

| | | | | | | | |
|---|---|---|---|---|---|---|---|
| ENSMUSG00000074796.11 | 0 | NA | NA | NA | NA | NA | #VALUE! |
| ENSMUSG00000074804.4 | 0 | NA | NA | NA | NA | NA | #VALUE! |
| ENSMUSG00000074805.6 | 0 | NA | NA | NA | NA | NA | #VALUE! |
| ENSMUSG00000074807.4 | 3.95837 | -1.44629 | 3.561439 | -0.4061 | 0.684672 | NA | #VALUE! |
| ENSMUSG00000074811.4 | 0.245976 | 1.027628 | 4.067126 | 0.252667 | 0.800526 | NA | #VALUE! |
| ENSMUSG00000074812.13 | 0 | NA | NA | NA | NA | NA | #VALUE! |
| ENSMUSG00000074813.15 | 1.492083 | 4.088114 | 3.936282 | 1.038572 | 0.299004 | NA | #VALUE! |
| ENSMUSG00000074817.3 | 0 | NA | NA | NA | NA | NA | #VALUE! |
| ENSMUSG00000074818.14 | 0 | NA | NA | NA | NA | NA | #VALUE! |
| ENSMUSG00000074824.4 | 3.513232 | 1.823777 | 3.80173 | 0.479723 | 0.631424 | NA | #VALUE! |
| ENSMUSG00000074830.5 | 0 | NA | NA | NA | NA | NA | #VALUE! |
| ENSMUSG00000074832.12 | 0 | NA | NA | NA | NA | NA | #VALUE! |
| ENSMUSG00000074837.3 | 0 | NA | NA | NA | NA | NA | #VALUE! |
| ENSMUSG00000074841.3 | 0 | NA | NA | NA | NA | NA | #VALUE! |
| ENSMUSG00000074846.4 | 0 | NA | NA | NA | NA | NA | #VALUE! |
| ENSMUSG00000074847.3 | 0 | NA | NA | NA | NA | NA | #VALUE! |
| ENSMUSG00000074849.3 | 0 | NA | NA | NA | NA | NA | #VALUE! |
| ENSMUSG00000074852.5 | 0 | NA | NA | NA | NA | NA | #VALUE! |
| ENSMUSG00000074862.12 | 0 | NA | NA | NA | NA | NA | #VALUE! |
| ENSMUSG00000074863.6 | 4.728045 | 5.732592 | 3.916255 | 1.463794 | 0.14325 | NA | #VALUE! |
| ENSMUSG00000074865.12 | 0 | NA | NA | NA | NA | NA | #VALUE! |
| ENSMUSG00000074867.5 | 0 | NA | NA | NA | NA | NA | #VALUE! |
| ENSMUSG00000074868.3 | 0 | NA | NA | NA | NA | NA | #VALUE! |
| ENSMUSG00000074870.5 | 0 | NA | NA | NA | NA | NA | #VALUE! |
| ENSMUSG00000074871.4 | 0 | NA | NA | NA | NA | NA | #VALUE! |
| ENSMUSG00000074872.4 | 0 | NA | NA | NA | NA | NA | #VALUE! |
| ENSMUSG00000074874.11 | 0 | NA | NA | NA | NA | NA | #VALUE! |
| ENSMUSG00000074877.4 | 0 | NA | NA | NA | NA | NA | #VALUE! |
| ENSMUSG00000074881.5 | 0 | NA | NA | NA | NA | NA | #VALUE! |
| ENSMUSG00000074885.5 | 0 | NA | NA | NA | NA | NA | #VALUE! |
| ENSMUSG00000074886.12 | 1.050131 | -3.6251 | 3.948379 | -0.91812 | 0.358554 | NA | #VALUE! |
| ENSMUSG00000074887.5 | 0 | NA | NA | NA | NA | NA | #VALUE! |
| ENSMUSG00000074892.10 | 0 | NA | NA | NA | NA | NA | #VALUE! |
| ENSMUSG00000074895.11 | 0 | NA | NA | NA | NA | NA | #VALUE! |
| ENSMUSG00000074899.7 | 0 | NA | NA | NA | NA | NA | #VALUE! |
| ENSMUSG00000074903.6 | 0 | NA | NA | NA | NA | NA | #VALUE! |
| ENSMUSG00000074912.11 | 0 | NA | NA | NA | NA | NA | #VALUE! |
| ENSMUSG00000074913.4 | 0 | NA | NA | NA | NA | NA | #VALUE! |
| ENSMUSG00000074916.6 | 6.520691 | -2.44561 | 3.779605 | -0.64706 | 0.517596 | NA | #VALUE! |
| ENSMUSG00000074919.3 | 0 | NA | NA | NA | NA | NA | #VALUE! |
| ENSMUSG00000074923.11 | 0 | NA | NA | NA | NA | NA | #VALUE! |
| ENSMUSG00000074928.5 | 0 | NA | NA | NA | NA | NA | #VALUE! |
| ENSMUSG00000074930.6 | 0 | NA | NA | NA | NA | NA | #VALUE! |
| ENSMUSG00000074934.4 | 0 | NA | NA | NA | NA | NA | #VALUE! |

| | | | | | | | |
|---|---|---|---|---|---|---|---|
| ENSMUSG00000074939.4 | 0 | NA | NA | NA | NA | NA | #VALUE! |
| ENSMUSG00000074945.4 | 0 | NA | NA | NA | NA | NA | #VALUE! |
| ENSMUSG00000074946.6 | 0 | NA | NA | NA | NA | NA | #VALUE! |
| ENSMUSG00000074947.4 | 0 | NA | NA | NA | NA | NA | #VALUE! |
| ENSMUSG00000074952.4 | 0 | NA | NA | NA | NA | NA | #VALUE! |
| ENSMUSG00000074955.4 | 0 | NA | NA | NA | NA | NA | #VALUE! |
| ENSMUSG00000074965.6 | 0 | NA | NA | NA | NA | NA | #VALUE! |
| ENSMUSG00000074966.6 | 0 | NA | NA | NA | NA | NA | #VALUE! |
| ENSMUSG00000074968.12 | 0 | NA | NA | NA | NA | NA | #VALUE! |
| ENSMUSG00000074970.2 | 0 | NA | NA | NA | NA | NA | #VALUE! |
| ENSMUSG00000074971.5 | 2.508557 | -3.90495 | 3.916787 | -0.99698 | 0.318775 | NA | #VALUE! |
| ENSMUSG00000074973.5 | 0 | NA | NA | NA | NA | NA | #VALUE! |
| ENSMUSG00000074978.5 | 0 | NA | NA | NA | NA | NA | #VALUE! |
| ENSMUSG00000074981.14 | 0 | NA | NA | NA | NA | NA | #VALUE! |
| ENSMUSG00000074987.10 | 0 | NA | NA | NA | NA | NA | #VALUE! |
| ENSMUSG00000074991.6 | 0 | NA | NA | NA | NA | NA | #VALUE! |
| ENSMUSG00000074995.3 | 0 | NA | NA | NA | NA | NA | #VALUE! |
| ENSMUSG00000074996.5 | 0 | NA | NA | NA | NA | NA | #VALUE! |
| ENSMUSG00000074997.4 | 0 | NA | NA | NA | NA | NA | #VALUE! |
| ENSMUSG00000074999.3 | 0 | NA | NA | NA | NA | NA | #VALUE! |
| ENSMUSG00000075000.12 | 0.664544 | 1.770909 | 3.978581 | 0.445111 | 0.65624 | NA | #VALUE! |
| ENSMUSG00000075002.3 | 0 | NA | NA | NA | NA | NA | #VALUE! |
| ENSMUSG00000075006.3 | 0 | NA | NA | NA | NA | NA | #VALUE! |
| ENSMUSG00000075012.5 | 0 | NA | NA | NA | NA | NA | #VALUE! |
| ENSMUSG00000075015.4 | 0.499071 | -1.3391 | 4.005362 | -0.33433 | 0.738133 | NA | #VALUE! |
| ENSMUSG00000075020.7 | 0 | NA | NA | NA | NA | NA | #VALUE! |
| ENSMUSG00000075023.12 | 0 | NA | NA | NA | NA | NA | #VALUE! |
| ENSMUSG00000075025.11 | 0 | NA | NA | NA | NA | NA | #VALUE! |
| ENSMUSG00000075027.11 | 0 | NA | NA | NA | NA | NA | #VALUE! |
| ENSMUSG00000075028.13 | 0 | NA | NA | NA | NA | NA | #VALUE! |
| ENSMUSG00000075029.8 | 0 | NA | NA | NA | NA | NA | #VALUE! |
| ENSMUSG00000075031.5 | 0 | NA | NA | NA | NA | NA | #VALUE! |
| ENSMUSG00000075033.5 | 19.75044 | -3.77209 | 2.734138 | -1.37963 | NA | NA | #VALUE! |
| ENSMUSG00000075036.2 | 0 | NA | NA | NA | NA | NA | #VALUE! |
| ENSMUSG00000075040.10 | 6.680079 | 1.309565 | 3.30244 | 0.396545 | 0.691703 | NA | #VALUE! |
| ENSMUSG00000075042.11 | 0 | NA | NA | NA | NA | NA | #VALUE! |
| ENSMUSG00000075044.15 | 7.13058 | 6.324018 | 3.091681 | 2.045495 | 0.040806 | NA | #VALUE! |
| ENSMUSG00000075045.5 | 0 | NA | NA | NA | NA | NA | #VALUE! |
| ENSMUSG00000075046.8 | 0 | NA | NA | NA | NA | NA | #VALUE! |
| ENSMUSG00000075053.4 | 0.204111 | -1.05581 | 4.073302 | -0.2592 | 0.795479 | NA | #VALUE! |
| ENSMUSG00000075061.3 | 0 | NA | NA | NA | NA | NA | #VALUE! |
| ENSMUSG00000075062.5 | 0 | NA | NA | NA | NA | NA | #VALUE! |
| ENSMUSG00000075063.3 | 0 | NA | NA | NA | NA | NA | #VALUE! |
| ENSMUSG00000075064.6 | 0 | NA | NA | NA | NA | NA | #VALUE! |

| | | | | | | | |
|---|---|---|---|---|---|---|---|
| ENSMUSG00000075065.6 | 0 | NA | NA | NA | NA | NA | #VALUE! |
| ENSMUSG00000075066.8 | 0 | NA | NA | NA | NA | NA | #VALUE! |
| ENSMUSG00000075067.5 | 0 | NA | NA | NA | NA | NA | #VALUE! |
| ENSMUSG00000075068.4 | 0 | NA | NA | NA | NA | NA | #VALUE! |
| ENSMUSG00000075069.4 | 0 | NA | NA | NA | NA | NA | #VALUE! |
| ENSMUSG00000075070.7 | 0 | NA | NA | NA | NA | NA | #VALUE! |
| ENSMUSG00000075072.4 | 0 | NA | NA | NA | NA | NA | #VALUE! |
| ENSMUSG00000075073.4 | 0 | NA | NA | NA | NA | NA | #VALUE! |
| ENSMUSG00000075074.4 | 0 | NA | NA | NA | NA | NA | #VALUE! |
| ENSMUSG00000075075.3 | 0 | NA | NA | NA | NA | NA | #VALUE! |
| ENSMUSG00000075078.6 | 0 | NA | NA | NA | NA | NA | #VALUE! |
| ENSMUSG00000075079.4 | 0 | NA | NA | NA | NA | NA | #VALUE! |
| ENSMUSG00000075081.7 | 0 | NA | NA | NA | NA | NA | #VALUE! |
| ENSMUSG00000075084.4 | 0 | NA | NA | NA | NA | NA | #VALUE! |
| ENSMUSG00000075085.7 | 0 | NA | NA | NA | NA | NA | #VALUE! |
| ENSMUSG00000075086.3 | 0 | NA | NA | NA | NA | NA | #VALUE! |
| ENSMUSG00000075088.3 | 0 | NA | NA | NA | NA | NA | #VALUE! |
| ENSMUSG00000075090.5 | 0 | NA | NA | NA | NA | NA | #VALUE! |
| ENSMUSG00000075091.6 | 0 | NA | NA | NA | NA | NA | #VALUE! |
| ENSMUSG00000075092.3 | 0 | NA | NA | NA | NA | NA | #VALUE! |
| ENSMUSG00000075093.4 | 0 | NA | NA | NA | NA | NA | #VALUE! |
| ENSMUSG00000075094.2 | 0 | NA | NA | NA | NA | NA | #VALUE! |
| ENSMUSG00000075095.6 | 0 | NA | NA | NA | NA | NA | #VALUE! |
| ENSMUSG00000075097.4 | 0 | NA | NA | NA | NA | NA | #VALUE! |
| ENSMUSG00000075099.3 | 0 | NA | NA | NA | NA | NA | #VALUE! |
| ENSMUSG00000075100.5 | 0 | NA | NA | NA | NA | NA | #VALUE! |
| ENSMUSG00000075101.5 | 0 | NA | NA | NA | NA | NA | #VALUE! |
| ENSMUSG00000075102.3 | 0 | NA | NA | NA | NA | NA | #VALUE! |
| ENSMUSG00000075104.4 | 0 | NA | NA | NA | NA | NA | #VALUE! |
| ENSMUSG00000075105.4 | 0 | NA | NA | NA | NA | NA | #VALUE! |
| ENSMUSG00000075107.4 | 0 | NA | NA | NA | NA | NA | #VALUE! |
| ENSMUSG00000075110.3 | 0 | NA | NA | NA | NA | NA | #VALUE! |
| ENSMUSG00000075111.12 | 0 | NA | NA | NA | NA | NA | #VALUE! |
| ENSMUSG00000075112.3 | 0 | NA | NA | NA | NA | NA | #VALUE! |
| ENSMUSG00000075113.3 | 0 | NA | NA | NA | NA | NA | #VALUE! |
| ENSMUSG00000075114.5 | 0 | NA | NA | NA | NA | NA | #VALUE! |
| ENSMUSG00000075115.4 | 0 | NA | NA | NA | NA | NA | #VALUE! |
| ENSMUSG00000075117.4 | 0.391578 | -2.1489 | 4.020883 | -0.53444 | 0.59304 | NA | #VALUE! |
| ENSMUSG00000075119.3 | 0 | NA | NA | NA | NA | NA | #VALUE! |
| ENSMUSG00000075120.5 | 0 | NA | NA | NA | NA | NA | #VALUE! |
| ENSMUSG00000075121.3 | 0 | NA | NA | NA | NA | NA | #VALUE! |
| ENSMUSG00000075122.6 | 0 | NA | NA | NA | NA | NA | #VALUE! |
| ENSMUSG00000075125.3 | 0 | NA | NA | NA | NA | NA | #VALUE! |
| ENSMUSG00000075126.5 | 0 | NA | NA | NA | NA | NA | #VALUE! |

| | | | | | | | |
|---|---|---|---|---|---|---|---|
| ENSMUSG00000075127.3 | 0 | NA | NA | NA | NA | NA | #VALUE! |
| ENSMUSG00000075128.8 | 0 | NA | NA | NA | NA | NA | #VALUE! |
| ENSMUSG00000075132.3 | 0 | NA | NA | NA | NA | NA | #VALUE! |
| ENSMUSG00000075133.3 | 0 | NA | NA | NA | NA | NA | #VALUE! |
| ENSMUSG00000075136.5 | 0 | NA | NA | NA | NA | NA | #VALUE! |
| ENSMUSG00000075137.5 | 0 | NA | NA | NA | NA | NA | #VALUE! |
| ENSMUSG00000075139.4 | 0 | NA | NA | NA | NA | NA | #VALUE! |
| ENSMUSG00000075140.5 | 0 | NA | NA | NA | NA | NA | #VALUE! |
| ENSMUSG00000075141.6 | 0 | NA | NA | NA | NA | NA | #VALUE! |
| ENSMUSG00000075142.2 | 0 | NA | NA | NA | NA | NA | #VALUE! |
| ENSMUSG00000075143.6 | 0 | NA | NA | NA | NA | NA | #VALUE! |
| ENSMUSG00000075144.6 | 0 | NA | NA | NA | NA | NA | #VALUE! |
| ENSMUSG00000075145.3 | 0 | NA | NA | NA | NA | NA | #VALUE! |
| ENSMUSG00000075146.3 | 0 | NA | NA | NA | NA | NA | #VALUE! |
| ENSMUSG00000075148.2 | 0 | NA | NA | NA | NA | NA | #VALUE! |
| ENSMUSG00000075149.3 | 0 | NA | NA | NA | NA | NA | #VALUE! |
| ENSMUSG00000075150.3 | 0 | NA | NA | NA | NA | NA | #VALUE! |
| ENSMUSG00000075151.7 | 0 | NA | NA | NA | NA | NA | #VALUE! |
| ENSMUSG00000075153.3 | 0 | NA | NA | NA | NA | NA | #VALUE! |
| ENSMUSG00000075155.3 | 0 | NA | NA | NA | NA | NA | #VALUE! |
| ENSMUSG00000075156.5 | 0 | NA | NA | NA | NA | NA | #VALUE! |
| ENSMUSG00000075158.6 | 0 | NA | NA | NA | NA | NA | #VALUE! |
| ENSMUSG00000075159.6 | 0 | NA | NA | NA | NA | NA | #VALUE! |
| ENSMUSG00000075160.4 | 0 | NA | NA | NA | NA | NA | #VALUE! |
| ENSMUSG00000075161.4 | 0 | NA | NA | NA | NA | NA | #VALUE! |
| ENSMUSG00000075162.4 | 0 | NA | NA | NA | NA | NA | #VALUE! |
| ENSMUSG00000075163.6 | 0 | NA | NA | NA | NA | NA | #VALUE! |
| ENSMUSG00000075164.3 | 0 | NA | NA | NA | NA | NA | #VALUE! |
| ENSMUSG00000075165.4 | 0 | NA | NA | NA | NA | NA | #VALUE! |
| ENSMUSG00000075166.3 | 0 | NA | NA | NA | NA | NA | #VALUE! |
| ENSMUSG00000075167.4 | 0 | NA | NA | NA | NA | NA | #VALUE! |
| ENSMUSG00000075168.5 | 0 | NA | NA | NA | NA | NA | #VALUE! |
| ENSMUSG00000075169.6 | 0 | NA | NA | NA | NA | NA | #VALUE! |
| ENSMUSG00000075170.5 | 0 | NA | NA | NA | NA | NA | #VALUE! |
| ENSMUSG00000075171.4 | 0 | NA | NA | NA | NA | NA | #VALUE! |
| ENSMUSG00000075172.4 | 0 | NA | NA | NA | NA | NA | #VALUE! |
| ENSMUSG00000075174.3 | 0 | NA | NA | NA | NA | NA | #VALUE! |
| ENSMUSG00000075175.3 | 0 | NA | NA | NA | NA | NA | #VALUE! |
| ENSMUSG00000075176.4 | 0 | NA | NA | NA | NA | NA | #VALUE! |
| ENSMUSG00000075179.4 | 0 | NA | NA | NA | NA | NA | #VALUE! |
| ENSMUSG00000075181.2 | 0 | NA | NA | NA | NA | NA | #VALUE! |
| ENSMUSG00000075184.4 | 0 | NA | NA | NA | NA | NA | #VALUE! |
| ENSMUSG00000075185.2 | 0 | NA | NA | NA | NA | NA | #VALUE! |
| ENSMUSG00000075186.5 | 0 | NA | NA | NA | NA | NA | #VALUE! |

| | | | | | | | |
|---|---|---|---|---|---|---|---|
| ENSMUSG00000075187.3 | 0 | NA | NA | NA | NA | NA | #VALUE! |
| ENSMUSG00000075188.5 | 0 | NA | NA | NA | NA | NA | #VALUE! |
| ENSMUSG00000075189.6 | 0 | NA | NA | NA | NA | NA | #VALUE! |
| ENSMUSG00000075190.3 | 0 | NA | NA | NA | NA | NA | #VALUE! |
| ENSMUSG00000075192.5 | 0 | NA | NA | NA | NA | NA | #VALUE! |
| ENSMUSG00000075193.3 | 0 | NA | NA | NA | NA | NA | #VALUE! |
| ENSMUSG00000075194.4 | 0 | NA | NA | NA | NA | NA | #VALUE! |
| ENSMUSG00000075195.4 | 0 | NA | NA | NA | NA | NA | #VALUE! |
| ENSMUSG00000075196.4 | 0 | NA | NA | NA | NA | NA | #VALUE! |
| ENSMUSG00000075197.4 | 0 | NA | NA | NA | NA | NA | #VALUE! |
| ENSMUSG00000075198.3 | 0 | NA | NA | NA | NA | NA | #VALUE! |
| ENSMUSG00000075199.4 | 0 | NA | NA | NA | NA | NA | #VALUE! |
| ENSMUSG00000075200.3 | 0 | NA | NA | NA | NA | NA | #VALUE! |
| ENSMUSG00000075201.3 | 0 | NA | NA | NA | NA | NA | #VALUE! |
| ENSMUSG00000075202.6 | 0 | NA | NA | NA | NA | NA | #VALUE! |
| ENSMUSG00000075203.2 | 0 | NA | NA | NA | NA | NA | #VALUE! |
| ENSMUSG00000075204.4 | 0 | NA | NA | NA | NA | NA | #VALUE! |
| ENSMUSG00000075205.6 | 0 | NA | NA | NA | NA | NA | #VALUE! |
| ENSMUSG00000075206.2 | 0 | NA | NA | NA | NA | NA | #VALUE! |
| ENSMUSG00000075208.7 | 0 | NA | NA | NA | NA | NA | #VALUE! |
| ENSMUSG00000075209.4 | 0 | NA | NA | NA | NA | NA | #VALUE! |
| ENSMUSG00000075210.5 | 0 | NA | NA | NA | NA | NA | #VALUE! |
| ENSMUSG00000075211.5 | 0 | NA | NA | NA | NA | NA | #VALUE! |
| ENSMUSG00000075212.6 | 0 | NA | NA | NA | NA | NA | #VALUE! |
| ENSMUSG00000075214.4 | 0 | NA | NA | NA | NA | NA | #VALUE! |
| ENSMUSG00000075215.3 | 0 | NA | NA | NA | NA | NA | #VALUE! |
| ENSMUSG00000075217.2 | 0 | NA | NA | NA | NA | NA | #VALUE! |
| ENSMUSG00000075218.4 | 0 | NA | NA | NA | NA | NA | #VALUE! |
| ENSMUSG00000075219.4 | 0 | NA | NA | NA | NA | NA | #VALUE! |
| ENSMUSG00000075220.2 | 0 | NA | NA | NA | NA | NA | #VALUE! |
| ENSMUSG00000075221.4 | 0 | NA | NA | NA | NA | NA | #VALUE! |
| ENSMUSG00000075222.6 | 0 | NA | NA | NA | NA | NA | #VALUE! |
| ENSMUSG00000075223.5 | 0 | NA | NA | NA | NA | NA | #VALUE! |
| ENSMUSG00000075224.11 | 0 | NA | NA | NA | NA | NA | #VALUE! |
| ENSMUSG00000075228.2 | 0 | NA | NA | NA | NA | NA | #VALUE! |
| ENSMUSG00000075245.5 | 0 | NA | NA | NA | NA | NA | #VALUE! |
| ENSMUSG00000075249.13 | 0 | NA | NA | NA | NA | NA | #VALUE! |
| ENSMUSG00000075256.10 | 0 | NA | NA | NA | NA | NA | #VALUE! |
| ENSMUSG00000075265.5 | 0 | NA | NA | NA | NA | NA | #VALUE! |
| ENSMUSG00000075267.10 | 0 | NA | NA | NA | NA | NA | #VALUE! |
| ENSMUSG00000075268.3 | 0 | NA | NA | NA | NA | NA | #VALUE! |
| ENSMUSG00000075269.6 | 0 | NA | NA | NA | NA | NA | #VALUE! |
| ENSMUSG00000075270.12 | 0 | NA | NA | NA | NA | NA | #VALUE! |
| ENSMUSG00000075271.5 | 0 | NA | NA | NA | NA | NA | #VALUE! |

| | | | | | | | |
|---|---|---|---|---|---|---|---|
| ENSMUSG00000075272.5 | 0 | NA | NA | NA | NA | NA | #VALUE! |
| ENSMUSG00000075276.6 | 0 | NA | NA | NA | NA | NA | #VALUE! |
| ENSMUSG00000075277.6 | 0 | NA | NA | NA | NA | NA | #VALUE! |
| ENSMUSG00000075286.5 | 0 | NA | NA | NA | NA | NA | #VALUE! |
| ENSMUSG00000075288.3 | 0 | NA | NA | NA | NA | NA | #VALUE! |
| ENSMUSG00000075289.6 | 2.793612 | -3.86483 | 3.909778 | -0.9885 | 0.322906 | NA | #VALUE! |
| ENSMUSG00000075296.6 | 0 | NA | NA | NA | NA | NA | #VALUE! |
| ENSMUSG00000075297.11 | 0 | NA | NA | NA | NA | NA | #VALUE! |
| ENSMUSG00000075302.11 | 0 | NA | NA | NA | NA | NA | #VALUE! |
| ENSMUSG00000075304.3 | 6.94605 | 0.231056 | 3.281296 | 0.070416 | 0.943862 | NA | #VALUE! |
| ENSMUSG00000075307.4 | 0 | NA | NA | NA | NA | NA | #VALUE! |
| ENSMUSG00000075312.5 | 0 | NA | NA | NA | NA | NA | #VALUE! |
| ENSMUSG00000075316.12 | 0 | NA | NA | NA | NA | NA | #VALUE! |
| ENSMUSG00000075318.14 | 0 | NA | NA | NA | NA | NA | #VALUE! |
| ENSMUSG00000075325.4 | 0 | NA | NA | NA | NA | NA | #VALUE! |
| ENSMUSG00000075330.5 | 0 | NA | NA | NA | NA | NA | #VALUE! |
| ENSMUSG00000075334.3 | 0 | NA | NA | NA | NA | NA | #VALUE! |
| ENSMUSG00000075335.4 | 0 | NA | NA | NA | NA | NA | #VALUE! |
| ENSMUSG00000075359.3 | 0 | NA | NA | NA | NA | NA | #VALUE! |
| ENSMUSG00000075370.13 | 0 | NA | NA | NA | NA | NA | #VALUE! |
| ENSMUSG00000075377.3 | 0 | NA | NA | NA | NA | NA | #VALUE! |
| ENSMUSG00000075378.3 | 0 | NA | NA | NA | NA | NA | #VALUE! |
| ENSMUSG00000075379.5 | 0 | NA | NA | NA | NA | NA | #VALUE! |
| ENSMUSG00000075380.3 | 0 | NA | NA | NA | NA | NA | #VALUE! |
| ENSMUSG00000075382.4 | 0 | NA | NA | NA | NA | NA | #VALUE! |
| ENSMUSG00000075383.5 | 0 | NA | NA | NA | NA | NA | #VALUE! |
| ENSMUSG00000075384.3 | 0 | NA | NA | NA | NA | NA | #VALUE! |
| ENSMUSG00000075387.3 | 0 | NA | NA | NA | NA | NA | #VALUE! |
| ENSMUSG00000075389.4 | 0 | NA | NA | NA | NA | NA | #VALUE! |
| ENSMUSG00000075390.6 | 0 | NA | NA | NA | NA | NA | #VALUE! |
| ENSMUSG00000075391.7 | 0.847396 | -0.59766 | 3.940051 | -0.15169 | 0.879432 | NA | #VALUE! |
| ENSMUSG00000075394.5 | 0 | NA | NA | NA | NA | NA | #VALUE! |
| ENSMUSG00000075395.12 | 0 | NA | NA | NA | NA | NA | #VALUE! |
| ENSMUSG00000075402.2 | 0 | NA | NA | NA | NA | NA | #VALUE! |
| ENSMUSG00000075405.4 | 0 | NA | NA | NA | NA | NA | #VALUE! |
| ENSMUSG00000075408.6 | 0 | NA | NA | NA | NA | NA | #VALUE! |
| ENSMUSG00000075410.14 | 0.420053 | -2.33493 | 4.007343 | -0.58266 | 0.56012 | NA | #VALUE! |
| ENSMUSG00000075416.6 | 0 | NA | NA | NA | NA | NA | #VALUE! |
| ENSMUSG00000075419.5 | 36.83676 | 0.20028 | 1.700088 | 0.117806 | NA | NA | #VALUE! |
| ENSMUSG00000075420.7 | 0.420053 | -2.33493 | 4.007343 | -0.58266 | 0.56012 | NA | #VALUE! |
| ENSMUSG00000075425.11 | 0 | NA | NA | NA | NA | NA | #VALUE! |
| ENSMUSG00000075427.16 | 0 | NA | NA | NA | NA | NA | #VALUE! |
| ENSMUSG00000075431.7 | 0 | NA | NA | NA | NA | NA | #VALUE! |
| ENSMUSG00000075437.5 | 0 | NA | NA | NA | NA | NA | #VALUE! |

| | | | | | | | |
|---|---|---|---|---|---|---|---|
| ENSMUSG00000075463.4 | 1.938301 | -4.49059 | 3.929692 | -1.14273 | 0.253149 | NA | #VALUE! |
| ENSMUSG00000075465.3 | 0.864345 | 1.839466 | 3.968067 | 0.463567 | 0.642958 | NA | #VALUE! |
| ENSMUSG00000075469.4 | 0 | NA | NA | NA | NA | NA | #VALUE! |
| ENSMUSG00000075478.4 | 0 | NA | NA | NA | NA | NA | #VALUE! |
| ENSMUSG00000075480.3 | 0 | NA | NA | NA | NA | NA | #VALUE! |
| ENSMUSG00000075502.3 | 2.908471 | 5.031766 | 3.922168 | 1.282904 | 0.199526 | NA | #VALUE! |
| ENSMUSG00000075510.4 | 0 | NA | NA | NA | NA | NA | #VALUE! |
| ENSMUSG00000075512.6 | 0 | NA | NA | NA | NA | NA | #VALUE! |
| ENSMUSG00000075514.5 | 0 | NA | NA | NA | NA | NA | #VALUE! |
| ENSMUSG00000075520.6 | 0 | NA | NA | NA | NA | NA | #VALUE! |
| ENSMUSG00000075524.4 | 0 | NA | NA | NA | NA | NA | #VALUE! |
| ENSMUSG00000075525.4 | 0 | NA | NA | NA | NA | NA | #VALUE! |
| ENSMUSG00000075528.14 | 0 | NA | NA | NA | NA | NA | #VALUE! |
| ENSMUSG00000075534.6 | 0 | NA | NA | NA | NA | NA | #VALUE! |
| ENSMUSG00000075538.3 | 0 | NA | NA | NA | NA | NA | #VALUE! |
| ENSMUSG00000075549.3 | 0 | NA | NA | NA | NA | NA | #VALUE! |
| ENSMUSG00000075551.5 | 0 | NA | NA | NA | NA | NA | #VALUE! |
| ENSMUSG00000075552.5 | 0 | NA | NA | NA | NA | NA | #VALUE! |
| ENSMUSG00000075553.3 | 0 | NA | NA | NA | NA | NA | #VALUE! |
| ENSMUSG00000075555.13 | 0 | NA | NA | NA | NA | NA | #VALUE! |
| ENSMUSG00000075558.4 | 0 | NA | NA | NA | NA | NA | #VALUE! |
| ENSMUSG00000075566.3 | 0 | NA | NA | NA | NA | NA | #VALUE! |
| ENSMUSG00000075567.5 | 0 | NA | NA | NA | NA | NA | #VALUE! |
| ENSMUSG00000075569.11 | 0 | NA | NA | NA | NA | NA | #VALUE! |
| ENSMUSG00000075570.3 | 0 | NA | NA | NA | NA | NA | #VALUE! |
| ENSMUSG00000075571.9 | 0 | NA | NA | NA | NA | NA | #VALUE! |
| ENSMUSG00000075572.2 | 0 | NA | NA | NA | NA | NA | #VALUE! |
| ENSMUSG00000075573.5 | 0 | NA | NA | NA | NA | NA | #VALUE! |
| ENSMUSG00000075574.3 | 0 | NA | NA | NA | NA | NA | #VALUE! |
| ENSMUSG00000075576.13 | 0 | NA | NA | NA | NA | NA | #VALUE! |
| ENSMUSG00000075581.6 | 0.600769 | 2.749741 | 3.980531 | 0.690798 | 0.489693 | NA | #VALUE! |
| ENSMUSG00000075585.4 | 4.91951 | 5.781601 | 3.915938 | 1.476428 | 0.139829 | NA | #VALUE! |
| ENSMUSG00000075586.5 | 0 | NA | NA | NA | NA | NA | #VALUE! |
| ENSMUSG00000075587.6 | 0 | NA | NA | NA | NA | NA | #VALUE! |
| ENSMUSG00000075588.7 | 0.673134 | 2.918386 | 3.972531 | 0.734641 | 0.462558 | NA | #VALUE! |
| ENSMUSG00000075589.4 | 0 | NA | NA | NA | NA | NA | #VALUE! |
| ENSMUSG00000075591.3 | 0 | NA | NA | NA | NA | NA | #VALUE! |
| ENSMUSG00000075592.11 | 2.545258 | -4.8732 | 3.92437 | -1.24178 | 0.214318 | NA | #VALUE! |
| ENSMUSG00000075593.11 | 0 | NA | NA | NA | NA | NA | #VALUE! |
| ENSMUSG00000075598.6 | 0 | NA | NA | NA | NA | NA | #VALUE! |
| ENSMUSG00000075599.5 | 0 | NA | NA | NA | NA | NA | #VALUE! |
| ENSMUSG00000075600.4 | 1.762102 | -4.34067 | 3.93219 | -1.10388 | 0.269645 | NA | #VALUE! |
| ENSMUSG00000075604.6 | 0 | NA | NA | NA | NA | NA | #VALUE! |
| ENSMUSG00000075605.4 | 0 | NA | NA | NA | NA | NA | #VALUE! |

| | | | | | | | |
|---|---|---|---|---|---|---|---|
| ENSMUSG00000075609.4 | 0 | NA | NA | NA | NA | NA | #VALUE! |
| ENSMUSG00000075610.10 | 0 | NA | NA | NA | NA | NA | #VALUE! |
| ENSMUSG00000075611.6 | 0 | NA | NA | NA | NA | NA | #VALUE! |
| ENSMUSG00000075702.11 | 0 | NA | NA | NA | NA | NA | #VALUE! |
| ENSMUSG00000075706.12 | 4.486506 | -0.23504 | 3.766678 | -0.0624 | 0.950244 | NA | #VALUE! |
| ENSMUSG00000075707.7 | 0 | NA | NA | NA | NA | NA | #VALUE! |
| ENSMUSG00000075719.4 | 0 | NA | NA | NA | NA | NA | #VALUE! |
| ENSMUSG00000075726.3 | 0 | NA | NA | NA | NA | NA | #VALUE! |
| ENSMUSG00000075770.3 | 0 | NA | NA | NA | NA | NA | #VALUE! |
| ENSMUSG00000075826.3 | 0 | NA | NA | NA | NA | NA | #VALUE! |
| ENSMUSG00000075829.4 | 0 | NA | NA | NA | NA | NA | #VALUE! |
| ENSMUSG00000075832.3 | 0 | NA | NA | NA | NA | NA | #VALUE! |
| ENSMUSG00000075837.3 | 0 | NA | NA | NA | NA | NA | #VALUE! |
| ENSMUSG00000075874.2 | 0 | NA | NA | NA | NA | NA | #VALUE! |
| ENSMUSG00000075903.3 | 0 | NA | NA | NA | NA | NA | #VALUE! |
| ENSMUSG00000075908.3 | 0 | NA | NA | NA | NA | NA | #VALUE! |
| ENSMUSG00000075934.3 | 0 | NA | NA | NA | NA | NA | #VALUE! |
| ENSMUSG00000075952.3 | 0 | NA | NA | NA | NA | NA | #VALUE! |
| ENSMUSG00000075958.2 | 0 | NA | NA | NA | NA | NA | #VALUE! |
| ENSMUSG00000075968.3 | 0 | NA | NA | NA | NA | NA | #VALUE! |
| ENSMUSG00000075983.3 | 0 | NA | NA | NA | NA | NA | #VALUE! |
| ENSMUSG00000076005.3 | 0 | NA | NA | NA | NA | NA | #VALUE! |
| ENSMUSG00000076006.4 | 0 | NA | NA | NA | NA | NA | #VALUE! |
| ENSMUSG00000076009.3 | 0 | NA | NA | NA | NA | NA | #VALUE! |
| ENSMUSG00000076010.3 | 0 | NA | NA | NA | NA | NA | #VALUE! |
| ENSMUSG00000076011.3 | 0 | NA | NA | NA | NA | NA | #VALUE! |
| ENSMUSG00000076012.3 | 0 | NA | NA | NA | NA | NA | #VALUE! |
| ENSMUSG00000076014.3 | 0 | NA | NA | NA | NA | NA | #VALUE! |
| ENSMUSG00000076028.3 | 0 | NA | NA | NA | NA | NA | #VALUE! |
| ENSMUSG00000076033.3 | 0 | NA | NA | NA | NA | NA | #VALUE! |
| ENSMUSG00000076041.3 | 0 | NA | NA | NA | NA | NA | #VALUE! |
| ENSMUSG00000076042.3 | 0 | NA | NA | NA | NA | NA | #VALUE! |
| ENSMUSG00000076049.6 | 0 | NA | NA | NA | NA | NA | #VALUE! |
| ENSMUSG00000076050.3 | 0 | NA | NA | NA | NA | NA | #VALUE! |
| ENSMUSG00000076051.3 | 0 | NA | NA | NA | NA | NA | #VALUE! |
| ENSMUSG00000076052.3 | 0 | NA | NA | NA | NA | NA | #VALUE! |
| ENSMUSG00000076060.3 | 0 | NA | NA | NA | NA | NA | #VALUE! |
| ENSMUSG00000076062.3 | 0 | NA | NA | NA | NA | NA | #VALUE! |
| ENSMUSG00000076063.3 | 0 | NA | NA | NA | NA | NA | #VALUE! |
| ENSMUSG00000076064.3 | 0 | NA | NA | NA | NA | NA | #VALUE! |
| ENSMUSG00000076065.3 | 0 | NA | NA | NA | NA | NA | #VALUE! |
| ENSMUSG00000076066.3 | 0 | NA | NA | NA | NA | NA | #VALUE! |
| ENSMUSG00000076067.3 | 0 | NA | NA | NA | NA | NA | #VALUE! |
| ENSMUSG00000076068.3 | 0 | NA | NA | NA | NA | NA | #VALUE! |

| | | | | | | | |
|---|---|---|---|---|---|---|---|
| ENSMUSG00000076118.3 | 0 | NA | NA | NA | NA | NA | #VALUE! |
| ENSMUSG00000076119.3 | 0 | NA | NA | NA | NA | NA | #VALUE! |
| ENSMUSG00000076120.3 | 0 | NA | NA | NA | NA | NA | #VALUE! |
| ENSMUSG00000076121.3 | 0 | NA | NA | NA | NA | NA | #VALUE! |
| ENSMUSG00000076122.3 | 0 | NA | NA | NA | NA | NA | #VALUE! |
| ENSMUSG00000076123.3 | 0 | NA | NA | NA | NA | NA | #VALUE! |
| ENSMUSG00000076126.3 | 0 | NA | NA | NA | NA | NA | #VALUE! |
| ENSMUSG00000076127.3 | 0 | NA | NA | NA | NA | NA | #VALUE! |
| ENSMUSG00000076128.3 | 0 | NA | NA | NA | NA | NA | #VALUE! |
| ENSMUSG00000076138.3 | 0 | NA | NA | NA | NA | NA | #VALUE! |
| ENSMUSG00000076140.3 | 0 | NA | NA | NA | NA | NA | #VALUE! |
| ENSMUSG00000076141.3 | 0 | NA | NA | NA | NA | NA | #VALUE! |
| ENSMUSG00000076142.3 | 0 | NA | NA | NA | NA | NA | #VALUE! |
| ENSMUSG00000076143.3 | 0 | NA | NA | NA | NA | NA | #VALUE! |
| ENSMUSG00000076144.3 | 0 | NA | NA | NA | NA | NA | #VALUE! |
| ENSMUSG00000076145.3 | 0 | NA | NA | NA | NA | NA | #VALUE! |
| ENSMUSG00000076146.3 | 0 | NA | NA | NA | NA | NA | #VALUE! |
| ENSMUSG00000076147.3 | 0 | NA | NA | NA | NA | NA | #VALUE! |
| ENSMUSG00000076163.3 | 0 | NA | NA | NA | NA | NA | #VALUE! |
| ENSMUSG00000076212.3 | 0 | NA | NA | NA | NA | NA | #VALUE! |
| ENSMUSG00000076214.3 | 0 | NA | NA | NA | NA | NA | #VALUE! |
| ENSMUSG00000076216.3 | 0 | NA | NA | NA | NA | NA | #VALUE! |
| ENSMUSG00000076217.3 | 0 | NA | NA | NA | NA | NA | #VALUE! |
| ENSMUSG00000076218.3 | 0 | NA | NA | NA | NA | NA | #VALUE! |
| ENSMUSG00000076219.3 | 0 | NA | NA | NA | NA | NA | #VALUE! |
| ENSMUSG00000076222.3 | 0 | NA | NA | NA | NA | NA | #VALUE! |
| ENSMUSG00000076236.3 | 0 | NA | NA | NA | NA | NA | #VALUE! |
| ENSMUSG00000076240.3 | 0 | NA | NA | NA | NA | NA | #VALUE! |
| ENSMUSG00000076241.4 | 0 | NA | NA | NA | NA | NA | #VALUE! |
| ENSMUSG00000076251.3 | 0 | NA | NA | NA | NA | NA | #VALUE! |
| ENSMUSG00000076252.3 | 0 | NA | NA | NA | NA | NA | #VALUE! |
| ENSMUSG00000076253.3 | 0 | NA | NA | NA | NA | NA | #VALUE! |
| ENSMUSG00000076255.4 | 0 | NA | NA | NA | NA | NA | #VALUE! |
| ENSMUSG00000076256.3 | 0 | NA | NA | NA | NA | NA | #VALUE! |
| ENSMUSG00000076269.3 | 0 | NA | NA | NA | NA | NA | #VALUE! |
| ENSMUSG00000076270.3 | 0 | NA | NA | NA | NA | NA | #VALUE! |
| ENSMUSG00000076271.3 | 0 | NA | NA | NA | NA | NA | #VALUE! |
| ENSMUSG00000076272.3 | 0 | NA | NA | NA | NA | NA | #VALUE! |
| ENSMUSG00000076275.3 | 0 | NA | NA | NA | NA | NA | #VALUE! |
| ENSMUSG00000076286.3 | 0 | NA | NA | NA | NA | NA | #VALUE! |
| ENSMUSG00000076288.3 | 0 | NA | NA | NA | NA | NA | #VALUE! |
| ENSMUSG00000076309.4 | 0 | NA | NA | NA | NA | NA | #VALUE! |
| ENSMUSG00000076312.3 | 0 | NA | NA | NA | NA | NA | #VALUE! |
| ENSMUSG00000076313.3 | 0 | NA | NA | NA | NA | NA | #VALUE! |

| | | | | | | | |
|---|---|---|---|---|---|---|---|
| ENSMUSG00000076315.4 | 0 | NA | NA | NA | NA | NA | #VALUE! |
| ENSMUSG00000076316.3 | 0 | NA | NA | NA | NA | NA | #VALUE! |
| ENSMUSG00000076318.3 | 0 | NA | NA | NA | NA | NA | #VALUE! |
| ENSMUSG00000076333.3 | 0 | NA | NA | NA | NA | NA | #VALUE! |
| ENSMUSG00000076335.3 | 0 | NA | NA | NA | NA | NA | #VALUE! |
| ENSMUSG00000076338.3 | 0 | NA | NA | NA | NA | NA | #VALUE! |
| ENSMUSG00000076357.3 | 0 | NA | NA | NA | NA | NA | #VALUE! |
| ENSMUSG00000076361.3 | 0 | NA | NA | NA | NA | NA | #VALUE! |
| ENSMUSG00000076372.4 | 0 | NA | NA | NA | NA | NA | #VALUE! |
| ENSMUSG00000076376.3 | 0 | NA | NA | NA | NA | NA | #VALUE! |
| ENSMUSG00000076377.4 | 0 | NA | NA | NA | NA | NA | #VALUE! |
| ENSMUSG00000076379.4 | 0 | NA | NA | NA | NA | NA | #VALUE! |
| ENSMUSG00000076387.3 | 0 | NA | NA | NA | NA | NA | #VALUE! |
| ENSMUSG00000076396.3 | 0 | NA | NA | NA | NA | NA | #VALUE! |
| ENSMUSG00000076398.3 | 0 | NA | NA | NA | NA | NA | #VALUE! |
| ENSMUSG00000076399.3 | 0 | NA | NA | NA | NA | NA | #VALUE! |
| ENSMUSG00000076424.4 | 0 | NA | NA | NA | NA | NA | #VALUE! |
| ENSMUSG00000076430.2 | 0 | NA | NA | NA | NA | NA | #VALUE! |
| ENSMUSG00000076431.5 | 9.679149 | 1.96155 | 2.605463 | 0.75286 | 0.451534 | NA | #VALUE! |
| ENSMUSG00000076433.5 | 0 | NA | NA | NA | NA | NA | #VALUE! |
| ENSMUSG00000076434.10 | 0 | NA | NA | NA | NA | NA | #VALUE! |
| ENSMUSG00000076436.2 | 0 | NA | NA | NA | NA | NA | #VALUE! |
| ENSMUSG00000076438.6 | 0 | NA | NA | NA | NA | NA | #VALUE! |
| ENSMUSG00000076439.13 | 0 | NA | NA | NA | NA | NA | #VALUE! |
| ENSMUSG00000076444.3 | 0 | NA | NA | NA | NA | NA | #VALUE! |
| ENSMUSG00000076448.3 | 0 | NA | NA | NA | NA | NA | #VALUE! |
| ENSMUSG00000076450.3 | 0 | NA | NA | NA | NA | NA | #VALUE! |
| ENSMUSG00000076451.3 | 0 | NA | NA | NA | NA | NA | #VALUE! |
| ENSMUSG00000076453.3 | 0 | NA | NA | NA | NA | NA | #VALUE! |
| ENSMUSG00000076454.3 | 0 | NA | NA | NA | NA | NA | #VALUE! |
| ENSMUSG00000076455.3 | 0 | NA | NA | NA | NA | NA | #VALUE! |
| ENSMUSG00000076456.3 | 0 | NA | NA | NA | NA | NA | #VALUE! |
| ENSMUSG00000076457.3 | 0 | NA | NA | NA | NA | NA | #VALUE! |
| ENSMUSG00000076458.3 | 0 | NA | NA | NA | NA | NA | #VALUE! |
| ENSMUSG00000076459.3 | 0 | NA | NA | NA | NA | NA | #VALUE! |
| ENSMUSG00000076460.3 | 0 | NA | NA | NA | NA | NA | #VALUE! |
| ENSMUSG00000076461.3 | 0 | NA | NA | NA | NA | NA | #VALUE! |
| ENSMUSG00000076462.3 | 0 | NA | NA | NA | NA | NA | #VALUE! |
| ENSMUSG00000076463.3 | 0 | NA | NA | NA | NA | NA | #VALUE! |
| ENSMUSG00000076465.3 | 0 | NA | NA | NA | NA | NA | #VALUE! |
| ENSMUSG00000076467.4 | 0 | NA | NA | NA | NA | NA | #VALUE! |
| ENSMUSG00000076469.4 | 0 | NA | NA | NA | NA | NA | #VALUE! |
| ENSMUSG00000076470.2 | 0 | NA | NA | NA | NA | NA | #VALUE! |
| ENSMUSG00000076471.4 | 0 | NA | NA | NA | NA | NA | #VALUE! |

| | | | | | | | |
|---|---|---|---|---|---|---|---|
| ENSMUSG00000076472.3 | 0 | NA | NA | NA | NA | NA | #VALUE! |
| ENSMUSG00000076473.4 | 0 | NA | NA | NA | NA | NA | #VALUE! |
| ENSMUSG00000076474.4 | 0 | NA | NA | NA | NA | NA | #VALUE! |
| ENSMUSG00000076475.3 | 0 | NA | NA | NA | NA | NA | #VALUE! |
| ENSMUSG00000076476.2 | 0 | NA | NA | NA | NA | NA | #VALUE! |
| ENSMUSG00000076477.4 | 0 | NA | NA | NA | NA | NA | #VALUE! |
| ENSMUSG00000076478.8 | 0 | NA | NA | NA | NA | NA | #VALUE! |
| ENSMUSG00000076479.4 | 0 | NA | NA | NA | NA | NA | #VALUE! |
| ENSMUSG00000076480.3 | 0 | NA | NA | NA | NA | NA | #VALUE! |
| ENSMUSG00000076481.8 | 0 | NA | NA | NA | NA | NA | #VALUE! |
| ENSMUSG00000076483.2 | 0 | NA | NA | NA | NA | NA | #VALUE! |
| ENSMUSG00000076484.2 | 0 | NA | NA | NA | NA | NA | #VALUE! |
| ENSMUSG00000076485.3 | 0 | NA | NA | NA | NA | NA | #VALUE! |
| ENSMUSG00000076486.2 | 0 | NA | NA | NA | NA | NA | #VALUE! |
| ENSMUSG00000076487.2 | 0 | NA | NA | NA | NA | NA | #VALUE! |
| ENSMUSG00000076488.2 | 0 | NA | NA | NA | NA | NA | #VALUE! |
| ENSMUSG00000076489.3 | 0 | NA | NA | NA | NA | NA | #VALUE! |
| ENSMUSG00000076490.7 | 2.062904 | -4.56996 | 3.92847 | -1.16329 | 0.244711 | NA | #VALUE! |
| ENSMUSG00000076492.2 | 0 | NA | NA | NA | NA | NA | #VALUE! |
| ENSMUSG00000076493.2 | 0 | NA | NA | NA | NA | NA | #VALUE! |
| ENSMUSG00000076494.2 | 0 | NA | NA | NA | NA | NA | #VALUE! |
| ENSMUSG00000076495.2 | 0 | NA | NA | NA | NA | NA | #VALUE! |
| ENSMUSG00000076496.2 | 0 | NA | NA | NA | NA | NA | #VALUE! |
| ENSMUSG00000076497.2 | 0 | NA | NA | NA | NA | NA | #VALUE! |
| ENSMUSG00000076498.3 | 1.890112 | 4.392518 | 3.930703 | 1.117489 | 0.263785 | NA | #VALUE! |
| ENSMUSG00000076499.4 | 0 | NA | NA | NA | NA | NA | #VALUE! |
| ENSMUSG00000076500.3 | 0 | NA | NA | NA | NA | NA | #VALUE! |
| ENSMUSG00000076501.3 | 0 | NA | NA | NA | NA | NA | #VALUE! |
| ENSMUSG00000076505.2 | 0 | NA | NA | NA | NA | NA | #VALUE! |
| ENSMUSG00000076508.4 | 0 | NA | NA | NA | NA | NA | #VALUE! |
| ENSMUSG00000076512.3 | 0 | NA | NA | NA | NA | NA | #VALUE! |
| ENSMUSG00000076514.4 | 0 | NA | NA | NA | NA | NA | #VALUE! |
| ENSMUSG00000076518.7 | 0 | NA | NA | NA | NA | NA | #VALUE! |
| ENSMUSG00000076522.3 | 0 | NA | NA | NA | NA | NA | #VALUE! |
| ENSMUSG00000076523.3 | 0 | NA | NA | NA | NA | NA | #VALUE! |
| ENSMUSG00000076525.4 | 0 | NA | NA | NA | NA | NA | #VALUE! |
| ENSMUSG00000076526.7 | 0 | NA | NA | NA | NA | NA | #VALUE! |
| ENSMUSG00000076530.5 | 0 | NA | NA | NA | NA | NA | #VALUE! |
| ENSMUSG00000076531.2 | 0 | NA | NA | NA | NA | NA | #VALUE! |
| ENSMUSG00000076532.4 | 0 | NA | NA | NA | NA | NA | #VALUE! |
| ENSMUSG00000076533.4 | 0 | NA | NA | NA | NA | NA | #VALUE! |
| ENSMUSG00000076534.7 | 0.373021 | 2.140748 | 4.018182 | 0.532765 | 0.594196 | NA | #VALUE! |
| ENSMUSG00000076535.2 | 0 | NA | NA | NA | NA | NA | #VALUE! |
| ENSMUSG00000076536.4 | 0 | NA | NA | NA | NA | NA | #VALUE! |

| | | | | | | | |
|---|---|---|---|---|---|---|---|
| ENSMUSG00000076538.2 | 0 | NA | NA | NA | NA | NA | #VALUE! |
| ENSMUSG00000076539.4 | 0 | NA | NA | NA | NA | NA | #VALUE! |
| ENSMUSG00000076540.4 | 0 | NA | NA | NA | NA | NA | #VALUE! |
| ENSMUSG00000076541.4 | 0 | NA | NA | NA | NA | NA | #VALUE! |
| ENSMUSG00000076543.3 | 0 | NA | NA | NA | NA | NA | #VALUE! |
| ENSMUSG00000076545.3 | 0 | NA | NA | NA | NA | NA | #VALUE! |
| ENSMUSG00000076547.3 | 0 | NA | NA | NA | NA | NA | #VALUE! |
| ENSMUSG00000076548.2 | 0 | NA | NA | NA | NA | NA | #VALUE! |
| ENSMUSG00000076549.3 | 0 | NA | NA | NA | NA | NA | #VALUE! |
| ENSMUSG00000076550.4 | 0 | NA | NA | NA | NA | NA | #VALUE! |
| ENSMUSG00000076552.4 | 0 | NA | NA | NA | NA | NA | #VALUE! |
| ENSMUSG00000076555.4 | 0 | NA | NA | NA | NA | NA | #VALUE! |
| ENSMUSG00000076556.4 | 0 | NA | NA | NA | NA | NA | #VALUE! |
| ENSMUSG00000076557.3 | 0 | NA | NA | NA | NA | NA | #VALUE! |
| ENSMUSG00000076562.2 | 0 | NA | NA | NA | NA | NA | #VALUE! |
| ENSMUSG00000076563.3 | 0 | NA | NA | NA | NA | NA | #VALUE! |
| ENSMUSG00000076564.3 | 0 | NA | NA | NA | NA | NA | #VALUE! |
| ENSMUSG00000076569.4 | 0 | NA | NA | NA | NA | NA | #VALUE! |
| ENSMUSG00000076570.7 | 0 | NA | NA | NA | NA | NA | #VALUE! |
| ENSMUSG00000076571.3 | 0 | NA | NA | NA | NA | NA | #VALUE! |
| ENSMUSG00000076572.7 | 0 | NA | NA | NA | NA | NA | #VALUE! |
| ENSMUSG00000076573.7 | 0 | NA | NA | NA | NA | NA | #VALUE! |
| ENSMUSG00000076575.7 | 0 | NA | NA | NA | NA | NA | #VALUE! |
| ENSMUSG00000076576.3 | 0 | NA | NA | NA | NA | NA | #VALUE! |
| ENSMUSG00000076577.3 | 0 | NA | NA | NA | NA | NA | #VALUE! |
| ENSMUSG00000076578.4 | 0 | NA | NA | NA | NA | NA | #VALUE! |
| ENSMUSG00000076580.4 | 0 | NA | NA | NA | NA | NA | #VALUE! |
| ENSMUSG00000076581.2 | 0 | NA | NA | NA | NA | NA | #VALUE! |
| ENSMUSG00000076583.2 | 0 | NA | NA | NA | NA | NA | #VALUE! |
| ENSMUSG00000076586.3 | 0 | NA | NA | NA | NA | NA | #VALUE! |
| ENSMUSG00000076587.4 | 0 | NA | NA | NA | NA | NA | #VALUE! |
| ENSMUSG00000076589.3 | 0 | NA | NA | NA | NA | NA | #VALUE! |
| ENSMUSG00000076591.4 | 0 | NA | NA | NA | NA | NA | #VALUE! |
| ENSMUSG00000076594.3 | 0 | NA | NA | NA | NA | NA | #VALUE! |
| ENSMUSG00000076596.4 | 0 | NA | NA | NA | NA | NA | #VALUE! |
| ENSMUSG00000076598.4 | 0 | NA | NA | NA | NA | NA | #VALUE! |
| ENSMUSG00000076604.2 | 0 | NA | NA | NA | NA | NA | #VALUE! |
| ENSMUSG00000076605.3 | 0 | NA | NA | NA | NA | NA | #VALUE! |
| ENSMUSG00000076606.4 | 0 | NA | NA | NA | NA | NA | #VALUE! |
| ENSMUSG00000076607.2 | 0 | NA | NA | NA | NA | NA | #VALUE! |
| ENSMUSG00000076608.3 | 0 | NA | NA | NA | NA | NA | #VALUE! |
| ENSMUSG00000076614.8 | 6.300788 | -6.18982 | 3.913848 | -1.58152 | 0.11376 | NA | #VALUE! |
| ENSMUSG00000076615.5 | 0 | NA | NA | NA | NA | NA | #VALUE! |
| ENSMUSG00000076618.2 | 0 | NA | NA | NA | NA | NA | #VALUE! |

| | | | | | | | |
|---|---|---|---|---|---|---|---|
| ENSMUSG00000076619.2 | 0 | NA | NA | NA | NA | NA | #VALUE! |
| ENSMUSG00000076620.3 | 0 | NA | NA | NA | NA | NA | #VALUE! |
| ENSMUSG00000076621.2 | 0 | NA | NA | NA | NA | NA | #VALUE! |
| ENSMUSG00000076630.2 | 0 | NA | NA | NA | NA | NA | #VALUE! |
| ENSMUSG00000076632.2 | 0 | NA | NA | NA | NA | NA | #VALUE! |
| ENSMUSG00000076633.8 | 0 | NA | NA | NA | NA | NA | #VALUE! |
| ENSMUSG00000076646.4 | 0 | NA | NA | NA | NA | NA | #VALUE! |
| ENSMUSG00000076652.5 | 0 | NA | NA | NA | NA | NA | #VALUE! |
| ENSMUSG00000076653.5 | 0 | NA | NA | NA | NA | NA | #VALUE! |
| ENSMUSG00000076655.3 | 0 | NA | NA | NA | NA | NA | #VALUE! |
| ENSMUSG00000076661.3 | 0 | NA | NA | NA | NA | NA | #VALUE! |
| ENSMUSG00000076665.5 | 0 | NA | NA | NA | NA | NA | #VALUE! |
| ENSMUSG00000076666.3 | 0 | NA | NA | NA | NA | NA | #VALUE! |
| ENSMUSG00000076668.5 | 0 | NA | NA | NA | NA | NA | #VALUE! |
| ENSMUSG00000076670.5 | 0 | NA | NA | NA | NA | NA | #VALUE! |
| ENSMUSG00000076671.6 | 0 | NA | NA | NA | NA | NA | #VALUE! |
| ENSMUSG00000076672.8 | 0 | NA | NA | NA | NA | NA | #VALUE! |
| ENSMUSG00000076674.3 | 0 | NA | NA | NA | NA | NA | #VALUE! |
| ENSMUSG00000076676.3 | 0 | NA | NA | NA | NA | NA | #VALUE! |
| ENSMUSG00000076677.2 | 0 | NA | NA | NA | NA | NA | #VALUE! |
| ENSMUSG00000076680.2 | 0 | NA | NA | NA | NA | NA | #VALUE! |
| ENSMUSG00000076688.3 | 0 | NA | NA | NA | NA | NA | #VALUE! |
| ENSMUSG00000076695.8 | 0 | NA | NA | NA | NA | NA | #VALUE! |
| ENSMUSG00000076709.3 | 0 | NA | NA | NA | NA | NA | #VALUE! |
| ENSMUSG00000076710.3 | 0 | NA | NA | NA | NA | NA | #VALUE! |
| ENSMUSG00000076731.4 | 0 | NA | NA | NA | NA | NA | #VALUE! |
| ENSMUSG00000076733.3 | 0 | NA | NA | NA | NA | NA | #VALUE! |
| ENSMUSG00000076744.3 | 0 | NA | NA | NA | NA | NA | #VALUE! |
| ENSMUSG00000076745.5 | 0 | NA | NA | NA | NA | NA | #VALUE! |
| ENSMUSG00000076746.4 | 0 | NA | NA | NA | NA | NA | #VALUE! |
| ENSMUSG00000076747.4 | 0 | NA | NA | NA | NA | NA | #VALUE! |
| ENSMUSG00000076749.3 | 0 | NA | NA | NA | NA | NA | #VALUE! |
| ENSMUSG00000076750.7 | 0 | NA | NA | NA | NA | NA | #VALUE! |
| ENSMUSG00000076752.3 | 2.86082 | 5.006323 | 3.922441 | 1.276328 | 0.201839 | NA | #VALUE! |
| ENSMUSG00000076754.3 | 0 | NA | NA | NA | NA | NA | #VALUE! |
| ENSMUSG00000076755.3 | 0 | NA | NA | NA | NA | NA | #VALUE! |
| ENSMUSG00000076757.10 | 0.204111 | -1.05581 | 4.073302 | -0.2592 | 0.795479 | NA | #VALUE! |
| ENSMUSG00000076758.7 | 0 | NA | NA | NA | NA | NA | #VALUE! |
| ENSMUSG00000076759.4 | 0 | NA | NA | NA | NA | NA | #VALUE! |
| ENSMUSG00000076760.3 | 0 | NA | NA | NA | NA | NA | #VALUE! |
| ENSMUSG00000076761.2 | 0 | NA | NA | NA | NA | NA | #VALUE! |
| ENSMUSG00000076762.2 | 0 | NA | NA | NA | NA | NA | #VALUE! |
| ENSMUSG00000076764.4 | 0 | NA | NA | NA | NA | NA | #VALUE! |
| ENSMUSG00000076770.4 | 0 | NA | NA | NA | NA | NA | #VALUE! |

| | | | | | | | |
|---|---|---|---|---|---|---|---|
| ENSMUSG00000076778.4 | 0 | NA | NA | NA | NA | NA | #VALUE! |
| ENSMUSG00000076795.7 | 0 | NA | NA | NA | NA | NA | #VALUE! |
| ENSMUSG00000076796.2 | 0 | NA | NA | NA | NA | NA | #VALUE! |
| ENSMUSG00000076800.2 | 0 | NA | NA | NA | NA | NA | #VALUE! |
| ENSMUSG00000076802.4 | 0 | NA | NA | NA | NA | NA | #VALUE! |
| ENSMUSG00000076821.4 | 0 | NA | NA | NA | NA | NA | #VALUE! |
| ENSMUSG00000076822.3 | 0 | NA | NA | NA | NA | NA | #VALUE! |
| ENSMUSG00000076823.4 | 0 | NA | NA | NA | NA | NA | #VALUE! |
| ENSMUSG00000076824.4 | 0 | NA | NA | NA | NA | NA | #VALUE! |
| ENSMUSG00000076826.6 | 0 | NA | NA | NA | NA | NA | #VALUE! |
| ENSMUSG00000076831.4 | 0 | NA | NA | NA | NA | NA | #VALUE! |
| ENSMUSG00000076839.4 | 0 | NA | NA | NA | NA | NA | #VALUE! |
| ENSMUSG00000076840.5 | 0 | NA | NA | NA | NA | NA | #VALUE! |
| ENSMUSG00000076846.4 | 0 | NA | NA | NA | NA | NA | #VALUE! |
| ENSMUSG00000076848.4 | 0 | NA | NA | NA | NA | NA | #VALUE! |
| ENSMUSG00000076855.6 | 0 | NA | NA | NA | NA | NA | #VALUE! |
| ENSMUSG00000076858.4 | 0 | NA | NA | NA | NA | NA | #VALUE! |
| ENSMUSG00000076861.11 | 0 | NA | NA | NA | NA | NA | #VALUE! |
| ENSMUSG00000076862.6 | 0 | NA | NA | NA | NA | NA | #VALUE! |
| ENSMUSG00000076863.9 | 0 | NA | NA | NA | NA | NA | #VALUE! |
| ENSMUSG00000076864.10 | 0 | NA | NA | NA | NA | NA | #VALUE! |
| ENSMUSG00000076865.7 | 0 | NA | NA | NA | NA | NA | #VALUE! |
| ENSMUSG00000076866.4 | 0 | NA | NA | NA | NA | NA | #VALUE! |
| ENSMUSG00000076867.3 | 0 | NA | NA | NA | NA | NA | #VALUE! |
| ENSMUSG00000076870.3 | 0 | NA | NA | NA | NA | NA | #VALUE! |
| ENSMUSG00000076871.2 | 0 | NA | NA | NA | NA | NA | #VALUE! |
| ENSMUSG00000076873.3 | 0 | NA | NA | NA | NA | NA | #VALUE! |
| ENSMUSG00000076874.3 | 0 | NA | NA | NA | NA | NA | #VALUE! |
| ENSMUSG00000076875.2 | 0 | NA | NA | NA | NA | NA | #VALUE! |
| ENSMUSG00000076876.4 | 0 | NA | NA | NA | NA | NA | #VALUE! |
| ENSMUSG00000076877.2 | 0 | NA | NA | NA | NA | NA | #VALUE! |
| ENSMUSG00000076878.2 | 0 | NA | NA | NA | NA | NA | #VALUE! |
| ENSMUSG00000076879.2 | 0 | NA | NA | NA | NA | NA | #VALUE! |
| ENSMUSG00000076880.3 | 0 | NA | NA | NA | NA | NA | #VALUE! |
| ENSMUSG00000076881.2 | 0 | NA | NA | NA | NA | NA | #VALUE! |
| ENSMUSG00000076882.4 | 0 | NA | NA | NA | NA | NA | #VALUE! |
| ENSMUSG00000076883.3 | 0 | NA | NA | NA | NA | NA | #VALUE! |
| ENSMUSG00000076884.2 | 0 | NA | NA | NA | NA | NA | #VALUE! |
| ENSMUSG00000076885.2 | 0 | NA | NA | NA | NA | NA | #VALUE! |
| ENSMUSG00000076886.2 | 0 | NA | NA | NA | NA | NA | #VALUE! |
| ENSMUSG00000076887.2 | 0 | NA | NA | NA | NA | NA | #VALUE! |
| ENSMUSG00000076888.2 | 0 | NA | NA | NA | NA | NA | #VALUE! |
| ENSMUSG00000076889.2 | 0 | NA | NA | NA | NA | NA | #VALUE! |
| ENSMUSG00000076890.2 | 0 | NA | NA | NA | NA | NA | #VALUE! |

| | | | | | | | |
|---|---|---|---|---|---|---|---|
| ENSMUSG00000076891.2 | 0 | NA | NA | NA | NA | NA | #VALUE! |
| ENSMUSG00000076892.2 | 0 | NA | NA | NA | NA | NA | #VALUE! |
| ENSMUSG00000076893.2 | 0 | NA | NA | NA | NA | NA | #VALUE! |
| ENSMUSG00000076894.2 | 0 | NA | NA | NA | NA | NA | #VALUE! |
| ENSMUSG00000076895.2 | 0 | NA | NA | NA | NA | NA | #VALUE! |
| ENSMUSG00000076896.2 | 0 | NA | NA | NA | NA | NA | #VALUE! |
| ENSMUSG00000076897.2 | 0 | NA | NA | NA | NA | NA | #VALUE! |
| ENSMUSG00000076898.2 | 0 | NA | NA | NA | NA | NA | #VALUE! |
| ENSMUSG00000076899.2 | 0 | NA | NA | NA | NA | NA | #VALUE! |
| ENSMUSG00000076900.2 | 0 | NA | NA | NA | NA | NA | #VALUE! |
| ENSMUSG00000076901.2 | 0 | NA | NA | NA | NA | NA | #VALUE! |
| ENSMUSG00000076902.2 | 0 | NA | NA | NA | NA | NA | #VALUE! |
| ENSMUSG00000076903.2 | 0 | NA | NA | NA | NA | NA | #VALUE! |
| ENSMUSG00000076904.3 | 0 | NA | NA | NA | NA | NA | #VALUE! |
| ENSMUSG00000076905.3 | 0 | NA | NA | NA | NA | NA | #VALUE! |
| ENSMUSG00000076906.2 | 0 | NA | NA | NA | NA | NA | #VALUE! |
| ENSMUSG00000076907.2 | 0 | NA | NA | NA | NA | NA | #VALUE! |
| ENSMUSG00000076908.2 | 0 | NA | NA | NA | NA | NA | #VALUE! |
| ENSMUSG00000076909.3 | 0 | NA | NA | NA | NA | NA | #VALUE! |
| ENSMUSG00000076910.2 | 0 | NA | NA | NA | NA | NA | #VALUE! |
| ENSMUSG00000076911.2 | 0 | NA | NA | NA | NA | NA | #VALUE! |
| ENSMUSG00000076912.2 | 0 | NA | NA | NA | NA | NA | #VALUE! |
| ENSMUSG00000076913.2 | 0 | NA | NA | NA | NA | NA | #VALUE! |
| ENSMUSG00000076914.2 | 0 | NA | NA | NA | NA | NA | #VALUE! |
| ENSMUSG00000076915.4 | 0 | NA | NA | NA | NA | NA | #VALUE! |
| ENSMUSG00000076916.2 | 0 | NA | NA | NA | NA | NA | #VALUE! |
| ENSMUSG00000076917.2 | 0 | NA | NA | NA | NA | NA | #VALUE! |
| ENSMUSG00000076918.2 | 0 | NA | NA | NA | NA | NA | #VALUE! |
| ENSMUSG00000076919.2 | 0 | NA | NA | NA | NA | NA | #VALUE! |
| ENSMUSG00000076920.2 | 0 | NA | NA | NA | NA | NA | #VALUE! |
| ENSMUSG00000076921.2 | 0 | NA | NA | NA | NA | NA | #VALUE! |
| ENSMUSG00000076922.2 | 0 | NA | NA | NA | NA | NA | #VALUE! |
| ENSMUSG00000076923.2 | 0 | NA | NA | NA | NA | NA | #VALUE! |
| ENSMUSG00000076924.2 | 0 | NA | NA | NA | NA | NA | #VALUE! |
| ENSMUSG00000076925.3 | 0 | NA | NA | NA | NA | NA | #VALUE! |
| ENSMUSG00000076926.2 | 0 | NA | NA | NA | NA | NA | #VALUE! |
| ENSMUSG00000076927.4 | 0 | NA | NA | NA | NA | NA | #VALUE! |
| ENSMUSG00000076928.6 | 0.336567 | 1.944925 | 4.029936 | 0.482619 | 0.629366 | NA | #VALUE! |
| ENSMUSG00000076934.3 | 3.702238 | 5.378281 | 3.918889 | 1.372399 | 0.169939 | NA | #VALUE! |
| ENSMUSG00000076937.4 | 9.379996 | -6.76345 | 3.208389 | -2.10805 | 0.035027 | NA | #VALUE! |
| ENSMUSG00000076939.3 | 0 | NA | NA | NA | NA | NA | #VALUE! |
| ENSMUSG00000076940.4 | 0 | NA | NA | NA | NA | NA | #VALUE! |
| ENSMUSG00000076946.4 | 0 | NA | NA | NA | NA | NA | #VALUE! |
| ENSMUSG00000076948.4 | 0 | NA | NA | NA | NA | NA | #VALUE! |

| | | | | | | | |
|---|---|---|---|---|---|---|---|
| ENSMUSG00000076956.4 | 0 | NA | NA | NA | NA | NA | #VALUE! |
| ENSMUSG00000076959.3 | 0 | NA | NA | NA | NA | NA | #VALUE! |
| ENSMUSG00000076972.4 | 0 | NA | NA | NA | NA | NA | #VALUE! |
| ENSMUSG00000076976.4 | 0 | NA | NA | NA | NA | NA | #VALUE! |
| ENSMUSG00000076983.4 | 0 | NA | NA | NA | NA | NA | #VALUE! |
| ENSMUSG00000076994.4 | 0 | NA | NA | NA | NA | NA | #VALUE! |
| ENSMUSG00000077001.4 | 0 | NA | NA | NA | NA | NA | #VALUE! |
| ENSMUSG00000077021.4 | 0 | NA | NA | NA | NA | NA | #VALUE! |
| ENSMUSG00000077025.4 | 0 | NA | NA | NA | NA | NA | #VALUE! |
| ENSMUSG00000077038.3 | 0 | NA | NA | NA | NA | NA | #VALUE! |
| ENSMUSG00000077042.4 | 0 | NA | NA | NA | NA | NA | #VALUE! |
| ENSMUSG00000077049.4 | 0 | NA | NA | NA | NA | NA | #VALUE! |
| ENSMUSG00000077086.3 | 0 | NA | NA | NA | NA | NA | #VALUE! |
| ENSMUSG00000077092.3 | 0 | NA | NA | NA | NA | NA | #VALUE! |
| ENSMUSG00000077107.3 | 0 | NA | NA | NA | NA | NA | #VALUE! |
| ENSMUSG00000077111.4 | 0 | NA | NA | NA | NA | NA | #VALUE! |
| ENSMUSG00000077113.4 | 0 | NA | NA | NA | NA | NA | #VALUE! |
| ENSMUSG00000077119.3 | 0 | NA | NA | NA | NA | NA | #VALUE! |
| ENSMUSG00000077127.3 | 0 | NA | NA | NA | NA | NA | #VALUE! |
| ENSMUSG00000077142.3 | 0 | NA | NA | NA | NA | NA | #VALUE! |
| ENSMUSG00000077143.3 | 0 | NA | NA | NA | NA | NA | #VALUE! |
| ENSMUSG00000077144.3 | 0 | NA | NA | NA | NA | NA | #VALUE! |
| ENSMUSG00000077146.3 | 0 | NA | NA | NA | NA | NA | #VALUE! |
| ENSMUSG00000077148.3 | 0 | NA | NA | NA | NA | NA | #VALUE! |
| ENSMUSG00000077149.3 | 0 | NA | NA | NA | NA | NA | #VALUE! |
| ENSMUSG00000077150.3 | 0 | NA | NA | NA | NA | NA | #VALUE! |
| ENSMUSG00000077152.4 | 0 | NA | NA | NA | NA | NA | #VALUE! |
| ENSMUSG00000077153.3 | 0 | NA | NA | NA | NA | NA | #VALUE! |
| ENSMUSG00000077155.3 | 0 | NA | NA | NA | NA | NA | #VALUE! |
| ENSMUSG00000077158.3 | 0 | NA | NA | NA | NA | NA | #VALUE! |
| ENSMUSG00000077160.3 | 0 | NA | NA | NA | NA | NA | #VALUE! |
| ENSMUSG00000077161.4 | 0 | NA | NA | NA | NA | NA | #VALUE! |
| ENSMUSG00000077163.4 | 0 | NA | NA | NA | NA | NA | #VALUE! |
| ENSMUSG00000077166.3 | 0 | NA | NA | NA | NA | NA | #VALUE! |
| ENSMUSG00000077167.3 | 0 | NA | NA | NA | NA | NA | #VALUE! |
| ENSMUSG00000077169.3 | 0 | NA | NA | NA | NA | NA | #VALUE! |
| ENSMUSG00000077171.3 | 0 | NA | NA | NA | NA | NA | #VALUE! |
| ENSMUSG00000077172.3 | 0 | NA | NA | NA | NA | NA | #VALUE! |
| ENSMUSG00000077174.3 | 0 | NA | NA | NA | NA | NA | #VALUE! |
| ENSMUSG00000077175.3 | 0 | NA | NA | NA | NA | NA | #VALUE! |
| ENSMUSG00000077176.3 | 0 | NA | NA | NA | NA | NA | #VALUE! |
| ENSMUSG00000077177.3 | 0 | NA | NA | NA | NA | NA | #VALUE! |
| ENSMUSG00000077178.3 | 0 | NA | NA | NA | NA | NA | #VALUE! |
| ENSMUSG00000077180.3 | 0 | NA | NA | NA | NA | NA | #VALUE! |

| | | | | | | | |
|---|---|---|---|---|---|---|---|
| ENSMUSG00000077181.3 | 0 | NA | NA | NA | NA | NA | #VALUE! |
| ENSMUSG00000077184.4 | 0 | NA | NA | NA | NA | NA | #VALUE! |
| ENSMUSG00000077185.3 | 0 | NA | NA | NA | NA | NA | #VALUE! |
| ENSMUSG00000077191.3 | 0 | NA | NA | NA | NA | NA | #VALUE! |
| ENSMUSG00000077192.5 | 0 | NA | NA | NA | NA | NA | #VALUE! |
| ENSMUSG00000077194.3 | 0 | NA | NA | NA | NA | NA | #VALUE! |
| ENSMUSG00000077196.3 | 0 | NA | NA | NA | NA | NA | #VALUE! |
| ENSMUSG00000077197.3 | 0 | NA | NA | NA | NA | NA | #VALUE! |
| ENSMUSG00000077199.5 | 0 | NA | NA | NA | NA | NA | #VALUE! |
| ENSMUSG00000077200.3 | 0 | NA | NA | NA | NA | NA | #VALUE! |
| ENSMUSG00000077201.3 | 0 | NA | NA | NA | NA | NA | #VALUE! |
| ENSMUSG00000077202.3 | 0 | NA | NA | NA | NA | NA | #VALUE! |
| ENSMUSG00000077204.3 | 0 | NA | NA | NA | NA | NA | #VALUE! |
| ENSMUSG00000077205.3 | 0 | NA | NA | NA | NA | NA | #VALUE! |
| ENSMUSG00000077207.3 | 0 | NA | NA | NA | NA | NA | #VALUE! |
| ENSMUSG00000077208.3 | 0 | NA | NA | NA | NA | NA | #VALUE! |
| ENSMUSG00000077209.3 | 0 | NA | NA | NA | NA | NA | #VALUE! |
| ENSMUSG00000077212.3 | 0 | NA | NA | NA | NA | NA | #VALUE! |
| ENSMUSG00000077214.3 | 0 | NA | NA | NA | NA | NA | #VALUE! |
| ENSMUSG00000077215.3 | 0 | NA | NA | NA | NA | NA | #VALUE! |
| ENSMUSG00000077217.3 | 0 | NA | NA | NA | NA | NA | #VALUE! |
| ENSMUSG00000077220.3 | 0 | NA | NA | NA | NA | NA | #VALUE! |
| ENSMUSG00000077221.3 | 0 | NA | NA | NA | NA | NA | #VALUE! |
| ENSMUSG00000077222.3 | 0 | NA | NA | NA | NA | NA | #VALUE! |
| ENSMUSG00000077223.3 | 0 | NA | NA | NA | NA | NA | #VALUE! |
| ENSMUSG00000077224.3 | 0 | NA | NA | NA | NA | NA | #VALUE! |
| ENSMUSG00000077227.3 | 0 | NA | NA | NA | NA | NA | #VALUE! |
| ENSMUSG00000077228.3 | 0 | NA | NA | NA | NA | NA | #VALUE! |
| ENSMUSG00000077230.3 | 0 | NA | NA | NA | NA | NA | #VALUE! |
| ENSMUSG00000077231.3 | 0 | NA | NA | NA | NA | NA | #VALUE! |
| ENSMUSG00000077235.3 | 0 | NA | NA | NA | NA | NA | #VALUE! |
| ENSMUSG00000077237.3 | 0 | NA | NA | NA | NA | NA | #VALUE! |
| ENSMUSG00000077239.3 | 0 | NA | NA | NA | NA | NA | #VALUE! |
| ENSMUSG00000077240.3 | 0 | NA | NA | NA | NA | NA | #VALUE! |
| ENSMUSG00000077241.4 | 0 | NA | NA | NA | NA | NA | #VALUE! |
| ENSMUSG00000077243.3 | 0 | NA | NA | NA | NA | NA | #VALUE! |
| ENSMUSG00000077244.3 | 0 | NA | NA | NA | NA | NA | #VALUE! |
| ENSMUSG00000077246.3 | 0 | NA | NA | NA | NA | NA | #VALUE! |
| ENSMUSG00000077248.2 | 0 | NA | NA | NA | NA | NA | #VALUE! |
| ENSMUSG00000077250.2 | 0 | NA | NA | NA | NA | NA | #VALUE! |
| ENSMUSG00000077251.3 | 0 | NA | NA | NA | NA | NA | #VALUE! |
| ENSMUSG00000077252.3 | 0 | NA | NA | NA | NA | NA | #VALUE! |
| ENSMUSG00000077254.3 | 0 | NA | NA | NA | NA | NA | #VALUE! |
| ENSMUSG00000077255.3 | 0 | NA | NA | NA | NA | NA | #VALUE! |

| | | | | | | | |
|---|---|---|---|---|---|---|---|
| ENSMUSG00000077256.3 | 0 | NA | NA | NA | NA | NA | #VALUE! |
| ENSMUSG00000077257.3 | 0 | NA | NA | NA | NA | NA | #VALUE! |
| ENSMUSG00000077258.4 | 0 | NA | NA | NA | NA | NA | #VALUE! |
| ENSMUSG00000077259.3 | 0 | NA | NA | NA | NA | NA | #VALUE! |
| ENSMUSG00000077260.3 | 0 | NA | NA | NA | NA | NA | #VALUE! |
| ENSMUSG00000077263.4 | 0 | NA | NA | NA | NA | NA | #VALUE! |
| ENSMUSG00000077264.3 | 0 | NA | NA | NA | NA | NA | #VALUE! |
| ENSMUSG00000077265.3 | 0 | NA | NA | NA | NA | NA | #VALUE! |
| ENSMUSG00000077268.3 | 0 | NA | NA | NA | NA | NA | #VALUE! |
| ENSMUSG00000077269.3 | 0 | NA | NA | NA | NA | NA | #VALUE! |
| ENSMUSG00000077271.2 | 0 | NA | NA | NA | NA | NA | #VALUE! |
| ENSMUSG00000077273.3 | 0 | NA | NA | NA | NA | NA | #VALUE! |
| ENSMUSG00000077274.3 | 0 | NA | NA | NA | NA | NA | #VALUE! |
| ENSMUSG00000077275.3 | 0 | NA | NA | NA | NA | NA | #VALUE! |
| ENSMUSG00000077276.3 | 0 | NA | NA | NA | NA | NA | #VALUE! |
| ENSMUSG00000077277.3 | 0 | NA | NA | NA | NA | NA | #VALUE! |
| ENSMUSG00000077278.3 | 0 | NA | NA | NA | NA | NA | #VALUE! |
| ENSMUSG00000077279.3 | 0 | NA | NA | NA | NA | NA | #VALUE! |
| ENSMUSG00000077280.3 | 0 | NA | NA | NA | NA | NA | #VALUE! |
| ENSMUSG00000077282.3 | 0 | NA | NA | NA | NA | NA | #VALUE! |
| ENSMUSG00000077287.3 | 0 | NA | NA | NA | NA | NA | #VALUE! |
| ENSMUSG00000077288.3 | 0 | NA | NA | NA | NA | NA | #VALUE! |
| ENSMUSG00000077289.3 | 0 | NA | NA | NA | NA | NA | #VALUE! |
| ENSMUSG00000077290.3 | 0 | NA | NA | NA | NA | NA | #VALUE! |
| ENSMUSG00000077291.3 | 0 | NA | NA | NA | NA | NA | #VALUE! |
| ENSMUSG00000077292.3 | 0 | NA | NA | NA | NA | NA | #VALUE! |
| ENSMUSG00000077293.2 | 0 | NA | NA | NA | NA | NA | #VALUE! |
| ENSMUSG00000077294.3 | 0 | NA | NA | NA | NA | NA | #VALUE! |
| ENSMUSG00000077295.3 | 0 | NA | NA | NA | NA | NA | #VALUE! |
| ENSMUSG00000077298.3 | 0 | NA | NA | NA | NA | NA | #VALUE! |
| ENSMUSG00000077299.3 | 0 | NA | NA | NA | NA | NA | #VALUE! |
| ENSMUSG00000077302.2 | 0 | NA | NA | NA | NA | NA | #VALUE! |
| ENSMUSG00000077303.3 | 0 | NA | NA | NA | NA | NA | #VALUE! |
| ENSMUSG00000077304.3 | 0 | NA | NA | NA | NA | NA | #VALUE! |
| ENSMUSG00000077305.3 | 0 | NA | NA | NA | NA | NA | #VALUE! |
| ENSMUSG00000077306.3 | 0 | NA | NA | NA | NA | NA | #VALUE! |
| ENSMUSG00000077312.3 | 0 | NA | NA | NA | NA | NA | #VALUE! |
| ENSMUSG00000077314.3 | 0 | NA | NA | NA | NA | NA | #VALUE! |
| ENSMUSG00000077315.4 | 0 | NA | NA | NA | NA | NA | #VALUE! |
| ENSMUSG00000077317.3 | 0 | NA | NA | NA | NA | NA | #VALUE! |
| ENSMUSG00000077318.3 | 0 | NA | NA | NA | NA | NA | #VALUE! |
| ENSMUSG00000077320.3 | 0 | NA | NA | NA | NA | NA | #VALUE! |
| ENSMUSG00000077323.3 | 0 | NA | NA | NA | NA | NA | #VALUE! |
| ENSMUSG00000077325.3 | 0 | NA | NA | NA | NA | NA | #VALUE! |

| | | | | | | | |
|---|---|---|---|---|---|---|---|
| ENSMUSG00000077327.3 | 0 | NA | NA | NA | NA | NA | #VALUE! |
| ENSMUSG00000077328.3 | 0 | NA | NA | NA | NA | NA | #VALUE! |
| ENSMUSG00000077332.3 | 0 | NA | NA | NA | NA | NA | #VALUE! |
| ENSMUSG00000077333.3 | 0 | NA | NA | NA | NA | NA | #VALUE! |
| ENSMUSG00000077334.4 | 0 | NA | NA | NA | NA | NA | #VALUE! |
| ENSMUSG00000077336.3 | 0 | NA | NA | NA | NA | NA | #VALUE! |
| ENSMUSG00000077337.4 | 0 | NA | NA | NA | NA | NA | #VALUE! |
| ENSMUSG00000077338.3 | 0 | NA | NA | NA | NA | NA | #VALUE! |
| ENSMUSG00000077339.3 | 0 | NA | NA | NA | NA | NA | #VALUE! |
| ENSMUSG00000077342.3 | 0 | NA | NA | NA | NA | NA | #VALUE! |
| ENSMUSG00000077344.3 | 0 | NA | NA | NA | NA | NA | #VALUE! |
| ENSMUSG00000077345.3 | 0 | NA | NA | NA | NA | NA | #VALUE! |
| ENSMUSG00000077347.4 | 0 | NA | NA | NA | NA | NA | #VALUE! |
| ENSMUSG00000077348.3 | 0 | NA | NA | NA | NA | NA | #VALUE! |
| ENSMUSG00000077349.4 | 0 | NA | NA | NA | NA | NA | #VALUE! |
| ENSMUSG00000077354.3 | 0 | NA | NA | NA | NA | NA | #VALUE! |
| ENSMUSG00000077357.3 | 0 | NA | NA | NA | NA | NA | #VALUE! |
| ENSMUSG00000077358.3 | 0 | NA | NA | NA | NA | NA | #VALUE! |
| ENSMUSG00000077359.3 | 0 | NA | NA | NA | NA | NA | #VALUE! |
| ENSMUSG00000077360.3 | 0 | NA | NA | NA | NA | NA | #VALUE! |
| ENSMUSG00000077361.3 | 0 | NA | NA | NA | NA | NA | #VALUE! |
| ENSMUSG00000077366.3 | 0 | NA | NA | NA | NA | NA | #VALUE! |
| ENSMUSG00000077367.3 | 0 | NA | NA | NA | NA | NA | #VALUE! |
| ENSMUSG00000077368.3 | 0 | NA | NA | NA | NA | NA | #VALUE! |
| ENSMUSG00000077369.4 | 0 | NA | NA | NA | NA | NA | #VALUE! |
| ENSMUSG00000077376.3 | 0 | NA | NA | NA | NA | NA | #VALUE! |
| ENSMUSG00000077377.3 | 0 | NA | NA | NA | NA | NA | #VALUE! |
| ENSMUSG00000077378.3 | 0 | NA | NA | NA | NA | NA | #VALUE! |
| ENSMUSG00000077379.3 | 0 | NA | NA | NA | NA | NA | #VALUE! |
| ENSMUSG00000077380.3 | 0 | NA | NA | NA | NA | NA | #VALUE! |
| ENSMUSG00000077381.3 | 0 | NA | NA | NA | NA | NA | #VALUE! |
| ENSMUSG00000077383.4 | 0 | NA | NA | NA | NA | NA | #VALUE! |
| ENSMUSG00000077384.3 | 0 | NA | NA | NA | NA | NA | #VALUE! |
| ENSMUSG00000077385.3 | 0 | NA | NA | NA | NA | NA | #VALUE! |
| ENSMUSG00000077387.3 | 0 | NA | NA | NA | NA | NA | #VALUE! |
| ENSMUSG00000077391.3 | 0.673134 | 2.918386 | 3.972531 | 0.734641 | 0.462558 | NA | #VALUE! |
| ENSMUSG00000077392.2 | 0 | NA | NA | NA | NA | NA | #VALUE! |
| ENSMUSG00000077393.3 | 0 | NA | NA | NA | NA | NA | #VALUE! |
| ENSMUSG00000077394.3 | 0 | NA | NA | NA | NA | NA | #VALUE! |
| ENSMUSG00000077396.3 | 0 | NA | NA | NA | NA | NA | #VALUE! |
| ENSMUSG00000077397.2 | 0 | NA | NA | NA | NA | NA | #VALUE! |
| ENSMUSG00000077398.3 | 0 | NA | NA | NA | NA | NA | #VALUE! |
| ENSMUSG00000077399.3 | 0 | NA | NA | NA | NA | NA | #VALUE! |
| ENSMUSG00000077400.3 | 0 | NA | NA | NA | NA | NA | #VALUE! |

| | | | | | | | |
|---|---|---|---|---|---|---|---|
| ENSMUSG00000077402.3 | 0 | NA | NA | NA | NA | NA | #VALUE! |
| ENSMUSG00000077403.3 | 0 | NA | NA | NA | NA | NA | #VALUE! |
| ENSMUSG00000077404.3 | 0 | NA | NA | NA | NA | NA | #VALUE! |
| ENSMUSG00000077406.3 | 0 | NA | NA | NA | NA | NA | #VALUE! |
| ENSMUSG00000077407.3 | 0 | NA | NA | NA | NA | NA | #VALUE! |
| ENSMUSG00000077408.3 | 0 | NA | NA | NA | NA | NA | #VALUE! |
| ENSMUSG00000077409.3 | 0 | NA | NA | NA | NA | NA | #VALUE! |
| ENSMUSG00000077410.3 | 0 | NA | NA | NA | NA | NA | #VALUE! |
| ENSMUSG00000077411.3 | 0 | NA | NA | NA | NA | NA | #VALUE! |
| ENSMUSG00000077413.3 | 0 | NA | NA | NA | NA | NA | #VALUE! |
| ENSMUSG00000077415.3 | 0 | NA | NA | NA | NA | NA | #VALUE! |
| ENSMUSG00000077416.3 | 0 | NA | NA | NA | NA | NA | #VALUE! |
| ENSMUSG00000077417.2 | 0 | NA | NA | NA | NA | NA | #VALUE! |
| ENSMUSG00000077418.2 | 0 | NA | NA | NA | NA | NA | #VALUE! |
| ENSMUSG00000077419.4 | 0 | NA | NA | NA | NA | NA | #VALUE! |
| ENSMUSG00000077422.3 | 0 | NA | NA | NA | NA | NA | #VALUE! |
| ENSMUSG00000077424.3 | 0 | NA | NA | NA | NA | NA | #VALUE! |
| ENSMUSG00000077425.3 | 0 | NA | NA | NA | NA | NA | #VALUE! |
| ENSMUSG00000077426.3 | 0 | NA | NA | NA | NA | NA | #VALUE! |
| ENSMUSG00000077427.3 | 0 | NA | NA | NA | NA | NA | #VALUE! |
| ENSMUSG00000077430.3 | 0 | NA | NA | NA | NA | NA | #VALUE! |
| ENSMUSG00000077431.3 | 0 | NA | NA | NA | NA | NA | #VALUE! |
| ENSMUSG00000077433.3 | 0 | NA | NA | NA | NA | NA | #VALUE! |
| ENSMUSG00000077435.3 | 0 | NA | NA | NA | NA | NA | #VALUE! |
| ENSMUSG00000077438.3 | 0 | NA | NA | NA | NA | NA | #VALUE! |
| ENSMUSG00000077439.3 | 0 | NA | NA | NA | NA | NA | #VALUE! |
| ENSMUSG00000077440.3 | 0 | NA | NA | NA | NA | NA | #VALUE! |
| ENSMUSG00000077441.3 | 0 | NA | NA | NA | NA | NA | #VALUE! |
| ENSMUSG00000077447.3 | 0 | NA | NA | NA | NA | NA | #VALUE! |
| ENSMUSG00000077448.3 | 0 | NA | NA | NA | NA | NA | #VALUE! |
| ENSMUSG00000077449.4 | 0 | NA | NA | NA | NA | NA | #VALUE! |
| ENSMUSG00000077451.3 | 0 | NA | NA | NA | NA | NA | #VALUE! |
| ENSMUSG00000077453.3 | 0 | NA | NA | NA | NA | NA | #VALUE! |
| ENSMUSG00000077454.3 | 0 | NA | NA | NA | NA | NA | #VALUE! |
| ENSMUSG00000077455.3 | 0 | NA | NA | NA | NA | NA | #VALUE! |
| ENSMUSG00000077456.3 | 0 | NA | NA | NA | NA | NA | #VALUE! |
| ENSMUSG00000077457.3 | 0 | NA | NA | NA | NA | NA | #VALUE! |
| ENSMUSG00000077459.3 | 0 | NA | NA | NA | NA | NA | #VALUE! |
| ENSMUSG00000077460.3 | 0 | NA | NA | NA | NA | NA | #VALUE! |
| ENSMUSG00000077461.3 | 0 | NA | NA | NA | NA | NA | #VALUE! |
| ENSMUSG00000077462.3 | 0 | NA | NA | NA | NA | NA | #VALUE! |
| ENSMUSG00000077463.3 | 0 | NA | NA | NA | NA | NA | #VALUE! |
| ENSMUSG00000077465.3 | 0 | NA | NA | NA | NA | NA | #VALUE! |
| ENSMUSG00000077468.3 | 0 | NA | NA | NA | NA | NA | #VALUE! |

| | | | | | | | |
|---|---|---|---|---|---|---|---|
| ENSMUSG00000077469.3 | 0 | NA | NA | NA | NA | NA | #VALUE! |
| ENSMUSG00000077470.3 | 0 | NA | NA | NA | NA | NA | #VALUE! |
| ENSMUSG00000077471.4 | 0 | NA | NA | NA | NA | NA | #VALUE! |
| ENSMUSG00000077472.3 | 0 | NA | NA | NA | NA | NA | #VALUE! |
| ENSMUSG00000077473.3 | 0 | NA | NA | NA | NA | NA | #VALUE! |
| ENSMUSG00000077474.3 | 0 | NA | NA | NA | NA | NA | #VALUE! |
| ENSMUSG00000077476.3 | 0 | NA | NA | NA | NA | NA | #VALUE! |
| ENSMUSG00000077477.3 | 0 | NA | NA | NA | NA | NA | #VALUE! |
| ENSMUSG00000077481.3 | 0 | NA | NA | NA | NA | NA | #VALUE! |
| ENSMUSG00000077484.3 | 0 | NA | NA | NA | NA | NA | #VALUE! |
| ENSMUSG00000077485.3 | 0 | NA | NA | NA | NA | NA | #VALUE! |
| ENSMUSG00000077488.3 | 0 | NA | NA | NA | NA | NA | #VALUE! |
| ENSMUSG00000077492.4 | 0 | NA | NA | NA | NA | NA | #VALUE! |
| ENSMUSG00000077493.3 | 0 | NA | NA | NA | NA | NA | #VALUE! |
| ENSMUSG00000077494.3 | 0 | NA | NA | NA | NA | NA | #VALUE! |
| ENSMUSG00000077495.3 | 0 | NA | NA | NA | NA | NA | #VALUE! |
| ENSMUSG00000077497.3 | 0 | NA | NA | NA | NA | NA | #VALUE! |
| ENSMUSG00000077501.3 | 0 | NA | NA | NA | NA | NA | #VALUE! |
| ENSMUSG00000077502.3 | 0 | NA | NA | NA | NA | NA | #VALUE! |
| ENSMUSG00000077503.4 | 0 | NA | NA | NA | NA | NA | #VALUE! |
| ENSMUSG00000077504.3 | 0 | NA | NA | NA | NA | NA | #VALUE! |
| ENSMUSG00000077505.3 | 0 | NA | NA | NA | NA | NA | #VALUE! |
| ENSMUSG00000077506.3 | 0 | NA | NA | NA | NA | NA | #VALUE! |
| ENSMUSG00000077507.3 | 0 | NA | NA | NA | NA | NA | #VALUE! |
| ENSMUSG00000077511.3 | 0 | NA | NA | NA | NA | NA | #VALUE! |
| ENSMUSG00000077514.3 | 0 | NA | NA | NA | NA | NA | #VALUE! |
| ENSMUSG00000077515.3 | 0 | NA | NA | NA | NA | NA | #VALUE! |
| ENSMUSG00000077518.3 | 0 | NA | NA | NA | NA | NA | #VALUE! |
| ENSMUSG00000077519.3 | 0 | NA | NA | NA | NA | NA | #VALUE! |
| ENSMUSG00000077521.2 | 0 | NA | NA | NA | NA | NA | #VALUE! |
| ENSMUSG00000077522.3 | 0 | NA | NA | NA | NA | NA | #VALUE! |
| ENSMUSG00000077526.3 | 0 | NA | NA | NA | NA | NA | #VALUE! |
| ENSMUSG00000077527.3 | 0 | NA | NA | NA | NA | NA | #VALUE! |
| ENSMUSG00000077528.3 | 0 | NA | NA | NA | NA | NA | #VALUE! |
| ENSMUSG00000077535.4 | 0 | NA | NA | NA | NA | NA | #VALUE! |
| ENSMUSG00000077537.3 | 0 | NA | NA | NA | NA | NA | #VALUE! |
| ENSMUSG00000077538.3 | 0 | NA | NA | NA | NA | NA | #VALUE! |
| ENSMUSG00000077539.3 | 0 | NA | NA | NA | NA | NA | #VALUE! |
| ENSMUSG00000077541.3 | 0 | NA | NA | NA | NA | NA | #VALUE! |
| ENSMUSG00000077542.3 | 0 | NA | NA | NA | NA | NA | #VALUE! |
| ENSMUSG00000077543.3 | 0 | NA | NA | NA | NA | NA | #VALUE! |
| ENSMUSG00000077545.3 | 0 | NA | NA | NA | NA | NA | #VALUE! |
| ENSMUSG00000077546.3 | 0 | NA | NA | NA | NA | NA | #VALUE! |
| ENSMUSG00000077547.3 | 0 | NA | NA | NA | NA | NA | #VALUE! |

| | | | | | | | |
|---|---|---|---|---|---|---|---|
| ENSMUSG00000077549.3 | 0 | NA | NA | NA | NA | NA | #VALUE! |
| ENSMUSG00000077552.2 | 0 | NA | NA | NA | NA | NA | #VALUE! |
| ENSMUSG00000077553.3 | 0 | NA | NA | NA | NA | NA | #VALUE! |
| ENSMUSG00000077554.3 | 0 | NA | NA | NA | NA | NA | #VALUE! |
| ENSMUSG00000077555.3 | 0 | NA | NA | NA | NA | NA | #VALUE! |
| ENSMUSG00000077557.3 | 0 | NA | NA | NA | NA | NA | #VALUE! |
| ENSMUSG00000077558.3 | 0 | NA | NA | NA | NA | NA | #VALUE! |
| ENSMUSG00000077561.2 | 0 | NA | NA | NA | NA | NA | #VALUE! |
| ENSMUSG00000077562.3 | 0 | NA | NA | NA | NA | NA | #VALUE! |
| ENSMUSG00000077563.3 | 0 | NA | NA | NA | NA | NA | #VALUE! |
| ENSMUSG00000077564.3 | 0 | NA | NA | NA | NA | NA | #VALUE! |
| ENSMUSG00000077565.3 | 0 | NA | NA | NA | NA | NA | #VALUE! |
| ENSMUSG00000077567.3 | 0 | NA | NA | NA | NA | NA | #VALUE! |
| ENSMUSG00000077569.3 | 0 | NA | NA | NA | NA | NA | #VALUE! |
| ENSMUSG00000077572.3 | 0 | NA | NA | NA | NA | NA | #VALUE! |
| ENSMUSG00000077574.3 | 0 | NA | NA | NA | NA | NA | #VALUE! |
| ENSMUSG00000077575.3 | 0 | NA | NA | NA | NA | NA | #VALUE! |
| ENSMUSG00000077576.3 | 0 | NA | NA | NA | NA | NA | #VALUE! |
| ENSMUSG00000077577.4 | 0 | NA | NA | NA | NA | NA | #VALUE! |
| ENSMUSG00000077578.3 | 0 | NA | NA | NA | NA | NA | #VALUE! |
| ENSMUSG00000077579.3 | 0 | NA | NA | NA | NA | NA | #VALUE! |
| ENSMUSG00000077581.3 | 0 | NA | NA | NA | NA | NA | #VALUE! |
| ENSMUSG00000077582.3 | 0 | NA | NA | NA | NA | NA | #VALUE! |
| ENSMUSG00000077585.3 | 0 | NA | NA | NA | NA | NA | #VALUE! |
| ENSMUSG00000077586.3 | 0 | NA | NA | NA | NA | NA | #VALUE! |
| ENSMUSG00000077587.2 | 0 | NA | NA | NA | NA | NA | #VALUE! |
| ENSMUSG00000077590.3 | 0 | NA | NA | NA | NA | NA | #VALUE! |
| ENSMUSG00000077591.3 | 0 | NA | NA | NA | NA | NA | #VALUE! |
| ENSMUSG00000077592.3 | 0 | NA | NA | NA | NA | NA | #VALUE! |
| ENSMUSG00000077593.3 | 0 | NA | NA | NA | NA | NA | #VALUE! |
| ENSMUSG00000077594.4 | 0 | NA | NA | NA | NA | NA | #VALUE! |
| ENSMUSG00000077595.3 | 0 | NA | NA | NA | NA | NA | #VALUE! |
| ENSMUSG00000077597.4 | 0 | NA | NA | NA | NA | NA | #VALUE! |
| ENSMUSG00000077598.4 | 0 | NA | NA | NA | NA | NA | #VALUE! |
| ENSMUSG00000077599.3 | 0 | NA | NA | NA | NA | NA | #VALUE! |
| ENSMUSG00000077601.3 | 0 | NA | NA | NA | NA | NA | #VALUE! |
| ENSMUSG00000077602.3 | 0 | NA | NA | NA | NA | NA | #VALUE! |
| ENSMUSG00000077603.3 | 0 | NA | NA | NA | NA | NA | #VALUE! |
| ENSMUSG00000077604.3 | 0 | NA | NA | NA | NA | NA | #VALUE! |
| ENSMUSG00000077607.3 | 0 | NA | NA | NA | NA | NA | #VALUE! |
| ENSMUSG00000077608.3 | 0 | NA | NA | NA | NA | NA | #VALUE! |
| ENSMUSG00000077609.3 | 0 | NA | NA | NA | NA | NA | #VALUE! |
| ENSMUSG00000077611.3 | 0 | NA | NA | NA | NA | NA | #VALUE! |
| ENSMUSG00000077612.3 | 0 | NA | NA | NA | NA | NA | #VALUE! |

| | | | | | | | |
|---|---|---|---|---|---|---|---|
| ENSMUSG00000077615.3 | 0 | NA | NA | NA | NA | NA | #VALUE! |
| ENSMUSG00000077616.3 | 0 | NA | NA | NA | NA | NA | #VALUE! |
| ENSMUSG00000077617.3 | 0 | NA | NA | NA | NA | NA | #VALUE! |
| ENSMUSG00000077618.3 | 0 | NA | NA | NA | NA | NA | #VALUE! |
| ENSMUSG00000077619.3 | 0 | NA | NA | NA | NA | NA | #VALUE! |
| ENSMUSG00000077624.3 | 0 | NA | NA | NA | NA | NA | #VALUE! |
| ENSMUSG00000077625.3 | 0 | NA | NA | NA | NA | NA | #VALUE! |
| ENSMUSG00000077627.3 | 0 | NA | NA | NA | NA | NA | #VALUE! |
| ENSMUSG00000077628.3 | 0 | NA | NA | NA | NA | NA | #VALUE! |
| ENSMUSG00000077629.3 | 0 | NA | NA | NA | NA | NA | #VALUE! |
| ENSMUSG00000077630.3 | 0 | NA | NA | NA | NA | NA | #VALUE! |
| ENSMUSG00000077633.3 | 0 | NA | NA | NA | NA | NA | #VALUE! |
| ENSMUSG00000077635.2 | 0 | NA | NA | NA | NA | NA | #VALUE! |
| ENSMUSG00000077637.3 | 0 | NA | NA | NA | NA | NA | #VALUE! |
| ENSMUSG00000077639.3 | 0 | NA | NA | NA | NA | NA | #VALUE! |
| ENSMUSG00000077641.3 | 0 | NA | NA | NA | NA | NA | #VALUE! |
| ENSMUSG00000077642.3 | 0 | NA | NA | NA | NA | NA | #VALUE! |
| ENSMUSG00000077643.3 | 0 | NA | NA | NA | NA | NA | #VALUE! |
| ENSMUSG00000077645.3 | 0 | NA | NA | NA | NA | NA | #VALUE! |
| ENSMUSG00000077647.3 | 0 | NA | NA | NA | NA | NA | #VALUE! |
| ENSMUSG00000077648.3 | 0 | NA | NA | NA | NA | NA | #VALUE! |
| ENSMUSG00000077649.3 | 0 | NA | NA | NA | NA | NA | #VALUE! |
| ENSMUSG00000077650.3 | 0 | NA | NA | NA | NA | NA | #VALUE! |
| ENSMUSG00000077652.3 | 0 | NA | NA | NA | NA | NA | #VALUE! |
| ENSMUSG00000077653.3 | 0 | NA | NA | NA | NA | NA | #VALUE! |
| ENSMUSG00000077655.3 | 0 | NA | NA | NA | NA | NA | #VALUE! |
| ENSMUSG00000077659.3 | 0 | NA | NA | NA | NA | NA | #VALUE! |
| ENSMUSG00000077660.3 | 0 | NA | NA | NA | NA | NA | #VALUE! |
| ENSMUSG00000077663.2 | 0 | NA | NA | NA | NA | NA | #VALUE! |
| ENSMUSG00000077664.3 | 0 | NA | NA | NA | NA | NA | #VALUE! |
| ENSMUSG00000077666.4 | 0 | NA | NA | NA | NA | NA | #VALUE! |
| ENSMUSG00000077667.3 | 0 | NA | NA | NA | NA | NA | #VALUE! |
| ENSMUSG00000077668.3 | 0 | NA | NA | NA | NA | NA | #VALUE! |
| ENSMUSG00000077669.3 | 0 | NA | NA | NA | NA | NA | #VALUE! |
| ENSMUSG00000077671.3 | 0 | NA | NA | NA | NA | NA | #VALUE! |
| ENSMUSG00000077672.3 | 0 | NA | NA | NA | NA | NA | #VALUE! |
| ENSMUSG00000077673.3 | 0 | NA | NA | NA | NA | NA | #VALUE! |
| ENSMUSG00000077674.3 | 0 | NA | NA | NA | NA | NA | #VALUE! |
| ENSMUSG00000077675.3 | 0 | NA | NA | NA | NA | NA | #VALUE! |
| ENSMUSG00000077677.3 | 0 | NA | NA | NA | NA | NA | #VALUE! |
| ENSMUSG00000077678.3 | 0 | NA | NA | NA | NA | NA | #VALUE! |
| ENSMUSG00000077680.3 | 0 | NA | NA | NA | NA | NA | #VALUE! |
| ENSMUSG00000077681.3 | 0 | NA | NA | NA | NA | NA | #VALUE! |
| ENSMUSG00000077682.3 | 0 | NA | NA | NA | NA | NA | #VALUE! |

| | | | | | | | |
|---|---|---|---|---|---|---|---|
| ENSMUSG00000077683.3 | 0 | NA | NA | NA | NA | NA | #VALUE! |
| ENSMUSG00000077685.3 | 0 | NA | NA | NA | NA | NA | #VALUE! |
| ENSMUSG00000077687.3 | 0 | NA | NA | NA | NA | NA | #VALUE! |
| ENSMUSG00000077689.3 | 0 | NA | NA | NA | NA | NA | #VALUE! |
| ENSMUSG00000077691.4 | 0 | NA | NA | NA | NA | NA | #VALUE! |
| ENSMUSG00000077693.3 | 0 | NA | NA | NA | NA | NA | #VALUE! |
| ENSMUSG00000077697.3 | 0 | NA | NA | NA | NA | NA | #VALUE! |
| ENSMUSG00000077698.3 | 0 | NA | NA | NA | NA | NA | #VALUE! |
| ENSMUSG00000077700.4 | 0 | NA | NA | NA | NA | NA | #VALUE! |
| ENSMUSG00000077701.2 | 0 | NA | NA | NA | NA | NA | #VALUE! |
| ENSMUSG00000077702.3 | 0 | NA | NA | NA | NA | NA | #VALUE! |
| ENSMUSG00000077703.3 | 0 | NA | NA | NA | NA | NA | #VALUE! |
| ENSMUSG00000077704.3 | 0 | NA | NA | NA | NA | NA | #VALUE! |
| ENSMUSG00000077707.3 | 0 | NA | NA | NA | NA | NA | #VALUE! |
| ENSMUSG00000077708.3 | 0 | NA | NA | NA | NA | NA | #VALUE! |
| ENSMUSG00000077709.3 | 0 | NA | NA | NA | NA | NA | #VALUE! |
| ENSMUSG00000077711.3 | 0 | NA | NA | NA | NA | NA | #VALUE! |
| ENSMUSG00000077713.4 | 0 | NA | NA | NA | NA | NA | #VALUE! |
| ENSMUSG00000077714.3 | 0 | NA | NA | NA | NA | NA | #VALUE! |
| ENSMUSG00000077716.3 | 0 | NA | NA | NA | NA | NA | #VALUE! |
| ENSMUSG00000077717.3 | 0 | NA | NA | NA | NA | NA | #VALUE! |
| ENSMUSG00000077718.3 | 0 | NA | NA | NA | NA | NA | #VALUE! |
| ENSMUSG00000077719.3 | 0 | NA | NA | NA | NA | NA | #VALUE! |
| ENSMUSG00000077730.3 | 0 | NA | NA | NA | NA | NA | #VALUE! |
| ENSMUSG00000077733.3 | 0 | NA | NA | NA | NA | NA | #VALUE! |
| ENSMUSG00000077734.3 | 0 | NA | NA | NA | NA | NA | #VALUE! |
| ENSMUSG00000077735.3 | 0 | NA | NA | NA | NA | NA | #VALUE! |
| ENSMUSG00000077736.3 | 0 | NA | NA | NA | NA | NA | #VALUE! |
| ENSMUSG00000077737.3 | 3.878104 | -5.48139 | 3.918329 | -1.39891 | 0.16184 | NA | #VALUE! |
| ENSMUSG00000077738.2 | 0 | NA | NA | NA | NA | NA | #VALUE! |
| ENSMUSG00000077744.3 | 0 | NA | NA | NA | NA | NA | #VALUE! |
| ENSMUSG00000077745.3 | 0 | NA | NA | NA | NA | NA | #VALUE! |
| ENSMUSG00000077749.3 | 0 | NA | NA | NA | NA | NA | #VALUE! |
| ENSMUSG00000077752.3 | 0 | NA | NA | NA | NA | NA | #VALUE! |
| ENSMUSG00000077756.3 | 0 | NA | NA | NA | NA | NA | #VALUE! |
| ENSMUSG00000077757.3 | 0 | NA | NA | NA | NA | NA | #VALUE! |
| ENSMUSG00000077758.3 | 0 | NA | NA | NA | NA | NA | #VALUE! |
| ENSMUSG00000077759.3 | 0 | NA | NA | NA | NA | NA | #VALUE! |
| ENSMUSG00000077761.3 | 0 | NA | NA | NA | NA | NA | #VALUE! |
| ENSMUSG00000077767.3 | 0 | NA | NA | NA | NA | NA | #VALUE! |
| ENSMUSG00000077768.3 | 0 | NA | NA | NA | NA | NA | #VALUE! |
| ENSMUSG00000077769.3 | 0 | NA | NA | NA | NA | NA | #VALUE! |
| ENSMUSG00000077772.3 | 0 | NA | NA | NA | NA | NA | #VALUE! |
| ENSMUSG00000077776.3 | 0 | NA | NA | NA | NA | NA | #VALUE! |

| | | | | | | | |
|---|---|---|---|---|---|---|---|
| ENSMUSG00000077777.3 | 0 | NA | NA | NA | NA | NA | #VALUE! |
| ENSMUSG00000077780.3 | 0 | NA | NA | NA | NA | NA | #VALUE! |
| ENSMUSG00000077782.3 | 0 | NA | NA | NA | NA | NA | #VALUE! |
| ENSMUSG00000077783.4 | 0 | NA | NA | NA | NA | NA | #VALUE! |
| ENSMUSG00000077784.3 | 0 | NA | NA | NA | NA | NA | #VALUE! |
| ENSMUSG00000077786.3 | 0 | NA | NA | NA | NA | NA | #VALUE! |
| ENSMUSG00000077787.3 | 0 | NA | NA | NA | NA | NA | #VALUE! |
| ENSMUSG00000077791.4 | 0 | NA | NA | NA | NA | NA | #VALUE! |
| ENSMUSG00000077793.4 | 0 | NA | NA | NA | NA | NA | #VALUE! |
| ENSMUSG00000077794.4 | 0 | NA | NA | NA | NA | NA | #VALUE! |
| ENSMUSG00000077795.3 | 0 | NA | NA | NA | NA | NA | #VALUE! |
| ENSMUSG00000077796.3 | 0 | NA | NA | NA | NA | NA | #VALUE! |
| ENSMUSG00000077797.3 | 0 | NA | NA | NA | NA | NA | #VALUE! |
| ENSMUSG00000077799.3 | 0 | NA | NA | NA | NA | NA | #VALUE! |
| ENSMUSG00000077804.3 | 0 | NA | NA | NA | NA | NA | #VALUE! |
| ENSMUSG00000077815.3 | 0 | NA | NA | NA | NA | NA | #VALUE! |
| ENSMUSG00000077834.3 | 0 | NA | NA | NA | NA | NA | #VALUE! |
| ENSMUSG00000077847.3 | 0 | NA | NA | NA | NA | NA | #VALUE! |
| ENSMUSG00000077851.2 | 0 | NA | NA | NA | NA | NA | #VALUE! |
| ENSMUSG00000077869.3 | 0 | NA | NA | NA | NA | NA | #VALUE! |
| ENSMUSG00000077872.3 | 0 | NA | NA | NA | NA | NA | #VALUE! |
| ENSMUSG00000077881.3 | 0 | NA | NA | NA | NA | NA | #VALUE! |
| ENSMUSG00000077886.3 | 0 | NA | NA | NA | NA | NA | #VALUE! |
| ENSMUSG00000077890.3 | 0 | NA | NA | NA | NA | NA | #VALUE! |
| ENSMUSG00000077896.3 | 0 | NA | NA | NA | NA | NA | #VALUE! |
| ENSMUSG00000077897.3 | 0 | NA | NA | NA | NA | NA | #VALUE! |
| ENSMUSG00000077903.3 | 0 | NA | NA | NA | NA | NA | #VALUE! |
| ENSMUSG00000077921.3 | 0 | NA | NA | NA | NA | NA | #VALUE! |
| ENSMUSG00000077925.3 | 0 | NA | NA | NA | NA | NA | #VALUE! |
| ENSMUSG00000077928.3 | 0 | NA | NA | NA | NA | NA | #VALUE! |
| ENSMUSG00000077931.3 | 0 | NA | NA | NA | NA | NA | #VALUE! |
| ENSMUSG00000077941.3 | 0 | NA | NA | NA | NA | NA | #VALUE! |
| ENSMUSG00000077954.3 | 0 | NA | NA | NA | NA | NA | #VALUE! |
| ENSMUSG00000077962.3 | 0 | NA | NA | NA | NA | NA | #VALUE! |
| ENSMUSG00000077972.3 | 0 | NA | NA | NA | NA | NA | #VALUE! |
| ENSMUSG00000077980.3 | 0 | NA | NA | NA | NA | NA | #VALUE! |
| ENSMUSG00000077981.3 | 0 | NA | NA | NA | NA | NA | #VALUE! |
| ENSMUSG00000077991.3 | 0 | NA | NA | NA | NA | NA | #VALUE! |
| ENSMUSG00000077996.3 | 0 | NA | NA | NA | NA | NA | #VALUE! |
| ENSMUSG00000077998.3 | 0 | NA | NA | NA | NA | NA | #VALUE! |
| ENSMUSG00000078002.3 | 0 | NA | NA | NA | NA | NA | #VALUE! |
| ENSMUSG00000078008.3 | 0 | NA | NA | NA | NA | NA | #VALUE! |
| ENSMUSG00000078017.3 | 0 | NA | NA | NA | NA | NA | #VALUE! |
| ENSMUSG00000078024.3 | 0 | NA | NA | NA | NA | NA | #VALUE! |

| | | | | | | | |
|---|---|---|---|---|---|---|---|
| ENSMUSG00000078025.3 | 0 | NA | NA | NA | NA | NA | #VALUE! |
| ENSMUSG00000078026.3 | 0 | NA | NA | NA | NA | NA | #VALUE! |
| ENSMUSG00000078027.3 | 0 | NA | NA | NA | NA | NA | #VALUE! |
| ENSMUSG00000078031.3 | 0 | NA | NA | NA | NA | NA | #VALUE! |
| ENSMUSG00000078032.3 | 0 | NA | NA | NA | NA | NA | #VALUE! |
| ENSMUSG00000078035.3 | 0 | NA | NA | NA | NA | NA | #VALUE! |
| ENSMUSG00000078038.3 | 0 | NA | NA | NA | NA | NA | #VALUE! |
| ENSMUSG00000078041.3 | 0 | NA | NA | NA | NA | NA | #VALUE! |
| ENSMUSG00000078057.2 | 0 | NA | NA | NA | NA | NA | #VALUE! |
| ENSMUSG00000078060.3 | 0 | NA | NA | NA | NA | NA | #VALUE! |
| ENSMUSG00000078080.7 | 0 | NA | NA | NA | NA | NA | #VALUE! |
| ENSMUSG00000078087.6 | 0.093255 | 0.868093 | 4.072464 | 0.213162 | 0.831201 | NA | #VALUE! |
| ENSMUSG00000078091.3 | 0 | NA | NA | NA | NA | NA | #VALUE! |
| ENSMUSG00000078097.4 | 0 | NA | NA | NA | NA | NA | #VALUE! |
| ENSMUSG00000078116.5 | 0 | NA | NA | NA | NA | NA | #VALUE! |
| ENSMUSG00000078117.3 | 0 | NA | NA | NA | NA | NA | #VALUE! |
| ENSMUSG00000078118.6 | 0 | NA | NA | NA | NA | NA | #VALUE! |
| ENSMUSG00000078122.5 | 1.575197 | -4.20222 | 3.934736 | -1.06798 | 0.285529 | NA | #VALUE! |
| ENSMUSG00000078123.4 | 0 | NA | NA | NA | NA | NA | #VALUE! |
| ENSMUSG00000078125.3 | 0 | NA | NA | NA | NA | NA | #VALUE! |
| ENSMUSG00000078126.5 | 0.900246 | 0.785241 | 3.941206 | 0.199239 | 0.842076 | NA | #VALUE! |
| ENSMUSG00000078127.4 | 0 | NA | NA | NA | NA | NA | #VALUE! |
| ENSMUSG00000078128.2 | 0 | NA | NA | NA | NA | NA | #VALUE! |
| ENSMUSG00000078129.4 | 0 | NA | NA | NA | NA | NA | #VALUE! |
| ENSMUSG00000078130.5 | 0 | NA | NA | NA | NA | NA | #VALUE! |
| ENSMUSG00000078131.2 | 0 | NA | NA | NA | NA | NA | #VALUE! |
| ENSMUSG00000078132.2 | 0 | NA | NA | NA | NA | NA | #VALUE! |
| ENSMUSG00000078137.2 | 0 | NA | NA | NA | NA | NA | #VALUE! |
| ENSMUSG00000078141.4 | 0 | NA | NA | NA | NA | NA | #VALUE! |
| ENSMUSG00000078143.4 | 0 | NA | NA | NA | NA | NA | #VALUE! |
| ENSMUSG00000078144.5 | 0 | NA | NA | NA | NA | NA | #VALUE! |
| ENSMUSG00000078151.4 | 0 | NA | NA | NA | NA | NA | #VALUE! |
| ENSMUSG00000078157.2 | 0 | NA | NA | NA | NA | NA | #VALUE! |
| ENSMUSG00000078161.9 | 0 | NA | NA | NA | NA | NA | #VALUE! |
| ENSMUSG00000078162.7 | 0 | NA | NA | NA | NA | NA | #VALUE! |
| ENSMUSG00000078173.4 | 0 | NA | NA | NA | NA | NA | #VALUE! |
| ENSMUSG00000078177.4 | 0 | NA | NA | NA | NA | NA | #VALUE! |
| ENSMUSG00000078179.2 | 0 | NA | NA | NA | NA | NA | #VALUE! |
| ENSMUSG00000078180.5 | 1.574768 | -0.06729 | 3.842406 | -0.01751 | 0.986027 | NA | #VALUE! |
| ENSMUSG00000078182.5 | 0 | NA | NA | NA | NA | NA | #VALUE! |
| ENSMUSG00000078183.4 | 0 | NA | NA | NA | NA | NA | #VALUE! |
| ENSMUSG00000078190.8 | 0 | NA | NA | NA | NA | NA | #VALUE! |
| ENSMUSG00000078192.6 | 0.168284 | 1.027628 | 4.067126 | 0.252667 | 0.800526 | NA | #VALUE! |
| ENSMUSG00000078197.3 | 0 | NA | NA | NA | NA | NA | #VALUE! |

| | | | | | | | |
|---|---|---|---|---|---|---|---|
| ENSMUSG00000078198.3 | 0 | NA | NA | NA | NA | NA | #VALUE! |
| ENSMUSG00000078202.4 | 5.883122 | -3.4436 | 2.988493 | -1.15229 | 0.249204 | NA | #VALUE! |
| ENSMUSG00000078206.2 | 0 | NA | NA | NA | NA | NA | #VALUE! |
| ENSMUSG00000078208.2 | 0 | NA | NA | NA | NA | NA | #VALUE! |
| ENSMUSG00000078210.4 | 0 | NA | NA | NA | NA | NA | #VALUE! |
| ENSMUSG00000078212.5 | 0 | NA | NA | NA | NA | NA | #VALUE! |
| ENSMUSG00000078213.4 | 0 | NA | NA | NA | NA | NA | #VALUE! |
| ENSMUSG00000078214.6 | 0 | NA | NA | NA | NA | NA | #VALUE! |
| ENSMUSG00000078218.3 | 0 | NA | NA | NA | NA | NA | #VALUE! |
| ENSMUSG00000078235.4 | 0 | NA | NA | NA | NA | NA | #VALUE! |
| ENSMUSG00000078238.5 | 0 | NA | NA | NA | NA | NA | #VALUE! |
| ENSMUSG00000078247.6 | 4.765512 | -5.77989 | 3.916172 | -1.4759 | 0.13997 | NA | #VALUE! |
| ENSMUSG00000078249.6 | 0 | NA | NA | NA | NA | NA | #VALUE! |
| ENSMUSG00000078251.3 | 0 | NA | NA | NA | NA | NA | #VALUE! |
| ENSMUSG00000078252.2 | 0 | NA | NA | NA | NA | NA | #VALUE! |
| ENSMUSG00000078253.4 | 0 | NA | NA | NA | NA | NA | #VALUE! |
| ENSMUSG00000078254.2 | 0 | NA | NA | NA | NA | NA | #VALUE! |
| ENSMUSG00000078255.2 | 0 | NA | NA | NA | NA | NA | #VALUE! |
| ENSMUSG00000078256.4 | 0 | NA | NA | NA | NA | NA | #VALUE! |
| ENSMUSG00000078257.3 | 0 | NA | NA | NA | NA | NA | #VALUE! |
| ENSMUSG00000078258.2 | 0 | NA | NA | NA | NA | NA | #VALUE! |
| ENSMUSG00000078259.4 | 0 | NA | NA | NA | NA | NA | #VALUE! |
| ENSMUSG00000078260.2 | 0 | NA | NA | NA | NA | NA | #VALUE! |
| ENSMUSG00000078261.2 | 0 | NA | NA | NA | NA | NA | #VALUE! |
| ENSMUSG00000078262.6 | 0 | NA | NA | NA | NA | NA | #VALUE! |
| ENSMUSG00000078263.2 | 0 | NA | NA | NA | NA | NA | #VALUE! |
| ENSMUSG00000078269.2 | 0 | NA | NA | NA | NA | NA | #VALUE! |
| ENSMUSG00000078270.4 | 0 | NA | NA | NA | NA | NA | #VALUE! |
| ENSMUSG00000078276.2 | 0 | NA | NA | NA | NA | NA | #VALUE! |
| ENSMUSG00000078280.2 | 0 | NA | NA | NA | NA | NA | #VALUE! |
| ENSMUSG00000078289.3 | 0 | NA | NA | NA | NA | NA | #VALUE! |
| ENSMUSG00000078291.2 | 0.105013 | -0.97324 | 4.076293 | -0.23876 | 0.811295 | NA | #VALUE! |
| ENSMUSG00000078300.4 | 0 | NA | NA | NA | NA | NA | #VALUE! |
| ENSMUSG00000078302.6 | 0 | NA | NA | NA | NA | NA | #VALUE! |
| ENSMUSG00000078307.4 | 0 | NA | NA | NA | NA | NA | #VALUE! |
| ENSMUSG00000078308.3 | 0 | NA | NA | NA | NA | NA | #VALUE! |
| ENSMUSG00000078314.3 | 0 | NA | NA | NA | NA | NA | #VALUE! |
| ENSMUSG00000078315.6 | 0 | NA | NA | NA | NA | NA | #VALUE! |
| ENSMUSG00000078320.5 | 0 | NA | NA | NA | NA | NA | #VALUE! |
| ENSMUSG00000078324.3 | 0 | NA | NA | NA | NA | NA | #VALUE! |
| ENSMUSG00000078346.4 | 0 | NA | NA | NA | NA | NA | #VALUE! |
| ENSMUSG00000078354.2 | 0 | NA | NA | NA | NA | NA | #VALUE! |
| ENSMUSG00000078355.2 | 0 | NA | NA | NA | NA | NA | #VALUE! |
| ENSMUSG00000078365.6 | 0 | NA | NA | NA | NA | NA | #VALUE! |

| | | | | | | | |
|---|---|---|---|---|---|---|---|
| ENSMUSG00000078370.2 | 0 | NA | NA | NA | NA | NA | #VALUE! |
| ENSMUSG00000078377.6 | 0.204111 | -1.05581 | 4.073302 | -0.2592 | 0.795479 | NA | #VALUE! |
| ENSMUSG00000078380.2 | 0 | NA | NA | NA | NA | NA | #VALUE! |
| ENSMUSG00000078420.7 | 0 | NA | NA | NA | NA | NA | #VALUE! |
| ENSMUSG00000078430.4 | 0 | NA | NA | NA | NA | NA | #VALUE! |
| ENSMUSG00000078439.10 | 0 | NA | NA | NA | NA | NA | #VALUE! |
| ENSMUSG00000078440.11 | 6.474041 | 1.425733 | 3.761546 | 0.379029 | 0.704667 | NA | #VALUE! |
| ENSMUSG00000078442.4 | 0 | NA | NA | NA | NA | NA | #VALUE! |
| ENSMUSG00000078444.4 | 0 | NA | NA | NA | NA | NA | #VALUE! |
| ENSMUSG00000078451.6 | 1.836997 | -4.39907 | 3.931186 | -1.11902 | 0.263133 | NA | #VALUE! |
| ENSMUSG00000078452.11 | 3.971596 | -0.97365 | 3.777281 | -0.25777 | 0.796588 | NA | #VALUE! |
| ENSMUSG00000078480.4 | 0.764599 | 0.032051 | 3.957652 | 0.008098 | 0.993538 | NA | #VALUE! |
| ENSMUSG00000078481.2 | 0 | NA | NA | NA | NA | NA | #VALUE! |
| ENSMUSG00000078485.5 | 0 | NA | NA | NA | NA | NA | #VALUE! |
| ENSMUSG00000078486.4 | 0 | NA | NA | NA | NA | NA | #VALUE! |
| ENSMUSG00000078487.9 | 0 | NA | NA | NA | NA | NA | #VALUE! |
| ENSMUSG00000078488.2 | 0 | NA | NA | NA | NA | NA | #VALUE! |
| ENSMUSG00000078489.3 | 0 | NA | NA | NA | NA | NA | #VALUE! |
| ENSMUSG00000078490.11 | 0 | NA | NA | NA | NA | NA | #VALUE! |
| ENSMUSG00000078491.3 | 0 | NA | NA | NA | NA | NA | #VALUE! |
| ENSMUSG00000078492.4 | 0 | NA | NA | NA | NA | NA | #VALUE! |
| ENSMUSG00000078494.6 | 0 | NA | NA | NA | NA | NA | #VALUE! |
| ENSMUSG00000078495.11 | 4.903871 | -1.6011 | 3.776351 | -0.42398 | 0.671579 | NA | #VALUE! |
| ENSMUSG00000078496.10 | 9.593045 | 6.747792 | 3.658637 | 1.844346 | 0.065133 | NA | #VALUE! |
| ENSMUSG00000078497.10 | 0 | NA | NA | NA | NA | NA | #VALUE! |
| ENSMUSG00000078498.6 | 0 | NA | NA | NA | NA | NA | #VALUE! |
| ENSMUSG00000078500.4 | 0 | NA | NA | NA | NA | NA | #VALUE! |
| ENSMUSG00000078502.11 | 0 | NA | NA | NA | NA | NA | #VALUE! |
| ENSMUSG00000078503.10 | 0 | NA | NA | NA | NA | NA | #VALUE! |
| ENSMUSG00000078504.3 | 0 | NA | NA | NA | NA | NA | #VALUE! |
| ENSMUSG00000078505.2 | 0 | NA | NA | NA | NA | NA | #VALUE! |
| ENSMUSG00000078506.2 | 0 | NA | NA | NA | NA | NA | #VALUE! |
| ENSMUSG00000078507.2 | 0 | NA | NA | NA | NA | NA | #VALUE! |
| ENSMUSG00000078508.2 | 0 | NA | NA | NA | NA | NA | #VALUE! |
| ENSMUSG00000078509.2 | 0 | NA | NA | NA | NA | NA | #VALUE! |
| ENSMUSG00000078510.4 | 0 | NA | NA | NA | NA | NA | #VALUE! |
| ENSMUSG00000078511.4 | 0 | NA | NA | NA | NA | NA | #VALUE! |
| ENSMUSG00000078512.9 | 0 | NA | NA | NA | NA | NA | #VALUE! |
| ENSMUSG00000078513.3 | 0 | NA | NA | NA | NA | NA | #VALUE! |
| ENSMUSG00000078518.2 | 0 | NA | NA | NA | NA | NA | #VALUE! |
| ENSMUSG00000078519.4 | 0 | NA | NA | NA | NA | NA | #VALUE! |
| ENSMUSG00000078520.4 | 0 | NA | NA | NA | NA | NA | #VALUE! |
| ENSMUSG00000078521.3 | 0 | NA | NA | NA | NA | NA | #VALUE! |
| ENSMUSG00000078528.3 | 0 | NA | NA | NA | NA | NA | #VALUE! |

| | | | | | | | |
|---|---|---|---|---|---|---|---|
| ENSMUSG00000078531.4 | 0 | NA | NA | NA | NA | NA | #VALUE! |
| ENSMUSG00000078532.10 | 0 | NA | NA | NA | NA | NA | #VALUE! |
| ENSMUSG00000078537.4 | 0 | NA | NA | NA | NA | NA | #VALUE! |
| ENSMUSG00000078546.9 | 6.504898 | -6.23597 | 3.713017 | -1.67949 | 0.093057 | NA | #VALUE! |
| ENSMUSG00000078552.11 | 3.689633 | 5.364344 | 3.919007 | 1.368802 | 0.171061 | NA | #VALUE! |
| ENSMUSG00000078554.3 | 0 | NA | NA | NA | NA | NA | #VALUE! |
| ENSMUSG00000078575.3 | 0 | NA | NA | NA | NA | NA | #VALUE! |
| ENSMUSG00000078576.2 | 0 | NA | NA | NA | NA | NA | #VALUE! |
| ENSMUSG00000078577.4 | 0 | NA | NA | NA | NA | NA | #VALUE! |
| ENSMUSG00000078580.4 | 19.29142 | -7.79813 | 2.742101 | -2.84385 | NA | NA | #VALUE! |
| ENSMUSG00000078586.4 | 0 | NA | NA | NA | NA | NA | #VALUE! |
| ENSMUSG00000078588.11 | 0 | NA | NA | NA | NA | NA | #VALUE! |
| ENSMUSG00000078590.2 | 0 | NA | NA | NA | NA | NA | #VALUE! |
| ENSMUSG00000078591.2 | 0 | NA | NA | NA | NA | NA | #VALUE! |
| ENSMUSG00000078592.3 | 0 | NA | NA | NA | NA | NA | #VALUE! |
| ENSMUSG00000078593.3 | 0 | NA | NA | NA | NA | NA | #VALUE! |
| ENSMUSG00000078598.12 | 0 | NA | NA | NA | NA | NA | #VALUE! |
| ENSMUSG00000078599.10 | 0 | NA | NA | NA | NA | NA | #VALUE! |
| ENSMUSG00000078600.2 | 0 | NA | NA | NA | NA | NA | #VALUE! |
| ENSMUSG00000078601.5 | 0 | NA | NA | NA | NA | NA | #VALUE! |
| ENSMUSG00000078604.2 | 0 | NA | NA | NA | NA | NA | #VALUE! |
| ENSMUSG00000078605.3 | 0 | NA | NA | NA | NA | NA | #VALUE! |
| ENSMUSG00000078606.9 | 0 | NA | NA | NA | NA | NA | #VALUE! |
| ENSMUSG00000078607.7 | 4.377437 | -3.05263 | 3.834493 | -0.7961 | 0.425976 | NA | #VALUE! |
| ENSMUSG00000078611.4 | 0 | NA | NA | NA | NA | NA | #VALUE! |
| ENSMUSG00000078616.3 | 0 | NA | NA | NA | NA | NA | #VALUE! |
| ENSMUSG00000078618.4 | 0 | NA | NA | NA | NA | NA | #VALUE! |
| ENSMUSG00000078620.2 | 0 | NA | NA | NA | NA | NA | #VALUE! |
| ENSMUSG00000078621.3 | 0 | NA | NA | NA | NA | NA | #VALUE! |
| ENSMUSG00000078624.5 | 0 | NA | NA | NA | NA | NA | #VALUE! |
| ENSMUSG00000078625.11 | 0 | NA | NA | NA | NA | NA | #VALUE! |
| ENSMUSG00000078626.3 | 0 | NA | NA | NA | NA | NA | #VALUE! |
| ENSMUSG00000078627.10 | 0 | NA | NA | NA | NA | NA | #VALUE! |
| ENSMUSG00000078630.9 | 0 | NA | NA | NA | NA | NA | #VALUE! |
| ENSMUSG00000078632.5 | 0 | NA | NA | NA | NA | NA | #VALUE! |
| ENSMUSG00000078635.4 | 0 | NA | NA | NA | NA | NA | #VALUE! |
| ENSMUSG00000078639.2 | 0 | NA | NA | NA | NA | NA | #VALUE! |
| ENSMUSG00000078640.10 | 0 | NA | NA | NA | NA | NA | #VALUE! |
| ENSMUSG00000078648.4 | 0 | NA | NA | NA | NA | NA | #VALUE! |
| ENSMUSG00000078651.9 | 0.093255 | 0.868093 | 4.072464 | 0.213162 | 0.831201 | NA | #VALUE! |
| ENSMUSG00000078653.5 | 37.80385 | 1.108449 | 3.730712 | 0.297115 | NA | NA | #VALUE! |
| ENSMUSG00000078657.7 | 0 | NA | NA | NA | NA | NA | #VALUE! |
| ENSMUSG00000078664.3 | 0 | NA | NA | NA | NA | NA | #VALUE! |
| ENSMUSG00000078668.4 | 0 | NA | NA | NA | NA | NA | #VALUE! |

| | | | | | | | |
|---|---|---|---|---|---|---|---|
| ENSMUSG00000078670.4 | 3.558688 | -4.4225 | 3.911258 | -1.13071 | 0.258177 | NA | #VALUE! |
| ENSMUSG00000078677.2 | 0 | NA | NA | NA | NA | NA | #VALUE! |
| ENSMUSG00000078680.4 | 0.980405 | 2.13809 | 3.955392 | 0.540551 | 0.588817 | NA | #VALUE! |
| ENSMUSG00000078684.9 | 0 | NA | NA | NA | NA | NA | #VALUE! |
| ENSMUSG00000078687.10 | 1.775321 | -1.52346 | 3.859187 | -0.39476 | 0.693019 | NA | #VALUE! |
| ENSMUSG00000078688.12 | 2.875494 | 3.098893 | 3.892382 | 0.796143 | 0.425949 | NA | #VALUE! |
| ENSMUSG00000078691.4 | 0 | NA | NA | NA | NA | NA | #VALUE! |
| ENSMUSG00000078698.4 | 0 | NA | NA | NA | NA | NA | #VALUE! |
| ENSMUSG00000078700.11 | 0 | NA | NA | NA | NA | NA | #VALUE! |
| ENSMUSG00000078706.5 | 0 | NA | NA | NA | NA | NA | #VALUE! |
| ENSMUSG00000078716.11 | 0.105013 | -0.97324 | 4.076293 | -0.23876 | 0.811295 | NA | #VALUE! |
| ENSMUSG00000078719.3 | 0 | NA | NA | NA | NA | NA | #VALUE! |
| ENSMUSG00000078721.2 | 0 | NA | NA | NA | NA | NA | #VALUE! |
| ENSMUSG00000078722.3 | 0 | NA | NA | NA | NA | NA | #VALUE! |
| ENSMUSG00000078735.5 | 0 | NA | NA | NA | NA | NA | #VALUE! |
| ENSMUSG00000078737.4 | 0 | NA | NA | NA | NA | NA | #VALUE! |
| ENSMUSG00000078742.2 | 0 | NA | NA | NA | NA | NA | #VALUE! |
| ENSMUSG00000078746.3 | 0 | NA | NA | NA | NA | NA | #VALUE! |
| ENSMUSG00000078747.10 | 0 | NA | NA | NA | NA | NA | #VALUE! |
| ENSMUSG00000078752.4 | 0 | NA | NA | NA | NA | NA | #VALUE! |
| ENSMUSG00000078753.4 | 0 | NA | NA | NA | NA | NA | #VALUE! |
| ENSMUSG00000078754.3 | 0 | NA | NA | NA | NA | NA | #VALUE! |
| ENSMUSG00000078757.5 | 0 | NA | NA | NA | NA | NA | #VALUE! |
| ENSMUSG00000078759.4 | 0 | NA | NA | NA | NA | NA | #VALUE! |
| ENSMUSG00000078763.3 | 0 | NA | NA | NA | NA | NA | #VALUE! |
| ENSMUSG00000078765.12 | 0 | NA | NA | NA | NA | NA | #VALUE! |
| ENSMUSG00000078768.3 | 0 | NA | NA | NA | NA | NA | #VALUE! |
| ENSMUSG00000078772.3 | 5.540457 | 5.959333 | 3.9129 | 1.522997 | 0.12776 | NA | #VALUE! |
| ENSMUSG00000078773.12 | 0 | NA | NA | NA | NA | NA | #VALUE! |
| ENSMUSG00000078776.10 | 0 | NA | NA | NA | NA | NA | #VALUE! |
| ENSMUSG00000078779.5 | 0 | NA | NA | NA | NA | NA | #VALUE! |
| ENSMUSG00000078780.8 | 5.899401 | 2.189356 | 3.778142 | 0.57948 | 0.562266 | NA | #VALUE! |
| ENSMUSG00000078783.2 | 0 | NA | NA | NA | NA | NA | #VALUE! |
| ENSMUSG00000078787.10 | 0 | NA | NA | NA | NA | NA | #VALUE! |
| ENSMUSG00000078789.10 | 0 | NA | NA | NA | NA | NA | #VALUE! |
| ENSMUSG00000078793.11 | 0 | NA | NA | NA | NA | NA | #VALUE! |
| ENSMUSG00000078794.5 | 0 | NA | NA | NA | NA | NA | #VALUE! |
| ENSMUSG00000078795.3 | 0 | NA | NA | NA | NA | NA | #VALUE! |
| ENSMUSG00000078796.5 | 0 | NA | NA | NA | NA | NA | #VALUE! |
| ENSMUSG00000078798.5 | 0 | NA | NA | NA | NA | NA | #VALUE! |
| ENSMUSG00000078799.6 | 0 | NA | NA | NA | NA | NA | #VALUE! |
| ENSMUSG00000078800.6 | 0 | NA | NA | NA | NA | NA | #VALUE! |
| ENSMUSG00000078808.6 | 0 | NA | NA | NA | NA | NA | #VALUE! |
| ENSMUSG00000078810.5 | 0 | NA | NA | NA | NA | NA | #VALUE! |

| | | | | | | | |
|---|---|---|---|---|---|---|---|
| ENSMUSG00000078815.9 | 0 | NA | NA | NA | NA | NA | #VALUE! |
| ENSMUSG00000078816.11 | 0 | NA | NA | NA | NA | NA | #VALUE! |
| ENSMUSG00000078838.4 | 0 | NA | NA | NA | NA | NA | #VALUE! |
| ENSMUSG00000078840.2 | 0 | NA | NA | NA | NA | NA | #VALUE! |
| ENSMUSG00000078847.2 | 0 | NA | NA | NA | NA | NA | #VALUE! |
| ENSMUSG00000078850.2 | 0 | NA | NA | NA | NA | NA | #VALUE! |
| ENSMUSG00000078862.11 | 3.533954 | 5.311169 | 3.919465 | 1.355075 | 0.175394 | NA | #VALUE! |
| ENSMUSG00000078864.11 | 1.514552 | 4.088766 | 3.936269 | 1.038742 | 0.298925 | NA | #VALUE! |
| ENSMUSG00000078865.9 | 0 | NA | NA | NA | NA | NA | #VALUE! |
| ENSMUSG00000078866.11 | 0 | NA | NA | NA | NA | NA | #VALUE! |
| ENSMUSG00000078867.10 | 0 | NA | NA | NA | NA | NA | #VALUE! |
| ENSMUSG00000078868.4 | 0 | NA | NA | NA | NA | NA | #VALUE! |
| ENSMUSG00000078869.3 | 0 | NA | NA | NA | NA | NA | #VALUE! |
| ENSMUSG00000078870.10 | 0 | NA | NA | NA | NA | NA | #VALUE! |
| ENSMUSG00000078872.10 | 0 | NA | NA | NA | NA | NA | #VALUE! |
| ENSMUSG00000078875.11 | 0 | NA | NA | NA | NA | NA | #VALUE! |
| ENSMUSG00000078876.11 | 0 | NA | NA | NA | NA | NA | #VALUE! |
| ENSMUSG00000078878.10 | 2.077289 | 4.552599 | 3.928203 | 1.158952 | 0.246476 | NA | #VALUE! |
| ENSMUSG00000078879.11 | 0 | NA | NA | NA | NA | NA | #VALUE! |
| ENSMUSG00000078880.11 | 0 | NA | NA | NA | NA | NA | #VALUE! |
| ENSMUSG00000078881.11 | 0 | NA | NA | NA | NA | NA | #VALUE! |
| ENSMUSG00000078886.13 | 0 | NA | NA | NA | NA | NA | #VALUE! |
| ENSMUSG00000078887.11 | 0 | NA | NA | NA | NA | NA | #VALUE! |
| ENSMUSG00000078889.12 | 0 | NA | NA | NA | NA | NA | #VALUE! |
| ENSMUSG00000078891.5 | 0 | NA | NA | NA | NA | NA | #VALUE! |
| ENSMUSG00000078894.10 | 0 | NA | NA | NA | NA | NA | #VALUE! |
| ENSMUSG00000078895.5 | 0 | NA | NA | NA | NA | NA | #VALUE! |
| ENSMUSG00000078896.12 | 0 | NA | NA | NA | NA | NA | #VALUE! |
| ENSMUSG00000078897.11 | 0 | NA | NA | NA | NA | NA | #VALUE! |
| ENSMUSG00000078898.10 | 0 | NA | NA | NA | NA | NA | #VALUE! |
| ENSMUSG00000078899.10 | 0 | NA | NA | NA | NA | NA | #VALUE! |
| ENSMUSG00000078901.9 | 0 | NA | NA | NA | NA | NA | #VALUE! |
| ENSMUSG00000078902.4 | 0 | NA | NA | NA | NA | NA | #VALUE! |
| ENSMUSG00000078903.11 | 0 | NA | NA | NA | NA | NA | #VALUE! |
| ENSMUSG00000078905.9 | 0.18651 | 1.401769 | 4.056354 | 0.345574 | 0.729663 | NA | #VALUE! |
| ENSMUSG00000078906.10 | 0 | NA | NA | NA | NA | NA | #VALUE! |
| ENSMUSG00000078907.3 | 0 | NA | NA | NA | NA | NA | #VALUE! |
| ENSMUSG00000078908.11 | 9.684765 | -3.09143 | 2.604363 | -1.18702 | 0.23522 | NA | #VALUE! |
| ENSMUSG00000078912.3 | 0 | NA | NA | NA | NA | NA | #VALUE! |
| ENSMUSG00000078915.3 | 0 | NA | NA | NA | NA | NA | #VALUE! |
| ENSMUSG00000078917.4 | 0 | NA | NA | NA | NA | NA | #VALUE! |
| ENSMUSG00000078921.4 | 6.49759 | -2.70493 | 3.275773 | -0.82574 | 0.408952 | NA | #VALUE! |
| ENSMUSG00000078922.10 | 3.438518 | -4.14407 | 3.750688 | -1.10488 | 0.269211 | NA | #VALUE! |
| ENSMUSG00000078923.11 | 0 | NA | NA | NA | NA | NA | #VALUE! |

| | | | | | | | |
|---|---|---|---|---|---|---|---|
| ENSMUSG00000078924.2 | 0 | NA | NA | NA | NA | NA | #VALUE! |
| ENSMUSG00000078925.12 | 0 | NA | NA | NA | NA | NA | #VALUE! |
| ENSMUSG00000078926.9 | 0 | NA | NA | NA | NA | NA | #VALUE! |
| ENSMUSG00000078931.5 | 0 | NA | NA | NA | NA | NA | #VALUE! |
| ENSMUSG00000078932.9 | 0 | NA | NA | NA | NA | NA | #VALUE! |
| ENSMUSG00000078934.5 | 0 | NA | NA | NA | NA | NA | #VALUE! |
| ENSMUSG00000078935.2 | 0 | NA | NA | NA | NA | NA | #VALUE! |
| ENSMUSG00000078937.9 | 0 | NA | NA | NA | NA | NA | #VALUE! |
| ENSMUSG00000078938.10 | 0 | NA | NA | NA | NA | NA | #VALUE! |
| ENSMUSG00000078940.9 | 0 | NA | NA | NA | NA | NA | #VALUE! |
| ENSMUSG00000078942.11 | 0.630079 | -2.90291 | 3.975047 | -0.73028 | 0.465217 | NA | #VALUE! |
| ENSMUSG00000078945.11 | 1.685563 | -4.2806 | 3.933265 | -1.08831 | 0.276459 | NA | #VALUE! |
| ENSMUSG00000078949.3 | 0 | NA | NA | NA | NA | NA | #VALUE! |
| ENSMUSG00000078952.10 | 0 | NA | NA | NA | NA | NA | #VALUE! |
| ENSMUSG00000078953.2 | 0 | NA | NA | NA | NA | NA | #VALUE! |
| ENSMUSG00000078954.10 | 0 | NA | NA | NA | NA | NA | #VALUE! |
| ENSMUSG00000078955.4 | 0 | NA | NA | NA | NA | NA | #VALUE! |
| ENSMUSG00000078956.4 | 3.775101 | -2.76677 | 3.833494 | -0.72174 | 0.470457 | NA | #VALUE! |
| ENSMUSG00000078957.4 | 0 | NA | NA | NA | NA | NA | #VALUE! |
| ENSMUSG00000078958.10 | 0 | NA | NA | NA | NA | NA | #VALUE! |
| ENSMUSG00000078962.5 | 0 | NA | NA | NA | NA | NA | #VALUE! |
| ENSMUSG00000078963.4 | 3.626001 | -3.26136 | 3.867737 | -0.84322 | 0.399104 | NA | #VALUE! |
| ENSMUSG00000078965.6 | 0 | NA | NA | NA | NA | NA | #VALUE! |
| ENSMUSG00000078967.6 | 0.204111 | -1.05581 | 4.073302 | -0.2592 | 0.795479 | NA | #VALUE! |
| ENSMUSG00000078972.2 | 0 | NA | NA | NA | NA | NA | #VALUE! |
| ENSMUSG00000078974.11 | 8.234198 | -1.01857 | 2.864665 | -0.35556 | 0.722168 | NA | #VALUE! |
| ENSMUSG00000078982.4 | 0 | NA | NA | NA | NA | NA | #VALUE! |
| ENSMUSG00000078984.2 | 0 | NA | NA | NA | NA | NA | #VALUE! |
| ENSMUSG00000078994.10 | 0.737927 | 3.007825 | 3.968641 | 0.757898 | 0.448512 | NA | #VALUE! |
| ENSMUSG00000078995.11 | 0.105013 | -0.97324 | 4.076293 | -0.23876 | 0.811295 | NA | #VALUE! |
| ENSMUSG00000078998.4 | 0 | NA | NA | NA | NA | NA | #VALUE! |
| ENSMUSG00000079000.4 | 0 | NA | NA | NA | NA | NA | #VALUE! |
| ENSMUSG00000079001.2 | 0 | NA | NA | NA | NA | NA | #VALUE! |
| ENSMUSG00000079003.4 | 0.606563 | -0.89296 | 3.98278 | -0.22421 | 0.822597 | NA | #VALUE! |
| ENSMUSG00000079005.4 | 0 | NA | NA | NA | NA | NA | #VALUE! |
| ENSMUSG00000079006.4 | 0 | NA | NA | NA | NA | NA | #VALUE! |
| ENSMUSG00000079007.5 | 0 | NA | NA | NA | NA | NA | #VALUE! |
| ENSMUSG00000079008.3 | 0 | NA | NA | NA | NA | NA | #VALUE! |
| ENSMUSG00000079009.9 | 0 | NA | NA | NA | NA | NA | #VALUE! |
| ENSMUSG00000079010.2 | 0 | NA | NA | NA | NA | NA | #VALUE! |
| ENSMUSG00000079011.10 | 0 | NA | NA | NA | NA | NA | #VALUE! |
| ENSMUSG00000079013.3 | 0 | NA | NA | NA | NA | NA | #VALUE! |
| ENSMUSG00000079014.5 | 0 | NA | NA | NA | NA | NA | #VALUE! |
| ENSMUSG00000079016.2 | 0 | NA | NA | NA | NA | NA | #VALUE! |

| | | | | | | | |
|---|---|---|---|---|---|---|---|
| ENSMUSG00000079018.11 | 0.195789 | -1.05581 | 4.073302 | -0.2592 | 0.795479 | NA | #VALUE! |
| ENSMUSG00000079019.3 | 0 | NA | NA | NA | NA | NA | #VALUE! |
| ENSMUSG00000079022.11 | 0 | NA | NA | NA | NA | NA | #VALUE! |
| ENSMUSG00000079024.3 | 0 | NA | NA | NA | NA | NA | #VALUE! |
| ENSMUSG00000079025.10 | 0 | NA | NA | NA | NA | NA | #VALUE! |
| ENSMUSG00000079029.3 | 0 | NA | NA | NA | NA | NA | #VALUE! |
| ENSMUSG00000079031.2 | 0 | NA | NA | NA | NA | NA | #VALUE! |
| ENSMUSG00000079033.11 | 0 | NA | NA | NA | NA | NA | #VALUE! |
| ENSMUSG00000079034.3 | 0 | NA | NA | NA | NA | NA | #VALUE! |
| ENSMUSG00000079038.4 | 0 | NA | NA | NA | NA | NA | #VALUE! |
| ENSMUSG00000079039.2 | 0 | NA | NA | NA | NA | NA | #VALUE! |
| ENSMUSG00000079042.11 | 0 | NA | NA | NA | NA | NA | #VALUE! |
| ENSMUSG00000079048.9 | 0 | NA | NA | NA | NA | NA | #VALUE! |
| ENSMUSG00000079049.10 | 0 | NA | NA | NA | NA | NA | #VALUE! |
| ENSMUSG00000079051.6 | 0 | NA | NA | NA | NA | NA | #VALUE! |
| ENSMUSG00000079053.4 | 0 | NA | NA | NA | NA | NA | #VALUE! |
| ENSMUSG00000079055.11 | 0 | NA | NA | NA | NA | NA | #VALUE! |
| ENSMUSG00000079056.13 | 2.625328 | -4.93268 | 3.923662 | -1.25716 | 0.208695 | NA | #VALUE! |
| ENSMUSG00000079058.4 | 0 | NA | NA | NA | NA | NA | #VALUE! |
| ENSMUSG00000079061.4 | 0 | NA | NA | NA | NA | NA | #VALUE! |
| ENSMUSG00000079064.3 | 0 | NA | NA | NA | NA | NA | #VALUE! |
| ENSMUSG00000079067.6 | 0 | NA | NA | NA | NA | NA | #VALUE! |
| ENSMUSG00000079069.2 | 0 | NA | NA | NA | NA | NA | #VALUE! |
| ENSMUSG00000079070.11 | 0 | NA | NA | NA | NA | NA | #VALUE! |
| ENSMUSG00000079071.3 | 0 | NA | NA | NA | NA | NA | #VALUE! |
| ENSMUSG00000079076.3 | 0 | NA | NA | NA | NA | NA | #VALUE! |
| ENSMUSG00000079084.13 | 7.207409 | 0.175537 | 2.852759 | 0.061532 | 0.950935 | NA | #VALUE! |
| ENSMUSG00000079091.2 | 0 | NA | NA | NA | NA | NA | #VALUE! |
| ENSMUSG00000079092.5 | 0 | NA | NA | NA | NA | NA | #VALUE! |
| ENSMUSG00000079093.3 | 0 | NA | NA | NA | NA | NA | #VALUE! |
| ENSMUSG00000079101.4 | 0 | NA | NA | NA | NA | NA | #VALUE! |
| ENSMUSG00000079103.4 | 0 | NA | NA | NA | NA | NA | #VALUE! |
| ENSMUSG00000079105.5 | 0 | NA | NA | NA | NA | NA | #VALUE! |
| ENSMUSG00000079108.7 | 0.210026 | -1.52494 | 4.058692 | -0.37572 | 0.707123 | NA | #VALUE! |
| ENSMUSG00000079109.12 | 10.46361 | -1.44374 | 3.239647 | -0.44565 | NA | NA | #VALUE! |
| ENSMUSG00000079110.13 | 4.082215 | -5.55558 | 3.91775 | -1.41805 | 0.156175 | NA | #VALUE! |
| ENSMUSG00000079112.2 | 0 | NA | NA | NA | NA | NA | #VALUE! |
| ENSMUSG00000079113.4 | 0 | NA | NA | NA | NA | NA | #VALUE! |
| ENSMUSG00000079114.4 | 0 | NA | NA | NA | NA | NA | #VALUE! |
| ENSMUSG00000079116.2 | 0 | NA | NA | NA | NA | NA | #VALUE! |
| ENSMUSG00000079120.4 | 0 | NA | NA | NA | NA | NA | #VALUE! |
| ENSMUSG00000079137.4 | 0.168284 | 1.027628 | 4.067126 | 0.252667 | 0.800526 | NA | #VALUE! |
| ENSMUSG00000079138.4 | 0 | NA | NA | NA | NA | NA | #VALUE! |
| ENSMUSG00000079139.5 | 6.007173 | 0.56388 | 3.079299 | 0.183119 | 0.854704 | NA | #VALUE! |

| | | | | | | | |
|---|---|---|---|---|---|---|---|
| ENSMUSG00000079140.4 | 0 | NA | NA | NA | NA | NA | #VALUE! |
| ENSMUSG00000079143.4 | 0 | NA | NA | NA | NA | NA | #VALUE! |
| ENSMUSG00000079146.3 | 0 | NA | NA | NA | NA | NA | #VALUE! |
| ENSMUSG00000079156.12 | 0 | NA | NA | NA | NA | NA | #VALUE! |
| ENSMUSG00000079157.5 | 0 | NA | NA | NA | NA | NA | #VALUE! |
| ENSMUSG00000079162.9 | 0 | NA | NA | NA | NA | NA | #VALUE! |
| ENSMUSG00000079163.4 | 0 | NA | NA | NA | NA | NA | #VALUE! |
| ENSMUSG00000079165.11 | 0 | NA | NA | NA | NA | NA | #VALUE! |
| ENSMUSG00000079168.5 | 0 | NA | NA | NA | NA | NA | #VALUE! |
| ENSMUSG00000079169.4 | 0 | NA | NA | NA | NA | NA | #VALUE! |
| ENSMUSG00000079170.4 | 0 | NA | NA | NA | NA | NA | #VALUE! |
| ENSMUSG00000079171.4 | 0 | NA | NA | NA | NA | NA | #VALUE! |
| ENSMUSG00000079173.12 | 0 | NA | NA | NA | NA | NA | #VALUE! |
| ENSMUSG00000079175.2 | 0 | NA | NA | NA | NA | NA | #VALUE! |
| ENSMUSG00000079177.5 | 0 | NA | NA | NA | NA | NA | #VALUE! |
| ENSMUSG00000079180.5 | 0 | NA | NA | NA | NA | NA | #VALUE! |
| ENSMUSG00000079183.3 | 0 | NA | NA | NA | NA | NA | #VALUE! |
| ENSMUSG00000079186.4 | 0 | NA | NA | NA | NA | NA | #VALUE! |
| ENSMUSG00000079190.4 | 0 | NA | NA | NA | NA | NA | #VALUE! |
| ENSMUSG00000079192.3 | 0 | NA | NA | NA | NA | NA | #VALUE! |
| ENSMUSG00000079222.3 | 0 | NA | NA | NA | NA | NA | #VALUE! |
| ENSMUSG00000079223.4 | 0 | NA | NA | NA | NA | NA | #VALUE! |
| ENSMUSG00000079224.6 | 0 | NA | NA | NA | NA | NA | #VALUE! |
| ENSMUSG00000079230.2 | 0 | NA | NA | NA | NA | NA | #VALUE! |
| ENSMUSG00000079235.11 | 0 | NA | NA | NA | NA | NA | #VALUE! |
| ENSMUSG00000079242.3 | 0 | NA | NA | NA | NA | NA | #VALUE! |
| ENSMUSG00000079243.6 | 0 | NA | NA | NA | NA | NA | #VALUE! |
| ENSMUSG00000079244.4 | 0 | NA | NA | NA | NA | NA | #VALUE! |
| ENSMUSG00000079247.3 | 0 | NA | NA | NA | NA | NA | #VALUE! |
| ENSMUSG00000079258.3 | 0 | NA | NA | NA | NA | NA | #VALUE! |
| ENSMUSG00000079259.3 | 0 | NA | NA | NA | NA | NA | #VALUE! |
| ENSMUSG00000079260.4 | 0 | NA | NA | NA | NA | NA | #VALUE! |
| ENSMUSG00000079261.3 | 0 | NA | NA | NA | NA | NA | #VALUE! |
| ENSMUSG00000079262.4 | 0 | NA | NA | NA | NA | NA | #VALUE! |
| ENSMUSG00000079263.9 | 0 | NA | NA | NA | NA | NA | #VALUE! |
| ENSMUSG00000079264.2 | 0 | NA | NA | NA | NA | NA | #VALUE! |
| ENSMUSG00000079265.11 | 0 | NA | NA | NA | NA | NA | #VALUE! |
| ENSMUSG00000079266.3 | 0 | NA | NA | NA | NA | NA | #VALUE! |
| ENSMUSG00000079267.10 | 0 | NA | NA | NA | NA | NA | #VALUE! |
| ENSMUSG00000079269.3 | 0 | NA | NA | NA | NA | NA | #VALUE! |
| ENSMUSG00000079271.5 | 0 | NA | NA | NA | NA | NA | #VALUE! |
| ENSMUSG00000079272.3 | 0 | NA | NA | NA | NA | NA | #VALUE! |
| ENSMUSG00000079273.3 | 0 | NA | NA | NA | NA | NA | #VALUE! |
| ENSMUSG00000079276.5 | 0 | NA | NA | NA | NA | NA | #VALUE! |

| | | | | | | | |
|---|---|---|---|---|---|---|---|
| ENSMUSG00000079277.10 | 0 | NA | NA | NA | NA | NA | #VALUE! |
| ENSMUSG00000079278.3 | 0 | NA | NA | NA | NA | NA | #VALUE! |
| ENSMUSG00000079286.2 | 0 | NA | NA | NA | NA | NA | #VALUE! |
| ENSMUSG00000079295.6 | 0 | NA | NA | NA | NA | NA | #VALUE! |
| ENSMUSG00000079297.5 | 0 | NA | NA | NA | NA | NA | #VALUE! |
| ENSMUSG00000079298.10 | 0 | NA | NA | NA | NA | NA | #VALUE! |
| ENSMUSG00000079299.4 | 0 | NA | NA | NA | NA | NA | #VALUE! |
| ENSMUSG00000079304.6 | 0 | NA | NA | NA | NA | NA | #VALUE! |
| ENSMUSG00000079311.2 | 0 | NA | NA | NA | NA | NA | #VALUE! |
| ENSMUSG00000079323.10 | 0 | NA | NA | NA | NA | NA | #VALUE! |
| ENSMUSG00000079324.12 | 0 | NA | NA | NA | NA | NA | #VALUE! |
| ENSMUSG00000079330.10 | 0 | NA | NA | NA | NA | NA | #VALUE! |
| ENSMUSG00000079333.4 | 0 | NA | NA | NA | NA | NA | #VALUE! |
| ENSMUSG00000079339.8 | 0.630079 | -2.90291 | 3.975047 | -0.73028 | 0.465217 | NA | #VALUE! |
| ENSMUSG00000079342.12 | 0 | NA | NA | NA | NA | NA | #VALUE! |
| ENSMUSG00000079343.5 | 0 | NA | NA | NA | NA | NA | #VALUE! |
| ENSMUSG00000079344.12 | 0 | NA | NA | NA | NA | NA | #VALUE! |
| ENSMUSG00000079346.5 | 0 | NA | NA | NA | NA | NA | #VALUE! |
| ENSMUSG00000079349.10 | 0 | NA | NA | NA | NA | NA | #VALUE! |
| ENSMUSG00000079350.3 | 0 | NA | NA | NA | NA | NA | #VALUE! |
| ENSMUSG00000079353.3 | 0 | NA | NA | NA | NA | NA | #VALUE! |
| ENSMUSG00000079355.6 | 2.203867 | -3.42547 | 3.915786 | -0.87479 | 0.381691 | NA | #VALUE! |
| ENSMUSG00000079356.11 | 0 | NA | NA | NA | NA | NA | #VALUE! |
| ENSMUSG00000079359.3 | 0 | NA | NA | NA | NA | NA | #VALUE! |
| ENSMUSG00000079362.6 | 0 | NA | NA | NA | NA | NA | #VALUE! |
| ENSMUSG00000079363.8 | 0.210026 | -1.52494 | 4.058692 | -0.37572 | 0.707123 | NA | #VALUE! |
| ENSMUSG00000079364.4 | 0 | NA | NA | NA | NA | NA | #VALUE! |
| ENSMUSG00000079371.10 | 0 | NA | NA | NA | NA | NA | #VALUE! |
| ENSMUSG00000079374.3 | 0 | NA | NA | NA | NA | NA | #VALUE! |
| ENSMUSG00000079378.9 | 0 | NA | NA | NA | NA | NA | #VALUE! |
| ENSMUSG00000079379.3 | 0 | NA | NA | NA | NA | NA | #VALUE! |
| ENSMUSG00000079380.5 | 0 | NA | NA | NA | NA | NA | #VALUE! |
| ENSMUSG00000079382.4 | 0 | NA | NA | NA | NA | NA | #VALUE! |
| ENSMUSG00000079383.10 | 0 | NA | NA | NA | NA | NA | #VALUE! |
| ENSMUSG00000079386.9 | 0 | NA | NA | NA | NA | NA | #VALUE! |
| ENSMUSG00000079387.11 | 0 | NA | NA | NA | NA | NA | #VALUE! |
| ENSMUSG00000079388.10 | 0 | NA | NA | NA | NA | NA | #VALUE! |
| ENSMUSG00000079389.11 | 0 | NA | NA | NA | NA | NA | #VALUE! |
| ENSMUSG00000079391.11 | 0 | NA | NA | NA | NA | NA | #VALUE! |
| ENSMUSG00000079392.4 | 0 | NA | NA | NA | NA | NA | #VALUE! |
| ENSMUSG00000079395.4 | 0 | NA | NA | NA | NA | NA | #VALUE! |
| ENSMUSG00000079396.5 | 0 | NA | NA | NA | NA | NA | #VALUE! |
| ENSMUSG00000079402.10 | 0 | NA | NA | NA | NA | NA | #VALUE! |
| ENSMUSG00000079407.10 | 0 | NA | NA | NA | NA | NA | #VALUE! |

| | | | | | | | |
|---|---|---|---|---|---|---|---|
| ENSMUSG00000079409.9 | 0 | NA | NA | NA | NA | NA | #VALUE! |
| ENSMUSG00000079410.5 | 0 | NA | NA | NA | NA | NA | #VALUE! |
| ENSMUSG00000079414.4 | 0 | NA | NA | NA | NA | NA | #VALUE! |
| ENSMUSG00000079415.3 | 0 | NA | NA | NA | NA | NA | #VALUE! |
| ENSMUSG00000079416.2 | 0 | NA | NA | NA | NA | NA | #VALUE! |
| ENSMUSG00000079421.9 | 0 | NA | NA | NA | NA | NA | #VALUE! |
| ENSMUSG00000079423.7 | 0 | NA | NA | NA | NA | NA | #VALUE! |
| ENSMUSG00000079424.9 | 0 | NA | NA | NA | NA | NA | #VALUE! |
| ENSMUSG00000079428.9 | 0 | NA | NA | NA | NA | NA | #VALUE! |
| ENSMUSG00000079432.2 | 0 | NA | NA | NA | NA | NA | #VALUE! |
| ENSMUSG00000079433.2 | 0 | NA | NA | NA | NA | NA | #VALUE! |
| ENSMUSG00000079434.9 | 0 | NA | NA | NA | NA | NA | #VALUE! |
| ENSMUSG00000079436.4 | 0 | NA | NA | NA | NA | NA | #VALUE! |
| ENSMUSG00000079438.2 | 0 | NA | NA | NA | NA | NA | #VALUE! |
| ENSMUSG00000079439.2 | 0 | NA | NA | NA | NA | NA | #VALUE! |
| ENSMUSG00000079440.3 | 0 | NA | NA | NA | NA | NA | #VALUE! |
| ENSMUSG00000079442.13 | 0 | NA | NA | NA | NA | NA | #VALUE! |
| ENSMUSG00000079444.10 | 0 | NA | NA | NA | NA | NA | #VALUE! |
| ENSMUSG00000079445.4 | 0 | NA | NA | NA | NA | NA | #VALUE! |
| ENSMUSG00000079450.12 | 0.630079 | -2.90291 | 3.975047 | -0.73028 | 0.465217 | NA | #VALUE! |
| ENSMUSG00000079451.10 | 0 | NA | NA | NA | NA | NA | #VALUE! |
| ENSMUSG00000079455.10 | 0 | NA | NA | NA | NA | NA | #VALUE! |
| ENSMUSG00000079457.12 | 0 | NA | NA | NA | NA | NA | #VALUE! |
| ENSMUSG00000079460.12 | 0 | NA | NA | NA | NA | NA | #VALUE! |
| ENSMUSG00000079462.2 | 0 | NA | NA | NA | NA | NA | #VALUE! |
| ENSMUSG00000079465.9 | 0.245976 | 1.027628 | 4.067126 | 0.252667 | 0.800526 | NA | #VALUE! |
| ENSMUSG00000079466.3 | 0 | NA | NA | NA | NA | NA | #VALUE! |
| ENSMUSG00000079467.9 | 0 | NA | NA | NA | NA | NA | #VALUE! |
| ENSMUSG00000079471.5 | 0 | NA | NA | NA | NA | NA | #VALUE! |
| ENSMUSG00000079476.2 | 0 | NA | NA | NA | NA | NA | #VALUE! |
| ENSMUSG00000079478.10 | 0 | NA | NA | NA | NA | NA | #VALUE! |
| ENSMUSG00000079479.3 | 0 | NA | NA | NA | NA | NA | #VALUE! |
| ENSMUSG00000079481.12 | 2.51789 | 4.834703 | 3.924413 | 1.231956 | 0.217966 | NA | #VALUE! |
| ENSMUSG00000079484.13 | 4.043916 | -5.54733 | 3.917813 | -1.41593 | 0.156797 | NA | #VALUE! |
| ENSMUSG00000079487.12 | 0 | NA | NA | NA | NA | NA | #VALUE! |
| ENSMUSG00000079491.10 | 0 | NA | NA | NA | NA | NA | #VALUE! |
| ENSMUSG00000079492.3 | 0 | NA | NA | NA | NA | NA | #VALUE! |
| ENSMUSG00000079495.3 | 1.858277 | 3.215098 | 3.926082 | 0.818907 | 0.412839 | NA | #VALUE! |
| ENSMUSG00000079497.5 | 0 | NA | NA | NA | NA | NA | #VALUE! |
| ENSMUSG00000079499.10 | 0 | NA | NA | NA | NA | NA | #VALUE! |
| ENSMUSG00000079501.4 | 0 | NA | NA | NA | NA | NA | #VALUE! |
| ENSMUSG00000079502.9 | 0 | NA | NA | NA | NA | NA | #VALUE! |
| ENSMUSG00000079505.3 | 5.216789 | 5.873026 | 3.915376 | 1.49999 | 0.133617 | NA | #VALUE! |
| ENSMUSG00000079507.11 | 4.97073 | 4.729945 | 3.665097 | 1.290538 | 0.196864 | NA | #VALUE! |

| | | | | | | | |
|---|---|---|---|---|---|---|---|
| ENSMUSG00000079511.9 | 0 | NA | NA | NA | NA | NA | #VALUE! |
| ENSMUSG00000079513.2 | 0 | NA | NA | NA | NA | NA | #VALUE! |
| ENSMUSG00000079516.3 | 0 | NA | NA | NA | NA | NA | #VALUE! |
| ENSMUSG00000079519.3 | 0 | NA | NA | NA | NA | NA | #VALUE! |
| ENSMUSG00000079521.3 | 0 | NA | NA | NA | NA | NA | #VALUE! |
| ENSMUSG00000079522.3 | 0 | NA | NA | NA | NA | NA | #VALUE! |
| ENSMUSG00000079523.9 | 3.431417 | 0.276291 | 3.779618 | 0.0731 | 0.941726 | NA | #VALUE! |
| ENSMUSG00000079524.3 | 0 | NA | NA | NA | NA | NA | #VALUE! |
| ENSMUSG00000079525.2 | 0 | NA | NA | NA | NA | NA | #VALUE! |
| ENSMUSG00000079528.10 | 0 | NA | NA | NA | NA | NA | #VALUE! |
| ENSMUSG00000079531.2 | 0 | NA | NA | NA | NA | NA | #VALUE! |
| ENSMUSG00000079532.2 | 0 | NA | NA | NA | NA | NA | #VALUE! |
| ENSMUSG00000079534.2 | 0 | NA | NA | NA | NA | NA | #VALUE! |
| ENSMUSG00000079536.2 | 0 | NA | NA | NA | NA | NA | #VALUE! |
| ENSMUSG00000079539.3 | 0 | NA | NA | NA | NA | NA | #VALUE! |
| ENSMUSG00000079542.4 | 0 | NA | NA | NA | NA | NA | #VALUE! |
| ENSMUSG00000079543.4 | 0 | NA | NA | NA | NA | NA | #VALUE! |
| ENSMUSG00000079544.3 | 0 | NA | NA | NA | NA | NA | #VALUE! |
| ENSMUSG00000079546.2 | 0 | NA | NA | NA | NA | NA | #VALUE! |
| ENSMUSG00000079547.5 | 3.197682 | 5.156529 | 3.920897 | 1.31514 | 0.188463 | NA | #VALUE! |
| ENSMUSG00000079550.10 | 5.636842 | 3.5783 | 3.839099 | 0.932068 | 0.351302 | NA | #VALUE! |
| ENSMUSG00000079553.12 | 0 | NA | NA | NA | NA | NA | #VALUE! |
| ENSMUSG00000079554.3 | 0 | NA | NA | NA | NA | NA | #VALUE! |
| ENSMUSG00000079559.6 | 0 | NA | NA | NA | NA | NA | #VALUE! |
| ENSMUSG00000079560.14 | 0 | NA | NA | NA | NA | NA | #VALUE! |
| ENSMUSG00000079564.4 | 0 | NA | NA | NA | NA | NA | #VALUE! |
| ENSMUSG00000079566.2 | 0 | NA | NA | NA | NA | NA | #VALUE! |
| ENSMUSG00000079575.4 | 0 | NA | NA | NA | NA | NA | #VALUE! |
| ENSMUSG00000079577.2 | 0 | NA | NA | NA | NA | NA | #VALUE! |
| ENSMUSG00000079578.2 | 0 | NA | NA | NA | NA | NA | #VALUE! |
| ENSMUSG00000079579.4 | 0 | NA | NA | NA | NA | NA | #VALUE! |
| ENSMUSG00000079580.10 | 0 | NA | NA | NA | NA | NA | #VALUE! |
| ENSMUSG00000079583.10 | 0 | NA | NA | NA | NA | NA | #VALUE! |
| ENSMUSG00000079584.3 | 0 | NA | NA | NA | NA | NA | #VALUE! |
| ENSMUSG00000079588.4 | 0 | NA | NA | NA | NA | NA | #VALUE! |
| ENSMUSG00000079592.10 | 0 | NA | NA | NA | NA | NA | #VALUE! |
| ENSMUSG00000079593.12 | 0 | NA | NA | NA | NA | NA | #VALUE! |
| ENSMUSG00000079594.3 | 0 | NA | NA | NA | NA | NA | #VALUE! |
| ENSMUSG00000079595.3 | 0 | NA | NA | NA | NA | NA | #VALUE! |
| ENSMUSG00000079597.3 | 0.504851 | 2.502843 | 3.993993 | 0.626652 | 0.530887 | NA | #VALUE! |
| ENSMUSG00000079598.5 | 0 | NA | NA | NA | NA | NA | #VALUE! |
| ENSMUSG00000079600.3 | 0 | NA | NA | NA | NA | NA | #VALUE! |
| ENSMUSG00000079602.3 | 0 | NA | NA | NA | NA | NA | #VALUE! |
| ENSMUSG00000079603.4 | 0 | NA | NA | NA | NA | NA | #VALUE! |

| | | | | | | | |
|---|---|---|---|---|---|---|---|
| ENSMUSG00000079604.10 | 0 | NA | NA | NA | NA | NA | #VALUE! |
| ENSMUSG00000079605.6 | 0 | NA | NA | NA | NA | NA | #VALUE! |
| ENSMUSG00000079606.2 | 0 | NA | NA | NA | NA | NA | #VALUE! |
| ENSMUSG00000079608.11 | 0 | NA | NA | NA | NA | NA | #VALUE! |
| ENSMUSG00000079610.10 | 4.490437 | -5.69342 | 3.916751 | -1.45361 | 0.146055 | NA | #VALUE! |
| ENSMUSG00000079619.3 | 0 | NA | NA | NA | NA | NA | #VALUE! |
| ENSMUSG00000079625.3 | 0 | NA | NA | NA | NA | NA | #VALUE! |
| ENSMUSG00000079626.3 | 0 | NA | NA | NA | NA | NA | #VALUE! |
| ENSMUSG00000079627.3 | 0 | NA | NA | NA | NA | NA | #VALUE! |
| ENSMUSG00000079628.4 | 0 | NA | NA | NA | NA | NA | #VALUE! |
| ENSMUSG00000079629.6 | 0 | NA | NA | NA | NA | NA | #VALUE! |
| ENSMUSG00000079630.3 | 0 | NA | NA | NA | NA | NA | #VALUE! |
| ENSMUSG00000079633.4 | 0 | NA | NA | NA | NA | NA | #VALUE! |
| ENSMUSG00000079635.3 | 0 | NA | NA | NA | NA | NA | #VALUE! |
| ENSMUSG00000079636.12 | 0 | NA | NA | NA | NA | NA | #VALUE! |
| ENSMUSG00000079637.11 | 0 | NA | NA | NA | NA | NA | #VALUE! |
| ENSMUSG00000079638.9 | 0 | NA | NA | NA | NA | NA | #VALUE! |
| ENSMUSG00000079639.5 | 0 | NA | NA | NA | NA | NA | #VALUE! |
| ENSMUSG00000079642.3 | 0 | NA | NA | NA | NA | NA | #VALUE! |
| ENSMUSG00000079644.3 | 0 | NA | NA | NA | NA | NA | #VALUE! |
| ENSMUSG00000079645.3 | 0 | NA | NA | NA | NA | NA | #VALUE! |
| ENSMUSG00000079649.4 | 0 | NA | NA | NA | NA | NA | #VALUE! |
| ENSMUSG00000079652.9 | 0 | NA | NA | NA | NA | NA | #VALUE! |
| ENSMUSG00000079654.4 | 0 | NA | NA | NA | NA | NA | #VALUE! |
| ENSMUSG00000079655.10 | 0 | NA | NA | NA | NA | NA | #VALUE! |
| ENSMUSG00000079657.11 | 0 | NA | NA | NA | NA | NA | #VALUE! |
| ENSMUSG00000079666.9 | 0 | NA | NA | NA | NA | NA | #VALUE! |
| ENSMUSG00000079669.3 | 0 | NA | NA | NA | NA | NA | #VALUE! |
| ENSMUSG00000079671.9 | 6.996491 | -0.9813 | 3.236506 | -0.3032 | 0.761739 | NA | #VALUE! |
| ENSMUSG00000079677.3 | 0 | NA | NA | NA | NA | NA | #VALUE! |
| ENSMUSG00000079679.4 | 0 | NA | NA | NA | NA | NA | #VALUE! |
| ENSMUSG00000079681.4 | 0 | NA | NA | NA | NA | NA | #VALUE! |
| ENSMUSG00000079685.11 | 0 | NA | NA | NA | NA | NA | #VALUE! |
| ENSMUSG00000079693.4 | 0 | NA | NA | NA | NA | NA | #VALUE! |
| ENSMUSG00000079694.3 | 0 | NA | NA | NA | NA | NA | #VALUE! |
| ENSMUSG00000079697.2 | 0 | NA | NA | NA | NA | NA | #VALUE! |
| ENSMUSG00000079698.8 | 0 | NA | NA | NA | NA | NA | #VALUE! |
| ENSMUSG00000079699.3 | 0 | NA | NA | NA | NA | NA | #VALUE! |
| ENSMUSG00000079700.8 | 0 | NA | NA | NA | NA | NA | #VALUE! |
| ENSMUSG00000079701.8 | 0 | NA | NA | NA | NA | NA | #VALUE! |
| ENSMUSG00000079702.6 | 0 | NA | NA | NA | NA | NA | #VALUE! |
| ENSMUSG00000079703.13 | 0 | NA | NA | NA | NA | NA | #VALUE! |
| ENSMUSG00000079704.3 | 0 | NA | NA | NA | NA | NA | #VALUE! |
| ENSMUSG00000079705.5 | 0 | NA | NA | NA | NA | NA | #VALUE! |

| | | | | | | | |
|---|---|---|---|---|---|---|---|
| ENSMUSG00000079707.11 | 0 | NA | NA | NA | NA | NA | #VALUE! |
| ENSMUSG00000079710.11 | 0 | NA | NA | NA | NA | NA | #VALUE! |
| ENSMUSG00000079711.5 | 0 | NA | NA | NA | NA | NA | #VALUE! |
| ENSMUSG00000079722.3 | 0 | NA | NA | NA | NA | NA | #VALUE! |
| ENSMUSG00000079737.5 | 3.935608 | 5.458007 | 3.918239 | 1.392975 | 0.163627 | NA | #VALUE! |
| ENSMUSG00000079740.2 | 0 | NA | NA | NA | NA | NA | #VALUE! |
| ENSMUSG00000079741.6 | 0 | NA | NA | NA | NA | NA | #VALUE! |
| ENSMUSG00000079794.3 | 0 | NA | NA | NA | NA | NA | #VALUE! |
| ENSMUSG00000079800.3 | 0 | NA | NA | NA | NA | NA | #VALUE! |
| ENSMUSG00000079806.3 | 0 | NA | NA | NA | NA | NA | #VALUE! |
| ENSMUSG00000079808.4 | 0 | NA | NA | NA | NA | NA | #VALUE! |
| ENSMUSG00000079842.2 | 0 | NA | NA | NA | NA | NA | #VALUE! |
| ENSMUSG00000079845.9 | 6.735655 | -6.27951 | 3.913422 | -1.60461 | 0.10858 | NA | #VALUE! |
| ENSMUSG00000079852.5 | 0.105013 | -0.97324 | 4.076293 | -0.23876 | 0.811295 | NA | #VALUE! |
| ENSMUSG00000079853.6 | 2.144869 | 4.605508 | 3.927435 | 1.17265 | 0.240936 | NA | #VALUE! |
| ENSMUSG00000079871.5 | 0 | NA | NA | NA | NA | NA | #VALUE! |
| ENSMUSG00000079875.6 | 0 | NA | NA | NA | NA | NA | #VALUE! |
| ENSMUSG00000079884.7 | 0 | NA | NA | NA | NA | NA | #VALUE! |
| ENSMUSG00000079889.5 | 0 | NA | NA | NA | NA | NA | #VALUE! |
| ENSMUSG00000079890.3 | 0 | NA | NA | NA | NA | NA | #VALUE! |
| ENSMUSG00000079892.3 | 0 | NA | NA | NA | NA | NA | #VALUE! |
| ENSMUSG00000079893.4 | 0 | NA | NA | NA | NA | NA | #VALUE! |
| ENSMUSG00000079906.3 | 0 | NA | NA | NA | NA | NA | #VALUE! |
| ENSMUSG00000079915.3 | 0 | NA | NA | NA | NA | NA | #VALUE! |
| ENSMUSG00000079962.3 | 0 | NA | NA | NA | NA | NA | #VALUE! |
| ENSMUSG00000079963.3 | 0 | NA | NA | NA | NA | NA | #VALUE! |
| ENSMUSG00000079965.3 | 0 | NA | NA | NA | NA | NA | #VALUE! |
| ENSMUSG00000079966.3 | 0 | NA | NA | NA | NA | NA | #VALUE! |
| ENSMUSG00000079988.3 | 0.670387 | 1.434266 | 3.989858 | 0.359478 | 0.719237 | NA | #VALUE! |
| ENSMUSG00000079993.3 | 0 | NA | NA | NA | NA | NA | #VALUE! |
| ENSMUSG00000079998.3 | 0 | NA | NA | NA | NA | NA | #VALUE! |
| ENSMUSG00000080002.3 | 0 | NA | NA | NA | NA | NA | #VALUE! |
| ENSMUSG00000080021.6 | 0 | NA | NA | NA | NA | NA | #VALUE! |
| ENSMUSG00000080046.3 | 0 | NA | NA | NA | NA | NA | #VALUE! |
| ENSMUSG00000080050.3 | 0 | NA | NA | NA | NA | NA | #VALUE! |
| ENSMUSG00000080051.3 | 0 | NA | NA | NA | NA | NA | #VALUE! |
| ENSMUSG00000080054.4 | 3.234572 | -1.15492 | 3.792145 | -0.30455 | 0.760705 | NA | #VALUE! |
| ENSMUSG00000080057.3 | 0 | NA | NA | NA | NA | NA | #VALUE! |
| ENSMUSG00000080058.3 | 0 | NA | NA | NA | NA | NA | #VALUE! |
| ENSMUSG00000080059.5 | 0.245976 | 1.027628 | 4.067126 | 0.252667 | 0.800526 | NA | #VALUE! |
| ENSMUSG00000080065.3 | 0 | NA | NA | NA | NA | NA | #VALUE! |
| ENSMUSG00000080069.3 | 0 | NA | NA | NA | NA | NA | #VALUE! |
| ENSMUSG00000080071.3 | 0 | NA | NA | NA | NA | NA | #VALUE! |
| ENSMUSG00000080072.3 | 0 | NA | NA | NA | NA | NA | #VALUE! |

| | | | | | | | |
|---|---|---|---|---|---|---|---|
| ENSMUSG00000080073.3 | 0 | NA | NA | NA | NA | NA | #VALUE! |
| ENSMUSG00000080076.5 | 0 | NA | NA | NA | NA | NA | #VALUE! |
| ENSMUSG00000080078.3 | 0 | NA | NA | NA | NA | NA | #VALUE! |
| ENSMUSG00000080115.4 | 0 | NA | NA | NA | NA | NA | #VALUE! |
| ENSMUSG00000080142.3 | 0 | NA | NA | NA | NA | NA | #VALUE! |
| ENSMUSG00000080152.6 | 0 | NA | NA | NA | NA | NA | #VALUE! |
| ENSMUSG00000080155.3 | 0 | NA | NA | NA | NA | NA | #VALUE! |
| ENSMUSG00000080158.3 | 0 | NA | NA | NA | NA | NA | #VALUE! |
| ENSMUSG00000080175.3 | 0 | NA | NA | NA | NA | NA | #VALUE! |
| ENSMUSG00000080180.4 | 0 | NA | NA | NA | NA | NA | #VALUE! |
| ENSMUSG00000080186.3 | 0 | NA | NA | NA | NA | NA | #VALUE! |
| ENSMUSG00000080193.3 | 0 | NA | NA | NA | NA | NA | #VALUE! |
| ENSMUSG00000080198.3 | 0.466276 | 2.449889 | 3.997179 | 0.612904 | 0.53994 | NA | #VALUE! |
| ENSMUSG00000080205.3 | 0 | NA | NA | NA | NA | NA | #VALUE! |
| ENSMUSG00000080222.3 | 0 | NA | NA | NA | NA | NA | #VALUE! |
| ENSMUSG00000080223.3 | 0 | NA | NA | NA | NA | NA | #VALUE! |
| ENSMUSG00000080227.3 | 0 | NA | NA | NA | NA | NA | #VALUE! |
| ENSMUSG00000080237.3 | 0 | NA | NA | NA | NA | NA | #VALUE! |
| ENSMUSG00000080241.4 | 0 | NA | NA | NA | NA | NA | #VALUE! |
| ENSMUSG00000080242.6 | 0 | NA | NA | NA | NA | NA | #VALUE! |
| ENSMUSG00000080250.3 | 0 | NA | NA | NA | NA | NA | #VALUE! |
| ENSMUSG00000080254.5 | 0 | NA | NA | NA | NA | NA | #VALUE! |
| ENSMUSG00000080266.3 | 0 | NA | NA | NA | NA | NA | #VALUE! |
| ENSMUSG00000080268.5 | 0 | NA | NA | NA | NA | NA | #VALUE! |
| ENSMUSG00000080283.3 | 0 | NA | NA | NA | NA | NA | #VALUE! |
| ENSMUSG00000080298.3 | 0 | NA | NA | NA | NA | NA | #VALUE! |
| ENSMUSG00000080299.3 | 0 | NA | NA | NA | NA | NA | #VALUE! |
| ENSMUSG00000080300.5 | 0 | NA | NA | NA | NA | NA | #VALUE! |
| ENSMUSG00000080303.3 | 0 | NA | NA | NA | NA | NA | #VALUE! |
| ENSMUSG00000080316.12 | 2.423806 | 3.136761 | 3.904571 | 0.803356 | 0.421769 | NA | #VALUE! |
| ENSMUSG00000080317.4 | 0 | NA | NA | NA | NA | NA | #VALUE! |
| ENSMUSG00000080321.3 | 0 | NA | NA | NA | NA | NA | #VALUE! |
| ENSMUSG00000080331.3 | 0 | NA | NA | NA | NA | NA | #VALUE! |
| ENSMUSG00000080352.3 | 0 | NA | NA | NA | NA | NA | #VALUE! |
| ENSMUSG00000080356.3 | 0 | NA | NA | NA | NA | NA | #VALUE! |
| ENSMUSG00000080358.3 | 0 | NA | NA | NA | NA | NA | #VALUE! |
| ENSMUSG00000080364.3 | 0 | NA | NA | NA | NA | NA | #VALUE! |
| ENSMUSG00000080365.3 | 0 | NA | NA | NA | NA | NA | #VALUE! |
| ENSMUSG00000080374.3 | 0 | NA | NA | NA | NA | NA | #VALUE! |
| ENSMUSG00000080378.4 | 0 | NA | NA | NA | NA | NA | #VALUE! |
| ENSMUSG00000080384.3 | 0 | NA | NA | NA | NA | NA | #VALUE! |
| ENSMUSG00000080396.3 | 0 | NA | NA | NA | NA | NA | #VALUE! |
| ENSMUSG00000080406.3 | 0 | NA | NA | NA | NA | NA | #VALUE! |
| ENSMUSG00000080409.3 | 0 | NA | NA | NA | NA | NA | #VALUE! |

| | | | | | | | |
|---|---|---|---|---|---|---|---|
| ENSMUSG00000080411.3 | 0 | NA | NA | NA | NA | NA | #VALUE! |
| ENSMUSG00000080413.3 | 0 | NA | NA | NA | NA | NA | #VALUE! |
| ENSMUSG00000080424.3 | 0 | NA | NA | NA | NA | NA | #VALUE! |
| ENSMUSG00000080425.3 | 0 | NA | NA | NA | NA | NA | #VALUE! |
| ENSMUSG00000080428.3 | 0 | NA | NA | NA | NA | NA | #VALUE! |
| ENSMUSG00000080437.3 | 0 | NA | NA | NA | NA | NA | #VALUE! |
| ENSMUSG00000080440.3 | 0 | NA | NA | NA | NA | NA | #VALUE! |
| ENSMUSG00000080441.3 | 0 | NA | NA | NA | NA | NA | #VALUE! |
| ENSMUSG00000080452.5 | 0 | NA | NA | NA | NA | NA | #VALUE! |
| ENSMUSG00000080454.6 | 0 | NA | NA | NA | NA | NA | #VALUE! |
| ENSMUSG00000080459.3 | 0 | NA | NA | NA | NA | NA | #VALUE! |
| ENSMUSG00000080461.3 | 0 | NA | NA | NA | NA | NA | #VALUE! |
| ENSMUSG00000080463.3 | 0 | NA | NA | NA | NA | NA | #VALUE! |
| ENSMUSG00000080465.3 | 0 | NA | NA | NA | NA | NA | #VALUE! |
| ENSMUSG00000080468.4 | 0 | NA | NA | NA | NA | NA | #VALUE! |
| ENSMUSG00000080469.3 | 0 | NA | NA | NA | NA | NA | #VALUE! |
| ENSMUSG00000080478.3 | 0 | NA | NA | NA | NA | NA | #VALUE! |
| ENSMUSG00000080480.3 | 0 | NA | NA | NA | NA | NA | #VALUE! |
| ENSMUSG00000080499.3 | 0 | NA | NA | NA | NA | NA | #VALUE! |
| ENSMUSG00000080504.3 | 0 | NA | NA | NA | NA | NA | #VALUE! |
| ENSMUSG00000080517.3 | 0 | NA | NA | NA | NA | NA | #VALUE! |
| ENSMUSG00000080518.3 | 0 | NA | NA | NA | NA | NA | #VALUE! |
| ENSMUSG00000080524.3 | 0 | NA | NA | NA | NA | NA | #VALUE! |
| ENSMUSG00000080526.3 | 0 | NA | NA | NA | NA | NA | #VALUE! |
| ENSMUSG00000080533.3 | 0 | NA | NA | NA | NA | NA | #VALUE! |
| ENSMUSG00000080536.3 | 0 | NA | NA | NA | NA | NA | #VALUE! |
| ENSMUSG00000080538.3 | 0 | NA | NA | NA | NA | NA | #VALUE! |
| ENSMUSG00000080539.4 | 0 | NA | NA | NA | NA | NA | #VALUE! |
| ENSMUSG00000080540.4 | 0 | NA | NA | NA | NA | NA | #VALUE! |
| ENSMUSG00000080542.3 | 0 | NA | NA | NA | NA | NA | #VALUE! |
| ENSMUSG00000080545.3 | 0 | NA | NA | NA | NA | NA | #VALUE! |
| ENSMUSG00000080566.3 | 0 | NA | NA | NA | NA | NA | #VALUE! |
| ENSMUSG00000080573.3 | 0 | NA | NA | NA | NA | NA | #VALUE! |
| ENSMUSG00000080574.3 | 0 | NA | NA | NA | NA | NA | #VALUE! |
| ENSMUSG00000080575.4 | 0 | NA | NA | NA | NA | NA | #VALUE! |
| ENSMUSG00000080577.3 | 0 | NA | NA | NA | NA | NA | #VALUE! |
| ENSMUSG00000080580.3 | 0 | NA | NA | NA | NA | NA | #VALUE! |
| ENSMUSG00000080582.3 | 0 | NA | NA | NA | NA | NA | #VALUE! |
| ENSMUSG00000080586.3 | 0 | NA | NA | NA | NA | NA | #VALUE! |
| ENSMUSG00000080594.3 | 0 | NA | NA | NA | NA | NA | #VALUE! |
| ENSMUSG00000080595.4 | 0 | NA | NA | NA | NA | NA | #VALUE! |
| ENSMUSG00000080607.3 | 0 | NA | NA | NA | NA | NA | #VALUE! |
| ENSMUSG00000080610.3 | 0 | NA | NA | NA | NA | NA | #VALUE! |
| ENSMUSG00000080615.3 | 0 | NA | NA | NA | NA | NA | #VALUE! |

| | | | | | | | |
|---|---|---|---|---|---|---|---|
| ENSMUSG00000080616.3 | 0 | NA | NA | NA | NA | NA | #VALUE! |
| ENSMUSG00000080622.3 | 0 | NA | NA | NA | NA | NA | #VALUE! |
| ENSMUSG00000080626.3 | 0 | NA | NA | NA | NA | NA | #VALUE! |
| ENSMUSG00000080645.3 | 0 | NA | NA | NA | NA | NA | #VALUE! |
| ENSMUSG00000080653.3 | 0 | NA | NA | NA | NA | NA | #VALUE! |
| ENSMUSG00000080657.3 | 0 | NA | NA | NA | NA | NA | #VALUE! |
| ENSMUSG00000080662.2 | 0 | NA | NA | NA | NA | NA | #VALUE! |
| ENSMUSG00000080665.3 | 0 | NA | NA | NA | NA | NA | #VALUE! |
| ENSMUSG00000080669.3 | 0 | NA | NA | NA | NA | NA | #VALUE! |
| ENSMUSG00000080673.3 | 0 | NA | NA | NA | NA | NA | #VALUE! |
| ENSMUSG00000080683.3 | 0 | NA | NA | NA | NA | NA | #VALUE! |
| ENSMUSG00000080687.3 | 0 | NA | NA | NA | NA | NA | #VALUE! |
| ENSMUSG00000080689.3 | 0 | NA | NA | NA | NA | NA | #VALUE! |
| ENSMUSG00000080694.2 | 0 | NA | NA | NA | NA | NA | #VALUE! |
| ENSMUSG00000080695.2 | 0 | NA | NA | NA | NA | NA | #VALUE! |
| ENSMUSG00000080696.2 | 0 | NA | NA | NA | NA | NA | #VALUE! |
| ENSMUSG00000080697.2 | 0 | NA | NA | NA | NA | NA | #VALUE! |
| ENSMUSG00000080698.2 | 0 | NA | NA | NA | NA | NA | #VALUE! |
| ENSMUSG00000080700.3 | 0 | NA | NA | NA | NA | NA | #VALUE! |
| ENSMUSG00000080701.2 | 0 | NA | NA | NA | NA | NA | #VALUE! |
| ENSMUSG00000080703.3 | 0 | NA | NA | NA | NA | NA | #VALUE! |
| ENSMUSG00000080704.2 | 0 | NA | NA | NA | NA | NA | #VALUE! |
| ENSMUSG00000080705.2 | 0 | NA | NA | NA | NA | NA | #VALUE! |
| ENSMUSG00000080706.2 | 0 | NA | NA | NA | NA | NA | #VALUE! |
| ENSMUSG00000080707.2 | 0 | NA | NA | NA | NA | NA | #VALUE! |
| ENSMUSG00000080708.2 | 0 | NA | NA | NA | NA | NA | #VALUE! |
| ENSMUSG00000080709.2 | 0 | NA | NA | NA | NA | NA | #VALUE! |
| ENSMUSG00000080710.2 | 0 | NA | NA | NA | NA | NA | #VALUE! |
| ENSMUSG00000080711.4 | 0 | NA | NA | NA | NA | NA | #VALUE! |
| ENSMUSG00000080712.5 | 0 | NA | NA | NA | NA | NA | #VALUE! |
| ENSMUSG00000080713.4 | 0 | NA | NA | NA | NA | NA | #VALUE! |
| ENSMUSG00000080714.2 | 0 | NA | NA | NA | NA | NA | #VALUE! |
| ENSMUSG00000080715.2 | 0 | NA | NA | NA | NA | NA | #VALUE! |
| ENSMUSG00000080716.2 | 0 | NA | NA | NA | NA | NA | #VALUE! |
| ENSMUSG00000080718.2 | 0 | NA | NA | NA | NA | NA | #VALUE! |
| ENSMUSG00000080719.2 | 0 | NA | NA | NA | NA | NA | #VALUE! |
| ENSMUSG00000080720.3 | 0 | NA | NA | NA | NA | NA | #VALUE! |
| ENSMUSG00000080722.4 | 0 | NA | NA | NA | NA | NA | #VALUE! |
| ENSMUSG00000080723.2 | 0 | NA | NA | NA | NA | NA | #VALUE! |
| ENSMUSG00000080725.10 | 0 | NA | NA | NA | NA | NA | #VALUE! |
| ENSMUSG00000080726.3 | 0 | NA | NA | NA | NA | NA | #VALUE! |
| ENSMUSG00000080728.2 | 0 | NA | NA | NA | NA | NA | #VALUE! |
| ENSMUSG00000080729.2 | 0 | NA | NA | NA | NA | NA | #VALUE! |
| ENSMUSG00000080730.2 | 0 | NA | NA | NA | NA | NA | #VALUE! |

| | | | | | | | |
|---|---|---|---|---|---|---|---|
| ENSMUSG00000080731.3 | 0.195789 | -1.05581 | 4.073302 | -0.2592 | 0.795479 | NA | #VALUE! |
| ENSMUSG00000080732.2 | 0 | NA | NA | NA | NA | NA | #VALUE! |
| ENSMUSG00000080733.3 | 0 | NA | NA | NA | NA | NA | #VALUE! |
| ENSMUSG00000080734.2 | 0 | NA | NA | NA | NA | NA | #VALUE! |
| ENSMUSG00000080735.2 | 0 | NA | NA | NA | NA | NA | #VALUE! |
| ENSMUSG00000080736.2 | 0 | NA | NA | NA | NA | NA | #VALUE! |
| ENSMUSG00000080737.2 | 0 | NA | NA | NA | NA | NA | #VALUE! |
| ENSMUSG00000080741.4 | 0 | NA | NA | NA | NA | NA | #VALUE! |
| ENSMUSG00000080742.2 | 0 | NA | NA | NA | NA | NA | #VALUE! |
| ENSMUSG00000080743.3 | 0 | NA | NA | NA | NA | NA | #VALUE! |
| ENSMUSG00000080744.2 | 0 | NA | NA | NA | NA | NA | #VALUE! |
| ENSMUSG00000080746.4 | 0.402379 | -0.92565 | 4.039907 | -0.22913 | 0.81877 | NA | #VALUE! |
| ENSMUSG00000080747.2 | 0 | NA | NA | NA | NA | NA | #VALUE! |
| ENSMUSG00000080748.2 | 0.168284 | 1.027628 | 4.067126 | 0.252667 | 0.800526 | NA | #VALUE! |
| ENSMUSG00000080749.2 | 0 | NA | NA | NA | NA | NA | #VALUE! |
| ENSMUSG00000080750.3 | 0 | NA | NA | NA | NA | NA | #VALUE! |
| ENSMUSG00000080751.2 | 0 | NA | NA | NA | NA | NA | #VALUE! |
| ENSMUSG00000080752.3 | 0 | NA | NA | NA | NA | NA | #VALUE! |
| ENSMUSG00000080753.2 | 0 | NA | NA | NA | NA | NA | #VALUE! |
| ENSMUSG00000080754.2 | 0 | NA | NA | NA | NA | NA | #VALUE! |
| ENSMUSG00000080755.2 | 0 | NA | NA | NA | NA | NA | #VALUE! |
| ENSMUSG00000080756.2 | 0 | NA | NA | NA | NA | NA | #VALUE! |
| ENSMUSG00000080757.2 | 0 | NA | NA | NA | NA | NA | #VALUE! |
| ENSMUSG00000080758.2 | 0 | NA | NA | NA | NA | NA | #VALUE! |
| ENSMUSG00000080759.2 | 0 | NA | NA | NA | NA | NA | #VALUE! |
| ENSMUSG00000080760.2 | 0 | NA | NA | NA | NA | NA | #VALUE! |
| ENSMUSG00000080762.2 | 0 | NA | NA | NA | NA | NA | #VALUE! |
| ENSMUSG00000080763.3 | 0 | NA | NA | NA | NA | NA | #VALUE! |
| ENSMUSG00000080764.2 | 0 | NA | NA | NA | NA | NA | #VALUE! |
| ENSMUSG00000080765.2 | 0 | NA | NA | NA | NA | NA | #VALUE! |
| ENSMUSG00000080766.2 | 0 | NA | NA | NA | NA | NA | #VALUE! |
| ENSMUSG00000080767.2 | 0 | NA | NA | NA | NA | NA | #VALUE! |
| ENSMUSG00000080768.2 | 0 | NA | NA | NA | NA | NA | #VALUE! |
| ENSMUSG00000080770.2 | 0 | NA | NA | NA | NA | NA | #VALUE! |
| ENSMUSG00000080771.2 | 0 | NA | NA | NA | NA | NA | #VALUE! |
| ENSMUSG00000080772.2 | 5.090517 | -5.87548 | 3.91557 | -1.50054 | 0.133474 | NA | #VALUE! |
| ENSMUSG00000080773.2 | 0 | NA | NA | NA | NA | NA | #VALUE! |
| ENSMUSG00000080774.2 | 0 | NA | NA | NA | NA | NA | #VALUE! |
| ENSMUSG00000080775.2 | 0.471565 | 0.203595 | 4.021039 | 0.050632 | 0.959618 | NA | #VALUE! |
| ENSMUSG00000080776.2 | 5.357352 | 1.29224 | 2.455456 | 0.526273 | 0.598699 | NA | #VALUE! |
| ENSMUSG00000080777.2 | 0 | NA | NA | NA | NA | NA | #VALUE! |
| ENSMUSG00000080778.2 | 0 | NA | NA | NA | NA | NA | #VALUE! |
| ENSMUSG00000080779.2 | 0.245976 | 1.027628 | 4.067126 | 0.252667 | 0.800526 | NA | #VALUE! |
| ENSMUSG00000080780.2 | 0 | NA | NA | NA | NA | NA | #VALUE! |

| | | | | | | | |
|---|---|---|---|---|---|---|---|
| ENSMUSG00000080781.4 | 0 | NA | NA | NA | NA | NA | #VALUE! |
| ENSMUSG00000080782.2 | 0 | NA | NA | NA | NA | NA | #VALUE! |
| ENSMUSG00000080783.2 | 0 | NA | NA | NA | NA | NA | #VALUE! |
| ENSMUSG00000080785.2 | 0 | NA | NA | NA | NA | NA | #VALUE! |
| ENSMUSG00000080788.2 | 0 | NA | NA | NA | NA | NA | #VALUE! |
| ENSMUSG00000080789.2 | 0 | NA | NA | NA | NA | NA | #VALUE! |
| ENSMUSG00000080790.2 | 0 | NA | NA | NA | NA | NA | #VALUE! |
| ENSMUSG00000080791.2 | 0 | NA | NA | NA | NA | NA | #VALUE! |
| ENSMUSG00000080792.2 | 0 | NA | NA | NA | NA | NA | #VALUE! |
| ENSMUSG00000080793.2 | 0 | NA | NA | NA | NA | NA | #VALUE! |
| ENSMUSG00000080794.2 | 0 | NA | NA | NA | NA | NA | #VALUE! |
| ENSMUSG00000080795.2 | 0 | NA | NA | NA | NA | NA | #VALUE! |
| ENSMUSG00000080796.2 | 0 | NA | NA | NA | NA | NA | #VALUE! |
| ENSMUSG00000080797.3 | 4.141803 | 5.546926 | 3.917555 | 1.415915 | 0.1568 | NA | #VALUE! |
| ENSMUSG00000080798.4 | 0 | NA | NA | NA | NA | NA | #VALUE! |
| ENSMUSG00000080799.2 | 0 | NA | NA | NA | NA | NA | #VALUE! |
| ENSMUSG00000080800.2 | 0 | NA | NA | NA | NA | NA | #VALUE! |
| ENSMUSG00000080801.3 | 0 | NA | NA | NA | NA | NA | #VALUE! |
| ENSMUSG00000080802.2 | 0 | NA | NA | NA | NA | NA | #VALUE! |
| ENSMUSG00000080803.2 | 0 | NA | NA | NA | NA | NA | #VALUE! |
| ENSMUSG00000080805.2 | 0 | NA | NA | NA | NA | NA | #VALUE! |
| ENSMUSG00000080806.2 | 0 | NA | NA | NA | NA | NA | #VALUE! |
| ENSMUSG00000080807.2 | 0 | NA | NA | NA | NA | NA | #VALUE! |
| ENSMUSG00000080808.2 | 0 | NA | NA | NA | NA | NA | #VALUE! |
| ENSMUSG00000080809.4 | 0 | NA | NA | NA | NA | NA | #VALUE! |
| ENSMUSG00000080810.2 | 1.009701 | 3.503687 | 3.950852 | 0.886818 | 0.375177 | NA | #VALUE! |
| ENSMUSG00000080811.2 | 0.105013 | -0.97324 | 4.076293 | -0.23876 | 0.811295 | NA | #VALUE! |
| ENSMUSG00000080812.2 | 0 | NA | NA | NA | NA | NA | #VALUE! |
| ENSMUSG00000080813.2 | 0 | NA | NA | NA | NA | NA | #VALUE! |
| ENSMUSG00000080814.2 | 0 | NA | NA | NA | NA | NA | #VALUE! |
| ENSMUSG00000080815.2 | 0 | NA | NA | NA | NA | NA | #VALUE! |
| ENSMUSG00000080816.2 | 0 | NA | NA | NA | NA | NA | #VALUE! |
| ENSMUSG00000080817.2 | 0 | NA | NA | NA | NA | NA | #VALUE! |
| ENSMUSG00000080818.5 | 0 | NA | NA | NA | NA | NA | #VALUE! |
| ENSMUSG00000080819.2 | 0 | NA | NA | NA | NA | NA | #VALUE! |
| ENSMUSG00000080820.2 | 0 | NA | NA | NA | NA | NA | #VALUE! |
| ENSMUSG00000080822.2 | 0 | NA | NA | NA | NA | NA | #VALUE! |
| ENSMUSG00000080823.2 | 2.27115 | 0.020388 | 3.470667 | 0.005874 | 0.995313 | NA | #VALUE! |
| ENSMUSG00000080824.2 | 0.105013 | -0.97324 | 4.076293 | -0.23876 | 0.811295 | NA | #VALUE! |
| ENSMUSG00000080825.2 | 0 | NA | NA | NA | NA | NA | #VALUE! |
| ENSMUSG00000080826.2 | 0 | NA | NA | NA | NA | NA | #VALUE! |
| ENSMUSG00000080827.2 | 0 | NA | NA | NA | NA | NA | #VALUE! |
| ENSMUSG00000080828.2 | 0 | NA | NA | NA | NA | NA | #VALUE! |
| ENSMUSG00000080829.2 | 0 | NA | NA | NA | NA | NA | #VALUE! |

| | | | | | | | |
|---|---|---|---|---|---|---|---|
| ENSMUSG00000080831.2 | 0 | NA | NA | NA | NA | NA | #VALUE! |
| ENSMUSG00000080832.5 | 0 | NA | NA | NA | NA | NA | #VALUE! |
| ENSMUSG00000080833.2 | 0 | NA | NA | NA | NA | NA | #VALUE! |
| ENSMUSG00000080834.2 | 0 | NA | NA | NA | NA | NA | #VALUE! |
| ENSMUSG00000080835.2 | 0 | NA | NA | NA | NA | NA | #VALUE! |
| ENSMUSG00000080836.2 | 6.894784 | -2.34362 | 3.670984 | -0.63842 | 0.523202 | NA | #VALUE! |
| ENSMUSG00000080839.2 | 0 | NA | NA | NA | NA | NA | #VALUE! |
| ENSMUSG00000080840.2 | 0 | NA | NA | NA | NA | NA | #VALUE! |
| ENSMUSG00000080841.2 | 0 | NA | NA | NA | NA | NA | #VALUE! |
| ENSMUSG00000080842.3 | 0 | NA | NA | NA | NA | NA | #VALUE! |
| ENSMUSG00000080843.2 | 0 | NA | NA | NA | NA | NA | #VALUE! |
| ENSMUSG00000080844.2 | 0 | NA | NA | NA | NA | NA | #VALUE! |
| ENSMUSG00000080845.2 | 0 | NA | NA | NA | NA | NA | #VALUE! |
| ENSMUSG00000080846.2 | 0 | NA | NA | NA | NA | NA | #VALUE! |
| ENSMUSG00000080847.2 | 0 | NA | NA | NA | NA | NA | #VALUE! |
| ENSMUSG00000080849.2 | 0 | NA | NA | NA | NA | NA | #VALUE! |
| ENSMUSG00000080850.2 | 0 | NA | NA | NA | NA | NA | #VALUE! |
| ENSMUSG00000080851.2 | 0 | NA | NA | NA | NA | NA | #VALUE! |
| ENSMUSG00000080852.2 | 0 | NA | NA | NA | NA | NA | #VALUE! |
| ENSMUSG00000080855.4 | 0 | NA | NA | NA | NA | NA | #VALUE! |
| ENSMUSG00000080857.2 | 0 | NA | NA | NA | NA | NA | #VALUE! |
| ENSMUSG00000080859.2 | 0 | NA | NA | NA | NA | NA | #VALUE! |
| ENSMUSG00000080860.2 | 0.168284 | 1.027628 | 4.067126 | 0.252667 | 0.800526 | NA | #VALUE! |
| ENSMUSG00000080861.2 | 0 | NA | NA | NA | NA | NA | #VALUE! |
| ENSMUSG00000080862.4 | 0 | NA | NA | NA | NA | NA | #VALUE! |
| ENSMUSG00000080863.2 | 0 | NA | NA | NA | NA | NA | #VALUE! |
| ENSMUSG00000080865.2 | 0.093255 | 0.868093 | 4.072464 | 0.213162 | 0.831201 | NA | #VALUE! |
| ENSMUSG00000080866.2 | 0.093255 | 0.868093 | 4.072464 | 0.213162 | 0.831201 | NA | #VALUE! |
| ENSMUSG00000080868.2 | 0.093255 | 0.868093 | 4.072464 | 0.213162 | 0.831201 | NA | #VALUE! |
| ENSMUSG00000080869.2 | 0 | NA | NA | NA | NA | NA | #VALUE! |
| ENSMUSG00000080870.2 | 0 | NA | NA | NA | NA | NA | #VALUE! |
| ENSMUSG00000080871.2 | 0 | NA | NA | NA | NA | NA | #VALUE! |
| ENSMUSG00000080872.2 | 0 | NA | NA | NA | NA | NA | #VALUE! |
| ENSMUSG00000080873.4 | 0.975827 | 2.089354 | 3.957291 | 0.527976 | 0.597516 | NA | #VALUE! |
| ENSMUSG00000080874.2 | 0 | NA | NA | NA | NA | NA | #VALUE! |
| ENSMUSG00000080875.5 | 0 | NA | NA | NA | NA | NA | #VALUE! |
| ENSMUSG00000080876.4 | 0 | NA | NA | NA | NA | NA | #VALUE! |
| ENSMUSG00000080878.2 | 0 | NA | NA | NA | NA | NA | #VALUE! |
| ENSMUSG00000080880.2 | 0 | NA | NA | NA | NA | NA | #VALUE! |
| ENSMUSG00000080884.2 | 0 | NA | NA | NA | NA | NA | #VALUE! |
| ENSMUSG00000080885.2 | 0 | NA | NA | NA | NA | NA | #VALUE! |
| ENSMUSG00000080886.2 | 0 | NA | NA | NA | NA | NA | #VALUE! |
| ENSMUSG00000080887.2 | 0 | NA | NA | NA | NA | NA | #VALUE! |
| ENSMUSG00000080888.4 | 0 | NA | NA | NA | NA | NA | #VALUE! |

| | | | | | | | |
|---|---|---|---|---|---|---|---|
| ENSMUSG00000080889.2 | 0 | NA | NA | NA | NA | NA | #VALUE! |
| ENSMUSG00000080890.2 | 0 | NA | NA | NA | NA | NA | #VALUE! |
| ENSMUSG00000080891.5 | 0 | NA | NA | NA | NA | NA | #VALUE! |
| ENSMUSG00000080892.2 | 0 | NA | NA | NA | NA | NA | #VALUE! |
| ENSMUSG00000080893.2 | 1.524519 | 1.061919 | 3.862425 | 0.274936 | 0.783365 | NA | #VALUE! |
| ENSMUSG00000080894.2 | 0 | NA | NA | NA | NA | NA | #VALUE! |
| ENSMUSG00000080895.2 | 0 | NA | NA | NA | NA | NA | #VALUE! |
| ENSMUSG00000080896.6 | 0.822481 | 0.012393 | 3.951017 | 0.003137 | 0.997497 | NA | #VALUE! |
| ENSMUSG00000080897.2 | 0 | NA | NA | NA | NA | NA | #VALUE! |
| ENSMUSG00000080898.2 | 0 | NA | NA | NA | NA | NA | #VALUE! |
| ENSMUSG00000080899.2 | 0 | NA | NA | NA | NA | NA | #VALUE! |
| ENSMUSG00000080902.2 | 0 | NA | NA | NA | NA | NA | #VALUE! |
| ENSMUSG00000080903.2 | 0 | NA | NA | NA | NA | NA | #VALUE! |
| ENSMUSG00000080906.2 | 0 | NA | NA | NA | NA | NA | #VALUE! |
| ENSMUSG00000080907.2 | 0 | NA | NA | NA | NA | NA | #VALUE! |
| ENSMUSG00000080910.2 | 0 | NA | NA | NA | NA | NA | #VALUE! |
| ENSMUSG00000080911.2 | 0 | NA | NA | NA | NA | NA | #VALUE! |
| ENSMUSG00000080912.2 | 0 | NA | NA | NA | NA | NA | #VALUE! |
| ENSMUSG00000080913.2 | 0 | NA | NA | NA | NA | NA | #VALUE! |
| ENSMUSG00000080914.2 | 0 | NA | NA | NA | NA | NA | #VALUE! |
| ENSMUSG00000080915.2 | 0 | NA | NA | NA | NA | NA | #VALUE! |
| ENSMUSG00000080916.4 | 0 | NA | NA | NA | NA | NA | #VALUE! |
| ENSMUSG00000080917.3 | 0 | NA | NA | NA | NA | NA | #VALUE! |
| ENSMUSG00000080918.2 | 0 | NA | NA | NA | NA | NA | #VALUE! |
| ENSMUSG00000080919.2 | 0 | NA | NA | NA | NA | NA | #VALUE! |
| ENSMUSG00000080921.5 | 2.611131 | -2.33559 | 3.852766 | -0.60621 | 0.544374 | NA | #VALUE! |
| ENSMUSG00000080922.2 | 0 | NA | NA | NA | NA | NA | #VALUE! |
| ENSMUSG00000080923.2 | 0 | NA | NA | NA | NA | NA | #VALUE! |
| ENSMUSG00000080924.2 | 0 | NA | NA | NA | NA | NA | #VALUE! |
| ENSMUSG00000080925.2 | 0 | NA | NA | NA | NA | NA | #VALUE! |
| ENSMUSG00000080927.2 | 0 | NA | NA | NA | NA | NA | #VALUE! |
| ENSMUSG00000080928.2 | 0.273297 | 0.094602 | 4.062933 | 0.023284 | 0.981424 | NA | #VALUE! |
| ENSMUSG00000080929.2 | 0 | NA | NA | NA | NA | NA | #VALUE! |
| ENSMUSG00000080930.3 | 0 | NA | NA | NA | NA | NA | #VALUE! |
| ENSMUSG00000080932.4 | 5.925524 | -2.10562 | 3.140976 | -0.67037 | 0.502621 | NA | #VALUE! |
| ENSMUSG00000080933.5 | 0 | NA | NA | NA | NA | NA | #VALUE! |
| ENSMUSG00000080934.2 | 0 | NA | NA | NA | NA | NA | #VALUE! |
| ENSMUSG00000080935.4 | 0 | NA | NA | NA | NA | NA | #VALUE! |
| ENSMUSG00000080936.3 | 0 | NA | NA | NA | NA | NA | #VALUE! |
| ENSMUSG00000080937.2 | 0 | NA | NA | NA | NA | NA | #VALUE! |
| ENSMUSG00000080938.2 | 0 | NA | NA | NA | NA | NA | #VALUE! |
| ENSMUSG00000080939.2 | 0 | NA | NA | NA | NA | NA | #VALUE! |
| ENSMUSG00000080941.2 | 0 | NA | NA | NA | NA | NA | #VALUE! |
| ENSMUSG00000080942.2 | 0 | NA | NA | NA | NA | NA | #VALUE! |

| | | | | | | | |
|---|---|---|---|---|---|---|---|
| ENSMUSG00000080943.2 | 0 | NA | NA | NA | NA | NA | #VALUE! |
| ENSMUSG00000080944.2 | 0 | NA | NA | NA | NA | NA | #VALUE! |
| ENSMUSG00000080946.2 | 0 | NA | NA | NA | NA | NA | #VALUE! |
| ENSMUSG00000080947.3 | 0 | NA | NA | NA | NA | NA | #VALUE! |
| ENSMUSG00000080948.2 | 0 | NA | NA | NA | NA | NA | #VALUE! |
| ENSMUSG00000080949.2 | 0 | NA | NA | NA | NA | NA | #VALUE! |
| ENSMUSG00000080950.2 | 0 | NA | NA | NA | NA | NA | #VALUE! |
| ENSMUSG00000080951.2 | 0 | NA | NA | NA | NA | NA | #VALUE! |
| ENSMUSG00000080952.2 | 0 | NA | NA | NA | NA | NA | #VALUE! |
| ENSMUSG00000080953.2 | 0 | NA | NA | NA | NA | NA | #VALUE! |
| ENSMUSG00000080954.2 | 0 | NA | NA | NA | NA | NA | #VALUE! |
| ENSMUSG00000080955.2 | 0 | NA | NA | NA | NA | NA | #VALUE! |
| ENSMUSG00000080956.2 | 0 | NA | NA | NA | NA | NA | #VALUE! |
| ENSMUSG00000080957.2 | 0 | NA | NA | NA | NA | NA | #VALUE! |
| ENSMUSG00000080958.2 | 0 | NA | NA | NA | NA | NA | #VALUE! |
| ENSMUSG00000080959.2 | 0 | NA | NA | NA | NA | NA | #VALUE! |
| ENSMUSG00000080960.2 | 0 | NA | NA | NA | NA | NA | #VALUE! |
| ENSMUSG00000080961.2 | 2.882611 | -1.25359 | 3.801558 | -0.32976 | 0.741585 | NA | #VALUE! |
| ENSMUSG00000080962.2 | 0 | NA | NA | NA | NA | NA | #VALUE! |
| ENSMUSG00000080964.2 | 0 | NA | NA | NA | NA | NA | #VALUE! |
| ENSMUSG00000080965.2 | 0 | NA | NA | NA | NA | NA | #VALUE! |
| ENSMUSG00000080966.2 | 0 | NA | NA | NA | NA | NA | #VALUE! |
| ENSMUSG00000080967.2 | 0 | NA | NA | NA | NA | NA | #VALUE! |
| ENSMUSG00000080968.4 | 0 | NA | NA | NA | NA | NA | #VALUE! |
| ENSMUSG00000080969.2 | 0 | NA | NA | NA | NA | NA | #VALUE! |
| ENSMUSG00000080970.2 | 0 | NA | NA | NA | NA | NA | #VALUE! |
| ENSMUSG00000080971.2 | 0 | NA | NA | NA | NA | NA | #VALUE! |
| ENSMUSG00000080972.2 | 0 | NA | NA | NA | NA | NA | #VALUE! |
| ENSMUSG00000080974.3 | 0 | NA | NA | NA | NA | NA | #VALUE! |
| ENSMUSG00000080976.2 | 0 | NA | NA | NA | NA | NA | #VALUE! |
| ENSMUSG00000080977.2 | 0.204111 | -1.05581 | 4.073302 | -0.2592 | 0.795479 | NA | #VALUE! |
| ENSMUSG00000080978.3 | 0 | NA | NA | NA | NA | NA | #VALUE! |
| ENSMUSG00000080979.4 | 0.372394 | 0.012025 | 4.059931 | 0.002962 | 0.997637 | NA | #VALUE! |
| ENSMUSG00000080980.2 | 0 | NA | NA | NA | NA | NA | #VALUE! |
| ENSMUSG00000080981.2 | 0 | NA | NA | NA | NA | NA | #VALUE! |
| ENSMUSG00000080982.2 | 0 | NA | NA | NA | NA | NA | #VALUE! |
| ENSMUSG00000080983.2 | 0 | NA | NA | NA | NA | NA | #VALUE! |
| ENSMUSG00000080984.2 | 0 | NA | NA | NA | NA | NA | #VALUE! |
| ENSMUSG00000080985.2 | 0 | NA | NA | NA | NA | NA | #VALUE! |
| ENSMUSG00000080986.2 | 0 | NA | NA | NA | NA | NA | #VALUE! |
| ENSMUSG00000080987.2 | 0 | NA | NA | NA | NA | NA | #VALUE! |
| ENSMUSG00000080989.2 | 0.792753 | 0.636055 | 3.969089 | 0.160252 | 0.872682 | NA | #VALUE! |
| ENSMUSG00000080990.3 | 0 | NA | NA | NA | NA | NA | #VALUE! |
| ENSMUSG00000080992.2 | 0 | NA | NA | NA | NA | NA | #VALUE! |

| | | | | | | | |
|---|---|---|---|---|---|---|---|
| ENSMUSG00000080993.2 | 0 | NA | NA | NA | NA | NA | #VALUE! |
| ENSMUSG00000080994.3 | 0 | NA | NA | NA | NA | NA | #VALUE! |
| ENSMUSG00000080995.2 | 0 | NA | NA | NA | NA | NA | #VALUE! |
| ENSMUSG00000080996.4 | 1.966531 | -0.9384 | 3.83004 | -0.24501 | 0.806448 | NA | #VALUE! |
| ENSMUSG00000080997.2 | 0 | NA | NA | NA | NA | NA | #VALUE! |
| ENSMUSG00000080998.2 | 0 | NA | NA | NA | NA | NA | #VALUE! |
| ENSMUSG00000080999.2 | 0 | NA | NA | NA | NA | NA | #VALUE! |
| ENSMUSG00000081000.2 | 0 | NA | NA | NA | NA | NA | #VALUE! |
| ENSMUSG00000081001.2 | 0 | NA | NA | NA | NA | NA | #VALUE! |
| ENSMUSG00000081002.2 | 0 | NA | NA | NA | NA | NA | #VALUE! |
| ENSMUSG00000081003.2 | 0 | NA | NA | NA | NA | NA | #VALUE! |
| ENSMUSG00000081004.2 | 0.195789 | -1.05581 | 4.073302 | -0.2592 | 0.795479 | NA | #VALUE! |
| ENSMUSG00000081005.2 | 0 | NA | NA | NA | NA | NA | #VALUE! |
| ENSMUSG00000081006.2 | 0 | NA | NA | NA | NA | NA | #VALUE! |
| ENSMUSG00000081007.2 | 0 | NA | NA | NA | NA | NA | #VALUE! |
| ENSMUSG00000081008.3 | 0 | NA | NA | NA | NA | NA | #VALUE! |
| ENSMUSG00000081009.2 | 0 | NA | NA | NA | NA | NA | #VALUE! |
| ENSMUSG00000081010.4 | 0 | NA | NA | NA | NA | NA | #VALUE! |
| ENSMUSG00000081011.2 | 0 | NA | NA | NA | NA | NA | #VALUE! |
| ENSMUSG00000081012.2 | 0 | NA | NA | NA | NA | NA | #VALUE! |
| ENSMUSG00000081013.2 | 0 | NA | NA | NA | NA | NA | #VALUE! |
| ENSMUSG00000081014.2 | 0 | NA | NA | NA | NA | NA | #VALUE! |
| ENSMUSG00000081016.2 | 0 | NA | NA | NA | NA | NA | #VALUE! |
| ENSMUSG00000081017.2 | 0 | NA | NA | NA | NA | NA | #VALUE! |
| ENSMUSG00000081018.2 | 0 | NA | NA | NA | NA | NA | #VALUE! |
| ENSMUSG00000081020.2 | 0 | NA | NA | NA | NA | NA | #VALUE! |
| ENSMUSG00000081021.2 | 0 | NA | NA | NA | NA | NA | #VALUE! |
| ENSMUSG00000081022.2 | 0 | NA | NA | NA | NA | NA | #VALUE! |
| ENSMUSG00000081023.2 | 0 | NA | NA | NA | NA | NA | #VALUE! |
| ENSMUSG00000081024.2 | 0 | NA | NA | NA | NA | NA | #VALUE! |
| ENSMUSG00000081025.2 | 0 | NA | NA | NA | NA | NA | #VALUE! |
| ENSMUSG00000081026.2 | 0 | NA | NA | NA | NA | NA | #VALUE! |
| ENSMUSG00000081029.2 | 0 | NA | NA | NA | NA | NA | #VALUE! |
| ENSMUSG00000081030.2 | 0 | NA | NA | NA | NA | NA | #VALUE! |
| ENSMUSG00000081033.2 | 0 | NA | NA | NA | NA | NA | #VALUE! |
| ENSMUSG00000081034.2 | 0.105013 | -0.97324 | 4.076293 | -0.23876 | 0.811295 | NA | #VALUE! |
| ENSMUSG00000081036.2 | 0 | NA | NA | NA | NA | NA | #VALUE! |
| ENSMUSG00000081037.4 | 0 | NA | NA | NA | NA | NA | #VALUE! |
| ENSMUSG00000081038.3 | 0 | NA | NA | NA | NA | NA | #VALUE! |
| ENSMUSG00000081041.2 | 0 | NA | NA | NA | NA | NA | #VALUE! |
| ENSMUSG00000081042.2 | 0 | NA | NA | NA | NA | NA | #VALUE! |
| ENSMUSG00000081043.2 | 0 | NA | NA | NA | NA | NA | #VALUE! |
| ENSMUSG00000081044.9 | 0 | NA | NA | NA | NA | NA | #VALUE! |
| ENSMUSG00000081045.2 | 0 | NA | NA | NA | NA | NA | #VALUE! |

| | | | | | | | |
|---|---|---|---|---|---|---|---|
| ENSMUSG00000081046.2 | 0 | NA | NA | NA | NA | NA | #VALUE! |
| ENSMUSG00000081047.3 | 0 | NA | NA | NA | NA | NA | #VALUE! |
| ENSMUSG00000081048.2 | 0 | NA | NA | NA | NA | NA | #VALUE! |
| ENSMUSG00000081050.2 | 0 | NA | NA | NA | NA | NA | #VALUE! |
| ENSMUSG00000081052.2 | 0 | NA | NA | NA | NA | NA | #VALUE! |
| ENSMUSG00000081053.2 | 0.195789 | -1.05581 | 4.073302 | -0.2592 | 0.795479 | NA | #VALUE! |
| ENSMUSG00000081054.2 | 0 | NA | NA | NA | NA | NA | #VALUE! |
| ENSMUSG00000081055.2 | 0 | NA | NA | NA | NA | NA | #VALUE! |
| ENSMUSG00000081056.2 | 0 | NA | NA | NA | NA | NA | #VALUE! |
| ENSMUSG00000081057.3 | 0 | NA | NA | NA | NA | NA | #VALUE! |
| ENSMUSG00000081059.4 | 1.356296 | 2.654574 | 3.938679 | 0.673976 | 0.500327 | NA | #VALUE! |
| ENSMUSG00000081060.2 | 0 | NA | NA | NA | NA | NA | #VALUE! |
| ENSMUSG00000081061.2 | 0 | NA | NA | NA | NA | NA | #VALUE! |
| ENSMUSG00000081062.2 | 0 | NA | NA | NA | NA | NA | #VALUE! |
| ENSMUSG00000081063.2 | 0 | NA | NA | NA | NA | NA | #VALUE! |
| ENSMUSG00000081064.2 | 0 | NA | NA | NA | NA | NA | #VALUE! |
| ENSMUSG00000081065.2 | 0 | NA | NA | NA | NA | NA | #VALUE! |
| ENSMUSG00000081066.2 | 0 | NA | NA | NA | NA | NA | #VALUE! |
| ENSMUSG00000081067.2 | 0 | NA | NA | NA | NA | NA | #VALUE! |
| ENSMUSG00000081068.2 | 0 | NA | NA | NA | NA | NA | #VALUE! |
| ENSMUSG00000081069.2 | 0 | NA | NA | NA | NA | NA | #VALUE! |
| ENSMUSG00000081070.2 | 0 | NA | NA | NA | NA | NA | #VALUE! |
| ENSMUSG00000081071.2 | 0 | NA | NA | NA | NA | NA | #VALUE! |
| ENSMUSG00000081072.2 | 0 | NA | NA | NA | NA | NA | #VALUE! |
| ENSMUSG00000081073.2 | 0 | NA | NA | NA | NA | NA | #VALUE! |
| ENSMUSG00000081076.4 | 0 | NA | NA | NA | NA | NA | #VALUE! |
| ENSMUSG00000081077.2 | 0 | NA | NA | NA | NA | NA | #VALUE! |
| ENSMUSG00000081078.2 | 0 | NA | NA | NA | NA | NA | #VALUE! |
| ENSMUSG00000081079.2 | 0 | NA | NA | NA | NA | NA | #VALUE! |
| ENSMUSG00000081080.2 | 0 | NA | NA | NA | NA | NA | #VALUE! |
| ENSMUSG00000081081.2 | 0 | NA | NA | NA | NA | NA | #VALUE! |
| ENSMUSG00000081082.2 | 0 | NA | NA | NA | NA | NA | #VALUE! |
| ENSMUSG00000081083.2 | 0 | NA | NA | NA | NA | NA | #VALUE! |
| ENSMUSG00000081084.2 | 0 | NA | NA | NA | NA | NA | #VALUE! |
| ENSMUSG00000081085.2 | 0 | NA | NA | NA | NA | NA | #VALUE! |
| ENSMUSG00000081086.2 | 0 | NA | NA | NA | NA | NA | #VALUE! |
| ENSMUSG00000081087.2 | 1.25256 | 1.029236 | 3.890651 | 0.264541 | 0.791363 | NA | #VALUE! |
| ENSMUSG00000081088.2 | 0 | NA | NA | NA | NA | NA | #VALUE! |
| ENSMUSG00000081089.2 | 0 | NA | NA | NA | NA | NA | #VALUE! |
| ENSMUSG00000081092.2 | 0 | NA | NA | NA | NA | NA | #VALUE! |
| ENSMUSG00000081093.2 | 0 | NA | NA | NA | NA | NA | #VALUE! |
| ENSMUSG00000081094.5 | 1.057606 | -2.17087 | 3.947268 | -0.54997 | 0.582342 | NA | #VALUE! |
| ENSMUSG00000081095.3 | 0 | NA | NA | NA | NA | NA | #VALUE! |
| ENSMUSG00000081096.3 | 0 | NA | NA | NA | NA | NA | #VALUE! |

| | | | | | | | |
|---|---|---|---|---|---|---|---|
| ENSMUSG00000081097.2 | 0 | NA | NA | NA | NA | NA | #VALUE! |
| ENSMUSG00000081098.2 | 0 | NA | NA | NA | NA | NA | #VALUE! |
| ENSMUSG00000081099.3 | 0 | NA | NA | NA | NA | NA | #VALUE! |
| ENSMUSG00000081100.2 | 0.384779 | 0.875002 | 4.032432 | 0.216991 | 0.828215 | NA | #VALUE! |
| ENSMUSG00000081101.2 | 0 | NA | NA | NA | NA | NA | #VALUE! |
| ENSMUSG00000081103.2 | 0 | NA | NA | NA | NA | NA | #VALUE! |
| ENSMUSG00000081104.2 | 0 | NA | NA | NA | NA | NA | #VALUE! |
| ENSMUSG00000081106.2 | 0 | NA | NA | NA | NA | NA | #VALUE! |
| ENSMUSG00000081107.2 | 0 | NA | NA | NA | NA | NA | #VALUE! |
| ENSMUSG00000081108.5 | 0 | NA | NA | NA | NA | NA | #VALUE! |
| ENSMUSG00000081110.2 | 0 | NA | NA | NA | NA | NA | #VALUE! |
| ENSMUSG00000081111.4 | 0 | NA | NA | NA | NA | NA | #VALUE! |
| ENSMUSG00000081112.2 | 0 | NA | NA | NA | NA | NA | #VALUE! |
| ENSMUSG00000081113.2 | 0.105013 | -0.97324 | 4.076293 | -0.23876 | 0.811295 | NA | #VALUE! |
| ENSMUSG00000081114.2 | 0 | NA | NA | NA | NA | NA | #VALUE! |
| ENSMUSG00000081115.3 | 0 | NA | NA | NA | NA | NA | #VALUE! |
| ENSMUSG00000081116.2 | 0 | NA | NA | NA | NA | NA | #VALUE! |
| ENSMUSG00000081117.2 | 0 | NA | NA | NA | NA | NA | #VALUE! |
| ENSMUSG00000081118.2 | 0 | NA | NA | NA | NA | NA | #VALUE! |
| ENSMUSG00000081119.2 | 0 | NA | NA | NA | NA | NA | #VALUE! |
| ENSMUSG00000081120.2 | 0 | NA | NA | NA | NA | NA | #VALUE! |
| ENSMUSG00000081121.2 | 0 | NA | NA | NA | NA | NA | #VALUE! |
| ENSMUSG00000081122.2 | 0 | NA | NA | NA | NA | NA | #VALUE! |
| ENSMUSG00000081124.2 | 0 | NA | NA | NA | NA | NA | #VALUE! |
| ENSMUSG00000081125.4 | 0 | NA | NA | NA | NA | NA | #VALUE! |
| ENSMUSG00000081126.2 | 0 | NA | NA | NA | NA | NA | #VALUE! |
| ENSMUSG00000081127.2 | 0 | NA | NA | NA | NA | NA | #VALUE! |
| ENSMUSG00000081128.2 | 0 | NA | NA | NA | NA | NA | #VALUE! |
| ENSMUSG00000081129.2 | 0 | NA | NA | NA | NA | NA | #VALUE! |
| ENSMUSG00000081130.2 | 0 | NA | NA | NA | NA | NA | #VALUE! |
| ENSMUSG00000081131.2 | 0 | NA | NA | NA | NA | NA | #VALUE! |
| ENSMUSG00000081132.2 | 0 | NA | NA | NA | NA | NA | #VALUE! |
| ENSMUSG00000081133.3 | 0 | NA | NA | NA | NA | NA | #VALUE! |
| ENSMUSG00000081134.2 | 0 | NA | NA | NA | NA | NA | #VALUE! |
| ENSMUSG00000081135.2 | 0 | NA | NA | NA | NA | NA | #VALUE! |
| ENSMUSG00000081136.2 | 0 | NA | NA | NA | NA | NA | #VALUE! |
| ENSMUSG00000081137.4 | 0 | NA | NA | NA | NA | NA | #VALUE! |
| ENSMUSG00000081138.2 | 0 | NA | NA | NA | NA | NA | #VALUE! |
| ENSMUSG00000081139.2 | 0 | NA | NA | NA | NA | NA | #VALUE! |
| ENSMUSG00000081141.2 | 0 | NA | NA | NA | NA | NA | #VALUE! |
| ENSMUSG00000081142.2 | 0 | NA | NA | NA | NA | NA | #VALUE! |
| ENSMUSG00000081143.3 | 0 | NA | NA | NA | NA | NA | #VALUE! |
| ENSMUSG00000081144.2 | 0 | NA | NA | NA | NA | NA | #VALUE! |
| ENSMUSG00000081145.3 | 0 | NA | NA | NA | NA | NA | #VALUE! |

| | | | | | | | |
|---|---|---|---|---|---|---|---|
| ENSMUSG00000081146.2 | 0 | NA | NA | NA | NA | NA | #VALUE! |
| ENSMUSG00000081147.2 | 0 | NA | NA | NA | NA | NA | #VALUE! |
| ENSMUSG00000081148.2 | 0 | NA | NA | NA | NA | NA | #VALUE! |
| ENSMUSG00000081150.2 | 0 | NA | NA | NA | NA | NA | #VALUE! |
| ENSMUSG00000081151.2 | 0 | NA | NA | NA | NA | NA | #VALUE! |
| ENSMUSG00000081152.2 | 0 | NA | NA | NA | NA | NA | #VALUE! |
| ENSMUSG00000081153.2 | 1.762102 | -4.34067 | 3.93219 | -1.10388 | 0.269645 | NA | #VALUE! |
| ENSMUSG00000081154.3 | 0 | NA | NA | NA | NA | NA | #VALUE! |
| ENSMUSG00000081155.2 | 0 | NA | NA | NA | NA | NA | #VALUE! |
| ENSMUSG00000081156.2 | 0 | NA | NA | NA | NA | NA | #VALUE! |
| ENSMUSG00000081157.2 | 0 | NA | NA | NA | NA | NA | #VALUE! |
| ENSMUSG00000081158.2 | 0 | NA | NA | NA | NA | NA | #VALUE! |
| ENSMUSG00000081159.2 | 0 | NA | NA | NA | NA | NA | #VALUE! |
| ENSMUSG00000081160.2 | 0 | NA | NA | NA | NA | NA | #VALUE! |
| ENSMUSG00000081161.2 | 0 | NA | NA | NA | NA | NA | #VALUE! |
| ENSMUSG00000081162.3 | 0 | NA | NA | NA | NA | NA | #VALUE! |
| ENSMUSG00000081163.3 | 0 | NA | NA | NA | NA | NA | #VALUE! |
| ENSMUSG00000081164.2 | 0.402379 | -0.92565 | 4.039907 | -0.22913 | 0.81877 | NA | #VALUE! |
| ENSMUSG00000081165.2 | 0 | NA | NA | NA | NA | NA | #VALUE! |
| ENSMUSG00000081166.2 | 0 | NA | NA | NA | NA | NA | #VALUE! |
| ENSMUSG00000081167.2 | 0 | NA | NA | NA | NA | NA | #VALUE! |
| ENSMUSG00000081168.2 | 0 | NA | NA | NA | NA | NA | #VALUE! |
| ENSMUSG00000081169.2 | 0 | NA | NA | NA | NA | NA | #VALUE! |
| ENSMUSG00000081170.2 | 0 | NA | NA | NA | NA | NA | #VALUE! |
| ENSMUSG00000081171.2 | 0 | NA | NA | NA | NA | NA | #VALUE! |
| ENSMUSG00000081172.2 | 0 | NA | NA | NA | NA | NA | #VALUE! |
| ENSMUSG00000081173.2 | 0 | NA | NA | NA | NA | NA | #VALUE! |
| ENSMUSG00000081174.2 | 0 | NA | NA | NA | NA | NA | #VALUE! |
| ENSMUSG00000081175.4 | 0 | NA | NA | NA | NA | NA | #VALUE! |
| ENSMUSG00000081176.2 | 0 | NA | NA | NA | NA | NA | #VALUE! |
| ENSMUSG00000081177.2 | 0 | NA | NA | NA | NA | NA | #VALUE! |
| ENSMUSG00000081179.2 | 0 | NA | NA | NA | NA | NA | #VALUE! |
| ENSMUSG00000081180.2 | 0 | NA | NA | NA | NA | NA | #VALUE! |
| ENSMUSG00000081181.2 | 0 | NA | NA | NA | NA | NA | #VALUE! |
| ENSMUSG00000081182.3 | 0 | NA | NA | NA | NA | NA | #VALUE! |
| ENSMUSG00000081183.2 | 0 | NA | NA | NA | NA | NA | #VALUE! |
| ENSMUSG00000081184.2 | 0 | NA | NA | NA | NA | NA | #VALUE! |
| ENSMUSG00000081185.2 | 0 | NA | NA | NA | NA | NA | #VALUE! |
| ENSMUSG00000081186.2 | 0 | NA | NA | NA | NA | NA | #VALUE! |
| ENSMUSG00000081187.2 | 0 | NA | NA | NA | NA | NA | #VALUE! |
| ENSMUSG00000081188.2 | 0 | NA | NA | NA | NA | NA | #VALUE! |
| ENSMUSG00000081189.2 | 0 | NA | NA | NA | NA | NA | #VALUE! |
| ENSMUSG00000081191.2 | 0 | NA | NA | NA | NA | NA | #VALUE! |
| ENSMUSG00000081194.2 | 0 | NA | NA | NA | NA | NA | #VALUE! |

| | | | | | | | |
|---|---|---|---|---|---|---|---|
| ENSMUSG00000081195.3 | 0 | NA | NA | NA | NA | NA | #VALUE! |
| ENSMUSG00000081196.2 | 0 | NA | NA | NA | NA | NA | #VALUE! |
| ENSMUSG00000081197.2 | 0 | NA | NA | NA | NA | NA | #VALUE! |
| ENSMUSG00000081199.4 | 0 | NA | NA | NA | NA | NA | #VALUE! |
| ENSMUSG00000081200.2 | 0 | NA | NA | NA | NA | NA | #VALUE! |
| ENSMUSG00000081201.2 | 0 | NA | NA | NA | NA | NA | #VALUE! |
| ENSMUSG00000081202.2 | 0 | NA | NA | NA | NA | NA | #VALUE! |
| ENSMUSG00000081205.5 | 0 | NA | NA | NA | NA | NA | #VALUE! |
| ENSMUSG00000081206.2 | 0 | NA | NA | NA | NA | NA | #VALUE! |
| ENSMUSG00000081208.2 | 0 | NA | NA | NA | NA | NA | #VALUE! |
| ENSMUSG00000081209.2 | 0 | NA | NA | NA | NA | NA | #VALUE! |
| ENSMUSG00000081210.2 | 0.195789 | -1.05581 | 4.073302 | -0.2592 | 0.795479 | NA | #VALUE! |
| ENSMUSG00000081211.2 | 0 | NA | NA | NA | NA | NA | #VALUE! |
| ENSMUSG00000081212.2 | 0 | NA | NA | NA | NA | NA | #VALUE! |
| ENSMUSG00000081215.2 | 0 | NA | NA | NA | NA | NA | #VALUE! |
| ENSMUSG00000081216.4 | 0 | NA | NA | NA | NA | NA | #VALUE! |
| ENSMUSG00000081217.2 | 0 | NA | NA | NA | NA | NA | #VALUE! |
| ENSMUSG00000081218.2 | 0 | NA | NA | NA | NA | NA | #VALUE! |
| ENSMUSG00000081219.2 | 0 | NA | NA | NA | NA | NA | #VALUE! |
| ENSMUSG00000081220.3 | 0 | NA | NA | NA | NA | NA | #VALUE! |
| ENSMUSG00000081221.5 | 0 | NA | NA | NA | NA | NA | #VALUE! |
| ENSMUSG00000081222.2 | 0 | NA | NA | NA | NA | NA | #VALUE! |
| ENSMUSG00000081223.2 | 0.673134 | 2.918386 | 3.972531 | 0.734641 | 0.462558 | NA | #VALUE! |
| ENSMUSG00000081224.2 | 0 | NA | NA | NA | NA | NA | #VALUE! |
| ENSMUSG00000081225.4 | 0 | NA | NA | NA | NA | NA | #VALUE! |
| ENSMUSG00000081226.2 | 0 | NA | NA | NA | NA | NA | #VALUE! |
| ENSMUSG00000081227.2 | 0 | NA | NA | NA | NA | NA | #VALUE! |
| ENSMUSG00000081228.2 | 5.31317 | 0.721986 | 2.426539 | 0.297537 | 0.766056 | NA | #VALUE! |
| ENSMUSG00000081229.2 | 0 | NA | NA | NA | NA | NA | #VALUE! |
| ENSMUSG00000081230.2 | 0 | NA | NA | NA | NA | NA | #VALUE! |
| ENSMUSG00000081231.2 | 0 | NA | NA | NA | NA | NA | #VALUE! |
| ENSMUSG00000081232.4 | 0 | NA | NA | NA | NA | NA | #VALUE! |
| ENSMUSG00000081233.2 | 0 | NA | NA | NA | NA | NA | #VALUE! |
| ENSMUSG00000081234.4 | 0 | NA | NA | NA | NA | NA | #VALUE! |
| ENSMUSG00000081235.2 | 0 | NA | NA | NA | NA | NA | #VALUE! |
| ENSMUSG00000081236.2 | 0 | NA | NA | NA | NA | NA | #VALUE! |
| ENSMUSG00000081237.4 | 0 | NA | NA | NA | NA | NA | #VALUE! |
| ENSMUSG00000081238.4 | 0 | NA | NA | NA | NA | NA | #VALUE! |
| ENSMUSG00000081239.4 | 0 | NA | NA | NA | NA | NA | #VALUE! |
| ENSMUSG00000081240.3 | 0 | NA | NA | NA | NA | NA | #VALUE! |
| ENSMUSG00000081242.2 | 0 | NA | NA | NA | NA | NA | #VALUE! |
| ENSMUSG00000081243.2 | 0 | NA | NA | NA | NA | NA | #VALUE! |
| ENSMUSG00000081244.4 | 0 | NA | NA | NA | NA | NA | #VALUE! |
| ENSMUSG00000081245.3 | 0 | NA | NA | NA | NA | NA | #VALUE! |

| | | | | | | | |
|---|---|---|---|---|---|---|---|
| ENSMUSG00000081248.2 | 0 | NA | NA | NA | NA | NA | #VALUE! |
| ENSMUSG00000081249.4 | 0 | NA | NA | NA | NA | NA | #VALUE! |
| ENSMUSG00000081250.4 | 0 | NA | NA | NA | NA | NA | #VALUE! |
| ENSMUSG00000081251.2 | 0 | NA | NA | NA | NA | NA | #VALUE! |
| ENSMUSG00000081252.2 | 0 | NA | NA | NA | NA | NA | #VALUE! |
| ENSMUSG00000081253.2 | 0 | NA | NA | NA | NA | NA | #VALUE! |
| ENSMUSG00000081254.2 | 0 | NA | NA | NA | NA | NA | #VALUE! |
| ENSMUSG00000081255.2 | 0 | NA | NA | NA | NA | NA | #VALUE! |
| ENSMUSG00000081256.2 | 0 | NA | NA | NA | NA | NA | #VALUE! |
| ENSMUSG00000081257.2 | 0 | NA | NA | NA | NA | NA | #VALUE! |
| ENSMUSG00000081258.2 | 0 | NA | NA | NA | NA | NA | #VALUE! |
| ENSMUSG00000081259.3 | 0 | NA | NA | NA | NA | NA | #VALUE! |
| ENSMUSG00000081260.2 | 0 | NA | NA | NA | NA | NA | #VALUE! |
| ENSMUSG00000081261.2 | 0 | NA | NA | NA | NA | NA | #VALUE! |
| ENSMUSG00000081262.2 | 0 | NA | NA | NA | NA | NA | #VALUE! |
| ENSMUSG00000081263.2 | 0 | NA | NA | NA | NA | NA | #VALUE! |
| ENSMUSG00000081264.2 | 0 | NA | NA | NA | NA | NA | #VALUE! |
| ENSMUSG00000081265.2 | 0 | NA | NA | NA | NA | NA | #VALUE! |
| ENSMUSG00000081268.2 | 0 | NA | NA | NA | NA | NA | #VALUE! |
| ENSMUSG00000081269.2 | 0 | NA | NA | NA | NA | NA | #VALUE! |
| ENSMUSG00000081270.2 | 0 | NA | NA | NA | NA | NA | #VALUE! |
| ENSMUSG00000081271.2 | 0 | NA | NA | NA | NA | NA | #VALUE! |
| ENSMUSG00000081272.2 | 0 | NA | NA | NA | NA | NA | #VALUE! |
| ENSMUSG00000081273.2 | 0 | NA | NA | NA | NA | NA | #VALUE! |
| ENSMUSG00000081274.3 | 0 | NA | NA | NA | NA | NA | #VALUE! |
| ENSMUSG00000081275.2 | 0 | NA | NA | NA | NA | NA | #VALUE! |
| ENSMUSG00000081276.2 | 0 | NA | NA | NA | NA | NA | #VALUE! |
| ENSMUSG00000081277.2 | 0.372394 | 0.012025 | 4.059931 | 0.002962 | 0.997637 | NA | #VALUE! |
| ENSMUSG00000081278.2 | 0.537499 | 1.33877 | 4.004498 | 0.334317 | 0.738141 | NA | #VALUE! |
| ENSMUSG00000081279.2 | 0 | NA | NA | NA | NA | NA | #VALUE! |
| ENSMUSG00000081280.2 | 0 | NA | NA | NA | NA | NA | #VALUE! |
| ENSMUSG00000081281.2 | 2.68903 | 0.050581 | 3.253631 | 0.015546 | 0.987597 | NA | #VALUE! |
| ENSMUSG00000081282.3 | 0 | NA | NA | NA | NA | NA | #VALUE! |
| ENSMUSG00000081283.2 | 0 | NA | NA | NA | NA | NA | #VALUE! |
| ENSMUSG00000081284.2 | 0 | NA | NA | NA | NA | NA | #VALUE! |
| ENSMUSG00000081285.3 | 0 | NA | NA | NA | NA | NA | #VALUE! |
| ENSMUSG00000081286.2 | 0 | NA | NA | NA | NA | NA | #VALUE! |
| ENSMUSG00000081287.4 | 0 | NA | NA | NA | NA | NA | #VALUE! |
| ENSMUSG00000081288.2 | 0 | NA | NA | NA | NA | NA | #VALUE! |
| ENSMUSG00000081289.2 | 0.105013 | -0.97324 | 4.076293 | -0.23876 | 0.811295 | NA | #VALUE! |
| ENSMUSG00000081290.2 | 0 | NA | NA | NA | NA | NA | #VALUE! |
| ENSMUSG00000081292.2 | 0 | NA | NA | NA | NA | NA | #VALUE! |
| ENSMUSG00000081293.2 | 0 | NA | NA | NA | NA | NA | #VALUE! |
| ENSMUSG00000081295.2 | 0 | NA | NA | NA | NA | NA | #VALUE! |

| | | | | | | | |
|---|---|---|---|---|---|---|---|
| ENSMUSG00000081296.2 | 0 | NA | NA | NA | NA | NA | #VALUE! |
| ENSMUSG00000081297.2 | 0 | NA | NA | NA | NA | NA | #VALUE! |
| ENSMUSG00000081298.2 | 0 | NA | NA | NA | NA | NA | #VALUE! |
| ENSMUSG00000081299.2 | 0 | NA | NA | NA | NA | NA | #VALUE! |
| ENSMUSG00000081300.2 | 0 | NA | NA | NA | NA | NA | #VALUE! |
| ENSMUSG00000081301.2 | 0 | NA | NA | NA | NA | NA | #VALUE! |
| ENSMUSG00000081302.2 | 0 | NA | NA | NA | NA | NA | #VALUE! |
| ENSMUSG00000081303.3 | 0 | NA | NA | NA | NA | NA | #VALUE! |
| ENSMUSG00000081304.2 | 0 | NA | NA | NA | NA | NA | #VALUE! |
| ENSMUSG00000081305.2 | 0 | NA | NA | NA | NA | NA | #VALUE! |
| ENSMUSG00000081306.2 | 0 | NA | NA | NA | NA | NA | #VALUE! |
| ENSMUSG00000081307.2 | 0 | NA | NA | NA | NA | NA | #VALUE! |
| ENSMUSG00000081308.2 | 0 | NA | NA | NA | NA | NA | #VALUE! |
| ENSMUSG00000081309.4 | 0 | NA | NA | NA | NA | NA | #VALUE! |
| ENSMUSG00000081310.2 | 0 | NA | NA | NA | NA | NA | #VALUE! |
| ENSMUSG00000081312.2 | 0 | NA | NA | NA | NA | NA | #VALUE! |
| ENSMUSG00000081313.2 | 0 | NA | NA | NA | NA | NA | #VALUE! |
| ENSMUSG00000081314.2 | 0 | NA | NA | NA | NA | NA | #VALUE! |
| ENSMUSG00000081315.3 | 0 | NA | NA | NA | NA | NA | #VALUE! |
| ENSMUSG00000081316.2 | 0 | NA | NA | NA | NA | NA | #VALUE! |
| ENSMUSG00000081317.2 | 0 | NA | NA | NA | NA | NA | #VALUE! |
| ENSMUSG00000081318.2 | 0 | NA | NA | NA | NA | NA | #VALUE! |
| ENSMUSG00000081319.2 | 0 | NA | NA | NA | NA | NA | #VALUE! |
| ENSMUSG00000081320.2 | 0 | NA | NA | NA | NA | NA | #VALUE! |
| ENSMUSG00000081321.2 | 0 | NA | NA | NA | NA | NA | #VALUE! |
| ENSMUSG00000081323.2 | 0 | NA | NA | NA | NA | NA | #VALUE! |
| ENSMUSG00000081324.2 | 0 | NA | NA | NA | NA | NA | #VALUE! |
| ENSMUSG00000081325.2 | 0.279766 | 1.808031 | 4.036658 | 0.447903 | 0.654223 | NA | #VALUE! |
| ENSMUSG00000081326.2 | 0 | NA | NA | NA | NA | NA | #VALUE! |
| ENSMUSG00000081327.2 | 0 | NA | NA | NA | NA | NA | #VALUE! |
| ENSMUSG00000081329.2 | 0 | NA | NA | NA | NA | NA | #VALUE! |
| ENSMUSG00000081330.2 | 0 | NA | NA | NA | NA | NA | #VALUE! |
| ENSMUSG00000081331.2 | 0 | NA | NA | NA | NA | NA | #VALUE! |
| ENSMUSG00000081332.2 | 0 | NA | NA | NA | NA | NA | #VALUE! |
| ENSMUSG00000081333.2 | 0 | NA | NA | NA | NA | NA | #VALUE! |
| ENSMUSG00000081334.2 | 0 | NA | NA | NA | NA | NA | #VALUE! |
| ENSMUSG00000081335.2 | 0 | NA | NA | NA | NA | NA | #VALUE! |
| ENSMUSG00000081336.2 | 0 | NA | NA | NA | NA | NA | #VALUE! |
| ENSMUSG00000081339.2 | 0 | NA | NA | NA | NA | NA | #VALUE! |
| ENSMUSG00000081340.2 | 0 | NA | NA | NA | NA | NA | #VALUE! |
| ENSMUSG00000081341.2 | 0 | NA | NA | NA | NA | NA | #VALUE! |
| ENSMUSG00000081342.2 | 0 | NA | NA | NA | NA | NA | #VALUE! |
| ENSMUSG00000081343.2 | 0 | NA | NA | NA | NA | NA | #VALUE! |
| ENSMUSG00000081345.3 | 0 | NA | NA | NA | NA | NA | #VALUE! |

| | | | | | | | |
|---|---|---|---|---|---|---|---|
| ENSMUSG00000081346.2 | 0 | NA | NA | NA | NA | NA | #VALUE! |
| ENSMUSG00000081348.2 | 0 | NA | NA | NA | NA | NA | #VALUE! |
| ENSMUSG00000081349.2 | 0 | NA | NA | NA | NA | NA | #VALUE! |
| ENSMUSG00000081350.2 | 0 | NA | NA | NA | NA | NA | #VALUE! |
| ENSMUSG00000081352.2 | 0 | NA | NA | NA | NA | NA | #VALUE! |
| ENSMUSG00000081353.3 | 0 | NA | NA | NA | NA | NA | #VALUE! |
| ENSMUSG00000081354.2 | 0 | NA | NA | NA | NA | NA | #VALUE! |
| ENSMUSG00000081355.2 | 0 | NA | NA | NA | NA | NA | #VALUE! |
| ENSMUSG00000081357.2 | 0 | NA | NA | NA | NA | NA | #VALUE! |
| ENSMUSG00000081358.2 | 0 | NA | NA | NA | NA | NA | #VALUE! |
| ENSMUSG00000081359.2 | 0.105013 | -0.97324 | 4.076293 | -0.23876 | 0.811295 | NA | #VALUE! |
| ENSMUSG00000081360.2 | 0 | NA | NA | NA | NA | NA | #VALUE! |
| ENSMUSG00000081361.2 | 0 | NA | NA | NA | NA | NA | #VALUE! |
| ENSMUSG00000081362.4 | 0 | NA | NA | NA | NA | NA | #VALUE! |
| ENSMUSG00000081363.3 | 0 | NA | NA | NA | NA | NA | #VALUE! |
| ENSMUSG00000081364.2 | 0 | NA | NA | NA | NA | NA | #VALUE! |
| ENSMUSG00000081365.2 | 0 | NA | NA | NA | NA | NA | #VALUE! |
| ENSMUSG00000081367.3 | 0 | NA | NA | NA | NA | NA | #VALUE! |
| ENSMUSG00000081369.2 | 0 | NA | NA | NA | NA | NA | #VALUE! |
| ENSMUSG00000081370.2 | 0 | NA | NA | NA | NA | NA | #VALUE! |
| ENSMUSG00000081372.2 | 0 | NA | NA | NA | NA | NA | #VALUE! |
| ENSMUSG00000081373.2 | 0 | NA | NA | NA | NA | NA | #VALUE! |
| ENSMUSG00000081374.2 | 0 | NA | NA | NA | NA | NA | #VALUE! |
| ENSMUSG00000081375.2 | 0 | NA | NA | NA | NA | NA | #VALUE! |
| ENSMUSG00000081376.2 | 0 | NA | NA | NA | NA | NA | #VALUE! |
| ENSMUSG00000081377.2 | 0 | NA | NA | NA | NA | NA | #VALUE! |
| ENSMUSG00000081378.2 | 5.391524 | 0.567885 | 2.451026 | 0.231693 | 0.816777 | NA | #VALUE! |
| ENSMUSG00000081379.2 | 0 | NA | NA | NA | NA | NA | #VALUE! |
| ENSMUSG00000081381.2 | 0 | NA | NA | NA | NA | NA | #VALUE! |
| ENSMUSG00000081383.2 | 0 | NA | NA | NA | NA | NA | #VALUE! |
| ENSMUSG00000081384.2 | 0 | NA | NA | NA | NA | NA | #VALUE! |
| ENSMUSG00000081386.2 | 0 | NA | NA | NA | NA | NA | #VALUE! |
| ENSMUSG00000081388.2 | 0 | NA | NA | NA | NA | NA | #VALUE! |
| ENSMUSG00000081389.3 | 0 | NA | NA | NA | NA | NA | #VALUE! |
| ENSMUSG00000081390.2 | 0 | NA | NA | NA | NA | NA | #VALUE! |
| ENSMUSG00000081391.2 | 0 | NA | NA | NA | NA | NA | #VALUE! |
| ENSMUSG00000081392.2 | 0 | NA | NA | NA | NA | NA | #VALUE! |
| ENSMUSG00000081394.2 | 0.598168 | -1.6263 | 3.986382 | -0.40796 | 0.6833 | NA | #VALUE! |
| ENSMUSG00000081395.2 | 0 | NA | NA | NA | NA | NA | #VALUE! |
| ENSMUSG00000081396.2 | 0 | NA | NA | NA | NA | NA | #VALUE! |
| ENSMUSG00000081397.2 | 0 | NA | NA | NA | NA | NA | #VALUE! |
| ENSMUSG00000081398.2 | 0 | NA | NA | NA | NA | NA | #VALUE! |
| ENSMUSG00000081399.3 | 0 | NA | NA | NA | NA | NA | #VALUE! |
| ENSMUSG00000081402.5 | 0 | NA | NA | NA | NA | NA | #VALUE! |

| | | | | | | | |
|---|---|---|---|---|---|---|---|
| ENSMUSG00000081404.2 | 0 | NA | NA | NA | NA | NA | #VALUE! |
| ENSMUSG00000081405.2 | 0 | NA | NA | NA | NA | NA | #VALUE! |
| ENSMUSG00000081407.2 | 0 | NA | NA | NA | NA | NA | #VALUE! |
| ENSMUSG00000081408.2 | 0 | NA | NA | NA | NA | NA | #VALUE! |
| ENSMUSG00000081409.2 | 0 | NA | NA | NA | NA | NA | #VALUE! |
| ENSMUSG00000081410.2 | 0 | NA | NA | NA | NA | NA | #VALUE! |
| ENSMUSG00000081412.2 | 0 | NA | NA | NA | NA | NA | #VALUE! |
| ENSMUSG00000081413.4 | 3.811295 | -0.32872 | 3.104192 | -0.1059 | 0.915665 | NA | #VALUE! |
| ENSMUSG00000081414.3 | 0 | NA | NA | NA | NA | NA | #VALUE! |
| ENSMUSG00000081415.2 | 0 | NA | NA | NA | NA | NA | #VALUE! |
| ENSMUSG00000081416.2 | 0 | NA | NA | NA | NA | NA | #VALUE! |
| ENSMUSG00000081417.2 | 0 | NA | NA | NA | NA | NA | #VALUE! |
| ENSMUSG00000081418.2 | 0.105013 | -0.97324 | 4.076293 | -0.23876 | 0.811295 | NA | #VALUE! |
| ENSMUSG00000081421.2 | 0 | NA | NA | NA | NA | NA | #VALUE! |
| ENSMUSG00000081422.2 | 0 | NA | NA | NA | NA | NA | #VALUE! |
| ENSMUSG00000081423.2 | 0.634559 | 2.867985 | 3.974828 | 0.721537 | 0.470579 | NA | #VALUE! |
| ENSMUSG00000081424.2 | 0 | NA | NA | NA | NA | NA | #VALUE! |
| ENSMUSG00000081427.2 | 0 | NA | NA | NA | NA | NA | #VALUE! |
| ENSMUSG00000081428.2 | 0 | NA | NA | NA | NA | NA | #VALUE! |
| ENSMUSG00000081429.2 | 0 | NA | NA | NA | NA | NA | #VALUE! |
| ENSMUSG00000081431.2 | 0 | NA | NA | NA | NA | NA | #VALUE! |
| ENSMUSG00000081432.2 | 0 | NA | NA | NA | NA | NA | #VALUE! |
| ENSMUSG00000081433.2 | 0 | NA | NA | NA | NA | NA | #VALUE! |
| ENSMUSG00000081435.2 | 0 | NA | NA | NA | NA | NA | #VALUE! |
| ENSMUSG00000081436.2 | 0 | NA | NA | NA | NA | NA | #VALUE! |
| ENSMUSG00000081437.2 | 0 | NA | NA | NA | NA | NA | #VALUE! |
| ENSMUSG00000081438.2 | 0 | NA | NA | NA | NA | NA | #VALUE! |
| ENSMUSG00000081439.2 | 0 | NA | NA | NA | NA | NA | #VALUE! |
| ENSMUSG00000081440.3 | 0 | NA | NA | NA | NA | NA | #VALUE! |
| ENSMUSG00000081441.4 | 0 | NA | NA | NA | NA | NA | #VALUE! |
| ENSMUSG00000081442.2 | 0 | NA | NA | NA | NA | NA | #VALUE! |
| ENSMUSG00000081443.3 | 0 | NA | NA | NA | NA | NA | #VALUE! |
| ENSMUSG00000081444.2 | 0 | NA | NA | NA | NA | NA | #VALUE! |
| ENSMUSG00000081445.2 | 0 | NA | NA | NA | NA | NA | #VALUE! |
| ENSMUSG00000081447.2 | 0 | NA | NA | NA | NA | NA | #VALUE! |
| ENSMUSG00000081449.2 | 0 | NA | NA | NA | NA | NA | #VALUE! |
| ENSMUSG00000081450.2 | 0 | NA | NA | NA | NA | NA | #VALUE! |
| ENSMUSG00000081451.2 | 0 | NA | NA | NA | NA | NA | #VALUE! |
| ENSMUSG00000081452.2 | 0 | NA | NA | NA | NA | NA | #VALUE! |
| ENSMUSG00000081453.2 | 0.273297 | 0.094602 | 4.062933 | 0.023284 | 0.981424 | NA | #VALUE! |
| ENSMUSG00000081455.2 | 1.448031 | -2.81141 | 3.927885 | -0.71576 | 0.474141 | NA | #VALUE! |
| ENSMUSG00000081456.2 | 0 | NA | NA | NA | NA | NA | #VALUE! |
| ENSMUSG00000081457.2 | 0 | NA | NA | NA | NA | NA | #VALUE! |
| ENSMUSG00000081458.3 | 0 | NA | NA | NA | NA | NA | #VALUE! |

| | | | | | | | |
|---|---|---|---|---|---|---|---|
| ENSMUSG00000081459.4 | 0 | NA | NA | NA | NA | NA | #VALUE! |
| ENSMUSG00000081460.2 | 0 | NA | NA | NA | NA | NA | #VALUE! |
| ENSMUSG00000081461.4 | 0 | NA | NA | NA | NA | NA | #VALUE! |
| ENSMUSG00000081463.2 | 0 | NA | NA | NA | NA | NA | #VALUE! |
| ENSMUSG00000081464.2 | 0 | NA | NA | NA | NA | NA | #VALUE! |
| ENSMUSG00000081465.2 | 0 | NA | NA | NA | NA | NA | #VALUE! |
| ENSMUSG00000081466.2 | 0 | NA | NA | NA | NA | NA | #VALUE! |
| ENSMUSG00000081467.2 | 0 | NA | NA | NA | NA | NA | #VALUE! |
| ENSMUSG00000081469.2 | 0 | NA | NA | NA | NA | NA | #VALUE! |
| ENSMUSG00000081470.2 | 0 | NA | NA | NA | NA | NA | #VALUE! |
| ENSMUSG00000081471.2 | 0 | NA | NA | NA | NA | NA | #VALUE! |
| ENSMUSG00000081473.4 | 0 | NA | NA | NA | NA | NA | #VALUE! |
| ENSMUSG00000081474.2 | 0 | NA | NA | NA | NA | NA | #VALUE! |
| ENSMUSG00000081475.2 | 0 | NA | NA | NA | NA | NA | #VALUE! |
| ENSMUSG00000081476.2 | 0 | NA | NA | NA | NA | NA | #VALUE! |
| ENSMUSG00000081477.2 | 0 | NA | NA | NA | NA | NA | #VALUE! |
| ENSMUSG00000081478.2 | 0 | NA | NA | NA | NA | NA | #VALUE! |
| ENSMUSG00000081480.2 | 0 | NA | NA | NA | NA | NA | #VALUE! |
| ENSMUSG00000081481.2 | 0 | NA | NA | NA | NA | NA | #VALUE! |
| ENSMUSG00000081483.2 | 0 | NA | NA | NA | NA | NA | #VALUE! |
| ENSMUSG00000081484.2 | 0 | NA | NA | NA | NA | NA | #VALUE! |
| ENSMUSG00000081485.6 | 4.430093 | 1.421957 | 2.694974 | 0.527633 | 0.597754 | NA | #VALUE! |
| ENSMUSG00000081486.2 | 0 | NA | NA | NA | NA | NA | #VALUE! |
| ENSMUSG00000081487.2 | 0 | NA | NA | NA | NA | NA | #VALUE! |
| ENSMUSG00000081488.2 | 0 | NA | NA | NA | NA | NA | #VALUE! |
| ENSMUSG00000081490.2 | 0 | NA | NA | NA | NA | NA | #VALUE! |
| ENSMUSG00000081491.2 | 0.195789 | -1.05581 | 4.073302 | -0.2592 | 0.795479 | NA | #VALUE! |
| ENSMUSG00000081492.2 | 0 | NA | NA | NA | NA | NA | #VALUE! |
| ENSMUSG00000081494.2 | 0 | NA | NA | NA | NA | NA | #VALUE! |
| ENSMUSG00000081495.2 | 0 | NA | NA | NA | NA | NA | #VALUE! |
| ENSMUSG00000081496.2 | 0 | NA | NA | NA | NA | NA | #VALUE! |
| ENSMUSG00000081497.2 | 0 | NA | NA | NA | NA | NA | #VALUE! |
| ENSMUSG00000081498.3 | 0 | NA | NA | NA | NA | NA | #VALUE! |
| ENSMUSG00000081499.2 | 0 | NA | NA | NA | NA | NA | #VALUE! |
| ENSMUSG00000081500.2 | 0 | NA | NA | NA | NA | NA | #VALUE! |
| ENSMUSG00000081501.6 | 0 | NA | NA | NA | NA | NA | #VALUE! |
| ENSMUSG00000081502.2 | 0 | NA | NA | NA | NA | NA | #VALUE! |
| ENSMUSG00000081504.4 | 0.210026 | -1.52494 | 4.058692 | -0.37572 | 0.707123 | NA | #VALUE! |
| ENSMUSG00000081505.2 | 0 | NA | NA | NA | NA | NA | #VALUE! |
| ENSMUSG00000081506.2 | 0 | NA | NA | NA | NA | NA | #VALUE! |
| ENSMUSG00000081507.2 | 0 | NA | NA | NA | NA | NA | #VALUE! |
| ENSMUSG00000081508.2 | 0 | NA | NA | NA | NA | NA | #VALUE! |
| ENSMUSG00000081510.2 | 0 | NA | NA | NA | NA | NA | #VALUE! |
| ENSMUSG00000081511.2 | 0 | NA | NA | NA | NA | NA | #VALUE! |

| | | | | | | | |
|---|---|---|---|---|---|---|---|
| ENSMUSG00000081512.2 | 0 | NA | NA | NA | NA | NA | #VALUE! |
| ENSMUSG00000081513.2 | 0 | NA | NA | NA | NA | NA | #VALUE! |
| ENSMUSG00000081514.3 | 0 | NA | NA | NA | NA | NA | #VALUE! |
| ENSMUSG00000081515.2 | 0 | NA | NA | NA | NA | NA | #VALUE! |
| ENSMUSG00000081516.2 | 0 | NA | NA | NA | NA | NA | #VALUE! |
| ENSMUSG00000081517.2 | 0 | NA | NA | NA | NA | NA | #VALUE! |
| ENSMUSG00000081519.2 | 0 | NA | NA | NA | NA | NA | #VALUE! |
| ENSMUSG00000081520.2 | 1.008389 | -2.241 | 3.944711 | -0.5681 | 0.569965 | NA | #VALUE! |
| ENSMUSG00000081522.2 | 0 | NA | NA | NA | NA | NA | #VALUE! |
| ENSMUSG00000081523.2 | 0 | NA | NA | NA | NA | NA | #VALUE! |
| ENSMUSG00000081524.4 | 0 | NA | NA | NA | NA | NA | #VALUE! |
| ENSMUSG00000081525.2 | 0 | NA | NA | NA | NA | NA | #VALUE! |
| ENSMUSG00000081526.2 | 0 | NA | NA | NA | NA | NA | #VALUE! |
| ENSMUSG00000081528.2 | 0 | NA | NA | NA | NA | NA | #VALUE! |
| ENSMUSG00000081529.2 | 0 | NA | NA | NA | NA | NA | #VALUE! |
| ENSMUSG00000081530.2 | 0 | NA | NA | NA | NA | NA | #VALUE! |
| ENSMUSG00000081531.2 | 0 | NA | NA | NA | NA | NA | #VALUE! |
| ENSMUSG00000081532.2 | 0 | NA | NA | NA | NA | NA | #VALUE! |
| ENSMUSG00000081533.2 | 0 | NA | NA | NA | NA | NA | #VALUE! |
| ENSMUSG00000081536.2 | 0 | NA | NA | NA | NA | NA | #VALUE! |
| ENSMUSG00000081537.3 | 0 | NA | NA | NA | NA | NA | #VALUE! |
| ENSMUSG00000081538.2 | 0 | NA | NA | NA | NA | NA | #VALUE! |
| ENSMUSG00000081539.2 | 0 | NA | NA | NA | NA | NA | #VALUE! |
| ENSMUSG00000081540.4 | 0 | NA | NA | NA | NA | NA | #VALUE! |
| ENSMUSG00000081541.2 | 0 | NA | NA | NA | NA | NA | #VALUE! |
| ENSMUSG00000081542.2 | 0 | NA | NA | NA | NA | NA | #VALUE! |
| ENSMUSG00000081543.2 | 0 | NA | NA | NA | NA | NA | #VALUE! |
| ENSMUSG00000081544.2 | 0 | NA | NA | NA | NA | NA | #VALUE! |
| ENSMUSG00000081546.2 | 0 | NA | NA | NA | NA | NA | #VALUE! |
| ENSMUSG00000081547.2 | 0 | NA | NA | NA | NA | NA | #VALUE! |
| ENSMUSG00000081548.2 | 0 | NA | NA | NA | NA | NA | #VALUE! |
| ENSMUSG00000081550.2 | 0 | NA | NA | NA | NA | NA | #VALUE! |
| ENSMUSG00000081551.2 | 0 | NA | NA | NA | NA | NA | #VALUE! |
| ENSMUSG00000081552.2 | 0 | NA | NA | NA | NA | NA | #VALUE! |
| ENSMUSG00000081553.2 | 0 | NA | NA | NA | NA | NA | #VALUE! |
| ENSMUSG00000081555.2 | 0 | NA | NA | NA | NA | NA | #VALUE! |
| ENSMUSG00000081556.2 | 0 | NA | NA | NA | NA | NA | #VALUE! |
| ENSMUSG00000081557.3 | 0 | NA | NA | NA | NA | NA | #VALUE! |
| ENSMUSG00000081559.4 | 0 | NA | NA | NA | NA | NA | #VALUE! |
| ENSMUSG00000081560.2 | 0 | NA | NA | NA | NA | NA | #VALUE! |
| ENSMUSG00000081561.2 | 0 | NA | NA | NA | NA | NA | #VALUE! |
| ENSMUSG00000081562.2 | 0 | NA | NA | NA | NA | NA | #VALUE! |
| ENSMUSG00000081563.2 | 0 | NA | NA | NA | NA | NA | #VALUE! |
| ENSMUSG00000081564.2 | 0.105013 | -0.97324 | 4.076293 | -0.23876 | 0.811295 | NA | #VALUE! |

| | | | | | | | |
|---|---|---|---|---|---|---|---|
| ENSMUSG00000081565.2 | 0 | NA | NA | NA | NA | NA | #VALUE! |
| ENSMUSG00000081566.2 | 0 | NA | NA | NA | NA | NA | #VALUE! |
| ENSMUSG00000081567.2 | 0 | NA | NA | NA | NA | NA | #VALUE! |
| ENSMUSG00000081568.2 | 0 | NA | NA | NA | NA | NA | #VALUE! |
| ENSMUSG00000081569.2 | 0 | NA | NA | NA | NA | NA | #VALUE! |
| ENSMUSG00000081570.2 | 0 | NA | NA | NA | NA | NA | #VALUE! |
| ENSMUSG00000081571.2 | 0 | NA | NA | NA | NA | NA | #VALUE! |
| ENSMUSG00000081572.4 | 0.998006 | 0.620307 | 3.917008 | 0.158362 | 0.874171 | NA | #VALUE! |
| ENSMUSG00000081573.2 | 0 | NA | NA | NA | NA | NA | #VALUE! |
| ENSMUSG00000081574.2 | 0 | NA | NA | NA | NA | NA | #VALUE! |
| ENSMUSG00000081575.2 | 0 | NA | NA | NA | NA | NA | #VALUE! |
| ENSMUSG00000081576.2 | 0 | NA | NA | NA | NA | NA | #VALUE! |
| ENSMUSG00000081577.2 | 0 | NA | NA | NA | NA | NA | #VALUE! |
| ENSMUSG00000081578.3 | 0 | NA | NA | NA | NA | NA | #VALUE! |
| ENSMUSG00000081579.2 | 0 | NA | NA | NA | NA | NA | #VALUE! |
| ENSMUSG00000081580.2 | 0 | NA | NA | NA | NA | NA | #VALUE! |
| ENSMUSG00000081583.2 | 0 | NA | NA | NA | NA | NA | #VALUE! |
| ENSMUSG00000081584.2 | 0 | NA | NA | NA | NA | NA | #VALUE! |
| ENSMUSG00000081585.2 | 0 | NA | NA | NA | NA | NA | #VALUE! |
| ENSMUSG00000081586.4 | 0 | NA | NA | NA | NA | NA | #VALUE! |
| ENSMUSG00000081587.2 | 0 | NA | NA | NA | NA | NA | #VALUE! |
| ENSMUSG00000081588.2 | 0 | NA | NA | NA | NA | NA | #VALUE! |
| ENSMUSG00000081589.2 | 0 | NA | NA | NA | NA | NA | #VALUE! |
| ENSMUSG00000081590.2 | 0 | NA | NA | NA | NA | NA | #VALUE! |
| ENSMUSG00000081591.2 | 0 | NA | NA | NA | NA | NA | #VALUE! |
| ENSMUSG00000081592.2 | 0 | NA | NA | NA | NA | NA | #VALUE! |
| ENSMUSG00000081593.2 | 0 | NA | NA | NA | NA | NA | #VALUE! |
| ENSMUSG00000081594.2 | 0 | NA | NA | NA | NA | NA | #VALUE! |
| ENSMUSG00000081595.2 | 0 | NA | NA | NA | NA | NA | #VALUE! |
| ENSMUSG00000081596.2 | 0 | NA | NA | NA | NA | NA | #VALUE! |
| ENSMUSG00000081597.2 | 0 | NA | NA | NA | NA | NA | #VALUE! |
| ENSMUSG00000081598.2 | 0 | NA | NA | NA | NA | NA | #VALUE! |
| ENSMUSG00000081599.2 | 0 | NA | NA | NA | NA | NA | #VALUE! |
| ENSMUSG00000081600.4 | 0 | NA | NA | NA | NA | NA | #VALUE! |
| ENSMUSG00000081601.2 | 0 | NA | NA | NA | NA | NA | #VALUE! |
| ENSMUSG00000081602.2 | 0 | NA | NA | NA | NA | NA | #VALUE! |
| ENSMUSG00000081603.2 | 0.600647 | -0.82995 | 3.987207 | -0.20815 | 0.835109 | NA | #VALUE! |
| ENSMUSG00000081604.6 | 8.683303 | 0.452631 | 2.701274 | 0.167562 | 0.866928 | NA | #VALUE! |
| ENSMUSG00000081605.2 | 0 | NA | NA | NA | NA | NA | #VALUE! |
| ENSMUSG00000081606.2 | 0 | NA | NA | NA | NA | NA | #VALUE! |
| ENSMUSG00000081607.4 | 0 | NA | NA | NA | NA | NA | #VALUE! |
| ENSMUSG00000081608.2 | 0 | NA | NA | NA | NA | NA | #VALUE! |
| ENSMUSG00000081609.2 | 0 | NA | NA | NA | NA | NA | #VALUE! |
| ENSMUSG00000081610.2 | 0 | NA | NA | NA | NA | NA | #VALUE! |

| | | | | | | | |
|---|---|---|---|---|---|---|---|
| ENSMUSG00000081611.2 | 0 | NA | NA | NA | NA | NA | #VALUE! |
| ENSMUSG00000081612.2 | 0 | NA | NA | NA | NA | NA | #VALUE! |
| ENSMUSG00000081613.2 | 0.372394 | 0.012025 | 4.059931 | 0.002962 | 0.997637 | NA | #VALUE! |
| ENSMUSG00000081614.2 | 0 | NA | NA | NA | NA | NA | #VALUE! |
| ENSMUSG00000081615.2 | 0 | NA | NA | NA | NA | NA | #VALUE! |
| ENSMUSG00000081616.2 | 0 | NA | NA | NA | NA | NA | #VALUE! |
| ENSMUSG00000081617.2 | 0 | NA | NA | NA | NA | NA | #VALUE! |
| ENSMUSG00000081618.3 | 0 | NA | NA | NA | NA | NA | #VALUE! |
| ENSMUSG00000081619.2 | 0 | NA | NA | NA | NA | NA | #VALUE! |
| ENSMUSG00000081620.2 | 0.105013 | -0.97324 | 4.076293 | -0.23876 | 0.811295 | NA | #VALUE! |
| ENSMUSG00000081621.2 | 0 | NA | NA | NA | NA | NA | #VALUE! |
| ENSMUSG00000081622.2 | 0 | NA | NA | NA | NA | NA | #VALUE! |
| ENSMUSG00000081623.2 | 0 | NA | NA | NA | NA | NA | #VALUE! |
| ENSMUSG00000081624.2 | 0 | NA | NA | NA | NA | NA | #VALUE! |
| ENSMUSG00000081625.2 | 0 | NA | NA | NA | NA | NA | #VALUE! |
| ENSMUSG00000081626.4 | 0 | NA | NA | NA | NA | NA | #VALUE! |
| ENSMUSG00000081627.2 | 0 | NA | NA | NA | NA | NA | #VALUE! |
| ENSMUSG00000081628.2 | 0 | NA | NA | NA | NA | NA | #VALUE! |
| ENSMUSG00000081629.2 | 0.457328 | 0.67272 | 4.035785 | 0.166689 | 0.867615 | NA | #VALUE! |
| ENSMUSG00000081630.2 | 0 | NA | NA | NA | NA | NA | #VALUE! |
| ENSMUSG00000081631.2 | 1.28187 | -2.53127 | 3.935335 | -0.64321 | 0.520085 | NA | #VALUE! |
| ENSMUSG00000081632.2 | 0 | NA | NA | NA | NA | NA | #VALUE! |
| ENSMUSG00000081633.2 | 0 | NA | NA | NA | NA | NA | #VALUE! |
| ENSMUSG00000081635.2 | 0 | NA | NA | NA | NA | NA | #VALUE! |
| ENSMUSG00000081636.3 | 0 | NA | NA | NA | NA | NA | #VALUE! |
| ENSMUSG00000081638.5 | 0 | NA | NA | NA | NA | NA | #VALUE! |
| ENSMUSG00000081639.2 | 0 | NA | NA | NA | NA | NA | #VALUE! |
| ENSMUSG00000081640.2 | 0.245976 | 1.027628 | 4.067126 | 0.252667 | 0.800526 | NA | #VALUE! |
| ENSMUSG00000081641.4 | 0 | NA | NA | NA | NA | NA | #VALUE! |
| ENSMUSG00000081642.2 | 3.357187 | 5.246639 | 3.920044 | 1.338413 | 0.180762 | NA | #VALUE! |
| ENSMUSG00000081643.2 | 0.198268 | -0.06494 | 4.068276 | -0.01596 | 0.987265 | NA | #VALUE! |
| ENSMUSG00000081646.2 | 0 | NA | NA | NA | NA | NA | #VALUE! |
| ENSMUSG00000081647.2 | 0 | NA | NA | NA | NA | NA | #VALUE! |
| ENSMUSG00000081648.2 | 3.411862 | 1.762436 | 3.802459 | 0.463499 | 0.643007 | NA | #VALUE! |
| ENSMUSG00000081649.3 | 0 | NA | NA | NA | NA | NA | #VALUE! |
| ENSMUSG00000081650.2 | 0 | NA | NA | NA | NA | NA | #VALUE! |
| ENSMUSG00000081651.2 | 0 | NA | NA | NA | NA | NA | #VALUE! |
| ENSMUSG00000081652.2 | 0 | NA | NA | NA | NA | NA | #VALUE! |
| ENSMUSG00000081653.2 | 0 | NA | NA | NA | NA | NA | #VALUE! |
| ENSMUSG00000081654.2 | 0 | NA | NA | NA | NA | NA | #VALUE! |
| ENSMUSG00000081656.2 | 0 | NA | NA | NA | NA | NA | #VALUE! |
| ENSMUSG00000081657.2 | 0 | NA | NA | NA | NA | NA | #VALUE! |
| ENSMUSG00000081658.2 | 0 | NA | NA | NA | NA | NA | #VALUE! |
| ENSMUSG00000081659.2 | 0 | NA | NA | NA | NA | NA | #VALUE! |

| | | | | | | | |
|---|---|---|---|---|---|---|---|
| ENSMUSG00000081660.2 | 0 | NA | NA | NA | NA | NA | #VALUE! |
| ENSMUSG00000081661.2 | 0.814159 | 0.038777 | 3.953041 | 0.009809 | 0.992173 | NA | #VALUE! |
| ENSMUSG00000081662.2 | 0 | NA | NA | NA | NA | NA | #VALUE! |
| ENSMUSG00000081663.2 | 0 | NA | NA | NA | NA | NA | #VALUE! |
| ENSMUSG00000081664.2 | 0 | NA | NA | NA | NA | NA | #VALUE! |
| ENSMUSG00000081665.3 | 0 | NA | NA | NA | NA | NA | #VALUE! |
| ENSMUSG00000081666.2 | 0 | NA | NA | NA | NA | NA | #VALUE! |
| ENSMUSG00000081667.2 | 0 | NA | NA | NA | NA | NA | #VALUE! |
| ENSMUSG00000081668.3 | 0 | NA | NA | NA | NA | NA | #VALUE! |
| ENSMUSG00000081669.2 | 0 | NA | NA | NA | NA | NA | #VALUE! |
| ENSMUSG00000081670.2 | 0.315039 | -1.93549 | 4.038621 | -0.47925 | 0.631763 | NA | #VALUE! |
| ENSMUSG00000081671.2 | 0 | NA | NA | NA | NA | NA | #VALUE! |
| ENSMUSG00000081672.2 | 0 | NA | NA | NA | NA | NA | #VALUE! |
| ENSMUSG00000081673.2 | 2.197128 | 0.281727 | 3.810649 | 0.073931 | 0.941065 | NA | #VALUE! |
| ENSMUSG00000081674.2 | 0 | NA | NA | NA | NA | NA | #VALUE! |
| ENSMUSG00000081675.2 | 0.429822 | 2.285488 | 4.007814 | 0.570258 | 0.568503 | NA | #VALUE! |
| ENSMUSG00000081676.2 | 0 | NA | NA | NA | NA | NA | #VALUE! |
| ENSMUSG00000081677.2 | 0 | NA | NA | NA | NA | NA | #VALUE! |
| ENSMUSG00000081678.2 | 0 | NA | NA | NA | NA | NA | #VALUE! |
| ENSMUSG00000081679.2 | 0 | NA | NA | NA | NA | NA | #VALUE! |
| ENSMUSG00000081680.2 | 0 | NA | NA | NA | NA | NA | #VALUE! |
| ENSMUSG00000081681.2 | 0 | NA | NA | NA | NA | NA | #VALUE! |
| ENSMUSG00000081683.6 | 0 | NA | NA | NA | NA | NA | #VALUE! |
| ENSMUSG00000081685.2 | 0 | NA | NA | NA | NA | NA | #VALUE! |
| ENSMUSG00000081686.2 | 0 | NA | NA | NA | NA | NA | #VALUE! |
| ENSMUSG00000081687.2 | 0 | NA | NA | NA | NA | NA | #VALUE! |
| ENSMUSG00000081688.2 | 0 | NA | NA | NA | NA | NA | #VALUE! |
| ENSMUSG00000081689.2 | 0 | NA | NA | NA | NA | NA | #VALUE! |
| ENSMUSG00000081690.2 | 0 | NA | NA | NA | NA | NA | #VALUE! |
| ENSMUSG00000081691.2 | 0 | NA | NA | NA | NA | NA | #VALUE! |
| ENSMUSG00000081692.2 | 0 | NA | NA | NA | NA | NA | #VALUE! |
| ENSMUSG00000081693.3 | 0 | NA | NA | NA | NA | NA | #VALUE! |
| ENSMUSG00000081694.2 | 0 | NA | NA | NA | NA | NA | #VALUE! |
| ENSMUSG00000081695.2 | 0 | NA | NA | NA | NA | NA | #VALUE! |
| ENSMUSG00000081696.3 | 0 | NA | NA | NA | NA | NA | #VALUE! |
| ENSMUSG00000081697.2 | 0 | NA | NA | NA | NA | NA | #VALUE! |
| ENSMUSG00000081698.2 | 0 | NA | NA | NA | NA | NA | #VALUE! |
| ENSMUSG00000081699.2 | 0 | NA | NA | NA | NA | NA | #VALUE! |
| ENSMUSG00000081700.2 | 0.933963 | 0.404526 | 3.925219 | 0.103058 | 0.917917 | NA | #VALUE! |
| ENSMUSG00000081701.2 | 0 | NA | NA | NA | NA | NA | #VALUE! |
| ENSMUSG00000081703.2 | 0 | NA | NA | NA | NA | NA | #VALUE! |
| ENSMUSG00000081704.2 | 0 | NA | NA | NA | NA | NA | #VALUE! |
| ENSMUSG00000081705.2 | 0 | NA | NA | NA | NA | NA | #VALUE! |
| ENSMUSG00000081706.2 | 0 | NA | NA | NA | NA | NA | #VALUE! |

| | | | | | | | |
|---|---|---|---|---|---|---|---|
| ENSMUSG00000081707.2 | 0 | NA | NA | NA | NA | NA | #VALUE! |
| ENSMUSG00000081709.2 | 0 | NA | NA | NA | NA | NA | #VALUE! |
| ENSMUSG00000081711.2 | 0 | NA | NA | NA | NA | NA | #VALUE! |
| ENSMUSG00000081712.2 | 0 | NA | NA | NA | NA | NA | #VALUE! |
| ENSMUSG00000081713.2 | 0 | NA | NA | NA | NA | NA | #VALUE! |
| ENSMUSG00000081714.2 | 0 | NA | NA | NA | NA | NA | #VALUE! |
| ENSMUSG00000081715.2 | 3.689633 | 5.364344 | 3.919007 | 1.368802 | 0.171061 | NA | #VALUE! |
| ENSMUSG00000081716.2 | 0 | NA | NA | NA | NA | NA | #VALUE! |
| ENSMUSG00000081717.2 | 0 | NA | NA | NA | NA | NA | #VALUE! |
| ENSMUSG00000081718.2 | 0 | NA | NA | NA | NA | NA | #VALUE! |
| ENSMUSG00000081719.2 | 0 | NA | NA | NA | NA | NA | #VALUE! |
| ENSMUSG00000081720.2 | 0 | NA | NA | NA | NA | NA | #VALUE! |
| ENSMUSG00000081721.4 | 0.168284 | 1.027628 | 4.067126 | 0.252667 | 0.800526 | NA | #VALUE! |
| ENSMUSG00000081722.2 | 0 | NA | NA | NA | NA | NA | #VALUE! |
| ENSMUSG00000081723.2 | 0 | NA | NA | NA | NA | NA | #VALUE! |
| ENSMUSG00000081724.5 | 0 | NA | NA | NA | NA | NA | #VALUE! |
| ENSMUSG00000081728.2 | 0 | NA | NA | NA | NA | NA | #VALUE! |
| ENSMUSG00000081729.2 | 0.18651 | 1.401769 | 4.056354 | 0.345574 | 0.729663 | NA | #VALUE! |
| ENSMUSG00000081730.2 | 0 | NA | NA | NA | NA | NA | #VALUE! |
| ENSMUSG00000081731.2 | 0 | NA | NA | NA | NA | NA | #VALUE! |
| ENSMUSG00000081732.2 | 0 | NA | NA | NA | NA | NA | #VALUE! |
| ENSMUSG00000081733.2 | 0 | NA | NA | NA | NA | NA | #VALUE! |
| ENSMUSG00000081734.2 | 0 | NA | NA | NA | NA | NA | #VALUE! |
| ENSMUSG00000081735.2 | 0.297366 | -0.14751 | 4.065279 | -0.03629 | 0.971054 | NA | #VALUE! |
| ENSMUSG00000081736.2 | 0 | NA | NA | NA | NA | NA | #VALUE! |
| ENSMUSG00000081737.3 | 0 | NA | NA | NA | NA | NA | #VALUE! |
| ENSMUSG00000081738.2 | 0 | NA | NA | NA | NA | NA | #VALUE! |
| ENSMUSG00000081739.4 | 0 | NA | NA | NA | NA | NA | #VALUE! |
| ENSMUSG00000081740.4 | 0 | NA | NA | NA | NA | NA | #VALUE! |
| ENSMUSG00000081741.2 | 0 | NA | NA | NA | NA | NA | #VALUE! |
| ENSMUSG00000081742.3 | 0 | NA | NA | NA | NA | NA | #VALUE! |
| ENSMUSG00000081743.4 | 0 | NA | NA | NA | NA | NA | #VALUE! |
| ENSMUSG00000081744.2 | 0 | NA | NA | NA | NA | NA | #VALUE! |
| ENSMUSG00000081745.2 | 0 | NA | NA | NA | NA | NA | #VALUE! |
| ENSMUSG00000081746.3 | 0 | NA | NA | NA | NA | NA | #VALUE! |
| ENSMUSG00000081747.2 | 0 | NA | NA | NA | NA | NA | #VALUE! |
| ENSMUSG00000081748.2 | 0 | NA | NA | NA | NA | NA | #VALUE! |
| ENSMUSG00000081749.4 | 0 | NA | NA | NA | NA | NA | #VALUE! |
| ENSMUSG00000081750.4 | 0 | NA | NA | NA | NA | NA | #VALUE! |
| ENSMUSG00000081751.2 | 0 | NA | NA | NA | NA | NA | #VALUE! |
| ENSMUSG00000081752.4 | 0 | NA | NA | NA | NA | NA | #VALUE! |
| ENSMUSG00000081755.2 | 0 | NA | NA | NA | NA | NA | #VALUE! |
| ENSMUSG00000081756.2 | 0 | NA | NA | NA | NA | NA | #VALUE! |
| ENSMUSG00000081757.2 | 0 | NA | NA | NA | NA | NA | #VALUE! |

| | | | | | | | |
|---|---|---|---|---|---|---|---|
| ENSMUSG00000081758.2 | 0 | NA | NA | NA | NA | NA | #VALUE! |
| ENSMUSG00000081761.3 | 0 | NA | NA | NA | NA | NA | #VALUE! |
| ENSMUSG00000081762.2 | 0 | NA | NA | NA | NA | NA | #VALUE! |
| ENSMUSG00000081763.2 | 0 | NA | NA | NA | NA | NA | #VALUE! |
| ENSMUSG00000081765.2 | 0 | NA | NA | NA | NA | NA | #VALUE! |
| ENSMUSG00000081766.4 | 0 | NA | NA | NA | NA | NA | #VALUE! |
| ENSMUSG00000081767.2 | 0 | NA | NA | NA | NA | NA | #VALUE! |
| ENSMUSG00000081768.3 | 0 | NA | NA | NA | NA | NA | #VALUE! |
| ENSMUSG00000081769.10 | 0 | NA | NA | NA | NA | NA | #VALUE! |
| ENSMUSG00000081770.2 | 0 | NA | NA | NA | NA | NA | #VALUE! |
| ENSMUSG00000081771.2 | 0 | NA | NA | NA | NA | NA | #VALUE! |
| ENSMUSG00000081772.2 | 0.598106 | 2.755755 | 3.98023 | 0.692361 | 0.488711 | NA | #VALUE! |
| ENSMUSG00000081773.2 | 0 | NA | NA | NA | NA | NA | #VALUE! |
| ENSMUSG00000081774.2 | 0 | NA | NA | NA | NA | NA | #VALUE! |
| ENSMUSG00000081775.2 | 0 | NA | NA | NA | NA | NA | #VALUE! |
| ENSMUSG00000081777.2 | 0 | NA | NA | NA | NA | NA | #VALUE! |
| ENSMUSG00000081778.2 | 0 | NA | NA | NA | NA | NA | #VALUE! |
| ENSMUSG00000081779.2 | 0 | NA | NA | NA | NA | NA | #VALUE! |
| ENSMUSG00000081781.2 | 0 | NA | NA | NA | NA | NA | #VALUE! |
| ENSMUSG00000081782.2 | 0 | NA | NA | NA | NA | NA | #VALUE! |
| ENSMUSG00000081783.3 | 0 | NA | NA | NA | NA | NA | #VALUE! |
| ENSMUSG00000081784.2 | 0 | NA | NA | NA | NA | NA | #VALUE! |
| ENSMUSG00000081786.2 | 0 | NA | NA | NA | NA | NA | #VALUE! |
| ENSMUSG00000081787.2 | 0 | NA | NA | NA | NA | NA | #VALUE! |
| ENSMUSG00000081788.3 | 0 | NA | NA | NA | NA | NA | #VALUE! |
| ENSMUSG00000081789.2 | 0 | NA | NA | NA | NA | NA | #VALUE! |
| ENSMUSG00000081791.2 | 0 | NA | NA | NA | NA | NA | #VALUE! |
| ENSMUSG00000081792.5 | 0 | NA | NA | NA | NA | NA | #VALUE! |
| ENSMUSG00000081793.2 | 0 | NA | NA | NA | NA | NA | #VALUE! |
| ENSMUSG00000081796.2 | 0 | NA | NA | NA | NA | NA | #VALUE! |
| ENSMUSG00000081797.2 | 0 | NA | NA | NA | NA | NA | #VALUE! |
| ENSMUSG00000081798.2 | 0 | NA | NA | NA | NA | NA | #VALUE! |
| ENSMUSG00000081799.2 | 0 | NA | NA | NA | NA | NA | #VALUE! |
| ENSMUSG00000081800.4 | 0 | NA | NA | NA | NA | NA | #VALUE! |
| ENSMUSG00000081801.2 | 0 | NA | NA | NA | NA | NA | #VALUE! |
| ENSMUSG00000081803.2 | 0 | NA | NA | NA | NA | NA | #VALUE! |
| ENSMUSG00000081804.2 | 0 | NA | NA | NA | NA | NA | #VALUE! |
| ENSMUSG00000081805.2 | 0 | NA | NA | NA | NA | NA | #VALUE! |
| ENSMUSG00000081806.2 | 0 | NA | NA | NA | NA | NA | #VALUE! |
| ENSMUSG00000081807.2 | 0 | NA | NA | NA | NA | NA | #VALUE! |
| ENSMUSG00000081809.3 | 0 | NA | NA | NA | NA | NA | #VALUE! |
| ENSMUSG00000081810.2 | 0 | NA | NA | NA | NA | NA | #VALUE! |
| ENSMUSG00000081811.2 | 0 | NA | NA | NA | NA | NA | #VALUE! |
| ENSMUSG00000081812.2 | 0 | NA | NA | NA | NA | NA | #VALUE! |

| | | | | | | | |
|---|---|---|---|---|---|---|---|
| ENSMUSG00000081813.2 | 0 | NA | NA | NA | NA | NA | #VALUE! |
| ENSMUSG00000081814.2 | 0 | NA | NA | NA | NA | NA | #VALUE! |
| ENSMUSG00000081816.2 | 0 | NA | NA | NA | NA | NA | #VALUE! |
| ENSMUSG00000081817.2 | 0 | NA | NA | NA | NA | NA | #VALUE! |
| ENSMUSG00000081819.2 | 0 | NA | NA | NA | NA | NA | #VALUE! |
| ENSMUSG00000081820.2 | 0 | NA | NA | NA | NA | NA | #VALUE! |
| ENSMUSG00000081821.2 | 0 | NA | NA | NA | NA | NA | #VALUE! |
| ENSMUSG00000081822.2 | 0 | NA | NA | NA | NA | NA | #VALUE! |
| ENSMUSG00000081823.2 | 0 | NA | NA | NA | NA | NA | #VALUE! |
| ENSMUSG00000081827.2 | 0 | NA | NA | NA | NA | NA | #VALUE! |
| ENSMUSG00000081828.2 | 0 | NA | NA | NA | NA | NA | #VALUE! |
| ENSMUSG00000081829.2 | 0 | NA | NA | NA | NA | NA | #VALUE! |
| ENSMUSG00000081830.2 | 0 | NA | NA | NA | NA | NA | #VALUE! |
| ENSMUSG00000081831.2 | 0 | NA | NA | NA | NA | NA | #VALUE! |
| ENSMUSG00000081832.2 | 0 | NA | NA | NA | NA | NA | #VALUE! |
| ENSMUSG00000081833.2 | 0 | NA | NA | NA | NA | NA | #VALUE! |
| ENSMUSG00000081834.2 | 0 | NA | NA | NA | NA | NA | #VALUE! |
| ENSMUSG00000081835.2 | 0 | NA | NA | NA | NA | NA | #VALUE! |
| ENSMUSG00000081836.4 | 0 | NA | NA | NA | NA | NA | #VALUE! |
| ENSMUSG00000081837.2 | 0 | NA | NA | NA | NA | NA | #VALUE! |
| ENSMUSG00000081838.2 | 0 | NA | NA | NA | NA | NA | #VALUE! |
| ENSMUSG00000081839.2 | 0 | NA | NA | NA | NA | NA | #VALUE! |
| ENSMUSG00000081840.2 | 0 | NA | NA | NA | NA | NA | #VALUE! |
| ENSMUSG00000081841.2 | 0 | NA | NA | NA | NA | NA | #VALUE! |
| ENSMUSG00000081842.2 | 0 | NA | NA | NA | NA | NA | #VALUE! |
| ENSMUSG00000081843.2 | 0 | NA | NA | NA | NA | NA | #VALUE! |
| ENSMUSG00000081844.2 | 0 | NA | NA | NA | NA | NA | #VALUE! |
| ENSMUSG00000081845.2 | 0 | NA | NA | NA | NA | NA | #VALUE! |
| ENSMUSG00000081847.2 | 0 | NA | NA | NA | NA | NA | #VALUE! |
| ENSMUSG00000081848.2 | 0 | NA | NA | NA | NA | NA | #VALUE! |
| ENSMUSG00000081849.3 | 0 | NA | NA | NA | NA | NA | #VALUE! |
| ENSMUSG00000081851.2 | 0 | NA | NA | NA | NA | NA | #VALUE! |
| ENSMUSG00000081852.2 | 0 | NA | NA | NA | NA | NA | #VALUE! |
| ENSMUSG00000081853.2 | 0 | NA | NA | NA | NA | NA | #VALUE! |
| ENSMUSG00000081854.2 | 0 | NA | NA | NA | NA | NA | #VALUE! |
| ENSMUSG00000081856.2 | 0 | NA | NA | NA | NA | NA | #VALUE! |
| ENSMUSG00000081857.2 | 0 | NA | NA | NA | NA | NA | #VALUE! |
| ENSMUSG00000081858.2 | 0 | NA | NA | NA | NA | NA | #VALUE! |
| ENSMUSG00000081859.4 | 0 | NA | NA | NA | NA | NA | #VALUE! |
| ENSMUSG00000081860.2 | 0 | NA | NA | NA | NA | NA | #VALUE! |
| ENSMUSG00000081861.2 | 0 | NA | NA | NA | NA | NA | #VALUE! |
| ENSMUSG00000081862.2 | 0 | NA | NA | NA | NA | NA | #VALUE! |
| ENSMUSG00000081863.2 | 0 | NA | NA | NA | NA | NA | #VALUE! |
| ENSMUSG00000081864.3 | 0 | NA | NA | NA | NA | NA | #VALUE! |

| | | | | | | | |
|---|---|---|---|---|---|---|---|
| ENSMUSG00000081865.2 | 0 | NA | NA | NA | NA | NA | #VALUE! |
| ENSMUSG00000081866.2 | 0 | NA | NA | NA | NA | NA | #VALUE! |
| ENSMUSG00000081868.2 | 0 | NA | NA | NA | NA | NA | #VALUE! |
| ENSMUSG00000081869.2 | 0 | NA | NA | NA | NA | NA | #VALUE! |
| ENSMUSG00000081870.2 | 0 | NA | NA | NA | NA | NA | #VALUE! |
| ENSMUSG00000081871.2 | 0 | NA | NA | NA | NA | NA | #VALUE! |
| ENSMUSG00000081872.2 | 0 | NA | NA | NA | NA | NA | #VALUE! |
| ENSMUSG00000081873.2 | 0 | NA | NA | NA | NA | NA | #VALUE! |
| ENSMUSG00000081874.2 | 0 | NA | NA | NA | NA | NA | #VALUE! |
| ENSMUSG00000081875.2 | 0 | NA | NA | NA | NA | NA | #VALUE! |
| ENSMUSG00000081876.2 | 0 | NA | NA | NA | NA | NA | #VALUE! |
| ENSMUSG00000081877.2 | 0 | NA | NA | NA | NA | NA | #VALUE! |
| ENSMUSG00000081878.2 | 0 | NA | NA | NA | NA | NA | #VALUE! |
| ENSMUSG00000081879.2 | 0 | NA | NA | NA | NA | NA | #VALUE! |
| ENSMUSG00000081880.2 | 0 | NA | NA | NA | NA | NA | #VALUE! |
| ENSMUSG00000081881.2 | 0 | NA | NA | NA | NA | NA | #VALUE! |
| ENSMUSG00000081882.2 | 0 | NA | NA | NA | NA | NA | #VALUE! |
| ENSMUSG00000081883.3 | 0 | NA | NA | NA | NA | NA | #VALUE! |
| ENSMUSG00000081884.2 | 0 | NA | NA | NA | NA | NA | #VALUE! |
| ENSMUSG00000081885.2 | 0 | NA | NA | NA | NA | NA | #VALUE! |
| ENSMUSG00000081886.2 | 0 | NA | NA | NA | NA | NA | #VALUE! |
| ENSMUSG00000081887.2 | 0 | NA | NA | NA | NA | NA | #VALUE! |
| ENSMUSG00000081888.2 | 0.82107 | 3.234732 | 3.959761 | 0.816901 | 0.413985 | NA | #VALUE! |
| ENSMUSG00000081889.2 | 0 | NA | NA | NA | NA | NA | #VALUE! |
| ENSMUSG00000081890.2 | 0 | NA | NA | NA | NA | NA | #VALUE! |
| ENSMUSG00000081891.2 | 0 | NA | NA | NA | NA | NA | #VALUE! |
| ENSMUSG00000081892.3 | 0 | NA | NA | NA | NA | NA | #VALUE! |
| ENSMUSG00000081893.2 | 0 | NA | NA | NA | NA | NA | #VALUE! |
| ENSMUSG00000081894.2 | 0 | NA | NA | NA | NA | NA | #VALUE! |
| ENSMUSG00000081895.4 | 0.469086 | -0.7385 | 4.036584 | -0.18295 | 0.854836 | NA | #VALUE! |
| ENSMUSG00000081896.2 | 0 | NA | NA | NA | NA | NA | #VALUE! |
| ENSMUSG00000081897.2 | 0 | NA | NA | NA | NA | NA | #VALUE! |
| ENSMUSG00000081898.2 | 0 | NA | NA | NA | NA | NA | #VALUE! |
| ENSMUSG00000081900.2 | 0 | NA | NA | NA | NA | NA | #VALUE! |
| ENSMUSG00000081901.2 | 0 | NA | NA | NA | NA | NA | #VALUE! |
| ENSMUSG00000081903.2 | 0 | NA | NA | NA | NA | NA | #VALUE! |
| ENSMUSG00000081904.3 | 0 | NA | NA | NA | NA | NA | #VALUE! |
| ENSMUSG00000081905.2 | 0 | NA | NA | NA | NA | NA | #VALUE! |
| ENSMUSG00000081906.4 | 0 | NA | NA | NA | NA | NA | #VALUE! |
| ENSMUSG00000081907.2 | 0 | NA | NA | NA | NA | NA | #VALUE! |
| ENSMUSG00000081908.3 | 0 | NA | NA | NA | NA | NA | #VALUE! |
| ENSMUSG00000081909.2 | 0 | NA | NA | NA | NA | NA | #VALUE! |
| ENSMUSG00000081910.2 | 0 | NA | NA | NA | NA | NA | #VALUE! |
| ENSMUSG00000081911.2 | 0 | NA | NA | NA | NA | NA | #VALUE! |

| | | | | | | | |
|---|---|---|---|---|---|---|---|
| ENSMUSG00000081912.2 | 0 | NA | NA | NA | NA | NA | #VALUE! |
| ENSMUSG00000081913.2 | 0 | NA | NA | NA | NA | NA | #VALUE! |
| ENSMUSG00000081914.2 | 0 | NA | NA | NA | NA | NA | #VALUE! |
| ENSMUSG00000081916.2 | 0 | NA | NA | NA | NA | NA | #VALUE! |
| ENSMUSG00000081917.2 | 0 | NA | NA | NA | NA | NA | #VALUE! |
| ENSMUSG00000081918.5 | 0 | NA | NA | NA | NA | NA | #VALUE! |
| ENSMUSG00000081919.3 | 0 | NA | NA | NA | NA | NA | #VALUE! |
| ENSMUSG00000081920.3 | 0 | NA | NA | NA | NA | NA | #VALUE! |
| ENSMUSG00000081921.2 | 0 | NA | NA | NA | NA | NA | #VALUE! |
| ENSMUSG00000081923.2 | 0 | NA | NA | NA | NA | NA | #VALUE! |
| ENSMUSG00000081924.2 | 0 | NA | NA | NA | NA | NA | #VALUE! |
| ENSMUSG00000081925.2 | 0 | NA | NA | NA | NA | NA | #VALUE! |
| ENSMUSG00000081928.2 | 0 | NA | NA | NA | NA | NA | #VALUE! |
| ENSMUSG00000081929.2 | 1.510663 | 1.958364 | 3.910318 | 0.50082 | 0.616498 | NA | #VALUE! |
| ENSMUSG00000081931.2 | 0 | NA | NA | NA | NA | NA | #VALUE! |
| ENSMUSG00000081932.2 | 0.093255 | 0.868093 | 4.072464 | 0.213162 | 0.831201 | NA | #VALUE! |
| ENSMUSG00000081933.2 | 0 | NA | NA | NA | NA | NA | #VALUE! |
| ENSMUSG00000081934.2 | 0 | NA | NA | NA | NA | NA | #VALUE! |
| ENSMUSG00000081935.2 | 0 | NA | NA | NA | NA | NA | #VALUE! |
| ENSMUSG00000081936.2 | 0 | NA | NA | NA | NA | NA | #VALUE! |
| ENSMUSG00000081937.2 | 0 | NA | NA | NA | NA | NA | #VALUE! |
| ENSMUSG00000081938.2 | 0 | NA | NA | NA | NA | NA | #VALUE! |
| ENSMUSG00000081939.3 | 0 | NA | NA | NA | NA | NA | #VALUE! |
| ENSMUSG00000081940.2 | 0 | NA | NA | NA | NA | NA | #VALUE! |
| ENSMUSG00000081941.2 | 0 | NA | NA | NA | NA | NA | #VALUE! |
| ENSMUSG00000081942.2 | 0 | NA | NA | NA | NA | NA | #VALUE! |
| ENSMUSG00000081943.2 | 0 | NA | NA | NA | NA | NA | #VALUE! |
| ENSMUSG00000081944.2 | 1.020554 | -3.54462 | 3.950745 | -0.8972 | 0.36961 | NA | #VALUE! |
| ENSMUSG00000081945.5 | 0 | NA | NA | NA | NA | NA | #VALUE! |
| ENSMUSG00000081946.2 | 0 | NA | NA | NA | NA | NA | #VALUE! |
| ENSMUSG00000081947.3 | 0 | NA | NA | NA | NA | NA | #VALUE! |
| ENSMUSG00000081948.5 | 0 | NA | NA | NA | NA | NA | #VALUE! |
| ENSMUSG00000081949.4 | 0 | NA | NA | NA | NA | NA | #VALUE! |
| ENSMUSG00000081950.2 | 0 | NA | NA | NA | NA | NA | #VALUE! |
| ENSMUSG00000081951.2 | 0 | NA | NA | NA | NA | NA | #VALUE! |
| ENSMUSG00000081952.2 | 0 | NA | NA | NA | NA | NA | #VALUE! |
| ENSMUSG00000081953.2 | 0 | NA | NA | NA | NA | NA | #VALUE! |
| ENSMUSG00000081954.2 | 0 | NA | NA | NA | NA | NA | #VALUE! |
| ENSMUSG00000081955.3 | 0 | NA | NA | NA | NA | NA | #VALUE! |
| ENSMUSG00000081957.6 | 0 | NA | NA | NA | NA | NA | #VALUE! |
| ENSMUSG00000081958.2 | 0 | NA | NA | NA | NA | NA | #VALUE! |
| ENSMUSG00000081959.2 | 0 | NA | NA | NA | NA | NA | #VALUE! |
| ENSMUSG00000081960.4 | 0 | NA | NA | NA | NA | NA | #VALUE! |
| ENSMUSG00000081961.2 | 0 | NA | NA | NA | NA | NA | #VALUE! |

| | | | | | | | |
|---|---|---|---|---|---|---|---|
| ENSMUSG00000081962.2 | 0 | NA | NA | NA | NA | NA | #VALUE! |
| ENSMUSG00000081964.2 | 0 | NA | NA | NA | NA | NA | #VALUE! |
| ENSMUSG00000081965.2 | 0 | NA | NA | NA | NA | NA | #VALUE! |
| ENSMUSG00000081966.2 | 0 | NA | NA | NA | NA | NA | #VALUE! |
| ENSMUSG00000081967.2 | 0.391578 | -2.1489 | 4.020883 | -0.53444 | 0.59304 | NA | #VALUE! |
| ENSMUSG00000081968.2 | 0 | NA | NA | NA | NA | NA | #VALUE! |
| ENSMUSG00000081969.2 | 0 | NA | NA | NA | NA | NA | #VALUE! |
| ENSMUSG00000081970.2 | 0 | NA | NA | NA | NA | NA | #VALUE! |
| ENSMUSG00000081971.2 | 0 | NA | NA | NA | NA | NA | #VALUE! |
| ENSMUSG00000081972.2 | 0 | NA | NA | NA | NA | NA | #VALUE! |
| ENSMUSG00000081973.2 | 0 | NA | NA | NA | NA | NA | #VALUE! |
| ENSMUSG00000081974.4 | 0 | NA | NA | NA | NA | NA | #VALUE! |
| ENSMUSG00000081975.2 | 0.857228 | -1.03306 | 3.943583 | -0.26196 | 0.793352 | NA | #VALUE! |
| ENSMUSG00000081976.2 | 0 | NA | NA | NA | NA | NA | #VALUE! |
| ENSMUSG00000081978.2 | 0 | NA | NA | NA | NA | NA | #VALUE! |
| ENSMUSG00000081979.2 | 0 | NA | NA | NA | NA | NA | #VALUE! |
| ENSMUSG00000081980.2 | 0 | NA | NA | NA | NA | NA | #VALUE! |
| ENSMUSG00000081982.6 | 0 | NA | NA | NA | NA | NA | #VALUE! |
| ENSMUSG00000081984.4 | 0 | NA | NA | NA | NA | NA | #VALUE! |
| ENSMUSG00000081985.2 | 0 | NA | NA | NA | NA | NA | #VALUE! |
| ENSMUSG00000081986.2 | 0 | NA | NA | NA | NA | NA | #VALUE! |
| ENSMUSG00000081987.2 | 0 | NA | NA | NA | NA | NA | #VALUE! |
| ENSMUSG00000081988.2 | 0 | NA | NA | NA | NA | NA | #VALUE! |
| ENSMUSG00000081991.2 | 0 | NA | NA | NA | NA | NA | #VALUE! |
| ENSMUSG00000081992.2 | 0 | NA | NA | NA | NA | NA | #VALUE! |
| ENSMUSG00000081993.2 | 0 | NA | NA | NA | NA | NA | #VALUE! |
| ENSMUSG00000081994.2 | 0 | NA | NA | NA | NA | NA | #VALUE! |
| ENSMUSG00000081995.2 | 0 | NA | NA | NA | NA | NA | #VALUE! |
| ENSMUSG00000081996.2 | 0 | NA | NA | NA | NA | NA | #VALUE! |
| ENSMUSG00000081998.2 | 0 | NA | NA | NA | NA | NA | #VALUE! |
| ENSMUSG00000081999.4 | 0.093255 | 0.868093 | 4.072464 | 0.213162 | 0.831201 | NA | #VALUE! |
| ENSMUSG00000082000.2 | 0 | NA | NA | NA | NA | NA | #VALUE! |
| ENSMUSG00000082001.4 | 0 | NA | NA | NA | NA | NA | #VALUE! |
| ENSMUSG00000082002.2 | 0 | NA | NA | NA | NA | NA | #VALUE! |
| ENSMUSG00000082005.2 | 0 | NA | NA | NA | NA | NA | #VALUE! |
| ENSMUSG00000082006.2 | 0 | NA | NA | NA | NA | NA | #VALUE! |
| ENSMUSG00000082007.2 | 0 | NA | NA | NA | NA | NA | #VALUE! |
| ENSMUSG00000082008.2 | 0 | NA | NA | NA | NA | NA | #VALUE! |
| ENSMUSG00000082009.2 | 0 | NA | NA | NA | NA | NA | #VALUE! |
| ENSMUSG00000082010.4 | 0 | NA | NA | NA | NA | NA | #VALUE! |
| ENSMUSG00000082011.2 | 0 | NA | NA | NA | NA | NA | #VALUE! |
| ENSMUSG00000082012.2 | 0 | NA | NA | NA | NA | NA | #VALUE! |
| ENSMUSG00000082013.2 | 0 | NA | NA | NA | NA | NA | #VALUE! |
| ENSMUSG00000082014.3 | 0 | NA | NA | NA | NA | NA | #VALUE! |

| | | | | | | | |
|---|---|---|---|---|---|---|---|
| ENSMUSG00000082015.2 | 0 | NA | NA | NA | NA | NA | #VALUE! |
| ENSMUSG00000082016.4 | 0 | NA | NA | NA | NA | NA | #VALUE! |
| ENSMUSG00000082017.2 | 0 | NA | NA | NA | NA | NA | #VALUE! |
| ENSMUSG00000082018.3 | 0 | NA | NA | NA | NA | NA | #VALUE! |
| ENSMUSG00000082019.2 | 0 | NA | NA | NA | NA | NA | #VALUE! |
| ENSMUSG00000082020.2 | 0 | NA | NA | NA | NA | NA | #VALUE! |
| ENSMUSG00000082021.2 | 0 | NA | NA | NA | NA | NA | #VALUE! |
| ENSMUSG00000082022.2 | 0 | NA | NA | NA | NA | NA | #VALUE! |
| ENSMUSG00000082023.4 | 0 | NA | NA | NA | NA | NA | #VALUE! |
| ENSMUSG00000082024.2 | 0 | NA | NA | NA | NA | NA | #VALUE! |
| ENSMUSG00000082025.2 | 0 | NA | NA | NA | NA | NA | #VALUE! |
| ENSMUSG00000082026.2 | 0 | NA | NA | NA | NA | NA | #VALUE! |
| ENSMUSG00000082027.2 | 0 | NA | NA | NA | NA | NA | #VALUE! |
| ENSMUSG00000082028.2 | 0 | NA | NA | NA | NA | NA | #VALUE! |
| ENSMUSG00000082029.2 | 0.168284 | 1.027628 | 4.067126 | 0.252667 | 0.800526 | NA | #VALUE! |
| ENSMUSG00000082030.2 | 0 | NA | NA | NA | NA | NA | #VALUE! |
| ENSMUSG00000082031.2 | 0 | NA | NA | NA | NA | NA | #VALUE! |
| ENSMUSG00000082032.2 | 0 | NA | NA | NA | NA | NA | #VALUE! |
| ENSMUSG00000082033.4 | 0 | NA | NA | NA | NA | NA | #VALUE! |
| ENSMUSG00000082035.2 | 3.858142 | -1.50614 | 2.809504 | -0.53609 | 0.591897 | NA | #VALUE! |
| ENSMUSG00000082036.2 | 0 | NA | NA | NA | NA | NA | #VALUE! |
| ENSMUSG00000082037.2 | 0 | NA | NA | NA | NA | NA | #VALUE! |
| ENSMUSG00000082038.2 | 0 | NA | NA | NA | NA | NA | #VALUE! |
| ENSMUSG00000082039.3 | 1.224665 | -3.81006 | 3.943408 | -0.96618 | 0.333952 | NA | #VALUE! |
| ENSMUSG00000082040.2 | 0 | NA | NA | NA | NA | NA | #VALUE! |
| ENSMUSG00000082042.2 | 0 | NA | NA | NA | NA | NA | #VALUE! |
| ENSMUSG00000082043.2 | 0 | NA | NA | NA | NA | NA | #VALUE! |
| ENSMUSG00000082044.2 | 2.629076 | 1.806633 | 3.826154 | 0.47218 | 0.636798 | NA | #VALUE! |
| ENSMUSG00000082045.5 | 0 | NA | NA | NA | NA | NA | #VALUE! |
| ENSMUSG00000082046.2 | 0 | NA | NA | NA | NA | NA | #VALUE! |
| ENSMUSG00000082047.2 | 0 | NA | NA | NA | NA | NA | #VALUE! |
| ENSMUSG00000082048.2 | 0 | NA | NA | NA | NA | NA | #VALUE! |
| ENSMUSG00000082049.2 | 0 | NA | NA | NA | NA | NA | #VALUE! |
| ENSMUSG00000082051.2 | 1.531821 | 2.880544 | 3.933017 | 0.7324 | 0.463924 | NA | #VALUE! |
| ENSMUSG00000082052.2 | 0 | NA | NA | NA | NA | NA | #VALUE! |
| ENSMUSG00000082053.2 | 0 | NA | NA | NA | NA | NA | #VALUE! |
| ENSMUSG00000082054.2 | 0 | NA | NA | NA | NA | NA | #VALUE! |
| ENSMUSG00000082056.2 | 0 | NA | NA | NA | NA | NA | #VALUE! |
| ENSMUSG00000082057.2 | 0 | NA | NA | NA | NA | NA | #VALUE! |
| ENSMUSG00000082058.2 | 0 | NA | NA | NA | NA | NA | #VALUE! |
| ENSMUSG00000082059.2 | 0 | NA | NA | NA | NA | NA | #VALUE! |
| ENSMUSG00000082060.2 | 0 | NA | NA | NA | NA | NA | #VALUE! |
| ENSMUSG00000082061.2 | 0 | NA | NA | NA | NA | NA | #VALUE! |
| ENSMUSG00000082062.2 | 0 | NA | NA | NA | NA | NA | #VALUE! |

| | | | | | | | |
|---|---|---|---|---|---|---|---|
| ENSMUSG00000082063.2 | 0 | NA | NA | NA | NA | NA | #VALUE! |
| ENSMUSG00000082064.4 | 0 | NA | NA | NA | NA | NA | #VALUE! |
| ENSMUSG00000082066.2 | 0 | NA | NA | NA | NA | NA | #VALUE! |
| ENSMUSG00000082068.2 | 0 | NA | NA | NA | NA | NA | #VALUE! |
| ENSMUSG00000082069.3 | 0 | NA | NA | NA | NA | NA | #VALUE! |
| ENSMUSG00000082070.2 | 0 | NA | NA | NA | NA | NA | #VALUE! |
| ENSMUSG00000082071.2 | 0 | NA | NA | NA | NA | NA | #VALUE! |
| ENSMUSG00000082072.2 | 0 | NA | NA | NA | NA | NA | #VALUE! |
| ENSMUSG00000082073.2 | 0 | NA | NA | NA | NA | NA | #VALUE! |
| ENSMUSG00000082074.2 | 0 | NA | NA | NA | NA | NA | #VALUE! |
| ENSMUSG00000082076.2 | 0 | NA | NA | NA | NA | NA | #VALUE! |
| ENSMUSG00000082077.2 | 0 | NA | NA | NA | NA | NA | #VALUE! |
| ENSMUSG00000082078.2 | 0 | NA | NA | NA | NA | NA | #VALUE! |
| ENSMUSG00000082079.3 | 0 | NA | NA | NA | NA | NA | #VALUE! |
| ENSMUSG00000082080.4 | 0 | NA | NA | NA | NA | NA | #VALUE! |
| ENSMUSG00000082081.2 | 0 | NA | NA | NA | NA | NA | #VALUE! |
| ENSMUSG00000082082.2 | 0 | NA | NA | NA | NA | NA | #VALUE! |
| ENSMUSG00000082083.3 | 0 | NA | NA | NA | NA | NA | #VALUE! |
| ENSMUSG00000082084.2 | 0 | NA | NA | NA | NA | NA | #VALUE! |
| ENSMUSG00000082085.2 | 0 | NA | NA | NA | NA | NA | #VALUE! |
| ENSMUSG00000082087.5 | 0 | NA | NA | NA | NA | NA | #VALUE! |
| ENSMUSG00000082088.2 | 0 | NA | NA | NA | NA | NA | #VALUE! |
| ENSMUSG00000082089.2 | 0 | NA | NA | NA | NA | NA | #VALUE! |
| ENSMUSG00000082090.2 | 0 | NA | NA | NA | NA | NA | #VALUE! |
| ENSMUSG00000082091.2 | 0 | NA | NA | NA | NA | NA | #VALUE! |
| ENSMUSG00000082092.2 | 0 | NA | NA | NA | NA | NA | #VALUE! |
| ENSMUSG00000082093.3 | 0 | NA | NA | NA | NA | NA | #VALUE! |
| ENSMUSG00000082095.2 | 0 | NA | NA | NA | NA | NA | #VALUE! |
| ENSMUSG00000082096.2 | 0 | NA | NA | NA | NA | NA | #VALUE! |
| ENSMUSG00000082097.2 | 0 | NA | NA | NA | NA | NA | #VALUE! |
| ENSMUSG00000082098.2 | 0 | NA | NA | NA | NA | NA | #VALUE! |
| ENSMUSG00000082099.2 | 0.18651 | 1.401769 | 4.056354 | 0.345574 | 0.729663 | NA | #VALUE! |
| ENSMUSG00000082100.2 | 0 | NA | NA | NA | NA | NA | #VALUE! |
| ENSMUSG00000082101.4 | 0 | NA | NA | NA | NA | NA | #VALUE! |
| ENSMUSG00000082103.3 | 0 | NA | NA | NA | NA | NA | #VALUE! |
| ENSMUSG00000082104.2 | 0 | NA | NA | NA | NA | NA | #VALUE! |
| ENSMUSG00000082106.4 | 0 | NA | NA | NA | NA | NA | #VALUE! |
| ENSMUSG00000082107.2 | 0 | NA | NA | NA | NA | NA | #VALUE! |
| ENSMUSG00000082108.4 | 0.168284 | 1.027628 | 4.067126 | 0.252667 | 0.800526 | NA | #VALUE! |
| ENSMUSG00000082109.2 | 0.198268 | -0.06494 | 4.068276 | -0.01596 | 0.987265 | NA | #VALUE! |
| ENSMUSG00000082110.2 | 1.019338 | 2.454816 | 3.950178 | 0.621444 | 0.534307 | NA | #VALUE! |
| ENSMUSG00000082111.2 | 0 | NA | NA | NA | NA | NA | #VALUE! |
| ENSMUSG00000082112.2 | 0 | NA | NA | NA | NA | NA | #VALUE! |
| ENSMUSG00000082113.2 | 0 | NA | NA | NA | NA | NA | #VALUE! |

| | | | | | | | |
|---|---|---|---|---|---|---|---|
| ENSMUSG00000082114.3 | 0 | NA | NA | NA | NA | NA | #VALUE! |
| ENSMUSG00000082115.3 | 0 | NA | NA | NA | NA | NA | #VALUE! |
| ENSMUSG00000082116.2 | 0 | NA | NA | NA | NA | NA | #VALUE! |
| ENSMUSG00000082117.2 | 0 | NA | NA | NA | NA | NA | #VALUE! |
| ENSMUSG00000082118.2 | 0 | NA | NA | NA | NA | NA | #VALUE! |
| ENSMUSG00000082119.2 | 0 | NA | NA | NA | NA | NA | #VALUE! |
| ENSMUSG00000082120.2 | 0 | NA | NA | NA | NA | NA | #VALUE! |
| ENSMUSG00000082121.2 | 0 | NA | NA | NA | NA | NA | #VALUE! |
| ENSMUSG00000082122.2 | 0 | NA | NA | NA | NA | NA | #VALUE! |
| ENSMUSG00000082123.2 | 0 | NA | NA | NA | NA | NA | #VALUE! |
| ENSMUSG00000082125.2 | 0 | NA | NA | NA | NA | NA | #VALUE! |
| ENSMUSG00000082126.2 | 0 | NA | NA | NA | NA | NA | #VALUE! |
| ENSMUSG00000082127.3 | 1.1982 | 0.311726 | 3.87758 | 0.080392 | 0.935926 | NA | #VALUE! |
| ENSMUSG00000082128.2 | 0.105013 | -0.97324 | 4.076293 | -0.23876 | 0.811295 | NA | #VALUE! |
| ENSMUSG00000082129.2 | 0 | NA | NA | NA | NA | NA | #VALUE! |
| ENSMUSG00000082130.2 | 0 | NA | NA | NA | NA | NA | #VALUE! |
| ENSMUSG00000082132.2 | 0 | NA | NA | NA | NA | NA | #VALUE! |
| ENSMUSG00000082134.2 | 0 | NA | NA | NA | NA | NA | #VALUE! |
| ENSMUSG00000082135.2 | 0 | NA | NA | NA | NA | NA | #VALUE! |
| ENSMUSG00000082136.2 | 0 | NA | NA | NA | NA | NA | #VALUE! |
| ENSMUSG00000082138.2 | 0 | NA | NA | NA | NA | NA | #VALUE! |
| ENSMUSG00000082139.2 | 0 | NA | NA | NA | NA | NA | #VALUE! |
| ENSMUSG00000082140.2 | 0 | NA | NA | NA | NA | NA | #VALUE! |
| ENSMUSG00000082141.2 | 0 | NA | NA | NA | NA | NA | #VALUE! |
| ENSMUSG00000082142.2 | 0 | NA | NA | NA | NA | NA | #VALUE! |
| ENSMUSG00000082143.2 | 0 | NA | NA | NA | NA | NA | #VALUE! |
| ENSMUSG00000082144.2 | 0 | NA | NA | NA | NA | NA | #VALUE! |
| ENSMUSG00000082145.2 | 0 | NA | NA | NA | NA | NA | #VALUE! |
| ENSMUSG00000082146.2 | 0 | NA | NA | NA | NA | NA | #VALUE! |
| ENSMUSG00000082147.2 | 0 | NA | NA | NA | NA | NA | #VALUE! |
| ENSMUSG00000082148.2 | 0 | NA | NA | NA | NA | NA | #VALUE! |
| ENSMUSG00000082149.2 | 0 | NA | NA | NA | NA | NA | #VALUE! |
| ENSMUSG00000082151.4 | 0 | NA | NA | NA | NA | NA | #VALUE! |
| ENSMUSG00000082152.2 | 0 | NA | NA | NA | NA | NA | #VALUE! |
| ENSMUSG00000082154.2 | 0 | NA | NA | NA | NA | NA | #VALUE! |
| ENSMUSG00000082155.2 | 0 | NA | NA | NA | NA | NA | #VALUE! |
| ENSMUSG00000082156.2 | 0 | NA | NA | NA | NA | NA | #VALUE! |
| ENSMUSG00000082157.2 | 0 | NA | NA | NA | NA | NA | #VALUE! |
| ENSMUSG00000082158.2 | 0 | NA | NA | NA | NA | NA | #VALUE! |
| ENSMUSG00000082159.3 | 0 | NA | NA | NA | NA | NA | #VALUE! |
| ENSMUSG00000082160.2 | 0 | NA | NA | NA | NA | NA | #VALUE! |
| ENSMUSG00000082161.2 | 0 | NA | NA | NA | NA | NA | #VALUE! |
| ENSMUSG00000082162.2 | 0 | NA | NA | NA | NA | NA | #VALUE! |
| ENSMUSG00000082163.3 | 0 | NA | NA | NA | NA | NA | #VALUE! |

| | | | | | | | |
|---|---|---|---|---|---|---|---|
| ENSMUSG00000082164.2 | 0 | NA | NA | NA | NA | NA | #VALUE! |
| ENSMUSG00000082165.2 | 0 | NA | NA | NA | NA | NA | #VALUE! |
| ENSMUSG00000082167.2 | 0 | NA | NA | NA | NA | NA | #VALUE! |
| ENSMUSG00000082168.2 | 0 | NA | NA | NA | NA | NA | #VALUE! |
| ENSMUSG00000082169.2 | 0 | NA | NA | NA | NA | NA | #VALUE! |
| ENSMUSG00000082170.2 | 0 | NA | NA | NA | NA | NA | #VALUE! |
| ENSMUSG00000082171.2 | 0 | NA | NA | NA | NA | NA | #VALUE! |
| ENSMUSG00000082172.2 | 0 | NA | NA | NA | NA | NA | #VALUE! |
| ENSMUSG00000082173.2 | 0.709588 | 3.017181 | 3.968248 | 0.760331 | 0.447057 | NA | #VALUE! |
| ENSMUSG00000082174.2 | 0 | NA | NA | NA | NA | NA | #VALUE! |
| ENSMUSG00000082175.2 | 0 | NA | NA | NA | NA | NA | #VALUE! |
| ENSMUSG00000082176.2 | 0 | NA | NA | NA | NA | NA | #VALUE! |
| ENSMUSG00000082177.3 | 0 | NA | NA | NA | NA | NA | #VALUE! |
| ENSMUSG00000082178.2 | 0 | NA | NA | NA | NA | NA | #VALUE! |
| ENSMUSG00000082179.2 | 0 | NA | NA | NA | NA | NA | #VALUE! |
| ENSMUSG00000082180.2 | 0 | NA | NA | NA | NA | NA | #VALUE! |
| ENSMUSG00000082181.2 | 0 | NA | NA | NA | NA | NA | #VALUE! |
| ENSMUSG00000082183.2 | 0 | NA | NA | NA | NA | NA | #VALUE! |
| ENSMUSG00000082185.2 | 0 | NA | NA | NA | NA | NA | #VALUE! |
| ENSMUSG00000082186.3 | 0 | NA | NA | NA | NA | NA | #VALUE! |
| ENSMUSG00000082187.2 | 0 | NA | NA | NA | NA | NA | #VALUE! |
| ENSMUSG00000082188.2 | 0 | NA | NA | NA | NA | NA | #VALUE! |
| ENSMUSG00000082189.2 | 0 | NA | NA | NA | NA | NA | #VALUE! |
| ENSMUSG00000082190.2 | 0 | NA | NA | NA | NA | NA | #VALUE! |
| ENSMUSG00000082191.2 | 0 | NA | NA | NA | NA | NA | #VALUE! |
| ENSMUSG00000082192.3 | 0 | NA | NA | NA | NA | NA | #VALUE! |
| ENSMUSG00000082194.4 | 0 | NA | NA | NA | NA | NA | #VALUE! |
| ENSMUSG00000082195.4 | 0 | NA | NA | NA | NA | NA | #VALUE! |
| ENSMUSG00000082196.4 | 0 | NA | NA | NA | NA | NA | #VALUE! |
| ENSMUSG00000082197.2 | 0 | NA | NA | NA | NA | NA | #VALUE! |
| ENSMUSG00000082198.2 | 0 | NA | NA | NA | NA | NA | #VALUE! |
| ENSMUSG00000082199.2 | 0 | NA | NA | NA | NA | NA | #VALUE! |
| ENSMUSG00000082200.2 | 0 | NA | NA | NA | NA | NA | #VALUE! |
| ENSMUSG00000082202.4 | 0 | NA | NA | NA | NA | NA | #VALUE! |
| ENSMUSG00000082203.2 | 0 | NA | NA | NA | NA | NA | #VALUE! |
| ENSMUSG00000082204.3 | 0 | NA | NA | NA | NA | NA | #VALUE! |
| ENSMUSG00000082205.2 | 0 | NA | NA | NA | NA | NA | #VALUE! |
| ENSMUSG00000082207.2 | 0 | NA | NA | NA | NA | NA | #VALUE! |
| ENSMUSG00000082208.2 | 0 | NA | NA | NA | NA | NA | #VALUE! |
| ENSMUSG00000082210.2 | 0 | NA | NA | NA | NA | NA | #VALUE! |
| ENSMUSG00000082211.4 | 0 | NA | NA | NA | NA | NA | #VALUE! |
| ENSMUSG00000082212.2 | 0 | NA | NA | NA | NA | NA | #VALUE! |
| ENSMUSG00000082213.2 | 0 | NA | NA | NA | NA | NA | #VALUE! |
| ENSMUSG00000082214.2 | 0 | NA | NA | NA | NA | NA | #VALUE! |

| | | | | | | | |
|---|---|---|---|---|---|---|---|
| ENSMUSG00000082215.4 | 0 | NA | NA | NA | NA | NA | #VALUE! |
| ENSMUSG00000082216.5 | 0 | NA | NA | NA | NA | NA | #VALUE! |
| ENSMUSG00000082217.2 | 0 | NA | NA | NA | NA | NA | #VALUE! |
| ENSMUSG00000082218.2 | 0 | NA | NA | NA | NA | NA | #VALUE! |
| ENSMUSG00000082220.4 | 3.807703 | -1.99193 | 3.800678 | -0.5241 | 0.60021 | NA | #VALUE! |
| ENSMUSG00000082222.2 | 0 | NA | NA | NA | NA | NA | #VALUE! |
| ENSMUSG00000082223.2 | 0 | NA | NA | NA | NA | NA | #VALUE! |
| ENSMUSG00000082225.2 | 0 | NA | NA | NA | NA | NA | #VALUE! |
| ENSMUSG00000082226.5 | 0 | NA | NA | NA | NA | NA | #VALUE! |
| ENSMUSG00000082227.5 | 0 | NA | NA | NA | NA | NA | #VALUE! |
| ENSMUSG00000082228.2 | 0 | NA | NA | NA | NA | NA | #VALUE! |
| ENSMUSG00000082229.2 | 0 | NA | NA | NA | NA | NA | #VALUE! |
| ENSMUSG00000082233.3 | 0 | NA | NA | NA | NA | NA | #VALUE! |
| ENSMUSG00000082234.2 | 0 | NA | NA | NA | NA | NA | #VALUE! |
| ENSMUSG00000082235.4 | 0 | NA | NA | NA | NA | NA | #VALUE! |
| ENSMUSG00000082236.2 | 0 | NA | NA | NA | NA | NA | #VALUE! |
| ENSMUSG00000082237.3 | 0 | NA | NA | NA | NA | NA | #VALUE! |
| ENSMUSG00000082238.2 | 0 | NA | NA | NA | NA | NA | #VALUE! |
| ENSMUSG00000082239.2 | 0 | NA | NA | NA | NA | NA | #VALUE! |
| ENSMUSG00000082240.2 | 0 | NA | NA | NA | NA | NA | #VALUE! |
| ENSMUSG00000082241.5 | 0.093255 | 0.868093 | 4.072464 | 0.213162 | 0.831201 | NA | #VALUE! |
| ENSMUSG00000082242.2 | 0 | NA | NA | NA | NA | NA | #VALUE! |
| ENSMUSG00000082243.2 | 0 | NA | NA | NA | NA | NA | #VALUE! |
| ENSMUSG00000082244.2 | 0 | NA | NA | NA | NA | NA | #VALUE! |
| ENSMUSG00000082245.2 | 0 | NA | NA | NA | NA | NA | #VALUE! |
| ENSMUSG00000082246.2 | 0 | NA | NA | NA | NA | NA | #VALUE! |
| ENSMUSG00000082247.2 | 0 | NA | NA | NA | NA | NA | #VALUE! |
| ENSMUSG00000082248.2 | 0 | NA | NA | NA | NA | NA | #VALUE! |
| ENSMUSG00000082249.2 | 0 | NA | NA | NA | NA | NA | #VALUE! |
| ENSMUSG00000082250.2 | 0 | NA | NA | NA | NA | NA | #VALUE! |
| ENSMUSG00000082251.2 | 0 | NA | NA | NA | NA | NA | #VALUE! |
| ENSMUSG00000082252.2 | 0 | NA | NA | NA | NA | NA | #VALUE! |
| ENSMUSG00000082253.4 | 0 | NA | NA | NA | NA | NA | #VALUE! |
| ENSMUSG00000082254.2 | 0 | NA | NA | NA | NA | NA | #VALUE! |
| ENSMUSG00000082255.2 | 0 | NA | NA | NA | NA | NA | #VALUE! |
| ENSMUSG00000082256.2 | 0 | NA | NA | NA | NA | NA | #VALUE! |
| ENSMUSG00000082258.2 | 0 | NA | NA | NA | NA | NA | #VALUE! |
| ENSMUSG00000082259.2 | 0 | NA | NA | NA | NA | NA | #VALUE! |
| ENSMUSG00000082260.4 | 0.300802 | -1.80634 | 4.050033 | -0.44601 | 0.655592 | NA | #VALUE! |
| ENSMUSG00000082261.2 | 0 | NA | NA | NA | NA | NA | #VALUE! |
| ENSMUSG00000082262.2 | 0 | NA | NA | NA | NA | NA | #VALUE! |
| ENSMUSG00000082263.2 | 0 | NA | NA | NA | NA | NA | #VALUE! |
| ENSMUSG00000082264.2 | 0.105013 | -0.97324 | 4.076293 | -0.23876 | 0.811295 | NA | #VALUE! |
| ENSMUSG00000082265.2 | 0 | NA | NA | NA | NA | NA | #VALUE! |

| | | | | | | | |
|---|---|---|---|---|---|---|---|
| ENSMUSG00000082266.2 | 0 | NA | NA | NA | NA | NA | #VALUE! |
| ENSMUSG00000082267.2 | 0 | NA | NA | NA | NA | NA | #VALUE! |
| ENSMUSG00000082268.2 | 0 | NA | NA | NA | NA | NA | #VALUE! |
| ENSMUSG00000082269.2 | 0 | NA | NA | NA | NA | NA | #VALUE! |
| ENSMUSG00000082270.2 | 0 | NA | NA | NA | NA | NA | #VALUE! |
| ENSMUSG00000082271.2 | 0 | NA | NA | NA | NA | NA | #VALUE! |
| ENSMUSG00000082272.2 | 4.954985 | 1.870707 | 3.603761 | 0.519099 | 0.603692 | NA | #VALUE! |
| ENSMUSG00000082274.2 | 0 | NA | NA | NA | NA | NA | #VALUE! |
| ENSMUSG00000082275.2 | 0 | NA | NA | NA | NA | NA | #VALUE! |
| ENSMUSG00000082276.2 | 0 | NA | NA | NA | NA | NA | #VALUE! |
| ENSMUSG00000082277.3 | 0 | NA | NA | NA | NA | NA | #VALUE! |
| ENSMUSG00000082278.2 | 0 | NA | NA | NA | NA | NA | #VALUE! |
| ENSMUSG00000082279.2 | 0 | NA | NA | NA | NA | NA | #VALUE! |
| ENSMUSG00000082280.5 | 0 | NA | NA | NA | NA | NA | #VALUE! |
| ENSMUSG00000082281.2 | 0 | NA | NA | NA | NA | NA | #VALUE! |
| ENSMUSG00000082283.2 | 0 | NA | NA | NA | NA | NA | #VALUE! |
| ENSMUSG00000082284.2 | 6.123323 | -6.14178 | 3.914088 | -1.56915 | 0.116614 | NA | #VALUE! |
| ENSMUSG00000082285.2 | 0 | NA | NA | NA | NA | NA | #VALUE! |
| ENSMUSG00000082286.12 | 0 | NA | NA | NA | NA | NA | #VALUE! |
| ENSMUSG00000082287.2 | 0 | NA | NA | NA | NA | NA | #VALUE! |
| ENSMUSG00000082288.2 | 0 | NA | NA | NA | NA | NA | #VALUE! |
| ENSMUSG00000082289.2 | 0.466276 | 2.449889 | 3.997179 | 0.612904 | 0.53994 | NA | #VALUE! |
| ENSMUSG00000082291.2 | 0 | NA | NA | NA | NA | NA | #VALUE! |
| ENSMUSG00000082292.4 | 0.093255 | 0.868093 | 4.072464 | 0.213162 | 0.831201 | NA | #VALUE! |
| ENSMUSG00000082294.3 | 0 | NA | NA | NA | NA | NA | #VALUE! |
| ENSMUSG00000082295.2 | 0 | NA | NA | NA | NA | NA | #VALUE! |
| ENSMUSG00000082296.2 | 0 | NA | NA | NA | NA | NA | #VALUE! |
| ENSMUSG00000082297.3 | 0 | NA | NA | NA | NA | NA | #VALUE! |
| ENSMUSG00000082301.2 | 0 | NA | NA | NA | NA | NA | #VALUE! |
| ENSMUSG00000082302.2 | 0 | NA | NA | NA | NA | NA | #VALUE! |
| ENSMUSG00000082303.2 | 0 | NA | NA | NA | NA | NA | #VALUE! |
| ENSMUSG00000082304.2 | 0 | NA | NA | NA | NA | NA | #VALUE! |
| ENSMUSG00000082305.5 | 0 | NA | NA | NA | NA | NA | #VALUE! |
| ENSMUSG00000082306.3 | 0 | NA | NA | NA | NA | NA | #VALUE! |
| ENSMUSG00000082307.4 | 0 | NA | NA | NA | NA | NA | #VALUE! |
| ENSMUSG00000082308.2 | 0 | NA | NA | NA | NA | NA | #VALUE! |
| ENSMUSG00000082309.4 | 0 | NA | NA | NA | NA | NA | #VALUE! |
| ENSMUSG00000082310.2 | 0 | NA | NA | NA | NA | NA | #VALUE! |
| ENSMUSG00000082311.2 | 0 | NA | NA | NA | NA | NA | #VALUE! |
| ENSMUSG00000082312.2 | 0.195789 | -1.05581 | 4.073302 | -0.2592 | 0.795479 | NA | #VALUE! |
| ENSMUSG00000082313.2 | 0 | NA | NA | NA | NA | NA | #VALUE! |
| ENSMUSG00000082314.3 | 0 | NA | NA | NA | NA | NA | #VALUE! |
| ENSMUSG00000082315.3 | 1.967804 | 4.449735 | 3.929778 | 1.132312 | 0.257503 | NA | #VALUE! |
| ENSMUSG00000082316.5 | 0 | NA | NA | NA | NA | NA | #VALUE! |

| | | | | | | | |
|---|---|---|---|---|---|---|---|
| ENSMUSG00000082317.3 | 0 | NA | NA | NA | NA | NA | #VALUE! |
| ENSMUSG00000082318.4 | 0 | NA | NA | NA | NA | NA | #VALUE! |
| ENSMUSG00000082319.4 | 3.858623 | 4.346749 | 3.911554 | 1.111259 | 0.266457 | NA | #VALUE! |
| ENSMUSG00000082320.2 | 0 | NA | NA | NA | NA | NA | #VALUE! |
| ENSMUSG00000082321.4 | 0 | NA | NA | NA | NA | NA | #VALUE! |
| ENSMUSG00000082322.3 | 0 | NA | NA | NA | NA | NA | #VALUE! |
| ENSMUSG00000082323.3 | 0 | NA | NA | NA | NA | NA | #VALUE! |
| ENSMUSG00000082324.2 | 0 | NA | NA | NA | NA | NA | #VALUE! |
| ENSMUSG00000082325.2 | 0 | NA | NA | NA | NA | NA | #VALUE! |
| ENSMUSG00000082326.2 | 0 | NA | NA | NA | NA | NA | #VALUE! |
| ENSMUSG00000082327.2 | 0 | NA | NA | NA | NA | NA | #VALUE! |
| ENSMUSG00000082328.2 | 0 | NA | NA | NA | NA | NA | #VALUE! |
| ENSMUSG00000082329.2 | 0 | NA | NA | NA | NA | NA | #VALUE! |
| ENSMUSG00000082330.2 | 0 | NA | NA | NA | NA | NA | #VALUE! |
| ENSMUSG00000082332.3 | 0 | NA | NA | NA | NA | NA | #VALUE! |
| ENSMUSG00000082333.2 | 0 | NA | NA | NA | NA | NA | #VALUE! |
| ENSMUSG00000082334.2 | 2.355969 | 4.72622 | 3.925784 | 1.203892 | 0.228631 | NA | #VALUE! |
| ENSMUSG00000082336.2 | 0 | NA | NA | NA | NA | NA | #VALUE! |
| ENSMUSG00000082337.2 | 0 | NA | NA | NA | NA | NA | #VALUE! |
| ENSMUSG00000082338.2 | 0 | NA | NA | NA | NA | NA | #VALUE! |
| ENSMUSG00000082339.2 | 0 | NA | NA | NA | NA | NA | #VALUE! |
| ENSMUSG00000082340.2 | 0 | NA | NA | NA | NA | NA | #VALUE! |
| ENSMUSG00000082341.2 | 0 | NA | NA | NA | NA | NA | #VALUE! |
| ENSMUSG00000082342.2 | 0 | NA | NA | NA | NA | NA | #VALUE! |
| ENSMUSG00000082343.2 | 0 | NA | NA | NA | NA | NA | #VALUE! |
| ENSMUSG00000082344.2 | 0 | NA | NA | NA | NA | NA | #VALUE! |
| ENSMUSG00000082345.2 | 2.664654 | -0.46094 | 3.266793 | -0.1411 | 0.887791 | NA | #VALUE! |
| ENSMUSG00000082346.2 | 0 | NA | NA | NA | NA | NA | #VALUE! |
| ENSMUSG00000082347.2 | 0 | NA | NA | NA | NA | NA | #VALUE! |
| ENSMUSG00000082348.2 | 0 | NA | NA | NA | NA | NA | #VALUE! |
| ENSMUSG00000082350.2 | 0 | NA | NA | NA | NA | NA | #VALUE! |
| ENSMUSG00000082351.2 | 0 | NA | NA | NA | NA | NA | #VALUE! |
| ENSMUSG00000082352.2 | 0 | NA | NA | NA | NA | NA | #VALUE! |
| ENSMUSG00000082353.2 | 0 | NA | NA | NA | NA | NA | #VALUE! |
| ENSMUSG00000082354.2 | 0 | NA | NA | NA | NA | NA | #VALUE! |
| ENSMUSG00000082356.2 | 0 | NA | NA | NA | NA | NA | #VALUE! |
| ENSMUSG00000082357.2 | 0 | NA | NA | NA | NA | NA | #VALUE! |
| ENSMUSG00000082358.2 | 0 | NA | NA | NA | NA | NA | #VALUE! |
| ENSMUSG00000082359.2 | 0 | NA | NA | NA | NA | NA | #VALUE! |
| ENSMUSG00000082360.2 | 0 | NA | NA | NA | NA | NA | #VALUE! |
| ENSMUSG00000082361.7 | 3.266746 | 3.144943 | 3.878516 | 0.810862 | 0.417445 | NA | #VALUE! |
| ENSMUSG00000082362.2 | 0 | NA | NA | NA | NA | NA | #VALUE! |
| ENSMUSG00000082364.2 | 0 | NA | NA | NA | NA | NA | #VALUE! |
| ENSMUSG00000082365.3 | 0 | NA | NA | NA | NA | NA | #VALUE! |

| | | | | | | | |
|---|---|---|---|---|---|---|---|
| ENSMUSG00000082367.2 | 0 | NA | NA | NA | NA | NA | #VALUE! |
| ENSMUSG00000082368.2 | 0 | NA | NA | NA | NA | NA | #VALUE! |
| ENSMUSG00000082369.2 | 0 | NA | NA | NA | NA | NA | #VALUE! |
| ENSMUSG00000082370.2 | 0 | NA | NA | NA | NA | NA | #VALUE! |
| ENSMUSG00000082371.2 | 0 | NA | NA | NA | NA | NA | #VALUE! |
| ENSMUSG00000082372.2 | 0 | NA | NA | NA | NA | NA | #VALUE! |
| ENSMUSG00000082373.3 | 0 | NA | NA | NA | NA | NA | #VALUE! |
| ENSMUSG00000082374.2 | 0 | NA | NA | NA | NA | NA | #VALUE! |
| ENSMUSG00000082375.2 | 0 | NA | NA | NA | NA | NA | #VALUE! |
| ENSMUSG00000082376.2 | 0 | NA | NA | NA | NA | NA | #VALUE! |
| ENSMUSG00000082377.2 | 0.195789 | -1.05581 | 4.073302 | -0.2592 | 0.795479 | NA | #VALUE! |
| ENSMUSG00000082378.3 | 0 | NA | NA | NA | NA | NA | #VALUE! |
| ENSMUSG00000082379.2 | 0 | NA | NA | NA | NA | NA | #VALUE! |
| ENSMUSG00000082380.2 | 0 | NA | NA | NA | NA | NA | #VALUE! |
| ENSMUSG00000082381.2 | 0 | NA | NA | NA | NA | NA | #VALUE! |
| ENSMUSG00000082382.2 | 0 | NA | NA | NA | NA | NA | #VALUE! |
| ENSMUSG00000082383.3 | 0 | NA | NA | NA | NA | NA | #VALUE! |
| ENSMUSG00000082388.2 | 0 | NA | NA | NA | NA | NA | #VALUE! |
| ENSMUSG00000082389.3 | 0 | NA | NA | NA | NA | NA | #VALUE! |
| ENSMUSG00000082390.2 | 0 | NA | NA | NA | NA | NA | #VALUE! |
| ENSMUSG00000082392.2 | 0 | NA | NA | NA | NA | NA | #VALUE! |
| ENSMUSG00000082393.3 | 0 | NA | NA | NA | NA | NA | #VALUE! |
| ENSMUSG00000082394.2 | 0.350989 | 0.094602 | 4.062933 | 0.023284 | 0.981424 | NA | #VALUE! |
| ENSMUSG00000082395.2 | 0 | NA | NA | NA | NA | NA | #VALUE! |
| ENSMUSG00000082396.5 | 0 | NA | NA | NA | NA | NA | #VALUE! |
| ENSMUSG00000082397.2 | 0 | NA | NA | NA | NA | NA | #VALUE! |
| ENSMUSG00000082398.2 | 0 | NA | NA | NA | NA | NA | #VALUE! |
| ENSMUSG00000082399.2 | 0 | NA | NA | NA | NA | NA | #VALUE! |
| ENSMUSG00000082400.2 | 0 | NA | NA | NA | NA | NA | #VALUE! |
| ENSMUSG00000082402.2 | 0 | NA | NA | NA | NA | NA | #VALUE! |
| ENSMUSG00000082403.2 | 0 | NA | NA | NA | NA | NA | #VALUE! |
| ENSMUSG00000082404.2 | 0 | NA | NA | NA | NA | NA | #VALUE! |
| ENSMUSG00000082405.2 | 0 | NA | NA | NA | NA | NA | #VALUE! |
| ENSMUSG00000082406.2 | 0 | NA | NA | NA | NA | NA | #VALUE! |
| ENSMUSG00000082407.2 | 0 | NA | NA | NA | NA | NA | #VALUE! |
| ENSMUSG00000082408.2 | 0 | NA | NA | NA | NA | NA | #VALUE! |
| ENSMUSG00000082410.2 | 0 | NA | NA | NA | NA | NA | #VALUE! |
| ENSMUSG00000082411.2 | 0 | NA | NA | NA | NA | NA | #VALUE! |
| ENSMUSG00000082412.2 | 0 | NA | NA | NA | NA | NA | #VALUE! |
| ENSMUSG00000082413.2 | 0 | NA | NA | NA | NA | NA | #VALUE! |
| ENSMUSG00000082414.3 | 0 | NA | NA | NA | NA | NA | #VALUE! |
| ENSMUSG00000082415.2 | 0 | NA | NA | NA | NA | NA | #VALUE! |
| ENSMUSG00000082416.2 | 0 | NA | NA | NA | NA | NA | #VALUE! |
| ENSMUSG00000082417.2 | 0 | NA | NA | NA | NA | NA | #VALUE! |

| | | | | | | | |
|---|---|---|---|---|---|---|---|
| ENSMUSG00000082418.2 | 0 | NA | NA | NA | NA | NA | #VALUE! |
| ENSMUSG00000082420.3 | 0 | NA | NA | NA | NA | NA | #VALUE! |
| ENSMUSG00000082421.2 | 0 | NA | NA | NA | NA | NA | #VALUE! |
| ENSMUSG00000082422.2 | 0 | NA | NA | NA | NA | NA | #VALUE! |
| ENSMUSG00000082423.2 | 0 | NA | NA | NA | NA | NA | #VALUE! |
| ENSMUSG00000082424.5 | 0 | NA | NA | NA | NA | NA | #VALUE! |
| ENSMUSG00000082426.2 | 0 | NA | NA | NA | NA | NA | #VALUE! |
| ENSMUSG00000082427.3 | 0 | NA | NA | NA | NA | NA | #VALUE! |
| ENSMUSG00000082428.2 | 0 | NA | NA | NA | NA | NA | #VALUE! |
| ENSMUSG00000082429.2 | 0 | NA | NA | NA | NA | NA | #VALUE! |
| ENSMUSG00000082431.4 | 0.245976 | 1.027628 | 4.067126 | 0.252667 | 0.800526 | NA | #VALUE! |
| ENSMUSG00000082432.2 | 0 | NA | NA | NA | NA | NA | #VALUE! |
| ENSMUSG00000082433.3 | 0 | NA | NA | NA | NA | NA | #VALUE! |
| ENSMUSG00000082435.2 | 0 | NA | NA | NA | NA | NA | #VALUE! |
| ENSMUSG00000082436.2 | 0 | NA | NA | NA | NA | NA | #VALUE! |
| ENSMUSG00000082437.2 | 0 | NA | NA | NA | NA | NA | #VALUE! |
| ENSMUSG00000082440.2 | 0 | NA | NA | NA | NA | NA | #VALUE! |
| ENSMUSG00000082442.2 | 0 | NA | NA | NA | NA | NA | #VALUE! |
| ENSMUSG00000082443.2 | 0 | NA | NA | NA | NA | NA | #VALUE! |
| ENSMUSG00000082444.2 | 0 | NA | NA | NA | NA | NA | #VALUE! |
| ENSMUSG00000082446.2 | 0 | NA | NA | NA | NA | NA | #VALUE! |
| ENSMUSG00000082447.2 | 0 | NA | NA | NA | NA | NA | #VALUE! |
| ENSMUSG00000082448.2 | 0 | NA | NA | NA | NA | NA | #VALUE! |
| ENSMUSG00000082449.2 | 0 | NA | NA | NA | NA | NA | #VALUE! |
| ENSMUSG00000082450.2 | 0 | NA | NA | NA | NA | NA | #VALUE! |
| ENSMUSG00000082451.2 | 0 | NA | NA | NA | NA | NA | #VALUE! |
| ENSMUSG00000082453.2 | 0 | NA | NA | NA | NA | NA | #VALUE! |
| ENSMUSG00000082454.4 | 0 | NA | NA | NA | NA | NA | #VALUE! |
| ENSMUSG00000082455.4 | 0 | NA | NA | NA | NA | NA | #VALUE! |
| ENSMUSG00000082456.2 | 0.391578 | -2.1489 | 4.020883 | -0.53444 | 0.59304 | NA | #VALUE! |
| ENSMUSG00000082457.2 | 0 | NA | NA | NA | NA | NA | #VALUE! |
| ENSMUSG00000082458.2 | 0 | NA | NA | NA | NA | NA | #VALUE! |
| ENSMUSG00000082459.2 | 0 | NA | NA | NA | NA | NA | #VALUE! |
| ENSMUSG00000082460.2 | 0 | NA | NA | NA | NA | NA | #VALUE! |
| ENSMUSG00000082461.2 | 0 | NA | NA | NA | NA | NA | #VALUE! |
| ENSMUSG00000082462.2 | 0 | NA | NA | NA | NA | NA | #VALUE! |
| ENSMUSG00000082463.2 | 0 | NA | NA | NA | NA | NA | #VALUE! |
| ENSMUSG00000082464.4 | 0 | NA | NA | NA | NA | NA | #VALUE! |
| ENSMUSG00000082467.2 | 0 | NA | NA | NA | NA | NA | #VALUE! |
| ENSMUSG00000082468.2 | 0 | NA | NA | NA | NA | NA | #VALUE! |
| ENSMUSG00000082472.2 | 0 | NA | NA | NA | NA | NA | #VALUE! |
| ENSMUSG00000082473.2 | 0 | NA | NA | NA | NA | NA | #VALUE! |
| ENSMUSG00000082474.4 | 7.756209 | -6.48334 | 3.912546 | -1.65706 | 0.097506 | NA | #VALUE! |
| ENSMUSG00000082475.3 | 0.491951 | 2.402185 | 4.000147 | 0.600524 | 0.548157 | NA | #VALUE! |

| | | | | | | | |
|---|---|---|---|---|---|---|---|
| ENSMUSG00000082476.2 | 0 | NA | NA | NA | NA | NA | #VALUE! |
| ENSMUSG00000082478.2 | 0.350989 | 0.094602 | 4.062933 | 0.023284 | 0.981424 | NA | #VALUE! |
| ENSMUSG00000082479.4 | 0 | NA | NA | NA | NA | NA | #VALUE! |
| ENSMUSG00000082480.2 | 0.195789 | -1.05581 | 4.073302 | -0.2592 | 0.795479 | NA | #VALUE! |
| ENSMUSG00000082481.2 | 0 | NA | NA | NA | NA | NA | #VALUE! |
| ENSMUSG00000082482.3 | 0 | NA | NA | NA | NA | NA | #VALUE! |
| ENSMUSG00000082483.2 | 0 | NA | NA | NA | NA | NA | #VALUE! |
| ENSMUSG00000082484.2 | 0 | NA | NA | NA | NA | NA | #VALUE! |
| ENSMUSG00000082485.2 | 0 | NA | NA | NA | NA | NA | #VALUE! |
| ENSMUSG00000082486.2 | 0 | NA | NA | NA | NA | NA | #VALUE! |
| ENSMUSG00000082487.2 | 0 | NA | NA | NA | NA | NA | #VALUE! |
| ENSMUSG00000082488.3 | 0 | NA | NA | NA | NA | NA | #VALUE! |
| ENSMUSG00000082489.2 | 0 | NA | NA | NA | NA | NA | #VALUE! |
| ENSMUSG00000082490.2 | 0 | NA | NA | NA | NA | NA | #VALUE! |
| ENSMUSG00000082491.2 | 8.872713 | 1.494633 | 2.310375 | 0.646922 | 0.517682 | NA | #VALUE! |
| ENSMUSG00000082492.2 | 0 | NA | NA | NA | NA | NA | #VALUE! |
| ENSMUSG00000082494.2 | 0 | NA | NA | NA | NA | NA | #VALUE! |
| ENSMUSG00000082495.3 | 0 | NA | NA | NA | NA | NA | #VALUE! |
| ENSMUSG00000082496.4 | 0 | NA | NA | NA | NA | NA | #VALUE! |
| ENSMUSG00000082497.2 | 0 | NA | NA | NA | NA | NA | #VALUE! |
| ENSMUSG00000082498.2 | 0 | NA | NA | NA | NA | NA | #VALUE! |
| ENSMUSG00000082499.2 | 0 | NA | NA | NA | NA | NA | #VALUE! |
| ENSMUSG00000082500.2 | 0 | NA | NA | NA | NA | NA | #VALUE! |
| ENSMUSG00000082501.2 | 0 | NA | NA | NA | NA | NA | #VALUE! |
| ENSMUSG00000082502.2 | 0 | NA | NA | NA | NA | NA | #VALUE! |
| ENSMUSG00000082503.2 | 0 | NA | NA | NA | NA | NA | #VALUE! |
| ENSMUSG00000082504.2 | 0 | NA | NA | NA | NA | NA | #VALUE! |
| ENSMUSG00000082505.2 | 0 | NA | NA | NA | NA | NA | #VALUE! |
| ENSMUSG00000082507.2 | 0 | NA | NA | NA | NA | NA | #VALUE! |
| ENSMUSG00000082508.2 | 0 | NA | NA | NA | NA | NA | #VALUE! |
| ENSMUSG00000082510.2 | 0 | NA | NA | NA | NA | NA | #VALUE! |
| ENSMUSG00000082511.2 | 0 | NA | NA | NA | NA | NA | #VALUE! |
| ENSMUSG00000082512.2 | 0 | NA | NA | NA | NA | NA | #VALUE! |
| ENSMUSG00000082513.2 | 0 | NA | NA | NA | NA | NA | #VALUE! |
| ENSMUSG00000082514.2 | 0 | NA | NA | NA | NA | NA | #VALUE! |
| ENSMUSG00000082515.2 | 0 | NA | NA | NA | NA | NA | #VALUE! |
| ENSMUSG00000082516.2 | 0.309124 | -1.83395 | 4.047982 | -0.45305 | 0.650511 | NA | #VALUE! |
| ENSMUSG00000082517.2 | 0 | NA | NA | NA | NA | NA | #VALUE! |
| ENSMUSG00000082519.2 | 0 | NA | NA | NA | NA | NA | #VALUE! |
| ENSMUSG00000082520.3 | 0 | NA | NA | NA | NA | NA | #VALUE! |
| ENSMUSG00000082523.2 | 0 | NA | NA | NA | NA | NA | #VALUE! |
| ENSMUSG00000082524.4 | 0 | NA | NA | NA | NA | NA | #VALUE! |
| ENSMUSG00000082526.2 | 0 | NA | NA | NA | NA | NA | #VALUE! |
| ENSMUSG00000082527.2 | 0 | NA | NA | NA | NA | NA | #VALUE! |

| | | | | | | | |
|---|---|---|---|---|---|---|---|
| ENSMUSG00000082528.2 | 0 | NA | NA | NA | NA | NA | #VALUE! |
| ENSMUSG00000082529.2 | 0 | NA | NA | NA | NA | NA | #VALUE! |
| ENSMUSG00000082530.2 | 0 | NA | NA | NA | NA | NA | #VALUE! |
| ENSMUSG00000082531.2 | 0 | NA | NA | NA | NA | NA | #VALUE! |
| ENSMUSG00000082532.2 | 0 | NA | NA | NA | NA | NA | #VALUE! |
| ENSMUSG00000082533.2 | 0 | NA | NA | NA | NA | NA | #VALUE! |
| ENSMUSG00000082534.2 | 0.093255 | 0.868093 | 4.072464 | 0.213162 | 0.831201 | NA | #VALUE! |
| ENSMUSG00000082535.2 | 0 | NA | NA | NA | NA | NA | #VALUE! |
| ENSMUSG00000082536.2 | 5.063928 | 0.100547 | 2.523667 | 0.039842 | 0.968219 | NA | #VALUE! |
| ENSMUSG00000082537.2 | 0 | NA | NA | NA | NA | NA | #VALUE! |
| ENSMUSG00000082538.2 | 0 | NA | NA | NA | NA | NA | #VALUE! |
| ENSMUSG00000082539.2 | 0 | NA | NA | NA | NA | NA | #VALUE! |
| ENSMUSG00000082540.2 | 0 | NA | NA | NA | NA | NA | #VALUE! |
| ENSMUSG00000082541.2 | 0 | NA | NA | NA | NA | NA | #VALUE! |
| ENSMUSG00000082542.3 | 0 | NA | NA | NA | NA | NA | #VALUE! |
| ENSMUSG00000082544.2 | 0 | NA | NA | NA | NA | NA | #VALUE! |
| ENSMUSG00000082545.2 | 0 | NA | NA | NA | NA | NA | #VALUE! |
| ENSMUSG00000082546.2 | 0 | NA | NA | NA | NA | NA | #VALUE! |
| ENSMUSG00000082547.3 | 0 | NA | NA | NA | NA | NA | #VALUE! |
| ENSMUSG00000082548.2 | 0 | NA | NA | NA | NA | NA | #VALUE! |
| ENSMUSG00000082549.2 | 0 | NA | NA | NA | NA | NA | #VALUE! |
| ENSMUSG00000082550.2 | 0 | NA | NA | NA | NA | NA | #VALUE! |
| ENSMUSG00000082551.2 | 0 | NA | NA | NA | NA | NA | #VALUE! |
| ENSMUSG00000082552.2 | 0 | NA | NA | NA | NA | NA | #VALUE! |
| ENSMUSG00000082553.2 | 0 | NA | NA | NA | NA | NA | #VALUE! |
| ENSMUSG00000082554.2 | 0 | NA | NA | NA | NA | NA | #VALUE! |
| ENSMUSG00000082555.2 | 0 | NA | NA | NA | NA | NA | #VALUE! |
| ENSMUSG00000082556.2 | 0 | NA | NA | NA | NA | NA | #VALUE! |
| ENSMUSG00000082558.2 | 0 | NA | NA | NA | NA | NA | #VALUE! |
| ENSMUSG00000082560.2 | 0 | NA | NA | NA | NA | NA | #VALUE! |
| ENSMUSG00000082561.2 | 0 | NA | NA | NA | NA | NA | #VALUE! |
| ENSMUSG00000082562.2 | 0 | NA | NA | NA | NA | NA | #VALUE! |
| ENSMUSG00000082563.2 | 0 | NA | NA | NA | NA | NA | #VALUE! |
| ENSMUSG00000082564.2 | 0 | NA | NA | NA | NA | NA | #VALUE! |
| ENSMUSG00000082565.2 | 0 | NA | NA | NA | NA | NA | #VALUE! |
| ENSMUSG00000082566.2 | 0 | NA | NA | NA | NA | NA | #VALUE! |
| ENSMUSG00000082567.2 | 0 | NA | NA | NA | NA | NA | #VALUE! |
| ENSMUSG00000082569.2 | 0 | NA | NA | NA | NA | NA | #VALUE! |
| ENSMUSG00000082570.2 | 0 | NA | NA | NA | NA | NA | #VALUE! |
| ENSMUSG00000082571.2 | 0 | NA | NA | NA | NA | NA | #VALUE! |
| ENSMUSG00000082573.2 | 0 | NA | NA | NA | NA | NA | #VALUE! |
| ENSMUSG00000082574.4 | 0 | NA | NA | NA | NA | NA | #VALUE! |
| ENSMUSG00000082575.2 | 0 | NA | NA | NA | NA | NA | #VALUE! |
| ENSMUSG00000082576.2 | 0 | NA | NA | NA | NA | NA | #VALUE! |

| | | | | | | | |
|---|---|---|---|---|---|---|---|
| ENSMUSG00000082577.3 | 0 | NA | NA | NA | NA | NA | #VALUE! |
| ENSMUSG00000082578.2 | 0 | NA | NA | NA | NA | NA | #VALUE! |
| ENSMUSG00000082579.2 | 0.168284 | 1.027628 | 4.067126 | 0.252667 | 0.800526 | NA | #VALUE! |
| ENSMUSG00000082580.4 | 0 | NA | NA | NA | NA | NA | #VALUE! |
| ENSMUSG00000082581.3 | 0 | NA | NA | NA | NA | NA | #VALUE! |
| ENSMUSG00000082583.3 | 0 | NA | NA | NA | NA | NA | #VALUE! |
| ENSMUSG00000082584.2 | 0 | NA | NA | NA | NA | NA | #VALUE! |
| ENSMUSG00000082585.4 | 0 | NA | NA | NA | NA | NA | #VALUE! |
| ENSMUSG00000082586.3 | 0 | NA | NA | NA | NA | NA | #VALUE! |
| ENSMUSG00000082587.2 | 0 | NA | NA | NA | NA | NA | #VALUE! |
| ENSMUSG00000082588.2 | 0 | NA | NA | NA | NA | NA | #VALUE! |
| ENSMUSG00000082589.2 | 0 | NA | NA | NA | NA | NA | #VALUE! |
| ENSMUSG00000082590.2 | 0 | NA | NA | NA | NA | NA | #VALUE! |
| ENSMUSG00000082591.2 | 0 | NA | NA | NA | NA | NA | #VALUE! |
| ENSMUSG00000082592.2 | 0 | NA | NA | NA | NA | NA | #VALUE! |
| ENSMUSG00000082593.2 | 0 | NA | NA | NA | NA | NA | #VALUE! |
| ENSMUSG00000082594.2 | 0 | NA | NA | NA | NA | NA | #VALUE! |
| ENSMUSG00000082595.2 | 0 | NA | NA | NA | NA | NA | #VALUE! |
| ENSMUSG00000082596.2 | 0 | NA | NA | NA | NA | NA | #VALUE! |
| ENSMUSG00000082597.2 | 0 | NA | NA | NA | NA | NA | #VALUE! |
| ENSMUSG00000082598.2 | 0 | NA | NA | NA | NA | NA | #VALUE! |
| ENSMUSG00000082599.4 | 0 | NA | NA | NA | NA | NA | #VALUE! |
| ENSMUSG00000082600.2 | 0 | NA | NA | NA | NA | NA | #VALUE! |
| ENSMUSG00000082601.2 | 0 | NA | NA | NA | NA | NA | #VALUE! |
| ENSMUSG00000082602.2 | 0 | NA | NA | NA | NA | NA | #VALUE! |
| ENSMUSG00000082604.4 | 0 | NA | NA | NA | NA | NA | #VALUE! |
| ENSMUSG00000082605.2 | 0 | NA | NA | NA | NA | NA | #VALUE! |
| ENSMUSG00000082606.2 | 0 | NA | NA | NA | NA | NA | #VALUE! |
| ENSMUSG00000082607.2 | 0 | NA | NA | NA | NA | NA | #VALUE! |
| ENSMUSG00000082608.2 | 0 | NA | NA | NA | NA | NA | #VALUE! |
| ENSMUSG00000082610.2 | 0 | NA | NA | NA | NA | NA | #VALUE! |
| ENSMUSG00000082611.2 | 0 | NA | NA | NA | NA | NA | #VALUE! |
| ENSMUSG00000082612.2 | 0 | NA | NA | NA | NA | NA | #VALUE! |
| ENSMUSG00000082613.2 | 0 | NA | NA | NA | NA | NA | #VALUE! |
| ENSMUSG00000082614.2 | 0 | NA | NA | NA | NA | NA | #VALUE! |
| ENSMUSG00000082615.2 | 0 | NA | NA | NA | NA | NA | #VALUE! |
| ENSMUSG00000082616.2 | 0 | NA | NA | NA | NA | NA | #VALUE! |
| ENSMUSG00000082617.4 | 0 | NA | NA | NA | NA | NA | #VALUE! |
| ENSMUSG00000082618.2 | 0 | NA | NA | NA | NA | NA | #VALUE! |
| ENSMUSG00000082619.2 | 0 | NA | NA | NA | NA | NA | #VALUE! |
| ENSMUSG00000082620.2 | 0 | NA | NA | NA | NA | NA | #VALUE! |
| ENSMUSG00000082621.2 | 0 | NA | NA | NA | NA | NA | #VALUE! |
| ENSMUSG00000082622.2 | 0 | NA | NA | NA | NA | NA | #VALUE! |
| ENSMUSG00000082623.2 | 0 | NA | NA | NA | NA | NA | #VALUE! |

| | | | | | | | |
|---|---|---|---|---|---|---|---|
| ENSMUSG00000082624.2 | 0 | NA | NA | NA | NA | NA | #VALUE! |
| ENSMUSG00000082625.2 | 0 | NA | NA | NA | NA | NA | #VALUE! |
| ENSMUSG00000082626.2 | 0 | NA | NA | NA | NA | NA | #VALUE! |
| ENSMUSG00000082628.2 | 0 | NA | NA | NA | NA | NA | #VALUE! |
| ENSMUSG00000082629.4 | 0 | NA | NA | NA | NA | NA | #VALUE! |
| ENSMUSG00000082630.2 | 0 | NA | NA | NA | NA | NA | #VALUE! |
| ENSMUSG00000082632.3 | 0 | NA | NA | NA | NA | NA | #VALUE! |
| ENSMUSG00000082633.2 | 0 | NA | NA | NA | NA | NA | #VALUE! |
| ENSMUSG00000082634.2 | 0 | NA | NA | NA | NA | NA | #VALUE! |
| ENSMUSG00000082635.3 | 0 | NA | NA | NA | NA | NA | #VALUE! |
| ENSMUSG00000082636.2 | 0 | NA | NA | NA | NA | NA | #VALUE! |
| ENSMUSG00000082637.3 | 0 | NA | NA | NA | NA | NA | #VALUE! |
| ENSMUSG00000082638.2 | 0 | NA | NA | NA | NA | NA | #VALUE! |
| ENSMUSG00000082639.9 | 0 | NA | NA | NA | NA | NA | #VALUE! |
| ENSMUSG00000082641.2 | 0 | NA | NA | NA | NA | NA | #VALUE! |
| ENSMUSG00000082644.2 | 0 | NA | NA | NA | NA | NA | #VALUE! |
| ENSMUSG00000082645.4 | 0 | NA | NA | NA | NA | NA | #VALUE! |
| ENSMUSG00000082646.2 | 0 | NA | NA | NA | NA | NA | #VALUE! |
| ENSMUSG00000082647.2 | 0 | NA | NA | NA | NA | NA | #VALUE! |
| ENSMUSG00000082648.2 | 0 | NA | NA | NA | NA | NA | #VALUE! |
| ENSMUSG00000082649.2 | 0 | NA | NA | NA | NA | NA | #VALUE! |
| ENSMUSG00000082650.2 | 0 | NA | NA | NA | NA | NA | #VALUE! |
| ENSMUSG00000082651.2 | 0 | NA | NA | NA | NA | NA | #VALUE! |
| ENSMUSG00000082652.2 | 0 | NA | NA | NA | NA | NA | #VALUE! |
| ENSMUSG00000082653.2 | 0 | NA | NA | NA | NA | NA | #VALUE! |
| ENSMUSG00000082655.2 | 0 | NA | NA | NA | NA | NA | #VALUE! |
| ENSMUSG00000082656.2 | 0 | NA | NA | NA | NA | NA | #VALUE! |
| ENSMUSG00000082657.2 | 0 | NA | NA | NA | NA | NA | #VALUE! |
| ENSMUSG00000082658.2 | 0 | NA | NA | NA | NA | NA | #VALUE! |
| ENSMUSG00000082660.2 | 0 | NA | NA | NA | NA | NA | #VALUE! |
| ENSMUSG00000082661.2 | 0 | NA | NA | NA | NA | NA | #VALUE! |
| ENSMUSG00000082662.2 | 0 | NA | NA | NA | NA | NA | #VALUE! |
| ENSMUSG00000082663.3 | 0 | NA | NA | NA | NA | NA | #VALUE! |
| ENSMUSG00000082664.2 | 0 | NA | NA | NA | NA | NA | #VALUE! |
| ENSMUSG00000082665.2 | 3.045381 | -5.14543 | 3.921352 | -1.31216 | 0.189467 | NA | #VALUE! |
| ENSMUSG00000082666.2 | 0 | NA | NA | NA | NA | NA | #VALUE! |
| ENSMUSG00000082667.2 | 0 | NA | NA | NA | NA | NA | #VALUE! |
| ENSMUSG00000082669.2 | 0 | NA | NA | NA | NA | NA | #VALUE! |
| ENSMUSG00000082670.2 | 0 | NA | NA | NA | NA | NA | #VALUE! |
| ENSMUSG00000082671.3 | 0 | NA | NA | NA | NA | NA | #VALUE! |
| ENSMUSG00000082672.2 | 0 | NA | NA | NA | NA | NA | #VALUE! |
| ENSMUSG00000082673.2 | 0 | NA | NA | NA | NA | NA | #VALUE! |
| ENSMUSG00000082674.2 | 6.701218 | -2.06538 | 2.599662 | -0.79448 | 0.426915 | NA | #VALUE! |
| ENSMUSG00000082675.2 | 0 | NA | NA | NA | NA | NA | #VALUE! |

| | | | | | | | |
|---|---|---|---|---|---|---|---|
| ENSMUSG00000082676.2 | 0 | NA | NA | NA | NA | NA | #VALUE! |
| ENSMUSG00000082677.2 | 0 | NA | NA | NA | NA | NA | #VALUE! |
| ENSMUSG00000082678.2 | 0 | NA | NA | NA | NA | NA | #VALUE! |
| ENSMUSG00000082679.2 | 0 | NA | NA | NA | NA | NA | #VALUE! |
| ENSMUSG00000082680.2 | 0 | NA | NA | NA | NA | NA | #VALUE! |
| ENSMUSG00000082681.2 | 0 | NA | NA | NA | NA | NA | #VALUE! |
| ENSMUSG00000082682.2 | 0 | NA | NA | NA | NA | NA | #VALUE! |
| ENSMUSG00000082683.2 | 0 | NA | NA | NA | NA | NA | #VALUE! |
| ENSMUSG00000082684.2 | 0 | NA | NA | NA | NA | NA | #VALUE! |
| ENSMUSG00000082686.2 | 0 | NA | NA | NA | NA | NA | #VALUE! |
| ENSMUSG00000082687.2 | 0 | NA | NA | NA | NA | NA | #VALUE! |
| ENSMUSG00000082688.2 | 0 | NA | NA | NA | NA | NA | #VALUE! |
| ENSMUSG00000082689.2 | 0 | NA | NA | NA | NA | NA | #VALUE! |
| ENSMUSG00000082690.2 | 0 | NA | NA | NA | NA | NA | #VALUE! |
| ENSMUSG00000082691.7 | 0 | NA | NA | NA | NA | NA | #VALUE! |
| ENSMUSG00000082692.3 | 0 | NA | NA | NA | NA | NA | #VALUE! |
| ENSMUSG00000082693.4 | 0 | NA | NA | NA | NA | NA | #VALUE! |
| ENSMUSG00000082696.2 | 0 | NA | NA | NA | NA | NA | #VALUE! |
| ENSMUSG00000082698.2 | 0 | NA | NA | NA | NA | NA | #VALUE! |
| ENSMUSG00000082699.2 | 0 | NA | NA | NA | NA | NA | #VALUE! |
| ENSMUSG00000082700.2 | 0 | NA | NA | NA | NA | NA | #VALUE! |
| ENSMUSG00000082701.2 | 0 | NA | NA | NA | NA | NA | #VALUE! |
| ENSMUSG00000082703.2 | 0 | NA | NA | NA | NA | NA | #VALUE! |
| ENSMUSG00000082704.2 | 0 | NA | NA | NA | NA | NA | #VALUE! |
| ENSMUSG00000082705.2 | 0 | NA | NA | NA | NA | NA | #VALUE! |
| ENSMUSG00000082706.2 | 0 | NA | NA | NA | NA | NA | #VALUE! |
| ENSMUSG00000082707.2 | 0 | NA | NA | NA | NA | NA | #VALUE! |
| ENSMUSG00000082708.2 | 0 | NA | NA | NA | NA | NA | #VALUE! |
| ENSMUSG00000082709.2 | 0 | NA | NA | NA | NA | NA | #VALUE! |
| ENSMUSG00000082710.2 | 0 | NA | NA | NA | NA | NA | #VALUE! |
| ENSMUSG00000082711.2 | 0 | NA | NA | NA | NA | NA | #VALUE! |
| ENSMUSG00000082712.3 | 0 | NA | NA | NA | NA | NA | #VALUE! |
| ENSMUSG00000082713.2 | 0 | NA | NA | NA | NA | NA | #VALUE! |
| ENSMUSG00000082714.2 | 0 | NA | NA | NA | NA | NA | #VALUE! |
| ENSMUSG00000082715.2 | 0.444244 | 0.890672 | 4.031633 | 0.220921 | 0.825154 | NA | #VALUE! |
| ENSMUSG00000082716.2 | 0 | NA | NA | NA | NA | NA | #VALUE! |
| ENSMUSG00000082717.2 | 0 | NA | NA | NA | NA | NA | #VALUE! |
| ENSMUSG00000082718.2 | 0 | NA | NA | NA | NA | NA | #VALUE! |
| ENSMUSG00000082720.2 | 0 | NA | NA | NA | NA | NA | #VALUE! |
| ENSMUSG00000082721.2 | 0 | NA | NA | NA | NA | NA | #VALUE! |
| ENSMUSG00000082722.2 | 0 | NA | NA | NA | NA | NA | #VALUE! |
| ENSMUSG00000082723.2 | 0 | NA | NA | NA | NA | NA | #VALUE! |
| ENSMUSG00000082724.4 | 0 | NA | NA | NA | NA | NA | #VALUE! |
| ENSMUSG00000082725.2 | 0 | NA | NA | NA | NA | NA | #VALUE! |

| | | | | | | | |
|---|---|---|---|---|---|---|---|
| ENSMUSG00000082726.2 | 0 | NA | NA | NA | NA | NA | #VALUE! |
| ENSMUSG00000082727.3 | 0 | NA | NA | NA | NA | NA | #VALUE! |
| ENSMUSG00000082728.2 | 0 | NA | NA | NA | NA | NA | #VALUE! |
| ENSMUSG00000082729.4 | 0 | NA | NA | NA | NA | NA | #VALUE! |
| ENSMUSG00000082730.3 | 0 | NA | NA | NA | NA | NA | #VALUE! |
| ENSMUSG00000082732.2 | 0 | NA | NA | NA | NA | NA | #VALUE! |
| ENSMUSG00000082733.2 | 0 | NA | NA | NA | NA | NA | #VALUE! |
| ENSMUSG00000082734.3 | 0 | NA | NA | NA | NA | NA | #VALUE! |
| ENSMUSG00000082735.2 | 0 | NA | NA | NA | NA | NA | #VALUE! |
| ENSMUSG00000082736.2 | 0 | NA | NA | NA | NA | NA | #VALUE! |
| ENSMUSG00000082737.4 | 0 | NA | NA | NA | NA | NA | #VALUE! |
| ENSMUSG00000082738.2 | 0 | NA | NA | NA | NA | NA | #VALUE! |
| ENSMUSG00000082739.4 | 0 | NA | NA | NA | NA | NA | #VALUE! |
| ENSMUSG00000082740.2 | 0 | NA | NA | NA | NA | NA | #VALUE! |
| ENSMUSG00000082743.2 | 0 | NA | NA | NA | NA | NA | #VALUE! |
| ENSMUSG00000082744.2 | 0 | NA | NA | NA | NA | NA | #VALUE! |
| ENSMUSG00000082745.2 | 0 | NA | NA | NA | NA | NA | #VALUE! |
| ENSMUSG00000082746.2 | 0 | NA | NA | NA | NA | NA | #VALUE! |
| ENSMUSG00000082749.2 | 0 | NA | NA | NA | NA | NA | #VALUE! |
| ENSMUSG00000082750.2 | 0 | NA | NA | NA | NA | NA | #VALUE! |
| ENSMUSG00000082751.2 | 0 | NA | NA | NA | NA | NA | #VALUE! |
| ENSMUSG00000082752.2 | 0 | NA | NA | NA | NA | NA | #VALUE! |
| ENSMUSG00000082754.3 | 0 | NA | NA | NA | NA | NA | #VALUE! |
| ENSMUSG00000082755.2 | 0.18651 | 1.401769 | 4.056354 | 0.345574 | 0.729663 | NA | #VALUE! |
| ENSMUSG00000082757.2 | 0 | NA | NA | NA | NA | NA | #VALUE! |
| ENSMUSG00000082758.2 | 0 | NA | NA | NA | NA | NA | #VALUE! |
| ENSMUSG00000082759.2 | 0 | NA | NA | NA | NA | NA | #VALUE! |
| ENSMUSG00000082760.3 | 0 | NA | NA | NA | NA | NA | #VALUE! |
| ENSMUSG00000082762.2 | 0.105013 | -0.97324 | 4.076293 | -0.23876 | 0.811295 | NA | #VALUE! |
| ENSMUSG00000082763.2 | 0 | NA | NA | NA | NA | NA | #VALUE! |
| ENSMUSG00000082765.4 | 0 | NA | NA | NA | NA | NA | #VALUE! |
| ENSMUSG00000082766.2 | 0 | NA | NA | NA | NA | NA | #VALUE! |
| ENSMUSG00000082767.2 | 0 | NA | NA | NA | NA | NA | #VALUE! |
| ENSMUSG00000082768.2 | 0 | NA | NA | NA | NA | NA | #VALUE! |
| ENSMUSG00000082769.4 | 0.245976 | 1.027628 | 4.067126 | 0.252667 | 0.800526 | NA | #VALUE! |
| ENSMUSG00000082770.2 | 0 | NA | NA | NA | NA | NA | #VALUE! |
| ENSMUSG00000082771.2 | 0 | NA | NA | NA | NA | NA | #VALUE! |
| ENSMUSG00000082772.4 | 0 | NA | NA | NA | NA | NA | #VALUE! |
| ENSMUSG00000082773.2 | 0 | NA | NA | NA | NA | NA | #VALUE! |
| ENSMUSG00000082774.2 | 0 | NA | NA | NA | NA | NA | #VALUE! |
| ENSMUSG00000082775.2 | 0 | NA | NA | NA | NA | NA | #VALUE! |
| ENSMUSG00000082776.4 | 0 | NA | NA | NA | NA | NA | #VALUE! |
| ENSMUSG00000082777.2 | 0 | NA | NA | NA | NA | NA | #VALUE! |
| ENSMUSG00000082778.2 | 0 | NA | NA | NA | NA | NA | #VALUE! |

| | | | | | | | |
|---|---|---|---|---|---|---|---|
| ENSMUSG00000082779.2 | 0 | NA | NA | NA | NA | NA | #VALUE! |
| ENSMUSG00000082780.3 | 0 | NA | NA | NA | NA | NA | #VALUE! |
| ENSMUSG00000082781.2 | 0 | NA | NA | NA | NA | NA | #VALUE! |
| ENSMUSG00000082782.2 | 0 | NA | NA | NA | NA | NA | #VALUE! |
| ENSMUSG00000082783.3 | 0 | NA | NA | NA | NA | NA | #VALUE! |
| ENSMUSG00000082784.2 | 0 | NA | NA | NA | NA | NA | #VALUE! |
| ENSMUSG00000082785.4 | 0 | NA | NA | NA | NA | NA | #VALUE! |
| ENSMUSG00000082786.2 | 0 | NA | NA | NA | NA | NA | #VALUE! |
| ENSMUSG00000082788.2 | 0 | NA | NA | NA | NA | NA | #VALUE! |
| ENSMUSG00000082789.4 | 0 | NA | NA | NA | NA | NA | #VALUE! |
| ENSMUSG00000082790.2 | 0 | NA | NA | NA | NA | NA | #VALUE! |
| ENSMUSG00000082791.4 | 0 | NA | NA | NA | NA | NA | #VALUE! |
| ENSMUSG00000082792.2 | 0 | NA | NA | NA | NA | NA | #VALUE! |
| ENSMUSG00000082793.3 | 0.491951 | 2.402185 | 4.000147 | 0.600524 | 0.548157 | NA | #VALUE! |
| ENSMUSG00000082794.2 | 0 | NA | NA | NA | NA | NA | #VALUE! |
| ENSMUSG00000082795.2 | 0 | NA | NA | NA | NA | NA | #VALUE! |
| ENSMUSG00000082796.2 | 0 | NA | NA | NA | NA | NA | #VALUE! |
| ENSMUSG00000082797.4 | 0 | NA | NA | NA | NA | NA | #VALUE! |
| ENSMUSG00000082798.2 | 0 | NA | NA | NA | NA | NA | #VALUE! |
| ENSMUSG00000082799.2 | 0 | NA | NA | NA | NA | NA | #VALUE! |
| ENSMUSG00000082800.2 | 0 | NA | NA | NA | NA | NA | #VALUE! |
| ENSMUSG00000082802.3 | 0 | NA | NA | NA | NA | NA | #VALUE! |
| ENSMUSG00000082804.4 | 0 | NA | NA | NA | NA | NA | #VALUE! |
| ENSMUSG00000082806.2 | 0 | NA | NA | NA | NA | NA | #VALUE! |
| ENSMUSG00000082807.2 | 0 | NA | NA | NA | NA | NA | #VALUE! |
| ENSMUSG00000082809.4 | 0 | NA | NA | NA | NA | NA | #VALUE! |
| ENSMUSG00000082810.2 | 0 | NA | NA | NA | NA | NA | #VALUE! |
| ENSMUSG00000082812.2 | 0 | NA | NA | NA | NA | NA | #VALUE! |
| ENSMUSG00000082814.2 | 0 | NA | NA | NA | NA | NA | #VALUE! |
| ENSMUSG00000082815.3 | 0 | NA | NA | NA | NA | NA | #VALUE! |
| ENSMUSG00000082816.2 | 0 | NA | NA | NA | NA | NA | #VALUE! |
| ENSMUSG00000082817.2 | 0 | NA | NA | NA | NA | NA | #VALUE! |
| ENSMUSG00000082818.2 | 0 | NA | NA | NA | NA | NA | #VALUE! |
| ENSMUSG00000082819.3 | 0 | NA | NA | NA | NA | NA | #VALUE! |
| ENSMUSG00000082820.4 | 8.173368 | 0.041075 | 2.137178 | 0.019219 | 0.984666 | NA | #VALUE! |
| ENSMUSG00000082821.2 | 0 | NA | NA | NA | NA | NA | #VALUE! |
| ENSMUSG00000082822.2 | 0 | NA | NA | NA | NA | NA | #VALUE! |
| ENSMUSG00000082823.4 | 0 | NA | NA | NA | NA | NA | #VALUE! |
| ENSMUSG00000082824.2 | 0 | NA | NA | NA | NA | NA | #VALUE! |
| ENSMUSG00000082825.2 | 0 | NA | NA | NA | NA | NA | #VALUE! |
| ENSMUSG00000082826.2 | 0 | NA | NA | NA | NA | NA | #VALUE! |
| ENSMUSG00000082827.4 | 0 | NA | NA | NA | NA | NA | #VALUE! |
| ENSMUSG00000082828.2 | 0 | NA | NA | NA | NA | NA | #VALUE! |
| ENSMUSG00000082829.2 | 0 | NA | NA | NA | NA | NA | #VALUE! |

| | | | | | | | |
|---|---|---|---|---|---|---|---|
| ENSMUSG00000082831.2 | 0 | NA | NA | NA | NA | NA | #VALUE! |
| ENSMUSG00000082832.2 | 0 | NA | NA | NA | NA | NA | #VALUE! |
| ENSMUSG00000082833.2 | 0 | NA | NA | NA | NA | NA | #VALUE! |
| ENSMUSG00000082835.2 | 0 | NA | NA | NA | NA | NA | #VALUE! |
| ENSMUSG00000082836.2 | 0 | NA | NA | NA | NA | NA | #VALUE! |
| ENSMUSG00000082837.2 | 0 | NA | NA | NA | NA | NA | #VALUE! |
| ENSMUSG00000082838.2 | 0 | NA | NA | NA | NA | NA | #VALUE! |
| ENSMUSG00000082839.2 | 0 | NA | NA | NA | NA | NA | #VALUE! |
| ENSMUSG00000082840.2 | 0 | NA | NA | NA | NA | NA | #VALUE! |
| ENSMUSG00000082841.2 | 0 | NA | NA | NA | NA | NA | #VALUE! |
| ENSMUSG00000082842.2 | 0 | NA | NA | NA | NA | NA | #VALUE! |
| ENSMUSG00000082843.3 | 0 | NA | NA | NA | NA | NA | #VALUE! |
| ENSMUSG00000082844.2 | 0 | NA | NA | NA | NA | NA | #VALUE! |
| ENSMUSG00000082845.2 | 0 | NA | NA | NA | NA | NA | #VALUE! |
| ENSMUSG00000082846.2 | 0 | NA | NA | NA | NA | NA | #VALUE! |
| ENSMUSG00000082847.2 | 0 | NA | NA | NA | NA | NA | #VALUE! |
| ENSMUSG00000082848.2 | 0 | NA | NA | NA | NA | NA | #VALUE! |
| ENSMUSG00000082849.2 | 0 | NA | NA | NA | NA | NA | #VALUE! |
| ENSMUSG00000082850.2 | 0 | NA | NA | NA | NA | NA | #VALUE! |
| ENSMUSG00000082851.2 | 0 | NA | NA | NA | NA | NA | #VALUE! |
| ENSMUSG00000082852.3 | 0 | NA | NA | NA | NA | NA | #VALUE! |
| ENSMUSG00000082853.2 | 0 | NA | NA | NA | NA | NA | #VALUE! |
| ENSMUSG00000082855.2 | 0 | NA | NA | NA | NA | NA | #VALUE! |
| ENSMUSG00000082856.2 | 0 | NA | NA | NA | NA | NA | #VALUE! |
| ENSMUSG00000082857.2 | 0 | NA | NA | NA | NA | NA | #VALUE! |
| ENSMUSG00000082858.2 | 0 | NA | NA | NA | NA | NA | #VALUE! |
| ENSMUSG00000082859.2 | 0 | NA | NA | NA | NA | NA | #VALUE! |
| ENSMUSG00000082860.2 | 0 | NA | NA | NA | NA | NA | #VALUE! |
| ENSMUSG00000082861.4 | 0 | NA | NA | NA | NA | NA | #VALUE! |
| ENSMUSG00000082862.2 | 0 | NA | NA | NA | NA | NA | #VALUE! |
| ENSMUSG00000082863.2 | 0 | NA | NA | NA | NA | NA | #VALUE! |
| ENSMUSG00000082864.2 | 0 | NA | NA | NA | NA | NA | #VALUE! |
| ENSMUSG00000082865.2 | 0 | NA | NA | NA | NA | NA | #VALUE! |
| ENSMUSG00000082867.2 | 0 | NA | NA | NA | NA | NA | #VALUE! |
| ENSMUSG00000082868.2 | 0 | NA | NA | NA | NA | NA | #VALUE! |
| ENSMUSG00000082869.2 | 0 | NA | NA | NA | NA | NA | #VALUE! |
| ENSMUSG00000082870.2 | 0 | NA | NA | NA | NA | NA | #VALUE! |
| ENSMUSG00000082871.3 | 0 | NA | NA | NA | NA | NA | #VALUE! |
| ENSMUSG00000082872.2 | 0 | NA | NA | NA | NA | NA | #VALUE! |
| ENSMUSG00000082873.2 | 0 | NA | NA | NA | NA | NA | #VALUE! |
| ENSMUSG00000082874.5 | 0 | NA | NA | NA | NA | NA | #VALUE! |
| ENSMUSG00000082875.2 | 0 | NA | NA | NA | NA | NA | #VALUE! |
| ENSMUSG00000082876.2 | 0 | NA | NA | NA | NA | NA | #VALUE! |
| ENSMUSG00000082877.2 | 0 | NA | NA | NA | NA | NA | #VALUE! |

| | | | | | | | |
|---|---|---|---|---|---|---|---|
| ENSMUSG00000082878.2 | 0 | NA | NA | NA | NA | NA | #VALUE! |
| ENSMUSG00000082879.2 | 0 | NA | NA | NA | NA | NA | #VALUE! |
| ENSMUSG00000082882.3 | 0 | NA | NA | NA | NA | NA | #VALUE! |
| ENSMUSG00000082886.2 | 0 | NA | NA | NA | NA | NA | #VALUE! |
| ENSMUSG00000082887.2 | 0 | NA | NA | NA | NA | NA | #VALUE! |
| ENSMUSG00000082888.2 | 0 | NA | NA | NA | NA | NA | #VALUE! |
| ENSMUSG00000082889.2 | 0 | NA | NA | NA | NA | NA | #VALUE! |
| ENSMUSG00000082890.2 | 0 | NA | NA | NA | NA | NA | #VALUE! |
| ENSMUSG00000082891.2 | 0.093255 | 0.868093 | 4.072464 | 0.213162 | 0.831201 | NA | #VALUE! |
| ENSMUSG00000082892.3 | 0 | NA | NA | NA | NA | NA | #VALUE! |
| ENSMUSG00000082893.2 | 0 | NA | NA | NA | NA | NA | #VALUE! |
| ENSMUSG00000082894.2 | 0 | NA | NA | NA | NA | NA | #VALUE! |
| ENSMUSG00000082896.2 | 2.943571 | 2.302405 | 3.514491 | 0.655118 | 0.512392 | NA | #VALUE! |
| ENSMUSG00000082897.2 | 0 | NA | NA | NA | NA | NA | #VALUE! |
| ENSMUSG00000082898.2 | 0 | NA | NA | NA | NA | NA | #VALUE! |
| ENSMUSG00000082899.2 | 0 | NA | NA | NA | NA | NA | #VALUE! |
| ENSMUSG00000082900.2 | 0 | NA | NA | NA | NA | NA | #VALUE! |
| ENSMUSG00000082901.2 | 0.309124 | -1.83395 | 4.047982 | -0.45305 | 0.650511 | NA | #VALUE! |
| ENSMUSG00000082902.4 | 0 | NA | NA | NA | NA | NA | #VALUE! |
| ENSMUSG00000082903.2 | 0 | NA | NA | NA | NA | NA | #VALUE! |
| ENSMUSG00000082905.2 | 0 | NA | NA | NA | NA | NA | #VALUE! |
| ENSMUSG00000082906.2 | 0 | NA | NA | NA | NA | NA | #VALUE! |
| ENSMUSG00000082907.2 | 0 | NA | NA | NA | NA | NA | #VALUE! |
| ENSMUSG00000082908.5 | 0.210026 | -1.52494 | 4.058692 | -0.37572 | 0.707123 | NA | #VALUE! |
| ENSMUSG00000082910.2 | 0 | NA | NA | NA | NA | NA | #VALUE! |
| ENSMUSG00000082911.2 | 0 | NA | NA | NA | NA | NA | #VALUE! |
| ENSMUSG00000082912.2 | 0 | NA | NA | NA | NA | NA | #VALUE! |
| ENSMUSG00000082913.2 | 0 | NA | NA | NA | NA | NA | #VALUE! |
| ENSMUSG00000082915.2 | 0 | NA | NA | NA | NA | NA | #VALUE! |
| ENSMUSG00000082917.2 | 0 | NA | NA | NA | NA | NA | #VALUE! |
| ENSMUSG00000082918.4 | 0 | NA | NA | NA | NA | NA | #VALUE! |
| ENSMUSG00000082919.2 | 0 | NA | NA | NA | NA | NA | #VALUE! |
| ENSMUSG00000082920.2 | 0.093255 | 0.868093 | 4.072464 | 0.213162 | 0.831201 | NA | #VALUE! |
| ENSMUSG00000082922.4 | 0 | NA | NA | NA | NA | NA | #VALUE! |
| ENSMUSG00000082924.2 | 0 | NA | NA | NA | NA | NA | #VALUE! |
| ENSMUSG00000082925.2 | 0 | NA | NA | NA | NA | NA | #VALUE! |
| ENSMUSG00000082926.4 | 0.105013 | -0.97324 | 4.076293 | -0.23876 | 0.811295 | NA | #VALUE! |
| ENSMUSG00000082927.4 | 0.093255 | 0.868093 | 4.072464 | 0.213162 | 0.831201 | NA | #VALUE! |
| ENSMUSG00000082928.2 | 0 | NA | NA | NA | NA | NA | #VALUE! |
| ENSMUSG00000082929.2 | 0 | NA | NA | NA | NA | NA | #VALUE! |
| ENSMUSG00000082930.2 | 0 | NA | NA | NA | NA | NA | #VALUE! |
| ENSMUSG00000082931.2 | 0 | NA | NA | NA | NA | NA | #VALUE! |
| ENSMUSG00000082932.4 | 0 | NA | NA | NA | NA | NA | #VALUE! |
| ENSMUSG00000082933.2 | 0 | NA | NA | NA | NA | NA | #VALUE! |

| | | | | | | | |
|---|---|---|---|---|---|---|---|
| ENSMUSG00000082934.2 | 0 | NA | NA | NA | NA | NA | #VALUE! |
| ENSMUSG00000082935.4 | 4.508121 | 1.47484 | 3.008125 | 0.490285 | 0.623932 | NA | #VALUE! |
| ENSMUSG00000082936.2 | 0 | NA | NA | NA | NA | NA | #VALUE! |
| ENSMUSG00000082937.2 | 0 | NA | NA | NA | NA | NA | #VALUE! |
| ENSMUSG00000082938.2 | 1.317993 | -2.26327 | 3.921202 | -0.57719 | 0.563812 | NA | #VALUE! |
| ENSMUSG00000082939.2 | 0 | NA | NA | NA | NA | NA | #VALUE! |
| ENSMUSG00000082940.2 | 0 | NA | NA | NA | NA | NA | #VALUE! |
| ENSMUSG00000082941.2 | 0 | NA | NA | NA | NA | NA | #VALUE! |
| ENSMUSG00000082943.2 | 0 | NA | NA | NA | NA | NA | #VALUE! |
| ENSMUSG00000082945.2 | 0 | NA | NA | NA | NA | NA | #VALUE! |
| ENSMUSG00000082947.2 | 0 | NA | NA | NA | NA | NA | #VALUE! |
| ENSMUSG00000082949.2 | 0 | NA | NA | NA | NA | NA | #VALUE! |
| ENSMUSG00000082950.2 | 0 | NA | NA | NA | NA | NA | #VALUE! |
| ENSMUSG00000082951.2 | 0 | NA | NA | NA | NA | NA | #VALUE! |
| ENSMUSG00000082952.2 | 0 | NA | NA | NA | NA | NA | #VALUE! |
| ENSMUSG00000082953.4 | 0 | NA | NA | NA | NA | NA | #VALUE! |
| ENSMUSG00000082954.2 | 0 | NA | NA | NA | NA | NA | #VALUE! |
| ENSMUSG00000082955.2 | 0 | NA | NA | NA | NA | NA | #VALUE! |
| ENSMUSG00000082956.5 | 0 | NA | NA | NA | NA | NA | #VALUE! |
| ENSMUSG00000082957.2 | 0 | NA | NA | NA | NA | NA | #VALUE! |
| ENSMUSG00000082958.2 | 0 | NA | NA | NA | NA | NA | #VALUE! |
| ENSMUSG00000082960.2 | 0 | NA | NA | NA | NA | NA | #VALUE! |
| ENSMUSG00000082961.2 | 0 | NA | NA | NA | NA | NA | #VALUE! |
| ENSMUSG00000082962.4 | 0 | NA | NA | NA | NA | NA | #VALUE! |
| ENSMUSG00000082963.2 | 0 | NA | NA | NA | NA | NA | #VALUE! |
| ENSMUSG00000082964.4 | 0 | NA | NA | NA | NA | NA | #VALUE! |
| ENSMUSG00000082965.2 | 0 | NA | NA | NA | NA | NA | #VALUE! |
| ENSMUSG00000082966.2 | 0 | NA | NA | NA | NA | NA | #VALUE! |
| ENSMUSG00000082967.3 | 0 | NA | NA | NA | NA | NA | #VALUE! |
| ENSMUSG00000082968.2 | 0 | NA | NA | NA | NA | NA | #VALUE! |
| ENSMUSG00000082969.2 | 0.093255 | 0.868093 | 4.072464 | 0.213162 | 0.831201 | NA | #VALUE! |
| ENSMUSG00000082970.2 | 0 | NA | NA | NA | NA | NA | #VALUE! |
| ENSMUSG00000082971.2 | 0 | NA | NA | NA | NA | NA | #VALUE! |
| ENSMUSG00000082973.2 | 0 | NA | NA | NA | NA | NA | #VALUE! |
| ENSMUSG00000082974.2 | 0 | NA | NA | NA | NA | NA | #VALUE! |
| ENSMUSG00000082975.2 | 0 | NA | NA | NA | NA | NA | #VALUE! |
| ENSMUSG00000082976.11 | 0 | NA | NA | NA | NA | NA | #VALUE! |
| ENSMUSG00000082977.2 | 0 | NA | NA | NA | NA | NA | #VALUE! |
| ENSMUSG00000082978.2 | 0.204111 | -1.05581 | 4.073302 | -0.2592 | 0.795479 | NA | #VALUE! |
| ENSMUSG00000082980.6 | 0 | NA | NA | NA | NA | NA | #VALUE! |
| ENSMUSG00000082981.2 | 0 | NA | NA | NA | NA | NA | #VALUE! |
| ENSMUSG00000082982.3 | 0 | NA | NA | NA | NA | NA | #VALUE! |
| ENSMUSG00000082983.2 | 0 | NA | NA | NA | NA | NA | #VALUE! |
| ENSMUSG00000082985.3 | 0 | NA | NA | NA | NA | NA | #VALUE! |

| | | | | | | | |
|---|---|---|---|---|---|---|---|
| ENSMUSG00000082986.2 | 0 | NA | NA | NA | NA | NA | #VALUE! |
| ENSMUSG00000082987.2 | 0 | NA | NA | NA | NA | NA | #VALUE! |
| ENSMUSG00000082988.4 | 0 | NA | NA | NA | NA | NA | #VALUE! |
| ENSMUSG00000082989.2 | 0 | NA | NA | NA | NA | NA | #VALUE! |
| ENSMUSG00000082990.2 | 0 | NA | NA | NA | NA | NA | #VALUE! |
| ENSMUSG00000082991.2 | 1.780461 | 3.124898 | 3.927799 | 0.795585 | 0.426273 | NA | #VALUE! |
| ENSMUSG00000082993.2 | 0 | NA | NA | NA | NA | NA | #VALUE! |
| ENSMUSG00000082994.2 | 0 | NA | NA | NA | NA | NA | #VALUE! |
| ENSMUSG00000082995.2 | 0 | NA | NA | NA | NA | NA | #VALUE! |
| ENSMUSG00000082996.2 | 0.289044 | -0.14751 | 4.065279 | -0.03629 | 0.971054 | NA | #VALUE! |
| ENSMUSG00000082997.2 | 0 | NA | NA | NA | NA | NA | #VALUE! |
| ENSMUSG00000082998.4 | 0.364073 | 0.012025 | 4.059931 | 0.002962 | 0.997637 | NA | #VALUE! |
| ENSMUSG00000083000.2 | 0 | NA | NA | NA | NA | NA | #VALUE! |
| ENSMUSG00000083001.2 | 0 | NA | NA | NA | NA | NA | #VALUE! |
| ENSMUSG00000083004.2 | 0 | NA | NA | NA | NA | NA | #VALUE! |
| ENSMUSG00000083005.2 | 0 | NA | NA | NA | NA | NA | #VALUE! |
| ENSMUSG00000083006.2 | 0 | NA | NA | NA | NA | NA | #VALUE! |
| ENSMUSG00000083007.2 | 0 | NA | NA | NA | NA | NA | #VALUE! |
| ENSMUSG00000083008.2 | 0 | NA | NA | NA | NA | NA | #VALUE! |
| ENSMUSG00000083009.2 | 0 | NA | NA | NA | NA | NA | #VALUE! |
| ENSMUSG00000083010.2 | 0 | NA | NA | NA | NA | NA | #VALUE! |
| ENSMUSG00000083011.2 | 0 | NA | NA | NA | NA | NA | #VALUE! |
| ENSMUSG00000083012.10 | 0 | NA | NA | NA | NA | NA | #VALUE! |
| ENSMUSG00000083013.2 | 0 | NA | NA | NA | NA | NA | #VALUE! |
| ENSMUSG00000083014.2 | 0 | NA | NA | NA | NA | NA | #VALUE! |
| ENSMUSG00000083015.2 | 0 | NA | NA | NA | NA | NA | #VALUE! |
| ENSMUSG00000083016.5 | 0.195789 | -1.05581 | 4.073302 | -0.2592 | 0.795479 | NA | #VALUE! |
| ENSMUSG00000083017.2 | 0.195789 | -1.05581 | 4.073302 | -0.2592 | 0.795479 | NA | #VALUE! |
| ENSMUSG00000083018.2 | 0 | NA | NA | NA | NA | NA | #VALUE! |
| ENSMUSG00000083019.3 | 1.506024 | 0.575669 | 3.851045 | 0.149484 | 0.881172 | NA | #VALUE! |
| ENSMUSG00000083020.2 | 0 | NA | NA | NA | NA | NA | #VALUE! |
| ENSMUSG00000083023.2 | 0 | NA | NA | NA | NA | NA | #VALUE! |
| ENSMUSG00000083024.2 | 0 | NA | NA | NA | NA | NA | #VALUE! |
| ENSMUSG00000083025.2 | 0 | NA | NA | NA | NA | NA | #VALUE! |
| ENSMUSG00000083026.2 | 0 | NA | NA | NA | NA | NA | #VALUE! |
| ENSMUSG00000083027.2 | 0 | NA | NA | NA | NA | NA | #VALUE! |
| ENSMUSG00000083029.2 | 0 | NA | NA | NA | NA | NA | #VALUE! |
| ENSMUSG00000083030.2 | 0 | NA | NA | NA | NA | NA | #VALUE! |
| ENSMUSG00000083031.2 | 0 | NA | NA | NA | NA | NA | #VALUE! |
| ENSMUSG00000083032.2 | 0 | NA | NA | NA | NA | NA | #VALUE! |
| ENSMUSG00000083033.2 | 0 | NA | NA | NA | NA | NA | #VALUE! |
| ENSMUSG00000083034.3 | 0 | NA | NA | NA | NA | NA | #VALUE! |
| ENSMUSG00000083036.2 | 0 | NA | NA | NA | NA | NA | #VALUE! |
| ENSMUSG00000083037.2 | 0 | NA | NA | NA | NA | NA | #VALUE! |

| | | | | | | | |
|---|---|---|---|---|---|---|---|
| ENSMUSG00000083038.2 | 0 | NA | NA | NA | NA | NA | #VALUE! |
| ENSMUSG00000083039.2 | 0 | NA | NA | NA | NA | NA | #VALUE! |
| ENSMUSG00000083041.3 | 0 | NA | NA | NA | NA | NA | #VALUE! |
| ENSMUSG00000083042.2 | 0 | NA | NA | NA | NA | NA | #VALUE! |
| ENSMUSG00000083043.2 | 0.3999 | -2.17566 | 4.018831 | -0.54137 | 0.588256 | NA | #VALUE! |
| ENSMUSG00000083044.2 | 0 | NA | NA | NA | NA | NA | #VALUE! |
| ENSMUSG00000083045.2 | 0 | NA | NA | NA | NA | NA | #VALUE! |
| ENSMUSG00000083046.2 | 0 | NA | NA | NA | NA | NA | #VALUE! |
| ENSMUSG00000083047.2 | 0 | NA | NA | NA | NA | NA | #VALUE! |
| ENSMUSG00000083049.2 | 0 | NA | NA | NA | NA | NA | #VALUE! |
| ENSMUSG00000083050.3 | 0 | NA | NA | NA | NA | NA | #VALUE! |
| ENSMUSG00000083052.2 | 0 | NA | NA | NA | NA | NA | #VALUE! |
| ENSMUSG00000083053.2 | 0 | NA | NA | NA | NA | NA | #VALUE! |
| ENSMUSG00000083054.2 | 0 | NA | NA | NA | NA | NA | #VALUE! |
| ENSMUSG00000083055.2 | 0 | NA | NA | NA | NA | NA | #VALUE! |
| ENSMUSG00000083057.2 | 0 | NA | NA | NA | NA | NA | #VALUE! |
| ENSMUSG00000083058.2 | 0 | NA | NA | NA | NA | NA | #VALUE! |
| ENSMUSG00000083059.4 | 0 | NA | NA | NA | NA | NA | #VALUE! |
| ENSMUSG00000083060.2 | 0 | NA | NA | NA | NA | NA | #VALUE! |
| ENSMUSG00000083061.2 | 0.195789 | -1.05581 | 4.073302 | -0.2592 | 0.795479 | NA | #VALUE! |
| ENSMUSG00000083062.2 | 0 | NA | NA | NA | NA | NA | #VALUE! |
| ENSMUSG00000083063.2 | 0 | NA | NA | NA | NA | NA | #VALUE! |
| ENSMUSG00000083064.2 | 0 | NA | NA | NA | NA | NA | #VALUE! |
| ENSMUSG00000083065.2 | 0 | NA | NA | NA | NA | NA | #VALUE! |
| ENSMUSG00000083066.2 | 0 | NA | NA | NA | NA | NA | #VALUE! |
| ENSMUSG00000083067.2 | 0 | NA | NA | NA | NA | NA | #VALUE! |
| ENSMUSG00000083068.2 | 0 | NA | NA | NA | NA | NA | #VALUE! |
| ENSMUSG00000083069.2 | 0 | NA | NA | NA | NA | NA | #VALUE! |
| ENSMUSG00000083070.3 | 0 | NA | NA | NA | NA | NA | #VALUE! |
| ENSMUSG00000083072.3 | 0 | NA | NA | NA | NA | NA | #VALUE! |
| ENSMUSG00000083073.2 | 0 | NA | NA | NA | NA | NA | #VALUE! |
| ENSMUSG00000083074.2 | 0 | NA | NA | NA | NA | NA | #VALUE! |
| ENSMUSG00000083075.2 | 0 | NA | NA | NA | NA | NA | #VALUE! |
| ENSMUSG00000083076.2 | 0 | NA | NA | NA | NA | NA | #VALUE! |
| ENSMUSG00000083077.2 | 0 | NA | NA | NA | NA | NA | #VALUE! |
| ENSMUSG00000083078.2 | 0 | NA | NA | NA | NA | NA | #VALUE! |
| ENSMUSG00000083079.5 | 0 | NA | NA | NA | NA | NA | #VALUE! |
| ENSMUSG00000083080.2 | 0 | NA | NA | NA | NA | NA | #VALUE! |
| ENSMUSG00000083081.2 | 0 | NA | NA | NA | NA | NA | #VALUE! |
| ENSMUSG00000083083.2 | 0 | NA | NA | NA | NA | NA | #VALUE! |
| ENSMUSG00000083084.2 | 0 | NA | NA | NA | NA | NA | #VALUE! |
| ENSMUSG00000083085.3 | 0 | NA | NA | NA | NA | NA | #VALUE! |
| ENSMUSG00000083086.2 | 0 | NA | NA | NA | NA | NA | #VALUE! |
| ENSMUSG00000083088.2 | 0 | NA | NA | NA | NA | NA | #VALUE! |

| | | | | | | | |
|---|---|---|---|---|---|---|---|
| ENSMUSG00000083089.2 | 0 | NA | NA | NA | NA | NA | #VALUE! |
| ENSMUSG00000083090.2 | 0 | NA | NA | NA | NA | NA | #VALUE! |
| ENSMUSG00000083091.2 | 0 | NA | NA | NA | NA | NA | #VALUE! |
| ENSMUSG00000083093.3 | 0 | NA | NA | NA | NA | NA | #VALUE! |
| ENSMUSG00000083094.3 | 0.093255 | 0.868093 | 4.072464 | 0.213162 | 0.831201 | NA | #VALUE! |
| ENSMUSG00000083096.3 | 0 | NA | NA | NA | NA | NA | #VALUE! |
| ENSMUSG00000083097.4 | 0 | NA | NA | NA | NA | NA | #VALUE! |
| ENSMUSG00000083098.2 | 0 | NA | NA | NA | NA | NA | #VALUE! |
| ENSMUSG00000083099.2 | 0 | NA | NA | NA | NA | NA | #VALUE! |
| ENSMUSG00000083100.2 | 0 | NA | NA | NA | NA | NA | #VALUE! |
| ENSMUSG00000083101.2 | 0 | NA | NA | NA | NA | NA | #VALUE! |
| ENSMUSG00000083103.2 | 0 | NA | NA | NA | NA | NA | #VALUE! |
| ENSMUSG00000083104.2 | 0 | NA | NA | NA | NA | NA | #VALUE! |
| ENSMUSG00000083105.2 | 0 | NA | NA | NA | NA | NA | #VALUE! |
| ENSMUSG00000083106.2 | 0 | NA | NA | NA | NA | NA | #VALUE! |
| ENSMUSG00000083107.2 | 0 | NA | NA | NA | NA | NA | #VALUE! |
| ENSMUSG00000083109.4 | 0 | NA | NA | NA | NA | NA | #VALUE! |
| ENSMUSG00000083110.2 | 0 | NA | NA | NA | NA | NA | #VALUE! |
| ENSMUSG00000083111.2 | 0.105013 | -0.97324 | 4.076293 | -0.23876 | 0.811295 | NA | #VALUE! |
| ENSMUSG00000083112.2 | 0 | NA | NA | NA | NA | NA | #VALUE! |
| ENSMUSG00000083113.2 | 0 | NA | NA | NA | NA | NA | #VALUE! |
| ENSMUSG00000083114.2 | 0 | NA | NA | NA | NA | NA | #VALUE! |
| ENSMUSG00000083115.2 | 0 | NA | NA | NA | NA | NA | #VALUE! |
| ENSMUSG00000083116.2 | 0 | NA | NA | NA | NA | NA | #VALUE! |
| ENSMUSG00000083117.2 | 0 | NA | NA | NA | NA | NA | #VALUE! |
| ENSMUSG00000083118.2 | 0 | NA | NA | NA | NA | NA | #VALUE! |
| ENSMUSG00000083119.2 | 0 | NA | NA | NA | NA | NA | #VALUE! |
| ENSMUSG00000083120.4 | 0 | NA | NA | NA | NA | NA | #VALUE! |
| ENSMUSG00000083121.4 | 0.469086 | -0.7385 | 4.036584 | -0.18295 | 0.854836 | NA | #VALUE! |
| ENSMUSG00000083122.2 | 0 | NA | NA | NA | NA | NA | #VALUE! |
| ENSMUSG00000083123.2 | 0 | NA | NA | NA | NA | NA | #VALUE! |
| ENSMUSG00000083124.2 | 0 | NA | NA | NA | NA | NA | #VALUE! |
| ENSMUSG00000083126.2 | 0 | NA | NA | NA | NA | NA | #VALUE! |
| ENSMUSG00000083127.2 | 0 | NA | NA | NA | NA | NA | #VALUE! |
| ENSMUSG00000083128.2 | 0 | NA | NA | NA | NA | NA | #VALUE! |
| ENSMUSG00000083129.2 | 0 | NA | NA | NA | NA | NA | #VALUE! |
| ENSMUSG00000083130.2 | 0 | NA | NA | NA | NA | NA | #VALUE! |
| ENSMUSG00000083135.2 | 0 | NA | NA | NA | NA | NA | #VALUE! |
| ENSMUSG00000083136.2 | 6.46104 | -6.21992 | 3.913702 | -1.58927 | 0.112 | NA | #VALUE! |
| ENSMUSG00000083137.2 | 0 | NA | NA | NA | NA | NA | #VALUE! |
| ENSMUSG00000083138.4 | 0 | NA | NA | NA | NA | NA | #VALUE! |
| ENSMUSG00000083139.2 | 0 | NA | NA | NA | NA | NA | #VALUE! |
| ENSMUSG00000083140.2 | 0 | NA | NA | NA | NA | NA | #VALUE! |
| ENSMUSG00000083141.2 | 0 | NA | NA | NA | NA | NA | #VALUE! |

| | | | | | | | |
|---|---|---|---|---|---|---|---|
| ENSMUSG00000083142.2 | 0 | NA | NA | NA | NA | NA | #VALUE! |
| ENSMUSG00000083143.3 | 0 | NA | NA | NA | NA | NA | #VALUE! |
| ENSMUSG00000083144.2 | 0 | NA | NA | NA | NA | NA | #VALUE! |
| ENSMUSG00000083145.2 | 0 | NA | NA | NA | NA | NA | #VALUE! |
| ENSMUSG00000083147.3 | 0 | NA | NA | NA | NA | NA | #VALUE! |
| ENSMUSG00000083149.3 | 0 | NA | NA | NA | NA | NA | #VALUE! |
| ENSMUSG00000083150.2 | 0 | NA | NA | NA | NA | NA | #VALUE! |
| ENSMUSG00000083151.2 | 0 | NA | NA | NA | NA | NA | #VALUE! |
| ENSMUSG00000083152.2 | 0 | NA | NA | NA | NA | NA | #VALUE! |
| ENSMUSG00000083153.2 | 0 | NA | NA | NA | NA | NA | #VALUE! |
| ENSMUSG00000083154.2 | 0 | NA | NA | NA | NA | NA | #VALUE! |
| ENSMUSG00000083155.2 | 0 | NA | NA | NA | NA | NA | #VALUE! |
| ENSMUSG00000083156.2 | 0 | NA | NA | NA | NA | NA | #VALUE! |
| ENSMUSG00000083157.2 | 0 | NA | NA | NA | NA | NA | #VALUE! |
| ENSMUSG00000083158.2 | 0 | NA | NA | NA | NA | NA | #VALUE! |
| ENSMUSG00000083159.2 | 0 | NA | NA | NA | NA | NA | #VALUE! |
| ENSMUSG00000083160.2 | 0 | NA | NA | NA | NA | NA | #VALUE! |
| ENSMUSG00000083161.3 | 0 | NA | NA | NA | NA | NA | #VALUE! |
| ENSMUSG00000083162.2 | 0 | NA | NA | NA | NA | NA | #VALUE! |
| ENSMUSG00000083163.2 | 0 | NA | NA | NA | NA | NA | #VALUE! |
| ENSMUSG00000083165.2 | 0 | NA | NA | NA | NA | NA | #VALUE! |
| ENSMUSG00000083166.2 | 0 | NA | NA | NA | NA | NA | #VALUE! |
| ENSMUSG00000083167.2 | 0 | NA | NA | NA | NA | NA | #VALUE! |
| ENSMUSG00000083168.2 | 0 | NA | NA | NA | NA | NA | #VALUE! |
| ENSMUSG00000083169.2 | 0 | NA | NA | NA | NA | NA | #VALUE! |
| ENSMUSG00000083170.2 | 0 | NA | NA | NA | NA | NA | #VALUE! |
| ENSMUSG00000083171.2 | 0 | NA | NA | NA | NA | NA | #VALUE! |
| ENSMUSG00000083172.2 | 0 | NA | NA | NA | NA | NA | #VALUE! |
| ENSMUSG00000083173.2 | 0 | NA | NA | NA | NA | NA | #VALUE! |
| ENSMUSG00000083174.2 | 0 | NA | NA | NA | NA | NA | #VALUE! |
| ENSMUSG00000083175.2 | 0 | NA | NA | NA | NA | NA | #VALUE! |
| ENSMUSG00000083176.2 | 0 | NA | NA | NA | NA | NA | #VALUE! |
| ENSMUSG00000083177.2 | 0 | NA | NA | NA | NA | NA | #VALUE! |
| ENSMUSG00000083178.2 | 0 | NA | NA | NA | NA | NA | #VALUE! |
| ENSMUSG00000083179.4 | 0.394057 | -0.89804 | 4.041963 | -0.22218 | 0.824174 | NA | #VALUE! |
| ENSMUSG00000083180.2 | 0 | NA | NA | NA | NA | NA | #VALUE! |
| ENSMUSG00000083181.2 | 0 | NA | NA | NA | NA | NA | #VALUE! |
| ENSMUSG00000083182.2 | 0 | NA | NA | NA | NA | NA | #VALUE! |
| ENSMUSG00000083183.2 | 0 | NA | NA | NA | NA | NA | #VALUE! |
| ENSMUSG00000083185.3 | 0 | NA | NA | NA | NA | NA | #VALUE! |
| ENSMUSG00000083186.2 | 1.229878 | 3.761857 | 3.943699 | 0.95389 | 0.340139 | NA | #VALUE! |
| ENSMUSG00000083188.3 | 0 | NA | NA | NA | NA | NA | #VALUE! |
| ENSMUSG00000083190.2 | 0 | NA | NA | NA | NA | NA | #VALUE! |
| ENSMUSG00000083191.2 | 0 | NA | NA | NA | NA | NA | #VALUE! |

| | | | | | | | |
|---|---|---|---|---|---|---|---|
| ENSMUSG00000083193.2 | 0 | NA | NA | NA | NA | NA | #VALUE! |
| ENSMUSG00000083195.2 | 0 | NA | NA | NA | NA | NA | #VALUE! |
| ENSMUSG00000083196.2 | 0 | NA | NA | NA | NA | NA | #VALUE! |
| ENSMUSG00000083197.2 | 0 | NA | NA | NA | NA | NA | #VALUE! |
| ENSMUSG00000083199.2 | 0 | NA | NA | NA | NA | NA | #VALUE! |
| ENSMUSG00000083200.2 | 0 | NA | NA | NA | NA | NA | #VALUE! |
| ENSMUSG00000083201.2 | 0 | NA | NA | NA | NA | NA | #VALUE! |
| ENSMUSG00000083202.2 | 0 | NA | NA | NA | NA | NA | #VALUE! |
| ENSMUSG00000083203.2 | 0 | NA | NA | NA | NA | NA | #VALUE! |
| ENSMUSG00000083204.2 | 0 | NA | NA | NA | NA | NA | #VALUE! |
| ENSMUSG00000083205.2 | 0 | NA | NA | NA | NA | NA | #VALUE! |
| ENSMUSG00000083206.2 | 0 | NA | NA | NA | NA | NA | #VALUE! |
| ENSMUSG00000083207.2 | 0 | NA | NA | NA | NA | NA | #VALUE! |
| ENSMUSG00000083208.2 | 0 | NA | NA | NA | NA | NA | #VALUE! |
| ENSMUSG00000083209.2 | 0 | NA | NA | NA | NA | NA | #VALUE! |
| ENSMUSG00000083210.2 | 0 | NA | NA | NA | NA | NA | #VALUE! |
| ENSMUSG00000083211.2 | 0 | NA | NA | NA | NA | NA | #VALUE! |
| ENSMUSG00000083212.4 | 0 | NA | NA | NA | NA | NA | #VALUE! |
| ENSMUSG00000083213.3 | 0 | NA | NA | NA | NA | NA | #VALUE! |
| ENSMUSG00000083214.2 | 0 | NA | NA | NA | NA | NA | #VALUE! |
| ENSMUSG00000083215.4 | 0 | NA | NA | NA | NA | NA | #VALUE! |
| ENSMUSG00000083216.3 | 0 | NA | NA | NA | NA | NA | #VALUE! |
| ENSMUSG00000083218.3 | 0 | NA | NA | NA | NA | NA | #VALUE! |
| ENSMUSG00000083219.2 | 9.308031 | -5.23492 | 2.511449 | -2.08442 | 0.037122 | NA | #VALUE! |
| ENSMUSG00000083220.2 | 0 | NA | NA | NA | NA | NA | #VALUE! |
| ENSMUSG00000083221.2 | 0 | NA | NA | NA | NA | NA | #VALUE! |
| ENSMUSG00000083222.2 | 0 | NA | NA | NA | NA | NA | #VALUE! |
| ENSMUSG00000083223.2 | 0 | NA | NA | NA | NA | NA | #VALUE! |
| ENSMUSG00000083224.2 | 0 | NA | NA | NA | NA | NA | #VALUE! |
| ENSMUSG00000083225.2 | 0 | NA | NA | NA | NA | NA | #VALUE! |
| ENSMUSG00000083226.3 | 0 | NA | NA | NA | NA | NA | #VALUE! |
| ENSMUSG00000083227.2 | 0 | NA | NA | NA | NA | NA | #VALUE! |
| ENSMUSG00000083228.2 | 0 | NA | NA | NA | NA | NA | #VALUE! |
| ENSMUSG00000083229.2 | 0 | NA | NA | NA | NA | NA | #VALUE! |
| ENSMUSG00000083230.2 | 0 | NA | NA | NA | NA | NA | #VALUE! |
| ENSMUSG00000083231.2 | 0 | NA | NA | NA | NA | NA | #VALUE! |
| ENSMUSG00000083232.2 | 0 | NA | NA | NA | NA | NA | #VALUE! |
| ENSMUSG00000083233.2 | 0 | NA | NA | NA | NA | NA | #VALUE! |
| ENSMUSG00000083234.2 | 0 | NA | NA | NA | NA | NA | #VALUE! |
| ENSMUSG00000083235.2 | 0 | NA | NA | NA | NA | NA | #VALUE! |
| ENSMUSG00000083236.2 | 0 | NA | NA | NA | NA | NA | #VALUE! |
| ENSMUSG00000083237.2 | 0 | NA | NA | NA | NA | NA | #VALUE! |
| ENSMUSG00000083238.2 | 0 | NA | NA | NA | NA | NA | #VALUE! |
| ENSMUSG00000083239.2 | 0 | NA | NA | NA | NA | NA | #VALUE! |

| | | | | | | | |
|---|---|---|---|---|---|---|---|
| ENSMUSG00000083240.4 | 0 | NA | NA | NA | NA | NA | #VALUE! |
| ENSMUSG00000083241.2 | 0 | NA | NA | NA | NA | NA | #VALUE! |
| ENSMUSG00000083242.2 | 0 | NA | NA | NA | NA | NA | #VALUE! |
| ENSMUSG00000083243.2 | 0 | NA | NA | NA | NA | NA | #VALUE! |
| ENSMUSG00000083244.2 | 0 | NA | NA | NA | NA | NA | #VALUE! |
| ENSMUSG00000083245.2 | 0 | NA | NA | NA | NA | NA | #VALUE! |
| ENSMUSG00000083246.2 | 0 | NA | NA | NA | NA | NA | #VALUE! |
| ENSMUSG00000083248.2 | 0 | NA | NA | NA | NA | NA | #VALUE! |
| ENSMUSG00000083249.2 | 0 | NA | NA | NA | NA | NA | #VALUE! |
| ENSMUSG00000083250.2 | 0 | NA | NA | NA | NA | NA | #VALUE! |
| ENSMUSG00000083251.2 | 0 | NA | NA | NA | NA | NA | #VALUE! |
| ENSMUSG00000083252.2 | 0 | NA | NA | NA | NA | NA | #VALUE! |
| ENSMUSG00000083253.2 | 0.168284 | 1.027628 | 4.067126 | 0.252667 | 0.800526 | NA | #VALUE! |
| ENSMUSG00000083254.2 | 0 | NA | NA | NA | NA | NA | #VALUE! |
| ENSMUSG00000083255.2 | 0 | NA | NA | NA | NA | NA | #VALUE! |
| ENSMUSG00000083256.2 | 0 | NA | NA | NA | NA | NA | #VALUE! |
| ENSMUSG00000083257.2 | 0 | NA | NA | NA | NA | NA | #VALUE! |
| ENSMUSG00000083258.4 | 0 | NA | NA | NA | NA | NA | #VALUE! |
| ENSMUSG00000083260.2 | 0 | NA | NA | NA | NA | NA | #VALUE! |
| ENSMUSG00000083261.5 | 0 | NA | NA | NA | NA | NA | #VALUE! |
| ENSMUSG00000083262.2 | 0 | NA | NA | NA | NA | NA | #VALUE! |
| ENSMUSG00000083263.2 | 0 | NA | NA | NA | NA | NA | #VALUE! |
| ENSMUSG00000083264.2 | 0 | NA | NA | NA | NA | NA | #VALUE! |
| ENSMUSG00000083267.3 | 0 | NA | NA | NA | NA | NA | #VALUE! |
| ENSMUSG00000083268.2 | 0 | NA | NA | NA | NA | NA | #VALUE! |
| ENSMUSG00000083269.2 | 0 | NA | NA | NA | NA | NA | #VALUE! |
| ENSMUSG00000083270.2 | 0 | NA | NA | NA | NA | NA | #VALUE! |
| ENSMUSG00000083271.2 | 0 | NA | NA | NA | NA | NA | #VALUE! |
| ENSMUSG00000083272.2 | 0 | NA | NA | NA | NA | NA | #VALUE! |
| ENSMUSG00000083274.2 | 0 | NA | NA | NA | NA | NA | #VALUE! |
| ENSMUSG00000083275.2 | 0 | NA | NA | NA | NA | NA | #VALUE! |
| ENSMUSG00000083276.2 | 0 | NA | NA | NA | NA | NA | #VALUE! |
| ENSMUSG00000083277.2 | 0 | NA | NA | NA | NA | NA | #VALUE! |
| ENSMUSG00000083278.2 | 0 | NA | NA | NA | NA | NA | #VALUE! |
| ENSMUSG00000083279.2 | 0 | NA | NA | NA | NA | NA | #VALUE! |
| ENSMUSG00000083280.2 | 0 | NA | NA | NA | NA | NA | #VALUE! |
| ENSMUSG00000083281.2 | 0 | NA | NA | NA | NA | NA | #VALUE! |
| ENSMUSG00000083283.4 | 0.210026 | -1.52494 | 4.058692 | -0.37572 | 0.707123 | NA | #VALUE! |
| ENSMUSG00000083284.2 | 0 | NA | NA | NA | NA | NA | #VALUE! |
| ENSMUSG00000083285.3 | 0 | NA | NA | NA | NA | NA | #VALUE! |
| ENSMUSG00000083287.5 | 0 | NA | NA | NA | NA | NA | #VALUE! |
| ENSMUSG00000083288.2 | 0 | NA | NA | NA | NA | NA | #VALUE! |
| ENSMUSG00000083289.2 | 0 | NA | NA | NA | NA | NA | #VALUE! |
| ENSMUSG00000083290.2 | 0 | NA | NA | NA | NA | NA | #VALUE! |

| | | | | | | | |
|---|---|---|---|---|---|---|---|
| ENSMUSG00000083291.4 | 0 | NA | NA | NA | NA | NA | #VALUE! |
| ENSMUSG00000083292.2 | 0 | NA | NA | NA | NA | NA | #VALUE! |
| ENSMUSG00000083293.2 | 0 | NA | NA | NA | NA | NA | #VALUE! |
| ENSMUSG00000083294.4 | 0 | NA | NA | NA | NA | NA | #VALUE! |
| ENSMUSG00000083296.2 | 0 | NA | NA | NA | NA | NA | #VALUE! |
| ENSMUSG00000083297.2 | 0 | NA | NA | NA | NA | NA | #VALUE! |
| ENSMUSG00000083298.2 | 0 | NA | NA | NA | NA | NA | #VALUE! |
| ENSMUSG00000083299.2 | 0 | NA | NA | NA | NA | NA | #VALUE! |
| ENSMUSG00000083300.2 | 0 | NA | NA | NA | NA | NA | #VALUE! |
| ENSMUSG00000083303.2 | 0 | NA | NA | NA | NA | NA | #VALUE! |
| ENSMUSG00000083304.2 | 0 | NA | NA | NA | NA | NA | #VALUE! |
| ENSMUSG00000083305.3 | 0 | NA | NA | NA | NA | NA | #VALUE! |
| ENSMUSG00000083306.2 | 0 | NA | NA | NA | NA | NA | #VALUE! |
| ENSMUSG00000083307.2 | 0 | NA | NA | NA | NA | NA | #VALUE! |
| ENSMUSG00000083309.2 | 0 | NA | NA | NA | NA | NA | #VALUE! |
| ENSMUSG00000083310.3 | 0 | NA | NA | NA | NA | NA | #VALUE! |
| ENSMUSG00000083311.4 | 0 | NA | NA | NA | NA | NA | #VALUE! |
| ENSMUSG00000083312.2 | 0 | NA | NA | NA | NA | NA | #VALUE! |
| ENSMUSG00000083314.2 | 0 | NA | NA | NA | NA | NA | #VALUE! |
| ENSMUSG00000083315.2 | 0 | NA | NA | NA | NA | NA | #VALUE! |
| ENSMUSG00000083316.2 | 0 | NA | NA | NA | NA | NA | #VALUE! |
| ENSMUSG00000083317.2 | 0 | NA | NA | NA | NA | NA | #VALUE! |
| ENSMUSG00000083318.2 | 0 | NA | NA | NA | NA | NA | #VALUE! |
| ENSMUSG00000083319.2 | 0 | NA | NA | NA | NA | NA | #VALUE! |
| ENSMUSG00000083320.2 | 0.195789 | -1.05581 | 4.073302 | -0.2592 | 0.795479 | NA | #VALUE! |
| ENSMUSG00000083321.2 | 0 | NA | NA | NA | NA | NA | #VALUE! |
| ENSMUSG00000083324.2 | 0 | NA | NA | NA | NA | NA | #VALUE! |
| ENSMUSG00000083325.4 | 0 | NA | NA | NA | NA | NA | #VALUE! |
| ENSMUSG00000083326.5 | 1.195132 | -0.09666 | 3.880247 | -0.02491 | 0.980126 | NA | #VALUE! |
| ENSMUSG00000083327.2 | 0.628091 | 1.641687 | 3.985568 | 0.411908 | 0.680407 | NA | #VALUE! |
| ENSMUSG00000083328.4 | 1.113108 | -0.13639 | 3.890512 | -0.03506 | 0.972034 | NA | #VALUE! |
| ENSMUSG00000083329.2 | 0 | NA | NA | NA | NA | NA | #VALUE! |
| ENSMUSG00000083330.2 | 0 | NA | NA | NA | NA | NA | #VALUE! |
| ENSMUSG00000083331.2 | 0 | NA | NA | NA | NA | NA | #VALUE! |
| ENSMUSG00000083332.2 | 0 | NA | NA | NA | NA | NA | #VALUE! |
| ENSMUSG00000083333.4 | 0 | NA | NA | NA | NA | NA | #VALUE! |
| ENSMUSG00000083334.2 | 0 | NA | NA | NA | NA | NA | #VALUE! |
| ENSMUSG00000083335.2 | 0 | NA | NA | NA | NA | NA | #VALUE! |
| ENSMUSG00000083336.3 | 0 | NA | NA | NA | NA | NA | #VALUE! |
| ENSMUSG00000083338.2 | 0 | NA | NA | NA | NA | NA | #VALUE! |
| ENSMUSG00000083339.2 | 0 | NA | NA | NA | NA | NA | #VALUE! |
| ENSMUSG00000083340.2 | 0 | NA | NA | NA | NA | NA | #VALUE! |
| ENSMUSG00000083341.2 | 0 | NA | NA | NA | NA | NA | #VALUE! |
| ENSMUSG00000083342.2 | 0 | NA | NA | NA | NA | NA | #VALUE! |

| | | | | | | | |
|---|---|---|---|---|---|---|---|
| ENSMUSG00000083343.2 | 0 | NA | NA | NA | NA | NA | #VALUE! |
| ENSMUSG00000083344.3 | 0.093255 | 0.868093 | 4.072464 | 0.213162 | 0.831201 | NA | #VALUE! |
| ENSMUSG00000083345.2 | 0 | NA | NA | NA | NA | NA | #VALUE! |
| ENSMUSG00000083346.4 | 0 | NA | NA | NA | NA | NA | #VALUE! |
| ENSMUSG00000083347.2 | 0 | NA | NA | NA | NA | NA | #VALUE! |
| ENSMUSG00000083348.2 | 0 | NA | NA | NA | NA | NA | #VALUE! |
| ENSMUSG00000083350.2 | 0 | NA | NA | NA | NA | NA | #VALUE! |
| ENSMUSG00000083351.2 | 0 | NA | NA | NA | NA | NA | #VALUE! |
| ENSMUSG00000083352.2 | 0 | NA | NA | NA | NA | NA | #VALUE! |
| ENSMUSG00000083353.2 | 0 | NA | NA | NA | NA | NA | #VALUE! |
| ENSMUSG00000083355.2 | 0 | NA | NA | NA | NA | NA | #VALUE! |
| ENSMUSG00000083356.2 | 0 | NA | NA | NA | NA | NA | #VALUE! |
| ENSMUSG00000083358.3 | 0 | NA | NA | NA | NA | NA | #VALUE! |
| ENSMUSG00000083359.2 | 0 | NA | NA | NA | NA | NA | #VALUE! |
| ENSMUSG00000083361.5 | 0 | NA | NA | NA | NA | NA | #VALUE! |
| ENSMUSG00000083362.2 | 0 | NA | NA | NA | NA | NA | #VALUE! |
| ENSMUSG00000083363.2 | 0 | NA | NA | NA | NA | NA | #VALUE! |
| ENSMUSG00000083364.5 | 0.696062 | 1.386576 | 3.99283 | 0.347267 | 0.728391 | NA | #VALUE! |
| ENSMUSG00000083365.2 | 0 | NA | NA | NA | NA | NA | #VALUE! |
| ENSMUSG00000083366.2 | 0 | NA | NA | NA | NA | NA | #VALUE! |
| ENSMUSG00000083367.2 | 0 | NA | NA | NA | NA | NA | #VALUE! |
| ENSMUSG00000083368.2 | 0 | NA | NA | NA | NA | NA | #VALUE! |
| ENSMUSG00000083369.2 | 0 | NA | NA | NA | NA | NA | #VALUE! |
| ENSMUSG00000083371.2 | 0 | NA | NA | NA | NA | NA | #VALUE! |
| ENSMUSG00000083372.2 | 0 | NA | NA | NA | NA | NA | #VALUE! |
| ENSMUSG00000083373.2 | 0 | NA | NA | NA | NA | NA | #VALUE! |
| ENSMUSG00000083374.2 | 0 | NA | NA | NA | NA | NA | #VALUE! |
| ENSMUSG00000083375.2 | 0 | NA | NA | NA | NA | NA | #VALUE! |
| ENSMUSG00000083376.2 | 0 | NA | NA | NA | NA | NA | #VALUE! |
| ENSMUSG00000083377.2 | 0 | NA | NA | NA | NA | NA | #VALUE! |
| ENSMUSG00000083378.4 | 0.198268 | -0.06494 | 4.068276 | -0.01596 | 0.987265 | NA | #VALUE! |
| ENSMUSG00000083379.2 | 0 | NA | NA | NA | NA | NA | #VALUE! |
| ENSMUSG00000083381.2 | 0 | NA | NA | NA | NA | NA | #VALUE! |
| ENSMUSG00000083382.2 | 0 | NA | NA | NA | NA | NA | #VALUE! |
| ENSMUSG00000083383.3 | 0 | NA | NA | NA | NA | NA | #VALUE! |
| ENSMUSG00000083384.2 | 0 | NA | NA | NA | NA | NA | #VALUE! |
| ENSMUSG00000083385.2 | 0 | NA | NA | NA | NA | NA | #VALUE! |
| ENSMUSG00000083386.2 | 0 | NA | NA | NA | NA | NA | #VALUE! |
| ENSMUSG00000083387.2 | 0 | NA | NA | NA | NA | NA | #VALUE! |
| ENSMUSG00000083388.2 | 0 | NA | NA | NA | NA | NA | #VALUE! |
| ENSMUSG00000083389.2 | 0 | NA | NA | NA | NA | NA | #VALUE! |
| ENSMUSG00000083390.2 | 0 | NA | NA | NA | NA | NA | #VALUE! |
| ENSMUSG00000083391.3 | 0 | NA | NA | NA | NA | NA | #VALUE! |
| ENSMUSG00000083392.2 | 0 | NA | NA | NA | NA | NA | #VALUE! |

| | | | | | | | |
|---|---|---|---|---|---|---|---|
| ENSMUSG00000083393.2 | 0 | NA | NA | NA | NA | NA | #VALUE! |
| ENSMUSG00000083394.3 | 0 | NA | NA | NA | NA | NA | #VALUE! |
| ENSMUSG00000083395.2 | 0 | NA | NA | NA | NA | NA | #VALUE! |
| ENSMUSG00000083396.2 | 3.170677 | 5.164728 | 3.920817 | 1.317258 | 0.187752 | NA | #VALUE! |
| ENSMUSG00000083397.2 | 0 | NA | NA | NA | NA | NA | #VALUE! |
| ENSMUSG00000083398.2 | 0 | NA | NA | NA | NA | NA | #VALUE! |
| ENSMUSG00000083399.2 | 0 | NA | NA | NA | NA | NA | #VALUE! |
| ENSMUSG00000083401.2 | 0 | NA | NA | NA | NA | NA | #VALUE! |
| ENSMUSG00000083402.2 | 0 | NA | NA | NA | NA | NA | #VALUE! |
| ENSMUSG00000083403.2 | 0 | NA | NA | NA | NA | NA | #VALUE! |
| ENSMUSG00000083404.2 | 0 | NA | NA | NA | NA | NA | #VALUE! |
| ENSMUSG00000083405.2 | 0 | NA | NA | NA | NA | NA | #VALUE! |
| ENSMUSG00000083406.2 | 0 | NA | NA | NA | NA | NA | #VALUE! |
| ENSMUSG00000083407.2 | 0 | NA | NA | NA | NA | NA | #VALUE! |
| ENSMUSG00000083409.2 | 0 | NA | NA | NA | NA | NA | #VALUE! |
| ENSMUSG00000083410.2 | 0 | NA | NA | NA | NA | NA | #VALUE! |
| ENSMUSG00000083411.4 | 0.18651 | 1.401769 | 4.056354 | 0.345574 | 0.729663 | NA | #VALUE! |
| ENSMUSG00000083412.2 | 0 | NA | NA | NA | NA | NA | #VALUE! |
| ENSMUSG00000083413.2 | 0 | NA | NA | NA | NA | NA | #VALUE! |
| ENSMUSG00000083414.2 | 0 | NA | NA | NA | NA | NA | #VALUE! |
| ENSMUSG00000083416.2 | 0 | NA | NA | NA | NA | NA | #VALUE! |
| ENSMUSG00000083417.2 | 0 | NA | NA | NA | NA | NA | #VALUE! |
| ENSMUSG00000083418.3 | 0 | NA | NA | NA | NA | NA | #VALUE! |
| ENSMUSG00000083419.2 | 0 | NA | NA | NA | NA | NA | #VALUE! |
| ENSMUSG00000083420.2 | 0 | NA | NA | NA | NA | NA | #VALUE! |
| ENSMUSG00000083421.2 | 0 | NA | NA | NA | NA | NA | #VALUE! |
| ENSMUSG00000083422.2 | 0 | NA | NA | NA | NA | NA | #VALUE! |
| ENSMUSG00000083423.2 | 0 | NA | NA | NA | NA | NA | #VALUE! |
| ENSMUSG00000083424.4 | 7.492163 | -0.57374 | 3.215039 | -0.17846 | 0.858365 | NA | #VALUE! |
| ENSMUSG00000083425.2 | 0 | NA | NA | NA | NA | NA | #VALUE! |
| ENSMUSG00000083427.2 | 0 | NA | NA | NA | NA | NA | #VALUE! |
| ENSMUSG00000083428.2 | 0 | NA | NA | NA | NA | NA | #VALUE! |
| ENSMUSG00000083429.3 | 0 | NA | NA | NA | NA | NA | #VALUE! |
| ENSMUSG00000083430.2 | 0 | NA | NA | NA | NA | NA | #VALUE! |
| ENSMUSG00000083431.2 | 0 | NA | NA | NA | NA | NA | #VALUE! |
| ENSMUSG00000083432.2 | 0 | NA | NA | NA | NA | NA | #VALUE! |
| ENSMUSG00000083434.2 | 0 | NA | NA | NA | NA | NA | #VALUE! |
| ENSMUSG00000083435.3 | 0 | NA | NA | NA | NA | NA | #VALUE! |
| ENSMUSG00000083436.2 | 0 | NA | NA | NA | NA | NA | #VALUE! |
| ENSMUSG00000083437.2 | 0 | NA | NA | NA | NA | NA | #VALUE! |
| ENSMUSG00000083438.2 | 0 | NA | NA | NA | NA | NA | #VALUE! |
| ENSMUSG00000083439.2 | 0.093255 | 0.868093 | 4.072464 | 0.213162 | 0.831201 | NA | #VALUE! |
| ENSMUSG00000083440.2 | 0.093255 | 0.868093 | 4.072464 | 0.213162 | 0.831201 | NA | #VALUE! |
| ENSMUSG00000083441.2 | 0 | NA | NA | NA | NA | NA | #VALUE! |

| | | | | | | | |
|---|---|---|---|---|---|---|---|
| ENSMUSG00000083443.2 | 0 | NA | NA | NA | NA | NA | #VALUE! |
| ENSMUSG00000083444.2 | 0 | NA | NA | NA | NA | NA | #VALUE! |
| ENSMUSG00000083445.2 | 0 | NA | NA | NA | NA | NA | #VALUE! |
| ENSMUSG00000083446.2 | 0 | NA | NA | NA | NA | NA | #VALUE! |
| ENSMUSG00000083447.2 | 0 | NA | NA | NA | NA | NA | #VALUE! |
| ENSMUSG00000083448.2 | 0 | NA | NA | NA | NA | NA | #VALUE! |
| ENSMUSG00000083449.2 | 0 | NA | NA | NA | NA | NA | #VALUE! |
| ENSMUSG00000083450.2 | 0 | NA | NA | NA | NA | NA | #VALUE! |
| ENSMUSG00000083451.2 | 0 | NA | NA | NA | NA | NA | #VALUE! |
| ENSMUSG00000083452.2 | 0 | NA | NA | NA | NA | NA | #VALUE! |
| ENSMUSG00000083453.2 | 0 | NA | NA | NA | NA | NA | #VALUE! |
| ENSMUSG00000083454.2 | 0.168284 | 1.027628 | 4.067126 | 0.252667 | 0.800526 | NA | #VALUE! |
| ENSMUSG00000083455.2 | 0 | NA | NA | NA | NA | NA | #VALUE! |
| ENSMUSG00000083456.2 | 0 | NA | NA | NA | NA | NA | #VALUE! |
| ENSMUSG00000083457.2 | 0 | NA | NA | NA | NA | NA | #VALUE! |
| ENSMUSG00000083458.2 | 0 | NA | NA | NA | NA | NA | #VALUE! |
| ENSMUSG00000083460.2 | 0 | NA | NA | NA | NA | NA | #VALUE! |
| ENSMUSG00000083461.2 | 0 | NA | NA | NA | NA | NA | #VALUE! |
| ENSMUSG00000083462.2 | 0.336567 | 1.944925 | 4.029936 | 0.482619 | 0.629366 | NA | #VALUE! |
| ENSMUSG00000083463.2 | 0 | NA | NA | NA | NA | NA | #VALUE! |
| ENSMUSG00000083465.2 | 0 | NA | NA | NA | NA | NA | #VALUE! |
| ENSMUSG00000083466.2 | 0 | NA | NA | NA | NA | NA | #VALUE! |
| ENSMUSG00000083467.2 | 0 | NA | NA | NA | NA | NA | #VALUE! |
| ENSMUSG00000083468.2 | 0 | NA | NA | NA | NA | NA | #VALUE! |
| ENSMUSG00000083469.2 | 0 | NA | NA | NA | NA | NA | #VALUE! |
| ENSMUSG00000083470.2 | 0 | NA | NA | NA | NA | NA | #VALUE! |
| ENSMUSG00000083471.2 | 0 | NA | NA | NA | NA | NA | #VALUE! |
| ENSMUSG00000083472.2 | 0 | NA | NA | NA | NA | NA | #VALUE! |
| ENSMUSG00000083473.2 | 0 | NA | NA | NA | NA | NA | #VALUE! |
| ENSMUSG00000083474.2 | 0 | NA | NA | NA | NA | NA | #VALUE! |
| ENSMUSG00000083475.2 | 0 | NA | NA | NA | NA | NA | #VALUE! |
| ENSMUSG00000083477.3 | 0 | NA | NA | NA | NA | NA | #VALUE! |
| ENSMUSG00000083478.2 | 0 | NA | NA | NA | NA | NA | #VALUE! |
| ENSMUSG00000083481.5 | 0 | NA | NA | NA | NA | NA | #VALUE! |
| ENSMUSG00000083482.2 | 0.922977 | 0.439266 | 3.926776 | 0.111864 | 0.910931 | NA | #VALUE! |
| ENSMUSG00000083483.4 | 0.391578 | -2.1489 | 4.020883 | -0.53444 | 0.59304 | NA | #VALUE! |
| ENSMUSG00000083484.2 | 0 | NA | NA | NA | NA | NA | #VALUE! |
| ENSMUSG00000083485.2 | 0 | NA | NA | NA | NA | NA | #VALUE! |
| ENSMUSG00000083486.2 | 0 | NA | NA | NA | NA | NA | #VALUE! |
| ENSMUSG00000083487.2 | 0 | NA | NA | NA | NA | NA | #VALUE! |
| ENSMUSG00000083488.2 | 0 | NA | NA | NA | NA | NA | #VALUE! |
| ENSMUSG00000083489.2 | 0 | NA | NA | NA | NA | NA | #VALUE! |
| ENSMUSG00000083490.2 | 0 | NA | NA | NA | NA | NA | #VALUE! |
| ENSMUSG00000083491.2 | 0 | NA | NA | NA | NA | NA | #VALUE! |

| | | | | | | | |
|---|---|---|---|---|---|---|---|
| ENSMUSG00000083492.2 | 0 | NA | NA | NA | NA | NA | #VALUE! |
| ENSMUSG00000083493.2 | 0 | NA | NA | NA | NA | NA | #VALUE! |
| ENSMUSG00000083494.2 | 0 | NA | NA | NA | NA | NA | #VALUE! |
| ENSMUSG00000083495.2 | 0 | NA | NA | NA | NA | NA | #VALUE! |
| ENSMUSG00000083496.2 | 1.978032 | -2.37278 | 3.887783 | -0.61032 | 0.541652 | NA | #VALUE! |
| ENSMUSG00000083498.2 | 0 | NA | NA | NA | NA | NA | #VALUE! |
| ENSMUSG00000083499.2 | 0 | NA | NA | NA | NA | NA | #VALUE! |
| ENSMUSG00000083500.2 | 0 | NA | NA | NA | NA | NA | #VALUE! |
| ENSMUSG00000083501.3 | 0 | NA | NA | NA | NA | NA | #VALUE! |
| ENSMUSG00000083502.2 | 0 | NA | NA | NA | NA | NA | #VALUE! |
| ENSMUSG00000083503.2 | 0 | NA | NA | NA | NA | NA | #VALUE! |
| ENSMUSG00000083505.2 | 0 | NA | NA | NA | NA | NA | #VALUE! |
| ENSMUSG00000083507.2 | 0 | NA | NA | NA | NA | NA | #VALUE! |
| ENSMUSG00000083508.2 | 0 | NA | NA | NA | NA | NA | #VALUE! |
| ENSMUSG00000083509.2 | 0 | NA | NA | NA | NA | NA | #VALUE! |
| ENSMUSG00000083510.2 | 0 | NA | NA | NA | NA | NA | #VALUE! |
| ENSMUSG00000083512.2 | 0 | NA | NA | NA | NA | NA | #VALUE! |
| ENSMUSG00000083513.3 | 0 | NA | NA | NA | NA | NA | #VALUE! |
| ENSMUSG00000083514.2 | 0 | NA | NA | NA | NA | NA | #VALUE! |
| ENSMUSG00000083515.2 | 0 | NA | NA | NA | NA | NA | #VALUE! |
| ENSMUSG00000083516.2 | 0 | NA | NA | NA | NA | NA | #VALUE! |
| ENSMUSG00000083517.2 | 0 | NA | NA | NA | NA | NA | #VALUE! |
| ENSMUSG00000083518.2 | 0 | NA | NA | NA | NA | NA | #VALUE! |
| ENSMUSG00000083519.2 | 0 | NA | NA | NA | NA | NA | #VALUE! |
| ENSMUSG00000083520.2 | 0 | NA | NA | NA | NA | NA | #VALUE! |
| ENSMUSG00000083522.2 | 0 | NA | NA | NA | NA | NA | #VALUE! |
| ENSMUSG00000083524.2 | 1.830687 | 3.187606 | 3.926595 | 0.811799 | 0.416907 | NA | #VALUE! |
| ENSMUSG00000083527.2 | 0 | NA | NA | NA | NA | NA | #VALUE! |
| ENSMUSG00000083528.2 | 0 | NA | NA | NA | NA | NA | #VALUE! |
| ENSMUSG00000083530.2 | 0 | NA | NA | NA | NA | NA | #VALUE! |
| ENSMUSG00000083533.3 | 0.245976 | 1.027628 | 4.067126 | 0.252667 | 0.800526 | NA | #VALUE! |
| ENSMUSG00000083534.2 | 0 | NA | NA | NA | NA | NA | #VALUE! |
| ENSMUSG00000083535.2 | 0 | NA | NA | NA | NA | NA | #VALUE! |
| ENSMUSG00000083536.2 | 0 | NA | NA | NA | NA | NA | #VALUE! |
| ENSMUSG00000083537.4 | 0 | NA | NA | NA | NA | NA | #VALUE! |
| ENSMUSG00000083538.2 | 0 | NA | NA | NA | NA | NA | #VALUE! |
| ENSMUSG00000083539.2 | 0 | NA | NA | NA | NA | NA | #VALUE! |
| ENSMUSG00000083541.2 | 0 | NA | NA | NA | NA | NA | #VALUE! |
| ENSMUSG00000083542.3 | 0 | NA | NA | NA | NA | NA | #VALUE! |
| ENSMUSG00000083543.2 | 0 | NA | NA | NA | NA | NA | #VALUE! |
| ENSMUSG00000083544.2 | 0 | NA | NA | NA | NA | NA | #VALUE! |
| ENSMUSG00000083545.3 | 0 | NA | NA | NA | NA | NA | #VALUE! |
| ENSMUSG00000083546.2 | 0 | NA | NA | NA | NA | NA | #VALUE! |
| ENSMUSG00000083547.3 | 0 | NA | NA | NA | NA | NA | #VALUE! |

| | | | | | | | |
|---|---|---|---|---|---|---|---|
| ENSMUSG00000083548.2 | 0 | NA | NA | NA | NA | NA | #VALUE! |
| ENSMUSG00000083549.2 | 0 | NA | NA | NA | NA | NA | #VALUE! |
| ENSMUSG00000083550.2 | 0 | NA | NA | NA | NA | NA | #VALUE! |
| ENSMUSG00000083551.2 | 0 | NA | NA | NA | NA | NA | #VALUE! |
| ENSMUSG00000083553.2 | 4.985574 | 0.243773 | 2.827107 | 0.086227 | 0.931286 | NA | #VALUE! |
| ENSMUSG00000083554.2 | 0 | NA | NA | NA | NA | NA | #VALUE! |
| ENSMUSG00000083555.2 | 0 | NA | NA | NA | NA | NA | #VALUE! |
| ENSMUSG00000083556.2 | 0 | NA | NA | NA | NA | NA | #VALUE! |
| ENSMUSG00000083557.2 | 0 | NA | NA | NA | NA | NA | #VALUE! |
| ENSMUSG00000083558.2 | 0 | NA | NA | NA | NA | NA | #VALUE! |
| ENSMUSG00000083560.2 | 0 | NA | NA | NA | NA | NA | #VALUE! |
| ENSMUSG00000083561.4 | 0 | NA | NA | NA | NA | NA | #VALUE! |
| ENSMUSG00000083562.2 | 0 | NA | NA | NA | NA | NA | #VALUE! |
| ENSMUSG00000083564.2 | 0 | NA | NA | NA | NA | NA | #VALUE! |
| ENSMUSG00000083565.2 | 0 | NA | NA | NA | NA | NA | #VALUE! |
| ENSMUSG00000083566.2 | 0 | NA | NA | NA | NA | NA | #VALUE! |
| ENSMUSG00000083567.4 | 8.800212 | -1.35116 | 2.758973 | -0.48973 | 0.624322 | NA | #VALUE! |
| ENSMUSG00000083568.4 | 0 | NA | NA | NA | NA | NA | #VALUE! |
| ENSMUSG00000083569.2 | 0 | NA | NA | NA | NA | NA | #VALUE! |
| ENSMUSG00000083570.2 | 0 | NA | NA | NA | NA | NA | #VALUE! |
| ENSMUSG00000083572.2 | 0 | NA | NA | NA | NA | NA | #VALUE! |
| ENSMUSG00000083573.2 | 0 | NA | NA | NA | NA | NA | #VALUE! |
| ENSMUSG00000083574.2 | 0 | NA | NA | NA | NA | NA | #VALUE! |
| ENSMUSG00000083576.2 | 0 | NA | NA | NA | NA | NA | #VALUE! |
| ENSMUSG00000083577.2 | 0 | NA | NA | NA | NA | NA | #VALUE! |
| ENSMUSG00000083579.3 | 0 | NA | NA | NA | NA | NA | #VALUE! |
| ENSMUSG00000083580.2 | 0 | NA | NA | NA | NA | NA | #VALUE! |
| ENSMUSG00000083581.2 | 0 | NA | NA | NA | NA | NA | #VALUE! |
| ENSMUSG00000083582.2 | 0 | NA | NA | NA | NA | NA | #VALUE! |
| ENSMUSG00000083584.2 | 0 | NA | NA | NA | NA | NA | #VALUE! |
| ENSMUSG00000083585.2 | 0 | NA | NA | NA | NA | NA | #VALUE! |
| ENSMUSG00000083586.3 | 0 | NA | NA | NA | NA | NA | #VALUE! |
| ENSMUSG00000083587.2 | 0 | NA | NA | NA | NA | NA | #VALUE! |
| ENSMUSG00000083588.2 | 0 | NA | NA | NA | NA | NA | #VALUE! |
| ENSMUSG00000083590.2 | 0 | NA | NA | NA | NA | NA | #VALUE! |
| ENSMUSG00000083592.2 | 0 | NA | NA | NA | NA | NA | #VALUE! |
| ENSMUSG00000083594.2 | 0 | NA | NA | NA | NA | NA | #VALUE! |
| ENSMUSG00000083595.2 | 0 | NA | NA | NA | NA | NA | #VALUE! |
| ENSMUSG00000083596.5 | 0 | NA | NA | NA | NA | NA | #VALUE! |
| ENSMUSG00000083597.2 | 0 | NA | NA | NA | NA | NA | #VALUE! |
| ENSMUSG00000083598.2 | 0 | NA | NA | NA | NA | NA | #VALUE! |
| ENSMUSG00000083599.2 | 0 | NA | NA | NA | NA | NA | #VALUE! |
| ENSMUSG00000083601.2 | 0 | NA | NA | NA | NA | NA | #VALUE! |
| ENSMUSG00000083602.2 | 0 | NA | NA | NA | NA | NA | #VALUE! |

| | | | | | | | |
|---|---|---|---|---|---|---|---|
| ENSMUSG00000083603.2 | 0 | NA | NA | NA | NA | NA | #VALUE! |
| ENSMUSG00000083605.2 | 0 | NA | NA | NA | NA | NA | #VALUE! |
| ENSMUSG00000083606.4 | 0 | NA | NA | NA | NA | NA | #VALUE! |
| ENSMUSG00000083607.2 | 0 | NA | NA | NA | NA | NA | #VALUE! |
| ENSMUSG00000083608.2 | 0 | NA | NA | NA | NA | NA | #VALUE! |
| ENSMUSG00000083609.2 | 0 | NA | NA | NA | NA | NA | #VALUE! |
| ENSMUSG00000083610.2 | 0 | NA | NA | NA | NA | NA | #VALUE! |
| ENSMUSG00000083611.2 | 0 | NA | NA | NA | NA | NA | #VALUE! |
| ENSMUSG00000083613.2 | 0 | NA | NA | NA | NA | NA | #VALUE! |
| ENSMUSG00000083614.2 | 0 | NA | NA | NA | NA | NA | #VALUE! |
| ENSMUSG00000083615.2 | 0 | NA | NA | NA | NA | NA | #VALUE! |
| ENSMUSG00000083616.4 | 0 | NA | NA | NA | NA | NA | #VALUE! |
| ENSMUSG00000083617.2 | 0 | NA | NA | NA | NA | NA | #VALUE! |
| ENSMUSG00000083618.2 | 0 | NA | NA | NA | NA | NA | #VALUE! |
| ENSMUSG00000083619.4 | 0 | NA | NA | NA | NA | NA | #VALUE! |
| ENSMUSG00000083620.2 | 0 | NA | NA | NA | NA | NA | #VALUE! |
| ENSMUSG00000083622.2 | 0 | NA | NA | NA | NA | NA | #VALUE! |
| ENSMUSG00000083623.2 | 0 | NA | NA | NA | NA | NA | #VALUE! |
| ENSMUSG00000083624.2 | 0.493155 | -1.26736 | 4.010698 | -0.31599 | 0.752007 | NA | #VALUE! |
| ENSMUSG00000083625.2 | 0 | NA | NA | NA | NA | NA | #VALUE! |
| ENSMUSG00000083626.2 | 0.757173 | 0.577595 | 3.968981 | 0.145527 | 0.884295 | NA | #VALUE! |
| ENSMUSG00000083627.2 | 0 | NA | NA | NA | NA | NA | #VALUE! |
| ENSMUSG00000083628.2 | 0 | NA | NA | NA | NA | NA | #VALUE! |
| ENSMUSG00000083629.2 | 0 | NA | NA | NA | NA | NA | #VALUE! |
| ENSMUSG00000083630.3 | 0 | NA | NA | NA | NA | NA | #VALUE! |
| ENSMUSG00000083631.3 | 0 | NA | NA | NA | NA | NA | #VALUE! |
| ENSMUSG00000083632.2 | 0 | NA | NA | NA | NA | NA | #VALUE! |
| ENSMUSG00000083633.3 | 0 | NA | NA | NA | NA | NA | #VALUE! |
| ENSMUSG00000083635.2 | 0 | NA | NA | NA | NA | NA | #VALUE! |
| ENSMUSG00000083636.2 | 0 | NA | NA | NA | NA | NA | #VALUE! |
| ENSMUSG00000083637.2 | 0 | NA | NA | NA | NA | NA | #VALUE! |
| ENSMUSG00000083640.2 | 0 | NA | NA | NA | NA | NA | #VALUE! |
| ENSMUSG00000083641.2 | 0 | NA | NA | NA | NA | NA | #VALUE! |
| ENSMUSG00000083642.2 | 0 | NA | NA | NA | NA | NA | #VALUE! |
| ENSMUSG00000083643.2 | 0 | NA | NA | NA | NA | NA | #VALUE! |
| ENSMUSG00000083644.2 | 0 | NA | NA | NA | NA | NA | #VALUE! |
| ENSMUSG00000083645.2 | 0 | NA | NA | NA | NA | NA | #VALUE! |
| ENSMUSG00000083646.2 | 0 | NA | NA | NA | NA | NA | #VALUE! |
| ENSMUSG00000083648.2 | 0 | NA | NA | NA | NA | NA | #VALUE! |
| ENSMUSG00000083649.6 | 0.420053 | -2.33493 | 4.007343 | -0.58266 | 0.56012 | NA | #VALUE! |
| ENSMUSG00000083650.2 | 0 | NA | NA | NA | NA | NA | #VALUE! |
| ENSMUSG00000083651.2 | 0.245976 | 1.027628 | 4.067126 | 0.252667 | 0.800526 | NA | #VALUE! |
| ENSMUSG00000083653.3 | 0 | NA | NA | NA | NA | NA | #VALUE! |
| ENSMUSG00000083654.2 | 0 | NA | NA | NA | NA | NA | #VALUE! |

| | | | | | | | |
|---|---|---|---|---|---|---|---|
| ENSMUSG00000083655.2 | 0 | NA | NA | NA | NA | NA | #VALUE! |
| ENSMUSG00000083657.2 | 0 | NA | NA | NA | NA | NA | #VALUE! |
| ENSMUSG00000083658.2 | 0 | NA | NA | NA | NA | NA | #VALUE! |
| ENSMUSG00000083659.2 | 0 | NA | NA | NA | NA | NA | #VALUE! |
| ENSMUSG00000083660.2 | 0 | NA | NA | NA | NA | NA | #VALUE! |
| ENSMUSG00000083661.2 | 0 | NA | NA | NA | NA | NA | #VALUE! |
| ENSMUSG00000083662.2 | 0 | NA | NA | NA | NA | NA | #VALUE! |
| ENSMUSG00000083663.2 | 0 | NA | NA | NA | NA | NA | #VALUE! |
| ENSMUSG00000083664.2 | 0 | NA | NA | NA | NA | NA | #VALUE! |
| ENSMUSG00000083665.2 | 0 | NA | NA | NA | NA | NA | #VALUE! |
| ENSMUSG00000083666.2 | 0 | NA | NA | NA | NA | NA | #VALUE! |
| ENSMUSG00000083667.2 | 0 | NA | NA | NA | NA | NA | #VALUE! |
| ENSMUSG00000083668.2 | 0 | NA | NA | NA | NA | NA | #VALUE! |
| ENSMUSG00000083669.6 | 0.382299 | 0.386159 | 4.04914 | 0.095368 | 0.924022 | NA | #VALUE! |
| ENSMUSG00000083670.2 | 0 | NA | NA | NA | NA | NA | #VALUE! |
| ENSMUSG00000083671.2 | 0 | NA | NA | NA | NA | NA | #VALUE! |
| ENSMUSG00000083672.2 | 0 | NA | NA | NA | NA | NA | #VALUE! |
| ENSMUSG00000083673.4 | 0 | NA | NA | NA | NA | NA | #VALUE! |
| ENSMUSG00000083674.4 | 0 | NA | NA | NA | NA | NA | #VALUE! |
| ENSMUSG00000083676.3 | 0 | NA | NA | NA | NA | NA | #VALUE! |
| ENSMUSG00000083677.2 | 0 | NA | NA | NA | NA | NA | #VALUE! |
| ENSMUSG00000083678.4 | 0.893128 | -1.5945 | 3.944282 | -0.40426 | 0.686024 | NA | #VALUE! |
| ENSMUSG00000083679.2 | 0 | NA | NA | NA | NA | NA | #VALUE! |
| ENSMUSG00000083680.2 | 0 | NA | NA | NA | NA | NA | #VALUE! |
| ENSMUSG00000083681.2 | 0 | NA | NA | NA | NA | NA | #VALUE! |
| ENSMUSG00000083682.2 | 0 | NA | NA | NA | NA | NA | #VALUE! |
| ENSMUSG00000083683.2 | 0 | NA | NA | NA | NA | NA | #VALUE! |
| ENSMUSG00000083684.2 | 0 | NA | NA | NA | NA | NA | #VALUE! |
| ENSMUSG00000083685.2 | 0 | NA | NA | NA | NA | NA | #VALUE! |
| ENSMUSG00000083686.3 | 0 | NA | NA | NA | NA | NA | #VALUE! |
| ENSMUSG00000083687.2 | 0 | NA | NA | NA | NA | NA | #VALUE! |
| ENSMUSG00000083688.2 | 0 | NA | NA | NA | NA | NA | #VALUE! |
| ENSMUSG00000083689.2 | 0.105013 | -0.97324 | 4.076293 | -0.23876 | 0.811295 | NA | #VALUE! |
| ENSMUSG00000083691.2 | 0 | NA | NA | NA | NA | NA | #VALUE! |
| ENSMUSG00000083692.2 | 0.168284 | 1.027628 | 4.067126 | 0.252667 | 0.800526 | NA | #VALUE! |
| ENSMUSG00000083693.2 | 0 | NA | NA | NA | NA | NA | #VALUE! |
| ENSMUSG00000083695.7 | 0 | NA | NA | NA | NA | NA | #VALUE! |
| ENSMUSG00000083697.2 | 0 | NA | NA | NA | NA | NA | #VALUE! |
| ENSMUSG00000083698.2 | 0 | NA | NA | NA | NA | NA | #VALUE! |
| ENSMUSG00000083699.2 | 0 | NA | NA | NA | NA | NA | #VALUE! |
| ENSMUSG00000083700.2 | 0 | NA | NA | NA | NA | NA | #VALUE! |
| ENSMUSG00000083701.4 | 0 | NA | NA | NA | NA | NA | #VALUE! |
| ENSMUSG00000083702.2 | 0 | NA | NA | NA | NA | NA | #VALUE! |
| ENSMUSG00000083704.2 | 0 | NA | NA | NA | NA | NA | #VALUE! |

| | | | | | | | |
|---|---|---|---|---|---|---|---|
| ENSMUSG00000083705.3 | 0.297366 | -0.14751 | 4.065279 | -0.03629 | 0.971054 | NA | #VALUE! |
| ENSMUSG00000083706.4 | 0 | NA | NA | NA | NA | NA | #VALUE! |
| ENSMUSG00000083707.2 | 0 | NA | NA | NA | NA | NA | #VALUE! |
| ENSMUSG00000083708.2 | 0 | NA | NA | NA | NA | NA | #VALUE! |
| ENSMUSG00000083709.2 | 0 | NA | NA | NA | NA | NA | #VALUE! |
| ENSMUSG00000083710.2 | 0 | NA | NA | NA | NA | NA | #VALUE! |
| ENSMUSG00000083711.2 | 0 | NA | NA | NA | NA | NA | #VALUE! |
| ENSMUSG00000083713.2 | 8.235669 | 4.978316 | 2.64632 | 1.881222 | 0.059942 | NA | #VALUE! |
| ENSMUSG00000083714.4 | 0 | NA | NA | NA | NA | NA | #VALUE! |
| ENSMUSG00000083715.2 | 0 | NA | NA | NA | NA | NA | #VALUE! |
| ENSMUSG00000083716.5 | 0.364073 | 0.012025 | 4.059931 | 0.002962 | 0.997637 | NA | #VALUE! |
| ENSMUSG00000083717.2 | 0 | NA | NA | NA | NA | NA | #VALUE! |
| ENSMUSG00000083718.3 | 0 | NA | NA | NA | NA | NA | #VALUE! |
| ENSMUSG00000083719.2 | 0 | NA | NA | NA | NA | NA | #VALUE! |
| ENSMUSG00000083720.2 | 0 | NA | NA | NA | NA | NA | #VALUE! |
| ENSMUSG00000083721.2 | 0 | NA | NA | NA | NA | NA | #VALUE! |
| ENSMUSG00000083722.3 | 0 | NA | NA | NA | NA | NA | #VALUE! |
| ENSMUSG00000083723.2 | 0 | NA | NA | NA | NA | NA | #VALUE! |
| ENSMUSG00000083725.2 | 0 | NA | NA | NA | NA | NA | #VALUE! |
| ENSMUSG00000083728.2 | 0 | NA | NA | NA | NA | NA | #VALUE! |
| ENSMUSG00000083729.2 | 0 | NA | NA | NA | NA | NA | #VALUE! |
| ENSMUSG00000083731.2 | 0 | NA | NA | NA | NA | NA | #VALUE! |
| ENSMUSG00000083732.2 | 0 | NA | NA | NA | NA | NA | #VALUE! |
| ENSMUSG00000083733.4 | 0 | NA | NA | NA | NA | NA | #VALUE! |
| ENSMUSG00000083734.2 | 0 | NA | NA | NA | NA | NA | #VALUE! |
| ENSMUSG00000083735.2 | 0 | NA | NA | NA | NA | NA | #VALUE! |
| ENSMUSG00000083736.2 | 0 | NA | NA | NA | NA | NA | #VALUE! |
| ENSMUSG00000083737.2 | 0 | NA | NA | NA | NA | NA | #VALUE! |
| ENSMUSG00000083738.2 | 0 | NA | NA | NA | NA | NA | #VALUE! |
| ENSMUSG00000083740.2 | 0 | NA | NA | NA | NA | NA | #VALUE! |
| ENSMUSG00000083741.2 | 0 | NA | NA | NA | NA | NA | #VALUE! |
| ENSMUSG00000083742.3 | 0 | NA | NA | NA | NA | NA | #VALUE! |
| ENSMUSG00000083743.2 | 0 | NA | NA | NA | NA | NA | #VALUE! |
| ENSMUSG00000083744.2 | 0 | NA | NA | NA | NA | NA | #VALUE! |
| ENSMUSG00000083745.2 | 0 | NA | NA | NA | NA | NA | #VALUE! |
| ENSMUSG00000083747.2 | 0 | NA | NA | NA | NA | NA | #VALUE! |
| ENSMUSG00000083748.2 | 0 | NA | NA | NA | NA | NA | #VALUE! |
| ENSMUSG00000083749.2 | 0 | NA | NA | NA | NA | NA | #VALUE! |
| ENSMUSG00000083750.2 | 0 | NA | NA | NA | NA | NA | #VALUE! |
| ENSMUSG00000083751.2 | 0 | NA | NA | NA | NA | NA | #VALUE! |
| ENSMUSG00000083752.2 | 0 | NA | NA | NA | NA | NA | #VALUE! |
| ENSMUSG00000083753.2 | 0 | NA | NA | NA | NA | NA | #VALUE! |
| ENSMUSG00000083754.4 | 0 | NA | NA | NA | NA | NA | #VALUE! |
| ENSMUSG00000083755.2 | 0 | NA | NA | NA | NA | NA | #VALUE! |

| | | | | | | | |
|---|---|---|---|---|---|---|---|
| ENSMUSG00000083757.3 | 0.245976 | 1.027628 | 4.067126 | 0.252667 | 0.800526 | NA | #VALUE! |
| ENSMUSG00000083760.2 | 0 | NA | NA | NA | NA | NA | #VALUE! |
| ENSMUSG00000083761.2 | 0 | NA | NA | NA | NA | NA | #VALUE! |
| ENSMUSG00000083763.2 | 0 | NA | NA | NA | NA | NA | #VALUE! |
| ENSMUSG00000083764.2 | 0 | NA | NA | NA | NA | NA | #VALUE! |
| ENSMUSG00000083766.2 | 0 | NA | NA | NA | NA | NA | #VALUE! |
| ENSMUSG00000083767.2 | 0 | NA | NA | NA | NA | NA | #VALUE! |
| ENSMUSG00000083768.2 | 0 | NA | NA | NA | NA | NA | #VALUE! |
| ENSMUSG00000083769.2 | 0 | NA | NA | NA | NA | NA | #VALUE! |
| ENSMUSG00000083771.2 | 0 | NA | NA | NA | NA | NA | #VALUE! |
| ENSMUSG00000083772.2 | 0 | NA | NA | NA | NA | NA | #VALUE! |
| ENSMUSG00000083773.5 | 0 | NA | NA | NA | NA | NA | #VALUE! |
| ENSMUSG00000083774.2 | 0 | NA | NA | NA | NA | NA | #VALUE! |
| ENSMUSG00000083775.2 | 0 | NA | NA | NA | NA | NA | #VALUE! |
| ENSMUSG00000083776.2 | 0 | NA | NA | NA | NA | NA | #VALUE! |
| ENSMUSG00000083777.2 | 0 | NA | NA | NA | NA | NA | #VALUE! |
| ENSMUSG00000083778.2 | 0 | NA | NA | NA | NA | NA | #VALUE! |
| ENSMUSG00000083779.2 | 0.168284 | 1.027628 | 4.067126 | 0.252667 | 0.800526 | NA | #VALUE! |
| ENSMUSG00000083780.10 | 0 | NA | NA | NA | NA | NA | #VALUE! |
| ENSMUSG00000083781.2 | 0 | NA | NA | NA | NA | NA | #VALUE! |
| ENSMUSG00000083782.2 | 0 | NA | NA | NA | NA | NA | #VALUE! |
| ENSMUSG00000083783.2 | 0 | NA | NA | NA | NA | NA | #VALUE! |
| ENSMUSG00000083784.2 | 0 | NA | NA | NA | NA | NA | #VALUE! |
| ENSMUSG00000083785.2 | 0 | NA | NA | NA | NA | NA | #VALUE! |
| ENSMUSG00000083786.2 | 0 | NA | NA | NA | NA | NA | #VALUE! |
| ENSMUSG00000083787.2 | 0 | NA | NA | NA | NA | NA | #VALUE! |
| ENSMUSG00000083789.2 | 0 | NA | NA | NA | NA | NA | #VALUE! |
| ENSMUSG00000083791.2 | 0 | NA | NA | NA | NA | NA | #VALUE! |
| ENSMUSG00000083792.2 | 0 | NA | NA | NA | NA | NA | #VALUE! |
| ENSMUSG00000083793.2 | 2.46116 | -4.82608 | 3.924953 | -1.22959 | 0.218851 | NA | #VALUE! |
| ENSMUSG00000083794.2 | 0 | NA | NA | NA | NA | NA | #VALUE! |
| ENSMUSG00000083795.2 | 0 | NA | NA | NA | NA | NA | #VALUE! |
| ENSMUSG00000083796.4 | 0 | NA | NA | NA | NA | NA | #VALUE! |
| ENSMUSG00000083798.2 | 0 | NA | NA | NA | NA | NA | #VALUE! |
| ENSMUSG00000083801.2 | 0 | NA | NA | NA | NA | NA | #VALUE! |
| ENSMUSG00000083803.2 | 0 | NA | NA | NA | NA | NA | #VALUE! |
| ENSMUSG00000083804.2 | 0 | NA | NA | NA | NA | NA | #VALUE! |
| ENSMUSG00000083805.3 | 0 | NA | NA | NA | NA | NA | #VALUE! |
| ENSMUSG00000083806.2 | 0 | NA | NA | NA | NA | NA | #VALUE! |
| ENSMUSG00000083807.2 | 0 | NA | NA | NA | NA | NA | #VALUE! |
| ENSMUSG00000083808.2 | 0 | NA | NA | NA | NA | NA | #VALUE! |
| ENSMUSG00000083809.4 | 0 | NA | NA | NA | NA | NA | #VALUE! |
| ENSMUSG00000083810.2 | 0 | NA | NA | NA | NA | NA | #VALUE! |
| ENSMUSG00000083811.2 | 0 | NA | NA | NA | NA | NA | #VALUE! |

| | | | | | | | |
|---|---|---|---|---|---|---|---|
| ENSMUSG00000083812.4 | 0 | NA | NA | NA | NA | NA | #VALUE! |
| ENSMUSG00000083813.2 | 0 | NA | NA | NA | NA | NA | #VALUE! |
| ENSMUSG00000083814.2 | 0 | NA | NA | NA | NA | NA | #VALUE! |
| ENSMUSG00000083815.2 | 0 | NA | NA | NA | NA | NA | #VALUE! |
| ENSMUSG00000083816.4 | 0.210026 | -1.52494 | 4.058692 | -0.37572 | 0.707123 | NA | #VALUE! |
| ENSMUSG00000083817.2 | 0 | NA | NA | NA | NA | NA | #VALUE! |
| ENSMUSG00000083818.2 | 0 | NA | NA | NA | NA | NA | #VALUE! |
| ENSMUSG00000083819.2 | 0 | NA | NA | NA | NA | NA | #VALUE! |
| ENSMUSG00000083820.2 | 0.456002 | -0.4571 | 4.045272 | -0.113 | 0.910033 | NA | #VALUE! |
| ENSMUSG00000083821.2 | 0 | NA | NA | NA | NA | NA | #VALUE! |
| ENSMUSG00000083822.2 | 0 | NA | NA | NA | NA | NA | #VALUE! |
| ENSMUSG00000083823.4 | 0 | NA | NA | NA | NA | NA | #VALUE! |
| ENSMUSG00000083825.3 | 0 | NA | NA | NA | NA | NA | #VALUE! |
| ENSMUSG00000083826.2 | 0 | NA | NA | NA | NA | NA | #VALUE! |
| ENSMUSG00000083827.2 | 0.510828 | -2.58279 | 3.991728 | -0.64704 | 0.517609 | NA | #VALUE! |
| ENSMUSG00000083828.2 | 0 | NA | NA | NA | NA | NA | #VALUE! |
| ENSMUSG00000083829.2 | 0.640033 | 0.057569 | 4.005078 | 0.014374 | 0.988532 | NA | #VALUE! |
| ENSMUSG00000083830.2 | 0 | NA | NA | NA | NA | NA | #VALUE! |
| ENSMUSG00000083831.2 | 0 | NA | NA | NA | NA | NA | #VALUE! |
| ENSMUSG00000083832.2 | 1.594197 | 1.56364 | 3.87877 | 0.403128 | 0.686854 | NA | #VALUE! |
| ENSMUSG00000083833.5 | 0 | NA | NA | NA | NA | NA | #VALUE! |
| ENSMUSG00000083834.2 | 0.673134 | 2.918386 | 3.972531 | 0.734641 | 0.462558 | NA | #VALUE! |
| ENSMUSG00000083835.2 | 0 | NA | NA | NA | NA | NA | #VALUE! |
| ENSMUSG00000083836.2 | 0 | NA | NA | NA | NA | NA | #VALUE! |
| ENSMUSG00000083837.2 | 0 | NA | NA | NA | NA | NA | #VALUE! |
| ENSMUSG00000083838.2 | 0 | NA | NA | NA | NA | NA | #VALUE! |
| ENSMUSG00000083839.2 | 0 | NA | NA | NA | NA | NA | #VALUE! |
| ENSMUSG00000083841.2 | 0 | NA | NA | NA | NA | NA | #VALUE! |
| ENSMUSG00000083842.2 | 0 | NA | NA | NA | NA | NA | #VALUE! |
| ENSMUSG00000083843.2 | 0 | NA | NA | NA | NA | NA | #VALUE! |
| ENSMUSG00000083845.2 | 0 | NA | NA | NA | NA | NA | #VALUE! |
| ENSMUSG00000083846.2 | 0 | NA | NA | NA | NA | NA | #VALUE! |
| ENSMUSG00000083847.2 | 0 | NA | NA | NA | NA | NA | #VALUE! |
| ENSMUSG00000083848.2 | 0 | NA | NA | NA | NA | NA | #VALUE! |
| ENSMUSG00000083849.2 | 0 | NA | NA | NA | NA | NA | #VALUE! |
| ENSMUSG00000083851.2 | 0 | NA | NA | NA | NA | NA | #VALUE! |
| ENSMUSG00000083852.2 | 0 | NA | NA | NA | NA | NA | #VALUE! |
| ENSMUSG00000083853.2 | 0 | NA | NA | NA | NA | NA | #VALUE! |
| ENSMUSG00000083854.4 | 4.468237 | 1.963438 | 2.715316 | 0.723097 | 0.46962 | NA | #VALUE! |
| ENSMUSG00000083855.7 | 0 | NA | NA | NA | NA | NA | #VALUE! |
| ENSMUSG00000083856.2 | 0 | NA | NA | NA | NA | NA | #VALUE! |
| ENSMUSG00000083857.2 | 0 | NA | NA | NA | NA | NA | #VALUE! |
| ENSMUSG00000083858.2 | 0 | NA | NA | NA | NA | NA | #VALUE! |
| ENSMUSG00000083859.2 | 0 | NA | NA | NA | NA | NA | #VALUE! |

| | | | | | | | |
|---|---|---|---|---|---|---|---|
| ENSMUSG00000083860.3 | 0 | NA | NA | NA | NA | NA | #VALUE! |
| ENSMUSG00000083861.2 | 0 | NA | NA | NA | NA | NA | #VALUE! |
| ENSMUSG00000083864.2 | 0 | NA | NA | NA | NA | NA | #VALUE! |
| ENSMUSG00000083866.2 | 0 | NA | NA | NA | NA | NA | #VALUE! |
| ENSMUSG00000083867.2 | 0 | NA | NA | NA | NA | NA | #VALUE! |
| ENSMUSG00000083868.2 | 0 | NA | NA | NA | NA | NA | #VALUE! |
| ENSMUSG00000083869.2 | 0 | NA | NA | NA | NA | NA | #VALUE! |
| ENSMUSG00000083870.2 | 0 | NA | NA | NA | NA | NA | #VALUE! |
| ENSMUSG00000083871.2 | 0 | NA | NA | NA | NA | NA | #VALUE! |
| ENSMUSG00000083872.2 | 0 | NA | NA | NA | NA | NA | #VALUE! |
| ENSMUSG00000083873.2 | 0 | NA | NA | NA | NA | NA | #VALUE! |
| ENSMUSG00000083874.2 | 0 | NA | NA | NA | NA | NA | #VALUE! |
| ENSMUSG00000083875.2 | 0 | NA | NA | NA | NA | NA | #VALUE! |
| ENSMUSG00000083876.2 | 0 | NA | NA | NA | NA | NA | #VALUE! |
| ENSMUSG00000083877.2 | 0 | NA | NA | NA | NA | NA | #VALUE! |
| ENSMUSG00000083878.4 | 0 | NA | NA | NA | NA | NA | #VALUE! |
| ENSMUSG00000083879.5 | 0 | NA | NA | NA | NA | NA | #VALUE! |
| ENSMUSG00000083880.2 | 0 | NA | NA | NA | NA | NA | #VALUE! |
| ENSMUSG00000083882.2 | 0 | NA | NA | NA | NA | NA | #VALUE! |
| ENSMUSG00000083884.2 | 0 | NA | NA | NA | NA | NA | #VALUE! |
| ENSMUSG00000083885.2 | 0 | NA | NA | NA | NA | NA | #VALUE! |
| ENSMUSG00000083886.3 | 0 | NA | NA | NA | NA | NA | #VALUE! |
| ENSMUSG00000083887.2 | 0 | NA | NA | NA | NA | NA | #VALUE! |
| ENSMUSG00000083888.2 | 0 | NA | NA | NA | NA | NA | #VALUE! |
| ENSMUSG00000083889.5 | 0 | NA | NA | NA | NA | NA | #VALUE! |
| ENSMUSG00000083890.2 | 0 | NA | NA | NA | NA | NA | #VALUE! |
| ENSMUSG00000083892.4 | 0 | NA | NA | NA | NA | NA | #VALUE! |
| ENSMUSG00000083893.2 | 0 | NA | NA | NA | NA | NA | #VALUE! |
| ENSMUSG00000083894.2 | 0 | NA | NA | NA | NA | NA | #VALUE! |
| ENSMUSG00000083895.2 | 0 | NA | NA | NA | NA | NA | #VALUE! |
| ENSMUSG00000083896.9 | 0 | NA | NA | NA | NA | NA | #VALUE! |
| ENSMUSG00000083897.2 | 0.336567 | 1.944925 | 4.029936 | 0.482619 | 0.629366 | NA | #VALUE! |
| ENSMUSG00000083898.3 | 0 | NA | NA | NA | NA | NA | #VALUE! |
| ENSMUSG00000083899.3 | 8.841664 | 1.314576 | 2.005404 | 0.655517 | 0.512135 | NA | #VALUE! |
| ENSMUSG00000083900.3 | 0 | NA | NA | NA | NA | NA | #VALUE! |
| ENSMUSG00000083901.2 | 0 | NA | NA | NA | NA | NA | #VALUE! |
| ENSMUSG00000083902.3 | 0 | NA | NA | NA | NA | NA | #VALUE! |
| ENSMUSG00000083903.2 | 0 | NA | NA | NA | NA | NA | #VALUE! |
| ENSMUSG00000083904.2 | 0 | NA | NA | NA | NA | NA | #VALUE! |
| ENSMUSG00000083905.2 | 0 | NA | NA | NA | NA | NA | #VALUE! |
| ENSMUSG00000083906.2 | 0.093255 | 0.868093 | 4.072464 | 0.213162 | 0.831201 | NA | #VALUE! |
| ENSMUSG00000083907.3 | 0 | NA | NA | NA | NA | NA | #VALUE! |
| ENSMUSG00000083908.2 | 0 | NA | NA | NA | NA | NA | #VALUE! |
| ENSMUSG00000083909.2 | 0 | NA | NA | NA | NA | NA | #VALUE! |

| | | | | | | | |
|---|---|---|---|---|---|---|---|
| ENSMUSG00000083910.2 | 0.195789 | -1.05581 | 4.073302 | -0.2592 | 0.795479 | NA | #VALUE! |
| ENSMUSG00000083911.2 | 0.336567 | 1.944925 | 4.029936 | 0.482619 | 0.629366 | NA | #VALUE! |
| ENSMUSG00000083912.2 | 0.300802 | -1.80634 | 4.050033 | -0.44601 | 0.655592 | NA | #VALUE! |
| ENSMUSG00000083913.2 | 0 | NA | NA | NA | NA | NA | #VALUE! |
| ENSMUSG00000083914.4 | 8.334157 | 2.885706 | 2.208422 | 1.306682 | 0.191321 | NA | #VALUE! |
| ENSMUSG00000083915.2 | 0 | NA | NA | NA | NA | NA | #VALUE! |
| ENSMUSG00000083916.3 | 0 | NA | NA | NA | NA | NA | #VALUE! |
| ENSMUSG00000083917.2 | 0 | NA | NA | NA | NA | NA | #VALUE! |
| ENSMUSG00000083918.2 | 0 | NA | NA | NA | NA | NA | #VALUE! |
| ENSMUSG00000083919.2 | 0 | NA | NA | NA | NA | NA | #VALUE! |
| ENSMUSG00000083920.2 | 0 | NA | NA | NA | NA | NA | #VALUE! |
| ENSMUSG00000083921.3 | 0 | NA | NA | NA | NA | NA | #VALUE! |
| ENSMUSG00000083922.2 | 0 | NA | NA | NA | NA | NA | #VALUE! |
| ENSMUSG00000083923.2 | 0 | NA | NA | NA | NA | NA | #VALUE! |
| ENSMUSG00000083924.2 | 0 | NA | NA | NA | NA | NA | #VALUE! |
| ENSMUSG00000083925.2 | 0 | NA | NA | NA | NA | NA | #VALUE! |
| ENSMUSG00000083926.2 | 0 | NA | NA | NA | NA | NA | #VALUE! |
| ENSMUSG00000083927.2 | 0 | NA | NA | NA | NA | NA | #VALUE! |
| ENSMUSG00000083928.2 | 0 | NA | NA | NA | NA | NA | #VALUE! |
| ENSMUSG00000083929.5 | 0 | NA | NA | NA | NA | NA | #VALUE! |
| ENSMUSG00000083930.2 | 0 | NA | NA | NA | NA | NA | #VALUE! |
| ENSMUSG00000083931.2 | 0 | NA | NA | NA | NA | NA | #VALUE! |
| ENSMUSG00000083932.2 | 0 | NA | NA | NA | NA | NA | #VALUE! |
| ENSMUSG00000083933.2 | 0 | NA | NA | NA | NA | NA | #VALUE! |
| ENSMUSG00000083934.2 | 0 | NA | NA | NA | NA | NA | #VALUE! |
| ENSMUSG00000083935.2 | 0 | NA | NA | NA | NA | NA | #VALUE! |
| ENSMUSG00000083936.2 | 0 | NA | NA | NA | NA | NA | #VALUE! |
| ENSMUSG00000083937.4 | 0 | NA | NA | NA | NA | NA | #VALUE! |
| ENSMUSG00000083938.3 | 0 | NA | NA | NA | NA | NA | #VALUE! |
| ENSMUSG00000083939.2 | 0 | NA | NA | NA | NA | NA | #VALUE! |
| ENSMUSG00000083940.4 | 0 | NA | NA | NA | NA | NA | #VALUE! |
| ENSMUSG00000083941.2 | 0 | NA | NA | NA | NA | NA | #VALUE! |
| ENSMUSG00000083942.5 | 0 | NA | NA | NA | NA | NA | #VALUE! |
| ENSMUSG00000083943.2 | 0 | NA | NA | NA | NA | NA | #VALUE! |
| ENSMUSG00000083944.2 | 0 | NA | NA | NA | NA | NA | #VALUE! |
| ENSMUSG00000083945.2 | 0 | NA | NA | NA | NA | NA | #VALUE! |
| ENSMUSG00000083946.2 | 0 | NA | NA | NA | NA | NA | #VALUE! |
| ENSMUSG00000083947.6 | 0 | NA | NA | NA | NA | NA | #VALUE! |
| ENSMUSG00000083948.2 | 0 | NA | NA | NA | NA | NA | #VALUE! |
| ENSMUSG00000083949.2 | 0 | NA | NA | NA | NA | NA | #VALUE! |
| ENSMUSG00000083950.2 | 0 | NA | NA | NA | NA | NA | #VALUE! |
| ENSMUSG00000083951.2 | 0 | NA | NA | NA | NA | NA | #VALUE! |
| ENSMUSG00000083953.2 | 0.391578 | -2.1489 | 4.020883 | -0.53444 | 0.59304 | NA | #VALUE! |
| ENSMUSG00000083954.2 | 0 | NA | NA | NA | NA | NA | #VALUE! |

| | | | | | | | |
|---|---|---|---|---|---|---|---|
| ENSMUSG00000083955.2 | 0 | NA | NA | NA | NA | NA | #VALUE! |
| ENSMUSG00000083956.2 | 0 | NA | NA | NA | NA | NA | #VALUE! |
| ENSMUSG00000083957.2 | 0 | NA | NA | NA | NA | NA | #VALUE! |
| ENSMUSG00000083958.2 | 0 | NA | NA | NA | NA | NA | #VALUE! |
| ENSMUSG00000083960.2 | 0 | NA | NA | NA | NA | NA | #VALUE! |
| ENSMUSG00000083961.2 | 0 | NA | NA | NA | NA | NA | #VALUE! |
| ENSMUSG00000083962.2 | 0 | NA | NA | NA | NA | NA | #VALUE! |
| ENSMUSG00000083963.2 | 0 | NA | NA | NA | NA | NA | #VALUE! |
| ENSMUSG00000083964.2 | 0 | NA | NA | NA | NA | NA | #VALUE! |
| ENSMUSG00000083966.2 | 0 | NA | NA | NA | NA | NA | #VALUE! |
| ENSMUSG00000083967.2 | 0 | NA | NA | NA | NA | NA | #VALUE! |
| ENSMUSG00000083969.2 | 0 | NA | NA | NA | NA | NA | #VALUE! |
| ENSMUSG00000083970.2 | 0 | NA | NA | NA | NA | NA | #VALUE! |
| ENSMUSG00000083971.4 | 0 | NA | NA | NA | NA | NA | #VALUE! |
| ENSMUSG00000083972.2 | 0 | NA | NA | NA | NA | NA | #VALUE! |
| ENSMUSG00000083973.2 | 0 | NA | NA | NA | NA | NA | #VALUE! |
| ENSMUSG00000083974.2 | 0 | NA | NA | NA | NA | NA | #VALUE! |
| ENSMUSG00000083975.2 | 0 | NA | NA | NA | NA | NA | #VALUE! |
| ENSMUSG00000083976.2 | 0 | NA | NA | NA | NA | NA | #VALUE! |
| ENSMUSG00000083977.4 | 0 | NA | NA | NA | NA | NA | #VALUE! |
| ENSMUSG00000083978.2 | 0 | NA | NA | NA | NA | NA | #VALUE! |
| ENSMUSG00000083979.2 | 0 | NA | NA | NA | NA | NA | #VALUE! |
| ENSMUSG00000083981.5 | 0 | NA | NA | NA | NA | NA | #VALUE! |
| ENSMUSG00000083982.2 | 0 | NA | NA | NA | NA | NA | #VALUE! |
| ENSMUSG00000083983.2 | 0 | NA | NA | NA | NA | NA | #VALUE! |
| ENSMUSG00000083985.4 | 0 | NA | NA | NA | NA | NA | #VALUE! |
| ENSMUSG00000083986.2 | 0 | NA | NA | NA | NA | NA | #VALUE! |
| ENSMUSG00000083987.2 | 0 | NA | NA | NA | NA | NA | #VALUE! |
| ENSMUSG00000083988.2 | 0 | NA | NA | NA | NA | NA | #VALUE! |
| ENSMUSG00000083989.3 | 0 | NA | NA | NA | NA | NA | #VALUE! |
| ENSMUSG00000083990.2 | 0 | NA | NA | NA | NA | NA | #VALUE! |
| ENSMUSG00000083991.2 | 0 | NA | NA | NA | NA | NA | #VALUE! |
| ENSMUSG00000083993.2 | 0 | NA | NA | NA | NA | NA | #VALUE! |
| ENSMUSG00000083994.2 | 0 | NA | NA | NA | NA | NA | #VALUE! |
| ENSMUSG00000083995.2 | 0 | NA | NA | NA | NA | NA | #VALUE! |
| ENSMUSG00000083996.2 | 0 | NA | NA | NA | NA | NA | #VALUE! |
| ENSMUSG00000083997.2 | 0 | NA | NA | NA | NA | NA | #VALUE! |
| ENSMUSG00000083998.2 | 0 | NA | NA | NA | NA | NA | #VALUE! |
| ENSMUSG00000083999.2 | 0 | NA | NA | NA | NA | NA | #VALUE! |
| ENSMUSG00000084000.2 | 0 | NA | NA | NA | NA | NA | #VALUE! |
| ENSMUSG00000084001.2 | 0 | NA | NA | NA | NA | NA | #VALUE! |
| ENSMUSG00000084004.2 | 0 | NA | NA | NA | NA | NA | #VALUE! |
| ENSMUSG00000084006.2 | 0 | NA | NA | NA | NA | NA | #VALUE! |
| ENSMUSG00000084007.4 | 0 | NA | NA | NA | NA | NA | #VALUE! |

| | | | | | | | |
|---|---|---|---|---|---|---|---|
| ENSMUSG00000084009.2 | 0 | NA | NA | NA | NA | NA | #VALUE! |
| ENSMUSG00000084010.2 | 0 | NA | NA | NA | NA | NA | #VALUE! |
| ENSMUSG00000084011.2 | 0 | NA | NA | NA | NA | NA | #VALUE! |
| ENSMUSG00000084012.2 | 4.384141 | -0.891 | 3.356282 | -0.26547 | 0.790645 | NA | #VALUE! |
| ENSMUSG00000084013.2 | 0.3999 | -2.17566 | 4.018831 | -0.54137 | 0.588256 | NA | #VALUE! |
| ENSMUSG00000084014.4 | 0 | NA | NA | NA | NA | NA | #VALUE! |
| ENSMUSG00000084015.2 | 0 | NA | NA | NA | NA | NA | #VALUE! |
| ENSMUSG00000084017.2 | 0 | NA | NA | NA | NA | NA | #VALUE! |
| ENSMUSG00000084018.2 | 0 | NA | NA | NA | NA | NA | #VALUE! |
| ENSMUSG00000084019.2 | 0 | NA | NA | NA | NA | NA | #VALUE! |
| ENSMUSG00000084020.4 | 0 | NA | NA | NA | NA | NA | #VALUE! |
| ENSMUSG00000084021.2 | 0 | NA | NA | NA | NA | NA | #VALUE! |
| ENSMUSG00000084022.2 | 0 | NA | NA | NA | NA | NA | #VALUE! |
| ENSMUSG00000084024.2 | 0.373021 | 2.140748 | 4.018182 | 0.532765 | 0.594196 | NA | #VALUE! |
| ENSMUSG00000084025.2 | 0 | NA | NA | NA | NA | NA | #VALUE! |
| ENSMUSG00000084026.2 | 0 | NA | NA | NA | NA | NA | #VALUE! |
| ENSMUSG00000084027.2 | 0 | NA | NA | NA | NA | NA | #VALUE! |
| ENSMUSG00000084028.2 | 0 | NA | NA | NA | NA | NA | #VALUE! |
| ENSMUSG00000084029.2 | 0 | NA | NA | NA | NA | NA | #VALUE! |
| ENSMUSG00000084030.2 | 0 | NA | NA | NA | NA | NA | #VALUE! |
| ENSMUSG00000084031.2 | 0 | NA | NA | NA | NA | NA | #VALUE! |
| ENSMUSG00000084032.2 | 0 | NA | NA | NA | NA | NA | #VALUE! |
| ENSMUSG00000084033.4 | 0 | NA | NA | NA | NA | NA | #VALUE! |
| ENSMUSG00000084034.2 | 0 | NA | NA | NA | NA | NA | #VALUE! |
| ENSMUSG00000084038.3 | 0 | NA | NA | NA | NA | NA | #VALUE! |
| ENSMUSG00000084039.2 | 0 | NA | NA | NA | NA | NA | #VALUE! |
| ENSMUSG00000084041.2 | 0 | NA | NA | NA | NA | NA | #VALUE! |
| ENSMUSG00000084043.2 | 0 | NA | NA | NA | NA | NA | #VALUE! |
| ENSMUSG00000084044.2 | 0 | NA | NA | NA | NA | NA | #VALUE! |
| ENSMUSG00000084045.2 | 0 | NA | NA | NA | NA | NA | #VALUE! |
| ENSMUSG00000084046.2 | 0 | NA | NA | NA | NA | NA | #VALUE! |
| ENSMUSG00000084047.2 | 0 | NA | NA | NA | NA | NA | #VALUE! |
| ENSMUSG00000084048.2 | 0 | NA | NA | NA | NA | NA | #VALUE! |
| ENSMUSG00000084050.4 | 0 | NA | NA | NA | NA | NA | #VALUE! |
| ENSMUSG00000084052.2 | 0 | NA | NA | NA | NA | NA | #VALUE! |
| ENSMUSG00000084053.2 | 0 | NA | NA | NA | NA | NA | #VALUE! |
| ENSMUSG00000084054.2 | 0 | NA | NA | NA | NA | NA | #VALUE! |
| ENSMUSG00000084055.2 | 0 | NA | NA | NA | NA | NA | #VALUE! |
| ENSMUSG00000084056.3 | 0 | NA | NA | NA | NA | NA | #VALUE! |
| ENSMUSG00000084057.2 | 0 | NA | NA | NA | NA | NA | #VALUE! |
| ENSMUSG00000084058.2 | 0 | NA | NA | NA | NA | NA | #VALUE! |
| ENSMUSG00000084059.2 | 0 | NA | NA | NA | NA | NA | #VALUE! |
| ENSMUSG00000084060.4 | 0 | NA | NA | NA | NA | NA | #VALUE! |
| ENSMUSG00000084061.2 | 0 | NA | NA | NA | NA | NA | #VALUE! |

| | | | | | | | |
|---|---|---|---|---|---|---|---|
| ENSMUSG00000084063.2 | 0 | NA | NA | NA | NA | NA | #VALUE! |
| ENSMUSG00000084064.2 | 0 | NA | NA | NA | NA | NA | #VALUE! |
| ENSMUSG00000084065.2 | 0.195789 | -1.05581 | 4.073302 | -0.2592 | 0.795479 | NA | #VALUE! |
| ENSMUSG00000084066.2 | 0 | NA | NA | NA | NA | NA | #VALUE! |
| ENSMUSG00000084067.4 | 0 | NA | NA | NA | NA | NA | #VALUE! |
| ENSMUSG00000084068.2 | 0 | NA | NA | NA | NA | NA | #VALUE! |
| ENSMUSG00000084069.2 | 0 | NA | NA | NA | NA | NA | #VALUE! |
| ENSMUSG00000084070.2 | 0 | NA | NA | NA | NA | NA | #VALUE! |
| ENSMUSG00000084071.2 | 0 | NA | NA | NA | NA | NA | #VALUE! |
| ENSMUSG00000084072.2 | 0 | NA | NA | NA | NA | NA | #VALUE! |
| ENSMUSG00000084073.2 | 0 | NA | NA | NA | NA | NA | #VALUE! |
| ENSMUSG00000084074.2 | 0 | NA | NA | NA | NA | NA | #VALUE! |
| ENSMUSG00000084076.2 | 0 | NA | NA | NA | NA | NA | #VALUE! |
| ENSMUSG00000084077.2 | 0 | NA | NA | NA | NA | NA | #VALUE! |
| ENSMUSG00000084078.3 | 0 | NA | NA | NA | NA | NA | #VALUE! |
| ENSMUSG00000084079.2 | 0 | NA | NA | NA | NA | NA | #VALUE! |
| ENSMUSG00000084080.2 | 0 | NA | NA | NA | NA | NA | #VALUE! |
| ENSMUSG00000084081.4 | 0 | NA | NA | NA | NA | NA | #VALUE! |
| ENSMUSG00000084082.2 | 0 | NA | NA | NA | NA | NA | #VALUE! |
| ENSMUSG00000084083.2 | 0 | NA | NA | NA | NA | NA | #VALUE! |
| ENSMUSG00000084085.3 | 0 | NA | NA | NA | NA | NA | #VALUE! |
| ENSMUSG00000084086.2 | 0 | NA | NA | NA | NA | NA | #VALUE! |
| ENSMUSG00000084087.2 | 0 | NA | NA | NA | NA | NA | #VALUE! |
| ENSMUSG00000084088.2 | 0 | NA | NA | NA | NA | NA | #VALUE! |
| ENSMUSG00000084089.2 | 0 | NA | NA | NA | NA | NA | #VALUE! |
| ENSMUSG00000084090.2 | 0 | NA | NA | NA | NA | NA | #VALUE! |
| ENSMUSG00000084092.2 | 0 | NA | NA | NA | NA | NA | #VALUE! |
| ENSMUSG00000084093.2 | 1.46314 | 0.933075 | 3.864476 | 0.241449 | 0.809207 | NA | #VALUE! |
| ENSMUSG00000084094.2 | 0 | NA | NA | NA | NA | NA | #VALUE! |
| ENSMUSG00000084095.2 | 0 | NA | NA | NA | NA | NA | #VALUE! |
| ENSMUSG00000084096.2 | 0 | NA | NA | NA | NA | NA | #VALUE! |
| ENSMUSG00000084097.2 | 0 | NA | NA | NA | NA | NA | #VALUE! |
| ENSMUSG00000084098.2 | 0 | NA | NA | NA | NA | NA | #VALUE! |
| ENSMUSG00000084099.2 | 0 | NA | NA | NA | NA | NA | #VALUE! |
| ENSMUSG00000084101.2 | 0 | NA | NA | NA | NA | NA | #VALUE! |
| ENSMUSG00000084102.3 | 0 | NA | NA | NA | NA | NA | #VALUE! |
| ENSMUSG00000084103.2 | 0 | NA | NA | NA | NA | NA | #VALUE! |
| ENSMUSG00000084104.2 | 0 | NA | NA | NA | NA | NA | #VALUE! |
| ENSMUSG00000084105.2 | 0 | NA | NA | NA | NA | NA | #VALUE! |
| ENSMUSG00000084107.2 | 0 | NA | NA | NA | NA | NA | #VALUE! |
| ENSMUSG00000084108.3 | 0 | NA | NA | NA | NA | NA | #VALUE! |
| ENSMUSG00000084110.5 | 0 | NA | NA | NA | NA | NA | #VALUE! |
| ENSMUSG00000084112.2 | 0 | NA | NA | NA | NA | NA | #VALUE! |
| ENSMUSG00000084113.2 | 0.168284 | 1.027628 | 4.067126 | 0.252667 | 0.800526 | NA | #VALUE! |

| | | | | | | | |
|---|---|---|---|---|---|---|---|
| ENSMUSG00000084114.2 | 0 | NA | NA | NA | NA | NA | #VALUE! |
| ENSMUSG00000084115.2 | 0 | NA | NA | NA | NA | NA | #VALUE! |
| ENSMUSG00000084116.2 | 0 | NA | NA | NA | NA | NA | #VALUE! |
| ENSMUSG00000084117.4 | 0 | NA | NA | NA | NA | NA | #VALUE! |
| ENSMUSG00000084118.2 | 0 | NA | NA | NA | NA | NA | #VALUE! |
| ENSMUSG00000084119.2 | 0 | NA | NA | NA | NA | NA | #VALUE! |
| ENSMUSG00000084120.2 | 0 | NA | NA | NA | NA | NA | #VALUE! |
| ENSMUSG00000084123.2 | 0 | NA | NA | NA | NA | NA | #VALUE! |
| ENSMUSG00000084124.2 | 0 | NA | NA | NA | NA | NA | #VALUE! |
| ENSMUSG00000084125.2 | 0 | NA | NA | NA | NA | NA | #VALUE! |
| ENSMUSG00000084126.2 | 0 | NA | NA | NA | NA | NA | #VALUE! |
| ENSMUSG00000084127.2 | 0.373021 | 2.140748 | 4.018182 | 0.532765 | 0.594196 | NA | #VALUE! |
| ENSMUSG00000084129.2 | 0 | NA | NA | NA | NA | NA | #VALUE! |
| ENSMUSG00000084130.2 | 0 | NA | NA | NA | NA | NA | #VALUE! |
| ENSMUSG00000084131.2 | 0.204111 | -1.05581 | 4.073302 | -0.2592 | 0.795479 | NA | #VALUE! |
| ENSMUSG00000084132.3 | 0 | NA | NA | NA | NA | NA | #VALUE! |
| ENSMUSG00000084133.2 | 0 | NA | NA | NA | NA | NA | #VALUE! |
| ENSMUSG00000084134.2 | 0 | NA | NA | NA | NA | NA | #VALUE! |
| ENSMUSG00000084135.5 | 0 | NA | NA | NA | NA | NA | #VALUE! |
| ENSMUSG00000084136.5 | 0 | NA | NA | NA | NA | NA | #VALUE! |
| ENSMUSG00000084137.2 | 0 | NA | NA | NA | NA | NA | #VALUE! |
| ENSMUSG00000084138.2 | 0 | NA | NA | NA | NA | NA | #VALUE! |
| ENSMUSG00000084139.2 | 0 | NA | NA | NA | NA | NA | #VALUE! |
| ENSMUSG00000084140.2 | 0 | NA | NA | NA | NA | NA | #VALUE! |
| ENSMUSG00000084142.2 | 0 | NA | NA | NA | NA | NA | #VALUE! |
| ENSMUSG00000084143.2 | 0 | NA | NA | NA | NA | NA | #VALUE! |
| ENSMUSG00000084144.2 | 0 | NA | NA | NA | NA | NA | #VALUE! |
| ENSMUSG00000084145.4 | 0 | NA | NA | NA | NA | NA | #VALUE! |
| ENSMUSG00000084146.2 | 0 | NA | NA | NA | NA | NA | #VALUE! |
| ENSMUSG00000084147.2 | 0 | NA | NA | NA | NA | NA | #VALUE! |
| ENSMUSG00000084148.2 | 0 | NA | NA | NA | NA | NA | #VALUE! |
| ENSMUSG00000084149.2 | 0.198268 | -0.06494 | 4.068276 | -0.01596 | 0.987265 | NA | #VALUE! |
| ENSMUSG00000084150.2 | 0 | NA | NA | NA | NA | NA | #VALUE! |
| ENSMUSG00000084151.2 | 0 | NA | NA | NA | NA | NA | #VALUE! |
| ENSMUSG00000084152.2 | 0 | NA | NA | NA | NA | NA | #VALUE! |
| ENSMUSG00000084153.3 | 0 | NA | NA | NA | NA | NA | #VALUE! |
| ENSMUSG00000084154.2 | 0 | NA | NA | NA | NA | NA | #VALUE! |
| ENSMUSG00000084155.2 | 0 | NA | NA | NA | NA | NA | #VALUE! |
| ENSMUSG00000084157.3 | 0 | NA | NA | NA | NA | NA | #VALUE! |
| ENSMUSG00000084158.2 | 0 | NA | NA | NA | NA | NA | #VALUE! |
| ENSMUSG00000084159.2 | 0 | NA | NA | NA | NA | NA | #VALUE! |
| ENSMUSG00000084160.2 | 0 | NA | NA | NA | NA | NA | #VALUE! |
| ENSMUSG00000084161.2 | 0 | NA | NA | NA | NA | NA | #VALUE! |
| ENSMUSG00000084162.2 | 0 | NA | NA | NA | NA | NA | #VALUE! |

| | | | | | | | |
|---|---|---|---|---|---|---|---|
| ENSMUSG00000084163.2 | 0 | NA | NA | NA | NA | NA | #VALUE! |
| ENSMUSG00000084164.2 | 0 | NA | NA | NA | NA | NA | #VALUE! |
| ENSMUSG00000084166.2 | 0.279766 | 1.808031 | 4.036658 | 0.447903 | 0.654223 | NA | #VALUE! |
| ENSMUSG00000084167.2 | 0 | NA | NA | NA | NA | NA | #VALUE! |
| ENSMUSG00000084168.4 | 0.198268 | -0.06494 | 4.068276 | -0.01596 | 0.987265 | NA | #VALUE! |
| ENSMUSG00000084169.2 | 0 | NA | NA | NA | NA | NA | #VALUE! |
| ENSMUSG00000084170.2 | 0 | NA | NA | NA | NA | NA | #VALUE! |
| ENSMUSG00000084171.2 | 0 | NA | NA | NA | NA | NA | #VALUE! |
| ENSMUSG00000084172.2 | 0 | NA | NA | NA | NA | NA | #VALUE! |
| ENSMUSG00000084173.2 | 0 | NA | NA | NA | NA | NA | #VALUE! |
| ENSMUSG00000084174.4 | 0 | NA | NA | NA | NA | NA | #VALUE! |
| ENSMUSG00000084175.2 | 0 | NA | NA | NA | NA | NA | #VALUE! |
| ENSMUSG00000084177.2 | 0 | NA | NA | NA | NA | NA | #VALUE! |
| ENSMUSG00000084178.2 | 0.504851 | 2.502843 | 3.993993 | 0.626652 | 0.530887 | NA | #VALUE! |
| ENSMUSG00000084179.2 | 0 | NA | NA | NA | NA | NA | #VALUE! |
| ENSMUSG00000084180.2 | 0 | NA | NA | NA | NA | NA | #VALUE! |
| ENSMUSG00000084181.2 | 0 | NA | NA | NA | NA | NA | #VALUE! |
| ENSMUSG00000084182.2 | 0 | NA | NA | NA | NA | NA | #VALUE! |
| ENSMUSG00000084183.2 | 0 | NA | NA | NA | NA | NA | #VALUE! |
| ENSMUSG00000084184.2 | 0 | NA | NA | NA | NA | NA | #VALUE! |
| ENSMUSG00000084185.2 | 0 | NA | NA | NA | NA | NA | #VALUE! |
| ENSMUSG00000084186.2 | 0 | NA | NA | NA | NA | NA | #VALUE! |
| ENSMUSG00000084188.2 | 0 | NA | NA | NA | NA | NA | #VALUE! |
| ENSMUSG00000084189.2 | 0 | NA | NA | NA | NA | NA | #VALUE! |
| ENSMUSG00000084190.2 | 0 | NA | NA | NA | NA | NA | #VALUE! |
| ENSMUSG00000084191.2 | 0 | NA | NA | NA | NA | NA | #VALUE! |
| ENSMUSG00000084192.2 | 0 | NA | NA | NA | NA | NA | #VALUE! |
| ENSMUSG00000084193.2 | 0 | NA | NA | NA | NA | NA | #VALUE! |
| ENSMUSG00000084195.3 | 0 | NA | NA | NA | NA | NA | #VALUE! |
| ENSMUSG00000084196.2 | 0 | NA | NA | NA | NA | NA | #VALUE! |
| ENSMUSG00000084197.2 | 0 | NA | NA | NA | NA | NA | #VALUE! |
| ENSMUSG00000084198.2 | 0 | NA | NA | NA | NA | NA | #VALUE! |
| ENSMUSG00000084199.2 | 0 | NA | NA | NA | NA | NA | #VALUE! |
| ENSMUSG00000084200.2 | 0 | NA | NA | NA | NA | NA | #VALUE! |
| ENSMUSG00000084201.2 | 0 | NA | NA | NA | NA | NA | #VALUE! |
| ENSMUSG00000084202.2 | 0 | NA | NA | NA | NA | NA | #VALUE! |
| ENSMUSG00000084203.2 | 0 | NA | NA | NA | NA | NA | #VALUE! |
| ENSMUSG00000084204.3 | 0 | NA | NA | NA | NA | NA | #VALUE! |
| ENSMUSG00000084205.2 | 0.093255 | 0.868093 | 4.072464 | 0.213162 | 0.831201 | NA | #VALUE! |
| ENSMUSG00000084206.2 | 0 | NA | NA | NA | NA | NA | #VALUE! |
| ENSMUSG00000084207.2 | 0 | NA | NA | NA | NA | NA | #VALUE! |
| ENSMUSG00000084208.2 | 0.168284 | 1.027628 | 4.067126 | 0.252667 | 0.800526 | NA | #VALUE! |
| ENSMUSG00000084209.2 | 0 | NA | NA | NA | NA | NA | #VALUE! |
| ENSMUSG00000084210.2 | 0 | NA | NA | NA | NA | NA | #VALUE! |

| | | | | | | | |
|---|---|---|---|---|---|---|---|
| ENSMUSG00000084211.2 | 0 | NA | NA | NA | NA | NA | #VALUE! |
| ENSMUSG00000084212.2 | 0 | NA | NA | NA | NA | NA | #VALUE! |
| ENSMUSG00000084213.2 | 0 | NA | NA | NA | NA | NA | #VALUE! |
| ENSMUSG00000084214.2 | 0 | NA | NA | NA | NA | NA | #VALUE! |
| ENSMUSG00000084215.4 | 0 | NA | NA | NA | NA | NA | #VALUE! |
| ENSMUSG00000084216.2 | 0 | NA | NA | NA | NA | NA | #VALUE! |
| ENSMUSG00000084217.3 | 0 | NA | NA | NA | NA | NA | #VALUE! |
| ENSMUSG00000084218.2 | 0 | NA | NA | NA | NA | NA | #VALUE! |
| ENSMUSG00000084219.2 | 0 | NA | NA | NA | NA | NA | #VALUE! |
| ENSMUSG00000084220.2 | 0 | NA | NA | NA | NA | NA | #VALUE! |
| ENSMUSG00000084221.2 | 0.18651 | 1.401769 | 4.056354 | 0.345574 | 0.729663 | NA | #VALUE! |
| ENSMUSG00000084222.2 | 0 | NA | NA | NA | NA | NA | #VALUE! |
| ENSMUSG00000084223.2 | 0.093255 | 0.868093 | 4.072464 | 0.213162 | 0.831201 | NA | #VALUE! |
| ENSMUSG00000084224.2 | 0 | NA | NA | NA | NA | NA | #VALUE! |
| ENSMUSG00000084226.2 | 0 | NA | NA | NA | NA | NA | #VALUE! |
| ENSMUSG00000084228.2 | 0 | NA | NA | NA | NA | NA | #VALUE! |
| ENSMUSG00000084230.2 | 0 | NA | NA | NA | NA | NA | #VALUE! |
| ENSMUSG00000084231.2 | 0 | NA | NA | NA | NA | NA | #VALUE! |
| ENSMUSG00000084234.3 | 0 | NA | NA | NA | NA | NA | #VALUE! |
| ENSMUSG00000084236.3 | 0 | NA | NA | NA | NA | NA | #VALUE! |
| ENSMUSG00000084237.2 | 0 | NA | NA | NA | NA | NA | #VALUE! |
| ENSMUSG00000084238.2 | 0 | NA | NA | NA | NA | NA | #VALUE! |
| ENSMUSG00000084239.2 | 0 | NA | NA | NA | NA | NA | #VALUE! |
| ENSMUSG00000084240.2 | 0 | NA | NA | NA | NA | NA | #VALUE! |
| ENSMUSG00000084241.2 | 0 | NA | NA | NA | NA | NA | #VALUE! |
| ENSMUSG00000084243.2 | 0 | NA | NA | NA | NA | NA | #VALUE! |
| ENSMUSG00000084244.2 | 0 | NA | NA | NA | NA | NA | #VALUE! |
| ENSMUSG00000084245.2 | 0 | NA | NA | NA | NA | NA | #VALUE! |
| ENSMUSG00000084246.2 | 0 | NA | NA | NA | NA | NA | #VALUE! |
| ENSMUSG00000084247.2 | 0 | NA | NA | NA | NA | NA | #VALUE! |
| ENSMUSG00000084248.2 | 0 | NA | NA | NA | NA | NA | #VALUE! |
| ENSMUSG00000084249.2 | 0 | NA | NA | NA | NA | NA | #VALUE! |
| ENSMUSG00000084250.2 | 0 | NA | NA | NA | NA | NA | #VALUE! |
| ENSMUSG00000084251.2 | 0 | NA | NA | NA | NA | NA | #VALUE! |
| ENSMUSG00000084252.2 | 2.611145 | 4.886731 | 3.92379 | 1.245411 | 0.212981 | NA | #VALUE! |
| ENSMUSG00000084253.2 | 0 | NA | NA | NA | NA | NA | #VALUE! |
| ENSMUSG00000084254.2 | 0 | NA | NA | NA | NA | NA | #VALUE! |
| ENSMUSG00000084256.2 | 0 | NA | NA | NA | NA | NA | #VALUE! |
| ENSMUSG00000084257.2 | 0 | NA | NA | NA | NA | NA | #VALUE! |
| ENSMUSG00000084259.2 | 0 | NA | NA | NA | NA | NA | #VALUE! |
| ENSMUSG00000084260.2 | 0 | NA | NA | NA | NA | NA | #VALUE! |
| ENSMUSG00000084261.2 | 0 | NA | NA | NA | NA | NA | #VALUE! |
| ENSMUSG00000084262.6 | 0 | NA | NA | NA | NA | NA | #VALUE! |
| ENSMUSG00000084264.2 | 0 | NA | NA | NA | NA | NA | #VALUE! |

| | | | | | | | |
|---|---|---|---|---|---|---|---|
| ENSMUSG00000084267.2 | 0 | NA | NA | NA | NA | NA | #VALUE! |
| ENSMUSG00000084269.2 | 0 | NA | NA | NA | NA | NA | #VALUE! |
| ENSMUSG00000084270.2 | 0 | NA | NA | NA | NA | NA | #VALUE! |
| ENSMUSG00000084271.2 | 0 | NA | NA | NA | NA | NA | #VALUE! |
| ENSMUSG00000084272.2 | 0 | NA | NA | NA | NA | NA | #VALUE! |
| ENSMUSG00000084273.2 | 0 | NA | NA | NA | NA | NA | #VALUE! |
| ENSMUSG00000084274.3 | 0.093255 | 0.868093 | 4.072464 | 0.213162 | 0.831201 | NA | #VALUE! |
| ENSMUSG00000084275.4 | 0 | NA | NA | NA | NA | NA | #VALUE! |
| ENSMUSG00000084276.2 | 0 | NA | NA | NA | NA | NA | #VALUE! |
| ENSMUSG00000084277.2 | 0 | NA | NA | NA | NA | NA | #VALUE! |
| ENSMUSG00000084278.2 | 0 | NA | NA | NA | NA | NA | #VALUE! |
| ENSMUSG00000084279.2 | 0 | NA | NA | NA | NA | NA | #VALUE! |
| ENSMUSG00000084280.2 | 0 | NA | NA | NA | NA | NA | #VALUE! |
| ENSMUSG00000084281.2 | 0 | NA | NA | NA | NA | NA | #VALUE! |
| ENSMUSG00000084282.2 | 0 | NA | NA | NA | NA | NA | #VALUE! |
| ENSMUSG00000084283.2 | 0 | NA | NA | NA | NA | NA | #VALUE! |
| ENSMUSG00000084284.2 | 0 | NA | NA | NA | NA | NA | #VALUE! |
| ENSMUSG00000084285.2 | 0 | NA | NA | NA | NA | NA | #VALUE! |
| ENSMUSG00000084286.2 | 0 | NA | NA | NA | NA | NA | #VALUE! |
| ENSMUSG00000084287.2 | 0 | NA | NA | NA | NA | NA | #VALUE! |
| ENSMUSG00000084288.2 | 0 | NA | NA | NA | NA | NA | #VALUE! |
| ENSMUSG00000084291.4 | 0 | NA | NA | NA | NA | NA | #VALUE! |
| ENSMUSG00000084292.2 | 0 | NA | NA | NA | NA | NA | #VALUE! |
| ENSMUSG00000084293.2 | 0 | NA | NA | NA | NA | NA | #VALUE! |
| ENSMUSG00000084294.2 | 0 | NA | NA | NA | NA | NA | #VALUE! |
| ENSMUSG00000084295.3 | 0 | NA | NA | NA | NA | NA | #VALUE! |
| ENSMUSG00000084296.2 | 0 | NA | NA | NA | NA | NA | #VALUE! |
| ENSMUSG00000084297.2 | 0 | NA | NA | NA | NA | NA | #VALUE! |
| ENSMUSG00000084298.2 | 0 | NA | NA | NA | NA | NA | #VALUE! |
| ENSMUSG00000084299.4 | 0 | NA | NA | NA | NA | NA | #VALUE! |
| ENSMUSG00000084300.2 | 0 | NA | NA | NA | NA | NA | #VALUE! |
| ENSMUSG00000084301.2 | 0 | NA | NA | NA | NA | NA | #VALUE! |
| ENSMUSG00000084302.2 | 0 | NA | NA | NA | NA | NA | #VALUE! |
| ENSMUSG00000084303.2 | 0 | NA | NA | NA | NA | NA | #VALUE! |
| ENSMUSG00000084304.2 | 1.384244 | 2.904951 | 3.937491 | 0.737767 | 0.460656 | NA | #VALUE! |
| ENSMUSG00000084305.2 | 0 | NA | NA | NA | NA | NA | #VALUE! |
| ENSMUSG00000084306.2 | 0 | NA | NA | NA | NA | NA | #VALUE! |
| ENSMUSG00000084307.2 | 0 | NA | NA | NA | NA | NA | #VALUE! |
| ENSMUSG00000084308.2 | 0 | NA | NA | NA | NA | NA | #VALUE! |
| ENSMUSG00000084310.2 | 0 | NA | NA | NA | NA | NA | #VALUE! |
| ENSMUSG00000084311.2 | 0 | NA | NA | NA | NA | NA | #VALUE! |
| ENSMUSG00000084312.2 | 0 | NA | NA | NA | NA | NA | #VALUE! |
| ENSMUSG00000084313.2 | 0 | NA | NA | NA | NA | NA | #VALUE! |
| ENSMUSG00000084314.2 | 0.297366 | -0.14751 | 4.065279 | -0.03629 | 0.971054 | NA | #VALUE! |

| | | | | | | | |
|---|---|---|---|---|---|---|---|
| ENSMUSG00000084315.3 | 0 | NA | NA | NA | NA | NA | #VALUE! |
| ENSMUSG00000084316.2 | 0 | NA | NA | NA | NA | NA | #VALUE! |
| ENSMUSG00000084318.2 | 0 | NA | NA | NA | NA | NA | #VALUE! |
| ENSMUSG00000084320.3 | 0 | NA | NA | NA | NA | NA | #VALUE! |
| ENSMUSG00000084323.5 | 0 | NA | NA | NA | NA | NA | #VALUE! |
| ENSMUSG00000084324.2 | 0 | NA | NA | NA | NA | NA | #VALUE! |
| ENSMUSG00000084325.3 | 0 | NA | NA | NA | NA | NA | #VALUE! |
| ENSMUSG00000084326.2 | 0 | NA | NA | NA | NA | NA | #VALUE! |
| ENSMUSG00000084327.2 | 0 | NA | NA | NA | NA | NA | #VALUE! |
| ENSMUSG00000084328.2 | 0 | NA | NA | NA | NA | NA | #VALUE! |
| ENSMUSG00000084329.2 | 0 | NA | NA | NA | NA | NA | #VALUE! |
| ENSMUSG00000084330.2 | 0 | NA | NA | NA | NA | NA | #VALUE! |
| ENSMUSG00000084331.2 | 0.204111 | -1.05581 | 4.073302 | -0.2592 | 0.795479 | NA | #VALUE! |
| ENSMUSG00000084332.2 | 0 | NA | NA | NA | NA | NA | #VALUE! |
| ENSMUSG00000084333.2 | 0.093255 | 0.868093 | 4.072464 | 0.213162 | 0.831201 | NA | #VALUE! |
| ENSMUSG00000084334.2 | 0 | NA | NA | NA | NA | NA | #VALUE! |
| ENSMUSG00000084335.2 | 0 | NA | NA | NA | NA | NA | #VALUE! |
| ENSMUSG00000084336.3 | 0 | NA | NA | NA | NA | NA | #VALUE! |
| ENSMUSG00000084337.2 | 0 | NA | NA | NA | NA | NA | #VALUE! |
| ENSMUSG00000084338.6 | 0 | NA | NA | NA | NA | NA | #VALUE! |
| ENSMUSG00000084339.2 | 0 | NA | NA | NA | NA | NA | #VALUE! |
| ENSMUSG00000084340.2 | 0 | NA | NA | NA | NA | NA | #VALUE! |
| ENSMUSG00000084341.2 | 0 | NA | NA | NA | NA | NA | #VALUE! |
| ENSMUSG00000084342.2 | 0.402379 | -0.92565 | 4.039907 | -0.22913 | 0.81877 | NA | #VALUE! |
| ENSMUSG00000084343.2 | 0 | NA | NA | NA | NA | NA | #VALUE! |
| ENSMUSG00000084344.2 | 0 | NA | NA | NA | NA | NA | #VALUE! |
| ENSMUSG00000084345.4 | 0.778332 | 1.671893 | 3.976372 | 0.420457 | 0.674152 | NA | #VALUE! |
| ENSMUSG00000084346.5 | 0 | NA | NA | NA | NA | NA | #VALUE! |
| ENSMUSG00000084347.4 | 0 | NA | NA | NA | NA | NA | #VALUE! |
| ENSMUSG00000084348.2 | 1.566313 | -4.16993 | 3.935366 | -1.0596 | 0.289325 | NA | #VALUE! |
| ENSMUSG00000084351.2 | 0 | NA | NA | NA | NA | NA | #VALUE! |
| ENSMUSG00000084352.2 | 0 | NA | NA | NA | NA | NA | #VALUE! |
| ENSMUSG00000084353.2 | 0.309124 | -1.83395 | 4.047982 | -0.45305 | 0.650511 | NA | #VALUE! |
| ENSMUSG00000084355.2 | 0 | NA | NA | NA | NA | NA | #VALUE! |
| ENSMUSG00000084356.3 | 0 | NA | NA | NA | NA | NA | #VALUE! |
| ENSMUSG00000084357.2 | 0 | NA | NA | NA | NA | NA | #VALUE! |
| ENSMUSG00000084358.2 | 0 | NA | NA | NA | NA | NA | #VALUE! |
| ENSMUSG00000084359.2 | 0 | NA | NA | NA | NA | NA | #VALUE! |
| ENSMUSG00000084360.2 | 0 | NA | NA | NA | NA | NA | #VALUE! |
| ENSMUSG00000084361.2 | 0 | NA | NA | NA | NA | NA | #VALUE! |
| ENSMUSG00000084362.2 | 0 | NA | NA | NA | NA | NA | #VALUE! |
| ENSMUSG00000084363.2 | 0 | NA | NA | NA | NA | NA | #VALUE! |
| ENSMUSG00000084366.2 | 0 | NA | NA | NA | NA | NA | #VALUE! |
| ENSMUSG00000084367.4 | 0 | NA | NA | NA | NA | NA | #VALUE! |

| | | | | | | | |
|---|---|---|---|---|---|---|---|
| ENSMUSG00000084368.2 | 0 | NA | NA | NA | NA | NA | #VALUE! |
| ENSMUSG00000084369.2 | 0 | NA | NA | NA | NA | NA | #VALUE! |
| ENSMUSG00000084370.4 | 0 | NA | NA | NA | NA | NA | #VALUE! |
| ENSMUSG00000084371.2 | 0 | NA | NA | NA | NA | NA | #VALUE! |
| ENSMUSG00000084372.2 | 0 | NA | NA | NA | NA | NA | #VALUE! |
| ENSMUSG00000084373.2 | 0 | NA | NA | NA | NA | NA | #VALUE! |
| ENSMUSG00000084374.2 | 0 | NA | NA | NA | NA | NA | #VALUE! |
| ENSMUSG00000084375.2 | 0 | NA | NA | NA | NA | NA | #VALUE! |
| ENSMUSG00000084376.2 | 0 | NA | NA | NA | NA | NA | #VALUE! |
| ENSMUSG00000084377.2 | 0 | NA | NA | NA | NA | NA | #VALUE! |
| ENSMUSG00000084378.2 | 0 | NA | NA | NA | NA | NA | #VALUE! |
| ENSMUSG00000084379.2 | 0 | NA | NA | NA | NA | NA | #VALUE! |
| ENSMUSG00000084380.2 | 0 | NA | NA | NA | NA | NA | #VALUE! |
| ENSMUSG00000084381.2 | 0 | NA | NA | NA | NA | NA | #VALUE! |
| ENSMUSG00000084382.2 | 0 | NA | NA | NA | NA | NA | #VALUE! |
| ENSMUSG00000084383.2 | 0 | NA | NA | NA | NA | NA | #VALUE! |
| ENSMUSG00000084385.2 | 0 | NA | NA | NA | NA | NA | #VALUE! |
| ENSMUSG00000084386.2 | 0 | NA | NA | NA | NA | NA | #VALUE! |
| ENSMUSG00000084387.5 | 0 | NA | NA | NA | NA | NA | #VALUE! |
| ENSMUSG00000084390.2 | 0 | NA | NA | NA | NA | NA | #VALUE! |
| ENSMUSG00000084391.2 | 0 | NA | NA | NA | NA | NA | #VALUE! |
| ENSMUSG00000084392.2 | 0 | NA | NA | NA | NA | NA | #VALUE! |
| ENSMUSG00000084393.2 | 0 | NA | NA | NA | NA | NA | #VALUE! |
| ENSMUSG00000084394.2 | 0 | NA | NA | NA | NA | NA | #VALUE! |
| ENSMUSG00000084395.2 | 0 | NA | NA | NA | NA | NA | #VALUE! |
| ENSMUSG00000084396.2 | 0 | NA | NA | NA | NA | NA | #VALUE! |
| ENSMUSG00000084397.2 | 0 | NA | NA | NA | NA | NA | #VALUE! |
| ENSMUSG00000084399.3 | 0 | NA | NA | NA | NA | NA | #VALUE! |
| ENSMUSG00000084400.2 | 0 | NA | NA | NA | NA | NA | #VALUE! |
| ENSMUSG00000084401.2 | 0 | NA | NA | NA | NA | NA | #VALUE! |
| ENSMUSG00000084403.2 | 0 | NA | NA | NA | NA | NA | #VALUE! |
| ENSMUSG00000084404.2 | 0 | NA | NA | NA | NA | NA | #VALUE! |
| ENSMUSG00000084405.3 | 0 | NA | NA | NA | NA | NA | #VALUE! |
| ENSMUSG00000084407.2 | 0 | NA | NA | NA | NA | NA | #VALUE! |
| ENSMUSG00000084408.2 | 0 | NA | NA | NA | NA | NA | #VALUE! |
| ENSMUSG00000084411.3 | 8.077609 | 6.503772 | 3.856709 | 1.686353 | 0.091728 | NA | #VALUE! |
| ENSMUSG00000084412.2 | 0 | NA | NA | NA | NA | NA | #VALUE! |
| ENSMUSG00000084413.2 | 0 | NA | NA | NA | NA | NA | #VALUE! |
| ENSMUSG00000084414.4 | 0 | NA | NA | NA | NA | NA | #VALUE! |
| ENSMUSG00000084415.2 | 0 | NA | NA | NA | NA | NA | #VALUE! |
| ENSMUSG00000084417.4 | 0 | NA | NA | NA | NA | NA | #VALUE! |
| ENSMUSG00000084421.3 | 0 | NA | NA | NA | NA | NA | #VALUE! |
| ENSMUSG00000084425.3 | 0 | NA | NA | NA | NA | NA | #VALUE! |
| ENSMUSG00000084428.3 | 0 | NA | NA | NA | NA | NA | #VALUE! |

| | | | | | | | |
|---|---|---|---|---|---|---|---|
| ENSMUSG00000084435.4 | 0 | NA | NA | NA | NA | NA | #VALUE! |
| ENSMUSG00000084437.3 | 0 | NA | NA | NA | NA | NA | #VALUE! |
| ENSMUSG00000084440.4 | 0 | NA | NA | NA | NA | NA | #VALUE! |
| ENSMUSG00000084441.3 | 0 | NA | NA | NA | NA | NA | #VALUE! |
| ENSMUSG00000084442.4 | 0 | NA | NA | NA | NA | NA | #VALUE! |
| ENSMUSG00000084447.3 | 0 | NA | NA | NA | NA | NA | #VALUE! |
| ENSMUSG00000084448.3 | 0 | NA | NA | NA | NA | NA | #VALUE! |
| ENSMUSG00000084449.3 | 0 | NA | NA | NA | NA | NA | #VALUE! |
| ENSMUSG00000084450.4 | 0 | NA | NA | NA | NA | NA | #VALUE! |
| ENSMUSG00000084453.3 | 0 | NA | NA | NA | NA | NA | #VALUE! |
| ENSMUSG00000084454.3 | 0 | NA | NA | NA | NA | NA | #VALUE! |
| ENSMUSG00000084456.4 | 0 | NA | NA | NA | NA | NA | #VALUE! |
| ENSMUSG00000084458.3 | 0 | NA | NA | NA | NA | NA | #VALUE! |
| ENSMUSG00000084459.3 | 0 | NA | NA | NA | NA | NA | #VALUE! |
| ENSMUSG00000084461.4 | 0 | NA | NA | NA | NA | NA | #VALUE! |
| ENSMUSG00000084466.4 | 0 | NA | NA | NA | NA | NA | #VALUE! |
| ENSMUSG00000084469.4 | 0 | NA | NA | NA | NA | NA | #VALUE! |
| ENSMUSG00000084473.4 | 0 | NA | NA | NA | NA | NA | #VALUE! |
| ENSMUSG00000084474.4 | 0 | NA | NA | NA | NA | NA | #VALUE! |
| ENSMUSG00000084478.4 | 0 | NA | NA | NA | NA | NA | #VALUE! |
| ENSMUSG00000084479.2 | 0 | NA | NA | NA | NA | NA | #VALUE! |
| ENSMUSG00000084484.4 | 0 | NA | NA | NA | NA | NA | #VALUE! |
| ENSMUSG00000084487.3 | 0 | NA | NA | NA | NA | NA | #VALUE! |
| ENSMUSG00000084493.3 | 0 | NA | NA | NA | NA | NA | #VALUE! |
| ENSMUSG00000084497.3 | 0 | NA | NA | NA | NA | NA | #VALUE! |
| ENSMUSG00000084504.3 | 0 | NA | NA | NA | NA | NA | #VALUE! |
| ENSMUSG00000084505.4 | 0 | NA | NA | NA | NA | NA | #VALUE! |
| ENSMUSG00000084506.3 | 0 | NA | NA | NA | NA | NA | #VALUE! |
| ENSMUSG00000084507.3 | 0 | NA | NA | NA | NA | NA | #VALUE! |
| ENSMUSG00000084514.2 | 0 | NA | NA | NA | NA | NA | #VALUE! |
| ENSMUSG00000084515.4 | 0 | NA | NA | NA | NA | NA | #VALUE! |
| ENSMUSG00000084516.3 | 0 | NA | NA | NA | NA | NA | #VALUE! |
| ENSMUSG00000084517.3 | 0 | NA | NA | NA | NA | NA | #VALUE! |
| ENSMUSG00000084521.3 | 0 | NA | NA | NA | NA | NA | #VALUE! |
| ENSMUSG00000084523.3 | 0 | NA | NA | NA | NA | NA | #VALUE! |
| ENSMUSG00000084527.3 | 0 | NA | NA | NA | NA | NA | #VALUE! |
| ENSMUSG00000084535.3 | 0 | NA | NA | NA | NA | NA | #VALUE! |
| ENSMUSG00000084549.3 | 0 | NA | NA | NA | NA | NA | #VALUE! |
| ENSMUSG00000084550.3 | 0 | NA | NA | NA | NA | NA | #VALUE! |
| ENSMUSG00000084551.3 | 0 | NA | NA | NA | NA | NA | #VALUE! |
| ENSMUSG00000084552.4 | 0 | NA | NA | NA | NA | NA | #VALUE! |
| ENSMUSG00000084555.3 | 0 | NA | NA | NA | NA | NA | #VALUE! |
| ENSMUSG00000084557.4 | 0 | NA | NA | NA | NA | NA | #VALUE! |
| ENSMUSG00000084558.3 | 0 | NA | NA | NA | NA | NA | #VALUE! |

| | | | | | | | |
|---|---|---|---|---|---|---|---|
| ENSMUSG00000084559.3 | 0 | NA | NA | NA | NA | NA | #VALUE! |
| ENSMUSG00000084560.3 | 0 | NA | NA | NA | NA | NA | #VALUE! |
| ENSMUSG00000084564.3 | 0 | NA | NA | NA | NA | NA | #VALUE! |
| ENSMUSG00000084565.3 | 0 | NA | NA | NA | NA | NA | #VALUE! |
| ENSMUSG00000084569.3 | 0 | NA | NA | NA | NA | NA | #VALUE! |
| ENSMUSG00000084575.3 | 0 | NA | NA | NA | NA | NA | #VALUE! |
| ENSMUSG00000084576.3 | 0 | NA | NA | NA | NA | NA | #VALUE! |
| ENSMUSG00000084579.4 | 0 | NA | NA | NA | NA | NA | #VALUE! |
| ENSMUSG00000084583.4 | 0 | NA | NA | NA | NA | NA | #VALUE! |
| ENSMUSG00000084589.4 | 0 | NA | NA | NA | NA | NA | #VALUE! |
| ENSMUSG00000084590.3 | 0 | NA | NA | NA | NA | NA | #VALUE! |
| ENSMUSG00000084592.3 | 0 | NA | NA | NA | NA | NA | #VALUE! |
| ENSMUSG00000084593.3 | 0 | NA | NA | NA | NA | NA | #VALUE! |
| ENSMUSG00000084601.4 | 0 | NA | NA | NA | NA | NA | #VALUE! |
| ENSMUSG00000084604.3 | 0 | NA | NA | NA | NA | NA | #VALUE! |
| ENSMUSG00000084606.3 | 0 | NA | NA | NA | NA | NA | #VALUE! |
| ENSMUSG00000084607.3 | 0 | NA | NA | NA | NA | NA | #VALUE! |
| ENSMUSG00000084609.4 | 0 | NA | NA | NA | NA | NA | #VALUE! |
| ENSMUSG00000084610.4 | 0 | NA | NA | NA | NA | NA | #VALUE! |
| ENSMUSG00000084617.3 | 0 | NA | NA | NA | NA | NA | #VALUE! |
| ENSMUSG00000084619.3 | 0 | NA | NA | NA | NA | NA | #VALUE! |
| ENSMUSG00000084621.4 | 0 | NA | NA | NA | NA | NA | #VALUE! |
| ENSMUSG00000084629.3 | 0 | NA | NA | NA | NA | NA | #VALUE! |
| ENSMUSG00000084632.3 | 0 | NA | NA | NA | NA | NA | #VALUE! |
| ENSMUSG00000084638.3 | 0 | NA | NA | NA | NA | NA | #VALUE! |
| ENSMUSG00000084644.3 | 0 | NA | NA | NA | NA | NA | #VALUE! |
| ENSMUSG00000084646.3 | 0 | NA | NA | NA | NA | NA | #VALUE! |
| ENSMUSG00000084647.3 | 0 | NA | NA | NA | NA | NA | #VALUE! |
| ENSMUSG00000084648.4 | 0 | NA | NA | NA | NA | NA | #VALUE! |
| ENSMUSG00000084651.3 | 0 | NA | NA | NA | NA | NA | #VALUE! |
| ENSMUSG00000084652.3 | 0 | NA | NA | NA | NA | NA | #VALUE! |
| ENSMUSG00000084659.3 | 0 | NA | NA | NA | NA | NA | #VALUE! |
| ENSMUSG00000084661.4 | 0 | NA | NA | NA | NA | NA | #VALUE! |
| ENSMUSG00000084663.3 | 0 | NA | NA | NA | NA | NA | #VALUE! |
| ENSMUSG00000084665.3 | 0 | NA | NA | NA | NA | NA | #VALUE! |
| ENSMUSG00000084667.3 | 0 | NA | NA | NA | NA | NA | #VALUE! |
| ENSMUSG00000084669.3 | 0 | NA | NA | NA | NA | NA | #VALUE! |
| ENSMUSG00000084678.3 | 0 | NA | NA | NA | NA | NA | #VALUE! |
| ENSMUSG00000084686.3 | 0 | NA | NA | NA | NA | NA | #VALUE! |
| ENSMUSG00000084690.2 | 0 | NA | NA | NA | NA | NA | #VALUE! |
| ENSMUSG00000084691.3 | 0 | NA | NA | NA | NA | NA | #VALUE! |
| ENSMUSG00000084696.3 | 0 | NA | NA | NA | NA | NA | #VALUE! |
| ENSMUSG00000084700.4 | 0 | NA | NA | NA | NA | NA | #VALUE! |
| ENSMUSG00000084703.4 | 0 | NA | NA | NA | NA | NA | #VALUE! |

| | | | | | | | |
|---|---|---|---|---|---|---|---|
| ENSMUSG00000084711.3 | 0 | NA | NA | NA | NA | NA | #VALUE! |
| ENSMUSG00000084713.5 | 0 | NA | NA | NA | NA | NA | #VALUE! |
| ENSMUSG00000084714.4 | 0 | NA | NA | NA | NA | NA | #VALUE! |
| ENSMUSG00000084717.4 | 0 | NA | NA | NA | NA | NA | #VALUE! |
| ENSMUSG00000084725.5 | 0 | NA | NA | NA | NA | NA | #VALUE! |
| ENSMUSG00000084728.3 | 0 | NA | NA | NA | NA | NA | #VALUE! |
| ENSMUSG00000084731.4 | 0 | NA | NA | NA | NA | NA | #VALUE! |
| ENSMUSG00000084737.3 | 0 | NA | NA | NA | NA | NA | #VALUE! |
| ENSMUSG00000084739.4 | 0 | NA | NA | NA | NA | NA | #VALUE! |
| ENSMUSG00000084740.3 | 0 | NA | NA | NA | NA | NA | #VALUE! |
| ENSMUSG00000084745.3 | 0 | NA | NA | NA | NA | NA | #VALUE! |
| ENSMUSG00000084747.3 | 0 | NA | NA | NA | NA | NA | #VALUE! |
| ENSMUSG00000084748.3 | 0 | NA | NA | NA | NA | NA | #VALUE! |
| ENSMUSG00000084751.3 | 0 | NA | NA | NA | NA | NA | #VALUE! |
| ENSMUSG00000084752.3 | 0 | NA | NA | NA | NA | NA | #VALUE! |
| ENSMUSG00000084753.3 | 0 | NA | NA | NA | NA | NA | #VALUE! |
| ENSMUSG00000084754.2 | 0 | NA | NA | NA | NA | NA | #VALUE! |
| ENSMUSG00000084755.2 | 0 | NA | NA | NA | NA | NA | #VALUE! |
| ENSMUSG00000084756.2 | 0 | NA | NA | NA | NA | NA | #VALUE! |
| ENSMUSG00000084757.8 | 0 | NA | NA | NA | NA | NA | #VALUE! |
| ENSMUSG00000084758.2 | 0 | NA | NA | NA | NA | NA | #VALUE! |
| ENSMUSG00000084760.2 | 0 | NA | NA | NA | NA | NA | #VALUE! |
| ENSMUSG00000084761.2 | 0 | NA | NA | NA | NA | NA | #VALUE! |
| ENSMUSG00000084762.2 | 0 | NA | NA | NA | NA | NA | #VALUE! |
| ENSMUSG00000084764.2 | 0 | NA | NA | NA | NA | NA | #VALUE! |
| ENSMUSG00000084766.2 | 0 | NA | NA | NA | NA | NA | #VALUE! |
| ENSMUSG00000084768.3 | 0 | NA | NA | NA | NA | NA | #VALUE! |
| ENSMUSG00000084769.2 | 0 | NA | NA | NA | NA | NA | #VALUE! |
| ENSMUSG00000084772.2 | 0 | NA | NA | NA | NA | NA | #VALUE! |
| ENSMUSG00000084773.2 | 0 | NA | NA | NA | NA | NA | #VALUE! |
| ENSMUSG00000084774.2 | 0 | NA | NA | NA | NA | NA | #VALUE! |
| ENSMUSG00000084775.3 | 0 | NA | NA | NA | NA | NA | #VALUE! |
| ENSMUSG00000084776.2 | 0 | NA | NA | NA | NA | NA | #VALUE! |
| ENSMUSG00000084780.8 | 0 | NA | NA | NA | NA | NA | #VALUE! |
| ENSMUSG00000084781.2 | 0 | NA | NA | NA | NA | NA | #VALUE! |
| ENSMUSG00000084782.3 | 0 | NA | NA | NA | NA | NA | #VALUE! |
| ENSMUSG00000084783.2 | 0 | NA | NA | NA | NA | NA | #VALUE! |
| ENSMUSG00000084785.2 | 0 | NA | NA | NA | NA | NA | #VALUE! |
| ENSMUSG00000084787.2 | 0 | NA | NA | NA | NA | NA | #VALUE! |
| ENSMUSG00000084788.3 | 0.408222 | -2.20205 | 4.016842 | -0.54821 | 0.583551 | NA | #VALUE! |
| ENSMUSG00000084789.2 | 0 | NA | NA | NA | NA | NA | #VALUE! |
| ENSMUSG00000084791.2 | 0 | NA | NA | NA | NA | NA | #VALUE! |
| ENSMUSG00000084792.2 | 1.177985 | 3.72614 | 3.944615 | 0.944614 | 0.344856 | NA | #VALUE! |
| ENSMUSG00000084793.2 | 0 | NA | NA | NA | NA | NA | #VALUE! |

| | | | | | | | |
|---|---|---|---|---|---|---|---|
| ENSMUSG00000084794.2 | 0 | NA | NA | NA | NA | NA | #VALUE! |
| ENSMUSG00000084795.2 | 0 | NA | NA | NA | NA | NA | #VALUE! |
| ENSMUSG00000084796.2 | 3.990499 | -5.53332 | 3.917921 | -1.41231 | 0.157859 | NA | #VALUE! |
| ENSMUSG00000084797.2 | 0 | NA | NA | NA | NA | NA | #VALUE! |
| ENSMUSG00000084798.2 | 0 | NA | NA | NA | NA | NA | #VALUE! |
| ENSMUSG00000084799.8 | 0 | NA | NA | NA | NA | NA | #VALUE! |
| ENSMUSG00000084800.2 | 0 | NA | NA | NA | NA | NA | #VALUE! |
| ENSMUSG00000084802.2 | 0 | NA | NA | NA | NA | NA | #VALUE! |
| ENSMUSG00000084803.9 | 6.320394 | -2.07342 | 3.77282 | -0.54957 | 0.582616 | NA | #VALUE! |
| ENSMUSG00000084804.2 | 0 | NA | NA | NA | NA | NA | #VALUE! |
| ENSMUSG00000084805.2 | 0 | NA | NA | NA | NA | NA | #VALUE! |
| ENSMUSG00000084806.2 | 0.105013 | -0.97324 | 4.076293 | -0.23876 | 0.811295 | NA | #VALUE! |
| ENSMUSG00000084807.2 | 0 | NA | NA | NA | NA | NA | #VALUE! |
| ENSMUSG00000084808.8 | 0 | NA | NA | NA | NA | NA | #VALUE! |
| ENSMUSG00000084809.2 | 0 | NA | NA | NA | NA | NA | #VALUE! |
| ENSMUSG00000084811.3 | 0 | NA | NA | NA | NA | NA | #VALUE! |
| ENSMUSG00000084816.2 | 0 | NA | NA | NA | NA | NA | #VALUE! |
| ENSMUSG00000084817.4 | 0.448049 | 2.371337 | 4.002117 | 0.592521 | 0.553502 | NA | #VALUE! |
| ENSMUSG00000084818.2 | 0 | NA | NA | NA | NA | NA | #VALUE! |
| ENSMUSG00000084821.2 | 0 | NA | NA | NA | NA | NA | #VALUE! |
| ENSMUSG00000084822.2 | 0 | NA | NA | NA | NA | NA | #VALUE! |
| ENSMUSG00000084823.3 | 0 | NA | NA | NA | NA | NA | #VALUE! |
| ENSMUSG00000084824.2 | 0 | NA | NA | NA | NA | NA | #VALUE! |
| ENSMUSG00000084825.2 | 0 | NA | NA | NA | NA | NA | #VALUE! |
| ENSMUSG00000084826.8 | 0 | NA | NA | NA | NA | NA | #VALUE! |
| ENSMUSG00000084828.2 | 0 | NA | NA | NA | NA | NA | #VALUE! |
| ENSMUSG00000084829.8 | 0 | NA | NA | NA | NA | NA | #VALUE! |
| ENSMUSG00000084830.3 | 9.036864 | 1.342651 | 1.972924 | 0.680539 | 0.496163 | NA | #VALUE! |
| ENSMUSG00000084831.2 | 0 | NA | NA | NA | NA | NA | #VALUE! |
| ENSMUSG00000084833.2 | 0 | NA | NA | NA | NA | NA | #VALUE! |
| ENSMUSG00000084834.2 | 0 | NA | NA | NA | NA | NA | #VALUE! |
| ENSMUSG00000084835.2 | 0 | NA | NA | NA | NA | NA | #VALUE! |
| ENSMUSG00000084836.2 | 0 | NA | NA | NA | NA | NA | #VALUE! |
| ENSMUSG00000084837.2 | 0 | NA | NA | NA | NA | NA | #VALUE! |
| ENSMUSG00000084838.4 | 0 | NA | NA | NA | NA | NA | #VALUE! |
| ENSMUSG00000084839.2 | 0 | NA | NA | NA | NA | NA | #VALUE! |
| ENSMUSG00000084840.2 | 0 | NA | NA | NA | NA | NA | #VALUE! |
| ENSMUSG00000084842.4 | 0 | NA | NA | NA | NA | NA | #VALUE! |
| ENSMUSG00000084843.2 | 0 | NA | NA | NA | NA | NA | #VALUE! |
| ENSMUSG00000084845.10 | 0 | NA | NA | NA | NA | NA | #VALUE! |
| ENSMUSG00000084846.2 | 0 | NA | NA | NA | NA | NA | #VALUE! |
| ENSMUSG00000084847.8 | 0 | NA | NA | NA | NA | NA | #VALUE! |
| ENSMUSG00000084848.2 | 0 | NA | NA | NA | NA | NA | #VALUE! |
| ENSMUSG00000084849.2 | 0 | NA | NA | NA | NA | NA | #VALUE! |

| | | | | | | | |
|---|---|---|---|---|---|---|---|
| ENSMUSG00000084850.2 | 0 | NA | NA | NA | NA | NA | #VALUE! |
| ENSMUSG00000084851.2 | 0 | NA | NA | NA | NA | NA | #VALUE! |
| ENSMUSG00000084852.2 | 0 | NA | NA | NA | NA | NA | #VALUE! |
| ENSMUSG00000084853.2 | 0 | NA | NA | NA | NA | NA | #VALUE! |
| ENSMUSG00000084854.2 | 0 | NA | NA | NA | NA | NA | #VALUE! |
| ENSMUSG00000084855.3 | 0 | NA | NA | NA | NA | NA | #VALUE! |
| ENSMUSG00000084856.2 | 0 | NA | NA | NA | NA | NA | #VALUE! |
| ENSMUSG00000084858.2 | 0 | NA | NA | NA | NA | NA | #VALUE! |
| ENSMUSG00000084859.8 | 0 | NA | NA | NA | NA | NA | #VALUE! |
| ENSMUSG00000084860.2 | 0 | NA | NA | NA | NA | NA | #VALUE! |
| ENSMUSG00000084861.2 | 0 | NA | NA | NA | NA | NA | #VALUE! |
| ENSMUSG00000084862.2 | 0 | NA | NA | NA | NA | NA | #VALUE! |
| ENSMUSG00000084863.2 | 0 | NA | NA | NA | NA | NA | #VALUE! |
| ENSMUSG00000084864.2 | 0 | NA | NA | NA | NA | NA | #VALUE! |
| ENSMUSG00000084865.2 | 0 | NA | NA | NA | NA | NA | #VALUE! |
| ENSMUSG00000084866.3 | 0 | NA | NA | NA | NA | NA | #VALUE! |
| ENSMUSG00000084869.2 | 0 | NA | NA | NA | NA | NA | #VALUE! |
| ENSMUSG00000084870.2 | 0 | NA | NA | NA | NA | NA | #VALUE! |
| ENSMUSG00000084871.2 | 0 | NA | NA | NA | NA | NA | #VALUE! |
| ENSMUSG00000084873.4 | 0 | NA | NA | NA | NA | NA | #VALUE! |
| ENSMUSG00000084874.5 | 0 | NA | NA | NA | NA | NA | #VALUE! |
| ENSMUSG00000084875.3 | 0 | NA | NA | NA | NA | NA | #VALUE! |
| ENSMUSG00000084876.8 | 0 | NA | NA | NA | NA | NA | #VALUE! |
| ENSMUSG00000084880.2 | 5.165486 | 5.852308 | 3.9155 | 1.494651 | 0.135005 | NA | #VALUE! |
| ENSMUSG00000084883.3 | 0.198268 | -0.06494 | 4.068276 | -0.01596 | 0.987265 | NA | #VALUE! |
| ENSMUSG00000084884.2 | 0 | NA | NA | NA | NA | NA | #VALUE! |
| ENSMUSG00000084885.8 | 1.174735 | -3.75246 | 3.944889 | -0.95122 | 0.341493 | NA | #VALUE! |
| ENSMUSG00000084887.2 | 0 | NA | NA | NA | NA | NA | #VALUE! |
| ENSMUSG00000084889.2 | 0 | NA | NA | NA | NA | NA | #VALUE! |
| ENSMUSG00000084890.2 | 0 | NA | NA | NA | NA | NA | #VALUE! |
| ENSMUSG00000084891.2 | 0 | NA | NA | NA | NA | NA | #VALUE! |
| ENSMUSG00000084892.2 | 0 | NA | NA | NA | NA | NA | #VALUE! |
| ENSMUSG00000084893.4 | 0 | NA | NA | NA | NA | NA | #VALUE! |
| ENSMUSG00000084895.2 | 0 | NA | NA | NA | NA | NA | #VALUE! |
| ENSMUSG00000084896.2 | 0 | NA | NA | NA | NA | NA | #VALUE! |
| ENSMUSG00000084897.2 | 0 | NA | NA | NA | NA | NA | #VALUE! |
| ENSMUSG00000084898.2 | 0 | NA | NA | NA | NA | NA | #VALUE! |
| ENSMUSG00000084899.2 | 0 | NA | NA | NA | NA | NA | #VALUE! |
| ENSMUSG00000084901.7 | 0 | NA | NA | NA | NA | NA | #VALUE! |
| ENSMUSG00000084902.3 | 0 | NA | NA | NA | NA | NA | #VALUE! |
| ENSMUSG00000084904.2 | 0 | NA | NA | NA | NA | NA | #VALUE! |
| ENSMUSG00000084905.2 | 0 | NA | NA | NA | NA | NA | #VALUE! |
| ENSMUSG00000084906.2 | 0 | NA | NA | NA | NA | NA | #VALUE! |
| ENSMUSG00000084907.2 | 0 | NA | NA | NA | NA | NA | #VALUE! |

| | | | | | | | |
|---|---|---|---|---|---|---|---|
| ENSMUSG00000084908.3 | 0 | NA | NA | NA | NA | NA | #VALUE! |
| ENSMUSG00000084909.2 | 0 | NA | NA | NA | NA | NA | #VALUE! |
| ENSMUSG00000084911.2 | 0 | NA | NA | NA | NA | NA | #VALUE! |
| ENSMUSG00000084913.4 | 0 | NA | NA | NA | NA | NA | #VALUE! |
| ENSMUSG00000084915.3 | 0.652786 | 2.919659 | 3.972474 | 0.734972 | 0.462356 | NA | #VALUE! |
| ENSMUSG00000084916.2 | 0 | NA | NA | NA | NA | NA | #VALUE! |
| ENSMUSG00000084917.4 | 0 | NA | NA | NA | NA | NA | #VALUE! |
| ENSMUSG00000084918.2 | 0 | NA | NA | NA | NA | NA | #VALUE! |
| ENSMUSG00000084919.2 | 0 | NA | NA | NA | NA | NA | #VALUE! |
| ENSMUSG00000084920.2 | 0 | NA | NA | NA | NA | NA | #VALUE! |
| ENSMUSG00000084921.2 | 0 | NA | NA | NA | NA | NA | #VALUE! |
| ENSMUSG00000084923.2 | 0 | NA | NA | NA | NA | NA | #VALUE! |
| ENSMUSG00000084924.4 | 0 | NA | NA | NA | NA | NA | #VALUE! |
| ENSMUSG00000084925.9 | 1.346268 | 3.918821 | 3.939925 | 0.994644 | 0.31991 | NA | #VALUE! |
| ENSMUSG00000084927.2 | 0 | NA | NA | NA | NA | NA | #VALUE! |
| ENSMUSG00000084929.2 | 0 | NA | NA | NA | NA | NA | #VALUE! |
| ENSMUSG00000084930.2 | 0 | NA | NA | NA | NA | NA | #VALUE! |
| ENSMUSG00000084932.2 | 0 | NA | NA | NA | NA | NA | #VALUE! |
| ENSMUSG00000084933.2 | 0 | NA | NA | NA | NA | NA | #VALUE! |
| ENSMUSG00000084934.2 | 0 | NA | NA | NA | NA | NA | #VALUE! |
| ENSMUSG00000084935.2 | 0 | NA | NA | NA | NA | NA | #VALUE! |
| ENSMUSG00000084937.2 | 0 | NA | NA | NA | NA | NA | #VALUE! |
| ENSMUSG00000084938.2 | 0 | NA | NA | NA | NA | NA | #VALUE! |
| ENSMUSG00000084939.3 | 0 | NA | NA | NA | NA | NA | #VALUE! |
| ENSMUSG00000084940.3 | 0 | NA | NA | NA | NA | NA | #VALUE! |
| ENSMUSG00000084941.2 | 0 | NA | NA | NA | NA | NA | #VALUE! |
| ENSMUSG00000084942.2 | 0 | NA | NA | NA | NA | NA | #VALUE! |
| ENSMUSG00000084944.2 | 0 | NA | NA | NA | NA | NA | #VALUE! |
| ENSMUSG00000084945.2 | 0 | NA | NA | NA | NA | NA | #VALUE! |
| ENSMUSG00000084946.3 | 0 | NA | NA | NA | NA | NA | #VALUE! |
| ENSMUSG00000084947.2 | 0 | NA | NA | NA | NA | NA | #VALUE! |
| ENSMUSG00000084948.2 | 0 | NA | NA | NA | NA | NA | #VALUE! |
| ENSMUSG00000084949.2 | 0 | NA | NA | NA | NA | NA | #VALUE! |
| ENSMUSG00000084950.8 | 0 | NA | NA | NA | NA | NA | #VALUE! |
| ENSMUSG00000084951.2 | 0 | NA | NA | NA | NA | NA | #VALUE! |
| ENSMUSG00000084954.3 | 0 | NA | NA | NA | NA | NA | #VALUE! |
| ENSMUSG00000084955.2 | 0 | NA | NA | NA | NA | NA | #VALUE! |
| ENSMUSG00000084959.2 | 0 | NA | NA | NA | NA | NA | #VALUE! |
| ENSMUSG00000084960.2 | 0 | NA | NA | NA | NA | NA | #VALUE! |
| ENSMUSG00000084962.2 | 0 | NA | NA | NA | NA | NA | #VALUE! |
| ENSMUSG00000084963.2 | 0 | NA | NA | NA | NA | NA | #VALUE! |
| ENSMUSG00000084964.3 | 0 | NA | NA | NA | NA | NA | #VALUE! |
| ENSMUSG00000084965.2 | 0 | NA | NA | NA | NA | NA | #VALUE! |
| ENSMUSG00000084966.2 | 0 | NA | NA | NA | NA | NA | #VALUE! |

| | | | | | | | |
|---|---|---|---|---|---|---|---|
| ENSMUSG00000084967.2 | 0 | NA | NA | NA | NA | NA | #VALUE! |
| ENSMUSG00000084968.2 | 0 | NA | NA | NA | NA | NA | #VALUE! |
| ENSMUSG00000084969.2 | 0 | NA | NA | NA | NA | NA | #VALUE! |
| ENSMUSG00000084970.2 | 0 | NA | NA | NA | NA | NA | #VALUE! |
| ENSMUSG00000084971.2 | 0 | NA | NA | NA | NA | NA | #VALUE! |
| ENSMUSG00000084973.3 | 0 | NA | NA | NA | NA | NA | #VALUE! |
| ENSMUSG00000084974.2 | 0 | NA | NA | NA | NA | NA | #VALUE! |
| ENSMUSG00000084978.2 | 0 | NA | NA | NA | NA | NA | #VALUE! |
| ENSMUSG00000084979.2 | 0 | NA | NA | NA | NA | NA | #VALUE! |
| ENSMUSG00000084980.2 | 0 | NA | NA | NA | NA | NA | #VALUE! |
| ENSMUSG00000084981.8 | 0 | NA | NA | NA | NA | NA | #VALUE! |
| ENSMUSG00000084982.2 | 0 | NA | NA | NA | NA | NA | #VALUE! |
| ENSMUSG00000084984.9 | 0 | NA | NA | NA | NA | NA | #VALUE! |
| ENSMUSG00000084985.2 | 0 | NA | NA | NA | NA | NA | #VALUE! |
| ENSMUSG00000084986.2 | 0 | NA | NA | NA | NA | NA | #VALUE! |
| ENSMUSG00000084987.2 | 0 | NA | NA | NA | NA | NA | #VALUE! |
| ENSMUSG00000084989.4 | 0 | NA | NA | NA | NA | NA | #VALUE! |
| ENSMUSG00000084990.2 | 0 | NA | NA | NA | NA | NA | #VALUE! |
| ENSMUSG00000084991.2 | 0 | NA | NA | NA | NA | NA | #VALUE! |
| ENSMUSG00000084992.2 | 0 | NA | NA | NA | NA | NA | #VALUE! |
| ENSMUSG00000084993.2 | 0 | NA | NA | NA | NA | NA | #VALUE! |
| ENSMUSG00000084995.9 | 0 | NA | NA | NA | NA | NA | #VALUE! |
| ENSMUSG00000084996.2 | 0 | NA | NA | NA | NA | NA | #VALUE! |
| ENSMUSG00000084997.8 | 0 | NA | NA | NA | NA | NA | #VALUE! |
| ENSMUSG00000084998.2 | 0 | NA | NA | NA | NA | NA | #VALUE! |
| ENSMUSG00000084999.3 | 0 | NA | NA | NA | NA | NA | #VALUE! |
| ENSMUSG00000085000.2 | 0 | NA | NA | NA | NA | NA | #VALUE! |
| ENSMUSG00000085001.2 | 0 | NA | NA | NA | NA | NA | #VALUE! |
| ENSMUSG00000085002.2 | 0 | NA | NA | NA | NA | NA | #VALUE! |
| ENSMUSG00000085003.2 | 0 | NA | NA | NA | NA | NA | #VALUE! |
| ENSMUSG00000085004.3 | 0 | NA | NA | NA | NA | NA | #VALUE! |
| ENSMUSG00000085006.8 | 0 | NA | NA | NA | NA | NA | #VALUE! |
| ENSMUSG00000085007.4 | 0 | NA | NA | NA | NA | NA | #VALUE! |
| ENSMUSG00000085008.2 | 0 | NA | NA | NA | NA | NA | #VALUE! |
| ENSMUSG00000085009.2 | 0 | NA | NA | NA | NA | NA | #VALUE! |
| ENSMUSG00000085010.2 | 0.373021 | 2.140748 | 4.018182 | 0.532765 | 0.594196 | NA | #VALUE! |
| ENSMUSG00000085012.2 | 0.587367 | -2.74297 | 3.982932 | -0.68868 | 0.491024 | NA | #VALUE! |
| ENSMUSG00000085014.2 | 0 | NA | NA | NA | NA | NA | #VALUE! |
| ENSMUSG00000085015.2 | 0 | NA | NA | NA | NA | NA | #VALUE! |
| ENSMUSG00000085016.2 | 0 | NA | NA | NA | NA | NA | #VALUE! |
| ENSMUSG00000085017.2 | 0 | NA | NA | NA | NA | NA | #VALUE! |
| ENSMUSG00000085019.3 | 0 | NA | NA | NA | NA | NA | #VALUE! |
| ENSMUSG00000085020.2 | 0 | NA | NA | NA | NA | NA | #VALUE! |
| ENSMUSG00000085021.2 | 0 | NA | NA | NA | NA | NA | #VALUE! |

| | | | | | | | |
|---|---|---|---|---|---|---|---|
| ENSMUSG00000085022.3 | 0 | NA | NA | NA | NA | NA | #VALUE! |
| ENSMUSG00000085024.2 | 0 | NA | NA | NA | NA | NA | #VALUE! |
| ENSMUSG00000085025.2 | 0 | NA | NA | NA | NA | NA | #VALUE! |
| ENSMUSG00000085026.2 | 0 | NA | NA | NA | NA | NA | #VALUE! |
| ENSMUSG00000085027.2 | 0 | NA | NA | NA | NA | NA | #VALUE! |
| ENSMUSG00000085029.2 | 0 | NA | NA | NA | NA | NA | #VALUE! |
| ENSMUSG00000085030.2 | 0 | NA | NA | NA | NA | NA | #VALUE! |
| ENSMUSG00000085032.2 | 0 | NA | NA | NA | NA | NA | #VALUE! |
| ENSMUSG00000085033.2 | 0 | NA | NA | NA | NA | NA | #VALUE! |
| ENSMUSG00000085034.2 | 0 | NA | NA | NA | NA | NA | #VALUE! |
| ENSMUSG00000085035.2 | 0 | NA | NA | NA | NA | NA | #VALUE! |
| ENSMUSG00000085036.2 | 0 | NA | NA | NA | NA | NA | #VALUE! |
| ENSMUSG00000085039.3 | 0 | NA | NA | NA | NA | NA | #VALUE! |
| ENSMUSG00000085041.2 | 0 | NA | NA | NA | NA | NA | #VALUE! |
| ENSMUSG00000085042.9 | 0 | NA | NA | NA | NA | NA | #VALUE! |
| ENSMUSG00000085043.2 | 0 | NA | NA | NA | NA | NA | #VALUE! |
| ENSMUSG00000085044.2 | 0 | NA | NA | NA | NA | NA | #VALUE! |
| ENSMUSG00000085046.2 | 0 | NA | NA | NA | NA | NA | #VALUE! |
| ENSMUSG00000085047.2 | 0 | NA | NA | NA | NA | NA | #VALUE! |
| ENSMUSG00000085048.2 | 0 | NA | NA | NA | NA | NA | #VALUE! |
| ENSMUSG00000085049.2 | 0 | NA | NA | NA | NA | NA | #VALUE! |
| ENSMUSG00000085051.2 | 0 | NA | NA | NA | NA | NA | #VALUE! |
| ENSMUSG00000085053.2 | 0 | NA | NA | NA | NA | NA | #VALUE! |
| ENSMUSG00000085054.3 | 0 | NA | NA | NA | NA | NA | #VALUE! |
| ENSMUSG00000085055.3 | 0 | NA | NA | NA | NA | NA | #VALUE! |
| ENSMUSG00000085057.8 | 0 | NA | NA | NA | NA | NA | #VALUE! |
| ENSMUSG00000085058.3 | 0 | NA | NA | NA | NA | NA | #VALUE! |
| ENSMUSG00000085059.2 | 0 | NA | NA | NA | NA | NA | #VALUE! |
| ENSMUSG00000085060.2 | 0 | NA | NA | NA | NA | NA | #VALUE! |
| ENSMUSG00000085063.2 | 0 | NA | NA | NA | NA | NA | #VALUE! |
| ENSMUSG00000085066.3 | 0 | NA | NA | NA | NA | NA | #VALUE! |
| ENSMUSG00000085067.3 | 0 | NA | NA | NA | NA | NA | #VALUE! |
| ENSMUSG00000085068.2 | 0 | NA | NA | NA | NA | NA | #VALUE! |
| ENSMUSG00000085069.3 | 0 | NA | NA | NA | NA | NA | #VALUE! |
| ENSMUSG00000085070.2 | 0 | NA | NA | NA | NA | NA | #VALUE! |
| ENSMUSG00000085071.2 | 0 | NA | NA | NA | NA | NA | #VALUE! |
| ENSMUSG00000085072.2 | 0 | NA | NA | NA | NA | NA | #VALUE! |
| ENSMUSG00000085073.4 | 0 | NA | NA | NA | NA | NA | #VALUE! |
| ENSMUSG00000085074.2 | 0 | NA | NA | NA | NA | NA | #VALUE! |
| ENSMUSG00000085075.2 | 0 | NA | NA | NA | NA | NA | #VALUE! |
| ENSMUSG00000085077.2 | 0 | NA | NA | NA | NA | NA | #VALUE! |
| ENSMUSG00000085078.2 | 0 | NA | NA | NA | NA | NA | #VALUE! |
| ENSMUSG00000085079.4 | 0 | NA | NA | NA | NA | NA | #VALUE! |
| ENSMUSG00000085080.2 | 0 | NA | NA | NA | NA | NA | #VALUE! |

| | | | | | | | |
|---|---|---|---|---|---|---|---|
| ENSMUSG00000085081.2 | 0 | NA | NA | NA | NA | NA | #VALUE! |
| ENSMUSG00000085082.2 | 0 | NA | NA | NA | NA | NA | #VALUE! |
| ENSMUSG00000085083.2 | 0 | NA | NA | NA | NA | NA | #VALUE! |
| ENSMUSG00000085084.8 | 0 | NA | NA | NA | NA | NA | #VALUE! |
| ENSMUSG00000085085.3 | 0 | NA | NA | NA | NA | NA | #VALUE! |
| ENSMUSG00000085087.2 | 0 | NA | NA | NA | NA | NA | #VALUE! |
| ENSMUSG00000085088.2 | 0 | NA | NA | NA | NA | NA | #VALUE! |
| ENSMUSG00000085089.2 | 0 | NA | NA | NA | NA | NA | #VALUE! |
| ENSMUSG00000085090.2 | 0 | NA | NA | NA | NA | NA | #VALUE! |
| ENSMUSG00000085091.2 | 0 | NA | NA | NA | NA | NA | #VALUE! |
| ENSMUSG00000085093.2 | 0 | NA | NA | NA | NA | NA | #VALUE! |
| ENSMUSG00000085094.3 | 0.105013 | -0.97324 | 4.076293 | -0.23876 | 0.811295 | NA | #VALUE! |
| ENSMUSG00000085095.3 | 0 | NA | NA | NA | NA | NA | #VALUE! |
| ENSMUSG00000085097.2 | 0 | NA | NA | NA | NA | NA | #VALUE! |
| ENSMUSG00000085099.2 | 0 | NA | NA | NA | NA | NA | #VALUE! |
| ENSMUSG00000085100.2 | 0 | NA | NA | NA | NA | NA | #VALUE! |
| ENSMUSG00000085101.8 | 0 | NA | NA | NA | NA | NA | #VALUE! |
| ENSMUSG00000085102.2 | 0 | NA | NA | NA | NA | NA | #VALUE! |
| ENSMUSG00000085103.4 | 0 | NA | NA | NA | NA | NA | #VALUE! |
| ENSMUSG00000085104.2 | 0 | NA | NA | NA | NA | NA | #VALUE! |
| ENSMUSG00000085105.3 | 0 | NA | NA | NA | NA | NA | #VALUE! |
| ENSMUSG00000085106.2 | 0 | NA | NA | NA | NA | NA | #VALUE! |
| ENSMUSG00000085107.4 | 0 | NA | NA | NA | NA | NA | #VALUE! |
| ENSMUSG00000085108.2 | 0 | NA | NA | NA | NA | NA | #VALUE! |
| ENSMUSG00000085109.2 | 0 | NA | NA | NA | NA | NA | #VALUE! |
| ENSMUSG00000085110.2 | 0 | NA | NA | NA | NA | NA | #VALUE! |
| ENSMUSG00000085111.4 | 0 | NA | NA | NA | NA | NA | #VALUE! |
| ENSMUSG00000085112.2 | 0 | NA | NA | NA | NA | NA | #VALUE! |
| ENSMUSG00000085113.2 | 0 | NA | NA | NA | NA | NA | #VALUE! |
| ENSMUSG00000085114.2 | 0 | NA | NA | NA | NA | NA | #VALUE! |
| ENSMUSG00000085115.2 | 0 | NA | NA | NA | NA | NA | #VALUE! |
| ENSMUSG00000085116.2 | 0.105013 | -0.97324 | 4.076293 | -0.23876 | 0.811295 | NA | #VALUE! |
| ENSMUSG00000085118.3 | 0 | NA | NA | NA | NA | NA | #VALUE! |
| ENSMUSG00000085120.2 | 0 | NA | NA | NA | NA | NA | #VALUE! |
| ENSMUSG00000085121.3 | 0 | NA | NA | NA | NA | NA | #VALUE! |
| ENSMUSG00000085122.2 | 0 | NA | NA | NA | NA | NA | #VALUE! |
| ENSMUSG00000085123.3 | 0 | NA | NA | NA | NA | NA | #VALUE! |
| ENSMUSG00000085124.2 | 0 | NA | NA | NA | NA | NA | #VALUE! |
| ENSMUSG00000085125.8 | 0 | NA | NA | NA | NA | NA | #VALUE! |
| ENSMUSG00000085126.2 | 0 | NA | NA | NA | NA | NA | #VALUE! |
| ENSMUSG00000085127.2 | 0 | NA | NA | NA | NA | NA | #VALUE! |
| ENSMUSG00000085129.2 | 3.730208 | 5.397701 | 3.918728 | 1.377412 | 0.168385 | NA | #VALUE! |
| ENSMUSG00000085130.2 | 0 | NA | NA | NA | NA | NA | #VALUE! |
| ENSMUSG00000085132.2 | 0 | NA | NA | NA | NA | NA | #VALUE! |

| | | | | | | | |
|---|---|---|---|---|---|---|---|
| ENSMUSG00000085133.3 | 4.152466 | -1.75853 | 3.78818 | -0.46421 | 0.642494 | NA | #VALUE! |
| ENSMUSG00000085134.2 | 0 | NA | NA | NA | NA | NA | #VALUE! |
| ENSMUSG00000085135.8 | 0 | NA | NA | NA | NA | NA | #VALUE! |
| ENSMUSG00000085139.2 | 0 | NA | NA | NA | NA | NA | #VALUE! |
| ENSMUSG00000085141.2 | 0 | NA | NA | NA | NA | NA | #VALUE! |
| ENSMUSG00000085142.2 | 0 | NA | NA | NA | NA | NA | #VALUE! |
| ENSMUSG00000085143.2 | 0 | NA | NA | NA | NA | NA | #VALUE! |
| ENSMUSG00000085145.2 | 0 | NA | NA | NA | NA | NA | #VALUE! |
| ENSMUSG00000085146.2 | 0 | NA | NA | NA | NA | NA | #VALUE! |
| ENSMUSG00000085147.2 | 0 | NA | NA | NA | NA | NA | #VALUE! |
| ENSMUSG00000085150.2 | 0 | NA | NA | NA | NA | NA | #VALUE! |
| ENSMUSG00000085151.8 | 0 | NA | NA | NA | NA | NA | #VALUE! |
| ENSMUSG00000085152.2 | 0 | NA | NA | NA | NA | NA | #VALUE! |
| ENSMUSG00000085154.8 | 0 | NA | NA | NA | NA | NA | #VALUE! |
| ENSMUSG00000085155.4 | 0 | NA | NA | NA | NA | NA | #VALUE! |
| ENSMUSG00000085157.2 | 0 | NA | NA | NA | NA | NA | #VALUE! |
| ENSMUSG00000085158.2 | 0 | NA | NA | NA | NA | NA | #VALUE! |
| ENSMUSG00000085160.2 | 0 | NA | NA | NA | NA | NA | #VALUE! |
| ENSMUSG00000085161.2 | 0 | NA | NA | NA | NA | NA | #VALUE! |
| ENSMUSG00000085162.8 | 0 | NA | NA | NA | NA | NA | #VALUE! |
| ENSMUSG00000085165.2 | 0 | NA | NA | NA | NA | NA | #VALUE! |
| ENSMUSG00000085166.2 | 0 | NA | NA | NA | NA | NA | #VALUE! |
| ENSMUSG00000085167.2 | 0 | NA | NA | NA | NA | NA | #VALUE! |
| ENSMUSG00000085168.3 | 0 | NA | NA | NA | NA | NA | #VALUE! |
| ENSMUSG00000085169.3 | 0.105013 | -0.97324 | 4.076293 | -0.23876 | 0.811295 | NA | #VALUE! |
| ENSMUSG00000085170.2 | 0 | NA | NA | NA | NA | NA | #VALUE! |
| ENSMUSG00000085171.8 | 0 | NA | NA | NA | NA | NA | #VALUE! |
| ENSMUSG00000085173.2 | 0 | NA | NA | NA | NA | NA | #VALUE! |
| ENSMUSG00000085174.2 | 0 | NA | NA | NA | NA | NA | #VALUE! |
| ENSMUSG00000085176.2 | 0 | NA | NA | NA | NA | NA | #VALUE! |
| ENSMUSG00000085177.2 | 0 | NA | NA | NA | NA | NA | #VALUE! |
| ENSMUSG00000085178.2 | 0 | NA | NA | NA | NA | NA | #VALUE! |
| ENSMUSG00000085179.2 | 0 | NA | NA | NA | NA | NA | #VALUE! |
| ENSMUSG00000085180.2 | 0 | NA | NA | NA | NA | NA | #VALUE! |
| ENSMUSG00000085181.2 | 0 | NA | NA | NA | NA | NA | #VALUE! |
| ENSMUSG00000085182.2 | 0 | NA | NA | NA | NA | NA | #VALUE! |
| ENSMUSG00000085183.2 | 0 | NA | NA | NA | NA | NA | #VALUE! |
| ENSMUSG00000085184.10 | 0.210026 | -1.52494 | 4.058692 | -0.37572 | 0.707123 | NA | #VALUE! |
| ENSMUSG00000085185.11 | 1.514552 | 4.088766 | 3.936269 | 1.038742 | 0.298925 | NA | #VALUE! |
| ENSMUSG00000085187.2 | 0 | NA | NA | NA | NA | NA | #VALUE! |
| ENSMUSG00000085188.2 | 0 | NA | NA | NA | NA | NA | #VALUE! |
| ENSMUSG00000085189.2 | 5.236419 | -5.92346 | 3.915282 | -1.51291 | 0.130303 | NA | #VALUE! |
| ENSMUSG00000085192.2 | 0 | NA | NA | NA | NA | NA | #VALUE! |
| ENSMUSG00000085193.2 | 0 | NA | NA | NA | NA | NA | #VALUE! |

| | | | | | | | |
|---|---|---|---|---|---|---|---|
| ENSMUSG00000085194.2 | 0 | NA | NA | NA | NA | NA | #VALUE! |
| ENSMUSG00000085196.2 | 0 | NA | NA | NA | NA | NA | #VALUE! |
| ENSMUSG00000085197.8 | 0 | NA | NA | NA | NA | NA | #VALUE! |
| ENSMUSG00000085198.2 | 0 | NA | NA | NA | NA | NA | #VALUE! |
| ENSMUSG00000085199.2 | 0 | NA | NA | NA | NA | NA | #VALUE! |
| ENSMUSG00000085201.2 | 0 | NA | NA | NA | NA | NA | #VALUE! |
| ENSMUSG00000085202.2 | 0 | NA | NA | NA | NA | NA | #VALUE! |
| ENSMUSG00000085203.2 | 0 | NA | NA | NA | NA | NA | #VALUE! |
| ENSMUSG00000085204.2 | 0 | NA | NA | NA | NA | NA | #VALUE! |
| ENSMUSG00000085205.2 | 0 | NA | NA | NA | NA | NA | #VALUE! |
| ENSMUSG00000085206.2 | 0 | NA | NA | NA | NA | NA | #VALUE! |
| ENSMUSG00000085207.2 | 0 | NA | NA | NA | NA | NA | #VALUE! |
| ENSMUSG00000085209.2 | 0 | NA | NA | NA | NA | NA | #VALUE! |
| ENSMUSG00000085211.3 | 1.009701 | 3.503687 | 3.950852 | 0.886818 | 0.375177 | NA | #VALUE! |
| ENSMUSG00000085213.2 | 0 | NA | NA | NA | NA | NA | #VALUE! |
| ENSMUSG00000085217.2 | 0 | NA | NA | NA | NA | NA | #VALUE! |
| ENSMUSG00000085218.8 | 5.838992 | -4.97043 | 3.900715 | -1.27424 | 0.20258 | NA | #VALUE! |
| ENSMUSG00000085221.3 | 0 | NA | NA | NA | NA | NA | #VALUE! |
| ENSMUSG00000085222.2 | 0 | NA | NA | NA | NA | NA | #VALUE! |
| ENSMUSG00000085223.2 | 0 | NA | NA | NA | NA | NA | #VALUE! |
| ENSMUSG00000085224.3 | 0 | NA | NA | NA | NA | NA | #VALUE! |
| ENSMUSG00000085225.2 | 0 | NA | NA | NA | NA | NA | #VALUE! |
| ENSMUSG00000085226.2 | 0 | NA | NA | NA | NA | NA | #VALUE! |
| ENSMUSG00000085227.2 | 0 | NA | NA | NA | NA | NA | #VALUE! |
| ENSMUSG00000085228.2 | 0 | NA | NA | NA | NA | NA | #VALUE! |
| ENSMUSG00000085229.2 | 0 | NA | NA | NA | NA | NA | #VALUE! |
| ENSMUSG00000085230.2 | 0 | NA | NA | NA | NA | NA | #VALUE! |
| ENSMUSG00000085232.4 | 0 | NA | NA | NA | NA | NA | #VALUE! |
| ENSMUSG00000085234.2 | 0 | NA | NA | NA | NA | NA | #VALUE! |
| ENSMUSG00000085235.2 | 0 | NA | NA | NA | NA | NA | #VALUE! |
| ENSMUSG00000085236.4 | 0 | NA | NA | NA | NA | NA | #VALUE! |
| ENSMUSG00000085237.3 | 0 | NA | NA | NA | NA | NA | #VALUE! |
| ENSMUSG00000085238.2 | 0 | NA | NA | NA | NA | NA | #VALUE! |
| ENSMUSG00000085239.2 | 0 | NA | NA | NA | NA | NA | #VALUE! |
| ENSMUSG00000085240.2 | 0 | NA | NA | NA | NA | NA | #VALUE! |
| ENSMUSG00000085242.2 | 0 | NA | NA | NA | NA | NA | #VALUE! |
| ENSMUSG00000085244.9 | 0 | NA | NA | NA | NA | NA | #VALUE! |
| ENSMUSG00000085245.2 | 0 | NA | NA | NA | NA | NA | #VALUE! |
| ENSMUSG00000085246.2 | 0 | NA | NA | NA | NA | NA | #VALUE! |
| ENSMUSG00000085247.2 | 0 | NA | NA | NA | NA | NA | #VALUE! |
| ENSMUSG00000085248.3 | 0 | NA | NA | NA | NA | NA | #VALUE! |
| ENSMUSG00000085250.2 | 0 | NA | NA | NA | NA | NA | #VALUE! |
| ENSMUSG00000085251.2 | 0 | NA | NA | NA | NA | NA | #VALUE! |
| ENSMUSG00000085252.2 | 0 | NA | NA | NA | NA | NA | #VALUE! |

| | | | | | | | |
|---|---|---|---|---|---|---|---|
| ENSMUSG00000085253.4 | 0 | NA | NA | NA | NA | NA | #VALUE! |
| ENSMUSG00000085254.2 | 0 | NA | NA | NA | NA | NA | #VALUE! |
| ENSMUSG00000085255.2 | 2.792643 | -5.01835 | 3.922691 | -1.27931 | 0.200787 | NA | #VALUE! |
| ENSMUSG00000085256.2 | 0 | NA | NA | NA | NA | NA | #VALUE! |
| ENSMUSG00000085257.3 | 0 | NA | NA | NA | NA | NA | #VALUE! |
| ENSMUSG00000085260.8 | 0 | NA | NA | NA | NA | NA | #VALUE! |
| ENSMUSG00000085261.2 | 0 | NA | NA | NA | NA | NA | #VALUE! |
| ENSMUSG00000085262.2 | 0 | NA | NA | NA | NA | NA | #VALUE! |
| ENSMUSG00000085263.2 | 0 | NA | NA | NA | NA | NA | #VALUE! |
| ENSMUSG00000085264.2 | 0 | NA | NA | NA | NA | NA | #VALUE! |
| ENSMUSG00000085265.2 | 0 | NA | NA | NA | NA | NA | #VALUE! |
| ENSMUSG00000085267.2 | 0 | NA | NA | NA | NA | NA | #VALUE! |
| ENSMUSG00000085269.2 | 0 | NA | NA | NA | NA | NA | #VALUE! |
| ENSMUSG00000085271.8 | 0 | NA | NA | NA | NA | NA | #VALUE! |
| ENSMUSG00000085272.8 | 0 | NA | NA | NA | NA | NA | #VALUE! |
| ENSMUSG00000085274.2 | 0 | NA | NA | NA | NA | NA | #VALUE! |
| ENSMUSG00000085275.2 | 0 | NA | NA | NA | NA | NA | #VALUE! |
| ENSMUSG00000085276.4 | 0 | NA | NA | NA | NA | NA | #VALUE! |
| ENSMUSG00000085277.2 | 0 | NA | NA | NA | NA | NA | #VALUE! |
| ENSMUSG00000085278.2 | 0 | NA | NA | NA | NA | NA | #VALUE! |
| ENSMUSG00000085279.2 | 0 | NA | NA | NA | NA | NA | #VALUE! |
| ENSMUSG00000085280.2 | 0 | NA | NA | NA | NA | NA | #VALUE! |
| ENSMUSG00000085281.4 | 0 | NA | NA | NA | NA | NA | #VALUE! |
| ENSMUSG00000085282.9 | 0 | NA | NA | NA | NA | NA | #VALUE! |
| ENSMUSG00000085284.2 | 0 | NA | NA | NA | NA | NA | #VALUE! |
| ENSMUSG00000085285.2 | 0 | NA | NA | NA | NA | NA | #VALUE! |
| ENSMUSG00000085286.2 | 0 | NA | NA | NA | NA | NA | #VALUE! |
| ENSMUSG00000085288.2 | 0 | NA | NA | NA | NA | NA | #VALUE! |
| ENSMUSG00000085289.2 | 0 | NA | NA | NA | NA | NA | #VALUE! |
| ENSMUSG00000085290.2 | 0 | NA | NA | NA | NA | NA | #VALUE! |
| ENSMUSG00000085292.2 | 0 | NA | NA | NA | NA | NA | #VALUE! |
| ENSMUSG00000085293.3 | 0 | NA | NA | NA | NA | NA | #VALUE! |
| ENSMUSG00000085295.8 | 0 | NA | NA | NA | NA | NA | #VALUE! |
| ENSMUSG00000085297.2 | 0 | NA | NA | NA | NA | NA | #VALUE! |
| ENSMUSG00000085298.3 | 0 | NA | NA | NA | NA | NA | #VALUE! |
| ENSMUSG00000085300.2 | 0 | NA | NA | NA | NA | NA | #VALUE! |
| ENSMUSG00000085301.2 | 0 | NA | NA | NA | NA | NA | #VALUE! |
| ENSMUSG00000085302.4 | 0 | NA | NA | NA | NA | NA | #VALUE! |
| ENSMUSG00000085303.4 | 6.360794 | -0.24784 | 3.754603 | -0.06601 | 0.94737 | NA | #VALUE! |
| ENSMUSG00000085304.2 | 4.181584 | 5.545962 | 3.917562 | 1.415667 | 0.156873 | NA | #VALUE! |
| ENSMUSG00000085305.2 | 0 | NA | NA | NA | NA | NA | #VALUE! |
| ENSMUSG00000085307.2 | 0 | NA | NA | NA | NA | NA | #VALUE! |
| ENSMUSG00000085310.8 | 0 | NA | NA | NA | NA | NA | #VALUE! |
| ENSMUSG00000085311.2 | 0 | NA | NA | NA | NA | NA | #VALUE! |

| | | | | | | | |
|---|---|---|---|---|---|---|---|
| ENSMUSG00000085312.2 | 0 | NA | NA | NA | NA | NA | #VALUE! |
| ENSMUSG00000085314.2 | 0 | NA | NA | NA | NA | NA | #VALUE! |
| ENSMUSG00000085315.8 | 4.541573 | -1.87056 | 3.776086 | -0.49537 | 0.62034 | NA | #VALUE! |
| ENSMUSG00000085316.2 | 0 | NA | NA | NA | NA | NA | #VALUE! |
| ENSMUSG00000085317.2 | 0 | NA | NA | NA | NA | NA | #VALUE! |
| ENSMUSG00000085318.2 | 0 | NA | NA | NA | NA | NA | #VALUE! |
| ENSMUSG00000085319.2 | 0 | NA | NA | NA | NA | NA | #VALUE! |
| ENSMUSG00000085320.2 | 0 | NA | NA | NA | NA | NA | #VALUE! |
| ENSMUSG00000085321.2 | 0 | NA | NA | NA | NA | NA | #VALUE! |
| ENSMUSG00000085322.3 | 0 | NA | NA | NA | NA | NA | #VALUE! |
| ENSMUSG00000085323.2 | 0 | NA | NA | NA | NA | NA | #VALUE! |
| ENSMUSG00000085324.2 | 0 | NA | NA | NA | NA | NA | #VALUE! |
| ENSMUSG00000085325.8 | 0 | NA | NA | NA | NA | NA | #VALUE! |
| ENSMUSG00000085326.2 | 0 | NA | NA | NA | NA | NA | #VALUE! |
| ENSMUSG00000085327.2 | 0 | NA | NA | NA | NA | NA | #VALUE! |
| ENSMUSG00000085328.3 | 0 | NA | NA | NA | NA | NA | #VALUE! |
| ENSMUSG00000085329.2 | 0 | NA | NA | NA | NA | NA | #VALUE! |
| ENSMUSG00000085330.2 | 0 | NA | NA | NA | NA | NA | #VALUE! |
| ENSMUSG00000085331.2 | 0.105013 | -0.97324 | 4.076293 | -0.23876 | 0.811295 | NA | #VALUE! |
| ENSMUSG00000085333.2 | 0 | NA | NA | NA | NA | NA | #VALUE! |
| ENSMUSG00000085334.8 | 2.308672 | 1.393092 | 3.823575 | 0.364343 | 0.715602 | NA | #VALUE! |
| ENSMUSG00000085335.2 | 0 | NA | NA | NA | NA | NA | #VALUE! |
| ENSMUSG00000085336.2 | 0 | NA | NA | NA | NA | NA | #VALUE! |
| ENSMUSG00000085337.2 | 0 | NA | NA | NA | NA | NA | #VALUE! |
| ENSMUSG00000085338.2 | 0 | NA | NA | NA | NA | NA | #VALUE! |
| ENSMUSG00000085340.2 | 0 | NA | NA | NA | NA | NA | #VALUE! |
| ENSMUSG00000085342.2 | 4.355912 | 0.442747 | 2.623833 | 0.16874 | 0.866001 | NA | #VALUE! |
| ENSMUSG00000085345.2 | 0 | NA | NA | NA | NA | NA | #VALUE! |
| ENSMUSG00000085347.2 | 0 | NA | NA | NA | NA | NA | #VALUE! |
| ENSMUSG00000085348.2 | 0 | NA | NA | NA | NA | NA | #VALUE! |
| ENSMUSG00000085350.2 | 0 | NA | NA | NA | NA | NA | #VALUE! |
| ENSMUSG00000085351.2 | 0 | NA | NA | NA | NA | NA | #VALUE! |
| ENSMUSG00000085352.2 | 0 | NA | NA | NA | NA | NA | #VALUE! |
| ENSMUSG00000085353.3 | 0 | NA | NA | NA | NA | NA | #VALUE! |
| ENSMUSG00000085354.2 | 0 | NA | NA | NA | NA | NA | #VALUE! |
| ENSMUSG00000085355.3 | 0 | NA | NA | NA | NA | NA | #VALUE! |
| ENSMUSG00000085357.2 | 0 | NA | NA | NA | NA | NA | #VALUE! |
| ENSMUSG00000085358.2 | 0 | NA | NA | NA | NA | NA | #VALUE! |
| ENSMUSG00000085359.2 | 0 | NA | NA | NA | NA | NA | #VALUE! |
| ENSMUSG00000085360.2 | 0 | NA | NA | NA | NA | NA | #VALUE! |
| ENSMUSG00000085361.3 | 0 | NA | NA | NA | NA | NA | #VALUE! |
| ENSMUSG00000085363.2 | 0 | NA | NA | NA | NA | NA | #VALUE! |
| ENSMUSG00000085366.2 | 0 | NA | NA | NA | NA | NA | #VALUE! |
| ENSMUSG00000085368.2 | 0 | NA | NA | NA | NA | NA | #VALUE! |

| | | | | | | | |
|---|---|---|---|---|---|---|---|
| ENSMUSG00000085369.2 | 0 | NA | NA | NA | NA | NA | #VALUE! |
| ENSMUSG00000085371.2 | 0 | NA | NA | NA | NA | NA | #VALUE! |
| ENSMUSG00000085372.2 | 0 | NA | NA | NA | NA | NA | #VALUE! |
| ENSMUSG00000085375.2 | 0 | NA | NA | NA | NA | NA | #VALUE! |
| ENSMUSG00000085376.2 | 0 | NA | NA | NA | NA | NA | #VALUE! |
| ENSMUSG00000085377.2 | 0 | NA | NA | NA | NA | NA | #VALUE! |
| ENSMUSG00000085378.2 | 0 | NA | NA | NA | NA | NA | #VALUE! |
| ENSMUSG00000085379.2 | 0.408222 | -2.20205 | 4.016842 | -0.54821 | 0.583551 | NA | #VALUE! |
| ENSMUSG00000085380.2 | 0 | NA | NA | NA | NA | NA | #VALUE! |
| ENSMUSG00000085382.2 | 0 | NA | NA | NA | NA | NA | #VALUE! |
| ENSMUSG00000085383.2 | 0 | NA | NA | NA | NA | NA | #VALUE! |
| ENSMUSG00000085386.2 | 0 | NA | NA | NA | NA | NA | #VALUE! |
| ENSMUSG00000085387.2 | 0 | NA | NA | NA | NA | NA | #VALUE! |
| ENSMUSG00000085388.2 | 0 | NA | NA | NA | NA | NA | #VALUE! |
| ENSMUSG00000085389.8 | 0 | NA | NA | NA | NA | NA | #VALUE! |
| ENSMUSG00000085390.2 | 0 | NA | NA | NA | NA | NA | #VALUE! |
| ENSMUSG00000085391.2 | 0 | NA | NA | NA | NA | NA | #VALUE! |
| ENSMUSG00000085393.2 | 0 | NA | NA | NA | NA | NA | #VALUE! |
| ENSMUSG00000085394.2 | 0 | NA | NA | NA | NA | NA | #VALUE! |
| ENSMUSG00000085395.2 | 0 | NA | NA | NA | NA | NA | #VALUE! |
| ENSMUSG00000085397.2 | 0 | NA | NA | NA | NA | NA | #VALUE! |
| ENSMUSG00000085398.2 | 0 | NA | NA | NA | NA | NA | #VALUE! |
| ENSMUSG00000085399.8 | 0.978946 | -3.48749 | 3.952505 | -0.88235 | 0.377588 | NA | #VALUE! |
| ENSMUSG00000085401.3 | 0 | NA | NA | NA | NA | NA | #VALUE! |
| ENSMUSG00000085402.2 | 0 | NA | NA | NA | NA | NA | #VALUE! |
| ENSMUSG00000085403.2 | 0 | NA | NA | NA | NA | NA | #VALUE! |
| ENSMUSG00000085405.2 | 0 | NA | NA | NA | NA | NA | #VALUE! |
| ENSMUSG00000085407.2 | 0 | NA | NA | NA | NA | NA | #VALUE! |
| ENSMUSG00000085408.2 | 0 | NA | NA | NA | NA | NA | #VALUE! |
| ENSMUSG00000085409.2 | 0 | NA | NA | NA | NA | NA | #VALUE! |
| ENSMUSG00000085411.2 | 0 | NA | NA | NA | NA | NA | #VALUE! |
| ENSMUSG00000085412.8 | 0 | NA | NA | NA | NA | NA | #VALUE! |
| ENSMUSG00000085413.2 | 0 | NA | NA | NA | NA | NA | #VALUE! |
| ENSMUSG00000085414.2 | 0 | NA | NA | NA | NA | NA | #VALUE! |
| ENSMUSG00000085415.2 | 0.204111 | -1.05581 | 4.073302 | -0.2592 | 0.795479 | NA | #VALUE! |
| ENSMUSG00000085416.2 | 0 | NA | NA | NA | NA | NA | #VALUE! |
| ENSMUSG00000085417.2 | 0 | NA | NA | NA | NA | NA | #VALUE! |
| ENSMUSG00000085418.2 | 0 | NA | NA | NA | NA | NA | #VALUE! |
| ENSMUSG00000085419.2 | 0 | NA | NA | NA | NA | NA | #VALUE! |
| ENSMUSG00000085420.2 | 0 | NA | NA | NA | NA | NA | #VALUE! |
| ENSMUSG00000085421.2 | 0 | NA | NA | NA | NA | NA | #VALUE! |
| ENSMUSG00000085423.2 | 0 | NA | NA | NA | NA | NA | #VALUE! |
| ENSMUSG00000085424.2 | 0 | NA | NA | NA | NA | NA | #VALUE! |
| ENSMUSG00000085426.2 | 0 | NA | NA | NA | NA | NA | #VALUE! |

| | | | | | | | |
|---|---|---|---|---|---|---|---|
| ENSMUSG00000085427.8 | 0.746042 | 3.107177 | 3.964586 | 0.783733 | 0.433197 | NA | #VALUE! |
| ENSMUSG00000085428.2 | 0 | NA | NA | NA | NA | NA | #VALUE! |
| ENSMUSG00000085429.2 | 0 | NA | NA | NA | NA | NA | #VALUE! |
| ENSMUSG00000085430.3 | 0 | NA | NA | NA | NA | NA | #VALUE! |
| ENSMUSG00000085431.8 | 0 | NA | NA | NA | NA | NA | #VALUE! |
| ENSMUSG00000085433.3 | 0 | NA | NA | NA | NA | NA | #VALUE! |
| ENSMUSG00000085434.2 | 0 | NA | NA | NA | NA | NA | #VALUE! |
| ENSMUSG00000085436.2 | 0 | NA | NA | NA | NA | NA | #VALUE! |
| ENSMUSG00000085437.9 | 0 | NA | NA | NA | NA | NA | #VALUE! |
| ENSMUSG00000085439.2 | 0 | NA | NA | NA | NA | NA | #VALUE! |
| ENSMUSG00000085440.2 | 0 | NA | NA | NA | NA | NA | #VALUE! |
| ENSMUSG00000085441.2 | 0 | NA | NA | NA | NA | NA | #VALUE! |
| ENSMUSG00000085443.2 | 0 | NA | NA | NA | NA | NA | #VALUE! |
| ENSMUSG00000085444.2 | 0 | NA | NA | NA | NA | NA | #VALUE! |
| ENSMUSG00000085445.2 | 0 | NA | NA | NA | NA | NA | #VALUE! |
| ENSMUSG00000085447.2 | 0 | NA | NA | NA | NA | NA | #VALUE! |
| ENSMUSG00000085448.2 | 0 | NA | NA | NA | NA | NA | #VALUE! |
| ENSMUSG00000085449.2 | 0 | NA | NA | NA | NA | NA | #VALUE! |
| ENSMUSG00000085450.2 | 0 | NA | NA | NA | NA | NA | #VALUE! |
| ENSMUSG00000085451.3 | 0 | NA | NA | NA | NA | NA | #VALUE! |
| ENSMUSG00000085452.2 | 0 | NA | NA | NA | NA | NA | #VALUE! |
| ENSMUSG00000085453.2 | 0 | NA | NA | NA | NA | NA | #VALUE! |
| ENSMUSG00000085454.2 | 0 | NA | NA | NA | NA | NA | #VALUE! |
| ENSMUSG00000085456.3 | 0 | NA | NA | NA | NA | NA | #VALUE! |
| ENSMUSG00000085457.4 | 0 | NA | NA | NA | NA | NA | #VALUE! |
| ENSMUSG00000085459.2 | 0 | NA | NA | NA | NA | NA | #VALUE! |
| ENSMUSG00000085463.2 | 0 | NA | NA | NA | NA | NA | #VALUE! |
| ENSMUSG00000085464.2 | 0 | NA | NA | NA | NA | NA | #VALUE! |
| ENSMUSG00000085465.2 | 0 | NA | NA | NA | NA | NA | #VALUE! |
| ENSMUSG00000085466.3 | 0 | NA | NA | NA | NA | NA | #VALUE! |
| ENSMUSG00000085467.2 | 0 | NA | NA | NA | NA | NA | #VALUE! |
| ENSMUSG00000085468.2 | 6.458635 | -1.08352 | 3.305466 | -0.3278 | 0.743067 | NA | #VALUE! |
| ENSMUSG00000085469.2 | 0 | NA | NA | NA | NA | NA | #VALUE! |
| ENSMUSG00000085470.2 | 0 | NA | NA | NA | NA | NA | #VALUE! |
| ENSMUSG00000085471.2 | 0 | NA | NA | NA | NA | NA | #VALUE! |
| ENSMUSG00000085472.2 | 0 | NA | NA | NA | NA | NA | #VALUE! |
| ENSMUSG00000085476.2 | 0 | NA | NA | NA | NA | NA | #VALUE! |
| ENSMUSG00000085477.8 | 0 | NA | NA | NA | NA | NA | #VALUE! |
| ENSMUSG00000085478.2 | 0 | NA | NA | NA | NA | NA | #VALUE! |
| ENSMUSG00000085479.2 | 0 | NA | NA | NA | NA | NA | #VALUE! |
| ENSMUSG00000085481.2 | 0 | NA | NA | NA | NA | NA | #VALUE! |
| ENSMUSG00000085482.2 | 0 | NA | NA | NA | NA | NA | #VALUE! |
| ENSMUSG00000085483.2 | 0 | NA | NA | NA | NA | NA | #VALUE! |
| ENSMUSG00000085484.3 | 0 | NA | NA | NA | NA | NA | #VALUE! |

| | | | | | | | |
|---|---|---|---|---|---|---|---|
| ENSMUSG00000085485.2 | 0 | NA | NA | NA | NA | NA | #VALUE! |
| ENSMUSG00000085486.3 | 0 | NA | NA | NA | NA | NA | #VALUE! |
| ENSMUSG00000085487.2 | 0 | NA | NA | NA | NA | NA | #VALUE! |
| ENSMUSG00000085488.2 | 0 | NA | NA | NA | NA | NA | #VALUE! |
| ENSMUSG00000085489.8 | 0 | NA | NA | NA | NA | NA | #VALUE! |
| ENSMUSG00000085491.2 | 0 | NA | NA | NA | NA | NA | #VALUE! |
| ENSMUSG00000085495.2 | 0 | NA | NA | NA | NA | NA | #VALUE! |
| ENSMUSG00000085496.2 | 0 | NA | NA | NA | NA | NA | #VALUE! |
| ENSMUSG00000085497.3 | 0 | NA | NA | NA | NA | NA | #VALUE! |
| ENSMUSG00000085498.2 | 2.63165 | -3.76871 | 3.910942 | -0.96363 | 0.335231 | NA | #VALUE! |
| ENSMUSG00000085499.2 | 0 | NA | NA | NA | NA | NA | #VALUE! |
| ENSMUSG00000085500.10 | 0 | NA | NA | NA | NA | NA | #VALUE! |
| ENSMUSG00000085501.2 | 2.805471 | -3.81467 | 3.910376 | -0.97553 | 0.3293 | NA | #VALUE! |
| ENSMUSG00000085502.2 | 0 | NA | NA | NA | NA | NA | #VALUE! |
| ENSMUSG00000085504.2 | 0 | NA | NA | NA | NA | NA | #VALUE! |
| ENSMUSG00000085505.2 | 0 | NA | NA | NA | NA | NA | #VALUE! |
| ENSMUSG00000085507.2 | 0 | NA | NA | NA | NA | NA | #VALUE! |
| ENSMUSG00000085509.2 | 0 | NA | NA | NA | NA | NA | #VALUE! |
| ENSMUSG00000085510.2 | 0 | NA | NA | NA | NA | NA | #VALUE! |
| ENSMUSG00000085511.2 | 0 | NA | NA | NA | NA | NA | #VALUE! |
| ENSMUSG00000085512.2 | 0 | NA | NA | NA | NA | NA | #VALUE! |
| ENSMUSG00000085513.2 | 0 | NA | NA | NA | NA | NA | #VALUE! |
| ENSMUSG00000085514.2 | 0 | NA | NA | NA | NA | NA | #VALUE! |
| ENSMUSG00000085515.2 | 0 | NA | NA | NA | NA | NA | #VALUE! |
| ENSMUSG00000085516.2 | 0 | NA | NA | NA | NA | NA | #VALUE! |
| ENSMUSG00000085517.8 | 0 | NA | NA | NA | NA | NA | #VALUE! |
| ENSMUSG00000085519.2 | 0 | NA | NA | NA | NA | NA | #VALUE! |
| ENSMUSG00000085521.5 | 0 | NA | NA | NA | NA | NA | #VALUE! |
| ENSMUSG00000085522.2 | 0 | NA | NA | NA | NA | NA | #VALUE! |
| ENSMUSG00000085523.2 | 0 | NA | NA | NA | NA | NA | #VALUE! |
| ENSMUSG00000085524.2 | 0 | NA | NA | NA | NA | NA | #VALUE! |
| ENSMUSG00000085526.2 | 0 | NA | NA | NA | NA | NA | #VALUE! |
| ENSMUSG00000085527.2 | 0 | NA | NA | NA | NA | NA | #VALUE! |
| ENSMUSG00000085528.2 | 0 | NA | NA | NA | NA | NA | #VALUE! |
| ENSMUSG00000085530.2 | 0 | NA | NA | NA | NA | NA | #VALUE! |
| ENSMUSG00000085531.2 | 0 | NA | NA | NA | NA | NA | #VALUE! |
| ENSMUSG00000085532.2 | 0 | NA | NA | NA | NA | NA | #VALUE! |
| ENSMUSG00000085533.2 | 0 | NA | NA | NA | NA | NA | #VALUE! |
| ENSMUSG00000085534.8 | 0 | NA | NA | NA | NA | NA | #VALUE! |
| ENSMUSG00000085536.2 | 0 | NA | NA | NA | NA | NA | #VALUE! |
| ENSMUSG00000085538.2 | 0 | NA | NA | NA | NA | NA | #VALUE! |
| ENSMUSG00000085539.2 | 0 | NA | NA | NA | NA | NA | #VALUE! |
| ENSMUSG00000085540.2 | 0 | NA | NA | NA | NA | NA | #VALUE! |
| ENSMUSG00000085541.2 | 0 | NA | NA | NA | NA | NA | #VALUE! |

| | | | | | | | |
|---|---|---|---|---|---|---|---|
| ENSMUSG00000085542.2 | 0 | NA | NA | NA | NA | NA | #VALUE! |
| ENSMUSG00000085543.2 | 3.702238 | 5.378281 | 3.918889 | 1.372399 | 0.169939 | NA | #VALUE! |
| ENSMUSG00000085545.2 | 0 | NA | NA | NA | NA | NA | #VALUE! |
| ENSMUSG00000085546.2 | 0 | NA | NA | NA | NA | NA | #VALUE! |
| ENSMUSG00000085547.5 | 0 | NA | NA | NA | NA | NA | #VALUE! |
| ENSMUSG00000085548.2 | 0 | NA | NA | NA | NA | NA | #VALUE! |
| ENSMUSG00000085549.8 | 0 | NA | NA | NA | NA | NA | #VALUE! |
| ENSMUSG00000085551.2 | 0 | NA | NA | NA | NA | NA | #VALUE! |
| ENSMUSG00000085552.2 | 0 | NA | NA | NA | NA | NA | #VALUE! |
| ENSMUSG00000085553.2 | 0 | NA | NA | NA | NA | NA | #VALUE! |
| ENSMUSG00000085555.10 | 0 | NA | NA | NA | NA | NA | #VALUE! |
| ENSMUSG00000085556.2 | 0 | NA | NA | NA | NA | NA | #VALUE! |
| ENSMUSG00000085558.8 | 1.737899 | -4.32008 | 3.932553 | -1.09854 | 0.271968 | NA | #VALUE! |
| ENSMUSG00000085559.2 | 0 | NA | NA | NA | NA | NA | #VALUE! |
| ENSMUSG00000085560.3 | 0 | NA | NA | NA | NA | NA | #VALUE! |
| ENSMUSG00000085561.2 | 0 | NA | NA | NA | NA | NA | #VALUE! |
| ENSMUSG00000085562.8 | 0 | NA | NA | NA | NA | NA | #VALUE! |
| ENSMUSG00000085563.3 | 0 | NA | NA | NA | NA | NA | #VALUE! |
| ENSMUSG00000085564.2 | 0 | NA | NA | NA | NA | NA | #VALUE! |
| ENSMUSG00000085565.2 | 0 | NA | NA | NA | NA | NA | #VALUE! |
| ENSMUSG00000085566.2 | 1.174735 | -3.75246 | 3.944889 | -0.95122 | 0.341493 | NA | #VALUE! |
| ENSMUSG00000085568.8 | 0 | NA | NA | NA | NA | NA | #VALUE! |
| ENSMUSG00000085569.2 | 0 | NA | NA | NA | NA | NA | #VALUE! |
| ENSMUSG00000085572.2 | 0 | NA | NA | NA | NA | NA | #VALUE! |
| ENSMUSG00000085573.2 | 0 | NA | NA | NA | NA | NA | #VALUE! |
| ENSMUSG00000085574.2 | 0 | NA | NA | NA | NA | NA | #VALUE! |
| ENSMUSG00000085575.2 | 0 | NA | NA | NA | NA | NA | #VALUE! |
| ENSMUSG00000085576.4 | 0 | NA | NA | NA | NA | NA | #VALUE! |
| ENSMUSG00000085577.2 | 0 | NA | NA | NA | NA | NA | #VALUE! |
| ENSMUSG00000085578.2 | 0 | NA | NA | NA | NA | NA | #VALUE! |
| ENSMUSG00000085579.2 | 0 | NA | NA | NA | NA | NA | #VALUE! |
| ENSMUSG00000085580.2 | 0 | NA | NA | NA | NA | NA | #VALUE! |
| ENSMUSG00000085581.2 | 0 | NA | NA | NA | NA | NA | #VALUE! |
| ENSMUSG00000085582.8 | 0 | NA | NA | NA | NA | NA | #VALUE! |
| ENSMUSG00000085584.4 | 0 | NA | NA | NA | NA | NA | #VALUE! |
| ENSMUSG00000085585.2 | 0 | NA | NA | NA | NA | NA | #VALUE! |
| ENSMUSG00000085586.2 | 0 | NA | NA | NA | NA | NA | #VALUE! |
| ENSMUSG00000085587.2 | 0 | NA | NA | NA | NA | NA | #VALUE! |
| ENSMUSG00000085588.2 | 0 | NA | NA | NA | NA | NA | #VALUE! |
| ENSMUSG00000085589.8 | 0 | NA | NA | NA | NA | NA | #VALUE! |
| ENSMUSG00000085591.2 | 0 | NA | NA | NA | NA | NA | #VALUE! |
| ENSMUSG00000085592.2 | 0 | NA | NA | NA | NA | NA | #VALUE! |
| ENSMUSG00000085593.2 | 0 | NA | NA | NA | NA | NA | #VALUE! |
| ENSMUSG00000085594.2 | 0 | NA | NA | NA | NA | NA | #VALUE! |

| | | | | | | | |
|---|---|---|---|---|---|---|---|
| ENSMUSG00000085595.2 | 0 | NA | NA | NA | NA | NA | #VALUE! |
| ENSMUSG00000085596.2 | 0 | NA | NA | NA | NA | NA | #VALUE! |
| ENSMUSG00000085598.2 | 0 | NA | NA | NA | NA | NA | #VALUE! |
| ENSMUSG00000085599.2 | 0 | NA | NA | NA | NA | NA | #VALUE! |
| ENSMUSG00000085600.8 | 0 | NA | NA | NA | NA | NA | #VALUE! |
| ENSMUSG00000085601.8 | 0 | NA | NA | NA | NA | NA | #VALUE! |
| ENSMUSG00000085603.4 | 0 | NA | NA | NA | NA | NA | #VALUE! |
| ENSMUSG00000085604.2 | 0 | NA | NA | NA | NA | NA | #VALUE! |
| ENSMUSG00000085606.2 | 0 | NA | NA | NA | NA | NA | #VALUE! |
| ENSMUSG00000085608.2 | 0 | NA | NA | NA | NA | NA | #VALUE! |
| ENSMUSG00000085609.2 | 0 | NA | NA | NA | NA | NA | #VALUE! |
| ENSMUSG00000085610.2 | 0 | NA | NA | NA | NA | NA | #VALUE! |
| ENSMUSG00000085611.4 | 0 | NA | NA | NA | NA | NA | #VALUE! |
| ENSMUSG00000085612.2 | 0 | NA | NA | NA | NA | NA | #VALUE! |
| ENSMUSG00000085614.2 | 0 | NA | NA | NA | NA | NA | #VALUE! |
| ENSMUSG00000085615.3 | 6.126873 | 6.107425 | 3.670344 | 1.663992 | 0.096114 | NA | #VALUE! |
| ENSMUSG00000085616.2 | 0 | NA | NA | NA | NA | NA | #VALUE! |
| ENSMUSG00000085618.2 | 0 | NA | NA | NA | NA | NA | #VALUE! |
| ENSMUSG00000085619.2 | 0 | NA | NA | NA | NA | NA | #VALUE! |
| ENSMUSG00000085620.3 | 0 | NA | NA | NA | NA | NA | #VALUE! |
| ENSMUSG00000085622.10 | 7.695772 | 2.788217 | 2.718324 | 1.025712 | 0.305028 | NA | #VALUE! |
| ENSMUSG00000085623.2 | 0 | NA | NA | NA | NA | NA | #VALUE! |
| ENSMUSG00000085624.2 | 0 | NA | NA | NA | NA | NA | #VALUE! |
| ENSMUSG00000085625.2 | 0 | NA | NA | NA | NA | NA | #VALUE! |
| ENSMUSG00000085626.2 | 0 | NA | NA | NA | NA | NA | #VALUE! |
| ENSMUSG00000085627.3 | 0 | NA | NA | NA | NA | NA | #VALUE! |
| ENSMUSG00000085628.5 | 6.082279 | -3.51435 | 3.467611 | -1.01348 | 0.310832 | NA | #VALUE! |
| ENSMUSG00000085629.2 | 0 | NA | NA | NA | NA | NA | #VALUE! |
| ENSMUSG00000085630.2 | 0 | NA | NA | NA | NA | NA | #VALUE! |
| ENSMUSG00000085631.2 | 0 | NA | NA | NA | NA | NA | #VALUE! |
| ENSMUSG00000085632.2 | 0 | NA | NA | NA | NA | NA | #VALUE! |
| ENSMUSG00000085633.2 | 0.336567 | 1.944925 | 4.029936 | 0.482619 | 0.629366 | NA | #VALUE! |
| ENSMUSG00000085634.3 | 0 | NA | NA | NA | NA | NA | #VALUE! |
| ENSMUSG00000085635.2 | 0 | NA | NA | NA | NA | NA | #VALUE! |
| ENSMUSG00000085636.2 | 0.735092 | -3.12016 | 3.965612 | -0.7868 | 0.431397 | NA | #VALUE! |
| ENSMUSG00000085637.3 | 0 | NA | NA | NA | NA | NA | #VALUE! |
| ENSMUSG00000085638.2 | 0 | NA | NA | NA | NA | NA | #VALUE! |
| ENSMUSG00000085639.2 | 0 | NA | NA | NA | NA | NA | #VALUE! |
| ENSMUSG00000085640.2 | 0 | NA | NA | NA | NA | NA | #VALUE! |
| ENSMUSG00000085641.2 | 0 | NA | NA | NA | NA | NA | #VALUE! |
| ENSMUSG00000085642.9 | 13.23113 | 6.268956 | 3.072552 | 2.040309 | NA | NA | #VALUE! |
| ENSMUSG00000085643.2 | 0 | NA | NA | NA | NA | NA | #VALUE! |
| ENSMUSG00000085645.2 | 0 | NA | NA | NA | NA | NA | #VALUE! |
| ENSMUSG00000085646.2 | 0 | NA | NA | NA | NA | NA | #VALUE! |

| | | | | | | | |
|---|---|---|---|---|---|---|---|
| ENSMUSG00000085648.2 | 0 | NA | NA | NA | NA | NA | #VALUE! |
| ENSMUSG00000085649.8 | 0 | NA | NA | NA | NA | NA | #VALUE! |
| ENSMUSG00000085651.2 | 0 | NA | NA | NA | NA | NA | #VALUE! |
| ENSMUSG00000085652.2 | 0 | NA | NA | NA | NA | NA | #VALUE! |
| ENSMUSG00000085653.8 | 0 | NA | NA | NA | NA | NA | #VALUE! |
| ENSMUSG00000085654.2 | 0 | NA | NA | NA | NA | NA | #VALUE! |
| ENSMUSG00000085656.2 | 0 | NA | NA | NA | NA | NA | #VALUE! |
| ENSMUSG00000085657.8 | 0 | NA | NA | NA | NA | NA | #VALUE! |
| ENSMUSG00000085658.2 | 0 | NA | NA | NA | NA | NA | #VALUE! |
| ENSMUSG00000085659.3 | 0 | NA | NA | NA | NA | NA | #VALUE! |
| ENSMUSG00000085660.2 | 0 | NA | NA | NA | NA | NA | #VALUE! |
| ENSMUSG00000085662.2 | 0 | NA | NA | NA | NA | NA | #VALUE! |
| ENSMUSG00000085663.2 | 0 | NA | NA | NA | NA | NA | #VALUE! |
| ENSMUSG00000085664.2 | 0 | NA | NA | NA | NA | NA | #VALUE! |
| ENSMUSG00000085665.2 | 6.206724 | -2.83665 | 3.79399 | -0.74767 | 0.45466 | NA | #VALUE! |
| ENSMUSG00000085666.4 | 0 | NA | NA | NA | NA | NA | #VALUE! |
| ENSMUSG00000085667.2 | 7.236192 | 6.345076 | 3.912976 | 1.621547 | 0.1049 | NA | #VALUE! |
| ENSMUSG00000085668.3 | 0 | NA | NA | NA | NA | NA | #VALUE! |
| ENSMUSG00000085669.2 | 0 | NA | NA | NA | NA | NA | #VALUE! |
| ENSMUSG00000085672.2 | 0 | NA | NA | NA | NA | NA | #VALUE! |
| ENSMUSG00000085673.2 | 0 | NA | NA | NA | NA | NA | #VALUE! |
| ENSMUSG00000085675.2 | 0 | NA | NA | NA | NA | NA | #VALUE! |
| ENSMUSG00000085676.2 | 0 | NA | NA | NA | NA | NA | #VALUE! |
| ENSMUSG00000085677.8 | 0 | NA | NA | NA | NA | NA | #VALUE! |
| ENSMUSG00000085678.2 | 0 | NA | NA | NA | NA | NA | #VALUE! |
| ENSMUSG00000085679.3 | 0 | NA | NA | NA | NA | NA | #VALUE! |
| ENSMUSG00000085680.2 | 0 | NA | NA | NA | NA | NA | #VALUE! |
| ENSMUSG00000085681.2 | 0 | NA | NA | NA | NA | NA | #VALUE! |
| ENSMUSG00000085682.3 | 0.093255 | 0.868093 | 4.072464 | 0.213162 | 0.831201 | NA | #VALUE! |
| ENSMUSG00000085683.3 | 0 | NA | NA | NA | NA | NA | #VALUE! |
| ENSMUSG00000085684.3 | 0 | NA | NA | NA | NA | NA | #VALUE! |
| ENSMUSG00000085686.2 | 0 | NA | NA | NA | NA | NA | #VALUE! |
| ENSMUSG00000085687.2 | 0 | NA | NA | NA | NA | NA | #VALUE! |
| ENSMUSG00000085688.2 | 0 | NA | NA | NA | NA | NA | #VALUE! |
| ENSMUSG00000085689.2 | 0 | NA | NA | NA | NA | NA | #VALUE! |
| ENSMUSG00000085690.2 | 0 | NA | NA | NA | NA | NA | #VALUE! |
| ENSMUSG00000085691.2 | 0 | NA | NA | NA | NA | NA | #VALUE! |
| ENSMUSG00000085693.2 | 0 | NA | NA | NA | NA | NA | #VALUE! |
| ENSMUSG00000085695.2 | 0 | NA | NA | NA | NA | NA | #VALUE! |
| ENSMUSG00000085696.9 | 0 | NA | NA | NA | NA | NA | #VALUE! |
| ENSMUSG00000085699.2 | 0 | NA | NA | NA | NA | NA | #VALUE! |
| ENSMUSG00000085700.2 | 0 | NA | NA | NA | NA | NA | #VALUE! |
| ENSMUSG00000085701.2 | 0 | NA | NA | NA | NA | NA | #VALUE! |
| ENSMUSG00000085702.9 | 0 | NA | NA | NA | NA | NA | #VALUE! |

| | | | | | | | |
|---|---|---|---|---|---|---|---|
| ENSMUSG00000085705.2 | 0 | NA | NA | NA | NA | NA | #VALUE! |
| ENSMUSG00000085706.2 | 0 | NA | NA | NA | NA | NA | #VALUE! |
| ENSMUSG00000085707.2 | 0 | NA | NA | NA | NA | NA | #VALUE! |
| ENSMUSG00000085708.2 | 0.587367 | -2.74297 | 3.982932 | -0.68868 | 0.491024 | NA | #VALUE! |
| ENSMUSG00000085709.2 | 0 | NA | NA | NA | NA | NA | #VALUE! |
| ENSMUSG00000085710.2 | 0 | NA | NA | NA | NA | NA | #VALUE! |
| ENSMUSG00000085711.5 | 0.273297 | 0.094602 | 4.062933 | 0.023284 | 0.981424 | NA | #VALUE! |
| ENSMUSG00000085712.2 | 0 | NA | NA | NA | NA | NA | #VALUE! |
| ENSMUSG00000085713.2 | 0 | NA | NA | NA | NA | NA | #VALUE! |
| ENSMUSG00000085714.2 | 0 | NA | NA | NA | NA | NA | #VALUE! |
| ENSMUSG00000085715.3 | 0 | NA | NA | NA | NA | NA | #VALUE! |
| ENSMUSG00000085716.2 | 0 | NA | NA | NA | NA | NA | #VALUE! |
| ENSMUSG00000085717.2 | 0 | NA | NA | NA | NA | NA | #VALUE! |
| ENSMUSG00000085718.3 | 0 | NA | NA | NA | NA | NA | #VALUE! |
| ENSMUSG00000085719.2 | 0 | NA | NA | NA | NA | NA | #VALUE! |
| ENSMUSG00000085720.9 | 0 | NA | NA | NA | NA | NA | #VALUE! |
| ENSMUSG00000085721.2 | 0 | NA | NA | NA | NA | NA | #VALUE! |
| ENSMUSG00000085722.5 | 0 | NA | NA | NA | NA | NA | #VALUE! |
| ENSMUSG00000085723.3 | 0 | NA | NA | NA | NA | NA | #VALUE! |
| ENSMUSG00000085724.2 | 0 | NA | NA | NA | NA | NA | #VALUE! |
| ENSMUSG00000085725.3 | 0 | NA | NA | NA | NA | NA | #VALUE! |
| ENSMUSG00000085728.2 | 0 | NA | NA | NA | NA | NA | #VALUE! |
| ENSMUSG00000085730.8 | 0 | NA | NA | NA | NA | NA | #VALUE! |
| ENSMUSG00000085731.2 | 0 | NA | NA | NA | NA | NA | #VALUE! |
| ENSMUSG00000085732.2 | 0 | NA | NA | NA | NA | NA | #VALUE! |
| ENSMUSG00000085733.2 | 0 | NA | NA | NA | NA | NA | #VALUE! |
| ENSMUSG00000085734.2 | 0 | NA | NA | NA | NA | NA | #VALUE! |
| ENSMUSG00000085737.2 | 0 | NA | NA | NA | NA | NA | #VALUE! |
| ENSMUSG00000085738.3 | 0 | NA | NA | NA | NA | NA | #VALUE! |
| ENSMUSG00000085739.2 | 0 | NA | NA | NA | NA | NA | #VALUE! |
| ENSMUSG00000085740.2 | 0 | NA | NA | NA | NA | NA | #VALUE! |
| ENSMUSG00000085741.9 | 25.01598 | -0.99734 | 1.970577 | -0.50611 | NA | NA | #VALUE! |
| ENSMUSG00000085742.2 | 0 | NA | NA | NA | NA | NA | #VALUE! |
| ENSMUSG00000085743.2 | 0 | NA | NA | NA | NA | NA | #VALUE! |
| ENSMUSG00000085744.2 | 0 | NA | NA | NA | NA | NA | #VALUE! |
| ENSMUSG00000085745.2 | 0 | NA | NA | NA | NA | NA | #VALUE! |
| ENSMUSG00000085747.2 | 0 | NA | NA | NA | NA | NA | #VALUE! |
| ENSMUSG00000085748.2 | 0 | NA | NA | NA | NA | NA | #VALUE! |
| ENSMUSG00000085749.2 | 0 | NA | NA | NA | NA | NA | #VALUE! |
| ENSMUSG00000085750.2 | 0 | NA | NA | NA | NA | NA | #VALUE! |
| ENSMUSG00000085751.2 | 0 | NA | NA | NA | NA | NA | #VALUE! |
| ENSMUSG00000085753.2 | 0 | NA | NA | NA | NA | NA | #VALUE! |
| ENSMUSG00000085754.8 | 0 | NA | NA | NA | NA | NA | #VALUE! |
| ENSMUSG00000085755.2 | 0 | NA | NA | NA | NA | NA | #VALUE! |

| | | | | | | | |
|---|---|---|---|---|---|---|---|
| ENSMUSG00000085756.2 | 0 | NA | NA | NA | NA | NA | #VALUE! |
| ENSMUSG00000085757.8 | 0 | NA | NA | NA | NA | NA | #VALUE! |
| ENSMUSG00000085759.2 | 0 | NA | NA | NA | NA | NA | #VALUE! |
| ENSMUSG00000085760.2 | 0 | NA | NA | NA | NA | NA | #VALUE! |
| ENSMUSG00000085761.2 | 1.771849 | 4.332864 | 3.931707 | 1.102031 | 0.270448 | NA | #VALUE! |
| ENSMUSG00000085762.2 | 0 | NA | NA | NA | NA | NA | #VALUE! |
| ENSMUSG00000085763.2 | 0 | NA | NA | NA | NA | NA | #VALUE! |
| ENSMUSG00000085765.2 | 0 | NA | NA | NA | NA | NA | #VALUE! |
| ENSMUSG00000085766.2 | 0 | NA | NA | NA | NA | NA | #VALUE! |
| ENSMUSG00000085767.2 | 0 | NA | NA | NA | NA | NA | #VALUE! |
| ENSMUSG00000085768.2 | 0 | NA | NA | NA | NA | NA | #VALUE! |
| ENSMUSG00000085770.2 | 0 | NA | NA | NA | NA | NA | #VALUE! |
| ENSMUSG00000085771.2 | 0 | NA | NA | NA | NA | NA | #VALUE! |
| ENSMUSG00000085772.8 | 7.552098 | -6.44481 | 3.912702 | -1.64715 | 0.099527 | NA | #VALUE! |
| ENSMUSG00000085773.4 | 0 | NA | NA | NA | NA | NA | #VALUE! |
| ENSMUSG00000085774.2 | 0 | NA | NA | NA | NA | NA | #VALUE! |
| ENSMUSG00000085775.2 | 0 | NA | NA | NA | NA | NA | #VALUE! |
| ENSMUSG00000085776.9 | 4.620578 | -5.74391 | 3.916409 | -1.46663 | 0.142478 | NA | #VALUE! |
| ENSMUSG00000085779.2 | 0 | NA | NA | NA | NA | NA | #VALUE! |
| ENSMUSG00000085781.2 | 0 | NA | NA | NA | NA | NA | #VALUE! |
| ENSMUSG00000085782.3 | 0 | NA | NA | NA | NA | NA | #VALUE! |
| ENSMUSG00000085783.4 | 0.204111 | -1.05581 | 4.073302 | -0.2592 | 0.795479 | NA | #VALUE! |
| ENSMUSG00000085784.2 | 0 | NA | NA | NA | NA | NA | #VALUE! |
| ENSMUSG00000085785.8 | 0 | NA | NA | NA | NA | NA | #VALUE! |
| ENSMUSG00000085786.2 | 0 | NA | NA | NA | NA | NA | #VALUE! |
| ENSMUSG00000085787.4 | 0.561015 | -0.86763 | 4.025136 | -0.21555 | 0.829336 | NA | #VALUE! |
| ENSMUSG00000085788.2 | 0 | NA | NA | NA | NA | NA | #VALUE! |
| ENSMUSG00000085790.2 | 0 | NA | NA | NA | NA | NA | #VALUE! |
| ENSMUSG00000085791.5 | 1.512824 | -0.50997 | 3.850174 | -0.13245 | 0.894625 | NA | #VALUE! |
| ENSMUSG00000085792.3 | 0 | NA | NA | NA | NA | NA | #VALUE! |
| ENSMUSG00000085794.6 | 0 | NA | NA | NA | NA | NA | #VALUE! |
| ENSMUSG00000085796.2 | 0 | NA | NA | NA | NA | NA | #VALUE! |
| ENSMUSG00000085797.2 | 0 | NA | NA | NA | NA | NA | #VALUE! |
| ENSMUSG00000085798.2 | 0 | NA | NA | NA | NA | NA | #VALUE! |
| ENSMUSG00000085799.2 | 0 | NA | NA | NA | NA | NA | #VALUE! |
| ENSMUSG00000085800.8 | 0 | NA | NA | NA | NA | NA | #VALUE! |
| ENSMUSG00000085801.2 | 0 | NA | NA | NA | NA | NA | #VALUE! |
| ENSMUSG00000085802.3 | 0 | NA | NA | NA | NA | NA | #VALUE! |
| ENSMUSG00000085803.2 | 0 | NA | NA | NA | NA | NA | #VALUE! |
| ENSMUSG00000085804.2 | 0 | NA | NA | NA | NA | NA | #VALUE! |
| ENSMUSG00000085805.2 | 0 | NA | NA | NA | NA | NA | #VALUE! |
| ENSMUSG00000085806.2 | 0 | NA | NA | NA | NA | NA | #VALUE! |
| ENSMUSG00000085807.2 | 0 | NA | NA | NA | NA | NA | #VALUE! |
| ENSMUSG00000085808.2 | 0 | NA | NA | NA | NA | NA | #VALUE! |

| | | | | | | | |
|---|---|---|---|---|---|---|---|
| ENSMUSG00000085810.8 | 0 | NA | NA | NA | NA | NA | #VALUE! |
| ENSMUSG00000085811.2 | 0 | NA | NA | NA | NA | NA | #VALUE! |
| ENSMUSG00000085812.2 | 8.275877 | 5.585607 | 3.20017 | 1.74541 | 0.080914 | NA | #VALUE! |
| ENSMUSG00000085813.2 | 0 | NA | NA | NA | NA | NA | #VALUE! |
| ENSMUSG00000085815.2 | 0 | NA | NA | NA | NA | NA | #VALUE! |
| ENSMUSG00000085816.2 | 0 | NA | NA | NA | NA | NA | #VALUE! |
| ENSMUSG00000085817.2 | 0 | NA | NA | NA | NA | NA | #VALUE! |
| ENSMUSG00000085818.8 | 0 | NA | NA | NA | NA | NA | #VALUE! |
| ENSMUSG00000085819.8 | 0 | NA | NA | NA | NA | NA | #VALUE! |
| ENSMUSG00000085820.8 | 0 | NA | NA | NA | NA | NA | #VALUE! |
| ENSMUSG00000085822.3 | 0 | NA | NA | NA | NA | NA | #VALUE! |
| ENSMUSG00000085823.8 | 0 | NA | NA | NA | NA | NA | #VALUE! |
| ENSMUSG00000085824.8 | 0 | NA | NA | NA | NA | NA | #VALUE! |
| ENSMUSG00000085827.9 | 0 | NA | NA | NA | NA | NA | #VALUE! |
| ENSMUSG00000085828.2 | 0 | NA | NA | NA | NA | NA | #VALUE! |
| ENSMUSG00000085829.2 | 17.27609 | -1.46694 | 2.833075 | -0.51779 | NA | NA | #VALUE! |
| ENSMUSG00000085830.2 | 0 | NA | NA | NA | NA | NA | #VALUE! |
| ENSMUSG00000085831.8 | 0 | NA | NA | NA | NA | NA | #VALUE! |
| ENSMUSG00000085833.3 | 0 | NA | NA | NA | NA | NA | #VALUE! |
| ENSMUSG00000085834.2 | 0.420053 | -2.33493 | 4.007343 | -0.58266 | 0.56012 | NA | #VALUE! |
| ENSMUSG00000085835.2 | 0 | NA | NA | NA | NA | NA | #VALUE! |
| ENSMUSG00000085836.2 | 0 | NA | NA | NA | NA | NA | #VALUE! |
| ENSMUSG00000085837.9 | 0 | NA | NA | NA | NA | NA | #VALUE! |
| ENSMUSG00000085838.2 | 0 | NA | NA | NA | NA | NA | #VALUE! |
| ENSMUSG00000085839.2 | 0 | NA | NA | NA | NA | NA | #VALUE! |
| ENSMUSG00000085840.2 | 0 | NA | NA | NA | NA | NA | #VALUE! |
| ENSMUSG00000085841.2 | 0 | NA | NA | NA | NA | NA | #VALUE! |
| ENSMUSG00000085842.8 | 0 | NA | NA | NA | NA | NA | #VALUE! |
| ENSMUSG00000085843.2 | 0 | NA | NA | NA | NA | NA | #VALUE! |
| ENSMUSG00000085844.2 | 0 | NA | NA | NA | NA | NA | #VALUE! |
| ENSMUSG00000085845.8 | 0 | NA | NA | NA | NA | NA | #VALUE! |
| ENSMUSG00000085846.2 | 0 | NA | NA | NA | NA | NA | #VALUE! |
| ENSMUSG00000085847.2 | 0 | NA | NA | NA | NA | NA | #VALUE! |
| ENSMUSG00000085851.2 | 0 | NA | NA | NA | NA | NA | #VALUE! |
| ENSMUSG00000085852.2 | 0 | NA | NA | NA | NA | NA | #VALUE! |
| ENSMUSG00000085853.2 | 0 | NA | NA | NA | NA | NA | #VALUE! |
| ENSMUSG00000085856.2 | 0 | NA | NA | NA | NA | NA | #VALUE! |
| ENSMUSG00000085857.2 | 0 | NA | NA | NA | NA | NA | #VALUE! |
| ENSMUSG00000085858.2 | 0 | NA | NA | NA | NA | NA | #VALUE! |
| ENSMUSG00000085860.2 | 0 | NA | NA | NA | NA | NA | #VALUE! |
| ENSMUSG00000085861.2 | 0 | NA | NA | NA | NA | NA | #VALUE! |
| ENSMUSG00000085862.3 | 0 | NA | NA | NA | NA | NA | #VALUE! |
| ENSMUSG00000085863.2 | 0 | NA | NA | NA | NA | NA | #VALUE! |
| ENSMUSG00000085864.2 | 0 | NA | NA | NA | NA | NA | #VALUE! |

| | | | | | | | |
|---|---|---|---|---|---|---|---|
| ENSMUSG00000085865.3 | 0 | NA | NA | NA | NA | NA | #VALUE! |
| ENSMUSG00000085866.2 | 0 | NA | NA | NA | NA | NA | #VALUE! |
| ENSMUSG00000085867.2 | 1.049323 | 1.788281 | 3.935934 | 0.454347 | 0.649579 | NA | #VALUE! |
| ENSMUSG00000085868.2 | 0 | NA | NA | NA | NA | NA | #VALUE! |
| ENSMUSG00000085869.2 | 0 | NA | NA | NA | NA | NA | #VALUE! |
| ENSMUSG00000085870.2 | 0 | NA | NA | NA | NA | NA | #VALUE! |
| ENSMUSG00000085871.2 | 0 | NA | NA | NA | NA | NA | #VALUE! |
| ENSMUSG00000085872.2 | 0 | NA | NA | NA | NA | NA | #VALUE! |
| ENSMUSG00000085873.2 | 0 | NA | NA | NA | NA | NA | #VALUE! |
| ENSMUSG00000085875.4 | 0 | NA | NA | NA | NA | NA | #VALUE! |
| ENSMUSG00000085876.2 | 0 | NA | NA | NA | NA | NA | #VALUE! |
| ENSMUSG00000085877.5 | 0 | NA | NA | NA | NA | NA | #VALUE! |
| ENSMUSG00000085878.2 | 0 | NA | NA | NA | NA | NA | #VALUE! |
| ENSMUSG00000085880.5 | 0 | NA | NA | NA | NA | NA | #VALUE! |
| ENSMUSG00000085881.3 | 0 | NA | NA | NA | NA | NA | #VALUE! |
| ENSMUSG00000085882.2 | 7.74547 | 4.91126 | 3.22508 | 1.522833 | NA | NA | #VALUE! |
| ENSMUSG00000085884.2 | 0 | NA | NA | NA | NA | NA | #VALUE! |
| ENSMUSG00000085885.2 | 0 | NA | NA | NA | NA | NA | #VALUE! |
| ENSMUSG00000085886.2 | 0 | NA | NA | NA | NA | NA | #VALUE! |
| ENSMUSG00000085887.3 | 0 | NA | NA | NA | NA | NA | #VALUE! |
| ENSMUSG00000085888.8 | 0 | NA | NA | NA | NA | NA | #VALUE! |
| ENSMUSG00000085889.2 | 0 | NA | NA | NA | NA | NA | #VALUE! |
| ENSMUSG00000085890.8 | 8.821103 | -6.67439 | 3.721088 | -1.79367 | NA | NA | #VALUE! |
| ENSMUSG00000085891.8 | 0 | NA | NA | NA | NA | NA | #VALUE! |
| ENSMUSG00000085892.2 | 0 | NA | NA | NA | NA | NA | #VALUE! |
| ENSMUSG00000085893.2 | 0 | NA | NA | NA | NA | NA | #VALUE! |
| ENSMUSG00000085895.2 | 0 | NA | NA | NA | NA | NA | #VALUE! |
| ENSMUSG00000085896.3 | 0 | NA | NA | NA | NA | NA | #VALUE! |
| ENSMUSG00000085897.3 | 0.501477 | -1.29375 | 4.008705 | -0.32274 | 0.746896 | NA | #VALUE! |
| ENSMUSG00000085898.2 | 0 | NA | NA | NA | NA | NA | #VALUE! |
| ENSMUSG00000085899.2 | 0 | NA | NA | NA | NA | NA | #VALUE! |
| ENSMUSG00000085900.2 | 0 | NA | NA | NA | NA | NA | #VALUE! |
| ENSMUSG00000085902.2 | 0 | NA | NA | NA | NA | NA | #VALUE! |
| ENSMUSG00000085903.2 | 0 | NA | NA | NA | NA | NA | #VALUE! |
| ENSMUSG00000085905.2 | 0 | NA | NA | NA | NA | NA | #VALUE! |
| ENSMUSG00000085908.2 | 0 | NA | NA | NA | NA | NA | #VALUE! |
| ENSMUSG00000085909.2 | 0 | NA | NA | NA | NA | NA | #VALUE! |
| ENSMUSG00000085910.2 | 0 | NA | NA | NA | NA | NA | #VALUE! |
| ENSMUSG00000085911.2 | 0 | NA | NA | NA | NA | NA | #VALUE! |
| ENSMUSG00000085912.3 | 0 | NA | NA | NA | NA | NA | #VALUE! |
| ENSMUSG00000085913.3 | 0 | NA | NA | NA | NA | NA | #VALUE! |
| ENSMUSG00000085914.2 | 0 | NA | NA | NA | NA | NA | #VALUE! |
| ENSMUSG00000085915.2 | 0 | NA | NA | NA | NA | NA | #VALUE! |
| ENSMUSG00000085917.2 | 0 | NA | NA | NA | NA | NA | #VALUE! |

| | | | | | | | |
|---|---|---|---|---|---|---|---|
| ENSMUSG00000085918.2 | 0 | NA | NA | NA | NA | NA | #VALUE! |
| ENSMUSG00000085919.2 | 0 | NA | NA | NA | NA | NA | #VALUE! |
| ENSMUSG00000085920.2 | 0 | NA | NA | NA | NA | NA | #VALUE! |
| ENSMUSG00000085921.2 | 0 | NA | NA | NA | NA | NA | #VALUE! |
| ENSMUSG00000085922.2 | 0 | NA | NA | NA | NA | NA | #VALUE! |
| ENSMUSG00000085923.2 | 6.144357 | -6.1469 | 3.713342 | -1.65536 | 0.097852 | NA | #VALUE! |
| ENSMUSG00000085924.2 | 0 | NA | NA | NA | NA | NA | #VALUE! |
| ENSMUSG00000085925.4 | 0 | NA | NA | NA | NA | NA | #VALUE! |
| ENSMUSG00000085926.2 | 0 | NA | NA | NA | NA | NA | #VALUE! |
| ENSMUSG00000085927.2 | 0 | NA | NA | NA | NA | NA | #VALUE! |
| ENSMUSG00000085928.2 | 0 | NA | NA | NA | NA | NA | #VALUE! |
| ENSMUSG00000085929.2 | 0 | NA | NA | NA | NA | NA | #VALUE! |
| ENSMUSG00000085931.2 | 0 | NA | NA | NA | NA | NA | #VALUE! |
| ENSMUSG00000085932.2 | 0 | NA | NA | NA | NA | NA | #VALUE! |
| ENSMUSG00000085933.4 | 0 | NA | NA | NA | NA | NA | #VALUE! |
| ENSMUSG00000085934.2 | 0 | NA | NA | NA | NA | NA | #VALUE! |
| ENSMUSG00000085936.8 | 0 | NA | NA | NA | NA | NA | #VALUE! |
| ENSMUSG00000085937.2 | 0 | NA | NA | NA | NA | NA | #VALUE! |
| ENSMUSG00000085938.2 | 0 | NA | NA | NA | NA | NA | #VALUE! |
| ENSMUSG00000085939.4 | 0 | NA | NA | NA | NA | NA | #VALUE! |
| ENSMUSG00000085940.2 | 0 | NA | NA | NA | NA | NA | #VALUE! |
| ENSMUSG00000085941.2 | 0 | NA | NA | NA | NA | NA | #VALUE! |
| ENSMUSG00000085944.2 | 0 | NA | NA | NA | NA | NA | #VALUE! |
| ENSMUSG00000085946.2 | 0 | NA | NA | NA | NA | NA | #VALUE! |
| ENSMUSG00000085949.2 | 0 | NA | NA | NA | NA | NA | #VALUE! |
| ENSMUSG00000085950.2 | 0 | NA | NA | NA | NA | NA | #VALUE! |
| ENSMUSG00000085951.2 | 0 | NA | NA | NA | NA | NA | #VALUE! |
| ENSMUSG00000085952.2 | 0 | NA | NA | NA | NA | NA | #VALUE! |
| ENSMUSG00000085953.3 | 0 | NA | NA | NA | NA | NA | #VALUE! |
| ENSMUSG00000085955.2 | 0 | NA | NA | NA | NA | NA | #VALUE! |
| ENSMUSG00000085956.2 | 0 | NA | NA | NA | NA | NA | #VALUE! |
| ENSMUSG00000085957.5 | 0 | NA | NA | NA | NA | NA | #VALUE! |
| ENSMUSG00000085958.2 | 0 | NA | NA | NA | NA | NA | #VALUE! |
| ENSMUSG00000085961.2 | 0 | NA | NA | NA | NA | NA | #VALUE! |
| ENSMUSG00000085962.9 | 0 | NA | NA | NA | NA | NA | #VALUE! |
| ENSMUSG00000085965.2 | 0 | NA | NA | NA | NA | NA | #VALUE! |
| ENSMUSG00000085966.2 | 0 | NA | NA | NA | NA | NA | #VALUE! |
| ENSMUSG00000085967.2 | 0 | NA | NA | NA | NA | NA | #VALUE! |
| ENSMUSG00000085968.2 | 0 | NA | NA | NA | NA | NA | #VALUE! |
| ENSMUSG00000085969.2 | 0 | NA | NA | NA | NA | NA | #VALUE! |
| ENSMUSG00000085970.2 | 0 | NA | NA | NA | NA | NA | #VALUE! |
| ENSMUSG00000085971.5 | 0 | NA | NA | NA | NA | NA | #VALUE! |
| ENSMUSG00000085973.2 | 0 | NA | NA | NA | NA | NA | #VALUE! |
| ENSMUSG00000085975.2 | 0 | NA | NA | NA | NA | NA | #VALUE! |

| | | | | | | | |
|---|---|---|---|---|---|---|---|
| ENSMUSG00000085976.2 | 0 | NA | NA | NA | NA | NA | #VALUE! |
| ENSMUSG00000085977.2 | 0 | NA | NA | NA | NA | NA | #VALUE! |
| ENSMUSG00000085978.2 | 0 | NA | NA | NA | NA | NA | #VALUE! |
| ENSMUSG00000085979.2 | 0 | NA | NA | NA | NA | NA | #VALUE! |
| ENSMUSG00000085980.3 | 0 | NA | NA | NA | NA | NA | #VALUE! |
| ENSMUSG00000085982.2 | 0 | NA | NA | NA | NA | NA | #VALUE! |
| ENSMUSG00000085984.8 | 0 | NA | NA | NA | NA | NA | #VALUE! |
| ENSMUSG00000085985.2 | 0 | NA | NA | NA | NA | NA | #VALUE! |
| ENSMUSG00000085987.3 | 0 | NA | NA | NA | NA | NA | #VALUE! |
| ENSMUSG00000085989.2 | 0 | NA | NA | NA | NA | NA | #VALUE! |
| ENSMUSG00000085990.2 | 4.503149 | -5.6983 | 3.916718 | -1.45487 | 0.145706 | NA | #VALUE! |
| ENSMUSG00000085991.2 | 0 | NA | NA | NA | NA | NA | #VALUE! |
| ENSMUSG00000085992.2 | 0 | NA | NA | NA | NA | NA | #VALUE! |
| ENSMUSG00000085993.3 | 0 | NA | NA | NA | NA | NA | #VALUE! |
| ENSMUSG00000085996.2 | 0 | NA | NA | NA | NA | NA | #VALUE! |
| ENSMUSG00000085998.9 | 0.245976 | 1.027628 | 4.067126 | 0.252667 | 0.800526 | NA | #VALUE! |
| ENSMUSG00000085999.2 | 0 | NA | NA | NA | NA | NA | #VALUE! |
| ENSMUSG00000086000.2 | 0 | NA | NA | NA | NA | NA | #VALUE! |
| ENSMUSG00000086001.2 | 0 | NA | NA | NA | NA | NA | #VALUE! |
| ENSMUSG00000086003.8 | 0 | NA | NA | NA | NA | NA | #VALUE! |
| ENSMUSG00000086004.2 | 0 | NA | NA | NA | NA | NA | #VALUE! |
| ENSMUSG00000086006.3 | 0 | NA | NA | NA | NA | NA | #VALUE! |
| ENSMUSG00000086007.2 | 0 | NA | NA | NA | NA | NA | #VALUE! |
| ENSMUSG00000086008.9 | 0 | NA | NA | NA | NA | NA | #VALUE! |
| ENSMUSG00000086009.2 | 0 | NA | NA | NA | NA | NA | #VALUE! |
| ENSMUSG00000086010.9 | 5.590276 | -6.01038 | 3.837573 | -1.56619 | 0.117303 | NA | #VALUE! |
| ENSMUSG00000086012.3 | 0 | NA | NA | NA | NA | NA | #VALUE! |
| ENSMUSG00000086013.2 | 0 | NA | NA | NA | NA | NA | #VALUE! |
| ENSMUSG00000086014.2 | 0 | NA | NA | NA | NA | NA | #VALUE! |
| ENSMUSG00000086015.2 | 0 | NA | NA | NA | NA | NA | #VALUE! |
| ENSMUSG00000086016.2 | 0 | NA | NA | NA | NA | NA | #VALUE! |
| ENSMUSG00000086017.2 | 0 | NA | NA | NA | NA | NA | #VALUE! |
| ENSMUSG00000086018.8 | 0 | NA | NA | NA | NA | NA | #VALUE! |
| ENSMUSG00000086019.2 | 0 | NA | NA | NA | NA | NA | #VALUE! |
| ENSMUSG00000086020.2 | 0 | NA | NA | NA | NA | NA | #VALUE! |
| ENSMUSG00000086021.2 | 0 | NA | NA | NA | NA | NA | #VALUE! |
| ENSMUSG00000086022.2 | 0 | NA | NA | NA | NA | NA | #VALUE! |
| ENSMUSG00000086025.2 | 0 | NA | NA | NA | NA | NA | #VALUE! |
| ENSMUSG00000086026.2 | 0 | NA | NA | NA | NA | NA | #VALUE! |
| ENSMUSG00000086027.2 | 0 | NA | NA | NA | NA | NA | #VALUE! |
| ENSMUSG00000086028.2 | 0 | NA | NA | NA | NA | NA | #VALUE! |
| ENSMUSG00000086029.8 | 0 | NA | NA | NA | NA | NA | #VALUE! |
| ENSMUSG00000086030.2 | 0 | NA | NA | NA | NA | NA | #VALUE! |
| ENSMUSG00000086031.2 | 0 | NA | NA | NA | NA | NA | #VALUE! |

| | | | | | | | |
|---|---|---|---|---|---|---|---|
| ENSMUSG00000086032.2 | 0 | NA | NA | NA | NA | NA | #VALUE! |
| ENSMUSG00000086034.2 | 0 | NA | NA | NA | NA | NA | #VALUE! |
| ENSMUSG00000086035.2 | 0 | NA | NA | NA | NA | NA | #VALUE! |
| ENSMUSG00000086036.2 | 0 | NA | NA | NA | NA | NA | #VALUE! |
| ENSMUSG00000086037.2 | 0 | NA | NA | NA | NA | NA | #VALUE! |
| ENSMUSG00000086039.2 | 0 | NA | NA | NA | NA | NA | #VALUE! |
| ENSMUSG00000086040.9 | 4.98015 | -5.84579 | 3.915752 | -1.49289 | 0.135466 | NA | #VALUE! |
| ENSMUSG00000086041.2 | 0 | NA | NA | NA | NA | NA | #VALUE! |
| ENSMUSG00000086042.2 | 0 | NA | NA | NA | NA | NA | #VALUE! |
| ENSMUSG00000086043.2 | 0 | NA | NA | NA | NA | NA | #VALUE! |
| ENSMUSG00000086044.2 | 0 | NA | NA | NA | NA | NA | #VALUE! |
| ENSMUSG00000086046.2 | 0 | NA | NA | NA | NA | NA | #VALUE! |
| ENSMUSG00000086047.2 | 0 | NA | NA | NA | NA | NA | #VALUE! |
| ENSMUSG00000086048.8 | 0 | NA | NA | NA | NA | NA | #VALUE! |
| ENSMUSG00000086049.2 | 0 | NA | NA | NA | NA | NA | #VALUE! |
| ENSMUSG00000086050.2 | 0 | NA | NA | NA | NA | NA | #VALUE! |
| ENSMUSG00000086052.2 | 0 | NA | NA | NA | NA | NA | #VALUE! |
| ENSMUSG00000086053.2 | 0 | NA | NA | NA | NA | NA | #VALUE! |
| ENSMUSG00000086055.2 | 0 | NA | NA | NA | NA | NA | #VALUE! |
| ENSMUSG00000086056.2 | 0 | NA | NA | NA | NA | NA | #VALUE! |
| ENSMUSG00000086057.2 | 0 | NA | NA | NA | NA | NA | #VALUE! |
| ENSMUSG00000086058.2 | 0 | NA | NA | NA | NA | NA | #VALUE! |
| ENSMUSG00000086060.2 | 0 | NA | NA | NA | NA | NA | #VALUE! |
| ENSMUSG00000086061.8 | 0 | NA | NA | NA | NA | NA | #VALUE! |
| ENSMUSG00000086063.2 | 0 | NA | NA | NA | NA | NA | #VALUE! |
| ENSMUSG00000086064.2 | 0 | NA | NA | NA | NA | NA | #VALUE! |
| ENSMUSG00000086065.2 | 0 | NA | NA | NA | NA | NA | #VALUE! |
| ENSMUSG00000086066.8 | 0 | NA | NA | NA | NA | NA | #VALUE! |
| ENSMUSG00000086067.2 | 0 | NA | NA | NA | NA | NA | #VALUE! |
| ENSMUSG00000086068.2 | 0 | NA | NA | NA | NA | NA | #VALUE! |
| ENSMUSG00000086069.2 | 0 | NA | NA | NA | NA | NA | #VALUE! |
| ENSMUSG00000086070.2 | 0 | NA | NA | NA | NA | NA | #VALUE! |
| ENSMUSG00000086071.8 | 8.064371 | -6.54006 | 2.878491 | -2.27204 | 0.023084 | NA | #VALUE! |
| ENSMUSG00000086072.2 | 0 | NA | NA | NA | NA | NA | #VALUE! |
| ENSMUSG00000086074.3 | 0 | NA | NA | NA | NA | NA | #VALUE! |
| ENSMUSG00000086075.2 | 0 | NA | NA | NA | NA | NA | #VALUE! |
| ENSMUSG00000086076.2 | 0 | NA | NA | NA | NA | NA | #VALUE! |
| ENSMUSG00000086077.8 | 0 | NA | NA | NA | NA | NA | #VALUE! |
| ENSMUSG00000086078.2 | 0 | NA | NA | NA | NA | NA | #VALUE! |
| ENSMUSG00000086080.8 | 0 | NA | NA | NA | NA | NA | #VALUE! |
| ENSMUSG00000086081.5 | 0 | NA | NA | NA | NA | NA | #VALUE! |
| ENSMUSG00000086083.2 | 0 | NA | NA | NA | NA | NA | #VALUE! |
| ENSMUSG00000086087.2 | 0 | NA | NA | NA | NA | NA | #VALUE! |
| ENSMUSG00000086089.2 | 0 | NA | NA | NA | NA | NA | #VALUE! |

| | | | | | | | |
|---|---|---|---|---|---|---|---|
| ENSMUSG00000086090.2 | 0 | NA | NA | NA | NA | NA | #VALUE! |
| ENSMUSG00000086092.2 | 0 | NA | NA | NA | NA | NA | #VALUE! |
| ENSMUSG00000086095.3 | 0 | NA | NA | NA | NA | NA | #VALUE! |
| ENSMUSG00000086096.2 | 0 | NA | NA | NA | NA | NA | #VALUE! |
| ENSMUSG00000086097.2 | 0 | NA | NA | NA | NA | NA | #VALUE! |
| ENSMUSG00000086098.2 | 0 | NA | NA | NA | NA | NA | #VALUE! |
| ENSMUSG00000086103.2 | 0 | NA | NA | NA | NA | NA | #VALUE! |
| ENSMUSG00000086104.2 | 0 | NA | NA | NA | NA | NA | #VALUE! |
| ENSMUSG00000086105.2 | 0 | NA | NA | NA | NA | NA | #VALUE! |
| ENSMUSG00000086107.2 | 0 | NA | NA | NA | NA | NA | #VALUE! |
| ENSMUSG00000086108.3 | 0 | NA | NA | NA | NA | NA | #VALUE! |
| ENSMUSG00000086109.3 | 0 | NA | NA | NA | NA | NA | #VALUE! |
| ENSMUSG00000086111.2 | 0 | NA | NA | NA | NA | NA | #VALUE! |
| ENSMUSG00000086112.2 | 0 | NA | NA | NA | NA | NA | #VALUE! |
| ENSMUSG00000086114.2 | 0 | NA | NA | NA | NA | NA | #VALUE! |
| ENSMUSG00000086115.2 | 0 | NA | NA | NA | NA | NA | #VALUE! |
| ENSMUSG00000086117.2 | 0 | NA | NA | NA | NA | NA | #VALUE! |
| ENSMUSG00000086118.2 | 0 | NA | NA | NA | NA | NA | #VALUE! |
| ENSMUSG00000086119.3 | 0 | NA | NA | NA | NA | NA | #VALUE! |
| ENSMUSG00000086121.2 | 0 | NA | NA | NA | NA | NA | #VALUE! |
| ENSMUSG00000086122.2 | 0 | NA | NA | NA | NA | NA | #VALUE! |
| ENSMUSG00000086123.4 | 0 | NA | NA | NA | NA | NA | #VALUE! |
| ENSMUSG00000086124.3 | 0 | NA | NA | NA | NA | NA | #VALUE! |
| ENSMUSG00000086125.2 | 0 | NA | NA | NA | NA | NA | #VALUE! |
| ENSMUSG00000086126.2 | 0 | NA | NA | NA | NA | NA | #VALUE! |
| ENSMUSG00000086127.2 | 0 | NA | NA | NA | NA | NA | #VALUE! |
| ENSMUSG00000086128.9 | 0 | NA | NA | NA | NA | NA | #VALUE! |
| ENSMUSG00000086129.2 | 0 | NA | NA | NA | NA | NA | #VALUE! |
| ENSMUSG00000086130.2 | 0 | NA | NA | NA | NA | NA | #VALUE! |
| ENSMUSG00000086132.2 | 0 | NA | NA | NA | NA | NA | #VALUE! |
| ENSMUSG00000086133.2 | 0 | NA | NA | NA | NA | NA | #VALUE! |
| ENSMUSG00000086134.2 | 0 | NA | NA | NA | NA | NA | #VALUE! |
| ENSMUSG00000086135.2 | 0 | NA | NA | NA | NA | NA | #VALUE! |
| ENSMUSG00000086137.2 | 0 | NA | NA | NA | NA | NA | #VALUE! |
| ENSMUSG00000086139.2 | 0 | NA | NA | NA | NA | NA | #VALUE! |
| ENSMUSG00000086140.2 | 0 | NA | NA | NA | NA | NA | #VALUE! |
| ENSMUSG00000086141.8 | 8.904587 | -1.27356 | 2.273686 | -0.56013 | 0.575392 | NA | #VALUE! |
| ENSMUSG00000086142.4 | 0 | NA | NA | NA | NA | NA | #VALUE! |
| ENSMUSG00000086144.3 | 0 | NA | NA | NA | NA | NA | #VALUE! |
| ENSMUSG00000086145.2 | 0 | NA | NA | NA | NA | NA | #VALUE! |
| ENSMUSG00000086146.2 | 0 | NA | NA | NA | NA | NA | #VALUE! |
| ENSMUSG00000086147.3 | 0 | NA | NA | NA | NA | NA | #VALUE! |
| ENSMUSG00000086148.2 | 0 | NA | NA | NA | NA | NA | #VALUE! |
| ENSMUSG00000086150.2 | 3.496942 | -1.47676 | 3.694185 | -0.39975 | 0.689339 | NA | #VALUE! |

| | | | | | | | |
|---|---|---|---|---|---|---|---|
| ENSMUSG00000086152.2 | 0 | NA | NA | NA | NA | NA | #VALUE! |
| ENSMUSG00000086153.2 | 0 | NA | NA | NA | NA | NA | #VALUE! |
| ENSMUSG00000086154.3 | 0 | NA | NA | NA | NA | NA | #VALUE! |
| ENSMUSG00000086155.8 | 0 | NA | NA | NA | NA | NA | #VALUE! |
| ENSMUSG00000086156.2 | 0 | NA | NA | NA | NA | NA | #VALUE! |
| ENSMUSG00000086157.2 | 0 | NA | NA | NA | NA | NA | #VALUE! |
| ENSMUSG00000086159.2 | 0 | NA | NA | NA | NA | NA | #VALUE! |
| ENSMUSG00000086160.2 | 0 | NA | NA | NA | NA | NA | #VALUE! |
| ENSMUSG00000086161.2 | 0 | NA | NA | NA | NA | NA | #VALUE! |
| ENSMUSG00000086162.2 | 0 | NA | NA | NA | NA | NA | #VALUE! |
| ENSMUSG00000086163.2 | 0 | NA | NA | NA | NA | NA | #VALUE! |
| ENSMUSG00000086164.2 | 0 | NA | NA | NA | NA | NA | #VALUE! |
| ENSMUSG00000086165.2 | 1.428775 | -4.03418 | 3.938169 | -1.02438 | 0.305656 | NA | #VALUE! |
| ENSMUSG00000086166.2 | 0 | NA | NA | NA | NA | NA | #VALUE! |
| ENSMUSG00000086167.4 | 0 | NA | NA | NA | NA | NA | #VALUE! |
| ENSMUSG00000086168.2 | 0 | NA | NA | NA | NA | NA | #VALUE! |
| ENSMUSG00000086170.2 | 0 | NA | NA | NA | NA | NA | #VALUE! |
| ENSMUSG00000086171.8 | 0 | NA | NA | NA | NA | NA | #VALUE! |
| ENSMUSG00000086172.2 | 0 | NA | NA | NA | NA | NA | #VALUE! |
| ENSMUSG00000086173.8 | 0 | NA | NA | NA | NA | NA | #VALUE! |
| ENSMUSG00000086174.2 | 0 | NA | NA | NA | NA | NA | #VALUE! |
| ENSMUSG00000086175.3 | 0 | NA | NA | NA | NA | NA | #VALUE! |
| ENSMUSG00000086176.5 | 0 | NA | NA | NA | NA | NA | #VALUE! |
| ENSMUSG00000086177.2 | 0 | NA | NA | NA | NA | NA | #VALUE! |
| ENSMUSG00000086179.2 | 0 | NA | NA | NA | NA | NA | #VALUE! |
| ENSMUSG00000086181.2 | 0 | NA | NA | NA | NA | NA | #VALUE! |
| ENSMUSG00000086184.2 | 0 | NA | NA | NA | NA | NA | #VALUE! |
| ENSMUSG00000086185.2 | 0 | NA | NA | NA | NA | NA | #VALUE! |
| ENSMUSG00000086186.2 | 0 | NA | NA | NA | NA | NA | #VALUE! |
| ENSMUSG00000086187.2 | 0 | NA | NA | NA | NA | NA | #VALUE! |
| ENSMUSG00000086188.2 | 0 | NA | NA | NA | NA | NA | #VALUE! |
| ENSMUSG00000086189.2 | 0 | NA | NA | NA | NA | NA | #VALUE! |
| ENSMUSG00000086190.2 | 0 | NA | NA | NA | NA | NA | #VALUE! |
| ENSMUSG00000086191.2 | 0 | NA | NA | NA | NA | NA | #VALUE! |
| ENSMUSG00000086192.2 | 0 | NA | NA | NA | NA | NA | #VALUE! |
| ENSMUSG00000086193.2 | 0 | NA | NA | NA | NA | NA | #VALUE! |
| ENSMUSG00000086194.2 | 0 | NA | NA | NA | NA | NA | #VALUE! |
| ENSMUSG00000086195.2 | 0 | NA | NA | NA | NA | NA | #VALUE! |
| ENSMUSG00000086196.8 | 0 | NA | NA | NA | NA | NA | #VALUE! |
| ENSMUSG00000086197.2 | 0 | NA | NA | NA | NA | NA | #VALUE! |
| ENSMUSG00000086199.8 | 0 | NA | NA | NA | NA | NA | #VALUE! |
| ENSMUSG00000086201.2 | 0 | NA | NA | NA | NA | NA | #VALUE! |
| ENSMUSG00000086202.2 | 0 | NA | NA | NA | NA | NA | #VALUE! |
| ENSMUSG00000086203.2 | 0 | NA | NA | NA | NA | NA | #VALUE! |

| | | | | | | | |
|---|---|---|---|---|---|---|---|
| ENSMUSG00000086204.2 | 0 | NA | NA | NA | NA | NA | #VALUE! |
| ENSMUSG00000086205.2 | 0 | NA | NA | NA | NA | NA | #VALUE! |
| ENSMUSG00000086206.2 | 0 | NA | NA | NA | NA | NA | #VALUE! |
| ENSMUSG00000086207.2 | 0 | NA | NA | NA | NA | NA | #VALUE! |
| ENSMUSG00000086209.2 | 0 | NA | NA | NA | NA | NA | #VALUE! |
| ENSMUSG00000086211.2 | 0 | NA | NA | NA | NA | NA | #VALUE! |
| ENSMUSG00000086212.8 | 0 | NA | NA | NA | NA | NA | #VALUE! |
| ENSMUSG00000086213.3 | 8.013406 | 6.488223 | 3.381866 | 1.918534 | 0.055043 | NA | #VALUE! |
| ENSMUSG00000086214.2 | 0 | NA | NA | NA | NA | NA | #VALUE! |
| ENSMUSG00000086216.2 | 0 | NA | NA | NA | NA | NA | #VALUE! |
| ENSMUSG00000086220.2 | 0 | NA | NA | NA | NA | NA | #VALUE! |
| ENSMUSG00000086221.2 | 0 | NA | NA | NA | NA | NA | #VALUE! |
| ENSMUSG00000086222.2 | 0 | NA | NA | NA | NA | NA | #VALUE! |
| ENSMUSG00000086224.9 | 0 | NA | NA | NA | NA | NA | #VALUE! |
| ENSMUSG00000086225.2 | 0 | NA | NA | NA | NA | NA | #VALUE! |
| ENSMUSG00000086226.2 | 0 | NA | NA | NA | NA | NA | #VALUE! |
| ENSMUSG00000086227.2 | 0 | NA | NA | NA | NA | NA | #VALUE! |
| ENSMUSG00000086228.4 | 0 | NA | NA | NA | NA | NA | #VALUE! |
| ENSMUSG00000086229.4 | 0 | NA | NA | NA | NA | NA | #VALUE! |
| ENSMUSG00000086231.2 | 0 | NA | NA | NA | NA | NA | #VALUE! |
| ENSMUSG00000086232.3 | 0 | NA | NA | NA | NA | NA | #VALUE! |
| ENSMUSG00000086233.2 | 0 | NA | NA | NA | NA | NA | #VALUE! |
| ENSMUSG00000086234.4 | 0 | NA | NA | NA | NA | NA | #VALUE! |
| ENSMUSG00000086235.2 | 0 | NA | NA | NA | NA | NA | #VALUE! |
| ENSMUSG00000086236.10 | 0 | NA | NA | NA | NA | NA | #VALUE! |
| ENSMUSG00000086237.2 | 0 | NA | NA | NA | NA | NA | #VALUE! |
| ENSMUSG00000086238.2 | 0 | NA | NA | NA | NA | NA | #VALUE! |
| ENSMUSG00000086239.2 | 0 | NA | NA | NA | NA | NA | #VALUE! |
| ENSMUSG00000086240.5 | 0.444244 | 0.890672 | 4.031633 | 0.220921 | 0.825154 | NA | #VALUE! |
| ENSMUSG00000086241.3 | 0 | NA | NA | NA | NA | NA | #VALUE! |
| ENSMUSG00000086243.2 | 0 | NA | NA | NA | NA | NA | #VALUE! |
| ENSMUSG00000086244.2 | 0 | NA | NA | NA | NA | NA | #VALUE! |
| ENSMUSG00000086245.2 | 0 | NA | NA | NA | NA | NA | #VALUE! |
| ENSMUSG00000086246.3 | 0 | NA | NA | NA | NA | NA | #VALUE! |
| ENSMUSG00000086247.8 | 0 | NA | NA | NA | NA | NA | #VALUE! |
| ENSMUSG00000086248.3 | 0 | NA | NA | NA | NA | NA | #VALUE! |
| ENSMUSG00000086249.2 | 0 | NA | NA | NA | NA | NA | #VALUE! |
| ENSMUSG00000086250.2 | 0 | NA | NA | NA | NA | NA | #VALUE! |
| ENSMUSG00000086251.2 | 0 | NA | NA | NA | NA | NA | #VALUE! |
| ENSMUSG00000086253.2 | 0 | NA | NA | NA | NA | NA | #VALUE! |
| ENSMUSG00000086254.2 | 0 | NA | NA | NA | NA | NA | #VALUE! |
| ENSMUSG00000086255.2 | 0 | NA | NA | NA | NA | NA | #VALUE! |
| ENSMUSG00000086256.2 | 0 | NA | NA | NA | NA | NA | #VALUE! |
| ENSMUSG00000086257.2 | 0 | NA | NA | NA | NA | NA | #VALUE! |

| | | | | | | | |
|---|---|---|---|---|---|---|---|
| ENSMUSG00000086258.2 | 0 | NA | NA | NA | NA | NA | #VALUE! |
| ENSMUSG00000086259.8 | 0 | NA | NA | NA | NA | NA | #VALUE! |
| ENSMUSG00000086260.2 | 0 | NA | NA | NA | NA | NA | #VALUE! |
| ENSMUSG00000086261.2 | 0 | NA | NA | NA | NA | NA | #VALUE! |
| ENSMUSG00000086262.8 | 0 | NA | NA | NA | NA | NA | #VALUE! |
| ENSMUSG00000086263.2 | 0 | NA | NA | NA | NA | NA | #VALUE! |
| ENSMUSG00000086264.10 | 0 | NA | NA | NA | NA | NA | #VALUE! |
| ENSMUSG00000086265.2 | 0 | NA | NA | NA | NA | NA | #VALUE! |
| ENSMUSG00000086266.2 | 0 | NA | NA | NA | NA | NA | #VALUE! |
| ENSMUSG00000086267.2 | 0 | NA | NA | NA | NA | NA | #VALUE! |
| ENSMUSG00000086268.2 | 0 | NA | NA | NA | NA | NA | #VALUE! |
| ENSMUSG00000086269.3 | 0 | NA | NA | NA | NA | NA | #VALUE! |
| ENSMUSG00000086272.2 | 0 | NA | NA | NA | NA | NA | #VALUE! |
| ENSMUSG00000086273.8 | 0 | NA | NA | NA | NA | NA | #VALUE! |
| ENSMUSG00000086275.2 | 0 | NA | NA | NA | NA | NA | #VALUE! |
| ENSMUSG00000086277.3 | 0 | NA | NA | NA | NA | NA | #VALUE! |
| ENSMUSG00000086278.6 | 0 | NA | NA | NA | NA | NA | #VALUE! |
| ENSMUSG00000086279.2 | 4.15549 | 5.541116 | 3.917598 | 1.414417 | 0.15724 | NA | #VALUE! |
| ENSMUSG00000086282.2 | 4.261361 | -5.61781 | 3.917288 | -1.43411 | 0.151541 | NA | #VALUE! |
| ENSMUSG00000086283.8 | 0 | NA | NA | NA | NA | NA | #VALUE! |
| ENSMUSG00000086284.2 | 0 | NA | NA | NA | NA | NA | #VALUE! |
| ENSMUSG00000086286.2 | 0 | NA | NA | NA | NA | NA | #VALUE! |
| ENSMUSG00000086287.2 | 0 | NA | NA | NA | NA | NA | #VALUE! |
| ENSMUSG00000086288.2 | 0 | NA | NA | NA | NA | NA | #VALUE! |
| ENSMUSG00000086289.4 | 0 | NA | NA | NA | NA | NA | #VALUE! |
| ENSMUSG00000086291.2 | 0 | NA | NA | NA | NA | NA | #VALUE! |
| ENSMUSG00000086293.2 | 0 | NA | NA | NA | NA | NA | #VALUE! |
| ENSMUSG00000086296.3 | 0 | NA | NA | NA | NA | NA | #VALUE! |
| ENSMUSG00000086297.2 | 0 | NA | NA | NA | NA | NA | #VALUE! |
| ENSMUSG00000086298.2 | 0 | NA | NA | NA | NA | NA | #VALUE! |
| ENSMUSG00000086302.2 | 0 | NA | NA | NA | NA | NA | #VALUE! |
| ENSMUSG00000086306.2 | 0 | NA | NA | NA | NA | NA | #VALUE! |
| ENSMUSG00000086308.2 | 0 | NA | NA | NA | NA | NA | #VALUE! |
| ENSMUSG00000086310.2 | 0 | NA | NA | NA | NA | NA | #VALUE! |
| ENSMUSG00000086311.2 | 0 | NA | NA | NA | NA | NA | #VALUE! |
| ENSMUSG00000086312.2 | 0 | NA | NA | NA | NA | NA | #VALUE! |
| ENSMUSG00000086313.2 | 0 | NA | NA | NA | NA | NA | #VALUE! |
| ENSMUSG00000086315.2 | 0 | NA | NA | NA | NA | NA | #VALUE! |
| ENSMUSG00000086319.2 | 0 | NA | NA | NA | NA | NA | #VALUE! |
| ENSMUSG00000086320.2 | 0 | NA | NA | NA | NA | NA | #VALUE! |
| ENSMUSG00000086321.2 | 0 | NA | NA | NA | NA | NA | #VALUE! |
| ENSMUSG00000086322.8 | 0 | NA | NA | NA | NA | NA | #VALUE! |
| ENSMUSG00000086324.9 | 0 | NA | NA | NA | NA | NA | #VALUE! |
| ENSMUSG00000086326.2 | 0 | NA | NA | NA | NA | NA | #VALUE! |

| | | | | | | | |
|---|---|---|---|---|---|---|---|
| ENSMUSG00000086327.3 | 0 | NA | NA | NA | NA | NA | #VALUE! |
| ENSMUSG00000086328.2 | 0 | NA | NA | NA | NA | NA | #VALUE! |
| ENSMUSG00000086329.2 | 0 | NA | NA | NA | NA | NA | #VALUE! |
| ENSMUSG00000086330.8 | 0 | NA | NA | NA | NA | NA | #VALUE! |
| ENSMUSG00000086331.2 | 0.105013 | -0.97324 | 4.076293 | -0.23876 | 0.811295 | NA | #VALUE! |
| ENSMUSG00000086333.2 | 0 | NA | NA | NA | NA | NA | #VALUE! |
| ENSMUSG00000086335.2 | 0 | NA | NA | NA | NA | NA | #VALUE! |
| ENSMUSG00000086336.2 | 0 | NA | NA | NA | NA | NA | #VALUE! |
| ENSMUSG00000086337.2 | 0 | NA | NA | NA | NA | NA | #VALUE! |
| ENSMUSG00000086338.3 | 0 | NA | NA | NA | NA | NA | #VALUE! |
| ENSMUSG00000086339.2 | 0 | NA | NA | NA | NA | NA | #VALUE! |
| ENSMUSG00000086340.2 | 0 | NA | NA | NA | NA | NA | #VALUE! |
| ENSMUSG00000086341.3 | 0 | NA | NA | NA | NA | NA | #VALUE! |
| ENSMUSG00000086342.2 | 0 | NA | NA | NA | NA | NA | #VALUE! |
| ENSMUSG00000086346.2 | 0 | NA | NA | NA | NA | NA | #VALUE! |
| ENSMUSG00000086347.2 | 0 | NA | NA | NA | NA | NA | #VALUE! |
| ENSMUSG00000086348.2 | 0 | NA | NA | NA | NA | NA | #VALUE! |
| ENSMUSG00000086349.2 | 0 | NA | NA | NA | NA | NA | #VALUE! |
| ENSMUSG00000086350.2 | 0 | NA | NA | NA | NA | NA | #VALUE! |
| ENSMUSG00000086352.2 | 0 | NA | NA | NA | NA | NA | #VALUE! |
| ENSMUSG00000086353.2 | 0 | NA | NA | NA | NA | NA | #VALUE! |
| ENSMUSG00000086354.3 | 0 | NA | NA | NA | NA | NA | #VALUE! |
| ENSMUSG00000086355.2 | 0 | NA | NA | NA | NA | NA | #VALUE! |
| ENSMUSG00000086356.2 | 0 | NA | NA | NA | NA | NA | #VALUE! |
| ENSMUSG00000086358.3 | 0 | NA | NA | NA | NA | NA | #VALUE! |
| ENSMUSG00000086359.2 | 0 | NA | NA | NA | NA | NA | #VALUE! |
| ENSMUSG00000086360.2 | 0 | NA | NA | NA | NA | NA | #VALUE! |
| ENSMUSG00000086361.2 | 0 | NA | NA | NA | NA | NA | #VALUE! |
| ENSMUSG00000086363.8 | 0 | NA | NA | NA | NA | NA | #VALUE! |
| ENSMUSG00000086364.2 | 0 | NA | NA | NA | NA | NA | #VALUE! |
| ENSMUSG00000086365.3 | 0 | NA | NA | NA | NA | NA | #VALUE! |
| ENSMUSG00000086366.2 | 0 | NA | NA | NA | NA | NA | #VALUE! |
| ENSMUSG00000086367.2 | 0 | NA | NA | NA | NA | NA | #VALUE! |
| ENSMUSG00000086368.3 | 0 | NA | NA | NA | NA | NA | #VALUE! |
| ENSMUSG00000086369.8 | 0 | NA | NA | NA | NA | NA | #VALUE! |
| ENSMUSG00000086371.2 | 0 | NA | NA | NA | NA | NA | #VALUE! |
| ENSMUSG00000086372.2 | 0 | NA | NA | NA | NA | NA | #VALUE! |
| ENSMUSG00000086373.9 | 0 | NA | NA | NA | NA | NA | #VALUE! |
| ENSMUSG00000086375.2 | 0 | NA | NA | NA | NA | NA | #VALUE! |
| ENSMUSG00000086376.3 | 0 | NA | NA | NA | NA | NA | #VALUE! |
| ENSMUSG00000086377.2 | 0 | NA | NA | NA | NA | NA | #VALUE! |
| ENSMUSG00000086379.8 | 0 | NA | NA | NA | NA | NA | #VALUE! |
| ENSMUSG00000086380.2 | 0 | NA | NA | NA | NA | NA | #VALUE! |
| ENSMUSG00000086381.2 | 0 | NA | NA | NA | NA | NA | #VALUE! |

| | | | | | | | |
|---|---|---|---|---|---|---|---|
| ENSMUSG00000086384.2 | 0 | NA | NA | NA | NA | NA | #VALUE! |
| ENSMUSG00000086385.2 | 0 | NA | NA | NA | NA | NA | #VALUE! |
| ENSMUSG00000086386.2 | 0 | NA | NA | NA | NA | NA | #VALUE! |
| ENSMUSG00000086387.2 | 0 | NA | NA | NA | NA | NA | #VALUE! |
| ENSMUSG00000086389.2 | 6.396754 | -6.20432 | 3.245419 | -1.91172 | 0.055912 | NA | #VALUE! |
| ENSMUSG00000086391.2 | 0 | NA | NA | NA | NA | NA | #VALUE! |
| ENSMUSG00000086392.10 | 0 | NA | NA | NA | NA | NA | #VALUE! |
| ENSMUSG00000086395.2 | 0 | NA | NA | NA | NA | NA | #VALUE! |
| ENSMUSG00000086396.2 | 0 | NA | NA | NA | NA | NA | #VALUE! |
| ENSMUSG00000086399.2 | 0 | NA | NA | NA | NA | NA | #VALUE! |
| ENSMUSG00000086401.2 | 0 | NA | NA | NA | NA | NA | #VALUE! |
| ENSMUSG00000086402.2 | 0 | NA | NA | NA | NA | NA | #VALUE! |
| ENSMUSG00000086404.2 | 0 | NA | NA | NA | NA | NA | #VALUE! |
| ENSMUSG00000086405.2 | 0 | NA | NA | NA | NA | NA | #VALUE! |
| ENSMUSG00000086407.2 | 0 | NA | NA | NA | NA | NA | #VALUE! |
| ENSMUSG00000086408.2 | 0 | NA | NA | NA | NA | NA | #VALUE! |
| ENSMUSG00000086409.4 | 0 | NA | NA | NA | NA | NA | #VALUE! |
| ENSMUSG00000086410.8 | 0 | NA | NA | NA | NA | NA | #VALUE! |
| ENSMUSG00000086411.2 | 0 | NA | NA | NA | NA | NA | #VALUE! |
| ENSMUSG00000086413.2 | 0 | NA | NA | NA | NA | NA | #VALUE! |
| ENSMUSG00000086414.3 | 0 | NA | NA | NA | NA | NA | #VALUE! |
| ENSMUSG00000086415.2 | 0 | NA | NA | NA | NA | NA | #VALUE! |
| ENSMUSG00000086416.2 | 0 | NA | NA | NA | NA | NA | #VALUE! |
| ENSMUSG00000086417.2 | 0 | NA | NA | NA | NA | NA | #VALUE! |
| ENSMUSG00000086418.2 | 0 | NA | NA | NA | NA | NA | #VALUE! |
| ENSMUSG00000086420.2 | 0 | NA | NA | NA | NA | NA | #VALUE! |
| ENSMUSG00000086421.2 | 0 | NA | NA | NA | NA | NA | #VALUE! |
| ENSMUSG00000086422.2 | 0 | NA | NA | NA | NA | NA | #VALUE! |
| ENSMUSG00000086423.2 | 0 | NA | NA | NA | NA | NA | #VALUE! |
| ENSMUSG00000086424.2 | 0 | NA | NA | NA | NA | NA | #VALUE! |
| ENSMUSG00000086425.8 | 0 | NA | NA | NA | NA | NA | #VALUE! |
| ENSMUSG00000086426.2 | 0 | NA | NA | NA | NA | NA | #VALUE! |
| ENSMUSG00000086427.9 | 0 | NA | NA | NA | NA | NA | #VALUE! |
| ENSMUSG00000086428.2 | 0 | NA | NA | NA | NA | NA | #VALUE! |
| ENSMUSG00000086430.2 | 0 | NA | NA | NA | NA | NA | #VALUE! |
| ENSMUSG00000086431.2 | 0 | NA | NA | NA | NA | NA | #VALUE! |
| ENSMUSG00000086433.2 | 0 | NA | NA | NA | NA | NA | #VALUE! |
| ENSMUSG00000086434.2 | 0 | NA | NA | NA | NA | NA | #VALUE! |
| ENSMUSG00000086436.3 | 0 | NA | NA | NA | NA | NA | #VALUE! |
| ENSMUSG00000086438.3 | 0 | NA | NA | NA | NA | NA | #VALUE! |
| ENSMUSG00000086439.2 | 0 | NA | NA | NA | NA | NA | #VALUE! |
| ENSMUSG00000086441.2 | 0 | NA | NA | NA | NA | NA | #VALUE! |
| ENSMUSG00000086442.2 | 0 | NA | NA | NA | NA | NA | #VALUE! |
| ENSMUSG00000086443.2 | 0 | NA | NA | NA | NA | NA | #VALUE! |

| | | | | | | | |
|---|---|---|---|---|---|---|---|
| ENSMUSG00000086444.3 | 0 | NA | NA | NA | NA | NA | #VALUE! |
| ENSMUSG00000086445.2 | 0 | NA | NA | NA | NA | NA | #VALUE! |
| ENSMUSG00000086446.2 | 0 | NA | NA | NA | NA | NA | #VALUE! |
| ENSMUSG00000086449.2 | 0 | NA | NA | NA | NA | NA | #VALUE! |
| ENSMUSG00000086450.2 | 0 | NA | NA | NA | NA | NA | #VALUE! |
| ENSMUSG00000086451.9 | 0 | NA | NA | NA | NA | NA | #VALUE! |
| ENSMUSG00000086452.2 | 0 | NA | NA | NA | NA | NA | #VALUE! |
| ENSMUSG00000086453.8 | 0 | NA | NA | NA | NA | NA | #VALUE! |
| ENSMUSG00000086454.8 | 0 | NA | NA | NA | NA | NA | #VALUE! |
| ENSMUSG00000086455.2 | 0 | NA | NA | NA | NA | NA | #VALUE! |
| ENSMUSG00000086458.2 | 0 | NA | NA | NA | NA | NA | #VALUE! |
| ENSMUSG00000086459.3 | 0 | NA | NA | NA | NA | NA | #VALUE! |
| ENSMUSG00000086460.2 | 0 | NA | NA | NA | NA | NA | #VALUE! |
| ENSMUSG00000086462.2 | 0 | NA | NA | NA | NA | NA | #VALUE! |
| ENSMUSG00000086464.3 | 0 | NA | NA | NA | NA | NA | #VALUE! |
| ENSMUSG00000086467.3 | 0 | NA | NA | NA | NA | NA | #VALUE! |
| ENSMUSG00000086468.2 | 0 | NA | NA | NA | NA | NA | #VALUE! |
| ENSMUSG00000086470.2 | 0 | NA | NA | NA | NA | NA | #VALUE! |
| ENSMUSG00000086472.2 | 0 | NA | NA | NA | NA | NA | #VALUE! |
| ENSMUSG00000086473.2 | 0.093255 | 0.868093 | 4.072464 | 0.213162 | 0.831201 | NA | #VALUE! |
| ENSMUSG00000086474.3 | 0 | NA | NA | NA | NA | NA | #VALUE! |
| ENSMUSG00000086475.2 | 0 | NA | NA | NA | NA | NA | #VALUE! |
| ENSMUSG00000086477.4 | 0.210026 | -1.52494 | 4.058692 | -0.37572 | 0.707123 | NA | #VALUE! |
| ENSMUSG00000086478.2 | 0.093255 | 0.868093 | 4.072464 | 0.213162 | 0.831201 | NA | #VALUE! |
| ENSMUSG00000086479.3 | 0 | NA | NA | NA | NA | NA | #VALUE! |
| ENSMUSG00000086480.2 | 0 | NA | NA | NA | NA | NA | #VALUE! |
| ENSMUSG00000086481.2 | 0 | NA | NA | NA | NA | NA | #VALUE! |
| ENSMUSG00000086482.2 | 0 | NA | NA | NA | NA | NA | #VALUE! |
| ENSMUSG00000086483.2 | 0 | NA | NA | NA | NA | NA | #VALUE! |
| ENSMUSG00000086484.2 | 0 | NA | NA | NA | NA | NA | #VALUE! |
| ENSMUSG00000086485.3 | 0 | NA | NA | NA | NA | NA | #VALUE! |
| ENSMUSG00000086486.2 | 0 | NA | NA | NA | NA | NA | #VALUE! |
| ENSMUSG00000086487.2 | 0 | NA | NA | NA | NA | NA | #VALUE! |
| ENSMUSG00000086489.2 | 0 | NA | NA | NA | NA | NA | #VALUE! |
| ENSMUSG00000086490.2 | 0 | NA | NA | NA | NA | NA | #VALUE! |
| ENSMUSG00000086491.2 | 0 | NA | NA | NA | NA | NA | #VALUE! |
| ENSMUSG00000086493.2 | 0 | NA | NA | NA | NA | NA | #VALUE! |
| ENSMUSG00000086495.8 | 0 | NA | NA | NA | NA | NA | #VALUE! |
| ENSMUSG00000086496.2 | 0 | NA | NA | NA | NA | NA | #VALUE! |
| ENSMUSG00000086497.2 | 0 | NA | NA | NA | NA | NA | #VALUE! |
| ENSMUSG00000086498.2 | 0 | NA | NA | NA | NA | NA | #VALUE! |
| ENSMUSG00000086499.2 | 0 | NA | NA | NA | NA | NA | #VALUE! |
| ENSMUSG00000086500.2 | 0 | NA | NA | NA | NA | NA | #VALUE! |
| ENSMUSG00000086501.2 | 0 | NA | NA | NA | NA | NA | #VALUE! |

| | | | | | | | |
|---|---|---|---|---|---|---|---|
| ENSMUSG00000086502.2 | 0 | NA | NA | NA | NA | NA | #VALUE! |
| ENSMUSG00000086503.5 | 0 | NA | NA | NA | NA | NA | #VALUE! |
| ENSMUSG00000086504.2 | 0 | NA | NA | NA | NA | NA | #VALUE! |
| ENSMUSG00000086505.2 | 0 | NA | NA | NA | NA | NA | #VALUE! |
| ENSMUSG00000086506.2 | 0 | NA | NA | NA | NA | NA | #VALUE! |
| ENSMUSG00000086507.2 | 0 | NA | NA | NA | NA | NA | #VALUE! |
| ENSMUSG00000086509.3 | 0 | NA | NA | NA | NA | NA | #VALUE! |
| ENSMUSG00000086512.2 | 0 | NA | NA | NA | NA | NA | #VALUE! |
| ENSMUSG00000086513.4 | 0 | NA | NA | NA | NA | NA | #VALUE! |
| ENSMUSG00000086514.2 | 0 | NA | NA | NA | NA | NA | #VALUE! |
| ENSMUSG00000086515.2 | 0 | NA | NA | NA | NA | NA | #VALUE! |
| ENSMUSG00000086517.2 | 0 | NA | NA | NA | NA | NA | #VALUE! |
| ENSMUSG00000086518.2 | 0 | NA | NA | NA | NA | NA | #VALUE! |
| ENSMUSG00000086519.2 | 0 | NA | NA | NA | NA | NA | #VALUE! |
| ENSMUSG00000086521.4 | 0 | NA | NA | NA | NA | NA | #VALUE! |
| ENSMUSG00000086522.2 | 0 | NA | NA | NA | NA | NA | #VALUE! |
| ENSMUSG00000086523.2 | 0 | NA | NA | NA | NA | NA | #VALUE! |
| ENSMUSG00000086524.4 | 0 | NA | NA | NA | NA | NA | #VALUE! |
| ENSMUSG00000086525.2 | 0 | NA | NA | NA | NA | NA | #VALUE! |
| ENSMUSG00000086526.3 | 0 | NA | NA | NA | NA | NA | #VALUE! |
| ENSMUSG00000086527.2 | 0 | NA | NA | NA | NA | NA | #VALUE! |
| ENSMUSG00000086528.2 | 0 | NA | NA | NA | NA | NA | #VALUE! |
| ENSMUSG00000086529.2 | 0 | NA | NA | NA | NA | NA | #VALUE! |
| ENSMUSG00000086530.2 | 0 | NA | NA | NA | NA | NA | #VALUE! |
| ENSMUSG00000086531.3 | 0 | NA | NA | NA | NA | NA | #VALUE! |
| ENSMUSG00000086533.3 | 0 | NA | NA | NA | NA | NA | #VALUE! |
| ENSMUSG00000086534.3 | 0 | NA | NA | NA | NA | NA | #VALUE! |
| ENSMUSG00000086536.2 | 0 | NA | NA | NA | NA | NA | #VALUE! |
| ENSMUSG00000086537.8 | 0 | NA | NA | NA | NA | NA | #VALUE! |
| ENSMUSG00000086539.9 | 0 | NA | NA | NA | NA | NA | #VALUE! |
| ENSMUSG00000086540.3 | 0 | NA | NA | NA | NA | NA | #VALUE! |
| ENSMUSG00000086541.9 | 0 | NA | NA | NA | NA | NA | #VALUE! |
| ENSMUSG00000086542.2 | 0 | NA | NA | NA | NA | NA | #VALUE! |
| ENSMUSG00000086544.2 | 4.143924 | 5.546101 | 3.917561 | 1.415702 | 0.156863 | NA | #VALUE! |
| ENSMUSG00000086545.5 | 0 | NA | NA | NA | NA | NA | #VALUE! |
| ENSMUSG00000086546.2 | 0 | NA | NA | NA | NA | NA | #VALUE! |
| ENSMUSG00000086547.2 | 0 | NA | NA | NA | NA | NA | #VALUE! |
| ENSMUSG00000086548.2 | 0 | NA | NA | NA | NA | NA | #VALUE! |
| ENSMUSG00000086549.2 | 0 | NA | NA | NA | NA | NA | #VALUE! |
| ENSMUSG00000086550.3 | 0 | NA | NA | NA | NA | NA | #VALUE! |
| ENSMUSG00000086552.4 | 0 | NA | NA | NA | NA | NA | #VALUE! |
| ENSMUSG00000086554.2 | 0.816443 | -3.21914 | 3.961753 | -0.81255 | 0.416474 | NA | #VALUE! |
| ENSMUSG00000086555.3 | 0 | NA | NA | NA | NA | NA | #VALUE! |
| ENSMUSG00000086556.3 | 0 | NA | NA | NA | NA | NA | #VALUE! |

| | | | | | | | |
|---|---|---|---|---|---|---|---|
| ENSMUSG00000086557.2 | 0 | NA | NA | NA | NA | NA | #VALUE! |
| ENSMUSG00000086558.2 | 0 | NA | NA | NA | NA | NA | #VALUE! |
| ENSMUSG00000086559.2 | 0 | NA | NA | NA | NA | NA | #VALUE! |
| ENSMUSG00000086560.2 | 0 | NA | NA | NA | NA | NA | #VALUE! |
| ENSMUSG00000086562.2 | 0 | NA | NA | NA | NA | NA | #VALUE! |
| ENSMUSG00000086563.2 | 0 | NA | NA | NA | NA | NA | #VALUE! |
| ENSMUSG00000086564.10 | 0 | NA | NA | NA | NA | NA | #VALUE! |
| ENSMUSG00000086565.2 | 0 | NA | NA | NA | NA | NA | #VALUE! |
| ENSMUSG00000086569.2 | 0 | NA | NA | NA | NA | NA | #VALUE! |
| ENSMUSG00000086573.2 | 0 | NA | NA | NA | NA | NA | #VALUE! |
| ENSMUSG00000086575.2 | 0 | NA | NA | NA | NA | NA | #VALUE! |
| ENSMUSG00000086576.2 | 0 | NA | NA | NA | NA | NA | #VALUE! |
| ENSMUSG00000086577.2 | 0 | NA | NA | NA | NA | NA | #VALUE! |
| ENSMUSG00000086578.3 | 0 | NA | NA | NA | NA | NA | #VALUE! |
| ENSMUSG00000086580.2 | 0 | NA | NA | NA | NA | NA | #VALUE! |
| ENSMUSG00000086582.2 | 0 | NA | NA | NA | NA | NA | #VALUE! |
| ENSMUSG00000086584.9 | 0 | NA | NA | NA | NA | NA | #VALUE! |
| ENSMUSG00000086585.2 | 0 | NA | NA | NA | NA | NA | #VALUE! |
| ENSMUSG00000086586.2 | 0 | NA | NA | NA | NA | NA | #VALUE! |
| ENSMUSG00000086588.2 | 0 | NA | NA | NA | NA | NA | #VALUE! |
| ENSMUSG00000086589.2 | 0 | NA | NA | NA | NA | NA | #VALUE! |
| ENSMUSG00000086591.2 | 0 | NA | NA | NA | NA | NA | #VALUE! |
| ENSMUSG00000086592.2 | 0 | NA | NA | NA | NA | NA | #VALUE! |
| ENSMUSG00000086594.3 | 0 | NA | NA | NA | NA | NA | #VALUE! |
| ENSMUSG00000086595.2 | 0 | NA | NA | NA | NA | NA | #VALUE! |
| ENSMUSG00000086596.3 | 0 | NA | NA | NA | NA | NA | #VALUE! |
| ENSMUSG00000086598.10 | 0 | NA | NA | NA | NA | NA | #VALUE! |
| ENSMUSG00000086600.9 | 0 | NA | NA | NA | NA | NA | #VALUE! |
| ENSMUSG00000086601.2 | 0 | NA | NA | NA | NA | NA | #VALUE! |
| ENSMUSG00000086602.2 | 0 | NA | NA | NA | NA | NA | #VALUE! |
| ENSMUSG00000086603.2 | 0 | NA | NA | NA | NA | NA | #VALUE! |
| ENSMUSG00000086604.3 | 0 | NA | NA | NA | NA | NA | #VALUE! |
| ENSMUSG00000086606.2 | 0 | NA | NA | NA | NA | NA | #VALUE! |
| ENSMUSG00000086607.8 | 0 | NA | NA | NA | NA | NA | #VALUE! |
| ENSMUSG00000086608.2 | 0 | NA | NA | NA | NA | NA | #VALUE! |
| ENSMUSG00000086610.2 | 2.915831 | 2.717058 | 3.862523 | 0.703441 | 0.481781 | NA | #VALUE! |
| ENSMUSG00000086612.2 | 0 | NA | NA | NA | NA | NA | #VALUE! |
| ENSMUSG00000086613.2 | 0 | NA | NA | NA | NA | NA | #VALUE! |
| ENSMUSG00000086614.3 | 0 | NA | NA | NA | NA | NA | #VALUE! |
| ENSMUSG00000086617.2 | 0 | NA | NA | NA | NA | NA | #VALUE! |
| ENSMUSG00000086618.2 | 0 | NA | NA | NA | NA | NA | #VALUE! |
| ENSMUSG00000086620.2 | 0 | NA | NA | NA | NA | NA | #VALUE! |
| ENSMUSG00000086621.2 | 0 | NA | NA | NA | NA | NA | #VALUE! |
| ENSMUSG00000086624.2 | 0 | NA | NA | NA | NA | NA | #VALUE! |

| | | | | | | | |
|---|---|---|---|---|---|---|---|
| ENSMUSG00000086625.2 | 0 | NA | NA | NA | NA | NA | #VALUE! |
| ENSMUSG00000086629.9 | 4.680645 | -3.16201 | 3.768809 | -0.83899 | 0.401473 | NA | #VALUE! |
| ENSMUSG00000086630.2 | 0 | NA | NA | NA | NA | NA | #VALUE! |
| ENSMUSG00000086631.3 | 0 | NA | NA | NA | NA | NA | #VALUE! |
| ENSMUSG00000086632.2 | 0 | NA | NA | NA | NA | NA | #VALUE! |
| ENSMUSG00000086633.2 | 0 | NA | NA | NA | NA | NA | #VALUE! |
| ENSMUSG00000086636.2 | 0 | NA | NA | NA | NA | NA | #VALUE! |
| ENSMUSG00000086637.2 | 0 | NA | NA | NA | NA | NA | #VALUE! |
| ENSMUSG00000086638.8 | 0 | NA | NA | NA | NA | NA | #VALUE! |
| ENSMUSG00000086641.2 | 0 | NA | NA | NA | NA | NA | #VALUE! |
| ENSMUSG00000086643.2 | 0 | NA | NA | NA | NA | NA | #VALUE! |
| ENSMUSG00000086644.2 | 0 | NA | NA | NA | NA | NA | #VALUE! |
| ENSMUSG00000086645.3 | 0 | NA | NA | NA | NA | NA | #VALUE! |
| ENSMUSG00000086646.8 | 0 | NA | NA | NA | NA | NA | #VALUE! |
| ENSMUSG00000086648.2 | 0 | NA | NA | NA | NA | NA | #VALUE! |
| ENSMUSG00000086649.2 | 0 | NA | NA | NA | NA | NA | #VALUE! |
| ENSMUSG00000086650.2 | 0 | NA | NA | NA | NA | NA | #VALUE! |
| ENSMUSG00000086652.2 | 0 | NA | NA | NA | NA | NA | #VALUE! |
| ENSMUSG00000086653.2 | 0 | NA | NA | NA | NA | NA | #VALUE! |
| ENSMUSG00000086654.2 | 0 | NA | NA | NA | NA | NA | #VALUE! |
| ENSMUSG00000086655.2 | 0 | NA | NA | NA | NA | NA | #VALUE! |
| ENSMUSG00000086656.2 | 0 | NA | NA | NA | NA | NA | #VALUE! |
| ENSMUSG00000086657.3 | 0 | NA | NA | NA | NA | NA | #VALUE! |
| ENSMUSG00000086658.2 | 0 | NA | NA | NA | NA | NA | #VALUE! |
| ENSMUSG00000086659.2 | 0 | NA | NA | NA | NA | NA | #VALUE! |
| ENSMUSG00000086660.2 | 0 | NA | NA | NA | NA | NA | #VALUE! |
| ENSMUSG00000086662.8 | 0 | NA | NA | NA | NA | NA | #VALUE! |
| ENSMUSG00000086665.2 | 0 | NA | NA | NA | NA | NA | #VALUE! |
| ENSMUSG00000086666.2 | 0 | NA | NA | NA | NA | NA | #VALUE! |
| ENSMUSG00000086668.2 | 0 | NA | NA | NA | NA | NA | #VALUE! |
| ENSMUSG00000086669.2 | 0.673134 | 2.918386 | 3.972531 | 0.734641 | 0.462558 | NA | #VALUE! |
| ENSMUSG00000086670.2 | 0.735092 | -3.12016 | 3.965612 | -0.7868 | 0.431397 | NA | #VALUE! |
| ENSMUSG00000086671.2 | 0 | NA | NA | NA | NA | NA | #VALUE! |
| ENSMUSG00000086674.2 | 0 | NA | NA | NA | NA | NA | #VALUE! |
| ENSMUSG00000086675.2 | 0 | NA | NA | NA | NA | NA | #VALUE! |
| ENSMUSG00000086676.3 | 0 | NA | NA | NA | NA | NA | #VALUE! |
| ENSMUSG00000086677.2 | 0.816443 | -3.21914 | 3.961753 | -0.81255 | 0.416474 | NA | #VALUE! |
| ENSMUSG00000086679.3 | 0 | NA | NA | NA | NA | NA | #VALUE! |
| ENSMUSG00000086680.3 | 0 | NA | NA | NA | NA | NA | #VALUE! |
| ENSMUSG00000086681.2 | 0 | NA | NA | NA | NA | NA | #VALUE! |
| ENSMUSG00000086682.2 | 0 | NA | NA | NA | NA | NA | #VALUE! |
| ENSMUSG00000086683.2 | 0 | NA | NA | NA | NA | NA | #VALUE! |
| ENSMUSG00000086684.2 | 0 | NA | NA | NA | NA | NA | #VALUE! |
| ENSMUSG00000086685.2 | 0 | NA | NA | NA | NA | NA | #VALUE! |

| | | | | | | | |
|---|---|---|---|---|---|---|---|
| ENSMUSG00000086686.2 | 0 | NA | NA | NA | NA | NA | #VALUE! |
| ENSMUSG00000086687.2 | 0 | NA | NA | NA | NA | NA | #VALUE! |
| ENSMUSG00000086688.5 | 0 | NA | NA | NA | NA | NA | #VALUE! |
| ENSMUSG00000086689.2 | 0 | NA | NA | NA | NA | NA | #VALUE! |
| ENSMUSG00000086690.2 | 0 | NA | NA | NA | NA | NA | #VALUE! |
| ENSMUSG00000086691.3 | 0.245976 | 1.027628 | 4.067126 | 0.252667 | 0.800526 | NA | #VALUE! |
| ENSMUSG00000086693.2 | 0 | NA | NA | NA | NA | NA | #VALUE! |
| ENSMUSG00000086694.2 | 0 | NA | NA | NA | NA | NA | #VALUE! |
| ENSMUSG00000086695.2 | 0.315039 | -1.93549 | 4.038621 | -0.47925 | 0.631763 | NA | #VALUE! |
| ENSMUSG00000086696.2 | 0 | NA | NA | NA | NA | NA | #VALUE! |
| ENSMUSG00000086697.3 | 0 | NA | NA | NA | NA | NA | #VALUE! |
| ENSMUSG00000086698.2 | 0 | NA | NA | NA | NA | NA | #VALUE! |
| ENSMUSG00000086699.4 | 0 | NA | NA | NA | NA | NA | #VALUE! |
| ENSMUSG00000086700.3 | 0 | NA | NA | NA | NA | NA | #VALUE! |
| ENSMUSG00000086701.2 | 0 | NA | NA | NA | NA | NA | #VALUE! |
| ENSMUSG00000086702.2 | 0 | NA | NA | NA | NA | NA | #VALUE! |
| ENSMUSG00000086703.2 | 0 | NA | NA | NA | NA | NA | #VALUE! |
| ENSMUSG00000086704.2 | 0 | NA | NA | NA | NA | NA | #VALUE! |
| ENSMUSG00000086706.8 | 0 | NA | NA | NA | NA | NA | #VALUE! |
| ENSMUSG00000086707.2 | 0 | NA | NA | NA | NA | NA | #VALUE! |
| ENSMUSG00000086708.2 | 0 | NA | NA | NA | NA | NA | #VALUE! |
| ENSMUSG00000086709.2 | 0 | NA | NA | NA | NA | NA | #VALUE! |
| ENSMUSG00000086710.2 | 0 | NA | NA | NA | NA | NA | #VALUE! |
| ENSMUSG00000086711.2 | 0 | NA | NA | NA | NA | NA | #VALUE! |
| ENSMUSG00000086712.3 | 0.279766 | 1.808031 | 4.036658 | 0.447903 | 0.654223 | NA | #VALUE! |
| ENSMUSG00000086713.2 | 0 | NA | NA | NA | NA | NA | #VALUE! |
| ENSMUSG00000086714.2 | 8.414176 | 6.562668 | 3.80431 | 1.725061 | 0.084516 | NA | #VALUE! |
| ENSMUSG00000086715.2 | 0 | NA | NA | NA | NA | NA | #VALUE! |
| ENSMUSG00000086716.2 | 0 | NA | NA | NA | NA | NA | #VALUE! |
| ENSMUSG00000086717.2 | 1.119062 | 3.679627 | 3.945843 | 0.932533 | 0.351061 | NA | #VALUE! |
| ENSMUSG00000086718.2 | 0 | NA | NA | NA | NA | NA | #VALUE! |
| ENSMUSG00000086719.2 | 0 | NA | NA | NA | NA | NA | #VALUE! |
| ENSMUSG00000086720.2 | 0 | NA | NA | NA | NA | NA | #VALUE! |
| ENSMUSG00000086721.3 | 0 | NA | NA | NA | NA | NA | #VALUE! |
| ENSMUSG00000086722.2 | 0 | NA | NA | NA | NA | NA | #VALUE! |
| ENSMUSG00000086724.2 | 0 | NA | NA | NA | NA | NA | #VALUE! |
| ENSMUSG00000086725.2 | 2.646266 | 4.883103 | 3.923833 | 1.244473 | 0.213326 | NA | #VALUE! |
| ENSMUSG00000086726.4 | 0 | NA | NA | NA | NA | NA | #VALUE! |
| ENSMUSG00000086727.9 | 1.346268 | 3.918821 | 3.939925 | 0.994644 | 0.31991 | NA | #VALUE! |
| ENSMUSG00000086728.3 | 1.232996 | 1.131842 | 3.89811 | 0.290357 | 0.771543 | NA | #VALUE! |
| ENSMUSG00000086729.2 | 0 | NA | NA | NA | NA | NA | #VALUE! |
| ENSMUSG00000086730.3 | 0 | NA | NA | NA | NA | NA | #VALUE! |
| ENSMUSG00000086731.2 | 0 | NA | NA | NA | NA | NA | #VALUE! |
| ENSMUSG00000086732.2 | 0 | NA | NA | NA | NA | NA | #VALUE! |

| | | | | | | | |
|---|---|---|---|---|---|---|---|
| ENSMUSG00000086733.2 | 0 | NA | NA | NA | NA | NA | #VALUE! |
| ENSMUSG00000086735.2 | 0 | NA | NA | NA | NA | NA | #VALUE! |
| ENSMUSG00000086736.2 | 0 | NA | NA | NA | NA | NA | #VALUE! |
| ENSMUSG00000086738.2 | 0 | NA | NA | NA | NA | NA | #VALUE! |
| ENSMUSG00000086739.2 | 0 | NA | NA | NA | NA | NA | #VALUE! |
| ENSMUSG00000086740.2 | 0 | NA | NA | NA | NA | NA | #VALUE! |
| ENSMUSG00000086741.2 | 0 | NA | NA | NA | NA | NA | #VALUE! |
| ENSMUSG00000086742.9 | 0 | NA | NA | NA | NA | NA | #VALUE! |
| ENSMUSG00000086743.3 | 0 | NA | NA | NA | NA | NA | #VALUE! |
| ENSMUSG00000086745.2 | 0 | NA | NA | NA | NA | NA | #VALUE! |
| ENSMUSG00000086746.3 | 0 | NA | NA | NA | NA | NA | #VALUE! |
| ENSMUSG00000086748.2 | 0 | NA | NA | NA | NA | NA | #VALUE! |
| ENSMUSG00000086749.2 | 0 | NA | NA | NA | NA | NA | #VALUE! |
| ENSMUSG00000086750.2 | 0 | NA | NA | NA | NA | NA | #VALUE! |
| ENSMUSG00000086752.2 | 0 | NA | NA | NA | NA | NA | #VALUE! |
| ENSMUSG00000086753.2 | 0 | NA | NA | NA | NA | NA | #VALUE! |
| ENSMUSG00000086754.2 | 0 | NA | NA | NA | NA | NA | #VALUE! |
| ENSMUSG00000086755.2 | 0 | NA | NA | NA | NA | NA | #VALUE! |
| ENSMUSG00000086756.2 | 0 | NA | NA | NA | NA | NA | #VALUE! |
| ENSMUSG00000086757.2 | 0 | NA | NA | NA | NA | NA | #VALUE! |
| ENSMUSG00000086758.2 | 0 | NA | NA | NA | NA | NA | #VALUE! |
| ENSMUSG00000086760.2 | 0 | NA | NA | NA | NA | NA | #VALUE! |
| ENSMUSG00000086761.2 | 0 | NA | NA | NA | NA | NA | #VALUE! |
| ENSMUSG00000086762.2 | 0 | NA | NA | NA | NA | NA | #VALUE! |
| ENSMUSG00000086763.2 | 0 | NA | NA | NA | NA | NA | #VALUE! |
| ENSMUSG00000086764.2 | 0 | NA | NA | NA | NA | NA | #VALUE! |
| ENSMUSG00000086765.8 | 0 | NA | NA | NA | NA | NA | #VALUE! |
| ENSMUSG00000086766.2 | 0 | NA | NA | NA | NA | NA | #VALUE! |
| ENSMUSG00000086767.2 | 0 | NA | NA | NA | NA | NA | #VALUE! |
| ENSMUSG00000086769.8 | 0 | NA | NA | NA | NA | NA | #VALUE! |
| ENSMUSG00000086770.2 | 0 | NA | NA | NA | NA | NA | #VALUE! |
| ENSMUSG00000086773.2 | 0 | NA | NA | NA | NA | NA | #VALUE! |
| ENSMUSG00000086774.2 | 0 | NA | NA | NA | NA | NA | #VALUE! |
| ENSMUSG00000086775.8 | 0 | NA | NA | NA | NA | NA | #VALUE! |
| ENSMUSG00000086777.4 | 0 | NA | NA | NA | NA | NA | #VALUE! |
| ENSMUSG00000086779.2 | 0 | NA | NA | NA | NA | NA | #VALUE! |
| ENSMUSG00000086781.2 | 0 | NA | NA | NA | NA | NA | #VALUE! |
| ENSMUSG00000086782.3 | 0.408222 | -2.20205 | 4.016842 | -0.54821 | 0.583551 | NA | #VALUE! |
| ENSMUSG00000086783.2 | 0 | NA | NA | NA | NA | NA | #VALUE! |
| ENSMUSG00000086785.2 | 0 | NA | NA | NA | NA | NA | #VALUE! |
| ENSMUSG00000086786.2 | 0 | NA | NA | NA | NA | NA | #VALUE! |
| ENSMUSG00000086788.3 | 0 | NA | NA | NA | NA | NA | #VALUE! |
| ENSMUSG00000086789.2 | 0 | NA | NA | NA | NA | NA | #VALUE! |
| ENSMUSG00000086790.2 | 0 | NA | NA | NA | NA | NA | #VALUE! |

| | | | | | | | |
|---|---|---|---|---|---|---|---|
| ENSMUSG00000086791.2 | 0 | NA | NA | NA | NA | NA | #VALUE! |
| ENSMUSG00000086792.2 | 0 | NA | NA | NA | NA | NA | #VALUE! |
| ENSMUSG00000086794.2 | 0 | NA | NA | NA | NA | NA | #VALUE! |
| ENSMUSG00000086795.2 | 0 | NA | NA | NA | NA | NA | #VALUE! |
| ENSMUSG00000086796.3 | 0 | NA | NA | NA | NA | NA | #VALUE! |
| ENSMUSG00000086798.2 | 0 | NA | NA | NA | NA | NA | #VALUE! |
| ENSMUSG00000086799.2 | 0 | NA | NA | NA | NA | NA | #VALUE! |
| ENSMUSG00000086800.2 | 0 | NA | NA | NA | NA | NA | #VALUE! |
| ENSMUSG00000086801.2 | 0 | NA | NA | NA | NA | NA | #VALUE! |
| ENSMUSG00000086804.9 | 0 | NA | NA | NA | NA | NA | #VALUE! |
| ENSMUSG00000086805.10 | 0 | NA | NA | NA | NA | NA | #VALUE! |
| ENSMUSG00000086806.2 | 0 | NA | NA | NA | NA | NA | #VALUE! |
| ENSMUSG00000086807.9 | 0 | NA | NA | NA | NA | NA | #VALUE! |
| ENSMUSG00000086808.2 | 0 | NA | NA | NA | NA | NA | #VALUE! |
| ENSMUSG00000086810.2 | 0 | NA | NA | NA | NA | NA | #VALUE! |
| ENSMUSG00000086812.3 | 0 | NA | NA | NA | NA | NA | #VALUE! |
| ENSMUSG00000086813.2 | 1.847869 | -0.79387 | 3.833129 | -0.20711 | 0.835927 | NA | #VALUE! |
| ENSMUSG00000086814.2 | 0 | NA | NA | NA | NA | NA | #VALUE! |
| ENSMUSG00000086815.5 | 0 | NA | NA | NA | NA | NA | #VALUE! |
| ENSMUSG00000086816.2 | 0 | NA | NA | NA | NA | NA | #VALUE! |
| ENSMUSG00000086817.2 | 0 | NA | NA | NA | NA | NA | #VALUE! |
| ENSMUSG00000086818.2 | 0 | NA | NA | NA | NA | NA | #VALUE! |
| ENSMUSG00000086819.2 | 0 | NA | NA | NA | NA | NA | #VALUE! |
| ENSMUSG00000086820.2 | 0 | NA | NA | NA | NA | NA | #VALUE! |
| ENSMUSG00000086821.4 | 0 | NA | NA | NA | NA | NA | #VALUE! |
| ENSMUSG00000086822.3 | 0 | NA | NA | NA | NA | NA | #VALUE! |
| ENSMUSG00000086823.8 | 0 | NA | NA | NA | NA | NA | #VALUE! |
| ENSMUSG00000086824.2 | 0 | NA | NA | NA | NA | NA | #VALUE! |
| ENSMUSG00000086825.2 | 7.980998 | -6.53008 | 3.849249 | -1.69645 | NA | NA | #VALUE! |
| ENSMUSG00000086826.2 | 0 | NA | NA | NA | NA | NA | #VALUE! |
| ENSMUSG00000086827.2 | 0 | NA | NA | NA | NA | NA | #VALUE! |
| ENSMUSG00000086828.2 | 0 | NA | NA | NA | NA | NA | #VALUE! |
| ENSMUSG00000086830.2 | 0 | NA | NA | NA | NA | NA | #VALUE! |
| ENSMUSG00000086831.2 | 0 | NA | NA | NA | NA | NA | #VALUE! |
| ENSMUSG00000086833.3 | 0 | NA | NA | NA | NA | NA | #VALUE! |
| ENSMUSG00000086834.2 | 0 | NA | NA | NA | NA | NA | #VALUE! |
| ENSMUSG00000086835.3 | 0 | NA | NA | NA | NA | NA | #VALUE! |
| ENSMUSG00000086836.2 | 0 | NA | NA | NA | NA | NA | #VALUE! |
| ENSMUSG00000086837.2 | 0 | NA | NA | NA | NA | NA | #VALUE! |
| ENSMUSG00000086838.2 | 0 | NA | NA | NA | NA | NA | #VALUE! |
| ENSMUSG00000086839.8 | 0 | NA | NA | NA | NA | NA | #VALUE! |
| ENSMUSG00000086840.2 | 0 | NA | NA | NA | NA | NA | #VALUE! |
| ENSMUSG00000086842.3 | 0 | NA | NA | NA | NA | NA | #VALUE! |
| ENSMUSG00000086843.4 | 0 | NA | NA | NA | NA | NA | #VALUE! |

| | | | | | | | |
|---|---|---|---|---|---|---|---|
| ENSMUSG00000086844.2 | 0 | NA | NA | NA | NA | NA | #VALUE! |
| ENSMUSG00000086845.2 | 0 | NA | NA | NA | NA | NA | #VALUE! |
| ENSMUSG00000086846.3 | 0 | NA | NA | NA | NA | NA | #VALUE! |
| ENSMUSG00000086847.2 | 0 | NA | NA | NA | NA | NA | #VALUE! |
| ENSMUSG00000086848.4 | 0 | NA | NA | NA | NA | NA | #VALUE! |
| ENSMUSG00000086849.2 | 5.060157 | 1.428317 | 3.771948 | 0.378668 | 0.704934 | NA | #VALUE! |
| ENSMUSG00000086850.2 | 0 | NA | NA | NA | NA | NA | #VALUE! |
| ENSMUSG00000086851.2 | 0 | NA | NA | NA | NA | NA | #VALUE! |
| ENSMUSG00000086854.2 | 0 | NA | NA | NA | NA | NA | #VALUE! |
| ENSMUSG00000086855.2 | 0 | NA | NA | NA | NA | NA | #VALUE! |
| ENSMUSG00000086856.2 | 0 | NA | NA | NA | NA | NA | #VALUE! |
| ENSMUSG00000086858.3 | 0 | NA | NA | NA | NA | NA | #VALUE! |
| ENSMUSG00000086860.2 | 0 | NA | NA | NA | NA | NA | #VALUE! |
| ENSMUSG00000086861.2 | 0 | NA | NA | NA | NA | NA | #VALUE! |
| ENSMUSG00000086862.2 | 0 | NA | NA | NA | NA | NA | #VALUE! |
| ENSMUSG00000086863.2 | 0 | NA | NA | NA | NA | NA | #VALUE! |
| ENSMUSG00000086866.3 | 0 | NA | NA | NA | NA | NA | #VALUE! |
| ENSMUSG00000086867.2 | 0 | NA | NA | NA | NA | NA | #VALUE! |
| ENSMUSG00000086868.2 | 6.41088 | 6.171344 | 3.636375 | 1.697115 | 0.089675 | NA | #VALUE! |
| ENSMUSG00000086870.2 | 0 | NA | NA | NA | NA | NA | #VALUE! |
| ENSMUSG00000086872.2 | 0 | NA | NA | NA | NA | NA | #VALUE! |
| ENSMUSG00000086873.2 | 0 | NA | NA | NA | NA | NA | #VALUE! |
| ENSMUSG00000086874.2 | 0 | NA | NA | NA | NA | NA | #VALUE! |
| ENSMUSG00000086875.3 | 0 | NA | NA | NA | NA | NA | #VALUE! |
| ENSMUSG00000086876.2 | 0 | NA | NA | NA | NA | NA | #VALUE! |
| ENSMUSG00000086877.8 | 1.305573 | 3.898287 | 3.940396 | 0.989314 | 0.32251 | NA | #VALUE! |
| ENSMUSG00000086881.2 | 0.168284 | 1.027628 | 4.067126 | 0.252667 | 0.800526 | NA | #VALUE! |
| ENSMUSG00000086882.5 | 0.168284 | 1.027628 | 4.067126 | 0.252667 | 0.800526 | NA | #VALUE! |
| ENSMUSG00000086884.2 | 0 | NA | NA | NA | NA | NA | #VALUE! |
| ENSMUSG00000086887.2 | 0 | NA | NA | NA | NA | NA | #VALUE! |
| ENSMUSG00000086889.2 | 0 | NA | NA | NA | NA | NA | #VALUE! |
| ENSMUSG00000086890.2 | 0 | NA | NA | NA | NA | NA | #VALUE! |
| ENSMUSG00000086891.3 | 0 | NA | NA | NA | NA | NA | #VALUE! |
| ENSMUSG00000086892.2 | 0 | NA | NA | NA | NA | NA | #VALUE! |
| ENSMUSG00000086893.2 | 0 | NA | NA | NA | NA | NA | #VALUE! |
| ENSMUSG00000086894.3 | 0 | NA | NA | NA | NA | NA | #VALUE! |
| ENSMUSG00000086896.2 | 0 | NA | NA | NA | NA | NA | #VALUE! |
| ENSMUSG00000086898.2 | 0 | NA | NA | NA | NA | NA | #VALUE! |
| ENSMUSG00000086900.2 | 0 | NA | NA | NA | NA | NA | #VALUE! |
| ENSMUSG00000086902.8 | 0 | NA | NA | NA | NA | NA | #VALUE! |
| ENSMUSG00000086903.5 | 0 | NA | NA | NA | NA | NA | #VALUE! |
| ENSMUSG00000086904.2 | 0 | NA | NA | NA | NA | NA | #VALUE! |
| ENSMUSG00000086905.2 | 0 | NA | NA | NA | NA | NA | #VALUE! |
| ENSMUSG00000086906.2 | 0 | NA | NA | NA | NA | NA | #VALUE! |

| | | | | | | | |
|---|---|---|---|---|---|---|---|
| ENSMUSG00000086907.2 | 0 | NA | NA | NA | NA | NA | #VALUE! |
| ENSMUSG00000086908.2 | 0 | NA | NA | NA | NA | NA | #VALUE! |
| ENSMUSG00000086909.2 | 0 | NA | NA | NA | NA | NA | #VALUE! |
| ENSMUSG00000086910.2 | 0 | NA | NA | NA | NA | NA | #VALUE! |
| ENSMUSG00000086911.2 | 0 | NA | NA | NA | NA | NA | #VALUE! |
| ENSMUSG00000086912.4 | 0 | NA | NA | NA | NA | NA | #VALUE! |
| ENSMUSG00000086913.2 | 0 | NA | NA | NA | NA | NA | #VALUE! |
| ENSMUSG00000086915.8 | 0 | NA | NA | NA | NA | NA | #VALUE! |
| ENSMUSG00000086916.3 | 2.692536 | 4.918862 | 3.923417 | 1.253719 | 0.209944 | NA | #VALUE! |
| ENSMUSG00000086917.2 | 0 | NA | NA | NA | NA | NA | #VALUE! |
| ENSMUSG00000086918.2 | 0.630079 | -2.90291 | 3.975047 | -0.73028 | 0.465217 | NA | #VALUE! |
| ENSMUSG00000086919.2 | 0 | NA | NA | NA | NA | NA | #VALUE! |
| ENSMUSG00000086920.2 | 0 | NA | NA | NA | NA | NA | #VALUE! |
| ENSMUSG00000086921.2 | 0 | NA | NA | NA | NA | NA | #VALUE! |
| ENSMUSG00000086922.3 | 3.979907 | 0.37767 | 2.802 | 0.134786 | 0.892781 | NA | #VALUE! |
| ENSMUSG00000086923.2 | 0 | NA | NA | NA | NA | NA | #VALUE! |
| ENSMUSG00000086924.2 | 0 | NA | NA | NA | NA | NA | #VALUE! |
| ENSMUSG00000086925.2 | 0 | NA | NA | NA | NA | NA | #VALUE! |
| ENSMUSG00000086926.3 | 0 | NA | NA | NA | NA | NA | #VALUE! |
| ENSMUSG00000086927.2 | 0 | NA | NA | NA | NA | NA | #VALUE! |
| ENSMUSG00000086928.2 | 0 | NA | NA | NA | NA | NA | #VALUE! |
| ENSMUSG00000086929.2 | 0 | NA | NA | NA | NA | NA | #VALUE! |
| ENSMUSG00000086930.2 | 0 | NA | NA | NA | NA | NA | #VALUE! |
| ENSMUSG00000086931.2 | 0 | NA | NA | NA | NA | NA | #VALUE! |
| ENSMUSG00000086932.2 | 0 | NA | NA | NA | NA | NA | #VALUE! |
| ENSMUSG00000086933.2 | 0 | NA | NA | NA | NA | NA | #VALUE! |
| ENSMUSG00000086935.2 | 5.915773 | 6.059096 | 3.734665 | 1.622393 | 0.104719 | NA | #VALUE! |
| ENSMUSG00000086936.2 | 0 | NA | NA | NA | NA | NA | #VALUE! |
| ENSMUSG00000086937.2 | 0 | NA | NA | NA | NA | NA | #VALUE! |
| ENSMUSG00000086939.2 | 0 | NA | NA | NA | NA | NA | #VALUE! |
| ENSMUSG00000086940.2 | 0 | NA | NA | NA | NA | NA | #VALUE! |
| ENSMUSG00000086941.2 | 0 | NA | NA | NA | NA | NA | #VALUE! |
| ENSMUSG00000086942.2 | 0 | NA | NA | NA | NA | NA | #VALUE! |
| ENSMUSG00000086943.3 | 0 | NA | NA | NA | NA | NA | #VALUE! |
| ENSMUSG00000086944.2 | 0 | NA | NA | NA | NA | NA | #VALUE! |
| ENSMUSG00000086945.2 | 0 | NA | NA | NA | NA | NA | #VALUE! |
| ENSMUSG00000086946.8 | 0 | NA | NA | NA | NA | NA | #VALUE! |
| ENSMUSG00000086947.2 | 0 | NA | NA | NA | NA | NA | #VALUE! |
| ENSMUSG00000086948.2 | 0 | NA | NA | NA | NA | NA | #VALUE! |
| ENSMUSG00000086949.2 | 0 | NA | NA | NA | NA | NA | #VALUE! |
| ENSMUSG00000086951.2 | 0 | NA | NA | NA | NA | NA | #VALUE! |
| ENSMUSG00000086952.2 | 0 | NA | NA | NA | NA | NA | #VALUE! |
| ENSMUSG00000086953.2 | 0 | NA | NA | NA | NA | NA | #VALUE! |
| ENSMUSG00000086954.2 | 0 | NA | NA | NA | NA | NA | #VALUE! |

| | | | | | | | |
|---|---|---|---|---|---|---|---|
| ENSMUSG00000086955.2 | 0 | NA | NA | NA | NA | NA | #VALUE! |
| ENSMUSG00000086957.2 | 0 | NA | NA | NA | NA | NA | #VALUE! |
| ENSMUSG00000086959.2 | 0 | NA | NA | NA | NA | NA | #VALUE! |
| ENSMUSG00000086960.2 | 0 | NA | NA | NA | NA | NA | #VALUE! |
| ENSMUSG00000086961.2 | 0 | NA | NA | NA | NA | NA | #VALUE! |
| ENSMUSG00000086962.5 | 0 | NA | NA | NA | NA | NA | #VALUE! |
| ENSMUSG00000086963.2 | 0 | NA | NA | NA | NA | NA | #VALUE! |
| ENSMUSG00000086964.2 | 0 | NA | NA | NA | NA | NA | #VALUE! |
| ENSMUSG00000086965.2 | 0 | NA | NA | NA | NA | NA | #VALUE! |
| ENSMUSG00000086969.2 | 0 | NA | NA | NA | NA | NA | #VALUE! |
| ENSMUSG00000086970.2 | 0 | NA | NA | NA | NA | NA | #VALUE! |
| ENSMUSG00000086971.2 | 0 | NA | NA | NA | NA | NA | #VALUE! |
| ENSMUSG00000086974.2 | 0 | NA | NA | NA | NA | NA | #VALUE! |
| ENSMUSG00000086975.2 | 0 | NA | NA | NA | NA | NA | #VALUE! |
| ENSMUSG00000086976.2 | 0 | NA | NA | NA | NA | NA | #VALUE! |
| ENSMUSG00000086978.2 | 0 | NA | NA | NA | NA | NA | #VALUE! |
| ENSMUSG00000086980.2 | 0 | NA | NA | NA | NA | NA | #VALUE! |
| ENSMUSG00000086981.4 | 0 | NA | NA | NA | NA | NA | #VALUE! |
| ENSMUSG00000086984.5 | 0 | NA | NA | NA | NA | NA | #VALUE! |
| ENSMUSG00000086986.4 | 0 | NA | NA | NA | NA | NA | #VALUE! |
| ENSMUSG00000086987.2 | 0 | NA | NA | NA | NA | NA | #VALUE! |
| ENSMUSG00000086989.4 | 0 | NA | NA | NA | NA | NA | #VALUE! |
| ENSMUSG00000086991.2 | 0 | NA | NA | NA | NA | NA | #VALUE! |
| ENSMUSG00000086992.8 | 0 | NA | NA | NA | NA | NA | #VALUE! |
| ENSMUSG00000086993.2 | 0 | NA | NA | NA | NA | NA | #VALUE! |
| ENSMUSG00000086994.2 | 0 | NA | NA | NA | NA | NA | #VALUE! |
| ENSMUSG00000086995.2 | 0 | NA | NA | NA | NA | NA | #VALUE! |
| ENSMUSG00000086996.2 | 0 | NA | NA | NA | NA | NA | #VALUE! |
| ENSMUSG00000086997.2 | 0 | NA | NA | NA | NA | NA | #VALUE! |
| ENSMUSG00000086998.2 | 0 | NA | NA | NA | NA | NA | #VALUE! |
| ENSMUSG00000086999.2 | 0 | NA | NA | NA | NA | NA | #VALUE! |
| ENSMUSG00000087001.3 | 0 | NA | NA | NA | NA | NA | #VALUE! |
| ENSMUSG00000087002.2 | 8.652712 | -0.96661 | 3.054113 | -0.3165 | 0.751627 | NA | #VALUE! |
| ENSMUSG00000087003.2 | 0 | NA | NA | NA | NA | NA | #VALUE! |
| ENSMUSG00000087004.2 | 0 | NA | NA | NA | NA | NA | #VALUE! |
| ENSMUSG00000087006.4 | 1.470184 | -4.10383 | 3.936698 | -1.04245 | 0.297201 | NA | #VALUE! |
| ENSMUSG00000087007.2 | 0 | NA | NA | NA | NA | NA | #VALUE! |
| ENSMUSG00000087008.5 | 0 | NA | NA | NA | NA | NA | #VALUE! |
| ENSMUSG00000087009.2 | 0 | NA | NA | NA | NA | NA | #VALUE! |
| ENSMUSG00000087010.2 | 0 | NA | NA | NA | NA | NA | #VALUE! |
| ENSMUSG00000087011.2 | 0 | NA | NA | NA | NA | NA | #VALUE! |
| ENSMUSG00000087013.8 | 0 | NA | NA | NA | NA | NA | #VALUE! |
| ENSMUSG00000087014.2 | 0 | NA | NA | NA | NA | NA | #VALUE! |
| ENSMUSG00000087015.2 | 0 | NA | NA | NA | NA | NA | #VALUE! |

| | | | | | | | |
|---|---|---|---|---|---|---|---|
| ENSMUSG00000087016.2 | 0 | NA | NA | NA | NA | NA | #VALUE! |
| ENSMUSG00000087017.8 | 2.96806 | 5.066512 | 3.921803 | 1.291883 | 0.196398 | NA | #VALUE! |
| ENSMUSG00000087018.2 | 0 | NA | NA | NA | NA | NA | #VALUE! |
| ENSMUSG00000087020.2 | 5.219613 | -5.91181 | 3.915351 | -1.5099 | 0.131068 | NA | #VALUE! |
| ENSMUSG00000087021.4 | 0 | NA | NA | NA | NA | NA | #VALUE! |
| ENSMUSG00000087022.5 | 0 | NA | NA | NA | NA | NA | #VALUE! |
| ENSMUSG00000087024.2 | 0 | NA | NA | NA | NA | NA | #VALUE! |
| ENSMUSG00000087026.8 | 0 | NA | NA | NA | NA | NA | #VALUE! |
| ENSMUSG00000087027.3 | 0 | NA | NA | NA | NA | NA | #VALUE! |
| ENSMUSG00000087028.8 | 0 | NA | NA | NA | NA | NA | #VALUE! |
| ENSMUSG00000087029.2 | 0 | NA | NA | NA | NA | NA | #VALUE! |
| ENSMUSG00000087030.2 | 0 | NA | NA | NA | NA | NA | #VALUE! |
| ENSMUSG00000087031.2 | 0 | NA | NA | NA | NA | NA | #VALUE! |
| ENSMUSG00000087032.2 | 0 | NA | NA | NA | NA | NA | #VALUE! |
| ENSMUSG00000087033.2 | 0 | NA | NA | NA | NA | NA | #VALUE! |
| ENSMUSG00000087034.5 | 0 | NA | NA | NA | NA | NA | #VALUE! |
| ENSMUSG00000087035.2 | 0 | NA | NA | NA | NA | NA | #VALUE! |
| ENSMUSG00000087036.2 | 0 | NA | NA | NA | NA | NA | #VALUE! |
| ENSMUSG00000087037.2 | 0 | NA | NA | NA | NA | NA | #VALUE! |
| ENSMUSG00000087038.10 | 0 | NA | NA | NA | NA | NA | #VALUE! |
| ENSMUSG00000087039.2 | 0 | NA | NA | NA | NA | NA | #VALUE! |
| ENSMUSG00000087040.2 | 0 | NA | NA | NA | NA | NA | #VALUE! |
| ENSMUSG00000087041.2 | 0 | NA | NA | NA | NA | NA | #VALUE! |
| ENSMUSG00000087042.2 | 0 | NA | NA | NA | NA | NA | #VALUE! |
| ENSMUSG00000087043.2 | 0 | NA | NA | NA | NA | NA | #VALUE! |
| ENSMUSG00000087044.2 | 0 | NA | NA | NA | NA | NA | #VALUE! |
| ENSMUSG00000087045.2 | 0 | NA | NA | NA | NA | NA | #VALUE! |
| ENSMUSG00000087047.9 | 0 | NA | NA | NA | NA | NA | #VALUE! |
| ENSMUSG00000087049.2 | 6.265251 | -6.17547 | 3.913919 | -1.57782 | 0.114607 | NA | #VALUE! |
| ENSMUSG00000087050.2 | 2.580136 | 1.69167 | 3.825459 | 0.442214 | 0.658335 | NA | #VALUE! |
| ENSMUSG00000087051.2 | 0 | NA | NA | NA | NA | NA | #VALUE! |
| ENSMUSG00000087052.2 | 0 | NA | NA | NA | NA | NA | #VALUE! |
| ENSMUSG00000087053.2 | 0 | NA | NA | NA | NA | NA | #VALUE! |
| ENSMUSG00000087054.2 | 0 | NA | NA | NA | NA | NA | #VALUE! |
| ENSMUSG00000087055.2 | 0 | NA | NA | NA | NA | NA | #VALUE! |
| ENSMUSG00000087056.2 | 0 | NA | NA | NA | NA | NA | #VALUE! |
| ENSMUSG00000087057.8 | 0 | NA | NA | NA | NA | NA | #VALUE! |
| ENSMUSG00000087058.2 | 0 | NA | NA | NA | NA | NA | #VALUE! |
| ENSMUSG00000087059.8 | 0 | NA | NA | NA | NA | NA | #VALUE! |
| ENSMUSG00000087060.8 | 0 | NA | NA | NA | NA | NA | #VALUE! |
| ENSMUSG00000087061.2 | 0 | NA | NA | NA | NA | NA | #VALUE! |
| ENSMUSG00000087063.2 | 0 | NA | NA | NA | NA | NA | #VALUE! |
| ENSMUSG00000087064.2 | 0 | NA | NA | NA | NA | NA | #VALUE! |
| ENSMUSG00000087066.2 | 0 | NA | NA | NA | NA | NA | #VALUE! |

| | | | | | | | |
|---|---|---|---|---|---|---|---|
| ENSMUSG00000087067.2 | 0 | NA | NA | NA | NA | NA | #VALUE! |
| ENSMUSG00000087069.2 | 0 | NA | NA | NA | NA | NA | #VALUE! |
| ENSMUSG00000087070.2 | 0 | NA | NA | NA | NA | NA | #VALUE! |
| ENSMUSG00000087071.2 | 0 | NA | NA | NA | NA | NA | #VALUE! |
| ENSMUSG00000087072.2 | 0 | NA | NA | NA | NA | NA | #VALUE! |
| ENSMUSG00000087074.2 | 0 | NA | NA | NA | NA | NA | #VALUE! |
| ENSMUSG00000087075.3 | 0 | NA | NA | NA | NA | NA | #VALUE! |
| ENSMUSG00000087076.3 | 0 | NA | NA | NA | NA | NA | #VALUE! |
| ENSMUSG00000087077.2 | 0 | NA | NA | NA | NA | NA | #VALUE! |
| ENSMUSG00000087079.2 | 0 | NA | NA | NA | NA | NA | #VALUE! |
| ENSMUSG00000087080.2 | 0 | NA | NA | NA | NA | NA | #VALUE! |
| ENSMUSG00000087082.2 | 0 | NA | NA | NA | NA | NA | #VALUE! |
| ENSMUSG00000087083.3 | 0 | NA | NA | NA | NA | NA | #VALUE! |
| ENSMUSG00000087084.2 | 0 | NA | NA | NA | NA | NA | #VALUE! |
| ENSMUSG00000087085.2 | 0 | NA | NA | NA | NA | NA | #VALUE! |
| ENSMUSG00000087086.8 | 0 | NA | NA | NA | NA | NA | #VALUE! |
| ENSMUSG00000087088.3 | 0 | NA | NA | NA | NA | NA | #VALUE! |
| ENSMUSG00000087089.2 | 0 | NA | NA | NA | NA | NA | #VALUE! |
| ENSMUSG00000087090.8 | 0 | NA | NA | NA | NA | NA | #VALUE! |
| ENSMUSG00000087093.3 | 0 | NA | NA | NA | NA | NA | #VALUE! |
| ENSMUSG00000087094.2 | 0 | NA | NA | NA | NA | NA | #VALUE! |
| ENSMUSG00000087095.3 | 0 | NA | NA | NA | NA | NA | #VALUE! |
| ENSMUSG00000087097.2 | 0 | NA | NA | NA | NA | NA | #VALUE! |
| ENSMUSG00000087098.2 | 0 | NA | NA | NA | NA | NA | #VALUE! |
| ENSMUSG00000087099.2 | 0 | NA | NA | NA | NA | NA | #VALUE! |
| ENSMUSG00000087100.2 | 0 | NA | NA | NA | NA | NA | #VALUE! |
| ENSMUSG00000087102.2 | 0 | NA | NA | NA | NA | NA | #VALUE! |
| ENSMUSG00000087103.2 | 0 | NA | NA | NA | NA | NA | #VALUE! |
| ENSMUSG00000087104.8 | 0 | NA | NA | NA | NA | NA | #VALUE! |
| ENSMUSG00000087106.2 | 0 | NA | NA | NA | NA | NA | #VALUE! |
| ENSMUSG00000087108.2 | 0 | NA | NA | NA | NA | NA | #VALUE! |
| ENSMUSG00000087109.2 | 0 | NA | NA | NA | NA | NA | #VALUE! |
| ENSMUSG00000087110.2 | 0 | NA | NA | NA | NA | NA | #VALUE! |
| ENSMUSG00000087111.2 | 0 | NA | NA | NA | NA | NA | #VALUE! |
| ENSMUSG00000087112.2 | 0 | NA | NA | NA | NA | NA | #VALUE! |
| ENSMUSG00000087113.2 | 6.99414 | 6.300596 | 3.91317 | 1.6101 | NA | NA | #VALUE! |
| ENSMUSG00000087114.2 | 0 | NA | NA | NA | NA | NA | #VALUE! |
| ENSMUSG00000087115.8 | 0 | NA | NA | NA | NA | NA | #VALUE! |
| ENSMUSG00000087116.2 | 0 | NA | NA | NA | NA | NA | #VALUE! |
| ENSMUSG00000087117.9 | 0 | NA | NA | NA | NA | NA | #VALUE! |
| ENSMUSG00000087119.5 | 3.288778 | 3.619338 | 3.898747 | 0.928334 | 0.353235 | NA | #VALUE! |
| ENSMUSG00000087120.2 | 0 | NA | NA | NA | NA | NA | #VALUE! |
| ENSMUSG00000087121.2 | 0 | NA | NA | NA | NA | NA | #VALUE! |
| ENSMUSG00000087122.2 | 0 | NA | NA | NA | NA | NA | #VALUE! |

| | | | | | | | |
|---|---|---|---|---|---|---|---|
| ENSMUSG00000087123.2 | 0 | NA | NA | NA | NA | NA | #VALUE! |
| ENSMUSG00000087124.2 | 0 | NA | NA | NA | NA | NA | #VALUE! |
| ENSMUSG00000087125.2 | 0 | NA | NA | NA | NA | NA | #VALUE! |
| ENSMUSG00000087127.2 | 0 | NA | NA | NA | NA | NA | #VALUE! |
| ENSMUSG00000087128.8 | 0 | NA | NA | NA | NA | NA | #VALUE! |
| ENSMUSG00000087129.2 | 0 | NA | NA | NA | NA | NA | #VALUE! |
| ENSMUSG00000087131.8 | 0 | NA | NA | NA | NA | NA | #VALUE! |
| ENSMUSG00000087133.3 | 0 | NA | NA | NA | NA | NA | #VALUE! |
| ENSMUSG00000087135.8 | 0 | NA | NA | NA | NA | NA | #VALUE! |
| ENSMUSG00000087136.2 | 0 | NA | NA | NA | NA | NA | #VALUE! |
| ENSMUSG00000087137.2 | 0 | NA | NA | NA | NA | NA | #VALUE! |
| ENSMUSG00000087138.2 | 6.590284 | 0.876465 | 3.301588 | 0.265468 | 0.790649 | NA | #VALUE! |
| ENSMUSG00000087139.2 | 0 | NA | NA | NA | NA | NA | #VALUE! |
| ENSMUSG00000087140.3 | 0 | NA | NA | NA | NA | NA | #VALUE! |
| ENSMUSG00000087142.8 | 0 | NA | NA | NA | NA | NA | #VALUE! |
| ENSMUSG00000087143.10 | 1.406238 | 2.267689 | 3.921398 | 0.578286 | 0.563071 | NA | #VALUE! |
| ENSMUSG00000087144.2 | 0 | NA | NA | NA | NA | NA | #VALUE! |
| ENSMUSG00000087145.8 | 0 | NA | NA | NA | NA | NA | #VALUE! |
| ENSMUSG00000087147.2 | 0 | NA | NA | NA | NA | NA | #VALUE! |
| ENSMUSG00000087148.2 | 0 | NA | NA | NA | NA | NA | #VALUE! |
| ENSMUSG00000087149.9 | 0 | NA | NA | NA | NA | NA | #VALUE! |
| ENSMUSG00000087150.8 | 0 | NA | NA | NA | NA | NA | #VALUE! |
| ENSMUSG00000087151.2 | 0 | NA | NA | NA | NA | NA | #VALUE! |
| ENSMUSG00000087152.4 | 0 | NA | NA | NA | NA | NA | #VALUE! |
| ENSMUSG00000087153.9 | 0 | NA | NA | NA | NA | NA | #VALUE! |
| ENSMUSG00000087155.2 | 0 | NA | NA | NA | NA | NA | #VALUE! |
| ENSMUSG00000087157.2 | 0 | NA | NA | NA | NA | NA | #VALUE! |
| ENSMUSG00000087158.2 | 0 | NA | NA | NA | NA | NA | #VALUE! |
| ENSMUSG00000087159.8 | 0 | NA | NA | NA | NA | NA | #VALUE! |
| ENSMUSG00000087160.2 | 0 | NA | NA | NA | NA | NA | #VALUE! |
| ENSMUSG00000087162.2 | 0 | NA | NA | NA | NA | NA | #VALUE! |
| ENSMUSG00000087164.2 | 0 | NA | NA | NA | NA | NA | #VALUE! |
| ENSMUSG00000087166.10 | 0 | NA | NA | NA | NA | NA | #VALUE! |
| ENSMUSG00000087167.2 | 0 | NA | NA | NA | NA | NA | #VALUE! |
| ENSMUSG00000087168.2 | 0 | NA | NA | NA | NA | NA | #VALUE! |
| ENSMUSG00000087170.8 | 0 | NA | NA | NA | NA | NA | #VALUE! |
| ENSMUSG00000087171.2 | 0 | NA | NA | NA | NA | NA | #VALUE! |
| ENSMUSG00000087172.2 | 0 | NA | NA | NA | NA | NA | #VALUE! |
| ENSMUSG00000087173.2 | 0 | NA | NA | NA | NA | NA | #VALUE! |
| ENSMUSG00000087175.8 | 0 | NA | NA | NA | NA | NA | #VALUE! |
| ENSMUSG00000087176.2 | 1.224665 | -3.81006 | 3.943408 | -0.96618 | 0.333952 | NA | #VALUE! |
| ENSMUSG00000087177.8 | 0.525066 | -2.64687 | 3.988095 | -0.66369 | 0.506887 | NA | #VALUE! |
| ENSMUSG00000087178.8 | 0 | NA | NA | NA | NA | NA | #VALUE! |
| ENSMUSG00000087179.9 | 0 | NA | NA | NA | NA | NA | #VALUE! |

| | | | | | | | |
|---|---|---|---|---|---|---|---|
| ENSMUSG00000087180.2 | 0 | NA | NA | NA | NA | NA | #VALUE! |
| ENSMUSG00000087181.4 | 0 | NA | NA | NA | NA | NA | #VALUE! |
| ENSMUSG00000087184.2 | 0 | NA | NA | NA | NA | NA | #VALUE! |
| ENSMUSG00000087185.2 | 0 | NA | NA | NA | NA | NA | #VALUE! |
| ENSMUSG00000087187.2 | 0 | NA | NA | NA | NA | NA | #VALUE! |
| ENSMUSG00000087190.3 | 0 | NA | NA | NA | NA | NA | #VALUE! |
| ENSMUSG00000087192.8 | 0 | NA | NA | NA | NA | NA | #VALUE! |
| ENSMUSG00000087193.8 | 0 | NA | NA | NA | NA | NA | #VALUE! |
| ENSMUSG00000087194.10 | 0 | NA | NA | NA | NA | NA | #VALUE! |
| ENSMUSG00000087195.2 | 0 | NA | NA | NA | NA | NA | #VALUE! |
| ENSMUSG00000087196.2 | 0 | NA | NA | NA | NA | NA | #VALUE! |
| ENSMUSG00000087197.2 | 0 | NA | NA | NA | NA | NA | #VALUE! |
| ENSMUSG00000087198.5 | 0 | NA | NA | NA | NA | NA | #VALUE! |
| ENSMUSG00000087199.2 | 0 | NA | NA | NA | NA | NA | #VALUE! |
| ENSMUSG00000087200.2 | 0 | NA | NA | NA | NA | NA | #VALUE! |
| ENSMUSG00000087201.2 | 0 | NA | NA | NA | NA | NA | #VALUE! |
| ENSMUSG00000087202.2 | 0 | NA | NA | NA | NA | NA | #VALUE! |
| ENSMUSG00000087203.8 | 0 | NA | NA | NA | NA | NA | #VALUE! |
| ENSMUSG00000087204.2 | 0 | NA | NA | NA | NA | NA | #VALUE! |
| ENSMUSG00000087205.2 | 0 | NA | NA | NA | NA | NA | #VALUE! |
| ENSMUSG00000087206.2 | 0 | NA | NA | NA | NA | NA | #VALUE! |
| ENSMUSG00000087207.9 | 0 | NA | NA | NA | NA | NA | #VALUE! |
| ENSMUSG00000087208.2 | 0 | NA | NA | NA | NA | NA | #VALUE! |
| ENSMUSG00000087210.2 | 0 | NA | NA | NA | NA | NA | #VALUE! |
| ENSMUSG00000087211.8 | 0 | NA | NA | NA | NA | NA | #VALUE! |
| ENSMUSG00000087214.3 | 0 | NA | NA | NA | NA | NA | #VALUE! |
| ENSMUSG00000087215.2 | 0 | NA | NA | NA | NA | NA | #VALUE! |
| ENSMUSG00000087216.2 | 0 | NA | NA | NA | NA | NA | #VALUE! |
| ENSMUSG00000087217.2 | 0 | NA | NA | NA | NA | NA | #VALUE! |
| ENSMUSG00000087219.2 | 0 | NA | NA | NA | NA | NA | #VALUE! |
| ENSMUSG00000087220.2 | 0 | NA | NA | NA | NA | NA | #VALUE! |
| ENSMUSG00000087221.4 | 0 | NA | NA | NA | NA | NA | #VALUE! |
| ENSMUSG00000087222.2 | 0 | NA | NA | NA | NA | NA | #VALUE! |
| ENSMUSG00000087223.3 | 0 | NA | NA | NA | NA | NA | #VALUE! |
| ENSMUSG00000087224.2 | 0 | NA | NA | NA | NA | NA | #VALUE! |
| ENSMUSG00000087225.2 | 0 | NA | NA | NA | NA | NA | #VALUE! |
| ENSMUSG00000087226.2 | 0 | NA | NA | NA | NA | NA | #VALUE! |
| ENSMUSG00000087228.2 | 0 | NA | NA | NA | NA | NA | #VALUE! |
| ENSMUSG00000087230.11 | 0 | NA | NA | NA | NA | NA | #VALUE! |
| ENSMUSG00000087231.8 | 2.100263 | -4.61322 | 3.927832 | -1.17449 | 0.240197 | NA | #VALUE! |
| ENSMUSG00000087232.2 | 0 | NA | NA | NA | NA | NA | #VALUE! |
| ENSMUSG00000087233.2 | 0 | NA | NA | NA | NA | NA | #VALUE! |
| ENSMUSG00000087235.2 | 0 | NA | NA | NA | NA | NA | #VALUE! |
| ENSMUSG00000087236.4 | 0 | NA | NA | NA | NA | NA | #VALUE! |

| | | | | | | | |
|---|---|---|---|---|---|---|---|
| ENSMUSG00000087238.2 | 0 | NA | NA | NA | NA | NA | #VALUE! |
| ENSMUSG00000087239.2 | 0 | NA | NA | NA | NA | NA | #VALUE! |
| ENSMUSG00000087240.2 | 0 | NA | NA | NA | NA | NA | #VALUE! |
| ENSMUSG00000087241.2 | 0 | NA | NA | NA | NA | NA | #VALUE! |
| ENSMUSG00000087242.3 | 0 | NA | NA | NA | NA | NA | #VALUE! |
| ENSMUSG00000087244.2 | 0 | NA | NA | NA | NA | NA | #VALUE! |
| ENSMUSG00000087245.2 | 0 | NA | NA | NA | NA | NA | #VALUE! |
| ENSMUSG00000087246.2 | 0 | NA | NA | NA | NA | NA | #VALUE! |
| ENSMUSG00000087247.4 | 0 | NA | NA | NA | NA | NA | #VALUE! |
| ENSMUSG00000087248.8 | 0 | NA | NA | NA | NA | NA | #VALUE! |
| ENSMUSG00000087249.2 | 9.39427 | -1.21878 | 2.925599 | -0.41659 | 0.676977 | NA | #VALUE! |
| ENSMUSG00000087250.3 | 0 | NA | NA | NA | NA | NA | #VALUE! |
| ENSMUSG00000087252.2 | 0 | NA | NA | NA | NA | NA | #VALUE! |
| ENSMUSG00000087253.2 | 0 | NA | NA | NA | NA | NA | #VALUE! |
| ENSMUSG00000087254.2 | 0 | NA | NA | NA | NA | NA | #VALUE! |
| ENSMUSG00000087255.3 | 0 | NA | NA | NA | NA | NA | #VALUE! |
| ENSMUSG00000087256.2 | 0 | NA | NA | NA | NA | NA | #VALUE! |
| ENSMUSG00000087258.2 | 0 | NA | NA | NA | NA | NA | #VALUE! |
| ENSMUSG00000087259.8 | 4.463969 | -5.69019 | 3.916774 | -1.45278 | 0.146286 | NA | #VALUE! |
| ENSMUSG00000087261.3 | 0 | NA | NA | NA | NA | NA | #VALUE! |
| ENSMUSG00000087263.2 | 0 | NA | NA | NA | NA | NA | #VALUE! |
| ENSMUSG00000087264.2 | 0 | NA | NA | NA | NA | NA | #VALUE! |
| ENSMUSG00000087265.2 | 0 | NA | NA | NA | NA | NA | #VALUE! |
| ENSMUSG00000087266.3 | 0 | NA | NA | NA | NA | NA | #VALUE! |
| ENSMUSG00000087267.2 | 0 | NA | NA | NA | NA | NA | #VALUE! |
| ENSMUSG00000087268.2 | 0 | NA | NA | NA | NA | NA | #VALUE! |
| ENSMUSG00000087270.3 | 0 | NA | NA | NA | NA | NA | #VALUE! |
| ENSMUSG00000087272.2 | 0 | NA | NA | NA | NA | NA | #VALUE! |
| ENSMUSG00000087273.2 | 0.630079 | -2.90291 | 3.975047 | -0.73028 | 0.465217 | NA | #VALUE! |
| ENSMUSG00000087276.2 | 0 | NA | NA | NA | NA | NA | #VALUE! |
| ENSMUSG00000087277.2 | 0 | NA | NA | NA | NA | NA | #VALUE! |
| ENSMUSG00000087278.2 | 0 | NA | NA | NA | NA | NA | #VALUE! |
| ENSMUSG00000087279.11 | 0 | NA | NA | NA | NA | NA | #VALUE! |
| ENSMUSG00000087281.2 | 0 | NA | NA | NA | NA | NA | #VALUE! |
| ENSMUSG00000087283.2 | 0 | NA | NA | NA | NA | NA | #VALUE! |
| ENSMUSG00000087286.2 | 0.093255 | 0.868093 | 4.072464 | 0.213162 | 0.831201 | NA | #VALUE! |
| ENSMUSG00000087288.2 | 0 | NA | NA | NA | NA | NA | #VALUE! |
| ENSMUSG00000087289.3 | 0 | NA | NA | NA | NA | NA | #VALUE! |
| ENSMUSG00000087290.3 | 0 | NA | NA | NA | NA | NA | #VALUE! |
| ENSMUSG00000087291.2 | 0 | NA | NA | NA | NA | NA | #VALUE! |
| ENSMUSG00000087292.2 | 0 | NA | NA | NA | NA | NA | #VALUE! |
| ENSMUSG00000087293.2 | 0 | NA | NA | NA | NA | NA | #VALUE! |
| ENSMUSG00000087294.3 | 0 | NA | NA | NA | NA | NA | #VALUE! |
| ENSMUSG00000087295.2 | 0 | NA | NA | NA | NA | NA | #VALUE! |

| | | | | | | | |
|---|---|---|---|---|---|---|---|
| ENSMUSG00000087296.2 | 0 | NA | NA | NA | NA | NA | #VALUE! |
| ENSMUSG00000087298.4 | 0 | NA | NA | NA | NA | NA | #VALUE! |
| ENSMUSG00000087299.2 | 0 | NA | NA | NA | NA | NA | #VALUE! |
| ENSMUSG00000087300.2 | 0 | NA | NA | NA | NA | NA | #VALUE! |
| ENSMUSG00000087301.8 | 0 | NA | NA | NA | NA | NA | #VALUE! |
| ENSMUSG00000087302.2 | 0 | NA | NA | NA | NA | NA | #VALUE! |
| ENSMUSG00000087303.4 | 0 | NA | NA | NA | NA | NA | #VALUE! |
| ENSMUSG00000087305.9 | 7.479856 | 0.222045 | 3.627787 | 0.061207 | 0.951195 | NA | #VALUE! |
| ENSMUSG00000087306.8 | 0 | NA | NA | NA | NA | NA | #VALUE! |
| ENSMUSG00000087307.2 | 6.930867 | -6.32693 | 3.913207 | -1.61681 | 0.105918 | NA | #VALUE! |
| ENSMUSG00000087308.2 | 0 | NA | NA | NA | NA | NA | #VALUE! |
| ENSMUSG00000087309.2 | 0 | NA | NA | NA | NA | NA | #VALUE! |
| ENSMUSG00000087312.2 | 0 | NA | NA | NA | NA | NA | #VALUE! |
| ENSMUSG00000087314.2 | 0 | NA | NA | NA | NA | NA | #VALUE! |
| ENSMUSG00000087316.2 | 0 | NA | NA | NA | NA | NA | #VALUE! |
| ENSMUSG00000087317.2 | 0 | NA | NA | NA | NA | NA | #VALUE! |
| ENSMUSG00000087318.2 | 0 | NA | NA | NA | NA | NA | #VALUE! |
| ENSMUSG00000087319.2 | 0 | NA | NA | NA | NA | NA | #VALUE! |
| ENSMUSG00000087321.2 | 0 | NA | NA | NA | NA | NA | #VALUE! |
| ENSMUSG00000087322.2 | 0 | NA | NA | NA | NA | NA | #VALUE! |
| ENSMUSG00000087323.2 | 0 | NA | NA | NA | NA | NA | #VALUE! |
| ENSMUSG00000087325.4 | 0 | NA | NA | NA | NA | NA | #VALUE! |
| ENSMUSG00000087326.2 | 0 | NA | NA | NA | NA | NA | #VALUE! |
| ENSMUSG00000087327.2 | 0 | NA | NA | NA | NA | NA | #VALUE! |
| ENSMUSG00000087328.2 | 0 | NA | NA | NA | NA | NA | #VALUE! |
| ENSMUSG00000087329.4 | 0 | NA | NA | NA | NA | NA | #VALUE! |
| ENSMUSG00000087330.2 | 0 | NA | NA | NA | NA | NA | #VALUE! |
| ENSMUSG00000087331.9 | 1.174735 | -3.75246 | 3.944889 | -0.95122 | 0.341493 | NA | #VALUE! |
| ENSMUSG00000087332.2 | 0 | NA | NA | NA | NA | NA | #VALUE! |
| ENSMUSG00000087333.2 | 0 | NA | NA | NA | NA | NA | #VALUE! |
| ENSMUSG00000087334.2 | 4.490437 | -5.69342 | 3.916751 | -1.45361 | 0.146055 | NA | #VALUE! |
| ENSMUSG00000087335.3 | 0 | NA | NA | NA | NA | NA | #VALUE! |
| ENSMUSG00000087336.3 | 3.077421 | 5.121959 | 3.921238 | 1.30621 | 0.191481 | NA | #VALUE! |
| ENSMUSG00000087337.2 | 0 | NA | NA | NA | NA | NA | #VALUE! |
| ENSMUSG00000087338.3 | 0 | NA | NA | NA | NA | NA | #VALUE! |
| ENSMUSG00000087339.2 | 0 | NA | NA | NA | NA | NA | #VALUE! |
| ENSMUSG00000087340.2 | 0 | NA | NA | NA | NA | NA | #VALUE! |
| ENSMUSG00000087341.5 | 0 | NA | NA | NA | NA | NA | #VALUE! |
| ENSMUSG00000087342.2 | 0 | NA | NA | NA | NA | NA | #VALUE! |
| ENSMUSG00000087343.2 | 0 | NA | NA | NA | NA | NA | #VALUE! |
| ENSMUSG00000087344.2 | 0 | NA | NA | NA | NA | NA | #VALUE! |
| ENSMUSG00000087345.2 | 0 | NA | NA | NA | NA | NA | #VALUE! |
| ENSMUSG00000087347.2 | 0 | NA | NA | NA | NA | NA | #VALUE! |
| ENSMUSG00000087348.2 | 0 | NA | NA | NA | NA | NA | #VALUE! |

| | | | | | | | |
|---|---|---|---|---|---|---|---|
| ENSMUSG00000087350.2 | 0 | NA | NA | NA | NA | NA | #VALUE! |
| ENSMUSG00000087351.2 | 0 | NA | NA | NA | NA | NA | #VALUE! |
| ENSMUSG00000087352.2 | 0 | NA | NA | NA | NA | NA | #VALUE! |
| ENSMUSG00000087353.2 | 0 | NA | NA | NA | NA | NA | #VALUE! |
| ENSMUSG00000087354.2 | 7.690654 | 6.434392 | 2.954036 | 2.17817 | 0.029393 | NA | #VALUE! |
| ENSMUSG00000087355.8 | 0 | NA | NA | NA | NA | NA | #VALUE! |
| ENSMUSG00000087356.2 | 0 | NA | NA | NA | NA | NA | #VALUE! |
| ENSMUSG00000087357.2 | 0 | NA | NA | NA | NA | NA | #VALUE! |
| ENSMUSG00000087358.9 | 0 | NA | NA | NA | NA | NA | #VALUE! |
| ENSMUSG00000087360.3 | 0 | NA | NA | NA | NA | NA | #VALUE! |
| ENSMUSG00000087362.8 | 0 | NA | NA | NA | NA | NA | #VALUE! |
| ENSMUSG00000087363.2 | 0 | NA | NA | NA | NA | NA | #VALUE! |
| ENSMUSG00000087364.2 | 0 | NA | NA | NA | NA | NA | #VALUE! |
| ENSMUSG00000087365.8 | 0 | NA | NA | NA | NA | NA | #VALUE! |
| ENSMUSG00000087366.8 | 5.832895 | -1.9106 | 3.772872 | -0.5064 | 0.612573 | NA | #VALUE! |
| ENSMUSG00000087367.2 | 1.585338 | 4.174397 | 3.934581 | 1.060951 | 0.288712 | NA | #VALUE! |
| ENSMUSG00000087368.8 | 0 | NA | NA | NA | NA | NA | #VALUE! |
| ENSMUSG00000087369.2 | 0 | NA | NA | NA | NA | NA | #VALUE! |
| ENSMUSG00000087371.2 | 0 | NA | NA | NA | NA | NA | #VALUE! |
| ENSMUSG00000087373.8 | 0 | NA | NA | NA | NA | NA | #VALUE! |
| ENSMUSG00000087374.2 | 0 | NA | NA | NA | NA | NA | #VALUE! |
| ENSMUSG00000087376.4 | 0 | NA | NA | NA | NA | NA | #VALUE! |
| ENSMUSG00000087377.2 | 0.315039 | -1.93549 | 4.038621 | -0.47925 | 0.631763 | NA | #VALUE! |
| ENSMUSG00000087378.2 | 0 | NA | NA | NA | NA | NA | #VALUE! |
| ENSMUSG00000087379.2 | 0 | NA | NA | NA | NA | NA | #VALUE! |
| ENSMUSG00000087381.9 | 0 | NA | NA | NA | NA | NA | #VALUE! |
| ENSMUSG00000087382.8 | 8.78638 | 1.461663 | 2.589843 | 0.564383 | 0.572494 | NA | #VALUE! |
| ENSMUSG00000087383.8 | 0 | NA | NA | NA | NA | NA | #VALUE! |
| ENSMUSG00000087384.2 | 0 | NA | NA | NA | NA | NA | #VALUE! |
| ENSMUSG00000087386.2 | 0 | NA | NA | NA | NA | NA | #VALUE! |
| ENSMUSG00000087387.3 | 0 | NA | NA | NA | NA | NA | #VALUE! |
| ENSMUSG00000087388.2 | 0 | NA | NA | NA | NA | NA | #VALUE! |
| ENSMUSG00000087389.2 | 0 | NA | NA | NA | NA | NA | #VALUE! |
| ENSMUSG00000087390.2 | 0 | NA | NA | NA | NA | NA | #VALUE! |
| ENSMUSG00000087393.2 | 0 | NA | NA | NA | NA | NA | #VALUE! |
| ENSMUSG00000087394.2 | 0 | NA | NA | NA | NA | NA | #VALUE! |
| ENSMUSG00000087395.2 | 0 | NA | NA | NA | NA | NA | #VALUE! |
| ENSMUSG00000087396.8 | 0 | NA | NA | NA | NA | NA | #VALUE! |
| ENSMUSG00000087397.2 | 0 | NA | NA | NA | NA | NA | #VALUE! |
| ENSMUSG00000087398.2 | 0 | NA | NA | NA | NA | NA | #VALUE! |
| ENSMUSG00000087399.2 | 0 | NA | NA | NA | NA | NA | #VALUE! |
| ENSMUSG00000087400.9 | 0 | NA | NA | NA | NA | NA | #VALUE! |
| ENSMUSG00000087401.2 | 0 | NA | NA | NA | NA | NA | #VALUE! |
| ENSMUSG00000087402.2 | 0 | NA | NA | NA | NA | NA | #VALUE! |

| | | | | | | | |
|---|---|---|---|---|---|---|---|
| ENSMUSG00000087403.10 | 4.194788 | -5.59536 | 3.917452 | -1.42832 | 0.153201 | NA | #VALUE! |
| ENSMUSG00000087404.2 | 2.33138 | 4.724655 | 3.925805 | 1.203487 | 0.228788 | NA | #VALUE! |
| ENSMUSG00000087405.2 | 0 | NA | NA | NA | NA | NA | #VALUE! |
| ENSMUSG00000087406.2 | 0 | NA | NA | NA | NA | NA | #VALUE! |
| ENSMUSG00000087407.2 | 0 | NA | NA | NA | NA | NA | #VALUE! |
| ENSMUSG00000087408.11 | 0 | NA | NA | NA | NA | NA | #VALUE! |
| ENSMUSG00000087409.2 | 0 | NA | NA | NA | NA | NA | #VALUE! |
| ENSMUSG00000087410.8 | 0 | NA | NA | NA | NA | NA | #VALUE! |
| ENSMUSG00000087411.2 | 0 | NA | NA | NA | NA | NA | #VALUE! |
| ENSMUSG00000087415.5 | 0 | NA | NA | NA | NA | NA | #VALUE! |
| ENSMUSG00000087416.2 | 0 | NA | NA | NA | NA | NA | #VALUE! |
| ENSMUSG00000087417.2 | 0 | NA | NA | NA | NA | NA | #VALUE! |
| ENSMUSG00000087422.2 | 0 | NA | NA | NA | NA | NA | #VALUE! |
| ENSMUSG00000087424.2 | 0 | NA | NA | NA | NA | NA | #VALUE! |
| ENSMUSG00000087425.2 | 0 | NA | NA | NA | NA | NA | #VALUE! |
| ENSMUSG00000087428.2 | 0 | NA | NA | NA | NA | NA | #VALUE! |
| ENSMUSG00000087429.2 | 0 | NA | NA | NA | NA | NA | #VALUE! |
| ENSMUSG00000087430.2 | 0 | NA | NA | NA | NA | NA | #VALUE! |
| ENSMUSG00000087431.2 | 0 | NA | NA | NA | NA | NA | #VALUE! |
| ENSMUSG00000087432.2 | 0 | NA | NA | NA | NA | NA | #VALUE! |
| ENSMUSG00000087433.2 | 0 | NA | NA | NA | NA | NA | #VALUE! |
| ENSMUSG00000087434.2 | 0 | NA | NA | NA | NA | NA | #VALUE! |
| ENSMUSG00000087435.2 | 0 | NA | NA | NA | NA | NA | #VALUE! |
| ENSMUSG00000087436.2 | 0 | NA | NA | NA | NA | NA | #VALUE! |
| ENSMUSG00000087437.8 | 0 | NA | NA | NA | NA | NA | #VALUE! |
| ENSMUSG00000087439.2 | 9.006509 | 6.658398 | 3.218264 | 2.068941 | 0.038552 | NA | #VALUE! |
| ENSMUSG00000087440.3 | 8.282346 | 6.540163 | 3.376665 | 1.93687 | 0.052761 | NA | #VALUE! |
| ENSMUSG00000087441.2 | 0 | NA | NA | NA | NA | NA | #VALUE! |
| ENSMUSG00000087442.2 | 0 | NA | NA | NA | NA | NA | #VALUE! |
| ENSMUSG00000087443.2 | 0 | NA | NA | NA | NA | NA | #VALUE! |
| ENSMUSG00000087444.2 | 0 | NA | NA | NA | NA | NA | #VALUE! |
| ENSMUSG00000087445.2 | 0 | NA | NA | NA | NA | NA | #VALUE! |
| ENSMUSG00000087449.2 | 0 | NA | NA | NA | NA | NA | #VALUE! |
| ENSMUSG00000087450.2 | 0 | NA | NA | NA | NA | NA | #VALUE! |
| ENSMUSG00000087451.2 | 0 | NA | NA | NA | NA | NA | #VALUE! |
| ENSMUSG00000087452.2 | 0 | NA | NA | NA | NA | NA | #VALUE! |
| ENSMUSG00000087454.2 | 0 | NA | NA | NA | NA | NA | #VALUE! |
| ENSMUSG00000087455.8 | 0 | NA | NA | NA | NA | NA | #VALUE! |
| ENSMUSG00000087456.2 | 0 | NA | NA | NA | NA | NA | #VALUE! |
| ENSMUSG00000087457.2 | 0 | NA | NA | NA | NA | NA | #VALUE! |
| ENSMUSG00000087458.2 | 0 | NA | NA | NA | NA | NA | #VALUE! |
| ENSMUSG00000087459.9 | 0 | NA | NA | NA | NA | NA | #VALUE! |
| ENSMUSG00000087460.3 | 0 | NA | NA | NA | NA | NA | #VALUE! |
| ENSMUSG00000087461.3 | 0.735092 | -3.12016 | 3.965612 | -0.7868 | 0.431397 | NA | #VALUE! |

| | | | | | | | |
|---|---|---|---|---|---|---|---|
| ENSMUSG00000087462.2 | 0 | NA | NA | NA | NA | NA | #VALUE! |
| ENSMUSG00000087464.2 | 0 | NA | NA | NA | NA | NA | #VALUE! |
| ENSMUSG00000087465.2 | 0 | NA | NA | NA | NA | NA | #VALUE! |
| ENSMUSG00000087466.4 | 0 | NA | NA | NA | NA | NA | #VALUE! |
| ENSMUSG00000087467.2 | 0 | NA | NA | NA | NA | NA | #VALUE! |
| ENSMUSG00000087468.2 | 0 | NA | NA | NA | NA | NA | #VALUE! |
| ENSMUSG00000087469.2 | 0 | NA | NA | NA | NA | NA | #VALUE! |
| ENSMUSG00000087473.8 | 0 | NA | NA | NA | NA | NA | #VALUE! |
| ENSMUSG00000087474.2 | 0 | NA | NA | NA | NA | NA | #VALUE! |
| ENSMUSG00000087475.4 | 0.673134 | 2.918386 | 3.972531 | 0.734641 | 0.462558 | NA | #VALUE! |
| ENSMUSG00000087476.2 | 0 | NA | NA | NA | NA | NA | #VALUE! |
| ENSMUSG00000087477.2 | 1.995249 | -4.53986 | 3.928926 | -1.1555 | 0.247887 | NA | #VALUE! |
| ENSMUSG00000087478.2 | 0.995589 | -3.51059 | 3.951785 | -0.88835 | 0.37435 | NA | #VALUE! |
| ENSMUSG00000087479.2 | 0 | NA | NA | NA | NA | NA | #VALUE! |
| ENSMUSG00000087480.2 | 0 | NA | NA | NA | NA | NA | #VALUE! |
| ENSMUSG00000087482.2 | 0 | NA | NA | NA | NA | NA | #VALUE! |
| ENSMUSG00000087484.2 | 0.805715 | -3.21534 | 3.961897 | -0.81156 | 0.417041 | NA | #VALUE! |
| ENSMUSG00000087485.2 | 0 | NA | NA | NA | NA | NA | #VALUE! |
| ENSMUSG00000087486.2 | 0 | NA | NA | NA | NA | NA | #VALUE! |
| ENSMUSG00000087487.2 | 0 | NA | NA | NA | NA | NA | #VALUE! |
| ENSMUSG00000087489.2 | 0 | NA | NA | NA | NA | NA | #VALUE! |
| ENSMUSG00000087491.8 | 0 | NA | NA | NA | NA | NA | #VALUE! |
| ENSMUSG00000087492.3 | 0 | NA | NA | NA | NA | NA | #VALUE! |
| ENSMUSG00000087495.8 | 0 | NA | NA | NA | NA | NA | #VALUE! |
| ENSMUSG00000087496.2 | 0 | NA | NA | NA | NA | NA | #VALUE! |
| ENSMUSG00000087500.2 | 0 | NA | NA | NA | NA | NA | #VALUE! |
| ENSMUSG00000087501.2 | 0 | NA | NA | NA | NA | NA | #VALUE! |
| ENSMUSG00000087502.2 | 0 | NA | NA | NA | NA | NA | #VALUE! |
| ENSMUSG00000087504.2 | 0 | NA | NA | NA | NA | NA | #VALUE! |
| ENSMUSG00000087505.2 | 0 | NA | NA | NA | NA | NA | #VALUE! |
| ENSMUSG00000087507.2 | 0 | NA | NA | NA | NA | NA | #VALUE! |
| ENSMUSG00000087508.2 | 0 | NA | NA | NA | NA | NA | #VALUE! |
| ENSMUSG00000087510.2 | 0 | NA | NA | NA | NA | NA | #VALUE! |
| ENSMUSG00000087512.3 | 0 | NA | NA | NA | NA | NA | #VALUE! |
| ENSMUSG00000087513.2 | 0 | NA | NA | NA | NA | NA | #VALUE! |
| ENSMUSG00000087514.2 | 0 | NA | NA | NA | NA | NA | #VALUE! |
| ENSMUSG00000087515.2 | 0 | NA | NA | NA | NA | NA | #VALUE! |
| ENSMUSG00000087516.8 | 0 | NA | NA | NA | NA | NA | #VALUE! |
| ENSMUSG00000087517.2 | 0 | NA | NA | NA | NA | NA | #VALUE! |
| ENSMUSG00000087518.2 | 0 | NA | NA | NA | NA | NA | #VALUE! |
| ENSMUSG00000087519.9 | 0 | NA | NA | NA | NA | NA | #VALUE! |
| ENSMUSG00000087520.3 | 0 | NA | NA | NA | NA | NA | #VALUE! |
| ENSMUSG00000087521.3 | 0 | NA | NA | NA | NA | NA | #VALUE! |
| ENSMUSG00000087522.4 | 0 | NA | NA | NA | NA | NA | #VALUE! |

| | | | | | | | |
|---|---|---|---|---|---|---|---|
| ENSMUSG00000087523.2 | 0 | NA | NA | NA | NA | NA | #VALUE! |
| ENSMUSG00000087524.2 | 0 | NA | NA | NA | NA | NA | #VALUE! |
| ENSMUSG00000087525.2 | 0 | NA | NA | NA | NA | NA | #VALUE! |
| ENSMUSG00000087526.3 | 0 | NA | NA | NA | NA | NA | #VALUE! |
| ENSMUSG00000087527.2 | 0 | NA | NA | NA | NA | NA | #VALUE! |
| ENSMUSG00000087528.2 | 0 | NA | NA | NA | NA | NA | #VALUE! |
| ENSMUSG00000087530.3 | 0 | NA | NA | NA | NA | NA | #VALUE! |
| ENSMUSG00000087531.8 | 0 | NA | NA | NA | NA | NA | #VALUE! |
| ENSMUSG00000087532.2 | 0 | NA | NA | NA | NA | NA | #VALUE! |
| ENSMUSG00000087534.2 | 0 | NA | NA | NA | NA | NA | #VALUE! |
| ENSMUSG00000087535.3 | 0 | NA | NA | NA | NA | NA | #VALUE! |
| ENSMUSG00000087536.2 | 0 | NA | NA | NA | NA | NA | #VALUE! |
| ENSMUSG00000087537.2 | 0 | NA | NA | NA | NA | NA | #VALUE! |
| ENSMUSG00000087538.2 | 0 | NA | NA | NA | NA | NA | #VALUE! |
| ENSMUSG00000087539.2 | 0 | NA | NA | NA | NA | NA | #VALUE! |
| ENSMUSG00000087540.2 | 0 | NA | NA | NA | NA | NA | #VALUE! |
| ENSMUSG00000087541.2 | 0.168284 | 1.027628 | 4.067126 | 0.252667 | 0.800526 | NA | #VALUE! |
| ENSMUSG00000087542.2 | 0 | NA | NA | NA | NA | NA | #VALUE! |
| ENSMUSG00000087543.2 | 0 | NA | NA | NA | NA | NA | #VALUE! |
| ENSMUSG00000087544.2 | 0 | NA | NA | NA | NA | NA | #VALUE! |
| ENSMUSG00000087545.2 | 0 | NA | NA | NA | NA | NA | #VALUE! |
| ENSMUSG00000087547.2 | 0 | NA | NA | NA | NA | NA | #VALUE! |
| ENSMUSG00000087549.2 | 0 | NA | NA | NA | NA | NA | #VALUE! |
| ENSMUSG00000087550.2 | 0 | NA | NA | NA | NA | NA | #VALUE! |
| ENSMUSG00000087552.3 | 0 | NA | NA | NA | NA | NA | #VALUE! |
| ENSMUSG00000087553.2 | 0 | NA | NA | NA | NA | NA | #VALUE! |
| ENSMUSG00000087555.2 | 0 | NA | NA | NA | NA | NA | #VALUE! |
| ENSMUSG00000087556.2 | 0 | NA | NA | NA | NA | NA | #VALUE! |
| ENSMUSG00000087559.2 | 0 | NA | NA | NA | NA | NA | #VALUE! |
| ENSMUSG00000087561.8 | 0 | NA | NA | NA | NA | NA | #VALUE! |
| ENSMUSG00000087563.2 | 0 | NA | NA | NA | NA | NA | #VALUE! |
| ENSMUSG00000087564.2 | 0 | NA | NA | NA | NA | NA | #VALUE! |
| ENSMUSG00000087565.2 | 0 | NA | NA | NA | NA | NA | #VALUE! |
| ENSMUSG00000087568.2 | 0 | NA | NA | NA | NA | NA | #VALUE! |
| ENSMUSG00000087569.2 | 0 | NA | NA | NA | NA | NA | #VALUE! |
| ENSMUSG00000087570.2 | 0 | NA | NA | NA | NA | NA | #VALUE! |
| ENSMUSG00000087571.2 | 0 | NA | NA | NA | NA | NA | #VALUE! |
| ENSMUSG00000087574.2 | 7.652584 | -1.71606 | 3.558368 | -0.48226 | 0.629622 | NA | #VALUE! |
| ENSMUSG00000087575.8 | 0 | NA | NA | NA | NA | NA | #VALUE! |
| ENSMUSG00000087577.2 | 0 | NA | NA | NA | NA | NA | #VALUE! |
| ENSMUSG00000087581.2 | 0 | NA | NA | NA | NA | NA | #VALUE! |
| ENSMUSG00000087582.7 | 0 | NA | NA | NA | NA | NA | #VALUE! |
| ENSMUSG00000087584.2 | 0 | NA | NA | NA | NA | NA | #VALUE! |
| ENSMUSG00000087585.2 | 0 | NA | NA | NA | NA | NA | #VALUE! |

| | | | | | | | |
|---|---|---|---|---|---|---|---|
| ENSMUSG00000087586.2 | 0 | NA | NA | NA | NA | NA | #VALUE! |
| ENSMUSG00000087587.2 | 0 | NA | NA | NA | NA | NA | #VALUE! |
| ENSMUSG00000087588.2 | 0 | NA | NA | NA | NA | NA | #VALUE! |
| ENSMUSG00000087589.4 | 0 | NA | NA | NA | NA | NA | #VALUE! |
| ENSMUSG00000087591.8 | 0 | NA | NA | NA | NA | NA | #VALUE! |
| ENSMUSG00000087592.2 | 0 | NA | NA | NA | NA | NA | #VALUE! |
| ENSMUSG00000087593.2 | 0 | NA | NA | NA | NA | NA | #VALUE! |
| ENSMUSG00000087595.2 | 0 | NA | NA | NA | NA | NA | #VALUE! |
| ENSMUSG00000087596.2 | 0 | NA | NA | NA | NA | NA | #VALUE! |
| ENSMUSG00000087597.2 | 0 | NA | NA | NA | NA | NA | #VALUE! |
| ENSMUSG00000087598.10 | 3.748932 | 4.30227 | 3.911937 | 1.09978 | 0.271428 | NA | #VALUE! |
| ENSMUSG00000087600.2 | 0 | NA | NA | NA | NA | NA | #VALUE! |
| ENSMUSG00000087601.4 | 0 | NA | NA | NA | NA | NA | #VALUE! |
| ENSMUSG00000087602.2 | 0 | NA | NA | NA | NA | NA | #VALUE! |
| ENSMUSG00000087603.2 | 0 | NA | NA | NA | NA | NA | #VALUE! |
| ENSMUSG00000087604.8 | 0 | NA | NA | NA | NA | NA | #VALUE! |
| ENSMUSG00000087605.2 | 0 | NA | NA | NA | NA | NA | #VALUE! |
| ENSMUSG00000087606.2 | 0.279766 | 1.808031 | 4.036658 | 0.447903 | 0.654223 | NA | #VALUE! |
| ENSMUSG00000087607.2 | 0 | NA | NA | NA | NA | NA | #VALUE! |
| ENSMUSG00000087608.2 | 0 | NA | NA | NA | NA | NA | #VALUE! |
| ENSMUSG00000087609.2 | 0 | NA | NA | NA | NA | NA | #VALUE! |
| ENSMUSG00000087610.2 | 0 | NA | NA | NA | NA | NA | #VALUE! |
| ENSMUSG00000087611.2 | 0 | NA | NA | NA | NA | NA | #VALUE! |
| ENSMUSG00000087612.2 | 0 | NA | NA | NA | NA | NA | #VALUE! |
| ENSMUSG00000087613.2 | 0 | NA | NA | NA | NA | NA | #VALUE! |
| ENSMUSG00000087614.2 | 0 | NA | NA | NA | NA | NA | #VALUE! |
| ENSMUSG00000087615.3 | 0 | NA | NA | NA | NA | NA | #VALUE! |
| ENSMUSG00000087616.2 | 0 | NA | NA | NA | NA | NA | #VALUE! |
| ENSMUSG00000087617.2 | 0 | NA | NA | NA | NA | NA | #VALUE! |
| ENSMUSG00000087618.2 | 0 | NA | NA | NA | NA | NA | #VALUE! |
| ENSMUSG00000087620.8 | 0 | NA | NA | NA | NA | NA | #VALUE! |
| ENSMUSG00000087621.8 | 0 | NA | NA | NA | NA | NA | #VALUE! |
| ENSMUSG00000087622.2 | 0 | NA | NA | NA | NA | NA | #VALUE! |
| ENSMUSG00000087623.2 | 0 | NA | NA | NA | NA | NA | #VALUE! |
| ENSMUSG00000087624.2 | 0 | NA | NA | NA | NA | NA | #VALUE! |
| ENSMUSG00000087625.2 | 0 | NA | NA | NA | NA | NA | #VALUE! |
| ENSMUSG00000087626.2 | 0 | NA | NA | NA | NA | NA | #VALUE! |
| ENSMUSG00000087627.3 | 0 | NA | NA | NA | NA | NA | #VALUE! |
| ENSMUSG00000087628.2 | 0 | NA | NA | NA | NA | NA | #VALUE! |
| ENSMUSG00000087629.2 | 0 | NA | NA | NA | NA | NA | #VALUE! |
| ENSMUSG00000087630.2 | 0 | NA | NA | NA | NA | NA | #VALUE! |
| ENSMUSG00000087632.2 | 0 | NA | NA | NA | NA | NA | #VALUE! |
| ENSMUSG00000087633.2 | 0.195789 | -1.05581 | 4.073302 | -0.2592 | 0.795479 | NA | #VALUE! |
| ENSMUSG00000087635.4 | 0 | NA | NA | NA | NA | NA | #VALUE! |

| | | | | | | | |
|---|---|---|---|---|---|---|---|
| ENSMUSG00000087636.2 | 0 | NA | NA | NA | NA | NA | #VALUE! |
| ENSMUSG00000087637.2 | 0 | NA | NA | NA | NA | NA | #VALUE! |
| ENSMUSG00000087638.2 | 4.880222 | 5.776815 | 3.915969 | 1.475194 | 0.14016 | NA | #VALUE! |
| ENSMUSG00000087639.2 | 0 | NA | NA | NA | NA | NA | #VALUE! |
| ENSMUSG00000087640.2 | 0 | NA | NA | NA | NA | NA | #VALUE! |
| ENSMUSG00000087641.2 | 0 | NA | NA | NA | NA | NA | #VALUE! |
| ENSMUSG00000087642.4 | 0 | NA | NA | NA | NA | NA | #VALUE! |
| ENSMUSG00000087643.2 | 0 | NA | NA | NA | NA | NA | #VALUE! |
| ENSMUSG00000087644.2 | 9.389159 | -0.53724 | 3.356872 | -0.16004 | 0.872848 | NA | #VALUE! |
| ENSMUSG00000087646.2 | 0 | NA | NA | NA | NA | NA | #VALUE! |
| ENSMUSG00000087647.3 | 0 | NA | NA | NA | NA | NA | #VALUE! |
| ENSMUSG00000087648.2 | 0 | NA | NA | NA | NA | NA | #VALUE! |
| ENSMUSG00000087649.3 | 0 | NA | NA | NA | NA | NA | #VALUE! |
| ENSMUSG00000087651.4 | 0 | NA | NA | NA | NA | NA | #VALUE! |
| ENSMUSG00000087652.2 | 0 | NA | NA | NA | NA | NA | #VALUE! |
| ENSMUSG00000087654.2 | 0 | NA | NA | NA | NA | NA | #VALUE! |
| ENSMUSG00000087657.9 | 0 | NA | NA | NA | NA | NA | #VALUE! |
| ENSMUSG00000087658.3 | 0 | NA | NA | NA | NA | NA | #VALUE! |
| ENSMUSG00000087659.8 | 0 | NA | NA | NA | NA | NA | #VALUE! |
| ENSMUSG00000087660.2 | 0 | NA | NA | NA | NA | NA | #VALUE! |
| ENSMUSG00000087662.3 | 0 | NA | NA | NA | NA | NA | #VALUE! |
| ENSMUSG00000087663.2 | 0 | NA | NA | NA | NA | NA | #VALUE! |
| ENSMUSG00000087666.4 | 0 | NA | NA | NA | NA | NA | #VALUE! |
| ENSMUSG00000087667.9 | 0 | NA | NA | NA | NA | NA | #VALUE! |
| ENSMUSG00000087668.2 | 0 | NA | NA | NA | NA | NA | #VALUE! |
| ENSMUSG00000087669.2 | 0 | NA | NA | NA | NA | NA | #VALUE! |
| ENSMUSG00000087670.2 | 0 | NA | NA | NA | NA | NA | #VALUE! |
| ENSMUSG00000087672.2 | 0 | NA | NA | NA | NA | NA | #VALUE! |
| ENSMUSG00000087674.8 | 0 | NA | NA | NA | NA | NA | #VALUE! |
| ENSMUSG00000087675.2 | 0 | NA | NA | NA | NA | NA | #VALUE! |
| ENSMUSG00000087677.2 | 0 | NA | NA | NA | NA | NA | #VALUE! |
| ENSMUSG00000087678.2 | 0 | NA | NA | NA | NA | NA | #VALUE! |
| ENSMUSG00000087681.2 | 0 | NA | NA | NA | NA | NA | #VALUE! |
| ENSMUSG00000087682.2 | 0 | NA | NA | NA | NA | NA | #VALUE! |
| ENSMUSG00000087683.2 | 0 | NA | NA | NA | NA | NA | #VALUE! |
| ENSMUSG00000087684.2 | 0 | NA | NA | NA | NA | NA | #VALUE! |
| ENSMUSG00000087685.2 | 0 | NA | NA | NA | NA | NA | #VALUE! |
| ENSMUSG00000087688.2 | 0 | NA | NA | NA | NA | NA | #VALUE! |
| ENSMUSG00000087689.2 | 0 | NA | NA | NA | NA | NA | #VALUE! |
| ENSMUSG00000087690.2 | 0 | NA | NA | NA | NA | NA | #VALUE! |
| ENSMUSG00000087691.2 | 0 | NA | NA | NA | NA | NA | #VALUE! |
| ENSMUSG00000087692.2 | 0 | NA | NA | NA | NA | NA | #VALUE! |
| ENSMUSG00000087693.2 | 0 | NA | NA | NA | NA | NA | #VALUE! |
| ENSMUSG00000087694.9 | 0 | NA | NA | NA | NA | NA | #VALUE! |

| | | | | | | | |
|---|---|---|---|---|---|---|---|
| ENSMUSG00000087695.2 | 0 | NA | NA | NA | NA | NA | #VALUE! |
| ENSMUSG00000087698.2 | 0 | NA | NA | NA | NA | NA | #VALUE! |
| ENSMUSG00000087700.3 | 6.939766 | -6.32265 | 3.913226 | -1.61571 | 0.106157 | NA | #VALUE! |
| ENSMUSG00000087701.5 | 0 | NA | NA | NA | NA | NA | #VALUE! |
| ENSMUSG00000087702.2 | 0 | NA | NA | NA | NA | NA | #VALUE! |
| ENSMUSG00000087703.2 | 0 | NA | NA | NA | NA | NA | #VALUE! |
| ENSMUSG00000087704.2 | 0 | NA | NA | NA | NA | NA | #VALUE! |
| ENSMUSG00000087707.4 | 0 | NA | NA | NA | NA | NA | #VALUE! |
| ENSMUSG00000087708.3 | 0 | NA | NA | NA | NA | NA | #VALUE! |
| ENSMUSG00000087712.4 | 0 | NA | NA | NA | NA | NA | #VALUE! |
| ENSMUSG00000087714.3 | 0 | NA | NA | NA | NA | NA | #VALUE! |
| ENSMUSG00000087715.3 | 0 | NA | NA | NA | NA | NA | #VALUE! |
| ENSMUSG00000087717.3 | 0 | NA | NA | NA | NA | NA | #VALUE! |
| ENSMUSG00000087718.3 | 0 | NA | NA | NA | NA | NA | #VALUE! |
| ENSMUSG00000087719.3 | 0 | NA | NA | NA | NA | NA | #VALUE! |
| ENSMUSG00000087720.4 | 0 | NA | NA | NA | NA | NA | #VALUE! |
| ENSMUSG00000087722.4 | 0 | NA | NA | NA | NA | NA | #VALUE! |
| ENSMUSG00000087723.3 | 0 | NA | NA | NA | NA | NA | #VALUE! |
| ENSMUSG00000087724.3 | 0 | NA | NA | NA | NA | NA | #VALUE! |
| ENSMUSG00000087734.3 | 0 | NA | NA | NA | NA | NA | #VALUE! |
| ENSMUSG00000087736.3 | 0 | NA | NA | NA | NA | NA | #VALUE! |
| ENSMUSG00000087739.3 | 0 | NA | NA | NA | NA | NA | #VALUE! |
| ENSMUSG00000087742.3 | 0 | NA | NA | NA | NA | NA | #VALUE! |
| ENSMUSG00000087746.4 | 0 | NA | NA | NA | NA | NA | #VALUE! |
| ENSMUSG00000087747.3 | 0 | NA | NA | NA | NA | NA | #VALUE! |
| ENSMUSG00000087752.3 | 0 | NA | NA | NA | NA | NA | #VALUE! |
| ENSMUSG00000087754.4 | 0 | NA | NA | NA | NA | NA | #VALUE! |
| ENSMUSG00000087760.4 | 0 | NA | NA | NA | NA | NA | #VALUE! |
| ENSMUSG00000087761.4 | 0 | NA | NA | NA | NA | NA | #VALUE! |
| ENSMUSG00000087764.3 | 0 | NA | NA | NA | NA | NA | #VALUE! |
| ENSMUSG00000087765.3 | 0 | NA | NA | NA | NA | NA | #VALUE! |
| ENSMUSG00000087769.3 | 0 | NA | NA | NA | NA | NA | #VALUE! |
| ENSMUSG00000087772.3 | 0 | NA | NA | NA | NA | NA | #VALUE! |
| ENSMUSG00000087775.3 | 0 | NA | NA | NA | NA | NA | #VALUE! |
| ENSMUSG00000087776.3 | 0 | NA | NA | NA | NA | NA | #VALUE! |
| ENSMUSG00000087777.3 | 0 | NA | NA | NA | NA | NA | #VALUE! |
| ENSMUSG00000087780.4 | 0 | NA | NA | NA | NA | NA | #VALUE! |
| ENSMUSG00000087781.3 | 0 | NA | NA | NA | NA | NA | #VALUE! |
| ENSMUSG00000087782.3 | 0 | NA | NA | NA | NA | NA | #VALUE! |
| ENSMUSG00000087790.3 | 0 | NA | NA | NA | NA | NA | #VALUE! |
| ENSMUSG00000087792.3 | 0 | NA | NA | NA | NA | NA | #VALUE! |
| ENSMUSG00000087793.3 | 0 | NA | NA | NA | NA | NA | #VALUE! |
| ENSMUSG00000087795.3 | 0 | NA | NA | NA | NA | NA | #VALUE! |
| ENSMUSG00000087796.3 | 0 | NA | NA | NA | NA | NA | #VALUE! |

| | | | | | | | |
|---|---|---|---|---|---|---|---|
| ENSMUSG00000087800.3 | 0 | NA | NA | NA | NA | NA | #VALUE! |
| ENSMUSG00000087802.3 | 0 | NA | NA | NA | NA | NA | #VALUE! |
| ENSMUSG00000087804.4 | 0 | NA | NA | NA | NA | NA | #VALUE! |
| ENSMUSG00000087805.4 | 0 | NA | NA | NA | NA | NA | #VALUE! |
| ENSMUSG00000087806.3 | 0 | NA | NA | NA | NA | NA | #VALUE! |
| ENSMUSG00000087808.4 | 0 | NA | NA | NA | NA | NA | #VALUE! |
| ENSMUSG00000087810.3 | 0 | NA | NA | NA | NA | NA | #VALUE! |
| ENSMUSG00000087812.3 | 0 | NA | NA | NA | NA | NA | #VALUE! |
| ENSMUSG00000087813.3 | 0 | NA | NA | NA | NA | NA | #VALUE! |
| ENSMUSG00000087815.3 | 0 | NA | NA | NA | NA | NA | #VALUE! |
| ENSMUSG00000087816.2 | 0 | NA | NA | NA | NA | NA | #VALUE! |
| ENSMUSG00000087819.3 | 0 | NA | NA | NA | NA | NA | #VALUE! |
| ENSMUSG00000087820.3 | 0 | NA | NA | NA | NA | NA | #VALUE! |
| ENSMUSG00000087823.3 | 0 | NA | NA | NA | NA | NA | #VALUE! |
| ENSMUSG00000087825.4 | 0 | NA | NA | NA | NA | NA | #VALUE! |
| ENSMUSG00000087831.3 | 0 | NA | NA | NA | NA | NA | #VALUE! |
| ENSMUSG00000087832.4 | 0 | NA | NA | NA | NA | NA | #VALUE! |
| ENSMUSG00000087835.4 | 0 | NA | NA | NA | NA | NA | #VALUE! |
| ENSMUSG00000087839.3 | 0 | NA | NA | NA | NA | NA | #VALUE! |
| ENSMUSG00000087841.2 | 0 | NA | NA | NA | NA | NA | #VALUE! |
| ENSMUSG00000087842.3 | 0 | NA | NA | NA | NA | NA | #VALUE! |
| ENSMUSG00000087844.2 | 0 | NA | NA | NA | NA | NA | #VALUE! |
| ENSMUSG00000087847.3 | 0 | NA | NA | NA | NA | NA | #VALUE! |
| ENSMUSG00000087850.3 | 0 | NA | NA | NA | NA | NA | #VALUE! |
| ENSMUSG00000087855.3 | 0 | NA | NA | NA | NA | NA | #VALUE! |
| ENSMUSG00000087857.3 | 0 | NA | NA | NA | NA | NA | #VALUE! |
| ENSMUSG00000087858.3 | 0 | NA | NA | NA | NA | NA | #VALUE! |
| ENSMUSG00000087861.3 | 0 | NA | NA | NA | NA | NA | #VALUE! |
| ENSMUSG00000087862.3 | 0 | NA | NA | NA | NA | NA | #VALUE! |
| ENSMUSG00000087867.3 | 0 | NA | NA | NA | NA | NA | #VALUE! |
| ENSMUSG00000087870.3 | 0 | NA | NA | NA | NA | NA | #VALUE! |
| ENSMUSG00000087871.3 | 0 | NA | NA | NA | NA | NA | #VALUE! |
| ENSMUSG00000087873.3 | 0 | NA | NA | NA | NA | NA | #VALUE! |
| ENSMUSG00000087875.4 | 0 | NA | NA | NA | NA | NA | #VALUE! |
| ENSMUSG00000087878.3 | 0 | NA | NA | NA | NA | NA | #VALUE! |
| ENSMUSG00000087880.3 | 0 | NA | NA | NA | NA | NA | #VALUE! |
| ENSMUSG00000087881.3 | 0 | NA | NA | NA | NA | NA | #VALUE! |
| ENSMUSG00000087883.3 | 0 | NA | NA | NA | NA | NA | #VALUE! |
| ENSMUSG00000087884.3 | 0 | NA | NA | NA | NA | NA | #VALUE! |
| ENSMUSG00000087887.3 | 0 | NA | NA | NA | NA | NA | #VALUE! |
| ENSMUSG00000087892.3 | 0 | NA | NA | NA | NA | NA | #VALUE! |
| ENSMUSG00000087893.3 | 0 | NA | NA | NA | NA | NA | #VALUE! |
| ENSMUSG00000087894.3 | 0 | NA | NA | NA | NA | NA | #VALUE! |
| ENSMUSG00000087897.3 | 0 | NA | NA | NA | NA | NA | #VALUE! |

| | | | | | | | |
|---|---|---|---|---|---|---|---|
| ENSMUSG00000087899.3 | 0 | NA | NA | NA | NA | NA | #VALUE! |
| ENSMUSG00000087903.3 | 0 | NA | NA | NA | NA | NA | #VALUE! |
| ENSMUSG00000087908.3 | 0 | NA | NA | NA | NA | NA | #VALUE! |
| ENSMUSG00000087912.3 | 0 | NA | NA | NA | NA | NA | #VALUE! |
| ENSMUSG00000087916.3 | 0 | NA | NA | NA | NA | NA | #VALUE! |
| ENSMUSG00000087917.3 | 0 | NA | NA | NA | NA | NA | #VALUE! |
| ENSMUSG00000087919.2 | 0 | NA | NA | NA | NA | NA | #VALUE! |
| ENSMUSG00000087922.3 | 0 | NA | NA | NA | NA | NA | #VALUE! |
| ENSMUSG00000087923.3 | 0 | NA | NA | NA | NA | NA | #VALUE! |
| ENSMUSG00000087924.3 | 0 | NA | NA | NA | NA | NA | #VALUE! |
| ENSMUSG00000087926.3 | 0 | NA | NA | NA | NA | NA | #VALUE! |
| ENSMUSG00000087929.3 | 0 | NA | NA | NA | NA | NA | #VALUE! |
| ENSMUSG00000087930.3 | 0 | NA | NA | NA | NA | NA | #VALUE! |
| ENSMUSG00000087932.3 | 0 | NA | NA | NA | NA | NA | #VALUE! |
| ENSMUSG00000087933.3 | 0 | NA | NA | NA | NA | NA | #VALUE! |
| ENSMUSG00000087935.3 | 0 | NA | NA | NA | NA | NA | #VALUE! |
| ENSMUSG00000087936.3 | 0 | NA | NA | NA | NA | NA | #VALUE! |
| ENSMUSG00000087937.3 | 0 | NA | NA | NA | NA | NA | #VALUE! |
| ENSMUSG00000087938.4 | 0 | NA | NA | NA | NA | NA | #VALUE! |
| ENSMUSG00000087939.3 | 0 | NA | NA | NA | NA | NA | #VALUE! |
| ENSMUSG00000087940.3 | 0 | NA | NA | NA | NA | NA | #VALUE! |
| ENSMUSG00000087942.3 | 0 | NA | NA | NA | NA | NA | #VALUE! |
| ENSMUSG00000087947.3 | 0 | NA | NA | NA | NA | NA | #VALUE! |
| ENSMUSG00000087949.3 | 0 | NA | NA | NA | NA | NA | #VALUE! |
| ENSMUSG00000087950.3 | 0 | NA | NA | NA | NA | NA | #VALUE! |
| ENSMUSG00000087954.3 | 0 | NA | NA | NA | NA | NA | #VALUE! |
| ENSMUSG00000087963.3 | 0 | NA | NA | NA | NA | NA | #VALUE! |
| ENSMUSG00000087964.4 | 0 | NA | NA | NA | NA | NA | #VALUE! |
| ENSMUSG00000087966.4 | 0 | NA | NA | NA | NA | NA | #VALUE! |
| ENSMUSG00000087967.3 | 0 | NA | NA | NA | NA | NA | #VALUE! |
| ENSMUSG00000087968.3 | 0 | NA | NA | NA | NA | NA | #VALUE! |
| ENSMUSG00000087973.3 | 0 | NA | NA | NA | NA | NA | #VALUE! |
| ENSMUSG00000087980.3 | 0 | NA | NA | NA | NA | NA | #VALUE! |
| ENSMUSG00000087985.3 | 0 | NA | NA | NA | NA | NA | #VALUE! |
| ENSMUSG00000087987.3 | 0 | NA | NA | NA | NA | NA | #VALUE! |
| ENSMUSG00000087990.4 | 0 | NA | NA | NA | NA | NA | #VALUE! |
| ENSMUSG00000087992.3 | 0 | NA | NA | NA | NA | NA | #VALUE! |
| ENSMUSG00000087993.3 | 0 | NA | NA | NA | NA | NA | #VALUE! |
| ENSMUSG00000087995.3 | 0 | NA | NA | NA | NA | NA | #VALUE! |
| ENSMUSG00000087997.3 | 0 | NA | NA | NA | NA | NA | #VALUE! |
| ENSMUSG00000088001.3 | 0 | NA | NA | NA | NA | NA | #VALUE! |
| ENSMUSG00000088002.3 | 0 | NA | NA | NA | NA | NA | #VALUE! |
| ENSMUSG00000088005.3 | 0 | NA | NA | NA | NA | NA | #VALUE! |
| ENSMUSG00000088008.3 | 0 | NA | NA | NA | NA | NA | #VALUE! |

| | | | | | | | |
|---|---|---|---|---|---|---|---|
| ENSMUSG00000088009.3 | 0 | NA | NA | NA | NA | NA | #VALUE! |
| ENSMUSG00000088015.3 | 0 | NA | NA | NA | NA | NA | #VALUE! |
| ENSMUSG00000088018.4 | 0 | NA | NA | NA | NA | NA | #VALUE! |
| ENSMUSG00000088019.3 | 0 | NA | NA | NA | NA | NA | #VALUE! |
| ENSMUSG00000088020.3 | 0 | NA | NA | NA | NA | NA | #VALUE! |
| ENSMUSG00000088021.3 | 0 | NA | NA | NA | NA | NA | #VALUE! |
| ENSMUSG00000088025.3 | 0 | NA | NA | NA | NA | NA | #VALUE! |
| ENSMUSG00000088027.3 | 0 | NA | NA | NA | NA | NA | #VALUE! |
| ENSMUSG00000088037.3 | 0 | NA | NA | NA | NA | NA | #VALUE! |
| ENSMUSG00000088039.3 | 0 | NA | NA | NA | NA | NA | #VALUE! |
| ENSMUSG00000088040.4 | 0 | NA | NA | NA | NA | NA | #VALUE! |
| ENSMUSG00000088041.3 | 0 | NA | NA | NA | NA | NA | #VALUE! |
| ENSMUSG00000088043.3 | 0 | NA | NA | NA | NA | NA | #VALUE! |
| ENSMUSG00000088044.3 | 0 | NA | NA | NA | NA | NA | #VALUE! |
| ENSMUSG00000088045.4 | 0 | NA | NA | NA | NA | NA | #VALUE! |
| ENSMUSG00000088046.3 | 0 | NA | NA | NA | NA | NA | #VALUE! |
| ENSMUSG00000088047.3 | 0 | NA | NA | NA | NA | NA | #VALUE! |
| ENSMUSG00000088048.3 | 0 | NA | NA | NA | NA | NA | #VALUE! |
| ENSMUSG00000088050.4 | 0 | NA | NA | NA | NA | NA | #VALUE! |
| ENSMUSG00000088051.3 | 0 | NA | NA | NA | NA | NA | #VALUE! |
| ENSMUSG00000088054.3 | 0 | NA | NA | NA | NA | NA | #VALUE! |
| ENSMUSG00000088055.4 | 0 | NA | NA | NA | NA | NA | #VALUE! |
| ENSMUSG00000088058.3 | 0 | NA | NA | NA | NA | NA | #VALUE! |
| ENSMUSG00000088060.3 | 0 | NA | NA | NA | NA | NA | #VALUE! |
| ENSMUSG00000088068.3 | 0 | NA | NA | NA | NA | NA | #VALUE! |
| ENSMUSG00000088071.3 | 0 | NA | NA | NA | NA | NA | #VALUE! |
| ENSMUSG00000088073.2 | 0 | NA | NA | NA | NA | NA | #VALUE! |
| ENSMUSG00000088076.3 | 0 | NA | NA | NA | NA | NA | #VALUE! |
| ENSMUSG00000088084.3 | 0 | NA | NA | NA | NA | NA | #VALUE! |
| ENSMUSG00000088088.2 | 0 | NA | NA | NA | NA | NA | #VALUE! |
| ENSMUSG00000088093.3 | 0 | NA | NA | NA | NA | NA | #VALUE! |
| ENSMUSG00000088094.3 | 0 | NA | NA | NA | NA | NA | #VALUE! |
| ENSMUSG00000088095.3 | 0 | NA | NA | NA | NA | NA | #VALUE! |
| ENSMUSG00000088098.3 | 0 | NA | NA | NA | NA | NA | #VALUE! |
| ENSMUSG00000088103.3 | 0 | NA | NA | NA | NA | NA | #VALUE! |
| ENSMUSG00000088104.3 | 0 | NA | NA | NA | NA | NA | #VALUE! |
| ENSMUSG00000088105.3 | 0 | NA | NA | NA | NA | NA | #VALUE! |
| ENSMUSG00000088106.3 | 0 | NA | NA | NA | NA | NA | #VALUE! |
| ENSMUSG00000088108.3 | 0 | NA | NA | NA | NA | NA | #VALUE! |
| ENSMUSG00000088111.3 | 0 | NA | NA | NA | NA | NA | #VALUE! |
| ENSMUSG00000088113.3 | 0 | NA | NA | NA | NA | NA | #VALUE! |
| ENSMUSG00000088114.3 | 0 | NA | NA | NA | NA | NA | #VALUE! |
| ENSMUSG00000088121.3 | 0 | NA | NA | NA | NA | NA | #VALUE! |
| ENSMUSG00000088125.3 | 0 | NA | NA | NA | NA | NA | #VALUE! |

| | | | | | | | |
|---|---|---|---|---|---|---|---|
| ENSMUSG00000088126.3 | 0 | NA | NA | NA | NA | NA | #VALUE! |
| ENSMUSG00000088134.3 | 0 | NA | NA | NA | NA | NA | #VALUE! |
| ENSMUSG00000088135.3 | 0 | NA | NA | NA | NA | NA | #VALUE! |
| ENSMUSG00000088139.4 | 0 | NA | NA | NA | NA | NA | #VALUE! |
| ENSMUSG00000088141.3 | 0 | NA | NA | NA | NA | NA | #VALUE! |
| ENSMUSG00000088145.3 | 0 | NA | NA | NA | NA | NA | #VALUE! |
| ENSMUSG00000088146.3 | 0 | NA | NA | NA | NA | NA | #VALUE! |
| ENSMUSG00000088147.3 | 0 | NA | NA | NA | NA | NA | #VALUE! |
| ENSMUSG00000088148.3 | 0 | NA | NA | NA | NA | NA | #VALUE! |
| ENSMUSG00000088149.3 | 0 | NA | NA | NA | NA | NA | #VALUE! |
| ENSMUSG00000088151.3 | 0 | NA | NA | NA | NA | NA | #VALUE! |
| ENSMUSG00000088152.4 | 0 | NA | NA | NA | NA | NA | #VALUE! |
| ENSMUSG00000088154.2 | 0 | NA | NA | NA | NA | NA | #VALUE! |
| ENSMUSG00000088155.3 | 0 | NA | NA | NA | NA | NA | #VALUE! |
| ENSMUSG00000088157.4 | 0 | NA | NA | NA | NA | NA | #VALUE! |
| ENSMUSG00000088159.3 | 0 | NA | NA | NA | NA | NA | #VALUE! |
| ENSMUSG00000088160.4 | 0 | NA | NA | NA | NA | NA | #VALUE! |
| ENSMUSG00000088164.4 | 0 | NA | NA | NA | NA | NA | #VALUE! |
| ENSMUSG00000088167.4 | 0 | NA | NA | NA | NA | NA | #VALUE! |
| ENSMUSG00000088168.3 | 0 | NA | NA | NA | NA | NA | #VALUE! |
| ENSMUSG00000088169.3 | 0 | NA | NA | NA | NA | NA | #VALUE! |
| ENSMUSG00000088172.3 | 0 | NA | NA | NA | NA | NA | #VALUE! |
| ENSMUSG00000088175.3 | 0 | NA | NA | NA | NA | NA | #VALUE! |
| ENSMUSG00000088180.3 | 0 | NA | NA | NA | NA | NA | #VALUE! |
| ENSMUSG00000088183.4 | 0 | NA | NA | NA | NA | NA | #VALUE! |
| ENSMUSG00000088185.3 | 0 | NA | NA | NA | NA | NA | #VALUE! |
| ENSMUSG00000088191.3 | 0 | NA | NA | NA | NA | NA | #VALUE! |
| ENSMUSG00000088192.3 | 0 | NA | NA | NA | NA | NA | #VALUE! |
| ENSMUSG00000088195.3 | 0 | NA | NA | NA | NA | NA | #VALUE! |
| ENSMUSG00000088196.3 | 0 | NA | NA | NA | NA | NA | #VALUE! |
| ENSMUSG00000088197.3 | 0 | NA | NA | NA | NA | NA | #VALUE! |
| ENSMUSG00000088198.3 | 0 | NA | NA | NA | NA | NA | #VALUE! |
| ENSMUSG00000088201.3 | 0 | NA | NA | NA | NA | NA | #VALUE! |
| ENSMUSG00000088208.3 | 0 | NA | NA | NA | NA | NA | #VALUE! |
| ENSMUSG00000088212.3 | 0 | NA | NA | NA | NA | NA | #VALUE! |
| ENSMUSG00000088217.4 | 0 | NA | NA | NA | NA | NA | #VALUE! |
| ENSMUSG00000088218.3 | 0 | NA | NA | NA | NA | NA | #VALUE! |
| ENSMUSG00000088224.3 | 0 | NA | NA | NA | NA | NA | #VALUE! |
| ENSMUSG00000088225.3 | 0 | NA | NA | NA | NA | NA | #VALUE! |
| ENSMUSG00000088227.3 | 0 | NA | NA | NA | NA | NA | #VALUE! |
| ENSMUSG00000088228.3 | 0 | NA | NA | NA | NA | NA | #VALUE! |
| ENSMUSG00000088232.3 | 0 | NA | NA | NA | NA | NA | #VALUE! |
| ENSMUSG00000088236.3 | 0 | NA | NA | NA | NA | NA | #VALUE! |
| ENSMUSG00000088240.3 | 0 | NA | NA | NA | NA | NA | #VALUE! |

| | | | | | | | |
|---|---|---|---|---|---|---|---|
| ENSMUSG00000088241.3 | 0 | NA | NA | NA | NA | NA | #VALUE! |
| ENSMUSG00000088242.3 | 0 | NA | NA | NA | NA | NA | #VALUE! |
| ENSMUSG00000088244.3 | 0 | NA | NA | NA | NA | NA | #VALUE! |
| ENSMUSG00000088245.3 | 0 | NA | NA | NA | NA | NA | #VALUE! |
| ENSMUSG00000088248.3 | 0 | NA | NA | NA | NA | NA | #VALUE! |
| ENSMUSG00000088252.3 | 15.46734 | -21.7006 | 3.156273 | -6.87538 | NA | NA | #VALUE! |
| ENSMUSG00000088253.4 | 0 | NA | NA | NA | NA | NA | #VALUE! |
| ENSMUSG00000088254.3 | 0 | NA | NA | NA | NA | NA | #VALUE! |
| ENSMUSG00000088255.3 | 0 | NA | NA | NA | NA | NA | #VALUE! |
| ENSMUSG00000088257.3 | 0 | NA | NA | NA | NA | NA | #VALUE! |
| ENSMUSG00000088258.3 | 0 | NA | NA | NA | NA | NA | #VALUE! |
| ENSMUSG00000088260.3 | 0 | NA | NA | NA | NA | NA | #VALUE! |
| ENSMUSG00000088261.3 | 0 | NA | NA | NA | NA | NA | #VALUE! |
| ENSMUSG00000088265.3 | 0 | NA | NA | NA | NA | NA | #VALUE! |
| ENSMUSG00000088266.3 | 0 | NA | NA | NA | NA | NA | #VALUE! |
| ENSMUSG00000088273.3 | 0 | NA | NA | NA | NA | NA | #VALUE! |
| ENSMUSG00000088275.3 | 0 | NA | NA | NA | NA | NA | #VALUE! |
| ENSMUSG00000088276.3 | 0 | NA | NA | NA | NA | NA | #VALUE! |
| ENSMUSG00000088281.3 | 0 | NA | NA | NA | NA | NA | #VALUE! |
| ENSMUSG00000088282.3 | 0 | NA | NA | NA | NA | NA | #VALUE! |
| ENSMUSG00000088293.3 | 0 | NA | NA | NA | NA | NA | #VALUE! |
| ENSMUSG00000088296.3 | 0 | NA | NA | NA | NA | NA | #VALUE! |
| ENSMUSG00000088297.4 | 0 | NA | NA | NA | NA | NA | #VALUE! |
| ENSMUSG00000088303.4 | 0 | NA | NA | NA | NA | NA | #VALUE! |
| ENSMUSG00000088304.3 | 0 | NA | NA | NA | NA | NA | #VALUE! |
| ENSMUSG00000088307.3 | 0 | NA | NA | NA | NA | NA | #VALUE! |
| ENSMUSG00000088308.3 | 0 | NA | NA | NA | NA | NA | #VALUE! |
| ENSMUSG00000088310.3 | 0 | NA | NA | NA | NA | NA | #VALUE! |
| ENSMUSG00000088311.3 | 0 | NA | NA | NA | NA | NA | #VALUE! |
| ENSMUSG00000088312.3 | 0 | NA | NA | NA | NA | NA | #VALUE! |
| ENSMUSG00000088313.4 | 0 | NA | NA | NA | NA | NA | #VALUE! |
| ENSMUSG00000088318.3 | 0 | NA | NA | NA | NA | NA | #VALUE! |
| ENSMUSG00000088320.3 | 0 | NA | NA | NA | NA | NA | #VALUE! |
| ENSMUSG00000088323.3 | 0 | NA | NA | NA | NA | NA | #VALUE! |
| ENSMUSG00000088325.4 | 0 | NA | NA | NA | NA | NA | #VALUE! |
| ENSMUSG00000088329.4 | 0 | NA | NA | NA | NA | NA | #VALUE! |
| ENSMUSG00000088332.4 | 0 | NA | NA | NA | NA | NA | #VALUE! |
| ENSMUSG00000088333.4 | 0 | NA | NA | NA | NA | NA | #VALUE! |
| ENSMUSG00000088337.3 | 0 | NA | NA | NA | NA | NA | #VALUE! |
| ENSMUSG00000088339.4 | 0 | NA | NA | NA | NA | NA | #VALUE! |
| ENSMUSG00000088343.3 | 0 | NA | NA | NA | NA | NA | #VALUE! |
| ENSMUSG00000088344.3 | 0 | NA | NA | NA | NA | NA | #VALUE! |
| ENSMUSG00000088346.3 | 0 | NA | NA | NA | NA | NA | #VALUE! |
| ENSMUSG00000088349.3 | 0 | NA | NA | NA | NA | NA | #VALUE! |

| | | | | | | | |
|---|---|---|---|---|---|---|---|
| ENSMUSG00000088351.2 | 0 | NA | NA | NA | NA | NA | #VALUE! |
| ENSMUSG00000088352.3 | 0 | NA | NA | NA | NA | NA | #VALUE! |
| ENSMUSG00000088358.3 | 0 | NA | NA | NA | NA | NA | #VALUE! |
| ENSMUSG00000088362.3 | 0 | NA | NA | NA | NA | NA | #VALUE! |
| ENSMUSG00000088363.3 | 0 | NA | NA | NA | NA | NA | #VALUE! |
| ENSMUSG00000088364.4 | 0 | NA | NA | NA | NA | NA | #VALUE! |
| ENSMUSG00000088370.3 | 0 | NA | NA | NA | NA | NA | #VALUE! |
| ENSMUSG00000088372.3 | 0 | NA | NA | NA | NA | NA | #VALUE! |
| ENSMUSG00000088373.5 | 0 | NA | NA | NA | NA | NA | #VALUE! |
| ENSMUSG00000088374.3 | 0 | NA | NA | NA | NA | NA | #VALUE! |
| ENSMUSG00000088377.3 | 0 | NA | NA | NA | NA | NA | #VALUE! |
| ENSMUSG00000088378.4 | 0 | NA | NA | NA | NA | NA | #VALUE! |
| ENSMUSG00000088379.3 | 0 | NA | NA | NA | NA | NA | #VALUE! |
| ENSMUSG00000088380.3 | 0 | NA | NA | NA | NA | NA | #VALUE! |
| ENSMUSG00000088383.4 | 0 | NA | NA | NA | NA | NA | #VALUE! |
| ENSMUSG00000088387.3 | 0 | NA | NA | NA | NA | NA | #VALUE! |
| ENSMUSG00000088391.3 | 0 | NA | NA | NA | NA | NA | #VALUE! |
| ENSMUSG00000088392.3 | 0 | NA | NA | NA | NA | NA | #VALUE! |
| ENSMUSG00000088394.4 | 0 | NA | NA | NA | NA | NA | #VALUE! |
| ENSMUSG00000088398.3 | 0 | NA | NA | NA | NA | NA | #VALUE! |
| ENSMUSG00000088402.3 | 0 | NA | NA | NA | NA | NA | #VALUE! |
| ENSMUSG00000088405.4 | 0 | NA | NA | NA | NA | NA | #VALUE! |
| ENSMUSG00000088407.3 | 0 | NA | NA | NA | NA | NA | #VALUE! |
| ENSMUSG00000088408.3 | 0 | NA | NA | NA | NA | NA | #VALUE! |
| ENSMUSG00000088410.3 | 0 | NA | NA | NA | NA | NA | #VALUE! |
| ENSMUSG00000088412.3 | 0 | NA | NA | NA | NA | NA | #VALUE! |
| ENSMUSG00000088414.3 | 0 | NA | NA | NA | NA | NA | #VALUE! |
| ENSMUSG00000088418.3 | 0 | NA | NA | NA | NA | NA | #VALUE! |
| ENSMUSG00000088419.3 | 0 | NA | NA | NA | NA | NA | #VALUE! |
| ENSMUSG00000088420.3 | 0 | NA | NA | NA | NA | NA | #VALUE! |
| ENSMUSG00000088423.3 | 0 | NA | NA | NA | NA | NA | #VALUE! |
| ENSMUSG00000088425.3 | 0 | NA | NA | NA | NA | NA | #VALUE! |
| ENSMUSG00000088429.4 | 0 | NA | NA | NA | NA | NA | #VALUE! |
| ENSMUSG00000088432.4 | 0 | NA | NA | NA | NA | NA | #VALUE! |
| ENSMUSG00000088435.3 | 0 | NA | NA | NA | NA | NA | #VALUE! |
| ENSMUSG00000088437.4 | 0 | NA | NA | NA | NA | NA | #VALUE! |
| ENSMUSG00000088440.3 | 0 | NA | NA | NA | NA | NA | #VALUE! |
| ENSMUSG00000088441.3 | 0 | NA | NA | NA | NA | NA | #VALUE! |
| ENSMUSG00000088443.3 | 0 | NA | NA | NA | NA | NA | #VALUE! |
| ENSMUSG00000088444.3 | 0 | NA | NA | NA | NA | NA | #VALUE! |
| ENSMUSG00000088445.3 | 0 | NA | NA | NA | NA | NA | #VALUE! |
| ENSMUSG00000088447.3 | 0 | NA | NA | NA | NA | NA | #VALUE! |
| ENSMUSG00000088450.3 | 0 | NA | NA | NA | NA | NA | #VALUE! |
| ENSMUSG00000088451.3 | 0 | NA | NA | NA | NA | NA | #VALUE! |

| | | | | | | | |
|---|---|---|---|---|---|---|---|
| ENSMUSG00000088454.3 | 0 | NA | NA | NA | NA | NA | #VALUE! |
| ENSMUSG00000088458.3 | 0 | NA | NA | NA | NA | NA | #VALUE! |
| ENSMUSG00000088459.3 | 0 | NA | NA | NA | NA | NA | #VALUE! |
| ENSMUSG00000088464.3 | 0 | NA | NA | NA | NA | NA | #VALUE! |
| ENSMUSG00000088465.3 | 0 | NA | NA | NA | NA | NA | #VALUE! |
| ENSMUSG00000088468.4 | 0 | NA | NA | NA | NA | NA | #VALUE! |
| ENSMUSG00000088469.3 | 0 | NA | NA | NA | NA | NA | #VALUE! |
| ENSMUSG00000088471.3 | 0 | NA | NA | NA | NA | NA | #VALUE! |
| ENSMUSG00000088472.3 | 0 | NA | NA | NA | NA | NA | #VALUE! |
| ENSMUSG00000088476.3 | 0 | NA | NA | NA | NA | NA | #VALUE! |
| ENSMUSG00000088477.3 | 0 | NA | NA | NA | NA | NA | #VALUE! |
| ENSMUSG00000088478.3 | 0 | NA | NA | NA | NA | NA | #VALUE! |
| ENSMUSG00000088479.3 | 0 | NA | NA | NA | NA | NA | #VALUE! |
| ENSMUSG00000088481.4 | 0 | NA | NA | NA | NA | NA | #VALUE! |
| ENSMUSG00000088482.3 | 0 | NA | NA | NA | NA | NA | #VALUE! |
| ENSMUSG00000088484.4 | 0 | NA | NA | NA | NA | NA | #VALUE! |
| ENSMUSG00000088490.3 | 0 | NA | NA | NA | NA | NA | #VALUE! |
| ENSMUSG00000088491.3 | 0 | NA | NA | NA | NA | NA | #VALUE! |
| ENSMUSG00000088493.3 | 0 | NA | NA | NA | NA | NA | #VALUE! |
| ENSMUSG00000088494.4 | 0 | NA | NA | NA | NA | NA | #VALUE! |
| ENSMUSG00000088495.3 | 0 | NA | NA | NA | NA | NA | #VALUE! |
| ENSMUSG00000088499.4 | 0 | NA | NA | NA | NA | NA | #VALUE! |
| ENSMUSG00000088501.3 | 0 | NA | NA | NA | NA | NA | #VALUE! |
| ENSMUSG00000088504.4 | 0 | NA | NA | NA | NA | NA | #VALUE! |
| ENSMUSG00000088505.3 | 0 | NA | NA | NA | NA | NA | #VALUE! |
| ENSMUSG00000088506.3 | 0 | NA | NA | NA | NA | NA | #VALUE! |
| ENSMUSG00000088509.3 | 0 | NA | NA | NA | NA | NA | #VALUE! |
| ENSMUSG00000088510.3 | 0 | NA | NA | NA | NA | NA | #VALUE! |
| ENSMUSG00000088511.4 | 0 | NA | NA | NA | NA | NA | #VALUE! |
| ENSMUSG00000088512.3 | 0 | NA | NA | NA | NA | NA | #VALUE! |
| ENSMUSG00000088515.4 | 0 | NA | NA | NA | NA | NA | #VALUE! |
| ENSMUSG00000088516.3 | 0 | NA | NA | NA | NA | NA | #VALUE! |
| ENSMUSG00000088517.3 | 0 | NA | NA | NA | NA | NA | #VALUE! |
| ENSMUSG00000088521.3 | 0 | NA | NA | NA | NA | NA | #VALUE! |
| ENSMUSG00000088524.3 | 0 | NA | NA | NA | NA | NA | #VALUE! |
| ENSMUSG00000088525.3 | 0 | NA | NA | NA | NA | NA | #VALUE! |
| ENSMUSG00000088526.2 | 0 | NA | NA | NA | NA | NA | #VALUE! |
| ENSMUSG00000088528.4 | 0 | NA | NA | NA | NA | NA | #VALUE! |
| ENSMUSG00000088530.4 | 0 | NA | NA | NA | NA | NA | #VALUE! |
| ENSMUSG00000088533.4 | 0 | NA | NA | NA | NA | NA | #VALUE! |
| ENSMUSG00000088536.3 | 0 | NA | NA | NA | NA | NA | #VALUE! |
| ENSMUSG00000088538.3 | 0 | NA | NA | NA | NA | NA | #VALUE! |
| ENSMUSG00000088539.3 | 0 | NA | NA | NA | NA | NA | #VALUE! |
| ENSMUSG00000088541.3 | 0 | NA | NA | NA | NA | NA | #VALUE! |

| | | | | | | | |
|---|---|---|---|---|---|---|---|
| ENSMUSG00000088544.3 | 0 | NA | NA | NA | NA | NA | #VALUE! |
| ENSMUSG00000088545.4 | 0 | NA | NA | NA | NA | NA | #VALUE! |
| ENSMUSG00000088547.4 | 0 | NA | NA | NA | NA | NA | #VALUE! |
| ENSMUSG00000088550.3 | 0 | NA | NA | NA | NA | NA | #VALUE! |
| ENSMUSG00000088551.3 | 0 | NA | NA | NA | NA | NA | #VALUE! |
| ENSMUSG00000088552.3 | 0 | NA | NA | NA | NA | NA | #VALUE! |
| ENSMUSG00000088558.3 | 0 | NA | NA | NA | NA | NA | #VALUE! |
| ENSMUSG00000088559.3 | 0 | NA | NA | NA | NA | NA | #VALUE! |
| ENSMUSG00000088563.3 | 0 | NA | NA | NA | NA | NA | #VALUE! |
| ENSMUSG00000088575.3 | 0 | NA | NA | NA | NA | NA | #VALUE! |
| ENSMUSG00000088577.3 | 0 | NA | NA | NA | NA | NA | #VALUE! |
| ENSMUSG00000088578.3 | 0 | NA | NA | NA | NA | NA | #VALUE! |
| ENSMUSG00000088579.3 | 0 | NA | NA | NA | NA | NA | #VALUE! |
| ENSMUSG00000088582.3 | 0 | NA | NA | NA | NA | NA | #VALUE! |
| ENSMUSG00000088583.3 | 0 | NA | NA | NA | NA | NA | #VALUE! |
| ENSMUSG00000088594.3 | 0 | NA | NA | NA | NA | NA | #VALUE! |
| ENSMUSG00000088603.4 | 0 | NA | NA | NA | NA | NA | #VALUE! |
| ENSMUSG00000088606.3 | 0 | NA | NA | NA | NA | NA | #VALUE! |
| ENSMUSG00000088616.3 | 0 | NA | NA | NA | NA | NA | #VALUE! |
| ENSMUSG00000088620.3 | 0 | NA | NA | NA | NA | NA | #VALUE! |
| ENSMUSG00000088621.3 | 0 | NA | NA | NA | NA | NA | #VALUE! |
| ENSMUSG00000088622.4 | 0 | NA | NA | NA | NA | NA | #VALUE! |
| ENSMUSG00000088623.3 | 0 | NA | NA | NA | NA | NA | #VALUE! |
| ENSMUSG00000088625.3 | 0 | NA | NA | NA | NA | NA | #VALUE! |
| ENSMUSG00000088626.5 | 0 | NA | NA | NA | NA | NA | #VALUE! |
| ENSMUSG00000088627.4 | 0 | NA | NA | NA | NA | NA | #VALUE! |
| ENSMUSG00000088628.4 | 0 | NA | NA | NA | NA | NA | #VALUE! |
| ENSMUSG00000088630.4 | 0 | NA | NA | NA | NA | NA | #VALUE! |
| ENSMUSG00000088633.3 | 0 | NA | NA | NA | NA | NA | #VALUE! |
| ENSMUSG00000088634.3 | 0 | NA | NA | NA | NA | NA | #VALUE! |
| ENSMUSG00000088635.3 | 0 | NA | NA | NA | NA | NA | #VALUE! |
| ENSMUSG00000088638.3 | 0 | NA | NA | NA | NA | NA | #VALUE! |
| ENSMUSG00000088639.3 | 0 | NA | NA | NA | NA | NA | #VALUE! |
| ENSMUSG00000088640.3 | 0 | NA | NA | NA | NA | NA | #VALUE! |
| ENSMUSG00000088644.4 | 0 | NA | NA | NA | NA | NA | #VALUE! |
| ENSMUSG00000088647.3 | 0 | NA | NA | NA | NA | NA | #VALUE! |
| ENSMUSG00000088649.3 | 0 | NA | NA | NA | NA | NA | #VALUE! |
| ENSMUSG00000088651.3 | 0 | NA | NA | NA | NA | NA | #VALUE! |
| ENSMUSG00000088653.3 | 0 | NA | NA | NA | NA | NA | #VALUE! |
| ENSMUSG00000088654.3 | 0 | NA | NA | NA | NA | NA | #VALUE! |
| ENSMUSG00000088656.3 | 0 | NA | NA | NA | NA | NA | #VALUE! |
| ENSMUSG00000088658.3 | 0 | NA | NA | NA | NA | NA | #VALUE! |
| ENSMUSG00000088668.3 | 0 | NA | NA | NA | NA | NA | #VALUE! |
| ENSMUSG00000088669.4 | 0 | NA | NA | NA | NA | NA | #VALUE! |

| | | | | | | | |
|---|---|---|---|---|---|---|---|
| ENSMUSG00000088672.3 | 0 | NA | NA | NA | NA | NA | #VALUE! |
| ENSMUSG00000088675.4 | 0 | NA | NA | NA | NA | NA | #VALUE! |
| ENSMUSG00000088678.3 | 0 | NA | NA | NA | NA | NA | #VALUE! |
| ENSMUSG00000088679.3 | 0 | NA | NA | NA | NA | NA | #VALUE! |
| ENSMUSG00000088684.3 | 0 | NA | NA | NA | NA | NA | #VALUE! |
| ENSMUSG00000088685.3 | 0 | NA | NA | NA | NA | NA | #VALUE! |
| ENSMUSG00000088687.3 | 0 | NA | NA | NA | NA | NA | #VALUE! |
| ENSMUSG00000088688.3 | 0 | NA | NA | NA | NA | NA | #VALUE! |
| ENSMUSG00000088689.3 | 0.735092 | -3.12016 | 3.965612 | -0.7868 | 0.431397 | NA | #VALUE! |
| ENSMUSG00000088691.3 | 0 | NA | NA | NA | NA | NA | #VALUE! |
| ENSMUSG00000088692.3 | 0 | NA | NA | NA | NA | NA | #VALUE! |
| ENSMUSG00000088693.3 | 0 | NA | NA | NA | NA | NA | #VALUE! |
| ENSMUSG00000088695.3 | 0 | NA | NA | NA | NA | NA | #VALUE! |
| ENSMUSG00000088696.3 | 0 | NA | NA | NA | NA | NA | #VALUE! |
| ENSMUSG00000088697.4 | 0 | NA | NA | NA | NA | NA | #VALUE! |
| ENSMUSG00000088704.4 | 0 | NA | NA | NA | NA | NA | #VALUE! |
| ENSMUSG00000088705.3 | 0 | NA | NA | NA | NA | NA | #VALUE! |
| ENSMUSG00000088706.4 | 0 | NA | NA | NA | NA | NA | #VALUE! |
| ENSMUSG00000088708.4 | 0 | NA | NA | NA | NA | NA | #VALUE! |
| ENSMUSG00000088712.3 | 0 | NA | NA | NA | NA | NA | #VALUE! |
| ENSMUSG00000088715.3 | 0 | NA | NA | NA | NA | NA | #VALUE! |
| ENSMUSG00000088716.3 | 0 | NA | NA | NA | NA | NA | #VALUE! |
| ENSMUSG00000088717.4 | 0 | NA | NA | NA | NA | NA | #VALUE! |
| ENSMUSG00000088718.4 | 0 | NA | NA | NA | NA | NA | #VALUE! |
| ENSMUSG00000088719.4 | 0 | NA | NA | NA | NA | NA | #VALUE! |
| ENSMUSG00000088721.5 | 0 | NA | NA | NA | NA | NA | #VALUE! |
| ENSMUSG00000088726.3 | 0 | NA | NA | NA | NA | NA | #VALUE! |
| ENSMUSG00000088730.3 | 0 | NA | NA | NA | NA | NA | #VALUE! |
| ENSMUSG00000088733.4 | 0 | NA | NA | NA | NA | NA | #VALUE! |
| ENSMUSG00000088734.3 | 0 | NA | NA | NA | NA | NA | #VALUE! |
| ENSMUSG00000088735.4 | 0 | NA | NA | NA | NA | NA | #VALUE! |
| ENSMUSG00000088738.5 | 0 | NA | NA | NA | NA | NA | #VALUE! |
| ENSMUSG00000088739.3 | 0 | NA | NA | NA | NA | NA | #VALUE! |
| ENSMUSG00000088740.3 | 0 | NA | NA | NA | NA | NA | #VALUE! |
| ENSMUSG00000088744.3 | 0 | NA | NA | NA | NA | NA | #VALUE! |
| ENSMUSG00000088746.3 | 0 | NA | NA | NA | NA | NA | #VALUE! |
| ENSMUSG00000088748.3 | 0 | NA | NA | NA | NA | NA | #VALUE! |
| ENSMUSG00000088750.3 | 0 | NA | NA | NA | NA | NA | #VALUE! |
| ENSMUSG00000088751.3 | 0 | NA | NA | NA | NA | NA | #VALUE! |
| ENSMUSG00000088752.3 | 0 | NA | NA | NA | NA | NA | #VALUE! |
| ENSMUSG00000088754.4 | 0 | NA | NA | NA | NA | NA | #VALUE! |
| ENSMUSG00000088756.3 | 0 | NA | NA | NA | NA | NA | #VALUE! |
| ENSMUSG00000088760.3 | 0 | NA | NA | NA | NA | NA | #VALUE! |
| ENSMUSG00000088761.3 | 0 | NA | NA | NA | NA | NA | #VALUE! |

| | | | | | | | |
|---|---|---|---|---|---|---|---|
| ENSMUSG00000088764.3 | 0 | NA | NA | NA | NA | NA | #VALUE! |
| ENSMUSG00000088766.3 | 0 | NA | NA | NA | NA | NA | #VALUE! |
| ENSMUSG00000088768.3 | 0 | NA | NA | NA | NA | NA | #VALUE! |
| ENSMUSG00000088772.3 | 0 | NA | NA | NA | NA | NA | #VALUE! |
| ENSMUSG00000088774.3 | 0 | NA | NA | NA | NA | NA | #VALUE! |
| ENSMUSG00000088776.3 | 0 | NA | NA | NA | NA | NA | #VALUE! |
| ENSMUSG00000088781.3 | 0 | NA | NA | NA | NA | NA | #VALUE! |
| ENSMUSG00000088782.3 | 0 | NA | NA | NA | NA | NA | #VALUE! |
| ENSMUSG00000088784.3 | 0 | NA | NA | NA | NA | NA | #VALUE! |
| ENSMUSG00000088791.3 | 0 | NA | NA | NA | NA | NA | #VALUE! |
| ENSMUSG00000088792.3 | 0 | NA | NA | NA | NA | NA | #VALUE! |
| ENSMUSG00000088793.4 | 0 | NA | NA | NA | NA | NA | #VALUE! |
| ENSMUSG00000088796.3 | 0 | NA | NA | NA | NA | NA | #VALUE! |
| ENSMUSG00000088798.3 | 0 | NA | NA | NA | NA | NA | #VALUE! |
| ENSMUSG00000088800.4 | 0 | NA | NA | NA | NA | NA | #VALUE! |
| ENSMUSG00000088810.3 | 0 | NA | NA | NA | NA | NA | #VALUE! |
| ENSMUSG00000088811.3 | 0 | NA | NA | NA | NA | NA | #VALUE! |
| ENSMUSG00000088816.3 | 0 | NA | NA | NA | NA | NA | #VALUE! |
| ENSMUSG00000088817.3 | 0 | NA | NA | NA | NA | NA | #VALUE! |
| ENSMUSG00000088820.3 | 0 | NA | NA | NA | NA | NA | #VALUE! |
| ENSMUSG00000088823.3 | 0 | NA | NA | NA | NA | NA | #VALUE! |
| ENSMUSG00000088824.3 | 0 | NA | NA | NA | NA | NA | #VALUE! |
| ENSMUSG00000088829.3 | 0 | NA | NA | NA | NA | NA | #VALUE! |
| ENSMUSG00000088830.4 | 0 | NA | NA | NA | NA | NA | #VALUE! |
| ENSMUSG00000088835.3 | 0 | NA | NA | NA | NA | NA | #VALUE! |
| ENSMUSG00000088836.3 | 0 | NA | NA | NA | NA | NA | #VALUE! |
| ENSMUSG00000088839.4 | 0 | NA | NA | NA | NA | NA | #VALUE! |
| ENSMUSG00000088840.3 | 0 | NA | NA | NA | NA | NA | #VALUE! |
| ENSMUSG00000088843.4 | 0 | NA | NA | NA | NA | NA | #VALUE! |
| ENSMUSG00000088846.3 | 0 | NA | NA | NA | NA | NA | #VALUE! |
| ENSMUSG00000088848.3 | 0 | NA | NA | NA | NA | NA | #VALUE! |
| ENSMUSG00000088850.3 | 0 | NA | NA | NA | NA | NA | #VALUE! |
| ENSMUSG00000088851.3 | 0 | NA | NA | NA | NA | NA | #VALUE! |
| ENSMUSG00000088852.3 | 0 | NA | NA | NA | NA | NA | #VALUE! |
| ENSMUSG00000088853.3 | 0 | NA | NA | NA | NA | NA | #VALUE! |
| ENSMUSG00000088854.4 | 0 | NA | NA | NA | NA | NA | #VALUE! |
| ENSMUSG00000088857.3 | 0 | NA | NA | NA | NA | NA | #VALUE! |
| ENSMUSG00000088858.3 | 0 | NA | NA | NA | NA | NA | #VALUE! |
| ENSMUSG00000088860.3 | 0 | NA | NA | NA | NA | NA | #VALUE! |
| ENSMUSG00000088863.3 | 0 | NA | NA | NA | NA | NA | #VALUE! |
| ENSMUSG00000088868.3 | 0 | NA | NA | NA | NA | NA | #VALUE! |
| ENSMUSG00000088870.3 | 0 | NA | NA | NA | NA | NA | #VALUE! |
| ENSMUSG00000088874.3 | 0 | NA | NA | NA | NA | NA | #VALUE! |
| ENSMUSG00000088875.3 | 0 | NA | NA | NA | NA | NA | #VALUE! |

| | | | | | | | |
|---|---|---|---|---|---|---|---|
| ENSMUSG00000088876.4 | 0 | NA | NA | NA | NA | NA | #VALUE! |
| ENSMUSG00000088877.3 | 0 | NA | NA | NA | NA | NA | #VALUE! |
| ENSMUSG00000088879.3 | 0 | NA | NA | NA | NA | NA | #VALUE! |
| ENSMUSG00000088883.3 | 0 | NA | NA | NA | NA | NA | #VALUE! |
| ENSMUSG00000088884.3 | 0 | NA | NA | NA | NA | NA | #VALUE! |
| ENSMUSG00000088888.3 | 0 | NA | NA | NA | NA | NA | #VALUE! |
| ENSMUSG00000088889.3 | 0 | NA | NA | NA | NA | NA | #VALUE! |
| ENSMUSG00000088891.4 | 0 | NA | NA | NA | NA | NA | #VALUE! |
| ENSMUSG00000088894.3 | 0 | NA | NA | NA | NA | NA | #VALUE! |
| ENSMUSG00000088895.3 | 0 | NA | NA | NA | NA | NA | #VALUE! |
| ENSMUSG00000088896.3 | 0 | NA | NA | NA | NA | NA | #VALUE! |
| ENSMUSG00000088897.3 | 0 | NA | NA | NA | NA | NA | #VALUE! |
| ENSMUSG00000088900.3 | 0 | NA | NA | NA | NA | NA | #VALUE! |
| ENSMUSG00000088901.3 | 0 | NA | NA | NA | NA | NA | #VALUE! |
| ENSMUSG00000088903.3 | 0 | NA | NA | NA | NA | NA | #VALUE! |
| ENSMUSG00000088905.3 | 0 | NA | NA | NA | NA | NA | #VALUE! |
| ENSMUSG00000088906.3 | 0 | NA | NA | NA | NA | NA | #VALUE! |
| ENSMUSG00000088907.3 | 0 | NA | NA | NA | NA | NA | #VALUE! |
| ENSMUSG00000088911.3 | 0 | NA | NA | NA | NA | NA | #VALUE! |
| ENSMUSG00000088914.3 | 0 | NA | NA | NA | NA | NA | #VALUE! |
| ENSMUSG00000088915.3 | 0 | NA | NA | NA | NA | NA | #VALUE! |
| ENSMUSG00000088922.3 | 0 | NA | NA | NA | NA | NA | #VALUE! |
| ENSMUSG00000088923.3 | 0 | NA | NA | NA | NA | NA | #VALUE! |
| ENSMUSG00000088924.4 | 0 | NA | NA | NA | NA | NA | #VALUE! |
| ENSMUSG00000088928.3 | 0 | NA | NA | NA | NA | NA | #VALUE! |
| ENSMUSG00000088929.3 | 0 | NA | NA | NA | NA | NA | #VALUE! |
| ENSMUSG00000088930.3 | 0 | NA | NA | NA | NA | NA | #VALUE! |
| ENSMUSG00000088935.3 | 0 | NA | NA | NA | NA | NA | #VALUE! |
| ENSMUSG00000088942.4 | 0 | NA | NA | NA | NA | NA | #VALUE! |
| ENSMUSG00000088944.3 | 0 | NA | NA | NA | NA | NA | #VALUE! |
| ENSMUSG00000088948.3 | 0 | NA | NA | NA | NA | NA | #VALUE! |
| ENSMUSG00000088951.4 | 0 | NA | NA | NA | NA | NA | #VALUE! |
| ENSMUSG00000088953.3 | 0 | NA | NA | NA | NA | NA | #VALUE! |
| ENSMUSG00000088954.4 | 0 | NA | NA | NA | NA | NA | #VALUE! |
| ENSMUSG00000088958.3 | 0 | NA | NA | NA | NA | NA | #VALUE! |
| ENSMUSG00000088959.3 | 0 | NA | NA | NA | NA | NA | #VALUE! |
| ENSMUSG00000088960.3 | 0 | NA | NA | NA | NA | NA | #VALUE! |
| ENSMUSG00000088961.3 | 0 | NA | NA | NA | NA | NA | #VALUE! |
| ENSMUSG00000088964.4 | 0 | NA | NA | NA | NA | NA | #VALUE! |
| ENSMUSG00000088969.3 | 0 | NA | NA | NA | NA | NA | #VALUE! |
| ENSMUSG00000088970.3 | 0 | NA | NA | NA | NA | NA | #VALUE! |
| ENSMUSG00000088975.3 | 0 | NA | NA | NA | NA | NA | #VALUE! |
| ENSMUSG00000088979.3 | 0 | NA | NA | NA | NA | NA | #VALUE! |
| ENSMUSG00000088980.3 | 0 | NA | NA | NA | NA | NA | #VALUE! |

| | | | | | | | |
|---|---|---|---|---|---|---|---|
| ENSMUSG00000088982.3 | 0 | NA | NA | NA | NA | NA | #VALUE! |
| ENSMUSG00000088983.3 | 0 | NA | NA | NA | NA | NA | #VALUE! |
| ENSMUSG00000088984.3 | 0 | NA | NA | NA | NA | NA | #VALUE! |
| ENSMUSG00000088985.3 | 0 | NA | NA | NA | NA | NA | #VALUE! |
| ENSMUSG00000088987.3 | 0 | NA | NA | NA | NA | NA | #VALUE! |
| ENSMUSG00000088990.3 | 0 | NA | NA | NA | NA | NA | #VALUE! |
| ENSMUSG00000088992.4 | 0 | NA | NA | NA | NA | NA | #VALUE! |
| ENSMUSG00000088995.3 | 0 | NA | NA | NA | NA | NA | #VALUE! |
| ENSMUSG00000088997.4 | 0 | NA | NA | NA | NA | NA | #VALUE! |
| ENSMUSG00000088999.3 | 0 | NA | NA | NA | NA | NA | #VALUE! |
| ENSMUSG00000089001.3 | 0 | NA | NA | NA | NA | NA | #VALUE! |
| ENSMUSG00000089002.3 | 0 | NA | NA | NA | NA | NA | #VALUE! |
| ENSMUSG00000089005.4 | 0 | NA | NA | NA | NA | NA | #VALUE! |
| ENSMUSG00000089007.3 | 0 | NA | NA | NA | NA | NA | #VALUE! |
| ENSMUSG00000089008.3 | 0 | NA | NA | NA | NA | NA | #VALUE! |
| ENSMUSG00000089009.3 | 0 | NA | NA | NA | NA | NA | #VALUE! |
| ENSMUSG00000089011.4 | 0 | NA | NA | NA | NA | NA | #VALUE! |
| ENSMUSG00000089014.3 | 0 | NA | NA | NA | NA | NA | #VALUE! |
| ENSMUSG00000089017.3 | 0 | NA | NA | NA | NA | NA | #VALUE! |
| ENSMUSG00000089018.3 | 0 | NA | NA | NA | NA | NA | #VALUE! |
| ENSMUSG00000089019.3 | 0 | NA | NA | NA | NA | NA | #VALUE! |
| ENSMUSG00000089024.3 | 0 | NA | NA | NA | NA | NA | #VALUE! |
| ENSMUSG00000089025.3 | 0 | NA | NA | NA | NA | NA | #VALUE! |
| ENSMUSG00000089026.5 | 0 | NA | NA | NA | NA | NA | #VALUE! |
| ENSMUSG00000089027.3 | 0 | NA | NA | NA | NA | NA | #VALUE! |
| ENSMUSG00000089028.4 | 0 | NA | NA | NA | NA | NA | #VALUE! |
| ENSMUSG00000089029.3 | 0 | NA | NA | NA | NA | NA | #VALUE! |
| ENSMUSG00000089033.3 | 0 | NA | NA | NA | NA | NA | #VALUE! |
| ENSMUSG00000089034.3 | 0 | NA | NA | NA | NA | NA | #VALUE! |
| ENSMUSG00000089035.3 | 0 | NA | NA | NA | NA | NA | #VALUE! |
| ENSMUSG00000089036.3 | 0 | NA | NA | NA | NA | NA | #VALUE! |
| ENSMUSG00000089037.3 | 0 | NA | NA | NA | NA | NA | #VALUE! |
| ENSMUSG00000089039.4 | 0 | NA | NA | NA | NA | NA | #VALUE! |
| ENSMUSG00000089040.4 | 0 | NA | NA | NA | NA | NA | #VALUE! |
| ENSMUSG00000089045.3 | 0 | NA | NA | NA | NA | NA | #VALUE! |
| ENSMUSG00000089046.4 | 0 | NA | NA | NA | NA | NA | #VALUE! |
| ENSMUSG00000089051.3 | 0 | NA | NA | NA | NA | NA | #VALUE! |
| ENSMUSG00000089052.4 | 0 | NA | NA | NA | NA | NA | #VALUE! |
| ENSMUSG00000089053.3 | 0 | NA | NA | NA | NA | NA | #VALUE! |
| ENSMUSG00000089054.2 | 0 | NA | NA | NA | NA | NA | #VALUE! |
| ENSMUSG00000089055.3 | 0 | NA | NA | NA | NA | NA | #VALUE! |
| ENSMUSG00000089057.4 | 0 | NA | NA | NA | NA | NA | #VALUE! |
| ENSMUSG00000089060.4 | 0 | NA | NA | NA | NA | NA | #VALUE! |
| ENSMUSG00000089066.3 | 0 | NA | NA | NA | NA | NA | #VALUE! |

| | | | | | | | |
|---|---|---|---|---|---|---|---|
| ENSMUSG00000089068.3 | 0 | NA | NA | NA | NA | NA | #VALUE! |
| ENSMUSG00000089069.3 | 0 | NA | NA | NA | NA | NA | #VALUE! |
| ENSMUSG00000089071.3 | 0 | NA | NA | NA | NA | NA | #VALUE! |
| ENSMUSG00000089074.3 | 0 | NA | NA | NA | NA | NA | #VALUE! |
| ENSMUSG00000089078.3 | 0 | NA | NA | NA | NA | NA | #VALUE! |
| ENSMUSG00000089079.3 | 0 | NA | NA | NA | NA | NA | #VALUE! |
| ENSMUSG00000089080.3 | 0 | NA | NA | NA | NA | NA | #VALUE! |
| ENSMUSG00000089081.3 | 0 | NA | NA | NA | NA | NA | #VALUE! |
| ENSMUSG00000089083.3 | 0 | NA | NA | NA | NA | NA | #VALUE! |
| ENSMUSG00000089085.3 | 0 | NA | NA | NA | NA | NA | #VALUE! |
| ENSMUSG00000089087.3 | 0 | NA | NA | NA | NA | NA | #VALUE! |
| ENSMUSG00000089091.3 | 0 | NA | NA | NA | NA | NA | #VALUE! |
| ENSMUSG00000089093.5 | 0 | NA | NA | NA | NA | NA | #VALUE! |
| ENSMUSG00000089096.3 | 0 | NA | NA | NA | NA | NA | #VALUE! |
| ENSMUSG00000089097.3 | 0 | NA | NA | NA | NA | NA | #VALUE! |
| ENSMUSG00000089098.3 | 0 | NA | NA | NA | NA | NA | #VALUE! |
| ENSMUSG00000089102.3 | 0 | NA | NA | NA | NA | NA | #VALUE! |
| ENSMUSG00000089106.3 | 0 | NA | NA | NA | NA | NA | #VALUE! |
| ENSMUSG00000089107.3 | 0 | NA | NA | NA | NA | NA | #VALUE! |
| ENSMUSG00000089109.4 | 0 | NA | NA | NA | NA | NA | #VALUE! |
| ENSMUSG00000089113.3 | 0 | NA | NA | NA | NA | NA | #VALUE! |
| ENSMUSG00000089114.3 | 0 | NA | NA | NA | NA | NA | #VALUE! |
| ENSMUSG00000089116.3 | 0 | NA | NA | NA | NA | NA | #VALUE! |
| ENSMUSG00000089117.3 | 0 | NA | NA | NA | NA | NA | #VALUE! |
| ENSMUSG00000089119.3 | 0 | NA | NA | NA | NA | NA | #VALUE! |
| ENSMUSG00000089123.3 | 0 | NA | NA | NA | NA | NA | #VALUE! |
| ENSMUSG00000089124.4 | 0 | NA | NA | NA | NA | NA | #VALUE! |
| ENSMUSG00000089125.3 | 0 | NA | NA | NA | NA | NA | #VALUE! |
| ENSMUSG00000089127.4 | 0 | NA | NA | NA | NA | NA | #VALUE! |
| ENSMUSG00000089128.3 | 0 | NA | NA | NA | NA | NA | #VALUE! |
| ENSMUSG00000089130.3 | 0 | NA | NA | NA | NA | NA | #VALUE! |
| ENSMUSG00000089133.4 | 0 | NA | NA | NA | NA | NA | #VALUE! |
| ENSMUSG00000089136.4 | 0 | NA | NA | NA | NA | NA | #VALUE! |
| ENSMUSG00000089139.3 | 0 | NA | NA | NA | NA | NA | #VALUE! |
| ENSMUSG00000089143.4 | 0 | NA | NA | NA | NA | NA | #VALUE! |
| ENSMUSG00000089144.3 | 0 | NA | NA | NA | NA | NA | #VALUE! |
| ENSMUSG00000089147.3 | 0 | NA | NA | NA | NA | NA | #VALUE! |
| ENSMUSG00000089151.3 | 0 | NA | NA | NA | NA | NA | #VALUE! |
| ENSMUSG00000089152.4 | 0 | NA | NA | NA | NA | NA | #VALUE! |
| ENSMUSG00000089155.3 | 0 | NA | NA | NA | NA | NA | #VALUE! |
| ENSMUSG00000089156.3 | 0 | NA | NA | NA | NA | NA | #VALUE! |
| ENSMUSG00000089157.3 | 0 | NA | NA | NA | NA | NA | #VALUE! |
| ENSMUSG00000089161.3 | 0 | NA | NA | NA | NA | NA | #VALUE! |
| ENSMUSG00000089162.4 | 0 | NA | NA | NA | NA | NA | #VALUE! |

| | | | | | | | |
|---|---|---|---|---|---|---|---|
| ENSMUSG00000089164.3 | 0 | NA | NA | NA | NA | NA | #VALUE! |
| ENSMUSG00000089168.3 | 0 | NA | NA | NA | NA | NA | #VALUE! |
| ENSMUSG00000089171.3 | 0 | NA | NA | NA | NA | NA | #VALUE! |
| ENSMUSG00000089174.3 | 0 | NA | NA | NA | NA | NA | #VALUE! |
| ENSMUSG00000089175.3 | 0 | NA | NA | NA | NA | NA | #VALUE! |
| ENSMUSG00000089178.3 | 0 | NA | NA | NA | NA | NA | #VALUE! |
| ENSMUSG00000089179.4 | 0 | NA | NA | NA | NA | NA | #VALUE! |
| ENSMUSG00000089182.3 | 0 | NA | NA | NA | NA | NA | #VALUE! |
| ENSMUSG00000089183.4 | 0 | NA | NA | NA | NA | NA | #VALUE! |
| ENSMUSG00000089184.3 | 0 | NA | NA | NA | NA | NA | #VALUE! |
| ENSMUSG00000089188.4 | 0 | NA | NA | NA | NA | NA | #VALUE! |
| ENSMUSG00000089190.3 | 0 | NA | NA | NA | NA | NA | #VALUE! |
| ENSMUSG00000089191.3 | 0 | NA | NA | NA | NA | NA | #VALUE! |
| ENSMUSG00000089194.3 | 0 | NA | NA | NA | NA | NA | #VALUE! |
| ENSMUSG00000089195.3 | 0 | NA | NA | NA | NA | NA | #VALUE! |
| ENSMUSG00000089197.3 | 0 | NA | NA | NA | NA | NA | #VALUE! |
| ENSMUSG00000089203.3 | 0 | NA | NA | NA | NA | NA | #VALUE! |
| ENSMUSG00000089204.3 | 0 | NA | NA | NA | NA | NA | #VALUE! |
| ENSMUSG00000089205.3 | 0 | NA | NA | NA | NA | NA | #VALUE! |
| ENSMUSG00000089210.3 | 0 | NA | NA | NA | NA | NA | #VALUE! |
| ENSMUSG00000089214.3 | 0 | NA | NA | NA | NA | NA | #VALUE! |
| ENSMUSG00000089219.3 | 0 | NA | NA | NA | NA | NA | #VALUE! |
| ENSMUSG00000089220.3 | 0 | NA | NA | NA | NA | NA | #VALUE! |
| ENSMUSG00000089222.4 | 0 | NA | NA | NA | NA | NA | #VALUE! |
| ENSMUSG00000089229.4 | 0 | NA | NA | NA | NA | NA | #VALUE! |
| ENSMUSG00000089230.3 | 0 | NA | NA | NA | NA | NA | #VALUE! |
| ENSMUSG00000089231.3 | 0 | NA | NA | NA | NA | NA | #VALUE! |
| ENSMUSG00000089237.3 | 0 | NA | NA | NA | NA | NA | #VALUE! |
| ENSMUSG00000089239.3 | 0 | NA | NA | NA | NA | NA | #VALUE! |
| ENSMUSG00000089242.3 | 0 | NA | NA | NA | NA | NA | #VALUE! |
| ENSMUSG00000089252.3 | 0 | NA | NA | NA | NA | NA | #VALUE! |
| ENSMUSG00000089255.3 | 0 | NA | NA | NA | NA | NA | #VALUE! |
| ENSMUSG00000089257.3 | 0 | NA | NA | NA | NA | NA | #VALUE! |
| ENSMUSG00000089258.3 | 0 | NA | NA | NA | NA | NA | #VALUE! |
| ENSMUSG00000089260.3 | 0 | NA | NA | NA | NA | NA | #VALUE! |
| ENSMUSG00000089261.3 | 0 | NA | NA | NA | NA | NA | #VALUE! |
| ENSMUSG00000089262.3 | 0 | NA | NA | NA | NA | NA | #VALUE! |
| ENSMUSG00000089264.4 | 0 | NA | NA | NA | NA | NA | #VALUE! |
| ENSMUSG00000089265.2 | 0 | NA | NA | NA | NA | NA | #VALUE! |
| ENSMUSG00000089267.3 | 0 | NA | NA | NA | NA | NA | #VALUE! |
| ENSMUSG00000089268.3 | 0 | NA | NA | NA | NA | NA | #VALUE! |
| ENSMUSG00000089269.3 | 0 | NA | NA | NA | NA | NA | #VALUE! |
| ENSMUSG00000089271.3 | 0 | NA | NA | NA | NA | NA | #VALUE! |
| ENSMUSG00000089272.3 | 0 | NA | NA | NA | NA | NA | #VALUE! |

| | | | | | | | |
|---|---|---|---|---|---|---|---|
| ENSMUSG00000089274.3 | 0 | NA | NA | NA | NA | NA | #VALUE! |
| ENSMUSG00000089275.3 | 0 | NA | NA | NA | NA | NA | #VALUE! |
| ENSMUSG00000089277.4 | 0 | NA | NA | NA | NA | NA | #VALUE! |
| ENSMUSG00000089278.3 | 0 | NA | NA | NA | NA | NA | #VALUE! |
| ENSMUSG00000089279.3 | 0 | NA | NA | NA | NA | NA | #VALUE! |
| ENSMUSG00000089281.3 | 0 | NA | NA | NA | NA | NA | #VALUE! |
| ENSMUSG00000089284.3 | 0 | NA | NA | NA | NA | NA | #VALUE! |
| ENSMUSG00000089287.3 | 0 | NA | NA | NA | NA | NA | #VALUE! |
| ENSMUSG00000089288.3 | 0 | NA | NA | NA | NA | NA | #VALUE! |
| ENSMUSG00000089289.3 | 0 | NA | NA | NA | NA | NA | #VALUE! |
| ENSMUSG00000089290.3 | 0 | NA | NA | NA | NA | NA | #VALUE! |
| ENSMUSG00000089293.4 | 0 | NA | NA | NA | NA | NA | #VALUE! |
| ENSMUSG00000089294.3 | 0 | NA | NA | NA | NA | NA | #VALUE! |
| ENSMUSG00000089296.4 | 0 | NA | NA | NA | NA | NA | #VALUE! |
| ENSMUSG00000089299.3 | 0 | NA | NA | NA | NA | NA | #VALUE! |
| ENSMUSG00000089302.3 | 0 | NA | NA | NA | NA | NA | #VALUE! |
| ENSMUSG00000089304.3 | 0 | NA | NA | NA | NA | NA | #VALUE! |
| ENSMUSG00000089307.4 | 0 | NA | NA | NA | NA | NA | #VALUE! |
| ENSMUSG00000089308.2 | 0 | NA | NA | NA | NA | NA | #VALUE! |
| ENSMUSG00000089309.3 | 0 | NA | NA | NA | NA | NA | #VALUE! |
| ENSMUSG00000089310.3 | 0 | NA | NA | NA | NA | NA | #VALUE! |
| ENSMUSG00000089311.3 | 0 | NA | NA | NA | NA | NA | #VALUE! |
| ENSMUSG00000089313.4 | 0 | NA | NA | NA | NA | NA | #VALUE! |
| ENSMUSG00000089314.3 | 0 | NA | NA | NA | NA | NA | #VALUE! |
| ENSMUSG00000089317.3 | 0 | NA | NA | NA | NA | NA | #VALUE! |
| ENSMUSG00000089320.3 | 0 | NA | NA | NA | NA | NA | #VALUE! |
| ENSMUSG00000089321.3 | 0 | NA | NA | NA | NA | NA | #VALUE! |
| ENSMUSG00000089322.4 | 0 | NA | NA | NA | NA | NA | #VALUE! |
| ENSMUSG00000089324.3 | 0 | NA | NA | NA | NA | NA | #VALUE! |
| ENSMUSG00000089327.2 | 0 | NA | NA | NA | NA | NA | #VALUE! |
| ENSMUSG00000089331.3 | 0 | NA | NA | NA | NA | NA | #VALUE! |
| ENSMUSG00000089333.3 | 0 | NA | NA | NA | NA | NA | #VALUE! |
| ENSMUSG00000089334.4 | 0 | NA | NA | NA | NA | NA | #VALUE! |
| ENSMUSG00000089335.4 | 0 | NA | NA | NA | NA | NA | #VALUE! |
| ENSMUSG00000089336.3 | 0 | NA | NA | NA | NA | NA | #VALUE! |
| ENSMUSG00000089346.4 | 0 | NA | NA | NA | NA | NA | #VALUE! |
| ENSMUSG00000089349.2 | 0 | NA | NA | NA | NA | NA | #VALUE! |
| ENSMUSG00000089350.3 | 0 | NA | NA | NA | NA | NA | #VALUE! |
| ENSMUSG00000089351.3 | 0 | NA | NA | NA | NA | NA | #VALUE! |
| ENSMUSG00000089352.3 | 0 | NA | NA | NA | NA | NA | #VALUE! |
| ENSMUSG00000089357.3 | 0 | NA | NA | NA | NA | NA | #VALUE! |
| ENSMUSG00000089359.3 | 0 | NA | NA | NA | NA | NA | #VALUE! |
| ENSMUSG00000089364.3 | 0 | NA | NA | NA | NA | NA | #VALUE! |
| ENSMUSG00000089369.3 | 0 | NA | NA | NA | NA | NA | #VALUE! |

| | | | | | | | |
|---|---|---|---|---|---|---|---|
| ENSMUSG00000089370.4 | 0 | NA | NA | NA | NA | NA | #VALUE! |
| ENSMUSG00000089371.3 | 0 | NA | NA | NA | NA | NA | #VALUE! |
| ENSMUSG00000089372.3 | 0 | NA | NA | NA | NA | NA | #VALUE! |
| ENSMUSG00000089373.3 | 0 | NA | NA | NA | NA | NA | #VALUE! |
| ENSMUSG00000089375.3 | 0 | NA | NA | NA | NA | NA | #VALUE! |
| ENSMUSG00000089380.4 | 0 | NA | NA | NA | NA | NA | #VALUE! |
| ENSMUSG00000089381.4 | 0 | NA | NA | NA | NA | NA | #VALUE! |
| ENSMUSG00000089384.3 | 0 | NA | NA | NA | NA | NA | #VALUE! |
| ENSMUSG00000089385.3 | 0 | NA | NA | NA | NA | NA | #VALUE! |
| ENSMUSG00000089387.3 | 0 | NA | NA | NA | NA | NA | #VALUE! |
| ENSMUSG00000089390.3 | 0 | NA | NA | NA | NA | NA | #VALUE! |
| ENSMUSG00000089393.3 | 0 | NA | NA | NA | NA | NA | #VALUE! |
| ENSMUSG00000089394.3 | 0 | NA | NA | NA | NA | NA | #VALUE! |
| ENSMUSG00000089399.4 | 0 | NA | NA | NA | NA | NA | #VALUE! |
| ENSMUSG00000089406.3 | 0 | NA | NA | NA | NA | NA | #VALUE! |
| ENSMUSG00000089407.3 | 0 | NA | NA | NA | NA | NA | #VALUE! |
| ENSMUSG00000089408.4 | 0 | NA | NA | NA | NA | NA | #VALUE! |
| ENSMUSG00000089409.3 | 0 | NA | NA | NA | NA | NA | #VALUE! |
| ENSMUSG00000089413.3 | 0 | NA | NA | NA | NA | NA | #VALUE! |
| ENSMUSG00000089415.3 | 0 | NA | NA | NA | NA | NA | #VALUE! |
| ENSMUSG00000089416.4 | 0 | NA | NA | NA | NA | NA | #VALUE! |
| ENSMUSG00000089417.3 | 0 | NA | NA | NA | NA | NA | #VALUE! |
| ENSMUSG00000089420.4 | 0 | NA | NA | NA | NA | NA | #VALUE! |
| ENSMUSG00000089422.3 | 0 | NA | NA | NA | NA | NA | #VALUE! |
| ENSMUSG00000089423.3 | 0 | NA | NA | NA | NA | NA | #VALUE! |
| ENSMUSG00000089424.3 | 0 | NA | NA | NA | NA | NA | #VALUE! |
| ENSMUSG00000089425.4 | 0 | NA | NA | NA | NA | NA | #VALUE! |
| ENSMUSG00000089427.3 | 0 | NA | NA | NA | NA | NA | #VALUE! |
| ENSMUSG00000089431.4 | 0 | NA | NA | NA | NA | NA | #VALUE! |
| ENSMUSG00000089432.3 | 0 | NA | NA | NA | NA | NA | #VALUE! |
| ENSMUSG00000089437.3 | 0 | NA | NA | NA | NA | NA | #VALUE! |
| ENSMUSG00000089438.3 | 0 | NA | NA | NA | NA | NA | #VALUE! |
| ENSMUSG00000089439.3 | 0 | NA | NA | NA | NA | NA | #VALUE! |
| ENSMUSG00000089442.4 | 0 | NA | NA | NA | NA | NA | #VALUE! |
| ENSMUSG00000089444.3 | 0 | NA | NA | NA | NA | NA | #VALUE! |
| ENSMUSG00000089445.3 | 0 | NA | NA | NA | NA | NA | #VALUE! |
| ENSMUSG00000089452.4 | 0 | NA | NA | NA | NA | NA | #VALUE! |
| ENSMUSG00000089454.3 | 0 | NA | NA | NA | NA | NA | #VALUE! |
| ENSMUSG00000089455.3 | 0 | NA | NA | NA | NA | NA | #VALUE! |
| ENSMUSG00000089458.4 | 0 | NA | NA | NA | NA | NA | #VALUE! |
| ENSMUSG00000089460.3 | 0 | NA | NA | NA | NA | NA | #VALUE! |
| ENSMUSG00000089461.3 | 0 | NA | NA | NA | NA | NA | #VALUE! |
| ENSMUSG00000089464.3 | 0 | NA | NA | NA | NA | NA | #VALUE! |
| ENSMUSG00000089468.4 | 0 | NA | NA | NA | NA | NA | #VALUE! |

| | | | | | | | |
|---|---|---|---|---|---|---|---|
| ENSMUSG00000089473.3 | 0 | NA | NA | NA | NA | NA | #VALUE! |
| ENSMUSG00000089476.4 | 0 | NA | NA | NA | NA | NA | #VALUE! |
| ENSMUSG00000089479.3 | 0 | NA | NA | NA | NA | NA | #VALUE! |
| ENSMUSG00000089484.3 | 0 | NA | NA | NA | NA | NA | #VALUE! |
| ENSMUSG00000089486.3 | 0 | NA | NA | NA | NA | NA | #VALUE! |
| ENSMUSG00000089489.3 | 0 | NA | NA | NA | NA | NA | #VALUE! |
| ENSMUSG00000089493.4 | 0 | NA | NA | NA | NA | NA | #VALUE! |
| ENSMUSG00000089498.3 | 0 | NA | NA | NA | NA | NA | #VALUE! |
| ENSMUSG00000089501.3 | 0 | NA | NA | NA | NA | NA | #VALUE! |
| ENSMUSG00000089504.3 | 0 | NA | NA | NA | NA | NA | #VALUE! |
| ENSMUSG00000089507.3 | 0 | NA | NA | NA | NA | NA | #VALUE! |
| ENSMUSG00000089512.5 | 0 | NA | NA | NA | NA | NA | #VALUE! |
| ENSMUSG00000089513.4 | 0 | NA | NA | NA | NA | NA | #VALUE! |
| ENSMUSG00000089514.3 | 0 | NA | NA | NA | NA | NA | #VALUE! |
| ENSMUSG00000089524.3 | 0 | NA | NA | NA | NA | NA | #VALUE! |
| ENSMUSG00000089528.3 | 0 | NA | NA | NA | NA | NA | #VALUE! |
| ENSMUSG00000089530.4 | 0 | NA | NA | NA | NA | NA | #VALUE! |
| ENSMUSG00000089531.4 | 0 | NA | NA | NA | NA | NA | #VALUE! |
| ENSMUSG00000089534.3 | 0 | NA | NA | NA | NA | NA | #VALUE! |
| ENSMUSG00000089538.3 | 0 | NA | NA | NA | NA | NA | #VALUE! |
| ENSMUSG00000089539.3 | 0 | NA | NA | NA | NA | NA | #VALUE! |
| ENSMUSG00000089541.3 | 0 | NA | NA | NA | NA | NA | #VALUE! |
| ENSMUSG00000089542.3 | 0 | NA | NA | NA | NA | NA | #VALUE! |
| ENSMUSG00000089543.3 | 0 | NA | NA | NA | NA | NA | #VALUE! |
| ENSMUSG00000089548.3 | 0 | NA | NA | NA | NA | NA | #VALUE! |
| ENSMUSG00000089549.3 | 0 | NA | NA | NA | NA | NA | #VALUE! |
| ENSMUSG00000089550.4 | 0 | NA | NA | NA | NA | NA | #VALUE! |
| ENSMUSG00000089551.3 | 0 | NA | NA | NA | NA | NA | #VALUE! |
| ENSMUSG00000089552.2 | 0 | NA | NA | NA | NA | NA | #VALUE! |
| ENSMUSG00000089553.3 | 0 | NA | NA | NA | NA | NA | #VALUE! |
| ENSMUSG00000089555.3 | 0 | NA | NA | NA | NA | NA | #VALUE! |
| ENSMUSG00000089556.3 | 0 | NA | NA | NA | NA | NA | #VALUE! |
| ENSMUSG00000089560.3 | 0 | NA | NA | NA | NA | NA | #VALUE! |
| ENSMUSG00000089561.3 | 0 | NA | NA | NA | NA | NA | #VALUE! |
| ENSMUSG00000089562.4 | 0 | NA | NA | NA | NA | NA | #VALUE! |
| ENSMUSG00000089563.3 | 0 | NA | NA | NA | NA | NA | #VALUE! |
| ENSMUSG00000089570.3 | 0 | NA | NA | NA | NA | NA | #VALUE! |
| ENSMUSG00000089571.3 | 0 | NA | NA | NA | NA | NA | #VALUE! |
| ENSMUSG00000089575.3 | 0 | NA | NA | NA | NA | NA | #VALUE! |
| ENSMUSG00000089580.3 | 0 | NA | NA | NA | NA | NA | #VALUE! |
| ENSMUSG00000089581.3 | 0 | NA | NA | NA | NA | NA | #VALUE! |
| ENSMUSG00000089584.4 | 0 | NA | NA | NA | NA | NA | #VALUE! |
| ENSMUSG00000089585.2 | 0 | NA | NA | NA | NA | NA | #VALUE! |
| ENSMUSG00000089587.4 | 0 | NA | NA | NA | NA | NA | #VALUE! |

| | | | | | | | |
|---|---|---|---|---|---|---|---|
| ENSMUSG00000089589.3 | 0 | NA | NA | NA | NA | NA | #VALUE! |
| ENSMUSG00000089590.3 | 0 | NA | NA | NA | NA | NA | #VALUE! |
| ENSMUSG00000089591.3 | 0 | NA | NA | NA | NA | NA | #VALUE! |
| ENSMUSG00000089593.3 | 0 | NA | NA | NA | NA | NA | #VALUE! |
| ENSMUSG00000089606.3 | 0 | NA | NA | NA | NA | NA | #VALUE! |
| ENSMUSG00000089607.3 | 0 | NA | NA | NA | NA | NA | #VALUE! |
| ENSMUSG00000089608.4 | 0 | NA | NA | NA | NA | NA | #VALUE! |
| ENSMUSG00000089612.3 | 0 | NA | NA | NA | NA | NA | #VALUE! |
| ENSMUSG00000089614.4 | 0 | NA | NA | NA | NA | NA | #VALUE! |
| ENSMUSG00000089615.3 | 0 | NA | NA | NA | NA | NA | #VALUE! |
| ENSMUSG00000089617.3 | 0 | NA | NA | NA | NA | NA | #VALUE! |
| ENSMUSG00000089625.3 | 0 | NA | NA | NA | NA | NA | #VALUE! |
| ENSMUSG00000089627.2 | 0 | NA | NA | NA | NA | NA | #VALUE! |
| ENSMUSG00000089628.2 | 0.168284 | 1.027628 | 4.067126 | 0.252667 | 0.800526 | NA | #VALUE! |
| ENSMUSG00000089630.2 | 0 | NA | NA | NA | NA | NA | #VALUE! |
| ENSMUSG00000089631.3 | 0 | NA | NA | NA | NA | NA | #VALUE! |
| ENSMUSG00000089633.3 | 0 | NA | NA | NA | NA | NA | #VALUE! |
| ENSMUSG00000089634.2 | 0 | NA | NA | NA | NA | NA | #VALUE! |
| ENSMUSG00000089635.2 | 1.470184 | -4.10383 | 3.936698 | -1.04245 | 0.297201 | NA | #VALUE! |
| ENSMUSG00000089636.2 | 0 | NA | NA | NA | NA | NA | #VALUE! |
| ENSMUSG00000089639.2 | 0 | NA | NA | NA | NA | NA | #VALUE! |
| ENSMUSG00000089640.2 | 0 | NA | NA | NA | NA | NA | #VALUE! |
| ENSMUSG00000089643.3 | 0 | NA | NA | NA | NA | NA | #VALUE! |
| ENSMUSG00000089645.4 | 0 | NA | NA | NA | NA | NA | #VALUE! |
| ENSMUSG00000089646.2 | 0.210026 | -1.52494 | 4.058692 | -0.37572 | 0.707123 | NA | #VALUE! |
| ENSMUSG00000089647.3 | 0 | NA | NA | NA | NA | NA | #VALUE! |
| ENSMUSG00000089648.2 | 0 | NA | NA | NA | NA | NA | #VALUE! |
| ENSMUSG00000089651.2 | 0 | NA | NA | NA | NA | NA | #VALUE! |
| ENSMUSG00000089652.2 | 0 | NA | NA | NA | NA | NA | #VALUE! |
| ENSMUSG00000089653.2 | 0 | NA | NA | NA | NA | NA | #VALUE! |
| ENSMUSG00000089654.2 | 0 | NA | NA | NA | NA | NA | #VALUE! |
| ENSMUSG00000089655.2 | 0 | NA | NA | NA | NA | NA | #VALUE! |
| ENSMUSG00000089656.2 | 0 | NA | NA | NA | NA | NA | #VALUE! |
| ENSMUSG00000089657.2 | 0 | NA | NA | NA | NA | NA | #VALUE! |
| ENSMUSG00000089659.3 | 0 | NA | NA | NA | NA | NA | #VALUE! |
| ENSMUSG00000089661.10 | 0 | NA | NA | NA | NA | NA | #VALUE! |
| ENSMUSG00000089662.2 | 0.930881 | -3.44601 | 3.953827 | -0.87156 | 0.383447 | NA | #VALUE! |
| ENSMUSG00000089663.2 | 0 | NA | NA | NA | NA | NA | #VALUE! |
| ENSMUSG00000089664.3 | 0 | NA | NA | NA | NA | NA | #VALUE! |
| ENSMUSG00000089666.2 | 0 | NA | NA | NA | NA | NA | #VALUE! |
| ENSMUSG00000089667.2 | 0 | NA | NA | NA | NA | NA | #VALUE! |
| ENSMUSG00000089669.8 | 0 | NA | NA | NA | NA | NA | #VALUE! |
| ENSMUSG00000089670.2 | 0 | NA | NA | NA | NA | NA | #VALUE! |
| ENSMUSG00000089671.2 | 0 | NA | NA | NA | NA | NA | #VALUE! |

| | | | | | | | |
|---|---|---|---|---|---|---|---|
| ENSMUSG00000089673.2 | 0 | NA | NA | NA | NA | NA | #VALUE! |
| ENSMUSG00000089675.2 | 0 | NA | NA | NA | NA | NA | #VALUE! |
| ENSMUSG00000089679.2 | 0 | NA | NA | NA | NA | NA | #VALUE! |
| ENSMUSG00000089680.2 | 0 | NA | NA | NA | NA | NA | #VALUE! |
| ENSMUSG00000089683.3 | 0 | NA | NA | NA | NA | NA | #VALUE! |
| ENSMUSG00000089684.2 | 0 | NA | NA | NA | NA | NA | #VALUE! |
| ENSMUSG00000089685.8 | 0 | NA | NA | NA | NA | NA | #VALUE! |
| ENSMUSG00000089686.3 | 3.8034 | 4.324703 | 3.533904 | 1.223775 | 0.221037 | NA | #VALUE! |
| ENSMUSG00000089687.3 | 0 | NA | NA | NA | NA | NA | #VALUE! |
| ENSMUSG00000089689.2 | 0 | NA | NA | NA | NA | NA | #VALUE! |
| ENSMUSG00000089690.2 | 0 | NA | NA | NA | NA | NA | #VALUE! |
| ENSMUSG00000089694.3 | 0 | NA | NA | NA | NA | NA | #VALUE! |
| ENSMUSG00000089695.3 | 7.186756 | 3.589179 | 2.808376 | 1.278027 | 0.20124 | NA | #VALUE! |
| ENSMUSG00000089696.4 | 0 | NA | NA | NA | NA | NA | #VALUE! |
| ENSMUSG00000089697.2 | 0 | NA | NA | NA | NA | NA | #VALUE! |
| ENSMUSG00000089698.2 | 0 | NA | NA | NA | NA | NA | #VALUE! |
| ENSMUSG00000089699.2 | 0 | NA | NA | NA | NA | NA | #VALUE! |
| ENSMUSG00000089700.2 | 0 | NA | NA | NA | NA | NA | #VALUE! |
| ENSMUSG00000089701.2 | 0 | NA | NA | NA | NA | NA | #VALUE! |
| ENSMUSG00000089702.2 | 0 | NA | NA | NA | NA | NA | #VALUE! |
| ENSMUSG00000089703.2 | 0 | NA | NA | NA | NA | NA | #VALUE! |
| ENSMUSG00000089705.2 | 0 | NA | NA | NA | NA | NA | #VALUE! |
| ENSMUSG00000089706.8 | 0 | NA | NA | NA | NA | NA | #VALUE! |
| ENSMUSG00000089707.2 | 0 | NA | NA | NA | NA | NA | #VALUE! |
| ENSMUSG00000089708.2 | 0 | NA | NA | NA | NA | NA | #VALUE! |
| ENSMUSG00000089709.2 | 0 | NA | NA | NA | NA | NA | #VALUE! |
| ENSMUSG00000089710.2 | 0 | NA | NA | NA | NA | NA | #VALUE! |
| ENSMUSG00000089711.2 | 0 | NA | NA | NA | NA | NA | #VALUE! |
| ENSMUSG00000089712.3 | 5.72164 | 6.006287 | 3.914617 | 1.534323 | 0.12495 | NA | #VALUE! |
| ENSMUSG00000089713.2 | 0 | NA | NA | NA | NA | NA | #VALUE! |
| ENSMUSG00000089714.2 | 0 | NA | NA | NA | NA | NA | #VALUE! |
| ENSMUSG00000089715.13 | 1.398828 | 3.99633 | 3.938205 | 1.014759 | 0.310221 | NA | #VALUE! |
| ENSMUSG00000089716.8 | 0 | NA | NA | NA | NA | NA | #VALUE! |
| ENSMUSG00000089717.3 | 0 | NA | NA | NA | NA | NA | #VALUE! |
| ENSMUSG00000089718.2 | 0 | NA | NA | NA | NA | NA | #VALUE! |
| ENSMUSG00000089719.2 | 0 | NA | NA | NA | NA | NA | #VALUE! |
| ENSMUSG00000089722.3 | 0 | NA | NA | NA | NA | NA | #VALUE! |
| ENSMUSG00000089723.2 | 0 | NA | NA | NA | NA | NA | #VALUE! |
| ENSMUSG00000089724.2 | 0 | NA | NA | NA | NA | NA | #VALUE! |
| ENSMUSG00000089726.3 | 9.770188 | 6.780302 | 3.103278 | 2.184884 | 0.028897 | NA | #VALUE! |
| ENSMUSG00000089727.3 | 0 | NA | NA | NA | NA | NA | #VALUE! |
| ENSMUSG00000089728.4 | 0 | NA | NA | NA | NA | NA | #VALUE! |
| ENSMUSG00000089729.2 | 0 | NA | NA | NA | NA | NA | #VALUE! |
| ENSMUSG00000089730.3 | 0 | NA | NA | NA | NA | NA | #VALUE! |

| | | | | | | | |
|---|---|---|---|---|---|---|---|
| ENSMUSG00000089731.4 | 0 | NA | NA | NA | NA | NA | #VALUE! |
| ENSMUSG00000089732.6 | 0 | NA | NA | NA | NA | NA | #VALUE! |
| ENSMUSG00000089733.2 | 0 | NA | NA | NA | NA | NA | #VALUE! |
| ENSMUSG00000089735.2 | 0 | NA | NA | NA | NA | NA | #VALUE! |
| ENSMUSG00000089736.3 | 0 | NA | NA | NA | NA | NA | #VALUE! |
| ENSMUSG00000089737.2 | 0 | NA | NA | NA | NA | NA | #VALUE! |
| ENSMUSG00000089739.3 | 0 | NA | NA | NA | NA | NA | #VALUE! |
| ENSMUSG00000089742.2 | 0 | NA | NA | NA | NA | NA | #VALUE! |
| ENSMUSG00000089743.8 | 0 | NA | NA | NA | NA | NA | #VALUE! |
| ENSMUSG00000089744.3 | 0 | NA | NA | NA | NA | NA | #VALUE! |
| ENSMUSG00000089745.2 | 0 | NA | NA | NA | NA | NA | #VALUE! |
| ENSMUSG00000089746.2 | 0 | NA | NA | NA | NA | NA | #VALUE! |
| ENSMUSG00000089748.3 | 0 | NA | NA | NA | NA | NA | #VALUE! |
| ENSMUSG00000089749.2 | 0 | NA | NA | NA | NA | NA | #VALUE! |
| ENSMUSG00000089753.10 | 0 | NA | NA | NA | NA | NA | #VALUE! |
| ENSMUSG00000089754.2 | 0 | NA | NA | NA | NA | NA | #VALUE! |
| ENSMUSG00000089755.2 | 0 | NA | NA | NA | NA | NA | #VALUE! |
| ENSMUSG00000089756.3 | 0 | NA | NA | NA | NA | NA | #VALUE! |
| ENSMUSG00000089759.8 | 0 | NA | NA | NA | NA | NA | #VALUE! |
| ENSMUSG00000089760.2 | 0 | NA | NA | NA | NA | NA | #VALUE! |
| ENSMUSG00000089761.4 | 0 | NA | NA | NA | NA | NA | #VALUE! |
| ENSMUSG00000089762.4 | 2.424635 | 4.780725 | 3.925082 | 1.217993 | 0.223226 | NA | #VALUE! |
| ENSMUSG00000089764.3 | 9.062632 | -1.41753 | 2.595556 | -0.54614 | 0.58497 | NA | #VALUE! |
| ENSMUSG00000089766.2 | 0 | NA | NA | NA | NA | NA | #VALUE! |
| ENSMUSG00000089767.2 | 0 | NA | NA | NA | NA | NA | #VALUE! |
| ENSMUSG00000089768.3 | 0.210026 | -1.52494 | 4.058692 | -0.37572 | 0.707123 | NA | #VALUE! |
| ENSMUSG00000089769.2 | 0 | NA | NA | NA | NA | NA | #VALUE! |
| ENSMUSG00000089771.2 | 0 | NA | NA | NA | NA | NA | #VALUE! |
| ENSMUSG00000089773.8 | 0 | NA | NA | NA | NA | NA | #VALUE! |
| ENSMUSG00000089775.2 | 0 | NA | NA | NA | NA | NA | #VALUE! |
| ENSMUSG00000089776.2 | 0 | NA | NA | NA | NA | NA | #VALUE! |
| ENSMUSG00000089779.2 | 0 | NA | NA | NA | NA | NA | #VALUE! |
| ENSMUSG00000089780.2 | 0 | NA | NA | NA | NA | NA | #VALUE! |
| ENSMUSG00000089781.3 | 0 | NA | NA | NA | NA | NA | #VALUE! |
| ENSMUSG00000089782.2 | 0.804831 | -0.79862 | 3.943253 | -0.20253 | 0.839504 | NA | #VALUE! |
| ENSMUSG00000089783.3 | 0 | NA | NA | NA | NA | NA | #VALUE! |
| ENSMUSG00000089785.3 | 0.913208 | -2.34381 | 3.952271 | -0.59303 | 0.553161 | NA | #VALUE! |
| ENSMUSG00000089787.3 | 0 | NA | NA | NA | NA | NA | #VALUE! |
| ENSMUSG00000089789.2 | 0 | NA | NA | NA | NA | NA | #VALUE! |
| ENSMUSG00000089790.2 | 0 | NA | NA | NA | NA | NA | #VALUE! |
| ENSMUSG00000089791.2 | 0.195789 | -1.05581 | 4.073302 | -0.2592 | 0.795479 | NA | #VALUE! |
| ENSMUSG00000089792.4 | 0 | NA | NA | NA | NA | NA | #VALUE! |
| ENSMUSG00000089793.2 | 0 | NA | NA | NA | NA | NA | #VALUE! |
| ENSMUSG00000089794.2 | 0 | NA | NA | NA | NA | NA | #VALUE! |

| | | | | | | | |
|---|---|---|---|---|---|---|---|
| ENSMUSG00000089795.2 | 0 | NA | NA | NA | NA | NA | #VALUE! |
| ENSMUSG00000089796.2 | 0 | NA | NA | NA | NA | NA | #VALUE! |
| ENSMUSG00000089797.2 | 0 | NA | NA | NA | NA | NA | #VALUE! |
| ENSMUSG00000089798.11 | 0 | NA | NA | NA | NA | NA | #VALUE! |
| ENSMUSG00000089800.2 | 0 | NA | NA | NA | NA | NA | #VALUE! |
| ENSMUSG00000089802.2 | 0 | NA | NA | NA | NA | NA | #VALUE! |
| ENSMUSG00000089803.4 | 0 | NA | NA | NA | NA | NA | #VALUE! |
| ENSMUSG00000089804.3 | 0 | NA | NA | NA | NA | NA | #VALUE! |
| ENSMUSG00000089806.2 | 0 | NA | NA | NA | NA | NA | #VALUE! |
| ENSMUSG00000089807.2 | 0 | NA | NA | NA | NA | NA | #VALUE! |
| ENSMUSG00000089808.4 | 0 | NA | NA | NA | NA | NA | #VALUE! |
| ENSMUSG00000089811.3 | 0 | NA | NA | NA | NA | NA | #VALUE! |
| ENSMUSG00000089812.3 | 0 | NA | NA | NA | NA | NA | #VALUE! |
| ENSMUSG00000089814.2 | 0 | NA | NA | NA | NA | NA | #VALUE! |
| ENSMUSG00000089815.8 | 0 | NA | NA | NA | NA | NA | #VALUE! |
| ENSMUSG00000089816.2 | 0 | NA | NA | NA | NA | NA | #VALUE! |
| ENSMUSG00000089817.2 | 0 | NA | NA | NA | NA | NA | #VALUE! |
| ENSMUSG00000089818.2 | 0 | NA | NA | NA | NA | NA | #VALUE! |
| ENSMUSG00000089819.2 | 0 | NA | NA | NA | NA | NA | #VALUE! |
| ENSMUSG00000089820.2 | 0 | NA | NA | NA | NA | NA | #VALUE! |
| ENSMUSG00000089821.2 | 0 | NA | NA | NA | NA | NA | #VALUE! |
| ENSMUSG00000089822.2 | 0 | NA | NA | NA | NA | NA | #VALUE! |
| ENSMUSG00000089826.2 | 0 | NA | NA | NA | NA | NA | #VALUE! |
| ENSMUSG00000089827.2 | 0 | NA | NA | NA | NA | NA | #VALUE! |
| ENSMUSG00000089828.3 | 0 | NA | NA | NA | NA | NA | #VALUE! |
| ENSMUSG00000089829.3 | 0 | NA | NA | NA | NA | NA | #VALUE! |
| ENSMUSG00000089830.2 | 0 | NA | NA | NA | NA | NA | #VALUE! |
| ENSMUSG00000089831.2 | 0 | NA | NA | NA | NA | NA | #VALUE! |
| ENSMUSG00000089832.9 | 0 | NA | NA | NA | NA | NA | #VALUE! |
| ENSMUSG00000089833.3 | 0 | NA | NA | NA | NA | NA | #VALUE! |
| ENSMUSG00000089834.3 | 0 | NA | NA | NA | NA | NA | #VALUE! |
| ENSMUSG00000089835.2 | 0 | NA | NA | NA | NA | NA | #VALUE! |
| ENSMUSG00000089836.3 | 0 | NA | NA | NA | NA | NA | #VALUE! |
| ENSMUSG00000089837.10 | 0 | NA | NA | NA | NA | NA | #VALUE! |
| ENSMUSG00000089838.2 | 1.039933 | -1.14338 | 3.924473 | -0.29135 | 0.770786 | NA | #VALUE! |
| ENSMUSG00000089839.2 | 0 | NA | NA | NA | NA | NA | #VALUE! |
| ENSMUSG00000089840.2 | 0 | NA | NA | NA | NA | NA | #VALUE! |
| ENSMUSG00000089842.3 | 0 | NA | NA | NA | NA | NA | #VALUE! |
| ENSMUSG00000089844.8 | 0 | NA | NA | NA | NA | NA | #VALUE! |
| ENSMUSG00000089849.2 | 0 | NA | NA | NA | NA | NA | #VALUE! |
| ENSMUSG00000089851.2 | 0 | NA | NA | NA | NA | NA | #VALUE! |
| ENSMUSG00000089854.3 | 0 | NA | NA | NA | NA | NA | #VALUE! |
| ENSMUSG00000089857.10 | 0 | NA | NA | NA | NA | NA | #VALUE! |
| ENSMUSG00000089859.6 | 0 | NA | NA | NA | NA | NA | #VALUE! |

| | | | | | | | |
|---|---|---|---|---|---|---|---|
| ENSMUSG00000089861.2 | 0 | NA | NA | NA | NA | NA | #VALUE! |
| ENSMUSG00000089865.3 | 0 | NA | NA | NA | NA | NA | #VALUE! |
| ENSMUSG00000089866.2 | 0 | NA | NA | NA | NA | NA | #VALUE! |
| ENSMUSG00000089868.2 | 0 | NA | NA | NA | NA | NA | #VALUE! |
| ENSMUSG00000089870.2 | 0 | NA | NA | NA | NA | NA | #VALUE! |
| ENSMUSG00000089871.2 | 0 | NA | NA | NA | NA | NA | #VALUE! |
| ENSMUSG00000089872.11 | 2.685572 | -1.58627 | 3.81654 | -0.41563 | 0.677681 | NA | #VALUE! |
| ENSMUSG00000089873.9 | 3.364093 | -0.35411 | 3.780639 | -0.09366 | 0.925376 | NA | #VALUE! |
| ENSMUSG00000089874.2 | 0 | NA | NA | NA | NA | NA | #VALUE! |
| ENSMUSG00000089875.3 | 2.83293 | 2.939438 | 3.880876 | 0.757416 | 0.448801 | NA | #VALUE! |
| ENSMUSG00000089876.4 | 3.357187 | 5.246639 | 3.920044 | 1.338413 | 0.180762 | NA | #VALUE! |
| ENSMUSG00000089877.2 | 0 | NA | NA | NA | NA | NA | #VALUE! |
| ENSMUSG00000089879.2 | 0 | NA | NA | NA | NA | NA | #VALUE! |
| ENSMUSG00000089881.2 | 0 | NA | NA | NA | NA | NA | #VALUE! |
| ENSMUSG00000089883.2 | 0 | NA | NA | NA | NA | NA | #VALUE! |
| ENSMUSG00000089885.3 | 0 | NA | NA | NA | NA | NA | #VALUE! |
| ENSMUSG00000089886.2 | 0 | NA | NA | NA | NA | NA | #VALUE! |
| ENSMUSG00000089887.3 | 0 | NA | NA | NA | NA | NA | #VALUE! |
| ENSMUSG00000089888.2 | 0 | NA | NA | NA | NA | NA | #VALUE! |
| ENSMUSG00000089889.3 | 4.379581 | -4.71786 | 3.908882 | -1.20696 | 0.227448 | NA | #VALUE! |
| ENSMUSG00000089890.2 | 0 | NA | NA | NA | NA | NA | #VALUE! |
| ENSMUSG00000089891.2 | 0 | NA | NA | NA | NA | NA | #VALUE! |
| ENSMUSG00000089892.4 | 0 | NA | NA | NA | NA | NA | #VALUE! |
| ENSMUSG00000089893.2 | 0 | NA | NA | NA | NA | NA | #VALUE! |
| ENSMUSG00000089894.3 | 0 | NA | NA | NA | NA | NA | #VALUE! |
| ENSMUSG00000089898.3 | 0 | NA | NA | NA | NA | NA | #VALUE! |
| ENSMUSG00000089900.3 | 0 | NA | NA | NA | NA | NA | #VALUE! |
| ENSMUSG00000089901.2 | 0 | NA | NA | NA | NA | NA | #VALUE! |
| ENSMUSG00000089904.2 | 0 | NA | NA | NA | NA | NA | #VALUE! |
| ENSMUSG00000089905.2 | 0 | NA | NA | NA | NA | NA | #VALUE! |
| ENSMUSG00000089906.2 | 0 | NA | NA | NA | NA | NA | #VALUE! |
| ENSMUSG00000089907.3 | 0 | NA | NA | NA | NA | NA | #VALUE! |
| ENSMUSG00000089908.2 | 0 | NA | NA | NA | NA | NA | #VALUE! |
| ENSMUSG00000089909.2 | 0 | NA | NA | NA | NA | NA | #VALUE! |
| ENSMUSG00000089910.2 | 0 | NA | NA | NA | NA | NA | #VALUE! |
| ENSMUSG00000089914.3 | 0 | NA | NA | NA | NA | NA | #VALUE! |
| ENSMUSG00000089916.2 | 0 | NA | NA | NA | NA | NA | #VALUE! |
| ENSMUSG00000089922.3 | 0 | NA | NA | NA | NA | NA | #VALUE! |
| ENSMUSG00000089924.2 | 0 | NA | NA | NA | NA | NA | #VALUE! |
| ENSMUSG00000089926.2 | 0 | NA | NA | NA | NA | NA | #VALUE! |
| ENSMUSG00000089927.2 | 0 | NA | NA | NA | NA | NA | #VALUE! |
| ENSMUSG00000089929.2 | 0 | NA | NA | NA | NA | NA | #VALUE! |
| ENSMUSG00000089931.2 | 0 | NA | NA | NA | NA | NA | #VALUE! |
| ENSMUSG00000089932.2 | 0 | NA | NA | NA | NA | NA | #VALUE! |

| | | | | | | | |
|---|---|---|---|---|---|---|---|
| ENSMUSG00000089933.2 | 0 | NA | NA | NA | NA | NA | #VALUE! |
| ENSMUSG00000089934.2 | 0 | NA | NA | NA | NA | NA | #VALUE! |
| ENSMUSG00000089935.2 | 0 | NA | NA | NA | NA | NA | #VALUE! |
| ENSMUSG00000089936.2 | 0 | NA | NA | NA | NA | NA | #VALUE! |
| ENSMUSG00000089937.2 | 0 | NA | NA | NA | NA | NA | #VALUE! |
| ENSMUSG00000089938.2 | 0.168284 | 1.027628 | 4.067126 | 0.252667 | 0.800526 | NA | #VALUE! |
| ENSMUSG00000089940.2 | 0 | NA | NA | NA | NA | NA | #VALUE! |
| ENSMUSG00000089941.2 | 0 | NA | NA | NA | NA | NA | #VALUE! |
| ENSMUSG00000089942.10 | 0 | NA | NA | NA | NA | NA | #VALUE! |
| ENSMUSG00000089943.2 | 0 | NA | NA | NA | NA | NA | #VALUE! |
| ENSMUSG00000089945.14 | 0 | NA | NA | NA | NA | NA | #VALUE! |
| ENSMUSG00000089947.2 | 0 | NA | NA | NA | NA | NA | #VALUE! |
| ENSMUSG00000089948.2 | 0 | NA | NA | NA | NA | NA | #VALUE! |
| ENSMUSG00000089949.3 | 0 | NA | NA | NA | NA | NA | #VALUE! |
| ENSMUSG00000089950.3 | 0 | NA | NA | NA | NA | NA | #VALUE! |
| ENSMUSG00000089951.9 | 0 | NA | NA | NA | NA | NA | #VALUE! |
| ENSMUSG00000089952.2 | 0 | NA | NA | NA | NA | NA | #VALUE! |
| ENSMUSG00000089953.4 | 0 | NA | NA | NA | NA | NA | #VALUE! |
| ENSMUSG00000089956.2 | 0 | NA | NA | NA | NA | NA | #VALUE! |
| ENSMUSG00000089957.3 | 0 | NA | NA | NA | NA | NA | #VALUE! |
| ENSMUSG00000089958.2 | 0 | NA | NA | NA | NA | NA | #VALUE! |
| ENSMUSG00000089959.2 | 0 | NA | NA | NA | NA | NA | #VALUE! |
| ENSMUSG00000089960.2 | 0 | NA | NA | NA | NA | NA | #VALUE! |
| ENSMUSG00000089961.2 | 0.168284 | 1.027628 | 4.067126 | 0.252667 | 0.800526 | NA | #VALUE! |
| ENSMUSG00000089963.4 | 0 | NA | NA | NA | NA | NA | #VALUE! |
| ENSMUSG00000089964.2 | 0 | NA | NA | NA | NA | NA | #VALUE! |
| ENSMUSG00000089965.2 | 0 | NA | NA | NA | NA | NA | #VALUE! |
| ENSMUSG00000089966.2 | 0 | NA | NA | NA | NA | NA | #VALUE! |
| ENSMUSG00000089967.2 | 0 | NA | NA | NA | NA | NA | #VALUE! |
| ENSMUSG00000089968.2 | 0 | NA | NA | NA | NA | NA | #VALUE! |
| ENSMUSG00000089972.4 | 0.168284 | 1.027628 | 4.067126 | 0.252667 | 0.800526 | NA | #VALUE! |
| ENSMUSG00000089973.4 | 0 | NA | NA | NA | NA | NA | #VALUE! |
| ENSMUSG00000089975.3 | 0 | NA | NA | NA | NA | NA | #VALUE! |
| ENSMUSG00000089976.2 | 0 | NA | NA | NA | NA | NA | #VALUE! |
| ENSMUSG00000089978.2 | 0 | NA | NA | NA | NA | NA | #VALUE! |
| ENSMUSG00000089982.8 | 0 | NA | NA | NA | NA | NA | #VALUE! |
| ENSMUSG00000089983.3 | 0 | NA | NA | NA | NA | NA | #VALUE! |
| ENSMUSG00000089984.8 | 0 | NA | NA | NA | NA | NA | #VALUE! |
| ENSMUSG00000089987.3 | 0 | NA | NA | NA | NA | NA | #VALUE! |
| ENSMUSG00000089988.3 | 0.660235 | 2.855092 | 3.975428 | 0.718185 | 0.472643 | NA | #VALUE! |
| ENSMUSG00000089989.11 | 0 | NA | NA | NA | NA | NA | #VALUE! |
| ENSMUSG00000089990.2 | 0 | NA | NA | NA | NA | NA | #VALUE! |
| ENSMUSG00000089992.4 | 0 | NA | NA | NA | NA | NA | #VALUE! |
| ENSMUSG00000089993.2 | 0.168284 | 1.027628 | 4.067126 | 0.252667 | 0.800526 | NA | #VALUE! |

| | | | | | | | |
|---|---|---|---|---|---|---|---|
| ENSMUSG00000089994.2 | 0 | NA | NA | NA | NA | NA | #VALUE! |
| ENSMUSG00000089995.3 | 0 | NA | NA | NA | NA | NA | #VALUE! |
| ENSMUSG00000089996.7 | 0 | NA | NA | NA | NA | NA | #VALUE! |
| ENSMUSG00000089997.3 | 0 | NA | NA | NA | NA | NA | #VALUE! |
| ENSMUSG00000089998.3 | 1.025807 | 3.556426 | 3.949287 | 0.900523 | 0.367842 | NA | #VALUE! |
| ENSMUSG00000089999.2 | 0.093255 | 0.868093 | 4.072464 | 0.213162 | 0.831201 | NA | #VALUE! |
| ENSMUSG00000090001.2 | 0 | NA | NA | NA | NA | NA | #VALUE! |
| ENSMUSG00000090002.2 | 0.504851 | 2.502843 | 3.993993 | 0.626652 | 0.530887 | NA | #VALUE! |
| ENSMUSG00000090003.2 | 0 | NA | NA | NA | NA | NA | #VALUE! |
| ENSMUSG00000090004.2 | 0 | NA | NA | NA | NA | NA | #VALUE! |
| ENSMUSG00000090005.2 | 0 | NA | NA | NA | NA | NA | #VALUE! |
| ENSMUSG00000090006.2 | 0 | NA | NA | NA | NA | NA | #VALUE! |
| ENSMUSG00000090007.2 | 0.168284 | 1.027628 | 4.067126 | 0.252667 | 0.800526 | NA | #VALUE! |
| ENSMUSG00000090008.2 | 0 | NA | NA | NA | NA | NA | #VALUE! |
| ENSMUSG00000090009.2 | 0 | NA | NA | NA | NA | NA | #VALUE! |
| ENSMUSG00000090012.2 | 0 | NA | NA | NA | NA | NA | #VALUE! |
| ENSMUSG00000090013.2 | 0 | NA | NA | NA | NA | NA | #VALUE! |
| ENSMUSG00000090016.2 | 0 | NA | NA | NA | NA | NA | #VALUE! |
| ENSMUSG00000090017.2 | 0 | NA | NA | NA | NA | NA | #VALUE! |
| ENSMUSG00000090019.6 | 1.224921 | -2.41965 | 3.938723 | -0.61432 | 0.539002 | NA | #VALUE! |
| ENSMUSG00000090021.3 | 1.762102 | -4.34067 | 3.93219 | -1.10388 | 0.269645 | NA | #VALUE! |
| ENSMUSG00000090023.2 | 0 | NA | NA | NA | NA | NA | #VALUE! |
| ENSMUSG00000090024.2 | 0 | NA | NA | NA | NA | NA | #VALUE! |
| ENSMUSG00000090026.2 | 0 | NA | NA | NA | NA | NA | #VALUE! |
| ENSMUSG00000090027.2 | 0 | NA | NA | NA | NA | NA | #VALUE! |
| ENSMUSG00000090030.2 | 0 | NA | NA | NA | NA | NA | #VALUE! |
| ENSMUSG00000090031.3 | 0 | NA | NA | NA | NA | NA | #VALUE! |
| ENSMUSG00000090033.2 | 0 | NA | NA | NA | NA | NA | #VALUE! |
| ENSMUSG00000090034.2 | 0 | NA | NA | NA | NA | NA | #VALUE! |
| ENSMUSG00000090036.2 | 0 | NA | NA | NA | NA | NA | #VALUE! |
| ENSMUSG00000090039.2 | 0 | NA | NA | NA | NA | NA | #VALUE! |
| ENSMUSG00000090040.2 | 0 | NA | NA | NA | NA | NA | #VALUE! |
| ENSMUSG00000090041.2 | 0 | NA | NA | NA | NA | NA | #VALUE! |
| ENSMUSG00000090042.2 | 0 | NA | NA | NA | NA | NA | #VALUE! |
| ENSMUSG00000090043.2 | 0 | NA | NA | NA | NA | NA | #VALUE! |
| ENSMUSG00000090044.2 | 0 | NA | NA | NA | NA | NA | #VALUE! |
| ENSMUSG00000090047.2 | 0 | NA | NA | NA | NA | NA | #VALUE! |
| ENSMUSG00000090048.2 | 0 | NA | NA | NA | NA | NA | #VALUE! |
| ENSMUSG00000090050.2 | 0 | NA | NA | NA | NA | NA | #VALUE! |
| ENSMUSG00000090051.2 | 0 | NA | NA | NA | NA | NA | #VALUE! |
| ENSMUSG00000090053.10 | 0 | NA | NA | NA | NA | NA | #VALUE! |
| ENSMUSG00000090054.2 | 0 | NA | NA | NA | NA | NA | #VALUE! |
| ENSMUSG00000090057.2 | 0 | NA | NA | NA | NA | NA | #VALUE! |
| ENSMUSG00000090059.5 | 0 | NA | NA | NA | NA | NA | #VALUE! |

| | | | | | | | |
|---|---|---|---|---|---|---|---|
| ENSMUSG00000090061.10 | 0 | NA | NA | NA | NA | NA | #VALUE! |
| ENSMUSG00000090062.3 | 0 | NA | NA | NA | NA | NA | #VALUE! |
| ENSMUSG00000090063.7 | 0 | NA | NA | NA | NA | NA | #VALUE! |
| ENSMUSG00000090066.3 | 0 | NA | NA | NA | NA | NA | #VALUE! |
| ENSMUSG00000090067.2 | 0 | NA | NA | NA | NA | NA | #VALUE! |
| ENSMUSG00000090069.3 | 0 | NA | NA | NA | NA | NA | #VALUE! |
| ENSMUSG00000090070.2 | 0 | NA | NA | NA | NA | NA | #VALUE! |
| ENSMUSG00000090071.5 | 0 | NA | NA | NA | NA | NA | #VALUE! |
| ENSMUSG00000090073.2 | 0 | NA | NA | NA | NA | NA | #VALUE! |
| ENSMUSG00000090074.2 | 0 | NA | NA | NA | NA | NA | #VALUE! |
| ENSMUSG00000090075.2 | 0 | NA | NA | NA | NA | NA | #VALUE! |
| ENSMUSG00000090077.10 | 0 | NA | NA | NA | NA | NA | #VALUE! |
| ENSMUSG00000090078.3 | 0 | NA | NA | NA | NA | NA | #VALUE! |
| ENSMUSG00000090079.2 | 0 | NA | NA | NA | NA | NA | #VALUE! |
| ENSMUSG00000090080.3 | 0 | NA | NA | NA | NA | NA | #VALUE! |
| ENSMUSG00000090084.8 | 0 | NA | NA | NA | NA | NA | #VALUE! |
| ENSMUSG00000090086.8 | 9.51203 | 6.743611 | 3.627857 | 1.858842 | NA | NA | #VALUE! |
| ENSMUSG00000090087.2 | 0 | NA | NA | NA | NA | NA | #VALUE! |
| ENSMUSG00000090088.2 | 0 | NA | NA | NA | NA | NA | #VALUE! |
| ENSMUSG00000090089.4 | 0 | NA | NA | NA | NA | NA | #VALUE! |
| ENSMUSG00000090090.2 | 1.177985 | 3.72614 | 3.944615 | 0.944614 | 0.344856 | NA | #VALUE! |
| ENSMUSG00000090093.9 | 5.968332 | 6.072569 | 3.914265 | 1.551394 | 0.120807 | NA | #VALUE! |
| ENSMUSG00000090095.2 | 0 | NA | NA | NA | NA | NA | #VALUE! |
| ENSMUSG00000090097.9 | 0 | NA | NA | NA | NA | NA | #VALUE! |
| ENSMUSG00000090098.8 | 0 | NA | NA | NA | NA | NA | #VALUE! |
| ENSMUSG00000090099.3 | 0 | NA | NA | NA | NA | NA | #VALUE! |
| ENSMUSG00000090100.8 | 1.808964 | 0.433931 | 3.828743 | 0.113335 | 0.909765 | NA | #VALUE! |
| ENSMUSG00000090101.2 | 0 | NA | NA | NA | NA | NA | #VALUE! |
| ENSMUSG00000090102.2 | 0 | NA | NA | NA | NA | NA | #VALUE! |
| ENSMUSG00000090103.2 | 0 | NA | NA | NA | NA | NA | #VALUE! |
| ENSMUSG00000090104.2 | 0 | NA | NA | NA | NA | NA | #VALUE! |
| ENSMUSG00000090105.3 | 0 | NA | NA | NA | NA | NA | #VALUE! |
| ENSMUSG00000090106.2 | 0 | NA | NA | NA | NA | NA | #VALUE! |
| ENSMUSG00000090109.2 | 0 | NA | NA | NA | NA | NA | #VALUE! |
| ENSMUSG00000090111.2 | 0 | NA | NA | NA | NA | NA | #VALUE! |
| ENSMUSG00000090113.8 | 0 | NA | NA | NA | NA | NA | #VALUE! |
| ENSMUSG00000090114.2 | 0.504851 | 2.502843 | 3.993993 | 0.626652 | 0.530887 | NA | #VALUE! |
| ENSMUSG00000090115.8 | 0.210026 | -1.52494 | 4.058692 | -0.37572 | 0.707123 | NA | #VALUE! |
| ENSMUSG00000090116.2 | 0 | NA | NA | NA | NA | NA | #VALUE! |
| ENSMUSG00000090117.8 | 0.093255 | 0.868093 | 4.072464 | 0.213162 | 0.831201 | NA | #VALUE! |
| ENSMUSG00000090118.2 | 0 | NA | NA | NA | NA | NA | #VALUE! |
| ENSMUSG00000090119.2 | 0.432486 | 2.271797 | 4.008753 | 0.566709 | 0.570912 | NA | #VALUE! |
| ENSMUSG00000090120.2 | 0 | NA | NA | NA | NA | NA | #VALUE! |
| ENSMUSG00000090121.11 | 0 | NA | NA | NA | NA | NA | #VALUE! |

| | | | | | | | |
|---|---|---|---|---|---|---|---|
| ENSMUSG00000090122.4 | 0 | NA | NA | NA | NA | NA | #VALUE! |
| ENSMUSG00000090124.8 | 0 | NA | NA | NA | NA | NA | #VALUE! |
| ENSMUSG00000090125.4 | 0 | NA | NA | NA | NA | NA | #VALUE! |
| ENSMUSG00000090126.2 | 0 | NA | NA | NA | NA | NA | #VALUE! |
| ENSMUSG00000090128.2 | 0 | NA | NA | NA | NA | NA | #VALUE! |
| ENSMUSG00000090129.11 | 0 | NA | NA | NA | NA | NA | #VALUE! |
| ENSMUSG00000090131.2 | 0 | NA | NA | NA | NA | NA | #VALUE! |
| ENSMUSG00000090132.3 | 0 | NA | NA | NA | NA | NA | #VALUE! |
| ENSMUSG00000090135.2 | 0 | NA | NA | NA | NA | NA | #VALUE! |
| ENSMUSG00000090137.9 | 0 | NA | NA | NA | NA | NA | #VALUE! |
| ENSMUSG00000090138.3 | 0 | NA | NA | NA | NA | NA | #VALUE! |
| ENSMUSG00000090141.2 | 0 | NA | NA | NA | NA | NA | #VALUE! |
| ENSMUSG00000090142.3 | 0 | NA | NA | NA | NA | NA | #VALUE! |
| ENSMUSG00000090145.8 | 0 | NA | NA | NA | NA | NA | #VALUE! |
| ENSMUSG00000090146.2 | 0 | NA | NA | NA | NA | NA | #VALUE! |
| ENSMUSG00000090147.2 | 0 | NA | NA | NA | NA | NA | #VALUE! |
| ENSMUSG00000090151.2 | 0 | NA | NA | NA | NA | NA | #VALUE! |
| ENSMUSG00000090152.2 | 0 | NA | NA | NA | NA | NA | #VALUE! |
| ENSMUSG00000090157.8 | 0 | NA | NA | NA | NA | NA | #VALUE! |
| ENSMUSG00000090159.2 | 0 | NA | NA | NA | NA | NA | #VALUE! |
| ENSMUSG00000090160.2 | 0 | NA | NA | NA | NA | NA | #VALUE! |
| ENSMUSG00000090162.2 | 0.300802 | -1.80634 | 4.050033 | -0.44601 | 0.655592 | NA | #VALUE! |
| ENSMUSG00000090163.2 | 0 | NA | NA | NA | NA | NA | #VALUE! |
| ENSMUSG00000090164.9 | 0 | NA | NA | NA | NA | NA | #VALUE! |
| ENSMUSG00000090165.9 | 0 | NA | NA | NA | NA | NA | #VALUE! |
| ENSMUSG00000090166.3 | 0 | NA | NA | NA | NA | NA | #VALUE! |
| ENSMUSG00000090168.2 | 0 | NA | NA | NA | NA | NA | #VALUE! |
| ENSMUSG00000090171.2 | 0 | NA | NA | NA | NA | NA | #VALUE! |
| ENSMUSG00000090173.9 | 0 | NA | NA | NA | NA | NA | #VALUE! |
| ENSMUSG00000090174.2 | 0 | NA | NA | NA | NA | NA | #VALUE! |
| ENSMUSG00000090175.2 | 0 | NA | NA | NA | NA | NA | #VALUE! |
| ENSMUSG00000090176.2 | 0 | NA | NA | NA | NA | NA | #VALUE! |
| ENSMUSG00000090179.2 | 0 | NA | NA | NA | NA | NA | #VALUE! |
| ENSMUSG00000090180.2 | 0 | NA | NA | NA | NA | NA | #VALUE! |
| ENSMUSG00000090185.2 | 0 | NA | NA | NA | NA | NA | #VALUE! |
| ENSMUSG00000090186.8 | 0 | NA | NA | NA | NA | NA | #VALUE! |
| ENSMUSG00000090188.2 | 0 | NA | NA | NA | NA | NA | #VALUE! |
| ENSMUSG00000090191.2 | 0 | NA | NA | NA | NA | NA | #VALUE! |
| ENSMUSG00000090192.2 | 0 | NA | NA | NA | NA | NA | #VALUE! |
| ENSMUSG00000090193.2 | 0 | NA | NA | NA | NA | NA | #VALUE! |
| ENSMUSG00000090194.2 | 0 | NA | NA | NA | NA | NA | #VALUE! |
| ENSMUSG00000090197.2 | 0 | NA | NA | NA | NA | NA | #VALUE! |
| ENSMUSG00000090199.2 | 0 | NA | NA | NA | NA | NA | #VALUE! |
| ENSMUSG00000090200.2 | 0 | NA | NA | NA | NA | NA | #VALUE! |

| | | | | | | | |
|---|---|---|---|---|---|---|---|
| ENSMUSG00000090202.2 | 0 | NA | NA | NA | NA | NA | #VALUE! |
| ENSMUSG00000090203.3 | 0 | NA | NA | NA | NA | NA | #VALUE! |
| ENSMUSG00000090206.9 | 0 | NA | NA | NA | NA | NA | #VALUE! |
| ENSMUSG00000090207.3 | 7.458522 | 0.016187 | 3.283318 | 0.00493 | 0.996066 | NA | #VALUE! |
| ENSMUSG00000090208.2 | 0 | NA | NA | NA | NA | NA | #VALUE! |
| ENSMUSG00000090210.8 | 0 | NA | NA | NA | NA | NA | #VALUE! |
| ENSMUSG00000090211.4 | 0 | NA | NA | NA | NA | NA | #VALUE! |
| ENSMUSG00000090212.2 | 0 | NA | NA | NA | NA | NA | #VALUE! |
| ENSMUSG00000090214.9 | 0 | NA | NA | NA | NA | NA | #VALUE! |
| ENSMUSG00000090215.10 | 0.204111 | -1.05581 | 4.073302 | -0.2592 | 0.795479 | NA | #VALUE! |
| ENSMUSG00000090216.2 | 0 | NA | NA | NA | NA | NA | #VALUE! |
| ENSMUSG00000090217.2 | 16.05471 | -21.746 | 3.123812 | -6.96135 | NA | NA | #VALUE! |
| ENSMUSG00000090219.4 | 0 | NA | NA | NA | NA | NA | #VALUE! |
| ENSMUSG00000090220.2 | 0 | NA | NA | NA | NA | NA | #VALUE! |
| ENSMUSG00000090222.3 | 1.370524 | -3.97623 | 3.939448 | -1.00934 | 0.312813 | NA | #VALUE! |
| ENSMUSG00000090223.3 | 0 | NA | NA | NA | NA | NA | #VALUE! |
| ENSMUSG00000090224.2 | 0 | NA | NA | NA | NA | NA | #VALUE! |
| ENSMUSG00000090225.2 | 0 | NA | NA | NA | NA | NA | #VALUE! |
| ENSMUSG00000090226.2 | 0 | NA | NA | NA | NA | NA | #VALUE! |
| ENSMUSG00000090227.2 | 0 | NA | NA | NA | NA | NA | #VALUE! |
| ENSMUSG00000090228.2 | 0 | NA | NA | NA | NA | NA | #VALUE! |
| ENSMUSG00000090230.2 | 0 | NA | NA | NA | NA | NA | #VALUE! |
| ENSMUSG00000090231.11 | 0 | NA | NA | NA | NA | NA | #VALUE! |
| ENSMUSG00000090232.2 | 0 | NA | NA | NA | NA | NA | #VALUE! |
| ENSMUSG00000090235.2 | 0 | NA | NA | NA | NA | NA | #VALUE! |
| ENSMUSG00000090236.4 | 0 | NA | NA | NA | NA | NA | #VALUE! |
| ENSMUSG00000090237.2 | 0 | NA | NA | NA | NA | NA | #VALUE! |
| ENSMUSG00000090242.2 | 0 | NA | NA | NA | NA | NA | #VALUE! |
| ENSMUSG00000090243.2 | 0 | NA | NA | NA | NA | NA | #VALUE! |
| ENSMUSG00000090244.2 | 0 | NA | NA | NA | NA | NA | #VALUE! |
| ENSMUSG00000090246.2 | 0 | NA | NA | NA | NA | NA | #VALUE! |
| ENSMUSG00000090247.8 | 0 | NA | NA | NA | NA | NA | #VALUE! |
| ENSMUSG00000090248.2 | 0 | NA | NA | NA | NA | NA | #VALUE! |
| ENSMUSG00000090249.10 | 0 | NA | NA | NA | NA | NA | #VALUE! |
| ENSMUSG00000090252.2 | 0 | NA | NA | NA | NA | NA | #VALUE! |
| ENSMUSG00000090254.8 | 0 | NA | NA | NA | NA | NA | #VALUE! |
| ENSMUSG00000090255.8 | 0 | NA | NA | NA | NA | NA | #VALUE! |
| ENSMUSG00000090256.2 | 0 | NA | NA | NA | NA | NA | #VALUE! |
| ENSMUSG00000090257.2 | 0 | NA | NA | NA | NA | NA | #VALUE! |
| ENSMUSG00000090259.2 | 0 | NA | NA | NA | NA | NA | #VALUE! |
| ENSMUSG00000090260.2 | 0 | NA | NA | NA | NA | NA | #VALUE! |
| ENSMUSG00000090263.2 | 0 | NA | NA | NA | NA | NA | #VALUE! |
| ENSMUSG00000090264.2 | 0 | NA | NA | NA | NA | NA | #VALUE! |
| ENSMUSG00000090265.2 | 0 | NA | NA | NA | NA | NA | #VALUE! |

| | | | | | | | |
|---|---|---|---|---|---|---|---|
| ENSMUSG00000090268.2 | 0 | NA | NA | NA | NA | NA | #VALUE! |
| ENSMUSG00000090270.2 | 0 | NA | NA | NA | NA | NA | #VALUE! |
| ENSMUSG00000090273.4 | 0 | NA | NA | NA | NA | NA | #VALUE! |
| ENSMUSG00000090277.2 | 0 | NA | NA | NA | NA | NA | #VALUE! |
| ENSMUSG00000090284.3 | 0 | NA | NA | NA | NA | NA | #VALUE! |
| ENSMUSG00000090291.4 | 0 | NA | NA | NA | NA | NA | #VALUE! |
| ENSMUSG00000090292.2 | 0 | NA | NA | NA | NA | NA | #VALUE! |
| ENSMUSG00000090293.2 | 0 | NA | NA | NA | NA | NA | #VALUE! |
| ENSMUSG00000090298.3 | 0 | NA | NA | NA | NA | NA | #VALUE! |
| ENSMUSG00000090300.2 | 0 | NA | NA | NA | NA | NA | #VALUE! |
| ENSMUSG00000090302.2 | 0 | NA | NA | NA | NA | NA | #VALUE! |
| ENSMUSG00000090303.4 | 0 | NA | NA | NA | NA | NA | #VALUE! |
| ENSMUSG00000090304.5 | 0 | NA | NA | NA | NA | NA | #VALUE! |
| ENSMUSG00000090305.3 | 0 | NA | NA | NA | NA | NA | #VALUE! |
| ENSMUSG00000090307.8 | 0 | NA | NA | NA | NA | NA | #VALUE! |
| ENSMUSG00000090308.3 | 0 | NA | NA | NA | NA | NA | #VALUE! |
| ENSMUSG00000090314.3 | 0 | NA | NA | NA | NA | NA | #VALUE! |
| ENSMUSG00000090315.3 | 0 | NA | NA | NA | NA | NA | #VALUE! |
| ENSMUSG00000090317.2 | 0 | NA | NA | NA | NA | NA | #VALUE! |
| ENSMUSG00000090318.2 | 0 | NA | NA | NA | NA | NA | #VALUE! |
| ENSMUSG00000090319.3 | 0 | NA | NA | NA | NA | NA | #VALUE! |
| ENSMUSG00000090322.2 | 0 | NA | NA | NA | NA | NA | #VALUE! |
| ENSMUSG00000090323.3 | 0 | NA | NA | NA | NA | NA | #VALUE! |
| ENSMUSG00000090326.3 | 0 | NA | NA | NA | NA | NA | #VALUE! |
| ENSMUSG00000090327.2 | 0 | NA | NA | NA | NA | NA | #VALUE! |
| ENSMUSG00000090328.2 | 0 | NA | NA | NA | NA | NA | #VALUE! |
| ENSMUSG00000090329.2 | 0 | NA | NA | NA | NA | NA | #VALUE! |
| ENSMUSG00000090331.2 | 0 | NA | NA | NA | NA | NA | #VALUE! |
| ENSMUSG00000090332.2 | 0 | NA | NA | NA | NA | NA | #VALUE! |
| ENSMUSG00000090333.2 | 0 | NA | NA | NA | NA | NA | #VALUE! |
| ENSMUSG00000090334.2 | 0 | NA | NA | NA | NA | NA | #VALUE! |
| ENSMUSG00000090336.6 | 0 | NA | NA | NA | NA | NA | #VALUE! |
| ENSMUSG00000090338.3 | 0 | NA | NA | NA | NA | NA | #VALUE! |
| ENSMUSG00000090341.2 | 0 | NA | NA | NA | NA | NA | #VALUE! |
| ENSMUSG00000090346.3 | 0 | NA | NA | NA | NA | NA | #VALUE! |
| ENSMUSG00000090350.2 | 0 | NA | NA | NA | NA | NA | #VALUE! |
| ENSMUSG00000090351.2 | 0 | NA | NA | NA | NA | NA | #VALUE! |
| ENSMUSG00000090352.4 | 0 | NA | NA | NA | NA | NA | #VALUE! |
| ENSMUSG00000090353.3 | 0 | NA | NA | NA | NA | NA | #VALUE! |
| ENSMUSG00000090355.4 | 0 | NA | NA | NA | NA | NA | #VALUE! |
| ENSMUSG00000090356.3 | 0 | NA | NA | NA | NA | NA | #VALUE! |
| ENSMUSG00000090357.2 | 0.210026 | -1.52494 | 4.058692 | -0.37572 | 0.707123 | NA | #VALUE! |
| ENSMUSG00000090358.2 | 0 | NA | NA | NA | NA | NA | #VALUE! |
| ENSMUSG00000090362.3 | 0 | NA | NA | NA | NA | NA | #VALUE! |

| | | | | | | | |
|---|---|---|---|---|---|---|---|
| ENSMUSG00000090363.9 | 0 | NA | NA | NA | NA | NA | #VALUE! |
| ENSMUSG00000090364.2 | 0 | NA | NA | NA | NA | NA | #VALUE! |
| ENSMUSG00000090365.2 | 0 | NA | NA | NA | NA | NA | #VALUE! |
| ENSMUSG00000090369.4 | 0 | NA | NA | NA | NA | NA | #VALUE! |
| ENSMUSG00000090370.5 | 0 | NA | NA | NA | NA | NA | #VALUE! |
| ENSMUSG00000090374.2 | 0 | NA | NA | NA | NA | NA | #VALUE! |
| ENSMUSG00000090379.2 | 0 | NA | NA | NA | NA | NA | #VALUE! |
| ENSMUSG00000090381.3 | 0 | NA | NA | NA | NA | NA | #VALUE! |
| ENSMUSG00000090383.3 | 0 | NA | NA | NA | NA | NA | #VALUE! |
| ENSMUSG00000090385.2 | 0 | NA | NA | NA | NA | NA | #VALUE! |
| ENSMUSG00000090387.2 | 0 | NA | NA | NA | NA | NA | #VALUE! |
| ENSMUSG00000090389.3 | 0 | NA | NA | NA | NA | NA | #VALUE! |
| ENSMUSG00000090395.2 | 0 | NA | NA | NA | NA | NA | #VALUE! |
| ENSMUSG00000090396.2 | 0 | NA | NA | NA | NA | NA | #VALUE! |
| ENSMUSG00000090397.2 | 0 | NA | NA | NA | NA | NA | #VALUE! |
| ENSMUSG00000090404.8 | 0 | NA | NA | NA | NA | NA | #VALUE! |
| ENSMUSG00000090406.2 | 0 | NA | NA | NA | NA | NA | #VALUE! |
| ENSMUSG00000090408.2 | 0 | NA | NA | NA | NA | NA | #VALUE! |
| ENSMUSG00000090411.3 | 0 | NA | NA | NA | NA | NA | #VALUE! |
| ENSMUSG00000090417.5 | 0 | NA | NA | NA | NA | NA | #VALUE! |
| ENSMUSG00000090427.2 | 0 | NA | NA | NA | NA | NA | #VALUE! |
| ENSMUSG00000090429.4 | 0 | NA | NA | NA | NA | NA | #VALUE! |
| ENSMUSG00000090436.3 | 0 | NA | NA | NA | NA | NA | #VALUE! |
| ENSMUSG00000090439.4 | 0 | NA | NA | NA | NA | NA | #VALUE! |
| ENSMUSG00000090440.3 | 0 | NA | NA | NA | NA | NA | #VALUE! |
| ENSMUSG00000090441.3 | 0 | NA | NA | NA | NA | NA | #VALUE! |
| ENSMUSG00000090448.2 | 0 | NA | NA | NA | NA | NA | #VALUE! |
| ENSMUSG00000090449.2 | 0 | NA | NA | NA | NA | NA | #VALUE! |
| ENSMUSG00000090451.2 | 1.825211 | -1.13053 | 3.842528 | -0.29421 | 0.768594 | NA | #VALUE! |
| ENSMUSG00000090455.3 | 0 | NA | NA | NA | NA | NA | #VALUE! |
| ENSMUSG00000090457.4 | 0 | NA | NA | NA | NA | NA | #VALUE! |
| ENSMUSG00000090458.2 | 0 | NA | NA | NA | NA | NA | #VALUE! |
| ENSMUSG00000090459.2 | 0 | NA | NA | NA | NA | NA | #VALUE! |
| ENSMUSG00000090460.4 | 0.562341 | -0.07781 | 4.012298 | -0.01939 | 0.984528 | NA | #VALUE! |
| ENSMUSG00000090461.3 | 0 | NA | NA | NA | NA | NA | #VALUE! |
| ENSMUSG00000090462.8 | 0 | NA | NA | NA | NA | NA | #VALUE! |
| ENSMUSG00000090464.2 | 0 | NA | NA | NA | NA | NA | #VALUE! |
| ENSMUSG00000090465.2 | 0 | NA | NA | NA | NA | NA | #VALUE! |
| ENSMUSG00000090466.2 | 0 | NA | NA | NA | NA | NA | #VALUE! |
| ENSMUSG00000090467.4 | 0 | NA | NA | NA | NA | NA | #VALUE! |
| ENSMUSG00000090468.4 | 0 | NA | NA | NA | NA | NA | #VALUE! |
| ENSMUSG00000090470.3 | 0 | NA | NA | NA | NA | NA | #VALUE! |
| ENSMUSG00000090471.3 | 0 | NA | NA | NA | NA | NA | #VALUE! |
| ENSMUSG00000090472.9 | 0 | NA | NA | NA | NA | NA | #VALUE! |

| | | | | | | | |
|---|---|---|---|---|---|---|---|
| ENSMUSG00000090474.4 | 0 | NA | NA | NA | NA | NA | #VALUE! |
| ENSMUSG00000090475.2 | 0.195789 | -1.05581 | 4.073302 | -0.2592 | 0.795479 | NA | #VALUE! |
| ENSMUSG00000090476.2 | 0 | NA | NA | NA | NA | NA | #VALUE! |
| ENSMUSG00000090478.2 | 0 | NA | NA | NA | NA | NA | #VALUE! |
| ENSMUSG00000090481.2 | 0 | NA | NA | NA | NA | NA | #VALUE! |
| ENSMUSG00000090482.4 | 0 | NA | NA | NA | NA | NA | #VALUE! |
| ENSMUSG00000090485.3 | 0 | NA | NA | NA | NA | NA | #VALUE! |
| ENSMUSG00000090486.3 | 0 | NA | NA | NA | NA | NA | #VALUE! |
| ENSMUSG00000090487.9 | 0 | NA | NA | NA | NA | NA | #VALUE! |
| ENSMUSG00000090489.4 | 0 | NA | NA | NA | NA | NA | #VALUE! |
| ENSMUSG00000090494.2 | 0 | NA | NA | NA | NA | NA | #VALUE! |
| ENSMUSG00000090497.3 | 0 | NA | NA | NA | NA | NA | #VALUE! |
| ENSMUSG00000090500.2 | 0 | NA | NA | NA | NA | NA | #VALUE! |
| ENSMUSG00000090505.3 | 0 | NA | NA | NA | NA | NA | #VALUE! |
| ENSMUSG00000090509.3 | 0 | NA | NA | NA | NA | NA | #VALUE! |
| ENSMUSG00000090512.9 | 0 | NA | NA | NA | NA | NA | #VALUE! |
| ENSMUSG00000090515.3 | 0 | NA | NA | NA | NA | NA | #VALUE! |
| ENSMUSG00000090516.2 | 9.503029 | -1.16048 | 2.055852 | -0.56447 | 0.572431 | NA | #VALUE! |
| ENSMUSG00000090519.2 | 0 | NA | NA | NA | NA | NA | #VALUE! |
| ENSMUSG00000090527.3 | 0 | NA | NA | NA | NA | NA | #VALUE! |
| ENSMUSG00000090533.3 | 0 | NA | NA | NA | NA | NA | #VALUE! |
| ENSMUSG00000090534.2 | 0 | NA | NA | NA | NA | NA | #VALUE! |
| ENSMUSG00000090535.2 | 0 | NA | NA | NA | NA | NA | #VALUE! |
| ENSMUSG00000090538.3 | 0 | NA | NA | NA | NA | NA | #VALUE! |
| ENSMUSG00000090539.9 | 0 | NA | NA | NA | NA | NA | #VALUE! |
| ENSMUSG00000090544.4 | 0.847396 | -0.59766 | 3.940051 | -0.15169 | 0.879432 | NA | #VALUE! |
| ENSMUSG00000090546.3 | 0 | NA | NA | NA | NA | NA | #VALUE! |
| ENSMUSG00000090547.2 | 0 | NA | NA | NA | NA | NA | #VALUE! |
| ENSMUSG00000090549.3 | 0 | NA | NA | NA | NA | NA | #VALUE! |
| ENSMUSG00000090550.3 | 0 | NA | NA | NA | NA | NA | #VALUE! |
| ENSMUSG00000090552.2 | 0 | NA | NA | NA | NA | NA | #VALUE! |
| ENSMUSG00000090556.6 | 0 | NA | NA | NA | NA | NA | #VALUE! |
| ENSMUSG00000090558.2 | 0 | NA | NA | NA | NA | NA | #VALUE! |
| ENSMUSG00000090559.2 | 0 | NA | NA | NA | NA | NA | #VALUE! |
| ENSMUSG00000090561.3 | 0 | NA | NA | NA | NA | NA | #VALUE! |
| ENSMUSG00000090564.10 | 0 | NA | NA | NA | NA | NA | #VALUE! |
| ENSMUSG00000090565.8 | 0 | NA | NA | NA | NA | NA | #VALUE! |
| ENSMUSG00000090570.2 | 0 | NA | NA | NA | NA | NA | #VALUE! |
| ENSMUSG00000090572.3 | 0 | NA | NA | NA | NA | NA | #VALUE! |
| ENSMUSG00000090574.2 | 0 | NA | NA | NA | NA | NA | #VALUE! |
| ENSMUSG00000090576.3 | 0 | NA | NA | NA | NA | NA | #VALUE! |
| ENSMUSG00000090577.2 | 0 | NA | NA | NA | NA | NA | #VALUE! |
| ENSMUSG00000090578.2 | 0 | NA | NA | NA | NA | NA | #VALUE! |
| ENSMUSG00000090581.10 | 0 | NA | NA | NA | NA | NA | #VALUE! |

| | | | | | | | |
|---|---|---|---|---|---|---|---|
| ENSMUSG00000090582.2 | 0 | NA | NA | NA | NA | NA | #VALUE! |
| ENSMUSG00000090584.2 | 0 | NA | NA | NA | NA | NA | #VALUE! |
| ENSMUSG00000090585.9 | 0 | NA | NA | NA | NA | NA | #VALUE! |
| ENSMUSG00000090588.3 | 0 | NA | NA | NA | NA | NA | #VALUE! |
| ENSMUSG00000090589.2 | 0 | NA | NA | NA | NA | NA | #VALUE! |
| ENSMUSG00000090592.3 | 0 | NA | NA | NA | NA | NA | #VALUE! |
| ENSMUSG00000090594.2 | 0 | NA | NA | NA | NA | NA | #VALUE! |
| ENSMUSG00000090598.2 | 0 | NA | NA | NA | NA | NA | #VALUE! |
| ENSMUSG00000090602.4 | 0 | NA | NA | NA | NA | NA | #VALUE! |
| ENSMUSG00000090603.2 | 0 | NA | NA | NA | NA | NA | #VALUE! |
| ENSMUSG00000090607.2 | 0 | NA | NA | NA | NA | NA | #VALUE! |
| ENSMUSG00000090608.2 | 0 | NA | NA | NA | NA | NA | #VALUE! |
| ENSMUSG00000090610.4 | 1.971109 | -0.87004 | 3.827538 | -0.22731 | 0.820181 | NA | #VALUE! |
| ENSMUSG00000090618.2 | 0 | NA | NA | NA | NA | NA | #VALUE! |
| ENSMUSG00000090619.3 | 0 | NA | NA | NA | NA | NA | #VALUE! |
| ENSMUSG00000090622.2 | 2.758453 | -4.9889 | 3.923018 | -1.2717 | 0.20348 | NA | #VALUE! |
| ENSMUSG00000090623.4 | 0 | NA | NA | NA | NA | NA | #VALUE! |
| ENSMUSG00000090624.2 | 0 | NA | NA | NA | NA | NA | #VALUE! |
| ENSMUSG00000090626.10 | 8.830002 | -6.67135 | 3.243238 | -2.057 | 0.039686 | NA | #VALUE! |
| ENSMUSG00000090627.9 | 0 | NA | NA | NA | NA | NA | #VALUE! |
| ENSMUSG00000090628.2 | 0 | NA | NA | NA | NA | NA | #VALUE! |
| ENSMUSG00000090629.3 | 0 | NA | NA | NA | NA | NA | #VALUE! |
| ENSMUSG00000090631.4 | 0 | NA | NA | NA | NA | NA | #VALUE! |
| ENSMUSG00000090634.2 | 0 | NA | NA | NA | NA | NA | #VALUE! |
| ENSMUSG00000090635.2 | 0 | NA | NA | NA | NA | NA | #VALUE! |
| ENSMUSG00000090636.2 | 0 | NA | NA | NA | NA | NA | #VALUE! |
| ENSMUSG00000090637.3 | 0 | NA | NA | NA | NA | NA | #VALUE! |
| ENSMUSG00000090638.2 | 0 | NA | NA | NA | NA | NA | #VALUE! |
| ENSMUSG00000090641.3 | 7.546886 | 6.406198 | 3.415308 | 1.875731 | 0.060692 | NA | #VALUE! |
| ENSMUSG00000090642.3 | 0 | NA | NA | NA | NA | NA | #VALUE! |
| ENSMUSG00000090643.9 | 0 | NA | NA | NA | NA | NA | #VALUE! |
| ENSMUSG00000090655.3 | 0 | NA | NA | NA | NA | NA | #VALUE! |
| ENSMUSG00000090657.2 | 0 | NA | NA | NA | NA | NA | #VALUE! |
| ENSMUSG00000090658.8 | 0 | NA | NA | NA | NA | NA | #VALUE! |
| ENSMUSG00000090660.2 | 0 | NA | NA | NA | NA | NA | #VALUE! |
| ENSMUSG00000090662.4 | 0 | NA | NA | NA | NA | NA | #VALUE! |
| ENSMUSG00000090665.3 | 0 | NA | NA | NA | NA | NA | #VALUE! |
| ENSMUSG00000090667.3 | 0 | NA | NA | NA | NA | NA | #VALUE! |
| ENSMUSG00000090671.2 | 0 | NA | NA | NA | NA | NA | #VALUE! |
| ENSMUSG00000090674.2 | 0 | NA | NA | NA | NA | NA | #VALUE! |
| ENSMUSG00000090675.3 | 0 | NA | NA | NA | NA | NA | #VALUE! |
| ENSMUSG00000090681.2 | 0 | NA | NA | NA | NA | NA | #VALUE! |
| ENSMUSG00000090682.5 | 0 | NA | NA | NA | NA | NA | #VALUE! |
| ENSMUSG00000090684.2 | 0 | NA | NA | NA | NA | NA | #VALUE! |

| | | | | | | | |
|---|---|---|---|---|---|---|---|
| ENSMUSG00000090685.2 | 0 | NA | NA | NA | NA | NA | #VALUE! |
| ENSMUSG00000090688.3 | 0 | NA | NA | NA | NA | NA | #VALUE! |
| ENSMUSG00000090690.2 | 0 | NA | NA | NA | NA | NA | #VALUE! |
| ENSMUSG00000090691.4 | 0 | NA | NA | NA | NA | NA | #VALUE! |
| ENSMUSG00000090692.2 | 0 | NA | NA | NA | NA | NA | #VALUE! |
| ENSMUSG00000090693.5 | 0 | NA | NA | NA | NA | NA | #VALUE! |
| ENSMUSG00000090698.3 | 3.636953 | 5.361394 | 3.919032 | 1.36804 | 0.171299 | NA | #VALUE! |
| ENSMUSG00000090699.2 | 0 | NA | NA | NA | NA | NA | #VALUE! |
| ENSMUSG00000090700.3 | 0 | NA | NA | NA | NA | NA | #VALUE! |
| ENSMUSG00000090701.2 | 0 | NA | NA | NA | NA | NA | #VALUE! |
| ENSMUSG00000090704.3 | 0 | NA | NA | NA | NA | NA | #VALUE! |
| ENSMUSG00000090706.2 | 0 | NA | NA | NA | NA | NA | #VALUE! |
| ENSMUSG00000090707.9 | 0 | NA | NA | NA | NA | NA | #VALUE! |
| ENSMUSG00000090708.2 | 2.051614 | 4.542018 | 3.92836 | 1.156212 | 0.247594 | NA | #VALUE! |
| ENSMUSG00000090709.3 | 0 | NA | NA | NA | NA | NA | #VALUE! |
| ENSMUSG00000090710.4 | 0 | NA | NA | NA | NA | NA | #VALUE! |
| ENSMUSG00000090713.3 | 0 | NA | NA | NA | NA | NA | #VALUE! |
| ENSMUSG00000090714.11 | 0 | NA | NA | NA | NA | NA | #VALUE! |
| ENSMUSG00000090715.3 | 0 | NA | NA | NA | NA | NA | #VALUE! |
| ENSMUSG00000090716.3 | 0 | NA | NA | NA | NA | NA | #VALUE! |
| ENSMUSG00000090718.2 | 0 | NA | NA | NA | NA | NA | #VALUE! |
| ENSMUSG00000090722.9 | 0 | NA | NA | NA | NA | NA | #VALUE! |
| ENSMUSG00000090724.2 | 0 | NA | NA | NA | NA | NA | #VALUE! |
| ENSMUSG00000090726.5 | 0 | NA | NA | NA | NA | NA | #VALUE! |
| ENSMUSG00000090727.3 | 0 | NA | NA | NA | NA | NA | #VALUE! |
| ENSMUSG00000090728.2 | 0 | NA | NA | NA | NA | NA | #VALUE! |
| ENSMUSG00000090731.2 | 0 | NA | NA | NA | NA | NA | #VALUE! |
| ENSMUSG00000090732.2 | 0 | NA | NA | NA | NA | NA | #VALUE! |
| ENSMUSG00000090736.2 | 0 | NA | NA | NA | NA | NA | #VALUE! |
| ENSMUSG00000090737.2 | 0 | NA | NA | NA | NA | NA | #VALUE! |
| ENSMUSG00000090738.3 | 0 | NA | NA | NA | NA | NA | #VALUE! |
| ENSMUSG00000090740.3 | 0 | NA | NA | NA | NA | NA | #VALUE! |
| ENSMUSG00000090742.2 | 0 | NA | NA | NA | NA | NA | #VALUE! |
| ENSMUSG00000090743.2 | 0 | NA | NA | NA | NA | NA | #VALUE! |
| ENSMUSG00000090744.8 | 0 | NA | NA | NA | NA | NA | #VALUE! |
| ENSMUSG00000090747.4 | 0 | NA | NA | NA | NA | NA | #VALUE! |
| ENSMUSG00000090750.2 | 0 | NA | NA | NA | NA | NA | #VALUE! |
| ENSMUSG00000090751.3 | 0 | NA | NA | NA | NA | NA | #VALUE! |
| ENSMUSG00000090753.2 | 0 | NA | NA | NA | NA | NA | #VALUE! |
| ENSMUSG00000090758.2 | 0 | NA | NA | NA | NA | NA | #VALUE! |
| ENSMUSG00000090761.2 | 0 | NA | NA | NA | NA | NA | #VALUE! |
| ENSMUSG00000090762.3 | 0 | NA | NA | NA | NA | NA | #VALUE! |
| ENSMUSG00000090764.9 | 0 | NA | NA | NA | NA | NA | #VALUE! |
| ENSMUSG00000090765.2 | 0 | NA | NA | NA | NA | NA | #VALUE! |

| | | | | | | | |
|---|---|---|---|---|---|---|---|
| ENSMUSG00000090772.2 | 0 | NA | NA | NA | NA | NA | #VALUE! |
| ENSMUSG00000090773.4 | 0 | NA | NA | NA | NA | NA | #VALUE! |
| ENSMUSG00000090774.3 | 0 | NA | NA | NA | NA | NA | #VALUE! |
| ENSMUSG00000090777.3 | 0 | NA | NA | NA | NA | NA | #VALUE! |
| ENSMUSG00000090778.2 | 0 | NA | NA | NA | NA | NA | #VALUE! |
| ENSMUSG00000090779.2 | 0 | NA | NA | NA | NA | NA | #VALUE! |
| ENSMUSG00000090781.3 | 0 | NA | NA | NA | NA | NA | #VALUE! |
| ENSMUSG00000090782.2 | 0 | NA | NA | NA | NA | NA | #VALUE! |
| ENSMUSG00000090785.2 | 0 | NA | NA | NA | NA | NA | #VALUE! |
| ENSMUSG00000090788.3 | 0 | NA | NA | NA | NA | NA | #VALUE! |
| ENSMUSG00000090790.3 | 0 | NA | NA | NA | NA | NA | #VALUE! |
| ENSMUSG00000090793.3 | 0 | NA | NA | NA | NA | NA | #VALUE! |
| ENSMUSG00000090795.2 | 0 | NA | NA | NA | NA | NA | #VALUE! |
| ENSMUSG00000090796.2 | 0 | NA | NA | NA | NA | NA | #VALUE! |
| ENSMUSG00000090798.2 | 0 | NA | NA | NA | NA | NA | #VALUE! |
| ENSMUSG00000090799.3 | 0 | NA | NA | NA | NA | NA | #VALUE! |
| ENSMUSG00000090801.2 | 0 | NA | NA | NA | NA | NA | #VALUE! |
| ENSMUSG00000090802.3 | 0 | NA | NA | NA | NA | NA | #VALUE! |
| ENSMUSG00000090806.3 | 0 | NA | NA | NA | NA | NA | #VALUE! |
| ENSMUSG00000090808.4 | 0 | NA | NA | NA | NA | NA | #VALUE! |
| ENSMUSG00000090812.9 | 0 | NA | NA | NA | NA | NA | #VALUE! |
| ENSMUSG00000090816.2 | 0 | NA | NA | NA | NA | NA | #VALUE! |
| ENSMUSG00000090817.8 | 0 | NA | NA | NA | NA | NA | #VALUE! |
| ENSMUSG00000090818.2 | 0 | NA | NA | NA | NA | NA | #VALUE! |
| ENSMUSG00000090821.2 | 0 | NA | NA | NA | NA | NA | #VALUE! |
| ENSMUSG00000090824.3 | 0 | NA | NA | NA | NA | NA | #VALUE! |
| ENSMUSG00000090827.9 | 0 | NA | NA | NA | NA | NA | #VALUE! |
| ENSMUSG00000090839.2 | 0 | NA | NA | NA | NA | NA | #VALUE! |
| ENSMUSG00000090840.3 | 0 | NA | NA | NA | NA | NA | #VALUE! |
| ENSMUSG00000090843.3 | 0 | NA | NA | NA | NA | NA | #VALUE! |
| ENSMUSG00000090849.4 | 0 | NA | NA | NA | NA | NA | #VALUE! |
| ENSMUSG00000090853.2 | 0 | NA | NA | NA | NA | NA | #VALUE! |
| ENSMUSG00000090854.10 | 0 | NA | NA | NA | NA | NA | #VALUE! |
| ENSMUSG00000090855.2 | 0 | NA | NA | NA | NA | NA | #VALUE! |
| ENSMUSG00000090860.2 | 0 | NA | NA | NA | NA | NA | #VALUE! |
| ENSMUSG00000090861.2 | 0 | NA | NA | NA | NA | NA | #VALUE! |
| ENSMUSG00000090863.2 | 2.844049 | 0.958724 | 3.79746 | 0.252464 | 0.800682 | NA | #VALUE! |
| ENSMUSG00000090864.12 | 0 | NA | NA | NA | NA | NA | #VALUE! |
| ENSMUSG00000090865.3 | 0 | NA | NA | NA | NA | NA | #VALUE! |
| ENSMUSG00000090868.3 | 0 | NA | NA | NA | NA | NA | #VALUE! |
| ENSMUSG00000090872.9 | 0 | NA | NA | NA | NA | NA | #VALUE! |
| ENSMUSG00000090873.2 | 0 | NA | NA | NA | NA | NA | #VALUE! |
| ENSMUSG00000090874.3 | 0 | NA | NA | NA | NA | NA | #VALUE! |
| ENSMUSG00000090881.4 | 0 | NA | NA | NA | NA | NA | #VALUE! |

| | | | | | | | |
|---|---|---|---|---|---|---|---|
| ENSMUSG00000090882.2 | 0 | NA | NA | NA | NA | NA | #VALUE! |
| ENSMUSG00000090889.2 | 0 | NA | NA | NA | NA | NA | #VALUE! |
| ENSMUSG00000090891.4 | 0 | NA | NA | NA | NA | NA | #VALUE! |
| ENSMUSG00000090892.3 | 0 | NA | NA | NA | NA | NA | #VALUE! |
| ENSMUSG00000090894.5 | 0 | NA | NA | NA | NA | NA | #VALUE! |
| ENSMUSG00000090897.3 | 0 | NA | NA | NA | NA | NA | #VALUE! |
| ENSMUSG00000090900.2 | 0 | NA | NA | NA | NA | NA | #VALUE! |
| ENSMUSG00000090907.2 | 0 | NA | NA | NA | NA | NA | #VALUE! |
| ENSMUSG00000090919.7 | 0 | NA | NA | NA | NA | NA | #VALUE! |
| ENSMUSG00000090927.2 | 0 | NA | NA | NA | NA | NA | #VALUE! |
| ENSMUSG00000090936.3 | 0 | NA | NA | NA | NA | NA | #VALUE! |
| ENSMUSG00000090939.2 | 0 | NA | NA | NA | NA | NA | #VALUE! |
| ENSMUSG00000090941.2 | 0 | NA | NA | NA | NA | NA | #VALUE! |
| ENSMUSG00000090947.2 | 0 | NA | NA | NA | NA | NA | #VALUE! |
| ENSMUSG00000090949.3 | 0 | NA | NA | NA | NA | NA | #VALUE! |
| ENSMUSG00000090951.5 | 0 | NA | NA | NA | NA | NA | #VALUE! |
| ENSMUSG00000090952.3 | 4.251271 | -2.66795 | 3.817154 | -0.69894 | 0.484592 | NA | #VALUE! |
| ENSMUSG00000090955.2 | 0 | NA | NA | NA | NA | NA | #VALUE! |
| ENSMUSG00000090957.3 | 0 | NA | NA | NA | NA | NA | #VALUE! |
| ENSMUSG00000090958.3 | 4.103229 | 5.534434 | 3.917649 | 1.412693 | 0.157746 | NA | #VALUE! |
| ENSMUSG00000090961.3 | 0 | NA | NA | NA | NA | NA | #VALUE! |
| ENSMUSG00000090963.3 | 4.29378 | -4.69869 | 3.909022 | -1.20201 | 0.229359 | NA | #VALUE! |
| ENSMUSG00000090964.2 | 0 | NA | NA | NA | NA | NA | #VALUE! |
| ENSMUSG00000090965.2 | 0 | NA | NA | NA | NA | NA | #VALUE! |
| ENSMUSG00000090966.3 | 0 | NA | NA | NA | NA | NA | #VALUE! |
| ENSMUSG00000090967.4 | 0 | NA | NA | NA | NA | NA | #VALUE! |
| ENSMUSG00000090968.3 | 0 | NA | NA | NA | NA | NA | #VALUE! |
| ENSMUSG00000090975.2 | 0 | NA | NA | NA | NA | NA | #VALUE! |
| ENSMUSG00000090981.7 | 0 | NA | NA | NA | NA | NA | #VALUE! |
| ENSMUSG00000090989.2 | 0 | NA | NA | NA | NA | NA | #VALUE! |
| ENSMUSG00000090990.2 | 0 | NA | NA | NA | NA | NA | #VALUE! |
| ENSMUSG00000090991.2 | 0 | NA | NA | NA | NA | NA | #VALUE! |
| ENSMUSG00000090994.2 | 0 | NA | NA | NA | NA | NA | #VALUE! |
| ENSMUSG00000090996.2 | 0 | NA | NA | NA | NA | NA | #VALUE! |
| ENSMUSG00000091002.3 | 0 | NA | NA | NA | NA | NA | #VALUE! |
| ENSMUSG00000091003.3 | 0 | NA | NA | NA | NA | NA | #VALUE! |
| ENSMUSG00000091004.3 | 0 | NA | NA | NA | NA | NA | #VALUE! |
| ENSMUSG00000091005.2 | 0 | NA | NA | NA | NA | NA | #VALUE! |
| ENSMUSG00000091006.4 | 0 | NA | NA | NA | NA | NA | #VALUE! |
| ENSMUSG00000091007.2 | 0 | NA | NA | NA | NA | NA | #VALUE! |
| ENSMUSG00000091008.2 | 0 | NA | NA | NA | NA | NA | #VALUE! |
| ENSMUSG00000091010.2 | 0 | NA | NA | NA | NA | NA | #VALUE! |
| ENSMUSG00000091011.2 | 0 | NA | NA | NA | NA | NA | #VALUE! |
| ENSMUSG00000091012.3 | 0 | NA | NA | NA | NA | NA | #VALUE! |

| | | | | | | | |
|---|---|---|---|---|---|---|---|
| ENSMUSG00000091013.4 | 0 | NA | NA | NA | NA | NA | #VALUE! |
| ENSMUSG00000091014.4 | 0 | NA | NA | NA | NA | NA | #VALUE! |
| ENSMUSG00000091016.2 | 0 | NA | NA | NA | NA | NA | #VALUE! |
| ENSMUSG00000091017.2 | 0 | NA | NA | NA | NA | NA | #VALUE! |
| ENSMUSG00000091019.2 | 0 | NA | NA | NA | NA | NA | #VALUE! |
| ENSMUSG00000091020.4 | 0 | NA | NA | NA | NA | NA | #VALUE! |
| ENSMUSG00000091021.2 | 2.388181 | 4.757222 | 3.925382 | 1.211913 | 0.225546 | NA | #VALUE! |
| ENSMUSG00000091022.9 | 0 | NA | NA | NA | NA | NA | #VALUE! |
| ENSMUSG00000091027.2 | 0 | NA | NA | NA | NA | NA | #VALUE! |
| ENSMUSG00000091028.2 | 0 | NA | NA | NA | NA | NA | #VALUE! |
| ENSMUSG00000091031.2 | 0 | NA | NA | NA | NA | NA | #VALUE! |
| ENSMUSG00000091034.9 | 0 | NA | NA | NA | NA | NA | #VALUE! |
| ENSMUSG00000091037.2 | 0 | NA | NA | NA | NA | NA | #VALUE! |
| ENSMUSG00000091039.3 | 0 | NA | NA | NA | NA | NA | #VALUE! |
| ENSMUSG00000091040.2 | 0 | NA | NA | NA | NA | NA | #VALUE! |
| ENSMUSG00000091041.2 | 0 | NA | NA | NA | NA | NA | #VALUE! |
| ENSMUSG00000091043.3 | 0 | NA | NA | NA | NA | NA | #VALUE! |
| ENSMUSG00000091045.10 | 0 | NA | NA | NA | NA | NA | #VALUE! |
| ENSMUSG00000091048.2 | 0 | NA | NA | NA | NA | NA | #VALUE! |
| ENSMUSG00000091049.9 | 0 | NA | NA | NA | NA | NA | #VALUE! |
| ENSMUSG00000091050.2 | 0 | NA | NA | NA | NA | NA | #VALUE! |
| ENSMUSG00000091055.3 | 0 | NA | NA | NA | NA | NA | #VALUE! |
| ENSMUSG00000091058.3 | 0 | NA | NA | NA | NA | NA | #VALUE! |
| ENSMUSG00000091059.3 | 0 | NA | NA | NA | NA | NA | #VALUE! |
| ENSMUSG00000091060.3 | 0 | NA | NA | NA | NA | NA | #VALUE! |
| ENSMUSG00000091061.2 | 0 | NA | NA | NA | NA | NA | #VALUE! |
| ENSMUSG00000091071.3 | 0 | NA | NA | NA | NA | NA | #VALUE! |
| ENSMUSG00000091076.3 | 0 | NA | NA | NA | NA | NA | #VALUE! |
| ENSMUSG00000091077.4 | 0 | NA | NA | NA | NA | NA | #VALUE! |
| ENSMUSG00000091078.2 | 0 | NA | NA | NA | NA | NA | #VALUE! |
| ENSMUSG00000091079.2 | 0 | NA | NA | NA | NA | NA | #VALUE! |
| ENSMUSG00000091080.3 | 0 | NA | NA | NA | NA | NA | #VALUE! |
| ENSMUSG00000091083.5 | 0 | NA | NA | NA | NA | NA | #VALUE! |
| ENSMUSG00000091087.3 | 0 | NA | NA | NA | NA | NA | #VALUE! |
| ENSMUSG00000091091.3 | 0 | NA | NA | NA | NA | NA | #VALUE! |
| ENSMUSG00000091096.2 | 0 | NA | NA | NA | NA | NA | #VALUE! |
| ENSMUSG00000091099.2 | 0 | NA | NA | NA | NA | NA | #VALUE! |
| ENSMUSG00000091101.3 | 0 | NA | NA | NA | NA | NA | #VALUE! |
| ENSMUSG00000091102.2 | 0 | NA | NA | NA | NA | NA | #VALUE! |
| ENSMUSG00000091105.2 | 0 | NA | NA | NA | NA | NA | #VALUE! |
| ENSMUSG00000091109.2 | 0 | NA | NA | NA | NA | NA | #VALUE! |
| ENSMUSG00000091110.2 | 0 | NA | NA | NA | NA | NA | #VALUE! |
| ENSMUSG00000091114.2 | 0 | NA | NA | NA | NA | NA | #VALUE! |
| ENSMUSG00000091116.2 | 0 | NA | NA | NA | NA | NA | #VALUE! |

| | | | | | | | |
|---|---|---|---|---|---|---|---|
| ENSMUSG00000091117.2 | 0 | NA | NA | NA | NA | NA | #VALUE! |
| ENSMUSG00000091122.9 | 0 | NA | NA | NA | NA | NA | #VALUE! |
| ENSMUSG00000091123.2 | 0 | NA | NA | NA | NA | NA | #VALUE! |
| ENSMUSG00000091129.4 | 0 | NA | NA | NA | NA | NA | #VALUE! |
| ENSMUSG00000091131.2 | 0 | NA | NA | NA | NA | NA | #VALUE! |
| ENSMUSG00000091132.2 | 0 | NA | NA | NA | NA | NA | #VALUE! |
| ENSMUSG00000091133.2 | 0 | NA | NA | NA | NA | NA | #VALUE! |
| ENSMUSG00000091135.2 | 0 | NA | NA | NA | NA | NA | #VALUE! |
| ENSMUSG00000091140.2 | 0 | NA | NA | NA | NA | NA | #VALUE! |
| ENSMUSG00000091142.2 | 0 | NA | NA | NA | NA | NA | #VALUE! |
| ENSMUSG00000091146.2 | 0 | NA | NA | NA | NA | NA | #VALUE! |
| ENSMUSG00000091147.2 | 0 | NA | NA | NA | NA | NA | #VALUE! |
| ENSMUSG00000091148.8 | 0 | NA | NA | NA | NA | NA | #VALUE! |
| ENSMUSG00000091149.2 | 0 | NA | NA | NA | NA | NA | #VALUE! |
| ENSMUSG00000091151.3 | 0 | NA | NA | NA | NA | NA | #VALUE! |
| ENSMUSG00000091154.3 | 0 | NA | NA | NA | NA | NA | #VALUE! |
| ENSMUSG00000091155.2 | 0 | NA | NA | NA | NA | NA | #VALUE! |
| ENSMUSG00000091157.2 | 0 | NA | NA | NA | NA | NA | #VALUE! |
| ENSMUSG00000091159.2 | 0 | NA | NA | NA | NA | NA | #VALUE! |
| ENSMUSG00000091160.2 | 0 | NA | NA | NA | NA | NA | #VALUE! |
| ENSMUSG00000091163.5 | 0 | NA | NA | NA | NA | NA | #VALUE! |
| ENSMUSG00000091165.2 | 0 | NA | NA | NA | NA | NA | #VALUE! |
| ENSMUSG00000091172.2 | 0 | NA | NA | NA | NA | NA | #VALUE! |
| ENSMUSG00000091174.2 | 0 | NA | NA | NA | NA | NA | #VALUE! |
| ENSMUSG00000091175.3 | 0 | NA | NA | NA | NA | NA | #VALUE! |
| ENSMUSG00000091177.3 | 0 | NA | NA | NA | NA | NA | #VALUE! |
| ENSMUSG00000091183.6 | 0 | NA | NA | NA | NA | NA | #VALUE! |
| ENSMUSG00000091184.2 | 0 | NA | NA | NA | NA | NA | #VALUE! |
| ENSMUSG00000091185.2 | 0 | NA | NA | NA | NA | NA | #VALUE! |
| ENSMUSG00000091189.2 | 0 | NA | NA | NA | NA | NA | #VALUE! |
| ENSMUSG00000091191.4 | 0 | NA | NA | NA | NA | NA | #VALUE! |
| ENSMUSG00000091192.2 | 0 | NA | NA | NA | NA | NA | #VALUE! |
| ENSMUSG00000091193.3 | 0 | NA | NA | NA | NA | NA | #VALUE! |
| ENSMUSG00000091194.2 | 0 | NA | NA | NA | NA | NA | #VALUE! |
| ENSMUSG00000091195.2 | 0 | NA | NA | NA | NA | NA | #VALUE! |
| ENSMUSG00000091198.3 | 0 | NA | NA | NA | NA | NA | #VALUE! |
| ENSMUSG00000091199.11 | 0 | NA | NA | NA | NA | NA | #VALUE! |
| ENSMUSG00000091204.9 | 0 | NA | NA | NA | NA | NA | #VALUE! |
| ENSMUSG00000091205.4 | 0 | NA | NA | NA | NA | NA | #VALUE! |
| ENSMUSG00000091206.3 | 0 | NA | NA | NA | NA | NA | #VALUE! |
| ENSMUSG00000091212.2 | 0 | NA | NA | NA | NA | NA | #VALUE! |
| ENSMUSG00000091215.4 | 0 | NA | NA | NA | NA | NA | #VALUE! |
| ENSMUSG00000091216.3 | 0 | NA | NA | NA | NA | NA | #VALUE! |
| ENSMUSG00000091221.2 | 0 | NA | NA | NA | NA | NA | #VALUE! |

| | | | | | | | |
|---|---|---|---|---|---|---|---|
| ENSMUSG00000091223.3 | 0 | NA | NA | NA | NA | NA | #VALUE! |
| ENSMUSG00000091227.2 | 0 | NA | NA | NA | NA | NA | #VALUE! |
| ENSMUSG00000091228.2 | 0 | NA | NA | NA | NA | NA | #VALUE! |
| ENSMUSG00000091230.2 | 0 | NA | NA | NA | NA | NA | #VALUE! |
| ENSMUSG00000091233.4 | 0 | NA | NA | NA | NA | NA | #VALUE! |
| ENSMUSG00000091237.2 | 0 | NA | NA | NA | NA | NA | #VALUE! |
| ENSMUSG00000091238.2 | 0 | NA | NA | NA | NA | NA | #VALUE! |
| ENSMUSG00000091239.3 | 0 | NA | NA | NA | NA | NA | #VALUE! |
| ENSMUSG00000091243.3 | 0 | NA | NA | NA | NA | NA | #VALUE! |
| ENSMUSG00000091248.2 | 0 | NA | NA | NA | NA | NA | #VALUE! |
| ENSMUSG00000091255.2 | 0 | NA | NA | NA | NA | NA | #VALUE! |
| ENSMUSG00000091259.3 | 0 | NA | NA | NA | NA | NA | #VALUE! |
| ENSMUSG00000091260.2 | 0 | NA | NA | NA | NA | NA | #VALUE! |
| ENSMUSG00000091268.3 | 0 | NA | NA | NA | NA | NA | #VALUE! |
| ENSMUSG00000091269.3 | 0.093255 | 0.868093 | 4.072464 | 0.213162 | 0.831201 | NA | #VALUE! |
| ENSMUSG00000091272.3 | 0 | NA | NA | NA | NA | NA | #VALUE! |
| ENSMUSG00000091273.2 | 0 | NA | NA | NA | NA | NA | #VALUE! |
| ENSMUSG00000091275.2 | 0 | NA | NA | NA | NA | NA | #VALUE! |
| ENSMUSG00000091277.3 | 0 | NA | NA | NA | NA | NA | #VALUE! |
| ENSMUSG00000091280.2 | 0 | NA | NA | NA | NA | NA | #VALUE! |
| ENSMUSG00000091281.2 | 0 | NA | NA | NA | NA | NA | #VALUE! |
| ENSMUSG00000091283.2 | 0 | NA | NA | NA | NA | NA | #VALUE! |
| ENSMUSG00000091285.3 | 0 | NA | NA | NA | NA | NA | #VALUE! |
| ENSMUSG00000091286.2 | 0 | NA | NA | NA | NA | NA | #VALUE! |
| ENSMUSG00000091288.2 | 0 | NA | NA | NA | NA | NA | #VALUE! |
| ENSMUSG00000091289.2 | 0 | NA | NA | NA | NA | NA | #VALUE! |
| ENSMUSG00000091290.2 | 0 | NA | NA | NA | NA | NA | #VALUE! |
| ENSMUSG00000091292.2 | 0 | NA | NA | NA | NA | NA | #VALUE! |
| ENSMUSG00000091293.2 | 0 | NA | NA | NA | NA | NA | #VALUE! |
| ENSMUSG00000091296.3 | 0 | NA | NA | NA | NA | NA | #VALUE! |
| ENSMUSG00000091297.3 | 0 | NA | NA | NA | NA | NA | #VALUE! |
| ENSMUSG00000091304.3 | 0 | NA | NA | NA | NA | NA | #VALUE! |
| ENSMUSG00000091305.2 | 0 | NA | NA | NA | NA | NA | #VALUE! |
| ENSMUSG00000091306.3 | 0 | NA | NA | NA | NA | NA | #VALUE! |
| ENSMUSG00000091307.2 | 0 | NA | NA | NA | NA | NA | #VALUE! |
| ENSMUSG00000091311.4 | 0 | NA | NA | NA | NA | NA | #VALUE! |
| ENSMUSG00000091312.2 | 0 | NA | NA | NA | NA | NA | #VALUE! |
| ENSMUSG00000091313.2 | 0 | NA | NA | NA | NA | NA | #VALUE! |
| ENSMUSG00000091315.2 | 0 | NA | NA | NA | NA | NA | #VALUE! |
| ENSMUSG00000091318.3 | 0 | NA | NA | NA | NA | NA | #VALUE! |
| ENSMUSG00000091325.3 | 0 | NA | NA | NA | NA | NA | #VALUE! |
| ENSMUSG00000091326.2 | 0 | NA | NA | NA | NA | NA | #VALUE! |
| ENSMUSG00000091329.3 | 0 | NA | NA | NA | NA | NA | #VALUE! |
| ENSMUSG00000091336.2 | 0 | NA | NA | NA | NA | NA | #VALUE! |

| | | | | | | | |
|---|---|---|---|---|---|---|---|
| ENSMUSG00000091345.10 | 0 | NA | NA | NA | NA | NA | #VALUE! |
| ENSMUSG00000091347.3 | 0 | NA | NA | NA | NA | NA | #VALUE! |
| ENSMUSG00000091350.3 | 0 | NA | NA | NA | NA | NA | #VALUE! |
| ENSMUSG00000091351.2 | 0 | NA | NA | NA | NA | NA | #VALUE! |
| ENSMUSG00000091355.2 | 0 | NA | NA | NA | NA | NA | #VALUE! |
| ENSMUSG00000091358.3 | 0 | NA | NA | NA | NA | NA | #VALUE! |
| ENSMUSG00000091361.3 | 0 | NA | NA | NA | NA | NA | #VALUE! |
| ENSMUSG00000091363.2 | 0 | NA | NA | NA | NA | NA | #VALUE! |
| ENSMUSG00000091366.2 | 0.336567 | 1.944925 | 4.029936 | 0.482619 | 0.629366 | NA | #VALUE! |
| ENSMUSG00000091367.2 | 0 | NA | NA | NA | NA | NA | #VALUE! |
| ENSMUSG00000091368.2 | 0 | NA | NA | NA | NA | NA | #VALUE! |
| ENSMUSG00000091370.2 | 1.009701 | 3.503687 | 3.950852 | 0.886818 | 0.375177 | NA | #VALUE! |
| ENSMUSG00000091373.5 | 0 | NA | NA | NA | NA | NA | #VALUE! |
| ENSMUSG00000091375.3 | 0 | NA | NA | NA | NA | NA | #VALUE! |
| ENSMUSG00000091376.2 | 0 | NA | NA | NA | NA | NA | #VALUE! |
| ENSMUSG00000091378.2 | 0 | NA | NA | NA | NA | NA | #VALUE! |
| ENSMUSG00000091381.4 | 0 | NA | NA | NA | NA | NA | #VALUE! |
| ENSMUSG00000091382.3 | 0 | NA | NA | NA | NA | NA | #VALUE! |
| ENSMUSG00000091383.3 | 0.391578 | -2.1489 | 4.020883 | -0.53444 | 0.59304 | NA | #VALUE! |
| ENSMUSG00000091386.3 | 0 | NA | NA | NA | NA | NA | #VALUE! |
| ENSMUSG00000091390.2 | 0 | NA | NA | NA | NA | NA | #VALUE! |
| ENSMUSG00000091393.2 | 0 | NA | NA | NA | NA | NA | #VALUE! |
| ENSMUSG00000091396.3 | 0 | NA | NA | NA | NA | NA | #VALUE! |
| ENSMUSG00000091400.2 | 0 | NA | NA | NA | NA | NA | #VALUE! |
| ENSMUSG00000091401.2 | 0 | NA | NA | NA | NA | NA | #VALUE! |
| ENSMUSG00000091402.8 | 0 | NA | NA | NA | NA | NA | #VALUE! |
| ENSMUSG00000091404.2 | 0 | NA | NA | NA | NA | NA | #VALUE! |
| ENSMUSG00000091405.3 | 0 | NA | NA | NA | NA | NA | #VALUE! |
| ENSMUSG00000091407.3 | 0 | NA | NA | NA | NA | NA | #VALUE! |
| ENSMUSG00000091408.4 | 0 | NA | NA | NA | NA | NA | #VALUE! |
| ENSMUSG00000091410.2 | 0 | NA | NA | NA | NA | NA | #VALUE! |
| ENSMUSG00000091411.2 | 0 | NA | NA | NA | NA | NA | #VALUE! |
| ENSMUSG00000091412.2 | 0 | NA | NA | NA | NA | NA | #VALUE! |
| ENSMUSG00000091415.6 | 0 | NA | NA | NA | NA | NA | #VALUE! |
| ENSMUSG00000091416.3 | 0 | NA | NA | NA | NA | NA | #VALUE! |
| ENSMUSG00000091418.3 | 0 | NA | NA | NA | NA | NA | #VALUE! |
| ENSMUSG00000091419.3 | 0 | NA | NA | NA | NA | NA | #VALUE! |
| ENSMUSG00000091421.4 | 0 | NA | NA | NA | NA | NA | #VALUE! |
| ENSMUSG00000091422.9 | 0 | NA | NA | NA | NA | NA | #VALUE! |
| ENSMUSG00000091423.3 | 0 | NA | NA | NA | NA | NA | #VALUE! |
| ENSMUSG00000091425.2 | 0 | NA | NA | NA | NA | NA | #VALUE! |
| ENSMUSG00000091426.2 | 0 | NA | NA | NA | NA | NA | #VALUE! |
| ENSMUSG00000091429.2 | 0 | NA | NA | NA | NA | NA | #VALUE! |
| ENSMUSG00000091430.2 | 0 | NA | NA | NA | NA | NA | #VALUE! |

| | | | | | | | |
|---|---|---|---|---|---|---|---|
| ENSMUSG00000091435.2 | 0 | NA | NA | NA | NA | NA | #VALUE! |
| ENSMUSG00000091438.2 | 0.168284 | 1.027628 | 4.067126 | 0.252667 | 0.800526 | NA | #VALUE! |
| ENSMUSG00000091441.2 | 0 | NA | NA | NA | NA | NA | #VALUE! |
| ENSMUSG00000091442.2 | 0 | NA | NA | NA | NA | NA | #VALUE! |
| ENSMUSG00000091443.2 | 0 | NA | NA | NA | NA | NA | #VALUE! |
| ENSMUSG00000091449.4 | 0.810796 | 1.265012 | 3.958199 | 0.319593 | 0.749277 | NA | #VALUE! |
| ENSMUSG00000091450.3 | 0 | NA | NA | NA | NA | NA | #VALUE! |
| ENSMUSG00000091453.3 | 0 | NA | NA | NA | NA | NA | #VALUE! |
| ENSMUSG00000091455.5 | 0 | NA | NA | NA | NA | NA | #VALUE! |
| ENSMUSG00000091457.2 | 0 | NA | NA | NA | NA | NA | #VALUE! |
| ENSMUSG00000091459.2 | 0 | NA | NA | NA | NA | NA | #VALUE! |
| ENSMUSG00000091462.3 | 0 | NA | NA | NA | NA | NA | #VALUE! |
| ENSMUSG00000091466.2 | 0 | NA | NA | NA | NA | NA | #VALUE! |
| ENSMUSG00000091468.4 | 0 | NA | NA | NA | NA | NA | #VALUE! |
| ENSMUSG00000091470.2 | 0 | NA | NA | NA | NA | NA | #VALUE! |
| ENSMUSG00000091471.2 | 0 | NA | NA | NA | NA | NA | #VALUE! |
| ENSMUSG00000091472.3 | 0 | NA | NA | NA | NA | NA | #VALUE! |
| ENSMUSG00000091475.5 | 0 | NA | NA | NA | NA | NA | #VALUE! |
| ENSMUSG00000091476.9 | 0 | NA | NA | NA | NA | NA | #VALUE! |
| ENSMUSG00000091477.2 | 0 | NA | NA | NA | NA | NA | #VALUE! |
| ENSMUSG00000091478.3 | 7.303383 | -3.56427 | 3.290873 | -1.08308 | 0.278773 | NA | #VALUE! |
| ENSMUSG00000091479.2 | 0 | NA | NA | NA | NA | NA | #VALUE! |
| ENSMUSG00000091481.2 | 0 | NA | NA | NA | NA | NA | #VALUE! |
| ENSMUSG00000091483.2 | 0 | NA | NA | NA | NA | NA | #VALUE! |
| ENSMUSG00000091486.2 | 0 | NA | NA | NA | NA | NA | #VALUE! |
| ENSMUSG00000091487.2 | 0 | NA | NA | NA | NA | NA | #VALUE! |
| ENSMUSG00000091490.4 | 0 | NA | NA | NA | NA | NA | #VALUE! |
| ENSMUSG00000091491.3 | 0 | NA | NA | NA | NA | NA | #VALUE! |
| ENSMUSG00000091492.2 | 1.044695 | -1.43417 | 3.925359 | -0.36536 | 0.714843 | NA | #VALUE! |
| ENSMUSG00000091494.9 | 0 | NA | NA | NA | NA | NA | #VALUE! |
| ENSMUSG00000091496.2 | 0 | NA | NA | NA | NA | NA | #VALUE! |
| ENSMUSG00000091497.2 | 0 | NA | NA | NA | NA | NA | #VALUE! |
| ENSMUSG00000091498.5 | 0.168284 | 1.027628 | 4.067126 | 0.252667 | 0.800526 | NA | #VALUE! |
| ENSMUSG00000091504.3 | 0 | NA | NA | NA | NA | NA | #VALUE! |
| ENSMUSG00000091507.2 | 0 | NA | NA | NA | NA | NA | #VALUE! |
| ENSMUSG00000091510.5 | 0 | NA | NA | NA | NA | NA | #VALUE! |
| ENSMUSG00000091511.4 | 0 | NA | NA | NA | NA | NA | #VALUE! |
| ENSMUSG00000091514.3 | 0 | NA | NA | NA | NA | NA | #VALUE! |
| ENSMUSG00000091515.6 | 0 | NA | NA | NA | NA | NA | #VALUE! |
| ENSMUSG00000091519.2 | 0 | NA | NA | NA | NA | NA | #VALUE! |
| ENSMUSG00000091523.2 | 0 | NA | NA | NA | NA | NA | #VALUE! |
| ENSMUSG00000091526.3 | 0 | NA | NA | NA | NA | NA | #VALUE! |
| ENSMUSG00000091528.10 | 0 | NA | NA | NA | NA | NA | #VALUE! |
| ENSMUSG00000091530.3 | 0 | NA | NA | NA | NA | NA | #VALUE! |

| | | | | | | | |
|---|---|---|---|---|---|---|---|
| ENSMUSG00000091531.3 | 0 | NA | NA | NA | NA | NA | #VALUE! |
| ENSMUSG00000091534.3 | 0 | NA | NA | NA | NA | NA | #VALUE! |
| ENSMUSG00000091539.3 | 0 | NA | NA | NA | NA | NA | #VALUE! |
| ENSMUSG00000091541.3 | 0 | NA | NA | NA | NA | NA | #VALUE! |
| ENSMUSG00000091542.2 | 0 | NA | NA | NA | NA | NA | #VALUE! |
| ENSMUSG00000091548.3 | 0 | NA | NA | NA | NA | NA | #VALUE! |
| ENSMUSG00000091549.2 | 3.804228 | -4.29217 | 3.675908 | -1.16765 | 0.242949 | NA | #VALUE! |
| ENSMUSG00000091550.2 | 0 | NA | NA | NA | NA | NA | #VALUE! |
| ENSMUSG00000091552.3 | 0 | NA | NA | NA | NA | NA | #VALUE! |
| ENSMUSG00000091553.2 | 0 | NA | NA | NA | NA | NA | #VALUE! |
| ENSMUSG00000091556.9 | 0 | NA | NA | NA | NA | NA | #VALUE! |
| ENSMUSG00000091557.3 | 0 | NA | NA | NA | NA | NA | #VALUE! |
| ENSMUSG00000091558.2 | 0 | NA | NA | NA | NA | NA | #VALUE! |
| ENSMUSG00000091561.4 | 0 | NA | NA | NA | NA | NA | #VALUE! |
| ENSMUSG00000091563.9 | 0 | NA | NA | NA | NA | NA | #VALUE! |
| ENSMUSG00000091568.9 | 0 | NA | NA | NA | NA | NA | #VALUE! |
| ENSMUSG00000091569.2 | 0 | NA | NA | NA | NA | NA | #VALUE! |
| ENSMUSG00000091570.2 | 0 | NA | NA | NA | NA | NA | #VALUE! |
| ENSMUSG00000091572.10 | 0 | NA | NA | NA | NA | NA | #VALUE! |
| ENSMUSG00000091573.2 | 0 | NA | NA | NA | NA | NA | #VALUE! |
| ENSMUSG00000091575.3 | 0 | NA | NA | NA | NA | NA | #VALUE! |
| ENSMUSG00000091577.3 | 0 | NA | NA | NA | NA | NA | #VALUE! |
| ENSMUSG00000091583.2 | 0 | NA | NA | NA | NA | NA | #VALUE! |
| ENSMUSG00000091584.2 | 0 | NA | NA | NA | NA | NA | #VALUE! |
| ENSMUSG00000091585.3 | 0 | NA | NA | NA | NA | NA | #VALUE! |
| ENSMUSG00000091587.2 | 0 | NA | NA | NA | NA | NA | #VALUE! |
| ENSMUSG00000091594.3 | 0 | NA | NA | NA | NA | NA | #VALUE! |
| ENSMUSG00000091595.2 | 0 | NA | NA | NA | NA | NA | #VALUE! |
| ENSMUSG00000091601.6 | 0 | NA | NA | NA | NA | NA | #VALUE! |
| ENSMUSG00000091604.3 | 0 | NA | NA | NA | NA | NA | #VALUE! |
| ENSMUSG00000091605.3 | 0 | NA | NA | NA | NA | NA | #VALUE! |
| ENSMUSG00000091606.3 | 0 | NA | NA | NA | NA | NA | #VALUE! |
| ENSMUSG00000091613.2 | 0 | NA | NA | NA | NA | NA | #VALUE! |
| ENSMUSG00000091614.3 | 0 | NA | NA | NA | NA | NA | #VALUE! |
| ENSMUSG00000091615.3 | 0 | NA | NA | NA | NA | NA | #VALUE! |
| ENSMUSG00000091617.9 | 0 | NA | NA | NA | NA | NA | #VALUE! |
| ENSMUSG00000091618.4 | 0 | NA | NA | NA | NA | NA | #VALUE! |
| ENSMUSG00000091620.4 | 0 | NA | NA | NA | NA | NA | #VALUE! |
| ENSMUSG00000091623.3 | 1.836997 | -4.39907 | 3.931186 | -1.11902 | 0.263133 | NA | #VALUE! |
| ENSMUSG00000091624.3 | 0 | NA | NA | NA | NA | NA | #VALUE! |
| ENSMUSG00000091627.2 | 0 | NA | NA | NA | NA | NA | #VALUE! |
| ENSMUSG00000091631.3 | 0 | NA | NA | NA | NA | NA | #VALUE! |
| ENSMUSG00000091635.3 | 0 | NA | NA | NA | NA | NA | #VALUE! |
| ENSMUSG00000091636.3 | 0 | NA | NA | NA | NA | NA | #VALUE! |

| | | | | | | | |
|---|---|---|---|---|---|---|---|
| ENSMUSG00000091638.3 | 0 | NA | NA | NA | NA | NA | #VALUE! |
| ENSMUSG00000091639.3 | 0 | NA | NA | NA | NA | NA | #VALUE! |
| ENSMUSG00000091648.3 | 3.967378 | -5.52066 | 3.918019 | -1.40904 | 0.158822 | NA | #VALUE! |
| ENSMUSG00000091649.4 | 0.18651 | 1.401769 | 4.056354 | 0.345574 | 0.729663 | NA | #VALUE! |
| ENSMUSG00000091650.3 | 0 | NA | NA | NA | NA | NA | #VALUE! |
| ENSMUSG00000091651.4 | 0 | NA | NA | NA | NA | NA | #VALUE! |
| ENSMUSG00000091652.3 | 0 | NA | NA | NA | NA | NA | #VALUE! |
| ENSMUSG00000091655.4 | 0 | NA | NA | NA | NA | NA | #VALUE! |
| ENSMUSG00000091656.3 | 0 | NA | NA | NA | NA | NA | #VALUE! |
| ENSMUSG00000091657.2 | 0 | NA | NA | NA | NA | NA | #VALUE! |
| ENSMUSG00000091659.2 | 0 | NA | NA | NA | NA | NA | #VALUE! |
| ENSMUSG00000091661.3 | 0 | NA | NA | NA | NA | NA | #VALUE! |
| ENSMUSG00000091662.10 | 0 | NA | NA | NA | NA | NA | #VALUE! |
| ENSMUSG00000091665.2 | 0 | NA | NA | NA | NA | NA | #VALUE! |
| ENSMUSG00000091668.4 | 0 | NA | NA | NA | NA | NA | #VALUE! |
| ENSMUSG00000091669.2 | 0 | NA | NA | NA | NA | NA | #VALUE! |
| ENSMUSG00000091670.4 | 0 | NA | NA | NA | NA | NA | #VALUE! |
| ENSMUSG00000091671.2 | 0 | NA | NA | NA | NA | NA | #VALUE! |
| ENSMUSG00000091672.3 | 0 | NA | NA | NA | NA | NA | #VALUE! |
| ENSMUSG00000091676.2 | 0 | NA | NA | NA | NA | NA | #VALUE! |
| ENSMUSG00000091679.10 | 0 | NA | NA | NA | NA | NA | #VALUE! |
| ENSMUSG00000091680.4 | 0 | NA | NA | NA | NA | NA | #VALUE! |
| ENSMUSG00000091682.3 | 0 | NA | NA | NA | NA | NA | #VALUE! |
| ENSMUSG00000091685.7 | 0 | NA | NA | NA | NA | NA | #VALUE! |
| ENSMUSG00000091686.2 | 0 | NA | NA | NA | NA | NA | #VALUE! |
| ENSMUSG00000091692.3 | 0 | NA | NA | NA | NA | NA | #VALUE! |
| ENSMUSG00000091694.10 | 0 | NA | NA | NA | NA | NA | #VALUE! |
| ENSMUSG00000091695.3 | 0.168284 | 1.027628 | 4.067126 | 0.252667 | 0.800526 | NA | #VALUE! |
| ENSMUSG00000091697.3 | 0.204111 | -1.05581 | 4.073302 | -0.2592 | 0.795479 | NA | #VALUE! |
| ENSMUSG00000091698.9 | 0 | NA | NA | NA | NA | NA | #VALUE! |
| ENSMUSG00000091700.8 | 0 | NA | NA | NA | NA | NA | #VALUE! |
| ENSMUSG00000091703.3 | 0 | NA | NA | NA | NA | NA | #VALUE! |
| ENSMUSG00000091705.9 | 0.840105 | -3.30885 | 3.958472 | -0.83589 | 0.403216 | NA | #VALUE! |
| ENSMUSG00000091707.2 | 0 | NA | NA | NA | NA | NA | #VALUE! |
| ENSMUSG00000091709.2 | 0 | NA | NA | NA | NA | NA | #VALUE! |
| ENSMUSG00000091712.3 | 0 | NA | NA | NA | NA | NA | #VALUE! |
| ENSMUSG00000091713.2 | 0 | NA | NA | NA | NA | NA | #VALUE! |
| ENSMUSG00000091718.2 | 0 | NA | NA | NA | NA | NA | #VALUE! |
| ENSMUSG00000091722.3 | 0 | NA | NA | NA | NA | NA | #VALUE! |
| ENSMUSG00000091723.3 | 0 | NA | NA | NA | NA | NA | #VALUE! |
| ENSMUSG00000091725.9 | 0 | NA | NA | NA | NA | NA | #VALUE! |
| ENSMUSG00000091730.2 | 0 | NA | NA | NA | NA | NA | #VALUE! |
| ENSMUSG00000091731.2 | 0 | NA | NA | NA | NA | NA | #VALUE! |
| ENSMUSG00000091732.2 | 0 | NA | NA | NA | NA | NA | #VALUE! |

| | | | | | | | |
|---|---|---|---|---|---|---|---|
| ENSMUSG00000091733.2 | 0 | NA | NA | NA | NA | NA | #VALUE! |
| ENSMUSG00000091734.3 | 0 | NA | NA | NA | NA | NA | #VALUE! |
| ENSMUSG00000091735.3 | 0 | NA | NA | NA | NA | NA | #VALUE! |
| ENSMUSG00000091736.4 | 0 | NA | NA | NA | NA | NA | #VALUE! |
| ENSMUSG00000091740.4 | 0 | NA | NA | NA | NA | NA | #VALUE! |
| ENSMUSG00000091741.2 | 0 | NA | NA | NA | NA | NA | #VALUE! |
| ENSMUSG00000091742.3 | 0 | NA | NA | NA | NA | NA | #VALUE! |
| ENSMUSG00000091743.3 | 0 | NA | NA | NA | NA | NA | #VALUE! |
| ENSMUSG00000091745.2 | 0 | NA | NA | NA | NA | NA | #VALUE! |
| ENSMUSG00000091749.8 | 0 | NA | NA | NA | NA | NA | #VALUE! |
| ENSMUSG00000091752.3 | 0 | NA | NA | NA | NA | NA | #VALUE! |
| ENSMUSG00000091754.9 | 0 | NA | NA | NA | NA | NA | #VALUE! |
| ENSMUSG00000091756.2 | 0 | NA | NA | NA | NA | NA | #VALUE! |
| ENSMUSG00000091757.2 | 0 | NA | NA | NA | NA | NA | #VALUE! |
| ENSMUSG00000091762.2 | 0 | NA | NA | NA | NA | NA | #VALUE! |
| ENSMUSG00000091768.2 | 0 | NA | NA | NA | NA | NA | #VALUE! |
| ENSMUSG00000091771.2 | 0 | NA | NA | NA | NA | NA | #VALUE! |
| ENSMUSG00000091775.3 | 0 | NA | NA | NA | NA | NA | #VALUE! |
| ENSMUSG00000091776.2 | 0 | NA | NA | NA | NA | NA | #VALUE! |
| ENSMUSG00000091779.9 | 0 | NA | NA | NA | NA | NA | #VALUE! |
| ENSMUSG00000091784.3 | 0 | NA | NA | NA | NA | NA | #VALUE! |
| ENSMUSG00000091785.3 | 0 | NA | NA | NA | NA | NA | #VALUE! |
| ENSMUSG00000091786.2 | 0 | NA | NA | NA | NA | NA | #VALUE! |
| ENSMUSG00000091788.4 | 0 | NA | NA | NA | NA | NA | #VALUE! |
| ENSMUSG00000091792.2 | 0 | NA | NA | NA | NA | NA | #VALUE! |
| ENSMUSG00000091794.4 | 0 | NA | NA | NA | NA | NA | #VALUE! |
| ENSMUSG00000091800.3 | 0 | NA | NA | NA | NA | NA | #VALUE! |
| ENSMUSG00000091803.8 | 8.234717 | 1.722593 | 2.63374 | 0.654048 | 0.513081 | NA | #VALUE! |
| ENSMUSG00000091805.3 | 0 | NA | NA | NA | NA | NA | #VALUE! |
| ENSMUSG00000091806.2 | 0 | NA | NA | NA | NA | NA | #VALUE! |
| ENSMUSG00000091808.2 | 0 | NA | NA | NA | NA | NA | #VALUE! |
| ENSMUSG00000091809.5 | 0 | NA | NA | NA | NA | NA | #VALUE! |
| ENSMUSG00000091812.2 | 0 | NA | NA | NA | NA | NA | #VALUE! |
| ENSMUSG00000091813.5 | 4.569505 | 5.688921 | 3.916546 | 1.452535 | 0.146353 | NA | #VALUE! |
| ENSMUSG00000091814.2 | 0 | NA | NA | NA | NA | NA | #VALUE! |
| ENSMUSG00000091815.2 | 0 | NA | NA | NA | NA | NA | #VALUE! |
| ENSMUSG00000091816.2 | 0 | NA | NA | NA | NA | NA | #VALUE! |
| ENSMUSG00000091817.2 | 0 | NA | NA | NA | NA | NA | #VALUE! |
| ENSMUSG00000091818.4 | 0.093255 | 0.868093 | 4.072464 | 0.213162 | 0.831201 | NA | #VALUE! |
| ENSMUSG00000091821.2 | 0 | NA | NA | NA | NA | NA | #VALUE! |
| ENSMUSG00000091825.2 | 0 | NA | NA | NA | NA | NA | #VALUE! |
| ENSMUSG00000091827.3 | 0 | NA | NA | NA | NA | NA | #VALUE! |
| ENSMUSG00000091831.7 | 0.645691 | 0.15118 | 3.997031 | 0.037823 | 0.969829 | NA | #VALUE! |
| ENSMUSG00000091832.2 | 0 | NA | NA | NA | NA | NA | #VALUE! |

| | | | | | | | |
|---|---|---|---|---|---|---|---|
| ENSMUSG00000091834.3 | 0 | NA | NA | NA | NA | NA | #VALUE! |
| ENSMUSG00000091842.3 | 2.235434 | 0.032433 | 3.522612 | 0.009207 | 0.992654 | NA | #VALUE! |
| ENSMUSG00000091844.3 | 0 | NA | NA | NA | NA | NA | #VALUE! |
| ENSMUSG00000091845.2 | 1.969254 | -3.54139 | 3.922032 | -0.90295 | 0.366554 | NA | #VALUE! |
| ENSMUSG00000091847.2 | 0 | NA | NA | NA | NA | NA | #VALUE! |
| ENSMUSG00000091849.8 | 0 | NA | NA | NA | NA | NA | #VALUE! |
| ENSMUSG00000091859.3 | 0 | NA | NA | NA | NA | NA | #VALUE! |
| ENSMUSG00000091863.3 | 0 | NA | NA | NA | NA | NA | #VALUE! |
| ENSMUSG00000091864.2 | 0.093255 | 0.868093 | 4.072464 | 0.213162 | 0.831201 | NA | #VALUE! |
| ENSMUSG00000091866.2 | 0 | NA | NA | NA | NA | NA | #VALUE! |
| ENSMUSG00000091867.3 | 4.613815 | 4.618433 | 3.734019 | 1.236853 | 0.216142 | NA | #VALUE! |
| ENSMUSG00000091873.3 | 0 | NA | NA | NA | NA | NA | #VALUE! |
| ENSMUSG00000091874.3 | 0 | NA | NA | NA | NA | NA | #VALUE! |
| ENSMUSG00000091876.2 | 0 | NA | NA | NA | NA | NA | #VALUE! |
| ENSMUSG00000091879.3 | 0.105013 | -0.97324 | 4.076293 | -0.23876 | 0.811295 | NA | #VALUE! |
| ENSMUSG00000091881.2 | 0 | NA | NA | NA | NA | NA | #VALUE! |
| ENSMUSG00000091882.9 | 0 | NA | NA | NA | NA | NA | #VALUE! |
| ENSMUSG00000091887.2 | 0 | NA | NA | NA | NA | NA | #VALUE! |
| ENSMUSG00000091888.4 | 0 | NA | NA | NA | NA | NA | #VALUE! |
| ENSMUSG00000091890.3 | 0 | NA | NA | NA | NA | NA | #VALUE! |
| ENSMUSG00000091894.2 | 0 | NA | NA | NA | NA | NA | #VALUE! |
| ENSMUSG00000091897.2 | 0 | NA | NA | NA | NA | NA | #VALUE! |
| ENSMUSG00000091898.9 | 0 | NA | NA | NA | NA | NA | #VALUE! |
| ENSMUSG00000091900.4 | 0 | NA | NA | NA | NA | NA | #VALUE! |
| ENSMUSG00000091902.2 | 0 | NA | NA | NA | NA | NA | #VALUE! |
| ENSMUSG00000091903.9 | 0 | NA | NA | NA | NA | NA | #VALUE! |
| ENSMUSG00000091905.4 | 0 | NA | NA | NA | NA | NA | #VALUE! |
| ENSMUSG00000091906.3 | 0 | NA | NA | NA | NA | NA | #VALUE! |
| ENSMUSG00000091908.8 | 0 | NA | NA | NA | NA | NA | #VALUE! |
| ENSMUSG00000091911.2 | 0 | NA | NA | NA | NA | NA | #VALUE! |
| ENSMUSG00000091922.2 | 0 | NA | NA | NA | NA | NA | #VALUE! |
| ENSMUSG00000091923.9 | 0 | NA | NA | NA | NA | NA | #VALUE! |
| ENSMUSG00000091924.3 | 0 | NA | NA | NA | NA | NA | #VALUE! |
| ENSMUSG00000091926.4 | 0 | NA | NA | NA | NA | NA | #VALUE! |
| ENSMUSG00000091929.3 | 0 | NA | NA | NA | NA | NA | #VALUE! |
| ENSMUSG00000091930.3 | 0 | NA | NA | NA | NA | NA | #VALUE! |
| ENSMUSG00000091933.3 | 0 | NA | NA | NA | NA | NA | #VALUE! |
| ENSMUSG00000091936.2 | 0 | NA | NA | NA | NA | NA | #VALUE! |
| ENSMUSG00000091937.4 | 0 | NA | NA | NA | NA | NA | #VALUE! |
| ENSMUSG00000091938.4 | 0 | NA | NA | NA | NA | NA | #VALUE! |
| ENSMUSG00000091941.2 | 0 | NA | NA | NA | NA | NA | #VALUE! |
| ENSMUSG00000091942.2 | 0 | NA | NA | NA | NA | NA | #VALUE! |
| ENSMUSG00000091945.3 | 0 | NA | NA | NA | NA | NA | #VALUE! |
| ENSMUSG00000091947.2 | 0 | NA | NA | NA | NA | NA | #VALUE! |

| | | | | | | | |
|---|---|---|---|---|---|---|---|
| ENSMUSG00000091950.4 | 0 | NA | NA | NA | NA | NA | #VALUE! |
| ENSMUSG00000091952.4 | 0 | NA | NA | NA | NA | NA | #VALUE! |
| ENSMUSG00000091955.3 | 0 | NA | NA | NA | NA | NA | #VALUE! |
| ENSMUSG00000091956.3 | 0 | NA | NA | NA | NA | NA | #VALUE! |
| ENSMUSG00000091960.2 | 0 | NA | NA | NA | NA | NA | #VALUE! |
| ENSMUSG00000091961.2 | 0 | NA | NA | NA | NA | NA | #VALUE! |
| ENSMUSG00000091962.4 | 0 | NA | NA | NA | NA | NA | #VALUE! |
| ENSMUSG00000091967.6 | 0 | NA | NA | NA | NA | NA | #VALUE! |
| ENSMUSG00000091968.2 | 1.395489 | -4.00133 | 3.938888 | -1.01585 | 0.3097 | NA | #VALUE! |
| ENSMUSG00000091970.4 | 0 | NA | NA | NA | NA | NA | #VALUE! |
| ENSMUSG00000091971.4 | 0.735092 | -3.12016 | 3.965612 | -0.7868 | 0.431397 | NA | #VALUE! |
| ENSMUSG00000091972.3 | 0 | NA | NA | NA | NA | NA | #VALUE! |
| ENSMUSG00000091983.4 | 0 | NA | NA | NA | NA | NA | #VALUE! |
| ENSMUSG00000091985.2 | 0.504851 | 2.502843 | 3.993993 | 0.626652 | 0.530887 | NA | #VALUE! |
| ENSMUSG00000091987.9 | 0 | NA | NA | NA | NA | NA | #VALUE! |
| ENSMUSG00000091989.2 | 0.168284 | 1.027628 | 4.067126 | 0.252667 | 0.800526 | NA | #VALUE! |
| ENSMUSG00000091990.2 | 0 | NA | NA | NA | NA | NA | #VALUE! |
| ENSMUSG00000091993.3 | 0 | NA | NA | NA | NA | NA | #VALUE! |
| ENSMUSG00000091994.3 | 0 | NA | NA | NA | NA | NA | #VALUE! |
| ENSMUSG00000091995.2 | 0 | NA | NA | NA | NA | NA | #VALUE! |
| ENSMUSG00000091996.9 | 0 | NA | NA | NA | NA | NA | #VALUE! |
| ENSMUSG00000091997.2 | 0 | NA | NA | NA | NA | NA | #VALUE! |
| ENSMUSG00000091998.3 | 0 | NA | NA | NA | NA | NA | #VALUE! |
| ENSMUSG00000092004.2 | 0 | NA | NA | NA | NA | NA | #VALUE! |
| ENSMUSG00000092006.2 | 0.737927 | 3.007825 | 3.968641 | 0.757898 | 0.448512 | NA | #VALUE! |
| ENSMUSG00000092008.3 | 0 | NA | NA | NA | NA | NA | #VALUE! |
| ENSMUSG00000092009.2 | 0 | NA | NA | NA | NA | NA | #VALUE! |
| ENSMUSG00000092014.2 | 0.540678 | 0.929314 | 4.022674 | 0.231019 | 0.8173 | NA | #VALUE! |
| ENSMUSG00000092019.5 | 0 | NA | NA | NA | NA | NA | #VALUE! |
| ENSMUSG00000092027.2 | 0 | NA | NA | NA | NA | NA | #VALUE! |
| ENSMUSG00000092029.2 | 0 | NA | NA | NA | NA | NA | #VALUE! |
| ENSMUSG00000092031.2 | 0 | NA | NA | NA | NA | NA | #VALUE! |
| ENSMUSG00000092032.3 | 0 | NA | NA | NA | NA | NA | #VALUE! |
| ENSMUSG00000092035.9 | 0 | NA | NA | NA | NA | NA | #VALUE! |
| ENSMUSG00000092036.3 | 0 | NA | NA | NA | NA | NA | #VALUE! |
| ENSMUSG00000092039.3 | 0 | NA | NA | NA | NA | NA | #VALUE! |
| ENSMUSG00000092040.2 | 0 | NA | NA | NA | NA | NA | #VALUE! |
| ENSMUSG00000092042.2 | 0 | NA | NA | NA | NA | NA | #VALUE! |
| ENSMUSG00000092043.3 | 0 | NA | NA | NA | NA | NA | #VALUE! |
| ENSMUSG00000092045.2 | 0 | NA | NA | NA | NA | NA | #VALUE! |
| ENSMUSG00000092047.2 | 0 | NA | NA | NA | NA | NA | #VALUE! |
| ENSMUSG00000092048.4 | 0 | NA | NA | NA | NA | NA | #VALUE! |
| ENSMUSG00000092049.10 | 0 | NA | NA | NA | NA | NA | #VALUE! |
| ENSMUSG00000092051.3 | 0 | NA | NA | NA | NA | NA | #VALUE! |

| | | | | | | | |
|---|---|---|---|---|---|---|---|
| ENSMUSG00000092054.4 | 0 | NA | NA | NA | NA | NA | #VALUE! |
| ENSMUSG00000092055.2 | 0 | NA | NA | NA | NA | NA | #VALUE! |
| ENSMUSG00000092056.3 | 0 | NA | NA | NA | NA | NA | #VALUE! |
| ENSMUSG00000092060.5 | 0.932552 | 3.421684 | 3.953399 | 0.865504 | 0.386762 | NA | #VALUE! |
| ENSMUSG00000092066.2 | 0 | NA | NA | NA | NA | NA | #VALUE! |
| ENSMUSG00000092071.2 | 0 | NA | NA | NA | NA | NA | #VALUE! |
| ENSMUSG00000092073.9 | 0 | NA | NA | NA | NA | NA | #VALUE! |
| ENSMUSG00000092074.3 | 0 | NA | NA | NA | NA | NA | #VALUE! |
| ENSMUSG00000092077.3 | 0 | NA | NA | NA | NA | NA | #VALUE! |
| ENSMUSG00000092080.3 | 0 | NA | NA | NA | NA | NA | #VALUE! |
| ENSMUSG00000092083.5 | 0 | NA | NA | NA | NA | NA | #VALUE! |
| ENSMUSG00000092085.2 | 0 | NA | NA | NA | NA | NA | #VALUE! |
| ENSMUSG00000092086.3 | 0 | NA | NA | NA | NA | NA | #VALUE! |
| ENSMUSG00000092090.3 | 0 | NA | NA | NA | NA | NA | #VALUE! |
| ENSMUSG00000092092.2 | 0 | NA | NA | NA | NA | NA | #VALUE! |
| ENSMUSG00000092093.2 | 0 | NA | NA | NA | NA | NA | #VALUE! |
| ENSMUSG00000092094.6 | 0 | NA | NA | NA | NA | NA | #VALUE! |
| ENSMUSG00000092097.3 | 0 | NA | NA | NA | NA | NA | #VALUE! |
| ENSMUSG00000092103.3 | 0 | NA | NA | NA | NA | NA | #VALUE! |
| ENSMUSG00000092107.2 | 0 | NA | NA | NA | NA | NA | #VALUE! |
| ENSMUSG00000092109.3 | 0 | NA | NA | NA | NA | NA | #VALUE! |
| ENSMUSG00000092111.3 | 0 | NA | NA | NA | NA | NA | #VALUE! |
| ENSMUSG00000092116.2 | 2.741747 | -2.14177 | 3.836871 | -0.55821 | 0.576703 | NA | #VALUE! |
| ENSMUSG00000092118.2 | 6.105797 | 0.143048 | 2.685536 | 0.053266 | 0.95752 | NA | #VALUE! |
| ENSMUSG00000092120.3 | 0 | NA | NA | NA | NA | NA | #VALUE! |
| ENSMUSG00000092123.2 | 0 | NA | NA | NA | NA | NA | #VALUE! |
| ENSMUSG00000092124.2 | 3.36567 | 5.240782 | 3.920098 | 1.336901 | 0.181255 | NA | #VALUE! |
| ENSMUSG00000092125.2 | 0 | NA | NA | NA | NA | NA | #VALUE! |
| ENSMUSG00000092130.3 | 0 | NA | NA | NA | NA | NA | #VALUE! |
| ENSMUSG00000092131.2 | 0 | NA | NA | NA | NA | NA | #VALUE! |
| ENSMUSG00000092134.2 | 0 | NA | NA | NA | NA | NA | #VALUE! |
| ENSMUSG00000092142.2 | 0 | NA | NA | NA | NA | NA | #VALUE! |
| ENSMUSG00000092144.2 | 0 | NA | NA | NA | NA | NA | #VALUE! |
| ENSMUSG00000092147.2 | 0 | NA | NA | NA | NA | NA | #VALUE! |
| ENSMUSG00000092148.2 | 0 | NA | NA | NA | NA | NA | #VALUE! |
| ENSMUSG00000092152.2 | 0 | NA | NA | NA | NA | NA | #VALUE! |
| ENSMUSG00000092157.2 | 0 | NA | NA | NA | NA | NA | #VALUE! |
| ENSMUSG00000092162.4 | 0 | NA | NA | NA | NA | NA | #VALUE! |
| ENSMUSG00000092164.3 | 0 | NA | NA | NA | NA | NA | #VALUE! |
| ENSMUSG00000092165.10 | 0 | NA | NA | NA | NA | NA | #VALUE! |
| ENSMUSG00000092166.3 | 0 | NA | NA | NA | NA | NA | #VALUE! |
| ENSMUSG00000092167.9 | 0 | NA | NA | NA | NA | NA | #VALUE! |
| ENSMUSG00000092168.2 | 0 | NA | NA | NA | NA | NA | #VALUE! |
| ENSMUSG00000092171.2 | 0 | NA | NA | NA | NA | NA | #VALUE! |

| | | | | | | | |
|---|---|---|---|---|---|---|---|
| ENSMUSG00000092172.2 | 0 | NA | NA | NA | NA | NA | #VALUE! |
| ENSMUSG00000092173.2 | 0 | NA | NA | NA | NA | NA | #VALUE! |
| ENSMUSG00000092175.2 | 0 | NA | NA | NA | NA | NA | #VALUE! |
| ENSMUSG00000092176.2 | 0 | NA | NA | NA | NA | NA | #VALUE! |
| ENSMUSG00000092178.3 | 0 | NA | NA | NA | NA | NA | #VALUE! |
| ENSMUSG00000092179.3 | 0 | NA | NA | NA | NA | NA | #VALUE! |
| ENSMUSG00000092181.2 | 0 | NA | NA | NA | NA | NA | #VALUE! |
| ENSMUSG00000092182.2 | 0 | NA | NA | NA | NA | NA | #VALUE! |
| ENSMUSG00000092183.2 | 3.533954 | 5.311169 | 3.919465 | 1.355075 | 0.175394 | NA | #VALUE! |
| ENSMUSG00000092184.2 | 0 | NA | NA | NA | NA | NA | #VALUE! |
| ENSMUSG00000092185.2 | 0 | NA | NA | NA | NA | NA | #VALUE! |
| ENSMUSG00000092186.2 | 0 | NA | NA | NA | NA | NA | #VALUE! |
| ENSMUSG00000092188.2 | 0 | NA | NA | NA | NA | NA | #VALUE! |
| ENSMUSG00000092190.2 | 0 | NA | NA | NA | NA | NA | #VALUE! |
| ENSMUSG00000092191.2 | 0 | NA | NA | NA | NA | NA | #VALUE! |
| ENSMUSG00000092192.8 | 0 | NA | NA | NA | NA | NA | #VALUE! |
| ENSMUSG00000092193.2 | 0 | NA | NA | NA | NA | NA | #VALUE! |
| ENSMUSG00000092195.3 | 0 | NA | NA | NA | NA | NA | #VALUE! |
| ENSMUSG00000092196.3 | 0 | NA | NA | NA | NA | NA | #VALUE! |
| ENSMUSG00000092197.2 | 0 | NA | NA | NA | NA | NA | #VALUE! |
| ENSMUSG00000092199.2 | 0 | NA | NA | NA | NA | NA | #VALUE! |
| ENSMUSG00000092200.3 | 0 | NA | NA | NA | NA | NA | #VALUE! |
| ENSMUSG00000092201.8 | 0 | NA | NA | NA | NA | NA | #VALUE! |
| ENSMUSG00000092202.2 | 0 | NA | NA | NA | NA | NA | #VALUE! |
| ENSMUSG00000092204.2 | 0 | NA | NA | NA | NA | NA | #VALUE! |
| ENSMUSG00000092208.2 | 0 | NA | NA | NA | NA | NA | #VALUE! |
| ENSMUSG00000092210.2 | 0 | NA | NA | NA | NA | NA | #VALUE! |
| ENSMUSG00000092211.2 | 0 | NA | NA | NA | NA | NA | #VALUE! |
| ENSMUSG00000092212.4 | 0 | NA | NA | NA | NA | NA | #VALUE! |
| ENSMUSG00000092213.2 | 0 | NA | NA | NA | NA | NA | #VALUE! |
| ENSMUSG00000092214.2 | 0 | NA | NA | NA | NA | NA | #VALUE! |
| ENSMUSG00000092216.4 | 0 | NA | NA | NA | NA | NA | #VALUE! |
| ENSMUSG00000092217.3 | 0 | NA | NA | NA | NA | NA | #VALUE! |
| ENSMUSG00000092218.2 | 0 | NA | NA | NA | NA | NA | #VALUE! |
| ENSMUSG00000092219.2 | 0 | NA | NA | NA | NA | NA | #VALUE! |
| ENSMUSG00000092221.2 | 0 | NA | NA | NA | NA | NA | #VALUE! |
| ENSMUSG00000092223.2 | 0 | NA | NA | NA | NA | NA | #VALUE! |
| ENSMUSG00000092224.3 | 0 | NA | NA | NA | NA | NA | #VALUE! |
| ENSMUSG00000092225.2 | 0 | NA | NA | NA | NA | NA | #VALUE! |
| ENSMUSG00000092227.2 | 0 | NA | NA | NA | NA | NA | #VALUE! |
| ENSMUSG00000092228.2 | 0 | NA | NA | NA | NA | NA | #VALUE! |
| ENSMUSG00000092229.4 | 0 | NA | NA | NA | NA | NA | #VALUE! |
| ENSMUSG00000092230.2 | 0 | NA | NA | NA | NA | NA | #VALUE! |
| ENSMUSG00000092231.2 | 0 | NA | NA | NA | NA | NA | #VALUE! |

| | | | | | | | |
|---|---|---|---|---|---|---|---|
| ENSMUSG00000092232.2 | 0 | NA | NA | NA | NA | NA | #VALUE! |
| ENSMUSG00000092233.2 | 0 | NA | NA | NA | NA | NA | #VALUE! |
| ENSMUSG00000092237.3 | 0 | NA | NA | NA | NA | NA | #VALUE! |
| ENSMUSG00000092238.2 | 0 | NA | NA | NA | NA | NA | #VALUE! |
| ENSMUSG00000092239.2 | 0 | NA | NA | NA | NA | NA | #VALUE! |
| ENSMUSG00000092240.2 | 0 | NA | NA | NA | NA | NA | #VALUE! |
| ENSMUSG00000092242.2 | 0 | NA | NA | NA | NA | NA | #VALUE! |
| ENSMUSG00000092243.9 | 0 | NA | NA | NA | NA | NA | #VALUE! |
| ENSMUSG00000092244.4 | 0 | NA | NA | NA | NA | NA | #VALUE! |
| ENSMUSG00000092245.2 | 0 | NA | NA | NA | NA | NA | #VALUE! |
| ENSMUSG00000092246.3 | 0 | NA | NA | NA | NA | NA | #VALUE! |
| ENSMUSG00000092247.3 | 0.630079 | -2.90291 | 3.975047 | -0.73028 | 0.465217 | NA | #VALUE! |
| ENSMUSG00000092248.2 | 0 | NA | NA | NA | NA | NA | #VALUE! |
| ENSMUSG00000092250.2 | 0 | NA | NA | NA | NA | NA | #VALUE! |
| ENSMUSG00000092251.4 | 0 | NA | NA | NA | NA | NA | #VALUE! |
| ENSMUSG00000092253.2 | 0 | NA | NA | NA | NA | NA | #VALUE! |
| ENSMUSG00000092255.2 | 0 | NA | NA | NA | NA | NA | #VALUE! |
| ENSMUSG00000092256.2 | 0 | NA | NA | NA | NA | NA | #VALUE! |
| ENSMUSG00000092257.2 | 0 | NA | NA | NA | NA | NA | #VALUE! |
| ENSMUSG00000092258.3 | 0 | NA | NA | NA | NA | NA | #VALUE! |
| ENSMUSG00000092259.2 | 0 | NA | NA | NA | NA | NA | #VALUE! |
| ENSMUSG00000092260.8 | 0.612332 | -2.79827 | 3.980109 | -0.70306 | 0.482016 | NA | #VALUE! |
| ENSMUSG00000092261.2 | 0 | NA | NA | NA | NA | NA | #VALUE! |
| ENSMUSG00000092263.2 | 0 | NA | NA | NA | NA | NA | #VALUE! |
| ENSMUSG00000092264.2 | 0 | NA | NA | NA | NA | NA | #VALUE! |
| ENSMUSG00000092265.3 | 0 | NA | NA | NA | NA | NA | #VALUE! |
| ENSMUSG00000092266.2 | 0 | NA | NA | NA | NA | NA | #VALUE! |
| ENSMUSG00000092268.2 | 1.174735 | -3.75246 | 3.944889 | -0.95122 | 0.341493 | NA | #VALUE! |
| ENSMUSG00000092269.2 | 0 | NA | NA | NA | NA | NA | #VALUE! |
| ENSMUSG00000092270.2 | 0 | NA | NA | NA | NA | NA | #VALUE! |
| ENSMUSG00000092271.2 | 0 | NA | NA | NA | NA | NA | #VALUE! |
| ENSMUSG00000092272.2 | 0 | NA | NA | NA | NA | NA | #VALUE! |
| ENSMUSG00000092275.2 | 0 | NA | NA | NA | NA | NA | #VALUE! |
| ENSMUSG00000092276.3 | 0 | NA | NA | NA | NA | NA | #VALUE! |
| ENSMUSG00000092277.2 | 0 | NA | NA | NA | NA | NA | #VALUE! |
| ENSMUSG00000092278.2 | 0 | NA | NA | NA | NA | NA | #VALUE! |
| ENSMUSG00000092281.2 | 3.465433 | -5.33079 | 3.919598 | -1.36004 | 0.173819 | NA | #VALUE! |
| ENSMUSG00000092283.2 | 0 | NA | NA | NA | NA | NA | #VALUE! |
| ENSMUSG00000092284.2 | 0 | NA | NA | NA | NA | NA | #VALUE! |
| ENSMUSG00000092285.2 | 0 | NA | NA | NA | NA | NA | #VALUE! |
| ENSMUSG00000092286.2 | 0 | NA | NA | NA | NA | NA | #VALUE! |
| ENSMUSG00000092287.3 | 0 | NA | NA | NA | NA | NA | #VALUE! |
| ENSMUSG00000092289.2 | 0 | NA | NA | NA | NA | NA | #VALUE! |
| ENSMUSG00000092290.2 | 0 | NA | NA | NA | NA | NA | #VALUE! |

| | | | | | | | |
|---|---|---|---|---|---|---|---|
| ENSMUSG00000092292.4 | 0 | NA | NA | NA | NA | NA | #VALUE! |
| ENSMUSG00000092293.2 | 0 | NA | NA | NA | NA | NA | #VALUE! |
| ENSMUSG00000092294.2 | 0 | NA | NA | NA | NA | NA | #VALUE! |
| ENSMUSG00000092295.2 | 0 | NA | NA | NA | NA | NA | #VALUE! |
| ENSMUSG00000092296.2 | 0 | NA | NA | NA | NA | NA | #VALUE! |
| ENSMUSG00000092297.2 | 0 | NA | NA | NA | NA | NA | #VALUE! |
| ENSMUSG00000092298.2 | 0 | NA | NA | NA | NA | NA | #VALUE! |
| ENSMUSG00000092300.8 | 0 | NA | NA | NA | NA | NA | #VALUE! |
| ENSMUSG00000092301.2 | 0 | NA | NA | NA | NA | NA | #VALUE! |
| ENSMUSG00000092303.2 | 0 | NA | NA | NA | NA | NA | #VALUE! |
| ENSMUSG00000092304.3 | 0 | NA | NA | NA | NA | NA | #VALUE! |
| ENSMUSG00000092305.2 | 0 | NA | NA | NA | NA | NA | #VALUE! |
| ENSMUSG00000092306.4 | 0 | NA | NA | NA | NA | NA | #VALUE! |
| ENSMUSG00000092307.2 | 0 | NA | NA | NA | NA | NA | #VALUE! |
| ENSMUSG00000092308.4 | 0 | NA | NA | NA | NA | NA | #VALUE! |
| ENSMUSG00000092309.3 | 0 | NA | NA | NA | NA | NA | #VALUE! |
| ENSMUSG00000092310.3 | 0 | NA | NA | NA | NA | NA | #VALUE! |
| ENSMUSG00000092311.2 | 0 | NA | NA | NA | NA | NA | #VALUE! |
| ENSMUSG00000092312.2 | 0 | NA | NA | NA | NA | NA | #VALUE! |
| ENSMUSG00000092313.2 | 0 | NA | NA | NA | NA | NA | #VALUE! |
| ENSMUSG00000092314.2 | 0 | NA | NA | NA | NA | NA | #VALUE! |
| ENSMUSG00000092316.2 | 0 | NA | NA | NA | NA | NA | #VALUE! |
| ENSMUSG00000092317.2 | 0 | NA | NA | NA | NA | NA | #VALUE! |
| ENSMUSG00000092318.2 | 0 | NA | NA | NA | NA | NA | #VALUE! |
| ENSMUSG00000092322.9 | 0 | NA | NA | NA | NA | NA | #VALUE! |
| ENSMUSG00000092323.2 | 0 | NA | NA | NA | NA | NA | #VALUE! |
| ENSMUSG00000092325.3 | 0 | NA | NA | NA | NA | NA | #VALUE! |
| ENSMUSG00000092326.2 | 0 | NA | NA | NA | NA | NA | #VALUE! |
| ENSMUSG00000092328.2 | 0 | NA | NA | NA | NA | NA | #VALUE! |
| ENSMUSG00000092330.3 | 0 | NA | NA | NA | NA | NA | #VALUE! |
| ENSMUSG00000092331.2 | 0 | NA | NA | NA | NA | NA | #VALUE! |
| ENSMUSG00000092332.2 | 0 | NA | NA | NA | NA | NA | #VALUE! |
| ENSMUSG00000092335.2 | 5.903413 | 6.04575 | 3.914406 | 1.544487 | 0.12247 | NA | #VALUE! |
| ENSMUSG00000092336.2 | 0 | NA | NA | NA | NA | NA | #VALUE! |
| ENSMUSG00000092338.2 | 0 | NA | NA | NA | NA | NA | #VALUE! |
| ENSMUSG00000092339.2 | 0 | NA | NA | NA | NA | NA | #VALUE! |
| ENSMUSG00000092340.4 | 0 | NA | NA | NA | NA | NA | #VALUE! |
| ENSMUSG00000092342.3 | 0 | NA | NA | NA | NA | NA | #VALUE! |
| ENSMUSG00000092343.2 | 0 | NA | NA | NA | NA | NA | #VALUE! |
| ENSMUSG00000092344.2 | 0 | NA | NA | NA | NA | NA | #VALUE! |
| ENSMUSG00000092345.2 | 0 | NA | NA | NA | NA | NA | #VALUE! |
| ENSMUSG00000092346.2 | 0 | NA | NA | NA | NA | NA | #VALUE! |
| ENSMUSG00000092349.2 | 0 | NA | NA | NA | NA | NA | #VALUE! |
| ENSMUSG00000092350.2 | 0 | NA | NA | NA | NA | NA | #VALUE! |

| | | | | | | | |
|---|---|---|---|---|---|---|---|
| ENSMUSG00000092351.2 | 0 | NA | NA | NA | NA | NA | #VALUE! |
| ENSMUSG00000092353.2 | 0 | NA | NA | NA | NA | NA | #VALUE! |
| ENSMUSG00000092354.8 | 0 | NA | NA | NA | NA | NA | #VALUE! |
| ENSMUSG00000092356.3 | 0 | NA | NA | NA | NA | NA | #VALUE! |
| ENSMUSG00000092357.2 | 0 | NA | NA | NA | NA | NA | #VALUE! |
| ENSMUSG00000092358.2 | 0 | NA | NA | NA | NA | NA | #VALUE! |
| ENSMUSG00000092359.2 | 0 | NA | NA | NA | NA | NA | #VALUE! |
| ENSMUSG00000092360.2 | 0 | NA | NA | NA | NA | NA | #VALUE! |
| ENSMUSG00000092362.4 | 0 | NA | NA | NA | NA | NA | #VALUE! |
| ENSMUSG00000092363.2 | 0 | NA | NA | NA | NA | NA | #VALUE! |
| ENSMUSG00000092364.3 | 0 | NA | NA | NA | NA | NA | #VALUE! |
| ENSMUSG00000092365.3 | 0 | NA | NA | NA | NA | NA | #VALUE! |
| ENSMUSG00000092366.3 | 0 | NA | NA | NA | NA | NA | #VALUE! |
| ENSMUSG00000092367.2 | 0 | NA | NA | NA | NA | NA | #VALUE! |
| ENSMUSG00000092368.2 | 0 | NA | NA | NA | NA | NA | #VALUE! |
| ENSMUSG00000092369.2 | 0 | NA | NA | NA | NA | NA | #VALUE! |
| ENSMUSG00000092370.2 | 0 | NA | NA | NA | NA | NA | #VALUE! |
| ENSMUSG00000092371.2 | 0 | NA | NA | NA | NA | NA | #VALUE! |
| ENSMUSG00000092372.4 | 0 | NA | NA | NA | NA | NA | #VALUE! |
| ENSMUSG00000092373.2 | 0 | NA | NA | NA | NA | NA | #VALUE! |
| ENSMUSG00000092374.2 | 0 | NA | NA | NA | NA | NA | #VALUE! |
| ENSMUSG00000092375.2 | 0 | NA | NA | NA | NA | NA | #VALUE! |
| ENSMUSG00000092376.2 | 0 | NA | NA | NA | NA | NA | #VALUE! |
| ENSMUSG00000092377.2 | 0 | NA | NA | NA | NA | NA | #VALUE! |
| ENSMUSG00000092381.2 | 0 | NA | NA | NA | NA | NA | #VALUE! |
| ENSMUSG00000092382.3 | 0 | NA | NA | NA | NA | NA | #VALUE! |
| ENSMUSG00000092383.3 | 0 | NA | NA | NA | NA | NA | #VALUE! |
| ENSMUSG00000092384.8 | 2.041108 | -4.55188 | 3.928743 | -1.15861 | 0.246615 | NA | #VALUE! |
| ENSMUSG00000092385.2 | 0 | NA | NA | NA | NA | NA | #VALUE! |
| ENSMUSG00000092386.2 | 0 | NA | NA | NA | NA | NA | #VALUE! |
| ENSMUSG00000092387.2 | 0 | NA | NA | NA | NA | NA | #VALUE! |
| ENSMUSG00000092388.3 | 0 | NA | NA | NA | NA | NA | #VALUE! |
| ENSMUSG00000092389.3 | 0 | NA | NA | NA | NA | NA | #VALUE! |
| ENSMUSG00000092390.2 | 0 | NA | NA | NA | NA | NA | #VALUE! |
| ENSMUSG00000092392.2 | 0 | NA | NA | NA | NA | NA | #VALUE! |
| ENSMUSG00000092393.3 | 0 | NA | NA | NA | NA | NA | #VALUE! |
| ENSMUSG00000092394.2 | 0 | NA | NA | NA | NA | NA | #VALUE! |
| ENSMUSG00000092395.2 | 9.109393 | -0.64615 | 3.38642 | -0.1908 | 0.848678 | NA | #VALUE! |
| ENSMUSG00000092396.3 | 0 | NA | NA | NA | NA | NA | #VALUE! |
| ENSMUSG00000092397.8 | 0 | NA | NA | NA | NA | NA | #VALUE! |
| ENSMUSG00000092398.2 | 0 | NA | NA | NA | NA | NA | #VALUE! |
| ENSMUSG00000092399.2 | 0 | NA | NA | NA | NA | NA | #VALUE! |
| ENSMUSG00000092401.4 | 0 | NA | NA | NA | NA | NA | #VALUE! |
| ENSMUSG00000092402.2 | 0 | NA | NA | NA | NA | NA | #VALUE! |

| | | | | | | | |
|---|---|---|---|---|---|---|---|
| ENSMUSG00000092404.3 | 0 | NA | NA | NA | NA | NA | #VALUE! |
| ENSMUSG00000092405.2 | 0 | NA | NA | NA | NA | NA | #VALUE! |
| ENSMUSG00000092406.2 | 0 | NA | NA | NA | NA | NA | #VALUE! |
| ENSMUSG00000092408.2 | 0 | NA | NA | NA | NA | NA | #VALUE! |
| ENSMUSG00000092409.2 | 0 | NA | NA | NA | NA | NA | #VALUE! |
| ENSMUSG00000092410.2 | 0 | NA | NA | NA | NA | NA | #VALUE! |
| ENSMUSG00000092411.2 | 0 | NA | NA | NA | NA | NA | #VALUE! |
| ENSMUSG00000092412.2 | 0 | NA | NA | NA | NA | NA | #VALUE! |
| ENSMUSG00000092413.7 | 0 | NA | NA | NA | NA | NA | #VALUE! |
| ENSMUSG00000092414.2 | 0 | NA | NA | NA | NA | NA | #VALUE! |
| ENSMUSG00000092415.3 | 0 | NA | NA | NA | NA | NA | #VALUE! |
| ENSMUSG00000092418.2 | 0 | NA | NA | NA | NA | NA | #VALUE! |
| ENSMUSG00000092419.2 | 0 | NA | NA | NA | NA | NA | #VALUE! |
| ENSMUSG00000092422.2 | 0 | NA | NA | NA | NA | NA | #VALUE! |
| ENSMUSG00000092425.2 | 0 | NA | NA | NA | NA | NA | #VALUE! |
| ENSMUSG00000092426.2 | 0 | NA | NA | NA | NA | NA | #VALUE! |
| ENSMUSG00000092427.2 | 0 | NA | NA | NA | NA | NA | #VALUE! |
| ENSMUSG00000092428.2 | 0 | NA | NA | NA | NA | NA | #VALUE! |
| ENSMUSG00000092429.2 | 0 | NA | NA | NA | NA | NA | #VALUE! |
| ENSMUSG00000092431.2 | 0 | NA | NA | NA | NA | NA | #VALUE! |
| ENSMUSG00000092433.2 | 0 | NA | NA | NA | NA | NA | #VALUE! |
| ENSMUSG00000092435.2 | 0 | NA | NA | NA | NA | NA | #VALUE! |
| ENSMUSG00000092437.2 | 0 | NA | NA | NA | NA | NA | #VALUE! |
| ENSMUSG00000092438.2 | 0 | NA | NA | NA | NA | NA | #VALUE! |
| ENSMUSG00000092439.3 | 0 | NA | NA | NA | NA | NA | #VALUE! |
| ENSMUSG00000092440.2 | 0 | NA | NA | NA | NA | NA | #VALUE! |
| ENSMUSG00000092441.2 | 0 | NA | NA | NA | NA | NA | #VALUE! |
| ENSMUSG00000092444.2 | 0 | NA | NA | NA | NA | NA | #VALUE! |
| ENSMUSG00000092447.2 | 0 | NA | NA | NA | NA | NA | #VALUE! |
| ENSMUSG00000092448.2 | 0 | NA | NA | NA | NA | NA | #VALUE! |
| ENSMUSG00000092450.2 | 0 | NA | NA | NA | NA | NA | #VALUE! |
| ENSMUSG00000092451.4 | 0 | NA | NA | NA | NA | NA | #VALUE! |
| ENSMUSG00000092452.2 | 0 | NA | NA | NA | NA | NA | #VALUE! |
| ENSMUSG00000092454.2 | 0 | NA | NA | NA | NA | NA | #VALUE! |
| ENSMUSG00000092455.2 | 0 | NA | NA | NA | NA | NA | #VALUE! |
| ENSMUSG00000092456.3 | 0 | NA | NA | NA | NA | NA | #VALUE! |
| ENSMUSG00000092457.4 | 0 | NA | NA | NA | NA | NA | #VALUE! |
| ENSMUSG00000092458.2 | 0 | NA | NA | NA | NA | NA | #VALUE! |
| ENSMUSG00000092460.2 | 0 | NA | NA | NA | NA | NA | #VALUE! |
| ENSMUSG00000092461.2 | 0 | NA | NA | NA | NA | NA | #VALUE! |
| ENSMUSG00000092462.2 | 0 | NA | NA | NA | NA | NA | #VALUE! |
| ENSMUSG00000092463.2 | 0 | NA | NA | NA | NA | NA | #VALUE! |
| ENSMUSG00000092464.2 | 0 | NA | NA | NA | NA | NA | #VALUE! |
| ENSMUSG00000092465.3 | 0 | NA | NA | NA | NA | NA | #VALUE! |

| | | | | | | | |
|---|---|---|---|---|---|---|---|
| ENSMUSG00000092466.2 | 0 | NA | NA | NA | NA | NA | #VALUE! |
| ENSMUSG00000092467.2 | 0 | NA | NA | NA | NA | NA | #VALUE! |
| ENSMUSG00000092468.2 | 0 | NA | NA | NA | NA | NA | #VALUE! |
| ENSMUSG00000092469.2 | 0 | NA | NA | NA | NA | NA | #VALUE! |
| ENSMUSG00000092470.2 | 0 | NA | NA | NA | NA | NA | #VALUE! |
| ENSMUSG00000092471.2 | 0 | NA | NA | NA | NA | NA | #VALUE! |
| ENSMUSG00000092472.2 | 0 | NA | NA | NA | NA | NA | #VALUE! |
| ENSMUSG00000092473.5 | 0 | NA | NA | NA | NA | NA | #VALUE! |
| ENSMUSG00000092474.2 | 0 | NA | NA | NA | NA | NA | #VALUE! |
| ENSMUSG00000092476.2 | 0 | NA | NA | NA | NA | NA | #VALUE! |
| ENSMUSG00000092477.2 | 0 | NA | NA | NA | NA | NA | #VALUE! |
| ENSMUSG00000092480.2 | 0 | NA | NA | NA | NA | NA | #VALUE! |
| ENSMUSG00000092482.2 | 0 | NA | NA | NA | NA | NA | #VALUE! |
| ENSMUSG00000092483.2 | 0 | NA | NA | NA | NA | NA | #VALUE! |
| ENSMUSG00000092485.2 | 0 | NA | NA | NA | NA | NA | #VALUE! |
| ENSMUSG00000092487.2 | 0 | NA | NA | NA | NA | NA | #VALUE! |
| ENSMUSG00000092488.2 | 0 | NA | NA | NA | NA | NA | #VALUE! |
| ENSMUSG00000092489.2 | 0 | NA | NA | NA | NA | NA | #VALUE! |
| ENSMUSG00000092490.8 | 0 | NA | NA | NA | NA | NA | #VALUE! |
| ENSMUSG00000092492.2 | 0 | NA | NA | NA | NA | NA | #VALUE! |
| ENSMUSG00000092493.2 | 0 | NA | NA | NA | NA | NA | #VALUE! |
| ENSMUSG00000092494.3 | 0 | NA | NA | NA | NA | NA | #VALUE! |
| ENSMUSG00000092495.2 | 0 | NA | NA | NA | NA | NA | #VALUE! |
| ENSMUSG00000092496.2 | 0 | NA | NA | NA | NA | NA | #VALUE! |
| ENSMUSG00000092499.2 | 0 | NA | NA | NA | NA | NA | #VALUE! |
| ENSMUSG00000092500.2 | 0 | NA | NA | NA | NA | NA | #VALUE! |
| ENSMUSG00000092502.2 | 0 | NA | NA | NA | NA | NA | #VALUE! |
| ENSMUSG00000092503.2 | 0 | NA | NA | NA | NA | NA | #VALUE! |
| ENSMUSG00000092504.2 | 0 | NA | NA | NA | NA | NA | #VALUE! |
| ENSMUSG00000092505.2 | 0 | NA | NA | NA | NA | NA | #VALUE! |
| ENSMUSG00000092506.2 | 0 | NA | NA | NA | NA | NA | #VALUE! |
| ENSMUSG00000092508.2 | 0 | NA | NA | NA | NA | NA | #VALUE! |
| ENSMUSG00000092509.2 | 0 | NA | NA | NA | NA | NA | #VALUE! |
| ENSMUSG00000092510.2 | 0 | NA | NA | NA | NA | NA | #VALUE! |
| ENSMUSG00000092511.8 | 0 | NA | NA | NA | NA | NA | #VALUE! |
| ENSMUSG00000092512.3 | 0 | NA | NA | NA | NA | NA | #VALUE! |
| ENSMUSG00000092513.2 | 0 | NA | NA | NA | NA | NA | #VALUE! |
| ENSMUSG00000092514.2 | 0 | NA | NA | NA | NA | NA | #VALUE! |
| ENSMUSG00000092515.9 | 0 | NA | NA | NA | NA | NA | #VALUE! |
| ENSMUSG00000092516.2 | 0 | NA | NA | NA | NA | NA | #VALUE! |
| ENSMUSG00000092517.2 | 0 | NA | NA | NA | NA | NA | #VALUE! |
| ENSMUSG00000092518.10 | 0 | NA | NA | NA | NA | NA | #VALUE! |
| ENSMUSG00000092519.3 | 0 | NA | NA | NA | NA | NA | #VALUE! |
| ENSMUSG00000092520.2 | 0 | NA | NA | NA | NA | NA | #VALUE! |

| | | | | | | | |
|---|---|---|---|---|---|---|---|
| ENSMUSG00000092521.3 | 0 | NA | NA | NA | NA | NA | #VALUE! |
| ENSMUSG00000092522.2 | 0 | NA | NA | NA | NA | NA | #VALUE! |
| ENSMUSG00000092523.2 | 0 | NA | NA | NA | NA | NA | #VALUE! |
| ENSMUSG00000092524.4 | 0 | NA | NA | NA | NA | NA | #VALUE! |
| ENSMUSG00000092525.2 | 0 | NA | NA | NA | NA | NA | #VALUE! |
| ENSMUSG00000092526.2 | 0 | NA | NA | NA | NA | NA | #VALUE! |
| ENSMUSG00000092528.10 | 0 | NA | NA | NA | NA | NA | #VALUE! |
| ENSMUSG00000092529.3 | 0 | NA | NA | NA | NA | NA | #VALUE! |
| ENSMUSG00000092530.2 | 0 | NA | NA | NA | NA | NA | #VALUE! |
| ENSMUSG00000092532.2 | 0 | NA | NA | NA | NA | NA | #VALUE! |
| ENSMUSG00000092533.2 | 0 | NA | NA | NA | NA | NA | #VALUE! |
| ENSMUSG00000092534.9 | 0 | NA | NA | NA | NA | NA | #VALUE! |
| ENSMUSG00000092535.2 | 0 | NA | NA | NA | NA | NA | #VALUE! |
| ENSMUSG00000092536.3 | 0 | NA | NA | NA | NA | NA | #VALUE! |
| ENSMUSG00000092538.2 | 0 | NA | NA | NA | NA | NA | #VALUE! |
| ENSMUSG00000092539.3 | 0 | NA | NA | NA | NA | NA | #VALUE! |
| ENSMUSG00000092540.2 | 0 | NA | NA | NA | NA | NA | #VALUE! |
| ENSMUSG00000092541.8 | 0 | NA | NA | NA | NA | NA | #VALUE! |
| ENSMUSG00000092542.3 | 0 | NA | NA | NA | NA | NA | #VALUE! |
| ENSMUSG00000092543.2 | 0 | NA | NA | NA | NA | NA | #VALUE! |
| ENSMUSG00000092544.9 | 0 | NA | NA | NA | NA | NA | #VALUE! |
| ENSMUSG00000092547.2 | 0 | NA | NA | NA | NA | NA | #VALUE! |
| ENSMUSG00000092548.2 | 0 | NA | NA | NA | NA | NA | #VALUE! |
| ENSMUSG00000092549.2 | 0 | NA | NA | NA | NA | NA | #VALUE! |
| ENSMUSG00000092550.3 | 0 | NA | NA | NA | NA | NA | #VALUE! |
| ENSMUSG00000092551.4 | 0 | NA | NA | NA | NA | NA | #VALUE! |
| ENSMUSG00000092552.2 | 0 | NA | NA | NA | NA | NA | #VALUE! |
| ENSMUSG00000092553.2 | 0 | NA | NA | NA | NA | NA | #VALUE! |
| ENSMUSG00000092554.2 | 0 | NA | NA | NA | NA | NA | #VALUE! |
| ENSMUSG00000092555.2 | 0 | NA | NA | NA | NA | NA | #VALUE! |
| ENSMUSG00000092556.3 | 0 | NA | NA | NA | NA | NA | #VALUE! |
| ENSMUSG00000092557.2 | 0 | NA | NA | NA | NA | NA | #VALUE! |
| ENSMUSG00000092559.2 | 0 | NA | NA | NA | NA | NA | #VALUE! |
| ENSMUSG00000092560.3 | 0 | NA | NA | NA | NA | NA | #VALUE! |
| ENSMUSG00000092561.2 | 0 | NA | NA | NA | NA | NA | #VALUE! |
| ENSMUSG00000092562.2 | 0 | NA | NA | NA | NA | NA | #VALUE! |
| ENSMUSG00000092563.2 | 0 | NA | NA | NA | NA | NA | #VALUE! |
| ENSMUSG00000092564.3 | 0 | NA | NA | NA | NA | NA | #VALUE! |
| ENSMUSG00000092565.2 | 0 | NA | NA | NA | NA | NA | #VALUE! |
| ENSMUSG00000092566.2 | 0 | NA | NA | NA | NA | NA | #VALUE! |
| ENSMUSG00000092567.3 | 0 | NA | NA | NA | NA | NA | #VALUE! |
| ENSMUSG00000092568.3 | 0 | NA | NA | NA | NA | NA | #VALUE! |
| ENSMUSG00000092569.2 | 0 | NA | NA | NA | NA | NA | #VALUE! |
| ENSMUSG00000092570.8 | 0 | NA | NA | NA | NA | NA | #VALUE! |

| | | | | | | | |
|---|---|---|---|---|---|---|---|
| ENSMUSG00000092571.2 | 0 | NA | NA | NA | NA | NA | #VALUE! |
| ENSMUSG00000092572.9 | 0 | NA | NA | NA | NA | NA | #VALUE! |
| ENSMUSG00000092573.2 | 0 | NA | NA | NA | NA | NA | #VALUE! |
| ENSMUSG00000092574.5 | 0 | NA | NA | NA | NA | NA | #VALUE! |
| ENSMUSG00000092575.3 | 0 | NA | NA | NA | NA | NA | #VALUE! |
| ENSMUSG00000092576.4 | 0 | NA | NA | NA | NA | NA | #VALUE! |
| ENSMUSG00000092577.2 | 0 | NA | NA | NA | NA | NA | #VALUE! |
| ENSMUSG00000092578.2 | 0 | NA | NA | NA | NA | NA | #VALUE! |
| ENSMUSG00000092579.3 | 0 | NA | NA | NA | NA | NA | #VALUE! |
| ENSMUSG00000092580.2 | 0 | NA | NA | NA | NA | NA | #VALUE! |
| ENSMUSG00000092581.2 | 0 | NA | NA | NA | NA | NA | #VALUE! |
| ENSMUSG00000092583.2 | 0 | NA | NA | NA | NA | NA | #VALUE! |
| ENSMUSG00000092584.2 | 0 | NA | NA | NA | NA | NA | #VALUE! |
| ENSMUSG00000092586.9 | 0 | NA | NA | NA | NA | NA | #VALUE! |
| ENSMUSG00000092587.2 | 0 | NA | NA | NA | NA | NA | #VALUE! |
| ENSMUSG00000092588.2 | 0 | NA | NA | NA | NA | NA | #VALUE! |
| ENSMUSG00000092591.2 | 0 | NA | NA | NA | NA | NA | #VALUE! |
| ENSMUSG00000092592.2 | 0 | NA | NA | NA | NA | NA | #VALUE! |
| ENSMUSG00000092593.2 | 1.370524 | -3.97623 | 3.939448 | -1.00934 | 0.312813 | NA | #VALUE! |
| ENSMUSG00000092595.3 | 0 | NA | NA | NA | NA | NA | #VALUE! |
| ENSMUSG00000092596.2 | 0 | NA | NA | NA | NA | NA | #VALUE! |
| ENSMUSG00000092597.3 | 0 | NA | NA | NA | NA | NA | #VALUE! |
| ENSMUSG00000092599.2 | 0 | NA | NA | NA | NA | NA | #VALUE! |
| ENSMUSG00000092600.2 | 0 | NA | NA | NA | NA | NA | #VALUE! |
| ENSMUSG00000092601.3 | 0 | NA | NA | NA | NA | NA | #VALUE! |
| ENSMUSG00000092602.2 | 0 | NA | NA | NA | NA | NA | #VALUE! |
| ENSMUSG00000092603.2 | 0 | NA | NA | NA | NA | NA | #VALUE! |
| ENSMUSG00000092604.2 | 0 | NA | NA | NA | NA | NA | #VALUE! |
| ENSMUSG00000092606.2 | 0 | NA | NA | NA | NA | NA | #VALUE! |
| ENSMUSG00000092608.3 | 0 | NA | NA | NA | NA | NA | #VALUE! |
| ENSMUSG00000092609.2 | 0 | NA | NA | NA | NA | NA | #VALUE! |
| ENSMUSG00000092610.2 | 0 | NA | NA | NA | NA | NA | #VALUE! |
| ENSMUSG00000092611.3 | 0 | NA | NA | NA | NA | NA | #VALUE! |
| ENSMUSG00000092612.3 | 0 | NA | NA | NA | NA | NA | #VALUE! |
| ENSMUSG00000092614.3 | 0 | NA | NA | NA | NA | NA | #VALUE! |
| ENSMUSG00000092615.3 | 0 | NA | NA | NA | NA | NA | #VALUE! |
| ENSMUSG00000092616.2 | 0 | NA | NA | NA | NA | NA | #VALUE! |
| ENSMUSG00000092617.2 | 0 | NA | NA | NA | NA | NA | #VALUE! |
| ENSMUSG00000092618.3 | 0 | NA | NA | NA | NA | NA | #VALUE! |
| ENSMUSG00000092619.2 | 0 | NA | NA | NA | NA | NA | #VALUE! |
| ENSMUSG00000092620.2 | 0 | NA | NA | NA | NA | NA | #VALUE! |
| ENSMUSG00000092621.2 | 0 | NA | NA | NA | NA | NA | #VALUE! |
| ENSMUSG00000092622.9 | 0 | NA | NA | NA | NA | NA | #VALUE! |
| ENSMUSG00000092624.3 | 0 | NA | NA | NA | NA | NA | #VALUE! |

| | | | | | | | |
|---|---|---|---|---|---|---|---|
| ENSMUSG00000092625.3 | 0 | NA | NA | NA | NA | NA | #VALUE! |
| ENSMUSG00000092627.3 | 0 | NA | NA | NA | NA | NA | #VALUE! |
| ENSMUSG00000092629.2 | 0 | NA | NA | NA | NA | NA | #VALUE! |
| ENSMUSG00000092633.3 | 0 | NA | NA | NA | NA | NA | #VALUE! |
| ENSMUSG00000092638.3 | 0 | NA | NA | NA | NA | NA | #VALUE! |
| ENSMUSG00000092639.3 | 0 | NA | NA | NA | NA | NA | #VALUE! |
| ENSMUSG00000092652.3 | 0 | NA | NA | NA | NA | NA | #VALUE! |
| ENSMUSG00000092659.3 | 0 | NA | NA | NA | NA | NA | #VALUE! |
| ENSMUSG00000092673.3 | 0 | NA | NA | NA | NA | NA | #VALUE! |
| ENSMUSG00000092674.5 | 0 | NA | NA | NA | NA | NA | #VALUE! |
| ENSMUSG00000092677.2 | 0 | NA | NA | NA | NA | NA | #VALUE! |
| ENSMUSG00000092679.3 | 0 | NA | NA | NA | NA | NA | #VALUE! |
| ENSMUSG00000092680.3 | 0 | NA | NA | NA | NA | NA | #VALUE! |
| ENSMUSG00000092687.3 | 0 | NA | NA | NA | NA | NA | #VALUE! |
| ENSMUSG00000092695.3 | 0 | NA | NA | NA | NA | NA | #VALUE! |
| ENSMUSG00000092696.3 | 0 | NA | NA | NA | NA | NA | #VALUE! |
| ENSMUSG00000092702.5 | 0 | NA | NA | NA | NA | NA | #VALUE! |
| ENSMUSG00000092704.3 | 0 | NA | NA | NA | NA | NA | #VALUE! |
| ENSMUSG00000092710.3 | 0 | NA | NA | NA | NA | NA | #VALUE! |
| ENSMUSG00000092713.3 | 0 | NA | NA | NA | NA | NA | #VALUE! |
| ENSMUSG00000092727.3 | 0 | NA | NA | NA | NA | NA | #VALUE! |
| ENSMUSG00000092730.3 | 0 | NA | NA | NA | NA | NA | #VALUE! |
| ENSMUSG00000092734.3 | 0 | NA | NA | NA | NA | NA | #VALUE! |
| ENSMUSG00000092738.3 | 0 | NA | NA | NA | NA | NA | #VALUE! |
| ENSMUSG00000092741.4 | 0 | NA | NA | NA | NA | NA | #VALUE! |
| ENSMUSG00000092743.3 | 0 | NA | NA | NA | NA | NA | #VALUE! |
| ENSMUSG00000092745.3 | 0 | NA | NA | NA | NA | NA | #VALUE! |
| ENSMUSG00000092746.5 | 0 | NA | NA | NA | NA | NA | #VALUE! |
| ENSMUSG00000092747.4 | 0 | NA | NA | NA | NA | NA | #VALUE! |
| ENSMUSG00000092751.3 | 0 | NA | NA | NA | NA | NA | #VALUE! |
| ENSMUSG00000092752.3 | 0 | NA | NA | NA | NA | NA | #VALUE! |
| ENSMUSG00000092761.2 | 0 | NA | NA | NA | NA | NA | #VALUE! |
| ENSMUSG00000092765.5 | 0 | NA | NA | NA | NA | NA | #VALUE! |
| ENSMUSG00000092770.3 | 0 | NA | NA | NA | NA | NA | #VALUE! |
| ENSMUSG00000092781.3 | 0 | NA | NA | NA | NA | NA | #VALUE! |
| ENSMUSG00000092784.3 | 0 | NA | NA | NA | NA | NA | #VALUE! |
| ENSMUSG00000092787.3 | 0 | NA | NA | NA | NA | NA | #VALUE! |
| ENSMUSG00000092793.3 | 0 | NA | NA | NA | NA | NA | #VALUE! |
| ENSMUSG00000092794.3 | 0 | NA | NA | NA | NA | NA | #VALUE! |
| ENSMUSG00000092805.4 | 0 | NA | NA | NA | NA | NA | #VALUE! |
| ENSMUSG00000092807.3 | 0 | NA | NA | NA | NA | NA | #VALUE! |
| ENSMUSG00000092813.3 | 0 | NA | NA | NA | NA | NA | #VALUE! |
| ENSMUSG00000092815.3 | 0 | NA | NA | NA | NA | NA | #VALUE! |
| ENSMUSG00000092819.3 | 0 | NA | NA | NA | NA | NA | #VALUE! |

| | | | | | | | |
|---|---|---|---|---|---|---|---|
| ENSMUSG00000092827.3 | 0 | NA | NA | NA | NA | NA | #VALUE! |
| ENSMUSG00000092830.3 | 0 | NA | NA | NA | NA | NA | #VALUE! |
| ENSMUSG00000092832.3 | 0 | NA | NA | NA | NA | NA | #VALUE! |
| ENSMUSG00000092837.3 | 0 | NA | NA | NA | NA | NA | #VALUE! |
| ENSMUSG00000092847.3 | 0 | NA | NA | NA | NA | NA | #VALUE! |
| ENSMUSG00000092850.3 | 0 | NA | NA | NA | NA | NA | #VALUE! |
| ENSMUSG00000092852.3 | 0 | NA | NA | NA | NA | NA | #VALUE! |
| ENSMUSG00000092862.2 | 0 | NA | NA | NA | NA | NA | #VALUE! |
| ENSMUSG00000092867.3 | 0 | NA | NA | NA | NA | NA | #VALUE! |
| ENSMUSG00000092868.3 | 0 | NA | NA | NA | NA | NA | #VALUE! |
| ENSMUSG00000092870.3 | 0 | NA | NA | NA | NA | NA | #VALUE! |
| ENSMUSG00000092875.4 | 0 | NA | NA | NA | NA | NA | #VALUE! |
| ENSMUSG00000092879.3 | 0 | NA | NA | NA | NA | NA | #VALUE! |
| ENSMUSG00000092880.2 | 0 | NA | NA | NA | NA | NA | #VALUE! |
| ENSMUSG00000092885.4 | 0 | NA | NA | NA | NA | NA | #VALUE! |
| ENSMUSG00000092887.3 | 0 | NA | NA | NA | NA | NA | #VALUE! |
| ENSMUSG00000092888.3 | 0 | NA | NA | NA | NA | NA | #VALUE! |
| ENSMUSG00000092901.3 | 0 | NA | NA | NA | NA | NA | #VALUE! |
| ENSMUSG00000092905.5 | 0 | NA | NA | NA | NA | NA | #VALUE! |
| ENSMUSG00000092907.3 | 0 | NA | NA | NA | NA | NA | #VALUE! |
| ENSMUSG00000092912.3 | 0 | NA | NA | NA | NA | NA | #VALUE! |
| ENSMUSG00000092921.3 | 0 | NA | NA | NA | NA | NA | #VALUE! |
| ENSMUSG00000092922.3 | 0 | NA | NA | NA | NA | NA | #VALUE! |
| ENSMUSG00000092952.3 | 0 | NA | NA | NA | NA | NA | #VALUE! |
| ENSMUSG00000092958.3 | 0 | NA | NA | NA | NA | NA | #VALUE! |
| ENSMUSG00000092972.3 | 0 | NA | NA | NA | NA | NA | #VALUE! |
| ENSMUSG00000092974.5 | 0 | NA | NA | NA | NA | NA | #VALUE! |
| ENSMUSG00000092981.3 | 0 | NA | NA | NA | NA | NA | #VALUE! |
| ENSMUSG00000092985.3 | 0 | NA | NA | NA | NA | NA | #VALUE! |
| ENSMUSG00000092986.5 | 0 | NA | NA | NA | NA | NA | #VALUE! |
| ENSMUSG00000092988.3 | 0 | NA | NA | NA | NA | NA | #VALUE! |
| ENSMUSG00000092990.3 | 0 | NA | NA | NA | NA | NA | #VALUE! |
| ENSMUSG00000092995.3 | 0 | NA | NA | NA | NA | NA | #VALUE! |
| ENSMUSG00000092998.3 | 0 | NA | NA | NA | NA | NA | #VALUE! |
| ENSMUSG00000092999.3 | 0 | NA | NA | NA | NA | NA | #VALUE! |
| ENSMUSG00000093002.3 | 0 | NA | NA | NA | NA | NA | #VALUE! |
| ENSMUSG00000093003.3 | 0 | NA | NA | NA | NA | NA | #VALUE! |
| ENSMUSG00000093007.3 | 0 | NA | NA | NA | NA | NA | #VALUE! |
| ENSMUSG00000093009.3 | 0 | NA | NA | NA | NA | NA | #VALUE! |
| ENSMUSG00000093011.3 | 0 | NA | NA | NA | NA | NA | #VALUE! |
| ENSMUSG00000093017.3 | 0 | NA | NA | NA | NA | NA | #VALUE! |
| ENSMUSG00000093036.3 | 0 | NA | NA | NA | NA | NA | #VALUE! |
| ENSMUSG00000093037.3 | 0 | NA | NA | NA | NA | NA | #VALUE! |
| ENSMUSG00000093039.3 | 0 | NA | NA | NA | NA | NA | #VALUE! |

| | | | | | | | |
|---|---|---|---|---|---|---|---|
| ENSMUSG00000093042.3 | 0 | NA | NA | NA | NA | NA | #VALUE! |
| ENSMUSG00000093044.3 | 0 | NA | NA | NA | NA | NA | #VALUE! |
| ENSMUSG00000093046.3 | 0 | NA | NA | NA | NA | NA | #VALUE! |
| ENSMUSG00000093056.5 | 0 | NA | NA | NA | NA | NA | #VALUE! |
| ENSMUSG00000093059.5 | 0 | NA | NA | NA | NA | NA | #VALUE! |
| ENSMUSG00000093060.3 | 0 | NA | NA | NA | NA | NA | #VALUE! |
| ENSMUSG00000093061.3 | 0 | NA | NA | NA | NA | NA | #VALUE! |
| ENSMUSG00000093064.5 | 0 | NA | NA | NA | NA | NA | #VALUE! |
| ENSMUSG00000093073.3 | 0 | NA | NA | NA | NA | NA | #VALUE! |
| ENSMUSG00000093076.3 | 0 | NA | NA | NA | NA | NA | #VALUE! |
| ENSMUSG00000093080.3 | 0 | NA | NA | NA | NA | NA | #VALUE! |
| ENSMUSG00000093081.3 | 0 | NA | NA | NA | NA | NA | #VALUE! |
| ENSMUSG00000093087.3 | 0 | NA | NA | NA | NA | NA | #VALUE! |
| ENSMUSG00000093090.3 | 0 | NA | NA | NA | NA | NA | #VALUE! |
| ENSMUSG00000093091.3 | 0 | NA | NA | NA | NA | NA | #VALUE! |
| ENSMUSG00000093093.5 | 0 | NA | NA | NA | NA | NA | #VALUE! |
| ENSMUSG00000093096.3 | 0 | NA | NA | NA | NA | NA | #VALUE! |
| ENSMUSG00000093097.3 | 0 | NA | NA | NA | NA | NA | #VALUE! |
| ENSMUSG00000093100.3 | 0 | NA | NA | NA | NA | NA | #VALUE! |
| ENSMUSG00000093102.3 | 0 | NA | NA | NA | NA | NA | #VALUE! |
| ENSMUSG00000093108.3 | 0 | NA | NA | NA | NA | NA | #VALUE! |
| ENSMUSG00000093112.4 | 0 | NA | NA | NA | NA | NA | #VALUE! |
| ENSMUSG00000093117.3 | 0 | NA | NA | NA | NA | NA | #VALUE! |
| ENSMUSG00000093118.3 | 0 | NA | NA | NA | NA | NA | #VALUE! |
| ENSMUSG00000093120.3 | 0 | NA | NA | NA | NA | NA | #VALUE! |
| ENSMUSG00000093122.3 | 0 | NA | NA | NA | NA | NA | #VALUE! |
| ENSMUSG00000093137.3 | 0 | NA | NA | NA | NA | NA | #VALUE! |
| ENSMUSG00000093139.3 | 0 | NA | NA | NA | NA | NA | #VALUE! |
| ENSMUSG00000093140.3 | 0 | NA | NA | NA | NA | NA | #VALUE! |
| ENSMUSG00000093142.3 | 0 | NA | NA | NA | NA | NA | #VALUE! |
| ENSMUSG00000093147.3 | 0 | NA | NA | NA | NA | NA | #VALUE! |
| ENSMUSG00000093156.3 | 0 | NA | NA | NA | NA | NA | #VALUE! |
| ENSMUSG00000093157.3 | 0 | NA | NA | NA | NA | NA | #VALUE! |
| ENSMUSG00000093167.3 | 0 | NA | NA | NA | NA | NA | #VALUE! |
| ENSMUSG00000093169.3 | 0 | NA | NA | NA | NA | NA | #VALUE! |
| ENSMUSG00000093172.3 | 0 | NA | NA | NA | NA | NA | #VALUE! |
| ENSMUSG00000093176.3 | 0 | NA | NA | NA | NA | NA | #VALUE! |
| ENSMUSG00000093178.3 | 0 | NA | NA | NA | NA | NA | #VALUE! |
| ENSMUSG00000093183.5 | 0 | NA | NA | NA | NA | NA | #VALUE! |
| ENSMUSG00000093187.3 | 0 | NA | NA | NA | NA | NA | #VALUE! |
| ENSMUSG00000093189.4 | 0 | NA | NA | NA | NA | NA | #VALUE! |
| ENSMUSG00000093191.5 | 0 | NA | NA | NA | NA | NA | #VALUE! |
| ENSMUSG00000093202.4 | 0 | NA | NA | NA | NA | NA | #VALUE! |
| ENSMUSG00000093215.3 | 0 | NA | NA | NA | NA | NA | #VALUE! |

| | | | | | | | |
|---|---|---|---|---|---|---|---|
| ENSMUSG00000093219.3 | 0 | NA | NA | NA | NA | NA | #VALUE! |
| ENSMUSG00000093221.3 | 0 | NA | NA | NA | NA | NA | #VALUE! |
| ENSMUSG00000093227.3 | 0 | NA | NA | NA | NA | NA | #VALUE! |
| ENSMUSG00000093234.3 | 0 | NA | NA | NA | NA | NA | #VALUE! |
| ENSMUSG00000093238.3 | 0 | NA | NA | NA | NA | NA | #VALUE! |
| ENSMUSG00000093241.3 | 0 | NA | NA | NA | NA | NA | #VALUE! |
| ENSMUSG00000093242.4 | 0 | NA | NA | NA | NA | NA | #VALUE! |
| ENSMUSG00000093246.3 | 0 | NA | NA | NA | NA | NA | #VALUE! |
| ENSMUSG00000093249.4 | 0 | NA | NA | NA | NA | NA | #VALUE! |
| ENSMUSG00000093254.3 | 0 | NA | NA | NA | NA | NA | #VALUE! |
| ENSMUSG00000093258.4 | 0 | NA | NA | NA | NA | NA | #VALUE! |
| ENSMUSG00000093269.3 | 0 | NA | NA | NA | NA | NA | #VALUE! |
| ENSMUSG00000093270.3 | 0 | NA | NA | NA | NA | NA | #VALUE! |
| ENSMUSG00000093273.4 | 0 | NA | NA | NA | NA | NA | #VALUE! |
| ENSMUSG00000093277.3 | 1.009701 | 3.503687 | 3.950852 | 0.886818 | 0.375177 | NA | #VALUE! |
| ENSMUSG00000093278.3 | 0 | NA | NA | NA | NA | NA | #VALUE! |
| ENSMUSG00000093280.3 | 0 | NA | NA | NA | NA | NA | #VALUE! |
| ENSMUSG00000093283.4 | 0 | NA | NA | NA | NA | NA | #VALUE! |
| ENSMUSG00000093289.3 | 0 | NA | NA | NA | NA | NA | #VALUE! |
| ENSMUSG00000093290.3 | 0 | NA | NA | NA | NA | NA | #VALUE! |
| ENSMUSG00000093291.4 | 0 | NA | NA | NA | NA | NA | #VALUE! |
| ENSMUSG00000093292.3 | 0 | NA | NA | NA | NA | NA | #VALUE! |
| ENSMUSG00000093293.2 | 0 | NA | NA | NA | NA | NA | #VALUE! |
| ENSMUSG00000093296.3 | 0 | NA | NA | NA | NA | NA | #VALUE! |
| ENSMUSG00000093298.3 | 0 | NA | NA | NA | NA | NA | #VALUE! |
| ENSMUSG00000093311.3 | 0 | NA | NA | NA | NA | NA | #VALUE! |
| ENSMUSG00000093314.3 | 0.978946 | -3.48749 | 3.952505 | -0.88235 | 0.377588 | NA | #VALUE! |
| ENSMUSG00000093315.3 | 0 | NA | NA | NA | NA | NA | #VALUE! |
| ENSMUSG00000093325.3 | 0 | NA | NA | NA | NA | NA | #VALUE! |
| ENSMUSG00000093327.3 | 0 | NA | NA | NA | NA | NA | #VALUE! |
| ENSMUSG00000093336.3 | 0 | NA | NA | NA | NA | NA | #VALUE! |
| ENSMUSG00000093338.3 | 0 | NA | NA | NA | NA | NA | #VALUE! |
| ENSMUSG00000093345.5 | 0 | NA | NA | NA | NA | NA | #VALUE! |
| ENSMUSG00000093348.3 | 0 | NA | NA | NA | NA | NA | #VALUE! |
| ENSMUSG00000093351.3 | 0 | NA | NA | NA | NA | NA | #VALUE! |
| ENSMUSG00000093354.3 | 0 | NA | NA | NA | NA | NA | #VALUE! |
| ENSMUSG00000093355.3 | 0 | NA | NA | NA | NA | NA | #VALUE! |
| ENSMUSG00000093358.3 | 0 | NA | NA | NA | NA | NA | #VALUE! |
| ENSMUSG00000093376.2 | 0 | NA | NA | NA | NA | NA | #VALUE! |
| ENSMUSG00000093378.3 | 0 | NA | NA | NA | NA | NA | #VALUE! |
| ENSMUSG00000093380.2 | 0 | NA | NA | NA | NA | NA | #VALUE! |
| ENSMUSG00000093381.2 | 0 | NA | NA | NA | NA | NA | #VALUE! |
| ENSMUSG00000093382.3 | 0 | NA | NA | NA | NA | NA | #VALUE! |
| ENSMUSG00000093383.2 | 0 | NA | NA | NA | NA | NA | #VALUE! |

| | | | | | | | |
|---|---|---|---|---|---|---|---|
| ENSMUSG00000093384.2 | 0 | NA | NA | NA | NA | NA | #VALUE! |
| ENSMUSG00000093385.4 | 0 | NA | NA | NA | NA | NA | #VALUE! |
| ENSMUSG00000093386.2 | 0 | NA | NA | NA | NA | NA | #VALUE! |
| ENSMUSG00000093387.2 | 0 | NA | NA | NA | NA | NA | #VALUE! |
| ENSMUSG00000093390.2 | 0 | NA | NA | NA | NA | NA | #VALUE! |
| ENSMUSG00000093392.2 | 0.195789 | -1.05581 | 4.073302 | -0.2592 | 0.795479 | NA | #VALUE! |
| ENSMUSG00000093394.2 | 2.713846 | 4.924714 | 3.92335 | 1.255232 | 0.209395 | NA | #VALUE! |
| ENSMUSG00000093395.2 | 0 | NA | NA | NA | NA | NA | #VALUE! |
| ENSMUSG00000093397.2 | 0 | NA | NA | NA | NA | NA | #VALUE! |
| ENSMUSG00000093398.2 | 0 | NA | NA | NA | NA | NA | #VALUE! |
| ENSMUSG00000093400.2 | 0 | NA | NA | NA | NA | NA | #VALUE! |
| ENSMUSG00000093401.3 | 0 | NA | NA | NA | NA | NA | #VALUE! |
| ENSMUSG00000093402.2 | 0 | NA | NA | NA | NA | NA | #VALUE! |
| ENSMUSG00000093403.3 | 0 | NA | NA | NA | NA | NA | #VALUE! |
| ENSMUSG00000093405.2 | 0 | NA | NA | NA | NA | NA | #VALUE! |
| ENSMUSG00000093406.2 | 0 | NA | NA | NA | NA | NA | #VALUE! |
| ENSMUSG00000093408.3 | 0 | NA | NA | NA | NA | NA | #VALUE! |
| ENSMUSG00000093410.2 | 0 | NA | NA | NA | NA | NA | #VALUE! |
| ENSMUSG00000093413.3 | 0 | NA | NA | NA | NA | NA | #VALUE! |
| ENSMUSG00000093416.2 | 0 | NA | NA | NA | NA | NA | #VALUE! |
| ENSMUSG00000093419.2 | 0 | NA | NA | NA | NA | NA | #VALUE! |
| ENSMUSG00000093420.2 | 0 | NA | NA | NA | NA | NA | #VALUE! |
| ENSMUSG00000093421.2 | 0 | NA | NA | NA | NA | NA | #VALUE! |
| ENSMUSG00000093424.4 | 0 | NA | NA | NA | NA | NA | #VALUE! |
| ENSMUSG00000093426.2 | 0 | NA | NA | NA | NA | NA | #VALUE! |
| ENSMUSG00000093427.2 | 0 | NA | NA | NA | NA | NA | #VALUE! |
| ENSMUSG00000093429.2 | 0 | NA | NA | NA | NA | NA | #VALUE! |
| ENSMUSG00000093433.2 | 0 | NA | NA | NA | NA | NA | #VALUE! |
| ENSMUSG00000093434.4 | 0 | NA | NA | NA | NA | NA | #VALUE! |
| ENSMUSG00000093436.7 | 0 | NA | NA | NA | NA | NA | #VALUE! |
| ENSMUSG00000093437.2 | 0 | NA | NA | NA | NA | NA | #VALUE! |
| ENSMUSG00000093441.3 | 0 | NA | NA | NA | NA | NA | #VALUE! |
| ENSMUSG00000093443.3 | 0 | NA | NA | NA | NA | NA | #VALUE! |
| ENSMUSG00000093444.2 | 0 | NA | NA | NA | NA | NA | #VALUE! |
| ENSMUSG00000093445.8 | 0 | NA | NA | NA | NA | NA | #VALUE! |
| ENSMUSG00000093446.2 | 0 | NA | NA | NA | NA | NA | #VALUE! |
| ENSMUSG00000093447.2 | 0 | NA | NA | NA | NA | NA | #VALUE! |
| ENSMUSG00000093449.2 | 0 | NA | NA | NA | NA | NA | #VALUE! |
| ENSMUSG00000093452.5 | 0 | NA | NA | NA | NA | NA | #VALUE! |
| ENSMUSG00000093453.2 | 0 | NA | NA | NA | NA | NA | #VALUE! |
| ENSMUSG00000093454.2 | 0 | NA | NA | NA | NA | NA | #VALUE! |
| ENSMUSG00000093456.8 | 0 | NA | NA | NA | NA | NA | #VALUE! |
| ENSMUSG00000093458.2 | 0 | NA | NA | NA | NA | NA | #VALUE! |
| ENSMUSG00000093460.10 | 0 | NA | NA | NA | NA | NA | #VALUE! |

| | | | | | | | |
|---|---|---|---|---|---|---|---|
| ENSMUSG00000093461.2 | 0 | NA | NA | NA | NA | NA | #VALUE! |
| ENSMUSG00000093465.2 | 0 | NA | NA | NA | NA | NA | #VALUE! |
| ENSMUSG00000093467.2 | 0.612332 | -2.79827 | 3.980109 | -0.70306 | 0.482016 | NA | #VALUE! |
| ENSMUSG00000093469.2 | 0 | NA | NA | NA | NA | NA | #VALUE! |
| ENSMUSG00000093472.2 | 0 | NA | NA | NA | NA | NA | #VALUE! |
| ENSMUSG00000093473.2 | 0 | NA | NA | NA | NA | NA | #VALUE! |
| ENSMUSG00000093474.2 | 0 | NA | NA | NA | NA | NA | #VALUE! |
| ENSMUSG00000093479.2 | 0 | NA | NA | NA | NA | NA | #VALUE! |
| ENSMUSG00000093480.3 | 0 | NA | NA | NA | NA | NA | #VALUE! |
| ENSMUSG00000093482.9 | 0 | NA | NA | NA | NA | NA | #VALUE! |
| ENSMUSG00000093483.2 | 0 | NA | NA | NA | NA | NA | #VALUE! |
| ENSMUSG00000093484.2 | 0 | NA | NA | NA | NA | NA | #VALUE! |
| ENSMUSG00000093485.2 | 0 | NA | NA | NA | NA | NA | #VALUE! |
| ENSMUSG00000093487.3 | 0 | NA | NA | NA | NA | NA | #VALUE! |
| ENSMUSG00000093489.2 | 0 | NA | NA | NA | NA | NA | #VALUE! |
| ENSMUSG00000093490.3 | 0 | NA | NA | NA | NA | NA | #VALUE! |
| ENSMUSG00000093492.2 | 0 | NA | NA | NA | NA | NA | #VALUE! |
| ENSMUSG00000093493.2 | 0 | NA | NA | NA | NA | NA | #VALUE! |
| ENSMUSG00000093497.2 | 0 | NA | NA | NA | NA | NA | #VALUE! |
| ENSMUSG00000093498.3 | 0 | NA | NA | NA | NA | NA | #VALUE! |
| ENSMUSG00000093502.3 | 0 | NA | NA | NA | NA | NA | #VALUE! |
| ENSMUSG00000093503.2 | 0 | NA | NA | NA | NA | NA | #VALUE! |
| ENSMUSG00000093504.3 | 0 | NA | NA | NA | NA | NA | #VALUE! |
| ENSMUSG00000093505.2 | 0 | NA | NA | NA | NA | NA | #VALUE! |
| ENSMUSG00000093507.2 | 1.962848 | -0.01381 | 3.820788 | -0.00361 | 0.997116 | NA | #VALUE! |
| ENSMUSG00000093508.3 | 0 | NA | NA | NA | NA | NA | #VALUE! |
| ENSMUSG00000093509.2 | 0 | NA | NA | NA | NA | NA | #VALUE! |
| ENSMUSG00000093510.2 | 0 | NA | NA | NA | NA | NA | #VALUE! |
| ENSMUSG00000093513.2 | 0 | NA | NA | NA | NA | NA | #VALUE! |
| ENSMUSG00000093515.3 | 0 | NA | NA | NA | NA | NA | #VALUE! |
| ENSMUSG00000093516.2 | 0 | NA | NA | NA | NA | NA | #VALUE! |
| ENSMUSG00000093518.2 | 0 | NA | NA | NA | NA | NA | #VALUE! |
| ENSMUSG00000093520.3 | 0 | NA | NA | NA | NA | NA | #VALUE! |
| ENSMUSG00000093522.2 | 0 | NA | NA | NA | NA | NA | #VALUE! |
| ENSMUSG00000093525.2 | 0 | NA | NA | NA | NA | NA | #VALUE! |
| ENSMUSG00000093528.2 | 0 | NA | NA | NA | NA | NA | #VALUE! |
| ENSMUSG00000093530.2 | 0 | NA | NA | NA | NA | NA | #VALUE! |
| ENSMUSG00000093531.4 | 0 | NA | NA | NA | NA | NA | #VALUE! |
| ENSMUSG00000093533.4 | 0 | NA | NA | NA | NA | NA | #VALUE! |
| ENSMUSG00000093534.2 | 0 | NA | NA | NA | NA | NA | #VALUE! |
| ENSMUSG00000093535.2 | 0 | NA | NA | NA | NA | NA | #VALUE! |
| ENSMUSG00000093536.8 | 0 | NA | NA | NA | NA | NA | #VALUE! |
| ENSMUSG00000093537.2 | 0 | NA | NA | NA | NA | NA | #VALUE! |
| ENSMUSG00000093538.3 | 0 | NA | NA | NA | NA | NA | #VALUE! |

| | | | | | | | |
|---|---|---|---|---|---|---|---|
| ENSMUSG00000093539.3 | 0 | NA | NA | NA | NA | NA | #VALUE! |
| ENSMUSG00000093540.3 | 0 | NA | NA | NA | NA | NA | #VALUE! |
| ENSMUSG00000093541.2 | 0 | NA | NA | NA | NA | NA | #VALUE! |
| ENSMUSG00000093543.2 | 0 | NA | NA | NA | NA | NA | #VALUE! |
| ENSMUSG00000093544.2 | 0 | NA | NA | NA | NA | NA | #VALUE! |
| ENSMUSG00000093545.2 | 0 | NA | NA | NA | NA | NA | #VALUE! |
| ENSMUSG00000093546.2 | 0 | NA | NA | NA | NA | NA | #VALUE! |
| ENSMUSG00000093547.2 | 0 | NA | NA | NA | NA | NA | #VALUE! |
| ENSMUSG00000093548.2 | 0 | NA | NA | NA | NA | NA | #VALUE! |
| ENSMUSG00000093549.3 | 0 | NA | NA | NA | NA | NA | #VALUE! |
| ENSMUSG00000093550.8 | 0 | NA | NA | NA | NA | NA | #VALUE! |
| ENSMUSG00000093551.2 | 0 | NA | NA | NA | NA | NA | #VALUE! |
| ENSMUSG00000093552.2 | 0 | NA | NA | NA | NA | NA | #VALUE! |
| ENSMUSG00000093553.2 | 0 | NA | NA | NA | NA | NA | #VALUE! |
| ENSMUSG00000093554.2 | 0 | NA | NA | NA | NA | NA | #VALUE! |
| ENSMUSG00000093556.2 | 0 | NA | NA | NA | NA | NA | #VALUE! |
| ENSMUSG00000093557.2 | 0 | NA | NA | NA | NA | NA | #VALUE! |
| ENSMUSG00000093559.2 | 0 | NA | NA | NA | NA | NA | #VALUE! |
| ENSMUSG00000093560.3 | 0 | NA | NA | NA | NA | NA | #VALUE! |
| ENSMUSG00000093562.2 | 0 | NA | NA | NA | NA | NA | #VALUE! |
| ENSMUSG00000093563.2 | 0 | NA | NA | NA | NA | NA | #VALUE! |
| ENSMUSG00000093564.2 | 0 | NA | NA | NA | NA | NA | #VALUE! |
| ENSMUSG00000093567.2 | 0 | NA | NA | NA | NA | NA | #VALUE! |
| ENSMUSG00000093568.2 | 0 | NA | NA | NA | NA | NA | #VALUE! |
| ENSMUSG00000093570.4 | 0 | NA | NA | NA | NA | NA | #VALUE! |
| ENSMUSG00000093573.2 | 0 | NA | NA | NA | NA | NA | #VALUE! |
| ENSMUSG00000093574.2 | 0 | NA | NA | NA | NA | NA | #VALUE! |
| ENSMUSG00000093575.2 | 0 | NA | NA | NA | NA | NA | #VALUE! |
| ENSMUSG00000093577.2 | 9.165172 | -6.72324 | 2.913589 | -2.30755 | 0.021024 | NA | #VALUE! |
| ENSMUSG00000093579.2 | 0 | NA | NA | NA | NA | NA | #VALUE! |
| ENSMUSG00000093580.2 | 0 | NA | NA | NA | NA | NA | #VALUE! |
| ENSMUSG00000093581.2 | 0 | NA | NA | NA | NA | NA | #VALUE! |
| ENSMUSG00000093584.2 | 0 | NA | NA | NA | NA | NA | #VALUE! |
| ENSMUSG00000093587.4 | 0 | NA | NA | NA | NA | NA | #VALUE! |
| ENSMUSG00000093588.2 | 0 | NA | NA | NA | NA | NA | #VALUE! |
| ENSMUSG00000093589.2 | 0 | NA | NA | NA | NA | NA | #VALUE! |
| ENSMUSG00000093590.2 | 0 | NA | NA | NA | NA | NA | #VALUE! |
| ENSMUSG00000093592.2 | 0 | NA | NA | NA | NA | NA | #VALUE! |
| ENSMUSG00000093593.2 | 0 | NA | NA | NA | NA | NA | #VALUE! |
| ENSMUSG00000093594.2 | 0 | NA | NA | NA | NA | NA | #VALUE! |
| ENSMUSG00000093595.2 | 0 | NA | NA | NA | NA | NA | #VALUE! |
| ENSMUSG00000093598.2 | 0.279766 | 1.808031 | 4.036658 | 0.447903 | 0.654223 | NA | #VALUE! |
| ENSMUSG00000093599.3 | 0 | NA | NA | NA | NA | NA | #VALUE! |
| ENSMUSG00000093601.3 | 0 | NA | NA | NA | NA | NA | #VALUE! |

| | | | | | | | |
|---|---|---|---|---|---|---|---|
| ENSMUSG00000093602.3 | 0 | NA | NA | NA | NA | NA | #VALUE! |
| ENSMUSG00000093606.2 | 0.210026 | -1.52494 | 4.058692 | -0.37572 | 0.707123 | NA | #VALUE! |
| ENSMUSG00000093607.2 | 0 | NA | NA | NA | NA | NA | #VALUE! |
| ENSMUSG00000093610.8 | 0 | NA | NA | NA | NA | NA | #VALUE! |
| ENSMUSG00000093614.2 | 0 | NA | NA | NA | NA | NA | #VALUE! |
| ENSMUSG00000093615.3 | 0 | NA | NA | NA | NA | NA | #VALUE! |
| ENSMUSG00000093616.2 | 0 | NA | NA | NA | NA | NA | #VALUE! |
| ENSMUSG00000093617.2 | 0 | NA | NA | NA | NA | NA | #VALUE! |
| ENSMUSG00000093618.3 | 0 | NA | NA | NA | NA | NA | #VALUE! |
| ENSMUSG00000093619.2 | 0 | NA | NA | NA | NA | NA | #VALUE! |
| ENSMUSG00000093621.2 | 0 | NA | NA | NA | NA | NA | #VALUE! |
| ENSMUSG00000093623.2 | 0 | NA | NA | NA | NA | NA | #VALUE! |
| ENSMUSG00000093624.2 | 0 | NA | NA | NA | NA | NA | #VALUE! |
| ENSMUSG00000093625.2 | 0 | NA | NA | NA | NA | NA | #VALUE! |
| ENSMUSG00000093626.2 | 0 | NA | NA | NA | NA | NA | #VALUE! |
| ENSMUSG00000093627.3 | 0 | NA | NA | NA | NA | NA | #VALUE! |
| ENSMUSG00000093628.2 | 0 | NA | NA | NA | NA | NA | #VALUE! |
| ENSMUSG00000093629.2 | 0 | NA | NA | NA | NA | NA | #VALUE! |
| ENSMUSG00000093634.2 | 0 | NA | NA | NA | NA | NA | #VALUE! |
| ENSMUSG00000093637.2 | 0 | NA | NA | NA | NA | NA | #VALUE! |
| ENSMUSG00000093639.5 | 0 | NA | NA | NA | NA | NA | #VALUE! |
| ENSMUSG00000093642.2 | 0 | NA | NA | NA | NA | NA | #VALUE! |
| ENSMUSG00000093644.2 | 0 | NA | NA | NA | NA | NA | #VALUE! |
| ENSMUSG00000093645.3 | 0 | NA | NA | NA | NA | NA | #VALUE! |
| ENSMUSG00000093648.2 | 0 | NA | NA | NA | NA | NA | #VALUE! |
| ENSMUSG00000093649.2 | 0 | NA | NA | NA | NA | NA | #VALUE! |
| ENSMUSG00000093650.3 | 0 | NA | NA | NA | NA | NA | #VALUE! |
| ENSMUSG00000093651.2 | 0.366552 | 0.755301 | 4.038805 | 0.187011 | 0.851652 | NA | #VALUE! |
| ENSMUSG00000093652.3 | 0 | NA | NA | NA | NA | NA | #VALUE! |
| ENSMUSG00000093656.2 | 0 | NA | NA | NA | NA | NA | #VALUE! |
| ENSMUSG00000093658.2 | 0 | NA | NA | NA | NA | NA | #VALUE! |
| ENSMUSG00000093660.3 | 0 | NA | NA | NA | NA | NA | #VALUE! |
| ENSMUSG00000093662.3 | 0 | NA | NA | NA | NA | NA | #VALUE! |
| ENSMUSG00000093663.2 | 0 | NA | NA | NA | NA | NA | #VALUE! |
| ENSMUSG00000093664.3 | 0 | NA | NA | NA | NA | NA | #VALUE! |
| ENSMUSG00000093665.3 | 0 | NA | NA | NA | NA | NA | #VALUE! |
| ENSMUSG00000093666.3 | 0 | NA | NA | NA | NA | NA | #VALUE! |
| ENSMUSG00000093668.4 | 8.368541 | -6.59301 | 3.815764 | -1.72783 | 0.084018 | NA | #VALUE! |
| ENSMUSG00000093669.2 | 0 | NA | NA | NA | NA | NA | #VALUE! |
| ENSMUSG00000093671.2 | 0 | NA | NA | NA | NA | NA | #VALUE! |
| ENSMUSG00000093672.3 | 0 | NA | NA | NA | NA | NA | #VALUE! |
| ENSMUSG00000093673.2 | 0 | NA | NA | NA | NA | NA | #VALUE! |
| ENSMUSG00000093675.8 | 0 | NA | NA | NA | NA | NA | #VALUE! |
| ENSMUSG00000093677.2 | 0.195789 | -1.05581 | 4.073302 | -0.2592 | 0.795479 | NA | #VALUE! |

| | | | | | | | |
|---|---|---|---|---|---|---|---|
| ENSMUSG00000093681.2 | 0 | NA | NA | NA | NA | NA | #VALUE! |
| ENSMUSG00000093682.2 | 0 | NA | NA | NA | NA | NA | #VALUE! |
| ENSMUSG00000093683.3 | 0 | NA | NA | NA | NA | NA | #VALUE! |
| ENSMUSG00000093685.2 | 0 | NA | NA | NA | NA | NA | #VALUE! |
| ENSMUSG00000093686.2 | 0.18651 | 1.401769 | 4.056354 | 0.345574 | 0.729663 | NA | #VALUE! |
| ENSMUSG00000093687.2 | 0 | NA | NA | NA | NA | NA | #VALUE! |
| ENSMUSG00000093688.2 | 0 | NA | NA | NA | NA | NA | #VALUE! |
| ENSMUSG00000093689.2 | 0 | NA | NA | NA | NA | NA | #VALUE! |
| ENSMUSG00000093690.3 | 0 | NA | NA | NA | NA | NA | #VALUE! |
| ENSMUSG00000093695.3 | 0 | NA | NA | NA | NA | NA | #VALUE! |
| ENSMUSG00000093696.2 | 0 | NA | NA | NA | NA | NA | #VALUE! |
| ENSMUSG00000093697.2 | 0 | NA | NA | NA | NA | NA | #VALUE! |
| ENSMUSG00000093699.2 | 0 | NA | NA | NA | NA | NA | #VALUE! |
| ENSMUSG00000093701.8 | 0 | NA | NA | NA | NA | NA | #VALUE! |
| ENSMUSG00000093702.2 | 0 | NA | NA | NA | NA | NA | #VALUE! |
| ENSMUSG00000093703.2 | 0 | NA | NA | NA | NA | NA | #VALUE! |
| ENSMUSG00000093705.2 | 0 | NA | NA | NA | NA | NA | #VALUE! |
| ENSMUSG00000093707.2 | 0 | NA | NA | NA | NA | NA | #VALUE! |
| ENSMUSG00000093708.2 | 0 | NA | NA | NA | NA | NA | #VALUE! |
| ENSMUSG00000093710.3 | 0 | NA | NA | NA | NA | NA | #VALUE! |
| ENSMUSG00000093711.2 | 0 | NA | NA | NA | NA | NA | #VALUE! |
| ENSMUSG00000093714.2 | 0 | NA | NA | NA | NA | NA | #VALUE! |
| ENSMUSG00000093716.2 | 0 | NA | NA | NA | NA | NA | #VALUE! |
| ENSMUSG00000093719.2 | 0 | NA | NA | NA | NA | NA | #VALUE! |
| ENSMUSG00000093720.2 | 0 | NA | NA | NA | NA | NA | #VALUE! |
| ENSMUSG00000093721.2 | 0 | NA | NA | NA | NA | NA | #VALUE! |
| ENSMUSG00000093722.2 | 0 | NA | NA | NA | NA | NA | #VALUE! |
| ENSMUSG00000093724.2 | 0 | NA | NA | NA | NA | NA | #VALUE! |
| ENSMUSG00000093725.2 | 0 | NA | NA | NA | NA | NA | #VALUE! |
| ENSMUSG00000093726.2 | 0 | NA | NA | NA | NA | NA | #VALUE! |
| ENSMUSG00000093727.2 | 0 | NA | NA | NA | NA | NA | #VALUE! |
| ENSMUSG00000093728.2 | 0 | NA | NA | NA | NA | NA | #VALUE! |
| ENSMUSG00000093729.2 | 0 | NA | NA | NA | NA | NA | #VALUE! |
| ENSMUSG00000093730.2 | 0 | NA | NA | NA | NA | NA | #VALUE! |
| ENSMUSG00000093731.2 | 0 | NA | NA | NA | NA | NA | #VALUE! |
| ENSMUSG00000093732.2 | 0 | NA | NA | NA | NA | NA | #VALUE! |
| ENSMUSG00000093736.2 | 0 | NA | NA | NA | NA | NA | #VALUE! |
| ENSMUSG00000093738.3 | 0 | NA | NA | NA | NA | NA | #VALUE! |
| ENSMUSG00000093739.2 | 0 | NA | NA | NA | NA | NA | #VALUE! |
| ENSMUSG00000093740.2 | 0 | NA | NA | NA | NA | NA | #VALUE! |
| ENSMUSG00000093741.2 | 0 | NA | NA | NA | NA | NA | #VALUE! |
| ENSMUSG00000093743.2 | 0 | NA | NA | NA | NA | NA | #VALUE! |
| ENSMUSG00000093744.2 | 0 | NA | NA | NA | NA | NA | #VALUE! |
| ENSMUSG00000093746.3 | 0 | NA | NA | NA | NA | NA | #VALUE! |

| | | | | | | | |
|---|---|---|---|---|---|---|---|
| ENSMUSG00000093747.2 | 0 | NA | NA | NA | NA | NA | #VALUE! |
| ENSMUSG00000093748.2 | 0 | NA | NA | NA | NA | NA | #VALUE! |
| ENSMUSG00000093749.2 | 0 | NA | NA | NA | NA | NA | #VALUE! |
| ENSMUSG00000093751.2 | 0 | NA | NA | NA | NA | NA | #VALUE! |
| ENSMUSG00000093752.2 | 0 | NA | NA | NA | NA | NA | #VALUE! |
| ENSMUSG00000093753.2 | 0 | NA | NA | NA | NA | NA | #VALUE! |
| ENSMUSG00000093755.3 | 0 | NA | NA | NA | NA | NA | #VALUE! |
| ENSMUSG00000093756.2 | 0 | NA | NA | NA | NA | NA | #VALUE! |
| ENSMUSG00000093757.3 | 0 | NA | NA | NA | NA | NA | #VALUE! |
| ENSMUSG00000093758.2 | 0 | NA | NA | NA | NA | NA | #VALUE! |
| ENSMUSG00000093759.2 | 0 | NA | NA | NA | NA | NA | #VALUE! |
| ENSMUSG00000093761.2 | 0 | NA | NA | NA | NA | NA | #VALUE! |
| ENSMUSG00000093763.3 | 0 | NA | NA | NA | NA | NA | #VALUE! |
| ENSMUSG00000093765.8 | 0 | NA | NA | NA | NA | NA | #VALUE! |
| ENSMUSG00000093766.2 | 0 | NA | NA | NA | NA | NA | #VALUE! |
| ENSMUSG00000093767.3 | 0 | NA | NA | NA | NA | NA | #VALUE! |
| ENSMUSG00000093768.2 | 0 | NA | NA | NA | NA | NA | #VALUE! |
| ENSMUSG00000093769.5 | 0 | NA | NA | NA | NA | NA | #VALUE! |
| ENSMUSG00000093770.2 | 0 | NA | NA | NA | NA | NA | #VALUE! |
| ENSMUSG00000093771.2 | 0 | NA | NA | NA | NA | NA | #VALUE! |
| ENSMUSG00000093772.2 | 0 | NA | NA | NA | NA | NA | #VALUE! |
| ENSMUSG00000093773.2 | 0 | NA | NA | NA | NA | NA | #VALUE! |
| ENSMUSG00000093774.2 | 0 | NA | NA | NA | NA | NA | #VALUE! |
| ENSMUSG00000093775.3 | 0 | NA | NA | NA | NA | NA | #VALUE! |
| ENSMUSG00000093776.2 | 0 | NA | NA | NA | NA | NA | #VALUE! |
| ENSMUSG00000093779.2 | 0 | NA | NA | NA | NA | NA | #VALUE! |
| ENSMUSG00000093781.3 | 0 | NA | NA | NA | NA | NA | #VALUE! |
| ENSMUSG00000093784.3 | 0 | NA | NA | NA | NA | NA | #VALUE! |
| ENSMUSG00000093788.3 | 0 | NA | NA | NA | NA | NA | #VALUE! |
| ENSMUSG00000093789.4 | 0 | NA | NA | NA | NA | NA | #VALUE! |
| ENSMUSG00000093796.2 | 0 | NA | NA | NA | NA | NA | #VALUE! |
| ENSMUSG00000093798.3 | 0.336567 | 1.944925 | 4.029936 | 0.482619 | 0.629366 | NA | #VALUE! |
| ENSMUSG00000093800.3 | 0 | NA | NA | NA | NA | NA | #VALUE! |
| ENSMUSG00000093801.3 | 0 | NA | NA | NA | NA | NA | #VALUE! |
| ENSMUSG00000093803.6 | 6.466396 | -1.38463 | 3.761062 | -0.36815 | 0.712762 | NA | #VALUE! |
| ENSMUSG00000093804.4 | 0 | NA | NA | NA | NA | NA | #VALUE! |
| ENSMUSG00000093805.8 | 0 | NA | NA | NA | NA | NA | #VALUE! |
| ENSMUSG00000093806.4 | 0 | NA | NA | NA | NA | NA | #VALUE! |
| ENSMUSG00000093812.2 | 0 | NA | NA | NA | NA | NA | #VALUE! |
| ENSMUSG00000093813.6 | 0 | NA | NA | NA | NA | NA | #VALUE! |
| ENSMUSG00000093814.2 | 0 | NA | NA | NA | NA | NA | #VALUE! |
| ENSMUSG00000093817.8 | 0 | NA | NA | NA | NA | NA | #VALUE! |
| ENSMUSG00000093818.2 | 0 | NA | NA | NA | NA | NA | #VALUE! |
| ENSMUSG00000093820.2 | 0 | NA | NA | NA | NA | NA | #VALUE! |

| | | | | | | | |
|---|---|---|---|---|---|---|---|
| ENSMUSG00000093825.3 | 0 | NA | NA | NA | NA | NA | #VALUE! |
| ENSMUSG00000093826.2 | 0.168284 | 1.027628 | 4.067126 | 0.252667 | 0.800526 | NA | #VALUE! |
| ENSMUSG00000093827.2 | 0 | NA | NA | NA | NA | NA | #VALUE! |
| ENSMUSG00000093833.2 | 0 | NA | NA | NA | NA | NA | #VALUE! |
| ENSMUSG00000093838.2 | 0 | NA | NA | NA | NA | NA | #VALUE! |
| ENSMUSG00000093839.3 | 0 | NA | NA | NA | NA | NA | #VALUE! |
| ENSMUSG00000093846.2 | 0 | NA | NA | NA | NA | NA | #VALUE! |
| ENSMUSG00000093847.3 | 0 | NA | NA | NA | NA | NA | #VALUE! |
| ENSMUSG00000093848.3 | 0 | NA | NA | NA | NA | NA | #VALUE! |
| ENSMUSG00000093849.2 | 0 | NA | NA | NA | NA | NA | #VALUE! |
| ENSMUSG00000093851.2 | 0 | NA | NA | NA | NA | NA | #VALUE! |
| ENSMUSG00000093853.2 | 0 | NA | NA | NA | NA | NA | #VALUE! |
| ENSMUSG00000093858.2 | 0 | NA | NA | NA | NA | NA | #VALUE! |
| ENSMUSG00000093859.3 | 0 | NA | NA | NA | NA | NA | #VALUE! |
| ENSMUSG00000093861.3 | 0 | NA | NA | NA | NA | NA | #VALUE! |
| ENSMUSG00000093862.3 | 0 | NA | NA | NA | NA | NA | #VALUE! |
| ENSMUSG00000093863.2 | 0 | NA | NA | NA | NA | NA | #VALUE! |
| ENSMUSG00000093865.8 | 0 | NA | NA | NA | NA | NA | #VALUE! |
| ENSMUSG00000093866.4 | 0 | NA | NA | NA | NA | NA | #VALUE! |
| ENSMUSG00000093867.8 | 0 | NA | NA | NA | NA | NA | #VALUE! |
| ENSMUSG00000093868.3 | 0 | NA | NA | NA | NA | NA | #VALUE! |
| ENSMUSG00000093871.2 | 0 | NA | NA | NA | NA | NA | #VALUE! |
| ENSMUSG00000093876.2 | 0 | NA | NA | NA | NA | NA | #VALUE! |
| ENSMUSG00000093877.5 | 0 | NA | NA | NA | NA | NA | #VALUE! |
| ENSMUSG00000093880.4 | 0 | NA | NA | NA | NA | NA | #VALUE! |
| ENSMUSG00000093883.2 | 0 | NA | NA | NA | NA | NA | #VALUE! |
| ENSMUSG00000093884.3 | 0 | NA | NA | NA | NA | NA | #VALUE! |
| ENSMUSG00000093885.3 | 3.170677 | 5.164728 | 3.920817 | 1.317258 | 0.187752 | NA | #VALUE! |
| ENSMUSG00000093887.10 | 0 | NA | NA | NA | NA | NA | #VALUE! |
| ENSMUSG00000093889.2 | 0 | NA | NA | NA | NA | NA | #VALUE! |
| ENSMUSG00000093890.2 | 0 | NA | NA | NA | NA | NA | #VALUE! |
| ENSMUSG00000093891.2 | 0 | NA | NA | NA | NA | NA | #VALUE! |
| ENSMUSG00000093892.2 | 0 | NA | NA | NA | NA | NA | #VALUE! |
| ENSMUSG00000093894.2 | 0 | NA | NA | NA | NA | NA | #VALUE! |
| ENSMUSG00000093895.8 | 0 | NA | NA | NA | NA | NA | #VALUE! |
| ENSMUSG00000093896.3 | 0 | NA | NA | NA | NA | NA | #VALUE! |
| ENSMUSG00000093898.3 | 0 | NA | NA | NA | NA | NA | #VALUE! |
| ENSMUSG00000093899.2 | 0 | NA | NA | NA | NA | NA | #VALUE! |
| ENSMUSG00000093901.4 | 0 | NA | NA | NA | NA | NA | #VALUE! |
| ENSMUSG00000093903.7 | 0 | NA | NA | NA | NA | NA | #VALUE! |
| ENSMUSG00000093906.6 | 0 | NA | NA | NA | NA | NA | #VALUE! |
| ENSMUSG00000093908.3 | 0 | NA | NA | NA | NA | NA | #VALUE! |
| ENSMUSG00000093909.3 | 0 | NA | NA | NA | NA | NA | #VALUE! |
| ENSMUSG00000093910.3 | 0 | NA | NA | NA | NA | NA | #VALUE! |

| | | | | | | | |
|---|---|---|---|---|---|---|---|
| ENSMUSG00000093913.2 | 0 | NA | NA | NA | NA | NA | #VALUE! |
| ENSMUSG00000093915.3 | 0 | NA | NA | NA | NA | NA | #VALUE! |
| ENSMUSG00000093916.3 | 5.971511 | -6.11264 | 3.823682 | -1.59863 | 0.109904 | NA | #VALUE! |
| ENSMUSG00000093917.2 | 0 | NA | NA | NA | NA | NA | #VALUE! |
| ENSMUSG00000093918.8 | 0 | NA | NA | NA | NA | NA | #VALUE! |
| ENSMUSG00000093919.2 | 0 | NA | NA | NA | NA | NA | #VALUE! |
| ENSMUSG00000093920.3 | 0 | NA | NA | NA | NA | NA | #VALUE! |
| ENSMUSG00000093922.2 | 0 | NA | NA | NA | NA | NA | #VALUE! |
| ENSMUSG00000093923.8 | 0 | NA | NA | NA | NA | NA | #VALUE! |
| ENSMUSG00000093924.2 | 0 | NA | NA | NA | NA | NA | #VALUE! |
| ENSMUSG00000093926.2 | 0 | NA | NA | NA | NA | NA | #VALUE! |
| ENSMUSG00000093927.3 | 0 | NA | NA | NA | NA | NA | #VALUE! |
| ENSMUSG00000093931.3 | 0 | NA | NA | NA | NA | NA | #VALUE! |
| ENSMUSG00000093934.3 | 0 | NA | NA | NA | NA | NA | #VALUE! |
| ENSMUSG00000093938.3 | 0 | NA | NA | NA | NA | NA | #VALUE! |
| ENSMUSG00000093939.2 | 0 | NA | NA | NA | NA | NA | #VALUE! |
| ENSMUSG00000093940.2 | 0 | NA | NA | NA | NA | NA | #VALUE! |
| ENSMUSG00000093941.2 | 0 | NA | NA | NA | NA | NA | #VALUE! |
| ENSMUSG00000093942.4 | 0 | NA | NA | NA | NA | NA | #VALUE! |
| ENSMUSG00000093945.2 | 0 | NA | NA | NA | NA | NA | #VALUE! |
| ENSMUSG00000093948.8 | 0 | NA | NA | NA | NA | NA | #VALUE! |
| ENSMUSG00000093949.3 | 0 | NA | NA | NA | NA | NA | #VALUE! |
| ENSMUSG00000093950.3 | 0 | NA | NA | NA | NA | NA | #VALUE! |
| ENSMUSG00000093953.3 | 0 | NA | NA | NA | NA | NA | #VALUE! |
| ENSMUSG00000093954.9 | 0 | NA | NA | NA | NA | NA | #VALUE! |
| ENSMUSG00000093955.3 | 0 | NA | NA | NA | NA | NA | #VALUE! |
| ENSMUSG00000093957.2 | 0 | NA | NA | NA | NA | NA | #VALUE! |
| ENSMUSG00000093961.2 | 0 | NA | NA | NA | NA | NA | #VALUE! |
| ENSMUSG00000093962.8 | 0 | NA | NA | NA | NA | NA | #VALUE! |
| ENSMUSG00000093963.2 | 0 | NA | NA | NA | NA | NA | #VALUE! |
| ENSMUSG00000093966.3 | 0 | NA | NA | NA | NA | NA | #VALUE! |
| ENSMUSG00000093968.2 | 0 | NA | NA | NA | NA | NA | #VALUE! |
| ENSMUSG00000093969.3 | 0 | NA | NA | NA | NA | NA | #VALUE! |
| ENSMUSG00000093973.2 | 0 | NA | NA | NA | NA | NA | #VALUE! |
| ENSMUSG00000093975.2 | 0 | NA | NA | NA | NA | NA | #VALUE! |
| ENSMUSG00000093976.3 | 0 | NA | NA | NA | NA | NA | #VALUE! |
| ENSMUSG00000093977.2 | 0 | NA | NA | NA | NA | NA | #VALUE! |
| ENSMUSG00000093978.2 | 0 | NA | NA | NA | NA | NA | #VALUE! |
| ENSMUSG00000093979.9 | 0 | NA | NA | NA | NA | NA | #VALUE! |
| ENSMUSG00000093980.2 | 0 | NA | NA | NA | NA | NA | #VALUE! |
| ENSMUSG00000093984.2 | 0 | NA | NA | NA | NA | NA | #VALUE! |
| ENSMUSG00000093985.8 | 0 | NA | NA | NA | NA | NA | #VALUE! |
| ENSMUSG00000093986.2 | 0 | NA | NA | NA | NA | NA | #VALUE! |
| ENSMUSG00000093987.8 | 0 | NA | NA | NA | NA | NA | #VALUE! |

| | | | | | | | |
|---|---|---|---|---|---|---|---|
| ENSMUSG00000093989.2 | 0 | NA | NA | NA | NA | NA | #VALUE! |
| ENSMUSG00000093990.2 | 0 | NA | NA | NA | NA | NA | #VALUE! |
| ENSMUSG00000093993.8 | 0 | NA | NA | NA | NA | NA | #VALUE! |
| ENSMUSG00000093996.3 | 0 | NA | NA | NA | NA | NA | #VALUE! |
| ENSMUSG00000094001.2 | 0 | NA | NA | NA | NA | NA | #VALUE! |
| ENSMUSG00000094002.4 | 0 | NA | NA | NA | NA | NA | #VALUE! |
| ENSMUSG00000094004.8 | 0 | NA | NA | NA | NA | NA | #VALUE! |
| ENSMUSG00000094005.2 | 0 | NA | NA | NA | NA | NA | #VALUE! |
| ENSMUSG00000094006.3 | 0 | NA | NA | NA | NA | NA | #VALUE! |
| ENSMUSG00000094010.2 | 0 | NA | NA | NA | NA | NA | #VALUE! |
| ENSMUSG00000094011.2 | 0 | NA | NA | NA | NA | NA | #VALUE! |
| ENSMUSG00000094012.2 | 0 | NA | NA | NA | NA | NA | #VALUE! |
| ENSMUSG00000094015.2 | 0 | NA | NA | NA | NA | NA | #VALUE! |
| ENSMUSG00000094016.3 | 0 | NA | NA | NA | NA | NA | #VALUE! |
| ENSMUSG00000094017.2 | 0 | NA | NA | NA | NA | NA | #VALUE! |
| ENSMUSG00000094018.3 | 0 | NA | NA | NA | NA | NA | #VALUE! |
| ENSMUSG00000094021.8 | 0 | NA | NA | NA | NA | NA | #VALUE! |
| ENSMUSG00000094023.4 | 0 | NA | NA | NA | NA | NA | #VALUE! |
| ENSMUSG00000094024.2 | 0 | NA | NA | NA | NA | NA | #VALUE! |
| ENSMUSG00000094025.8 | 0 | NA | NA | NA | NA | NA | #VALUE! |
| ENSMUSG00000094027.3 | 0 | NA | NA | NA | NA | NA | #VALUE! |
| ENSMUSG00000094028.2 | 0 | NA | NA | NA | NA | NA | #VALUE! |
| ENSMUSG00000094029.3 | 0 | NA | NA | NA | NA | NA | #VALUE! |
| ENSMUSG00000094030.2 | 0 | NA | NA | NA | NA | NA | #VALUE! |
| ENSMUSG00000094035.4 | 0 | NA | NA | NA | NA | NA | #VALUE! |
| ENSMUSG00000094036.2 | 0 | NA | NA | NA | NA | NA | #VALUE! |
| ENSMUSG00000094043.3 | 0 | NA | NA | NA | NA | NA | #VALUE! |
| ENSMUSG00000094051.7 | 0 | NA | NA | NA | NA | NA | #VALUE! |
| ENSMUSG00000094052.3 | 0 | NA | NA | NA | NA | NA | #VALUE! |
| ENSMUSG00000094053.3 | 0 | NA | NA | NA | NA | NA | #VALUE! |
| ENSMUSG00000094054.2 | 0 | NA | NA | NA | NA | NA | #VALUE! |
| ENSMUSG00000094057.2 | 0 | NA | NA | NA | NA | NA | #VALUE! |
| ENSMUSG00000094063.5 | 0 | NA | NA | NA | NA | NA | #VALUE! |
| ENSMUSG00000094064.3 | 0 | NA | NA | NA | NA | NA | #VALUE! |
| ENSMUSG00000094065.2 | 0 | NA | NA | NA | NA | NA | #VALUE! |
| ENSMUSG00000094066.2 | 0 | NA | NA | NA | NA | NA | #VALUE! |
| ENSMUSG00000094071.3 | 0 | NA | NA | NA | NA | NA | #VALUE! |
| ENSMUSG00000094072.2 | 0 | NA | NA | NA | NA | NA | #VALUE! |
| ENSMUSG00000094074.2 | 0 | NA | NA | NA | NA | NA | #VALUE! |
| ENSMUSG00000094075.2 | 0 | NA | NA | NA | NA | NA | #VALUE! |
| ENSMUSG00000094076.8 | 0 | NA | NA | NA | NA | NA | #VALUE! |
| ENSMUSG00000094079.2 | 0 | NA | NA | NA | NA | NA | #VALUE! |
| ENSMUSG00000094080.4 | 0 | NA | NA | NA | NA | NA | #VALUE! |
| ENSMUSG00000094081.3 | 0 | NA | NA | NA | NA | NA | #VALUE! |

| | | | | | | | |
|---|---|---|---|---|---|---|---|
| ENSMUSG00000094082.2 | 0 | NA | NA | NA | NA | NA | #VALUE! |
| ENSMUSG00000094083.3 | 0 | NA | NA | NA | NA | NA | #VALUE! |
| ENSMUSG00000094084.3 | 0 | NA | NA | NA | NA | NA | #VALUE! |
| ENSMUSG00000094085.2 | 0 | NA | NA | NA | NA | NA | #VALUE! |
| ENSMUSG00000094087.2 | 0 | NA | NA | NA | NA | NA | #VALUE! |
| ENSMUSG00000094088.3 | 0 | NA | NA | NA | NA | NA | #VALUE! |
| ENSMUSG00000094090.3 | 0.793073 | -0.15398 | 3.943895 | -0.03904 | 0.968856 | NA | #VALUE! |
| ENSMUSG00000094091.4 | 2.745096 | 4.95268 | 3.923033 | 1.262462 | 0.206783 | NA | #VALUE! |
| ENSMUSG00000094092.2 | 0 | NA | NA | NA | NA | NA | #VALUE! |
| ENSMUSG00000094094.3 | 0 | NA | NA | NA | NA | NA | #VALUE! |
| ENSMUSG00000094098.4 | 0 | NA | NA | NA | NA | NA | #VALUE! |
| ENSMUSG00000094102.2 | 0 | NA | NA | NA | NA | NA | #VALUE! |
| ENSMUSG00000094103.3 | 0 | NA | NA | NA | NA | NA | #VALUE! |
| ENSMUSG00000094105.8 | 0 | NA | NA | NA | NA | NA | #VALUE! |
| ENSMUSG00000094108.2 | 0 | NA | NA | NA | NA | NA | #VALUE! |
| ENSMUSG00000094111.3 | 0 | NA | NA | NA | NA | NA | #VALUE! |
| ENSMUSG00000094112.3 | 0 | NA | NA | NA | NA | NA | #VALUE! |
| ENSMUSG00000094113.8 | 0 | NA | NA | NA | NA | NA | #VALUE! |
| ENSMUSG00000094115.3 | 0 | NA | NA | NA | NA | NA | #VALUE! |
| ENSMUSG00000094116.2 | 0 | NA | NA | NA | NA | NA | #VALUE! |
| ENSMUSG00000094117.2 | 0 | NA | NA | NA | NA | NA | #VALUE! |
| ENSMUSG00000094118.3 | 0 | NA | NA | NA | NA | NA | #VALUE! |
| ENSMUSG00000094119.4 | 0 | NA | NA | NA | NA | NA | #VALUE! |
| ENSMUSG00000094120.2 | 0 | NA | NA | NA | NA | NA | #VALUE! |
| ENSMUSG00000094122.2 | 0.408295 | -1.02718 | 4.03053 | -0.25485 | 0.79884 | NA | #VALUE! |
| ENSMUSG00000094124.6 | 0 | NA | NA | NA | NA | NA | #VALUE! |
| ENSMUSG00000094125.2 | 0 | NA | NA | NA | NA | NA | #VALUE! |
| ENSMUSG00000094127.2 | 0 | NA | NA | NA | NA | NA | #VALUE! |
| ENSMUSG00000094132.8 | 0 | NA | NA | NA | NA | NA | #VALUE! |
| ENSMUSG00000094133.3 | 0 | NA | NA | NA | NA | NA | #VALUE! |
| ENSMUSG00000094134.3 | 0 | NA | NA | NA | NA | NA | #VALUE! |
| ENSMUSG00000094135.2 | 0 | NA | NA | NA | NA | NA | #VALUE! |
| ENSMUSG00000094136.3 | 0 | NA | NA | NA | NA | NA | #VALUE! |
| ENSMUSG00000094140.3 | 0 | NA | NA | NA | NA | NA | #VALUE! |
| ENSMUSG00000094144.9 | 0 | NA | NA | NA | NA | NA | #VALUE! |
| ENSMUSG00000094145.3 | 0 | NA | NA | NA | NA | NA | #VALUE! |
| ENSMUSG00000094146.2 | 0 | NA | NA | NA | NA | NA | #VALUE! |
| ENSMUSG00000094149.2 | 0 | NA | NA | NA | NA | NA | #VALUE! |
| ENSMUSG00000094151.2 | 0 | NA | NA | NA | NA | NA | #VALUE! |
| ENSMUSG00000094152.6 | 0 | NA | NA | NA | NA | NA | #VALUE! |
| ENSMUSG00000094153.3 | 0 | NA | NA | NA | NA | NA | #VALUE! |
| ENSMUSG00000094157.8 | 0 | NA | NA | NA | NA | NA | #VALUE! |
| ENSMUSG00000094160.3 | 0 | NA | NA | NA | NA | NA | #VALUE! |
| ENSMUSG00000094161.2 | 0 | NA | NA | NA | NA | NA | #VALUE! |

| | | | | | | | |
|---|---|---|---|---|---|---|---|
| ENSMUSG00000094162.2 | 0 | NA | NA | NA | NA | NA | #VALUE! |
| ENSMUSG00000094163.2 | 0 | NA | NA | NA | NA | NA | #VALUE! |
| ENSMUSG00000094164.3 | 0 | NA | NA | NA | NA | NA | #VALUE! |
| ENSMUSG00000094167.2 | 0 | NA | NA | NA | NA | NA | #VALUE! |
| ENSMUSG00000094168.2 | 0 | NA | NA | NA | NA | NA | #VALUE! |
| ENSMUSG00000094172.2 | 0 | NA | NA | NA | NA | NA | #VALUE! |
| ENSMUSG00000094174.3 | 0 | NA | NA | NA | NA | NA | #VALUE! |
| ENSMUSG00000094176.7 | 0 | NA | NA | NA | NA | NA | #VALUE! |
| ENSMUSG00000094177.2 | 0 | NA | NA | NA | NA | NA | #VALUE! |
| ENSMUSG00000094180.2 | 0 | NA | NA | NA | NA | NA | #VALUE! |
| ENSMUSG00000094181.8 | 0 | NA | NA | NA | NA | NA | #VALUE! |
| ENSMUSG00000094182.3 | 0 | NA | NA | NA | NA | NA | #VALUE! |
| ENSMUSG00000094186.3 | 0 | NA | NA | NA | NA | NA | #VALUE! |
| ENSMUSG00000094187.2 | 0 | NA | NA | NA | NA | NA | #VALUE! |
| ENSMUSG00000094189.2 | 0 | NA | NA | NA | NA | NA | #VALUE! |
| ENSMUSG00000094191.2 | 0 | NA | NA | NA | NA | NA | #VALUE! |
| ENSMUSG00000094192.6 | 0 | NA | NA | NA | NA | NA | #VALUE! |
| ENSMUSG00000094194.3 | 0 | NA | NA | NA | NA | NA | #VALUE! |
| ENSMUSG00000094195.3 | 0 | NA | NA | NA | NA | NA | #VALUE! |
| ENSMUSG00000094196.3 | 0 | NA | NA | NA | NA | NA | #VALUE! |
| ENSMUSG00000094197.2 | 0 | NA | NA | NA | NA | NA | #VALUE! |
| ENSMUSG00000094198.3 | 0 | NA | NA | NA | NA | NA | #VALUE! |
| ENSMUSG00000094200.3 | 0 | NA | NA | NA | NA | NA | #VALUE! |
| ENSMUSG00000094202.2 | 0 | NA | NA | NA | NA | NA | #VALUE! |
| ENSMUSG00000094204.2 | 0 | NA | NA | NA | NA | NA | #VALUE! |
| ENSMUSG00000094205.8 | 0 | NA | NA | NA | NA | NA | #VALUE! |
| ENSMUSG00000094208.2 | 0 | NA | NA | NA | NA | NA | #VALUE! |
| ENSMUSG00000094210.3 | 0 | NA | NA | NA | NA | NA | #VALUE! |
| ENSMUSG00000094212.4 | 0 | NA | NA | NA | NA | NA | #VALUE! |
| ENSMUSG00000094215.3 | 0 | NA | NA | NA | NA | NA | #VALUE! |
| ENSMUSG00000094217.3 | 0 | NA | NA | NA | NA | NA | #VALUE! |
| ENSMUSG00000094219.2 | 0 | NA | NA | NA | NA | NA | #VALUE! |
| ENSMUSG00000094220.6 | 0 | NA | NA | NA | NA | NA | #VALUE! |
| ENSMUSG00000094221.2 | 0 | NA | NA | NA | NA | NA | #VALUE! |
| ENSMUSG00000094222.2 | 0 | NA | NA | NA | NA | NA | #VALUE! |
| ENSMUSG00000094226.4 | 0 | NA | NA | NA | NA | NA | #VALUE! |
| ENSMUSG00000094227.2 | 0 | NA | NA | NA | NA | NA | #VALUE! |
| ENSMUSG00000094230.6 | 0 | NA | NA | NA | NA | NA | #VALUE! |
| ENSMUSG00000094231.3 | 0 | NA | NA | NA | NA | NA | #VALUE! |
| ENSMUSG00000094232.3 | 0 | NA | NA | NA | NA | NA | #VALUE! |
| ENSMUSG00000094233.3 | 0 | NA | NA | NA | NA | NA | #VALUE! |
| ENSMUSG00000094235.2 | 0 | NA | NA | NA | NA | NA | #VALUE! |
| ENSMUSG00000094237.3 | 0 | NA | NA | NA | NA | NA | #VALUE! |
| ENSMUSG00000094238.2 | 0 | NA | NA | NA | NA | NA | #VALUE! |

| | | | | | | | |
|---|---|---|---|---|---|---|---|
| ENSMUSG00000094240.2 | 0 | NA | NA | NA | NA | NA | #VALUE! |
| ENSMUSG00000094241.3 | 0 | NA | NA | NA | NA | NA | #VALUE! |
| ENSMUSG00000094242.3 | 0 | NA | NA | NA | NA | NA | #VALUE! |
| ENSMUSG00000094243.2 | 0 | NA | NA | NA | NA | NA | #VALUE! |
| ENSMUSG00000094248.2 | 0 | NA | NA | NA | NA | NA | #VALUE! |
| ENSMUSG00000094249.3 | 0 | NA | NA | NA | NA | NA | #VALUE! |
| ENSMUSG00000094250.2 | 0 | NA | NA | NA | NA | NA | #VALUE! |
| ENSMUSG00000094253.2 | 0 | NA | NA | NA | NA | NA | #VALUE! |
| ENSMUSG00000094254.3 | 0 | NA | NA | NA | NA | NA | #VALUE! |
| ENSMUSG00000094256.3 | 0 | NA | NA | NA | NA | NA | #VALUE! |
| ENSMUSG00000094257.3 | 0 | NA | NA | NA | NA | NA | #VALUE! |
| ENSMUSG00000094258.8 | 0 | NA | NA | NA | NA | NA | #VALUE! |
| ENSMUSG00000094259.2 | 0 | NA | NA | NA | NA | NA | #VALUE! |
| ENSMUSG00000094262.3 | 0 | NA | NA | NA | NA | NA | #VALUE! |
| ENSMUSG00000094264.3 | 0 | NA | NA | NA | NA | NA | #VALUE! |
| ENSMUSG00000094266.2 | 0 | NA | NA | NA | NA | NA | #VALUE! |
| ENSMUSG00000094268.2 | 0 | NA | NA | NA | NA | NA | #VALUE! |
| ENSMUSG00000094269.3 | 0 | NA | NA | NA | NA | NA | #VALUE! |
| ENSMUSG00000094271.2 | 0 | NA | NA | NA | NA | NA | #VALUE! |
| ENSMUSG00000094273.2 | 0 | NA | NA | NA | NA | NA | #VALUE! |
| ENSMUSG00000094276.2 | 0 | NA | NA | NA | NA | NA | #VALUE! |
| ENSMUSG00000094277.3 | 0 | NA | NA | NA | NA | NA | #VALUE! |
| ENSMUSG00000094278.2 | 0 | NA | NA | NA | NA | NA | #VALUE! |
| ENSMUSG00000094280.2 | 0 | NA | NA | NA | NA | NA | #VALUE! |
| ENSMUSG00000094281.2 | 0 | NA | NA | NA | NA | NA | #VALUE! |
| ENSMUSG00000094282.2 | 0 | NA | NA | NA | NA | NA | #VALUE! |
| ENSMUSG00000094284.2 | 0 | NA | NA | NA | NA | NA | #VALUE! |
| ENSMUSG00000094285.3 | 0 | NA | NA | NA | NA | NA | #VALUE! |
| ENSMUSG00000094286.2 | 0 | NA | NA | NA | NA | NA | #VALUE! |
| ENSMUSG00000094288.2 | 0 | NA | NA | NA | NA | NA | #VALUE! |
| ENSMUSG00000094293.2 | 0 | NA | NA | NA | NA | NA | #VALUE! |
| ENSMUSG00000094294.3 | 0 | NA | NA | NA | NA | NA | #VALUE! |
| ENSMUSG00000094295.4 | 0 | NA | NA | NA | NA | NA | #VALUE! |
| ENSMUSG00000094296.2 | 0 | NA | NA | NA | NA | NA | #VALUE! |
| ENSMUSG00000094298.2 | 0 | NA | NA | NA | NA | NA | #VALUE! |
| ENSMUSG00000094303.2 | 0 | NA | NA | NA | NA | NA | #VALUE! |
| ENSMUSG00000094305.2 | 0 | NA | NA | NA | NA | NA | #VALUE! |
| ENSMUSG00000094306.3 | 0 | NA | NA | NA | NA | NA | #VALUE! |
| ENSMUSG00000094307.2 | 0 | NA | NA | NA | NA | NA | #VALUE! |
| ENSMUSG00000094311.2 | 0 | NA | NA | NA | NA | NA | #VALUE! |
| ENSMUSG00000094314.4 | 0 | NA | NA | NA | NA | NA | #VALUE! |
| ENSMUSG00000094315.3 | 0 | NA | NA | NA | NA | NA | #VALUE! |
| ENSMUSG00000094317.2 | 0 | NA | NA | NA | NA | NA | #VALUE! |
| ENSMUSG00000094318.2 | 0 | NA | NA | NA | NA | NA | #VALUE! |

| | | | | | | | |
|---|---|---|---|---|---|---|---|
| ENSMUSG00000094319.3 | 0 | NA | NA | NA | NA | NA | #VALUE! |
| ENSMUSG00000094320.2 | 0 | NA | NA | NA | NA | NA | #VALUE! |
| ENSMUSG00000094321.2 | 0 | NA | NA | NA | NA | NA | #VALUE! |
| ENSMUSG00000094322.3 | 0 | NA | NA | NA | NA | NA | #VALUE! |
| ENSMUSG00000094323.2 | 0 | NA | NA | NA | NA | NA | #VALUE! |
| ENSMUSG00000094325.8 | 0 | NA | NA | NA | NA | NA | #VALUE! |
| ENSMUSG00000094326.2 | 0 | NA | NA | NA | NA | NA | #VALUE! |
| ENSMUSG00000094328.3 | 0 | NA | NA | NA | NA | NA | #VALUE! |
| ENSMUSG00000094334.3 | 0 | NA | NA | NA | NA | NA | #VALUE! |
| ENSMUSG00000094335.3 | 0 | NA | NA | NA | NA | NA | #VALUE! |
| ENSMUSG00000094336.2 | 0 | NA | NA | NA | NA | NA | #VALUE! |
| ENSMUSG00000094337.8 | 0 | NA | NA | NA | NA | NA | #VALUE! |
| ENSMUSG00000094338.2 | 0 | NA | NA | NA | NA | NA | #VALUE! |
| ENSMUSG00000094342.4 | 0 | NA | NA | NA | NA | NA | #VALUE! |
| ENSMUSG00000094345.3 | 0 | NA | NA | NA | NA | NA | #VALUE! |
| ENSMUSG00000094347.3 | 0 | NA | NA | NA | NA | NA | #VALUE! |
| ENSMUSG00000094349.2 | 0 | NA | NA | NA | NA | NA | #VALUE! |
| ENSMUSG00000094353.3 | 0 | NA | NA | NA | NA | NA | #VALUE! |
| ENSMUSG00000094354.2 | 0 | NA | NA | NA | NA | NA | #VALUE! |
| ENSMUSG00000094356.3 | 0 | NA | NA | NA | NA | NA | #VALUE! |
| ENSMUSG00000094358.2 | 0 | NA | NA | NA | NA | NA | #VALUE! |
| ENSMUSG00000094360.2 | 0 | NA | NA | NA | NA | NA | #VALUE! |
| ENSMUSG00000094362.3 | 0 | NA | NA | NA | NA | NA | #VALUE! |
| ENSMUSG00000094365.2 | 0 | NA | NA | NA | NA | NA | #VALUE! |
| ENSMUSG00000094370.8 | 0 | NA | NA | NA | NA | NA | #VALUE! |
| ENSMUSG00000094371.2 | 0 | NA | NA | NA | NA | NA | #VALUE! |
| ENSMUSG00000094372.2 | 0 | NA | NA | NA | NA | NA | #VALUE! |
| ENSMUSG00000094374.3 | 0 | NA | NA | NA | NA | NA | #VALUE! |
| ENSMUSG00000094378.8 | 0 | NA | NA | NA | NA | NA | #VALUE! |
| ENSMUSG00000094379.3 | 0 | NA | NA | NA | NA | NA | #VALUE! |
| ENSMUSG00000094380.4 | 0 | NA | NA | NA | NA | NA | #VALUE! |
| ENSMUSG00000094383.2 | 0 | NA | NA | NA | NA | NA | #VALUE! |
| ENSMUSG00000094385.2 | 0 | NA | NA | NA | NA | NA | #VALUE! |
| ENSMUSG00000094388.2 | 0 | NA | NA | NA | NA | NA | #VALUE! |
| ENSMUSG00000094391.2 | 0 | NA | NA | NA | NA | NA | #VALUE! |
| ENSMUSG00000094392.2 | 0 | NA | NA | NA | NA | NA | #VALUE! |
| ENSMUSG00000094396.4 | 0 | NA | NA | NA | NA | NA | #VALUE! |
| ENSMUSG00000094399.8 | 0 | NA | NA | NA | NA | NA | #VALUE! |
| ENSMUSG00000094401.2 | 0 | NA | NA | NA | NA | NA | #VALUE! |
| ENSMUSG00000094404.2 | 0 | NA | NA | NA | NA | NA | #VALUE! |
| ENSMUSG00000094409.2 | 0 | NA | NA | NA | NA | NA | #VALUE! |
| ENSMUSG00000094412.2 | 0 | NA | NA | NA | NA | NA | #VALUE! |
| ENSMUSG00000094413.6 | 0 | NA | NA | NA | NA | NA | #VALUE! |
| ENSMUSG00000094416.2 | 0 | NA | NA | NA | NA | NA | #VALUE! |

| | | | | | | | |
|---|---|---|---|---|---|---|---|
| ENSMUSG00000094417.2 | 0 | NA | NA | NA | NA | NA | #VALUE! |
| ENSMUSG00000094420.3 | 0 | NA | NA | NA | NA | NA | #VALUE! |
| ENSMUSG00000094421.3 | 0 | NA | NA | NA | NA | NA | #VALUE! |
| ENSMUSG00000094422.4 | 0 | NA | NA | NA | NA | NA | #VALUE! |
| ENSMUSG00000094423.2 | 0 | NA | NA | NA | NA | NA | #VALUE! |
| ENSMUSG00000094424.2 | 0 | NA | NA | NA | NA | NA | #VALUE! |
| ENSMUSG00000094426.4 | 0 | NA | NA | NA | NA | NA | #VALUE! |
| ENSMUSG00000094427.2 | 0 | NA | NA | NA | NA | NA | #VALUE! |
| ENSMUSG00000094429.3 | 0 | NA | NA | NA | NA | NA | #VALUE! |
| ENSMUSG00000094430.3 | 0 | NA | NA | NA | NA | NA | #VALUE! |
| ENSMUSG00000094433.3 | 0 | NA | NA | NA | NA | NA | #VALUE! |
| ENSMUSG00000094436.2 | 0 | NA | NA | NA | NA | NA | #VALUE! |
| ENSMUSG00000094437.3 | 0 | NA | NA | NA | NA | NA | #VALUE! |
| ENSMUSG00000094439.2 | 0 | NA | NA | NA | NA | NA | #VALUE! |
| ENSMUSG00000094441.3 | 6.919608 | 5.232067 | 3.251831 | 1.60896 | 0.107625 | NA | #VALUE! |
| ENSMUSG00000094443.3 | 0 | NA | NA | NA | NA | NA | #VALUE! |
| ENSMUSG00000094444.3 | 0 | NA | NA | NA | NA | NA | #VALUE! |
| ENSMUSG00000094445.8 | 0 | NA | NA | NA | NA | NA | #VALUE! |
| ENSMUSG00000094446.2 | 0 | NA | NA | NA | NA | NA | #VALUE! |
| ENSMUSG00000094447.3 | 0 | NA | NA | NA | NA | NA | #VALUE! |
| ENSMUSG00000094449.4 | 0 | NA | NA | NA | NA | NA | #VALUE! |
| ENSMUSG00000094460.8 | 0 | NA | NA | NA | NA | NA | #VALUE! |
| ENSMUSG00000094461.2 | 0 | NA | NA | NA | NA | NA | #VALUE! |
| ENSMUSG00000094462.2 | 0 | NA | NA | NA | NA | NA | #VALUE! |
| ENSMUSG00000094463.3 | 0 | NA | NA | NA | NA | NA | #VALUE! |
| ENSMUSG00000094464.3 | 0 | NA | NA | NA | NA | NA | #VALUE! |
| ENSMUSG00000094468.6 | 0 | NA | NA | NA | NA | NA | #VALUE! |
| ENSMUSG00000094472.2 | 0 | NA | NA | NA | NA | NA | #VALUE! |
| ENSMUSG00000094474.2 | 0 | NA | NA | NA | NA | NA | #VALUE! |
| ENSMUSG00000094475.8 | 0 | NA | NA | NA | NA | NA | #VALUE! |
| ENSMUSG00000094478.2 | 0 | NA | NA | NA | NA | NA | #VALUE! |
| ENSMUSG00000094482.2 | 0 | NA | NA | NA | NA | NA | #VALUE! |
| ENSMUSG00000094484.3 | 0 | NA | NA | NA | NA | NA | #VALUE! |
| ENSMUSG00000094485.2 | 0 | NA | NA | NA | NA | NA | #VALUE! |
| ENSMUSG00000094487.2 | 0 | NA | NA | NA | NA | NA | #VALUE! |
| ENSMUSG00000094488.3 | 0 | NA | NA | NA | NA | NA | #VALUE! |
| ENSMUSG00000094491.3 | 0 | NA | NA | NA | NA | NA | #VALUE! |
| ENSMUSG00000094493.3 | 0 | NA | NA | NA | NA | NA | #VALUE! |
| ENSMUSG00000094496.3 | 0 | NA | NA | NA | NA | NA | #VALUE! |
| ENSMUSG00000094497.2 | 0 | NA | NA | NA | NA | NA | #VALUE! |
| ENSMUSG00000094498.2 | 0 | NA | NA | NA | NA | NA | #VALUE! |
| ENSMUSG00000094499.4 | 0 | NA | NA | NA | NA | NA | #VALUE! |
| ENSMUSG00000094500.3 | 0 | NA | NA | NA | NA | NA | #VALUE! |
| ENSMUSG00000094501.4 | 0 | NA | NA | NA | NA | NA | #VALUE! |

| | | | | | | | |
|---|---|---|---|---|---|---|---|
| ENSMUSG00000094502.6 | 0 | NA | NA | NA | NA | NA | #VALUE! |
| ENSMUSG00000094504.4 | 0 | NA | NA | NA | NA | NA | #VALUE! |
| ENSMUSG00000094505.3 | 0 | NA | NA | NA | NA | NA | #VALUE! |
| ENSMUSG00000094507.2 | 0 | NA | NA | NA | NA | NA | #VALUE! |
| ENSMUSG00000094509.2 | 0 | NA | NA | NA | NA | NA | #VALUE! |
| ENSMUSG00000094511.8 | 0 | NA | NA | NA | NA | NA | #VALUE! |
| ENSMUSG00000094514.2 | 0 | NA | NA | NA | NA | NA | #VALUE! |
| ENSMUSG00000094520.5 | 0 | NA | NA | NA | NA | NA | #VALUE! |
| ENSMUSG00000094525.3 | 0 | NA | NA | NA | NA | NA | #VALUE! |
| ENSMUSG00000094526.2 | 0 | NA | NA | NA | NA | NA | #VALUE! |
| ENSMUSG00000094529.3 | 0 | NA | NA | NA | NA | NA | #VALUE! |
| ENSMUSG00000094530.3 | 1.089935 | 0.682304 | 3.902097 | 0.174856 | 0.861193 | NA | #VALUE! |
| ENSMUSG00000094531.4 | 0 | NA | NA | NA | NA | NA | #VALUE! |
| ENSMUSG00000094532.2 | 0 | NA | NA | NA | NA | NA | #VALUE! |
| ENSMUSG00000094533.2 | 0 | NA | NA | NA | NA | NA | #VALUE! |
| ENSMUSG00000094534.2 | 0 | NA | NA | NA | NA | NA | #VALUE! |
| ENSMUSG00000094535.3 | 0 | NA | NA | NA | NA | NA | #VALUE! |
| ENSMUSG00000094539.4 | 0 | NA | NA | NA | NA | NA | #VALUE! |
| ENSMUSG00000094540.2 | 0 | NA | NA | NA | NA | NA | #VALUE! |
| ENSMUSG00000094542.2 | 0 | NA | NA | NA | NA | NA | #VALUE! |
| ENSMUSG00000094543.2 | 0 | NA | NA | NA | NA | NA | #VALUE! |
| ENSMUSG00000094545.2 | 0 | NA | NA | NA | NA | NA | #VALUE! |
| ENSMUSG00000094546.2 | 0 | NA | NA | NA | NA | NA | #VALUE! |
| ENSMUSG00000094552.2 | 0 | NA | NA | NA | NA | NA | #VALUE! |
| ENSMUSG00000094553.4 | 0 | NA | NA | NA | NA | NA | #VALUE! |
| ENSMUSG00000094555.2 | 0 | NA | NA | NA | NA | NA | #VALUE! |
| ENSMUSG00000094556.3 | 0 | NA | NA | NA | NA | NA | #VALUE! |
| ENSMUSG00000094558.2 | 0 | NA | NA | NA | NA | NA | #VALUE! |
| ENSMUSG00000094559.3 | 0 | NA | NA | NA | NA | NA | #VALUE! |
| ENSMUSG00000094560.2 | 0 | NA | NA | NA | NA | NA | #VALUE! |
| ENSMUSG00000094561.2 | 0 | NA | NA | NA | NA | NA | #VALUE! |
| ENSMUSG00000094562.6 | 0 | NA | NA | NA | NA | NA | #VALUE! |
| ENSMUSG00000094566.2 | 0 | NA | NA | NA | NA | NA | #VALUE! |
| ENSMUSG00000094568.2 | 0 | NA | NA | NA | NA | NA | #VALUE! |
| ENSMUSG00000094569.2 | 0 | NA | NA | NA | NA | NA | #VALUE! |
| ENSMUSG00000094570.8 | 0 | NA | NA | NA | NA | NA | #VALUE! |
| ENSMUSG00000094574.2 | 0 | NA | NA | NA | NA | NA | #VALUE! |
| ENSMUSG00000094575.3 | 0 | NA | NA | NA | NA | NA | #VALUE! |
| ENSMUSG00000094577.8 | 0 | NA | NA | NA | NA | NA | #VALUE! |
| ENSMUSG00000094578.10 | 0 | NA | NA | NA | NA | NA | #VALUE! |
| ENSMUSG00000094582.2 | 0 | NA | NA | NA | NA | NA | #VALUE! |
| ENSMUSG00000094584.3 | 0 | NA | NA | NA | NA | NA | #VALUE! |
| ENSMUSG00000094586.5 | 0 | NA | NA | NA | NA | NA | #VALUE! |
| ENSMUSG00000094588.3 | 0 | NA | NA | NA | NA | NA | #VALUE! |

| | | | | | | | |
|---|---|---|---|---|---|---|---|
| ENSMUSG00000094589.2 | 0 | NA | NA | NA | NA | NA | #VALUE! |
| ENSMUSG00000094590.3 | 0 | NA | NA | NA | NA | NA | #VALUE! |
| ENSMUSG00000094592.2 | 0 | NA | NA | NA | NA | NA | #VALUE! |
| ENSMUSG00000094594.3 | 0 | NA | NA | NA | NA | NA | #VALUE! |
| ENSMUSG00000094595.2 | 0 | NA | NA | NA | NA | NA | #VALUE! |
| ENSMUSG00000094596.2 | 0 | NA | NA | NA | NA | NA | #VALUE! |
| ENSMUSG00000094597.3 | 0 | NA | NA | NA | NA | NA | #VALUE! |
| ENSMUSG00000094601.3 | 0 | NA | NA | NA | NA | NA | #VALUE! |
| ENSMUSG00000094602.3 | 0 | NA | NA | NA | NA | NA | #VALUE! |
| ENSMUSG00000094606.9 | 0 | NA | NA | NA | NA | NA | #VALUE! |
| ENSMUSG00000094607.3 | 0 | NA | NA | NA | NA | NA | #VALUE! |
| ENSMUSG00000094612.4 | 0 | NA | NA | NA | NA | NA | #VALUE! |
| ENSMUSG00000094613.3 | 0 | NA | NA | NA | NA | NA | #VALUE! |
| ENSMUSG00000094615.3 | 0 | NA | NA | NA | NA | NA | #VALUE! |
| ENSMUSG00000094616.8 | 0 | NA | NA | NA | NA | NA | #VALUE! |
| ENSMUSG00000094618.2 | 0 | NA | NA | NA | NA | NA | #VALUE! |
| ENSMUSG00000094619.3 | 0 | NA | NA | NA | NA | NA | #VALUE! |
| ENSMUSG00000094622.3 | 0 | NA | NA | NA | NA | NA | #VALUE! |
| ENSMUSG00000094623.3 | 0 | NA | NA | NA | NA | NA | #VALUE! |
| ENSMUSG00000094624.2 | 0 | NA | NA | NA | NA | NA | #VALUE! |
| ENSMUSG00000094626.2 | 0 | NA | NA | NA | NA | NA | #VALUE! |
| ENSMUSG00000094627.3 | 0 | NA | NA | NA | NA | NA | #VALUE! |
| ENSMUSG00000094628.8 | 0 | NA | NA | NA | NA | NA | #VALUE! |
| ENSMUSG00000094632.4 | 0 | NA | NA | NA | NA | NA | #VALUE! |
| ENSMUSG00000094634.9 | 0 | NA | NA | NA | NA | NA | #VALUE! |
| ENSMUSG00000094635.3 | 0 | NA | NA | NA | NA | NA | #VALUE! |
| ENSMUSG00000094637.3 | 0 | NA | NA | NA | NA | NA | #VALUE! |
| ENSMUSG00000094638.2 | 0 | NA | NA | NA | NA | NA | #VALUE! |
| ENSMUSG00000094643.2 | 0 | NA | NA | NA | NA | NA | #VALUE! |
| ENSMUSG00000094647.8 | 0 | NA | NA | NA | NA | NA | #VALUE! |
| ENSMUSG00000094648.2 | 0 | NA | NA | NA | NA | NA | #VALUE! |
| ENSMUSG00000094649.2 | 0 | NA | NA | NA | NA | NA | #VALUE! |
| ENSMUSG00000094651.3 | 0 | NA | NA | NA | NA | NA | #VALUE! |
| ENSMUSG00000094652.3 | 0 | NA | NA | NA | NA | NA | #VALUE! |
| ENSMUSG00000094658.8 | 0 | NA | NA | NA | NA | NA | #VALUE! |
| ENSMUSG00000094660.3 | 0 | NA | NA | NA | NA | NA | #VALUE! |
| ENSMUSG00000094662.3 | 0 | NA | NA | NA | NA | NA | #VALUE! |
| ENSMUSG00000094664.2 | 0 | NA | NA | NA | NA | NA | #VALUE! |
| ENSMUSG00000094669.4 | 0 | NA | NA | NA | NA | NA | #VALUE! |
| ENSMUSG00000094672.3 | 0 | NA | NA | NA | NA | NA | #VALUE! |
| ENSMUSG00000094673.5 | 0 | NA | NA | NA | NA | NA | #VALUE! |
| ENSMUSG00000094675.2 | 0 | NA | NA | NA | NA | NA | #VALUE! |
| ENSMUSG00000094676.4 | 0 | NA | NA | NA | NA | NA | #VALUE! |
| ENSMUSG00000094678.6 | 0 | NA | NA | NA | NA | NA | #VALUE! |

| | | | | | | | |
|---|---|---|---|---|---|---|---|
| ENSMUSG00000094679.3 | 0 | NA | NA | NA | NA | NA | #VALUE! |
| ENSMUSG00000094680.2 | 0 | NA | NA | NA | NA | NA | #VALUE! |
| ENSMUSG00000094682.3 | 0 | NA | NA | NA | NA | NA | #VALUE! |
| ENSMUSG00000094683.3 | 0 | NA | NA | NA | NA | NA | #VALUE! |
| ENSMUSG00000094684.2 | 0 | NA | NA | NA | NA | NA | #VALUE! |
| ENSMUSG00000094685.3 | 0 | NA | NA | NA | NA | NA | #VALUE! |
| ENSMUSG00000094686.2 | 0 | NA | NA | NA | NA | NA | #VALUE! |
| ENSMUSG00000094687.2 | 0 | NA | NA | NA | NA | NA | #VALUE! |
| ENSMUSG00000094689.2 | 0 | NA | NA | NA | NA | NA | #VALUE! |
| ENSMUSG00000094690.3 | 0 | NA | NA | NA | NA | NA | #VALUE! |
| ENSMUSG00000094692.3 | 0 | NA | NA | NA | NA | NA | #VALUE! |
| ENSMUSG00000094693.2 | 0 | NA | NA | NA | NA | NA | #VALUE! |
| ENSMUSG00000094694.7 | 0 | NA | NA | NA | NA | NA | #VALUE! |
| ENSMUSG00000094695.3 | 0 | NA | NA | NA | NA | NA | #VALUE! |
| ENSMUSG00000094696.2 | 0 | NA | NA | NA | NA | NA | #VALUE! |
| ENSMUSG00000094698.2 | 0 | NA | NA | NA | NA | NA | #VALUE! |
| ENSMUSG00000094699.3 | 0 | NA | NA | NA | NA | NA | #VALUE! |
| ENSMUSG00000094700.2 | 0 | NA | NA | NA | NA | NA | #VALUE! |
| ENSMUSG00000094701.4 | 0 | NA | NA | NA | NA | NA | #VALUE! |
| ENSMUSG00000094706.9 | 0 | NA | NA | NA | NA | NA | #VALUE! |
| ENSMUSG00000094707.3 | 0 | NA | NA | NA | NA | NA | #VALUE! |
| ENSMUSG00000094708.2 | 0 | NA | NA | NA | NA | NA | #VALUE! |
| ENSMUSG00000094712.2 | 0 | NA | NA | NA | NA | NA | #VALUE! |
| ENSMUSG00000094714.8 | 0 | NA | NA | NA | NA | NA | #VALUE! |
| ENSMUSG00000094715.4 | 0 | NA | NA | NA | NA | NA | #VALUE! |
| ENSMUSG00000094719.5 | 0 | NA | NA | NA | NA | NA | #VALUE! |
| ENSMUSG00000094721.4 | 0 | NA | NA | NA | NA | NA | #VALUE! |
| ENSMUSG00000094722.2 | 0 | NA | NA | NA | NA | NA | #VALUE! |
| ENSMUSG00000094724.9 | 2.524253 | 4.825754 | 3.924522 | 1.229641 | 0.218831 | NA | #VALUE! |
| ENSMUSG00000094725.2 | 0 | NA | NA | NA | NA | NA | #VALUE! |
| ENSMUSG00000094728.2 | 0 | NA | NA | NA | NA | NA | #VALUE! |
| ENSMUSG00000094729.3 | 0 | NA | NA | NA | NA | NA | #VALUE! |
| ENSMUSG00000094732.3 | 0 | NA | NA | NA | NA | NA | #VALUE! |
| ENSMUSG00000094733.2 | 0 | NA | NA | NA | NA | NA | #VALUE! |
| ENSMUSG00000094734.3 | 0 | NA | NA | NA | NA | NA | #VALUE! |
| ENSMUSG00000094735.2 | 0 | NA | NA | NA | NA | NA | #VALUE! |
| ENSMUSG00000094736.3 | 0 | NA | NA | NA | NA | NA | #VALUE! |
| ENSMUSG00000094737.2 | 0 | NA | NA | NA | NA | NA | #VALUE! |
| ENSMUSG00000094739.3 | 0 | NA | NA | NA | NA | NA | #VALUE! |
| ENSMUSG00000094741.2 | 0 | NA | NA | NA | NA | NA | #VALUE! |
| ENSMUSG00000094743.3 | 6.147607 | -2.0736 | 3.39327 | -0.61109 | 0.541138 | NA | #VALUE! |
| ENSMUSG00000094745.4 | 0 | NA | NA | NA | NA | NA | #VALUE! |
| ENSMUSG00000094746.8 | 0 | NA | NA | NA | NA | NA | #VALUE! |
| ENSMUSG00000094747.4 | 0 | NA | NA | NA | NA | NA | #VALUE! |

| | | | | | | | |
|---|---|---|---|---|---|---|---|
| ENSMUSG00000094748.2 | 0 | NA | NA | NA | NA | NA | #VALUE! |
| ENSMUSG00000094752.2 | 0 | NA | NA | NA | NA | NA | #VALUE! |
| ENSMUSG00000094755.2 | 0 | NA | NA | NA | NA | NA | #VALUE! |
| ENSMUSG00000094757.2 | 0 | NA | NA | NA | NA | NA | #VALUE! |
| ENSMUSG00000094758.3 | 0 | NA | NA | NA | NA | NA | #VALUE! |
| ENSMUSG00000094759.8 | 0 | NA | NA | NA | NA | NA | #VALUE! |
| ENSMUSG00000094762.2 | 0 | NA | NA | NA | NA | NA | #VALUE! |
| ENSMUSG00000094763.4 | 0 | NA | NA | NA | NA | NA | #VALUE! |
| ENSMUSG00000094764.3 | 0 | NA | NA | NA | NA | NA | #VALUE! |
| ENSMUSG00000094766.4 | 0 | NA | NA | NA | NA | NA | #VALUE! |
| ENSMUSG00000094767.2 | 0 | NA | NA | NA | NA | NA | #VALUE! |
| ENSMUSG00000094771.2 | 0 | NA | NA | NA | NA | NA | #VALUE! |
| ENSMUSG00000094772.3 | 0 | NA | NA | NA | NA | NA | #VALUE! |
| ENSMUSG00000094773.3 | 0 | NA | NA | NA | NA | NA | #VALUE! |
| ENSMUSG00000094775.2 | 0 | NA | NA | NA | NA | NA | #VALUE! |
| ENSMUSG00000094777.3 | 0 | NA | NA | NA | NA | NA | #VALUE! |
| ENSMUSG00000094778.4 | 0 | NA | NA | NA | NA | NA | #VALUE! |
| ENSMUSG00000094779.3 | 0 | NA | NA | NA | NA | NA | #VALUE! |
| ENSMUSG00000094782.8 | 0 | NA | NA | NA | NA | NA | #VALUE! |
| ENSMUSG00000094784.2 | 0 | NA | NA | NA | NA | NA | #VALUE! |
| ENSMUSG00000094787.3 | 0 | NA | NA | NA | NA | NA | #VALUE! |
| ENSMUSG00000094789.8 | 0 | NA | NA | NA | NA | NA | #VALUE! |
| ENSMUSG00000094790.3 | 0 | NA | NA | NA | NA | NA | #VALUE! |
| ENSMUSG00000094791.2 | 0 | NA | NA | NA | NA | NA | #VALUE! |
| ENSMUSG00000094792.4 | 0 | NA | NA | NA | NA | NA | #VALUE! |
| ENSMUSG00000094794.2 | 0 | NA | NA | NA | NA | NA | #VALUE! |
| ENSMUSG00000094796.3 | 0 | NA | NA | NA | NA | NA | #VALUE! |
| ENSMUSG00000094797.2 | 0 | NA | NA | NA | NA | NA | #VALUE! |
| ENSMUSG00000094799.2 | 0 | NA | NA | NA | NA | NA | #VALUE! |
| ENSMUSG00000094802.8 | 0 | NA | NA | NA | NA | NA | #VALUE! |
| ENSMUSG00000094804.2 | 0 | NA | NA | NA | NA | NA | #VALUE! |
| ENSMUSG00000094805.3 | 0 | NA | NA | NA | NA | NA | #VALUE! |
| ENSMUSG00000094808.3 | 0 | NA | NA | NA | NA | NA | #VALUE! |
| ENSMUSG00000094810.4 | 0 | NA | NA | NA | NA | NA | #VALUE! |
| ENSMUSG00000094811.8 | 0 | NA | NA | NA | NA | NA | #VALUE! |
| ENSMUSG00000094813.3 | 0 | NA | NA | NA | NA | NA | #VALUE! |
| ENSMUSG00000094814.2 | 0 | NA | NA | NA | NA | NA | #VALUE! |
| ENSMUSG00000094818.2 | 0 | NA | NA | NA | NA | NA | #VALUE! |
| ENSMUSG00000094819.4 | 0 | NA | NA | NA | NA | NA | #VALUE! |
| ENSMUSG00000094821.8 | 0 | NA | NA | NA | NA | NA | #VALUE! |
| ENSMUSG00000094822.3 | 0 | NA | NA | NA | NA | NA | #VALUE! |
| ENSMUSG00000094825.9 | 0 | NA | NA | NA | NA | NA | #VALUE! |
| ENSMUSG00000094827.4 | 0 | NA | NA | NA | NA | NA | #VALUE! |
| ENSMUSG00000094828.6 | 0 | NA | NA | NA | NA | NA | #VALUE! |

| | | | | | | | |
|---|---|---|---|---|---|---|---|
| ENSMUSG00000094829.3 | 0 | NA | NA | NA | NA | NA | #VALUE! |
| ENSMUSG00000094832.2 | 0 | NA | NA | NA | NA | NA | #VALUE! |
| ENSMUSG00000094836.2 | 0 | NA | NA | NA | NA | NA | #VALUE! |
| ENSMUSG00000094838.3 | 0 | NA | NA | NA | NA | NA | #VALUE! |
| ENSMUSG00000094840.6 | 0 | NA | NA | NA | NA | NA | #VALUE! |
| ENSMUSG00000094841.3 | 0 | NA | NA | NA | NA | NA | #VALUE! |
| ENSMUSG00000094843.3 | 0 | NA | NA | NA | NA | NA | #VALUE! |
| ENSMUSG00000094845.3 | 0 | NA | NA | NA | NA | NA | #VALUE! |
| ENSMUSG00000094846.5 | 0 | NA | NA | NA | NA | NA | #VALUE! |
| ENSMUSG00000094849.2 | 0 | NA | NA | NA | NA | NA | #VALUE! |
| ENSMUSG00000094852.4 | 0 | NA | NA | NA | NA | NA | #VALUE! |
| ENSMUSG00000094854.4 | 0 | NA | NA | NA | NA | NA | #VALUE! |
| ENSMUSG00000094855.2 | 0 | NA | NA | NA | NA | NA | #VALUE! |
| ENSMUSG00000094856.2 | 0 | NA | NA | NA | NA | NA | #VALUE! |
| ENSMUSG00000094858.4 | 0 | NA | NA | NA | NA | NA | #VALUE! |
| ENSMUSG00000094860.2 | 0 | NA | NA | NA | NA | NA | #VALUE! |
| ENSMUSG00000094861.2 | 0 | NA | NA | NA | NA | NA | #VALUE! |
| ENSMUSG00000094862.3 | 0 | NA | NA | NA | NA | NA | #VALUE! |
| ENSMUSG00000094864.3 | 0 | NA | NA | NA | NA | NA | #VALUE! |
| ENSMUSG00000094865.3 | 0 | NA | NA | NA | NA | NA | #VALUE! |
| ENSMUSG00000094868.2 | 0 | NA | NA | NA | NA | NA | #VALUE! |
| ENSMUSG00000094872.2 | 0 | NA | NA | NA | NA | NA | #VALUE! |
| ENSMUSG00000094873.2 | 0 | NA | NA | NA | NA | NA | #VALUE! |
| ENSMUSG00000094874.2 | 0 | NA | NA | NA | NA | NA | #VALUE! |
| ENSMUSG00000094876.2 | 0 | NA | NA | NA | NA | NA | #VALUE! |
| ENSMUSG00000094877.3 | 0 | NA | NA | NA | NA | NA | #VALUE! |
| ENSMUSG00000094878.3 | 0 | NA | NA | NA | NA | NA | #VALUE! |
| ENSMUSG00000094879.2 | 0 | NA | NA | NA | NA | NA | #VALUE! |
| ENSMUSG00000094880.2 | 0 | NA | NA | NA | NA | NA | #VALUE! |
| ENSMUSG00000094881.4 | 0 | NA | NA | NA | NA | NA | #VALUE! |
| ENSMUSG00000094884.4 | 0 | NA | NA | NA | NA | NA | #VALUE! |
| ENSMUSG00000094885.8 | 0 | NA | NA | NA | NA | NA | #VALUE! |
| ENSMUSG00000094886.3 | 0 | NA | NA | NA | NA | NA | #VALUE! |
| ENSMUSG00000094887.2 | 0 | NA | NA | NA | NA | NA | #VALUE! |
| ENSMUSG00000094891.3 | 0 | NA | NA | NA | NA | NA | #VALUE! |
| ENSMUSG00000094892.3 | 0 | NA | NA | NA | NA | NA | #VALUE! |
| ENSMUSG00000094893.3 | 0 | NA | NA | NA | NA | NA | #VALUE! |
| ENSMUSG00000094898.3 | 0 | NA | NA | NA | NA | NA | #VALUE! |
| ENSMUSG00000094902.3 | 0 | NA | NA | NA | NA | NA | #VALUE! |
| ENSMUSG00000094904.2 | 0 | NA | NA | NA | NA | NA | #VALUE! |
| ENSMUSG00000094905.2 | 0 | NA | NA | NA | NA | NA | #VALUE! |
| ENSMUSG00000094910.2 | 5.57535 | -5.07427 | 3.906591 | -1.2989 | 0.193979 | NA | #VALUE! |
| ENSMUSG00000094911.2 | 0 | NA | NA | NA | NA | NA | #VALUE! |
| ENSMUSG00000094913.4 | 0 | NA | NA | NA | NA | NA | #VALUE! |

| | | | | | | | |
|---|---|---|---|---|---|---|---|
| ENSMUSG00000094914.6 | 0 | NA | NA | NA | NA | NA | #VALUE! |
| ENSMUSG00000094915.2 | 0 | NA | NA | NA | NA | NA | #VALUE! |
| ENSMUSG00000094918.4 | 0 | NA | NA | NA | NA | NA | #VALUE! |
| ENSMUSG00000094920.3 | 0 | NA | NA | NA | NA | NA | #VALUE! |
| ENSMUSG00000094921.3 | 0 | NA | NA | NA | NA | NA | #VALUE! |
| ENSMUSG00000094925.8 | 0 | NA | NA | NA | NA | NA | #VALUE! |
| ENSMUSG00000094926.2 | 0 | NA | NA | NA | NA | NA | #VALUE! |
| ENSMUSG00000094928.2 | 0 | NA | NA | NA | NA | NA | #VALUE! |
| ENSMUSG00000094929.2 | 0 | NA | NA | NA | NA | NA | #VALUE! |
| ENSMUSG00000094930.3 | 0 | NA | NA | NA | NA | NA | #VALUE! |
| ENSMUSG00000094931.2 | 0 | NA | NA | NA | NA | NA | #VALUE! |
| ENSMUSG00000094932.8 | 0 | NA | NA | NA | NA | NA | #VALUE! |
| ENSMUSG00000094934.2 | 0 | NA | NA | NA | NA | NA | #VALUE! |
| ENSMUSG00000094935.3 | 0 | NA | NA | NA | NA | NA | #VALUE! |
| ENSMUSG00000094939.2 | 0 | NA | NA | NA | NA | NA | #VALUE! |
| ENSMUSG00000094940.2 | 0 | NA | NA | NA | NA | NA | #VALUE! |
| ENSMUSG00000094941.2 | 0 | NA | NA | NA | NA | NA | #VALUE! |
| ENSMUSG00000094942.8 | 0.315039 | -1.93549 | 4.038621 | -0.47925 | 0.631763 | NA | #VALUE! |
| ENSMUSG00000094945.4 | 0 | NA | NA | NA | NA | NA | #VALUE! |
| ENSMUSG00000094948.3 | 0 | NA | NA | NA | NA | NA | #VALUE! |
| ENSMUSG00000094950.3 | 0 | NA | NA | NA | NA | NA | #VALUE! |
| ENSMUSG00000094951.2 | 0 | NA | NA | NA | NA | NA | #VALUE! |
| ENSMUSG00000094954.2 | 0 | NA | NA | NA | NA | NA | #VALUE! |
| ENSMUSG00000094955.3 | 0 | NA | NA | NA | NA | NA | #VALUE! |
| ENSMUSG00000094957.2 | 0 | NA | NA | NA | NA | NA | #VALUE! |
| ENSMUSG00000094958.2 | 0 | NA | NA | NA | NA | NA | #VALUE! |
| ENSMUSG00000094962.2 | 0 | NA | NA | NA | NA | NA | #VALUE! |
| ENSMUSG00000094964.2 | 0 | NA | NA | NA | NA | NA | #VALUE! |
| ENSMUSG00000094966.3 | 0 | NA | NA | NA | NA | NA | #VALUE! |
| ENSMUSG00000094967.2 | 0 | NA | NA | NA | NA | NA | #VALUE! |
| ENSMUSG00000094970.4 | 0 | NA | NA | NA | NA | NA | #VALUE! |
| ENSMUSG00000094973.3 | 0 | NA | NA | NA | NA | NA | #VALUE! |
| ENSMUSG00000094974.2 | 0 | NA | NA | NA | NA | NA | #VALUE! |
| ENSMUSG00000094975.2 | 0 | NA | NA | NA | NA | NA | #VALUE! |
| ENSMUSG00000094978.3 | 0 | NA | NA | NA | NA | NA | #VALUE! |
| ENSMUSG00000094981.2 | 0 | NA | NA | NA | NA | NA | #VALUE! |
| ENSMUSG00000094985.3 | 0 | NA | NA | NA | NA | NA | #VALUE! |
| ENSMUSG00000094986.4 | 0 | NA | NA | NA | NA | NA | #VALUE! |
| ENSMUSG00000094989.2 | 0 | NA | NA | NA | NA | NA | #VALUE! |
| ENSMUSG00000094991.3 | 0 | NA | NA | NA | NA | NA | #VALUE! |
| ENSMUSG00000094992.2 | 0 | NA | NA | NA | NA | NA | #VALUE! |
| ENSMUSG00000094993.3 | 0 | NA | NA | NA | NA | NA | #VALUE! |
| ENSMUSG00000094995.3 | 0 | NA | NA | NA | NA | NA | #VALUE! |
| ENSMUSG00000094999.3 | 0 | NA | NA | NA | NA | NA | #VALUE! |

| | | | | | | | |
|---|---|---|---|---|---|---|---|
| ENSMUSG00000095002.4 | 0 | NA | NA | NA | NA | NA | #VALUE! |
| ENSMUSG00000095003.2 | 0 | NA | NA | NA | NA | NA | #VALUE! |
| ENSMUSG00000095005.2 | 0 | NA | NA | NA | NA | NA | #VALUE! |
| ENSMUSG00000095007.2 | 0 | NA | NA | NA | NA | NA | #VALUE! |
| ENSMUSG00000095008.2 | 0 | NA | NA | NA | NA | NA | #VALUE! |
| ENSMUSG00000095011.8 | 0 | NA | NA | NA | NA | NA | #VALUE! |
| ENSMUSG00000095015.8 | 0 | NA | NA | NA | NA | NA | #VALUE! |
| ENSMUSG00000095017.2 | 0 | NA | NA | NA | NA | NA | #VALUE! |
| ENSMUSG00000095019.2 | 0 | NA | NA | NA | NA | NA | #VALUE! |
| ENSMUSG00000095022.9 | 0 | NA | NA | NA | NA | NA | #VALUE! |
| ENSMUSG00000095024.9 | 0 | NA | NA | NA | NA | NA | #VALUE! |
| ENSMUSG00000095026.3 | 0 | NA | NA | NA | NA | NA | #VALUE! |
| ENSMUSG00000095028.7 | 0 | NA | NA | NA | NA | NA | #VALUE! |
| ENSMUSG00000095029.2 | 0 | NA | NA | NA | NA | NA | #VALUE! |
| ENSMUSG00000095030.3 | 0 | NA | NA | NA | NA | NA | #VALUE! |
| ENSMUSG00000095032.3 | 0 | NA | NA | NA | NA | NA | #VALUE! |
| ENSMUSG00000095034.3 | 0 | NA | NA | NA | NA | NA | #VALUE! |
| ENSMUSG00000095040.3 | 0 | NA | NA | NA | NA | NA | #VALUE! |
| ENSMUSG00000095042.2 | 0 | NA | NA | NA | NA | NA | #VALUE! |
| ENSMUSG00000095044.8 | 0 | NA | NA | NA | NA | NA | #VALUE! |
| ENSMUSG00000095048.8 | 0 | NA | NA | NA | NA | NA | #VALUE! |
| ENSMUSG00000095049.3 | 0 | NA | NA | NA | NA | NA | #VALUE! |
| ENSMUSG00000095054.2 | 0 | NA | NA | NA | NA | NA | #VALUE! |
| ENSMUSG00000095056.8 | 0 | NA | NA | NA | NA | NA | #VALUE! |
| ENSMUSG00000095057.2 | 0 | NA | NA | NA | NA | NA | #VALUE! |
| ENSMUSG00000095061.2 | 1.177985 | 3.72614 | 3.944615 | 0.944614 | 0.344856 | NA | #VALUE! |
| ENSMUSG00000095063.2 | 0 | NA | NA | NA | NA | NA | #VALUE! |
| ENSMUSG00000095064.2 | 0 | NA | NA | NA | NA | NA | #VALUE! |
| ENSMUSG00000095065.2 | 0 | NA | NA | NA | NA | NA | #VALUE! |
| ENSMUSG00000095066.2 | 0 | NA | NA | NA | NA | NA | #VALUE! |
| ENSMUSG00000095071.3 | 0 | NA | NA | NA | NA | NA | #VALUE! |
| ENSMUSG00000095074.5 | 0 | NA | NA | NA | NA | NA | #VALUE! |
| ENSMUSG00000095075.3 | 0 | NA | NA | NA | NA | NA | #VALUE! |
| ENSMUSG00000095076.2 | 0 | NA | NA | NA | NA | NA | #VALUE! |
| ENSMUSG00000095078.3 | 0 | NA | NA | NA | NA | NA | #VALUE! |
| ENSMUSG00000095079.7 | 1.492083 | 4.088114 | 3.936282 | 1.038572 | 0.299004 | NA | #VALUE! |
| ENSMUSG00000095081.2 | 0 | NA | NA | NA | NA | NA | #VALUE! |
| ENSMUSG00000095082.2 | 0 | NA | NA | NA | NA | NA | #VALUE! |
| ENSMUSG00000095088.4 | 0 | NA | NA | NA | NA | NA | #VALUE! |
| ENSMUSG00000095089.2 | 0 | NA | NA | NA | NA | NA | #VALUE! |
| ENSMUSG00000095092.2 | 0 | NA | NA | NA | NA | NA | #VALUE! |
| ENSMUSG00000095093.3 | 0 | NA | NA | NA | NA | NA | #VALUE! |
| ENSMUSG00000095095.3 | 0 | NA | NA | NA | NA | NA | #VALUE! |
| ENSMUSG00000095096.2 | 0 | NA | NA | NA | NA | NA | #VALUE! |

| | | | | | | | |
|---|---|---|---|---|---|---|---|
| ENSMUSG00000095101.3 | 0 | NA | NA | NA | NA | NA | #VALUE! |
| ENSMUSG00000095103.3 | 0 | NA | NA | NA | NA | NA | #VALUE! |
| ENSMUSG00000095104.3 | 0 | NA | NA | NA | NA | NA | #VALUE! |
| ENSMUSG00000095105.3 | 0 | NA | NA | NA | NA | NA | #VALUE! |
| ENSMUSG00000095107.2 | 0 | NA | NA | NA | NA | NA | #VALUE! |
| ENSMUSG00000095108.3 | 0 | NA | NA | NA | NA | NA | #VALUE! |
| ENSMUSG00000095110.2 | 0 | NA | NA | NA | NA | NA | #VALUE! |
| ENSMUSG00000095113.2 | 0 | NA | NA | NA | NA | NA | #VALUE! |
| ENSMUSG00000095115.3 | 8.22132 | -6.57092 | 3.302286 | -1.98981 | 0.046612 | NA | #VALUE! |
| ENSMUSG00000095117.3 | 0 | NA | NA | NA | NA | NA | #VALUE! |
| ENSMUSG00000095123.2 | 2.33138 | 4.724655 | 3.925805 | 1.203487 | 0.228788 | NA | #VALUE! |
| ENSMUSG00000095125.3 | 0 | NA | NA | NA | NA | NA | #VALUE! |
| ENSMUSG00000095126.4 | 0 | NA | NA | NA | NA | NA | #VALUE! |
| ENSMUSG00000095127.3 | 0 | NA | NA | NA | NA | NA | #VALUE! |
| ENSMUSG00000095128.3 | 0 | NA | NA | NA | NA | NA | #VALUE! |
| ENSMUSG00000095130.2 | 0 | NA | NA | NA | NA | NA | #VALUE! |
| ENSMUSG00000095134.3 | 0 | NA | NA | NA | NA | NA | #VALUE! |
| ENSMUSG00000095135.8 | 0 | NA | NA | NA | NA | NA | #VALUE! |
| ENSMUSG00000095136.2 | 0 | NA | NA | NA | NA | NA | #VALUE! |
| ENSMUSG00000095137.2 | 0 | NA | NA | NA | NA | NA | #VALUE! |
| ENSMUSG00000095138.4 | 0 | NA | NA | NA | NA | NA | #VALUE! |
| ENSMUSG00000095139.3 | 0 | NA | NA | NA | NA | NA | #VALUE! |
| ENSMUSG00000095141.8 | 0 | NA | NA | NA | NA | NA | #VALUE! |
| ENSMUSG00000095143.6 | 0 | NA | NA | NA | NA | NA | #VALUE! |
| ENSMUSG00000095148.3 | 0 | NA | NA | NA | NA | NA | #VALUE! |
| ENSMUSG00000095150.4 | 0 | NA | NA | NA | NA | NA | #VALUE! |
| ENSMUSG00000095152.2 | 0 | NA | NA | NA | NA | NA | #VALUE! |
| ENSMUSG00000095153.8 | 0 | NA | NA | NA | NA | NA | #VALUE! |
| ENSMUSG00000095156.3 | 0 | NA | NA | NA | NA | NA | #VALUE! |
| ENSMUSG00000095159.3 | 0 | NA | NA | NA | NA | NA | #VALUE! |
| ENSMUSG00000095163.2 | 0 | NA | NA | NA | NA | NA | #VALUE! |
| ENSMUSG00000095168.2 | 0 | NA | NA | NA | NA | NA | #VALUE! |
| ENSMUSG00000095170.3 | 0 | NA | NA | NA | NA | NA | #VALUE! |
| ENSMUSG00000095171.2 | 0 | NA | NA | NA | NA | NA | #VALUE! |
| ENSMUSG00000095172.3 | 0 | NA | NA | NA | NA | NA | #VALUE! |
| ENSMUSG00000095174.2 | 0 | NA | NA | NA | NA | NA | #VALUE! |
| ENSMUSG00000095179.3 | 0 | NA | NA | NA | NA | NA | #VALUE! |
| ENSMUSG00000095185.2 | 0 | NA | NA | NA | NA | NA | #VALUE! |
| ENSMUSG00000095186.2 | 3.570446 | -5.37363 | 3.919223 | -1.3711 | 0.170345 | NA | #VALUE! |
| ENSMUSG00000095187.3 | 0 | NA | NA | NA | NA | NA | #VALUE! |
| ENSMUSG00000095189.3 | 0 | NA | NA | NA | NA | NA | #VALUE! |
| ENSMUSG00000095190.4 | 0 | NA | NA | NA | NA | NA | #VALUE! |
| ENSMUSG00000095191.2 | 0 | NA | NA | NA | NA | NA | #VALUE! |
| ENSMUSG00000095192.3 | 0 | NA | NA | NA | NA | NA | #VALUE! |

| | | | | | | | |
|---|---|---|---|---|---|---|---|
| ENSMUSG00000095193.3 | 7.316007 | 0.73527 | 2.593968 | 0.283454 | 0.776829 | NA | #VALUE! |
| ENSMUSG00000095194.3 | 0 | NA | NA | NA | NA | NA | #VALUE! |
| ENSMUSG00000095195.8 | 0 | NA | NA | NA | NA | NA | #VALUE! |
| ENSMUSG00000095196.2 | 0 | NA | NA | NA | NA | NA | #VALUE! |
| ENSMUSG00000095197.7 | 0 | NA | NA | NA | NA | NA | #VALUE! |
| ENSMUSG00000095199.9 | 0 | NA | NA | NA | NA | NA | #VALUE! |
| ENSMUSG00000095200.3 | 0 | NA | NA | NA | NA | NA | #VALUE! |
| ENSMUSG00000095201.3 | 0 | NA | NA | NA | NA | NA | #VALUE! |
| ENSMUSG00000095202.3 | 0 | NA | NA | NA | NA | NA | #VALUE! |
| ENSMUSG00000095203.2 | 0 | NA | NA | NA | NA | NA | #VALUE! |
| ENSMUSG00000095204.7 | 0 | NA | NA | NA | NA | NA | #VALUE! |
| ENSMUSG00000095210.3 | 0 | NA | NA | NA | NA | NA | #VALUE! |
| ENSMUSG00000095212.5 | 0 | NA | NA | NA | NA | NA | #VALUE! |
| ENSMUSG00000095213.3 | 0 | NA | NA | NA | NA | NA | #VALUE! |
| ENSMUSG00000095217.2 | 0 | NA | NA | NA | NA | NA | #VALUE! |
| ENSMUSG00000095218.4 | 0 | NA | NA | NA | NA | NA | #VALUE! |
| ENSMUSG00000095226.2 | 0 | NA | NA | NA | NA | NA | #VALUE! |
| ENSMUSG00000095231.2 | 0 | NA | NA | NA | NA | NA | #VALUE! |
| ENSMUSG00000095232.3 | 0 | NA | NA | NA | NA | NA | #VALUE! |
| ENSMUSG00000095234.3 | 0 | NA | NA | NA | NA | NA | #VALUE! |
| ENSMUSG00000095236.3 | 0 | NA | NA | NA | NA | NA | #VALUE! |
| ENSMUSG00000095238.3 | 0 | NA | NA | NA | NA | NA | #VALUE! |
| ENSMUSG00000095239.5 | 0 | NA | NA | NA | NA | NA | #VALUE! |
| ENSMUSG00000095240.2 | 0 | NA | NA | NA | NA | NA | #VALUE! |
| ENSMUSG00000095241.3 | 0 | NA | NA | NA | NA | NA | #VALUE! |
| ENSMUSG00000095242.3 | 0 | NA | NA | NA | NA | NA | #VALUE! |
| ENSMUSG00000095248.4 | 0 | NA | NA | NA | NA | NA | #VALUE! |
| ENSMUSG00000095250.2 | 0 | NA | NA | NA | NA | NA | #VALUE! |
| ENSMUSG00000095251.2 | 0 | NA | NA | NA | NA | NA | #VALUE! |
| ENSMUSG00000095257.2 | 0 | NA | NA | NA | NA | NA | #VALUE! |
| ENSMUSG00000095258.3 | 0 | NA | NA | NA | NA | NA | #VALUE! |
| ENSMUSG00000095259.2 | 0 | NA | NA | NA | NA | NA | #VALUE! |
| ENSMUSG00000095261.3 | 0 | NA | NA | NA | NA | NA | #VALUE! |
| ENSMUSG00000095262.3 | 0 | NA | NA | NA | NA | NA | #VALUE! |
| ENSMUSG00000095263.8 | 0 | NA | NA | NA | NA | NA | #VALUE! |
| ENSMUSG00000095266.3 | 0 | NA | NA | NA | NA | NA | #VALUE! |
| ENSMUSG00000095267.3 | 0 | NA | NA | NA | NA | NA | #VALUE! |
| ENSMUSG00000095269.3 | 0 | NA | NA | NA | NA | NA | #VALUE! |
| ENSMUSG00000095270.2 | 0 | NA | NA | NA | NA | NA | #VALUE! |
| ENSMUSG00000095273.2 | 0 | NA | NA | NA | NA | NA | #VALUE! |
| ENSMUSG00000095274.2 | 0 | NA | NA | NA | NA | NA | #VALUE! |
| ENSMUSG00000095275.2 | 0 | NA | NA | NA | NA | NA | #VALUE! |
| ENSMUSG00000095276.3 | 0 | NA | NA | NA | NA | NA | #VALUE! |
| ENSMUSG00000095277.2 | 0.093255 | 0.868093 | 4.072464 | 0.213162 | 0.831201 | NA | #VALUE! |

| | | | | | | | |
|---|---|---|---|---|---|---|---|
| ENSMUSG00000095279.3 | 0 | NA | NA | NA | NA | NA | #VALUE! |
| ENSMUSG00000095283.2 | 0 | NA | NA | NA | NA | NA | #VALUE! |
| ENSMUSG00000095285.3 | 0 | NA | NA | NA | NA | NA | #VALUE! |
| ENSMUSG00000095286.4 | 0 | NA | NA | NA | NA | NA | #VALUE! |
| ENSMUSG00000095287.2 | 0 | NA | NA | NA | NA | NA | #VALUE! |
| ENSMUSG00000095288.2 | 0 | NA | NA | NA | NA | NA | #VALUE! |
| ENSMUSG00000095291.2 | 0 | NA | NA | NA | NA | NA | #VALUE! |
| ENSMUSG00000095293.8 | 0 | NA | NA | NA | NA | NA | #VALUE! |
| ENSMUSG00000095294.3 | 0 | NA | NA | NA | NA | NA | #VALUE! |
| ENSMUSG00000095295.9 | 0 | NA | NA | NA | NA | NA | #VALUE! |
| ENSMUSG00000095296.8 | 0 | NA | NA | NA | NA | NA | #VALUE! |
| ENSMUSG00000095298.3 | 0 | NA | NA | NA | NA | NA | #VALUE! |
| ENSMUSG00000095300.2 | 0 | NA | NA | NA | NA | NA | #VALUE! |
| ENSMUSG00000095301.3 | 0 | NA | NA | NA | NA | NA | #VALUE! |
| ENSMUSG00000095302.3 | 0 | NA | NA | NA | NA | NA | #VALUE! |
| ENSMUSG00000095307.2 | 0 | NA | NA | NA | NA | NA | #VALUE! |
| ENSMUSG00000095309.2 | 0 | NA | NA | NA | NA | NA | #VALUE! |
| ENSMUSG00000095312.2 | 0 | NA | NA | NA | NA | NA | #VALUE! |
| ENSMUSG00000095315.3 | 9.34707 | 6.710288 | 3.689587 | 1.81871 | 0.068956 | NA | #VALUE! |
| ENSMUSG00000095316.2 | 0 | NA | NA | NA | NA | NA | #VALUE! |
| ENSMUSG00000095318.2 | 0 | NA | NA | NA | NA | NA | #VALUE! |
| ENSMUSG00000095322.3 | 0 | NA | NA | NA | NA | NA | #VALUE! |
| ENSMUSG00000095324.3 | 0 | NA | NA | NA | NA | NA | #VALUE! |
| ENSMUSG00000095327.2 | 0 | NA | NA | NA | NA | NA | #VALUE! |
| ENSMUSG00000095328.2 | 0 | NA | NA | NA | NA | NA | #VALUE! |
| ENSMUSG00000095330.2 | 0 | NA | NA | NA | NA | NA | #VALUE! |
| ENSMUSG00000095331.4 | 0 | NA | NA | NA | NA | NA | #VALUE! |
| ENSMUSG00000095332.3 | 0 | NA | NA | NA | NA | NA | #VALUE! |
| ENSMUSG00000095334.3 | 0 | NA | NA | NA | NA | NA | #VALUE! |
| ENSMUSG00000095335.3 | 0 | NA | NA | NA | NA | NA | #VALUE! |
| ENSMUSG00000095338.2 | 0 | NA | NA | NA | NA | NA | #VALUE! |
| ENSMUSG00000095339.3 | 0 | NA | NA | NA | NA | NA | #VALUE! |
| ENSMUSG00000095341.2 | 0 | NA | NA | NA | NA | NA | #VALUE! |
| ENSMUSG00000095344.3 | 0 | NA | NA | NA | NA | NA | #VALUE! |
| ENSMUSG00000095345.3 | 0 | NA | NA | NA | NA | NA | #VALUE! |
| ENSMUSG00000095346.2 | 0 | NA | NA | NA | NA | NA | #VALUE! |
| ENSMUSG00000095348.2 | 0 | NA | NA | NA | NA | NA | #VALUE! |
| ENSMUSG00000095349.3 | 0 | NA | NA | NA | NA | NA | #VALUE! |
| ENSMUSG00000095351.3 | 0 | NA | NA | NA | NA | NA | #VALUE! |
| ENSMUSG00000095353.3 | 0 | NA | NA | NA | NA | NA | #VALUE! |
| ENSMUSG00000095356.2 | 0 | NA | NA | NA | NA | NA | #VALUE! |
| ENSMUSG00000095358.2 | 0 | NA | NA | NA | NA | NA | #VALUE! |
| ENSMUSG00000095360.8 | 0 | NA | NA | NA | NA | NA | #VALUE! |
| ENSMUSG00000095363.3 | 0 | NA | NA | NA | NA | NA | #VALUE! |

| | | | | | | | |
|---|---|---|---|---|---|---|---|
| ENSMUSG00000095365.4 | 0 | NA | NA | NA | NA | NA | #VALUE! |
| ENSMUSG00000095366.3 | 0 | NA | NA | NA | NA | NA | #VALUE! |
| ENSMUSG00000095368.8 | 0 | NA | NA | NA | NA | NA | #VALUE! |
| ENSMUSG00000095369.3 | 0 | NA | NA | NA | NA | NA | #VALUE! |
| ENSMUSG00000095370.3 | 0 | NA | NA | NA | NA | NA | #VALUE! |
| ENSMUSG00000095371.2 | 0 | NA | NA | NA | NA | NA | #VALUE! |
| ENSMUSG00000095377.3 | 0 | NA | NA | NA | NA | NA | #VALUE! |
| ENSMUSG00000095383.2 | 0 | NA | NA | NA | NA | NA | #VALUE! |
| ENSMUSG00000095384.2 | 0 | NA | NA | NA | NA | NA | #VALUE! |
| ENSMUSG00000095386.2 | 0 | NA | NA | NA | NA | NA | #VALUE! |
| ENSMUSG00000095387.6 | 0 | NA | NA | NA | NA | NA | #VALUE! |
| ENSMUSG00000095388.8 | 0 | NA | NA | NA | NA | NA | #VALUE! |
| ENSMUSG00000095390.3 | 0 | NA | NA | NA | NA | NA | #VALUE! |
| ENSMUSG00000095392.2 | 0 | NA | NA | NA | NA | NA | #VALUE! |
| ENSMUSG00000095394.2 | 0 | NA | NA | NA | NA | NA | #VALUE! |
| ENSMUSG00000095398.3 | 0 | NA | NA | NA | NA | NA | #VALUE! |
| ENSMUSG00000095399.2 | 0 | NA | NA | NA | NA | NA | #VALUE! |
| ENSMUSG00000095401.3 | 0 | NA | NA | NA | NA | NA | #VALUE! |
| ENSMUSG00000095403.9 | 0 | NA | NA | NA | NA | NA | #VALUE! |
| ENSMUSG00000095407.3 | 0 | NA | NA | NA | NA | NA | #VALUE! |
| ENSMUSG00000095408.2 | 0 | NA | NA | NA | NA | NA | #VALUE! |
| ENSMUSG00000095409.2 | 0 | NA | NA | NA | NA | NA | #VALUE! |
| ENSMUSG00000095411.2 | 0 | NA | NA | NA | NA | NA | #VALUE! |
| ENSMUSG00000095412.4 | 0 | NA | NA | NA | NA | NA | #VALUE! |
| ENSMUSG00000095413.2 | 0 | NA | NA | NA | NA | NA | #VALUE! |
| ENSMUSG00000095416.7 | 0 | NA | NA | NA | NA | NA | #VALUE! |
| ENSMUSG00000095419.2 | 0 | NA | NA | NA | NA | NA | #VALUE! |
| ENSMUSG00000095420.2 | 0 | NA | NA | NA | NA | NA | #VALUE! |
| ENSMUSG00000095423.2 | 0 | NA | NA | NA | NA | NA | #VALUE! |
| ENSMUSG00000095426.7 | 0 | NA | NA | NA | NA | NA | #VALUE! |
| ENSMUSG00000095427.3 | 0 | NA | NA | NA | NA | NA | #VALUE! |
| ENSMUSG00000095429.2 | 0 | NA | NA | NA | NA | NA | #VALUE! |
| ENSMUSG00000095430.4 | 0 | NA | NA | NA | NA | NA | #VALUE! |
| ENSMUSG00000095433.3 | 0 | NA | NA | NA | NA | NA | #VALUE! |
| ENSMUSG00000095436.2 | 0 | NA | NA | NA | NA | NA | #VALUE! |
| ENSMUSG00000095438.2 | 0 | NA | NA | NA | NA | NA | #VALUE! |
| ENSMUSG00000095442.2 | 0 | NA | NA | NA | NA | NA | #VALUE! |
| ENSMUSG00000095444.2 | 0 | NA | NA | NA | NA | NA | #VALUE! |
| ENSMUSG00000095445.2 | 0 | NA | NA | NA | NA | NA | #VALUE! |
| ENSMUSG00000095448.4 | 0 | NA | NA | NA | NA | NA | #VALUE! |
| ENSMUSG00000095450.2 | 0 | NA | NA | NA | NA | NA | #VALUE! |
| ENSMUSG00000095452.3 | 0 | NA | NA | NA | NA | NA | #VALUE! |
| ENSMUSG00000095455.2 | 0 | NA | NA | NA | NA | NA | #VALUE! |
| ENSMUSG00000095457.4 | 0 | NA | NA | NA | NA | NA | #VALUE! |

| | | | | | | | |
|---|---|---|---|---|---|---|---|
| ENSMUSG00000095460.2 | 0 | NA | NA | NA | NA | NA | #VALUE! |
| ENSMUSG00000095462.2 | 0 | NA | NA | NA | NA | NA | #VALUE! |
| ENSMUSG00000095463.9 | 0 | NA | NA | NA | NA | NA | #VALUE! |
| ENSMUSG00000095464.2 | 0 | NA | NA | NA | NA | NA | #VALUE! |
| ENSMUSG00000095465.3 | 0 | NA | NA | NA | NA | NA | #VALUE! |
| ENSMUSG00000095466.8 | 0 | NA | NA | NA | NA | NA | #VALUE! |
| ENSMUSG00000095469.2 | 0 | NA | NA | NA | NA | NA | #VALUE! |
| ENSMUSG00000095470.2 | 0 | NA | NA | NA | NA | NA | #VALUE! |
| ENSMUSG00000095474.3 | 0 | NA | NA | NA | NA | NA | #VALUE! |
| ENSMUSG00000095475.2 | 0 | NA | NA | NA | NA | NA | #VALUE! |
| ENSMUSG00000095478.3 | 0 | NA | NA | NA | NA | NA | #VALUE! |
| ENSMUSG00000095483.3 | 0 | NA | NA | NA | NA | NA | #VALUE! |
| ENSMUSG00000095484.5 | 0 | NA | NA | NA | NA | NA | #VALUE! |
| ENSMUSG00000095486.3 | 0 | NA | NA | NA | NA | NA | #VALUE! |
| ENSMUSG00000095487.4 | 0 | NA | NA | NA | NA | NA | #VALUE! |
| ENSMUSG00000095490.2 | 0 | NA | NA | NA | NA | NA | #VALUE! |
| ENSMUSG00000095492.2 | 0 | NA | NA | NA | NA | NA | #VALUE! |
| ENSMUSG00000095493.3 | 0 | NA | NA | NA | NA | NA | #VALUE! |
| ENSMUSG00000095494.2 | 0 | NA | NA | NA | NA | NA | #VALUE! |
| ENSMUSG00000095495.3 | 0 | NA | NA | NA | NA | NA | #VALUE! |
| ENSMUSG00000095497.3 | 0 | NA | NA | NA | NA | NA | #VALUE! |
| ENSMUSG00000095498.2 | 0 | NA | NA | NA | NA | NA | #VALUE! |
| ENSMUSG00000095499.2 | 0 | NA | NA | NA | NA | NA | #VALUE! |
| ENSMUSG00000095500.2 | 0 | NA | NA | NA | NA | NA | #VALUE! |
| ENSMUSG00000095503.5 | 0 | NA | NA | NA | NA | NA | #VALUE! |
| ENSMUSG00000095505.2 | 0 | NA | NA | NA | NA | NA | #VALUE! |
| ENSMUSG00000095508.3 | 0 | NA | NA | NA | NA | NA | #VALUE! |
| ENSMUSG00000095510.9 | 0 | NA | NA | NA | NA | NA | #VALUE! |
| ENSMUSG00000095512.2 | 0 | NA | NA | NA | NA | NA | #VALUE! |
| ENSMUSG00000095514.3 | 0 | NA | NA | NA | NA | NA | #VALUE! |
| ENSMUSG00000095517.2 | 0 | NA | NA | NA | NA | NA | #VALUE! |
| ENSMUSG00000095518.2 | 0 | NA | NA | NA | NA | NA | #VALUE! |
| ENSMUSG00000095519.3 | 0 | NA | NA | NA | NA | NA | #VALUE! |
| ENSMUSG00000095520.3 | 0 | NA | NA | NA | NA | NA | #VALUE! |
| ENSMUSG00000095521.2 | 0 | NA | NA | NA | NA | NA | #VALUE! |
| ENSMUSG00000095522.3 | 0 | NA | NA | NA | NA | NA | #VALUE! |
| ENSMUSG00000095523.2 | 0 | NA | NA | NA | NA | NA | #VALUE! |
| ENSMUSG00000095524.2 | 0 | NA | NA | NA | NA | NA | #VALUE! |
| ENSMUSG00000095525.2 | 0 | NA | NA | NA | NA | NA | #VALUE! |
| ENSMUSG00000095527.4 | 0 | NA | NA | NA | NA | NA | #VALUE! |
| ENSMUSG00000095528.9 | 0 | NA | NA | NA | NA | NA | #VALUE! |
| ENSMUSG00000095533.8 | 0 | NA | NA | NA | NA | NA | #VALUE! |
| ENSMUSG00000095534.2 | 0 | NA | NA | NA | NA | NA | #VALUE! |
| ENSMUSG00000095538.2 | 0 | NA | NA | NA | NA | NA | #VALUE! |

| | | | | | | | |
|---|---|---|---|---|---|---|---|
| ENSMUSG00000095540.2 | 0 | NA | NA | NA | NA | NA | #VALUE! |
| ENSMUSG00000095543.3 | 0 | NA | NA | NA | NA | NA | #VALUE! |
| ENSMUSG00000095544.2 | 0 | NA | NA | NA | NA | NA | #VALUE! |
| ENSMUSG00000095545.8 | 0 | NA | NA | NA | NA | NA | #VALUE! |
| ENSMUSG00000095546.8 | 0 | NA | NA | NA | NA | NA | #VALUE! |
| ENSMUSG00000095547.2 | 0.093255 | 0.868093 | 4.072464 | 0.213162 | 0.831201 | NA | #VALUE! |
| ENSMUSG00000095549.2 | 0 | NA | NA | NA | NA | NA | #VALUE! |
| ENSMUSG00000095550.3 | 0 | NA | NA | NA | NA | NA | #VALUE! |
| ENSMUSG00000095551.2 | 0 | NA | NA | NA | NA | NA | #VALUE! |
| ENSMUSG00000095553.3 | 0 | NA | NA | NA | NA | NA | #VALUE! |
| ENSMUSG00000095554.3 | 0 | NA | NA | NA | NA | NA | #VALUE! |
| ENSMUSG00000095557.8 | 0 | NA | NA | NA | NA | NA | #VALUE! |
| ENSMUSG00000095560.3 | 0 | NA | NA | NA | NA | NA | #VALUE! |
| ENSMUSG00000095561.2 | 0 | NA | NA | NA | NA | NA | #VALUE! |
| ENSMUSG00000095562.9 | 0 | NA | NA | NA | NA | NA | #VALUE! |
| ENSMUSG00000095563.2 | 0 | NA | NA | NA | NA | NA | #VALUE! |
| ENSMUSG00000095564.3 | 0 | NA | NA | NA | NA | NA | #VALUE! |
| ENSMUSG00000095565.3 | 0 | NA | NA | NA | NA | NA | #VALUE! |
| ENSMUSG00000095569.2 | 0 | NA | NA | NA | NA | NA | #VALUE! |
| ENSMUSG00000095570.2 | 0 | NA | NA | NA | NA | NA | #VALUE! |
| ENSMUSG00000095571.3 | 0 | NA | NA | NA | NA | NA | #VALUE! |
| ENSMUSG00000095572.2 | 0 | NA | NA | NA | NA | NA | #VALUE! |
| ENSMUSG00000095573.4 | 0 | NA | NA | NA | NA | NA | #VALUE! |
| ENSMUSG00000095574.3 | 0 | NA | NA | NA | NA | NA | #VALUE! |
| ENSMUSG00000095576.6 | 0 | NA | NA | NA | NA | NA | #VALUE! |
| ENSMUSG00000095577.3 | 0 | NA | NA | NA | NA | NA | #VALUE! |
| ENSMUSG00000095578.8 | 0 | NA | NA | NA | NA | NA | #VALUE! |
| ENSMUSG00000095583.2 | 0 | NA | NA | NA | NA | NA | #VALUE! |
| ENSMUSG00000095586.2 | 0 | NA | NA | NA | NA | NA | #VALUE! |
| ENSMUSG00000095588.2 | 1.821948 | -0.94106 | 3.837053 | -0.24526 | 0.806257 | NA | #VALUE! |
| ENSMUSG00000095589.3 | 0 | NA | NA | NA | NA | NA | #VALUE! |
| ENSMUSG00000095591.2 | 0 | NA | NA | NA | NA | NA | #VALUE! |
| ENSMUSG00000095592.2 | 0 | NA | NA | NA | NA | NA | #VALUE! |
| ENSMUSG00000095593.2 | 0 | NA | NA | NA | NA | NA | #VALUE! |
| ENSMUSG00000095595.3 | 0 | NA | NA | NA | NA | NA | #VALUE! |
| ENSMUSG00000095599.2 | 0 | NA | NA | NA | NA | NA | #VALUE! |
| ENSMUSG00000095600.3 | 0 | NA | NA | NA | NA | NA | #VALUE! |
| ENSMUSG00000095601.9 | 0 | NA | NA | NA | NA | NA | #VALUE! |
| ENSMUSG00000095603.5 | 0 | NA | NA | NA | NA | NA | #VALUE! |
| ENSMUSG00000095605.2 | 0 | NA | NA | NA | NA | NA | #VALUE! |
| ENSMUSG00000095606.8 | 0 | NA | NA | NA | NA | NA | #VALUE! |
| ENSMUSG00000095607.4 | 0 | NA | NA | NA | NA | NA | #VALUE! |
| ENSMUSG00000095608.4 | 0 | NA | NA | NA | NA | NA | #VALUE! |
| ENSMUSG00000095609.3 | 0 | NA | NA | NA | NA | NA | #VALUE! |

| | | | | | | | |
|---|---|---|---|---|---|---|---|
| ENSMUSG00000095612.3 | 0 | NA | NA | NA | NA | NA | #VALUE! |
| ENSMUSG00000095614.2 | 0 | NA | NA | NA | NA | NA | #VALUE! |
| ENSMUSG00000095616.2 | 6.254401 | 5.171616 | 3.864462 | 1.33825 | 0.180815 | NA | #VALUE! |
| ENSMUSG00000095619.2 | 0 | NA | NA | NA | NA | NA | #VALUE! |
| ENSMUSG00000095620.8 | 0 | NA | NA | NA | NA | NA | #VALUE! |
| ENSMUSG00000095621.2 | 0 | NA | NA | NA | NA | NA | #VALUE! |
| ENSMUSG00000095625.3 | 0 | NA | NA | NA | NA | NA | #VALUE! |
| ENSMUSG00000095626.3 | 0 | NA | NA | NA | NA | NA | #VALUE! |
| ENSMUSG00000095627.2 | 0 | NA | NA | NA | NA | NA | #VALUE! |
| ENSMUSG00000095629.3 | 0 | NA | NA | NA | NA | NA | #VALUE! |
| ENSMUSG00000095630.3 | 0 | NA | NA | NA | NA | NA | #VALUE! |
| ENSMUSG00000095632.3 | 0 | NA | NA | NA | NA | NA | #VALUE! |
| ENSMUSG00000095633.3 | 0 | NA | NA | NA | NA | NA | #VALUE! |
| ENSMUSG00000095634.3 | 0 | NA | NA | NA | NA | NA | #VALUE! |
| ENSMUSG00000095638.3 | 0 | NA | NA | NA | NA | NA | #VALUE! |
| ENSMUSG00000095640.5 | 0 | NA | NA | NA | NA | NA | #VALUE! |
| ENSMUSG00000095642.4 | 0 | NA | NA | NA | NA | NA | #VALUE! |
| ENSMUSG00000095643.3 | 0 | NA | NA | NA | NA | NA | #VALUE! |
| ENSMUSG00000095646.2 | 0 | NA | NA | NA | NA | NA | #VALUE! |
| ENSMUSG00000095647.2 | 0 | NA | NA | NA | NA | NA | #VALUE! |
| ENSMUSG00000095648.8 | 0 | NA | NA | NA | NA | NA | #VALUE! |
| ENSMUSG00000095649.4 | 0 | NA | NA | NA | NA | NA | #VALUE! |
| ENSMUSG00000095650.3 | 0 | NA | NA | NA | NA | NA | #VALUE! |
| ENSMUSG00000095651.2 | 0 | NA | NA | NA | NA | NA | #VALUE! |
| ENSMUSG00000095653.3 | 0 | NA | NA | NA | NA | NA | #VALUE! |
| ENSMUSG00000095654.3 | 0 | NA | NA | NA | NA | NA | #VALUE! |
| ENSMUSG00000095655.2 | 0 | NA | NA | NA | NA | NA | #VALUE! |
| ENSMUSG00000095656.2 | 0 | NA | NA | NA | NA | NA | #VALUE! |
| ENSMUSG00000095657.2 | 0 | NA | NA | NA | NA | NA | #VALUE! |
| ENSMUSG00000095658.3 | 0 | NA | NA | NA | NA | NA | #VALUE! |
| ENSMUSG00000095659.3 | 0 | NA | NA | NA | NA | NA | #VALUE! |
| ENSMUSG00000095661.2 | 0 | NA | NA | NA | NA | NA | #VALUE! |
| ENSMUSG00000095662.2 | 0 | NA | NA | NA | NA | NA | #VALUE! |
| ENSMUSG00000095663.3 | 0 | NA | NA | NA | NA | NA | #VALUE! |
| ENSMUSG00000095664.3 | 0 | NA | NA | NA | NA | NA | #VALUE! |
| ENSMUSG00000095666.2 | 0 | NA | NA | NA | NA | NA | #VALUE! |
| ENSMUSG00000095667.3 | 0 | NA | NA | NA | NA | NA | #VALUE! |
| ENSMUSG00000095668.2 | 0 | NA | NA | NA | NA | NA | #VALUE! |
| ENSMUSG00000095670.3 | 0 | NA | NA | NA | NA | NA | #VALUE! |
| ENSMUSG00000095672.2 | 0 | NA | NA | NA | NA | NA | #VALUE! |
| ENSMUSG00000095673.2 | 0 | NA | NA | NA | NA | NA | #VALUE! |
| ENSMUSG00000095675.3 | 0 | NA | NA | NA | NA | NA | #VALUE! |
| ENSMUSG00000095677.3 | 0 | NA | NA | NA | NA | NA | #VALUE! |
| ENSMUSG00000095680.2 | 0 | NA | NA | NA | NA | NA | #VALUE! |

| | | | | | | | |
|---|---|---|---|---|---|---|---|
| ENSMUSG00000095681.8 | 0 | NA | NA | NA | NA | NA | #VALUE! |
| ENSMUSG00000095682.2 | 0 | NA | NA | NA | NA | NA | #VALUE! |
| ENSMUSG00000095683.3 | 0 | NA | NA | NA | NA | NA | #VALUE! |
| ENSMUSG00000095686.8 | 0 | NA | NA | NA | NA | NA | #VALUE! |
| ENSMUSG00000095687.3 | 0 | NA | NA | NA | NA | NA | #VALUE! |
| ENSMUSG00000095689.6 | 0 | NA | NA | NA | NA | NA | #VALUE! |
| ENSMUSG00000095690.3 | 0 | NA | NA | NA | NA | NA | #VALUE! |
| ENSMUSG00000095693.2 | 0 | NA | NA | NA | NA | NA | #VALUE! |
| ENSMUSG00000095696.4 | 0 | NA | NA | NA | NA | NA | #VALUE! |
| ENSMUSG00000095698.3 | 0 | NA | NA | NA | NA | NA | #VALUE! |
| ENSMUSG00000095700.3 | 0 | NA | NA | NA | NA | NA | #VALUE! |
| ENSMUSG00000095702.2 | 0 | NA | NA | NA | NA | NA | #VALUE! |
| ENSMUSG00000095704.3 | 0 | NA | NA | NA | NA | NA | #VALUE! |
| ENSMUSG00000095706.4 | 0 | NA | NA | NA | NA | NA | #VALUE! |
| ENSMUSG00000095710.8 | 0 | NA | NA | NA | NA | NA | #VALUE! |
| ENSMUSG00000095711.6 | 0 | NA | NA | NA | NA | NA | #VALUE! |
| ENSMUSG00000095716.2 | 0 | NA | NA | NA | NA | NA | #VALUE! |
| ENSMUSG00000095717.2 | 0 | NA | NA | NA | NA | NA | #VALUE! |
| ENSMUSG00000095718.3 | 0 | NA | NA | NA | NA | NA | #VALUE! |
| ENSMUSG00000095721.2 | 0 | NA | NA | NA | NA | NA | #VALUE! |
| ENSMUSG00000095724.2 | 0 | NA | NA | NA | NA | NA | #VALUE! |
| ENSMUSG00000095725.3 | 0 | NA | NA | NA | NA | NA | #VALUE! |
| ENSMUSG00000095728.2 | 0 | NA | NA | NA | NA | NA | #VALUE! |
| ENSMUSG00000095730.4 | 0 | NA | NA | NA | NA | NA | #VALUE! |
| ENSMUSG00000095736.2 | 0 | NA | NA | NA | NA | NA | #VALUE! |
| ENSMUSG00000095737.6 | 0 | NA | NA | NA | NA | NA | #VALUE! |
| ENSMUSG00000095741.4 | 0 | NA | NA | NA | NA | NA | #VALUE! |
| ENSMUSG00000095742.2 | 0 | NA | NA | NA | NA | NA | #VALUE! |
| ENSMUSG00000095743.2 | 0 | NA | NA | NA | NA | NA | #VALUE! |
| ENSMUSG00000095745.2 | 0 | NA | NA | NA | NA | NA | #VALUE! |
| ENSMUSG00000095753.3 | 0 | NA | NA | NA | NA | NA | #VALUE! |
| ENSMUSG00000095754.3 | 0 | NA | NA | NA | NA | NA | #VALUE! |
| ENSMUSG00000095756.3 | 0 | NA | NA | NA | NA | NA | #VALUE! |
| ENSMUSG00000095758.2 | 0 | NA | NA | NA | NA | NA | #VALUE! |
| ENSMUSG00000095759.2 | 0 | NA | NA | NA | NA | NA | #VALUE! |
| ENSMUSG00000095761.7 | 0 | NA | NA | NA | NA | NA | #VALUE! |
| ENSMUSG00000095762.2 | 0.204111 | -1.05581 | 4.073302 | -0.2592 | 0.795479 | NA | #VALUE! |
| ENSMUSG00000095763.2 | 0 | NA | NA | NA | NA | NA | #VALUE! |
| ENSMUSG00000095765.4 | 0 | NA | NA | NA | NA | NA | #VALUE! |
| ENSMUSG00000095766.3 | 0 | NA | NA | NA | NA | NA | #VALUE! |
| ENSMUSG00000095768.2 | 0 | NA | NA | NA | NA | NA | #VALUE! |
| ENSMUSG00000095769.2 | 0 | NA | NA | NA | NA | NA | #VALUE! |
| ENSMUSG00000095770.2 | 0 | NA | NA | NA | NA | NA | #VALUE! |
| ENSMUSG00000095771.3 | 0 | NA | NA | NA | NA | NA | #VALUE! |

| | | | | | | | |
|---|---|---|---|---|---|---|---|
| ENSMUSG00000095773.4 | 0 | NA | NA | NA | NA | NA | #VALUE! |
| ENSMUSG00000095774.3 | 0 | NA | NA | NA | NA | NA | #VALUE! |
| ENSMUSG00000095779.3 | 0 | NA | NA | NA | NA | NA | #VALUE! |
| ENSMUSG00000095785.3 | 0 | NA | NA | NA | NA | NA | #VALUE! |
| ENSMUSG00000095787.2 | 0 | NA | NA | NA | NA | NA | #VALUE! |
| ENSMUSG00000095788.7 | 0.525066 | -2.64687 | 3.988095 | -0.66369 | 0.506887 | NA | #VALUE! |
| ENSMUSG00000095789.7 | 0 | NA | NA | NA | NA | NA | #VALUE! |
| ENSMUSG00000095793.8 | 0 | NA | NA | NA | NA | NA | #VALUE! |
| ENSMUSG00000095794.3 | 0 | NA | NA | NA | NA | NA | #VALUE! |
| ENSMUSG00000095797.9 | 0 | NA | NA | NA | NA | NA | #VALUE! |
| ENSMUSG00000095799.2 | 0 | NA | NA | NA | NA | NA | #VALUE! |
| ENSMUSG00000095802.2 | 0 | NA | NA | NA | NA | NA | #VALUE! |
| ENSMUSG00000095804.3 | 0 | NA | NA | NA | NA | NA | #VALUE! |
| ENSMUSG00000095806.2 | 0 | NA | NA | NA | NA | NA | #VALUE! |
| ENSMUSG00000095807.3 | 0 | NA | NA | NA | NA | NA | #VALUE! |
| ENSMUSG00000095809.4 | 0 | NA | NA | NA | NA | NA | #VALUE! |
| ENSMUSG00000095812.2 | 0 | NA | NA | NA | NA | NA | #VALUE! |
| ENSMUSG00000095813.5 | 0 | NA | NA | NA | NA | NA | #VALUE! |
| ENSMUSG00000095814.2 | 0 | NA | NA | NA | NA | NA | #VALUE! |
| ENSMUSG00000095816.3 | 0 | NA | NA | NA | NA | NA | #VALUE! |
| ENSMUSG00000095817.2 | 0 | NA | NA | NA | NA | NA | #VALUE! |
| ENSMUSG00000095820.2 | 0 | NA | NA | NA | NA | NA | #VALUE! |
| ENSMUSG00000095821.3 | 0 | NA | NA | NA | NA | NA | #VALUE! |
| ENSMUSG00000095822.3 | 0 | NA | NA | NA | NA | NA | #VALUE! |
| ENSMUSG00000095824.2 | 0 | NA | NA | NA | NA | NA | #VALUE! |
| ENSMUSG00000095826.2 | 0 | NA | NA | NA | NA | NA | #VALUE! |
| ENSMUSG00000095827.3 | 0 | NA | NA | NA | NA | NA | #VALUE! |
| ENSMUSG00000095829.9 | 0 | NA | NA | NA | NA | NA | #VALUE! |
| ENSMUSG00000095831.3 | 0 | NA | NA | NA | NA | NA | #VALUE! |
| ENSMUSG00000095834.3 | 0 | NA | NA | NA | NA | NA | #VALUE! |
| ENSMUSG00000095836.4 | 0 | NA | NA | NA | NA | NA | #VALUE! |
| ENSMUSG00000095837.2 | 0 | NA | NA | NA | NA | NA | #VALUE! |
| ENSMUSG00000095839.4 | 0 | NA | NA | NA | NA | NA | #VALUE! |
| ENSMUSG00000095841.3 | 0 | NA | NA | NA | NA | NA | #VALUE! |
| ENSMUSG00000095845.4 | 0 | NA | NA | NA | NA | NA | #VALUE! |
| ENSMUSG00000095847.2 | 0 | NA | NA | NA | NA | NA | #VALUE! |
| ENSMUSG00000095852.8 | 0 | NA | NA | NA | NA | NA | #VALUE! |
| ENSMUSG00000095854.5 | 0 | NA | NA | NA | NA | NA | #VALUE! |
| ENSMUSG00000095858.5 | 0 | NA | NA | NA | NA | NA | #VALUE! |
| ENSMUSG00000095859.3 | 0 | NA | NA | NA | NA | NA | #VALUE! |
| ENSMUSG00000095862.2 | 0 | NA | NA | NA | NA | NA | #VALUE! |
| ENSMUSG00000095863.3 | 0 | NA | NA | NA | NA | NA | #VALUE! |
| ENSMUSG00000095864.3 | 0 | NA | NA | NA | NA | NA | #VALUE! |
| ENSMUSG00000095865.2 | 0 | NA | NA | NA | NA | NA | #VALUE! |

| | | | | | | | |
|---|---|---|---|---|---|---|---|
| ENSMUSG00000095866.2 | 0 | NA | NA | NA | NA | NA | #VALUE! |
| ENSMUSG00000095867.3 | 0 | NA | NA | NA | NA | NA | #VALUE! |
| ENSMUSG00000095869.2 | 0 | NA | NA | NA | NA | NA | #VALUE! |
| ENSMUSG00000095870.2 | 0 | NA | NA | NA | NA | NA | #VALUE! |
| ENSMUSG00000095872.8 | 0 | NA | NA | NA | NA | NA | #VALUE! |
| ENSMUSG00000095873.2 | 0 | NA | NA | NA | NA | NA | #VALUE! |
| ENSMUSG00000095874.3 | 0 | NA | NA | NA | NA | NA | #VALUE! |
| ENSMUSG00000095879.2 | 0 | NA | NA | NA | NA | NA | #VALUE! |
| ENSMUSG00000095886.3 | 0 | NA | NA | NA | NA | NA | #VALUE! |
| ENSMUSG00000095887.8 | 0 | NA | NA | NA | NA | NA | #VALUE! |
| ENSMUSG00000095889.2 | 0 | NA | NA | NA | NA | NA | #VALUE! |
| ENSMUSG00000095890.2 | 0 | NA | NA | NA | NA | NA | #VALUE! |
| ENSMUSG00000095891.8 | 8.577905 | -2.84196 | 3.439452 | -0.82628 | 0.408643 | NA | #VALUE! |
| ENSMUSG00000095893.5 | 0 | NA | NA | NA | NA | NA | #VALUE! |
| ENSMUSG00000095895.3 | 0 | NA | NA | NA | NA | NA | #VALUE! |
| ENSMUSG00000095896.2 | 0 | NA | NA | NA | NA | NA | #VALUE! |
| ENSMUSG00000095897.2 | 0 | NA | NA | NA | NA | NA | #VALUE! |
| ENSMUSG00000095899.2 | 0 | NA | NA | NA | NA | NA | #VALUE! |
| ENSMUSG00000095900.3 | 0 | NA | NA | NA | NA | NA | #VALUE! |
| ENSMUSG00000095901.3 | 0 | NA | NA | NA | NA | NA | #VALUE! |
| ENSMUSG00000095903.4 | 0 | NA | NA | NA | NA | NA | #VALUE! |
| ENSMUSG00000095905.3 | 0 | NA | NA | NA | NA | NA | #VALUE! |
| ENSMUSG00000095906.2 | 0 | NA | NA | NA | NA | NA | #VALUE! |
| ENSMUSG00000095907.3 | 0 | NA | NA | NA | NA | NA | #VALUE! |
| ENSMUSG00000095909.3 | 0 | NA | NA | NA | NA | NA | #VALUE! |
| ENSMUSG00000095910.3 | 0 | NA | NA | NA | NA | NA | #VALUE! |
| ENSMUSG00000095912.2 | 0 | NA | NA | NA | NA | NA | #VALUE! |
| ENSMUSG00000095913.2 | 0 | NA | NA | NA | NA | NA | #VALUE! |
| ENSMUSG00000095914.2 | 0 | NA | NA | NA | NA | NA | #VALUE! |
| ENSMUSG00000095916.3 | 0 | NA | NA | NA | NA | NA | #VALUE! |
| ENSMUSG00000095917.3 | 0 | NA | NA | NA | NA | NA | #VALUE! |
| ENSMUSG00000095918.8 | 0 | NA | NA | NA | NA | NA | #VALUE! |
| ENSMUSG00000095920.3 | 0 | NA | NA | NA | NA | NA | #VALUE! |
| ENSMUSG00000095922.2 | 0 | NA | NA | NA | NA | NA | #VALUE! |
| ENSMUSG00000095923.7 | 0 | NA | NA | NA | NA | NA | #VALUE! |
| ENSMUSG00000095925.2 | 0 | NA | NA | NA | NA | NA | #VALUE! |
| ENSMUSG00000095927.2 | 0 | NA | NA | NA | NA | NA | #VALUE! |
| ENSMUSG00000095928.4 | 0 | NA | NA | NA | NA | NA | #VALUE! |
| ENSMUSG00000095929.3 | 0 | NA | NA | NA | NA | NA | #VALUE! |
| ENSMUSG00000095931.2 | 0 | NA | NA | NA | NA | NA | #VALUE! |
| ENSMUSG00000095932.4 | 0 | NA | NA | NA | NA | NA | #VALUE! |
| ENSMUSG00000095934.2 | 0 | NA | NA | NA | NA | NA | #VALUE! |
| ENSMUSG00000095935.2 | 0 | NA | NA | NA | NA | NA | #VALUE! |
| ENSMUSG00000095936.4 | 0 | NA | NA | NA | NA | NA | #VALUE! |

| | | | | | | | |
|---|---|---|---|---|---|---|---|
| ENSMUSG00000095937.2 | 0 | NA | NA | NA | NA | NA | #VALUE! |
| ENSMUSG00000095940.2 | 0.093255 | 0.868093 | 4.072464 | 0.213162 | 0.831201 | NA | #VALUE! |
| ENSMUSG00000095941.4 | 0 | NA | NA | NA | NA | NA | #VALUE! |
| ENSMUSG00000095948.8 | 0 | NA | NA | NA | NA | NA | #VALUE! |
| ENSMUSG00000095950.3 | 0 | NA | NA | NA | NA | NA | #VALUE! |
| ENSMUSG00000095952.2 | 0 | NA | NA | NA | NA | NA | #VALUE! |
| ENSMUSG00000095953.4 | 0 | NA | NA | NA | NA | NA | #VALUE! |
| ENSMUSG00000095954.5 | 0 | NA | NA | NA | NA | NA | #VALUE! |
| ENSMUSG00000095956.3 | 0 | NA | NA | NA | NA | NA | #VALUE! |
| ENSMUSG00000095957.3 | 0 | NA | NA | NA | NA | NA | #VALUE! |
| ENSMUSG00000095958.4 | 0 | NA | NA | NA | NA | NA | #VALUE! |
| ENSMUSG00000095959.3 | 0 | NA | NA | NA | NA | NA | #VALUE! |
| ENSMUSG00000095960.2 | 0 | NA | NA | NA | NA | NA | #VALUE! |
| ENSMUSG00000095961.3 | 0 | NA | NA | NA | NA | NA | #VALUE! |
| ENSMUSG00000095962.2 | 0 | NA | NA | NA | NA | NA | #VALUE! |
| ENSMUSG00000095963.2 | 0 | NA | NA | NA | NA | NA | #VALUE! |
| ENSMUSG00000095964.2 | 0 | NA | NA | NA | NA | NA | #VALUE! |
| ENSMUSG00000095968.3 | 0 | NA | NA | NA | NA | NA | #VALUE! |
| ENSMUSG00000095970.2 | 0 | NA | NA | NA | NA | NA | #VALUE! |
| ENSMUSG00000095971.2 | 0 | NA | NA | NA | NA | NA | #VALUE! |
| ENSMUSG00000095973.2 | 0 | NA | NA | NA | NA | NA | #VALUE! |
| ENSMUSG00000095974.2 | 0 | NA | NA | NA | NA | NA | #VALUE! |
| ENSMUSG00000095975.4 | 0 | NA | NA | NA | NA | NA | #VALUE! |
| ENSMUSG00000095976.3 | 0 | NA | NA | NA | NA | NA | #VALUE! |
| ENSMUSG00000095978.2 | 0 | NA | NA | NA | NA | NA | #VALUE! |
| ENSMUSG00000095979.8 | 0 | NA | NA | NA | NA | NA | #VALUE! |
| ENSMUSG00000095981.2 | 0 | NA | NA | NA | NA | NA | #VALUE! |
| ENSMUSG00000095984.2 | 0 | NA | NA | NA | NA | NA | #VALUE! |
| ENSMUSG00000095987.2 | 0 | NA | NA | NA | NA | NA | #VALUE! |
| ENSMUSG00000095988.2 | 0 | NA | NA | NA | NA | NA | #VALUE! |
| ENSMUSG00000095990.2 | 0.491951 | 2.402185 | 4.000147 | 0.600524 | 0.548157 | NA | #VALUE! |
| ENSMUSG00000095991.2 | 0 | NA | NA | NA | NA | NA | #VALUE! |
| ENSMUSG00000095992.3 | 0 | NA | NA | NA | NA | NA | #VALUE! |
| ENSMUSG00000095993.2 | 0 | NA | NA | NA | NA | NA | #VALUE! |
| ENSMUSG00000095994.2 | 0 | NA | NA | NA | NA | NA | #VALUE! |
| ENSMUSG00000095996.5 | 0 | NA | NA | NA | NA | NA | #VALUE! |
| ENSMUSG00000096000.2 | 0 | NA | NA | NA | NA | NA | #VALUE! |
| ENSMUSG00000096001.3 | 0 | NA | NA | NA | NA | NA | #VALUE! |
| ENSMUSG00000096002.3 | 0 | NA | NA | NA | NA | NA | #VALUE! |
| ENSMUSG00000096003.8 | 0 | NA | NA | NA | NA | NA | #VALUE! |
| ENSMUSG00000096006.2 | 5.790616 | -2.2505 | 3.068564 | -0.7334 | 0.463311 | NA | #VALUE! |
| ENSMUSG00000096008.2 | 0 | NA | NA | NA | NA | NA | #VALUE! |
| ENSMUSG00000096009.3 | 0 | NA | NA | NA | NA | NA | #VALUE! |
| ENSMUSG00000096010.3 | 0 | NA | NA | NA | NA | NA | #VALUE! |

| | | | | | | | |
|---|---|---|---|---|---|---|---|
| ENSMUSG00000096011.2 | 0 | NA | NA | NA | NA | NA | #VALUE! |
| ENSMUSG00000096014.2 | 0 | NA | NA | NA | NA | NA | #VALUE! |
| ENSMUSG00000096016.8 | 0 | NA | NA | NA | NA | NA | #VALUE! |
| ENSMUSG00000096020.3 | 0 | NA | NA | NA | NA | NA | #VALUE! |
| ENSMUSG00000096023.2 | 0 | NA | NA | NA | NA | NA | #VALUE! |
| ENSMUSG00000096024.4 | 0 | NA | NA | NA | NA | NA | #VALUE! |
| ENSMUSG00000096029.5 | 0 | NA | NA | NA | NA | NA | #VALUE! |
| ENSMUSG00000096033.3 | 0 | NA | NA | NA | NA | NA | #VALUE! |
| ENSMUSG00000096035.3 | 0 | NA | NA | NA | NA | NA | #VALUE! |
| ENSMUSG00000096036.3 | 0 | NA | NA | NA | NA | NA | #VALUE! |
| ENSMUSG00000096038.3 | 0 | NA | NA | NA | NA | NA | #VALUE! |
| ENSMUSG00000096039.9 | 0 | NA | NA | NA | NA | NA | #VALUE! |
| ENSMUSG00000096042.4 | 0 | NA | NA | NA | NA | NA | #VALUE! |
| ENSMUSG00000096044.5 | 0 | NA | NA | NA | NA | NA | #VALUE! |
| ENSMUSG00000096045.8 | 0 | NA | NA | NA | NA | NA | #VALUE! |
| ENSMUSG00000096049.2 | 0 | NA | NA | NA | NA | NA | #VALUE! |
| ENSMUSG00000096051.3 | 0 | NA | NA | NA | NA | NA | #VALUE! |
| ENSMUSG00000096052.2 | 0 | NA | NA | NA | NA | NA | #VALUE! |
| ENSMUSG00000096054.4 | 8.329378 | 4.167358 | 3.467608 | 1.201796 | 0.229443 | NA | #VALUE! |
| ENSMUSG00000096055.3 | 0 | NA | NA | NA | NA | NA | #VALUE! |
| ENSMUSG00000096056.2 | 0 | NA | NA | NA | NA | NA | #VALUE! |
| ENSMUSG00000096065.2 | 0 | NA | NA | NA | NA | NA | #VALUE! |
| ENSMUSG00000096066.5 | 0 | NA | NA | NA | NA | NA | #VALUE! |
| ENSMUSG00000096068.2 | 0 | NA | NA | NA | NA | NA | #VALUE! |
| ENSMUSG00000096069.5 | 0 | NA | NA | NA | NA | NA | #VALUE! |
| ENSMUSG00000096071.2 | 0 | NA | NA | NA | NA | NA | #VALUE! |
| ENSMUSG00000096072.2 | 0 | NA | NA | NA | NA | NA | #VALUE! |
| ENSMUSG00000096073.2 | 0 | NA | NA | NA | NA | NA | #VALUE! |
| ENSMUSG00000096074.3 | 0 | NA | NA | NA | NA | NA | #VALUE! |
| ENSMUSG00000096078.3 | 0 | NA | NA | NA | NA | NA | #VALUE! |
| ENSMUSG00000096081.2 | 0 | NA | NA | NA | NA | NA | #VALUE! |
| ENSMUSG00000096083.2 | 0 | NA | NA | NA | NA | NA | #VALUE! |
| ENSMUSG00000096084.3 | 0 | NA | NA | NA | NA | NA | #VALUE! |
| ENSMUSG00000096086.2 | 0 | NA | NA | NA | NA | NA | #VALUE! |
| ENSMUSG00000096088.2 | 0 | NA | NA | NA | NA | NA | #VALUE! |
| ENSMUSG00000096094.3 | 0 | NA | NA | NA | NA | NA | #VALUE! |
| ENSMUSG00000096096.4 | 0 | NA | NA | NA | NA | NA | #VALUE! |
| ENSMUSG00000096097.2 | 0 | NA | NA | NA | NA | NA | #VALUE! |
| ENSMUSG00000096099.3 | 0 | NA | NA | NA | NA | NA | #VALUE! |
| ENSMUSG00000096100.2 | 0 | NA | NA | NA | NA | NA | #VALUE! |
| ENSMUSG00000096102.4 | 0 | NA | NA | NA | NA | NA | #VALUE! |
| ENSMUSG00000096105.2 | 0.105013 | -0.97324 | 4.076293 | -0.23876 | 0.811295 | NA | #VALUE! |
| ENSMUSG00000096106.2 | 0 | NA | NA | NA | NA | NA | #VALUE! |
| ENSMUSG00000096107.3 | 0 | NA | NA | NA | NA | NA | #VALUE! |

| | | | | | | | |
|---|---|---|---|---|---|---|---|
| ENSMUSG00000096108.3 | 0 | NA | NA | NA | NA | NA | #VALUE! |
| ENSMUSG00000096109.3 | 0 | NA | NA | NA | NA | NA | #VALUE! |
| ENSMUSG00000096111.8 | 0.840105 | -3.30885 | 3.958472 | -0.83589 | 0.403216 | NA | #VALUE! |
| ENSMUSG00000096119.2 | 0 | NA | NA | NA | NA | NA | #VALUE! |
| ENSMUSG00000096120.2 | 0 | NA | NA | NA | NA | NA | #VALUE! |
| ENSMUSG00000096122.3 | 0 | NA | NA | NA | NA | NA | #VALUE! |
| ENSMUSG00000096127.3 | 0 | NA | NA | NA | NA | NA | #VALUE! |
| ENSMUSG00000096129.3 | 0 | NA | NA | NA | NA | NA | #VALUE! |
| ENSMUSG00000096131.4 | 0 | NA | NA | NA | NA | NA | #VALUE! |
| ENSMUSG00000096135.2 | 0 | NA | NA | NA | NA | NA | #VALUE! |
| ENSMUSG00000096137.2 | 0 | NA | NA | NA | NA | NA | #VALUE! |
| ENSMUSG00000096138.6 | 0 | NA | NA | NA | NA | NA | #VALUE! |
| ENSMUSG00000096139.5 | 0 | NA | NA | NA | NA | NA | #VALUE! |
| ENSMUSG00000096140.3 | 0 | NA | NA | NA | NA | NA | #VALUE! |
| ENSMUSG00000096141.3 | 0 | NA | NA | NA | NA | NA | #VALUE! |
| ENSMUSG00000096144.4 | 0 | NA | NA | NA | NA | NA | #VALUE! |
| ENSMUSG00000096146.3 | 0 | NA | NA | NA | NA | NA | #VALUE! |
| ENSMUSG00000096149.6 | 0 | NA | NA | NA | NA | NA | #VALUE! |
| ENSMUSG00000096150.2 | 0 | NA | NA | NA | NA | NA | #VALUE! |
| ENSMUSG00000096151.3 | 0 | NA | NA | NA | NA | NA | #VALUE! |
| ENSMUSG00000096152.2 | 0 | NA | NA | NA | NA | NA | #VALUE! |
| ENSMUSG00000096153.2 | 0 | NA | NA | NA | NA | NA | #VALUE! |
| ENSMUSG00000096154.3 | 0 | NA | NA | NA | NA | NA | #VALUE! |
| ENSMUSG00000096156.2 | 0 | NA | NA | NA | NA | NA | #VALUE! |
| ENSMUSG00000096157.3 | 0 | NA | NA | NA | NA | NA | #VALUE! |
| ENSMUSG00000096158.2 | 0 | NA | NA | NA | NA | NA | #VALUE! |
| ENSMUSG00000096160.3 | 0 | NA | NA | NA | NA | NA | #VALUE! |
| ENSMUSG00000096164.2 | 0 | NA | NA | NA | NA | NA | #VALUE! |
| ENSMUSG00000096167.5 | 0 | NA | NA | NA | NA | NA | #VALUE! |
| ENSMUSG00000096168.2 | 0 | NA | NA | NA | NA | NA | #VALUE! |
| ENSMUSG00000096169.4 | 0 | NA | NA | NA | NA | NA | #VALUE! |
| ENSMUSG00000096171.2 | 0 | NA | NA | NA | NA | NA | #VALUE! |
| ENSMUSG00000096173.3 | 0.89433 | 0.886815 | 3.950796 | 0.224465 | 0.822395 | NA | #VALUE! |
| ENSMUSG00000096175.4 | 0 | NA | NA | NA | NA | NA | #VALUE! |
| ENSMUSG00000096176.2 | 0 | NA | NA | NA | NA | NA | #VALUE! |
| ENSMUSG00000096177.3 | 0.198268 | -0.06494 | 4.068276 | -0.01596 | 0.987265 | NA | #VALUE! |
| ENSMUSG00000096178.8 | 0 | NA | NA | NA | NA | NA | #VALUE! |
| ENSMUSG00000096179.2 | 0 | NA | NA | NA | NA | NA | #VALUE! |
| ENSMUSG00000096180.3 | 0 | NA | NA | NA | NA | NA | #VALUE! |
| ENSMUSG00000096183.8 | 0 | NA | NA | NA | NA | NA | #VALUE! |
| ENSMUSG00000096192.3 | 0 | NA | NA | NA | NA | NA | #VALUE! |
| ENSMUSG00000096194.3 | 0 | NA | NA | NA | NA | NA | #VALUE! |
| ENSMUSG00000096195.2 | 0 | NA | NA | NA | NA | NA | #VALUE! |
| ENSMUSG00000096197.4 | 0 | NA | NA | NA | NA | NA | #VALUE! |

| | | | | | | | |
|---|---|---|---|---|---|---|---|
| ENSMUSG00000096201.2 | 0 | NA | NA | NA | NA | NA | #VALUE! |
| ENSMUSG00000096209.5 | 0 | NA | NA | NA | NA | NA | #VALUE! |
| ENSMUSG00000096213.2 | 0 | NA | NA | NA | NA | NA | #VALUE! |
| ENSMUSG00000096216.3 | 0 | NA | NA | NA | NA | NA | #VALUE! |
| ENSMUSG00000096217.2 | 0 | NA | NA | NA | NA | NA | #VALUE! |
| ENSMUSG00000096218.3 | 0 | NA | NA | NA | NA | NA | #VALUE! |
| ENSMUSG00000096220.3 | 0 | NA | NA | NA | NA | NA | #VALUE! |
| ENSMUSG00000096221.4 | 0 | NA | NA | NA | NA | NA | #VALUE! |
| ENSMUSG00000096223.3 | 0 | NA | NA | NA | NA | NA | #VALUE! |
| ENSMUSG00000096225.8 | 0 | NA | NA | NA | NA | NA | #VALUE! |
| ENSMUSG00000096228.6 | 0 | NA | NA | NA | NA | NA | #VALUE! |
| ENSMUSG00000096229.4 | 0 | NA | NA | NA | NA | NA | #VALUE! |
| ENSMUSG00000096230.5 | 0 | NA | NA | NA | NA | NA | #VALUE! |
| ENSMUSG00000096231.3 | 0 | NA | NA | NA | NA | NA | #VALUE! |
| ENSMUSG00000096233.2 | 0 | NA | NA | NA | NA | NA | #VALUE! |
| ENSMUSG00000096235.2 | 0 | NA | NA | NA | NA | NA | #VALUE! |
| ENSMUSG00000096236.2 | 0 | NA | NA | NA | NA | NA | #VALUE! |
| ENSMUSG00000096240.2 | 0 | NA | NA | NA | NA | NA | #VALUE! |
| ENSMUSG00000096244.2 | 0 | NA | NA | NA | NA | NA | #VALUE! |
| ENSMUSG00000096250.2 | 0 | NA | NA | NA | NA | NA | #VALUE! |
| ENSMUSG00000096254.3 | 0 | NA | NA | NA | NA | NA | #VALUE! |
| ENSMUSG00000096255.3 | 0 | NA | NA | NA | NA | NA | #VALUE! |
| ENSMUSG00000096256.3 | 0 | NA | NA | NA | NA | NA | #VALUE! |
| ENSMUSG00000096257.3 | 0 | NA | NA | NA | NA | NA | #VALUE! |
| ENSMUSG00000096258.3 | 0 | NA | NA | NA | NA | NA | #VALUE! |
| ENSMUSG00000096259.3 | 0 | NA | NA | NA | NA | NA | #VALUE! |
| ENSMUSG00000096265.3 | 0 | NA | NA | NA | NA | NA | #VALUE! |
| ENSMUSG00000096268.3 | 0 | NA | NA | NA | NA | NA | #VALUE! |
| ENSMUSG00000096269.4 | 0 | NA | NA | NA | NA | NA | #VALUE! |
| ENSMUSG00000096270.2 | 0 | NA | NA | NA | NA | NA | #VALUE! |
| ENSMUSG00000096272.3 | 0 | NA | NA | NA | NA | NA | #VALUE! |
| ENSMUSG00000096273.4 | 0 | NA | NA | NA | NA | NA | #VALUE! |
| ENSMUSG00000096275.3 | 0 | NA | NA | NA | NA | NA | #VALUE! |
| ENSMUSG00000096276.10 | 0 | NA | NA | NA | NA | NA | #VALUE! |
| ENSMUSG00000096277.3 | 0 | NA | NA | NA | NA | NA | #VALUE! |
| ENSMUSG00000096278.9 | 0 | NA | NA | NA | NA | NA | #VALUE! |
| ENSMUSG00000096283.3 | 0 | NA | NA | NA | NA | NA | #VALUE! |
| ENSMUSG00000096284.3 | 0 | NA | NA | NA | NA | NA | #VALUE! |
| ENSMUSG00000096286.2 | 0 | NA | NA | NA | NA | NA | #VALUE! |
| ENSMUSG00000096289.5 | 0 | NA | NA | NA | NA | NA | #VALUE! |
| ENSMUSG00000096294.2 | 0 | NA | NA | NA | NA | NA | #VALUE! |
| ENSMUSG00000096295.3 | 0 | NA | NA | NA | NA | NA | #VALUE! |
| ENSMUSG00000096299.4 | 0 | NA | NA | NA | NA | NA | #VALUE! |
| ENSMUSG00000096301.2 | 0 | NA | NA | NA | NA | NA | #VALUE! |

| | | | | | | | |
|---|---|---|---|---|---|---|---|
| ENSMUSG00000096304.2 | 0 | NA | NA | NA | NA | NA | #VALUE! |
| ENSMUSG00000096308.2 | 0 | NA | NA | NA | NA | NA | #VALUE! |
| ENSMUSG00000096311.2 | 0 | NA | NA | NA | NA | NA | #VALUE! |
| ENSMUSG00000096316.8 | 0 | NA | NA | NA | NA | NA | #VALUE! |
| ENSMUSG00000096318.2 | 0 | NA | NA | NA | NA | NA | #VALUE! |
| ENSMUSG00000096320.3 | 0 | NA | NA | NA | NA | NA | #VALUE! |
| ENSMUSG00000096323.3 | 0 | NA | NA | NA | NA | NA | #VALUE! |
| ENSMUSG00000096326.3 | 0 | NA | NA | NA | NA | NA | #VALUE! |
| ENSMUSG00000096327.2 | 0 | NA | NA | NA | NA | NA | #VALUE! |
| ENSMUSG00000096329.3 | 0 | NA | NA | NA | NA | NA | #VALUE! |
| ENSMUSG00000096330.9 | 0 | NA | NA | NA | NA | NA | #VALUE! |
| ENSMUSG00000096332.3 | 0 | NA | NA | NA | NA | NA | #VALUE! |
| ENSMUSG00000096333.8 | 0 | NA | NA | NA | NA | NA | #VALUE! |
| ENSMUSG00000096336.3 | 0 | NA | NA | NA | NA | NA | #VALUE! |
| ENSMUSG00000096337.2 | 0 | NA | NA | NA | NA | NA | #VALUE! |
| ENSMUSG00000096338.2 | 0 | NA | NA | NA | NA | NA | #VALUE! |
| ENSMUSG00000096340.2 | 0 | NA | NA | NA | NA | NA | #VALUE! |
| ENSMUSG00000096344.3 | 0 | NA | NA | NA | NA | NA | #VALUE! |
| ENSMUSG00000096345.2 | 0 | NA | NA | NA | NA | NA | #VALUE! |
| ENSMUSG00000096347.3 | 0 | NA | NA | NA | NA | NA | #VALUE! |
| ENSMUSG00000096348.2 | 0 | NA | NA | NA | NA | NA | #VALUE! |
| ENSMUSG00000096351.3 | 0 | NA | NA | NA | NA | NA | #VALUE! |
| ENSMUSG00000096354.2 | 0 | NA | NA | NA | NA | NA | #VALUE! |
| ENSMUSG00000096355.3 | 0 | NA | NA | NA | NA | NA | #VALUE! |
| ENSMUSG00000096356.4 | 0 | NA | NA | NA | NA | NA | #VALUE! |
| ENSMUSG00000096361.3 | 0 | NA | NA | NA | NA | NA | #VALUE! |
| ENSMUSG00000096362.2 | 0 | NA | NA | NA | NA | NA | #VALUE! |
| ENSMUSG00000096363.2 | 0 | NA | NA | NA | NA | NA | #VALUE! |
| ENSMUSG00000096365.5 | 0 | NA | NA | NA | NA | NA | #VALUE! |
| ENSMUSG00000096366.2 | 0 | NA | NA | NA | NA | NA | #VALUE! |
| ENSMUSG00000096370.9 | 8.368541 | -6.59301 | 3.815764 | -1.72783 | 0.084018 | NA | #VALUE! |
| ENSMUSG00000096372.2 | 0 | NA | NA | NA | NA | NA | #VALUE! |
| ENSMUSG00000096373.4 | 0 | NA | NA | NA | NA | NA | #VALUE! |
| ENSMUSG00000096374.2 | 0 | NA | NA | NA | NA | NA | #VALUE! |
| ENSMUSG00000096375.3 | 0 | NA | NA | NA | NA | NA | #VALUE! |
| ENSMUSG00000096376.2 | 0 | NA | NA | NA | NA | NA | #VALUE! |
| ENSMUSG00000096377.2 | 0 | NA | NA | NA | NA | NA | #VALUE! |
| ENSMUSG00000096380.2 | 0 | NA | NA | NA | NA | NA | #VALUE! |
| ENSMUSG00000096385.8 | 0.204111 | -1.05581 | 4.073302 | -0.2592 | 0.795479 | NA | #VALUE! |
| ENSMUSG00000096386.2 | 0 | NA | NA | NA | NA | NA | #VALUE! |
| ENSMUSG00000096393.2 | 0 | NA | NA | NA | NA | NA | #VALUE! |
| ENSMUSG00000096396.2 | 0 | NA | NA | NA | NA | NA | #VALUE! |
| ENSMUSG00000096397.8 | 0 | NA | NA | NA | NA | NA | #VALUE! |
| ENSMUSG00000096399.2 | 0 | NA | NA | NA | NA | NA | #VALUE! |

| | | | | | | | |
|---|---|---|---|---|---|---|---|
| ENSMUSG00000096400.3 | 0 | NA | NA | NA | NA | NA | #VALUE! |
| ENSMUSG00000096401.3 | 0 | NA | NA | NA | NA | NA | #VALUE! |
| ENSMUSG00000096403.3 | 0 | NA | NA | NA | NA | NA | #VALUE! |
| ENSMUSG00000096405.4 | 0 | NA | NA | NA | NA | NA | #VALUE! |
| ENSMUSG00000096407.3 | 0 | NA | NA | NA | NA | NA | #VALUE! |
| ENSMUSG00000096409.4 | 0 | NA | NA | NA | NA | NA | #VALUE! |
| ENSMUSG00000096410.3 | 0 | NA | NA | NA | NA | NA | #VALUE! |
| ENSMUSG00000096413.3 | 0 | NA | NA | NA | NA | NA | #VALUE! |
| ENSMUSG00000096414.2 | 0 | NA | NA | NA | NA | NA | #VALUE! |
| ENSMUSG00000096417.3 | 0 | NA | NA | NA | NA | NA | #VALUE! |
| ENSMUSG00000096420.3 | 0 | NA | NA | NA | NA | NA | #VALUE! |
| ENSMUSG00000096421.2 | 0 | NA | NA | NA | NA | NA | #VALUE! |
| ENSMUSG00000096422.3 | 0 | NA | NA | NA | NA | NA | #VALUE! |
| ENSMUSG00000096423.2 | 0 | NA | NA | NA | NA | NA | #VALUE! |
| ENSMUSG00000096424.2 | 0 | NA | NA | NA | NA | NA | #VALUE! |
| ENSMUSG00000096426.3 | 0 | NA | NA | NA | NA | NA | #VALUE! |
| ENSMUSG00000096427.4 | 0 | NA | NA | NA | NA | NA | #VALUE! |
| ENSMUSG00000096429.2 | 0 | NA | NA | NA | NA | NA | #VALUE! |
| ENSMUSG00000096436.4 | 0 | NA | NA | NA | NA | NA | #VALUE! |
| ENSMUSG00000096437.2 | 0 | NA | NA | NA | NA | NA | #VALUE! |
| ENSMUSG00000096438.4 | 0 | NA | NA | NA | NA | NA | #VALUE! |
| ENSMUSG00000096439.2 | 0 | NA | NA | NA | NA | NA | #VALUE! |
| ENSMUSG00000096440.2 | 0 | NA | NA | NA | NA | NA | #VALUE! |
| ENSMUSG00000096442.2 | 0 | NA | NA | NA | NA | NA | #VALUE! |
| ENSMUSG00000096445.2 | 0 | NA | NA | NA | NA | NA | #VALUE! |
| ENSMUSG00000096446.2 | 0 | NA | NA | NA | NA | NA | #VALUE! |
| ENSMUSG00000096448.3 | 0 | NA | NA | NA | NA | NA | #VALUE! |
| ENSMUSG00000096449.3 | 1.367983 | -2.71367 | 3.930324 | -0.69044 | 0.489915 | NA | #VALUE! |
| ENSMUSG00000096450.2 | 0 | NA | NA | NA | NA | NA | #VALUE! |
| ENSMUSG00000096452.3 | 0 | NA | NA | NA | NA | NA | #VALUE! |
| ENSMUSG00000096453.2 | 0 | NA | NA | NA | NA | NA | #VALUE! |
| ENSMUSG00000096457.8 | 0 | NA | NA | NA | NA | NA | #VALUE! |
| ENSMUSG00000096458.8 | 0 | NA | NA | NA | NA | NA | #VALUE! |
| ENSMUSG00000096459.2 | 0 | NA | NA | NA | NA | NA | #VALUE! |
| ENSMUSG00000096461.6 | 0 | NA | NA | NA | NA | NA | #VALUE! |
| ENSMUSG00000096463.2 | 0 | NA | NA | NA | NA | NA | #VALUE! |
| ENSMUSG00000096464.3 | 0 | NA | NA | NA | NA | NA | #VALUE! |
| ENSMUSG00000096465.4 | 0 | NA | NA | NA | NA | NA | #VALUE! |
| ENSMUSG00000096467.3 | 0 | NA | NA | NA | NA | NA | #VALUE! |
| ENSMUSG00000096468.8 | 0 | NA | NA | NA | NA | NA | #VALUE! |
| ENSMUSG00000096470.2 | 0 | NA | NA | NA | NA | NA | #VALUE! |
| ENSMUSG00000096472.3 | 4.435079 | 1.901692 | 3.362133 | 0.565621 | 0.571652 | NA | #VALUE! |
| ENSMUSG00000096474.3 | 0 | NA | NA | NA | NA | NA | #VALUE! |
| ENSMUSG00000096476.2 | 0 | NA | NA | NA | NA | NA | #VALUE! |

| | | | | | | | |
|---|---|---|---|---|---|---|---|
| ENSMUSG00000096477.3 | 0 | NA | NA | NA | NA | NA | #VALUE! |
| ENSMUSG00000096478.4 | 0 | NA | NA | NA | NA | NA | #VALUE! |
| ENSMUSG00000096481.2 | 0 | NA | NA | NA | NA | NA | #VALUE! |
| ENSMUSG00000096483.2 | 0 | NA | NA | NA | NA | NA | #VALUE! |
| ENSMUSG00000096484.2 | 0 | NA | NA | NA | NA | NA | #VALUE! |
| ENSMUSG00000096485.4 | 0 | NA | NA | NA | NA | NA | #VALUE! |
| ENSMUSG00000096486.2 | 0 | NA | NA | NA | NA | NA | #VALUE! |
| ENSMUSG00000096488.2 | 0 | NA | NA | NA | NA | NA | #VALUE! |
| ENSMUSG00000096490.2 | 0 | NA | NA | NA | NA | NA | #VALUE! |
| ENSMUSG00000096497.3 | 0 | NA | NA | NA | NA | NA | #VALUE! |
| ENSMUSG00000096498.4 | 0 | NA | NA | NA | NA | NA | #VALUE! |
| ENSMUSG00000096499.3 | 0 | NA | NA | NA | NA | NA | #VALUE! |
| ENSMUSG00000096505.3 | 0 | NA | NA | NA | NA | NA | #VALUE! |
| ENSMUSG00000096507.2 | 0 | NA | NA | NA | NA | NA | #VALUE! |
| ENSMUSG00000096508.8 | 0 | NA | NA | NA | NA | NA | #VALUE! |
| ENSMUSG00000096513.2 | 0 | NA | NA | NA | NA | NA | #VALUE! |
| ENSMUSG00000096515.6 | 0 | NA | NA | NA | NA | NA | #VALUE! |
| ENSMUSG00000096516.4 | 0 | NA | NA | NA | NA | NA | #VALUE! |
| ENSMUSG00000096517.2 | 0 | NA | NA | NA | NA | NA | #VALUE! |
| ENSMUSG00000096518.3 | 0 | NA | NA | NA | NA | NA | #VALUE! |
| ENSMUSG00000096519.2 | 9.34707 | 6.710288 | 3.689587 | 1.81871 | 0.068956 | NA | #VALUE! |
| ENSMUSG00000096520.8 | 0 | NA | NA | NA | NA | NA | #VALUE! |
| ENSMUSG00000096521.2 | 0 | NA | NA | NA | NA | NA | #VALUE! |
| ENSMUSG00000096524.2 | 0 | NA | NA | NA | NA | NA | #VALUE! |
| ENSMUSG00000096525.3 | 0 | NA | NA | NA | NA | NA | #VALUE! |
| ENSMUSG00000096527.6 | 0 | NA | NA | NA | NA | NA | #VALUE! |
| ENSMUSG00000096528.2 | 0 | NA | NA | NA | NA | NA | #VALUE! |
| ENSMUSG00000096530.8 | 0 | NA | NA | NA | NA | NA | #VALUE! |
| ENSMUSG00000096531.3 | 0 | NA | NA | NA | NA | NA | #VALUE! |
| ENSMUSG00000096532.3 | 0 | NA | NA | NA | NA | NA | #VALUE! |
| ENSMUSG00000096534.3 | 0 | NA | NA | NA | NA | NA | #VALUE! |
| ENSMUSG00000096537.7 | 0 | NA | NA | NA | NA | NA | #VALUE! |
| ENSMUSG00000096538.3 | 0 | NA | NA | NA | NA | NA | #VALUE! |
| ENSMUSG00000096539.2 | 0 | NA | NA | NA | NA | NA | #VALUE! |
| ENSMUSG00000096541.3 | 0 | NA | NA | NA | NA | NA | #VALUE! |
| ENSMUSG00000096544.3 | 0.168284 | 1.027628 | 4.067126 | 0.252667 | 0.800526 | NA | #VALUE! |
| ENSMUSG00000096549.10 | 0 | NA | NA | NA | NA | NA | #VALUE! |
| ENSMUSG00000096550.2 | 0 | NA | NA | NA | NA | NA | #VALUE! |
| ENSMUSG00000096551.3 | 0 | NA | NA | NA | NA | NA | #VALUE! |
| ENSMUSG00000096553.3 | 0 | NA | NA | NA | NA | NA | #VALUE! |
| ENSMUSG00000096554.2 | 0 | NA | NA | NA | NA | NA | #VALUE! |
| ENSMUSG00000096555.4 | 0 | NA | NA | NA | NA | NA | #VALUE! |
| ENSMUSG00000096557.3 | 0 | NA | NA | NA | NA | NA | #VALUE! |
| ENSMUSG00000096566.3 | 0 | NA | NA | NA | NA | NA | #VALUE! |

| | | | | | | | |
|---|---|---|---|---|---|---|---|
| ENSMUSG00000096568.2 | 0 | NA | NA | NA | NA | NA | #VALUE! |
| ENSMUSG00000096569.3 | 0 | NA | NA | NA | NA | NA | #VALUE! |
| ENSMUSG00000096573.3 | 0 | NA | NA | NA | NA | NA | #VALUE! |
| ENSMUSG00000096574.8 | 0 | NA | NA | NA | NA | NA | #VALUE! |
| ENSMUSG00000096576.3 | 0 | NA | NA | NA | NA | NA | #VALUE! |
| ENSMUSG00000096577.3 | 0 | NA | NA | NA | NA | NA | #VALUE! |
| ENSMUSG00000096578.3 | 0 | NA | NA | NA | NA | NA | #VALUE! |
| ENSMUSG00000096579.2 | 0 | NA | NA | NA | NA | NA | #VALUE! |
| ENSMUSG00000096580.2 | 0 | NA | NA | NA | NA | NA | #VALUE! |
| ENSMUSG00000096582.2 | 0 | NA | NA | NA | NA | NA | #VALUE! |
| ENSMUSG00000096584.3 | 0 | NA | NA | NA | NA | NA | #VALUE! |
| ENSMUSG00000096589.2 | 0 | NA | NA | NA | NA | NA | #VALUE! |
| ENSMUSG00000096591.2 | 0 | NA | NA | NA | NA | NA | #VALUE! |
| ENSMUSG00000096592.2 | 0 | NA | NA | NA | NA | NA | #VALUE! |
| ENSMUSG00000096593.3 | 0 | NA | NA | NA | NA | NA | #VALUE! |
| ENSMUSG00000096594.3 | 0 | NA | NA | NA | NA | NA | #VALUE! |
| ENSMUSG00000096596.3 | 0 | NA | NA | NA | NA | NA | #VALUE! |
| ENSMUSG00000096597.2 | 0 | NA | NA | NA | NA | NA | #VALUE! |
| ENSMUSG00000096600.6 | 0 | NA | NA | NA | NA | NA | #VALUE! |
| ENSMUSG00000096601.2 | 0 | NA | NA | NA | NA | NA | #VALUE! |
| ENSMUSG00000096606.3 | 14.20827 | -0.89464 | 2.622761 | -0.34111 | NA | NA | #VALUE! |
| ENSMUSG00000096607.2 | 0 | NA | NA | NA | NA | NA | #VALUE! |
| ENSMUSG00000096614.2 | 0 | NA | NA | NA | NA | NA | #VALUE! |
| ENSMUSG00000096615.2 | 0 | NA | NA | NA | NA | NA | #VALUE! |
| ENSMUSG00000096617.4 | 0 | NA | NA | NA | NA | NA | #VALUE! |
| ENSMUSG00000096618.3 | 0 | NA | NA | NA | NA | NA | #VALUE! |
| ENSMUSG00000096619.2 | 0 | NA | NA | NA | NA | NA | #VALUE! |
| ENSMUSG00000096620.3 | 0 | NA | NA | NA | NA | NA | #VALUE! |
| ENSMUSG00000096621.3 | 0 | NA | NA | NA | NA | NA | #VALUE! |
| ENSMUSG00000096626.8 | 0 | NA | NA | NA | NA | NA | #VALUE! |
| ENSMUSG00000096629.8 | 0 | NA | NA | NA | NA | NA | #VALUE! |
| ENSMUSG00000096630.3 | 0 | NA | NA | NA | NA | NA | #VALUE! |
| ENSMUSG00000096632.3 | 0 | NA | NA | NA | NA | NA | #VALUE! |
| ENSMUSG00000096638.3 | 0 | NA | NA | NA | NA | NA | #VALUE! |
| ENSMUSG00000096640.4 | 0 | NA | NA | NA | NA | NA | #VALUE! |
| ENSMUSG00000096641.8 | 0 | NA | NA | NA | NA | NA | #VALUE! |
| ENSMUSG00000096644.2 | 0 | NA | NA | NA | NA | NA | #VALUE! |
| ENSMUSG00000096645.3 | 0 | NA | NA | NA | NA | NA | #VALUE! |
| ENSMUSG00000096647.3 | 0 | NA | NA | NA | NA | NA | #VALUE! |
| ENSMUSG00000096649.3 | 0 | NA | NA | NA | NA | NA | #VALUE! |
| ENSMUSG00000096650.8 | 0 | NA | NA | NA | NA | NA | #VALUE! |
| ENSMUSG00000096652.2 | 0 | NA | NA | NA | NA | NA | #VALUE! |
| ENSMUSG00000096653.5 | 0 | NA | NA | NA | NA | NA | #VALUE! |
| ENSMUSG00000096654.2 | 0 | NA | NA | NA | NA | NA | #VALUE! |

| | | | | | | | |
|---|---|---|---|---|---|---|---|
| ENSMUSG00000096655.3 | 0 | NA | NA | NA | NA | NA | #VALUE! |
| ENSMUSG00000096656.7 | 0 | NA | NA | NA | NA | NA | #VALUE! |
| ENSMUSG00000096658.4 | 0 | NA | NA | NA | NA | NA | #VALUE! |
| ENSMUSG00000096661.2 | 0 | NA | NA | NA | NA | NA | #VALUE! |
| ENSMUSG00000096663.2 | 0 | NA | NA | NA | NA | NA | #VALUE! |
| ENSMUSG00000096664.8 | 0 | NA | NA | NA | NA | NA | #VALUE! |
| ENSMUSG00000096666.3 | 0 | NA | NA | NA | NA | NA | #VALUE! |
| ENSMUSG00000096668.2 | 0 | NA | NA | NA | NA | NA | #VALUE! |
| ENSMUSG00000096670.2 | 0 | NA | NA | NA | NA | NA | #VALUE! |
| ENSMUSG00000096672.2 | 0 | NA | NA | NA | NA | NA | #VALUE! |
| ENSMUSG00000096678.4 | 0 | NA | NA | NA | NA | NA | #VALUE! |
| ENSMUSG00000096679.4 | 0 | NA | NA | NA | NA | NA | #VALUE! |
| ENSMUSG00000096680.2 | 0 | NA | NA | NA | NA | NA | #VALUE! |
| ENSMUSG00000096681.2 | 0 | NA | NA | NA | NA | NA | #VALUE! |
| ENSMUSG00000096682.2 | 0 | NA | NA | NA | NA | NA | #VALUE! |
| ENSMUSG00000096685.8 | 0 | NA | NA | NA | NA | NA | #VALUE! |
| ENSMUSG00000096686.3 | 0.195789 | -1.05581 | 4.073302 | -0.2592 | 0.795479 | NA | #VALUE! |
| ENSMUSG00000096687.9 | 2.071295 | 4.531036 | 3.928524 | 1.153369 | 0.248759 | NA | #VALUE! |
| ENSMUSG00000096690.2 | 0 | NA | NA | NA | NA | NA | #VALUE! |
| ENSMUSG00000096691.2 | 0 | NA | NA | NA | NA | NA | #VALUE! |
| ENSMUSG00000096695.4 | 0 | NA | NA | NA | NA | NA | #VALUE! |
| ENSMUSG00000096696.2 | 0 | NA | NA | NA | NA | NA | #VALUE! |
| ENSMUSG00000096697.2 | 0 | NA | NA | NA | NA | NA | #VALUE! |
| ENSMUSG00000096698.3 | 0 | NA | NA | NA | NA | NA | #VALUE! |
| ENSMUSG00000096699.2 | 0.245976 | 1.027628 | 4.067126 | 0.252667 | 0.800526 | NA | #VALUE! |
| ENSMUSG00000096700.2 | 0 | NA | NA | NA | NA | NA | #VALUE! |
| ENSMUSG00000096701.3 | 0 | NA | NA | NA | NA | NA | #VALUE! |
| ENSMUSG00000096703.4 | 0 | NA | NA | NA | NA | NA | #VALUE! |
| ENSMUSG00000096705.2 | 0 | NA | NA | NA | NA | NA | #VALUE! |
| ENSMUSG00000096706.3 | 0 | NA | NA | NA | NA | NA | #VALUE! |
| ENSMUSG00000096707.3 | 0 | NA | NA | NA | NA | NA | #VALUE! |
| ENSMUSG00000096708.2 | 0 | NA | NA | NA | NA | NA | #VALUE! |
| ENSMUSG00000096709.2 | 0 | NA | NA | NA | NA | NA | #VALUE! |
| ENSMUSG00000096711.3 | 0 | NA | NA | NA | NA | NA | #VALUE! |
| ENSMUSG00000096712.2 | 0 | NA | NA | NA | NA | NA | #VALUE! |
| ENSMUSG00000096714.3 | 0 | NA | NA | NA | NA | NA | #VALUE! |
| ENSMUSG00000096715.3 | 0 | NA | NA | NA | NA | NA | #VALUE! |
| ENSMUSG00000096716.2 | 0 | NA | NA | NA | NA | NA | #VALUE! |
| ENSMUSG00000096717.3 | 0 | NA | NA | NA | NA | NA | #VALUE! |
| ENSMUSG00000096718.3 | 0 | NA | NA | NA | NA | NA | #VALUE! |
| ENSMUSG00000096719.2 | 0 | NA | NA | NA | NA | NA | #VALUE! |
| ENSMUSG00000096722.3 | 0 | NA | NA | NA | NA | NA | #VALUE! |
| ENSMUSG00000096724.2 | 0 | NA | NA | NA | NA | NA | #VALUE! |
| ENSMUSG00000096726.3 | 0.093255 | 0.868093 | 4.072464 | 0.213162 | 0.831201 | NA | #VALUE! |

| | | | | | | | |
|---|---|---|---|---|---|---|---|
| ENSMUSG00000096728.2 | 0 | NA | NA | NA | NA | NA | #VALUE! |
| ENSMUSG00000096729.2 | 0 | NA | NA | NA | NA | NA | #VALUE! |
| ENSMUSG00000096732.8 | 0 | NA | NA | NA | NA | NA | #VALUE! |
| ENSMUSG00000096733.2 | 0 | NA | NA | NA | NA | NA | #VALUE! |
| ENSMUSG00000096734.2 | 0 | NA | NA | NA | NA | NA | #VALUE! |
| ENSMUSG00000096735.4 | 0 | NA | NA | NA | NA | NA | #VALUE! |
| ENSMUSG00000096736.2 | 1.020554 | -3.54462 | 3.950745 | -0.8972 | 0.36961 | NA | #VALUE! |
| ENSMUSG00000096737.2 | 0 | NA | NA | NA | NA | NA | #VALUE! |
| ENSMUSG00000096738.2 | 0 | NA | NA | NA | NA | NA | #VALUE! |
| ENSMUSG00000096740.9 | 0.877871 | 3.316761 | 3.95687 | 0.838228 | 0.401902 | NA | #VALUE! |
| ENSMUSG00000096741.2 | 0 | NA | NA | NA | NA | NA | #VALUE! |
| ENSMUSG00000096742.2 | 0 | NA | NA | NA | NA | NA | #VALUE! |
| ENSMUSG00000096743.4 | 0 | NA | NA | NA | NA | NA | #VALUE! |
| ENSMUSG00000096744.2 | 0 | NA | NA | NA | NA | NA | #VALUE! |
| ENSMUSG00000096745.2 | 0 | NA | NA | NA | NA | NA | #VALUE! |
| ENSMUSG00000096746.4 | 0 | NA | NA | NA | NA | NA | #VALUE! |
| ENSMUSG00000096747.3 | 0 | NA | NA | NA | NA | NA | #VALUE! |
| ENSMUSG00000096749.3 | 0 | NA | NA | NA | NA | NA | #VALUE! |
| ENSMUSG00000096750.8 | 0 | NA | NA | NA | NA | NA | #VALUE! |
| ENSMUSG00000096751.4 | 0 | NA | NA | NA | NA | NA | #VALUE! |
| ENSMUSG00000096753.8 | 0 | NA | NA | NA | NA | NA | #VALUE! |
| ENSMUSG00000096755.3 | 0 | NA | NA | NA | NA | NA | #VALUE! |
| ENSMUSG00000096756.2 | 0 | NA | NA | NA | NA | NA | #VALUE! |
| ENSMUSG00000096757.3 | 0 | NA | NA | NA | NA | NA | #VALUE! |
| ENSMUSG00000096760.2 | 0 | NA | NA | NA | NA | NA | #VALUE! |
| ENSMUSG00000096761.2 | 0 | NA | NA | NA | NA | NA | #VALUE! |
| ENSMUSG00000096763.2 | 0 | NA | NA | NA | NA | NA | #VALUE! |
| ENSMUSG00000096764.2 | 0 | NA | NA | NA | NA | NA | #VALUE! |
| ENSMUSG00000096766.3 | 0 | NA | NA | NA | NA | NA | #VALUE! |
| ENSMUSG00000096767.3 | 0 | NA | NA | NA | NA | NA | #VALUE! |
| ENSMUSG00000096769.8 | 0 | NA | NA | NA | NA | NA | #VALUE! |
| ENSMUSG00000096770.3 | 0 | NA | NA | NA | NA | NA | #VALUE! |
| ENSMUSG00000096773.5 | 0 | NA | NA | NA | NA | NA | #VALUE! |
| ENSMUSG00000096775.2 | 0 | NA | NA | NA | NA | NA | #VALUE! |
| ENSMUSG00000096776.2 | 0 | NA | NA | NA | NA | NA | #VALUE! |
| ENSMUSG00000096777.3 | 0 | NA | NA | NA | NA | NA | #VALUE! |
| ENSMUSG00000096780.8 | 0 | NA | NA | NA | NA | NA | #VALUE! |
| ENSMUSG00000096788.9 | 0 | NA | NA | NA | NA | NA | #VALUE! |
| ENSMUSG00000096792.3 | 0 | NA | NA | NA | NA | NA | #VALUE! |
| ENSMUSG00000096793.8 | 0 | NA | NA | NA | NA | NA | #VALUE! |
| ENSMUSG00000096794.3 | 0 | NA | NA | NA | NA | NA | #VALUE! |
| ENSMUSG00000096795.5 | 0 | NA | NA | NA | NA | NA | #VALUE! |
| ENSMUSG00000096796.2 | 0 | NA | NA | NA | NA | NA | #VALUE! |
| ENSMUSG00000096797.2 | 0 | NA | NA | NA | NA | NA | #VALUE! |

| | | | | | | | |
|---|---|---|---|---|---|---|---|
| ENSMUSG00000096798.2 | 0 | NA | NA | NA | NA | NA | #VALUE! |
| ENSMUSG00000096800.2 | 0 | NA | NA | NA | NA | NA | #VALUE! |
| ENSMUSG00000096803.2 | 0 | NA | NA | NA | NA | NA | #VALUE! |
| ENSMUSG00000096805.3 | 0 | NA | NA | NA | NA | NA | #VALUE! |
| ENSMUSG00000096806.3 | 0 | NA | NA | NA | NA | NA | #VALUE! |
| ENSMUSG00000096808.2 | 0 | NA | NA | NA | NA | NA | #VALUE! |
| ENSMUSG00000096810.2 | 0 | NA | NA | NA | NA | NA | #VALUE! |
| ENSMUSG00000096813.2 | 0 | NA | NA | NA | NA | NA | #VALUE! |
| ENSMUSG00000096818.3 | 0 | NA | NA | NA | NA | NA | #VALUE! |
| ENSMUSG00000096819.2 | 0 | NA | NA | NA | NA | NA | #VALUE! |
| ENSMUSG00000096820.3 | 0 | NA | NA | NA | NA | NA | #VALUE! |
| ENSMUSG00000096822.3 | 0 | NA | NA | NA | NA | NA | #VALUE! |
| ENSMUSG00000096824.3 | 0 | NA | NA | NA | NA | NA | #VALUE! |
| ENSMUSG00000096825.3 | 0 | NA | NA | NA | NA | NA | #VALUE! |
| ENSMUSG00000096826.3 | 0 | NA | NA | NA | NA | NA | #VALUE! |
| ENSMUSG00000096827.3 | 0 | NA | NA | NA | NA | NA | #VALUE! |
| ENSMUSG00000096828.3 | 0 | NA | NA | NA | NA | NA | #VALUE! |
| ENSMUSG00000096831.2 | 0 | NA | NA | NA | NA | NA | #VALUE! |
| ENSMUSG00000096833.3 | 0 | NA | NA | NA | NA | NA | #VALUE! |
| ENSMUSG00000096834.8 | 0 | NA | NA | NA | NA | NA | #VALUE! |
| ENSMUSG00000096839.8 | 0 | NA | NA | NA | NA | NA | #VALUE! |
| ENSMUSG00000096840.4 | 0 | NA | NA | NA | NA | NA | #VALUE! |
| ENSMUSG00000096842.2 | 0.291524 | 0.468739 | 4.052149 | 0.115677 | 0.907909 | NA | #VALUE! |
| ENSMUSG00000096844.2 | 0 | NA | NA | NA | NA | NA | #VALUE! |
| ENSMUSG00000096846.8 | 0 | NA | NA | NA | NA | NA | #VALUE! |
| ENSMUSG00000096847.3 | 0 | NA | NA | NA | NA | NA | #VALUE! |
| ENSMUSG00000096850.2 | 0 | NA | NA | NA | NA | NA | #VALUE! |
| ENSMUSG00000096852.3 | 5.960786 | -3.48739 | 3.380975 | -1.03147 | 0.302318 | NA | #VALUE! |
| ENSMUSG00000096854.2 | 0 | NA | NA | NA | NA | NA | #VALUE! |
| ENSMUSG00000096856.8 | 0 | NA | NA | NA | NA | NA | #VALUE! |
| ENSMUSG00000096857.2 | 0 | NA | NA | NA | NA | NA | #VALUE! |
| ENSMUSG00000096858.5 | 0 | NA | NA | NA | NA | NA | #VALUE! |
| ENSMUSG00000096859.3 | 0 | NA | NA | NA | NA | NA | #VALUE! |
| ENSMUSG00000096861.3 | 0 | NA | NA | NA | NA | NA | #VALUE! |
| ENSMUSG00000096862.4 | 0 | NA | NA | NA | NA | NA | #VALUE! |
| ENSMUSG00000096864.2 | 0 | NA | NA | NA | NA | NA | #VALUE! |
| ENSMUSG00000096867.2 | 0 | NA | NA | NA | NA | NA | #VALUE! |
| ENSMUSG00000096869.8 | 0 | NA | NA | NA | NA | NA | #VALUE! |
| ENSMUSG00000096870.3 | 0 | NA | NA | NA | NA | NA | #VALUE! |
| ENSMUSG00000096871.2 | 0 | NA | NA | NA | NA | NA | #VALUE! |
| ENSMUSG00000096872.3 | 0 | NA | NA | NA | NA | NA | #VALUE! |
| ENSMUSG00000096874.3 | 0 | NA | NA | NA | NA | NA | #VALUE! |
| ENSMUSG00000096875.3 | 0 | NA | NA | NA | NA | NA | #VALUE! |
| ENSMUSG00000096878.3 | 0 | NA | NA | NA | NA | NA | #VALUE! |

| | | | | | | | |
|---|---|---|---|---|---|---|---|
| ENSMUSG00000096879.3 | 0 | NA | NA | NA | NA | NA | #VALUE! |
| ENSMUSG00000096881.3 | 0 | NA | NA | NA | NA | NA | #VALUE! |
| ENSMUSG00000096883.3 | 0 | NA | NA | NA | NA | NA | #VALUE! |
| ENSMUSG00000096884.2 | 0 | NA | NA | NA | NA | NA | #VALUE! |
| ENSMUSG00000096885.2 | 0 | NA | NA | NA | NA | NA | #VALUE! |
| ENSMUSG00000096887.3 | 0 | NA | NA | NA | NA | NA | #VALUE! |
| ENSMUSG00000096890.2 | 0 | NA | NA | NA | NA | NA | #VALUE! |
| ENSMUSG00000096891.2 | 0 | NA | NA | NA | NA | NA | #VALUE! |
| ENSMUSG00000096896.3 | 0 | NA | NA | NA | NA | NA | #VALUE! |
| ENSMUSG00000096897.2 | 0 | NA | NA | NA | NA | NA | #VALUE! |
| ENSMUSG00000096898.3 | 0 | NA | NA | NA | NA | NA | #VALUE! |
| ENSMUSG00000096899.2 | 0 | NA | NA | NA | NA | NA | #VALUE! |
| ENSMUSG00000096900.4 | 0 | NA | NA | NA | NA | NA | #VALUE! |
| ENSMUSG00000096901.8 | 0 | NA | NA | NA | NA | NA | #VALUE! |
| ENSMUSG00000096902.8 | 0 | NA | NA | NA | NA | NA | #VALUE! |
| ENSMUSG00000096903.2 | 0 | NA | NA | NA | NA | NA | #VALUE! |
| ENSMUSG00000096904.8 | 0 | NA | NA | NA | NA | NA | #VALUE! |
| ENSMUSG00000096905.2 | 0 | NA | NA | NA | NA | NA | #VALUE! |
| ENSMUSG00000096907.2 | 0 | NA | NA | NA | NA | NA | #VALUE! |
| ENSMUSG00000096908.4 | 0 | NA | NA | NA | NA | NA | #VALUE! |
| ENSMUSG00000096911.8 | 0 | NA | NA | NA | NA | NA | #VALUE! |
| ENSMUSG00000096914.4 | 0 | NA | NA | NA | NA | NA | #VALUE! |
| ENSMUSG00000096915.2 | 0 | NA | NA | NA | NA | NA | #VALUE! |
| ENSMUSG00000096916.8 | 1.365171 | -3.99822 | 3.938957 | -1.01505 | 0.310084 | NA | #VALUE! |
| ENSMUSG00000096918.3 | 0 | NA | NA | NA | NA | NA | #VALUE! |
| ENSMUSG00000096920.2 | 0 | NA | NA | NA | NA | NA | #VALUE! |
| ENSMUSG00000096921.2 | 0 | NA | NA | NA | NA | NA | #VALUE! |
| ENSMUSG00000096922.2 | 0 | NA | NA | NA | NA | NA | #VALUE! |
| ENSMUSG00000096923.2 | 0 | NA | NA | NA | NA | NA | #VALUE! |
| ENSMUSG00000096927.2 | 0 | NA | NA | NA | NA | NA | #VALUE! |
| ENSMUSG00000096930.2 | 0 | NA | NA | NA | NA | NA | #VALUE! |
| ENSMUSG00000096932.3 | 0 | NA | NA | NA | NA | NA | #VALUE! |
| ENSMUSG00000096934.2 | 0 | NA | NA | NA | NA | NA | #VALUE! |
| ENSMUSG00000096936.2 | 0 | NA | NA | NA | NA | NA | #VALUE! |
| ENSMUSG00000096937.2 | 0 | NA | NA | NA | NA | NA | #VALUE! |
| ENSMUSG00000096938.8 | 0 | NA | NA | NA | NA | NA | #VALUE! |
| ENSMUSG00000096939.2 | 0 | NA | NA | NA | NA | NA | #VALUE! |
| ENSMUSG00000096940.2 | 0.466276 | 2.449889 | 3.997179 | 0.612904 | 0.53994 | NA | #VALUE! |
| ENSMUSG00000096942.2 | 9.770128 | 2.268837 | 1.996673 | 1.136308 | 0.255828 | NA | #VALUE! |
| ENSMUSG00000096948.3 | 0 | NA | NA | NA | NA | NA | #VALUE! |
| ENSMUSG00000096949.2 | 0 | NA | NA | NA | NA | NA | #VALUE! |
| ENSMUSG00000096950.3 | 0 | NA | NA | NA | NA | NA | #VALUE! |
| ENSMUSG00000096953.3 | 3.450442 | 5.285914 | 3.919688 | 1.348555 | 0.17748 | NA | #VALUE! |
| ENSMUSG00000096955.2 | 0 | NA | NA | NA | NA | NA | #VALUE! |

| | | | | | | | |
|---|---|---|---|---|---|---|---|
| ENSMUSG00000096957.2 | 3.543697 | 5.324148 | 3.919351 | 1.358426 | 0.174329 | NA | #VALUE! |
| ENSMUSG00000096959.2 | 0 | NA | NA | NA | NA | NA | #VALUE! |
| ENSMUSG00000096960.2 | 0 | NA | NA | NA | NA | NA | #VALUE! |
| ENSMUSG00000096962.2 | 0 | NA | NA | NA | NA | NA | #VALUE! |
| ENSMUSG00000096963.2 | 0 | NA | NA | NA | NA | NA | #VALUE! |
| ENSMUSG00000096964.2 | 3.357187 | 5.246639 | 3.920044 | 1.338413 | 0.180762 | NA | #VALUE! |
| ENSMUSG00000096965.8 | 0 | NA | NA | NA | NA | NA | #VALUE! |
| ENSMUSG00000096966.3 | 0 | NA | NA | NA | NA | NA | #VALUE! |
| ENSMUSG00000096967.4 | 2.705731 | 4.913666 | 3.923477 | 1.252375 | 0.210433 | NA | #VALUE! |
| ENSMUSG00000096968.2 | 0.245976 | 1.027628 | 4.067126 | 0.252667 | 0.800526 | NA | #VALUE! |
| ENSMUSG00000096973.2 | 0 | NA | NA | NA | NA | NA | #VALUE! |
| ENSMUSG00000096974.3 | 9.365616 | 3.472749 | 3.362845 | 1.032682 | 0.301753 | NA | #VALUE! |
| ENSMUSG00000096977.3 | 0 | NA | NA | NA | NA | NA | #VALUE! |
| ENSMUSG00000096981.3 | 0 | NA | NA | NA | NA | NA | #VALUE! |
| ENSMUSG00000096982.3 | 0 | NA | NA | NA | NA | NA | #VALUE! |
| ENSMUSG00000096986.3 | 0 | NA | NA | NA | NA | NA | #VALUE! |
| ENSMUSG00000096988.2 | 0 | NA | NA | NA | NA | NA | #VALUE! |
| ENSMUSG00000096991.2 | 0 | NA | NA | NA | NA | NA | #VALUE! |
| ENSMUSG00000096992.4 | 0 | NA | NA | NA | NA | NA | #VALUE! |
| ENSMUSG00000096993.4 | 0 | NA | NA | NA | NA | NA | #VALUE! |
| ENSMUSG00000096995.3 | 0 | NA | NA | NA | NA | NA | #VALUE! |
| ENSMUSG00000096997.2 | 0 | NA | NA | NA | NA | NA | #VALUE! |
| ENSMUSG00000096999.2 | 0 | NA | NA | NA | NA | NA | #VALUE! |
| ENSMUSG00000097000.3 | 0.466276 | 2.449889 | 3.997179 | 0.612904 | 0.53994 | NA | #VALUE! |
| ENSMUSG00000097001.2 | 0 | NA | NA | NA | NA | NA | #VALUE! |
| ENSMUSG00000097002.2 | 0 | NA | NA | NA | NA | NA | #VALUE! |
| ENSMUSG00000097003.2 | 0 | NA | NA | NA | NA | NA | #VALUE! |
| ENSMUSG00000097004.2 | 0 | NA | NA | NA | NA | NA | #VALUE! |
| ENSMUSG00000097006.8 | 1.851119 | 4.378291 | 3.930938 | 1.113803 | 0.265364 | NA | #VALUE! |
| ENSMUSG00000097007.4 | 0 | NA | NA | NA | NA | NA | #VALUE! |
| ENSMUSG00000097009.3 | 0 | NA | NA | NA | NA | NA | #VALUE! |
| ENSMUSG00000097010.3 | 0 | NA | NA | NA | NA | NA | #VALUE! |
| ENSMUSG00000097011.3 | 0 | NA | NA | NA | NA | NA | #VALUE! |
| ENSMUSG00000097012.2 | 0 | NA | NA | NA | NA | NA | #VALUE! |
| ENSMUSG00000097015.3 | 0 | NA | NA | NA | NA | NA | #VALUE! |
| ENSMUSG00000097016.3 | 4.620578 | -5.74391 | 3.916409 | -1.46663 | 0.142478 | NA | #VALUE! |
| ENSMUSG00000097017.2 | 0 | NA | NA | NA | NA | NA | #VALUE! |
| ENSMUSG00000097018.3 | 0 | NA | NA | NA | NA | NA | #VALUE! |
| ENSMUSG00000097021.8 | 0 | NA | NA | NA | NA | NA | #VALUE! |
| ENSMUSG00000097022.3 | 0 | NA | NA | NA | NA | NA | #VALUE! |
| ENSMUSG00000097023.9 | 0 | NA | NA | NA | NA | NA | #VALUE! |
| ENSMUSG00000097024.2 | 0 | NA | NA | NA | NA | NA | #VALUE! |
| ENSMUSG00000097025.3 | 3.719993 | -5.42213 | 3.918812 | -1.38361 | 0.166476 | NA | #VALUE! |
| ENSMUSG00000097027.3 | 0 | NA | NA | NA | NA | NA | #VALUE! |

| | | | | | | | |
|---|---|---|---|---|---|---|---|
| ENSMUSG00000097028.3 | 0 | NA | NA | NA | NA | NA | #VALUE! |
| ENSMUSG00000097029.3 | 0 | NA | NA | NA | NA | NA | #VALUE! |
| ENSMUSG00000097030.2 | 0 | NA | NA | NA | NA | NA | #VALUE! |
| ENSMUSG00000097031.3 | 0 | NA | NA | NA | NA | NA | #VALUE! |
| ENSMUSG00000097032.3 | 1.771849 | 4.332864 | 3.931707 | 1.102031 | 0.270448 | NA | #VALUE! |
| ENSMUSG00000097035.3 | 0 | NA | NA | NA | NA | NA | #VALUE! |
| ENSMUSG00000097040.7 | 0 | NA | NA | NA | NA | NA | #VALUE! |
| ENSMUSG00000097041.2 | 0 | NA | NA | NA | NA | NA | #VALUE! |
| ENSMUSG00000097043.2 | 0 | NA | NA | NA | NA | NA | #VALUE! |
| ENSMUSG00000097044.2 | 0 | NA | NA | NA | NA | NA | #VALUE! |
| ENSMUSG00000097047.3 | 3.333626 | 2.165543 | 3.818869 | 0.567064 | 0.570671 | NA | #VALUE! |
| ENSMUSG00000097049.2 | 0 | NA | NA | NA | NA | NA | #VALUE! |
| ENSMUSG00000097050.2 | 0 | NA | NA | NA | NA | NA | #VALUE! |
| ENSMUSG00000097052.4 | 0 | NA | NA | NA | NA | NA | #VALUE! |
| ENSMUSG00000097053.2 | 0 | NA | NA | NA | NA | NA | #VALUE! |
| ENSMUSG00000097054.2 | 0 | NA | NA | NA | NA | NA | #VALUE! |
| ENSMUSG00000097055.3 | 0 | NA | NA | NA | NA | NA | #VALUE! |
| ENSMUSG00000097057.2 | 0 | NA | NA | NA | NA | NA | #VALUE! |
| ENSMUSG00000097060.2 | 9.424172 | 6.724361 | 3.196071 | 2.103946 | 0.035383 | NA | #VALUE! |
| ENSMUSG00000097063.4 | 0 | NA | NA | NA | NA | NA | #VALUE! |
| ENSMUSG00000097067.2 | 0 | NA | NA | NA | NA | NA | #VALUE! |
| ENSMUSG00000097068.3 | 0 | NA | NA | NA | NA | NA | #VALUE! |
| ENSMUSG00000097069.2 | 0 | NA | NA | NA | NA | NA | #VALUE! |
| ENSMUSG00000097071.3 | 0 | NA | NA | NA | NA | NA | #VALUE! |
| ENSMUSG00000097072.8 | 0 | NA | NA | NA | NA | NA | #VALUE! |
| ENSMUSG00000097073.8 | 1.470184 | -4.10383 | 3.936698 | -1.04245 | 0.297201 | NA | #VALUE! |
| ENSMUSG00000097074.3 | 0 | NA | NA | NA | NA | NA | #VALUE! |
| ENSMUSG00000097075.4 | 0 | NA | NA | NA | NA | NA | #VALUE! |
| ENSMUSG00000097076.2 | 0 | NA | NA | NA | NA | NA | #VALUE! |
| ENSMUSG00000097077.2 | 0 | NA | NA | NA | NA | NA | #VALUE! |
| ENSMUSG00000097078.2 | 0 | NA | NA | NA | NA | NA | #VALUE! |
| ENSMUSG00000097079.3 | 0 | NA | NA | NA | NA | NA | #VALUE! |
| ENSMUSG00000097081.2 | 0 | NA | NA | NA | NA | NA | #VALUE! |
| ENSMUSG00000097082.2 | 0 | NA | NA | NA | NA | NA | #VALUE! |
| ENSMUSG00000097083.2 | 0 | NA | NA | NA | NA | NA | #VALUE! |
| ENSMUSG00000097084.2 | 0 | NA | NA | NA | NA | NA | #VALUE! |
| ENSMUSG00000097085.3 | 0 | NA | NA | NA | NA | NA | #VALUE! |
| ENSMUSG00000097086.2 | 0 | NA | NA | NA | NA | NA | #VALUE! |
| ENSMUSG00000097087.2 | 0 | NA | NA | NA | NA | NA | #VALUE! |
| ENSMUSG00000097089.3 | 0 | NA | NA | NA | NA | NA | #VALUE! |
| ENSMUSG00000097090.2 | 0 | NA | NA | NA | NA | NA | #VALUE! |
| ENSMUSG00000097091.2 | 0 | NA | NA | NA | NA | NA | #VALUE! |
| ENSMUSG00000097092.3 | 0 | NA | NA | NA | NA | NA | #VALUE! |
| ENSMUSG00000097097.3 | 0 | NA | NA | NA | NA | NA | #VALUE! |

| | | | | | | | |
|---|---|---|---|---|---|---|---|
| ENSMUSG00000097098.9 | 0 | NA | NA | NA | NA | NA | #VALUE! |
| ENSMUSG00000097099.3 | 1.68021 | -4.29432 | 3.933015 | -1.09187 | 0.274892 | NA | #VALUE! |
| ENSMUSG00000097100.2 | 0 | NA | NA | NA | NA | NA | #VALUE! |
| ENSMUSG00000097101.4 | 18.48816 | -0.32588 | 2.222577 | -0.14662 | NA | NA | #VALUE! |
| ENSMUSG00000097104.2 | 0 | NA | NA | NA | NA | NA | #VALUE! |
| ENSMUSG00000097106.2 | 0 | NA | NA | NA | NA | NA | #VALUE! |
| ENSMUSG00000097107.2 | 0 | NA | NA | NA | NA | NA | #VALUE! |
| ENSMUSG00000097108.2 | 0 | NA | NA | NA | NA | NA | #VALUE! |
| ENSMUSG00000097110.2 | 0 | NA | NA | NA | NA | NA | #VALUE! |
| ENSMUSG00000097111.2 | 0 | NA | NA | NA | NA | NA | #VALUE! |
| ENSMUSG00000097113.8 | 5.978686 | -6.10788 | 3.847895 | -1.58733 | 0.112438 | NA | #VALUE! |
| ENSMUSG00000097115.3 | 0 | NA | NA | NA | NA | NA | #VALUE! |
| ENSMUSG00000097116.2 | 0 | NA | NA | NA | NA | NA | #VALUE! |
| ENSMUSG00000097120.2 | 0 | NA | NA | NA | NA | NA | #VALUE! |
| ENSMUSG00000097124.3 | 0 | NA | NA | NA | NA | NA | #VALUE! |
| ENSMUSG00000097125.3 | 0 | NA | NA | NA | NA | NA | #VALUE! |
| ENSMUSG00000097127.8 | 0 | NA | NA | NA | NA | NA | #VALUE! |
| ENSMUSG00000097129.3 | 0 | NA | NA | NA | NA | NA | #VALUE! |
| ENSMUSG00000097130.2 | 0 | NA | NA | NA | NA | NA | #VALUE! |
| ENSMUSG00000097133.2 | 0 | NA | NA | NA | NA | NA | #VALUE! |
| ENSMUSG00000097134.3 | 0 | NA | NA | NA | NA | NA | #VALUE! |
| ENSMUSG00000097137.2 | 0 | NA | NA | NA | NA | NA | #VALUE! |
| ENSMUSG00000097139.4 | 0 | NA | NA | NA | NA | NA | #VALUE! |
| ENSMUSG00000097140.2 | 0 | NA | NA | NA | NA | NA | #VALUE! |
| ENSMUSG00000097141.3 | 0 | NA | NA | NA | NA | NA | #VALUE! |
| ENSMUSG00000097145.6 | 8.272813 | 0.666444 | 2.736899 | 0.243503 | 0.807616 | NA | #VALUE! |
| ENSMUSG00000097146.2 | 0 | NA | NA | NA | NA | NA | #VALUE! |
| ENSMUSG00000097149.2 | 0 | NA | NA | NA | NA | NA | #VALUE! |
| ENSMUSG00000097150.2 | 0.466276 | 2.449889 | 3.997179 | 0.612904 | 0.53994 | NA | #VALUE! |
| ENSMUSG00000097154.3 | 4.207088 | 5.562698 | 3.917438 | 1.419984 | 0.155612 | NA | #VALUE! |
| ENSMUSG00000097155.2 | 0 | NA | NA | NA | NA | NA | #VALUE! |
| ENSMUSG00000097156.8 | 0 | NA | NA | NA | NA | NA | #VALUE! |
| ENSMUSG00000097157.3 | 0 | NA | NA | NA | NA | NA | #VALUE! |
| ENSMUSG00000097160.3 | 0 | NA | NA | NA | NA | NA | #VALUE! |
| ENSMUSG00000097163.3 | 0 | NA | NA | NA | NA | NA | #VALUE! |
| ENSMUSG00000097164.3 | 1.589447 | 0.988526 | 3.852621 | 0.256585 | 0.797499 | NA | #VALUE! |
| ENSMUSG00000097166.2 | 0 | NA | NA | NA | NA | NA | #VALUE! |
| ENSMUSG00000097169.3 | 0 | NA | NA | NA | NA | NA | #VALUE! |
| ENSMUSG00000097170.2 | 0 | NA | NA | NA | NA | NA | #VALUE! |
| ENSMUSG00000097171.2 | 0 | NA | NA | NA | NA | NA | #VALUE! |
| ENSMUSG00000097174.3 | 0 | NA | NA | NA | NA | NA | #VALUE! |
| ENSMUSG00000097175.2 | 0 | NA | NA | NA | NA | NA | #VALUE! |
| ENSMUSG00000097178.8 | 0 | NA | NA | NA | NA | NA | #VALUE! |
| ENSMUSG00000097180.6 | 0 | NA | NA | NA | NA | NA | #VALUE! |

| | | | | | | | |
|---|---|---|---|---|---|---|---|
| ENSMUSG00000097181.2 | 0 | NA | NA | NA | NA | NA | #VALUE! |
| ENSMUSG00000097182.3 | 0 | NA | NA | NA | NA | NA | #VALUE! |
| ENSMUSG00000097183.3 | 0 | NA | NA | NA | NA | NA | #VALUE! |
| ENSMUSG00000097184.3 | 0 | NA | NA | NA | NA | NA | #VALUE! |
| ENSMUSG00000097185.2 | 0 | NA | NA | NA | NA | NA | #VALUE! |
| ENSMUSG00000097186.2 | 0 | NA | NA | NA | NA | NA | #VALUE! |
| ENSMUSG00000097189.2 | 0 | NA | NA | NA | NA | NA | #VALUE! |
| ENSMUSG00000097190.2 | 0 | NA | NA | NA | NA | NA | #VALUE! |
| ENSMUSG00000097191.3 | 0 | NA | NA | NA | NA | NA | #VALUE! |
| ENSMUSG00000097196.3 | 0 | NA | NA | NA | NA | NA | #VALUE! |
| ENSMUSG00000097199.2 | 0 | NA | NA | NA | NA | NA | #VALUE! |
| ENSMUSG00000097202.2 | 0 | NA | NA | NA | NA | NA | #VALUE! |
| ENSMUSG00000097204.3 | 0.105013 | -0.97324 | 4.076293 | -0.23876 | 0.811295 | NA | #VALUE! |
| ENSMUSG00000097205.2 | 0 | NA | NA | NA | NA | NA | #VALUE! |
| ENSMUSG00000097207.8 | 0 | NA | NA | NA | NA | NA | #VALUE! |
| ENSMUSG00000097209.3 | 0 | NA | NA | NA | NA | NA | #VALUE! |
| ENSMUSG00000097211.3 | 0 | NA | NA | NA | NA | NA | #VALUE! |
| ENSMUSG00000097214.2 | 0 | NA | NA | NA | NA | NA | #VALUE! |
| ENSMUSG00000097215.2 | 0 | NA | NA | NA | NA | NA | #VALUE! |
| ENSMUSG00000097216.4 | 0 | NA | NA | NA | NA | NA | #VALUE! |
| ENSMUSG00000097217.3 | 0 | NA | NA | NA | NA | NA | #VALUE! |
| ENSMUSG00000097218.8 | 0 | NA | NA | NA | NA | NA | #VALUE! |
| ENSMUSG00000097221.2 | 0 | NA | NA | NA | NA | NA | #VALUE! |
| ENSMUSG00000097222.3 | 0 | NA | NA | NA | NA | NA | #VALUE! |
| ENSMUSG00000097223.3 | 0 | NA | NA | NA | NA | NA | #VALUE! |
| ENSMUSG00000097224.2 | 23.85963 | 21.45502 | 2.83687 | 7.562921 | NA | NA | #VALUE! |
| ENSMUSG00000097226.2 | 0 | NA | NA | NA | NA | NA | #VALUE! |
| ENSMUSG00000097228.2 | 0 | NA | NA | NA | NA | NA | #VALUE! |
| ENSMUSG00000097229.2 | 0 | NA | NA | NA | NA | NA | #VALUE! |
| ENSMUSG00000097231.3 | 0 | NA | NA | NA | NA | NA | #VALUE! |
| ENSMUSG00000097232.2 | 0 | NA | NA | NA | NA | NA | #VALUE! |
| ENSMUSG00000097233.2 | 0 | NA | NA | NA | NA | NA | #VALUE! |
| ENSMUSG00000097237.2 | 0 | NA | NA | NA | NA | NA | #VALUE! |
| ENSMUSG00000097239.8 | 3.039465 | -5.14172 | 3.921389 | -1.3112 | 0.189791 | NA | #VALUE! |
| ENSMUSG00000097240.2 | 0 | NA | NA | NA | NA | NA | #VALUE! |
| ENSMUSG00000097241.3 | 0 | NA | NA | NA | NA | NA | #VALUE! |
| ENSMUSG00000097244.2 | 0 | NA | NA | NA | NA | NA | #VALUE! |
| ENSMUSG00000097245.2 | 0 | NA | NA | NA | NA | NA | #VALUE! |
| ENSMUSG00000097247.3 | 0 | NA | NA | NA | NA | NA | #VALUE! |
| ENSMUSG00000097248.10 | 0 | NA | NA | NA | NA | NA | #VALUE! |
| ENSMUSG00000097250.2 | 0 | NA | NA | NA | NA | NA | #VALUE! |
| ENSMUSG00000097251.3 | 1.155144 | -3.7605 | 3.944679 | -0.95331 | 0.340433 | NA | #VALUE! |
| ENSMUSG00000097253.3 | 0 | NA | NA | NA | NA | NA | #VALUE! |
| ENSMUSG00000097254.3 | 0 | NA | NA | NA | NA | NA | #VALUE! |

| | | | | | | | |
|---|---|---|---|---|---|---|---|
| ENSMUSG00000097255.2 | 0 | NA | NA | NA | NA | NA | #VALUE! |
| ENSMUSG00000097256.2 | 0 | NA | NA | NA | NA | NA | #VALUE! |
| ENSMUSG00000097259.3 | 0 | NA | NA | NA | NA | NA | #VALUE! |
| ENSMUSG00000097260.2 | 0 | NA | NA | NA | NA | NA | #VALUE! |
| ENSMUSG00000097261.3 | 1.009701 | 3.503687 | 3.950852 | 0.886818 | 0.375177 | NA | #VALUE! |
| ENSMUSG00000097262.3 | 0 | NA | NA | NA | NA | NA | #VALUE! |
| ENSMUSG00000097264.2 | 0 | NA | NA | NA | NA | NA | #VALUE! |
| ENSMUSG00000097265.2 | 0 | NA | NA | NA | NA | NA | #VALUE! |
| ENSMUSG00000097266.2 | 0 | NA | NA | NA | NA | NA | #VALUE! |
| ENSMUSG00000097267.3 | 0 | NA | NA | NA | NA | NA | #VALUE! |
| ENSMUSG00000097268.2 | 0 | NA | NA | NA | NA | NA | #VALUE! |
| ENSMUSG00000097271.2 | 0 | NA | NA | NA | NA | NA | #VALUE! |
| ENSMUSG00000097274.3 | 0 | NA | NA | NA | NA | NA | #VALUE! |
| ENSMUSG00000097275.2 | 0 | NA | NA | NA | NA | NA | #VALUE! |
| ENSMUSG00000097276.2 | 0 | NA | NA | NA | NA | NA | #VALUE! |
| ENSMUSG00000097278.8 | 0 | NA | NA | NA | NA | NA | #VALUE! |
| ENSMUSG00000097279.2 | 0 | NA | NA | NA | NA | NA | #VALUE! |
| ENSMUSG00000097280.2 | 0 | NA | NA | NA | NA | NA | #VALUE! |
| ENSMUSG00000097281.3 | 0 | NA | NA | NA | NA | NA | #VALUE! |
| ENSMUSG00000097282.2 | 0 | NA | NA | NA | NA | NA | #VALUE! |
| ENSMUSG00000097283.2 | 0 | NA | NA | NA | NA | NA | #VALUE! |
| ENSMUSG00000097286.3 | 2.977324 | 0.28073 | 3.56961 | 0.078645 | 0.937315 | NA | #VALUE! |
| ENSMUSG00000097287.4 | 7.434633 | -6.42344 | 3.449352 | -1.86222 | 0.062573 | NA | #VALUE! |
| ENSMUSG00000097288.2 | 0 | NA | NA | NA | NA | NA | #VALUE! |
| ENSMUSG00000097291.2 | 0 | NA | NA | NA | NA | NA | #VALUE! |
| ENSMUSG00000097292.3 | 1.025807 | 3.556426 | 3.949287 | 0.900523 | 0.367842 | NA | #VALUE! |
| ENSMUSG00000097293.2 | 0 | NA | NA | NA | NA | NA | #VALUE! |
| ENSMUSG00000097295.2 | 1.260158 | -3.88425 | 3.941584 | -0.98545 | 0.324401 | NA | #VALUE! |
| ENSMUSG00000097296.2 | 5.565696 | -6.01141 | 3.914779 | -1.53557 | 0.124644 | NA | #VALUE! |
| ENSMUSG00000097299.2 | 0 | NA | NA | NA | NA | NA | #VALUE! |
| ENSMUSG00000097302.2 | 0 | NA | NA | NA | NA | NA | #VALUE! |
| ENSMUSG00000097303.3 | 0 | NA | NA | NA | NA | NA | #VALUE! |
| ENSMUSG00000097304.2 | 0.195789 | -1.05581 | 4.073302 | -0.2592 | 0.795479 | NA | #VALUE! |
| ENSMUSG00000097305.3 | 0 | NA | NA | NA | NA | NA | #VALUE! |
| ENSMUSG00000097307.2 | 0 | NA | NA | NA | NA | NA | #VALUE! |
| ENSMUSG00000097308.2 | 0 | NA | NA | NA | NA | NA | #VALUE! |
| ENSMUSG00000097309.3 | 2.984166 | 5.077883 | 3.921685 | 1.294822 | 0.195382 | NA | #VALUE! |
| ENSMUSG00000097310.3 | 8.217789 | -6.56783 | 3.333139 | -1.97046 | 0.048785 | NA | #VALUE! |
| ENSMUSG00000097311.8 | 4.111571 | -5.56681 | 3.917665 | -1.42095 | 0.155331 | NA | #VALUE! |
| ENSMUSG00000097313.3 | 1.68021 | -4.29432 | 3.933015 | -1.09187 | 0.274892 | NA | #VALUE! |
| ENSMUSG00000097315.2 | 0 | NA | NA | NA | NA | NA | #VALUE! |
| ENSMUSG00000097316.3 | 0 | NA | NA | NA | NA | NA | #VALUE! |
| ENSMUSG00000097318.3 | 3.789673 | 5.419286 | 3.91855 | 1.382982 | 0.16667 | NA | #VALUE! |
| ENSMUSG00000097320.9 | 0 | NA | NA | NA | NA | NA | #VALUE! |

| | | | | | | | |
|---|---|---|---|---|---|---|---|
| ENSMUSG00000097322.9 | 0 | NA | NA | NA | NA | NA | #VALUE! |
| ENSMUSG00000097323.3 | 0 | NA | NA | NA | NA | NA | #VALUE! |
| ENSMUSG00000097324.9 | 0 | NA | NA | NA | NA | NA | #VALUE! |
| ENSMUSG00000097325.3 | 0 | NA | NA | NA | NA | NA | #VALUE! |
| ENSMUSG00000097326.3 | 0 | NA | NA | NA | NA | NA | #VALUE! |
| ENSMUSG00000097330.3 | 0 | NA | NA | NA | NA | NA | #VALUE! |
| ENSMUSG00000097331.3 | 0 | NA | NA | NA | NA | NA | #VALUE! |
| ENSMUSG00000097336.9 | 5.657437 | 5.984105 | 3.914739 | 1.528609 | 0.126361 | NA | #VALUE! |
| ENSMUSG00000097339.6 | 0 | NA | NA | NA | NA | NA | #VALUE! |
| ENSMUSG00000097342.2 | 0 | NA | NA | NA | NA | NA | #VALUE! |
| ENSMUSG00000097343.3 | 4.686473 | -0.62447 | 3.766616 | -0.16579 | 0.868321 | NA | #VALUE! |
| ENSMUSG00000097346.2 | 0 | NA | NA | NA | NA | NA | #VALUE! |
| ENSMUSG00000097349.2 | 0 | NA | NA | NA | NA | NA | #VALUE! |
| ENSMUSG00000097350.3 | 3.34744 | -0.12606 | 3.780578 | -0.03334 | 0.9734 | NA | #VALUE! |
| ENSMUSG00000097352.8 | 0 | NA | NA | NA | NA | NA | #VALUE! |
| ENSMUSG00000097356.2 | 0 | NA | NA | NA | NA | NA | #VALUE! |
| ENSMUSG00000097357.3 | 1.346268 | 3.918821 | 3.939925 | 0.994644 | 0.31991 | NA | #VALUE! |
| ENSMUSG00000097360.9 | 1.650289 | -2.128 | 3.894045 | -0.54648 | 0.584739 | NA | #VALUE! |
| ENSMUSG00000097361.2 | 0 | NA | NA | NA | NA | NA | #VALUE! |
| ENSMUSG00000097362.2 | 0 | NA | NA | NA | NA | NA | #VALUE! |
| ENSMUSG00000097363.2 | 0 | NA | NA | NA | NA | NA | #VALUE! |
| ENSMUSG00000097365.8 | 0 | NA | NA | NA | NA | NA | #VALUE! |
| ENSMUSG00000097366.2 | 0 | NA | NA | NA | NA | NA | #VALUE! |
| ENSMUSG00000097368.3 | 0.336567 | 1.944925 | 4.029936 | 0.482619 | 0.629366 | NA | #VALUE! |
| ENSMUSG00000097370.3 | 0 | NA | NA | NA | NA | NA | #VALUE! |
| ENSMUSG00000097371.2 | 0 | NA | NA | NA | NA | NA | #VALUE! |
| ENSMUSG00000097372.2 | 0 | NA | NA | NA | NA | NA | #VALUE! |
| ENSMUSG00000097373.2 | 0 | NA | NA | NA | NA | NA | #VALUE! |
| ENSMUSG00000097379.3 | 0 | NA | NA | NA | NA | NA | #VALUE! |
| ENSMUSG00000097381.3 | 0.105013 | -0.97324 | 4.076293 | -0.23876 | 0.811295 | NA | #VALUE! |
| ENSMUSG00000097382.2 | 0 | NA | NA | NA | NA | NA | #VALUE! |
| ENSMUSG00000097383.2 | 7.697255 | 1.292692 | 3.156181 | 0.409575 | 0.682118 | NA | #VALUE! |
| ENSMUSG00000097384.2 | 0 | NA | NA | NA | NA | NA | #VALUE! |
| ENSMUSG00000097385.2 | 0 | NA | NA | NA | NA | NA | #VALUE! |
| ENSMUSG00000097386.2 | 0 | NA | NA | NA | NA | NA | #VALUE! |
| ENSMUSG00000097387.2 | 0 | NA | NA | NA | NA | NA | #VALUE! |
| ENSMUSG00000097388.2 | 0 | NA | NA | NA | NA | NA | #VALUE! |
| ENSMUSG00000097391.9 | 0 | NA | NA | NA | NA | NA | #VALUE! |
| ENSMUSG00000097393.9 | 0 | NA | NA | NA | NA | NA | #VALUE! |
| ENSMUSG00000097396.2 | 0 | NA | NA | NA | NA | NA | #VALUE! |
| ENSMUSG00000097397.3 | 0 | NA | NA | NA | NA | NA | #VALUE! |
| ENSMUSG00000097398.3 | 0 | NA | NA | NA | NA | NA | #VALUE! |
| ENSMUSG00000097399.8 | 0 | NA | NA | NA | NA | NA | #VALUE! |
| ENSMUSG00000097401.2 | 0 | NA | NA | NA | NA | NA | #VALUE! |

| | | | | | | | |
|---|---|---|---|---|---|---|---|
| ENSMUSG00000097402.2 | 0 | NA | NA | NA | NA | NA | #VALUE! |
| ENSMUSG00000097403.3 | 0.932552 | 3.421684 | 3.953399 | 0.865504 | 0.386762 | NA | #VALUE! |
| ENSMUSG00000097404.3 | 0 | NA | NA | NA | NA | NA | #VALUE! |
| ENSMUSG00000097405.2 | 0 | NA | NA | NA | NA | NA | #VALUE! |
| ENSMUSG00000097406.2 | 0 | NA | NA | NA | NA | NA | #VALUE! |
| ENSMUSG00000097407.2 | 0 | NA | NA | NA | NA | NA | #VALUE! |
| ENSMUSG00000097408.2 | 0 | NA | NA | NA | NA | NA | #VALUE! |
| ENSMUSG00000097409.2 | 0 | NA | NA | NA | NA | NA | #VALUE! |
| ENSMUSG00000097411.3 | 0.840105 | -3.30885 | 3.958472 | -0.83589 | 0.403216 | NA | #VALUE! |
| ENSMUSG00000097413.3 | 4.701929 | -5.76743 | 3.916253 | -1.47269 | 0.140834 | NA | #VALUE! |
| ENSMUSG00000097414.3 | 0 | NA | NA | NA | NA | NA | #VALUE! |
| ENSMUSG00000097415.3 | 2.07945 | 1.385352 | 3.835073 | 0.361232 | 0.717926 | NA | #VALUE! |
| ENSMUSG00000097416.4 | 2.595015 | -0.80823 | 3.80147 | -0.21261 | 0.831632 | NA | #VALUE! |
| ENSMUSG00000097418.8 | 0 | NA | NA | NA | NA | NA | #VALUE! |
| ENSMUSG00000097419.2 | 0 | NA | NA | NA | NA | NA | #VALUE! |
| ENSMUSG00000097421.3 | 0 | NA | NA | NA | NA | NA | #VALUE! |
| ENSMUSG00000097424.3 | 0 | NA | NA | NA | NA | NA | #VALUE! |
| ENSMUSG00000097425.4 | 0 | NA | NA | NA | NA | NA | #VALUE! |
| ENSMUSG00000097426.2 | 0 | NA | NA | NA | NA | NA | #VALUE! |
| ENSMUSG00000097427.6 | 0 | NA | NA | NA | NA | NA | #VALUE! |
| ENSMUSG00000097428.2 | 0 | NA | NA | NA | NA | NA | #VALUE! |
| ENSMUSG00000097429.2 | 0 | NA | NA | NA | NA | NA | #VALUE! |
| ENSMUSG00000097430.3 | 0 | NA | NA | NA | NA | NA | #VALUE! |
| ENSMUSG00000097431.3 | 4.45919 | 4.084885 | 3.810418 | 1.072031 | 0.283706 | NA | #VALUE! |
| ENSMUSG00000097433.2 | 0 | NA | NA | NA | NA | NA | #VALUE! |
| ENSMUSG00000097434.3 | 0 | NA | NA | NA | NA | NA | #VALUE! |
| ENSMUSG00000097435.2 | 0 | NA | NA | NA | NA | NA | #VALUE! |
| ENSMUSG00000097438.2 | 0 | NA | NA | NA | NA | NA | #VALUE! |
| ENSMUSG00000097439.7 | 0 | NA | NA | NA | NA | NA | #VALUE! |
| ENSMUSG00000097440.3 | 7.344146 | 6.364389 | 3.454431 | 1.842384 | 0.065419 | NA | #VALUE! |
| ENSMUSG00000097441.2 | 0 | NA | NA | NA | NA | NA | #VALUE! |
| ENSMUSG00000097442.2 | 0 | NA | NA | NA | NA | NA | #VALUE! |
| ENSMUSG00000097443.3 | 0 | NA | NA | NA | NA | NA | #VALUE! |
| ENSMUSG00000097446.2 | 0 | NA | NA | NA | NA | NA | #VALUE! |
| ENSMUSG00000097447.8 | 0 | NA | NA | NA | NA | NA | #VALUE! |
| ENSMUSG00000097448.2 | 0 | NA | NA | NA | NA | NA | #VALUE! |
| ENSMUSG00000097450.8 | 0 | NA | NA | NA | NA | NA | #VALUE! |
| ENSMUSG00000097451.12 | 0 | NA | NA | NA | NA | NA | #VALUE! |
| ENSMUSG00000097452.3 | 0 | NA | NA | NA | NA | NA | #VALUE! |
| ENSMUSG00000097455.2 | 0 | NA | NA | NA | NA | NA | #VALUE! |
| ENSMUSG00000097456.8 | 0 | NA | NA | NA | NA | NA | #VALUE! |
| ENSMUSG00000097462.8 | 4.427559 | 5.628857 | 3.916961 | 1.437047 | 0.150705 | NA | #VALUE! |
| ENSMUSG00000097463.3 | 0 | NA | NA | NA | NA | NA | #VALUE! |
| ENSMUSG00000097464.2 | 0 | NA | NA | NA | NA | NA | #VALUE! |

| | | | | | | | |
|---|---|---|---|---|---|---|---|
| ENSMUSG00000097466.9 | 0 | NA | NA | NA | NA | NA | #VALUE! |
| ENSMUSG00000097467.3 | 0 | NA | NA | NA | NA | NA | #VALUE! |
| ENSMUSG00000097470.2 | 0 | NA | NA | NA | NA | NA | #VALUE! |
| ENSMUSG00000097471.9 | 0 | NA | NA | NA | NA | NA | #VALUE! |
| ENSMUSG00000097472.2 | 0 | NA | NA | NA | NA | NA | #VALUE! |
| ENSMUSG00000097473.2 | 0 | NA | NA | NA | NA | NA | #VALUE! |
| ENSMUSG00000097476.2 | 0 | NA | NA | NA | NA | NA | #VALUE! |
| ENSMUSG00000097480.2 | 0 | NA | NA | NA | NA | NA | #VALUE! |
| ENSMUSG00000097483.3 | 0 | NA | NA | NA | NA | NA | #VALUE! |
| ENSMUSG00000097484.2 | 0 | NA | NA | NA | NA | NA | #VALUE! |
| ENSMUSG00000097486.3 | 0 | NA | NA | NA | NA | NA | #VALUE! |
| ENSMUSG00000097487.8 | 0.408222 | -2.20205 | 4.016842 | -0.54821 | 0.583551 | NA | #VALUE! |
| ENSMUSG00000097488.2 | 0 | NA | NA | NA | NA | NA | #VALUE! |
| ENSMUSG00000097491.2 | 0 | NA | NA | NA | NA | NA | #VALUE! |
| ENSMUSG00000097493.4 | 0 | NA | NA | NA | NA | NA | #VALUE! |
| ENSMUSG00000097494.8 | 0 | NA | NA | NA | NA | NA | #VALUE! |
| ENSMUSG00000097495.2 | 0 | NA | NA | NA | NA | NA | #VALUE! |
| ENSMUSG00000097496.2 | 0 | NA | NA | NA | NA | NA | #VALUE! |
| ENSMUSG00000097497.2 | 0 | NA | NA | NA | NA | NA | #VALUE! |
| ENSMUSG00000097500.2 | 0 | NA | NA | NA | NA | NA | #VALUE! |
| ENSMUSG00000097502.3 | 0 | NA | NA | NA | NA | NA | #VALUE! |
| ENSMUSG00000097504.2 | 0 | NA | NA | NA | NA | NA | #VALUE! |
| ENSMUSG00000097509.2 | 1.566313 | -4.16993 | 3.935366 | -1.0596 | 0.289325 | NA | #VALUE! |
| ENSMUSG00000097511.3 | 0 | NA | NA | NA | NA | NA | #VALUE! |
| ENSMUSG00000097513.2 | 0 | NA | NA | NA | NA | NA | #VALUE! |
| ENSMUSG00000097515.3 | 0 | NA | NA | NA | NA | NA | #VALUE! |
| ENSMUSG00000097516.8 | 0 | NA | NA | NA | NA | NA | #VALUE! |
| ENSMUSG00000097519.3 | 0 | NA | NA | NA | NA | NA | #VALUE! |
| ENSMUSG00000097520.3 | 0 | NA | NA | NA | NA | NA | #VALUE! |
| ENSMUSG00000097522.2 | 0 | NA | NA | NA | NA | NA | #VALUE! |
| ENSMUSG00000097523.3 | 1.851119 | 4.378291 | 3.930938 | 1.113803 | 0.265364 | NA | #VALUE! |
| ENSMUSG00000097525.3 | 0 | NA | NA | NA | NA | NA | #VALUE! |
| ENSMUSG00000097527.3 | 1.365171 | -3.99822 | 3.938957 | -1.01505 | 0.310084 | NA | #VALUE! |
| ENSMUSG00000097529.2 | 0 | NA | NA | NA | NA | NA | #VALUE! |
| ENSMUSG00000097530.2 | 0 | NA | NA | NA | NA | NA | #VALUE! |
| ENSMUSG00000097532.3 | 0 | NA | NA | NA | NA | NA | #VALUE! |
| ENSMUSG00000097533.3 | 6.300788 | -6.18982 | 3.913848 | -1.58152 | 0.11376 | NA | #VALUE! |
| ENSMUSG00000097534.4 | 0 | NA | NA | NA | NA | NA | #VALUE! |
| ENSMUSG00000097535.2 | 0 | NA | NA | NA | NA | NA | #VALUE! |
| ENSMUSG00000097536.3 | 3.36042 | -5.28663 | 3.919996 | -1.34863 | 0.177455 | NA | #VALUE! |
| ENSMUSG00000097537.3 | 0 | NA | NA | NA | NA | NA | #VALUE! |
| ENSMUSG00000097538.2 | 0 | NA | NA | NA | NA | NA | #VALUE! |
| ENSMUSG00000097539.2 | 0 | NA | NA | NA | NA | NA | #VALUE! |
| ENSMUSG00000097543.2 | 0 | NA | NA | NA | NA | NA | #VALUE! |

| | | | | | | | |
|---|---|---|---|---|---|---|---|
| ENSMUSG00000097545.8 | 0 | NA | NA | NA | NA | NA | #VALUE! |
| ENSMUSG00000097546.2 | 0 | NA | NA | NA | NA | NA | #VALUE! |
| ENSMUSG00000097547.9 | 0 | NA | NA | NA | NA | NA | #VALUE! |
| ENSMUSG00000097548.2 | 0 | NA | NA | NA | NA | NA | #VALUE! |
| ENSMUSG00000097550.8 | 0 | NA | NA | NA | NA | NA | #VALUE! |
| ENSMUSG00000097552.3 | 0 | NA | NA | NA | NA | NA | #VALUE! |
| ENSMUSG00000097557.3 | 0 | NA | NA | NA | NA | NA | #VALUE! |
| ENSMUSG00000097558.2 | 0 | NA | NA | NA | NA | NA | #VALUE! |
| ENSMUSG00000097559.2 | 0 | NA | NA | NA | NA | NA | #VALUE! |
| ENSMUSG00000097561.2 | 0 | NA | NA | NA | NA | NA | #VALUE! |
| ENSMUSG00000097562.4 | 0 | NA | NA | NA | NA | NA | #VALUE! |
| ENSMUSG00000097563.2 | 0 | NA | NA | NA | NA | NA | #VALUE! |
| ENSMUSG00000097564.3 | 0.105013 | -0.97324 | 4.076293 | -0.23876 | 0.811295 | NA | #VALUE! |
| ENSMUSG00000097565.2 | 0 | NA | NA | NA | NA | NA | #VALUE! |
| ENSMUSG00000097569.3 | 0 | NA | NA | NA | NA | NA | #VALUE! |
| ENSMUSG00000097572.3 | 0 | NA | NA | NA | NA | NA | #VALUE! |
| ENSMUSG00000097573.3 | 2.019402 | 4.503826 | 3.928936 | 1.146322 | 0.251662 | NA | #VALUE! |
| ENSMUSG00000097575.3 | 0 | NA | NA | NA | NA | NA | #VALUE! |
| ENSMUSG00000097576.3 | 0 | NA | NA | NA | NA | NA | #VALUE! |
| ENSMUSG00000097577.3 | 0 | NA | NA | NA | NA | NA | #VALUE! |
| ENSMUSG00000097580.2 | 0 | NA | NA | NA | NA | NA | #VALUE! |
| ENSMUSG00000097583.3 | 0.924903 | 0.208114 | 3.919247 | 0.053101 | 0.957652 | NA | #VALUE! |
| ENSMUSG00000097584.2 | 0 | NA | NA | NA | NA | NA | #VALUE! |
| ENSMUSG00000097585.3 | 0.966646 | -0.91778 | 3.92315 | -0.23394 | 0.815031 | NA | #VALUE! |
| ENSMUSG00000097586.2 | 0 | NA | NA | NA | NA | NA | #VALUE! |
| ENSMUSG00000097587.3 | 0 | NA | NA | NA | NA | NA | #VALUE! |
| ENSMUSG00000097591.2 | 0 | NA | NA | NA | NA | NA | #VALUE! |
| ENSMUSG00000097592.3 | 0 | NA | NA | NA | NA | NA | #VALUE! |
| ENSMUSG00000097593.3 | 0 | NA | NA | NA | NA | NA | #VALUE! |
| ENSMUSG00000097594.2 | 0 | NA | NA | NA | NA | NA | #VALUE! |
| ENSMUSG00000097595.3 | 0 | NA | NA | NA | NA | NA | #VALUE! |
| ENSMUSG00000097596.2 | 0 | NA | NA | NA | NA | NA | #VALUE! |
| ENSMUSG00000097597.3 | 0 | NA | NA | NA | NA | NA | #VALUE! |
| ENSMUSG00000097601.3 | 0.504851 | 2.502843 | 3.993993 | 0.626652 | 0.530887 | NA | #VALUE! |
| ENSMUSG00000097602.3 | 0 | NA | NA | NA | NA | NA | #VALUE! |
| ENSMUSG00000097603.4 | 0 | NA | NA | NA | NA | NA | #VALUE! |
| ENSMUSG00000097604.2 | 0 | NA | NA | NA | NA | NA | #VALUE! |
| ENSMUSG00000097608.2 | 0 | NA | NA | NA | NA | NA | #VALUE! |
| ENSMUSG00000097610.3 | 0 | NA | NA | NA | NA | NA | #VALUE! |
| ENSMUSG00000097612.2 | 0 | NA | NA | NA | NA | NA | #VALUE! |
| ENSMUSG00000097613.2 | 0.168284 | 1.027628 | 4.067126 | 0.252667 | 0.800526 | NA | #VALUE! |
| ENSMUSG00000097615.2 | 5.129036 | 5.854808 | 3.915485 | 1.495296 | 0.134837 | NA | #VALUE! |
| ENSMUSG00000097617.3 | 0 | NA | NA | NA | NA | NA | #VALUE! |
| ENSMUSG00000097619.9 | 0 | NA | NA | NA | NA | NA | #VALUE! |

| | | | | | | | |
|---|---|---|---|---|---|---|---|
| ENSMUSG00000097620.8 | 0 | NA | NA | NA | NA | NA | #VALUE! |
| ENSMUSG00000097621.2 | 0 | NA | NA | NA | NA | NA | #VALUE! |
| ENSMUSG00000097622.3 | 0 | NA | NA | NA | NA | NA | #VALUE! |
| ENSMUSG00000097623.8 | 0 | NA | NA | NA | NA | NA | #VALUE! |
| ENSMUSG00000097624.3 | 0 | NA | NA | NA | NA | NA | #VALUE! |
| ENSMUSG00000097626.6 | 0 | NA | NA | NA | NA | NA | #VALUE! |
| ENSMUSG00000097627.3 | 0 | NA | NA | NA | NA | NA | #VALUE! |
| ENSMUSG00000097628.2 | 0 | NA | NA | NA | NA | NA | #VALUE! |
| ENSMUSG00000097630.3 | 0 | NA | NA | NA | NA | NA | #VALUE! |
| ENSMUSG00000097633.3 | 0 | NA | NA | NA | NA | NA | #VALUE! |
| ENSMUSG00000097634.3 | 0 | NA | NA | NA | NA | NA | #VALUE! |
| ENSMUSG00000097635.3 | 0 | NA | NA | NA | NA | NA | #VALUE! |
| ENSMUSG00000097636.9 | 0 | NA | NA | NA | NA | NA | #VALUE! |
| ENSMUSG00000097637.2 | 0 | NA | NA | NA | NA | NA | #VALUE! |
| ENSMUSG00000097638.2 | 0 | NA | NA | NA | NA | NA | #VALUE! |
| ENSMUSG00000097639.3 | 5.216789 | 5.873026 | 3.915376 | 1.49999 | 0.133617 | NA | #VALUE! |
| ENSMUSG00000097640.8 | 0 | NA | NA | NA | NA | NA | #VALUE! |
| ENSMUSG00000097641.3 | 0 | NA | NA | NA | NA | NA | #VALUE! |
| ENSMUSG00000097642.2 | 0 | NA | NA | NA | NA | NA | #VALUE! |
| ENSMUSG00000097643.2 | 0 | NA | NA | NA | NA | NA | #VALUE! |
| ENSMUSG00000097644.2 | 0 | NA | NA | NA | NA | NA | #VALUE! |
| ENSMUSG00000097646.2 | 0 | NA | NA | NA | NA | NA | #VALUE! |
| ENSMUSG00000097648.2 | 0 | NA | NA | NA | NA | NA | #VALUE! |
| ENSMUSG00000097649.3 | 0 | NA | NA | NA | NA | NA | #VALUE! |
| ENSMUSG00000097650.2 | 0 | NA | NA | NA | NA | NA | #VALUE! |
| ENSMUSG00000097651.3 | 0 | NA | NA | NA | NA | NA | #VALUE! |
| ENSMUSG00000097652.3 | 0 | NA | NA | NA | NA | NA | #VALUE! |
| ENSMUSG00000097653.3 | 0 | NA | NA | NA | NA | NA | #VALUE! |
| ENSMUSG00000097654.2 | 0 | NA | NA | NA | NA | NA | #VALUE! |
| ENSMUSG00000097655.4 | 0 | NA | NA | NA | NA | NA | #VALUE! |
| ENSMUSG00000097656.2 | 0 | NA | NA | NA | NA | NA | #VALUE! |
| ENSMUSG00000097657.3 | 0 | NA | NA | NA | NA | NA | #VALUE! |
| ENSMUSG00000097662.2 | 0 | NA | NA | NA | NA | NA | #VALUE! |
| ENSMUSG00000097665.2 | 0 | NA | NA | NA | NA | NA | #VALUE! |
| ENSMUSG00000097666.3 | 0 | NA | NA | NA | NA | NA | #VALUE! |
| ENSMUSG00000097668.3 | 0 | NA | NA | NA | NA | NA | #VALUE! |
| ENSMUSG00000097669.2 | 0 | NA | NA | NA | NA | NA | #VALUE! |
| ENSMUSG00000097670.2 | 0 | NA | NA | NA | NA | NA | #VALUE! |
| ENSMUSG00000097671.3 | 0 | NA | NA | NA | NA | NA | #VALUE! |
| ENSMUSG00000097673.2 | 2.811839 | -3.49071 | 3.898572 | -0.89538 | 0.370582 | NA | #VALUE! |
| ENSMUSG00000097675.3 | 0 | NA | NA | NA | NA | NA | #VALUE! |
| ENSMUSG00000097676.2 | 0 | NA | NA | NA | NA | NA | #VALUE! |
| ENSMUSG00000097677.2 | 0 | NA | NA | NA | NA | NA | #VALUE! |
| ENSMUSG00000097679.2 | 0 | NA | NA | NA | NA | NA | #VALUE! |

| | | | | | | | |
|---|---|---|---|---|---|---|---|
| ENSMUSG00000097680.2 | 0 | NA | NA | NA | NA | NA | #VALUE! |
| ENSMUSG00000097682.2 | 0 | NA | NA | NA | NA | NA | #VALUE! |
| ENSMUSG00000097683.3 | 0 | NA | NA | NA | NA | NA | #VALUE! |
| ENSMUSG00000097684.2 | 0 | NA | NA | NA | NA | NA | #VALUE! |
| ENSMUSG00000097685.3 | 0 | NA | NA | NA | NA | NA | #VALUE! |
| ENSMUSG00000097686.2 | 0 | NA | NA | NA | NA | NA | #VALUE! |
| ENSMUSG00000097689.2 | 0 | NA | NA | NA | NA | NA | #VALUE! |
| ENSMUSG00000097690.3 | 0 | NA | NA | NA | NA | NA | #VALUE! |
| ENSMUSG00000097692.3 | 0 | NA | NA | NA | NA | NA | #VALUE! |
| ENSMUSG00000097693.4 | 0 | NA | NA | NA | NA | NA | #VALUE! |
| ENSMUSG00000097694.9 | 0 | NA | NA | NA | NA | NA | #VALUE! |
| ENSMUSG00000097697.3 | 0 | NA | NA | NA | NA | NA | #VALUE! |
| ENSMUSG00000097699.3 | 0 | NA | NA | NA | NA | NA | #VALUE! |
| ENSMUSG00000097700.2 | 0 | NA | NA | NA | NA | NA | #VALUE! |
| ENSMUSG00000097701.2 | 0 | NA | NA | NA | NA | NA | #VALUE! |
| ENSMUSG00000097702.2 | 0 | NA | NA | NA | NA | NA | #VALUE! |
| ENSMUSG00000097704.2 | 0 | NA | NA | NA | NA | NA | #VALUE! |
| ENSMUSG00000097705.2 | 6.486717 | 2.711709 | 3.392676 | 0.799283 | 0.424126 | NA | #VALUE! |
| ENSMUSG00000097709.9 | 0 | NA | NA | NA | NA | NA | #VALUE! |
| ENSMUSG00000097710.3 | 0 | NA | NA | NA | NA | NA | #VALUE! |
| ENSMUSG00000097711.2 | 0 | NA | NA | NA | NA | NA | #VALUE! |
| ENSMUSG00000097712.2 | 0 | NA | NA | NA | NA | NA | #VALUE! |
| ENSMUSG00000097713.2 | 0 | NA | NA | NA | NA | NA | #VALUE! |
| ENSMUSG00000097714.2 | 0 | NA | NA | NA | NA | NA | #VALUE! |
| ENSMUSG00000097715.3 | 7.496934 | -6.435 | 3.45986 | -1.8599 | 0.062899 | NA | #VALUE! |
| ENSMUSG00000097716.2 | 0 | NA | NA | NA | NA | NA | #VALUE! |
| ENSMUSG00000097717.3 | 0 | NA | NA | NA | NA | NA | #VALUE! |
| ENSMUSG00000097719.2 | 0 | NA | NA | NA | NA | NA | #VALUE! |
| ENSMUSG00000097724.3 | 3.132626 | -5.17361 | 3.921071 | -1.31944 | 0.187023 | NA | #VALUE! |
| ENSMUSG00000097725.5 | 0 | NA | NA | NA | NA | NA | #VALUE! |
| ENSMUSG00000097726.8 | 0 | NA | NA | NA | NA | NA | #VALUE! |
| ENSMUSG00000097727.9 | 0 | NA | NA | NA | NA | NA | #VALUE! |
| ENSMUSG00000097729.3 | 0 | NA | NA | NA | NA | NA | #VALUE! |
| ENSMUSG00000097735.8 | 0 | NA | NA | NA | NA | NA | #VALUE! |
| ENSMUSG00000097736.9 | 0 | NA | NA | NA | NA | NA | #VALUE! |
| ENSMUSG00000097738.3 | 0 | NA | NA | NA | NA | NA | #VALUE! |
| ENSMUSG00000097739.2 | 0 | NA | NA | NA | NA | NA | #VALUE! |
| ENSMUSG00000097740.3 | 0 | NA | NA | NA | NA | NA | #VALUE! |
| ENSMUSG00000097742.2 | 0 | NA | NA | NA | NA | NA | #VALUE! |
| ENSMUSG00000097743.2 | 2.882264 | -0.69973 | 3.793011 | -0.18448 | 0.853637 | NA | #VALUE! |
| ENSMUSG00000097744.2 | 0 | NA | NA | NA | NA | NA | #VALUE! |
| ENSMUSG00000097745.2 | 0 | NA | NA | NA | NA | NA | #VALUE! |
| ENSMUSG00000097746.3 | 2.424635 | 4.780725 | 3.925082 | 1.217993 | 0.223226 | NA | #VALUE! |
| ENSMUSG00000097748.2 | 0 | NA | NA | NA | NA | NA | #VALUE! |

| | | | | | | | |
|---|---|---|---|---|---|---|---|
| ENSMUSG00000097749.3 | 0 | NA | NA | NA | NA | NA | #VALUE! |
| ENSMUSG00000097752.3 | 0 | NA | NA | NA | NA | NA | #VALUE! |
| ENSMUSG00000097754.2 | 0 | NA | NA | NA | NA | NA | #VALUE! |
| ENSMUSG00000097755.3 | 0 | NA | NA | NA | NA | NA | #VALUE! |
| ENSMUSG00000097756.3 | 0 | NA | NA | NA | NA | NA | #VALUE! |
| ENSMUSG00000097760.3 | 0 | NA | NA | NA | NA | NA | #VALUE! |
| ENSMUSG00000097761.2 | 0 | NA | NA | NA | NA | NA | #VALUE! |
| ENSMUSG00000097762.2 | 1.585338 | 4.174397 | 3.934581 | 1.060951 | 0.288712 | NA | #VALUE! |
| ENSMUSG00000097763.2 | 0 | NA | NA | NA | NA | NA | #VALUE! |
| ENSMUSG00000097766.2 | 0 | NA | NA | NA | NA | NA | #VALUE! |
| ENSMUSG00000097767.9 | 0 | NA | NA | NA | NA | NA | #VALUE! |
| ENSMUSG00000097768.3 | 0 | NA | NA | NA | NA | NA | #VALUE! |
| ENSMUSG00000097773.2 | 4.543655 | 5.673725 | 3.916649 | 1.448617 | 0.147445 | NA | #VALUE! |
| ENSMUSG00000097776.2 | 0 | NA | NA | NA | NA | NA | #VALUE! |
| ENSMUSG00000097777.8 | 3.061661 | -5.13931 | 3.921414 | -1.31058 | 0.190001 | NA | #VALUE! |
| ENSMUSG00000097779.3 | 0 | NA | NA | NA | NA | NA | #VALUE! |
| ENSMUSG00000097780.2 | 0 | NA | NA | NA | NA | NA | #VALUE! |
| ENSMUSG00000097781.2 | 0 | NA | NA | NA | NA | NA | #VALUE! |
| ENSMUSG00000097783.3 | 0 | NA | NA | NA | NA | NA | #VALUE! |
| ENSMUSG00000097784.2 | 0 | NA | NA | NA | NA | NA | #VALUE! |
| ENSMUSG00000097785.3 | 0 | NA | NA | NA | NA | NA | #VALUE! |
| ENSMUSG00000097786.4 | 2.33138 | 4.724655 | 3.925805 | 1.203487 | 0.228788 | NA | #VALUE! |
| ENSMUSG00000097787.3 | 5.451618 | 5.940245 | 3.837345 | 1.548009 | 0.12162 | NA | #VALUE! |
| ENSMUSG00000097788.3 | 0 | NA | NA | NA | NA | NA | #VALUE! |
| ENSMUSG00000097789.3 | 0 | NA | NA | NA | NA | NA | #VALUE! |
| ENSMUSG00000097790.2 | 1.084019 | 0.752172 | 3.907339 | 0.192502 | 0.847349 | NA | #VALUE! |
| ENSMUSG00000097792.3 | 0 | NA | NA | NA | NA | NA | #VALUE! |
| ENSMUSG00000097793.2 | 0 | NA | NA | NA | NA | NA | #VALUE! |
| ENSMUSG00000097794.3 | 0 | NA | NA | NA | NA | NA | #VALUE! |
| ENSMUSG00000097795.2 | 0 | NA | NA | NA | NA | NA | #VALUE! |
| ENSMUSG00000097797.7 | 0 | NA | NA | NA | NA | NA | #VALUE! |
| ENSMUSG00000097798.2 | 0 | NA | NA | NA | NA | NA | #VALUE! |
| ENSMUSG00000097801.2 | 0 | NA | NA | NA | NA | NA | #VALUE! |
| ENSMUSG00000097803.2 | 3.409991 | 5.253872 | 3.919978 | 1.340281 | 0.180154 | NA | #VALUE! |
| ENSMUSG00000097804.2 | 0 | NA | NA | NA | NA | NA | #VALUE! |
| ENSMUSG00000097805.3 | 0 | NA | NA | NA | NA | NA | #VALUE! |
| ENSMUSG00000097806.4 | 0 | NA | NA | NA | NA | NA | #VALUE! |
| ENSMUSG00000097807.2 | 0 | NA | NA | NA | NA | NA | #VALUE! |
| ENSMUSG00000097809.3 | 2.449329 | -4.81615 | 3.925078 | -1.22702 | 0.219815 | NA | #VALUE! |
| ENSMUSG00000097812.2 | 0 | NA | NA | NA | NA | NA | #VALUE! |
| ENSMUSG00000097814.6 | 0 | NA | NA | NA | NA | NA | #VALUE! |
| ENSMUSG00000097816.2 | 0 | NA | NA | NA | NA | NA | #VALUE! |
| ENSMUSG00000097817.3 | 0 | NA | NA | NA | NA | NA | #VALUE! |
| ENSMUSG00000097819.2 | 0 | NA | NA | NA | NA | NA | #VALUE! |

| | | | | | | | |
|---|---|---|---|---|---|---|---|
| ENSMUSG00000097820.4 | 1.212318 | 3.793111 | 3.942915 | 0.962007 | 0.336046 | NA | #VALUE! |
| ENSMUSG00000097821.3 | 0 | NA | NA | NA | NA | NA | #VALUE! |
| ENSMUSG00000097822.8 | 0 | NA | NA | NA | NA | NA | #VALUE! |
| ENSMUSG00000097823.3 | 0 | NA | NA | NA | NA | NA | #VALUE! |
| ENSMUSG00000097825.6 | 0 | NA | NA | NA | NA | NA | #VALUE! |
| ENSMUSG00000097827.2 | 0 | NA | NA | NA | NA | NA | #VALUE! |
| ENSMUSG00000097828.3 | 0.390621 | 0.386159 | 4.04914 | 0.095368 | 0.924022 | NA | #VALUE! |
| ENSMUSG00000097831.2 | 0 | NA | NA | NA | NA | NA | #VALUE! |
| ENSMUSG00000097832.2 | 0 | NA | NA | NA | NA | NA | #VALUE! |
| ENSMUSG00000097833.2 | 0 | NA | NA | NA | NA | NA | #VALUE! |
| ENSMUSG00000097834.3 | 0 | NA | NA | NA | NA | NA | #VALUE! |
| ENSMUSG00000097835.2 | 3.757723 | -2.79033 | 3.836821 | -0.72725 | 0.467072 | NA | #VALUE! |
| ENSMUSG00000097836.2 | 0.466276 | 2.449889 | 3.997179 | 0.612904 | 0.53994 | NA | #VALUE! |
| ENSMUSG00000097840.2 | 0 | NA | NA | NA | NA | NA | #VALUE! |
| ENSMUSG00000097841.3 | 0 | NA | NA | NA | NA | NA | #VALUE! |
| ENSMUSG00000097842.3 | 0 | NA | NA | NA | NA | NA | #VALUE! |
| ENSMUSG00000097845.3 | 0 | NA | NA | NA | NA | NA | #VALUE! |
| ENSMUSG00000097847.2 | 0 | NA | NA | NA | NA | NA | #VALUE! |
| ENSMUSG00000097848.3 | 0 | NA | NA | NA | NA | NA | #VALUE! |
| ENSMUSG00000097850.3 | 0 | NA | NA | NA | NA | NA | #VALUE! |
| ENSMUSG00000097851.6 | 0 | NA | NA | NA | NA | NA | #VALUE! |
| ENSMUSG00000097852.6 | 0 | NA | NA | NA | NA | NA | #VALUE! |
| ENSMUSG00000097853.3 | 0 | NA | NA | NA | NA | NA | #VALUE! |
| ENSMUSG00000097854.2 | 0 | NA | NA | NA | NA | NA | #VALUE! |
| ENSMUSG00000097855.9 | 0 | NA | NA | NA | NA | NA | #VALUE! |
| ENSMUSG00000097858.3 | 0 | NA | NA | NA | NA | NA | #VALUE! |
| ENSMUSG00000097863.2 | 1.771849 | 4.332864 | 3.931707 | 1.102031 | 0.270448 | NA | #VALUE! |
| ENSMUSG00000097870.3 | 0 | NA | NA | NA | NA | NA | #VALUE! |
| ENSMUSG00000097871.2 | 0 | NA | NA | NA | NA | NA | #VALUE! |
| ENSMUSG00000097874.4 | 0 | NA | NA | NA | NA | NA | #VALUE! |
| ENSMUSG00000097875.3 | 0 | NA | NA | NA | NA | NA | #VALUE! |
| ENSMUSG00000097876.2 | 0 | NA | NA | NA | NA | NA | #VALUE! |
| ENSMUSG00000097877.3 | 0 | NA | NA | NA | NA | NA | #VALUE! |
| ENSMUSG00000097878.6 | 0 | NA | NA | NA | NA | NA | #VALUE! |
| ENSMUSG00000097881.8 | 0 | NA | NA | NA | NA | NA | #VALUE! |
| ENSMUSG00000097882.3 | 0.210026 | -1.52494 | 4.058692 | -0.37572 | 0.707123 | NA | #VALUE! |
| ENSMUSG00000097883.2 | 0 | NA | NA | NA | NA | NA | #VALUE! |
| ENSMUSG00000097884.2 | 0 | NA | NA | NA | NA | NA | #VALUE! |
| ENSMUSG00000097885.8 | 0 | NA | NA | NA | NA | NA | #VALUE! |
| ENSMUSG00000097886.3 | 0 | NA | NA | NA | NA | NA | #VALUE! |
| ENSMUSG00000097887.3 | 0 | NA | NA | NA | NA | NA | #VALUE! |
| ENSMUSG00000097888.3 | 0 | NA | NA | NA | NA | NA | #VALUE! |
| ENSMUSG00000097890.3 | 0 | NA | NA | NA | NA | NA | #VALUE! |
| ENSMUSG00000097891.2 | 0 | NA | NA | NA | NA | NA | #VALUE! |

| | | | | | | | |
|---|---|---|---|---|---|---|---|
| ENSMUSG00000097892.3 | 0 | NA | NA | NA | NA | NA | #VALUE! |
| ENSMUSG00000097893.9 | 2.100263 | -4.61322 | 3.927832 | -1.17449 | 0.240197 | NA | #VALUE! |
| ENSMUSG00000097895.2 | 0 | NA | NA | NA | NA | NA | #VALUE! |
| ENSMUSG00000097896.2 | 0 | NA | NA | NA | NA | NA | #VALUE! |
| ENSMUSG00000097897.2 | 0 | NA | NA | NA | NA | NA | #VALUE! |
| ENSMUSG00000097898.9 | 0 | NA | NA | NA | NA | NA | #VALUE! |
| ENSMUSG00000097899.3 | 0 | NA | NA | NA | NA | NA | #VALUE! |
| ENSMUSG00000097900.2 | 0 | NA | NA | NA | NA | NA | #VALUE! |
| ENSMUSG00000097901.8 | 0 | NA | NA | NA | NA | NA | #VALUE! |
| ENSMUSG00000097902.3 | 0 | NA | NA | NA | NA | NA | #VALUE! |
| ENSMUSG00000097903.3 | 0 | NA | NA | NA | NA | NA | #VALUE! |
| ENSMUSG00000097907.3 | 0 | NA | NA | NA | NA | NA | #VALUE! |
| ENSMUSG00000097910.3 | 5.553356 | 5.96322 | 3.914855 | 1.523229 | 0.127701 | NA | #VALUE! |
| ENSMUSG00000097911.3 | 0 | NA | NA | NA | NA | NA | #VALUE! |
| ENSMUSG00000097914.2 | 0 | NA | NA | NA | NA | NA | #VALUE! |
| ENSMUSG00000097915.3 | 0 | NA | NA | NA | NA | NA | #VALUE! |
| ENSMUSG00000097917.3 | 0 | NA | NA | NA | NA | NA | #VALUE! |
| ENSMUSG00000097918.2 | 0 | NA | NA | NA | NA | NA | #VALUE! |
| ENSMUSG00000097919.2 | 0 | NA | NA | NA | NA | NA | #VALUE! |
| ENSMUSG00000097920.2 | 0 | NA | NA | NA | NA | NA | #VALUE! |
| ENSMUSG00000097921.3 | 0 | NA | NA | NA | NA | NA | #VALUE! |
| ENSMUSG00000097923.2 | 0 | NA | NA | NA | NA | NA | #VALUE! |
| ENSMUSG00000097924.3 | 0 | NA | NA | NA | NA | NA | #VALUE! |
| ENSMUSG00000097925.2 | 0 | NA | NA | NA | NA | NA | #VALUE! |
| ENSMUSG00000097926.2 | 0 | NA | NA | NA | NA | NA | #VALUE! |
| ENSMUSG00000097927.3 | 0 | NA | NA | NA | NA | NA | #VALUE! |
| ENSMUSG00000097929.10 | 0 | NA | NA | NA | NA | NA | #VALUE! |
| ENSMUSG00000097930.3 | 0 | NA | NA | NA | NA | NA | #VALUE! |
| ENSMUSG00000097931.2 | 0 | NA | NA | NA | NA | NA | #VALUE! |
| ENSMUSG00000097932.3 | 0 | NA | NA | NA | NA | NA | #VALUE! |
| ENSMUSG00000097933.3 | 0 | NA | NA | NA | NA | NA | #VALUE! |
| ENSMUSG00000097934.2 | 0 | NA | NA | NA | NA | NA | #VALUE! |
| ENSMUSG00000097935.2 | 0.105013 | -0.97324 | 4.076293 | -0.23876 | 0.811295 | NA | #VALUE! |
| ENSMUSG00000097936.2 | 0.093255 | 0.868093 | 4.072464 | 0.213162 | 0.831201 | NA | #VALUE! |
| ENSMUSG00000097937.2 | 0 | NA | NA | NA | NA | NA | #VALUE! |
| ENSMUSG00000097938.2 | 0 | NA | NA | NA | NA | NA | #VALUE! |
| ENSMUSG00000097939.2 | 0 | NA | NA | NA | NA | NA | #VALUE! |
| ENSMUSG00000097940.3 | 0 | NA | NA | NA | NA | NA | #VALUE! |
| ENSMUSG00000097941.2 | 0 | NA | NA | NA | NA | NA | #VALUE! |
| ENSMUSG00000097942.2 | 0 | NA | NA | NA | NA | NA | #VALUE! |
| ENSMUSG00000097943.2 | 0 | NA | NA | NA | NA | NA | #VALUE! |
| ENSMUSG00000097945.2 | 0 | NA | NA | NA | NA | NA | #VALUE! |
| ENSMUSG00000097946.2 | 0 | NA | NA | NA | NA | NA | #VALUE! |
| ENSMUSG00000097947.2 | 0 | NA | NA | NA | NA | NA | #VALUE! |

| | | | | | | | |
|---|---|---|---|---|---|---|---|
| ENSMUSG00000097948.2 | 0 | NA | NA | NA | NA | NA | #VALUE! |
| ENSMUSG00000097949.2 | 0 | NA | NA | NA | NA | NA | #VALUE! |
| ENSMUSG00000097950.2 | 0 | NA | NA | NA | NA | NA | #VALUE! |
| ENSMUSG00000097951.2 | 0 | NA | NA | NA | NA | NA | #VALUE! |
| ENSMUSG00000097952.2 | 0 | NA | NA | NA | NA | NA | #VALUE! |
| ENSMUSG00000097953.2 | 0 | NA | NA | NA | NA | NA | #VALUE! |
| ENSMUSG00000097954.2 | 0 | NA | NA | NA | NA | NA | #VALUE! |
| ENSMUSG00000097955.2 | 0 | NA | NA | NA | NA | NA | #VALUE! |
| ENSMUSG00000097957.3 | 0 | NA | NA | NA | NA | NA | #VALUE! |
| ENSMUSG00000097958.2 | 0 | NA | NA | NA | NA | NA | #VALUE! |
| ENSMUSG00000097959.3 | 0 | NA | NA | NA | NA | NA | #VALUE! |
| ENSMUSG00000097960.2 | 0 | NA | NA | NA | NA | NA | #VALUE! |
| ENSMUSG00000097961.2 | 0 | NA | NA | NA | NA | NA | #VALUE! |
| ENSMUSG00000097962.8 | 0 | NA | NA | NA | NA | NA | #VALUE! |
| ENSMUSG00000097963.2 | 0 | NA | NA | NA | NA | NA | #VALUE! |
| ENSMUSG00000097964.2 | 0 | NA | NA | NA | NA | NA | #VALUE! |
| ENSMUSG00000097965.2 | 0 | NA | NA | NA | NA | NA | #VALUE! |
| ENSMUSG00000097966.2 | 0 | NA | NA | NA | NA | NA | #VALUE! |
| ENSMUSG00000097967.2 | 0 | NA | NA | NA | NA | NA | #VALUE! |
| ENSMUSG00000097968.2 | 0 | NA | NA | NA | NA | NA | #VALUE! |
| ENSMUSG00000097970.2 | 0 | NA | NA | NA | NA | NA | #VALUE! |
| ENSMUSG00000097972.2 | 0 | NA | NA | NA | NA | NA | #VALUE! |
| ENSMUSG00000097973.2 | 0.496591 | -2.51334 | 3.995845 | -0.62899 | 0.529356 | NA | #VALUE! |
| ENSMUSG00000097974.2 | 0 | NA | NA | NA | NA | NA | #VALUE! |
| ENSMUSG00000097975.2 | 0 | NA | NA | NA | NA | NA | #VALUE! |
| ENSMUSG00000097976.2 | 0 | NA | NA | NA | NA | NA | #VALUE! |
| ENSMUSG00000097977.2 | 0 | NA | NA | NA | NA | NA | #VALUE! |
| ENSMUSG00000097979.3 | 0 | NA | NA | NA | NA | NA | #VALUE! |
| ENSMUSG00000097980.3 | 0 | NA | NA | NA | NA | NA | #VALUE! |
| ENSMUSG00000097981.2 | 0 | NA | NA | NA | NA | NA | #VALUE! |
| ENSMUSG00000097982.2 | 0 | NA | NA | NA | NA | NA | #VALUE! |
| ENSMUSG00000097983.2 | 0 | NA | NA | NA | NA | NA | #VALUE! |
| ENSMUSG00000097984.2 | 0 | NA | NA | NA | NA | NA | #VALUE! |
| ENSMUSG00000097985.2 | 0 | NA | NA | NA | NA | NA | #VALUE! |
| ENSMUSG00000097986.3 | 0 | NA | NA | NA | NA | NA | #VALUE! |
| ENSMUSG00000097987.2 | 0 | NA | NA | NA | NA | NA | #VALUE! |
| ENSMUSG00000097988.2 | 0 | NA | NA | NA | NA | NA | #VALUE! |
| ENSMUSG00000097989.2 | 0.18651 | 1.401769 | 4.056354 | 0.345574 | 0.729663 | NA | #VALUE! |
| ENSMUSG00000097990.2 | 0 | NA | NA | NA | NA | NA | #VALUE! |
| ENSMUSG00000097991.2 | 0 | NA | NA | NA | NA | NA | #VALUE! |
| ENSMUSG00000097992.2 | 0 | NA | NA | NA | NA | NA | #VALUE! |
| ENSMUSG00000097993.8 | 0 | NA | NA | NA | NA | NA | #VALUE! |
| ENSMUSG00000097994.2 | 1.742512 | -4.33906 | 3.932218 | -1.10346 | 0.269826 | NA | #VALUE! |
| ENSMUSG00000097995.2 | 0 | NA | NA | NA | NA | NA | #VALUE! |

| | | | | | | | |
|---|---|---|---|---|---|---|---|
| ENSMUSG00000097996.2 | 0 | NA | NA | NA | NA | NA | #VALUE! |
| ENSMUSG00000097997.2 | 0 | NA | NA | NA | NA | NA | #VALUE! |
| ENSMUSG00000097998.2 | 0 | NA | NA | NA | NA | NA | #VALUE! |
| ENSMUSG00000097999.2 | 0 | NA | NA | NA | NA | NA | #VALUE! |
| ENSMUSG00000098000.2 | 0 | NA | NA | NA | NA | NA | #VALUE! |
| ENSMUSG00000098001.2 | 0 | NA | NA | NA | NA | NA | #VALUE! |
| ENSMUSG00000098002.2 | 0 | NA | NA | NA | NA | NA | #VALUE! |
| ENSMUSG00000098003.3 | 0 | NA | NA | NA | NA | NA | #VALUE! |
| ENSMUSG00000098004.2 | 0 | NA | NA | NA | NA | NA | #VALUE! |
| ENSMUSG00000098005.2 | 0 | NA | NA | NA | NA | NA | #VALUE! |
| ENSMUSG00000098006.2 | 0 | NA | NA | NA | NA | NA | #VALUE! |
| ENSMUSG00000098007.2 | 0 | NA | NA | NA | NA | NA | #VALUE! |
| ENSMUSG00000098008.2 | 0 | NA | NA | NA | NA | NA | #VALUE! |
| ENSMUSG00000098009.2 | 0 | NA | NA | NA | NA | NA | #VALUE! |
| ENSMUSG00000098010.2 | 0 | NA | NA | NA | NA | NA | #VALUE! |
| ENSMUSG00000098011.2 | 0 | NA | NA | NA | NA | NA | #VALUE! |
| ENSMUSG00000098012.2 | 0 | NA | NA | NA | NA | NA | #VALUE! |
| ENSMUSG00000098013.2 | 0 | NA | NA | NA | NA | NA | #VALUE! |
| ENSMUSG00000098014.2 | 1.942129 | 4.438743 | 3.929952 | 1.129465 | 0.258702 | NA | #VALUE! |
| ENSMUSG00000098015.4 | 0 | NA | NA | NA | NA | NA | #VALUE! |
| ENSMUSG00000098016.2 | 0 | NA | NA | NA | NA | NA | #VALUE! |
| ENSMUSG00000098017.2 | 0 | NA | NA | NA | NA | NA | #VALUE! |
| ENSMUSG00000098019.2 | 0.300802 | -1.80634 | 4.050033 | -0.44601 | 0.655592 | NA | #VALUE! |
| ENSMUSG00000098021.3 | 0.168284 | 1.027628 | 4.067126 | 0.252667 | 0.800526 | NA | #VALUE! |
| ENSMUSG00000098022.8 | 0 | NA | NA | NA | NA | NA | #VALUE! |
| ENSMUSG00000098023.2 | 0 | NA | NA | NA | NA | NA | #VALUE! |
| ENSMUSG00000098024.2 | 0 | NA | NA | NA | NA | NA | #VALUE! |
| ENSMUSG00000098025.3 | 0 | NA | NA | NA | NA | NA | #VALUE! |
| ENSMUSG00000098026.3 | 0 | NA | NA | NA | NA | NA | #VALUE! |
| ENSMUSG00000098027.2 | 0 | NA | NA | NA | NA | NA | #VALUE! |
| ENSMUSG00000098028.2 | 0 | NA | NA | NA | NA | NA | #VALUE! |
| ENSMUSG00000098029.3 | 0 | NA | NA | NA | NA | NA | #VALUE! |
| ENSMUSG00000098030.2 | 0 | NA | NA | NA | NA | NA | #VALUE! |
| ENSMUSG00000098031.2 | 0 | NA | NA | NA | NA | NA | #VALUE! |
| ENSMUSG00000098033.3 | 0 | NA | NA | NA | NA | NA | #VALUE! |
| ENSMUSG00000098035.2 | 0 | NA | NA | NA | NA | NA | #VALUE! |
| ENSMUSG00000098036.2 | 0 | NA | NA | NA | NA | NA | #VALUE! |
| ENSMUSG00000098037.2 | 0 | NA | NA | NA | NA | NA | #VALUE! |
| ENSMUSG00000098038.2 | 0 | NA | NA | NA | NA | NA | #VALUE! |
| ENSMUSG00000098039.2 | 0 | NA | NA | NA | NA | NA | #VALUE! |
| ENSMUSG00000098040.2 | 0 | NA | NA | NA | NA | NA | #VALUE! |
| ENSMUSG00000098041.2 | 0 | NA | NA | NA | NA | NA | #VALUE! |
| ENSMUSG00000098042.2 | 0 | NA | NA | NA | NA | NA | #VALUE! |
| ENSMUSG00000098043.2 | 0 | NA | NA | NA | NA | NA | #VALUE! |

| | | | | | | | |
|---|---|---|---|---|---|---|---|
| ENSMUSG00000098044.2 | 0 | NA | NA | NA | NA | NA | #VALUE! |
| ENSMUSG00000098045.2 | 0 | NA | NA | NA | NA | NA | #VALUE! |
| ENSMUSG00000098046.2 | 0 | NA | NA | NA | NA | NA | #VALUE! |
| ENSMUSG00000098047.2 | 0 | NA | NA | NA | NA | NA | #VALUE! |
| ENSMUSG00000098048.2 | 0 | NA | NA | NA | NA | NA | #VALUE! |
| ENSMUSG00000098050.2 | 0 | NA | NA | NA | NA | NA | #VALUE! |
| ENSMUSG00000098051.2 | 0 | NA | NA | NA | NA | NA | #VALUE! |
| ENSMUSG00000098052.2 | 0 | NA | NA | NA | NA | NA | #VALUE! |
| ENSMUSG00000098053.2 | 0 | NA | NA | NA | NA | NA | #VALUE! |
| ENSMUSG00000098054.2 | 0 | NA | NA | NA | NA | NA | #VALUE! |
| ENSMUSG00000098055.2 | 0 | NA | NA | NA | NA | NA | #VALUE! |
| ENSMUSG00000098056.2 | 0 | NA | NA | NA | NA | NA | #VALUE! |
| ENSMUSG00000098057.2 | 0 | NA | NA | NA | NA | NA | #VALUE! |
| ENSMUSG00000098058.2 | 0 | NA | NA | NA | NA | NA | #VALUE! |
| ENSMUSG00000098059.2 | 0 | NA | NA | NA | NA | NA | #VALUE! |
| ENSMUSG00000098060.2 | 0 | NA | NA | NA | NA | NA | #VALUE! |
| ENSMUSG00000098062.2 | 0 | NA | NA | NA | NA | NA | #VALUE! |
| ENSMUSG00000098064.2 | 0 | NA | NA | NA | NA | NA | #VALUE! |
| ENSMUSG00000098065.2 | 0.105013 | -0.97324 | 4.076293 | -0.23876 | 0.811295 | NA | #VALUE! |
| ENSMUSG00000098066.3 | 0 | NA | NA | NA | NA | NA | #VALUE! |
| ENSMUSG00000098067.2 | 0 | NA | NA | NA | NA | NA | #VALUE! |
| ENSMUSG00000098068.2 | 0 | NA | NA | NA | NA | NA | #VALUE! |
| ENSMUSG00000098069.2 | 0 | NA | NA | NA | NA | NA | #VALUE! |
| ENSMUSG00000098070.2 | 0 | NA | NA | NA | NA | NA | #VALUE! |
| ENSMUSG00000098071.3 | 0 | NA | NA | NA | NA | NA | #VALUE! |
| ENSMUSG00000098072.2 | 0 | NA | NA | NA | NA | NA | #VALUE! |
| ENSMUSG00000098073.2 | 0 | NA | NA | NA | NA | NA | #VALUE! |
| ENSMUSG00000098074.2 | 0 | NA | NA | NA | NA | NA | #VALUE! |
| ENSMUSG00000098075.2 | 0 | NA | NA | NA | NA | NA | #VALUE! |
| ENSMUSG00000098076.2 | 0 | NA | NA | NA | NA | NA | #VALUE! |
| ENSMUSG00000098077.2 | 0 | NA | NA | NA | NA | NA | #VALUE! |
| ENSMUSG00000098078.2 | 0 | NA | NA | NA | NA | NA | #VALUE! |
| ENSMUSG00000098079.2 | 0 | NA | NA | NA | NA | NA | #VALUE! |
| ENSMUSG00000098080.2 | 0 | NA | NA | NA | NA | NA | #VALUE! |
| ENSMUSG00000098081.2 | 0 | NA | NA | NA | NA | NA | #VALUE! |
| ENSMUSG00000098082.2 | 0 | NA | NA | NA | NA | NA | #VALUE! |
| ENSMUSG00000098083.2 | 0 | NA | NA | NA | NA | NA | #VALUE! |
| ENSMUSG00000098084.2 | 0 | NA | NA | NA | NA | NA | #VALUE! |
| ENSMUSG00000098085.2 | 0 | NA | NA | NA | NA | NA | #VALUE! |
| ENSMUSG00000098086.2 | 0 | NA | NA | NA | NA | NA | #VALUE! |
| ENSMUSG00000098087.2 | 0 | NA | NA | NA | NA | NA | #VALUE! |
| ENSMUSG00000098088.2 | 0 | NA | NA | NA | NA | NA | #VALUE! |
| ENSMUSG00000098089.3 | 0 | NA | NA | NA | NA | NA | #VALUE! |
| ENSMUSG00000098090.9 | 0 | NA | NA | NA | NA | NA | #VALUE! |

| | | | | | | | |
|---|---|---|---|---|---|---|---|
| ENSMUSG00000098091.2 | 0 | NA | NA | NA | NA | NA | #VALUE! |
| ENSMUSG00000098092.2 | 0 | NA | NA | NA | NA | NA | #VALUE! |
| ENSMUSG00000098093.3 | 0 | NA | NA | NA | NA | NA | #VALUE! |
| ENSMUSG00000098094.8 | 0 | NA | NA | NA | NA | NA | #VALUE! |
| ENSMUSG00000098095.2 | 0 | NA | NA | NA | NA | NA | #VALUE! |
| ENSMUSG00000098096.2 | 0 | NA | NA | NA | NA | NA | #VALUE! |
| ENSMUSG00000098097.8 | 0 | NA | NA | NA | NA | NA | #VALUE! |
| ENSMUSG00000098099.2 | 0 | NA | NA | NA | NA | NA | #VALUE! |
| ENSMUSG00000098100.2 | 0 | NA | NA | NA | NA | NA | #VALUE! |
| ENSMUSG00000098101.2 | 0 | NA | NA | NA | NA | NA | #VALUE! |
| ENSMUSG00000098102.2 | 0 | NA | NA | NA | NA | NA | #VALUE! |
| ENSMUSG00000098103.2 | 0 | NA | NA | NA | NA | NA | #VALUE! |
| ENSMUSG00000098105.2 | 0.195789 | -1.05581 | 4.073302 | -0.2592 | 0.795479 | NA | #VALUE! |
| ENSMUSG00000098106.2 | 0 | NA | NA | NA | NA | NA | #VALUE! |
| ENSMUSG00000098107.2 | 0 | NA | NA | NA | NA | NA | #VALUE! |
| ENSMUSG00000098108.8 | 0 | NA | NA | NA | NA | NA | #VALUE! |
| ENSMUSG00000098109.2 | 0 | NA | NA | NA | NA | NA | #VALUE! |
| ENSMUSG00000098110.2 | 0 | NA | NA | NA | NA | NA | #VALUE! |
| ENSMUSG00000098111.2 | 0 | NA | NA | NA | NA | NA | #VALUE! |
| ENSMUSG00000098112.9 | 7.874105 | -3.37806 | 2.627089 | -1.28586 | 0.198493 | NA | #VALUE! |
| ENSMUSG00000098113.2 | 0 | NA | NA | NA | NA | NA | #VALUE! |
| ENSMUSG00000098114.2 | 0 | NA | NA | NA | NA | NA | #VALUE! |
| ENSMUSG00000098115.2 | 0 | NA | NA | NA | NA | NA | #VALUE! |
| ENSMUSG00000098116.2 | 0 | NA | NA | NA | NA | NA | #VALUE! |
| ENSMUSG00000098118.2 | 0 | NA | NA | NA | NA | NA | #VALUE! |
| ENSMUSG00000098119.2 | 0 | NA | NA | NA | NA | NA | #VALUE! |
| ENSMUSG00000098120.2 | 0.420053 | -2.33493 | 4.007343 | -0.58266 | 0.56012 | NA | #VALUE! |
| ENSMUSG00000098121.2 | 0 | NA | NA | NA | NA | NA | #VALUE! |
| ENSMUSG00000098122.2 | 0 | NA | NA | NA | NA | NA | #VALUE! |
| ENSMUSG00000098123.2 | 0 | NA | NA | NA | NA | NA | #VALUE! |
| ENSMUSG00000098124.2 | 0 | NA | NA | NA | NA | NA | #VALUE! |
| ENSMUSG00000098125.2 | 0 | NA | NA | NA | NA | NA | #VALUE! |
| ENSMUSG00000098126.2 | 0 | NA | NA | NA | NA | NA | #VALUE! |
| ENSMUSG00000098127.2 | 0 | NA | NA | NA | NA | NA | #VALUE! |
| ENSMUSG00000098128.2 | 0 | NA | NA | NA | NA | NA | #VALUE! |
| ENSMUSG00000098129.2 | 0 | NA | NA | NA | NA | NA | #VALUE! |
| ENSMUSG00000098130.2 | 0 | NA | NA | NA | NA | NA | #VALUE! |
| ENSMUSG00000098131.2 | 0 | NA | NA | NA | NA | NA | #VALUE! |
| ENSMUSG00000098132.2 | 0 | NA | NA | NA | NA | NA | #VALUE! |
| ENSMUSG00000098135.2 | 0 | NA | NA | NA | NA | NA | #VALUE! |
| ENSMUSG00000098136.2 | 0 | NA | NA | NA | NA | NA | #VALUE! |
| ENSMUSG00000098137.4 | 0 | NA | NA | NA | NA | NA | #VALUE! |
| ENSMUSG00000098138.2 | 0 | NA | NA | NA | NA | NA | #VALUE! |
| ENSMUSG00000098139.2 | 0 | NA | NA | NA | NA | NA | #VALUE! |

| | | | | | | | |
|---|---|---|---|---|---|---|---|
| ENSMUSG00000098140.2 | 0 | NA | NA | NA | NA | NA | #VALUE! |
| ENSMUSG00000098141.2 | 0 | NA | NA | NA | NA | NA | #VALUE! |
| ENSMUSG00000098142.2 | 0 | NA | NA | NA | NA | NA | #VALUE! |
| ENSMUSG00000098143.2 | 0 | NA | NA | NA | NA | NA | #VALUE! |
| ENSMUSG00000098144.9 | 0 | NA | NA | NA | NA | NA | #VALUE! |
| ENSMUSG00000098145.2 | 0 | NA | NA | NA | NA | NA | #VALUE! |
| ENSMUSG00000098146.2 | 0 | NA | NA | NA | NA | NA | #VALUE! |
| ENSMUSG00000098147.2 | 0 | NA | NA | NA | NA | NA | #VALUE! |
| ENSMUSG00000098149.2 | 0 | NA | NA | NA | NA | NA | #VALUE! |
| ENSMUSG00000098150.3 | 0 | NA | NA | NA | NA | NA | #VALUE! |
| ENSMUSG00000098152.2 | 0 | NA | NA | NA | NA | NA | #VALUE! |
| ENSMUSG00000098153.2 | 0 | NA | NA | NA | NA | NA | #VALUE! |
| ENSMUSG00000098154.3 | 0 | NA | NA | NA | NA | NA | #VALUE! |
| ENSMUSG00000098155.2 | 0 | NA | NA | NA | NA | NA | #VALUE! |
| ENSMUSG00000098156.2 | 0 | NA | NA | NA | NA | NA | #VALUE! |
| ENSMUSG00000098157.2 | 0 | NA | NA | NA | NA | NA | #VALUE! |
| ENSMUSG00000098158.3 | 0 | NA | NA | NA | NA | NA | #VALUE! |
| ENSMUSG00000098159.2 | 0 | NA | NA | NA | NA | NA | #VALUE! |
| ENSMUSG00000098160.4 | 0 | NA | NA | NA | NA | NA | #VALUE! |
| ENSMUSG00000098161.2 | 0 | NA | NA | NA | NA | NA | #VALUE! |
| ENSMUSG00000098162.2 | 0 | NA | NA | NA | NA | NA | #VALUE! |
| ENSMUSG00000098163.2 | 0 | NA | NA | NA | NA | NA | #VALUE! |
| ENSMUSG00000098164.3 | 0 | NA | NA | NA | NA | NA | #VALUE! |
| ENSMUSG00000098165.2 | 0 | NA | NA | NA | NA | NA | #VALUE! |
| ENSMUSG00000098166.2 | 0 | NA | NA | NA | NA | NA | #VALUE! |
| ENSMUSG00000098167.2 | 0 | NA | NA | NA | NA | NA | #VALUE! |
| ENSMUSG00000098169.2 | 0 | NA | NA | NA | NA | NA | #VALUE! |
| ENSMUSG00000098170.2 | 0 | NA | NA | NA | NA | NA | #VALUE! |
| ENSMUSG00000098171.2 | 0 | NA | NA | NA | NA | NA | #VALUE! |
| ENSMUSG00000098172.8 | 0 | NA | NA | NA | NA | NA | #VALUE! |
| ENSMUSG00000098173.2 | 0 | NA | NA | NA | NA | NA | #VALUE! |
| ENSMUSG00000098174.2 | 0 | NA | NA | NA | NA | NA | #VALUE! |
| ENSMUSG00000098175.2 | 0 | NA | NA | NA | NA | NA | #VALUE! |
| ENSMUSG00000098176.3 | 4.914886 | 3.562785 | 3.663286 | 0.972565 | 0.330769 | NA | #VALUE! |
| ENSMUSG00000098177.2 | 0 | NA | NA | NA | NA | NA | #VALUE! |
| ENSMUSG00000098179.2 | 0 | NA | NA | NA | NA | NA | #VALUE! |
| ENSMUSG00000098180.2 | 0.093255 | 0.868093 | 4.072464 | 0.213162 | 0.831201 | NA | #VALUE! |
| ENSMUSG00000098181.2 | 6.079188 | 5.034707 | 3.86756 | 1.301779 | 0.192992 | NA | #VALUE! |
| ENSMUSG00000098183.2 | 0 | NA | NA | NA | NA | NA | #VALUE! |
| ENSMUSG00000098184.2 | 0 | NA | NA | NA | NA | NA | #VALUE! |
| ENSMUSG00000098185.2 | 0 | NA | NA | NA | NA | NA | #VALUE! |
| ENSMUSG00000098186.2 | 0.093255 | 0.868093 | 4.072464 | 0.213162 | 0.831201 | NA | #VALUE! |
| ENSMUSG00000098187.2 | 0 | NA | NA | NA | NA | NA | #VALUE! |
| ENSMUSG00000098189.2 | 0 | NA | NA | NA | NA | NA | #VALUE! |

| | | | | | | | |
|---|---|---|---|---|---|---|---|
| ENSMUSG00000098190.2 | 0 | NA | NA | NA | NA | NA | #VALUE! |
| ENSMUSG00000098191.2 | 0 | NA | NA | NA | NA | NA | #VALUE! |
| ENSMUSG00000098192.2 | 0 | NA | NA | NA | NA | NA | #VALUE! |
| ENSMUSG00000098193.2 | 0 | NA | NA | NA | NA | NA | #VALUE! |
| ENSMUSG00000098194.2 | 0 | NA | NA | NA | NA | NA | #VALUE! |
| ENSMUSG00000098195.2 | 0 | NA | NA | NA | NA | NA | #VALUE! |
| ENSMUSG00000098196.2 | 0 | NA | NA | NA | NA | NA | #VALUE! |
| ENSMUSG00000098197.3 | 0 | NA | NA | NA | NA | NA | #VALUE! |
| ENSMUSG00000098198.2 | 0 | NA | NA | NA | NA | NA | #VALUE! |
| ENSMUSG00000098199.2 | 0 | NA | NA | NA | NA | NA | #VALUE! |
| ENSMUSG00000098200.2 | 0 | NA | NA | NA | NA | NA | #VALUE! |
| ENSMUSG00000098201.2 | 0 | NA | NA | NA | NA | NA | #VALUE! |
| ENSMUSG00000098202.3 | 0 | NA | NA | NA | NA | NA | #VALUE! |
| ENSMUSG00000098203.2 | 0 | NA | NA | NA | NA | NA | #VALUE! |
| ENSMUSG00000098204.2 | 0 | NA | NA | NA | NA | NA | #VALUE! |
| ENSMUSG00000098205.2 | 0 | NA | NA | NA | NA | NA | #VALUE! |
| ENSMUSG00000098206.2 | 0 | NA | NA | NA | NA | NA | #VALUE! |
| ENSMUSG00000098207.2 | 0.735092 | -3.12016 | 3.965612 | -0.7868 | 0.431397 | NA | #VALUE! |
| ENSMUSG00000098208.2 | 0 | NA | NA | NA | NA | NA | #VALUE! |
| ENSMUSG00000098209.3 | 0 | NA | NA | NA | NA | NA | #VALUE! |
| ENSMUSG00000098210.2 | 0 | NA | NA | NA | NA | NA | #VALUE! |
| ENSMUSG00000098211.3 | 0 | NA | NA | NA | NA | NA | #VALUE! |
| ENSMUSG00000098212.3 | 0 | NA | NA | NA | NA | NA | #VALUE! |
| ENSMUSG00000098213.2 | 0 | NA | NA | NA | NA | NA | #VALUE! |
| ENSMUSG00000098214.3 | 0 | NA | NA | NA | NA | NA | #VALUE! |
| ENSMUSG00000098215.2 | 0 | NA | NA | NA | NA | NA | #VALUE! |
| ENSMUSG00000098216.2 | 0 | NA | NA | NA | NA | NA | #VALUE! |
| ENSMUSG00000098217.2 | 0 | NA | NA | NA | NA | NA | #VALUE! |
| ENSMUSG00000098218.2 | 0 | NA | NA | NA | NA | NA | #VALUE! |
| ENSMUSG00000098219.2 | 0 | NA | NA | NA | NA | NA | #VALUE! |
| ENSMUSG00000098221.2 | 0 | NA | NA | NA | NA | NA | #VALUE! |
| ENSMUSG00000098222.3 | 0.093255 | 0.868093 | 4.072464 | 0.213162 | 0.831201 | NA | #VALUE! |
| ENSMUSG00000098223.2 | 0 | NA | NA | NA | NA | NA | #VALUE! |
| ENSMUSG00000098224.2 | 0 | NA | NA | NA | NA | NA | #VALUE! |
| ENSMUSG00000098225.2 | 0 | NA | NA | NA | NA | NA | #VALUE! |
| ENSMUSG00000098226.2 | 0 | NA | NA | NA | NA | NA | #VALUE! |
| ENSMUSG00000098227.2 | 0 | NA | NA | NA | NA | NA | #VALUE! |
| ENSMUSG00000098228.2 | 0 | NA | NA | NA | NA | NA | #VALUE! |
| ENSMUSG00000098229.2 | 0 | NA | NA | NA | NA | NA | #VALUE! |
| ENSMUSG00000098230.8 | 0 | NA | NA | NA | NA | NA | #VALUE! |
| ENSMUSG00000098231.2 | 0 | NA | NA | NA | NA | NA | #VALUE! |
| ENSMUSG00000098232.2 | 0 | NA | NA | NA | NA | NA | #VALUE! |
| ENSMUSG00000098233.2 | 0 | NA | NA | NA | NA | NA | #VALUE! |
| ENSMUSG00000098235.3 | 0 | NA | NA | NA | NA | NA | #VALUE! |

| | | | | | | | |
|---|---|---|---|---|---|---|---|
| ENSMUSG00000098236.2 | 0 | NA | NA | NA | NA | NA | #VALUE! |
| ENSMUSG00000098237.2 | 0 | NA | NA | NA | NA | NA | #VALUE! |
| ENSMUSG00000098238.2 | 0 | NA | NA | NA | NA | NA | #VALUE! |
| ENSMUSG00000098240.2 | 0 | NA | NA | NA | NA | NA | #VALUE! |
| ENSMUSG00000098241.3 | 0 | NA | NA | NA | NA | NA | #VALUE! |
| ENSMUSG00000098242.2 | 0 | NA | NA | NA | NA | NA | #VALUE! |
| ENSMUSG00000098243.4 | 0 | NA | NA | NA | NA | NA | #VALUE! |
| ENSMUSG00000098244.2 | 0 | NA | NA | NA | NA | NA | #VALUE! |
| ENSMUSG00000098245.2 | 0 | NA | NA | NA | NA | NA | #VALUE! |
| ENSMUSG00000098246.2 | 0 | NA | NA | NA | NA | NA | #VALUE! |
| ENSMUSG00000098247.3 | 0 | NA | NA | NA | NA | NA | #VALUE! |
| ENSMUSG00000098248.2 | 0 | NA | NA | NA | NA | NA | #VALUE! |
| ENSMUSG00000098251.3 | 0 | NA | NA | NA | NA | NA | #VALUE! |
| ENSMUSG00000098252.3 | 0 | NA | NA | NA | NA | NA | #VALUE! |
| ENSMUSG00000098253.3 | 0 | NA | NA | NA | NA | NA | #VALUE! |
| ENSMUSG00000098255.2 | 0 | NA | NA | NA | NA | NA | #VALUE! |
| ENSMUSG00000098257.2 | 0 | NA | NA | NA | NA | NA | #VALUE! |
| ENSMUSG00000098258.3 | 0 | NA | NA | NA | NA | NA | #VALUE! |
| ENSMUSG00000098259.3 | 0 | NA | NA | NA | NA | NA | #VALUE! |
| ENSMUSG00000098261.3 | 0 | NA | NA | NA | NA | NA | #VALUE! |
| ENSMUSG00000098263.3 | 0 | NA | NA | NA | NA | NA | #VALUE! |
| ENSMUSG00000098264.3 | 0 | NA | NA | NA | NA | NA | #VALUE! |
| ENSMUSG00000098267.3 | 0 | NA | NA | NA | NA | NA | #VALUE! |
| ENSMUSG00000098269.3 | 0 | NA | NA | NA | NA | NA | #VALUE! |
| ENSMUSG00000098270.2 | 0 | NA | NA | NA | NA | NA | #VALUE! |
| ENSMUSG00000098272.2 | 0 | NA | NA | NA | NA | NA | #VALUE! |
| ENSMUSG00000098273.3 | 0 | NA | NA | NA | NA | NA | #VALUE! |
| ENSMUSG00000098274.8 | 0.093255 | 0.868093 | 4.072464 | 0.213162 | 0.831201 | NA | #VALUE! |
| ENSMUSG00000098275.2 | 0 | NA | NA | NA | NA | NA | #VALUE! |
| ENSMUSG00000098276.3 | 0 | NA | NA | NA | NA | NA | #VALUE! |
| ENSMUSG00000098278.2 | 0 | NA | NA | NA | NA | NA | #VALUE! |
| ENSMUSG00000098279.3 | 0 | NA | NA | NA | NA | NA | #VALUE! |
| ENSMUSG00000098280.3 | 0 | NA | NA | NA | NA | NA | #VALUE! |
| ENSMUSG00000098282.3 | 0 | NA | NA | NA | NA | NA | #VALUE! |
| ENSMUSG00000098284.9 | 0.168284 | 1.027628 | 4.067126 | 0.252667 | 0.800526 | NA | #VALUE! |
| ENSMUSG00000098285.3 | 0 | NA | NA | NA | NA | NA | #VALUE! |
| ENSMUSG00000098286.3 | 0 | NA | NA | NA | NA | NA | #VALUE! |
| ENSMUSG00000098287.3 | 0 | NA | NA | NA | NA | NA | #VALUE! |
| ENSMUSG00000098288.3 | 0 | NA | NA | NA | NA | NA | #VALUE! |
| ENSMUSG00000098289.3 | 0 | NA | NA | NA | NA | NA | #VALUE! |
| ENSMUSG00000098290.2 | 0 | NA | NA | NA | NA | NA | #VALUE! |
| ENSMUSG00000098291.3 | 0 | NA | NA | NA | NA | NA | #VALUE! |
| ENSMUSG00000098292.2 | 0 | NA | NA | NA | NA | NA | #VALUE! |
| ENSMUSG00000098293.3 | 0 | NA | NA | NA | NA | NA | #VALUE! |

| | | | | | | | |
|---|---|---|---|---|---|---|---|
| ENSMUSG00000098294.2 | 0 | NA | NA | NA | NA | NA | #VALUE! |
| ENSMUSG00000098295.2 | 0 | NA | NA | NA | NA | NA | #VALUE! |
| ENSMUSG00000098296.3 | 0 | NA | NA | NA | NA | NA | #VALUE! |
| ENSMUSG00000098297.3 | 0 | NA | NA | NA | NA | NA | #VALUE! |
| ENSMUSG00000098299.3 | 0 | NA | NA | NA | NA | NA | #VALUE! |
| ENSMUSG00000098300.3 | 0 | NA | NA | NA | NA | NA | #VALUE! |
| ENSMUSG00000098302.2 | 0 | NA | NA | NA | NA | NA | #VALUE! |
| ENSMUSG00000098304.3 | 0 | NA | NA | NA | NA | NA | #VALUE! |
| ENSMUSG00000098306.10 | 0 | NA | NA | NA | NA | NA | #VALUE! |
| ENSMUSG00000098309.2 | 0 | NA | NA | NA | NA | NA | #VALUE! |
| ENSMUSG00000098310.2 | 0 | NA | NA | NA | NA | NA | #VALUE! |
| ENSMUSG00000098311.2 | 0 | NA | NA | NA | NA | NA | #VALUE! |
| ENSMUSG00000098313.2 | 0 | NA | NA | NA | NA | NA | #VALUE! |
| ENSMUSG00000098315.3 | 0 | NA | NA | NA | NA | NA | #VALUE! |
| ENSMUSG00000098316.3 | 0 | NA | NA | NA | NA | NA | #VALUE! |
| ENSMUSG00000098317.3 | 0 | NA | NA | NA | NA | NA | #VALUE! |
| ENSMUSG00000098318.8 | 0.204111 | -1.05581 | 4.073302 | -0.2592 | 0.795479 | NA | #VALUE! |
| ENSMUSG00000098319.3 | 0 | NA | NA | NA | NA | NA | #VALUE! |
| ENSMUSG00000098322.2 | 0 | NA | NA | NA | NA | NA | #VALUE! |
| ENSMUSG00000098325.3 | 0 | NA | NA | NA | NA | NA | #VALUE! |
| ENSMUSG00000098327.3 | 0 | NA | NA | NA | NA | NA | #VALUE! |
| ENSMUSG00000098329.3 | 0 | NA | NA | NA | NA | NA | #VALUE! |
| ENSMUSG00000098330.2 | 1.260158 | -3.88425 | 3.941584 | -0.98545 | 0.324401 | NA | #VALUE! |
| ENSMUSG00000098331.3 | 0 | NA | NA | NA | NA | NA | #VALUE! |
| ENSMUSG00000098333.2 | 0 | NA | NA | NA | NA | NA | #VALUE! |
| ENSMUSG00000098334.3 | 0 | NA | NA | NA | NA | NA | #VALUE! |
| ENSMUSG00000098335.2 | 0 | NA | NA | NA | NA | NA | #VALUE! |
| ENSMUSG00000098336.3 | 0 | NA | NA | NA | NA | NA | #VALUE! |
| ENSMUSG00000098338.3 | 0 | NA | NA | NA | NA | NA | #VALUE! |
| ENSMUSG00000098339.3 | 0 | NA | NA | NA | NA | NA | #VALUE! |
| ENSMUSG00000098340.2 | 0 | NA | NA | NA | NA | NA | #VALUE! |
| ENSMUSG00000098341.3 | 0 | NA | NA | NA | NA | NA | #VALUE! |
| ENSMUSG00000098343.3 | 0 | NA | NA | NA | NA | NA | #VALUE! |
| ENSMUSG00000098344.3 | 0 | NA | NA | NA | NA | NA | #VALUE! |
| ENSMUSG00000098346.3 | 0 | NA | NA | NA | NA | NA | #VALUE! |
| ENSMUSG00000098347.3 | 0 | NA | NA | NA | NA | NA | #VALUE! |
| ENSMUSG00000098348.8 | 0 | NA | NA | NA | NA | NA | #VALUE! |
| ENSMUSG00000098353.3 | 0 | NA | NA | NA | NA | NA | #VALUE! |
| ENSMUSG00000098354.2 | 0 | NA | NA | NA | NA | NA | #VALUE! |
| ENSMUSG00000098358.2 | 0 | NA | NA | NA | NA | NA | #VALUE! |
| ENSMUSG00000098360.2 | 0 | NA | NA | NA | NA | NA | #VALUE! |
| ENSMUSG00000098361.3 | 0 | NA | NA | NA | NA | NA | #VALUE! |
| ENSMUSG00000098362.3 | 0 | NA | NA | NA | NA | NA | #VALUE! |
| ENSMUSG00000098363.3 | 0 | NA | NA | NA | NA | NA | #VALUE! |

| | | | | | | | |
|---|---|---|---|---|---|---|---|
| ENSMUSG00000098366.3 | 0 | NA | NA | NA | NA | NA | #VALUE! |
| ENSMUSG00000098370.2 | 0 | NA | NA | NA | NA | NA | #VALUE! |
| ENSMUSG00000098371.2 | 0 | NA | NA | NA | NA | NA | #VALUE! |
| ENSMUSG00000098374.2 | 0 | NA | NA | NA | NA | NA | #VALUE! |
| ENSMUSG00000098376.3 | 0 | NA | NA | NA | NA | NA | #VALUE! |
| ENSMUSG00000098379.3 | 0 | NA | NA | NA | NA | NA | #VALUE! |
| ENSMUSG00000098380.3 | 0 | NA | NA | NA | NA | NA | #VALUE! |
| ENSMUSG00000098381.3 | 0 | NA | NA | NA | NA | NA | #VALUE! |
| ENSMUSG00000098382.2 | 0 | NA | NA | NA | NA | NA | #VALUE! |
| ENSMUSG00000098383.2 | 0 | NA | NA | NA | NA | NA | #VALUE! |
| ENSMUSG00000098386.2 | 0 | NA | NA | NA | NA | NA | #VALUE! |
| ENSMUSG00000098387.3 | 0 | NA | NA | NA | NA | NA | #VALUE! |
| ENSMUSG00000098388.3 | 0 | NA | NA | NA | NA | NA | #VALUE! |
| ENSMUSG00000098389.3 | 0 | NA | NA | NA | NA | NA | #VALUE! |
| ENSMUSG00000098390.8 | 0 | NA | NA | NA | NA | NA | #VALUE! |
| ENSMUSG00000098391.3 | 0 | NA | NA | NA | NA | NA | #VALUE! |
| ENSMUSG00000098397.3 | 0 | NA | NA | NA | NA | NA | #VALUE! |
| ENSMUSG00000098400.3 | 0 | NA | NA | NA | NA | NA | #VALUE! |
| ENSMUSG00000098401.3 | 0 | NA | NA | NA | NA | NA | #VALUE! |
| ENSMUSG00000098404.2 | 0 | NA | NA | NA | NA | NA | #VALUE! |
| ENSMUSG00000098405.3 | 0 | NA | NA | NA | NA | NA | #VALUE! |
| ENSMUSG00000098406.3 | 0 | NA | NA | NA | NA | NA | #VALUE! |
| ENSMUSG00000098407.3 | 0 | NA | NA | NA | NA | NA | #VALUE! |
| ENSMUSG00000098408.3 | 0 | NA | NA | NA | NA | NA | #VALUE! |
| ENSMUSG00000098410.3 | 0 | NA | NA | NA | NA | NA | #VALUE! |
| ENSMUSG00000098412.2 | 0.261539 | 1.68833 | 4.043024 | 0.417591 | 0.676246 | NA | #VALUE! |
| ENSMUSG00000098413.3 | 0 | NA | NA | NA | NA | NA | #VALUE! |
| ENSMUSG00000098414.3 | 0 | NA | NA | NA | NA | NA | #VALUE! |
| ENSMUSG00000098415.3 | 0 | NA | NA | NA | NA | NA | #VALUE! |
| ENSMUSG00000098416.3 | 0 | NA | NA | NA | NA | NA | #VALUE! |
| ENSMUSG00000098417.3 | 0 | NA | NA | NA | NA | NA | #VALUE! |
| ENSMUSG00000098419.3 | 0 | NA | NA | NA | NA | NA | #VALUE! |
| ENSMUSG00000098420.2 | 0 | NA | NA | NA | NA | NA | #VALUE! |
| ENSMUSG00000098421.2 | 0 | NA | NA | NA | NA | NA | #VALUE! |
| ENSMUSG00000098422.3 | 0 | NA | NA | NA | NA | NA | #VALUE! |
| ENSMUSG00000098424.2 | 0 | NA | NA | NA | NA | NA | #VALUE! |
| ENSMUSG00000098426.2 | 0 | NA | NA | NA | NA | NA | #VALUE! |
| ENSMUSG00000098429.2 | 0 | NA | NA | NA | NA | NA | #VALUE! |
| ENSMUSG00000098431.3 | 0 | NA | NA | NA | NA | NA | #VALUE! |
| ENSMUSG00000098433.3 | 0 | NA | NA | NA | NA | NA | #VALUE! |
| ENSMUSG00000098434.2 | 0 | NA | NA | NA | NA | NA | #VALUE! |
| ENSMUSG00000098437.3 | 0 | NA | NA | NA | NA | NA | #VALUE! |
| ENSMUSG00000098438.2 | 0 | NA | NA | NA | NA | NA | #VALUE! |
| ENSMUSG00000098439.2 | 0 | NA | NA | NA | NA | NA | #VALUE! |

| | | | | | | | |
|---|---|---|---|---|---|---|---|
| ENSMUSG00000098441.3 | 0 | NA | NA | NA | NA | NA | #VALUE! |
| ENSMUSG00000098442.3 | 0 | NA | NA | NA | NA | NA | #VALUE! |
| ENSMUSG00000098443.3 | 0 | NA | NA | NA | NA | NA | #VALUE! |
| ENSMUSG00000098444.3 | 0 | NA | NA | NA | NA | NA | #VALUE! |
| ENSMUSG00000098445.3 | 0 | NA | NA | NA | NA | NA | #VALUE! |
| ENSMUSG00000098446.3 | 0 | NA | NA | NA | NA | NA | #VALUE! |
| ENSMUSG00000098447.3 | 12.13152 | 0.973756 | 3.140908 | 0.310024 | NA | NA | #VALUE! |
| ENSMUSG00000098448.3 | 0 | NA | NA | NA | NA | NA | #VALUE! |
| ENSMUSG00000098449.2 | 0 | NA | NA | NA | NA | NA | #VALUE! |
| ENSMUSG00000098451.3 | 0 | NA | NA | NA | NA | NA | #VALUE! |
| ENSMUSG00000098453.3 | 0 | NA | NA | NA | NA | NA | #VALUE! |
| ENSMUSG00000098454.3 | 0 | NA | NA | NA | NA | NA | #VALUE! |
| ENSMUSG00000098456.3 | 0 | NA | NA | NA | NA | NA | #VALUE! |
| ENSMUSG00000098457.3 | 0 | NA | NA | NA | NA | NA | #VALUE! |
| ENSMUSG00000098458.3 | 0 | NA | NA | NA | NA | NA | #VALUE! |
| ENSMUSG00000098464.3 | 0 | NA | NA | NA | NA | NA | #VALUE! |
| ENSMUSG00000098465.3 | 0 | NA | NA | NA | NA | NA | #VALUE! |
| ENSMUSG00000098468.2 | 0 | NA | NA | NA | NA | NA | #VALUE! |
| ENSMUSG00000098469.2 | 0 | NA | NA | NA | NA | NA | #VALUE! |
| ENSMUSG00000098470.2 | 0 | NA | NA | NA | NA | NA | #VALUE! |
| ENSMUSG00000098471.2 | 0 | NA | NA | NA | NA | NA | #VALUE! |
| ENSMUSG00000098473.3 | 0 | NA | NA | NA | NA | NA | #VALUE! |
| ENSMUSG00000098474.3 | 0 | NA | NA | NA | NA | NA | #VALUE! |
| ENSMUSG00000098475.3 | 0 | NA | NA | NA | NA | NA | #VALUE! |
| ENSMUSG00000098480.3 | 0 | NA | NA | NA | NA | NA | #VALUE! |
| ENSMUSG00000098482.3 | 0 | NA | NA | NA | NA | NA | #VALUE! |
| ENSMUSG00000098483.3 | 0 | NA | NA | NA | NA | NA | #VALUE! |
| ENSMUSG00000098485.2 | 0 | NA | NA | NA | NA | NA | #VALUE! |
| ENSMUSG00000098487.3 | 0 | NA | NA | NA | NA | NA | #VALUE! |
| ENSMUSG00000098488.8 | 0 | NA | NA | NA | NA | NA | #VALUE! |
| ENSMUSG00000098489.3 | 0 | NA | NA | NA | NA | NA | #VALUE! |
| ENSMUSG00000098490.3 | 0 | NA | NA | NA | NA | NA | #VALUE! |
| ENSMUSG00000098492.2 | 0 | NA | NA | NA | NA | NA | #VALUE! |
| ENSMUSG00000098493.3 | 0 | NA | NA | NA | NA | NA | #VALUE! |
| ENSMUSG00000098497.3 | 0 | NA | NA | NA | NA | NA | #VALUE! |
| ENSMUSG00000098498.3 | 0 | NA | NA | NA | NA | NA | #VALUE! |
| ENSMUSG00000098499.3 | 0 | NA | NA | NA | NA | NA | #VALUE! |
| ENSMUSG00000098500.3 | 0 | NA | NA | NA | NA | NA | #VALUE! |
| ENSMUSG00000098501.8 | 0 | NA | NA | NA | NA | NA | #VALUE! |
| ENSMUSG00000098504.3 | 0 | NA | NA | NA | NA | NA | #VALUE! |
| ENSMUSG00000098505.4 | 0 | NA | NA | NA | NA | NA | #VALUE! |
| ENSMUSG00000098506.3 | 0 | NA | NA | NA | NA | NA | #VALUE! |
| ENSMUSG00000098508.2 | 0 | NA | NA | NA | NA | NA | #VALUE! |
| ENSMUSG00000098509.3 | 0 | NA | NA | NA | NA | NA | #VALUE! |

| | | | | | | | |
|---|---|---|---|---|---|---|---|
| ENSMUSG00000098510.3 | 0 | NA | NA | NA | NA | NA | #VALUE! |
| ENSMUSG00000098511.3 | 0 | NA | NA | NA | NA | NA | #VALUE! |
| ENSMUSG00000098513.3 | 0 | NA | NA | NA | NA | NA | #VALUE! |
| ENSMUSG00000098515.2 | 0 | NA | NA | NA | NA | NA | #VALUE! |
| ENSMUSG00000098516.3 | 0 | NA | NA | NA | NA | NA | #VALUE! |
| ENSMUSG00000098519.2 | 0 | NA | NA | NA | NA | NA | #VALUE! |
| ENSMUSG00000098520.2 | 0 | NA | NA | NA | NA | NA | #VALUE! |
| ENSMUSG00000098521.3 | 0 | NA | NA | NA | NA | NA | #VALUE! |
| ENSMUSG00000098522.3 | 0 | NA | NA | NA | NA | NA | #VALUE! |
| ENSMUSG00000098527.3 | 0 | NA | NA | NA | NA | NA | #VALUE! |
| ENSMUSG00000098528.2 | 0 | NA | NA | NA | NA | NA | #VALUE! |
| ENSMUSG00000098530.2 | 0 | NA | NA | NA | NA | NA | #VALUE! |
| ENSMUSG00000098531.3 | 0 | NA | NA | NA | NA | NA | #VALUE! |
| ENSMUSG00000098532.3 | 0 | NA | NA | NA | NA | NA | #VALUE! |
| ENSMUSG00000098533.3 | 0 | NA | NA | NA | NA | NA | #VALUE! |
| ENSMUSG00000098534.2 | 0 | NA | NA | NA | NA | NA | #VALUE! |
| ENSMUSG00000098539.3 | 0 | NA | NA | NA | NA | NA | #VALUE! |
| ENSMUSG00000098540.3 | 0 | NA | NA | NA | NA | NA | #VALUE! |
| ENSMUSG00000098544.3 | 0 | NA | NA | NA | NA | NA | #VALUE! |
| ENSMUSG00000098547.3 | 0 | NA | NA | NA | NA | NA | #VALUE! |
| ENSMUSG00000098548.3 | 0 | NA | NA | NA | NA | NA | #VALUE! |
| ENSMUSG00000098549.3 | 0 | NA | NA | NA | NA | NA | #VALUE! |
| ENSMUSG00000098550.3 | 0 | NA | NA | NA | NA | NA | #VALUE! |
| ENSMUSG00000098552.2 | 0 | NA | NA | NA | NA | NA | #VALUE! |
| ENSMUSG00000098553.3 | 0 | NA | NA | NA | NA | NA | #VALUE! |
| ENSMUSG00000098555.3 | 0 | NA | NA | NA | NA | NA | #VALUE! |
| ENSMUSG00000098558.3 | 0 | NA | NA | NA | NA | NA | #VALUE! |
| ENSMUSG00000098559.3 | 0 | NA | NA | NA | NA | NA | #VALUE! |
| ENSMUSG00000098561.2 | 0 | NA | NA | NA | NA | NA | #VALUE! |
| ENSMUSG00000098562.3 | 0 | NA | NA | NA | NA | NA | #VALUE! |
| ENSMUSG00000098563.3 | 0 | NA | NA | NA | NA | NA | #VALUE! |
| ENSMUSG00000098564.2 | 0 | NA | NA | NA | NA | NA | #VALUE! |
| ENSMUSG00000098568.3 | 0 | NA | NA | NA | NA | NA | #VALUE! |
| ENSMUSG00000098569.3 | 0 | NA | NA | NA | NA | NA | #VALUE! |
| ENSMUSG00000098573.2 | 0 | NA | NA | NA | NA | NA | #VALUE! |
| ENSMUSG00000098574.3 | 0 | NA | NA | NA | NA | NA | #VALUE! |
| ENSMUSG00000098578.4 | 0 | NA | NA | NA | NA | NA | #VALUE! |
| ENSMUSG00000098579.3 | 0 | NA | NA | NA | NA | NA | #VALUE! |
| ENSMUSG00000098580.3 | 0 | NA | NA | NA | NA | NA | #VALUE! |
| ENSMUSG00000098582.3 | 0 | NA | NA | NA | NA | NA | #VALUE! |
| ENSMUSG00000098583.3 | 0 | NA | NA | NA | NA | NA | #VALUE! |
| ENSMUSG00000098588.3 | 0 | NA | NA | NA | NA | NA | #VALUE! |
| ENSMUSG00000098590.5 | 0 | NA | NA | NA | NA | NA | #VALUE! |
| ENSMUSG00000098592.3 | 0 | NA | NA | NA | NA | NA | #VALUE! |

| | | | | | | | |
|---|---|---|---|---|---|---|---|
| ENSMUSG00000098593.3 | 0 | NA | NA | NA | NA | NA | #VALUE! |
| ENSMUSG00000098596.3 | 0 | NA | NA | NA | NA | NA | #VALUE! |
| ENSMUSG00000098597.3 | 0 | NA | NA | NA | NA | NA | #VALUE! |
| ENSMUSG00000098600.3 | 0 | NA | NA | NA | NA | NA | #VALUE! |
| ENSMUSG00000098602.3 | 0 | NA | NA | NA | NA | NA | #VALUE! |
| ENSMUSG00000098606.3 | 0 | NA | NA | NA | NA | NA | #VALUE! |
| ENSMUSG00000098607.3 | 0 | NA | NA | NA | NA | NA | #VALUE! |
| ENSMUSG00000098608.3 | 0 | NA | NA | NA | NA | NA | #VALUE! |
| ENSMUSG00000098609.2 | 0 | NA | NA | NA | NA | NA | #VALUE! |
| ENSMUSG00000098611.3 | 0 | NA | NA | NA | NA | NA | #VALUE! |
| ENSMUSG00000098612.3 | 0 | NA | NA | NA | NA | NA | #VALUE! |
| ENSMUSG00000098613.3 | 0 | NA | NA | NA | NA | NA | #VALUE! |
| ENSMUSG00000098619.3 | 0 | NA | NA | NA | NA | NA | #VALUE! |
| ENSMUSG00000098620.2 | 0 | NA | NA | NA | NA | NA | #VALUE! |
| ENSMUSG00000098623.3 | 0 | NA | NA | NA | NA | NA | #VALUE! |
| ENSMUSG00000098624.3 | 0 | NA | NA | NA | NA | NA | #VALUE! |
| ENSMUSG00000098625.3 | 0 | NA | NA | NA | NA | NA | #VALUE! |
| ENSMUSG00000098627.8 | 0 | NA | NA | NA | NA | NA | #VALUE! |
| ENSMUSG00000098630.2 | 0 | NA | NA | NA | NA | NA | #VALUE! |
| ENSMUSG00000098631.3 | 0 | NA | NA | NA | NA | NA | #VALUE! |
| ENSMUSG00000098632.3 | 0 | NA | NA | NA | NA | NA | #VALUE! |
| ENSMUSG00000098634.3 | 0 | NA | NA | NA | NA | NA | #VALUE! |
| ENSMUSG00000098640.3 | 0 | NA | NA | NA | NA | NA | #VALUE! |
| ENSMUSG00000098644.2 | 0 | NA | NA | NA | NA | NA | #VALUE! |
| ENSMUSG00000098645.2 | 0 | NA | NA | NA | NA | NA | #VALUE! |
| ENSMUSG00000098646.3 | 0 | NA | NA | NA | NA | NA | #VALUE! |
| ENSMUSG00000098648.3 | 0 | NA | NA | NA | NA | NA | #VALUE! |
| ENSMUSG00000098649.3 | 0 | NA | NA | NA | NA | NA | #VALUE! |
| ENSMUSG00000098650.2 | 0 | NA | NA | NA | NA | NA | #VALUE! |
| ENSMUSG00000098651.3 | 0 | NA | NA | NA | NA | NA | #VALUE! |
| ENSMUSG00000098655.3 | 0 | NA | NA | NA | NA | NA | #VALUE! |
| ENSMUSG00000098656.3 | 0 | NA | NA | NA | NA | NA | #VALUE! |
| ENSMUSG00000098658.3 | 0 | NA | NA | NA | NA | NA | #VALUE! |
| ENSMUSG00000098659.8 | 0 | NA | NA | NA | NA | NA | #VALUE! |
| ENSMUSG00000098660.3 | 0 | NA | NA | NA | NA | NA | #VALUE! |
| ENSMUSG00000098661.3 | 0 | NA | NA | NA | NA | NA | #VALUE! |
| ENSMUSG00000098663.3 | 0 | NA | NA | NA | NA | NA | #VALUE! |
| ENSMUSG00000098666.3 | 0 | NA | NA | NA | NA | NA | #VALUE! |
| ENSMUSG00000098668.2 | 0 | NA | NA | NA | NA | NA | #VALUE! |
| ENSMUSG00000098671.3 | 0 | NA | NA | NA | NA | NA | #VALUE! |
| ENSMUSG00000098672.3 | 0 | NA | NA | NA | NA | NA | #VALUE! |
| ENSMUSG00000098675.3 | 0 | NA | NA | NA | NA | NA | #VALUE! |
| ENSMUSG00000098676.3 | 0 | NA | NA | NA | NA | NA | #VALUE! |
| ENSMUSG00000098678.2 | 0 | NA | NA | NA | NA | NA | #VALUE! |

| | | | | | | | |
|---|---|---|---|---|---|---|---|
| ENSMUSG00000098679.3 | 0 | NA | NA | NA | NA | NA | #VALUE! |
| ENSMUSG00000098680.3 | 0 | NA | NA | NA | NA | NA | #VALUE! |
| ENSMUSG00000098682.9 | 0 | NA | NA | NA | NA | NA | #VALUE! |
| ENSMUSG00000098684.2 | 0 | NA | NA | NA | NA | NA | #VALUE! |
| ENSMUSG00000098685.2 | 0 | NA | NA | NA | NA | NA | #VALUE! |
| ENSMUSG00000098686.3 | 0 | NA | NA | NA | NA | NA | #VALUE! |
| ENSMUSG00000098687.3 | 0 | NA | NA | NA | NA | NA | #VALUE! |
| ENSMUSG00000098689.3 | 0 | NA | NA | NA | NA | NA | #VALUE! |
| ENSMUSG00000098692.2 | 0 | NA | NA | NA | NA | NA | #VALUE! |
| ENSMUSG00000098698.3 | 0 | NA | NA | NA | NA | NA | #VALUE! |
| ENSMUSG00000098701.3 | 0 | NA | NA | NA | NA | NA | #VALUE! |
| ENSMUSG00000098702.2 | 0.839297 | 3.273019 | 3.958392 | 0.826856 | 0.408319 | NA | #VALUE! |
| ENSMUSG00000098704.3 | 0 | NA | NA | NA | NA | NA | #VALUE! |
| ENSMUSG00000098705.9 | 0 | NA | NA | NA | NA | NA | #VALUE! |
| ENSMUSG00000098706.3 | 0 | NA | NA | NA | NA | NA | #VALUE! |
| ENSMUSG00000098707.2 | 0 | NA | NA | NA | NA | NA | #VALUE! |
| ENSMUSG00000098708.8 | 0 | NA | NA | NA | NA | NA | #VALUE! |
| ENSMUSG00000098710.3 | 0 | NA | NA | NA | NA | NA | #VALUE! |
| ENSMUSG00000098712.3 | 0 | NA | NA | NA | NA | NA | #VALUE! |
| ENSMUSG00000098713.2 | 0 | NA | NA | NA | NA | NA | #VALUE! |
| ENSMUSG00000098714.3 | 0 | NA | NA | NA | NA | NA | #VALUE! |
| ENSMUSG00000098715.7 | 0 | NA | NA | NA | NA | NA | #VALUE! |
| ENSMUSG00000098718.3 | 0 | NA | NA | NA | NA | NA | #VALUE! |
| ENSMUSG00000098719.3 | 0 | NA | NA | NA | NA | NA | #VALUE! |
| ENSMUSG00000098720.2 | 0 | NA | NA | NA | NA | NA | #VALUE! |
| ENSMUSG00000098721.3 | 0 | NA | NA | NA | NA | NA | #VALUE! |
| ENSMUSG00000098726.3 | 0 | NA | NA | NA | NA | NA | #VALUE! |
| ENSMUSG00000098727.2 | 0 | NA | NA | NA | NA | NA | #VALUE! |
| ENSMUSG00000098729.2 | 0 | NA | NA | NA | NA | NA | #VALUE! |
| ENSMUSG00000098730.3 | 0 | NA | NA | NA | NA | NA | #VALUE! |
| ENSMUSG00000098731.3 | 0 | NA | NA | NA | NA | NA | #VALUE! |
| ENSMUSG00000098734.3 | 0 | NA | NA | NA | NA | NA | #VALUE! |
| ENSMUSG00000098735.3 | 0 | NA | NA | NA | NA | NA | #VALUE! |
| ENSMUSG00000098737.3 | 0 | NA | NA | NA | NA | NA | #VALUE! |
| ENSMUSG00000098739.2 | 0 | NA | NA | NA | NA | NA | #VALUE! |
| ENSMUSG00000098741.2 | 0 | NA | NA | NA | NA | NA | #VALUE! |
| ENSMUSG00000098744.2 | 0 | NA | NA | NA | NA | NA | #VALUE! |
| ENSMUSG00000098748.2 | 0 | NA | NA | NA | NA | NA | #VALUE! |
| ENSMUSG00000098751.3 | 0 | NA | NA | NA | NA | NA | #VALUE! |
| ENSMUSG00000098752.3 | 0 | NA | NA | NA | NA | NA | #VALUE! |
| ENSMUSG00000098754.8 | 0 | NA | NA | NA | NA | NA | #VALUE! |
| ENSMUSG00000098756.3 | 0 | NA | NA | NA | NA | NA | #VALUE! |
| ENSMUSG00000098758.2 | 0 | NA | NA | NA | NA | NA | #VALUE! |
| ENSMUSG00000098759.3 | 0 | NA | NA | NA | NA | NA | #VALUE! |

| | | | | | | | |
|---|---|---|---|---|---|---|---|
| ENSMUSG00000098760.2 | 0 | NA | NA | NA | NA | NA | #VALUE! |
| ENSMUSG00000098761.2 | 0 | NA | NA | NA | NA | NA | #VALUE! |
| ENSMUSG00000098762.3 | 0 | NA | NA | NA | NA | NA | #VALUE! |
| ENSMUSG00000098764.3 | 0 | NA | NA | NA | NA | NA | #VALUE! |
| ENSMUSG00000098765.3 | 0 | NA | NA | NA | NA | NA | #VALUE! |
| ENSMUSG00000098766.3 | 0 | NA | NA | NA | NA | NA | #VALUE! |
| ENSMUSG00000098768.2 | 0 | NA | NA | NA | NA | NA | #VALUE! |
| ENSMUSG00000098769.3 | 0 | NA | NA | NA | NA | NA | #VALUE! |
| ENSMUSG00000098771.3 | 0 | NA | NA | NA | NA | NA | #VALUE! |
| ENSMUSG00000098773.2 | 0 | NA | NA | NA | NA | NA | #VALUE! |
| ENSMUSG00000098774.3 | 0 | NA | NA | NA | NA | NA | #VALUE! |
| ENSMUSG00000098778.2 | 0 | NA | NA | NA | NA | NA | #VALUE! |
| ENSMUSG00000098779.3 | 0 | NA | NA | NA | NA | NA | #VALUE! |
| ENSMUSG00000098780.3 | 0 | NA | NA | NA | NA | NA | #VALUE! |
| ENSMUSG00000098781.2 | 0 | NA | NA | NA | NA | NA | #VALUE! |
| ENSMUSG00000098782.3 | 0 | NA | NA | NA | NA | NA | #VALUE! |
| ENSMUSG00000098783.3 | 0 | NA | NA | NA | NA | NA | #VALUE! |
| ENSMUSG00000098784.2 | 0 | NA | NA | NA | NA | NA | #VALUE! |
| ENSMUSG00000098785.3 | 0 | NA | NA | NA | NA | NA | #VALUE! |
| ENSMUSG00000098786.3 | 0 | NA | NA | NA | NA | NA | #VALUE! |
| ENSMUSG00000098787.3 | 0 | NA | NA | NA | NA | NA | #VALUE! |
| ENSMUSG00000098789.8 | 3.004514 | 5.085263 | 3.92161 | 1.296728 | 0.194725 | NA | #VALUE! |
| ENSMUSG00000098790.3 | 0 | NA | NA | NA | NA | NA | #VALUE! |
| ENSMUSG00000098794.2 | 0 | NA | NA | NA | NA | NA | #VALUE! |
| ENSMUSG00000098796.3 | 0 | NA | NA | NA | NA | NA | #VALUE! |
| ENSMUSG00000098797.3 | 0 | NA | NA | NA | NA | NA | #VALUE! |
| ENSMUSG00000098799.2 | 0 | NA | NA | NA | NA | NA | #VALUE! |
| ENSMUSG00000098800.2 | 0 | NA | NA | NA | NA | NA | #VALUE! |
| ENSMUSG00000098801.2 | 0 | NA | NA | NA | NA | NA | #VALUE! |
| ENSMUSG00000098802.3 | 0 | NA | NA | NA | NA | NA | #VALUE! |
| ENSMUSG00000098807.4 | 0 | NA | NA | NA | NA | NA | #VALUE! |
| ENSMUSG00000098808.3 | 0 | NA | NA | NA | NA | NA | #VALUE! |
| ENSMUSG00000098811.3 | 0 | NA | NA | NA | NA | NA | #VALUE! |
| ENSMUSG00000098812.3 | 0 | NA | NA | NA | NA | NA | #VALUE! |
| ENSMUSG00000098814.3 | 0 | NA | NA | NA | NA | NA | #VALUE! |
| ENSMUSG00000098817.3 | 0 | NA | NA | NA | NA | NA | #VALUE! |
| ENSMUSG00000098822.3 | 0 | NA | NA | NA | NA | NA | #VALUE! |
| ENSMUSG00000098823.3 | 0 | NA | NA | NA | NA | NA | #VALUE! |
| ENSMUSG00000098824.3 | 0 | NA | NA | NA | NA | NA | #VALUE! |
| ENSMUSG00000098826.3 | 0 | NA | NA | NA | NA | NA | #VALUE! |
| ENSMUSG00000098829.3 | 0 | NA | NA | NA | NA | NA | #VALUE! |
| ENSMUSG00000098830.3 | 0 | NA | NA | NA | NA | NA | #VALUE! |
| ENSMUSG00000098831.3 | 0 | NA | NA | NA | NA | NA | #VALUE! |
| ENSMUSG00000098832.3 | 0 | NA | NA | NA | NA | NA | #VALUE! |

| | | | | | | | |
|---|---|---|---|---|---|---|---|
| ENSMUSG00000098833.3 | 0 | NA | NA | NA | NA | NA | #VALUE! |
| ENSMUSG00000098837.2 | 0 | NA | NA | NA | NA | NA | #VALUE! |
| ENSMUSG00000098838.4 | 0 | NA | NA | NA | NA | NA | #VALUE! |
| ENSMUSG00000098839.3 | 0 | NA | NA | NA | NA | NA | #VALUE! |
| ENSMUSG00000098843.2 | 0 | NA | NA | NA | NA | NA | #VALUE! |
| ENSMUSG00000098844.3 | 0 | NA | NA | NA | NA | NA | #VALUE! |
| ENSMUSG00000098847.2 | 0 | NA | NA | NA | NA | NA | #VALUE! |
| ENSMUSG00000098851.3 | 0 | NA | NA | NA | NA | NA | #VALUE! |
| ENSMUSG00000098853.3 | 0 | NA | NA | NA | NA | NA | #VALUE! |
| ENSMUSG00000098854.2 | 0 | NA | NA | NA | NA | NA | #VALUE! |
| ENSMUSG00000098855.3 | 0 | NA | NA | NA | NA | NA | #VALUE! |
| ENSMUSG00000098856.3 | 0 | NA | NA | NA | NA | NA | #VALUE! |
| ENSMUSG00000098858.3 | 0 | NA | NA | NA | NA | NA | #VALUE! |
| ENSMUSG00000098861.2 | 0 | NA | NA | NA | NA | NA | #VALUE! |
| ENSMUSG00000098862.3 | 0 | NA | NA | NA | NA | NA | #VALUE! |
| ENSMUSG00000098863.3 | 0 | NA | NA | NA | NA | NA | #VALUE! |
| ENSMUSG00000098866.3 | 0 | NA | NA | NA | NA | NA | #VALUE! |
| ENSMUSG00000098871.3 | 0 | NA | NA | NA | NA | NA | #VALUE! |
| ENSMUSG00000098873.3 | 0 | NA | NA | NA | NA | NA | #VALUE! |
| ENSMUSG00000098875.2 | 0 | NA | NA | NA | NA | NA | #VALUE! |
| ENSMUSG00000098876.3 | 0 | NA | NA | NA | NA | NA | #VALUE! |
| ENSMUSG00000098879.3 | 0 | NA | NA | NA | NA | NA | #VALUE! |
| ENSMUSG00000098881.3 | 0 | NA | NA | NA | NA | NA | #VALUE! |
| ENSMUSG00000098882.3 | 0 | NA | NA | NA | NA | NA | #VALUE! |
| ENSMUSG00000098884.3 | 0 | NA | NA | NA | NA | NA | #VALUE! |
| ENSMUSG00000098885.3 | 0 | NA | NA | NA | NA | NA | #VALUE! |
| ENSMUSG00000098886.3 | 0 | NA | NA | NA | NA | NA | #VALUE! |
| ENSMUSG00000098887.3 | 0 | NA | NA | NA | NA | NA | #VALUE! |
| ENSMUSG00000098888.2 | 0 | NA | NA | NA | NA | NA | #VALUE! |
| ENSMUSG00000098889.2 | 0 | NA | NA | NA | NA | NA | #VALUE! |
| ENSMUSG00000098890.3 | 0 | NA | NA | NA | NA | NA | #VALUE! |
| ENSMUSG00000098893.2 | 0 | NA | NA | NA | NA | NA | #VALUE! |
| ENSMUSG00000098894.2 | 0 | NA | NA | NA | NA | NA | #VALUE! |
| ENSMUSG00000098895.2 | 0 | NA | NA | NA | NA | NA | #VALUE! |
| ENSMUSG00000098896.3 | 0 | NA | NA | NA | NA | NA | #VALUE! |
| ENSMUSG00000098897.2 | 0 | NA | NA | NA | NA | NA | #VALUE! |
| ENSMUSG00000098900.3 | 0 | NA | NA | NA | NA | NA | #VALUE! |
| ENSMUSG00000098904.3 | 0 | NA | NA | NA | NA | NA | #VALUE! |
| ENSMUSG00000098905.3 | 8.941755 | -0.26431 | 2.776439 | -0.0952 | 0.924157 | NA | #VALUE! |
| ENSMUSG00000098906.3 | 0 | NA | NA | NA | NA | NA | #VALUE! |
| ENSMUSG00000098907.3 | 0 | NA | NA | NA | NA | NA | #VALUE! |
| ENSMUSG00000098909.2 | 0 | NA | NA | NA | NA | NA | #VALUE! |
| ENSMUSG00000098910.3 | 0 | NA | NA | NA | NA | NA | #VALUE! |
| ENSMUSG00000098912.8 | 0 | NA | NA | NA | NA | NA | #VALUE! |

| | | | | | | | |
|---|---|---|---|---|---|---|---|
| ENSMUSG00000098915.2 | 0 | NA | NA | NA | NA | NA | #VALUE! |
| ENSMUSG00000098917.3 | 0 | NA | NA | NA | NA | NA | #VALUE! |
| ENSMUSG00000098918.3 | 0 | NA | NA | NA | NA | NA | #VALUE! |
| ENSMUSG00000098919.3 | 0 | NA | NA | NA | NA | NA | #VALUE! |
| ENSMUSG00000098922.3 | 0 | NA | NA | NA | NA | NA | #VALUE! |
| ENSMUSG00000098924.2 | 0 | NA | NA | NA | NA | NA | #VALUE! |
| ENSMUSG00000098926.3 | 0 | NA | NA | NA | NA | NA | #VALUE! |
| ENSMUSG00000098928.3 | 0 | NA | NA | NA | NA | NA | #VALUE! |
| ENSMUSG00000098929.2 | 0 | NA | NA | NA | NA | NA | #VALUE! |
| ENSMUSG00000098931.3 | 0 | NA | NA | NA | NA | NA | #VALUE! |
| ENSMUSG00000098932.3 | 0 | NA | NA | NA | NA | NA | #VALUE! |
| ENSMUSG00000098934.2 | 0 | NA | NA | NA | NA | NA | #VALUE! |
| ENSMUSG00000098935.2 | 0 | NA | NA | NA | NA | NA | #VALUE! |
| ENSMUSG00000098937.3 | 0 | NA | NA | NA | NA | NA | #VALUE! |
| ENSMUSG00000098938.3 | 0 | NA | NA | NA | NA | NA | #VALUE! |
| ENSMUSG00000098942.3 | 0 | NA | NA | NA | NA | NA | #VALUE! |
| ENSMUSG00000098944.3 | 0 | NA | NA | NA | NA | NA | #VALUE! |
| ENSMUSG00000098945.2 | 0 | NA | NA | NA | NA | NA | #VALUE! |
| ENSMUSG00000098947.3 | 0 | NA | NA | NA | NA | NA | #VALUE! |
| ENSMUSG00000098949.2 | 0 | NA | NA | NA | NA | NA | #VALUE! |
| ENSMUSG00000098950.8 | 0 | NA | NA | NA | NA | NA | #VALUE! |
| ENSMUSG00000098951.3 | 0 | NA | NA | NA | NA | NA | #VALUE! |
| ENSMUSG00000098952.3 | 0 | NA | NA | NA | NA | NA | #VALUE! |
| ENSMUSG00000098954.3 | 0 | NA | NA | NA | NA | NA | #VALUE! |
| ENSMUSG00000098955.3 | 0 | NA | NA | NA | NA | NA | #VALUE! |
| ENSMUSG00000098957.3 | 0 | NA | NA | NA | NA | NA | #VALUE! |
| ENSMUSG00000098958.3 | 0 | NA | NA | NA | NA | NA | #VALUE! |
| ENSMUSG00000098959.2 | 0 | NA | NA | NA | NA | NA | #VALUE! |
| ENSMUSG00000098960.3 | 0 | NA | NA | NA | NA | NA | #VALUE! |
| ENSMUSG00000098961.3 | 0 | NA | NA | NA | NA | NA | #VALUE! |
| ENSMUSG00000098970.3 | 0 | NA | NA | NA | NA | NA | #VALUE! |
| ENSMUSG00000098972.3 | 0 | NA | NA | NA | NA | NA | #VALUE! |
| ENSMUSG00000098973.3 | 0 | NA | NA | NA | NA | NA | #VALUE! |
| ENSMUSG00000098975.8 | 0 | NA | NA | NA | NA | NA | #VALUE! |
| ENSMUSG00000098976.3 | 0 | NA | NA | NA | NA | NA | #VALUE! |
| ENSMUSG00000098977.2 | 0 | NA | NA | NA | NA | NA | #VALUE! |
| ENSMUSG00000098980.3 | 0 | NA | NA | NA | NA | NA | #VALUE! |
| ENSMUSG00000098981.3 | 0 | NA | NA | NA | NA | NA | #VALUE! |
| ENSMUSG00000098984.3 | 0 | NA | NA | NA | NA | NA | #VALUE! |
| ENSMUSG00000098985.2 | 0 | NA | NA | NA | NA | NA | #VALUE! |
| ENSMUSG00000098987.2 | 0 | NA | NA | NA | NA | NA | #VALUE! |
| ENSMUSG00000098989.3 | 0 | NA | NA | NA | NA | NA | #VALUE! |
| ENSMUSG00000098997.3 | 0 | NA | NA | NA | NA | NA | #VALUE! |
| ENSMUSG00000099003.3 | 0 | NA | NA | NA | NA | NA | #VALUE! |

| | | | | | | | |
|---|---|---|---|---|---|---|---|
| ENSMUSG00000099005.3 | 0 | NA | NA | NA | NA | NA | #VALUE! |
| ENSMUSG00000099006.3 | 0 | NA | NA | NA | NA | NA | #VALUE! |
| ENSMUSG00000099008.2 | 0 | NA | NA | NA | NA | NA | #VALUE! |
| ENSMUSG00000099009.2 | 0 | NA | NA | NA | NA | NA | #VALUE! |
| ENSMUSG00000099011.3 | 0 | NA | NA | NA | NA | NA | #VALUE! |
| ENSMUSG00000099013.2 | 0 | NA | NA | NA | NA | NA | #VALUE! |
| ENSMUSG00000099015.3 | 0 | NA | NA | NA | NA | NA | #VALUE! |
| ENSMUSG00000099016.2 | 0 | NA | NA | NA | NA | NA | #VALUE! |
| ENSMUSG00000099017.3 | 0 | NA | NA | NA | NA | NA | #VALUE! |
| ENSMUSG00000099019.2 | 0.105013 | -0.97324 | 4.076293 | -0.23876 | 0.811295 | NA | #VALUE! |
| ENSMUSG00000099020.2 | 0 | NA | NA | NA | NA | NA | #VALUE! |
| ENSMUSG00000099025.2 | 0 | NA | NA | NA | NA | NA | #VALUE! |
| ENSMUSG00000099027.2 | 0 | NA | NA | NA | NA | NA | #VALUE! |
| ENSMUSG00000099030.3 | 0 | NA | NA | NA | NA | NA | #VALUE! |
| ENSMUSG00000099031.3 | 0 | NA | NA | NA | NA | NA | #VALUE! |
| ENSMUSG00000099032.3 | 1.221596 | 2.540809 | 3.941876 | 0.644569 | 0.519207 | NA | #VALUE! |
| ENSMUSG00000099033.3 | 0 | NA | NA | NA | NA | NA | #VALUE! |
| ENSMUSG00000099034.8 | 0 | NA | NA | NA | NA | NA | #VALUE! |
| ENSMUSG00000099035.3 | 0 | NA | NA | NA | NA | NA | #VALUE! |
| ENSMUSG00000099036.3 | 0 | NA | NA | NA | NA | NA | #VALUE! |
| ENSMUSG00000099041.2 | 0 | NA | NA | NA | NA | NA | #VALUE! |
| ENSMUSG00000099042.3 | 0 | NA | NA | NA | NA | NA | #VALUE! |
| ENSMUSG00000099043.3 | 0 | NA | NA | NA | NA | NA | #VALUE! |
| ENSMUSG00000099045.3 | 0 | NA | NA | NA | NA | NA | #VALUE! |
| ENSMUSG00000099047.3 | 0 | NA | NA | NA | NA | NA | #VALUE! |
| ENSMUSG00000099050.3 | 0 | NA | NA | NA | NA | NA | #VALUE! |
| ENSMUSG00000099051.2 | 0 | NA | NA | NA | NA | NA | #VALUE! |
| ENSMUSG00000099054.2 | 0 | NA | NA | NA | NA | NA | #VALUE! |
| ENSMUSG00000099055.2 | 0 | NA | NA | NA | NA | NA | #VALUE! |
| ENSMUSG00000099056.2 | 0 | NA | NA | NA | NA | NA | #VALUE! |
| ENSMUSG00000099057.2 | 0 | NA | NA | NA | NA | NA | #VALUE! |
| ENSMUSG00000099058.3 | 0 | NA | NA | NA | NA | NA | #VALUE! |
| ENSMUSG00000099060.3 | 3.465433 | -5.33079 | 3.919598 | -1.36004 | 0.173819 | NA | #VALUE! |
| ENSMUSG00000099061.2 | 0 | NA | NA | NA | NA | NA | #VALUE! |
| ENSMUSG00000099062.3 | 0 | NA | NA | NA | NA | NA | #VALUE! |
| ENSMUSG00000099063.3 | 0 | NA | NA | NA | NA | NA | #VALUE! |
| ENSMUSG00000099064.2 | 0 | NA | NA | NA | NA | NA | #VALUE! |
| ENSMUSG00000099066.3 | 0 | NA | NA | NA | NA | NA | #VALUE! |
| ENSMUSG00000099069.3 | 0 | NA | NA | NA | NA | NA | #VALUE! |
| ENSMUSG00000099070.3 | 0 | NA | NA | NA | NA | NA | #VALUE! |
| ENSMUSG00000099071.3 | 0 | NA | NA | NA | NA | NA | #VALUE! |
| ENSMUSG00000099072.3 | 0 | NA | NA | NA | NA | NA | #VALUE! |
| ENSMUSG00000099075.3 | 0 | NA | NA | NA | NA | NA | #VALUE! |
| ENSMUSG00000099076.3 | 0 | NA | NA | NA | NA | NA | #VALUE! |

| | | | | | | | |
|---|---|---|---|---|---|---|---|
| ENSMUSG00000099078.3 | 0 | NA | NA | NA | NA | NA | #VALUE! |
| ENSMUSG00000099079.3 | 0 | NA | NA | NA | NA | NA | #VALUE! |
| ENSMUSG00000099080.3 | 0 | NA | NA | NA | NA | NA | #VALUE! |
| ENSMUSG00000099082.3 | 0 | NA | NA | NA | NA | NA | #VALUE! |
| ENSMUSG00000099085.2 | 0 | NA | NA | NA | NA | NA | #VALUE! |
| ENSMUSG00000099087.2 | 0 | NA | NA | NA | NA | NA | #VALUE! |
| ENSMUSG00000099091.3 | 0 | NA | NA | NA | NA | NA | #VALUE! |
| ENSMUSG00000099092.3 | 0 | NA | NA | NA | NA | NA | #VALUE! |
| ENSMUSG00000099093.3 | 0 | NA | NA | NA | NA | NA | #VALUE! |
| ENSMUSG00000099094.2 | 0 | NA | NA | NA | NA | NA | #VALUE! |
| ENSMUSG00000099095.3 | 0 | NA | NA | NA | NA | NA | #VALUE! |
| ENSMUSG00000099096.3 | 0 | NA | NA | NA | NA | NA | #VALUE! |
| ENSMUSG00000099098.5 | 0 | NA | NA | NA | NA | NA | #VALUE! |
| ENSMUSG00000099101.3 | 0 | NA | NA | NA | NA | NA | #VALUE! |
| ENSMUSG00000099102.3 | 0 | NA | NA | NA | NA | NA | #VALUE! |
| ENSMUSG00000099104.3 | 0 | NA | NA | NA | NA | NA | #VALUE! |
| ENSMUSG00000099105.3 | 0 | NA | NA | NA | NA | NA | #VALUE! |
| ENSMUSG00000099109.3 | 0 | NA | NA | NA | NA | NA | #VALUE! |
| ENSMUSG00000099110.3 | 0 | NA | NA | NA | NA | NA | #VALUE! |
| ENSMUSG00000099115.3 | 0 | NA | NA | NA | NA | NA | #VALUE! |
| ENSMUSG00000099118.3 | 0 | NA | NA | NA | NA | NA | #VALUE! |
| ENSMUSG00000099120.2 | 0 | NA | NA | NA | NA | NA | #VALUE! |
| ENSMUSG00000099123.3 | 0 | NA | NA | NA | NA | NA | #VALUE! |
| ENSMUSG00000099124.3 | 0 | NA | NA | NA | NA | NA | #VALUE! |
| ENSMUSG00000099129.3 | 0 | NA | NA | NA | NA | NA | #VALUE! |
| ENSMUSG00000099131.3 | 0 | NA | NA | NA | NA | NA | #VALUE! |
| ENSMUSG00000099134.3 | 0 | NA | NA | NA | NA | NA | #VALUE! |
| ENSMUSG00000099137.2 | 0 | NA | NA | NA | NA | NA | #VALUE! |
| ENSMUSG00000099138.2 | 0 | NA | NA | NA | NA | NA | #VALUE! |
| ENSMUSG00000099139.2 | 0 | NA | NA | NA | NA | NA | #VALUE! |
| ENSMUSG00000099140.3 | 0 | NA | NA | NA | NA | NA | #VALUE! |
| ENSMUSG00000099141.3 | 0 | NA | NA | NA | NA | NA | #VALUE! |
| ENSMUSG00000099142.3 | 0 | NA | NA | NA | NA | NA | #VALUE! |
| ENSMUSG00000099145.3 | 0 | NA | NA | NA | NA | NA | #VALUE! |
| ENSMUSG00000099147.3 | 0 | NA | NA | NA | NA | NA | #VALUE! |
| ENSMUSG00000099148.2 | 0 | NA | NA | NA | NA | NA | #VALUE! |
| ENSMUSG00000099149.3 | 0 | NA | NA | NA | NA | NA | #VALUE! |
| ENSMUSG00000099150.2 | 0 | NA | NA | NA | NA | NA | #VALUE! |
| ENSMUSG00000099152.3 | 0 | NA | NA | NA | NA | NA | #VALUE! |
| ENSMUSG00000099153.3 | 0 | NA | NA | NA | NA | NA | #VALUE! |
| ENSMUSG00000099158.3 | 0 | NA | NA | NA | NA | NA | #VALUE! |
| ENSMUSG00000099160.3 | 0 | NA | NA | NA | NA | NA | #VALUE! |
| ENSMUSG00000099161.3 | 0 | NA | NA | NA | NA | NA | #VALUE! |
| ENSMUSG00000099162.2 | 0 | NA | NA | NA | NA | NA | #VALUE! |

| | | | | | | | |
|---|---|---|---|---|---|---|---|
| ENSMUSG00000099163.3 | 0 | NA | NA | NA | NA | NA | #VALUE! |
| ENSMUSG00000099165.3 | 0 | NA | NA | NA | NA | NA | #VALUE! |
| ENSMUSG00000099169.3 | 0 | NA | NA | NA | NA | NA | #VALUE! |
| ENSMUSG00000099170.2 | 0 | NA | NA | NA | NA | NA | #VALUE! |
| ENSMUSG00000099172.3 | 0 | NA | NA | NA | NA | NA | #VALUE! |
| ENSMUSG00000099174.3 | 0 | NA | NA | NA | NA | NA | #VALUE! |
| ENSMUSG00000099176.3 | 0 | NA | NA | NA | NA | NA | #VALUE! |
| ENSMUSG00000099178.2 | 0 | NA | NA | NA | NA | NA | #VALUE! |
| ENSMUSG00000099180.3 | 0 | NA | NA | NA | NA | NA | #VALUE! |
| ENSMUSG00000099181.3 | 0 | NA | NA | NA | NA | NA | #VALUE! |
| ENSMUSG00000099182.2 | 0 | NA | NA | NA | NA | NA | #VALUE! |
| ENSMUSG00000099184.3 | 0 | NA | NA | NA | NA | NA | #VALUE! |
| ENSMUSG00000099185.3 | 0 | NA | NA | NA | NA | NA | #VALUE! |
| ENSMUSG00000099188.2 | 0 | NA | NA | NA | NA | NA | #VALUE! |
| ENSMUSG00000099190.2 | 0 | NA | NA | NA | NA | NA | #VALUE! |
| ENSMUSG00000099196.3 | 0 | NA | NA | NA | NA | NA | #VALUE! |
| ENSMUSG00000099197.3 | 0 | NA | NA | NA | NA | NA | #VALUE! |
| ENSMUSG00000099198.2 | 0 | NA | NA | NA | NA | NA | #VALUE! |
| ENSMUSG00000099199.3 | 0 | NA | NA | NA | NA | NA | #VALUE! |
| ENSMUSG00000099200.3 | 0 | NA | NA | NA | NA | NA | #VALUE! |
| ENSMUSG00000099201.9 | 0 | NA | NA | NA | NA | NA | #VALUE! |
| ENSMUSG00000099202.3 | 0 | NA | NA | NA | NA | NA | #VALUE! |
| ENSMUSG00000099203.3 | 0 | NA | NA | NA | NA | NA | #VALUE! |
| ENSMUSG00000099207.8 | 0 | NA | NA | NA | NA | NA | #VALUE! |
| ENSMUSG00000099210.3 | 0 | NA | NA | NA | NA | NA | #VALUE! |
| ENSMUSG00000099212.3 | 0 | NA | NA | NA | NA | NA | #VALUE! |
| ENSMUSG00000099214.3 | 0 | NA | NA | NA | NA | NA | #VALUE! |
| ENSMUSG00000099216.2 | 0 | NA | NA | NA | NA | NA | #VALUE! |
| ENSMUSG00000099218.3 | 0 | NA | NA | NA | NA | NA | #VALUE! |
| ENSMUSG00000099220.2 | 0 | NA | NA | NA | NA | NA | #VALUE! |
| ENSMUSG00000099221.3 | 0 | NA | NA | NA | NA | NA | #VALUE! |
| ENSMUSG00000099222.3 | 0 | NA | NA | NA | NA | NA | #VALUE! |
| ENSMUSG00000099223.3 | 0 | NA | NA | NA | NA | NA | #VALUE! |
| ENSMUSG00000099225.3 | 0 | NA | NA | NA | NA | NA | #VALUE! |
| ENSMUSG00000099226.3 | 0 | NA | NA | NA | NA | NA | #VALUE! |
| ENSMUSG00000099227.3 | 0 | NA | NA | NA | NA | NA | #VALUE! |
| ENSMUSG00000099230.3 | 0 | NA | NA | NA | NA | NA | #VALUE! |
| ENSMUSG00000099236.4 | 0 | NA | NA | NA | NA | NA | #VALUE! |
| ENSMUSG00000099237.3 | 0 | NA | NA | NA | NA | NA | #VALUE! |
| ENSMUSG00000099238.3 | 0 | NA | NA | NA | NA | NA | #VALUE! |
| ENSMUSG00000099239.2 | 0 | NA | NA | NA | NA | NA | #VALUE! |
| ENSMUSG00000099241.3 | 0 | NA | NA | NA | NA | NA | #VALUE! |
| ENSMUSG00000099242.2 | 0 | NA | NA | NA | NA | NA | #VALUE! |
| ENSMUSG00000099245.3 | 0 | NA | NA | NA | NA | NA | #VALUE! |

| | | | | | | | |
|---|---|---|---|---|---|---|---|
| ENSMUSG00000099253.2 | 0 | NA | NA | NA | NA | NA | #VALUE! |
| ENSMUSG00000099254.3 | 0 | NA | NA | NA | NA | NA | #VALUE! |
| ENSMUSG00000099256.2 | 0 | NA | NA | NA | NA | NA | #VALUE! |
| ENSMUSG00000099257.3 | 0 | NA | NA | NA | NA | NA | #VALUE! |
| ENSMUSG00000099258.3 | 0 | NA | NA | NA | NA | NA | #VALUE! |
| ENSMUSG00000099259.3 | 0 | NA | NA | NA | NA | NA | #VALUE! |
| ENSMUSG00000099260.3 | 0 | NA | NA | NA | NA | NA | #VALUE! |
| ENSMUSG00000099261.3 | 0 | NA | NA | NA | NA | NA | #VALUE! |
| ENSMUSG00000099264.3 | 0 | NA | NA | NA | NA | NA | #VALUE! |
| ENSMUSG00000099266.2 | 0 | NA | NA | NA | NA | NA | #VALUE! |
| ENSMUSG00000099267.3 | 0 | NA | NA | NA | NA | NA | #VALUE! |
| ENSMUSG00000099269.4 | 0 | NA | NA | NA | NA | NA | #VALUE! |
| ENSMUSG00000099272.3 | 0 | NA | NA | NA | NA | NA | #VALUE! |
| ENSMUSG00000099274.2 | 0 | NA | NA | NA | NA | NA | #VALUE! |
| ENSMUSG00000099275.3 | 0 | NA | NA | NA | NA | NA | #VALUE! |
| ENSMUSG00000099276.2 | 0 | NA | NA | NA | NA | NA | #VALUE! |
| ENSMUSG00000099279.3 | 0 | NA | NA | NA | NA | NA | #VALUE! |
| ENSMUSG00000099280.4 | 0 | NA | NA | NA | NA | NA | #VALUE! |
| ENSMUSG00000099283.3 | 0 | NA | NA | NA | NA | NA | #VALUE! |
| ENSMUSG00000099284.3 | 0 | NA | NA | NA | NA | NA | #VALUE! |
| ENSMUSG00000099286.3 | 0 | NA | NA | NA | NA | NA | #VALUE! |
| ENSMUSG00000099289.3 | 0 | NA | NA | NA | NA | NA | #VALUE! |
| ENSMUSG00000099290.3 | 0 | NA | NA | NA | NA | NA | #VALUE! |
| ENSMUSG00000099292.3 | 0 | NA | NA | NA | NA | NA | #VALUE! |
| ENSMUSG00000099293.3 | 0 | NA | NA | NA | NA | NA | #VALUE! |
| ENSMUSG00000099294.3 | 0 | NA | NA | NA | NA | NA | #VALUE! |
| ENSMUSG00000099298.2 | 0 | NA | NA | NA | NA | NA | #VALUE! |
| ENSMUSG00000099300.3 | 0 | NA | NA | NA | NA | NA | #VALUE! |
| ENSMUSG00000099301.3 | 0 | NA | NA | NA | NA | NA | #VALUE! |
| ENSMUSG00000099308.3 | 0 | NA | NA | NA | NA | NA | #VALUE! |
| ENSMUSG00000099309.3 | 0 | NA | NA | NA | NA | NA | #VALUE! |
| ENSMUSG00000099310.2 | 0.587367 | -2.74297 | 3.982932 | -0.68868 | 0.491024 | NA | #VALUE! |
| ENSMUSG00000099311.3 | 0 | NA | NA | NA | NA | NA | #VALUE! |
| ENSMUSG00000099313.2 | 0 | NA | NA | NA | NA | NA | #VALUE! |
| ENSMUSG00000099315.2 | 0 | NA | NA | NA | NA | NA | #VALUE! |
| ENSMUSG00000099319.3 | 0 | NA | NA | NA | NA | NA | #VALUE! |
| ENSMUSG00000099320.3 | 0 | NA | NA | NA | NA | NA | #VALUE! |
| ENSMUSG00000099321.3 | 0 | NA | NA | NA | NA | NA | #VALUE! |
| ENSMUSG00000099324.3 | 0 | NA | NA | NA | NA | NA | #VALUE! |
| ENSMUSG00000099326.3 | 0 | NA | NA | NA | NA | NA | #VALUE! |
| ENSMUSG00000099329.3 | 0 | NA | NA | NA | NA | NA | #VALUE! |
| ENSMUSG00000099330.3 | 0 | NA | NA | NA | NA | NA | #VALUE! |
| ENSMUSG00000099331.3 | 0 | NA | NA | NA | NA | NA | #VALUE! |
| ENSMUSG00000099335.2 | 0 | NA | NA | NA | NA | NA | #VALUE! |

| | | | | | | | |
|---|---|---|---|---|---|---|---|
| ENSMUSG00000099336.2 | 0 | NA | NA | NA | NA | NA | #VALUE! |
| ENSMUSG00000099337.2 | 0 | NA | NA | NA | NA | NA | #VALUE! |
| ENSMUSG00000099338.2 | 1.575197 | -4.20222 | 3.934736 | -1.06798 | 0.285529 | NA | #VALUE! |
| ENSMUSG00000099339.2 | 0 | NA | NA | NA | NA | NA | #VALUE! |
| ENSMUSG00000099340.2 | 0 | NA | NA | NA | NA | NA | #VALUE! |
| ENSMUSG00000099341.2 | 0 | NA | NA | NA | NA | NA | #VALUE! |
| ENSMUSG00000099342.2 | 0 | NA | NA | NA | NA | NA | #VALUE! |
| ENSMUSG00000099343.2 | 0 | NA | NA | NA | NA | NA | #VALUE! |
| ENSMUSG00000099344.2 | 0 | NA | NA | NA | NA | NA | #VALUE! |
| ENSMUSG00000099346.2 | 0 | NA | NA | NA | NA | NA | #VALUE! |
| ENSMUSG00000099347.2 | 0 | NA | NA | NA | NA | NA | #VALUE! |
| ENSMUSG00000099348.2 | 0 | NA | NA | NA | NA | NA | #VALUE! |
| ENSMUSG00000099349.8 | 0 | NA | NA | NA | NA | NA | #VALUE! |
| ENSMUSG00000099350.2 | 0 | NA | NA | NA | NA | NA | #VALUE! |
| ENSMUSG00000099351.2 | 0 | NA | NA | NA | NA | NA | #VALUE! |
| ENSMUSG00000099352.2 | 0 | NA | NA | NA | NA | NA | #VALUE! |
| ENSMUSG00000099353.8 | 0 | NA | NA | NA | NA | NA | #VALUE! |
| ENSMUSG00000099354.7 | 0 | NA | NA | NA | NA | NA | #VALUE! |
| ENSMUSG00000099355.2 | 0 | NA | NA | NA | NA | NA | #VALUE! |
| ENSMUSG00000099357.2 | 0 | NA | NA | NA | NA | NA | #VALUE! |
| ENSMUSG00000099359.2 | 0 | NA | NA | NA | NA | NA | #VALUE! |
| ENSMUSG00000099360.2 | 0 | NA | NA | NA | NA | NA | #VALUE! |
| ENSMUSG00000099361.2 | 0 | NA | NA | NA | NA | NA | #VALUE! |
| ENSMUSG00000099362.2 | 0 | NA | NA | NA | NA | NA | #VALUE! |
| ENSMUSG00000099363.2 | 0 | NA | NA | NA | NA | NA | #VALUE! |
| ENSMUSG00000099364.3 | 0 | NA | NA | NA | NA | NA | #VALUE! |
| ENSMUSG00000099366.2 | 0 | NA | NA | NA | NA | NA | #VALUE! |
| ENSMUSG00000099367.2 | 0 | NA | NA | NA | NA | NA | #VALUE! |
| ENSMUSG00000099368.2 | 0 | NA | NA | NA | NA | NA | #VALUE! |
| ENSMUSG00000099369.2 | 0 | NA | NA | NA | NA | NA | #VALUE! |
| ENSMUSG00000099370.2 | 0 | NA | NA | NA | NA | NA | #VALUE! |
| ENSMUSG00000099371.2 | 0 | NA | NA | NA | NA | NA | #VALUE! |
| ENSMUSG00000099372.2 | 0 | NA | NA | NA | NA | NA | #VALUE! |
| ENSMUSG00000099373.2 | 0 | NA | NA | NA | NA | NA | #VALUE! |
| ENSMUSG00000099374.2 | 0 | NA | NA | NA | NA | NA | #VALUE! |
| ENSMUSG00000099375.2 | 0 | NA | NA | NA | NA | NA | #VALUE! |
| ENSMUSG00000099376.2 | 0 | NA | NA | NA | NA | NA | #VALUE! |
| ENSMUSG00000099377.2 | 0.336567 | 1.944925 | 4.029936 | 0.482619 | 0.629366 | NA | #VALUE! |
| ENSMUSG00000099378.2 | 0 | NA | NA | NA | NA | NA | #VALUE! |
| ENSMUSG00000099379.2 | 0 | NA | NA | NA | NA | NA | #VALUE! |
| ENSMUSG00000099380.7 | 0 | NA | NA | NA | NA | NA | #VALUE! |
| ENSMUSG00000099381.2 | 0 | NA | NA | NA | NA | NA | #VALUE! |
| ENSMUSG00000099382.2 | 0 | NA | NA | NA | NA | NA | #VALUE! |
| ENSMUSG00000099383.2 | 0 | NA | NA | NA | NA | NA | #VALUE! |

| | | | | | | | |
|---|---|---|---|---|---|---|---|
| ENSMUSG00000099384.7 | 0 | NA | NA | NA | NA | NA | #VALUE! |
| ENSMUSG00000099385.2 | 0 | NA | NA | NA | NA | NA | #VALUE! |
| ENSMUSG00000099386.2 | 0 | NA | NA | NA | NA | NA | #VALUE! |
| ENSMUSG00000099387.2 | 0 | NA | NA | NA | NA | NA | #VALUE! |
| ENSMUSG00000099388.2 | 0 | NA | NA | NA | NA | NA | #VALUE! |
| ENSMUSG00000099389.2 | 0 | NA | NA | NA | NA | NA | #VALUE! |
| ENSMUSG00000099390.7 | 0 | NA | NA | NA | NA | NA | #VALUE! |
| ENSMUSG00000099391.3 | 0 | NA | NA | NA | NA | NA | #VALUE! |
| ENSMUSG00000099392.2 | 0 | NA | NA | NA | NA | NA | #VALUE! |
| ENSMUSG00000099393.7 | 0 | NA | NA | NA | NA | NA | #VALUE! |
| ENSMUSG00000099394.2 | 0 | NA | NA | NA | NA | NA | #VALUE! |
| ENSMUSG00000099395.2 | 0 | NA | NA | NA | NA | NA | #VALUE! |
| ENSMUSG00000099396.2 | 0 | NA | NA | NA | NA | NA | #VALUE! |
| ENSMUSG00000099397.2 | 0 | NA | NA | NA | NA | NA | #VALUE! |
| ENSMUSG00000099398.2 | 0 | NA | NA | NA | NA | NA | #VALUE! |
| ENSMUSG00000099399.7 | 0 | NA | NA | NA | NA | NA | #VALUE! |
| ENSMUSG00000099400.2 | 0 | NA | NA | NA | NA | NA | #VALUE! |
| ENSMUSG00000099401.2 | 0 | NA | NA | NA | NA | NA | #VALUE! |
| ENSMUSG00000099403.2 | 0 | NA | NA | NA | NA | NA | #VALUE! |
| ENSMUSG00000099404.2 | 0 | NA | NA | NA | NA | NA | #VALUE! |
| ENSMUSG00000099405.2 | 0 | NA | NA | NA | NA | NA | #VALUE! |
| ENSMUSG00000099406.2 | 0 | NA | NA | NA | NA | NA | #VALUE! |
| ENSMUSG00000099407.2 | 0 | NA | NA | NA | NA | NA | #VALUE! |
| ENSMUSG00000099408.2 | 0 | NA | NA | NA | NA | NA | #VALUE! |
| ENSMUSG00000099409.2 | 0 | NA | NA | NA | NA | NA | #VALUE! |
| ENSMUSG00000099410.2 | 0 | NA | NA | NA | NA | NA | #VALUE! |
| ENSMUSG00000099411.2 | 0 | NA | NA | NA | NA | NA | #VALUE! |
| ENSMUSG00000099412.2 | 0 | NA | NA | NA | NA | NA | #VALUE! |
| ENSMUSG00000099413.2 | 0 | NA | NA | NA | NA | NA | #VALUE! |
| ENSMUSG00000099414.2 | 0 | NA | NA | NA | NA | NA | #VALUE! |
| ENSMUSG00000099415.2 | 0 | NA | NA | NA | NA | NA | #VALUE! |
| ENSMUSG00000099416.2 | 0 | NA | NA | NA | NA | NA | #VALUE! |
| ENSMUSG00000099417.2 | 0 | NA | NA | NA | NA | NA | #VALUE! |
| ENSMUSG00000099418.4 | 0 | NA | NA | NA | NA | NA | #VALUE! |
| ENSMUSG00000099419.2 | 0 | NA | NA | NA | NA | NA | #VALUE! |
| ENSMUSG00000099420.2 | 0 | NA | NA | NA | NA | NA | #VALUE! |
| ENSMUSG00000099421.2 | 0 | NA | NA | NA | NA | NA | #VALUE! |
| ENSMUSG00000099422.5 | 0 | NA | NA | NA | NA | NA | #VALUE! |
| ENSMUSG00000099423.2 | 0 | NA | NA | NA | NA | NA | #VALUE! |
| ENSMUSG00000099424.2 | 0 | NA | NA | NA | NA | NA | #VALUE! |
| ENSMUSG00000099425.2 | 0 | NA | NA | NA | NA | NA | #VALUE! |
| ENSMUSG00000099426.7 | 0 | NA | NA | NA | NA | NA | #VALUE! |
| ENSMUSG00000099427.2 | 0 | NA | NA | NA | NA | NA | #VALUE! |
| ENSMUSG00000099428.2 | 0 | NA | NA | NA | NA | NA | #VALUE! |

| | | | | | | | |
|---|---|---|---|---|---|---|---|
| ENSMUSG00000099429.2 | 0 | NA | NA | NA | NA | NA | #VALUE! |
| ENSMUSG00000099430.2 | 0 | NA | NA | NA | NA | NA | #VALUE! |
| ENSMUSG00000099431.2 | 0 | NA | NA | NA | NA | NA | #VALUE! |
| ENSMUSG00000099432.2 | 0 | NA | NA | NA | NA | NA | #VALUE! |
| ENSMUSG00000099433.2 | 0 | NA | NA | NA | NA | NA | #VALUE! |
| ENSMUSG00000099434.2 | 0 | NA | NA | NA | NA | NA | #VALUE! |
| ENSMUSG00000099435.7 | 0 | NA | NA | NA | NA | NA | #VALUE! |
| ENSMUSG00000099436.2 | 0 | NA | NA | NA | NA | NA | #VALUE! |
| ENSMUSG00000099437.2 | 0 | NA | NA | NA | NA | NA | #VALUE! |
| ENSMUSG00000099438.2 | 0 | NA | NA | NA | NA | NA | #VALUE! |
| ENSMUSG00000099439.2 | 0 | NA | NA | NA | NA | NA | #VALUE! |
| ENSMUSG00000099441.2 | 0 | NA | NA | NA | NA | NA | #VALUE! |
| ENSMUSG00000099442.2 | 0 | NA | NA | NA | NA | NA | #VALUE! |
| ENSMUSG00000099443.2 | 0 | NA | NA | NA | NA | NA | #VALUE! |
| ENSMUSG00000099444.2 | 0 | NA | NA | NA | NA | NA | #VALUE! |
| ENSMUSG00000099445.2 | 0 | NA | NA | NA | NA | NA | #VALUE! |
| ENSMUSG00000099446.2 | 0 | NA | NA | NA | NA | NA | #VALUE! |
| ENSMUSG00000099447.2 | 0 | NA | NA | NA | NA | NA | #VALUE! |
| ENSMUSG00000099448.2 | 0 | NA | NA | NA | NA | NA | #VALUE! |
| ENSMUSG00000099449.2 | 0 | NA | NA | NA | NA | NA | #VALUE! |
| ENSMUSG00000099450.2 | 0 | NA | NA | NA | NA | NA | #VALUE! |
| ENSMUSG00000099451.2 | 0 | NA | NA | NA | NA | NA | #VALUE! |
| ENSMUSG00000099452.2 | 0 | NA | NA | NA | NA | NA | #VALUE! |
| ENSMUSG00000099453.2 | 0 | NA | NA | NA | NA | NA | #VALUE! |
| ENSMUSG00000099454.2 | 0 | NA | NA | NA | NA | NA | #VALUE! |
| ENSMUSG00000099455.2 | 0 | NA | NA | NA | NA | NA | #VALUE! |
| ENSMUSG00000099456.7 | 0 | NA | NA | NA | NA | NA | #VALUE! |
| ENSMUSG00000099457.2 | 0 | NA | NA | NA | NA | NA | #VALUE! |
| ENSMUSG00000099458.2 | 0 | NA | NA | NA | NA | NA | #VALUE! |
| ENSMUSG00000099459.3 | 0 | NA | NA | NA | NA | NA | #VALUE! |
| ENSMUSG00000099460.2 | 0 | NA | NA | NA | NA | NA | #VALUE! |
| ENSMUSG00000099461.7 | 0 | NA | NA | NA | NA | NA | #VALUE! |
| ENSMUSG00000099462.2 | 0 | NA | NA | NA | NA | NA | #VALUE! |
| ENSMUSG00000099463.7 | 0 | NA | NA | NA | NA | NA | #VALUE! |
| ENSMUSG00000099464.2 | 0 | NA | NA | NA | NA | NA | #VALUE! |
| ENSMUSG00000099465.2 | 0 | NA | NA | NA | NA | NA | #VALUE! |
| ENSMUSG00000099466.2 | 0 | NA | NA | NA | NA | NA | #VALUE! |
| ENSMUSG00000099467.2 | 0 | NA | NA | NA | NA | NA | #VALUE! |
| ENSMUSG00000099468.2 | 0 | NA | NA | NA | NA | NA | #VALUE! |
| ENSMUSG00000099469.2 | 0 | NA | NA | NA | NA | NA | #VALUE! |
| ENSMUSG00000099470.2 | 0 | NA | NA | NA | NA | NA | #VALUE! |
| ENSMUSG00000099472.2 | 0 | NA | NA | NA | NA | NA | #VALUE! |
| ENSMUSG00000099473.2 | 0 | NA | NA | NA | NA | NA | #VALUE! |
| ENSMUSG00000099474.2 | 0 | NA | NA | NA | NA | NA | #VALUE! |

| | | | | | | | |
|---|---|---|---|---|---|---|---|
| ENSMUSG00000099476.2 | 0 | NA | NA | NA | NA | NA | #VALUE! |
| ENSMUSG00000099477.2 | 0 | NA | NA | NA | NA | NA | #VALUE! |
| ENSMUSG00000099478.2 | 0 | NA | NA | NA | NA | NA | #VALUE! |
| ENSMUSG00000099479.2 | 0 | NA | NA | NA | NA | NA | #VALUE! |
| ENSMUSG00000099480.2 | 0 | NA | NA | NA | NA | NA | #VALUE! |
| ENSMUSG00000099481.7 | 6.030107 | 5.11899 | 3.023399 | 1.693124 | 0.090432 | NA | #VALUE! |
| ENSMUSG00000099482.2 | 0 | NA | NA | NA | NA | NA | #VALUE! |
| ENSMUSG00000099483.2 | 0 | NA | NA | NA | NA | NA | #VALUE! |
| ENSMUSG00000099484.2 | 0 | NA | NA | NA | NA | NA | #VALUE! |
| ENSMUSG00000099485.2 | 0 | NA | NA | NA | NA | NA | #VALUE! |
| ENSMUSG00000099486.4 | 0 | NA | NA | NA | NA | NA | #VALUE! |
| ENSMUSG00000099487.2 | 0 | NA | NA | NA | NA | NA | #VALUE! |
| ENSMUSG00000099488.2 | 0 | NA | NA | NA | NA | NA | #VALUE! |
| ENSMUSG00000099489.2 | 0 | NA | NA | NA | NA | NA | #VALUE! |
| ENSMUSG00000099490.2 | 0 | NA | NA | NA | NA | NA | #VALUE! |
| ENSMUSG00000099491.2 | 0 | NA | NA | NA | NA | NA | #VALUE! |
| ENSMUSG00000099492.2 | 0 | NA | NA | NA | NA | NA | #VALUE! |
| ENSMUSG00000099493.2 | 0 | NA | NA | NA | NA | NA | #VALUE! |
| ENSMUSG00000099494.7 | 0 | NA | NA | NA | NA | NA | #VALUE! |
| ENSMUSG00000099495.2 | 0 | NA | NA | NA | NA | NA | #VALUE! |
| ENSMUSG00000099496.2 | 0 | NA | NA | NA | NA | NA | #VALUE! |
| ENSMUSG00000099497.2 | 0 | NA | NA | NA | NA | NA | #VALUE! |
| ENSMUSG00000099498.2 | 0 | NA | NA | NA | NA | NA | #VALUE! |
| ENSMUSG00000099499.2 | 0 | NA | NA | NA | NA | NA | #VALUE! |
| ENSMUSG00000099500.2 | 0 | NA | NA | NA | NA | NA | #VALUE! |
| ENSMUSG00000099501.2 | 0 | NA | NA | NA | NA | NA | #VALUE! |
| ENSMUSG00000099502.2 | 0 | NA | NA | NA | NA | NA | #VALUE! |
| ENSMUSG00000099503.2 | 0 | NA | NA | NA | NA | NA | #VALUE! |
| ENSMUSG00000099504.2 | 0 | NA | NA | NA | NA | NA | #VALUE! |
| ENSMUSG00000099505.2 | 0 | NA | NA | NA | NA | NA | #VALUE! |
| ENSMUSG00000099506.2 | 0 | NA | NA | NA | NA | NA | #VALUE! |
| ENSMUSG00000099507.2 | 0 | NA | NA | NA | NA | NA | #VALUE! |
| ENSMUSG00000099508.2 | 0 | NA | NA | NA | NA | NA | #VALUE! |
| ENSMUSG00000099509.2 | 0 | NA | NA | NA | NA | NA | #VALUE! |
| ENSMUSG00000099510.3 | 0 | NA | NA | NA | NA | NA | #VALUE! |
| ENSMUSG00000099511.2 | 0 | NA | NA | NA | NA | NA | #VALUE! |
| ENSMUSG00000099512.2 | 0 | NA | NA | NA | NA | NA | #VALUE! |
| ENSMUSG00000099513.2 | 0 | NA | NA | NA | NA | NA | #VALUE! |
| ENSMUSG00000099514.2 | 0 | NA | NA | NA | NA | NA | #VALUE! |
| ENSMUSG00000099516.2 | 0.105013 | -0.97324 | 4.076293 | -0.23876 | 0.811295 | NA | #VALUE! |
| ENSMUSG00000099517.3 | 0 | NA | NA | NA | NA | NA | #VALUE! |
| ENSMUSG00000099518.2 | 0 | NA | NA | NA | NA | NA | #VALUE! |
| ENSMUSG00000099519.2 | 0 | NA | NA | NA | NA | NA | #VALUE! |
| ENSMUSG00000099520.2 | 0 | NA | NA | NA | NA | NA | #VALUE! |

| | | | | | | | |
|---|---|---|---|---|---|---|---|
| ENSMUSG00000099521.2 | 0 | NA | NA | NA | NA | NA | #VALUE! |
| ENSMUSG00000099522.2 | 0 | NA | NA | NA | NA | NA | #VALUE! |
| ENSMUSG00000099523.2 | 0 | NA | NA | NA | NA | NA | #VALUE! |
| ENSMUSG00000099524.2 | 0 | NA | NA | NA | NA | NA | #VALUE! |
| ENSMUSG00000099525.2 | 0 | NA | NA | NA | NA | NA | #VALUE! |
| ENSMUSG00000099526.2 | 0 | NA | NA | NA | NA | NA | #VALUE! |
| ENSMUSG00000099527.2 | 0 | NA | NA | NA | NA | NA | #VALUE! |
| ENSMUSG00000099528.2 | 0 | NA | NA | NA | NA | NA | #VALUE! |
| ENSMUSG00000099529.2 | 0 | NA | NA | NA | NA | NA | #VALUE! |
| ENSMUSG00000099530.2 | 0 | NA | NA | NA | NA | NA | #VALUE! |
| ENSMUSG00000099531.2 | 0 | NA | NA | NA | NA | NA | #VALUE! |
| ENSMUSG00000099532.2 | 0 | NA | NA | NA | NA | NA | #VALUE! |
| ENSMUSG00000099534.2 | 0 | NA | NA | NA | NA | NA | #VALUE! |
| ENSMUSG00000099535.2 | 0 | NA | NA | NA | NA | NA | #VALUE! |
| ENSMUSG00000099536.2 | 0 | NA | NA | NA | NA | NA | #VALUE! |
| ENSMUSG00000099537.7 | 0 | NA | NA | NA | NA | NA | #VALUE! |
| ENSMUSG00000099539.2 | 0 | NA | NA | NA | NA | NA | #VALUE! |
| ENSMUSG00000099541.2 | 0 | NA | NA | NA | NA | NA | #VALUE! |
| ENSMUSG00000099542.2 | 0 | NA | NA | NA | NA | NA | #VALUE! |
| ENSMUSG00000099543.2 | 0 | NA | NA | NA | NA | NA | #VALUE! |
| ENSMUSG00000099544.2 | 0 | NA | NA | NA | NA | NA | #VALUE! |
| ENSMUSG00000099545.2 | 0 | NA | NA | NA | NA | NA | #VALUE! |
| ENSMUSG00000099546.2 | 0 | NA | NA | NA | NA | NA | #VALUE! |
| ENSMUSG00000099547.2 | 0 | NA | NA | NA | NA | NA | #VALUE! |
| ENSMUSG00000099548.2 | 0 | NA | NA | NA | NA | NA | #VALUE! |
| ENSMUSG00000099549.2 | 0 | NA | NA | NA | NA | NA | #VALUE! |
| ENSMUSG00000099550.7 | 0 | NA | NA | NA | NA | NA | #VALUE! |
| ENSMUSG00000099551.7 | 0 | NA | NA | NA | NA | NA | #VALUE! |
| ENSMUSG00000099552.2 | 0 | NA | NA | NA | NA | NA | #VALUE! |
| ENSMUSG00000099553.2 | 0 | NA | NA | NA | NA | NA | #VALUE! |
| ENSMUSG00000099554.2 | 0 | NA | NA | NA | NA | NA | #VALUE! |
| ENSMUSG00000099555.2 | 0 | NA | NA | NA | NA | NA | #VALUE! |
| ENSMUSG00000099556.2 | 0 | NA | NA | NA | NA | NA | #VALUE! |
| ENSMUSG00000099558.2 | 0 | NA | NA | NA | NA | NA | #VALUE! |
| ENSMUSG00000099559.2 | 0 | NA | NA | NA | NA | NA | #VALUE! |
| ENSMUSG00000099560.2 | 0 | NA | NA | NA | NA | NA | #VALUE! |
| ENSMUSG00000099561.2 | 0 | NA | NA | NA | NA | NA | #VALUE! |
| ENSMUSG00000099562.2 | 0 | NA | NA | NA | NA | NA | #VALUE! |
| ENSMUSG00000099563.2 | 0 | NA | NA | NA | NA | NA | #VALUE! |
| ENSMUSG00000099564.3 | 0 | NA | NA | NA | NA | NA | #VALUE! |
| ENSMUSG00000099565.2 | 0 | NA | NA | NA | NA | NA | #VALUE! |
| ENSMUSG00000099566.2 | 0 | NA | NA | NA | NA | NA | #VALUE! |
| ENSMUSG00000099568.2 | 0 | NA | NA | NA | NA | NA | #VALUE! |
| ENSMUSG00000099569.2 | 0 | NA | NA | NA | NA | NA | #VALUE! |

| | | | | | | | |
|---|---|---|---|---|---|---|---|
| ENSMUSG00000099570.7 | 0 | NA | NA | NA | NA | NA | #VALUE! |
| ENSMUSG00000099571.2 | 0 | NA | NA | NA | NA | NA | #VALUE! |
| ENSMUSG00000099572.2 | 0 | NA | NA | NA | NA | NA | #VALUE! |
| ENSMUSG00000099573.2 | 0 | NA | NA | NA | NA | NA | #VALUE! |
| ENSMUSG00000099574.2 | 0 | NA | NA | NA | NA | NA | #VALUE! |
| ENSMUSG00000099575.2 | 0 | NA | NA | NA | NA | NA | #VALUE! |
| ENSMUSG00000099576.2 | 0 | NA | NA | NA | NA | NA | #VALUE! |
| ENSMUSG00000099577.2 | 0 | NA | NA | NA | NA | NA | #VALUE! |
| ENSMUSG00000099578.2 | 0 | NA | NA | NA | NA | NA | #VALUE! |
| ENSMUSG00000099579.2 | 0 | NA | NA | NA | NA | NA | #VALUE! |
| ENSMUSG00000099581.2 | 0 | NA | NA | NA | NA | NA | #VALUE! |
| ENSMUSG00000099582.2 | 0 | NA | NA | NA | NA | NA | #VALUE! |
| ENSMUSG00000099583.3 | 0 | NA | NA | NA | NA | NA | #VALUE! |
| ENSMUSG00000099584.2 | 0 | NA | NA | NA | NA | NA | #VALUE! |
| ENSMUSG00000099585.2 | 0 | NA | NA | NA | NA | NA | #VALUE! |
| ENSMUSG00000099586.2 | 0 | NA | NA | NA | NA | NA | #VALUE! |
| ENSMUSG00000099587.2 | 0 | NA | NA | NA | NA | NA | #VALUE! |
| ENSMUSG00000099588.2 | 0 | NA | NA | NA | NA | NA | #VALUE! |
| ENSMUSG00000099589.2 | 0 | NA | NA | NA | NA | NA | #VALUE! |
| ENSMUSG00000099591.2 | 0 | NA | NA | NA | NA | NA | #VALUE! |
| ENSMUSG00000099592.2 | 0 | NA | NA | NA | NA | NA | #VALUE! |
| ENSMUSG00000099593.2 | 0 | NA | NA | NA | NA | NA | #VALUE! |
| ENSMUSG00000099594.2 | 0 | NA | NA | NA | NA | NA | #VALUE! |
| ENSMUSG00000099595.2 | 0 | NA | NA | NA | NA | NA | #VALUE! |
| ENSMUSG00000099596.2 | 0 | NA | NA | NA | NA | NA | #VALUE! |
| ENSMUSG00000099598.2 | 0 | NA | NA | NA | NA | NA | #VALUE! |
| ENSMUSG00000099599.2 | 3.36567 | 5.240782 | 3.920098 | 1.336901 | 0.181255 | NA | #VALUE! |
| ENSMUSG00000099600.2 | 0 | NA | NA | NA | NA | NA | #VALUE! |
| ENSMUSG00000099601.2 | 0 | NA | NA | NA | NA | NA | #VALUE! |
| ENSMUSG00000099603.2 | 0 | NA | NA | NA | NA | NA | #VALUE! |
| ENSMUSG00000099604.2 | 0 | NA | NA | NA | NA | NA | #VALUE! |
| ENSMUSG00000099605.2 | 0 | NA | NA | NA | NA | NA | #VALUE! |
| ENSMUSG00000099606.2 | 0 | NA | NA | NA | NA | NA | #VALUE! |
| ENSMUSG00000099607.2 | 0 | NA | NA | NA | NA | NA | #VALUE! |
| ENSMUSG00000099608.2 | 0 | NA | NA | NA | NA | NA | #VALUE! |
| ENSMUSG00000099609.2 | 0 | NA | NA | NA | NA | NA | #VALUE! |
| ENSMUSG00000099610.2 | 0 | NA | NA | NA | NA | NA | #VALUE! |
| ENSMUSG00000099611.3 | 0 | NA | NA | NA | NA | NA | #VALUE! |
| ENSMUSG00000099612.2 | 0 | NA | NA | NA | NA | NA | #VALUE! |
| ENSMUSG00000099613.2 | 0 | NA | NA | NA | NA | NA | #VALUE! |
| ENSMUSG00000099614.2 | 0 | NA | NA | NA | NA | NA | #VALUE! |
| ENSMUSG00000099615.2 | 0 | NA | NA | NA | NA | NA | #VALUE! |
| ENSMUSG00000099616.2 | 0 | NA | NA | NA | NA | NA | #VALUE! |
| ENSMUSG00000099617.2 | 0 | NA | NA | NA | NA | NA | #VALUE! |

| | | | | | | | |
|---|---|---|---|---|---|---|---|
| ENSMUSG00000099618.2 | 0 | NA | NA | NA | NA | NA | #VALUE! |
| ENSMUSG00000099619.7 | 0 | NA | NA | NA | NA | NA | #VALUE! |
| ENSMUSG00000099620.2 | 0 | NA | NA | NA | NA | NA | #VALUE! |
| ENSMUSG00000099621.2 | 0 | NA | NA | NA | NA | NA | #VALUE! |
| ENSMUSG00000099622.2 | 0 | NA | NA | NA | NA | NA | #VALUE! |
| ENSMUSG00000099623.3 | 0 | NA | NA | NA | NA | NA | #VALUE! |
| ENSMUSG00000099624.2 | 0 | NA | NA | NA | NA | NA | #VALUE! |
| ENSMUSG00000099625.2 | 0 | NA | NA | NA | NA | NA | #VALUE! |
| ENSMUSG00000099626.2 | 0 | NA | NA | NA | NA | NA | #VALUE! |
| ENSMUSG00000099627.2 | 0 | NA | NA | NA | NA | NA | #VALUE! |
| ENSMUSG00000099628.2 | 0 | NA | NA | NA | NA | NA | #VALUE! |
| ENSMUSG00000099629.2 | 0 | NA | NA | NA | NA | NA | #VALUE! |
| ENSMUSG00000099630.2 | 0 | NA | NA | NA | NA | NA | #VALUE! |
| ENSMUSG00000099631.2 | 0 | NA | NA | NA | NA | NA | #VALUE! |
| ENSMUSG00000099632.2 | 2.832857 | 2.618883 | 3.858887 | 0.678663 | 0.497352 | NA | #VALUE! |
| ENSMUSG00000099633.2 | 0 | NA | NA | NA | NA | NA | #VALUE! |
| ENSMUSG00000099634.2 | 0 | NA | NA | NA | NA | NA | #VALUE! |
| ENSMUSG00000099635.2 | 0 | NA | NA | NA | NA | NA | #VALUE! |
| ENSMUSG00000099636.2 | 0 | NA | NA | NA | NA | NA | #VALUE! |
| ENSMUSG00000099637.2 | 0 | NA | NA | NA | NA | NA | #VALUE! |
| ENSMUSG00000099638.2 | 0 | NA | NA | NA | NA | NA | #VALUE! |
| ENSMUSG00000099639.7 | 0 | NA | NA | NA | NA | NA | #VALUE! |
| ENSMUSG00000099640.2 | 0 | NA | NA | NA | NA | NA | #VALUE! |
| ENSMUSG00000099641.2 | 0 | NA | NA | NA | NA | NA | #VALUE! |
| ENSMUSG00000099642.2 | 0 | NA | NA | NA | NA | NA | #VALUE! |
| ENSMUSG00000099643.2 | 0 | NA | NA | NA | NA | NA | #VALUE! |
| ENSMUSG00000099644.2 | 0 | NA | NA | NA | NA | NA | #VALUE! |
| ENSMUSG00000099645.2 | 0 | NA | NA | NA | NA | NA | #VALUE! |
| ENSMUSG00000099646.2 | 0 | NA | NA | NA | NA | NA | #VALUE! |
| ENSMUSG00000099647.2 | 0.204111 | -1.05581 | 4.073302 | -0.2592 | 0.795479 | NA | #VALUE! |
| ENSMUSG00000099648.2 | 0 | NA | NA | NA | NA | NA | #VALUE! |
| ENSMUSG00000099649.2 | 0 | NA | NA | NA | NA | NA | #VALUE! |
| ENSMUSG00000099650.7 | 0 | NA | NA | NA | NA | NA | #VALUE! |
| ENSMUSG00000099651.2 | 0 | NA | NA | NA | NA | NA | #VALUE! |
| ENSMUSG00000099652.2 | 0 | NA | NA | NA | NA | NA | #VALUE! |
| ENSMUSG00000099653.2 | 0 | NA | NA | NA | NA | NA | #VALUE! |
| ENSMUSG00000099654.7 | 0 | NA | NA | NA | NA | NA | #VALUE! |
| ENSMUSG00000099655.3 | 0 | NA | NA | NA | NA | NA | #VALUE! |
| ENSMUSG00000099656.3 | 0 | NA | NA | NA | NA | NA | #VALUE! |
| ENSMUSG00000099657.3 | 0 | NA | NA | NA | NA | NA | #VALUE! |
| ENSMUSG00000099658.2 | 0 | NA | NA | NA | NA | NA | #VALUE! |
| ENSMUSG00000099659.2 | 0 | NA | NA | NA | NA | NA | #VALUE! |
| ENSMUSG00000099660.2 | 0 | NA | NA | NA | NA | NA | #VALUE! |
| ENSMUSG00000099661.2 | 0 | NA | NA | NA | NA | NA | #VALUE! |

| | | | | | | | |
|---|---|---|---|---|---|---|---|
| ENSMUSG00000099662.2 | 0 | NA | NA | NA | NA | NA | #VALUE! |
| ENSMUSG00000099663.2 | 0 | NA | NA | NA | NA | NA | #VALUE! |
| ENSMUSG00000099664.2 | 0 | NA | NA | NA | NA | NA | #VALUE! |
| ENSMUSG00000099665.2 | 0 | NA | NA | NA | NA | NA | #VALUE! |
| ENSMUSG00000099666.7 | 0 | NA | NA | NA | NA | NA | #VALUE! |
| ENSMUSG00000099667.2 | 0 | NA | NA | NA | NA | NA | #VALUE! |
| ENSMUSG00000099668.2 | 0 | NA | NA | NA | NA | NA | #VALUE! |
| ENSMUSG00000099669.2 | 0 | NA | NA | NA | NA | NA | #VALUE! |
| ENSMUSG00000099670.2 | 0 | NA | NA | NA | NA | NA | #VALUE! |
| ENSMUSG00000099671.3 | 0 | NA | NA | NA | NA | NA | #VALUE! |
| ENSMUSG00000099672.2 | 0 | NA | NA | NA | NA | NA | #VALUE! |
| ENSMUSG00000099673.2 | 0 | NA | NA | NA | NA | NA | #VALUE! |
| ENSMUSG00000099674.7 | 0 | NA | NA | NA | NA | NA | #VALUE! |
| ENSMUSG00000099675.2 | 0 | NA | NA | NA | NA | NA | #VALUE! |
| ENSMUSG00000099676.2 | 0 | NA | NA | NA | NA | NA | #VALUE! |
| ENSMUSG00000099677.2 | 0 | NA | NA | NA | NA | NA | #VALUE! |
| ENSMUSG00000099678.2 | 0 | NA | NA | NA | NA | NA | #VALUE! |
| ENSMUSG00000099679.2 | 0 | NA | NA | NA | NA | NA | #VALUE! |
| ENSMUSG00000099681.2 | 0.94431 | 2.339985 | 3.954083 | 0.59179 | 0.553991 | NA | #VALUE! |
| ENSMUSG00000099682.2 | 0 | NA | NA | NA | NA | NA | #VALUE! |
| ENSMUSG00000099683.2 | 0 | NA | NA | NA | NA | NA | #VALUE! |
| ENSMUSG00000099684.7 | 0 | NA | NA | NA | NA | NA | #VALUE! |
| ENSMUSG00000099685.7 | 0 | NA | NA | NA | NA | NA | #VALUE! |
| ENSMUSG00000099686.2 | 0 | NA | NA | NA | NA | NA | #VALUE! |
| ENSMUSG00000099687.4 | 0 | NA | NA | NA | NA | NA | #VALUE! |
| ENSMUSG00000099688.2 | 0 | NA | NA | NA | NA | NA | #VALUE! |
| ENSMUSG00000099689.7 | 9.505151 | 4.561973 | 2.51109 | 1.81673 | 0.069258 | NA | #VALUE! |
| ENSMUSG00000099690.2 | 0 | NA | NA | NA | NA | NA | #VALUE! |
| ENSMUSG00000099691.2 | 0 | NA | NA | NA | NA | NA | #VALUE! |
| ENSMUSG00000099692.2 | 0 | NA | NA | NA | NA | NA | #VALUE! |
| ENSMUSG00000099693.2 | 0 | NA | NA | NA | NA | NA | #VALUE! |
| ENSMUSG00000099694.2 | 0 | NA | NA | NA | NA | NA | #VALUE! |
| ENSMUSG00000099695.7 | 0 | NA | NA | NA | NA | NA | #VALUE! |
| ENSMUSG00000099696.7 | 0 | NA | NA | NA | NA | NA | #VALUE! |
| ENSMUSG00000099698.2 | 0 | NA | NA | NA | NA | NA | #VALUE! |
| ENSMUSG00000099699.2 | 0 | NA | NA | NA | NA | NA | #VALUE! |
| ENSMUSG00000099700.2 | 0 | NA | NA | NA | NA | NA | #VALUE! |
| ENSMUSG00000099701.2 | 0 | NA | NA | NA | NA | NA | #VALUE! |
| ENSMUSG00000099702.2 | 0 | NA | NA | NA | NA | NA | #VALUE! |
| ENSMUSG00000099704.2 | 0 | NA | NA | NA | NA | NA | #VALUE! |
| ENSMUSG00000099706.2 | 0 | NA | NA | NA | NA | NA | #VALUE! |
| ENSMUSG00000099708.2 | 0 | NA | NA | NA | NA | NA | #VALUE! |
| ENSMUSG00000099709.2 | 0 | NA | NA | NA | NA | NA | #VALUE! |
| ENSMUSG00000099711.2 | 0 | NA | NA | NA | NA | NA | #VALUE! |

| | | | | | | | |
|---|---|---|---|---|---|---|---|
| ENSMUSG00000099712.2 | 0 | NA | NA | NA | NA | NA | #VALUE! |
| ENSMUSG00000099713.2 | 0 | NA | NA | NA | NA | NA | #VALUE! |
| ENSMUSG00000099714.2 | 0 | NA | NA | NA | NA | NA | #VALUE! |
| ENSMUSG00000099715.3 | 0 | NA | NA | NA | NA | NA | #VALUE! |
| ENSMUSG00000099716.2 | 0 | NA | NA | NA | NA | NA | #VALUE! |
| ENSMUSG00000099717.2 | 0 | NA | NA | NA | NA | NA | #VALUE! |
| ENSMUSG00000099718.2 | 0 | NA | NA | NA | NA | NA | #VALUE! |
| ENSMUSG00000099719.2 | 0 | NA | NA | NA | NA | NA | #VALUE! |
| ENSMUSG00000099721.2 | 0 | NA | NA | NA | NA | NA | #VALUE! |
| ENSMUSG00000099722.4 | 0 | NA | NA | NA | NA | NA | #VALUE! |
| ENSMUSG00000099723.2 | 0 | NA | NA | NA | NA | NA | #VALUE! |
| ENSMUSG00000099724.3 | 0 | NA | NA | NA | NA | NA | #VALUE! |
| ENSMUSG00000099725.2 | 0 | NA | NA | NA | NA | NA | #VALUE! |
| ENSMUSG00000099726.2 | 0 | NA | NA | NA | NA | NA | #VALUE! |
| ENSMUSG00000099727.2 | 0 | NA | NA | NA | NA | NA | #VALUE! |
| ENSMUSG00000099728.2 | 0 | NA | NA | NA | NA | NA | #VALUE! |
| ENSMUSG00000099729.2 | 0 | NA | NA | NA | NA | NA | #VALUE! |
| ENSMUSG00000099730.2 | 0 | NA | NA | NA | NA | NA | #VALUE! |
| ENSMUSG00000099731.2 | 0 | NA | NA | NA | NA | NA | #VALUE! |
| ENSMUSG00000099732.3 | 0 | NA | NA | NA | NA | NA | #VALUE! |
| ENSMUSG00000099733.2 | 0 | NA | NA | NA | NA | NA | #VALUE! |
| ENSMUSG00000099734.2 | 0 | NA | NA | NA | NA | NA | #VALUE! |
| ENSMUSG00000099735.2 | 0 | NA | NA | NA | NA | NA | #VALUE! |
| ENSMUSG00000099736.2 | 0 | NA | NA | NA | NA | NA | #VALUE! |
| ENSMUSG00000099737.7 | 0 | NA | NA | NA | NA | NA | #VALUE! |
| ENSMUSG00000099738.2 | 0 | NA | NA | NA | NA | NA | #VALUE! |
| ENSMUSG00000099739.2 | 0 | NA | NA | NA | NA | NA | #VALUE! |
| ENSMUSG00000099740.2 | 0 | NA | NA | NA | NA | NA | #VALUE! |
| ENSMUSG00000099741.2 | 0 | NA | NA | NA | NA | NA | #VALUE! |
| ENSMUSG00000099742.2 | 0 | NA | NA | NA | NA | NA | #VALUE! |
| ENSMUSG00000099744.2 | 0 | NA | NA | NA | NA | NA | #VALUE! |
| ENSMUSG00000099745.2 | 0 | NA | NA | NA | NA | NA | #VALUE! |
| ENSMUSG00000099746.2 | 0 | NA | NA | NA | NA | NA | #VALUE! |
| ENSMUSG00000099747.2 | 0 | NA | NA | NA | NA | NA | #VALUE! |
| ENSMUSG00000099748.2 | 0 | NA | NA | NA | NA | NA | #VALUE! |
| ENSMUSG00000099749.2 | 0 | NA | NA | NA | NA | NA | #VALUE! |
| ENSMUSG00000099750.2 | 0 | NA | NA | NA | NA | NA | #VALUE! |
| ENSMUSG00000099751.7 | 0 | NA | NA | NA | NA | NA | #VALUE! |
| ENSMUSG00000099752.2 | 0 | NA | NA | NA | NA | NA | #VALUE! |
| ENSMUSG00000099753.2 | 0 | NA | NA | NA | NA | NA | #VALUE! |
| ENSMUSG00000099754.2 | 0 | NA | NA | NA | NA | NA | #VALUE! |
| ENSMUSG00000099755.2 | 0 | NA | NA | NA | NA | NA | #VALUE! |
| ENSMUSG00000099756.2 | 0 | NA | NA | NA | NA | NA | #VALUE! |
| ENSMUSG00000099757.2 | 0 | NA | NA | NA | NA | NA | #VALUE! |

| | | | | | | | |
|---|---|---|---|---|---|---|---|
| ENSMUSG00000099758.2 | 0 | NA | NA | NA | NA | NA | #VALUE! |
| ENSMUSG00000099759.2 | 8.644421 | 2.197888 | 2.175335 | 1.010368 | 0.312319 | NA | #VALUE! |
| ENSMUSG00000099760.2 | 0 | NA | NA | NA | NA | NA | #VALUE! |
| ENSMUSG00000099761.2 | 0 | NA | NA | NA | NA | NA | #VALUE! |
| ENSMUSG00000099762.4 | 0 | NA | NA | NA | NA | NA | #VALUE! |
| ENSMUSG00000099765.2 | 0 | NA | NA | NA | NA | NA | #VALUE! |
| ENSMUSG00000099766.2 | 0 | NA | NA | NA | NA | NA | #VALUE! |
| ENSMUSG00000099767.2 | 0 | NA | NA | NA | NA | NA | #VALUE! |
| ENSMUSG00000099768.2 | 0 | NA | NA | NA | NA | NA | #VALUE! |
| ENSMUSG00000099770.2 | 0 | NA | NA | NA | NA | NA | #VALUE! |
| ENSMUSG00000099771.2 | 0 | NA | NA | NA | NA | NA | #VALUE! |
| ENSMUSG00000099772.2 | 0 | NA | NA | NA | NA | NA | #VALUE! |
| ENSMUSG00000099773.2 | 0 | NA | NA | NA | NA | NA | #VALUE! |
| ENSMUSG00000099774.2 | 0 | NA | NA | NA | NA | NA | #VALUE! |
| ENSMUSG00000099775.2 | 0 | NA | NA | NA | NA | NA | #VALUE! |
| ENSMUSG00000099776.2 | 0 | NA | NA | NA | NA | NA | #VALUE! |
| ENSMUSG00000099777.2 | 0 | NA | NA | NA | NA | NA | #VALUE! |
| ENSMUSG00000099778.2 | 0 | NA | NA | NA | NA | NA | #VALUE! |
| ENSMUSG00000099779.2 | 0 | NA | NA | NA | NA | NA | #VALUE! |
| ENSMUSG00000099780.2 | 0 | NA | NA | NA | NA | NA | #VALUE! |
| ENSMUSG00000099781.2 | 0 | NA | NA | NA | NA | NA | #VALUE! |
| ENSMUSG00000099782.2 | 0 | NA | NA | NA | NA | NA | #VALUE! |
| ENSMUSG00000099783.2 | 0 | NA | NA | NA | NA | NA | #VALUE! |
| ENSMUSG00000099784.2 | 0 | NA | NA | NA | NA | NA | #VALUE! |
| ENSMUSG00000099785.7 | 0 | NA | NA | NA | NA | NA | #VALUE! |
| ENSMUSG00000099786.7 | 0 | NA | NA | NA | NA | NA | #VALUE! |
| ENSMUSG00000099787.4 | 0 | NA | NA | NA | NA | NA | #VALUE! |
| ENSMUSG00000099789.7 | 0 | NA | NA | NA | NA | NA | #VALUE! |
| ENSMUSG00000099790.2 | 0 | NA | NA | NA | NA | NA | #VALUE! |
| ENSMUSG00000099791.2 | 0 | NA | NA | NA | NA | NA | #VALUE! |
| ENSMUSG00000099792.2 | 0 | NA | NA | NA | NA | NA | #VALUE! |
| ENSMUSG00000099793.2 | 0 | NA | NA | NA | NA | NA | #VALUE! |
| ENSMUSG00000099794.2 | 0 | NA | NA | NA | NA | NA | #VALUE! |
| ENSMUSG00000099795.2 | 0 | NA | NA | NA | NA | NA | #VALUE! |
| ENSMUSG00000099796.2 | 0 | NA | NA | NA | NA | NA | #VALUE! |
| ENSMUSG00000099797.2 | 0 | NA | NA | NA | NA | NA | #VALUE! |
| ENSMUSG00000099798.2 | 0 | NA | NA | NA | NA | NA | #VALUE! |
| ENSMUSG00000099799.7 | 0 | NA | NA | NA | NA | NA | #VALUE! |
| ENSMUSG00000099800.2 | 0 | NA | NA | NA | NA | NA | #VALUE! |
| ENSMUSG00000099801.2 | 0 | NA | NA | NA | NA | NA | #VALUE! |
| ENSMUSG00000099802.2 | 0 | NA | NA | NA | NA | NA | #VALUE! |
| ENSMUSG00000099803.2 | 0 | NA | NA | NA | NA | NA | #VALUE! |
| ENSMUSG00000099804.2 | 0 | NA | NA | NA | NA | NA | #VALUE! |
| ENSMUSG00000099805.2 | 0 | NA | NA | NA | NA | NA | #VALUE! |

| | | | | | | | |
|---|---|---|---|---|---|---|---|
| ENSMUSG00000099806.2 | 0 | NA | NA | NA | NA | NA | #VALUE! |
| ENSMUSG00000099807.2 | 0 | NA | NA | NA | NA | NA | #VALUE! |
| ENSMUSG00000099808.3 | 0 | NA | NA | NA | NA | NA | #VALUE! |
| ENSMUSG00000099809.2 | 0 | NA | NA | NA | NA | NA | #VALUE! |
| ENSMUSG00000099810.2 | 0 | NA | NA | NA | NA | NA | #VALUE! |
| ENSMUSG00000099811.7 | 0 | NA | NA | NA | NA | NA | #VALUE! |
| ENSMUSG00000099812.2 | 0 | NA | NA | NA | NA | NA | #VALUE! |
| ENSMUSG00000099814.7 | 0 | NA | NA | NA | NA | NA | #VALUE! |
| ENSMUSG00000099815.2 | 0 | NA | NA | NA | NA | NA | #VALUE! |
| ENSMUSG00000099816.2 | 0 | NA | NA | NA | NA | NA | #VALUE! |
| ENSMUSG00000099817.2 | 0 | NA | NA | NA | NA | NA | #VALUE! |
| ENSMUSG00000099818.2 | 0 | NA | NA | NA | NA | NA | #VALUE! |
| ENSMUSG00000099819.2 | 0 | NA | NA | NA | NA | NA | #VALUE! |
| ENSMUSG00000099820.3 | 0 | NA | NA | NA | NA | NA | #VALUE! |
| ENSMUSG00000099821.2 | 0 | NA | NA | NA | NA | NA | #VALUE! |
| ENSMUSG00000099822.7 | 0 | NA | NA | NA | NA | NA | #VALUE! |
| ENSMUSG00000099823.2 | 0 | NA | NA | NA | NA | NA | #VALUE! |
| ENSMUSG00000099824.2 | 0 | NA | NA | NA | NA | NA | #VALUE! |
| ENSMUSG00000099825.2 | 0 | NA | NA | NA | NA | NA | #VALUE! |
| ENSMUSG00000099826.2 | 0 | NA | NA | NA | NA | NA | #VALUE! |
| ENSMUSG00000099827.2 | 0 | NA | NA | NA | NA | NA | #VALUE! |
| ENSMUSG00000099828.2 | 0 | NA | NA | NA | NA | NA | #VALUE! |
| ENSMUSG00000099829.2 | 0 | NA | NA | NA | NA | NA | #VALUE! |
| ENSMUSG00000099831.2 | 0 | NA | NA | NA | NA | NA | #VALUE! |
| ENSMUSG00000099832.2 | 0 | NA | NA | NA | NA | NA | #VALUE! |
| ENSMUSG00000099833.2 | 0 | NA | NA | NA | NA | NA | #VALUE! |
| ENSMUSG00000099834.2 | 0 | NA | NA | NA | NA | NA | #VALUE! |
| ENSMUSG00000099835.2 | 0 | NA | NA | NA | NA | NA | #VALUE! |
| ENSMUSG00000099836.2 | 0 | NA | NA | NA | NA | NA | #VALUE! |
| ENSMUSG00000099838.2 | 0 | NA | NA | NA | NA | NA | #VALUE! |
| ENSMUSG00000099839.2 | 0 | NA | NA | NA | NA | NA | #VALUE! |
| ENSMUSG00000099840.2 | 0 | NA | NA | NA | NA | NA | #VALUE! |
| ENSMUSG00000099841.2 | 0 | NA | NA | NA | NA | NA | #VALUE! |
| ENSMUSG00000099842.2 | 0 | NA | NA | NA | NA | NA | #VALUE! |
| ENSMUSG00000099843.2 | 9.73805 | 3.125645 | 2.610495 | 1.197338 | 0.231175 | NA | #VALUE! |
| ENSMUSG00000099845.2 | 0 | NA | NA | NA | NA | NA | #VALUE! |
| ENSMUSG00000099846.2 | 0 | NA | NA | NA | NA | NA | #VALUE! |
| ENSMUSG00000099847.2 | 0 | NA | NA | NA | NA | NA | #VALUE! |
| ENSMUSG00000099848.2 | 0 | NA | NA | NA | NA | NA | #VALUE! |
| ENSMUSG00000099849.2 | 0 | NA | NA | NA | NA | NA | #VALUE! |
| ENSMUSG00000099850.2 | 0 | NA | NA | NA | NA | NA | #VALUE! |
| ENSMUSG00000099851.2 | 0 | NA | NA | NA | NA | NA | #VALUE! |
| ENSMUSG00000099852.2 | 0 | NA | NA | NA | NA | NA | #VALUE! |
| ENSMUSG00000099853.4 | 0 | NA | NA | NA | NA | NA | #VALUE! |

| | | | | | | | |
|---|---|---|---|---|---|---|---|
| ENSMUSG00000099854.2 | 0 | NA | NA | NA | NA | NA | #VALUE! |
| ENSMUSG00000099855.7 | 0 | NA | NA | NA | NA | NA | #VALUE! |
| ENSMUSG00000099856.2 | 0 | NA | NA | NA | NA | NA | #VALUE! |
| ENSMUSG00000099858.2 | 0 | NA | NA | NA | NA | NA | #VALUE! |
| ENSMUSG00000099860.2 | 0 | NA | NA | NA | NA | NA | #VALUE! |
| ENSMUSG00000099861.2 | 0 | NA | NA | NA | NA | NA | #VALUE! |
| ENSMUSG00000099862.2 | 0 | NA | NA | NA | NA | NA | #VALUE! |
| ENSMUSG00000099863.2 | 0 | NA | NA | NA | NA | NA | #VALUE! |
| ENSMUSG00000099864.2 | 0 | NA | NA | NA | NA | NA | #VALUE! |
| ENSMUSG00000099865.2 | 0 | NA | NA | NA | NA | NA | #VALUE! |
| ENSMUSG00000099866.7 | 0 | NA | NA | NA | NA | NA | #VALUE! |
| ENSMUSG00000099867.2 | 0 | NA | NA | NA | NA | NA | #VALUE! |
| ENSMUSG00000099868.2 | 0 | NA | NA | NA | NA | NA | #VALUE! |
| ENSMUSG00000099869.7 | 0 | NA | NA | NA | NA | NA | #VALUE! |
| ENSMUSG00000099870.2 | 0 | NA | NA | NA | NA | NA | #VALUE! |
| ENSMUSG00000099871.2 | 0 | NA | NA | NA | NA | NA | #VALUE! |
| ENSMUSG00000099872.2 | 0 | NA | NA | NA | NA | NA | #VALUE! |
| ENSMUSG00000099873.2 | 0 | NA | NA | NA | NA | NA | #VALUE! |
| ENSMUSG00000099874.2 | 5.968332 | 6.072569 | 3.914265 | 1.551394 | 0.120807 | NA | #VALUE! |
| ENSMUSG00000099875.2 | 0 | NA | NA | NA | NA | NA | #VALUE! |
| ENSMUSG00000099876.2 | 0 | NA | NA | NA | NA | NA | #VALUE! |
| ENSMUSG00000099877.7 | 0 | NA | NA | NA | NA | NA | #VALUE! |
| ENSMUSG00000099878.2 | 0 | NA | NA | NA | NA | NA | #VALUE! |
| ENSMUSG00000099879.2 | 0 | NA | NA | NA | NA | NA | #VALUE! |
| ENSMUSG00000099880.2 | 0 | NA | NA | NA | NA | NA | #VALUE! |
| ENSMUSG00000099882.2 | 0 | NA | NA | NA | NA | NA | #VALUE! |
| ENSMUSG00000099883.2 | 0 | NA | NA | NA | NA | NA | #VALUE! |
| ENSMUSG00000099884.2 | 0 | NA | NA | NA | NA | NA | #VALUE! |
| ENSMUSG00000099886.2 | 0 | NA | NA | NA | NA | NA | #VALUE! |
| ENSMUSG00000099888.2 | 0 | NA | NA | NA | NA | NA | #VALUE! |
| ENSMUSG00000099889.2 | 0 | NA | NA | NA | NA | NA | #VALUE! |
| ENSMUSG00000099890.3 | 0 | NA | NA | NA | NA | NA | #VALUE! |
| ENSMUSG00000099891.2 | 0 | NA | NA | NA | NA | NA | #VALUE! |
| ENSMUSG00000099892.2 | 0 | NA | NA | NA | NA | NA | #VALUE! |
| ENSMUSG00000099893.2 | 0 | NA | NA | NA | NA | NA | #VALUE! |
| ENSMUSG00000099894.2 | 0 | NA | NA | NA | NA | NA | #VALUE! |
| ENSMUSG00000099895.2 | 0 | NA | NA | NA | NA | NA | #VALUE! |
| ENSMUSG00000099896.2 | 0 | NA | NA | NA | NA | NA | #VALUE! |
| ENSMUSG00000099897.7 | 0 | NA | NA | NA | NA | NA | #VALUE! |
| ENSMUSG00000099898.2 | 0 | NA | NA | NA | NA | NA | #VALUE! |
| ENSMUSG00000099899.2 | 0 | NA | NA | NA | NA | NA | #VALUE! |
| ENSMUSG00000099900.2 | 0 | NA | NA | NA | NA | NA | #VALUE! |
| ENSMUSG00000099901.2 | 0 | NA | NA | NA | NA | NA | #VALUE! |
| ENSMUSG00000099902.2 | 0.336567 | 1.944925 | 4.029936 | 0.482619 | 0.629366 | NA | #VALUE! |

| | | | | | | | |
|---|---|---|---|---|---|---|---|
| ENSMUSG00000099903.2 | 0 | NA | NA | NA | NA | NA | #VALUE! |
| ENSMUSG00000099904.2 | 0 | NA | NA | NA | NA | NA | #VALUE! |
| ENSMUSG00000099905.2 | 0 | NA | NA | NA | NA | NA | #VALUE! |
| ENSMUSG00000099906.3 | 0 | NA | NA | NA | NA | NA | #VALUE! |
| ENSMUSG00000099907.2 | 0 | NA | NA | NA | NA | NA | #VALUE! |
| ENSMUSG00000099908.2 | 0 | NA | NA | NA | NA | NA | #VALUE! |
| ENSMUSG00000099910.2 | 0 | NA | NA | NA | NA | NA | #VALUE! |
| ENSMUSG00000099911.2 | 0 | NA | NA | NA | NA | NA | #VALUE! |
| ENSMUSG00000099912.2 | 0 | NA | NA | NA | NA | NA | #VALUE! |
| ENSMUSG00000099913.2 | 0.491951 | 2.402185 | 4.000147 | 0.600524 | 0.548157 | NA | #VALUE! |
| ENSMUSG00000099914.2 | 0 | NA | NA | NA | NA | NA | #VALUE! |
| ENSMUSG00000099915.2 | 0 | NA | NA | NA | NA | NA | #VALUE! |
| ENSMUSG00000099916.7 | 0 | NA | NA | NA | NA | NA | #VALUE! |
| ENSMUSG00000099917.3 | 0 | NA | NA | NA | NA | NA | #VALUE! |
| ENSMUSG00000099918.2 | 0 | NA | NA | NA | NA | NA | #VALUE! |
| ENSMUSG00000099920.2 | 0 | NA | NA | NA | NA | NA | #VALUE! |
| ENSMUSG00000099921.2 | 0 | NA | NA | NA | NA | NA | #VALUE! |
| ENSMUSG00000099922.2 | 0 | NA | NA | NA | NA | NA | #VALUE! |
| ENSMUSG00000099923.3 | 0 | NA | NA | NA | NA | NA | #VALUE! |
| ENSMUSG00000099924.2 | 5.286306 | -5.93001 | 3.915244 | -1.5146 | 0.129875 | NA | #VALUE! |
| ENSMUSG00000099925.2 | 0 | NA | NA | NA | NA | NA | #VALUE! |
| ENSMUSG00000099926.2 | 0 | NA | NA | NA | NA | NA | #VALUE! |
| ENSMUSG00000099927.2 | 0.105013 | -0.97324 | 4.076293 | -0.23876 | 0.811295 | NA | #VALUE! |
| ENSMUSG00000099928.2 | 0 | NA | NA | NA | NA | NA | #VALUE! |
| ENSMUSG00000099929.2 | 0 | NA | NA | NA | NA | NA | #VALUE! |
| ENSMUSG00000099930.2 | 0 | NA | NA | NA | NA | NA | #VALUE! |
| ENSMUSG00000099931.2 | 0 | NA | NA | NA | NA | NA | #VALUE! |
| ENSMUSG00000099932.2 | 0 | NA | NA | NA | NA | NA | #VALUE! |
| ENSMUSG00000099933.2 | 0 | NA | NA | NA | NA | NA | #VALUE! |
| ENSMUSG00000099934.2 | 0 | NA | NA | NA | NA | NA | #VALUE! |
| ENSMUSG00000099935.2 | 0 | NA | NA | NA | NA | NA | #VALUE! |
| ENSMUSG00000099936.2 | 0 | NA | NA | NA | NA | NA | #VALUE! |
| ENSMUSG00000099937.2 | 0 | NA | NA | NA | NA | NA | #VALUE! |
| ENSMUSG00000099938.2 | 0 | NA | NA | NA | NA | NA | #VALUE! |
| ENSMUSG00000099939.2 | 0 | NA | NA | NA | NA | NA | #VALUE! |
| ENSMUSG00000099940.2 | 0 | NA | NA | NA | NA | NA | #VALUE! |
| ENSMUSG00000099941.2 | 0 | NA | NA | NA | NA | NA | #VALUE! |
| ENSMUSG00000099942.2 | 0 | NA | NA | NA | NA | NA | #VALUE! |
| ENSMUSG00000099943.2 | 0 | NA | NA | NA | NA | NA | #VALUE! |
| ENSMUSG00000099944.2 | 0 | NA | NA | NA | NA | NA | #VALUE! |
| ENSMUSG00000099945.2 | 0 | NA | NA | NA | NA | NA | #VALUE! |
| ENSMUSG00000099946.2 | 0 | NA | NA | NA | NA | NA | #VALUE! |
| ENSMUSG00000099947.2 | 0 | NA | NA | NA | NA | NA | #VALUE! |
| ENSMUSG00000099948.2 | 0 | NA | NA | NA | NA | NA | #VALUE! |

| | | | | | | | |
|---|---|---|---|---|---|---|---|
| ENSMUSG00000099949.2 | 0 | NA | NA | NA | NA | NA | #VALUE! |
| ENSMUSG00000099950.2 | 0 | NA | NA | NA | NA | NA | #VALUE! |
| ENSMUSG00000099951.2 | 0 | NA | NA | NA | NA | NA | #VALUE! |
| ENSMUSG00000099952.2 | 0 | NA | NA | NA | NA | NA | #VALUE! |
| ENSMUSG00000099953.2 | 0 | NA | NA | NA | NA | NA | #VALUE! |
| ENSMUSG00000099954.2 | 0 | NA | NA | NA | NA | NA | #VALUE! |
| ENSMUSG00000099955.2 | 0 | NA | NA | NA | NA | NA | #VALUE! |
| ENSMUSG00000099956.2 | 0 | NA | NA | NA | NA | NA | #VALUE! |
| ENSMUSG00000099957.5 | 0 | NA | NA | NA | NA | NA | #VALUE! |
| ENSMUSG00000099958.2 | 0 | NA | NA | NA | NA | NA | #VALUE! |
| ENSMUSG00000099959.2 | 0 | NA | NA | NA | NA | NA | #VALUE! |
| ENSMUSG00000099960.2 | 0 | NA | NA | NA | NA | NA | #VALUE! |
| ENSMUSG00000099961.2 | 0 | NA | NA | NA | NA | NA | #VALUE! |
| ENSMUSG00000099962.2 | 0 | NA | NA | NA | NA | NA | #VALUE! |
| ENSMUSG00000099963.7 | 0 | NA | NA | NA | NA | NA | #VALUE! |
| ENSMUSG00000099964.7 | 0 | NA | NA | NA | NA | NA | #VALUE! |
| ENSMUSG00000099965.2 | 0 | NA | NA | NA | NA | NA | #VALUE! |
| ENSMUSG00000099967.2 | 0 | NA | NA | NA | NA | NA | #VALUE! |
| ENSMUSG00000099968.8 | 0 | NA | NA | NA | NA | NA | #VALUE! |
| ENSMUSG00000099969.2 | 0 | NA | NA | NA | NA | NA | #VALUE! |
| ENSMUSG00000099970.2 | 0 | NA | NA | NA | NA | NA | #VALUE! |
| ENSMUSG00000099971.2 | 0 | NA | NA | NA | NA | NA | #VALUE! |
| ENSMUSG00000099972.2 | 0 | NA | NA | NA | NA | NA | #VALUE! |
| ENSMUSG00000099973.2 | 0 | NA | NA | NA | NA | NA | #VALUE! |
| ENSMUSG00000099974.2 | 0.630079 | -2.90291 | 3.975047 | -0.73028 | 0.465217 | NA | #VALUE! |
| ENSMUSG00000099975.2 | 0 | NA | NA | NA | NA | NA | #VALUE! |
| ENSMUSG00000099976.2 | 0 | NA | NA | NA | NA | NA | #VALUE! |
| ENSMUSG00000099977.7 | 0 | NA | NA | NA | NA | NA | #VALUE! |
| ENSMUSG00000099978.2 | 0 | NA | NA | NA | NA | NA | #VALUE! |
| ENSMUSG00000099979.2 | 0 | NA | NA | NA | NA | NA | #VALUE! |
| ENSMUSG00000099980.2 | 0 | NA | NA | NA | NA | NA | #VALUE! |
| ENSMUSG00000099981.2 | 0 | NA | NA | NA | NA | NA | #VALUE! |
| ENSMUSG00000099982.2 | 0 | NA | NA | NA | NA | NA | #VALUE! |
| ENSMUSG00000099983.2 | 0 | NA | NA | NA | NA | NA | #VALUE! |
| ENSMUSG00000099985.2 | 0 | NA | NA | NA | NA | NA | #VALUE! |
| ENSMUSG00000099986.2 | 0 | NA | NA | NA | NA | NA | #VALUE! |
| ENSMUSG00000099988.8 | 0 | NA | NA | NA | NA | NA | #VALUE! |
| ENSMUSG00000099989.2 | 0.245976 | 1.027628 | 4.067126 | 0.252667 | 0.800526 | NA | #VALUE! |
| ENSMUSG00000099990.7 | 0 | NA | NA | NA | NA | NA | #VALUE! |
| ENSMUSG00000099991.2 | 0 | NA | NA | NA | NA | NA | #VALUE! |
| ENSMUSG00000099992.2 | 0 | NA | NA | NA | NA | NA | #VALUE! |
| ENSMUSG00000099993.2 | 0 | NA | NA | NA | NA | NA | #VALUE! |
| ENSMUSG00000099994.2 | 0 | NA | NA | NA | NA | NA | #VALUE! |
| ENSMUSG00000099995.2 | 0 | NA | NA | NA | NA | NA | #VALUE! |

| | | | | | | | |
|---|---|---|---|---|---|---|---|
| ENSMUSG00000099996.2 | 0 | NA | NA | NA | NA | NA | #VALUE! |
| ENSMUSG00000099997.2 | 0 | NA | NA | NA | NA | NA | #VALUE! |
| ENSMUSG00000099998.2 | 0 | NA | NA | NA | NA | NA | #VALUE! |
| ENSMUSG00000099999.2 | 0 | NA | NA | NA | NA | NA | #VALUE! |
| ENSMUSG00000100000.7 | 0 | NA | NA | NA | NA | NA | #VALUE! |
| ENSMUSG00000100001.2 | 0 | NA | NA | NA | NA | NA | #VALUE! |
| ENSMUSG00000100002.3 | 5.222291 | 5.88069 | 3.915331 | 1.501965 | 0.133106 | NA | #VALUE! |
| ENSMUSG00000100003.2 | 0 | NA | NA | NA | NA | NA | #VALUE! |
| ENSMUSG00000100004.2 | 0 | NA | NA | NA | NA | NA | #VALUE! |
| ENSMUSG00000100005.8 | 1.305573 | 3.898287 | 3.940396 | 0.989314 | 0.32251 | NA | #VALUE! |
| ENSMUSG00000100006.2 | 0 | NA | NA | NA | NA | NA | #VALUE! |
| ENSMUSG00000100007.2 | 0 | NA | NA | NA | NA | NA | #VALUE! |
| ENSMUSG00000100008.2 | 0 | NA | NA | NA | NA | NA | #VALUE! |
| ENSMUSG00000100009.7 | 0 | NA | NA | NA | NA | NA | #VALUE! |
| ENSMUSG00000100010.6 | 0 | NA | NA | NA | NA | NA | #VALUE! |
| ENSMUSG00000100011.2 | 0 | NA | NA | NA | NA | NA | #VALUE! |
| ENSMUSG00000100012.2 | 0 | NA | NA | NA | NA | NA | #VALUE! |
| ENSMUSG00000100013.2 | 0 | NA | NA | NA | NA | NA | #VALUE! |
| ENSMUSG00000100014.2 | 0 | NA | NA | NA | NA | NA | #VALUE! |
| ENSMUSG00000100015.2 | 0 | NA | NA | NA | NA | NA | #VALUE! |
| ENSMUSG00000100016.4 | 0 | NA | NA | NA | NA | NA | #VALUE! |
| ENSMUSG00000100017.3 | 8.866517 | 6.640196 | 2.804106 | 2.368026 | 0.017883 | NA | #VALUE! |
| ENSMUSG00000100018.2 | 0.300802 | -1.80634 | 4.050033 | -0.44601 | 0.655592 | NA | #VALUE! |
| ENSMUSG00000100019.2 | 0 | NA | NA | NA | NA | NA | #VALUE! |
| ENSMUSG00000100020.2 | 0 | NA | NA | NA | NA | NA | #VALUE! |
| ENSMUSG00000100021.2 | 0 | NA | NA | NA | NA | NA | #VALUE! |
| ENSMUSG00000100022.3 | 0 | NA | NA | NA | NA | NA | #VALUE! |
| ENSMUSG00000100023.2 | 0 | NA | NA | NA | NA | NA | #VALUE! |
| ENSMUSG00000100024.2 | 0 | NA | NA | NA | NA | NA | #VALUE! |
| ENSMUSG00000100025.2 | 0 | NA | NA | NA | NA | NA | #VALUE! |
| ENSMUSG00000100026.2 | 0 | NA | NA | NA | NA | NA | #VALUE! |
| ENSMUSG00000100027.2 | 0 | NA | NA | NA | NA | NA | #VALUE! |
| ENSMUSG00000100028.2 | 0 | NA | NA | NA | NA | NA | #VALUE! |
| ENSMUSG00000100029.2 | 0 | NA | NA | NA | NA | NA | #VALUE! |
| ENSMUSG00000100031.2 | 0 | NA | NA | NA | NA | NA | #VALUE! |
| ENSMUSG00000100032.2 | 0 | NA | NA | NA | NA | NA | #VALUE! |
| ENSMUSG00000100033.2 | 0 | NA | NA | NA | NA | NA | #VALUE! |
| ENSMUSG00000100034.2 | 0 | NA | NA | NA | NA | NA | #VALUE! |
| ENSMUSG00000100035.2 | 0 | NA | NA | NA | NA | NA | #VALUE! |
| ENSMUSG00000100036.2 | 0 | NA | NA | NA | NA | NA | #VALUE! |
| ENSMUSG00000100037.2 | 0 | NA | NA | NA | NA | NA | #VALUE! |
| ENSMUSG00000100038.2 | 0 | NA | NA | NA | NA | NA | #VALUE! |
| ENSMUSG00000100039.2 | 18.00502 | 4.298392 | 2.762329 | 1.556075 | NA | NA | #VALUE! |
| ENSMUSG00000100040.2 | 0 | NA | NA | NA | NA | NA | #VALUE! |

| | | | | | | | |
|---|---|---|---|---|---|---|---|
| ENSMUSG00000100041.2 | 0 | NA | NA | NA | NA | NA | #VALUE! |
| ENSMUSG00000100042.2 | 0 | NA | NA | NA | NA | NA | #VALUE! |
| ENSMUSG00000100043.2 | 0 | NA | NA | NA | NA | NA | #VALUE! |
| ENSMUSG00000100044.2 | 0 | NA | NA | NA | NA | NA | #VALUE! |
| ENSMUSG00000100045.2 | 0 | NA | NA | NA | NA | NA | #VALUE! |
| ENSMUSG00000100046.2 | 0 | NA | NA | NA | NA | NA | #VALUE! |
| ENSMUSG00000100047.2 | 0 | NA | NA | NA | NA | NA | #VALUE! |
| ENSMUSG00000100048.2 | 0 | NA | NA | NA | NA | NA | #VALUE! |
| ENSMUSG00000100049.2 | 0 | NA | NA | NA | NA | NA | #VALUE! |
| ENSMUSG00000100050.2 | 0 | NA | NA | NA | NA | NA | #VALUE! |
| ENSMUSG00000100051.2 | 0 | NA | NA | NA | NA | NA | #VALUE! |
| ENSMUSG00000100052.2 | 0 | NA | NA | NA | NA | NA | #VALUE! |
| ENSMUSG00000100053.2 | 0 | NA | NA | NA | NA | NA | #VALUE! |
| ENSMUSG00000100054.7 | 0 | NA | NA | NA | NA | NA | #VALUE! |
| ENSMUSG00000100055.2 | 0 | NA | NA | NA | NA | NA | #VALUE! |
| ENSMUSG00000100056.2 | 0 | NA | NA | NA | NA | NA | #VALUE! |
| ENSMUSG00000100057.7 | 0 | NA | NA | NA | NA | NA | #VALUE! |
| ENSMUSG00000100058.2 | 0 | NA | NA | NA | NA | NA | #VALUE! |
| ENSMUSG00000100059.2 | 0 | NA | NA | NA | NA | NA | #VALUE! |
| ENSMUSG00000100060.2 | 0 | NA | NA | NA | NA | NA | #VALUE! |
| ENSMUSG00000100061.2 | 0 | NA | NA | NA | NA | NA | #VALUE! |
| ENSMUSG00000100062.2 | 0 | NA | NA | NA | NA | NA | #VALUE! |
| ENSMUSG00000100063.2 | 0 | NA | NA | NA | NA | NA | #VALUE! |
| ENSMUSG00000100064.2 | 0 | NA | NA | NA | NA | NA | #VALUE! |
| ENSMUSG00000100065.2 | 0 | NA | NA | NA | NA | NA | #VALUE! |
| ENSMUSG00000100066.2 | 0 | NA | NA | NA | NA | NA | #VALUE! |
| ENSMUSG00000100067.2 | 0 | NA | NA | NA | NA | NA | #VALUE! |
| ENSMUSG00000100068.2 | 0 | NA | NA | NA | NA | NA | #VALUE! |
| ENSMUSG00000100069.2 | 0 | NA | NA | NA | NA | NA | #VALUE! |
| ENSMUSG00000100070.7 | 0 | NA | NA | NA | NA | NA | #VALUE! |
| ENSMUSG00000100071.2 | 0 | NA | NA | NA | NA | NA | #VALUE! |
| ENSMUSG00000100072.2 | 0 | NA | NA | NA | NA | NA | #VALUE! |
| ENSMUSG00000100073.2 | 0 | NA | NA | NA | NA | NA | #VALUE! |
| ENSMUSG00000100074.2 | 0 | NA | NA | NA | NA | NA | #VALUE! |
| ENSMUSG00000100076.2 | 0 | NA | NA | NA | NA | NA | #VALUE! |
| ENSMUSG00000100077.2 | 0 | NA | NA | NA | NA | NA | #VALUE! |
| ENSMUSG00000100078.2 | 0.093255 | 0.868093 | 4.072464 | 0.213162 | 0.831201 | NA | #VALUE! |
| ENSMUSG00000100079.2 | 0 | NA | NA | NA | NA | NA | #VALUE! |
| ENSMUSG00000100080.2 | 0 | NA | NA | NA | NA | NA | #VALUE! |
| ENSMUSG00000100081.2 | 0 | NA | NA | NA | NA | NA | #VALUE! |
| ENSMUSG00000100082.2 | 0 | NA | NA | NA | NA | NA | #VALUE! |
| ENSMUSG00000100083.7 | 0 | NA | NA | NA | NA | NA | #VALUE! |
| ENSMUSG00000100084.3 | 0 | NA | NA | NA | NA | NA | #VALUE! |
| ENSMUSG00000100085.2 | 0 | NA | NA | NA | NA | NA | #VALUE! |

| | | | | | | | |
|---|---|---|---|---|---|---|---|
| ENSMUSG00000100086.2 | 0 | NA | NA | NA | NA | NA | #VALUE! |
| ENSMUSG00000100087.2 | 0 | NA | NA | NA | NA | NA | #VALUE! |
| ENSMUSG00000100088.2 | 0 | NA | NA | NA | NA | NA | #VALUE! |
| ENSMUSG00000100089.2 | 0 | NA | NA | NA | NA | NA | #VALUE! |
| ENSMUSG00000100090.2 | 0 | NA | NA | NA | NA | NA | #VALUE! |
| ENSMUSG00000100091.2 | 0 | NA | NA | NA | NA | NA | #VALUE! |
| ENSMUSG00000100092.2 | 0 | NA | NA | NA | NA | NA | #VALUE! |
| ENSMUSG00000100093.2 | 0 | NA | NA | NA | NA | NA | #VALUE! |
| ENSMUSG00000100095.2 | 0 | NA | NA | NA | NA | NA | #VALUE! |
| ENSMUSG00000100096.2 | 0 | NA | NA | NA | NA | NA | #VALUE! |
| ENSMUSG00000100097.7 | 0 | NA | NA | NA | NA | NA | #VALUE! |
| ENSMUSG00000100098.2 | 0 | NA | NA | NA | NA | NA | #VALUE! |
| ENSMUSG00000100099.2 | 0 | NA | NA | NA | NA | NA | #VALUE! |
| ENSMUSG00000100100.2 | 0 | NA | NA | NA | NA | NA | #VALUE! |
| ENSMUSG00000100101.2 | 0 | NA | NA | NA | NA | NA | #VALUE! |
| ENSMUSG00000100103.2 | 0 | NA | NA | NA | NA | NA | #VALUE! |
| ENSMUSG00000100104.2 | 0 | NA | NA | NA | NA | NA | #VALUE! |
| ENSMUSG00000100105.2 | 0 | NA | NA | NA | NA | NA | #VALUE! |
| ENSMUSG00000100106.2 | 0 | NA | NA | NA | NA | NA | #VALUE! |
| ENSMUSG00000100107.2 | 0 | NA | NA | NA | NA | NA | #VALUE! |
| ENSMUSG00000100108.2 | 0 | NA | NA | NA | NA | NA | #VALUE! |
| ENSMUSG00000100109.2 | 0 | NA | NA | NA | NA | NA | #VALUE! |
| ENSMUSG00000100110.2 | 0 | NA | NA | NA | NA | NA | #VALUE! |
| ENSMUSG00000100111.3 | 0 | NA | NA | NA | NA | NA | #VALUE! |
| ENSMUSG00000100112.2 | 0 | NA | NA | NA | NA | NA | #VALUE! |
| ENSMUSG00000100113.2 | 4.711939 | 5.72619 | 3.916297 | 1.462144 | 0.143702 | NA | #VALUE! |
| ENSMUSG00000100115.2 | 0 | NA | NA | NA | NA | NA | #VALUE! |
| ENSMUSG00000100116.2 | 0 | NA | NA | NA | NA | NA | #VALUE! |
| ENSMUSG00000100117.2 | 0 | NA | NA | NA | NA | NA | #VALUE! |
| ENSMUSG00000100118.2 | 0 | NA | NA | NA | NA | NA | #VALUE! |
| ENSMUSG00000100119.2 | 0 | NA | NA | NA | NA | NA | #VALUE! |
| ENSMUSG00000100120.2 | 0 | NA | NA | NA | NA | NA | #VALUE! |
| ENSMUSG00000100121.2 | 0 | NA | NA | NA | NA | NA | #VALUE! |
| ENSMUSG00000100122.2 | 0 | NA | NA | NA | NA | NA | #VALUE! |
| ENSMUSG00000100123.2 | 0 | NA | NA | NA | NA | NA | #VALUE! |
| ENSMUSG00000100124.2 | 0 | NA | NA | NA | NA | NA | #VALUE! |
| ENSMUSG00000100125.2 | 0 | NA | NA | NA | NA | NA | #VALUE! |
| ENSMUSG00000100126.2 | 0 | NA | NA | NA | NA | NA | #VALUE! |
| ENSMUSG00000100127.7 | 0 | NA | NA | NA | NA | NA | #VALUE! |
| ENSMUSG00000100128.2 | 0 | NA | NA | NA | NA | NA | #VALUE! |
| ENSMUSG00000100129.2 | 0 | NA | NA | NA | NA | NA | #VALUE! |
| ENSMUSG00000100130.2 | 0 | NA | NA | NA | NA | NA | #VALUE! |
| ENSMUSG00000100131.2 | 2.189124 | 2.602928 | 3.89689 | 0.66795 | 0.504165 | NA | #VALUE! |
| ENSMUSG00000100132.2 | 0 | NA | NA | NA | NA | NA | #VALUE! |

| | | | | | | | |
|---|---|---|---|---|---|---|---|
| ENSMUSG00000100133.2 | 0 | NA | NA | NA | NA | NA | #VALUE! |
| ENSMUSG00000100134.2 | 0 | NA | NA | NA | NA | NA | #VALUE! |
| ENSMUSG00000100135.2 | 0 | NA | NA | NA | NA | NA | #VALUE! |
| ENSMUSG00000100136.2 | 0 | NA | NA | NA | NA | NA | #VALUE! |
| ENSMUSG00000100137.2 | 0 | NA | NA | NA | NA | NA | #VALUE! |
| ENSMUSG00000100138.2 | 0 | NA | NA | NA | NA | NA | #VALUE! |
| ENSMUSG00000100140.2 | 0 | NA | NA | NA | NA | NA | #VALUE! |
| ENSMUSG00000100141.2 | 1.009701 | 3.503687 | 3.950852 | 0.886818 | 0.375177 | NA | #VALUE! |
| ENSMUSG00000100142.2 | 0 | NA | NA | NA | NA | NA | #VALUE! |
| ENSMUSG00000100144.7 | 0 | NA | NA | NA | NA | NA | #VALUE! |
| ENSMUSG00000100145.2 | 0 | NA | NA | NA | NA | NA | #VALUE! |
| ENSMUSG00000100146.2 | 0 | NA | NA | NA | NA | NA | #VALUE! |
| ENSMUSG00000100147.3 | 0 | NA | NA | NA | NA | NA | #VALUE! |
| ENSMUSG00000100148.2 | 0 | NA | NA | NA | NA | NA | #VALUE! |
| ENSMUSG00000100149.2 | 0 | NA | NA | NA | NA | NA | #VALUE! |
| ENSMUSG00000100150.3 | 2.310289 | -4.7496 | 3.925939 | -1.2098 | 0.226355 | NA | #VALUE! |
| ENSMUSG00000100151.2 | 0 | NA | NA | NA | NA | NA | #VALUE! |
| ENSMUSG00000100152.2 | 0 | NA | NA | NA | NA | NA | #VALUE! |
| ENSMUSG00000100154.2 | 0 | NA | NA | NA | NA | NA | #VALUE! |
| ENSMUSG00000100155.2 | 0 | NA | NA | NA | NA | NA | #VALUE! |
| ENSMUSG00000100156.2 | 0 | NA | NA | NA | NA | NA | #VALUE! |
| ENSMUSG00000100157.7 | 0 | NA | NA | NA | NA | NA | #VALUE! |
| ENSMUSG00000100158.2 | 0 | NA | NA | NA | NA | NA | #VALUE! |
| ENSMUSG00000100159.2 | 0 | NA | NA | NA | NA | NA | #VALUE! |
| ENSMUSG00000100160.2 | 0 | NA | NA | NA | NA | NA | #VALUE! |
| ENSMUSG00000100161.2 | 0 | NA | NA | NA | NA | NA | #VALUE! |
| ENSMUSG00000100162.3 | 0 | NA | NA | NA | NA | NA | #VALUE! |
| ENSMUSG00000100165.2 | 0 | NA | NA | NA | NA | NA | #VALUE! |
| ENSMUSG00000100166.2 | 0 | NA | NA | NA | NA | NA | #VALUE! |
| ENSMUSG00000100167.7 | 0 | NA | NA | NA | NA | NA | #VALUE! |
| ENSMUSG00000100168.3 | 0 | NA | NA | NA | NA | NA | #VALUE! |
| ENSMUSG00000100169.2 | 0 | NA | NA | NA | NA | NA | #VALUE! |
| ENSMUSG00000100170.2 | 0 | NA | NA | NA | NA | NA | #VALUE! |
| ENSMUSG00000100171.2 | 0 | NA | NA | NA | NA | NA | #VALUE! |
| ENSMUSG00000100172.2 | 0 | NA | NA | NA | NA | NA | #VALUE! |
| ENSMUSG00000100174.2 | 0 | NA | NA | NA | NA | NA | #VALUE! |
| ENSMUSG00000100175.7 | 0 | NA | NA | NA | NA | NA | #VALUE! |
| ENSMUSG00000100176.2 | 0 | NA | NA | NA | NA | NA | #VALUE! |
| ENSMUSG00000100177.2 | 0 | NA | NA | NA | NA | NA | #VALUE! |
| ENSMUSG00000100178.2 | 0 | NA | NA | NA | NA | NA | #VALUE! |
| ENSMUSG00000100179.2 | 0 | NA | NA | NA | NA | NA | #VALUE! |
| ENSMUSG00000100180.2 | 0 | NA | NA | NA | NA | NA | #VALUE! |
| ENSMUSG00000100182.7 | 0 | NA | NA | NA | NA | NA | #VALUE! |
| ENSMUSG00000100183.2 | 0 | NA | NA | NA | NA | NA | #VALUE! |

| | | | | | | | |
|---|---|---|---|---|---|---|---|
| ENSMUSG00000100184.2 | 0 | NA | NA | NA | NA | NA | #VALUE! |
| ENSMUSG00000100185.2 | 0 | NA | NA | NA | NA | NA | #VALUE! |
| ENSMUSG00000100186.2 | 0 | NA | NA | NA | NA | NA | #VALUE! |
| ENSMUSG00000100187.2 | 0 | NA | NA | NA | NA | NA | #VALUE! |
| ENSMUSG00000100188.2 | 0 | NA | NA | NA | NA | NA | #VALUE! |
| ENSMUSG00000100189.2 | 0 | NA | NA | NA | NA | NA | #VALUE! |
| ENSMUSG00000100190.2 | 0 | NA | NA | NA | NA | NA | #VALUE! |
| ENSMUSG00000100191.2 | 0 | NA | NA | NA | NA | NA | #VALUE! |
| ENSMUSG00000100192.2 | 0 | NA | NA | NA | NA | NA | #VALUE! |
| ENSMUSG00000100193.2 | 0 | NA | NA | NA | NA | NA | #VALUE! |
| ENSMUSG00000100194.2 | 0 | NA | NA | NA | NA | NA | #VALUE! |
| ENSMUSG00000100195.2 | 0 | NA | NA | NA | NA | NA | #VALUE! |
| ENSMUSG00000100196.2 | 0 | NA | NA | NA | NA | NA | #VALUE! |
| ENSMUSG00000100197.2 | 0 | NA | NA | NA | NA | NA | #VALUE! |
| ENSMUSG00000100198.2 | 0 | NA | NA | NA | NA | NA | #VALUE! |
| ENSMUSG00000100199.2 | 4.506076 | 2.185416 | 3.795237 | 0.575831 | 0.564729 | NA | #VALUE! |
| ENSMUSG00000100200.2 | 0 | NA | NA | NA | NA | NA | #VALUE! |
| ENSMUSG00000100201.2 | 0 | NA | NA | NA | NA | NA | #VALUE! |
| ENSMUSG00000100202.2 | 0 | NA | NA | NA | NA | NA | #VALUE! |
| ENSMUSG00000100203.2 | 0 | NA | NA | NA | NA | NA | #VALUE! |
| ENSMUSG00000100204.2 | 0 | NA | NA | NA | NA | NA | #VALUE! |
| ENSMUSG00000100205.2 | 0 | NA | NA | NA | NA | NA | #VALUE! |
| ENSMUSG00000100206.2 | 0 | NA | NA | NA | NA | NA | #VALUE! |
| ENSMUSG00000100207.2 | 0 | NA | NA | NA | NA | NA | #VALUE! |
| ENSMUSG00000100209.2 | 0 | NA | NA | NA | NA | NA | #VALUE! |
| ENSMUSG00000100210.3 | 0 | NA | NA | NA | NA | NA | #VALUE! |
| ENSMUSG00000100211.2 | 0 | NA | NA | NA | NA | NA | #VALUE! |
| ENSMUSG00000100212.2 | 0 | NA | NA | NA | NA | NA | #VALUE! |
| ENSMUSG00000100213.2 | 0 | NA | NA | NA | NA | NA | #VALUE! |
| ENSMUSG00000100215.2 | 0.934036 | 0.912864 | 3.93471 | 0.232003 | 0.816536 | NA | #VALUE! |
| ENSMUSG00000100216.2 | 0 | NA | NA | NA | NA | NA | #VALUE! |
| ENSMUSG00000100217.2 | 0 | NA | NA | NA | NA | NA | #VALUE! |
| ENSMUSG00000100218.2 | 0 | NA | NA | NA | NA | NA | #VALUE! |
| ENSMUSG00000100219.2 | 0 | NA | NA | NA | NA | NA | #VALUE! |
| ENSMUSG00000100220.2 | 0 | NA | NA | NA | NA | NA | #VALUE! |
| ENSMUSG00000100221.7 | 0 | NA | NA | NA | NA | NA | #VALUE! |
| ENSMUSG00000100222.2 | 0 | NA | NA | NA | NA | NA | #VALUE! |
| ENSMUSG00000100223.2 | 0 | NA | NA | NA | NA | NA | #VALUE! |
| ENSMUSG00000100225.2 | 0 | NA | NA | NA | NA | NA | #VALUE! |
| ENSMUSG00000100226.2 | 0 | NA | NA | NA | NA | NA | #VALUE! |
| ENSMUSG00000100227.7 | 0 | NA | NA | NA | NA | NA | #VALUE! |
| ENSMUSG00000100228.2 | 0.168284 | 1.027628 | 4.067126 | 0.252667 | 0.800526 | NA | #VALUE! |
| ENSMUSG00000100229.2 | 0 | NA | NA | NA | NA | NA | #VALUE! |
| ENSMUSG00000100230.2 | 0 | NA | NA | NA | NA | NA | #VALUE! |

| | | | | | | | |
|---|---|---|---|---|---|---|---|
| ENSMUSG00000100231.2 | 0 | NA | NA | NA | NA | NA | #VALUE! |
| ENSMUSG00000100232.7 | 0 | NA | NA | NA | NA | NA | #VALUE! |
| ENSMUSG00000100233.2 | 0 | NA | NA | NA | NA | NA | #VALUE! |
| ENSMUSG00000100234.2 | 0 | NA | NA | NA | NA | NA | #VALUE! |
| ENSMUSG00000100236.2 | 0 | NA | NA | NA | NA | NA | #VALUE! |
| ENSMUSG00000100237.2 | 6.394774 | 6.166745 | 3.913792 | 1.575645 | 0.115108 | NA | #VALUE! |
| ENSMUSG00000100238.2 | 0 | NA | NA | NA | NA | NA | #VALUE! |
| ENSMUSG00000100239.2 | 0 | NA | NA | NA | NA | NA | #VALUE! |
| ENSMUSG00000100240.2 | 0 | NA | NA | NA | NA | NA | #VALUE! |
| ENSMUSG00000100241.2 | 0 | NA | NA | NA | NA | NA | #VALUE! |
| ENSMUSG00000100242.2 | 0 | NA | NA | NA | NA | NA | #VALUE! |
| ENSMUSG00000100243.2 | 0 | NA | NA | NA | NA | NA | #VALUE! |
| ENSMUSG00000100244.2 | 0 | NA | NA | NA | NA | NA | #VALUE! |
| ENSMUSG00000100245.2 | 0 | NA | NA | NA | NA | NA | #VALUE! |
| ENSMUSG00000100246.2 | 0 | NA | NA | NA | NA | NA | #VALUE! |
| ENSMUSG00000100247.2 | 0 | NA | NA | NA | NA | NA | #VALUE! |
| ENSMUSG00000100248.7 | 0 | NA | NA | NA | NA | NA | #VALUE! |
| ENSMUSG00000100249.2 | 0 | NA | NA | NA | NA | NA | #VALUE! |
| ENSMUSG00000100250.2 | 0 | NA | NA | NA | NA | NA | #VALUE! |
| ENSMUSG00000100251.2 | 0 | NA | NA | NA | NA | NA | #VALUE! |
| ENSMUSG00000100252.7 | 0 | NA | NA | NA | NA | NA | #VALUE! |
| ENSMUSG00000100253.2 | 4.830604 | -5.80779 | 3.915992 | -1.4831 | 0.138049 | NA | #VALUE! |
| ENSMUSG00000100254.2 | 0 | NA | NA | NA | NA | NA | #VALUE! |
| ENSMUSG00000100255.2 | 0 | NA | NA | NA | NA | NA | #VALUE! |
| ENSMUSG00000100256.2 | 0 | NA | NA | NA | NA | NA | #VALUE! |
| ENSMUSG00000100257.2 | 0 | NA | NA | NA | NA | NA | #VALUE! |
| ENSMUSG00000100258.2 | 0 | NA | NA | NA | NA | NA | #VALUE! |
| ENSMUSG00000100260.2 | 0 | NA | NA | NA | NA | NA | #VALUE! |
| ENSMUSG00000100261.2 | 0 | NA | NA | NA | NA | NA | #VALUE! |
| ENSMUSG00000100262.2 | 0 | NA | NA | NA | NA | NA | #VALUE! |
| ENSMUSG00000100263.2 | 0 | NA | NA | NA | NA | NA | #VALUE! |
| ENSMUSG00000100264.2 | 0 | NA | NA | NA | NA | NA | #VALUE! |
| ENSMUSG00000100265.2 | 0 | NA | NA | NA | NA | NA | #VALUE! |
| ENSMUSG00000100266.3 | 0 | NA | NA | NA | NA | NA | #VALUE! |
| ENSMUSG00000100267.2 | 0 | NA | NA | NA | NA | NA | #VALUE! |
| ENSMUSG00000100268.2 | 0 | NA | NA | NA | NA | NA | #VALUE! |
| ENSMUSG00000100269.2 | 0 | NA | NA | NA | NA | NA | #VALUE! |
| ENSMUSG00000100270.2 | 0 | NA | NA | NA | NA | NA | #VALUE! |
| ENSMUSG00000100271.2 | 0 | NA | NA | NA | NA | NA | #VALUE! |
| ENSMUSG00000100272.2 | 0 | NA | NA | NA | NA | NA | #VALUE! |
| ENSMUSG00000100273.2 | 0 | NA | NA | NA | NA | NA | #VALUE! |
| ENSMUSG00000100274.2 | 0 | NA | NA | NA | NA | NA | #VALUE! |
| ENSMUSG00000100275.2 | 0 | NA | NA | NA | NA | NA | #VALUE! |
| ENSMUSG00000100276.2 | 0 | NA | NA | NA | NA | NA | #VALUE! |

| | | | | | | | |
|---|---|---|---|---|---|---|---|
| ENSMUSG00000100277.9 | 0.093255 | 0.868093 | 4.072464 | 0.213162 | 0.831201 | NA | #VALUE! |
| ENSMUSG00000100278.2 | 0 | NA | NA | NA | NA | NA | #VALUE! |
| ENSMUSG00000100279.2 | 0 | NA | NA | NA | NA | NA | #VALUE! |
| ENSMUSG00000100280.3 | 0 | NA | NA | NA | NA | NA | #VALUE! |
| ENSMUSG00000100281.2 | 0 | NA | NA | NA | NA | NA | #VALUE! |
| ENSMUSG00000100282.2 | 0 | NA | NA | NA | NA | NA | #VALUE! |
| ENSMUSG00000100283.2 | 0 | NA | NA | NA | NA | NA | #VALUE! |
| ENSMUSG00000100284.2 | 0 | NA | NA | NA | NA | NA | #VALUE! |
| ENSMUSG00000100285.2 | 0 | NA | NA | NA | NA | NA | #VALUE! |
| ENSMUSG00000100286.2 | 0 | NA | NA | NA | NA | NA | #VALUE! |
| ENSMUSG00000100287.2 | 0 | NA | NA | NA | NA | NA | #VALUE! |
| ENSMUSG00000100288.2 | 0 | NA | NA | NA | NA | NA | #VALUE! |
| ENSMUSG00000100289.2 | 0 | NA | NA | NA | NA | NA | #VALUE! |
| ENSMUSG00000100290.2 | 0 | NA | NA | NA | NA | NA | #VALUE! |
| ENSMUSG00000100291.7 | 0 | NA | NA | NA | NA | NA | #VALUE! |
| ENSMUSG00000100292.2 | 0 | NA | NA | NA | NA | NA | #VALUE! |
| ENSMUSG00000100294.7 | 0 | NA | NA | NA | NA | NA | #VALUE! |
| ENSMUSG00000100295.2 | 0 | NA | NA | NA | NA | NA | #VALUE! |
| ENSMUSG00000100296.3 | 0 | NA | NA | NA | NA | NA | #VALUE! |
| ENSMUSG00000100297.2 | 0 | NA | NA | NA | NA | NA | #VALUE! |
| ENSMUSG00000100298.2 | 0 | NA | NA | NA | NA | NA | #VALUE! |
| ENSMUSG00000100299.2 | 0 | NA | NA | NA | NA | NA | #VALUE! |
| ENSMUSG00000100300.2 | 0 | NA | NA | NA | NA | NA | #VALUE! |
| ENSMUSG00000100301.7 | 0 | NA | NA | NA | NA | NA | #VALUE! |
| ENSMUSG00000100302.2 | 0 | NA | NA | NA | NA | NA | #VALUE! |
| ENSMUSG00000100303.4 | 0 | NA | NA | NA | NA | NA | #VALUE! |
| ENSMUSG00000100304.2 | 0 | NA | NA | NA | NA | NA | #VALUE! |
| ENSMUSG00000100305.2 | 0 | NA | NA | NA | NA | NA | #VALUE! |
| ENSMUSG00000100306.2 | 0 | NA | NA | NA | NA | NA | #VALUE! |
| ENSMUSG00000100307.2 | 0 | NA | NA | NA | NA | NA | #VALUE! |
| ENSMUSG00000100308.2 | 0 | NA | NA | NA | NA | NA | #VALUE! |
| ENSMUSG00000100309.2 | 0 | NA | NA | NA | NA | NA | #VALUE! |
| ENSMUSG00000100310.2 | 0 | NA | NA | NA | NA | NA | #VALUE! |
| ENSMUSG00000100311.7 | 0 | NA | NA | NA | NA | NA | #VALUE! |
| ENSMUSG00000100312.2 | 0 | NA | NA | NA | NA | NA | #VALUE! |
| ENSMUSG00000100313.2 | 0 | NA | NA | NA | NA | NA | #VALUE! |
| ENSMUSG00000100314.7 | 0 | NA | NA | NA | NA | NA | #VALUE! |
| ENSMUSG00000100315.7 | 0 | NA | NA | NA | NA | NA | #VALUE! |
| ENSMUSG00000100316.2 | 0 | NA | NA | NA | NA | NA | #VALUE! |
| ENSMUSG00000100317.2 | 0 | NA | NA | NA | NA | NA | #VALUE! |
| ENSMUSG00000100318.3 | 0 | NA | NA | NA | NA | NA | #VALUE! |
| ENSMUSG00000100319.2 | 0 | NA | NA | NA | NA | NA | #VALUE! |
| ENSMUSG00000100320.2 | 0 | NA | NA | NA | NA | NA | #VALUE! |
| ENSMUSG00000100321.2 | 0 | NA | NA | NA | NA | NA | #VALUE! |

| | | | | | | | |
|---|---|---|---|---|---|---|---|
| ENSMUSG00000100322.7 | 0 | NA | NA | NA | NA | NA | #VALUE! |
| ENSMUSG00000100324.2 | 0 | NA | NA | NA | NA | NA | #VALUE! |
| ENSMUSG00000100325.2 | 0 | NA | NA | NA | NA | NA | #VALUE! |
| ENSMUSG00000100326.2 | 0 | NA | NA | NA | NA | NA | #VALUE! |
| ENSMUSG00000100328.7 | 0 | NA | NA | NA | NA | NA | #VALUE! |
| ENSMUSG00000100329.2 | 0 | NA | NA | NA | NA | NA | #VALUE! |
| ENSMUSG00000100330.2 | 0 | NA | NA | NA | NA | NA | #VALUE! |
| ENSMUSG00000100331.2 | 0 | NA | NA | NA | NA | NA | #VALUE! |
| ENSMUSG00000100332.2 | 0 | NA | NA | NA | NA | NA | #VALUE! |
| ENSMUSG00000100333.2 | 0 | NA | NA | NA | NA | NA | #VALUE! |
| ENSMUSG00000100334.2 | 0 | NA | NA | NA | NA | NA | #VALUE! |
| ENSMUSG00000100335.3 | 0 | NA | NA | NA | NA | NA | #VALUE! |
| ENSMUSG00000100336.2 | 0 | NA | NA | NA | NA | NA | #VALUE! |
| ENSMUSG00000100337.2 | 0 | NA | NA | NA | NA | NA | #VALUE! |
| ENSMUSG00000100338.3 | 0 | NA | NA | NA | NA | NA | #VALUE! |
| ENSMUSG00000100339.2 | 0 | NA | NA | NA | NA | NA | #VALUE! |
| ENSMUSG00000100340.2 | 0 | NA | NA | NA | NA | NA | #VALUE! |
| ENSMUSG00000100341.2 | 0 | NA | NA | NA | NA | NA | #VALUE! |
| ENSMUSG00000100343.2 | 0 | NA | NA | NA | NA | NA | #VALUE! |
| ENSMUSG00000100344.2 | 0 | NA | NA | NA | NA | NA | #VALUE! |
| ENSMUSG00000100345.2 | 0 | NA | NA | NA | NA | NA | #VALUE! |
| ENSMUSG00000100346.3 | 0 | NA | NA | NA | NA | NA | #VALUE! |
| ENSMUSG00000100347.2 | 0 | NA | NA | NA | NA | NA | #VALUE! |
| ENSMUSG00000100349.2 | 0 | NA | NA | NA | NA | NA | #VALUE! |
| ENSMUSG00000100350.2 | 0 | NA | NA | NA | NA | NA | #VALUE! |
| ENSMUSG00000100351.2 | 0.105013 | -0.97324 | 4.076293 | -0.23876 | 0.811295 | NA | #VALUE! |
| ENSMUSG00000100352.2 | 0 | NA | NA | NA | NA | NA | #VALUE! |
| ENSMUSG00000100353.2 | 0 | NA | NA | NA | NA | NA | #VALUE! |
| ENSMUSG00000100354.2 | 0 | NA | NA | NA | NA | NA | #VALUE! |
| ENSMUSG00000100355.2 | 0 | NA | NA | NA | NA | NA | #VALUE! |
| ENSMUSG00000100356.2 | 0 | NA | NA | NA | NA | NA | #VALUE! |
| ENSMUSG00000100357.2 | 0 | NA | NA | NA | NA | NA | #VALUE! |
| ENSMUSG00000100358.2 | 0 | NA | NA | NA | NA | NA | #VALUE! |
| ENSMUSG00000100359.2 | 0 | NA | NA | NA | NA | NA | #VALUE! |
| ENSMUSG00000100360.2 | 0 | NA | NA | NA | NA | NA | #VALUE! |
| ENSMUSG00000100361.2 | 0 | NA | NA | NA | NA | NA | #VALUE! |
| ENSMUSG00000100362.2 | 0 | NA | NA | NA | NA | NA | #VALUE! |
| ENSMUSG00000100363.2 | 0 | NA | NA | NA | NA | NA | #VALUE! |
| ENSMUSG00000100364.2 | 0 | NA | NA | NA | NA | NA | #VALUE! |
| ENSMUSG00000100365.2 | 0 | NA | NA | NA | NA | NA | #VALUE! |
| ENSMUSG00000100366.2 | 0 | NA | NA | NA | NA | NA | #VALUE! |
| ENSMUSG00000100367.2 | 0 | NA | NA | NA | NA | NA | #VALUE! |
| ENSMUSG00000100368.2 | 0 | NA | NA | NA | NA | NA | #VALUE! |
| ENSMUSG00000100369.2 | 0 | NA | NA | NA | NA | NA | #VALUE! |

| | | | | | | | |
|---|---|---|---|---|---|---|---|
| ENSMUSG00000100372.2 | 0 | NA | NA | NA | NA | NA | #VALUE! |
| ENSMUSG00000100373.7 | 0 | NA | NA | NA | NA | NA | #VALUE! |
| ENSMUSG00000100374.2 | 0 | NA | NA | NA | NA | NA | #VALUE! |
| ENSMUSG00000100375.2 | 0 | NA | NA | NA | NA | NA | #VALUE! |
| ENSMUSG00000100376.2 | 0 | NA | NA | NA | NA | NA | #VALUE! |
| ENSMUSG00000100377.2 | 0 | NA | NA | NA | NA | NA | #VALUE! |
| ENSMUSG00000100378.2 | 0 | NA | NA | NA | NA | NA | #VALUE! |
| ENSMUSG00000100379.7 | 0 | NA | NA | NA | NA | NA | #VALUE! |
| ENSMUSG00000100380.2 | 0 | NA | NA | NA | NA | NA | #VALUE! |
| ENSMUSG00000100381.2 | 0 | NA | NA | NA | NA | NA | #VALUE! |
| ENSMUSG00000100382.2 | 0 | NA | NA | NA | NA | NA | #VALUE! |
| ENSMUSG00000100383.2 | 0 | NA | NA | NA | NA | NA | #VALUE! |
| ENSMUSG00000100384.2 | 0 | NA | NA | NA | NA | NA | #VALUE! |
| ENSMUSG00000100385.2 | 0 | NA | NA | NA | NA | NA | #VALUE! |
| ENSMUSG00000100386.2 | 0 | NA | NA | NA | NA | NA | #VALUE! |
| ENSMUSG00000100387.2 | 0 | NA | NA | NA | NA | NA | #VALUE! |
| ENSMUSG00000100388.2 | 0 | NA | NA | NA | NA | NA | #VALUE! |
| ENSMUSG00000100389.2 | 0 | NA | NA | NA | NA | NA | #VALUE! |
| ENSMUSG00000100390.2 | 0 | NA | NA | NA | NA | NA | #VALUE! |
| ENSMUSG00000100391.2 | 0 | NA | NA | NA | NA | NA | #VALUE! |
| ENSMUSG00000100392.2 | 0.279766 | 1.808031 | 4.036658 | 0.447903 | 0.654223 | NA | #VALUE! |
| ENSMUSG00000100393.2 | 0 | NA | NA | NA | NA | NA | #VALUE! |
| ENSMUSG00000100394.2 | 0 | NA | NA | NA | NA | NA | #VALUE! |
| ENSMUSG00000100395.2 | 0 | NA | NA | NA | NA | NA | #VALUE! |
| ENSMUSG00000100396.2 | 0.204111 | -1.05581 | 4.073302 | -0.2592 | 0.795479 | NA | #VALUE! |
| ENSMUSG00000100397.2 | 0 | NA | NA | NA | NA | NA | #VALUE! |
| ENSMUSG00000100398.2 | 0 | NA | NA | NA | NA | NA | #VALUE! |
| ENSMUSG00000100399.7 | 0 | NA | NA | NA | NA | NA | #VALUE! |
| ENSMUSG00000100400.2 | 0 | NA | NA | NA | NA | NA | #VALUE! |
| ENSMUSG00000100401.7 | 0 | NA | NA | NA | NA | NA | #VALUE! |
| ENSMUSG00000100402.2 | 0 | NA | NA | NA | NA | NA | #VALUE! |
| ENSMUSG00000100403.2 | 0 | NA | NA | NA | NA | NA | #VALUE! |
| ENSMUSG00000100404.2 | 0 | NA | NA | NA | NA | NA | #VALUE! |
| ENSMUSG00000100405.2 | 0 | NA | NA | NA | NA | NA | #VALUE! |
| ENSMUSG00000100406.2 | 0 | NA | NA | NA | NA | NA | #VALUE! |
| ENSMUSG00000100407.2 | 0 | NA | NA | NA | NA | NA | #VALUE! |
| ENSMUSG00000100408.2 | 0 | NA | NA | NA | NA | NA | #VALUE! |
| ENSMUSG00000100410.2 | 0 | NA | NA | NA | NA | NA | #VALUE! |
| ENSMUSG00000100411.2 | 0 | NA | NA | NA | NA | NA | #VALUE! |
| ENSMUSG00000100412.2 | 0 | NA | NA | NA | NA | NA | #VALUE! |
| ENSMUSG00000100413.2 | 0 | NA | NA | NA | NA | NA | #VALUE! |
| ENSMUSG00000100415.7 | 0 | NA | NA | NA | NA | NA | #VALUE! |
| ENSMUSG00000100416.2 | 0 | NA | NA | NA | NA | NA | #VALUE! |
| ENSMUSG00000100417.3 | 0 | NA | NA | NA | NA | NA | #VALUE! |

| | | | | | | | |
|---|---|---|---|---|---|---|---|
| ENSMUSG00000100418.2 | 0 | NA | NA | NA | NA | NA | #VALUE! |
| ENSMUSG00000100420.2 | 0 | NA | NA | NA | NA | NA | #VALUE! |
| ENSMUSG00000100421.2 | 0 | NA | NA | NA | NA | NA | #VALUE! |
| ENSMUSG00000100422.2 | 0 | NA | NA | NA | NA | NA | #VALUE! |
| ENSMUSG00000100423.2 | 0 | NA | NA | NA | NA | NA | #VALUE! |
| ENSMUSG00000100424.2 | 0 | NA | NA | NA | NA | NA | #VALUE! |
| ENSMUSG00000100425.2 | 0 | NA | NA | NA | NA | NA | #VALUE! |
| ENSMUSG00000100426.2 | 0 | NA | NA | NA | NA | NA | #VALUE! |
| ENSMUSG00000100428.2 | 0 | NA | NA | NA | NA | NA | #VALUE! |
| ENSMUSG00000100429.2 | 0 | NA | NA | NA | NA | NA | #VALUE! |
| ENSMUSG00000100430.3 | 0 | NA | NA | NA | NA | NA | #VALUE! |
| ENSMUSG00000100431.3 | 0 | NA | NA | NA | NA | NA | #VALUE! |
| ENSMUSG00000100432.2 | 0 | NA | NA | NA | NA | NA | #VALUE! |
| ENSMUSG00000100433.2 | 0 | NA | NA | NA | NA | NA | #VALUE! |
| ENSMUSG00000100434.2 | 0 | NA | NA | NA | NA | NA | #VALUE! |
| ENSMUSG00000100436.2 | 0 | NA | NA | NA | NA | NA | #VALUE! |
| ENSMUSG00000100437.2 | 0 | NA | NA | NA | NA | NA | #VALUE! |
| ENSMUSG00000100438.2 | 0 | NA | NA | NA | NA | NA | #VALUE! |
| ENSMUSG00000100439.2 | 0 | NA | NA | NA | NA | NA | #VALUE! |
| ENSMUSG00000100440.2 | 0 | NA | NA | NA | NA | NA | #VALUE! |
| ENSMUSG00000100441.2 | 0 | NA | NA | NA | NA | NA | #VALUE! |
| ENSMUSG00000100442.2 | 0 | NA | NA | NA | NA | NA | #VALUE! |
| ENSMUSG00000100443.2 | 0 | NA | NA | NA | NA | NA | #VALUE! |
| ENSMUSG00000100444.2 | 0 | NA | NA | NA | NA | NA | #VALUE! |
| ENSMUSG00000100445.2 | 0 | NA | NA | NA | NA | NA | #VALUE! |
| ENSMUSG00000100446.2 | 0 | NA | NA | NA | NA | NA | #VALUE! |
| ENSMUSG00000100447.2 | 0 | NA | NA | NA | NA | NA | #VALUE! |
| ENSMUSG00000100448.2 | 0 | NA | NA | NA | NA | NA | #VALUE! |
| ENSMUSG00000100449.2 | 0 | NA | NA | NA | NA | NA | #VALUE! |
| ENSMUSG00000100450.2 | 0 | NA | NA | NA | NA | NA | #VALUE! |
| ENSMUSG00000100451.2 | 0 | NA | NA | NA | NA | NA | #VALUE! |
| ENSMUSG00000100452.2 | 0 | NA | NA | NA | NA | NA | #VALUE! |
| ENSMUSG00000100453.2 | 0 | NA | NA | NA | NA | NA | #VALUE! |
| ENSMUSG00000100454.2 | 0 | NA | NA | NA | NA | NA | #VALUE! |
| ENSMUSG00000100455.2 | 2.459755 | 4.774963 | 3.925155 | 1.216503 | 0.223793 | NA | #VALUE! |
| ENSMUSG00000100456.2 | 0 | NA | NA | NA | NA | NA | #VALUE! |
| ENSMUSG00000100457.7 | 0 | NA | NA | NA | NA | NA | #VALUE! |
| ENSMUSG00000100458.3 | 0 | NA | NA | NA | NA | NA | #VALUE! |
| ENSMUSG00000100459.7 | 0 | NA | NA | NA | NA | NA | #VALUE! |
| ENSMUSG00000100460.2 | 0 | NA | NA | NA | NA | NA | #VALUE! |
| ENSMUSG00000100461.2 | 0 | NA | NA | NA | NA | NA | #VALUE! |
| ENSMUSG00000100462.2 | 0 | NA | NA | NA | NA | NA | #VALUE! |
| ENSMUSG00000100463.2 | 0 | NA | NA | NA | NA | NA | #VALUE! |
| ENSMUSG00000100464.2 | 0 | NA | NA | NA | NA | NA | #VALUE! |

| | | | | | | | |
|---|---|---|---|---|---|---|---|
| ENSMUSG00000100465.2 | 0 | NA | NA | NA | NA | NA | #VALUE! |
| ENSMUSG00000100466.2 | 0 | NA | NA | NA | NA | NA | #VALUE! |
| ENSMUSG00000100467.2 | 0 | NA | NA | NA | NA | NA | #VALUE! |
| ENSMUSG00000100468.2 | 0 | NA | NA | NA | NA | NA | #VALUE! |
| ENSMUSG00000100469.2 | 0 | NA | NA | NA | NA | NA | #VALUE! |
| ENSMUSG00000100470.2 | 0 | NA | NA | NA | NA | NA | #VALUE! |
| ENSMUSG00000100471.2 | 0 | NA | NA | NA | NA | NA | #VALUE! |
| ENSMUSG00000100472.2 | 0 | NA | NA | NA | NA | NA | #VALUE! |
| ENSMUSG00000100473.2 | 0 | NA | NA | NA | NA | NA | #VALUE! |
| ENSMUSG00000100474.2 | 0 | NA | NA | NA | NA | NA | #VALUE! |
| ENSMUSG00000100475.2 | 0 | NA | NA | NA | NA | NA | #VALUE! |
| ENSMUSG00000100476.2 | 0 | NA | NA | NA | NA | NA | #VALUE! |
| ENSMUSG00000100477.2 | 0 | NA | NA | NA | NA | NA | #VALUE! |
| ENSMUSG00000100478.2 | 0 | NA | NA | NA | NA | NA | #VALUE! |
| ENSMUSG00000100479.2 | 0 | NA | NA | NA | NA | NA | #VALUE! |
| ENSMUSG00000100480.2 | 0 | NA | NA | NA | NA | NA | #VALUE! |
| ENSMUSG00000100481.2 | 0 | NA | NA | NA | NA | NA | #VALUE! |
| ENSMUSG00000100483.2 | 0 | NA | NA | NA | NA | NA | #VALUE! |
| ENSMUSG00000100484.2 | 0 | NA | NA | NA | NA | NA | #VALUE! |
| ENSMUSG00000100485.2 | 0 | NA | NA | NA | NA | NA | #VALUE! |
| ENSMUSG00000100486.8 | 0 | NA | NA | NA | NA | NA | #VALUE! |
| ENSMUSG00000100487.2 | 0 | NA | NA | NA | NA | NA | #VALUE! |
| ENSMUSG00000100488.2 | 0 | NA | NA | NA | NA | NA | #VALUE! |
| ENSMUSG00000100489.2 | 0 | NA | NA | NA | NA | NA | #VALUE! |
| ENSMUSG00000100490.2 | 0 | NA | NA | NA | NA | NA | #VALUE! |
| ENSMUSG00000100491.2 | 0 | NA | NA | NA | NA | NA | #VALUE! |
| ENSMUSG00000100492.2 | 0 | NA | NA | NA | NA | NA | #VALUE! |
| ENSMUSG00000100494.2 | 0 | NA | NA | NA | NA | NA | #VALUE! |
| ENSMUSG00000100495.2 | 0 | NA | NA | NA | NA | NA | #VALUE! |
| ENSMUSG00000100496.2 | 0 | NA | NA | NA | NA | NA | #VALUE! |
| ENSMUSG00000100497.2 | 0 | NA | NA | NA | NA | NA | #VALUE! |
| ENSMUSG00000100499.7 | 0 | NA | NA | NA | NA | NA | #VALUE! |
| ENSMUSG00000100500.2 | 0 | NA | NA | NA | NA | NA | #VALUE! |
| ENSMUSG00000100501.2 | 0 | NA | NA | NA | NA | NA | #VALUE! |
| ENSMUSG00000100502.2 | 0 | NA | NA | NA | NA | NA | #VALUE! |
| ENSMUSG00000100503.2 | 0 | NA | NA | NA | NA | NA | #VALUE! |
| ENSMUSG00000100504.2 | 0 | NA | NA | NA | NA | NA | #VALUE! |
| ENSMUSG00000100505.2 | 0 | NA | NA | NA | NA | NA | #VALUE! |
| ENSMUSG00000100506.2 | 0 | NA | NA | NA | NA | NA | #VALUE! |
| ENSMUSG00000100507.2 | 0 | NA | NA | NA | NA | NA | #VALUE! |
| ENSMUSG00000100508.7 | 0 | NA | NA | NA | NA | NA | #VALUE! |
| ENSMUSG00000100509.2 | 0 | NA | NA | NA | NA | NA | #VALUE! |
| ENSMUSG00000100511.2 | 0 | NA | NA | NA | NA | NA | #VALUE! |
| ENSMUSG00000100512.2 | 5.275125 | 3.896772 | 3.546615 | 1.09873 | 0.271886 | NA | #VALUE! |

| | | | | | | | |
|---|---|---|---|---|---|---|---|
| ENSMUSG00000100513.2 | 0 | NA | NA | NA | NA | NA | #VALUE! |
| ENSMUSG00000100514.2 | 0 | NA | NA | NA | NA | NA | #VALUE! |
| ENSMUSG00000100515.7 | 0 | NA | NA | NA | NA | NA | #VALUE! |
| ENSMUSG00000100516.2 | 0 | NA | NA | NA | NA | NA | #VALUE! |
| ENSMUSG00000100517.2 | 0 | NA | NA | NA | NA | NA | #VALUE! |
| ENSMUSG00000100518.2 | 0 | NA | NA | NA | NA | NA | #VALUE! |
| ENSMUSG00000100519.2 | 0 | NA | NA | NA | NA | NA | #VALUE! |
| ENSMUSG00000100520.2 | 0 | NA | NA | NA | NA | NA | #VALUE! |
| ENSMUSG00000100521.2 | 0 | NA | NA | NA | NA | NA | #VALUE! |
| ENSMUSG00000100522.2 | 0 | NA | NA | NA | NA | NA | #VALUE! |
| ENSMUSG00000100523.2 | 0 | NA | NA | NA | NA | NA | #VALUE! |
| ENSMUSG00000100524.2 | 0 | NA | NA | NA | NA | NA | #VALUE! |
| ENSMUSG00000100525.2 | 0 | NA | NA | NA | NA | NA | #VALUE! |
| ENSMUSG00000100526.2 | 0 | NA | NA | NA | NA | NA | #VALUE! |
| ENSMUSG00000100527.2 | 0 | NA | NA | NA | NA | NA | #VALUE! |
| ENSMUSG00000100528.2 | 0 | NA | NA | NA | NA | NA | #VALUE! |
| ENSMUSG00000100529.2 | 0 | NA | NA | NA | NA | NA | #VALUE! |
| ENSMUSG00000100530.2 | 0 | NA | NA | NA | NA | NA | #VALUE! |
| ENSMUSG00000100531.2 | 0 | NA | NA | NA | NA | NA | #VALUE! |
| ENSMUSG00000100532.2 | 0 | NA | NA | NA | NA | NA | #VALUE! |
| ENSMUSG00000100533.7 | 0 | NA | NA | NA | NA | NA | #VALUE! |
| ENSMUSG00000100534.2 | 0 | NA | NA | NA | NA | NA | #VALUE! |
| ENSMUSG00000100535.2 | 0 | NA | NA | NA | NA | NA | #VALUE! |
| ENSMUSG00000100537.2 | 0 | NA | NA | NA | NA | NA | #VALUE! |
| ENSMUSG00000100538.2 | 0 | NA | NA | NA | NA | NA | #VALUE! |
| ENSMUSG00000100539.2 | 0 | NA | NA | NA | NA | NA | #VALUE! |
| ENSMUSG00000100540.2 | 0 | NA | NA | NA | NA | NA | #VALUE! |
| ENSMUSG00000100541.2 | 0 | NA | NA | NA | NA | NA | #VALUE! |
| ENSMUSG00000100542.2 | 0 | NA | NA | NA | NA | NA | #VALUE! |
| ENSMUSG00000100543.2 | 0 | NA | NA | NA | NA | NA | #VALUE! |
| ENSMUSG00000100544.2 | 0 | NA | NA | NA | NA | NA | #VALUE! |
| ENSMUSG00000100546.2 | 0 | NA | NA | NA | NA | NA | #VALUE! |
| ENSMUSG00000100547.2 | 0 | NA | NA | NA | NA | NA | #VALUE! |
| ENSMUSG00000100548.2 | 0 | NA | NA | NA | NA | NA | #VALUE! |
| ENSMUSG00000100549.2 | 0 | NA | NA | NA | NA | NA | #VALUE! |
| ENSMUSG00000100550.3 | 0 | NA | NA | NA | NA | NA | #VALUE! |
| ENSMUSG00000100551.2 | 0.834312 | -0.12048 | 3.94136 | -0.03057 | 0.975613 | NA | #VALUE! |
| ENSMUSG00000100552.2 | 0 | NA | NA | NA | NA | NA | #VALUE! |
| ENSMUSG00000100553.7 | 0 | NA | NA | NA | NA | NA | #VALUE! |
| ENSMUSG00000100554.2 | 0 | NA | NA | NA | NA | NA | #VALUE! |
| ENSMUSG00000100555.2 | 0 | NA | NA | NA | NA | NA | #VALUE! |
| ENSMUSG00000100556.2 | 0 | NA | NA | NA | NA | NA | #VALUE! |
| ENSMUSG00000100557.2 | 0 | NA | NA | NA | NA | NA | #VALUE! |
| ENSMUSG00000100558.2 | 0 | NA | NA | NA | NA | NA | #VALUE! |

| | | | | | | | |
|---|---|---|---|---|---|---|---|
| ENSMUSG00000100559.2 | 0 | NA | NA | NA | NA | NA | #VALUE! |
| ENSMUSG00000100560.2 | 0 | NA | NA | NA | NA | NA | #VALUE! |
| ENSMUSG00000100561.2 | 0 | NA | NA | NA | NA | NA | #VALUE! |
| ENSMUSG00000100562.2 | 0 | NA | NA | NA | NA | NA | #VALUE! |
| ENSMUSG00000100563.2 | 0 | NA | NA | NA | NA | NA | #VALUE! |
| ENSMUSG00000100564.2 | 0 | NA | NA | NA | NA | NA | #VALUE! |
| ENSMUSG00000100565.7 | 0 | NA | NA | NA | NA | NA | #VALUE! |
| ENSMUSG00000100566.2 | 0 | NA | NA | NA | NA | NA | #VALUE! |
| ENSMUSG00000100567.2 | 0 | NA | NA | NA | NA | NA | #VALUE! |
| ENSMUSG00000100568.2 | 0 | NA | NA | NA | NA | NA | #VALUE! |
| ENSMUSG00000100569.2 | 0 | NA | NA | NA | NA | NA | #VALUE! |
| ENSMUSG00000100570.7 | 0 | NA | NA | NA | NA | NA | #VALUE! |
| ENSMUSG00000100571.2 | 0 | NA | NA | NA | NA | NA | #VALUE! |
| ENSMUSG00000100572.2 | 0 | NA | NA | NA | NA | NA | #VALUE! |
| ENSMUSG00000100573.2 | 0 | NA | NA | NA | NA | NA | #VALUE! |
| ENSMUSG00000100574.2 | 0 | NA | NA | NA | NA | NA | #VALUE! |
| ENSMUSG00000100575.2 | 0 | NA | NA | NA | NA | NA | #VALUE! |
| ENSMUSG00000100576.2 | 0 | NA | NA | NA | NA | NA | #VALUE! |
| ENSMUSG00000100577.2 | 0 | NA | NA | NA | NA | NA | #VALUE! |
| ENSMUSG00000100578.2 | 0 | NA | NA | NA | NA | NA | #VALUE! |
| ENSMUSG00000100579.7 | 0 | NA | NA | NA | NA | NA | #VALUE! |
| ENSMUSG00000100580.7 | 0 | NA | NA | NA | NA | NA | #VALUE! |
| ENSMUSG00000100581.2 | 0 | NA | NA | NA | NA | NA | #VALUE! |
| ENSMUSG00000100582.2 | 0 | NA | NA | NA | NA | NA | #VALUE! |
| ENSMUSG00000100583.7 | 0 | NA | NA | NA | NA | NA | #VALUE! |
| ENSMUSG00000100584.7 | 0 | NA | NA | NA | NA | NA | #VALUE! |
| ENSMUSG00000100585.2 | 0 | NA | NA | NA | NA | NA | #VALUE! |
| ENSMUSG00000100586.3 | 0 | NA | NA | NA | NA | NA | #VALUE! |
| ENSMUSG00000100587.2 | 0 | NA | NA | NA | NA | NA | #VALUE! |
| ENSMUSG00000100588.2 | 0 | NA | NA | NA | NA | NA | #VALUE! |
| ENSMUSG00000100589.2 | 0 | NA | NA | NA | NA | NA | #VALUE! |
| ENSMUSG00000100590.2 | 0 | NA | NA | NA | NA | NA | #VALUE! |
| ENSMUSG00000100591.2 | 0 | NA | NA | NA | NA | NA | #VALUE! |
| ENSMUSG00000100592.2 | 0 | NA | NA | NA | NA | NA | #VALUE! |
| ENSMUSG00000100593.2 | 0 | NA | NA | NA | NA | NA | #VALUE! |
| ENSMUSG00000100595.2 | 0 | NA | NA | NA | NA | NA | #VALUE! |
| ENSMUSG00000100596.2 | 0 | NA | NA | NA | NA | NA | #VALUE! |
| ENSMUSG00000100597.2 | 0 | NA | NA | NA | NA | NA | #VALUE! |
| ENSMUSG00000100598.2 | 0 | NA | NA | NA | NA | NA | #VALUE! |
| ENSMUSG00000100599.3 | 0 | NA | NA | NA | NA | NA | #VALUE! |
| ENSMUSG00000100600.7 | 0 | NA | NA | NA | NA | NA | #VALUE! |
| ENSMUSG00000100602.2 | 0 | NA | NA | NA | NA | NA | #VALUE! |
| ENSMUSG00000100603.7 | 0 | NA | NA | NA | NA | NA | #VALUE! |
| ENSMUSG00000100604.2 | 0 | NA | NA | NA | NA | NA | #VALUE! |

| | | | | | | | |
|---|---|---|---|---|---|---|---|
| ENSMUSG00000100605.2 | 1.224665 | -3.81006 | 3.943408 | -0.96618 | 0.333952 | NA | #VALUE! |
| ENSMUSG00000100606.2 | 0 | NA | NA | NA | NA | NA | #VALUE! |
| ENSMUSG00000100607.2 | 0 | NA | NA | NA | NA | NA | #VALUE! |
| ENSMUSG00000100608.2 | 0 | NA | NA | NA | NA | NA | #VALUE! |
| ENSMUSG00000100609.7 | 0 | NA | NA | NA | NA | NA | #VALUE! |
| ENSMUSG00000100610.2 | 0 | NA | NA | NA | NA | NA | #VALUE! |
| ENSMUSG00000100611.2 | 0 | NA | NA | NA | NA | NA | #VALUE! |
| ENSMUSG00000100613.2 | 0 | NA | NA | NA | NA | NA | #VALUE! |
| ENSMUSG00000100614.2 | 0 | NA | NA | NA | NA | NA | #VALUE! |
| ENSMUSG00000100615.2 | 0.093255 | 0.868093 | 4.072464 | 0.213162 | 0.831201 | NA | #VALUE! |
| ENSMUSG00000100616.2 | 0 | NA | NA | NA | NA | NA | #VALUE! |
| ENSMUSG00000100617.2 | 0 | NA | NA | NA | NA | NA | #VALUE! |
| ENSMUSG00000100619.2 | 0 | NA | NA | NA | NA | NA | #VALUE! |
| ENSMUSG00000100620.2 | 0 | NA | NA | NA | NA | NA | #VALUE! |
| ENSMUSG00000100621.3 | 0 | NA | NA | NA | NA | NA | #VALUE! |
| ENSMUSG00000100622.2 | 0 | NA | NA | NA | NA | NA | #VALUE! |
| ENSMUSG00000100623.2 | 0 | NA | NA | NA | NA | NA | #VALUE! |
| ENSMUSG00000100624.2 | 0 | NA | NA | NA | NA | NA | #VALUE! |
| ENSMUSG00000100625.8 | 0 | NA | NA | NA | NA | NA | #VALUE! |
| ENSMUSG00000100626.2 | 0 | NA | NA | NA | NA | NA | #VALUE! |
| ENSMUSG00000100627.7 | 0 | NA | NA | NA | NA | NA | #VALUE! |
| ENSMUSG00000100628.2 | 0 | NA | NA | NA | NA | NA | #VALUE! |
| ENSMUSG00000100629.2 | 0 | NA | NA | NA | NA | NA | #VALUE! |
| ENSMUSG00000100630.2 | 0 | NA | NA | NA | NA | NA | #VALUE! |
| ENSMUSG00000100632.2 | 0 | NA | NA | NA | NA | NA | #VALUE! |
| ENSMUSG00000100634.2 | 0 | NA | NA | NA | NA | NA | #VALUE! |
| ENSMUSG00000100635.2 | 0 | NA | NA | NA | NA | NA | #VALUE! |
| ENSMUSG00000100636.2 | 0 | NA | NA | NA | NA | NA | #VALUE! |
| ENSMUSG00000100637.2 | 0 | NA | NA | NA | NA | NA | #VALUE! |
| ENSMUSG00000100639.2 | 0 | NA | NA | NA | NA | NA | #VALUE! |
| ENSMUSG00000100640.2 | 0 | NA | NA | NA | NA | NA | #VALUE! |
| ENSMUSG00000100641.2 | 0 | NA | NA | NA | NA | NA | #VALUE! |
| ENSMUSG00000100642.2 | 0 | NA | NA | NA | NA | NA | #VALUE! |
| ENSMUSG00000100643.2 | 0 | NA | NA | NA | NA | NA | #VALUE! |
| ENSMUSG00000100644.2 | 0 | NA | NA | NA | NA | NA | #VALUE! |
| ENSMUSG00000100645.2 | 0 | NA | NA | NA | NA | NA | #VALUE! |
| ENSMUSG00000100646.2 | 0 | NA | NA | NA | NA | NA | #VALUE! |
| ENSMUSG00000100647.7 | 0 | NA | NA | NA | NA | NA | #VALUE! |
| ENSMUSG00000100648.2 | 0 | NA | NA | NA | NA | NA | #VALUE! |
| ENSMUSG00000100649.2 | 0 | NA | NA | NA | NA | NA | #VALUE! |
| ENSMUSG00000100650.2 | 0 | NA | NA | NA | NA | NA | #VALUE! |
| ENSMUSG00000100651.2 | 0 | NA | NA | NA | NA | NA | #VALUE! |
| ENSMUSG00000100652.2 | 0 | NA | NA | NA | NA | NA | #VALUE! |
| ENSMUSG00000100653.2 | 0 | NA | NA | NA | NA | NA | #VALUE! |

| | | | | | | | |
|---|---|---|---|---|---|---|---|
| ENSMUSG00000100654.2 | 0 | NA | NA | NA | NA | NA | #VALUE! |
| ENSMUSG00000100655.2 | 0 | NA | NA | NA | NA | NA | #VALUE! |
| ENSMUSG00000100656.2 | 0 | NA | NA | NA | NA | NA | #VALUE! |
| ENSMUSG00000100657.2 | 0 | NA | NA | NA | NA | NA | #VALUE! |
| ENSMUSG00000100658.2 | 0 | NA | NA | NA | NA | NA | #VALUE! |
| ENSMUSG00000100659.2 | 0 | NA | NA | NA | NA | NA | #VALUE! |
| ENSMUSG00000100660.2 | 0 | NA | NA | NA | NA | NA | #VALUE! |
| ENSMUSG00000100661.2 | 0 | NA | NA | NA | NA | NA | #VALUE! |
| ENSMUSG00000100662.2 | 0 | NA | NA | NA | NA | NA | #VALUE! |
| ENSMUSG00000100663.2 | 0 | NA | NA | NA | NA | NA | #VALUE! |
| ENSMUSG00000100664.2 | 0 | NA | NA | NA | NA | NA | #VALUE! |
| ENSMUSG00000100665.2 | 0 | NA | NA | NA | NA | NA | #VALUE! |
| ENSMUSG00000100666.2 | 0 | NA | NA | NA | NA | NA | #VALUE! |
| ENSMUSG00000100667.2 | 0 | NA | NA | NA | NA | NA | #VALUE! |
| ENSMUSG00000100668.2 | 0 | NA | NA | NA | NA | NA | #VALUE! |
| ENSMUSG00000100669.2 | 0 | NA | NA | NA | NA | NA | #VALUE! |
| ENSMUSG00000100670.2 | 0 | NA | NA | NA | NA | NA | #VALUE! |
| ENSMUSG00000100671.2 | 0 | NA | NA | NA | NA | NA | #VALUE! |
| ENSMUSG00000100672.2 | 0 | NA | NA | NA | NA | NA | #VALUE! |
| ENSMUSG00000100673.2 | 0 | NA | NA | NA | NA | NA | #VALUE! |
| ENSMUSG00000100674.2 | 0 | NA | NA | NA | NA | NA | #VALUE! |
| ENSMUSG00000100675.2 | 0 | NA | NA | NA | NA | NA | #VALUE! |
| ENSMUSG00000100676.2 | 0 | NA | NA | NA | NA | NA | #VALUE! |
| ENSMUSG00000100677.2 | 0 | NA | NA | NA | NA | NA | #VALUE! |
| ENSMUSG00000100678.2 | 0 | NA | NA | NA | NA | NA | #VALUE! |
| ENSMUSG00000100679.2 | 0 | NA | NA | NA | NA | NA | #VALUE! |
| ENSMUSG00000100681.2 | 0 | NA | NA | NA | NA | NA | #VALUE! |
| ENSMUSG00000100682.2 | 0 | NA | NA | NA | NA | NA | #VALUE! |
| ENSMUSG00000100683.2 | 0 | NA | NA | NA | NA | NA | #VALUE! |
| ENSMUSG00000100684.2 | 0 | NA | NA | NA | NA | NA | #VALUE! |
| ENSMUSG00000100685.2 | 0 | NA | NA | NA | NA | NA | #VALUE! |
| ENSMUSG00000100687.2 | 0 | NA | NA | NA | NA | NA | #VALUE! |
| ENSMUSG00000100689.2 | 0 | NA | NA | NA | NA | NA | #VALUE! |
| ENSMUSG00000100690.2 | 0 | NA | NA | NA | NA | NA | #VALUE! |
| ENSMUSG00000100692.2 | 0 | NA | NA | NA | NA | NA | #VALUE! |
| ENSMUSG00000100693.2 | 0 | NA | NA | NA | NA | NA | #VALUE! |
| ENSMUSG00000100694.2 | 0 | NA | NA | NA | NA | NA | #VALUE! |
| ENSMUSG00000100695.2 | 0 | NA | NA | NA | NA | NA | #VALUE! |
| ENSMUSG00000100696.2 | 0 | NA | NA | NA | NA | NA | #VALUE! |
| ENSMUSG00000100697.2 | 0 | NA | NA | NA | NA | NA | #VALUE! |
| ENSMUSG00000100698.2 | 0 | NA | NA | NA | NA | NA | #VALUE! |
| ENSMUSG00000100699.2 | 0 | NA | NA | NA | NA | NA | #VALUE! |
| ENSMUSG00000100700.2 | 0 | NA | NA | NA | NA | NA | #VALUE! |
| ENSMUSG00000100701.2 | 0 | NA | NA | NA | NA | NA | #VALUE! |

| | | | | | | | |
|---|---|---|---|---|---|---|---|
| ENSMUSG00000100702.2 | 0.093255 | 0.868093 | 4.072464 | 0.213162 | 0.831201 | NA | #VALUE! |
| ENSMUSG00000100703.7 | 0 | NA | NA | NA | NA | NA | #VALUE! |
| ENSMUSG00000100704.2 | 0 | NA | NA | NA | NA | NA | #VALUE! |
| ENSMUSG00000100705.2 | 0 | NA | NA | NA | NA | NA | #VALUE! |
| ENSMUSG00000100706.7 | 0 | NA | NA | NA | NA | NA | #VALUE! |
| ENSMUSG00000100707.2 | 3.870521 | 5.442409 | 3.918364 | 1.388949 | 0.164848 | NA | #VALUE! |
| ENSMUSG00000100708.2 | 0 | NA | NA | NA | NA | NA | #VALUE! |
| ENSMUSG00000100709.7 | 0 | NA | NA | NA | NA | NA | #VALUE! |
| ENSMUSG00000100710.2 | 0 | NA | NA | NA | NA | NA | #VALUE! |
| ENSMUSG00000100711.2 | 0 | NA | NA | NA | NA | NA | #VALUE! |
| ENSMUSG00000100712.2 | 0 | NA | NA | NA | NA | NA | #VALUE! |
| ENSMUSG00000100713.2 | 0 | NA | NA | NA | NA | NA | #VALUE! |
| ENSMUSG00000100714.7 | 0 | NA | NA | NA | NA | NA | #VALUE! |
| ENSMUSG00000100715.7 | 0 | NA | NA | NA | NA | NA | #VALUE! |
| ENSMUSG00000100716.7 | 0 | NA | NA | NA | NA | NA | #VALUE! |
| ENSMUSG00000100717.7 | 0 | NA | NA | NA | NA | NA | #VALUE! |
| ENSMUSG00000100718.2 | 0 | NA | NA | NA | NA | NA | #VALUE! |
| ENSMUSG00000100719.2 | 0 | NA | NA | NA | NA | NA | #VALUE! |
| ENSMUSG00000100720.2 | 0.093255 | 0.868093 | 4.072464 | 0.213162 | 0.831201 | NA | #VALUE! |
| ENSMUSG00000100721.2 | 0 | NA | NA | NA | NA | NA | #VALUE! |
| ENSMUSG00000100722.2 | 0 | NA | NA | NA | NA | NA | #VALUE! |
| ENSMUSG00000100723.2 | 0 | NA | NA | NA | NA | NA | #VALUE! |
| ENSMUSG00000100724.2 | 0 | NA | NA | NA | NA | NA | #VALUE! |
| ENSMUSG00000100726.2 | 0 | NA | NA | NA | NA | NA | #VALUE! |
| ENSMUSG00000100727.7 | 0 | NA | NA | NA | NA | NA | #VALUE! |
| ENSMUSG00000100728.2 | 0 | NA | NA | NA | NA | NA | #VALUE! |
| ENSMUSG00000100730.2 | 0 | NA | NA | NA | NA | NA | #VALUE! |
| ENSMUSG00000100731.2 | 0 | NA | NA | NA | NA | NA | #VALUE! |
| ENSMUSG00000100732.2 | 0 | NA | NA | NA | NA | NA | #VALUE! |
| ENSMUSG00000100733.3 | 0 | NA | NA | NA | NA | NA | #VALUE! |
| ENSMUSG00000100734.2 | 0 | NA | NA | NA | NA | NA | #VALUE! |
| ENSMUSG00000100736.2 | 0.168284 | 1.027628 | 4.067126 | 0.252667 | 0.800526 | NA | #VALUE! |
| ENSMUSG00000100737.2 | 0 | NA | NA | NA | NA | NA | #VALUE! |
| ENSMUSG00000100738.2 | 0 | NA | NA | NA | NA | NA | #VALUE! |
| ENSMUSG00000100739.7 | 0 | NA | NA | NA | NA | NA | #VALUE! |
| ENSMUSG00000100740.2 | 0 | NA | NA | NA | NA | NA | #VALUE! |
| ENSMUSG00000100741.2 | 0 | NA | NA | NA | NA | NA | #VALUE! |
| ENSMUSG00000100742.2 | 0 | NA | NA | NA | NA | NA | #VALUE! |
| ENSMUSG00000100743.2 | 0 | NA | NA | NA | NA | NA | #VALUE! |
| ENSMUSG00000100744.2 | 0 | NA | NA | NA | NA | NA | #VALUE! |
| ENSMUSG00000100745.2 | 0 | NA | NA | NA | NA | NA | #VALUE! |
| ENSMUSG00000100746.2 | 0 | NA | NA | NA | NA | NA | #VALUE! |
| ENSMUSG00000100747.5 | 14.60461 | 7.362655 | 2.861039 | 2.57342 | NA | NA | #VALUE! |
| ENSMUSG00000100749.2 | 0 | NA | NA | NA | NA | NA | #VALUE! |

| | | | | | | | |
|---|---|---|---|---|---|---|---|
| ENSMUSG00000100750.2 | 0 | NA | NA | NA | NA | NA | #VALUE! |
| ENSMUSG00000100751.2 | 0 | NA | NA | NA | NA | NA | #VALUE! |
| ENSMUSG00000100752.2 | 0 | NA | NA | NA | NA | NA | #VALUE! |
| ENSMUSG00000100753.2 | 0 | NA | NA | NA | NA | NA | #VALUE! |
| ENSMUSG00000100754.2 | 0 | NA | NA | NA | NA | NA | #VALUE! |
| ENSMUSG00000100755.2 | 0.879786 | -1.10645 | 3.93987 | -0.28083 | 0.778837 | NA | #VALUE! |
| ENSMUSG00000100756.2 | 0 | NA | NA | NA | NA | NA | #VALUE! |
| ENSMUSG00000100757.2 | 0 | NA | NA | NA | NA | NA | #VALUE! |
| ENSMUSG00000100758.2 | 0 | NA | NA | NA | NA | NA | #VALUE! |
| ENSMUSG00000100759.7 | 0 | NA | NA | NA | NA | NA | #VALUE! |
| ENSMUSG00000100760.2 | 0 | NA | NA | NA | NA | NA | #VALUE! |
| ENSMUSG00000100761.2 | 0 | NA | NA | NA | NA | NA | #VALUE! |
| ENSMUSG00000100762.2 | 0 | NA | NA | NA | NA | NA | #VALUE! |
| ENSMUSG00000100763.2 | 0 | NA | NA | NA | NA | NA | #VALUE! |
| ENSMUSG00000100764.2 | 5.715101 | -6.04207 | 3.914611 | -1.54347 | 0.122718 | NA | #VALUE! |
| ENSMUSG00000100765.2 | 0 | NA | NA | NA | NA | NA | #VALUE! |
| ENSMUSG00000100767.2 | 0 | NA | NA | NA | NA | NA | #VALUE! |
| ENSMUSG00000100768.2 | 2.471029 | 0.640827 | 3.803464 | 0.168485 | 0.866202 | NA | #VALUE! |
| ENSMUSG00000100769.2 | 0 | NA | NA | NA | NA | NA | #VALUE! |
| ENSMUSG00000100770.2 | 0 | NA | NA | NA | NA | NA | #VALUE! |
| ENSMUSG00000100771.2 | 0 | NA | NA | NA | NA | NA | #VALUE! |
| ENSMUSG00000100772.2 | 0 | NA | NA | NA | NA | NA | #VALUE! |
| ENSMUSG00000100773.2 | 0 | NA | NA | NA | NA | NA | #VALUE! |
| ENSMUSG00000100774.2 | 0 | NA | NA | NA | NA | NA | #VALUE! |
| ENSMUSG00000100775.2 | 4.969622 | -5.84048 | 3.915785 | -1.49152 | 0.135824 | NA | #VALUE! |
| ENSMUSG00000100776.2 | 0 | NA | NA | NA | NA | NA | #VALUE! |
| ENSMUSG00000100777.2 | 0 | NA | NA | NA | NA | NA | #VALUE! |
| ENSMUSG00000100778.2 | 0 | NA | NA | NA | NA | NA | #VALUE! |
| ENSMUSG00000100779.2 | 0 | NA | NA | NA | NA | NA | #VALUE! |
| ENSMUSG00000100780.2 | 0 | NA | NA | NA | NA | NA | #VALUE! |
| ENSMUSG00000100781.2 | 0 | NA | NA | NA | NA | NA | #VALUE! |
| ENSMUSG00000100782.2 | 0 | NA | NA | NA | NA | NA | #VALUE! |
| ENSMUSG00000100783.2 | 0 | NA | NA | NA | NA | NA | #VALUE! |
| ENSMUSG00000100784.2 | 0 | NA | NA | NA | NA | NA | #VALUE! |
| ENSMUSG00000100785.2 | 0 | NA | NA | NA | NA | NA | #VALUE! |
| ENSMUSG00000100786.2 | 0 | NA | NA | NA | NA | NA | #VALUE! |
| ENSMUSG00000100787.7 | 0 | NA | NA | NA | NA | NA | #VALUE! |
| ENSMUSG00000100788.2 | 0 | NA | NA | NA | NA | NA | #VALUE! |
| ENSMUSG00000100789.7 | 0 | NA | NA | NA | NA | NA | #VALUE! |
| ENSMUSG00000100790.7 | 0 | NA | NA | NA | NA | NA | #VALUE! |
| ENSMUSG00000100791.2 | 0 | NA | NA | NA | NA | NA | #VALUE! |
| ENSMUSG00000100792.2 | 0 | NA | NA | NA | NA | NA | #VALUE! |
| ENSMUSG00000100794.2 | 0 | NA | NA | NA | NA | NA | #VALUE! |
| ENSMUSG00000100795.3 | 0 | NA | NA | NA | NA | NA | #VALUE! |

| | | | | | | | |
|---|---|---|---|---|---|---|---|
| ENSMUSG00000100796.2 | 0 | NA | NA | NA | NA | NA | #VALUE! |
| ENSMUSG00000100797.2 | 0 | NA | NA | NA | NA | NA | #VALUE! |
| ENSMUSG00000100798.2 | 0 | NA | NA | NA | NA | NA | #VALUE! |
| ENSMUSG00000100799.3 | 0 | NA | NA | NA | NA | NA | #VALUE! |
| ENSMUSG00000100800.2 | 0 | NA | NA | NA | NA | NA | #VALUE! |
| ENSMUSG00000100802.2 | 0 | NA | NA | NA | NA | NA | #VALUE! |
| ENSMUSG00000100803.2 | 0 | NA | NA | NA | NA | NA | #VALUE! |
| ENSMUSG00000100804.2 | 0 | NA | NA | NA | NA | NA | #VALUE! |
| ENSMUSG00000100805.2 | 0 | NA | NA | NA | NA | NA | #VALUE! |
| ENSMUSG00000100806.2 | 0 | NA | NA | NA | NA | NA | #VALUE! |
| ENSMUSG00000100807.3 | 0 | NA | NA | NA | NA | NA | #VALUE! |
| ENSMUSG00000100808.2 | 0 | NA | NA | NA | NA | NA | #VALUE! |
| ENSMUSG00000100809.2 | 0 | NA | NA | NA | NA | NA | #VALUE! |
| ENSMUSG00000100810.2 | 0 | NA | NA | NA | NA | NA | #VALUE! |
| ENSMUSG00000100811.2 | 0 | NA | NA | NA | NA | NA | #VALUE! |
| ENSMUSG00000100812.2 | 0 | NA | NA | NA | NA | NA | #VALUE! |
| ENSMUSG00000100813.2 | 0 | NA | NA | NA | NA | NA | #VALUE! |
| ENSMUSG00000100814.2 | 0 | NA | NA | NA | NA | NA | #VALUE! |
| ENSMUSG00000100815.2 | 0 | NA | NA | NA | NA | NA | #VALUE! |
| ENSMUSG00000100816.2 | 0 | NA | NA | NA | NA | NA | #VALUE! |
| ENSMUSG00000100817.2 | 0 | NA | NA | NA | NA | NA | #VALUE! |
| ENSMUSG00000100818.2 | 0 | NA | NA | NA | NA | NA | #VALUE! |
| ENSMUSG00000100819.2 | 0 | NA | NA | NA | NA | NA | #VALUE! |
| ENSMUSG00000100820.2 | 0 | NA | NA | NA | NA | NA | #VALUE! |
| ENSMUSG00000100821.2 | 0 | NA | NA | NA | NA | NA | #VALUE! |
| ENSMUSG00000100822.2 | 0 | NA | NA | NA | NA | NA | #VALUE! |
| ENSMUSG00000100823.2 | 0 | NA | NA | NA | NA | NA | #VALUE! |
| ENSMUSG00000100825.2 | 0 | NA | NA | NA | NA | NA | #VALUE! |
| ENSMUSG00000100827.2 | 0 | NA | NA | NA | NA | NA | #VALUE! |
| ENSMUSG00000100830.2 | 0 | NA | NA | NA | NA | NA | #VALUE! |
| ENSMUSG00000100831.2 | 0 | NA | NA | NA | NA | NA | #VALUE! |
| ENSMUSG00000100832.2 | 0 | NA | NA | NA | NA | NA | #VALUE! |
| ENSMUSG00000100833.2 | 0 | NA | NA | NA | NA | NA | #VALUE! |
| ENSMUSG00000100834.2 | 0 | NA | NA | NA | NA | NA | #VALUE! |
| ENSMUSG00000100835.2 | 0 | NA | NA | NA | NA | NA | #VALUE! |
| ENSMUSG00000100836.2 | 0 | NA | NA | NA | NA | NA | #VALUE! |
| ENSMUSG00000100837.8 | 0 | NA | NA | NA | NA | NA | #VALUE! |
| ENSMUSG00000100838.2 | 0 | NA | NA | NA | NA | NA | #VALUE! |
| ENSMUSG00000100839.2 | 0 | NA | NA | NA | NA | NA | #VALUE! |
| ENSMUSG00000100841.2 | 0 | NA | NA | NA | NA | NA | #VALUE! |
| ENSMUSG00000100842.2 | 0 | NA | NA | NA | NA | NA | #VALUE! |
| ENSMUSG00000100843.2 | 0 | NA | NA | NA | NA | NA | #VALUE! |
| ENSMUSG00000100844.2 | 0 | NA | NA | NA | NA | NA | #VALUE! |
| ENSMUSG00000100845.2 | 0 | NA | NA | NA | NA | NA | #VALUE! |

| | | | | | | | |
|---|---|---|---|---|---|---|---|
| ENSMUSG00000100846.2 | 0 | NA | NA | NA | NA | NA | #VALUE! |
| ENSMUSG00000100847.2 | 0 | NA | NA | NA | NA | NA | #VALUE! |
| ENSMUSG00000100848.2 | 0 | NA | NA | NA | NA | NA | #VALUE! |
| ENSMUSG00000100849.2 | 0 | NA | NA | NA | NA | NA | #VALUE! |
| ENSMUSG00000100850.2 | 0 | NA | NA | NA | NA | NA | #VALUE! |
| ENSMUSG00000100852.2 | 0 | NA | NA | NA | NA | NA | #VALUE! |
| ENSMUSG00000100853.2 | 0 | NA | NA | NA | NA | NA | #VALUE! |
| ENSMUSG00000100854.2 | 0 | NA | NA | NA | NA | NA | #VALUE! |
| ENSMUSG00000100856.2 | 0 | NA | NA | NA | NA | NA | #VALUE! |
| ENSMUSG00000100857.3 | 0 | NA | NA | NA | NA | NA | #VALUE! |
| ENSMUSG00000100858.2 | 0 | NA | NA | NA | NA | NA | #VALUE! |
| ENSMUSG00000100859.7 | 0 | NA | NA | NA | NA | NA | #VALUE! |
| ENSMUSG00000100860.2 | 0 | NA | NA | NA | NA | NA | #VALUE! |
| ENSMUSG00000100861.2 | 0 | NA | NA | NA | NA | NA | #VALUE! |
| ENSMUSG00000100862.2 | 5.135223 | 0.927812 | 2.828768 | 0.327992 | 0.742918 | NA | #VALUE! |
| ENSMUSG00000100863.2 | 1.737971 | -2.02686 | 3.88627 | -0.52154 | 0.601988 | NA | #VALUE! |
| ENSMUSG00000100864.2 | 0 | NA | NA | NA | NA | NA | #VALUE! |
| ENSMUSG00000100866.2 | 0 | NA | NA | NA | NA | NA | #VALUE! |
| ENSMUSG00000100867.2 | 0 | NA | NA | NA | NA | NA | #VALUE! |
| ENSMUSG00000100868.2 | 0 | NA | NA | NA | NA | NA | #VALUE! |
| ENSMUSG00000100869.2 | 0 | NA | NA | NA | NA | NA | #VALUE! |
| ENSMUSG00000100870.2 | 0 | NA | NA | NA | NA | NA | #VALUE! |
| ENSMUSG00000100871.2 | 0 | NA | NA | NA | NA | NA | #VALUE! |
| ENSMUSG00000100872.3 | 0 | NA | NA | NA | NA | NA | #VALUE! |
| ENSMUSG00000100873.2 | 0 | NA | NA | NA | NA | NA | #VALUE! |
| ENSMUSG00000100874.2 | 0 | NA | NA | NA | NA | NA | #VALUE! |
| ENSMUSG00000100875.7 | 0 | NA | NA | NA | NA | NA | #VALUE! |
| ENSMUSG00000100876.2 | 0 | NA | NA | NA | NA | NA | #VALUE! |
| ENSMUSG00000100877.2 | 0 | NA | NA | NA | NA | NA | #VALUE! |
| ENSMUSG00000100878.2 | 0 | NA | NA | NA | NA | NA | #VALUE! |
| ENSMUSG00000100879.2 | 0 | NA | NA | NA | NA | NA | #VALUE! |
| ENSMUSG00000100880.3 | 0 | NA | NA | NA | NA | NA | #VALUE! |
| ENSMUSG00000100881.2 | 0 | NA | NA | NA | NA | NA | #VALUE! |
| ENSMUSG00000100882.2 | 0 | NA | NA | NA | NA | NA | #VALUE! |
| ENSMUSG00000100883.2 | 0 | NA | NA | NA | NA | NA | #VALUE! |
| ENSMUSG00000100884.2 | 0 | NA | NA | NA | NA | NA | #VALUE! |
| ENSMUSG00000100885.2 | 0 | NA | NA | NA | NA | NA | #VALUE! |
| ENSMUSG00000100886.2 | 0 | NA | NA | NA | NA | NA | #VALUE! |
| ENSMUSG00000100887.2 | 0 | NA | NA | NA | NA | NA | #VALUE! |
| ENSMUSG00000100888.2 | 0 | NA | NA | NA | NA | NA | #VALUE! |
| ENSMUSG00000100889.2 | 0 | NA | NA | NA | NA | NA | #VALUE! |
| ENSMUSG00000100890.2 | 0 | NA | NA | NA | NA | NA | #VALUE! |
| ENSMUSG00000100891.2 | 0 | NA | NA | NA | NA | NA | #VALUE! |
| ENSMUSG00000100892.2 | 0 | NA | NA | NA | NA | NA | #VALUE! |

| | | | | | | | |
|---|---|---|---|---|---|---|---|
| ENSMUSG00000100893.2 | 0 | NA | NA | NA | NA | NA | #VALUE! |
| ENSMUSG00000100894.2 | 0 | NA | NA | NA | NA | NA | #VALUE! |
| ENSMUSG00000100895.2 | 0 | NA | NA | NA | NA | NA | #VALUE! |
| ENSMUSG00000100896.2 | 0 | NA | NA | NA | NA | NA | #VALUE! |
| ENSMUSG00000100897.2 | 0 | NA | NA | NA | NA | NA | #VALUE! |
| ENSMUSG00000100898.7 | 0 | NA | NA | NA | NA | NA | #VALUE! |
| ENSMUSG00000100899.3 | 0 | NA | NA | NA | NA | NA | #VALUE! |
| ENSMUSG00000100900.7 | 0 | NA | NA | NA | NA | NA | #VALUE! |
| ENSMUSG00000100901.2 | 0 | NA | NA | NA | NA | NA | #VALUE! |
| ENSMUSG00000100902.2 | 0 | NA | NA | NA | NA | NA | #VALUE! |
| ENSMUSG00000100903.2 | 0 | NA | NA | NA | NA | NA | #VALUE! |
| ENSMUSG00000100904.2 | 0 | NA | NA | NA | NA | NA | #VALUE! |
| ENSMUSG00000100905.2 | 0 | NA | NA | NA | NA | NA | #VALUE! |
| ENSMUSG00000100906.2 | 0 | NA | NA | NA | NA | NA | #VALUE! |
| ENSMUSG00000100907.2 | 0 | NA | NA | NA | NA | NA | #VALUE! |
| ENSMUSG00000100908.2 | 0 | NA | NA | NA | NA | NA | #VALUE! |
| ENSMUSG00000100910.2 | 0 | NA | NA | NA | NA | NA | #VALUE! |
| ENSMUSG00000100911.3 | 0 | NA | NA | NA | NA | NA | #VALUE! |
| ENSMUSG00000100912.2 | 0 | NA | NA | NA | NA | NA | #VALUE! |
| ENSMUSG00000100913.2 | 0 | NA | NA | NA | NA | NA | #VALUE! |
| ENSMUSG00000100914.2 | 0 | NA | NA | NA | NA | NA | #VALUE! |
| ENSMUSG00000100915.7 | 0 | NA | NA | NA | NA | NA | #VALUE! |
| ENSMUSG00000100916.4 | 0 | NA | NA | NA | NA | NA | #VALUE! |
| ENSMUSG00000100917.2 | 0 | NA | NA | NA | NA | NA | #VALUE! |
| ENSMUSG00000100918.2 | 0 | NA | NA | NA | NA | NA | #VALUE! |
| ENSMUSG00000100919.2 | 0 | NA | NA | NA | NA | NA | #VALUE! |
| ENSMUSG00000100920.2 | 0 | NA | NA | NA | NA | NA | #VALUE! |
| ENSMUSG00000100922.2 | 0 | NA | NA | NA | NA | NA | #VALUE! |
| ENSMUSG00000100923.3 | 0 | NA | NA | NA | NA | NA | #VALUE! |
| ENSMUSG00000100925.2 | 0 | NA | NA | NA | NA | NA | #VALUE! |
| ENSMUSG00000100926.2 | 0 | NA | NA | NA | NA | NA | #VALUE! |
| ENSMUSG00000100927.2 | 0 | NA | NA | NA | NA | NA | #VALUE! |
| ENSMUSG00000100928.7 | 0 | NA | NA | NA | NA | NA | #VALUE! |
| ENSMUSG00000100929.2 | 0 | NA | NA | NA | NA | NA | #VALUE! |
| ENSMUSG00000100930.2 | 0 | NA | NA | NA | NA | NA | #VALUE! |
| ENSMUSG00000100931.2 | 0 | NA | NA | NA | NA | NA | #VALUE! |
| ENSMUSG00000100932.2 | 0 | NA | NA | NA | NA | NA | #VALUE! |
| ENSMUSG00000100933.2 | 0 | NA | NA | NA | NA | NA | #VALUE! |
| ENSMUSG00000100935.2 | 0 | NA | NA | NA | NA | NA | #VALUE! |
| ENSMUSG00000100936.2 | 0 | NA | NA | NA | NA | NA | #VALUE! |
| ENSMUSG00000100937.2 | 0 | NA | NA | NA | NA | NA | #VALUE! |
| ENSMUSG00000100938.2 | 0 | NA | NA | NA | NA | NA | #VALUE! |
| ENSMUSG00000100939.2 | 0 | NA | NA | NA | NA | NA | #VALUE! |
| ENSMUSG00000100940.2 | 0 | NA | NA | NA | NA | NA | #VALUE! |

| | | | | | | | |
|---|---|---|---|---|---|---|---|
| ENSMUSG00000100941.2 | 0 | NA | NA | NA | NA | NA | #VALUE! |
| ENSMUSG00000100942.2 | 0 | NA | NA | NA | NA | NA | #VALUE! |
| ENSMUSG00000100943.2 | 0 | NA | NA | NA | NA | NA | #VALUE! |
| ENSMUSG00000100944.2 | 0 | NA | NA | NA | NA | NA | #VALUE! |
| ENSMUSG00000100945.2 | 0 | NA | NA | NA | NA | NA | #VALUE! |
| ENSMUSG00000100946.2 | 0 | NA | NA | NA | NA | NA | #VALUE! |
| ENSMUSG00000100947.2 | 0 | NA | NA | NA | NA | NA | #VALUE! |
| ENSMUSG00000100948.2 | 0 | NA | NA | NA | NA | NA | #VALUE! |
| ENSMUSG00000100949.2 | 0 | NA | NA | NA | NA | NA | #VALUE! |
| ENSMUSG00000100950.2 | 0 | NA | NA | NA | NA | NA | #VALUE! |
| ENSMUSG00000100951.2 | 0 | NA | NA | NA | NA | NA | #VALUE! |
| ENSMUSG00000100952.2 | 0 | NA | NA | NA | NA | NA | #VALUE! |
| ENSMUSG00000100953.7 | 0 | NA | NA | NA | NA | NA | #VALUE! |
| ENSMUSG00000100954.6 | 4.503149 | -5.6983 | 3.916718 | -1.45487 | 0.145706 | NA | #VALUE! |
| ENSMUSG00000100955.2 | 0 | NA | NA | NA | NA | NA | #VALUE! |
| ENSMUSG00000100956.2 | 0 | NA | NA | NA | NA | NA | #VALUE! |
| ENSMUSG00000100957.7 | 0 | NA | NA | NA | NA | NA | #VALUE! |
| ENSMUSG00000100958.2 | 0 | NA | NA | NA | NA | NA | #VALUE! |
| ENSMUSG00000100959.2 | 0 | NA | NA | NA | NA | NA | #VALUE! |
| ENSMUSG00000100960.2 | 0 | NA | NA | NA | NA | NA | #VALUE! |
| ENSMUSG00000100961.2 | 0 | NA | NA | NA | NA | NA | #VALUE! |
| ENSMUSG00000100962.4 | 0 | NA | NA | NA | NA | NA | #VALUE! |
| ENSMUSG00000100963.2 | 0 | NA | NA | NA | NA | NA | #VALUE! |
| ENSMUSG00000100964.2 | 0 | NA | NA | NA | NA | NA | #VALUE! |
| ENSMUSG00000100965.2 | 0 | NA | NA | NA | NA | NA | #VALUE! |
| ENSMUSG00000100966.2 | 0 | NA | NA | NA | NA | NA | #VALUE! |
| ENSMUSG00000100967.2 | 2.524253 | 4.825754 | 3.924522 | 1.229641 | 0.218831 | NA | #VALUE! |
| ENSMUSG00000100968.2 | 0 | NA | NA | NA | NA | NA | #VALUE! |
| ENSMUSG00000100969.8 | 0 | NA | NA | NA | NA | NA | #VALUE! |
| ENSMUSG00000100970.2 | 0 | NA | NA | NA | NA | NA | #VALUE! |
| ENSMUSG00000100971.2 | 0 | NA | NA | NA | NA | NA | #VALUE! |
| ENSMUSG00000100972.2 | 0 | NA | NA | NA | NA | NA | #VALUE! |
| ENSMUSG00000100973.2 | 0 | NA | NA | NA | NA | NA | #VALUE! |
| ENSMUSG00000100974.2 | 0 | NA | NA | NA | NA | NA | #VALUE! |
| ENSMUSG00000100976.2 | 0 | NA | NA | NA | NA | NA | #VALUE! |
| ENSMUSG00000100977.2 | 0 | NA | NA | NA | NA | NA | #VALUE! |
| ENSMUSG00000100978.2 | 0 | NA | NA | NA | NA | NA | #VALUE! |
| ENSMUSG00000100979.2 | 0 | NA | NA | NA | NA | NA | #VALUE! |
| ENSMUSG00000100980.2 | 0 | NA | NA | NA | NA | NA | #VALUE! |
| ENSMUSG00000100981.2 | 0 | NA | NA | NA | NA | NA | #VALUE! |
| ENSMUSG00000100982.2 | 0 | NA | NA | NA | NA | NA | #VALUE! |
| ENSMUSG00000100983.2 | 0 | NA | NA | NA | NA | NA | #VALUE! |
| ENSMUSG00000100984.2 | 0 | NA | NA | NA | NA | NA | #VALUE! |
| ENSMUSG00000100985.2 | 0 | NA | NA | NA | NA | NA | #VALUE! |

| | | | | | | | |
|---|---|---|---|---|---|---|---|
| ENSMUSG00000100986.2 | 0 | NA | NA | NA | NA | NA | #VALUE! |
| ENSMUSG00000100987.2 | 1.365171 | -3.99822 | 3.938957 | -1.01505 | 0.310084 | NA | #VALUE! |
| ENSMUSG00000100988.7 | 0 | NA | NA | NA | NA | NA | #VALUE! |
| ENSMUSG00000100989.2 | 0 | NA | NA | NA | NA | NA | #VALUE! |
| ENSMUSG00000100990.2 | 0 | NA | NA | NA | NA | NA | #VALUE! |
| ENSMUSG00000100991.2 | 0 | NA | NA | NA | NA | NA | #VALUE! |
| ENSMUSG00000100992.2 | 0 | NA | NA | NA | NA | NA | #VALUE! |
| ENSMUSG00000100993.2 | 0 | NA | NA | NA | NA | NA | #VALUE! |
| ENSMUSG00000100994.2 | 0 | NA | NA | NA | NA | NA | #VALUE! |
| ENSMUSG00000100995.2 | 0 | NA | NA | NA | NA | NA | #VALUE! |
| ENSMUSG00000100996.2 | 0 | NA | NA | NA | NA | NA | #VALUE! |
| ENSMUSG00000100998.2 | 0 | NA | NA | NA | NA | NA | #VALUE! |
| ENSMUSG00000100999.2 | 0 | NA | NA | NA | NA | NA | #VALUE! |
| ENSMUSG00000101000.2 | 0 | NA | NA | NA | NA | NA | #VALUE! |
| ENSMUSG00000101001.2 | 0 | NA | NA | NA | NA | NA | #VALUE! |
| ENSMUSG00000101002.2 | 0 | NA | NA | NA | NA | NA | #VALUE! |
| ENSMUSG00000101003.7 | 0 | NA | NA | NA | NA | NA | #VALUE! |
| ENSMUSG00000101004.2 | 0 | NA | NA | NA | NA | NA | #VALUE! |
| ENSMUSG00000101005.2 | 0 | NA | NA | NA | NA | NA | #VALUE! |
| ENSMUSG00000101006.2 | 0 | NA | NA | NA | NA | NA | #VALUE! |
| ENSMUSG00000101007.2 | 0 | NA | NA | NA | NA | NA | #VALUE! |
| ENSMUSG00000101008.2 | 0 | NA | NA | NA | NA | NA | #VALUE! |
| ENSMUSG00000101009.7 | 0 | NA | NA | NA | NA | NA | #VALUE! |
| ENSMUSG00000101010.2 | 0 | NA | NA | NA | NA | NA | #VALUE! |
| ENSMUSG00000101011.7 | 0 | NA | NA | NA | NA | NA | #VALUE! |
| ENSMUSG00000101012.7 | 0 | NA | NA | NA | NA | NA | #VALUE! |
| ENSMUSG00000101014.2 | 0 | NA | NA | NA | NA | NA | #VALUE! |
| ENSMUSG00000101015.2 | 0 | NA | NA | NA | NA | NA | #VALUE! |
| ENSMUSG00000101016.2 | 0 | NA | NA | NA | NA | NA | #VALUE! |
| ENSMUSG00000101017.2 | 0 | NA | NA | NA | NA | NA | #VALUE! |
| ENSMUSG00000101018.2 | 0 | NA | NA | NA | NA | NA | #VALUE! |
| ENSMUSG00000101019.2 | 0 | NA | NA | NA | NA | NA | #VALUE! |
| ENSMUSG00000101020.2 | 0 | NA | NA | NA | NA | NA | #VALUE! |
| ENSMUSG00000101021.2 | 0 | NA | NA | NA | NA | NA | #VALUE! |
| ENSMUSG00000101022.2 | 0 | NA | NA | NA | NA | NA | #VALUE! |
| ENSMUSG00000101023.2 | 0 | NA | NA | NA | NA | NA | #VALUE! |
| ENSMUSG00000101024.2 | 0 | NA | NA | NA | NA | NA | #VALUE! |
| ENSMUSG00000101025.2 | 0 | NA | NA | NA | NA | NA | #VALUE! |
| ENSMUSG00000101026.2 | 0 | NA | NA | NA | NA | NA | #VALUE! |
| ENSMUSG00000101027.2 | 0 | NA | NA | NA | NA | NA | #VALUE! |
| ENSMUSG00000101028.2 | 0 | NA | NA | NA | NA | NA | #VALUE! |
| ENSMUSG00000101029.2 | 0 | NA | NA | NA | NA | NA | #VALUE! |
| ENSMUSG00000101030.2 | 0 | NA | NA | NA | NA | NA | #VALUE! |
| ENSMUSG00000101031.3 | 0 | NA | NA | NA | NA | NA | #VALUE! |

| | | | | | | | |
|---|---|---|---|---|---|---|---|
| ENSMUSG00000101032.7 | 0 | NA | NA | NA | NA | NA | #VALUE! |
| ENSMUSG00000101033.2 | 0 | NA | NA | NA | NA | NA | #VALUE! |
| ENSMUSG00000101035.2 | 0 | NA | NA | NA | NA | NA | #VALUE! |
| ENSMUSG00000101036.2 | 0 | NA | NA | NA | NA | NA | #VALUE! |
| ENSMUSG00000101037.2 | 0 | NA | NA | NA | NA | NA | #VALUE! |
| ENSMUSG00000101038.2 | 0 | NA | NA | NA | NA | NA | #VALUE! |
| ENSMUSG00000101039.2 | 0 | NA | NA | NA | NA | NA | #VALUE! |
| ENSMUSG00000101040.7 | 0 | NA | NA | NA | NA | NA | #VALUE! |
| ENSMUSG00000101041.2 | 0 | NA | NA | NA | NA | NA | #VALUE! |
| ENSMUSG00000101042.2 | 0 | NA | NA | NA | NA | NA | #VALUE! |
| ENSMUSG00000101043.2 | 0 | NA | NA | NA | NA | NA | #VALUE! |
| ENSMUSG00000101044.2 | 0 | NA | NA | NA | NA | NA | #VALUE! |
| ENSMUSG00000101045.2 | 0 | NA | NA | NA | NA | NA | #VALUE! |
| ENSMUSG00000101046.2 | 0 | NA | NA | NA | NA | NA | #VALUE! |
| ENSMUSG00000101049.2 | 0 | NA | NA | NA | NA | NA | #VALUE! |
| ENSMUSG00000101050.2 | 0 | NA | NA | NA | NA | NA | #VALUE! |
| ENSMUSG00000101051.7 | 0 | NA | NA | NA | NA | NA | #VALUE! |
| ENSMUSG00000101052.2 | 0 | NA | NA | NA | NA | NA | #VALUE! |
| ENSMUSG00000101053.2 | 0 | NA | NA | NA | NA | NA | #VALUE! |
| ENSMUSG00000101054.2 | 0 | NA | NA | NA | NA | NA | #VALUE! |
| ENSMUSG00000101055.2 | 0 | NA | NA | NA | NA | NA | #VALUE! |
| ENSMUSG00000101056.2 | 0 | NA | NA | NA | NA | NA | #VALUE! |
| ENSMUSG00000101057.2 | 0 | NA | NA | NA | NA | NA | #VALUE! |
| ENSMUSG00000101058.2 | 0 | NA | NA | NA | NA | NA | #VALUE! |
| ENSMUSG00000101059.2 | 0.168284 | 1.027628 | 4.067126 | 0.252667 | 0.800526 | NA | #VALUE! |
| ENSMUSG00000101060.2 | 0 | NA | NA | NA | NA | NA | #VALUE! |
| ENSMUSG00000101061.2 | 0 | NA | NA | NA | NA | NA | #VALUE! |
| ENSMUSG00000101062.2 | 0 | NA | NA | NA | NA | NA | #VALUE! |
| ENSMUSG00000101063.2 | 0 | NA | NA | NA | NA | NA | #VALUE! |
| ENSMUSG00000101064.2 | 0 | NA | NA | NA | NA | NA | #VALUE! |
| ENSMUSG00000101065.2 | 0 | NA | NA | NA | NA | NA | #VALUE! |
| ENSMUSG00000101066.2 | 0 | NA | NA | NA | NA | NA | #VALUE! |
| ENSMUSG00000101067.2 | 0 | NA | NA | NA | NA | NA | #VALUE! |
| ENSMUSG00000101068.2 | 0 | NA | NA | NA | NA | NA | #VALUE! |
| ENSMUSG00000101069.2 | 0 | NA | NA | NA | NA | NA | #VALUE! |
| ENSMUSG00000101070.2 | 0 | NA | NA | NA | NA | NA | #VALUE! |
| ENSMUSG00000101071.2 | 0 | NA | NA | NA | NA | NA | #VALUE! |
| ENSMUSG00000101072.2 | 0 | NA | NA | NA | NA | NA | #VALUE! |
| ENSMUSG00000101073.3 | 0 | NA | NA | NA | NA | NA | #VALUE! |
| ENSMUSG00000101074.2 | 0 | NA | NA | NA | NA | NA | #VALUE! |
| ENSMUSG00000101075.2 | 0 | NA | NA | NA | NA | NA | #VALUE! |
| ENSMUSG00000101076.3 | 0 | NA | NA | NA | NA | NA | #VALUE! |
| ENSMUSG00000101077.2 | 0 | NA | NA | NA | NA | NA | #VALUE! |
| ENSMUSG00000101078.4 | 0 | NA | NA | NA | NA | NA | #VALUE! |

| | | | | | | | |
|---|---|---|---|---|---|---|---|
| ENSMUSG00000101079.2 | 0 | NA | NA | NA | NA | NA | #VALUE! |
| ENSMUSG00000101080.2 | 0 | NA | NA | NA | NA | NA | #VALUE! |
| ENSMUSG00000101081.2 | 0 | NA | NA | NA | NA | NA | #VALUE! |
| ENSMUSG00000101082.2 | 0 | NA | NA | NA | NA | NA | #VALUE! |
| ENSMUSG00000101083.2 | 0 | NA | NA | NA | NA | NA | #VALUE! |
| ENSMUSG00000101084.2 | 0 | NA | NA | NA | NA | NA | #VALUE! |
| ENSMUSG00000101085.2 | 0 | NA | NA | NA | NA | NA | #VALUE! |
| ENSMUSG00000101086.3 | 0 | NA | NA | NA | NA | NA | #VALUE! |
| ENSMUSG00000101087.2 | 0 | NA | NA | NA | NA | NA | #VALUE! |
| ENSMUSG00000101088.7 | 0 | NA | NA | NA | NA | NA | #VALUE! |
| ENSMUSG00000101089.2 | 0 | NA | NA | NA | NA | NA | #VALUE! |
| ENSMUSG00000101090.7 | 0 | NA | NA | NA | NA | NA | #VALUE! |
| ENSMUSG00000101091.2 | 0 | NA | NA | NA | NA | NA | #VALUE! |
| ENSMUSG00000101092.2 | 0 | NA | NA | NA | NA | NA | #VALUE! |
| ENSMUSG00000101093.2 | 0 | NA | NA | NA | NA | NA | #VALUE! |
| ENSMUSG00000101094.2 | 0 | NA | NA | NA | NA | NA | #VALUE! |
| ENSMUSG00000101095.2 | 0 | NA | NA | NA | NA | NA | #VALUE! |
| ENSMUSG00000101096.2 | 0 | NA | NA | NA | NA | NA | #VALUE! |
| ENSMUSG00000101097.2 | 0 | NA | NA | NA | NA | NA | #VALUE! |
| ENSMUSG00000101098.2 | 0 | NA | NA | NA | NA | NA | #VALUE! |
| ENSMUSG00000101099.7 | 0 | NA | NA | NA | NA | NA | #VALUE! |
| ENSMUSG00000101100.7 | 0 | NA | NA | NA | NA | NA | #VALUE! |
| ENSMUSG00000101101.2 | 0 | NA | NA | NA | NA | NA | #VALUE! |
| ENSMUSG00000101102.2 | 0 | NA | NA | NA | NA | NA | #VALUE! |
| ENSMUSG00000101103.2 | 0 | NA | NA | NA | NA | NA | #VALUE! |
| ENSMUSG00000101104.7 | 0 | NA | NA | NA | NA | NA | #VALUE! |
| ENSMUSG00000101105.2 | 0 | NA | NA | NA | NA | NA | #VALUE! |
| ENSMUSG00000101106.7 | 0 | NA | NA | NA | NA | NA | #VALUE! |
| ENSMUSG00000101107.2 | 0 | NA | NA | NA | NA | NA | #VALUE! |
| ENSMUSG00000101108.2 | 0 | NA | NA | NA | NA | NA | #VALUE! |
| ENSMUSG00000101109.7 | 0 | NA | NA | NA | NA | NA | #VALUE! |
| ENSMUSG00000101110.2 | 0 | NA | NA | NA | NA | NA | #VALUE! |
| ENSMUSG00000101112.2 | 0 | NA | NA | NA | NA | NA | #VALUE! |
| ENSMUSG00000101113.2 | 0 | NA | NA | NA | NA | NA | #VALUE! |
| ENSMUSG00000101114.2 | 0 | NA | NA | NA | NA | NA | #VALUE! |
| ENSMUSG00000101115.2 | 0 | NA | NA | NA | NA | NA | #VALUE! |
| ENSMUSG00000101116.2 | 0 | NA | NA | NA | NA | NA | #VALUE! |
| ENSMUSG00000101117.2 | 0 | NA | NA | NA | NA | NA | #VALUE! |
| ENSMUSG00000101118.2 | 0 | NA | NA | NA | NA | NA | #VALUE! |
| ENSMUSG00000101119.2 | 0 | NA | NA | NA | NA | NA | #VALUE! |
| ENSMUSG00000101120.2 | 0 | NA | NA | NA | NA | NA | #VALUE! |
| ENSMUSG00000101122.2 | 1.995249 | -4.53986 | 3.928926 | -1.1555 | 0.247887 | NA | #VALUE! |
| ENSMUSG00000101123.2 | 0 | NA | NA | NA | NA | NA | #VALUE! |
| ENSMUSG00000101124.2 | 0 | NA | NA | NA | NA | NA | #VALUE! |

| | | | | | | | |
|---|---|---|---|---|---|---|---|
| ENSMUSG00000101125.2 | 0 | NA | NA | NA | NA | NA | #VALUE! |
| ENSMUSG00000101126.2 | 0 | NA | NA | NA | NA | NA | #VALUE! |
| ENSMUSG00000101127.2 | 0 | NA | NA | NA | NA | NA | #VALUE! |
| ENSMUSG00000101128.2 | 0 | NA | NA | NA | NA | NA | #VALUE! |
| ENSMUSG00000101129.2 | 0 | NA | NA | NA | NA | NA | #VALUE! |
| ENSMUSG00000101130.2 | 0 | NA | NA | NA | NA | NA | #VALUE! |
| ENSMUSG00000101131.2 | 0 | NA | NA | NA | NA | NA | #VALUE! |
| ENSMUSG00000101132.3 | 0.652786 | 2.919659 | 3.972474 | 0.734972 | 0.462356 | NA | #VALUE! |
| ENSMUSG00000101133.2 | 0 | NA | NA | NA | NA | NA | #VALUE! |
| ENSMUSG00000101134.2 | 0 | NA | NA | NA | NA | NA | #VALUE! |
| ENSMUSG00000101135.2 | 0 | NA | NA | NA | NA | NA | #VALUE! |
| ENSMUSG00000101136.2 | 0 | NA | NA | NA | NA | NA | #VALUE! |
| ENSMUSG00000101137.2 | 0 | NA | NA | NA | NA | NA | #VALUE! |
| ENSMUSG00000101138.7 | 0 | NA | NA | NA | NA | NA | #VALUE! |
| ENSMUSG00000101139.2 | 0 | NA | NA | NA | NA | NA | #VALUE! |
| ENSMUSG00000101140.2 | 0 | NA | NA | NA | NA | NA | #VALUE! |
| ENSMUSG00000101141.2 | 0 | NA | NA | NA | NA | NA | #VALUE! |
| ENSMUSG00000101142.2 | 0 | NA | NA | NA | NA | NA | #VALUE! |
| ENSMUSG00000101143.2 | 0 | NA | NA | NA | NA | NA | #VALUE! |
| ENSMUSG00000101144.3 | 0 | NA | NA | NA | NA | NA | #VALUE! |
| ENSMUSG00000101145.2 | 0 | NA | NA | NA | NA | NA | #VALUE! |
| ENSMUSG00000101146.2 | 0 | NA | NA | NA | NA | NA | #VALUE! |
| ENSMUSG00000101147.2 | 0 | NA | NA | NA | NA | NA | #VALUE! |
| ENSMUSG00000101148.2 | 0 | NA | NA | NA | NA | NA | #VALUE! |
| ENSMUSG00000101149.2 | 0 | NA | NA | NA | NA | NA | #VALUE! |
| ENSMUSG00000101150.2 | 0 | NA | NA | NA | NA | NA | #VALUE! |
| ENSMUSG00000101151.2 | 0 | NA | NA | NA | NA | NA | #VALUE! |
| ENSMUSG00000101152.2 | 0 | NA | NA | NA | NA | NA | #VALUE! |
| ENSMUSG00000101153.2 | 0 | NA | NA | NA | NA | NA | #VALUE! |
| ENSMUSG00000101154.2 | 0 | NA | NA | NA | NA | NA | #VALUE! |
| ENSMUSG00000101155.7 | 0 | NA | NA | NA | NA | NA | #VALUE! |
| ENSMUSG00000101156.2 | 0 | NA | NA | NA | NA | NA | #VALUE! |
| ENSMUSG00000101157.2 | 0 | NA | NA | NA | NA | NA | #VALUE! |
| ENSMUSG00000101158.2 | 0 | NA | NA | NA | NA | NA | #VALUE! |
| ENSMUSG00000101159.2 | 0 | NA | NA | NA | NA | NA | #VALUE! |
| ENSMUSG00000101160.2 | 0 | NA | NA | NA | NA | NA | #VALUE! |
| ENSMUSG00000101161.2 | 0 | NA | NA | NA | NA | NA | #VALUE! |
| ENSMUSG00000101162.3 | 0 | NA | NA | NA | NA | NA | #VALUE! |
| ENSMUSG00000101163.2 | 0 | NA | NA | NA | NA | NA | #VALUE! |
| ENSMUSG00000101164.7 | 0 | NA | NA | NA | NA | NA | #VALUE! |
| ENSMUSG00000101166.7 | 0 | NA | NA | NA | NA | NA | #VALUE! |
| ENSMUSG00000101167.2 | 0 | NA | NA | NA | NA | NA | #VALUE! |
| ENSMUSG00000101168.2 | 0 | NA | NA | NA | NA | NA | #VALUE! |
| ENSMUSG00000101170.2 | 0 | NA | NA | NA | NA | NA | #VALUE! |

| | | | | | | | |
|---|---|---|---|---|---|---|---|
| ENSMUSG00000101171.2 | 0 | NA | NA | NA | NA | NA | #VALUE! |
| ENSMUSG00000101172.2 | 0 | NA | NA | NA | NA | NA | #VALUE! |
| ENSMUSG00000101173.2 | 0 | NA | NA | NA | NA | NA | #VALUE! |
| ENSMUSG00000101174.9 | 0 | NA | NA | NA | NA | NA | #VALUE! |
| ENSMUSG00000101177.7 | 0 | NA | NA | NA | NA | NA | #VALUE! |
| ENSMUSG00000101178.2 | 0 | NA | NA | NA | NA | NA | #VALUE! |
| ENSMUSG00000101179.2 | 0 | NA | NA | NA | NA | NA | #VALUE! |
| ENSMUSG00000101180.2 | 0 | NA | NA | NA | NA | NA | #VALUE! |
| ENSMUSG00000101181.3 | 0 | NA | NA | NA | NA | NA | #VALUE! |
| ENSMUSG00000101182.2 | 0 | NA | NA | NA | NA | NA | #VALUE! |
| ENSMUSG00000101183.2 | 0 | NA | NA | NA | NA | NA | #VALUE! |
| ENSMUSG00000101184.2 | 0 | NA | NA | NA | NA | NA | #VALUE! |
| ENSMUSG00000101185.2 | 0 | NA | NA | NA | NA | NA | #VALUE! |
| ENSMUSG00000101186.2 | 0 | NA | NA | NA | NA | NA | #VALUE! |
| ENSMUSG00000101188.2 | 0 | NA | NA | NA | NA | NA | #VALUE! |
| ENSMUSG00000101189.2 | 0 | NA | NA | NA | NA | NA | #VALUE! |
| ENSMUSG00000101190.7 | 0 | NA | NA | NA | NA | NA | #VALUE! |
| ENSMUSG00000101191.7 | 0 | NA | NA | NA | NA | NA | #VALUE! |
| ENSMUSG00000101192.7 | 0 | NA | NA | NA | NA | NA | #VALUE! |
| ENSMUSG00000101193.2 | 0 | NA | NA | NA | NA | NA | #VALUE! |
| ENSMUSG00000101194.2 | 0 | NA | NA | NA | NA | NA | #VALUE! |
| ENSMUSG00000101195.2 | 0 | NA | NA | NA | NA | NA | #VALUE! |
| ENSMUSG00000101196.3 | 0 | NA | NA | NA | NA | NA | #VALUE! |
| ENSMUSG00000101197.2 | 0 | NA | NA | NA | NA | NA | #VALUE! |
| ENSMUSG00000101198.7 | 0 | NA | NA | NA | NA | NA | #VALUE! |
| ENSMUSG00000101199.2 | 0.540678 | 0.929314 | 4.022674 | 0.231019 | 0.8173 | NA | #VALUE! |
| ENSMUSG00000101200.2 | 0 | NA | NA | NA | NA | NA | #VALUE! |
| ENSMUSG00000101201.2 | 0 | NA | NA | NA | NA | NA | #VALUE! |
| ENSMUSG00000101202.7 | 0 | NA | NA | NA | NA | NA | #VALUE! |
| ENSMUSG00000101203.2 | 0 | NA | NA | NA | NA | NA | #VALUE! |
| ENSMUSG00000101204.7 | 0 | NA | NA | NA | NA | NA | #VALUE! |
| ENSMUSG00000101205.2 | 0 | NA | NA | NA | NA | NA | #VALUE! |
| ENSMUSG00000101206.2 | 0 | NA | NA | NA | NA | NA | #VALUE! |
| ENSMUSG00000101207.2 | 0 | NA | NA | NA | NA | NA | #VALUE! |
| ENSMUSG00000101208.2 | 0 | NA | NA | NA | NA | NA | #VALUE! |
| ENSMUSG00000101209.2 | 0 | NA | NA | NA | NA | NA | #VALUE! |
| ENSMUSG00000101210.2 | 0 | NA | NA | NA | NA | NA | #VALUE! |
| ENSMUSG00000101211.2 | 0 | NA | NA | NA | NA | NA | #VALUE! |
| ENSMUSG00000101212.2 | 0 | NA | NA | NA | NA | NA | #VALUE! |
| ENSMUSG00000101213.2 | 0 | NA | NA | NA | NA | NA | #VALUE! |
| ENSMUSG00000101214.2 | 0 | NA | NA | NA | NA | NA | #VALUE! |
| ENSMUSG00000101215.2 | 0 | NA | NA | NA | NA | NA | #VALUE! |
| ENSMUSG00000101216.2 | 0 | NA | NA | NA | NA | NA | #VALUE! |
| ENSMUSG00000101218.2 | 0 | NA | NA | NA | NA | NA | #VALUE! |

| | | | | | | | |
|---|---|---|---|---|---|---|---|
| ENSMUSG00000101219.2 | 0 | NA | NA | NA | NA | NA | #VALUE! |
| ENSMUSG00000101220.2 | 0 | NA | NA | NA | NA | NA | #VALUE! |
| ENSMUSG00000101221.2 | 0 | NA | NA | NA | NA | NA | #VALUE! |
| ENSMUSG00000101222.2 | 0 | NA | NA | NA | NA | NA | #VALUE! |
| ENSMUSG00000101223.7 | 0 | NA | NA | NA | NA | NA | #VALUE! |
| ENSMUSG00000101224.2 | 0 | NA | NA | NA | NA | NA | #VALUE! |
| ENSMUSG00000101226.2 | 0 | NA | NA | NA | NA | NA | #VALUE! |
| ENSMUSG00000101227.2 | 0 | NA | NA | NA | NA | NA | #VALUE! |
| ENSMUSG00000101228.2 | 0 | NA | NA | NA | NA | NA | #VALUE! |
| ENSMUSG00000101229.2 | 0 | NA | NA | NA | NA | NA | #VALUE! |
| ENSMUSG00000101230.2 | 0 | NA | NA | NA | NA | NA | #VALUE! |
| ENSMUSG00000101231.2 | 0 | NA | NA | NA | NA | NA | #VALUE! |
| ENSMUSG00000101232.2 | 0 | NA | NA | NA | NA | NA | #VALUE! |
| ENSMUSG00000101233.2 | 0 | NA | NA | NA | NA | NA | #VALUE! |
| ENSMUSG00000101234.2 | 0 | NA | NA | NA | NA | NA | #VALUE! |
| ENSMUSG00000101235.2 | 0 | NA | NA | NA | NA | NA | #VALUE! |
| ENSMUSG00000101236.2 | 0 | NA | NA | NA | NA | NA | #VALUE! |
| ENSMUSG00000101237.2 | 0 | NA | NA | NA | NA | NA | #VALUE! |
| ENSMUSG00000101238.2 | 0 | NA | NA | NA | NA | NA | #VALUE! |
| ENSMUSG00000101239.2 | 0 | NA | NA | NA | NA | NA | #VALUE! |
| ENSMUSG00000101240.2 | 0 | NA | NA | NA | NA | NA | #VALUE! |
| ENSMUSG00000101241.2 | 0 | NA | NA | NA | NA | NA | #VALUE! |
| ENSMUSG00000101242.2 | 0 | NA | NA | NA | NA | NA | #VALUE! |
| ENSMUSG00000101243.2 | 0 | NA | NA | NA | NA | NA | #VALUE! |
| ENSMUSG00000101244.7 | 0 | NA | NA | NA | NA | NA | #VALUE! |
| ENSMUSG00000101245.2 | 0 | NA | NA | NA | NA | NA | #VALUE! |
| ENSMUSG00000101246.2 | 0 | NA | NA | NA | NA | NA | #VALUE! |
| ENSMUSG00000101247.2 | 0 | NA | NA | NA | NA | NA | #VALUE! |
| ENSMUSG00000101248.2 | 0 | NA | NA | NA | NA | NA | #VALUE! |
| ENSMUSG00000101250.2 | 0 | NA | NA | NA | NA | NA | #VALUE! |
| ENSMUSG00000101251.2 | 0 | NA | NA | NA | NA | NA | #VALUE! |
| ENSMUSG00000101252.2 | 0 | NA | NA | NA | NA | NA | #VALUE! |
| ENSMUSG00000101253.2 | 0 | NA | NA | NA | NA | NA | #VALUE! |
| ENSMUSG00000101254.2 | 0 | NA | NA | NA | NA | NA | #VALUE! |
| ENSMUSG00000101255.2 | 0 | NA | NA | NA | NA | NA | #VALUE! |
| ENSMUSG00000101256.2 | 0 | NA | NA | NA | NA | NA | #VALUE! |
| ENSMUSG00000101257.2 | 0 | NA | NA | NA | NA | NA | #VALUE! |
| ENSMUSG00000101258.2 | 0 | NA | NA | NA | NA | NA | #VALUE! |
| ENSMUSG00000101259.2 | 0 | NA | NA | NA | NA | NA | #VALUE! |
| ENSMUSG00000101260.2 | 0 | NA | NA | NA | NA | NA | #VALUE! |
| ENSMUSG00000101261.2 | 0 | NA | NA | NA | NA | NA | #VALUE! |
| ENSMUSG00000101262.2 | 0 | NA | NA | NA | NA | NA | #VALUE! |
| ENSMUSG00000101263.2 | 0 | NA | NA | NA | NA | NA | #VALUE! |
| ENSMUSG00000101264.2 | 1.514552 | 4.088766 | 3.936269 | 1.038742 | 0.298925 | NA | #VALUE! |

| | | | | | | | |
|---|---|---|---|---|---|---|---|
| ENSMUSG00000101265.2 | 0 | NA | NA | NA | NA | NA | #VALUE! |
| ENSMUSG00000101266.2 | 0 | NA | NA | NA | NA | NA | #VALUE! |
| ENSMUSG00000101267.2 | 0 | NA | NA | NA | NA | NA | #VALUE! |
| ENSMUSG00000101268.7 | 0 | NA | NA | NA | NA | NA | #VALUE! |
| ENSMUSG00000101270.2 | 0 | NA | NA | NA | NA | NA | #VALUE! |
| ENSMUSG00000101271.2 | 0 | NA | NA | NA | NA | NA | #VALUE! |
| ENSMUSG00000101272.2 | 0 | NA | NA | NA | NA | NA | #VALUE! |
| ENSMUSG00000101273.2 | 0 | NA | NA | NA | NA | NA | #VALUE! |
| ENSMUSG00000101275.2 | 0 | NA | NA | NA | NA | NA | #VALUE! |
| ENSMUSG00000101276.2 | 0 | NA | NA | NA | NA | NA | #VALUE! |
| ENSMUSG00000101277.2 | 0 | NA | NA | NA | NA | NA | #VALUE! |
| ENSMUSG00000101278.2 | 0 | NA | NA | NA | NA | NA | #VALUE! |
| ENSMUSG00000101279.2 | 0 | NA | NA | NA | NA | NA | #VALUE! |
| ENSMUSG00000101280.2 | 0 | NA | NA | NA | NA | NA | #VALUE! |
| ENSMUSG00000101281.2 | 0 | NA | NA | NA | NA | NA | #VALUE! |
| ENSMUSG00000101282.2 | 0 | NA | NA | NA | NA | NA | #VALUE! |
| ENSMUSG00000101283.2 | 0 | NA | NA | NA | NA | NA | #VALUE! |
| ENSMUSG00000101284.2 | 0 | NA | NA | NA | NA | NA | #VALUE! |
| ENSMUSG00000101285.7 | 0 | NA | NA | NA | NA | NA | #VALUE! |
| ENSMUSG00000101286.2 | 0 | NA | NA | NA | NA | NA | #VALUE! |
| ENSMUSG00000101287.2 | 0 | NA | NA | NA | NA | NA | #VALUE! |
| ENSMUSG00000101288.2 | 0 | NA | NA | NA | NA | NA | #VALUE! |
| ENSMUSG00000101289.2 | 0 | NA | NA | NA | NA | NA | #VALUE! |
| ENSMUSG00000101290.2 | 0 | NA | NA | NA | NA | NA | #VALUE! |
| ENSMUSG00000101291.2 | 0 | NA | NA | NA | NA | NA | #VALUE! |
| ENSMUSG00000101292.2 | 0 | NA | NA | NA | NA | NA | #VALUE! |
| ENSMUSG00000101294.2 | 0 | NA | NA | NA | NA | NA | #VALUE! |
| ENSMUSG00000101295.2 | 0 | NA | NA | NA | NA | NA | #VALUE! |
| ENSMUSG00000101296.2 | 0 | NA | NA | NA | NA | NA | #VALUE! |
| ENSMUSG00000101297.2 | 0 | NA | NA | NA | NA | NA | #VALUE! |
| ENSMUSG00000101298.2 | 0 | NA | NA | NA | NA | NA | #VALUE! |
| ENSMUSG00000101299.3 | 0 | NA | NA | NA | NA | NA | #VALUE! |
| ENSMUSG00000101300.7 | 0 | NA | NA | NA | NA | NA | #VALUE! |
| ENSMUSG00000101301.2 | 0 | NA | NA | NA | NA | NA | #VALUE! |
| ENSMUSG00000101302.7 | 0 | NA | NA | NA | NA | NA | #VALUE! |
| ENSMUSG00000101303.2 | 0 | NA | NA | NA | NA | NA | #VALUE! |
| ENSMUSG00000101304.2 | 6.583272 | 3.480387 | 3.743751 | 0.929652 | 0.352551 | NA | #VALUE! |
| ENSMUSG00000101305.2 | 0 | NA | NA | NA | NA | NA | #VALUE! |
| ENSMUSG00000101306.2 | 0 | NA | NA | NA | NA | NA | #VALUE! |
| ENSMUSG00000101307.3 | 0 | NA | NA | NA | NA | NA | #VALUE! |
| ENSMUSG00000101308.2 | 0 | NA | NA | NA | NA | NA | #VALUE! |
| ENSMUSG00000101309.2 | 0 | NA | NA | NA | NA | NA | #VALUE! |
| ENSMUSG00000101310.2 | 0 | NA | NA | NA | NA | NA | #VALUE! |
| ENSMUSG00000101311.2 | 0 | NA | NA | NA | NA | NA | #VALUE! |

| | | | | | | | |
|---|---|---|---|---|---|---|---|
| ENSMUSG00000101312.2 | 0 | NA | NA | NA | NA | NA | #VALUE! |
| ENSMUSG00000101313.2 | 0 | NA | NA | NA | NA | NA | #VALUE! |
| ENSMUSG00000101314.2 | 0 | NA | NA | NA | NA | NA | #VALUE! |
| ENSMUSG00000101315.2 | 0 | NA | NA | NA | NA | NA | #VALUE! |
| ENSMUSG00000101316.2 | 0 | NA | NA | NA | NA | NA | #VALUE! |
| ENSMUSG00000101317.2 | 0 | NA | NA | NA | NA | NA | #VALUE! |
| ENSMUSG00000101318.2 | 0 | NA | NA | NA | NA | NA | #VALUE! |
| ENSMUSG00000101319.2 | 0 | NA | NA | NA | NA | NA | #VALUE! |
| ENSMUSG00000101320.2 | 0 | NA | NA | NA | NA | NA | #VALUE! |
| ENSMUSG00000101321.2 | 0 | NA | NA | NA | NA | NA | #VALUE! |
| ENSMUSG00000101323.2 | 0 | NA | NA | NA | NA | NA | #VALUE! |
| ENSMUSG00000101324.2 | 0 | NA | NA | NA | NA | NA | #VALUE! |
| ENSMUSG00000101325.2 | 0 | NA | NA | NA | NA | NA | #VALUE! |
| ENSMUSG00000101326.7 | 0 | NA | NA | NA | NA | NA | #VALUE! |
| ENSMUSG00000101327.2 | 0 | NA | NA | NA | NA | NA | #VALUE! |
| ENSMUSG00000101328.2 | 0 | NA | NA | NA | NA | NA | #VALUE! |
| ENSMUSG00000101329.2 | 0 | NA | NA | NA | NA | NA | #VALUE! |
| ENSMUSG00000101330.2 | 0 | NA | NA | NA | NA | NA | #VALUE! |
| ENSMUSG00000101331.2 | 0 | NA | NA | NA | NA | NA | #VALUE! |
| ENSMUSG00000101332.2 | 0 | NA | NA | NA | NA | NA | #VALUE! |
| ENSMUSG00000101333.2 | 0 | NA | NA | NA | NA | NA | #VALUE! |
| ENSMUSG00000101334.2 | 0 | NA | NA | NA | NA | NA | #VALUE! |
| ENSMUSG00000101335.2 | 31.10303 | -2.25433 | 2.522583 | -0.89366 | NA | NA | #VALUE! |
| ENSMUSG00000101336.2 | 0 | NA | NA | NA | NA | NA | #VALUE! |
| ENSMUSG00000101337.7 | 0 | NA | NA | NA | NA | NA | #VALUE! |
| ENSMUSG00000101338.2 | 0 | NA | NA | NA | NA | NA | #VALUE! |
| ENSMUSG00000101339.2 | 0 | NA | NA | NA | NA | NA | #VALUE! |
| ENSMUSG00000101340.2 | 0 | NA | NA | NA | NA | NA | #VALUE! |
| ENSMUSG00000101341.2 | 0 | NA | NA | NA | NA | NA | #VALUE! |
| ENSMUSG00000101342.2 | 0 | NA | NA | NA | NA | NA | #VALUE! |
| ENSMUSG00000101343.2 | 0 | NA | NA | NA | NA | NA | #VALUE! |
| ENSMUSG00000101344.2 | 0 | NA | NA | NA | NA | NA | #VALUE! |
| ENSMUSG00000101345.2 | 0 | NA | NA | NA | NA | NA | #VALUE! |
| ENSMUSG00000101346.2 | 0 | NA | NA | NA | NA | NA | #VALUE! |
| ENSMUSG00000101347.2 | 0 | NA | NA | NA | NA | NA | #VALUE! |
| ENSMUSG00000101348.7 | 0 | NA | NA | NA | NA | NA | #VALUE! |
| ENSMUSG00000101349.2 | 0 | NA | NA | NA | NA | NA | #VALUE! |
| ENSMUSG00000101350.7 | 0 | NA | NA | NA | NA | NA | #VALUE! |
| ENSMUSG00000101351.2 | 0 | NA | NA | NA | NA | NA | #VALUE! |
| ENSMUSG00000101352.2 | 0 | NA | NA | NA | NA | NA | #VALUE! |
| ENSMUSG00000101353.7 | 0 | NA | NA | NA | NA | NA | #VALUE! |
| ENSMUSG00000101354.2 | 0 | NA | NA | NA | NA | NA | #VALUE! |
| ENSMUSG00000101355.2 | 0 | NA | NA | NA | NA | NA | #VALUE! |
| ENSMUSG00000101356.2 | 0 | NA | NA | NA | NA | NA | #VALUE! |

| | | | | | | | |
|---|---|---|---|---|---|---|---|
| ENSMUSG00000101357.2 | 0 | NA | NA | NA | NA | NA | #VALUE! |
| ENSMUSG00000101358.2 | 0 | NA | NA | NA | NA | NA | #VALUE! |
| ENSMUSG00000101359.2 | 0 | NA | NA | NA | NA | NA | #VALUE! |
| ENSMUSG00000101360.3 | 0 | NA | NA | NA | NA | NA | #VALUE! |
| ENSMUSG00000101361.3 | 0 | NA | NA | NA | NA | NA | #VALUE! |
| ENSMUSG00000101362.2 | 0 | NA | NA | NA | NA | NA | #VALUE! |
| ENSMUSG00000101363.2 | 0 | NA | NA | NA | NA | NA | #VALUE! |
| ENSMUSG00000101364.2 | 0 | NA | NA | NA | NA | NA | #VALUE! |
| ENSMUSG00000101365.2 | 0 | NA | NA | NA | NA | NA | #VALUE! |
| ENSMUSG00000101366.2 | 0 | NA | NA | NA | NA | NA | #VALUE! |
| ENSMUSG00000101367.2 | 0 | NA | NA | NA | NA | NA | #VALUE! |
| ENSMUSG00000101368.2 | 0 | NA | NA | NA | NA | NA | #VALUE! |
| ENSMUSG00000101369.2 | 0 | NA | NA | NA | NA | NA | #VALUE! |
| ENSMUSG00000101370.2 | 0 | NA | NA | NA | NA | NA | #VALUE! |
| ENSMUSG00000101371.2 | 0 | NA | NA | NA | NA | NA | #VALUE! |
| ENSMUSG00000101372.2 | 0 | NA | NA | NA | NA | NA | #VALUE! |
| ENSMUSG00000101373.3 | 0 | NA | NA | NA | NA | NA | #VALUE! |
| ENSMUSG00000101374.7 | 0 | NA | NA | NA | NA | NA | #VALUE! |
| ENSMUSG00000101375.2 | 0 | NA | NA | NA | NA | NA | #VALUE! |
| ENSMUSG00000101376.2 | 0 | NA | NA | NA | NA | NA | #VALUE! |
| ENSMUSG00000101377.2 | 0 | NA | NA | NA | NA | NA | #VALUE! |
| ENSMUSG00000101378.2 | 1.865104 | 4.40601 | 3.930481 | 1.120985 | 0.262294 | NA | #VALUE! |
| ENSMUSG00000101379.2 | 0 | NA | NA | NA | NA | NA | #VALUE! |
| ENSMUSG00000101380.2 | 0 | NA | NA | NA | NA | NA | #VALUE! |
| ENSMUSG00000101381.2 | 0 | NA | NA | NA | NA | NA | #VALUE! |
| ENSMUSG00000101382.2 | 0 | NA | NA | NA | NA | NA | #VALUE! |
| ENSMUSG00000101383.2 | 0 | NA | NA | NA | NA | NA | #VALUE! |
| ENSMUSG00000101385.2 | 0 | NA | NA | NA | NA | NA | #VALUE! |
| ENSMUSG00000101386.2 | 0 | NA | NA | NA | NA | NA | #VALUE! |
| ENSMUSG00000101387.2 | 0 | NA | NA | NA | NA | NA | #VALUE! |
| ENSMUSG00000101388.2 | 0 | NA | NA | NA | NA | NA | #VALUE! |
| ENSMUSG00000101389.2 | 0 | NA | NA | NA | NA | NA | #VALUE! |
| ENSMUSG00000101390.2 | 0 | NA | NA | NA | NA | NA | #VALUE! |
| ENSMUSG00000101391.4 | 0 | NA | NA | NA | NA | NA | #VALUE! |
| ENSMUSG00000101393.2 | 0 | NA | NA | NA | NA | NA | #VALUE! |
| ENSMUSG00000101394.2 | 0 | NA | NA | NA | NA | NA | #VALUE! |
| ENSMUSG00000101395.2 | 0 | NA | NA | NA | NA | NA | #VALUE! |
| ENSMUSG00000101396.2 | 0 | NA | NA | NA | NA | NA | #VALUE! |
| ENSMUSG00000101398.2 | 0 | NA | NA | NA | NA | NA | #VALUE! |
| ENSMUSG00000101399.2 | 0 | NA | NA | NA | NA | NA | #VALUE! |
| ENSMUSG00000101400.2 | 0 | NA | NA | NA | NA | NA | #VALUE! |
| ENSMUSG00000101401.2 | 0 | NA | NA | NA | NA | NA | #VALUE! |
| ENSMUSG00000101402.2 | 0 | NA | NA | NA | NA | NA | #VALUE! |
| ENSMUSG00000101403.2 | 0 | NA | NA | NA | NA | NA | #VALUE! |

| | | | | | | | |
|---|---|---|---|---|---|---|---|
| ENSMUSG00000101404.2 | 0 | NA | NA | NA | NA | NA | #VALUE! |
| ENSMUSG00000101405.2 | 0 | NA | NA | NA | NA | NA | #VALUE! |
| ENSMUSG00000101406.2 | 0 | NA | NA | NA | NA | NA | #VALUE! |
| ENSMUSG00000101407.7 | 0 | NA | NA | NA | NA | NA | #VALUE! |
| ENSMUSG00000101408.2 | 0 | NA | NA | NA | NA | NA | #VALUE! |
| ENSMUSG00000101409.2 | 0 | NA | NA | NA | NA | NA | #VALUE! |
| ENSMUSG00000101410.2 | 0 | NA | NA | NA | NA | NA | #VALUE! |
| ENSMUSG00000101411.2 | 0 | NA | NA | NA | NA | NA | #VALUE! |
| ENSMUSG00000101412.7 | 0 | NA | NA | NA | NA | NA | #VALUE! |
| ENSMUSG00000101413.2 | 0 | NA | NA | NA | NA | NA | #VALUE! |
| ENSMUSG00000101414.2 | 0 | NA | NA | NA | NA | NA | #VALUE! |
| ENSMUSG00000101415.2 | 0 | NA | NA | NA | NA | NA | #VALUE! |
| ENSMUSG00000101416.2 | 0 | NA | NA | NA | NA | NA | #VALUE! |
| ENSMUSG00000101417.2 | 0 | NA | NA | NA | NA | NA | #VALUE! |
| ENSMUSG00000101418.2 | 0 | NA | NA | NA | NA | NA | #VALUE! |
| ENSMUSG00000101419.2 | 0 | NA | NA | NA | NA | NA | #VALUE! |
| ENSMUSG00000101420.2 | 0 | NA | NA | NA | NA | NA | #VALUE! |
| ENSMUSG00000101421.2 | 0 | NA | NA | NA | NA | NA | #VALUE! |
| ENSMUSG00000101422.7 | 0 | NA | NA | NA | NA | NA | #VALUE! |
| ENSMUSG00000101423.7 | 0 | NA | NA | NA | NA | NA | #VALUE! |
| ENSMUSG00000101424.7 | 0 | NA | NA | NA | NA | NA | #VALUE! |
| ENSMUSG00000101425.2 | 0 | NA | NA | NA | NA | NA | #VALUE! |
| ENSMUSG00000101426.2 | 0 | NA | NA | NA | NA | NA | #VALUE! |
| ENSMUSG00000101427.2 | 0 | NA | NA | NA | NA | NA | #VALUE! |
| ENSMUSG00000101428.2 | 0 | NA | NA | NA | NA | NA | #VALUE! |
| ENSMUSG00000101429.2 | 0 | NA | NA | NA | NA | NA | #VALUE! |
| ENSMUSG00000101431.2 | 0 | NA | NA | NA | NA | NA | #VALUE! |
| ENSMUSG00000101433.2 | 0 | NA | NA | NA | NA | NA | #VALUE! |
| ENSMUSG00000101434.2 | 0 | NA | NA | NA | NA | NA | #VALUE! |
| ENSMUSG00000101435.2 | 0 | NA | NA | NA | NA | NA | #VALUE! |
| ENSMUSG00000101436.2 | 0 | NA | NA | NA | NA | NA | #VALUE! |
| ENSMUSG00000101437.2 | 0 | NA | NA | NA | NA | NA | #VALUE! |
| ENSMUSG00000101438.3 | 0 | NA | NA | NA | NA | NA | #VALUE! |
| ENSMUSG00000101439.2 | 0 | NA | NA | NA | NA | NA | #VALUE! |
| ENSMUSG00000101440.2 | 0 | NA | NA | NA | NA | NA | #VALUE! |
| ENSMUSG00000101441.2 | 0 | NA | NA | NA | NA | NA | #VALUE! |
| ENSMUSG00000101442.2 | 0 | NA | NA | NA | NA | NA | #VALUE! |
| ENSMUSG00000101443.2 | 0 | NA | NA | NA | NA | NA | #VALUE! |
| ENSMUSG00000101444.7 | 0 | NA | NA | NA | NA | NA | #VALUE! |
| ENSMUSG00000101445.2 | 0 | NA | NA | NA | NA | NA | #VALUE! |
| ENSMUSG00000101446.2 | 0 | NA | NA | NA | NA | NA | #VALUE! |
| ENSMUSG00000101447.2 | 0 | NA | NA | NA | NA | NA | #VALUE! |
| ENSMUSG00000101448.2 | 0 | NA | NA | NA | NA | NA | #VALUE! |
| ENSMUSG00000101449.2 | 0 | NA | NA | NA | NA | NA | #VALUE! |

| | | | | | | | |
|---|---|---|---|---|---|---|---|
| ENSMUSG00000101450.2 | 0 | NA | NA | NA | NA | NA | #VALUE! |
| ENSMUSG00000101451.2 | 0 | NA | NA | NA | NA | NA | #VALUE! |
| ENSMUSG00000101452.2 | 0.245976 | 1.027628 | 4.067126 | 0.252667 | 0.800526 | NA | #VALUE! |
| ENSMUSG00000101453.2 | 0 | NA | NA | NA | NA | NA | #VALUE! |
| ENSMUSG00000101454.2 | 0 | NA | NA | NA | NA | NA | #VALUE! |
| ENSMUSG00000101455.2 | 0 | NA | NA | NA | NA | NA | #VALUE! |
| ENSMUSG00000101456.2 | 0 | NA | NA | NA | NA | NA | #VALUE! |
| ENSMUSG00000101457.2 | 0 | NA | NA | NA | NA | NA | #VALUE! |
| ENSMUSG00000101459.2 | 0 | NA | NA | NA | NA | NA | #VALUE! |
| ENSMUSG00000101462.3 | 0 | NA | NA | NA | NA | NA | #VALUE! |
| ENSMUSG00000101463.2 | 0 | NA | NA | NA | NA | NA | #VALUE! |
| ENSMUSG00000101464.2 | 0.093255 | 0.868093 | 4.072464 | 0.213162 | 0.831201 | NA | #VALUE! |
| ENSMUSG00000101465.2 | 0 | NA | NA | NA | NA | NA | #VALUE! |
| ENSMUSG00000101466.2 | 0 | NA | NA | NA | NA | NA | #VALUE! |
| ENSMUSG00000101467.2 | 0 | NA | NA | NA | NA | NA | #VALUE! |
| ENSMUSG00000101468.2 | 0 | NA | NA | NA | NA | NA | #VALUE! |
| ENSMUSG00000101469.2 | 0 | NA | NA | NA | NA | NA | #VALUE! |
| ENSMUSG00000101470.2 | 0 | NA | NA | NA | NA | NA | #VALUE! |
| ENSMUSG00000101471.2 | 0 | NA | NA | NA | NA | NA | #VALUE! |
| ENSMUSG00000101472.2 | 0 | NA | NA | NA | NA | NA | #VALUE! |
| ENSMUSG00000101473.2 | 0 | NA | NA | NA | NA | NA | #VALUE! |
| ENSMUSG00000101474.2 | 0 | NA | NA | NA | NA | NA | #VALUE! |
| ENSMUSG00000101475.2 | 0 | NA | NA | NA | NA | NA | #VALUE! |
| ENSMUSG00000101476.2 | 0 | NA | NA | NA | NA | NA | #VALUE! |
| ENSMUSG00000101477.2 | 0 | NA | NA | NA | NA | NA | #VALUE! |
| ENSMUSG00000101478.2 | 0 | NA | NA | NA | NA | NA | #VALUE! |
| ENSMUSG00000101479.2 | 0 | NA | NA | NA | NA | NA | #VALUE! |
| ENSMUSG00000101480.4 | 0 | NA | NA | NA | NA | NA | #VALUE! |
| ENSMUSG00000101481.2 | 0 | NA | NA | NA | NA | NA | #VALUE! |
| ENSMUSG00000101482.2 | 0 | NA | NA | NA | NA | NA | #VALUE! |
| ENSMUSG00000101483.2 | 0 | NA | NA | NA | NA | NA | #VALUE! |
| ENSMUSG00000101484.2 | 0 | NA | NA | NA | NA | NA | #VALUE! |
| ENSMUSG00000101486.2 | 0 | NA | NA | NA | NA | NA | #VALUE! |
| ENSMUSG00000101487.2 | 0 | NA | NA | NA | NA | NA | #VALUE! |
| ENSMUSG00000101489.2 | 0 | NA | NA | NA | NA | NA | #VALUE! |
| ENSMUSG00000101490.2 | 0 | NA | NA | NA | NA | NA | #VALUE! |
| ENSMUSG00000101491.2 | 0 | NA | NA | NA | NA | NA | #VALUE! |
| ENSMUSG00000101492.2 | 0 | NA | NA | NA | NA | NA | #VALUE! |
| ENSMUSG00000101493.2 | 1.475853 | 4.029233 | 3.937502 | 1.023297 | 0.306167 | NA | #VALUE! |
| ENSMUSG00000101494.2 | 0 | NA | NA | NA | NA | NA | #VALUE! |
| ENSMUSG00000101495.2 | 0 | NA | NA | NA | NA | NA | #VALUE! |
| ENSMUSG00000101496.2 | 0 | NA | NA | NA | NA | NA | #VALUE! |
| ENSMUSG00000101497.2 | 0 | NA | NA | NA | NA | NA | #VALUE! |
| ENSMUSG00000101498.2 | 0 | NA | NA | NA | NA | NA | #VALUE! |

| | | | | | | | |
|---|---|---|---|---|---|---|---|
| ENSMUSG00000101499.2 | 0 | NA | NA | NA | NA | NA | #VALUE! |
| ENSMUSG00000101500.2 | 0 | NA | NA | NA | NA | NA | #VALUE! |
| ENSMUSG00000101501.2 | 0 | NA | NA | NA | NA | NA | #VALUE! |
| ENSMUSG00000101502.2 | 0.198268 | -0.06494 | 4.068276 | -0.01596 | 0.987265 | NA | #VALUE! |
| ENSMUSG00000101503.2 | 0 | NA | NA | NA | NA | NA | #VALUE! |
| ENSMUSG00000101504.2 | 0 | NA | NA | NA | NA | NA | #VALUE! |
| ENSMUSG00000101505.2 | 0 | NA | NA | NA | NA | NA | #VALUE! |
| ENSMUSG00000101506.2 | 0 | NA | NA | NA | NA | NA | #VALUE! |
| ENSMUSG00000101507.2 | 0 | NA | NA | NA | NA | NA | #VALUE! |
| ENSMUSG00000101508.2 | 0 | NA | NA | NA | NA | NA | #VALUE! |
| ENSMUSG00000101509.2 | 0 | NA | NA | NA | NA | NA | #VALUE! |
| ENSMUSG00000101510.2 | 0 | NA | NA | NA | NA | NA | #VALUE! |
| ENSMUSG00000101511.2 | 0 | NA | NA | NA | NA | NA | #VALUE! |
| ENSMUSG00000101512.2 | 0 | NA | NA | NA | NA | NA | #VALUE! |
| ENSMUSG00000101513.7 | 0 | NA | NA | NA | NA | NA | #VALUE! |
| ENSMUSG00000101515.2 | 0 | NA | NA | NA | NA | NA | #VALUE! |
| ENSMUSG00000101516.7 | 0 | NA | NA | NA | NA | NA | #VALUE! |
| ENSMUSG00000101518.2 | 0 | NA | NA | NA | NA | NA | #VALUE! |
| ENSMUSG00000101519.2 | 0 | NA | NA | NA | NA | NA | #VALUE! |
| ENSMUSG00000101520.2 | 0 | NA | NA | NA | NA | NA | #VALUE! |
| ENSMUSG00000101521.2 | 0 | NA | NA | NA | NA | NA | #VALUE! |
| ENSMUSG00000101522.2 | 0 | NA | NA | NA | NA | NA | #VALUE! |
| ENSMUSG00000101523.3 | 9.311139 | 1.414805 | 2.437164 | 0.580513 | 0.561569 | NA | #VALUE! |
| ENSMUSG00000101525.2 | 0 | NA | NA | NA | NA | NA | #VALUE! |
| ENSMUSG00000101526.2 | 0 | NA | NA | NA | NA | NA | #VALUE! |
| ENSMUSG00000101527.2 | 0 | NA | NA | NA | NA | NA | #VALUE! |
| ENSMUSG00000101528.2 | 0 | NA | NA | NA | NA | NA | #VALUE! |
| ENSMUSG00000101529.2 | 0 | NA | NA | NA | NA | NA | #VALUE! |
| ENSMUSG00000101530.2 | 0 | NA | NA | NA | NA | NA | #VALUE! |
| ENSMUSG00000101531.7 | 0 | NA | NA | NA | NA | NA | #VALUE! |
| ENSMUSG00000101532.2 | 0 | NA | NA | NA | NA | NA | #VALUE! |
| ENSMUSG00000101533.2 | 0 | NA | NA | NA | NA | NA | #VALUE! |
| ENSMUSG00000101534.2 | 0 | NA | NA | NA | NA | NA | #VALUE! |
| ENSMUSG00000101535.2 | 0 | NA | NA | NA | NA | NA | #VALUE! |
| ENSMUSG00000101536.7 | 0 | NA | NA | NA | NA | NA | #VALUE! |
| ENSMUSG00000101537.2 | 0.587367 | -2.74297 | 3.982932 | -0.68868 | 0.491024 | NA | #VALUE! |
| ENSMUSG00000101538.2 | 0 | NA | NA | NA | NA | NA | #VALUE! |
| ENSMUSG00000101539.2 | 0 | NA | NA | NA | NA | NA | #VALUE! |
| ENSMUSG00000101540.3 | 0 | NA | NA | NA | NA | NA | #VALUE! |
| ENSMUSG00000101541.2 | 0 | NA | NA | NA | NA | NA | #VALUE! |
| ENSMUSG00000101542.7 | 0 | NA | NA | NA | NA | NA | #VALUE! |
| ENSMUSG00000101543.2 | 0 | NA | NA | NA | NA | NA | #VALUE! |
| ENSMUSG00000101544.2 | 0 | NA | NA | NA | NA | NA | #VALUE! |
| ENSMUSG00000101545.2 | 0 | NA | NA | NA | NA | NA | #VALUE! |

| | | | | | | | |
|---|---|---|---|---|---|---|---|
| ENSMUSG00000101546.2 | 0 | NA | NA | NA | NA | NA | #VALUE! |
| ENSMUSG00000101547.2 | 0 | NA | NA | NA | NA | NA | #VALUE! |
| ENSMUSG00000101548.2 | 0 | NA | NA | NA | NA | NA | #VALUE! |
| ENSMUSG00000101549.2 | 0 | NA | NA | NA | NA | NA | #VALUE! |
| ENSMUSG00000101550.2 | 0 | NA | NA | NA | NA | NA | #VALUE! |
| ENSMUSG00000101551.2 | 0 | NA | NA | NA | NA | NA | #VALUE! |
| ENSMUSG00000101552.2 | 0 | NA | NA | NA | NA | NA | #VALUE! |
| ENSMUSG00000101553.2 | 0 | NA | NA | NA | NA | NA | #VALUE! |
| ENSMUSG00000101554.3 | 0 | NA | NA | NA | NA | NA | #VALUE! |
| ENSMUSG00000101555.2 | 0 | NA | NA | NA | NA | NA | #VALUE! |
| ENSMUSG00000101557.2 | 0 | NA | NA | NA | NA | NA | #VALUE! |
| ENSMUSG00000101558.2 | 0 | NA | NA | NA | NA | NA | #VALUE! |
| ENSMUSG00000101559.2 | 0 | NA | NA | NA | NA | NA | #VALUE! |
| ENSMUSG00000101560.7 | 0 | NA | NA | NA | NA | NA | #VALUE! |
| ENSMUSG00000101562.2 | 0 | NA | NA | NA | NA | NA | #VALUE! |
| ENSMUSG00000101563.2 | 0 | NA | NA | NA | NA | NA | #VALUE! |
| ENSMUSG00000101564.2 | 0 | NA | NA | NA | NA | NA | #VALUE! |
| ENSMUSG00000101565.3 | 0 | NA | NA | NA | NA | NA | #VALUE! |
| ENSMUSG00000101566.2 | 0 | NA | NA | NA | NA | NA | #VALUE! |
| ENSMUSG00000101567.2 | 0.855902 | -1.74958 | 3.965467 | -0.4412 | 0.659065 | NA | #VALUE! |
| ENSMUSG00000101568.2 | 0 | NA | NA | NA | NA | NA | #VALUE! |
| ENSMUSG00000101569.2 | 0 | NA | NA | NA | NA | NA | #VALUE! |
| ENSMUSG00000101570.2 | 0 | NA | NA | NA | NA | NA | #VALUE! |
| ENSMUSG00000101571.3 | 0 | NA | NA | NA | NA | NA | #VALUE! |
| ENSMUSG00000101573.2 | 0 | NA | NA | NA | NA | NA | #VALUE! |
| ENSMUSG00000101574.2 | 0 | NA | NA | NA | NA | NA | #VALUE! |
| ENSMUSG00000101575.2 | 0 | NA | NA | NA | NA | NA | #VALUE! |
| ENSMUSG00000101576.2 | 0 | NA | NA | NA | NA | NA | #VALUE! |
| ENSMUSG00000101578.3 | 0 | NA | NA | NA | NA | NA | #VALUE! |
| ENSMUSG00000101579.2 | 0 | NA | NA | NA | NA | NA | #VALUE! |
| ENSMUSG00000101580.2 | 0 | NA | NA | NA | NA | NA | #VALUE! |
| ENSMUSG00000101581.7 | 0 | NA | NA | NA | NA | NA | #VALUE! |
| ENSMUSG00000101582.2 | 0 | NA | NA | NA | NA | NA | #VALUE! |
| ENSMUSG00000101583.2 | 0 | NA | NA | NA | NA | NA | #VALUE! |
| ENSMUSG00000101584.3 | 0 | NA | NA | NA | NA | NA | #VALUE! |
| ENSMUSG00000101585.7 | 0 | NA | NA | NA | NA | NA | #VALUE! |
| ENSMUSG00000101586.2 | 0 | NA | NA | NA | NA | NA | #VALUE! |
| ENSMUSG00000101587.3 | 0 | NA | NA | NA | NA | NA | #VALUE! |
| ENSMUSG00000101588.2 | 0 | NA | NA | NA | NA | NA | #VALUE! |
| ENSMUSG00000101589.2 | 0 | NA | NA | NA | NA | NA | #VALUE! |
| ENSMUSG00000101590.2 | 0 | NA | NA | NA | NA | NA | #VALUE! |
| ENSMUSG00000101592.2 | 0 | NA | NA | NA | NA | NA | #VALUE! |
| ENSMUSG00000101593.2 | 0 | NA | NA | NA | NA | NA | #VALUE! |
| ENSMUSG00000101594.2 | 0 | NA | NA | NA | NA | NA | #VALUE! |

| | | | | | | | |
|---|---|---|---|---|---|---|---|
| ENSMUSG00000101595.2 | 0 | NA | NA | NA | NA | NA | #VALUE! |
| ENSMUSG00000101596.2 | 0 | NA | NA | NA | NA | NA | #VALUE! |
| ENSMUSG00000101597.3 | 0.261539 | 1.68833 | 4.043024 | 0.417591 | 0.676246 | NA | #VALUE! |
| ENSMUSG00000101598.2 | 0 | NA | NA | NA | NA | NA | #VALUE! |
| ENSMUSG00000101599.2 | 4.330482 | -5.64551 | 3.917088 | -1.44125 | 0.149514 | NA | #VALUE! |
| ENSMUSG00000101600.2 | 0 | NA | NA | NA | NA | NA | #VALUE! |
| ENSMUSG00000101601.2 | 0 | NA | NA | NA | NA | NA | #VALUE! |
| ENSMUSG00000101602.2 | 0 | NA | NA | NA | NA | NA | #VALUE! |
| ENSMUSG00000101603.2 | 0 | NA | NA | NA | NA | NA | #VALUE! |
| ENSMUSG00000101605.2 | 0 | NA | NA | NA | NA | NA | #VALUE! |
| ENSMUSG00000101606.2 | 0 | NA | NA | NA | NA | NA | #VALUE! |
| ENSMUSG00000101607.2 | 0 | NA | NA | NA | NA | NA | #VALUE! |
| ENSMUSG00000101608.2 | 0 | NA | NA | NA | NA | NA | #VALUE! |
| ENSMUSG00000101610.2 | 0 | NA | NA | NA | NA | NA | #VALUE! |
| ENSMUSG00000101611.2 | 0 | NA | NA | NA | NA | NA | #VALUE! |
| ENSMUSG00000101612.3 | 0 | NA | NA | NA | NA | NA | #VALUE! |
| ENSMUSG00000101613.2 | 0 | NA | NA | NA | NA | NA | #VALUE! |
| ENSMUSG00000101614.2 | 0 | NA | NA | NA | NA | NA | #VALUE! |
| ENSMUSG00000101615.2 | 0 | NA | NA | NA | NA | NA | #VALUE! |
| ENSMUSG00000101616.2 | 0 | NA | NA | NA | NA | NA | #VALUE! |
| ENSMUSG00000101617.2 | 0 | NA | NA | NA | NA | NA | #VALUE! |
| ENSMUSG00000101618.2 | 0 | NA | NA | NA | NA | NA | #VALUE! |
| ENSMUSG00000101619.2 | 0 | NA | NA | NA | NA | NA | #VALUE! |
| ENSMUSG00000101621.3 | 0 | NA | NA | NA | NA | NA | #VALUE! |
| ENSMUSG00000101622.2 | 0 | NA | NA | NA | NA | NA | #VALUE! |
| ENSMUSG00000101623.2 | 0 | NA | NA | NA | NA | NA | #VALUE! |
| ENSMUSG00000101624.2 | 0 | NA | NA | NA | NA | NA | #VALUE! |
| ENSMUSG00000101625.2 | 0 | NA | NA | NA | NA | NA | #VALUE! |
| ENSMUSG00000101626.7 | 0 | NA | NA | NA | NA | NA | #VALUE! |
| ENSMUSG00000101627.2 | 0 | NA | NA | NA | NA | NA | #VALUE! |
| ENSMUSG00000101628.2 | 0 | NA | NA | NA | NA | NA | #VALUE! |
| ENSMUSG00000101629.2 | 0 | NA | NA | NA | NA | NA | #VALUE! |
| ENSMUSG00000101630.2 | 0 | NA | NA | NA | NA | NA | #VALUE! |
| ENSMUSG00000101631.2 | 0 | NA | NA | NA | NA | NA | #VALUE! |
| ENSMUSG00000101632.2 | 0 | NA | NA | NA | NA | NA | #VALUE! |
| ENSMUSG00000101633.2 | 0 | NA | NA | NA | NA | NA | #VALUE! |
| ENSMUSG00000101634.7 | 0 | NA | NA | NA | NA | NA | #VALUE! |
| ENSMUSG00000101636.2 | 0 | NA | NA | NA | NA | NA | #VALUE! |
| ENSMUSG00000101637.2 | 0 | NA | NA | NA | NA | NA | #VALUE! |
| ENSMUSG00000101638.2 | 0 | NA | NA | NA | NA | NA | #VALUE! |
| ENSMUSG00000101639.2 | 0 | NA | NA | NA | NA | NA | #VALUE! |
| ENSMUSG00000101640.2 | 0 | NA | NA | NA | NA | NA | #VALUE! |
| ENSMUSG00000101641.2 | 0 | NA | NA | NA | NA | NA | #VALUE! |
| ENSMUSG00000101642.7 | 0 | NA | NA | NA | NA | NA | #VALUE! |

| | | | | | | | |
|---|---|---|---|---|---|---|---|
| ENSMUSG00000101643.2 | 0 | NA | NA | NA | NA | NA | #VALUE! |
| ENSMUSG00000101644.2 | 0 | NA | NA | NA | NA | NA | #VALUE! |
| ENSMUSG00000101645.2 | 0 | NA | NA | NA | NA | NA | #VALUE! |
| ENSMUSG00000101646.2 | 0 | NA | NA | NA | NA | NA | #VALUE! |
| ENSMUSG00000101647.2 | 0 | NA | NA | NA | NA | NA | #VALUE! |
| ENSMUSG00000101648.2 | 0 | NA | NA | NA | NA | NA | #VALUE! |
| ENSMUSG00000101649.2 | 0 | NA | NA | NA | NA | NA | #VALUE! |
| ENSMUSG00000101650.2 | 0 | NA | NA | NA | NA | NA | #VALUE! |
| ENSMUSG00000101651.2 | 0 | NA | NA | NA | NA | NA | #VALUE! |
| ENSMUSG00000101652.2 | 0 | NA | NA | NA | NA | NA | #VALUE! |
| ENSMUSG00000101653.2 | 0 | NA | NA | NA | NA | NA | #VALUE! |
| ENSMUSG00000101654.2 | 0 | NA | NA | NA | NA | NA | #VALUE! |
| ENSMUSG00000101656.2 | 0 | NA | NA | NA | NA | NA | #VALUE! |
| ENSMUSG00000101657.2 | 0 | NA | NA | NA | NA | NA | #VALUE! |
| ENSMUSG00000101658.2 | 0 | NA | NA | NA | NA | NA | #VALUE! |
| ENSMUSG00000101659.2 | 0 | NA | NA | NA | NA | NA | #VALUE! |
| ENSMUSG00000101660.2 | 0 | NA | NA | NA | NA | NA | #VALUE! |
| ENSMUSG00000101661.2 | 0 | NA | NA | NA | NA | NA | #VALUE! |
| ENSMUSG00000101662.2 | 0 | NA | NA | NA | NA | NA | #VALUE! |
| ENSMUSG00000101664.2 | 0 | NA | NA | NA | NA | NA | #VALUE! |
| ENSMUSG00000101665.2 | 0 | NA | NA | NA | NA | NA | #VALUE! |
| ENSMUSG00000101666.2 | 0 | NA | NA | NA | NA | NA | #VALUE! |
| ENSMUSG00000101667.2 | 0 | NA | NA | NA | NA | NA | #VALUE! |
| ENSMUSG00000101668.2 | 0 | NA | NA | NA | NA | NA | #VALUE! |
| ENSMUSG00000101669.2 | 0 | NA | NA | NA | NA | NA | #VALUE! |
| ENSMUSG00000101670.2 | 0 | NA | NA | NA | NA | NA | #VALUE! |
| ENSMUSG00000101671.2 | 0 | NA | NA | NA | NA | NA | #VALUE! |
| ENSMUSG00000101672.2 | 0 | NA | NA | NA | NA | NA | #VALUE! |
| ENSMUSG00000101673.2 | 0 | NA | NA | NA | NA | NA | #VALUE! |
| ENSMUSG00000101674.7 | 0 | NA | NA | NA | NA | NA | #VALUE! |
| ENSMUSG00000101675.2 | 0 | NA | NA | NA | NA | NA | #VALUE! |
| ENSMUSG00000101676.2 | 0 | NA | NA | NA | NA | NA | #VALUE! |
| ENSMUSG00000101678.4 | 0 | NA | NA | NA | NA | NA | #VALUE! |
| ENSMUSG00000101679.2 | 0 | NA | NA | NA | NA | NA | #VALUE! |
| ENSMUSG00000101680.2 | 0 | NA | NA | NA | NA | NA | #VALUE! |
| ENSMUSG00000101681.7 | 0 | NA | NA | NA | NA | NA | #VALUE! |
| ENSMUSG00000101682.2 | 0.372394 | 0.012025 | 4.059931 | 0.002962 | 0.997637 | NA | #VALUE! |
| ENSMUSG00000101683.7 | 0 | NA | NA | NA | NA | NA | #VALUE! |
| ENSMUSG00000101684.2 | 0 | NA | NA | NA | NA | NA | #VALUE! |
| ENSMUSG00000101685.2 | 0 | NA | NA | NA | NA | NA | #VALUE! |
| ENSMUSG00000101686.7 | 0 | NA | NA | NA | NA | NA | #VALUE! |
| ENSMUSG00000101687.2 | 0 | NA | NA | NA | NA | NA | #VALUE! |
| ENSMUSG00000101688.2 | 0 | NA | NA | NA | NA | NA | #VALUE! |
| ENSMUSG00000101689.2 | 0 | NA | NA | NA | NA | NA | #VALUE! |

| | | | | | | | |
|---|---|---|---|---|---|---|---|
| ENSMUSG00000101690.2 | 0 | NA | NA | NA | NA | NA | #VALUE! |
| ENSMUSG00000101691.2 | 0 | NA | NA | NA | NA | NA | #VALUE! |
| ENSMUSG00000101692.2 | 0 | NA | NA | NA | NA | NA | #VALUE! |
| ENSMUSG00000101693.2 | 0 | NA | NA | NA | NA | NA | #VALUE! |
| ENSMUSG00000101694.2 | 0 | NA | NA | NA | NA | NA | #VALUE! |
| ENSMUSG00000101695.2 | 0 | NA | NA | NA | NA | NA | #VALUE! |
| ENSMUSG00000101698.2 | 0 | NA | NA | NA | NA | NA | #VALUE! |
| ENSMUSG00000101699.2 | 0 | NA | NA | NA | NA | NA | #VALUE! |
| ENSMUSG00000101700.2 | 0 | NA | NA | NA | NA | NA | #VALUE! |
| ENSMUSG00000101701.2 | 0 | NA | NA | NA | NA | NA | #VALUE! |
| ENSMUSG00000101702.2 | 0 | NA | NA | NA | NA | NA | #VALUE! |
| ENSMUSG00000101703.2 | 0 | NA | NA | NA | NA | NA | #VALUE! |
| ENSMUSG00000101704.2 | 0 | NA | NA | NA | NA | NA | #VALUE! |
| ENSMUSG00000101705.2 | 0 | NA | NA | NA | NA | NA | #VALUE! |
| ENSMUSG00000101706.2 | 0 | NA | NA | NA | NA | NA | #VALUE! |
| ENSMUSG00000101707.2 | 0 | NA | NA | NA | NA | NA | #VALUE! |
| ENSMUSG00000101708.7 | 0 | NA | NA | NA | NA | NA | #VALUE! |
| ENSMUSG00000101709.2 | 0 | NA | NA | NA | NA | NA | #VALUE! |
| ENSMUSG00000101710.2 | 0 | NA | NA | NA | NA | NA | #VALUE! |
| ENSMUSG00000101711.2 | 0.093255 | 0.868093 | 4.072464 | 0.213162 | 0.831201 | NA | #VALUE! |
| ENSMUSG00000101712.2 | 0 | NA | NA | NA | NA | NA | #VALUE! |
| ENSMUSG00000101713.2 | 0 | NA | NA | NA | NA | NA | #VALUE! |
| ENSMUSG00000101714.2 | 0 | NA | NA | NA | NA | NA | #VALUE! |
| ENSMUSG00000101715.2 | 0 | NA | NA | NA | NA | NA | #VALUE! |
| ENSMUSG00000101717.3 | 0 | NA | NA | NA | NA | NA | #VALUE! |
| ENSMUSG00000101718.2 | 0 | NA | NA | NA | NA | NA | #VALUE! |
| ENSMUSG00000101719.2 | 0 | NA | NA | NA | NA | NA | #VALUE! |
| ENSMUSG00000101720.2 | 0 | NA | NA | NA | NA | NA | #VALUE! |
| ENSMUSG00000101721.2 | 0 | NA | NA | NA | NA | NA | #VALUE! |
| ENSMUSG00000101722.2 | 0 | NA | NA | NA | NA | NA | #VALUE! |
| ENSMUSG00000101723.2 | 0 | NA | NA | NA | NA | NA | #VALUE! |
| ENSMUSG00000101724.2 | 0 | NA | NA | NA | NA | NA | #VALUE! |
| ENSMUSG00000101725.7 | 0 | NA | NA | NA | NA | NA | #VALUE! |
| ENSMUSG00000101726.2 | 0 | NA | NA | NA | NA | NA | #VALUE! |
| ENSMUSG00000101727.2 | 0 | NA | NA | NA | NA | NA | #VALUE! |
| ENSMUSG00000101728.7 | 0 | NA | NA | NA | NA | NA | #VALUE! |
| ENSMUSG00000101729.2 | 0 | NA | NA | NA | NA | NA | #VALUE! |
| ENSMUSG00000101730.2 | 0 | NA | NA | NA | NA | NA | #VALUE! |
| ENSMUSG00000101731.2 | 0 | NA | NA | NA | NA | NA | #VALUE! |
| ENSMUSG00000101732.2 | 0 | NA | NA | NA | NA | NA | #VALUE! |
| ENSMUSG00000101733.2 | 0 | NA | NA | NA | NA | NA | #VALUE! |
| ENSMUSG00000101734.2 | 0 | NA | NA | NA | NA | NA | #VALUE! |
| ENSMUSG00000101735.2 | 0 | NA | NA | NA | NA | NA | #VALUE! |
| ENSMUSG00000101736.2 | 0 | NA | NA | NA | NA | NA | #VALUE! |

| | | | | | | | |
|---|---|---|---|---|---|---|---|
| ENSMUSG00000101737.2 | 0 | NA | NA | NA | NA | NA | #VALUE! |
| ENSMUSG00000101739.2 | 0 | NA | NA | NA | NA | NA | #VALUE! |
| ENSMUSG00000101740.7 | 0 | NA | NA | NA | NA | NA | #VALUE! |
| ENSMUSG00000101741.2 | 0 | NA | NA | NA | NA | NA | #VALUE! |
| ENSMUSG00000101742.2 | 0 | NA | NA | NA | NA | NA | #VALUE! |
| ENSMUSG00000101743.7 | 0 | NA | NA | NA | NA | NA | #VALUE! |
| ENSMUSG00000101744.7 | 0 | NA | NA | NA | NA | NA | #VALUE! |
| ENSMUSG00000101745.7 | 0 | NA | NA | NA | NA | NA | #VALUE! |
| ENSMUSG00000101746.2 | 0 | NA | NA | NA | NA | NA | #VALUE! |
| ENSMUSG00000101747.2 | 0 | NA | NA | NA | NA | NA | #VALUE! |
| ENSMUSG00000101748.2 | 0 | NA | NA | NA | NA | NA | #VALUE! |
| ENSMUSG00000101750.4 | 0 | NA | NA | NA | NA | NA | #VALUE! |
| ENSMUSG00000101751.2 | 0 | NA | NA | NA | NA | NA | #VALUE! |
| ENSMUSG00000101752.2 | 0 | NA | NA | NA | NA | NA | #VALUE! |
| ENSMUSG00000101753.2 | 0 | NA | NA | NA | NA | NA | #VALUE! |
| ENSMUSG00000101754.3 | 0 | NA | NA | NA | NA | NA | #VALUE! |
| ENSMUSG00000101755.2 | 0 | NA | NA | NA | NA | NA | #VALUE! |
| ENSMUSG00000101756.2 | 0 | NA | NA | NA | NA | NA | #VALUE! |
| ENSMUSG00000101757.2 | 0 | NA | NA | NA | NA | NA | #VALUE! |
| ENSMUSG00000101758.2 | 0 | NA | NA | NA | NA | NA | #VALUE! |
| ENSMUSG00000101759.2 | 0 | NA | NA | NA | NA | NA | #VALUE! |
| ENSMUSG00000101760.2 | 0 | NA | NA | NA | NA | NA | #VALUE! |
| ENSMUSG00000101761.7 | 0 | NA | NA | NA | NA | NA | #VALUE! |
| ENSMUSG00000101762.2 | 0 | NA | NA | NA | NA | NA | #VALUE! |
| ENSMUSG00000101763.2 | 0 | NA | NA | NA | NA | NA | #VALUE! |
| ENSMUSG00000101764.2 | 0 | NA | NA | NA | NA | NA | #VALUE! |
| ENSMUSG00000101765.2 | 0 | NA | NA | NA | NA | NA | #VALUE! |
| ENSMUSG00000101766.2 | 0 | NA | NA | NA | NA | NA | #VALUE! |
| ENSMUSG00000101768.2 | 0 | NA | NA | NA | NA | NA | #VALUE! |
| ENSMUSG00000101769.2 | 0 | NA | NA | NA | NA | NA | #VALUE! |
| ENSMUSG00000101770.2 | 0 | NA | NA | NA | NA | NA | #VALUE! |
| ENSMUSG00000101771.7 | 0 | NA | NA | NA | NA | NA | #VALUE! |
| ENSMUSG00000101772.2 | 0 | NA | NA | NA | NA | NA | #VALUE! |
| ENSMUSG00000101773.2 | 0 | NA | NA | NA | NA | NA | #VALUE! |
| ENSMUSG00000101774.7 | 0 | NA | NA | NA | NA | NA | #VALUE! |
| ENSMUSG00000101775.7 | 0 | NA | NA | NA | NA | NA | #VALUE! |
| ENSMUSG00000101776.2 | 0 | NA | NA | NA | NA | NA | #VALUE! |
| ENSMUSG00000101777.2 | 0 | NA | NA | NA | NA | NA | #VALUE! |
| ENSMUSG00000101778.2 | 0 | NA | NA | NA | NA | NA | #VALUE! |
| ENSMUSG00000101779.7 | 0 | NA | NA | NA | NA | NA | #VALUE! |
| ENSMUSG00000101780.2 | 0 | NA | NA | NA | NA | NA | #VALUE! |
| ENSMUSG00000101781.2 | 0 | NA | NA | NA | NA | NA | #VALUE! |
| ENSMUSG00000101782.2 | 0 | NA | NA | NA | NA | NA | #VALUE! |
| ENSMUSG00000101783.2 | 0 | NA | NA | NA | NA | NA | #VALUE! |

| | | | | | | | |
|---|---|---|---|---|---|---|---|
| ENSMUSG00000101784.2 | 0 | NA | NA | NA | NA | NA | #VALUE! |
| ENSMUSG00000101785.2 | 0 | NA | NA | NA | NA | NA | #VALUE! |
| ENSMUSG00000101786.7 | 0 | NA | NA | NA | NA | NA | #VALUE! |
| ENSMUSG00000101787.2 | 0 | NA | NA | NA | NA | NA | #VALUE! |
| ENSMUSG00000101788.2 | 0 | NA | NA | NA | NA | NA | #VALUE! |
| ENSMUSG00000101789.2 | 0 | NA | NA | NA | NA | NA | #VALUE! |
| ENSMUSG00000101790.2 | 0 | NA | NA | NA | NA | NA | #VALUE! |
| ENSMUSG00000101791.2 | 0 | NA | NA | NA | NA | NA | #VALUE! |
| ENSMUSG00000101792.2 | 0 | NA | NA | NA | NA | NA | #VALUE! |
| ENSMUSG00000101793.2 | 0 | NA | NA | NA | NA | NA | #VALUE! |
| ENSMUSG00000101794.2 | 0 | NA | NA | NA | NA | NA | #VALUE! |
| ENSMUSG00000101796.2 | 0 | NA | NA | NA | NA | NA | #VALUE! |
| ENSMUSG00000101797.2 | 0 | NA | NA | NA | NA | NA | #VALUE! |
| ENSMUSG00000101798.2 | 0 | NA | NA | NA | NA | NA | #VALUE! |
| ENSMUSG00000101799.2 | 0 | NA | NA | NA | NA | NA | #VALUE! |
| ENSMUSG00000101800.2 | 0 | NA | NA | NA | NA | NA | #VALUE! |
| ENSMUSG00000101801.7 | 0 | NA | NA | NA | NA | NA | #VALUE! |
| ENSMUSG00000101802.2 | 0 | NA | NA | NA | NA | NA | #VALUE! |
| ENSMUSG00000101803.2 | 0 | NA | NA | NA | NA | NA | #VALUE! |
| ENSMUSG00000101804.2 | 0 | NA | NA | NA | NA | NA | #VALUE! |
| ENSMUSG00000101805.2 | 0 | NA | NA | NA | NA | NA | #VALUE! |
| ENSMUSG00000101806.2 | 0 | NA | NA | NA | NA | NA | #VALUE! |
| ENSMUSG00000101807.2 | 0 | NA | NA | NA | NA | NA | #VALUE! |
| ENSMUSG00000101808.2 | 0 | NA | NA | NA | NA | NA | #VALUE! |
| ENSMUSG00000101809.2 | 0 | NA | NA | NA | NA | NA | #VALUE! |
| ENSMUSG00000101810.2 | 0 | NA | NA | NA | NA | NA | #VALUE! |
| ENSMUSG00000101811.2 | 0 | NA | NA | NA | NA | NA | #VALUE! |
| ENSMUSG00000101812.2 | 0 | NA | NA | NA | NA | NA | #VALUE! |
| ENSMUSG00000101813.2 | 0 | NA | NA | NA | NA | NA | #VALUE! |
| ENSMUSG00000101814.2 | 0 | NA | NA | NA | NA | NA | #VALUE! |
| ENSMUSG00000101815.2 | 0 | NA | NA | NA | NA | NA | #VALUE! |
| ENSMUSG00000101816.2 | 0 | NA | NA | NA | NA | NA | #VALUE! |
| ENSMUSG00000101817.2 | 0 | NA | NA | NA | NA | NA | #VALUE! |
| ENSMUSG00000101818.2 | 0 | NA | NA | NA | NA | NA | #VALUE! |
| ENSMUSG00000101819.2 | 0 | NA | NA | NA | NA | NA | #VALUE! |
| ENSMUSG00000101820.2 | 0 | NA | NA | NA | NA | NA | #VALUE! |
| ENSMUSG00000101821.2 | 0 | NA | NA | NA | NA | NA | #VALUE! |
| ENSMUSG00000101822.2 | 0 | NA | NA | NA | NA | NA | #VALUE! |
| ENSMUSG00000101823.2 | 0 | NA | NA | NA | NA | NA | #VALUE! |
| ENSMUSG00000101824.2 | 0 | NA | NA | NA | NA | NA | #VALUE! |
| ENSMUSG00000101825.2 | 0 | NA | NA | NA | NA | NA | #VALUE! |
| ENSMUSG00000101826.2 | 0 | NA | NA | NA | NA | NA | #VALUE! |
| ENSMUSG00000101827.2 | 0 | NA | NA | NA | NA | NA | #VALUE! |
| ENSMUSG00000101829.2 | 0 | NA | NA | NA | NA | NA | #VALUE! |

| | | | | | | | |
|---|---|---|---|---|---|---|---|
| ENSMUSG00000101830.2 | 0 | NA | NA | NA | NA | NA | #VALUE! |
| ENSMUSG00000101831.2 | 0 | NA | NA | NA | NA | NA | #VALUE! |
| ENSMUSG00000101832.2 | 0 | NA | NA | NA | NA | NA | #VALUE! |
| ENSMUSG00000101833.2 | 0 | NA | NA | NA | NA | NA | #VALUE! |
| ENSMUSG00000101834.2 | 0 | NA | NA | NA | NA | NA | #VALUE! |
| ENSMUSG00000101835.2 | 0 | NA | NA | NA | NA | NA | #VALUE! |
| ENSMUSG00000101836.2 | 0 | NA | NA | NA | NA | NA | #VALUE! |
| ENSMUSG00000101837.2 | 0 | NA | NA | NA | NA | NA | #VALUE! |
| ENSMUSG00000101838.7 | 0 | NA | NA | NA | NA | NA | #VALUE! |
| ENSMUSG00000101839.2 | 0 | NA | NA | NA | NA | NA | #VALUE! |
| ENSMUSG00000101840.2 | 0 | NA | NA | NA | NA | NA | #VALUE! |
| ENSMUSG00000101841.2 | 0 | NA | NA | NA | NA | NA | #VALUE! |
| ENSMUSG00000101842.2 | 0 | NA | NA | NA | NA | NA | #VALUE! |
| ENSMUSG00000101843.2 | 0 | NA | NA | NA | NA | NA | #VALUE! |
| ENSMUSG00000101844.2 | 0 | NA | NA | NA | NA | NA | #VALUE! |
| ENSMUSG00000101845.2 | 0 | NA | NA | NA | NA | NA | #VALUE! |
| ENSMUSG00000101846.2 | 0 | NA | NA | NA | NA | NA | #VALUE! |
| ENSMUSG00000101847.2 | 0 | NA | NA | NA | NA | NA | #VALUE! |
| ENSMUSG00000101848.2 | 0 | NA | NA | NA | NA | NA | #VALUE! |
| ENSMUSG00000101849.2 | 0 | NA | NA | NA | NA | NA | #VALUE! |
| ENSMUSG00000101850.2 | 0 | NA | NA | NA | NA | NA | #VALUE! |
| ENSMUSG00000101851.2 | 0 | NA | NA | NA | NA | NA | #VALUE! |
| ENSMUSG00000101852.2 | 0 | NA | NA | NA | NA | NA | #VALUE! |
| ENSMUSG00000101853.2 | 0 | NA | NA | NA | NA | NA | #VALUE! |
| ENSMUSG00000101854.3 | 0 | NA | NA | NA | NA | NA | #VALUE! |
| ENSMUSG00000101855.2 | 0 | NA | NA | NA | NA | NA | #VALUE! |
| ENSMUSG00000101856.2 | 7.486434 | -0.57521 | 3.208038 | -0.1793 | 0.857699 | NA | #VALUE! |
| ENSMUSG00000101857.2 | 0 | NA | NA | NA | NA | NA | #VALUE! |
| ENSMUSG00000101858.2 | 0 | NA | NA | NA | NA | NA | #VALUE! |
| ENSMUSG00000101859.2 | 0 | NA | NA | NA | NA | NA | #VALUE! |
| ENSMUSG00000101860.5 | 0 | NA | NA | NA | NA | NA | #VALUE! |
| ENSMUSG00000101861.2 | 0 | NA | NA | NA | NA | NA | #VALUE! |
| ENSMUSG00000101862.2 | 0 | NA | NA | NA | NA | NA | #VALUE! |
| ENSMUSG00000101863.2 | 0 | NA | NA | NA | NA | NA | #VALUE! |
| ENSMUSG00000101864.2 | 0 | NA | NA | NA | NA | NA | #VALUE! |
| ENSMUSG00000101865.2 | 0 | NA | NA | NA | NA | NA | #VALUE! |
| ENSMUSG00000101866.2 | 0 | NA | NA | NA | NA | NA | #VALUE! |
| ENSMUSG00000101867.2 | 0 | NA | NA | NA | NA | NA | #VALUE! |
| ENSMUSG00000101868.2 | 0 | NA | NA | NA | NA | NA | #VALUE! |
| ENSMUSG00000101869.2 | 0 | NA | NA | NA | NA | NA | #VALUE! |
| ENSMUSG00000101870.2 | 0 | NA | NA | NA | NA | NA | #VALUE! |
| ENSMUSG00000101871.2 | 0 | NA | NA | NA | NA | NA | #VALUE! |
| ENSMUSG00000101872.2 | 0 | NA | NA | NA | NA | NA | #VALUE! |
| ENSMUSG00000101873.2 | 0 | NA | NA | NA | NA | NA | #VALUE! |

| | | | | | | | |
|---|---|---|---|---|---|---|---|
| ENSMUSG00000101874.3 | 0 | NA | NA | NA | NA | NA | #VALUE! |
| ENSMUSG00000101875.2 | 0.105013 | -0.97324 | 4.076293 | -0.23876 | 0.811295 | NA | #VALUE! |
| ENSMUSG00000101877.2 | 0 | NA | NA | NA | NA | NA | #VALUE! |
| ENSMUSG00000101878.2 | 0.093255 | 0.868093 | 4.072464 | 0.213162 | 0.831201 | NA | #VALUE! |
| ENSMUSG00000101879.2 | 0 | NA | NA | NA | NA | NA | #VALUE! |
| ENSMUSG00000101880.2 | 0 | NA | NA | NA | NA | NA | #VALUE! |
| ENSMUSG00000101881.2 | 0 | NA | NA | NA | NA | NA | #VALUE! |
| ENSMUSG00000101882.2 | 0 | NA | NA | NA | NA | NA | #VALUE! |
| ENSMUSG00000101883.2 | 0 | NA | NA | NA | NA | NA | #VALUE! |
| ENSMUSG00000101884.2 | 0 | NA | NA | NA | NA | NA | #VALUE! |
| ENSMUSG00000101885.2 | 0 | NA | NA | NA | NA | NA | #VALUE! |
| ENSMUSG00000101886.2 | 0 | NA | NA | NA | NA | NA | #VALUE! |
| ENSMUSG00000101887.2 | 0 | NA | NA | NA | NA | NA | #VALUE! |
| ENSMUSG00000101888.2 | 0 | NA | NA | NA | NA | NA | #VALUE! |
| ENSMUSG00000101890.2 | 0 | NA | NA | NA | NA | NA | #VALUE! |
| ENSMUSG00000101891.7 | 0 | NA | NA | NA | NA | NA | #VALUE! |
| ENSMUSG00000101893.2 | 0 | NA | NA | NA | NA | NA | #VALUE! |
| ENSMUSG00000101894.2 | 0 | NA | NA | NA | NA | NA | #VALUE! |
| ENSMUSG00000101895.2 | 0 | NA | NA | NA | NA | NA | #VALUE! |
| ENSMUSG00000101896.2 | 0 | NA | NA | NA | NA | NA | #VALUE! |
| ENSMUSG00000101897.2 | 0 | NA | NA | NA | NA | NA | #VALUE! |
| ENSMUSG00000101898.2 | 0 | NA | NA | NA | NA | NA | #VALUE! |
| ENSMUSG00000101899.2 | 0 | NA | NA | NA | NA | NA | #VALUE! |
| ENSMUSG00000101900.2 | 0 | NA | NA | NA | NA | NA | #VALUE! |
| ENSMUSG00000101901.2 | 0 | NA | NA | NA | NA | NA | #VALUE! |
| ENSMUSG00000101902.2 | 0 | NA | NA | NA | NA | NA | #VALUE! |
| ENSMUSG00000101903.2 | 0 | NA | NA | NA | NA | NA | #VALUE! |
| ENSMUSG00000101904.2 | 0.652786 | 2.919659 | 3.972474 | 0.734972 | 0.462356 | NA | #VALUE! |
| ENSMUSG00000101905.2 | 0 | NA | NA | NA | NA | NA | #VALUE! |
| ENSMUSG00000101906.2 | 0 | NA | NA | NA | NA | NA | #VALUE! |
| ENSMUSG00000101907.2 | 0 | NA | NA | NA | NA | NA | #VALUE! |
| ENSMUSG00000101908.2 | 0 | NA | NA | NA | NA | NA | #VALUE! |
| ENSMUSG00000101909.3 | 0 | NA | NA | NA | NA | NA | #VALUE! |
| ENSMUSG00000101910.2 | 0 | NA | NA | NA | NA | NA | #VALUE! |
| ENSMUSG00000101911.2 | 0 | NA | NA | NA | NA | NA | #VALUE! |
| ENSMUSG00000101912.2 | 0 | NA | NA | NA | NA | NA | #VALUE! |
| ENSMUSG00000101913.2 | 0 | NA | NA | NA | NA | NA | #VALUE! |
| ENSMUSG00000101914.2 | 0 | NA | NA | NA | NA | NA | #VALUE! |
| ENSMUSG00000101915.2 | 0 | NA | NA | NA | NA | NA | #VALUE! |
| ENSMUSG00000101916.2 | 0 | NA | NA | NA | NA | NA | #VALUE! |
| ENSMUSG00000101917.2 | 0 | NA | NA | NA | NA | NA | #VALUE! |
| ENSMUSG00000101918.3 | 0 | NA | NA | NA | NA | NA | #VALUE! |
| ENSMUSG00000101919.2 | 0 | NA | NA | NA | NA | NA | #VALUE! |
| ENSMUSG00000101920.7 | 0 | NA | NA | NA | NA | NA | #VALUE! |

| | | | | | | | |
|---|---|---|---|---|---|---|---|
| ENSMUSG00000101921.2 | 0 | NA | NA | NA | NA | NA | #VALUE! |
| ENSMUSG00000101922.2 | 0 | NA | NA | NA | NA | NA | #VALUE! |
| ENSMUSG00000101923.2 | 0 | NA | NA | NA | NA | NA | #VALUE! |
| ENSMUSG00000101925.2 | 0 | NA | NA | NA | NA | NA | #VALUE! |
| ENSMUSG00000101927.2 | 0 | NA | NA | NA | NA | NA | #VALUE! |
| ENSMUSG00000101928.2 | 0 | NA | NA | NA | NA | NA | #VALUE! |
| ENSMUSG00000101929.7 | 0 | NA | NA | NA | NA | NA | #VALUE! |
| ENSMUSG00000101930.2 | 0 | NA | NA | NA | NA | NA | #VALUE! |
| ENSMUSG00000101931.2 | 0 | NA | NA | NA | NA | NA | #VALUE! |
| ENSMUSG00000101932.2 | 0 | NA | NA | NA | NA | NA | #VALUE! |
| ENSMUSG00000101933.2 | 0 | NA | NA | NA | NA | NA | #VALUE! |
| ENSMUSG00000101934.2 | 0 | NA | NA | NA | NA | NA | #VALUE! |
| ENSMUSG00000101935.2 | 0.336567 | 1.944925 | 4.029936 | 0.482619 | 0.629366 | NA | #VALUE! |
| ENSMUSG00000101936.2 | 0 | NA | NA | NA | NA | NA | #VALUE! |
| ENSMUSG00000101937.2 | 0 | NA | NA | NA | NA | NA | #VALUE! |
| ENSMUSG00000101938.2 | 0 | NA | NA | NA | NA | NA | #VALUE! |
| ENSMUSG00000101940.3 | 0 | NA | NA | NA | NA | NA | #VALUE! |
| ENSMUSG00000101941.2 | 0 | NA | NA | NA | NA | NA | #VALUE! |
| ENSMUSG00000101942.2 | 0 | NA | NA | NA | NA | NA | #VALUE! |
| ENSMUSG00000101943.2 | 0 | NA | NA | NA | NA | NA | #VALUE! |
| ENSMUSG00000101944.2 | 0 | NA | NA | NA | NA | NA | #VALUE! |
| ENSMUSG00000101945.2 | 0 | NA | NA | NA | NA | NA | #VALUE! |
| ENSMUSG00000101947.2 | 0 | NA | NA | NA | NA | NA | #VALUE! |
| ENSMUSG00000101948.2 | 0 | NA | NA | NA | NA | NA | #VALUE! |
| ENSMUSG00000101949.2 | 0 | NA | NA | NA | NA | NA | #VALUE! |
| ENSMUSG00000101950.2 | 0 | NA | NA | NA | NA | NA | #VALUE! |
| ENSMUSG00000101951.2 | 0 | NA | NA | NA | NA | NA | #VALUE! |
| ENSMUSG00000101952.2 | 0 | NA | NA | NA | NA | NA | #VALUE! |
| ENSMUSG00000101953.2 | 0 | NA | NA | NA | NA | NA | #VALUE! |
| ENSMUSG00000101954.2 | 0 | NA | NA | NA | NA | NA | #VALUE! |
| ENSMUSG00000101955.2 | 0 | NA | NA | NA | NA | NA | #VALUE! |
| ENSMUSG00000101956.2 | 0 | NA | NA | NA | NA | NA | #VALUE! |
| ENSMUSG00000101957.2 | 0 | NA | NA | NA | NA | NA | #VALUE! |
| ENSMUSG00000101958.2 | 0 | NA | NA | NA | NA | NA | #VALUE! |
| ENSMUSG00000101959.2 | 0 | NA | NA | NA | NA | NA | #VALUE! |
| ENSMUSG00000101960.2 | 0 | NA | NA | NA | NA | NA | #VALUE! |
| ENSMUSG00000101961.2 | 0 | NA | NA | NA | NA | NA | #VALUE! |
| ENSMUSG00000101962.2 | 0 | NA | NA | NA | NA | NA | #VALUE! |
| ENSMUSG00000101963.3 | 0 | NA | NA | NA | NA | NA | #VALUE! |
| ENSMUSG00000101964.2 | 0 | NA | NA | NA | NA | NA | #VALUE! |
| ENSMUSG00000101965.2 | 0 | NA | NA | NA | NA | NA | #VALUE! |
| ENSMUSG00000101966.3 | 0 | NA | NA | NA | NA | NA | #VALUE! |
| ENSMUSG00000101967.7 | 0 | NA | NA | NA | NA | NA | #VALUE! |
| ENSMUSG00000101968.7 | 0 | NA | NA | NA | NA | NA | #VALUE! |

| | | | | | | | |
|---|---|---|---|---|---|---|---|
| ENSMUSG00000101969.3 | 0 | NA | NA | NA | NA | NA | #VALUE! |
| ENSMUSG00000101971.2 | 0 | NA | NA | NA | NA | NA | #VALUE! |
| ENSMUSG00000101972.2 | 0.168284 | 1.027628 | 4.067126 | 0.252667 | 0.800526 | NA | #VALUE! |
| ENSMUSG00000101973.2 | 0 | NA | NA | NA | NA | NA | #VALUE! |
| ENSMUSG00000101974.2 | 0 | NA | NA | NA | NA | NA | #VALUE! |
| ENSMUSG00000101975.2 | 0 | NA | NA | NA | NA | NA | #VALUE! |
| ENSMUSG00000101977.2 | 0 | NA | NA | NA | NA | NA | #VALUE! |
| ENSMUSG00000101978.2 | 0 | NA | NA | NA | NA | NA | #VALUE! |
| ENSMUSG00000101980.2 | 0 | NA | NA | NA | NA | NA | #VALUE! |
| ENSMUSG00000101981.2 | 0 | NA | NA | NA | NA | NA | #VALUE! |
| ENSMUSG00000101982.7 | 0 | NA | NA | NA | NA | NA | #VALUE! |
| ENSMUSG00000101983.2 | 0 | NA | NA | NA | NA | NA | #VALUE! |
| ENSMUSG00000101984.2 | 0 | NA | NA | NA | NA | NA | #VALUE! |
| ENSMUSG00000101985.2 | 0 | NA | NA | NA | NA | NA | #VALUE! |
| ENSMUSG00000101986.2 | 0.195789 | -1.05581 | 4.073302 | -0.2592 | 0.795479 | NA | #VALUE! |
| ENSMUSG00000101987.2 | 0 | NA | NA | NA | NA | NA | #VALUE! |
| ENSMUSG00000101988.2 | 0 | NA | NA | NA | NA | NA | #VALUE! |
| ENSMUSG00000101989.2 | 0 | NA | NA | NA | NA | NA | #VALUE! |
| ENSMUSG00000101990.2 | 0 | NA | NA | NA | NA | NA | #VALUE! |
| ENSMUSG00000101991.2 | 0 | NA | NA | NA | NA | NA | #VALUE! |
| ENSMUSG00000101992.2 | 0 | NA | NA | NA | NA | NA | #VALUE! |
| ENSMUSG00000101993.2 | 0 | NA | NA | NA | NA | NA | #VALUE! |
| ENSMUSG00000101994.2 | 0 | NA | NA | NA | NA | NA | #VALUE! |
| ENSMUSG00000101995.2 | 0 | NA | NA | NA | NA | NA | #VALUE! |
| ENSMUSG00000101996.2 | 0 | NA | NA | NA | NA | NA | #VALUE! |
| ENSMUSG00000101997.2 | 0 | NA | NA | NA | NA | NA | #VALUE! |
| ENSMUSG00000101998.2 | 0 | NA | NA | NA | NA | NA | #VALUE! |
| ENSMUSG00000102000.7 | 0 | NA | NA | NA | NA | NA | #VALUE! |
| ENSMUSG00000102001.2 | 0 | NA | NA | NA | NA | NA | #VALUE! |
| ENSMUSG00000102002.2 | 0 | NA | NA | NA | NA | NA | #VALUE! |
| ENSMUSG00000102003.2 | 0 | NA | NA | NA | NA | NA | #VALUE! |
| ENSMUSG00000102004.4 | 0 | NA | NA | NA | NA | NA | #VALUE! |
| ENSMUSG00000102005.7 | 0 | NA | NA | NA | NA | NA | #VALUE! |
| ENSMUSG00000102006.2 | 0 | NA | NA | NA | NA | NA | #VALUE! |
| ENSMUSG00000102008.2 | 0 | NA | NA | NA | NA | NA | #VALUE! |
| ENSMUSG00000102009.7 | 0 | NA | NA | NA | NA | NA | #VALUE! |
| ENSMUSG00000102010.2 | 0 | NA | NA | NA | NA | NA | #VALUE! |
| ENSMUSG00000102011.2 | 0 | NA | NA | NA | NA | NA | #VALUE! |
| ENSMUSG00000102013.2 | 0 | NA | NA | NA | NA | NA | #VALUE! |
| ENSMUSG00000102015.2 | 0 | NA | NA | NA | NA | NA | #VALUE! |
| ENSMUSG00000102016.7 | 0 | NA | NA | NA | NA | NA | #VALUE! |
| ENSMUSG00000102017.2 | 0 | NA | NA | NA | NA | NA | #VALUE! |
| ENSMUSG00000102018.9 | 0 | NA | NA | NA | NA | NA | #VALUE! |
| ENSMUSG00000102019.2 | 0 | NA | NA | NA | NA | NA | #VALUE! |

| | | | | | | | |
|---|---|---|---|---|---|---|---|
| ENSMUSG00000102020.2 | 0 | NA | NA | NA | NA | NA | #VALUE! |
| ENSMUSG00000102022.7 | 0 | NA | NA | NA | NA | NA | #VALUE! |
| ENSMUSG00000102023.2 | 0 | NA | NA | NA | NA | NA | #VALUE! |
| ENSMUSG00000102025.2 | 0 | NA | NA | NA | NA | NA | #VALUE! |
| ENSMUSG00000102026.2 | 0 | NA | NA | NA | NA | NA | #VALUE! |
| ENSMUSG00000102027.2 | 0 | NA | NA | NA | NA | NA | #VALUE! |
| ENSMUSG00000102028.2 | 0 | NA | NA | NA | NA | NA | #VALUE! |
| ENSMUSG00000102029.2 | 0 | NA | NA | NA | NA | NA | #VALUE! |
| ENSMUSG00000102030.2 | 0 | NA | NA | NA | NA | NA | #VALUE! |
| ENSMUSG00000102031.7 | 0 | NA | NA | NA | NA | NA | #VALUE! |
| ENSMUSG00000102032.2 | 0 | NA | NA | NA | NA | NA | #VALUE! |
| ENSMUSG00000102033.2 | 0 | NA | NA | NA | NA | NA | #VALUE! |
| ENSMUSG00000102035.2 | 0 | NA | NA | NA | NA | NA | #VALUE! |
| ENSMUSG00000102036.2 | 0 | NA | NA | NA | NA | NA | #VALUE! |
| ENSMUSG00000102037.2 | 5.11037 | -3.95956 | 3.870313 | -1.02306 | 0.30628 | NA | #VALUE! |
| ENSMUSG00000102038.2 | 0 | NA | NA | NA | NA | NA | #VALUE! |
| ENSMUSG00000102039.2 | 0 | NA | NA | NA | NA | NA | #VALUE! |
| ENSMUSG00000102041.2 | 0 | NA | NA | NA | NA | NA | #VALUE! |
| ENSMUSG00000102042.2 | 0 | NA | NA | NA | NA | NA | #VALUE! |
| ENSMUSG00000102043.2 | 0 | NA | NA | NA | NA | NA | #VALUE! |
| ENSMUSG00000102044.2 | 0 | NA | NA | NA | NA | NA | #VALUE! |
| ENSMUSG00000102045.2 | 0 | NA | NA | NA | NA | NA | #VALUE! |
| ENSMUSG00000102046.2 | 0 | NA | NA | NA | NA | NA | #VALUE! |
| ENSMUSG00000102047.2 | 0 | NA | NA | NA | NA | NA | #VALUE! |
| ENSMUSG00000102048.2 | 0 | NA | NA | NA | NA | NA | #VALUE! |
| ENSMUSG00000102050.2 | 0 | NA | NA | NA | NA | NA | #VALUE! |
| ENSMUSG00000102051.2 | 0 | NA | NA | NA | NA | NA | #VALUE! |
| ENSMUSG00000102052.2 | 0 | NA | NA | NA | NA | NA | #VALUE! |
| ENSMUSG00000102053.2 | 0 | NA | NA | NA | NA | NA | #VALUE! |
| ENSMUSG00000102054.7 | 0 | NA | NA | NA | NA | NA | #VALUE! |
| ENSMUSG00000102055.2 | 0 | NA | NA | NA | NA | NA | #VALUE! |
| ENSMUSG00000102056.2 | 0 | NA | NA | NA | NA | NA | #VALUE! |
| ENSMUSG00000102057.2 | 0 | NA | NA | NA | NA | NA | #VALUE! |
| ENSMUSG00000102058.7 | 0 | NA | NA | NA | NA | NA | #VALUE! |
| ENSMUSG00000102060.2 | 0 | NA | NA | NA | NA | NA | #VALUE! |
| ENSMUSG00000102061.2 | 0 | NA | NA | NA | NA | NA | #VALUE! |
| ENSMUSG00000102062.2 | 0 | NA | NA | NA | NA | NA | #VALUE! |
| ENSMUSG00000102063.2 | 0 | NA | NA | NA | NA | NA | #VALUE! |
| ENSMUSG00000102064.2 | 0 | NA | NA | NA | NA | NA | #VALUE! |
| ENSMUSG00000102065.2 | 0 | NA | NA | NA | NA | NA | #VALUE! |
| ENSMUSG00000102067.2 | 0 | NA | NA | NA | NA | NA | #VALUE! |
| ENSMUSG00000102068.2 | 0 | NA | NA | NA | NA | NA | #VALUE! |
| ENSMUSG00000102069.2 | 0 | NA | NA | NA | NA | NA | #VALUE! |
| ENSMUSG00000102071.2 | 0 | NA | NA | NA | NA | NA | #VALUE! |

| | | | | | | | |
|---|---|---|---|---|---|---|---|
| ENSMUSG00000102072.2 | 0 | NA | NA | NA | NA | NA | #VALUE! |
| ENSMUSG00000102073.2 | 0 | NA | NA | NA | NA | NA | #VALUE! |
| ENSMUSG00000102074.2 | 0 | NA | NA | NA | NA | NA | #VALUE! |
| ENSMUSG00000102075.3 | 0 | NA | NA | NA | NA | NA | #VALUE! |
| ENSMUSG00000102076.2 | 0 | NA | NA | NA | NA | NA | #VALUE! |
| ENSMUSG00000102077.2 | 0 | NA | NA | NA | NA | NA | #VALUE! |
| ENSMUSG00000102079.2 | 0 | NA | NA | NA | NA | NA | #VALUE! |
| ENSMUSG00000102080.3 | 0 | NA | NA | NA | NA | NA | #VALUE! |
| ENSMUSG00000102081.2 | 0 | NA | NA | NA | NA | NA | #VALUE! |
| ENSMUSG00000102082.2 | 0 | NA | NA | NA | NA | NA | #VALUE! |
| ENSMUSG00000102083.2 | 0 | NA | NA | NA | NA | NA | #VALUE! |
| ENSMUSG00000102084.2 | 0 | NA | NA | NA | NA | NA | #VALUE! |
| ENSMUSG00000102085.3 | 0 | NA | NA | NA | NA | NA | #VALUE! |
| ENSMUSG00000102087.2 | 0 | NA | NA | NA | NA | NA | #VALUE! |
| ENSMUSG00000102088.7 | 0 | NA | NA | NA | NA | NA | #VALUE! |
| ENSMUSG00000102090.2 | 0 | NA | NA | NA | NA | NA | #VALUE! |
| ENSMUSG00000102092.2 | 0 | NA | NA | NA | NA | NA | #VALUE! |
| ENSMUSG00000102093.2 | 0 | NA | NA | NA | NA | NA | #VALUE! |
| ENSMUSG00000102096.2 | 0 | NA | NA | NA | NA | NA | #VALUE! |
| ENSMUSG00000102097.2 | 0 | NA | NA | NA | NA | NA | #VALUE! |
| ENSMUSG00000102098.2 | 0 | NA | NA | NA | NA | NA | #VALUE! |
| ENSMUSG00000102099.3 | 0 | NA | NA | NA | NA | NA | #VALUE! |
| ENSMUSG00000102100.2 | 0 | NA | NA | NA | NA | NA | #VALUE! |
| ENSMUSG00000102102.2 | 0 | NA | NA | NA | NA | NA | #VALUE! |
| ENSMUSG00000102103.2 | 0 | NA | NA | NA | NA | NA | #VALUE! |
| ENSMUSG00000102104.2 | 0 | NA | NA | NA | NA | NA | #VALUE! |
| ENSMUSG00000102105.2 | 0 | NA | NA | NA | NA | NA | #VALUE! |
| ENSMUSG00000102106.3 | 0 | NA | NA | NA | NA | NA | #VALUE! |
| ENSMUSG00000102107.2 | 0 | NA | NA | NA | NA | NA | #VALUE! |
| ENSMUSG00000102108.2 | 0 | NA | NA | NA | NA | NA | #VALUE! |
| ENSMUSG00000102109.2 | 0 | NA | NA | NA | NA | NA | #VALUE! |
| ENSMUSG00000102110.2 | 0 | NA | NA | NA | NA | NA | #VALUE! |
| ENSMUSG00000102111.2 | 0 | NA | NA | NA | NA | NA | #VALUE! |
| ENSMUSG00000102112.2 | 8.263326 | -0.18941 | 3.130539 | -0.0605 | 0.951755 | NA | #VALUE! |
| ENSMUSG00000102113.2 | 0 | NA | NA | NA | NA | NA | #VALUE! |
| ENSMUSG00000102114.3 | 0 | NA | NA | NA | NA | NA | #VALUE! |
| ENSMUSG00000102115.2 | 0 | NA | NA | NA | NA | NA | #VALUE! |
| ENSMUSG00000102116.2 | 0 | NA | NA | NA | NA | NA | #VALUE! |
| ENSMUSG00000102117.2 | 0 | NA | NA | NA | NA | NA | #VALUE! |
| ENSMUSG00000102118.2 | 0 | NA | NA | NA | NA | NA | #VALUE! |
| ENSMUSG00000102119.7 | 0 | NA | NA | NA | NA | NA | #VALUE! |
| ENSMUSG00000102120.2 | 0 | NA | NA | NA | NA | NA | #VALUE! |
| ENSMUSG00000102121.2 | 0 | NA | NA | NA | NA | NA | #VALUE! |
| ENSMUSG00000102122.2 | 0 | NA | NA | NA | NA | NA | #VALUE! |

| | | | | | | | |
|---|---|---|---|---|---|---|---|
| ENSMUSG00000102123.2 | 0 | NA | NA | NA | NA | NA | #VALUE! |
| ENSMUSG00000102124.2 | 0 | NA | NA | NA | NA | NA | #VALUE! |
| ENSMUSG00000102125.7 | 0 | NA | NA | NA | NA | NA | #VALUE! |
| ENSMUSG00000102126.2 | 0 | NA | NA | NA | NA | NA | #VALUE! |
| ENSMUSG00000102127.2 | 0 | NA | NA | NA | NA | NA | #VALUE! |
| ENSMUSG00000102128.2 | 0 | NA | NA | NA | NA | NA | #VALUE! |
| ENSMUSG00000102129.7 | 0 | NA | NA | NA | NA | NA | #VALUE! |
| ENSMUSG00000102131.2 | 0 | NA | NA | NA | NA | NA | #VALUE! |
| ENSMUSG00000102132.2 | 0 | NA | NA | NA | NA | NA | #VALUE! |
| ENSMUSG00000102133.2 | 0 | NA | NA | NA | NA | NA | #VALUE! |
| ENSMUSG00000102134.2 | 0 | NA | NA | NA | NA | NA | #VALUE! |
| ENSMUSG00000102135.2 | 0.366552 | 0.755301 | 4.038805 | 0.187011 | 0.851652 | NA | #VALUE! |
| ENSMUSG00000102136.2 | 0 | NA | NA | NA | NA | NA | #VALUE! |
| ENSMUSG00000102137.2 | 0 | NA | NA | NA | NA | NA | #VALUE! |
| ENSMUSG00000102138.2 | 0 | NA | NA | NA | NA | NA | #VALUE! |
| ENSMUSG00000102139.2 | 0 | NA | NA | NA | NA | NA | #VALUE! |
| ENSMUSG00000102140.2 | 0 | NA | NA | NA | NA | NA | #VALUE! |
| ENSMUSG00000102141.2 | 0 | NA | NA | NA | NA | NA | #VALUE! |
| ENSMUSG00000102142.2 | 0 | NA | NA | NA | NA | NA | #VALUE! |
| ENSMUSG00000102143.2 | 0 | NA | NA | NA | NA | NA | #VALUE! |
| ENSMUSG00000102144.2 | 0 | NA | NA | NA | NA | NA | #VALUE! |
| ENSMUSG00000102145.2 | 0 | NA | NA | NA | NA | NA | #VALUE! |
| ENSMUSG00000102146.2 | 0 | NA | NA | NA | NA | NA | #VALUE! |
| ENSMUSG00000102147.2 | 0 | NA | NA | NA | NA | NA | #VALUE! |
| ENSMUSG00000102148.2 | 0 | NA | NA | NA | NA | NA | #VALUE! |
| ENSMUSG00000102149.2 | 0 | NA | NA | NA | NA | NA | #VALUE! |
| ENSMUSG00000102150.2 | 0 | NA | NA | NA | NA | NA | #VALUE! |
| ENSMUSG00000102151.2 | 0 | NA | NA | NA | NA | NA | #VALUE! |
| ENSMUSG00000102152.2 | 0 | NA | NA | NA | NA | NA | #VALUE! |
| ENSMUSG00000102153.2 | 6.135817 | 1.559203 | 3.765124 | 0.414117 | 0.678788 | NA | #VALUE! |
| ENSMUSG00000102154.2 | 0 | NA | NA | NA | NA | NA | #VALUE! |
| ENSMUSG00000102155.2 | 0 | NA | NA | NA | NA | NA | #VALUE! |
| ENSMUSG00000102157.2 | 0 | NA | NA | NA | NA | NA | #VALUE! |
| ENSMUSG00000102158.2 | 0 | NA | NA | NA | NA | NA | #VALUE! |
| ENSMUSG00000102159.2 | 0 | NA | NA | NA | NA | NA | #VALUE! |
| ENSMUSG00000102160.2 | 0.093255 | 0.868093 | 4.072464 | 0.213162 | 0.831201 | NA | #VALUE! |
| ENSMUSG00000102162.2 | 0 | NA | NA | NA | NA | NA | #VALUE! |
| ENSMUSG00000102163.2 | 0 | NA | NA | NA | NA | NA | #VALUE! |
| ENSMUSG00000102164.2 | 0 | NA | NA | NA | NA | NA | #VALUE! |
| ENSMUSG00000102166.2 | 0 | NA | NA | NA | NA | NA | #VALUE! |
| ENSMUSG00000102167.2 | 0 | NA | NA | NA | NA | NA | #VALUE! |
| ENSMUSG00000102168.2 | 0 | NA | NA | NA | NA | NA | #VALUE! |
| ENSMUSG00000102169.2 | 0 | NA | NA | NA | NA | NA | #VALUE! |
| ENSMUSG00000102170.2 | 0 | NA | NA | NA | NA | NA | #VALUE! |

| | | | | | | | |
|---|---|---|---|---|---|---|---|
| ENSMUSG00000102172.2 | 0 | NA | NA | NA | NA | NA | #VALUE! |
| ENSMUSG00000102173.2 | 0 | NA | NA | NA | NA | NA | #VALUE! |
| ENSMUSG00000102174.2 | 0 | NA | NA | NA | NA | NA | #VALUE! |
| ENSMUSG00000102175.2 | 0 | NA | NA | NA | NA | NA | #VALUE! |
| ENSMUSG00000102176.2 | 0 | NA | NA | NA | NA | NA | #VALUE! |
| ENSMUSG00000102177.6 | 0 | NA | NA | NA | NA | NA | #VALUE! |
| ENSMUSG00000102178.2 | 0 | NA | NA | NA | NA | NA | #VALUE! |
| ENSMUSG00000102179.2 | 0 | NA | NA | NA | NA | NA | #VALUE! |
| ENSMUSG00000102180.2 | 0 | NA | NA | NA | NA | NA | #VALUE! |
| ENSMUSG00000102181.2 | 0 | NA | NA | NA | NA | NA | #VALUE! |
| ENSMUSG00000102182.2 | 0 | NA | NA | NA | NA | NA | #VALUE! |
| ENSMUSG00000102184.2 | 0 | NA | NA | NA | NA | NA | #VALUE! |
| ENSMUSG00000102185.2 | 0 | NA | NA | NA | NA | NA | #VALUE! |
| ENSMUSG00000102186.2 | 0 | NA | NA | NA | NA | NA | #VALUE! |
| ENSMUSG00000102187.2 | 0 | NA | NA | NA | NA | NA | #VALUE! |
| ENSMUSG00000102188.2 | 0 | NA | NA | NA | NA | NA | #VALUE! |
| ENSMUSG00000102189.2 | 0 | NA | NA | NA | NA | NA | #VALUE! |
| ENSMUSG00000102190.2 | 0 | NA | NA | NA | NA | NA | #VALUE! |
| ENSMUSG00000102191.2 | 0 | NA | NA | NA | NA | NA | #VALUE! |
| ENSMUSG00000102192.2 | 0 | NA | NA | NA | NA | NA | #VALUE! |
| ENSMUSG00000102193.2 | 0 | NA | NA | NA | NA | NA | #VALUE! |
| ENSMUSG00000102194.2 | 0 | NA | NA | NA | NA | NA | #VALUE! |
| ENSMUSG00000102195.6 | 0 | NA | NA | NA | NA | NA | #VALUE! |
| ENSMUSG00000102196.2 | 0 | NA | NA | NA | NA | NA | #VALUE! |
| ENSMUSG00000102197.2 | 0 | NA | NA | NA | NA | NA | #VALUE! |
| ENSMUSG00000102198.2 | 0 | NA | NA | NA | NA | NA | #VALUE! |
| ENSMUSG00000102199.2 | 0 | NA | NA | NA | NA | NA | #VALUE! |
| ENSMUSG00000102200.2 | 0 | NA | NA | NA | NA | NA | #VALUE! |
| ENSMUSG00000102201.2 | 0 | NA | NA | NA | NA | NA | #VALUE! |
| ENSMUSG00000102202.2 | 0 | NA | NA | NA | NA | NA | #VALUE! |
| ENSMUSG00000102203.2 | 0 | NA | NA | NA | NA | NA | #VALUE! |
| ENSMUSG00000102204.2 | 0 | NA | NA | NA | NA | NA | #VALUE! |
| ENSMUSG00000102205.2 | 0 | NA | NA | NA | NA | NA | #VALUE! |
| ENSMUSG00000102206.6 | 0 | NA | NA | NA | NA | NA | #VALUE! |
| ENSMUSG00000102207.2 | 0 | NA | NA | NA | NA | NA | #VALUE! |
| ENSMUSG00000102208.2 | 0 | NA | NA | NA | NA | NA | #VALUE! |
| ENSMUSG00000102211.2 | 2.7437 | -0.30759 | 3.793146 | -0.08109 | 0.935369 | NA | #VALUE! |
| ENSMUSG00000102212.2 | 0 | NA | NA | NA | NA | NA | #VALUE! |
| ENSMUSG00000102213.2 | 0 | NA | NA | NA | NA | NA | #VALUE! |
| ENSMUSG00000102214.2 | 0 | NA | NA | NA | NA | NA | #VALUE! |
| ENSMUSG00000102215.2 | 0 | NA | NA | NA | NA | NA | #VALUE! |
| ENSMUSG00000102216.2 | 0 | NA | NA | NA | NA | NA | #VALUE! |
| ENSMUSG00000102217.2 | 0 | NA | NA | NA | NA | NA | #VALUE! |
| ENSMUSG00000102218.2 | 0 | NA | NA | NA | NA | NA | #VALUE! |

| | | | | | | | |
|---|---|---|---|---|---|---|---|
| ENSMUSG00000102219.2 | 0 | NA | NA | NA | NA | NA | #VALUE! |
| ENSMUSG00000102220.2 | 0 | NA | NA | NA | NA | NA | #VALUE! |
| ENSMUSG00000102221.2 | 0 | NA | NA | NA | NA | NA | #VALUE! |
| ENSMUSG00000102222.2 | 0 | NA | NA | NA | NA | NA | #VALUE! |
| ENSMUSG00000102223.2 | 0 | NA | NA | NA | NA | NA | #VALUE! |
| ENSMUSG00000102224.6 | 0 | NA | NA | NA | NA | NA | #VALUE! |
| ENSMUSG00000102225.2 | 0 | NA | NA | NA | NA | NA | #VALUE! |
| ENSMUSG00000102227.2 | 0 | NA | NA | NA | NA | NA | #VALUE! |
| ENSMUSG00000102229.2 | 0 | NA | NA | NA | NA | NA | #VALUE! |
| ENSMUSG00000102230.2 | 0 | NA | NA | NA | NA | NA | #VALUE! |
| ENSMUSG00000102231.2 | 0 | NA | NA | NA | NA | NA | #VALUE! |
| ENSMUSG00000102232.2 | 0 | NA | NA | NA | NA | NA | #VALUE! |
| ENSMUSG00000102233.2 | 0 | NA | NA | NA | NA | NA | #VALUE! |
| ENSMUSG00000102234.2 | 0 | NA | NA | NA | NA | NA | #VALUE! |
| ENSMUSG00000102235.2 | 0 | NA | NA | NA | NA | NA | #VALUE! |
| ENSMUSG00000102236.2 | 0 | NA | NA | NA | NA | NA | #VALUE! |
| ENSMUSG00000102238.2 | 0 | NA | NA | NA | NA | NA | #VALUE! |
| ENSMUSG00000102239.2 | 0 | NA | NA | NA | NA | NA | #VALUE! |
| ENSMUSG00000102240.2 | 0 | NA | NA | NA | NA | NA | #VALUE! |
| ENSMUSG00000102241.2 | 0 | NA | NA | NA | NA | NA | #VALUE! |
| ENSMUSG00000102242.2 | 0 | NA | NA | NA | NA | NA | #VALUE! |
| ENSMUSG00000102243.2 | 0.420053 | -2.33493 | 4.007343 | -0.58266 | 0.56012 | NA | #VALUE! |
| ENSMUSG00000102244.2 | 0 | NA | NA | NA | NA | NA | #VALUE! |
| ENSMUSG00000102245.2 | 0 | NA | NA | NA | NA | NA | #VALUE! |
| ENSMUSG00000102246.2 | 0 | NA | NA | NA | NA | NA | #VALUE! |
| ENSMUSG00000102247.2 | 0 | NA | NA | NA | NA | NA | #VALUE! |
| ENSMUSG00000102248.2 | 0 | NA | NA | NA | NA | NA | #VALUE! |
| ENSMUSG00000102249.2 | 0 | NA | NA | NA | NA | NA | #VALUE! |
| ENSMUSG00000102250.2 | 0 | NA | NA | NA | NA | NA | #VALUE! |
| ENSMUSG00000102251.6 | 0 | NA | NA | NA | NA | NA | #VALUE! |
| ENSMUSG00000102252.6 | 1.830771 | 4.368534 | 3.931101 | 1.111275 | 0.26645 | NA | #VALUE! |
| ENSMUSG00000102253.2 | 0 | NA | NA | NA | NA | NA | #VALUE! |
| ENSMUSG00000102254.2 | 0 | NA | NA | NA | NA | NA | #VALUE! |
| ENSMUSG00000102255.2 | 0 | NA | NA | NA | NA | NA | #VALUE! |
| ENSMUSG00000102256.2 | 0 | NA | NA | NA | NA | NA | #VALUE! |
| ENSMUSG00000102257.2 | 0 | NA | NA | NA | NA | NA | #VALUE! |
| ENSMUSG00000102258.2 | 0 | NA | NA | NA | NA | NA | #VALUE! |
| ENSMUSG00000102259.2 | 0 | NA | NA | NA | NA | NA | #VALUE! |
| ENSMUSG00000102260.2 | 0 | NA | NA | NA | NA | NA | #VALUE! |
| ENSMUSG00000102261.2 | 0 | NA | NA | NA | NA | NA | #VALUE! |
| ENSMUSG00000102262.2 | 0 | NA | NA | NA | NA | NA | #VALUE! |
| ENSMUSG00000102263.2 | 0 | NA | NA | NA | NA | NA | #VALUE! |
| ENSMUSG00000102264.2 | 0 | NA | NA | NA | NA | NA | #VALUE! |
| ENSMUSG00000102265.2 | 0 | NA | NA | NA | NA | NA | #VALUE! |

| | | | | | | | |
|---|---|---|---|---|---|---|---|
| ENSMUSG00000102266.2 | 0 | NA | NA | NA | NA | NA | #VALUE! |
| ENSMUSG00000102267.2 | 0 | NA | NA | NA | NA | NA | #VALUE! |
| ENSMUSG00000102268.2 | 0 | NA | NA | NA | NA | NA | #VALUE! |
| ENSMUSG00000102269.2 | 0 | NA | NA | NA | NA | NA | #VALUE! |
| ENSMUSG00000102270.2 | 0 | NA | NA | NA | NA | NA | #VALUE! |
| ENSMUSG00000102271.2 | 0 | NA | NA | NA | NA | NA | #VALUE! |
| ENSMUSG00000102272.2 | 0 | NA | NA | NA | NA | NA | #VALUE! |
| ENSMUSG00000102273.6 | 0 | NA | NA | NA | NA | NA | #VALUE! |
| ENSMUSG00000102274.2 | 0 | NA | NA | NA | NA | NA | #VALUE! |
| ENSMUSG00000102275.2 | 0.3999 | -2.17566 | 4.018831 | -0.54137 | 0.588256 | NA | #VALUE! |
| ENSMUSG00000102276.2 | 0 | NA | NA | NA | NA | NA | #VALUE! |
| ENSMUSG00000102277.2 | 0 | NA | NA | NA | NA | NA | #VALUE! |
| ENSMUSG00000102278.2 | 0 | NA | NA | NA | NA | NA | #VALUE! |
| ENSMUSG00000102279.2 | 0 | NA | NA | NA | NA | NA | #VALUE! |
| ENSMUSG00000102280.2 | 0 | NA | NA | NA | NA | NA | #VALUE! |
| ENSMUSG00000102281.2 | 0 | NA | NA | NA | NA | NA | #VALUE! |
| ENSMUSG00000102282.2 | 0 | NA | NA | NA | NA | NA | #VALUE! |
| ENSMUSG00000102283.2 | 0 | NA | NA | NA | NA | NA | #VALUE! |
| ENSMUSG00000102284.2 | 0 | NA | NA | NA | NA | NA | #VALUE! |
| ENSMUSG00000102285.2 | 0 | NA | NA | NA | NA | NA | #VALUE! |
| ENSMUSG00000102286.2 | 0 | NA | NA | NA | NA | NA | #VALUE! |
| ENSMUSG00000102287.2 | 0 | NA | NA | NA | NA | NA | #VALUE! |
| ENSMUSG00000102288.2 | 0 | NA | NA | NA | NA | NA | #VALUE! |
| ENSMUSG00000102289.2 | 0 | NA | NA | NA | NA | NA | #VALUE! |
| ENSMUSG00000102290.2 | 0 | NA | NA | NA | NA | NA | #VALUE! |
| ENSMUSG00000102291.2 | 0 | NA | NA | NA | NA | NA | #VALUE! |
| ENSMUSG00000102293.2 | 0 | NA | NA | NA | NA | NA | #VALUE! |
| ENSMUSG00000102294.2 | 0 | NA | NA | NA | NA | NA | #VALUE! |
| ENSMUSG00000102295.2 | 0 | NA | NA | NA | NA | NA | #VALUE! |
| ENSMUSG00000102296.2 | 0 | NA | NA | NA | NA | NA | #VALUE! |
| ENSMUSG00000102297.2 | 0 | NA | NA | NA | NA | NA | #VALUE! |
| ENSMUSG00000102298.2 | 0 | NA | NA | NA | NA | NA | #VALUE! |
| ENSMUSG00000102299.2 | 0 | NA | NA | NA | NA | NA | #VALUE! |
| ENSMUSG00000102300.2 | 0 | NA | NA | NA | NA | NA | #VALUE! |
| ENSMUSG00000102301.2 | 0 | NA | NA | NA | NA | NA | #VALUE! |
| ENSMUSG00000102302.2 | 0 | NA | NA | NA | NA | NA | #VALUE! |
| ENSMUSG00000102303.2 | 0 | NA | NA | NA | NA | NA | #VALUE! |
| ENSMUSG00000102304.2 | 0 | NA | NA | NA | NA | NA | #VALUE! |
| ENSMUSG00000102305.2 | 0 | NA | NA | NA | NA | NA | #VALUE! |
| ENSMUSG00000102307.2 | 0 | NA | NA | NA | NA | NA | #VALUE! |
| ENSMUSG00000102308.2 | 0 | NA | NA | NA | NA | NA | #VALUE! |
| ENSMUSG00000102309.2 | 0 | NA | NA | NA | NA | NA | #VALUE! |
| ENSMUSG00000102310.2 | 0 | NA | NA | NA | NA | NA | #VALUE! |
| ENSMUSG00000102311.2 | 0 | NA | NA | NA | NA | NA | #VALUE! |

| | | | | | | | |
|---|---|---|---|---|---|---|---|
| ENSMUSG00000102312.2 | 0 | NA | NA | NA | NA | NA | #VALUE! |
| ENSMUSG00000102313.2 | 0 | NA | NA | NA | NA | NA | #VALUE! |
| ENSMUSG00000102314.2 | 0 | NA | NA | NA | NA | NA | #VALUE! |
| ENSMUSG00000102315.2 | 0 | NA | NA | NA | NA | NA | #VALUE! |
| ENSMUSG00000102316.2 | 7.646926 | 6.428952 | 3.90533 | 1.6462 | NA | NA | #VALUE! |
| ENSMUSG00000102317.2 | 0 | NA | NA | NA | NA | NA | #VALUE! |
| ENSMUSG00000102318.2 | 0 | NA | NA | NA | NA | NA | #VALUE! |
| ENSMUSG00000102319.2 | 0 | NA | NA | NA | NA | NA | #VALUE! |
| ENSMUSG00000102320.2 | 0 | NA | NA | NA | NA | NA | #VALUE! |
| ENSMUSG00000102321.2 | 0 | NA | NA | NA | NA | NA | #VALUE! |
| ENSMUSG00000102322.2 | 0 | NA | NA | NA | NA | NA | #VALUE! |
| ENSMUSG00000102323.2 | 0 | NA | NA | NA | NA | NA | #VALUE! |
| ENSMUSG00000102324.2 | 0 | NA | NA | NA | NA | NA | #VALUE! |
| ENSMUSG00000102325.2 | 0 | NA | NA | NA | NA | NA | #VALUE! |
| ENSMUSG00000102326.2 | 0.587367 | -2.74297 | 3.982932 | -0.68868 | 0.491024 | NA | #VALUE! |
| ENSMUSG00000102327.2 | 0 | NA | NA | NA | NA | NA | #VALUE! |
| ENSMUSG00000102328.2 | 0 | NA | NA | NA | NA | NA | #VALUE! |
| ENSMUSG00000102329.2 | 0 | NA | NA | NA | NA | NA | #VALUE! |
| ENSMUSG00000102330.2 | 0 | NA | NA | NA | NA | NA | #VALUE! |
| ENSMUSG00000102331.2 | 0 | NA | NA | NA | NA | NA | #VALUE! |
| ENSMUSG00000102332.2 | 0 | NA | NA | NA | NA | NA | #VALUE! |
| ENSMUSG00000102333.2 | 0 | NA | NA | NA | NA | NA | #VALUE! |
| ENSMUSG00000102334.2 | 0 | NA | NA | NA | NA | NA | #VALUE! |
| ENSMUSG00000102335.6 | 0 | NA | NA | NA | NA | NA | #VALUE! |
| ENSMUSG00000102336.2 | 0 | NA | NA | NA | NA | NA | #VALUE! |
| ENSMUSG00000102337.2 | 0 | NA | NA | NA | NA | NA | #VALUE! |
| ENSMUSG00000102338.2 | 0 | NA | NA | NA | NA | NA | #VALUE! |
| ENSMUSG00000102339.2 | 0 | NA | NA | NA | NA | NA | #VALUE! |
| ENSMUSG00000102341.2 | 0 | NA | NA | NA | NA | NA | #VALUE! |
| ENSMUSG00000102342.2 | 0 | NA | NA | NA | NA | NA | #VALUE! |
| ENSMUSG00000102343.2 | 0 | NA | NA | NA | NA | NA | #VALUE! |
| ENSMUSG00000102344.2 | 0 | NA | NA | NA | NA | NA | #VALUE! |
| ENSMUSG00000102345.2 | 0 | NA | NA | NA | NA | NA | #VALUE! |
| ENSMUSG00000102346.6 | 0 | NA | NA | NA | NA | NA | #VALUE! |
| ENSMUSG00000102347.2 | 0 | NA | NA | NA | NA | NA | #VALUE! |
| ENSMUSG00000102348.2 | 0 | NA | NA | NA | NA | NA | #VALUE! |
| ENSMUSG00000102350.6 | 0 | NA | NA | NA | NA | NA | #VALUE! |
| ENSMUSG00000102352.2 | 0 | NA | NA | NA | NA | NA | #VALUE! |
| ENSMUSG00000102353.2 | 9.40235 | 4.757884 | 3.352327 | 1.419278 | 0.155818 | NA | #VALUE! |
| ENSMUSG00000102354.2 | 0 | NA | NA | NA | NA | NA | #VALUE! |
| ENSMUSG00000102355.2 | 0 | NA | NA | NA | NA | NA | #VALUE! |
| ENSMUSG00000102356.2 | 0 | NA | NA | NA | NA | NA | #VALUE! |
| ENSMUSG00000102357.2 | 0 | NA | NA | NA | NA | NA | #VALUE! |
| ENSMUSG00000102358.2 | 0 | NA | NA | NA | NA | NA | #VALUE! |

| | | | | | | | |
|---|---|---|---|---|---|---|---|
| ENSMUSG00000102359.2 | 0 | NA | NA | NA | NA | NA | #VALUE! |
| ENSMUSG00000102360.2 | 0 | NA | NA | NA | NA | NA | #VALUE! |
| ENSMUSG00000102362.2 | 0 | NA | NA | NA | NA | NA | #VALUE! |
| ENSMUSG00000102363.2 | 0 | NA | NA | NA | NA | NA | #VALUE! |
| ENSMUSG00000102364.2 | 0 | NA | NA | NA | NA | NA | #VALUE! |
| ENSMUSG00000102365.2 | 0 | NA | NA | NA | NA | NA | #VALUE! |
| ENSMUSG00000102366.2 | 0 | NA | NA | NA | NA | NA | #VALUE! |
| ENSMUSG00000102368.2 | 0 | NA | NA | NA | NA | NA | #VALUE! |
| ENSMUSG00000102369.2 | 0 | NA | NA | NA | NA | NA | #VALUE! |
| ENSMUSG00000102370.2 | 0 | NA | NA | NA | NA | NA | #VALUE! |
| ENSMUSG00000102371.5 | 0 | NA | NA | NA | NA | NA | #VALUE! |
| ENSMUSG00000102372.2 | 0 | NA | NA | NA | NA | NA | #VALUE! |
| ENSMUSG00000102373.2 | 0 | NA | NA | NA | NA | NA | #VALUE! |
| ENSMUSG00000102374.2 | 0 | NA | NA | NA | NA | NA | #VALUE! |
| ENSMUSG00000102375.2 | 0 | NA | NA | NA | NA | NA | #VALUE! |
| ENSMUSG00000102376.2 | 0 | NA | NA | NA | NA | NA | #VALUE! |
| ENSMUSG00000102377.2 | 0 | NA | NA | NA | NA | NA | #VALUE! |
| ENSMUSG00000102378.2 | 0 | NA | NA | NA | NA | NA | #VALUE! |
| ENSMUSG00000102379.2 | 0 | NA | NA | NA | NA | NA | #VALUE! |
| ENSMUSG00000102380.2 | 0 | NA | NA | NA | NA | NA | #VALUE! |
| ENSMUSG00000102381.2 | 0 | NA | NA | NA | NA | NA | #VALUE! |
| ENSMUSG00000102382.2 | 0 | NA | NA | NA | NA | NA | #VALUE! |
| ENSMUSG00000102383.2 | 0 | NA | NA | NA | NA | NA | #VALUE! |
| ENSMUSG00000102384.2 | 0 | NA | NA | NA | NA | NA | #VALUE! |
| ENSMUSG00000102385.2 | 0 | NA | NA | NA | NA | NA | #VALUE! |
| ENSMUSG00000102386.2 | 0 | NA | NA | NA | NA | NA | #VALUE! |
| ENSMUSG00000102387.2 | 0 | NA | NA | NA | NA | NA | #VALUE! |
| ENSMUSG00000102388.6 | 0 | NA | NA | NA | NA | NA | #VALUE! |
| ENSMUSG00000102389.2 | 0 | NA | NA | NA | NA | NA | #VALUE! |
| ENSMUSG00000102390.2 | 0 | NA | NA | NA | NA | NA | #VALUE! |
| ENSMUSG00000102391.2 | 0 | NA | NA | NA | NA | NA | #VALUE! |
| ENSMUSG00000102392.6 | 0 | NA | NA | NA | NA | NA | #VALUE! |
| ENSMUSG00000102393.2 | 0 | NA | NA | NA | NA | NA | #VALUE! |
| ENSMUSG00000102394.2 | 0 | NA | NA | NA | NA | NA | #VALUE! |
| ENSMUSG00000102396.2 | 0 | NA | NA | NA | NA | NA | #VALUE! |
| ENSMUSG00000102397.2 | 0 | NA | NA | NA | NA | NA | #VALUE! |
| ENSMUSG00000102398.2 | 0 | NA | NA | NA | NA | NA | #VALUE! |
| ENSMUSG00000102399.2 | 0 | NA | NA | NA | NA | NA | #VALUE! |
| ENSMUSG00000102400.2 | 0 | NA | NA | NA | NA | NA | #VALUE! |
| ENSMUSG00000102401.2 | 0 | NA | NA | NA | NA | NA | #VALUE! |
| ENSMUSG00000102402.2 | 0 | NA | NA | NA | NA | NA | #VALUE! |
| ENSMUSG00000102403.2 | 0 | NA | NA | NA | NA | NA | #VALUE! |
| ENSMUSG00000102404.2 | 0 | NA | NA | NA | NA | NA | #VALUE! |
| ENSMUSG00000102405.2 | 0 | NA | NA | NA | NA | NA | #VALUE! |

| | | | | | | | |
|---|---|---|---|---|---|---|---|
| ENSMUSG00000102406.2 | 0 | NA | NA | NA | NA | NA | #VALUE! |
| ENSMUSG00000102407.2 | 0 | NA | NA | NA | NA | NA | #VALUE! |
| ENSMUSG00000102408.2 | 0 | NA | NA | NA | NA | NA | #VALUE! |
| ENSMUSG00000102409.2 | 0 | NA | NA | NA | NA | NA | #VALUE! |
| ENSMUSG00000102410.2 | 0 | NA | NA | NA | NA | NA | #VALUE! |
| ENSMUSG00000102411.2 | 0 | NA | NA | NA | NA | NA | #VALUE! |
| ENSMUSG00000102413.2 | 0 | NA | NA | NA | NA | NA | #VALUE! |
| ENSMUSG00000102414.2 | 0 | NA | NA | NA | NA | NA | #VALUE! |
| ENSMUSG00000102415.2 | 0 | NA | NA | NA | NA | NA | #VALUE! |
| ENSMUSG00000102416.5 | 0 | NA | NA | NA | NA | NA | #VALUE! |
| ENSMUSG00000102417.2 | 0 | NA | NA | NA | NA | NA | #VALUE! |
| ENSMUSG00000102418.2 | 0 | NA | NA | NA | NA | NA | #VALUE! |
| ENSMUSG00000102419.2 | 0 | NA | NA | NA | NA | NA | #VALUE! |
| ENSMUSG00000102420.2 | 0 | NA | NA | NA | NA | NA | #VALUE! |
| ENSMUSG00000102421.2 | 0 | NA | NA | NA | NA | NA | #VALUE! |
| ENSMUSG00000102422.2 | 0 | NA | NA | NA | NA | NA | #VALUE! |
| ENSMUSG00000102423.2 | 0 | NA | NA | NA | NA | NA | #VALUE! |
| ENSMUSG00000102424.3 | 0 | NA | NA | NA | NA | NA | #VALUE! |
| ENSMUSG00000102425.2 | 0.841418 | 3.240539 | 3.959551 | 0.818411 | 0.413123 | NA | #VALUE! |
| ENSMUSG00000102427.2 | 0 | NA | NA | NA | NA | NA | #VALUE! |
| ENSMUSG00000102428.2 | 0 | NA | NA | NA | NA | NA | #VALUE! |
| ENSMUSG00000102429.6 | 0 | NA | NA | NA | NA | NA | #VALUE! |
| ENSMUSG00000102430.2 | 0 | NA | NA | NA | NA | NA | #VALUE! |
| ENSMUSG00000102432.2 | 0.420053 | -2.33493 | 4.007343 | -0.58266 | 0.56012 | NA | #VALUE! |
| ENSMUSG00000102433.2 | 0 | NA | NA | NA | NA | NA | #VALUE! |
| ENSMUSG00000102434.2 | 0 | NA | NA | NA | NA | NA | #VALUE! |
| ENSMUSG00000102436.2 | 0 | NA | NA | NA | NA | NA | #VALUE! |
| ENSMUSG00000102438.2 | 0 | NA | NA | NA | NA | NA | #VALUE! |
| ENSMUSG00000102439.5 | 0 | NA | NA | NA | NA | NA | #VALUE! |
| ENSMUSG00000102440.2 | 0 | NA | NA | NA | NA | NA | #VALUE! |
| ENSMUSG00000102441.2 | 0 | NA | NA | NA | NA | NA | #VALUE! |
| ENSMUSG00000102442.2 | 0 | NA | NA | NA | NA | NA | #VALUE! |
| ENSMUSG00000102443.2 | 0 | NA | NA | NA | NA | NA | #VALUE! |
| ENSMUSG00000102445.2 | 0 | NA | NA | NA | NA | NA | #VALUE! |
| ENSMUSG00000102446.2 | 0 | NA | NA | NA | NA | NA | #VALUE! |
| ENSMUSG00000102447.2 | 0 | NA | NA | NA | NA | NA | #VALUE! |
| ENSMUSG00000102448.2 | 0 | NA | NA | NA | NA | NA | #VALUE! |
| ENSMUSG00000102449.2 | 0 | NA | NA | NA | NA | NA | #VALUE! |
| ENSMUSG00000102450.2 | 0 | NA | NA | NA | NA | NA | #VALUE! |
| ENSMUSG00000102451.2 | 0 | NA | NA | NA | NA | NA | #VALUE! |
| ENSMUSG00000102452.2 | 0 | NA | NA | NA | NA | NA | #VALUE! |
| ENSMUSG00000102453.2 | 0 | NA | NA | NA | NA | NA | #VALUE! |
| ENSMUSG00000102456.2 | 0 | NA | NA | NA | NA | NA | #VALUE! |
| ENSMUSG00000102457.2 | 0 | NA | NA | NA | NA | NA | #VALUE! |

| | | | | | | | |
|---|---|---|---|---|---|---|---|
| ENSMUSG00000102458.2 | 0 | NA | NA | NA | NA | NA | #VALUE! |
| ENSMUSG00000102459.2 | 0 | NA | NA | NA | NA | NA | #VALUE! |
| ENSMUSG00000102460.2 | 0 | NA | NA | NA | NA | NA | #VALUE! |
| ENSMUSG00000102461.2 | 0 | NA | NA | NA | NA | NA | #VALUE! |
| ENSMUSG00000102462.2 | 0 | NA | NA | NA | NA | NA | #VALUE! |
| ENSMUSG00000102463.2 | 0 | NA | NA | NA | NA | NA | #VALUE! |
| ENSMUSG00000102464.4 | 0 | NA | NA | NA | NA | NA | #VALUE! |
| ENSMUSG00000102465.2 | 0 | NA | NA | NA | NA | NA | #VALUE! |
| ENSMUSG00000102466.2 | 0 | NA | NA | NA | NA | NA | #VALUE! |
| ENSMUSG00000102467.2 | 0 | NA | NA | NA | NA | NA | #VALUE! |
| ENSMUSG00000102469.2 | 0 | NA | NA | NA | NA | NA | #VALUE! |
| ENSMUSG00000102470.2 | 0 | NA | NA | NA | NA | NA | #VALUE! |
| ENSMUSG00000102471.2 | 0 | NA | NA | NA | NA | NA | #VALUE! |
| ENSMUSG00000102473.2 | 0 | NA | NA | NA | NA | NA | #VALUE! |
| ENSMUSG00000102474.2 | 0 | NA | NA | NA | NA | NA | #VALUE! |
| ENSMUSG00000102475.2 | 0 | NA | NA | NA | NA | NA | #VALUE! |
| ENSMUSG00000102476.2 | 0 | NA | NA | NA | NA | NA | #VALUE! |
| ENSMUSG00000102478.2 | 0 | NA | NA | NA | NA | NA | #VALUE! |
| ENSMUSG00000102479.2 | 0 | NA | NA | NA | NA | NA | #VALUE! |
| ENSMUSG00000102480.2 | 0 | NA | NA | NA | NA | NA | #VALUE! |
| ENSMUSG00000102481.2 | 0 | NA | NA | NA | NA | NA | #VALUE! |
| ENSMUSG00000102482.2 | 0 | NA | NA | NA | NA | NA | #VALUE! |
| ENSMUSG00000102483.3 | 0 | NA | NA | NA | NA | NA | #VALUE! |
| ENSMUSG00000102484.2 | 0 | NA | NA | NA | NA | NA | #VALUE! |
| ENSMUSG00000102485.2 | 0 | NA | NA | NA | NA | NA | #VALUE! |
| ENSMUSG00000102486.2 | 0 | NA | NA | NA | NA | NA | #VALUE! |
| ENSMUSG00000102487.2 | 0 | NA | NA | NA | NA | NA | #VALUE! |
| ENSMUSG00000102488.2 | 0 | NA | NA | NA | NA | NA | #VALUE! |
| ENSMUSG00000102489.2 | 0 | NA | NA | NA | NA | NA | #VALUE! |
| ENSMUSG00000102490.2 | 0 | NA | NA | NA | NA | NA | #VALUE! |
| ENSMUSG00000102491.2 | 0 | NA | NA | NA | NA | NA | #VALUE! |
| ENSMUSG00000102492.2 | 0 | NA | NA | NA | NA | NA | #VALUE! |
| ENSMUSG00000102493.2 | 0 | NA | NA | NA | NA | NA | #VALUE! |
| ENSMUSG00000102494.2 | 0 | NA | NA | NA | NA | NA | #VALUE! |
| ENSMUSG00000102495.2 | 0 | NA | NA | NA | NA | NA | #VALUE! |
| ENSMUSG00000102496.2 | 0 | NA | NA | NA | NA | NA | #VALUE! |
| ENSMUSG00000102497.2 | 0 | NA | NA | NA | NA | NA | #VALUE! |
| ENSMUSG00000102498.2 | 0 | NA | NA | NA | NA | NA | #VALUE! |
| ENSMUSG00000102499.2 | 0 | NA | NA | NA | NA | NA | #VALUE! |
| ENSMUSG00000102500.2 | 0 | NA | NA | NA | NA | NA | #VALUE! |
| ENSMUSG00000102501.2 | 0 | NA | NA | NA | NA | NA | #VALUE! |
| ENSMUSG00000102502.2 | 0 | NA | NA | NA | NA | NA | #VALUE! |
| ENSMUSG00000102503.2 | 0 | NA | NA | NA | NA | NA | #VALUE! |
| ENSMUSG00000102504.2 | 0 | NA | NA | NA | NA | NA | #VALUE! |

| | | | | | | | |
|---|---|---|---|---|---|---|---|
| ENSMUSG00000102505.2 | 0 | NA | NA | NA | NA | NA | #VALUE! |
| ENSMUSG00000102506.2 | 0 | NA | NA | NA | NA | NA | #VALUE! |
| ENSMUSG00000102507.2 | 0 | NA | NA | NA | NA | NA | #VALUE! |
| ENSMUSG00000102508.2 | 0 | NA | NA | NA | NA | NA | #VALUE! |
| ENSMUSG00000102509.2 | 0 | NA | NA | NA | NA | NA | #VALUE! |
| ENSMUSG00000102510.3 | 0 | NA | NA | NA | NA | NA | #VALUE! |
| ENSMUSG00000102511.2 | 0 | NA | NA | NA | NA | NA | #VALUE! |
| ENSMUSG00000102512.2 | 0 | NA | NA | NA | NA | NA | #VALUE! |
| ENSMUSG00000102513.2 | 0 | NA | NA | NA | NA | NA | #VALUE! |
| ENSMUSG00000102514.2 | 0 | NA | NA | NA | NA | NA | #VALUE! |
| ENSMUSG00000102515.2 | 0 | NA | NA | NA | NA | NA | #VALUE! |
| ENSMUSG00000102516.2 | 0 | NA | NA | NA | NA | NA | #VALUE! |
| ENSMUSG00000102517.2 | 0 | NA | NA | NA | NA | NA | #VALUE! |
| ENSMUSG00000102518.2 | 0 | NA | NA | NA | NA | NA | #VALUE! |
| ENSMUSG00000102520.6 | 0 | NA | NA | NA | NA | NA | #VALUE! |
| ENSMUSG00000102521.2 | 0 | NA | NA | NA | NA | NA | #VALUE! |
| ENSMUSG00000102522.2 | 0 | NA | NA | NA | NA | NA | #VALUE! |
| ENSMUSG00000102523.2 | 0 | NA | NA | NA | NA | NA | #VALUE! |
| ENSMUSG00000102524.2 | 0 | NA | NA | NA | NA | NA | #VALUE! |
| ENSMUSG00000102525.2 | 0 | NA | NA | NA | NA | NA | #VALUE! |
| ENSMUSG00000102526.2 | 0 | NA | NA | NA | NA | NA | #VALUE! |
| ENSMUSG00000102528.2 | 0 | NA | NA | NA | NA | NA | #VALUE! |
| ENSMUSG00000102529.2 | 0 | NA | NA | NA | NA | NA | #VALUE! |
| ENSMUSG00000102530.2 | 0 | NA | NA | NA | NA | NA | #VALUE! |
| ENSMUSG00000102531.2 | 0 | NA | NA | NA | NA | NA | #VALUE! |
| ENSMUSG00000102532.2 | 0 | NA | NA | NA | NA | NA | #VALUE! |
| ENSMUSG00000102533.2 | 0 | NA | NA | NA | NA | NA | #VALUE! |
| ENSMUSG00000102534.2 | 0 | NA | NA | NA | NA | NA | #VALUE! |
| ENSMUSG00000102535.2 | 0 | NA | NA | NA | NA | NA | #VALUE! |
| ENSMUSG00000102536.2 | 0 | NA | NA | NA | NA | NA | #VALUE! |
| ENSMUSG00000102537.2 | 0 | NA | NA | NA | NA | NA | #VALUE! |
| ENSMUSG00000102538.2 | 0 | NA | NA | NA | NA | NA | #VALUE! |
| ENSMUSG00000102539.2 | 0 | NA | NA | NA | NA | NA | #VALUE! |
| ENSMUSG00000102540.2 | 0 | NA | NA | NA | NA | NA | #VALUE! |
| ENSMUSG00000102541.2 | 0 | NA | NA | NA | NA | NA | #VALUE! |
| ENSMUSG00000102542.2 | 0 | NA | NA | NA | NA | NA | #VALUE! |
| ENSMUSG00000102543.6 | 0 | NA | NA | NA | NA | NA | #VALUE! |
| ENSMUSG00000102544.2 | 0 | NA | NA | NA | NA | NA | #VALUE! |
| ENSMUSG00000102545.2 | 6.692366 | -6.27687 | 3.638115 | -1.72531 | 0.084472 | NA | #VALUE! |
| ENSMUSG00000102546.2 | 0.364073 | 0.012025 | 4.059931 | 0.002962 | 0.997637 | NA | #VALUE! |
| ENSMUSG00000102547.2 | 0 | NA | NA | NA | NA | NA | #VALUE! |
| ENSMUSG00000102548.2 | 0 | NA | NA | NA | NA | NA | #VALUE! |
| ENSMUSG00000102549.2 | 0 | NA | NA | NA | NA | NA | #VALUE! |
| ENSMUSG00000102550.2 | 0.372394 | 0.012025 | 4.059931 | 0.002962 | 0.997637 | NA | #VALUE! |

| | | | | | | | |
|---|---|---|---|---|---|---|---|
| ENSMUSG00000102551.2 | 0 | NA | NA | NA | NA | NA | #VALUE! |
| ENSMUSG00000102552.2 | 0 | NA | NA | NA | NA | NA | #VALUE! |
| ENSMUSG00000102553.2 | 5.670709 | -6.03829 | 3.914631 | -1.54249 | 0.122954 | NA | #VALUE! |
| ENSMUSG00000102554.2 | 0 | NA | NA | NA | NA | NA | #VALUE! |
| ENSMUSG00000102555.2 | 0 | NA | NA | NA | NA | NA | #VALUE! |
| ENSMUSG00000102556.2 | 0 | NA | NA | NA | NA | NA | #VALUE! |
| ENSMUSG00000102558.2 | 0 | NA | NA | NA | NA | NA | #VALUE! |
| ENSMUSG00000102559.2 | 0 | NA | NA | NA | NA | NA | #VALUE! |
| ENSMUSG00000102561.2 | 0 | NA | NA | NA | NA | NA | #VALUE! |
| ENSMUSG00000102563.2 | 0 | NA | NA | NA | NA | NA | #VALUE! |
| ENSMUSG00000102564.2 | 0 | NA | NA | NA | NA | NA | #VALUE! |
| ENSMUSG00000102565.2 | 0 | NA | NA | NA | NA | NA | #VALUE! |
| ENSMUSG00000102566.2 | 0 | NA | NA | NA | NA | NA | #VALUE! |
| ENSMUSG00000102568.2 | 0 | NA | NA | NA | NA | NA | #VALUE! |
| ENSMUSG00000102569.2 | 0 | NA | NA | NA | NA | NA | #VALUE! |
| ENSMUSG00000102570.2 | 0 | NA | NA | NA | NA | NA | #VALUE! |
| ENSMUSG00000102571.2 | 0.816443 | -3.21914 | 3.961753 | -0.81255 | 0.416474 | NA | #VALUE! |
| ENSMUSG00000102572.2 | 0 | NA | NA | NA | NA | NA | #VALUE! |
| ENSMUSG00000102573.2 | 0 | NA | NA | NA | NA | NA | #VALUE! |
| ENSMUSG00000102574.2 | 0 | NA | NA | NA | NA | NA | #VALUE! |
| ENSMUSG00000102575.2 | 0 | NA | NA | NA | NA | NA | #VALUE! |
| ENSMUSG00000102576.6 | 0 | NA | NA | NA | NA | NA | #VALUE! |
| ENSMUSG00000102577.2 | 0 | NA | NA | NA | NA | NA | #VALUE! |
| ENSMUSG00000102578.2 | 0 | NA | NA | NA | NA | NA | #VALUE! |
| ENSMUSG00000102579.2 | 0 | NA | NA | NA | NA | NA | #VALUE! |
| ENSMUSG00000102580.2 | 0 | NA | NA | NA | NA | NA | #VALUE! |
| ENSMUSG00000102581.2 | 0 | NA | NA | NA | NA | NA | #VALUE! |
| ENSMUSG00000102582.2 | 0 | NA | NA | NA | NA | NA | #VALUE! |
| ENSMUSG00000102583.2 | 0 | NA | NA | NA | NA | NA | #VALUE! |
| ENSMUSG00000102584.2 | 0 | NA | NA | NA | NA | NA | #VALUE! |
| ENSMUSG00000102585.2 | 0 | NA | NA | NA | NA | NA | #VALUE! |
| ENSMUSG00000102586.6 | 0 | NA | NA | NA | NA | NA | #VALUE! |
| ENSMUSG00000102587.2 | 0 | NA | NA | NA | NA | NA | #VALUE! |
| ENSMUSG00000102588.2 | 0 | NA | NA | NA | NA | NA | #VALUE! |
| ENSMUSG00000102589.2 | 0 | NA | NA | NA | NA | NA | #VALUE! |
| ENSMUSG00000102590.2 | 0 | NA | NA | NA | NA | NA | #VALUE! |
| ENSMUSG00000102591.2 | 0 | NA | NA | NA | NA | NA | #VALUE! |
| ENSMUSG00000102592.2 | 0 | NA | NA | NA | NA | NA | #VALUE! |
| ENSMUSG00000102593.2 | 0 | NA | NA | NA | NA | NA | #VALUE! |
| ENSMUSG00000102594.2 | 0 | NA | NA | NA | NA | NA | #VALUE! |
| ENSMUSG00000102595.2 | 13.64928 | -2.1285 | 2.993137 | -0.71113 | NA | NA | #VALUE! |
| ENSMUSG00000102596.2 | 0 | NA | NA | NA | NA | NA | #VALUE! |
| ENSMUSG00000102597.2 | 0 | NA | NA | NA | NA | NA | #VALUE! |
| ENSMUSG00000102598.2 | 0 | NA | NA | NA | NA | NA | #VALUE! |

| | | | | | | | |
|---|---|---|---|---|---|---|---|
| ENSMUSG00000102599.2 | 0 | NA | NA | NA | NA | NA | #VALUE! |
| ENSMUSG00000102600.2 | 0 | NA | NA | NA | NA | NA | #VALUE! |
| ENSMUSG00000102601.2 | 0 | NA | NA | NA | NA | NA | #VALUE! |
| ENSMUSG00000102602.2 | 0 | NA | NA | NA | NA | NA | #VALUE! |
| ENSMUSG00000102603.3 | 0 | NA | NA | NA | NA | NA | #VALUE! |
| ENSMUSG00000102605.2 | 0 | NA | NA | NA | NA | NA | #VALUE! |
| ENSMUSG00000102606.2 | 0 | NA | NA | NA | NA | NA | #VALUE! |
| ENSMUSG00000102607.2 | 0 | NA | NA | NA | NA | NA | #VALUE! |
| ENSMUSG00000102608.2 | 0 | NA | NA | NA | NA | NA | #VALUE! |
| ENSMUSG00000102609.2 | 0 | NA | NA | NA | NA | NA | #VALUE! |
| ENSMUSG00000102610.2 | 0 | NA | NA | NA | NA | NA | #VALUE! |
| ENSMUSG00000102611.2 | 0 | NA | NA | NA | NA | NA | #VALUE! |
| ENSMUSG00000102612.2 | 0 | NA | NA | NA | NA | NA | #VALUE! |
| ENSMUSG00000102613.2 | 0 | NA | NA | NA | NA | NA | #VALUE! |
| ENSMUSG00000102614.2 | 0 | NA | NA | NA | NA | NA | #VALUE! |
| ENSMUSG00000102615.2 | 0 | NA | NA | NA | NA | NA | #VALUE! |
| ENSMUSG00000102616.2 | 0 | NA | NA | NA | NA | NA | #VALUE! |
| ENSMUSG00000102617.2 | 0 | NA | NA | NA | NA | NA | #VALUE! |
| ENSMUSG00000102618.2 | 0 | NA | NA | NA | NA | NA | #VALUE! |
| ENSMUSG00000102619.2 | 0 | NA | NA | NA | NA | NA | #VALUE! |
| ENSMUSG00000102620.2 | 0 | NA | NA | NA | NA | NA | #VALUE! |
| ENSMUSG00000102621.2 | 0 | NA | NA | NA | NA | NA | #VALUE! |
| ENSMUSG00000102622.2 | 0 | NA | NA | NA | NA | NA | #VALUE! |
| ENSMUSG00000102623.2 | 0 | NA | NA | NA | NA | NA | #VALUE! |
| ENSMUSG00000102624.2 | 0 | NA | NA | NA | NA | NA | #VALUE! |
| ENSMUSG00000102625.2 | 0 | NA | NA | NA | NA | NA | #VALUE! |
| ENSMUSG00000102626.2 | 0 | NA | NA | NA | NA | NA | #VALUE! |
| ENSMUSG00000102627.2 | 0 | NA | NA | NA | NA | NA | #VALUE! |
| ENSMUSG00000102628.2 | 0 | NA | NA | NA | NA | NA | #VALUE! |
| ENSMUSG00000102629.2 | 0 | NA | NA | NA | NA | NA | #VALUE! |
| ENSMUSG00000102630.2 | 0 | NA | NA | NA | NA | NA | #VALUE! |
| ENSMUSG00000102632.2 | 0 | NA | NA | NA | NA | NA | #VALUE! |
| ENSMUSG00000102633.2 | 0 | NA | NA | NA | NA | NA | #VALUE! |
| ENSMUSG00000102634.2 | 0 | NA | NA | NA | NA | NA | #VALUE! |
| ENSMUSG00000102635.2 | 0 | NA | NA | NA | NA | NA | #VALUE! |
| ENSMUSG00000102636.2 | 0 | NA | NA | NA | NA | NA | #VALUE! |
| ENSMUSG00000102637.2 | 0 | NA | NA | NA | NA | NA | #VALUE! |
| ENSMUSG00000102639.2 | 0 | NA | NA | NA | NA | NA | #VALUE! |
| ENSMUSG00000102641.2 | 0 | NA | NA | NA | NA | NA | #VALUE! |
| ENSMUSG00000102642.2 | 0 | NA | NA | NA | NA | NA | #VALUE! |
| ENSMUSG00000102643.2 | 0 | NA | NA | NA | NA | NA | #VALUE! |
| ENSMUSG00000102645.2 | 0 | NA | NA | NA | NA | NA | #VALUE! |
| ENSMUSG00000102646.2 | 0 | NA | NA | NA | NA | NA | #VALUE! |
| ENSMUSG00000102647.2 | 0 | NA | NA | NA | NA | NA | #VALUE! |

| | | | | | | | |
|---|---|---|---|---|---|---|---|
| ENSMUSG00000102648.2 | 0 | NA | NA | NA | NA | NA | #VALUE! |
| ENSMUSG00000102649.2 | 4.207088 | 5.562698 | 3.917438 | 1.419984 | 0.155612 | NA | #VALUE! |
| ENSMUSG00000102650.2 | 0 | NA | NA | NA | NA | NA | #VALUE! |
| ENSMUSG00000102651.2 | 0 | NA | NA | NA | NA | NA | #VALUE! |
| ENSMUSG00000102652.2 | 0 | NA | NA | NA | NA | NA | #VALUE! |
| ENSMUSG00000102653.2 | 0 | NA | NA | NA | NA | NA | #VALUE! |
| ENSMUSG00000102654.2 | 0 | NA | NA | NA | NA | NA | #VALUE! |
| ENSMUSG00000102655.2 | 0 | NA | NA | NA | NA | NA | #VALUE! |
| ENSMUSG00000102656.2 | 0 | NA | NA | NA | NA | NA | #VALUE! |
| ENSMUSG00000102657.2 | 0 | NA | NA | NA | NA | NA | #VALUE! |
| ENSMUSG00000102658.2 | 0 | NA | NA | NA | NA | NA | #VALUE! |
| ENSMUSG00000102659.2 | 0 | NA | NA | NA | NA | NA | #VALUE! |
| ENSMUSG00000102660.2 | 0 | NA | NA | NA | NA | NA | #VALUE! |
| ENSMUSG00000102661.2 | 0 | NA | NA | NA | NA | NA | #VALUE! |
| ENSMUSG00000102662.2 | 0 | NA | NA | NA | NA | NA | #VALUE! |
| ENSMUSG00000102663.2 | 0 | NA | NA | NA | NA | NA | #VALUE! |
| ENSMUSG00000102664.2 | 1.514552 | 4.088766 | 3.936269 | 1.038742 | 0.298925 | NA | #VALUE! |
| ENSMUSG00000102665.2 | 0 | NA | NA | NA | NA | NA | #VALUE! |
| ENSMUSG00000102666.2 | 0.198268 | -0.06494 | 4.068276 | -0.01596 | 0.987265 | NA | #VALUE! |
| ENSMUSG00000102667.2 | 0 | NA | NA | NA | NA | NA | #VALUE! |
| ENSMUSG00000102668.2 | 0 | NA | NA | NA | NA | NA | #VALUE! |
| ENSMUSG00000102670.2 | 0 | NA | NA | NA | NA | NA | #VALUE! |
| ENSMUSG00000102672.2 | 0 | NA | NA | NA | NA | NA | #VALUE! |
| ENSMUSG00000102673.2 | 0 | NA | NA | NA | NA | NA | #VALUE! |
| ENSMUSG00000102674.2 | 0 | NA | NA | NA | NA | NA | #VALUE! |
| ENSMUSG00000102676.2 | 0 | NA | NA | NA | NA | NA | #VALUE! |
| ENSMUSG00000102677.2 | 0 | NA | NA | NA | NA | NA | #VALUE! |
| ENSMUSG00000102678.2 | 0 | NA | NA | NA | NA | NA | #VALUE! |
| ENSMUSG00000102679.2 | 0 | NA | NA | NA | NA | NA | #VALUE! |
| ENSMUSG00000102680.2 | 0 | NA | NA | NA | NA | NA | #VALUE! |
| ENSMUSG00000102681.2 | 0 | NA | NA | NA | NA | NA | #VALUE! |
| ENSMUSG00000102682.6 | 0 | NA | NA | NA | NA | NA | #VALUE! |
| ENSMUSG00000102683.2 | 0 | NA | NA | NA | NA | NA | #VALUE! |
| ENSMUSG00000102684.2 | 0 | NA | NA | NA | NA | NA | #VALUE! |
| ENSMUSG00000102685.2 | 0 | NA | NA | NA | NA | NA | #VALUE! |
| ENSMUSG00000102686.2 | 0 | NA | NA | NA | NA | NA | #VALUE! |
| ENSMUSG00000102687.2 | 0 | NA | NA | NA | NA | NA | #VALUE! |
| ENSMUSG00000102688.2 | 0 | NA | NA | NA | NA | NA | #VALUE! |
| ENSMUSG00000102689.2 | 0 | NA | NA | NA | NA | NA | #VALUE! |
| ENSMUSG00000102690.2 | 0 | NA | NA | NA | NA | NA | #VALUE! |
| ENSMUSG00000102691.2 | 8.40105 | -6.60404 | 3.782709 | -1.74585 | NA | NA | #VALUE! |
| ENSMUSG00000102692.2 | 0 | NA | NA | NA | NA | NA | #VALUE! |
| ENSMUSG00000102693.2 | 0 | NA | NA | NA | NA | NA | #VALUE! |
| ENSMUSG00000102694.2 | 0 | NA | NA | NA | NA | NA | #VALUE! |

| | | | | | | | |
|---|---|---|---|---|---|---|---|
| ENSMUSG00000102696.2 | 0 | NA | NA | NA | NA | NA | #VALUE! |
| ENSMUSG00000102697.2 | 0 | NA | NA | NA | NA | NA | #VALUE! |
| ENSMUSG00000102698.2 | 0 | NA | NA | NA | NA | NA | #VALUE! |
| ENSMUSG00000102699.2 | 0 | NA | NA | NA | NA | NA | #VALUE! |
| ENSMUSG00000102700.2 | 0 | NA | NA | NA | NA | NA | #VALUE! |
| ENSMUSG00000102701.2 | 2.801898 | 4.980858 | 3.922719 | 1.269746 | 0.204175 | NA | #VALUE! |
| ENSMUSG00000102702.2 | 0 | NA | NA | NA | NA | NA | #VALUE! |
| ENSMUSG00000102703.2 | 0 | NA | NA | NA | NA | NA | #VALUE! |
| ENSMUSG00000102704.2 | 0 | NA | NA | NA | NA | NA | #VALUE! |
| ENSMUSG00000102705.3 | 0 | NA | NA | NA | NA | NA | #VALUE! |
| ENSMUSG00000102706.2 | 0 | NA | NA | NA | NA | NA | #VALUE! |
| ENSMUSG00000102707.2 | 0 | NA | NA | NA | NA | NA | #VALUE! |
| ENSMUSG00000102709.2 | 0 | NA | NA | NA | NA | NA | #VALUE! |
| ENSMUSG00000102710.2 | 0 | NA | NA | NA | NA | NA | #VALUE! |
| ENSMUSG00000102712.2 | 0 | NA | NA | NA | NA | NA | #VALUE! |
| ENSMUSG00000102713.2 | 0 | NA | NA | NA | NA | NA | #VALUE! |
| ENSMUSG00000102714.2 | 0 | NA | NA | NA | NA | NA | #VALUE! |
| ENSMUSG00000102715.2 | 0 | NA | NA | NA | NA | NA | #VALUE! |
| ENSMUSG00000102716.2 | 0 | NA | NA | NA | NA | NA | #VALUE! |
| ENSMUSG00000102717.2 | 0 | NA | NA | NA | NA | NA | #VALUE! |
| ENSMUSG00000102718.2 | 0 | NA | NA | NA | NA | NA | #VALUE! |
| ENSMUSG00000102719.2 | 2.187686 | 4.619304 | 3.927239 | 1.176222 | 0.239506 | NA | #VALUE! |
| ENSMUSG00000102721.2 | 0 | NA | NA | NA | NA | NA | #VALUE! |
| ENSMUSG00000102722.2 | 0 | NA | NA | NA | NA | NA | #VALUE! |
| ENSMUSG00000102723.2 | 0 | NA | NA | NA | NA | NA | #VALUE! |
| ENSMUSG00000102725.3 | 0 | NA | NA | NA | NA | NA | #VALUE! |
| ENSMUSG00000102726.2 | 0 | NA | NA | NA | NA | NA | #VALUE! |
| ENSMUSG00000102728.2 | 0 | NA | NA | NA | NA | NA | #VALUE! |
| ENSMUSG00000102729.2 | 0 | NA | NA | NA | NA | NA | #VALUE! |
| ENSMUSG00000102730.2 | 0 | NA | NA | NA | NA | NA | #VALUE! |
| ENSMUSG00000102731.2 | 0 | NA | NA | NA | NA | NA | #VALUE! |
| ENSMUSG00000102732.2 | 0 | NA | NA | NA | NA | NA | #VALUE! |
| ENSMUSG00000102733.2 | 0 | NA | NA | NA | NA | NA | #VALUE! |
| ENSMUSG00000102734.2 | 0 | NA | NA | NA | NA | NA | #VALUE! |
| ENSMUSG00000102735.2 | 0 | NA | NA | NA | NA | NA | #VALUE! |
| ENSMUSG00000102737.2 | 0 | NA | NA | NA | NA | NA | #VALUE! |
| ENSMUSG00000102738.2 | 0 | NA | NA | NA | NA | NA | #VALUE! |
| ENSMUSG00000102739.2 | 0 | NA | NA | NA | NA | NA | #VALUE! |
| ENSMUSG00000102740.2 | 0 | NA | NA | NA | NA | NA | #VALUE! |
| ENSMUSG00000102741.2 | 0 | NA | NA | NA | NA | NA | #VALUE! |
| ENSMUSG00000102742.2 | 0 | NA | NA | NA | NA | NA | #VALUE! |
| ENSMUSG00000102743.2 | 0 | NA | NA | NA | NA | NA | #VALUE! |
| ENSMUSG00000102744.2 | 0 | NA | NA | NA | NA | NA | #VALUE! |
| ENSMUSG00000102745.2 | 0 | NA | NA | NA | NA | NA | #VALUE! |

| | | | | | | | |
|---|---|---|---|---|---|---|---|
| ENSMUSG00000102746.2 | 0 | NA | NA | NA | NA | NA | #VALUE! |
| ENSMUSG00000102747.2 | 0 | NA | NA | NA | NA | NA | #VALUE! |
| ENSMUSG00000102748.2 | 0 | NA | NA | NA | NA | NA | #VALUE! |
| ENSMUSG00000102749.2 | 0 | NA | NA | NA | NA | NA | #VALUE! |
| ENSMUSG00000102750.2 | 0 | NA | NA | NA | NA | NA | #VALUE! |
| ENSMUSG00000102751.2 | 0 | NA | NA | NA | NA | NA | #VALUE! |
| ENSMUSG00000102752.2 | 4.579762 | -0.17141 | 3.742865 | -0.0458 | 0.963472 | NA | #VALUE! |
| ENSMUSG00000102753.2 | 0 | NA | NA | NA | NA | NA | #VALUE! |
| ENSMUSG00000102754.2 | 0 | NA | NA | NA | NA | NA | #VALUE! |
| ENSMUSG00000102755.2 | 0 | NA | NA | NA | NA | NA | #VALUE! |
| ENSMUSG00000102756.2 | 0 | NA | NA | NA | NA | NA | #VALUE! |
| ENSMUSG00000102757.2 | 0 | NA | NA | NA | NA | NA | #VALUE! |
| ENSMUSG00000102759.2 | 0 | NA | NA | NA | NA | NA | #VALUE! |
| ENSMUSG00000102760.2 | 0 | NA | NA | NA | NA | NA | #VALUE! |
| ENSMUSG00000102761.2 | 0 | NA | NA | NA | NA | NA | #VALUE! |
| ENSMUSG00000102762.2 | 0 | NA | NA | NA | NA | NA | #VALUE! |
| ENSMUSG00000102763.2 | 0 | NA | NA | NA | NA | NA | #VALUE! |
| ENSMUSG00000102764.2 | 0 | NA | NA | NA | NA | NA | #VALUE! |
| ENSMUSG00000102765.2 | 0 | NA | NA | NA | NA | NA | #VALUE! |
| ENSMUSG00000102766.2 | 0 | NA | NA | NA | NA | NA | #VALUE! |
| ENSMUSG00000102768.2 | 0 | NA | NA | NA | NA | NA | #VALUE! |
| ENSMUSG00000102769.2 | 0 | NA | NA | NA | NA | NA | #VALUE! |
| ENSMUSG00000102770.2 | 0 | NA | NA | NA | NA | NA | #VALUE! |
| ENSMUSG00000102771.2 | 0 | NA | NA | NA | NA | NA | #VALUE! |
| ENSMUSG00000102772.6 | 0 | NA | NA | NA | NA | NA | #VALUE! |
| ENSMUSG00000102773.2 | 0 | NA | NA | NA | NA | NA | #VALUE! |
| ENSMUSG00000102774.2 | 0 | NA | NA | NA | NA | NA | #VALUE! |
| ENSMUSG00000102775.2 | 0 | NA | NA | NA | NA | NA | #VALUE! |
| ENSMUSG00000102776.2 | 0 | NA | NA | NA | NA | NA | #VALUE! |
| ENSMUSG00000102777.2 | 0 | NA | NA | NA | NA | NA | #VALUE! |
| ENSMUSG00000102778.2 | 0 | NA | NA | NA | NA | NA | #VALUE! |
| ENSMUSG00000102779.2 | 0 | NA | NA | NA | NA | NA | #VALUE! |
| ENSMUSG00000102780.2 | 0 | NA | NA | NA | NA | NA | #VALUE! |
| ENSMUSG00000102782.2 | 0 | NA | NA | NA | NA | NA | #VALUE! |
| ENSMUSG00000102783.2 | 0 | NA | NA | NA | NA | NA | #VALUE! |
| ENSMUSG00000102784.2 | 0 | NA | NA | NA | NA | NA | #VALUE! |
| ENSMUSG00000102785.6 | 0 | NA | NA | NA | NA | NA | #VALUE! |
| ENSMUSG00000102786.2 | 0 | NA | NA | NA | NA | NA | #VALUE! |
| ENSMUSG00000102787.6 | 0 | NA | NA | NA | NA | NA | #VALUE! |
| ENSMUSG00000102788.2 | 0 | NA | NA | NA | NA | NA | #VALUE! |
| ENSMUSG00000102789.2 | 0 | NA | NA | NA | NA | NA | #VALUE! |
| ENSMUSG00000102790.2 | 0 | NA | NA | NA | NA | NA | #VALUE! |
| ENSMUSG00000102791.2 | 0 | NA | NA | NA | NA | NA | #VALUE! |
| ENSMUSG00000102792.2 | 0 | NA | NA | NA | NA | NA | #VALUE! |

| | | | | | | | |
|---|---|---|---|---|---|---|---|
| ENSMUSG00000102793.2 | 0 | NA | NA | NA | NA | NA | #VALUE! |
| ENSMUSG00000102794.2 | 0 | NA | NA | NA | NA | NA | #VALUE! |
| ENSMUSG00000102795.2 | 0 | NA | NA | NA | NA | NA | #VALUE! |
| ENSMUSG00000102796.2 | 0 | NA | NA | NA | NA | NA | #VALUE! |
| ENSMUSG00000102797.2 | 0 | NA | NA | NA | NA | NA | #VALUE! |
| ENSMUSG00000102799.2 | 0 | NA | NA | NA | NA | NA | #VALUE! |
| ENSMUSG00000102800.2 | 0 | NA | NA | NA | NA | NA | #VALUE! |
| ENSMUSG00000102801.2 | 0 | NA | NA | NA | NA | NA | #VALUE! |
| ENSMUSG00000102802.4 | 0 | NA | NA | NA | NA | NA | #VALUE! |
| ENSMUSG00000102803.2 | 0 | NA | NA | NA | NA | NA | #VALUE! |
| ENSMUSG00000102804.2 | 0 | NA | NA | NA | NA | NA | #VALUE! |
| ENSMUSG00000102805.6 | 0 | NA | NA | NA | NA | NA | #VALUE! |
| ENSMUSG00000102806.2 | 0 | NA | NA | NA | NA | NA | #VALUE! |
| ENSMUSG00000102807.2 | 0 | NA | NA | NA | NA | NA | #VALUE! |
| ENSMUSG00000102808.2 | 0 | NA | NA | NA | NA | NA | #VALUE! |
| ENSMUSG00000102809.2 | 0 | NA | NA | NA | NA | NA | #VALUE! |
| ENSMUSG00000102810.2 | 0 | NA | NA | NA | NA | NA | #VALUE! |
| ENSMUSG00000102813.2 | 0 | NA | NA | NA | NA | NA | #VALUE! |
| ENSMUSG00000102814.2 | 0 | NA | NA | NA | NA | NA | #VALUE! |
| ENSMUSG00000102815.2 | 0 | NA | NA | NA | NA | NA | #VALUE! |
| ENSMUSG00000102816.2 | 0 | NA | NA | NA | NA | NA | #VALUE! |
| ENSMUSG00000102817.3 | 0 | NA | NA | NA | NA | NA | #VALUE! |
| ENSMUSG00000102818.2 | 0 | NA | NA | NA | NA | NA | #VALUE! |
| ENSMUSG00000102819.2 | 0 | NA | NA | NA | NA | NA | #VALUE! |
| ENSMUSG00000102820.2 | 0 | NA | NA | NA | NA | NA | #VALUE! |
| ENSMUSG00000102821.2 | 0 | NA | NA | NA | NA | NA | #VALUE! |
| ENSMUSG00000102822.2 | 0 | NA | NA | NA | NA | NA | #VALUE! |
| ENSMUSG00000102824.2 | 2.221661 | 1.071399 | 3.820982 | 0.280399 | 0.779172 | NA | #VALUE! |
| ENSMUSG00000102825.2 | 0 | NA | NA | NA | NA | NA | #VALUE! |
| ENSMUSG00000102826.2 | 0 | NA | NA | NA | NA | NA | #VALUE! |
| ENSMUSG00000102828.6 | 0 | NA | NA | NA | NA | NA | #VALUE! |
| ENSMUSG00000102830.2 | 0 | NA | NA | NA | NA | NA | #VALUE! |
| ENSMUSG00000102831.2 | 0 | NA | NA | NA | NA | NA | #VALUE! |
| ENSMUSG00000102832.2 | 0 | NA | NA | NA | NA | NA | #VALUE! |
| ENSMUSG00000102833.2 | 0 | NA | NA | NA | NA | NA | #VALUE! |
| ENSMUSG00000102834.3 | 0 | NA | NA | NA | NA | NA | #VALUE! |
| ENSMUSG00000102835.6 | 0 | NA | NA | NA | NA | NA | #VALUE! |
| ENSMUSG00000102836.2 | 0 | NA | NA | NA | NA | NA | #VALUE! |
| ENSMUSG00000102837.2 | 0 | NA | NA | NA | NA | NA | #VALUE! |
| ENSMUSG00000102838.2 | 0 | NA | NA | NA | NA | NA | #VALUE! |
| ENSMUSG00000102839.2 | 0 | NA | NA | NA | NA | NA | #VALUE! |
| ENSMUSG00000102840.2 | 0 | NA | NA | NA | NA | NA | #VALUE! |
| ENSMUSG00000102841.2 | 0 | NA | NA | NA | NA | NA | #VALUE! |
| ENSMUSG00000102842.2 | 0 | NA | NA | NA | NA | NA | #VALUE! |

| | | | | | | | |
|---|---|---|---|---|---|---|---|
| ENSMUSG00000102844.2 | 0 | NA | NA | NA | NA | NA | #VALUE! |
| ENSMUSG00000102845.2 | 0 | NA | NA | NA | NA | NA | #VALUE! |
| ENSMUSG00000102846.2 | 0 | NA | NA | NA | NA | NA | #VALUE! |
| ENSMUSG00000102847.2 | 0 | NA | NA | NA | NA | NA | #VALUE! |
| ENSMUSG00000102848.2 | 0 | NA | NA | NA | NA | NA | #VALUE! |
| ENSMUSG00000102849.2 | 0 | NA | NA | NA | NA | NA | #VALUE! |
| ENSMUSG00000102850.2 | 0 | NA | NA | NA | NA | NA | #VALUE! |
| ENSMUSG00000102851.2 | 0 | NA | NA | NA | NA | NA | #VALUE! |
| ENSMUSG00000102852.2 | 0 | NA | NA | NA | NA | NA | #VALUE! |
| ENSMUSG00000102853.2 | 0 | NA | NA | NA | NA | NA | #VALUE! |
| ENSMUSG00000102854.2 | 3.734863 | 1.917715 | 3.800671 | 0.504573 | 0.613859 | NA | #VALUE! |
| ENSMUSG00000102855.2 | 0 | NA | NA | NA | NA | NA | #VALUE! |
| ENSMUSG00000102856.2 | 0 | NA | NA | NA | NA | NA | #VALUE! |
| ENSMUSG00000102857.2 | 0 | NA | NA | NA | NA | NA | #VALUE! |
| ENSMUSG00000102858.2 | 0 | NA | NA | NA | NA | NA | #VALUE! |
| ENSMUSG00000102859.2 | 0 | NA | NA | NA | NA | NA | #VALUE! |
| ENSMUSG00000102860.2 | 0 | NA | NA | NA | NA | NA | #VALUE! |
| ENSMUSG00000102861.2 | 0 | NA | NA | NA | NA | NA | #VALUE! |
| ENSMUSG00000102862.2 | 0 | NA | NA | NA | NA | NA | #VALUE! |
| ENSMUSG00000102863.2 | 0 | NA | NA | NA | NA | NA | #VALUE! |
| ENSMUSG00000102864.2 | 0 | NA | NA | NA | NA | NA | #VALUE! |
| ENSMUSG00000102865.2 | 0 | NA | NA | NA | NA | NA | #VALUE! |
| ENSMUSG00000102866.2 | 0 | NA | NA | NA | NA | NA | #VALUE! |
| ENSMUSG00000102867.2 | 0 | NA | NA | NA | NA | NA | #VALUE! |
| ENSMUSG00000102868.2 | 0 | NA | NA | NA | NA | NA | #VALUE! |
| ENSMUSG00000102870.2 | 0 | NA | NA | NA | NA | NA | #VALUE! |
| ENSMUSG00000102871.2 | 0 | NA | NA | NA | NA | NA | #VALUE! |
| ENSMUSG00000102872.2 | 0 | NA | NA | NA | NA | NA | #VALUE! |
| ENSMUSG00000102874.2 | 0 | NA | NA | NA | NA | NA | #VALUE! |
| ENSMUSG00000102875.2 | 0 | NA | NA | NA | NA | NA | #VALUE! |
| ENSMUSG00000102876.2 | 0 | NA | NA | NA | NA | NA | #VALUE! |
| ENSMUSG00000102877.2 | 0 | NA | NA | NA | NA | NA | #VALUE! |
| ENSMUSG00000102878.2 | 0 | NA | NA | NA | NA | NA | #VALUE! |
| ENSMUSG00000102879.2 | 0 | NA | NA | NA | NA | NA | #VALUE! |
| ENSMUSG00000102880.2 | 0 | NA | NA | NA | NA | NA | #VALUE! |
| ENSMUSG00000102881.2 | 0 | NA | NA | NA | NA | NA | #VALUE! |
| ENSMUSG00000102882.2 | 0 | NA | NA | NA | NA | NA | #VALUE! |
| ENSMUSG00000102883.2 | 0 | NA | NA | NA | NA | NA | #VALUE! |
| ENSMUSG00000102884.2 | 0 | NA | NA | NA | NA | NA | #VALUE! |
| ENSMUSG00000102885.2 | 0 | NA | NA | NA | NA | NA | #VALUE! |
| ENSMUSG00000102886.2 | 0 | NA | NA | NA | NA | NA | #VALUE! |
| ENSMUSG00000102887.2 | 0.093255 | 0.868093 | 4.072464 | 0.213162 | 0.831201 | NA | #VALUE! |
| ENSMUSG00000102888.2 | 0 | NA | NA | NA | NA | NA | #VALUE! |
| ENSMUSG00000102889.2 | 0 | NA | NA | NA | NA | NA | #VALUE! |

| | | | | | | | |
|---|---|---|---|---|---|---|---|
| ENSMUSG00000102890.2 | 0 | NA | NA | NA | NA | NA | #VALUE! |
| ENSMUSG00000102891.2 | 0 | NA | NA | NA | NA | NA | #VALUE! |
| ENSMUSG00000102892.2 | 1.015028 | 3.501945 | 3.950905 | 0.886365 | 0.375421 | NA | #VALUE! |
| ENSMUSG00000102893.2 | 0 | NA | NA | NA | NA | NA | #VALUE! |
| ENSMUSG00000102894.2 | 0 | NA | NA | NA | NA | NA | #VALUE! |
| ENSMUSG00000102895.2 | 0 | NA | NA | NA | NA | NA | #VALUE! |
| ENSMUSG00000102896.2 | 0 | NA | NA | NA | NA | NA | #VALUE! |
| ENSMUSG00000102897.2 | 0 | NA | NA | NA | NA | NA | #VALUE! |
| ENSMUSG00000102899.2 | 0 | NA | NA | NA | NA | NA | #VALUE! |
| ENSMUSG00000102900.2 | 0 | NA | NA | NA | NA | NA | #VALUE! |
| ENSMUSG00000102901.2 | 0 | NA | NA | NA | NA | NA | #VALUE! |
| ENSMUSG00000102902.2 | 0 | NA | NA | NA | NA | NA | #VALUE! |
| ENSMUSG00000102903.2 | 0 | NA | NA | NA | NA | NA | #VALUE! |
| ENSMUSG00000102904.2 | 0 | NA | NA | NA | NA | NA | #VALUE! |
| ENSMUSG00000102905.2 | 0 | NA | NA | NA | NA | NA | #VALUE! |
| ENSMUSG00000102906.2 | 0 | NA | NA | NA | NA | NA | #VALUE! |
| ENSMUSG00000102907.2 | 0 | NA | NA | NA | NA | NA | #VALUE! |
| ENSMUSG00000102908.2 | 0 | NA | NA | NA | NA | NA | #VALUE! |
| ENSMUSG00000102909.2 | 0 | NA | NA | NA | NA | NA | #VALUE! |
| ENSMUSG00000102910.2 | 0 | NA | NA | NA | NA | NA | #VALUE! |
| ENSMUSG00000102911.2 | 0 | NA | NA | NA | NA | NA | #VALUE! |
| ENSMUSG00000102912.2 | 0 | NA | NA | NA | NA | NA | #VALUE! |
| ENSMUSG00000102913.2 | 0 | NA | NA | NA | NA | NA | #VALUE! |
| ENSMUSG00000102914.2 | 0.459807 | 1.105905 | 4.021405 | 0.275005 | 0.783313 | NA | #VALUE! |
| ENSMUSG00000102915.2 | 0 | NA | NA | NA | NA | NA | #VALUE! |
| ENSMUSG00000102916.2 | 0 | NA | NA | NA | NA | NA | #VALUE! |
| ENSMUSG00000102917.2 | 0 | NA | NA | NA | NA | NA | #VALUE! |
| ENSMUSG00000102918.2 | 0 | NA | NA | NA | NA | NA | #VALUE! |
| ENSMUSG00000102919.2 | 0 | NA | NA | NA | NA | NA | #VALUE! |
| ENSMUSG00000102920.2 | 0 | NA | NA | NA | NA | NA | #VALUE! |
| ENSMUSG00000102921.2 | 0 | NA | NA | NA | NA | NA | #VALUE! |
| ENSMUSG00000102922.2 | 0 | NA | NA | NA | NA | NA | #VALUE! |
| ENSMUSG00000102923.2 | 0 | NA | NA | NA | NA | NA | #VALUE! |
| ENSMUSG00000102925.2 | 0 | NA | NA | NA | NA | NA | #VALUE! |
| ENSMUSG00000102926.2 | 0 | NA | NA | NA | NA | NA | #VALUE! |
| ENSMUSG00000102927.2 | 0 | NA | NA | NA | NA | NA | #VALUE! |
| ENSMUSG00000102928.2 | 0 | NA | NA | NA | NA | NA | #VALUE! |
| ENSMUSG00000102929.2 | 0 | NA | NA | NA | NA | NA | #VALUE! |
| ENSMUSG00000102930.2 | 0 | NA | NA | NA | NA | NA | #VALUE! |
| ENSMUSG00000102931.2 | 0.210026 | -1.52494 | 4.058692 | -0.37572 | 0.707123 | NA | #VALUE! |
| ENSMUSG00000102932.2 | 0 | NA | NA | NA | NA | NA | #VALUE! |
| ENSMUSG00000102933.2 | 0 | NA | NA | NA | NA | NA | #VALUE! |
| ENSMUSG00000102934.2 | 0 | NA | NA | NA | NA | NA | #VALUE! |
| ENSMUSG00000102935.2 | 0 | NA | NA | NA | NA | NA | #VALUE! |

| | | | | | | | |
|---|---|---|---|---|---|---|---|
| ENSMUSG00000102936.2 | 0 | NA | NA | NA | NA | NA | #VALUE! |
| ENSMUSG00000102937.2 | 0 | NA | NA | NA | NA | NA | #VALUE! |
| ENSMUSG00000102939.2 | 0 | NA | NA | NA | NA | NA | #VALUE! |
| ENSMUSG00000102940.2 | 0 | NA | NA | NA | NA | NA | #VALUE! |
| ENSMUSG00000102941.2 | 0 | NA | NA | NA | NA | NA | #VALUE! |
| ENSMUSG00000102942.2 | 0 | NA | NA | NA | NA | NA | #VALUE! |
| ENSMUSG00000102943.2 | 0 | NA | NA | NA | NA | NA | #VALUE! |
| ENSMUSG00000102945.2 | 0 | NA | NA | NA | NA | NA | #VALUE! |
| ENSMUSG00000102946.3 | 0 | NA | NA | NA | NA | NA | #VALUE! |
| ENSMUSG00000102948.2 | 0 | NA | NA | NA | NA | NA | #VALUE! |
| ENSMUSG00000102949.2 | 0 | NA | NA | NA | NA | NA | #VALUE! |
| ENSMUSG00000102950.2 | 0 | NA | NA | NA | NA | NA | #VALUE! |
| ENSMUSG00000102951.2 | 0 | NA | NA | NA | NA | NA | #VALUE! |
| ENSMUSG00000102952.2 | 0 | NA | NA | NA | NA | NA | #VALUE! |
| ENSMUSG00000102953.2 | 0 | NA | NA | NA | NA | NA | #VALUE! |
| ENSMUSG00000102955.2 | 0 | NA | NA | NA | NA | NA | #VALUE! |
| ENSMUSG00000102956.2 | 0 | NA | NA | NA | NA | NA | #VALUE! |
| ENSMUSG00000102957.2 | 0 | NA | NA | NA | NA | NA | #VALUE! |
| ENSMUSG00000102958.2 | 0 | NA | NA | NA | NA | NA | #VALUE! |
| ENSMUSG00000102959.2 | 0 | NA | NA | NA | NA | NA | #VALUE! |
| ENSMUSG00000102960.2 | 0 | NA | NA | NA | NA | NA | #VALUE! |
| ENSMUSG00000102961.2 | 0 | NA | NA | NA | NA | NA | #VALUE! |
| ENSMUSG00000102962.2 | 0 | NA | NA | NA | NA | NA | #VALUE! |
| ENSMUSG00000102963.2 | 0 | NA | NA | NA | NA | NA | #VALUE! |
| ENSMUSG00000102964.2 | 0 | NA | NA | NA | NA | NA | #VALUE! |
| ENSMUSG00000102965.2 | 0 | NA | NA | NA | NA | NA | #VALUE! |
| ENSMUSG00000102966.2 | 0 | NA | NA | NA | NA | NA | #VALUE! |
| ENSMUSG00000102967.2 | 0 | NA | NA | NA | NA | NA | #VALUE! |
| ENSMUSG00000102968.6 | 0 | NA | NA | NA | NA | NA | #VALUE! |
| ENSMUSG00000102969.2 | 0 | NA | NA | NA | NA | NA | #VALUE! |
| ENSMUSG00000102970.2 | 0 | NA | NA | NA | NA | NA | #VALUE! |
| ENSMUSG00000102971.2 | 0 | NA | NA | NA | NA | NA | #VALUE! |
| ENSMUSG00000102972.2 | 0 | NA | NA | NA | NA | NA | #VALUE! |
| ENSMUSG00000102973.2 | 0 | NA | NA | NA | NA | NA | #VALUE! |
| ENSMUSG00000102975.2 | 0 | NA | NA | NA | NA | NA | #VALUE! |
| ENSMUSG00000102976.7 | 0.391578 | -2.1489 | 4.020883 | -0.53444 | 0.59304 | NA | #VALUE! |
| ENSMUSG00000102977.6 | 0 | NA | NA | NA | NA | NA | #VALUE! |
| ENSMUSG00000102979.2 | 0 | NA | NA | NA | NA | NA | #VALUE! |
| ENSMUSG00000102980.2 | 0 | NA | NA | NA | NA | NA | #VALUE! |
| ENSMUSG00000102981.2 | 0 | NA | NA | NA | NA | NA | #VALUE! |
| ENSMUSG00000102982.2 | 0 | NA | NA | NA | NA | NA | #VALUE! |
| ENSMUSG00000102983.2 | 0 | NA | NA | NA | NA | NA | #VALUE! |
| ENSMUSG00000102984.2 | 0 | NA | NA | NA | NA | NA | #VALUE! |
| ENSMUSG00000102985.2 | 0 | NA | NA | NA | NA | NA | #VALUE! |

| | | | | | | | |
|---|---|---|---|---|---|---|---|
| ENSMUSG00000102986.2 | 0 | NA | NA | NA | NA | NA | #VALUE! |
| ENSMUSG00000102987.2 | 0 | NA | NA | NA | NA | NA | #VALUE! |
| ENSMUSG00000102988.2 | 0 | NA | NA | NA | NA | NA | #VALUE! |
| ENSMUSG00000102989.2 | 0 | NA | NA | NA | NA | NA | #VALUE! |
| ENSMUSG00000102990.2 | 0 | NA | NA | NA | NA | NA | #VALUE! |
| ENSMUSG00000102991.2 | 0 | NA | NA | NA | NA | NA | #VALUE! |
| ENSMUSG00000102992.2 | 0 | NA | NA | NA | NA | NA | #VALUE! |
| ENSMUSG00000102993.2 | 0 | NA | NA | NA | NA | NA | #VALUE! |
| ENSMUSG00000102994.2 | 0 | NA | NA | NA | NA | NA | #VALUE! |
| ENSMUSG00000102995.2 | 0 | NA | NA | NA | NA | NA | #VALUE! |
| ENSMUSG00000102997.2 | 0 | NA | NA | NA | NA | NA | #VALUE! |
| ENSMUSG00000102998.2 | 0 | NA | NA | NA | NA | NA | #VALUE! |
| ENSMUSG00000102999.2 | 0 | NA | NA | NA | NA | NA | #VALUE! |
| ENSMUSG00000103000.2 | 0 | NA | NA | NA | NA | NA | #VALUE! |
| ENSMUSG00000103001.2 | 0 | NA | NA | NA | NA | NA | #VALUE! |
| ENSMUSG00000103002.2 | 0 | NA | NA | NA | NA | NA | #VALUE! |
| ENSMUSG00000103003.2 | 0 | NA | NA | NA | NA | NA | #VALUE! |
| ENSMUSG00000103004.2 | 0 | NA | NA | NA | NA | NA | #VALUE! |
| ENSMUSG00000103005.2 | 0 | NA | NA | NA | NA | NA | #VALUE! |
| ENSMUSG00000103006.2 | 0 | NA | NA | NA | NA | NA | #VALUE! |
| ENSMUSG00000103007.2 | 0 | NA | NA | NA | NA | NA | #VALUE! |
| ENSMUSG00000103008.2 | 0 | NA | NA | NA | NA | NA | #VALUE! |
| ENSMUSG00000103009.2 | 1.155144 | -3.7605 | 3.944679 | -0.95331 | 0.340433 | NA | #VALUE! |
| ENSMUSG00000103010.2 | 0 | NA | NA | NA | NA | NA | #VALUE! |
| ENSMUSG00000103011.2 | 0 | NA | NA | NA | NA | NA | #VALUE! |
| ENSMUSG00000103012.2 | 0 | NA | NA | NA | NA | NA | #VALUE! |
| ENSMUSG00000103013.2 | 0 | NA | NA | NA | NA | NA | #VALUE! |
| ENSMUSG00000103015.2 | 0 | NA | NA | NA | NA | NA | #VALUE! |
| ENSMUSG00000103016.2 | 0 | NA | NA | NA | NA | NA | #VALUE! |
| ENSMUSG00000103017.2 | 0 | NA | NA | NA | NA | NA | #VALUE! |
| ENSMUSG00000103018.2 | 0 | NA | NA | NA | NA | NA | #VALUE! |
| ENSMUSG00000103020.2 | 0 | NA | NA | NA | NA | NA | #VALUE! |
| ENSMUSG00000103021.2 | 0 | NA | NA | NA | NA | NA | #VALUE! |
| ENSMUSG00000103023.2 | 0 | NA | NA | NA | NA | NA | #VALUE! |
| ENSMUSG00000103024.2 | 0 | NA | NA | NA | NA | NA | #VALUE! |
| ENSMUSG00000103025.2 | 0 | NA | NA | NA | NA | NA | #VALUE! |
| ENSMUSG00000103026.2 | 0 | NA | NA | NA | NA | NA | #VALUE! |
| ENSMUSG00000103027.2 | 0 | NA | NA | NA | NA | NA | #VALUE! |
| ENSMUSG00000103028.2 | 0 | NA | NA | NA | NA | NA | #VALUE! |
| ENSMUSG00000103029.6 | 0 | NA | NA | NA | NA | NA | #VALUE! |
| ENSMUSG00000103030.2 | 0 | NA | NA | NA | NA | NA | #VALUE! |
| ENSMUSG00000103031.2 | 0 | NA | NA | NA | NA | NA | #VALUE! |
| ENSMUSG00000103032.2 | 0 | NA | NA | NA | NA | NA | #VALUE! |
| ENSMUSG00000103033.2 | 0 | NA | NA | NA | NA | NA | #VALUE! |

| | | | | | | | |
|---|---|---|---|---|---|---|---|
| ENSMUSG00000103034.2 | 0.093255 | 0.868093 | 4.072464 | 0.213162 | 0.831201 | NA | #VALUE! |
| ENSMUSG00000103035.2 | 0 | NA | NA | NA | NA | NA | #VALUE! |
| ENSMUSG00000103037.2 | 0 | NA | NA | NA | NA | NA | #VALUE! |
| ENSMUSG00000103038.2 | 0 | NA | NA | NA | NA | NA | #VALUE! |
| ENSMUSG00000103039.2 | 5.42419 | 0.888804 | 3.762702 | 0.236214 | 0.813266 | NA | #VALUE! |
| ENSMUSG00000103041.2 | 2.30602 | -1.26954 | 3.82217 | -0.33215 | 0.739775 | NA | #VALUE! |
| ENSMUSG00000103042.2 | 0 | NA | NA | NA | NA | NA | #VALUE! |
| ENSMUSG00000103043.2 | 0 | NA | NA | NA | NA | NA | #VALUE! |
| ENSMUSG00000103044.2 | 0 | NA | NA | NA | NA | NA | #VALUE! |
| ENSMUSG00000103045.2 | 0 | NA | NA | NA | NA | NA | #VALUE! |
| ENSMUSG00000103046.2 | 0 | NA | NA | NA | NA | NA | #VALUE! |
| ENSMUSG00000103047.2 | 0 | NA | NA | NA | NA | NA | #VALUE! |
| ENSMUSG00000103049.2 | 0 | NA | NA | NA | NA | NA | #VALUE! |
| ENSMUSG00000103050.2 | 0 | NA | NA | NA | NA | NA | #VALUE! |
| ENSMUSG00000103051.2 | 0 | NA | NA | NA | NA | NA | #VALUE! |
| ENSMUSG00000103052.2 | 0 | NA | NA | NA | NA | NA | #VALUE! |
| ENSMUSG00000103053.2 | 0 | NA | NA | NA | NA | NA | #VALUE! |
| ENSMUSG00000103054.2 | 0 | NA | NA | NA | NA | NA | #VALUE! |
| ENSMUSG00000103055.2 | 0 | NA | NA | NA | NA | NA | #VALUE! |
| ENSMUSG00000103056.2 | 0 | NA | NA | NA | NA | NA | #VALUE! |
| ENSMUSG00000103058.2 | 0 | NA | NA | NA | NA | NA | #VALUE! |
| ENSMUSG00000103059.2 | 0 | NA | NA | NA | NA | NA | #VALUE! |
| ENSMUSG00000103060.2 | 0 | NA | NA | NA | NA | NA | #VALUE! |
| ENSMUSG00000103061.2 | 0 | NA | NA | NA | NA | NA | #VALUE! |
| ENSMUSG00000103062.2 | 0 | NA | NA | NA | NA | NA | #VALUE! |
| ENSMUSG00000103063.2 | 0 | NA | NA | NA | NA | NA | #VALUE! |
| ENSMUSG00000103064.2 | 0 | NA | NA | NA | NA | NA | #VALUE! |
| ENSMUSG00000103065.2 | 0 | NA | NA | NA | NA | NA | #VALUE! |
| ENSMUSG00000103066.2 | 0 | NA | NA | NA | NA | NA | #VALUE! |
| ENSMUSG00000103067.2 | 0 | NA | NA | NA | NA | NA | #VALUE! |
| ENSMUSG00000103069.2 | 0 | NA | NA | NA | NA | NA | #VALUE! |
| ENSMUSG00000103070.2 | 0 | NA | NA | NA | NA | NA | #VALUE! |
| ENSMUSG00000103071.2 | 0 | NA | NA | NA | NA | NA | #VALUE! |
| ENSMUSG00000103072.2 | 0 | NA | NA | NA | NA | NA | #VALUE! |
| ENSMUSG00000103073.2 | 0 | NA | NA | NA | NA | NA | #VALUE! |
| ENSMUSG00000103074.2 | 0 | NA | NA | NA | NA | NA | #VALUE! |
| ENSMUSG00000103075.2 | 0 | NA | NA | NA | NA | NA | #VALUE! |
| ENSMUSG00000103076.2 | 0.315039 | -1.93549 | 4.038621 | -0.47925 | 0.631763 | NA | #VALUE! |
| ENSMUSG00000103077.6 | 0 | NA | NA | NA | NA | NA | #VALUE! |
| ENSMUSG00000103078.2 | 0 | NA | NA | NA | NA | NA | #VALUE! |
| ENSMUSG00000103079.2 | 0 | NA | NA | NA | NA | NA | #VALUE! |
| ENSMUSG00000103080.2 | 0 | NA | NA | NA | NA | NA | #VALUE! |
| ENSMUSG00000103081.4 | 0 | NA | NA | NA | NA | NA | #VALUE! |
| ENSMUSG00000103082.2 | 0 | NA | NA | NA | NA | NA | #VALUE! |

| | | | | | | | |
|---|---|---|---|---|---|---|---|
| ENSMUSG00000103083.2 | 0 | NA | NA | NA | NA | NA | #VALUE! |
| ENSMUSG00000103084.2 | 0 | NA | NA | NA | NA | NA | #VALUE! |
| ENSMUSG00000103085.2 | 0 | NA | NA | NA | NA | NA | #VALUE! |
| ENSMUSG00000103086.2 | 0 | NA | NA | NA | NA | NA | #VALUE! |
| ENSMUSG00000103087.2 | 0 | NA | NA | NA | NA | NA | #VALUE! |
| ENSMUSG00000103088.2 | 0 | NA | NA | NA | NA | NA | #VALUE! |
| ENSMUSG00000103089.2 | 0 | NA | NA | NA | NA | NA | #VALUE! |
| ENSMUSG00000103090.2 | 0 | NA | NA | NA | NA | NA | #VALUE! |
| ENSMUSG00000103091.2 | 0 | NA | NA | NA | NA | NA | #VALUE! |
| ENSMUSG00000103092.2 | 0 | NA | NA | NA | NA | NA | #VALUE! |
| ENSMUSG00000103094.2 | 0 | NA | NA | NA | NA | NA | #VALUE! |
| ENSMUSG00000103095.2 | 0 | NA | NA | NA | NA | NA | #VALUE! |
| ENSMUSG00000103096.2 | 0 | NA | NA | NA | NA | NA | #VALUE! |
| ENSMUSG00000103097.2 | 0 | NA | NA | NA | NA | NA | #VALUE! |
| ENSMUSG00000103098.2 | 0 | NA | NA | NA | NA | NA | #VALUE! |
| ENSMUSG00000103099.2 | 0 | NA | NA | NA | NA | NA | #VALUE! |
| ENSMUSG00000103100.2 | 0 | NA | NA | NA | NA | NA | #VALUE! |
| ENSMUSG00000103101.2 | 0 | NA | NA | NA | NA | NA | #VALUE! |
| ENSMUSG00000103102.2 | 0 | NA | NA | NA | NA | NA | #VALUE! |
| ENSMUSG00000103103.2 | 9.08731 | 6.673702 | 3.708455 | 1.799591 | 0.071925 | NA | #VALUE! |
| ENSMUSG00000103105.6 | 0 | NA | NA | NA | NA | NA | #VALUE! |
| ENSMUSG00000103106.2 | 0 | NA | NA | NA | NA | NA | #VALUE! |
| ENSMUSG00000103107.2 | 0 | NA | NA | NA | NA | NA | #VALUE! |
| ENSMUSG00000103108.3 | 0 | NA | NA | NA | NA | NA | #VALUE! |
| ENSMUSG00000103110.2 | 0 | NA | NA | NA | NA | NA | #VALUE! |
| ENSMUSG00000103111.2 | 0 | NA | NA | NA | NA | NA | #VALUE! |
| ENSMUSG00000103112.2 | 0 | NA | NA | NA | NA | NA | #VALUE! |
| ENSMUSG00000103113.2 | 0 | NA | NA | NA | NA | NA | #VALUE! |
| ENSMUSG00000103114.2 | 0 | NA | NA | NA | NA | NA | #VALUE! |
| ENSMUSG00000103115.2 | 0 | NA | NA | NA | NA | NA | #VALUE! |
| ENSMUSG00000103116.2 | 0 | NA | NA | NA | NA | NA | #VALUE! |
| ENSMUSG00000103118.2 | 0 | NA | NA | NA | NA | NA | #VALUE! |
| ENSMUSG00000103119.2 | 0 | NA | NA | NA | NA | NA | #VALUE! |
| ENSMUSG00000103120.2 | 0 | NA | NA | NA | NA | NA | #VALUE! |
| ENSMUSG00000103121.2 | 0 | NA | NA | NA | NA | NA | #VALUE! |
| ENSMUSG00000103122.2 | 0 | NA | NA | NA | NA | NA | #VALUE! |
| ENSMUSG00000103123.2 | 0 | NA | NA | NA | NA | NA | #VALUE! |
| ENSMUSG00000103124.2 | 0 | NA | NA | NA | NA | NA | #VALUE! |
| ENSMUSG00000103125.2 | 0 | NA | NA | NA | NA | NA | #VALUE! |
| ENSMUSG00000103127.2 | 0 | NA | NA | NA | NA | NA | #VALUE! |
| ENSMUSG00000103129.2 | 0 | NA | NA | NA | NA | NA | #VALUE! |
| ENSMUSG00000103130.2 | 0 | NA | NA | NA | NA | NA | #VALUE! |
| ENSMUSG00000103131.2 | 0 | NA | NA | NA | NA | NA | #VALUE! |
| ENSMUSG00000103133.2 | 0 | NA | NA | NA | NA | NA | #VALUE! |

| | | | | | | | |
|---|---|---|---|---|---|---|---|
| ENSMUSG00000103135.2 | 0 | NA | NA | NA | NA | NA | #VALUE! |
| ENSMUSG00000103136.2 | 0 | NA | NA | NA | NA | NA | #VALUE! |
| ENSMUSG00000103137.2 | 0 | NA | NA | NA | NA | NA | #VALUE! |
| ENSMUSG00000103138.2 | 0 | NA | NA | NA | NA | NA | #VALUE! |
| ENSMUSG00000103139.2 | 0 | NA | NA | NA | NA | NA | #VALUE! |
| ENSMUSG00000103140.2 | 0 | NA | NA | NA | NA | NA | #VALUE! |
| ENSMUSG00000103141.2 | 0 | NA | NA | NA | NA | NA | #VALUE! |
| ENSMUSG00000103142.2 | 0 | NA | NA | NA | NA | NA | #VALUE! |
| ENSMUSG00000103143.2 | 7.960319 | -6.52083 | 3.880481 | -1.68042 | 0.092876 | NA | #VALUE! |
| ENSMUSG00000103144.2 | 0 | NA | NA | NA | NA | NA | #VALUE! |
| ENSMUSG00000103145.2 | 0 | NA | NA | NA | NA | NA | #VALUE! |
| ENSMUSG00000103146.2 | 0 | NA | NA | NA | NA | NA | #VALUE! |
| ENSMUSG00000103147.2 | 0 | NA | NA | NA | NA | NA | #VALUE! |
| ENSMUSG00000103148.2 | 0 | NA | NA | NA | NA | NA | #VALUE! |
| ENSMUSG00000103149.2 | 0 | NA | NA | NA | NA | NA | #VALUE! |
| ENSMUSG00000103150.2 | 0 | NA | NA | NA | NA | NA | #VALUE! |
| ENSMUSG00000103151.2 | 0 | NA | NA | NA | NA | NA | #VALUE! |
| ENSMUSG00000103152.2 | 0 | NA | NA | NA | NA | NA | #VALUE! |
| ENSMUSG00000103153.2 | 0 | NA | NA | NA | NA | NA | #VALUE! |
| ENSMUSG00000103154.2 | 0 | NA | NA | NA | NA | NA | #VALUE! |
| ENSMUSG00000103155.2 | 0 | NA | NA | NA | NA | NA | #VALUE! |
| ENSMUSG00000103156.2 | 0 | NA | NA | NA | NA | NA | #VALUE! |
| ENSMUSG00000103157.2 | 0 | NA | NA | NA | NA | NA | #VALUE! |
| ENSMUSG00000103158.6 | 0 | NA | NA | NA | NA | NA | #VALUE! |
| ENSMUSG00000103159.2 | 0 | NA | NA | NA | NA | NA | #VALUE! |
| ENSMUSG00000103160.2 | 0 | NA | NA | NA | NA | NA | #VALUE! |
| ENSMUSG00000103161.2 | 0 | NA | NA | NA | NA | NA | #VALUE! |
| ENSMUSG00000103162.2 | 0 | NA | NA | NA | NA | NA | #VALUE! |
| ENSMUSG00000103163.2 | 0 | NA | NA | NA | NA | NA | #VALUE! |
| ENSMUSG00000103164.2 | 0 | NA | NA | NA | NA | NA | #VALUE! |
| ENSMUSG00000103165.2 | 0 | NA | NA | NA | NA | NA | #VALUE! |
| ENSMUSG00000103166.2 | 0 | NA | NA | NA | NA | NA | #VALUE! |
| ENSMUSG00000103167.2 | 0 | NA | NA | NA | NA | NA | #VALUE! |
| ENSMUSG00000103168.2 | 0 | NA | NA | NA | NA | NA | #VALUE! |
| ENSMUSG00000103170.2 | 0 | NA | NA | NA | NA | NA | #VALUE! |
| ENSMUSG00000103172.2 | 0 | NA | NA | NA | NA | NA | #VALUE! |
| ENSMUSG00000103174.2 | 0 | NA | NA | NA | NA | NA | #VALUE! |
| ENSMUSG00000103175.2 | 0 | NA | NA | NA | NA | NA | #VALUE! |
| ENSMUSG00000103176.2 | 0 | NA | NA | NA | NA | NA | #VALUE! |
| ENSMUSG00000103177.2 | 0 | NA | NA | NA | NA | NA | #VALUE! |
| ENSMUSG00000103178.2 | 0 | NA | NA | NA | NA | NA | #VALUE! |
| ENSMUSG00000103179.2 | 0 | NA | NA | NA | NA | NA | #VALUE! |
| ENSMUSG00000103180.2 | 0 | NA | NA | NA | NA | NA | #VALUE! |
| ENSMUSG00000103181.2 | 0 | NA | NA | NA | NA | NA | #VALUE! |

| | | | | | | | |
|---|---|---|---|---|---|---|---|
| ENSMUSG00000103182.2 | 0 | NA | NA | NA | NA | NA | #VALUE! |
| ENSMUSG00000103183.2 | 0 | NA | NA | NA | NA | NA | #VALUE! |
| ENSMUSG00000103184.2 | 0 | NA | NA | NA | NA | NA | #VALUE! |
| ENSMUSG00000103185.2 | 0 | NA | NA | NA | NA | NA | #VALUE! |
| ENSMUSG00000103186.2 | 0 | NA | NA | NA | NA | NA | #VALUE! |
| ENSMUSG00000103187.2 | 0 | NA | NA | NA | NA | NA | #VALUE! |
| ENSMUSG00000103188.2 | 0 | NA | NA | NA | NA | NA | #VALUE! |
| ENSMUSG00000103189.2 | 0 | NA | NA | NA | NA | NA | #VALUE! |
| ENSMUSG00000103190.6 | 0 | NA | NA | NA | NA | NA | #VALUE! |
| ENSMUSG00000103191.2 | 0 | NA | NA | NA | NA | NA | #VALUE! |
| ENSMUSG00000103192.2 | 0 | NA | NA | NA | NA | NA | #VALUE! |
| ENSMUSG00000103193.2 | 0 | NA | NA | NA | NA | NA | #VALUE! |
| ENSMUSG00000103194.2 | 0 | NA | NA | NA | NA | NA | #VALUE! |
| ENSMUSG00000103195.2 | 0 | NA | NA | NA | NA | NA | #VALUE! |
| ENSMUSG00000103196.2 | 0 | NA | NA | NA | NA | NA | #VALUE! |
| ENSMUSG00000103197.2 | 0 | NA | NA | NA | NA | NA | #VALUE! |
| ENSMUSG00000103198.2 | 0 | NA | NA | NA | NA | NA | #VALUE! |
| ENSMUSG00000103199.2 | 0 | NA | NA | NA | NA | NA | #VALUE! |
| ENSMUSG00000103200.2 | 0 | NA | NA | NA | NA | NA | #VALUE! |
| ENSMUSG00000103201.2 | 0 | NA | NA | NA | NA | NA | #VALUE! |
| ENSMUSG00000103202.2 | 0 | NA | NA | NA | NA | NA | #VALUE! |
| ENSMUSG00000103203.2 | 0 | NA | NA | NA | NA | NA | #VALUE! |
| ENSMUSG00000103204.2 | 0 | NA | NA | NA | NA | NA | #VALUE! |
| ENSMUSG00000103206.2 | 0 | NA | NA | NA | NA | NA | #VALUE! |
| ENSMUSG00000103207.2 | 0 | NA | NA | NA | NA | NA | #VALUE! |
| ENSMUSG00000103208.2 | 0 | NA | NA | NA | NA | NA | #VALUE! |
| ENSMUSG00000103209.2 | 0 | NA | NA | NA | NA | NA | #VALUE! |
| ENSMUSG00000103210.2 | 0 | NA | NA | NA | NA | NA | #VALUE! |
| ENSMUSG00000103211.2 | 0 | NA | NA | NA | NA | NA | #VALUE! |
| ENSMUSG00000103213.2 | 0 | NA | NA | NA | NA | NA | #VALUE! |
| ENSMUSG00000103214.2 | 0 | NA | NA | NA | NA | NA | #VALUE! |
| ENSMUSG00000103215.2 | 0.105013 | -0.97324 | 4.076293 | -0.23876 | 0.811295 | NA | #VALUE! |
| ENSMUSG00000103216.2 | 0 | NA | NA | NA | NA | NA | #VALUE! |
| ENSMUSG00000103217.2 | 2.625328 | -4.93268 | 3.923662 | -1.25716 | 0.208695 | NA | #VALUE! |
| ENSMUSG00000103218.2 | 0 | NA | NA | NA | NA | NA | #VALUE! |
| ENSMUSG00000103219.2 | 0 | NA | NA | NA | NA | NA | #VALUE! |
| ENSMUSG00000103220.2 | 0.559531 | 2.703958 | 3.982863 | 0.678898 | 0.497202 | NA | #VALUE! |
| ENSMUSG00000103221.2 | 0 | NA | NA | NA | NA | NA | #VALUE! |
| ENSMUSG00000103222.2 | 0.093255 | 0.868093 | 4.072464 | 0.213162 | 0.831201 | NA | #VALUE! |
| ENSMUSG00000103223.2 | 0 | NA | NA | NA | NA | NA | #VALUE! |
| ENSMUSG00000103224.2 | 0 | NA | NA | NA | NA | NA | #VALUE! |
| ENSMUSG00000103225.2 | 0 | NA | NA | NA | NA | NA | #VALUE! |
| ENSMUSG00000103226.2 | 0 | NA | NA | NA | NA | NA | #VALUE! |
| ENSMUSG00000103227.2 | 0.168284 | 1.027628 | 4.067126 | 0.252667 | 0.800526 | NA | #VALUE! |

| | | | | | | | |
|---|---|---|---|---|---|---|---|
| ENSMUSG00000103228.2 | 0 | NA | NA | NA | NA | NA | #VALUE! |
| ENSMUSG00000103230.2 | 0 | NA | NA | NA | NA | NA | #VALUE! |
| ENSMUSG00000103231.2 | 0 | NA | NA | NA | NA | NA | #VALUE! |
| ENSMUSG00000103232.2 | 0 | NA | NA | NA | NA | NA | #VALUE! |
| ENSMUSG00000103233.2 | 0 | NA | NA | NA | NA | NA | #VALUE! |
| ENSMUSG00000103234.2 | 0 | NA | NA | NA | NA | NA | #VALUE! |
| ENSMUSG00000103236.2 | 0 | NA | NA | NA | NA | NA | #VALUE! |
| ENSMUSG00000103237.2 | 0 | NA | NA | NA | NA | NA | #VALUE! |
| ENSMUSG00000103238.2 | 0 | NA | NA | NA | NA | NA | #VALUE! |
| ENSMUSG00000103239.2 | 0 | NA | NA | NA | NA | NA | #VALUE! |
| ENSMUSG00000103240.2 | 0 | NA | NA | NA | NA | NA | #VALUE! |
| ENSMUSG00000103241.2 | 0 | NA | NA | NA | NA | NA | #VALUE! |
| ENSMUSG00000103242.2 | 0 | NA | NA | NA | NA | NA | #VALUE! |
| ENSMUSG00000103243.2 | 0 | NA | NA | NA | NA | NA | #VALUE! |
| ENSMUSG00000103244.2 | 0 | NA | NA | NA | NA | NA | #VALUE! |
| ENSMUSG00000103245.2 | 0 | NA | NA | NA | NA | NA | #VALUE! |
| ENSMUSG00000103246.2 | 0 | NA | NA | NA | NA | NA | #VALUE! |
| ENSMUSG00000103247.2 | 0 | NA | NA | NA | NA | NA | #VALUE! |
| ENSMUSG00000103248.2 | 0 | NA | NA | NA | NA | NA | #VALUE! |
| ENSMUSG00000103249.2 | 0 | NA | NA | NA | NA | NA | #VALUE! |
| ENSMUSG00000103250.2 | 0 | NA | NA | NA | NA | NA | #VALUE! |
| ENSMUSG00000103251.2 | 0 | NA | NA | NA | NA | NA | #VALUE! |
| ENSMUSG00000103252.2 | 0 | NA | NA | NA | NA | NA | #VALUE! |
| ENSMUSG00000103253.2 | 0.245976 | 1.027628 | 4.067126 | 0.252667 | 0.800526 | NA | #VALUE! |
| ENSMUSG00000103254.2 | 0 | NA | NA | NA | NA | NA | #VALUE! |
| ENSMUSG00000103255.2 | 0 | NA | NA | NA | NA | NA | #VALUE! |
| ENSMUSG00000103256.2 | 0 | NA | NA | NA | NA | NA | #VALUE! |
| ENSMUSG00000103257.2 | 0 | NA | NA | NA | NA | NA | #VALUE! |
| ENSMUSG00000103258.2 | 0 | NA | NA | NA | NA | NA | #VALUE! |
| ENSMUSG00000103259.2 | 0 | NA | NA | NA | NA | NA | #VALUE! |
| ENSMUSG00000103260.2 | 0.093255 | 0.868093 | 4.072464 | 0.213162 | 0.831201 | NA | #VALUE! |
| ENSMUSG00000103261.2 | 0 | NA | NA | NA | NA | NA | #VALUE! |
| ENSMUSG00000103262.2 | 0 | NA | NA | NA | NA | NA | #VALUE! |
| ENSMUSG00000103263.2 | 0 | NA | NA | NA | NA | NA | #VALUE! |
| ENSMUSG00000103264.2 | 0 | NA | NA | NA | NA | NA | #VALUE! |
| ENSMUSG00000103265.2 | 0 | NA | NA | NA | NA | NA | #VALUE! |
| ENSMUSG00000103266.2 | 0 | NA | NA | NA | NA | NA | #VALUE! |
| ENSMUSG00000103267.2 | 0 | NA | NA | NA | NA | NA | #VALUE! |
| ENSMUSG00000103269.2 | 0 | NA | NA | NA | NA | NA | #VALUE! |
| ENSMUSG00000103270.2 | 0 | NA | NA | NA | NA | NA | #VALUE! |
| ENSMUSG00000103271.2 | 0 | NA | NA | NA | NA | NA | #VALUE! |
| ENSMUSG00000103272.2 | 0 | NA | NA | NA | NA | NA | #VALUE! |
| ENSMUSG00000103273.2 | 0 | NA | NA | NA | NA | NA | #VALUE! |
| ENSMUSG00000103274.2 | 0 | NA | NA | NA | NA | NA | #VALUE! |

| | | | | | | | |
|---|---|---|---|---|---|---|---|
| ENSMUSG00000103275.2 | 0 | NA | NA | NA | NA | NA | #VALUE! |
| ENSMUSG00000103276.2 | 0 | NA | NA | NA | NA | NA | #VALUE! |
| ENSMUSG00000103277.2 | 0 | NA | NA | NA | NA | NA | #VALUE! |
| ENSMUSG00000103279.2 | 0 | NA | NA | NA | NA | NA | #VALUE! |
| ENSMUSG00000103280.2 | 0 | NA | NA | NA | NA | NA | #VALUE! |
| ENSMUSG00000103281.2 | 0 | NA | NA | NA | NA | NA | #VALUE! |
| ENSMUSG00000103282.2 | 0 | NA | NA | NA | NA | NA | #VALUE! |
| ENSMUSG00000103284.2 | 0 | NA | NA | NA | NA | NA | #VALUE! |
| ENSMUSG00000103285.2 | 0 | NA | NA | NA | NA | NA | #VALUE! |
| ENSMUSG00000103286.2 | 0 | NA | NA | NA | NA | NA | #VALUE! |
| ENSMUSG00000103287.2 | 0 | NA | NA | NA | NA | NA | #VALUE! |
| ENSMUSG00000103288.2 | 0 | NA | NA | NA | NA | NA | #VALUE! |
| ENSMUSG00000103289.2 | 0 | NA | NA | NA | NA | NA | #VALUE! |
| ENSMUSG00000103290.2 | 0 | NA | NA | NA | NA | NA | #VALUE! |
| ENSMUSG00000103291.2 | 0 | NA | NA | NA | NA | NA | #VALUE! |
| ENSMUSG00000103292.2 | 0 | NA | NA | NA | NA | NA | #VALUE! |
| ENSMUSG00000103293.2 | 0 | NA | NA | NA | NA | NA | #VALUE! |
| ENSMUSG00000103294.2 | 0 | NA | NA | NA | NA | NA | #VALUE! |
| ENSMUSG00000103295.2 | 0 | NA | NA | NA | NA | NA | #VALUE! |
| ENSMUSG00000103296.2 | 0 | NA | NA | NA | NA | NA | #VALUE! |
| ENSMUSG00000103297.2 | 0 | NA | NA | NA | NA | NA | #VALUE! |
| ENSMUSG00000103298.2 | 0 | NA | NA | NA | NA | NA | #VALUE! |
| ENSMUSG00000103299.2 | 0 | NA | NA | NA | NA | NA | #VALUE! |
| ENSMUSG00000103300.2 | 0 | NA | NA | NA | NA | NA | #VALUE! |
| ENSMUSG00000103301.2 | 0 | NA | NA | NA | NA | NA | #VALUE! |
| ENSMUSG00000103302.2 | 0 | NA | NA | NA | NA | NA | #VALUE! |
| ENSMUSG00000103303.2 | 0 | NA | NA | NA | NA | NA | #VALUE! |
| ENSMUSG00000103304.2 | 0 | NA | NA | NA | NA | NA | #VALUE! |
| ENSMUSG00000103305.2 | 0 | NA | NA | NA | NA | NA | #VALUE! |
| ENSMUSG00000103306.2 | 0 | NA | NA | NA | NA | NA | #VALUE! |
| ENSMUSG00000103307.2 | 0 | NA | NA | NA | NA | NA | #VALUE! |
| ENSMUSG00000103308.6 | 0 | NA | NA | NA | NA | NA | #VALUE! |
| ENSMUSG00000103309.2 | 0 | NA | NA | NA | NA | NA | #VALUE! |
| ENSMUSG00000103310.2 | 0 | NA | NA | NA | NA | NA | #VALUE! |
| ENSMUSG00000103311.2 | 0 | NA | NA | NA | NA | NA | #VALUE! |
| ENSMUSG00000103312.2 | 0 | NA | NA | NA | NA | NA | #VALUE! |
| ENSMUSG00000103313.2 | 0 | NA | NA | NA | NA | NA | #VALUE! |
| ENSMUSG00000103314.2 | 0 | NA | NA | NA | NA | NA | #VALUE! |
| ENSMUSG00000103315.2 | 0 | NA | NA | NA | NA | NA | #VALUE! |
| ENSMUSG00000103317.2 | 0 | NA | NA | NA | NA | NA | #VALUE! |
| ENSMUSG00000103318.2 | 0 | NA | NA | NA | NA | NA | #VALUE! |
| ENSMUSG00000103320.2 | 0 | NA | NA | NA | NA | NA | #VALUE! |
| ENSMUSG00000103321.2 | 0 | NA | NA | NA | NA | NA | #VALUE! |
| ENSMUSG00000103322.2 | 0 | NA | NA | NA | NA | NA | #VALUE! |

| | | | | | | | |
|---|---|---|---|---|---|---|---|
| ENSMUSG00000103323.2 | 0 | NA | NA | NA | NA | NA | #VALUE! |
| ENSMUSG00000103324.2 | 0 | NA | NA | NA | NA | NA | #VALUE! |
| ENSMUSG00000103325.2 | 0 | NA | NA | NA | NA | NA | #VALUE! |
| ENSMUSG00000103327.2 | 0 | NA | NA | NA | NA | NA | #VALUE! |
| ENSMUSG00000103328.2 | 0 | NA | NA | NA | NA | NA | #VALUE! |
| ENSMUSG00000103329.3 | 0 | NA | NA | NA | NA | NA | #VALUE! |
| ENSMUSG00000103330.2 | 0 | NA | NA | NA | NA | NA | #VALUE! |
| ENSMUSG00000103331.2 | 0 | NA | NA | NA | NA | NA | #VALUE! |
| ENSMUSG00000103332.2 | 0 | NA | NA | NA | NA | NA | #VALUE! |
| ENSMUSG00000103333.2 | 0 | NA | NA | NA | NA | NA | #VALUE! |
| ENSMUSG00000103334.2 | 0 | NA | NA | NA | NA | NA | #VALUE! |
| ENSMUSG00000103336.2 | 0 | NA | NA | NA | NA | NA | #VALUE! |
| ENSMUSG00000103338.2 | 0 | NA | NA | NA | NA | NA | #VALUE! |
| ENSMUSG00000103339.2 | 0 | NA | NA | NA | NA | NA | #VALUE! |
| ENSMUSG00000103340.2 | 0 | NA | NA | NA | NA | NA | #VALUE! |
| ENSMUSG00000103341.2 | 0 | NA | NA | NA | NA | NA | #VALUE! |
| ENSMUSG00000103343.2 | 0 | NA | NA | NA | NA | NA | #VALUE! |
| ENSMUSG00000103345.2 | 0 | NA | NA | NA | NA | NA | #VALUE! |
| ENSMUSG00000103346.2 | 0 | NA | NA | NA | NA | NA | #VALUE! |
| ENSMUSG00000103347.2 | 0 | NA | NA | NA | NA | NA | #VALUE! |
| ENSMUSG00000103348.2 | 0 | NA | NA | NA | NA | NA | #VALUE! |
| ENSMUSG00000103349.2 | 0 | NA | NA | NA | NA | NA | #VALUE! |
| ENSMUSG00000103350.2 | 0 | NA | NA | NA | NA | NA | #VALUE! |
| ENSMUSG00000103351.2 | 0 | NA | NA | NA | NA | NA | #VALUE! |
| ENSMUSG00000103352.2 | 0 | NA | NA | NA | NA | NA | #VALUE! |
| ENSMUSG00000103353.2 | 0 | NA | NA | NA | NA | NA | #VALUE! |
| ENSMUSG00000103354.2 | 0 | NA | NA | NA | NA | NA | #VALUE! |
| ENSMUSG00000103355.2 | 0 | NA | NA | NA | NA | NA | #VALUE! |
| ENSMUSG00000103356.2 | 0 | NA | NA | NA | NA | NA | #VALUE! |
| ENSMUSG00000103357.2 | 0 | NA | NA | NA | NA | NA | #VALUE! |
| ENSMUSG00000103358.2 | 0 | NA | NA | NA | NA | NA | #VALUE! |
| ENSMUSG00000103360.2 | 0 | NA | NA | NA | NA | NA | #VALUE! |
| ENSMUSG00000103361.2 | 0 | NA | NA | NA | NA | NA | #VALUE! |
| ENSMUSG00000103362.2 | 0 | NA | NA | NA | NA | NA | #VALUE! |
| ENSMUSG00000103364.2 | 0 | NA | NA | NA | NA | NA | #VALUE! |
| ENSMUSG00000103365.2 | 0 | NA | NA | NA | NA | NA | #VALUE! |
| ENSMUSG00000103366.2 | 0 | NA | NA | NA | NA | NA | #VALUE! |
| ENSMUSG00000103367.2 | 0 | NA | NA | NA | NA | NA | #VALUE! |
| ENSMUSG00000103368.2 | 0 | NA | NA | NA | NA | NA | #VALUE! |
| ENSMUSG00000103369.2 | 0 | NA | NA | NA | NA | NA | #VALUE! |
| ENSMUSG00000103370.2 | 0 | NA | NA | NA | NA | NA | #VALUE! |
| ENSMUSG00000103371.2 | 0 | NA | NA | NA | NA | NA | #VALUE! |
| ENSMUSG00000103372.2 | 0 | NA | NA | NA | NA | NA | #VALUE! |
| ENSMUSG00000103373.2 | 0 | NA | NA | NA | NA | NA | #VALUE! |

| | | | | | | | |
|---|---|---|---|---|---|---|---|
| ENSMUSG00000103374.2 | 0 | NA | NA | NA | NA | NA | #VALUE! |
| ENSMUSG00000103375.2 | 0 | NA | NA | NA | NA | NA | #VALUE! |
| ENSMUSG00000103376.2 | 0 | NA | NA | NA | NA | NA | #VALUE! |
| ENSMUSG00000103377.2 | 0 | NA | NA | NA | NA | NA | #VALUE! |
| ENSMUSG00000103378.2 | 0 | NA | NA | NA | NA | NA | #VALUE! |
| ENSMUSG00000103379.2 | 0 | NA | NA | NA | NA | NA | #VALUE! |
| ENSMUSG00000103380.2 | 0 | NA | NA | NA | NA | NA | #VALUE! |
| ENSMUSG00000103382.2 | 0 | NA | NA | NA | NA | NA | #VALUE! |
| ENSMUSG00000103383.2 | 0 | NA | NA | NA | NA | NA | #VALUE! |
| ENSMUSG00000103384.2 | 0 | NA | NA | NA | NA | NA | #VALUE! |
| ENSMUSG00000103385.2 | 0 | NA | NA | NA | NA | NA | #VALUE! |
| ENSMUSG00000103386.2 | 0 | NA | NA | NA | NA | NA | #VALUE! |
| ENSMUSG00000103387.2 | 0 | NA | NA | NA | NA | NA | #VALUE! |
| ENSMUSG00000103388.2 | 0 | NA | NA | NA | NA | NA | #VALUE! |
| ENSMUSG00000103389.2 | 0 | NA | NA | NA | NA | NA | #VALUE! |
| ENSMUSG00000103390.2 | 0 | NA | NA | NA | NA | NA | #VALUE! |
| ENSMUSG00000103391.2 | 0 | NA | NA | NA | NA | NA | #VALUE! |
| ENSMUSG00000103392.2 | 0 | NA | NA | NA | NA | NA | #VALUE! |
| ENSMUSG00000103393.2 | 0 | NA | NA | NA | NA | NA | #VALUE! |
| ENSMUSG00000103394.2 | 0 | NA | NA | NA | NA | NA | #VALUE! |
| ENSMUSG00000103395.2 | 0 | NA | NA | NA | NA | NA | #VALUE! |
| ENSMUSG00000103396.2 | 0 | NA | NA | NA | NA | NA | #VALUE! |
| ENSMUSG00000103397.2 | 0 | NA | NA | NA | NA | NA | #VALUE! |
| ENSMUSG00000103398.2 | 1.346268 | 3.918821 | 3.939925 | 0.994644 | 0.31991 | NA | #VALUE! |
| ENSMUSG00000103399.2 | 0 | NA | NA | NA | NA | NA | #VALUE! |
| ENSMUSG00000103400.3 | 0 | NA | NA | NA | NA | NA | #VALUE! |
| ENSMUSG00000103401.2 | 0 | NA | NA | NA | NA | NA | #VALUE! |
| ENSMUSG00000103402.2 | 0 | NA | NA | NA | NA | NA | #VALUE! |
| ENSMUSG00000103403.2 | 0 | NA | NA | NA | NA | NA | #VALUE! |
| ENSMUSG00000103404.2 | 0 | NA | NA | NA | NA | NA | #VALUE! |
| ENSMUSG00000103406.2 | 0 | NA | NA | NA | NA | NA | #VALUE! |
| ENSMUSG00000103407.2 | 0 | NA | NA | NA | NA | NA | #VALUE! |
| ENSMUSG00000103408.2 | 0 | NA | NA | NA | NA | NA | #VALUE! |
| ENSMUSG00000103409.3 | 0 | NA | NA | NA | NA | NA | #VALUE! |
| ENSMUSG00000103410.2 | 0 | NA | NA | NA | NA | NA | #VALUE! |
| ENSMUSG00000103411.2 | 0 | NA | NA | NA | NA | NA | #VALUE! |
| ENSMUSG00000103412.2 | 0 | NA | NA | NA | NA | NA | #VALUE! |
| ENSMUSG00000103413.2 | 0 | NA | NA | NA | NA | NA | #VALUE! |
| ENSMUSG00000103415.2 | 5.566071 | -4.08178 | 3.869556 | -1.05485 | 0.291496 | NA | #VALUE! |
| ENSMUSG00000103416.2 | 0 | NA | NA | NA | NA | NA | #VALUE! |
| ENSMUSG00000103417.2 | 0 | NA | NA | NA | NA | NA | #VALUE! |
| ENSMUSG00000103419.2 | 0 | NA | NA | NA | NA | NA | #VALUE! |
| ENSMUSG00000103420.2 | 0 | NA | NA | NA | NA | NA | #VALUE! |
| ENSMUSG00000103421.2 | 1.917618 | 3.239121 | 3.925643 | 0.825119 | 0.409304 | NA | #VALUE! |

| | | | | | | | |
|---|---|---|---|---|---|---|---|
| ENSMUSG00000103422.2 | 0 | NA | NA | NA | NA | NA | #VALUE! |
| ENSMUSG00000103423.2 | 0.245976 | 1.027628 | 4.067126 | 0.252667 | 0.800526 | NA | #VALUE! |
| ENSMUSG00000103424.2 | 0 | NA | NA | NA | NA | NA | #VALUE! |
| ENSMUSG00000103425.2 | 0 | NA | NA | NA | NA | NA | #VALUE! |
| ENSMUSG00000103426.2 | 0 | NA | NA | NA | NA | NA | #VALUE! |
| ENSMUSG00000103427.2 | 0 | NA | NA | NA | NA | NA | #VALUE! |
| ENSMUSG00000103428.2 | 0 | NA | NA | NA | NA | NA | #VALUE! |
| ENSMUSG00000103430.2 | 0 | NA | NA | NA | NA | NA | #VALUE! |
| ENSMUSG00000103431.6 | 0 | NA | NA | NA | NA | NA | #VALUE! |
| ENSMUSG00000103432.2 | 5.670709 | -6.03829 | 3.914631 | -1.54249 | 0.122954 | NA | #VALUE! |
| ENSMUSG00000103433.2 | 0 | NA | NA | NA | NA | NA | #VALUE! |
| ENSMUSG00000103434.2 | 0 | NA | NA | NA | NA | NA | #VALUE! |
| ENSMUSG00000103435.2 | 0 | NA | NA | NA | NA | NA | #VALUE! |
| ENSMUSG00000103436.2 | 0 | NA | NA | NA | NA | NA | #VALUE! |
| ENSMUSG00000103437.2 | 0 | NA | NA | NA | NA | NA | #VALUE! |
| ENSMUSG00000103438.2 | 0 | NA | NA | NA | NA | NA | #VALUE! |
| ENSMUSG00000103439.2 | 0 | NA | NA | NA | NA | NA | #VALUE! |
| ENSMUSG00000103440.2 | 0 | NA | NA | NA | NA | NA | #VALUE! |
| ENSMUSG00000103441.2 | 0 | NA | NA | NA | NA | NA | #VALUE! |
| ENSMUSG00000103442.6 | 0 | NA | NA | NA | NA | NA | #VALUE! |
| ENSMUSG00000103443.2 | 0 | NA | NA | NA | NA | NA | #VALUE! |
| ENSMUSG00000103444.2 | 0 | NA | NA | NA | NA | NA | #VALUE! |
| ENSMUSG00000103445.2 | 0 | NA | NA | NA | NA | NA | #VALUE! |
| ENSMUSG00000103446.2 | 0 | NA | NA | NA | NA | NA | #VALUE! |
| ENSMUSG00000103448.2 | 0 | NA | NA | NA | NA | NA | #VALUE! |
| ENSMUSG00000103449.2 | 0 | NA | NA | NA | NA | NA | #VALUE! |
| ENSMUSG00000103451.2 | 0 | NA | NA | NA | NA | NA | #VALUE! |
| ENSMUSG00000103452.2 | 0 | NA | NA | NA | NA | NA | #VALUE! |
| ENSMUSG00000103454.2 | 0 | NA | NA | NA | NA | NA | #VALUE! |
| ENSMUSG00000103456.2 | 0 | NA | NA | NA | NA | NA | #VALUE! |
| ENSMUSG00000103457.2 | 0 | NA | NA | NA | NA | NA | #VALUE! |
| ENSMUSG00000103458.2 | 0 | NA | NA | NA | NA | NA | #VALUE! |
| ENSMUSG00000103459.2 | 0 | NA | NA | NA | NA | NA | #VALUE! |
| ENSMUSG00000103460.2 | 0 | NA | NA | NA | NA | NA | #VALUE! |
| ENSMUSG00000103461.2 | 0 | NA | NA | NA | NA | NA | #VALUE! |
| ENSMUSG00000103462.6 | 0 | NA | NA | NA | NA | NA | #VALUE! |
| ENSMUSG00000103463.2 | 0 | NA | NA | NA | NA | NA | #VALUE! |
| ENSMUSG00000103464.2 | 0 | NA | NA | NA | NA | NA | #VALUE! |
| ENSMUSG00000103465.2 | 0 | NA | NA | NA | NA | NA | #VALUE! |
| ENSMUSG00000103466.2 | 0 | NA | NA | NA | NA | NA | #VALUE! |
| ENSMUSG00000103467.2 | 0 | NA | NA | NA | NA | NA | #VALUE! |
| ENSMUSG00000103468.2 | 0 | NA | NA | NA | NA | NA | #VALUE! |
| ENSMUSG00000103469.2 | 0 | NA | NA | NA | NA | NA | #VALUE! |
| ENSMUSG00000103470.2 | 0 | NA | NA | NA | NA | NA | #VALUE! |

| | | | | | | | |
|---|---|---|---|---|---|---|---|
| ENSMUSG00000103471.2 | 0 | NA | NA | NA | NA | NA | #VALUE! |
| ENSMUSG00000103472.2 | 0 | NA | NA | NA | NA | NA | #VALUE! |
| ENSMUSG00000103473.2 | 0 | NA | NA | NA | NA | NA | #VALUE! |
| ENSMUSG00000103474.2 | 0 | NA | NA | NA | NA | NA | #VALUE! |
| ENSMUSG00000103475.2 | 0 | NA | NA | NA | NA | NA | #VALUE! |
| ENSMUSG00000103476.2 | 0 | NA | NA | NA | NA | NA | #VALUE! |
| ENSMUSG00000103477.2 | 0 | NA | NA | NA | NA | NA | #VALUE! |
| ENSMUSG00000103478.2 | 0 | NA | NA | NA | NA | NA | #VALUE! |
| ENSMUSG00000103479.2 | 0 | NA | NA | NA | NA | NA | #VALUE! |
| ENSMUSG00000103481.2 | 0 | NA | NA | NA | NA | NA | #VALUE! |
| ENSMUSG00000103482.2 | 0 | NA | NA | NA | NA | NA | #VALUE! |
| ENSMUSG00000103483.2 | 0 | NA | NA | NA | NA | NA | #VALUE! |
| ENSMUSG00000103484.2 | 0 | NA | NA | NA | NA | NA | #VALUE! |
| ENSMUSG00000103485.2 | 0 | NA | NA | NA | NA | NA | #VALUE! |
| ENSMUSG00000103486.2 | 0 | NA | NA | NA | NA | NA | #VALUE! |
| ENSMUSG00000103487.2 | 0 | NA | NA | NA | NA | NA | #VALUE! |
| ENSMUSG00000103488.2 | 0 | NA | NA | NA | NA | NA | #VALUE! |
| ENSMUSG00000103489.2 | 0 | NA | NA | NA | NA | NA | #VALUE! |
| ENSMUSG00000103491.2 | 0 | NA | NA | NA | NA | NA | #VALUE! |
| ENSMUSG00000103492.2 | 0 | NA | NA | NA | NA | NA | #VALUE! |
| ENSMUSG00000103493.2 | 0 | NA | NA | NA | NA | NA | #VALUE! |
| ENSMUSG00000103494.2 | 0 | NA | NA | NA | NA | NA | #VALUE! |
| ENSMUSG00000103495.2 | 0 | NA | NA | NA | NA | NA | #VALUE! |
| ENSMUSG00000103497.2 | 6.069462 | -6.1296 | 3.91415 | -1.56601 | 0.117346 | NA | #VALUE! |
| ENSMUSG00000103498.2 | 0 | NA | NA | NA | NA | NA | #VALUE! |
| ENSMUSG00000103499.6 | 0 | NA | NA | NA | NA | NA | #VALUE! |
| ENSMUSG00000103500.2 | 0 | NA | NA | NA | NA | NA | #VALUE! |
| ENSMUSG00000103501.2 | 0 | NA | NA | NA | NA | NA | #VALUE! |
| ENSMUSG00000103503.2 | 0 | NA | NA | NA | NA | NA | #VALUE! |
| ENSMUSG00000103504.2 | 0 | NA | NA | NA | NA | NA | #VALUE! |
| ENSMUSG00000103505.2 | 0 | NA | NA | NA | NA | NA | #VALUE! |
| ENSMUSG00000103506.2 | 0 | NA | NA | NA | NA | NA | #VALUE! |
| ENSMUSG00000103507.2 | 0 | NA | NA | NA | NA | NA | #VALUE! |
| ENSMUSG00000103509.2 | 0 | NA | NA | NA | NA | NA | #VALUE! |
| ENSMUSG00000103510.2 | 0 | NA | NA | NA | NA | NA | #VALUE! |
| ENSMUSG00000103511.2 | 0 | NA | NA | NA | NA | NA | #VALUE! |
| ENSMUSG00000103512.2 | 0 | NA | NA | NA | NA | NA | #VALUE! |
| ENSMUSG00000103513.2 | 0 | NA | NA | NA | NA | NA | #VALUE! |
| ENSMUSG00000103514.2 | 0 | NA | NA | NA | NA | NA | #VALUE! |
| ENSMUSG00000103516.2 | 0 | NA | NA | NA | NA | NA | #VALUE! |
| ENSMUSG00000103517.2 | 0 | NA | NA | NA | NA | NA | #VALUE! |
| ENSMUSG00000103518.2 | 0 | NA | NA | NA | NA | NA | #VALUE! |
| ENSMUSG00000103519.2 | 0 | NA | NA | NA | NA | NA | #VALUE! |
| ENSMUSG00000103520.2 | 0 | NA | NA | NA | NA | NA | #VALUE! |

| | | | | | | | |
|---|---|---|---|---|---|---|---|
| ENSMUSG00000103521.2 | 0 | NA | NA | NA | NA | NA | #VALUE! |
| ENSMUSG00000103522.2 | 0 | NA | NA | NA | NA | NA | #VALUE! |
| ENSMUSG00000103523.2 | 0 | NA | NA | NA | NA | NA | #VALUE! |
| ENSMUSG00000103524.2 | 0 | NA | NA | NA | NA | NA | #VALUE! |
| ENSMUSG00000103525.2 | 0 | NA | NA | NA | NA | NA | #VALUE! |
| ENSMUSG00000103526.2 | 0 | NA | NA | NA | NA | NA | #VALUE! |
| ENSMUSG00000103527.2 | 0 | NA | NA | NA | NA | NA | #VALUE! |
| ENSMUSG00000103528.2 | 0 | NA | NA | NA | NA | NA | #VALUE! |
| ENSMUSG00000103529.2 | 0 | NA | NA | NA | NA | NA | #VALUE! |
| ENSMUSG00000103530.2 | 0 | NA | NA | NA | NA | NA | #VALUE! |
| ENSMUSG00000103531.2 | 0 | NA | NA | NA | NA | NA | #VALUE! |
| ENSMUSG00000103532.2 | 0 | NA | NA | NA | NA | NA | #VALUE! |
| ENSMUSG00000103533.2 | 0 | NA | NA | NA | NA | NA | #VALUE! |
| ENSMUSG00000103534.2 | 0 | NA | NA | NA | NA | NA | #VALUE! |
| ENSMUSG00000103535.2 | 0 | NA | NA | NA | NA | NA | #VALUE! |
| ENSMUSG00000103536.2 | 0 | NA | NA | NA | NA | NA | #VALUE! |
| ENSMUSG00000103537.2 | 0 | NA | NA | NA | NA | NA | #VALUE! |
| ENSMUSG00000103538.4 | 0 | NA | NA | NA | NA | NA | #VALUE! |
| ENSMUSG00000103539.2 | 0 | NA | NA | NA | NA | NA | #VALUE! |
| ENSMUSG00000103540.2 | 0 | NA | NA | NA | NA | NA | #VALUE! |
| ENSMUSG00000103541.2 | 0 | NA | NA | NA | NA | NA | #VALUE! |
| ENSMUSG00000103542.2 | 0 | NA | NA | NA | NA | NA | #VALUE! |
| ENSMUSG00000103543.2 | 0 | NA | NA | NA | NA | NA | #VALUE! |
| ENSMUSG00000103544.3 | 0 | NA | NA | NA | NA | NA | #VALUE! |
| ENSMUSG00000103545.2 | 0 | NA | NA | NA | NA | NA | #VALUE! |
| ENSMUSG00000103546.2 | 0 | NA | NA | NA | NA | NA | #VALUE! |
| ENSMUSG00000103547.2 | 0 | NA | NA | NA | NA | NA | #VALUE! |
| ENSMUSG00000103548.2 | 0 | NA | NA | NA | NA | NA | #VALUE! |
| ENSMUSG00000103549.2 | 0 | NA | NA | NA | NA | NA | #VALUE! |
| ENSMUSG00000103552.2 | 0 | NA | NA | NA | NA | NA | #VALUE! |
| ENSMUSG00000103553.2 | 0 | NA | NA | NA | NA | NA | #VALUE! |
| ENSMUSG00000103554.2 | 0 | NA | NA | NA | NA | NA | #VALUE! |
| ENSMUSG00000103555.2 | 0 | NA | NA | NA | NA | NA | #VALUE! |
| ENSMUSG00000103556.2 | 0 | NA | NA | NA | NA | NA | #VALUE! |
| ENSMUSG00000103557.2 | 0 | NA | NA | NA | NA | NA | #VALUE! |
| ENSMUSG00000103558.2 | 0 | NA | NA | NA | NA | NA | #VALUE! |
| ENSMUSG00000103559.2 | 0 | NA | NA | NA | NA | NA | #VALUE! |
| ENSMUSG00000103560.2 | 1.575197 | -4.20222 | 3.934736 | -1.06798 | 0.285529 | NA | #VALUE! |
| ENSMUSG00000103561.2 | 0 | NA | NA | NA | NA | NA | #VALUE! |
| ENSMUSG00000103562.2 | 0 | NA | NA | NA | NA | NA | #VALUE! |
| ENSMUSG00000103563.2 | 0 | NA | NA | NA | NA | NA | #VALUE! |
| ENSMUSG00000103564.2 | 0 | NA | NA | NA | NA | NA | #VALUE! |
| ENSMUSG00000103566.2 | 0 | NA | NA | NA | NA | NA | #VALUE! |
| ENSMUSG00000103567.2 | 0 | NA | NA | NA | NA | NA | #VALUE! |

| | | | | | | | |
|---|---|---|---|---|---|---|---|
| ENSMUSG00000103568.2 | 0 | NA | NA | NA | NA | NA | #VALUE! |
| ENSMUSG00000103569.2 | 0 | NA | NA | NA | NA | NA | #VALUE! |
| ENSMUSG00000103570.2 | 0 | NA | NA | NA | NA | NA | #VALUE! |
| ENSMUSG00000103571.2 | 0 | NA | NA | NA | NA | NA | #VALUE! |
| ENSMUSG00000103572.2 | 0 | NA | NA | NA | NA | NA | #VALUE! |
| ENSMUSG00000103573.2 | 0 | NA | NA | NA | NA | NA | #VALUE! |
| ENSMUSG00000103574.2 | 0 | NA | NA | NA | NA | NA | #VALUE! |
| ENSMUSG00000103575.2 | 0 | NA | NA | NA | NA | NA | #VALUE! |
| ENSMUSG00000103576.2 | 0 | NA | NA | NA | NA | NA | #VALUE! |
| ENSMUSG00000103577.2 | 0 | NA | NA | NA | NA | NA | #VALUE! |
| ENSMUSG00000103579.2 | 0 | NA | NA | NA | NA | NA | #VALUE! |
| ENSMUSG00000103580.2 | 0 | NA | NA | NA | NA | NA | #VALUE! |
| ENSMUSG00000103581.2 | 0 | NA | NA | NA | NA | NA | #VALUE! |
| ENSMUSG00000103583.2 | 0 | NA | NA | NA | NA | NA | #VALUE! |
| ENSMUSG00000103584.2 | 0 | NA | NA | NA | NA | NA | #VALUE! |
| ENSMUSG00000103585.2 | 0 | NA | NA | NA | NA | NA | #VALUE! |
| ENSMUSG00000103586.6 | 0 | NA | NA | NA | NA | NA | #VALUE! |
| ENSMUSG00000103587.2 | 0 | NA | NA | NA | NA | NA | #VALUE! |
| ENSMUSG00000103588.2 | 0 | NA | NA | NA | NA | NA | #VALUE! |
| ENSMUSG00000103589.6 | 0 | NA | NA | NA | NA | NA | #VALUE! |
| ENSMUSG00000103590.2 | 0 | NA | NA | NA | NA | NA | #VALUE! |
| ENSMUSG00000103591.2 | 0 | NA | NA | NA | NA | NA | #VALUE! |
| ENSMUSG00000103592.2 | 0 | NA | NA | NA | NA | NA | #VALUE! |
| ENSMUSG00000103593.2 | 0 | NA | NA | NA | NA | NA | #VALUE! |
| ENSMUSG00000103594.2 | 0 | NA | NA | NA | NA | NA | #VALUE! |
| ENSMUSG00000103595.2 | 0 | NA | NA | NA | NA | NA | #VALUE! |
| ENSMUSG00000103596.2 | 0 | NA | NA | NA | NA | NA | #VALUE! |
| ENSMUSG00000103597.2 | 0 | NA | NA | NA | NA | NA | #VALUE! |
| ENSMUSG00000103598.2 | 0 | NA | NA | NA | NA | NA | #VALUE! |
| ENSMUSG00000103599.2 | 0 | NA | NA | NA | NA | NA | #VALUE! |
| ENSMUSG00000103600.2 | 0 | NA | NA | NA | NA | NA | #VALUE! |
| ENSMUSG00000103601.2 | 0 | NA | NA | NA | NA | NA | #VALUE! |
| ENSMUSG00000103602.2 | 0 | NA | NA | NA | NA | NA | #VALUE! |
| ENSMUSG00000103603.2 | 0 | NA | NA | NA | NA | NA | #VALUE! |
| ENSMUSG00000103604.2 | 0 | NA | NA | NA | NA | NA | #VALUE! |
| ENSMUSG00000103606.2 | 0 | NA | NA | NA | NA | NA | #VALUE! |
| ENSMUSG00000103607.2 | 0 | NA | NA | NA | NA | NA | #VALUE! |
| ENSMUSG00000103608.2 | 0 | NA | NA | NA | NA | NA | #VALUE! |
| ENSMUSG00000103609.2 | 0 | NA | NA | NA | NA | NA | #VALUE! |
| ENSMUSG00000103610.2 | 0 | NA | NA | NA | NA | NA | #VALUE! |
| ENSMUSG00000103611.2 | 0 | NA | NA | NA | NA | NA | #VALUE! |
| ENSMUSG00000103612.2 | 0 | NA | NA | NA | NA | NA | #VALUE! |
| ENSMUSG00000103613.2 | 0 | NA | NA | NA | NA | NA | #VALUE! |
| ENSMUSG00000103614.2 | 0 | NA | NA | NA | NA | NA | #VALUE! |

| | | | | | | | |
|---|---|---|---|---|---|---|---|
| ENSMUSG00000103615.6 | 0 | NA | NA | NA | NA | NA | #VALUE! |
| ENSMUSG00000103616.2 | 0 | NA | NA | NA | NA | NA | #VALUE! |
| ENSMUSG00000103617.2 | 0 | NA | NA | NA | NA | NA | #VALUE! |
| ENSMUSG00000103618.2 | 0 | NA | NA | NA | NA | NA | #VALUE! |
| ENSMUSG00000103620.2 | 0 | NA | NA | NA | NA | NA | #VALUE! |
| ENSMUSG00000103621.2 | 0 | NA | NA | NA | NA | NA | #VALUE! |
| ENSMUSG00000103622.2 | 0 | NA | NA | NA | NA | NA | #VALUE! |
| ENSMUSG00000103624.2 | 0 | NA | NA | NA | NA | NA | #VALUE! |
| ENSMUSG00000103625.2 | 0 | NA | NA | NA | NA | NA | #VALUE! |
| ENSMUSG00000103626.2 | 0 | NA | NA | NA | NA | NA | #VALUE! |
| ENSMUSG00000103628.2 | 0 | NA | NA | NA | NA | NA | #VALUE! |
| ENSMUSG00000103629.2 | 0 | NA | NA | NA | NA | NA | #VALUE! |
| ENSMUSG00000103630.2 | 0 | NA | NA | NA | NA | NA | #VALUE! |
| ENSMUSG00000103631.2 | 0 | NA | NA | NA | NA | NA | #VALUE! |
| ENSMUSG00000103632.2 | 0.105013 | -0.97324 | 4.076293 | -0.23876 | 0.811295 | NA | #VALUE! |
| ENSMUSG00000103633.2 | 0 | NA | NA | NA | NA | NA | #VALUE! |
| ENSMUSG00000103634.2 | 0 | NA | NA | NA | NA | NA | #VALUE! |
| ENSMUSG00000103635.2 | 0 | NA | NA | NA | NA | NA | #VALUE! |
| ENSMUSG00000103636.2 | 0 | NA | NA | NA | NA | NA | #VALUE! |
| ENSMUSG00000103637.2 | 0 | NA | NA | NA | NA | NA | #VALUE! |
| ENSMUSG00000103638.2 | 0 | NA | NA | NA | NA | NA | #VALUE! |
| ENSMUSG00000103639.2 | 0 | NA | NA | NA | NA | NA | #VALUE! |
| ENSMUSG00000103640.2 | 0 | NA | NA | NA | NA | NA | #VALUE! |
| ENSMUSG00000103641.2 | 0 | NA | NA | NA | NA | NA | #VALUE! |
| ENSMUSG00000103642.2 | 0 | NA | NA | NA | NA | NA | #VALUE! |
| ENSMUSG00000103643.2 | 0 | NA | NA | NA | NA | NA | #VALUE! |
| ENSMUSG00000103644.2 | 0 | NA | NA | NA | NA | NA | #VALUE! |
| ENSMUSG00000103646.2 | 0 | NA | NA | NA | NA | NA | #VALUE! |
| ENSMUSG00000103647.2 | 0 | NA | NA | NA | NA | NA | #VALUE! |
| ENSMUSG00000103649.2 | 0 | NA | NA | NA | NA | NA | #VALUE! |
| ENSMUSG00000103650.2 | 0 | NA | NA | NA | NA | NA | #VALUE! |
| ENSMUSG00000103651.2 | 0 | NA | NA | NA | NA | NA | #VALUE! |
| ENSMUSG00000103652.2 | 0 | NA | NA | NA | NA | NA | #VALUE! |
| ENSMUSG00000103654.2 | 0 | NA | NA | NA | NA | NA | #VALUE! |
| ENSMUSG00000103655.2 | 0 | NA | NA | NA | NA | NA | #VALUE! |
| ENSMUSG00000103656.2 | 0 | NA | NA | NA | NA | NA | #VALUE! |
| ENSMUSG00000103657.2 | 0 | NA | NA | NA | NA | NA | #VALUE! |
| ENSMUSG00000103658.2 | 0 | NA | NA | NA | NA | NA | #VALUE! |
| ENSMUSG00000103659.2 | 0 | NA | NA | NA | NA | NA | #VALUE! |
| ENSMUSG00000103660.6 | 0 | NA | NA | NA | NA | NA | #VALUE! |
| ENSMUSG00000103662.2 | 0 | NA | NA | NA | NA | NA | #VALUE! |
| ENSMUSG00000103663.2 | 0 | NA | NA | NA | NA | NA | #VALUE! |
| ENSMUSG00000103664.6 | 0 | NA | NA | NA | NA | NA | #VALUE! |
| ENSMUSG00000103665.2 | 0 | NA | NA | NA | NA | NA | #VALUE! |

| | | | | | | | |
|---|---|---|---|---|---|---|---|
| ENSMUSG00000103666.2 | 0 | NA | NA | NA | NA | NA | #VALUE! |
| ENSMUSG00000103667.2 | 0 | NA | NA | NA | NA | NA | #VALUE! |
| ENSMUSG00000103668.2 | 0 | NA | NA | NA | NA | NA | #VALUE! |
| ENSMUSG00000103669.2 | 0 | NA | NA | NA | NA | NA | #VALUE! |
| ENSMUSG00000103670.2 | 0 | NA | NA | NA | NA | NA | #VALUE! |
| ENSMUSG00000103672.2 | 0 | NA | NA | NA | NA | NA | #VALUE! |
| ENSMUSG00000103673.2 | 0 | NA | NA | NA | NA | NA | #VALUE! |
| ENSMUSG00000103675.2 | 0 | NA | NA | NA | NA | NA | #VALUE! |
| ENSMUSG00000103676.2 | 0 | NA | NA | NA | NA | NA | #VALUE! |
| ENSMUSG00000103677.2 | 0 | NA | NA | NA | NA | NA | #VALUE! |
| ENSMUSG00000103678.2 | 0 | NA | NA | NA | NA | NA | #VALUE! |
| ENSMUSG00000103679.2 | 0 | NA | NA | NA | NA | NA | #VALUE! |
| ENSMUSG00000103680.2 | 0 | NA | NA | NA | NA | NA | #VALUE! |
| ENSMUSG00000103681.6 | 0 | NA | NA | NA | NA | NA | #VALUE! |
| ENSMUSG00000103684.2 | 0 | NA | NA | NA | NA | NA | #VALUE! |
| ENSMUSG00000103685.2 | 0 | NA | NA | NA | NA | NA | #VALUE! |
| ENSMUSG00000103686.2 | 0 | NA | NA | NA | NA | NA | #VALUE! |
| ENSMUSG00000103688.2 | 0 | NA | NA | NA | NA | NA | #VALUE! |
| ENSMUSG00000103689.2 | 0 | NA | NA | NA | NA | NA | #VALUE! |
| ENSMUSG00000103690.2 | 0 | NA | NA | NA | NA | NA | #VALUE! |
| ENSMUSG00000103691.2 | 7.085498 | 1.133505 | 3.679497 | 0.30806 | 0.758037 | NA | #VALUE! |
| ENSMUSG00000103692.2 | 0 | NA | NA | NA | NA | NA | #VALUE! |
| ENSMUSG00000103693.2 | 6.149388 | 6.104864 | 3.9141 | 1.559711 | 0.118828 | NA | #VALUE! |
| ENSMUSG00000103694.2 | 0 | NA | NA | NA | NA | NA | #VALUE! |
| ENSMUSG00000103695.2 | 0 | NA | NA | NA | NA | NA | #VALUE! |
| ENSMUSG00000103696.2 | 0 | NA | NA | NA | NA | NA | #VALUE! |
| ENSMUSG00000103697.2 | 0 | NA | NA | NA | NA | NA | #VALUE! |
| ENSMUSG00000103698.2 | 0 | NA | NA | NA | NA | NA | #VALUE! |
| ENSMUSG00000103699.2 | 0 | NA | NA | NA | NA | NA | #VALUE! |
| ENSMUSG00000103700.2 | 0 | NA | NA | NA | NA | NA | #VALUE! |
| ENSMUSG00000103701.2 | 0 | NA | NA | NA | NA | NA | #VALUE! |
| ENSMUSG00000103702.2 | 0 | NA | NA | NA | NA | NA | #VALUE! |
| ENSMUSG00000103703.3 | 2.338473 | -3.78176 | 3.918419 | -0.96513 | 0.334482 | NA | #VALUE! |
| ENSMUSG00000103704.2 | 0 | NA | NA | NA | NA | NA | #VALUE! |
| ENSMUSG00000103705.2 | 0 | NA | NA | NA | NA | NA | #VALUE! |
| ENSMUSG00000103706.2 | 0.841418 | 3.240539 | 3.959551 | 0.818411 | 0.413123 | NA | #VALUE! |
| ENSMUSG00000103707.6 | 0 | NA | NA | NA | NA | NA | #VALUE! |
| ENSMUSG00000103708.2 | 0 | NA | NA | NA | NA | NA | #VALUE! |
| ENSMUSG00000103709.2 | 0 | NA | NA | NA | NA | NA | #VALUE! |
| ENSMUSG00000103710.2 | 0 | NA | NA | NA | NA | NA | #VALUE! |
| ENSMUSG00000103712.2 | 0 | NA | NA | NA | NA | NA | #VALUE! |
| ENSMUSG00000103713.2 | 0 | NA | NA | NA | NA | NA | #VALUE! |
| ENSMUSG00000103714.2 | 0 | NA | NA | NA | NA | NA | #VALUE! |
| ENSMUSG00000103715.2 | 0.105013 | -0.97324 | 4.076293 | -0.23876 | 0.811295 | NA | #VALUE! |

| | | | | | | | |
|---|---|---|---|---|---|---|---|
| ENSMUSG00000103716.2 | 0 | NA | NA | NA | NA | NA | #VALUE! |
| ENSMUSG00000103717.2 | 0 | NA | NA | NA | NA | NA | #VALUE! |
| ENSMUSG00000103718.2 | 0 | NA | NA | NA | NA | NA | #VALUE! |
| ENSMUSG00000103719.2 | 0 | NA | NA | NA | NA | NA | #VALUE! |
| ENSMUSG00000103720.2 | 0 | NA | NA | NA | NA | NA | #VALUE! |
| ENSMUSG00000103722.2 | 0 | NA | NA | NA | NA | NA | #VALUE! |
| ENSMUSG00000103723.2 | 0 | NA | NA | NA | NA | NA | #VALUE! |
| ENSMUSG00000103724.3 | 0 | NA | NA | NA | NA | NA | #VALUE! |
| ENSMUSG00000103725.2 | 0 | NA | NA | NA | NA | NA | #VALUE! |
| ENSMUSG00000103726.2 | 0 | NA | NA | NA | NA | NA | #VALUE! |
| ENSMUSG00000103727.2 | 0 | NA | NA | NA | NA | NA | #VALUE! |
| ENSMUSG00000103728.2 | 0 | NA | NA | NA | NA | NA | #VALUE! |
| ENSMUSG00000103729.2 | 0 | NA | NA | NA | NA | NA | #VALUE! |
| ENSMUSG00000103730.2 | 0.816443 | -3.21914 | 3.961753 | -0.81255 | 0.416474 | NA | #VALUE! |
| ENSMUSG00000103731.2 | 1.492083 | 4.088114 | 3.936282 | 1.038572 | 0.299004 | NA | #VALUE! |
| ENSMUSG00000103732.2 | 0 | NA | NA | NA | NA | NA | #VALUE! |
| ENSMUSG00000103733.2 | 0 | NA | NA | NA | NA | NA | #VALUE! |
| ENSMUSG00000103734.2 | 0 | NA | NA | NA | NA | NA | #VALUE! |
| ENSMUSG00000103735.2 | 0 | NA | NA | NA | NA | NA | #VALUE! |
| ENSMUSG00000103736.2 | 0 | NA | NA | NA | NA | NA | #VALUE! |
| ENSMUSG00000103737.2 | 0 | NA | NA | NA | NA | NA | #VALUE! |
| ENSMUSG00000103738.2 | 0 | NA | NA | NA | NA | NA | #VALUE! |
| ENSMUSG00000103739.2 | 0 | NA | NA | NA | NA | NA | #VALUE! |
| ENSMUSG00000103740.2 | 0 | NA | NA | NA | NA | NA | #VALUE! |
| ENSMUSG00000103741.2 | 0 | NA | NA | NA | NA | NA | #VALUE! |
| ENSMUSG00000103742.2 | 0 | NA | NA | NA | NA | NA | #VALUE! |
| ENSMUSG00000103744.2 | 0 | NA | NA | NA | NA | NA | #VALUE! |
| ENSMUSG00000103745.2 | 0 | NA | NA | NA | NA | NA | #VALUE! |
| ENSMUSG00000103746.2 | 0 | NA | NA | NA | NA | NA | #VALUE! |
| ENSMUSG00000103747.2 | 0 | NA | NA | NA | NA | NA | #VALUE! |
| ENSMUSG00000103748.2 | 0 | NA | NA | NA | NA | NA | #VALUE! |
| ENSMUSG00000103749.2 | 0 | NA | NA | NA | NA | NA | #VALUE! |
| ENSMUSG00000103751.2 | 2.15368 | -4.63136 | 3.92757 | -1.17919 | 0.238321 | NA | #VALUE! |
| ENSMUSG00000103752.6 | 0 | NA | NA | NA | NA | NA | #VALUE! |
| ENSMUSG00000103753.2 | 0 | NA | NA | NA | NA | NA | #VALUE! |
| ENSMUSG00000103754.2 | 0.093255 | 0.868093 | 4.072464 | 0.213162 | 0.831201 | NA | #VALUE! |
| ENSMUSG00000103755.2 | 4.942525 | 5.801604 | 3.915812 | 1.481584 | 0.138451 | NA | #VALUE! |
| ENSMUSG00000103756.2 | 0 | NA | NA | NA | NA | NA | #VALUE! |
| ENSMUSG00000103757.2 | 0 | NA | NA | NA | NA | NA | #VALUE! |
| ENSMUSG00000103758.2 | 0 | NA | NA | NA | NA | NA | #VALUE! |
| ENSMUSG00000103759.2 | 0 | NA | NA | NA | NA | NA | #VALUE! |
| ENSMUSG00000103760.2 | 0 | NA | NA | NA | NA | NA | #VALUE! |
| ENSMUSG00000103761.2 | 0 | NA | NA | NA | NA | NA | #VALUE! |
| ENSMUSG00000103762.2 | 0 | NA | NA | NA | NA | NA | #VALUE! |

| | | | | | | | |
|---|---|---|---|---|---|---|---|
| ENSMUSG00000103763.2 | 0 | NA | NA | NA | NA | NA | #VALUE! |
| ENSMUSG00000103764.2 | 0 | NA | NA | NA | NA | NA | #VALUE! |
| ENSMUSG00000103765.2 | 0 | NA | NA | NA | NA | NA | #VALUE! |
| ENSMUSG00000103766.6 | 0 | NA | NA | NA | NA | NA | #VALUE! |
| ENSMUSG00000103767.2 | 0 | NA | NA | NA | NA | NA | #VALUE! |
| ENSMUSG00000103768.2 | 0 | NA | NA | NA | NA | NA | #VALUE! |
| ENSMUSG00000103769.2 | 0 | NA | NA | NA | NA | NA | #VALUE! |
| ENSMUSG00000103770.2 | 0 | NA | NA | NA | NA | NA | #VALUE! |
| ENSMUSG00000103771.2 | 0 | NA | NA | NA | NA | NA | #VALUE! |
| ENSMUSG00000103772.2 | 0 | NA | NA | NA | NA | NA | #VALUE! |
| ENSMUSG00000103773.2 | 0 | NA | NA | NA | NA | NA | #VALUE! |
| ENSMUSG00000103774.2 | 0 | NA | NA | NA | NA | NA | #VALUE! |
| ENSMUSG00000103775.2 | 0 | NA | NA | NA | NA | NA | #VALUE! |
| ENSMUSG00000103776.2 | 0 | NA | NA | NA | NA | NA | #VALUE! |
| ENSMUSG00000103778.2 | 0 | NA | NA | NA | NA | NA | #VALUE! |
| ENSMUSG00000103779.2 | 0 | NA | NA | NA | NA | NA | #VALUE! |
| ENSMUSG00000103780.2 | 0 | NA | NA | NA | NA | NA | #VALUE! |
| ENSMUSG00000103781.2 | 0 | NA | NA | NA | NA | NA | #VALUE! |
| ENSMUSG00000103782.2 | 0 | NA | NA | NA | NA | NA | #VALUE! |
| ENSMUSG00000103783.2 | 0 | NA | NA | NA | NA | NA | #VALUE! |
| ENSMUSG00000103784.2 | 0 | NA | NA | NA | NA | NA | #VALUE! |
| ENSMUSG00000103785.2 | 0 | NA | NA | NA | NA | NA | #VALUE! |
| ENSMUSG00000103786.2 | 0 | NA | NA | NA | NA | NA | #VALUE! |
| ENSMUSG00000103787.2 | 0 | NA | NA | NA | NA | NA | #VALUE! |
| ENSMUSG00000103788.2 | 0 | NA | NA | NA | NA | NA | #VALUE! |
| ENSMUSG00000103789.2 | 0 | NA | NA | NA | NA | NA | #VALUE! |
| ENSMUSG00000103790.2 | 0 | NA | NA | NA | NA | NA | #VALUE! |
| ENSMUSG00000103791.2 | 0 | NA | NA | NA | NA | NA | #VALUE! |
| ENSMUSG00000103792.2 | 0 | NA | NA | NA | NA | NA | #VALUE! |
| ENSMUSG00000103793.2 | 1.632886 | -4.22813 | 3.934241 | -1.0747 | 0.282509 | NA | #VALUE! |
| ENSMUSG00000103794.2 | 0 | NA | NA | NA | NA | NA | #VALUE! |
| ENSMUSG00000103795.2 | 0 | NA | NA | NA | NA | NA | #VALUE! |
| ENSMUSG00000103796.2 | 0 | NA | NA | NA | NA | NA | #VALUE! |
| ENSMUSG00000103797.2 | 0 | NA | NA | NA | NA | NA | #VALUE! |
| ENSMUSG00000103798.2 | 0 | NA | NA | NA | NA | NA | #VALUE! |
| ENSMUSG00000103799.2 | 0 | NA | NA | NA | NA | NA | #VALUE! |
| ENSMUSG00000103800.2 | 0 | NA | NA | NA | NA | NA | #VALUE! |
| ENSMUSG00000103801.2 | 0 | NA | NA | NA | NA | NA | #VALUE! |
| ENSMUSG00000103802.2 | 0 | NA | NA | NA | NA | NA | #VALUE! |
| ENSMUSG00000103803.2 | 0 | NA | NA | NA | NA | NA | #VALUE! |
| ENSMUSG00000103804.2 | 0 | NA | NA | NA | NA | NA | #VALUE! |
| ENSMUSG00000103805.2 | 0 | NA | NA | NA | NA | NA | #VALUE! |
| ENSMUSG00000103807.2 | 0 | NA | NA | NA | NA | NA | #VALUE! |
| ENSMUSG00000103808.2 | 0 | NA | NA | NA | NA | NA | #VALUE! |

| | | | | | | | |
|---|---|---|---|---|---|---|---|
| ENSMUSG00000103809.2 | 0 | NA | NA | NA | NA | NA | #VALUE! |
| ENSMUSG00000103810.2 | 0 | NA | NA | NA | NA | NA | #VALUE! |
| ENSMUSG00000103811.2 | 0 | NA | NA | NA | NA | NA | #VALUE! |
| ENSMUSG00000103812.6 | 0 | NA | NA | NA | NA | NA | #VALUE! |
| ENSMUSG00000103813.6 | 0 | NA | NA | NA | NA | NA | #VALUE! |
| ENSMUSG00000103814.2 | 0 | NA | NA | NA | NA | NA | #VALUE! |
| ENSMUSG00000103815.2 | 0 | NA | NA | NA | NA | NA | #VALUE! |
| ENSMUSG00000103816.2 | 0 | NA | NA | NA | NA | NA | #VALUE! |
| ENSMUSG00000103817.2 | 0 | NA | NA | NA | NA | NA | #VALUE! |
| ENSMUSG00000103818.2 | 0.945118 | -3.47564 | 3.952879 | -0.87927 | 0.379256 | NA | #VALUE! |
| ENSMUSG00000103819.2 | 0 | NA | NA | NA | NA | NA | #VALUE! |
| ENSMUSG00000103821.2 | 0 | NA | NA | NA | NA | NA | #VALUE! |
| ENSMUSG00000103822.2 | 0 | NA | NA | NA | NA | NA | #VALUE! |
| ENSMUSG00000103823.2 | 0 | NA | NA | NA | NA | NA | #VALUE! |
| ENSMUSG00000103824.2 | 0 | NA | NA | NA | NA | NA | #VALUE! |
| ENSMUSG00000103825.2 | 0 | NA | NA | NA | NA | NA | #VALUE! |
| ENSMUSG00000103827.2 | 0 | NA | NA | NA | NA | NA | #VALUE! |
| ENSMUSG00000103828.2 | 0 | NA | NA | NA | NA | NA | #VALUE! |
| ENSMUSG00000103829.6 | 0 | NA | NA | NA | NA | NA | #VALUE! |
| ENSMUSG00000103830.2 | 0 | NA | NA | NA | NA | NA | #VALUE! |
| ENSMUSG00000103831.2 | 0 | NA | NA | NA | NA | NA | #VALUE! |
| ENSMUSG00000103832.2 | 0 | NA | NA | NA | NA | NA | #VALUE! |
| ENSMUSG00000103833.2 | 0 | NA | NA | NA | NA | NA | #VALUE! |
| ENSMUSG00000103834.2 | 0 | NA | NA | NA | NA | NA | #VALUE! |
| ENSMUSG00000103835.2 | 0 | NA | NA | NA | NA | NA | #VALUE! |
| ENSMUSG00000103837.2 | 0 | NA | NA | NA | NA | NA | #VALUE! |
| ENSMUSG00000103839.2 | 6.735655 | -6.27951 | 3.913422 | -1.60461 | 0.10858 | NA | #VALUE! |
| ENSMUSG00000103840.2 | 0 | NA | NA | NA | NA | NA | #VALUE! |
| ENSMUSG00000103841.2 | 0 | NA | NA | NA | NA | NA | #VALUE! |
| ENSMUSG00000103842.2 | 0 | NA | NA | NA | NA | NA | #VALUE! |
| ENSMUSG00000103843.2 | 0 | NA | NA | NA | NA | NA | #VALUE! |
| ENSMUSG00000103844.2 | 0 | NA | NA | NA | NA | NA | #VALUE! |
| ENSMUSG00000103845.2 | 0 | NA | NA | NA | NA | NA | #VALUE! |
| ENSMUSG00000103846.2 | 0 | NA | NA | NA | NA | NA | #VALUE! |
| ENSMUSG00000103847.2 | 0 | NA | NA | NA | NA | NA | #VALUE! |
| ENSMUSG00000103848.2 | 0 | NA | NA | NA | NA | NA | #VALUE! |
| ENSMUSG00000103849.2 | 0 | NA | NA | NA | NA | NA | #VALUE! |
| ENSMUSG00000103850.2 | 0 | NA | NA | NA | NA | NA | #VALUE! |
| ENSMUSG00000103851.2 | 0.198268 | -0.06494 | 4.068276 | -0.01596 | 0.987265 | NA | #VALUE! |
| ENSMUSG00000103852.2 | 0 | NA | NA | NA | NA | NA | #VALUE! |
| ENSMUSG00000103854.2 | 0 | NA | NA | NA | NA | NA | #VALUE! |
| ENSMUSG00000103856.2 | 0 | NA | NA | NA | NA | NA | #VALUE! |
| ENSMUSG00000103857.2 | 0 | NA | NA | NA | NA | NA | #VALUE! |
| ENSMUSG00000103858.2 | 0 | NA | NA | NA | NA | NA | #VALUE! |

| | | | | | | | |
|---|---|---|---|---|---|---|---|
| ENSMUSG00000103859.2 | 0 | NA | NA | NA | NA | NA | #VALUE! |
| ENSMUSG00000103860.2 | 0 | NA | NA | NA | NA | NA | #VALUE! |
| ENSMUSG00000103862.4 | 0 | NA | NA | NA | NA | NA | #VALUE! |
| ENSMUSG00000103863.3 | 0 | NA | NA | NA | NA | NA | #VALUE! |
| ENSMUSG00000103864.2 | 0 | NA | NA | NA | NA | NA | #VALUE! |
| ENSMUSG00000103865.2 | 0 | NA | NA | NA | NA | NA | #VALUE! |
| ENSMUSG00000103867.2 | 0 | NA | NA | NA | NA | NA | #VALUE! |
| ENSMUSG00000103869.2 | 3.524204 | -5.34395 | 3.919482 | -1.36343 | 0.172746 | NA | #VALUE! |
| ENSMUSG00000103870.2 | 0 | NA | NA | NA | NA | NA | #VALUE! |
| ENSMUSG00000103871.2 | 0 | NA | NA | NA | NA | NA | #VALUE! |
| ENSMUSG00000103872.6 | 0 | NA | NA | NA | NA | NA | #VALUE! |
| ENSMUSG00000103873.2 | 0 | NA | NA | NA | NA | NA | #VALUE! |
| ENSMUSG00000103874.2 | 0 | NA | NA | NA | NA | NA | #VALUE! |
| ENSMUSG00000103875.2 | 0 | NA | NA | NA | NA | NA | #VALUE! |
| ENSMUSG00000103876.2 | 0 | NA | NA | NA | NA | NA | #VALUE! |
| ENSMUSG00000103877.3 | 0 | NA | NA | NA | NA | NA | #VALUE! |
| ENSMUSG00000103878.2 | 0 | NA | NA | NA | NA | NA | #VALUE! |
| ENSMUSG00000103880.2 | 0 | NA | NA | NA | NA | NA | #VALUE! |
| ENSMUSG00000103881.2 | 0 | NA | NA | NA | NA | NA | #VALUE! |
| ENSMUSG00000103882.2 | 0 | NA | NA | NA | NA | NA | #VALUE! |
| ENSMUSG00000103883.2 | 0 | NA | NA | NA | NA | NA | #VALUE! |
| ENSMUSG00000103884.2 | 0 | NA | NA | NA | NA | NA | #VALUE! |
| ENSMUSG00000103885.2 | 0 | NA | NA | NA | NA | NA | #VALUE! |
| ENSMUSG00000103886.2 | 0 | NA | NA | NA | NA | NA | #VALUE! |
| ENSMUSG00000103887.2 | 0 | NA | NA | NA | NA | NA | #VALUE! |
| ENSMUSG00000103889.2 | 0 | NA | NA | NA | NA | NA | #VALUE! |
| ENSMUSG00000103890.2 | 0 | NA | NA | NA | NA | NA | #VALUE! |
| ENSMUSG00000103891.2 | 0 | NA | NA | NA | NA | NA | #VALUE! |
| ENSMUSG00000103892.2 | 0 | NA | NA | NA | NA | NA | #VALUE! |
| ENSMUSG00000103893.2 | 0 | NA | NA | NA | NA | NA | #VALUE! |
| ENSMUSG00000103894.2 | 0 | NA | NA | NA | NA | NA | #VALUE! |
| ENSMUSG00000103895.2 | 0 | NA | NA | NA | NA | NA | #VALUE! |
| ENSMUSG00000103896.2 | 0 | NA | NA | NA | NA | NA | #VALUE! |
| ENSMUSG00000103897.2 | 0 | NA | NA | NA | NA | NA | #VALUE! |
| ENSMUSG00000103898.2 | 0 | NA | NA | NA | NA | NA | #VALUE! |
| ENSMUSG00000103899.2 | 0 | NA | NA | NA | NA | NA | #VALUE! |
| ENSMUSG00000103900.2 | 0 | NA | NA | NA | NA | NA | #VALUE! |
| ENSMUSG00000103901.2 | 0 | NA | NA | NA | NA | NA | #VALUE! |
| ENSMUSG00000103902.2 | 0 | NA | NA | NA | NA | NA | #VALUE! |
| ENSMUSG00000103903.2 | 0.105013 | -0.97324 | 4.076293 | -0.23876 | 0.811295 | NA | #VALUE! |
| ENSMUSG00000103904.2 | 0 | NA | NA | NA | NA | NA | #VALUE! |
| ENSMUSG00000103905.2 | 0 | NA | NA | NA | NA | NA | #VALUE! |
| ENSMUSG00000103906.2 | 0 | NA | NA | NA | NA | NA | #VALUE! |
| ENSMUSG00000103907.2 | 0 | NA | NA | NA | NA | NA | #VALUE! |

| | | | | | | | |
|---|---|---|---|---|---|---|---|
| ENSMUSG00000103908.2 | 0 | NA | NA | NA | NA | NA | #VALUE! |
| ENSMUSG00000103909.2 | 0 | NA | NA | NA | NA | NA | #VALUE! |
| ENSMUSG00000103911.2 | 0 | NA | NA | NA | NA | NA | #VALUE! |
| ENSMUSG00000103912.2 | 0 | NA | NA | NA | NA | NA | #VALUE! |
| ENSMUSG00000103913.2 | 0 | NA | NA | NA | NA | NA | #VALUE! |
| ENSMUSG00000103914.2 | 0 | NA | NA | NA | NA | NA | #VALUE! |
| ENSMUSG00000103915.2 | 0 | NA | NA | NA | NA | NA | #VALUE! |
| ENSMUSG00000103916.2 | 3.946036 | 4.501295 | 3.914075 | 1.150028 | 0.250132 | NA | #VALUE! |
| ENSMUSG00000103917.2 | 0 | NA | NA | NA | NA | NA | #VALUE! |
| ENSMUSG00000103918.2 | 0 | NA | NA | NA | NA | NA | #VALUE! |
| ENSMUSG00000103919.2 | 0 | NA | NA | NA | NA | NA | #VALUE! |
| ENSMUSG00000103920.2 | 0 | NA | NA | NA | NA | NA | #VALUE! |
| ENSMUSG00000103921.2 | 0.195789 | -1.05581 | 4.073302 | -0.2592 | 0.795479 | NA | #VALUE! |
| ENSMUSG00000103922.2 | 0 | NA | NA | NA | NA | NA | #VALUE! |
| ENSMUSG00000103923.2 | 0 | NA | NA | NA | NA | NA | #VALUE! |
| ENSMUSG00000103924.2 | 0 | NA | NA | NA | NA | NA | #VALUE! |
| ENSMUSG00000103925.2 | 0 | NA | NA | NA | NA | NA | #VALUE! |
| ENSMUSG00000103926.2 | 0 | NA | NA | NA | NA | NA | #VALUE! |
| ENSMUSG00000103927.2 | 0 | NA | NA | NA | NA | NA | #VALUE! |
| ENSMUSG00000103928.2 | 0 | NA | NA | NA | NA | NA | #VALUE! |
| ENSMUSG00000103929.2 | 0 | NA | NA | NA | NA | NA | #VALUE! |
| ENSMUSG00000103930.2 | 0 | NA | NA | NA | NA | NA | #VALUE! |
| ENSMUSG00000103931.2 | 0 | NA | NA | NA | NA | NA | #VALUE! |
| ENSMUSG00000103932.2 | 5.825721 | 6.026672 | 3.896315 | 1.546762 | 0.121921 | NA | #VALUE! |
| ENSMUSG00000103933.2 | 0.105013 | -0.97324 | 4.076293 | -0.23876 | 0.811295 | NA | #VALUE! |
| ENSMUSG00000103934.2 | 0 | NA | NA | NA | NA | NA | #VALUE! |
| ENSMUSG00000103935.2 | 0 | NA | NA | NA | NA | NA | #VALUE! |
| ENSMUSG00000103936.2 | 0 | NA | NA | NA | NA | NA | #VALUE! |
| ENSMUSG00000103937.2 | 0 | NA | NA | NA | NA | NA | #VALUE! |
| ENSMUSG00000103938.2 | 0 | NA | NA | NA | NA | NA | #VALUE! |
| ENSMUSG00000103939.2 | 0 | NA | NA | NA | NA | NA | #VALUE! |
| ENSMUSG00000103940.2 | 0 | NA | NA | NA | NA | NA | #VALUE! |
| ENSMUSG00000103942.2 | 0 | NA | NA | NA | NA | NA | #VALUE! |
| ENSMUSG00000103943.6 | 0 | NA | NA | NA | NA | NA | #VALUE! |
| ENSMUSG00000103944.2 | 0 | NA | NA | NA | NA | NA | #VALUE! |
| ENSMUSG00000103945.2 | 0 | NA | NA | NA | NA | NA | #VALUE! |
| ENSMUSG00000103947.2 | 0 | NA | NA | NA | NA | NA | #VALUE! |
| ENSMUSG00000103948.2 | 0 | NA | NA | NA | NA | NA | #VALUE! |
| ENSMUSG00000103949.2 | 0 | NA | NA | NA | NA | NA | #VALUE! |
| ENSMUSG00000103950.2 | 0 | NA | NA | NA | NA | NA | #VALUE! |
| ENSMUSG00000103951.2 | 0 | NA | NA | NA | NA | NA | #VALUE! |
| ENSMUSG00000103952.2 | 0 | NA | NA | NA | NA | NA | #VALUE! |
| ENSMUSG00000103953.2 | 0 | NA | NA | NA | NA | NA | #VALUE! |
| ENSMUSG00000103954.6 | 0 | NA | NA | NA | NA | NA | #VALUE! |

| | | | | | | | |
|---|---|---|---|---|---|---|---|
| ENSMUSG00000103955.2 | 0 | NA | NA | NA | NA | NA | #VALUE! |
| ENSMUSG00000103956.2 | 0 | NA | NA | NA | NA | NA | #VALUE! |
| ENSMUSG00000103957.2 | 0 | NA | NA | NA | NA | NA | #VALUE! |
| ENSMUSG00000103958.2 | 0 | NA | NA | NA | NA | NA | #VALUE! |
| ENSMUSG00000103959.3 | 0 | NA | NA | NA | NA | NA | #VALUE! |
| ENSMUSG00000103960.2 | 0 | NA | NA | NA | NA | NA | #VALUE! |
| ENSMUSG00000103961.2 | 0 | NA | NA | NA | NA | NA | #VALUE! |
| ENSMUSG00000103962.2 | 0 | NA | NA | NA | NA | NA | #VALUE! |
| ENSMUSG00000103963.2 | 0 | NA | NA | NA | NA | NA | #VALUE! |
| ENSMUSG00000103964.2 | 0 | NA | NA | NA | NA | NA | #VALUE! |
| ENSMUSG00000103965.2 | 0 | NA | NA | NA | NA | NA | #VALUE! |
| ENSMUSG00000103966.2 | 0 | NA | NA | NA | NA | NA | #VALUE! |
| ENSMUSG00000103967.2 | 0 | NA | NA | NA | NA | NA | #VALUE! |
| ENSMUSG00000103968.2 | 0 | NA | NA | NA | NA | NA | #VALUE! |
| ENSMUSG00000103969.2 | 0 | NA | NA | NA | NA | NA | #VALUE! |
| ENSMUSG00000103970.2 | 0 | NA | NA | NA | NA | NA | #VALUE! |
| ENSMUSG00000103971.2 | 0 | NA | NA | NA | NA | NA | #VALUE! |
| ENSMUSG00000103972.2 | 3.328415 | -5.26129 | 3.920229 | -1.34209 | 0.179567 | NA | #VALUE! |
| ENSMUSG00000103973.2 | 0 | NA | NA | NA | NA | NA | #VALUE! |
| ENSMUSG00000103975.2 | 3.870521 | 5.442409 | 3.918364 | 1.388949 | 0.164848 | NA | #VALUE! |
| ENSMUSG00000103976.2 | 1.009701 | 3.503687 | 3.950852 | 0.886818 | 0.375177 | NA | #VALUE! |
| ENSMUSG00000103977.6 | 0 | NA | NA | NA | NA | NA | #VALUE! |
| ENSMUSG00000103978.2 | 0 | NA | NA | NA | NA | NA | #VALUE! |
| ENSMUSG00000103979.2 | 0 | NA | NA | NA | NA | NA | #VALUE! |
| ENSMUSG00000103980.2 | 0 | NA | NA | NA | NA | NA | #VALUE! |
| ENSMUSG00000103981.2 | 0 | NA | NA | NA | NA | NA | #VALUE! |
| ENSMUSG00000103982.2 | 0 | NA | NA | NA | NA | NA | #VALUE! |
| ENSMUSG00000103983.2 | 0 | NA | NA | NA | NA | NA | #VALUE! |
| ENSMUSG00000103984.2 | 0.630079 | -2.90291 | 3.975047 | -0.73028 | 0.465217 | NA | #VALUE! |
| ENSMUSG00000103985.2 | 0 | NA | NA | NA | NA | NA | #VALUE! |
| ENSMUSG00000103986.2 | 0 | NA | NA | NA | NA | NA | #VALUE! |
| ENSMUSG00000103987.2 | 0 | NA | NA | NA | NA | NA | #VALUE! |
| ENSMUSG00000103989.2 | 0 | NA | NA | NA | NA | NA | #VALUE! |
| ENSMUSG00000103990.2 | 0 | NA | NA | NA | NA | NA | #VALUE! |
| ENSMUSG00000103991.2 | 0 | NA | NA | NA | NA | NA | #VALUE! |
| ENSMUSG00000103992.2 | 0 | NA | NA | NA | NA | NA | #VALUE! |
| ENSMUSG00000103993.2 | 0 | NA | NA | NA | NA | NA | #VALUE! |
| ENSMUSG00000103994.2 | 0 | NA | NA | NA | NA | NA | #VALUE! |
| ENSMUSG00000103995.2 | 0 | NA | NA | NA | NA | NA | #VALUE! |
| ENSMUSG00000103996.2 | 0 | NA | NA | NA | NA | NA | #VALUE! |
| ENSMUSG00000103997.2 | 0 | NA | NA | NA | NA | NA | #VALUE! |
| ENSMUSG00000103998.2 | 0 | NA | NA | NA | NA | NA | #VALUE! |
| ENSMUSG00000103999.2 | 0 | NA | NA | NA | NA | NA | #VALUE! |
| ENSMUSG00000104000.2 | 0 | NA | NA | NA | NA | NA | #VALUE! |

| | | | | | | | |
|---|---|---|---|---|---|---|---|
| ENSMUSG00000104001.2 | 0 | NA | NA | NA | NA | NA | #VALUE! |
| ENSMUSG00000104002.3 | 0 | NA | NA | NA | NA | NA | #VALUE! |
| ENSMUSG00000104004.2 | 0 | NA | NA | NA | NA | NA | #VALUE! |
| ENSMUSG00000104006.2 | 0 | NA | NA | NA | NA | NA | #VALUE! |
| ENSMUSG00000104007.2 | 0 | NA | NA | NA | NA | NA | #VALUE! |
| ENSMUSG00000104009.2 | 0 | NA | NA | NA | NA | NA | #VALUE! |
| ENSMUSG00000104010.2 | 0 | NA | NA | NA | NA | NA | #VALUE! |
| ENSMUSG00000104011.2 | 0 | NA | NA | NA | NA | NA | #VALUE! |
| ENSMUSG00000104012.2 | 0 | NA | NA | NA | NA | NA | #VALUE! |
| ENSMUSG00000104013.2 | 0 | NA | NA | NA | NA | NA | #VALUE! |
| ENSMUSG00000104014.2 | 0 | NA | NA | NA | NA | NA | #VALUE! |
| ENSMUSG00000104015.2 | 0 | NA | NA | NA | NA | NA | #VALUE! |
| ENSMUSG00000104016.2 | 0 | NA | NA | NA | NA | NA | #VALUE! |
| ENSMUSG00000104017.2 | 0 | NA | NA | NA | NA | NA | #VALUE! |
| ENSMUSG00000104018.2 | 0.737927 | 3.007825 | 3.968641 | 0.757898 | 0.448512 | NA | #VALUE! |
| ENSMUSG00000104020.2 | 0.354794 | 2.038933 | 4.025642 | 0.506487 | 0.612515 | NA | #VALUE! |
| ENSMUSG00000104021.2 | 0 | NA | NA | NA | NA | NA | #VALUE! |
| ENSMUSG00000104022.2 | 0 | NA | NA | NA | NA | NA | #VALUE! |
| ENSMUSG00000104023.2 | 0 | NA | NA | NA | NA | NA | #VALUE! |
| ENSMUSG00000104024.2 | 0 | NA | NA | NA | NA | NA | #VALUE! |
| ENSMUSG00000104025.2 | 0 | NA | NA | NA | NA | NA | #VALUE! |
| ENSMUSG00000104026.2 | 0 | NA | NA | NA | NA | NA | #VALUE! |
| ENSMUSG00000104027.2 | 0 | NA | NA | NA | NA | NA | #VALUE! |
| ENSMUSG00000104028.2 | 0 | NA | NA | NA | NA | NA | #VALUE! |
| ENSMUSG00000104029.2 | 0 | NA | NA | NA | NA | NA | #VALUE! |
| ENSMUSG00000104030.2 | 7.914342 | 0.524708 | 3.215676 | 0.163172 | 0.870383 | NA | #VALUE! |
| ENSMUSG00000104031.2 | 0 | NA | NA | NA | NA | NA | #VALUE! |
| ENSMUSG00000104032.2 | 0 | NA | NA | NA | NA | NA | #VALUE! |
| ENSMUSG00000104033.2 | 0 | NA | NA | NA | NA | NA | #VALUE! |
| ENSMUSG00000104034.2 | 0 | NA | NA | NA | NA | NA | #VALUE! |
| ENSMUSG00000104035.2 | 0 | NA | NA | NA | NA | NA | #VALUE! |
| ENSMUSG00000104036.2 | 0 | NA | NA | NA | NA | NA | #VALUE! |
| ENSMUSG00000104037.2 | 0 | NA | NA | NA | NA | NA | #VALUE! |
| ENSMUSG00000104038.2 | 0 | NA | NA | NA | NA | NA | #VALUE! |
| ENSMUSG00000104039.2 | 0 | NA | NA | NA | NA | NA | #VALUE! |
| ENSMUSG00000104040.2 | 0 | NA | NA | NA | NA | NA | #VALUE! |
| ENSMUSG00000104041.2 | 0 | NA | NA | NA | NA | NA | #VALUE! |
| ENSMUSG00000104042.2 | 0 | NA | NA | NA | NA | NA | #VALUE! |
| ENSMUSG00000104043.2 | 0.105013 | -0.97324 | 4.076293 | -0.23876 | 0.811295 | NA | #VALUE! |
| ENSMUSG00000104044.2 | 4.711939 | 5.72619 | 3.916297 | 1.462144 | 0.143702 | NA | #VALUE! |
| ENSMUSG00000104045.2 | 0 | NA | NA | NA | NA | NA | #VALUE! |
| ENSMUSG00000104046.2 | 0 | NA | NA | NA | NA | NA | #VALUE! |
| ENSMUSG00000104048.2 | 0 | NA | NA | NA | NA | NA | #VALUE! |
| ENSMUSG00000104050.2 | 0 | NA | NA | NA | NA | NA | #VALUE! |

| | | | | | | | |
|---|---|---|---|---|---|---|---|
| ENSMUSG00000104051.2 | 0 | NA | NA | NA | NA | NA | #VALUE! |
| ENSMUSG00000104052.2 | 0 | NA | NA | NA | NA | NA | #VALUE! |
| ENSMUSG00000104053.2 | 0 | NA | NA | NA | NA | NA | #VALUE! |
| ENSMUSG00000104054.2 | 0 | NA | NA | NA | NA | NA | #VALUE! |
| ENSMUSG00000104055.2 | 0 | NA | NA | NA | NA | NA | #VALUE! |
| ENSMUSG00000104057.2 | 0 | NA | NA | NA | NA | NA | #VALUE! |
| ENSMUSG00000104059.2 | 0 | NA | NA | NA | NA | NA | #VALUE! |
| ENSMUSG00000104060.2 | 0 | NA | NA | NA | NA | NA | #VALUE! |
| ENSMUSG00000104061.2 | 0 | NA | NA | NA | NA | NA | #VALUE! |
| ENSMUSG00000104062.6 | 0 | NA | NA | NA | NA | NA | #VALUE! |
| ENSMUSG00000104063.2 | 0 | NA | NA | NA | NA | NA | #VALUE! |
| ENSMUSG00000104064.2 | 0 | NA | NA | NA | NA | NA | #VALUE! |
| ENSMUSG00000104065.2 | 0 | NA | NA | NA | NA | NA | #VALUE! |
| ENSMUSG00000104066.2 | 0 | NA | NA | NA | NA | NA | #VALUE! |
| ENSMUSG00000104067.2 | 0 | NA | NA | NA | NA | NA | #VALUE! |
| ENSMUSG00000104068.2 | 0 | NA | NA | NA | NA | NA | #VALUE! |
| ENSMUSG00000104069.2 | 0 | NA | NA | NA | NA | NA | #VALUE! |
| ENSMUSG00000104070.2 | 0 | NA | NA | NA | NA | NA | #VALUE! |
| ENSMUSG00000104071.2 | 0 | NA | NA | NA | NA | NA | #VALUE! |
| ENSMUSG00000104072.2 | 0 | NA | NA | NA | NA | NA | #VALUE! |
| ENSMUSG00000104073.2 | 0 | NA | NA | NA | NA | NA | #VALUE! |
| ENSMUSG00000104074.2 | 0 | NA | NA | NA | NA | NA | #VALUE! |
| ENSMUSG00000104075.2 | 0 | NA | NA | NA | NA | NA | #VALUE! |
| ENSMUSG00000104076.2 | 0 | NA | NA | NA | NA | NA | #VALUE! |
| ENSMUSG00000104077.2 | 0 | NA | NA | NA | NA | NA | #VALUE! |
| ENSMUSG00000104078.2 | 0 | NA | NA | NA | NA | NA | #VALUE! |
| ENSMUSG00000104079.2 | 0 | NA | NA | NA | NA | NA | #VALUE! |
| ENSMUSG00000104080.2 | 0 | NA | NA | NA | NA | NA | #VALUE! |
| ENSMUSG00000104081.2 | 0 | NA | NA | NA | NA | NA | #VALUE! |
| ENSMUSG00000104082.3 | 0 | NA | NA | NA | NA | NA | #VALUE! |
| ENSMUSG00000104083.2 | 0 | NA | NA | NA | NA | NA | #VALUE! |
| ENSMUSG00000104084.2 | 0 | NA | NA | NA | NA | NA | #VALUE! |
| ENSMUSG00000104085.2 | 0 | NA | NA | NA | NA | NA | #VALUE! |
| ENSMUSG00000104086.2 | 0 | NA | NA | NA | NA | NA | #VALUE! |
| ENSMUSG00000104087.2 | 0 | NA | NA | NA | NA | NA | #VALUE! |
| ENSMUSG00000104088.2 | 0 | NA | NA | NA | NA | NA | #VALUE! |
| ENSMUSG00000104089.2 | 0 | NA | NA | NA | NA | NA | #VALUE! |
| ENSMUSG00000104091.2 | 0 | NA | NA | NA | NA | NA | #VALUE! |
| ENSMUSG00000104092.3 | 0 | NA | NA | NA | NA | NA | #VALUE! |
| ENSMUSG00000104093.2 | 0 | NA | NA | NA | NA | NA | #VALUE! |
| ENSMUSG00000104094.2 | 0 | NA | NA | NA | NA | NA | #VALUE! |
| ENSMUSG00000104095.2 | 0 | NA | NA | NA | NA | NA | #VALUE! |
| ENSMUSG00000104096.2 | 0 | NA | NA | NA | NA | NA | #VALUE! |
| ENSMUSG00000104097.2 | 0 | NA | NA | NA | NA | NA | #VALUE! |

| | | | | | | | |
|---|---|---|---|---|---|---|---|
| ENSMUSG00000104098.2 | 0 | NA | NA | NA | NA | NA | #VALUE! |
| ENSMUSG00000104099.2 | 0 | NA | NA | NA | NA | NA | #VALUE! |
| ENSMUSG00000104100.2 | 0 | NA | NA | NA | NA | NA | #VALUE! |
| ENSMUSG00000104101.2 | 0 | NA | NA | NA | NA | NA | #VALUE! |
| ENSMUSG00000104102.2 | 0 | NA | NA | NA | NA | NA | #VALUE! |
| ENSMUSG00000104103.2 | 0.195789 | -1.05581 | 4.073302 | -0.2592 | 0.795479 | NA | #VALUE! |
| ENSMUSG00000104104.2 | 0 | NA | NA | NA | NA | NA | #VALUE! |
| ENSMUSG00000104105.2 | 0 | NA | NA | NA | NA | NA | #VALUE! |
| ENSMUSG00000104106.2 | 0 | NA | NA | NA | NA | NA | #VALUE! |
| ENSMUSG00000104107.2 | 0 | NA | NA | NA | NA | NA | #VALUE! |
| ENSMUSG00000104108.2 | 0 | NA | NA | NA | NA | NA | #VALUE! |
| ENSMUSG00000104109.2 | 0 | NA | NA | NA | NA | NA | #VALUE! |
| ENSMUSG00000104110.2 | 0 | NA | NA | NA | NA | NA | #VALUE! |
| ENSMUSG00000104111.2 | 0 | NA | NA | NA | NA | NA | #VALUE! |
| ENSMUSG00000104112.2 | 0 | NA | NA | NA | NA | NA | #VALUE! |
| ENSMUSG00000104113.2 | 0 | NA | NA | NA | NA | NA | #VALUE! |
| ENSMUSG00000104114.2 | 0 | NA | NA | NA | NA | NA | #VALUE! |
| ENSMUSG00000104115.2 | 0 | NA | NA | NA | NA | NA | #VALUE! |
| ENSMUSG00000104116.2 | 2.65344 | -4.9321 | 3.923668 | -1.25701 | 0.208749 | NA | #VALUE! |
| ENSMUSG00000104117.6 | 0 | NA | NA | NA | NA | NA | #VALUE! |
| ENSMUSG00000104118.2 | 0 | NA | NA | NA | NA | NA | #VALUE! |
| ENSMUSG00000104120.2 | 0 | NA | NA | NA | NA | NA | #VALUE! |
| ENSMUSG00000104121.2 | 0 | NA | NA | NA | NA | NA | #VALUE! |
| ENSMUSG00000104122.2 | 0 | NA | NA | NA | NA | NA | #VALUE! |
| ENSMUSG00000104123.2 | 0 | NA | NA | NA | NA | NA | #VALUE! |
| ENSMUSG00000104125.2 | 0 | NA | NA | NA | NA | NA | #VALUE! |
| ENSMUSG00000104126.2 | 0 | NA | NA | NA | NA | NA | #VALUE! |
| ENSMUSG00000104127.2 | 0 | NA | NA | NA | NA | NA | #VALUE! |
| ENSMUSG00000104129.2 | 0 | NA | NA | NA | NA | NA | #VALUE! |
| ENSMUSG00000104130.2 | 0 | NA | NA | NA | NA | NA | #VALUE! |
| ENSMUSG00000104131.6 | 0 | NA | NA | NA | NA | NA | #VALUE! |
| ENSMUSG00000104132.2 | 0 | NA | NA | NA | NA | NA | #VALUE! |
| ENSMUSG00000104133.2 | 0 | NA | NA | NA | NA | NA | #VALUE! |
| ENSMUSG00000104134.2 | 0 | NA | NA | NA | NA | NA | #VALUE! |
| ENSMUSG00000104135.2 | 0 | NA | NA | NA | NA | NA | #VALUE! |
| ENSMUSG00000104136.2 | 0 | NA | NA | NA | NA | NA | #VALUE! |
| ENSMUSG00000104137.2 | 0 | NA | NA | NA | NA | NA | #VALUE! |
| ENSMUSG00000104138.2 | 0 | NA | NA | NA | NA | NA | #VALUE! |
| ENSMUSG00000104140.2 | 0 | NA | NA | NA | NA | NA | #VALUE! |
| ENSMUSG00000104141.2 | 0 | NA | NA | NA | NA | NA | #VALUE! |
| ENSMUSG00000104142.2 | 0 | NA | NA | NA | NA | NA | #VALUE! |
| ENSMUSG00000104144.2 | 0 | NA | NA | NA | NA | NA | #VALUE! |
| ENSMUSG00000104145.2 | 0 | NA | NA | NA | NA | NA | #VALUE! |
| ENSMUSG00000104146.2 | 0 | NA | NA | NA | NA | NA | #VALUE! |

| | | | | | | | |
|---|---|---|---|---|---|---|---|
| ENSMUSG00000104147.2 | 0 | NA | NA | NA | NA | NA | #VALUE! |
| ENSMUSG00000104148.6 | 0 | NA | NA | NA | NA | NA | #VALUE! |
| ENSMUSG00000104149.2 | 1.305573 | 3.898287 | 3.940396 | 0.989314 | 0.32251 | NA | #VALUE! |
| ENSMUSG00000104150.3 | 0 | NA | NA | NA | NA | NA | #VALUE! |
| ENSMUSG00000104152.2 | 0 | NA | NA | NA | NA | NA | #VALUE! |
| ENSMUSG00000104153.2 | 0 | NA | NA | NA | NA | NA | #VALUE! |
| ENSMUSG00000104154.2 | 0 | NA | NA | NA | NA | NA | #VALUE! |
| ENSMUSG00000104155.2 | 0 | NA | NA | NA | NA | NA | #VALUE! |
| ENSMUSG00000104156.3 | 0 | NA | NA | NA | NA | NA | #VALUE! |
| ENSMUSG00000104157.2 | 0 | NA | NA | NA | NA | NA | #VALUE! |
| ENSMUSG00000104158.3 | 0 | NA | NA | NA | NA | NA | #VALUE! |
| ENSMUSG00000104159.2 | 0 | NA | NA | NA | NA | NA | #VALUE! |
| ENSMUSG00000104161.2 | 0 | NA | NA | NA | NA | NA | #VALUE! |
| ENSMUSG00000104162.2 | 0 | NA | NA | NA | NA | NA | #VALUE! |
| ENSMUSG00000104163.2 | 0 | NA | NA | NA | NA | NA | #VALUE! |
| ENSMUSG00000104164.2 | 0 | NA | NA | NA | NA | NA | #VALUE! |
| ENSMUSG00000104165.2 | 0 | NA | NA | NA | NA | NA | #VALUE! |
| ENSMUSG00000104166.2 | 0 | NA | NA | NA | NA | NA | #VALUE! |
| ENSMUSG00000104167.2 | 0 | NA | NA | NA | NA | NA | #VALUE! |
| ENSMUSG00000104168.2 | 0 | NA | NA | NA | NA | NA | #VALUE! |
| ENSMUSG00000104169.2 | 0 | NA | NA | NA | NA | NA | #VALUE! |
| ENSMUSG00000104170.2 | 0 | NA | NA | NA | NA | NA | #VALUE! |
| ENSMUSG00000104171.2 | 0 | NA | NA | NA | NA | NA | #VALUE! |
| ENSMUSG00000104172.2 | 0 | NA | NA | NA | NA | NA | #VALUE! |
| ENSMUSG00000104173.2 | 0 | NA | NA | NA | NA | NA | #VALUE! |
| ENSMUSG00000104174.2 | 0 | NA | NA | NA | NA | NA | #VALUE! |
| ENSMUSG00000104175.2 | 0 | NA | NA | NA | NA | NA | #VALUE! |
| ENSMUSG00000104176.2 | 0 | NA | NA | NA | NA | NA | #VALUE! |
| ENSMUSG00000104177.2 | 0 | NA | NA | NA | NA | NA | #VALUE! |
| ENSMUSG00000104178.2 | 0 | NA | NA | NA | NA | NA | #VALUE! |
| ENSMUSG00000104180.2 | 0 | NA | NA | NA | NA | NA | #VALUE! |
| ENSMUSG00000104181.2 | 0 | NA | NA | NA | NA | NA | #VALUE! |
| ENSMUSG00000104182.2 | 0 | NA | NA | NA | NA | NA | #VALUE! |
| ENSMUSG00000104183.2 | 0 | NA | NA | NA | NA | NA | #VALUE! |
| ENSMUSG00000104184.2 | 0 | NA | NA | NA | NA | NA | #VALUE! |
| ENSMUSG00000104185.2 | 0 | NA | NA | NA | NA | NA | #VALUE! |
| ENSMUSG00000104186.2 | 0 | NA | NA | NA | NA | NA | #VALUE! |
| ENSMUSG00000104187.2 | 0 | NA | NA | NA | NA | NA | #VALUE! |
| ENSMUSG00000104188.2 | 0 | NA | NA | NA | NA | NA | #VALUE! |
| ENSMUSG00000104189.2 | 0 | NA | NA | NA | NA | NA | #VALUE! |
| ENSMUSG00000104190.2 | 0 | NA | NA | NA | NA | NA | #VALUE! |
| ENSMUSG00000104191.2 | 0 | NA | NA | NA | NA | NA | #VALUE! |
| ENSMUSG00000104192.2 | 0 | NA | NA | NA | NA | NA | #VALUE! |
| ENSMUSG00000104194.2 | 0 | NA | NA | NA | NA | NA | #VALUE! |

| | | | | | | | |
|---|---|---|---|---|---|---|---|
| ENSMUSG00000104195.2 | 0.210026 | -1.52494 | 4.058692 | -0.37572 | 0.707123 | NA | #VALUE! |
| ENSMUSG00000104197.2 | 0 | NA | NA | NA | NA | NA | #VALUE! |
| ENSMUSG00000104198.2 | 0 | NA | NA | NA | NA | NA | #VALUE! |
| ENSMUSG00000104199.2 | 0 | NA | NA | NA | NA | NA | #VALUE! |
| ENSMUSG00000104200.2 | 0 | NA | NA | NA | NA | NA | #VALUE! |
| ENSMUSG00000104201.2 | 0 | NA | NA | NA | NA | NA | #VALUE! |
| ENSMUSG00000104202.2 | 0 | NA | NA | NA | NA | NA | #VALUE! |
| ENSMUSG00000104203.2 | 0 | NA | NA | NA | NA | NA | #VALUE! |
| ENSMUSG00000104205.2 | 0 | NA | NA | NA | NA | NA | #VALUE! |
| ENSMUSG00000104206.2 | 0 | NA | NA | NA | NA | NA | #VALUE! |
| ENSMUSG00000104207.2 | 0 | NA | NA | NA | NA | NA | #VALUE! |
| ENSMUSG00000104209.2 | 0 | NA | NA | NA | NA | NA | #VALUE! |
| ENSMUSG00000104210.2 | 0 | NA | NA | NA | NA | NA | #VALUE! |
| ENSMUSG00000104211.2 | 0 | NA | NA | NA | NA | NA | #VALUE! |
| ENSMUSG00000104212.2 | 0 | NA | NA | NA | NA | NA | #VALUE! |
| ENSMUSG00000104213.6 | 2.310289 | -4.7496 | 3.925939 | -1.2098 | 0.226355 | NA | #VALUE! |
| ENSMUSG00000104214.2 | 0 | NA | NA | NA | NA | NA | #VALUE! |
| ENSMUSG00000104215.2 | 0 | NA | NA | NA | NA | NA | #VALUE! |
| ENSMUSG00000104216.2 | 0 | NA | NA | NA | NA | NA | #VALUE! |
| ENSMUSG00000104217.2 | 0 | NA | NA | NA | NA | NA | #VALUE! |
| ENSMUSG00000104218.2 | 0.105013 | -0.97324 | 4.076293 | -0.23876 | 0.811295 | NA | #VALUE! |
| ENSMUSG00000104219.2 | 0 | NA | NA | NA | NA | NA | #VALUE! |
| ENSMUSG00000104220.2 | 0 | NA | NA | NA | NA | NA | #VALUE! |
| ENSMUSG00000104221.2 | 0 | NA | NA | NA | NA | NA | #VALUE! |
| ENSMUSG00000104222.2 | 15.48208 | 20.85669 | 3.152387 | 6.61616 | NA | NA | #VALUE! |
| ENSMUSG00000104223.2 | 0 | NA | NA | NA | NA | NA | #VALUE! |
| ENSMUSG00000104224.2 | 0 | NA | NA | NA | NA | NA | #VALUE! |
| ENSMUSG00000104225.2 | 0 | NA | NA | NA | NA | NA | #VALUE! |
| ENSMUSG00000104226.2 | 0 | NA | NA | NA | NA | NA | #VALUE! |
| ENSMUSG00000104227.2 | 0 | NA | NA | NA | NA | NA | #VALUE! |
| ENSMUSG00000104228.2 | 0 | NA | NA | NA | NA | NA | #VALUE! |
| ENSMUSG00000104229.2 | 0 | NA | NA | NA | NA | NA | #VALUE! |
| ENSMUSG00000104230.2 | 0 | NA | NA | NA | NA | NA | #VALUE! |
| ENSMUSG00000104231.2 | 0 | NA | NA | NA | NA | NA | #VALUE! |
| ENSMUSG00000104232.2 | 0 | NA | NA | NA | NA | NA | #VALUE! |
| ENSMUSG00000104233.2 | 0 | NA | NA | NA | NA | NA | #VALUE! |
| ENSMUSG00000104234.2 | 0 | NA | NA | NA | NA | NA | #VALUE! |
| ENSMUSG00000104235.2 | 0 | NA | NA | NA | NA | NA | #VALUE! |
| ENSMUSG00000104236.2 | 0 | NA | NA | NA | NA | NA | #VALUE! |
| ENSMUSG00000104237.2 | 0 | NA | NA | NA | NA | NA | #VALUE! |
| ENSMUSG00000104238.2 | 0 | NA | NA | NA | NA | NA | #VALUE! |
| ENSMUSG00000104239.2 | 0 | NA | NA | NA | NA | NA | #VALUE! |
| ENSMUSG00000104240.2 | 0 | NA | NA | NA | NA | NA | #VALUE! |
| ENSMUSG00000104241.2 | 0 | NA | NA | NA | NA | NA | #VALUE! |

| | | | | | | | |
|---|---|---|---|---|---|---|---|
| ENSMUSG00000104242.2 | 0 | NA | NA | NA | NA | NA | #VALUE! |
| ENSMUSG00000104243.2 | 0 | NA | NA | NA | NA | NA | #VALUE! |
| ENSMUSG00000104244.2 | 0 | NA | NA | NA | NA | NA | #VALUE! |
| ENSMUSG00000104245.2 | 0 | NA | NA | NA | NA | NA | #VALUE! |
| ENSMUSG00000104247.2 | 0 | NA | NA | NA | NA | NA | #VALUE! |
| ENSMUSG00000104248.2 | 0.105013 | -0.97324 | 4.076293 | -0.23876 | 0.811295 | NA | #VALUE! |
| ENSMUSG00000104249.2 | 0 | NA | NA | NA | NA | NA | #VALUE! |
| ENSMUSG00000104251.2 | 0 | NA | NA | NA | NA | NA | #VALUE! |
| ENSMUSG00000104252.6 | 0 | NA | NA | NA | NA | NA | #VALUE! |
| ENSMUSG00000104253.2 | 8.150982 | 6.511708 | 3.425924 | 1.900716 | 0.057339 | NA | #VALUE! |
| ENSMUSG00000104254.2 | 0 | NA | NA | NA | NA | NA | #VALUE! |
| ENSMUSG00000104255.2 | 0 | NA | NA | NA | NA | NA | #VALUE! |
| ENSMUSG00000104256.3 | 0 | NA | NA | NA | NA | NA | #VALUE! |
| ENSMUSG00000104257.2 | 0 | NA | NA | NA | NA | NA | #VALUE! |
| ENSMUSG00000104258.2 | 0 | NA | NA | NA | NA | NA | #VALUE! |
| ENSMUSG00000104259.2 | 0 | NA | NA | NA | NA | NA | #VALUE! |
| ENSMUSG00000104260.2 | 0 | NA | NA | NA | NA | NA | #VALUE! |
| ENSMUSG00000104261.2 | 0 | NA | NA | NA | NA | NA | #VALUE! |
| ENSMUSG00000104262.2 | 0 | NA | NA | NA | NA | NA | #VALUE! |
| ENSMUSG00000104263.2 | 2.310289 | -4.7496 | 3.925939 | -1.2098 | 0.226355 | NA | #VALUE! |
| ENSMUSG00000104264.2 | 0 | NA | NA | NA | NA | NA | #VALUE! |
| ENSMUSG00000104265.2 | 0 | NA | NA | NA | NA | NA | #VALUE! |
| ENSMUSG00000104266.2 | 0 | NA | NA | NA | NA | NA | #VALUE! |
| ENSMUSG00000104267.2 | 0 | NA | NA | NA | NA | NA | #VALUE! |
| ENSMUSG00000104268.2 | 0 | NA | NA | NA | NA | NA | #VALUE! |
| ENSMUSG00000104269.2 | 0 | NA | NA | NA | NA | NA | #VALUE! |
| ENSMUSG00000104270.2 | 0 | NA | NA | NA | NA | NA | #VALUE! |
| ENSMUSG00000104271.2 | 0 | NA | NA | NA | NA | NA | #VALUE! |
| ENSMUSG00000104272.2 | 0 | NA | NA | NA | NA | NA | #VALUE! |
| ENSMUSG00000104273.2 | 0 | NA | NA | NA | NA | NA | #VALUE! |
| ENSMUSG00000104274.2 | 0 | NA | NA | NA | NA | NA | #VALUE! |
| ENSMUSG00000104275.2 | 0 | NA | NA | NA | NA | NA | #VALUE! |
| ENSMUSG00000104276.2 | 0 | NA | NA | NA | NA | NA | #VALUE! |
| ENSMUSG00000104277.2 | 0 | NA | NA | NA | NA | NA | #VALUE! |
| ENSMUSG00000104278.2 | 0 | NA | NA | NA | NA | NA | #VALUE! |
| ENSMUSG00000104279.2 | 3.050109 | -2.33015 | 3.834368 | -0.6077 | 0.543386 | NA | #VALUE! |
| ENSMUSG00000104280.2 | 0.168284 | 1.027628 | 4.067126 | 0.252667 | 0.800526 | NA | #VALUE! |
| ENSMUSG00000104281.2 | 0 | NA | NA | NA | NA | NA | #VALUE! |
| ENSMUSG00000104282.2 | 0 | NA | NA | NA | NA | NA | #VALUE! |
| ENSMUSG00000104283.2 | 0 | NA | NA | NA | NA | NA | #VALUE! |
| ENSMUSG00000104284.2 | 0 | NA | NA | NA | NA | NA | #VALUE! |
| ENSMUSG00000104285.2 | 0 | NA | NA | NA | NA | NA | #VALUE! |
| ENSMUSG00000104286.2 | 0 | NA | NA | NA | NA | NA | #VALUE! |
| ENSMUSG00000104287.2 | 0 | NA | NA | NA | NA | NA | #VALUE! |

| | | | | | | | |
|---|---|---|---|---|---|---|---|
| ENSMUSG00000104288.2 | 0 | NA | NA | NA | NA | NA | #VALUE! |
| ENSMUSG00000104289.6 | 0 | NA | NA | NA | NA | NA | #VALUE! |
| ENSMUSG00000104290.2 | 0 | NA | NA | NA | NA | NA | #VALUE! |
| ENSMUSG00000104291.2 | 0.391578 | -2.1489 | 4.020883 | -0.53444 | 0.59304 | NA | #VALUE! |
| ENSMUSG00000104292.2 | 2.741047 | -4.98044 | 3.923113 | -1.26951 | 0.204259 | NA | #VALUE! |
| ENSMUSG00000104293.2 | 0 | NA | NA | NA | NA | NA | #VALUE! |
| ENSMUSG00000104294.2 | 0 | NA | NA | NA | NA | NA | #VALUE! |
| ENSMUSG00000104295.2 | 0 | NA | NA | NA | NA | NA | #VALUE! |
| ENSMUSG00000104296.2 | 0 | NA | NA | NA | NA | NA | #VALUE! |
| ENSMUSG00000104297.2 | 0 | NA | NA | NA | NA | NA | #VALUE! |
| ENSMUSG00000104298.2 | 0 | NA | NA | NA | NA | NA | #VALUE! |
| ENSMUSG00000104299.2 | 0 | NA | NA | NA | NA | NA | #VALUE! |
| ENSMUSG00000104301.4 | 0 | NA | NA | NA | NA | NA | #VALUE! |
| ENSMUSG00000104302.2 | 0 | NA | NA | NA | NA | NA | #VALUE! |
| ENSMUSG00000104303.2 | 0 | NA | NA | NA | NA | NA | #VALUE! |
| ENSMUSG00000104304.2 | 0 | NA | NA | NA | NA | NA | #VALUE! |
| ENSMUSG00000104306.2 | 0 | NA | NA | NA | NA | NA | #VALUE! |
| ENSMUSG00000104307.2 | 0 | NA | NA | NA | NA | NA | #VALUE! |
| ENSMUSG00000104308.2 | 0 | NA | NA | NA | NA | NA | #VALUE! |
| ENSMUSG00000104309.2 | 0 | NA | NA | NA | NA | NA | #VALUE! |
| ENSMUSG00000104310.2 | 0 | NA | NA | NA | NA | NA | #VALUE! |
| ENSMUSG00000104311.2 | 0 | NA | NA | NA | NA | NA | #VALUE! |
| ENSMUSG00000104312.2 | 0 | NA | NA | NA | NA | NA | #VALUE! |
| ENSMUSG00000104313.2 | 0 | NA | NA | NA | NA | NA | #VALUE! |
| ENSMUSG00000104314.2 | 0 | NA | NA | NA | NA | NA | #VALUE! |
| ENSMUSG00000104315.2 | 0 | NA | NA | NA | NA | NA | #VALUE! |
| ENSMUSG00000104316.2 | 0 | NA | NA | NA | NA | NA | #VALUE! |
| ENSMUSG00000104317.2 | 0 | NA | NA | NA | NA | NA | #VALUE! |
| ENSMUSG00000104318.2 | 0 | NA | NA | NA | NA | NA | #VALUE! |
| ENSMUSG00000104319.2 | 0 | NA | NA | NA | NA | NA | #VALUE! |
| ENSMUSG00000104320.2 | 0 | NA | NA | NA | NA | NA | #VALUE! |
| ENSMUSG00000104321.2 | 0 | NA | NA | NA | NA | NA | #VALUE! |
| ENSMUSG00000104323.2 | 0 | NA | NA | NA | NA | NA | #VALUE! |
| ENSMUSG00000104324.2 | 0 | NA | NA | NA | NA | NA | #VALUE! |
| ENSMUSG00000104325.2 | 0 | NA | NA | NA | NA | NA | #VALUE! |
| ENSMUSG00000104326.2 | 0 | NA | NA | NA | NA | NA | #VALUE! |
| ENSMUSG00000104327.2 | 0 | NA | NA | NA | NA | NA | #VALUE! |
| ENSMUSG00000104328.2 | 0 | NA | NA | NA | NA | NA | #VALUE! |
| ENSMUSG00000104329.2 | 0 | NA | NA | NA | NA | NA | #VALUE! |
| ENSMUSG00000104330.2 | 0 | NA | NA | NA | NA | NA | #VALUE! |
| ENSMUSG00000104331.2 | 0 | NA | NA | NA | NA | NA | #VALUE! |
| ENSMUSG00000104332.2 | 0 | NA | NA | NA | NA | NA | #VALUE! |
| ENSMUSG00000104333.2 | 0 | NA | NA | NA | NA | NA | #VALUE! |
| ENSMUSG00000104334.2 | 0 | NA | NA | NA | NA | NA | #VALUE! |

| | | | | | | | |
|---|---|---|---|---|---|---|---|
| ENSMUSG00000104335.2 | 0 | NA | NA | NA | NA | NA | #VALUE! |
| ENSMUSG00000104336.2 | 0 | NA | NA | NA | NA | NA | #VALUE! |
| ENSMUSG00000104337.2 | 0 | NA | NA | NA | NA | NA | #VALUE! |
| ENSMUSG00000104338.2 | 0 | NA | NA | NA | NA | NA | #VALUE! |
| ENSMUSG00000104339.2 | 0 | NA | NA | NA | NA | NA | #VALUE! |
| ENSMUSG00000104340.2 | 0 | NA | NA | NA | NA | NA | #VALUE! |
| ENSMUSG00000104341.2 | 0 | NA | NA | NA | NA | NA | #VALUE! |
| ENSMUSG00000104342.2 | 0 | NA | NA | NA | NA | NA | #VALUE! |
| ENSMUSG00000104343.2 | 0 | NA | NA | NA | NA | NA | #VALUE! |
| ENSMUSG00000104344.2 | 0 | NA | NA | NA | NA | NA | #VALUE! |
| ENSMUSG00000104345.6 | 0 | NA | NA | NA | NA | NA | #VALUE! |
| ENSMUSG00000104346.2 | 0 | NA | NA | NA | NA | NA | #VALUE! |
| ENSMUSG00000104347.2 | 0 | NA | NA | NA | NA | NA | #VALUE! |
| ENSMUSG00000104348.2 | 0 | NA | NA | NA | NA | NA | #VALUE! |
| ENSMUSG00000104349.2 | 0 | NA | NA | NA | NA | NA | #VALUE! |
| ENSMUSG00000104350.2 | 0 | NA | NA | NA | NA | NA | #VALUE! |
| ENSMUSG00000104351.2 | 0 | NA | NA | NA | NA | NA | #VALUE! |
| ENSMUSG00000104352.2 | 0 | NA | NA | NA | NA | NA | #VALUE! |
| ENSMUSG00000104354.2 | 0 | NA | NA | NA | NA | NA | #VALUE! |
| ENSMUSG00000104355.2 | 0 | NA | NA | NA | NA | NA | #VALUE! |
| ENSMUSG00000104356.2 | 0 | NA | NA | NA | NA | NA | #VALUE! |
| ENSMUSG00000104358.2 | 0 | NA | NA | NA | NA | NA | #VALUE! |
| ENSMUSG00000104360.2 | 0 | NA | NA | NA | NA | NA | #VALUE! |
| ENSMUSG00000104361.2 | 0 | NA | NA | NA | NA | NA | #VALUE! |
| ENSMUSG00000104362.2 | 0 | NA | NA | NA | NA | NA | #VALUE! |
| ENSMUSG00000104363.2 | 0 | NA | NA | NA | NA | NA | #VALUE! |
| ENSMUSG00000104364.2 | 0 | NA | NA | NA | NA | NA | #VALUE! |
| ENSMUSG00000104365.2 | 0 | NA | NA | NA | NA | NA | #VALUE! |
| ENSMUSG00000104366.2 | 0 | NA | NA | NA | NA | NA | #VALUE! |
| ENSMUSG00000104368.2 | 0 | NA | NA | NA | NA | NA | #VALUE! |
| ENSMUSG00000104369.2 | 0 | NA | NA | NA | NA | NA | #VALUE! |
| ENSMUSG00000104370.2 | 0 | NA | NA | NA | NA | NA | #VALUE! |
| ENSMUSG00000104371.2 | 0 | NA | NA | NA | NA | NA | #VALUE! |
| ENSMUSG00000104373.2 | 0 | NA | NA | NA | NA | NA | #VALUE! |
| ENSMUSG00000104374.2 | 0 | NA | NA | NA | NA | NA | #VALUE! |
| ENSMUSG00000104375.2 | 0 | NA | NA | NA | NA | NA | #VALUE! |
| ENSMUSG00000104376.2 | 0 | NA | NA | NA | NA | NA | #VALUE! |
| ENSMUSG00000104377.2 | 0 | NA | NA | NA | NA | NA | #VALUE! |
| ENSMUSG00000104378.2 | 0 | NA | NA | NA | NA | NA | #VALUE! |
| ENSMUSG00000104379.2 | 0 | NA | NA | NA | NA | NA | #VALUE! |
| ENSMUSG00000104380.2 | 0 | NA | NA | NA | NA | NA | #VALUE! |
| ENSMUSG00000104381.2 | 0 | NA | NA | NA | NA | NA | #VALUE! |
| ENSMUSG00000104382.2 | 0 | NA | NA | NA | NA | NA | #VALUE! |
| ENSMUSG00000104383.2 | 0 | NA | NA | NA | NA | NA | #VALUE! |

| | | | | | | | |
|---|---|---|---|---|---|---|---|
| ENSMUSG00000104384.2 | 0 | NA | NA | NA | NA | NA | #VALUE! |
| ENSMUSG00000104385.2 | 0.195789 | -1.05581 | 4.073302 | -0.2592 | 0.795479 | NA | #VALUE! |
| ENSMUSG00000104386.2 | 0 | NA | NA | NA | NA | NA | #VALUE! |
| ENSMUSG00000104388.2 | 0 | NA | NA | NA | NA | NA | #VALUE! |
| ENSMUSG00000104390.2 | 0 | NA | NA | NA | NA | NA | #VALUE! |
| ENSMUSG00000104391.2 | 0 | NA | NA | NA | NA | NA | #VALUE! |
| ENSMUSG00000104392.2 | 0 | NA | NA | NA | NA | NA | #VALUE! |
| ENSMUSG00000104393.2 | 0 | NA | NA | NA | NA | NA | #VALUE! |
| ENSMUSG00000104395.2 | 0 | NA | NA | NA | NA | NA | #VALUE! |
| ENSMUSG00000104396.2 | 0 | NA | NA | NA | NA | NA | #VALUE! |
| ENSMUSG00000104397.2 | 0 | NA | NA | NA | NA | NA | #VALUE! |
| ENSMUSG00000104398.2 | 0 | NA | NA | NA | NA | NA | #VALUE! |
| ENSMUSG00000104399.2 | 0 | NA | NA | NA | NA | NA | #VALUE! |
| ENSMUSG00000104400.2 | 0 | NA | NA | NA | NA | NA | #VALUE! |
| ENSMUSG00000104401.2 | 0 | NA | NA | NA | NA | NA | #VALUE! |
| ENSMUSG00000104402.2 | 0 | NA | NA | NA | NA | NA | #VALUE! |
| ENSMUSG00000104403.2 | 0 | NA | NA | NA | NA | NA | #VALUE! |
| ENSMUSG00000104404.2 | 0 | NA | NA | NA | NA | NA | #VALUE! |
| ENSMUSG00000104405.2 | 0 | NA | NA | NA | NA | NA | #VALUE! |
| ENSMUSG00000104406.2 | 3.570446 | -5.37363 | 3.919223 | -1.3711 | 0.170345 | NA | #VALUE! |
| ENSMUSG00000104407.6 | 0 | NA | NA | NA | NA | NA | #VALUE! |
| ENSMUSG00000104408.2 | 0 | NA | NA | NA | NA | NA | #VALUE! |
| ENSMUSG00000104409.6 | 0 | NA | NA | NA | NA | NA | #VALUE! |
| ENSMUSG00000104410.2 | 0 | NA | NA | NA | NA | NA | #VALUE! |
| ENSMUSG00000104411.2 | 0 | NA | NA | NA | NA | NA | #VALUE! |
| ENSMUSG00000104412.2 | 0 | NA | NA | NA | NA | NA | #VALUE! |
| ENSMUSG00000104413.2 | 0 | NA | NA | NA | NA | NA | #VALUE! |
| ENSMUSG00000104414.2 | 0 | NA | NA | NA | NA | NA | #VALUE! |
| ENSMUSG00000104415.2 | 0 | NA | NA | NA | NA | NA | #VALUE! |
| ENSMUSG00000104416.2 | 0 | NA | NA | NA | NA | NA | #VALUE! |
| ENSMUSG00000104417.2 | 0 | NA | NA | NA | NA | NA | #VALUE! |
| ENSMUSG00000104418.2 | 1.740388 | 0.220476 | 3.831756 | 0.057539 | 0.954116 | NA | #VALUE! |
| ENSMUSG00000104419.2 | 0 | NA | NA | NA | NA | NA | #VALUE! |
| ENSMUSG00000104420.2 | 0 | NA | NA | NA | NA | NA | #VALUE! |
| ENSMUSG00000104422.2 | 0 | NA | NA | NA | NA | NA | #VALUE! |
| ENSMUSG00000104423.2 | 0 | NA | NA | NA | NA | NA | #VALUE! |
| ENSMUSG00000104424.2 | 0 | NA | NA | NA | NA | NA | #VALUE! |
| ENSMUSG00000104425.6 | 0 | NA | NA | NA | NA | NA | #VALUE! |
| ENSMUSG00000104426.2 | 0 | NA | NA | NA | NA | NA | #VALUE! |
| ENSMUSG00000104427.2 | 0 | NA | NA | NA | NA | NA | #VALUE! |
| ENSMUSG00000104428.2 | 0 | NA | NA | NA | NA | NA | #VALUE! |
| ENSMUSG00000104429.2 | 0 | NA | NA | NA | NA | NA | #VALUE! |
| ENSMUSG00000104430.2 | 0 | NA | NA | NA | NA | NA | #VALUE! |
| ENSMUSG00000104431.2 | 0 | NA | NA | NA | NA | NA | #VALUE! |

| | | | | | | | |
|---|---|---|---|---|---|---|---|
| ENSMUSG00000104432.2 | 0.210026 | -1.52494 | 4.058692 | -0.37572 | 0.707123 | NA | #VALUE! |
| ENSMUSG00000104433.2 | 0 | NA | NA | NA | NA | NA | #VALUE! |
| ENSMUSG00000104434.2 | 0 | NA | NA | NA | NA | NA | #VALUE! |
| ENSMUSG00000104435.2 | 2.869182 | -5.05543 | 3.922288 | -1.2889 | 0.197433 | NA | #VALUE! |
| ENSMUSG00000104436.2 | 0 | NA | NA | NA | NA | NA | #VALUE! |
| ENSMUSG00000104437.2 | 0 | NA | NA | NA | NA | NA | #VALUE! |
| ENSMUSG00000104438.2 | 0 | NA | NA | NA | NA | NA | #VALUE! |
| ENSMUSG00000104439.2 | 0 | NA | NA | NA | NA | NA | #VALUE! |
| ENSMUSG00000104440.2 | 0 | NA | NA | NA | NA | NA | #VALUE! |
| ENSMUSG00000104441.2 | 0 | NA | NA | NA | NA | NA | #VALUE! |
| ENSMUSG00000104442.2 | 0 | NA | NA | NA | NA | NA | #VALUE! |
| ENSMUSG00000104443.2 | 0 | NA | NA | NA | NA | NA | #VALUE! |
| ENSMUSG00000104444.2 | 6.722249 | -3.16774 | 3.718478 | -0.85189 | 0.394274 | NA | #VALUE! |
| ENSMUSG00000104446.2 | 0 | NA | NA | NA | NA | NA | #VALUE! |
| ENSMUSG00000104448.2 | 0 | NA | NA | NA | NA | NA | #VALUE! |
| ENSMUSG00000104449.2 | 0 | NA | NA | NA | NA | NA | #VALUE! |
| ENSMUSG00000104450.2 | 0 | NA | NA | NA | NA | NA | #VALUE! |
| ENSMUSG00000104451.2 | 0 | NA | NA | NA | NA | NA | #VALUE! |
| ENSMUSG00000104452.2 | 0 | NA | NA | NA | NA | NA | #VALUE! |
| ENSMUSG00000104453.2 | 0 | NA | NA | NA | NA | NA | #VALUE! |
| ENSMUSG00000104454.2 | 0 | NA | NA | NA | NA | NA | #VALUE! |
| ENSMUSG00000104455.2 | 0 | NA | NA | NA | NA | NA | #VALUE! |
| ENSMUSG00000104456.2 | 0 | NA | NA | NA | NA | NA | #VALUE! |
| ENSMUSG00000104457.2 | 0 | NA | NA | NA | NA | NA | #VALUE! |
| ENSMUSG00000104460.2 | 0 | NA | NA | NA | NA | NA | #VALUE! |
| ENSMUSG00000104461.2 | 0 | NA | NA | NA | NA | NA | #VALUE! |
| ENSMUSG00000104462.2 | 0 | NA | NA | NA | NA | NA | #VALUE! |
| ENSMUSG00000104464.2 | 0 | NA | NA | NA | NA | NA | #VALUE! |
| ENSMUSG00000104465.2 | 0 | NA | NA | NA | NA | NA | #VALUE! |
| ENSMUSG00000104466.2 | 0 | NA | NA | NA | NA | NA | #VALUE! |
| ENSMUSG00000104467.2 | 0.210026 | -1.52494 | 4.058692 | -0.37572 | 0.707123 | NA | #VALUE! |
| ENSMUSG00000104468.2 | 0 | NA | NA | NA | NA | NA | #VALUE! |
| ENSMUSG00000104469.2 | 0 | NA | NA | NA | NA | NA | #VALUE! |
| ENSMUSG00000104470.2 | 0 | NA | NA | NA | NA | NA | #VALUE! |
| ENSMUSG00000104471.2 | 0 | NA | NA | NA | NA | NA | #VALUE! |
| ENSMUSG00000104472.2 | 0 | NA | NA | NA | NA | NA | #VALUE! |
| ENSMUSG00000104473.2 | 0 | NA | NA | NA | NA | NA | #VALUE! |
| ENSMUSG00000104474.2 | 0 | NA | NA | NA | NA | NA | #VALUE! |
| ENSMUSG00000104476.2 | 0 | NA | NA | NA | NA | NA | #VALUE! |
| ENSMUSG00000104477.2 | 0 | NA | NA | NA | NA | NA | #VALUE! |
| ENSMUSG00000104478.2 | 0 | NA | NA | NA | NA | NA | #VALUE! |
| ENSMUSG00000104479.2 | 0 | NA | NA | NA | NA | NA | #VALUE! |
| ENSMUSG00000104480.2 | 0 | NA | NA | NA | NA | NA | #VALUE! |
| ENSMUSG00000104481.2 | 0 | NA | NA | NA | NA | NA | #VALUE! |

| | | | | | | | |
|---|---|---|---|---|---|---|---|
| ENSMUSG00000104482.2 | 0 | NA | NA | NA | NA | NA | #VALUE! |
| ENSMUSG00000104484.2 | 0 | NA | NA | NA | NA | NA | #VALUE! |
| ENSMUSG00000104485.2 | 0 | NA | NA | NA | NA | NA | #VALUE! |
| ENSMUSG00000104486.2 | 0 | NA | NA | NA | NA | NA | #VALUE! |
| ENSMUSG00000104487.2 | 0 | NA | NA | NA | NA | NA | #VALUE! |
| ENSMUSG00000104488.2 | 0 | NA | NA | NA | NA | NA | #VALUE! |
| ENSMUSG00000104489.2 | 0 | NA | NA | NA | NA | NA | #VALUE! |
| ENSMUSG00000104490.2 | 0 | NA | NA | NA | NA | NA | #VALUE! |
| ENSMUSG00000104491.2 | 0 | NA | NA | NA | NA | NA | #VALUE! |
| ENSMUSG00000104493.2 | 0.783156 | -3.1627 | 3.963922 | -0.79787 | 0.424945 | NA | #VALUE! |
| ENSMUSG00000104494.2 | 0 | NA | NA | NA | NA | NA | #VALUE! |
| ENSMUSG00000104496.2 | 0 | NA | NA | NA | NA | NA | #VALUE! |
| ENSMUSG00000104497.2 | 0 | NA | NA | NA | NA | NA | #VALUE! |
| ENSMUSG00000104498.2 | 0 | NA | NA | NA | NA | NA | #VALUE! |
| ENSMUSG00000104499.6 | 0 | NA | NA | NA | NA | NA | #VALUE! |
| ENSMUSG00000104500.6 | 0 | NA | NA | NA | NA | NA | #VALUE! |
| ENSMUSG00000104501.2 | 0 | NA | NA | NA | NA | NA | #VALUE! |
| ENSMUSG00000104502.2 | 0 | NA | NA | NA | NA | NA | #VALUE! |
| ENSMUSG00000104503.2 | 0 | NA | NA | NA | NA | NA | #VALUE! |
| ENSMUSG00000104504.2 | 0 | NA | NA | NA | NA | NA | #VALUE! |
| ENSMUSG00000104505.6 | 0 | NA | NA | NA | NA | NA | #VALUE! |
| ENSMUSG00000104506.6 | 0 | NA | NA | NA | NA | NA | #VALUE! |
| ENSMUSG00000104507.2 | 0 | NA | NA | NA | NA | NA | #VALUE! |
| ENSMUSG00000104508.2 | 0 | NA | NA | NA | NA | NA | #VALUE! |
| ENSMUSG00000104509.2 | 0 | NA | NA | NA | NA | NA | #VALUE! |
| ENSMUSG00000104510.2 | 0 | NA | NA | NA | NA | NA | #VALUE! |
| ENSMUSG00000104511.2 | 0 | NA | NA | NA | NA | NA | #VALUE! |
| ENSMUSG00000104512.2 | 0 | NA | NA | NA | NA | NA | #VALUE! |
| ENSMUSG00000104513.2 | 0 | NA | NA | NA | NA | NA | #VALUE! |
| ENSMUSG00000104514.2 | 0 | NA | NA | NA | NA | NA | #VALUE! |
| ENSMUSG00000104515.2 | 0 | NA | NA | NA | NA | NA | #VALUE! |
| ENSMUSG00000104516.2 | 0 | NA | NA | NA | NA | NA | #VALUE! |
| ENSMUSG00000104517.2 | 0 | NA | NA | NA | NA | NA | #VALUE! |
| ENSMUSG00000104518.2 | 0 | NA | NA | NA | NA | NA | #VALUE! |
| ENSMUSG00000104519.2 | 0 | NA | NA | NA | NA | NA | #VALUE! |
| ENSMUSG00000104520.2 | 0.616333 | 2.813477 | 3.977401 | 0.707366 | 0.479339 | NA | #VALUE! |
| ENSMUSG00000104521.2 | 0 | NA | NA | NA | NA | NA | #VALUE! |
| ENSMUSG00000104522.6 | 0 | NA | NA | NA | NA | NA | #VALUE! |
| ENSMUSG00000104523.2 | 0 | NA | NA | NA | NA | NA | #VALUE! |
| ENSMUSG00000104524.2 | 0 | NA | NA | NA | NA | NA | #VALUE! |
| ENSMUSG00000104525.2 | 0 | NA | NA | NA | NA | NA | #VALUE! |
| ENSMUSG00000104526.2 | 0 | NA | NA | NA | NA | NA | #VALUE! |
| ENSMUSG00000104527.2 | 0 | NA | NA | NA | NA | NA | #VALUE! |
| ENSMUSG00000104528.2 | 1.91379 | 4.427138 | 3.930139 | 1.126459 | 0.259971 | NA | #VALUE! |

| | | | | | | | |
|---|---|---|---|---|---|---|---|
| ENSMUSG00000104529.5 | 0 | NA | NA | NA | NA | NA | #VALUE! |
| ENSMUSG00000104530.2 | 0 | NA | NA | NA | NA | NA | #VALUE! |
| ENSMUSG00000104531.2 | 0 | NA | NA | NA | NA | NA | #VALUE! |
| ENSMUSG00000104532.2 | 0 | NA | NA | NA | NA | NA | #VALUE! |
| ENSMUSG00000104533.2 | 0 | NA | NA | NA | NA | NA | #VALUE! |
| ENSMUSG00000104534.2 | 0 | NA | NA | NA | NA | NA | #VALUE! |
| ENSMUSG00000104535.2 | 0 | NA | NA | NA | NA | NA | #VALUE! |
| ENSMUSG00000104536.3 | 0 | NA | NA | NA | NA | NA | #VALUE! |
| ENSMUSG00000104538.2 | 0 | NA | NA | NA | NA | NA | #VALUE! |
| ENSMUSG00000104539.2 | 0 | NA | NA | NA | NA | NA | #VALUE! |
| ENSMUSG00000104540.2 | 0 | NA | NA | NA | NA | NA | #VALUE! |
| ENSMUSG00000104541.2 | 0 | NA | NA | NA | NA | NA | #VALUE! |
| ENSMUSG00000104542.2 | 0 | NA | NA | NA | NA | NA | #VALUE! |
| ENSMUSG00000104543.2 | 0 | NA | NA | NA | NA | NA | #VALUE! |
| ENSMUSG00000104545.2 | 0 | NA | NA | NA | NA | NA | #VALUE! |
| ENSMUSG00000104546.2 | 0 | NA | NA | NA | NA | NA | #VALUE! |
| ENSMUSG00000104547.2 | 0 | NA | NA | NA | NA | NA | #VALUE! |
| ENSMUSG00000104549.2 | 0 | NA | NA | NA | NA | NA | #VALUE! |
| ENSMUSG00000104550.2 | 0 | NA | NA | NA | NA | NA | #VALUE! |
| ENSMUSG00000104551.2 | 0 | NA | NA | NA | NA | NA | #VALUE! |
| ENSMUSG00000104553.2 | 0 | NA | NA | NA | NA | NA | #VALUE! |
| ENSMUSG00000104554.2 | 0 | NA | NA | NA | NA | NA | #VALUE! |
| ENSMUSG00000104555.2 | 0 | NA | NA | NA | NA | NA | #VALUE! |
| ENSMUSG00000104556.2 | 0 | NA | NA | NA | NA | NA | #VALUE! |
| ENSMUSG00000104557.2 | 0 | NA | NA | NA | NA | NA | #VALUE! |
| ENSMUSG00000104559.2 | 13.31153 | -2.59515 | 3.001331 | -0.86467 | NA | NA | #VALUE! |
| ENSMUSG00000104560.2 | 0 | NA | NA | NA | NA | NA | #VALUE! |
| ENSMUSG00000104562.2 | 0 | NA | NA | NA | NA | NA | #VALUE! |
| ENSMUSG00000104563.2 | 0 | NA | NA | NA | NA | NA | #VALUE! |
| ENSMUSG00000104564.2 | 0 | NA | NA | NA | NA | NA | #VALUE! |
| ENSMUSG00000104566.3 | 0 | NA | NA | NA | NA | NA | #VALUE! |
| ENSMUSG00000104567.2 | 0 | NA | NA | NA | NA | NA | #VALUE! |
| ENSMUSG00000104568.2 | 0 | NA | NA | NA | NA | NA | #VALUE! |
| ENSMUSG00000104569.2 | 0 | NA | NA | NA | NA | NA | #VALUE! |
| ENSMUSG00000104571.2 | 0 | NA | NA | NA | NA | NA | #VALUE! |
| ENSMUSG00000104572.6 | 0 | NA | NA | NA | NA | NA | #VALUE! |
| ENSMUSG00000104573.2 | 0 | NA | NA | NA | NA | NA | #VALUE! |
| ENSMUSG00000104574.2 | 0 | NA | NA | NA | NA | NA | #VALUE! |
| ENSMUSG00000104575.2 | 0 | NA | NA | NA | NA | NA | #VALUE! |
| ENSMUSG00000104576.2 | 1.177985 | 3.72614 | 3.944615 | 0.944614 | 0.344856 | NA | #VALUE! |
| ENSMUSG00000104577.2 | 0 | NA | NA | NA | NA | NA | #VALUE! |
| ENSMUSG00000104578.2 | 0 | NA | NA | NA | NA | NA | #VALUE! |
| ENSMUSG00000104579.2 | 0 | NA | NA | NA | NA | NA | #VALUE! |
| ENSMUSG00000104580.2 | 0 | NA | NA | NA | NA | NA | #VALUE! |

| | | | | | | | |
|---|---|---|---|---|---|---|---|
| ENSMUSG00000104581.2 | 0 | NA | NA | NA | NA | NA | #VALUE! |
| ENSMUSG00000104582.3 | 0 | NA | NA | NA | NA | NA | #VALUE! |
| ENSMUSG00000104583.2 | 0.195789 | -1.05581 | 4.073302 | -0.2592 | 0.795479 | NA | #VALUE! |
| ENSMUSG00000104584.2 | 0 | NA | NA | NA | NA | NA | #VALUE! |
| ENSMUSG00000104585.2 | 0 | NA | NA | NA | NA | NA | #VALUE! |
| ENSMUSG00000104586.2 | 0 | NA | NA | NA | NA | NA | #VALUE! |
| ENSMUSG00000104587.2 | 0 | NA | NA | NA | NA | NA | #VALUE! |
| ENSMUSG00000104588.2 | 0 | NA | NA | NA | NA | NA | #VALUE! |
| ENSMUSG00000104590.3 | 0 | NA | NA | NA | NA | NA | #VALUE! |
| ENSMUSG00000104591.2 | 0 | NA | NA | NA | NA | NA | #VALUE! |
| ENSMUSG00000104592.2 | 0 | NA | NA | NA | NA | NA | #VALUE! |
| ENSMUSG00000104593.2 | 0 | NA | NA | NA | NA | NA | #VALUE! |
| ENSMUSG00000104595.2 | 0 | NA | NA | NA | NA | NA | #VALUE! |
| ENSMUSG00000104596.2 | 0 | NA | NA | NA | NA | NA | #VALUE! |
| ENSMUSG00000104597.2 | 0.587367 | -2.74297 | 3.982932 | -0.68868 | 0.491024 | NA | #VALUE! |
| ENSMUSG00000104598.2 | 0 | NA | NA | NA | NA | NA | #VALUE! |
| ENSMUSG00000104600.5 | 0 | NA | NA | NA | NA | NA | #VALUE! |
| ENSMUSG00000104601.2 | 0 | NA | NA | NA | NA | NA | #VALUE! |
| ENSMUSG00000104602.2 | 0 | NA | NA | NA | NA | NA | #VALUE! |
| ENSMUSG00000104604.2 | 0 | NA | NA | NA | NA | NA | #VALUE! |
| ENSMUSG00000104605.2 | 0 | NA | NA | NA | NA | NA | #VALUE! |
| ENSMUSG00000104606.2 | 0 | NA | NA | NA | NA | NA | #VALUE! |
| ENSMUSG00000104607.2 | 0 | NA | NA | NA | NA | NA | #VALUE! |
| ENSMUSG00000104609.2 | 0 | NA | NA | NA | NA | NA | #VALUE! |
| ENSMUSG00000104610.2 | 0 | NA | NA | NA | NA | NA | #VALUE! |
| ENSMUSG00000104612.2 | 0 | NA | NA | NA | NA | NA | #VALUE! |
| ENSMUSG00000104613.2 | 0 | NA | NA | NA | NA | NA | #VALUE! |
| ENSMUSG00000104614.6 | 0 | NA | NA | NA | NA | NA | #VALUE! |
| ENSMUSG00000104615.2 | 0 | NA | NA | NA | NA | NA | #VALUE! |
| ENSMUSG00000104616.2 | 0 | NA | NA | NA | NA | NA | #VALUE! |
| ENSMUSG00000104617.2 | 0 | NA | NA | NA | NA | NA | #VALUE! |
| ENSMUSG00000104618.3 | 0 | NA | NA | NA | NA | NA | #VALUE! |
| ENSMUSG00000104619.2 | 0 | NA | NA | NA | NA | NA | #VALUE! |
| ENSMUSG00000104620.2 | 0 | NA | NA | NA | NA | NA | #VALUE! |
| ENSMUSG00000104621.2 | 0 | NA | NA | NA | NA | NA | #VALUE! |
| ENSMUSG00000104622.2 | 0 | NA | NA | NA | NA | NA | #VALUE! |
| ENSMUSG00000104624.2 | 0 | NA | NA | NA | NA | NA | #VALUE! |
| ENSMUSG00000104625.2 | 0 | NA | NA | NA | NA | NA | #VALUE! |
| ENSMUSG00000104626.2 | 0 | NA | NA | NA | NA | NA | #VALUE! |
| ENSMUSG00000104627.3 | 0 | NA | NA | NA | NA | NA | #VALUE! |
| ENSMUSG00000104629.2 | 0 | NA | NA | NA | NA | NA | #VALUE! |
| ENSMUSG00000104630.2 | 0 | NA | NA | NA | NA | NA | #VALUE! |
| ENSMUSG00000104631.2 | 6.641339 | 6.216291 | 3.913555 | 1.5884 | 0.112196 | NA | #VALUE! |
| ENSMUSG00000104632.2 | 0 | NA | NA | NA | NA | NA | #VALUE! |

| | | | | | | | |
|---|---|---|---|---|---|---|---|
| ENSMUSG00000104633.2 | 0 | NA | NA | NA | NA | NA | #VALUE! |
| ENSMUSG00000104634.2 | 0 | NA | NA | NA | NA | NA | #VALUE! |
| ENSMUSG00000104636.2 | 0 | NA | NA | NA | NA | NA | #VALUE! |
| ENSMUSG00000104637.2 | 0 | NA | NA | NA | NA | NA | #VALUE! |
| ENSMUSG00000104638.2 | 0 | NA | NA | NA | NA | NA | #VALUE! |
| ENSMUSG00000104639.2 | 0 | NA | NA | NA | NA | NA | #VALUE! |
| ENSMUSG00000104640.2 | 0 | NA | NA | NA | NA | NA | #VALUE! |
| ENSMUSG00000104641.2 | 0 | NA | NA | NA | NA | NA | #VALUE! |
| ENSMUSG00000104642.2 | 0 | NA | NA | NA | NA | NA | #VALUE! |
| ENSMUSG00000104643.2 | 0 | NA | NA | NA | NA | NA | #VALUE! |
| ENSMUSG00000104644.2 | 0 | NA | NA | NA | NA | NA | #VALUE! |
| ENSMUSG00000104645.2 | 0 | NA | NA | NA | NA | NA | #VALUE! |
| ENSMUSG00000104646.2 | 0 | NA | NA | NA | NA | NA | #VALUE! |
| ENSMUSG00000104647.2 | 0 | NA | NA | NA | NA | NA | #VALUE! |
| ENSMUSG00000104648.2 | 0 | NA | NA | NA | NA | NA | #VALUE! |
| ENSMUSG00000104650.2 | 0 | NA | NA | NA | NA | NA | #VALUE! |
| ENSMUSG00000104651.2 | 0 | NA | NA | NA | NA | NA | #VALUE! |
| ENSMUSG00000104653.2 | 0 | NA | NA | NA | NA | NA | #VALUE! |
| ENSMUSG00000104654.2 | 0.606563 | -0.89296 | 3.98278 | -0.22421 | 0.822597 | NA | #VALUE! |
| ENSMUSG00000104655.2 | 0 | NA | NA | NA | NA | NA | #VALUE! |
| ENSMUSG00000104658.5 | 0 | NA | NA | NA | NA | NA | #VALUE! |
| ENSMUSG00000104659.2 | 0 | NA | NA | NA | NA | NA | #VALUE! |
| ENSMUSG00000104660.2 | 0 | NA | NA | NA | NA | NA | #VALUE! |
| ENSMUSG00000104661.2 | 0 | NA | NA | NA | NA | NA | #VALUE! |
| ENSMUSG00000104662.2 | 0 | NA | NA | NA | NA | NA | #VALUE! |
| ENSMUSG00000104663.2 | 0 | NA | NA | NA | NA | NA | #VALUE! |
| ENSMUSG00000104664.2 | 0 | NA | NA | NA | NA | NA | #VALUE! |
| ENSMUSG00000104665.2 | 0 | NA | NA | NA | NA | NA | #VALUE! |
| ENSMUSG00000104667.2 | 0 | NA | NA | NA | NA | NA | #VALUE! |
| ENSMUSG00000104668.2 | 0 | NA | NA | NA | NA | NA | #VALUE! |
| ENSMUSG00000104669.2 | 0 | NA | NA | NA | NA | NA | #VALUE! |
| ENSMUSG00000104670.2 | 0 | NA | NA | NA | NA | NA | #VALUE! |
| ENSMUSG00000104671.2 | 0.408222 | -2.20205 | 4.016842 | -0.54821 | 0.583551 | NA | #VALUE! |
| ENSMUSG00000104672.2 | 0 | NA | NA | NA | NA | NA | #VALUE! |
| ENSMUSG00000104673.2 | 0 | NA | NA | NA | NA | NA | #VALUE! |
| ENSMUSG00000104674.2 | 0 | NA | NA | NA | NA | NA | #VALUE! |
| ENSMUSG00000104675.2 | 0 | NA | NA | NA | NA | NA | #VALUE! |
| ENSMUSG00000104676.2 | 0 | NA | NA | NA | NA | NA | #VALUE! |
| ENSMUSG00000104677.2 | 0 | NA | NA | NA | NA | NA | #VALUE! |
| ENSMUSG00000104678.2 | 0 | NA | NA | NA | NA | NA | #VALUE! |
| ENSMUSG00000104679.2 | 0 | NA | NA | NA | NA | NA | #VALUE! |
| ENSMUSG00000104680.2 | 0 | NA | NA | NA | NA | NA | #VALUE! |
| ENSMUSG00000104681.2 | 0 | NA | NA | NA | NA | NA | #VALUE! |
| ENSMUSG00000104682.2 | 0 | NA | NA | NA | NA | NA | #VALUE! |

| | | | | | | | |
|---|---|---|---|---|---|---|---|
| ENSMUSG00000104684.2 | 0 | NA | NA | NA | NA | NA | #VALUE! |
| ENSMUSG00000104685.2 | 0 | NA | NA | NA | NA | NA | #VALUE! |
| ENSMUSG00000104686.2 | 4.426611 | -5.67416 | 3.916885 | -1.44864 | 0.147438 | NA | #VALUE! |
| ENSMUSG00000104687.2 | 0 | NA | NA | NA | NA | NA | #VALUE! |
| ENSMUSG00000104688.2 | 0 | NA | NA | NA | NA | NA | #VALUE! |
| ENSMUSG00000104691.2 | 0 | NA | NA | NA | NA | NA | #VALUE! |
| ENSMUSG00000104692.2 | 0 | NA | NA | NA | NA | NA | #VALUE! |
| ENSMUSG00000104693.2 | 0 | NA | NA | NA | NA | NA | #VALUE! |
| ENSMUSG00000104694.2 | 0 | NA | NA | NA | NA | NA | #VALUE! |
| ENSMUSG00000104695.2 | 0 | NA | NA | NA | NA | NA | #VALUE! |
| ENSMUSG00000104696.2 | 0 | NA | NA | NA | NA | NA | #VALUE! |
| ENSMUSG00000104697.2 | 0 | NA | NA | NA | NA | NA | #VALUE! |
| ENSMUSG00000104698.2 | 0 | NA | NA | NA | NA | NA | #VALUE! |
| ENSMUSG00000104699.2 | 4.872318 | 0.334895 | 3.71753 | 0.090085 | 0.928219 | NA | #VALUE! |
| ENSMUSG00000104700.3 | 0 | NA | NA | NA | NA | NA | #VALUE! |
| ENSMUSG00000104701.2 | 0 | NA | NA | NA | NA | NA | #VALUE! |
| ENSMUSG00000104703.2 | 0 | NA | NA | NA | NA | NA | #VALUE! |
| ENSMUSG00000104704.2 | 0 | NA | NA | NA | NA | NA | #VALUE! |
| ENSMUSG00000104705.2 | 0 | NA | NA | NA | NA | NA | #VALUE! |
| ENSMUSG00000104706.2 | 0 | NA | NA | NA | NA | NA | #VALUE! |
| ENSMUSG00000104707.2 | 0 | NA | NA | NA | NA | NA | #VALUE! |
| ENSMUSG00000104708.2 | 0 | NA | NA | NA | NA | NA | #VALUE! |
| ENSMUSG00000104709.2 | 0 | NA | NA | NA | NA | NA | #VALUE! |
| ENSMUSG00000104710.2 | 0 | NA | NA | NA | NA | NA | #VALUE! |
| ENSMUSG00000104711.2 | 0 | NA | NA | NA | NA | NA | #VALUE! |
| ENSMUSG00000104712.2 | 0 | NA | NA | NA | NA | NA | #VALUE! |
| ENSMUSG00000104713.5 | 3.730731 | -2.37765 | 3.816641 | -0.62297 | 0.533304 | NA | #VALUE! |
| ENSMUSG00000104714.2 | 0 | NA | NA | NA | NA | NA | #VALUE! |
| ENSMUSG00000104715.2 | 0 | NA | NA | NA | NA | NA | #VALUE! |
| ENSMUSG00000104716.2 | 0 | NA | NA | NA | NA | NA | #VALUE! |
| ENSMUSG00000104717.2 | 0 | NA | NA | NA | NA | NA | #VALUE! |
| ENSMUSG00000104718.2 | 0 | NA | NA | NA | NA | NA | #VALUE! |
| ENSMUSG00000104719.2 | 0.195789 | -1.05581 | 4.073302 | -0.2592 | 0.795479 | NA | #VALUE! |
| ENSMUSG00000104720.2 | 0 | NA | NA | NA | NA | NA | #VALUE! |
| ENSMUSG00000104721.2 | 0 | NA | NA | NA | NA | NA | #VALUE! |
| ENSMUSG00000104722.2 | 0 | NA | NA | NA | NA | NA | #VALUE! |
| ENSMUSG00000104723.2 | 0 | NA | NA | NA | NA | NA | #VALUE! |
| ENSMUSG00000104724.2 | 2.249883 | 3.672485 | 3.923091 | 0.93612 | 0.349211 | NA | #VALUE! |
| ENSMUSG00000104725.2 | 0 | NA | NA | NA | NA | NA | #VALUE! |
| ENSMUSG00000104726.2 | 0 | NA | NA | NA | NA | NA | #VALUE! |
| ENSMUSG00000104727.2 | 0 | NA | NA | NA | NA | NA | #VALUE! |
| ENSMUSG00000104728.2 | 3.265772 | -5.23273 | 3.920498 | -1.33471 | 0.181971 | NA | #VALUE! |
| ENSMUSG00000104730.2 | 0 | NA | NA | NA | NA | NA | #VALUE! |
| ENSMUSG00000104731.2 | 0 | NA | NA | NA | NA | NA | #VALUE! |

| | | | | | | | |
|---|---|---|---|---|---|---|---|
| ENSMUSG00000104732.2 | 0 | NA | NA | NA | NA | NA | #VALUE! |
| ENSMUSG00000104733.2 | 0 | NA | NA | NA | NA | NA | #VALUE! |
| ENSMUSG00000104735.2 | 0 | NA | NA | NA | NA | NA | #VALUE! |
| ENSMUSG00000104736.2 | 0 | NA | NA | NA | NA | NA | #VALUE! |
| ENSMUSG00000104737.2 | 0 | NA | NA | NA | NA | NA | #VALUE! |
| ENSMUSG00000104738.2 | 0 | NA | NA | NA | NA | NA | #VALUE! |
| ENSMUSG00000104739.2 | 0 | NA | NA | NA | NA | NA | #VALUE! |
| ENSMUSG00000104740.3 | 0 | NA | NA | NA | NA | NA | #VALUE! |
| ENSMUSG00000104741.3 | 0 | NA | NA | NA | NA | NA | #VALUE! |
| ENSMUSG00000104742.2 | 0 | NA | NA | NA | NA | NA | #VALUE! |
| ENSMUSG00000104744.2 | 0 | NA | NA | NA | NA | NA | #VALUE! |
| ENSMUSG00000104745.2 | 0 | NA | NA | NA | NA | NA | #VALUE! |
| ENSMUSG00000104747.3 | 0 | NA | NA | NA | NA | NA | #VALUE! |
| ENSMUSG00000104748.2 | 0.18651 | 1.401769 | 4.056354 | 0.345574 | 0.729663 | NA | #VALUE! |
| ENSMUSG00000104749.2 | 0 | NA | NA | NA | NA | NA | #VALUE! |
| ENSMUSG00000104751.2 | 0 | NA | NA | NA | NA | NA | #VALUE! |
| ENSMUSG00000104752.2 | 0 | NA | NA | NA | NA | NA | #VALUE! |
| ENSMUSG00000104753.2 | 0 | NA | NA | NA | NA | NA | #VALUE! |
| ENSMUSG00000104754.2 | 0 | NA | NA | NA | NA | NA | #VALUE! |
| ENSMUSG00000104755.3 | 0 | NA | NA | NA | NA | NA | #VALUE! |
| ENSMUSG00000104756.2 | 0 | NA | NA | NA | NA | NA | #VALUE! |
| ENSMUSG00000104757.4 | 0 | NA | NA | NA | NA | NA | #VALUE! |
| ENSMUSG00000104758.3 | 0 | NA | NA | NA | NA | NA | #VALUE! |
| ENSMUSG00000104759.3 | 0 | NA | NA | NA | NA | NA | #VALUE! |
| ENSMUSG00000104760.2 | 0 | NA | NA | NA | NA | NA | #VALUE! |
| ENSMUSG00000104761.2 | 0 | NA | NA | NA | NA | NA | #VALUE! |
| ENSMUSG00000104763.2 | 0 | NA | NA | NA | NA | NA | #VALUE! |
| ENSMUSG00000104764.2 | 0 | NA | NA | NA | NA | NA | #VALUE! |
| ENSMUSG00000104765.2 | 0 | NA | NA | NA | NA | NA | #VALUE! |
| ENSMUSG00000104766.2 | 0 | NA | NA | NA | NA | NA | #VALUE! |
| ENSMUSG00000104768.2 | 0 | NA | NA | NA | NA | NA | #VALUE! |
| ENSMUSG00000104769.2 | 0 | NA | NA | NA | NA | NA | #VALUE! |
| ENSMUSG00000104770.2 | 0 | NA | NA | NA | NA | NA | #VALUE! |
| ENSMUSG00000104772.2 | 0 | NA | NA | NA | NA | NA | #VALUE! |
| ENSMUSG00000104773.3 | 0 | NA | NA | NA | NA | NA | #VALUE! |
| ENSMUSG00000104774.2 | 0 | NA | NA | NA | NA | NA | #VALUE! |
| ENSMUSG00000104775.2 | 0 | NA | NA | NA | NA | NA | #VALUE! |
| ENSMUSG00000104776.2 | 0 | NA | NA | NA | NA | NA | #VALUE! |
| ENSMUSG00000104777.2 | 0 | NA | NA | NA | NA | NA | #VALUE! |
| ENSMUSG00000104778.2 | 0 | NA | NA | NA | NA | NA | #VALUE! |
| ENSMUSG00000104779.2 | 0 | NA | NA | NA | NA | NA | #VALUE! |
| ENSMUSG00000104780.2 | 0 | NA | NA | NA | NA | NA | #VALUE! |
| ENSMUSG00000104781.2 | 0 | NA | NA | NA | NA | NA | #VALUE! |
| ENSMUSG00000104782.2 | 0 | NA | NA | NA | NA | NA | #VALUE! |

| | | | | | | | |
|---|---|---|---|---|---|---|---|
| ENSMUSG00000104784.2 | 0 | NA | NA | NA | NA | NA | #VALUE! |
| ENSMUSG00000104785.2 | 3.138636 | -4.23713 | 3.913016 | -1.08283 | 0.278884 | NA | #VALUE! |
| ENSMUSG00000104787.2 | 0 | NA | NA | NA | NA | NA | #VALUE! |
| ENSMUSG00000104788.2 | 0 | NA | NA | NA | NA | NA | #VALUE! |
| ENSMUSG00000104789.2 | 0 | NA | NA | NA | NA | NA | #VALUE! |
| ENSMUSG00000104790.2 | 0 | NA | NA | NA | NA | NA | #VALUE! |
| ENSMUSG00000104791.2 | 0 | NA | NA | NA | NA | NA | #VALUE! |
| ENSMUSG00000104792.2 | 0 | NA | NA | NA | NA | NA | #VALUE! |
| ENSMUSG00000104793.2 | 1.721829 | 4.254739 | 3.933085 | 1.081782 | 0.27935 | NA | #VALUE! |
| ENSMUSG00000104794.3 | 0 | NA | NA | NA | NA | NA | #VALUE! |
| ENSMUSG00000104795.2 | 0 | NA | NA | NA | NA | NA | #VALUE! |
| ENSMUSG00000104797.3 | 0 | NA | NA | NA | NA | NA | #VALUE! |
| ENSMUSG00000104798.2 | 0 | NA | NA | NA | NA | NA | #VALUE! |
| ENSMUSG00000104799.2 | 0 | NA | NA | NA | NA | NA | #VALUE! |
| ENSMUSG00000104800.2 | 0 | NA | NA | NA | NA | NA | #VALUE! |
| ENSMUSG00000104801.2 | 0 | NA | NA | NA | NA | NA | #VALUE! |
| ENSMUSG00000104802.2 | 0.300802 | -1.80634 | 4.050033 | -0.44601 | 0.655592 | NA | #VALUE! |
| ENSMUSG00000104803.2 | 0 | NA | NA | NA | NA | NA | #VALUE! |
| ENSMUSG00000104804.3 | 0 | NA | NA | NA | NA | NA | #VALUE! |
| ENSMUSG00000104805.2 | 0 | NA | NA | NA | NA | NA | #VALUE! |
| ENSMUSG00000104806.2 | 0 | NA | NA | NA | NA | NA | #VALUE! |
| ENSMUSG00000104807.2 | 0 | NA | NA | NA | NA | NA | #VALUE! |
| ENSMUSG00000104808.2 | 0 | NA | NA | NA | NA | NA | #VALUE! |
| ENSMUSG00000104810.2 | 0 | NA | NA | NA | NA | NA | #VALUE! |
| ENSMUSG00000104811.2 | 0 | NA | NA | NA | NA | NA | #VALUE! |
| ENSMUSG00000104814.2 | 8.722902 | -6.65261 | 3.277612 | -2.02971 | 0.042386 | NA | #VALUE! |
| ENSMUSG00000104815.2 | 0 | NA | NA | NA | NA | NA | #VALUE! |
| ENSMUSG00000104816.2 | 0 | NA | NA | NA | NA | NA | #VALUE! |
| ENSMUSG00000104818.2 | 0 | NA | NA | NA | NA | NA | #VALUE! |
| ENSMUSG00000104819.2 | 0 | NA | NA | NA | NA | NA | #VALUE! |
| ENSMUSG00000104821.2 | 0 | NA | NA | NA | NA | NA | #VALUE! |
| ENSMUSG00000104822.2 | 0 | NA | NA | NA | NA | NA | #VALUE! |
| ENSMUSG00000104823.2 | 0 | NA | NA | NA | NA | NA | #VALUE! |
| ENSMUSG00000104824.2 | 0 | NA | NA | NA | NA | NA | #VALUE! |
| ENSMUSG00000104825.2 | 0 | NA | NA | NA | NA | NA | #VALUE! |
| ENSMUSG00000104826.2 | 0 | NA | NA | NA | NA | NA | #VALUE! |
| ENSMUSG00000104828.2 | 0 | NA | NA | NA | NA | NA | #VALUE! |
| ENSMUSG00000104829.2 | 0 | NA | NA | NA | NA | NA | #VALUE! |
| ENSMUSG00000104830.2 | 0 | NA | NA | NA | NA | NA | #VALUE! |
| ENSMUSG00000104831.2 | 0 | NA | NA | NA | NA | NA | #VALUE! |
| ENSMUSG00000104832.2 | 0 | NA | NA | NA | NA | NA | #VALUE! |
| ENSMUSG00000104833.2 | 0 | NA | NA | NA | NA | NA | #VALUE! |
| ENSMUSG00000104834.2 | 0 | NA | NA | NA | NA | NA | #VALUE! |
| ENSMUSG00000104835.5 | 0.105013 | -0.97324 | 4.076293 | -0.23876 | 0.811295 | NA | #VALUE! |

| | | | | | | | |
|---|---|---|---|---|---|---|---|
| ENSMUSG00000104837.2 | 0 | NA | NA | NA | NA | NA | #VALUE! |
| ENSMUSG00000104838.2 | 0 | NA | NA | NA | NA | NA | #VALUE! |
| ENSMUSG00000104839.2 | 0 | NA | NA | NA | NA | NA | #VALUE! |
| ENSMUSG00000104840.3 | 0 | NA | NA | NA | NA | NA | #VALUE! |
| ENSMUSG00000104841.2 | 0 | NA | NA | NA | NA | NA | #VALUE! |
| ENSMUSG00000104842.2 | 0 | NA | NA | NA | NA | NA | #VALUE! |
| ENSMUSG00000104843.2 | 0 | NA | NA | NA | NA | NA | #VALUE! |
| ENSMUSG00000104845.2 | 0 | NA | NA | NA | NA | NA | #VALUE! |
| ENSMUSG00000104846.2 | 0 | NA | NA | NA | NA | NA | #VALUE! |
| ENSMUSG00000104847.2 | 0 | NA | NA | NA | NA | NA | #VALUE! |
| ENSMUSG00000104849.2 | 0 | NA | NA | NA | NA | NA | #VALUE! |
| ENSMUSG00000104850.2 | 0 | NA | NA | NA | NA | NA | #VALUE! |
| ENSMUSG00000104851.2 | 0 | NA | NA | NA | NA | NA | #VALUE! |
| ENSMUSG00000104852.2 | 4.009973 | 5.501447 | 3.9179 | 1.404183 | 0.160264 | NA | #VALUE! |
| ENSMUSG00000104853.3 | 0 | NA | NA | NA | NA | NA | #VALUE! |
| ENSMUSG00000104854.3 | 0 | NA | NA | NA | NA | NA | #VALUE! |
| ENSMUSG00000104855.2 | 0 | NA | NA | NA | NA | NA | #VALUE! |
| ENSMUSG00000104858.3 | 0 | NA | NA | NA | NA | NA | #VALUE! |
| ENSMUSG00000104859.2 | 0 | NA | NA | NA | NA | NA | #VALUE! |
| ENSMUSG00000104860.2 | 0 | NA | NA | NA | NA | NA | #VALUE! |
| ENSMUSG00000104861.2 | 0 | NA | NA | NA | NA | NA | #VALUE! |
| ENSMUSG00000104862.2 | 0 | NA | NA | NA | NA | NA | #VALUE! |
| ENSMUSG00000104864.2 | 0 | NA | NA | NA | NA | NA | #VALUE! |
| ENSMUSG00000104865.2 | 0 | NA | NA | NA | NA | NA | #VALUE! |
| ENSMUSG00000104866.2 | 0 | NA | NA | NA | NA | NA | #VALUE! |
| ENSMUSG00000104867.2 | 0 | NA | NA | NA | NA | NA | #VALUE! |
| ENSMUSG00000104868.2 | 0 | NA | NA | NA | NA | NA | #VALUE! |
| ENSMUSG00000104869.2 | 0 | NA | NA | NA | NA | NA | #VALUE! |
| ENSMUSG00000104870.2 | 0 | NA | NA | NA | NA | NA | #VALUE! |
| ENSMUSG00000104871.2 | 0 | NA | NA | NA | NA | NA | #VALUE! |
| ENSMUSG00000104872.2 | 0 | NA | NA | NA | NA | NA | #VALUE! |
| ENSMUSG00000104873.2 | 2.705731 | 4.913666 | 3.923477 | 1.252375 | 0.210433 | NA | #VALUE! |
| ENSMUSG00000104874.2 | 0 | NA | NA | NA | NA | NA | #VALUE! |
| ENSMUSG00000104875.2 | 0 | NA | NA | NA | NA | NA | #VALUE! |
| ENSMUSG00000104876.2 | 0 | NA | NA | NA | NA | NA | #VALUE! |
| ENSMUSG00000104877.2 | 0 | NA | NA | NA | NA | NA | #VALUE! |
| ENSMUSG00000104878.2 | 0 | NA | NA | NA | NA | NA | #VALUE! |
| ENSMUSG00000104881.2 | 0 | NA | NA | NA | NA | NA | #VALUE! |
| ENSMUSG00000104882.2 | 0 | NA | NA | NA | NA | NA | #VALUE! |
| ENSMUSG00000104883.2 | 0 | NA | NA | NA | NA | NA | #VALUE! |
| ENSMUSG00000104884.3 | 0 | NA | NA | NA | NA | NA | #VALUE! |
| ENSMUSG00000104885.2 | 0 | NA | NA | NA | NA | NA | #VALUE! |
| ENSMUSG00000104886.2 | 0 | NA | NA | NA | NA | NA | #VALUE! |
| ENSMUSG00000104887.2 | 5.04063 | -5.86896 | 3.915609 | -1.49886 | 0.133909 | NA | #VALUE! |

| | | | | | | | |
|---|---|---|---|---|---|---|---|
| ENSMUSG00000104888.2 | 0 | NA | NA | NA | NA | NA | #VALUE! |
| ENSMUSG00000104889.2 | 0 | NA | NA | NA | NA | NA | #VALUE! |
| ENSMUSG00000104891.2 | 1.632886 | -4.22813 | 3.934241 | -1.0747 | 0.282509 | NA | #VALUE! |
| ENSMUSG00000104892.2 | 0 | NA | NA | NA | NA | NA | #VALUE! |
| ENSMUSG00000104893.2 | 0 | NA | NA | NA | NA | NA | #VALUE! |
| ENSMUSG00000104894.5 | 0 | NA | NA | NA | NA | NA | #VALUE! |
| ENSMUSG00000104895.2 | 0 | NA | NA | NA | NA | NA | #VALUE! |
| ENSMUSG00000104897.2 | 0 | NA | NA | NA | NA | NA | #VALUE! |
| ENSMUSG00000104898.2 | 0 | NA | NA | NA | NA | NA | #VALUE! |
| ENSMUSG00000104899.2 | 0 | NA | NA | NA | NA | NA | #VALUE! |
| ENSMUSG00000104900.2 | 0 | NA | NA | NA | NA | NA | #VALUE! |
| ENSMUSG00000104901.2 | 0 | NA | NA | NA | NA | NA | #VALUE! |
| ENSMUSG00000104902.2 | 0 | NA | NA | NA | NA | NA | #VALUE! |
| ENSMUSG00000104903.2 | 0 | NA | NA | NA | NA | NA | #VALUE! |
| ENSMUSG00000104904.2 | 0 | NA | NA | NA | NA | NA | #VALUE! |
| ENSMUSG00000104905.2 | 0 | NA | NA | NA | NA | NA | #VALUE! |
| ENSMUSG00000104906.2 | 0 | NA | NA | NA | NA | NA | #VALUE! |
| ENSMUSG00000104908.2 | 0 | NA | NA | NA | NA | NA | #VALUE! |
| ENSMUSG00000104909.2 | 0 | NA | NA | NA | NA | NA | #VALUE! |
| ENSMUSG00000104910.2 | 0 | NA | NA | NA | NA | NA | #VALUE! |
| ENSMUSG00000104912.2 | 0 | NA | NA | NA | NA | NA | #VALUE! |
| ENSMUSG00000104913.2 | 0 | NA | NA | NA | NA | NA | #VALUE! |
| ENSMUSG00000104914.3 | 0 | NA | NA | NA | NA | NA | #VALUE! |
| ENSMUSG00000104915.2 | 0 | NA | NA | NA | NA | NA | #VALUE! |
| ENSMUSG00000104916.2 | 0 | NA | NA | NA | NA | NA | #VALUE! |
| ENSMUSG00000104917.2 | 5.250656 | -5.92764 | 3.915257 | -1.51398 | 0.13003 | NA | #VALUE! |
| ENSMUSG00000104918.2 | 0 | NA | NA | NA | NA | NA | #VALUE! |
| ENSMUSG00000104919.2 | 0 | NA | NA | NA | NA | NA | #VALUE! |
| ENSMUSG00000104920.2 | 0 | NA | NA | NA | NA | NA | #VALUE! |
| ENSMUSG00000104921.3 | 0 | NA | NA | NA | NA | NA | #VALUE! |
| ENSMUSG00000104922.2 | 0 | NA | NA | NA | NA | NA | #VALUE! |
| ENSMUSG00000104923.2 | 0.18651 | 1.401769 | 4.056354 | 0.345574 | 0.729663 | NA | #VALUE! |
| ENSMUSG00000104924.2 | 0 | NA | NA | NA | NA | NA | #VALUE! |
| ENSMUSG00000104925.2 | 0 | NA | NA | NA | NA | NA | #VALUE! |
| ENSMUSG00000104926.3 | 0 | NA | NA | NA | NA | NA | #VALUE! |
| ENSMUSG00000104927.2 | 0 | NA | NA | NA | NA | NA | #VALUE! |
| ENSMUSG00000104928.2 | 0 | NA | NA | NA | NA | NA | #VALUE! |
| ENSMUSG00000104929.2 | 0 | NA | NA | NA | NA | NA | #VALUE! |
| ENSMUSG00000104930.2 | 0 | NA | NA | NA | NA | NA | #VALUE! |
| ENSMUSG00000104931.2 | 0 | NA | NA | NA | NA | NA | #VALUE! |
| ENSMUSG00000104932.2 | 0 | NA | NA | NA | NA | NA | #VALUE! |
| ENSMUSG00000104934.2 | 0 | NA | NA | NA | NA | NA | #VALUE! |
| ENSMUSG00000104935.2 | 0 | NA | NA | NA | NA | NA | #VALUE! |
| ENSMUSG00000104936.2 | 0 | NA | NA | NA | NA | NA | #VALUE! |

| | | | | | | | |
|---|---|---|---|---|---|---|---|
| ENSMUSG00000104937.2 | 0 | NA | NA | NA | NA | NA | #VALUE! |
| ENSMUSG00000104938.2 | 0 | NA | NA | NA | NA | NA | #VALUE! |
| ENSMUSG00000104939.2 | 0 | NA | NA | NA | NA | NA | #VALUE! |
| ENSMUSG00000104940.2 | 0 | NA | NA | NA | NA | NA | #VALUE! |
| ENSMUSG00000104941.2 | 0 | NA | NA | NA | NA | NA | #VALUE! |
| ENSMUSG00000104942.2 | 0 | NA | NA | NA | NA | NA | #VALUE! |
| ENSMUSG00000104943.2 | 0 | NA | NA | NA | NA | NA | #VALUE! |
| ENSMUSG00000104944.2 | 0 | NA | NA | NA | NA | NA | #VALUE! |
| ENSMUSG00000104945.2 | 0 | NA | NA | NA | NA | NA | #VALUE! |
| ENSMUSG00000104946.2 | 0 | NA | NA | NA | NA | NA | #VALUE! |
| ENSMUSG00000104947.2 | 0 | NA | NA | NA | NA | NA | #VALUE! |
| ENSMUSG00000104949.2 | 0 | NA | NA | NA | NA | NA | #VALUE! |
| ENSMUSG00000104951.2 | 0 | NA | NA | NA | NA | NA | #VALUE! |
| ENSMUSG00000104952.2 | 0 | NA | NA | NA | NA | NA | #VALUE! |
| ENSMUSG00000104954.2 | 0 | NA | NA | NA | NA | NA | #VALUE! |
| ENSMUSG00000104955.2 | 0 | NA | NA | NA | NA | NA | #VALUE! |
| ENSMUSG00000104956.2 | 0 | NA | NA | NA | NA | NA | #VALUE! |
| ENSMUSG00000104957.2 | 0 | NA | NA | NA | NA | NA | #VALUE! |
| ENSMUSG00000104958.3 | 0 | NA | NA | NA | NA | NA | #VALUE! |
| ENSMUSG00000104959.2 | 0 | NA | NA | NA | NA | NA | #VALUE! |
| ENSMUSG00000104961.2 | 0 | NA | NA | NA | NA | NA | #VALUE! |
| ENSMUSG00000104962.2 | 0 | NA | NA | NA | NA | NA | #VALUE! |
| ENSMUSG00000104963.2 | 0.093255 | 0.868093 | 4.072464 | 0.213162 | 0.831201 | NA | #VALUE! |
| ENSMUSG00000104964.2 | 0 | NA | NA | NA | NA | NA | #VALUE! |
| ENSMUSG00000104965.2 | 0 | NA | NA | NA | NA | NA | #VALUE! |
| ENSMUSG00000104966.2 | 0 | NA | NA | NA | NA | NA | #VALUE! |
| ENSMUSG00000104967.2 | 0 | NA | NA | NA | NA | NA | #VALUE! |
| ENSMUSG00000104968.2 | 0 | NA | NA | NA | NA | NA | #VALUE! |
| ENSMUSG00000104969.2 | 0 | NA | NA | NA | NA | NA | #VALUE! |
| ENSMUSG00000104970.2 | 0 | NA | NA | NA | NA | NA | #VALUE! |
| ENSMUSG00000104971.2 | 0 | NA | NA | NA | NA | NA | #VALUE! |
| ENSMUSG00000104972.2 | 0 | NA | NA | NA | NA | NA | #VALUE! |
| ENSMUSG00000104973.2 | 6.595688 | 6.212383 | 3.102263 | 2.002533 | 0.045227 | NA | #VALUE! |
| ENSMUSG00000104974.2 | 0 | NA | NA | NA | NA | NA | #VALUE! |
| ENSMUSG00000104975.2 | 0 | NA | NA | NA | NA | NA | #VALUE! |
| ENSMUSG00000104976.2 | 0 | NA | NA | NA | NA | NA | #VALUE! |
| ENSMUSG00000104978.2 | 0 | NA | NA | NA | NA | NA | #VALUE! |
| ENSMUSG00000104979.3 | 0 | NA | NA | NA | NA | NA | #VALUE! |
| ENSMUSG00000104980.2 | 0 | NA | NA | NA | NA | NA | #VALUE! |
| ENSMUSG00000104981.2 | 0 | NA | NA | NA | NA | NA | #VALUE! |
| ENSMUSG00000104982.2 | 0 | NA | NA | NA | NA | NA | #VALUE! |
| ENSMUSG00000104983.2 | 0 | NA | NA | NA | NA | NA | #VALUE! |
| ENSMUSG00000104984.2 | 0 | NA | NA | NA | NA | NA | #VALUE! |
| ENSMUSG00000104985.2 | 0 | NA | NA | NA | NA | NA | #VALUE! |

| | | | | | | | |
|---|---|---|---|---|---|---|---|
| ENSMUSG00000104986.2 | 0 | NA | NA | NA | NA | NA | #VALUE! |
| ENSMUSG00000104987.2 | 0 | NA | NA | NA | NA | NA | #VALUE! |
| ENSMUSG00000104988.2 | 0 | NA | NA | NA | NA | NA | #VALUE! |
| ENSMUSG00000104990.2 | 0 | NA | NA | NA | NA | NA | #VALUE! |
| ENSMUSG00000104991.4 | 0 | NA | NA | NA | NA | NA | #VALUE! |
| ENSMUSG00000104992.2 | 0 | NA | NA | NA | NA | NA | #VALUE! |
| ENSMUSG00000104993.2 | 0 | NA | NA | NA | NA | NA | #VALUE! |
| ENSMUSG00000104994.2 | 0 | NA | NA | NA | NA | NA | #VALUE! |
| ENSMUSG00000104995.2 | 0 | NA | NA | NA | NA | NA | #VALUE! |
| ENSMUSG00000104996.3 | 0 | NA | NA | NA | NA | NA | #VALUE! |
| ENSMUSG00000104998.2 | 0 | NA | NA | NA | NA | NA | #VALUE! |
| ENSMUSG00000104999.2 | 0 | NA | NA | NA | NA | NA | #VALUE! |
| ENSMUSG00000105000.2 | 0 | NA | NA | NA | NA | NA | #VALUE! |
| ENSMUSG00000105003.2 | 0 | NA | NA | NA | NA | NA | #VALUE! |
| ENSMUSG00000105004.2 | 0 | NA | NA | NA | NA | NA | #VALUE! |
| ENSMUSG00000105005.2 | 0 | NA | NA | NA | NA | NA | #VALUE! |
| ENSMUSG00000105006.2 | 5.460683 | -5.98402 | 3.914932 | -1.52851 | 0.126385 | NA | #VALUE! |
| ENSMUSG00000105007.2 | 0 | NA | NA | NA | NA | NA | #VALUE! |
| ENSMUSG00000105008.2 | 0 | NA | NA | NA | NA | NA | #VALUE! |
| ENSMUSG00000105009.2 | 0 | NA | NA | NA | NA | NA | #VALUE! |
| ENSMUSG00000105010.3 | 0 | NA | NA | NA | NA | NA | #VALUE! |
| ENSMUSG00000105011.2 | 0 | NA | NA | NA | NA | NA | #VALUE! |
| ENSMUSG00000105012.2 | 0 | NA | NA | NA | NA | NA | #VALUE! |
| ENSMUSG00000105013.2 | 0 | NA | NA | NA | NA | NA | #VALUE! |
| ENSMUSG00000105015.2 | 0 | NA | NA | NA | NA | NA | #VALUE! |
| ENSMUSG00000105016.2 | 0 | NA | NA | NA | NA | NA | #VALUE! |
| ENSMUSG00000105017.2 | 0 | NA | NA | NA | NA | NA | #VALUE! |
| ENSMUSG00000105018.2 | 0 | NA | NA | NA | NA | NA | #VALUE! |
| ENSMUSG00000105019.2 | 0 | NA | NA | NA | NA | NA | #VALUE! |
| ENSMUSG00000105020.2 | 0 | NA | NA | NA | NA | NA | #VALUE! |
| ENSMUSG00000105021.2 | 0 | NA | NA | NA | NA | NA | #VALUE! |
| ENSMUSG00000105023.2 | 0 | NA | NA | NA | NA | NA | #VALUE! |
| ENSMUSG00000105024.2 | 0 | NA | NA | NA | NA | NA | #VALUE! |
| ENSMUSG00000105026.2 | 0 | NA | NA | NA | NA | NA | #VALUE! |
| ENSMUSG00000105027.2 | 0 | NA | NA | NA | NA | NA | #VALUE! |
| ENSMUSG00000105028.2 | 0 | NA | NA | NA | NA | NA | #VALUE! |
| ENSMUSG00000105030.3 | 0 | NA | NA | NA | NA | NA | #VALUE! |
| ENSMUSG00000105031.2 | 31.5871 | -3.17243 | 2.499329 | -1.26931 | NA | NA | #VALUE! |
| ENSMUSG00000105032.2 | 0 | NA | NA | NA | NA | NA | #VALUE! |
| ENSMUSG00000105033.2 | 0 | NA | NA | NA | NA | NA | #VALUE! |
| ENSMUSG00000105034.2 | 0 | NA | NA | NA | NA | NA | #VALUE! |
| ENSMUSG00000105035.2 | 0 | NA | NA | NA | NA | NA | #VALUE! |
| ENSMUSG00000105036.2 | 0 | NA | NA | NA | NA | NA | #VALUE! |
| ENSMUSG00000105037.3 | 0 | NA | NA | NA | NA | NA | #VALUE! |

| | | | | | | | |
|---|---|---|---|---|---|---|---|
| ENSMUSG00000105038.2 | 0 | NA | NA | NA | NA | NA | #VALUE! |
| ENSMUSG00000105039.2 | 0 | NA | NA | NA | NA | NA | #VALUE! |
| ENSMUSG00000105040.2 | 0 | NA | NA | NA | NA | NA | #VALUE! |
| ENSMUSG00000105041.3 | 0 | NA | NA | NA | NA | NA | #VALUE! |
| ENSMUSG00000105042.2 | 0 | NA | NA | NA | NA | NA | #VALUE! |
| ENSMUSG00000105043.2 | 0 | NA | NA | NA | NA | NA | #VALUE! |
| ENSMUSG00000105044.2 | 0 | NA | NA | NA | NA | NA | #VALUE! |
| ENSMUSG00000105045.2 | 0 | NA | NA | NA | NA | NA | #VALUE! |
| ENSMUSG00000105047.2 | 0 | NA | NA | NA | NA | NA | #VALUE! |
| ENSMUSG00000105048.2 | 0 | NA | NA | NA | NA | NA | #VALUE! |
| ENSMUSG00000105049.2 | 0 | NA | NA | NA | NA | NA | #VALUE! |
| ENSMUSG00000105050.2 | 0 | NA | NA | NA | NA | NA | #VALUE! |
| ENSMUSG00000105051.2 | 0 | NA | NA | NA | NA | NA | #VALUE! |
| ENSMUSG00000105052.5 | 0 | NA | NA | NA | NA | NA | #VALUE! |
| ENSMUSG00000105053.2 | 0 | NA | NA | NA | NA | NA | #VALUE! |
| ENSMUSG00000105054.2 | 0 | NA | NA | NA | NA | NA | #VALUE! |
| ENSMUSG00000105055.2 | 0 | NA | NA | NA | NA | NA | #VALUE! |
| ENSMUSG00000105056.2 | 0 | NA | NA | NA | NA | NA | #VALUE! |
| ENSMUSG00000105057.2 | 0 | NA | NA | NA | NA | NA | #VALUE! |
| ENSMUSG00000105058.2 | 0 | NA | NA | NA | NA | NA | #VALUE! |
| ENSMUSG00000105059.2 | 0 | NA | NA | NA | NA | NA | #VALUE! |
| ENSMUSG00000105061.2 | 0 | NA | NA | NA | NA | NA | #VALUE! |
| ENSMUSG00000105062.2 | 0 | NA | NA | NA | NA | NA | #VALUE! |
| ENSMUSG00000105063.2 | 0 | NA | NA | NA | NA | NA | #VALUE! |
| ENSMUSG00000105065.2 | 0 | NA | NA | NA | NA | NA | #VALUE! |
| ENSMUSG00000105066.2 | 0 | NA | NA | NA | NA | NA | #VALUE! |
| ENSMUSG00000105067.2 | 0 | NA | NA | NA | NA | NA | #VALUE! |
| ENSMUSG00000105068.2 | 0 | NA | NA | NA | NA | NA | #VALUE! |
| ENSMUSG00000105070.2 | 0 | NA | NA | NA | NA | NA | #VALUE! |
| ENSMUSG00000105071.2 | 0 | NA | NA | NA | NA | NA | #VALUE! |
| ENSMUSG00000105073.2 | 0 | NA | NA | NA | NA | NA | #VALUE! |
| ENSMUSG00000105074.2 | 0 | NA | NA | NA | NA | NA | #VALUE! |
| ENSMUSG00000105075.2 | 0 | NA | NA | NA | NA | NA | #VALUE! |
| ENSMUSG00000105076.2 | 0 | NA | NA | NA | NA | NA | #VALUE! |
| ENSMUSG00000105077.2 | 0 | NA | NA | NA | NA | NA | #VALUE! |
| ENSMUSG00000105078.3 | 0 | NA | NA | NA | NA | NA | #VALUE! |
| ENSMUSG00000105079.2 | 0 | NA | NA | NA | NA | NA | #VALUE! |
| ENSMUSG00000105080.2 | 0 | NA | NA | NA | NA | NA | #VALUE! |
| ENSMUSG00000105081.2 | 0 | NA | NA | NA | NA | NA | #VALUE! |
| ENSMUSG00000105082.3 | 0 | NA | NA | NA | NA | NA | #VALUE! |
| ENSMUSG00000105083.2 | 0 | NA | NA | NA | NA | NA | #VALUE! |
| ENSMUSG00000105084.2 | 0 | NA | NA | NA | NA | NA | #VALUE! |
| ENSMUSG00000105085.2 | 0 | NA | NA | NA | NA | NA | #VALUE! |
| ENSMUSG00000105086.2 | 0 | NA | NA | NA | NA | NA | #VALUE! |

| | | | | | | | |
|---|---|---|---|---|---|---|---|
| ENSMUSG00000105087.2 | 0 | NA | NA | NA | NA | NA | #VALUE! |
| ENSMUSG00000105088.2 | 0 | NA | NA | NA | NA | NA | #VALUE! |
| ENSMUSG00000105089.3 | 0 | NA | NA | NA | NA | NA | #VALUE! |
| ENSMUSG00000105090.2 | 0 | NA | NA | NA | NA | NA | #VALUE! |
| ENSMUSG00000105091.2 | 0 | NA | NA | NA | NA | NA | #VALUE! |
| ENSMUSG00000105092.2 | 0 | NA | NA | NA | NA | NA | #VALUE! |
| ENSMUSG00000105093.2 | 0 | NA | NA | NA | NA | NA | #VALUE! |
| ENSMUSG00000105094.2 | 0 | NA | NA | NA | NA | NA | #VALUE! |
| ENSMUSG00000105095.2 | 0 | NA | NA | NA | NA | NA | #VALUE! |
| ENSMUSG00000105096.2 | 0 | NA | NA | NA | NA | NA | #VALUE! |
| ENSMUSG00000105097.2 | 0 | NA | NA | NA | NA | NA | #VALUE! |
| ENSMUSG00000105099.2 | 0 | NA | NA | NA | NA | NA | #VALUE! |
| ENSMUSG00000105101.2 | 0 | NA | NA | NA | NA | NA | #VALUE! |
| ENSMUSG00000105102.2 | 0 | NA | NA | NA | NA | NA | #VALUE! |
| ENSMUSG00000105103.2 | 0 | NA | NA | NA | NA | NA | #VALUE! |
| ENSMUSG00000105104.2 | 0 | NA | NA | NA | NA | NA | #VALUE! |
| ENSMUSG00000105105.2 | 0 | NA | NA | NA | NA | NA | #VALUE! |
| ENSMUSG00000105106.2 | 0 | NA | NA | NA | NA | NA | #VALUE! |
| ENSMUSG00000105107.2 | 0 | NA | NA | NA | NA | NA | #VALUE! |
| ENSMUSG00000105108.2 | 0 | NA | NA | NA | NA | NA | #VALUE! |
| ENSMUSG00000105110.2 | 0 | NA | NA | NA | NA | NA | #VALUE! |
| ENSMUSG00000105111.2 | 0 | NA | NA | NA | NA | NA | #VALUE! |
| ENSMUSG00000105112.2 | 0 | NA | NA | NA | NA | NA | #VALUE! |
| ENSMUSG00000105113.2 | 0 | NA | NA | NA | NA | NA | #VALUE! |
| ENSMUSG00000105114.2 | 0 | NA | NA | NA | NA | NA | #VALUE! |
| ENSMUSG00000105117.2 | 0 | NA | NA | NA | NA | NA | #VALUE! |
| ENSMUSG00000105119.2 | 0 | NA | NA | NA | NA | NA | #VALUE! |
| ENSMUSG00000105120.3 | 0 | NA | NA | NA | NA | NA | #VALUE! |
| ENSMUSG00000105121.2 | 0 | NA | NA | NA | NA | NA | #VALUE! |
| ENSMUSG00000105123.2 | 0 | NA | NA | NA | NA | NA | #VALUE! |
| ENSMUSG00000105124.3 | 0.942037 | 1.992212 | 3.961267 | 0.502923 | 0.615018 | NA | #VALUE! |
| ENSMUSG00000105125.2 | 0 | NA | NA | NA | NA | NA | #VALUE! |
| ENSMUSG00000105126.2 | 0 | NA | NA | NA | NA | NA | #VALUE! |
| ENSMUSG00000105127.2 | 0 | NA | NA | NA | NA | NA | #VALUE! |
| ENSMUSG00000105128.2 | 0 | NA | NA | NA | NA | NA | #VALUE! |
| ENSMUSG00000105129.2 | 0 | NA | NA | NA | NA | NA | #VALUE! |
| ENSMUSG00000105130.2 | 6.825853 | -6.30496 | 3.913306 | -1.61116 | 0.107145 | NA | #VALUE! |
| ENSMUSG00000105132.2 | 0 | NA | NA | NA | NA | NA | #VALUE! |
| ENSMUSG00000105134.2 | 0 | NA | NA | NA | NA | NA | #VALUE! |
| ENSMUSG00000105136.2 | 2.310289 | -4.7496 | 3.925939 | -1.2098 | 0.226355 | NA | #VALUE! |
| ENSMUSG00000105137.2 | 0 | NA | NA | NA | NA | NA | #VALUE! |
| ENSMUSG00000105138.3 | 0 | NA | NA | NA | NA | NA | #VALUE! |
| ENSMUSG00000105139.2 | 0 | NA | NA | NA | NA | NA | #VALUE! |
| ENSMUSG00000105140.2 | 0 | NA | NA | NA | NA | NA | #VALUE! |

| | | | | | | | |
|---|---|---|---|---|---|---|---|
| ENSMUSG00000105141.2 | 0 | NA | NA | NA | NA | NA | #VALUE! |
| ENSMUSG00000105142.3 | 0 | NA | NA | NA | NA | NA | #VALUE! |
| ENSMUSG00000105143.2 | 0 | NA | NA | NA | NA | NA | #VALUE! |
| ENSMUSG00000105144.2 | 0 | NA | NA | NA | NA | NA | #VALUE! |
| ENSMUSG00000105146.2 | 0 | NA | NA | NA | NA | NA | #VALUE! |
| ENSMUSG00000105147.2 | 0 | NA | NA | NA | NA | NA | #VALUE! |
| ENSMUSG00000105148.2 | 0 | NA | NA | NA | NA | NA | #VALUE! |
| ENSMUSG00000105149.2 | 0 | NA | NA | NA | NA | NA | #VALUE! |
| ENSMUSG00000105150.2 | 0 | NA | NA | NA | NA | NA | #VALUE! |
| ENSMUSG00000105151.2 | 0 | NA | NA | NA | NA | NA | #VALUE! |
| ENSMUSG00000105152.2 | 0 | NA | NA | NA | NA | NA | #VALUE! |
| ENSMUSG00000105153.5 | 0 | NA | NA | NA | NA | NA | #VALUE! |
| ENSMUSG00000105154.2 | 0 | NA | NA | NA | NA | NA | #VALUE! |
| ENSMUSG00000105155.2 | 0 | NA | NA | NA | NA | NA | #VALUE! |
| ENSMUSG00000105156.2 | 0 | NA | NA | NA | NA | NA | #VALUE! |
| ENSMUSG00000105157.2 | 0 | NA | NA | NA | NA | NA | #VALUE! |
| ENSMUSG00000105158.2 | 0 | NA | NA | NA | NA | NA | #VALUE! |
| ENSMUSG00000105159.2 | 0 | NA | NA | NA | NA | NA | #VALUE! |
| ENSMUSG00000105160.2 | 0.816443 | -3.21914 | 3.961753 | -0.81255 | 0.416474 | NA | #VALUE! |
| ENSMUSG00000105161.2 | 5.688567 | 6.00357 | 3.914632 | 1.533623 | 0.125122 | NA | #VALUE! |
| ENSMUSG00000105162.2 | 0 | NA | NA | NA | NA | NA | #VALUE! |
| ENSMUSG00000105163.2 | 0 | NA | NA | NA | NA | NA | #VALUE! |
| ENSMUSG00000105165.3 | 0 | NA | NA | NA | NA | NA | #VALUE! |
| ENSMUSG00000105166.2 | 0 | NA | NA | NA | NA | NA | #VALUE! |
| ENSMUSG00000105167.4 | 0 | NA | NA | NA | NA | NA | #VALUE! |
| ENSMUSG00000105168.2 | 0 | NA | NA | NA | NA | NA | #VALUE! |
| ENSMUSG00000105170.2 | 0 | NA | NA | NA | NA | NA | #VALUE! |
| ENSMUSG00000105171.2 | 0 | NA | NA | NA | NA | NA | #VALUE! |
| ENSMUSG00000105172.2 | 0 | NA | NA | NA | NA | NA | #VALUE! |
| ENSMUSG00000105173.2 | 0 | NA | NA | NA | NA | NA | #VALUE! |
| ENSMUSG00000105174.2 | 0 | NA | NA | NA | NA | NA | #VALUE! |
| ENSMUSG00000105175.2 | 0 | NA | NA | NA | NA | NA | #VALUE! |
| ENSMUSG00000105176.2 | 8.78815 | 0.05339 | 3.00118 | 0.01779 | 0.985807 | NA | #VALUE! |
| ENSMUSG00000105177.2 | 0 | NA | NA | NA | NA | NA | #VALUE! |
| ENSMUSG00000105178.3 | 0 | NA | NA | NA | NA | NA | #VALUE! |
| ENSMUSG00000105179.2 | 0.408222 | -2.20205 | 4.016842 | -0.54821 | 0.583551 | NA | #VALUE! |
| ENSMUSG00000105180.2 | 0 | NA | NA | NA | NA | NA | #VALUE! |
| ENSMUSG00000105181.2 | 0.195789 | -1.05581 | 4.073302 | -0.2592 | 0.795479 | NA | #VALUE! |
| ENSMUSG00000105183.2 | 0 | NA | NA | NA | NA | NA | #VALUE! |
| ENSMUSG00000105184.2 | 0 | NA | NA | NA | NA | NA | #VALUE! |
| ENSMUSG00000105185.5 | 0 | NA | NA | NA | NA | NA | #VALUE! |
| ENSMUSG00000105186.2 | 0 | NA | NA | NA | NA | NA | #VALUE! |
| ENSMUSG00000105187.3 | 0 | NA | NA | NA | NA | NA | #VALUE! |
| ENSMUSG00000105188.2 | 0 | NA | NA | NA | NA | NA | #VALUE! |

| | | | | | | | |
|---|---|---|---|---|---|---|---|
| ENSMUSG00000105189.2 | 0 | NA | NA | NA | NA | NA | #VALUE! |
| ENSMUSG00000105190.2 | 0 | NA | NA | NA | NA | NA | #VALUE! |
| ENSMUSG00000105191.2 | 0 | NA | NA | NA | NA | NA | #VALUE! |
| ENSMUSG00000105192.2 | 0 | NA | NA | NA | NA | NA | #VALUE! |
| ENSMUSG00000105193.2 | 0 | NA | NA | NA | NA | NA | #VALUE! |
| ENSMUSG00000105194.2 | 0 | NA | NA | NA | NA | NA | #VALUE! |
| ENSMUSG00000105195.2 | 0 | NA | NA | NA | NA | NA | #VALUE! |
| ENSMUSG00000105196.3 | 0 | NA | NA | NA | NA | NA | #VALUE! |
| ENSMUSG00000105197.2 | 0 | NA | NA | NA | NA | NA | #VALUE! |
| ENSMUSG00000105198.2 | 0 | NA | NA | NA | NA | NA | #VALUE! |
| ENSMUSG00000105199.2 | 0 | NA | NA | NA | NA | NA | #VALUE! |
| ENSMUSG00000105200.3 | 0 | NA | NA | NA | NA | NA | #VALUE! |
| ENSMUSG00000105201.2 | 4.894209 | -5.82407 | 3.915888 | -1.48729 | 0.136937 | NA | #VALUE! |
| ENSMUSG00000105202.2 | 0 | NA | NA | NA | NA | NA | #VALUE! |
| ENSMUSG00000105203.2 | 0 | NA | NA | NA | NA | NA | #VALUE! |
| ENSMUSG00000105204.2 | 0 | NA | NA | NA | NA | NA | #VALUE! |
| ENSMUSG00000105206.2 | 0 | NA | NA | NA | NA | NA | #VALUE! |
| ENSMUSG00000105207.2 | 0 | NA | NA | NA | NA | NA | #VALUE! |
| ENSMUSG00000105208.2 | 0 | NA | NA | NA | NA | NA | #VALUE! |
| ENSMUSG00000105209.2 | 0 | NA | NA | NA | NA | NA | #VALUE! |
| ENSMUSG00000105210.3 | 0 | NA | NA | NA | NA | NA | #VALUE! |
| ENSMUSG00000105212.2 | 0 | NA | NA | NA | NA | NA | #VALUE! |
| ENSMUSG00000105213.2 | 0 | NA | NA | NA | NA | NA | #VALUE! |
| ENSMUSG00000105216.2 | 0 | NA | NA | NA | NA | NA | #VALUE! |
| ENSMUSG00000105217.2 | 0 | NA | NA | NA | NA | NA | #VALUE! |
| ENSMUSG00000105218.5 | 0 | NA | NA | NA | NA | NA | #VALUE! |
| ENSMUSG00000105219.2 | 0 | NA | NA | NA | NA | NA | #VALUE! |
| ENSMUSG00000105220.3 | 0 | NA | NA | NA | NA | NA | #VALUE! |
| ENSMUSG00000105221.2 | 0 | NA | NA | NA | NA | NA | #VALUE! |
| ENSMUSG00000105222.2 | 0 | NA | NA | NA | NA | NA | #VALUE! |
| ENSMUSG00000105223.2 | 0 | NA | NA | NA | NA | NA | #VALUE! |
| ENSMUSG00000105224.2 | 0 | NA | NA | NA | NA | NA | #VALUE! |
| ENSMUSG00000105225.2 | 0 | NA | NA | NA | NA | NA | #VALUE! |
| ENSMUSG00000105226.2 | 0 | NA | NA | NA | NA | NA | #VALUE! |
| ENSMUSG00000105227.2 | 0 | NA | NA | NA | NA | NA | #VALUE! |
| ENSMUSG00000105228.2 | 0 | NA | NA | NA | NA | NA | #VALUE! |
| ENSMUSG00000105229.2 | 0 | NA | NA | NA | NA | NA | #VALUE! |
| ENSMUSG00000105230.2 | 0 | NA | NA | NA | NA | NA | #VALUE! |
| ENSMUSG00000105231.2 | 0 | NA | NA | NA | NA | NA | #VALUE! |
| ENSMUSG00000105233.2 | 7.327856 | 2.268078 | 2.869048 | 0.790533 | 0.429216 | NA | #VALUE! |
| ENSMUSG00000105234.2 | 0 | NA | NA | NA | NA | NA | #VALUE! |
| ENSMUSG00000105235.2 | 0 | NA | NA | NA | NA | NA | #VALUE! |
| ENSMUSG00000105237.2 | 0 | NA | NA | NA | NA | NA | #VALUE! |
| ENSMUSG00000105239.2 | 0 | NA | NA | NA | NA | NA | #VALUE! |

| | | | | | | | |
|---|---|---|---|---|---|---|---|
| ENSMUSG00000105240.2 | 0 | NA | NA | NA | NA | NA | #VALUE! |
| ENSMUSG00000105241.2 | 0 | NA | NA | NA | NA | NA | #VALUE! |
| ENSMUSG00000105242.2 | 0 | NA | NA | NA | NA | NA | #VALUE! |
| ENSMUSG00000105243.2 | 0.195789 | -1.05581 | 4.073302 | -0.2592 | 0.795479 | NA | #VALUE! |
| ENSMUSG00000105244.2 | 0 | NA | NA | NA | NA | NA | #VALUE! |
| ENSMUSG00000105245.2 | 0 | NA | NA | NA | NA | NA | #VALUE! |
| ENSMUSG00000105246.2 | 0 | NA | NA | NA | NA | NA | #VALUE! |
| ENSMUSG00000105247.2 | 0 | NA | NA | NA | NA | NA | #VALUE! |
| ENSMUSG00000105248.2 | 0 | NA | NA | NA | NA | NA | #VALUE! |
| ENSMUSG00000105250.2 | 0 | NA | NA | NA | NA | NA | #VALUE! |
| ENSMUSG00000105251.2 | 0 | NA | NA | NA | NA | NA | #VALUE! |
| ENSMUSG00000105252.2 | 0 | NA | NA | NA | NA | NA | #VALUE! |
| ENSMUSG00000105253.2 | 0 | NA | NA | NA | NA | NA | #VALUE! |
| ENSMUSG00000105254.2 | 0 | NA | NA | NA | NA | NA | #VALUE! |
| ENSMUSG00000105257.2 | 0 | NA | NA | NA | NA | NA | #VALUE! |
| ENSMUSG00000105258.2 | 0 | NA | NA | NA | NA | NA | #VALUE! |
| ENSMUSG00000105259.2 | 0 | NA | NA | NA | NA | NA | #VALUE! |
| ENSMUSG00000105260.2 | 0.391578 | -2.1489 | 4.020883 | -0.53444 | 0.59304 | NA | #VALUE! |
| ENSMUSG00000105261.2 | 0 | NA | NA | NA | NA | NA | #VALUE! |
| ENSMUSG00000105262.2 | 0 | NA | NA | NA | NA | NA | #VALUE! |
| ENSMUSG00000105263.2 | 0 | NA | NA | NA | NA | NA | #VALUE! |
| ENSMUSG00000105264.2 | 0 | NA | NA | NA | NA | NA | #VALUE! |
| ENSMUSG00000105265.6 | 0 | NA | NA | NA | NA | NA | #VALUE! |
| ENSMUSG00000105266.2 | 0 | NA | NA | NA | NA | NA | #VALUE! |
| ENSMUSG00000105267.2 | 0 | NA | NA | NA | NA | NA | #VALUE! |
| ENSMUSG00000105269.2 | 0 | NA | NA | NA | NA | NA | #VALUE! |
| ENSMUSG00000105270.2 | 0 | NA | NA | NA | NA | NA | #VALUE! |
| ENSMUSG00000105271.2 | 0 | NA | NA | NA | NA | NA | #VALUE! |
| ENSMUSG00000105272.2 | 0 | NA | NA | NA | NA | NA | #VALUE! |
| ENSMUSG00000105273.2 | 0 | NA | NA | NA | NA | NA | #VALUE! |
| ENSMUSG00000105274.2 | 0 | NA | NA | NA | NA | NA | #VALUE! |
| ENSMUSG00000105275.2 | 0 | NA | NA | NA | NA | NA | #VALUE! |
| ENSMUSG00000105277.2 | 0 | NA | NA | NA | NA | NA | #VALUE! |
| ENSMUSG00000105278.2 | 0 | NA | NA | NA | NA | NA | #VALUE! |
| ENSMUSG00000105279.2 | 0 | NA | NA | NA | NA | NA | #VALUE! |
| ENSMUSG00000105280.2 | 0 | NA | NA | NA | NA | NA | #VALUE! |
| ENSMUSG00000105281.2 | 0 | NA | NA | NA | NA | NA | #VALUE! |
| ENSMUSG00000105282.2 | 0 | NA | NA | NA | NA | NA | #VALUE! |
| ENSMUSG00000105283.2 | 0 | NA | NA | NA | NA | NA | #VALUE! |
| ENSMUSG00000105284.2 | 0 | NA | NA | NA | NA | NA | #VALUE! |
| ENSMUSG00000105285.2 | 0 | NA | NA | NA | NA | NA | #VALUE! |
| ENSMUSG00000105286.2 | 0 | NA | NA | NA | NA | NA | #VALUE! |
| ENSMUSG00000105287.2 | 0 | NA | NA | NA | NA | NA | #VALUE! |
| ENSMUSG00000105289.2 | 0 | NA | NA | NA | NA | NA | #VALUE! |

| | | | | | | | |
|---|---|---|---|---|---|---|---|
| ENSMUSG00000105290.2 | 0 | NA | NA | NA | NA | NA | #VALUE! |
| ENSMUSG00000105291.2 | 0 | NA | NA | NA | NA | NA | #VALUE! |
| ENSMUSG00000105293.2 | 0 | NA | NA | NA | NA | NA | #VALUE! |
| ENSMUSG00000105294.2 | 0 | NA | NA | NA | NA | NA | #VALUE! |
| ENSMUSG00000105295.2 | 0 | NA | NA | NA | NA | NA | #VALUE! |
| ENSMUSG00000105296.2 | 0 | NA | NA | NA | NA | NA | #VALUE! |
| ENSMUSG00000105297.2 | 0 | NA | NA | NA | NA | NA | #VALUE! |
| ENSMUSG00000105298.2 | 0 | NA | NA | NA | NA | NA | #VALUE! |
| ENSMUSG00000105299.2 | 0 | NA | NA | NA | NA | NA | #VALUE! |
| ENSMUSG00000105300.2 | 0 | NA | NA | NA | NA | NA | #VALUE! |
| ENSMUSG00000105301.2 | 0 | NA | NA | NA | NA | NA | #VALUE! |
| ENSMUSG00000105302.2 | 0 | NA | NA | NA | NA | NA | #VALUE! |
| ENSMUSG00000105303.2 | 0 | NA | NA | NA | NA | NA | #VALUE! |
| ENSMUSG00000105304.2 | 3.197682 | 5.156529 | 3.920897 | 1.31514 | 0.188463 | NA | #VALUE! |
| ENSMUSG00000105305.2 | 0 | NA | NA | NA | NA | NA | #VALUE! |
| ENSMUSG00000105306.2 | 0 | NA | NA | NA | NA | NA | #VALUE! |
| ENSMUSG00000105307.2 | 0 | NA | NA | NA | NA | NA | #VALUE! |
| ENSMUSG00000105308.5 | 0 | NA | NA | NA | NA | NA | #VALUE! |
| ENSMUSG00000105309.2 | 0 | NA | NA | NA | NA | NA | #VALUE! |
| ENSMUSG00000105310.2 | 0 | NA | NA | NA | NA | NA | #VALUE! |
| ENSMUSG00000105311.2 | 0 | NA | NA | NA | NA | NA | #VALUE! |
| ENSMUSG00000105313.2 | 0 | NA | NA | NA | NA | NA | #VALUE! |
| ENSMUSG00000105314.2 | 0 | NA | NA | NA | NA | NA | #VALUE! |
| ENSMUSG00000105316.2 | 0 | NA | NA | NA | NA | NA | #VALUE! |
| ENSMUSG00000105317.3 | 0 | NA | NA | NA | NA | NA | #VALUE! |
| ENSMUSG00000105319.2 | 0 | NA | NA | NA | NA | NA | #VALUE! |
| ENSMUSG00000105320.2 | 0 | NA | NA | NA | NA | NA | #VALUE! |
| ENSMUSG00000105322.2 | 0 | NA | NA | NA | NA | NA | #VALUE! |
| ENSMUSG00000105323.2 | 0 | NA | NA | NA | NA | NA | #VALUE! |
| ENSMUSG00000105324.2 | 0 | NA | NA | NA | NA | NA | #VALUE! |
| ENSMUSG00000105325.2 | 0 | NA | NA | NA | NA | NA | #VALUE! |
| ENSMUSG00000105326.2 | 0.932552 | 3.421684 | 3.953399 | 0.865504 | 0.386762 | NA | #VALUE! |
| ENSMUSG00000105327.2 | 0.204111 | -1.05581 | 4.073302 | -0.2592 | 0.795479 | NA | #VALUE! |
| ENSMUSG00000105328.2 | 0 | NA | NA | NA | NA | NA | #VALUE! |
| ENSMUSG00000105329.3 | 0 | NA | NA | NA | NA | NA | #VALUE! |
| ENSMUSG00000105331.2 | 0 | NA | NA | NA | NA | NA | #VALUE! |
| ENSMUSG00000105332.2 | 0 | NA | NA | NA | NA | NA | #VALUE! |
| ENSMUSG00000105333.2 | 0 | NA | NA | NA | NA | NA | #VALUE! |
| ENSMUSG00000105334.2 | 0 | NA | NA | NA | NA | NA | #VALUE! |
| ENSMUSG00000105335.2 | 0 | NA | NA | NA | NA | NA | #VALUE! |
| ENSMUSG00000105336.2 | 0 | NA | NA | NA | NA | NA | #VALUE! |
| ENSMUSG00000105337.5 | 0 | NA | NA | NA | NA | NA | #VALUE! |
| ENSMUSG00000105338.2 | 0 | NA | NA | NA | NA | NA | #VALUE! |
| ENSMUSG00000105339.2 | 12.16969 | -7.13907 | 3.011576 | -2.37054 | NA | NA | #VALUE! |

| | | | | | | | |
|---|---|---|---|---|---|---|---|
| ENSMUSG00000105340.6 | 0 | NA | NA | NA | NA | NA | #VALUE! |
| ENSMUSG00000105341.2 | 0 | NA | NA | NA | NA | NA | #VALUE! |
| ENSMUSG00000105342.2 | 0 | NA | NA | NA | NA | NA | #VALUE! |
| ENSMUSG00000105343.2 | 0 | NA | NA | NA | NA | NA | #VALUE! |
| ENSMUSG00000105344.2 | 0 | NA | NA | NA | NA | NA | #VALUE! |
| ENSMUSG00000105345.2 | 0 | NA | NA | NA | NA | NA | #VALUE! |
| ENSMUSG00000105346.2 | 0 | NA | NA | NA | NA | NA | #VALUE! |
| ENSMUSG00000105347.2 | 0 | NA | NA | NA | NA | NA | #VALUE! |
| ENSMUSG00000105348.2 | 0 | NA | NA | NA | NA | NA | #VALUE! |
| ENSMUSG00000105349.2 | 0 | NA | NA | NA | NA | NA | #VALUE! |
| ENSMUSG00000105350.2 | 0 | NA | NA | NA | NA | NA | #VALUE! |
| ENSMUSG00000105351.2 | 0 | NA | NA | NA | NA | NA | #VALUE! |
| ENSMUSG00000105352.5 | 0 | NA | NA | NA | NA | NA | #VALUE! |
| ENSMUSG00000105353.2 | 0 | NA | NA | NA | NA | NA | #VALUE! |
| ENSMUSG00000105355.2 | 0 | NA | NA | NA | NA | NA | #VALUE! |
| ENSMUSG00000105356.2 | 0 | NA | NA | NA | NA | NA | #VALUE! |
| ENSMUSG00000105357.2 | 0 | NA | NA | NA | NA | NA | #VALUE! |
| ENSMUSG00000105358.2 | 0 | NA | NA | NA | NA | NA | #VALUE! |
| ENSMUSG00000105359.2 | 0.204111 | -1.05581 | 4.073302 | -0.2592 | 0.795479 | NA | #VALUE! |
| ENSMUSG00000105360.2 | 0 | NA | NA | NA | NA | NA | #VALUE! |
| ENSMUSG00000105362.2 | 0 | NA | NA | NA | NA | NA | #VALUE! |
| ENSMUSG00000105363.2 | 0 | NA | NA | NA | NA | NA | #VALUE! |
| ENSMUSG00000105364.2 | 0 | NA | NA | NA | NA | NA | #VALUE! |
| ENSMUSG00000105366.2 | 0 | NA | NA | NA | NA | NA | #VALUE! |
| ENSMUSG00000105368.2 | 0 | NA | NA | NA | NA | NA | #VALUE! |
| ENSMUSG00000105369.2 | 0 | NA | NA | NA | NA | NA | #VALUE! |
| ENSMUSG00000105370.2 | 0 | NA | NA | NA | NA | NA | #VALUE! |
| ENSMUSG00000105372.3 | 0 | NA | NA | NA | NA | NA | #VALUE! |
| ENSMUSG00000105373.2 | 0 | NA | NA | NA | NA | NA | #VALUE! |
| ENSMUSG00000105374.2 | 0 | NA | NA | NA | NA | NA | #VALUE! |
| ENSMUSG00000105375.2 | 0 | NA | NA | NA | NA | NA | #VALUE! |
| ENSMUSG00000105376.5 | 0 | NA | NA | NA | NA | NA | #VALUE! |
| ENSMUSG00000105377.2 | 0 | NA | NA | NA | NA | NA | #VALUE! |
| ENSMUSG00000105378.2 | 0 | NA | NA | NA | NA | NA | #VALUE! |
| ENSMUSG00000105379.3 | 0 | NA | NA | NA | NA | NA | #VALUE! |
| ENSMUSG00000105380.2 | 0 | NA | NA | NA | NA | NA | #VALUE! |
| ENSMUSG00000105381.3 | 0 | NA | NA | NA | NA | NA | #VALUE! |
| ENSMUSG00000105382.2 | 0 | NA | NA | NA | NA | NA | #VALUE! |
| ENSMUSG00000105384.2 | 0 | NA | NA | NA | NA | NA | #VALUE! |
| ENSMUSG00000105385.2 | 0 | NA | NA | NA | NA | NA | #VALUE! |
| ENSMUSG00000105386.2 | 0 | NA | NA | NA | NA | NA | #VALUE! |
| ENSMUSG00000105387.2 | 0 | NA | NA | NA | NA | NA | #VALUE! |
| ENSMUSG00000105389.2 | 0 | NA | NA | NA | NA | NA | #VALUE! |
| ENSMUSG00000105390.2 | 0 | NA | NA | NA | NA | NA | #VALUE! |

| | | | | | | | |
|---|---|---|---|---|---|---|---|
| ENSMUSG00000105391.2 | 0 | NA | NA | NA | NA | NA | #VALUE! |
| ENSMUSG00000105392.2 | 0 | NA | NA | NA | NA | NA | #VALUE! |
| ENSMUSG00000105393.2 | 0 | NA | NA | NA | NA | NA | #VALUE! |
| ENSMUSG00000105394.2 | 0 | NA | NA | NA | NA | NA | #VALUE! |
| ENSMUSG00000105395.2 | 0 | NA | NA | NA | NA | NA | #VALUE! |
| ENSMUSG00000105396.2 | 0 | NA | NA | NA | NA | NA | #VALUE! |
| ENSMUSG00000105397.2 | 0 | NA | NA | NA | NA | NA | #VALUE! |
| ENSMUSG00000105399.3 | 0 | NA | NA | NA | NA | NA | #VALUE! |
| ENSMUSG00000105400.3 | 0 | NA | NA | NA | NA | NA | #VALUE! |
| ENSMUSG00000105401.2 | 0 | NA | NA | NA | NA | NA | #VALUE! |
| ENSMUSG00000105402.5 | 0 | NA | NA | NA | NA | NA | #VALUE! |
| ENSMUSG00000105403.2 | 0 | NA | NA | NA | NA | NA | #VALUE! |
| ENSMUSG00000105404.2 | 0 | NA | NA | NA | NA | NA | #VALUE! |
| ENSMUSG00000105405.2 | 0 | NA | NA | NA | NA | NA | #VALUE! |
| ENSMUSG00000105406.2 | 0 | NA | NA | NA | NA | NA | #VALUE! |
| ENSMUSG00000105407.2 | 0 | NA | NA | NA | NA | NA | #VALUE! |
| ENSMUSG00000105408.2 | 0.951758 | 2.303402 | 3.955391 | 0.582345 | 0.560334 | NA | #VALUE! |
| ENSMUSG00000105410.2 | 0 | NA | NA | NA | NA | NA | #VALUE! |
| ENSMUSG00000105412.2 | 0 | NA | NA | NA | NA | NA | #VALUE! |
| ENSMUSG00000105416.2 | 0 | NA | NA | NA | NA | NA | #VALUE! |
| ENSMUSG00000105418.2 | 0 | NA | NA | NA | NA | NA | #VALUE! |
| ENSMUSG00000105419.2 | 0 | NA | NA | NA | NA | NA | #VALUE! |
| ENSMUSG00000105421.2 | 0 | NA | NA | NA | NA | NA | #VALUE! |
| ENSMUSG00000105422.2 | 0 | NA | NA | NA | NA | NA | #VALUE! |
| ENSMUSG00000105423.5 | 0 | NA | NA | NA | NA | NA | #VALUE! |
| ENSMUSG00000105425.2 | 0 | NA | NA | NA | NA | NA | #VALUE! |
| ENSMUSG00000105426.2 | 0 | NA | NA | NA | NA | NA | #VALUE! |
| ENSMUSG00000105427.2 | 0 | NA | NA | NA | NA | NA | #VALUE! |
| ENSMUSG00000105428.3 | 0 | NA | NA | NA | NA | NA | #VALUE! |
| ENSMUSG00000105429.2 | 0.093255 | 0.868093 | 4.072464 | 0.213162 | 0.831201 | NA | #VALUE! |
| ENSMUSG00000105430.2 | 0 | NA | NA | NA | NA | NA | #VALUE! |
| ENSMUSG00000105431.2 | 0 | NA | NA | NA | NA | NA | #VALUE! |
| ENSMUSG00000105432.2 | 0 | NA | NA | NA | NA | NA | #VALUE! |
| ENSMUSG00000105434.2 | 3.163872 | -1.89186 | 3.812296 | -0.49625 | 0.619716 | NA | #VALUE! |
| ENSMUSG00000105436.2 | 0 | NA | NA | NA | NA | NA | #VALUE! |
| ENSMUSG00000105437.2 | 0 | NA | NA | NA | NA | NA | #VALUE! |
| ENSMUSG00000105439.2 | 0 | NA | NA | NA | NA | NA | #VALUE! |
| ENSMUSG00000105440.2 | 0 | NA | NA | NA | NA | NA | #VALUE! |
| ENSMUSG00000105441.2 | 0.336567 | 1.944925 | 4.029936 | 0.482619 | 0.629366 | NA | #VALUE! |
| ENSMUSG00000105442.2 | 0.245976 | 1.027628 | 4.067126 | 0.252667 | 0.800526 | NA | #VALUE! |
| ENSMUSG00000105443.2 | 0 | NA | NA | NA | NA | NA | #VALUE! |
| ENSMUSG00000105444.2 | 0 | NA | NA | NA | NA | NA | #VALUE! |
| ENSMUSG00000105445.2 | 0 | NA | NA | NA | NA | NA | #VALUE! |
| ENSMUSG00000105446.2 | 0 | NA | NA | NA | NA | NA | #VALUE! |

| | | | | | | | |
|---|---|---|---|---|---|---|---|
| ENSMUSG00000105447.2 | 0 | NA | NA | NA | NA | NA | #VALUE! |
| ENSMUSG00000105448.3 | 0 | NA | NA | NA | NA | NA | #VALUE! |
| ENSMUSG00000105449.2 | 0 | NA | NA | NA | NA | NA | #VALUE! |
| ENSMUSG00000105451.2 | 0 | NA | NA | NA | NA | NA | #VALUE! |
| ENSMUSG00000105452.2 | 0 | NA | NA | NA | NA | NA | #VALUE! |
| ENSMUSG00000105453.2 | 0 | NA | NA | NA | NA | NA | #VALUE! |
| ENSMUSG00000105454.2 | 0 | NA | NA | NA | NA | NA | #VALUE! |
| ENSMUSG00000105456.2 | 0 | NA | NA | NA | NA | NA | #VALUE! |
| ENSMUSG00000105457.2 | 0 | NA | NA | NA | NA | NA | #VALUE! |
| ENSMUSG00000105458.3 | 0 | NA | NA | NA | NA | NA | #VALUE! |
| ENSMUSG00000105459.2 | 0 | NA | NA | NA | NA | NA | #VALUE! |
| ENSMUSG00000105460.2 | 0 | NA | NA | NA | NA | NA | #VALUE! |
| ENSMUSG00000105461.2 | 0 | NA | NA | NA | NA | NA | #VALUE! |
| ENSMUSG00000105462.2 | 0 | NA | NA | NA | NA | NA | #VALUE! |
| ENSMUSG00000105463.2 | 0 | NA | NA | NA | NA | NA | #VALUE! |
| ENSMUSG00000105464.2 | 0 | NA | NA | NA | NA | NA | #VALUE! |
| ENSMUSG00000105466.2 | 0 | NA | NA | NA | NA | NA | #VALUE! |
| ENSMUSG00000105467.2 | 0 | NA | NA | NA | NA | NA | #VALUE! |
| ENSMUSG00000105468.3 | 0 | NA | NA | NA | NA | NA | #VALUE! |
| ENSMUSG00000105469.2 | 0 | NA | NA | NA | NA | NA | #VALUE! |
| ENSMUSG00000105470.2 | 0 | NA | NA | NA | NA | NA | #VALUE! |
| ENSMUSG00000105471.2 | 0 | NA | NA | NA | NA | NA | #VALUE! |
| ENSMUSG00000105472.2 | 0 | NA | NA | NA | NA | NA | #VALUE! |
| ENSMUSG00000105473.2 | 0 | NA | NA | NA | NA | NA | #VALUE! |
| ENSMUSG00000105474.2 | 0 | NA | NA | NA | NA | NA | #VALUE! |
| ENSMUSG00000105475.2 | 0 | NA | NA | NA | NA | NA | #VALUE! |
| ENSMUSG00000105476.2 | 0 | NA | NA | NA | NA | NA | #VALUE! |
| ENSMUSG00000105478.2 | 0 | NA | NA | NA | NA | NA | #VALUE! |
| ENSMUSG00000105479.2 | 0 | NA | NA | NA | NA | NA | #VALUE! |
| ENSMUSG00000105480.2 | 0 | NA | NA | NA | NA | NA | #VALUE! |
| ENSMUSG00000105481.2 | 0 | NA | NA | NA | NA | NA | #VALUE! |
| ENSMUSG00000105482.2 | 0 | NA | NA | NA | NA | NA | #VALUE! |
| ENSMUSG00000105483.2 | 0 | NA | NA | NA | NA | NA | #VALUE! |
| ENSMUSG00000105484.2 | 0 | NA | NA | NA | NA | NA | #VALUE! |
| ENSMUSG00000105485.2 | 0 | NA | NA | NA | NA | NA | #VALUE! |
| ENSMUSG00000105486.2 | 0 | NA | NA | NA | NA | NA | #VALUE! |
| ENSMUSG00000105487.2 | 0 | NA | NA | NA | NA | NA | #VALUE! |
| ENSMUSG00000105489.2 | 0 | NA | NA | NA | NA | NA | #VALUE! |
| ENSMUSG00000105490.2 | 0 | NA | NA | NA | NA | NA | #VALUE! |
| ENSMUSG00000105491.2 | 0 | NA | NA | NA | NA | NA | #VALUE! |
| ENSMUSG00000105492.2 | 0 | NA | NA | NA | NA | NA | #VALUE! |
| ENSMUSG00000105493.2 | 0 | NA | NA | NA | NA | NA | #VALUE! |
| ENSMUSG00000105495.2 | 0 | NA | NA | NA | NA | NA | #VALUE! |
| ENSMUSG00000105496.3 | 0 | NA | NA | NA | NA | NA | #VALUE! |

| | | | | | | | |
|---|---|---|---|---|---|---|---|
| ENSMUSG00000105497.3 | 0 | NA | NA | NA | NA | NA | #VALUE! |
| ENSMUSG00000105498.2 | 0 | NA | NA | NA | NA | NA | #VALUE! |
| ENSMUSG00000105499.2 | 0 | NA | NA | NA | NA | NA | #VALUE! |
| ENSMUSG00000105500.2 | 0 | NA | NA | NA | NA | NA | #VALUE! |
| ENSMUSG00000105501.2 | 0 | NA | NA | NA | NA | NA | #VALUE! |
| ENSMUSG00000105502.2 | 0 | NA | NA | NA | NA | NA | #VALUE! |
| ENSMUSG00000105503.2 | 0 | NA | NA | NA | NA | NA | #VALUE! |
| ENSMUSG00000105504.5 | 0 | NA | NA | NA | NA | NA | #VALUE! |
| ENSMUSG00000105505.2 | 0 | NA | NA | NA | NA | NA | #VALUE! |
| ENSMUSG00000105506.2 | 0.309124 | -1.83395 | 4.047982 | -0.45305 | 0.650511 | NA | #VALUE! |
| ENSMUSG00000105507.2 | 0 | NA | NA | NA | NA | NA | #VALUE! |
| ENSMUSG00000105508.2 | 0 | NA | NA | NA | NA | NA | #VALUE! |
| ENSMUSG00000105510.2 | 0 | NA | NA | NA | NA | NA | #VALUE! |
| ENSMUSG00000105511.2 | 0 | NA | NA | NA | NA | NA | #VALUE! |
| ENSMUSG00000105512.2 | 0.673134 | 2.918386 | 3.972531 | 0.734641 | 0.462558 | NA | #VALUE! |
| ENSMUSG00000105514.2 | 0 | NA | NA | NA | NA | NA | #VALUE! |
| ENSMUSG00000105515.2 | 0 | NA | NA | NA | NA | NA | #VALUE! |
| ENSMUSG00000105516.5 | 0 | NA | NA | NA | NA | NA | #VALUE! |
| ENSMUSG00000105517.2 | 0 | NA | NA | NA | NA | NA | #VALUE! |
| ENSMUSG00000105518.2 | 0 | NA | NA | NA | NA | NA | #VALUE! |
| ENSMUSG00000105519.2 | 0.245976 | 1.027628 | 4.067126 | 0.252667 | 0.800526 | NA | #VALUE! |
| ENSMUSG00000105520.2 | 0 | NA | NA | NA | NA | NA | #VALUE! |
| ENSMUSG00000105521.2 | 0 | NA | NA | NA | NA | NA | #VALUE! |
| ENSMUSG00000105522.2 | 0 | NA | NA | NA | NA | NA | #VALUE! |
| ENSMUSG00000105523.2 | 0 | NA | NA | NA | NA | NA | #VALUE! |
| ENSMUSG00000105524.3 | 0 | NA | NA | NA | NA | NA | #VALUE! |
| ENSMUSG00000105525.2 | 0 | NA | NA | NA | NA | NA | #VALUE! |
| ENSMUSG00000105526.2 | 0 | NA | NA | NA | NA | NA | #VALUE! |
| ENSMUSG00000105527.2 | 0 | NA | NA | NA | NA | NA | #VALUE! |
| ENSMUSG00000105528.2 | 0 | NA | NA | NA | NA | NA | #VALUE! |
| ENSMUSG00000105529.2 | 0.105013 | -0.97324 | 4.076293 | -0.23876 | 0.811295 | NA | #VALUE! |
| ENSMUSG00000105530.2 | 0 | NA | NA | NA | NA | NA | #VALUE! |
| ENSMUSG00000105531.2 | 0 | NA | NA | NA | NA | NA | #VALUE! |
| ENSMUSG00000105532.2 | 0 | NA | NA | NA | NA | NA | #VALUE! |
| ENSMUSG00000105533.2 | 0 | NA | NA | NA | NA | NA | #VALUE! |
| ENSMUSG00000105534.2 | 0 | NA | NA | NA | NA | NA | #VALUE! |
| ENSMUSG00000105535.2 | 0 | NA | NA | NA | NA | NA | #VALUE! |
| ENSMUSG00000105536.2 | 0 | NA | NA | NA | NA | NA | #VALUE! |
| ENSMUSG00000105538.2 | 0 | NA | NA | NA | NA | NA | #VALUE! |
| ENSMUSG00000105539.2 | 0 | NA | NA | NA | NA | NA | #VALUE! |
| ENSMUSG00000105540.2 | 0 | NA | NA | NA | NA | NA | #VALUE! |
| ENSMUSG00000105541.2 | 1.890236 | -4.46257 | 3.93014 | -1.13547 | 0.256177 | NA | #VALUE! |
| ENSMUSG00000105542.2 | 0 | NA | NA | NA | NA | NA | #VALUE! |
| ENSMUSG00000105543.2 | 0 | NA | NA | NA | NA | NA | #VALUE! |

| | | | | | | | |
|---|---|---|---|---|---|---|---|
| ENSMUSG00000105545.2 | 0 | NA | NA | NA | NA | NA | #VALUE! |
| ENSMUSG00000105547.2 | 0.525066 | -2.64687 | 3.988095 | -0.66369 | 0.506887 | NA | #VALUE! |
| ENSMUSG00000105548.2 | 0 | NA | NA | NA | NA | NA | #VALUE! |
| ENSMUSG00000105549.2 | 0 | NA | NA | NA | NA | NA | #VALUE! |
| ENSMUSG00000105550.2 | 0 | NA | NA | NA | NA | NA | #VALUE! |
| ENSMUSG00000105551.2 | 0 | NA | NA | NA | NA | NA | #VALUE! |
| ENSMUSG00000105552.2 | 0.651606 | 0.049656 | 3.987553 | 0.012453 | 0.990064 | NA | #VALUE! |
| ENSMUSG00000105553.2 | 0 | NA | NA | NA | NA | NA | #VALUE! |
| ENSMUSG00000105554.2 | 0 | NA | NA | NA | NA | NA | #VALUE! |
| ENSMUSG00000105555.2 | 0 | NA | NA | NA | NA | NA | #VALUE! |
| ENSMUSG00000105556.2 | 0 | NA | NA | NA | NA | NA | #VALUE! |
| ENSMUSG00000105557.3 | 0 | NA | NA | NA | NA | NA | #VALUE! |
| ENSMUSG00000105558.2 | 0 | NA | NA | NA | NA | NA | #VALUE! |
| ENSMUSG00000105559.2 | 0 | NA | NA | NA | NA | NA | #VALUE! |
| ENSMUSG00000105560.2 | 0 | NA | NA | NA | NA | NA | #VALUE! |
| ENSMUSG00000105561.2 | 0 | NA | NA | NA | NA | NA | #VALUE! |
| ENSMUSG00000105562.2 | 0 | NA | NA | NA | NA | NA | #VALUE! |
| ENSMUSG00000105564.2 | 0 | NA | NA | NA | NA | NA | #VALUE! |
| ENSMUSG00000105565.2 | 0 | NA | NA | NA | NA | NA | #VALUE! |
| ENSMUSG00000105566.2 | 0 | NA | NA | NA | NA | NA | #VALUE! |
| ENSMUSG00000105568.2 | 0 | NA | NA | NA | NA | NA | #VALUE! |
| ENSMUSG00000105570.2 | 0 | NA | NA | NA | NA | NA | #VALUE! |
| ENSMUSG00000105571.2 | 0 | NA | NA | NA | NA | NA | #VALUE! |
| ENSMUSG00000105572.2 | 0 | NA | NA | NA | NA | NA | #VALUE! |
| ENSMUSG00000105573.3 | 0 | NA | NA | NA | NA | NA | #VALUE! |
| ENSMUSG00000105574.2 | 0 | NA | NA | NA | NA | NA | #VALUE! |
| ENSMUSG00000105575.2 | 0 | NA | NA | NA | NA | NA | #VALUE! |
| ENSMUSG00000105576.2 | 0 | NA | NA | NA | NA | NA | #VALUE! |
| ENSMUSG00000105577.2 | 0 | NA | NA | NA | NA | NA | #VALUE! |
| ENSMUSG00000105578.2 | 0 | NA | NA | NA | NA | NA | #VALUE! |
| ENSMUSG00000105579.3 | 0 | NA | NA | NA | NA | NA | #VALUE! |
| ENSMUSG00000105582.2 | 0 | NA | NA | NA | NA | NA | #VALUE! |
| ENSMUSG00000105583.2 | 0 | NA | NA | NA | NA | NA | #VALUE! |
| ENSMUSG00000105584.2 | 0 | NA | NA | NA | NA | NA | #VALUE! |
| ENSMUSG00000105586.2 | 0 | NA | NA | NA | NA | NA | #VALUE! |
| ENSMUSG00000105588.2 | 0 | NA | NA | NA | NA | NA | #VALUE! |
| ENSMUSG00000105589.5 | 0 | NA | NA | NA | NA | NA | #VALUE! |
| ENSMUSG00000105590.2 | 0 | NA | NA | NA | NA | NA | #VALUE! |
| ENSMUSG00000105591.2 | 0 | NA | NA | NA | NA | NA | #VALUE! |
| ENSMUSG00000105592.2 | 0 | NA | NA | NA | NA | NA | #VALUE! |
| ENSMUSG00000105593.5 | 0 | NA | NA | NA | NA | NA | #VALUE! |
| ENSMUSG00000105594.2 | 0 | NA | NA | NA | NA | NA | #VALUE! |
| ENSMUSG00000105595.2 | 0 | NA | NA | NA | NA | NA | #VALUE! |
| ENSMUSG00000105596.2 | 0 | NA | NA | NA | NA | NA | #VALUE! |

| | | | | | | | |
|---|---|---|---|---|---|---|---|
| ENSMUSG00000105597.2 | 0 | NA | NA | NA | NA | NA | #VALUE! |
| ENSMUSG00000105598.2 | 0 | NA | NA | NA | NA | NA | #VALUE! |
| ENSMUSG00000105600.2 | 0 | NA | NA | NA | NA | NA | #VALUE! |
| ENSMUSG00000105601.2 | 0 | NA | NA | NA | NA | NA | #VALUE! |
| ENSMUSG00000105602.2 | 0 | NA | NA | NA | NA | NA | #VALUE! |
| ENSMUSG00000105603.2 | 0 | NA | NA | NA | NA | NA | #VALUE! |
| ENSMUSG00000105604.2 | 0 | NA | NA | NA | NA | NA | #VALUE! |
| ENSMUSG00000105605.2 | 0 | NA | NA | NA | NA | NA | #VALUE! |
| ENSMUSG00000105606.2 | 0 | NA | NA | NA | NA | NA | #VALUE! |
| ENSMUSG00000105607.2 | 3.936159 | -2.83038 | 3.832901 | -0.73844 | 0.460245 | NA | #VALUE! |
| ENSMUSG00000105608.2 | 0 | NA | NA | NA | NA | NA | #VALUE! |
| ENSMUSG00000105609.2 | 0 | NA | NA | NA | NA | NA | #VALUE! |
| ENSMUSG00000105611.2 | 0 | NA | NA | NA | NA | NA | #VALUE! |
| ENSMUSG00000105613.2 | 0 | NA | NA | NA | NA | NA | #VALUE! |
| ENSMUSG00000105614.2 | 0 | NA | NA | NA | NA | NA | #VALUE! |
| ENSMUSG00000105615.2 | 0 | NA | NA | NA | NA | NA | #VALUE! |
| ENSMUSG00000105616.2 | 0 | NA | NA | NA | NA | NA | #VALUE! |
| ENSMUSG00000105617.5 | 0 | NA | NA | NA | NA | NA | #VALUE! |
| ENSMUSG00000105618.2 | 0 | NA | NA | NA | NA | NA | #VALUE! |
| ENSMUSG00000105619.2 | 0 | NA | NA | NA | NA | NA | #VALUE! |
| ENSMUSG00000105620.2 | 0 | NA | NA | NA | NA | NA | #VALUE! |
| ENSMUSG00000105621.3 | 0 | NA | NA | NA | NA | NA | #VALUE! |
| ENSMUSG00000105622.2 | 0 | NA | NA | NA | NA | NA | #VALUE! |
| ENSMUSG00000105624.2 | 0 | NA | NA | NA | NA | NA | #VALUE! |
| ENSMUSG00000105625.2 | 0 | NA | NA | NA | NA | NA | #VALUE! |
| ENSMUSG00000105627.2 | 0 | NA | NA | NA | NA | NA | #VALUE! |
| ENSMUSG00000105628.2 | 0 | NA | NA | NA | NA | NA | #VALUE! |
| ENSMUSG00000105629.2 | 0 | NA | NA | NA | NA | NA | #VALUE! |
| ENSMUSG00000105630.2 | 0 | NA | NA | NA | NA | NA | #VALUE! |
| ENSMUSG00000105632.2 | 0 | NA | NA | NA | NA | NA | #VALUE! |
| ENSMUSG00000105633.2 | 0 | NA | NA | NA | NA | NA | #VALUE! |
| ENSMUSG00000105634.2 | 0 | NA | NA | NA | NA | NA | #VALUE! |
| ENSMUSG00000105636.2 | 0 | NA | NA | NA | NA | NA | #VALUE! |
| ENSMUSG00000105637.2 | 0 | NA | NA | NA | NA | NA | #VALUE! |
| ENSMUSG00000105638.2 | 0 | NA | NA | NA | NA | NA | #VALUE! |
| ENSMUSG00000105639.2 | 0 | NA | NA | NA | NA | NA | #VALUE! |
| ENSMUSG00000105640.2 | 0 | NA | NA | NA | NA | NA | #VALUE! |
| ENSMUSG00000105641.2 | 0 | NA | NA | NA | NA | NA | #VALUE! |
| ENSMUSG00000105642.2 | 0 | NA | NA | NA | NA | NA | #VALUE! |
| ENSMUSG00000105643.2 | 0 | NA | NA | NA | NA | NA | #VALUE! |
| ENSMUSG00000105644.2 | 0 | NA | NA | NA | NA | NA | #VALUE! |
| ENSMUSG00000105645.2 | 0 | NA | NA | NA | NA | NA | #VALUE! |
| ENSMUSG00000105646.2 | 0 | NA | NA | NA | NA | NA | #VALUE! |
| ENSMUSG00000105647.2 | 0 | NA | NA | NA | NA | NA | #VALUE! |

| | | | | | | | |
|---|---|---|---|---|---|---|---|
| ENSMUSG00000105648.2 | 0 | NA | NA | NA | NA | NA | #VALUE! |
| ENSMUSG00000105649.2 | 0 | NA | NA | NA | NA | NA | #VALUE! |
| ENSMUSG00000105650.2 | 0 | NA | NA | NA | NA | NA | #VALUE! |
| ENSMUSG00000105651.2 | 0 | NA | NA | NA | NA | NA | #VALUE! |
| ENSMUSG00000105652.2 | 0 | NA | NA | NA | NA | NA | #VALUE! |
| ENSMUSG00000105653.2 | 0 | NA | NA | NA | NA | NA | #VALUE! |
| ENSMUSG00000105654.2 | 0 | NA | NA | NA | NA | NA | #VALUE! |
| ENSMUSG00000105655.2 | 3.150394 | -5.19405 | 3.92087 | -1.32472 | 0.185265 | NA | #VALUE! |
| ENSMUSG00000105656.2 | 0 | NA | NA | NA | NA | NA | #VALUE! |
| ENSMUSG00000105657.2 | 0 | NA | NA | NA | NA | NA | #VALUE! |
| ENSMUSG00000105658.2 | 0 | NA | NA | NA | NA | NA | #VALUE! |
| ENSMUSG00000105659.2 | 0 | NA | NA | NA | NA | NA | #VALUE! |
| ENSMUSG00000105660.2 | 0 | NA | NA | NA | NA | NA | #VALUE! |
| ENSMUSG00000105661.2 | 0 | NA | NA | NA | NA | NA | #VALUE! |
| ENSMUSG00000105662.5 | 0 | NA | NA | NA | NA | NA | #VALUE! |
| ENSMUSG00000105663.2 | 0 | NA | NA | NA | NA | NA | #VALUE! |
| ENSMUSG00000105664.2 | 0 | NA | NA | NA | NA | NA | #VALUE! |
| ENSMUSG00000105665.2 | 0 | NA | NA | NA | NA | NA | #VALUE! |
| ENSMUSG00000105666.2 | 0 | NA | NA | NA | NA | NA | #VALUE! |
| ENSMUSG00000105667.2 | 0 | NA | NA | NA | NA | NA | #VALUE! |
| ENSMUSG00000105668.2 | 0 | NA | NA | NA | NA | NA | #VALUE! |
| ENSMUSG00000105669.2 | 0 | NA | NA | NA | NA | NA | #VALUE! |
| ENSMUSG00000105671.2 | 0 | NA | NA | NA | NA | NA | #VALUE! |
| ENSMUSG00000105672.2 | 0 | NA | NA | NA | NA | NA | #VALUE! |
| ENSMUSG00000105673.2 | 0 | NA | NA | NA | NA | NA | #VALUE! |
| ENSMUSG00000105674.2 | 0 | NA | NA | NA | NA | NA | #VALUE! |
| ENSMUSG00000105675.2 | 0 | NA | NA | NA | NA | NA | #VALUE! |
| ENSMUSG00000105676.2 | 0 | NA | NA | NA | NA | NA | #VALUE! |
| ENSMUSG00000105677.2 | 0 | NA | NA | NA | NA | NA | #VALUE! |
| ENSMUSG00000105678.2 | 0 | NA | NA | NA | NA | NA | #VALUE! |
| ENSMUSG00000105679.2 | 0 | NA | NA | NA | NA | NA | #VALUE! |
| ENSMUSG00000105680.2 | 0 | NA | NA | NA | NA | NA | #VALUE! |
| ENSMUSG00000105681.2 | 0 | NA | NA | NA | NA | NA | #VALUE! |
| ENSMUSG00000105683.2 | 0 | NA | NA | NA | NA | NA | #VALUE! |
| ENSMUSG00000105684.2 | 0 | NA | NA | NA | NA | NA | #VALUE! |
| ENSMUSG00000105685.2 | 0 | NA | NA | NA | NA | NA | #VALUE! |
| ENSMUSG00000105686.2 | 0 | NA | NA | NA | NA | NA | #VALUE! |
| ENSMUSG00000105687.2 | 0 | NA | NA | NA | NA | NA | #VALUE! |
| ENSMUSG00000105688.2 | 0 | NA | NA | NA | NA | NA | #VALUE! |
| ENSMUSG00000105689.2 | 0 | NA | NA | NA | NA | NA | #VALUE! |
| ENSMUSG00000105690.3 | 0 | NA | NA | NA | NA | NA | #VALUE! |
| ENSMUSG00000105691.2 | 0 | NA | NA | NA | NA | NA | #VALUE! |
| ENSMUSG00000105692.2 | 0.408222 | -2.20205 | 4.016842 | -0.54821 | 0.583551 | NA | #VALUE! |
| ENSMUSG00000105693.2 | 0 | NA | NA | NA | NA | NA | #VALUE! |

| | | | | | | | |
|---|---|---|---|---|---|---|---|
| ENSMUSG00000105694.2 | 0 | NA | NA | NA | NA | NA | #VALUE! |
| ENSMUSG00000105695.2 | 0 | NA | NA | NA | NA | NA | #VALUE! |
| ENSMUSG00000105696.2 | 0 | NA | NA | NA | NA | NA | #VALUE! |
| ENSMUSG00000105697.3 | 0 | NA | NA | NA | NA | NA | #VALUE! |
| ENSMUSG00000105698.2 | 0 | NA | NA | NA | NA | NA | #VALUE! |
| ENSMUSG00000105699.2 | 0 | NA | NA | NA | NA | NA | #VALUE! |
| ENSMUSG00000105700.2 | 0 | NA | NA | NA | NA | NA | #VALUE! |
| ENSMUSG00000105701.2 | 0 | NA | NA | NA | NA | NA | #VALUE! |
| ENSMUSG00000105702.2 | 0 | NA | NA | NA | NA | NA | #VALUE! |
| ENSMUSG00000105703.3 | 0 | NA | NA | NA | NA | NA | #VALUE! |
| ENSMUSG00000105704.2 | 0 | NA | NA | NA | NA | NA | #VALUE! |
| ENSMUSG00000105705.2 | 0 | NA | NA | NA | NA | NA | #VALUE! |
| ENSMUSG00000105706.2 | 0 | NA | NA | NA | NA | NA | #VALUE! |
| ENSMUSG00000105707.2 | 0 | NA | NA | NA | NA | NA | #VALUE! |
| ENSMUSG00000105708.2 | 0 | NA | NA | NA | NA | NA | #VALUE! |
| ENSMUSG00000105709.2 | 0 | NA | NA | NA | NA | NA | #VALUE! |
| ENSMUSG00000105710.2 | 0 | NA | NA | NA | NA | NA | #VALUE! |
| ENSMUSG00000105711.2 | 0 | NA | NA | NA | NA | NA | #VALUE! |
| ENSMUSG00000105712.2 | 0 | NA | NA | NA | NA | NA | #VALUE! |
| ENSMUSG00000105713.2 | 0 | NA | NA | NA | NA | NA | #VALUE! |
| ENSMUSG00000105714.2 | 0 | NA | NA | NA | NA | NA | #VALUE! |
| ENSMUSG00000105715.2 | 0 | NA | NA | NA | NA | NA | #VALUE! |
| ENSMUSG00000105716.2 | 0 | NA | NA | NA | NA | NA | #VALUE! |
| ENSMUSG00000105717.2 | 0 | NA | NA | NA | NA | NA | #VALUE! |
| ENSMUSG00000105718.2 | 0 | NA | NA | NA | NA | NA | #VALUE! |
| ENSMUSG00000105719.2 | 0 | NA | NA | NA | NA | NA | #VALUE! |
| ENSMUSG00000105720.2 | 0 | NA | NA | NA | NA | NA | #VALUE! |
| ENSMUSG00000105721.2 | 0 | NA | NA | NA | NA | NA | #VALUE! |
| ENSMUSG00000105722.2 | 0 | NA | NA | NA | NA | NA | #VALUE! |
| ENSMUSG00000105723.3 | 0 | NA | NA | NA | NA | NA | #VALUE! |
| ENSMUSG00000105725.2 | 0 | NA | NA | NA | NA | NA | #VALUE! |
| ENSMUSG00000105726.2 | 0 | NA | NA | NA | NA | NA | #VALUE! |
| ENSMUSG00000105727.2 | 0 | NA | NA | NA | NA | NA | #VALUE! |
| ENSMUSG00000105728.2 | 0 | NA | NA | NA | NA | NA | #VALUE! |
| ENSMUSG00000105729.2 | 0 | NA | NA | NA | NA | NA | #VALUE! |
| ENSMUSG00000105730.2 | 0 | NA | NA | NA | NA | NA | #VALUE! |
| ENSMUSG00000105731.2 | 0 | NA | NA | NA | NA | NA | #VALUE! |
| ENSMUSG00000105732.2 | 0 | NA | NA | NA | NA | NA | #VALUE! |
| ENSMUSG00000105733.2 | 0 | NA | NA | NA | NA | NA | #VALUE! |
| ENSMUSG00000105734.2 | 0 | NA | NA | NA | NA | NA | #VALUE! |
| ENSMUSG00000105735.2 | 0 | NA | NA | NA | NA | NA | #VALUE! |
| ENSMUSG00000105736.2 | 0 | NA | NA | NA | NA | NA | #VALUE! |
| ENSMUSG00000105737.2 | 0 | NA | NA | NA | NA | NA | #VALUE! |
| ENSMUSG00000105738.3 | 0 | NA | NA | NA | NA | NA | #VALUE! |

| | | | | | | | |
|---|---|---|---|---|---|---|---|
| ENSMUSG00000105739.2 | 0 | NA | NA | NA | NA | NA | #VALUE! |
| ENSMUSG00000105740.2 | 0 | NA | NA | NA | NA | NA | #VALUE! |
| ENSMUSG00000105741.2 | 0 | NA | NA | NA | NA | NA | #VALUE! |
| ENSMUSG00000105742.2 | 0 | NA | NA | NA | NA | NA | #VALUE! |
| ENSMUSG00000105743.3 | 0 | NA | NA | NA | NA | NA | #VALUE! |
| ENSMUSG00000105744.2 | 0 | NA | NA | NA | NA | NA | #VALUE! |
| ENSMUSG00000105745.2 | 0 | NA | NA | NA | NA | NA | #VALUE! |
| ENSMUSG00000105746.2 | 0 | NA | NA | NA | NA | NA | #VALUE! |
| ENSMUSG00000105747.2 | 0 | NA | NA | NA | NA | NA | #VALUE! |
| ENSMUSG00000105748.2 | 3.870521 | 5.442409 | 3.918364 | 1.388949 | 0.164848 | NA | #VALUE! |
| ENSMUSG00000105749.2 | 0 | NA | NA | NA | NA | NA | #VALUE! |
| ENSMUSG00000105750.2 | 0 | NA | NA | NA | NA | NA | #VALUE! |
| ENSMUSG00000105751.2 | 0 | NA | NA | NA | NA | NA | #VALUE! |
| ENSMUSG00000105752.2 | 0 | NA | NA | NA | NA | NA | #VALUE! |
| ENSMUSG00000105753.2 | 0 | NA | NA | NA | NA | NA | #VALUE! |
| ENSMUSG00000105755.2 | 0 | NA | NA | NA | NA | NA | #VALUE! |
| ENSMUSG00000105757.2 | 0 | NA | NA | NA | NA | NA | #VALUE! |
| ENSMUSG00000105758.3 | 0 | NA | NA | NA | NA | NA | #VALUE! |
| ENSMUSG00000105759.2 | 0 | NA | NA | NA | NA | NA | #VALUE! |
| ENSMUSG00000105760.2 | 0 | NA | NA | NA | NA | NA | #VALUE! |
| ENSMUSG00000105761.2 | 0 | NA | NA | NA | NA | NA | #VALUE! |
| ENSMUSG00000105762.2 | 0 | NA | NA | NA | NA | NA | #VALUE! |
| ENSMUSG00000105763.2 | 0 | NA | NA | NA | NA | NA | #VALUE! |
| ENSMUSG00000105764.2 | 0 | NA | NA | NA | NA | NA | #VALUE! |
| ENSMUSG00000105765.2 | 0 | NA | NA | NA | NA | NA | #VALUE! |
| ENSMUSG00000105766.2 | 0 | NA | NA | NA | NA | NA | #VALUE! |
| ENSMUSG00000105767.2 | 0 | NA | NA | NA | NA | NA | #VALUE! |
| ENSMUSG00000105769.2 | 0 | NA | NA | NA | NA | NA | #VALUE! |
| ENSMUSG00000105770.3 | 0 | NA | NA | NA | NA | NA | #VALUE! |
| ENSMUSG00000105771.2 | 0 | NA | NA | NA | NA | NA | #VALUE! |
| ENSMUSG00000105772.2 | 0 | NA | NA | NA | NA | NA | #VALUE! |
| ENSMUSG00000105773.2 | 0 | NA | NA | NA | NA | NA | #VALUE! |
| ENSMUSG00000105774.2 | 0 | NA | NA | NA | NA | NA | #VALUE! |
| ENSMUSG00000105775.2 | 0.093255 | 0.868093 | 4.072464 | 0.213162 | 0.831201 | NA | #VALUE! |
| ENSMUSG00000105776.2 | 0 | NA | NA | NA | NA | NA | #VALUE! |
| ENSMUSG00000105777.2 | 0 | NA | NA | NA | NA | NA | #VALUE! |
| ENSMUSG00000105778.3 | 0 | NA | NA | NA | NA | NA | #VALUE! |
| ENSMUSG00000105779.2 | 0 | NA | NA | NA | NA | NA | #VALUE! |
| ENSMUSG00000105781.2 | 0 | NA | NA | NA | NA | NA | #VALUE! |
| ENSMUSG00000105782.2 | 0 | NA | NA | NA | NA | NA | #VALUE! |
| ENSMUSG00000105784.2 | 0 | NA | NA | NA | NA | NA | #VALUE! |
| ENSMUSG00000105785.2 | 0 | NA | NA | NA | NA | NA | #VALUE! |
| ENSMUSG00000105786.2 | 0 | NA | NA | NA | NA | NA | #VALUE! |
| ENSMUSG00000105787.2 | 0 | NA | NA | NA | NA | NA | #VALUE! |

| | | | | | | | |
|---|---|---|---|---|---|---|---|
| ENSMUSG00000105789.2 | 0 | NA | NA | NA | NA | NA | #VALUE! |
| ENSMUSG00000105791.2 | 0 | NA | NA | NA | NA | NA | #VALUE! |
| ENSMUSG00000105792.2 | 0 | NA | NA | NA | NA | NA | #VALUE! |
| ENSMUSG00000105793.2 | 0 | NA | NA | NA | NA | NA | #VALUE! |
| ENSMUSG00000105794.2 | 0 | NA | NA | NA | NA | NA | #VALUE! |
| ENSMUSG00000105795.2 | 0 | NA | NA | NA | NA | NA | #VALUE! |
| ENSMUSG00000105796.2 | 0 | NA | NA | NA | NA | NA | #VALUE! |
| ENSMUSG00000105797.2 | 0 | NA | NA | NA | NA | NA | #VALUE! |
| ENSMUSG00000105798.2 | 0 | NA | NA | NA | NA | NA | #VALUE! |
| ENSMUSG00000105799.3 | 0 | NA | NA | NA | NA | NA | #VALUE! |
| ENSMUSG00000105800.2 | 0 | NA | NA | NA | NA | NA | #VALUE! |
| ENSMUSG00000105801.2 | 0 | NA | NA | NA | NA | NA | #VALUE! |
| ENSMUSG00000105802.2 | 0 | NA | NA | NA | NA | NA | #VALUE! |
| ENSMUSG00000105803.2 | 0 | NA | NA | NA | NA | NA | #VALUE! |
| ENSMUSG00000105804.2 | 0 | NA | NA | NA | NA | NA | #VALUE! |
| ENSMUSG00000105806.2 | 0 | NA | NA | NA | NA | NA | #VALUE! |
| ENSMUSG00000105807.2 | 0 | NA | NA | NA | NA | NA | #VALUE! |
| ENSMUSG00000105808.2 | 0 | NA | NA | NA | NA | NA | #VALUE! |
| ENSMUSG00000105809.2 | 0 | NA | NA | NA | NA | NA | #VALUE! |
| ENSMUSG00000105811.2 | 0 | NA | NA | NA | NA | NA | #VALUE! |
| ENSMUSG00000105813.2 | 0 | NA | NA | NA | NA | NA | #VALUE! |
| ENSMUSG00000105815.2 | 0 | NA | NA | NA | NA | NA | #VALUE! |
| ENSMUSG00000105816.5 | 0 | NA | NA | NA | NA | NA | #VALUE! |
| ENSMUSG00000105817.2 | 0 | NA | NA | NA | NA | NA | #VALUE! |
| ENSMUSG00000105818.2 | 0 | NA | NA | NA | NA | NA | #VALUE! |
| ENSMUSG00000105819.2 | 0 | NA | NA | NA | NA | NA | #VALUE! |
| ENSMUSG00000105820.2 | 2.984166 | 5.077883 | 3.921685 | 1.294822 | 0.195382 | NA | #VALUE! |
| ENSMUSG00000105822.2 | 0 | NA | NA | NA | NA | NA | #VALUE! |
| ENSMUSG00000105823.2 | 0 | NA | NA | NA | NA | NA | #VALUE! |
| ENSMUSG00000105824.2 | 0 | NA | NA | NA | NA | NA | #VALUE! |
| ENSMUSG00000105826.2 | 4.673535 | 5.707254 | 3.916423 | 1.457262 | 0.145044 | NA | #VALUE! |
| ENSMUSG00000105827.2 | 0 | NA | NA | NA | NA | NA | #VALUE! |
| ENSMUSG00000105828.2 | 0 | NA | NA | NA | NA | NA | #VALUE! |
| ENSMUSG00000105829.2 | 0 | NA | NA | NA | NA | NA | #VALUE! |
| ENSMUSG00000105830.2 | 0 | NA | NA | NA | NA | NA | #VALUE! |
| ENSMUSG00000105831.2 | 0 | NA | NA | NA | NA | NA | #VALUE! |
| ENSMUSG00000105832.2 | 7.121836 | -0.12586 | 2.201432 | -0.05717 | 0.954407 | NA | #VALUE! |
| ENSMUSG00000105833.2 | 0 | NA | NA | NA | NA | NA | #VALUE! |
| ENSMUSG00000105834.2 | 3.990499 | -5.53332 | 3.917921 | -1.41231 | 0.157859 | NA | #VALUE! |
| ENSMUSG00000105835.2 | 0 | NA | NA | NA | NA | NA | #VALUE! |
| ENSMUSG00000105836.2 | 0 | NA | NA | NA | NA | NA | #VALUE! |
| ENSMUSG00000105838.2 | 0 | NA | NA | NA | NA | NA | #VALUE! |
| ENSMUSG00000105840.2 | 0 | NA | NA | NA | NA | NA | #VALUE! |
| ENSMUSG00000105841.3 | 0 | NA | NA | NA | NA | NA | #VALUE! |

| | | | | | | | |
|---|---|---|---|---|---|---|---|
| ENSMUSG00000105842.2 | 0 | NA | NA | NA | NA | NA | #VALUE! |
| ENSMUSG00000105843.2 | 0 | NA | NA | NA | NA | NA | #VALUE! |
| ENSMUSG00000105845.2 | 0 | NA | NA | NA | NA | NA | #VALUE! |
| ENSMUSG00000105846.2 | 0 | NA | NA | NA | NA | NA | #VALUE! |
| ENSMUSG00000105847.3 | 0 | NA | NA | NA | NA | NA | #VALUE! |
| ENSMUSG00000105848.2 | 0 | NA | NA | NA | NA | NA | #VALUE! |
| ENSMUSG00000105849.3 | 0 | NA | NA | NA | NA | NA | #VALUE! |
| ENSMUSG00000105850.2 | 0 | NA | NA | NA | NA | NA | #VALUE! |
| ENSMUSG00000105851.2 | 0 | NA | NA | NA | NA | NA | #VALUE! |
| ENSMUSG00000105852.5 | 0 | NA | NA | NA | NA | NA | #VALUE! |
| ENSMUSG00000105853.3 | 0 | NA | NA | NA | NA | NA | #VALUE! |
| ENSMUSG00000105854.2 | 0.245976 | 1.027628 | 4.067126 | 0.252667 | 0.800526 | NA | #VALUE! |
| ENSMUSG00000105855.2 | 0 | NA | NA | NA | NA | NA | #VALUE! |
| ENSMUSG00000105856.2 | 0 | NA | NA | NA | NA | NA | #VALUE! |
| ENSMUSG00000105857.2 | 0 | NA | NA | NA | NA | NA | #VALUE! |
| ENSMUSG00000105858.2 | 1.177985 | 3.72614 | 3.944615 | 0.944614 | 0.344856 | NA | #VALUE! |
| ENSMUSG00000105859.2 | 0 | NA | NA | NA | NA | NA | #VALUE! |
| ENSMUSG00000105860.2 | 0 | NA | NA | NA | NA | NA | #VALUE! |
| ENSMUSG00000105861.2 | 0 | NA | NA | NA | NA | NA | #VALUE! |
| ENSMUSG00000105862.2 | 0 | NA | NA | NA | NA | NA | #VALUE! |
| ENSMUSG00000105864.2 | 0 | NA | NA | NA | NA | NA | #VALUE! |
| ENSMUSG00000105865.3 | 0 | NA | NA | NA | NA | NA | #VALUE! |
| ENSMUSG00000105866.2 | 0 | NA | NA | NA | NA | NA | #VALUE! |
| ENSMUSG00000105867.3 | 2.951706 | 5.040208 | 3.922079 | 1.285086 | 0.198762 | NA | #VALUE! |
| ENSMUSG00000105868.2 | 0 | NA | NA | NA | NA | NA | #VALUE! |
| ENSMUSG00000105869.2 | 0 | NA | NA | NA | NA | NA | #VALUE! |
| ENSMUSG00000105870.2 | 0 | NA | NA | NA | NA | NA | #VALUE! |
| ENSMUSG00000105871.2 | 0 | NA | NA | NA | NA | NA | #VALUE! |
| ENSMUSG00000105872.2 | 0 | NA | NA | NA | NA | NA | #VALUE! |
| ENSMUSG00000105873.2 | 0 | NA | NA | NA | NA | NA | #VALUE! |
| ENSMUSG00000105875.2 | 0 | NA | NA | NA | NA | NA | #VALUE! |
| ENSMUSG00000105876.2 | 0 | NA | NA | NA | NA | NA | #VALUE! |
| ENSMUSG00000105877.2 | 0 | NA | NA | NA | NA | NA | #VALUE! |
| ENSMUSG00000105880.2 | 0 | NA | NA | NA | NA | NA | #VALUE! |
| ENSMUSG00000105881.2 | 1.009701 | 3.503687 | 3.950852 | 0.886818 | 0.375177 | NA | #VALUE! |
| ENSMUSG00000105882.2 | 0 | NA | NA | NA | NA | NA | #VALUE! |
| ENSMUSG00000105883.2 | 0 | NA | NA | NA | NA | NA | #VALUE! |
| ENSMUSG00000105885.2 | 0 | NA | NA | NA | NA | NA | #VALUE! |
| ENSMUSG00000105886.3 | 0 | NA | NA | NA | NA | NA | #VALUE! |
| ENSMUSG00000105887.2 | 0 | NA | NA | NA | NA | NA | #VALUE! |
| ENSMUSG00000105888.2 | 0 | NA | NA | NA | NA | NA | #VALUE! |
| ENSMUSG00000105889.2 | 0 | NA | NA | NA | NA | NA | #VALUE! |
| ENSMUSG00000105890.2 | 0 | NA | NA | NA | NA | NA | #VALUE! |
| ENSMUSG00000105891.5 | 0 | NA | NA | NA | NA | NA | #VALUE! |

| | | | | | | | |
|---|---|---|---|---|---|---|---|
| ENSMUSG00000105892.2 | 0 | NA | NA | NA | NA | NA | #VALUE! |
| ENSMUSG00000105893.2 | 0 | NA | NA | NA | NA | NA | #VALUE! |
| ENSMUSG00000105894.2 | 0 | NA | NA | NA | NA | NA | #VALUE! |
| ENSMUSG00000105895.2 | 0 | NA | NA | NA | NA | NA | #VALUE! |
| ENSMUSG00000105896.2 | 0.559531 | 2.703958 | 3.982863 | 0.678898 | 0.497202 | NA | #VALUE! |
| ENSMUSG00000105897.3 | 0 | NA | NA | NA | NA | NA | #VALUE! |
| ENSMUSG00000105898.2 | 0 | NA | NA | NA | NA | NA | #VALUE! |
| ENSMUSG00000105900.2 | 0 | NA | NA | NA | NA | NA | #VALUE! |
| ENSMUSG00000105901.2 | 0 | NA | NA | NA | NA | NA | #VALUE! |
| ENSMUSG00000105903.2 | 0 | NA | NA | NA | NA | NA | #VALUE! |
| ENSMUSG00000105904.3 | 0 | NA | NA | NA | NA | NA | #VALUE! |
| ENSMUSG00000105905.2 | 0 | NA | NA | NA | NA | NA | #VALUE! |
| ENSMUSG00000105906.2 | 5.03578 | 5.828452 | 3.915646 | 1.488503 | 0.136618 | NA | #VALUE! |
| ENSMUSG00000105907.2 | 5.319587 | -1.88715 | 3.776925 | -0.49965 | 0.617319 | NA | #VALUE! |
| ENSMUSG00000105908.3 | 0 | NA | NA | NA | NA | NA | #VALUE! |
| ENSMUSG00000105909.2 | 1.398828 | 3.99633 | 3.938205 | 1.014759 | 0.310221 | NA | #VALUE! |
| ENSMUSG00000105910.2 | 0 | NA | NA | NA | NA | NA | #VALUE! |
| ENSMUSG00000105912.2 | 0 | NA | NA | NA | NA | NA | #VALUE! |
| ENSMUSG00000105913.3 | 0 | NA | NA | NA | NA | NA | #VALUE! |
| ENSMUSG00000105914.2 | 0 | NA | NA | NA | NA | NA | #VALUE! |
| ENSMUSG00000105915.2 | 0 | NA | NA | NA | NA | NA | #VALUE! |
| ENSMUSG00000105916.2 | 0 | NA | NA | NA | NA | NA | #VALUE! |
| ENSMUSG00000105917.2 | 0 | NA | NA | NA | NA | NA | #VALUE! |
| ENSMUSG00000105919.2 | 0 | NA | NA | NA | NA | NA | #VALUE! |
| ENSMUSG00000105920.2 | 0 | NA | NA | NA | NA | NA | #VALUE! |
| ENSMUSG00000105921.2 | 0 | NA | NA | NA | NA | NA | #VALUE! |
| ENSMUSG00000105922.2 | 0 | NA | NA | NA | NA | NA | #VALUE! |
| ENSMUSG00000105923.5 | 0 | NA | NA | NA | NA | NA | #VALUE! |
| ENSMUSG00000105925.2 | 0 | NA | NA | NA | NA | NA | #VALUE! |
| ENSMUSG00000105926.2 | 0 | NA | NA | NA | NA | NA | #VALUE! |
| ENSMUSG00000105928.2 | 0 | NA | NA | NA | NA | NA | #VALUE! |
| ENSMUSG00000105929.2 | 0 | NA | NA | NA | NA | NA | #VALUE! |
| ENSMUSG00000105930.2 | 0 | NA | NA | NA | NA | NA | #VALUE! |
| ENSMUSG00000105931.2 | 0 | NA | NA | NA | NA | NA | #VALUE! |
| ENSMUSG00000105932.3 | 0 | NA | NA | NA | NA | NA | #VALUE! |
| ENSMUSG00000105933.2 | 0 | NA | NA | NA | NA | NA | #VALUE! |
| ENSMUSG00000105934.2 | 0 | NA | NA | NA | NA | NA | #VALUE! |
| ENSMUSG00000105936.2 | 0 | NA | NA | NA | NA | NA | #VALUE! |
| ENSMUSG00000105937.2 | 0 | NA | NA | NA | NA | NA | #VALUE! |
| ENSMUSG00000105938.2 | 0 | NA | NA | NA | NA | NA | #VALUE! |
| ENSMUSG00000105939.2 | 0 | NA | NA | NA | NA | NA | #VALUE! |
| ENSMUSG00000105940.2 | 0 | NA | NA | NA | NA | NA | #VALUE! |
| ENSMUSG00000105941.2 | 0 | NA | NA | NA | NA | NA | #VALUE! |
| ENSMUSG00000105942.2 | 0 | NA | NA | NA | NA | NA | #VALUE! |

| | | | | | | | |
|---|---|---|---|---|---|---|---|
| ENSMUSG00000105943.2 | 0.168284 | 1.027628 | 4.067126 | 0.252667 | 0.800526 | NA | #VALUE! |
| ENSMUSG00000105944.2 | 0 | NA | NA | NA | NA | NA | #VALUE! |
| ENSMUSG00000105945.2 | 0 | NA | NA | NA | NA | NA | #VALUE! |
| ENSMUSG00000105946.2 | 0 | NA | NA | NA | NA | NA | #VALUE! |
| ENSMUSG00000105947.2 | 0 | NA | NA | NA | NA | NA | #VALUE! |
| ENSMUSG00000105948.2 | 0 | NA | NA | NA | NA | NA | #VALUE! |
| ENSMUSG00000105949.2 | 0 | NA | NA | NA | NA | NA | #VALUE! |
| ENSMUSG00000105950.2 | 0 | NA | NA | NA | NA | NA | #VALUE! |
| ENSMUSG00000105952.2 | 0 | NA | NA | NA | NA | NA | #VALUE! |
| ENSMUSG00000105953.2 | 0 | NA | NA | NA | NA | NA | #VALUE! |
| ENSMUSG00000105954.2 | 0 | NA | NA | NA | NA | NA | #VALUE! |
| ENSMUSG00000105955.2 | 0 | NA | NA | NA | NA | NA | #VALUE! |
| ENSMUSG00000105956.2 | 0 | NA | NA | NA | NA | NA | #VALUE! |
| ENSMUSG00000105957.2 | 3.942659 | -1.15557 | 3.779732 | -0.30573 | 0.759813 | NA | #VALUE! |
| ENSMUSG00000105958.2 | 0 | NA | NA | NA | NA | NA | #VALUE! |
| ENSMUSG00000105959.2 | 0 | NA | NA | NA | NA | NA | #VALUE! |
| ENSMUSG00000105960.2 | 0 | NA | NA | NA | NA | NA | #VALUE! |
| ENSMUSG00000105961.2 | 0 | NA | NA | NA | NA | NA | #VALUE! |
| ENSMUSG00000105962.2 | 0 | NA | NA | NA | NA | NA | #VALUE! |
| ENSMUSG00000105963.2 | 0 | NA | NA | NA | NA | NA | #VALUE! |
| ENSMUSG00000105964.3 | 0 | NA | NA | NA | NA | NA | #VALUE! |
| ENSMUSG00000105965.2 | 0.093255 | 0.868093 | 4.072464 | 0.213162 | 0.831201 | NA | #VALUE! |
| ENSMUSG00000105966.2 | 0 | NA | NA | NA | NA | NA | #VALUE! |
| ENSMUSG00000105968.2 | 0 | NA | NA | NA | NA | NA | #VALUE! |
| ENSMUSG00000105969.2 | 0 | NA | NA | NA | NA | NA | #VALUE! |
| ENSMUSG00000105970.2 | 0 | NA | NA | NA | NA | NA | #VALUE! |
| ENSMUSG00000105971.2 | 0 | NA | NA | NA | NA | NA | #VALUE! |
| ENSMUSG00000105972.3 | 0 | NA | NA | NA | NA | NA | #VALUE! |
| ENSMUSG00000105973.2 | 0 | NA | NA | NA | NA | NA | #VALUE! |
| ENSMUSG00000105974.2 | 0 | NA | NA | NA | NA | NA | #VALUE! |
| ENSMUSG00000105975.5 | 0 | NA | NA | NA | NA | NA | #VALUE! |
| ENSMUSG00000105976.2 | 0 | NA | NA | NA | NA | NA | #VALUE! |
| ENSMUSG00000105977.2 | 0 | NA | NA | NA | NA | NA | #VALUE! |
| ENSMUSG00000105978.2 | 0 | NA | NA | NA | NA | NA | #VALUE! |
| ENSMUSG00000105979.2 | 0 | NA | NA | NA | NA | NA | #VALUE! |
| ENSMUSG00000105980.2 | 0 | NA | NA | NA | NA | NA | #VALUE! |
| ENSMUSG00000105981.2 | 0 | NA | NA | NA | NA | NA | #VALUE! |
| ENSMUSG00000105982.3 | 0 | NA | NA | NA | NA | NA | #VALUE! |
| ENSMUSG00000105983.2 | 0 | NA | NA | NA | NA | NA | #VALUE! |
| ENSMUSG00000105985.2 | 0 | NA | NA | NA | NA | NA | #VALUE! |
| ENSMUSG00000105986.2 | 0 | NA | NA | NA | NA | NA | #VALUE! |
| ENSMUSG00000105987.5 | 0.105013 | -0.97324 | 4.076293 | -0.23876 | 0.811295 | NA | #VALUE! |
| ENSMUSG00000105989.2 | 0 | NA | NA | NA | NA | NA | #VALUE! |
| ENSMUSG00000105990.2 | 0 | NA | NA | NA | NA | NA | #VALUE! |

| | | | | | | | |
|---|---|---|---|---|---|---|---|
| ENSMUSG00000105991.2 | 0 | NA | NA | NA | NA | NA | #VALUE! |
| ENSMUSG00000105992.2 | 0 | NA | NA | NA | NA | NA | #VALUE! |
| ENSMUSG00000105993.2 | 0 | NA | NA | NA | NA | NA | #VALUE! |
| ENSMUSG00000105994.2 | 0 | NA | NA | NA | NA | NA | #VALUE! |
| ENSMUSG00000105995.2 | 0 | NA | NA | NA | NA | NA | #VALUE! |
| ENSMUSG00000105996.2 | 0 | NA | NA | NA | NA | NA | #VALUE! |
| ENSMUSG00000105999.2 | 0 | NA | NA | NA | NA | NA | #VALUE! |
| ENSMUSG00000106000.2 | 0 | NA | NA | NA | NA | NA | #VALUE! |
| ENSMUSG00000106001.2 | 0 | NA | NA | NA | NA | NA | #VALUE! |
| ENSMUSG00000106002.2 | 0 | NA | NA | NA | NA | NA | #VALUE! |
| ENSMUSG00000106003.2 | 0 | NA | NA | NA | NA | NA | #VALUE! |
| ENSMUSG00000106004.5 | 0 | NA | NA | NA | NA | NA | #VALUE! |
| ENSMUSG00000106005.2 | 0 | NA | NA | NA | NA | NA | #VALUE! |
| ENSMUSG00000106006.2 | 0 | NA | NA | NA | NA | NA | #VALUE! |
| ENSMUSG00000106008.2 | 0 | NA | NA | NA | NA | NA | #VALUE! |
| ENSMUSG00000106009.2 | 0 | NA | NA | NA | NA | NA | #VALUE! |
| ENSMUSG00000106010.2 | 0 | NA | NA | NA | NA | NA | #VALUE! |
| ENSMUSG00000106011.2 | 0 | NA | NA | NA | NA | NA | #VALUE! |
| ENSMUSG00000106012.2 | 0 | NA | NA | NA | NA | NA | #VALUE! |
| ENSMUSG00000106013.5 | 0 | NA | NA | NA | NA | NA | #VALUE! |
| ENSMUSG00000106015.2 | 0 | NA | NA | NA | NA | NA | #VALUE! |
| ENSMUSG00000106016.2 | 0 | NA | NA | NA | NA | NA | #VALUE! |
| ENSMUSG00000106018.2 | 0 | NA | NA | NA | NA | NA | #VALUE! |
| ENSMUSG00000106019.2 | 2.21378 | 4.621487 | 3.927208 | 1.176787 | 0.239281 | NA | #VALUE! |
| ENSMUSG00000106020.2 | 0 | NA | NA | NA | NA | NA | #VALUE! |
| ENSMUSG00000106022.2 | 0 | NA | NA | NA | NA | NA | #VALUE! |
| ENSMUSG00000106023.2 | 0 | NA | NA | NA | NA | NA | #VALUE! |
| ENSMUSG00000106024.2 | 0 | NA | NA | NA | NA | NA | #VALUE! |
| ENSMUSG00000106025.2 | 0 | NA | NA | NA | NA | NA | #VALUE! |
| ENSMUSG00000106026.2 | 0 | NA | NA | NA | NA | NA | #VALUE! |
| ENSMUSG00000106028.2 | 0 | NA | NA | NA | NA | NA | #VALUE! |
| ENSMUSG00000106029.2 | 0 | NA | NA | NA | NA | NA | #VALUE! |
| ENSMUSG00000106030.2 | 0 | NA | NA | NA | NA | NA | #VALUE! |
| ENSMUSG00000106031.2 | 0 | NA | NA | NA | NA | NA | #VALUE! |
| ENSMUSG00000106032.2 | 0 | NA | NA | NA | NA | NA | #VALUE! |
| ENSMUSG00000106033.2 | 0 | NA | NA | NA | NA | NA | #VALUE! |
| ENSMUSG00000106034.2 | 0 | NA | NA | NA | NA | NA | #VALUE! |
| ENSMUSG00000106035.2 | 0 | NA | NA | NA | NA | NA | #VALUE! |
| ENSMUSG00000106036.2 | 0 | NA | NA | NA | NA | NA | #VALUE! |
| ENSMUSG00000106037.2 | 7.273315 | 0.476095 | 2.205634 | 0.215854 | 0.829101 | NA | #VALUE! |
| ENSMUSG00000106038.2 | 0 | NA | NA | NA | NA | NA | #VALUE! |
| ENSMUSG00000106039.2 | 0 | NA | NA | NA | NA | NA | #VALUE! |
| ENSMUSG00000106040.2 | 0.093255 | 0.868093 | 4.072464 | 0.213162 | 0.831201 | NA | #VALUE! |
| ENSMUSG00000106041.2 | 0 | NA | NA | NA | NA | NA | #VALUE! |

| | | | | | | | |
|---|---|---|---|---|---|---|---|
| ENSMUSG00000106042.2 | 0 | NA | NA | NA | NA | NA | #VALUE! |
| ENSMUSG00000106043.2 | 0 | NA | NA | NA | NA | NA | #VALUE! |
| ENSMUSG00000106044.3 | 0 | NA | NA | NA | NA | NA | #VALUE! |
| ENSMUSG00000106045.2 | 0 | NA | NA | NA | NA | NA | #VALUE! |
| ENSMUSG00000106047.2 | 0 | NA | NA | NA | NA | NA | #VALUE! |
| ENSMUSG00000106048.2 | 0 | NA | NA | NA | NA | NA | #VALUE! |
| ENSMUSG00000106049.2 | 0 | NA | NA | NA | NA | NA | #VALUE! |
| ENSMUSG00000106050.2 | 0 | NA | NA | NA | NA | NA | #VALUE! |
| ENSMUSG00000106051.2 | 0 | NA | NA | NA | NA | NA | #VALUE! |
| ENSMUSG00000106052.2 | 0 | NA | NA | NA | NA | NA | #VALUE! |
| ENSMUSG00000106053.2 | 0 | NA | NA | NA | NA | NA | #VALUE! |
| ENSMUSG00000106054.2 | 0 | NA | NA | NA | NA | NA | #VALUE! |
| ENSMUSG00000106056.2 | 0 | NA | NA | NA | NA | NA | #VALUE! |
| ENSMUSG00000106057.2 | 0 | NA | NA | NA | NA | NA | #VALUE! |
| ENSMUSG00000106058.2 | 0 | NA | NA | NA | NA | NA | #VALUE! |
| ENSMUSG00000106059.2 | 0 | NA | NA | NA | NA | NA | #VALUE! |
| ENSMUSG00000106060.2 | 0 | NA | NA | NA | NA | NA | #VALUE! |
| ENSMUSG00000106061.2 | 1.678593 | 4.255795 | 3.933066 | 1.082055 | 0.279228 | NA | #VALUE! |
| ENSMUSG00000106062.2 | 0 | NA | NA | NA | NA | NA | #VALUE! |
| ENSMUSG00000106063.2 | 0 | NA | NA | NA | NA | NA | #VALUE! |
| ENSMUSG00000106064.2 | 0 | NA | NA | NA | NA | NA | #VALUE! |
| ENSMUSG00000106065.2 | 0 | NA | NA | NA | NA | NA | #VALUE! |
| ENSMUSG00000106066.2 | 0 | NA | NA | NA | NA | NA | #VALUE! |
| ENSMUSG00000106067.2 | 0 | NA | NA | NA | NA | NA | #VALUE! |
| ENSMUSG00000106068.2 | 0 | NA | NA | NA | NA | NA | #VALUE! |
| ENSMUSG00000106069.2 | 2.238933 | 0.727689 | 3.811651 | 0.190912 | 0.848595 | NA | #VALUE! |
| ENSMUSG00000106070.2 | 0 | NA | NA | NA | NA | NA | #VALUE! |
| ENSMUSG00000106071.2 | 0 | NA | NA | NA | NA | NA | #VALUE! |
| ENSMUSG00000106072.2 | 0 | NA | NA | NA | NA | NA | #VALUE! |
| ENSMUSG00000106073.2 | 7.327403 | -4.9237 | 3.62028 | -1.36003 | 0.173819 | NA | #VALUE! |
| ENSMUSG00000106074.3 | 0 | NA | NA | NA | NA | NA | #VALUE! |
| ENSMUSG00000106075.2 | 0 | NA | NA | NA | NA | NA | #VALUE! |
| ENSMUSG00000106076.2 | 0 | NA | NA | NA | NA | NA | #VALUE! |
| ENSMUSG00000106077.2 | 0 | NA | NA | NA | NA | NA | #VALUE! |
| ENSMUSG00000106078.2 | 0 | NA | NA | NA | NA | NA | #VALUE! |
| ENSMUSG00000106079.2 | 0 | NA | NA | NA | NA | NA | #VALUE! |
| ENSMUSG00000106080.2 | 0 | NA | NA | NA | NA | NA | #VALUE! |
| ENSMUSG00000106082.5 | 0 | NA | NA | NA | NA | NA | #VALUE! |
| ENSMUSG00000106083.2 | 0 | NA | NA | NA | NA | NA | #VALUE! |
| ENSMUSG00000106084.2 | 0 | NA | NA | NA | NA | NA | #VALUE! |
| ENSMUSG00000106085.2 | 0 | NA | NA | NA | NA | NA | #VALUE! |
| ENSMUSG00000106086.2 | 0 | NA | NA | NA | NA | NA | #VALUE! |
| ENSMUSG00000106089.2 | 0 | NA | NA | NA | NA | NA | #VALUE! |
| ENSMUSG00000106090.2 | 0 | NA | NA | NA | NA | NA | #VALUE! |

| | | | | | | | |
|---|---|---|---|---|---|---|---|
| ENSMUSG00000106092.2 | 0 | NA | NA | NA | NA | NA | #VALUE! |
| ENSMUSG00000106093.2 | 0 | NA | NA | NA | NA | NA | #VALUE! |
| ENSMUSG00000106094.2 | 0 | NA | NA | NA | NA | NA | #VALUE! |
| ENSMUSG00000106095.3 | 0 | NA | NA | NA | NA | NA | #VALUE! |
| ENSMUSG00000106096.2 | 0 | NA | NA | NA | NA | NA | #VALUE! |
| ENSMUSG00000106097.2 | 0 | NA | NA | NA | NA | NA | #VALUE! |
| ENSMUSG00000106098.2 | 0 | NA | NA | NA | NA | NA | #VALUE! |
| ENSMUSG00000106099.2 | 0 | NA | NA | NA | NA | NA | #VALUE! |
| ENSMUSG00000106100.2 | 0 | NA | NA | NA | NA | NA | #VALUE! |
| ENSMUSG00000106101.2 | 0 | NA | NA | NA | NA | NA | #VALUE! |
| ENSMUSG00000106102.2 | 0 | NA | NA | NA | NA | NA | #VALUE! |
| ENSMUSG00000106103.2 | 0 | NA | NA | NA | NA | NA | #VALUE! |
| ENSMUSG00000106104.2 | 0 | NA | NA | NA | NA | NA | #VALUE! |
| ENSMUSG00000106105.2 | 0 | NA | NA | NA | NA | NA | #VALUE! |
| ENSMUSG00000106107.2 | 0 | NA | NA | NA | NA | NA | #VALUE! |
| ENSMUSG00000106108.2 | 0 | NA | NA | NA | NA | NA | #VALUE! |
| ENSMUSG00000106109.2 | 0 | NA | NA | NA | NA | NA | #VALUE! |
| ENSMUSG00000106110.2 | 0 | NA | NA | NA | NA | NA | #VALUE! |
| ENSMUSG00000106111.2 | 0 | NA | NA | NA | NA | NA | #VALUE! |
| ENSMUSG00000106112.2 | 0 | NA | NA | NA | NA | NA | #VALUE! |
| ENSMUSG00000106113.2 | 0 | NA | NA | NA | NA | NA | #VALUE! |
| ENSMUSG00000106114.2 | 0 | NA | NA | NA | NA | NA | #VALUE! |
| ENSMUSG00000106115.2 | 0 | NA | NA | NA | NA | NA | #VALUE! |
| ENSMUSG00000106116.2 | 0 | NA | NA | NA | NA | NA | #VALUE! |
| ENSMUSG00000106117.2 | 0 | NA | NA | NA | NA | NA | #VALUE! |
| ENSMUSG00000106118.2 | 0 | NA | NA | NA | NA | NA | #VALUE! |
| ENSMUSG00000106119.2 | 0 | NA | NA | NA | NA | NA | #VALUE! |
| ENSMUSG00000106120.2 | 0 | NA | NA | NA | NA | NA | #VALUE! |
| ENSMUSG00000106121.2 | 4.66276 | 5.717927 | 3.916352 | 1.460014 | 0.144286 | NA | #VALUE! |
| ENSMUSG00000106122.2 | 0 | NA | NA | NA | NA | NA | #VALUE! |
| ENSMUSG00000106123.2 | 0 | NA | NA | NA | NA | NA | #VALUE! |
| ENSMUSG00000106124.2 | 0 | NA | NA | NA | NA | NA | #VALUE! |
| ENSMUSG00000106125.2 | 0 | NA | NA | NA | NA | NA | #VALUE! |
| ENSMUSG00000106126.2 | 0 | NA | NA | NA | NA | NA | #VALUE! |
| ENSMUSG00000106127.2 | 0 | NA | NA | NA | NA | NA | #VALUE! |
| ENSMUSG00000106128.2 | 0 | NA | NA | NA | NA | NA | #VALUE! |
| ENSMUSG00000106129.2 | 0 | NA | NA | NA | NA | NA | #VALUE! |
| ENSMUSG00000106131.3 | 0 | NA | NA | NA | NA | NA | #VALUE! |
| ENSMUSG00000106132.2 | 0 | NA | NA | NA | NA | NA | #VALUE! |
| ENSMUSG00000106133.2 | 0 | NA | NA | NA | NA | NA | #VALUE! |
| ENSMUSG00000106134.2 | 0 | NA | NA | NA | NA | NA | #VALUE! |
| ENSMUSG00000106135.2 | 4.419071 | -1.8164 | 2.833801 | -0.64098 | 0.521538 | NA | #VALUE! |
| ENSMUSG00000106136.3 | 0 | NA | NA | NA | NA | NA | #VALUE! |
| ENSMUSG00000106137.2 | 0 | NA | NA | NA | NA | NA | #VALUE! |

| | | | | | | | |
|---|---|---|---|---|---|---|---|
| ENSMUSG00000106139.2 | 0 | NA | NA | NA | NA | NA | #VALUE! |
| ENSMUSG00000106140.2 | 0 | NA | NA | NA | NA | NA | #VALUE! |
| ENSMUSG00000106141.2 | 0 | NA | NA | NA | NA | NA | #VALUE! |
| ENSMUSG00000106142.2 | 0 | NA | NA | NA | NA | NA | #VALUE! |
| ENSMUSG00000106143.2 | 0 | NA | NA | NA | NA | NA | #VALUE! |
| ENSMUSG00000106144.2 | 0 | NA | NA | NA | NA | NA | #VALUE! |
| ENSMUSG00000106145.3 | 0 | NA | NA | NA | NA | NA | #VALUE! |
| ENSMUSG00000106146.2 | 0 | NA | NA | NA | NA | NA | #VALUE! |
| ENSMUSG00000106147.3 | 0 | NA | NA | NA | NA | NA | #VALUE! |
| ENSMUSG00000106148.2 | 0 | NA | NA | NA | NA | NA | #VALUE! |
| ENSMUSG00000106149.2 | 0 | NA | NA | NA | NA | NA | #VALUE! |
| ENSMUSG00000106150.3 | 0 | NA | NA | NA | NA | NA | #VALUE! |
| ENSMUSG00000106153.2 | 0 | NA | NA | NA | NA | NA | #VALUE! |
| ENSMUSG00000106154.3 | 0 | NA | NA | NA | NA | NA | #VALUE! |
| ENSMUSG00000106155.2 | 0 | NA | NA | NA | NA | NA | #VALUE! |
| ENSMUSG00000106156.2 | 0 | NA | NA | NA | NA | NA | #VALUE! |
| ENSMUSG00000106157.2 | 0 | NA | NA | NA | NA | NA | #VALUE! |
| ENSMUSG00000106158.2 | 0 | NA | NA | NA | NA | NA | #VALUE! |
| ENSMUSG00000106159.2 | 0 | NA | NA | NA | NA | NA | #VALUE! |
| ENSMUSG00000106160.2 | 0 | NA | NA | NA | NA | NA | #VALUE! |
| ENSMUSG00000106161.2 | 0 | NA | NA | NA | NA | NA | #VALUE! |
| ENSMUSG00000106162.2 | 0 | NA | NA | NA | NA | NA | #VALUE! |
| ENSMUSG00000106164.2 | 0.746042 | 3.107177 | 3.964586 | 0.783733 | 0.433197 | NA | #VALUE! |
| ENSMUSG00000106165.2 | 0 | NA | NA | NA | NA | NA | #VALUE! |
| ENSMUSG00000106166.2 | 0 | NA | NA | NA | NA | NA | #VALUE! |
| ENSMUSG00000106168.2 | 0 | NA | NA | NA | NA | NA | #VALUE! |
| ENSMUSG00000106169.3 | 0 | NA | NA | NA | NA | NA | #VALUE! |
| ENSMUSG00000106170.2 | 0 | NA | NA | NA | NA | NA | #VALUE! |
| ENSMUSG00000106172.2 | 0 | NA | NA | NA | NA | NA | #VALUE! |
| ENSMUSG00000106173.2 | 0 | NA | NA | NA | NA | NA | #VALUE! |
| ENSMUSG00000106174.2 | 0 | NA | NA | NA | NA | NA | #VALUE! |
| ENSMUSG00000106176.2 | 0 | NA | NA | NA | NA | NA | #VALUE! |
| ENSMUSG00000106177.2 | 0 | NA | NA | NA | NA | NA | #VALUE! |
| ENSMUSG00000106178.2 | 0 | NA | NA | NA | NA | NA | #VALUE! |
| ENSMUSG00000106180.2 | 0 | NA | NA | NA | NA | NA | #VALUE! |
| ENSMUSG00000106181.2 | 0 | NA | NA | NA | NA | NA | #VALUE! |
| ENSMUSG00000106182.2 | 0 | NA | NA | NA | NA | NA | #VALUE! |
| ENSMUSG00000106183.2 | 0 | NA | NA | NA | NA | NA | #VALUE! |
| ENSMUSG00000106184.2 | 0 | NA | NA | NA | NA | NA | #VALUE! |
| ENSMUSG00000106185.2 | 0 | NA | NA | NA | NA | NA | #VALUE! |
| ENSMUSG00000106186.2 | 0 | NA | NA | NA | NA | NA | #VALUE! |
| ENSMUSG00000106187.2 | 0 | NA | NA | NA | NA | NA | #VALUE! |
| ENSMUSG00000106188.2 | 0 | NA | NA | NA | NA | NA | #VALUE! |
| ENSMUSG00000106189.2 | 5.940979 | -0.90981 | 2.688106 | -0.33846 | 0.735019 | NA | #VALUE! |

| | | | | | | | |
|---|---|---|---|---|---|---|---|
| ENSMUSG00000106190.3 | 1.217102 | 3.777985 | 3.943292 | 0.958079 | 0.338023 | NA | #VALUE! |
| ENSMUSG00000106191.2 | 0 | NA | NA | NA | NA | NA | #VALUE! |
| ENSMUSG00000106192.2 | 0 | NA | NA | NA | NA | NA | #VALUE! |
| ENSMUSG00000106193.3 | 0 | NA | NA | NA | NA | NA | #VALUE! |
| ENSMUSG00000106194.2 | 0 | NA | NA | NA | NA | NA | #VALUE! |
| ENSMUSG00000106197.2 | 0 | NA | NA | NA | NA | NA | #VALUE! |
| ENSMUSG00000106199.3 | 0 | NA | NA | NA | NA | NA | #VALUE! |
| ENSMUSG00000106201.2 | 0 | NA | NA | NA | NA | NA | #VALUE! |
| ENSMUSG00000106202.2 | 0 | NA | NA | NA | NA | NA | #VALUE! |
| ENSMUSG00000106204.2 | 0 | NA | NA | NA | NA | NA | #VALUE! |
| ENSMUSG00000106205.2 | 3.0485 | -2.44513 | 3.840605 | -0.63665 | 0.524351 | NA | #VALUE! |
| ENSMUSG00000106206.2 | 0 | NA | NA | NA | NA | NA | #VALUE! |
| ENSMUSG00000106207.2 | 0 | NA | NA | NA | NA | NA | #VALUE! |
| ENSMUSG00000106208.2 | 0.093255 | 0.868093 | 4.072464 | 0.213162 | 0.831201 | NA | #VALUE! |
| ENSMUSG00000106209.2 | 0 | NA | NA | NA | NA | NA | #VALUE! |
| ENSMUSG00000106210.2 | 0 | NA | NA | NA | NA | NA | #VALUE! |
| ENSMUSG00000106211.2 | 0 | NA | NA | NA | NA | NA | #VALUE! |
| ENSMUSG00000106212.2 | 0 | NA | NA | NA | NA | NA | #VALUE! |
| ENSMUSG00000106213.2 | 0 | NA | NA | NA | NA | NA | #VALUE! |
| ENSMUSG00000106214.2 | 0 | NA | NA | NA | NA | NA | #VALUE! |
| ENSMUSG00000106215.3 | 0 | NA | NA | NA | NA | NA | #VALUE! |
| ENSMUSG00000106216.2 | 0 | NA | NA | NA | NA | NA | #VALUE! |
| ENSMUSG00000106217.2 | 0 | NA | NA | NA | NA | NA | #VALUE! |
| ENSMUSG00000106218.2 | 0 | NA | NA | NA | NA | NA | #VALUE! |
| ENSMUSG00000106219.2 | 0 | NA | NA | NA | NA | NA | #VALUE! |
| ENSMUSG00000106220.2 | 0 | NA | NA | NA | NA | NA | #VALUE! |
| ENSMUSG00000106222.2 | 0 | NA | NA | NA | NA | NA | #VALUE! |
| ENSMUSG00000106223.2 | 0 | NA | NA | NA | NA | NA | #VALUE! |
| ENSMUSG00000106224.2 | 0 | NA | NA | NA | NA | NA | #VALUE! |
| ENSMUSG00000106225.2 | 0 | NA | NA | NA | NA | NA | #VALUE! |
| ENSMUSG00000106226.2 | 0 | NA | NA | NA | NA | NA | #VALUE! |
| ENSMUSG00000106227.2 | 0 | NA | NA | NA | NA | NA | #VALUE! |
| ENSMUSG00000106228.3 | 0 | NA | NA | NA | NA | NA | #VALUE! |
| ENSMUSG00000106229.2 | 0.945118 | -3.47564 | 3.952879 | -0.87927 | 0.379256 | NA | #VALUE! |
| ENSMUSG00000106230.2 | 0 | NA | NA | NA | NA | NA | #VALUE! |
| ENSMUSG00000106231.2 | 0 | NA | NA | NA | NA | NA | #VALUE! |
| ENSMUSG00000106232.2 | 0 | NA | NA | NA | NA | NA | #VALUE! |
| ENSMUSG00000106233.3 | 0 | NA | NA | NA | NA | NA | #VALUE! |
| ENSMUSG00000106234.2 | 0 | NA | NA | NA | NA | NA | #VALUE! |
| ENSMUSG00000106235.2 | 0 | NA | NA | NA | NA | NA | #VALUE! |
| ENSMUSG00000106236.3 | 0 | NA | NA | NA | NA | NA | #VALUE! |
| ENSMUSG00000106237.2 | 0 | NA | NA | NA | NA | NA | #VALUE! |
| ENSMUSG00000106238.2 | 0 | NA | NA | NA | NA | NA | #VALUE! |
| ENSMUSG00000106239.2 | 0 | NA | NA | NA | NA | NA | #VALUE! |

| | | | | | | | |
|---|---|---|---|---|---|---|---|
| ENSMUSG00000106240.2 | 0 | NA | NA | NA | NA | NA | #VALUE! |
| ENSMUSG00000106241.2 | 0 | NA | NA | NA | NA | NA | #VALUE! |
| ENSMUSG00000106243.2 | 0 | NA | NA | NA | NA | NA | #VALUE! |
| ENSMUSG00000106244.2 | 0 | NA | NA | NA | NA | NA | #VALUE! |
| ENSMUSG00000106245.2 | 0 | NA | NA | NA | NA | NA | #VALUE! |
| ENSMUSG00000106246.2 | 0.105013 | -0.97324 | 4.076293 | -0.23876 | 0.811295 | NA | #VALUE! |
| ENSMUSG00000106247.2 | 0 | NA | NA | NA | NA | NA | #VALUE! |
| ENSMUSG00000106248.2 | 0 | NA | NA | NA | NA | NA | #VALUE! |
| ENSMUSG00000106249.2 | 0 | NA | NA | NA | NA | NA | #VALUE! |
| ENSMUSG00000106250.2 | 0 | NA | NA | NA | NA | NA | #VALUE! |
| ENSMUSG00000106251.2 | 6.656829 | -6.26304 | 3.913498 | -1.60037 | 0.109517 | NA | #VALUE! |
| ENSMUSG00000106252.3 | 0.168284 | 1.027628 | 4.067126 | 0.252667 | 0.800526 | NA | #VALUE! |
| ENSMUSG00000106253.3 | 0 | NA | NA | NA | NA | NA | #VALUE! |
| ENSMUSG00000106254.2 | 0 | NA | NA | NA | NA | NA | #VALUE! |
| ENSMUSG00000106255.2 | 0 | NA | NA | NA | NA | NA | #VALUE! |
| ENSMUSG00000106256.2 | 0 | NA | NA | NA | NA | NA | #VALUE! |
| ENSMUSG00000106257.2 | 0 | NA | NA | NA | NA | NA | #VALUE! |
| ENSMUSG00000106259.2 | 0 | NA | NA | NA | NA | NA | #VALUE! |
| ENSMUSG00000106260.2 | 0 | NA | NA | NA | NA | NA | #VALUE! |
| ENSMUSG00000106261.2 | 0 | NA | NA | NA | NA | NA | #VALUE! |
| ENSMUSG00000106262.2 | 0 | NA | NA | NA | NA | NA | #VALUE! |
| ENSMUSG00000106263.2 | 0 | NA | NA | NA | NA | NA | #VALUE! |
| ENSMUSG00000106264.3 | 0 | NA | NA | NA | NA | NA | #VALUE! |
| ENSMUSG00000106265.2 | 0 | NA | NA | NA | NA | NA | #VALUE! |
| ENSMUSG00000106266.2 | 0 | NA | NA | NA | NA | NA | #VALUE! |
| ENSMUSG00000106267.2 | 0 | NA | NA | NA | NA | NA | #VALUE! |
| ENSMUSG00000106268.2 | 0 | NA | NA | NA | NA | NA | #VALUE! |
| ENSMUSG00000106269.2 | 0 | NA | NA | NA | NA | NA | #VALUE! |
| ENSMUSG00000106270.2 | 0 | NA | NA | NA | NA | NA | #VALUE! |
| ENSMUSG00000106271.2 | 0 | NA | NA | NA | NA | NA | #VALUE! |
| ENSMUSG00000106273.2 | 0 | NA | NA | NA | NA | NA | #VALUE! |
| ENSMUSG00000106274.2 | 0 | NA | NA | NA | NA | NA | #VALUE! |
| ENSMUSG00000106275.2 | 0 | NA | NA | NA | NA | NA | #VALUE! |
| ENSMUSG00000106276.2 | 0 | NA | NA | NA | NA | NA | #VALUE! |
| ENSMUSG00000106278.3 | 0 | NA | NA | NA | NA | NA | #VALUE! |
| ENSMUSG00000106280.2 | 0 | NA | NA | NA | NA | NA | #VALUE! |
| ENSMUSG00000106282.2 | 0 | NA | NA | NA | NA | NA | #VALUE! |
| ENSMUSG00000106283.2 | 0 | NA | NA | NA | NA | NA | #VALUE! |
| ENSMUSG00000106284.2 | 0 | NA | NA | NA | NA | NA | #VALUE! |
| ENSMUSG00000106285.2 | 0 | NA | NA | NA | NA | NA | #VALUE! |
| ENSMUSG00000106286.2 | 0 | NA | NA | NA | NA | NA | #VALUE! |
| ENSMUSG00000106287.2 | 0 | NA | NA | NA | NA | NA | #VALUE! |
| ENSMUSG00000106288.2 | 1.146898 | 0.001197 | 3.882869 | 0.000308 | 0.999754 | NA | #VALUE! |
| ENSMUSG00000106289.2 | 0 | NA | NA | NA | NA | NA | #VALUE! |

| | | | | | | | |
|---|---|---|---|---|---|---|---|
| ENSMUSG00000106290.2 | 0.612332 | -2.79827 | 3.980109 | -0.70306 | 0.482016 | NA | #VALUE! |
| ENSMUSG00000106291.2 | 0 | NA | NA | NA | NA | NA | #VALUE! |
| ENSMUSG00000106292.2 | 0 | NA | NA | NA | NA | NA | #VALUE! |
| ENSMUSG00000106293.2 | 0 | NA | NA | NA | NA | NA | #VALUE! |
| ENSMUSG00000106294.2 | 1.694325 | 0.843492 | 3.844053 | 0.219428 | 0.826317 | NA | #VALUE! |
| ENSMUSG00000106295.2 | 0 | NA | NA | NA | NA | NA | #VALUE! |
| ENSMUSG00000106296.2 | 0 | NA | NA | NA | NA | NA | #VALUE! |
| ENSMUSG00000106298.2 | 0 | NA | NA | NA | NA | NA | #VALUE! |
| ENSMUSG00000106300.2 | 0 | NA | NA | NA | NA | NA | #VALUE! |
| ENSMUSG00000106301.2 | 0 | NA | NA | NA | NA | NA | #VALUE! |
| ENSMUSG00000106302.3 | 0 | NA | NA | NA | NA | NA | #VALUE! |
| ENSMUSG00000106303.2 | 0 | NA | NA | NA | NA | NA | #VALUE! |
| ENSMUSG00000106304.2 | 0 | NA | NA | NA | NA | NA | #VALUE! |
| ENSMUSG00000106305.2 | 0 | NA | NA | NA | NA | NA | #VALUE! |
| ENSMUSG00000106306.5 | 0 | NA | NA | NA | NA | NA | #VALUE! |
| ENSMUSG00000106307.3 | 0 | NA | NA | NA | NA | NA | #VALUE! |
| ENSMUSG00000106308.2 | 0 | NA | NA | NA | NA | NA | #VALUE! |
| ENSMUSG00000106309.2 | 0 | NA | NA | NA | NA | NA | #VALUE! |
| ENSMUSG00000106310.2 | 0 | NA | NA | NA | NA | NA | #VALUE! |
| ENSMUSG00000106311.2 | 0 | NA | NA | NA | NA | NA | #VALUE! |
| ENSMUSG00000106313.2 | 0 | NA | NA | NA | NA | NA | #VALUE! |
| ENSMUSG00000106314.2 | 0 | NA | NA | NA | NA | NA | #VALUE! |
| ENSMUSG00000106315.2 | 0 | NA | NA | NA | NA | NA | #VALUE! |
| ENSMUSG00000106316.2 | 0 | NA | NA | NA | NA | NA | #VALUE! |
| ENSMUSG00000106317.2 | 1.836997 | -4.39907 | 3.931186 | -1.11902 | 0.263133 | NA | #VALUE! |
| ENSMUSG00000106318.2 | 0 | NA | NA | NA | NA | NA | #VALUE! |
| ENSMUSG00000106319.2 | 0 | NA | NA | NA | NA | NA | #VALUE! |
| ENSMUSG00000106320.2 | 0 | NA | NA | NA | NA | NA | #VALUE! |
| ENSMUSG00000106321.2 | 0 | NA | NA | NA | NA | NA | #VALUE! |
| ENSMUSG00000106322.2 | 0 | NA | NA | NA | NA | NA | #VALUE! |
| ENSMUSG00000106323.3 | 0 | NA | NA | NA | NA | NA | #VALUE! |
| ENSMUSG00000106324.2 | 0 | NA | NA | NA | NA | NA | #VALUE! |
| ENSMUSG00000106325.2 | 0 | NA | NA | NA | NA | NA | #VALUE! |
| ENSMUSG00000106326.2 | 0 | NA | NA | NA | NA | NA | #VALUE! |
| ENSMUSG00000106327.2 | 0 | NA | NA | NA | NA | NA | #VALUE! |
| ENSMUSG00000106328.3 | 0 | NA | NA | NA | NA | NA | #VALUE! |
| ENSMUSG00000106329.2 | 0 | NA | NA | NA | NA | NA | #VALUE! |
| ENSMUSG00000106330.2 | 0 | NA | NA | NA | NA | NA | #VALUE! |
| ENSMUSG00000106331.2 | 0 | NA | NA | NA | NA | NA | #VALUE! |
| ENSMUSG00000106332.2 | 0 | NA | NA | NA | NA | NA | #VALUE! |
| ENSMUSG00000106333.2 | 0 | NA | NA | NA | NA | NA | #VALUE! |
| ENSMUSG00000106334.2 | 0 | NA | NA | NA | NA | NA | #VALUE! |
| ENSMUSG00000106335.2 | 0 | NA | NA | NA | NA | NA | #VALUE! |
| ENSMUSG00000106336.3 | 0 | NA | NA | NA | NA | NA | #VALUE! |

| | | | | | | | |
|---|---|---|---|---|---|---|---|
| ENSMUSG00000106337.2 | 0 | NA | NA | NA | NA | NA | #VALUE! |
| ENSMUSG00000106338.2 | 0 | NA | NA | NA | NA | NA | #VALUE! |
| ENSMUSG00000106339.2 | 0 | NA | NA | NA | NA | NA | #VALUE! |
| ENSMUSG00000106340.3 | 0 | NA | NA | NA | NA | NA | #VALUE! |
| ENSMUSG00000106341.2 | 0 | NA | NA | NA | NA | NA | #VALUE! |
| ENSMUSG00000106342.2 | 0 | NA | NA | NA | NA | NA | #VALUE! |
| ENSMUSG00000106343.2 | 0 | NA | NA | NA | NA | NA | #VALUE! |
| ENSMUSG00000106344.3 | 0 | NA | NA | NA | NA | NA | #VALUE! |
| ENSMUSG00000106345.2 | 0 | NA | NA | NA | NA | NA | #VALUE! |
| ENSMUSG00000106346.3 | 0 | NA | NA | NA | NA | NA | #VALUE! |
| ENSMUSG00000106347.3 | 0 | NA | NA | NA | NA | NA | #VALUE! |
| ENSMUSG00000106348.2 | 0 | NA | NA | NA | NA | NA | #VALUE! |
| ENSMUSG00000106350.3 | 0 | NA | NA | NA | NA | NA | #VALUE! |
| ENSMUSG00000106351.2 | 0 | NA | NA | NA | NA | NA | #VALUE! |
| ENSMUSG00000106353.2 | 0 | NA | NA | NA | NA | NA | #VALUE! |
| ENSMUSG00000106354.2 | 0 | NA | NA | NA | NA | NA | #VALUE! |
| ENSMUSG00000106356.2 | 0 | NA | NA | NA | NA | NA | #VALUE! |
| ENSMUSG00000106357.2 | 0 | NA | NA | NA | NA | NA | #VALUE! |
| ENSMUSG00000106358.2 | 0 | NA | NA | NA | NA | NA | #VALUE! |
| ENSMUSG00000106359.2 | 0 | NA | NA | NA | NA | NA | #VALUE! |
| ENSMUSG00000106361.2 | 0 | NA | NA | NA | NA | NA | #VALUE! |
| ENSMUSG00000106362.4 | 0 | NA | NA | NA | NA | NA | #VALUE! |
| ENSMUSG00000106363.2 | 0 | NA | NA | NA | NA | NA | #VALUE! |
| ENSMUSG00000106364.2 | 0 | NA | NA | NA | NA | NA | #VALUE! |
| ENSMUSG00000106365.2 | 0.491951 | 2.402185 | 4.000147 | 0.600524 | 0.548157 | NA | #VALUE! |
| ENSMUSG00000106366.3 | 0 | NA | NA | NA | NA | NA | #VALUE! |
| ENSMUSG00000106367.2 | 0 | NA | NA | NA | NA | NA | #VALUE! |
| ENSMUSG00000106369.2 | 0 | NA | NA | NA | NA | NA | #VALUE! |
| ENSMUSG00000106370.3 | 0 | NA | NA | NA | NA | NA | #VALUE! |
| ENSMUSG00000106371.2 | 0 | NA | NA | NA | NA | NA | #VALUE! |
| ENSMUSG00000106372.2 | 0 | NA | NA | NA | NA | NA | #VALUE! |
| ENSMUSG00000106373.2 | 0 | NA | NA | NA | NA | NA | #VALUE! |
| ENSMUSG00000106374.2 | 0 | NA | NA | NA | NA | NA | #VALUE! |
| ENSMUSG00000106375.2 | 0 | NA | NA | NA | NA | NA | #VALUE! |
| ENSMUSG00000106376.2 | 0 | NA | NA | NA | NA | NA | #VALUE! |
| ENSMUSG00000106378.2 | 0 | NA | NA | NA | NA | NA | #VALUE! |
| ENSMUSG00000106379.2 | 0 | NA | NA | NA | NA | NA | #VALUE! |
| ENSMUSG00000106380.2 | 0 | NA | NA | NA | NA | NA | #VALUE! |
| ENSMUSG00000106381.2 | 0 | NA | NA | NA | NA | NA | #VALUE! |
| ENSMUSG00000106382.2 | 0 | NA | NA | NA | NA | NA | #VALUE! |
| ENSMUSG00000106383.2 | 0 | NA | NA | NA | NA | NA | #VALUE! |
| ENSMUSG00000106384.2 | 0 | NA | NA | NA | NA | NA | #VALUE! |
| ENSMUSG00000106385.2 | 0 | NA | NA | NA | NA | NA | #VALUE! |
| ENSMUSG00000106386.2 | 0 | NA | NA | NA | NA | NA | #VALUE! |

| | | | | | | | |
|---|---|---|---|---|---|---|---|
| ENSMUSG00000106387.2 | 0 | NA | NA | NA | NA | NA | #VALUE! |
| ENSMUSG00000106388.2 | 0 | NA | NA | NA | NA | NA | #VALUE! |
| ENSMUSG00000106389.2 | 0 | NA | NA | NA | NA | NA | #VALUE! |
| ENSMUSG00000106390.2 | 0 | NA | NA | NA | NA | NA | #VALUE! |
| ENSMUSG00000106391.2 | 0 | NA | NA | NA | NA | NA | #VALUE! |
| ENSMUSG00000106392.2 | 0 | NA | NA | NA | NA | NA | #VALUE! |
| ENSMUSG00000106393.2 | 0 | NA | NA | NA | NA | NA | #VALUE! |
| ENSMUSG00000106394.2 | 0 | NA | NA | NA | NA | NA | #VALUE! |
| ENSMUSG00000106395.2 | 0 | NA | NA | NA | NA | NA | #VALUE! |
| ENSMUSG00000106396.2 | 0 | NA | NA | NA | NA | NA | #VALUE! |
| ENSMUSG00000106397.2 | 0.168284 | 1.027628 | 4.067126 | 0.252667 | 0.800526 | NA | #VALUE! |
| ENSMUSG00000106398.2 | 0 | NA | NA | NA | NA | NA | #VALUE! |
| ENSMUSG00000106399.2 | 0 | NA | NA | NA | NA | NA | #VALUE! |
| ENSMUSG00000106400.3 | 0 | NA | NA | NA | NA | NA | #VALUE! |
| ENSMUSG00000106401.2 | 0 | NA | NA | NA | NA | NA | #VALUE! |
| ENSMUSG00000106402.2 | 0 | NA | NA | NA | NA | NA | #VALUE! |
| ENSMUSG00000106403.2 | 0 | NA | NA | NA | NA | NA | #VALUE! |
| ENSMUSG00000106404.2 | 0 | NA | NA | NA | NA | NA | #VALUE! |
| ENSMUSG00000106405.2 | 0 | NA | NA | NA | NA | NA | #VALUE! |
| ENSMUSG00000106406.3 | 0 | NA | NA | NA | NA | NA | #VALUE! |
| ENSMUSG00000106407.2 | 0 | NA | NA | NA | NA | NA | #VALUE! |
| ENSMUSG00000106408.2 | 0 | NA | NA | NA | NA | NA | #VALUE! |
| ENSMUSG00000106409.3 | 0 | NA | NA | NA | NA | NA | #VALUE! |
| ENSMUSG00000106410.2 | 0 | NA | NA | NA | NA | NA | #VALUE! |
| ENSMUSG00000106411.2 | 0 | NA | NA | NA | NA | NA | #VALUE! |
| ENSMUSG00000106413.2 | 0 | NA | NA | NA | NA | NA | #VALUE! |
| ENSMUSG00000106414.2 | 0 | NA | NA | NA | NA | NA | #VALUE! |
| ENSMUSG00000106415.2 | 0 | NA | NA | NA | NA | NA | #VALUE! |
| ENSMUSG00000106416.2 | 0.289044 | -0.14751 | 4.065279 | -0.03629 | 0.971054 | NA | #VALUE! |
| ENSMUSG00000106417.2 | 0 | NA | NA | NA | NA | NA | #VALUE! |
| ENSMUSG00000106419.2 | 0 | NA | NA | NA | NA | NA | #VALUE! |
| ENSMUSG00000106420.2 | 0 | NA | NA | NA | NA | NA | #VALUE! |
| ENSMUSG00000106421.2 | 0 | NA | NA | NA | NA | NA | #VALUE! |
| ENSMUSG00000106422.2 | 0 | NA | NA | NA | NA | NA | #VALUE! |
| ENSMUSG00000106423.2 | 0 | NA | NA | NA | NA | NA | #VALUE! |
| ENSMUSG00000106424.2 | 0 | NA | NA | NA | NA | NA | #VALUE! |
| ENSMUSG00000106425.2 | 0 | NA | NA | NA | NA | NA | #VALUE! |
| ENSMUSG00000106426.2 | 0 | NA | NA | NA | NA | NA | #VALUE! |
| ENSMUSG00000106427.2 | 0 | NA | NA | NA | NA | NA | #VALUE! |
| ENSMUSG00000106428.2 | 0 | NA | NA | NA | NA | NA | #VALUE! |
| ENSMUSG00000106429.2 | 0 | NA | NA | NA | NA | NA | #VALUE! |
| ENSMUSG00000106430.2 | 0 | NA | NA | NA | NA | NA | #VALUE! |
| ENSMUSG00000106432.2 | 0.315039 | -1.93549 | 4.038621 | -0.47925 | 0.631763 | NA | #VALUE! |
| ENSMUSG00000106433.2 | 0 | NA | NA | NA | NA | NA | #VALUE! |

| | | | | | | | |
|---|---|---|---|---|---|---|---|
| ENSMUSG00000106434.2 | 0 | NA | NA | NA | NA | NA | #VALUE! |
| ENSMUSG00000106435.2 | 0 | NA | NA | NA | NA | NA | #VALUE! |
| ENSMUSG00000106436.3 | 0 | NA | NA | NA | NA | NA | #VALUE! |
| ENSMUSG00000106437.2 | 0 | NA | NA | NA | NA | NA | #VALUE! |
| ENSMUSG00000106438.2 | 0 | NA | NA | NA | NA | NA | #VALUE! |
| ENSMUSG00000106441.2 | 0 | NA | NA | NA | NA | NA | #VALUE! |
| ENSMUSG00000106442.2 | 0 | NA | NA | NA | NA | NA | #VALUE! |
| ENSMUSG00000106443.2 | 6.090761 | -6.14106 | 3.914091 | -1.56896 | 0.116657 | NA | #VALUE! |
| ENSMUSG00000106444.2 | 0 | NA | NA | NA | NA | NA | #VALUE! |
| ENSMUSG00000106445.2 | 0 | NA | NA | NA | NA | NA | #VALUE! |
| ENSMUSG00000106446.2 | 0 | NA | NA | NA | NA | NA | #VALUE! |
| ENSMUSG00000106447.2 | 0 | NA | NA | NA | NA | NA | #VALUE! |
| ENSMUSG00000106448.2 | 0 | NA | NA | NA | NA | NA | #VALUE! |
| ENSMUSG00000106450.2 | 0 | NA | NA | NA | NA | NA | #VALUE! |
| ENSMUSG00000106451.2 | 0 | NA | NA | NA | NA | NA | #VALUE! |
| ENSMUSG00000106452.2 | 0 | NA | NA | NA | NA | NA | #VALUE! |
| ENSMUSG00000106453.2 | 0 | NA | NA | NA | NA | NA | #VALUE! |
| ENSMUSG00000106454.2 | 0 | NA | NA | NA | NA | NA | #VALUE! |
| ENSMUSG00000106455.2 | 0 | NA | NA | NA | NA | NA | #VALUE! |
| ENSMUSG00000106456.2 | 0 | NA | NA | NA | NA | NA | #VALUE! |
| ENSMUSG00000106457.2 | 0 | NA | NA | NA | NA | NA | #VALUE! |
| ENSMUSG00000106458.2 | 0 | NA | NA | NA | NA | NA | #VALUE! |
| ENSMUSG00000106459.2 | 0 | NA | NA | NA | NA | NA | #VALUE! |
| ENSMUSG00000106460.2 | 0 | NA | NA | NA | NA | NA | #VALUE! |
| ENSMUSG00000106461.2 | 0 | NA | NA | NA | NA | NA | #VALUE! |
| ENSMUSG00000106462.2 | 0 | NA | NA | NA | NA | NA | #VALUE! |
| ENSMUSG00000106463.3 | 0 | NA | NA | NA | NA | NA | #VALUE! |
| ENSMUSG00000106464.2 | 0 | NA | NA | NA | NA | NA | #VALUE! |
| ENSMUSG00000106465.3 | 0 | NA | NA | NA | NA | NA | #VALUE! |
| ENSMUSG00000106466.2 | 0 | NA | NA | NA | NA | NA | #VALUE! |
| ENSMUSG00000106467.2 | 0 | NA | NA | NA | NA | NA | #VALUE! |
| ENSMUSG00000106469.2 | 0 | NA | NA | NA | NA | NA | #VALUE! |
| ENSMUSG00000106470.3 | 0 | NA | NA | NA | NA | NA | #VALUE! |
| ENSMUSG00000106471.3 | 0 | NA | NA | NA | NA | NA | #VALUE! |
| ENSMUSG00000106472.2 | 0 | NA | NA | NA | NA | NA | #VALUE! |
| ENSMUSG00000106473.2 | 0 | NA | NA | NA | NA | NA | #VALUE! |
| ENSMUSG00000106475.2 | 0 | NA | NA | NA | NA | NA | #VALUE! |
| ENSMUSG00000106476.2 | 0 | NA | NA | NA | NA | NA | #VALUE! |
| ENSMUSG00000106477.2 | 0 | NA | NA | NA | NA | NA | #VALUE! |
| ENSMUSG00000106478.2 | 0 | NA | NA | NA | NA | NA | #VALUE! |
| ENSMUSG00000106479.3 | 0 | NA | NA | NA | NA | NA | #VALUE! |
| ENSMUSG00000106480.2 | 0 | NA | NA | NA | NA | NA | #VALUE! |
| ENSMUSG00000106481.2 | 0 | NA | NA | NA | NA | NA | #VALUE! |
| ENSMUSG00000106482.2 | 0 | NA | NA | NA | NA | NA | #VALUE! |

| | | | | | | | |
|---|---|---|---|---|---|---|---|
| ENSMUSG00000106483.2 | 0 | NA | NA | NA | NA | NA | #VALUE! |
| ENSMUSG00000106484.2 | 0 | NA | NA | NA | NA | NA | #VALUE! |
| ENSMUSG00000106485.2 | 0 | NA | NA | NA | NA | NA | #VALUE! |
| ENSMUSG00000106486.2 | 0 | NA | NA | NA | NA | NA | #VALUE! |
| ENSMUSG00000106487.2 | 0 | NA | NA | NA | NA | NA | #VALUE! |
| ENSMUSG00000106490.2 | 0 | NA | NA | NA | NA | NA | #VALUE! |
| ENSMUSG00000106491.2 | 0 | NA | NA | NA | NA | NA | #VALUE! |
| ENSMUSG00000106492.2 | 0 | NA | NA | NA | NA | NA | #VALUE! |
| ENSMUSG00000106493.3 | 0 | NA | NA | NA | NA | NA | #VALUE! |
| ENSMUSG00000106494.2 | 0 | NA | NA | NA | NA | NA | #VALUE! |
| ENSMUSG00000106495.2 | 0 | NA | NA | NA | NA | NA | #VALUE! |
| ENSMUSG00000106496.2 | 0 | NA | NA | NA | NA | NA | #VALUE! |
| ENSMUSG00000106497.2 | 0 | NA | NA | NA | NA | NA | #VALUE! |
| ENSMUSG00000106498.2 | 0 | NA | NA | NA | NA | NA | #VALUE! |
| ENSMUSG00000106499.2 | 0 | NA | NA | NA | NA | NA | #VALUE! |
| ENSMUSG00000106500.2 | 0 | NA | NA | NA | NA | NA | #VALUE! |
| ENSMUSG00000106501.2 | 0 | NA | NA | NA | NA | NA | #VALUE! |
| ENSMUSG00000106502.2 | 0 | NA | NA | NA | NA | NA | #VALUE! |
| ENSMUSG00000106503.2 | 0 | NA | NA | NA | NA | NA | #VALUE! |
| ENSMUSG00000106504.2 | 0 | NA | NA | NA | NA | NA | #VALUE! |
| ENSMUSG00000106505.2 | 0 | NA | NA | NA | NA | NA | #VALUE! |
| ENSMUSG00000106506.2 | 0 | NA | NA | NA | NA | NA | #VALUE! |
| ENSMUSG00000106507.2 | 0 | NA | NA | NA | NA | NA | #VALUE! |
| ENSMUSG00000106508.2 | 0 | NA | NA | NA | NA | NA | #VALUE! |
| ENSMUSG00000106509.2 | 0 | NA | NA | NA | NA | NA | #VALUE! |
| ENSMUSG00000106510.2 | 0 | NA | NA | NA | NA | NA | #VALUE! |
| ENSMUSG00000106511.2 | 0 | NA | NA | NA | NA | NA | #VALUE! |
| ENSMUSG00000106512.2 | 0 | NA | NA | NA | NA | NA | #VALUE! |
| ENSMUSG00000106514.3 | 0 | NA | NA | NA | NA | NA | #VALUE! |
| ENSMUSG00000106515.5 | 0 | NA | NA | NA | NA | NA | #VALUE! |
| ENSMUSG00000106516.2 | 0 | NA | NA | NA | NA | NA | #VALUE! |
| ENSMUSG00000106517.2 | 0 | NA | NA | NA | NA | NA | #VALUE! |
| ENSMUSG00000106518.2 | 0.210026 | -1.52494 | 4.058692 | -0.37572 | 0.707123 | NA | #VALUE! |
| ENSMUSG00000106519.2 | 0 | NA | NA | NA | NA | NA | #VALUE! |
| ENSMUSG00000106520.2 | 0 | NA | NA | NA | NA | NA | #VALUE! |
| ENSMUSG00000106522.2 | 0 | NA | NA | NA | NA | NA | #VALUE! |
| ENSMUSG00000106524.2 | 0 | NA | NA | NA | NA | NA | #VALUE! |
| ENSMUSG00000106525.2 | 0 | NA | NA | NA | NA | NA | #VALUE! |
| ENSMUSG00000106526.2 | 1.470184 | -4.10383 | 3.936698 | -1.04245 | 0.297201 | NA | #VALUE! |
| ENSMUSG00000106527.2 | 0 | NA | NA | NA | NA | NA | #VALUE! |
| ENSMUSG00000106528.2 | 0 | NA | NA | NA | NA | NA | #VALUE! |
| ENSMUSG00000106529.2 | 0 | NA | NA | NA | NA | NA | #VALUE! |
| ENSMUSG00000106530.3 | 0 | NA | NA | NA | NA | NA | #VALUE! |
| ENSMUSG00000106531.2 | 0 | NA | NA | NA | NA | NA | #VALUE! |

| | | | | | | | |
|---|---|---|---|---|---|---|---|
| ENSMUSG00000106532.2 | 0 | NA | NA | NA | NA | NA | #VALUE! |
| ENSMUSG00000106533.2 | 0 | NA | NA | NA | NA | NA | #VALUE! |
| ENSMUSG00000106534.2 | 0 | NA | NA | NA | NA | NA | #VALUE! |
| ENSMUSG00000106535.2 | 0 | NA | NA | NA | NA | NA | #VALUE! |
| ENSMUSG00000106536.2 | 0 | NA | NA | NA | NA | NA | #VALUE! |
| ENSMUSG00000106537.2 | 0 | NA | NA | NA | NA | NA | #VALUE! |
| ENSMUSG00000106538.2 | 0 | NA | NA | NA | NA | NA | #VALUE! |
| ENSMUSG00000106540.2 | 0 | NA | NA | NA | NA | NA | #VALUE! |
| ENSMUSG00000106541.2 | 0 | NA | NA | NA | NA | NA | #VALUE! |
| ENSMUSG00000106542.2 | 0.405815 | -2.2474 | 4.013505 | -0.55996 | 0.575507 | NA | #VALUE! |
| ENSMUSG00000106543.2 | 0 | NA | NA | NA | NA | NA | #VALUE! |
| ENSMUSG00000106545.2 | 0 | NA | NA | NA | NA | NA | #VALUE! |
| ENSMUSG00000106546.2 | 0 | NA | NA | NA | NA | NA | #VALUE! |
| ENSMUSG00000106547.2 | 0 | NA | NA | NA | NA | NA | #VALUE! |
| ENSMUSG00000106548.2 | 0 | NA | NA | NA | NA | NA | #VALUE! |
| ENSMUSG00000106549.2 | 0 | NA | NA | NA | NA | NA | #VALUE! |
| ENSMUSG00000106550.2 | 0 | NA | NA | NA | NA | NA | #VALUE! |
| ENSMUSG00000106551.2 | 0 | NA | NA | NA | NA | NA | #VALUE! |
| ENSMUSG00000106552.2 | 0 | NA | NA | NA | NA | NA | #VALUE! |
| ENSMUSG00000106553.2 | 0 | NA | NA | NA | NA | NA | #VALUE! |
| ENSMUSG00000106554.2 | 0 | NA | NA | NA | NA | NA | #VALUE! |
| ENSMUSG00000106555.5 | 0 | NA | NA | NA | NA | NA | #VALUE! |
| ENSMUSG00000106557.2 | 0 | NA | NA | NA | NA | NA | #VALUE! |
| ENSMUSG00000106558.2 | 0 | NA | NA | NA | NA | NA | #VALUE! |
| ENSMUSG00000106559.2 | 0 | NA | NA | NA | NA | NA | #VALUE! |
| ENSMUSG00000106560.2 | 0 | NA | NA | NA | NA | NA | #VALUE! |
| ENSMUSG00000106562.2 | 0.245976 | 1.027628 | 4.067126 | 0.252667 | 0.800526 | NA | #VALUE! |
| ENSMUSG00000106563.3 | 0 | NA | NA | NA | NA | NA | #VALUE! |
| ENSMUSG00000106564.2 | 0 | NA | NA | NA | NA | NA | #VALUE! |
| ENSMUSG00000106565.2 | 0 | NA | NA | NA | NA | NA | #VALUE! |
| ENSMUSG00000106566.2 | 0 | NA | NA | NA | NA | NA | #VALUE! |
| ENSMUSG00000106567.2 | 0 | NA | NA | NA | NA | NA | #VALUE! |
| ENSMUSG00000106569.2 | 0 | NA | NA | NA | NA | NA | #VALUE! |
| ENSMUSG00000106570.2 | 0 | NA | NA | NA | NA | NA | #VALUE! |
| ENSMUSG00000106572.2 | 0 | NA | NA | NA | NA | NA | #VALUE! |
| ENSMUSG00000106573.2 | 0 | NA | NA | NA | NA | NA | #VALUE! |
| ENSMUSG00000106574.2 | 0 | NA | NA | NA | NA | NA | #VALUE! |
| ENSMUSG00000106575.2 | 0 | NA | NA | NA | NA | NA | #VALUE! |
| ENSMUSG00000106576.2 | 0 | NA | NA | NA | NA | NA | #VALUE! |
| ENSMUSG00000106577.2 | 0 | NA | NA | NA | NA | NA | #VALUE! |
| ENSMUSG00000106578.3 | 0 | NA | NA | NA | NA | NA | #VALUE! |
| ENSMUSG00000106579.2 | 0 | NA | NA | NA | NA | NA | #VALUE! |
| ENSMUSG00000106580.3 | 0 | NA | NA | NA | NA | NA | #VALUE! |
| ENSMUSG00000106581.3 | 0 | NA | NA | NA | NA | NA | #VALUE! |

| | | | | | | | |
|---|---|---|---|---|---|---|---|
| ENSMUSG00000106582.2 | 0 | NA | NA | NA | NA | NA | #VALUE! |
| ENSMUSG00000106583.2 | 0 | NA | NA | NA | NA | NA | #VALUE! |
| ENSMUSG00000106584.2 | 0 | NA | NA | NA | NA | NA | #VALUE! |
| ENSMUSG00000106585.2 | 0 | NA | NA | NA | NA | NA | #VALUE! |
| ENSMUSG00000106586.2 | 0 | NA | NA | NA | NA | NA | #VALUE! |
| ENSMUSG00000106588.2 | 0 | NA | NA | NA | NA | NA | #VALUE! |
| ENSMUSG00000106589.2 | 0 | NA | NA | NA | NA | NA | #VALUE! |
| ENSMUSG00000106590.2 | 2.890911 | 5.03241 | 3.922161 | 1.283071 | 0.199467 | NA | #VALUE! |
| ENSMUSG00000106592.2 | 0 | NA | NA | NA | NA | NA | #VALUE! |
| ENSMUSG00000106593.2 | 0 | NA | NA | NA | NA | NA | #VALUE! |
| ENSMUSG00000106594.2 | 0 | NA | NA | NA | NA | NA | #VALUE! |
| ENSMUSG00000106596.2 | 0 | NA | NA | NA | NA | NA | #VALUE! |
| ENSMUSG00000106597.3 | 0 | NA | NA | NA | NA | NA | #VALUE! |
| ENSMUSG00000106598.2 | 0 | NA | NA | NA | NA | NA | #VALUE! |
| ENSMUSG00000106599.2 | 0 | NA | NA | NA | NA | NA | #VALUE! |
| ENSMUSG00000106600.3 | 0 | NA | NA | NA | NA | NA | #VALUE! |
| ENSMUSG00000106601.2 | 0 | NA | NA | NA | NA | NA | #VALUE! |
| ENSMUSG00000106602.2 | 0 | NA | NA | NA | NA | NA | #VALUE! |
| ENSMUSG00000106603.2 | 0 | NA | NA | NA | NA | NA | #VALUE! |
| ENSMUSG00000106604.3 | 0 | NA | NA | NA | NA | NA | #VALUE! |
| ENSMUSG00000106606.2 | 0 | NA | NA | NA | NA | NA | #VALUE! |
| ENSMUSG00000106607.2 | 0 | NA | NA | NA | NA | NA | #VALUE! |
| ENSMUSG00000106609.2 | 0 | NA | NA | NA | NA | NA | #VALUE! |
| ENSMUSG00000106610.2 | 0 | NA | NA | NA | NA | NA | #VALUE! |
| ENSMUSG00000106612.2 | 0 | NA | NA | NA | NA | NA | #VALUE! |
| ENSMUSG00000106613.2 | 0 | NA | NA | NA | NA | NA | #VALUE! |
| ENSMUSG00000106614.2 | 0 | NA | NA | NA | NA | NA | #VALUE! |
| ENSMUSG00000106615.2 | 0 | NA | NA | NA | NA | NA | #VALUE! |
| ENSMUSG00000106616.2 | 0 | NA | NA | NA | NA | NA | #VALUE! |
| ENSMUSG00000106617.2 | 0 | NA | NA | NA | NA | NA | #VALUE! |
| ENSMUSG00000106619.2 | 0 | NA | NA | NA | NA | NA | #VALUE! |
| ENSMUSG00000106620.2 | 0 | NA | NA | NA | NA | NA | #VALUE! |
| ENSMUSG00000106621.2 | 0 | NA | NA | NA | NA | NA | #VALUE! |
| ENSMUSG00000106622.3 | 0 | NA | NA | NA | NA | NA | #VALUE! |
| ENSMUSG00000106623.2 | 0.491951 | 2.402185 | 4.000147 | 0.600524 | 0.548157 | NA | #VALUE! |
| ENSMUSG00000106624.2 | 0 | NA | NA | NA | NA | NA | #VALUE! |
| ENSMUSG00000106625.2 | 0 | NA | NA | NA | NA | NA | #VALUE! |
| ENSMUSG00000106627.2 | 0 | NA | NA | NA | NA | NA | #VALUE! |
| ENSMUSG00000106628.2 | 0 | NA | NA | NA | NA | NA | #VALUE! |
| ENSMUSG00000106629.2 | 0 | NA | NA | NA | NA | NA | #VALUE! |
| ENSMUSG00000106630.2 | 0 | NA | NA | NA | NA | NA | #VALUE! |
| ENSMUSG00000106631.3 | 0 | NA | NA | NA | NA | NA | #VALUE! |
| ENSMUSG00000106632.2 | 0 | NA | NA | NA | NA | NA | #VALUE! |
| ENSMUSG00000106633.2 | 0 | NA | NA | NA | NA | NA | #VALUE! |

| | | | | | | | |
|---|---|---|---|---|---|---|---|
| ENSMUSG00000106634.2 | 0 | NA | NA | NA | NA | NA | #VALUE! |
| ENSMUSG00000106635.2 | 0 | NA | NA | NA | NA | NA | #VALUE! |
| ENSMUSG00000106636.2 | 0.595689 | -2.7616 | 3.981969 | -0.69353 | 0.48798 | NA | #VALUE! |
| ENSMUSG00000106637.2 | 0 | NA | NA | NA | NA | NA | #VALUE! |
| ENSMUSG00000106639.2 | 0 | NA | NA | NA | NA | NA | #VALUE! |
| ENSMUSG00000106640.2 | 0 | NA | NA | NA | NA | NA | #VALUE! |
| ENSMUSG00000106641.2 | 0 | NA | NA | NA | NA | NA | #VALUE! |
| ENSMUSG00000106642.3 | 0 | NA | NA | NA | NA | NA | #VALUE! |
| ENSMUSG00000106643.2 | 0 | NA | NA | NA | NA | NA | #VALUE! |
| ENSMUSG00000106644.2 | 0 | NA | NA | NA | NA | NA | #VALUE! |
| ENSMUSG00000106645.2 | 0 | NA | NA | NA | NA | NA | #VALUE! |
| ENSMUSG00000106646.2 | 0 | NA | NA | NA | NA | NA | #VALUE! |
| ENSMUSG00000106647.2 | 0 | NA | NA | NA | NA | NA | #VALUE! |
| ENSMUSG00000106648.2 | 0 | NA | NA | NA | NA | NA | #VALUE! |
| ENSMUSG00000106650.2 | 0 | NA | NA | NA | NA | NA | #VALUE! |
| ENSMUSG00000106651.2 | 0 | NA | NA | NA | NA | NA | #VALUE! |
| ENSMUSG00000106652.2 | 0 | NA | NA | NA | NA | NA | #VALUE! |
| ENSMUSG00000106653.3 | 0 | NA | NA | NA | NA | NA | #VALUE! |
| ENSMUSG00000106654.2 | 0 | NA | NA | NA | NA | NA | #VALUE! |
| ENSMUSG00000106655.3 | 0 | NA | NA | NA | NA | NA | #VALUE! |
| ENSMUSG00000106656.2 | 0 | NA | NA | NA | NA | NA | #VALUE! |
| ENSMUSG00000106658.2 | 0 | NA | NA | NA | NA | NA | #VALUE! |
| ENSMUSG00000106659.2 | 0 | NA | NA | NA | NA | NA | #VALUE! |
| ENSMUSG00000106660.2 | 0 | NA | NA | NA | NA | NA | #VALUE! |
| ENSMUSG00000106661.2 | 0 | NA | NA | NA | NA | NA | #VALUE! |
| ENSMUSG00000106662.2 | 0 | NA | NA | NA | NA | NA | #VALUE! |
| ENSMUSG00000106663.2 | 0 | NA | NA | NA | NA | NA | #VALUE! |
| ENSMUSG00000106664.2 | 0 | NA | NA | NA | NA | NA | #VALUE! |
| ENSMUSG00000106665.2 | 0 | NA | NA | NA | NA | NA | #VALUE! |
| ENSMUSG00000106666.2 | 0 | NA | NA | NA | NA | NA | #VALUE! |
| ENSMUSG00000106667.2 | 0 | NA | NA | NA | NA | NA | #VALUE! |
| ENSMUSG00000106668.2 | 0 | NA | NA | NA | NA | NA | #VALUE! |
| ENSMUSG00000106669.2 | 0 | NA | NA | NA | NA | NA | #VALUE! |
| ENSMUSG00000106671.2 | 0 | NA | NA | NA | NA | NA | #VALUE! |
| ENSMUSG00000106673.2 | 0 | NA | NA | NA | NA | NA | #VALUE! |
| ENSMUSG00000106674.2 | 0 | NA | NA | NA | NA | NA | #VALUE! |
| ENSMUSG00000106675.2 | 0 | NA | NA | NA | NA | NA | #VALUE! |
| ENSMUSG00000106676.2 | 4.293145 | -5.63431 | 3.70783 | -1.51957 | 0.128619 | NA | #VALUE! |
| ENSMUSG00000106677.2 | 0 | NA | NA | NA | NA | NA | #VALUE! |
| ENSMUSG00000106678.2 | 0 | NA | NA | NA | NA | NA | #VALUE! |
| ENSMUSG00000106679.2 | 0 | NA | NA | NA | NA | NA | #VALUE! |
| ENSMUSG00000106680.2 | 0 | NA | NA | NA | NA | NA | #VALUE! |
| ENSMUSG00000106681.2 | 0 | NA | NA | NA | NA | NA | #VALUE! |
| ENSMUSG00000106682.2 | 0 | NA | NA | NA | NA | NA | #VALUE! |

| | | | | | | | |
|---|---|---|---|---|---|---|---|
| ENSMUSG00000106683.2 | 0 | NA | NA | NA | NA | NA | #VALUE! |
| ENSMUSG00000106688.2 | 1.865104 | 4.40601 | 3.930481 | 1.120985 | 0.262294 | NA | #VALUE! |
| ENSMUSG00000106689.2 | 0 | NA | NA | NA | NA | NA | #VALUE! |
| ENSMUSG00000106690.2 | 0 | NA | NA | NA | NA | NA | #VALUE! |
| ENSMUSG00000106691.2 | 0 | NA | NA | NA | NA | NA | #VALUE! |
| ENSMUSG00000106692.2 | 0 | NA | NA | NA | NA | NA | #VALUE! |
| ENSMUSG00000106694.2 | 0 | NA | NA | NA | NA | NA | #VALUE! |
| ENSMUSG00000106696.2 | 0 | NA | NA | NA | NA | NA | #VALUE! |
| ENSMUSG00000106698.2 | 0 | NA | NA | NA | NA | NA | #VALUE! |
| ENSMUSG00000106699.2 | 0 | NA | NA | NA | NA | NA | #VALUE! |
| ENSMUSG00000106701.2 | 0 | NA | NA | NA | NA | NA | #VALUE! |
| ENSMUSG00000106702.2 | 0 | NA | NA | NA | NA | NA | #VALUE! |
| ENSMUSG00000106703.2 | 0 | NA | NA | NA | NA | NA | #VALUE! |
| ENSMUSG00000106704.2 | 0 | NA | NA | NA | NA | NA | #VALUE! |
| ENSMUSG00000106705.2 | 0 | NA | NA | NA | NA | NA | #VALUE! |
| ENSMUSG00000106706.2 | 0 | NA | NA | NA | NA | NA | #VALUE! |
| ENSMUSG00000106707.2 | 0 | NA | NA | NA | NA | NA | #VALUE! |
| ENSMUSG00000106708.2 | 0 | NA | NA | NA | NA | NA | #VALUE! |
| ENSMUSG00000106709.2 | 3.916718 | 5.467688 | 3.918163 | 1.395472 | 0.162873 | NA | #VALUE! |
| ENSMUSG00000106710.2 | 0 | NA | NA | NA | NA | NA | #VALUE! |
| ENSMUSG00000106712.2 | 0 | NA | NA | NA | NA | NA | #VALUE! |
| ENSMUSG00000106713.2 | 0 | NA | NA | NA | NA | NA | #VALUE! |
| ENSMUSG00000106714.2 | 0 | NA | NA | NA | NA | NA | #VALUE! |
| ENSMUSG00000106715.4 | 0 | NA | NA | NA | NA | NA | #VALUE! |
| ENSMUSG00000106718.2 | 0 | NA | NA | NA | NA | NA | #VALUE! |
| ENSMUSG00000106719.2 | 0 | NA | NA | NA | NA | NA | #VALUE! |
| ENSMUSG00000106720.2 | 0 | NA | NA | NA | NA | NA | #VALUE! |
| ENSMUSG00000106721.2 | 0.195789 | -1.05581 | 4.073302 | -0.2592 | 0.795479 | NA | #VALUE! |
| ENSMUSG00000106725.2 | 0 | NA | NA | NA | NA | NA | #VALUE! |
| ENSMUSG00000106726.2 | 0 | NA | NA | NA | NA | NA | #VALUE! |
| ENSMUSG00000106729.2 | 0 | NA | NA | NA | NA | NA | #VALUE! |
| ENSMUSG00000106730.2 | 0 | NA | NA | NA | NA | NA | #VALUE! |
| ENSMUSG00000106732.2 | 0 | NA | NA | NA | NA | NA | #VALUE! |
| ENSMUSG00000106733.2 | 0 | NA | NA | NA | NA | NA | #VALUE! |
| ENSMUSG00000106734.4 | 0 | NA | NA | NA | NA | NA | #VALUE! |
| ENSMUSG00000106735.2 | 0 | NA | NA | NA | NA | NA | #VALUE! |
| ENSMUSG00000106736.2 | 0 | NA | NA | NA | NA | NA | #VALUE! |
| ENSMUSG00000106737.2 | 0 | NA | NA | NA | NA | NA | #VALUE! |
| ENSMUSG00000106738.2 | 0 | NA | NA | NA | NA | NA | #VALUE! |
| ENSMUSG00000106739.2 | 0.279766 | 1.808031 | 4.036658 | 0.447903 | 0.654223 | NA | #VALUE! |
| ENSMUSG00000106740.2 | 0 | NA | NA | NA | NA | NA | #VALUE! |
| ENSMUSG00000106741.4 | 0 | NA | NA | NA | NA | NA | #VALUE! |
| ENSMUSG00000106742.2 | 0 | NA | NA | NA | NA | NA | #VALUE! |
| ENSMUSG00000106743.2 | 0 | NA | NA | NA | NA | NA | #VALUE! |

| | | | | | | | |
|---|---|---|---|---|---|---|---|
| ENSMUSG00000106745.2 | 0 | NA | NA | NA | NA | NA | #VALUE! |
| ENSMUSG00000106746.2 | 0 | NA | NA | NA | NA | NA | #VALUE! |
| ENSMUSG00000106747.2 | 0 | NA | NA | NA | NA | NA | #VALUE! |
| ENSMUSG00000106749.2 | 0 | NA | NA | NA | NA | NA | #VALUE! |
| ENSMUSG00000106750.2 | 0 | NA | NA | NA | NA | NA | #VALUE! |
| ENSMUSG00000106751.2 | 0 | NA | NA | NA | NA | NA | #VALUE! |
| ENSMUSG00000106752.2 | 0 | NA | NA | NA | NA | NA | #VALUE! |
| ENSMUSG00000106753.2 | 0 | NA | NA | NA | NA | NA | #VALUE! |
| ENSMUSG00000106755.2 | 0 | NA | NA | NA | NA | NA | #VALUE! |
| ENSMUSG00000106756.2 | 0 | NA | NA | NA | NA | NA | #VALUE! |
| ENSMUSG00000106757.2 | 0 | NA | NA | NA | NA | NA | #VALUE! |
| ENSMUSG00000106758.2 | 0 | NA | NA | NA | NA | NA | #VALUE! |
| ENSMUSG00000106759.2 | 0 | NA | NA | NA | NA | NA | #VALUE! |
| ENSMUSG00000106760.2 | 0 | NA | NA | NA | NA | NA | #VALUE! |
| ENSMUSG00000106762.4 | 0 | NA | NA | NA | NA | NA | #VALUE! |
| ENSMUSG00000106763.2 | 0 | NA | NA | NA | NA | NA | #VALUE! |
| ENSMUSG00000106764.2 | 0 | NA | NA | NA | NA | NA | #VALUE! |
| ENSMUSG00000106766.2 | 0 | NA | NA | NA | NA | NA | #VALUE! |
| ENSMUSG00000106767.2 | 2.936837 | -5.08026 | 3.922024 | -1.29532 | 0.195212 | NA | #VALUE! |
| ENSMUSG00000106768.2 | 0 | NA | NA | NA | NA | NA | #VALUE! |
| ENSMUSG00000106771.2 | 0 | NA | NA | NA | NA | NA | #VALUE! |
| ENSMUSG00000106772.2 | 1.224665 | -3.81006 | 3.943408 | -0.96618 | 0.333952 | NA | #VALUE! |
| ENSMUSG00000106773.2 | 0 | NA | NA | NA | NA | NA | #VALUE! |
| ENSMUSG00000106775.2 | 0 | NA | NA | NA | NA | NA | #VALUE! |
| ENSMUSG00000106776.2 | 0 | NA | NA | NA | NA | NA | #VALUE! |
| ENSMUSG00000106777.2 | 0 | NA | NA | NA | NA | NA | #VALUE! |
| ENSMUSG00000106779.2 | 0 | NA | NA | NA | NA | NA | #VALUE! |
| ENSMUSG00000106780.2 | 0 | NA | NA | NA | NA | NA | #VALUE! |
| ENSMUSG00000106783.2 | 0 | NA | NA | NA | NA | NA | #VALUE! |
| ENSMUSG00000106784.2 | 0 | NA | NA | NA | NA | NA | #VALUE! |
| ENSMUSG00000106785.2 | 0 | NA | NA | NA | NA | NA | #VALUE! |
| ENSMUSG00000106786.2 | 0 | NA | NA | NA | NA | NA | #VALUE! |
| ENSMUSG00000106791.2 | 0 | NA | NA | NA | NA | NA | #VALUE! |
| ENSMUSG00000106794.2 | 0 | NA | NA | NA | NA | NA | #VALUE! |
| ENSMUSG00000106795.4 | 0 | NA | NA | NA | NA | NA | #VALUE! |
| ENSMUSG00000106797.2 | 0 | NA | NA | NA | NA | NA | #VALUE! |
| ENSMUSG00000106798.2 | 0 | NA | NA | NA | NA | NA | #VALUE! |
| ENSMUSG00000106799.2 | 0 | NA | NA | NA | NA | NA | #VALUE! |
| ENSMUSG00000106800.2 | 0 | NA | NA | NA | NA | NA | #VALUE! |
| ENSMUSG00000106801.2 | 0 | NA | NA | NA | NA | NA | #VALUE! |
| ENSMUSG00000106802.2 | 0 | NA | NA | NA | NA | NA | #VALUE! |
| ENSMUSG00000106803.2 | 0 | NA | NA | NA | NA | NA | #VALUE! |
| ENSMUSG00000106805.2 | 0 | NA | NA | NA | NA | NA | #VALUE! |
| ENSMUSG00000106806.2 | 0 | NA | NA | NA | NA | NA | #VALUE! |

| | | | | | | | |
|---|---|---|---|---|---|---|---|
| ENSMUSG00000106807.2 | 0 | NA | NA | NA | NA | NA | #VALUE! |
| ENSMUSG00000106808.2 | 0 | NA | NA | NA | NA | NA | #VALUE! |
| ENSMUSG00000106810.2 | 0 | NA | NA | NA | NA | NA | #VALUE! |
| ENSMUSG00000106812.2 | 0 | NA | NA | NA | NA | NA | #VALUE! |
| ENSMUSG00000106813.2 | 0 | NA | NA | NA | NA | NA | #VALUE! |
| ENSMUSG00000106814.2 | 0 | NA | NA | NA | NA | NA | #VALUE! |
| ENSMUSG00000106815.2 | 0 | NA | NA | NA | NA | NA | #VALUE! |
| ENSMUSG00000106817.2 | 0 | NA | NA | NA | NA | NA | #VALUE! |
| ENSMUSG00000106818.2 | 0 | NA | NA | NA | NA | NA | #VALUE! |
| ENSMUSG00000106819.2 | 0 | NA | NA | NA | NA | NA | #VALUE! |
| ENSMUSG00000106820.2 | 0 | NA | NA | NA | NA | NA | #VALUE! |
| ENSMUSG00000106821.2 | 0 | NA | NA | NA | NA | NA | #VALUE! |
| ENSMUSG00000106822.2 | 0 | NA | NA | NA | NA | NA | #VALUE! |
| ENSMUSG00000106824.2 | 0 | NA | NA | NA | NA | NA | #VALUE! |
| ENSMUSG00000106825.2 | 0.839297 | 3.273019 | 3.958392 | 0.826856 | 0.408319 | NA | #VALUE! |
| ENSMUSG00000106826.2 | 0 | NA | NA | NA | NA | NA | #VALUE! |
| ENSMUSG00000106827.2 | 0 | NA | NA | NA | NA | NA | #VALUE! |
| ENSMUSG00000106828.2 | 0 | NA | NA | NA | NA | NA | #VALUE! |
| ENSMUSG00000106829.2 | 0 | NA | NA | NA | NA | NA | #VALUE! |
| ENSMUSG00000106831.2 | 0 | NA | NA | NA | NA | NA | #VALUE! |
| ENSMUSG00000106832.2 | 0 | NA | NA | NA | NA | NA | #VALUE! |
| ENSMUSG00000106833.2 | 0 | NA | NA | NA | NA | NA | #VALUE! |
| ENSMUSG00000106834.2 | 0 | NA | NA | NA | NA | NA | #VALUE! |
| ENSMUSG00000106835.4 | 0 | NA | NA | NA | NA | NA | #VALUE! |
| ENSMUSG00000106836.2 | 0 | NA | NA | NA | NA | NA | #VALUE! |
| ENSMUSG00000106837.2 | 0 | NA | NA | NA | NA | NA | #VALUE! |
| ENSMUSG00000106838.2 | 0 | NA | NA | NA | NA | NA | #VALUE! |
| ENSMUSG00000106839.2 | 0 | NA | NA | NA | NA | NA | #VALUE! |
| ENSMUSG00000106840.2 | 0 | NA | NA | NA | NA | NA | #VALUE! |
| ENSMUSG00000106841.2 | 0 | NA | NA | NA | NA | NA | #VALUE! |
| ENSMUSG00000106843.2 | 0 | NA | NA | NA | NA | NA | #VALUE! |
| ENSMUSG00000106845.2 | 0 | NA | NA | NA | NA | NA | #VALUE! |
| ENSMUSG00000106846.2 | 0 | NA | NA | NA | NA | NA | #VALUE! |
| ENSMUSG00000106847.2 | 8.28726 | -1.29186 | 2.837314 | -0.45531 | 0.648887 | NA | #VALUE! |
| ENSMUSG00000106849.2 | 0 | NA | NA | NA | NA | NA | #VALUE! |
| ENSMUSG00000106851.2 | 0 | NA | NA | NA | NA | NA | #VALUE! |
| ENSMUSG00000106853.2 | 0 | NA | NA | NA | NA | NA | #VALUE! |
| ENSMUSG00000106854.2 | 0 | NA | NA | NA | NA | NA | #VALUE! |
| ENSMUSG00000106855.2 | 0 | NA | NA | NA | NA | NA | #VALUE! |
| ENSMUSG00000106856.2 | 0 | NA | NA | NA | NA | NA | #VALUE! |
| ENSMUSG00000106858.2 | 0 | NA | NA | NA | NA | NA | #VALUE! |
| ENSMUSG00000106859.2 | 0 | NA | NA | NA | NA | NA | #VALUE! |
| ENSMUSG00000106860.2 | 0 | NA | NA | NA | NA | NA | #VALUE! |
| ENSMUSG00000106862.2 | 0 | NA | NA | NA | NA | NA | #VALUE! |

| | | | | | | | |
|---|---|---|---|---|---|---|---|
| ENSMUSG00000106863.4 | 1.265119 | 0.161593 | 3.869308 | 0.041763 | 0.966688 | NA | #VALUE! |
| ENSMUSG00000106865.2 | 0 | NA | NA | NA | NA | NA | #VALUE! |
| ENSMUSG00000106866.2 | 0 | NA | NA | NA | NA | NA | #VALUE! |
| ENSMUSG00000106867.2 | 0 | NA | NA | NA | NA | NA | #VALUE! |
| ENSMUSG00000106868.2 | 0 | NA | NA | NA | NA | NA | #VALUE! |
| ENSMUSG00000106869.2 | 0 | NA | NA | NA | NA | NA | #VALUE! |
| ENSMUSG00000106870.2 | 2.125844 | -1.42579 | 3.834062 | -0.37187 | 0.709987 | NA | #VALUE! |
| ENSMUSG00000106871.2 | 0 | NA | NA | NA | NA | NA | #VALUE! |
| ENSMUSG00000106872.2 | 0 | NA | NA | NA | NA | NA | #VALUE! |
| ENSMUSG00000106873.2 | 0 | NA | NA | NA | NA | NA | #VALUE! |
| ENSMUSG00000106874.2 | 0 | NA | NA | NA | NA | NA | #VALUE! |
| ENSMUSG00000106875.2 | 0 | NA | NA | NA | NA | NA | #VALUE! |
| ENSMUSG00000106876.2 | 0 | NA | NA | NA | NA | NA | #VALUE! |
| ENSMUSG00000106878.2 | 0 | NA | NA | NA | NA | NA | #VALUE! |
| ENSMUSG00000106879.2 | 0 | NA | NA | NA | NA | NA | #VALUE! |
| ENSMUSG00000106880.2 | 0 | NA | NA | NA | NA | NA | #VALUE! |
| ENSMUSG00000106883.2 | 0 | NA | NA | NA | NA | NA | #VALUE! |
| ENSMUSG00000106884.2 | 0 | NA | NA | NA | NA | NA | #VALUE! |
| ENSMUSG00000106885.2 | 0 | NA | NA | NA | NA | NA | #VALUE! |
| ENSMUSG00000106887.2 | 0 | NA | NA | NA | NA | NA | #VALUE! |
| ENSMUSG00000106888.2 | 0 | NA | NA | NA | NA | NA | #VALUE! |
| ENSMUSG00000106889.2 | 0 | NA | NA | NA | NA | NA | #VALUE! |
| ENSMUSG00000106891.2 | 0 | NA | NA | NA | NA | NA | #VALUE! |
| ENSMUSG00000106892.2 | 1.958359 | 4.475621 | 3.929371 | 1.139017 | 0.254696 | NA | #VALUE! |
| ENSMUSG00000106893.2 | 0 | NA | NA | NA | NA | NA | #VALUE! |
| ENSMUSG00000106894.2 | 2.349469 | -4.75734 | 3.925837 | -1.2118 | 0.225588 | NA | #VALUE! |
| ENSMUSG00000106896.2 | 0 | NA | NA | NA | NA | NA | #VALUE! |
| ENSMUSG00000106897.2 | 0 | NA | NA | NA | NA | NA | #VALUE! |
| ENSMUSG00000106898.2 | 0.168284 | 1.027628 | 4.067126 | 0.252667 | 0.800526 | NA | #VALUE! |
| ENSMUSG00000106899.2 | 0 | NA | NA | NA | NA | NA | #VALUE! |
| ENSMUSG00000106900.2 | 0 | NA | NA | NA | NA | NA | #VALUE! |
| ENSMUSG00000106902.2 | 0 | NA | NA | NA | NA | NA | #VALUE! |
| ENSMUSG00000106904.2 | 0 | NA | NA | NA | NA | NA | #VALUE! |
| ENSMUSG00000106905.2 | 0 | NA | NA | NA | NA | NA | #VALUE! |
| ENSMUSG00000106906.2 | 0 | NA | NA | NA | NA | NA | #VALUE! |
| ENSMUSG00000106909.2 | 0 | NA | NA | NA | NA | NA | #VALUE! |
| ENSMUSG00000106910.2 | 0 | NA | NA | NA | NA | NA | #VALUE! |
| ENSMUSG00000106912.2 | 0 | NA | NA | NA | NA | NA | #VALUE! |
| ENSMUSG00000106913.2 | 0 | NA | NA | NA | NA | NA | #VALUE! |
| ENSMUSG00000106914.2 | 0 | NA | NA | NA | NA | NA | #VALUE! |
| ENSMUSG00000106915.2 | 0 | NA | NA | NA | NA | NA | #VALUE! |
| ENSMUSG00000106916.2 | 0 | NA | NA | NA | NA | NA | #VALUE! |
| ENSMUSG00000106917.2 | 0.261539 | 1.68833 | 4.043024 | 0.417591 | 0.676246 | NA | #VALUE! |
| ENSMUSG00000106924.2 | 0 | NA | NA | NA | NA | NA | #VALUE! |

| | | | | | | | |
|---|---|---|---|---|---|---|---|
| ENSMUSG00000106925.2 | 0.093255 | 0.868093 | 4.072464 | 0.213162 | 0.831201 | NA | #VALUE! |
| ENSMUSG00000106927.2 | 0 | NA | NA | NA | NA | NA | #VALUE! |
| ENSMUSG00000106928.2 | 0 | NA | NA | NA | NA | NA | #VALUE! |
| ENSMUSG00000106930.2 | 0 | NA | NA | NA | NA | NA | #VALUE! |
| ENSMUSG00000106932.2 | 0 | NA | NA | NA | NA | NA | #VALUE! |
| ENSMUSG00000106933.2 | 0 | NA | NA | NA | NA | NA | #VALUE! |
| ENSMUSG00000106934.2 | 0 | NA | NA | NA | NA | NA | #VALUE! |
| ENSMUSG00000106935.2 | 0 | NA | NA | NA | NA | NA | #VALUE! |
| ENSMUSG00000106936.2 | 0 | NA | NA | NA | NA | NA | #VALUE! |
| ENSMUSG00000106938.2 | 0 | NA | NA | NA | NA | NA | #VALUE! |
| ENSMUSG00000106939.2 | 0 | NA | NA | NA | NA | NA | #VALUE! |
| ENSMUSG00000106940.2 | 0 | NA | NA | NA | NA | NA | #VALUE! |
| ENSMUSG00000106944.2 | 0 | NA | NA | NA | NA | NA | #VALUE! |
| ENSMUSG00000106945.2 | 0 | NA | NA | NA | NA | NA | #VALUE! |
| ENSMUSG00000106946.2 | 0 | NA | NA | NA | NA | NA | #VALUE! |
| ENSMUSG00000106947.2 | 0 | NA | NA | NA | NA | NA | #VALUE! |
| ENSMUSG00000106948.2 | 0 | NA | NA | NA | NA | NA | #VALUE! |
| ENSMUSG00000106950.2 | 0 | NA | NA | NA | NA | NA | #VALUE! |
| ENSMUSG00000106951.2 | 23.098 | 2.889076 | 2.359324 | 1.224535 | NA | NA | #VALUE! |
| ENSMUSG00000106954.2 | 0 | NA | NA | NA | NA | NA | #VALUE! |
| ENSMUSG00000106955.2 | 0 | NA | NA | NA | NA | NA | #VALUE! |
| ENSMUSG00000106956.2 | 0 | NA | NA | NA | NA | NA | #VALUE! |
| ENSMUSG00000106957.2 | 0 | NA | NA | NA | NA | NA | #VALUE! |
| ENSMUSG00000106958.2 | 0 | NA | NA | NA | NA | NA | #VALUE! |
| ENSMUSG00000106959.2 | 0 | NA | NA | NA | NA | NA | #VALUE! |
| ENSMUSG00000106960.2 | 0 | NA | NA | NA | NA | NA | #VALUE! |
| ENSMUSG00000106961.2 | 0 | NA | NA | NA | NA | NA | #VALUE! |
| ENSMUSG00000106962.2 | 0 | NA | NA | NA | NA | NA | #VALUE! |
| ENSMUSG00000106963.2 | 0 | NA | NA | NA | NA | NA | #VALUE! |
| ENSMUSG00000106965.2 | 0 | NA | NA | NA | NA | NA | #VALUE! |
| ENSMUSG00000106967.2 | 0 | NA | NA | NA | NA | NA | #VALUE! |
| ENSMUSG00000106968.2 | 0 | NA | NA | NA | NA | NA | #VALUE! |
| ENSMUSG00000106969.2 | 0 | NA | NA | NA | NA | NA | #VALUE! |
| ENSMUSG00000106970.2 | 0 | NA | NA | NA | NA | NA | #VALUE! |
| ENSMUSG00000106971.2 | 0 | NA | NA | NA | NA | NA | #VALUE! |
| ENSMUSG00000106973.2 | 0 | NA | NA | NA | NA | NA | #VALUE! |
| ENSMUSG00000106974.2 | 0 | NA | NA | NA | NA | NA | #VALUE! |
| ENSMUSG00000106975.2 | 0 | NA | NA | NA | NA | NA | #VALUE! |
| ENSMUSG00000106976.2 | 0 | NA | NA | NA | NA | NA | #VALUE! |
| ENSMUSG00000106978.2 | 7.723511 | 0.759833 | 2.915814 | 0.26059 | 0.794409 | NA | #VALUE! |
| ENSMUSG00000106980.2 | 0 | NA | NA | NA | NA | NA | #VALUE! |
| ENSMUSG00000106981.2 | 1.575197 | -4.20222 | 3.934736 | -1.06798 | 0.285529 | NA | #VALUE! |
| ENSMUSG00000106982.2 | 0 | NA | NA | NA | NA | NA | #VALUE! |
| ENSMUSG00000106985.2 | 0 | NA | NA | NA | NA | NA | #VALUE! |

| | | | | | | | |
|---|---|---|---|---|---|---|---|
| ENSMUSG00000106986.2 | 0 | NA | NA | NA | NA | NA | #VALUE! |
| ENSMUSG00000106987.2 | 0 | NA | NA | NA | NA | NA | #VALUE! |
| ENSMUSG00000106988.2 | 0 | NA | NA | NA | NA | NA | #VALUE! |
| ENSMUSG00000106989.2 | 0 | NA | NA | NA | NA | NA | #VALUE! |
| ENSMUSG00000106991.2 | 0 | NA | NA | NA | NA | NA | #VALUE! |
| ENSMUSG00000106992.2 | 0 | NA | NA | NA | NA | NA | #VALUE! |
| ENSMUSG00000106993.2 | 7.741042 | 6.442369 | 3.912571 | 1.646582 | 0.099644 | NA | #VALUE! |
| ENSMUSG00000106994.2 | 0 | NA | NA | NA | NA | NA | #VALUE! |
| ENSMUSG00000106995.2 | 0 | NA | NA | NA | NA | NA | #VALUE! |
| ENSMUSG00000106996.2 | 0 | NA | NA | NA | NA | NA | #VALUE! |
| ENSMUSG00000106999.2 | 0 | NA | NA | NA | NA | NA | #VALUE! |
| ENSMUSG00000107000.2 | 0 | NA | NA | NA | NA | NA | #VALUE! |
| ENSMUSG00000107004.2 | 0 | NA | NA | NA | NA | NA | #VALUE! |
| ENSMUSG00000107005.2 | 0 | NA | NA | NA | NA | NA | #VALUE! |
| ENSMUSG00000107006.2 | 0.204111 | -1.05581 | 4.073302 | -0.2592 | 0.795479 | NA | #VALUE! |
| ENSMUSG00000107008.4 | 0 | NA | NA | NA | NA | NA | #VALUE! |
| ENSMUSG00000107009.2 | 0 | NA | NA | NA | NA | NA | #VALUE! |
| ENSMUSG00000107010.2 | 0 | NA | NA | NA | NA | NA | #VALUE! |
| ENSMUSG00000107011.2 | 0 | NA | NA | NA | NA | NA | #VALUE! |
| ENSMUSG00000107013.2 | 0 | NA | NA | NA | NA | NA | #VALUE! |
| ENSMUSG00000107014.2 | 0 | NA | NA | NA | NA | NA | #VALUE! |
| ENSMUSG00000107015.2 | 0 | NA | NA | NA | NA | NA | #VALUE! |
| ENSMUSG00000107017.2 | 0 | NA | NA | NA | NA | NA | #VALUE! |
| ENSMUSG00000107018.2 | 0 | NA | NA | NA | NA | NA | #VALUE! |
| ENSMUSG00000107019.2 | 0 | NA | NA | NA | NA | NA | #VALUE! |
| ENSMUSG00000107020.2 | 0 | NA | NA | NA | NA | NA | #VALUE! |
| ENSMUSG00000107021.2 | 0 | NA | NA | NA | NA | NA | #VALUE! |
| ENSMUSG00000107022.2 | 0 | NA | NA | NA | NA | NA | #VALUE! |
| ENSMUSG00000107023.3 | 0 | NA | NA | NA | NA | NA | #VALUE! |
| ENSMUSG00000107026.2 | 0 | NA | NA | NA | NA | NA | #VALUE! |
| ENSMUSG00000107027.2 | 0 | NA | NA | NA | NA | NA | #VALUE! |
| ENSMUSG00000107028.2 | 0 | NA | NA | NA | NA | NA | #VALUE! |
| ENSMUSG00000107029.2 | 0 | NA | NA | NA | NA | NA | #VALUE! |
| ENSMUSG00000107030.2 | 0 | NA | NA | NA | NA | NA | #VALUE! |
| ENSMUSG00000107031.2 | 0 | NA | NA | NA | NA | NA | #VALUE! |
| ENSMUSG00000107032.3 | 0 | NA | NA | NA | NA | NA | #VALUE! |
| ENSMUSG00000107033.2 | 0 | NA | NA | NA | NA | NA | #VALUE! |
| ENSMUSG00000107034.2 | 0 | NA | NA | NA | NA | NA | #VALUE! |
| ENSMUSG00000107035.2 | 0 | NA | NA | NA | NA | NA | #VALUE! |
| ENSMUSG00000107036.2 | 0 | NA | NA | NA | NA | NA | #VALUE! |
| ENSMUSG00000107037.2 | 0 | NA | NA | NA | NA | NA | #VALUE! |
| ENSMUSG00000107040.2 | 0 | NA | NA | NA | NA | NA | #VALUE! |
| ENSMUSG00000107041.2 | 0 | NA | NA | NA | NA | NA | #VALUE! |
| ENSMUSG00000107042.2 | 0 | NA | NA | NA | NA | NA | #VALUE! |

| | | | | | | | |
|---|---|---|---|---|---|---|---|
| ENSMUSG00000107043.2 | 0 | NA | NA | NA | NA | NA | #VALUE! |
| ENSMUSG00000107044.2 | 0 | NA | NA | NA | NA | NA | #VALUE! |
| ENSMUSG00000107045.2 | 0 | NA | NA | NA | NA | NA | #VALUE! |
| ENSMUSG00000107046.2 | 0 | NA | NA | NA | NA | NA | #VALUE! |
| ENSMUSG00000107047.2 | 0 | NA | NA | NA | NA | NA | #VALUE! |
| ENSMUSG00000107048.2 | 0 | NA | NA | NA | NA | NA | #VALUE! |
| ENSMUSG00000107049.2 | 0 | NA | NA | NA | NA | NA | #VALUE! |
| ENSMUSG00000107050.2 | 0 | NA | NA | NA | NA | NA | #VALUE! |
| ENSMUSG00000107053.2 | 0 | NA | NA | NA | NA | NA | #VALUE! |
| ENSMUSG00000107054.2 | 0.983902 | 3.433476 | 3.953024 | 0.868569 | 0.385083 | NA | #VALUE! |
| ENSMUSG00000107056.2 | 0 | NA | NA | NA | NA | NA | #VALUE! |
| ENSMUSG00000107058.2 | 0 | NA | NA | NA | NA | NA | #VALUE! |
| ENSMUSG00000107059.2 | 0 | NA | NA | NA | NA | NA | #VALUE! |
| ENSMUSG00000107060.2 | 0 | NA | NA | NA | NA | NA | #VALUE! |
| ENSMUSG00000107061.2 | 0 | NA | NA | NA | NA | NA | #VALUE! |
| ENSMUSG00000107062.2 | 0 | NA | NA | NA | NA | NA | #VALUE! |
| ENSMUSG00000107063.2 | 0 | NA | NA | NA | NA | NA | #VALUE! |
| ENSMUSG00000107064.2 | 0 | NA | NA | NA | NA | NA | #VALUE! |
| ENSMUSG00000107065.2 | 0 | NA | NA | NA | NA | NA | #VALUE! |
| ENSMUSG00000107066.3 | 0 | NA | NA | NA | NA | NA | #VALUE! |
| ENSMUSG00000107067.2 | 0 | NA | NA | NA | NA | NA | #VALUE! |
| ENSMUSG00000107068.2 | 0 | NA | NA | NA | NA | NA | #VALUE! |
| ENSMUSG00000107069.2 | 0 | NA | NA | NA | NA | NA | #VALUE! |
| ENSMUSG00000107070.2 | 0 | NA | NA | NA | NA | NA | #VALUE! |
| ENSMUSG00000107071.2 | 0 | NA | NA | NA | NA | NA | #VALUE! |
| ENSMUSG00000107072.2 | 0 | NA | NA | NA | NA | NA | #VALUE! |
| ENSMUSG00000107073.2 | 0 | NA | NA | NA | NA | NA | #VALUE! |
| ENSMUSG00000107074.2 | 0 | NA | NA | NA | NA | NA | #VALUE! |
| ENSMUSG00000107075.2 | 0 | NA | NA | NA | NA | NA | #VALUE! |
| ENSMUSG00000107076.2 | 0 | NA | NA | NA | NA | NA | #VALUE! |
| ENSMUSG00000107078.2 | 0 | NA | NA | NA | NA | NA | #VALUE! |
| ENSMUSG00000107079.2 | 0 | NA | NA | NA | NA | NA | #VALUE! |
| ENSMUSG00000107081.2 | 0 | NA | NA | NA | NA | NA | #VALUE! |
| ENSMUSG00000107082.2 | 0 | NA | NA | NA | NA | NA | #VALUE! |
| ENSMUSG00000107083.2 | 0 | NA | NA | NA | NA | NA | #VALUE! |
| ENSMUSG00000107084.2 | 0 | NA | NA | NA | NA | NA | #VALUE! |
| ENSMUSG00000107085.2 | 0 | NA | NA | NA | NA | NA | #VALUE! |
| ENSMUSG00000107086.2 | 0 | NA | NA | NA | NA | NA | #VALUE! |
| ENSMUSG00000107087.2 | 0 | NA | NA | NA | NA | NA | #VALUE! |
| ENSMUSG00000107089.2 | 0 | NA | NA | NA | NA | NA | #VALUE! |
| ENSMUSG00000107090.2 | 0 | NA | NA | NA | NA | NA | #VALUE! |
| ENSMUSG00000107092.2 | 0.578089 | -0.70693 | 3.996403 | -0.17689 | 0.859593 | NA | #VALUE! |
| ENSMUSG00000107093.2 | 0 | NA | NA | NA | NA | NA | #VALUE! |
| ENSMUSG00000107094.2 | 0 | NA | NA | NA | NA | NA | #VALUE! |

| | | | | | | | |
|---|---|---|---|---|---|---|---|
| ENSMUSG00000107096.4 | 0 | NA | NA | NA | NA | NA | #VALUE! |
| ENSMUSG00000107097.2 | 0 | NA | NA | NA | NA | NA | #VALUE! |
| ENSMUSG00000107098.2 | 0 | NA | NA | NA | NA | NA | #VALUE! |
| ENSMUSG00000107100.2 | 0 | NA | NA | NA | NA | NA | #VALUE! |
| ENSMUSG00000107102.2 | 0 | NA | NA | NA | NA | NA | #VALUE! |
| ENSMUSG00000107103.2 | 0 | NA | NA | NA | NA | NA | #VALUE! |
| ENSMUSG00000107105.2 | 0 | NA | NA | NA | NA | NA | #VALUE! |
| ENSMUSG00000107106.2 | 0 | NA | NA | NA | NA | NA | #VALUE! |
| ENSMUSG00000107108.3 | 0 | NA | NA | NA | NA | NA | #VALUE! |
| ENSMUSG00000107109.2 | 0 | NA | NA | NA | NA | NA | #VALUE! |
| ENSMUSG00000107110.2 | 0 | NA | NA | NA | NA | NA | #VALUE! |
| ENSMUSG00000107111.2 | 0 | NA | NA | NA | NA | NA | #VALUE! |
| ENSMUSG00000107112.2 | 0 | NA | NA | NA | NA | NA | #VALUE! |
| ENSMUSG00000107113.2 | 0 | NA | NA | NA | NA | NA | #VALUE! |
| ENSMUSG00000107114.2 | 0 | NA | NA | NA | NA | NA | #VALUE! |
| ENSMUSG00000107115.2 | 0 | NA | NA | NA | NA | NA | #VALUE! |
| ENSMUSG00000107116.2 | 2.545258 | -4.8732 | 3.92437 | -1.24178 | 0.214318 | NA | #VALUE! |
| ENSMUSG00000107117.2 | 0 | NA | NA | NA | NA | NA | #VALUE! |
| ENSMUSG00000107118.2 | 0 | NA | NA | NA | NA | NA | #VALUE! |
| ENSMUSG00000107120.2 | 0 | NA | NA | NA | NA | NA | #VALUE! |
| ENSMUSG00000107121.2 | 0 | NA | NA | NA | NA | NA | #VALUE! |
| ENSMUSG00000107122.2 | 0 | NA | NA | NA | NA | NA | #VALUE! |
| ENSMUSG00000107123.2 | 0 | NA | NA | NA | NA | NA | #VALUE! |
| ENSMUSG00000107125.2 | 0 | NA | NA | NA | NA | NA | #VALUE! |
| ENSMUSG00000107126.2 | 0 | NA | NA | NA | NA | NA | #VALUE! |
| ENSMUSG00000107127.2 | 0 | NA | NA | NA | NA | NA | #VALUE! |
| ENSMUSG00000107128.2 | 0 | NA | NA | NA | NA | NA | #VALUE! |
| ENSMUSG00000107129.2 | 0 | NA | NA | NA | NA | NA | #VALUE! |
| ENSMUSG00000107130.2 | 0 | NA | NA | NA | NA | NA | #VALUE! |
| ENSMUSG00000107131.2 | 0 | NA | NA | NA | NA | NA | #VALUE! |
| ENSMUSG00000107132.2 | 0 | NA | NA | NA | NA | NA | #VALUE! |
| ENSMUSG00000107133.2 | 0 | NA | NA | NA | NA | NA | #VALUE! |
| ENSMUSG00000107134.2 | 0 | NA | NA | NA | NA | NA | #VALUE! |
| ENSMUSG00000107135.2 | 0 | NA | NA | NA | NA | NA | #VALUE! |
| ENSMUSG00000107136.2 | 0 | NA | NA | NA | NA | NA | #VALUE! |
| ENSMUSG00000107139.2 | 0 | NA | NA | NA | NA | NA | #VALUE! |
| ENSMUSG00000107140.2 | 0 | NA | NA | NA | NA | NA | #VALUE! |
| ENSMUSG00000107141.2 | 0 | NA | NA | NA | NA | NA | #VALUE! |
| ENSMUSG00000107142.2 | 0 | NA | NA | NA | NA | NA | #VALUE! |
| ENSMUSG00000107143.2 | 2.154315 | 4.584472 | 3.927737 | 1.167204 | 0.243128 | NA | #VALUE! |
| ENSMUSG00000107144.2 | 0 | NA | NA | NA | NA | NA | #VALUE! |
| ENSMUSG00000107145.2 | 0 | NA | NA | NA | NA | NA | #VALUE! |
| ENSMUSG00000107148.2 | 0 | NA | NA | NA | NA | NA | #VALUE! |
| ENSMUSG00000107149.2 | 0 | NA | NA | NA | NA | NA | #VALUE! |

| | | | | | | | |
|---|---|---|---|---|---|---|---|
| ENSMUSG00000107151.2 | 0 | NA | NA | NA | NA | NA | #VALUE! |
| ENSMUSG00000107152.2 | 0 | NA | NA | NA | NA | NA | #VALUE! |
| ENSMUSG00000107153.2 | 0 | NA | NA | NA | NA | NA | #VALUE! |
| ENSMUSG00000107154.2 | 0 | NA | NA | NA | NA | NA | #VALUE! |
| ENSMUSG00000107155.2 | 0 | NA | NA | NA | NA | NA | #VALUE! |
| ENSMUSG00000107157.2 | 0 | NA | NA | NA | NA | NA | #VALUE! |
| ENSMUSG00000107158.2 | 0 | NA | NA | NA | NA | NA | #VALUE! |
| ENSMUSG00000107159.2 | 0 | NA | NA | NA | NA | NA | #VALUE! |
| ENSMUSG00000107161.2 | 0 | NA | NA | NA | NA | NA | #VALUE! |
| ENSMUSG00000107162.2 | 0 | NA | NA | NA | NA | NA | #VALUE! |
| ENSMUSG00000107163.2 | 0 | NA | NA | NA | NA | NA | #VALUE! |
| ENSMUSG00000107164.2 | 0 | NA | NA | NA | NA | NA | #VALUE! |
| ENSMUSG00000107165.2 | 0 | NA | NA | NA | NA | NA | #VALUE! |
| ENSMUSG00000107167.4 | 0 | NA | NA | NA | NA | NA | #VALUE! |
| ENSMUSG00000107168.2 | 0 | NA | NA | NA | NA | NA | #VALUE! |
| ENSMUSG00000107171.2 | 0 | NA | NA | NA | NA | NA | #VALUE! |
| ENSMUSG00000107172.2 | 0 | NA | NA | NA | NA | NA | #VALUE! |
| ENSMUSG00000107173.2 | 0 | NA | NA | NA | NA | NA | #VALUE! |
| ENSMUSG00000107177.2 | 0 | NA | NA | NA | NA | NA | #VALUE! |
| ENSMUSG00000107178.2 | 0 | NA | NA | NA | NA | NA | #VALUE! |
| ENSMUSG00000107179.2 | 0 | NA | NA | NA | NA | NA | #VALUE! |
| ENSMUSG00000107180.2 | 0 | NA | NA | NA | NA | NA | #VALUE! |
| ENSMUSG00000107181.2 | 0 | NA | NA | NA | NA | NA | #VALUE! |
| ENSMUSG00000107182.2 | 0 | NA | NA | NA | NA | NA | #VALUE! |
| ENSMUSG00000107183.2 | 0 | NA | NA | NA | NA | NA | #VALUE! |
| ENSMUSG00000107184.2 | 0 | NA | NA | NA | NA | NA | #VALUE! |
| ENSMUSG00000107185.2 | 0 | NA | NA | NA | NA | NA | #VALUE! |
| ENSMUSG00000107187.2 | 0 | NA | NA | NA | NA | NA | #VALUE! |
| ENSMUSG00000107189.2 | 0.105013 | -0.97324 | 4.076293 | -0.23876 | 0.811295 | NA | #VALUE! |
| ENSMUSG00000107191.2 | 0 | NA | NA | NA | NA | NA | #VALUE! |
| ENSMUSG00000107192.2 | 0 | NA | NA | NA | NA | NA | #VALUE! |
| ENSMUSG00000107193.2 | 0 | NA | NA | NA | NA | NA | #VALUE! |
| ENSMUSG00000107194.2 | 0 | NA | NA | NA | NA | NA | #VALUE! |
| ENSMUSG00000107195.2 | 0 | NA | NA | NA | NA | NA | #VALUE! |
| ENSMUSG00000107197.2 | 0 | NA | NA | NA | NA | NA | #VALUE! |
| ENSMUSG00000107199.2 | 0 | NA | NA | NA | NA | NA | #VALUE! |
| ENSMUSG00000107200.2 | 0 | NA | NA | NA | NA | NA | #VALUE! |
| ENSMUSG00000107201.2 | 0 | NA | NA | NA | NA | NA | #VALUE! |
| ENSMUSG00000107203.2 | 0 | NA | NA | NA | NA | NA | #VALUE! |
| ENSMUSG00000107204.2 | 0 | NA | NA | NA | NA | NA | #VALUE! |
| ENSMUSG00000107205.2 | 8.81051 | -6.66773 | 3.74925 | -1.77842 | 0.075335 | NA | #VALUE! |
| ENSMUSG00000107206.2 | 0 | NA | NA | NA | NA | NA | #VALUE! |
| ENSMUSG00000107207.2 | 0 | NA | NA | NA | NA | NA | #VALUE! |
| ENSMUSG00000107208.2 | 0 | NA | NA | NA | NA | NA | #VALUE! |

| | | | | | | | |
|---|---|---|---|---|---|---|---|
| ENSMUSG00000107209.2 | 0 | NA | NA | NA | NA | NA | #VALUE! |
| ENSMUSG00000107211.2 | 0 | NA | NA | NA | NA | NA | #VALUE! |
| ENSMUSG00000107212.2 | 0 | NA | NA | NA | NA | NA | #VALUE! |
| ENSMUSG00000107213.2 | 0 | NA | NA | NA | NA | NA | #VALUE! |
| ENSMUSG00000107214.4 | 0 | NA | NA | NA | NA | NA | #VALUE! |
| ENSMUSG00000107215.2 | 0 | NA | NA | NA | NA | NA | #VALUE! |
| ENSMUSG00000107216.2 | 0 | NA | NA | NA | NA | NA | #VALUE! |
| ENSMUSG00000107217.2 | 0 | NA | NA | NA | NA | NA | #VALUE! |
| ENSMUSG00000107218.2 | 0 | NA | NA | NA | NA | NA | #VALUE! |
| ENSMUSG00000107219.2 | 0 | NA | NA | NA | NA | NA | #VALUE! |
| ENSMUSG00000107220.2 | 0 | NA | NA | NA | NA | NA | #VALUE! |
| ENSMUSG00000107222.2 | 0 | NA | NA | NA | NA | NA | #VALUE! |
| ENSMUSG00000107223.2 | 0 | NA | NA | NA | NA | NA | #VALUE! |
| ENSMUSG00000107225.2 | 0 | NA | NA | NA | NA | NA | #VALUE! |
| ENSMUSG00000107226.2 | 0 | NA | NA | NA | NA | NA | #VALUE! |
| ENSMUSG00000107227.2 | 0 | NA | NA | NA | NA | NA | #VALUE! |
| ENSMUSG00000107230.2 | 0 | NA | NA | NA | NA | NA | #VALUE! |
| ENSMUSG00000107232.2 | 0 | NA | NA | NA | NA | NA | #VALUE! |
| ENSMUSG00000107233.2 | 0 | NA | NA | NA | NA | NA | #VALUE! |
| ENSMUSG00000107234.2 | 0 | NA | NA | NA | NA | NA | #VALUE! |
| ENSMUSG00000107237.2 | 0 | NA | NA | NA | NA | NA | #VALUE! |
| ENSMUSG00000107238.2 | 0 | NA | NA | NA | NA | NA | #VALUE! |
| ENSMUSG00000107240.2 | 0 | NA | NA | NA | NA | NA | #VALUE! |
| ENSMUSG00000107241.2 | 0 | NA | NA | NA | NA | NA | #VALUE! |
| ENSMUSG00000107243.2 | 0 | NA | NA | NA | NA | NA | #VALUE! |
| ENSMUSG00000107244.2 | 0 | NA | NA | NA | NA | NA | #VALUE! |
| ENSMUSG00000107245.2 | 0 | NA | NA | NA | NA | NA | #VALUE! |
| ENSMUSG00000107246.2 | 0 | NA | NA | NA | NA | NA | #VALUE! |
| ENSMUSG00000107247.2 | 0 | NA | NA | NA | NA | NA | #VALUE! |
| ENSMUSG00000107248.2 | 0 | NA | NA | NA | NA | NA | #VALUE! |
| ENSMUSG00000107251.2 | 0 | NA | NA | NA | NA | NA | #VALUE! |
| ENSMUSG00000107252.2 | 0 | NA | NA | NA | NA | NA | #VALUE! |
| ENSMUSG00000107253.2 | 0 | NA | NA | NA | NA | NA | #VALUE! |
| ENSMUSG00000107254.2 | 0 | NA | NA | NA | NA | NA | #VALUE! |
| ENSMUSG00000107255.2 | 0 | NA | NA | NA | NA | NA | #VALUE! |
| ENSMUSG00000107256.2 | 0 | NA | NA | NA | NA | NA | #VALUE! |
| ENSMUSG00000107257.2 | 0.168284 | 1.027628 | 4.067126 | 0.252667 | 0.800526 | NA | #VALUE! |
| ENSMUSG00000107258.3 | 0 | NA | NA | NA | NA | NA | #VALUE! |
| ENSMUSG00000107259.2 | 0 | NA | NA | NA | NA | NA | #VALUE! |
| ENSMUSG00000107261.2 | 0 | NA | NA | NA | NA | NA | #VALUE! |
| ENSMUSG00000107262.2 | 0 | NA | NA | NA | NA | NA | #VALUE! |
| ENSMUSG00000107263.2 | 0 | NA | NA | NA | NA | NA | #VALUE! |
| ENSMUSG00000107265.2 | 0 | NA | NA | NA | NA | NA | #VALUE! |
| ENSMUSG00000107266.2 | 0 | NA | NA | NA | NA | NA | #VALUE! |

| | | | | | | | |
|---|---|---|---|---|---|---|---|
| ENSMUSG00000107270.2 | 0 | NA | NA | NA | NA | NA | #VALUE! |
| ENSMUSG00000107271.4 | 0 | NA | NA | NA | NA | NA | #VALUE! |
| ENSMUSG00000107272.2 | 0 | NA | NA | NA | NA | NA | #VALUE! |
| ENSMUSG00000107276.2 | 0 | NA | NA | NA | NA | NA | #VALUE! |
| ENSMUSG00000107277.2 | 1.797524 | 4.343839 | 3.931519 | 1.104876 | 0.269213 | NA | #VALUE! |
| ENSMUSG00000107278.2 | 1.678593 | 4.255795 | 3.933066 | 1.082055 | 0.279228 | NA | #VALUE! |
| ENSMUSG00000107279.2 | 0 | NA | NA | NA | NA | NA | #VALUE! |
| ENSMUSG00000107280.2 | 0 | NA | NA | NA | NA | NA | #VALUE! |
| ENSMUSG00000107282.2 | 0 | NA | NA | NA | NA | NA | #VALUE! |
| ENSMUSG00000107284.2 | 0 | NA | NA | NA | NA | NA | #VALUE! |
| ENSMUSG00000107285.2 | 0 | NA | NA | NA | NA | NA | #VALUE! |
| ENSMUSG00000107286.2 | 0 | NA | NA | NA | NA | NA | #VALUE! |
| ENSMUSG00000107287.2 | 0 | NA | NA | NA | NA | NA | #VALUE! |
| ENSMUSG00000107288.2 | 3.915782 | -5.49628 | 3.91821 | -1.40275 | 0.16069 | NA | #VALUE! |
| ENSMUSG00000107289.2 | 0 | NA | NA | NA | NA | NA | #VALUE! |
| ENSMUSG00000107290.2 | 0 | NA | NA | NA | NA | NA | #VALUE! |
| ENSMUSG00000107291.2 | 0 | NA | NA | NA | NA | NA | #VALUE! |
| ENSMUSG00000107292.2 | 0 | NA | NA | NA | NA | NA | #VALUE! |
| ENSMUSG00000107295.2 | 0 | NA | NA | NA | NA | NA | #VALUE! |
| ENSMUSG00000107296.2 | 0 | NA | NA | NA | NA | NA | #VALUE! |
| ENSMUSG00000107299.2 | 0 | NA | NA | NA | NA | NA | #VALUE! |
| ENSMUSG00000107300.2 | 0 | NA | NA | NA | NA | NA | #VALUE! |
| ENSMUSG00000107301.2 | 0 | NA | NA | NA | NA | NA | #VALUE! |
| ENSMUSG00000107303.2 | 0 | NA | NA | NA | NA | NA | #VALUE! |
| ENSMUSG00000107304.2 | 4.755547 | 0.532354 | 3.765527 | 0.141376 | 0.887573 | NA | #VALUE! |
| ENSMUSG00000107305.2 | 0 | NA | NA | NA | NA | NA | #VALUE! |
| ENSMUSG00000107306.2 | 0 | NA | NA | NA | NA | NA | #VALUE! |
| ENSMUSG00000107307.2 | 0 | NA | NA | NA | NA | NA | #VALUE! |
| ENSMUSG00000107308.2 | 0 | NA | NA | NA | NA | NA | #VALUE! |
| ENSMUSG00000107309.2 | 0 | NA | NA | NA | NA | NA | #VALUE! |
| ENSMUSG00000107310.2 | 0 | NA | NA | NA | NA | NA | #VALUE! |
| ENSMUSG00000107312.2 | 0 | NA | NA | NA | NA | NA | #VALUE! |
| ENSMUSG00000107313.2 | 0 | NA | NA | NA | NA | NA | #VALUE! |
| ENSMUSG00000107314.2 | 0 | NA | NA | NA | NA | NA | #VALUE! |
| ENSMUSG00000107315.2 | 0 | NA | NA | NA | NA | NA | #VALUE! |
| ENSMUSG00000107316.2 | 0 | NA | NA | NA | NA | NA | #VALUE! |
| ENSMUSG00000107317.2 | 0 | NA | NA | NA | NA | NA | #VALUE! |
| ENSMUSG00000107318.2 | 0 | NA | NA | NA | NA | NA | #VALUE! |
| ENSMUSG00000107319.2 | 6.893887 | 0.95816 | 3.381349 | 0.283366 | 0.776896 | NA | #VALUE! |
| ENSMUSG00000107320.2 | 5.250656 | -5.92764 | 3.915257 | -1.51398 | 0.13003 | NA | #VALUE! |
| ENSMUSG00000107322.2 | 0 | NA | NA | NA | NA | NA | #VALUE! |
| ENSMUSG00000107325.2 | 0 | NA | NA | NA | NA | NA | #VALUE! |
| ENSMUSG00000107326.2 | 0 | NA | NA | NA | NA | NA | #VALUE! |
| ENSMUSG00000107327.2 | 0 | NA | NA | NA | NA | NA | #VALUE! |

| | | | | | | | |
|---|---|---|---|---|---|---|---|
| ENSMUSG00000107331.2 | 0 | NA | NA | NA | NA | NA | #VALUE! |
| ENSMUSG00000107333.2 | 0 | NA | NA | NA | NA | NA | #VALUE! |
| ENSMUSG00000107334.2 | 0 | NA | NA | NA | NA | NA | #VALUE! |
| ENSMUSG00000107335.2 | 0 | NA | NA | NA | NA | NA | #VALUE! |
| ENSMUSG00000107336.2 | 0 | NA | NA | NA | NA | NA | #VALUE! |
| ENSMUSG00000107337.2 | 0 | NA | NA | NA | NA | NA | #VALUE! |
| ENSMUSG00000107340.2 | 0 | NA | NA | NA | NA | NA | #VALUE! |
| ENSMUSG00000107341.2 | 0 | NA | NA | NA | NA | NA | #VALUE! |
| ENSMUSG00000107344.2 | 0 | NA | NA | NA | NA | NA | #VALUE! |
| ENSMUSG00000107346.2 | 0 | NA | NA | NA | NA | NA | #VALUE! |
| ENSMUSG00000107348.2 | 0 | NA | NA | NA | NA | NA | #VALUE! |
| ENSMUSG00000107349.2 | 0 | NA | NA | NA | NA | NA | #VALUE! |
| ENSMUSG00000107350.2 | 0 | NA | NA | NA | NA | NA | #VALUE! |
| ENSMUSG00000107351.2 | 0 | NA | NA | NA | NA | NA | #VALUE! |
| ENSMUSG00000107352.2 | 0 | NA | NA | NA | NA | NA | #VALUE! |
| ENSMUSG00000107353.2 | 0 | NA | NA | NA | NA | NA | #VALUE! |
| ENSMUSG00000107354.2 | 0 | NA | NA | NA | NA | NA | #VALUE! |
| ENSMUSG00000107355.2 | 0 | NA | NA | NA | NA | NA | #VALUE! |
| ENSMUSG00000107356.2 | 0 | NA | NA | NA | NA | NA | #VALUE! |
| ENSMUSG00000107357.2 | 0 | NA | NA | NA | NA | NA | #VALUE! |
| ENSMUSG00000107358.2 | 0 | NA | NA | NA | NA | NA | #VALUE! |
| ENSMUSG00000107359.2 | 0 | NA | NA | NA | NA | NA | #VALUE! |
| ENSMUSG00000107360.2 | 0 | NA | NA | NA | NA | NA | #VALUE! |
| ENSMUSG00000107362.2 | 0 | NA | NA | NA | NA | NA | #VALUE! |
| ENSMUSG00000107363.2 | 0 | NA | NA | NA | NA | NA | #VALUE! |
| ENSMUSG00000107364.2 | 0 | NA | NA | NA | NA | NA | #VALUE! |
| ENSMUSG00000107365.2 | 0 | NA | NA | NA | NA | NA | #VALUE! |
| ENSMUSG00000107368.2 | 0 | NA | NA | NA | NA | NA | #VALUE! |
| ENSMUSG00000107369.2 | 0 | NA | NA | NA | NA | NA | #VALUE! |
| ENSMUSG00000107370.2 | 0 | NA | NA | NA | NA | NA | #VALUE! |
| ENSMUSG00000107371.2 | 0 | NA | NA | NA | NA | NA | #VALUE! |
| ENSMUSG00000107373.2 | 0 | NA | NA | NA | NA | NA | #VALUE! |
| ENSMUSG00000107374.2 | 0 | NA | NA | NA | NA | NA | #VALUE! |
| ENSMUSG00000107375.2 | 0 | NA | NA | NA | NA | NA | #VALUE! |
| ENSMUSG00000107376.2 | 0 | NA | NA | NA | NA | NA | #VALUE! |
| ENSMUSG00000107377.2 | 0 | NA | NA | NA | NA | NA | #VALUE! |
| ENSMUSG00000107378.2 | 0 | NA | NA | NA | NA | NA | #VALUE! |
| ENSMUSG00000107379.2 | 0 | NA | NA | NA | NA | NA | #VALUE! |
| ENSMUSG00000107380.2 | 0 | NA | NA | NA | NA | NA | #VALUE! |
| ENSMUSG00000107384.2 | 0 | NA | NA | NA | NA | NA | #VALUE! |
| ENSMUSG00000107385.4 | 0 | NA | NA | NA | NA | NA | #VALUE! |
| ENSMUSG00000107386.2 | 0 | NA | NA | NA | NA | NA | #VALUE! |
| ENSMUSG00000107387.2 | 0 | NA | NA | NA | NA | NA | #VALUE! |
| ENSMUSG00000107388.2 | 0 | NA | NA | NA | NA | NA | #VALUE! |

| | | | | | | | |
|---|---|---|---|---|---|---|---|
| ENSMUSG00000107389.2 | 0 | NA | NA | NA | NA | NA | #VALUE! |
| ENSMUSG00000107392.2 | 0 | NA | NA | NA | NA | NA | #VALUE! |
| ENSMUSG00000107394.2 | 0 | NA | NA | NA | NA | NA | #VALUE! |
| ENSMUSG00000107395.2 | 0 | NA | NA | NA | NA | NA | #VALUE! |
| ENSMUSG00000107396.3 | 0 | NA | NA | NA | NA | NA | #VALUE! |
| ENSMUSG00000107397.2 | 0 | NA | NA | NA | NA | NA | #VALUE! |
| ENSMUSG00000107398.2 | 0 | NA | NA | NA | NA | NA | #VALUE! |
| ENSMUSG00000107399.3 | 0 | NA | NA | NA | NA | NA | #VALUE! |
| ENSMUSG00000107400.2 | 0 | NA | NA | NA | NA | NA | #VALUE! |
| ENSMUSG00000107401.2 | 0 | NA | NA | NA | NA | NA | #VALUE! |
| ENSMUSG00000107402.2 | 3.066908 | -3.00376 | 3.870424 | -0.77608 | 0.437702 | NA | #VALUE! |
| ENSMUSG00000107403.2 | 0 | NA | NA | NA | NA | NA | #VALUE! |
| ENSMUSG00000107404.2 | 0 | NA | NA | NA | NA | NA | #VALUE! |
| ENSMUSG00000107405.2 | 0 | NA | NA | NA | NA | NA | #VALUE! |
| ENSMUSG00000107406.2 | 0 | NA | NA | NA | NA | NA | #VALUE! |
| ENSMUSG00000107407.2 | 0 | NA | NA | NA | NA | NA | #VALUE! |
| ENSMUSG00000107408.2 | 0 | NA | NA | NA | NA | NA | #VALUE! |
| ENSMUSG00000107409.2 | 0 | NA | NA | NA | NA | NA | #VALUE! |
| ENSMUSG00000107410.2 | 0 | NA | NA | NA | NA | NA | #VALUE! |
| ENSMUSG00000107411.2 | 0 | NA | NA | NA | NA | NA | #VALUE! |
| ENSMUSG00000107412.2 | 0 | NA | NA | NA | NA | NA | #VALUE! |
| ENSMUSG00000107413.2 | 0 | NA | NA | NA | NA | NA | #VALUE! |
| ENSMUSG00000107414.2 | 0 | NA | NA | NA | NA | NA | #VALUE! |
| ENSMUSG00000107415.2 | 0 | NA | NA | NA | NA | NA | #VALUE! |
| ENSMUSG00000107416.2 | 0 | NA | NA | NA | NA | NA | #VALUE! |
| ENSMUSG00000107417.3 | 0 | NA | NA | NA | NA | NA | #VALUE! |
| ENSMUSG00000107418.2 | 0 | NA | NA | NA | NA | NA | #VALUE! |
| ENSMUSG00000107419.2 | 0 | NA | NA | NA | NA | NA | #VALUE! |
| ENSMUSG00000107420.2 | 0 | NA | NA | NA | NA | NA | #VALUE! |
| ENSMUSG00000107421.2 | 0 | NA | NA | NA | NA | NA | #VALUE! |
| ENSMUSG00000107422.2 | 0 | NA | NA | NA | NA | NA | #VALUE! |
| ENSMUSG00000107423.2 | 0 | NA | NA | NA | NA | NA | #VALUE! |
| ENSMUSG00000107424.2 | 0 | NA | NA | NA | NA | NA | #VALUE! |
| ENSMUSG00000107425.2 | 0 | NA | NA | NA | NA | NA | #VALUE! |
| ENSMUSG00000107427.2 | 0 | NA | NA | NA | NA | NA | #VALUE! |
| ENSMUSG00000107428.2 | 0 | NA | NA | NA | NA | NA | #VALUE! |
| ENSMUSG00000107429.3 | 0 | NA | NA | NA | NA | NA | #VALUE! |
| ENSMUSG00000107430.2 | 0 | NA | NA | NA | NA | NA | #VALUE! |
| ENSMUSG00000107431.2 | 0 | NA | NA | NA | NA | NA | #VALUE! |
| ENSMUSG00000107432.2 | 0 | NA | NA | NA | NA | NA | #VALUE! |
| ENSMUSG00000107433.2 | 0 | NA | NA | NA | NA | NA | #VALUE! |
| ENSMUSG00000107434.2 | 0 | NA | NA | NA | NA | NA | #VALUE! |
| ENSMUSG00000107435.2 | 0 | NA | NA | NA | NA | NA | #VALUE! |
| ENSMUSG00000107436.2 | 0 | NA | NA | NA | NA | NA | #VALUE! |

| | | | | | | | |
|---|---|---|---|---|---|---|---|
| ENSMUSG00000107438.2 | 0 | NA | NA | NA | NA | NA | #VALUE! |
| ENSMUSG00000107439.2 | 0 | NA | NA | NA | NA | NA | #VALUE! |
| ENSMUSG00000107440.2 | 0 | NA | NA | NA | NA | NA | #VALUE! |
| ENSMUSG00000107441.2 | 0 | NA | NA | NA | NA | NA | #VALUE! |
| ENSMUSG00000107442.2 | 0 | NA | NA | NA | NA | NA | #VALUE! |
| ENSMUSG00000107443.2 | 0 | NA | NA | NA | NA | NA | #VALUE! |
| ENSMUSG00000107445.2 | 0 | NA | NA | NA | NA | NA | #VALUE! |
| ENSMUSG00000107446.2 | 0 | NA | NA | NA | NA | NA | #VALUE! |
| ENSMUSG00000107447.2 | 0 | NA | NA | NA | NA | NA | #VALUE! |
| ENSMUSG00000107448.2 | 0 | NA | NA | NA | NA | NA | #VALUE! |
| ENSMUSG00000107449.2 | 0 | NA | NA | NA | NA | NA | #VALUE! |
| ENSMUSG00000107450.2 | 0 | NA | NA | NA | NA | NA | #VALUE! |
| ENSMUSG00000107452.2 | 0 | NA | NA | NA | NA | NA | #VALUE! |
| ENSMUSG00000107453.2 | 0 | NA | NA | NA | NA | NA | #VALUE! |
| ENSMUSG00000107454.2 | 0 | NA | NA | NA | NA | NA | #VALUE! |
| ENSMUSG00000107455.2 | 0 | NA | NA | NA | NA | NA | #VALUE! |
| ENSMUSG00000107456.2 | 0 | NA | NA | NA | NA | NA | #VALUE! |
| ENSMUSG00000107457.2 | 0 | NA | NA | NA | NA | NA | #VALUE! |
| ENSMUSG00000107458.2 | 0 | NA | NA | NA | NA | NA | #VALUE! |
| ENSMUSG00000107459.2 | 0 | NA | NA | NA | NA | NA | #VALUE! |
| ENSMUSG00000107460.2 | 0 | NA | NA | NA | NA | NA | #VALUE! |
| ENSMUSG00000107461.2 | 0 | NA | NA | NA | NA | NA | #VALUE! |
| ENSMUSG00000107462.2 | 0 | NA | NA | NA | NA | NA | #VALUE! |
| ENSMUSG00000107463.2 | 0 | NA | NA | NA | NA | NA | #VALUE! |
| ENSMUSG00000107464.2 | 0 | NA | NA | NA | NA | NA | #VALUE! |
| ENSMUSG00000107465.2 | 0 | NA | NA | NA | NA | NA | #VALUE! |
| ENSMUSG00000107466.2 | 0 | NA | NA | NA | NA | NA | #VALUE! |
| ENSMUSG00000107467.2 | 0 | NA | NA | NA | NA | NA | #VALUE! |
| ENSMUSG00000107468.2 | 0 | NA | NA | NA | NA | NA | #VALUE! |
| ENSMUSG00000107469.2 | 0 | NA | NA | NA | NA | NA | #VALUE! |
| ENSMUSG00000107471.2 | 0 | NA | NA | NA | NA | NA | #VALUE! |
| ENSMUSG00000107472.2 | 0 | NA | NA | NA | NA | NA | #VALUE! |
| ENSMUSG00000107473.2 | 0 | NA | NA | NA | NA | NA | #VALUE! |
| ENSMUSG00000107474.2 | 0 | NA | NA | NA | NA | NA | #VALUE! |
| ENSMUSG00000107475.2 | 0 | NA | NA | NA | NA | NA | #VALUE! |
| ENSMUSG00000107476.4 | 0 | NA | NA | NA | NA | NA | #VALUE! |
| ENSMUSG00000107477.2 | 0 | NA | NA | NA | NA | NA | #VALUE! |
| ENSMUSG00000107478.2 | 0 | NA | NA | NA | NA | NA | #VALUE! |
| ENSMUSG00000107479.2 | 0 | NA | NA | NA | NA | NA | #VALUE! |
| ENSMUSG00000107480.2 | 0 | NA | NA | NA | NA | NA | #VALUE! |
| ENSMUSG00000107481.2 | 0 | NA | NA | NA | NA | NA | #VALUE! |
| ENSMUSG00000107482.3 | 0 | NA | NA | NA | NA | NA | #VALUE! |
| ENSMUSG00000107483.2 | 0 | NA | NA | NA | NA | NA | #VALUE! |
| ENSMUSG00000107484.2 | 0 | NA | NA | NA | NA | NA | #VALUE! |

| | | | | | | | |
|---|---|---|---|---|---|---|---|
| ENSMUSG00000107486.2 | 0 | NA | NA | NA | NA | NA | #VALUE! |
| ENSMUSG00000107487.2 | 0 | NA | NA | NA | NA | NA | #VALUE! |
| ENSMUSG00000107488.2 | 0 | NA | NA | NA | NA | NA | #VALUE! |
| ENSMUSG00000107489.2 | 0 | NA | NA | NA | NA | NA | #VALUE! |
| ENSMUSG00000107490.2 | 0 | NA | NA | NA | NA | NA | #VALUE! |
| ENSMUSG00000107491.2 | 0 | NA | NA | NA | NA | NA | #VALUE! |
| ENSMUSG00000107492.2 | 0 | NA | NA | NA | NA | NA | #VALUE! |
| ENSMUSG00000107493.2 | 0 | NA | NA | NA | NA | NA | #VALUE! |
| ENSMUSG00000107495.2 | 0 | NA | NA | NA | NA | NA | #VALUE! |
| ENSMUSG00000107496.2 | 0 | NA | NA | NA | NA | NA | #VALUE! |
| ENSMUSG00000107497.2 | 0 | NA | NA | NA | NA | NA | #VALUE! |
| ENSMUSG00000107498.2 | 0 | NA | NA | NA | NA | NA | #VALUE! |
| ENSMUSG00000107499.2 | 0 | NA | NA | NA | NA | NA | #VALUE! |
| ENSMUSG00000107500.2 | 0 | NA | NA | NA | NA | NA | #VALUE! |
| ENSMUSG00000107501.2 | 0 | NA | NA | NA | NA | NA | #VALUE! |
| ENSMUSG00000107502.2 | 0 | NA | NA | NA | NA | NA | #VALUE! |
| ENSMUSG00000107503.2 | 0 | NA | NA | NA | NA | NA | #VALUE! |
| ENSMUSG00000107504.4 | 0 | NA | NA | NA | NA | NA | #VALUE! |
| ENSMUSG00000107505.2 | 0 | NA | NA | NA | NA | NA | #VALUE! |
| ENSMUSG00000107506.2 | 0 | NA | NA | NA | NA | NA | #VALUE! |
| ENSMUSG00000107508.2 | 1.711717 | 4.260588 | 3.932979 | 1.083298 | 0.278676 | NA | #VALUE! |
| ENSMUSG00000107509.2 | 0 | NA | NA | NA | NA | NA | #VALUE! |
| ENSMUSG00000107510.2 | 0 | NA | NA | NA | NA | NA | #VALUE! |
| ENSMUSG00000107511.2 | 0 | NA | NA | NA | NA | NA | #VALUE! |
| ENSMUSG00000107512.2 | 0 | NA | NA | NA | NA | NA | #VALUE! |
| ENSMUSG00000107513.2 | 0 | NA | NA | NA | NA | NA | #VALUE! |
| ENSMUSG00000107514.2 | 0 | NA | NA | NA | NA | NA | #VALUE! |
| ENSMUSG00000107515.2 | 0 | NA | NA | NA | NA | NA | #VALUE! |
| ENSMUSG00000107516.2 | 3.569781 | 4.225161 | 3.912631 | 1.079877 | 0.280197 | NA | #VALUE! |
| ENSMUSG00000107517.2 | 0 | NA | NA | NA | NA | NA | #VALUE! |
| ENSMUSG00000107518.2 | 0 | NA | NA | NA | NA | NA | #VALUE! |
| ENSMUSG00000107519.2 | 0 | NA | NA | NA | NA | NA | #VALUE! |
| ENSMUSG00000107520.2 | 0 | NA | NA | NA | NA | NA | #VALUE! |
| ENSMUSG00000107521.2 | 0 | NA | NA | NA | NA | NA | #VALUE! |
| ENSMUSG00000107522.2 | 0 | NA | NA | NA | NA | NA | #VALUE! |
| ENSMUSG00000107523.2 | 0 | NA | NA | NA | NA | NA | #VALUE! |
| ENSMUSG00000107524.2 | 0 | NA | NA | NA | NA | NA | #VALUE! |
| ENSMUSG00000107525.2 | 0 | NA | NA | NA | NA | NA | #VALUE! |
| ENSMUSG00000107526.2 | 0 | NA | NA | NA | NA | NA | #VALUE! |
| ENSMUSG00000107527.2 | 0 | NA | NA | NA | NA | NA | #VALUE! |
| ENSMUSG00000107528.2 | 0 | NA | NA | NA | NA | NA | #VALUE! |
| ENSMUSG00000107529.2 | 0 | NA | NA | NA | NA | NA | #VALUE! |
| ENSMUSG00000107530.2 | 0 | NA | NA | NA | NA | NA | #VALUE! |
| ENSMUSG00000107531.2 | 1.020554 | -3.54462 | 3.950745 | -0.8972 | 0.36961 | NA | #VALUE! |

| | | | | | | | |
|---|---|---|---|---|---|---|---|
| ENSMUSG00000107532.2 | 0 | NA | NA | NA | NA | NA | #VALUE! |
| ENSMUSG00000107533.2 | 0 | NA | NA | NA | NA | NA | #VALUE! |
| ENSMUSG00000107534.2 | 0 | NA | NA | NA | NA | NA | #VALUE! |
| ENSMUSG00000107535.2 | 0 | NA | NA | NA | NA | NA | #VALUE! |
| ENSMUSG00000107536.2 | 0 | NA | NA | NA | NA | NA | #VALUE! |
| ENSMUSG00000107538.2 | 0 | NA | NA | NA | NA | NA | #VALUE! |
| ENSMUSG00000107539.2 | 0 | NA | NA | NA | NA | NA | #VALUE! |
| ENSMUSG00000107540.2 | 0 | NA | NA | NA | NA | NA | #VALUE! |
| ENSMUSG00000107541.2 | 0.289044 | -0.14751 | 4.065279 | -0.03629 | 0.971054 | NA | #VALUE! |
| ENSMUSG00000107542.2 | 0 | NA | NA | NA | NA | NA | #VALUE! |
| ENSMUSG00000107543.2 | 0 | NA | NA | NA | NA | NA | #VALUE! |
| ENSMUSG00000107544.2 | 0 | NA | NA | NA | NA | NA | #VALUE! |
| ENSMUSG00000107545.2 | 0 | NA | NA | NA | NA | NA | #VALUE! |
| ENSMUSG00000107546.2 | 0 | NA | NA | NA | NA | NA | #VALUE! |
| ENSMUSG00000107548.2 | 0 | NA | NA | NA | NA | NA | #VALUE! |
| ENSMUSG00000107549.2 | 0 | NA | NA | NA | NA | NA | #VALUE! |
| ENSMUSG00000107550.2 | 0 | NA | NA | NA | NA | NA | #VALUE! |
| ENSMUSG00000107551.2 | 0 | NA | NA | NA | NA | NA | #VALUE! |
| ENSMUSG00000107553.2 | 0 | NA | NA | NA | NA | NA | #VALUE! |
| ENSMUSG00000107555.2 | 0 | NA | NA | NA | NA | NA | #VALUE! |
| ENSMUSG00000107556.2 | 0 | NA | NA | NA | NA | NA | #VALUE! |
| ENSMUSG00000107557.2 | 0 | NA | NA | NA | NA | NA | #VALUE! |
| ENSMUSG00000107558.2 | 0 | NA | NA | NA | NA | NA | #VALUE! |
| ENSMUSG00000107559.2 | 0 | NA | NA | NA | NA | NA | #VALUE! |
| ENSMUSG00000107560.2 | 0 | NA | NA | NA | NA | NA | #VALUE! |
| ENSMUSG00000107561.2 | 0 | NA | NA | NA | NA | NA | #VALUE! |
| ENSMUSG00000107562.2 | 0 | NA | NA | NA | NA | NA | #VALUE! |
| ENSMUSG00000107563.3 | 0 | NA | NA | NA | NA | NA | #VALUE! |
| ENSMUSG00000107564.2 | 0 | NA | NA | NA | NA | NA | #VALUE! |
| ENSMUSG00000107565.2 | 0 | NA | NA | NA | NA | NA | #VALUE! |
| ENSMUSG00000107566.2 | 0 | NA | NA | NA | NA | NA | #VALUE! |
| ENSMUSG00000107567.2 | 0 | NA | NA | NA | NA | NA | #VALUE! |
| ENSMUSG00000107568.2 | 0 | NA | NA | NA | NA | NA | #VALUE! |
| ENSMUSG00000107569.2 | 0 | NA | NA | NA | NA | NA | #VALUE! |
| ENSMUSG00000107570.2 | 0 | NA | NA | NA | NA | NA | #VALUE! |
| ENSMUSG00000107571.2 | 0 | NA | NA | NA | NA | NA | #VALUE! |
| ENSMUSG00000107572.2 | 0 | NA | NA | NA | NA | NA | #VALUE! |
| ENSMUSG00000107573.3 | 0 | NA | NA | NA | NA | NA | #VALUE! |
| ENSMUSG00000107574.2 | 0 | NA | NA | NA | NA | NA | #VALUE! |
| ENSMUSG00000107575.2 | 0 | NA | NA | NA | NA | NA | #VALUE! |
| ENSMUSG00000107576.2 | 0 | NA | NA | NA | NA | NA | #VALUE! |
| ENSMUSG00000107577.2 | 1.514552 | 4.088766 | 3.936269 | 1.038742 | 0.298925 | NA | #VALUE! |
| ENSMUSG00000107578.2 | 0 | NA | NA | NA | NA | NA | #VALUE! |
| ENSMUSG00000107579.2 | 0 | NA | NA | NA | NA | NA | #VALUE! |

| | | | | | | | |
|---|---|---|---|---|---|---|---|
| ENSMUSG00000107580.2 | 0 | NA | NA | NA | NA | NA | #VALUE! |
| ENSMUSG00000107581.2 | 0 | NA | NA | NA | NA | NA | #VALUE! |
| ENSMUSG00000107582.3 | 0 | NA | NA | NA | NA | NA | #VALUE! |
| ENSMUSG00000107583.2 | 0 | NA | NA | NA | NA | NA | #VALUE! |
| ENSMUSG00000107584.2 | 0 | NA | NA | NA | NA | NA | #VALUE! |
| ENSMUSG00000107585.2 | 0 | NA | NA | NA | NA | NA | #VALUE! |
| ENSMUSG00000107586.2 | 0 | NA | NA | NA | NA | NA | #VALUE! |
| ENSMUSG00000107587.2 | 0 | NA | NA | NA | NA | NA | #VALUE! |
| ENSMUSG00000107588.2 | 0 | NA | NA | NA | NA | NA | #VALUE! |
| ENSMUSG00000107590.2 | 0 | NA | NA | NA | NA | NA | #VALUE! |
| ENSMUSG00000107591.2 | 0 | NA | NA | NA | NA | NA | #VALUE! |
| ENSMUSG00000107592.2 | 0 | NA | NA | NA | NA | NA | #VALUE! |
| ENSMUSG00000107593.2 | 0 | NA | NA | NA | NA | NA | #VALUE! |
| ENSMUSG00000107594.2 | 0 | NA | NA | NA | NA | NA | #VALUE! |
| ENSMUSG00000107595.2 | 0 | NA | NA | NA | NA | NA | #VALUE! |
| ENSMUSG00000107596.2 | 0.303281 | -0.61664 | 4.050639 | -0.15223 | 0.879003 | NA | #VALUE! |
| ENSMUSG00000107597.2 | 0 | NA | NA | NA | NA | NA | #VALUE! |
| ENSMUSG00000107598.2 | 0 | NA | NA | NA | NA | NA | #VALUE! |
| ENSMUSG00000107599.2 | 0 | NA | NA | NA | NA | NA | #VALUE! |
| ENSMUSG00000107600.2 | 0 | NA | NA | NA | NA | NA | #VALUE! |
| ENSMUSG00000107601.2 | 0 | NA | NA | NA | NA | NA | #VALUE! |
| ENSMUSG00000107602.2 | 0 | NA | NA | NA | NA | NA | #VALUE! |
| ENSMUSG00000107603.2 | 0 | NA | NA | NA | NA | NA | #VALUE! |
| ENSMUSG00000107604.2 | 0 | NA | NA | NA | NA | NA | #VALUE! |
| ENSMUSG00000107605.2 | 6.99414 | 6.300596 | 3.91317 | 1.6101 | NA | NA | #VALUE! |
| ENSMUSG00000107606.2 | 0 | NA | NA | NA | NA | NA | #VALUE! |
| ENSMUSG00000107608.2 | 0 | NA | NA | NA | NA | NA | #VALUE! |
| ENSMUSG00000107609.2 | 0 | NA | NA | NA | NA | NA | #VALUE! |
| ENSMUSG00000107610.2 | 0 | NA | NA | NA | NA | NA | #VALUE! |
| ENSMUSG00000107611.2 | 0 | NA | NA | NA | NA | NA | #VALUE! |
| ENSMUSG00000107612.2 | 0 | NA | NA | NA | NA | NA | #VALUE! |
| ENSMUSG00000107613.2 | 0 | NA | NA | NA | NA | NA | #VALUE! |
| ENSMUSG00000107614.2 | 0 | NA | NA | NA | NA | NA | #VALUE! |
| ENSMUSG00000107615.2 | 0 | NA | NA | NA | NA | NA | #VALUE! |
| ENSMUSG00000107616.2 | 0 | NA | NA | NA | NA | NA | #VALUE! |
| ENSMUSG00000107618.2 | 0 | NA | NA | NA | NA | NA | #VALUE! |
| ENSMUSG00000107619.2 | 0 | NA | NA | NA | NA | NA | #VALUE! |
| ENSMUSG00000107620.2 | 0 | NA | NA | NA | NA | NA | #VALUE! |
| ENSMUSG00000107622.2 | 0 | NA | NA | NA | NA | NA | #VALUE! |
| ENSMUSG00000107623.2 | 0 | NA | NA | NA | NA | NA | #VALUE! |
| ENSMUSG00000107624.2 | 0 | NA | NA | NA | NA | NA | #VALUE! |
| ENSMUSG00000107625.2 | 0 | NA | NA | NA | NA | NA | #VALUE! |
| ENSMUSG00000107626.2 | 0 | NA | NA | NA | NA | NA | #VALUE! |
| ENSMUSG00000107627.2 | 0 | NA | NA | NA | NA | NA | #VALUE! |

| | | | | | | | |
|---|---|---|---|---|---|---|---|
| ENSMUSG00000107628.2 | 0 | NA | NA | NA | NA | NA | #VALUE! |
| ENSMUSG00000107629.2 | 0 | NA | NA | NA | NA | NA | #VALUE! |
| ENSMUSG00000107630.2 | 0 | NA | NA | NA | NA | NA | #VALUE! |
| ENSMUSG00000107631.2 | 0 | NA | NA | NA | NA | NA | #VALUE! |
| ENSMUSG00000107632.2 | 0 | NA | NA | NA | NA | NA | #VALUE! |
| ENSMUSG00000107633.2 | 0 | NA | NA | NA | NA | NA | #VALUE! |
| ENSMUSG00000107634.2 | 0 | NA | NA | NA | NA | NA | #VALUE! |
| ENSMUSG00000107635.2 | 0 | NA | NA | NA | NA | NA | #VALUE! |
| ENSMUSG00000107637.2 | 0 | NA | NA | NA | NA | NA | #VALUE! |
| ENSMUSG00000107638.3 | 0 | NA | NA | NA | NA | NA | #VALUE! |
| ENSMUSG00000107639.2 | 0 | NA | NA | NA | NA | NA | #VALUE! |
| ENSMUSG00000107640.2 | 0 | NA | NA | NA | NA | NA | #VALUE! |
| ENSMUSG00000107641.2 | 0 | NA | NA | NA | NA | NA | #VALUE! |
| ENSMUSG00000107642.2 | 0 | NA | NA | NA | NA | NA | #VALUE! |
| ENSMUSG00000107643.2 | 0 | NA | NA | NA | NA | NA | #VALUE! |
| ENSMUSG00000107644.3 | 0 | NA | NA | NA | NA | NA | #VALUE! |
| ENSMUSG00000107645.3 | 0 | NA | NA | NA | NA | NA | #VALUE! |
| ENSMUSG00000107646.2 | 0 | NA | NA | NA | NA | NA | #VALUE! |
| ENSMUSG00000107647.2 | 0 | NA | NA | NA | NA | NA | #VALUE! |
| ENSMUSG00000107648.5 | 0 | NA | NA | NA | NA | NA | #VALUE! |
| ENSMUSG00000107649.2 | 0 | NA | NA | NA | NA | NA | #VALUE! |
| ENSMUSG00000107650.2 | 0 | NA | NA | NA | NA | NA | #VALUE! |
| ENSMUSG00000107651.2 | 0 | NA | NA | NA | NA | NA | #VALUE! |
| ENSMUSG00000107652.2 | 0 | NA | NA | NA | NA | NA | #VALUE! |
| ENSMUSG00000107653.2 | 0 | NA | NA | NA | NA | NA | #VALUE! |
| ENSMUSG00000107655.2 | 6.123323 | -6.14178 | 3.914088 | -1.56915 | 0.116614 | NA | #VALUE! |
| ENSMUSG00000107656.2 | 0 | NA | NA | NA | NA | NA | #VALUE! |
| ENSMUSG00000107657.2 | 0 | NA | NA | NA | NA | NA | #VALUE! |
| ENSMUSG00000107658.2 | 0 | NA | NA | NA | NA | NA | #VALUE! |
| ENSMUSG00000107659.2 | 27.58623 | -1.46084 | 2.619378 | -0.55771 | NA | NA | #VALUE! |
| ENSMUSG00000107660.2 | 0 | NA | NA | NA | NA | NA | #VALUE! |
| ENSMUSG00000107661.2 | 0 | NA | NA | NA | NA | NA | #VALUE! |
| ENSMUSG00000107662.3 | 0 | NA | NA | NA | NA | NA | #VALUE! |
| ENSMUSG00000107663.2 | 0 | NA | NA | NA | NA | NA | #VALUE! |
| ENSMUSG00000107664.2 | 0 | NA | NA | NA | NA | NA | #VALUE! |
| ENSMUSG00000107666.2 | 0 | NA | NA | NA | NA | NA | #VALUE! |
| ENSMUSG00000107667.3 | 0 | NA | NA | NA | NA | NA | #VALUE! |
| ENSMUSG00000107668.2 | 0 | NA | NA | NA | NA | NA | #VALUE! |
| ENSMUSG00000107669.2 | 0 | NA | NA | NA | NA | NA | #VALUE! |
| ENSMUSG00000107671.2 | 0 | NA | NA | NA | NA | NA | #VALUE! |
| ENSMUSG00000107672.2 | 0 | NA | NA | NA | NA | NA | #VALUE! |
| ENSMUSG00000107673.2 | 0 | NA | NA | NA | NA | NA | #VALUE! |
| ENSMUSG00000107674.2 | 0 | NA | NA | NA | NA | NA | #VALUE! |
| ENSMUSG00000107675.2 | 0 | NA | NA | NA | NA | NA | #VALUE! |

| | | | | | | | |
|---|---|---|---|---|---|---|---|
| ENSMUSG00000107676.2 | 0 | NA | NA | NA | NA | NA | #VALUE! |
| ENSMUSG00000107677.3 | 0 | NA | NA | NA | NA | NA | #VALUE! |
| ENSMUSG00000107678.2 | 0 | NA | NA | NA | NA | NA | #VALUE! |
| ENSMUSG00000107679.2 | 0 | NA | NA | NA | NA | NA | #VALUE! |
| ENSMUSG00000107680.2 | 0 | NA | NA | NA | NA | NA | #VALUE! |
| ENSMUSG00000107681.2 | 0 | NA | NA | NA | NA | NA | #VALUE! |
| ENSMUSG00000107682.2 | 0 | NA | NA | NA | NA | NA | #VALUE! |
| ENSMUSG00000107683.2 | 0 | NA | NA | NA | NA | NA | #VALUE! |
| ENSMUSG00000107684.2 | 0 | NA | NA | NA | NA | NA | #VALUE! |
| ENSMUSG00000107685.2 | 0 | NA | NA | NA | NA | NA | #VALUE! |
| ENSMUSG00000107686.2 | 0 | NA | NA | NA | NA | NA | #VALUE! |
| ENSMUSG00000107687.2 | 0 | NA | NA | NA | NA | NA | #VALUE! |
| ENSMUSG00000107688.2 | 0 | NA | NA | NA | NA | NA | #VALUE! |
| ENSMUSG00000107689.2 | 0.198268 | -0.06494 | 4.068276 | -0.01596 | 0.987265 | NA | #VALUE! |
| ENSMUSG00000107690.2 | 0 | NA | NA | NA | NA | NA | #VALUE! |
| ENSMUSG00000107691.2 | 0 | NA | NA | NA | NA | NA | #VALUE! |
| ENSMUSG00000107692.2 | 0 | NA | NA | NA | NA | NA | #VALUE! |
| ENSMUSG00000107693.2 | 0 | NA | NA | NA | NA | NA | #VALUE! |
| ENSMUSG00000107694.2 | 0 | NA | NA | NA | NA | NA | #VALUE! |
| ENSMUSG00000107695.2 | 0 | NA | NA | NA | NA | NA | #VALUE! |
| ENSMUSG00000107696.2 | 0 | NA | NA | NA | NA | NA | #VALUE! |
| ENSMUSG00000107697.2 | 0 | NA | NA | NA | NA | NA | #VALUE! |
| ENSMUSG00000107698.2 | 0 | NA | NA | NA | NA | NA | #VALUE! |
| ENSMUSG00000107699.2 | 0 | NA | NA | NA | NA | NA | #VALUE! |
| ENSMUSG00000107700.2 | 0 | NA | NA | NA | NA | NA | #VALUE! |
| ENSMUSG00000107701.2 | 0.245976 | 1.027628 | 4.067126 | 0.252667 | 0.800526 | NA | #VALUE! |
| ENSMUSG00000107702.2 | 0 | NA | NA | NA | NA | NA | #VALUE! |
| ENSMUSG00000107703.2 | 0 | NA | NA | NA | NA | NA | #VALUE! |
| ENSMUSG00000107704.2 | 0 | NA | NA | NA | NA | NA | #VALUE! |
| ENSMUSG00000107705.2 | 0 | NA | NA | NA | NA | NA | #VALUE! |
| ENSMUSG00000107706.2 | 0 | NA | NA | NA | NA | NA | #VALUE! |
| ENSMUSG00000107707.2 | 0 | NA | NA | NA | NA | NA | #VALUE! |
| ENSMUSG00000107708.2 | 0 | NA | NA | NA | NA | NA | #VALUE! |
| ENSMUSG00000107709.2 | 0 | NA | NA | NA | NA | NA | #VALUE! |
| ENSMUSG00000107710.2 | 0 | NA | NA | NA | NA | NA | #VALUE! |
| ENSMUSG00000107711.3 | 0 | NA | NA | NA | NA | NA | #VALUE! |
| ENSMUSG00000107712.2 | 0 | NA | NA | NA | NA | NA | #VALUE! |
| ENSMUSG00000107713.2 | 0 | NA | NA | NA | NA | NA | #VALUE! |
| ENSMUSG00000107714.2 | 0 | NA | NA | NA | NA | NA | #VALUE! |
| ENSMUSG00000107715.2 | 0 | NA | NA | NA | NA | NA | #VALUE! |
| ENSMUSG00000107716.2 | 0 | NA | NA | NA | NA | NA | #VALUE! |
| ENSMUSG00000107718.3 | 0 | NA | NA | NA | NA | NA | #VALUE! |
| ENSMUSG00000107719.2 | 0 | NA | NA | NA | NA | NA | #VALUE! |
| ENSMUSG00000107720.2 | 0 | NA | NA | NA | NA | NA | #VALUE! |

| | | | | | | | |
|---|---|---|---|---|---|---|---|
| ENSMUSG00000107721.2 | 0.245976 | 1.027628 | 4.067126 | 0.252667 | 0.800526 | NA | #VALUE! |
| ENSMUSG00000107722.2 | 0 | NA | NA | NA | NA | NA | #VALUE! |
| ENSMUSG00000107723.2 | 0 | NA | NA | NA | NA | NA | #VALUE! |
| ENSMUSG00000107724.2 | 0 | NA | NA | NA | NA | NA | #VALUE! |
| ENSMUSG00000107725.2 | 0 | NA | NA | NA | NA | NA | #VALUE! |
| ENSMUSG00000107726.2 | 0 | NA | NA | NA | NA | NA | #VALUE! |
| ENSMUSG00000107727.2 | 0 | NA | NA | NA | NA | NA | #VALUE! |
| ENSMUSG00000107728.2 | 0 | NA | NA | NA | NA | NA | #VALUE! |
| ENSMUSG00000107729.2 | 0 | NA | NA | NA | NA | NA | #VALUE! |
| ENSMUSG00000107730.2 | 0 | NA | NA | NA | NA | NA | #VALUE! |
| ENSMUSG00000107731.2 | 0 | NA | NA | NA | NA | NA | #VALUE! |
| ENSMUSG00000107732.2 | 0 | NA | NA | NA | NA | NA | #VALUE! |
| ENSMUSG00000107733.2 | 0 | NA | NA | NA | NA | NA | #VALUE! |
| ENSMUSG00000107734.2 | 0 | NA | NA | NA | NA | NA | #VALUE! |
| ENSMUSG00000107735.2 | 0 | NA | NA | NA | NA | NA | #VALUE! |
| ENSMUSG00000107736.2 | 0 | NA | NA | NA | NA | NA | #VALUE! |
| ENSMUSG00000107737.2 | 0 | NA | NA | NA | NA | NA | #VALUE! |
| ENSMUSG00000107738.2 | 0 | NA | NA | NA | NA | NA | #VALUE! |
| ENSMUSG00000107739.2 | 0 | NA | NA | NA | NA | NA | #VALUE! |
| ENSMUSG00000107740.2 | 0 | NA | NA | NA | NA | NA | #VALUE! |
| ENSMUSG00000107741.3 | 5.026698 | 2.454595 | 3.649186 | 0.672642 | 0.501175 | NA | #VALUE! |
| ENSMUSG00000107742.2 | 0 | NA | NA | NA | NA | NA | #VALUE! |
| ENSMUSG00000107743.2 | 0 | NA | NA | NA | NA | NA | #VALUE! |
| ENSMUSG00000107744.2 | 0 | NA | NA | NA | NA | NA | #VALUE! |
| ENSMUSG00000107745.2 | 0 | NA | NA | NA | NA | NA | #VALUE! |
| ENSMUSG00000107746.2 | 0 | NA | NA | NA | NA | NA | #VALUE! |
| ENSMUSG00000107747.2 | 0 | NA | NA | NA | NA | NA | #VALUE! |
| ENSMUSG00000107748.5 | 0 | NA | NA | NA | NA | NA | #VALUE! |
| ENSMUSG00000107749.2 | 0 | NA | NA | NA | NA | NA | #VALUE! |
| ENSMUSG00000107750.2 | 0 | NA | NA | NA | NA | NA | #VALUE! |
| ENSMUSG00000107751.2 | 0 | NA | NA | NA | NA | NA | #VALUE! |
| ENSMUSG00000107752.2 | 0 | NA | NA | NA | NA | NA | #VALUE! |
| ENSMUSG00000107753.2 | 0 | NA | NA | NA | NA | NA | #VALUE! |
| ENSMUSG00000107754.2 | 0 | NA | NA | NA | NA | NA | #VALUE! |
| ENSMUSG00000107755.2 | 0 | NA | NA | NA | NA | NA | #VALUE! |
| ENSMUSG00000107756.2 | 0 | NA | NA | NA | NA | NA | #VALUE! |
| ENSMUSG00000107757.2 | 0 | NA | NA | NA | NA | NA | #VALUE! |
| ENSMUSG00000107758.2 | 0 | NA | NA | NA | NA | NA | #VALUE! |
| ENSMUSG00000107759.2 | 0 | NA | NA | NA | NA | NA | #VALUE! |
| ENSMUSG00000107760.2 | 0 | NA | NA | NA | NA | NA | #VALUE! |
| ENSMUSG00000107761.2 | 0 | NA | NA | NA | NA | NA | #VALUE! |
| ENSMUSG00000107762.2 | 0 | NA | NA | NA | NA | NA | #VALUE! |
| ENSMUSG00000107763.2 | 0 | NA | NA | NA | NA | NA | #VALUE! |
| ENSMUSG00000107764.2 | 0 | NA | NA | NA | NA | NA | #VALUE! |

| | | | | | | | |
|---|---|---|---|---|---|---|---|
| ENSMUSG00000107765.2 | 0 | NA | NA | NA | NA | NA | #VALUE! |
| ENSMUSG00000107766.2 | 0 | NA | NA | NA | NA | NA | #VALUE! |
| ENSMUSG00000107767.2 | 0 | NA | NA | NA | NA | NA | #VALUE! |
| ENSMUSG00000107768.2 | 0 | NA | NA | NA | NA | NA | #VALUE! |
| ENSMUSG00000107769.2 | 0 | NA | NA | NA | NA | NA | #VALUE! |
| ENSMUSG00000107770.2 | 0 | NA | NA | NA | NA | NA | #VALUE! |
| ENSMUSG00000107771.2 | 0 | NA | NA | NA | NA | NA | #VALUE! |
| ENSMUSG00000107772.2 | 0 | NA | NA | NA | NA | NA | #VALUE! |
| ENSMUSG00000107773.2 | 0 | NA | NA | NA | NA | NA | #VALUE! |
| ENSMUSG00000107774.2 | 0 | NA | NA | NA | NA | NA | #VALUE! |
| ENSMUSG00000107775.2 | 0 | NA | NA | NA | NA | NA | #VALUE! |
| ENSMUSG00000107776.2 | 0 | NA | NA | NA | NA | NA | #VALUE! |
| ENSMUSG00000107777.2 | 0 | NA | NA | NA | NA | NA | #VALUE! |
| ENSMUSG00000107778.2 | 0 | NA | NA | NA | NA | NA | #VALUE! |
| ENSMUSG00000107779.2 | 0 | NA | NA | NA | NA | NA | #VALUE! |
| ENSMUSG00000107780.2 | 0 | NA | NA | NA | NA | NA | #VALUE! |
| ENSMUSG00000107781.2 | 0 | NA | NA | NA | NA | NA | #VALUE! |
| ENSMUSG00000107782.2 | 0 | NA | NA | NA | NA | NA | #VALUE! |
| ENSMUSG00000107783.2 | 0 | NA | NA | NA | NA | NA | #VALUE! |
| ENSMUSG00000107784.2 | 0 | NA | NA | NA | NA | NA | #VALUE! |
| ENSMUSG00000107785.2 | 0.420053 | -2.33493 | 4.007343 | -0.58266 | 0.56012 | NA | #VALUE! |
| ENSMUSG00000107786.2 | 0 | NA | NA | NA | NA | NA | #VALUE! |
| ENSMUSG00000107787.2 | 0 | NA | NA | NA | NA | NA | #VALUE! |
| ENSMUSG00000107788.2 | 0.210026 | -1.52494 | 4.058692 | -0.37572 | 0.707123 | NA | #VALUE! |
| ENSMUSG00000107789.2 | 0 | NA | NA | NA | NA | NA | #VALUE! |
| ENSMUSG00000107790.2 | 0 | NA | NA | NA | NA | NA | #VALUE! |
| ENSMUSG00000107791.2 | 0 | NA | NA | NA | NA | NA | #VALUE! |
| ENSMUSG00000107792.2 | 0 | NA | NA | NA | NA | NA | #VALUE! |
| ENSMUSG00000107793.2 | 0 | NA | NA | NA | NA | NA | #VALUE! |
| ENSMUSG00000107794.2 | 0 | NA | NA | NA | NA | NA | #VALUE! |
| ENSMUSG00000107795.2 | 0 | NA | NA | NA | NA | NA | #VALUE! |
| ENSMUSG00000107796.2 | 0 | NA | NA | NA | NA | NA | #VALUE! |
| ENSMUSG00000107797.2 | 0 | NA | NA | NA | NA | NA | #VALUE! |
| ENSMUSG00000107798.2 | 0 | NA | NA | NA | NA | NA | #VALUE! |
| ENSMUSG00000107799.2 | 0 | NA | NA | NA | NA | NA | #VALUE! |
| ENSMUSG00000107800.2 | 0 | NA | NA | NA | NA | NA | #VALUE! |
| ENSMUSG00000107801.2 | 0 | NA | NA | NA | NA | NA | #VALUE! |
| ENSMUSG00000107802.2 | 0 | NA | NA | NA | NA | NA | #VALUE! |
| ENSMUSG00000107803.2 | 0 | NA | NA | NA | NA | NA | #VALUE! |
| ENSMUSG00000107804.2 | 0 | NA | NA | NA | NA | NA | #VALUE! |
| ENSMUSG00000107805.2 | 0 | NA | NA | NA | NA | NA | #VALUE! |
| ENSMUSG00000107806.2 | 0.18651 | 1.401769 | 4.056354 | 0.345574 | 0.729663 | NA | #VALUE! |
| ENSMUSG00000107808.2 | 0 | NA | NA | NA | NA | NA | #VALUE! |
| ENSMUSG00000107809.2 | 0 | NA | NA | NA | NA | NA | #VALUE! |

| | | | | | | | |
|---|---|---|---|---|---|---|---|
| ENSMUSG00000107810.2 | 0 | NA | NA | NA | NA | NA | #VALUE! |
| ENSMUSG00000107811.2 | 0 | NA | NA | NA | NA | NA | #VALUE! |
| ENSMUSG00000107812.2 | 0 | NA | NA | NA | NA | NA | #VALUE! |
| ENSMUSG00000107813.2 | 0 | NA | NA | NA | NA | NA | #VALUE! |
| ENSMUSG00000107814.2 | 0 | NA | NA | NA | NA | NA | #VALUE! |
| ENSMUSG00000107815.2 | 0 | NA | NA | NA | NA | NA | #VALUE! |
| ENSMUSG00000107816.2 | 0 | NA | NA | NA | NA | NA | #VALUE! |
| ENSMUSG00000107817.2 | 0 | NA | NA | NA | NA | NA | #VALUE! |
| ENSMUSG00000107818.2 | 0 | NA | NA | NA | NA | NA | #VALUE! |
| ENSMUSG00000107819.2 | 0 | NA | NA | NA | NA | NA | #VALUE! |
| ENSMUSG00000107821.2 | 0 | NA | NA | NA | NA | NA | #VALUE! |
| ENSMUSG00000107822.5 | 0 | NA | NA | NA | NA | NA | #VALUE! |
| ENSMUSG00000107823.2 | 0 | NA | NA | NA | NA | NA | #VALUE! |
| ENSMUSG00000107824.2 | 0 | NA | NA | NA | NA | NA | #VALUE! |
| ENSMUSG00000107825.2 | 0 | NA | NA | NA | NA | NA | #VALUE! |
| ENSMUSG00000107826.2 | 0 | NA | NA | NA | NA | NA | #VALUE! |
| ENSMUSG00000107828.2 | 0 | NA | NA | NA | NA | NA | #VALUE! |
| ENSMUSG00000107829.2 | 0 | NA | NA | NA | NA | NA | #VALUE! |
| ENSMUSG00000107830.2 | 0 | NA | NA | NA | NA | NA | #VALUE! |
| ENSMUSG00000107831.2 | 0 | NA | NA | NA | NA | NA | #VALUE! |
| ENSMUSG00000107832.2 | 0 | NA | NA | NA | NA | NA | #VALUE! |
| ENSMUSG00000107833.2 | 0 | NA | NA | NA | NA | NA | #VALUE! |
| ENSMUSG00000107834.2 | 0 | NA | NA | NA | NA | NA | #VALUE! |
| ENSMUSG00000107835.2 | 0 | NA | NA | NA | NA | NA | #VALUE! |
| ENSMUSG00000107836.2 | 0 | NA | NA | NA | NA | NA | #VALUE! |
| ENSMUSG00000107837.2 | 0 | NA | NA | NA | NA | NA | #VALUE! |
| ENSMUSG00000107839.2 | 0 | NA | NA | NA | NA | NA | #VALUE! |
| ENSMUSG00000107840.2 | 0 | NA | NA | NA | NA | NA | #VALUE! |
| ENSMUSG00000107841.2 | 0 | NA | NA | NA | NA | NA | #VALUE! |
| ENSMUSG00000107842.2 | 0 | NA | NA | NA | NA | NA | #VALUE! |
| ENSMUSG00000107844.2 | 0 | NA | NA | NA | NA | NA | #VALUE! |
| ENSMUSG00000107845.2 | 0 | NA | NA | NA | NA | NA | #VALUE! |
| ENSMUSG00000107846.2 | 0 | NA | NA | NA | NA | NA | #VALUE! |
| ENSMUSG00000107847.2 | 0 | NA | NA | NA | NA | NA | #VALUE! |
| ENSMUSG00000107848.2 | 0 | NA | NA | NA | NA | NA | #VALUE! |
| ENSMUSG00000107849.2 | 0 | NA | NA | NA | NA | NA | #VALUE! |
| ENSMUSG00000107850.2 | 0 | NA | NA | NA | NA | NA | #VALUE! |
| ENSMUSG00000107851.2 | 0 | NA | NA | NA | NA | NA | #VALUE! |
| ENSMUSG00000107852.2 | 0 | NA | NA | NA | NA | NA | #VALUE! |
| ENSMUSG00000107853.2 | 0 | NA | NA | NA | NA | NA | #VALUE! |
| ENSMUSG00000107854.2 | 0 | NA | NA | NA | NA | NA | #VALUE! |
| ENSMUSG00000107855.2 | 0 | NA | NA | NA | NA | NA | #VALUE! |
| ENSMUSG00000107856.2 | 0 | NA | NA | NA | NA | NA | #VALUE! |
| ENSMUSG00000107857.2 | 0 | NA | NA | NA | NA | NA | #VALUE! |

| | | | | | | | |
|---|---|---|---|---|---|---|---|
| ENSMUSG00000107858.2 | 0 | NA | NA | NA | NA | NA | #VALUE! |
| ENSMUSG00000107859.2 | 0 | NA | NA | NA | NA | NA | #VALUE! |
| ENSMUSG00000107861.2 | 0 | NA | NA | NA | NA | NA | #VALUE! |
| ENSMUSG00000107862.2 | 0 | NA | NA | NA | NA | NA | #VALUE! |
| ENSMUSG00000107863.2 | 0 | NA | NA | NA | NA | NA | #VALUE! |
| ENSMUSG00000107864.2 | 0 | NA | NA | NA | NA | NA | #VALUE! |
| ENSMUSG00000107865.2 | 0 | NA | NA | NA | NA | NA | #VALUE! |
| ENSMUSG00000107866.2 | 0 | NA | NA | NA | NA | NA | #VALUE! |
| ENSMUSG00000107867.2 | 0 | NA | NA | NA | NA | NA | #VALUE! |
| ENSMUSG00000107868.2 | 0 | NA | NA | NA | NA | NA | #VALUE! |
| ENSMUSG00000107869.2 | 0 | NA | NA | NA | NA | NA | #VALUE! |
| ENSMUSG00000107870.2 | 0 | NA | NA | NA | NA | NA | #VALUE! |
| ENSMUSG00000107872.3 | 0 | NA | NA | NA | NA | NA | #VALUE! |
| ENSMUSG00000107873.2 | 0 | NA | NA | NA | NA | NA | #VALUE! |
| ENSMUSG00000107874.2 | 0 | NA | NA | NA | NA | NA | #VALUE! |
| ENSMUSG00000107875.2 | 0 | NA | NA | NA | NA | NA | #VALUE! |
| ENSMUSG00000107876.2 | 0 | NA | NA | NA | NA | NA | #VALUE! |
| ENSMUSG00000107877.3 | 0 | NA | NA | NA | NA | NA | #VALUE! |
| ENSMUSG00000107878.2 | 0 | NA | NA | NA | NA | NA | #VALUE! |
| ENSMUSG00000107879.3 | 0 | NA | NA | NA | NA | NA | #VALUE! |
| ENSMUSG00000107880.2 | 0 | NA | NA | NA | NA | NA | #VALUE! |
| ENSMUSG00000107882.2 | 0 | NA | NA | NA | NA | NA | #VALUE! |
| ENSMUSG00000107883.2 | 0 | NA | NA | NA | NA | NA | #VALUE! |
| ENSMUSG00000107884.2 | 0 | NA | NA | NA | NA | NA | #VALUE! |
| ENSMUSG00000107885.2 | 0 | NA | NA | NA | NA | NA | #VALUE! |
| ENSMUSG00000107886.2 | 0 | NA | NA | NA | NA | NA | #VALUE! |
| ENSMUSG00000107887.2 | 0 | NA | NA | NA | NA | NA | #VALUE! |
| ENSMUSG00000107888.2 | 0 | NA | NA | NA | NA | NA | #VALUE! |
| ENSMUSG00000107889.2 | 0 | NA | NA | NA | NA | NA | #VALUE! |
| ENSMUSG00000107890.2 | 0 | NA | NA | NA | NA | NA | #VALUE! |
| ENSMUSG00000107891.2 | 0 | NA | NA | NA | NA | NA | #VALUE! |
| ENSMUSG00000107892.2 | 0 | NA | NA | NA | NA | NA | #VALUE! |
| ENSMUSG00000107893.2 | 0 | NA | NA | NA | NA | NA | #VALUE! |
| ENSMUSG00000107894.2 | 0 | NA | NA | NA | NA | NA | #VALUE! |
| ENSMUSG00000107895.2 | 0 | NA | NA | NA | NA | NA | #VALUE! |
| ENSMUSG00000107896.2 | 0 | NA | NA | NA | NA | NA | #VALUE! |
| ENSMUSG00000107898.2 | 0 | NA | NA | NA | NA | NA | #VALUE! |
| ENSMUSG00000107899.2 | 0 | NA | NA | NA | NA | NA | #VALUE! |
| ENSMUSG00000107900.2 | 0 | NA | NA | NA | NA | NA | #VALUE! |
| ENSMUSG00000107901.3 | 0 | NA | NA | NA | NA | NA | #VALUE! |
| ENSMUSG00000107902.2 | 0 | NA | NA | NA | NA | NA | #VALUE! |
| ENSMUSG00000107903.2 | 0 | NA | NA | NA | NA | NA | #VALUE! |
| ENSMUSG00000107904.2 | 0 | NA | NA | NA | NA | NA | #VALUE! |
| ENSMUSG00000107905.2 | 0 | NA | NA | NA | NA | NA | #VALUE! |

| | | | | | | | |
|---|---|---|---|---|---|---|---|
| ENSMUSG00000107906.2 | 0 | NA | NA | NA | NA | NA | #VALUE! |
| ENSMUSG00000107907.2 | 0 | NA | NA | NA | NA | NA | #VALUE! |
| ENSMUSG00000107908.2 | 0 | NA | NA | NA | NA | NA | #VALUE! |
| ENSMUSG00000107909.2 | 0 | NA | NA | NA | NA | NA | #VALUE! |
| ENSMUSG00000107911.2 | 0.261539 | 1.68833 | 4.043024 | 0.417591 | 0.676246 | NA | #VALUE! |
| ENSMUSG00000107913.2 | 0 | NA | NA | NA | NA | NA | #VALUE! |
| ENSMUSG00000107914.2 | 0 | NA | NA | NA | NA | NA | #VALUE! |
| ENSMUSG00000107915.2 | 0 | NA | NA | NA | NA | NA | #VALUE! |
| ENSMUSG00000107916.2 | 0 | NA | NA | NA | NA | NA | #VALUE! |
| ENSMUSG00000107917.2 | 0 | NA | NA | NA | NA | NA | #VALUE! |
| ENSMUSG00000107918.2 | 0 | NA | NA | NA | NA | NA | #VALUE! |
| ENSMUSG00000107919.2 | 0 | NA | NA | NA | NA | NA | #VALUE! |
| ENSMUSG00000107920.2 | 0 | NA | NA | NA | NA | NA | #VALUE! |
| ENSMUSG00000107921.2 | 0 | NA | NA | NA | NA | NA | #VALUE! |
| ENSMUSG00000107922.2 | 0 | NA | NA | NA | NA | NA | #VALUE! |
| ENSMUSG00000107923.2 | 0 | NA | NA | NA | NA | NA | #VALUE! |
| ENSMUSG00000107924.2 | 0 | NA | NA | NA | NA | NA | #VALUE! |
| ENSMUSG00000107925.2 | 0 | NA | NA | NA | NA | NA | #VALUE! |
| ENSMUSG00000107926.2 | 0 | NA | NA | NA | NA | NA | #VALUE! |
| ENSMUSG00000107927.2 | 0.204111 | -1.05581 | 4.073302 | -0.2592 | 0.795479 | NA | #VALUE! |
| ENSMUSG00000107928.2 | 0 | NA | NA | NA | NA | NA | #VALUE! |
| ENSMUSG00000107929.2 | 0 | NA | NA | NA | NA | NA | #VALUE! |
| ENSMUSG00000107930.2 | 0 | NA | NA | NA | NA | NA | #VALUE! |
| ENSMUSG00000107931.2 | 0 | NA | NA | NA | NA | NA | #VALUE! |
| ENSMUSG00000107932.2 | 3.133984 | 1.954705 | 3.814891 | 0.512388 | 0.608379 | NA | #VALUE! |
| ENSMUSG00000107933.2 | 0 | NA | NA | NA | NA | NA | #VALUE! |
| ENSMUSG00000107934.2 | 0 | NA | NA | NA | NA | NA | #VALUE! |
| ENSMUSG00000107935.2 | 0 | NA | NA | NA | NA | NA | #VALUE! |
| ENSMUSG00000107936.2 | 0 | NA | NA | NA | NA | NA | #VALUE! |
| ENSMUSG00000107937.2 | 0 | NA | NA | NA | NA | NA | #VALUE! |
| ENSMUSG00000107938.2 | 0 | NA | NA | NA | NA | NA | #VALUE! |
| ENSMUSG00000107940.2 | 0 | NA | NA | NA | NA | NA | #VALUE! |
| ENSMUSG00000107941.2 | 0 | NA | NA | NA | NA | NA | #VALUE! |
| ENSMUSG00000107942.2 | 0 | NA | NA | NA | NA | NA | #VALUE! |
| ENSMUSG00000107943.2 | 0 | NA | NA | NA | NA | NA | #VALUE! |
| ENSMUSG00000107944.2 | 0 | NA | NA | NA | NA | NA | #VALUE! |
| ENSMUSG00000107945.2 | 0 | NA | NA | NA | NA | NA | #VALUE! |
| ENSMUSG00000107946.2 | 0 | NA | NA | NA | NA | NA | #VALUE! |
| ENSMUSG00000107947.2 | 0 | NA | NA | NA | NA | NA | #VALUE! |
| ENSMUSG00000107948.3 | 0 | NA | NA | NA | NA | NA | #VALUE! |
| ENSMUSG00000107949.2 | 0 | NA | NA | NA | NA | NA | #VALUE! |
| ENSMUSG00000107950.2 | 0 | NA | NA | NA | NA | NA | #VALUE! |
| ENSMUSG00000107951.2 | 0 | NA | NA | NA | NA | NA | #VALUE! |
| ENSMUSG00000107952.2 | 0 | NA | NA | NA | NA | NA | #VALUE! |

| | | | | | | | |
|---|---|---|---|---|---|---|---|
| ENSMUSG00000107953.2 | 0 | NA | NA | NA | NA | NA | #VALUE! |
| ENSMUSG00000107954.2 | 0 | NA | NA | NA | NA | NA | #VALUE! |
| ENSMUSG00000107955.2 | 0 | NA | NA | NA | NA | NA | #VALUE! |
| ENSMUSG00000107956.2 | 0 | NA | NA | NA | NA | NA | #VALUE! |
| ENSMUSG00000107957.2 | 0 | NA | NA | NA | NA | NA | #VALUE! |
| ENSMUSG00000107959.2 | 0 | NA | NA | NA | NA | NA | #VALUE! |
| ENSMUSG00000107960.2 | 0 | NA | NA | NA | NA | NA | #VALUE! |
| ENSMUSG00000107961.2 | 5.315546 | 5.90612 | 3.915181 | 1.508518 | 0.131422 | NA | #VALUE! |
| ENSMUSG00000107962.2 | 0 | NA | NA | NA | NA | NA | #VALUE! |
| ENSMUSG00000107964.2 | 0 | NA | NA | NA | NA | NA | #VALUE! |
| ENSMUSG00000107965.2 | 0 | NA | NA | NA | NA | NA | #VALUE! |
| ENSMUSG00000107966.2 | 0 | NA | NA | NA | NA | NA | #VALUE! |
| ENSMUSG00000107967.2 | 0 | NA | NA | NA | NA | NA | #VALUE! |
| ENSMUSG00000107968.2 | 0 | NA | NA | NA | NA | NA | #VALUE! |
| ENSMUSG00000107969.2 | 0 | NA | NA | NA | NA | NA | #VALUE! |
| ENSMUSG00000107970.2 | 0.336567 | 1.944925 | 4.029936 | 0.482619 | 0.629366 | NA | #VALUE! |
| ENSMUSG00000107971.2 | 0 | NA | NA | NA | NA | NA | #VALUE! |
| ENSMUSG00000107972.2 | 0 | NA | NA | NA | NA | NA | #VALUE! |
| ENSMUSG00000107973.2 | 0 | NA | NA | NA | NA | NA | #VALUE! |
| ENSMUSG00000107975.2 | 0 | NA | NA | NA | NA | NA | #VALUE! |
| ENSMUSG00000107976.2 | 0 | NA | NA | NA | NA | NA | #VALUE! |
| ENSMUSG00000107977.2 | 0 | NA | NA | NA | NA | NA | #VALUE! |
| ENSMUSG00000107978.2 | 0 | NA | NA | NA | NA | NA | #VALUE! |
| ENSMUSG00000107979.2 | 0 | NA | NA | NA | NA | NA | #VALUE! |
| ENSMUSG00000107980.2 | 0 | NA | NA | NA | NA | NA | #VALUE! |
| ENSMUSG00000107981.2 | 0 | NA | NA | NA | NA | NA | #VALUE! |
| ENSMUSG00000107982.2 | 0 | NA | NA | NA | NA | NA | #VALUE! |
| ENSMUSG00000107983.2 | 0 | NA | NA | NA | NA | NA | #VALUE! |
| ENSMUSG00000107984.2 | 0 | NA | NA | NA | NA | NA | #VALUE! |
| ENSMUSG00000107985.2 | 0 | NA | NA | NA | NA | NA | #VALUE! |
| ENSMUSG00000107986.2 | 0 | NA | NA | NA | NA | NA | #VALUE! |
| ENSMUSG00000107987.2 | 0 | NA | NA | NA | NA | NA | #VALUE! |
| ENSMUSG00000107988.2 | 0 | NA | NA | NA | NA | NA | #VALUE! |
| ENSMUSG00000107989.2 | 0 | NA | NA | NA | NA | NA | #VALUE! |
| ENSMUSG00000107990.2 | 0 | NA | NA | NA | NA | NA | #VALUE! |
| ENSMUSG00000107991.2 | 0 | NA | NA | NA | NA | NA | #VALUE! |
| ENSMUSG00000107992.2 | 0 | NA | NA | NA | NA | NA | #VALUE! |
| ENSMUSG00000107993.4 | 0 | NA | NA | NA | NA | NA | #VALUE! |
| ENSMUSG00000107995.2 | 0 | NA | NA | NA | NA | NA | #VALUE! |
| ENSMUSG00000107996.2 | 0 | NA | NA | NA | NA | NA | #VALUE! |
| ENSMUSG00000107997.2 | 0 | NA | NA | NA | NA | NA | #VALUE! |
| ENSMUSG00000107998.2 | 0 | NA | NA | NA | NA | NA | #VALUE! |
| ENSMUSG00000107999.2 | 0 | NA | NA | NA | NA | NA | #VALUE! |
| ENSMUSG00000108000.2 | 0 | NA | NA | NA | NA | NA | #VALUE! |

| | | | | | | | |
|---|---|---|---|---|---|---|---|
| ENSMUSG00000108002.2 | 0 | NA | NA | NA | NA | NA | #VALUE! |
| ENSMUSG00000108003.2 | 0 | NA | NA | NA | NA | NA | #VALUE! |
| ENSMUSG00000108004.2 | 0 | NA | NA | NA | NA | NA | #VALUE! |
| ENSMUSG00000108005.2 | 0 | NA | NA | NA | NA | NA | #VALUE! |
| ENSMUSG00000108006.2 | 0 | NA | NA | NA | NA | NA | #VALUE! |
| ENSMUSG00000108007.2 | 0 | NA | NA | NA | NA | NA | #VALUE! |
| ENSMUSG00000108008.2 | 0 | NA | NA | NA | NA | NA | #VALUE! |
| ENSMUSG00000108009.2 | 0 | NA | NA | NA | NA | NA | #VALUE! |
| ENSMUSG00000108010.2 | 0 | NA | NA | NA | NA | NA | #VALUE! |
| ENSMUSG00000108011.2 | 0 | NA | NA | NA | NA | NA | #VALUE! |
| ENSMUSG00000108012.2 | 0 | NA | NA | NA | NA | NA | #VALUE! |
| ENSMUSG00000108013.2 | 0 | NA | NA | NA | NA | NA | #VALUE! |
| ENSMUSG00000108014.2 | 0 | NA | NA | NA | NA | NA | #VALUE! |
| ENSMUSG00000108015.4 | 0 | NA | NA | NA | NA | NA | #VALUE! |
| ENSMUSG00000108016.2 | 0 | NA | NA | NA | NA | NA | #VALUE! |
| ENSMUSG00000108017.2 | 0 | NA | NA | NA | NA | NA | #VALUE! |
| ENSMUSG00000108018.2 | 0 | NA | NA | NA | NA | NA | #VALUE! |
| ENSMUSG00000108019.2 | 0 | NA | NA | NA | NA | NA | #VALUE! |
| ENSMUSG00000108020.2 | 0 | NA | NA | NA | NA | NA | #VALUE! |
| ENSMUSG00000108021.2 | 0 | NA | NA | NA | NA | NA | #VALUE! |
| ENSMUSG00000108022.2 | 0.105013 | -0.97324 | 4.076293 | -0.23876 | 0.811295 | NA | #VALUE! |
| ENSMUSG00000108023.2 | 0 | NA | NA | NA | NA | NA | #VALUE! |
| ENSMUSG00000108024.2 | 0 | NA | NA | NA | NA | NA | #VALUE! |
| ENSMUSG00000108025.2 | 0 | NA | NA | NA | NA | NA | #VALUE! |
| ENSMUSG00000108026.2 | 0 | NA | NA | NA | NA | NA | #VALUE! |
| ENSMUSG00000108027.2 | 0 | NA | NA | NA | NA | NA | #VALUE! |
| ENSMUSG00000108028.2 | 0 | NA | NA | NA | NA | NA | #VALUE! |
| ENSMUSG00000108029.2 | 0 | NA | NA | NA | NA | NA | #VALUE! |
| ENSMUSG00000108030.2 | 2.310289 | -4.7496 | 3.925939 | -1.2098 | 0.226355 | NA | #VALUE! |
| ENSMUSG00000108032.2 | 0 | NA | NA | NA | NA | NA | #VALUE! |
| ENSMUSG00000108033.2 | 0 | NA | NA | NA | NA | NA | #VALUE! |
| ENSMUSG00000108034.2 | 0 | NA | NA | NA | NA | NA | #VALUE! |
| ENSMUSG00000108035.2 | 0 | NA | NA | NA | NA | NA | #VALUE! |
| ENSMUSG00000108036.2 | 0 | NA | NA | NA | NA | NA | #VALUE! |
| ENSMUSG00000108037.3 | 0 | NA | NA | NA | NA | NA | #VALUE! |
| ENSMUSG00000108038.2 | 0 | NA | NA | NA | NA | NA | #VALUE! |
| ENSMUSG00000108039.2 | 0 | NA | NA | NA | NA | NA | #VALUE! |
| ENSMUSG00000108040.2 | 0 | NA | NA | NA | NA | NA | #VALUE! |
| ENSMUSG00000108041.2 | 0 | NA | NA | NA | NA | NA | #VALUE! |
| ENSMUSG00000108042.2 | 0 | NA | NA | NA | NA | NA | #VALUE! |
| ENSMUSG00000108043.3 | 0 | NA | NA | NA | NA | NA | #VALUE! |
| ENSMUSG00000108044.2 | 0 | NA | NA | NA | NA | NA | #VALUE! |
| ENSMUSG00000108045.2 | 0 | NA | NA | NA | NA | NA | #VALUE! |
| ENSMUSG00000108046.2 | 0 | NA | NA | NA | NA | NA | #VALUE! |

| | | | | | | | |
|---|---|---|---|---|---|---|---|
| ENSMUSG00000108047.2 | 0 | NA | NA | NA | NA | NA | #VALUE! |
| ENSMUSG00000108048.2 | 0 | NA | NA | NA | NA | NA | #VALUE! |
| ENSMUSG00000108049.2 | 0 | NA | NA | NA | NA | NA | #VALUE! |
| ENSMUSG00000108050.2 | 0 | NA | NA | NA | NA | NA | #VALUE! |
| ENSMUSG00000108052.2 | 0 | NA | NA | NA | NA | NA | #VALUE! |
| ENSMUSG00000108053.2 | 0 | NA | NA | NA | NA | NA | #VALUE! |
| ENSMUSG00000108054.2 | 0 | NA | NA | NA | NA | NA | #VALUE! |
| ENSMUSG00000108055.2 | 0 | NA | NA | NA | NA | NA | #VALUE! |
| ENSMUSG00000108056.2 | 0 | NA | NA | NA | NA | NA | #VALUE! |
| ENSMUSG00000108057.2 | 0 | NA | NA | NA | NA | NA | #VALUE! |
| ENSMUSG00000108058.2 | 0 | NA | NA | NA | NA | NA | #VALUE! |
| ENSMUSG00000108059.2 | 0 | NA | NA | NA | NA | NA | #VALUE! |
| ENSMUSG00000108060.3 | 0 | NA | NA | NA | NA | NA | #VALUE! |
| ENSMUSG00000108062.2 | 0 | NA | NA | NA | NA | NA | #VALUE! |
| ENSMUSG00000108063.2 | 0 | NA | NA | NA | NA | NA | #VALUE! |
| ENSMUSG00000108064.2 | 0 | NA | NA | NA | NA | NA | #VALUE! |
| ENSMUSG00000108065.2 | 0 | NA | NA | NA | NA | NA | #VALUE! |
| ENSMUSG00000108066.3 | 0 | NA | NA | NA | NA | NA | #VALUE! |
| ENSMUSG00000108067.2 | 0 | NA | NA | NA | NA | NA | #VALUE! |
| ENSMUSG00000108068.2 | 0 | NA | NA | NA | NA | NA | #VALUE! |
| ENSMUSG00000108069.2 | 0 | NA | NA | NA | NA | NA | #VALUE! |
| ENSMUSG00000108070.2 | 0 | NA | NA | NA | NA | NA | #VALUE! |
| ENSMUSG00000108071.2 | 0 | NA | NA | NA | NA | NA | #VALUE! |
| ENSMUSG00000108072.2 | 0 | NA | NA | NA | NA | NA | #VALUE! |
| ENSMUSG00000108073.2 | 0 | NA | NA | NA | NA | NA | #VALUE! |
| ENSMUSG00000108074.2 | 0 | NA | NA | NA | NA | NA | #VALUE! |
| ENSMUSG00000108075.2 | 0 | NA | NA | NA | NA | NA | #VALUE! |
| ENSMUSG00000108076.2 | 0 | NA | NA | NA | NA | NA | #VALUE! |
| ENSMUSG00000108077.2 | 0 | NA | NA | NA | NA | NA | #VALUE! |
| ENSMUSG00000108078.2 | 0 | NA | NA | NA | NA | NA | #VALUE! |
| ENSMUSG00000108079.2 | 0 | NA | NA | NA | NA | NA | #VALUE! |
| ENSMUSG00000108080.2 | 0 | NA | NA | NA | NA | NA | #VALUE! |
| ENSMUSG00000108081.2 | 0 | NA | NA | NA | NA | NA | #VALUE! |
| ENSMUSG00000108082.2 | 0 | NA | NA | NA | NA | NA | #VALUE! |
| ENSMUSG00000108083.2 | 2.039864 | -0.30045 | 3.816883 | -0.07872 | 0.937259 | NA | #VALUE! |
| ENSMUSG00000108084.2 | 0 | NA | NA | NA | NA | NA | #VALUE! |
| ENSMUSG00000108085.2 | 0 | NA | NA | NA | NA | NA | #VALUE! |
| ENSMUSG00000108086.2 | 0.093255 | 0.868093 | 4.072464 | 0.213162 | 0.831201 | NA | #VALUE! |
| ENSMUSG00000108087.2 | 0 | NA | NA | NA | NA | NA | #VALUE! |
| ENSMUSG00000108088.2 | 0 | NA | NA | NA | NA | NA | #VALUE! |
| ENSMUSG00000108089.2 | 0 | NA | NA | NA | NA | NA | #VALUE! |
| ENSMUSG00000108090.2 | 0 | NA | NA | NA | NA | NA | #VALUE! |
| ENSMUSG00000108091.2 | 0 | NA | NA | NA | NA | NA | #VALUE! |
| ENSMUSG00000108092.2 | 0 | NA | NA | NA | NA | NA | #VALUE! |

| | | | | | | | |
|---|---|---|---|---|---|---|---|
| ENSMUSG00000108093.2 | 0 | NA | NA | NA | NA | NA | #VALUE! |
| ENSMUSG00000108094.2 | 0 | NA | NA | NA | NA | NA | #VALUE! |
| ENSMUSG00000108095.2 | 0 | NA | NA | NA | NA | NA | #VALUE! |
| ENSMUSG00000108096.2 | 0 | NA | NA | NA | NA | NA | #VALUE! |
| ENSMUSG00000108097.2 | 0 | NA | NA | NA | NA | NA | #VALUE! |
| ENSMUSG00000108098.2 | 0 | NA | NA | NA | NA | NA | #VALUE! |
| ENSMUSG00000108099.2 | 0 | NA | NA | NA | NA | NA | #VALUE! |
| ENSMUSG00000108100.2 | 0 | NA | NA | NA | NA | NA | #VALUE! |
| ENSMUSG00000108101.2 | 0 | NA | NA | NA | NA | NA | #VALUE! |
| ENSMUSG00000108102.2 | 0 | NA | NA | NA | NA | NA | #VALUE! |
| ENSMUSG00000108103.2 | 0 | NA | NA | NA | NA | NA | #VALUE! |
| ENSMUSG00000108104.2 | 0 | NA | NA | NA | NA | NA | #VALUE! |
| ENSMUSG00000108105.2 | 0 | NA | NA | NA | NA | NA | #VALUE! |
| ENSMUSG00000108106.3 | 0 | NA | NA | NA | NA | NA | #VALUE! |
| ENSMUSG00000108107.3 | 0 | NA | NA | NA | NA | NA | #VALUE! |
| ENSMUSG00000108108.2 | 0 | NA | NA | NA | NA | NA | #VALUE! |
| ENSMUSG00000108109.2 | 0 | NA | NA | NA | NA | NA | #VALUE! |
| ENSMUSG00000108110.2 | 0 | NA | NA | NA | NA | NA | #VALUE! |
| ENSMUSG00000108111.2 | 0 | NA | NA | NA | NA | NA | #VALUE! |
| ENSMUSG00000108112.2 | 0 | NA | NA | NA | NA | NA | #VALUE! |
| ENSMUSG00000108113.3 | 0 | NA | NA | NA | NA | NA | #VALUE! |
| ENSMUSG00000108114.3 | 0 | NA | NA | NA | NA | NA | #VALUE! |
| ENSMUSG00000108115.2 | 0 | NA | NA | NA | NA | NA | #VALUE! |
| ENSMUSG00000108117.2 | 0 | NA | NA | NA | NA | NA | #VALUE! |
| ENSMUSG00000108118.2 | 0 | NA | NA | NA | NA | NA | #VALUE! |
| ENSMUSG00000108119.2 | 0 | NA | NA | NA | NA | NA | #VALUE! |
| ENSMUSG00000108120.2 | 0 | NA | NA | NA | NA | NA | #VALUE! |
| ENSMUSG00000108121.2 | 0 | NA | NA | NA | NA | NA | #VALUE! |
| ENSMUSG00000108122.2 | 0 | NA | NA | NA | NA | NA | #VALUE! |
| ENSMUSG00000108123.2 | 0 | NA | NA | NA | NA | NA | #VALUE! |
| ENSMUSG00000108124.2 | 0 | NA | NA | NA | NA | NA | #VALUE! |
| ENSMUSG00000108125.2 | 0 | NA | NA | NA | NA | NA | #VALUE! |
| ENSMUSG00000108126.2 | 0 | NA | NA | NA | NA | NA | #VALUE! |
| ENSMUSG00000108127.2 | 0 | NA | NA | NA | NA | NA | #VALUE! |
| ENSMUSG00000108128.2 | 0 | NA | NA | NA | NA | NA | #VALUE! |
| ENSMUSG00000108129.3 | 0 | NA | NA | NA | NA | NA | #VALUE! |
| ENSMUSG00000108130.2 | 0 | NA | NA | NA | NA | NA | #VALUE! |
| ENSMUSG00000108131.2 | 0 | NA | NA | NA | NA | NA | #VALUE! |
| ENSMUSG00000108132.2 | 0 | NA | NA | NA | NA | NA | #VALUE! |
| ENSMUSG00000108134.2 | 0 | NA | NA | NA | NA | NA | #VALUE! |
| ENSMUSG00000108135.2 | 0 | NA | NA | NA | NA | NA | #VALUE! |
| ENSMUSG00000108137.2 | 0 | NA | NA | NA | NA | NA | #VALUE! |
| ENSMUSG00000108138.2 | 0 | NA | NA | NA | NA | NA | #VALUE! |
| ENSMUSG00000108139.2 | 0 | NA | NA | NA | NA | NA | #VALUE! |

| | | | | | | | |
|---|---|---|---|---|---|---|---|
| ENSMUSG00000108140.2 | 0 | NA | NA | NA | NA | NA | #VALUE! |
| ENSMUSG00000108141.2 | 0 | NA | NA | NA | NA | NA | #VALUE! |
| ENSMUSG00000108142.2 | 0 | NA | NA | NA | NA | NA | #VALUE! |
| ENSMUSG00000108143.2 | 0 | NA | NA | NA | NA | NA | #VALUE! |
| ENSMUSG00000108144.2 | 0 | NA | NA | NA | NA | NA | #VALUE! |
| ENSMUSG00000108145.2 | 0 | NA | NA | NA | NA | NA | #VALUE! |
| ENSMUSG00000108146.2 | 0 | NA | NA | NA | NA | NA | #VALUE! |
| ENSMUSG00000108147.2 | 0 | NA | NA | NA | NA | NA | #VALUE! |
| ENSMUSG00000108148.2 | 0 | NA | NA | NA | NA | NA | #VALUE! |
| ENSMUSG00000108149.2 | 0 | NA | NA | NA | NA | NA | #VALUE! |
| ENSMUSG00000108150.2 | 0 | NA | NA | NA | NA | NA | #VALUE! |
| ENSMUSG00000108151.2 | 0 | NA | NA | NA | NA | NA | #VALUE! |
| ENSMUSG00000108152.2 | 0 | NA | NA | NA | NA | NA | #VALUE! |
| ENSMUSG00000108153.2 | 0 | NA | NA | NA | NA | NA | #VALUE! |
| ENSMUSG00000108154.2 | 0 | NA | NA | NA | NA | NA | #VALUE! |
| ENSMUSG00000108155.2 | 0 | NA | NA | NA | NA | NA | #VALUE! |
| ENSMUSG00000108156.2 | 0 | NA | NA | NA | NA | NA | #VALUE! |
| ENSMUSG00000108157.2 | 0 | NA | NA | NA | NA | NA | #VALUE! |
| ENSMUSG00000108158.2 | 0 | NA | NA | NA | NA | NA | #VALUE! |
| ENSMUSG00000108160.2 | 0 | NA | NA | NA | NA | NA | #VALUE! |
| ENSMUSG00000108161.2 | 0 | NA | NA | NA | NA | NA | #VALUE! |
| ENSMUSG00000108162.2 | 0 | NA | NA | NA | NA | NA | #VALUE! |
| ENSMUSG00000108163.2 | 0 | NA | NA | NA | NA | NA | #VALUE! |
| ENSMUSG00000108164.2 | 0 | NA | NA | NA | NA | NA | #VALUE! |
| ENSMUSG00000108166.2 | 0 | NA | NA | NA | NA | NA | #VALUE! |
| ENSMUSG00000108167.3 | 0 | NA | NA | NA | NA | NA | #VALUE! |
| ENSMUSG00000108168.2 | 0 | NA | NA | NA | NA | NA | #VALUE! |
| ENSMUSG00000108169.2 | 0 | NA | NA | NA | NA | NA | #VALUE! |
| ENSMUSG00000108170.2 | 1.370524 | -3.97623 | 3.939448 | -1.00934 | 0.312813 | NA | #VALUE! |
| ENSMUSG00000108171.2 | 0 | NA | NA | NA | NA | NA | #VALUE! |
| ENSMUSG00000108172.2 | 0 | NA | NA | NA | NA | NA | #VALUE! |
| ENSMUSG00000108173.2 | 0 | NA | NA | NA | NA | NA | #VALUE! |
| ENSMUSG00000108174.2 | 0 | NA | NA | NA | NA | NA | #VALUE! |
| ENSMUSG00000108175.2 | 0 | NA | NA | NA | NA | NA | #VALUE! |
| ENSMUSG00000108176.2 | 0 | NA | NA | NA | NA | NA | #VALUE! |
| ENSMUSG00000108177.2 | 0 | NA | NA | NA | NA | NA | #VALUE! |
| ENSMUSG00000108178.2 | 0 | NA | NA | NA | NA | NA | #VALUE! |
| ENSMUSG00000108179.2 | 0 | NA | NA | NA | NA | NA | #VALUE! |
| ENSMUSG00000108180.2 | 0 | NA | NA | NA | NA | NA | #VALUE! |
| ENSMUSG00000108181.2 | 3.02641 | -0.93405 | 3.792645 | -0.24628 | 0.805466 | NA | #VALUE! |
| ENSMUSG00000108183.2 | 0 | NA | NA | NA | NA | NA | #VALUE! |
| ENSMUSG00000108184.2 | 0 | NA | NA | NA | NA | NA | #VALUE! |
| ENSMUSG00000108185.2 | 0 | NA | NA | NA | NA | NA | #VALUE! |
| ENSMUSG00000108186.2 | 0 | NA | NA | NA | NA | NA | #VALUE! |

| | | | | | | | |
|---|---|---|---|---|---|---|---|
| ENSMUSG00000108187.2 | 0 | NA | NA | NA | NA | NA | #VALUE! |
| ENSMUSG00000108188.2 | 0 | NA | NA | NA | NA | NA | #VALUE! |
| ENSMUSG00000108189.2 | 0 | NA | NA | NA | NA | NA | #VALUE! |
| ENSMUSG00000108190.2 | 0 | NA | NA | NA | NA | NA | #VALUE! |
| ENSMUSG00000108191.2 | 0 | NA | NA | NA | NA | NA | #VALUE! |
| ENSMUSG00000108192.2 | 0 | NA | NA | NA | NA | NA | #VALUE! |
| ENSMUSG00000108193.2 | 0 | NA | NA | NA | NA | NA | #VALUE! |
| ENSMUSG00000108194.2 | 0 | NA | NA | NA | NA | NA | #VALUE! |
| ENSMUSG00000108195.2 | 0 | NA | NA | NA | NA | NA | #VALUE! |
| ENSMUSG00000108196.2 | 0 | NA | NA | NA | NA | NA | #VALUE! |
| ENSMUSG00000108197.2 | 0 | NA | NA | NA | NA | NA | #VALUE! |
| ENSMUSG00000108198.2 | 0 | NA | NA | NA | NA | NA | #VALUE! |
| ENSMUSG00000108199.2 | 0 | NA | NA | NA | NA | NA | #VALUE! |
| ENSMUSG00000108200.2 | 0 | NA | NA | NA | NA | NA | #VALUE! |
| ENSMUSG00000108201.2 | 0 | NA | NA | NA | NA | NA | #VALUE! |
| ENSMUSG00000108202.2 | 0 | NA | NA | NA | NA | NA | #VALUE! |
| ENSMUSG00000108203.2 | 0 | NA | NA | NA | NA | NA | #VALUE! |
| ENSMUSG00000108204.2 | 0 | NA | NA | NA | NA | NA | #VALUE! |
| ENSMUSG00000108205.2 | 0 | NA | NA | NA | NA | NA | #VALUE! |
| ENSMUSG00000108206.2 | 0 | NA | NA | NA | NA | NA | #VALUE! |
| ENSMUSG00000108207.3 | 8.689929 | -4.33395 | 2.568654 | -1.68724 | 0.091556 | NA | #VALUE! |
| ENSMUSG00000108208.2 | 0 | NA | NA | NA | NA | NA | #VALUE! |
| ENSMUSG00000108209.2 | 0 | NA | NA | NA | NA | NA | #VALUE! |
| ENSMUSG00000108210.2 | 0 | NA | NA | NA | NA | NA | #VALUE! |
| ENSMUSG00000108211.2 | 0.559531 | 2.703958 | 3.982863 | 0.678898 | 0.497202 | NA | #VALUE! |
| ENSMUSG00000108212.2 | 0 | NA | NA | NA | NA | NA | #VALUE! |
| ENSMUSG00000108213.2 | 0 | NA | NA | NA | NA | NA | #VALUE! |
| ENSMUSG00000108214.2 | 0 | NA | NA | NA | NA | NA | #VALUE! |
| ENSMUSG00000108215.2 | 0 | NA | NA | NA | NA | NA | #VALUE! |
| ENSMUSG00000108216.2 | 0 | NA | NA | NA | NA | NA | #VALUE! |
| ENSMUSG00000108217.2 | 0 | NA | NA | NA | NA | NA | #VALUE! |
| ENSMUSG00000108218.3 | 0 | NA | NA | NA | NA | NA | #VALUE! |
| ENSMUSG00000108219.2 | 0 | NA | NA | NA | NA | NA | #VALUE! |
| ENSMUSG00000108220.2 | 0 | NA | NA | NA | NA | NA | #VALUE! |
| ENSMUSG00000108221.2 | 0 | NA | NA | NA | NA | NA | #VALUE! |
| ENSMUSG00000108222.2 | 0 | NA | NA | NA | NA | NA | #VALUE! |
| ENSMUSG00000108223.2 | 0 | NA | NA | NA | NA | NA | #VALUE! |
| ENSMUSG00000108224.2 | 0 | NA | NA | NA | NA | NA | #VALUE! |
| ENSMUSG00000108225.2 | 0 | NA | NA | NA | NA | NA | #VALUE! |
| ENSMUSG00000108226.2 | 0 | NA | NA | NA | NA | NA | #VALUE! |
| ENSMUSG00000108227.2 | 0 | NA | NA | NA | NA | NA | #VALUE! |
| ENSMUSG00000108228.3 | 0 | NA | NA | NA | NA | NA | #VALUE! |
| ENSMUSG00000108229.2 | 0 | NA | NA | NA | NA | NA | #VALUE! |
| ENSMUSG00000108230.2 | 0 | NA | NA | NA | NA | NA | #VALUE! |

| | | | | | | | |
|---|---|---|---|---|---|---|---|
| ENSMUSG00000108232.2 | 0 | NA | NA | NA | NA | NA | #VALUE! |
| ENSMUSG00000108233.2 | 0 | NA | NA | NA | NA | NA | #VALUE! |
| ENSMUSG00000108234.2 | 0 | NA | NA | NA | NA | NA | #VALUE! |
| ENSMUSG00000108235.2 | 0 | NA | NA | NA | NA | NA | #VALUE! |
| ENSMUSG00000108236.2 | 0 | NA | NA | NA | NA | NA | #VALUE! |
| ENSMUSG00000108237.2 | 0 | NA | NA | NA | NA | NA | #VALUE! |
| ENSMUSG00000108238.2 | 0.168284 | 1.027628 | 4.067126 | 0.252667 | 0.800526 | NA | #VALUE! |
| ENSMUSG00000108239.2 | 0 | NA | NA | NA | NA | NA | #VALUE! |
| ENSMUSG00000108240.2 | 0 | NA | NA | NA | NA | NA | #VALUE! |
| ENSMUSG00000108241.2 | 0 | NA | NA | NA | NA | NA | #VALUE! |
| ENSMUSG00000108242.2 | 0 | NA | NA | NA | NA | NA | #VALUE! |
| ENSMUSG00000108244.2 | 0 | NA | NA | NA | NA | NA | #VALUE! |
| ENSMUSG00000108245.2 | 0 | NA | NA | NA | NA | NA | #VALUE! |
| ENSMUSG00000108246.2 | 0 | NA | NA | NA | NA | NA | #VALUE! |
| ENSMUSG00000108247.2 | 0 | NA | NA | NA | NA | NA | #VALUE! |
| ENSMUSG00000108248.2 | 0 | NA | NA | NA | NA | NA | #VALUE! |
| ENSMUSG00000108249.2 | 0 | NA | NA | NA | NA | NA | #VALUE! |
| ENSMUSG00000108250.2 | 0 | NA | NA | NA | NA | NA | #VALUE! |
| ENSMUSG00000108251.2 | 0 | NA | NA | NA | NA | NA | #VALUE! |
| ENSMUSG00000108252.2 | 0 | NA | NA | NA | NA | NA | #VALUE! |
| ENSMUSG00000108253.2 | 0 | NA | NA | NA | NA | NA | #VALUE! |
| ENSMUSG00000108254.2 | 0 | NA | NA | NA | NA | NA | #VALUE! |
| ENSMUSG00000108255.2 | 0 | NA | NA | NA | NA | NA | #VALUE! |
| ENSMUSG00000108256.2 | 0 | NA | NA | NA | NA | NA | #VALUE! |
| ENSMUSG00000108257.2 | 0 | NA | NA | NA | NA | NA | #VALUE! |
| ENSMUSG00000108258.2 | 0 | NA | NA | NA | NA | NA | #VALUE! |
| ENSMUSG00000108259.2 | 0 | NA | NA | NA | NA | NA | #VALUE! |
| ENSMUSG00000108260.2 | 0 | NA | NA | NA | NA | NA | #VALUE! |
| ENSMUSG00000108261.2 | 0 | NA | NA | NA | NA | NA | #VALUE! |
| ENSMUSG00000108262.2 | 0 | NA | NA | NA | NA | NA | #VALUE! |
| ENSMUSG00000108263.2 | 0 | NA | NA | NA | NA | NA | #VALUE! |
| ENSMUSG00000108264.2 | 0 | NA | NA | NA | NA | NA | #VALUE! |
| ENSMUSG00000108265.3 | 0 | NA | NA | NA | NA | NA | #VALUE! |
| ENSMUSG00000108266.2 | 0 | NA | NA | NA | NA | NA | #VALUE! |
| ENSMUSG00000108267.2 | 0 | NA | NA | NA | NA | NA | #VALUE! |
| ENSMUSG00000108268.2 | 0 | NA | NA | NA | NA | NA | #VALUE! |
| ENSMUSG00000108269.2 | 0 | NA | NA | NA | NA | NA | #VALUE! |
| ENSMUSG00000108270.2 | 0 | NA | NA | NA | NA | NA | #VALUE! |
| ENSMUSG00000108271.2 | 0 | NA | NA | NA | NA | NA | #VALUE! |
| ENSMUSG00000108272.2 | 0 | NA | NA | NA | NA | NA | #VALUE! |
| ENSMUSG00000108274.2 | 0 | NA | NA | NA | NA | NA | #VALUE! |
| ENSMUSG00000108275.2 | 0 | NA | NA | NA | NA | NA | #VALUE! |
| ENSMUSG00000108276.2 | 0 | NA | NA | NA | NA | NA | #VALUE! |
| ENSMUSG00000108278.2 | 0 | NA | NA | NA | NA | NA | #VALUE! |

| | | | | | | | |
|---|---|---|---|---|---|---|---|
| ENSMUSG00000108279.2 | 0 | NA | NA | NA | NA | NA | #VALUE! |
| ENSMUSG00000108280.2 | 0 | NA | NA | NA | NA | NA | #VALUE! |
| ENSMUSG00000108282.2 | 0 | NA | NA | NA | NA | NA | #VALUE! |
| ENSMUSG00000108283.2 | 0 | NA | NA | NA | NA | NA | #VALUE! |
| ENSMUSG00000108284.2 | 0 | NA | NA | NA | NA | NA | #VALUE! |
| ENSMUSG00000108285.2 | 0 | NA | NA | NA | NA | NA | #VALUE! |
| ENSMUSG00000108286.2 | 0 | NA | NA | NA | NA | NA | #VALUE! |
| ENSMUSG00000108287.2 | 0 | NA | NA | NA | NA | NA | #VALUE! |
| ENSMUSG00000108288.2 | 0 | NA | NA | NA | NA | NA | #VALUE! |
| ENSMUSG00000108289.2 | 0 | NA | NA | NA | NA | NA | #VALUE! |
| ENSMUSG00000108290.2 | 0 | NA | NA | NA | NA | NA | #VALUE! |
| ENSMUSG00000108291.2 | 0 | NA | NA | NA | NA | NA | #VALUE! |
| ENSMUSG00000108292.2 | 0 | NA | NA | NA | NA | NA | #VALUE! |
| ENSMUSG00000108293.2 | 0 | NA | NA | NA | NA | NA | #VALUE! |
| ENSMUSG00000108294.2 | 0 | NA | NA | NA | NA | NA | #VALUE! |
| ENSMUSG00000108295.3 | 0 | NA | NA | NA | NA | NA | #VALUE! |
| ENSMUSG00000108296.2 | 0 | NA | NA | NA | NA | NA | #VALUE! |
| ENSMUSG00000108297.2 | 0 | NA | NA | NA | NA | NA | #VALUE! |
| ENSMUSG00000108298.2 | 0 | NA | NA | NA | NA | NA | #VALUE! |
| ENSMUSG00000108299.2 | 0 | NA | NA | NA | NA | NA | #VALUE! |
| ENSMUSG00000108300.2 | 0 | NA | NA | NA | NA | NA | #VALUE! |
| ENSMUSG00000108301.2 | 0 | NA | NA | NA | NA | NA | #VALUE! |
| ENSMUSG00000108302.2 | 0 | NA | NA | NA | NA | NA | #VALUE! |
| ENSMUSG00000108303.2 | 0 | NA | NA | NA | NA | NA | #VALUE! |
| ENSMUSG00000108304.2 | 0 | NA | NA | NA | NA | NA | #VALUE! |
| ENSMUSG00000108305.2 | 0 | NA | NA | NA | NA | NA | #VALUE! |
| ENSMUSG00000108306.2 | 0 | NA | NA | NA | NA | NA | #VALUE! |
| ENSMUSG00000108307.2 | 0 | NA | NA | NA | NA | NA | #VALUE! |
| ENSMUSG00000108308.2 | 0 | NA | NA | NA | NA | NA | #VALUE! |
| ENSMUSG00000108309.2 | 0 | NA | NA | NA | NA | NA | #VALUE! |
| ENSMUSG00000108310.2 | 0 | NA | NA | NA | NA | NA | #VALUE! |
| ENSMUSG00000108311.2 | 0 | NA | NA | NA | NA | NA | #VALUE! |
| ENSMUSG00000108312.2 | 0 | NA | NA | NA | NA | NA | #VALUE! |
| ENSMUSG00000108313.2 | 0 | NA | NA | NA | NA | NA | #VALUE! |
| ENSMUSG00000108314.2 | 0.198268 | -0.06494 | 4.068276 | -0.01596 | 0.987265 | NA | #VALUE! |
| ENSMUSG00000108315.2 | 0 | NA | NA | NA | NA | NA | #VALUE! |
| ENSMUSG00000108316.2 | 0 | NA | NA | NA | NA | NA | #VALUE! |
| ENSMUSG00000108317.2 | 0 | NA | NA | NA | NA | NA | #VALUE! |
| ENSMUSG00000108318.2 | 0 | NA | NA | NA | NA | NA | #VALUE! |
| ENSMUSG00000108319.2 | 0 | NA | NA | NA | NA | NA | #VALUE! |
| ENSMUSG00000108320.2 | 0 | NA | NA | NA | NA | NA | #VALUE! |
| ENSMUSG00000108321.2 | 0 | NA | NA | NA | NA | NA | #VALUE! |
| ENSMUSG00000108322.3 | 0 | NA | NA | NA | NA | NA | #VALUE! |
| ENSMUSG00000108323.2 | 0 | NA | NA | NA | NA | NA | #VALUE! |

| | | | | | | | |
|---|---|---|---|---|---|---|---|
| ENSMUSG00000108324.2 | 0 | NA | NA | NA | NA | NA | #VALUE! |
| ENSMUSG00000108325.2 | 0 | NA | NA | NA | NA | NA | #VALUE! |
| ENSMUSG00000108326.2 | 0 | NA | NA | NA | NA | NA | #VALUE! |
| ENSMUSG00000108327.2 | 0 | NA | NA | NA | NA | NA | #VALUE! |
| ENSMUSG00000108328.2 | 0 | NA | NA | NA | NA | NA | #VALUE! |
| ENSMUSG00000108329.2 | 0 | NA | NA | NA | NA | NA | #VALUE! |
| ENSMUSG00000108330.2 | 0 | NA | NA | NA | NA | NA | #VALUE! |
| ENSMUSG00000108331.2 | 0 | NA | NA | NA | NA | NA | #VALUE! |
| ENSMUSG00000108332.2 | 8.821103 | -6.67439 | 3.721088 | -1.79367 | NA | NA | #VALUE! |
| ENSMUSG00000108333.2 | 0 | NA | NA | NA | NA | NA | #VALUE! |
| ENSMUSG00000108334.2 | 0 | NA | NA | NA | NA | NA | #VALUE! |
| ENSMUSG00000108335.2 | 0 | NA | NA | NA | NA | NA | #VALUE! |
| ENSMUSG00000108336.2 | 0 | NA | NA | NA | NA | NA | #VALUE! |
| ENSMUSG00000108337.2 | 0 | NA | NA | NA | NA | NA | #VALUE! |
| ENSMUSG00000108338.2 | 0 | NA | NA | NA | NA | NA | #VALUE! |
| ENSMUSG00000108339.2 | 0 | NA | NA | NA | NA | NA | #VALUE! |
| ENSMUSG00000108340.2 | 0 | NA | NA | NA | NA | NA | #VALUE! |
| ENSMUSG00000108341.2 | 0 | NA | NA | NA | NA | NA | #VALUE! |
| ENSMUSG00000108342.2 | 0 | NA | NA | NA | NA | NA | #VALUE! |
| ENSMUSG00000108343.2 | 0 | NA | NA | NA | NA | NA | #VALUE! |
| ENSMUSG00000108344.2 | 0 | NA | NA | NA | NA | NA | #VALUE! |
| ENSMUSG00000108345.2 | 0 | NA | NA | NA | NA | NA | #VALUE! |
| ENSMUSG00000108346.2 | 0 | NA | NA | NA | NA | NA | #VALUE! |
| ENSMUSG00000108347.2 | 0 | NA | NA | NA | NA | NA | #VALUE! |
| ENSMUSG00000108348.3 | 0 | NA | NA | NA | NA | NA | #VALUE! |
| ENSMUSG00000108349.2 | 0 | NA | NA | NA | NA | NA | #VALUE! |
| ENSMUSG00000108351.2 | 0 | NA | NA | NA | NA | NA | #VALUE! |
| ENSMUSG00000108352.2 | 0 | NA | NA | NA | NA | NA | #VALUE! |
| ENSMUSG00000108353.2 | 0 | NA | NA | NA | NA | NA | #VALUE! |
| ENSMUSG00000108354.2 | 0 | NA | NA | NA | NA | NA | #VALUE! |
| ENSMUSG00000108355.2 | 0 | NA | NA | NA | NA | NA | #VALUE! |
| ENSMUSG00000108356.2 | 0 | NA | NA | NA | NA | NA | #VALUE! |
| ENSMUSG00000108357.2 | 0 | NA | NA | NA | NA | NA | #VALUE! |
| ENSMUSG00000108358.2 | 4.327187 | 3.349311 | 3.856139 | 0.868566 | 0.385085 | NA | #VALUE! |
| ENSMUSG00000108359.2 | 56.38726 | -23.4552 | 3.90756 | -6.00253 | NA | NA | #VALUE! |
| ENSMUSG00000108360.2 | 0 | NA | NA | NA | NA | NA | #VALUE! |
| ENSMUSG00000108361.2 | 0 | NA | NA | NA | NA | NA | #VALUE! |
| ENSMUSG00000108362.2 | 0 | NA | NA | NA | NA | NA | #VALUE! |
| ENSMUSG00000108363.2 | 0 | NA | NA | NA | NA | NA | #VALUE! |
| ENSMUSG00000108364.2 | 0 | NA | NA | NA | NA | NA | #VALUE! |
| ENSMUSG00000108365.2 | 0 | NA | NA | NA | NA | NA | #VALUE! |
| ENSMUSG00000108367.3 | 0 | NA | NA | NA | NA | NA | #VALUE! |
| ENSMUSG00000108368.2 | 0 | NA | NA | NA | NA | NA | #VALUE! |
| ENSMUSG00000108370.2 | 0 | NA | NA | NA | NA | NA | #VALUE! |

| | | | | | | | |
|---|---|---|---|---|---|---|---|
| ENSMUSG00000108371.2 | 2.019402 | 4.503826 | 3.928936 | 1.146322 | 0.251662 | NA | #VALUE! |
| ENSMUSG00000108372.2 | 0 | NA | NA | NA | NA | NA | #VALUE! |
| ENSMUSG00000108373.2 | 0 | NA | NA | NA | NA | NA | #VALUE! |
| ENSMUSG00000108374.2 | 0 | NA | NA | NA | NA | NA | #VALUE! |
| ENSMUSG00000108375.2 | 0 | NA | NA | NA | NA | NA | #VALUE! |
| ENSMUSG00000108376.2 | 0 | NA | NA | NA | NA | NA | #VALUE! |
| ENSMUSG00000108377.2 | 0 | NA | NA | NA | NA | NA | #VALUE! |
| ENSMUSG00000108378.2 | 0 | NA | NA | NA | NA | NA | #VALUE! |
| ENSMUSG00000108379.2 | 0 | NA | NA | NA | NA | NA | #VALUE! |
| ENSMUSG00000108380.2 | 0 | NA | NA | NA | NA | NA | #VALUE! |
| ENSMUSG00000108381.2 | 0 | NA | NA | NA | NA | NA | #VALUE! |
| ENSMUSG00000108382.2 | 0 | NA | NA | NA | NA | NA | #VALUE! |
| ENSMUSG00000108383.2 | 0 | NA | NA | NA | NA | NA | #VALUE! |
| ENSMUSG00000108384.2 | 0 | NA | NA | NA | NA | NA | #VALUE! |
| ENSMUSG00000108385.2 | 0 | NA | NA | NA | NA | NA | #VALUE! |
| ENSMUSG00000108387.2 | 0 | NA | NA | NA | NA | NA | #VALUE! |
| ENSMUSG00000108388.2 | 0 | NA | NA | NA | NA | NA | #VALUE! |
| ENSMUSG00000108389.2 | 0.336567 | 1.944925 | 4.029936 | 0.482619 | 0.629366 | NA | #VALUE! |
| ENSMUSG00000108390.2 | 0 | NA | NA | NA | NA | NA | #VALUE! |
| ENSMUSG00000108391.2 | 0 | NA | NA | NA | NA | NA | #VALUE! |
| ENSMUSG00000108392.2 | 0 | NA | NA | NA | NA | NA | #VALUE! |
| ENSMUSG00000108393.2 | 0 | NA | NA | NA | NA | NA | #VALUE! |
| ENSMUSG00000108394.2 | 0 | NA | NA | NA | NA | NA | #VALUE! |
| ENSMUSG00000108395.2 | 0 | NA | NA | NA | NA | NA | #VALUE! |
| ENSMUSG00000108396.2 | 0 | NA | NA | NA | NA | NA | #VALUE! |
| ENSMUSG00000108397.2 | 0 | NA | NA | NA | NA | NA | #VALUE! |
| ENSMUSG00000108398.2 | 0 | NA | NA | NA | NA | NA | #VALUE! |
| ENSMUSG00000108400.2 | 0 | NA | NA | NA | NA | NA | #VALUE! |
| ENSMUSG00000108401.2 | 0 | NA | NA | NA | NA | NA | #VALUE! |
| ENSMUSG00000108402.2 | 0 | NA | NA | NA | NA | NA | #VALUE! |
| ENSMUSG00000108403.2 | 0 | NA | NA | NA | NA | NA | #VALUE! |
| ENSMUSG00000108404.2 | 0 | NA | NA | NA | NA | NA | #VALUE! |
| ENSMUSG00000108405.2 | 0 | NA | NA | NA | NA | NA | #VALUE! |
| ENSMUSG00000108406.2 | 0 | NA | NA | NA | NA | NA | #VALUE! |
| ENSMUSG00000108407.2 | 0 | NA | NA | NA | NA | NA | #VALUE! |
| ENSMUSG00000108409.2 | 0 | NA | NA | NA | NA | NA | #VALUE! |
| ENSMUSG00000108410.2 | 0 | NA | NA | NA | NA | NA | #VALUE! |
| ENSMUSG00000108411.2 | 0 | NA | NA | NA | NA | NA | #VALUE! |
| ENSMUSG00000108412.2 | 0 | NA | NA | NA | NA | NA | #VALUE! |
| ENSMUSG00000108413.2 | 0 | NA | NA | NA | NA | NA | #VALUE! |
| ENSMUSG00000108415.2 | 0 | NA | NA | NA | NA | NA | #VALUE! |
| ENSMUSG00000108416.2 | 0 | NA | NA | NA | NA | NA | #VALUE! |
| ENSMUSG00000108417.2 | 0 | NA | NA | NA | NA | NA | #VALUE! |
| ENSMUSG00000108418.2 | 0 | NA | NA | NA | NA | NA | #VALUE! |

| | | | | | | | |
|---|---|---|---|---|---|---|---|
| ENSMUSG00000108419.2 | 0 | NA | NA | NA | NA | NA | #VALUE! |
| ENSMUSG00000108420.2 | 0 | NA | NA | NA | NA | NA | #VALUE! |
| ENSMUSG00000108421.2 | 0 | NA | NA | NA | NA | NA | #VALUE! |
| ENSMUSG00000108422.2 | 0 | NA | NA | NA | NA | NA | #VALUE! |
| ENSMUSG00000108423.2 | 0 | NA | NA | NA | NA | NA | #VALUE! |
| ENSMUSG00000108424.2 | 0 | NA | NA | NA | NA | NA | #VALUE! |
| ENSMUSG00000108425.2 | 0 | NA | NA | NA | NA | NA | #VALUE! |
| ENSMUSG00000108426.3 | 0 | NA | NA | NA | NA | NA | #VALUE! |
| ENSMUSG00000108427.2 | 0 | NA | NA | NA | NA | NA | #VALUE! |
| ENSMUSG00000108428.2 | 0 | NA | NA | NA | NA | NA | #VALUE! |
| ENSMUSG00000108429.2 | 0 | NA | NA | NA | NA | NA | #VALUE! |
| ENSMUSG00000108430.2 | 0 | NA | NA | NA | NA | NA | #VALUE! |
| ENSMUSG00000108431.2 | 0 | NA | NA | NA | NA | NA | #VALUE! |
| ENSMUSG00000108432.2 | 0 | NA | NA | NA | NA | NA | #VALUE! |
| ENSMUSG00000108433.2 | 0 | NA | NA | NA | NA | NA | #VALUE! |
| ENSMUSG00000108436.2 | 0 | NA | NA | NA | NA | NA | #VALUE! |
| ENSMUSG00000108437.2 | 0 | NA | NA | NA | NA | NA | #VALUE! |
| ENSMUSG00000108438.2 | 0 | NA | NA | NA | NA | NA | #VALUE! |
| ENSMUSG00000108440.2 | 0 | NA | NA | NA | NA | NA | #VALUE! |
| ENSMUSG00000108441.2 | 0 | NA | NA | NA | NA | NA | #VALUE! |
| ENSMUSG00000108442.2 | 2.611145 | 4.886731 | 3.92379 | 1.245411 | 0.212981 | NA | #VALUE! |
| ENSMUSG00000108443.2 | 0 | NA | NA | NA | NA | NA | #VALUE! |
| ENSMUSG00000108444.2 | 0 | NA | NA | NA | NA | NA | #VALUE! |
| ENSMUSG00000108445.2 | 0 | NA | NA | NA | NA | NA | #VALUE! |
| ENSMUSG00000108446.2 | 0.396537 | -0.08297 | 4.034442 | -0.02056 | 0.983593 | NA | #VALUE! |
| ENSMUSG00000108447.2 | 9.456435 | 1.068753 | 3.394624 | 0.314837 | 0.752886 | NA | #VALUE! |
| ENSMUSG00000108448.3 | 0 | NA | NA | NA | NA | NA | #VALUE! |
| ENSMUSG00000108449.2 | 0 | NA | NA | NA | NA | NA | #VALUE! |
| ENSMUSG00000108450.2 | 0 | NA | NA | NA | NA | NA | #VALUE! |
| ENSMUSG00000108451.2 | 0 | NA | NA | NA | NA | NA | #VALUE! |
| ENSMUSG00000108452.2 | 0 | NA | NA | NA | NA | NA | #VALUE! |
| ENSMUSG00000108454.2 | 0 | NA | NA | NA | NA | NA | #VALUE! |
| ENSMUSG00000108455.2 | 0 | NA | NA | NA | NA | NA | #VALUE! |
| ENSMUSG00000108456.2 | 0 | NA | NA | NA | NA | NA | #VALUE! |
| ENSMUSG00000108457.2 | 0 | NA | NA | NA | NA | NA | #VALUE! |
| ENSMUSG00000108459.2 | 0 | NA | NA | NA | NA | NA | #VALUE! |
| ENSMUSG00000108460.2 | 0 | NA | NA | NA | NA | NA | #VALUE! |
| ENSMUSG00000108461.2 | 0 | NA | NA | NA | NA | NA | #VALUE! |
| ENSMUSG00000108462.2 | 0 | NA | NA | NA | NA | NA | #VALUE! |
| ENSMUSG00000108464.6 | 0 | NA | NA | NA | NA | NA | #VALUE! |
| ENSMUSG00000108465.2 | 0 | NA | NA | NA | NA | NA | #VALUE! |
| ENSMUSG00000108466.2 | 0 | NA | NA | NA | NA | NA | #VALUE! |
| ENSMUSG00000108467.2 | 0 | NA | NA | NA | NA | NA | #VALUE! |
| ENSMUSG00000108468.2 | 0 | NA | NA | NA | NA | NA | #VALUE! |

| | | | | | | | |
|---|---|---|---|---|---|---|---|
| ENSMUSG00000108469.2 | 0 | NA | NA | NA | NA | NA | #VALUE! |
| ENSMUSG00000108470.2 | 0 | NA | NA | NA | NA | NA | #VALUE! |
| ENSMUSG00000108471.2 | 0 | NA | NA | NA | NA | NA | #VALUE! |
| ENSMUSG00000108472.2 | 0 | NA | NA | NA | NA | NA | #VALUE! |
| ENSMUSG00000108473.2 | 0.195789 | -1.05581 | 4.073302 | -0.2592 | 0.795479 | NA | #VALUE! |
| ENSMUSG00000108474.2 | 0 | NA | NA | NA | NA | NA | #VALUE! |
| ENSMUSG00000108476.2 | 0 | NA | NA | NA | NA | NA | #VALUE! |
| ENSMUSG00000108477.2 | 6.270208 | -2.45633 | 3.782229 | -0.64944 | 0.516054 | NA | #VALUE! |
| ENSMUSG00000108478.2 | 0 | NA | NA | NA | NA | NA | #VALUE! |
| ENSMUSG00000108479.2 | 0 | NA | NA | NA | NA | NA | #VALUE! |
| ENSMUSG00000108480.2 | 0 | NA | NA | NA | NA | NA | #VALUE! |
| ENSMUSG00000108481.2 | 0 | NA | NA | NA | NA | NA | #VALUE! |
| ENSMUSG00000108482.2 | 0 | NA | NA | NA | NA | NA | #VALUE! |
| ENSMUSG00000108483.2 | 0 | NA | NA | NA | NA | NA | #VALUE! |
| ENSMUSG00000108484.2 | 0 | NA | NA | NA | NA | NA | #VALUE! |
| ENSMUSG00000108485.2 | 0 | NA | NA | NA | NA | NA | #VALUE! |
| ENSMUSG00000108486.2 | 0 | NA | NA | NA | NA | NA | #VALUE! |
| ENSMUSG00000108487.2 | 0 | NA | NA | NA | NA | NA | #VALUE! |
| ENSMUSG00000108488.2 | 0 | NA | NA | NA | NA | NA | #VALUE! |
| ENSMUSG00000108489.2 | 0 | NA | NA | NA | NA | NA | #VALUE! |
| ENSMUSG00000108490.2 | 0 | NA | NA | NA | NA | NA | #VALUE! |
| ENSMUSG00000108491.2 | 0 | NA | NA | NA | NA | NA | #VALUE! |
| ENSMUSG00000108492.2 | 0 | NA | NA | NA | NA | NA | #VALUE! |
| ENSMUSG00000108493.2 | 0 | NA | NA | NA | NA | NA | #VALUE! |
| ENSMUSG00000108494.2 | 0 | NA | NA | NA | NA | NA | #VALUE! |
| ENSMUSG00000108495.2 | 0 | NA | NA | NA | NA | NA | #VALUE! |
| ENSMUSG00000108496.2 | 0 | NA | NA | NA | NA | NA | #VALUE! |
| ENSMUSG00000108497.2 | 0 | NA | NA | NA | NA | NA | #VALUE! |
| ENSMUSG00000108498.2 | 0 | NA | NA | NA | NA | NA | #VALUE! |
| ENSMUSG00000108500.2 | 0 | NA | NA | NA | NA | NA | #VALUE! |
| ENSMUSG00000108502.2 | 0 | NA | NA | NA | NA | NA | #VALUE! |
| ENSMUSG00000108503.2 | 0 | NA | NA | NA | NA | NA | #VALUE! |
| ENSMUSG00000108504.2 | 0 | NA | NA | NA | NA | NA | #VALUE! |
| ENSMUSG00000108505.2 | 0 | NA | NA | NA | NA | NA | #VALUE! |
| ENSMUSG00000108506.2 | 0 | NA | NA | NA | NA | NA | #VALUE! |
| ENSMUSG00000108507.2 | 0 | NA | NA | NA | NA | NA | #VALUE! |
| ENSMUSG00000108508.2 | 0 | NA | NA | NA | NA | NA | #VALUE! |
| ENSMUSG00000108509.2 | 0 | NA | NA | NA | NA | NA | #VALUE! |
| ENSMUSG00000108510.2 | 0 | NA | NA | NA | NA | NA | #VALUE! |
| ENSMUSG00000108511.2 | 0 | NA | NA | NA | NA | NA | #VALUE! |
| ENSMUSG00000108512.2 | 0 | NA | NA | NA | NA | NA | #VALUE! |
| ENSMUSG00000108513.2 | 0 | NA | NA | NA | NA | NA | #VALUE! |
| ENSMUSG00000108514.2 | 0 | NA | NA | NA | NA | NA | #VALUE! |
| ENSMUSG00000108515.2 | 0 | NA | NA | NA | NA | NA | #VALUE! |

| | | | | | | | |
|---|---|---|---|---|---|---|---|
| ENSMUSG00000108516.2 | 0 | NA | NA | NA | NA | NA | #VALUE! |
| ENSMUSG00000108517.2 | 0 | NA | NA | NA | NA | NA | #VALUE! |
| ENSMUSG00000108518.2 | 0 | NA | NA | NA | NA | NA | #VALUE! |
| ENSMUSG00000108519.2 | 0 | NA | NA | NA | NA | NA | #VALUE! |
| ENSMUSG00000108520.2 | 0 | NA | NA | NA | NA | NA | #VALUE! |
| ENSMUSG00000108521.2 | 0 | NA | NA | NA | NA | NA | #VALUE! |
| ENSMUSG00000108522.2 | 0 | NA | NA | NA | NA | NA | #VALUE! |
| ENSMUSG00000108523.2 | 0 | NA | NA | NA | NA | NA | #VALUE! |
| ENSMUSG00000108524.2 | 0 | NA | NA | NA | NA | NA | #VALUE! |
| ENSMUSG00000108525.2 | 0 | NA | NA | NA | NA | NA | #VALUE! |
| ENSMUSG00000108526.2 | 0 | NA | NA | NA | NA | NA | #VALUE! |
| ENSMUSG00000108527.2 | 0 | NA | NA | NA | NA | NA | #VALUE! |
| ENSMUSG00000108528.2 | 0 | NA | NA | NA | NA | NA | #VALUE! |
| ENSMUSG00000108529.2 | 0 | NA | NA | NA | NA | NA | #VALUE! |
| ENSMUSG00000108530.2 | 0 | NA | NA | NA | NA | NA | #VALUE! |
| ENSMUSG00000108531.2 | 0 | NA | NA | NA | NA | NA | #VALUE! |
| ENSMUSG00000108532.2 | 0 | NA | NA | NA | NA | NA | #VALUE! |
| ENSMUSG00000108533.2 | 0 | NA | NA | NA | NA | NA | #VALUE! |
| ENSMUSG00000108534.3 | 0 | NA | NA | NA | NA | NA | #VALUE! |
| ENSMUSG00000108535.2 | 0 | NA | NA | NA | NA | NA | #VALUE! |
| ENSMUSG00000108536.2 | 0 | NA | NA | NA | NA | NA | #VALUE! |
| ENSMUSG00000108537.2 | 0 | NA | NA | NA | NA | NA | #VALUE! |
| ENSMUSG00000108538.2 | 0 | NA | NA | NA | NA | NA | #VALUE! |
| ENSMUSG00000108539.2 | 0 | NA | NA | NA | NA | NA | #VALUE! |
| ENSMUSG00000108540.2 | 0 | NA | NA | NA | NA | NA | #VALUE! |
| ENSMUSG00000108541.2 | 0 | NA | NA | NA | NA | NA | #VALUE! |
| ENSMUSG00000108542.2 | 0 | NA | NA | NA | NA | NA | #VALUE! |
| ENSMUSG00000108543.2 | 0 | NA | NA | NA | NA | NA | #VALUE! |
| ENSMUSG00000108544.2 | 0 | NA | NA | NA | NA | NA | #VALUE! |
| ENSMUSG00000108545.2 | 0 | NA | NA | NA | NA | NA | #VALUE! |
| ENSMUSG00000108546.2 | 0 | NA | NA | NA | NA | NA | #VALUE! |
| ENSMUSG00000108547.2 | 0 | NA | NA | NA | NA | NA | #VALUE! |
| ENSMUSG00000108548.2 | 0 | NA | NA | NA | NA | NA | #VALUE! |
| ENSMUSG00000108550.2 | 0 | NA | NA | NA | NA | NA | #VALUE! |
| ENSMUSG00000108551.2 | 0 | NA | NA | NA | NA | NA | #VALUE! |
| ENSMUSG00000108552.2 | 0 | NA | NA | NA | NA | NA | #VALUE! |
| ENSMUSG00000108553.2 | 0 | NA | NA | NA | NA | NA | #VALUE! |
| ENSMUSG00000108554.2 | 0 | NA | NA | NA | NA | NA | #VALUE! |
| ENSMUSG00000108556.2 | 0 | NA | NA | NA | NA | NA | #VALUE! |
| ENSMUSG00000108557.2 | 0 | NA | NA | NA | NA | NA | #VALUE! |
| ENSMUSG00000108558.2 | 0 | NA | NA | NA | NA | NA | #VALUE! |
| ENSMUSG00000108559.2 | 0 | NA | NA | NA | NA | NA | #VALUE! |
| ENSMUSG00000108560.2 | 0 | NA | NA | NA | NA | NA | #VALUE! |
| ENSMUSG00000108561.2 | 0 | NA | NA | NA | NA | NA | #VALUE! |

| | | | | | | | |
|---|---|---|---|---|---|---|---|
| ENSMUSG00000108562.2 | 0 | NA | NA | NA | NA | NA | #VALUE! |
| ENSMUSG00000108563.2 | 0 | NA | NA | NA | NA | NA | #VALUE! |
| ENSMUSG00000108565.2 | 0 | NA | NA | NA | NA | NA | #VALUE! |
| ENSMUSG00000108566.2 | 0 | NA | NA | NA | NA | NA | #VALUE! |
| ENSMUSG00000108567.2 | 0 | NA | NA | NA | NA | NA | #VALUE! |
| ENSMUSG00000108568.2 | 0 | NA | NA | NA | NA | NA | #VALUE! |
| ENSMUSG00000108570.2 | 0 | NA | NA | NA | NA | NA | #VALUE! |
| ENSMUSG00000108571.2 | 0 | NA | NA | NA | NA | NA | #VALUE! |
| ENSMUSG00000108572.2 | 0 | NA | NA | NA | NA | NA | #VALUE! |
| ENSMUSG00000108573.2 | 0 | NA | NA | NA | NA | NA | #VALUE! |
| ENSMUSG00000108574.2 | 0 | NA | NA | NA | NA | NA | #VALUE! |
| ENSMUSG00000108575.2 | 0 | NA | NA | NA | NA | NA | #VALUE! |
| ENSMUSG00000108576.2 | 0 | NA | NA | NA | NA | NA | #VALUE! |
| ENSMUSG00000108577.2 | 0 | NA | NA | NA | NA | NA | #VALUE! |
| ENSMUSG00000108578.2 | 0 | NA | NA | NA | NA | NA | #VALUE! |
| ENSMUSG00000108579.2 | 0 | NA | NA | NA | NA | NA | #VALUE! |
| ENSMUSG00000108580.2 | 0 | NA | NA | NA | NA | NA | #VALUE! |
| ENSMUSG00000108581.2 | 0 | NA | NA | NA | NA | NA | #VALUE! |
| ENSMUSG00000108582.2 | 0 | NA | NA | NA | NA | NA | #VALUE! |
| ENSMUSG00000108583.2 | 0 | NA | NA | NA | NA | NA | #VALUE! |
| ENSMUSG00000108584.2 | 0 | NA | NA | NA | NA | NA | #VALUE! |
| ENSMUSG00000108586.2 | 0 | NA | NA | NA | NA | NA | #VALUE! |
| ENSMUSG00000108587.2 | 0 | NA | NA | NA | NA | NA | #VALUE! |
| ENSMUSG00000108588.2 | 0 | NA | NA | NA | NA | NA | #VALUE! |
| ENSMUSG00000108589.2 | 0 | NA | NA | NA | NA | NA | #VALUE! |
| ENSMUSG00000108590.2 | 0 | NA | NA | NA | NA | NA | #VALUE! |
| ENSMUSG00000108591.2 | 0 | NA | NA | NA | NA | NA | #VALUE! |
| ENSMUSG00000108592.3 | 0.261539 | 1.68833 | 4.043024 | 0.417591 | 0.676246 | NA | #VALUE! |
| ENSMUSG00000108593.2 | 0 | NA | NA | NA | NA | NA | #VALUE! |
| ENSMUSG00000108594.2 | 0 | NA | NA | NA | NA | NA | #VALUE! |
| ENSMUSG00000108596.2 | 0 | NA | NA | NA | NA | NA | #VALUE! |
| ENSMUSG00000108597.2 | 0 | NA | NA | NA | NA | NA | #VALUE! |
| ENSMUSG00000108598.2 | 0 | NA | NA | NA | NA | NA | #VALUE! |
| ENSMUSG00000108599.2 | 0 | NA | NA | NA | NA | NA | #VALUE! |
| ENSMUSG00000108600.2 | 0 | NA | NA | NA | NA | NA | #VALUE! |
| ENSMUSG00000108601.2 | 6.22649 | 6.128273 | 3.913982 | 1.565739 | 0.11741 | NA | #VALUE! |
| ENSMUSG00000108602.2 | 0 | NA | NA | NA | NA | NA | #VALUE! |
| ENSMUSG00000108603.2 | 0 | NA | NA | NA | NA | NA | #VALUE! |
| ENSMUSG00000108604.2 | 0 | NA | NA | NA | NA | NA | #VALUE! |
| ENSMUSG00000108605.3 | 0 | NA | NA | NA | NA | NA | #VALUE! |
| ENSMUSG00000108606.2 | 0 | NA | NA | NA | NA | NA | #VALUE! |
| ENSMUSG00000108607.3 | 0 | NA | NA | NA | NA | NA | #VALUE! |
| ENSMUSG00000108608.2 | 0 | NA | NA | NA | NA | NA | #VALUE! |
| ENSMUSG00000108609.2 | 0 | NA | NA | NA | NA | NA | #VALUE! |

| | | | | | | | |
|---|---|---|---|---|---|---|---|
| ENSMUSG00000108610.2 | 0 | NA | NA | NA | NA | NA | #VALUE! |
| ENSMUSG00000108611.2 | 0 | NA | NA | NA | NA | NA | #VALUE! |
| ENSMUSG00000108612.3 | 0 | NA | NA | NA | NA | NA | #VALUE! |
| ENSMUSG00000108613.2 | 0 | NA | NA | NA | NA | NA | #VALUE! |
| ENSMUSG00000108614.2 | 0 | NA | NA | NA | NA | NA | #VALUE! |
| ENSMUSG00000108615.2 | 0 | NA | NA | NA | NA | NA | #VALUE! |
| ENSMUSG00000108616.2 | 0 | NA | NA | NA | NA | NA | #VALUE! |
| ENSMUSG00000108617.2 | 0 | NA | NA | NA | NA | NA | #VALUE! |
| ENSMUSG00000108618.2 | 0 | NA | NA | NA | NA | NA | #VALUE! |
| ENSMUSG00000108619.2 | 0 | NA | NA | NA | NA | NA | #VALUE! |
| ENSMUSG00000108620.2 | 0 | NA | NA | NA | NA | NA | #VALUE! |
| ENSMUSG00000108622.3 | 0 | NA | NA | NA | NA | NA | #VALUE! |
| ENSMUSG00000108623.2 | 0 | NA | NA | NA | NA | NA | #VALUE! |
| ENSMUSG00000108624.2 | 0 | NA | NA | NA | NA | NA | #VALUE! |
| ENSMUSG00000108625.2 | 0 | NA | NA | NA | NA | NA | #VALUE! |
| ENSMUSG00000108626.2 | 5.889923 | 6.048106 | 3.914393 | 1.545094 | 0.122324 | NA | #VALUE! |
| ENSMUSG00000108627.2 | 0 | NA | NA | NA | NA | NA | #VALUE! |
| ENSMUSG00000108628.2 | 0.105013 | -0.97324 | 4.076293 | -0.23876 | 0.811295 | NA | #VALUE! |
| ENSMUSG00000108629.2 | 0 | NA | NA | NA | NA | NA | #VALUE! |
| ENSMUSG00000108630.2 | 0 | NA | NA | NA | NA | NA | #VALUE! |
| ENSMUSG00000108633.2 | 0.18651 | 1.401769 | 4.056354 | 0.345574 | 0.729663 | NA | #VALUE! |
| ENSMUSG00000108634.2 | 0 | NA | NA | NA | NA | NA | #VALUE! |
| ENSMUSG00000108635.2 | 0 | NA | NA | NA | NA | NA | #VALUE! |
| ENSMUSG00000108636.2 | 0 | NA | NA | NA | NA | NA | #VALUE! |
| ENSMUSG00000108637.2 | 0 | NA | NA | NA | NA | NA | #VALUE! |
| ENSMUSG00000108638.2 | 0 | NA | NA | NA | NA | NA | #VALUE! |
| ENSMUSG00000108639.2 | 0 | NA | NA | NA | NA | NA | #VALUE! |
| ENSMUSG00000108640.2 | 0 | NA | NA | NA | NA | NA | #VALUE! |
| ENSMUSG00000108641.2 | 0 | NA | NA | NA | NA | NA | #VALUE! |
| ENSMUSG00000108642.2 | 0 | NA | NA | NA | NA | NA | #VALUE! |
| ENSMUSG00000108643.2 | 0 | NA | NA | NA | NA | NA | #VALUE! |
| ENSMUSG00000108644.2 | 0 | NA | NA | NA | NA | NA | #VALUE! |
| ENSMUSG00000108645.2 | 0 | NA | NA | NA | NA | NA | #VALUE! |
| ENSMUSG00000108646.2 | 0 | NA | NA | NA | NA | NA | #VALUE! |
| ENSMUSG00000108647.2 | 0 | NA | NA | NA | NA | NA | #VALUE! |
| ENSMUSG00000108648.2 | 0 | NA | NA | NA | NA | NA | #VALUE! |
| ENSMUSG00000108649.2 | 0 | NA | NA | NA | NA | NA | #VALUE! |
| ENSMUSG00000108650.2 | 0 | NA | NA | NA | NA | NA | #VALUE! |
| ENSMUSG00000108651.2 | 0 | NA | NA | NA | NA | NA | #VALUE! |
| ENSMUSG00000108652.2 | 0 | NA | NA | NA | NA | NA | #VALUE! |
| ENSMUSG00000108654.2 | 0 | NA | NA | NA | NA | NA | #VALUE! |
| ENSMUSG00000108655.2 | 0 | NA | NA | NA | NA | NA | #VALUE! |
| ENSMUSG00000108657.2 | 0 | NA | NA | NA | NA | NA | #VALUE! |
| ENSMUSG00000108660.2 | 0 | NA | NA | NA | NA | NA | #VALUE! |

| | | | | | | | |
|---|---|---|---|---|---|---|---|
| ENSMUSG00000108661.2 | 0 | NA | NA | NA | NA | NA | #VALUE! |
| ENSMUSG00000108662.4 | 0 | NA | NA | NA | NA | NA | #VALUE! |
| ENSMUSG00000108663.2 | 0 | NA | NA | NA | NA | NA | #VALUE! |
| ENSMUSG00000108665.2 | 0 | NA | NA | NA | NA | NA | #VALUE! |
| ENSMUSG00000108666.2 | 0 | NA | NA | NA | NA | NA | #VALUE! |
| ENSMUSG00000108667.2 | 0 | NA | NA | NA | NA | NA | #VALUE! |
| ENSMUSG00000108668.2 | 0 | NA | NA | NA | NA | NA | #VALUE! |
| ENSMUSG00000108669.2 | 0 | NA | NA | NA | NA | NA | #VALUE! |
| ENSMUSG00000108670.2 | 0 | NA | NA | NA | NA | NA | #VALUE! |
| ENSMUSG00000108671.2 | 0 | NA | NA | NA | NA | NA | #VALUE! |
| ENSMUSG00000108672.2 | 0 | NA | NA | NA | NA | NA | #VALUE! |
| ENSMUSG00000108673.2 | 0 | NA | NA | NA | NA | NA | #VALUE! |
| ENSMUSG00000108674.3 | 0 | NA | NA | NA | NA | NA | #VALUE! |
| ENSMUSG00000108675.2 | 0 | NA | NA | NA | NA | NA | #VALUE! |
| ENSMUSG00000108676.2 | 0 | NA | NA | NA | NA | NA | #VALUE! |
| ENSMUSG00000108677.2 | 0 | NA | NA | NA | NA | NA | #VALUE! |
| ENSMUSG00000108678.2 | 0 | NA | NA | NA | NA | NA | #VALUE! |
| ENSMUSG00000108679.2 | 0 | NA | NA | NA | NA | NA | #VALUE! |
| ENSMUSG00000108681.2 | 0 | NA | NA | NA | NA | NA | #VALUE! |
| ENSMUSG00000108682.2 | 0 | NA | NA | NA | NA | NA | #VALUE! |
| ENSMUSG00000108683.2 | 0 | NA | NA | NA | NA | NA | #VALUE! |
| ENSMUSG00000108685.2 | 0 | NA | NA | NA | NA | NA | #VALUE! |
| ENSMUSG00000108686.2 | 0 | NA | NA | NA | NA | NA | #VALUE! |
| ENSMUSG00000108687.2 | 0 | NA | NA | NA | NA | NA | #VALUE! |
| ENSMUSG00000108688.2 | 0 | NA | NA | NA | NA | NA | #VALUE! |
| ENSMUSG00000108689.2 | 0 | NA | NA | NA | NA | NA | #VALUE! |
| ENSMUSG00000108691.2 | 0 | NA | NA | NA | NA | NA | #VALUE! |
| ENSMUSG00000108692.2 | 0 | NA | NA | NA | NA | NA | #VALUE! |
| ENSMUSG00000108693.2 | 0 | NA | NA | NA | NA | NA | #VALUE! |
| ENSMUSG00000108694.2 | 0.519272 | 1.241215 | 4.011446 | 0.309418 | 0.757003 | NA | #VALUE! |
| ENSMUSG00000108695.2 | 0 | NA | NA | NA | NA | NA | #VALUE! |
| ENSMUSG00000108696.2 | 0 | NA | NA | NA | NA | NA | #VALUE! |
| ENSMUSG00000108697.2 | 0 | NA | NA | NA | NA | NA | #VALUE! |
| ENSMUSG00000108698.3 | 0 | NA | NA | NA | NA | NA | #VALUE! |
| ENSMUSG00000108699.2 | 0 | NA | NA | NA | NA | NA | #VALUE! |
| ENSMUSG00000108700.2 | 0 | NA | NA | NA | NA | NA | #VALUE! |
| ENSMUSG00000108701.2 | 1.967804 | 4.449735 | 3.929778 | 1.132312 | 0.257503 | NA | #VALUE! |
| ENSMUSG00000108702.2 | 0 | NA | NA | NA | NA | NA | #VALUE! |
| ENSMUSG00000108703.2 | 0 | NA | NA | NA | NA | NA | #VALUE! |
| ENSMUSG00000108704.2 | 0 | NA | NA | NA | NA | NA | #VALUE! |
| ENSMUSG00000108705.2 | 0 | NA | NA | NA | NA | NA | #VALUE! |
| ENSMUSG00000108707.2 | 0 | NA | NA | NA | NA | NA | #VALUE! |
| ENSMUSG00000108708.2 | 0 | NA | NA | NA | NA | NA | #VALUE! |
| ENSMUSG00000108709.2 | 0 | NA | NA | NA | NA | NA | #VALUE! |

| | | | | | | | |
|---|---|---|---|---|---|---|---|
| ENSMUSG00000108711.2 | 0 | NA | NA | NA | NA | NA | #VALUE! |
| ENSMUSG00000108712.2 | 0 | NA | NA | NA | NA | NA | #VALUE! |
| ENSMUSG00000108713.2 | 0 | NA | NA | NA | NA | NA | #VALUE! |
| ENSMUSG00000108714.2 | 0 | NA | NA | NA | NA | NA | #VALUE! |
| ENSMUSG00000108716.2 | 0 | NA | NA | NA | NA | NA | #VALUE! |
| ENSMUSG00000108717.2 | 0 | NA | NA | NA | NA | NA | #VALUE! |
| ENSMUSG00000108718.2 | 0 | NA | NA | NA | NA | NA | #VALUE! |
| ENSMUSG00000108719.2 | 0 | NA | NA | NA | NA | NA | #VALUE! |
| ENSMUSG00000108720.2 | 0 | NA | NA | NA | NA | NA | #VALUE! |
| ENSMUSG00000108721.2 | 0 | NA | NA | NA | NA | NA | #VALUE! |
| ENSMUSG00000108722.2 | 0 | NA | NA | NA | NA | NA | #VALUE! |
| ENSMUSG00000108723.2 | 0 | NA | NA | NA | NA | NA | #VALUE! |
| ENSMUSG00000108724.2 | 0 | NA | NA | NA | NA | NA | #VALUE! |
| ENSMUSG00000108725.2 | 0 | NA | NA | NA | NA | NA | #VALUE! |
| ENSMUSG00000108726.2 | 0 | NA | NA | NA | NA | NA | #VALUE! |
| ENSMUSG00000108727.2 | 0 | NA | NA | NA | NA | NA | #VALUE! |
| ENSMUSG00000108728.5 | 0 | NA | NA | NA | NA | NA | #VALUE! |
| ENSMUSG00000108729.2 | 0 | NA | NA | NA | NA | NA | #VALUE! |
| ENSMUSG00000108730.2 | 0 | NA | NA | NA | NA | NA | #VALUE! |
| ENSMUSG00000108731.2 | 0 | NA | NA | NA | NA | NA | #VALUE! |
| ENSMUSG00000108732.2 | 0 | NA | NA | NA | NA | NA | #VALUE! |
| ENSMUSG00000108734.2 | 0 | NA | NA | NA | NA | NA | #VALUE! |
| ENSMUSG00000108735.2 | 0 | NA | NA | NA | NA | NA | #VALUE! |
| ENSMUSG00000108736.2 | 0 | NA | NA | NA | NA | NA | #VALUE! |
| ENSMUSG00000108737.2 | 0 | NA | NA | NA | NA | NA | #VALUE! |
| ENSMUSG00000108739.2 | 0 | NA | NA | NA | NA | NA | #VALUE! |
| ENSMUSG00000108740.2 | 0 | NA | NA | NA | NA | NA | #VALUE! |
| ENSMUSG00000108741.2 | 0 | NA | NA | NA | NA | NA | #VALUE! |
| ENSMUSG00000108742.2 | 0 | NA | NA | NA | NA | NA | #VALUE! |
| ENSMUSG00000108743.2 | 0 | NA | NA | NA | NA | NA | #VALUE! |
| ENSMUSG00000108744.2 | 0 | NA | NA | NA | NA | NA | #VALUE! |
| ENSMUSG00000108745.2 | 0 | NA | NA | NA | NA | NA | #VALUE! |
| ENSMUSG00000108746.2 | 0 | NA | NA | NA | NA | NA | #VALUE! |
| ENSMUSG00000108748.2 | 0 | NA | NA | NA | NA | NA | #VALUE! |
| ENSMUSG00000108749.2 | 0 | NA | NA | NA | NA | NA | #VALUE! |
| ENSMUSG00000108750.2 | 0 | NA | NA | NA | NA | NA | #VALUE! |
| ENSMUSG00000108751.2 | 0 | NA | NA | NA | NA | NA | #VALUE! |
| ENSMUSG00000108752.2 | 0 | NA | NA | NA | NA | NA | #VALUE! |
| ENSMUSG00000108753.2 | 0 | NA | NA | NA | NA | NA | #VALUE! |
| ENSMUSG00000108754.2 | 0 | NA | NA | NA | NA | NA | #VALUE! |
| ENSMUSG00000108756.2 | 0 | NA | NA | NA | NA | NA | #VALUE! |
| ENSMUSG00000108757.2 | 0 | NA | NA | NA | NA | NA | #VALUE! |
| ENSMUSG00000108758.2 | 0 | NA | NA | NA | NA | NA | #VALUE! |
| ENSMUSG00000108759.2 | 0 | NA | NA | NA | NA | NA | #VALUE! |

| | | | | | | | |
|---|---|---|---|---|---|---|---|
| ENSMUSG00000108760.2 | 0 | NA | NA | NA | NA | NA | #VALUE! |
| ENSMUSG00000108761.2 | 0 | NA | NA | NA | NA | NA | #VALUE! |
| ENSMUSG00000108764.2 | 0 | NA | NA | NA | NA | NA | #VALUE! |
| ENSMUSG00000108765.2 | 0 | NA | NA | NA | NA | NA | #VALUE! |
| ENSMUSG00000108766.2 | 0 | NA | NA | NA | NA | NA | #VALUE! |
| ENSMUSG00000108767.2 | 0 | NA | NA | NA | NA | NA | #VALUE! |
| ENSMUSG00000108768.2 | 0 | NA | NA | NA | NA | NA | #VALUE! |
| ENSMUSG00000108770.2 | 0 | NA | NA | NA | NA | NA | #VALUE! |
| ENSMUSG00000108771.2 | 0 | NA | NA | NA | NA | NA | #VALUE! |
| ENSMUSG00000108772.2 | 0.168284 | 1.027628 | 4.067126 | 0.252667 | 0.800526 | NA | #VALUE! |
| ENSMUSG00000108773.2 | 0 | NA | NA | NA | NA | NA | #VALUE! |
| ENSMUSG00000108774.2 | 0 | NA | NA | NA | NA | NA | #VALUE! |
| ENSMUSG00000108775.2 | 0 | NA | NA | NA | NA | NA | #VALUE! |
| ENSMUSG00000108776.2 | 0 | NA | NA | NA | NA | NA | #VALUE! |
| ENSMUSG00000108777.2 | 0 | NA | NA | NA | NA | NA | #VALUE! |
| ENSMUSG00000108778.2 | 0 | NA | NA | NA | NA | NA | #VALUE! |
| ENSMUSG00000108779.2 | 0 | NA | NA | NA | NA | NA | #VALUE! |
| ENSMUSG00000108780.2 | 0 | NA | NA | NA | NA | NA | #VALUE! |
| ENSMUSG00000108781.2 | 0 | NA | NA | NA | NA | NA | #VALUE! |
| ENSMUSG00000108782.2 | 0 | NA | NA | NA | NA | NA | #VALUE! |
| ENSMUSG00000108783.2 | 0 | NA | NA | NA | NA | NA | #VALUE! |
| ENSMUSG00000108784.2 | 0 | NA | NA | NA | NA | NA | #VALUE! |
| ENSMUSG00000108785.2 | 0 | NA | NA | NA | NA | NA | #VALUE! |
| ENSMUSG00000108786.2 | 0 | NA | NA | NA | NA | NA | #VALUE! |
| ENSMUSG00000108787.2 | 0 | NA | NA | NA | NA | NA | #VALUE! |
| ENSMUSG00000108788.2 | 0 | NA | NA | NA | NA | NA | #VALUE! |
| ENSMUSG00000108790.2 | 0 | NA | NA | NA | NA | NA | #VALUE! |
| ENSMUSG00000108791.2 | 0 | NA | NA | NA | NA | NA | #VALUE! |
| ENSMUSG00000108792.2 | 0 | NA | NA | NA | NA | NA | #VALUE! |
| ENSMUSG00000108793.2 | 0 | NA | NA | NA | NA | NA | #VALUE! |
| ENSMUSG00000108794.2 | 0 | NA | NA | NA | NA | NA | #VALUE! |
| ENSMUSG00000108795.2 | 0 | NA | NA | NA | NA | NA | #VALUE! |
| ENSMUSG00000108796.2 | 0 | NA | NA | NA | NA | NA | #VALUE! |
| ENSMUSG00000108797.2 | 0 | NA | NA | NA | NA | NA | #VALUE! |
| ENSMUSG00000108798.2 | 0 | NA | NA | NA | NA | NA | #VALUE! |
| ENSMUSG00000108799.2 | 0 | NA | NA | NA | NA | NA | #VALUE! |
| ENSMUSG00000108800.2 | 0 | NA | NA | NA | NA | NA | #VALUE! |
| ENSMUSG00000108801.2 | 0 | NA | NA | NA | NA | NA | #VALUE! |
| ENSMUSG00000108802.2 | 0 | NA | NA | NA | NA | NA | #VALUE! |
| ENSMUSG00000108803.2 | 0 | NA | NA | NA | NA | NA | #VALUE! |
| ENSMUSG00000108804.2 | 0 | NA | NA | NA | NA | NA | #VALUE! |
| ENSMUSG00000108805.2 | 0 | NA | NA | NA | NA | NA | #VALUE! |
| ENSMUSG00000108806.2 | 0 | NA | NA | NA | NA | NA | #VALUE! |
| ENSMUSG00000108807.2 | 0 | NA | NA | NA | NA | NA | #VALUE! |

| | | | | | | | |
|---|---|---|---|---|---|---|---|
| ENSMUSG00000108808.2 | 0 | NA | NA | NA | NA | NA | #VALUE! |
| ENSMUSG00000108809.2 | 0 | NA | NA | NA | NA | NA | #VALUE! |
| ENSMUSG00000108810.2 | 0 | NA | NA | NA | NA | NA | #VALUE! |
| ENSMUSG00000108811.2 | 0 | NA | NA | NA | NA | NA | #VALUE! |
| ENSMUSG00000108812.2 | 0 | NA | NA | NA | NA | NA | #VALUE! |
| ENSMUSG00000108813.2 | 0.297366 | -0.14751 | 4.065279 | -0.03629 | 0.971054 | NA | #VALUE! |
| ENSMUSG00000108814.2 | 0 | NA | NA | NA | NA | NA | #VALUE! |
| ENSMUSG00000108815.2 | 0 | NA | NA | NA | NA | NA | #VALUE! |
| ENSMUSG00000108816.2 | 0 | NA | NA | NA | NA | NA | #VALUE! |
| ENSMUSG00000108818.2 | 0 | NA | NA | NA | NA | NA | #VALUE! |
| ENSMUSG00000108819.2 | 0 | NA | NA | NA | NA | NA | #VALUE! |
| ENSMUSG00000108820.2 | 0 | NA | NA | NA | NA | NA | #VALUE! |
| ENSMUSG00000108821.2 | 0 | NA | NA | NA | NA | NA | #VALUE! |
| ENSMUSG00000108822.2 | 0.673134 | 2.918386 | 3.972531 | 0.734641 | 0.462558 | NA | #VALUE! |
| ENSMUSG00000108823.2 | 0.105013 | -0.97324 | 4.076293 | -0.23876 | 0.811295 | NA | #VALUE! |
| ENSMUSG00000108825.2 | 0 | NA | NA | NA | NA | NA | #VALUE! |
| ENSMUSG00000108826.2 | 0 | NA | NA | NA | NA | NA | #VALUE! |
| ENSMUSG00000108827.4 | 0 | NA | NA | NA | NA | NA | #VALUE! |
| ENSMUSG00000108828.2 | 0 | NA | NA | NA | NA | NA | #VALUE! |
| ENSMUSG00000108829.2 | 0 | NA | NA | NA | NA | NA | #VALUE! |
| ENSMUSG00000108830.2 | 0 | NA | NA | NA | NA | NA | #VALUE! |
| ENSMUSG00000108831.2 | 0 | NA | NA | NA | NA | NA | #VALUE! |
| ENSMUSG00000108832.2 | 0 | NA | NA | NA | NA | NA | #VALUE! |
| ENSMUSG00000108833.2 | 0.093255 | 0.868093 | 4.072464 | 0.213162 | 0.831201 | NA | #VALUE! |
| ENSMUSG00000108834.2 | 0 | NA | NA | NA | NA | NA | #VALUE! |
| ENSMUSG00000108835.2 | 0 | NA | NA | NA | NA | NA | #VALUE! |
| ENSMUSG00000108836.2 | 0 | NA | NA | NA | NA | NA | #VALUE! |
| ENSMUSG00000108837.2 | 0 | NA | NA | NA | NA | NA | #VALUE! |
| ENSMUSG00000108838.2 | 0 | NA | NA | NA | NA | NA | #VALUE! |
| ENSMUSG00000108839.2 | 0 | NA | NA | NA | NA | NA | #VALUE! |
| ENSMUSG00000108840.2 | 0 | NA | NA | NA | NA | NA | #VALUE! |
| ENSMUSG00000108841.2 | 0 | NA | NA | NA | NA | NA | #VALUE! |
| ENSMUSG00000108842.2 | 0 | NA | NA | NA | NA | NA | #VALUE! |
| ENSMUSG00000108843.2 | 0 | NA | NA | NA | NA | NA | #VALUE! |
| ENSMUSG00000108844.2 | 0 | NA | NA | NA | NA | NA | #VALUE! |
| ENSMUSG00000108845.2 | 0 | NA | NA | NA | NA | NA | #VALUE! |
| ENSMUSG00000108846.2 | 0 | NA | NA | NA | NA | NA | #VALUE! |
| ENSMUSG00000108847.2 | 0 | NA | NA | NA | NA | NA | #VALUE! |
| ENSMUSG00000108848.2 | 0 | NA | NA | NA | NA | NA | #VALUE! |
| ENSMUSG00000108849.2 | 0 | NA | NA | NA | NA | NA | #VALUE! |
| ENSMUSG00000108850.2 | 0 | NA | NA | NA | NA | NA | #VALUE! |
| ENSMUSG00000108851.2 | 0 | NA | NA | NA | NA | NA | #VALUE! |
| ENSMUSG00000108852.2 | 0 | NA | NA | NA | NA | NA | #VALUE! |
| ENSMUSG00000108853.2 | 0 | NA | NA | NA | NA | NA | #VALUE! |

| | | | | | | | |
|---|---|---|---|---|---|---|---|
| ENSMUSG00000108854.2 | 0 | NA | NA | NA | NA | NA | #VALUE! |
| ENSMUSG00000108855.2 | 0 | NA | NA | NA | NA | NA | #VALUE! |
| ENSMUSG00000108856.2 | 0 | NA | NA | NA | NA | NA | #VALUE! |
| ENSMUSG00000108857.2 | 0 | NA | NA | NA | NA | NA | #VALUE! |
| ENSMUSG00000108858.2 | 0 | NA | NA | NA | NA | NA | #VALUE! |
| ENSMUSG00000108859.2 | 0 | NA | NA | NA | NA | NA | #VALUE! |
| ENSMUSG00000108860.2 | 0 | NA | NA | NA | NA | NA | #VALUE! |
| ENSMUSG00000108861.2 | 0 | NA | NA | NA | NA | NA | #VALUE! |
| ENSMUSG00000108862.2 | 0 | NA | NA | NA | NA | NA | #VALUE! |
| ENSMUSG00000108863.3 | 0 | NA | NA | NA | NA | NA | #VALUE! |
| ENSMUSG00000108864.2 | 0 | NA | NA | NA | NA | NA | #VALUE! |
| ENSMUSG00000108865.2 | 0 | NA | NA | NA | NA | NA | #VALUE! |
| ENSMUSG00000108866.2 | 0 | NA | NA | NA | NA | NA | #VALUE! |
| ENSMUSG00000108867.2 | 0 | NA | NA | NA | NA | NA | #VALUE! |
| ENSMUSG00000108868.2 | 0 | NA | NA | NA | NA | NA | #VALUE! |
| ENSMUSG00000108869.2 | 0 | NA | NA | NA | NA | NA | #VALUE! |
| ENSMUSG00000108870.2 | 0 | NA | NA | NA | NA | NA | #VALUE! |
| ENSMUSG00000108871.2 | 0 | NA | NA | NA | NA | NA | #VALUE! |
| ENSMUSG00000108873.2 | 0 | NA | NA | NA | NA | NA | #VALUE! |
| ENSMUSG00000108874.2 | 0 | NA | NA | NA | NA | NA | #VALUE! |
| ENSMUSG00000108875.2 | 0 | NA | NA | NA | NA | NA | #VALUE! |
| ENSMUSG00000108876.2 | 0 | NA | NA | NA | NA | NA | #VALUE! |
| ENSMUSG00000108877.2 | 0 | NA | NA | NA | NA | NA | #VALUE! |
| ENSMUSG00000108878.2 | 0 | NA | NA | NA | NA | NA | #VALUE! |
| ENSMUSG00000108879.2 | 0 | NA | NA | NA | NA | NA | #VALUE! |
| ENSMUSG00000108880.2 | 0 | NA | NA | NA | NA | NA | #VALUE! |
| ENSMUSG00000108881.2 | 0 | NA | NA | NA | NA | NA | #VALUE! |
| ENSMUSG00000108882.2 | 0 | NA | NA | NA | NA | NA | #VALUE! |
| ENSMUSG00000108883.2 | 0 | NA | NA | NA | NA | NA | #VALUE! |
| ENSMUSG00000108885.2 | 0 | NA | NA | NA | NA | NA | #VALUE! |
| ENSMUSG00000108886.2 | 0 | NA | NA | NA | NA | NA | #VALUE! |
| ENSMUSG00000108887.2 | 0 | NA | NA | NA | NA | NA | #VALUE! |
| ENSMUSG00000108888.2 | 0 | NA | NA | NA | NA | NA | #VALUE! |
| ENSMUSG00000108889.3 | 0 | NA | NA | NA | NA | NA | #VALUE! |
| ENSMUSG00000108890.2 | 0 | NA | NA | NA | NA | NA | #VALUE! |
| ENSMUSG00000108891.5 | 0 | NA | NA | NA | NA | NA | #VALUE! |
| ENSMUSG00000108892.2 | 0 | NA | NA | NA | NA | NA | #VALUE! |
| ENSMUSG00000108893.2 | 0 | NA | NA | NA | NA | NA | #VALUE! |
| ENSMUSG00000108894.2 | 0 | NA | NA | NA | NA | NA | #VALUE! |
| ENSMUSG00000108895.2 | 0 | NA | NA | NA | NA | NA | #VALUE! |
| ENSMUSG00000108896.2 | 0 | NA | NA | NA | NA | NA | #VALUE! |
| ENSMUSG00000108897.2 | 0 | NA | NA | NA | NA | NA | #VALUE! |
| ENSMUSG00000108899.2 | 0 | NA | NA | NA | NA | NA | #VALUE! |
| ENSMUSG00000108900.2 | 0 | NA | NA | NA | NA | NA | #VALUE! |

| | | | | | | | |
|---|---|---|---|---|---|---|---|
| ENSMUSG00000108902.2 | 0 | NA | NA | NA | NA | NA | #VALUE! |
| ENSMUSG00000108903.2 | 0 | NA | NA | NA | NA | NA | #VALUE! |
| ENSMUSG00000108904.2 | 0 | NA | NA | NA | NA | NA | #VALUE! |
| ENSMUSG00000108905.2 | 0 | NA | NA | NA | NA | NA | #VALUE! |
| ENSMUSG00000108906.2 | 0 | NA | NA | NA | NA | NA | #VALUE! |
| ENSMUSG00000108907.2 | 0 | NA | NA | NA | NA | NA | #VALUE! |
| ENSMUSG00000108908.5 | 0 | NA | NA | NA | NA | NA | #VALUE! |
| ENSMUSG00000108909.2 | 0 | NA | NA | NA | NA | NA | #VALUE! |
| ENSMUSG00000108910.2 | 0 | NA | NA | NA | NA | NA | #VALUE! |
| ENSMUSG00000108911.2 | 0.245976 | 1.027628 | 4.067126 | 0.252667 | 0.800526 | NA | #VALUE! |
| ENSMUSG00000108912.2 | 0 | NA | NA | NA | NA | NA | #VALUE! |
| ENSMUSG00000108913.2 | 0 | NA | NA | NA | NA | NA | #VALUE! |
| ENSMUSG00000108914.2 | 0 | NA | NA | NA | NA | NA | #VALUE! |
| ENSMUSG00000108915.2 | 0 | NA | NA | NA | NA | NA | #VALUE! |
| ENSMUSG00000108916.2 | 0 | NA | NA | NA | NA | NA | #VALUE! |
| ENSMUSG00000108917.2 | 0 | NA | NA | NA | NA | NA | #VALUE! |
| ENSMUSG00000108918.2 | 0 | NA | NA | NA | NA | NA | #VALUE! |
| ENSMUSG00000108919.4 | 0 | NA | NA | NA | NA | NA | #VALUE! |
| ENSMUSG00000108920.2 | 0 | NA | NA | NA | NA | NA | #VALUE! |
| ENSMUSG00000108921.2 | 0 | NA | NA | NA | NA | NA | #VALUE! |
| ENSMUSG00000108922.2 | 0 | NA | NA | NA | NA | NA | #VALUE! |
| ENSMUSG00000108923.2 | 0 | NA | NA | NA | NA | NA | #VALUE! |
| ENSMUSG00000108924.2 | 0 | NA | NA | NA | NA | NA | #VALUE! |
| ENSMUSG00000108925.2 | 0 | NA | NA | NA | NA | NA | #VALUE! |
| ENSMUSG00000108926.2 | 0 | NA | NA | NA | NA | NA | #VALUE! |
| ENSMUSG00000108928.2 | 0 | NA | NA | NA | NA | NA | #VALUE! |
| ENSMUSG00000108929.3 | 0 | NA | NA | NA | NA | NA | #VALUE! |
| ENSMUSG00000108930.2 | 0 | NA | NA | NA | NA | NA | #VALUE! |
| ENSMUSG00000108931.3 | 0 | NA | NA | NA | NA | NA | #VALUE! |
| ENSMUSG00000108932.2 | 0 | NA | NA | NA | NA | NA | #VALUE! |
| ENSMUSG00000108933.2 | 0 | NA | NA | NA | NA | NA | #VALUE! |
| ENSMUSG00000108934.2 | 0 | NA | NA | NA | NA | NA | #VALUE! |
| ENSMUSG00000108935.2 | 0 | NA | NA | NA | NA | NA | #VALUE! |
| ENSMUSG00000108936.2 | 0 | NA | NA | NA | NA | NA | #VALUE! |
| ENSMUSG00000108937.2 | 0 | NA | NA | NA | NA | NA | #VALUE! |
| ENSMUSG00000108938.2 | 0 | NA | NA | NA | NA | NA | #VALUE! |
| ENSMUSG00000108939.3 | 0 | NA | NA | NA | NA | NA | #VALUE! |
| ENSMUSG00000108940.2 | 0 | NA | NA | NA | NA | NA | #VALUE! |
| ENSMUSG00000108941.2 | 0 | NA | NA | NA | NA | NA | #VALUE! |
| ENSMUSG00000108942.2 | 0 | NA | NA | NA | NA | NA | #VALUE! |
| ENSMUSG00000108943.2 | 0 | NA | NA | NA | NA | NA | #VALUE! |
| ENSMUSG00000108944.2 | 0 | NA | NA | NA | NA | NA | #VALUE! |
| ENSMUSG00000108945.2 | 0 | NA | NA | NA | NA | NA | #VALUE! |
| ENSMUSG00000108946.2 | 0 | NA | NA | NA | NA | NA | #VALUE! |

| | | | | | | | |
|---|---|---|---|---|---|---|---|
| ENSMUSG00000108947.2 | 0 | NA | NA | NA | NA | NA | #VALUE! |
| ENSMUSG00000108948.4 | 0 | NA | NA | NA | NA | NA | #VALUE! |
| ENSMUSG00000108949.2 | 0 | NA | NA | NA | NA | NA | #VALUE! |
| ENSMUSG00000108950.2 | 0 | NA | NA | NA | NA | NA | #VALUE! |
| ENSMUSG00000108952.2 | 0 | NA | NA | NA | NA | NA | #VALUE! |
| ENSMUSG00000108953.2 | 0 | NA | NA | NA | NA | NA | #VALUE! |
| ENSMUSG00000108954.2 | 0 | NA | NA | NA | NA | NA | #VALUE! |
| ENSMUSG00000108955.2 | 0 | NA | NA | NA | NA | NA | #VALUE! |
| ENSMUSG00000108956.2 | 0 | NA | NA | NA | NA | NA | #VALUE! |
| ENSMUSG00000108957.2 | 0 | NA | NA | NA | NA | NA | #VALUE! |
| ENSMUSG00000108958.2 | 0 | NA | NA | NA | NA | NA | #VALUE! |
| ENSMUSG00000108959.2 | 0 | NA | NA | NA | NA | NA | #VALUE! |
| ENSMUSG00000108960.2 | 0 | NA | NA | NA | NA | NA | #VALUE! |
| ENSMUSG00000108961.2 | 0 | NA | NA | NA | NA | NA | #VALUE! |
| ENSMUSG00000108962.2 | 0 | NA | NA | NA | NA | NA | #VALUE! |
| ENSMUSG00000108963.2 | 0 | NA | NA | NA | NA | NA | #VALUE! |
| ENSMUSG00000108964.2 | 0 | NA | NA | NA | NA | NA | #VALUE! |
| ENSMUSG00000108965.2 | 0 | NA | NA | NA | NA | NA | #VALUE! |
| ENSMUSG00000108966.2 | 0 | NA | NA | NA | NA | NA | #VALUE! |
| ENSMUSG00000108967.2 | 0 | NA | NA | NA | NA | NA | #VALUE! |
| ENSMUSG00000108968.2 | 0 | NA | NA | NA | NA | NA | #VALUE! |
| ENSMUSG00000108969.2 | 0 | NA | NA | NA | NA | NA | #VALUE! |
| ENSMUSG00000108971.2 | 0 | NA | NA | NA | NA | NA | #VALUE! |
| ENSMUSG00000108972.2 | 0 | NA | NA | NA | NA | NA | #VALUE! |
| ENSMUSG00000108973.2 | 0 | NA | NA | NA | NA | NA | #VALUE! |
| ENSMUSG00000108974.2 | 0 | NA | NA | NA | NA | NA | #VALUE! |
| ENSMUSG00000108975.2 | 0 | NA | NA | NA | NA | NA | #VALUE! |
| ENSMUSG00000108976.2 | 0 | NA | NA | NA | NA | NA | #VALUE! |
| ENSMUSG00000108977.2 | 0 | NA | NA | NA | NA | NA | #VALUE! |
| ENSMUSG00000108978.2 | 0 | NA | NA | NA | NA | NA | #VALUE! |
| ENSMUSG00000108979.2 | 0 | NA | NA | NA | NA | NA | #VALUE! |
| ENSMUSG00000108980.2 | 0 | NA | NA | NA | NA | NA | #VALUE! |
| ENSMUSG00000108981.2 | 0 | NA | NA | NA | NA | NA | #VALUE! |
| ENSMUSG00000108982.2 | 0 | NA | NA | NA | NA | NA | #VALUE! |
| ENSMUSG00000108983.2 | 0 | NA | NA | NA | NA | NA | #VALUE! |
| ENSMUSG00000108984.2 | 0 | NA | NA | NA | NA | NA | #VALUE! |
| ENSMUSG00000108985.2 | 0 | NA | NA | NA | NA | NA | #VALUE! |
| ENSMUSG00000108986.2 | 0 | NA | NA | NA | NA | NA | #VALUE! |
| ENSMUSG00000108987.2 | 0 | NA | NA | NA | NA | NA | #VALUE! |
| ENSMUSG00000108988.2 | 0 | NA | NA | NA | NA | NA | #VALUE! |
| ENSMUSG00000108989.2 | 0 | NA | NA | NA | NA | NA | #VALUE! |
| ENSMUSG00000108990.2 | 0 | NA | NA | NA | NA | NA | #VALUE! |
| ENSMUSG00000108991.2 | 0 | NA | NA | NA | NA | NA | #VALUE! |
| ENSMUSG00000108992.2 | 0 | NA | NA | NA | NA | NA | #VALUE! |

| | | | | | | | |
|---|---|---|---|---|---|---|---|
| ENSMUSG00000108993.2 | 0 | NA | NA | NA | NA | NA | #VALUE! |
| ENSMUSG00000108994.2 | 0 | NA | NA | NA | NA | NA | #VALUE! |
| ENSMUSG00000108995.2 | 0 | NA | NA | NA | NA | NA | #VALUE! |
| ENSMUSG00000108997.2 | 0 | NA | NA | NA | NA | NA | #VALUE! |
| ENSMUSG00000108998.3 | 0 | NA | NA | NA | NA | NA | #VALUE! |
| ENSMUSG00000109000.2 | 0 | NA | NA | NA | NA | NA | #VALUE! |
| ENSMUSG00000109001.2 | 0 | NA | NA | NA | NA | NA | #VALUE! |
| ENSMUSG00000109002.2 | 0 | NA | NA | NA | NA | NA | #VALUE! |
| ENSMUSG00000109003.2 | 0 | NA | NA | NA | NA | NA | #VALUE! |
| ENSMUSG00000109004.2 | 0 | NA | NA | NA | NA | NA | #VALUE! |
| ENSMUSG00000109005.2 | 0 | NA | NA | NA | NA | NA | #VALUE! |
| ENSMUSG00000109006.3 | 0 | NA | NA | NA | NA | NA | #VALUE! |
| ENSMUSG00000109007.2 | 0 | NA | NA | NA | NA | NA | #VALUE! |
| ENSMUSG00000109008.3 | 0 | NA | NA | NA | NA | NA | #VALUE! |
| ENSMUSG00000109009.2 | 0 | NA | NA | NA | NA | NA | #VALUE! |
| ENSMUSG00000109010.2 | 0 | NA | NA | NA | NA | NA | #VALUE! |
| ENSMUSG00000109011.2 | 0 | NA | NA | NA | NA | NA | #VALUE! |
| ENSMUSG00000109012.2 | 0 | NA | NA | NA | NA | NA | #VALUE! |
| ENSMUSG00000109013.2 | 0 | NA | NA | NA | NA | NA | #VALUE! |
| ENSMUSG00000109014.2 | 0 | NA | NA | NA | NA | NA | #VALUE! |
| ENSMUSG00000109015.2 | 0 | NA | NA | NA | NA | NA | #VALUE! |
| ENSMUSG00000109016.2 | 0 | NA | NA | NA | NA | NA | #VALUE! |
| ENSMUSG00000109017.2 | 0 | NA | NA | NA | NA | NA | #VALUE! |
| ENSMUSG00000109018.2 | 0 | NA | NA | NA | NA | NA | #VALUE! |
| ENSMUSG00000109019.2 | 0 | NA | NA | NA | NA | NA | #VALUE! |
| ENSMUSG00000109020.5 | 0 | NA | NA | NA | NA | NA | #VALUE! |
| ENSMUSG00000109021.2 | 0 | NA | NA | NA | NA | NA | #VALUE! |
| ENSMUSG00000109022.3 | 0 | NA | NA | NA | NA | NA | #VALUE! |
| ENSMUSG00000109023.2 | 0 | NA | NA | NA | NA | NA | #VALUE! |
| ENSMUSG00000109024.2 | 0 | NA | NA | NA | NA | NA | #VALUE! |
| ENSMUSG00000109025.2 | 0 | NA | NA | NA | NA | NA | #VALUE! |
| ENSMUSG00000109026.2 | 0 | NA | NA | NA | NA | NA | #VALUE! |
| ENSMUSG00000109027.2 | 0 | NA | NA | NA | NA | NA | #VALUE! |
| ENSMUSG00000109028.2 | 0 | NA | NA | NA | NA | NA | #VALUE! |
| ENSMUSG00000109029.2 | 0 | NA | NA | NA | NA | NA | #VALUE! |
| ENSMUSG00000109030.2 | 0 | NA | NA | NA | NA | NA | #VALUE! |
| ENSMUSG00000109031.2 | 0 | NA | NA | NA | NA | NA | #VALUE! |
| ENSMUSG00000109032.2 | 0.652786 | 2.919659 | 3.972474 | 0.734972 | 0.462356 | NA | #VALUE! |
| ENSMUSG00000109033.3 | 0 | NA | NA | NA | NA | NA | #VALUE! |
| ENSMUSG00000109034.2 | 0 | NA | NA | NA | NA | NA | #VALUE! |
| ENSMUSG00000109035.2 | 0 | NA | NA | NA | NA | NA | #VALUE! |
| ENSMUSG00000109036.2 | 0 | NA | NA | NA | NA | NA | #VALUE! |
| ENSMUSG00000109037.2 | 0 | NA | NA | NA | NA | NA | #VALUE! |
| ENSMUSG00000109038.2 | 1.632886 | -4.22813 | 3.934241 | -1.0747 | 0.282509 | NA | #VALUE! |

| | | | | | | | |
|---|---|---|---|---|---|---|---|
| ENSMUSG00000109039.2 | 0 | NA | NA | NA | NA | NA | #VALUE! |
| ENSMUSG00000109040.2 | 0 | NA | NA | NA | NA | NA | #VALUE! |
| ENSMUSG00000109041.2 | 0 | NA | NA | NA | NA | NA | #VALUE! |
| ENSMUSG00000109042.2 | 0 | NA | NA | NA | NA | NA | #VALUE! |
| ENSMUSG00000109043.2 | 0 | NA | NA | NA | NA | NA | #VALUE! |
| ENSMUSG00000109044.2 | 0 | NA | NA | NA | NA | NA | #VALUE! |
| ENSMUSG00000109045.2 | 0 | NA | NA | NA | NA | NA | #VALUE! |
| ENSMUSG00000109046.2 | 0 | NA | NA | NA | NA | NA | #VALUE! |
| ENSMUSG00000109047.2 | 0 | NA | NA | NA | NA | NA | #VALUE! |
| ENSMUSG00000109049.2 | 0 | NA | NA | NA | NA | NA | #VALUE! |
| ENSMUSG00000109050.2 | 0 | NA | NA | NA | NA | NA | #VALUE! |
| ENSMUSG00000109051.2 | 0 | NA | NA | NA | NA | NA | #VALUE! |
| ENSMUSG00000109052.2 | 0 | NA | NA | NA | NA | NA | #VALUE! |
| ENSMUSG00000109053.2 | 0 | NA | NA | NA | NA | NA | #VALUE! |
| ENSMUSG00000109054.2 | 0 | NA | NA | NA | NA | NA | #VALUE! |
| ENSMUSG00000109055.2 | 0 | NA | NA | NA | NA | NA | #VALUE! |
| ENSMUSG00000109056.2 | 0 | NA | NA | NA | NA | NA | #VALUE! |
| ENSMUSG00000109057.2 | 0 | NA | NA | NA | NA | NA | #VALUE! |
| ENSMUSG00000109058.4 | 0 | NA | NA | NA | NA | NA | #VALUE! |
| ENSMUSG00000109059.2 | 2.540421 | 0.903755 | 3.750508 | 0.240969 | 0.809579 | NA | #VALUE! |
| ENSMUSG00000109060.2 | 0 | NA | NA | NA | NA | NA | #VALUE! |
| ENSMUSG00000109061.2 | 0 | NA | NA | NA | NA | NA | #VALUE! |
| ENSMUSG00000109062.2 | 0 | NA | NA | NA | NA | NA | #VALUE! |
| ENSMUSG00000109063.2 | 0 | NA | NA | NA | NA | NA | #VALUE! |
| ENSMUSG00000109064.2 | 0 | NA | NA | NA | NA | NA | #VALUE! |
| ENSMUSG00000109065.2 | 0 | NA | NA | NA | NA | NA | #VALUE! |
| ENSMUSG00000109066.2 | 0 | NA | NA | NA | NA | NA | #VALUE! |
| ENSMUSG00000109067.2 | 0 | NA | NA | NA | NA | NA | #VALUE! |
| ENSMUSG00000109068.2 | 0 | NA | NA | NA | NA | NA | #VALUE! |
| ENSMUSG00000109069.2 | 0 | NA | NA | NA | NA | NA | #VALUE! |
| ENSMUSG00000109070.3 | 0 | NA | NA | NA | NA | NA | #VALUE! |
| ENSMUSG00000109071.2 | 0 | NA | NA | NA | NA | NA | #VALUE! |
| ENSMUSG00000109072.2 | 0 | NA | NA | NA | NA | NA | #VALUE! |
| ENSMUSG00000109073.2 | 0 | NA | NA | NA | NA | NA | #VALUE! |
| ENSMUSG00000109074.2 | 0 | NA | NA | NA | NA | NA | #VALUE! |
| ENSMUSG00000109076.2 | 0 | NA | NA | NA | NA | NA | #VALUE! |
| ENSMUSG00000109077.2 | 0 | NA | NA | NA | NA | NA | #VALUE! |
| ENSMUSG00000109078.2 | 0 | NA | NA | NA | NA | NA | #VALUE! |
| ENSMUSG00000109079.2 | 0 | NA | NA | NA | NA | NA | #VALUE! |
| ENSMUSG00000109080.2 | 0 | NA | NA | NA | NA | NA | #VALUE! |
| ENSMUSG00000109081.2 | 0 | NA | NA | NA | NA | NA | #VALUE! |
| ENSMUSG00000109082.2 | 0 | NA | NA | NA | NA | NA | #VALUE! |
| ENSMUSG00000109083.3 | 0 | NA | NA | NA | NA | NA | #VALUE! |
| ENSMUSG00000109084.2 | 0 | NA | NA | NA | NA | NA | #VALUE! |

| | | | | | | | |
|---|---|---|---|---|---|---|---|
| ENSMUSG00000109085.2 | 0 | NA | NA | NA | NA | NA | #VALUE! |
| ENSMUSG00000109086.2 | 0 | NA | NA | NA | NA | NA | #VALUE! |
| ENSMUSG00000109087.2 | 0 | NA | NA | NA | NA | NA | #VALUE! |
| ENSMUSG00000109088.2 | 0 | NA | NA | NA | NA | NA | #VALUE! |
| ENSMUSG00000109090.2 | 0 | NA | NA | NA | NA | NA | #VALUE! |
| ENSMUSG00000109091.2 | 0 | NA | NA | NA | NA | NA | #VALUE! |
| ENSMUSG00000109092.2 | 0 | NA | NA | NA | NA | NA | #VALUE! |
| ENSMUSG00000109094.2 | 0 | NA | NA | NA | NA | NA | #VALUE! |
| ENSMUSG00000109095.2 | 0 | NA | NA | NA | NA | NA | #VALUE! |
| ENSMUSG00000109096.2 | 0 | NA | NA | NA | NA | NA | #VALUE! |
| ENSMUSG00000109097.2 | 0 | NA | NA | NA | NA | NA | #VALUE! |
| ENSMUSG00000109098.2 | 0 | NA | NA | NA | NA | NA | #VALUE! |
| ENSMUSG00000109099.2 | 0 | NA | NA | NA | NA | NA | #VALUE! |
| ENSMUSG00000109100.2 | 0 | NA | NA | NA | NA | NA | #VALUE! |
| ENSMUSG00000109101.2 | 0 | NA | NA | NA | NA | NA | #VALUE! |
| ENSMUSG00000109102.2 | 0 | NA | NA | NA | NA | NA | #VALUE! |
| ENSMUSG00000109103.2 | 0 | NA | NA | NA | NA | NA | #VALUE! |
| ENSMUSG00000109104.2 | 0 | NA | NA | NA | NA | NA | #VALUE! |
| ENSMUSG00000109105.2 | 0 | NA | NA | NA | NA | NA | #VALUE! |
| ENSMUSG00000109106.2 | 0 | NA | NA | NA | NA | NA | #VALUE! |
| ENSMUSG00000109107.2 | 0 | NA | NA | NA | NA | NA | #VALUE! |
| ENSMUSG00000109108.2 | 6.123323 | -6.14178 | 3.914088 | -1.56915 | 0.116614 | NA | #VALUE! |
| ENSMUSG00000109109.2 | 0 | NA | NA | NA | NA | NA | #VALUE! |
| ENSMUSG00000109110.2 | 0 | NA | NA | NA | NA | NA | #VALUE! |
| ENSMUSG00000109111.2 | 0 | NA | NA | NA | NA | NA | #VALUE! |
| ENSMUSG00000109112.2 | 0 | NA | NA | NA | NA | NA | #VALUE! |
| ENSMUSG00000109113.2 | 0 | NA | NA | NA | NA | NA | #VALUE! |
| ENSMUSG00000109114.2 | 0 | NA | NA | NA | NA | NA | #VALUE! |
| ENSMUSG00000109115.2 | 0 | NA | NA | NA | NA | NA | #VALUE! |
| ENSMUSG00000109116.2 | 0 | NA | NA | NA | NA | NA | #VALUE! |
| ENSMUSG00000109117.2 | 0 | NA | NA | NA | NA | NA | #VALUE! |
| ENSMUSG00000109119.2 | 0 | NA | NA | NA | NA | NA | #VALUE! |
| ENSMUSG00000109121.2 | 0 | NA | NA | NA | NA | NA | #VALUE! |
| ENSMUSG00000109122.2 | 0 | NA | NA | NA | NA | NA | #VALUE! |
| ENSMUSG00000109123.2 | 0 | NA | NA | NA | NA | NA | #VALUE! |
| ENSMUSG00000109124.2 | 0 | NA | NA | NA | NA | NA | #VALUE! |
| ENSMUSG00000109125.2 | 0 | NA | NA | NA | NA | NA | #VALUE! |
| ENSMUSG00000109126.2 | 0 | NA | NA | NA | NA | NA | #VALUE! |
| ENSMUSG00000109127.2 | 0 | NA | NA | NA | NA | NA | #VALUE! |
| ENSMUSG00000109128.2 | 0 | NA | NA | NA | NA | NA | #VALUE! |
| ENSMUSG00000109129.2 | 0 | NA | NA | NA | NA | NA | #VALUE! |
| ENSMUSG00000109130.2 | 0 | NA | NA | NA | NA | NA | #VALUE! |
| ENSMUSG00000109131.2 | 0 | NA | NA | NA | NA | NA | #VALUE! |
| ENSMUSG00000109132.2 | 0 | NA | NA | NA | NA | NA | #VALUE! |

| | | | | | | | |
|---|---|---|---|---|---|---|---|
| ENSMUSG00000109133.2 | 0 | NA | NA | NA | NA | NA | #VALUE! |
| ENSMUSG00000109134.2 | 0 | NA | NA | NA | NA | NA | #VALUE! |
| ENSMUSG00000109135.2 | 0 | NA | NA | NA | NA | NA | #VALUE! |
| ENSMUSG00000109136.2 | 0 | NA | NA | NA | NA | NA | #VALUE! |
| ENSMUSG00000109137.2 | 0 | NA | NA | NA | NA | NA | #VALUE! |
| ENSMUSG00000109138.2 | 0 | NA | NA | NA | NA | NA | #VALUE! |
| ENSMUSG00000109139.2 | 0 | NA | NA | NA | NA | NA | #VALUE! |
| ENSMUSG00000109140.2 | 0 | NA | NA | NA | NA | NA | #VALUE! |
| ENSMUSG00000109141.2 | 0 | NA | NA | NA | NA | NA | #VALUE! |
| ENSMUSG00000109142.2 | 0 | NA | NA | NA | NA | NA | #VALUE! |
| ENSMUSG00000109143.2 | 0 | NA | NA | NA | NA | NA | #VALUE! |
| ENSMUSG00000109145.2 | 0 | NA | NA | NA | NA | NA | #VALUE! |
| ENSMUSG00000109147.2 | 1.933292 | -0.92566 | 3.829728 | -0.2417 | 0.809009 | NA | #VALUE! |
| ENSMUSG00000109148.2 | 0 | NA | NA | NA | NA | NA | #VALUE! |
| ENSMUSG00000109149.2 | 0 | NA | NA | NA | NA | NA | #VALUE! |
| ENSMUSG00000109150.2 | 0 | NA | NA | NA | NA | NA | #VALUE! |
| ENSMUSG00000109151.2 | 0 | NA | NA | NA | NA | NA | #VALUE! |
| ENSMUSG00000109152.2 | 0 | NA | NA | NA | NA | NA | #VALUE! |
| ENSMUSG00000109153.2 | 0 | NA | NA | NA | NA | NA | #VALUE! |
| ENSMUSG00000109154.2 | 0 | NA | NA | NA | NA | NA | #VALUE! |
| ENSMUSG00000109155.2 | 0 | NA | NA | NA | NA | NA | #VALUE! |
| ENSMUSG00000109156.2 | 0 | NA | NA | NA | NA | NA | #VALUE! |
| ENSMUSG00000109157.2 | 0.105013 | -0.97324 | 4.076293 | -0.23876 | 0.811295 | NA | #VALUE! |
| ENSMUSG00000109158.2 | 0 | NA | NA | NA | NA | NA | #VALUE! |
| ENSMUSG00000109159.2 | 0 | NA | NA | NA | NA | NA | #VALUE! |
| ENSMUSG00000109160.2 | 0 | NA | NA | NA | NA | NA | #VALUE! |
| ENSMUSG00000109161.2 | 0 | NA | NA | NA | NA | NA | #VALUE! |
| ENSMUSG00000109162.2 | 0 | NA | NA | NA | NA | NA | #VALUE! |
| ENSMUSG00000109163.2 | 0 | NA | NA | NA | NA | NA | #VALUE! |
| ENSMUSG00000109164.2 | 0 | NA | NA | NA | NA | NA | #VALUE! |
| ENSMUSG00000109165.2 | 0 | NA | NA | NA | NA | NA | #VALUE! |
| ENSMUSG00000109166.2 | 0 | NA | NA | NA | NA | NA | #VALUE! |
| ENSMUSG00000109167.2 | 0 | NA | NA | NA | NA | NA | #VALUE! |
| ENSMUSG00000109168.2 | 0 | NA | NA | NA | NA | NA | #VALUE! |
| ENSMUSG00000109169.2 | 0 | NA | NA | NA | NA | NA | #VALUE! |
| ENSMUSG00000109170.2 | 0 | NA | NA | NA | NA | NA | #VALUE! |
| ENSMUSG00000109171.2 | 0 | NA | NA | NA | NA | NA | #VALUE! |
| ENSMUSG00000109172.2 | 0 | NA | NA | NA | NA | NA | #VALUE! |
| ENSMUSG00000109173.2 | 0 | NA | NA | NA | NA | NA | #VALUE! |
| ENSMUSG00000109174.2 | 0 | NA | NA | NA | NA | NA | #VALUE! |
| ENSMUSG00000109175.2 | 0 | NA | NA | NA | NA | NA | #VALUE! |
| ENSMUSG00000109176.2 | 0 | NA | NA | NA | NA | NA | #VALUE! |
| ENSMUSG00000109177.2 | 0 | NA | NA | NA | NA | NA | #VALUE! |
| ENSMUSG00000109178.2 | 0 | NA | NA | NA | NA | NA | #VALUE! |

| | | | | | | | |
|---|---|---|---|---|---|---|---|
| ENSMUSG00000109179.3 | 0 | NA | NA | NA | NA | NA | #VALUE! |
| ENSMUSG00000109180.2 | 0 | NA | NA | NA | NA | NA | #VALUE! |
| ENSMUSG00000109181.2 | 5.642491 | 0.761391 | 3.136419 | 0.242758 | 0.808193 | NA | #VALUE! |
| ENSMUSG00000109182.2 | 0 | NA | NA | NA | NA | NA | #VALUE! |
| ENSMUSG00000109183.2 | 0 | NA | NA | NA | NA | NA | #VALUE! |
| ENSMUSG00000109184.2 | 0 | NA | NA | NA | NA | NA | #VALUE! |
| ENSMUSG00000109185.2 | 0 | NA | NA | NA | NA | NA | #VALUE! |
| ENSMUSG00000109186.2 | 0 | NA | NA | NA | NA | NA | #VALUE! |
| ENSMUSG00000109188.2 | 0 | NA | NA | NA | NA | NA | #VALUE! |
| ENSMUSG00000109189.2 | 0 | NA | NA | NA | NA | NA | #VALUE! |
| ENSMUSG00000109190.2 | 0 | NA | NA | NA | NA | NA | #VALUE! |
| ENSMUSG00000109191.2 | 0 | NA | NA | NA | NA | NA | #VALUE! |
| ENSMUSG00000109192.3 | 0 | NA | NA | NA | NA | NA | #VALUE! |
| ENSMUSG00000109193.2 | 0 | NA | NA | NA | NA | NA | #VALUE! |
| ENSMUSG00000109194.2 | 0 | NA | NA | NA | NA | NA | #VALUE! |
| ENSMUSG00000109195.2 | 0 | NA | NA | NA | NA | NA | #VALUE! |
| ENSMUSG00000109196.2 | 0 | NA | NA | NA | NA | NA | #VALUE! |
| ENSMUSG00000109197.2 | 0 | NA | NA | NA | NA | NA | #VALUE! |
| ENSMUSG00000109198.2 | 0 | NA | NA | NA | NA | NA | #VALUE! |
| ENSMUSG00000109199.2 | 3.327096 | 5.225506 | 3.920239 | 1.332956 | 0.182546 | NA | #VALUE! |
| ENSMUSG00000109200.2 | 0 | NA | NA | NA | NA | NA | #VALUE! |
| ENSMUSG00000109201.2 | 0 | NA | NA | NA | NA | NA | #VALUE! |
| ENSMUSG00000109202.2 | 0 | NA | NA | NA | NA | NA | #VALUE! |
| ENSMUSG00000109203.2 | 0 | NA | NA | NA | NA | NA | #VALUE! |
| ENSMUSG00000109204.2 | 0 | NA | NA | NA | NA | NA | #VALUE! |
| ENSMUSG00000109205.2 | 0 | NA | NA | NA | NA | NA | #VALUE! |
| ENSMUSG00000109206.2 | 0 | NA | NA | NA | NA | NA | #VALUE! |
| ENSMUSG00000109207.2 | 0 | NA | NA | NA | NA | NA | #VALUE! |
| ENSMUSG00000109209.2 | 0 | NA | NA | NA | NA | NA | #VALUE! |
| ENSMUSG00000109210.2 | 0 | NA | NA | NA | NA | NA | #VALUE! |
| ENSMUSG00000109211.2 | 0 | NA | NA | NA | NA | NA | #VALUE! |
| ENSMUSG00000109212.4 | 0 | NA | NA | NA | NA | NA | #VALUE! |
| ENSMUSG00000109213.3 | 0 | NA | NA | NA | NA | NA | #VALUE! |
| ENSMUSG00000109214.2 | 0 | NA | NA | NA | NA | NA | #VALUE! |
| ENSMUSG00000109215.2 | 0 | NA | NA | NA | NA | NA | #VALUE! |
| ENSMUSG00000109216.2 | 0 | NA | NA | NA | NA | NA | #VALUE! |
| ENSMUSG00000109217.2 | 0 | NA | NA | NA | NA | NA | #VALUE! |
| ENSMUSG00000109218.2 | 0 | NA | NA | NA | NA | NA | #VALUE! |
| ENSMUSG00000109219.3 | 0 | NA | NA | NA | NA | NA | #VALUE! |
| ENSMUSG00000109220.2 | 0 | NA | NA | NA | NA | NA | #VALUE! |
| ENSMUSG00000109221.2 | 0 | NA | NA | NA | NA | NA | #VALUE! |
| ENSMUSG00000109222.2 | 0 | NA | NA | NA | NA | NA | #VALUE! |
| ENSMUSG00000109223.2 | 0 | NA | NA | NA | NA | NA | #VALUE! |
| ENSMUSG00000109224.2 | 0 | NA | NA | NA | NA | NA | #VALUE! |

| | | | | | | | |
|---|---|---|---|---|---|---|---|
| ENSMUSG00000109225.2 | 0 | NA | NA | NA | NA | NA | #VALUE! |
| ENSMUSG00000109226.2 | 0 | NA | NA | NA | NA | NA | #VALUE! |
| ENSMUSG00000109227.2 | 0 | NA | NA | NA | NA | NA | #VALUE! |
| ENSMUSG00000109228.3 | 0 | NA | NA | NA | NA | NA | #VALUE! |
| ENSMUSG00000109229.2 | 0 | NA | NA | NA | NA | NA | #VALUE! |
| ENSMUSG00000109230.2 | 0 | NA | NA | NA | NA | NA | #VALUE! |
| ENSMUSG00000109231.2 | 0 | NA | NA | NA | NA | NA | #VALUE! |
| ENSMUSG00000109232.2 | 0 | NA | NA | NA | NA | NA | #VALUE! |
| ENSMUSG00000109233.2 | 0 | NA | NA | NA | NA | NA | #VALUE! |
| ENSMUSG00000109234.2 | 0 | NA | NA | NA | NA | NA | #VALUE! |
| ENSMUSG00000109235.2 | 0 | NA | NA | NA | NA | NA | #VALUE! |
| ENSMUSG00000109236.2 | 0 | NA | NA | NA | NA | NA | #VALUE! |
| ENSMUSG00000109237.2 | 0 | NA | NA | NA | NA | NA | #VALUE! |
| ENSMUSG00000109238.3 | 0 | NA | NA | NA | NA | NA | #VALUE! |
| ENSMUSG00000109239.2 | 0 | NA | NA | NA | NA | NA | #VALUE! |
| ENSMUSG00000109240.2 | 0 | NA | NA | NA | NA | NA | #VALUE! |
| ENSMUSG00000109241.2 | 1.584478 | -2.05773 | 3.895684 | -0.52821 | 0.597355 | NA | #VALUE! |
| ENSMUSG00000109242.2 | 0 | NA | NA | NA | NA | NA | #VALUE! |
| ENSMUSG00000109243.2 | 0 | NA | NA | NA | NA | NA | #VALUE! |
| ENSMUSG00000109244.2 | 0 | NA | NA | NA | NA | NA | #VALUE! |
| ENSMUSG00000109245.2 | 0 | NA | NA | NA | NA | NA | #VALUE! |
| ENSMUSG00000109246.3 | 0 | NA | NA | NA | NA | NA | #VALUE! |
| ENSMUSG00000109247.2 | 0 | NA | NA | NA | NA | NA | #VALUE! |
| ENSMUSG00000109248.2 | 0 | NA | NA | NA | NA | NA | #VALUE! |
| ENSMUSG00000109249.2 | 0 | NA | NA | NA | NA | NA | #VALUE! |
| ENSMUSG00000109250.2 | 0 | NA | NA | NA | NA | NA | #VALUE! |
| ENSMUSG00000109251.2 | 0 | NA | NA | NA | NA | NA | #VALUE! |
| ENSMUSG00000109252.2 | 0 | NA | NA | NA | NA | NA | #VALUE! |
| ENSMUSG00000109253.2 | 0 | NA | NA | NA | NA | NA | #VALUE! |
| ENSMUSG00000109254.2 | 0 | NA | NA | NA | NA | NA | #VALUE! |
| ENSMUSG00000109255.2 | 0 | NA | NA | NA | NA | NA | #VALUE! |
| ENSMUSG00000109256.2 | 0 | NA | NA | NA | NA | NA | #VALUE! |
| ENSMUSG00000109257.2 | 0 | NA | NA | NA | NA | NA | #VALUE! |
| ENSMUSG00000109258.2 | 0 | NA | NA | NA | NA | NA | #VALUE! |
| ENSMUSG00000109259.2 | 0 | NA | NA | NA | NA | NA | #VALUE! |
| ENSMUSG00000109260.2 | 0 | NA | NA | NA | NA | NA | #VALUE! |
| ENSMUSG00000109261.2 | 0 | NA | NA | NA | NA | NA | #VALUE! |
| ENSMUSG00000109262.2 | 4.744151 | 5.738953 | 3.916213 | 1.465434 | 0.142802 | NA | #VALUE! |
| ENSMUSG00000109263.2 | 0 | NA | NA | NA | NA | NA | #VALUE! |
| ENSMUSG00000109264.2 | 0 | NA | NA | NA | NA | NA | #VALUE! |
| ENSMUSG00000109266.2 | 0 | NA | NA | NA | NA | NA | #VALUE! |
| ENSMUSG00000109267.2 | 0 | NA | NA | NA | NA | NA | #VALUE! |
| ENSMUSG00000109268.2 | 0 | NA | NA | NA | NA | NA | #VALUE! |
| ENSMUSG00000109269.2 | 0 | NA | NA | NA | NA | NA | #VALUE! |

| | | | | | | | |
|---|---|---|---|---|---|---|---|
| ENSMUSG00000109270.2 | 0 | NA | NA | NA | NA | NA | #VALUE! |
| ENSMUSG00000109271.2 | 0 | NA | NA | NA | NA | NA | #VALUE! |
| ENSMUSG00000109272.2 | 0 | NA | NA | NA | NA | NA | #VALUE! |
| ENSMUSG00000109273.2 | 0 | NA | NA | NA | NA | NA | #VALUE! |
| ENSMUSG00000109274.2 | 8.223142 | -6.56817 | 3.836124 | -1.71219 | 0.086862 | NA | #VALUE! |
| ENSMUSG00000109275.2 | 0 | NA | NA | NA | NA | NA | #VALUE! |
| ENSMUSG00000109276.2 | 0 | NA | NA | NA | NA | NA | #VALUE! |
| ENSMUSG00000109277.2 | 0 | NA | NA | NA | NA | NA | #VALUE! |
| ENSMUSG00000109278.2 | 0 | NA | NA | NA | NA | NA | #VALUE! |
| ENSMUSG00000109279.2 | 5.87863 | -2.34512 | 3.782931 | -0.61992 | 0.535309 | NA | #VALUE! |
| ENSMUSG00000109280.2 | 0 | NA | NA | NA | NA | NA | #VALUE! |
| ENSMUSG00000109281.2 | 0 | NA | NA | NA | NA | NA | #VALUE! |
| ENSMUSG00000109282.2 | 0 | NA | NA | NA | NA | NA | #VALUE! |
| ENSMUSG00000109283.2 | 0 | NA | NA | NA | NA | NA | #VALUE! |
| ENSMUSG00000109284.2 | 0 | NA | NA | NA | NA | NA | #VALUE! |
| ENSMUSG00000109285.2 | 0 | NA | NA | NA | NA | NA | #VALUE! |
| ENSMUSG00000109286.2 | 0 | NA | NA | NA | NA | NA | #VALUE! |
| ENSMUSG00000109287.2 | 0 | NA | NA | NA | NA | NA | #VALUE! |
| ENSMUSG00000109289.2 | 0 | NA | NA | NA | NA | NA | #VALUE! |
| ENSMUSG00000109290.2 | 0 | NA | NA | NA | NA | NA | #VALUE! |
| ENSMUSG00000109292.2 | 0 | NA | NA | NA | NA | NA | #VALUE! |
| ENSMUSG00000109293.2 | 0 | NA | NA | NA | NA | NA | #VALUE! |
| ENSMUSG00000109294.2 | 0 | NA | NA | NA | NA | NA | #VALUE! |
| ENSMUSG00000109295.2 | 0 | NA | NA | NA | NA | NA | #VALUE! |
| ENSMUSG00000109297.2 | 0 | NA | NA | NA | NA | NA | #VALUE! |
| ENSMUSG00000109298.2 | 0 | NA | NA | NA | NA | NA | #VALUE! |
| ENSMUSG00000109299.2 | 0 | NA | NA | NA | NA | NA | #VALUE! |
| ENSMUSG00000109300.2 | 0 | NA | NA | NA | NA | NA | #VALUE! |
| ENSMUSG00000109301.2 | 0 | NA | NA | NA | NA | NA | #VALUE! |
| ENSMUSG00000109302.2 | 0 | NA | NA | NA | NA | NA | #VALUE! |
| ENSMUSG00000109303.2 | 0 | NA | NA | NA | NA | NA | #VALUE! |
| ENSMUSG00000109304.2 | 0 | NA | NA | NA | NA | NA | #VALUE! |
| ENSMUSG00000109305.2 | 0 | NA | NA | NA | NA | NA | #VALUE! |
| ENSMUSG00000109306.2 | 0 | NA | NA | NA | NA | NA | #VALUE! |
| ENSMUSG00000109307.2 | 0 | NA | NA | NA | NA | NA | #VALUE! |
| ENSMUSG00000109308.2 | 0 | NA | NA | NA | NA | NA | #VALUE! |
| ENSMUSG00000109309.2 | 0 | NA | NA | NA | NA | NA | #VALUE! |
| ENSMUSG00000109310.2 | 0 | NA | NA | NA | NA | NA | #VALUE! |
| ENSMUSG00000109311.2 | 0 | NA | NA | NA | NA | NA | #VALUE! |
| ENSMUSG00000109312.2 | 0 | NA | NA | NA | NA | NA | #VALUE! |
| ENSMUSG00000109313.2 | 0 | NA | NA | NA | NA | NA | #VALUE! |
| ENSMUSG00000109314.2 | 0 | NA | NA | NA | NA | NA | #VALUE! |
| ENSMUSG00000109315.2 | 0 | NA | NA | NA | NA | NA | #VALUE! |
| ENSMUSG00000109316.2 | 0 | NA | NA | NA | NA | NA | #VALUE! |

| | | | | | | | |
|---|---|---|---|---|---|---|---|
| ENSMUSG00000109317.2 | 0 | NA | NA | NA | NA | NA | #VALUE! |
| ENSMUSG00000109318.2 | 0 | NA | NA | NA | NA | NA | #VALUE! |
| ENSMUSG00000109319.2 | 0 | NA | NA | NA | NA | NA | #VALUE! |
| ENSMUSG00000109320.2 | 0 | NA | NA | NA | NA | NA | #VALUE! |
| ENSMUSG00000109321.2 | 0 | NA | NA | NA | NA | NA | #VALUE! |
| ENSMUSG00000109322.3 | 0 | NA | NA | NA | NA | NA | #VALUE! |
| ENSMUSG00000109323.2 | 0 | NA | NA | NA | NA | NA | #VALUE! |
| ENSMUSG00000109324.3 | 6.062599 | -0.43404 | 3.096402 | -0.14018 | 0.888522 | NA | #VALUE! |
| ENSMUSG00000109325.2 | 0 | NA | NA | NA | NA | NA | #VALUE! |
| ENSMUSG00000109326.2 | 0 | NA | NA | NA | NA | NA | #VALUE! |
| ENSMUSG00000109327.2 | 0 | NA | NA | NA | NA | NA | #VALUE! |
| ENSMUSG00000109328.3 | 0 | NA | NA | NA | NA | NA | #VALUE! |
| ENSMUSG00000109329.2 | 0 | NA | NA | NA | NA | NA | #VALUE! |
| ENSMUSG00000109330.2 | 0 | NA | NA | NA | NA | NA | #VALUE! |
| ENSMUSG00000109331.2 | 0 | NA | NA | NA | NA | NA | #VALUE! |
| ENSMUSG00000109332.2 | 1.365171 | -3.99822 | 3.938957 | -1.01505 | 0.310084 | NA | #VALUE! |
| ENSMUSG00000109333.2 | 0 | NA | NA | NA | NA | NA | #VALUE! |
| ENSMUSG00000109334.2 | 0 | NA | NA | NA | NA | NA | #VALUE! |
| ENSMUSG00000109335.2 | 0 | NA | NA | NA | NA | NA | #VALUE! |
| ENSMUSG00000109337.2 | 0 | NA | NA | NA | NA | NA | #VALUE! |
| ENSMUSG00000109338.3 | 0 | NA | NA | NA | NA | NA | #VALUE! |
| ENSMUSG00000109339.2 | 0 | NA | NA | NA | NA | NA | #VALUE! |
| ENSMUSG00000109340.2 | 0 | NA | NA | NA | NA | NA | #VALUE! |
| ENSMUSG00000109341.2 | 0 | NA | NA | NA | NA | NA | #VALUE! |
| ENSMUSG00000109342.2 | 0 | NA | NA | NA | NA | NA | #VALUE! |
| ENSMUSG00000109343.2 | 0 | NA | NA | NA | NA | NA | #VALUE! |
| ENSMUSG00000109344.2 | 0 | NA | NA | NA | NA | NA | #VALUE! |
| ENSMUSG00000109345.2 | 0 | NA | NA | NA | NA | NA | #VALUE! |
| ENSMUSG00000109346.2 | 0 | NA | NA | NA | NA | NA | #VALUE! |
| ENSMUSG00000109347.2 | 0 | NA | NA | NA | NA | NA | #VALUE! |
| ENSMUSG00000109348.2 | 0 | NA | NA | NA | NA | NA | #VALUE! |
| ENSMUSG00000109350.2 | 0 | NA | NA | NA | NA | NA | #VALUE! |
| ENSMUSG00000109351.2 | 0 | NA | NA | NA | NA | NA | #VALUE! |
| ENSMUSG00000109352.2 | 0 | NA | NA | NA | NA | NA | #VALUE! |
| ENSMUSG00000109353.2 | 0 | NA | NA | NA | NA | NA | #VALUE! |
| ENSMUSG00000109354.3 | 0 | NA | NA | NA | NA | NA | #VALUE! |
| ENSMUSG00000109355.2 | 0 | NA | NA | NA | NA | NA | #VALUE! |
| ENSMUSG00000109356.2 | 0 | NA | NA | NA | NA | NA | #VALUE! |
| ENSMUSG00000109357.2 | 0.737927 | 3.007825 | 3.968641 | 0.757898 | 0.448512 | NA | #VALUE! |
| ENSMUSG00000109358.2 | 0 | NA | NA | NA | NA | NA | #VALUE! |
| ENSMUSG00000109359.2 | 0 | NA | NA | NA | NA | NA | #VALUE! |
| ENSMUSG00000109360.2 | 0 | NA | NA | NA | NA | NA | #VALUE! |
| ENSMUSG00000109361.2 | 0 | NA | NA | NA | NA | NA | #VALUE! |
| ENSMUSG00000109362.2 | 0 | NA | NA | NA | NA | NA | #VALUE! |

| | | | | | | | |
|---|---|---|---|---|---|---|---|
| ENSMUSG00000109363.2 | 0 | NA | NA | NA | NA | NA | #VALUE! |
| ENSMUSG00000109364.2 | 0 | NA | NA | NA | NA | NA | #VALUE! |
| ENSMUSG00000109365.3 | 0 | NA | NA | NA | NA | NA | #VALUE! |
| ENSMUSG00000109366.2 | 0 | NA | NA | NA | NA | NA | #VALUE! |
| ENSMUSG00000109367.2 | 0 | NA | NA | NA | NA | NA | #VALUE! |
| ENSMUSG00000109368.2 | 0 | NA | NA | NA | NA | NA | #VALUE! |
| ENSMUSG00000109369.3 | 0 | NA | NA | NA | NA | NA | #VALUE! |
| ENSMUSG00000109370.2 | 0 | NA | NA | NA | NA | NA | #VALUE! |
| ENSMUSG00000109371.2 | 0 | NA | NA | NA | NA | NA | #VALUE! |
| ENSMUSG00000109372.2 | 0 | NA | NA | NA | NA | NA | #VALUE! |
| ENSMUSG00000109373.2 | 0 | NA | NA | NA | NA | NA | #VALUE! |
| ENSMUSG00000109374.2 | 0 | NA | NA | NA | NA | NA | #VALUE! |
| ENSMUSG00000109375.2 | 0 | NA | NA | NA | NA | NA | #VALUE! |
| ENSMUSG00000109376.2 | 0 | NA | NA | NA | NA | NA | #VALUE! |
| ENSMUSG00000109377.2 | 0 | NA | NA | NA | NA | NA | #VALUE! |
| ENSMUSG00000109378.2 | 0 | NA | NA | NA | NA | NA | #VALUE! |
| ENSMUSG00000109379.2 | 1.566313 | -4.16993 | 3.935366 | -1.0596 | 0.289325 | NA | #VALUE! |
| ENSMUSG00000109380.2 | 0 | NA | NA | NA | NA | NA | #VALUE! |
| ENSMUSG00000109381.2 | 0 | NA | NA | NA | NA | NA | #VALUE! |
| ENSMUSG00000109382.2 | 0 | NA | NA | NA | NA | NA | #VALUE! |
| ENSMUSG00000109383.2 | 0 | NA | NA | NA | NA | NA | #VALUE! |
| ENSMUSG00000109384.2 | 0 | NA | NA | NA | NA | NA | #VALUE! |
| ENSMUSG00000109385.2 | 0 | NA | NA | NA | NA | NA | #VALUE! |
| ENSMUSG00000109387.2 | 0 | NA | NA | NA | NA | NA | #VALUE! |
| ENSMUSG00000109388.2 | 0 | NA | NA | NA | NA | NA | #VALUE! |
| ENSMUSG00000109389.2 | 0 | NA | NA | NA | NA | NA | #VALUE! |
| ENSMUSG00000109390.2 | 0 | NA | NA | NA | NA | NA | #VALUE! |
| ENSMUSG00000109391.2 | 0 | NA | NA | NA | NA | NA | #VALUE! |
| ENSMUSG00000109392.2 | 0 | NA | NA | NA | NA | NA | #VALUE! |
| ENSMUSG00000109393.2 | 0 | NA | NA | NA | NA | NA | #VALUE! |
| ENSMUSG00000109394.3 | 0 | NA | NA | NA | NA | NA | #VALUE! |
| ENSMUSG00000109395.2 | 0 | NA | NA | NA | NA | NA | #VALUE! |
| ENSMUSG00000109396.2 | 0 | NA | NA | NA | NA | NA | #VALUE! |
| ENSMUSG00000109397.2 | 0 | NA | NA | NA | NA | NA | #VALUE! |
| ENSMUSG00000109398.3 | 0 | NA | NA | NA | NA | NA | #VALUE! |
| ENSMUSG00000109399.2 | 0 | NA | NA | NA | NA | NA | #VALUE! |
| ENSMUSG00000109400.2 | 0 | NA | NA | NA | NA | NA | #VALUE! |
| ENSMUSG00000109401.2 | 0 | NA | NA | NA | NA | NA | #VALUE! |
| ENSMUSG00000109402.2 | 0 | NA | NA | NA | NA | NA | #VALUE! |
| ENSMUSG00000109403.3 | 0 | NA | NA | NA | NA | NA | #VALUE! |
| ENSMUSG00000109404.2 | 0 | NA | NA | NA | NA | NA | #VALUE! |
| ENSMUSG00000109405.2 | 0 | NA | NA | NA | NA | NA | #VALUE! |
| ENSMUSG00000109406.3 | 0 | NA | NA | NA | NA | NA | #VALUE! |
| ENSMUSG00000109407.2 | 0 | NA | NA | NA | NA | NA | #VALUE! |

| | | | | | | | |
|---|---|---|---|---|---|---|---|
| ENSMUSG00000109408.2 | 2.355969 | 4.72622 | 3.925784 | 1.203892 | 0.228631 | NA | #VALUE! |
| ENSMUSG00000109409.2 | 0 | NA | NA | NA | NA | NA | #VALUE! |
| ENSMUSG00000109410.2 | 0 | NA | NA | NA | NA | NA | #VALUE! |
| ENSMUSG00000109411.2 | 0 | NA | NA | NA | NA | NA | #VALUE! |
| ENSMUSG00000109412.2 | 0 | NA | NA | NA | NA | NA | #VALUE! |
| ENSMUSG00000109413.2 | 0 | NA | NA | NA | NA | NA | #VALUE! |
| ENSMUSG00000109414.2 | 0 | NA | NA | NA | NA | NA | #VALUE! |
| ENSMUSG00000109415.2 | 0 | NA | NA | NA | NA | NA | #VALUE! |
| ENSMUSG00000109416.2 | 0 | NA | NA | NA | NA | NA | #VALUE! |
| ENSMUSG00000109417.2 | 0 | NA | NA | NA | NA | NA | #VALUE! |
| ENSMUSG00000109418.2 | 0 | NA | NA | NA | NA | NA | #VALUE! |
| ENSMUSG00000109419.2 | 0 | NA | NA | NA | NA | NA | #VALUE! |
| ENSMUSG00000109420.2 | 0 | NA | NA | NA | NA | NA | #VALUE! |
| ENSMUSG00000109421.2 | 0 | NA | NA | NA | NA | NA | #VALUE! |
| ENSMUSG00000109422.2 | 0 | NA | NA | NA | NA | NA | #VALUE! |
| ENSMUSG00000109423.2 | 0 | NA | NA | NA | NA | NA | #VALUE! |
| ENSMUSG00000109424.2 | 0 | NA | NA | NA | NA | NA | #VALUE! |
| ENSMUSG00000109425.2 | 0 | NA | NA | NA | NA | NA | #VALUE! |
| ENSMUSG00000109426.3 | 0 | NA | NA | NA | NA | NA | #VALUE! |
| ENSMUSG00000109427.2 | 0 | NA | NA | NA | NA | NA | #VALUE! |
| ENSMUSG00000109428.2 | 0 | NA | NA | NA | NA | NA | #VALUE! |
| ENSMUSG00000109429.2 | 0.618321 | -1.73856 | 3.9799 | -0.43683 | 0.662231 | NA | #VALUE! |
| ENSMUSG00000109430.2 | 0 | NA | NA | NA | NA | NA | #VALUE! |
| ENSMUSG00000109431.2 | 0 | NA | NA | NA | NA | NA | #VALUE! |
| ENSMUSG00000109432.2 | 0 | NA | NA | NA | NA | NA | #VALUE! |
| ENSMUSG00000109433.2 | 0 | NA | NA | NA | NA | NA | #VALUE! |
| ENSMUSG00000109434.2 | 0 | NA | NA | NA | NA | NA | #VALUE! |
| ENSMUSG00000109435.2 | 0 | NA | NA | NA | NA | NA | #VALUE! |
| ENSMUSG00000109436.2 | 0 | NA | NA | NA | NA | NA | #VALUE! |
| ENSMUSG00000109437.2 | 0 | NA | NA | NA | NA | NA | #VALUE! |
| ENSMUSG00000109438.2 | 0 | NA | NA | NA | NA | NA | #VALUE! |
| ENSMUSG00000109439.2 | 0 | NA | NA | NA | NA | NA | #VALUE! |
| ENSMUSG00000109440.3 | 0 | NA | NA | NA | NA | NA | #VALUE! |
| ENSMUSG00000109441.2 | 0 | NA | NA | NA | NA | NA | #VALUE! |
| ENSMUSG00000109442.2 | 0 | NA | NA | NA | NA | NA | #VALUE! |
| ENSMUSG00000109443.2 | 0 | NA | NA | NA | NA | NA | #VALUE! |
| ENSMUSG00000109444.2 | 0 | NA | NA | NA | NA | NA | #VALUE! |
| ENSMUSG00000109445.2 | 0 | NA | NA | NA | NA | NA | #VALUE! |
| ENSMUSG00000109446.2 | 0 | NA | NA | NA | NA | NA | #VALUE! |
| ENSMUSG00000109447.2 | 0 | NA | NA | NA | NA | NA | #VALUE! |
| ENSMUSG00000109448.2 | 0 | NA | NA | NA | NA | NA | #VALUE! |
| ENSMUSG00000109449.4 | 0 | NA | NA | NA | NA | NA | #VALUE! |
| ENSMUSG00000109450.2 | 0 | NA | NA | NA | NA | NA | #VALUE! |
| ENSMUSG00000109451.2 | 0 | NA | NA | NA | NA | NA | #VALUE! |

| | | | | | | | |
|---|---|---|---|---|---|---|---|
| ENSMUSG00000109452.2 | 0 | NA | NA | NA | NA | NA | #VALUE! |
| ENSMUSG00000109453.2 | 0 | NA | NA | NA | NA | NA | #VALUE! |
| ENSMUSG00000109454.2 | 0 | NA | NA | NA | NA | NA | #VALUE! |
| ENSMUSG00000109455.2 | 0 | NA | NA | NA | NA | NA | #VALUE! |
| ENSMUSG00000109456.2 | 0 | NA | NA | NA | NA | NA | #VALUE! |
| ENSMUSG00000109457.2 | 0 | NA | NA | NA | NA | NA | #VALUE! |
| ENSMUSG00000109458.2 | 0 | NA | NA | NA | NA | NA | #VALUE! |
| ENSMUSG00000109459.2 | 0 | NA | NA | NA | NA | NA | #VALUE! |
| ENSMUSG00000109460.2 | 0 | NA | NA | NA | NA | NA | #VALUE! |
| ENSMUSG00000109461.2 | 0 | NA | NA | NA | NA | NA | #VALUE! |
| ENSMUSG00000109462.2 | 0 | NA | NA | NA | NA | NA | #VALUE! |
| ENSMUSG00000109463.2 | 0 | NA | NA | NA | NA | NA | #VALUE! |
| ENSMUSG00000109464.2 | 0 | NA | NA | NA | NA | NA | #VALUE! |
| ENSMUSG00000109465.2 | 0 | NA | NA | NA | NA | NA | #VALUE! |
| ENSMUSG00000109466.2 | 0 | NA | NA | NA | NA | NA | #VALUE! |
| ENSMUSG00000109467.2 | 0 | NA | NA | NA | NA | NA | #VALUE! |
| ENSMUSG00000109468.2 | 0 | NA | NA | NA | NA | NA | #VALUE! |
| ENSMUSG00000109469.2 | 0 | NA | NA | NA | NA | NA | #VALUE! |
| ENSMUSG00000109471.2 | 0 | NA | NA | NA | NA | NA | #VALUE! |
| ENSMUSG00000109472.2 | 0 | NA | NA | NA | NA | NA | #VALUE! |
| ENSMUSG00000109473.2 | 1.877451 | -0.8502 | 3.831546 | -0.22189 | 0.824397 | NA | #VALUE! |
| ENSMUSG00000109475.2 | 0 | NA | NA | NA | NA | NA | #VALUE! |
| ENSMUSG00000109476.2 | 0 | NA | NA | NA | NA | NA | #VALUE! |
| ENSMUSG00000109477.2 | 0 | NA | NA | NA | NA | NA | #VALUE! |
| ENSMUSG00000109478.2 | 0 | NA | NA | NA | NA | NA | #VALUE! |
| ENSMUSG00000109479.2 | 0 | NA | NA | NA | NA | NA | #VALUE! |
| ENSMUSG00000109480.2 | 0 | NA | NA | NA | NA | NA | #VALUE! |
| ENSMUSG00000109481.2 | 0 | NA | NA | NA | NA | NA | #VALUE! |
| ENSMUSG00000109484.2 | 0 | NA | NA | NA | NA | NA | #VALUE! |
| ENSMUSG00000109485.2 | 0 | NA | NA | NA | NA | NA | #VALUE! |
| ENSMUSG00000109487.4 | 0 | NA | NA | NA | NA | NA | #VALUE! |
| ENSMUSG00000109488.2 | 0 | NA | NA | NA | NA | NA | #VALUE! |
| ENSMUSG00000109489.2 | 0 | NA | NA | NA | NA | NA | #VALUE! |
| ENSMUSG00000109490.2 | 0 | NA | NA | NA | NA | NA | #VALUE! |
| ENSMUSG00000109491.2 | 0 | NA | NA | NA | NA | NA | #VALUE! |
| ENSMUSG00000109492.2 | 0.168284 | 1.027628 | 4.067126 | 0.252667 | 0.800526 | NA | #VALUE! |
| ENSMUSG00000109493.2 | 0 | NA | NA | NA | NA | NA | #VALUE! |
| ENSMUSG00000109494.2 | 0 | NA | NA | NA | NA | NA | #VALUE! |
| ENSMUSG00000109495.2 | 0 | NA | NA | NA | NA | NA | #VALUE! |
| ENSMUSG00000109496.2 | 0 | NA | NA | NA | NA | NA | #VALUE! |
| ENSMUSG00000109497.2 | 0 | NA | NA | NA | NA | NA | #VALUE! |
| ENSMUSG00000109498.2 | 5.172669 | -1.95989 | 3.680246 | -0.53254 | 0.59435 | NA | #VALUE! |
| ENSMUSG00000109499.2 | 0 | NA | NA | NA | NA | NA | #VALUE! |
| ENSMUSG00000109500.2 | 0 | NA | NA | NA | NA | NA | #VALUE! |

| | | | | | | | |
|---|---|---|---|---|---|---|---|
| ENSMUSG00000109501.2 | 0 | NA | NA | NA | NA | NA | #VALUE! |
| ENSMUSG00000109502.2 | 0 | NA | NA | NA | NA | NA | #VALUE! |
| ENSMUSG00000109503.2 | 0 | NA | NA | NA | NA | NA | #VALUE! |
| ENSMUSG00000109504.2 | 0 | NA | NA | NA | NA | NA | #VALUE! |
| ENSMUSG00000109505.2 | 0 | NA | NA | NA | NA | NA | #VALUE! |
| ENSMUSG00000109506.2 | 0 | NA | NA | NA | NA | NA | #VALUE! |
| ENSMUSG00000109507.2 | 0 | NA | NA | NA | NA | NA | #VALUE! |
| ENSMUSG00000109508.2 | 0 | NA | NA | NA | NA | NA | #VALUE! |
| ENSMUSG00000109510.2 | 0 | NA | NA | NA | NA | NA | #VALUE! |
| ENSMUSG00000109512.2 | 0 | NA | NA | NA | NA | NA | #VALUE! |
| ENSMUSG00000109513.2 | 0 | NA | NA | NA | NA | NA | #VALUE! |
| ENSMUSG00000109514.2 | 0 | NA | NA | NA | NA | NA | #VALUE! |
| ENSMUSG00000109515.2 | 0 | NA | NA | NA | NA | NA | #VALUE! |
| ENSMUSG00000109516.2 | 0 | NA | NA | NA | NA | NA | #VALUE! |
| ENSMUSG00000109517.2 | 0 | NA | NA | NA | NA | NA | #VALUE! |
| ENSMUSG00000109518.2 | 0 | NA | NA | NA | NA | NA | #VALUE! |
| ENSMUSG00000109519.2 | 0 | NA | NA | NA | NA | NA | #VALUE! |
| ENSMUSG00000109520.3 | 0 | NA | NA | NA | NA | NA | #VALUE! |
| ENSMUSG00000109521.4 | 0 | NA | NA | NA | NA | NA | #VALUE! |
| ENSMUSG00000109522.2 | 0 | NA | NA | NA | NA | NA | #VALUE! |
| ENSMUSG00000109523.2 | 0 | NA | NA | NA | NA | NA | #VALUE! |
| ENSMUSG00000109524.2 | 0 | NA | NA | NA | NA | NA | #VALUE! |
| ENSMUSG00000109525.2 | 0 | NA | NA | NA | NA | NA | #VALUE! |
| ENSMUSG00000109526.2 | 0 | NA | NA | NA | NA | NA | #VALUE! |
| ENSMUSG00000109527.2 | 0 | NA | NA | NA | NA | NA | #VALUE! |
| ENSMUSG00000109528.3 | 0 | NA | NA | NA | NA | NA | #VALUE! |
| ENSMUSG00000109529.2 | 0 | NA | NA | NA | NA | NA | #VALUE! |
| ENSMUSG00000109530.2 | 0 | NA | NA | NA | NA | NA | #VALUE! |
| ENSMUSG00000109532.2 | 0 | NA | NA | NA | NA | NA | #VALUE! |
| ENSMUSG00000109533.2 | 0 | NA | NA | NA | NA | NA | #VALUE! |
| ENSMUSG00000109534.2 | 0 | NA | NA | NA | NA | NA | #VALUE! |
| ENSMUSG00000109535.2 | 0 | NA | NA | NA | NA | NA | #VALUE! |
| ENSMUSG00000109536.2 | 0.408222 | -2.20205 | 4.016842 | -0.54821 | 0.583551 | NA | #VALUE! |
| ENSMUSG00000109537.2 | 0 | NA | NA | NA | NA | NA | #VALUE! |
| ENSMUSG00000109538.2 | 0 | NA | NA | NA | NA | NA | #VALUE! |
| ENSMUSG00000109539.2 | 7.665958 | -6.47201 | 3.903408 | -1.65804 | NA | NA | #VALUE! |
| ENSMUSG00000109540.2 | 0 | NA | NA | NA | NA | NA | #VALUE! |
| ENSMUSG00000109541.2 | 0 | NA | NA | NA | NA | NA | #VALUE! |
| ENSMUSG00000109542.3 | 0 | NA | NA | NA | NA | NA | #VALUE! |
| ENSMUSG00000109543.2 | 0 | NA | NA | NA | NA | NA | #VALUE! |
| ENSMUSG00000109544.3 | 0 | NA | NA | NA | NA | NA | #VALUE! |
| ENSMUSG00000109545.2 | 0 | NA | NA | NA | NA | NA | #VALUE! |
| ENSMUSG00000109546.2 | 0 | NA | NA | NA | NA | NA | #VALUE! |
| ENSMUSG00000109547.3 | 0 | NA | NA | NA | NA | NA | #VALUE! |

| | | | | | | | |
|---|---|---|---|---|---|---|---|
| ENSMUSG00000109548.2 | 0 | NA | NA | NA | NA | NA | #VALUE! |
| ENSMUSG00000109549.2 | 0 | NA | NA | NA | NA | NA | #VALUE! |
| ENSMUSG00000109550.2 | 0 | NA | NA | NA | NA | NA | #VALUE! |
| ENSMUSG00000109551.2 | 0 | NA | NA | NA | NA | NA | #VALUE! |
| ENSMUSG00000109552.2 | 0 | NA | NA | NA | NA | NA | #VALUE! |
| ENSMUSG00000109553.2 | 0 | NA | NA | NA | NA | NA | #VALUE! |
| ENSMUSG00000109555.2 | 0.839297 | 3.273019 | 3.958392 | 0.826856 | 0.408319 | NA | #VALUE! |
| ENSMUSG00000109556.2 | 0 | NA | NA | NA | NA | NA | #VALUE! |
| ENSMUSG00000109557.2 | 0 | NA | NA | NA | NA | NA | #VALUE! |
| ENSMUSG00000109558.2 | 0 | NA | NA | NA | NA | NA | #VALUE! |
| ENSMUSG00000109559.2 | 0 | NA | NA | NA | NA | NA | #VALUE! |
| ENSMUSG00000109560.2 | 0.105013 | -0.97324 | 4.076293 | -0.23876 | 0.811295 | NA | #VALUE! |
| ENSMUSG00000109561.2 | 0 | NA | NA | NA | NA | NA | #VALUE! |
| ENSMUSG00000109562.2 | 0 | NA | NA | NA | NA | NA | #VALUE! |
| ENSMUSG00000109564.2 | 0 | NA | NA | NA | NA | NA | #VALUE! |
| ENSMUSG00000109565.2 | 0 | NA | NA | NA | NA | NA | #VALUE! |
| ENSMUSG00000109566.2 | 0 | NA | NA | NA | NA | NA | #VALUE! |
| ENSMUSG00000109567.2 | 0 | NA | NA | NA | NA | NA | #VALUE! |
| ENSMUSG00000109568.2 | 0 | NA | NA | NA | NA | NA | #VALUE! |
| ENSMUSG00000109569.2 | 0 | NA | NA | NA | NA | NA | #VALUE! |
| ENSMUSG00000109570.2 | 0 | NA | NA | NA | NA | NA | #VALUE! |
| ENSMUSG00000109571.2 | 0 | NA | NA | NA | NA | NA | #VALUE! |
| ENSMUSG00000109572.2 | 0 | NA | NA | NA | NA | NA | #VALUE! |
| ENSMUSG00000109573.2 | 0 | NA | NA | NA | NA | NA | #VALUE! |
| ENSMUSG00000109574.2 | 0 | NA | NA | NA | NA | NA | #VALUE! |
| ENSMUSG00000109575.2 | 0 | NA | NA | NA | NA | NA | #VALUE! |
| ENSMUSG00000109576.2 | 0 | NA | NA | NA | NA | NA | #VALUE! |
| ENSMUSG00000109577.2 | 0 | NA | NA | NA | NA | NA | #VALUE! |
| ENSMUSG00000109578.2 | 0 | NA | NA | NA | NA | NA | #VALUE! |
| ENSMUSG00000109579.2 | 0 | NA | NA | NA | NA | NA | #VALUE! |
| ENSMUSG00000109580.2 | 0 | NA | NA | NA | NA | NA | #VALUE! |
| ENSMUSG00000109581.2 | 0 | NA | NA | NA | NA | NA | #VALUE! |
| ENSMUSG00000109582.2 | 0 | NA | NA | NA | NA | NA | #VALUE! |
| ENSMUSG00000109583.2 | 0 | NA | NA | NA | NA | NA | #VALUE! |
| ENSMUSG00000109584.2 | 0 | NA | NA | NA | NA | NA | #VALUE! |
| ENSMUSG00000109585.2 | 0 | NA | NA | NA | NA | NA | #VALUE! |
| ENSMUSG00000109586.2 | 0 | NA | NA | NA | NA | NA | #VALUE! |
| ENSMUSG00000109587.2 | 0.709588 | 3.017181 | 3.968248 | 0.760331 | 0.447057 | NA | #VALUE! |
| ENSMUSG00000109588.2 | 0 | NA | NA | NA | NA | NA | #VALUE! |
| ENSMUSG00000109589.2 | 0 | NA | NA | NA | NA | NA | #VALUE! |
| ENSMUSG00000109590.2 | 0 | NA | NA | NA | NA | NA | #VALUE! |
| ENSMUSG00000109591.2 | 0 | NA | NA | NA | NA | NA | #VALUE! |
| ENSMUSG00000109592.2 | 0 | NA | NA | NA | NA | NA | #VALUE! |
| ENSMUSG00000109593.2 | 0 | NA | NA | NA | NA | NA | #VALUE! |

| | | | | | | | |
|---|---|---|---|---|---|---|---|
| ENSMUSG00000109594.2 | 0 | NA | NA | NA | NA | NA | #VALUE! |
| ENSMUSG00000109595.2 | 0 | NA | NA | NA | NA | NA | #VALUE! |
| ENSMUSG00000109596.2 | 0 | NA | NA | NA | NA | NA | #VALUE! |
| ENSMUSG00000109597.2 | 0 | NA | NA | NA | NA | NA | #VALUE! |
| ENSMUSG00000109598.2 | 0 | NA | NA | NA | NA | NA | #VALUE! |
| ENSMUSG00000109600.2 | 0 | NA | NA | NA | NA | NA | #VALUE! |
| ENSMUSG00000109601.2 | 0 | NA | NA | NA | NA | NA | #VALUE! |
| ENSMUSG00000109602.2 | 0 | NA | NA | NA | NA | NA | #VALUE! |
| ENSMUSG00000109603.2 | 0 | NA | NA | NA | NA | NA | #VALUE! |
| ENSMUSG00000109604.2 | 0 | NA | NA | NA | NA | NA | #VALUE! |
| ENSMUSG00000109605.2 | 0 | NA | NA | NA | NA | NA | #VALUE! |
| ENSMUSG00000109606.2 | 0 | NA | NA | NA | NA | NA | #VALUE! |
| ENSMUSG00000109607.2 | 0 | NA | NA | NA | NA | NA | #VALUE! |
| ENSMUSG00000109608.2 | 0 | NA | NA | NA | NA | NA | #VALUE! |
| ENSMUSG00000109609.2 | 0 | NA | NA | NA | NA | NA | #VALUE! |
| ENSMUSG00000109610.2 | 0 | NA | NA | NA | NA | NA | #VALUE! |
| ENSMUSG00000109611.2 | 0.195789 | -1.05581 | 4.073302 | -0.2592 | 0.795479 | NA | #VALUE! |
| ENSMUSG00000109612.2 | 0 | NA | NA | NA | NA | NA | #VALUE! |
| ENSMUSG00000109613.2 | 0 | NA | NA | NA | NA | NA | #VALUE! |
| ENSMUSG00000109614.2 | 0 | NA | NA | NA | NA | NA | #VALUE! |
| ENSMUSG00000109615.2 | 0 | NA | NA | NA | NA | NA | #VALUE! |
| ENSMUSG00000109616.2 | 0 | NA | NA | NA | NA | NA | #VALUE! |
| ENSMUSG00000109617.2 | 0 | NA | NA | NA | NA | NA | #VALUE! |
| ENSMUSG00000109618.2 | 0 | NA | NA | NA | NA | NA | #VALUE! |
| ENSMUSG00000109619.2 | 0 | NA | NA | NA | NA | NA | #VALUE! |
| ENSMUSG00000109620.2 | 0 | NA | NA | NA | NA | NA | #VALUE! |
| ENSMUSG00000109621.2 | 0 | NA | NA | NA | NA | NA | #VALUE! |
| ENSMUSG00000109622.2 | 0 | NA | NA | NA | NA | NA | #VALUE! |
| ENSMUSG00000109623.2 | 0 | NA | NA | NA | NA | NA | #VALUE! |
| ENSMUSG00000109624.2 | 0 | NA | NA | NA | NA | NA | #VALUE! |
| ENSMUSG00000109625.2 | 0 | NA | NA | NA | NA | NA | #VALUE! |
| ENSMUSG00000109626.2 | 0 | NA | NA | NA | NA | NA | #VALUE! |
| ENSMUSG00000109627.2 | 0 | NA | NA | NA | NA | NA | #VALUE! |
| ENSMUSG00000109629.2 | 0 | NA | NA | NA | NA | NA | #VALUE! |
| ENSMUSG00000109630.2 | 0 | NA | NA | NA | NA | NA | #VALUE! |
| ENSMUSG00000109631.2 | 0 | NA | NA | NA | NA | NA | #VALUE! |
| ENSMUSG00000109632.2 | 0 | NA | NA | NA | NA | NA | #VALUE! |
| ENSMUSG00000109633.2 | 0 | NA | NA | NA | NA | NA | #VALUE! |
| ENSMUSG00000109634.2 | 0 | NA | NA | NA | NA | NA | #VALUE! |
| ENSMUSG00000109635.2 | 0 | NA | NA | NA | NA | NA | #VALUE! |
| ENSMUSG00000109636.2 | 0 | NA | NA | NA | NA | NA | #VALUE! |
| ENSMUSG00000109637.2 | 0 | NA | NA | NA | NA | NA | #VALUE! |
| ENSMUSG00000109638.2 | 0 | NA | NA | NA | NA | NA | #VALUE! |
| ENSMUSG00000109639.2 | 0 | NA | NA | NA | NA | NA | #VALUE! |

| | | | | | | | |
|---|---|---|---|---|---|---|---|
| ENSMUSG00000109640.2 | 0 | NA | NA | NA | NA | NA | #VALUE! |
| ENSMUSG00000109641.2 | 0 | NA | NA | NA | NA | NA | #VALUE! |
| ENSMUSG00000109642.2 | 0 | NA | NA | NA | NA | NA | #VALUE! |
| ENSMUSG00000109643.2 | 0 | NA | NA | NA | NA | NA | #VALUE! |
| ENSMUSG00000109645.2 | 0 | NA | NA | NA | NA | NA | #VALUE! |
| ENSMUSG00000109646.2 | 0 | NA | NA | NA | NA | NA | #VALUE! |
| ENSMUSG00000109647.2 | 0 | NA | NA | NA | NA | NA | #VALUE! |
| ENSMUSG00000109648.2 | 0 | NA | NA | NA | NA | NA | #VALUE! |
| ENSMUSG00000109649.2 | 0 | NA | NA | NA | NA | NA | #VALUE! |
| ENSMUSG00000109650.2 | 0 | NA | NA | NA | NA | NA | #VALUE! |
| ENSMUSG00000109651.2 | 0 | NA | NA | NA | NA | NA | #VALUE! |
| ENSMUSG00000109652.2 | 0 | NA | NA | NA | NA | NA | #VALUE! |
| ENSMUSG00000109653.2 | 0 | NA | NA | NA | NA | NA | #VALUE! |
| ENSMUSG00000109654.2 | 0 | NA | NA | NA | NA | NA | #VALUE! |
| ENSMUSG00000109655.2 | 0 | NA | NA | NA | NA | NA | #VALUE! |
| ENSMUSG00000109656.2 | 0 | NA | NA | NA | NA | NA | #VALUE! |
| ENSMUSG00000109657.2 | 0 | NA | NA | NA | NA | NA | #VALUE! |
| ENSMUSG00000109658.2 | 0.525066 | -2.64687 | 3.988095 | -0.66369 | 0.506887 | NA | #VALUE! |
| ENSMUSG00000109659.4 | 0 | NA | NA | NA | NA | NA | #VALUE! |
| ENSMUSG00000109662.2 | 0 | NA | NA | NA | NA | NA | #VALUE! |
| ENSMUSG00000109663.2 | 0 | NA | NA | NA | NA | NA | #VALUE! |
| ENSMUSG00000109664.2 | 0 | NA | NA | NA | NA | NA | #VALUE! |
| ENSMUSG00000109665.2 | 0 | NA | NA | NA | NA | NA | #VALUE! |
| ENSMUSG00000109666.2 | 0 | NA | NA | NA | NA | NA | #VALUE! |
| ENSMUSG00000109667.2 | 0 | NA | NA | NA | NA | NA | #VALUE! |
| ENSMUSG00000109668.2 | 0 | NA | NA | NA | NA | NA | #VALUE! |
| ENSMUSG00000109669.2 | 0 | NA | NA | NA | NA | NA | #VALUE! |
| ENSMUSG00000109670.2 | 0 | NA | NA | NA | NA | NA | #VALUE! |
| ENSMUSG00000109671.2 | 0 | NA | NA | NA | NA | NA | #VALUE! |
| ENSMUSG00000109672.2 | 0 | NA | NA | NA | NA | NA | #VALUE! |
| ENSMUSG00000109673.2 | 0.245976 | 1.027628 | 4.067126 | 0.252667 | 0.800526 | NA | #VALUE! |
| ENSMUSG00000109674.2 | 0 | NA | NA | NA | NA | NA | #VALUE! |
| ENSMUSG00000109675.3 | 0 | NA | NA | NA | NA | NA | #VALUE! |
| ENSMUSG00000109676.2 | 0 | NA | NA | NA | NA | NA | #VALUE! |
| ENSMUSG00000109677.2 | 0 | NA | NA | NA | NA | NA | #VALUE! |
| ENSMUSG00000109678.2 | 0 | NA | NA | NA | NA | NA | #VALUE! |
| ENSMUSG00000109679.2 | 0 | NA | NA | NA | NA | NA | #VALUE! |
| ENSMUSG00000109680.2 | 0 | NA | NA | NA | NA | NA | #VALUE! |
| ENSMUSG00000109681.2 | 0 | NA | NA | NA | NA | NA | #VALUE! |
| ENSMUSG00000109682.2 | 0 | NA | NA | NA | NA | NA | #VALUE! |
| ENSMUSG00000109683.2 | 0 | NA | NA | NA | NA | NA | #VALUE! |
| ENSMUSG00000109684.2 | 0 | NA | NA | NA | NA | NA | #VALUE! |
| ENSMUSG00000109685.2 | 0 | NA | NA | NA | NA | NA | #VALUE! |
| ENSMUSG00000109686.2 | 0 | NA | NA | NA | NA | NA | #VALUE! |

| | | | | | | | |
|---|---|---|---|---|---|---|---|
| ENSMUSG00000109687.2 | 0 | NA | NA | NA | NA | NA | #VALUE! |
| ENSMUSG00000109688.2 | 0 | NA | NA | NA | NA | NA | #VALUE! |
| ENSMUSG00000109689.2 | 0 | NA | NA | NA | NA | NA | #VALUE! |
| ENSMUSG00000109690.2 | 0 | NA | NA | NA | NA | NA | #VALUE! |
| ENSMUSG00000109691.2 | 0 | NA | NA | NA | NA | NA | #VALUE! |
| ENSMUSG00000109692.2 | 0 | NA | NA | NA | NA | NA | #VALUE! |
| ENSMUSG00000109693.2 | 0 | NA | NA | NA | NA | NA | #VALUE! |
| ENSMUSG00000109694.2 | 0 | NA | NA | NA | NA | NA | #VALUE! |
| ENSMUSG00000109695.2 | 0 | NA | NA | NA | NA | NA | #VALUE! |
| ENSMUSG00000109696.2 | 0 | NA | NA | NA | NA | NA | #VALUE! |
| ENSMUSG00000109697.2 | 0 | NA | NA | NA | NA | NA | #VALUE! |
| ENSMUSG00000109698.2 | 6.852619 | -6.30491 | 3.913306 | -1.61115 | 0.107148 | NA | #VALUE! |
| ENSMUSG00000109699.2 | 0 | NA | NA | NA | NA | NA | #VALUE! |
| ENSMUSG00000109700.2 | 0 | NA | NA | NA | NA | NA | #VALUE! |
| ENSMUSG00000109701.2 | 0 | NA | NA | NA | NA | NA | #VALUE! |
| ENSMUSG00000109702.2 | 0 | NA | NA | NA | NA | NA | #VALUE! |
| ENSMUSG00000109704.2 | 0 | NA | NA | NA | NA | NA | #VALUE! |
| ENSMUSG00000109705.2 | 0 | NA | NA | NA | NA | NA | #VALUE! |
| ENSMUSG00000109706.2 | 0 | NA | NA | NA | NA | NA | #VALUE! |
| ENSMUSG00000109707.2 | 0 | NA | NA | NA | NA | NA | #VALUE! |
| ENSMUSG00000109708.3 | 0 | NA | NA | NA | NA | NA | #VALUE! |
| ENSMUSG00000109709.2 | 0 | NA | NA | NA | NA | NA | #VALUE! |
| ENSMUSG00000109710.2 | 0 | NA | NA | NA | NA | NA | #VALUE! |
| ENSMUSG00000109711.2 | 0 | NA | NA | NA | NA | NA | #VALUE! |
| ENSMUSG00000109712.2 | 0 | NA | NA | NA | NA | NA | #VALUE! |
| ENSMUSG00000109713.2 | 0 | NA | NA | NA | NA | NA | #VALUE! |
| ENSMUSG00000109714.2 | 8.776762 | -6.66175 | 3.755766 | -1.77374 | 0.076106 | NA | #VALUE! |
| ENSMUSG00000109715.2 | 6.900884 | 6.28129 | 3.913256 | 1.605131 | NA | NA | #VALUE! |
| ENSMUSG00000109716.2 | 0 | NA | NA | NA | NA | NA | #VALUE! |
| ENSMUSG00000109717.2 | 0 | NA | NA | NA | NA | NA | #VALUE! |
| ENSMUSG00000109718.2 | 0 | NA | NA | NA | NA | NA | #VALUE! |
| ENSMUSG00000109719.2 | 0 | NA | NA | NA | NA | NA | #VALUE! |
| ENSMUSG00000109720.2 | 0 | NA | NA | NA | NA | NA | #VALUE! |
| ENSMUSG00000109721.2 | 0 | NA | NA | NA | NA | NA | #VALUE! |
| ENSMUSG00000109722.2 | 0 | NA | NA | NA | NA | NA | #VALUE! |
| ENSMUSG00000109723.2 | 0 | NA | NA | NA | NA | NA | #VALUE! |
| ENSMUSG00000109724.2 | 0 | NA | NA | NA | NA | NA | #VALUE! |
| ENSMUSG00000109725.2 | 0 | NA | NA | NA | NA | NA | #VALUE! |
| ENSMUSG00000109726.2 | 0 | NA | NA | NA | NA | NA | #VALUE! |
| ENSMUSG00000109727.2 | 0 | NA | NA | NA | NA | NA | #VALUE! |
| ENSMUSG00000109728.2 | 0 | NA | NA | NA | NA | NA | #VALUE! |
| ENSMUSG00000109729.2 | 0 | NA | NA | NA | NA | NA | #VALUE! |
| ENSMUSG00000109730.2 | 0 | NA | NA | NA | NA | NA | #VALUE! |
| ENSMUSG00000109731.2 | 0 | NA | NA | NA | NA | NA | #VALUE! |

| | | | | | | | |
|---|---|---|---|---|---|---|---|
| ENSMUSG00000109732.2 | 0 | NA | NA | NA | NA | NA | #VALUE! |
| ENSMUSG00000109733.2 | 0 | NA | NA | NA | NA | NA | #VALUE! |
| ENSMUSG00000109734.2 | 0 | NA | NA | NA | NA | NA | #VALUE! |
| ENSMUSG00000109735.2 | 0 | NA | NA | NA | NA | NA | #VALUE! |
| ENSMUSG00000109736.2 | 0 | NA | NA | NA | NA | NA | #VALUE! |
| ENSMUSG00000109737.2 | 0 | NA | NA | NA | NA | NA | #VALUE! |
| ENSMUSG00000109738.2 | 0 | NA | NA | NA | NA | NA | #VALUE! |
| ENSMUSG00000109739.2 | 0 | NA | NA | NA | NA | NA | #VALUE! |
| ENSMUSG00000109740.2 | 0.093255 | 0.868093 | 4.072464 | 0.213162 | 0.831201 | NA | #VALUE! |
| ENSMUSG00000109741.2 | 0 | NA | NA | NA | NA | NA | #VALUE! |
| ENSMUSG00000109742.2 | 0 | NA | NA | NA | NA | NA | #VALUE! |
| ENSMUSG00000109743.2 | 0 | NA | NA | NA | NA | NA | #VALUE! |
| ENSMUSG00000109744.2 | 0 | NA | NA | NA | NA | NA | #VALUE! |
| ENSMUSG00000109745.2 | 0 | NA | NA | NA | NA | NA | #VALUE! |
| ENSMUSG00000109746.2 | 0 | NA | NA | NA | NA | NA | #VALUE! |
| ENSMUSG00000109747.2 | 0 | NA | NA | NA | NA | NA | #VALUE! |
| ENSMUSG00000109748.2 | 0 | NA | NA | NA | NA | NA | #VALUE! |
| ENSMUSG00000109749.2 | 0 | NA | NA | NA | NA | NA | #VALUE! |
| ENSMUSG00000109750.2 | 0 | NA | NA | NA | NA | NA | #VALUE! |
| ENSMUSG00000109751.2 | 0 | NA | NA | NA | NA | NA | #VALUE! |
| ENSMUSG00000109752.2 | 0 | NA | NA | NA | NA | NA | #VALUE! |
| ENSMUSG00000109753.2 | 0 | NA | NA | NA | NA | NA | #VALUE! |
| ENSMUSG00000109754.2 | 0 | NA | NA | NA | NA | NA | #VALUE! |
| ENSMUSG00000109755.2 | 0 | NA | NA | NA | NA | NA | #VALUE! |
| ENSMUSG00000109756.2 | 0 | NA | NA | NA | NA | NA | #VALUE! |
| ENSMUSG00000109757.2 | 0 | NA | NA | NA | NA | NA | #VALUE! |
| ENSMUSG00000109758.2 | 0 | NA | NA | NA | NA | NA | #VALUE! |
| ENSMUSG00000109759.2 | 0 | NA | NA | NA | NA | NA | #VALUE! |
| ENSMUSG00000109760.2 | 0 | NA | NA | NA | NA | NA | #VALUE! |
| ENSMUSG00000109761.2 | 0 | NA | NA | NA | NA | NA | #VALUE! |
| ENSMUSG00000109762.2 | 0 | NA | NA | NA | NA | NA | #VALUE! |
| ENSMUSG00000109763.2 | 0 | NA | NA | NA | NA | NA | #VALUE! |
| ENSMUSG00000109765.3 | 0 | NA | NA | NA | NA | NA | #VALUE! |
| ENSMUSG00000109766.2 | 0 | NA | NA | NA | NA | NA | #VALUE! |
| ENSMUSG00000109767.2 | 0 | NA | NA | NA | NA | NA | #VALUE! |
| ENSMUSG00000109768.2 | 0 | NA | NA | NA | NA | NA | #VALUE! |
| ENSMUSG00000109769.2 | 0 | NA | NA | NA | NA | NA | #VALUE! |
| ENSMUSG00000109770.2 | 0 | NA | NA | NA | NA | NA | #VALUE! |
| ENSMUSG00000109771.2 | 7.432864 | 6.386473 | 2.979976 | 2.143129 | 0.032103 | NA | #VALUE! |
| ENSMUSG00000109772.2 | 0 | NA | NA | NA | NA | NA | #VALUE! |
| ENSMUSG00000109773.2 | 4.407584 | 5.633257 | 3.91693 | 1.438182 | 0.150382 | NA | #VALUE! |
| ENSMUSG00000109774.2 | 0 | NA | NA | NA | NA | NA | #VALUE! |
| ENSMUSG00000109775.2 | 0 | NA | NA | NA | NA | NA | #VALUE! |
| ENSMUSG00000109776.2 | 0 | NA | NA | NA | NA | NA | #VALUE! |

| | | | | | | | |
|---|---|---|---|---|---|---|---|
| ENSMUSG00000109777.2 | 0 | NA | NA | NA | NA | NA | #VALUE! |
| ENSMUSG00000109778.2 | 0 | NA | NA | NA | NA | NA | #VALUE! |
| ENSMUSG00000109779.2 | 0 | NA | NA | NA | NA | NA | #VALUE! |
| ENSMUSG00000109780.2 | 0 | NA | NA | NA | NA | NA | #VALUE! |
| ENSMUSG00000109781.2 | 0 | NA | NA | NA | NA | NA | #VALUE! |
| ENSMUSG00000109784.2 | 0 | NA | NA | NA | NA | NA | #VALUE! |
| ENSMUSG00000109785.2 | 0 | NA | NA | NA | NA | NA | #VALUE! |
| ENSMUSG00000109786.2 | 0 | NA | NA | NA | NA | NA | #VALUE! |
| ENSMUSG00000109787.2 | 0 | NA | NA | NA | NA | NA | #VALUE! |
| ENSMUSG00000109788.2 | 0 | NA | NA | NA | NA | NA | #VALUE! |
| ENSMUSG00000109789.2 | 0 | NA | NA | NA | NA | NA | #VALUE! |
| ENSMUSG00000109790.2 | 0 | NA | NA | NA | NA | NA | #VALUE! |
| ENSMUSG00000109791.2 | 0 | NA | NA | NA | NA | NA | #VALUE! |
| ENSMUSG00000109792.2 | 0 | NA | NA | NA | NA | NA | #VALUE! |
| ENSMUSG00000109793.2 | 0 | NA | NA | NA | NA | NA | #VALUE! |
| ENSMUSG00000109794.2 | 0 | NA | NA | NA | NA | NA | #VALUE! |
| ENSMUSG00000109795.2 | 0 | NA | NA | NA | NA | NA | #VALUE! |
| ENSMUSG00000109796.2 | 0 | NA | NA | NA | NA | NA | #VALUE! |
| ENSMUSG00000109797.2 | 0 | NA | NA | NA | NA | NA | #VALUE! |
| ENSMUSG00000109798.2 | 0 | NA | NA | NA | NA | NA | #VALUE! |
| ENSMUSG00000109799.2 | 0 | NA | NA | NA | NA | NA | #VALUE! |
| ENSMUSG00000109800.2 | 0 | NA | NA | NA | NA | NA | #VALUE! |
| ENSMUSG00000109801.3 | 0 | NA | NA | NA | NA | NA | #VALUE! |
| ENSMUSG00000109802.2 | 0 | NA | NA | NA | NA | NA | #VALUE! |
| ENSMUSG00000109803.2 | 0 | NA | NA | NA | NA | NA | #VALUE! |
| ENSMUSG00000109805.2 | 0 | NA | NA | NA | NA | NA | #VALUE! |
| ENSMUSG00000109806.2 | 0 | NA | NA | NA | NA | NA | #VALUE! |
| ENSMUSG00000109807.2 | 0 | NA | NA | NA | NA | NA | #VALUE! |
| ENSMUSG00000109808.2 | 0 | NA | NA | NA | NA | NA | #VALUE! |
| ENSMUSG00000109809.2 | 0 | NA | NA | NA | NA | NA | #VALUE! |
| ENSMUSG00000109810.2 | 0 | NA | NA | NA | NA | NA | #VALUE! |
| ENSMUSG00000109811.2 | 0 | NA | NA | NA | NA | NA | #VALUE! |
| ENSMUSG00000109812.2 | 31.74655 | 0.841321 | 2.30213 | 0.365453 | NA | NA | #VALUE! |
| ENSMUSG00000109813.3 | 0 | NA | NA | NA | NA | NA | #VALUE! |
| ENSMUSG00000109814.2 | 0 | NA | NA | NA | NA | NA | #VALUE! |
| ENSMUSG00000109815.2 | 0 | NA | NA | NA | NA | NA | #VALUE! |
| ENSMUSG00000109816.2 | 0 | NA | NA | NA | NA | NA | #VALUE! |
| ENSMUSG00000109817.2 | 0 | NA | NA | NA | NA | NA | #VALUE! |
| ENSMUSG00000109818.2 | 0 | NA | NA | NA | NA | NA | #VALUE! |
| ENSMUSG00000109819.2 | 0 | NA | NA | NA | NA | NA | #VALUE! |
| ENSMUSG00000109820.2 | 0 | NA | NA | NA | NA | NA | #VALUE! |
| ENSMUSG00000109821.2 | 0 | NA | NA | NA | NA | NA | #VALUE! |
| ENSMUSG00000109822.2 | 0 | NA | NA | NA | NA | NA | #VALUE! |
| ENSMUSG00000109823.2 | 0 | NA | NA | NA | NA | NA | #VALUE! |

| | | | | | | | |
|---|---|---|---|---|---|---|---|
| ENSMUSG00000109824.5 | 0 | NA | NA | NA | NA | NA | #VALUE! |
| ENSMUSG00000109825.2 | 0 | NA | NA | NA | NA | NA | #VALUE! |
| ENSMUSG00000109826.2 | 0 | NA | NA | NA | NA | NA | #VALUE! |
| ENSMUSG00000109827.2 | 0 | NA | NA | NA | NA | NA | #VALUE! |
| ENSMUSG00000109828.2 | 0 | NA | NA | NA | NA | NA | #VALUE! |
| ENSMUSG00000109829.2 | 0 | NA | NA | NA | NA | NA | #VALUE! |
| ENSMUSG00000109830.2 | 0 | NA | NA | NA | NA | NA | #VALUE! |
| ENSMUSG00000109831.2 | 0 | NA | NA | NA | NA | NA | #VALUE! |
| ENSMUSG00000109832.2 | 0 | NA | NA | NA | NA | NA | #VALUE! |
| ENSMUSG00000109833.2 | 0 | NA | NA | NA | NA | NA | #VALUE! |
| ENSMUSG00000109834.2 | 0 | NA | NA | NA | NA | NA | #VALUE! |
| ENSMUSG00000109835.3 | 0 | NA | NA | NA | NA | NA | #VALUE! |
| ENSMUSG00000109837.2 | 0 | NA | NA | NA | NA | NA | #VALUE! |
| ENSMUSG00000109838.2 | 0 | NA | NA | NA | NA | NA | #VALUE! |
| ENSMUSG00000109839.2 | 0 | NA | NA | NA | NA | NA | #VALUE! |
| ENSMUSG00000109840.2 | 0 | NA | NA | NA | NA | NA | #VALUE! |
| ENSMUSG00000109842.2 | 0 | NA | NA | NA | NA | NA | #VALUE! |
| ENSMUSG00000109843.2 | 0 | NA | NA | NA | NA | NA | #VALUE! |
| ENSMUSG00000109844.3 | 0 | NA | NA | NA | NA | NA | #VALUE! |
| ENSMUSG00000109845.2 | 0 | NA | NA | NA | NA | NA | #VALUE! |
| ENSMUSG00000109846.2 | 0 | NA | NA | NA | NA | NA | #VALUE! |
| ENSMUSG00000109847.2 | 0 | NA | NA | NA | NA | NA | #VALUE! |
| ENSMUSG00000109848.2 | 0 | NA | NA | NA | NA | NA | #VALUE! |
| ENSMUSG00000109849.2 | 0 | NA | NA | NA | NA | NA | #VALUE! |
| ENSMUSG00000109850.2 | 0 | NA | NA | NA | NA | NA | #VALUE! |
| ENSMUSG00000109851.2 | 0 | NA | NA | NA | NA | NA | #VALUE! |
| ENSMUSG00000109852.2 | 0 | NA | NA | NA | NA | NA | #VALUE! |
| ENSMUSG00000109853.2 | 0 | NA | NA | NA | NA | NA | #VALUE! |
| ENSMUSG00000109854.2 | 0 | NA | NA | NA | NA | NA | #VALUE! |
| ENSMUSG00000109855.2 | 0 | NA | NA | NA | NA | NA | #VALUE! |
| ENSMUSG00000109856.2 | 0 | NA | NA | NA | NA | NA | #VALUE! |
| ENSMUSG00000109857.2 | 0 | NA | NA | NA | NA | NA | #VALUE! |
| ENSMUSG00000109858.2 | 0 | NA | NA | NA | NA | NA | #VALUE! |
| ENSMUSG00000109859.2 | 0 | NA | NA | NA | NA | NA | #VALUE! |
| ENSMUSG00000109860.2 | 0 | NA | NA | NA | NA | NA | #VALUE! |
| ENSMUSG00000109861.2 | 0 | NA | NA | NA | NA | NA | #VALUE! |
| ENSMUSG00000109863.2 | 0 | NA | NA | NA | NA | NA | #VALUE! |
| ENSMUSG00000109864.2 | 0.841418 | 3.240539 | 3.959551 | 0.818411 | 0.413123 | NA | #VALUE! |
| ENSMUSG00000109866.2 | 0.279766 | 1.808031 | 4.036658 | 0.447903 | 0.654223 | NA | #VALUE! |
| ENSMUSG00000109867.2 | 0 | NA | NA | NA | NA | NA | #VALUE! |
| ENSMUSG00000109868.2 | 0 | NA | NA | NA | NA | NA | #VALUE! |
| ENSMUSG00000109869.2 | 0 | NA | NA | NA | NA | NA | #VALUE! |
| ENSMUSG00000109870.2 | 0 | NA | NA | NA | NA | NA | #VALUE! |
| ENSMUSG00000109871.2 | 0 | NA | NA | NA | NA | NA | #VALUE! |

| | | | | | | | |
|---|---|---|---|---|---|---|---|
| ENSMUSG00000109872.2 | 0 | NA | NA | NA | NA | NA | #VALUE! |
| ENSMUSG00000109873.2 | 0 | NA | NA | NA | NA | NA | #VALUE! |
| ENSMUSG00000109874.2 | 0 | NA | NA | NA | NA | NA | #VALUE! |
| ENSMUSG00000109875.2 | 0 | NA | NA | NA | NA | NA | #VALUE! |
| ENSMUSG00000109876.2 | 0 | NA | NA | NA | NA | NA | #VALUE! |
| ENSMUSG00000109877.2 | 7.987419 | 1.766906 | 2.691731 | 0.65642 | 0.511554 | NA | #VALUE! |
| ENSMUSG00000109878.2 | 0 | NA | NA | NA | NA | NA | #VALUE! |
| ENSMUSG00000109879.2 | 0 | NA | NA | NA | NA | NA | #VALUE! |
| ENSMUSG00000109880.2 | 0 | NA | NA | NA | NA | NA | #VALUE! |
| ENSMUSG00000109882.2 | 0 | NA | NA | NA | NA | NA | #VALUE! |
| ENSMUSG00000109883.2 | 0 | NA | NA | NA | NA | NA | #VALUE! |
| ENSMUSG00000109884.4 | 0 | NA | NA | NA | NA | NA | #VALUE! |
| ENSMUSG00000109885.2 | 0 | NA | NA | NA | NA | NA | #VALUE! |
| ENSMUSG00000109886.2 | 0 | NA | NA | NA | NA | NA | #VALUE! |
| ENSMUSG00000109887.2 | 0 | NA | NA | NA | NA | NA | #VALUE! |
| ENSMUSG00000109888.2 | 0 | NA | NA | NA | NA | NA | #VALUE! |
| ENSMUSG00000109889.2 | 0 | NA | NA | NA | NA | NA | #VALUE! |
| ENSMUSG00000109890.3 | 0 | NA | NA | NA | NA | NA | #VALUE! |
| ENSMUSG00000109891.2 | 0 | NA | NA | NA | NA | NA | #VALUE! |
| ENSMUSG00000109892.2 | 0 | NA | NA | NA | NA | NA | #VALUE! |
| ENSMUSG00000109893.2 | 0 | NA | NA | NA | NA | NA | #VALUE! |
| ENSMUSG00000109894.2 | 0 | NA | NA | NA | NA | NA | #VALUE! |
| ENSMUSG00000109895.2 | 0 | NA | NA | NA | NA | NA | #VALUE! |
| ENSMUSG00000109896.2 | 0 | NA | NA | NA | NA | NA | #VALUE! |
| ENSMUSG00000109897.2 | 0 | NA | NA | NA | NA | NA | #VALUE! |
| ENSMUSG00000109898.2 | 0 | NA | NA | NA | NA | NA | #VALUE! |
| ENSMUSG00000109899.2 | 0 | NA | NA | NA | NA | NA | #VALUE! |
| ENSMUSG00000109900.2 | 0 | NA | NA | NA | NA | NA | #VALUE! |
| ENSMUSG00000109902.2 | 0 | NA | NA | NA | NA | NA | #VALUE! |
| ENSMUSG00000109903.2 | 0 | NA | NA | NA | NA | NA | #VALUE! |
| ENSMUSG00000109904.2 | 0 | NA | NA | NA | NA | NA | #VALUE! |
| ENSMUSG00000109905.2 | 0 | NA | NA | NA | NA | NA | #VALUE! |
| ENSMUSG00000109906.2 | 0 | NA | NA | NA | NA | NA | #VALUE! |
| ENSMUSG00000109907.2 | 0 | NA | NA | NA | NA | NA | #VALUE! |
| ENSMUSG00000109909.2 | 0 | NA | NA | NA | NA | NA | #VALUE! |
| ENSMUSG00000109910.2 | 0 | NA | NA | NA | NA | NA | #VALUE! |
| ENSMUSG00000109911.2 | 0 | NA | NA | NA | NA | NA | #VALUE! |
| ENSMUSG00000109912.2 | 0 | NA | NA | NA | NA | NA | #VALUE! |
| ENSMUSG00000109913.2 | 0 | NA | NA | NA | NA | NA | #VALUE! |
| ENSMUSG00000109914.2 | 0 | NA | NA | NA | NA | NA | #VALUE! |
| ENSMUSG00000109915.2 | 0 | NA | NA | NA | NA | NA | #VALUE! |
| ENSMUSG00000109916.2 | 0 | NA | NA | NA | NA | NA | #VALUE! |
| ENSMUSG00000109917.2 | 0 | NA | NA | NA | NA | NA | #VALUE! |
| ENSMUSG00000109918.2 | 0 | NA | NA | NA | NA | NA | #VALUE! |

| | | | | | | | |
|---|---|---|---|---|---|---|---|
| ENSMUSG00000109919.2 | 0 | NA | NA | NA | NA | NA | #VALUE! |
| ENSMUSG00000109920.2 | 0 | NA | NA | NA | NA | NA | #VALUE! |
| ENSMUSG00000109921.2 | 0 | NA | NA | NA | NA | NA | #VALUE! |
| ENSMUSG00000109922.2 | 0 | NA | NA | NA | NA | NA | #VALUE! |
| ENSMUSG00000109923.2 | 0 | NA | NA | NA | NA | NA | #VALUE! |
| ENSMUSG00000109924.2 | 0 | NA | NA | NA | NA | NA | #VALUE! |
| ENSMUSG00000109925.2 | 0.210026 | -1.52494 | 4.058692 | -0.37572 | 0.707123 | NA | #VALUE! |
| ENSMUSG00000109926.2 | 0 | NA | NA | NA | NA | NA | #VALUE! |
| ENSMUSG00000109927.2 | 0 | NA | NA | NA | NA | NA | #VALUE! |
| ENSMUSG00000109928.2 | 0 | NA | NA | NA | NA | NA | #VALUE! |
| ENSMUSG00000109929.2 | 0 | NA | NA | NA | NA | NA | #VALUE! |
| ENSMUSG00000109930.2 | 0 | NA | NA | NA | NA | NA | #VALUE! |
| ENSMUSG00000109931.2 | 0 | NA | NA | NA | NA | NA | #VALUE! |
| ENSMUSG00000109932.2 | 0 | NA | NA | NA | NA | NA | #VALUE! |
| ENSMUSG00000109934.2 | 0 | NA | NA | NA | NA | NA | #VALUE! |
| ENSMUSG00000109935.2 | 0 | NA | NA | NA | NA | NA | #VALUE! |
| ENSMUSG00000109936.2 | 0 | NA | NA | NA | NA | NA | #VALUE! |
| ENSMUSG00000109937.2 | 0 | NA | NA | NA | NA | NA | #VALUE! |
| ENSMUSG00000109938.2 | 0 | NA | NA | NA | NA | NA | #VALUE! |
| ENSMUSG00000109939.2 | 0 | NA | NA | NA | NA | NA | #VALUE! |
| ENSMUSG00000109940.2 | 0 | NA | NA | NA | NA | NA | #VALUE! |
| ENSMUSG00000109941.2 | 0 | NA | NA | NA | NA | NA | #VALUE! |
| ENSMUSG00000109942.2 | 0 | NA | NA | NA | NA | NA | #VALUE! |
| ENSMUSG00000109943.2 | 0 | NA | NA | NA | NA | NA | #VALUE! |
| ENSMUSG00000109944.2 | 0 | NA | NA | NA | NA | NA | #VALUE! |
| ENSMUSG00000109945.2 | 0 | NA | NA | NA | NA | NA | #VALUE! |
| ENSMUSG00000109946.2 | 0 | NA | NA | NA | NA | NA | #VALUE! |
| ENSMUSG00000109947.2 | 0 | NA | NA | NA | NA | NA | #VALUE! |
| ENSMUSG00000109948.2 | 0 | NA | NA | NA | NA | NA | #VALUE! |
| ENSMUSG00000109949.2 | 0 | NA | NA | NA | NA | NA | #VALUE! |
| ENSMUSG00000109950.2 | 0 | NA | NA | NA | NA | NA | #VALUE! |
| ENSMUSG00000109951.3 | 0 | NA | NA | NA | NA | NA | #VALUE! |
| ENSMUSG00000109952.2 | 0 | NA | NA | NA | NA | NA | #VALUE! |
| ENSMUSG00000109953.2 | 0 | NA | NA | NA | NA | NA | #VALUE! |
| ENSMUSG00000109954.2 | 0 | NA | NA | NA | NA | NA | #VALUE! |
| ENSMUSG00000109955.2 | 0 | NA | NA | NA | NA | NA | #VALUE! |
| ENSMUSG00000109956.2 | 0 | NA | NA | NA | NA | NA | #VALUE! |
| ENSMUSG00000109957.2 | 16.64207 | -21.794 | 3.093228 | -7.04572 | NA | NA | #VALUE! |
| ENSMUSG00000109958.2 | 0 | NA | NA | NA | NA | NA | #VALUE! |
| ENSMUSG00000109959.2 | 0 | NA | NA | NA | NA | NA | #VALUE! |
| ENSMUSG00000109960.2 | 0 | NA | NA | NA | NA | NA | #VALUE! |
| ENSMUSG00000109961.2 | 0 | NA | NA | NA | NA | NA | #VALUE! |
| ENSMUSG00000109962.2 | 0 | NA | NA | NA | NA | NA | #VALUE! |
| ENSMUSG00000109963.2 | 0 | NA | NA | NA | NA | NA | #VALUE! |

| | | | | | | | |
|---|---|---|---|---|---|---|---|
| ENSMUSG00000109964.2 | 0 | NA | NA | NA | NA | NA | #VALUE! |
| ENSMUSG00000109965.2 | 0 | NA | NA | NA | NA | NA | #VALUE! |
| ENSMUSG00000109966.2 | 0 | NA | NA | NA | NA | NA | #VALUE! |
| ENSMUSG00000109967.2 | 0 | NA | NA | NA | NA | NA | #VALUE! |
| ENSMUSG00000109968.2 | 0 | NA | NA | NA | NA | NA | #VALUE! |
| ENSMUSG00000109969.2 | 0 | NA | NA | NA | NA | NA | #VALUE! |
| ENSMUSG00000109970.2 | 0 | NA | NA | NA | NA | NA | #VALUE! |
| ENSMUSG00000109971.2 | 0 | NA | NA | NA | NA | NA | #VALUE! |
| ENSMUSG00000109972.2 | 0 | NA | NA | NA | NA | NA | #VALUE! |
| ENSMUSG00000109973.2 | 0 | NA | NA | NA | NA | NA | #VALUE! |
| ENSMUSG00000109974.2 | 0 | NA | NA | NA | NA | NA | #VALUE! |
| ENSMUSG00000109975.2 | 0 | NA | NA | NA | NA | NA | #VALUE! |
| ENSMUSG00000109976.2 | 0 | NA | NA | NA | NA | NA | #VALUE! |
| ENSMUSG00000109977.2 | 0 | NA | NA | NA | NA | NA | #VALUE! |
| ENSMUSG00000109978.2 | 0 | NA | NA | NA | NA | NA | #VALUE! |
| ENSMUSG00000109979.2 | 0 | NA | NA | NA | NA | NA | #VALUE! |
| ENSMUSG00000109980.2 | 0 | NA | NA | NA | NA | NA | #VALUE! |
| ENSMUSG00000109981.2 | 0 | NA | NA | NA | NA | NA | #VALUE! |
| ENSMUSG00000109982.2 | 0 | NA | NA | NA | NA | NA | #VALUE! |
| ENSMUSG00000109983.2 | 0 | NA | NA | NA | NA | NA | #VALUE! |
| ENSMUSG00000109984.2 | 0 | NA | NA | NA | NA | NA | #VALUE! |
| ENSMUSG00000109985.2 | 0 | NA | NA | NA | NA | NA | #VALUE! |
| ENSMUSG00000109986.2 | 0 | NA | NA | NA | NA | NA | #VALUE! |
| ENSMUSG00000109987.2 | 0 | NA | NA | NA | NA | NA | #VALUE! |
| ENSMUSG00000109988.2 | 0 | NA | NA | NA | NA | NA | #VALUE! |
| ENSMUSG00000109990.2 | 0 | NA | NA | NA | NA | NA | #VALUE! |
| ENSMUSG00000109991.2 | 0 | NA | NA | NA | NA | NA | #VALUE! |
| ENSMUSG00000109992.2 | 0 | NA | NA | NA | NA | NA | #VALUE! |
| ENSMUSG00000109993.2 | 0 | NA | NA | NA | NA | NA | #VALUE! |
| ENSMUSG00000109994.2 | 0 | NA | NA | NA | NA | NA | #VALUE! |
| ENSMUSG00000109995.2 | 0 | NA | NA | NA | NA | NA | #VALUE! |
| ENSMUSG00000109996.2 | 0 | NA | NA | NA | NA | NA | #VALUE! |
| ENSMUSG00000109997.2 | 0 | NA | NA | NA | NA | NA | #VALUE! |
| ENSMUSG00000109998.2 | 0 | NA | NA | NA | NA | NA | #VALUE! |
| ENSMUSG00000109999.2 | 0 | NA | NA | NA | NA | NA | #VALUE! |
| ENSMUSG00000110000.2 | 0 | NA | NA | NA | NA | NA | #VALUE! |
| ENSMUSG00000110001.2 | 0 | NA | NA | NA | NA | NA | #VALUE! |
| ENSMUSG00000110002.2 | 0 | NA | NA | NA | NA | NA | #VALUE! |
| ENSMUSG00000110003.2 | 0 | NA | NA | NA | NA | NA | #VALUE! |
| ENSMUSG00000110004.2 | 0 | NA | NA | NA | NA | NA | #VALUE! |
| ENSMUSG00000110005.2 | 0 | NA | NA | NA | NA | NA | #VALUE! |
| ENSMUSG00000110006.2 | 0 | NA | NA | NA | NA | NA | #VALUE! |
| ENSMUSG00000110007.3 | 0 | NA | NA | NA | NA | NA | #VALUE! |
| ENSMUSG00000110008.6 | 0 | NA | NA | NA | NA | NA | #VALUE! |

| | | | | | | | |
|---|---|---|---|---|---|---|---|
| ENSMUSG00000110009.2 | 0 | NA | NA | NA | NA | NA | #VALUE! |
| ENSMUSG00000110010.2 | 0 | NA | NA | NA | NA | NA | #VALUE! |
| ENSMUSG00000110011.2 | 0 | NA | NA | NA | NA | NA | #VALUE! |
| ENSMUSG00000110012.3 | 0 | NA | NA | NA | NA | NA | #VALUE! |
| ENSMUSG00000110013.2 | 0 | NA | NA | NA | NA | NA | #VALUE! |
| ENSMUSG00000110014.2 | 0 | NA | NA | NA | NA | NA | #VALUE! |
| ENSMUSG00000110015.2 | 0 | NA | NA | NA | NA | NA | #VALUE! |
| ENSMUSG00000110016.2 | 0 | NA | NA | NA | NA | NA | #VALUE! |
| ENSMUSG00000110017.2 | 0 | NA | NA | NA | NA | NA | #VALUE! |
| ENSMUSG00000110018.3 | 0 | NA | NA | NA | NA | NA | #VALUE! |
| ENSMUSG00000110019.2 | 0 | NA | NA | NA | NA | NA | #VALUE! |
| ENSMUSG00000110020.2 | 0 | NA | NA | NA | NA | NA | #VALUE! |
| ENSMUSG00000110022.2 | 0 | NA | NA | NA | NA | NA | #VALUE! |
| ENSMUSG00000110023.2 | 0 | NA | NA | NA | NA | NA | #VALUE! |
| ENSMUSG00000110024.2 | 0 | NA | NA | NA | NA | NA | #VALUE! |
| ENSMUSG00000110025.2 | 0 | NA | NA | NA | NA | NA | #VALUE! |
| ENSMUSG00000110026.2 | 0 | NA | NA | NA | NA | NA | #VALUE! |
| ENSMUSG00000110027.2 | 0 | NA | NA | NA | NA | NA | #VALUE! |
| ENSMUSG00000110028.2 | 0 | NA | NA | NA | NA | NA | #VALUE! |
| ENSMUSG00000110030.2 | 0 | NA | NA | NA | NA | NA | #VALUE! |
| ENSMUSG00000110031.2 | 0 | NA | NA | NA | NA | NA | #VALUE! |
| ENSMUSG00000110032.2 | 0 | NA | NA | NA | NA | NA | #VALUE! |
| ENSMUSG00000110033.2 | 0 | NA | NA | NA | NA | NA | #VALUE! |
| ENSMUSG00000110034.2 | 0 | NA | NA | NA | NA | NA | #VALUE! |
| ENSMUSG00000110036.2 | 0 | NA | NA | NA | NA | NA | #VALUE! |
| ENSMUSG00000110037.2 | 0 | NA | NA | NA | NA | NA | #VALUE! |
| ENSMUSG00000110038.2 | 0 | NA | NA | NA | NA | NA | #VALUE! |
| ENSMUSG00000110039.2 | 0 | NA | NA | NA | NA | NA | #VALUE! |
| ENSMUSG00000110040.2 | 0 | NA | NA | NA | NA | NA | #VALUE! |
| ENSMUSG00000110041.2 | 0 | NA | NA | NA | NA | NA | #VALUE! |
| ENSMUSG00000110042.2 | 0 | NA | NA | NA | NA | NA | #VALUE! |
| ENSMUSG00000110043.2 | 0 | NA | NA | NA | NA | NA | #VALUE! |
| ENSMUSG00000110044.2 | 0 | NA | NA | NA | NA | NA | #VALUE! |
| ENSMUSG00000110045.2 | 0 | NA | NA | NA | NA | NA | #VALUE! |
| ENSMUSG00000110046.2 | 0 | NA | NA | NA | NA | NA | #VALUE! |
| ENSMUSG00000110047.2 | 0 | NA | NA | NA | NA | NA | #VALUE! |
| ENSMUSG00000110048.2 | 0 | NA | NA | NA | NA | NA | #VALUE! |
| ENSMUSG00000110049.2 | 0 | NA | NA | NA | NA | NA | #VALUE! |
| ENSMUSG00000110050.2 | 0 | NA | NA | NA | NA | NA | #VALUE! |
| ENSMUSG00000110051.2 | 0 | NA | NA | NA | NA | NA | #VALUE! |
| ENSMUSG00000110052.2 | 0 | NA | NA | NA | NA | NA | #VALUE! |
| ENSMUSG00000110053.2 | 0 | NA | NA | NA | NA | NA | #VALUE! |
| ENSMUSG00000110054.2 | 0 | NA | NA | NA | NA | NA | #VALUE! |
| ENSMUSG00000110055.2 | 0 | NA | NA | NA | NA | NA | #VALUE! |

| | | | | | | | |
|---|---|---|---|---|---|---|---|
| ENSMUSG00000110056.2 | 0 | NA | NA | NA | NA | NA | #VALUE! |
| ENSMUSG00000110058.2 | 0 | NA | NA | NA | NA | NA | #VALUE! |
| ENSMUSG00000110059.2 | 0 | NA | NA | NA | NA | NA | #VALUE! |
| ENSMUSG00000110060.2 | 0 | NA | NA | NA | NA | NA | #VALUE! |
| ENSMUSG00000110061.2 | 0 | NA | NA | NA | NA | NA | #VALUE! |
| ENSMUSG00000110062.2 | 0 | NA | NA | NA | NA | NA | #VALUE! |
| ENSMUSG00000110063.2 | 0 | NA | NA | NA | NA | NA | #VALUE! |
| ENSMUSG00000110064.2 | 0 | NA | NA | NA | NA | NA | #VALUE! |
| ENSMUSG00000110065.2 | 0 | NA | NA | NA | NA | NA | #VALUE! |
| ENSMUSG00000110066.2 | 0 | NA | NA | NA | NA | NA | #VALUE! |
| ENSMUSG00000110067.3 | 0 | NA | NA | NA | NA | NA | #VALUE! |
| ENSMUSG00000110068.2 | 0 | NA | NA | NA | NA | NA | #VALUE! |
| ENSMUSG00000110069.2 | 0 | NA | NA | NA | NA | NA | #VALUE! |
| ENSMUSG00000110070.2 | 0 | NA | NA | NA | NA | NA | #VALUE! |
| ENSMUSG00000110071.2 | 0 | NA | NA | NA | NA | NA | #VALUE! |
| ENSMUSG00000110072.2 | 0 | NA | NA | NA | NA | NA | #VALUE! |
| ENSMUSG00000110073.2 | 0 | NA | NA | NA | NA | NA | #VALUE! |
| ENSMUSG00000110074.2 | 0 | NA | NA | NA | NA | NA | #VALUE! |
| ENSMUSG00000110075.2 | 0 | NA | NA | NA | NA | NA | #VALUE! |
| ENSMUSG00000110076.2 | 0 | NA | NA | NA | NA | NA | #VALUE! |
| ENSMUSG00000110077.2 | 0 | NA | NA | NA | NA | NA | #VALUE! |
| ENSMUSG00000110078.2 | 0 | NA | NA | NA | NA | NA | #VALUE! |
| ENSMUSG00000110079.2 | 0 | NA | NA | NA | NA | NA | #VALUE! |
| ENSMUSG00000110080.2 | 0 | NA | NA | NA | NA | NA | #VALUE! |
| ENSMUSG00000110081.2 | 0 | NA | NA | NA | NA | NA | #VALUE! |
| ENSMUSG00000110082.2 | 0 | NA | NA | NA | NA | NA | #VALUE! |
| ENSMUSG00000110083.2 | 0 | NA | NA | NA | NA | NA | #VALUE! |
| ENSMUSG00000110084.2 | 0 | NA | NA | NA | NA | NA | #VALUE! |
| ENSMUSG00000110085.2 | 0 | NA | NA | NA | NA | NA | #VALUE! |
| ENSMUSG00000110086.2 | 0 | NA | NA | NA | NA | NA | #VALUE! |
| ENSMUSG00000110087.2 | 0 | NA | NA | NA | NA | NA | #VALUE! |
| ENSMUSG00000110088.2 | 1.771849 | 4.332864 | 3.931707 | 1.102031 | 0.270448 | NA | #VALUE! |
| ENSMUSG00000110089.2 | 0 | NA | NA | NA | NA | NA | #VALUE! |
| ENSMUSG00000110090.2 | 0 | NA | NA | NA | NA | NA | #VALUE! |
| ENSMUSG00000110091.2 | 0 | NA | NA | NA | NA | NA | #VALUE! |
| ENSMUSG00000110092.3 | 0 | NA | NA | NA | NA | NA | #VALUE! |
| ENSMUSG00000110093.2 | 0 | NA | NA | NA | NA | NA | #VALUE! |
| ENSMUSG00000110094.2 | 0 | NA | NA | NA | NA | NA | #VALUE! |
| ENSMUSG00000110095.2 | 0 | NA | NA | NA | NA | NA | #VALUE! |
| ENSMUSG00000110096.2 | 0 | NA | NA | NA | NA | NA | #VALUE! |
| ENSMUSG00000110097.2 | 0 | NA | NA | NA | NA | NA | #VALUE! |
| ENSMUSG00000110098.2 | 0 | NA | NA | NA | NA | NA | #VALUE! |
| ENSMUSG00000110099.2 | 0 | NA | NA | NA | NA | NA | #VALUE! |
| ENSMUSG00000110100.2 | 0 | NA | NA | NA | NA | NA | #VALUE! |

| | | | | | | | |
|---|---|---|---|---|---|---|---|
| ENSMUSG00000110101.2 | 0 | NA | NA | NA | NA | NA | #VALUE! |
| ENSMUSG00000110102.2 | 0 | NA | NA | NA | NA | NA | #VALUE! |
| ENSMUSG00000110103.2 | 0 | NA | NA | NA | NA | NA | #VALUE! |
| ENSMUSG00000110104.2 | 0 | NA | NA | NA | NA | NA | #VALUE! |
| ENSMUSG00000110105.2 | 19.11591 | 1.180822 | 2.797548 | 0.422092 | NA | NA | #VALUE! |
| ENSMUSG00000110106.2 | 0 | NA | NA | NA | NA | NA | #VALUE! |
| ENSMUSG00000110108.2 | 0 | NA | NA | NA | NA | NA | #VALUE! |
| ENSMUSG00000110109.2 | 0 | NA | NA | NA | NA | NA | #VALUE! |
| ENSMUSG00000110110.2 | 0 | NA | NA | NA | NA | NA | #VALUE! |
| ENSMUSG00000110111.2 | 0 | NA | NA | NA | NA | NA | #VALUE! |
| ENSMUSG00000110112.2 | 0 | NA | NA | NA | NA | NA | #VALUE! |
| ENSMUSG00000110113.2 | 0 | NA | NA | NA | NA | NA | #VALUE! |
| ENSMUSG00000110114.2 | 0 | NA | NA | NA | NA | NA | #VALUE! |
| ENSMUSG00000110115.2 | 0 | NA | NA | NA | NA | NA | #VALUE! |
| ENSMUSG00000110116.2 | 0 | NA | NA | NA | NA | NA | #VALUE! |
| ENSMUSG00000110117.2 | 0 | NA | NA | NA | NA | NA | #VALUE! |
| ENSMUSG00000110118.2 | 0 | NA | NA | NA | NA | NA | #VALUE! |
| ENSMUSG00000110119.2 | 0 | NA | NA | NA | NA | NA | #VALUE! |
| ENSMUSG00000110120.2 | 0 | NA | NA | NA | NA | NA | #VALUE! |
| ENSMUSG00000110121.2 | 0 | NA | NA | NA | NA | NA | #VALUE! |
| ENSMUSG00000110122.2 | 0 | NA | NA | NA | NA | NA | #VALUE! |
| ENSMUSG00000110123.2 | 0 | NA | NA | NA | NA | NA | #VALUE! |
| ENSMUSG00000110124.2 | 0 | NA | NA | NA | NA | NA | #VALUE! |
| ENSMUSG00000110125.2 | 0 | NA | NA | NA | NA | NA | #VALUE! |
| ENSMUSG00000110126.2 | 0 | NA | NA | NA | NA | NA | #VALUE! |
| ENSMUSG00000110127.2 | 0 | NA | NA | NA | NA | NA | #VALUE! |
| ENSMUSG00000110128.2 | 0 | NA | NA | NA | NA | NA | #VALUE! |
| ENSMUSG00000110129.2 | 0 | NA | NA | NA | NA | NA | #VALUE! |
| ENSMUSG00000110130.2 | 0 | NA | NA | NA | NA | NA | #VALUE! |
| ENSMUSG00000110131.2 | 0 | NA | NA | NA | NA | NA | #VALUE! |
| ENSMUSG00000110132.2 | 0 | NA | NA | NA | NA | NA | #VALUE! |
| ENSMUSG00000110133.2 | 0 | NA | NA | NA | NA | NA | #VALUE! |
| ENSMUSG00000110134.2 | 0 | NA | NA | NA | NA | NA | #VALUE! |
| ENSMUSG00000110135.2 | 0 | NA | NA | NA | NA | NA | #VALUE! |
| ENSMUSG00000110136.2 | 0 | NA | NA | NA | NA | NA | #VALUE! |
| ENSMUSG00000110137.2 | 0 | NA | NA | NA | NA | NA | #VALUE! |
| ENSMUSG00000110138.2 | 0 | NA | NA | NA | NA | NA | #VALUE! |
| ENSMUSG00000110139.2 | 0 | NA | NA | NA | NA | NA | #VALUE! |
| ENSMUSG00000110140.2 | 0 | NA | NA | NA | NA | NA | #VALUE! |
| ENSMUSG00000110141.2 | 0 | NA | NA | NA | NA | NA | #VALUE! |
| ENSMUSG00000110142.2 | 0 | NA | NA | NA | NA | NA | #VALUE! |
| ENSMUSG00000110143.2 | 0 | NA | NA | NA | NA | NA | #VALUE! |
| ENSMUSG00000110144.2 | 0 | NA | NA | NA | NA | NA | #VALUE! |
| ENSMUSG00000110145.2 | 0 | NA | NA | NA | NA | NA | #VALUE! |

| | | | | | | | |
|---|---|---|---|---|---|---|---|
| ENSMUSG00000110146.2 | 0 | NA | NA | NA | NA | NA | #VALUE! |
| ENSMUSG00000110147.2 | 0 | NA | NA | NA | NA | NA | #VALUE! |
| ENSMUSG00000110148.2 | 0 | NA | NA | NA | NA | NA | #VALUE! |
| ENSMUSG00000110149.2 | 0 | NA | NA | NA | NA | NA | #VALUE! |
| ENSMUSG00000110150.2 | 0 | NA | NA | NA | NA | NA | #VALUE! |
| ENSMUSG00000110151.3 | 2.835354 | -5.04297 | 3.922422 | -1.28568 | 0.198556 | NA | #VALUE! |
| ENSMUSG00000110152.2 | 0 | NA | NA | NA | NA | NA | #VALUE! |
| ENSMUSG00000110153.2 | 0 | NA | NA | NA | NA | NA | #VALUE! |
| ENSMUSG00000110154.2 | 0.168284 | 1.027628 | 4.067126 | 0.252667 | 0.800526 | NA | #VALUE! |
| ENSMUSG00000110155.2 | 0 | NA | NA | NA | NA | NA | #VALUE! |
| ENSMUSG00000110156.3 | 3.918335 | 0.412188 | 3.773118 | 0.109243 | 0.913009 | NA | #VALUE! |
| ENSMUSG00000110157.2 | 0 | NA | NA | NA | NA | NA | #VALUE! |
| ENSMUSG00000110158.2 | 0 | NA | NA | NA | NA | NA | #VALUE! |
| ENSMUSG00000110159.2 | 0 | NA | NA | NA | NA | NA | #VALUE! |
| ENSMUSG00000110160.2 | 0 | NA | NA | NA | NA | NA | #VALUE! |
| ENSMUSG00000110161.2 | 0 | NA | NA | NA | NA | NA | #VALUE! |
| ENSMUSG00000110162.2 | 0 | NA | NA | NA | NA | NA | #VALUE! |
| ENSMUSG00000110163.2 | 0 | NA | NA | NA | NA | NA | #VALUE! |
| ENSMUSG00000110164.2 | 0 | NA | NA | NA | NA | NA | #VALUE! |
| ENSMUSG00000110166.2 | 0 | NA | NA | NA | NA | NA | #VALUE! |
| ENSMUSG00000110167.2 | 0 | NA | NA | NA | NA | NA | #VALUE! |
| ENSMUSG00000110168.2 | 0 | NA | NA | NA | NA | NA | #VALUE! |
| ENSMUSG00000110169.2 | 0 | NA | NA | NA | NA | NA | #VALUE! |
| ENSMUSG00000110170.2 | 0 | NA | NA | NA | NA | NA | #VALUE! |
| ENSMUSG00000110171.3 | 0 | NA | NA | NA | NA | NA | #VALUE! |
| ENSMUSG00000110172.2 | 0 | NA | NA | NA | NA | NA | #VALUE! |
| ENSMUSG00000110173.2 | 0 | NA | NA | NA | NA | NA | #VALUE! |
| ENSMUSG00000110174.2 | 0 | NA | NA | NA | NA | NA | #VALUE! |
| ENSMUSG00000110175.2 | 0 | NA | NA | NA | NA | NA | #VALUE! |
| ENSMUSG00000110176.2 | 0 | NA | NA | NA | NA | NA | #VALUE! |
| ENSMUSG00000110177.2 | 0 | NA | NA | NA | NA | NA | #VALUE! |
| ENSMUSG00000110178.2 | 0 | NA | NA | NA | NA | NA | #VALUE! |
| ENSMUSG00000110179.2 | 0 | NA | NA | NA | NA | NA | #VALUE! |
| ENSMUSG00000110180.2 | 0 | NA | NA | NA | NA | NA | #VALUE! |
| ENSMUSG00000110181.2 | 0 | NA | NA | NA | NA | NA | #VALUE! |
| ENSMUSG00000110182.2 | 0 | NA | NA | NA | NA | NA | #VALUE! |
| ENSMUSG00000110183.2 | 0 | NA | NA | NA | NA | NA | #VALUE! |
| ENSMUSG00000110184.2 | 0 | NA | NA | NA | NA | NA | #VALUE! |
| ENSMUSG00000110186.2 | 0 | NA | NA | NA | NA | NA | #VALUE! |
| ENSMUSG00000110187.2 | 0 | NA | NA | NA | NA | NA | #VALUE! |
| ENSMUSG00000110188.2 | 0 | NA | NA | NA | NA | NA | #VALUE! |
| ENSMUSG00000110189.2 | 0 | NA | NA | NA | NA | NA | #VALUE! |
| ENSMUSG00000110190.2 | 0 | NA | NA | NA | NA | NA | #VALUE! |
| ENSMUSG00000110192.2 | 0 | NA | NA | NA | NA | NA | #VALUE! |

| | | | | | | | |
|---|---|---|---|---|---|---|---|
| ENSMUSG00000110193.2 | 0 | NA | NA | NA | NA | NA | #VALUE! |
| ENSMUSG00000110194.2 | 0 | NA | NA | NA | NA | NA | #VALUE! |
| ENSMUSG00000110195.2 | 0 | NA | NA | NA | NA | NA | #VALUE! |
| ENSMUSG00000110196.2 | 0 | NA | NA | NA | NA | NA | #VALUE! |
| ENSMUSG00000110197.2 | 0 | NA | NA | NA | NA | NA | #VALUE! |
| ENSMUSG00000110198.2 | 0 | NA | NA | NA | NA | NA | #VALUE! |
| ENSMUSG00000110199.3 | 0 | NA | NA | NA | NA | NA | #VALUE! |
| ENSMUSG00000110200.2 | 0 | NA | NA | NA | NA | NA | #VALUE! |
| ENSMUSG00000110201.2 | 0 | NA | NA | NA | NA | NA | #VALUE! |
| ENSMUSG00000110202.2 | 0 | NA | NA | NA | NA | NA | #VALUE! |
| ENSMUSG00000110203.2 | 0 | NA | NA | NA | NA | NA | #VALUE! |
| ENSMUSG00000110204.2 | 0 | NA | NA | NA | NA | NA | #VALUE! |
| ENSMUSG00000110205.2 | 0 | NA | NA | NA | NA | NA | #VALUE! |
| ENSMUSG00000110207.2 | 0 | NA | NA | NA | NA | NA | #VALUE! |
| ENSMUSG00000110208.2 | 0 | NA | NA | NA | NA | NA | #VALUE! |
| ENSMUSG00000110209.2 | 0 | NA | NA | NA | NA | NA | #VALUE! |
| ENSMUSG00000110210.2 | 0 | NA | NA | NA | NA | NA | #VALUE! |
| ENSMUSG00000110211.2 | 0 | NA | NA | NA | NA | NA | #VALUE! |
| ENSMUSG00000110212.2 | 0 | NA | NA | NA | NA | NA | #VALUE! |
| ENSMUSG00000110213.2 | 0 | NA | NA | NA | NA | NA | #VALUE! |
| ENSMUSG00000110214.2 | 0 | NA | NA | NA | NA | NA | #VALUE! |
| ENSMUSG00000110215.2 | 0 | NA | NA | NA | NA | NA | #VALUE! |
| ENSMUSG00000110216.2 | 0 | NA | NA | NA | NA | NA | #VALUE! |
| ENSMUSG00000110217.2 | 0 | NA | NA | NA | NA | NA | #VALUE! |
| ENSMUSG00000110218.2 | 0 | NA | NA | NA | NA | NA | #VALUE! |
| ENSMUSG00000110219.2 | 0 | NA | NA | NA | NA | NA | #VALUE! |
| ENSMUSG00000110220.2 | 0 | NA | NA | NA | NA | NA | #VALUE! |
| ENSMUSG00000110221.2 | 0 | NA | NA | NA | NA | NA | #VALUE! |
| ENSMUSG00000110222.2 | 0 | NA | NA | NA | NA | NA | #VALUE! |
| ENSMUSG00000110223.2 | 0 | NA | NA | NA | NA | NA | #VALUE! |
| ENSMUSG00000110224.2 | 0 | NA | NA | NA | NA | NA | #VALUE! |
| ENSMUSG00000110225.2 | 0 | NA | NA | NA | NA | NA | #VALUE! |
| ENSMUSG00000110226.2 | 0 | NA | NA | NA | NA | NA | #VALUE! |
| ENSMUSG00000110227.2 | 0 | NA | NA | NA | NA | NA | #VALUE! |
| ENSMUSG00000110228.2 | 0 | NA | NA | NA | NA | NA | #VALUE! |
| ENSMUSG00000110229.2 | 0 | NA | NA | NA | NA | NA | #VALUE! |
| ENSMUSG00000110230.2 | 0 | NA | NA | NA | NA | NA | #VALUE! |
| ENSMUSG00000110231.2 | 0 | NA | NA | NA | NA | NA | #VALUE! |
| ENSMUSG00000110232.2 | 0 | NA | NA | NA | NA | NA | #VALUE! |
| ENSMUSG00000110233.2 | 0 | NA | NA | NA | NA | NA | #VALUE! |
| ENSMUSG00000110235.2 | 2.355969 | 4.72622 | 3.925784 | 1.203892 | 0.228631 | NA | #VALUE! |
| ENSMUSG00000110236.2 | 0 | NA | NA | NA | NA | NA | #VALUE! |
| ENSMUSG00000110237.2 | 0 | NA | NA | NA | NA | NA | #VALUE! |
| ENSMUSG00000110238.2 | 0 | NA | NA | NA | NA | NA | #VALUE! |

| | | | | | | | |
|---|---|---|---|---|---|---|---|
| ENSMUSG00000110239.2 | 0 | NA | NA | NA | NA | NA | #VALUE! |
| ENSMUSG00000110240.2 | 0 | NA | NA | NA | NA | NA | #VALUE! |
| ENSMUSG00000110241.2 | 0 | NA | NA | NA | NA | NA | #VALUE! |
| ENSMUSG00000110242.2 | 0 | NA | NA | NA | NA | NA | #VALUE! |
| ENSMUSG00000110243.2 | 0 | NA | NA | NA | NA | NA | #VALUE! |
| ENSMUSG00000110244.2 | 0 | NA | NA | NA | NA | NA | #VALUE! |
| ENSMUSG00000110245.2 | 0 | NA | NA | NA | NA | NA | #VALUE! |
| ENSMUSG00000110246.2 | 0 | NA | NA | NA | NA | NA | #VALUE! |
| ENSMUSG00000110247.2 | 0 | NA | NA | NA | NA | NA | #VALUE! |
| ENSMUSG00000110248.2 | 0 | NA | NA | NA | NA | NA | #VALUE! |
| ENSMUSG00000110249.2 | 0 | NA | NA | NA | NA | NA | #VALUE! |
| ENSMUSG00000110250.2 | 0 | NA | NA | NA | NA | NA | #VALUE! |
| ENSMUSG00000110251.2 | 0 | NA | NA | NA | NA | NA | #VALUE! |
| ENSMUSG00000110252.2 | 0 | NA | NA | NA | NA | NA | #VALUE! |
| ENSMUSG00000110253.3 | 0 | NA | NA | NA | NA | NA | #VALUE! |
| ENSMUSG00000110254.2 | 0 | NA | NA | NA | NA | NA | #VALUE! |
| ENSMUSG00000110255.2 | 0 | NA | NA | NA | NA | NA | #VALUE! |
| ENSMUSG00000110256.2 | 0 | NA | NA | NA | NA | NA | #VALUE! |
| ENSMUSG00000110257.2 | 0 | NA | NA | NA | NA | NA | #VALUE! |
| ENSMUSG00000110258.2 | 0 | NA | NA | NA | NA | NA | #VALUE! |
| ENSMUSG00000110259.3 | 0 | NA | NA | NA | NA | NA | #VALUE! |
| ENSMUSG00000110260.2 | 0 | NA | NA | NA | NA | NA | #VALUE! |
| ENSMUSG00000110261.2 | 0 | NA | NA | NA | NA | NA | #VALUE! |
| ENSMUSG00000110262.2 | 0 | NA | NA | NA | NA | NA | #VALUE! |
| ENSMUSG00000110263.2 | 0 | NA | NA | NA | NA | NA | #VALUE! |
| ENSMUSG00000110264.2 | 0 | NA | NA | NA | NA | NA | #VALUE! |
| ENSMUSG00000110265.2 | 0 | NA | NA | NA | NA | NA | #VALUE! |
| ENSMUSG00000110266.2 | 0 | NA | NA | NA | NA | NA | #VALUE! |
| ENSMUSG00000110267.2 | 0 | NA | NA | NA | NA | NA | #VALUE! |
| ENSMUSG00000110268.2 | 0 | NA | NA | NA | NA | NA | #VALUE! |
| ENSMUSG00000110269.2 | 0 | NA | NA | NA | NA | NA | #VALUE! |
| ENSMUSG00000110270.2 | 0 | NA | NA | NA | NA | NA | #VALUE! |
| ENSMUSG00000110271.2 | 0 | NA | NA | NA | NA | NA | #VALUE! |
| ENSMUSG00000110272.2 | 0 | NA | NA | NA | NA | NA | #VALUE! |
| ENSMUSG00000110273.2 | 0.793073 | -0.15398 | 3.943895 | -0.03904 | 0.968856 | NA | #VALUE! |
| ENSMUSG00000110274.2 | 0 | NA | NA | NA | NA | NA | #VALUE! |
| ENSMUSG00000110276.2 | 0 | NA | NA | NA | NA | NA | #VALUE! |
| ENSMUSG00000110278.2 | 0 | NA | NA | NA | NA | NA | #VALUE! |
| ENSMUSG00000110279.2 | 0 | NA | NA | NA | NA | NA | #VALUE! |
| ENSMUSG00000110280.2 | 0 | NA | NA | NA | NA | NA | #VALUE! |
| ENSMUSG00000110281.2 | 0 | NA | NA | NA | NA | NA | #VALUE! |
| ENSMUSG00000110282.2 | 0 | NA | NA | NA | NA | NA | #VALUE! |
| ENSMUSG00000110283.2 | 0 | NA | NA | NA | NA | NA | #VALUE! |
| ENSMUSG00000110284.2 | 0 | NA | NA | NA | NA | NA | #VALUE! |

| | | | | | | | |
|---|---|---|---|---|---|---|---|
| ENSMUSG00000110285.2 | 0 | NA | NA | NA | NA | NA | #VALUE! |
| ENSMUSG00000110286.2 | 0 | NA | NA | NA | NA | NA | #VALUE! |
| ENSMUSG00000110287.2 | 0 | NA | NA | NA | NA | NA | #VALUE! |
| ENSMUSG00000110288.2 | 0 | NA | NA | NA | NA | NA | #VALUE! |
| ENSMUSG00000110289.2 | 0 | NA | NA | NA | NA | NA | #VALUE! |
| ENSMUSG00000110290.2 | 0 | NA | NA | NA | NA | NA | #VALUE! |
| ENSMUSG00000110291.2 | 0 | NA | NA | NA | NA | NA | #VALUE! |
| ENSMUSG00000110292.2 | 0 | NA | NA | NA | NA | NA | #VALUE! |
| ENSMUSG00000110293.2 | 0 | NA | NA | NA | NA | NA | #VALUE! |
| ENSMUSG00000110294.2 | 0 | NA | NA | NA | NA | NA | #VALUE! |
| ENSMUSG00000110295.2 | 0 | NA | NA | NA | NA | NA | #VALUE! |
| ENSMUSG00000110296.2 | 0 | NA | NA | NA | NA | NA | #VALUE! |
| ENSMUSG00000110297.2 | 0 | NA | NA | NA | NA | NA | #VALUE! |
| ENSMUSG00000110298.2 | 0 | NA | NA | NA | NA | NA | #VALUE! |
| ENSMUSG00000110299.2 | 0 | NA | NA | NA | NA | NA | #VALUE! |
| ENSMUSG00000110300.2 | 0 | NA | NA | NA | NA | NA | #VALUE! |
| ENSMUSG00000110301.2 | 0 | NA | NA | NA | NA | NA | #VALUE! |
| ENSMUSG00000110302.2 | 0 | NA | NA | NA | NA | NA | #VALUE! |
| ENSMUSG00000110303.2 | 0 | NA | NA | NA | NA | NA | #VALUE! |
| ENSMUSG00000110304.2 | 0 | NA | NA | NA | NA | NA | #VALUE! |
| ENSMUSG00000110305.2 | 0 | NA | NA | NA | NA | NA | #VALUE! |
| ENSMUSG00000110307.2 | 0 | NA | NA | NA | NA | NA | #VALUE! |
| ENSMUSG00000110308.2 | 0 | NA | NA | NA | NA | NA | #VALUE! |
| ENSMUSG00000110309.2 | 0 | NA | NA | NA | NA | NA | #VALUE! |
| ENSMUSG00000110310.2 | 0 | NA | NA | NA | NA | NA | #VALUE! |
| ENSMUSG00000110311.2 | 0 | NA | NA | NA | NA | NA | #VALUE! |
| ENSMUSG00000110312.2 | 0 | NA | NA | NA | NA | NA | #VALUE! |
| ENSMUSG00000110313.2 | 0 | NA | NA | NA | NA | NA | #VALUE! |
| ENSMUSG00000110314.2 | 0 | NA | NA | NA | NA | NA | #VALUE! |
| ENSMUSG00000110315.2 | 0 | NA | NA | NA | NA | NA | #VALUE! |
| ENSMUSG00000110316.2 | 0 | NA | NA | NA | NA | NA | #VALUE! |
| ENSMUSG00000110317.2 | 0 | NA | NA | NA | NA | NA | #VALUE! |
| ENSMUSG00000110318.2 | 0 | NA | NA | NA | NA | NA | #VALUE! |
| ENSMUSG00000110319.2 | 0 | NA | NA | NA | NA | NA | #VALUE! |
| ENSMUSG00000110320.2 | 0 | NA | NA | NA | NA | NA | #VALUE! |
| ENSMUSG00000110321.2 | 0 | NA | NA | NA | NA | NA | #VALUE! |
| ENSMUSG00000110322.2 | 0 | NA | NA | NA | NA | NA | #VALUE! |
| ENSMUSG00000110323.2 | 0 | NA | NA | NA | NA | NA | #VALUE! |
| ENSMUSG00000110324.2 | 0 | NA | NA | NA | NA | NA | #VALUE! |
| ENSMUSG00000110325.2 | 0 | NA | NA | NA | NA | NA | #VALUE! |
| ENSMUSG00000110326.2 | 0 | NA | NA | NA | NA | NA | #VALUE! |
| ENSMUSG00000110327.2 | 0 | NA | NA | NA | NA | NA | #VALUE! |
| ENSMUSG00000110328.2 | 0 | NA | NA | NA | NA | NA | #VALUE! |
| ENSMUSG00000110329.2 | 0 | NA | NA | NA | NA | NA | #VALUE! |

| | | | | | | | |
|---|---|---|---|---|---|---|---|
| ENSMUSG00000110330.2 | 0 | NA | NA | NA | NA | NA | #VALUE! |
| ENSMUSG00000110331.2 | 0 | NA | NA | NA | NA | NA | #VALUE! |
| ENSMUSG00000110332.2 | 0 | NA | NA | NA | NA | NA | #VALUE! |
| ENSMUSG00000110333.2 | 0 | NA | NA | NA | NA | NA | #VALUE! |
| ENSMUSG00000110334.2 | 0 | NA | NA | NA | NA | NA | #VALUE! |
| ENSMUSG00000110335.2 | 0 | NA | NA | NA | NA | NA | #VALUE! |
| ENSMUSG00000110336.2 | 0 | NA | NA | NA | NA | NA | #VALUE! |
| ENSMUSG00000110337.2 | 0 | NA | NA | NA | NA | NA | #VALUE! |
| ENSMUSG00000110338.2 | 0 | NA | NA | NA | NA | NA | #VALUE! |
| ENSMUSG00000110339.2 | 0 | NA | NA | NA | NA | NA | #VALUE! |
| ENSMUSG00000110340.2 | 0 | NA | NA | NA | NA | NA | #VALUE! |
| ENSMUSG00000110341.2 | 0 | NA | NA | NA | NA | NA | #VALUE! |
| ENSMUSG00000110342.2 | 0 | NA | NA | NA | NA | NA | #VALUE! |
| ENSMUSG00000110343.2 | 0 | NA | NA | NA | NA | NA | #VALUE! |
| ENSMUSG00000110345.2 | 0 | NA | NA | NA | NA | NA | #VALUE! |
| ENSMUSG00000110346.2 | 0 | NA | NA | NA | NA | NA | #VALUE! |
| ENSMUSG00000110347.2 | 0 | NA | NA | NA | NA | NA | #VALUE! |
| ENSMUSG00000110348.2 | 0 | NA | NA | NA | NA | NA | #VALUE! |
| ENSMUSG00000110349.2 | 0 | NA | NA | NA | NA | NA | #VALUE! |
| ENSMUSG00000110350.2 | 0 | NA | NA | NA | NA | NA | #VALUE! |
| ENSMUSG00000110351.2 | 0 | NA | NA | NA | NA | NA | #VALUE! |
| ENSMUSG00000110352.2 | 0 | NA | NA | NA | NA | NA | #VALUE! |
| ENSMUSG00000110354.2 | 0 | NA | NA | NA | NA | NA | #VALUE! |
| ENSMUSG00000110355.2 | 0.168284 | 1.027628 | 4.067126 | 0.252667 | 0.800526 | NA | #VALUE! |
| ENSMUSG00000110356.2 | 0 | NA | NA | NA | NA | NA | #VALUE! |
| ENSMUSG00000110357.2 | 0 | NA | NA | NA | NA | NA | #VALUE! |
| ENSMUSG00000110358.2 | 0 | NA | NA | NA | NA | NA | #VALUE! |
| ENSMUSG00000110359.2 | 0 | NA | NA | NA | NA | NA | #VALUE! |
| ENSMUSG00000110360.2 | 4.196484 | 5.566682 | 3.917409 | 1.421011 | 0.155313 | NA | #VALUE! |
| ENSMUSG00000110361.2 | 0 | NA | NA | NA | NA | NA | #VALUE! |
| ENSMUSG00000110362.2 | 0 | NA | NA | NA | NA | NA | #VALUE! |
| ENSMUSG00000110363.2 | 0 | NA | NA | NA | NA | NA | #VALUE! |
| ENSMUSG00000110364.2 | 0 | NA | NA | NA | NA | NA | #VALUE! |
| ENSMUSG00000110365.2 | 0 | NA | NA | NA | NA | NA | #VALUE! |
| ENSMUSG00000110366.2 | 0 | NA | NA | NA | NA | NA | #VALUE! |
| ENSMUSG00000110367.2 | 0 | NA | NA | NA | NA | NA | #VALUE! |
| ENSMUSG00000110368.2 | 0 | NA | NA | NA | NA | NA | #VALUE! |
| ENSMUSG00000110369.2 | 0.816443 | -3.21914 | 3.961753 | -0.81255 | 0.416474 | NA | #VALUE! |
| ENSMUSG00000110370.2 | 0 | NA | NA | NA | NA | NA | #VALUE! |
| ENSMUSG00000110371.2 | 0.18651 | 1.401769 | 4.056354 | 0.345574 | 0.729663 | NA | #VALUE! |
| ENSMUSG00000110372.2 | 0 | NA | NA | NA | NA | NA | #VALUE! |
| ENSMUSG00000110373.2 | 0.168284 | 1.027628 | 4.067126 | 0.252667 | 0.800526 | NA | #VALUE! |
| ENSMUSG00000110374.2 | 0 | NA | NA | NA | NA | NA | #VALUE! |
| ENSMUSG00000110375.2 | 0 | NA | NA | NA | NA | NA | #VALUE! |

| | | | | | | | |
|---|---|---|---|---|---|---|---|
| ENSMUSG00000110376.2 | 0 | NA | NA | NA | NA | NA | #VALUE! |
| ENSMUSG00000110377.2 | 0 | NA | NA | NA | NA | NA | #VALUE! |
| ENSMUSG00000110378.2 | 0 | NA | NA | NA | NA | NA | #VALUE! |
| ENSMUSG00000110379.2 | 0 | NA | NA | NA | NA | NA | #VALUE! |
| ENSMUSG00000110380.2 | 0 | NA | NA | NA | NA | NA | #VALUE! |
| ENSMUSG00000110381.2 | 0 | NA | NA | NA | NA | NA | #VALUE! |
| ENSMUSG00000110382.2 | 0 | NA | NA | NA | NA | NA | #VALUE! |
| ENSMUSG00000110383.2 | 0 | NA | NA | NA | NA | NA | #VALUE! |
| ENSMUSG00000110384.2 | 0 | NA | NA | NA | NA | NA | #VALUE! |
| ENSMUSG00000110385.2 | 0 | NA | NA | NA | NA | NA | #VALUE! |
| ENSMUSG00000110386.2 | 0 | NA | NA | NA | NA | NA | #VALUE! |
| ENSMUSG00000110387.2 | 0 | NA | NA | NA | NA | NA | #VALUE! |
| ENSMUSG00000110388.2 | 0 | NA | NA | NA | NA | NA | #VALUE! |
| ENSMUSG00000110389.2 | 0 | NA | NA | NA | NA | NA | #VALUE! |
| ENSMUSG00000110390.2 | 0 | NA | NA | NA | NA | NA | #VALUE! |
| ENSMUSG00000110391.3 | 0 | NA | NA | NA | NA | NA | #VALUE! |
| ENSMUSG00000110392.2 | 0 | NA | NA | NA | NA | NA | #VALUE! |
| ENSMUSG00000110393.3 | 0.373021 | 2.140748 | 4.018182 | 0.532765 | 0.594196 | NA | #VALUE! |
| ENSMUSG00000110394.2 | 0 | NA | NA | NA | NA | NA | #VALUE! |
| ENSMUSG00000110395.2 | 0 | NA | NA | NA | NA | NA | #VALUE! |
| ENSMUSG00000110396.2 | 0 | NA | NA | NA | NA | NA | #VALUE! |
| ENSMUSG00000110398.2 | 0 | NA | NA | NA | NA | NA | #VALUE! |
| ENSMUSG00000110399.2 | 0 | NA | NA | NA | NA | NA | #VALUE! |
| ENSMUSG00000110400.2 | 0 | NA | NA | NA | NA | NA | #VALUE! |
| ENSMUSG00000110401.2 | 0 | NA | NA | NA | NA | NA | #VALUE! |
| ENSMUSG00000110402.2 | 0 | NA | NA | NA | NA | NA | #VALUE! |
| ENSMUSG00000110403.2 | 0 | NA | NA | NA | NA | NA | #VALUE! |
| ENSMUSG00000110404.2 | 0 | NA | NA | NA | NA | NA | #VALUE! |
| ENSMUSG00000110405.2 | 0 | NA | NA | NA | NA | NA | #VALUE! |
| ENSMUSG00000110406.2 | 0 | NA | NA | NA | NA | NA | #VALUE! |
| ENSMUSG00000110407.2 | 0 | NA | NA | NA | NA | NA | #VALUE! |
| ENSMUSG00000110409.2 | 0 | NA | NA | NA | NA | NA | #VALUE! |
| ENSMUSG00000110410.2 | 0 | NA | NA | NA | NA | NA | #VALUE! |
| ENSMUSG00000110411.2 | 0 | NA | NA | NA | NA | NA | #VALUE! |
| ENSMUSG00000110412.2 | 0 | NA | NA | NA | NA | NA | #VALUE! |
| ENSMUSG00000110413.2 | 0 | NA | NA | NA | NA | NA | #VALUE! |
| ENSMUSG00000110414.2 | 0 | NA | NA | NA | NA | NA | #VALUE! |
| ENSMUSG00000110415.2 | 0 | NA | NA | NA | NA | NA | #VALUE! |
| ENSMUSG00000110416.2 | 0 | NA | NA | NA | NA | NA | #VALUE! |
| ENSMUSG00000110417.2 | 0 | NA | NA | NA | NA | NA | #VALUE! |
| ENSMUSG00000110419.2 | 0 | NA | NA | NA | NA | NA | #VALUE! |
| ENSMUSG00000110420.2 | 0 | NA | NA | NA | NA | NA | #VALUE! |
| ENSMUSG00000110421.2 | 0 | NA | NA | NA | NA | NA | #VALUE! |
| ENSMUSG00000110422.2 | 0 | NA | NA | NA | NA | NA | #VALUE! |

| | | | | | | | |
|---|---|---|---|---|---|---|---|
| ENSMUSG00000110424.2 | 0 | NA | NA | NA | NA | NA | #VALUE! |
| ENSMUSG00000110425.2 | 0 | NA | NA | NA | NA | NA | #VALUE! |
| ENSMUSG00000110426.2 | 0 | NA | NA | NA | NA | NA | #VALUE! |
| ENSMUSG00000110427.2 | 0 | NA | NA | NA | NA | NA | #VALUE! |
| ENSMUSG00000110428.2 | 0 | NA | NA | NA | NA | NA | #VALUE! |
| ENSMUSG00000110429.2 | 0 | NA | NA | NA | NA | NA | #VALUE! |
| ENSMUSG00000110430.3 | 0 | NA | NA | NA | NA | NA | #VALUE! |
| ENSMUSG00000110431.2 | 0 | NA | NA | NA | NA | NA | #VALUE! |
| ENSMUSG00000110432.2 | 0 | NA | NA | NA | NA | NA | #VALUE! |
| ENSMUSG00000110433.2 | 0 | NA | NA | NA | NA | NA | #VALUE! |
| ENSMUSG00000110434.2 | 0 | NA | NA | NA | NA | NA | #VALUE! |
| ENSMUSG00000110435.2 | 0 | NA | NA | NA | NA | NA | #VALUE! |
| ENSMUSG00000110437.2 | 0 | NA | NA | NA | NA | NA | #VALUE! |
| ENSMUSG00000110438.2 | 0 | NA | NA | NA | NA | NA | #VALUE! |
| ENSMUSG00000110439.2 | 0.105013 | -0.97324 | 4.076293 | -0.23876 | 0.811295 | NA | #VALUE! |
| ENSMUSG00000110440.2 | 0 | NA | NA | NA | NA | NA | #VALUE! |
| ENSMUSG00000110441.2 | 0 | NA | NA | NA | NA | NA | #VALUE! |
| ENSMUSG00000110442.2 | 0 | NA | NA | NA | NA | NA | #VALUE! |
| ENSMUSG00000110443.2 | 0 | NA | NA | NA | NA | NA | #VALUE! |
| ENSMUSG00000110445.2 | 0 | NA | NA | NA | NA | NA | #VALUE! |
| ENSMUSG00000110446.2 | 0 | NA | NA | NA | NA | NA | #VALUE! |
| ENSMUSG00000110447.2 | 0 | NA | NA | NA | NA | NA | #VALUE! |
| ENSMUSG00000110448.2 | 0 | NA | NA | NA | NA | NA | #VALUE! |
| ENSMUSG00000110449.2 | 0 | NA | NA | NA | NA | NA | #VALUE! |
| ENSMUSG00000110450.2 | 0 | NA | NA | NA | NA | NA | #VALUE! |
| ENSMUSG00000110452.2 | 0 | NA | NA | NA | NA | NA | #VALUE! |
| ENSMUSG00000110453.2 | 0 | NA | NA | NA | NA | NA | #VALUE! |
| ENSMUSG00000110454.2 | 0 | NA | NA | NA | NA | NA | #VALUE! |
| ENSMUSG00000110455.2 | 0 | NA | NA | NA | NA | NA | #VALUE! |
| ENSMUSG00000110456.2 | 0 | NA | NA | NA | NA | NA | #VALUE! |
| ENSMUSG00000110457.3 | 0 | NA | NA | NA | NA | NA | #VALUE! |
| ENSMUSG00000110458.2 | 0 | NA | NA | NA | NA | NA | #VALUE! |
| ENSMUSG00000110459.2 | 2.335612 | -0.13763 | 3.805268 | -0.03617 | 0.971148 | NA | #VALUE! |
| ENSMUSG00000110460.2 | 0 | NA | NA | NA | NA | NA | #VALUE! |
| ENSMUSG00000110461.2 | 0.093255 | 0.868093 | 4.072464 | 0.213162 | 0.831201 | NA | #VALUE! |
| ENSMUSG00000110462.2 | 0 | NA | NA | NA | NA | NA | #VALUE! |
| ENSMUSG00000110463.2 | 0 | NA | NA | NA | NA | NA | #VALUE! |
| ENSMUSG00000110464.2 | 0 | NA | NA | NA | NA | NA | #VALUE! |
| ENSMUSG00000110465.2 | 0 | NA | NA | NA | NA | NA | #VALUE! |
| ENSMUSG00000110466.4 | 0 | NA | NA | NA | NA | NA | #VALUE! |
| ENSMUSG00000110467.2 | 0 | NA | NA | NA | NA | NA | #VALUE! |
| ENSMUSG00000110468.2 | 0 | NA | NA | NA | NA | NA | #VALUE! |
| ENSMUSG00000110469.2 | 0 | NA | NA | NA | NA | NA | #VALUE! |
| ENSMUSG00000110470.2 | 0 | NA | NA | NA | NA | NA | #VALUE! |

| | | | | | | | |
|---|---|---|---|---|---|---|---|
| ENSMUSG00000110471.2 | 0 | NA | NA | NA | NA | NA | #VALUE! |
| ENSMUSG00000110472.2 | 0 | NA | NA | NA | NA | NA | #VALUE! |
| ENSMUSG00000110473.2 | 0 | NA | NA | NA | NA | NA | #VALUE! |
| ENSMUSG00000110474.2 | 0 | NA | NA | NA | NA | NA | #VALUE! |
| ENSMUSG00000110475.2 | 0 | NA | NA | NA | NA | NA | #VALUE! |
| ENSMUSG00000110476.2 | 0 | NA | NA | NA | NA | NA | #VALUE! |
| ENSMUSG00000110477.2 | 0 | NA | NA | NA | NA | NA | #VALUE! |
| ENSMUSG00000110478.2 | 0 | NA | NA | NA | NA | NA | #VALUE! |
| ENSMUSG00000110479.2 | 0 | NA | NA | NA | NA | NA | #VALUE! |
| ENSMUSG00000110480.2 | 0 | NA | NA | NA | NA | NA | #VALUE! |
| ENSMUSG00000110481.2 | 0 | NA | NA | NA | NA | NA | #VALUE! |
| ENSMUSG00000110482.2 | 0 | NA | NA | NA | NA | NA | #VALUE! |
| ENSMUSG00000110483.2 | 0 | NA | NA | NA | NA | NA | #VALUE! |
| ENSMUSG00000110484.2 | 0 | NA | NA | NA | NA | NA | #VALUE! |
| ENSMUSG00000110485.2 | 0 | NA | NA | NA | NA | NA | #VALUE! |
| ENSMUSG00000110486.2 | 0 | NA | NA | NA | NA | NA | #VALUE! |
| ENSMUSG00000110487.2 | 0 | NA | NA | NA | NA | NA | #VALUE! |
| ENSMUSG00000110488.2 | 0 | NA | NA | NA | NA | NA | #VALUE! |
| ENSMUSG00000110489.2 | 0 | NA | NA | NA | NA | NA | #VALUE! |
| ENSMUSG00000110490.2 | 0 | NA | NA | NA | NA | NA | #VALUE! |
| ENSMUSG00000110491.2 | 0 | NA | NA | NA | NA | NA | #VALUE! |
| ENSMUSG00000110492.2 | 0 | NA | NA | NA | NA | NA | #VALUE! |
| ENSMUSG00000110493.2 | 0 | NA | NA | NA | NA | NA | #VALUE! |
| ENSMUSG00000110494.2 | 0 | NA | NA | NA | NA | NA | #VALUE! |
| ENSMUSG00000110495.2 | 0 | NA | NA | NA | NA | NA | #VALUE! |
| ENSMUSG00000110496.2 | 0 | NA | NA | NA | NA | NA | #VALUE! |
| ENSMUSG00000110497.2 | 0 | NA | NA | NA | NA | NA | #VALUE! |
| ENSMUSG00000110498.2 | 0 | NA | NA | NA | NA | NA | #VALUE! |
| ENSMUSG00000110499.2 | 0 | NA | NA | NA | NA | NA | #VALUE! |
| ENSMUSG00000110500.2 | 0 | NA | NA | NA | NA | NA | #VALUE! |
| ENSMUSG00000110501.2 | 0 | NA | NA | NA | NA | NA | #VALUE! |
| ENSMUSG00000110502.2 | 0 | NA | NA | NA | NA | NA | #VALUE! |
| ENSMUSG00000110503.2 | 0 | NA | NA | NA | NA | NA | #VALUE! |
| ENSMUSG00000110504.2 | 8.919027 | 6.646734 | 3.731324 | 1.781334 | 0.074858 | NA | #VALUE! |
| ENSMUSG00000110505.2 | 0 | NA | NA | NA | NA | NA | #VALUE! |
| ENSMUSG00000110506.2 | 0 | NA | NA | NA | NA | NA | #VALUE! |
| ENSMUSG00000110507.2 | 0 | NA | NA | NA | NA | NA | #VALUE! |
| ENSMUSG00000110508.2 | 0 | NA | NA | NA | NA | NA | #VALUE! |
| ENSMUSG00000110509.2 | 0 | NA | NA | NA | NA | NA | #VALUE! |
| ENSMUSG00000110510.2 | 0 | NA | NA | NA | NA | NA | #VALUE! |
| ENSMUSG00000110511.2 | 0 | NA | NA | NA | NA | NA | #VALUE! |
| ENSMUSG00000110512.2 | 0 | NA | NA | NA | NA | NA | #VALUE! |
| ENSMUSG00000110513.2 | 0 | NA | NA | NA | NA | NA | #VALUE! |
| ENSMUSG00000110514.2 | 0 | NA | NA | NA | NA | NA | #VALUE! |

| | | | | | | | |
|---|---|---|---|---|---|---|---|
| ENSMUSG00000110515.2 | 0.093255 | 0.868093 | 4.072464 | 0.213162 | 0.831201 | NA | #VALUE! |
| ENSMUSG00000110516.2 | 0 | NA | NA | NA | NA | NA | #VALUE! |
| ENSMUSG00000110517.2 | 0 | NA | NA | NA | NA | NA | #VALUE! |
| ENSMUSG00000110518.2 | 0 | NA | NA | NA | NA | NA | #VALUE! |
| ENSMUSG00000110519.3 | 0 | NA | NA | NA | NA | NA | #VALUE! |
| ENSMUSG00000110520.2 | 0 | NA | NA | NA | NA | NA | #VALUE! |
| ENSMUSG00000110521.2 | 0 | NA | NA | NA | NA | NA | #VALUE! |
| ENSMUSG00000110522.2 | 0 | NA | NA | NA | NA | NA | #VALUE! |
| ENSMUSG00000110523.2 | 0 | NA | NA | NA | NA | NA | #VALUE! |
| ENSMUSG00000110524.2 | 0 | NA | NA | NA | NA | NA | #VALUE! |
| ENSMUSG00000110525.2 | 0 | NA | NA | NA | NA | NA | #VALUE! |
| ENSMUSG00000110526.4 | 0 | NA | NA | NA | NA | NA | #VALUE! |
| ENSMUSG00000110527.2 | 0 | NA | NA | NA | NA | NA | #VALUE! |
| ENSMUSG00000110528.2 | 0 | NA | NA | NA | NA | NA | #VALUE! |
| ENSMUSG00000110529.2 | 0 | NA | NA | NA | NA | NA | #VALUE! |
| ENSMUSG00000110530.2 | 0 | NA | NA | NA | NA | NA | #VALUE! |
| ENSMUSG00000110531.2 | 0 | NA | NA | NA | NA | NA | #VALUE! |
| ENSMUSG00000110532.2 | 0.093255 | 0.868093 | 4.072464 | 0.213162 | 0.831201 | NA | #VALUE! |
| ENSMUSG00000110533.2 | 0 | NA | NA | NA | NA | NA | #VALUE! |
| ENSMUSG00000110534.2 | 0 | NA | NA | NA | NA | NA | #VALUE! |
| ENSMUSG00000110535.2 | 0 | NA | NA | NA | NA | NA | #VALUE! |
| ENSMUSG00000110536.2 | 0 | NA | NA | NA | NA | NA | #VALUE! |
| ENSMUSG00000110537.2 | 0 | NA | NA | NA | NA | NA | #VALUE! |
| ENSMUSG00000110538.2 | 0.105013 | -0.97324 | 4.076293 | -0.23876 | 0.811295 | NA | #VALUE! |
| ENSMUSG00000110539.2 | 0 | NA | NA | NA | NA | NA | #VALUE! |
| ENSMUSG00000110540.2 | 0 | NA | NA | NA | NA | NA | #VALUE! |
| ENSMUSG00000110541.2 | 0 | NA | NA | NA | NA | NA | #VALUE! |
| ENSMUSG00000110542.2 | 0 | NA | NA | NA | NA | NA | #VALUE! |
| ENSMUSG00000110543.2 | 0 | NA | NA | NA | NA | NA | #VALUE! |
| ENSMUSG00000110544.2 | 0 | NA | NA | NA | NA | NA | #VALUE! |
| ENSMUSG00000110545.2 | 0 | NA | NA | NA | NA | NA | #VALUE! |
| ENSMUSG00000110546.2 | 0 | NA | NA | NA | NA | NA | #VALUE! |
| ENSMUSG00000110547.2 | 2.43801 | -1.42103 | 3.819282 | -0.37207 | 0.709843 | NA | #VALUE! |
| ENSMUSG00000110548.2 | 0 | NA | NA | NA | NA | NA | #VALUE! |
| ENSMUSG00000110549.2 | 0 | NA | NA | NA | NA | NA | #VALUE! |
| ENSMUSG00000110550.2 | 0 | NA | NA | NA | NA | NA | #VALUE! |
| ENSMUSG00000110551.2 | 0 | NA | NA | NA | NA | NA | #VALUE! |
| ENSMUSG00000110552.2 | 0 | NA | NA | NA | NA | NA | #VALUE! |
| ENSMUSG00000110553.2 | 0 | NA | NA | NA | NA | NA | #VALUE! |
| ENSMUSG00000110554.2 | 0 | NA | NA | NA | NA | NA | #VALUE! |
| ENSMUSG00000110555.2 | 0 | NA | NA | NA | NA | NA | #VALUE! |
| ENSMUSG00000110556.2 | 1.68021 | -4.29432 | 3.933015 | -1.09187 | 0.274892 | NA | #VALUE! |
| ENSMUSG00000110557.3 | 2.65344 | -4.9321 | 3.923668 | -1.25701 | 0.208749 | NA | #VALUE! |
| ENSMUSG00000110558.2 | 0 | NA | NA | NA | NA | NA | #VALUE! |

| | | | | | | | |
|---|---|---|---|---|---|---|---|
| ENSMUSG00000110559.2 | 0 | NA | NA | NA | NA | NA | #VALUE! |
| ENSMUSG00000110560.2 | 0 | NA | NA | NA | NA | NA | #VALUE! |
| ENSMUSG00000110561.2 | 0 | NA | NA | NA | NA | NA | #VALUE! |
| ENSMUSG00000110562.2 | 0 | NA | NA | NA | NA | NA | #VALUE! |
| ENSMUSG00000110563.2 | 0 | NA | NA | NA | NA | NA | #VALUE! |
| ENSMUSG00000110564.2 | 0 | NA | NA | NA | NA | NA | #VALUE! |
| ENSMUSG00000110565.2 | 0 | NA | NA | NA | NA | NA | #VALUE! |
| ENSMUSG00000110566.2 | 0 | NA | NA | NA | NA | NA | #VALUE! |
| ENSMUSG00000110567.2 | 0 | NA | NA | NA | NA | NA | #VALUE! |
| ENSMUSG00000110568.2 | 0 | NA | NA | NA | NA | NA | #VALUE! |
| ENSMUSG00000110569.2 | 0 | NA | NA | NA | NA | NA | #VALUE! |
| ENSMUSG00000110570.2 | 0 | NA | NA | NA | NA | NA | #VALUE! |
| ENSMUSG00000110571.2 | 0 | NA | NA | NA | NA | NA | #VALUE! |
| ENSMUSG00000110572.2 | 0 | NA | NA | NA | NA | NA | #VALUE! |
| ENSMUSG00000110573.2 | 0 | NA | NA | NA | NA | NA | #VALUE! |
| ENSMUSG00000110574.2 | 0 | NA | NA | NA | NA | NA | #VALUE! |
| ENSMUSG00000110575.2 | 0 | NA | NA | NA | NA | NA | #VALUE! |
| ENSMUSG00000110576.3 | 0 | NA | NA | NA | NA | NA | #VALUE! |
| ENSMUSG00000110577.2 | 0 | NA | NA | NA | NA | NA | #VALUE! |
| ENSMUSG00000110578.2 | 0 | NA | NA | NA | NA | NA | #VALUE! |
| ENSMUSG00000110580.2 | 0 | NA | NA | NA | NA | NA | #VALUE! |
| ENSMUSG00000110581.2 | 0 | NA | NA | NA | NA | NA | #VALUE! |
| ENSMUSG00000110582.2 | 0 | NA | NA | NA | NA | NA | #VALUE! |
| ENSMUSG00000110583.2 | 0 | NA | NA | NA | NA | NA | #VALUE! |
| ENSMUSG00000110585.2 | 0 | NA | NA | NA | NA | NA | #VALUE! |
| ENSMUSG00000110586.2 | 0 | NA | NA | NA | NA | NA | #VALUE! |
| ENSMUSG00000110587.2 | 0 | NA | NA | NA | NA | NA | #VALUE! |
| ENSMUSG00000110588.2 | 9.854957 | -1.17706 | 3.325872 | -0.35391 | 0.723407 | NA | #VALUE! |
| ENSMUSG00000110589.2 | 0 | NA | NA | NA | NA | NA | #VALUE! |
| ENSMUSG00000110590.2 | 0 | NA | NA | NA | NA | NA | #VALUE! |
| ENSMUSG00000110591.2 | 0 | NA | NA | NA | NA | NA | #VALUE! |
| ENSMUSG00000110592.2 | 0 | NA | NA | NA | NA | NA | #VALUE! |
| ENSMUSG00000110593.2 | 0 | NA | NA | NA | NA | NA | #VALUE! |
| ENSMUSG00000110594.2 | 0 | NA | NA | NA | NA | NA | #VALUE! |
| ENSMUSG00000110595.2 | 0 | NA | NA | NA | NA | NA | #VALUE! |
| ENSMUSG00000110596.2 | 0 | NA | NA | NA | NA | NA | #VALUE! |
| ENSMUSG00000110597.2 | 0 | NA | NA | NA | NA | NA | #VALUE! |
| ENSMUSG00000110598.2 | 0 | NA | NA | NA | NA | NA | #VALUE! |
| ENSMUSG00000110599.2 | 0 | NA | NA | NA | NA | NA | #VALUE! |
| ENSMUSG00000110600.2 | 0 | NA | NA | NA | NA | NA | #VALUE! |
| ENSMUSG00000110601.2 | 0 | NA | NA | NA | NA | NA | #VALUE! |
| ENSMUSG00000110602.2 | 0 | NA | NA | NA | NA | NA | #VALUE! |
| ENSMUSG00000110603.2 | 0 | NA | NA | NA | NA | NA | #VALUE! |
| ENSMUSG00000110604.2 | 0 | NA | NA | NA | NA | NA | #VALUE! |

| | | | | | | | |
|---|---|---|---|---|---|---|---|
| ENSMUSG00000110605.2 | 3.915782 | -5.49628 | 3.91821 | -1.40275 | 0.16069 | NA | #VALUE! |
| ENSMUSG00000110606.2 | 0 | NA | NA | NA | NA | NA | #VALUE! |
| ENSMUSG00000110607.2 | 0 | NA | NA | NA | NA | NA | #VALUE! |
| ENSMUSG00000110608.2 | 0 | NA | NA | NA | NA | NA | #VALUE! |
| ENSMUSG00000110609.2 | 0 | NA | NA | NA | NA | NA | #VALUE! |
| ENSMUSG00000110610.2 | 0 | NA | NA | NA | NA | NA | #VALUE! |
| ENSMUSG00000110611.2 | 0 | NA | NA | NA | NA | NA | #VALUE! |
| ENSMUSG00000110612.2 | 0 | NA | NA | NA | NA | NA | #VALUE! |
| ENSMUSG00000110614.2 | 0 | NA | NA | NA | NA | NA | #VALUE! |
| ENSMUSG00000110615.2 | 6.759624 | -2.59202 | 3.731347 | -0.69466 | 0.487268 | NA | #VALUE! |
| ENSMUSG00000110616.2 | 0 | NA | NA | NA | NA | NA | #VALUE! |
| ENSMUSG00000110617.2 | 0 | NA | NA | NA | NA | NA | #VALUE! |
| ENSMUSG00000110618.2 | 0 | NA | NA | NA | NA | NA | #VALUE! |
| ENSMUSG00000110619.2 | 0 | NA | NA | NA | NA | NA | #VALUE! |
| ENSMUSG00000110620.2 | 5.179485 | -0.29843 | 2.444241 | -0.12209 | 0.902824 | NA | #VALUE! |
| ENSMUSG00000110621.4 | 0 | NA | NA | NA | NA | NA | #VALUE! |
| ENSMUSG00000110622.2 | 0 | NA | NA | NA | NA | NA | #VALUE! |
| ENSMUSG00000110623.2 | 0 | NA | NA | NA | NA | NA | #VALUE! |
| ENSMUSG00000110624.2 | 0 | NA | NA | NA | NA | NA | #VALUE! |
| ENSMUSG00000110625.2 | 0 | NA | NA | NA | NA | NA | #VALUE! |
| ENSMUSG00000110626.2 | 0 | NA | NA | NA | NA | NA | #VALUE! |
| ENSMUSG00000110627.2 | 0 | NA | NA | NA | NA | NA | #VALUE! |
| ENSMUSG00000110628.2 | 0 | NA | NA | NA | NA | NA | #VALUE! |
| ENSMUSG00000110629.2 | 0 | NA | NA | NA | NA | NA | #VALUE! |
| ENSMUSG00000110630.2 | 2.40029 | 4.741481 | 3.925585 | 1.207841 | 0.227109 | NA | #VALUE! |
| ENSMUSG00000110631.2 | 0 | NA | NA | NA | NA | NA | #VALUE! |
| ENSMUSG00000110632.2 | 0 | NA | NA | NA | NA | NA | #VALUE! |
| ENSMUSG00000110633.2 | 0 | NA | NA | NA | NA | NA | #VALUE! |
| ENSMUSG00000110634.2 | 0 | NA | NA | NA | NA | NA | #VALUE! |
| ENSMUSG00000110635.2 | 0 | NA | NA | NA | NA | NA | #VALUE! |
| ENSMUSG00000110636.2 | 0 | NA | NA | NA | NA | NA | #VALUE! |
| ENSMUSG00000110637.2 | 0 | NA | NA | NA | NA | NA | #VALUE! |
| ENSMUSG00000110638.2 | 0 | NA | NA | NA | NA | NA | #VALUE! |
| ENSMUSG00000110639.2 | 0 | NA | NA | NA | NA | NA | #VALUE! |
| ENSMUSG00000110640.2 | 0 | NA | NA | NA | NA | NA | #VALUE! |
| ENSMUSG00000110641.2 | 0 | NA | NA | NA | NA | NA | #VALUE! |
| ENSMUSG00000110642.2 | 0 | NA | NA | NA | NA | NA | #VALUE! |
| ENSMUSG00000110643.2 | 0 | NA | NA | NA | NA | NA | #VALUE! |
| ENSMUSG00000110644.2 | 0 | NA | NA | NA | NA | NA | #VALUE! |
| ENSMUSG00000110645.2 | 0 | NA | NA | NA | NA | NA | #VALUE! |
| ENSMUSG00000110646.2 | 0 | NA | NA | NA | NA | NA | #VALUE! |
| ENSMUSG00000110647.3 | 0 | NA | NA | NA | NA | NA | #VALUE! |
| ENSMUSG00000110648.2 | 0.978957 | -0.79066 | 3.920325 | -0.20168 | 0.840166 | NA | #VALUE! |
| ENSMUSG00000110649.2 | 6.004783 | 6.071557 | 3.748679 | 1.619652 | 0.105307 | NA | #VALUE! |

| | | | | | | | |
|---|---|---|---|---|---|---|---|
| ENSMUSG00000110650.2 | 0 | NA | NA | NA | NA | NA | #VALUE! |
| ENSMUSG00000110651.2 | 0 | NA | NA | NA | NA | NA | #VALUE! |
| ENSMUSG00000110652.2 | 0 | NA | NA | NA | NA | NA | #VALUE! |
| ENSMUSG00000110653.2 | 0 | NA | NA | NA | NA | NA | #VALUE! |
| ENSMUSG00000110654.2 | 0.204111 | -1.05581 | 4.073302 | -0.2592 | 0.795479 | NA | #VALUE! |
| ENSMUSG00000110655.2 | 0 | NA | NA | NA | NA | NA | #VALUE! |
| ENSMUSG00000110656.2 | 0 | NA | NA | NA | NA | NA | #VALUE! |
| ENSMUSG00000110657.2 | 0 | NA | NA | NA | NA | NA | #VALUE! |
| ENSMUSG00000110658.2 | 0 | NA | NA | NA | NA | NA | #VALUE! |
| ENSMUSG00000110659.2 | 2.741047 | -4.98044 | 3.923113 | -1.26951 | 0.204259 | NA | #VALUE! |
| ENSMUSG00000110660.2 | 0 | NA | NA | NA | NA | NA | #VALUE! |
| ENSMUSG00000110661.2 | 0 | NA | NA | NA | NA | NA | #VALUE! |
| ENSMUSG00000110662.2 | 0 | NA | NA | NA | NA | NA | #VALUE! |
| ENSMUSG00000110663.2 | 7.154305 | 6.32307 | 3.113078 | 2.031131 | 0.042242 | NA | #VALUE! |
| ENSMUSG00000110664.2 | 0 | NA | NA | NA | NA | NA | #VALUE! |
| ENSMUSG00000110665.2 | 0 | NA | NA | NA | NA | NA | #VALUE! |
| ENSMUSG00000110666.2 | 0 | NA | NA | NA | NA | NA | #VALUE! |
| ENSMUSG00000110667.2 | 0 | NA | NA | NA | NA | NA | #VALUE! |
| ENSMUSG00000110668.2 | 0 | NA | NA | NA | NA | NA | #VALUE! |
| ENSMUSG00000110669.2 | 0 | NA | NA | NA | NA | NA | #VALUE! |
| ENSMUSG00000110670.2 | 0 | NA | NA | NA | NA | NA | #VALUE! |
| ENSMUSG00000110671.2 | 0 | NA | NA | NA | NA | NA | #VALUE! |
| ENSMUSG00000110672.2 | 0 | NA | NA | NA | NA | NA | #VALUE! |
| ENSMUSG00000110673.2 | 0 | NA | NA | NA | NA | NA | #VALUE! |
| ENSMUSG00000110674.2 | 0 | NA | NA | NA | NA | NA | #VALUE! |
| ENSMUSG00000110675.2 | 0 | NA | NA | NA | NA | NA | #VALUE! |
| ENSMUSG00000110676.2 | 0 | NA | NA | NA | NA | NA | #VALUE! |
| ENSMUSG00000110677.2 | 0 | NA | NA | NA | NA | NA | #VALUE! |
| ENSMUSG00000110678.2 | 0 | NA | NA | NA | NA | NA | #VALUE! |
| ENSMUSG00000110679.2 | 0 | NA | NA | NA | NA | NA | #VALUE! |
| ENSMUSG00000110680.2 | 0 | NA | NA | NA | NA | NA | #VALUE! |
| ENSMUSG00000110681.2 | 0 | NA | NA | NA | NA | NA | #VALUE! |
| ENSMUSG00000110682.2 | 0 | NA | NA | NA | NA | NA | #VALUE! |
| ENSMUSG00000110683.2 | 0 | NA | NA | NA | NA | NA | #VALUE! |
| ENSMUSG00000110684.2 | 0.093255 | 0.868093 | 4.072464 | 0.213162 | 0.831201 | NA | #VALUE! |
| ENSMUSG00000110685.2 | 0 | NA | NA | NA | NA | NA | #VALUE! |
| ENSMUSG00000110687.2 | 0 | NA | NA | NA | NA | NA | #VALUE! |
| ENSMUSG00000110688.2 | 0 | NA | NA | NA | NA | NA | #VALUE! |
| ENSMUSG00000110689.2 | 0 | NA | NA | NA | NA | NA | #VALUE! |
| ENSMUSG00000110690.2 | 0 | NA | NA | NA | NA | NA | #VALUE! |
| ENSMUSG00000110691.2 | 0 | NA | NA | NA | NA | NA | #VALUE! |
| ENSMUSG00000110692.2 | 0 | NA | NA | NA | NA | NA | #VALUE! |
| ENSMUSG00000110693.2 | 0 | NA | NA | NA | NA | NA | #VALUE! |
| ENSMUSG00000110694.2 | 0 | NA | NA | NA | NA | NA | #VALUE! |

| | | | | | | | |
|---|---|---|---|---|---|---|---|
| ENSMUSG00000110695.2 | 0 | NA | NA | NA | NA | NA | #VALUE! |
| ENSMUSG00000110696.2 | 0 | NA | NA | NA | NA | NA | #VALUE! |
| ENSMUSG00000110697.2 | 11.89326 | -1.15022 | 3.154 | -0.36468 | NA | NA | #VALUE! |
| ENSMUSG00000110698.2 | 0 | NA | NA | NA | NA | NA | #VALUE! |
| ENSMUSG00000110699.2 | 0 | NA | NA | NA | NA | NA | #VALUE! |
| ENSMUSG00000110700.2 | 0 | NA | NA | NA | NA | NA | #VALUE! |
| ENSMUSG00000110701.2 | 0 | NA | NA | NA | NA | NA | #VALUE! |
| ENSMUSG00000110702.2 | 0 | NA | NA | NA | NA | NA | #VALUE! |
| ENSMUSG00000110703.2 | 0 | NA | NA | NA | NA | NA | #VALUE! |
| ENSMUSG00000110704.2 | 0 | NA | NA | NA | NA | NA | #VALUE! |
| ENSMUSG00000110705.2 | 0 | NA | NA | NA | NA | NA | #VALUE! |
| ENSMUSG00000110706.2 | 0 | NA | NA | NA | NA | NA | #VALUE! |
| ENSMUSG00000110707.2 | 0 | NA | NA | NA | NA | NA | #VALUE! |
| ENSMUSG00000110708.2 | 0 | NA | NA | NA | NA | NA | #VALUE! |
| ENSMUSG00000110709.2 | 0 | NA | NA | NA | NA | NA | #VALUE! |
| ENSMUSG00000110710.2 | 0 | NA | NA | NA | NA | NA | #VALUE! |
| ENSMUSG00000110711.2 | 0 | NA | NA | NA | NA | NA | #VALUE! |
| ENSMUSG00000110712.2 | 0 | NA | NA | NA | NA | NA | #VALUE! |
| ENSMUSG00000110713.2 | 0 | NA | NA | NA | NA | NA | #VALUE! |
| ENSMUSG00000110714.2 | 0 | NA | NA | NA | NA | NA | #VALUE! |
| ENSMUSG00000110715.2 | 0.105013 | -0.97324 | 4.076293 | -0.23876 | 0.811295 | NA | #VALUE! |
| ENSMUSG00000110716.2 | 0 | NA | NA | NA | NA | NA | #VALUE! |
| ENSMUSG00000110717.2 | 0 | NA | NA | NA | NA | NA | #VALUE! |
| ENSMUSG00000110718.2 | 0 | NA | NA | NA | NA | NA | #VALUE! |
| ENSMUSG00000110719.2 | 0 | NA | NA | NA | NA | NA | #VALUE! |
| ENSMUSG00000110720.2 | 0 | NA | NA | NA | NA | NA | #VALUE! |
| ENSMUSG00000110721.2 | 0 | NA | NA | NA | NA | NA | #VALUE! |
| ENSMUSG00000110722.2 | 0 | NA | NA | NA | NA | NA | #VALUE! |
| ENSMUSG00000110723.2 | 0 | NA | NA | NA | NA | NA | #VALUE! |
| ENSMUSG00000110724.2 | 0 | NA | NA | NA | NA | NA | #VALUE! |
| ENSMUSG00000110725.2 | 0 | NA | NA | NA | NA | NA | #VALUE! |
| ENSMUSG00000110726.2 | 0 | NA | NA | NA | NA | NA | #VALUE! |
| ENSMUSG00000110727.2 | 0 | NA | NA | NA | NA | NA | #VALUE! |
| ENSMUSG00000110729.2 | 0 | NA | NA | NA | NA | NA | #VALUE! |
| ENSMUSG00000110730.2 | 0 | NA | NA | NA | NA | NA | #VALUE! |
| ENSMUSG00000110731.2 | 0 | NA | NA | NA | NA | NA | #VALUE! |
| ENSMUSG00000110732.2 | 0 | NA | NA | NA | NA | NA | #VALUE! |
| ENSMUSG00000110734.2 | 0 | NA | NA | NA | NA | NA | #VALUE! |
| ENSMUSG00000110735.2 | 0 | NA | NA | NA | NA | NA | #VALUE! |
| ENSMUSG00000110736.2 | 0 | NA | NA | NA | NA | NA | #VALUE! |
| ENSMUSG00000110737.2 | 0 | NA | NA | NA | NA | NA | #VALUE! |
| ENSMUSG00000110738.2 | 0 | NA | NA | NA | NA | NA | #VALUE! |
| ENSMUSG00000110739.2 | 0 | NA | NA | NA | NA | NA | #VALUE! |
| ENSMUSG00000110740.2 | 0 | NA | NA | NA | NA | NA | #VALUE! |

| | | | | | | | |
|---|---|---|---|---|---|---|---|
| ENSMUSG00000110741.2 | 0 | NA | NA | NA | NA | NA | #VALUE! |
| ENSMUSG00000110744.2 | 0 | NA | NA | NA | NA | NA | #VALUE! |
| ENSMUSG00000110745.2 | 0 | NA | NA | NA | NA | NA | #VALUE! |
| ENSMUSG00000110747.2 | 0 | NA | NA | NA | NA | NA | #VALUE! |
| ENSMUSG00000110748.2 | 0 | NA | NA | NA | NA | NA | #VALUE! |
| ENSMUSG00000110749.2 | 0 | NA | NA | NA | NA | NA | #VALUE! |
| ENSMUSG00000110750.2 | 2.100263 | -4.61322 | 3.927832 | -1.17449 | 0.240197 | NA | #VALUE! |
| ENSMUSG00000110751.2 | 0 | NA | NA | NA | NA | NA | #VALUE! |
| ENSMUSG00000110752.2 | 0 | NA | NA | NA | NA | NA | #VALUE! |
| ENSMUSG00000110753.2 | 0 | NA | NA | NA | NA | NA | #VALUE! |
| ENSMUSG00000110754.2 | 0 | NA | NA | NA | NA | NA | #VALUE! |
| ENSMUSG00000110756.2 | 0 | NA | NA | NA | NA | NA | #VALUE! |
| ENSMUSG00000110757.2 | 0 | NA | NA | NA | NA | NA | #VALUE! |
| ENSMUSG00000110758.2 | 0 | NA | NA | NA | NA | NA | #VALUE! |
| ENSMUSG00000110759.2 | 0 | NA | NA | NA | NA | NA | #VALUE! |
| ENSMUSG00000110760.2 | 0 | NA | NA | NA | NA | NA | #VALUE! |
| ENSMUSG00000110761.2 | 0 | NA | NA | NA | NA | NA | #VALUE! |
| ENSMUSG00000110762.2 | 0 | NA | NA | NA | NA | NA | #VALUE! |
| ENSMUSG00000110765.2 | 0 | NA | NA | NA | NA | NA | #VALUE! |
| ENSMUSG00000110766.2 | 0 | NA | NA | NA | NA | NA | #VALUE! |
| ENSMUSG00000110767.2 | 0 | NA | NA | NA | NA | NA | #VALUE! |
| ENSMUSG00000110768.2 | 0 | NA | NA | NA | NA | NA | #VALUE! |
| ENSMUSG00000110769.2 | 0 | NA | NA | NA | NA | NA | #VALUE! |
| ENSMUSG00000110770.2 | 0 | NA | NA | NA | NA | NA | #VALUE! |
| ENSMUSG00000110771.2 | 0 | NA | NA | NA | NA | NA | #VALUE! |
| ENSMUSG00000110772.2 | 0 | NA | NA | NA | NA | NA | #VALUE! |
| ENSMUSG00000110773.2 | 0 | NA | NA | NA | NA | NA | #VALUE! |
| ENSMUSG00000110774.2 | 0 | NA | NA | NA | NA | NA | #VALUE! |
| ENSMUSG00000110776.2 | 0 | NA | NA | NA | NA | NA | #VALUE! |
| ENSMUSG00000110778.2 | 0 | NA | NA | NA | NA | NA | #VALUE! |
| ENSMUSG00000110779.2 | 0 | NA | NA | NA | NA | NA | #VALUE! |
| ENSMUSG00000110780.2 | 0 | NA | NA | NA | NA | NA | #VALUE! |
| ENSMUSG00000110781.2 | 1.678593 | 4.255795 | 3.933066 | 1.082055 | 0.279228 | NA | #VALUE! |
| ENSMUSG00000110782.2 | 0 | NA | NA | NA | NA | NA | #VALUE! |
| ENSMUSG00000110783.2 | 0 | NA | NA | NA | NA | NA | #VALUE! |
| ENSMUSG00000110784.2 | 0 | NA | NA | NA | NA | NA | #VALUE! |
| ENSMUSG00000110786.2 | 0 | NA | NA | NA | NA | NA | #VALUE! |
| ENSMUSG00000110787.2 | 0 | NA | NA | NA | NA | NA | #VALUE! |
| ENSMUSG00000110788.2 | 0 | NA | NA | NA | NA | NA | #VALUE! |
| ENSMUSG00000110789.2 | 0 | NA | NA | NA | NA | NA | #VALUE! |
| ENSMUSG00000110790.2 | 0 | NA | NA | NA | NA | NA | #VALUE! |
| ENSMUSG00000110791.2 | 0.210026 | -1.52494 | 4.058692 | -0.37572 | 0.707123 | NA | #VALUE! |
| ENSMUSG00000110792.2 | 0 | NA | NA | NA | NA | NA | #VALUE! |
| ENSMUSG00000110793.2 | 0 | NA | NA | NA | NA | NA | #VALUE! |

| | | | | | | | |
|---|---|---|---|---|---|---|---|
| ENSMUSG00000110794.2 | 0 | NA | NA | NA | NA | NA | #VALUE! |
| ENSMUSG00000110795.2 | 0 | NA | NA | NA | NA | NA | #VALUE! |
| ENSMUSG00000110799.2 | 0 | NA | NA | NA | NA | NA | #VALUE! |
| ENSMUSG00000110801.2 | 0 | NA | NA | NA | NA | NA | #VALUE! |
| ENSMUSG00000110802.2 | 0 | NA | NA | NA | NA | NA | #VALUE! |
| ENSMUSG00000110803.2 | 0 | NA | NA | NA | NA | NA | #VALUE! |
| ENSMUSG00000110804.2 | 0 | NA | NA | NA | NA | NA | #VALUE! |
| ENSMUSG00000110808.2 | 0 | NA | NA | NA | NA | NA | #VALUE! |
| ENSMUSG00000110809.2 | 0 | NA | NA | NA | NA | NA | #VALUE! |
| ENSMUSG00000110810.2 | 0 | NA | NA | NA | NA | NA | #VALUE! |
| ENSMUSG00000110811.2 | 0 | NA | NA | NA | NA | NA | #VALUE! |
| ENSMUSG00000110812.2 | 0 | NA | NA | NA | NA | NA | #VALUE! |
| ENSMUSG00000110813.2 | 0 | NA | NA | NA | NA | NA | #VALUE! |
| ENSMUSG00000110814.2 | 0 | NA | NA | NA | NA | NA | #VALUE! |
| ENSMUSG00000110815.2 | 0 | NA | NA | NA | NA | NA | #VALUE! |
| ENSMUSG00000110816.2 | 0 | NA | NA | NA | NA | NA | #VALUE! |
| ENSMUSG00000110817.2 | 0 | NA | NA | NA | NA | NA | #VALUE! |
| ENSMUSG00000110819.2 | 0 | NA | NA | NA | NA | NA | #VALUE! |
| ENSMUSG00000110821.2 | 0 | NA | NA | NA | NA | NA | #VALUE! |
| ENSMUSG00000110822.2 | 0 | NA | NA | NA | NA | NA | #VALUE! |
| ENSMUSG00000110823.2 | 0 | NA | NA | NA | NA | NA | #VALUE! |
| ENSMUSG00000110824.2 | 0 | NA | NA | NA | NA | NA | #VALUE! |
| ENSMUSG00000110825.2 | 0 | NA | NA | NA | NA | NA | #VALUE! |
| ENSMUSG00000110826.2 | 0 | NA | NA | NA | NA | NA | #VALUE! |
| ENSMUSG00000110827.2 | 0 | NA | NA | NA | NA | NA | #VALUE! |
| ENSMUSG00000110828.2 | 0 | NA | NA | NA | NA | NA | #VALUE! |
| ENSMUSG00000110829.2 | 0 | NA | NA | NA | NA | NA | #VALUE! |
| ENSMUSG00000110830.2 | 0 | NA | NA | NA | NA | NA | #VALUE! |
| ENSMUSG00000110831.2 | 0 | NA | NA | NA | NA | NA | #VALUE! |
| ENSMUSG00000110832.2 | 0 | NA | NA | NA | NA | NA | #VALUE! |
| ENSMUSG00000110833.2 | 0 | NA | NA | NA | NA | NA | #VALUE! |
| ENSMUSG00000110835.2 | 0 | NA | NA | NA | NA | NA | #VALUE! |
| ENSMUSG00000110836.2 | 0 | NA | NA | NA | NA | NA | #VALUE! |
| ENSMUSG00000110837.2 | 0 | NA | NA | NA | NA | NA | #VALUE! |
| ENSMUSG00000110839.2 | 0 | NA | NA | NA | NA | NA | #VALUE! |
| ENSMUSG00000110840.2 | 0 | NA | NA | NA | NA | NA | #VALUE! |
| ENSMUSG00000110843.2 | 0 | NA | NA | NA | NA | NA | #VALUE! |
| ENSMUSG00000110844.2 | 0 | NA | NA | NA | NA | NA | #VALUE! |
| ENSMUSG00000110845.2 | 0 | NA | NA | NA | NA | NA | #VALUE! |
| ENSMUSG00000110846.2 | 0 | NA | NA | NA | NA | NA | #VALUE! |
| ENSMUSG00000110847.2 | 0 | NA | NA | NA | NA | NA | #VALUE! |
| ENSMUSG00000110848.2 | 0 | NA | NA | NA | NA | NA | #VALUE! |
| ENSMUSG00000110849.2 | 0 | NA | NA | NA | NA | NA | #VALUE! |
| ENSMUSG00000110851.2 | 0 | NA | NA | NA | NA | NA | #VALUE! |

| | | | | | | | |
|---|---|---|---|---|---|---|---|
| ENSMUSG00000110853.2 | 0 | NA | NA | NA | NA | NA | #VALUE! |
| ENSMUSG00000110854.2 | 0 | NA | NA | NA | NA | NA | #VALUE! |
| ENSMUSG00000110855.2 | 0 | NA | NA | NA | NA | NA | #VALUE! |
| ENSMUSG00000110856.2 | 0 | NA | NA | NA | NA | NA | #VALUE! |
| ENSMUSG00000110857.2 | 0 | NA | NA | NA | NA | NA | #VALUE! |
| ENSMUSG00000110858.2 | 0 | NA | NA | NA | NA | NA | #VALUE! |
| ENSMUSG00000110862.2 | 0 | NA | NA | NA | NA | NA | #VALUE! |
| ENSMUSG00000110863.2 | 0 | NA | NA | NA | NA | NA | #VALUE! |
| ENSMUSG00000110864.2 | 0 | NA | NA | NA | NA | NA | #VALUE! |
| ENSMUSG00000110865.2 | 0 | NA | NA | NA | NA | NA | #VALUE! |
| ENSMUSG00000110866.2 | 0 | NA | NA | NA | NA | NA | #VALUE! |
| ENSMUSG00000110867.2 | 0 | NA | NA | NA | NA | NA | #VALUE! |
| ENSMUSG00000110869.2 | 0 | NA | NA | NA | NA | NA | #VALUE! |
| ENSMUSG00000110870.2 | 0 | NA | NA | NA | NA | NA | #VALUE! |
| ENSMUSG00000110872.2 | 0 | NA | NA | NA | NA | NA | #VALUE! |
| ENSMUSG00000110873.2 | 0 | NA | NA | NA | NA | NA | #VALUE! |
| ENSMUSG00000110874.2 | 0 | NA | NA | NA | NA | NA | #VALUE! |
| ENSMUSG00000110875.2 | 0 | NA | NA | NA | NA | NA | #VALUE! |
| ENSMUSG00000110876.2 | 0 | NA | NA | NA | NA | NA | #VALUE! |
| ENSMUSG00000110877.2 | 0 | NA | NA | NA | NA | NA | #VALUE! |
| ENSMUSG00000110878.2 | 0 | NA | NA | NA | NA | NA | #VALUE! |
| ENSMUSG00000110879.3 | 0 | NA | NA | NA | NA | NA | #VALUE! |
| ENSMUSG00000110880.2 | 0 | NA | NA | NA | NA | NA | #VALUE! |
| ENSMUSG00000110881.2 | 0 | NA | NA | NA | NA | NA | #VALUE! |
| ENSMUSG00000110882.2 | 0 | NA | NA | NA | NA | NA | #VALUE! |
| ENSMUSG00000110884.2 | 0 | NA | NA | NA | NA | NA | #VALUE! |
| ENSMUSG00000110885.2 | 0 | NA | NA | NA | NA | NA | #VALUE! |
| ENSMUSG00000110886.2 | 0 | NA | NA | NA | NA | NA | #VALUE! |
| ENSMUSG00000110887.2 | 0 | NA | NA | NA | NA | NA | #VALUE! |
| ENSMUSG00000110888.2 | 0 | NA | NA | NA | NA | NA | #VALUE! |
| ENSMUSG00000110889.2 | 0 | NA | NA | NA | NA | NA | #VALUE! |
| ENSMUSG00000110891.2 | 0 | NA | NA | NA | NA | NA | #VALUE! |
| ENSMUSG00000110892.2 | 0 | NA | NA | NA | NA | NA | #VALUE! |
| ENSMUSG00000110893.2 | 0 | NA | NA | NA | NA | NA | #VALUE! |
| ENSMUSG00000110894.2 | 0 | NA | NA | NA | NA | NA | #VALUE! |
| ENSMUSG00000110895.2 | 0 | NA | NA | NA | NA | NA | #VALUE! |
| ENSMUSG00000110896.2 | 0 | NA | NA | NA | NA | NA | #VALUE! |
| ENSMUSG00000110897.2 | 0 | NA | NA | NA | NA | NA | #VALUE! |
| ENSMUSG00000110898.2 | 0 | NA | NA | NA | NA | NA | #VALUE! |
| ENSMUSG00000110901.2 | 0 | NA | NA | NA | NA | NA | #VALUE! |
| ENSMUSG00000110902.2 | 0 | NA | NA | NA | NA | NA | #VALUE! |
| ENSMUSG00000110903.2 | 0 | NA | NA | NA | NA | NA | #VALUE! |
| ENSMUSG00000110905.2 | 0 | NA | NA | NA | NA | NA | #VALUE! |
| ENSMUSG00000110908.2 | 0 | NA | NA | NA | NA | NA | #VALUE! |

| | | | | | | | |
|---|---|---|---|---|---|---|---|
| ENSMUSG00000110909.2 | 0 | NA | NA | NA | NA | NA | #VALUE! |
| ENSMUSG00000110910.2 | 0 | NA | NA | NA | NA | NA | #VALUE! |
| ENSMUSG00000110911.2 | 0 | NA | NA | NA | NA | NA | #VALUE! |
| ENSMUSG00000110912.2 | 0 | NA | NA | NA | NA | NA | #VALUE! |
| ENSMUSG00000110913.2 | 0 | NA | NA | NA | NA | NA | #VALUE! |
| ENSMUSG00000110914.2 | 0 | NA | NA | NA | NA | NA | #VALUE! |
| ENSMUSG00000110915.2 | 0 | NA | NA | NA | NA | NA | #VALUE! |
| ENSMUSG00000110916.2 | 0 | NA | NA | NA | NA | NA | #VALUE! |
| ENSMUSG00000110917.2 | 0 | NA | NA | NA | NA | NA | #VALUE! |
| ENSMUSG00000110918.2 | 0 | NA | NA | NA | NA | NA | #VALUE! |
| ENSMUSG00000110919.2 | 0 | NA | NA | NA | NA | NA | #VALUE! |
| ENSMUSG00000110920.2 | 0 | NA | NA | NA | NA | NA | #VALUE! |
| ENSMUSG00000110922.2 | 0 | NA | NA | NA | NA | NA | #VALUE! |
| ENSMUSG00000110923.2 | 0 | NA | NA | NA | NA | NA | #VALUE! |
| ENSMUSG00000110924.2 | 0 | NA | NA | NA | NA | NA | #VALUE! |
| ENSMUSG00000110925.2 | 0 | NA | NA | NA | NA | NA | #VALUE! |
| ENSMUSG00000110926.2 | 0 | NA | NA | NA | NA | NA | #VALUE! |
| ENSMUSG00000110927.2 | 0 | NA | NA | NA | NA | NA | #VALUE! |
| ENSMUSG00000110928.2 | 0 | NA | NA | NA | NA | NA | #VALUE! |
| ENSMUSG00000110929.2 | 0 | NA | NA | NA | NA | NA | #VALUE! |
| ENSMUSG00000110930.2 | 0 | NA | NA | NA | NA | NA | #VALUE! |
| ENSMUSG00000110932.2 | 0 | NA | NA | NA | NA | NA | #VALUE! |
| ENSMUSG00000110935.2 | 0 | NA | NA | NA | NA | NA | #VALUE! |
| ENSMUSG00000110936.2 | 0 | NA | NA | NA | NA | NA | #VALUE! |
| ENSMUSG00000110937.2 | 0 | NA | NA | NA | NA | NA | #VALUE! |
| ENSMUSG00000110938.2 | 0 | NA | NA | NA | NA | NA | #VALUE! |
| ENSMUSG00000110940.2 | 0 | NA | NA | NA | NA | NA | #VALUE! |
| ENSMUSG00000110941.2 | 0 | NA | NA | NA | NA | NA | #VALUE! |
| ENSMUSG00000110942.2 | 0 | NA | NA | NA | NA | NA | #VALUE! |
| ENSMUSG00000110943.2 | 0 | NA | NA | NA | NA | NA | #VALUE! |
| ENSMUSG00000110944.2 | 0 | NA | NA | NA | NA | NA | #VALUE! |
| ENSMUSG00000110945.2 | 0 | NA | NA | NA | NA | NA | #VALUE! |
| ENSMUSG00000110946.2 | 0 | NA | NA | NA | NA | NA | #VALUE! |
| ENSMUSG00000110947.2 | 0 | NA | NA | NA | NA | NA | #VALUE! |
| ENSMUSG00000110949.2 | 0 | NA | NA | NA | NA | NA | #VALUE! |
| ENSMUSG00000110950.2 | 0 | NA | NA | NA | NA | NA | #VALUE! |
| ENSMUSG00000110951.2 | 0 | NA | NA | NA | NA | NA | #VALUE! |
| ENSMUSG00000110952.2 | 0 | NA | NA | NA | NA | NA | #VALUE! |
| ENSMUSG00000110953.2 | 0 | NA | NA | NA | NA | NA | #VALUE! |
| ENSMUSG00000110956.2 | 0 | NA | NA | NA | NA | NA | #VALUE! |
| ENSMUSG00000110957.2 | 0 | NA | NA | NA | NA | NA | #VALUE! |
| ENSMUSG00000110958.2 | 0 | NA | NA | NA | NA | NA | #VALUE! |
| ENSMUSG00000110959.2 | 0 | NA | NA | NA | NA | NA | #VALUE! |
| ENSMUSG00000110960.2 | 0 | NA | NA | NA | NA | NA | #VALUE! |

| | | | | | | | |
|---|---|---|---|---|---|---|---|
| ENSMUSG00000110962.2 | 0 | NA | NA | NA | NA | NA | #VALUE! |
| ENSMUSG00000110964.2 | 0 | NA | NA | NA | NA | NA | #VALUE! |
| ENSMUSG00000110966.2 | 0 | NA | NA | NA | NA | NA | #VALUE! |
| ENSMUSG00000110967.2 | 0 | NA | NA | NA | NA | NA | #VALUE! |
| ENSMUSG00000110968.2 | 0 | NA | NA | NA | NA | NA | #VALUE! |
| ENSMUSG00000110969.2 | 0 | NA | NA | NA | NA | NA | #VALUE! |
| ENSMUSG00000110970.3 | 0 | NA | NA | NA | NA | NA | #VALUE! |
| ENSMUSG00000110972.2 | 0 | NA | NA | NA | NA | NA | #VALUE! |
| ENSMUSG00000110973.2 | 0 | NA | NA | NA | NA | NA | #VALUE! |
| ENSMUSG00000110976.2 | 0 | NA | NA | NA | NA | NA | #VALUE! |
| ENSMUSG00000110977.2 | 0 | NA | NA | NA | NA | NA | #VALUE! |
| ENSMUSG00000110978.2 | 0 | NA | NA | NA | NA | NA | #VALUE! |
| ENSMUSG00000110980.2 | 0 | NA | NA | NA | NA | NA | #VALUE! |
| ENSMUSG00000110981.2 | 0 | NA | NA | NA | NA | NA | #VALUE! |
| ENSMUSG00000110982.2 | 0 | NA | NA | NA | NA | NA | #VALUE! |
| ENSMUSG00000110983.2 | 0 | NA | NA | NA | NA | NA | #VALUE! |
| ENSMUSG00000110984.2 | 0 | NA | NA | NA | NA | NA | #VALUE! |
| ENSMUSG00000110985.2 | 0 | NA | NA | NA | NA | NA | #VALUE! |
| ENSMUSG00000110986.2 | 0 | NA | NA | NA | NA | NA | #VALUE! |
| ENSMUSG00000110987.2 | 0 | NA | NA | NA | NA | NA | #VALUE! |
| ENSMUSG00000110988.2 | 0 | NA | NA | NA | NA | NA | #VALUE! |
| ENSMUSG00000110989.2 | 0 | NA | NA | NA | NA | NA | #VALUE! |
| ENSMUSG00000110991.2 | 0 | NA | NA | NA | NA | NA | #VALUE! |
| ENSMUSG00000110992.2 | 0 | NA | NA | NA | NA | NA | #VALUE! |
| ENSMUSG00000110993.2 | 0 | NA | NA | NA | NA | NA | #VALUE! |
| ENSMUSG00000110995.2 | 0 | NA | NA | NA | NA | NA | #VALUE! |
| ENSMUSG00000110996.2 | 1.119062 | 3.679627 | 3.945843 | 0.932533 | 0.351061 | NA | #VALUE! |
| ENSMUSG00000110997.2 | 0 | NA | NA | NA | NA | NA | #VALUE! |
| ENSMUSG00000110998.2 | 0 | NA | NA | NA | NA | NA | #VALUE! |
| ENSMUSG00000110999.2 | 0 | NA | NA | NA | NA | NA | #VALUE! |
| ENSMUSG00000111000.2 | 0 | NA | NA | NA | NA | NA | #VALUE! |
| ENSMUSG00000111001.2 | 0 | NA | NA | NA | NA | NA | #VALUE! |
| ENSMUSG00000111002.2 | 0 | NA | NA | NA | NA | NA | #VALUE! |
| ENSMUSG00000111003.2 | 0 | NA | NA | NA | NA | NA | #VALUE! |
| ENSMUSG00000111004.2 | 0 | NA | NA | NA | NA | NA | #VALUE! |
| ENSMUSG00000111005.2 | 0 | NA | NA | NA | NA | NA | #VALUE! |
| ENSMUSG00000111006.2 | 0 | NA | NA | NA | NA | NA | #VALUE! |
| ENSMUSG00000111008.2 | 0 | NA | NA | NA | NA | NA | #VALUE! |
| ENSMUSG00000111010.2 | 0 | NA | NA | NA | NA | NA | #VALUE! |
| ENSMUSG00000111012.2 | 0 | NA | NA | NA | NA | NA | #VALUE! |
| ENSMUSG00000111014.2 | 0 | NA | NA | NA | NA | NA | #VALUE! |
| ENSMUSG00000111015.2 | 0 | NA | NA | NA | NA | NA | #VALUE! |
| ENSMUSG00000111016.2 | 0 | NA | NA | NA | NA | NA | #VALUE! |
| ENSMUSG00000111017.2 | 0 | NA | NA | NA | NA | NA | #VALUE! |

| | | | | | | | |
|---|---|---|---|---|---|---|---|
| ENSMUSG00000111018.2 | 0 | NA | NA | NA | NA | NA | #VALUE! |
| ENSMUSG00000111019.2 | 0 | NA | NA | NA | NA | NA | #VALUE! |
| ENSMUSG00000111020.2 | 0 | NA | NA | NA | NA | NA | #VALUE! |
| ENSMUSG00000111021.3 | 0 | NA | NA | NA | NA | NA | #VALUE! |
| ENSMUSG00000111022.2 | 0 | NA | NA | NA | NA | NA | #VALUE! |
| ENSMUSG00000111023.2 | 0 | NA | NA | NA | NA | NA | #VALUE! |
| ENSMUSG00000111025.2 | 0 | NA | NA | NA | NA | NA | #VALUE! |
| ENSMUSG00000111027.2 | 0 | NA | NA | NA | NA | NA | #VALUE! |
| ENSMUSG00000111028.2 | 0 | NA | NA | NA | NA | NA | #VALUE! |
| ENSMUSG00000111029.2 | 0 | NA | NA | NA | NA | NA | #VALUE! |
| ENSMUSG00000111030.2 | 0 | NA | NA | NA | NA | NA | #VALUE! |
| ENSMUSG00000111031.2 | 0 | NA | NA | NA | NA | NA | #VALUE! |
| ENSMUSG00000111032.2 | 0 | NA | NA | NA | NA | NA | #VALUE! |
| ENSMUSG00000111033.2 | 0 | NA | NA | NA | NA | NA | #VALUE! |
| ENSMUSG00000111034.2 | 0 | NA | NA | NA | NA | NA | #VALUE! |
| ENSMUSG00000111035.2 | 0 | NA | NA | NA | NA | NA | #VALUE! |
| ENSMUSG00000111036.2 | 0 | NA | NA | NA | NA | NA | #VALUE! |
| ENSMUSG00000111037.2 | 0 | NA | NA | NA | NA | NA | #VALUE! |
| ENSMUSG00000111039.2 | 0.737927 | 3.007825 | 3.968641 | 0.757898 | 0.448512 | NA | #VALUE! |
| ENSMUSG00000111041.2 | 0 | NA | NA | NA | NA | NA | #VALUE! |
| ENSMUSG00000111042.2 | 0 | NA | NA | NA | NA | NA | #VALUE! |
| ENSMUSG00000111043.2 | 0 | NA | NA | NA | NA | NA | #VALUE! |
| ENSMUSG00000111044.2 | 0 | NA | NA | NA | NA | NA | #VALUE! |
| ENSMUSG00000111045.2 | 0 | NA | NA | NA | NA | NA | #VALUE! |
| ENSMUSG00000111046.2 | 0 | NA | NA | NA | NA | NA | #VALUE! |
| ENSMUSG00000111047.2 | 0 | NA | NA | NA | NA | NA | #VALUE! |
| ENSMUSG00000111048.2 | 0 | NA | NA | NA | NA | NA | #VALUE! |
| ENSMUSG00000111049.2 | 0 | NA | NA | NA | NA | NA | #VALUE! |
| ENSMUSG00000111050.2 | 0 | NA | NA | NA | NA | NA | #VALUE! |
| ENSMUSG00000111051.2 | 0 | NA | NA | NA | NA | NA | #VALUE! |
| ENSMUSG00000111052.2 | 0 | NA | NA | NA | NA | NA | #VALUE! |
| ENSMUSG00000111053.2 | 0 | NA | NA | NA | NA | NA | #VALUE! |
| ENSMUSG00000111054.2 | 0 | NA | NA | NA | NA | NA | #VALUE! |
| ENSMUSG00000111056.2 | 0 | NA | NA | NA | NA | NA | #VALUE! |
| ENSMUSG00000111057.2 | 0 | NA | NA | NA | NA | NA | #VALUE! |
| ENSMUSG00000111058.2 | 0 | NA | NA | NA | NA | NA | #VALUE! |
| ENSMUSG00000111059.2 | 0 | NA | NA | NA | NA | NA | #VALUE! |
| ENSMUSG00000111060.2 | 0 | NA | NA | NA | NA | NA | #VALUE! |
| ENSMUSG00000111061.2 | 0 | NA | NA | NA | NA | NA | #VALUE! |
| ENSMUSG00000111062.2 | 0 | NA | NA | NA | NA | NA | #VALUE! |
| ENSMUSG00000111063.2 | 1.174735 | -3.75246 | 3.944889 | -0.95122 | 0.341493 | NA | #VALUE! |
| ENSMUSG00000111064.2 | 0.105013 | -0.97324 | 4.076293 | -0.23876 | 0.811295 | NA | #VALUE! |
| ENSMUSG00000111065.2 | 0 | NA | NA | NA | NA | NA | #VALUE! |
| ENSMUSG00000111066.2 | 0 | NA | NA | NA | NA | NA | #VALUE! |

| | | | | | | | |
|---|---|---|---|---|---|---|---|
| ENSMUSG00000111067.2 | 0 | NA | NA | NA | NA | NA | #VALUE! |
| ENSMUSG00000111068.2 | 0 | NA | NA | NA | NA | NA | #VALUE! |
| ENSMUSG00000111069.2 | 0 | NA | NA | NA | NA | NA | #VALUE! |
| ENSMUSG00000111070.2 | 0 | NA | NA | NA | NA | NA | #VALUE! |
| ENSMUSG00000111071.2 | 0 | NA | NA | NA | NA | NA | #VALUE! |
| ENSMUSG00000111072.2 | 0 | NA | NA | NA | NA | NA | #VALUE! |
| ENSMUSG00000111073.2 | 0 | NA | NA | NA | NA | NA | #VALUE! |
| ENSMUSG00000111075.2 | 0 | NA | NA | NA | NA | NA | #VALUE! |
| ENSMUSG00000111077.2 | 0 | NA | NA | NA | NA | NA | #VALUE! |
| ENSMUSG00000111078.2 | 0 | NA | NA | NA | NA | NA | #VALUE! |
| ENSMUSG00000111079.2 | 0 | NA | NA | NA | NA | NA | #VALUE! |
| ENSMUSG00000111081.2 | 0 | NA | NA | NA | NA | NA | #VALUE! |
| ENSMUSG00000111082.2 | 0.841418 | 3.240539 | 3.959551 | 0.818411 | 0.413123 | NA | #VALUE! |
| ENSMUSG00000111083.2 | 0 | NA | NA | NA | NA | NA | #VALUE! |
| ENSMUSG00000111084.2 | 0 | NA | NA | NA | NA | NA | #VALUE! |
| ENSMUSG00000111085.2 | 0 | NA | NA | NA | NA | NA | #VALUE! |
| ENSMUSG00000111086.2 | 0 | NA | NA | NA | NA | NA | #VALUE! |
| ENSMUSG00000111087.2 | 0 | NA | NA | NA | NA | NA | #VALUE! |
| ENSMUSG00000111088.2 | 4.077922 | 0.196938 | 3.7708 | 0.052227 | 0.958348 | NA | #VALUE! |
| ENSMUSG00000111089.2 | 0 | NA | NA | NA | NA | NA | #VALUE! |
| ENSMUSG00000111090.2 | 0 | NA | NA | NA | NA | NA | #VALUE! |
| ENSMUSG00000111091.2 | 0 | NA | NA | NA | NA | NA | #VALUE! |
| ENSMUSG00000111092.2 | 0 | NA | NA | NA | NA | NA | #VALUE! |
| ENSMUSG00000111093.2 | 0 | NA | NA | NA | NA | NA | #VALUE! |
| ENSMUSG00000111094.2 | 0 | NA | NA | NA | NA | NA | #VALUE! |
| ENSMUSG00000111095.2 | 0 | NA | NA | NA | NA | NA | #VALUE! |
| ENSMUSG00000111096.2 | 0 | NA | NA | NA | NA | NA | #VALUE! |
| ENSMUSG00000111097.2 | 0 | NA | NA | NA | NA | NA | #VALUE! |
| ENSMUSG00000111098.2 | 0 | NA | NA | NA | NA | NA | #VALUE! |
| ENSMUSG00000111100.2 | 0 | NA | NA | NA | NA | NA | #VALUE! |
| ENSMUSG00000111102.2 | 0 | NA | NA | NA | NA | NA | #VALUE! |
| ENSMUSG00000111103.2 | 0 | NA | NA | NA | NA | NA | #VALUE! |
| ENSMUSG00000111104.2 | 0 | NA | NA | NA | NA | NA | #VALUE! |
| ENSMUSG00000111105.2 | 0 | NA | NA | NA | NA | NA | #VALUE! |
| ENSMUSG00000111106.2 | 0 | NA | NA | NA | NA | NA | #VALUE! |
| ENSMUSG00000111107.2 | 0 | NA | NA | NA | NA | NA | #VALUE! |
| ENSMUSG00000111108.2 | 0 | NA | NA | NA | NA | NA | #VALUE! |
| ENSMUSG00000111109.2 | 0 | NA | NA | NA | NA | NA | #VALUE! |
| ENSMUSG00000111110.2 | 0 | NA | NA | NA | NA | NA | #VALUE! |
| ENSMUSG00000111111.2 | 0 | NA | NA | NA | NA | NA | #VALUE! |
| ENSMUSG00000111112.2 | 0 | NA | NA | NA | NA | NA | #VALUE! |
| ENSMUSG00000111114.2 | 0 | NA | NA | NA | NA | NA | #VALUE! |
| ENSMUSG00000111115.2 | 0 | NA | NA | NA | NA | NA | #VALUE! |
| ENSMUSG00000111116.2 | 0 | NA | NA | NA | NA | NA | #VALUE! |

| | | | | | | | |
|---|---|---|---|---|---|---|---|
| ENSMUSG00000111117.2 | 0 | NA | NA | NA | NA | NA | #VALUE! |
| ENSMUSG00000111118.2 | 0 | NA | NA | NA | NA | NA | #VALUE! |
| ENSMUSG00000111119.2 | 0 | NA | NA | NA | NA | NA | #VALUE! |
| ENSMUSG00000111120.2 | 0 | NA | NA | NA | NA | NA | #VALUE! |
| ENSMUSG00000111121.2 | 0 | NA | NA | NA | NA | NA | #VALUE! |
| ENSMUSG00000111122.2 | 0 | NA | NA | NA | NA | NA | #VALUE! |
| ENSMUSG00000111124.2 | 0 | NA | NA | NA | NA | NA | #VALUE! |
| ENSMUSG00000111126.2 | 0 | NA | NA | NA | NA | NA | #VALUE! |
| ENSMUSG00000111127.2 | 0 | NA | NA | NA | NA | NA | #VALUE! |
| ENSMUSG00000111128.2 | 8.746963 | 2.211192 | 3.428456 | 0.644953 | NA | NA | #VALUE! |
| ENSMUSG00000111129.2 | 6.714374 | 6.241894 | 3.913436 | 1.594991 | NA | NA | #VALUE! |
| ENSMUSG00000111131.2 | 0 | NA | NA | NA | NA | NA | #VALUE! |
| ENSMUSG00000111132.2 | 0 | NA | NA | NA | NA | NA | #VALUE! |
| ENSMUSG00000111133.2 | 0 | NA | NA | NA | NA | NA | #VALUE! |
| ENSMUSG00000111135.2 | 0 | NA | NA | NA | NA | NA | #VALUE! |
| ENSMUSG00000111137.2 | 0 | NA | NA | NA | NA | NA | #VALUE! |
| ENSMUSG00000111139.2 | 0 | NA | NA | NA | NA | NA | #VALUE! |
| ENSMUSG00000111141.2 | 0 | NA | NA | NA | NA | NA | #VALUE! |
| ENSMUSG00000111142.2 | 0 | NA | NA | NA | NA | NA | #VALUE! |
| ENSMUSG00000111143.2 | 0 | NA | NA | NA | NA | NA | #VALUE! |
| ENSMUSG00000111144.2 | 0 | NA | NA | NA | NA | NA | #VALUE! |
| ENSMUSG00000111146.2 | 0 | NA | NA | NA | NA | NA | #VALUE! |
| ENSMUSG00000111147.2 | 0 | NA | NA | NA | NA | NA | #VALUE! |
| ENSMUSG00000111148.2 | 0 | NA | NA | NA | NA | NA | #VALUE! |
| ENSMUSG00000111149.2 | 0 | NA | NA | NA | NA | NA | #VALUE! |
| ENSMUSG00000111151.2 | 0 | NA | NA | NA | NA | NA | #VALUE! |
| ENSMUSG00000111152.2 | 0 | NA | NA | NA | NA | NA | #VALUE! |
| ENSMUSG00000111153.2 | 0 | NA | NA | NA | NA | NA | #VALUE! |
| ENSMUSG00000111154.2 | 0 | NA | NA | NA | NA | NA | #VALUE! |
| ENSMUSG00000111155.2 | 0 | NA | NA | NA | NA | NA | #VALUE! |
| ENSMUSG00000111156.2 | 0 | NA | NA | NA | NA | NA | #VALUE! |
| ENSMUSG00000111157.2 | 0 | NA | NA | NA | NA | NA | #VALUE! |
| ENSMUSG00000111159.2 | 0 | NA | NA | NA | NA | NA | #VALUE! |
| ENSMUSG00000111160.2 | 0 | NA | NA | NA | NA | NA | #VALUE! |
| ENSMUSG00000111161.2 | 0 | NA | NA | NA | NA | NA | #VALUE! |
| ENSMUSG00000111162.2 | 0 | NA | NA | NA | NA | NA | #VALUE! |
| ENSMUSG00000111163.2 | 9.862026 | -1.74265 | 2.195395 | -0.79377 | 0.427327 | NA | #VALUE! |
| ENSMUSG00000111164.2 | 0 | NA | NA | NA | NA | NA | #VALUE! |
| ENSMUSG00000111165.2 | 0 | NA | NA | NA | NA | NA | #VALUE! |
| ENSMUSG00000111166.2 | 0 | NA | NA | NA | NA | NA | #VALUE! |
| ENSMUSG00000111167.2 | 0 | NA | NA | NA | NA | NA | #VALUE! |
| ENSMUSG00000111168.2 | 0 | NA | NA | NA | NA | NA | #VALUE! |
| ENSMUSG00000111169.2 | 0 | NA | NA | NA | NA | NA | #VALUE! |
| ENSMUSG00000111171.2 | 0 | NA | NA | NA | NA | NA | #VALUE! |

| | | | | | | | |
|---|---|---|---|---|---|---|---|
| ENSMUSG00000111172.2 | 0 | NA | NA | NA | NA | NA | #VALUE! |
| ENSMUSG00000111173.2 | 0 | NA | NA | NA | NA | NA | #VALUE! |
| ENSMUSG00000111174.2 | 0 | NA | NA | NA | NA | NA | #VALUE! |
| ENSMUSG00000111175.2 | 0 | NA | NA | NA | NA | NA | #VALUE! |
| ENSMUSG00000111176.2 | 0 | NA | NA | NA | NA | NA | #VALUE! |
| ENSMUSG00000111177.2 | 0 | NA | NA | NA | NA | NA | #VALUE! |
| ENSMUSG00000111179.2 | 0 | NA | NA | NA | NA | NA | #VALUE! |
| ENSMUSG00000111180.2 | 0 | NA | NA | NA | NA | NA | #VALUE! |
| ENSMUSG00000111181.2 | 0 | NA | NA | NA | NA | NA | #VALUE! |
| ENSMUSG00000111182.2 | 0 | NA | NA | NA | NA | NA | #VALUE! |
| ENSMUSG00000111183.2 | 0 | NA | NA | NA | NA | NA | #VALUE! |
| ENSMUSG00000111184.2 | 0 | NA | NA | NA | NA | NA | #VALUE! |
| ENSMUSG00000111186.2 | 0 | NA | NA | NA | NA | NA | #VALUE! |
| ENSMUSG00000111188.2 | 0 | NA | NA | NA | NA | NA | #VALUE! |
| ENSMUSG00000111189.2 | 0 | NA | NA | NA | NA | NA | #VALUE! |
| ENSMUSG00000111190.3 | 0 | NA | NA | NA | NA | NA | #VALUE! |
| ENSMUSG00000111191.2 | 0 | NA | NA | NA | NA | NA | #VALUE! |
| ENSMUSG00000111192.2 | 0 | NA | NA | NA | NA | NA | #VALUE! |
| ENSMUSG00000111193.2 | 0 | NA | NA | NA | NA | NA | #VALUE! |
| ENSMUSG00000111194.2 | 0 | NA | NA | NA | NA | NA | #VALUE! |
| ENSMUSG00000111195.2 | 0 | NA | NA | NA | NA | NA | #VALUE! |
| ENSMUSG00000111196.2 | 0 | NA | NA | NA | NA | NA | #VALUE! |
| ENSMUSG00000111197.2 | 0 | NA | NA | NA | NA | NA | #VALUE! |
| ENSMUSG00000111198.2 | 0 | NA | NA | NA | NA | NA | #VALUE! |
| ENSMUSG00000111199.2 | 0 | NA | NA | NA | NA | NA | #VALUE! |
| ENSMUSG00000111200.2 | 0 | NA | NA | NA | NA | NA | #VALUE! |
| ENSMUSG00000111201.2 | 0 | NA | NA | NA | NA | NA | #VALUE! |
| ENSMUSG00000111202.2 | 0 | NA | NA | NA | NA | NA | #VALUE! |
| ENSMUSG00000111203.2 | 0 | NA | NA | NA | NA | NA | #VALUE! |
| ENSMUSG00000111204.2 | 0 | NA | NA | NA | NA | NA | #VALUE! |
| ENSMUSG00000111205.2 | 0 | NA | NA | NA | NA | NA | #VALUE! |
| ENSMUSG00000111207.2 | 0 | NA | NA | NA | NA | NA | #VALUE! |
| ENSMUSG00000111208.2 | 0 | NA | NA | NA | NA | NA | #VALUE! |
| ENSMUSG00000111209.2 | 0 | NA | NA | NA | NA | NA | #VALUE! |
| ENSMUSG00000111210.2 | 0 | NA | NA | NA | NA | NA | #VALUE! |
| ENSMUSG00000111211.2 | 0 | NA | NA | NA | NA | NA | #VALUE! |
| ENSMUSG00000111212.2 | 0 | NA | NA | NA | NA | NA | #VALUE! |
| ENSMUSG00000111213.2 | 0 | NA | NA | NA | NA | NA | #VALUE! |
| ENSMUSG00000111215.2 | 0 | NA | NA | NA | NA | NA | #VALUE! |
| ENSMUSG00000111216.2 | 0 | NA | NA | NA | NA | NA | #VALUE! |
| ENSMUSG00000111217.2 | 0 | NA | NA | NA | NA | NA | #VALUE! |
| ENSMUSG00000111218.2 | 0 | NA | NA | NA | NA | NA | #VALUE! |
| ENSMUSG00000111219.2 | 0 | NA | NA | NA | NA | NA | #VALUE! |
| ENSMUSG00000111221.2 | 0 | NA | NA | NA | NA | NA | #VALUE! |

| | | | | | | | |
|---|---|---|---|---|---|---|---|
| ENSMUSG00000111222.2 | 0 | NA | NA | NA | NA | NA | #VALUE! |
| ENSMUSG00000111223.2 | 0 | NA | NA | NA | NA | NA | #VALUE! |
| ENSMUSG00000111224.2 | 0 | NA | NA | NA | NA | NA | #VALUE! |
| ENSMUSG00000111225.2 | 0 | NA | NA | NA | NA | NA | #VALUE! |
| ENSMUSG00000111227.2 | 0 | NA | NA | NA | NA | NA | #VALUE! |
| ENSMUSG00000111228.2 | 0 | NA | NA | NA | NA | NA | #VALUE! |
| ENSMUSG00000111229.2 | 0 | NA | NA | NA | NA | NA | #VALUE! |
| ENSMUSG00000111231.2 | 0 | NA | NA | NA | NA | NA | #VALUE! |
| ENSMUSG00000111232.2 | 0 | NA | NA | NA | NA | NA | #VALUE! |
| ENSMUSG00000111233.2 | 0 | NA | NA | NA | NA | NA | #VALUE! |
| ENSMUSG00000111234.2 | 0 | NA | NA | NA | NA | NA | #VALUE! |
| ENSMUSG00000111235.2 | 0 | NA | NA | NA | NA | NA | #VALUE! |
| ENSMUSG00000111236.2 | 0 | NA | NA | NA | NA | NA | #VALUE! |
| ENSMUSG00000111238.2 | 0 | NA | NA | NA | NA | NA | #VALUE! |
| ENSMUSG00000111239.2 | 0 | NA | NA | NA | NA | NA | #VALUE! |
| ENSMUSG00000111240.2 | 0 | NA | NA | NA | NA | NA | #VALUE! |
| ENSMUSG00000111241.2 | 0 | NA | NA | NA | NA | NA | #VALUE! |
| ENSMUSG00000111242.2 | 0 | NA | NA | NA | NA | NA | #VALUE! |
| ENSMUSG00000111243.2 | 0 | NA | NA | NA | NA | NA | #VALUE! |
| ENSMUSG00000111244.2 | 0 | NA | NA | NA | NA | NA | #VALUE! |
| ENSMUSG00000111245.2 | 0 | NA | NA | NA | NA | NA | #VALUE! |
| ENSMUSG00000111246.2 | 0 | NA | NA | NA | NA | NA | #VALUE! |
| ENSMUSG00000111248.2 | 0 | NA | NA | NA | NA | NA | #VALUE! |
| ENSMUSG00000111249.2 | 0 | NA | NA | NA | NA | NA | #VALUE! |
| ENSMUSG00000111250.2 | 0 | NA | NA | NA | NA | NA | #VALUE! |
| ENSMUSG00000111251.2 | 0 | NA | NA | NA | NA | NA | #VALUE! |
| ENSMUSG00000111252.2 | 0 | NA | NA | NA | NA | NA | #VALUE! |
| ENSMUSG00000111253.2 | 0 | NA | NA | NA | NA | NA | #VALUE! |
| ENSMUSG00000111254.2 | 0 | NA | NA | NA | NA | NA | #VALUE! |
| ENSMUSG00000111256.2 | 0 | NA | NA | NA | NA | NA | #VALUE! |
| ENSMUSG00000111257.2 | 0 | NA | NA | NA | NA | NA | #VALUE! |
| ENSMUSG00000111258.2 | 0 | NA | NA | NA | NA | NA | #VALUE! |
| ENSMUSG00000111259.3 | 0 | NA | NA | NA | NA | NA | #VALUE! |
| ENSMUSG00000111260.2 | 0 | NA | NA | NA | NA | NA | #VALUE! |
| ENSMUSG00000111262.2 | 0 | NA | NA | NA | NA | NA | #VALUE! |
| ENSMUSG00000111263.2 | 0 | NA | NA | NA | NA | NA | #VALUE! |
| ENSMUSG00000111264.2 | 0 | NA | NA | NA | NA | NA | #VALUE! |
| ENSMUSG00000111265.2 | 0 | NA | NA | NA | NA | NA | #VALUE! |
| ENSMUSG00000111266.2 | 0 | NA | NA | NA | NA | NA | #VALUE! |
| ENSMUSG00000111267.2 | 0 | NA | NA | NA | NA | NA | #VALUE! |
| ENSMUSG00000111268.2 | 0 | NA | NA | NA | NA | NA | #VALUE! |
| ENSMUSG00000111269.2 | 0 | NA | NA | NA | NA | NA | #VALUE! |
| ENSMUSG00000111270.2 | 0 | NA | NA | NA | NA | NA | #VALUE! |
| ENSMUSG00000111271.2 | 0 | NA | NA | NA | NA | NA | #VALUE! |

| | | | | | | | |
|---|---|---|---|---|---|---|---|
| ENSMUSG00000111272.2 | 0 | NA | NA | NA | NA | NA | #VALUE! |
| ENSMUSG00000111273.2 | 0 | NA | NA | NA | NA | NA | #VALUE! |
| ENSMUSG00000111274.2 | 0 | NA | NA | NA | NA | NA | #VALUE! |
| ENSMUSG00000111275.2 | 0 | NA | NA | NA | NA | NA | #VALUE! |
| ENSMUSG00000111276.2 | 0 | NA | NA | NA | NA | NA | #VALUE! |
| ENSMUSG00000111278.2 | 0 | NA | NA | NA | NA | NA | #VALUE! |
| ENSMUSG00000111280.2 | 0 | NA | NA | NA | NA | NA | #VALUE! |
| ENSMUSG00000111281.2 | 0 | NA | NA | NA | NA | NA | #VALUE! |
| ENSMUSG00000111282.2 | 0 | NA | NA | NA | NA | NA | #VALUE! |
| ENSMUSG00000111283.2 | 0 | NA | NA | NA | NA | NA | #VALUE! |
| ENSMUSG00000111284.2 | 0 | NA | NA | NA | NA | NA | #VALUE! |
| ENSMUSG00000111285.2 | 0 | NA | NA | NA | NA | NA | #VALUE! |
| ENSMUSG00000111286.2 | 0 | NA | NA | NA | NA | NA | #VALUE! |
| ENSMUSG00000111290.2 | 0 | NA | NA | NA | NA | NA | #VALUE! |
| ENSMUSG00000111291.2 | 2.310289 | -4.7496 | 3.925939 | -1.2098 | 0.226355 | NA | #VALUE! |
| ENSMUSG00000111292.2 | 0 | NA | NA | NA | NA | NA | #VALUE! |
| ENSMUSG00000111293.2 | 0 | NA | NA | NA | NA | NA | #VALUE! |
| ENSMUSG00000111294.2 | 0 | NA | NA | NA | NA | NA | #VALUE! |
| ENSMUSG00000111295.2 | 0 | NA | NA | NA | NA | NA | #VALUE! |
| ENSMUSG00000111296.2 | 0 | NA | NA | NA | NA | NA | #VALUE! |
| ENSMUSG00000111297.2 | 0 | NA | NA | NA | NA | NA | #VALUE! |
| ENSMUSG00000111298.2 | 0 | NA | NA | NA | NA | NA | #VALUE! |
| ENSMUSG00000111299.2 | 0 | NA | NA | NA | NA | NA | #VALUE! |
| ENSMUSG00000111300.2 | 0 | NA | NA | NA | NA | NA | #VALUE! |
| ENSMUSG00000111301.2 | 0 | NA | NA | NA | NA | NA | #VALUE! |
| ENSMUSG00000111302.2 | 0 | NA | NA | NA | NA | NA | #VALUE! |
| ENSMUSG00000111303.2 | 0 | NA | NA | NA | NA | NA | #VALUE! |
| ENSMUSG00000111304.2 | 0 | NA | NA | NA | NA | NA | #VALUE! |
| ENSMUSG00000111305.2 | 0 | NA | NA | NA | NA | NA | #VALUE! |
| ENSMUSG00000111306.3 | 0 | NA | NA | NA | NA | NA | #VALUE! |
| ENSMUSG00000111308.2 | 0 | NA | NA | NA | NA | NA | #VALUE! |
| ENSMUSG00000111309.2 | 0 | NA | NA | NA | NA | NA | #VALUE! |
| ENSMUSG00000111310.2 | 0 | NA | NA | NA | NA | NA | #VALUE! |
| ENSMUSG00000111312.2 | 0 | NA | NA | NA | NA | NA | #VALUE! |
| ENSMUSG00000111313.2 | 0 | NA | NA | NA | NA | NA | #VALUE! |
| ENSMUSG00000111315.2 | 0 | NA | NA | NA | NA | NA | #VALUE! |
| ENSMUSG00000111316.2 | 0 | NA | NA | NA | NA | NA | #VALUE! |
| ENSMUSG00000111317.2 | 0 | NA | NA | NA | NA | NA | #VALUE! |
| ENSMUSG00000111318.2 | 0 | NA | NA | NA | NA | NA | #VALUE! |
| ENSMUSG00000111320.2 | 0 | NA | NA | NA | NA | NA | #VALUE! |
| ENSMUSG00000111322.2 | 0 | NA | NA | NA | NA | NA | #VALUE! |
| ENSMUSG00000111323.2 | 0 | NA | NA | NA | NA | NA | #VALUE! |
| ENSMUSG00000111324.2 | 0 | NA | NA | NA | NA | NA | #VALUE! |
| ENSMUSG00000111325.2 | 0 | NA | NA | NA | NA | NA | #VALUE! |

| | | | | | | | |
|---|---|---|---|---|---|---|---|
| ENSMUSG00000111326.2 | 1.549202 | -3.17824 | 3.928755 | -0.80897 | 0.418533 | NA | #VALUE! |
| ENSMUSG00000111327.2 | 0 | NA | NA | NA | NA | NA | #VALUE! |
| ENSMUSG00000111329.2 | 0 | NA | NA | NA | NA | NA | #VALUE! |
| ENSMUSG00000111330.2 | 0 | NA | NA | NA | NA | NA | #VALUE! |
| ENSMUSG00000111333.2 | 0 | NA | NA | NA | NA | NA | #VALUE! |
| ENSMUSG00000111334.2 | 0 | NA | NA | NA | NA | NA | #VALUE! |
| ENSMUSG00000111335.2 | 0 | NA | NA | NA | NA | NA | #VALUE! |
| ENSMUSG00000111336.2 | 0 | NA | NA | NA | NA | NA | #VALUE! |
| ENSMUSG00000111337.2 | 0 | NA | NA | NA | NA | NA | #VALUE! |
| ENSMUSG00000111338.2 | 0 | NA | NA | NA | NA | NA | #VALUE! |
| ENSMUSG00000111339.2 | 0 | NA | NA | NA | NA | NA | #VALUE! |
| ENSMUSG00000111340.2 | 0 | NA | NA | NA | NA | NA | #VALUE! |
| ENSMUSG00000111341.2 | 0 | NA | NA | NA | NA | NA | #VALUE! |
| ENSMUSG00000111342.2 | 0 | NA | NA | NA | NA | NA | #VALUE! |
| ENSMUSG00000111343.2 | 0 | NA | NA | NA | NA | NA | #VALUE! |
| ENSMUSG00000111344.2 | 0 | NA | NA | NA | NA | NA | #VALUE! |
| ENSMUSG00000111345.2 | 4.503149 | -5.6983 | 3.916718 | -1.45487 | 0.145706 | NA | #VALUE! |
| ENSMUSG00000111346.2 | 0 | NA | NA | NA | NA | NA | #VALUE! |
| ENSMUSG00000111348.2 | 0 | NA | NA | NA | NA | NA | #VALUE! |
| ENSMUSG00000111349.2 | 0 | NA | NA | NA | NA | NA | #VALUE! |
| ENSMUSG00000111350.2 | 0 | NA | NA | NA | NA | NA | #VALUE! |
| ENSMUSG00000111351.2 | 0 | NA | NA | NA | NA | NA | #VALUE! |
| ENSMUSG00000111352.2 | 0 | NA | NA | NA | NA | NA | #VALUE! |
| ENSMUSG00000111353.2 | 0 | NA | NA | NA | NA | NA | #VALUE! |
| ENSMUSG00000111354.2 | 0 | NA | NA | NA | NA | NA | #VALUE! |
| ENSMUSG00000111355.2 | 0 | NA | NA | NA | NA | NA | #VALUE! |
| ENSMUSG00000111356.2 | 4.779179 | -0.29439 | 2.963971 | -0.09932 | 0.920882 | NA | #VALUE! |
| ENSMUSG00000111357.2 | 0 | NA | NA | NA | NA | NA | #VALUE! |
| ENSMUSG00000111358.2 | 0 | NA | NA | NA | NA | NA | #VALUE! |
| ENSMUSG00000111359.2 | 0 | NA | NA | NA | NA | NA | #VALUE! |
| ENSMUSG00000111360.2 | 0 | NA | NA | NA | NA | NA | #VALUE! |
| ENSMUSG00000111362.2 | 0 | NA | NA | NA | NA | NA | #VALUE! |
| ENSMUSG00000111363.2 | 0 | NA | NA | NA | NA | NA | #VALUE! |
| ENSMUSG00000111364.2 | 0 | NA | NA | NA | NA | NA | #VALUE! |
| ENSMUSG00000111365.2 | 0 | NA | NA | NA | NA | NA | #VALUE! |
| ENSMUSG00000111366.2 | 0 | NA | NA | NA | NA | NA | #VALUE! |
| ENSMUSG00000111368.2 | 0.630079 | -2.90291 | 3.975047 | -0.73028 | 0.465217 | NA | #VALUE! |
| ENSMUSG00000111369.2 | 0 | NA | NA | NA | NA | NA | #VALUE! |
| ENSMUSG00000111371.2 | 3.820133 | 5.426457 | 3.897428 | 1.392317 | 0.163826 | NA | #VALUE! |
| ENSMUSG00000111373.2 | 0 | NA | NA | NA | NA | NA | #VALUE! |
| ENSMUSG00000111374.2 | 0 | NA | NA | NA | NA | NA | #VALUE! |
| ENSMUSG00000111376.2 | 0 | NA | NA | NA | NA | NA | #VALUE! |
| ENSMUSG00000111377.2 | 7.833436 | 6.463695 | 3.872704 | 1.669039 | NA | NA | #VALUE! |
| ENSMUSG00000111378.2 | 0 | NA | NA | NA | NA | NA | #VALUE! |

| | | | | | | | |
|---|---|---|---|---|---|---|---|
| ENSMUSG00000111379.2 | 0 | NA | NA | NA | NA | NA | #VALUE! |
| ENSMUSG00000111380.2 | 0.245976 | 1.027628 | 4.067126 | 0.252667 | 0.800526 | NA | #VALUE! |
| ENSMUSG00000111381.2 | 0 | NA | NA | NA | NA | NA | #VALUE! |
| ENSMUSG00000111382.2 | 0 | NA | NA | NA | NA | NA | #VALUE! |
| ENSMUSG00000111383.2 | 0 | NA | NA | NA | NA | NA | #VALUE! |
| ENSMUSG00000111384.2 | 0 | NA | NA | NA | NA | NA | #VALUE! |
| ENSMUSG00000111386.2 | 0 | NA | NA | NA | NA | NA | #VALUE! |
| ENSMUSG00000111387.2 | 0 | NA | NA | NA | NA | NA | #VALUE! |
| ENSMUSG00000111388.2 | 0 | NA | NA | NA | NA | NA | #VALUE! |
| ENSMUSG00000111389.2 | 0 | NA | NA | NA | NA | NA | #VALUE! |
| ENSMUSG00000111390.2 | 0 | NA | NA | NA | NA | NA | #VALUE! |
| ENSMUSG00000111394.2 | 0 | NA | NA | NA | NA | NA | #VALUE! |
| ENSMUSG00000111395.2 | 0 | NA | NA | NA | NA | NA | #VALUE! |
| ENSMUSG00000111396.2 | 0 | NA | NA | NA | NA | NA | #VALUE! |
| ENSMUSG00000111397.2 | 0 | NA | NA | NA | NA | NA | #VALUE! |
| ENSMUSG00000111398.2 | 0 | NA | NA | NA | NA | NA | #VALUE! |
| ENSMUSG00000111399.2 | 0 | NA | NA | NA | NA | NA | #VALUE! |
| ENSMUSG00000111400.2 | 0 | NA | NA | NA | NA | NA | #VALUE! |
| ENSMUSG00000111404.2 | 0 | NA | NA | NA | NA | NA | #VALUE! |
| ENSMUSG00000111405.2 | 0 | NA | NA | NA | NA | NA | #VALUE! |
| ENSMUSG00000111406.2 | 0 | NA | NA | NA | NA | NA | #VALUE! |
| ENSMUSG00000111409.2 | 0 | NA | NA | NA | NA | NA | #VALUE! |
| ENSMUSG00000111410.2 | 0 | NA | NA | NA | NA | NA | #VALUE! |
| ENSMUSG00000111411.2 | 0 | NA | NA | NA | NA | NA | #VALUE! |
| ENSMUSG00000111412.2 | 0 | NA | NA | NA | NA | NA | #VALUE! |
| ENSMUSG00000111413.2 | 0 | NA | NA | NA | NA | NA | #VALUE! |
| ENSMUSG00000111414.2 | 0 | NA | NA | NA | NA | NA | #VALUE! |
| ENSMUSG00000111415.2 | 0 | NA | NA | NA | NA | NA | #VALUE! |
| ENSMUSG00000111416.2 | 0 | NA | NA | NA | NA | NA | #VALUE! |
| ENSMUSG00000111417.2 | 0 | NA | NA | NA | NA | NA | #VALUE! |
| ENSMUSG00000111418.2 | 0 | NA | NA | NA | NA | NA | #VALUE! |
| ENSMUSG00000111419.2 | 0 | NA | NA | NA | NA | NA | #VALUE! |
| ENSMUSG00000111420.2 | 0 | NA | NA | NA | NA | NA | #VALUE! |
| ENSMUSG00000111421.2 | 0.504851 | 2.502843 | 3.993993 | 0.626652 | 0.530887 | NA | #VALUE! |
| ENSMUSG00000111422.2 | 0 | NA | NA | NA | NA | NA | #VALUE! |
| ENSMUSG00000111423.2 | 0 | NA | NA | NA | NA | NA | #VALUE! |
| ENSMUSG00000111424.2 | 4.711939 | 5.72619 | 3.916297 | 1.462144 | 0.143702 | NA | #VALUE! |
| ENSMUSG00000111425.2 | 0 | NA | NA | NA | NA | NA | #VALUE! |
| ENSMUSG00000111426.2 | 0 | NA | NA | NA | NA | NA | #VALUE! |
| ENSMUSG00000111427.2 | 0 | NA | NA | NA | NA | NA | #VALUE! |
| ENSMUSG00000111428.2 | 0 | NA | NA | NA | NA | NA | #VALUE! |
| ENSMUSG00000111429.2 | 2.545258 | -4.8732 | 3.92437 | -1.24178 | 0.214318 | NA | #VALUE! |
| ENSMUSG00000111430.2 | 0 | NA | NA | NA | NA | NA | #VALUE! |
| ENSMUSG00000111431.2 | 0 | NA | NA | NA | NA | NA | #VALUE! |

| | | | | | | | |
|---|---|---|---|---|---|---|---|
| ENSMUSG00000111432.2 | 0 | NA | NA | NA | NA | NA | #VALUE! |
| ENSMUSG00000111433.2 | 0 | NA | NA | NA | NA | NA | #VALUE! |
| ENSMUSG00000111434.2 | 1.762102 | -4.34067 | 3.93219 | -1.10388 | 0.269645 | NA | #VALUE! |
| ENSMUSG00000111435.2 | 0 | NA | NA | NA | NA | NA | #VALUE! |
| ENSMUSG00000111436.2 | 0 | NA | NA | NA | NA | NA | #VALUE! |
| ENSMUSG00000111437.2 | 0 | NA | NA | NA | NA | NA | #VALUE! |
| ENSMUSG00000111438.2 | 0 | NA | NA | NA | NA | NA | #VALUE! |
| ENSMUSG00000111439.2 | 0 | NA | NA | NA | NA | NA | #VALUE! |
| ENSMUSG00000111440.2 | 0 | NA | NA | NA | NA | NA | #VALUE! |
| ENSMUSG00000111441.2 | 0 | NA | NA | NA | NA | NA | #VALUE! |
| ENSMUSG00000111442.2 | 3.132626 | -5.17361 | 3.921071 | -1.31944 | 0.187023 | NA | #VALUE! |
| ENSMUSG00000111443.2 | 0 | NA | NA | NA | NA | NA | #VALUE! |
| ENSMUSG00000111444.2 | 0 | NA | NA | NA | NA | NA | #VALUE! |
| ENSMUSG00000111445.2 | 0 | NA | NA | NA | NA | NA | #VALUE! |
| ENSMUSG00000111446.2 | 0.291524 | 0.468739 | 4.052149 | 0.115677 | 0.907909 | NA | #VALUE! |
| ENSMUSG00000111447.2 | 0 | NA | NA | NA | NA | NA | #VALUE! |
| ENSMUSG00000111448.3 | 0 | NA | NA | NA | NA | NA | #VALUE! |
| ENSMUSG00000111449.2 | 0 | NA | NA | NA | NA | NA | #VALUE! |
| ENSMUSG00000111450.2 | 0 | NA | NA | NA | NA | NA | #VALUE! |
| ENSMUSG00000111451.2 | 0 | NA | NA | NA | NA | NA | #VALUE! |
| ENSMUSG00000111453.2 | 0 | NA | NA | NA | NA | NA | #VALUE! |
| ENSMUSG00000111454.3 | 0 | NA | NA | NA | NA | NA | #VALUE! |
| ENSMUSG00000111455.2 | 0 | NA | NA | NA | NA | NA | #VALUE! |
| ENSMUSG00000111456.2 | 0 | NA | NA | NA | NA | NA | #VALUE! |
| ENSMUSG00000111457.2 | 0 | NA | NA | NA | NA | NA | #VALUE! |
| ENSMUSG00000111458.2 | 0 | NA | NA | NA | NA | NA | #VALUE! |
| ENSMUSG00000111459.2 | 0 | NA | NA | NA | NA | NA | #VALUE! |
| ENSMUSG00000111460.2 | 0 | NA | NA | NA | NA | NA | #VALUE! |
| ENSMUSG00000111462.3 | 0 | NA | NA | NA | NA | NA | #VALUE! |
| ENSMUSG00000111463.2 | 0 | NA | NA | NA | NA | NA | #VALUE! |
| ENSMUSG00000111464.2 | 0 | NA | NA | NA | NA | NA | #VALUE! |
| ENSMUSG00000111465.2 | 0 | NA | NA | NA | NA | NA | #VALUE! |
| ENSMUSG00000111466.2 | 0 | NA | NA | NA | NA | NA | #VALUE! |
| ENSMUSG00000111467.2 | 0 | NA | NA | NA | NA | NA | #VALUE! |
| ENSMUSG00000111468.2 | 0 | NA | NA | NA | NA | NA | #VALUE! |
| ENSMUSG00000111470.2 | 0 | NA | NA | NA | NA | NA | #VALUE! |
| ENSMUSG00000111471.2 | 0 | NA | NA | NA | NA | NA | #VALUE! |
| ENSMUSG00000111472.2 | 0 | NA | NA | NA | NA | NA | #VALUE! |
| ENSMUSG00000111473.2 | 0 | NA | NA | NA | NA | NA | #VALUE! |
| ENSMUSG00000111474.2 | 0 | NA | NA | NA | NA | NA | #VALUE! |
| ENSMUSG00000111477.2 | 0 | NA | NA | NA | NA | NA | #VALUE! |
| ENSMUSG00000111478.2 | 0 | NA | NA | NA | NA | NA | #VALUE! |
| ENSMUSG00000111479.2 | 0 | NA | NA | NA | NA | NA | #VALUE! |
| ENSMUSG00000111480.2 | 0 | NA | NA | NA | NA | NA | #VALUE! |

| | | | | | | | |
|---|---|---|---|---|---|---|---|
| ENSMUSG00000111481.2 | 0 | NA | NA | NA | NA | NA | #VALUE! |
| ENSMUSG00000111482.2 | 0 | NA | NA | NA | NA | NA | #VALUE! |
| ENSMUSG00000111483.2 | 0 | NA | NA | NA | NA | NA | #VALUE! |
| ENSMUSG00000111484.2 | 0 | NA | NA | NA | NA | NA | #VALUE! |
| ENSMUSG00000111485.2 | 0 | NA | NA | NA | NA | NA | #VALUE! |
| ENSMUSG00000111486.2 | 0 | NA | NA | NA | NA | NA | #VALUE! |
| ENSMUSG00000111487.2 | 0 | NA | NA | NA | NA | NA | #VALUE! |
| ENSMUSG00000111488.2 | 0 | NA | NA | NA | NA | NA | #VALUE! |
| ENSMUSG00000111489.2 | 0 | NA | NA | NA | NA | NA | #VALUE! |
| ENSMUSG00000111490.2 | 0 | NA | NA | NA | NA | NA | #VALUE! |
| ENSMUSG00000111491.2 | 0 | NA | NA | NA | NA | NA | #VALUE! |
| ENSMUSG00000111492.2 | 0 | NA | NA | NA | NA | NA | #VALUE! |
| ENSMUSG00000111493.2 | 0 | NA | NA | NA | NA | NA | #VALUE! |
| ENSMUSG00000111494.2 | 0 | NA | NA | NA | NA | NA | #VALUE! |
| ENSMUSG00000111495.2 | 0 | NA | NA | NA | NA | NA | #VALUE! |
| ENSMUSG00000111496.2 | 2.741047 | -4.98044 | 3.923113 | -1.26951 | 0.204259 | NA | #VALUE! |
| ENSMUSG00000111497.2 | 0 | NA | NA | NA | NA | NA | #VALUE! |
| ENSMUSG00000111498.2 | 0 | NA | NA | NA | NA | NA | #VALUE! |
| ENSMUSG00000111499.2 | 0 | NA | NA | NA | NA | NA | #VALUE! |
| ENSMUSG00000111500.2 | 0 | NA | NA | NA | NA | NA | #VALUE! |
| ENSMUSG00000111503.2 | 0 | NA | NA | NA | NA | NA | #VALUE! |
| ENSMUSG00000111504.2 | 0 | NA | NA | NA | NA | NA | #VALUE! |
| ENSMUSG00000111505.2 | 0 | NA | NA | NA | NA | NA | #VALUE! |
| ENSMUSG00000111506.2 | 0 | NA | NA | NA | NA | NA | #VALUE! |
| ENSMUSG00000111507.2 | 0 | NA | NA | NA | NA | NA | #VALUE! |
| ENSMUSG00000111508.2 | 0 | NA | NA | NA | NA | NA | #VALUE! |
| ENSMUSG00000111509.2 | 0 | NA | NA | NA | NA | NA | #VALUE! |
| ENSMUSG00000111510.3 | 0 | NA | NA | NA | NA | NA | #VALUE! |
| ENSMUSG00000111511.2 | 0 | NA | NA | NA | NA | NA | #VALUE! |
| ENSMUSG00000111512.2 | 0 | NA | NA | NA | NA | NA | #VALUE! |
| ENSMUSG00000111514.2 | 0 | NA | NA | NA | NA | NA | #VALUE! |
| ENSMUSG00000111515.2 | 0 | NA | NA | NA | NA | NA | #VALUE! |
| ENSMUSG00000111517.2 | 0 | NA | NA | NA | NA | NA | #VALUE! |
| ENSMUSG00000111518.2 | 0 | NA | NA | NA | NA | NA | #VALUE! |
| ENSMUSG00000111520.2 | 0 | NA | NA | NA | NA | NA | #VALUE! |
| ENSMUSG00000111521.2 | 9.461682 | 1.289748 | 3.364033 | 0.383393 | 0.701428 | NA | #VALUE! |
| ENSMUSG00000111523.2 | 0 | NA | NA | NA | NA | NA | #VALUE! |
| ENSMUSG00000111524.2 | 0 | NA | NA | NA | NA | NA | #VALUE! |
| ENSMUSG00000111525.2 | 0 | NA | NA | NA | NA | NA | #VALUE! |
| ENSMUSG00000111526.2 | 0 | NA | NA | NA | NA | NA | #VALUE! |
| ENSMUSG00000111528.2 | 4.095512 | -5.57062 | 3.917637 | -1.42193 | 0.155045 | NA | #VALUE! |
| ENSMUSG00000111529.2 | 0 | NA | NA | NA | NA | NA | #VALUE! |
| ENSMUSG00000111530.2 | 0 | NA | NA | NA | NA | NA | #VALUE! |
| ENSMUSG00000111531.2 | 0 | NA | NA | NA | NA | NA | #VALUE! |

| | | | | | | | |
|---|---|---|---|---|---|---|---|
| ENSMUSG00000111532.2 | 0 | NA | NA | NA | NA | NA | #VALUE! |
| ENSMUSG00000111533.2 | 0 | NA | NA | NA | NA | NA | #VALUE! |
| ENSMUSG00000111534.2 | 0 | NA | NA | NA | NA | NA | #VALUE! |
| ENSMUSG00000111535.2 | 0 | NA | NA | NA | NA | NA | #VALUE! |
| ENSMUSG00000111537.2 | 0 | NA | NA | NA | NA | NA | #VALUE! |
| ENSMUSG00000111539.2 | 0 | NA | NA | NA | NA | NA | #VALUE! |
| ENSMUSG00000111540.2 | 0 | NA | NA | NA | NA | NA | #VALUE! |
| ENSMUSG00000111541.2 | 0 | NA | NA | NA | NA | NA | #VALUE! |
| ENSMUSG00000111542.2 | 0 | NA | NA | NA | NA | NA | #VALUE! |
| ENSMUSG00000111543.2 | 0 | NA | NA | NA | NA | NA | #VALUE! |
| ENSMUSG00000111544.2 | 0.105013 | -0.97324 | 4.076293 | -0.23876 | 0.811295 | NA | #VALUE! |
| ENSMUSG00000111545.2 | 0 | NA | NA | NA | NA | NA | #VALUE! |
| ENSMUSG00000111546.2 | 0 | NA | NA | NA | NA | NA | #VALUE! |
| ENSMUSG00000111547.2 | 0 | NA | NA | NA | NA | NA | #VALUE! |
| ENSMUSG00000111548.2 | 0 | NA | NA | NA | NA | NA | #VALUE! |
| ENSMUSG00000111549.2 | 0 | NA | NA | NA | NA | NA | #VALUE! |
| ENSMUSG00000111550.2 | 0 | NA | NA | NA | NA | NA | #VALUE! |
| ENSMUSG00000111551.2 | 0 | NA | NA | NA | NA | NA | #VALUE! |
| ENSMUSG00000111552.2 | 0 | NA | NA | NA | NA | NA | #VALUE! |
| ENSMUSG00000111553.2 | 0 | NA | NA | NA | NA | NA | #VALUE! |
| ENSMUSG00000111554.2 | 0 | NA | NA | NA | NA | NA | #VALUE! |
| ENSMUSG00000111555.2 | 0 | NA | NA | NA | NA | NA | #VALUE! |
| ENSMUSG00000111556.2 | 0 | NA | NA | NA | NA | NA | #VALUE! |
| ENSMUSG00000111557.2 | 0 | NA | NA | NA | NA | NA | #VALUE! |
| ENSMUSG00000111558.2 | 0 | NA | NA | NA | NA | NA | #VALUE! |
| ENSMUSG00000111560.2 | 0 | NA | NA | NA | NA | NA | #VALUE! |
| ENSMUSG00000111561.2 | 0 | NA | NA | NA | NA | NA | #VALUE! |
| ENSMUSG00000111562.2 | 0 | NA | NA | NA | NA | NA | #VALUE! |
| ENSMUSG00000111563.2 | 0 | NA | NA | NA | NA | NA | #VALUE! |
| ENSMUSG00000111564.2 | 0 | NA | NA | NA | NA | NA | #VALUE! |
| ENSMUSG00000111565.2 | 0 | NA | NA | NA | NA | NA | #VALUE! |
| ENSMUSG00000111566.2 | 0 | NA | NA | NA | NA | NA | #VALUE! |
| ENSMUSG00000111567.2 | 0 | NA | NA | NA | NA | NA | #VALUE! |
| ENSMUSG00000111568.2 | 0 | NA | NA | NA | NA | NA | #VALUE! |
| ENSMUSG00000111569.2 | 0 | NA | NA | NA | NA | NA | #VALUE! |
| ENSMUSG00000111570.2 | 0 | NA | NA | NA | NA | NA | #VALUE! |
| ENSMUSG00000111572.2 | 0 | NA | NA | NA | NA | NA | #VALUE! |
| ENSMUSG00000111573.2 | 0 | NA | NA | NA | NA | NA | #VALUE! |
| ENSMUSG00000111574.2 | 0 | NA | NA | NA | NA | NA | #VALUE! |
| ENSMUSG00000111575.2 | 0 | NA | NA | NA | NA | NA | #VALUE! |
| ENSMUSG00000111576.2 | 0 | NA | NA | NA | NA | NA | #VALUE! |
| ENSMUSG00000111577.2 | 0 | NA | NA | NA | NA | NA | #VALUE! |
| ENSMUSG00000111578.2 | 0 | NA | NA | NA | NA | NA | #VALUE! |
| ENSMUSG00000111580.2 | 0 | NA | NA | NA | NA | NA | #VALUE! |

| | | | | | | | |
|---|---|---|---|---|---|---|---|
| ENSMUSG00000111581.2 | 0 | NA | NA | NA | NA | NA | #VALUE! |
| ENSMUSG00000111582.2 | 0 | NA | NA | NA | NA | NA | #VALUE! |
| ENSMUSG00000111583.2 | 0 | NA | NA | NA | NA | NA | #VALUE! |
| ENSMUSG00000111585.2 | 0 | NA | NA | NA | NA | NA | #VALUE! |
| ENSMUSG00000111587.2 | 0 | NA | NA | NA | NA | NA | #VALUE! |
| ENSMUSG00000111589.2 | 0 | NA | NA | NA | NA | NA | #VALUE! |
| ENSMUSG00000111590.3 | 0 | NA | NA | NA | NA | NA | #VALUE! |
| ENSMUSG00000111591.2 | 0 | NA | NA | NA | NA | NA | #VALUE! |
| ENSMUSG00000111593.2 | 0 | NA | NA | NA | NA | NA | #VALUE! |
| ENSMUSG00000111594.2 | 0 | NA | NA | NA | NA | NA | #VALUE! |
| ENSMUSG00000111596.2 | 0 | NA | NA | NA | NA | NA | #VALUE! |
| ENSMUSG00000111597.2 | 0 | NA | NA | NA | NA | NA | #VALUE! |
| ENSMUSG00000111598.2 | 0 | NA | NA | NA | NA | NA | #VALUE! |
| ENSMUSG00000111600.2 | 0 | NA | NA | NA | NA | NA | #VALUE! |
| ENSMUSG00000111601.2 | 0 | NA | NA | NA | NA | NA | #VALUE! |
| ENSMUSG00000111602.2 | 0 | NA | NA | NA | NA | NA | #VALUE! |
| ENSMUSG00000111604.2 | 0 | NA | NA | NA | NA | NA | #VALUE! |
| ENSMUSG00000111605.2 | 0 | NA | NA | NA | NA | NA | #VALUE! |
| ENSMUSG00000111606.2 | 0 | NA | NA | NA | NA | NA | #VALUE! |
| ENSMUSG00000111608.2 | 0 | NA | NA | NA | NA | NA | #VALUE! |
| ENSMUSG00000111609.2 | 0 | NA | NA | NA | NA | NA | #VALUE! |
| ENSMUSG00000111610.2 | 0 | NA | NA | NA | NA | NA | #VALUE! |
| ENSMUSG00000111611.2 | 0 | NA | NA | NA | NA | NA | #VALUE! |
| ENSMUSG00000111612.2 | 0 | NA | NA | NA | NA | NA | #VALUE! |
| ENSMUSG00000111613.2 | 0 | NA | NA | NA | NA | NA | #VALUE! |
| ENSMUSG00000111615.2 | 0 | NA | NA | NA | NA | NA | #VALUE! |
| ENSMUSG00000111616.2 | 0 | NA | NA | NA | NA | NA | #VALUE! |
| ENSMUSG00000111617.2 | 0 | NA | NA | NA | NA | NA | #VALUE! |
| ENSMUSG00000111618.2 | 0 | NA | NA | NA | NA | NA | #VALUE! |
| ENSMUSG00000111619.2 | 0 | NA | NA | NA | NA | NA | #VALUE! |
| ENSMUSG00000111620.2 | 0 | NA | NA | NA | NA | NA | #VALUE! |
| ENSMUSG00000111621.2 | 0 | NA | NA | NA | NA | NA | #VALUE! |
| ENSMUSG00000111622.2 | 0 | NA | NA | NA | NA | NA | #VALUE! |
| ENSMUSG00000111623.2 | 0 | NA | NA | NA | NA | NA | #VALUE! |
| ENSMUSG00000111625.2 | 0 | NA | NA | NA | NA | NA | #VALUE! |
| ENSMUSG00000111626.2 | 0 | NA | NA | NA | NA | NA | #VALUE! |
| ENSMUSG00000111627.2 | 0 | NA | NA | NA | NA | NA | #VALUE! |
| ENSMUSG00000111629.2 | 0 | NA | NA | NA | NA | NA | #VALUE! |
| ENSMUSG00000111630.2 | 0 | NA | NA | NA | NA | NA | #VALUE! |
| ENSMUSG00000111631.2 | 0 | NA | NA | NA | NA | NA | #VALUE! |
| ENSMUSG00000111632.2 | 0 | NA | NA | NA | NA | NA | #VALUE! |
| ENSMUSG00000111633.2 | 0 | NA | NA | NA | NA | NA | #VALUE! |
| ENSMUSG00000111635.2 | 0 | NA | NA | NA | NA | NA | #VALUE! |
| ENSMUSG00000111637.2 | 0 | NA | NA | NA | NA | NA | #VALUE! |

| | | | | | | | |
|---|---|---|---|---|---|---|---|
| ENSMUSG00000111638.2 | 0 | NA | NA | NA | NA | NA | #VALUE! |
| ENSMUSG00000111639.2 | 0 | NA | NA | NA | NA | NA | #VALUE! |
| ENSMUSG00000111641.2 | 0 | NA | NA | NA | NA | NA | #VALUE! |
| ENSMUSG00000111643.2 | 0 | NA | NA | NA | NA | NA | #VALUE! |
| ENSMUSG00000111645.2 | 0 | NA | NA | NA | NA | NA | #VALUE! |
| ENSMUSG00000111646.2 | 0 | NA | NA | NA | NA | NA | #VALUE! |
| ENSMUSG00000111648.2 | 0 | NA | NA | NA | NA | NA | #VALUE! |
| ENSMUSG00000111649.2 | 0 | NA | NA | NA | NA | NA | #VALUE! |
| ENSMUSG00000111651.2 | 0 | NA | NA | NA | NA | NA | #VALUE! |
| ENSMUSG00000111652.2 | 0 | NA | NA | NA | NA | NA | #VALUE! |
| ENSMUSG00000111654.2 | 0 | NA | NA | NA | NA | NA | #VALUE! |
| ENSMUSG00000111655.2 | 0 | NA | NA | NA | NA | NA | #VALUE! |
| ENSMUSG00000111656.2 | 0 | NA | NA | NA | NA | NA | #VALUE! |
| ENSMUSG00000111657.2 | 0 | NA | NA | NA | NA | NA | #VALUE! |
| ENSMUSG00000111658.2 | 0 | NA | NA | NA | NA | NA | #VALUE! |
| ENSMUSG00000111660.2 | 0 | NA | NA | NA | NA | NA | #VALUE! |
| ENSMUSG00000111661.2 | 0.093255 | 0.868093 | 4.072464 | 0.213162 | 0.831201 | NA | #VALUE! |
| ENSMUSG00000111662.2 | 0 | NA | NA | NA | NA | NA | #VALUE! |
| ENSMUSG00000111663.2 | 0 | NA | NA | NA | NA | NA | #VALUE! |
| ENSMUSG00000111664.2 | 0 | NA | NA | NA | NA | NA | #VALUE! |
| ENSMUSG00000111665.2 | 0 | NA | NA | NA | NA | NA | #VALUE! |
| ENSMUSG00000111666.2 | 8.872122 | 1.648895 | 3.093925 | 0.532946 | 0.594071 | NA | #VALUE! |
| ENSMUSG00000111667.2 | 0 | NA | NA | NA | NA | NA | #VALUE! |
| ENSMUSG00000111668.2 | 0 | NA | NA | NA | NA | NA | #VALUE! |
| ENSMUSG00000111669.2 | 0 | NA | NA | NA | NA | NA | #VALUE! |
| ENSMUSG00000111670.2 | 0 | NA | NA | NA | NA | NA | #VALUE! |
| ENSMUSG00000111671.2 | 0 | NA | NA | NA | NA | NA | #VALUE! |
| ENSMUSG00000111672.2 | 0 | NA | NA | NA | NA | NA | #VALUE! |
| ENSMUSG00000111673.2 | 0 | NA | NA | NA | NA | NA | #VALUE! |
| ENSMUSG00000111674.2 | 0 | NA | NA | NA | NA | NA | #VALUE! |
| ENSMUSG00000111678.3 | 0 | NA | NA | NA | NA | NA | #VALUE! |
| ENSMUSG00000111679.2 | 0 | NA | NA | NA | NA | NA | #VALUE! |
| ENSMUSG00000111680.2 | 0 | NA | NA | NA | NA | NA | #VALUE! |
| ENSMUSG00000111681.2 | 0 | NA | NA | NA | NA | NA | #VALUE! |
| ENSMUSG00000111682.2 | 0 | NA | NA | NA | NA | NA | #VALUE! |
| ENSMUSG00000111683.2 | 0 | NA | NA | NA | NA | NA | #VALUE! |
| ENSMUSG00000111684.2 | 0 | NA | NA | NA | NA | NA | #VALUE! |
| ENSMUSG00000111685.2 | 0 | NA | NA | NA | NA | NA | #VALUE! |
| ENSMUSG00000111686.2 | 0 | NA | NA | NA | NA | NA | #VALUE! |
| ENSMUSG00000111687.2 | 0 | NA | NA | NA | NA | NA | #VALUE! |
| ENSMUSG00000111689.2 | 0 | NA | NA | NA | NA | NA | #VALUE! |
| ENSMUSG00000111690.2 | 0 | NA | NA | NA | NA | NA | #VALUE! |
| ENSMUSG00000111691.2 | 0 | NA | NA | NA | NA | NA | #VALUE! |
| ENSMUSG00000111692.2 | 0 | NA | NA | NA | NA | NA | #VALUE! |

| | | | | | | | |
|---|---|---|---|---|---|---|---|
| ENSMUSG00000111693.2 | 0 | NA | NA | NA | NA | NA | #VALUE! |
| ENSMUSG00000111694.2 | 0 | NA | NA | NA | NA | NA | #VALUE! |
| ENSMUSG00000111695.2 | 0 | NA | NA | NA | NA | NA | #VALUE! |
| ENSMUSG00000111696.2 | 0 | NA | NA | NA | NA | NA | #VALUE! |
| ENSMUSG00000111697.2 | 0 | NA | NA | NA | NA | NA | #VALUE! |
| ENSMUSG00000111698.2 | 0 | NA | NA | NA | NA | NA | #VALUE! |
| ENSMUSG00000111699.2 | 0 | NA | NA | NA | NA | NA | #VALUE! |
| ENSMUSG00000111700.2 | 0 | NA | NA | NA | NA | NA | #VALUE! |
| ENSMUSG00000111702.2 | 0 | NA | NA | NA | NA | NA | #VALUE! |
| ENSMUSG00000111703.2 | 0 | NA | NA | NA | NA | NA | #VALUE! |
| ENSMUSG00000111704.2 | 0 | NA | NA | NA | NA | NA | #VALUE! |
| ENSMUSG00000111707.2 | 0 | NA | NA | NA | NA | NA | #VALUE! |
| ENSMUSG00000111708.2 | 0 | NA | NA | NA | NA | NA | #VALUE! |
| ENSMUSG00000111710.2 | 0 | NA | NA | NA | NA | NA | #VALUE! |
| ENSMUSG00000111711.2 | 0 | NA | NA | NA | NA | NA | #VALUE! |
| ENSMUSG00000111712.2 | 0 | NA | NA | NA | NA | NA | #VALUE! |
| ENSMUSG00000111713.2 | 0 | NA | NA | NA | NA | NA | #VALUE! |
| ENSMUSG00000111715.2 | 0 | NA | NA | NA | NA | NA | #VALUE! |
| ENSMUSG00000111716.2 | 0 | NA | NA | NA | NA | NA | #VALUE! |
| ENSMUSG00000111717.2 | 0 | NA | NA | NA | NA | NA | #VALUE! |
| ENSMUSG00000111719.2 | 0 | NA | NA | NA | NA | NA | #VALUE! |
| ENSMUSG00000111720.2 | 0 | NA | NA | NA | NA | NA | #VALUE! |
| ENSMUSG00000111722.2 | 0 | NA | NA | NA | NA | NA | #VALUE! |
| ENSMUSG00000111725.2 | 0 | NA | NA | NA | NA | NA | #VALUE! |
| ENSMUSG00000111726.2 | 0 | NA | NA | NA | NA | NA | #VALUE! |
| ENSMUSG00000111728.2 | 0 | NA | NA | NA | NA | NA | #VALUE! |
| ENSMUSG00000111729.2 | 0 | NA | NA | NA | NA | NA | #VALUE! |
| ENSMUSG00000111730.2 | 0 | NA | NA | NA | NA | NA | #VALUE! |
| ENSMUSG00000111731.2 | 0.945118 | -3.47564 | 3.952879 | -0.87927 | 0.379256 | NA | #VALUE! |
| ENSMUSG00000111732.2 | 0 | NA | NA | NA | NA | NA | #VALUE! |
| ENSMUSG00000111733.2 | 0 | NA | NA | NA | NA | NA | #VALUE! |
| ENSMUSG00000111734.2 | 0 | NA | NA | NA | NA | NA | #VALUE! |
| ENSMUSG00000111736.2 | 0 | NA | NA | NA | NA | NA | #VALUE! |
| ENSMUSG00000111737.2 | 1.836997 | -4.39907 | 3.931186 | -1.11902 | 0.263133 | NA | #VALUE! |
| ENSMUSG00000111738.2 | 0 | NA | NA | NA | NA | NA | #VALUE! |
| ENSMUSG00000111739.2 | 0 | NA | NA | NA | NA | NA | #VALUE! |
| ENSMUSG00000111740.2 | 0 | NA | NA | NA | NA | NA | #VALUE! |
| ENSMUSG00000111741.2 | 0 | NA | NA | NA | NA | NA | #VALUE! |
| ENSMUSG00000111743.2 | 0.279766 | 1.808031 | 4.036658 | 0.447903 | 0.654223 | NA | #VALUE! |
| ENSMUSG00000111744.2 | 0.204111 | -1.05581 | 4.073302 | -0.2592 | 0.795479 | NA | #VALUE! |
| ENSMUSG00000111746.2 | 0 | NA | NA | NA | NA | NA | #VALUE! |
| ENSMUSG00000111747.3 | 0 | NA | NA | NA | NA | NA | #VALUE! |
| ENSMUSG00000111748.2 | 0 | NA | NA | NA | NA | NA | #VALUE! |
| ENSMUSG00000111749.2 | 0 | NA | NA | NA | NA | NA | #VALUE! |

| | | | | | | | |
|---|---|---|---|---|---|---|---|
| ENSMUSG00000111750.2 | 0 | NA | NA | NA | NA | NA | #VALUE! |
| ENSMUSG00000111751.2 | 0 | NA | NA | NA | NA | NA | #VALUE! |
| ENSMUSG00000111752.2 | 0 | NA | NA | NA | NA | NA | #VALUE! |
| ENSMUSG00000111753.2 | 0 | NA | NA | NA | NA | NA | #VALUE! |
| ENSMUSG00000111754.2 | 0 | NA | NA | NA | NA | NA | #VALUE! |
| ENSMUSG00000111755.2 | 0 | NA | NA | NA | NA | NA | #VALUE! |
| ENSMUSG00000111756.2 | 0 | NA | NA | NA | NA | NA | #VALUE! |
| ENSMUSG00000111757.2 | 0 | NA | NA | NA | NA | NA | #VALUE! |
| ENSMUSG00000111758.2 | 0 | NA | NA | NA | NA | NA | #VALUE! |
| ENSMUSG00000111759.2 | 0 | NA | NA | NA | NA | NA | #VALUE! |
| ENSMUSG00000111760.2 | 0 | NA | NA | NA | NA | NA | #VALUE! |
| ENSMUSG00000111761.2 | 0 | NA | NA | NA | NA | NA | #VALUE! |
| ENSMUSG00000111762.2 | 0 | NA | NA | NA | NA | NA | #VALUE! |
| ENSMUSG00000111763.2 | 0 | NA | NA | NA | NA | NA | #VALUE! |
| ENSMUSG00000111764.2 | 0 | NA | NA | NA | NA | NA | #VALUE! |
| ENSMUSG00000111765.2 | 0 | NA | NA | NA | NA | NA | #VALUE! |
| ENSMUSG00000111766.2 | 0 | NA | NA | NA | NA | NA | #VALUE! |
| ENSMUSG00000111767.2 | 0 | NA | NA | NA | NA | NA | #VALUE! |
| ENSMUSG00000111769.2 | 0 | NA | NA | NA | NA | NA | #VALUE! |
| ENSMUSG00000111770.2 | 0 | NA | NA | NA | NA | NA | #VALUE! |
| ENSMUSG00000111771.2 | 0 | NA | NA | NA | NA | NA | #VALUE! |
| ENSMUSG00000111772.3 | 0 | NA | NA | NA | NA | NA | #VALUE! |
| ENSMUSG00000111774.2 | 0.093255 | 0.868093 | 4.072464 | 0.213162 | 0.831201 | NA | #VALUE! |
| ENSMUSG00000111776.2 | 0 | NA | NA | NA | NA | NA | #VALUE! |
| ENSMUSG00000111777.2 | 0 | NA | NA | NA | NA | NA | #VALUE! |
| ENSMUSG00000111778.2 | 0 | NA | NA | NA | NA | NA | #VALUE! |
| ENSMUSG00000111780.2 | 0 | NA | NA | NA | NA | NA | #VALUE! |
| ENSMUSG00000111781.2 | 0 | NA | NA | NA | NA | NA | #VALUE! |
| ENSMUSG00000111782.2 | 0 | NA | NA | NA | NA | NA | #VALUE! |
| ENSMUSG00000111783.2 | 0 | NA | NA | NA | NA | NA | #VALUE! |
| ENSMUSG00000111785.2 | 0 | NA | NA | NA | NA | NA | #VALUE! |
| ENSMUSG00000111786.2 | 0 | NA | NA | NA | NA | NA | #VALUE! |
| ENSMUSG00000111787.2 | 0 | NA | NA | NA | NA | NA | #VALUE! |
| ENSMUSG00000111788.2 | 0 | NA | NA | NA | NA | NA | #VALUE! |
| ENSMUSG00000111789.2 | 0 | NA | NA | NA | NA | NA | #VALUE! |
| ENSMUSG00000111792.2 | 0 | NA | NA | NA | NA | NA | #VALUE! |
| ENSMUSG00000111793.2 | 0 | NA | NA | NA | NA | NA | #VALUE! |
| ENSMUSG00000111794.2 | 0 | NA | NA | NA | NA | NA | #VALUE! |
| ENSMUSG00000111795.2 | 0 | NA | NA | NA | NA | NA | #VALUE! |
| ENSMUSG00000111796.2 | 0 | NA | NA | NA | NA | NA | #VALUE! |
| ENSMUSG00000111797.2 | 0 | NA | NA | NA | NA | NA | #VALUE! |
| ENSMUSG00000111799.2 | 0 | NA | NA | NA | NA | NA | #VALUE! |
| ENSMUSG00000111800.2 | 0 | NA | NA | NA | NA | NA | #VALUE! |
| ENSMUSG00000111801.2 | 0 | NA | NA | NA | NA | NA | #VALUE! |

| | | | | | | | |
|---|---|---|---|---|---|---|---|
| ENSMUSG00000111802.2 | 0 | NA | NA | NA | NA | NA | #VALUE! |
| ENSMUSG00000111803.2 | 0 | NA | NA | NA | NA | NA | #VALUE! |
| ENSMUSG00000111805.2 | 0 | NA | NA | NA | NA | NA | #VALUE! |
| ENSMUSG00000111806.2 | 0 | NA | NA | NA | NA | NA | #VALUE! |
| ENSMUSG00000111807.2 | 0 | NA | NA | NA | NA | NA | #VALUE! |
| ENSMUSG00000111808.2 | 0 | NA | NA | NA | NA | NA | #VALUE! |
| ENSMUSG00000111809.2 | 0 | NA | NA | NA | NA | NA | #VALUE! |
| ENSMUSG00000111810.2 | 0 | NA | NA | NA | NA | NA | #VALUE! |
| ENSMUSG00000111811.2 | 0 | NA | NA | NA | NA | NA | #VALUE! |
| ENSMUSG00000111812.2 | 0 | NA | NA | NA | NA | NA | #VALUE! |
| ENSMUSG00000111814.2 | 0 | NA | NA | NA | NA | NA | #VALUE! |
| ENSMUSG00000111815.2 | 0 | NA | NA | NA | NA | NA | #VALUE! |
| ENSMUSG00000111816.2 | 0 | NA | NA | NA | NA | NA | #VALUE! |
| ENSMUSG00000111817.2 | 0.195789 | -1.05581 | 4.073302 | -0.2592 | 0.795479 | NA | #VALUE! |
| ENSMUSG00000111818.2 | 0 | NA | NA | NA | NA | NA | #VALUE! |
| ENSMUSG00000111819.2 | 0 | NA | NA | NA | NA | NA | #VALUE! |
| ENSMUSG00000111820.2 | 0 | NA | NA | NA | NA | NA | #VALUE! |
| ENSMUSG00000111821.2 | 0 | NA | NA | NA | NA | NA | #VALUE! |
| ENSMUSG00000111822.2 | 0 | NA | NA | NA | NA | NA | #VALUE! |
| ENSMUSG00000111823.2 | 0 | NA | NA | NA | NA | NA | #VALUE! |
| ENSMUSG00000111824.2 | 0 | NA | NA | NA | NA | NA | #VALUE! |
| ENSMUSG00000111825.2 | 0 | NA | NA | NA | NA | NA | #VALUE! |
| ENSMUSG00000111827.2 | 0 | NA | NA | NA | NA | NA | #VALUE! |
| ENSMUSG00000111828.2 | 8.867611 | -5.74618 | 2.83378 | -2.02774 | 0.042586 | NA | #VALUE! |
| ENSMUSG00000111829.2 | 0 | NA | NA | NA | NA | NA | #VALUE! |
| ENSMUSG00000111830.2 | 0 | NA | NA | NA | NA | NA | #VALUE! |
| ENSMUSG00000111831.2 | 0 | NA | NA | NA | NA | NA | #VALUE! |
| ENSMUSG00000111833.2 | 0 | NA | NA | NA | NA | NA | #VALUE! |
| ENSMUSG00000111834.2 | 0 | NA | NA | NA | NA | NA | #VALUE! |
| ENSMUSG00000111836.2 | 0 | NA | NA | NA | NA | NA | #VALUE! |
| ENSMUSG00000111837.2 | 0 | NA | NA | NA | NA | NA | #VALUE! |
| ENSMUSG00000111839.2 | 0 | NA | NA | NA | NA | NA | #VALUE! |
| ENSMUSG00000111840.2 | 0 | NA | NA | NA | NA | NA | #VALUE! |
| ENSMUSG00000111841.2 | 0 | NA | NA | NA | NA | NA | #VALUE! |
| ENSMUSG00000111842.2 | 0 | NA | NA | NA | NA | NA | #VALUE! |
| ENSMUSG00000111843.2 | 0 | NA | NA | NA | NA | NA | #VALUE! |
| ENSMUSG00000111844.2 | 0 | NA | NA | NA | NA | NA | #VALUE! |
| ENSMUSG00000111845.2 | 0 | NA | NA | NA | NA | NA | #VALUE! |
| ENSMUSG00000111846.2 | 0 | NA | NA | NA | NA | NA | #VALUE! |
| ENSMUSG00000111847.2 | 0 | NA | NA | NA | NA | NA | #VALUE! |
| ENSMUSG00000111849.2 | 0 | NA | NA | NA | NA | NA | #VALUE! |
| ENSMUSG00000111850.2 | 0 | NA | NA | NA | NA | NA | #VALUE! |
| ENSMUSG00000111852.2 | 0 | NA | NA | NA | NA | NA | #VALUE! |
| ENSMUSG00000111853.2 | 0 | NA | NA | NA | NA | NA | #VALUE! |

| | | | | | | | |
|---|---|---|---|---|---|---|---|
| ENSMUSG00000111854.2 | 0 | NA | NA | NA | NA | NA | #VALUE! |
| ENSMUSG00000111856.2 | 0 | NA | NA | NA | NA | NA | #VALUE! |
| ENSMUSG00000111857.2 | 0 | NA | NA | NA | NA | NA | #VALUE! |
| ENSMUSG00000111858.2 | 0 | NA | NA | NA | NA | NA | #VALUE! |
| ENSMUSG00000111859.2 | 0 | NA | NA | NA | NA | NA | #VALUE! |
| ENSMUSG00000111862.2 | 0 | NA | NA | NA | NA | NA | #VALUE! |
| ENSMUSG00000111863.3 | 0 | NA | NA | NA | NA | NA | #VALUE! |
| ENSMUSG00000111864.2 | 0 | NA | NA | NA | NA | NA | #VALUE! |
| ENSMUSG00000111865.2 | 0 | NA | NA | NA | NA | NA | #VALUE! |
| ENSMUSG00000111866.2 | 0 | NA | NA | NA | NA | NA | #VALUE! |
| ENSMUSG00000111867.2 | 0 | NA | NA | NA | NA | NA | #VALUE! |
| ENSMUSG00000111868.2 | 0 | NA | NA | NA | NA | NA | #VALUE! |
| ENSMUSG00000111869.2 | 0 | NA | NA | NA | NA | NA | #VALUE! |
| ENSMUSG00000111870.2 | 0 | NA | NA | NA | NA | NA | #VALUE! |
| ENSMUSG00000111871.2 | 0 | NA | NA | NA | NA | NA | #VALUE! |
| ENSMUSG00000111873.2 | 0 | NA | NA | NA | NA | NA | #VALUE! |
| ENSMUSG00000111874.2 | 0 | NA | NA | NA | NA | NA | #VALUE! |
| ENSMUSG00000111875.2 | 0.559862 | -1.08106 | 4.007336 | -0.26977 | 0.787337 | NA | #VALUE! |
| ENSMUSG00000111876.2 | 0 | NA | NA | NA | NA | NA | #VALUE! |
| ENSMUSG00000111877.2 | 0.198268 | -0.06494 | 4.068276 | -0.01596 | 0.987265 | NA | #VALUE! |
| ENSMUSG00000111878.2 | 0 | NA | NA | NA | NA | NA | #VALUE! |
| ENSMUSG00000111880.2 | 0 | NA | NA | NA | NA | NA | #VALUE! |
| ENSMUSG00000111881.2 | 0 | NA | NA | NA | NA | NA | #VALUE! |
| ENSMUSG00000111882.3 | 0 | NA | NA | NA | NA | NA | #VALUE! |
| ENSMUSG00000111883.2 | 0.168284 | 1.027628 | 4.067126 | 0.252667 | 0.800526 | NA | #VALUE! |
| ENSMUSG00000111884.2 | 0 | NA | NA | NA | NA | NA | #VALUE! |
| ENSMUSG00000111885.2 | 0 | NA | NA | NA | NA | NA | #VALUE! |
| ENSMUSG00000111886.2 | 0 | NA | NA | NA | NA | NA | #VALUE! |
| ENSMUSG00000111887.2 | 0 | NA | NA | NA | NA | NA | #VALUE! |
| ENSMUSG00000111888.2 | 0 | NA | NA | NA | NA | NA | #VALUE! |
| ENSMUSG00000111889.2 | 0 | NA | NA | NA | NA | NA | #VALUE! |
| ENSMUSG00000111890.2 | 0 | NA | NA | NA | NA | NA | #VALUE! |
| ENSMUSG00000111891.2 | 0 | NA | NA | NA | NA | NA | #VALUE! |
| ENSMUSG00000111892.2 | 0 | NA | NA | NA | NA | NA | #VALUE! |
| ENSMUSG00000111893.2 | 0 | NA | NA | NA | NA | NA | #VALUE! |
| ENSMUSG00000111894.2 | 0 | NA | NA | NA | NA | NA | #VALUE! |
| ENSMUSG00000111895.2 | 0 | NA | NA | NA | NA | NA | #VALUE! |
| ENSMUSG00000111896.2 | 0 | NA | NA | NA | NA | NA | #VALUE! |
| ENSMUSG00000111897.2 | 2.913525 | 2.226075 | 3.834907 | 0.580477 | 0.561593 | NA | #VALUE! |
| ENSMUSG00000111898.2 | 0 | NA | NA | NA | NA | NA | #VALUE! |
| ENSMUSG00000111899.2 | 0 | NA | NA | NA | NA | NA | #VALUE! |
| ENSMUSG00000111900.2 | 0 | NA | NA | NA | NA | NA | #VALUE! |
| ENSMUSG00000111901.2 | 0 | NA | NA | NA | NA | NA | #VALUE! |
| ENSMUSG00000111902.2 | 0 | NA | NA | NA | NA | NA | #VALUE! |

| | | | | | | | |
|---|---|---|---|---|---|---|---|
| ENSMUSG00000111903.2 | 0 | NA | NA | NA | NA | NA | #VALUE! |
| ENSMUSG00000111904.2 | 0 | NA | NA | NA | NA | NA | #VALUE! |
| ENSMUSG00000111905.2 | 1.68021 | -4.29432 | 3.933015 | -1.09187 | 0.274892 | NA | #VALUE! |
| ENSMUSG00000111906.2 | 0 | NA | NA | NA | NA | NA | #VALUE! |
| ENSMUSG00000111907.2 | 0 | NA | NA | NA | NA | NA | #VALUE! |
| ENSMUSG00000111908.2 | 0 | NA | NA | NA | NA | NA | #VALUE! |
| ENSMUSG00000111909.2 | 0 | NA | NA | NA | NA | NA | #VALUE! |
| ENSMUSG00000111910.2 | 0 | NA | NA | NA | NA | NA | #VALUE! |
| ENSMUSG00000111911.2 | 0 | NA | NA | NA | NA | NA | #VALUE! |
| ENSMUSG00000111912.2 | 0 | NA | NA | NA | NA | NA | #VALUE! |
| ENSMUSG00000111913.2 | 0 | NA | NA | NA | NA | NA | #VALUE! |
| ENSMUSG00000111914.2 | 0 | NA | NA | NA | NA | NA | #VALUE! |
| ENSMUSG00000111915.2 | 0 | NA | NA | NA | NA | NA | #VALUE! |
| ENSMUSG00000111916.2 | 0 | NA | NA | NA | NA | NA | #VALUE! |
| ENSMUSG00000111917.2 | 0 | NA | NA | NA | NA | NA | #VALUE! |
| ENSMUSG00000111918.2 | 0 | NA | NA | NA | NA | NA | #VALUE! |
| ENSMUSG00000111919.2 | 0 | NA | NA | NA | NA | NA | #VALUE! |
| ENSMUSG00000111920.2 | 0 | NA | NA | NA | NA | NA | #VALUE! |
| ENSMUSG00000111921.2 | 0 | NA | NA | NA | NA | NA | #VALUE! |
| ENSMUSG00000111922.2 | 0 | NA | NA | NA | NA | NA | #VALUE! |
| ENSMUSG00000111923.2 | 7.831564 | -6.49776 | 3.90065 | -1.66581 | 0.095751 | NA | #VALUE! |
| ENSMUSG00000111924.2 | 0 | NA | NA | NA | NA | NA | #VALUE! |
| ENSMUSG00000111925.2 | 0 | NA | NA | NA | NA | NA | #VALUE! |
| ENSMUSG00000111926.2 | 0 | NA | NA | NA | NA | NA | #VALUE! |
| ENSMUSG00000111927.2 | 0 | NA | NA | NA | NA | NA | #VALUE! |
| ENSMUSG00000111928.2 | 0 | NA | NA | NA | NA | NA | #VALUE! |
| ENSMUSG00000111929.2 | 0 | NA | NA | NA | NA | NA | #VALUE! |
| ENSMUSG00000111930.2 | 0 | NA | NA | NA | NA | NA | #VALUE! |
| ENSMUSG00000111931.2 | 0 | NA | NA | NA | NA | NA | #VALUE! |
| ENSMUSG00000111932.2 | 0.568183 | -1.10781 | 4.005278 | -0.27659 | 0.782096 | NA | #VALUE! |
| ENSMUSG00000111933.2 | 0 | NA | NA | NA | NA | NA | #VALUE! |
| ENSMUSG00000111934.2 | 0 | NA | NA | NA | NA | NA | #VALUE! |
| ENSMUSG00000111935.2 | 0 | NA | NA | NA | NA | NA | #VALUE! |
| ENSMUSG00000111936.2 | 0 | NA | NA | NA | NA | NA | #VALUE! |
| ENSMUSG00000111937.2 | 0 | NA | NA | NA | NA | NA | #VALUE! |
| ENSMUSG00000111938.2 | 0 | NA | NA | NA | NA | NA | #VALUE! |
| ENSMUSG00000111939.2 | 0 | NA | NA | NA | NA | NA | #VALUE! |
| ENSMUSG00000111940.2 | 0 | NA | NA | NA | NA | NA | #VALUE! |
| ENSMUSG00000111941.2 | 0 | NA | NA | NA | NA | NA | #VALUE! |
| ENSMUSG00000111942.2 | 0 | NA | NA | NA | NA | NA | #VALUE! |
| ENSMUSG00000111943.2 | 0 | NA | NA | NA | NA | NA | #VALUE! |
| ENSMUSG00000111944.2 | 0 | NA | NA | NA | NA | NA | #VALUE! |
| ENSMUSG00000111945.2 | 0 | NA | NA | NA | NA | NA | #VALUE! |
| ENSMUSG00000111946.2 | 0 | NA | NA | NA | NA | NA | #VALUE! |

| | | | | | | | |
|---|---|---|---|---|---|---|---|
| ENSMUSG00000111947.2 | 0 | NA | NA | NA | NA | NA | #VALUE! |
| ENSMUSG00000111948.2 | 0 | NA | NA | NA | NA | NA | #VALUE! |
| ENSMUSG00000111949.2 | 0 | NA | NA | NA | NA | NA | #VALUE! |
| ENSMUSG00000111950.2 | 0 | NA | NA | NA | NA | NA | #VALUE! |
| ENSMUSG00000111951.2 | 0 | NA | NA | NA | NA | NA | #VALUE! |
| ENSMUSG00000111952.2 | 0 | NA | NA | NA | NA | NA | #VALUE! |
| ENSMUSG00000111953.2 | 0 | NA | NA | NA | NA | NA | #VALUE! |
| ENSMUSG00000111954.2 | 0 | NA | NA | NA | NA | NA | #VALUE! |
| ENSMUSG00000111955.2 | 0 | NA | NA | NA | NA | NA | #VALUE! |
| ENSMUSG00000111956.2 | 0 | NA | NA | NA | NA | NA | #VALUE! |
| ENSMUSG00000111957.2 | 0 | NA | NA | NA | NA | NA | #VALUE! |
| ENSMUSG00000111958.2 | 0 | NA | NA | NA | NA | NA | #VALUE! |
| ENSMUSG00000111959.2 | 0 | NA | NA | NA | NA | NA | #VALUE! |
| ENSMUSG00000111960.2 | 0 | NA | NA | NA | NA | NA | #VALUE! |
| ENSMUSG00000111961.2 | 0 | NA | NA | NA | NA | NA | #VALUE! |
| ENSMUSG00000111962.2 | 0 | NA | NA | NA | NA | NA | #VALUE! |
| ENSMUSG00000111963.2 | 5.869576 | 6.043788 | 3.772802 | 1.601936 | 0.10917 | NA | #VALUE! |
| ENSMUSG00000111964.2 | 0.168284 | 1.027628 | 4.067126 | 0.252667 | 0.800526 | NA | #VALUE! |
| ENSMUSG00000111965.2 | 0 | NA | NA | NA | NA | NA | #VALUE! |
| ENSMUSG00000111967.2 | 0 | NA | NA | NA | NA | NA | #VALUE! |
| ENSMUSG00000111968.2 | 0 | NA | NA | NA | NA | NA | #VALUE! |
| ENSMUSG00000111969.2 | 0 | NA | NA | NA | NA | NA | #VALUE! |
| ENSMUSG00000111970.2 | 0 | NA | NA | NA | NA | NA | #VALUE! |
| ENSMUSG00000111971.2 | 2.730341 | -4.98888 | 3.923018 | -1.27169 | 0.203482 | NA | #VALUE! |
| ENSMUSG00000111972.2 | 0 | NA | NA | NA | NA | NA | #VALUE! |
| ENSMUSG00000111973.2 | 0 | NA | NA | NA | NA | NA | #VALUE! |
| ENSMUSG00000111974.2 | 0.093255 | 0.868093 | 4.072464 | 0.213162 | 0.831201 | NA | #VALUE! |
| ENSMUSG00000111975.2 | 0 | NA | NA | NA | NA | NA | #VALUE! |
| ENSMUSG00000111976.2 | 0 | NA | NA | NA | NA | NA | #VALUE! |
| ENSMUSG00000111977.2 | 0 | NA | NA | NA | NA | NA | #VALUE! |
| ENSMUSG00000111978.2 | 0 | NA | NA | NA | NA | NA | #VALUE! |
| ENSMUSG00000111979.2 | 0 | NA | NA | NA | NA | NA | #VALUE! |
| ENSMUSG00000111980.2 | 0 | NA | NA | NA | NA | NA | #VALUE! |
| ENSMUSG00000111981.2 | 0 | NA | NA | NA | NA | NA | #VALUE! |
| ENSMUSG00000111982.2 | 0 | NA | NA | NA | NA | NA | #VALUE! |
| ENSMUSG00000111983.2 | 0 | NA | NA | NA | NA | NA | #VALUE! |
| ENSMUSG00000111984.2 | 0 | NA | NA | NA | NA | NA | #VALUE! |
| ENSMUSG00000111985.2 | 0 | NA | NA | NA | NA | NA | #VALUE! |
| ENSMUSG00000111986.2 | 0 | NA | NA | NA | NA | NA | #VALUE! |
| ENSMUSG00000111987.2 | 0 | NA | NA | NA | NA | NA | #VALUE! |
| ENSMUSG00000111988.2 | 0 | NA | NA | NA | NA | NA | #VALUE! |
| ENSMUSG00000111989.2 | 0 | NA | NA | NA | NA | NA | #VALUE! |
| ENSMUSG00000111990.2 | 0 | NA | NA | NA | NA | NA | #VALUE! |
| ENSMUSG00000111991.2 | 0 | NA | NA | NA | NA | NA | #VALUE! |

| | | | | | | | |
|---|---|---|---|---|---|---|---|
| ENSMUSG00000111992.2 | 0 | NA | NA | NA | NA | NA | #VALUE! |
| ENSMUSG00000111993.2 | 0 | NA | NA | NA | NA | NA | #VALUE! |
| ENSMUSG00000111994.2 | 0 | NA | NA | NA | NA | NA | #VALUE! |
| ENSMUSG00000111995.2 | 0 | NA | NA | NA | NA | NA | #VALUE! |
| ENSMUSG00000111996.2 | 0 | NA | NA | NA | NA | NA | #VALUE! |
| ENSMUSG00000111997.2 | 0 | NA | NA | NA | NA | NA | #VALUE! |
| ENSMUSG00000111998.2 | 0 | NA | NA | NA | NA | NA | #VALUE! |
| ENSMUSG00000111999.2 | 0 | NA | NA | NA | NA | NA | #VALUE! |
| ENSMUSG00000112000.2 | 0 | NA | NA | NA | NA | NA | #VALUE! |
| ENSMUSG00000112001.2 | 0 | NA | NA | NA | NA | NA | #VALUE! |
| ENSMUSG00000112002.2 | 0 | NA | NA | NA | NA | NA | #VALUE! |
| ENSMUSG00000112003.2 | 0 | NA | NA | NA | NA | NA | #VALUE! |
| ENSMUSG00000112004.2 | 0 | NA | NA | NA | NA | NA | #VALUE! |
| ENSMUSG00000112005.2 | 0 | NA | NA | NA | NA | NA | #VALUE! |
| ENSMUSG00000112007.2 | 0 | NA | NA | NA | NA | NA | #VALUE! |
| ENSMUSG00000112008.2 | 0 | NA | NA | NA | NA | NA | #VALUE! |
| ENSMUSG00000112009.2 | 0 | NA | NA | NA | NA | NA | #VALUE! |
| ENSMUSG00000112010.2 | 0 | NA | NA | NA | NA | NA | #VALUE! |
| ENSMUSG00000112011.2 | 0 | NA | NA | NA | NA | NA | #VALUE! |
| ENSMUSG00000112012.2 | 0 | NA | NA | NA | NA | NA | #VALUE! |
| ENSMUSG00000112013.2 | 0 | NA | NA | NA | NA | NA | #VALUE! |
| ENSMUSG00000112014.2 | 0 | NA | NA | NA | NA | NA | #VALUE! |
| ENSMUSG00000112015.2 | 0 | NA | NA | NA | NA | NA | #VALUE! |
| ENSMUSG00000112016.2 | 0 | NA | NA | NA | NA | NA | #VALUE! |
| ENSMUSG00000112017.2 | 0 | NA | NA | NA | NA | NA | #VALUE! |
| ENSMUSG00000112018.2 | 0 | NA | NA | NA | NA | NA | #VALUE! |
| ENSMUSG00000112019.2 | 0 | NA | NA | NA | NA | NA | #VALUE! |
| ENSMUSG00000112020.2 | 0 | NA | NA | NA | NA | NA | #VALUE! |
| ENSMUSG00000112021.2 | 0 | NA | NA | NA | NA | NA | #VALUE! |
| ENSMUSG00000112022.3 | 0 | NA | NA | NA | NA | NA | #VALUE! |
| ENSMUSG00000112024.2 | 0 | NA | NA | NA | NA | NA | #VALUE! |
| ENSMUSG00000112025.2 | 0 | NA | NA | NA | NA | NA | #VALUE! |
| ENSMUSG00000112026.2 | 0 | NA | NA | NA | NA | NA | #VALUE! |
| ENSMUSG00000112027.2 | 0 | NA | NA | NA | NA | NA | #VALUE! |
| ENSMUSG00000112028.2 | 0 | NA | NA | NA | NA | NA | #VALUE! |
| ENSMUSG00000112029.2 | 0 | NA | NA | NA | NA | NA | #VALUE! |
| ENSMUSG00000112030.2 | 0 | NA | NA | NA | NA | NA | #VALUE! |
| ENSMUSG00000112031.2 | 0 | NA | NA | NA | NA | NA | #VALUE! |
| ENSMUSG00000112032.2 | 0 | NA | NA | NA | NA | NA | #VALUE! |
| ENSMUSG00000112033.2 | 0 | NA | NA | NA | NA | NA | #VALUE! |
| ENSMUSG00000112034.2 | 0 | NA | NA | NA | NA | NA | #VALUE! |
| ENSMUSG00000112035.2 | 0 | NA | NA | NA | NA | NA | #VALUE! |
| ENSMUSG00000112036.2 | 0 | NA | NA | NA | NA | NA | #VALUE! |
| ENSMUSG00000112037.2 | 0 | NA | NA | NA | NA | NA | #VALUE! |

| | | | | | | | |
|---|---|---|---|---|---|---|---|
| ENSMUSG00000112038.2 | 0 | NA | NA | NA | NA | NA | #VALUE! |
| ENSMUSG00000112039.2 | 0 | NA | NA | NA | NA | NA | #VALUE! |
| ENSMUSG00000112040.2 | 0 | NA | NA | NA | NA | NA | #VALUE! |
| ENSMUSG00000112041.2 | 0 | NA | NA | NA | NA | NA | #VALUE! |
| ENSMUSG00000112042.2 | 0 | NA | NA | NA | NA | NA | #VALUE! |
| ENSMUSG00000112043.2 | 0 | NA | NA | NA | NA | NA | #VALUE! |
| ENSMUSG00000112044.2 | 0 | NA | NA | NA | NA | NA | #VALUE! |
| ENSMUSG00000112045.2 | 0 | NA | NA | NA | NA | NA | #VALUE! |
| ENSMUSG00000112046.2 | 0 | NA | NA | NA | NA | NA | #VALUE! |
| ENSMUSG00000112047.2 | 0 | NA | NA | NA | NA | NA | #VALUE! |
| ENSMUSG00000112048.2 | 0 | NA | NA | NA | NA | NA | #VALUE! |
| ENSMUSG00000112049.2 | 0 | NA | NA | NA | NA | NA | #VALUE! |
| ENSMUSG00000112050.2 | 0 | NA | NA | NA | NA | NA | #VALUE! |
| ENSMUSG00000112051.2 | 0 | NA | NA | NA | NA | NA | #VALUE! |
| ENSMUSG00000112052.2 | 0 | NA | NA | NA | NA | NA | #VALUE! |
| ENSMUSG00000112053.2 | 0 | NA | NA | NA | NA | NA | #VALUE! |
| ENSMUSG00000112054.2 | 0 | NA | NA | NA | NA | NA | #VALUE! |
| ENSMUSG00000112055.2 | 0 | NA | NA | NA | NA | NA | #VALUE! |
| ENSMUSG00000112056.2 | 0 | NA | NA | NA | NA | NA | #VALUE! |
| ENSMUSG00000112057.2 | 0 | NA | NA | NA | NA | NA | #VALUE! |
| ENSMUSG00000112058.2 | 0 | NA | NA | NA | NA | NA | #VALUE! |
| ENSMUSG00000112059.2 | 0 | NA | NA | NA | NA | NA | #VALUE! |
| ENSMUSG00000112060.2 | 0 | NA | NA | NA | NA | NA | #VALUE! |
| ENSMUSG00000112061.2 | 0 | NA | NA | NA | NA | NA | #VALUE! |
| ENSMUSG00000112062.2 | 0 | NA | NA | NA | NA | NA | #VALUE! |
| ENSMUSG00000112063.2 | 0 | NA | NA | NA | NA | NA | #VALUE! |
| ENSMUSG00000112064.2 | 0 | NA | NA | NA | NA | NA | #VALUE! |
| ENSMUSG00000112065.2 | 0 | NA | NA | NA | NA | NA | #VALUE! |
| ENSMUSG00000112066.2 | 0 | NA | NA | NA | NA | NA | #VALUE! |
| ENSMUSG00000112067.2 | 0 | NA | NA | NA | NA | NA | #VALUE! |
| ENSMUSG00000112068.2 | 0 | NA | NA | NA | NA | NA | #VALUE! |
| ENSMUSG00000112069.2 | 0 | NA | NA | NA | NA | NA | #VALUE! |
| ENSMUSG00000112070.2 | 0 | NA | NA | NA | NA | NA | #VALUE! |
| ENSMUSG00000112071.2 | 0 | NA | NA | NA | NA | NA | #VALUE! |
| ENSMUSG00000112072.2 | 0 | NA | NA | NA | NA | NA | #VALUE! |
| ENSMUSG00000112073.2 | 0 | NA | NA | NA | NA | NA | #VALUE! |
| ENSMUSG00000112074.2 | 0 | NA | NA | NA | NA | NA | #VALUE! |
| ENSMUSG00000112075.2 | 0 | NA | NA | NA | NA | NA | #VALUE! |
| ENSMUSG00000112076.2 | 0 | NA | NA | NA | NA | NA | #VALUE! |
| ENSMUSG00000112077.2 | 0 | NA | NA | NA | NA | NA | #VALUE! |
| ENSMUSG00000112078.2 | 0 | NA | NA | NA | NA | NA | #VALUE! |
| ENSMUSG00000112079.2 | 0 | NA | NA | NA | NA | NA | #VALUE! |
| ENSMUSG00000112080.2 | 0 | NA | NA | NA | NA | NA | #VALUE! |
| ENSMUSG00000112081.2 | 0 | NA | NA | NA | NA | NA | #VALUE! |

| | | | | | | | |
|---|---|---|---|---|---|---|---|
| ENSMUSG00000112082.2 | 0 | NA | NA | NA | NA | NA | #VALUE! |
| ENSMUSG00000112083.2 | 0 | NA | NA | NA | NA | NA | #VALUE! |
| ENSMUSG00000112084.2 | 0 | NA | NA | NA | NA | NA | #VALUE! |
| ENSMUSG00000112085.2 | 0 | NA | NA | NA | NA | NA | #VALUE! |
| ENSMUSG00000112086.2 | 0 | NA | NA | NA | NA | NA | #VALUE! |
| ENSMUSG00000112087.2 | 0 | NA | NA | NA | NA | NA | #VALUE! |
| ENSMUSG00000112088.2 | 0 | NA | NA | NA | NA | NA | #VALUE! |
| ENSMUSG00000112089.2 | 0 | NA | NA | NA | NA | NA | #VALUE! |
| ENSMUSG00000112091.2 | 0 | NA | NA | NA | NA | NA | #VALUE! |
| ENSMUSG00000112092.2 | 0 | NA | NA | NA | NA | NA | #VALUE! |
| ENSMUSG00000112093.2 | 0 | NA | NA | NA | NA | NA | #VALUE! |
| ENSMUSG00000112094.2 | 0 | NA | NA | NA | NA | NA | #VALUE! |
| ENSMUSG00000112095.2 | 0 | NA | NA | NA | NA | NA | #VALUE! |
| ENSMUSG00000112096.2 | 0.839297 | 3.273019 | 3.958392 | 0.826856 | 0.408319 | NA | #VALUE! |
| ENSMUSG00000112097.2 | 0 | NA | NA | NA | NA | NA | #VALUE! |
| ENSMUSG00000112098.2 | 0 | NA | NA | NA | NA | NA | #VALUE! |
| ENSMUSG00000112100.2 | 0 | NA | NA | NA | NA | NA | #VALUE! |
| ENSMUSG00000112101.2 | 0 | NA | NA | NA | NA | NA | #VALUE! |
| ENSMUSG00000112102.2 | 0 | NA | NA | NA | NA | NA | #VALUE! |
| ENSMUSG00000112103.2 | 0 | NA | NA | NA | NA | NA | #VALUE! |
| ENSMUSG00000112104.2 | 0 | NA | NA | NA | NA | NA | #VALUE! |
| ENSMUSG00000112105.2 | 0 | NA | NA | NA | NA | NA | #VALUE! |
| ENSMUSG00000112106.2 | 0 | NA | NA | NA | NA | NA | #VALUE! |
| ENSMUSG00000112107.2 | 0 | NA | NA | NA | NA | NA | #VALUE! |
| ENSMUSG00000112108.2 | 0 | NA | NA | NA | NA | NA | #VALUE! |
| ENSMUSG00000112109.2 | 0 | NA | NA | NA | NA | NA | #VALUE! |
| ENSMUSG00000112110.2 | 0 | NA | NA | NA | NA | NA | #VALUE! |
| ENSMUSG00000112111.2 | 0 | NA | NA | NA | NA | NA | #VALUE! |
| ENSMUSG00000112112.2 | 0 | NA | NA | NA | NA | NA | #VALUE! |
| ENSMUSG00000112113.2 | 0 | NA | NA | NA | NA | NA | #VALUE! |
| ENSMUSG00000112114.2 | 0 | NA | NA | NA | NA | NA | #VALUE! |
| ENSMUSG00000112115.2 | 0 | NA | NA | NA | NA | NA | #VALUE! |
| ENSMUSG00000112116.2 | 0 | NA | NA | NA | NA | NA | #VALUE! |
| ENSMUSG00000112117.3 | 0 | NA | NA | NA | NA | NA | #VALUE! |
| ENSMUSG00000112118.2 | 0 | NA | NA | NA | NA | NA | #VALUE! |
| ENSMUSG00000112119.2 | 0 | NA | NA | NA | NA | NA | #VALUE! |
| ENSMUSG00000112120.2 | 0 | NA | NA | NA | NA | NA | #VALUE! |
| ENSMUSG00000112121.2 | 0 | NA | NA | NA | NA | NA | #VALUE! |
| ENSMUSG00000112122.2 | 0 | NA | NA | NA | NA | NA | #VALUE! |
| ENSMUSG00000112123.2 | 0 | NA | NA | NA | NA | NA | #VALUE! |
| ENSMUSG00000112124.2 | 0 | NA | NA | NA | NA | NA | #VALUE! |
| ENSMUSG00000112125.2 | 0 | NA | NA | NA | NA | NA | #VALUE! |
| ENSMUSG00000112126.2 | 0 | NA | NA | NA | NA | NA | #VALUE! |
| ENSMUSG00000112127.2 | 0 | NA | NA | NA | NA | NA | #VALUE! |

| | | | | | | | |
|---|---|---|---|---|---|---|---|
| ENSMUSG00000112128.2 | 0 | NA | NA | NA | NA | NA | #VALUE! |
| ENSMUSG00000112130.2 | 0 | NA | NA | NA | NA | NA | #VALUE! |
| ENSMUSG00000112131.2 | 0 | NA | NA | NA | NA | NA | #VALUE! |
| ENSMUSG00000112132.2 | 0 | NA | NA | NA | NA | NA | #VALUE! |
| ENSMUSG00000112133.2 | 0 | NA | NA | NA | NA | NA | #VALUE! |
| ENSMUSG00000112134.2 | 0 | NA | NA | NA | NA | NA | #VALUE! |
| ENSMUSG00000112135.2 | 0 | NA | NA | NA | NA | NA | #VALUE! |
| ENSMUSG00000112136.2 | 0 | NA | NA | NA | NA | NA | #VALUE! |
| ENSMUSG00000112137.2 | 0 | NA | NA | NA | NA | NA | #VALUE! |
| ENSMUSG00000112138.2 | 0 | NA | NA | NA | NA | NA | #VALUE! |
| ENSMUSG00000112139.2 | 0 | NA | NA | NA | NA | NA | #VALUE! |
| ENSMUSG00000112140.2 | 0 | NA | NA | NA | NA | NA | #VALUE! |
| ENSMUSG00000112141.2 | 0 | NA | NA | NA | NA | NA | #VALUE! |
| ENSMUSG00000112142.2 | 0 | NA | NA | NA | NA | NA | #VALUE! |
| ENSMUSG00000112143.2 | 0 | NA | NA | NA | NA | NA | #VALUE! |
| ENSMUSG00000112144.2 | 0 | NA | NA | NA | NA | NA | #VALUE! |
| ENSMUSG00000112145.2 | 0 | NA | NA | NA | NA | NA | #VALUE! |
| ENSMUSG00000112146.2 | 0 | NA | NA | NA | NA | NA | #VALUE! |
| ENSMUSG00000112147.2 | 0 | NA | NA | NA | NA | NA | #VALUE! |
| ENSMUSG00000112148.2 | 0 | NA | NA | NA | NA | NA | #VALUE! |
| ENSMUSG00000112149.2 | 0 | NA | NA | NA | NA | NA | #VALUE! |
| ENSMUSG00000112150.2 | 0.210026 | -1.52494 | 4.058692 | -0.37572 | 0.707123 | NA | #VALUE! |
| ENSMUSG00000112151.2 | 0 | NA | NA | NA | NA | NA | #VALUE! |
| ENSMUSG00000112152.2 | 0 | NA | NA | NA | NA | NA | #VALUE! |
| ENSMUSG00000112153.2 | 0 | NA | NA | NA | NA | NA | #VALUE! |
| ENSMUSG00000112154.2 | 0 | NA | NA | NA | NA | NA | #VALUE! |
| ENSMUSG00000112155.2 | 0 | NA | NA | NA | NA | NA | #VALUE! |
| ENSMUSG00000112156.2 | 0 | NA | NA | NA | NA | NA | #VALUE! |
| ENSMUSG00000112157.2 | 0 | NA | NA | NA | NA | NA | #VALUE! |
| ENSMUSG00000112158.2 | 0 | NA | NA | NA | NA | NA | #VALUE! |
| ENSMUSG00000112159.2 | 0 | NA | NA | NA | NA | NA | #VALUE! |
| ENSMUSG00000112160.2 | 7.97215 | -6.52275 | 3.424492 | -1.90474 | 0.056814 | NA | #VALUE! |
| ENSMUSG00000112161.2 | 0 | NA | NA | NA | NA | NA | #VALUE! |
| ENSMUSG00000112162.2 | 0 | NA | NA | NA | NA | NA | #VALUE! |
| ENSMUSG00000112163.2 | 0 | NA | NA | NA | NA | NA | #VALUE! |
| ENSMUSG00000112164.2 | 0 | NA | NA | NA | NA | NA | #VALUE! |
| ENSMUSG00000112165.2 | 0 | NA | NA | NA | NA | NA | #VALUE! |
| ENSMUSG00000112166.2 | 0 | NA | NA | NA | NA | NA | #VALUE! |
| ENSMUSG00000112167.2 | 0 | NA | NA | NA | NA | NA | #VALUE! |
| ENSMUSG00000112168.2 | 0 | NA | NA | NA | NA | NA | #VALUE! |
| ENSMUSG00000112169.2 | 0 | NA | NA | NA | NA | NA | #VALUE! |
| ENSMUSG00000112170.2 | 0 | NA | NA | NA | NA | NA | #VALUE! |
| ENSMUSG00000112171.2 | 0 | NA | NA | NA | NA | NA | #VALUE! |
| ENSMUSG00000112172.2 | 0 | NA | NA | NA | NA | NA | #VALUE! |

| | | | | | | | |
|---|---|---|---|---|---|---|---|
| ENSMUSG00000112173.2 | 0 | NA | NA | NA | NA | NA | #VALUE! |
| ENSMUSG00000112174.2 | 0 | NA | NA | NA | NA | NA | #VALUE! |
| ENSMUSG00000112175.2 | 0 | NA | NA | NA | NA | NA | #VALUE! |
| ENSMUSG00000112176.2 | 0 | NA | NA | NA | NA | NA | #VALUE! |
| ENSMUSG00000112177.2 | 0 | NA | NA | NA | NA | NA | #VALUE! |
| ENSMUSG00000112178.2 | 0 | NA | NA | NA | NA | NA | #VALUE! |
| ENSMUSG00000112179.2 | 0 | NA | NA | NA | NA | NA | #VALUE! |
| ENSMUSG00000112180.2 | 0 | NA | NA | NA | NA | NA | #VALUE! |
| ENSMUSG00000112181.2 | 0 | NA | NA | NA | NA | NA | #VALUE! |
| ENSMUSG00000112182.2 | 0 | NA | NA | NA | NA | NA | #VALUE! |
| ENSMUSG00000112183.2 | 0 | NA | NA | NA | NA | NA | #VALUE! |
| ENSMUSG00000112184.2 | 0 | NA | NA | NA | NA | NA | #VALUE! |
| ENSMUSG00000112185.2 | 0 | NA | NA | NA | NA | NA | #VALUE! |
| ENSMUSG00000112186.2 | 0 | NA | NA | NA | NA | NA | #VALUE! |
| ENSMUSG00000112187.2 | 0 | NA | NA | NA | NA | NA | #VALUE! |
| ENSMUSG00000112188.2 | 0 | NA | NA | NA | NA | NA | #VALUE! |
| ENSMUSG00000112190.2 | 0 | NA | NA | NA | NA | NA | #VALUE! |
| ENSMUSG00000112191.2 | 0 | NA | NA | NA | NA | NA | #VALUE! |
| ENSMUSG00000112192.2 | 0 | NA | NA | NA | NA | NA | #VALUE! |
| ENSMUSG00000112193.2 | 0 | NA | NA | NA | NA | NA | #VALUE! |
| ENSMUSG00000112194.2 | 0 | NA | NA | NA | NA | NA | #VALUE! |
| ENSMUSG00000112195.2 | 0.105013 | -0.97324 | 4.076293 | -0.23876 | 0.811295 | NA | #VALUE! |
| ENSMUSG00000112196.2 | 0 | NA | NA | NA | NA | NA | #VALUE! |
| ENSMUSG00000112197.2 | 0 | NA | NA | NA | NA | NA | #VALUE! |
| ENSMUSG00000112198.2 | 0 | NA | NA | NA | NA | NA | #VALUE! |
| ENSMUSG00000112199.2 | 0 | NA | NA | NA | NA | NA | #VALUE! |
| ENSMUSG00000112200.2 | 0 | NA | NA | NA | NA | NA | #VALUE! |
| ENSMUSG00000112201.2 | 0 | NA | NA | NA | NA | NA | #VALUE! |
| ENSMUSG00000112202.2 | 0 | NA | NA | NA | NA | NA | #VALUE! |
| ENSMUSG00000112203.2 | 0 | NA | NA | NA | NA | NA | #VALUE! |
| ENSMUSG00000112204.2 | 0 | NA | NA | NA | NA | NA | #VALUE! |
| ENSMUSG00000112205.2 | 0 | NA | NA | NA | NA | NA | #VALUE! |
| ENSMUSG00000112206.2 | 0 | NA | NA | NA | NA | NA | #VALUE! |
| ENSMUSG00000112207.2 | 0 | NA | NA | NA | NA | NA | #VALUE! |
| ENSMUSG00000112208.2 | 0 | NA | NA | NA | NA | NA | #VALUE! |
| ENSMUSG00000112209.2 | 0 | NA | NA | NA | NA | NA | #VALUE! |
| ENSMUSG00000112210.2 | 0 | NA | NA | NA | NA | NA | #VALUE! |
| ENSMUSG00000112211.2 | 0 | NA | NA | NA | NA | NA | #VALUE! |
| ENSMUSG00000112212.2 | 0 | NA | NA | NA | NA | NA | #VALUE! |
| ENSMUSG00000112213.2 | 0 | NA | NA | NA | NA | NA | #VALUE! |
| ENSMUSG00000112214.2 | 0 | NA | NA | NA | NA | NA | #VALUE! |
| ENSMUSG00000112215.2 | 0 | NA | NA | NA | NA | NA | #VALUE! |
| ENSMUSG00000112216.2 | 0 | NA | NA | NA | NA | NA | #VALUE! |
| ENSMUSG00000112217.2 | 0 | NA | NA | NA | NA | NA | #VALUE! |

| | | | | | | | |
|---|---|---|---|---|---|---|---|
| ENSMUSG00000112218.2 | 0 | NA | NA | NA | NA | NA | #VALUE! |
| ENSMUSG00000112219.2 | 0 | NA | NA | NA | NA | NA | #VALUE! |
| ENSMUSG00000112220.2 | 0 | NA | NA | NA | NA | NA | #VALUE! |
| ENSMUSG00000112221.2 | 0 | NA | NA | NA | NA | NA | #VALUE! |
| ENSMUSG00000112222.2 | 0 | NA | NA | NA | NA | NA | #VALUE! |
| ENSMUSG00000112223.2 | 0 | NA | NA | NA | NA | NA | #VALUE! |
| ENSMUSG00000112224.2 | 0 | NA | NA | NA | NA | NA | #VALUE! |
| ENSMUSG00000112225.2 | 0 | NA | NA | NA | NA | NA | #VALUE! |
| ENSMUSG00000112226.2 | 0 | NA | NA | NA | NA | NA | #VALUE! |
| ENSMUSG00000112227.2 | 0 | NA | NA | NA | NA | NA | #VALUE! |
| ENSMUSG00000112228.2 | 0 | NA | NA | NA | NA | NA | #VALUE! |
| ENSMUSG00000112229.2 | 0 | NA | NA | NA | NA | NA | #VALUE! |
| ENSMUSG00000112230.3 | 0 | NA | NA | NA | NA | NA | #VALUE! |
| ENSMUSG00000112231.2 | 0 | NA | NA | NA | NA | NA | #VALUE! |
| ENSMUSG00000112232.2 | 0 | NA | NA | NA | NA | NA | #VALUE! |
| ENSMUSG00000112233.2 | 0 | NA | NA | NA | NA | NA | #VALUE! |
| ENSMUSG00000112234.2 | 0 | NA | NA | NA | NA | NA | #VALUE! |
| ENSMUSG00000112235.2 | 0 | NA | NA | NA | NA | NA | #VALUE! |
| ENSMUSG00000112236.2 | 0 | NA | NA | NA | NA | NA | #VALUE! |
| ENSMUSG00000112237.2 | 0 | NA | NA | NA | NA | NA | #VALUE! |
| ENSMUSG00000112238.2 | 0 | NA | NA | NA | NA | NA | #VALUE! |
| ENSMUSG00000112239.2 | 5.072671 | 0.622304 | 3.763282 | 0.165362 | 0.868659 | NA | #VALUE! |
| ENSMUSG00000112240.2 | 0 | NA | NA | NA | NA | NA | #VALUE! |
| ENSMUSG00000112241.2 | 0 | NA | NA | NA | NA | NA | #VALUE! |
| ENSMUSG00000112242.2 | 0 | NA | NA | NA | NA | NA | #VALUE! |
| ENSMUSG00000112243.2 | 0 | NA | NA | NA | NA | NA | #VALUE! |
| ENSMUSG00000112244.2 | 0 | NA | NA | NA | NA | NA | #VALUE! |
| ENSMUSG00000112245.2 | 0 | NA | NA | NA | NA | NA | #VALUE! |
| ENSMUSG00000112246.2 | 0 | NA | NA | NA | NA | NA | #VALUE! |
| ENSMUSG00000112247.2 | 0 | NA | NA | NA | NA | NA | #VALUE! |
| ENSMUSG00000112248.2 | 0 | NA | NA | NA | NA | NA | #VALUE! |
| ENSMUSG00000112250.2 | 0 | NA | NA | NA | NA | NA | #VALUE! |
| ENSMUSG00000112251.2 | 0 | NA | NA | NA | NA | NA | #VALUE! |
| ENSMUSG00000112252.2 | 0 | NA | NA | NA | NA | NA | #VALUE! |
| ENSMUSG00000112253.2 | 0 | NA | NA | NA | NA | NA | #VALUE! |
| ENSMUSG00000112254.2 | 0 | NA | NA | NA | NA | NA | #VALUE! |
| ENSMUSG00000112255.2 | 0 | NA | NA | NA | NA | NA | #VALUE! |
| ENSMUSG00000112256.2 | 0 | NA | NA | NA | NA | NA | #VALUE! |
| ENSMUSG00000112257.2 | 0 | NA | NA | NA | NA | NA | #VALUE! |
| ENSMUSG00000112258.2 | 0 | NA | NA | NA | NA | NA | #VALUE! |
| ENSMUSG00000112259.2 | 0 | NA | NA | NA | NA | NA | #VALUE! |
| ENSMUSG00000112260.2 | 0 | NA | NA | NA | NA | NA | #VALUE! |
| ENSMUSG00000112261.2 | 0 | NA | NA | NA | NA | NA | #VALUE! |
| ENSMUSG00000112262.2 | 0 | NA | NA | NA | NA | NA | #VALUE! |

| | | | | | | | |
|---|---|---|---|---|---|---|---|
| ENSMUSG00000112263.2 | 0 | NA | NA | NA | NA | NA | #VALUE! |
| ENSMUSG00000112264.2 | 0 | NA | NA | NA | NA | NA | #VALUE! |
| ENSMUSG00000112265.2 | 0.841418 | 3.240539 | 3.959551 | 0.818411 | 0.413123 | NA | #VALUE! |
| ENSMUSG00000112266.2 | 0 | NA | NA | NA | NA | NA | #VALUE! |
| ENSMUSG00000112267.2 | 0 | NA | NA | NA | NA | NA | #VALUE! |
| ENSMUSG00000112268.2 | 0 | NA | NA | NA | NA | NA | #VALUE! |
| ENSMUSG00000112269.2 | 0 | NA | NA | NA | NA | NA | #VALUE! |
| ENSMUSG00000112270.2 | 0 | NA | NA | NA | NA | NA | #VALUE! |
| ENSMUSG00000112271.2 | 0 | NA | NA | NA | NA | NA | #VALUE! |
| ENSMUSG00000112272.2 | 0 | NA | NA | NA | NA | NA | #VALUE! |
| ENSMUSG00000112273.2 | 0 | NA | NA | NA | NA | NA | #VALUE! |
| ENSMUSG00000112274.2 | 0 | NA | NA | NA | NA | NA | #VALUE! |
| ENSMUSG00000112275.2 | 31.13245 | 21.82905 | 2.655308 | 8.22091 | NA | NA | #VALUE! |
| ENSMUSG00000112277.2 | 0 | NA | NA | NA | NA | NA | #VALUE! |
| ENSMUSG00000112278.2 | 0 | NA | NA | NA | NA | NA | #VALUE! |
| ENSMUSG00000112279.2 | 0 | NA | NA | NA | NA | NA | #VALUE! |
| ENSMUSG00000112280.2 | 1.119062 | 3.679627 | 3.945843 | 0.932533 | 0.351061 | NA | #VALUE! |
| ENSMUSG00000112281.2 | 0 | NA | NA | NA | NA | NA | #VALUE! |
| ENSMUSG00000112282.2 | 0 | NA | NA | NA | NA | NA | #VALUE! |
| ENSMUSG00000112283.2 | 0 | NA | NA | NA | NA | NA | #VALUE! |
| ENSMUSG00000112284.2 | 0 | NA | NA | NA | NA | NA | #VALUE! |
| ENSMUSG00000112285.2 | 0 | NA | NA | NA | NA | NA | #VALUE! |
| ENSMUSG00000112286.2 | 0 | NA | NA | NA | NA | NA | #VALUE! |
| ENSMUSG00000112287.2 | 0.093255 | 0.868093 | 4.072464 | 0.213162 | 0.831201 | NA | #VALUE! |
| ENSMUSG00000112288.2 | 0 | NA | NA | NA | NA | NA | #VALUE! |
| ENSMUSG00000112289.2 | 0 | NA | NA | NA | NA | NA | #VALUE! |
| ENSMUSG00000112290.2 | 0 | NA | NA | NA | NA | NA | #VALUE! |
| ENSMUSG00000112291.2 | 0 | NA | NA | NA | NA | NA | #VALUE! |
| ENSMUSG00000112292.2 | 0 | NA | NA | NA | NA | NA | #VALUE! |
| ENSMUSG00000112293.2 | 0 | NA | NA | NA | NA | NA | #VALUE! |
| ENSMUSG00000112295.2 | 0 | NA | NA | NA | NA | NA | #VALUE! |
| ENSMUSG00000112296.2 | 0 | NA | NA | NA | NA | NA | #VALUE! |
| ENSMUSG00000112297.2 | 0.105013 | -0.97324 | 4.076293 | -0.23876 | 0.811295 | NA | #VALUE! |
| ENSMUSG00000112298.2 | 0 | NA | NA | NA | NA | NA | #VALUE! |
| ENSMUSG00000112299.2 | 0 | NA | NA | NA | NA | NA | #VALUE! |
| ENSMUSG00000112300.2 | 0 | NA | NA | NA | NA | NA | #VALUE! |
| ENSMUSG00000112301.2 | 0 | NA | NA | NA | NA | NA | #VALUE! |
| ENSMUSG00000112303.2 | 0 | NA | NA | NA | NA | NA | #VALUE! |
| ENSMUSG00000112304.2 | 0 | NA | NA | NA | NA | NA | #VALUE! |
| ENSMUSG00000112305.2 | 0 | NA | NA | NA | NA | NA | #VALUE! |
| ENSMUSG00000112306.2 | 0 | NA | NA | NA | NA | NA | #VALUE! |
| ENSMUSG00000112307.2 | 0 | NA | NA | NA | NA | NA | #VALUE! |
| ENSMUSG00000112308.2 | 0 | NA | NA | NA | NA | NA | #VALUE! |
| ENSMUSG00000112309.2 | 0 | NA | NA | NA | NA | NA | #VALUE! |

| | | | | | | | |
|---|---|---|---|---|---|---|---|
| ENSMUSG00000112311.2 | 0 | NA | NA | NA | NA | NA | #VALUE! |
| ENSMUSG00000112312.2 | 0 | NA | NA | NA | NA | NA | #VALUE! |
| ENSMUSG00000112313.2 | 0 | NA | NA | NA | NA | NA | #VALUE! |
| ENSMUSG00000112314.2 | 0 | NA | NA | NA | NA | NA | #VALUE! |
| ENSMUSG00000112315.2 | 0 | NA | NA | NA | NA | NA | #VALUE! |
| ENSMUSG00000112316.2 | 0 | NA | NA | NA | NA | NA | #VALUE! |
| ENSMUSG00000112317.2 | 0 | NA | NA | NA | NA | NA | #VALUE! |
| ENSMUSG00000112318.2 | 0 | NA | NA | NA | NA | NA | #VALUE! |
| ENSMUSG00000112319.2 | 0 | NA | NA | NA | NA | NA | #VALUE! |
| ENSMUSG00000112320.2 | 0 | NA | NA | NA | NA | NA | #VALUE! |
| ENSMUSG00000112321.2 | 0 | NA | NA | NA | NA | NA | #VALUE! |
| ENSMUSG00000112322.2 | 0 | NA | NA | NA | NA | NA | #VALUE! |
| ENSMUSG00000112323.2 | 0 | NA | NA | NA | NA | NA | #VALUE! |
| ENSMUSG00000112324.2 | 0 | NA | NA | NA | NA | NA | #VALUE! |
| ENSMUSG00000112325.2 | 0 | NA | NA | NA | NA | NA | #VALUE! |
| ENSMUSG00000112326.2 | 0 | NA | NA | NA | NA | NA | #VALUE! |
| ENSMUSG00000112327.2 | 0 | NA | NA | NA | NA | NA | #VALUE! |
| ENSMUSG00000112328.2 | 0 | NA | NA | NA | NA | NA | #VALUE! |
| ENSMUSG00000112329.2 | 0 | NA | NA | NA | NA | NA | #VALUE! |
| ENSMUSG00000112330.2 | 0 | NA | NA | NA | NA | NA | #VALUE! |
| ENSMUSG00000112331.2 | 0 | NA | NA | NA | NA | NA | #VALUE! |
| ENSMUSG00000112332.2 | 7.067908 | 6.31113 | 3.913123 | 1.612811 | 0.106786 | NA | #VALUE! |
| ENSMUSG00000112333.2 | 0 | NA | NA | NA | NA | NA | #VALUE! |
| ENSMUSG00000112334.2 | 0 | NA | NA | NA | NA | NA | #VALUE! |
| ENSMUSG00000112335.2 | 0 | NA | NA | NA | NA | NA | #VALUE! |
| ENSMUSG00000112336.2 | 0 | NA | NA | NA | NA | NA | #VALUE! |
| ENSMUSG00000112337.2 | 0 | NA | NA | NA | NA | NA | #VALUE! |
| ENSMUSG00000112338.2 | 0 | NA | NA | NA | NA | NA | #VALUE! |
| ENSMUSG00000112339.2 | 0 | NA | NA | NA | NA | NA | #VALUE! |
| ENSMUSG00000112340.2 | 0 | NA | NA | NA | NA | NA | #VALUE! |
| ENSMUSG00000112341.2 | 0 | NA | NA | NA | NA | NA | #VALUE! |
| ENSMUSG00000112342.2 | 0 | NA | NA | NA | NA | NA | #VALUE! |
| ENSMUSG00000112343.2 | 0 | NA | NA | NA | NA | NA | #VALUE! |
| ENSMUSG00000112344.2 | 0 | NA | NA | NA | NA | NA | #VALUE! |
| ENSMUSG00000112345.2 | 0 | NA | NA | NA | NA | NA | #VALUE! |
| ENSMUSG00000112346.2 | 6.902392 | -6.32144 | 3.599843 | -1.75603 | 0.079083 | NA | #VALUE! |
| ENSMUSG00000112347.2 | 0 | NA | NA | NA | NA | NA | #VALUE! |
| ENSMUSG00000112348.2 | 0 | NA | NA | NA | NA | NA | #VALUE! |
| ENSMUSG00000112349.2 | 0 | NA | NA | NA | NA | NA | #VALUE! |
| ENSMUSG00000112350.2 | 0 | NA | NA | NA | NA | NA | #VALUE! |
| ENSMUSG00000112351.2 | 0 | NA | NA | NA | NA | NA | #VALUE! |
| ENSMUSG00000112352.2 | 0 | NA | NA | NA | NA | NA | #VALUE! |
| ENSMUSG00000112353.2 | 0 | NA | NA | NA | NA | NA | #VALUE! |
| ENSMUSG00000112354.2 | 0 | NA | NA | NA | NA | NA | #VALUE! |

| | | | | | | | |
|---|---|---|---|---|---|---|---|
| ENSMUSG00000112355.2 | 0 | NA | NA | NA | NA | NA | #VALUE! |
| ENSMUSG00000112357.2 | 0 | NA | NA | NA | NA | NA | #VALUE! |
| ENSMUSG00000112358.2 | 0 | NA | NA | NA | NA | NA | #VALUE! |
| ENSMUSG00000112359.2 | 0 | NA | NA | NA | NA | NA | #VALUE! |
| ENSMUSG00000112360.2 | 0 | NA | NA | NA | NA | NA | #VALUE! |
| ENSMUSG00000112361.2 | 0 | NA | NA | NA | NA | NA | #VALUE! |
| ENSMUSG00000112362.2 | 0 | NA | NA | NA | NA | NA | #VALUE! |
| ENSMUSG00000112363.2 | 0 | NA | NA | NA | NA | NA | #VALUE! |
| ENSMUSG00000112364.2 | 0 | NA | NA | NA | NA | NA | #VALUE! |
| ENSMUSG00000112366.2 | 3.039838 | 0.693761 | 3.789533 | 0.183073 | 0.854741 | NA | #VALUE! |
| ENSMUSG00000112367.2 | 0 | NA | NA | NA | NA | NA | #VALUE! |
| ENSMUSG00000112368.2 | 0.886328 | 0.15516 | 3.922494 | 0.039557 | 0.968447 | NA | #VALUE! |
| ENSMUSG00000112369.2 | 0 | NA | NA | NA | NA | NA | #VALUE! |
| ENSMUSG00000112370.2 | 0 | NA | NA | NA | NA | NA | #VALUE! |
| ENSMUSG00000112371.2 | 0 | NA | NA | NA | NA | NA | #VALUE! |
| ENSMUSG00000112372.2 | 0 | NA | NA | NA | NA | NA | #VALUE! |
| ENSMUSG00000112373.2 | 0 | NA | NA | NA | NA | NA | #VALUE! |
| ENSMUSG00000112374.2 | 6.615827 | -6.26 | 3.913512 | -1.59959 | 0.10969 | NA | #VALUE! |
| ENSMUSG00000112375.2 | 0 | NA | NA | NA | NA | NA | #VALUE! |
| ENSMUSG00000112376.2 | 0 | NA | NA | NA | NA | NA | #VALUE! |
| ENSMUSG00000112377.2 | 0 | NA | NA | NA | NA | NA | #VALUE! |
| ENSMUSG00000112378.2 | 0 | NA | NA | NA | NA | NA | #VALUE! |
| ENSMUSG00000112379.2 | 0 | NA | NA | NA | NA | NA | #VALUE! |
| ENSMUSG00000112380.2 | 0 | NA | NA | NA | NA | NA | #VALUE! |
| ENSMUSG00000112381.2 | 0 | NA | NA | NA | NA | NA | #VALUE! |
| ENSMUSG00000112382.2 | 0 | NA | NA | NA | NA | NA | #VALUE! |
| ENSMUSG00000112383.2 | 0 | NA | NA | NA | NA | NA | #VALUE! |
| ENSMUSG00000112384.2 | 3.443657 | 5.264174 | 3.919884 | 1.342941 | 0.179291 | NA | #VALUE! |
| ENSMUSG00000112385.2 | 0 | NA | NA | NA | NA | NA | #VALUE! |
| ENSMUSG00000112386.2 | 0 | NA | NA | NA | NA | NA | #VALUE! |
| ENSMUSG00000112387.2 | 0 | NA | NA | NA | NA | NA | #VALUE! |
| ENSMUSG00000112389.2 | 0 | NA | NA | NA | NA | NA | #VALUE! |
| ENSMUSG00000112390.2 | 0 | NA | NA | NA | NA | NA | #VALUE! |
| ENSMUSG00000112391.2 | 0 | NA | NA | NA | NA | NA | #VALUE! |
| ENSMUSG00000112392.2 | 0 | NA | NA | NA | NA | NA | #VALUE! |
| ENSMUSG00000112393.2 | 0 | NA | NA | NA | NA | NA | #VALUE! |
| ENSMUSG00000112394.2 | 0 | NA | NA | NA | NA | NA | #VALUE! |
| ENSMUSG00000112395.2 | 0 | NA | NA | NA | NA | NA | #VALUE! |
| ENSMUSG00000112396.2 | 0 | NA | NA | NA | NA | NA | #VALUE! |
| ENSMUSG00000112397.2 | 0 | NA | NA | NA | NA | NA | #VALUE! |
| ENSMUSG00000112398.2 | 0 | NA | NA | NA | NA | NA | #VALUE! |
| ENSMUSG00000112399.2 | 0 | NA | NA | NA | NA | NA | #VALUE! |
| ENSMUSG00000112400.2 | 0 | NA | NA | NA | NA | NA | #VALUE! |
| ENSMUSG00000112401.2 | 0 | NA | NA | NA | NA | NA | #VALUE! |

| | | | | | | | |
|---|---|---|---|---|---|---|---|
| ENSMUSG00000112402.2 | 0 | NA | NA | NA | NA | NA | #VALUE! |
| ENSMUSG00000112403.2 | 0 | NA | NA | NA | NA | NA | #VALUE! |
| ENSMUSG00000112404.2 | 0 | NA | NA | NA | NA | NA | #VALUE! |
| ENSMUSG00000112405.2 | 0 | NA | NA | NA | NA | NA | #VALUE! |
| ENSMUSG00000112406.2 | 0 | NA | NA | NA | NA | NA | #VALUE! |
| ENSMUSG00000112407.2 | 0 | NA | NA | NA | NA | NA | #VALUE! |
| ENSMUSG00000112408.2 | 0 | NA | NA | NA | NA | NA | #VALUE! |
| ENSMUSG00000112409.2 | 0 | NA | NA | NA | NA | NA | #VALUE! |
| ENSMUSG00000112410.2 | 0 | NA | NA | NA | NA | NA | #VALUE! |
| ENSMUSG00000112411.2 | 0 | NA | NA | NA | NA | NA | #VALUE! |
| ENSMUSG00000112412.2 | 0 | NA | NA | NA | NA | NA | #VALUE! |
| ENSMUSG00000112413.2 | 0.093255 | 0.868093 | 4.072464 | 0.213162 | 0.831201 | NA | #VALUE! |
| ENSMUSG00000112414.2 | 0 | NA | NA | NA | NA | NA | #VALUE! |
| ENSMUSG00000112415.2 | 0 | NA | NA | NA | NA | NA | #VALUE! |
| ENSMUSG00000112416.2 | 0 | NA | NA | NA | NA | NA | #VALUE! |
| ENSMUSG00000112417.2 | 0 | NA | NA | NA | NA | NA | #VALUE! |
| ENSMUSG00000112418.2 | 0 | NA | NA | NA | NA | NA | #VALUE! |
| ENSMUSG00000112419.2 | 0 | NA | NA | NA | NA | NA | #VALUE! |
| ENSMUSG00000112420.2 | 0 | NA | NA | NA | NA | NA | #VALUE! |
| ENSMUSG00000112421.2 | 0 | NA | NA | NA | NA | NA | #VALUE! |
| ENSMUSG00000112422.2 | 0 | NA | NA | NA | NA | NA | #VALUE! |
| ENSMUSG00000112423.2 | 0 | NA | NA | NA | NA | NA | #VALUE! |
| ENSMUSG00000112424.2 | 0 | NA | NA | NA | NA | NA | #VALUE! |
| ENSMUSG00000112425.2 | 0 | NA | NA | NA | NA | NA | #VALUE! |
| ENSMUSG00000112426.2 | 0 | NA | NA | NA | NA | NA | #VALUE! |
| ENSMUSG00000112427.2 | 0 | NA | NA | NA | NA | NA | #VALUE! |
| ENSMUSG00000112428.2 | 0 | NA | NA | NA | NA | NA | #VALUE! |
| ENSMUSG00000112429.2 | 0 | NA | NA | NA | NA | NA | #VALUE! |
| ENSMUSG00000112430.2 | 0 | NA | NA | NA | NA | NA | #VALUE! |
| ENSMUSG00000112431.2 | 0 | NA | NA | NA | NA | NA | #VALUE! |
| ENSMUSG00000112432.2 | 0 | NA | NA | NA | NA | NA | #VALUE! |
| ENSMUSG00000112433.2 | 0 | NA | NA | NA | NA | NA | #VALUE! |
| ENSMUSG00000112434.2 | 0 | NA | NA | NA | NA | NA | #VALUE! |
| ENSMUSG00000112435.2 | 0 | NA | NA | NA | NA | NA | #VALUE! |
| ENSMUSG00000112436.2 | 0 | NA | NA | NA | NA | NA | #VALUE! |
| ENSMUSG00000112437.2 | 0 | NA | NA | NA | NA | NA | #VALUE! |
| ENSMUSG00000112438.2 | 0 | NA | NA | NA | NA | NA | #VALUE! |
| ENSMUSG00000112439.2 | 0 | NA | NA | NA | NA | NA | #VALUE! |
| ENSMUSG00000112440.2 | 0 | NA | NA | NA | NA | NA | #VALUE! |
| ENSMUSG00000112441.2 | 0 | NA | NA | NA | NA | NA | #VALUE! |
| ENSMUSG00000112442.2 | 0 | NA | NA | NA | NA | NA | #VALUE! |
| ENSMUSG00000112443.2 | 0 | NA | NA | NA | NA | NA | #VALUE! |
| ENSMUSG00000112444.2 | 0 | NA | NA | NA | NA | NA | #VALUE! |
| ENSMUSG00000112445.2 | 0 | NA | NA | NA | NA | NA | #VALUE! |

| | | | | | | | |
|---|---|---|---|---|---|---|---|
| ENSMUSG00000112446.2 | 0 | NA | NA | NA | NA | NA | #VALUE! |
| ENSMUSG00000112447.2 | 0 | NA | NA | NA | NA | NA | #VALUE! |
| ENSMUSG00000112448.2 | 0 | NA | NA | NA | NA | NA | #VALUE! |
| ENSMUSG00000112449.2 | 0 | NA | NA | NA | NA | NA | #VALUE! |
| ENSMUSG00000112450.2 | 0 | NA | NA | NA | NA | NA | #VALUE! |
| ENSMUSG00000112451.2 | 0 | NA | NA | NA | NA | NA | #VALUE! |
| ENSMUSG00000112452.2 | 0 | NA | NA | NA | NA | NA | #VALUE! |
| ENSMUSG00000112453.2 | 0 | NA | NA | NA | NA | NA | #VALUE! |
| ENSMUSG00000112454.2 | 0 | NA | NA | NA | NA | NA | #VALUE! |
| ENSMUSG00000112455.2 | 0 | NA | NA | NA | NA | NA | #VALUE! |
| ENSMUSG00000112456.2 | 0 | NA | NA | NA | NA | NA | #VALUE! |
| ENSMUSG00000112457.2 | 0 | NA | NA | NA | NA | NA | #VALUE! |
| ENSMUSG00000112458.2 | 0 | NA | NA | NA | NA | NA | #VALUE! |
| ENSMUSG00000112459.2 | 0 | NA | NA | NA | NA | NA | #VALUE! |
| ENSMUSG00000112460.2 | 0 | NA | NA | NA | NA | NA | #VALUE! |
| ENSMUSG00000112461.2 | 1.260158 | -3.88425 | 3.941584 | -0.98545 | 0.324401 | NA | #VALUE! |
| ENSMUSG00000112462.2 | 0 | NA | NA | NA | NA | NA | #VALUE! |
| ENSMUSG00000112463.2 | 0.245976 | 1.027628 | 4.067126 | 0.252667 | 0.800526 | NA | #VALUE! |
| ENSMUSG00000112464.2 | 0 | NA | NA | NA | NA | NA | #VALUE! |
| ENSMUSG00000112465.2 | 0 | NA | NA | NA | NA | NA | #VALUE! |
| ENSMUSG00000112466.2 | 0 | NA | NA | NA | NA | NA | #VALUE! |
| ENSMUSG00000112467.2 | 0 | NA | NA | NA | NA | NA | #VALUE! |
| ENSMUSG00000112468.2 | 0 | NA | NA | NA | NA | NA | #VALUE! |
| ENSMUSG00000112469.2 | 0 | NA | NA | NA | NA | NA | #VALUE! |
| ENSMUSG00000112470.2 | 0 | NA | NA | NA | NA | NA | #VALUE! |
| ENSMUSG00000112471.2 | 0 | NA | NA | NA | NA | NA | #VALUE! |
| ENSMUSG00000112472.2 | 0 | NA | NA | NA | NA | NA | #VALUE! |
| ENSMUSG00000112473.2 | 0 | NA | NA | NA | NA | NA | #VALUE! |
| ENSMUSG00000112474.2 | 0.168284 | 1.027628 | 4.067126 | 0.252667 | 0.800526 | NA | #VALUE! |
| ENSMUSG00000112475.2 | 0 | NA | NA | NA | NA | NA | #VALUE! |
| ENSMUSG00000112476.2 | 0 | NA | NA | NA | NA | NA | #VALUE! |
| ENSMUSG00000112477.2 | 0 | NA | NA | NA | NA | NA | #VALUE! |
| ENSMUSG00000112478.2 | 0 | NA | NA | NA | NA | NA | #VALUE! |
| ENSMUSG00000112479.2 | 0 | NA | NA | NA | NA | NA | #VALUE! |
| ENSMUSG00000112480.2 | 0 | NA | NA | NA | NA | NA | #VALUE! |
| ENSMUSG00000112481.2 | 0 | NA | NA | NA | NA | NA | #VALUE! |
| ENSMUSG00000112482.2 | 0 | NA | NA | NA | NA | NA | #VALUE! |
| ENSMUSG00000112483.2 | 0 | NA | NA | NA | NA | NA | #VALUE! |
| ENSMUSG00000112484.2 | 0 | NA | NA | NA | NA | NA | #VALUE! |
| ENSMUSG00000112485.2 | 0 | NA | NA | NA | NA | NA | #VALUE! |
| ENSMUSG00000112486.2 | 0 | NA | NA | NA | NA | NA | #VALUE! |
| ENSMUSG00000112487.2 | 0 | NA | NA | NA | NA | NA | #VALUE! |
| ENSMUSG00000112488.2 | 0 | NA | NA | NA | NA | NA | #VALUE! |
| ENSMUSG00000112489.2 | 0 | NA | NA | NA | NA | NA | #VALUE! |

| | | | | | | | |
|---|---|---|---|---|---|---|---|
| ENSMUSG00000112490.2 | 0 | NA | NA | NA | NA | NA | #VALUE! |
| ENSMUSG00000112491.2 | 0 | NA | NA | NA | NA | NA | #VALUE! |
| ENSMUSG00000112492.2 | 0 | NA | NA | NA | NA | NA | #VALUE! |
| ENSMUSG00000112493.2 | 0 | NA | NA | NA | NA | NA | #VALUE! |
| ENSMUSG00000112494.2 | 0 | NA | NA | NA | NA | NA | #VALUE! |
| ENSMUSG00000112495.2 | 0 | NA | NA | NA | NA | NA | #VALUE! |
| ENSMUSG00000112496.3 | 0 | NA | NA | NA | NA | NA | #VALUE! |
| ENSMUSG00000112497.2 | 0 | NA | NA | NA | NA | NA | #VALUE! |
| ENSMUSG00000112498.2 | 0 | NA | NA | NA | NA | NA | #VALUE! |
| ENSMUSG00000112499.2 | 0 | NA | NA | NA | NA | NA | #VALUE! |
| ENSMUSG00000112500.2 | 0 | NA | NA | NA | NA | NA | #VALUE! |
| ENSMUSG00000112501.2 | 0 | NA | NA | NA | NA | NA | #VALUE! |
| ENSMUSG00000112502.2 | 0 | NA | NA | NA | NA | NA | #VALUE! |
| ENSMUSG00000112503.2 | 0 | NA | NA | NA | NA | NA | #VALUE! |
| ENSMUSG00000112504.2 | 0 | NA | NA | NA | NA | NA | #VALUE! |
| ENSMUSG00000112505.2 | 0.297366 | -0.14751 | 4.065279 | -0.03629 | 0.971054 | NA | #VALUE! |
| ENSMUSG00000112506.2 | 0 | NA | NA | NA | NA | NA | #VALUE! |
| ENSMUSG00000112507.2 | 0 | NA | NA | NA | NA | NA | #VALUE! |
| ENSMUSG00000112509.2 | 0 | NA | NA | NA | NA | NA | #VALUE! |
| ENSMUSG00000112510.2 | 0 | NA | NA | NA | NA | NA | #VALUE! |
| ENSMUSG00000112511.2 | 0 | NA | NA | NA | NA | NA | #VALUE! |
| ENSMUSG00000112512.2 | 0 | NA | NA | NA | NA | NA | #VALUE! |
| ENSMUSG00000112513.2 | 0 | NA | NA | NA | NA | NA | #VALUE! |
| ENSMUSG00000112514.2 | 0 | NA | NA | NA | NA | NA | #VALUE! |
| ENSMUSG00000112515.2 | 0.210026 | -1.52494 | 4.058692 | -0.37572 | 0.707123 | NA | #VALUE! |
| ENSMUSG00000112516.2 | 0 | NA | NA | NA | NA | NA | #VALUE! |
| ENSMUSG00000112517.2 | 0 | NA | NA | NA | NA | NA | #VALUE! |
| ENSMUSG00000112518.2 | 0 | NA | NA | NA | NA | NA | #VALUE! |
| ENSMUSG00000112519.2 | 0 | NA | NA | NA | NA | NA | #VALUE! |
| ENSMUSG00000112520.2 | 0 | NA | NA | NA | NA | NA | #VALUE! |
| ENSMUSG00000112521.2 | 0 | NA | NA | NA | NA | NA | #VALUE! |
| ENSMUSG00000112522.2 | 0 | NA | NA | NA | NA | NA | #VALUE! |
| ENSMUSG00000112523.2 | 0 | NA | NA | NA | NA | NA | #VALUE! |
| ENSMUSG00000112524.2 | 0 | NA | NA | NA | NA | NA | #VALUE! |
| ENSMUSG00000112525.2 | 0 | NA | NA | NA | NA | NA | #VALUE! |
| ENSMUSG00000112526.2 | 0 | NA | NA | NA | NA | NA | #VALUE! |
| ENSMUSG00000112527.2 | 3.885486 | -5.49502 | 3.91822 | -1.40243 | 0.160787 | NA | #VALUE! |
| ENSMUSG00000112528.2 | 0 | NA | NA | NA | NA | NA | #VALUE! |
| ENSMUSG00000112529.2 | 0 | NA | NA | NA | NA | NA | #VALUE! |
| ENSMUSG00000112530.2 | 0 | NA | NA | NA | NA | NA | #VALUE! |
| ENSMUSG00000112531.2 | 0 | NA | NA | NA | NA | NA | #VALUE! |
| ENSMUSG00000112532.2 | 2.238125 | 4.666312 | 3.926586 | 1.188389 | 0.23468 | NA | #VALUE! |
| ENSMUSG00000112533.2 | 0 | NA | NA | NA | NA | NA | #VALUE! |
| ENSMUSG00000112534.2 | 0 | NA | NA | NA | NA | NA | #VALUE! |

| | | | | | | | |
|---|---|---|---|---|---|---|---|
| ENSMUSG00000112535.2 | 0 | NA | NA | NA | NA | NA | #VALUE! |
| ENSMUSG00000112536.2 | 0 | NA | NA | NA | NA | NA | #VALUE! |
| ENSMUSG00000112537.2 | 0 | NA | NA | NA | NA | NA | #VALUE! |
| ENSMUSG00000112538.2 | 0 | NA | NA | NA | NA | NA | #VALUE! |
| ENSMUSG00000112540.2 | 0 | NA | NA | NA | NA | NA | #VALUE! |
| ENSMUSG00000112541.2 | 0 | NA | NA | NA | NA | NA | #VALUE! |
| ENSMUSG00000112542.2 | 0 | NA | NA | NA | NA | NA | #VALUE! |
| ENSMUSG00000112543.2 | 0 | NA | NA | NA | NA | NA | #VALUE! |
| ENSMUSG00000112544.2 | 0 | NA | NA | NA | NA | NA | #VALUE! |
| ENSMUSG00000112545.2 | 2.349469 | -4.75734 | 3.925837 | -1.2118 | 0.225588 | NA | #VALUE! |
| ENSMUSG00000112546.2 | 0 | NA | NA | NA | NA | NA | #VALUE! |
| ENSMUSG00000112547.2 | 0 | NA | NA | NA | NA | NA | #VALUE! |
| ENSMUSG00000112548.2 | 0 | NA | NA | NA | NA | NA | #VALUE! |
| ENSMUSG00000112549.2 | 0 | NA | NA | NA | NA | NA | #VALUE! |
| ENSMUSG00000112550.2 | 0 | NA | NA | NA | NA | NA | #VALUE! |
| ENSMUSG00000112551.2 | 0 | NA | NA | NA | NA | NA | #VALUE! |
| ENSMUSG00000112552.2 | 0 | NA | NA | NA | NA | NA | #VALUE! |
| ENSMUSG00000112553.2 | 0 | NA | NA | NA | NA | NA | #VALUE! |
| ENSMUSG00000112554.2 | 0 | NA | NA | NA | NA | NA | #VALUE! |
| ENSMUSG00000112555.2 | 0 | NA | NA | NA | NA | NA | #VALUE! |
| ENSMUSG00000112556.2 | 0 | NA | NA | NA | NA | NA | #VALUE! |
| ENSMUSG00000112557.2 | 0 | NA | NA | NA | NA | NA | #VALUE! |
| ENSMUSG00000112558.2 | 0 | NA | NA | NA | NA | NA | #VALUE! |
| ENSMUSG00000112559.2 | 0 | NA | NA | NA | NA | NA | #VALUE! |
| ENSMUSG00000112560.2 | 0 | NA | NA | NA | NA | NA | #VALUE! |
| ENSMUSG00000112561.2 | 0 | NA | NA | NA | NA | NA | #VALUE! |
| ENSMUSG00000112562.2 | 0.093255 | 0.868093 | 4.072464 | 0.213162 | 0.831201 | NA | #VALUE! |
| ENSMUSG00000112563.2 | 0 | NA | NA | NA | NA | NA | #VALUE! |
| ENSMUSG00000112564.2 | 0 | NA | NA | NA | NA | NA | #VALUE! |
| ENSMUSG00000112565.2 | 0 | NA | NA | NA | NA | NA | #VALUE! |
| ENSMUSG00000112566.2 | 0 | NA | NA | NA | NA | NA | #VALUE! |
| ENSMUSG00000112568.2 | 0 | NA | NA | NA | NA | NA | #VALUE! |
| ENSMUSG00000112569.3 | 0 | NA | NA | NA | NA | NA | #VALUE! |
| ENSMUSG00000112570.2 | 0 | NA | NA | NA | NA | NA | #VALUE! |
| ENSMUSG00000112571.2 | 2.415302 | -4.81324 | 3.925115 | -1.22627 | 0.220098 | NA | #VALUE! |
| ENSMUSG00000112572.2 | 0 | NA | NA | NA | NA | NA | #VALUE! |
| ENSMUSG00000112573.2 | 0 | NA | NA | NA | NA | NA | #VALUE! |
| ENSMUSG00000112574.2 | 0 | NA | NA | NA | NA | NA | #VALUE! |
| ENSMUSG00000112575.2 | 0 | NA | NA | NA | NA | NA | #VALUE! |
| ENSMUSG00000112576.2 | 0 | NA | NA | NA | NA | NA | #VALUE! |
| ENSMUSG00000112577.2 | 0 | NA | NA | NA | NA | NA | #VALUE! |
| ENSMUSG00000112578.2 | 0 | NA | NA | NA | NA | NA | #VALUE! |
| ENSMUSG00000112579.2 | 0 | NA | NA | NA | NA | NA | #VALUE! |
| ENSMUSG00000112580.2 | 0 | NA | NA | NA | NA | NA | #VALUE! |

| | | | | | | | |
|---|---|---|---|---|---|---|---|
| ENSMUSG00000112581.2 | 0 | NA | NA | NA | NA | NA | #VALUE! |
| ENSMUSG00000112582.2 | 0 | NA | NA | NA | NA | NA | #VALUE! |
| ENSMUSG00000112583.2 | 0 | NA | NA | NA | NA | NA | #VALUE! |
| ENSMUSG00000112584.2 | 0 | NA | NA | NA | NA | NA | #VALUE! |
| ENSMUSG00000112585.2 | 0 | NA | NA | NA | NA | NA | #VALUE! |
| ENSMUSG00000112586.2 | 0 | NA | NA | NA | NA | NA | #VALUE! |
| ENSMUSG00000112587.2 | 0 | NA | NA | NA | NA | NA | #VALUE! |
| ENSMUSG00000112588.2 | 0 | NA | NA | NA | NA | NA | #VALUE! |
| ENSMUSG00000112589.2 | 0 | NA | NA | NA | NA | NA | #VALUE! |
| ENSMUSG00000112590.2 | 0 | NA | NA | NA | NA | NA | #VALUE! |
| ENSMUSG00000112591.2 | 0 | NA | NA | NA | NA | NA | #VALUE! |
| ENSMUSG00000112592.2 | 0 | NA | NA | NA | NA | NA | #VALUE! |
| ENSMUSG00000112593.2 | 0 | NA | NA | NA | NA | NA | #VALUE! |
| ENSMUSG00000112594.2 | 0 | NA | NA | NA | NA | NA | #VALUE! |
| ENSMUSG00000112595.2 | 0 | NA | NA | NA | NA | NA | #VALUE! |
| ENSMUSG00000112596.2 | 0 | NA | NA | NA | NA | NA | #VALUE! |
| ENSMUSG00000112597.2 | 0 | NA | NA | NA | NA | NA | #VALUE! |
| ENSMUSG00000112598.2 | 0 | NA | NA | NA | NA | NA | #VALUE! |
| ENSMUSG00000112599.2 | 0 | NA | NA | NA | NA | NA | #VALUE! |
| ENSMUSG00000112600.2 | 0 | NA | NA | NA | NA | NA | #VALUE! |
| ENSMUSG00000112601.2 | 0 | NA | NA | NA | NA | NA | #VALUE! |
| ENSMUSG00000112602.2 | 0 | NA | NA | NA | NA | NA | #VALUE! |
| ENSMUSG00000112603.2 | 0 | NA | NA | NA | NA | NA | #VALUE! |
| ENSMUSG00000112604.2 | 0 | NA | NA | NA | NA | NA | #VALUE! |
| ENSMUSG00000112605.2 | 0.195789 | -1.05581 | 4.073302 | -0.2592 | 0.795479 | NA | #VALUE! |
| ENSMUSG00000112606.2 | 0 | NA | NA | NA | NA | NA | #VALUE! |
| ENSMUSG00000112607.2 | 0 | NA | NA | NA | NA | NA | #VALUE! |
| ENSMUSG00000112608.2 | 0 | NA | NA | NA | NA | NA | #VALUE! |
| ENSMUSG00000112609.2 | 0 | NA | NA | NA | NA | NA | #VALUE! |
| ENSMUSG00000112610.2 | 0 | NA | NA | NA | NA | NA | #VALUE! |
| ENSMUSG00000112611.2 | 0 | NA | NA | NA | NA | NA | #VALUE! |
| ENSMUSG00000112612.2 | 0 | NA | NA | NA | NA | NA | #VALUE! |
| ENSMUSG00000112613.2 | 0 | NA | NA | NA | NA | NA | #VALUE! |
| ENSMUSG00000112614.2 | 0 | NA | NA | NA | NA | NA | #VALUE! |
| ENSMUSG00000112615.2 | 0 | NA | NA | NA | NA | NA | #VALUE! |
| ENSMUSG00000112616.2 | 0 | NA | NA | NA | NA | NA | #VALUE! |
| ENSMUSG00000112617.2 | 0 | NA | NA | NA | NA | NA | #VALUE! |
| ENSMUSG00000112618.2 | 0 | NA | NA | NA | NA | NA | #VALUE! |
| ENSMUSG00000112619.2 | 0 | NA | NA | NA | NA | NA | #VALUE! |
| ENSMUSG00000112620.2 | 0 | NA | NA | NA | NA | NA | #VALUE! |
| ENSMUSG00000112621.2 | 0 | NA | NA | NA | NA | NA | #VALUE! |
| ENSMUSG00000112622.2 | 0 | NA | NA | NA | NA | NA | #VALUE! |
| ENSMUSG00000112623.2 | 0 | NA | NA | NA | NA | NA | #VALUE! |
| ENSMUSG00000112624.2 | 0 | NA | NA | NA | NA | NA | #VALUE! |

| | | | | | | | |
|---|---|---|---|---|---|---|---|
| ENSMUSG00000112625.2 | 0 | NA | NA | NA | NA | NA | #VALUE! |
| ENSMUSG00000112626.2 | 0 | NA | NA | NA | NA | NA | #VALUE! |
| ENSMUSG00000112627.2 | 0 | NA | NA | NA | NA | NA | #VALUE! |
| ENSMUSG00000112628.2 | 0 | NA | NA | NA | NA | NA | #VALUE! |
| ENSMUSG00000112629.2 | 0 | NA | NA | NA | NA | NA | #VALUE! |
| ENSMUSG00000112630.2 | 0 | NA | NA | NA | NA | NA | #VALUE! |
| ENSMUSG00000112631.2 | 0 | NA | NA | NA | NA | NA | #VALUE! |
| ENSMUSG00000112632.2 | 0 | NA | NA | NA | NA | NA | #VALUE! |
| ENSMUSG00000112633.2 | 0 | NA | NA | NA | NA | NA | #VALUE! |
| ENSMUSG00000112634.2 | 0 | NA | NA | NA | NA | NA | #VALUE! |
| ENSMUSG00000112635.2 | 0 | NA | NA | NA | NA | NA | #VALUE! |
| ENSMUSG00000112636.2 | 0 | NA | NA | NA | NA | NA | #VALUE! |
| ENSMUSG00000112637.2 | 0 | NA | NA | NA | NA | NA | #VALUE! |
| ENSMUSG00000112638.2 | 0 | NA | NA | NA | NA | NA | #VALUE! |
| ENSMUSG00000112639.2 | 7.404201 | 0.035537 | 3.638412 | 0.009767 | 0.992207 | NA | #VALUE! |
| ENSMUSG00000112640.2 | 0 | NA | NA | NA | NA | NA | #VALUE! |
| ENSMUSG00000112641.2 | 0 | NA | NA | NA | NA | NA | #VALUE! |
| ENSMUSG00000112642.2 | 0 | NA | NA | NA | NA | NA | #VALUE! |
| ENSMUSG00000112643.2 | 0 | NA | NA | NA | NA | NA | #VALUE! |
| ENSMUSG00000112644.2 | 0 | NA | NA | NA | NA | NA | #VALUE! |
| ENSMUSG00000112645.2 | 0 | NA | NA | NA | NA | NA | #VALUE! |
| ENSMUSG00000112646.2 | 0 | NA | NA | NA | NA | NA | #VALUE! |
| ENSMUSG00000112647.2 | 0 | NA | NA | NA | NA | NA | #VALUE! |
| ENSMUSG00000112648.2 | 0 | NA | NA | NA | NA | NA | #VALUE! |
| ENSMUSG00000112649.2 | 0 | NA | NA | NA | NA | NA | #VALUE! |
| ENSMUSG00000112650.2 | 0 | NA | NA | NA | NA | NA | #VALUE! |
| ENSMUSG00000112651.2 | 0 | NA | NA | NA | NA | NA | #VALUE! |
| ENSMUSG00000112652.2 | 0 | NA | NA | NA | NA | NA | #VALUE! |
| ENSMUSG00000112653.2 | 0 | NA | NA | NA | NA | NA | #VALUE! |
| ENSMUSG00000112654.2 | 0 | NA | NA | NA | NA | NA | #VALUE! |
| ENSMUSG00000112655.2 | 0 | NA | NA | NA | NA | NA | #VALUE! |
| ENSMUSG00000112656.2 | 0 | NA | NA | NA | NA | NA | #VALUE! |
| ENSMUSG00000112657.2 | 0 | NA | NA | NA | NA | NA | #VALUE! |
| ENSMUSG00000112658.2 | 0 | NA | NA | NA | NA | NA | #VALUE! |
| ENSMUSG00000112659.2 | 0 | NA | NA | NA | NA | NA | #VALUE! |
| ENSMUSG00000112660.2 | 0 | NA | NA | NA | NA | NA | #VALUE! |
| ENSMUSG00000112661.2 | 0 | NA | NA | NA | NA | NA | #VALUE! |
| ENSMUSG00000112662.2 | 0 | NA | NA | NA | NA | NA | #VALUE! |
| ENSMUSG00000112663.2 | 0 | NA | NA | NA | NA | NA | #VALUE! |
| ENSMUSG00000112664.2 | 0 | NA | NA | NA | NA | NA | #VALUE! |
| ENSMUSG00000112665.2 | 0 | NA | NA | NA | NA | NA | #VALUE! |
| ENSMUSG00000112666.2 | 0 | NA | NA | NA | NA | NA | #VALUE! |
| ENSMUSG00000112667.2 | 0 | NA | NA | NA | NA | NA | #VALUE! |
| ENSMUSG00000112668.2 | 0 | NA | NA | NA | NA | NA | #VALUE! |

| | | | | | | | |
|---|---|---|---|---|---|---|---|
| ENSMUSG00000112669.2 | 0 | NA | NA | NA | NA | NA | #VALUE! |
| ENSMUSG00000112670.2 | 0 | NA | NA | NA | NA | NA | #VALUE! |
| ENSMUSG00000112671.2 | 0 | NA | NA | NA | NA | NA | #VALUE! |
| ENSMUSG00000112672.2 | 0 | NA | NA | NA | NA | NA | #VALUE! |
| ENSMUSG00000112673.2 | 0.168284 | 1.027628 | 4.067126 | 0.252667 | 0.800526 | NA | #VALUE! |
| ENSMUSG00000112674.2 | 0 | NA | NA | NA | NA | NA | #VALUE! |
| ENSMUSG00000112675.2 | 0 | NA | NA | NA | NA | NA | #VALUE! |
| ENSMUSG00000112676.2 | 0 | NA | NA | NA | NA | NA | #VALUE! |
| ENSMUSG00000112677.2 | 0 | NA | NA | NA | NA | NA | #VALUE! |
| ENSMUSG00000112678.2 | 0 | NA | NA | NA | NA | NA | #VALUE! |
| ENSMUSG00000112679.2 | 0 | NA | NA | NA | NA | NA | #VALUE! |
| ENSMUSG00000112680.2 | 0 | NA | NA | NA | NA | NA | #VALUE! |
| ENSMUSG00000112681.2 | 0 | NA | NA | NA | NA | NA | #VALUE! |
| ENSMUSG00000112682.2 | 0 | NA | NA | NA | NA | NA | #VALUE! |
| ENSMUSG00000112683.2 | 0 | NA | NA | NA | NA | NA | #VALUE! |
| ENSMUSG00000112684.2 | 0 | NA | NA | NA | NA | NA | #VALUE! |
| ENSMUSG00000112685.2 | 0 | NA | NA | NA | NA | NA | #VALUE! |
| ENSMUSG00000112686.2 | 0 | NA | NA | NA | NA | NA | #VALUE! |
| ENSMUSG00000112687.2 | 0 | NA | NA | NA | NA | NA | #VALUE! |
| ENSMUSG00000112688.2 | 0 | NA | NA | NA | NA | NA | #VALUE! |
| ENSMUSG00000112689.2 | 0 | NA | NA | NA | NA | NA | #VALUE! |
| ENSMUSG00000112690.3 | 0 | NA | NA | NA | NA | NA | #VALUE! |
| ENSMUSG00000112691.2 | 0 | NA | NA | NA | NA | NA | #VALUE! |
| ENSMUSG00000112692.2 | 0 | NA | NA | NA | NA | NA | #VALUE! |
| ENSMUSG00000112693.2 | 0.273297 | 0.094602 | 4.062933 | 0.023284 | 0.981424 | NA | #VALUE! |
| ENSMUSG00000112694.2 | 0 | NA | NA | NA | NA | NA | #VALUE! |
| ENSMUSG00000112696.2 | 0 | NA | NA | NA | NA | NA | #VALUE! |
| ENSMUSG00000112697.2 | 0 | NA | NA | NA | NA | NA | #VALUE! |
| ENSMUSG00000112698.2 | 0 | NA | NA | NA | NA | NA | #VALUE! |
| ENSMUSG00000112699.2 | 0 | NA | NA | NA | NA | NA | #VALUE! |
| ENSMUSG00000112700.2 | 0 | NA | NA | NA | NA | NA | #VALUE! |
| ENSMUSG00000112701.2 | 0 | NA | NA | NA | NA | NA | #VALUE! |
| ENSMUSG00000112702.2 | 0 | NA | NA | NA | NA | NA | #VALUE! |
| ENSMUSG00000112703.2 | 0 | NA | NA | NA | NA | NA | #VALUE! |
| ENSMUSG00000112704.2 | 0 | NA | NA | NA | NA | NA | #VALUE! |
| ENSMUSG00000112705.2 | 0 | NA | NA | NA | NA | NA | #VALUE! |
| ENSMUSG00000112706.2 | 0 | NA | NA | NA | NA | NA | #VALUE! |
| ENSMUSG00000112707.2 | 0 | NA | NA | NA | NA | NA | #VALUE! |
| ENSMUSG00000112708.2 | 0 | NA | NA | NA | NA | NA | #VALUE! |
| ENSMUSG00000112709.2 | 0 | NA | NA | NA | NA | NA | #VALUE! |
| ENSMUSG00000112710.2 | 0 | NA | NA | NA | NA | NA | #VALUE! |
| ENSMUSG00000112711.2 | 0 | NA | NA | NA | NA | NA | #VALUE! |
| ENSMUSG00000112712.2 | 0 | NA | NA | NA | NA | NA | #VALUE! |
| ENSMUSG00000112713.2 | 0 | NA | NA | NA | NA | NA | #VALUE! |

| | | | | | | | |
|---|---|---|---|---|---|---|---|
| ENSMUSG00000112714.2 | 0 | NA | NA | NA | NA | NA | #VALUE! |
| ENSMUSG00000112715.2 | 0 | NA | NA | NA | NA | NA | #VALUE! |
| ENSMUSG00000112716.2 | 0 | NA | NA | NA | NA | NA | #VALUE! |
| ENSMUSG00000112717.2 | 0 | NA | NA | NA | NA | NA | #VALUE! |
| ENSMUSG00000112718.2 | 0 | NA | NA | NA | NA | NA | #VALUE! |
| ENSMUSG00000112719.2 | 0 | NA | NA | NA | NA | NA | #VALUE! |
| ENSMUSG00000112720.2 | 0 | NA | NA | NA | NA | NA | #VALUE! |
| ENSMUSG00000112721.2 | 0 | NA | NA | NA | NA | NA | #VALUE! |
| ENSMUSG00000112722.2 | 0 | NA | NA | NA | NA | NA | #VALUE! |
| ENSMUSG00000112723.2 | 0 | NA | NA | NA | NA | NA | #VALUE! |
| ENSMUSG00000112724.2 | 0 | NA | NA | NA | NA | NA | #VALUE! |
| ENSMUSG00000112725.2 | 0 | NA | NA | NA | NA | NA | #VALUE! |
| ENSMUSG00000112726.2 | 0 | NA | NA | NA | NA | NA | #VALUE! |
| ENSMUSG00000112727.2 | 0.195789 | -1.05581 | 4.073302 | -0.2592 | 0.795479 | NA | #VALUE! |
| ENSMUSG00000112728.2 | 0 | NA | NA | NA | NA | NA | #VALUE! |
| ENSMUSG00000112729.2 | 0 | NA | NA | NA | NA | NA | #VALUE! |
| ENSMUSG00000112730.2 | 0 | NA | NA | NA | NA | NA | #VALUE! |
| ENSMUSG00000112731.2 | 0 | NA | NA | NA | NA | NA | #VALUE! |
| ENSMUSG00000112732.2 | 0 | NA | NA | NA | NA | NA | #VALUE! |
| ENSMUSG00000112733.2 | 0 | NA | NA | NA | NA | NA | #VALUE! |
| ENSMUSG00000112734.2 | 0.195789 | -1.05581 | 4.073302 | -0.2592 | 0.795479 | NA | #VALUE! |
| ENSMUSG00000112735.2 | 0 | NA | NA | NA | NA | NA | #VALUE! |
| ENSMUSG00000112736.2 | 0 | NA | NA | NA | NA | NA | #VALUE! |
| ENSMUSG00000112737.2 | 0 | NA | NA | NA | NA | NA | #VALUE! |
| ENSMUSG00000112738.2 | 0 | NA | NA | NA | NA | NA | #VALUE! |
| ENSMUSG00000112739.2 | 0 | NA | NA | NA | NA | NA | #VALUE! |
| ENSMUSG00000112740.2 | 0 | NA | NA | NA | NA | NA | #VALUE! |
| ENSMUSG00000112741.2 | 0 | NA | NA | NA | NA | NA | #VALUE! |
| ENSMUSG00000112742.2 | 0 | NA | NA | NA | NA | NA | #VALUE! |
| ENSMUSG00000112743.2 | 0 | NA | NA | NA | NA | NA | #VALUE! |
| ENSMUSG00000112744.2 | 0 | NA | NA | NA | NA | NA | #VALUE! |
| ENSMUSG00000112745.2 | 0 | NA | NA | NA | NA | NA | #VALUE! |
| ENSMUSG00000112746.2 | 0 | NA | NA | NA | NA | NA | #VALUE! |
| ENSMUSG00000112747.2 | 0 | NA | NA | NA | NA | NA | #VALUE! |
| ENSMUSG00000112748.2 | 0 | NA | NA | NA | NA | NA | #VALUE! |
| ENSMUSG00000112749.2 | 0 | NA | NA | NA | NA | NA | #VALUE! |
| ENSMUSG00000112750.2 | 0 | NA | NA | NA | NA | NA | #VALUE! |
| ENSMUSG00000112751.2 | 0 | NA | NA | NA | NA | NA | #VALUE! |
| ENSMUSG00000112752.2 | 0 | NA | NA | NA | NA | NA | #VALUE! |
| ENSMUSG00000112753.2 | 0 | NA | NA | NA | NA | NA | #VALUE! |
| ENSMUSG00000112754.2 | 0 | NA | NA | NA | NA | NA | #VALUE! |
| ENSMUSG00000112755.2 | 0 | NA | NA | NA | NA | NA | #VALUE! |
| ENSMUSG00000112756.2 | 0 | NA | NA | NA | NA | NA | #VALUE! |
| ENSMUSG00000112757.2 | 0 | NA | NA | NA | NA | NA | #VALUE! |

| | | | | | | | |
|---|---|---|---|---|---|---|---|
| ENSMUSG00000112759.2 | 2.940368 | -5.09511 | 3.921868 | -1.29915 | 0.193891 | NA | #VALUE! |
| ENSMUSG00000112760.2 | 0 | NA | NA | NA | NA | NA | #VALUE! |
| ENSMUSG00000112761.2 | 0 | NA | NA | NA | NA | NA | #VALUE! |
| ENSMUSG00000112762.2 | 0 | NA | NA | NA | NA | NA | #VALUE! |
| ENSMUSG00000112763.2 | 0 | NA | NA | NA | NA | NA | #VALUE! |
| ENSMUSG00000112764.2 | 0 | NA | NA | NA | NA | NA | #VALUE! |
| ENSMUSG00000112765.2 | 0 | NA | NA | NA | NA | NA | #VALUE! |
| ENSMUSG00000112766.2 | 0 | NA | NA | NA | NA | NA | #VALUE! |
| ENSMUSG00000112767.2 | 0 | NA | NA | NA | NA | NA | #VALUE! |
| ENSMUSG00000112768.2 | 0 | NA | NA | NA | NA | NA | #VALUE! |
| ENSMUSG00000112769.2 | 0 | NA | NA | NA | NA | NA | #VALUE! |
| ENSMUSG00000112770.2 | 0 | NA | NA | NA | NA | NA | #VALUE! |
| ENSMUSG00000112771.2 | 0.559531 | 2.703958 | 3.982863 | 0.678898 | 0.497202 | NA | #VALUE! |
| ENSMUSG00000112772.2 | 0 | NA | NA | NA | NA | NA | #VALUE! |
| ENSMUSG00000112773.2 | 0 | NA | NA | NA | NA | NA | #VALUE! |
| ENSMUSG00000112774.2 | 6.022644 | 0.589654 | 3.000069 | 0.196547 | 0.844182 | NA | #VALUE! |
| ENSMUSG00000112775.2 | 0 | NA | NA | NA | NA | NA | #VALUE! |
| ENSMUSG00000112776.2 | 0 | NA | NA | NA | NA | NA | #VALUE! |
| ENSMUSG00000112777.2 | 0 | NA | NA | NA | NA | NA | #VALUE! |
| ENSMUSG00000112778.2 | 0 | NA | NA | NA | NA | NA | #VALUE! |
| ENSMUSG00000112779.2 | 0 | NA | NA | NA | NA | NA | #VALUE! |
| ENSMUSG00000112780.2 | 0 | NA | NA | NA | NA | NA | #VALUE! |
| ENSMUSG00000112781.2 | 0 | NA | NA | NA | NA | NA | #VALUE! |
| ENSMUSG00000112782.2 | 0 | NA | NA | NA | NA | NA | #VALUE! |
| ENSMUSG00000112783.2 | 0 | NA | NA | NA | NA | NA | #VALUE! |
| ENSMUSG00000112784.2 | 0 | NA | NA | NA | NA | NA | #VALUE! |
| ENSMUSG00000112785.2 | 0 | NA | NA | NA | NA | NA | #VALUE! |
| ENSMUSG00000112786.2 | 0 | NA | NA | NA | NA | NA | #VALUE! |
| ENSMUSG00000112787.2 | 0 | NA | NA | NA | NA | NA | #VALUE! |
| ENSMUSG00000112788.2 | 0 | NA | NA | NA | NA | NA | #VALUE! |
| ENSMUSG00000112789.2 | 0.364073 | 0.012025 | 4.059931 | 0.002962 | 0.997637 | NA | #VALUE! |
| ENSMUSG00000112790.2 | 2.139047 | -0.28682 | 3.812691 | -0.07523 | 0.940034 | NA | #VALUE! |
| ENSMUSG00000112791.2 | 0 | NA | NA | NA | NA | NA | #VALUE! |
| ENSMUSG00000112792.2 | 0 | NA | NA | NA | NA | NA | #VALUE! |
| ENSMUSG00000112795.2 | 0 | NA | NA | NA | NA | NA | #VALUE! |
| ENSMUSG00000112797.2 | 0 | NA | NA | NA | NA | NA | #VALUE! |
| ENSMUSG00000112798.2 | 0 | NA | NA | NA | NA | NA | #VALUE! |
| ENSMUSG00000112799.2 | 0 | NA | NA | NA | NA | NA | #VALUE! |
| ENSMUSG00000112800.2 | 0.093255 | 0.868093 | 4.072464 | 0.213162 | 0.831201 | NA | #VALUE! |
| ENSMUSG00000112801.2 | 0 | NA | NA | NA | NA | NA | #VALUE! |
| ENSMUSG00000112802.2 | 0 | NA | NA | NA | NA | NA | #VALUE! |
| ENSMUSG00000112803.2 | 0 | NA | NA | NA | NA | NA | #VALUE! |
| ENSMUSG00000112804.2 | 0 | NA | NA | NA | NA | NA | #VALUE! |
| ENSMUSG00000112805.2 | 0 | NA | NA | NA | NA | NA | #VALUE! |

| | | | | | | | |
|---|---|---|---|---|---|---|---|
| ENSMUSG00000112806.2 | 4.375058 | -5.66277 | 3.916965 | -1.4457 | 0.14826 | NA | #VALUE! |
| ENSMUSG00000112807.2 | 0 | NA | NA | NA | NA | NA | #VALUE! |
| ENSMUSG00000112808.2 | 0 | NA | NA | NA | NA | NA | #VALUE! |
| ENSMUSG00000112809.2 | 0 | NA | NA | NA | NA | NA | #VALUE! |
| ENSMUSG00000112810.2 | 0 | NA | NA | NA | NA | NA | #VALUE! |
| ENSMUSG00000112811.2 | 0 | NA | NA | NA | NA | NA | #VALUE! |
| ENSMUSG00000112812.2 | 0 | NA | NA | NA | NA | NA | #VALUE! |
| ENSMUSG00000112813.2 | 0 | NA | NA | NA | NA | NA | #VALUE! |
| ENSMUSG00000112814.2 | 0 | NA | NA | NA | NA | NA | #VALUE! |
| ENSMUSG00000112815.2 | 0 | NA | NA | NA | NA | NA | #VALUE! |
| ENSMUSG00000112816.2 | 0 | NA | NA | NA | NA | NA | #VALUE! |
| ENSMUSG00000112817.2 | 0 | NA | NA | NA | NA | NA | #VALUE! |
| ENSMUSG00000112818.2 | 0 | NA | NA | NA | NA | NA | #VALUE! |
| ENSMUSG00000112819.2 | 1.346268 | 3.918821 | 3.939925 | 0.994644 | 0.31991 | NA | #VALUE! |
| ENSMUSG00000112820.2 | 0 | NA | NA | NA | NA | NA | #VALUE! |
| ENSMUSG00000112821.2 | 0 | NA | NA | NA | NA | NA | #VALUE! |
| ENSMUSG00000112822.2 | 0 | NA | NA | NA | NA | NA | #VALUE! |
| ENSMUSG00000112823.2 | 0 | NA | NA | NA | NA | NA | #VALUE! |
| ENSMUSG00000112824.2 | 0 | NA | NA | NA | NA | NA | #VALUE! |
| ENSMUSG00000112825.2 | 0.372394 | 0.012025 | 4.059931 | 0.002962 | 0.997637 | NA | #VALUE! |
| ENSMUSG00000112826.2 | 0 | NA | NA | NA | NA | NA | #VALUE! |
| ENSMUSG00000112827.2 | 0 | NA | NA | NA | NA | NA | #VALUE! |
| ENSMUSG00000112828.2 | 0 | NA | NA | NA | NA | NA | #VALUE! |
| ENSMUSG00000112829.2 | 0 | NA | NA | NA | NA | NA | #VALUE! |
| ENSMUSG00000112830.2 | 0 | NA | NA | NA | NA | NA | #VALUE! |
| ENSMUSG00000112831.2 | 0 | NA | NA | NA | NA | NA | #VALUE! |
| ENSMUSG00000112832.2 | 0 | NA | NA | NA | NA | NA | #VALUE! |
| ENSMUSG00000112833.2 | 0 | NA | NA | NA | NA | NA | #VALUE! |
| ENSMUSG00000112834.2 | 0 | NA | NA | NA | NA | NA | #VALUE! |
| ENSMUSG00000112835.2 | 0 | NA | NA | NA | NA | NA | #VALUE! |
| ENSMUSG00000112836.2 | 0 | NA | NA | NA | NA | NA | #VALUE! |
| ENSMUSG00000112837.2 | 0 | NA | NA | NA | NA | NA | #VALUE! |
| ENSMUSG00000112838.2 | 0 | NA | NA | NA | NA | NA | #VALUE! |
| ENSMUSG00000112839.2 | 0 | NA | NA | NA | NA | NA | #VALUE! |
| ENSMUSG00000112840.2 | 0 | NA | NA | NA | NA | NA | #VALUE! |
| ENSMUSG00000112841.2 | 0 | NA | NA | NA | NA | NA | #VALUE! |
| ENSMUSG00000112842.3 | 0 | NA | NA | NA | NA | NA | #VALUE! |
| ENSMUSG00000112843.2 | 2.205276 | -4.68302 | 3.926841 | -1.19257 | 0.233039 | NA | #VALUE! |
| ENSMUSG00000112844.2 | 0 | NA | NA | NA | NA | NA | #VALUE! |
| ENSMUSG00000112845.2 | 0 | NA | NA | NA | NA | NA | #VALUE! |
| ENSMUSG00000112846.2 | 0 | NA | NA | NA | NA | NA | #VALUE! |
| ENSMUSG00000112847.2 | 0 | NA | NA | NA | NA | NA | #VALUE! |
| ENSMUSG00000112848.2 | 0 | NA | NA | NA | NA | NA | #VALUE! |
| ENSMUSG00000112849.2 | 0 | NA | NA | NA | NA | NA | #VALUE! |

| | | | | | | | |
|---|---|---|---|---|---|---|---|
| ENSMUSG00000112850.2 | 0 | NA | NA | NA | NA | NA | #VALUE! |
| ENSMUSG00000112851.2 | 0 | NA | NA | NA | NA | NA | #VALUE! |
| ENSMUSG00000112852.2 | 0 | NA | NA | NA | NA | NA | #VALUE! |
| ENSMUSG00000112853.2 | 0 | NA | NA | NA | NA | NA | #VALUE! |
| ENSMUSG00000112854.2 | 0 | NA | NA | NA | NA | NA | #VALUE! |
| ENSMUSG00000112855.2 | 0 | NA | NA | NA | NA | NA | #VALUE! |
| ENSMUSG00000112856.2 | 0 | NA | NA | NA | NA | NA | #VALUE! |
| ENSMUSG00000112857.2 | 0 | NA | NA | NA | NA | NA | #VALUE! |
| ENSMUSG00000112858.3 | 0 | NA | NA | NA | NA | NA | #VALUE! |
| ENSMUSG00000112859.2 | 0 | NA | NA | NA | NA | NA | #VALUE! |
| ENSMUSG00000112860.2 | 0 | NA | NA | NA | NA | NA | #VALUE! |
| ENSMUSG00000112861.2 | 0 | NA | NA | NA | NA | NA | #VALUE! |
| ENSMUSG00000112862.2 | 0 | NA | NA | NA | NA | NA | #VALUE! |
| ENSMUSG00000112863.2 | 0 | NA | NA | NA | NA | NA | #VALUE! |
| ENSMUSG00000112864.2 | 0 | NA | NA | NA | NA | NA | #VALUE! |
| ENSMUSG00000112865.2 | 0 | NA | NA | NA | NA | NA | #VALUE! |
| ENSMUSG00000112866.2 | 0 | NA | NA | NA | NA | NA | #VALUE! |
| ENSMUSG00000112867.2 | 0 | NA | NA | NA | NA | NA | #VALUE! |
| ENSMUSG00000112868.2 | 0 | NA | NA | NA | NA | NA | #VALUE! |
| ENSMUSG00000112869.2 | 0 | NA | NA | NA | NA | NA | #VALUE! |
| ENSMUSG00000112870.2 | 0 | NA | NA | NA | NA | NA | #VALUE! |
| ENSMUSG00000112871.2 | 0 | NA | NA | NA | NA | NA | #VALUE! |
| ENSMUSG00000112872.2 | 0 | NA | NA | NA | NA | NA | #VALUE! |
| ENSMUSG00000112873.2 | 0 | NA | NA | NA | NA | NA | #VALUE! |
| ENSMUSG00000112874.2 | 0 | NA | NA | NA | NA | NA | #VALUE! |
| ENSMUSG00000112875.2 | 0 | NA | NA | NA | NA | NA | #VALUE! |
| ENSMUSG00000112876.2 | 0 | NA | NA | NA | NA | NA | #VALUE! |
| ENSMUSG00000112877.2 | 0 | NA | NA | NA | NA | NA | #VALUE! |
| ENSMUSG00000112878.2 | 0 | NA | NA | NA | NA | NA | #VALUE! |
| ENSMUSG00000112879.2 | 0 | NA | NA | NA | NA | NA | #VALUE! |
| ENSMUSG00000112880.2 | 4.778587 | -0.69156 | 3.747553 | -0.18454 | 0.853593 | NA | #VALUE! |
| ENSMUSG00000112881.2 | 0 | NA | NA | NA | NA | NA | #VALUE! |
| ENSMUSG00000112882.2 | 0 | NA | NA | NA | NA | NA | #VALUE! |
| ENSMUSG00000112883.2 | 0 | NA | NA | NA | NA | NA | #VALUE! |
| ENSMUSG00000112884.2 | 0 | NA | NA | NA | NA | NA | #VALUE! |
| ENSMUSG00000112885.2 | 0 | NA | NA | NA | NA | NA | #VALUE! |
| ENSMUSG00000112886.2 | 0 | NA | NA | NA | NA | NA | #VALUE! |
| ENSMUSG00000112887.2 | 0 | NA | NA | NA | NA | NA | #VALUE! |
| ENSMUSG00000112888.2 | 0 | NA | NA | NA | NA | NA | #VALUE! |
| ENSMUSG00000112889.2 | 2.238125 | 4.666312 | 3.926586 | 1.188389 | 0.23468 | NA | #VALUE! |
| ENSMUSG00000112890.2 | 0 | NA | NA | NA | NA | NA | #VALUE! |
| ENSMUSG00000112891.2 | 0 | NA | NA | NA | NA | NA | #VALUE! |
| ENSMUSG00000112892.2 | 0 | NA | NA | NA | NA | NA | #VALUE! |
| ENSMUSG00000112893.2 | 0 | NA | NA | NA | NA | NA | #VALUE! |

| | | | | | | | |
|---|---|---|---|---|---|---|---|
| ENSMUSG00000112894.2 | 0 | NA | NA | NA | NA | NA | #VALUE! |
| ENSMUSG00000112895.2 | 3.885486 | -5.49502 | 3.91822 | -1.40243 | 0.160787 | NA | #VALUE! |
| ENSMUSG00000112896.2 | 0 | NA | NA | NA | NA | NA | #VALUE! |
| ENSMUSG00000112897.2 | 0 | NA | NA | NA | NA | NA | #VALUE! |
| ENSMUSG00000112898.2 | 0 | NA | NA | NA | NA | NA | #VALUE! |
| ENSMUSG00000112899.2 | 0 | NA | NA | NA | NA | NA | #VALUE! |
| ENSMUSG00000112900.2 | 0 | NA | NA | NA | NA | NA | #VALUE! |
| ENSMUSG00000112901.2 | 0 | NA | NA | NA | NA | NA | #VALUE! |
| ENSMUSG00000112902.2 | 0 | NA | NA | NA | NA | NA | #VALUE! |
| ENSMUSG00000112903.2 | 0 | NA | NA | NA | NA | NA | #VALUE! |
| ENSMUSG00000112904.2 | 0 | NA | NA | NA | NA | NA | #VALUE! |
| ENSMUSG00000112905.2 | 0 | NA | NA | NA | NA | NA | #VALUE! |
| ENSMUSG00000112906.2 | 0 | NA | NA | NA | NA | NA | #VALUE! |
| ENSMUSG00000112907.2 | 0 | NA | NA | NA | NA | NA | #VALUE! |
| ENSMUSG00000112908.2 | 3.979423 | -1.89831 | 3.2515 | -0.58383 | 0.559338 | NA | #VALUE! |
| ENSMUSG00000112910.2 | 0 | NA | NA | NA | NA | NA | #VALUE! |
| ENSMUSG00000112911.2 | 0 | NA | NA | NA | NA | NA | #VALUE! |
| ENSMUSG00000112912.2 | 0 | NA | NA | NA | NA | NA | #VALUE! |
| ENSMUSG00000112913.2 | 0 | NA | NA | NA | NA | NA | #VALUE! |
| ENSMUSG00000112914.2 | 0 | NA | NA | NA | NA | NA | #VALUE! |
| ENSMUSG00000112915.2 | 0 | NA | NA | NA | NA | NA | #VALUE! |
| ENSMUSG00000112916.2 | 0 | NA | NA | NA | NA | NA | #VALUE! |
| ENSMUSG00000112917.2 | 0 | NA | NA | NA | NA | NA | #VALUE! |
| ENSMUSG00000112918.2 | 0 | NA | NA | NA | NA | NA | #VALUE! |
| ENSMUSG00000112919.2 | 0 | NA | NA | NA | NA | NA | #VALUE! |
| ENSMUSG00000112920.2 | 0 | NA | NA | NA | NA | NA | #VALUE! |
| ENSMUSG00000112921.2 | 0 | NA | NA | NA | NA | NA | #VALUE! |
| ENSMUSG00000112922.2 | 0.559531 | 2.703958 | 3.982863 | 0.678898 | 0.497202 | NA | #VALUE! |
| ENSMUSG00000112923.2 | 0 | NA | NA | NA | NA | NA | #VALUE! |
| ENSMUSG00000112924.2 | 0 | NA | NA | NA | NA | NA | #VALUE! |
| ENSMUSG00000112925.2 | 0 | NA | NA | NA | NA | NA | #VALUE! |
| ENSMUSG00000112926.2 | 0 | NA | NA | NA | NA | NA | #VALUE! |
| ENSMUSG00000112927.2 | 0 | NA | NA | NA | NA | NA | #VALUE! |
| ENSMUSG00000112928.2 | 0 | NA | NA | NA | NA | NA | #VALUE! |
| ENSMUSG00000112929.2 | 0 | NA | NA | NA | NA | NA | #VALUE! |
| ENSMUSG00000112930.3 | 0 | NA | NA | NA | NA | NA | #VALUE! |
| ENSMUSG00000112931.2 | 0 | NA | NA | NA | NA | NA | #VALUE! |
| ENSMUSG00000112932.2 | 0 | NA | NA | NA | NA | NA | #VALUE! |
| ENSMUSG00000112933.2 | 0 | NA | NA | NA | NA | NA | #VALUE! |
| ENSMUSG00000112934.2 | 0 | NA | NA | NA | NA | NA | #VALUE! |
| ENSMUSG00000112935.2 | 4.537092 | -3.62186 | 3.864695 | -0.93717 | 0.348673 | NA | #VALUE! |
| ENSMUSG00000112936.2 | 0 | NA | NA | NA | NA | NA | #VALUE! |
| ENSMUSG00000112937.2 | 0 | NA | NA | NA | NA | NA | #VALUE! |
| ENSMUSG00000112938.2 | 0 | NA | NA | NA | NA | NA | #VALUE! |

| | | | | | | | |
|---|---|---|---|---|---|---|---|
| ENSMUSG00000112939.2 | 0 | NA | NA | NA | NA | NA | #VALUE! |
| ENSMUSG00000112940.2 | 0 | NA | NA | NA | NA | NA | #VALUE! |
| ENSMUSG00000112941.2 | 0 | NA | NA | NA | NA | NA | #VALUE! |
| ENSMUSG00000112942.2 | 0 | NA | NA | NA | NA | NA | #VALUE! |
| ENSMUSG00000112943.2 | 0 | NA | NA | NA | NA | NA | #VALUE! |
| ENSMUSG00000112944.2 | 0 | NA | NA | NA | NA | NA | #VALUE! |
| ENSMUSG00000112945.2 | 0 | NA | NA | NA | NA | NA | #VALUE! |
| ENSMUSG00000112946.2 | 0 | NA | NA | NA | NA | NA | #VALUE! |
| ENSMUSG00000112947.2 | 0 | NA | NA | NA | NA | NA | #VALUE! |
| ENSMUSG00000112948.2 | 0 | NA | NA | NA | NA | NA | #VALUE! |
| ENSMUSG00000112949.2 | 0 | NA | NA | NA | NA | NA | #VALUE! |
| ENSMUSG00000112950.2 | 0 | NA | NA | NA | NA | NA | #VALUE! |
| ENSMUSG00000112951.3 | 0 | NA | NA | NA | NA | NA | #VALUE! |
| ENSMUSG00000112952.2 | 0 | NA | NA | NA | NA | NA | #VALUE! |
| ENSMUSG00000112953.2 | 0 | NA | NA | NA | NA | NA | #VALUE! |
| ENSMUSG00000112954.2 | 0 | NA | NA | NA | NA | NA | #VALUE! |
| ENSMUSG00000112955.2 | 0 | NA | NA | NA | NA | NA | #VALUE! |
| ENSMUSG00000112956.2 | 0 | NA | NA | NA | NA | NA | #VALUE! |
| ENSMUSG00000112957.2 | 0 | NA | NA | NA | NA | NA | #VALUE! |
| ENSMUSG00000112958.2 | 0 | NA | NA | NA | NA | NA | #VALUE! |
| ENSMUSG00000112959.2 | 0 | NA | NA | NA | NA | NA | #VALUE! |
| ENSMUSG00000112960.2 | 0 | NA | NA | NA | NA | NA | #VALUE! |
| ENSMUSG00000112961.2 | 0 | NA | NA | NA | NA | NA | #VALUE! |
| ENSMUSG00000112962.2 | 0 | NA | NA | NA | NA | NA | #VALUE! |
| ENSMUSG00000112963.2 | 0 | NA | NA | NA | NA | NA | #VALUE! |
| ENSMUSG00000112964.2 | 1.518248 | -4.13698 | 3.936022 | -1.05106 | 0.293233 | NA | #VALUE! |
| ENSMUSG00000112965.2 | 0 | NA | NA | NA | NA | NA | #VALUE! |
| ENSMUSG00000112966.2 | 0 | NA | NA | NA | NA | NA | #VALUE! |
| ENSMUSG00000112967.2 | 0 | NA | NA | NA | NA | NA | #VALUE! |
| ENSMUSG00000112968.2 | 0 | NA | NA | NA | NA | NA | #VALUE! |
| ENSMUSG00000112969.2 | 0 | NA | NA | NA | NA | NA | #VALUE! |
| ENSMUSG00000112970.2 | 0 | NA | NA | NA | NA | NA | #VALUE! |
| ENSMUSG00000112971.2 | 0 | NA | NA | NA | NA | NA | #VALUE! |
| ENSMUSG00000112972.2 | 0 | NA | NA | NA | NA | NA | #VALUE! |
| ENSMUSG00000112973.2 | 0 | NA | NA | NA | NA | NA | #VALUE! |
| ENSMUSG00000112974.2 | 0 | NA | NA | NA | NA | NA | #VALUE! |
| ENSMUSG00000112975.2 | 0 | NA | NA | NA | NA | NA | #VALUE! |
| ENSMUSG00000112976.2 | 0 | NA | NA | NA | NA | NA | #VALUE! |
| ENSMUSG00000112977.2 | 0 | NA | NA | NA | NA | NA | #VALUE! |
| ENSMUSG00000112978.2 | 0 | NA | NA | NA | NA | NA | #VALUE! |
| ENSMUSG00000112979.2 | 0 | NA | NA | NA | NA | NA | #VALUE! |
| ENSMUSG00000112980.2 | 0 | NA | NA | NA | NA | NA | #VALUE! |
| ENSMUSG00000112981.2 | 0 | NA | NA | NA | NA | NA | #VALUE! |
| ENSMUSG00000112982.2 | 0 | NA | NA | NA | NA | NA | #VALUE! |

| | | | | | | | |
|---|---|---|---|---|---|---|---|
| ENSMUSG00000112983.2 | 0 | NA | NA | NA | NA | NA | #VALUE! |
| ENSMUSG00000112984.2 | 0 | NA | NA | NA | NA | NA | #VALUE! |
| ENSMUSG00000112985.2 | 0 | NA | NA | NA | NA | NA | #VALUE! |
| ENSMUSG00000112986.2 | 0 | NA | NA | NA | NA | NA | #VALUE! |
| ENSMUSG00000112987.2 | 0 | NA | NA | NA | NA | NA | #VALUE! |
| ENSMUSG00000112988.2 | 0 | NA | NA | NA | NA | NA | #VALUE! |
| ENSMUSG00000112989.2 | 0 | NA | NA | NA | NA | NA | #VALUE! |
| ENSMUSG00000112990.2 | 0 | NA | NA | NA | NA | NA | #VALUE! |
| ENSMUSG00000112991.2 | 0 | NA | NA | NA | NA | NA | #VALUE! |
| ENSMUSG00000112992.2 | 0 | NA | NA | NA | NA | NA | #VALUE! |
| ENSMUSG00000112993.2 | 0 | NA | NA | NA | NA | NA | #VALUE! |
| ENSMUSG00000112994.2 | 0 | NA | NA | NA | NA | NA | #VALUE! |
| ENSMUSG00000112995.2 | 0 | NA | NA | NA | NA | NA | #VALUE! |
| ENSMUSG00000112996.2 | 0 | NA | NA | NA | NA | NA | #VALUE! |
| ENSMUSG00000112997.2 | 0 | NA | NA | NA | NA | NA | #VALUE! |
| ENSMUSG00000112998.2 | 0 | NA | NA | NA | NA | NA | #VALUE! |
| ENSMUSG00000112999.2 | 0 | NA | NA | NA | NA | NA | #VALUE! |
| ENSMUSG00000113000.2 | 0 | NA | NA | NA | NA | NA | #VALUE! |
| ENSMUSG00000113001.2 | 0 | NA | NA | NA | NA | NA | #VALUE! |
| ENSMUSG00000113002.2 | 0 | NA | NA | NA | NA | NA | #VALUE! |
| ENSMUSG00000113003.2 | 1.682835 | 4.240781 | 3.933339 | 1.078163 | 0.280961 | NA | #VALUE! |
| ENSMUSG00000113004.2 | 0 | NA | NA | NA | NA | NA | #VALUE! |
| ENSMUSG00000113005.2 | 0 | NA | NA | NA | NA | NA | #VALUE! |
| ENSMUSG00000113006.2 | 0 | NA | NA | NA | NA | NA | #VALUE! |
| ENSMUSG00000113007.2 | 0 | NA | NA | NA | NA | NA | #VALUE! |
| ENSMUSG00000113008.2 | 0 | NA | NA | NA | NA | NA | #VALUE! |
| ENSMUSG00000113009.2 | 0 | NA | NA | NA | NA | NA | #VALUE! |
| ENSMUSG00000113010.2 | 0 | NA | NA | NA | NA | NA | #VALUE! |
| ENSMUSG00000113011.2 | 0 | NA | NA | NA | NA | NA | #VALUE! |
| ENSMUSG00000113012.2 | 0 | NA | NA | NA | NA | NA | #VALUE! |
| ENSMUSG00000113013.2 | 0 | NA | NA | NA | NA | NA | #VALUE! |
| ENSMUSG00000113014.2 | 0 | NA | NA | NA | NA | NA | #VALUE! |
| ENSMUSG00000113015.2 | 0 | NA | NA | NA | NA | NA | #VALUE! |
| ENSMUSG00000113016.2 | 0 | NA | NA | NA | NA | NA | #VALUE! |
| ENSMUSG00000113017.2 | 0 | NA | NA | NA | NA | NA | #VALUE! |
| ENSMUSG00000113018.2 | 0 | NA | NA | NA | NA | NA | #VALUE! |
| ENSMUSG00000113019.2 | 7.770972 | -6.49162 | 3.884908 | -1.67099 | NA | NA | #VALUE! |
| ENSMUSG00000113020.2 | 0 | NA | NA | NA | NA | NA | #VALUE! |
| ENSMUSG00000113021.2 | 0 | NA | NA | NA | NA | NA | #VALUE! |
| ENSMUSG00000113022.2 | 0 | NA | NA | NA | NA | NA | #VALUE! |
| ENSMUSG00000113023.2 | 0 | NA | NA | NA | NA | NA | #VALUE! |
| ENSMUSG00000113024.2 | 0 | NA | NA | NA | NA | NA | #VALUE! |
| ENSMUSG00000113025.2 | 0 | NA | NA | NA | NA | NA | #VALUE! |
| ENSMUSG00000113026.2 | 0 | NA | NA | NA | NA | NA | #VALUE! |

| | | | | | | | |
|---|---|---|---|---|---|---|---|
| ENSMUSG00000113027.2 | 0 | NA | NA | NA | NA | NA | #VALUE! |
| ENSMUSG00000113028.2 | 0 | NA | NA | NA | NA | NA | #VALUE! |
| ENSMUSG00000113029.2 | 0 | NA | NA | NA | NA | NA | #VALUE! |
| ENSMUSG00000113030.2 | 0 | NA | NA | NA | NA | NA | #VALUE! |
| ENSMUSG00000113031.2 | 1.836997 | -4.39907 | 3.931186 | -1.11902 | 0.263133 | NA | #VALUE! |
| ENSMUSG00000113032.2 | 0 | NA | NA | NA | NA | NA | #VALUE! |
| ENSMUSG00000113033.2 | 0 | NA | NA | NA | NA | NA | #VALUE! |
| ENSMUSG00000113034.2 | 0 | NA | NA | NA | NA | NA | #VALUE! |
| ENSMUSG00000113035.2 | 0 | NA | NA | NA | NA | NA | #VALUE! |
| ENSMUSG00000113036.2 | 0 | NA | NA | NA | NA | NA | #VALUE! |
| ENSMUSG00000113037.2 | 0 | NA | NA | NA | NA | NA | #VALUE! |
| ENSMUSG00000113038.2 | 0 | NA | NA | NA | NA | NA | #VALUE! |
| ENSMUSG00000113039.2 | 0 | NA | NA | NA | NA | NA | #VALUE! |
| ENSMUSG00000113040.2 | 0 | NA | NA | NA | NA | NA | #VALUE! |
| ENSMUSG00000113041.2 | 0 | NA | NA | NA | NA | NA | #VALUE! |
| ENSMUSG00000113042.2 | 0 | NA | NA | NA | NA | NA | #VALUE! |
| ENSMUSG00000113043.2 | 0 | NA | NA | NA | NA | NA | #VALUE! |
| ENSMUSG00000113044.2 | 0 | NA | NA | NA | NA | NA | #VALUE! |
| ENSMUSG00000113045.2 | 0 | NA | NA | NA | NA | NA | #VALUE! |
| ENSMUSG00000113046.2 | 0 | NA | NA | NA | NA | NA | #VALUE! |
| ENSMUSG00000113047.2 | 0 | NA | NA | NA | NA | NA | #VALUE! |
| ENSMUSG00000113048.2 | 0 | NA | NA | NA | NA | NA | #VALUE! |
| ENSMUSG00000113049.2 | 0 | NA | NA | NA | NA | NA | #VALUE! |
| ENSMUSG00000113050.2 | 0 | NA | NA | NA | NA | NA | #VALUE! |
| ENSMUSG00000113051.2 | 0 | NA | NA | NA | NA | NA | #VALUE! |
| ENSMUSG00000113052.2 | 0.391578 | -2.1489 | 4.020883 | -0.53444 | 0.59304 | NA | #VALUE! |
| ENSMUSG00000113053.2 | 0 | NA | NA | NA | NA | NA | #VALUE! |
| ENSMUSG00000113054.2 | 0 | NA | NA | NA | NA | NA | #VALUE! |
| ENSMUSG00000113055.2 | 0 | NA | NA | NA | NA | NA | #VALUE! |
| ENSMUSG00000113056.2 | 0 | NA | NA | NA | NA | NA | #VALUE! |
| ENSMUSG00000113057.2 | 0 | NA | NA | NA | NA | NA | #VALUE! |
| ENSMUSG00000113058.2 | 0 | NA | NA | NA | NA | NA | #VALUE! |
| ENSMUSG00000113059.2 | 0 | NA | NA | NA | NA | NA | #VALUE! |
| ENSMUSG00000113060.2 | 0 | NA | NA | NA | NA | NA | #VALUE! |
| ENSMUSG00000113062.2 | 0 | NA | NA | NA | NA | NA | #VALUE! |
| ENSMUSG00000113064.2 | 0 | NA | NA | NA | NA | NA | #VALUE! |
| ENSMUSG00000113065.2 | 0 | NA | NA | NA | NA | NA | #VALUE! |
| ENSMUSG00000113066.2 | 0 | NA | NA | NA | NA | NA | #VALUE! |
| ENSMUSG00000113067.2 | 0 | NA | NA | NA | NA | NA | #VALUE! |
| ENSMUSG00000113068.2 | 0 | NA | NA | NA | NA | NA | #VALUE! |
| ENSMUSG00000113069.2 | 0 | NA | NA | NA | NA | NA | #VALUE! |
| ENSMUSG00000113070.2 | 1.575197 | -4.20222 | 3.934736 | -1.06798 | 0.285529 | NA | #VALUE! |
| ENSMUSG00000113071.2 | 0 | NA | NA | NA | NA | NA | #VALUE! |
| ENSMUSG00000113072.2 | 0 | NA | NA | NA | NA | NA | #VALUE! |

| | | | | | | | |
|---|---|---|---|---|---|---|---|
| ENSMUSG00000113073.2 | 0 | NA | NA | NA | NA | NA | #VALUE! |
| ENSMUSG00000113074.2 | 0 | NA | NA | NA | NA | NA | #VALUE! |
| ENSMUSG00000113075.2 | 0 | NA | NA | NA | NA | NA | #VALUE! |
| ENSMUSG00000113076.2 | 0 | NA | NA | NA | NA | NA | #VALUE! |
| ENSMUSG00000113077.2 | 0 | NA | NA | NA | NA | NA | #VALUE! |
| ENSMUSG00000113078.2 | 0 | NA | NA | NA | NA | NA | #VALUE! |
| ENSMUSG00000113079.2 | 0 | NA | NA | NA | NA | NA | #VALUE! |
| ENSMUSG00000113080.2 | 0 | NA | NA | NA | NA | NA | #VALUE! |
| ENSMUSG00000113081.2 | 0 | NA | NA | NA | NA | NA | #VALUE! |
| ENSMUSG00000113082.2 | 0 | NA | NA | NA | NA | NA | #VALUE! |
| ENSMUSG00000113083.2 | 0 | NA | NA | NA | NA | NA | #VALUE! |
| ENSMUSG00000113084.2 | 0 | NA | NA | NA | NA | NA | #VALUE! |
| ENSMUSG00000113085.2 | 0 | NA | NA | NA | NA | NA | #VALUE! |
| ENSMUSG00000113086.2 | 0 | NA | NA | NA | NA | NA | #VALUE! |
| ENSMUSG00000113087.2 | 0 | NA | NA | NA | NA | NA | #VALUE! |
| ENSMUSG00000113088.2 | 0 | NA | NA | NA | NA | NA | #VALUE! |
| ENSMUSG00000113089.2 | 0 | NA | NA | NA | NA | NA | #VALUE! |
| ENSMUSG00000113090.2 | 0 | NA | NA | NA | NA | NA | #VALUE! |
| ENSMUSG00000113091.2 | 0 | NA | NA | NA | NA | NA | #VALUE! |
| ENSMUSG00000113092.2 | 0 | NA | NA | NA | NA | NA | #VALUE! |
| ENSMUSG00000113093.2 | 0 | NA | NA | NA | NA | NA | #VALUE! |
| ENSMUSG00000113094.2 | 0 | NA | NA | NA | NA | NA | #VALUE! |
| ENSMUSG00000113095.2 | 0 | NA | NA | NA | NA | NA | #VALUE! |
| ENSMUSG00000113096.2 | 0 | NA | NA | NA | NA | NA | #VALUE! |
| ENSMUSG00000113097.2 | 0 | NA | NA | NA | NA | NA | #VALUE! |
| ENSMUSG00000113098.2 | 0 | NA | NA | NA | NA | NA | #VALUE! |
| ENSMUSG00000113099.2 | 0 | NA | NA | NA | NA | NA | #VALUE! |
| ENSMUSG00000113100.2 | 0 | NA | NA | NA | NA | NA | #VALUE! |
| ENSMUSG00000113101.2 | 0 | NA | NA | NA | NA | NA | #VALUE! |
| ENSMUSG00000113102.2 | 0 | NA | NA | NA | NA | NA | #VALUE! |
| ENSMUSG00000113103.2 | 0 | NA | NA | NA | NA | NA | #VALUE! |
| ENSMUSG00000113104.2 | 0 | NA | NA | NA | NA | NA | #VALUE! |
| ENSMUSG00000113105.2 | 0 | NA | NA | NA | NA | NA | #VALUE! |
| ENSMUSG00000113106.2 | 0 | NA | NA | NA | NA | NA | #VALUE! |
| ENSMUSG00000113107.2 | 0 | NA | NA | NA | NA | NA | #VALUE! |
| ENSMUSG00000113108.2 | 0 | NA | NA | NA | NA | NA | #VALUE! |
| ENSMUSG00000113109.2 | 0 | NA | NA | NA | NA | NA | #VALUE! |
| ENSMUSG00000113110.2 | 0 | NA | NA | NA | NA | NA | #VALUE! |
| ENSMUSG00000113111.2 | 0 | NA | NA | NA | NA | NA | #VALUE! |
| ENSMUSG00000113112.2 | 0 | NA | NA | NA | NA | NA | #VALUE! |
| ENSMUSG00000113113.2 | 1.739873 | 0.537277 | 3.834779 | 0.140106 | 0.888576 | NA | #VALUE! |
| ENSMUSG00000113114.2 | 0 | NA | NA | NA | NA | NA | #VALUE! |
| ENSMUSG00000113115.2 | 0 | NA | NA | NA | NA | NA | #VALUE! |
| ENSMUSG00000113116.2 | 0 | NA | NA | NA | NA | NA | #VALUE! |

| | | | | | | | |
|---|---|---|---|---|---|---|---|
| ENSMUSG00000113117.2 | 0 | NA | NA | NA | NA | NA | #VALUE! |
| ENSMUSG00000113118.2 | 0 | NA | NA | NA | NA | NA | #VALUE! |
| ENSMUSG00000113119.2 | 0 | NA | NA | NA | NA | NA | #VALUE! |
| ENSMUSG00000113120.2 | 0 | NA | NA | NA | NA | NA | #VALUE! |
| ENSMUSG00000113121.2 | 0 | NA | NA | NA | NA | NA | #VALUE! |
| ENSMUSG00000113122.2 | 0 | NA | NA | NA | NA | NA | #VALUE! |
| ENSMUSG00000113123.3 | 0 | NA | NA | NA | NA | NA | #VALUE! |
| ENSMUSG00000113124.2 | 0 | NA | NA | NA | NA | NA | #VALUE! |
| ENSMUSG00000113125.2 | 0 | NA | NA | NA | NA | NA | #VALUE! |
| ENSMUSG00000113126.2 | 0 | NA | NA | NA | NA | NA | #VALUE! |
| ENSMUSG00000113127.2 | 0 | NA | NA | NA | NA | NA | #VALUE! |
| ENSMUSG00000113128.2 | 0 | NA | NA | NA | NA | NA | #VALUE! |
| ENSMUSG00000113129.2 | 0 | NA | NA | NA | NA | NA | #VALUE! |
| ENSMUSG00000113130.2 | 0 | NA | NA | NA | NA | NA | #VALUE! |
| ENSMUSG00000113131.2 | 0 | NA | NA | NA | NA | NA | #VALUE! |
| ENSMUSG00000113132.2 | 0 | NA | NA | NA | NA | NA | #VALUE! |
| ENSMUSG00000113133.2 | 0 | NA | NA | NA | NA | NA | #VALUE! |
| ENSMUSG00000113134.3 | 0 | NA | NA | NA | NA | NA | #VALUE! |
| ENSMUSG00000113135.2 | 0 | NA | NA | NA | NA | NA | #VALUE! |
| ENSMUSG00000113136.2 | 0 | NA | NA | NA | NA | NA | #VALUE! |
| ENSMUSG00000113137.2 | 0 | NA | NA | NA | NA | NA | #VALUE! |
| ENSMUSG00000113138.2 | 0 | NA | NA | NA | NA | NA | #VALUE! |
| ENSMUSG00000113139.2 | 0 | NA | NA | NA | NA | NA | #VALUE! |
| ENSMUSG00000113140.2 | 0 | NA | NA | NA | NA | NA | #VALUE! |
| ENSMUSG00000113141.2 | 0 | NA | NA | NA | NA | NA | #VALUE! |
| ENSMUSG00000113142.2 | 0 | NA | NA | NA | NA | NA | #VALUE! |
| ENSMUSG00000113143.2 | 0 | NA | NA | NA | NA | NA | #VALUE! |
| ENSMUSG00000113144.2 | 0 | NA | NA | NA | NA | NA | #VALUE! |
| ENSMUSG00000113145.2 | 0 | NA | NA | NA | NA | NA | #VALUE! |
| ENSMUSG00000113146.2 | 0 | NA | NA | NA | NA | NA | #VALUE! |
| ENSMUSG00000113147.2 | 0 | NA | NA | NA | NA | NA | #VALUE! |
| ENSMUSG00000113148.2 | 0 | NA | NA | NA | NA | NA | #VALUE! |
| ENSMUSG00000113149.2 | 1.155144 | -3.7605 | 3.944679 | -0.95331 | 0.340433 | NA | #VALUE! |
| ENSMUSG00000113150.2 | 0 | NA | NA | NA | NA | NA | #VALUE! |
| ENSMUSG00000113151.2 | 0 | NA | NA | NA | NA | NA | #VALUE! |
| ENSMUSG00000113152.2 | 0 | NA | NA | NA | NA | NA | #VALUE! |
| ENSMUSG00000113153.2 | 0 | NA | NA | NA | NA | NA | #VALUE! |
| ENSMUSG00000113154.2 | 0 | NA | NA | NA | NA | NA | #VALUE! |
| ENSMUSG00000113155.2 | 0 | NA | NA | NA | NA | NA | #VALUE! |
| ENSMUSG00000113156.2 | 0 | NA | NA | NA | NA | NA | #VALUE! |
| ENSMUSG00000113157.3 | 0 | NA | NA | NA | NA | NA | #VALUE! |
| ENSMUSG00000113158.2 | 0 | NA | NA | NA | NA | NA | #VALUE! |
| ENSMUSG00000113159.2 | 1.32413 | 1.312987 | 3.891861 | 0.337367 | 0.73584 | NA | #VALUE! |
| ENSMUSG00000113160.2 | 0 | NA | NA | NA | NA | NA | #VALUE! |

| | | | | | | | |
|---|---|---|---|---|---|---|---|
| ENSMUSG00000113161.2 | 0 | NA | NA | NA | NA | NA | #VALUE! |
| ENSMUSG00000113162.2 | 0 | NA | NA | NA | NA | NA | #VALUE! |
| ENSMUSG00000113163.2 | 0 | NA | NA | NA | NA | NA | #VALUE! |
| ENSMUSG00000113164.2 | 0 | NA | NA | NA | NA | NA | #VALUE! |
| ENSMUSG00000113165.2 | 0 | NA | NA | NA | NA | NA | #VALUE! |
| ENSMUSG00000113166.2 | 0 | NA | NA | NA | NA | NA | #VALUE! |
| ENSMUSG00000113167.2 | 0 | NA | NA | NA | NA | NA | #VALUE! |
| ENSMUSG00000113168.2 | 0 | NA | NA | NA | NA | NA | #VALUE! |
| ENSMUSG00000113169.2 | 0 | NA | NA | NA | NA | NA | #VALUE! |
| ENSMUSG00000113170.2 | 0 | NA | NA | NA | NA | NA | #VALUE! |
| ENSMUSG00000113171.2 | 0 | NA | NA | NA | NA | NA | #VALUE! |
| ENSMUSG00000113172.2 | 0 | NA | NA | NA | NA | NA | #VALUE! |
| ENSMUSG00000113173.2 | 0 | NA | NA | NA | NA | NA | #VALUE! |
| ENSMUSG00000113174.2 | 0 | NA | NA | NA | NA | NA | #VALUE! |
| ENSMUSG00000113175.2 | 0 | NA | NA | NA | NA | NA | #VALUE! |
| ENSMUSG00000113177.2 | 0 | NA | NA | NA | NA | NA | #VALUE! |
| ENSMUSG00000113178.2 | 0 | NA | NA | NA | NA | NA | #VALUE! |
| ENSMUSG00000113179.2 | 0 | NA | NA | NA | NA | NA | #VALUE! |
| ENSMUSG00000113180.2 | 0 | NA | NA | NA | NA | NA | #VALUE! |
| ENSMUSG00000113181.2 | 0 | NA | NA | NA | NA | NA | #VALUE! |
| ENSMUSG00000113182.2 | 0 | NA | NA | NA | NA | NA | #VALUE! |
| ENSMUSG00000113183.2 | 0 | NA | NA | NA | NA | NA | #VALUE! |
| ENSMUSG00000113184.2 | 0 | NA | NA | NA | NA | NA | #VALUE! |
| ENSMUSG00000113185.2 | 0.391578 | -2.1489 | 4.020883 | -0.53444 | 0.59304 | NA | #VALUE! |
| ENSMUSG00000113186.2 | 0 | NA | NA | NA | NA | NA | #VALUE! |
| ENSMUSG00000113187.2 | 0 | NA | NA | NA | NA | NA | #VALUE! |
| ENSMUSG00000113188.2 | 0 | NA | NA | NA | NA | NA | #VALUE! |
| ENSMUSG00000113189.2 | 0 | NA | NA | NA | NA | NA | #VALUE! |
| ENSMUSG00000113190.2 | 0 | NA | NA | NA | NA | NA | #VALUE! |
| ENSMUSG00000113191.2 | 0 | NA | NA | NA | NA | NA | #VALUE! |
| ENSMUSG00000113192.2 | 0 | NA | NA | NA | NA | NA | #VALUE! |
| ENSMUSG00000113193.2 | 0 | NA | NA | NA | NA | NA | #VALUE! |
| ENSMUSG00000113194.2 | 0 | NA | NA | NA | NA | NA | #VALUE! |
| ENSMUSG00000113195.2 | 0 | NA | NA | NA | NA | NA | #VALUE! |
| ENSMUSG00000113196.2 | 0 | NA | NA | NA | NA | NA | #VALUE! |
| ENSMUSG00000113197.2 | 0 | NA | NA | NA | NA | NA | #VALUE! |
| ENSMUSG00000113199.2 | 0 | NA | NA | NA | NA | NA | #VALUE! |
| ENSMUSG00000113200.2 | 0 | NA | NA | NA | NA | NA | #VALUE! |
| ENSMUSG00000113201.2 | 0 | NA | NA | NA | NA | NA | #VALUE! |
| ENSMUSG00000113202.2 | 0 | NA | NA | NA | NA | NA | #VALUE! |
| ENSMUSG00000113203.2 | 0 | NA | NA | NA | NA | NA | #VALUE! |
| ENSMUSG00000113205.2 | 0 | NA | NA | NA | NA | NA | #VALUE! |
| ENSMUSG00000113206.2 | 0 | NA | NA | NA | NA | NA | #VALUE! |
| ENSMUSG00000113207.2 | 0 | NA | NA | NA | NA | NA | #VALUE! |

| | | | | | | | |
|---|---|---|---|---|---|---|---|
| ENSMUSG00000113208.2 | 0 | NA | NA | NA | NA | NA | #VALUE! |
| ENSMUSG00000113209.2 | 0 | NA | NA | NA | NA | NA | #VALUE! |
| ENSMUSG00000113210.2 | 0 | NA | NA | NA | NA | NA | #VALUE! |
| ENSMUSG00000113211.2 | 0 | NA | NA | NA | NA | NA | #VALUE! |
| ENSMUSG00000113212.2 | 0 | NA | NA | NA | NA | NA | #VALUE! |
| ENSMUSG00000113213.2 | 0 | NA | NA | NA | NA | NA | #VALUE! |
| ENSMUSG00000113214.2 | 0 | NA | NA | NA | NA | NA | #VALUE! |
| ENSMUSG00000113215.2 | 0 | NA | NA | NA | NA | NA | #VALUE! |
| ENSMUSG00000113216.2 | 0 | NA | NA | NA | NA | NA | #VALUE! |
| ENSMUSG00000113217.2 | 0 | NA | NA | NA | NA | NA | #VALUE! |
| ENSMUSG00000113218.2 | 0 | NA | NA | NA | NA | NA | #VALUE! |
| ENSMUSG00000113219.2 | 0 | NA | NA | NA | NA | NA | #VALUE! |
| ENSMUSG00000113220.2 | 0 | NA | NA | NA | NA | NA | #VALUE! |
| ENSMUSG00000113221.2 | 0 | NA | NA | NA | NA | NA | #VALUE! |
| ENSMUSG00000113222.2 | 0.168284 | 1.027628 | 4.067126 | 0.252667 | 0.800526 | NA | #VALUE! |
| ENSMUSG00000113223.2 | 0 | NA | NA | NA | NA | NA | #VALUE! |
| ENSMUSG00000113224.2 | 0 | NA | NA | NA | NA | NA | #VALUE! |
| ENSMUSG00000113225.2 | 0 | NA | NA | NA | NA | NA | #VALUE! |
| ENSMUSG00000113226.2 | 0 | NA | NA | NA | NA | NA | #VALUE! |
| ENSMUSG00000113227.2 | 0 | NA | NA | NA | NA | NA | #VALUE! |
| ENSMUSG00000113228.2 | 0 | NA | NA | NA | NA | NA | #VALUE! |
| ENSMUSG00000113229.2 | 0 | NA | NA | NA | NA | NA | #VALUE! |
| ENSMUSG00000113230.2 | 0 | NA | NA | NA | NA | NA | #VALUE! |
| ENSMUSG00000113231.2 | 0 | NA | NA | NA | NA | NA | #VALUE! |
| ENSMUSG00000113232.2 | 0 | NA | NA | NA | NA | NA | #VALUE! |
| ENSMUSG00000113233.2 | 0 | NA | NA | NA | NA | NA | #VALUE! |
| ENSMUSG00000113234.2 | 0 | NA | NA | NA | NA | NA | #VALUE! |
| ENSMUSG00000113235.2 | 0 | NA | NA | NA | NA | NA | #VALUE! |
| ENSMUSG00000113236.2 | 0 | NA | NA | NA | NA | NA | #VALUE! |
| ENSMUSG00000113237.2 | 3.581349 | 1.038448 | 3.320148 | 0.312772 | 0.754454 | NA | #VALUE! |
| ENSMUSG00000113238.2 | 0 | NA | NA | NA | NA | NA | #VALUE! |
| ENSMUSG00000113239.2 | 0 | NA | NA | NA | NA | NA | #VALUE! |
| ENSMUSG00000113240.2 | 0 | NA | NA | NA | NA | NA | #VALUE! |
| ENSMUSG00000113241.2 | 0 | NA | NA | NA | NA | NA | #VALUE! |
| ENSMUSG00000113242.2 | 0 | NA | NA | NA | NA | NA | #VALUE! |
| ENSMUSG00000113243.2 | 0 | NA | NA | NA | NA | NA | #VALUE! |
| ENSMUSG00000113244.2 | 0 | NA | NA | NA | NA | NA | #VALUE! |
| ENSMUSG00000113245.2 | 0 | NA | NA | NA | NA | NA | #VALUE! |
| ENSMUSG00000113246.2 | 0 | NA | NA | NA | NA | NA | #VALUE! |
| ENSMUSG00000113247.2 | 0.204111 | -1.05581 | 4.073302 | -0.2592 | 0.795479 | NA | #VALUE! |
| ENSMUSG00000113248.2 | 0 | NA | NA | NA | NA | NA | #VALUE! |
| ENSMUSG00000113250.2 | 0 | NA | NA | NA | NA | NA | #VALUE! |
| ENSMUSG00000113251.2 | 0 | NA | NA | NA | NA | NA | #VALUE! |
| ENSMUSG00000113252.2 | 0 | NA | NA | NA | NA | NA | #VALUE! |

| | | | | | | | |
|---|---|---|---|---|---|---|---|
| ENSMUSG00000113253.2 | 0 | NA | NA | NA | NA | NA | #VALUE! |
| ENSMUSG00000113254.2 | 0 | NA | NA | NA | NA | NA | #VALUE! |
| ENSMUSG00000113255.2 | 3.111401 | -1.43751 | 3.800534 | -0.37824 | 0.705254 | NA | #VALUE! |
| ENSMUSG00000113256.2 | 0 | NA | NA | NA | NA | NA | #VALUE! |
| ENSMUSG00000113257.2 | 0 | NA | NA | NA | NA | NA | #VALUE! |
| ENSMUSG00000113258.2 | 0 | NA | NA | NA | NA | NA | #VALUE! |
| ENSMUSG00000113259.2 | 0 | NA | NA | NA | NA | NA | #VALUE! |
| ENSMUSG00000113260.2 | 0 | NA | NA | NA | NA | NA | #VALUE! |
| ENSMUSG00000113261.2 | 0 | NA | NA | NA | NA | NA | #VALUE! |
| ENSMUSG00000113262.2 | 0 | NA | NA | NA | NA | NA | #VALUE! |
| ENSMUSG00000113263.2 | 0.168284 | 1.027628 | 4.067126 | 0.252667 | 0.800526 | NA | #VALUE! |
| ENSMUSG00000113264.2 | 0 | NA | NA | NA | NA | NA | #VALUE! |
| ENSMUSG00000113265.2 | 0 | NA | NA | NA | NA | NA | #VALUE! |
| ENSMUSG00000113266.2 | 0 | NA | NA | NA | NA | NA | #VALUE! |
| ENSMUSG00000113267.2 | 0 | NA | NA | NA | NA | NA | #VALUE! |
| ENSMUSG00000113268.2 | 0 | NA | NA | NA | NA | NA | #VALUE! |
| ENSMUSG00000113269.2 | 0 | NA | NA | NA | NA | NA | #VALUE! |
| ENSMUSG00000113270.2 | 0 | NA | NA | NA | NA | NA | #VALUE! |
| ENSMUSG00000113271.2 | 0 | NA | NA | NA | NA | NA | #VALUE! |
| ENSMUSG00000113272.2 | 0 | NA | NA | NA | NA | NA | #VALUE! |
| ENSMUSG00000113273.2 | 0 | NA | NA | NA | NA | NA | #VALUE! |
| ENSMUSG00000113274.2 | 0 | NA | NA | NA | NA | NA | #VALUE! |
| ENSMUSG00000113275.2 | 0 | NA | NA | NA | NA | NA | #VALUE! |
| ENSMUSG00000113276.2 | 0 | NA | NA | NA | NA | NA | #VALUE! |
| ENSMUSG00000113277.2 | 0 | NA | NA | NA | NA | NA | #VALUE! |
| ENSMUSG00000113278.2 | 0 | NA | NA | NA | NA | NA | #VALUE! |
| ENSMUSG00000113279.2 | 0 | NA | NA | NA | NA | NA | #VALUE! |
| ENSMUSG00000113280.2 | 0 | NA | NA | NA | NA | NA | #VALUE! |
| ENSMUSG00000113281.2 | 0 | NA | NA | NA | NA | NA | #VALUE! |
| ENSMUSG00000113282.2 | 0 | NA | NA | NA | NA | NA | #VALUE! |
| ENSMUSG00000113283.2 | 0 | NA | NA | NA | NA | NA | #VALUE! |
| ENSMUSG00000113284.2 | 0 | NA | NA | NA | NA | NA | #VALUE! |
| ENSMUSG00000113285.2 | 0 | NA | NA | NA | NA | NA | #VALUE! |
| ENSMUSG00000113286.2 | 0 | NA | NA | NA | NA | NA | #VALUE! |
| ENSMUSG00000113287.2 | 0 | NA | NA | NA | NA | NA | #VALUE! |
| ENSMUSG00000113288.2 | 0 | NA | NA | NA | NA | NA | #VALUE! |
| ENSMUSG00000113289.2 | 0 | NA | NA | NA | NA | NA | #VALUE! |
| ENSMUSG00000113290.2 | 0 | NA | NA | NA | NA | NA | #VALUE! |
| ENSMUSG00000113291.2 | 0 | NA | NA | NA | NA | NA | #VALUE! |
| ENSMUSG00000113292.2 | 0 | NA | NA | NA | NA | NA | #VALUE! |
| ENSMUSG00000113293.2 | 0 | NA | NA | NA | NA | NA | #VALUE! |
| ENSMUSG00000113294.2 | 0 | NA | NA | NA | NA | NA | #VALUE! |
| ENSMUSG00000113295.2 | 0 | NA | NA | NA | NA | NA | #VALUE! |
| ENSMUSG00000113296.2 | 0 | NA | NA | NA | NA | NA | #VALUE! |

| | | | | | | | |
|---|---|---|---|---|---|---|---|
| ENSMUSG00000113297.2 | 0 | NA | NA | NA | NA | NA | #VALUE! |
| ENSMUSG00000113298.2 | 0 | NA | NA | NA | NA | NA | #VALUE! |
| ENSMUSG00000113299.2 | 0 | NA | NA | NA | NA | NA | #VALUE! |
| ENSMUSG00000113300.2 | 0 | NA | NA | NA | NA | NA | #VALUE! |
| ENSMUSG00000113301.2 | 0 | NA | NA | NA | NA | NA | #VALUE! |
| ENSMUSG00000113302.2 | 0 | NA | NA | NA | NA | NA | #VALUE! |
| ENSMUSG00000113303.2 | 0 | NA | NA | NA | NA | NA | #VALUE! |
| ENSMUSG00000113304.2 | 0 | NA | NA | NA | NA | NA | #VALUE! |
| ENSMUSG00000113305.2 | 0 | NA | NA | NA | NA | NA | #VALUE! |
| ENSMUSG00000113306.2 | 0 | NA | NA | NA | NA | NA | #VALUE! |
| ENSMUSG00000113307.2 | 0 | NA | NA | NA | NA | NA | #VALUE! |
| ENSMUSG00000113308.2 | 0 | NA | NA | NA | NA | NA | #VALUE! |
| ENSMUSG00000113309.2 | 0 | NA | NA | NA | NA | NA | #VALUE! |
| ENSMUSG00000113310.2 | 0 | NA | NA | NA | NA | NA | #VALUE! |
| ENSMUSG00000113311.2 | 0 | NA | NA | NA | NA | NA | #VALUE! |
| ENSMUSG00000113312.2 | 0 | NA | NA | NA | NA | NA | #VALUE! |
| ENSMUSG00000113313.2 | 0 | NA | NA | NA | NA | NA | #VALUE! |
| ENSMUSG00000113314.2 | 0 | NA | NA | NA | NA | NA | #VALUE! |
| ENSMUSG00000113315.2 | 0 | NA | NA | NA | NA | NA | #VALUE! |
| ENSMUSG00000113316.2 | 0 | NA | NA | NA | NA | NA | #VALUE! |
| ENSMUSG00000113317.2 | 0 | NA | NA | NA | NA | NA | #VALUE! |
| ENSMUSG00000113319.2 | 0 | NA | NA | NA | NA | NA | #VALUE! |
| ENSMUSG00000113320.2 | 0 | NA | NA | NA | NA | NA | #VALUE! |
| ENSMUSG00000113321.2 | 0 | NA | NA | NA | NA | NA | #VALUE! |
| ENSMUSG00000113322.2 | 0 | NA | NA | NA | NA | NA | #VALUE! |
| ENSMUSG00000113323.2 | 0 | NA | NA | NA | NA | NA | #VALUE! |
| ENSMUSG00000113324.2 | 0 | NA | NA | NA | NA | NA | #VALUE! |
| ENSMUSG00000113325.2 | 0 | NA | NA | NA | NA | NA | #VALUE! |
| ENSMUSG00000113326.2 | 2.545258 | -4.8732 | 3.92437 | -1.24178 | 0.214318 | NA | #VALUE! |
| ENSMUSG00000113327.2 | 0 | NA | NA | NA | NA | NA | #VALUE! |
| ENSMUSG00000113328.2 | 0 | NA | NA | NA | NA | NA | #VALUE! |
| ENSMUSG00000113329.2 | 0 | NA | NA | NA | NA | NA | #VALUE! |
| ENSMUSG00000113330.2 | 0 | NA | NA | NA | NA | NA | #VALUE! |
| ENSMUSG00000113331.2 | 0 | NA | NA | NA | NA | NA | #VALUE! |
| ENSMUSG00000113332.2 | 0 | NA | NA | NA | NA | NA | #VALUE! |
| ENSMUSG00000113333.2 | 0 | NA | NA | NA | NA | NA | #VALUE! |
| ENSMUSG00000113334.2 | 0 | NA | NA | NA | NA | NA | #VALUE! |
| ENSMUSG00000113335.2 | 0 | NA | NA | NA | NA | NA | #VALUE! |
| ENSMUSG00000113336.2 | 0 | NA | NA | NA | NA | NA | #VALUE! |
| ENSMUSG00000113337.2 | 0 | NA | NA | NA | NA | NA | #VALUE! |
| ENSMUSG00000113338.2 | 0 | NA | NA | NA | NA | NA | #VALUE! |
| ENSMUSG00000113339.2 | 0 | NA | NA | NA | NA | NA | #VALUE! |
| ENSMUSG00000113340.2 | 0 | NA | NA | NA | NA | NA | #VALUE! |
| ENSMUSG00000113341.2 | 0 | NA | NA | NA | NA | NA | #VALUE! |

| | | | | | | | |
|---|---|---|---|---|---|---|---|
| ENSMUSG00000113342.2 | 0 | NA | NA | NA | NA | NA | #VALUE! |
| ENSMUSG00000113343.3 | 0 | NA | NA | NA | NA | NA | #VALUE! |
| ENSMUSG00000113344.2 | 0 | NA | NA | NA | NA | NA | #VALUE! |
| ENSMUSG00000113345.2 | 0 | NA | NA | NA | NA | NA | #VALUE! |
| ENSMUSG00000113346.3 | 0 | NA | NA | NA | NA | NA | #VALUE! |
| ENSMUSG00000113347.2 | 0 | NA | NA | NA | NA | NA | #VALUE! |
| ENSMUSG00000113348.2 | 0 | NA | NA | NA | NA | NA | #VALUE! |
| ENSMUSG00000113349.2 | 0 | NA | NA | NA | NA | NA | #VALUE! |
| ENSMUSG00000113350.3 | 0 | NA | NA | NA | NA | NA | #VALUE! |
| ENSMUSG00000113351.2 | 0 | NA | NA | NA | NA | NA | #VALUE! |
| ENSMUSG00000113352.2 | 0 | NA | NA | NA | NA | NA | #VALUE! |
| ENSMUSG00000113353.2 | 0 | NA | NA | NA | NA | NA | #VALUE! |
| ENSMUSG00000113354.2 | 0 | NA | NA | NA | NA | NA | #VALUE! |
| ENSMUSG00000113355.2 | 0 | NA | NA | NA | NA | NA | #VALUE! |
| ENSMUSG00000113356.2 | 0 | NA | NA | NA | NA | NA | #VALUE! |
| ENSMUSG00000113357.2 | 0 | NA | NA | NA | NA | NA | #VALUE! |
| ENSMUSG00000113358.2 | 0 | NA | NA | NA | NA | NA | #VALUE! |
| ENSMUSG00000113359.2 | 0 | NA | NA | NA | NA | NA | #VALUE! |
| ENSMUSG00000113360.2 | 0 | NA | NA | NA | NA | NA | #VALUE! |
| ENSMUSG00000113362.2 | 0 | NA | NA | NA | NA | NA | #VALUE! |
| ENSMUSG00000113363.2 | 0 | NA | NA | NA | NA | NA | #VALUE! |
| ENSMUSG00000113364.2 | 0 | NA | NA | NA | NA | NA | #VALUE! |
| ENSMUSG00000113365.2 | 0 | NA | NA | NA | NA | NA | #VALUE! |
| ENSMUSG00000113366.2 | 0 | NA | NA | NA | NA | NA | #VALUE! |
| ENSMUSG00000113367.2 | 0 | NA | NA | NA | NA | NA | #VALUE! |
| ENSMUSG00000113368.2 | 0.303281 | -0.61664 | 4.050639 | -0.15223 | 0.879003 | NA | #VALUE! |
| ENSMUSG00000113369.2 | 0 | NA | NA | NA | NA | NA | #VALUE! |
| ENSMUSG00000113370.2 | 0 | NA | NA | NA | NA | NA | #VALUE! |
| ENSMUSG00000113371.2 | 0 | NA | NA | NA | NA | NA | #VALUE! |
| ENSMUSG00000113372.2 | 0 | NA | NA | NA | NA | NA | #VALUE! |
| ENSMUSG00000113373.2 | 0 | NA | NA | NA | NA | NA | #VALUE! |
| ENSMUSG00000113374.2 | 0 | NA | NA | NA | NA | NA | #VALUE! |
| ENSMUSG00000113375.2 | 0 | NA | NA | NA | NA | NA | #VALUE! |
| ENSMUSG00000113376.2 | 0.168284 | 1.027628 | 4.067126 | 0.252667 | 0.800526 | NA | #VALUE! |
| ENSMUSG00000113377.2 | 0 | NA | NA | NA | NA | NA | #VALUE! |
| ENSMUSG00000113378.2 | 0 | NA | NA | NA | NA | NA | #VALUE! |
| ENSMUSG00000113379.2 | 0 | NA | NA | NA | NA | NA | #VALUE! |
| ENSMUSG00000113380.2 | 0 | NA | NA | NA | NA | NA | #VALUE! |
| ENSMUSG00000113381.3 | 0 | NA | NA | NA | NA | NA | #VALUE! |
| ENSMUSG00000113382.2 | 0 | NA | NA | NA | NA | NA | #VALUE! |
| ENSMUSG00000113383.2 | 0 | NA | NA | NA | NA | NA | #VALUE! |
| ENSMUSG00000113384.2 | 0 | NA | NA | NA | NA | NA | #VALUE! |
| ENSMUSG00000113385.2 | 0 | NA | NA | NA | NA | NA | #VALUE! |
| ENSMUSG00000113386.2 | 0 | NA | NA | NA | NA | NA | #VALUE! |

| | | | | | | | |
|---|---|---|---|---|---|---|---|
| ENSMUSG00000113387.2 | 0 | NA | NA | NA | NA | NA | #VALUE! |
| ENSMUSG00000113388.2 | 0 | NA | NA | NA | NA | NA | #VALUE! |
| ENSMUSG00000113389.2 | 0.105013 | -0.97324 | 4.076293 | -0.23876 | 0.811295 | NA | #VALUE! |
| ENSMUSG00000113390.2 | 0 | NA | NA | NA | NA | NA | #VALUE! |
| ENSMUSG00000113391.2 | 0 | NA | NA | NA | NA | NA | #VALUE! |
| ENSMUSG00000113392.2 | 0 | NA | NA | NA | NA | NA | #VALUE! |
| ENSMUSG00000113393.2 | 0 | NA | NA | NA | NA | NA | #VALUE! |
| ENSMUSG00000113394.2 | 0 | NA | NA | NA | NA | NA | #VALUE! |
| ENSMUSG00000113395.2 | 0 | NA | NA | NA | NA | NA | #VALUE! |
| ENSMUSG00000113396.2 | 0 | NA | NA | NA | NA | NA | #VALUE! |
| ENSMUSG00000113397.2 | 0 | NA | NA | NA | NA | NA | #VALUE! |
| ENSMUSG00000113398.2 | 0 | NA | NA | NA | NA | NA | #VALUE! |
| ENSMUSG00000113399.2 | 0 | NA | NA | NA | NA | NA | #VALUE! |
| ENSMUSG00000113400.2 | 0 | NA | NA | NA | NA | NA | #VALUE! |
| ENSMUSG00000113401.2 | 0 | NA | NA | NA | NA | NA | #VALUE! |
| ENSMUSG00000113402.2 | 0 | NA | NA | NA | NA | NA | #VALUE! |
| ENSMUSG00000113403.2 | 0 | NA | NA | NA | NA | NA | #VALUE! |
| ENSMUSG00000113404.2 | 0 | NA | NA | NA | NA | NA | #VALUE! |
| ENSMUSG00000113405.2 | 0.297366 | -0.14751 | 4.065279 | -0.03629 | 0.971054 | NA | #VALUE! |
| ENSMUSG00000113406.2 | 0 | NA | NA | NA | NA | NA | #VALUE! |
| ENSMUSG00000113407.2 | 0 | NA | NA | NA | NA | NA | #VALUE! |
| ENSMUSG00000113408.2 | 0 | NA | NA | NA | NA | NA | #VALUE! |
| ENSMUSG00000113409.2 | 4.158846 | -3.27286 | 3.845694 | -0.85105 | 0.394744 | NA | #VALUE! |
| ENSMUSG00000113410.2 | 0 | NA | NA | NA | NA | NA | #VALUE! |
| ENSMUSG00000113411.2 | 0 | NA | NA | NA | NA | NA | #VALUE! |
| ENSMUSG00000113412.2 | 0 | NA | NA | NA | NA | NA | #VALUE! |
| ENSMUSG00000113413.2 | 0 | NA | NA | NA | NA | NA | #VALUE! |
| ENSMUSG00000113414.2 | 0 | NA | NA | NA | NA | NA | #VALUE! |
| ENSMUSG00000113415.2 | 0 | NA | NA | NA | NA | NA | #VALUE! |
| ENSMUSG00000113416.2 | 0 | NA | NA | NA | NA | NA | #VALUE! |
| ENSMUSG00000113417.2 | 0 | NA | NA | NA | NA | NA | #VALUE! |
| ENSMUSG00000113418.2 | 0 | NA | NA | NA | NA | NA | #VALUE! |
| ENSMUSG00000113419.2 | 0 | NA | NA | NA | NA | NA | #VALUE! |
| ENSMUSG00000113420.2 | 0 | NA | NA | NA | NA | NA | #VALUE! |
| ENSMUSG00000113421.2 | 0 | NA | NA | NA | NA | NA | #VALUE! |
| ENSMUSG00000113422.2 | 0 | NA | NA | NA | NA | NA | #VALUE! |
| ENSMUSG00000113423.2 | 0 | NA | NA | NA | NA | NA | #VALUE! |
| ENSMUSG00000113424.2 | 0 | NA | NA | NA | NA | NA | #VALUE! |
| ENSMUSG00000113425.2 | 1.020554 | -3.54462 | 3.950745 | -0.8972 | 0.36961 | NA | #VALUE! |
| ENSMUSG00000113426.2 | 0 | NA | NA | NA | NA | NA | #VALUE! |
| ENSMUSG00000113427.2 | 0 | NA | NA | NA | NA | NA | #VALUE! |
| ENSMUSG00000113428.2 | 0 | NA | NA | NA | NA | NA | #VALUE! |
| ENSMUSG00000113429.2 | 0 | NA | NA | NA | NA | NA | #VALUE! |
| ENSMUSG00000113430.2 | 0 | NA | NA | NA | NA | NA | #VALUE! |

| | | | | | | | |
|---|---|---|---|---|---|---|---|
| ENSMUSG00000113431.2 | 0 | NA | NA | NA | NA | NA | #VALUE! |
| ENSMUSG00000113432.2 | 0 | NA | NA | NA | NA | NA | #VALUE! |
| ENSMUSG00000113433.2 | 0 | NA | NA | NA | NA | NA | #VALUE! |
| ENSMUSG00000113434.2 | 0 | NA | NA | NA | NA | NA | #VALUE! |
| ENSMUSG00000113435.2 | 0 | NA | NA | NA | NA | NA | #VALUE! |
| ENSMUSG00000113436.2 | 0 | NA | NA | NA | NA | NA | #VALUE! |
| ENSMUSG00000113437.2 | 0 | NA | NA | NA | NA | NA | #VALUE! |
| ENSMUSG00000113438.2 | 0 | NA | NA | NA | NA | NA | #VALUE! |
| ENSMUSG00000113439.2 | 0 | NA | NA | NA | NA | NA | #VALUE! |
| ENSMUSG00000113440.2 | 0.210026 | -1.52494 | 4.058692 | -0.37572 | 0.707123 | NA | #VALUE! |
| ENSMUSG00000113441.2 | 4.9682 | 5.807757 | 3.915774 | 1.48317 | 0.138029 | NA | #VALUE! |
| ENSMUSG00000113442.2 | 0 | NA | NA | NA | NA | NA | #VALUE! |
| ENSMUSG00000113443.2 | 0 | NA | NA | NA | NA | NA | #VALUE! |
| ENSMUSG00000113444.2 | 0 | NA | NA | NA | NA | NA | #VALUE! |
| ENSMUSG00000113445.2 | 0 | NA | NA | NA | NA | NA | #VALUE! |
| ENSMUSG00000113446.2 | 0 | NA | NA | NA | NA | NA | #VALUE! |
| ENSMUSG00000113447.2 | 0 | NA | NA | NA | NA | NA | #VALUE! |
| ENSMUSG00000113448.2 | 0 | NA | NA | NA | NA | NA | #VALUE! |
| ENSMUSG00000113449.2 | 0 | NA | NA | NA | NA | NA | #VALUE! |
| ENSMUSG00000113451.2 | 0 | NA | NA | NA | NA | NA | #VALUE! |
| ENSMUSG00000113452.2 | 0 | NA | NA | NA | NA | NA | #VALUE! |
| ENSMUSG00000113453.2 | 0 | NA | NA | NA | NA | NA | #VALUE! |
| ENSMUSG00000113454.2 | 0 | NA | NA | NA | NA | NA | #VALUE! |
| ENSMUSG00000113455.2 | 0 | NA | NA | NA | NA | NA | #VALUE! |
| ENSMUSG00000113456.2 | 0 | NA | NA | NA | NA | NA | #VALUE! |
| ENSMUSG00000113457.2 | 0 | NA | NA | NA | NA | NA | #VALUE! |
| ENSMUSG00000113458.2 | 0 | NA | NA | NA | NA | NA | #VALUE! |
| ENSMUSG00000113459.2 | 0 | NA | NA | NA | NA | NA | #VALUE! |
| ENSMUSG00000113460.2 | 0 | NA | NA | NA | NA | NA | #VALUE! |
| ENSMUSG00000113461.2 | 0 | NA | NA | NA | NA | NA | #VALUE! |
| ENSMUSG00000113462.2 | 0 | NA | NA | NA | NA | NA | #VALUE! |
| ENSMUSG00000113463.2 | 0 | NA | NA | NA | NA | NA | #VALUE! |
| ENSMUSG00000113464.2 | 0 | NA | NA | NA | NA | NA | #VALUE! |
| ENSMUSG00000113465.2 | 0 | NA | NA | NA | NA | NA | #VALUE! |
| ENSMUSG00000113466.2 | 0 | NA | NA | NA | NA | NA | #VALUE! |
| ENSMUSG00000113467.2 | 0.466276 | 2.449889 | 3.997179 | 0.612904 | 0.53994 | NA | #VALUE! |
| ENSMUSG00000113468.2 | 0 | NA | NA | NA | NA | NA | #VALUE! |
| ENSMUSG00000113469.2 | 0 | NA | NA | NA | NA | NA | #VALUE! |
| ENSMUSG00000113470.2 | 0 | NA | NA | NA | NA | NA | #VALUE! |
| ENSMUSG00000113471.2 | 0 | NA | NA | NA | NA | NA | #VALUE! |
| ENSMUSG00000113472.2 | 0 | NA | NA | NA | NA | NA | #VALUE! |
| ENSMUSG00000113473.2 | 0 | NA | NA | NA | NA | NA | #VALUE! |
| ENSMUSG00000113474.2 | 0 | NA | NA | NA | NA | NA | #VALUE! |
| ENSMUSG00000113475.2 | 0 | NA | NA | NA | NA | NA | #VALUE! |

| | | | | | | | |
|---|---|---|---|---|---|---|---|
| ENSMUSG00000113476.2 | 0 | NA | NA | NA | NA | NA | #VALUE! |
| ENSMUSG00000113477.2 | 0 | NA | NA | NA | NA | NA | #VALUE! |
| ENSMUSG00000113478.2 | 0 | NA | NA | NA | NA | NA | #VALUE! |
| ENSMUSG00000113479.2 | 0 | NA | NA | NA | NA | NA | #VALUE! |
| ENSMUSG00000113480.2 | 0 | NA | NA | NA | NA | NA | #VALUE! |
| ENSMUSG00000113481.2 | 0 | NA | NA | NA | NA | NA | #VALUE! |
| ENSMUSG00000113482.2 | 0 | NA | NA | NA | NA | NA | #VALUE! |
| ENSMUSG00000113483.2 | 0 | NA | NA | NA | NA | NA | #VALUE! |
| ENSMUSG00000113484.2 | 0 | NA | NA | NA | NA | NA | #VALUE! |
| ENSMUSG00000113485.2 | 0 | NA | NA | NA | NA | NA | #VALUE! |
| ENSMUSG00000113486.2 | 0 | NA | NA | NA | NA | NA | #VALUE! |
| ENSMUSG00000113487.2 | 0 | NA | NA | NA | NA | NA | #VALUE! |
| ENSMUSG00000113489.2 | 0 | NA | NA | NA | NA | NA | #VALUE! |
| ENSMUSG00000113490.2 | 0 | NA | NA | NA | NA | NA | #VALUE! |
| ENSMUSG00000113491.2 | 0 | NA | NA | NA | NA | NA | #VALUE! |
| ENSMUSG00000113492.2 | 0 | NA | NA | NA | NA | NA | #VALUE! |
| ENSMUSG00000113493.2 | 0 | NA | NA | NA | NA | NA | #VALUE! |
| ENSMUSG00000113494.2 | 0 | NA | NA | NA | NA | NA | #VALUE! |
| ENSMUSG00000113495.2 | 0 | NA | NA | NA | NA | NA | #VALUE! |
| ENSMUSG00000113496.2 | 0 | NA | NA | NA | NA | NA | #VALUE! |
| ENSMUSG00000113497.2 | 0 | NA | NA | NA | NA | NA | #VALUE! |
| ENSMUSG00000113498.2 | 0 | NA | NA | NA | NA | NA | #VALUE! |
| ENSMUSG00000113499.2 | 0 | NA | NA | NA | NA | NA | #VALUE! |
| ENSMUSG00000113500.2 | 0 | NA | NA | NA | NA | NA | #VALUE! |
| ENSMUSG00000113501.2 | 0 | NA | NA | NA | NA | NA | #VALUE! |
| ENSMUSG00000113502.2 | 0 | NA | NA | NA | NA | NA | #VALUE! |
| ENSMUSG00000113503.2 | 0 | NA | NA | NA | NA | NA | #VALUE! |
| ENSMUSG00000113504.2 | 0 | NA | NA | NA | NA | NA | #VALUE! |
| ENSMUSG00000113505.2 | 0 | NA | NA | NA | NA | NA | #VALUE! |
| ENSMUSG00000113506.2 | 0 | NA | NA | NA | NA | NA | #VALUE! |
| ENSMUSG00000113507.2 | 0 | NA | NA | NA | NA | NA | #VALUE! |
| ENSMUSG00000113508.2 | 0 | NA | NA | NA | NA | NA | #VALUE! |
| ENSMUSG00000113509.2 | 0 | NA | NA | NA | NA | NA | #VALUE! |
| ENSMUSG00000113510.2 | 0 | NA | NA | NA | NA | NA | #VALUE! |
| ENSMUSG00000113511.2 | 0 | NA | NA | NA | NA | NA | #VALUE! |
| ENSMUSG00000113512.2 | 0 | NA | NA | NA | NA | NA | #VALUE! |
| ENSMUSG00000113513.2 | 0 | NA | NA | NA | NA | NA | #VALUE! |
| ENSMUSG00000113514.2 | 0 | NA | NA | NA | NA | NA | #VALUE! |
| ENSMUSG00000113515.2 | 0 | NA | NA | NA | NA | NA | #VALUE! |
| ENSMUSG00000113516.2 | 0 | NA | NA | NA | NA | NA | #VALUE! |
| ENSMUSG00000113517.2 | 0 | NA | NA | NA | NA | NA | #VALUE! |
| ENSMUSG00000113518.2 | 0 | NA | NA | NA | NA | NA | #VALUE! |
| ENSMUSG00000113519.2 | 0 | NA | NA | NA | NA | NA | #VALUE! |
| ENSMUSG00000113520.2 | 0 | NA | NA | NA | NA | NA | #VALUE! |

| | | | | | | | |
|---|---|---|---|---|---|---|---|
| ENSMUSG00000113521.2 | 0 | NA | NA | NA | NA | NA | #VALUE! |
| ENSMUSG00000113522.2 | 0 | NA | NA | NA | NA | NA | #VALUE! |
| ENSMUSG00000113523.2 | 0 | NA | NA | NA | NA | NA | #VALUE! |
| ENSMUSG00000113524.2 | 0 | NA | NA | NA | NA | NA | #VALUE! |
| ENSMUSG00000113526.2 | 0 | NA | NA | NA | NA | NA | #VALUE! |
| ENSMUSG00000113527.2 | 0 | NA | NA | NA | NA | NA | #VALUE! |
| ENSMUSG00000113528.2 | 0 | NA | NA | NA | NA | NA | #VALUE! |
| ENSMUSG00000113529.2 | 0.491951 | 2.402185 | 4.000147 | 0.600524 | 0.548157 | NA | #VALUE! |
| ENSMUSG00000113530.2 | 0 | NA | NA | NA | NA | NA | #VALUE! |
| ENSMUSG00000113531.2 | 0 | NA | NA | NA | NA | NA | #VALUE! |
| ENSMUSG00000113532.2 | 0 | NA | NA | NA | NA | NA | #VALUE! |
| ENSMUSG00000113533.2 | 0 | NA | NA | NA | NA | NA | #VALUE! |
| ENSMUSG00000113534.2 | 0 | NA | NA | NA | NA | NA | #VALUE! |
| ENSMUSG00000113535.2 | 0 | NA | NA | NA | NA | NA | #VALUE! |
| ENSMUSG00000113536.2 | 0 | NA | NA | NA | NA | NA | #VALUE! |
| ENSMUSG00000113537.2 | 0 | NA | NA | NA | NA | NA | #VALUE! |
| ENSMUSG00000113538.2 | 0 | NA | NA | NA | NA | NA | #VALUE! |
| ENSMUSG00000113539.2 | 0 | NA | NA | NA | NA | NA | #VALUE! |
| ENSMUSG00000113540.2 | 0 | NA | NA | NA | NA | NA | #VALUE! |
| ENSMUSG00000113541.2 | 0 | NA | NA | NA | NA | NA | #VALUE! |
| ENSMUSG00000113542.2 | 0 | NA | NA | NA | NA | NA | #VALUE! |
| ENSMUSG00000113544.2 | 0 | NA | NA | NA | NA | NA | #VALUE! |
| ENSMUSG00000113545.2 | 0 | NA | NA | NA | NA | NA | #VALUE! |
| ENSMUSG00000113546.2 | 0 | NA | NA | NA | NA | NA | #VALUE! |
| ENSMUSG00000113547.2 | 0 | NA | NA | NA | NA | NA | #VALUE! |
| ENSMUSG00000113548.2 | 0 | NA | NA | NA | NA | NA | #VALUE! |
| ENSMUSG00000113549.2 | 0 | NA | NA | NA | NA | NA | #VALUE! |
| ENSMUSG00000113550.2 | 0 | NA | NA | NA | NA | NA | #VALUE! |
| ENSMUSG00000113551.2 | 0 | NA | NA | NA | NA | NA | #VALUE! |
| ENSMUSG00000113552.2 | 0 | NA | NA | NA | NA | NA | #VALUE! |
| ENSMUSG00000113553.2 | 0 | NA | NA | NA | NA | NA | #VALUE! |
| ENSMUSG00000113554.2 | 0 | NA | NA | NA | NA | NA | #VALUE! |
| ENSMUSG00000113556.2 | 0 | NA | NA | NA | NA | NA | #VALUE! |
| ENSMUSG00000113557.2 | 0 | NA | NA | NA | NA | NA | #VALUE! |
| ENSMUSG00000113558.2 | 0 | NA | NA | NA | NA | NA | #VALUE! |
| ENSMUSG00000113559.2 | 0 | NA | NA | NA | NA | NA | #VALUE! |
| ENSMUSG00000113560.2 | 0 | NA | NA | NA | NA | NA | #VALUE! |
| ENSMUSG00000113561.2 | 0 | NA | NA | NA | NA | NA | #VALUE! |
| ENSMUSG00000113562.2 | 0 | NA | NA | NA | NA | NA | #VALUE! |
| ENSMUSG00000113563.2 | 0 | NA | NA | NA | NA | NA | #VALUE! |
| ENSMUSG00000113564.2 | 0 | NA | NA | NA | NA | NA | #VALUE! |
| ENSMUSG00000113565.2 | 0 | NA | NA | NA | NA | NA | #VALUE! |
| ENSMUSG00000113566.2 | 0 | NA | NA | NA | NA | NA | #VALUE! |
| ENSMUSG00000113567.2 | 0 | NA | NA | NA | NA | NA | #VALUE! |

| | | | | | | | |
|---|---|---|---|---|---|---|---|
| ENSMUSG00000113568.2 | 0 | NA | NA | NA | NA | NA | #VALUE! |
| ENSMUSG00000113569.2 | 0 | NA | NA | NA | NA | NA | #VALUE! |
| ENSMUSG00000113570.2 | 0 | NA | NA | NA | NA | NA | #VALUE! |
| ENSMUSG00000113571.2 | 0 | NA | NA | NA | NA | NA | #VALUE! |
| ENSMUSG00000113572.2 | 0 | NA | NA | NA | NA | NA | #VALUE! |
| ENSMUSG00000113573.2 | 0 | NA | NA | NA | NA | NA | #VALUE! |
| ENSMUSG00000113574.2 | 0 | NA | NA | NA | NA | NA | #VALUE! |
| ENSMUSG00000113575.2 | 0 | NA | NA | NA | NA | NA | #VALUE! |
| ENSMUSG00000113576.2 | 0 | NA | NA | NA | NA | NA | #VALUE! |
| ENSMUSG00000113577.2 | 0 | NA | NA | NA | NA | NA | #VALUE! |
| ENSMUSG00000113578.2 | 0 | NA | NA | NA | NA | NA | #VALUE! |
| ENSMUSG00000113579.2 | 0 | NA | NA | NA | NA | NA | #VALUE! |
| ENSMUSG00000113580.2 | 0 | NA | NA | NA | NA | NA | #VALUE! |
| ENSMUSG00000113581.2 | 0 | NA | NA | NA | NA | NA | #VALUE! |
| ENSMUSG00000113582.2 | 0 | NA | NA | NA | NA | NA | #VALUE! |
| ENSMUSG00000113583.2 | 0 | NA | NA | NA | NA | NA | #VALUE! |
| ENSMUSG00000113584.2 | 0 | NA | NA | NA | NA | NA | #VALUE! |
| ENSMUSG00000113585.2 | 1.154263 | 0.945335 | 3.899062 | 0.242452 | 0.80843 | NA | #VALUE! |
| ENSMUSG00000113586.2 | 0 | NA | NA | NA | NA | NA | #VALUE! |
| ENSMUSG00000113587.2 | 0 | NA | NA | NA | NA | NA | #VALUE! |
| ENSMUSG00000113588.2 | 0 | NA | NA | NA | NA | NA | #VALUE! |
| ENSMUSG00000113589.2 | 0 | NA | NA | NA | NA | NA | #VALUE! |
| ENSMUSG00000113590.2 | 0 | NA | NA | NA | NA | NA | #VALUE! |
| ENSMUSG00000113591.2 | 0 | NA | NA | NA | NA | NA | #VALUE! |
| ENSMUSG00000113592.2 | 0 | NA | NA | NA | NA | NA | #VALUE! |
| ENSMUSG00000113593.2 | 0 | NA | NA | NA | NA | NA | #VALUE! |
| ENSMUSG00000113594.2 | 0 | NA | NA | NA | NA | NA | #VALUE! |
| ENSMUSG00000113595.2 | 0 | NA | NA | NA | NA | NA | #VALUE! |
| ENSMUSG00000113596.2 | 0 | NA | NA | NA | NA | NA | #VALUE! |
| ENSMUSG00000113597.2 | 0 | NA | NA | NA | NA | NA | #VALUE! |
| ENSMUSG00000113598.2 | 0 | NA | NA | NA | NA | NA | #VALUE! |
| ENSMUSG00000113599.2 | 0 | NA | NA | NA | NA | NA | #VALUE! |
| ENSMUSG00000113600.2 | 0 | NA | NA | NA | NA | NA | #VALUE! |
| ENSMUSG00000113601.2 | 0 | NA | NA | NA | NA | NA | #VALUE! |
| ENSMUSG00000113602.2 | 0 | NA | NA | NA | NA | NA | #VALUE! |
| ENSMUSG00000113603.2 | 0 | NA | NA | NA | NA | NA | #VALUE! |
| ENSMUSG00000113604.2 | 0 | NA | NA | NA | NA | NA | #VALUE! |
| ENSMUSG00000113605.2 | 0 | NA | NA | NA | NA | NA | #VALUE! |
| ENSMUSG00000113606.2 | 0 | NA | NA | NA | NA | NA | #VALUE! |
| ENSMUSG00000113607.2 | 0 | NA | NA | NA | NA | NA | #VALUE! |
| ENSMUSG00000113608.2 | 0 | NA | NA | NA | NA | NA | #VALUE! |
| ENSMUSG00000113609.2 | 0 | NA | NA | NA | NA | NA | #VALUE! |
| ENSMUSG00000113610.2 | 0 | NA | NA | NA | NA | NA | #VALUE! |
| ENSMUSG00000113611.2 | 0.093255 | 0.868093 | 4.072464 | 0.213162 | 0.831201 | NA | #VALUE! |

| | | | | | | | |
|---|---|---|---|---|---|---|---|
| ENSMUSG00000113612.2 | 0 | NA | NA | NA | NA | NA | #VALUE! |
| ENSMUSG00000113613.2 | 0 | NA | NA | NA | NA | NA | #VALUE! |
| ENSMUSG00000113614.2 | 0 | NA | NA | NA | NA | NA | #VALUE! |
| ENSMUSG00000113615.2 | 0 | NA | NA | NA | NA | NA | #VALUE! |
| ENSMUSG00000113616.2 | 0.967826 | 1.024393 | 3.929472 | 0.260695 | 0.794328 | NA | #VALUE! |
| ENSMUSG00000113617.2 | 0 | NA | NA | NA | NA | NA | #VALUE! |
| ENSMUSG00000113618.2 | 0 | NA | NA | NA | NA | NA | #VALUE! |
| ENSMUSG00000113619.2 | 0 | NA | NA | NA | NA | NA | #VALUE! |
| ENSMUSG00000113620.2 | 0 | NA | NA | NA | NA | NA | #VALUE! |
| ENSMUSG00000113621.2 | 0 | NA | NA | NA | NA | NA | #VALUE! |
| ENSMUSG00000113622.2 | 0 | NA | NA | NA | NA | NA | #VALUE! |
| ENSMUSG00000113623.2 | 0 | NA | NA | NA | NA | NA | #VALUE! |
| ENSMUSG00000113624.2 | 0 | NA | NA | NA | NA | NA | #VALUE! |
| ENSMUSG00000113626.2 | 0 | NA | NA | NA | NA | NA | #VALUE! |
| ENSMUSG00000113627.2 | 0 | NA | NA | NA | NA | NA | #VALUE! |
| ENSMUSG00000113628.2 | 0 | NA | NA | NA | NA | NA | #VALUE! |
| ENSMUSG00000113629.2 | 0 | NA | NA | NA | NA | NA | #VALUE! |
| ENSMUSG00000113630.2 | 2.126884 | 1.330701 | 3.832128 | 0.347249 | 0.728405 | NA | #VALUE! |
| ENSMUSG00000113631.2 | 0 | NA | NA | NA | NA | NA | #VALUE! |
| ENSMUSG00000113632.2 | 0 | NA | NA | NA | NA | NA | #VALUE! |
| ENSMUSG00000113633.2 | 0 | NA | NA | NA | NA | NA | #VALUE! |
| ENSMUSG00000113634.2 | 0 | NA | NA | NA | NA | NA | #VALUE! |
| ENSMUSG00000113635.2 | 0 | NA | NA | NA | NA | NA | #VALUE! |
| ENSMUSG00000113636.2 | 0 | NA | NA | NA | NA | NA | #VALUE! |
| ENSMUSG00000113637.2 | 0 | NA | NA | NA | NA | NA | #VALUE! |
| ENSMUSG00000113638.2 | 0 | NA | NA | NA | NA | NA | #VALUE! |
| ENSMUSG00000113639.2 | 0 | NA | NA | NA | NA | NA | #VALUE! |
| ENSMUSG00000113640.2 | 0 | NA | NA | NA | NA | NA | #VALUE! |
| ENSMUSG00000113641.2 | 0 | NA | NA | NA | NA | NA | #VALUE! |
| ENSMUSG00000113642.2 | 0 | NA | NA | NA | NA | NA | #VALUE! |
| ENSMUSG00000113643.2 | 0 | NA | NA | NA | NA | NA | #VALUE! |
| ENSMUSG00000113644.2 | 0 | NA | NA | NA | NA | NA | #VALUE! |
| ENSMUSG00000113645.2 | 0 | NA | NA | NA | NA | NA | #VALUE! |
| ENSMUSG00000113646.2 | 0 | NA | NA | NA | NA | NA | #VALUE! |
| ENSMUSG00000113647.2 | 0 | NA | NA | NA | NA | NA | #VALUE! |
| ENSMUSG00000113648.2 | 0 | NA | NA | NA | NA | NA | #VALUE! |
| ENSMUSG00000113649.2 | 0 | NA | NA | NA | NA | NA | #VALUE! |
| ENSMUSG00000113650.2 | 0 | NA | NA | NA | NA | NA | #VALUE! |
| ENSMUSG00000113651.2 | 0 | NA | NA | NA | NA | NA | #VALUE! |
| ENSMUSG00000113652.2 | 0 | NA | NA | NA | NA | NA | #VALUE! |
| ENSMUSG00000113653.2 | 0 | NA | NA | NA | NA | NA | #VALUE! |
| ENSMUSG00000113654.2 | 0 | NA | NA | NA | NA | NA | #VALUE! |
| ENSMUSG00000113655.2 | 0 | NA | NA | NA | NA | NA | #VALUE! |
| ENSMUSG00000113656.2 | 0 | NA | NA | NA | NA | NA | #VALUE! |

| | | | | | | | |
|---|---|---|---|---|---|---|---|
| ENSMUSG00000113657.2 | 0 | NA | NA | NA | NA | NA | #VALUE! |
| ENSMUSG00000113658.2 | 0 | NA | NA | NA | NA | NA | #VALUE! |
| ENSMUSG00000113659.2 | 0 | NA | NA | NA | NA | NA | #VALUE! |
| ENSMUSG00000113661.2 | 0 | NA | NA | NA | NA | NA | #VALUE! |
| ENSMUSG00000113662.2 | 0 | NA | NA | NA | NA | NA | #VALUE! |
| ENSMUSG00000113663.2 | 0 | NA | NA | NA | NA | NA | #VALUE! |
| ENSMUSG00000113664.2 | 0 | NA | NA | NA | NA | NA | #VALUE! |
| ENSMUSG00000113665.2 | 0 | NA | NA | NA | NA | NA | #VALUE! |
| ENSMUSG00000113666.2 | 0 | NA | NA | NA | NA | NA | #VALUE! |
| ENSMUSG00000113667.2 | 0 | NA | NA | NA | NA | NA | #VALUE! |
| ENSMUSG00000113668.2 | 0.562341 | -0.07781 | 4.012298 | -0.01939 | 0.984528 | NA | #VALUE! |
| ENSMUSG00000113669.2 | 0 | NA | NA | NA | NA | NA | #VALUE! |
| ENSMUSG00000113670.2 | 0 | NA | NA | NA | NA | NA | #VALUE! |
| ENSMUSG00000113671.2 | 0 | NA | NA | NA | NA | NA | #VALUE! |
| ENSMUSG00000113672.2 | 0 | NA | NA | NA | NA | NA | #VALUE! |
| ENSMUSG00000113673.2 | 0 | NA | NA | NA | NA | NA | #VALUE! |
| ENSMUSG00000113674.2 | 0 | NA | NA | NA | NA | NA | #VALUE! |
| ENSMUSG00000113675.2 | 0 | NA | NA | NA | NA | NA | #VALUE! |
| ENSMUSG00000113676.2 | 0 | NA | NA | NA | NA | NA | #VALUE! |
| ENSMUSG00000113677.2 | 0 | NA | NA | NA | NA | NA | #VALUE! |
| ENSMUSG00000113678.2 | 0 | NA | NA | NA | NA | NA | #VALUE! |
| ENSMUSG00000113679.2 | 8.609143 | 6.591549 | 3.791181 | 1.738653 | 0.082096 | NA | #VALUE! |
| ENSMUSG00000113680.2 | 0 | NA | NA | NA | NA | NA | #VALUE! |
| ENSMUSG00000113681.2 | 0 | NA | NA | NA | NA | NA | #VALUE! |
| ENSMUSG00000113682.2 | 0 | NA | NA | NA | NA | NA | #VALUE! |
| ENSMUSG00000113683.2 | 1.428775 | -4.03418 | 3.938169 | -1.02438 | 0.305656 | NA | #VALUE! |
| ENSMUSG00000113684.2 | 0 | NA | NA | NA | NA | NA | #VALUE! |
| ENSMUSG00000113685.2 | 0 | NA | NA | NA | NA | NA | #VALUE! |
| ENSMUSG00000113686.2 | 0 | NA | NA | NA | NA | NA | #VALUE! |
| ENSMUSG00000113687.2 | 0.168284 | 1.027628 | 4.067126 | 0.252667 | 0.800526 | NA | #VALUE! |
| ENSMUSG00000113688.2 | 0 | NA | NA | NA | NA | NA | #VALUE! |
| ENSMUSG00000113689.2 | 0 | NA | NA | NA | NA | NA | #VALUE! |
| ENSMUSG00000113690.2 | 0 | NA | NA | NA | NA | NA | #VALUE! |
| ENSMUSG00000113691.2 | 0 | NA | NA | NA | NA | NA | #VALUE! |
| ENSMUSG00000113692.2 | 0 | NA | NA | NA | NA | NA | #VALUE! |
| ENSMUSG00000113693.2 | 0 | NA | NA | NA | NA | NA | #VALUE! |
| ENSMUSG00000113694.2 | 0 | NA | NA | NA | NA | NA | #VALUE! |
| ENSMUSG00000113695.2 | 0 | NA | NA | NA | NA | NA | #VALUE! |
| ENSMUSG00000113696.2 | 0 | NA | NA | NA | NA | NA | #VALUE! |
| ENSMUSG00000113697.2 | 0 | NA | NA | NA | NA | NA | #VALUE! |
| ENSMUSG00000113698.2 | 0 | NA | NA | NA | NA | NA | #VALUE! |
| ENSMUSG00000113699.2 | 0 | NA | NA | NA | NA | NA | #VALUE! |
| ENSMUSG00000113700.2 | 0 | NA | NA | NA | NA | NA | #VALUE! |
| ENSMUSG00000113701.2 | 0 | NA | NA | NA | NA | NA | #VALUE! |

| | | | | | | | |
|---|---|---|---|---|---|---|---|
| ENSMUSG00000113702.2 | 0 | NA | NA | NA | NA | NA | #VALUE! |
| ENSMUSG00000113703.2 | 0 | NA | NA | NA | NA | NA | #VALUE! |
| ENSMUSG00000113704.2 | 0 | NA | NA | NA | NA | NA | #VALUE! |
| ENSMUSG00000113705.2 | 0 | NA | NA | NA | NA | NA | #VALUE! |
| ENSMUSG00000113706.2 | 0 | NA | NA | NA | NA | NA | #VALUE! |
| ENSMUSG00000113707.2 | 0 | NA | NA | NA | NA | NA | #VALUE! |
| ENSMUSG00000113708.2 | 0 | NA | NA | NA | NA | NA | #VALUE! |
| ENSMUSG00000113709.2 | 0 | NA | NA | NA | NA | NA | #VALUE! |
| ENSMUSG00000113710.2 | 0 | NA | NA | NA | NA | NA | #VALUE! |
| ENSMUSG00000113711.2 | 0 | NA | NA | NA | NA | NA | #VALUE! |
| ENSMUSG00000113712.2 | 0 | NA | NA | NA | NA | NA | #VALUE! |
| ENSMUSG00000113713.2 | 0 | NA | NA | NA | NA | NA | #VALUE! |
| ENSMUSG00000113714.2 | 0 | NA | NA | NA | NA | NA | #VALUE! |
| ENSMUSG00000113715.2 | 0 | NA | NA | NA | NA | NA | #VALUE! |
| ENSMUSG00000113716.2 | 0 | NA | NA | NA | NA | NA | #VALUE! |
| ENSMUSG00000113717.2 | 0 | NA | NA | NA | NA | NA | #VALUE! |
| ENSMUSG00000113718.2 | 0 | NA | NA | NA | NA | NA | #VALUE! |
| ENSMUSG00000113719.2 | 0 | NA | NA | NA | NA | NA | #VALUE! |
| ENSMUSG00000113720.2 | 0 | NA | NA | NA | NA | NA | #VALUE! |
| ENSMUSG00000113721.2 | 0 | NA | NA | NA | NA | NA | #VALUE! |
| ENSMUSG00000113722.2 | 0.737927 | 3.007825 | 3.968641 | 0.757898 | 0.448512 | NA | #VALUE! |
| ENSMUSG00000113723.3 | 0 | NA | NA | NA | NA | NA | #VALUE! |
| ENSMUSG00000113724.2 | 0 | NA | NA | NA | NA | NA | #VALUE! |
| ENSMUSG00000113725.2 | 0 | NA | NA | NA | NA | NA | #VALUE! |
| ENSMUSG00000113726.2 | 0 | NA | NA | NA | NA | NA | #VALUE! |
| ENSMUSG00000113727.2 | 0 | NA | NA | NA | NA | NA | #VALUE! |
| ENSMUSG00000113728.2 | 0 | NA | NA | NA | NA | NA | #VALUE! |
| ENSMUSG00000113729.2 | 0 | NA | NA | NA | NA | NA | #VALUE! |
| ENSMUSG00000113730.2 | 0 | NA | NA | NA | NA | NA | #VALUE! |
| ENSMUSG00000113731.2 | 0 | NA | NA | NA | NA | NA | #VALUE! |
| ENSMUSG00000113732.2 | 0 | NA | NA | NA | NA | NA | #VALUE! |
| ENSMUSG00000113733.2 | 0 | NA | NA | NA | NA | NA | #VALUE! |
| ENSMUSG00000113734.2 | 0 | NA | NA | NA | NA | NA | #VALUE! |
| ENSMUSG00000113735.2 | 0 | NA | NA | NA | NA | NA | #VALUE! |
| ENSMUSG00000113736.2 | 0 | NA | NA | NA | NA | NA | #VALUE! |
| ENSMUSG00000113737.2 | 0 | NA | NA | NA | NA | NA | #VALUE! |
| ENSMUSG00000113738.2 | 0 | NA | NA | NA | NA | NA | #VALUE! |
| ENSMUSG00000113739.2 | 0 | NA | NA | NA | NA | NA | #VALUE! |
| ENSMUSG00000113740.2 | 0 | NA | NA | NA | NA | NA | #VALUE! |
| ENSMUSG00000113741.2 | 0 | NA | NA | NA | NA | NA | #VALUE! |
| ENSMUSG00000113742.2 | 0 | NA | NA | NA | NA | NA | #VALUE! |
| ENSMUSG00000113743.2 | 3.132547 | 0.144227 | 3.784609 | 0.038109 | 0.969601 | NA | #VALUE! |
| ENSMUSG00000113744.2 | 0 | NA | NA | NA | NA | NA | #VALUE! |
| ENSMUSG00000113745.2 | 0 | NA | NA | NA | NA | NA | #VALUE! |

| | | | | | | | |
|---|---|---|---|---|---|---|---|
| ENSMUSG00000113746.2 | 0 | NA | NA | NA | NA | NA | #VALUE! |
| ENSMUSG00000113747.2 | 0 | NA | NA | NA | NA | NA | #VALUE! |
| ENSMUSG00000113748.2 | 0 | NA | NA | NA | NA | NA | #VALUE! |
| ENSMUSG00000113749.2 | 0 | NA | NA | NA | NA | NA | #VALUE! |
| ENSMUSG00000113750.2 | 0 | NA | NA | NA | NA | NA | #VALUE! |
| ENSMUSG00000113751.2 | 0 | NA | NA | NA | NA | NA | #VALUE! |
| ENSMUSG00000113752.2 | 1.212318 | 3.793111 | 3.942915 | 0.962007 | 0.336046 | NA | #VALUE! |
| ENSMUSG00000113753.2 | 0.261539 | 1.68833 | 4.043024 | 0.417591 | 0.676246 | NA | #VALUE! |
| ENSMUSG00000113754.2 | 0 | NA | NA | NA | NA | NA | #VALUE! |
| ENSMUSG00000113755.2 | 0 | NA | NA | NA | NA | NA | #VALUE! |
| ENSMUSG00000113756.2 | 0 | NA | NA | NA | NA | NA | #VALUE! |
| ENSMUSG00000113757.2 | 0 | NA | NA | NA | NA | NA | #VALUE! |
| ENSMUSG00000113758.2 | 0 | NA | NA | NA | NA | NA | #VALUE! |
| ENSMUSG00000113759.2 | 0 | NA | NA | NA | NA | NA | #VALUE! |
| ENSMUSG00000113760.2 | 0 | NA | NA | NA | NA | NA | #VALUE! |
| ENSMUSG00000113761.2 | 0 | NA | NA | NA | NA | NA | #VALUE! |
| ENSMUSG00000113762.2 | 0 | NA | NA | NA | NA | NA | #VALUE! |
| ENSMUSG00000113763.2 | 0 | NA | NA | NA | NA | NA | #VALUE! |
| ENSMUSG00000113764.2 | 0 | NA | NA | NA | NA | NA | #VALUE! |
| ENSMUSG00000113765.2 | 0 | NA | NA | NA | NA | NA | #VALUE! |
| ENSMUSG00000113766.2 | 0 | NA | NA | NA | NA | NA | #VALUE! |
| ENSMUSG00000113767.2 | 0 | NA | NA | NA | NA | NA | #VALUE! |
| ENSMUSG00000113768.2 | 0 | NA | NA | NA | NA | NA | #VALUE! |
| ENSMUSG00000113770.2 | 0 | NA | NA | NA | NA | NA | #VALUE! |
| ENSMUSG00000113771.2 | 0 | NA | NA | NA | NA | NA | #VALUE! |
| ENSMUSG00000113772.2 | 0 | NA | NA | NA | NA | NA | #VALUE! |
| ENSMUSG00000113773.2 | 0 | NA | NA | NA | NA | NA | #VALUE! |
| ENSMUSG00000113775.2 | 0 | NA | NA | NA | NA | NA | #VALUE! |
| ENSMUSG00000113776.2 | 0 | NA | NA | NA | NA | NA | #VALUE! |
| ENSMUSG00000113777.2 | 0 | NA | NA | NA | NA | NA | #VALUE! |
| ENSMUSG00000113778.2 | 0 | NA | NA | NA | NA | NA | #VALUE! |
| ENSMUSG00000113779.2 | 0 | NA | NA | NA | NA | NA | #VALUE! |
| ENSMUSG00000113780.2 | 0 | NA | NA | NA | NA | NA | #VALUE! |
| ENSMUSG00000113781.2 | 0 | NA | NA | NA | NA | NA | #VALUE! |
| ENSMUSG00000113782.2 | 0 | NA | NA | NA | NA | NA | #VALUE! |
| ENSMUSG00000113783.2 | 0 | NA | NA | NA | NA | NA | #VALUE! |
| ENSMUSG00000113784.2 | 0 | NA | NA | NA | NA | NA | #VALUE! |
| ENSMUSG00000113785.2 | 0 | NA | NA | NA | NA | NA | #VALUE! |
| ENSMUSG00000113786.2 | 0 | NA | NA | NA | NA | NA | #VALUE! |
| ENSMUSG00000113787.2 | 0 | NA | NA | NA | NA | NA | #VALUE! |
| ENSMUSG00000113788.2 | 0 | NA | NA | NA | NA | NA | #VALUE! |
| ENSMUSG00000113789.2 | 0 | NA | NA | NA | NA | NA | #VALUE! |
| ENSMUSG00000113790.2 | 0 | NA | NA | NA | NA | NA | #VALUE! |
| ENSMUSG00000113791.2 | 0 | NA | NA | NA | NA | NA | #VALUE! |

| | | | | | | | |
|---|---|---|---|---|---|---|---|
| ENSMUSG00000113792.2 | 0 | NA | NA | NA | NA | NA | #VALUE! |
| ENSMUSG00000113793.2 | 0 | NA | NA | NA | NA | NA | #VALUE! |
| ENSMUSG00000113794.2 | 0 | NA | NA | NA | NA | NA | #VALUE! |
| ENSMUSG00000113795.2 | 0 | NA | NA | NA | NA | NA | #VALUE! |
| ENSMUSG00000113796.2 | 0 | NA | NA | NA | NA | NA | #VALUE! |
| ENSMUSG00000113797.2 | 0 | NA | NA | NA | NA | NA | #VALUE! |
| ENSMUSG00000113798.2 | 0 | NA | NA | NA | NA | NA | #VALUE! |
| ENSMUSG00000113799.2 | 0 | NA | NA | NA | NA | NA | #VALUE! |
| ENSMUSG00000113800.2 | 2.165756 | -1.46557 | 3.835 | -0.38216 | 0.702345 | NA | #VALUE! |
| ENSMUSG00000113801.2 | 0 | NA | NA | NA | NA | NA | #VALUE! |
| ENSMUSG00000113802.2 | 0 | NA | NA | NA | NA | NA | #VALUE! |
| ENSMUSG00000113803.2 | 0 | NA | NA | NA | NA | NA | #VALUE! |
| ENSMUSG00000113804.2 | 0 | NA | NA | NA | NA | NA | #VALUE! |
| ENSMUSG00000113805.2 | 0 | NA | NA | NA | NA | NA | #VALUE! |
| ENSMUSG00000113806.2 | 0 | NA | NA | NA | NA | NA | #VALUE! |
| ENSMUSG00000113807.2 | 0 | NA | NA | NA | NA | NA | #VALUE! |
| ENSMUSG00000113808.2 | 0 | NA | NA | NA | NA | NA | #VALUE! |
| ENSMUSG00000113809.2 | 0 | NA | NA | NA | NA | NA | #VALUE! |
| ENSMUSG00000113810.2 | 0 | NA | NA | NA | NA | NA | #VALUE! |
| ENSMUSG00000113811.2 | 0 | NA | NA | NA | NA | NA | #VALUE! |
| ENSMUSG00000113812.2 | 0 | NA | NA | NA | NA | NA | #VALUE! |
| ENSMUSG00000113813.2 | 0 | NA | NA | NA | NA | NA | #VALUE! |
| ENSMUSG00000113814.2 | 0 | NA | NA | NA | NA | NA | #VALUE! |
| ENSMUSG00000113815.2 | 0 | NA | NA | NA | NA | NA | #VALUE! |
| ENSMUSG00000113816.2 | 0 | NA | NA | NA | NA | NA | #VALUE! |
| ENSMUSG00000113817.2 | 0 | NA | NA | NA | NA | NA | #VALUE! |
| ENSMUSG00000113818.2 | 0 | NA | NA | NA | NA | NA | #VALUE! |
| ENSMUSG00000113819.2 | 0 | NA | NA | NA | NA | NA | #VALUE! |
| ENSMUSG00000113820.3 | 0 | NA | NA | NA | NA | NA | #VALUE! |
| ENSMUSG00000113821.2 | 0 | NA | NA | NA | NA | NA | #VALUE! |
| ENSMUSG00000113822.2 | 0 | NA | NA | NA | NA | NA | #VALUE! |
| ENSMUSG00000113823.2 | 0 | NA | NA | NA | NA | NA | #VALUE! |
| ENSMUSG00000113824.2 | 0 | NA | NA | NA | NA | NA | #VALUE! |
| ENSMUSG00000113825.2 | 0 | NA | NA | NA | NA | NA | #VALUE! |
| ENSMUSG00000113826.2 | 0 | NA | NA | NA | NA | NA | #VALUE! |
| ENSMUSG00000113827.2 | 0 | NA | NA | NA | NA | NA | #VALUE! |
| ENSMUSG00000113828.2 | 0 | NA | NA | NA | NA | NA | #VALUE! |
| ENSMUSG00000113829.2 | 0 | NA | NA | NA | NA | NA | #VALUE! |
| ENSMUSG00000113830.2 | 0 | NA | NA | NA | NA | NA | #VALUE! |
| ENSMUSG00000113832.2 | 0 | NA | NA | NA | NA | NA | #VALUE! |
| ENSMUSG00000113833.2 | 0 | NA | NA | NA | NA | NA | #VALUE! |
| ENSMUSG00000113834.2 | 0 | NA | NA | NA | NA | NA | #VALUE! |
| ENSMUSG00000113835.2 | 0 | NA | NA | NA | NA | NA | #VALUE! |
| ENSMUSG00000113836.2 | 0 | NA | NA | NA | NA | NA | #VALUE! |

| | | | | | | | |
|---|---|---|---|---|---|---|---|
| ENSMUSG00000113837.2 | 0 | NA | NA | NA | NA | NA | #VALUE! |
| ENSMUSG00000113838.2 | 0 | NA | NA | NA | NA | NA | #VALUE! |
| ENSMUSG00000113840.2 | 0 | NA | NA | NA | NA | NA | #VALUE! |
| ENSMUSG00000113841.2 | 0 | NA | NA | NA | NA | NA | #VALUE! |
| ENSMUSG00000113842.2 | 0.839297 | 3.273019 | 3.958392 | 0.826856 | 0.408319 | NA | #VALUE! |
| ENSMUSG00000113843.2 | 0 | NA | NA | NA | NA | NA | #VALUE! |
| ENSMUSG00000113844.2 | 0 | NA | NA | NA | NA | NA | #VALUE! |
| ENSMUSG00000113845.2 | 0 | NA | NA | NA | NA | NA | #VALUE! |
| ENSMUSG00000113846.2 | 0 | NA | NA | NA | NA | NA | #VALUE! |
| ENSMUSG00000113847.2 | 0 | NA | NA | NA | NA | NA | #VALUE! |
| ENSMUSG00000113848.2 | 0 | NA | NA | NA | NA | NA | #VALUE! |
| ENSMUSG00000113849.2 | 0 | NA | NA | NA | NA | NA | #VALUE! |
| ENSMUSG00000113850.2 | 0 | NA | NA | NA | NA | NA | #VALUE! |
| ENSMUSG00000113851.2 | 0 | NA | NA | NA | NA | NA | #VALUE! |
| ENSMUSG00000113852.2 | 0 | NA | NA | NA | NA | NA | #VALUE! |
| ENSMUSG00000113853.2 | 0 | NA | NA | NA | NA | NA | #VALUE! |
| ENSMUSG00000113854.2 | 2.094849 | 4.546621 | 3.928291 | 1.157404 | 0.247107 | NA | #VALUE! |
| ENSMUSG00000113855.2 | 0 | NA | NA | NA | NA | NA | #VALUE! |
| ENSMUSG00000113856.2 | 0 | NA | NA | NA | NA | NA | #VALUE! |
| ENSMUSG00000113857.2 | 0 | NA | NA | NA | NA | NA | #VALUE! |
| ENSMUSG00000113858.2 | 0 | NA | NA | NA | NA | NA | #VALUE! |
| ENSMUSG00000113859.2 | 0 | NA | NA | NA | NA | NA | #VALUE! |
| ENSMUSG00000113860.2 | 0 | NA | NA | NA | NA | NA | #VALUE! |
| ENSMUSG00000113861.2 | 0 | NA | NA | NA | NA | NA | #VALUE! |
| ENSMUSG00000113862.2 | 0 | NA | NA | NA | NA | NA | #VALUE! |
| ENSMUSG00000113863.2 | 0 | NA | NA | NA | NA | NA | #VALUE! |
| ENSMUSG00000113864.2 | 0 | NA | NA | NA | NA | NA | #VALUE! |
| ENSMUSG00000113865.2 | 0 | NA | NA | NA | NA | NA | #VALUE! |
| ENSMUSG00000113866.2 | 0 | NA | NA | NA | NA | NA | #VALUE! |
| ENSMUSG00000113867.2 | 0 | NA | NA | NA | NA | NA | #VALUE! |
| ENSMUSG00000113868.2 | 0 | NA | NA | NA | NA | NA | #VALUE! |
| ENSMUSG00000113869.2 | 0 | NA | NA | NA | NA | NA | #VALUE! |
| ENSMUSG00000113870.2 | 0 | NA | NA | NA | NA | NA | #VALUE! |
| ENSMUSG00000113871.2 | 0 | NA | NA | NA | NA | NA | #VALUE! |
| ENSMUSG00000113872.2 | 0 | NA | NA | NA | NA | NA | #VALUE! |
| ENSMUSG00000113873.2 | 0 | NA | NA | NA | NA | NA | #VALUE! |
| ENSMUSG00000113874.2 | 0 | NA | NA | NA | NA | NA | #VALUE! |
| ENSMUSG00000113875.2 | 0 | NA | NA | NA | NA | NA | #VALUE! |
| ENSMUSG00000113876.2 | 0 | NA | NA | NA | NA | NA | #VALUE! |
| ENSMUSG00000113877.2 | 0 | NA | NA | NA | NA | NA | #VALUE! |
| ENSMUSG00000113878.2 | 0 | NA | NA | NA | NA | NA | #VALUE! |
| ENSMUSG00000113879.2 | 0 | NA | NA | NA | NA | NA | #VALUE! |
| ENSMUSG00000113880.2 | 0 | NA | NA | NA | NA | NA | #VALUE! |
| ENSMUSG00000113881.2 | 0 | NA | NA | NA | NA | NA | #VALUE! |

| | | | | | | | |
|---|---|---|---|---|---|---|---|
| ENSMUSG00000113882.2 | 0 | NA | NA | NA | NA | NA | #VALUE! |
| ENSMUSG00000113883.2 | 0 | NA | NA | NA | NA | NA | #VALUE! |
| ENSMUSG00000113884.2 | 0 | NA | NA | NA | NA | NA | #VALUE! |
| ENSMUSG00000113885.2 | 0 | NA | NA | NA | NA | NA | #VALUE! |
| ENSMUSG00000113886.2 | 0 | NA | NA | NA | NA | NA | #VALUE! |
| ENSMUSG00000113887.2 | 0 | NA | NA | NA | NA | NA | #VALUE! |
| ENSMUSG00000113888.2 | 0 | NA | NA | NA | NA | NA | #VALUE! |
| ENSMUSG00000113889.2 | 0 | NA | NA | NA | NA | NA | #VALUE! |
| ENSMUSG00000113890.2 | 0 | NA | NA | NA | NA | NA | #VALUE! |
| ENSMUSG00000113891.2 | 0 | NA | NA | NA | NA | NA | #VALUE! |
| ENSMUSG00000113892.2 | 0 | NA | NA | NA | NA | NA | #VALUE! |
| ENSMUSG00000113893.2 | 0 | NA | NA | NA | NA | NA | #VALUE! |
| ENSMUSG00000113894.2 | 0 | NA | NA | NA | NA | NA | #VALUE! |
| ENSMUSG00000113895.2 | 0 | NA | NA | NA | NA | NA | #VALUE! |
| ENSMUSG00000113896.2 | 0 | NA | NA | NA | NA | NA | #VALUE! |
| ENSMUSG00000113897.2 | 0 | NA | NA | NA | NA | NA | #VALUE! |
| ENSMUSG00000113898.2 | 0 | NA | NA | NA | NA | NA | #VALUE! |
| ENSMUSG00000113899.2 | 0 | NA | NA | NA | NA | NA | #VALUE! |
| ENSMUSG00000113900.2 | 0 | NA | NA | NA | NA | NA | #VALUE! |
| ENSMUSG00000113901.2 | 0 | NA | NA | NA | NA | NA | #VALUE! |
| ENSMUSG00000113903.2 | 0 | NA | NA | NA | NA | NA | #VALUE! |
| ENSMUSG00000113904.2 | 0 | NA | NA | NA | NA | NA | #VALUE! |
| ENSMUSG00000113905.2 | 0 | NA | NA | NA | NA | NA | #VALUE! |
| ENSMUSG00000113906.2 | 0 | NA | NA | NA | NA | NA | #VALUE! |
| ENSMUSG00000113907.2 | 0 | NA | NA | NA | NA | NA | #VALUE! |
| ENSMUSG00000113908.2 | 0 | NA | NA | NA | NA | NA | #VALUE! |
| ENSMUSG00000113909.2 | 0 | NA | NA | NA | NA | NA | #VALUE! |
| ENSMUSG00000113910.2 | 0.839297 | 3.273019 | 3.958392 | 0.826856 | 0.408319 | NA | #VALUE! |
| ENSMUSG00000113911.2 | 0 | NA | NA | NA | NA | NA | #VALUE! |
| ENSMUSG00000113912.2 | 0 | NA | NA | NA | NA | NA | #VALUE! |
| ENSMUSG00000113913.2 | 0 | NA | NA | NA | NA | NA | #VALUE! |
| ENSMUSG00000113914.2 | 0 | NA | NA | NA | NA | NA | #VALUE! |
| ENSMUSG00000113915.2 | 0 | NA | NA | NA | NA | NA | #VALUE! |
| ENSMUSG00000113916.3 | 0 | NA | NA | NA | NA | NA | #VALUE! |
| ENSMUSG00000113917.2 | 0 | NA | NA | NA | NA | NA | #VALUE! |
| ENSMUSG00000113918.2 | 0.466276 | 2.449889 | 3.997179 | 0.612904 | 0.53994 | NA | #VALUE! |
| ENSMUSG00000113919.2 | 0 | NA | NA | NA | NA | NA | #VALUE! |
| ENSMUSG00000113920.2 | 0 | NA | NA | NA | NA | NA | #VALUE! |
| ENSMUSG00000113921.2 | 0 | NA | NA | NA | NA | NA | #VALUE! |
| ENSMUSG00000113922.2 | 0 | NA | NA | NA | NA | NA | #VALUE! |
| ENSMUSG00000113923.2 | 0 | NA | NA | NA | NA | NA | #VALUE! |
| ENSMUSG00000113924.2 | 0 | NA | NA | NA | NA | NA | #VALUE! |
| ENSMUSG00000113925.2 | 0 | NA | NA | NA | NA | NA | #VALUE! |
| ENSMUSG00000113926.2 | 0 | NA | NA | NA | NA | NA | #VALUE! |

| | | | | | | | |
|---|---|---|---|---|---|---|---|
| ENSMUSG00000113927.2 | 0 | NA | NA | NA | NA | NA | #VALUE! |
| ENSMUSG00000113928.2 | 0 | NA | NA | NA | NA | NA | #VALUE! |
| ENSMUSG00000113929.2 | 0 | NA | NA | NA | NA | NA | #VALUE! |
| ENSMUSG00000113930.2 | 0 | NA | NA | NA | NA | NA | #VALUE! |
| ENSMUSG00000113931.2 | 0.3999 | -2.17566 | 4.018831 | -0.54137 | 0.588256 | NA | #VALUE! |
| ENSMUSG00000113932.2 | 0 | NA | NA | NA | NA | NA | #VALUE! |
| ENSMUSG00000113933.2 | 0 | NA | NA | NA | NA | NA | #VALUE! |
| ENSMUSG00000113934.2 | 0 | NA | NA | NA | NA | NA | #VALUE! |
| ENSMUSG00000113935.2 | 0 | NA | NA | NA | NA | NA | #VALUE! |
| ENSMUSG00000113936.2 | 0 | NA | NA | NA | NA | NA | #VALUE! |
| ENSMUSG00000113937.2 | 0 | NA | NA | NA | NA | NA | #VALUE! |
| ENSMUSG00000113938.2 | 0 | NA | NA | NA | NA | NA | #VALUE! |
| ENSMUSG00000113939.2 | 0 | NA | NA | NA | NA | NA | #VALUE! |
| ENSMUSG00000113940.2 | 0 | NA | NA | NA | NA | NA | #VALUE! |
| ENSMUSG00000113941.2 | 0 | NA | NA | NA | NA | NA | #VALUE! |
| ENSMUSG00000113942.2 | 0.093255 | 0.868093 | 4.072464 | 0.213162 | 0.831201 | NA | #VALUE! |
| ENSMUSG00000113943.2 | 0 | NA | NA | NA | NA | NA | #VALUE! |
| ENSMUSG00000113944.2 | 0 | NA | NA | NA | NA | NA | #VALUE! |
| ENSMUSG00000113945.2 | 0 | NA | NA | NA | NA | NA | #VALUE! |
| ENSMUSG00000113946.2 | 0 | NA | NA | NA | NA | NA | #VALUE! |
| ENSMUSG00000113947.2 | 0 | NA | NA | NA | NA | NA | #VALUE! |
| ENSMUSG00000113949.2 | 0 | NA | NA | NA | NA | NA | #VALUE! |
| ENSMUSG00000113950.2 | 0 | NA | NA | NA | NA | NA | #VALUE! |
| ENSMUSG00000113951.2 | 0 | NA | NA | NA | NA | NA | #VALUE! |
| ENSMUSG00000113952.2 | 0 | NA | NA | NA | NA | NA | #VALUE! |
| ENSMUSG00000113953.2 | 0 | NA | NA | NA | NA | NA | #VALUE! |
| ENSMUSG00000113954.2 | 0 | NA | NA | NA | NA | NA | #VALUE! |
| ENSMUSG00000113955.2 | 0 | NA | NA | NA | NA | NA | #VALUE! |
| ENSMUSG00000113956.2 | 0 | NA | NA | NA | NA | NA | #VALUE! |
| ENSMUSG00000113957.2 | 0 | NA | NA | NA | NA | NA | #VALUE! |
| ENSMUSG00000113958.2 | 0 | NA | NA | NA | NA | NA | #VALUE! |
| ENSMUSG00000113959.2 | 0 | NA | NA | NA | NA | NA | #VALUE! |
| ENSMUSG00000113960.2 | 0 | NA | NA | NA | NA | NA | #VALUE! |
| ENSMUSG00000113961.2 | 0 | NA | NA | NA | NA | NA | #VALUE! |
| ENSMUSG00000113962.2 | 0 | NA | NA | NA | NA | NA | #VALUE! |
| ENSMUSG00000113963.2 | 0 | NA | NA | NA | NA | NA | #VALUE! |
| ENSMUSG00000113964.2 | 0 | NA | NA | NA | NA | NA | #VALUE! |
| ENSMUSG00000113965.2 | 0 | NA | NA | NA | NA | NA | #VALUE! |
| ENSMUSG00000113966.2 | 0 | NA | NA | NA | NA | NA | #VALUE! |
| ENSMUSG00000113967.2 | 0 | NA | NA | NA | NA | NA | #VALUE! |
| ENSMUSG00000113968.2 | 0 | NA | NA | NA | NA | NA | #VALUE! |
| ENSMUSG00000113969.2 | 0 | NA | NA | NA | NA | NA | #VALUE! |
| ENSMUSG00000113970.2 | 0 | NA | NA | NA | NA | NA | #VALUE! |
| ENSMUSG00000113971.2 | 0 | NA | NA | NA | NA | NA | #VALUE! |

| | | | | | | | |
|---|---|---|---|---|---|---|---|
| ENSMUSG00000113972.2 | 0 | NA | NA | NA | NA | NA | #VALUE! |
| ENSMUSG00000113973.2 | 0 | NA | NA | NA | NA | NA | #VALUE! |
| ENSMUSG00000113974.2 | 0 | NA | NA | NA | NA | NA | #VALUE! |
| ENSMUSG00000113975.2 | 0 | NA | NA | NA | NA | NA | #VALUE! |
| ENSMUSG00000113976.2 | 0 | NA | NA | NA | NA | NA | #VALUE! |
| ENSMUSG00000113977.2 | 0 | NA | NA | NA | NA | NA | #VALUE! |
| ENSMUSG00000113978.2 | 0 | NA | NA | NA | NA | NA | #VALUE! |
| ENSMUSG00000113979.2 | 0 | NA | NA | NA | NA | NA | #VALUE! |
| ENSMUSG00000113980.2 | 0 | NA | NA | NA | NA | NA | #VALUE! |
| ENSMUSG00000113981.2 | 0 | NA | NA | NA | NA | NA | #VALUE! |
| ENSMUSG00000113982.2 | 0 | NA | NA | NA | NA | NA | #VALUE! |
| ENSMUSG00000113983.2 | 0 | NA | NA | NA | NA | NA | #VALUE! |
| ENSMUSG00000113984.2 | 0 | NA | NA | NA | NA | NA | #VALUE! |
| ENSMUSG00000113985.2 | 0 | NA | NA | NA | NA | NA | #VALUE! |
| ENSMUSG00000113986.2 | 0 | NA | NA | NA | NA | NA | #VALUE! |
| ENSMUSG00000113987.2 | 0 | NA | NA | NA | NA | NA | #VALUE! |
| ENSMUSG00000113989.2 | 0 | NA | NA | NA | NA | NA | #VALUE! |
| ENSMUSG00000113990.2 | 0 | NA | NA | NA | NA | NA | #VALUE! |
| ENSMUSG00000113991.2 | 0 | NA | NA | NA | NA | NA | #VALUE! |
| ENSMUSG00000113992.2 | 0 | NA | NA | NA | NA | NA | #VALUE! |
| ENSMUSG00000113993.2 | 0 | NA | NA | NA | NA | NA | #VALUE! |
| ENSMUSG00000113994.2 | 0 | NA | NA | NA | NA | NA | #VALUE! |
| ENSMUSG00000113995.2 | 0 | NA | NA | NA | NA | NA | #VALUE! |
| ENSMUSG00000113996.2 | 0 | NA | NA | NA | NA | NA | #VALUE! |
| ENSMUSG00000113997.2 | 0 | NA | NA | NA | NA | NA | #VALUE! |
| ENSMUSG00000113998.2 | 0 | NA | NA | NA | NA | NA | #VALUE! |
| ENSMUSG00000113999.2 | 0 | NA | NA | NA | NA | NA | #VALUE! |
| ENSMUSG00000114000.2 | 0 | NA | NA | NA | NA | NA | #VALUE! |
| ENSMUSG00000114001.2 | 0 | NA | NA | NA | NA | NA | #VALUE! |
| ENSMUSG00000114002.2 | 0 | NA | NA | NA | NA | NA | #VALUE! |
| ENSMUSG00000114003.2 | 1.606028 | 1.430762 | 3.868933 | 0.369808 | 0.711526 | NA | #VALUE! |
| ENSMUSG00000114004.2 | 0 | NA | NA | NA | NA | NA | #VALUE! |
| ENSMUSG00000114005.2 | 0 | NA | NA | NA | NA | NA | #VALUE! |
| ENSMUSG00000114006.2 | 0 | NA | NA | NA | NA | NA | #VALUE! |
| ENSMUSG00000114007.2 | 0 | NA | NA | NA | NA | NA | #VALUE! |
| ENSMUSG00000114008.2 | 0 | NA | NA | NA | NA | NA | #VALUE! |
| ENSMUSG00000114009.2 | 0 | NA | NA | NA | NA | NA | #VALUE! |
| ENSMUSG00000114010.2 | 0 | NA | NA | NA | NA | NA | #VALUE! |
| ENSMUSG00000114011.2 | 0 | NA | NA | NA | NA | NA | #VALUE! |
| ENSMUSG00000114012.2 | 0 | NA | NA | NA | NA | NA | #VALUE! |
| ENSMUSG00000114013.2 | 0 | NA | NA | NA | NA | NA | #VALUE! |
| ENSMUSG00000114014.2 | 0 | NA | NA | NA | NA | NA | #VALUE! |
| ENSMUSG00000114015.2 | 0 | NA | NA | NA | NA | NA | #VALUE! |
| ENSMUSG00000114016.2 | 0 | NA | NA | NA | NA | NA | #VALUE! |

| | | | | | | | |
|---|---|---|---|---|---|---|---|
| ENSMUSG00000114017.2 | 0 | NA | NA | NA | NA | NA | #VALUE! |
| ENSMUSG00000114018.2 | 0 | NA | NA | NA | NA | NA | #VALUE! |
| ENSMUSG00000114020.2 | 0 | NA | NA | NA | NA | NA | #VALUE! |
| ENSMUSG00000114021.2 | 0 | NA | NA | NA | NA | NA | #VALUE! |
| ENSMUSG00000114022.2 | 0 | NA | NA | NA | NA | NA | #VALUE! |
| ENSMUSG00000114023.2 | 0 | NA | NA | NA | NA | NA | #VALUE! |
| ENSMUSG00000114024.2 | 0 | NA | NA | NA | NA | NA | #VALUE! |
| ENSMUSG00000114025.2 | 0.816443 | -3.21914 | 3.961753 | -0.81255 | 0.416474 | NA | #VALUE! |
| ENSMUSG00000114026.2 | 0 | NA | NA | NA | NA | NA | #VALUE! |
| ENSMUSG00000114027.2 | 0 | NA | NA | NA | NA | NA | #VALUE! |
| ENSMUSG00000114028.2 | 0 | NA | NA | NA | NA | NA | #VALUE! |
| ENSMUSG00000114029.2 | 0 | NA | NA | NA | NA | NA | #VALUE! |
| ENSMUSG00000114030.2 | 0 | NA | NA | NA | NA | NA | #VALUE! |
| ENSMUSG00000114031.2 | 0 | NA | NA | NA | NA | NA | #VALUE! |
| ENSMUSG00000114032.2 | 0 | NA | NA | NA | NA | NA | #VALUE! |
| ENSMUSG00000114033.2 | 0 | NA | NA | NA | NA | NA | #VALUE! |
| ENSMUSG00000114034.2 | 0 | NA | NA | NA | NA | NA | #VALUE! |
| ENSMUSG00000114035.2 | 0 | NA | NA | NA | NA | NA | #VALUE! |
| ENSMUSG00000114036.2 | 0 | NA | NA | NA | NA | NA | #VALUE! |
| ENSMUSG00000114037.2 | 0 | NA | NA | NA | NA | NA | #VALUE! |
| ENSMUSG00000114038.2 | 0 | NA | NA | NA | NA | NA | #VALUE! |
| ENSMUSG00000114039.2 | 0 | NA | NA | NA | NA | NA | #VALUE! |
| ENSMUSG00000114040.2 | 0 | NA | NA | NA | NA | NA | #VALUE! |
| ENSMUSG00000114041.3 | 0 | NA | NA | NA | NA | NA | #VALUE! |
| ENSMUSG00000114042.2 | 0 | NA | NA | NA | NA | NA | #VALUE! |
| ENSMUSG00000114043.2 | 0 | NA | NA | NA | NA | NA | #VALUE! |
| ENSMUSG00000114044.2 | 0 | NA | NA | NA | NA | NA | #VALUE! |
| ENSMUSG00000114045.2 | 0.816443 | -3.21914 | 3.961753 | -0.81255 | 0.416474 | NA | #VALUE! |
| ENSMUSG00000114046.2 | 0 | NA | NA | NA | NA | NA | #VALUE! |
| ENSMUSG00000114047.2 | 0 | NA | NA | NA | NA | NA | #VALUE! |
| ENSMUSG00000114048.2 | 0 | NA | NA | NA | NA | NA | #VALUE! |
| ENSMUSG00000114049.2 | 0 | NA | NA | NA | NA | NA | #VALUE! |
| ENSMUSG00000114050.2 | 0 | NA | NA | NA | NA | NA | #VALUE! |
| ENSMUSG00000114051.2 | 0 | NA | NA | NA | NA | NA | #VALUE! |
| ENSMUSG00000114052.2 | 0 | NA | NA | NA | NA | NA | #VALUE! |
| ENSMUSG00000114053.2 | 0 | NA | NA | NA | NA | NA | #VALUE! |
| ENSMUSG00000114054.2 | 0 | NA | NA | NA | NA | NA | #VALUE! |
| ENSMUSG00000114055.2 | 0 | NA | NA | NA | NA | NA | #VALUE! |
| ENSMUSG00000114056.2 | 0 | NA | NA | NA | NA | NA | #VALUE! |
| ENSMUSG00000114057.2 | 0 | NA | NA | NA | NA | NA | #VALUE! |
| ENSMUSG00000114058.2 | 0 | NA | NA | NA | NA | NA | #VALUE! |
| ENSMUSG00000114059.2 | 0 | NA | NA | NA | NA | NA | #VALUE! |
| ENSMUSG00000114060.2 | 0 | NA | NA | NA | NA | NA | #VALUE! |
| ENSMUSG00000114061.2 | 0 | NA | NA | NA | NA | NA | #VALUE! |

| | | | | | | | |
|---|---|---|---|---|---|---|---|
| ENSMUSG00000114062.2 | 0 | NA | NA | NA | NA | NA | #VALUE! |
| ENSMUSG00000114063.2 | 0 | NA | NA | NA | NA | NA | #VALUE! |
| ENSMUSG00000114064.2 | 0.198268 | -0.06494 | 4.068276 | -0.01596 | 0.987265 | NA | #VALUE! |
| ENSMUSG00000114065.2 | 0 | NA | NA | NA | NA | NA | #VALUE! |
| ENSMUSG00000114066.2 | 0.105013 | -0.97324 | 4.076293 | -0.23876 | 0.811295 | NA | #VALUE! |
| ENSMUSG00000114067.2 | 0 | NA | NA | NA | NA | NA | #VALUE! |
| ENSMUSG00000114068.2 | 0 | NA | NA | NA | NA | NA | #VALUE! |
| ENSMUSG00000114069.2 | 0 | NA | NA | NA | NA | NA | #VALUE! |
| ENSMUSG00000114070.2 | 0 | NA | NA | NA | NA | NA | #VALUE! |
| ENSMUSG00000114071.2 | 0 | NA | NA | NA | NA | NA | #VALUE! |
| ENSMUSG00000114072.2 | 0 | NA | NA | NA | NA | NA | #VALUE! |
| ENSMUSG00000114073.2 | 0 | NA | NA | NA | NA | NA | #VALUE! |
| ENSMUSG00000114074.2 | 0 | NA | NA | NA | NA | NA | #VALUE! |
| ENSMUSG00000114075.2 | 0 | NA | NA | NA | NA | NA | #VALUE! |
| ENSMUSG00000114076.2 | 0 | NA | NA | NA | NA | NA | #VALUE! |
| ENSMUSG00000114077.2 | 0 | NA | NA | NA | NA | NA | #VALUE! |
| ENSMUSG00000114078.2 | 0 | NA | NA | NA | NA | NA | #VALUE! |
| ENSMUSG00000114079.2 | 0 | NA | NA | NA | NA | NA | #VALUE! |
| ENSMUSG00000114080.2 | 0 | NA | NA | NA | NA | NA | #VALUE! |
| ENSMUSG00000114081.2 | 0 | NA | NA | NA | NA | NA | #VALUE! |
| ENSMUSG00000114082.2 | 0 | NA | NA | NA | NA | NA | #VALUE! |
| ENSMUSG00000114083.2 | 0 | NA | NA | NA | NA | NA | #VALUE! |
| ENSMUSG00000114084.2 | 0 | NA | NA | NA | NA | NA | #VALUE! |
| ENSMUSG00000114085.2 | 0 | NA | NA | NA | NA | NA | #VALUE! |
| ENSMUSG00000114086.2 | 0 | NA | NA | NA | NA | NA | #VALUE! |
| ENSMUSG00000114087.2 | 0 | NA | NA | NA | NA | NA | #VALUE! |
| ENSMUSG00000114088.2 | 0 | NA | NA | NA | NA | NA | #VALUE! |
| ENSMUSG00000114089.2 | 0 | NA | NA | NA | NA | NA | #VALUE! |
| ENSMUSG00000114090.2 | 0 | NA | NA | NA | NA | NA | #VALUE! |
| ENSMUSG00000114091.2 | 0 | NA | NA | NA | NA | NA | #VALUE! |
| ENSMUSG00000114092.2 | 0 | NA | NA | NA | NA | NA | #VALUE! |
| ENSMUSG00000114093.2 | 0 | NA | NA | NA | NA | NA | #VALUE! |
| ENSMUSG00000114094.2 | 0.105013 | -0.97324 | 4.076293 | -0.23876 | 0.811295 | NA | #VALUE! |
| ENSMUSG00000114095.2 | 0 | NA | NA | NA | NA | NA | #VALUE! |
| ENSMUSG00000114096.2 | 0 | NA | NA | NA | NA | NA | #VALUE! |
| ENSMUSG00000114097.2 | 0 | NA | NA | NA | NA | NA | #VALUE! |
| ENSMUSG00000114098.2 | 0 | NA | NA | NA | NA | NA | #VALUE! |
| ENSMUSG00000114099.2 | 0 | NA | NA | NA | NA | NA | #VALUE! |
| ENSMUSG00000114100.2 | 0 | NA | NA | NA | NA | NA | #VALUE! |
| ENSMUSG00000114101.2 | 0 | NA | NA | NA | NA | NA | #VALUE! |
| ENSMUSG00000114102.2 | 0 | NA | NA | NA | NA | NA | #VALUE! |
| ENSMUSG00000114104.2 | 0 | NA | NA | NA | NA | NA | #VALUE! |
| ENSMUSG00000114105.2 | 0 | NA | NA | NA | NA | NA | #VALUE! |
| ENSMUSG00000114106.2 | 0 | NA | NA | NA | NA | NA | #VALUE! |

| | | | | | | | |
|---|---|---|---|---|---|---|---|
| ENSMUSG00000114107.2 | 0 | NA | NA | NA | NA | NA | #VALUE! |
| ENSMUSG00000114108.2 | 0 | NA | NA | NA | NA | NA | #VALUE! |
| ENSMUSG00000114109.2 | 0 | NA | NA | NA | NA | NA | #VALUE! |
| ENSMUSG00000114110.2 | 0 | NA | NA | NA | NA | NA | #VALUE! |
| ENSMUSG00000114111.2 | 0 | NA | NA | NA | NA | NA | #VALUE! |
| ENSMUSG00000114112.2 | 0 | NA | NA | NA | NA | NA | #VALUE! |
| ENSMUSG00000114113.2 | 0 | NA | NA | NA | NA | NA | #VALUE! |
| ENSMUSG00000114114.2 | 0 | NA | NA | NA | NA | NA | #VALUE! |
| ENSMUSG00000114115.2 | 0 | NA | NA | NA | NA | NA | #VALUE! |
| ENSMUSG00000114117.2 | 0 | NA | NA | NA | NA | NA | #VALUE! |
| ENSMUSG00000114118.2 | 0 | NA | NA | NA | NA | NA | #VALUE! |
| ENSMUSG00000114119.2 | 0 | NA | NA | NA | NA | NA | #VALUE! |
| ENSMUSG00000114120.2 | 0 | NA | NA | NA | NA | NA | #VALUE! |
| ENSMUSG00000114121.2 | 0 | NA | NA | NA | NA | NA | #VALUE! |
| ENSMUSG00000114122.2 | 0 | NA | NA | NA | NA | NA | #VALUE! |
| ENSMUSG00000114123.2 | 0 | NA | NA | NA | NA | NA | #VALUE! |
| ENSMUSG00000114124.2 | 0 | NA | NA | NA | NA | NA | #VALUE! |
| ENSMUSG00000114125.2 | 0 | NA | NA | NA | NA | NA | #VALUE! |
| ENSMUSG00000114126.2 | 0 | NA | NA | NA | NA | NA | #VALUE! |
| ENSMUSG00000114127.2 | 0 | NA | NA | NA | NA | NA | #VALUE! |
| ENSMUSG00000114128.2 | 0 | NA | NA | NA | NA | NA | #VALUE! |
| ENSMUSG00000114129.2 | 0 | NA | NA | NA | NA | NA | #VALUE! |
| ENSMUSG00000114130.2 | 0 | NA | NA | NA | NA | NA | #VALUE! |
| ENSMUSG00000114131.2 | 0 | NA | NA | NA | NA | NA | #VALUE! |
| ENSMUSG00000114132.2 | 0 | NA | NA | NA | NA | NA | #VALUE! |
| ENSMUSG00000114133.2 | 0.600647 | -0.82995 | 3.987207 | -0.20815 | 0.835109 | NA | #VALUE! |
| ENSMUSG00000114134.2 | 0 | NA | NA | NA | NA | NA | #VALUE! |
| ENSMUSG00000114135.2 | 0 | NA | NA | NA | NA | NA | #VALUE! |
| ENSMUSG00000114136.2 | 0 | NA | NA | NA | NA | NA | #VALUE! |
| ENSMUSG00000114137.2 | 0 | NA | NA | NA | NA | NA | #VALUE! |
| ENSMUSG00000114138.2 | 5.522084 | 0.542408 | 3.135738 | 0.172976 | 0.86267 | NA | #VALUE! |
| ENSMUSG00000114139.2 | 0 | NA | NA | NA | NA | NA | #VALUE! |
| ENSMUSG00000114140.2 | 0 | NA | NA | NA | NA | NA | #VALUE! |
| ENSMUSG00000114141.2 | 0 | NA | NA | NA | NA | NA | #VALUE! |
| ENSMUSG00000114142.2 | 0 | NA | NA | NA | NA | NA | #VALUE! |
| ENSMUSG00000114143.2 | 0 | NA | NA | NA | NA | NA | #VALUE! |
| ENSMUSG00000114144.2 | 0 | NA | NA | NA | NA | NA | #VALUE! |
| ENSMUSG00000114145.2 | 0 | NA | NA | NA | NA | NA | #VALUE! |
| ENSMUSG00000114147.2 | 0 | NA | NA | NA | NA | NA | #VALUE! |
| ENSMUSG00000114148.2 | 0 | NA | NA | NA | NA | NA | #VALUE! |
| ENSMUSG00000114149.2 | 0 | NA | NA | NA | NA | NA | #VALUE! |
| ENSMUSG00000114150.2 | 0 | NA | NA | NA | NA | NA | #VALUE! |
| ENSMUSG00000114151.2 | 0 | NA | NA | NA | NA | NA | #VALUE! |
| ENSMUSG00000114152.2 | 0 | NA | NA | NA | NA | NA | #VALUE! |

| | | | | | | | |
|---|---|---|---|---|---|---|---|
| ENSMUSG00000114153.2 | 0 | NA | NA | NA | NA | NA | #VALUE! |
| ENSMUSG00000114155.2 | 0 | NA | NA | NA | NA | NA | #VALUE! |
| ENSMUSG00000114156.2 | 0 | NA | NA | NA | NA | NA | #VALUE! |
| ENSMUSG00000114157.2 | 0 | NA | NA | NA | NA | NA | #VALUE! |
| ENSMUSG00000114158.2 | 0 | NA | NA | NA | NA | NA | #VALUE! |
| ENSMUSG00000114159.2 | 0 | NA | NA | NA | NA | NA | #VALUE! |
| ENSMUSG00000114160.2 | 0 | NA | NA | NA | NA | NA | #VALUE! |
| ENSMUSG00000114161.2 | 0 | NA | NA | NA | NA | NA | #VALUE! |
| ENSMUSG00000114162.2 | 0 | NA | NA | NA | NA | NA | #VALUE! |
| ENSMUSG00000114163.2 | 0 | NA | NA | NA | NA | NA | #VALUE! |
| ENSMUSG00000114164.2 | 0 | NA | NA | NA | NA | NA | #VALUE! |
| ENSMUSG00000114165.2 | 0 | NA | NA | NA | NA | NA | #VALUE! |
| ENSMUSG00000114166.2 | 0 | NA | NA | NA | NA | NA | #VALUE! |
| ENSMUSG00000114167.2 | 0 | NA | NA | NA | NA | NA | #VALUE! |
| ENSMUSG00000114168.2 | 0 | NA | NA | NA | NA | NA | #VALUE! |
| ENSMUSG00000114169.2 | 0 | NA | NA | NA | NA | NA | #VALUE! |
| ENSMUSG00000114170.2 | 0 | NA | NA | NA | NA | NA | #VALUE! |
| ENSMUSG00000114171.2 | 0 | NA | NA | NA | NA | NA | #VALUE! |
| ENSMUSG00000114172.2 | 0.336567 | 1.944925 | 4.029936 | 0.482619 | 0.629366 | NA | #VALUE! |
| ENSMUSG00000114173.2 | 0 | NA | NA | NA | NA | NA | #VALUE! |
| ENSMUSG00000114174.2 | 0 | NA | NA | NA | NA | NA | #VALUE! |
| ENSMUSG00000114175.2 | 0 | NA | NA | NA | NA | NA | #VALUE! |
| ENSMUSG00000114176.2 | 0 | NA | NA | NA | NA | NA | #VALUE! |
| ENSMUSG00000114177.2 | 0 | NA | NA | NA | NA | NA | #VALUE! |
| ENSMUSG00000114178.2 | 0 | NA | NA | NA | NA | NA | #VALUE! |
| ENSMUSG00000114179.2 | 0 | NA | NA | NA | NA | NA | #VALUE! |
| ENSMUSG00000114180.2 | 0 | NA | NA | NA | NA | NA | #VALUE! |
| ENSMUSG00000114181.2 | 0 | NA | NA | NA | NA | NA | #VALUE! |
| ENSMUSG00000114182.2 | 0 | NA | NA | NA | NA | NA | #VALUE! |
| ENSMUSG00000114183.2 | 0 | NA | NA | NA | NA | NA | #VALUE! |
| ENSMUSG00000114184.2 | 0.093255 | 0.868093 | 4.072464 | 0.213162 | 0.831201 | NA | #VALUE! |
| ENSMUSG00000114185.2 | 0 | NA | NA | NA | NA | NA | #VALUE! |
| ENSMUSG00000114186.2 | 0 | NA | NA | NA | NA | NA | #VALUE! |
| ENSMUSG00000114187.2 | 0 | NA | NA | NA | NA | NA | #VALUE! |
| ENSMUSG00000114188.2 | 0 | NA | NA | NA | NA | NA | #VALUE! |
| ENSMUSG00000114189.2 | 0 | NA | NA | NA | NA | NA | #VALUE! |
| ENSMUSG00000114190.2 | 0 | NA | NA | NA | NA | NA | #VALUE! |
| ENSMUSG00000114191.2 | 0 | NA | NA | NA | NA | NA | #VALUE! |
| ENSMUSG00000114192.2 | 0 | NA | NA | NA | NA | NA | #VALUE! |
| ENSMUSG00000114193.2 | 0 | NA | NA | NA | NA | NA | #VALUE! |
| ENSMUSG00000114194.2 | 0 | NA | NA | NA | NA | NA | #VALUE! |
| ENSMUSG00000114195.2 | 0 | NA | NA | NA | NA | NA | #VALUE! |
| ENSMUSG00000114196.2 | 0 | NA | NA | NA | NA | NA | #VALUE! |
| ENSMUSG00000114197.2 | 0 | NA | NA | NA | NA | NA | #VALUE! |

| | | | | | | | |
|---|---|---|---|---|---|---|---|
| ENSMUSG00000114198.2 | 0 | NA | NA | NA | NA | NA | #VALUE! |
| ENSMUSG00000114199.2 | 0 | NA | NA | NA | NA | NA | #VALUE! |
| ENSMUSG00000114200.2 | 0 | NA | NA | NA | NA | NA | #VALUE! |
| ENSMUSG00000114201.2 | 0 | NA | NA | NA | NA | NA | #VALUE! |
| ENSMUSG00000114202.3 | 0 | NA | NA | NA | NA | NA | #VALUE! |
| ENSMUSG00000114203.2 | 0 | NA | NA | NA | NA | NA | #VALUE! |
| ENSMUSG00000114204.2 | 0 | NA | NA | NA | NA | NA | #VALUE! |
| ENSMUSG00000114205.2 | 0 | NA | NA | NA | NA | NA | #VALUE! |
| ENSMUSG00000114206.2 | 0 | NA | NA | NA | NA | NA | #VALUE! |
| ENSMUSG00000114207.2 | 0 | NA | NA | NA | NA | NA | #VALUE! |
| ENSMUSG00000114208.2 | 0 | NA | NA | NA | NA | NA | #VALUE! |
| ENSMUSG00000114209.2 | 0 | NA | NA | NA | NA | NA | #VALUE! |
| ENSMUSG00000114210.3 | 0 | NA | NA | NA | NA | NA | #VALUE! |
| ENSMUSG00000114211.2 | 0 | NA | NA | NA | NA | NA | #VALUE! |
| ENSMUSG00000114212.2 | 0 | NA | NA | NA | NA | NA | #VALUE! |
| ENSMUSG00000114213.2 | 0 | NA | NA | NA | NA | NA | #VALUE! |
| ENSMUSG00000114214.2 | 0 | NA | NA | NA | NA | NA | #VALUE! |
| ENSMUSG00000114215.2 | 0 | NA | NA | NA | NA | NA | #VALUE! |
| ENSMUSG00000114216.2 | 0 | NA | NA | NA | NA | NA | #VALUE! |
| ENSMUSG00000114217.2 | 0 | NA | NA | NA | NA | NA | #VALUE! |
| ENSMUSG00000114218.2 | 0 | NA | NA | NA | NA | NA | #VALUE! |
| ENSMUSG00000114219.2 | 0 | NA | NA | NA | NA | NA | #VALUE! |
| ENSMUSG00000114220.2 | 0 | NA | NA | NA | NA | NA | #VALUE! |
| ENSMUSG00000114221.2 | 0 | NA | NA | NA | NA | NA | #VALUE! |
| ENSMUSG00000114222.2 | 0 | NA | NA | NA | NA | NA | #VALUE! |
| ENSMUSG00000114223.2 | 0 | NA | NA | NA | NA | NA | #VALUE! |
| ENSMUSG00000114225.2 | 0 | NA | NA | NA | NA | NA | #VALUE! |
| ENSMUSG00000114226.2 | 0 | NA | NA | NA | NA | NA | #VALUE! |
| ENSMUSG00000114227.2 | 0 | NA | NA | NA | NA | NA | #VALUE! |
| ENSMUSG00000114228.2 | 0 | NA | NA | NA | NA | NA | #VALUE! |
| ENSMUSG00000114229.2 | 0 | NA | NA | NA | NA | NA | #VALUE! |
| ENSMUSG00000114230.2 | 0 | NA | NA | NA | NA | NA | #VALUE! |
| ENSMUSG00000114231.2 | 0 | NA | NA | NA | NA | NA | #VALUE! |
| ENSMUSG00000114232.2 | 0 | NA | NA | NA | NA | NA | #VALUE! |
| ENSMUSG00000114233.2 | 0 | NA | NA | NA | NA | NA | #VALUE! |
| ENSMUSG00000114234.2 | 0 | NA | NA | NA | NA | NA | #VALUE! |
| ENSMUSG00000114235.2 | 0 | NA | NA | NA | NA | NA | #VALUE! |
| ENSMUSG00000114236.2 | 0 | NA | NA | NA | NA | NA | #VALUE! |
| ENSMUSG00000114237.2 | 0 | NA | NA | NA | NA | NA | #VALUE! |
| ENSMUSG00000114238.2 | 0 | NA | NA | NA | NA | NA | #VALUE! |
| ENSMUSG00000114239.2 | 0 | NA | NA | NA | NA | NA | #VALUE! |
| ENSMUSG00000114240.2 | 0 | NA | NA | NA | NA | NA | #VALUE! |
| ENSMUSG00000114241.2 | 0 | NA | NA | NA | NA | NA | #VALUE! |
| ENSMUSG00000114242.2 | 0 | NA | NA | NA | NA | NA | #VALUE! |

| | | | | | | | |
|---|---|---|---|---|---|---|---|
| ENSMUSG00000114243.2 | 7.487337 | -2.18708 | 3.190188 | -0.68556 | 0.492988 | NA | #VALUE! |
| ENSMUSG00000114244.2 | 0 | NA | NA | NA | NA | NA | #VALUE! |
| ENSMUSG00000114245.2 | 0 | NA | NA | NA | NA | NA | #VALUE! |
| ENSMUSG00000114246.2 | 0 | NA | NA | NA | NA | NA | #VALUE! |
| ENSMUSG00000114247.2 | 3.287895 | 4.100863 | 3.913828 | 1.047788 | 0.294736 | NA | #VALUE! |
| ENSMUSG00000114248.2 | 0 | NA | NA | NA | NA | NA | #VALUE! |
| ENSMUSG00000114249.2 | 0 | NA | NA | NA | NA | NA | #VALUE! |
| ENSMUSG00000114250.3 | 0 | NA | NA | NA | NA | NA | #VALUE! |
| ENSMUSG00000114251.2 | 0 | NA | NA | NA | NA | NA | #VALUE! |
| ENSMUSG00000114252.2 | 0 | NA | NA | NA | NA | NA | #VALUE! |
| ENSMUSG00000114253.3 | 0 | NA | NA | NA | NA | NA | #VALUE! |
| ENSMUSG00000114254.2 | 0.18651 | 1.401769 | 4.056354 | 0.345574 | 0.729663 | NA | #VALUE! |
| ENSMUSG00000114256.2 | 5.482095 | -5.98256 | 3.914941 | -1.52813 | 0.126479 | NA | #VALUE! |
| ENSMUSG00000114258.2 | 0 | NA | NA | NA | NA | NA | #VALUE! |
| ENSMUSG00000114259.2 | 0 | NA | NA | NA | NA | NA | #VALUE! |
| ENSMUSG00000114260.2 | 0 | NA | NA | NA | NA | NA | #VALUE! |
| ENSMUSG00000114261.2 | 0 | NA | NA | NA | NA | NA | #VALUE! |
| ENSMUSG00000114262.2 | 0 | NA | NA | NA | NA | NA | #VALUE! |
| ENSMUSG00000114263.2 | 0 | NA | NA | NA | NA | NA | #VALUE! |
| ENSMUSG00000114264.2 | 3.823463 | 5.43312 | 3.918438 | 1.386552 | 0.165578 | NA | #VALUE! |
| ENSMUSG00000114265.2 | 0 | NA | NA | NA | NA | NA | #VALUE! |
| ENSMUSG00000114266.2 | 0 | NA | NA | NA | NA | NA | #VALUE! |
| ENSMUSG00000114267.2 | 0 | NA | NA | NA | NA | NA | #VALUE! |
| ENSMUSG00000114270.2 | 0 | NA | NA | NA | NA | NA | #VALUE! |
| ENSMUSG00000114271.2 | 0 | NA | NA | NA | NA | NA | #VALUE! |
| ENSMUSG00000114272.2 | 0 | NA | NA | NA | NA | NA | #VALUE! |
| ENSMUSG00000114273.2 | 0 | NA | NA | NA | NA | NA | #VALUE! |
| ENSMUSG00000114274.2 | 0 | NA | NA | NA | NA | NA | #VALUE! |
| ENSMUSG00000114275.2 | 0 | NA | NA | NA | NA | NA | #VALUE! |
| ENSMUSG00000114276.2 | 0 | NA | NA | NA | NA | NA | #VALUE! |
| ENSMUSG00000114277.2 | 0 | NA | NA | NA | NA | NA | #VALUE! |
| ENSMUSG00000114278.3 | 0 | NA | NA | NA | NA | NA | #VALUE! |
| ENSMUSG00000114279.2 | 0 | NA | NA | NA | NA | NA | #VALUE! |
| ENSMUSG00000114280.2 | 0 | NA | NA | NA | NA | NA | #VALUE! |
| ENSMUSG00000114281.2 | 0 | NA | NA | NA | NA | NA | #VALUE! |
| ENSMUSG00000114282.2 | 0 | NA | NA | NA | NA | NA | #VALUE! |
| ENSMUSG00000114283.2 | 0 | NA | NA | NA | NA | NA | #VALUE! |
| ENSMUSG00000114284.2 | 0 | NA | NA | NA | NA | NA | #VALUE! |
| ENSMUSG00000114285.2 | 0 | NA | NA | NA | NA | NA | #VALUE! |
| ENSMUSG00000114286.2 | 0 | NA | NA | NA | NA | NA | #VALUE! |
| ENSMUSG00000114287.2 | 0 | NA | NA | NA | NA | NA | #VALUE! |
| ENSMUSG00000114288.2 | 0 | NA | NA | NA | NA | NA | #VALUE! |
| ENSMUSG00000114289.2 | 0 | NA | NA | NA | NA | NA | #VALUE! |
| ENSMUSG00000114290.2 | 0 | NA | NA | NA | NA | NA | #VALUE! |

| | | | | | | | |
|---|---|---|---|---|---|---|---|
| ENSMUSG00000114291.2 | 0 | NA | NA | NA | NA | NA | #VALUE! |
| ENSMUSG00000114292.2 | 0 | NA | NA | NA | NA | NA | #VALUE! |
| ENSMUSG00000114293.2 | 0 | NA | NA | NA | NA | NA | #VALUE! |
| ENSMUSG00000114294.2 | 0 | NA | NA | NA | NA | NA | #VALUE! |
| ENSMUSG00000114295.2 | 0.093255 | 0.868093 | 4.072464 | 0.213162 | 0.831201 | NA | #VALUE! |
| ENSMUSG00000114296.2 | 0 | NA | NA | NA | NA | NA | #VALUE! |
| ENSMUSG00000114297.2 | 0 | NA | NA | NA | NA | NA | #VALUE! |
| ENSMUSG00000114298.2 | 0 | NA | NA | NA | NA | NA | #VALUE! |
| ENSMUSG00000114299.2 | 0 | NA | NA | NA | NA | NA | #VALUE! |
| ENSMUSG00000114300.2 | 0 | NA | NA | NA | NA | NA | #VALUE! |
| ENSMUSG00000114301.2 | 0 | NA | NA | NA | NA | NA | #VALUE! |
| ENSMUSG00000114302.2 | 0 | NA | NA | NA | NA | NA | #VALUE! |
| ENSMUSG00000114303.2 | 0 | NA | NA | NA | NA | NA | #VALUE! |
| ENSMUSG00000114304.2 | 0 | NA | NA | NA | NA | NA | #VALUE! |
| ENSMUSG00000114305.2 | 0 | NA | NA | NA | NA | NA | #VALUE! |
| ENSMUSG00000114306.2 | 0 | NA | NA | NA | NA | NA | #VALUE! |
| ENSMUSG00000114307.2 | 0 | NA | NA | NA | NA | NA | #VALUE! |
| ENSMUSG00000114308.2 | 0 | NA | NA | NA | NA | NA | #VALUE! |
| ENSMUSG00000114309.2 | 0 | NA | NA | NA | NA | NA | #VALUE! |
| ENSMUSG00000114310.2 | 0 | NA | NA | NA | NA | NA | #VALUE! |
| ENSMUSG00000114311.2 | 0 | NA | NA | NA | NA | NA | #VALUE! |
| ENSMUSG00000114312.2 | 0 | NA | NA | NA | NA | NA | #VALUE! |
| ENSMUSG00000114313.2 | 0 | NA | NA | NA | NA | NA | #VALUE! |
| ENSMUSG00000114314.2 | 0 | NA | NA | NA | NA | NA | #VALUE! |
| ENSMUSG00000114315.2 | 0 | NA | NA | NA | NA | NA | #VALUE! |
| ENSMUSG00000114316.2 | 0 | NA | NA | NA | NA | NA | #VALUE! |
| ENSMUSG00000114317.2 | 0 | NA | NA | NA | NA | NA | #VALUE! |
| ENSMUSG00000114318.2 | 0 | NA | NA | NA | NA | NA | #VALUE! |
| ENSMUSG00000114319.2 | 0 | NA | NA | NA | NA | NA | #VALUE! |
| ENSMUSG00000114320.2 | 0 | NA | NA | NA | NA | NA | #VALUE! |
| ENSMUSG00000114321.2 | 0.093255 | 0.868093 | 4.072464 | 0.213162 | 0.831201 | NA | #VALUE! |
| ENSMUSG00000114322.2 | 0 | NA | NA | NA | NA | NA | #VALUE! |
| ENSMUSG00000114323.2 | 0 | NA | NA | NA | NA | NA | #VALUE! |
| ENSMUSG00000114324.2 | 0 | NA | NA | NA | NA | NA | #VALUE! |
| ENSMUSG00000114325.2 | 0 | NA | NA | NA | NA | NA | #VALUE! |
| ENSMUSG00000114326.2 | 0 | NA | NA | NA | NA | NA | #VALUE! |
| ENSMUSG00000114327.2 | 0 | NA | NA | NA | NA | NA | #VALUE! |
| ENSMUSG00000114328.2 | 0 | NA | NA | NA | NA | NA | #VALUE! |
| ENSMUSG00000114329.2 | 0 | NA | NA | NA | NA | NA | #VALUE! |
| ENSMUSG00000114330.2 | 0 | NA | NA | NA | NA | NA | #VALUE! |
| ENSMUSG00000114331.2 | 0 | NA | NA | NA | NA | NA | #VALUE! |
| ENSMUSG00000114332.2 | 0 | NA | NA | NA | NA | NA | #VALUE! |
| ENSMUSG00000114333.2 | 0 | NA | NA | NA | NA | NA | #VALUE! |
| ENSMUSG00000114334.2 | 0 | NA | NA | NA | NA | NA | #VALUE! |

| | | | | | | | |
|---|---|---|---|---|---|---|---|
| ENSMUSG00000114335.2 | 0 | NA | NA | NA | NA | NA | #VALUE! |
| ENSMUSG00000114336.2 | 0 | NA | NA | NA | NA | NA | #VALUE! |
| ENSMUSG00000114337.2 | 0 | NA | NA | NA | NA | NA | #VALUE! |
| ENSMUSG00000114338.2 | 0 | NA | NA | NA | NA | NA | #VALUE! |
| ENSMUSG00000114339.2 | 0 | NA | NA | NA | NA | NA | #VALUE! |
| ENSMUSG00000114340.2 | 0 | NA | NA | NA | NA | NA | #VALUE! |
| ENSMUSG00000114341.2 | 0 | NA | NA | NA | NA | NA | #VALUE! |
| ENSMUSG00000114342.2 | 0 | NA | NA | NA | NA | NA | #VALUE! |
| ENSMUSG00000114343.2 | 0 | NA | NA | NA | NA | NA | #VALUE! |
| ENSMUSG00000114344.2 | 0 | NA | NA | NA | NA | NA | #VALUE! |
| ENSMUSG00000114345.2 | 0 | NA | NA | NA | NA | NA | #VALUE! |
| ENSMUSG00000114346.2 | 0 | NA | NA | NA | NA | NA | #VALUE! |
| ENSMUSG00000114347.2 | 0 | NA | NA | NA | NA | NA | #VALUE! |
| ENSMUSG00000114348.2 | 0 | NA | NA | NA | NA | NA | #VALUE! |
| ENSMUSG00000114349.2 | 0 | NA | NA | NA | NA | NA | #VALUE! |
| ENSMUSG00000114350.2 | 0 | NA | NA | NA | NA | NA | #VALUE! |
| ENSMUSG00000114352.2 | 0 | NA | NA | NA | NA | NA | #VALUE! |
| ENSMUSG00000114353.2 | 0 | NA | NA | NA | NA | NA | #VALUE! |
| ENSMUSG00000114354.2 | 0 | NA | NA | NA | NA | NA | #VALUE! |
| ENSMUSG00000114355.2 | 0 | NA | NA | NA | NA | NA | #VALUE! |
| ENSMUSG00000114356.2 | 0 | NA | NA | NA | NA | NA | #VALUE! |
| ENSMUSG00000114357.2 | 0 | NA | NA | NA | NA | NA | #VALUE! |
| ENSMUSG00000114359.2 | 0 | NA | NA | NA | NA | NA | #VALUE! |
| ENSMUSG00000114360.2 | 0 | NA | NA | NA | NA | NA | #VALUE! |
| ENSMUSG00000114361.2 | 0 | NA | NA | NA | NA | NA | #VALUE! |
| ENSMUSG00000114362.2 | 0 | NA | NA | NA | NA | NA | #VALUE! |
| ENSMUSG00000114363.2 | 0 | NA | NA | NA | NA | NA | #VALUE! |
| ENSMUSG00000114364.2 | 0 | NA | NA | NA | NA | NA | #VALUE! |
| ENSMUSG00000114365.2 | 0 | NA | NA | NA | NA | NA | #VALUE! |
| ENSMUSG00000114366.2 | 0 | NA | NA | NA | NA | NA | #VALUE! |
| ENSMUSG00000114367.2 | 0 | NA | NA | NA | NA | NA | #VALUE! |
| ENSMUSG00000114368.2 | 0 | NA | NA | NA | NA | NA | #VALUE! |
| ENSMUSG00000114369.2 | 0 | NA | NA | NA | NA | NA | #VALUE! |
| ENSMUSG00000114370.2 | 0.195789 | -1.05581 | 4.073302 | -0.2592 | 0.795479 | NA | #VALUE! |
| ENSMUSG00000114371.2 | 0 | NA | NA | NA | NA | NA | #VALUE! |
| ENSMUSG00000114372.2 | 0 | NA | NA | NA | NA | NA | #VALUE! |
| ENSMUSG00000114373.2 | 0 | NA | NA | NA | NA | NA | #VALUE! |
| ENSMUSG00000114374.2 | 0 | NA | NA | NA | NA | NA | #VALUE! |
| ENSMUSG00000114375.2 | 0 | NA | NA | NA | NA | NA | #VALUE! |
| ENSMUSG00000114376.2 | 0.279766 | 1.808031 | 4.036658 | 0.447903 | 0.654223 | NA | #VALUE! |
| ENSMUSG00000114377.2 | 0 | NA | NA | NA | NA | NA | #VALUE! |
| ENSMUSG00000114378.2 | 0 | NA | NA | NA | NA | NA | #VALUE! |
| ENSMUSG00000114379.2 | 0 | NA | NA | NA | NA | NA | #VALUE! |
| ENSMUSG00000114380.2 | 0 | NA | NA | NA | NA | NA | #VALUE! |

| | | | | | | | |
|---|---|---|---|---|---|---|---|
| ENSMUSG00000114381.2 | 0 | NA | NA | NA | NA | NA | #VALUE! |
| ENSMUSG00000114382.2 | 0 | NA | NA | NA | NA | NA | #VALUE! |
| ENSMUSG00000114383.2 | 0 | NA | NA | NA | NA | NA | #VALUE! |
| ENSMUSG00000114384.2 | 0 | NA | NA | NA | NA | NA | #VALUE! |
| ENSMUSG00000114385.2 | 0 | NA | NA | NA | NA | NA | #VALUE! |
| ENSMUSG00000114386.2 | 0 | NA | NA | NA | NA | NA | #VALUE! |
| ENSMUSG00000114387.2 | 0 | NA | NA | NA | NA | NA | #VALUE! |
| ENSMUSG00000114388.2 | 0 | NA | NA | NA | NA | NA | #VALUE! |
| ENSMUSG00000114389.2 | 0 | NA | NA | NA | NA | NA | #VALUE! |
| ENSMUSG00000114390.2 | 0 | NA | NA | NA | NA | NA | #VALUE! |
| ENSMUSG00000114391.2 | 0 | NA | NA | NA | NA | NA | #VALUE! |
| ENSMUSG00000114392.2 | 0 | NA | NA | NA | NA | NA | #VALUE! |
| ENSMUSG00000114393.2 | 0 | NA | NA | NA | NA | NA | #VALUE! |
| ENSMUSG00000114394.2 | 0 | NA | NA | NA | NA | NA | #VALUE! |
| ENSMUSG00000114395.2 | 0 | NA | NA | NA | NA | NA | #VALUE! |
| ENSMUSG00000114396.2 | 0 | NA | NA | NA | NA | NA | #VALUE! |
| ENSMUSG00000114397.2 | 0 | NA | NA | NA | NA | NA | #VALUE! |
| ENSMUSG00000114398.2 | 0 | NA | NA | NA | NA | NA | #VALUE! |
| ENSMUSG00000114399.2 | 0 | NA | NA | NA | NA | NA | #VALUE! |
| ENSMUSG00000114400.2 | 0 | NA | NA | NA | NA | NA | #VALUE! |
| ENSMUSG00000114401.2 | 0 | NA | NA | NA | NA | NA | #VALUE! |
| ENSMUSG00000114402.2 | 0 | NA | NA | NA | NA | NA | #VALUE! |
| ENSMUSG00000114403.2 | 0 | NA | NA | NA | NA | NA | #VALUE! |
| ENSMUSG00000114404.2 | 0 | NA | NA | NA | NA | NA | #VALUE! |
| ENSMUSG00000114405.2 | 0 | NA | NA | NA | NA | NA | #VALUE! |
| ENSMUSG00000114406.2 | 0 | NA | NA | NA | NA | NA | #VALUE! |
| ENSMUSG00000114407.2 | 0 | NA | NA | NA | NA | NA | #VALUE! |
| ENSMUSG00000114408.2 | 0 | NA | NA | NA | NA | NA | #VALUE! |
| ENSMUSG00000114409.2 | 0.245976 | 1.027628 | 4.067126 | 0.252667 | 0.800526 | NA | #VALUE! |
| ENSMUSG00000114410.2 | 0 | NA | NA | NA | NA | NA | #VALUE! |
| ENSMUSG00000114411.2 | 0 | NA | NA | NA | NA | NA | #VALUE! |
| ENSMUSG00000114412.2 | 0 | NA | NA | NA | NA | NA | #VALUE! |
| ENSMUSG00000114414.2 | 0 | NA | NA | NA | NA | NA | #VALUE! |
| ENSMUSG00000114415.2 | 0 | NA | NA | NA | NA | NA | #VALUE! |
| ENSMUSG00000114416.2 | 0 | NA | NA | NA | NA | NA | #VALUE! |
| ENSMUSG00000114417.2 | 0 | NA | NA | NA | NA | NA | #VALUE! |
| ENSMUSG00000114418.2 | 0 | NA | NA | NA | NA | NA | #VALUE! |
| ENSMUSG00000114419.2 | 0 | NA | NA | NA | NA | NA | #VALUE! |
| ENSMUSG00000114420.2 | 0 | NA | NA | NA | NA | NA | #VALUE! |
| ENSMUSG00000114421.2 | 0 | NA | NA | NA | NA | NA | #VALUE! |
| ENSMUSG00000114422.2 | 0 | NA | NA | NA | NA | NA | #VALUE! |
| ENSMUSG00000114423.2 | 0 | NA | NA | NA | NA | NA | #VALUE! |
| ENSMUSG00000114424.2 | 0 | NA | NA | NA | NA | NA | #VALUE! |
| ENSMUSG00000114425.2 | 1.428775 | -4.03418 | 3.938169 | -1.02438 | 0.305656 | NA | #VALUE! |

| | | | | | | | |
|---|---|---|---|---|---|---|---|
| ENSMUSG00000114426.2 | 0 | NA | NA | NA | NA | NA | #VALUE! |
| ENSMUSG00000114427.2 | 0 | NA | NA | NA | NA | NA | #VALUE! |
| ENSMUSG00000114428.2 | 0 | NA | NA | NA | NA | NA | #VALUE! |
| ENSMUSG00000114429.2 | 0 | NA | NA | NA | NA | NA | #VALUE! |
| ENSMUSG00000114430.2 | 4.560749 | -0.83733 | 3.769443 | -0.22214 | 0.824207 | NA | #VALUE! |
| ENSMUSG00000114431.2 | 0 | NA | NA | NA | NA | NA | #VALUE! |
| ENSMUSG00000114432.2 | 0 | NA | NA | NA | NA | NA | #VALUE! |
| ENSMUSG00000114433.2 | 0 | NA | NA | NA | NA | NA | #VALUE! |
| ENSMUSG00000114434.2 | 0 | NA | NA | NA | NA | NA | #VALUE! |
| ENSMUSG00000114436.2 | 0 | NA | NA | NA | NA | NA | #VALUE! |
| ENSMUSG00000114437.2 | 0 | NA | NA | NA | NA | NA | #VALUE! |
| ENSMUSG00000114438.2 | 0 | NA | NA | NA | NA | NA | #VALUE! |
| ENSMUSG00000114439.2 | 0 | NA | NA | NA | NA | NA | #VALUE! |
| ENSMUSG00000114440.2 | 0.105013 | -0.97324 | 4.076293 | -0.23876 | 0.811295 | NA | #VALUE! |
| ENSMUSG00000114441.2 | 0 | NA | NA | NA | NA | NA | #VALUE! |
| ENSMUSG00000114442.2 | 0 | NA | NA | NA | NA | NA | #VALUE! |
| ENSMUSG00000114443.2 | 0 | NA | NA | NA | NA | NA | #VALUE! |
| ENSMUSG00000114444.2 | 0 | NA | NA | NA | NA | NA | #VALUE! |
| ENSMUSG00000114445.2 | 0 | NA | NA | NA | NA | NA | #VALUE! |
| ENSMUSG00000114446.2 | 0 | NA | NA | NA | NA | NA | #VALUE! |
| ENSMUSG00000114447.2 | 0 | NA | NA | NA | NA | NA | #VALUE! |
| ENSMUSG00000114448.2 | 0 | NA | NA | NA | NA | NA | #VALUE! |
| ENSMUSG00000114449.2 | 0 | NA | NA | NA | NA | NA | #VALUE! |
| ENSMUSG00000114450.2 | 0 | NA | NA | NA | NA | NA | #VALUE! |
| ENSMUSG00000114451.2 | 0 | NA | NA | NA | NA | NA | #VALUE! |
| ENSMUSG00000114452.2 | 0.746042 | 3.107177 | 3.964586 | 0.783733 | 0.433197 | NA | #VALUE! |
| ENSMUSG00000114453.2 | 0 | NA | NA | NA | NA | NA | #VALUE! |
| ENSMUSG00000114454.2 | 0 | NA | NA | NA | NA | NA | #VALUE! |
| ENSMUSG00000114456.2 | 0.279766 | 1.808031 | 4.036658 | 0.447903 | 0.654223 | NA | #VALUE! |
| ENSMUSG00000114457.2 | 0 | NA | NA | NA | NA | NA | #VALUE! |
| ENSMUSG00000114458.2 | 0 | NA | NA | NA | NA | NA | #VALUE! |
| ENSMUSG00000114459.2 | 0 | NA | NA | NA | NA | NA | #VALUE! |
| ENSMUSG00000114460.2 | 0 | NA | NA | NA | NA | NA | #VALUE! |
| ENSMUSG00000114461.2 | 0 | NA | NA | NA | NA | NA | #VALUE! |
| ENSMUSG00000114462.2 | 0 | NA | NA | NA | NA | NA | #VALUE! |
| ENSMUSG00000114463.2 | 0 | NA | NA | NA | NA | NA | #VALUE! |
| ENSMUSG00000114464.2 | 0 | NA | NA | NA | NA | NA | #VALUE! |
| ENSMUSG00000114465.2 | 0 | NA | NA | NA | NA | NA | #VALUE! |
| ENSMUSG00000114466.2 | 0 | NA | NA | NA | NA | NA | #VALUE! |
| ENSMUSG00000114467.2 | 0 | NA | NA | NA | NA | NA | #VALUE! |
| ENSMUSG00000114468.2 | 0 | NA | NA | NA | NA | NA | #VALUE! |
| ENSMUSG00000114469.3 | 8.333621 | -0.35269 | 3.105153 | -0.11358 | 0.909568 | NA | #VALUE! |
| ENSMUSG00000114470.2 | 2.984166 | 5.077883 | 3.921685 | 1.294822 | 0.195382 | NA | #VALUE! |
| ENSMUSG00000114471.2 | 0 | NA | NA | NA | NA | NA | #VALUE! |

| | | | | | | | |
|---|---|---|---|---|---|---|---|
| ENSMUSG00000114472.2 | 0 | NA | NA | NA | NA | NA | #VALUE! |
| ENSMUSG00000114473.2 | 0 | NA | NA | NA | NA | NA | #VALUE! |
| ENSMUSG00000114474.2 | 0 | NA | NA | NA | NA | NA | #VALUE! |
| ENSMUSG00000114475.2 | 0 | NA | NA | NA | NA | NA | #VALUE! |
| ENSMUSG00000114476.2 | 0 | NA | NA | NA | NA | NA | #VALUE! |
| ENSMUSG00000114477.2 | 0 | NA | NA | NA | NA | NA | #VALUE! |
| ENSMUSG00000114478.2 | 0 | NA | NA | NA | NA | NA | #VALUE! |
| ENSMUSG00000114479.2 | 0 | NA | NA | NA | NA | NA | #VALUE! |
| ENSMUSG00000114480.2 | 0 | NA | NA | NA | NA | NA | #VALUE! |
| ENSMUSG00000114481.2 | 0 | NA | NA | NA | NA | NA | #VALUE! |
| ENSMUSG00000114482.2 | 0 | NA | NA | NA | NA | NA | #VALUE! |
| ENSMUSG00000114483.2 | 0 | NA | NA | NA | NA | NA | #VALUE! |
| ENSMUSG00000114484.2 | 0 | NA | NA | NA | NA | NA | #VALUE! |
| ENSMUSG00000114485.2 | 0 | NA | NA | NA | NA | NA | #VALUE! |
| ENSMUSG00000114486.2 | 0 | NA | NA | NA | NA | NA | #VALUE! |
| ENSMUSG00000114487.2 | 0 | NA | NA | NA | NA | NA | #VALUE! |
| ENSMUSG00000114488.2 | 0 | NA | NA | NA | NA | NA | #VALUE! |
| ENSMUSG00000114489.3 | 0 | NA | NA | NA | NA | NA | #VALUE! |
| ENSMUSG00000114490.2 | 0 | NA | NA | NA | NA | NA | #VALUE! |
| ENSMUSG00000114491.2 | 0 | NA | NA | NA | NA | NA | #VALUE! |
| ENSMUSG00000114492.2 | 0 | NA | NA | NA | NA | NA | #VALUE! |
| ENSMUSG00000114493.3 | 0.746042 | 3.107177 | 3.964586 | 0.783733 | 0.433197 | NA | #VALUE! |
| ENSMUSG00000114494.2 | 0 | NA | NA | NA | NA | NA | #VALUE! |
| ENSMUSG00000114495.2 | 0 | NA | NA | NA | NA | NA | #VALUE! |
| ENSMUSG00000114496.2 | 0 | NA | NA | NA | NA | NA | #VALUE! |
| ENSMUSG00000114498.2 | 0 | NA | NA | NA | NA | NA | #VALUE! |
| ENSMUSG00000114499.2 | 0 | NA | NA | NA | NA | NA | #VALUE! |
| ENSMUSG00000114500.2 | 0 | NA | NA | NA | NA | NA | #VALUE! |
| ENSMUSG00000114501.2 | 12.30908 | 0.905061 | 2.856842 | 0.316805 | NA | NA | #VALUE! |
| ENSMUSG00000114502.2 | 0 | NA | NA | NA | NA | NA | #VALUE! |
| ENSMUSG00000114503.2 | 0 | NA | NA | NA | NA | NA | #VALUE! |
| ENSMUSG00000114504.2 | 0 | NA | NA | NA | NA | NA | #VALUE! |
| ENSMUSG00000114505.2 | 0 | NA | NA | NA | NA | NA | #VALUE! |
| ENSMUSG00000114506.2 | 0 | NA | NA | NA | NA | NA | #VALUE! |
| ENSMUSG00000114507.2 | 0 | NA | NA | NA | NA | NA | #VALUE! |
| ENSMUSG00000114508.2 | 0 | NA | NA | NA | NA | NA | #VALUE! |
| ENSMUSG00000114509.2 | 0 | NA | NA | NA | NA | NA | #VALUE! |
| ENSMUSG00000114510.2 | 0 | NA | NA | NA | NA | NA | #VALUE! |
| ENSMUSG00000114511.2 | 0 | NA | NA | NA | NA | NA | #VALUE! |
| ENSMUSG00000114512.2 | 0 | NA | NA | NA | NA | NA | #VALUE! |
| ENSMUSG00000114513.2 | 0 | NA | NA | NA | NA | NA | #VALUE! |
| ENSMUSG00000114514.2 | 0 | NA | NA | NA | NA | NA | #VALUE! |
| ENSMUSG00000114515.3 | 0 | NA | NA | NA | NA | NA | #VALUE! |
| ENSMUSG00000114516.2 | 1.958359 | 4.475621 | 3.929371 | 1.139017 | 0.254696 | NA | #VALUE! |

| | | | | | | | |
|---|---|---|---|---|---|---|---|
| ENSMUSG00000114517.2 | 0 | NA | NA | NA | NA | NA | #VALUE! |
| ENSMUSG00000114518.2 | 0 | NA | NA | NA | NA | NA | #VALUE! |
| ENSMUSG00000114519.2 | 0 | NA | NA | NA | NA | NA | #VALUE! |
| ENSMUSG00000114520.2 | 0.195789 | -1.05581 | 4.073302 | -0.2592 | 0.795479 | NA | #VALUE! |
| ENSMUSG00000114521.2 | 0 | NA | NA | NA | NA | NA | #VALUE! |
| ENSMUSG00000114522.2 | 0 | NA | NA | NA | NA | NA | #VALUE! |
| ENSMUSG00000114523.2 | 0 | NA | NA | NA | NA | NA | #VALUE! |
| ENSMUSG00000114524.2 | 0.18651 | 1.401769 | 4.056354 | 0.345574 | 0.729663 | NA | #VALUE! |
| ENSMUSG00000114525.2 | 0 | NA | NA | NA | NA | NA | #VALUE! |
| ENSMUSG00000114526.2 | 0 | NA | NA | NA | NA | NA | #VALUE! |
| ENSMUSG00000114527.3 | 0 | NA | NA | NA | NA | NA | #VALUE! |
| ENSMUSG00000114528.2 | 0 | NA | NA | NA | NA | NA | #VALUE! |
| ENSMUSG00000114529.2 | 0 | NA | NA | NA | NA | NA | #VALUE! |
| ENSMUSG00000114530.2 | 0 | NA | NA | NA | NA | NA | #VALUE! |
| ENSMUSG00000114531.2 | 0 | NA | NA | NA | NA | NA | #VALUE! |
| ENSMUSG00000114532.2 | 0 | NA | NA | NA | NA | NA | #VALUE! |
| ENSMUSG00000114533.2 | 0 | NA | NA | NA | NA | NA | #VALUE! |
| ENSMUSG00000114534.2 | 0 | NA | NA | NA | NA | NA | #VALUE! |
| ENSMUSG00000114535.2 | 4.84927 | 5.774248 | 3.915985 | 1.474533 | 0.140338 | NA | #VALUE! |
| ENSMUSG00000114536.2 | 0 | NA | NA | NA | NA | NA | #VALUE! |
| ENSMUSG00000114537.2 | 0 | NA | NA | NA | NA | NA | #VALUE! |
| ENSMUSG00000114538.2 | 0 | NA | NA | NA | NA | NA | #VALUE! |
| ENSMUSG00000114539.2 | 0 | NA | NA | NA | NA | NA | #VALUE! |
| ENSMUSG00000114540.2 | 0.195789 | -1.05581 | 4.073302 | -0.2592 | 0.795479 | NA | #VALUE! |
| ENSMUSG00000114541.2 | 0 | NA | NA | NA | NA | NA | #VALUE! |
| ENSMUSG00000114542.2 | 0 | NA | NA | NA | NA | NA | #VALUE! |
| ENSMUSG00000114543.2 | 0 | NA | NA | NA | NA | NA | #VALUE! |
| ENSMUSG00000114544.2 | 0 | NA | NA | NA | NA | NA | #VALUE! |
| ENSMUSG00000114545.2 | 0 | NA | NA | NA | NA | NA | #VALUE! |
| ENSMUSG00000114546.2 | 0 | NA | NA | NA | NA | NA | #VALUE! |
| ENSMUSG00000114547.2 | 0 | NA | NA | NA | NA | NA | #VALUE! |
| ENSMUSG00000114549.2 | 0 | NA | NA | NA | NA | NA | #VALUE! |
| ENSMUSG00000114551.2 | 0.18651 | 1.401769 | 4.056354 | 0.345574 | 0.729663 | NA | #VALUE! |
| ENSMUSG00000114552.2 | 0 | NA | NA | NA | NA | NA | #VALUE! |
| ENSMUSG00000114553.2 | 0 | NA | NA | NA | NA | NA | #VALUE! |
| ENSMUSG00000114554.2 | 0 | NA | NA | NA | NA | NA | #VALUE! |
| ENSMUSG00000114555.2 | 0 | NA | NA | NA | NA | NA | #VALUE! |
| ENSMUSG00000114556.2 | 0 | NA | NA | NA | NA | NA | #VALUE! |
| ENSMUSG00000114557.2 | 0 | NA | NA | NA | NA | NA | #VALUE! |
| ENSMUSG00000114558.2 | 0 | NA | NA | NA | NA | NA | #VALUE! |
| ENSMUSG00000114559.2 | 0 | NA | NA | NA | NA | NA | #VALUE! |
| ENSMUSG00000114560.2 | 0 | NA | NA | NA | NA | NA | #VALUE! |
| ENSMUSG00000114561.2 | 0 | NA | NA | NA | NA | NA | #VALUE! |
| ENSMUSG00000114562.2 | 0 | NA | NA | NA | NA | NA | #VALUE! |

| | | | | | | | |
|---|---|---|---|---|---|---|---|
| ENSMUSG00000114563.2 | 0 | NA | NA | NA | NA | NA | #VALUE! |
| ENSMUSG00000114564.2 | 0 | NA | NA | NA | NA | NA | #VALUE! |
| ENSMUSG00000114565.2 | 0 | NA | NA | NA | NA | NA | #VALUE! |
| ENSMUSG00000114566.2 | 0 | NA | NA | NA | NA | NA | #VALUE! |
| ENSMUSG00000114567.2 | 0 | NA | NA | NA | NA | NA | #VALUE! |
| ENSMUSG00000114568.2 | 0 | NA | NA | NA | NA | NA | #VALUE! |
| ENSMUSG00000114569.2 | 0 | NA | NA | NA | NA | NA | #VALUE! |
| ENSMUSG00000114570.2 | 0 | NA | NA | NA | NA | NA | #VALUE! |
| ENSMUSG00000114571.2 | 0 | NA | NA | NA | NA | NA | #VALUE! |
| ENSMUSG00000114572.2 | 0 | NA | NA | NA | NA | NA | #VALUE! |
| ENSMUSG00000114573.2 | 0 | NA | NA | NA | NA | NA | #VALUE! |
| ENSMUSG00000114575.2 | 0.210026 | -1.52494 | 4.058692 | -0.37572 | 0.707123 | NA | #VALUE! |
| ENSMUSG00000114576.2 | 0 | NA | NA | NA | NA | NA | #VALUE! |
| ENSMUSG00000114577.2 | 0 | NA | NA | NA | NA | NA | #VALUE! |
| ENSMUSG00000114578.2 | 0 | NA | NA | NA | NA | NA | #VALUE! |
| ENSMUSG00000114579.2 | 0 | NA | NA | NA | NA | NA | #VALUE! |
| ENSMUSG00000114580.2 | 0 | NA | NA | NA | NA | NA | #VALUE! |
| ENSMUSG00000114581.2 | 0.105013 | -0.97324 | 4.076293 | -0.23876 | 0.811295 | NA | #VALUE! |
| ENSMUSG00000114582.2 | 0 | NA | NA | NA | NA | NA | #VALUE! |
| ENSMUSG00000114583.2 | 0 | NA | NA | NA | NA | NA | #VALUE! |
| ENSMUSG00000114584.2 | 0 | NA | NA | NA | NA | NA | #VALUE! |
| ENSMUSG00000114585.2 | 0 | NA | NA | NA | NA | NA | #VALUE! |
| ENSMUSG00000114586.2 | 0 | NA | NA | NA | NA | NA | #VALUE! |
| ENSMUSG00000114587.2 | 0 | NA | NA | NA | NA | NA | #VALUE! |
| ENSMUSG00000114588.2 | 0 | NA | NA | NA | NA | NA | #VALUE! |
| ENSMUSG00000114589.2 | 0 | NA | NA | NA | NA | NA | #VALUE! |
| ENSMUSG00000114590.2 | 0 | NA | NA | NA | NA | NA | #VALUE! |
| ENSMUSG00000114591.2 | 0 | NA | NA | NA | NA | NA | #VALUE! |
| ENSMUSG00000114592.2 | 0 | NA | NA | NA | NA | NA | #VALUE! |
| ENSMUSG00000114593.2 | 0 | NA | NA | NA | NA | NA | #VALUE! |
| ENSMUSG00000114594.2 | 0 | NA | NA | NA | NA | NA | #VALUE! |
| ENSMUSG00000114595.2 | 0 | NA | NA | NA | NA | NA | #VALUE! |
| ENSMUSG00000114596.2 | 0 | NA | NA | NA | NA | NA | #VALUE! |
| ENSMUSG00000114597.2 | 0 | NA | NA | NA | NA | NA | #VALUE! |
| ENSMUSG00000114598.2 | 0 | NA | NA | NA | NA | NA | #VALUE! |
| ENSMUSG00000114599.2 | 0 | NA | NA | NA | NA | NA | #VALUE! |
| ENSMUSG00000114600.2 | 0 | NA | NA | NA | NA | NA | #VALUE! |
| ENSMUSG00000114601.2 | 0 | NA | NA | NA | NA | NA | #VALUE! |
| ENSMUSG00000114602.2 | 0 | NA | NA | NA | NA | NA | #VALUE! |
| ENSMUSG00000114603.2 | 0 | NA | NA | NA | NA | NA | #VALUE! |
| ENSMUSG00000114604.2 | 0 | NA | NA | NA | NA | NA | #VALUE! |
| ENSMUSG00000114605.2 | 0 | NA | NA | NA | NA | NA | #VALUE! |
| ENSMUSG00000114606.2 | 0 | NA | NA | NA | NA | NA | #VALUE! |
| ENSMUSG00000114607.2 | 0 | NA | NA | NA | NA | NA | #VALUE! |

| | | | | | | | |
|---|---|---|---|---|---|---|---|
| ENSMUSG00000114608.2 | 1.23058 | -3.82293 | 3.943085 | -0.96953 | 0.332282 | NA | #VALUE! |
| ENSMUSG00000114609.2 | 0 | NA | NA | NA | NA | NA | #VALUE! |
| ENSMUSG00000114610.2 | 0 | NA | NA | NA | NA | NA | #VALUE! |
| ENSMUSG00000114611.2 | 0 | NA | NA | NA | NA | NA | #VALUE! |
| ENSMUSG00000114612.2 | 0 | NA | NA | NA | NA | NA | #VALUE! |
| ENSMUSG00000114613.2 | 0 | NA | NA | NA | NA | NA | #VALUE! |
| ENSMUSG00000114614.2 | 0 | NA | NA | NA | NA | NA | #VALUE! |
| ENSMUSG00000114615.2 | 0 | NA | NA | NA | NA | NA | #VALUE! |
| ENSMUSG00000114616.2 | 0 | NA | NA | NA | NA | NA | #VALUE! |
| ENSMUSG00000114617.2 | 0 | NA | NA | NA | NA | NA | #VALUE! |
| ENSMUSG00000114618.2 | 0 | NA | NA | NA | NA | NA | #VALUE! |
| ENSMUSG00000114619.2 | 0 | NA | NA | NA | NA | NA | #VALUE! |
| ENSMUSG00000114620.3 | 0 | NA | NA | NA | NA | NA | #VALUE! |
| ENSMUSG00000114621.2 | 0 | NA | NA | NA | NA | NA | #VALUE! |
| ENSMUSG00000114622.2 | 0 | NA | NA | NA | NA | NA | #VALUE! |
| ENSMUSG00000114623.2 | 0 | NA | NA | NA | NA | NA | #VALUE! |
| ENSMUSG00000114624.2 | 0 | NA | NA | NA | NA | NA | #VALUE! |
| ENSMUSG00000114625.2 | 0 | NA | NA | NA | NA | NA | #VALUE! |
| ENSMUSG00000114626.2 | 0 | NA | NA | NA | NA | NA | #VALUE! |
| ENSMUSG00000114627.2 | 0 | NA | NA | NA | NA | NA | #VALUE! |
| ENSMUSG00000114628.2 | 0 | NA | NA | NA | NA | NA | #VALUE! |
| ENSMUSG00000114629.2 | 0 | NA | NA | NA | NA | NA | #VALUE! |
| ENSMUSG00000114630.2 | 0 | NA | NA | NA | NA | NA | #VALUE! |
| ENSMUSG00000114631.2 | 0 | NA | NA | NA | NA | NA | #VALUE! |
| ENSMUSG00000114632.2 | 0 | NA | NA | NA | NA | NA | #VALUE! |
| ENSMUSG00000114633.2 | 0 | NA | NA | NA | NA | NA | #VALUE! |
| ENSMUSG00000114634.2 | 0 | NA | NA | NA | NA | NA | #VALUE! |
| ENSMUSG00000114635.2 | 0 | NA | NA | NA | NA | NA | #VALUE! |
| ENSMUSG00000114636.2 | 0 | NA | NA | NA | NA | NA | #VALUE! |
| ENSMUSG00000114637.2 | 0 | NA | NA | NA | NA | NA | #VALUE! |
| ENSMUSG00000114639.2 | 0 | NA | NA | NA | NA | NA | #VALUE! |
| ENSMUSG00000114640.2 | 0 | NA | NA | NA | NA | NA | #VALUE! |
| ENSMUSG00000114641.2 | 0 | NA | NA | NA | NA | NA | #VALUE! |
| ENSMUSG00000114642.2 | 0 | NA | NA | NA | NA | NA | #VALUE! |
| ENSMUSG00000114643.2 | 0 | NA | NA | NA | NA | NA | #VALUE! |
| ENSMUSG00000114644.2 | 0 | NA | NA | NA | NA | NA | #VALUE! |
| ENSMUSG00000114645.2 | 0 | NA | NA | NA | NA | NA | #VALUE! |
| ENSMUSG00000114646.2 | 0 | NA | NA | NA | NA | NA | #VALUE! |
| ENSMUSG00000114647.2 | 0 | NA | NA | NA | NA | NA | #VALUE! |
| ENSMUSG00000114648.2 | 0 | NA | NA | NA | NA | NA | #VALUE! |
| ENSMUSG00000114649.2 | 0 | NA | NA | NA | NA | NA | #VALUE! |
| ENSMUSG00000114650.2 | 0 | NA | NA | NA | NA | NA | #VALUE! |
| ENSMUSG00000114651.2 | 0 | NA | NA | NA | NA | NA | #VALUE! |
| ENSMUSG00000114652.2 | 0 | NA | NA | NA | NA | NA | #VALUE! |

| | | | | | | | |
|---|---|---|---|---|---|---|---|
| ENSMUSG00000114653.2 | 0 | NA | NA | NA | NA | NA | #VALUE! |
| ENSMUSG00000114654.2 | 0 | NA | NA | NA | NA | NA | #VALUE! |
| ENSMUSG00000114655.2 | 0 | NA | NA | NA | NA | NA | #VALUE! |
| ENSMUSG00000114656.2 | 0 | NA | NA | NA | NA | NA | #VALUE! |
| ENSMUSG00000114657.2 | 0 | NA | NA | NA | NA | NA | #VALUE! |
| ENSMUSG00000114658.2 | 0 | NA | NA | NA | NA | NA | #VALUE! |
| ENSMUSG00000114659.2 | 0 | NA | NA | NA | NA | NA | #VALUE! |
| ENSMUSG00000114660.2 | 0 | NA | NA | NA | NA | NA | #VALUE! |
| ENSMUSG00000114661.2 | 0 | NA | NA | NA | NA | NA | #VALUE! |
| ENSMUSG00000114662.2 | 0 | NA | NA | NA | NA | NA | #VALUE! |
| ENSMUSG00000114663.2 | 0 | NA | NA | NA | NA | NA | #VALUE! |
| ENSMUSG00000114664.2 | 0.783156 | -3.1627 | 3.963922 | -0.79787 | 0.424945 | NA | #VALUE! |
| ENSMUSG00000114665.2 | 0 | NA | NA | NA | NA | NA | #VALUE! |
| ENSMUSG00000114666.2 | 0 | NA | NA | NA | NA | NA | #VALUE! |
| ENSMUSG00000114667.2 | 0 | NA | NA | NA | NA | NA | #VALUE! |
| ENSMUSG00000114668.2 | 0 | NA | NA | NA | NA | NA | #VALUE! |
| ENSMUSG00000114669.2 | 0 | NA | NA | NA | NA | NA | #VALUE! |
| ENSMUSG00000114670.2 | 0 | NA | NA | NA | NA | NA | #VALUE! |
| ENSMUSG00000114671.2 | 0 | NA | NA | NA | NA | NA | #VALUE! |
| ENSMUSG00000114672.2 | 0 | NA | NA | NA | NA | NA | #VALUE! |
| ENSMUSG00000114673.2 | 0 | NA | NA | NA | NA | NA | #VALUE! |
| ENSMUSG00000114674.2 | 0 | NA | NA | NA | NA | NA | #VALUE! |
| ENSMUSG00000114675.2 | 0 | NA | NA | NA | NA | NA | #VALUE! |
| ENSMUSG00000114677.2 | 0 | NA | NA | NA | NA | NA | #VALUE! |
| ENSMUSG00000114678.2 | 0 | NA | NA | NA | NA | NA | #VALUE! |
| ENSMUSG00000114679.2 | 0 | NA | NA | NA | NA | NA | #VALUE! |
| ENSMUSG00000114680.2 | 0 | NA | NA | NA | NA | NA | #VALUE! |
| ENSMUSG00000114681.2 | 0 | NA | NA | NA | NA | NA | #VALUE! |
| ENSMUSG00000114682.2 | 0 | NA | NA | NA | NA | NA | #VALUE! |
| ENSMUSG00000114683.2 | 0 | NA | NA | NA | NA | NA | #VALUE! |
| ENSMUSG00000114684.3 | 0 | NA | NA | NA | NA | NA | #VALUE! |
| ENSMUSG00000114685.2 | 0 | NA | NA | NA | NA | NA | #VALUE! |
| ENSMUSG00000114686.2 | 0 | NA | NA | NA | NA | NA | #VALUE! |
| ENSMUSG00000114687.2 | 0 | NA | NA | NA | NA | NA | #VALUE! |
| ENSMUSG00000114688.2 | 0 | NA | NA | NA | NA | NA | #VALUE! |
| ENSMUSG00000114689.2 | 0 | NA | NA | NA | NA | NA | #VALUE! |
| ENSMUSG00000114690.2 | 0 | NA | NA | NA | NA | NA | #VALUE! |
| ENSMUSG00000114691.2 | 0 | NA | NA | NA | NA | NA | #VALUE! |
| ENSMUSG00000114692.2 | 0 | NA | NA | NA | NA | NA | #VALUE! |
| ENSMUSG00000114693.2 | 0 | NA | NA | NA | NA | NA | #VALUE! |
| ENSMUSG00000114694.2 | 0 | NA | NA | NA | NA | NA | #VALUE! |
| ENSMUSG00000114695.2 | 0 | NA | NA | NA | NA | NA | #VALUE! |
| ENSMUSG00000114696.2 | 0 | NA | NA | NA | NA | NA | #VALUE! |
| ENSMUSG00000114697.2 | 0 | NA | NA | NA | NA | NA | #VALUE! |

| | | | | | | | |
|---|---|---|---|---|---|---|---|
| ENSMUSG00000114698.2 | 0 | NA | NA | NA | NA | NA | #VALUE! |
| ENSMUSG00000114699.2 | 0 | NA | NA | NA | NA | NA | #VALUE! |
| ENSMUSG00000114700.2 | 0 | NA | NA | NA | NA | NA | #VALUE! |
| ENSMUSG00000114701.2 | 0 | NA | NA | NA | NA | NA | #VALUE! |
| ENSMUSG00000114702.3 | 0 | NA | NA | NA | NA | NA | #VALUE! |
| ENSMUSG00000114703.2 | 0 | NA | NA | NA | NA | NA | #VALUE! |
| ENSMUSG00000114704.2 | 0 | NA | NA | NA | NA | NA | #VALUE! |
| ENSMUSG00000114705.2 | 0 | NA | NA | NA | NA | NA | #VALUE! |
| ENSMUSG00000114706.2 | 0 | NA | NA | NA | NA | NA | #VALUE! |
| ENSMUSG00000114707.2 | 5.408801 | 5.931107 | 3.915037 | 1.514955 | 0.129784 | NA | #VALUE! |
| ENSMUSG00000114708.2 | 0 | NA | NA | NA | NA | NA | #VALUE! |
| ENSMUSG00000114709.2 | 0 | NA | NA | NA | NA | NA | #VALUE! |
| ENSMUSG00000114710.2 | 0 | NA | NA | NA | NA | NA | #VALUE! |
| ENSMUSG00000114711.2 | 0 | NA | NA | NA | NA | NA | #VALUE! |
| ENSMUSG00000114712.2 | 0 | NA | NA | NA | NA | NA | #VALUE! |
| ENSMUSG00000114713.2 | 0 | NA | NA | NA | NA | NA | #VALUE! |
| ENSMUSG00000114714.2 | 0 | NA | NA | NA | NA | NA | #VALUE! |
| ENSMUSG00000114715.2 | 0 | NA | NA | NA | NA | NA | #VALUE! |
| ENSMUSG00000114717.2 | 0 | NA | NA | NA | NA | NA | #VALUE! |
| ENSMUSG00000114718.2 | 0 | NA | NA | NA | NA | NA | #VALUE! |
| ENSMUSG00000114719.2 | 0 | NA | NA | NA | NA | NA | #VALUE! |
| ENSMUSG00000114720.2 | 0 | NA | NA | NA | NA | NA | #VALUE! |
| ENSMUSG00000114721.2 | 0 | NA | NA | NA | NA | NA | #VALUE! |
| ENSMUSG00000114722.2 | 0 | NA | NA | NA | NA | NA | #VALUE! |
| ENSMUSG00000114723.2 | 0 | NA | NA | NA | NA | NA | #VALUE! |
| ENSMUSG00000114724.2 | 0 | NA | NA | NA | NA | NA | #VALUE! |
| ENSMUSG00000114725.2 | 0 | NA | NA | NA | NA | NA | #VALUE! |
| ENSMUSG00000114726.2 | 0 | NA | NA | NA | NA | NA | #VALUE! |
| ENSMUSG00000114727.2 | 0 | NA | NA | NA | NA | NA | #VALUE! |
| ENSMUSG00000114728.2 | 0 | NA | NA | NA | NA | NA | #VALUE! |
| ENSMUSG00000114729.2 | 0 | NA | NA | NA | NA | NA | #VALUE! |
| ENSMUSG00000114730.2 | 0 | NA | NA | NA | NA | NA | #VALUE! |
| ENSMUSG00000114731.2 | 0 | NA | NA | NA | NA | NA | #VALUE! |
| ENSMUSG00000114732.2 | 0 | NA | NA | NA | NA | NA | #VALUE! |
| ENSMUSG00000114733.2 | 0 | NA | NA | NA | NA | NA | #VALUE! |
| ENSMUSG00000114734.2 | 0 | NA | NA | NA | NA | NA | #VALUE! |
| ENSMUSG00000114735.2 | 0 | NA | NA | NA | NA | NA | #VALUE! |
| ENSMUSG00000114736.2 | 0 | NA | NA | NA | NA | NA | #VALUE! |
| ENSMUSG00000114737.2 | 0 | NA | NA | NA | NA | NA | #VALUE! |
| ENSMUSG00000114738.2 | 0 | NA | NA | NA | NA | NA | #VALUE! |
| ENSMUSG00000114739.2 | 0 | NA | NA | NA | NA | NA | #VALUE! |
| ENSMUSG00000114740.2 | 0 | NA | NA | NA | NA | NA | #VALUE! |
| ENSMUSG00000114741.2 | 0 | NA | NA | NA | NA | NA | #VALUE! |
| ENSMUSG00000114742.2 | 0 | NA | NA | NA | NA | NA | #VALUE! |

| | | | | | | | |
|---|---|---|---|---|---|---|---|
| ENSMUSG00000114743.2 | 0 | NA | NA | NA | NA | NA | #VALUE! |
| ENSMUSG00000114744.2 | 0 | NA | NA | NA | NA | NA | #VALUE! |
| ENSMUSG00000114745.2 | 0 | NA | NA | NA | NA | NA | #VALUE! |
| ENSMUSG00000114746.2 | 0 | NA | NA | NA | NA | NA | #VALUE! |
| ENSMUSG00000114747.2 | 0 | NA | NA | NA | NA | NA | #VALUE! |
| ENSMUSG00000114748.2 | 0 | NA | NA | NA | NA | NA | #VALUE! |
| ENSMUSG00000114749.2 | 0 | NA | NA | NA | NA | NA | #VALUE! |
| ENSMUSG00000114750.2 | 0 | NA | NA | NA | NA | NA | #VALUE! |
| ENSMUSG00000114751.2 | 0 | NA | NA | NA | NA | NA | #VALUE! |
| ENSMUSG00000114752.2 | 0 | NA | NA | NA | NA | NA | #VALUE! |
| ENSMUSG00000114753.2 | 0 | NA | NA | NA | NA | NA | #VALUE! |
| ENSMUSG00000114754.2 | 0.105013 | -0.97324 | 4.076293 | -0.23876 | 0.811295 | NA | #VALUE! |
| ENSMUSG00000114755.2 | 0 | NA | NA | NA | NA | NA | #VALUE! |
| ENSMUSG00000114756.2 | 0 | NA | NA | NA | NA | NA | #VALUE! |
| ENSMUSG00000114757.2 | 0 | NA | NA | NA | NA | NA | #VALUE! |
| ENSMUSG00000114758.2 | 0 | NA | NA | NA | NA | NA | #VALUE! |
| ENSMUSG00000114759.2 | 0 | NA | NA | NA | NA | NA | #VALUE! |
| ENSMUSG00000114760.2 | 0 | NA | NA | NA | NA | NA | #VALUE! |
| ENSMUSG00000114761.2 | 0 | NA | NA | NA | NA | NA | #VALUE! |
| ENSMUSG00000114762.2 | 0 | NA | NA | NA | NA | NA | #VALUE! |
| ENSMUSG00000114763.2 | 0 | NA | NA | NA | NA | NA | #VALUE! |
| ENSMUSG00000114764.2 | 0 | NA | NA | NA | NA | NA | #VALUE! |
| ENSMUSG00000114765.2 | 0 | NA | NA | NA | NA | NA | #VALUE! |
| ENSMUSG00000114766.2 | 0 | NA | NA | NA | NA | NA | #VALUE! |
| ENSMUSG00000114767.2 | 0 | NA | NA | NA | NA | NA | #VALUE! |
| ENSMUSG00000114768.2 | 4.898658 | -5.81923 | 3.915919 | -1.48605 | 0.137267 | NA | #VALUE! |
| ENSMUSG00000114769.2 | 0 | NA | NA | NA | NA | NA | #VALUE! |
| ENSMUSG00000114770.2 | 0 | NA | NA | NA | NA | NA | #VALUE! |
| ENSMUSG00000114771.2 | 0 | NA | NA | NA | NA | NA | #VALUE! |
| ENSMUSG00000114772.2 | 0 | NA | NA | NA | NA | NA | #VALUE! |
| ENSMUSG00000114773.2 | 0 | NA | NA | NA | NA | NA | #VALUE! |
| ENSMUSG00000114774.2 | 0 | NA | NA | NA | NA | NA | #VALUE! |
| ENSMUSG00000114775.2 | 0 | NA | NA | NA | NA | NA | #VALUE! |
| ENSMUSG00000114776.2 | 0 | NA | NA | NA | NA | NA | #VALUE! |
| ENSMUSG00000114777.2 | 0 | NA | NA | NA | NA | NA | #VALUE! |
| ENSMUSG00000114778.2 | 0 | NA | NA | NA | NA | NA | #VALUE! |
| ENSMUSG00000114779.2 | 0 | NA | NA | NA | NA | NA | #VALUE! |
| ENSMUSG00000114780.2 | 0 | NA | NA | NA | NA | NA | #VALUE! |
| ENSMUSG00000114781.2 | 0 | NA | NA | NA | NA | NA | #VALUE! |
| ENSMUSG00000114782.2 | 0 | NA | NA | NA | NA | NA | #VALUE! |
| ENSMUSG00000114783.2 | 0 | NA | NA | NA | NA | NA | #VALUE! |
| ENSMUSG00000114784.2 | 0 | NA | NA | NA | NA | NA | #VALUE! |
| ENSMUSG00000114785.2 | 0 | NA | NA | NA | NA | NA | #VALUE! |
| ENSMUSG00000114786.2 | 0 | NA | NA | NA | NA | NA | #VALUE! |

| | | | | | | | |
|---|---|---|---|---|---|---|---|
| ENSMUSG00000114787.2 | 0 | NA | NA | NA | NA | NA | #VALUE! |
| ENSMUSG00000114788.2 | 0.839297 | 3.273019 | 3.958392 | 0.826856 | 0.408319 | NA | #VALUE! |
| ENSMUSG00000114789.2 | 0 | NA | NA | NA | NA | NA | #VALUE! |
| ENSMUSG00000114790.2 | 0 | NA | NA | NA | NA | NA | #VALUE! |
| ENSMUSG00000114791.2 | 0 | NA | NA | NA | NA | NA | #VALUE! |
| ENSMUSG00000114792.2 | 0 | NA | NA | NA | NA | NA | #VALUE! |
| ENSMUSG00000114793.2 | 0 | NA | NA | NA | NA | NA | #VALUE! |
| ENSMUSG00000114794.2 | 0 | NA | NA | NA | NA | NA | #VALUE! |
| ENSMUSG00000114795.2 | 0 | NA | NA | NA | NA | NA | #VALUE! |
| ENSMUSG00000114796.2 | 0.559531 | 2.703958 | 3.982863 | 0.678898 | 0.497202 | NA | #VALUE! |
| ENSMUSG00000114798.2 | 0 | NA | NA | NA | NA | NA | #VALUE! |
| ENSMUSG00000114799.2 | 0 | NA | NA | NA | NA | NA | #VALUE! |
| ENSMUSG00000114800.2 | 0 | NA | NA | NA | NA | NA | #VALUE! |
| ENSMUSG00000114801.2 | 0 | NA | NA | NA | NA | NA | #VALUE! |
| ENSMUSG00000114802.2 | 0 | NA | NA | NA | NA | NA | #VALUE! |
| ENSMUSG00000114803.2 | 0 | NA | NA | NA | NA | NA | #VALUE! |
| ENSMUSG00000114804.2 | 0 | NA | NA | NA | NA | NA | #VALUE! |
| ENSMUSG00000114805.2 | 0 | NA | NA | NA | NA | NA | #VALUE! |
| ENSMUSG00000114806.2 | 0 | NA | NA | NA | NA | NA | #VALUE! |
| ENSMUSG00000114807.2 | 0 | NA | NA | NA | NA | NA | #VALUE! |
| ENSMUSG00000114808.2 | 0 | NA | NA | NA | NA | NA | #VALUE! |
| ENSMUSG00000114809.2 | 0.408222 | -2.20205 | 4.016842 | -0.54821 | 0.583551 | NA | #VALUE! |
| ENSMUSG00000114810.2 | 0 | NA | NA | NA | NA | NA | #VALUE! |
| ENSMUSG00000114811.2 | 0 | NA | NA | NA | NA | NA | #VALUE! |
| ENSMUSG00000114812.2 | 0 | NA | NA | NA | NA | NA | #VALUE! |
| ENSMUSG00000114813.2 | 0 | NA | NA | NA | NA | NA | #VALUE! |
| ENSMUSG00000114814.2 | 0 | NA | NA | NA | NA | NA | #VALUE! |
| ENSMUSG00000114815.2 | 0 | NA | NA | NA | NA | NA | #VALUE! |
| ENSMUSG00000114816.2 | 0 | NA | NA | NA | NA | NA | #VALUE! |
| ENSMUSG00000114817.2 | 0 | NA | NA | NA | NA | NA | #VALUE! |
| ENSMUSG00000114818.2 | 5.822612 | -0.90898 | 3.479662 | -0.26123 | 0.793918 | NA | #VALUE! |
| ENSMUSG00000114819.2 | 0 | NA | NA | NA | NA | NA | #VALUE! |
| ENSMUSG00000114820.2 | 0 | NA | NA | NA | NA | NA | #VALUE! |
| ENSMUSG00000114821.2 | 0 | NA | NA | NA | NA | NA | #VALUE! |
| ENSMUSG00000114822.2 | 0 | NA | NA | NA | NA | NA | #VALUE! |
| ENSMUSG00000114823.2 | 0 | NA | NA | NA | NA | NA | #VALUE! |
| ENSMUSG00000114824.2 | 0 | NA | NA | NA | NA | NA | #VALUE! |
| ENSMUSG00000114825.2 | 0 | NA | NA | NA | NA | NA | #VALUE! |
| ENSMUSG00000114826.3 | 0 | NA | NA | NA | NA | NA | #VALUE! |
| ENSMUSG00000114827.2 | 0 | NA | NA | NA | NA | NA | #VALUE! |
| ENSMUSG00000114829.2 | 0 | NA | NA | NA | NA | NA | #VALUE! |
| ENSMUSG00000114830.2 | 0 | NA | NA | NA | NA | NA | #VALUE! |
| ENSMUSG00000114831.2 | 0 | NA | NA | NA | NA | NA | #VALUE! |
| ENSMUSG00000114832.3 | 0 | NA | NA | NA | NA | NA | #VALUE! |

| | | | | | | | |
|---|---|---|---|---|---|---|---|
| ENSMUSG00000114833.2 | 0 | NA | NA | NA | NA | NA | #VALUE! |
| ENSMUSG00000114834.2 | 0 | NA | NA | NA | NA | NA | #VALUE! |
| ENSMUSG00000114835.2 | 8.716868 | -1.06464 | 3.453878 | -0.30824 | 0.757896 | NA | #VALUE! |
| ENSMUSG00000114836.2 | 0 | NA | NA | NA | NA | NA | #VALUE! |
| ENSMUSG00000114837.2 | 0 | NA | NA | NA | NA | NA | #VALUE! |
| ENSMUSG00000114838.2 | 0 | NA | NA | NA | NA | NA | #VALUE! |
| ENSMUSG00000114839.2 | 0 | NA | NA | NA | NA | NA | #VALUE! |
| ENSMUSG00000114840.2 | 0 | NA | NA | NA | NA | NA | #VALUE! |
| ENSMUSG00000114841.2 | 0 | NA | NA | NA | NA | NA | #VALUE! |
| ENSMUSG00000114842.2 | 4.924298 | 5.796077 | 3.915847 | 1.480159 | 0.138831 | NA | #VALUE! |
| ENSMUSG00000114843.2 | 0 | NA | NA | NA | NA | NA | #VALUE! |
| ENSMUSG00000114844.2 | 0 | NA | NA | NA | NA | NA | #VALUE! |
| ENSMUSG00000114845.2 | 0 | NA | NA | NA | NA | NA | #VALUE! |
| ENSMUSG00000114846.2 | 0 | NA | NA | NA | NA | NA | #VALUE! |
| ENSMUSG00000114847.2 | 0 | NA | NA | NA | NA | NA | #VALUE! |
| ENSMUSG00000114848.2 | 0 | NA | NA | NA | NA | NA | #VALUE! |
| ENSMUSG00000114849.2 | 0 | NA | NA | NA | NA | NA | #VALUE! |
| ENSMUSG00000114850.2 | 0 | NA | NA | NA | NA | NA | #VALUE! |
| ENSMUSG00000114851.2 | 0 | NA | NA | NA | NA | NA | #VALUE! |
| ENSMUSG00000114852.2 | 0 | NA | NA | NA | NA | NA | #VALUE! |
| ENSMUSG00000114853.2 | 0 | NA | NA | NA | NA | NA | #VALUE! |
| ENSMUSG00000114854.2 | 0 | NA | NA | NA | NA | NA | #VALUE! |
| ENSMUSG00000114855.2 | 0 | NA | NA | NA | NA | NA | #VALUE! |
| ENSMUSG00000114856.2 | 0 | NA | NA | NA | NA | NA | #VALUE! |
| ENSMUSG00000114857.2 | 0 | NA | NA | NA | NA | NA | #VALUE! |
| ENSMUSG00000114858.2 | 0 | NA | NA | NA | NA | NA | #VALUE! |
| ENSMUSG00000114859.2 | 0 | NA | NA | NA | NA | NA | #VALUE! |
| ENSMUSG00000114860.2 | 0 | NA | NA | NA | NA | NA | #VALUE! |
| ENSMUSG00000114861.2 | 0 | NA | NA | NA | NA | NA | #VALUE! |
| ENSMUSG00000114862.2 | 0 | NA | NA | NA | NA | NA | #VALUE! |
| ENSMUSG00000114863.2 | 0 | NA | NA | NA | NA | NA | #VALUE! |
| ENSMUSG00000114864.2 | 0 | NA | NA | NA | NA | NA | #VALUE! |
| ENSMUSG00000114865.3 | 0 | NA | NA | NA | NA | NA | #VALUE! |
| ENSMUSG00000114866.2 | 0 | NA | NA | NA | NA | NA | #VALUE! |
| ENSMUSG00000114867.2 | 0 | NA | NA | NA | NA | NA | #VALUE! |
| ENSMUSG00000114868.2 | 0 | NA | NA | NA | NA | NA | #VALUE! |
| ENSMUSG00000114869.2 | 0 | NA | NA | NA | NA | NA | #VALUE! |
| ENSMUSG00000114870.2 | 0 | NA | NA | NA | NA | NA | #VALUE! |
| ENSMUSG00000114871.2 | 0 | NA | NA | NA | NA | NA | #VALUE! |
| ENSMUSG00000114872.2 | 0 | NA | NA | NA | NA | NA | #VALUE! |
| ENSMUSG00000114873.2 | 0 | NA | NA | NA | NA | NA | #VALUE! |
| ENSMUSG00000114874.2 | 0 | NA | NA | NA | NA | NA | #VALUE! |
| ENSMUSG00000114875.2 | 0 | NA | NA | NA | NA | NA | #VALUE! |
| ENSMUSG00000114876.2 | 0 | NA | NA | NA | NA | NA | #VALUE! |

| | | | | | | | |
|---|---|---|---|---|---|---|---|
| ENSMUSG00000114877.2 | 0 | NA | NA | NA | NA | NA | #VALUE! |
| ENSMUSG00000114878.2 | 0 | NA | NA | NA | NA | NA | #VALUE! |
| ENSMUSG00000114879.2 | 0 | NA | NA | NA | NA | NA | #VALUE! |
| ENSMUSG00000114880.2 | 0 | NA | NA | NA | NA | NA | #VALUE! |
| ENSMUSG00000114881.2 | 0 | NA | NA | NA | NA | NA | #VALUE! |
| ENSMUSG00000114882.2 | 0 | NA | NA | NA | NA | NA | #VALUE! |
| ENSMUSG00000114883.2 | 0 | NA | NA | NA | NA | NA | #VALUE! |
| ENSMUSG00000114884.2 | 0 | NA | NA | NA | NA | NA | #VALUE! |
| ENSMUSG00000114885.2 | 0 | NA | NA | NA | NA | NA | #VALUE! |
| ENSMUSG00000114886.2 | 0.245976 | 1.027628 | 4.067126 | 0.252667 | 0.800526 | NA | #VALUE! |
| ENSMUSG00000114887.2 | 0 | NA | NA | NA | NA | NA | #VALUE! |
| ENSMUSG00000114888.2 | 0 | NA | NA | NA | NA | NA | #VALUE! |
| ENSMUSG00000114889.2 | 0 | NA | NA | NA | NA | NA | #VALUE! |
| ENSMUSG00000114890.2 | 0 | NA | NA | NA | NA | NA | #VALUE! |
| ENSMUSG00000114891.2 | 0 | NA | NA | NA | NA | NA | #VALUE! |
| ENSMUSG00000114892.2 | 0 | NA | NA | NA | NA | NA | #VALUE! |
| ENSMUSG00000114893.2 | 0 | NA | NA | NA | NA | NA | #VALUE! |
| ENSMUSG00000114894.2 | 0 | NA | NA | NA | NA | NA | #VALUE! |
| ENSMUSG00000114895.2 | 0 | NA | NA | NA | NA | NA | #VALUE! |
| ENSMUSG00000114896.2 | 0 | NA | NA | NA | NA | NA | #VALUE! |
| ENSMUSG00000114897.2 | 0 | NA | NA | NA | NA | NA | #VALUE! |
| ENSMUSG00000114898.2 | 0 | NA | NA | NA | NA | NA | #VALUE! |
| ENSMUSG00000114899.2 | 0 | NA | NA | NA | NA | NA | #VALUE! |
| ENSMUSG00000114900.2 | 0 | NA | NA | NA | NA | NA | #VALUE! |
| ENSMUSG00000114901.2 | 0 | NA | NA | NA | NA | NA | #VALUE! |
| ENSMUSG00000114902.2 | 0 | NA | NA | NA | NA | NA | #VALUE! |
| ENSMUSG00000114903.2 | 0 | NA | NA | NA | NA | NA | #VALUE! |
| ENSMUSG00000114904.2 | 0 | NA | NA | NA | NA | NA | #VALUE! |
| ENSMUSG00000114905.2 | 0 | NA | NA | NA | NA | NA | #VALUE! |
| ENSMUSG00000114906.2 | 0 | NA | NA | NA | NA | NA | #VALUE! |
| ENSMUSG00000114907.2 | 0 | NA | NA | NA | NA | NA | #VALUE! |
| ENSMUSG00000114908.2 | 0 | NA | NA | NA | NA | NA | #VALUE! |
| ENSMUSG00000114909.2 | 0 | NA | NA | NA | NA | NA | #VALUE! |
| ENSMUSG00000114910.2 | 0 | NA | NA | NA | NA | NA | #VALUE! |
| ENSMUSG00000114911.2 | 0 | NA | NA | NA | NA | NA | #VALUE! |
| ENSMUSG00000114912.2 | 0 | NA | NA | NA | NA | NA | #VALUE! |
| ENSMUSG00000114913.2 | 0 | NA | NA | NA | NA | NA | #VALUE! |
| ENSMUSG00000114914.2 | 0 | NA | NA | NA | NA | NA | #VALUE! |
| ENSMUSG00000114915.2 | 0 | NA | NA | NA | NA | NA | #VALUE! |
| ENSMUSG00000114916.2 | 0 | NA | NA | NA | NA | NA | #VALUE! |
| ENSMUSG00000114917.2 | 0 | NA | NA | NA | NA | NA | #VALUE! |
| ENSMUSG00000114918.2 | 0 | NA | NA | NA | NA | NA | #VALUE! |
| ENSMUSG00000114919.2 | 0 | NA | NA | NA | NA | NA | #VALUE! |
| ENSMUSG00000114920.2 | 0 | NA | NA | NA | NA | NA | #VALUE! |

| | | | | | | | |
|---|---|---|---|---|---|---|---|
| ENSMUSG00000114921.2 | 0 | NA | NA | NA | NA | NA | #VALUE! |
| ENSMUSG00000114922.2 | 0 | NA | NA | NA | NA | NA | #VALUE! |
| ENSMUSG00000114923.2 | 0 | NA | NA | NA | NA | NA | #VALUE! |
| ENSMUSG00000114924.2 | 0 | NA | NA | NA | NA | NA | #VALUE! |
| ENSMUSG00000114925.2 | 0 | NA | NA | NA | NA | NA | #VALUE! |
| ENSMUSG00000114926.2 | 0 | NA | NA | NA | NA | NA | #VALUE! |
| ENSMUSG00000114927.2 | 0 | NA | NA | NA | NA | NA | #VALUE! |
| ENSMUSG00000114928.2 | 0 | NA | NA | NA | NA | NA | #VALUE! |
| ENSMUSG00000114929.2 | 0 | NA | NA | NA | NA | NA | #VALUE! |
| ENSMUSG00000114930.2 | 0 | NA | NA | NA | NA | NA | #VALUE! |
| ENSMUSG00000114931.2 | 0 | NA | NA | NA | NA | NA | #VALUE! |
| ENSMUSG00000114932.2 | 0 | NA | NA | NA | NA | NA | #VALUE! |
| ENSMUSG00000114933.2 | 0 | NA | NA | NA | NA | NA | #VALUE! |
| ENSMUSG00000114934.2 | 0 | NA | NA | NA | NA | NA | #VALUE! |
| ENSMUSG00000114935.2 | 0 | NA | NA | NA | NA | NA | #VALUE! |
| ENSMUSG00000114936.2 | 0 | NA | NA | NA | NA | NA | #VALUE! |
| ENSMUSG00000114937.2 | 0.624163 | -2.86979 | 3.97661 | -0.72167 | 0.470499 | NA | #VALUE! |
| ENSMUSG00000114938.2 | 0 | NA | NA | NA | NA | NA | #VALUE! |
| ENSMUSG00000114939.2 | 0 | NA | NA | NA | NA | NA | #VALUE! |
| ENSMUSG00000114940.2 | 0 | NA | NA | NA | NA | NA | #VALUE! |
| ENSMUSG00000114941.2 | 0 | NA | NA | NA | NA | NA | #VALUE! |
| ENSMUSG00000114942.2 | 0 | NA | NA | NA | NA | NA | #VALUE! |
| ENSMUSG00000114943.2 | 0 | NA | NA | NA | NA | NA | #VALUE! |
| ENSMUSG00000114944.2 | 0 | NA | NA | NA | NA | NA | #VALUE! |
| ENSMUSG00000114945.2 | 0 | NA | NA | NA | NA | NA | #VALUE! |
| ENSMUSG00000114946.2 | 0 | NA | NA | NA | NA | NA | #VALUE! |
| ENSMUSG00000114947.2 | 0 | NA | NA | NA | NA | NA | #VALUE! |
| ENSMUSG00000114948.2 | 0 | NA | NA | NA | NA | NA | #VALUE! |
| ENSMUSG00000114949.2 | 0 | NA | NA | NA | NA | NA | #VALUE! |
| ENSMUSG00000114950.2 | 0 | NA | NA | NA | NA | NA | #VALUE! |
| ENSMUSG00000114951.2 | 0 | NA | NA | NA | NA | NA | #VALUE! |
| ENSMUSG00000114952.2 | 0 | NA | NA | NA | NA | NA | #VALUE! |
| ENSMUSG00000114953.2 | 0 | NA | NA | NA | NA | NA | #VALUE! |
| ENSMUSG00000114954.2 | 0 | NA | NA | NA | NA | NA | #VALUE! |
| ENSMUSG00000114955.2 | 0 | NA | NA | NA | NA | NA | #VALUE! |
| ENSMUSG00000114956.2 | 0 | NA | NA | NA | NA | NA | #VALUE! |
| ENSMUSG00000114957.2 | 0 | NA | NA | NA | NA | NA | #VALUE! |
| ENSMUSG00000114958.2 | 0 | NA | NA | NA | NA | NA | #VALUE! |
| ENSMUSG00000114959.2 | 0 | NA | NA | NA | NA | NA | #VALUE! |
| ENSMUSG00000114960.2 | 0 | NA | NA | NA | NA | NA | #VALUE! |
| ENSMUSG00000114961.2 | 0 | NA | NA | NA | NA | NA | #VALUE! |
| ENSMUSG00000114962.2 | 0 | NA | NA | NA | NA | NA | #VALUE! |
| ENSMUSG00000114963.2 | 0 | NA | NA | NA | NA | NA | #VALUE! |
| ENSMUSG00000114964.2 | 0 | NA | NA | NA | NA | NA | #VALUE! |

| | | | | | | | |
|---|---|---|---|---|---|---|---|
| ENSMUSG00000114965.2 | 0 | NA | NA | NA | NA | NA | #VALUE! |
| ENSMUSG00000114966.2 | 0 | NA | NA | NA | NA | NA | #VALUE! |
| ENSMUSG00000114967.2 | 0 | NA | NA | NA | NA | NA | #VALUE! |
| ENSMUSG00000114968.2 | 0 | NA | NA | NA | NA | NA | #VALUE! |
| ENSMUSG00000114969.2 | 0 | NA | NA | NA | NA | NA | #VALUE! |
| ENSMUSG00000114970.2 | 0 | NA | NA | NA | NA | NA | #VALUE! |
| ENSMUSG00000114971.2 | 0 | NA | NA | NA | NA | NA | #VALUE! |
| ENSMUSG00000114972.2 | 0 | NA | NA | NA | NA | NA | #VALUE! |
| ENSMUSG00000114973.2 | 0 | NA | NA | NA | NA | NA | #VALUE! |
| ENSMUSG00000114974.2 | 0 | NA | NA | NA | NA | NA | #VALUE! |
| ENSMUSG00000114975.2 | 0 | NA | NA | NA | NA | NA | #VALUE! |
| ENSMUSG00000114976.2 | 0 | NA | NA | NA | NA | NA | #VALUE! |
| ENSMUSG00000114977.2 | 0 | NA | NA | NA | NA | NA | #VALUE! |
| ENSMUSG00000114978.2 | 0 | NA | NA | NA | NA | NA | #VALUE! |
| ENSMUSG00000114979.2 | 0 | NA | NA | NA | NA | NA | #VALUE! |
| ENSMUSG00000114980.2 | 0 | NA | NA | NA | NA | NA | #VALUE! |
| ENSMUSG00000114981.2 | 0 | NA | NA | NA | NA | NA | #VALUE! |
| ENSMUSG00000114982.2 | 0 | NA | NA | NA | NA | NA | #VALUE! |
| ENSMUSG00000114983.2 | 0 | NA | NA | NA | NA | NA | #VALUE! |
| ENSMUSG00000114984.2 | 0 | NA | NA | NA | NA | NA | #VALUE! |
| ENSMUSG00000114985.2 | 0 | NA | NA | NA | NA | NA | #VALUE! |
| ENSMUSG00000114986.2 | 0 | NA | NA | NA | NA | NA | #VALUE! |
| ENSMUSG00000114987.2 | 0 | NA | NA | NA | NA | NA | #VALUE! |
| ENSMUSG00000114988.2 | 0 | NA | NA | NA | NA | NA | #VALUE! |
| ENSMUSG00000114989.2 | 0 | NA | NA | NA | NA | NA | #VALUE! |
| ENSMUSG00000114990.2 | 0 | NA | NA | NA | NA | NA | #VALUE! |
| ENSMUSG00000114991.2 | 0 | NA | NA | NA | NA | NA | #VALUE! |
| ENSMUSG00000114992.2 | 0 | NA | NA | NA | NA | NA | #VALUE! |
| ENSMUSG00000114993.2 | 0.289044 | -0.14751 | 4.065279 | -0.03629 | 0.971054 | NA | #VALUE! |
| ENSMUSG00000114994.2 | 0 | NA | NA | NA | NA | NA | #VALUE! |
| ENSMUSG00000114995.2 | 0 | NA | NA | NA | NA | NA | #VALUE! |
| ENSMUSG00000114996.2 | 0 | NA | NA | NA | NA | NA | #VALUE! |
| ENSMUSG00000114997.2 | 0 | NA | NA | NA | NA | NA | #VALUE! |
| ENSMUSG00000114998.2 | 0 | NA | NA | NA | NA | NA | #VALUE! |
| ENSMUSG00000114999.2 | 0.210026 | -1.52494 | 4.058692 | -0.37572 | 0.707123 | NA | #VALUE! |
| ENSMUSG00000115000.2 | 0 | NA | NA | NA | NA | NA | #VALUE! |
| ENSMUSG00000115001.2 | 0 | NA | NA | NA | NA | NA | #VALUE! |
| ENSMUSG00000115002.2 | 0 | NA | NA | NA | NA | NA | #VALUE! |
| ENSMUSG00000115003.2 | 0.210026 | -1.52494 | 4.058692 | -0.37572 | 0.707123 | NA | #VALUE! |
| ENSMUSG00000115004.2 | 0 | NA | NA | NA | NA | NA | #VALUE! |
| ENSMUSG00000115005.2 | 0 | NA | NA | NA | NA | NA | #VALUE! |
| ENSMUSG00000115006.2 | 0 | NA | NA | NA | NA | NA | #VALUE! |
| ENSMUSG00000115007.2 | 5.599523 | -6.0191 | 3.831887 | -1.57079 | 0.116231 | NA | #VALUE! |
| ENSMUSG00000115008.2 | 0 | NA | NA | NA | NA | NA | #VALUE! |

| | | | | | | | |
|---|---|---|---|---|---|---|---|
| ENSMUSG00000115009.2 | 0 | NA | NA | NA | NA | NA | #VALUE! |
| ENSMUSG00000115010.2 | 0 | NA | NA | NA | NA | NA | #VALUE! |
| ENSMUSG00000115011.2 | 0 | NA | NA | NA | NA | NA | #VALUE! |
| ENSMUSG00000115012.2 | 0 | NA | NA | NA | NA | NA | #VALUE! |
| ENSMUSG00000115013.2 | 0 | NA | NA | NA | NA | NA | #VALUE! |
| ENSMUSG00000115014.2 | 0 | NA | NA | NA | NA | NA | #VALUE! |
| ENSMUSG00000115015.2 | 0 | NA | NA | NA | NA | NA | #VALUE! |
| ENSMUSG00000115016.3 | 0 | NA | NA | NA | NA | NA | #VALUE! |
| ENSMUSG00000115017.2 | 0.105013 | -0.97324 | 4.076293 | -0.23876 | 0.811295 | NA | #VALUE! |
| ENSMUSG00000115018.3 | 0 | NA | NA | NA | NA | NA | #VALUE! |
| ENSMUSG00000115019.2 | 0 | NA | NA | NA | NA | NA | #VALUE! |
| ENSMUSG00000115020.2 | 0 | NA | NA | NA | NA | NA | #VALUE! |
| ENSMUSG00000115021.2 | 0 | NA | NA | NA | NA | NA | #VALUE! |
| ENSMUSG00000115022.2 | 0 | NA | NA | NA | NA | NA | #VALUE! |
| ENSMUSG00000115023.2 | 0 | NA | NA | NA | NA | NA | #VALUE! |
| ENSMUSG00000115024.2 | 0 | NA | NA | NA | NA | NA | #VALUE! |
| ENSMUSG00000115025.2 | 0 | NA | NA | NA | NA | NA | #VALUE! |
| ENSMUSG00000115026.2 | 0.466276 | 2.449889 | 3.997179 | 0.612904 | 0.53994 | NA | #VALUE! |
| ENSMUSG00000115027.2 | 0 | NA | NA | NA | NA | NA | #VALUE! |
| ENSMUSG00000115028.2 | 0 | NA | NA | NA | NA | NA | #VALUE! |
| ENSMUSG00000115029.2 | 0 | NA | NA | NA | NA | NA | #VALUE! |
| ENSMUSG00000115030.2 | 0 | NA | NA | NA | NA | NA | #VALUE! |
| ENSMUSG00000115031.2 | 0 | NA | NA | NA | NA | NA | #VALUE! |
| ENSMUSG00000115032.2 | 0 | NA | NA | NA | NA | NA | #VALUE! |
| ENSMUSG00000115033.2 | 0 | NA | NA | NA | NA | NA | #VALUE! |
| ENSMUSG00000115034.2 | 0 | NA | NA | NA | NA | NA | #VALUE! |
| ENSMUSG00000115035.2 | 0 | NA | NA | NA | NA | NA | #VALUE! |
| ENSMUSG00000115036.2 | 0 | NA | NA | NA | NA | NA | #VALUE! |
| ENSMUSG00000115037.2 | 0 | NA | NA | NA | NA | NA | #VALUE! |
| ENSMUSG00000115038.2 | 0 | NA | NA | NA | NA | NA | #VALUE! |
| ENSMUSG00000115039.2 | 0 | NA | NA | NA | NA | NA | #VALUE! |
| ENSMUSG00000115040.2 | 0 | NA | NA | NA | NA | NA | #VALUE! |
| ENSMUSG00000115041.2 | 0 | NA | NA | NA | NA | NA | #VALUE! |
| ENSMUSG00000115042.2 | 0 | NA | NA | NA | NA | NA | #VALUE! |
| ENSMUSG00000115043.2 | 0 | NA | NA | NA | NA | NA | #VALUE! |
| ENSMUSG00000115044.2 | 0 | NA | NA | NA | NA | NA | #VALUE! |
| ENSMUSG00000115045.2 | 0 | NA | NA | NA | NA | NA | #VALUE! |
| ENSMUSG00000115046.2 | 0 | NA | NA | NA | NA | NA | #VALUE! |
| ENSMUSG00000115047.2 | 0 | NA | NA | NA | NA | NA | #VALUE! |
| ENSMUSG00000115048.2 | 0 | NA | NA | NA | NA | NA | #VALUE! |
| ENSMUSG00000115049.2 | 0 | NA | NA | NA | NA | NA | #VALUE! |
| ENSMUSG00000115050.2 | 0 | NA | NA | NA | NA | NA | #VALUE! |
| ENSMUSG00000115051.2 | 0 | NA | NA | NA | NA | NA | #VALUE! |
| ENSMUSG00000115052.2 | 0 | NA | NA | NA | NA | NA | #VALUE! |

| | | | | | | | |
|---|---|---|---|---|---|---|---|
| ENSMUSG00000115053.2 | 0 | NA | NA | NA | NA | NA | #VALUE! |
| ENSMUSG00000115054.2 | 0 | NA | NA | NA | NA | NA | #VALUE! |
| ENSMUSG00000115056.2 | 0 | NA | NA | NA | NA | NA | #VALUE! |
| ENSMUSG00000115057.2 | 0 | NA | NA | NA | NA | NA | #VALUE! |
| ENSMUSG00000115058.2 | 0 | NA | NA | NA | NA | NA | #VALUE! |
| ENSMUSG00000115059.2 | 0 | NA | NA | NA | NA | NA | #VALUE! |
| ENSMUSG00000115060.2 | 0 | NA | NA | NA | NA | NA | #VALUE! |
| ENSMUSG00000115061.2 | 0 | NA | NA | NA | NA | NA | #VALUE! |
| ENSMUSG00000115062.2 | 0 | NA | NA | NA | NA | NA | #VALUE! |
| ENSMUSG00000115063.2 | 0 | NA | NA | NA | NA | NA | #VALUE! |
| ENSMUSG00000115064.2 | 0 | NA | NA | NA | NA | NA | #VALUE! |
| ENSMUSG00000115065.2 | 0 | NA | NA | NA | NA | NA | #VALUE! |
| ENSMUSG00000115066.2 | 0 | NA | NA | NA | NA | NA | #VALUE! |
| ENSMUSG00000115067.2 | 0 | NA | NA | NA | NA | NA | #VALUE! |
| ENSMUSG00000115068.2 | 0 | NA | NA | NA | NA | NA | #VALUE! |
| ENSMUSG00000115069.2 | 0 | NA | NA | NA | NA | NA | #VALUE! |
| ENSMUSG00000115070.2 | 0 | NA | NA | NA | NA | NA | #VALUE! |
| ENSMUSG00000115071.2 | 0 | NA | NA | NA | NA | NA | #VALUE! |
| ENSMUSG00000115072.2 | 0 | NA | NA | NA | NA | NA | #VALUE! |
| ENSMUSG00000115073.2 | 0 | NA | NA | NA | NA | NA | #VALUE! |
| ENSMUSG00000115074.2 | 0 | NA | NA | NA | NA | NA | #VALUE! |
| ENSMUSG00000115075.2 | 0 | NA | NA | NA | NA | NA | #VALUE! |
| ENSMUSG00000115076.2 | 0 | NA | NA | NA | NA | NA | #VALUE! |
| ENSMUSG00000115077.2 | 0 | NA | NA | NA | NA | NA | #VALUE! |
| ENSMUSG00000115079.2 | 0 | NA | NA | NA | NA | NA | #VALUE! |
| ENSMUSG00000115080.2 | 0 | NA | NA | NA | NA | NA | #VALUE! |
| ENSMUSG00000115081.2 | 0 | NA | NA | NA | NA | NA | #VALUE! |
| ENSMUSG00000115082.2 | 0 | NA | NA | NA | NA | NA | #VALUE! |
| ENSMUSG00000115083.2 | 0 | NA | NA | NA | NA | NA | #VALUE! |
| ENSMUSG00000115084.2 | 0 | NA | NA | NA | NA | NA | #VALUE! |
| ENSMUSG00000115085.2 | 0 | NA | NA | NA | NA | NA | #VALUE! |
| ENSMUSG00000115086.2 | 0 | NA | NA | NA | NA | NA | #VALUE! |
| ENSMUSG00000115087.2 | 0 | NA | NA | NA | NA | NA | #VALUE! |
| ENSMUSG00000115088.2 | 0 | NA | NA | NA | NA | NA | #VALUE! |
| ENSMUSG00000115089.2 | 0 | NA | NA | NA | NA | NA | #VALUE! |
| ENSMUSG00000115090.2 | 0 | NA | NA | NA | NA | NA | #VALUE! |
| ENSMUSG00000115091.2 | 0 | NA | NA | NA | NA | NA | #VALUE! |
| ENSMUSG00000115092.2 | 0 | NA | NA | NA | NA | NA | #VALUE! |
| ENSMUSG00000115093.2 | 0 | NA | NA | NA | NA | NA | #VALUE! |
| ENSMUSG00000115094.2 | 0 | NA | NA | NA | NA | NA | #VALUE! |
| ENSMUSG00000115095.2 | 0 | NA | NA | NA | NA | NA | #VALUE! |
| ENSMUSG00000115096.2 | 0 | NA | NA | NA | NA | NA | #VALUE! |
| ENSMUSG00000115097.2 | 0 | NA | NA | NA | NA | NA | #VALUE! |
| ENSMUSG00000115098.2 | 0 | NA | NA | NA | NA | NA | #VALUE! |

| | | | | | | | |
|---|---|---|---|---|---|---|---|
| ENSMUSG00000115099.2 | 0 | NA | NA | NA | NA | NA | #VALUE! |
| ENSMUSG00000115101.2 | 0 | NA | NA | NA | NA | NA | #VALUE! |
| ENSMUSG00000115102.2 | 0 | NA | NA | NA | NA | NA | #VALUE! |
| ENSMUSG00000115103.2 | 0 | NA | NA | NA | NA | NA | #VALUE! |
| ENSMUSG00000115104.2 | 0 | NA | NA | NA | NA | NA | #VALUE! |
| ENSMUSG00000115105.2 | 0 | NA | NA | NA | NA | NA | #VALUE! |
| ENSMUSG00000115106.2 | 0 | NA | NA | NA | NA | NA | #VALUE! |
| ENSMUSG00000115107.2 | 0 | NA | NA | NA | NA | NA | #VALUE! |
| ENSMUSG00000115108.2 | 0 | NA | NA | NA | NA | NA | #VALUE! |
| ENSMUSG00000115109.2 | 0 | NA | NA | NA | NA | NA | #VALUE! |
| ENSMUSG00000115110.2 | 0 | NA | NA | NA | NA | NA | #VALUE! |
| ENSMUSG00000115111.2 | 0 | NA | NA | NA | NA | NA | #VALUE! |
| ENSMUSG00000115112.2 | 0 | NA | NA | NA | NA | NA | #VALUE! |
| ENSMUSG00000115113.2 | 0 | NA | NA | NA | NA | NA | #VALUE! |
| ENSMUSG00000115114.2 | 0 | NA | NA | NA | NA | NA | #VALUE! |
| ENSMUSG00000115115.2 | 0 | NA | NA | NA | NA | NA | #VALUE! |
| ENSMUSG00000115116.2 | 0.093255 | 0.868093 | 4.072464 | 0.213162 | 0.831201 | NA | #VALUE! |
| ENSMUSG00000115117.2 | 0 | NA | NA | NA | NA | NA | #VALUE! |
| ENSMUSG00000115118.2 | 0 | NA | NA | NA | NA | NA | #VALUE! |
| ENSMUSG00000115119.2 | 0 | NA | NA | NA | NA | NA | #VALUE! |
| ENSMUSG00000115120.2 | 0 | NA | NA | NA | NA | NA | #VALUE! |
| ENSMUSG00000115121.2 | 0 | NA | NA | NA | NA | NA | #VALUE! |
| ENSMUSG00000115122.2 | 0 | NA | NA | NA | NA | NA | #VALUE! |
| ENSMUSG00000115123.2 | 0 | NA | NA | NA | NA | NA | #VALUE! |
| ENSMUSG00000115125.2 | 0 | NA | NA | NA | NA | NA | #VALUE! |
| ENSMUSG00000115126.2 | 0 | NA | NA | NA | NA | NA | #VALUE! |
| ENSMUSG00000115127.2 | 0 | NA | NA | NA | NA | NA | #VALUE! |
| ENSMUSG00000115128.2 | 0 | NA | NA | NA | NA | NA | #VALUE! |
| ENSMUSG00000115129.2 | 0 | NA | NA | NA | NA | NA | #VALUE! |
| ENSMUSG00000115130.2 | 0 | NA | NA | NA | NA | NA | #VALUE! |
| ENSMUSG00000115131.2 | 0 | NA | NA | NA | NA | NA | #VALUE! |
| ENSMUSG00000115132.2 | 0 | NA | NA | NA | NA | NA | #VALUE! |
| ENSMUSG00000115133.2 | 0 | NA | NA | NA | NA | NA | #VALUE! |
| ENSMUSG00000115134.2 | 0 | NA | NA | NA | NA | NA | #VALUE! |
| ENSMUSG00000115135.2 | 0 | NA | NA | NA | NA | NA | #VALUE! |
| ENSMUSG00000115136.2 | 0 | NA | NA | NA | NA | NA | #VALUE! |
| ENSMUSG00000115137.2 | 0 | NA | NA | NA | NA | NA | #VALUE! |
| ENSMUSG00000115138.2 | 0 | NA | NA | NA | NA | NA | #VALUE! |
| ENSMUSG00000115139.2 | 0 | NA | NA | NA | NA | NA | #VALUE! |
| ENSMUSG00000115140.2 | 0 | NA | NA | NA | NA | NA | #VALUE! |
| ENSMUSG00000115141.2 | 0 | NA | NA | NA | NA | NA | #VALUE! |
| ENSMUSG00000115142.2 | 0 | NA | NA | NA | NA | NA | #VALUE! |
| ENSMUSG00000115143.2 | 0 | NA | NA | NA | NA | NA | #VALUE! |
| ENSMUSG00000115144.2 | 0 | NA | NA | NA | NA | NA | #VALUE! |

| | | | | | | | |
|---|---|---|---|---|---|---|---|
| ENSMUSG00000115145.2 | 0 | NA | NA | NA | NA | NA | #VALUE! |
| ENSMUSG00000115146.2 | 0 | NA | NA | NA | NA | NA | #VALUE! |
| ENSMUSG00000115147.2 | 0 | NA | NA | NA | NA | NA | #VALUE! |
| ENSMUSG00000115148.2 | 0 | NA | NA | NA | NA | NA | #VALUE! |
| ENSMUSG00000115149.2 | 0 | NA | NA | NA | NA | NA | #VALUE! |
| ENSMUSG00000115150.2 | 0 | NA | NA | NA | NA | NA | #VALUE! |
| ENSMUSG00000115151.2 | 0 | NA | NA | NA | NA | NA | #VALUE! |
| ENSMUSG00000115152.2 | 0 | NA | NA | NA | NA | NA | #VALUE! |
| ENSMUSG00000115153.2 | 0 | NA | NA | NA | NA | NA | #VALUE! |
| ENSMUSG00000115154.2 | 0 | NA | NA | NA | NA | NA | #VALUE! |
| ENSMUSG00000115155.2 | 0 | NA | NA | NA | NA | NA | #VALUE! |
| ENSMUSG00000115156.2 | 0 | NA | NA | NA | NA | NA | #VALUE! |
| ENSMUSG00000115157.2 | 0 | NA | NA | NA | NA | NA | #VALUE! |
| ENSMUSG00000115158.2 | 0 | NA | NA | NA | NA | NA | #VALUE! |
| ENSMUSG00000115159.2 | 0 | NA | NA | NA | NA | NA | #VALUE! |
| ENSMUSG00000115160.2 | 0 | NA | NA | NA | NA | NA | #VALUE! |
| ENSMUSG00000115161.2 | 0 | NA | NA | NA | NA | NA | #VALUE! |
| ENSMUSG00000115162.2 | 0 | NA | NA | NA | NA | NA | #VALUE! |
| ENSMUSG00000115163.2 | 0 | NA | NA | NA | NA | NA | #VALUE! |
| ENSMUSG00000115164.3 | 0 | NA | NA | NA | NA | NA | #VALUE! |
| ENSMUSG00000115165.2 | 0 | NA | NA | NA | NA | NA | #VALUE! |
| ENSMUSG00000115166.2 | 0.093255 | 0.868093 | 4.072464 | 0.213162 | 0.831201 | NA | #VALUE! |
| ENSMUSG00000115167.2 | 0 | NA | NA | NA | NA | NA | #VALUE! |
| ENSMUSG00000115168.2 | 0 | NA | NA | NA | NA | NA | #VALUE! |
| ENSMUSG00000115169.2 | 0 | NA | NA | NA | NA | NA | #VALUE! |
| ENSMUSG00000115170.2 | 0 | NA | NA | NA | NA | NA | #VALUE! |
| ENSMUSG00000115171.2 | 0 | NA | NA | NA | NA | NA | #VALUE! |
| ENSMUSG00000115172.2 | 0 | NA | NA | NA | NA | NA | #VALUE! |
| ENSMUSG00000115173.2 | 0 | NA | NA | NA | NA | NA | #VALUE! |
| ENSMUSG00000115174.2 | 0 | NA | NA | NA | NA | NA | #VALUE! |
| ENSMUSG00000115175.2 | 0 | NA | NA | NA | NA | NA | #VALUE! |
| ENSMUSG00000115176.2 | 0 | NA | NA | NA | NA | NA | #VALUE! |
| ENSMUSG00000115177.2 | 0 | NA | NA | NA | NA | NA | #VALUE! |
| ENSMUSG00000115178.2 | 0 | NA | NA | NA | NA | NA | #VALUE! |
| ENSMUSG00000115179.2 | 0 | NA | NA | NA | NA | NA | #VALUE! |
| ENSMUSG00000115180.2 | 0 | NA | NA | NA | NA | NA | #VALUE! |
| ENSMUSG00000115181.2 | 0 | NA | NA | NA | NA | NA | #VALUE! |
| ENSMUSG00000115182.2 | 0 | NA | NA | NA | NA | NA | #VALUE! |
| ENSMUSG00000115183.2 | 0 | NA | NA | NA | NA | NA | #VALUE! |
| ENSMUSG00000115184.2 | 0 | NA | NA | NA | NA | NA | #VALUE! |
| ENSMUSG00000115185.2 | 0 | NA | NA | NA | NA | NA | #VALUE! |
| ENSMUSG00000115186.2 | 4.382994 | 5.629097 | 3.916959 | 1.437109 | 0.150687 | NA | #VALUE! |
| ENSMUSG00000115187.2 | 0 | NA | NA | NA | NA | NA | #VALUE! |
| ENSMUSG00000115188.2 | 0 | NA | NA | NA | NA | NA | #VALUE! |

| | | | | | | | |
|---|---|---|---|---|---|---|---|
| ENSMUSG00000115190.2 | 0 | NA | NA | NA | NA | NA | #VALUE! |
| ENSMUSG00000115191.2 | 0 | NA | NA | NA | NA | NA | #VALUE! |
| ENSMUSG00000115192.2 | 0 | NA | NA | NA | NA | NA | #VALUE! |
| ENSMUSG00000115193.2 | 0 | NA | NA | NA | NA | NA | #VALUE! |
| ENSMUSG00000115194.2 | 0.105013 | -0.97324 | 4.076293 | -0.23876 | 0.811295 | NA | #VALUE! |
| ENSMUSG00000115195.2 | 0 | NA | NA | NA | NA | NA | #VALUE! |
| ENSMUSG00000115196.2 | 0 | NA | NA | NA | NA | NA | #VALUE! |
| ENSMUSG00000115197.2 | 0 | NA | NA | NA | NA | NA | #VALUE! |
| ENSMUSG00000115198.2 | 0 | NA | NA | NA | NA | NA | #VALUE! |
| ENSMUSG00000115199.2 | 0 | NA | NA | NA | NA | NA | #VALUE! |
| ENSMUSG00000115201.2 | 0 | NA | NA | NA | NA | NA | #VALUE! |
| ENSMUSG00000115202.2 | 0 | NA | NA | NA | NA | NA | #VALUE! |
| ENSMUSG00000115203.2 | 0 | NA | NA | NA | NA | NA | #VALUE! |
| ENSMUSG00000115204.2 | 0 | NA | NA | NA | NA | NA | #VALUE! |
| ENSMUSG00000115205.2 | 0.198268 | -0.06494 | 4.068276 | -0.01596 | 0.987265 | NA | #VALUE! |
| ENSMUSG00000115206.2 | 0 | NA | NA | NA | NA | NA | #VALUE! |
| ENSMUSG00000115207.2 | 0 | NA | NA | NA | NA | NA | #VALUE! |
| ENSMUSG00000115208.2 | 0 | NA | NA | NA | NA | NA | #VALUE! |
| ENSMUSG00000115209.2 | 0 | NA | NA | NA | NA | NA | #VALUE! |
| ENSMUSG00000115210.2 | 0 | NA | NA | NA | NA | NA | #VALUE! |
| ENSMUSG00000115211.2 | 0 | NA | NA | NA | NA | NA | #VALUE! |
| ENSMUSG00000115212.2 | 0 | NA | NA | NA | NA | NA | #VALUE! |
| ENSMUSG00000115213.2 | 0 | NA | NA | NA | NA | NA | #VALUE! |
| ENSMUSG00000115214.2 | 0 | NA | NA | NA | NA | NA | #VALUE! |
| ENSMUSG00000115215.2 | 0 | NA | NA | NA | NA | NA | #VALUE! |
| ENSMUSG00000115216.2 | 0 | NA | NA | NA | NA | NA | #VALUE! |
| ENSMUSG00000115217.2 | 0 | NA | NA | NA | NA | NA | #VALUE! |
| ENSMUSG00000115218.2 | 0 | NA | NA | NA | NA | NA | #VALUE! |
| ENSMUSG00000115219.3 | 3.097374 | -3.18253 | 3.876953 | -0.82089 | 0.411712 | NA | #VALUE! |
| ENSMUSG00000115220.2 | 0 | NA | NA | NA | NA | NA | #VALUE! |
| ENSMUSG00000115221.2 | 0 | NA | NA | NA | NA | NA | #VALUE! |
| ENSMUSG00000115222.2 | 0 | NA | NA | NA | NA | NA | #VALUE! |
| ENSMUSG00000115223.2 | 0 | NA | NA | NA | NA | NA | #VALUE! |
| ENSMUSG00000115224.2 | 0 | NA | NA | NA | NA | NA | #VALUE! |
| ENSMUSG00000115225.2 | 0 | NA | NA | NA | NA | NA | #VALUE! |
| ENSMUSG00000115226.2 | 0 | NA | NA | NA | NA | NA | #VALUE! |
| ENSMUSG00000115227.2 | 0 | NA | NA | NA | NA | NA | #VALUE! |
| ENSMUSG00000115228.2 | 0 | NA | NA | NA | NA | NA | #VALUE! |
| ENSMUSG00000115229.2 | 0 | NA | NA | NA | NA | NA | #VALUE! |
| ENSMUSG00000115230.2 | 0 | NA | NA | NA | NA | NA | #VALUE! |
| ENSMUSG00000115231.2 | 0 | NA | NA | NA | NA | NA | #VALUE! |
| ENSMUSG00000115232.2 | 0 | NA | NA | NA | NA | NA | #VALUE! |
| ENSMUSG00000115233.2 | 0 | NA | NA | NA | NA | NA | #VALUE! |
| ENSMUSG00000115234.2 | 0 | NA | NA | NA | NA | NA | #VALUE! |

| | | | | | | | |
|---|---|---|---|---|---|---|---|
| ENSMUSG00000115235.2 | 0 | NA | NA | NA | NA | NA | #VALUE! |
| ENSMUSG00000115236.2 | 0 | NA | NA | NA | NA | NA | #VALUE! |
| ENSMUSG00000115237.2 | 0 | NA | NA | NA | NA | NA | #VALUE! |
| ENSMUSG00000115238.2 | 0 | NA | NA | NA | NA | NA | #VALUE! |
| ENSMUSG00000115239.2 | 0 | NA | NA | NA | NA | NA | #VALUE! |
| ENSMUSG00000115240.2 | 0 | NA | NA | NA | NA | NA | #VALUE! |
| ENSMUSG00000115241.2 | 0 | NA | NA | NA | NA | NA | #VALUE! |
| ENSMUSG00000115242.2 | 0 | NA | NA | NA | NA | NA | #VALUE! |
| ENSMUSG00000115243.2 | 0 | NA | NA | NA | NA | NA | #VALUE! |
| ENSMUSG00000115244.2 | 0 | NA | NA | NA | NA | NA | #VALUE! |
| ENSMUSG00000115245.2 | 0 | NA | NA | NA | NA | NA | #VALUE! |
| ENSMUSG00000115246.2 | 0 | NA | NA | NA | NA | NA | #VALUE! |
| ENSMUSG00000115247.2 | 0 | NA | NA | NA | NA | NA | #VALUE! |
| ENSMUSG00000115248.2 | 0 | NA | NA | NA | NA | NA | #VALUE! |
| ENSMUSG00000115249.2 | 0 | NA | NA | NA | NA | NA | #VALUE! |
| ENSMUSG00000115250.2 | 0 | NA | NA | NA | NA | NA | #VALUE! |
| ENSMUSG00000115251.2 | 0 | NA | NA | NA | NA | NA | #VALUE! |
| ENSMUSG00000115252.2 | 0 | NA | NA | NA | NA | NA | #VALUE! |
| ENSMUSG00000115253.2 | 0 | NA | NA | NA | NA | NA | #VALUE! |
| ENSMUSG00000115254.2 | 0 | NA | NA | NA | NA | NA | #VALUE! |
| ENSMUSG00000115256.2 | 0 | NA | NA | NA | NA | NA | #VALUE! |
| ENSMUSG00000115257.2 | 0 | NA | NA | NA | NA | NA | #VALUE! |
| ENSMUSG00000115258.2 | 0 | NA | NA | NA | NA | NA | #VALUE! |
| ENSMUSG00000115259.2 | 0 | NA | NA | NA | NA | NA | #VALUE! |
| ENSMUSG00000115260.2 | 0 | NA | NA | NA | NA | NA | #VALUE! |
| ENSMUSG00000115261.2 | 0 | NA | NA | NA | NA | NA | #VALUE! |
| ENSMUSG00000115262.2 | 0 | NA | NA | NA | NA | NA | #VALUE! |
| ENSMUSG00000115263.2 | 0 | NA | NA | NA | NA | NA | #VALUE! |
| ENSMUSG00000115264.2 | 0 | NA | NA | NA | NA | NA | #VALUE! |
| ENSMUSG00000115265.2 | 0 | NA | NA | NA | NA | NA | #VALUE! |
| ENSMUSG00000115266.2 | 0 | NA | NA | NA | NA | NA | #VALUE! |
| ENSMUSG00000115267.2 | 0 | NA | NA | NA | NA | NA | #VALUE! |
| ENSMUSG00000115268.2 | 0 | NA | NA | NA | NA | NA | #VALUE! |
| ENSMUSG00000115269.2 | 0 | NA | NA | NA | NA | NA | #VALUE! |
| ENSMUSG00000115270.2 | 0 | NA | NA | NA | NA | NA | #VALUE! |
| ENSMUSG00000115271.2 | 0 | NA | NA | NA | NA | NA | #VALUE! |
| ENSMUSG00000115272.2 | 0 | NA | NA | NA | NA | NA | #VALUE! |
| ENSMUSG00000115273.2 | 0 | NA | NA | NA | NA | NA | #VALUE! |
| ENSMUSG00000115274.2 | 0 | NA | NA | NA | NA | NA | #VALUE! |
| ENSMUSG00000115275.2 | 8.16443 | -6.55737 | 3.847188 | -1.70446 | 0.088296 | NA | #VALUE! |
| ENSMUSG00000115276.2 | 6.99399 | 2.176975 | 2.973873 | 0.732034 | 0.464148 | NA | #VALUE! |
| ENSMUSG00000115277.2 | 0 | NA | NA | NA | NA | NA | #VALUE! |
| ENSMUSG00000115278.2 | 0 | NA | NA | NA | NA | NA | #VALUE! |
| ENSMUSG00000115279.2 | 0 | NA | NA | NA | NA | NA | #VALUE! |

| | | | | | | | |
|---|---|---|---|---|---|---|---|
| ENSMUSG00000115280.2 | 0 | NA | NA | NA | NA | NA | #VALUE! |
| ENSMUSG00000115281.2 | 0 | NA | NA | NA | NA | NA | #VALUE! |
| ENSMUSG00000115282.2 | 0 | NA | NA | NA | NA | NA | #VALUE! |
| ENSMUSG00000115283.2 | 0 | NA | NA | NA | NA | NA | #VALUE! |
| ENSMUSG00000115284.2 | 0 | NA | NA | NA | NA | NA | #VALUE! |
| ENSMUSG00000115285.2 | 0 | NA | NA | NA | NA | NA | #VALUE! |
| ENSMUSG00000115286.2 | 0 | NA | NA | NA | NA | NA | #VALUE! |
| ENSMUSG00000115287.2 | 0 | NA | NA | NA | NA | NA | #VALUE! |
| ENSMUSG00000115288.2 | 0 | NA | NA | NA | NA | NA | #VALUE! |
| ENSMUSG00000115289.2 | 0 | NA | NA | NA | NA | NA | #VALUE! |
| ENSMUSG00000115290.2 | 0 | NA | NA | NA | NA | NA | #VALUE! |
| ENSMUSG00000115291.2 | 0 | NA | NA | NA | NA | NA | #VALUE! |
| ENSMUSG00000115292.2 | 0 | NA | NA | NA | NA | NA | #VALUE! |
| ENSMUSG00000115293.3 | 0 | NA | NA | NA | NA | NA | #VALUE! |
| ENSMUSG00000115294.2 | 0 | NA | NA | NA | NA | NA | #VALUE! |
| ENSMUSG00000115295.2 | 0 | NA | NA | NA | NA | NA | #VALUE! |
| ENSMUSG00000115296.2 | 0 | NA | NA | NA | NA | NA | #VALUE! |
| ENSMUSG00000115297.2 | 0 | NA | NA | NA | NA | NA | #VALUE! |
| ENSMUSG00000115298.2 | 0 | NA | NA | NA | NA | NA | #VALUE! |
| ENSMUSG00000115299.2 | 0 | NA | NA | NA | NA | NA | #VALUE! |
| ENSMUSG00000115300.2 | 0 | NA | NA | NA | NA | NA | #VALUE! |
| ENSMUSG00000115301.2 | 3.081686 | 1.204336 | 3.796868 | 0.317192 | 0.751098 | NA | #VALUE! |
| ENSMUSG00000115302.2 | 0 | NA | NA | NA | NA | NA | #VALUE! |
| ENSMUSG00000115303.2 | 0 | NA | NA | NA | NA | NA | #VALUE! |
| ENSMUSG00000115304.2 | 0 | NA | NA | NA | NA | NA | #VALUE! |
| ENSMUSG00000115305.2 | 0 | NA | NA | NA | NA | NA | #VALUE! |
| ENSMUSG00000115306.2 | 0 | NA | NA | NA | NA | NA | #VALUE! |
| ENSMUSG00000115307.2 | 0 | NA | NA | NA | NA | NA | #VALUE! |
| ENSMUSG00000115308.2 | 0 | NA | NA | NA | NA | NA | #VALUE! |
| ENSMUSG00000115309.2 | 0 | NA | NA | NA | NA | NA | #VALUE! |
| ENSMUSG00000115310.2 | 0 | NA | NA | NA | NA | NA | #VALUE! |
| ENSMUSG00000115311.2 | 0 | NA | NA | NA | NA | NA | #VALUE! |
| ENSMUSG00000115312.2 | 0 | NA | NA | NA | NA | NA | #VALUE! |
| ENSMUSG00000115313.2 | 0 | NA | NA | NA | NA | NA | #VALUE! |
| ENSMUSG00000115314.2 | 0 | NA | NA | NA | NA | NA | #VALUE! |
| ENSMUSG00000115315.2 | 0 | NA | NA | NA | NA | NA | #VALUE! |
| ENSMUSG00000115316.2 | 0 | NA | NA | NA | NA | NA | #VALUE! |
| ENSMUSG00000115317.2 | 0 | NA | NA | NA | NA | NA | #VALUE! |
| ENSMUSG00000115318.2 | 0 | NA | NA | NA | NA | NA | #VALUE! |
| ENSMUSG00000115319.2 | 0 | NA | NA | NA | NA | NA | #VALUE! |
| ENSMUSG00000115320.2 | 0 | NA | NA | NA | NA | NA | #VALUE! |
| ENSMUSG00000115321.2 | 0 | NA | NA | NA | NA | NA | #VALUE! |
| ENSMUSG00000115322.2 | 0 | NA | NA | NA | NA | NA | #VALUE! |
| ENSMUSG00000115323.2 | 0 | NA | NA | NA | NA | NA | #VALUE! |

| | | | | | | | |
|---|---|---|---|---|---|---|---|
| ENSMUSG00000115324.2 | 0 | NA | NA | NA | NA | NA | #VALUE! |
| ENSMUSG00000115325.3 | 0 | NA | NA | NA | NA | NA | #VALUE! |
| ENSMUSG00000115326.2 | 0 | NA | NA | NA | NA | NA | #VALUE! |
| ENSMUSG00000115327.2 | 0 | NA | NA | NA | NA | NA | #VALUE! |
| ENSMUSG00000115328.2 | 0 | NA | NA | NA | NA | NA | #VALUE! |
| ENSMUSG00000115329.2 | 0 | NA | NA | NA | NA | NA | #VALUE! |
| ENSMUSG00000115330.2 | 0 | NA | NA | NA | NA | NA | #VALUE! |
| ENSMUSG00000115331.2 | 0 | NA | NA | NA | NA | NA | #VALUE! |
| ENSMUSG00000115332.2 | 0 | NA | NA | NA | NA | NA | #VALUE! |
| ENSMUSG00000115333.2 | 0 | NA | NA | NA | NA | NA | #VALUE! |
| ENSMUSG00000115334.2 | 0 | NA | NA | NA | NA | NA | #VALUE! |
| ENSMUSG00000115335.2 | 0 | NA | NA | NA | NA | NA | #VALUE! |
| ENSMUSG00000115336.2 | 0 | NA | NA | NA | NA | NA | #VALUE! |
| ENSMUSG00000115337.2 | 0 | NA | NA | NA | NA | NA | #VALUE! |
| ENSMUSG00000115339.2 | 0 | NA | NA | NA | NA | NA | #VALUE! |
| ENSMUSG00000115340.2 | 0 | NA | NA | NA | NA | NA | #VALUE! |
| ENSMUSG00000115341.2 | 0 | NA | NA | NA | NA | NA | #VALUE! |
| ENSMUSG00000115342.2 | 0 | NA | NA | NA | NA | NA | #VALUE! |
| ENSMUSG00000115343.2 | 0 | NA | NA | NA | NA | NA | #VALUE! |
| ENSMUSG00000115344.2 | 0 | NA | NA | NA | NA | NA | #VALUE! |
| ENSMUSG00000115345.2 | 0 | NA | NA | NA | NA | NA | #VALUE! |
| ENSMUSG00000115346.2 | 0 | NA | NA | NA | NA | NA | #VALUE! |
| ENSMUSG00000115347.2 | 0.450086 | 0.012025 | 4.059931 | 0.002962 | 0.997637 | NA | #VALUE! |
| ENSMUSG00000115348.2 | 0 | NA | NA | NA | NA | NA | #VALUE! |
| ENSMUSG00000115349.2 | 0 | NA | NA | NA | NA | NA | #VALUE! |
| ENSMUSG00000115350.2 | 0 | NA | NA | NA | NA | NA | #VALUE! |
| ENSMUSG00000115351.2 | 0 | NA | NA | NA | NA | NA | #VALUE! |
| ENSMUSG00000115352.2 | 0 | NA | NA | NA | NA | NA | #VALUE! |
| ENSMUSG00000115353.2 | 0 | NA | NA | NA | NA | NA | #VALUE! |
| ENSMUSG00000115354.2 | 0 | NA | NA | NA | NA | NA | #VALUE! |
| ENSMUSG00000115355.2 | 0 | NA | NA | NA | NA | NA | #VALUE! |
| ENSMUSG00000115356.2 | 0 | NA | NA | NA | NA | NA | #VALUE! |
| ENSMUSG00000115357.2 | 0 | NA | NA | NA | NA | NA | #VALUE! |
| ENSMUSG00000115359.2 | 0 | NA | NA | NA | NA | NA | #VALUE! |
| ENSMUSG00000115360.2 | 0 | NA | NA | NA | NA | NA | #VALUE! |
| ENSMUSG00000115361.2 | 0 | NA | NA | NA | NA | NA | #VALUE! |
| ENSMUSG00000115362.2 | 0 | NA | NA | NA | NA | NA | #VALUE! |
| ENSMUSG00000115363.2 | 0 | NA | NA | NA | NA | NA | #VALUE! |
| ENSMUSG00000115365.2 | 0 | NA | NA | NA | NA | NA | #VALUE! |
| ENSMUSG00000115366.2 | 0 | NA | NA | NA | NA | NA | #VALUE! |
| ENSMUSG00000115367.2 | 0 | NA | NA | NA | NA | NA | #VALUE! |
| ENSMUSG00000115368.2 | 0 | NA | NA | NA | NA | NA | #VALUE! |
| ENSMUSG00000115369.2 | 0 | NA | NA | NA | NA | NA | #VALUE! |
| ENSMUSG00000115370.2 | 0 | NA | NA | NA | NA | NA | #VALUE! |

| | | | | | | | |
|---|---|---|---|---|---|---|---|
| ENSMUSG00000115371.2 | 0 | NA | NA | NA | NA | NA | #VALUE! |
| ENSMUSG00000115372.2 | 0 | NA | NA | NA | NA | NA | #VALUE! |
| ENSMUSG00000115373.2 | 0 | NA | NA | NA | NA | NA | #VALUE! |
| ENSMUSG00000115374.2 | 0 | NA | NA | NA | NA | NA | #VALUE! |
| ENSMUSG00000115375.2 | 0 | NA | NA | NA | NA | NA | #VALUE! |
| ENSMUSG00000115376.2 | 0 | NA | NA | NA | NA | NA | #VALUE! |
| ENSMUSG00000115377.2 | 0 | NA | NA | NA | NA | NA | #VALUE! |
| ENSMUSG00000115378.2 | 0 | NA | NA | NA | NA | NA | #VALUE! |
| ENSMUSG00000115379.2 | 0 | NA | NA | NA | NA | NA | #VALUE! |
| ENSMUSG00000115380.2 | 0 | NA | NA | NA | NA | NA | #VALUE! |
| ENSMUSG00000115381.2 | 0 | NA | NA | NA | NA | NA | #VALUE! |
| ENSMUSG00000115382.2 | 0 | NA | NA | NA | NA | NA | #VALUE! |
| ENSMUSG00000115383.2 | 0 | NA | NA | NA | NA | NA | #VALUE! |
| ENSMUSG00000115384.2 | 0 | NA | NA | NA | NA | NA | #VALUE! |
| ENSMUSG00000115385.2 | 0 | NA | NA | NA | NA | NA | #VALUE! |
| ENSMUSG00000115386.2 | 0 | NA | NA | NA | NA | NA | #VALUE! |
| ENSMUSG00000115387.2 | 0 | NA | NA | NA | NA | NA | #VALUE! |
| ENSMUSG00000115389.2 | 0 | NA | NA | NA | NA | NA | #VALUE! |
| ENSMUSG00000115390.2 | 0 | NA | NA | NA | NA | NA | #VALUE! |
| ENSMUSG00000115391.2 | 0 | NA | NA | NA | NA | NA | #VALUE! |
| ENSMUSG00000115392.2 | 0 | NA | NA | NA | NA | NA | #VALUE! |
| ENSMUSG00000115393.2 | 0 | NA | NA | NA | NA | NA | #VALUE! |
| ENSMUSG00000115394.2 | 0 | NA | NA | NA | NA | NA | #VALUE! |
| ENSMUSG00000115395.2 | 0 | NA | NA | NA | NA | NA | #VALUE! |
| ENSMUSG00000115396.2 | 0 | NA | NA | NA | NA | NA | #VALUE! |
| ENSMUSG00000115397.2 | 0 | NA | NA | NA | NA | NA | #VALUE! |
| ENSMUSG00000115398.2 | 0 | NA | NA | NA | NA | NA | #VALUE! |
| ENSMUSG00000115399.2 | 0 | NA | NA | NA | NA | NA | #VALUE! |
| ENSMUSG00000115400.2 | 0 | NA | NA | NA | NA | NA | #VALUE! |
| ENSMUSG00000115401.2 | 0 | NA | NA | NA | NA | NA | #VALUE! |
| ENSMUSG00000115402.2 | 0 | NA | NA | NA | NA | NA | #VALUE! |
| ENSMUSG00000115403.2 | 0 | NA | NA | NA | NA | NA | #VALUE! |
| ENSMUSG00000115404.2 | 0 | NA | NA | NA | NA | NA | #VALUE! |
| ENSMUSG00000115405.2 | 0 | NA | NA | NA | NA | NA | #VALUE! |
| ENSMUSG00000115406.2 | 0 | NA | NA | NA | NA | NA | #VALUE! |
| ENSMUSG00000115407.2 | 0 | NA | NA | NA | NA | NA | #VALUE! |
| ENSMUSG00000115408.2 | 0 | NA | NA | NA | NA | NA | #VALUE! |
| ENSMUSG00000115409.2 | 0 | NA | NA | NA | NA | NA | #VALUE! |
| ENSMUSG00000115410.2 | 0 | NA | NA | NA | NA | NA | #VALUE! |
| ENSMUSG00000115411.2 | 0 | NA | NA | NA | NA | NA | #VALUE! |
| ENSMUSG00000115412.2 | 0 | NA | NA | NA | NA | NA | #VALUE! |
| ENSMUSG00000115413.2 | 0 | NA | NA | NA | NA | NA | #VALUE! |
| ENSMUSG00000115414.2 | 0 | NA | NA | NA | NA | NA | #VALUE! |
| ENSMUSG00000115415.2 | 0 | NA | NA | NA | NA | NA | #VALUE! |

| | | | | | | | |
|---|---|---|---|---|---|---|---|
| ENSMUSG00000115416.2 | 0 | NA | NA | NA | NA | NA | #VALUE! |
| ENSMUSG00000115417.2 | 0 | NA | NA | NA | NA | NA | #VALUE! |
| ENSMUSG00000115418.2 | 0 | NA | NA | NA | NA | NA | #VALUE! |
| ENSMUSG00000115419.2 | 0 | NA | NA | NA | NA | NA | #VALUE! |
| ENSMUSG00000115420.2 | 0 | NA | NA | NA | NA | NA | #VALUE! |
| ENSMUSG00000115421.2 | 0 | NA | NA | NA | NA | NA | #VALUE! |
| ENSMUSG00000115422.2 | 0 | NA | NA | NA | NA | NA | #VALUE! |
| ENSMUSG00000115423.3 | 0 | NA | NA | NA | NA | NA | #VALUE! |
| ENSMUSG00000115424.2 | 0 | NA | NA | NA | NA | NA | #VALUE! |
| ENSMUSG00000115425.2 | 0 | NA | NA | NA | NA | NA | #VALUE! |
| ENSMUSG00000115426.2 | 0 | NA | NA | NA | NA | NA | #VALUE! |
| ENSMUSG00000115427.2 | 0 | NA | NA | NA | NA | NA | #VALUE! |
| ENSMUSG00000115428.2 | 0 | NA | NA | NA | NA | NA | #VALUE! |
| ENSMUSG00000115429.2 | 0 | NA | NA | NA | NA | NA | #VALUE! |
| ENSMUSG00000115430.2 | 0 | NA | NA | NA | NA | NA | #VALUE! |
| ENSMUSG00000115431.2 | 7.569917 | -5.51878 | 3.170206 | -1.74083 | 0.081714 | NA | #VALUE! |
| ENSMUSG00000115432.2 | 0 | NA | NA | NA | NA | NA | #VALUE! |
| ENSMUSG00000115433.2 | 0 | NA | NA | NA | NA | NA | #VALUE! |
| ENSMUSG00000115434.2 | 0 | NA | NA | NA | NA | NA | #VALUE! |
| ENSMUSG00000115435.2 | 0 | NA | NA | NA | NA | NA | #VALUE! |
| ENSMUSG00000115436.2 | 0 | NA | NA | NA | NA | NA | #VALUE! |
| ENSMUSG00000115437.2 | 0 | NA | NA | NA | NA | NA | #VALUE! |
| ENSMUSG00000115438.2 | 0 | NA | NA | NA | NA | NA | #VALUE! |
| ENSMUSG00000115439.2 | 0 | NA | NA | NA | NA | NA | #VALUE! |
| ENSMUSG00000115440.2 | 0 | NA | NA | NA | NA | NA | #VALUE! |
| ENSMUSG00000115441.2 | 0 | NA | NA | NA | NA | NA | #VALUE! |
| ENSMUSG00000115443.2 | 0 | NA | NA | NA | NA | NA | #VALUE! |
| ENSMUSG00000115444.2 | 0 | NA | NA | NA | NA | NA | #VALUE! |
| ENSMUSG00000115445.2 | 0 | NA | NA | NA | NA | NA | #VALUE! |
| ENSMUSG00000115446.2 | 0 | NA | NA | NA | NA | NA | #VALUE! |
| ENSMUSG00000115447.2 | 0 | NA | NA | NA | NA | NA | #VALUE! |
| ENSMUSG00000115448.2 | 0 | NA | NA | NA | NA | NA | #VALUE! |
| ENSMUSG00000115449.2 | 0 | NA | NA | NA | NA | NA | #VALUE! |
| ENSMUSG00000115450.2 | 0.093255 | 0.868093 | 4.072464 | 0.213162 | 0.831201 | NA | #VALUE! |
| ENSMUSG00000115451.2 | 0 | NA | NA | NA | NA | NA | #VALUE! |
| ENSMUSG00000115452.2 | 0 | NA | NA | NA | NA | NA | #VALUE! |
| ENSMUSG00000115453.2 | 0 | NA | NA | NA | NA | NA | #VALUE! |
| ENSMUSG00000115454.2 | 0 | NA | NA | NA | NA | NA | #VALUE! |
| ENSMUSG00000115455.2 | 0 | NA | NA | NA | NA | NA | #VALUE! |
| ENSMUSG00000115456.2 | 0 | NA | NA | NA | NA | NA | #VALUE! |
| ENSMUSG00000115457.2 | 0 | NA | NA | NA | NA | NA | #VALUE! |
| ENSMUSG00000115458.2 | 0 | NA | NA | NA | NA | NA | #VALUE! |
| ENSMUSG00000115459.2 | 0 | NA | NA | NA | NA | NA | #VALUE! |
| ENSMUSG00000115460.2 | 0 | NA | NA | NA | NA | NA | #VALUE! |

| | | | | | | | |
|---|---|---|---|---|---|---|---|
| ENSMUSG00000115461.2 | 0 | NA | NA | NA | NA | NA | #VALUE! |
| ENSMUSG00000115462.2 | 0 | NA | NA | NA | NA | NA | #VALUE! |
| ENSMUSG00000115463.2 | 0 | NA | NA | NA | NA | NA | #VALUE! |
| ENSMUSG00000115464.2 | 0 | NA | NA | NA | NA | NA | #VALUE! |
| ENSMUSG00000115465.2 | 0 | NA | NA | NA | NA | NA | #VALUE! |
| ENSMUSG00000115466.2 | 0 | NA | NA | NA | NA | NA | #VALUE! |
| ENSMUSG00000115467.2 | 0 | NA | NA | NA | NA | NA | #VALUE! |
| ENSMUSG00000115468.2 | 0 | NA | NA | NA | NA | NA | #VALUE! |
| ENSMUSG00000115469.2 | 0 | NA | NA | NA | NA | NA | #VALUE! |
| ENSMUSG00000115470.2 | 0 | NA | NA | NA | NA | NA | #VALUE! |
| ENSMUSG00000115471.2 | 0 | NA | NA | NA | NA | NA | #VALUE! |
| ENSMUSG00000115472.2 | 0 | NA | NA | NA | NA | NA | #VALUE! |
| ENSMUSG00000115473.2 | 0 | NA | NA | NA | NA | NA | #VALUE! |
| ENSMUSG00000115474.2 | 0 | NA | NA | NA | NA | NA | #VALUE! |
| ENSMUSG00000115475.2 | 0 | NA | NA | NA | NA | NA | #VALUE! |
| ENSMUSG00000115476.2 | 0 | NA | NA | NA | NA | NA | #VALUE! |
| ENSMUSG00000115477.2 | 0 | NA | NA | NA | NA | NA | #VALUE! |
| ENSMUSG00000115478.2 | 0 | NA | NA | NA | NA | NA | #VALUE! |
| ENSMUSG00000115479.2 | 0 | NA | NA | NA | NA | NA | #VALUE! |
| ENSMUSG00000115480.2 | 0 | NA | NA | NA | NA | NA | #VALUE! |
| ENSMUSG00000115481.2 | 0 | NA | NA | NA | NA | NA | #VALUE! |
| ENSMUSG00000115482.2 | 0 | NA | NA | NA | NA | NA | #VALUE! |
| ENSMUSG00000115483.2 | 0 | NA | NA | NA | NA | NA | #VALUE! |
| ENSMUSG00000115484.2 | 0 | NA | NA | NA | NA | NA | #VALUE! |
| ENSMUSG00000115485.2 | 0 | NA | NA | NA | NA | NA | #VALUE! |
| ENSMUSG00000115486.2 | 0.105013 | -0.97324 | 4.076293 | -0.23876 | 0.811295 | NA | #VALUE! |
| ENSMUSG00000115487.2 | 0 | NA | NA | NA | NA | NA | #VALUE! |
| ENSMUSG00000115488.2 | 0 | NA | NA | NA | NA | NA | #VALUE! |
| ENSMUSG00000115489.2 | 0 | NA | NA | NA | NA | NA | #VALUE! |
| ENSMUSG00000115490.2 | 0 | NA | NA | NA | NA | NA | #VALUE! |
| ENSMUSG00000115491.2 | 0 | NA | NA | NA | NA | NA | #VALUE! |
| ENSMUSG00000115492.2 | 0 | NA | NA | NA | NA | NA | #VALUE! |
| ENSMUSG00000115493.2 | 0 | NA | NA | NA | NA | NA | #VALUE! |
| ENSMUSG00000115494.2 | 0 | NA | NA | NA | NA | NA | #VALUE! |
| ENSMUSG00000115495.2 | 0 | NA | NA | NA | NA | NA | #VALUE! |
| ENSMUSG00000115498.2 | 0 | NA | NA | NA | NA | NA | #VALUE! |
| ENSMUSG00000115499.2 | 0 | NA | NA | NA | NA | NA | #VALUE! |
| ENSMUSG00000115500.2 | 0 | NA | NA | NA | NA | NA | #VALUE! |
| ENSMUSG00000115501.2 | 0 | NA | NA | NA | NA | NA | #VALUE! |
| ENSMUSG00000115502.2 | 0 | NA | NA | NA | NA | NA | #VALUE! |
| ENSMUSG00000115503.2 | 0 | NA | NA | NA | NA | NA | #VALUE! |
| ENSMUSG00000115504.2 | 0 | NA | NA | NA | NA | NA | #VALUE! |
| ENSMUSG00000115505.2 | 0 | NA | NA | NA | NA | NA | #VALUE! |
| ENSMUSG00000115506.2 | 0 | NA | NA | NA | NA | NA | #VALUE! |

| | | | | | | | |
|---|---|---|---|---|---|---|---|
| ENSMUSG00000115507.2 | 0 | NA | NA | NA | NA | NA | #VALUE! |
| ENSMUSG00000115508.2 | 0 | NA | NA | NA | NA | NA | #VALUE! |
| ENSMUSG00000115509.2 | 0 | NA | NA | NA | NA | NA | #VALUE! |
| ENSMUSG00000115510.2 | 0 | NA | NA | NA | NA | NA | #VALUE! |
| ENSMUSG00000115511.2 | 0 | NA | NA | NA | NA | NA | #VALUE! |
| ENSMUSG00000115512.2 | 0.093255 | 0.868093 | 4.072464 | 0.213162 | 0.831201 | NA | #VALUE! |
| ENSMUSG00000115513.2 | 0 | NA | NA | NA | NA | NA | #VALUE! |
| ENSMUSG00000115514.2 | 0 | NA | NA | NA | NA | NA | #VALUE! |
| ENSMUSG00000115515.2 | 0 | NA | NA | NA | NA | NA | #VALUE! |
| ENSMUSG00000115516.2 | 0 | NA | NA | NA | NA | NA | #VALUE! |
| ENSMUSG00000115517.2 | 0 | NA | NA | NA | NA | NA | #VALUE! |
| ENSMUSG00000115518.2 | 0 | NA | NA | NA | NA | NA | #VALUE! |
| ENSMUSG00000115519.2 | 0 | NA | NA | NA | NA | NA | #VALUE! |
| ENSMUSG00000115520.3 | 0 | NA | NA | NA | NA | NA | #VALUE! |
| ENSMUSG00000115521.2 | 0 | NA | NA | NA | NA | NA | #VALUE! |
| ENSMUSG00000115522.2 | 0 | NA | NA | NA | NA | NA | #VALUE! |
| ENSMUSG00000115523.2 | 0 | NA | NA | NA | NA | NA | #VALUE! |
| ENSMUSG00000115524.2 | 0 | NA | NA | NA | NA | NA | #VALUE! |
| ENSMUSG00000115525.2 | 0 | NA | NA | NA | NA | NA | #VALUE! |
| ENSMUSG00000115526.2 | 0 | NA | NA | NA | NA | NA | #VALUE! |
| ENSMUSG00000115527.2 | 0 | NA | NA | NA | NA | NA | #VALUE! |
| ENSMUSG00000115528.2 | 0 | NA | NA | NA | NA | NA | #VALUE! |
| ENSMUSG00000115529.2 | 0 | NA | NA | NA | NA | NA | #VALUE! |
| ENSMUSG00000115530.2 | 0 | NA | NA | NA | NA | NA | #VALUE! |
| ENSMUSG00000115531.2 | 0 | NA | NA | NA | NA | NA | #VALUE! |
| ENSMUSG00000115532.2 | 0 | NA | NA | NA | NA | NA | #VALUE! |
| ENSMUSG00000115533.2 | 0 | NA | NA | NA | NA | NA | #VALUE! |
| ENSMUSG00000115534.2 | 0 | NA | NA | NA | NA | NA | #VALUE! |
| ENSMUSG00000115535.2 | 0 | NA | NA | NA | NA | NA | #VALUE! |
| ENSMUSG00000115536.2 | 0 | NA | NA | NA | NA | NA | #VALUE! |
| ENSMUSG00000115537.2 | 0 | NA | NA | NA | NA | NA | #VALUE! |
| ENSMUSG00000115538.2 | 0 | NA | NA | NA | NA | NA | #VALUE! |
| ENSMUSG00000115539.2 | 0.391578 | -2.1489 | 4.020883 | -0.53444 | 0.59304 | NA | #VALUE! |
| ENSMUSG00000115540.2 | 0 | NA | NA | NA | NA | NA | #VALUE! |
| ENSMUSG00000115541.2 | 0 | NA | NA | NA | NA | NA | #VALUE! |
| ENSMUSG00000115542.2 | 0 | NA | NA | NA | NA | NA | #VALUE! |
| ENSMUSG00000115543.2 | 0 | NA | NA | NA | NA | NA | #VALUE! |
| ENSMUSG00000115544.2 | 0 | NA | NA | NA | NA | NA | #VALUE! |
| ENSMUSG00000115545.2 | 0 | NA | NA | NA | NA | NA | #VALUE! |
| ENSMUSG00000115546.2 | 0 | NA | NA | NA | NA | NA | #VALUE! |
| ENSMUSG00000115547.2 | 0 | NA | NA | NA | NA | NA | #VALUE! |
| ENSMUSG00000115548.2 | 0 | NA | NA | NA | NA | NA | #VALUE! |
| ENSMUSG00000115549.2 | 0 | NA | NA | NA | NA | NA | #VALUE! |
| ENSMUSG00000115550.2 | 0 | NA | NA | NA | NA | NA | #VALUE! |

| | | | | | | | |
|---|---|---|---|---|---|---|---|
| ENSMUSG00000115551.2 | 0 | NA | NA | NA | NA | NA | #VALUE! |
| ENSMUSG00000115552.2 | 0 | NA | NA | NA | NA | NA | #VALUE! |
| ENSMUSG00000115553.2 | 0 | NA | NA | NA | NA | NA | #VALUE! |
| ENSMUSG00000115554.2 | 0 | NA | NA | NA | NA | NA | #VALUE! |
| ENSMUSG00000115555.2 | 1.305573 | 3.898287 | 3.940396 | 0.989314 | 0.32251 | NA | #VALUE! |
| ENSMUSG00000115556.2 | 0 | NA | NA | NA | NA | NA | #VALUE! |
| ENSMUSG00000115557.2 | 0 | NA | NA | NA | NA | NA | #VALUE! |
| ENSMUSG00000115558.2 | 0 | NA | NA | NA | NA | NA | #VALUE! |
| ENSMUSG00000115559.2 | 0 | NA | NA | NA | NA | NA | #VALUE! |
| ENSMUSG00000115560.2 | 0 | NA | NA | NA | NA | NA | #VALUE! |
| ENSMUSG00000115561.2 | 0 | NA | NA | NA | NA | NA | #VALUE! |
| ENSMUSG00000115562.2 | 0 | NA | NA | NA | NA | NA | #VALUE! |
| ENSMUSG00000115563.2 | 0 | NA | NA | NA | NA | NA | #VALUE! |
| ENSMUSG00000115564.2 | 0 | NA | NA | NA | NA | NA | #VALUE! |
| ENSMUSG00000115565.2 | 0 | NA | NA | NA | NA | NA | #VALUE! |
| ENSMUSG00000115566.2 | 0 | NA | NA | NA | NA | NA | #VALUE! |
| ENSMUSG00000115567.2 | 0 | NA | NA | NA | NA | NA | #VALUE! |
| ENSMUSG00000115568.2 | 0 | NA | NA | NA | NA | NA | #VALUE! |
| ENSMUSG00000115569.2 | 0 | NA | NA | NA | NA | NA | #VALUE! |
| ENSMUSG00000115570.2 | 0 | NA | NA | NA | NA | NA | #VALUE! |
| ENSMUSG00000115571.2 | 0 | NA | NA | NA | NA | NA | #VALUE! |
| ENSMUSG00000115572.2 | 0 | NA | NA | NA | NA | NA | #VALUE! |
| ENSMUSG00000115573.2 | 0 | NA | NA | NA | NA | NA | #VALUE! |
| ENSMUSG00000115574.2 | 0 | NA | NA | NA | NA | NA | #VALUE! |
| ENSMUSG00000115575.2 | 0 | NA | NA | NA | NA | NA | #VALUE! |
| ENSMUSG00000115576.2 | 0 | NA | NA | NA | NA | NA | #VALUE! |
| ENSMUSG00000115577.2 | 0 | NA | NA | NA | NA | NA | #VALUE! |
| ENSMUSG00000115578.2 | 0 | NA | NA | NA | NA | NA | #VALUE! |
| ENSMUSG00000115579.2 | 0 | NA | NA | NA | NA | NA | #VALUE! |
| ENSMUSG00000115580.2 | 0 | NA | NA | NA | NA | NA | #VALUE! |
| ENSMUSG00000115581.2 | 0 | NA | NA | NA | NA | NA | #VALUE! |
| ENSMUSG00000115582.2 | 0 | NA | NA | NA | NA | NA | #VALUE! |
| ENSMUSG00000115584.2 | 0 | NA | NA | NA | NA | NA | #VALUE! |
| ENSMUSG00000115585.2 | 0 | NA | NA | NA | NA | NA | #VALUE! |
| ENSMUSG00000115586.2 | 0 | NA | NA | NA | NA | NA | #VALUE! |
| ENSMUSG00000115587.2 | 0 | NA | NA | NA | NA | NA | #VALUE! |
| ENSMUSG00000115588.2 | 0 | NA | NA | NA | NA | NA | #VALUE! |
| ENSMUSG00000115589.2 | 0 | NA | NA | NA | NA | NA | #VALUE! |
| ENSMUSG00000115590.2 | 0 | NA | NA | NA | NA | NA | #VALUE! |
| ENSMUSG00000115591.2 | 0 | NA | NA | NA | NA | NA | #VALUE! |
| ENSMUSG00000115592.2 | 0 | NA | NA | NA | NA | NA | #VALUE! |
| ENSMUSG00000115593.2 | 0.093255 | 0.868093 | 4.072464 | 0.213162 | 0.831201 | NA | #VALUE! |
| ENSMUSG00000115594.2 | 0 | NA | NA | NA | NA | NA | #VALUE! |
| ENSMUSG00000115595.2 | 1.229878 | 3.761857 | 3.943699 | 0.95389 | 0.340139 | NA | #VALUE! |

| | | | | | | | |
|---|---|---|---|---|---|---|---|
| ENSMUSG00000115596.2 | 0 | NA | NA | NA | NA | NA | #VALUE! |
| ENSMUSG00000115597.2 | 0 | NA | NA | NA | NA | NA | #VALUE! |
| ENSMUSG00000115598.2 | 0 | NA | NA | NA | NA | NA | #VALUE! |
| ENSMUSG00000115599.2 | 0 | NA | NA | NA | NA | NA | #VALUE! |
| ENSMUSG00000115600.2 | 0 | NA | NA | NA | NA | NA | #VALUE! |
| ENSMUSG00000115601.2 | 0 | NA | NA | NA | NA | NA | #VALUE! |
| ENSMUSG00000115602.2 | 0.493155 | -1.26736 | 4.010698 | -0.31599 | 0.752007 | NA | #VALUE! |
| ENSMUSG00000115603.2 | 0 | NA | NA | NA | NA | NA | #VALUE! |
| ENSMUSG00000115604.2 | 0.105013 | -0.97324 | 4.076293 | -0.23876 | 0.811295 | NA | #VALUE! |
| ENSMUSG00000115605.2 | 0 | NA | NA | NA | NA | NA | #VALUE! |
| ENSMUSG00000115606.2 | 0 | NA | NA | NA | NA | NA | #VALUE! |
| ENSMUSG00000115607.2 | 0 | NA | NA | NA | NA | NA | #VALUE! |
| ENSMUSG00000115608.2 | 0 | NA | NA | NA | NA | NA | #VALUE! |
| ENSMUSG00000115609.2 | 0 | NA | NA | NA | NA | NA | #VALUE! |
| ENSMUSG00000115610.2 | 0 | NA | NA | NA | NA | NA | #VALUE! |
| ENSMUSG00000115611.2 | 0 | NA | NA | NA | NA | NA | #VALUE! |
| ENSMUSG00000115612.2 | 0 | NA | NA | NA | NA | NA | #VALUE! |
| ENSMUSG00000115613.2 | 0 | NA | NA | NA | NA | NA | #VALUE! |
| ENSMUSG00000115614.2 | 0 | NA | NA | NA | NA | NA | #VALUE! |
| ENSMUSG00000115615.2 | 0 | NA | NA | NA | NA | NA | #VALUE! |
| ENSMUSG00000115616.2 | 0 | NA | NA | NA | NA | NA | #VALUE! |
| ENSMUSG00000115617.2 | 0 | NA | NA | NA | NA | NA | #VALUE! |
| ENSMUSG00000115618.2 | 0 | NA | NA | NA | NA | NA | #VALUE! |
| ENSMUSG00000115619.2 | 0 | NA | NA | NA | NA | NA | #VALUE! |
| ENSMUSG00000115620.2 | 0 | NA | NA | NA | NA | NA | #VALUE! |
| ENSMUSG00000115621.2 | 0 | NA | NA | NA | NA | NA | #VALUE! |
| ENSMUSG00000115622.2 | 0 | NA | NA | NA | NA | NA | #VALUE! |
| ENSMUSG00000115623.2 | 0 | NA | NA | NA | NA | NA | #VALUE! |
| ENSMUSG00000115624.2 | 0 | NA | NA | NA | NA | NA | #VALUE! |
| ENSMUSG00000115625.2 | 0 | NA | NA | NA | NA | NA | #VALUE! |
| ENSMUSG00000115626.2 | 0 | NA | NA | NA | NA | NA | #VALUE! |
| ENSMUSG00000115627.2 | 0 | NA | NA | NA | NA | NA | #VALUE! |
| ENSMUSG00000115628.2 | 0 | NA | NA | NA | NA | NA | #VALUE! |
| ENSMUSG00000115629.2 | 0 | NA | NA | NA | NA | NA | #VALUE! |
| ENSMUSG00000115630.2 | 0 | NA | NA | NA | NA | NA | #VALUE! |
| ENSMUSG00000115631.2 | 0 | NA | NA | NA | NA | NA | #VALUE! |
| ENSMUSG00000115632.2 | 0 | NA | NA | NA | NA | NA | #VALUE! |
| ENSMUSG00000115633.2 | 0 | NA | NA | NA | NA | NA | #VALUE! |
| ENSMUSG00000115635.2 | 0 | NA | NA | NA | NA | NA | #VALUE! |
| ENSMUSG00000115636.2 | 0 | NA | NA | NA | NA | NA | #VALUE! |
| ENSMUSG00000115637.2 | 0 | NA | NA | NA | NA | NA | #VALUE! |
| ENSMUSG00000115638.2 | 0 | NA | NA | NA | NA | NA | #VALUE! |
| ENSMUSG00000115639.2 | 0 | NA | NA | NA | NA | NA | #VALUE! |
| ENSMUSG00000115640.2 | 0 | NA | NA | NA | NA | NA | #VALUE! |

| | | | | | | | |
|---|---|---|---|---|---|---|---|
| ENSMUSG00000115641.2 | 0 | NA | NA | NA | NA | NA | #VALUE! |
| ENSMUSG00000115642.2 | 0 | NA | NA | NA | NA | NA | #VALUE! |
| ENSMUSG00000115643.2 | 0 | NA | NA | NA | NA | NA | #VALUE! |
| ENSMUSG00000115644.2 | 0 | NA | NA | NA | NA | NA | #VALUE! |
| ENSMUSG00000115645.2 | 0 | NA | NA | NA | NA | NA | #VALUE! |
| ENSMUSG00000115646.2 | 0 | NA | NA | NA | NA | NA | #VALUE! |
| ENSMUSG00000115647.2 | 7.36716 | 6.375323 | 3.912847 | 1.629331 | NA | NA | #VALUE! |
| ENSMUSG00000115648.2 | 0 | NA | NA | NA | NA | NA | #VALUE! |
| ENSMUSG00000115649.2 | 0 | NA | NA | NA | NA | NA | #VALUE! |
| ENSMUSG00000115650.2 | 0 | NA | NA | NA | NA | NA | #VALUE! |
| ENSMUSG00000115651.2 | 0 | NA | NA | NA | NA | NA | #VALUE! |
| ENSMUSG00000115652.2 | 0 | NA | NA | NA | NA | NA | #VALUE! |
| ENSMUSG00000115653.2 | 0 | NA | NA | NA | NA | NA | #VALUE! |
| ENSMUSG00000115654.2 | 0 | NA | NA | NA | NA | NA | #VALUE! |
| ENSMUSG00000115656.2 | 0 | NA | NA | NA | NA | NA | #VALUE! |
| ENSMUSG00000115657.2 | 0 | NA | NA | NA | NA | NA | #VALUE! |
| ENSMUSG00000115658.2 | 0 | NA | NA | NA | NA | NA | #VALUE! |
| ENSMUSG00000115659.2 | 0 | NA | NA | NA | NA | NA | #VALUE! |
| ENSMUSG00000115660.2 | 0 | NA | NA | NA | NA | NA | #VALUE! |
| ENSMUSG00000115661.2 | 0 | NA | NA | NA | NA | NA | #VALUE! |
| ENSMUSG00000115662.2 | 0.679665 | -0.32742 | 3.974335 | -0.08238 | 0.934342 | NA | #VALUE! |
| ENSMUSG00000115663.2 | 0 | NA | NA | NA | NA | NA | #VALUE! |
| ENSMUSG00000115664.2 | 0 | NA | NA | NA | NA | NA | #VALUE! |
| ENSMUSG00000115665.2 | 0 | NA | NA | NA | NA | NA | #VALUE! |
| ENSMUSG00000115666.2 | 0 | NA | NA | NA | NA | NA | #VALUE! |
| ENSMUSG00000115667.2 | 0 | NA | NA | NA | NA | NA | #VALUE! |
| ENSMUSG00000115668.2 | 0 | NA | NA | NA | NA | NA | #VALUE! |
| ENSMUSG00000115669.2 | 0 | NA | NA | NA | NA | NA | #VALUE! |
| ENSMUSG00000115670.2 | 0 | NA | NA | NA | NA | NA | #VALUE! |
| ENSMUSG00000115671.2 | 0 | NA | NA | NA | NA | NA | #VALUE! |
| ENSMUSG00000115672.2 | 0 | NA | NA | NA | NA | NA | #VALUE! |
| ENSMUSG00000115673.2 | 0 | NA | NA | NA | NA | NA | #VALUE! |
| ENSMUSG00000115674.2 | 0 | NA | NA | NA | NA | NA | #VALUE! |
| ENSMUSG00000115675.2 | 0 | NA | NA | NA | NA | NA | #VALUE! |
| ENSMUSG00000115676.2 | 0 | NA | NA | NA | NA | NA | #VALUE! |
| ENSMUSG00000115677.2 | 0 | NA | NA | NA | NA | NA | #VALUE! |
| ENSMUSG00000115678.2 | 0 | NA | NA | NA | NA | NA | #VALUE! |
| ENSMUSG00000115679.2 | 0 | NA | NA | NA | NA | NA | #VALUE! |
| ENSMUSG00000115680.2 | 0 | NA | NA | NA | NA | NA | #VALUE! |
| ENSMUSG00000115681.2 | 0 | NA | NA | NA | NA | NA | #VALUE! |
| ENSMUSG00000115682.2 | 0 | NA | NA | NA | NA | NA | #VALUE! |
| ENSMUSG00000115683.2 | 0 | NA | NA | NA | NA | NA | #VALUE! |
| ENSMUSG00000115684.2 | 0 | NA | NA | NA | NA | NA | #VALUE! |
| ENSMUSG00000115685.2 | 0.660235 | 2.855092 | 3.975428 | 0.718185 | 0.472643 | NA | #VALUE! |

| | | | | | | | |
|---|---|---|---|---|---|---|---|
| ENSMUSG00000115686.2 | 0 | NA | NA | NA | NA | NA | #VALUE! |
| ENSMUSG00000115687.2 | 0 | NA | NA | NA | NA | NA | #VALUE! |
| ENSMUSG00000115688.2 | 0 | NA | NA | NA | NA | NA | #VALUE! |
| ENSMUSG00000115689.2 | 0 | NA | NA | NA | NA | NA | #VALUE! |
| ENSMUSG00000115690.2 | 0 | NA | NA | NA | NA | NA | #VALUE! |
| ENSMUSG00000115691.2 | 0 | NA | NA | NA | NA | NA | #VALUE! |
| ENSMUSG00000115692.2 | 0 | NA | NA | NA | NA | NA | #VALUE! |
| ENSMUSG00000115693.2 | 0 | NA | NA | NA | NA | NA | #VALUE! |
| ENSMUSG00000115694.2 | 0 | NA | NA | NA | NA | NA | #VALUE! |
| ENSMUSG00000115695.2 | 0 | NA | NA | NA | NA | NA | #VALUE! |
| ENSMUSG00000115696.2 | 0 | NA | NA | NA | NA | NA | #VALUE! |
| ENSMUSG00000115697.3 | 0 | NA | NA | NA | NA | NA | #VALUE! |
| ENSMUSG00000115698.2 | 0 | NA | NA | NA | NA | NA | #VALUE! |
| ENSMUSG00000115699.2 | 0 | NA | NA | NA | NA | NA | #VALUE! |
| ENSMUSG00000115700.2 | 0 | NA | NA | NA | NA | NA | #VALUE! |
| ENSMUSG00000115701.2 | 0 | NA | NA | NA | NA | NA | #VALUE! |
| ENSMUSG00000115702.2 | 0 | NA | NA | NA | NA | NA | #VALUE! |
| ENSMUSG00000115703.2 | 0 | NA | NA | NA | NA | NA | #VALUE! |
| ENSMUSG00000115704.2 | 0 | NA | NA | NA | NA | NA | #VALUE! |
| ENSMUSG00000115705.2 | 0 | NA | NA | NA | NA | NA | #VALUE! |
| ENSMUSG00000115706.2 | 0 | NA | NA | NA | NA | NA | #VALUE! |
| ENSMUSG00000115707.2 | 0 | NA | NA | NA | NA | NA | #VALUE! |
| ENSMUSG00000115709.2 | 0 | NA | NA | NA | NA | NA | #VALUE! |
| ENSMUSG00000115710.2 | 0 | NA | NA | NA | NA | NA | #VALUE! |
| ENSMUSG00000115711.2 | 0 | NA | NA | NA | NA | NA | #VALUE! |
| ENSMUSG00000115712.2 | 0 | NA | NA | NA | NA | NA | #VALUE! |
| ENSMUSG00000115713.2 | 0 | NA | NA | NA | NA | NA | #VALUE! |
| ENSMUSG00000115714.2 | 0 | NA | NA | NA | NA | NA | #VALUE! |
| ENSMUSG00000115715.2 | 0 | NA | NA | NA | NA | NA | #VALUE! |
| ENSMUSG00000115716.2 | 0 | NA | NA | NA | NA | NA | #VALUE! |
| ENSMUSG00000115717.2 | 0 | NA | NA | NA | NA | NA | #VALUE! |
| ENSMUSG00000115718.2 | 0 | NA | NA | NA | NA | NA | #VALUE! |
| ENSMUSG00000115719.2 | 0 | NA | NA | NA | NA | NA | #VALUE! |
| ENSMUSG00000115720.2 | 0 | NA | NA | NA | NA | NA | #VALUE! |
| ENSMUSG00000115721.2 | 0 | NA | NA | NA | NA | NA | #VALUE! |
| ENSMUSG00000115722.2 | 0 | NA | NA | NA | NA | NA | #VALUE! |
| ENSMUSG00000115723.2 | 0 | NA | NA | NA | NA | NA | #VALUE! |
| ENSMUSG00000115724.2 | 0 | NA | NA | NA | NA | NA | #VALUE! |
| ENSMUSG00000115725.2 | 0 | NA | NA | NA | NA | NA | #VALUE! |
| ENSMUSG00000115726.2 | 0 | NA | NA | NA | NA | NA | #VALUE! |
| ENSMUSG00000115727.2 | 0 | NA | NA | NA | NA | NA | #VALUE! |
| ENSMUSG00000115728.2 | 0 | NA | NA | NA | NA | NA | #VALUE! |
| ENSMUSG00000115729.2 | 0 | NA | NA | NA | NA | NA | #VALUE! |
| ENSMUSG00000115730.2 | 0 | NA | NA | NA | NA | NA | #VALUE! |

| | | | | | | | |
|---|---|---|---|---|---|---|---|
| ENSMUSG00000115731.2 | 0 | NA | NA | NA | NA | NA | #VALUE! |
| ENSMUSG00000115732.2 | 0 | NA | NA | NA | NA | NA | #VALUE! |
| ENSMUSG00000115733.2 | 0 | NA | NA | NA | NA | NA | #VALUE! |
| ENSMUSG00000115734.2 | 0 | NA | NA | NA | NA | NA | #VALUE! |
| ENSMUSG00000115735.2 | 0 | NA | NA | NA | NA | NA | #VALUE! |
| ENSMUSG00000115736.2 | 0.105013 | -0.97324 | 4.076293 | -0.23876 | 0.811295 | NA | #VALUE! |
| ENSMUSG00000115737.2 | 0 | NA | NA | NA | NA | NA | #VALUE! |
| ENSMUSG00000115738.2 | 0 | NA | NA | NA | NA | NA | #VALUE! |
| ENSMUSG00000115739.2 | 0 | NA | NA | NA | NA | NA | #VALUE! |
| ENSMUSG00000115740.2 | 0 | NA | NA | NA | NA | NA | #VALUE! |
| ENSMUSG00000115741.2 | 0 | NA | NA | NA | NA | NA | #VALUE! |
| ENSMUSG00000115742.2 | 0 | NA | NA | NA | NA | NA | #VALUE! |
| ENSMUSG00000115743.2 | 0 | NA | NA | NA | NA | NA | #VALUE! |
| ENSMUSG00000115744.2 | 0 | NA | NA | NA | NA | NA | #VALUE! |
| ENSMUSG00000115745.2 | 0 | NA | NA | NA | NA | NA | #VALUE! |
| ENSMUSG00000115746.2 | 0 | NA | NA | NA | NA | NA | #VALUE! |
| ENSMUSG00000115747.2 | 0 | NA | NA | NA | NA | NA | #VALUE! |
| ENSMUSG00000115748.2 | 0 | NA | NA | NA | NA | NA | #VALUE! |
| ENSMUSG00000115749.2 | 0 | NA | NA | NA | NA | NA | #VALUE! |
| ENSMUSG00000115750.2 | 0 | NA | NA | NA | NA | NA | #VALUE! |
| ENSMUSG00000115751.2 | 0 | NA | NA | NA | NA | NA | #VALUE! |
| ENSMUSG00000115752.2 | 0 | NA | NA | NA | NA | NA | #VALUE! |
| ENSMUSG00000115754.2 | 0.093255 | 0.868093 | 4.072464 | 0.213162 | 0.831201 | NA | #VALUE! |
| ENSMUSG00000115755.2 | 0 | NA | NA | NA | NA | NA | #VALUE! |
| ENSMUSG00000115756.2 | 0 | NA | NA | NA | NA | NA | #VALUE! |
| ENSMUSG00000115757.2 | 0 | NA | NA | NA | NA | NA | #VALUE! |
| ENSMUSG00000115758.2 | 0 | NA | NA | NA | NA | NA | #VALUE! |
| ENSMUSG00000115759.2 | 0 | NA | NA | NA | NA | NA | #VALUE! |
| ENSMUSG00000115760.2 | 0 | NA | NA | NA | NA | NA | #VALUE! |
| ENSMUSG00000115761.2 | 0 | NA | NA | NA | NA | NA | #VALUE! |
| ENSMUSG00000115762.2 | 0 | NA | NA | NA | NA | NA | #VALUE! |
| ENSMUSG00000115763.2 | 0 | NA | NA | NA | NA | NA | #VALUE! |
| ENSMUSG00000115764.2 | 0 | NA | NA | NA | NA | NA | #VALUE! |
| ENSMUSG00000115765.2 | 0 | NA | NA | NA | NA | NA | #VALUE! |
| ENSMUSG00000115766.2 | 0 | NA | NA | NA | NA | NA | #VALUE! |
| ENSMUSG00000115767.2 | 0 | NA | NA | NA | NA | NA | #VALUE! |
| ENSMUSG00000115768.2 | 0 | NA | NA | NA | NA | NA | #VALUE! |
| ENSMUSG00000115769.2 | 0 | NA | NA | NA | NA | NA | #VALUE! |
| ENSMUSG00000115770.2 | 0 | NA | NA | NA | NA | NA | #VALUE! |
| ENSMUSG00000115771.2 | 0 | NA | NA | NA | NA | NA | #VALUE! |
| ENSMUSG00000115772.2 | 0 | NA | NA | NA | NA | NA | #VALUE! |
| ENSMUSG00000115773.2 | 0 | NA | NA | NA | NA | NA | #VALUE! |
| ENSMUSG00000115774.2 | 0 | NA | NA | NA | NA | NA | #VALUE! |
| ENSMUSG00000115775.2 | 0 | NA | NA | NA | NA | NA | #VALUE! |

| | | | | | | | |
|---|---|---|---|---|---|---|---|
| ENSMUSG00000115776.2 | 0 | NA | NA | NA | NA | NA | #VALUE! |
| ENSMUSG00000115777.2 | 0 | NA | NA | NA | NA | NA | #VALUE! |
| ENSMUSG00000115778.2 | 0 | NA | NA | NA | NA | NA | #VALUE! |
| ENSMUSG00000115779.2 | 4.618153 | 0.261692 | 3.765449 | 0.069498 | 0.944593 | NA | #VALUE! |
| ENSMUSG00000115780.2 | 0 | NA | NA | NA | NA | NA | #VALUE! |
| ENSMUSG00000115781.2 | 0 | NA | NA | NA | NA | NA | #VALUE! |
| ENSMUSG00000115782.2 | 0 | NA | NA | NA | NA | NA | #VALUE! |
| ENSMUSG00000115783.2 | 0 | NA | NA | NA | NA | NA | #VALUE! |
| ENSMUSG00000115784.2 | 0 | NA | NA | NA | NA | NA | #VALUE! |
| ENSMUSG00000115785.2 | 0 | NA | NA | NA | NA | NA | #VALUE! |
| ENSMUSG00000115786.2 | 0 | NA | NA | NA | NA | NA | #VALUE! |
| ENSMUSG00000115787.2 | 0 | NA | NA | NA | NA | NA | #VALUE! |
| ENSMUSG00000115788.2 | 0 | NA | NA | NA | NA | NA | #VALUE! |
| ENSMUSG00000115789.2 | 0 | NA | NA | NA | NA | NA | #VALUE! |
| ENSMUSG00000115790.2 | 0 | NA | NA | NA | NA | NA | #VALUE! |
| ENSMUSG00000115791.2 | 0 | NA | NA | NA | NA | NA | #VALUE! |
| ENSMUSG00000115792.2 | 0 | NA | NA | NA | NA | NA | #VALUE! |
| ENSMUSG00000115793.2 | 0 | NA | NA | NA | NA | NA | #VALUE! |
| ENSMUSG00000115794.2 | 0 | NA | NA | NA | NA | NA | #VALUE! |
| ENSMUSG00000115795.2 | 0 | NA | NA | NA | NA | NA | #VALUE! |
| ENSMUSG00000115796.2 | 0 | NA | NA | NA | NA | NA | #VALUE! |
| ENSMUSG00000115797.2 | 0 | NA | NA | NA | NA | NA | #VALUE! |
| ENSMUSG00000115798.2 | 0 | NA | NA | NA | NA | NA | #VALUE! |
| ENSMUSG00000115799.2 | 0 | NA | NA | NA | NA | NA | #VALUE! |
| ENSMUSG00000115800.2 | 0 | NA | NA | NA | NA | NA | #VALUE! |
| ENSMUSG00000115801.2 | 0.093255 | 0.868093 | 4.072464 | 0.213162 | 0.831201 | NA | #VALUE! |
| ENSMUSG00000115802.2 | 0 | NA | NA | NA | NA | NA | #VALUE! |
| ENSMUSG00000115803.2 | 0 | NA | NA | NA | NA | NA | #VALUE! |
| ENSMUSG00000115804.2 | 0 | NA | NA | NA | NA | NA | #VALUE! |
| ENSMUSG00000115805.2 | 0 | NA | NA | NA | NA | NA | #VALUE! |
| ENSMUSG00000115806.2 | 0 | NA | NA | NA | NA | NA | #VALUE! |
| ENSMUSG00000115807.2 | 0 | NA | NA | NA | NA | NA | #VALUE! |
| ENSMUSG00000115808.2 | 0 | NA | NA | NA | NA | NA | #VALUE! |
| ENSMUSG00000115809.2 | 0 | NA | NA | NA | NA | NA | #VALUE! |
| ENSMUSG00000115810.2 | 0 | NA | NA | NA | NA | NA | #VALUE! |
| ENSMUSG00000115811.2 | 0 | NA | NA | NA | NA | NA | #VALUE! |
| ENSMUSG00000115812.2 | 0 | NA | NA | NA | NA | NA | #VALUE! |
| ENSMUSG00000115813.2 | 0 | NA | NA | NA | NA | NA | #VALUE! |
| ENSMUSG00000115814.2 | 0 | NA | NA | NA | NA | NA | #VALUE! |
| ENSMUSG00000115815.2 | 0 | NA | NA | NA | NA | NA | #VALUE! |
| ENSMUSG00000115816.2 | 0 | NA | NA | NA | NA | NA | #VALUE! |
| ENSMUSG00000115817.2 | 0 | NA | NA | NA | NA | NA | #VALUE! |
| ENSMUSG00000115818.2 | 0 | NA | NA | NA | NA | NA | #VALUE! |
| ENSMUSG00000115819.2 | 0 | NA | NA | NA | NA | NA | #VALUE! |

| | | | | | | | |
|---|---|---|---|---|---|---|---|
| ENSMUSG00000115820.2 | 0 | NA | NA | NA | NA | NA | #VALUE! |
| ENSMUSG00000115821.2 | 0 | NA | NA | NA | NA | NA | #VALUE! |
| ENSMUSG00000115822.2 | 0 | NA | NA | NA | NA | NA | #VALUE! |
| ENSMUSG00000115823.2 | 0 | NA | NA | NA | NA | NA | #VALUE! |
| ENSMUSG00000115824.2 | 0 | NA | NA | NA | NA | NA | #VALUE! |
| ENSMUSG00000115825.2 | 0 | NA | NA | NA | NA | NA | #VALUE! |
| ENSMUSG00000115826.2 | 0 | NA | NA | NA | NA | NA | #VALUE! |
| ENSMUSG00000115827.2 | 0 | NA | NA | NA | NA | NA | #VALUE! |
| ENSMUSG00000115828.2 | 0 | NA | NA | NA | NA | NA | #VALUE! |
| ENSMUSG00000115829.2 | 0 | NA | NA | NA | NA | NA | #VALUE! |
| ENSMUSG00000115830.2 | 0 | NA | NA | NA | NA | NA | #VALUE! |
| ENSMUSG00000115831.2 | 0 | NA | NA | NA | NA | NA | #VALUE! |
| ENSMUSG00000115832.2 | 0 | NA | NA | NA | NA | NA | #VALUE! |
| ENSMUSG00000115833.2 | 0 | NA | NA | NA | NA | NA | #VALUE! |
| ENSMUSG00000115834.2 | 0 | NA | NA | NA | NA | NA | #VALUE! |
| ENSMUSG00000115835.2 | 0 | NA | NA | NA | NA | NA | #VALUE! |
| ENSMUSG00000115836.2 | 0 | NA | NA | NA | NA | NA | #VALUE! |
| ENSMUSG00000115837.2 | 5.601645 | -4.88825 | 3.901177 | -1.25302 | 0.210199 | NA | #VALUE! |
| ENSMUSG00000115838.2 | 0 | NA | NA | NA | NA | NA | #VALUE! |
| ENSMUSG00000115839.2 | 0 | NA | NA | NA | NA | NA | #VALUE! |
| ENSMUSG00000115840.2 | 0 | NA | NA | NA | NA | NA | #VALUE! |
| ENSMUSG00000115841.2 | 0 | NA | NA | NA | NA | NA | #VALUE! |
| ENSMUSG00000115842.2 | 0 | NA | NA | NA | NA | NA | #VALUE! |
| ENSMUSG00000115843.2 | 0 | NA | NA | NA | NA | NA | #VALUE! |
| ENSMUSG00000115844.2 | 0 | NA | NA | NA | NA | NA | #VALUE! |
| ENSMUSG00000115845.2 | 0 | NA | NA | NA | NA | NA | #VALUE! |
| ENSMUSG00000115846.2 | 0 | NA | NA | NA | NA | NA | #VALUE! |
| ENSMUSG00000115847.2 | 0 | NA | NA | NA | NA | NA | #VALUE! |
| ENSMUSG00000115848.2 | 0 | NA | NA | NA | NA | NA | #VALUE! |
| ENSMUSG00000115849.2 | 0 | NA | NA | NA | NA | NA | #VALUE! |
| ENSMUSG00000115850.2 | 0 | NA | NA | NA | NA | NA | #VALUE! |
| ENSMUSG00000115854.2 | 0 | NA | NA | NA | NA | NA | #VALUE! |
| ENSMUSG00000115855.2 | 0 | NA | NA | NA | NA | NA | #VALUE! |
| ENSMUSG00000115856.2 | 0 | NA | NA | NA | NA | NA | #VALUE! |
| ENSMUSG00000115858.2 | 0 | NA | NA | NA | NA | NA | #VALUE! |
| ENSMUSG00000115860.2 | 0 | NA | NA | NA | NA | NA | #VALUE! |
| ENSMUSG00000115862.2 | 0 | NA | NA | NA | NA | NA | #VALUE! |
| ENSMUSG00000115863.2 | 0 | NA | NA | NA | NA | NA | #VALUE! |
| ENSMUSG00000115866.2 | 0 | NA | NA | NA | NA | NA | #VALUE! |
| ENSMUSG00000115868.2 | 0 | NA | NA | NA | NA | NA | #VALUE! |
| ENSMUSG00000115869.2 | 3.474766 | 1.195832 | 3.78757 | 0.315725 | 0.752211 | NA | #VALUE! |
| ENSMUSG00000115871.2 | 0 | NA | NA | NA | NA | NA | #VALUE! |
| ENSMUSG00000115875.2 | 0 | NA | NA | NA | NA | NA | #VALUE! |
| ENSMUSG00000115877.2 | 0 | NA | NA | NA | NA | NA | #VALUE! |

| | | | | | | | |
|---|---|---|---|---|---|---|---|
| ENSMUSG00000115880.2 | 0 | NA | NA | NA | NA | NA | #VALUE! |
| ENSMUSG00000115882.2 | 0 | NA | NA | NA | NA | NA | #VALUE! |
| ENSMUSG00000115885.2 | 0 | NA | NA | NA | NA | NA | #VALUE! |
| ENSMUSG00000115887.2 | 0 | NA | NA | NA | NA | NA | #VALUE! |
| ENSMUSG00000115888.2 | 0 | NA | NA | NA | NA | NA | #VALUE! |
| ENSMUSG00000115894.2 | 0 | NA | NA | NA | NA | NA | #VALUE! |
| ENSMUSG00000115896.2 | 0 | NA | NA | NA | NA | NA | #VALUE! |
| ENSMUSG00000115898.2 | 0 | NA | NA | NA | NA | NA | #VALUE! |
| ENSMUSG00000115902.2 | 0 | NA | NA | NA | NA | NA | #VALUE! |
| ENSMUSG00000115903.2 | 0 | NA | NA | NA | NA | NA | #VALUE! |
| ENSMUSG00000115907.2 | 0 | NA | NA | NA | NA | NA | #VALUE! |
| ENSMUSG00000115908.2 | 0 | NA | NA | NA | NA | NA | #VALUE! |
| ENSMUSG00000115911.2 | 0 | NA | NA | NA | NA | NA | #VALUE! |
| ENSMUSG00000115912.2 | 0 | NA | NA | NA | NA | NA | #VALUE! |
| ENSMUSG00000115914.2 | 0 | NA | NA | NA | NA | NA | #VALUE! |
| ENSMUSG00000115916.2 | 0.291524 | 0.468739 | 4.052149 | 0.115677 | 0.907909 | NA | #VALUE! |
| ENSMUSG00000115917.2 | 0 | NA | NA | NA | NA | NA | #VALUE! |
| ENSMUSG00000115918.2 | 0 | NA | NA | NA | NA | NA | #VALUE! |
| ENSMUSG00000115920.2 | 0 | NA | NA | NA | NA | NA | #VALUE! |
| ENSMUSG00000115921.2 | 0 | NA | NA | NA | NA | NA | #VALUE! |
| ENSMUSG00000115923.2 | 0 | NA | NA | NA | NA | NA | #VALUE! |
| ENSMUSG00000115924.2 | 0 | NA | NA | NA | NA | NA | #VALUE! |
| ENSMUSG00000115925.2 | 0 | NA | NA | NA | NA | NA | #VALUE! |
| ENSMUSG00000115926.2 | 0 | NA | NA | NA | NA | NA | #VALUE! |
| ENSMUSG00000115928.2 | 0 | NA | NA | NA | NA | NA | #VALUE! |
| ENSMUSG00000115930.2 | 0 | NA | NA | NA | NA | NA | #VALUE! |
| ENSMUSG00000115931.2 | 0 | NA | NA | NA | NA | NA | #VALUE! |
| ENSMUSG00000115935.2 | 0 | NA | NA | NA | NA | NA | #VALUE! |
| ENSMUSG00000115936.2 | 0 | NA | NA | NA | NA | NA | #VALUE! |
| ENSMUSG00000115943.2 | 0 | NA | NA | NA | NA | NA | #VALUE! |
| ENSMUSG00000115944.2 | 0 | NA | NA | NA | NA | NA | #VALUE! |
| ENSMUSG00000115948.2 | 0 | NA | NA | NA | NA | NA | #VALUE! |
| ENSMUSG00000115952.2 | 0 | NA | NA | NA | NA | NA | #VALUE! |
| ENSMUSG00000115955.2 | 0 | NA | NA | NA | NA | NA | #VALUE! |
| ENSMUSG00000115957.2 | 0 | NA | NA | NA | NA | NA | #VALUE! |
| ENSMUSG00000115958.2 | 0 | NA | NA | NA | NA | NA | #VALUE! |
| ENSMUSG00000115959.2 | 0 | NA | NA | NA | NA | NA | #VALUE! |
| ENSMUSG00000115961.2 | 0 | NA | NA | NA | NA | NA | #VALUE! |
| ENSMUSG00000115962.2 | 0 | NA | NA | NA | NA | NA | #VALUE! |
| ENSMUSG00000115964.2 | 0 | NA | NA | NA | NA | NA | #VALUE! |
| ENSMUSG00000115965.2 | 0 | NA | NA | NA | NA | NA | #VALUE! |
| ENSMUSG00000115968.2 | 0 | NA | NA | NA | NA | NA | #VALUE! |
| ENSMUSG00000115969.2 | 0 | NA | NA | NA | NA | NA | #VALUE! |
| ENSMUSG00000115970.2 | 0 | NA | NA | NA | NA | NA | #VALUE! |

| | | | | | | | |
|---|---|---|---|---|---|---|---|
| ENSMUSG00000115971.2 | 0 | NA | NA | NA | NA | NA | #VALUE! |
| ENSMUSG00000115973.2 | 0 | NA | NA | NA | NA | NA | #VALUE! |
| ENSMUSG00000115975.2 | 0 | NA | NA | NA | NA | NA | #VALUE! |
| ENSMUSG00000115976.2 | 0 | NA | NA | NA | NA | NA | #VALUE! |
| ENSMUSG00000115979.2 | 0 | NA | NA | NA | NA | NA | #VALUE! |
| ENSMUSG00000115980.2 | 0 | NA | NA | NA | NA | NA | #VALUE! |
| ENSMUSG00000115983.2 | 0 | NA | NA | NA | NA | NA | #VALUE! |
| ENSMUSG00000115985.2 | 0 | NA | NA | NA | NA | NA | #VALUE! |
| ENSMUSG00000115986.2 | 0 | NA | NA | NA | NA | NA | #VALUE! |
| ENSMUSG00000115988.2 | 0 | NA | NA | NA | NA | NA | #VALUE! |
| ENSMUSG00000115989.2 | 0 | NA | NA | NA | NA | NA | #VALUE! |
| ENSMUSG00000115992.2 | 0 | NA | NA | NA | NA | NA | #VALUE! |
| ENSMUSG00000115993.2 | 0 | NA | NA | NA | NA | NA | #VALUE! |
| ENSMUSG00000115996.2 | 0 | NA | NA | NA | NA | NA | #VALUE! |
| ENSMUSG00000115997.2 | 0 | NA | NA | NA | NA | NA | #VALUE! |
| ENSMUSG00000116000.2 | 0 | NA | NA | NA | NA | NA | #VALUE! |
| ENSMUSG00000116004.2 | 0 | NA | NA | NA | NA | NA | #VALUE! |
| ENSMUSG00000116007.2 | 0 | NA | NA | NA | NA | NA | #VALUE! |
| ENSMUSG00000116008.2 | 0 | NA | NA | NA | NA | NA | #VALUE! |
| ENSMUSG00000116009.2 | 0 | NA | NA | NA | NA | NA | #VALUE! |
| ENSMUSG00000116010.2 | 0 | NA | NA | NA | NA | NA | #VALUE! |
| ENSMUSG00000116011.2 | 0 | NA | NA | NA | NA | NA | #VALUE! |
| ENSMUSG00000116013.2 | 0 | NA | NA | NA | NA | NA | #VALUE! |
| ENSMUSG00000116016.2 | 0 | NA | NA | NA | NA | NA | #VALUE! |
| ENSMUSG00000116017.2 | 0 | NA | NA | NA | NA | NA | #VALUE! |
| ENSMUSG00000116021.2 | 0 | NA | NA | NA | NA | NA | #VALUE! |
| ENSMUSG00000116024.2 | 0 | NA | NA | NA | NA | NA | #VALUE! |
| ENSMUSG00000116025.2 | 0 | NA | NA | NA | NA | NA | #VALUE! |
| ENSMUSG00000116029.2 | 0 | NA | NA | NA | NA | NA | #VALUE! |
| ENSMUSG00000116030.2 | 0 | NA | NA | NA | NA | NA | #VALUE! |
| ENSMUSG00000116031.2 | 0 | NA | NA | NA | NA | NA | #VALUE! |
| ENSMUSG00000116033.2 | 0 | NA | NA | NA | NA | NA | #VALUE! |
| ENSMUSG00000116038.2 | 0 | NA | NA | NA | NA | NA | #VALUE! |
| ENSMUSG00000116039.2 | 0 | NA | NA | NA | NA | NA | #VALUE! |
| ENSMUSG00000116040.2 | 0 | NA | NA | NA | NA | NA | #VALUE! |
| ENSMUSG00000116041.2 | 0 | NA | NA | NA | NA | NA | #VALUE! |
| ENSMUSG00000116044.2 | 0 | NA | NA | NA | NA | NA | #VALUE! |
| ENSMUSG00000116048.2 | 0 | NA | NA | NA | NA | NA | #VALUE! |
| ENSMUSG00000116056.2 | 0 | NA | NA | NA | NA | NA | #VALUE! |
| ENSMUSG00000116057.2 | 0 | NA | NA | NA | NA | NA | #VALUE! |
| ENSMUSG00000116058.2 | 0 | NA | NA | NA | NA | NA | #VALUE! |
| ENSMUSG00000116061.2 | 0 | NA | NA | NA | NA | NA | #VALUE! |
| ENSMUSG00000116066.2 | 0 | NA | NA | NA | NA | NA | #VALUE! |
| ENSMUSG00000116068.2 | 0 | NA | NA | NA | NA | NA | #VALUE! |

| | | | | | | | |
|---|---|---|---|---|---|---|---|
| ENSMUSG00000116069.2 | 0 | NA | NA | NA | NA | NA | #VALUE! |
| ENSMUSG00000116071.2 | 0 | NA | NA | NA | NA | NA | #VALUE! |
| ENSMUSG00000116072.2 | 0 | NA | NA | NA | NA | NA | #VALUE! |
| ENSMUSG00000116074.2 | 0 | NA | NA | NA | NA | NA | #VALUE! |
| ENSMUSG00000116075.2 | 0 | NA | NA | NA | NA | NA | #VALUE! |
| ENSMUSG00000116079.2 | 0 | NA | NA | NA | NA | NA | #VALUE! |
| ENSMUSG00000116081.2 | 0 | NA | NA | NA | NA | NA | #VALUE! |
| ENSMUSG00000116083.2 | 0 | NA | NA | NA | NA | NA | #VALUE! |
| ENSMUSG00000116088.2 | 0 | NA | NA | NA | NA | NA | #VALUE! |
| ENSMUSG00000116089.2 | 0.195789 | -1.05581 | 4.073302 | -0.2592 | 0.795479 | NA | #VALUE! |
| ENSMUSG00000116090.2 | 0.105013 | -0.97324 | 4.076293 | -0.23876 | 0.811295 | NA | #VALUE! |
| ENSMUSG00000116091.2 | 0 | NA | NA | NA | NA | NA | #VALUE! |
| ENSMUSG00000116092.2 | 0 | NA | NA | NA | NA | NA | #VALUE! |
| ENSMUSG00000116093.2 | 0 | NA | NA | NA | NA | NA | #VALUE! |
| ENSMUSG00000116095.2 | 0 | NA | NA | NA | NA | NA | #VALUE! |
| ENSMUSG00000116096.2 | 0 | NA | NA | NA | NA | NA | #VALUE! |
| ENSMUSG00000116102.2 | 0 | NA | NA | NA | NA | NA | #VALUE! |
| ENSMUSG00000116104.2 | 0 | NA | NA | NA | NA | NA | #VALUE! |
| ENSMUSG00000116107.2 | 0 | NA | NA | NA | NA | NA | #VALUE! |
| ENSMUSG00000116109.2 | 0 | NA | NA | NA | NA | NA | #VALUE! |
| ENSMUSG00000116113.2 | 0 | NA | NA | NA | NA | NA | #VALUE! |
| ENSMUSG00000116114.2 | 0 | NA | NA | NA | NA | NA | #VALUE! |
| ENSMUSG00000116116.2 | 0 | NA | NA | NA | NA | NA | #VALUE! |
| ENSMUSG00000116117.2 | 0 | NA | NA | NA | NA | NA | #VALUE! |
| ENSMUSG00000116118.2 | 0 | NA | NA | NA | NA | NA | #VALUE! |
| ENSMUSG00000116120.2 | 0 | NA | NA | NA | NA | NA | #VALUE! |
| ENSMUSG00000116121.2 | 0 | NA | NA | NA | NA | NA | #VALUE! |
| ENSMUSG00000116122.2 | 0 | NA | NA | NA | NA | NA | #VALUE! |
| ENSMUSG00000116123.2 | 0 | NA | NA | NA | NA | NA | #VALUE! |
| ENSMUSG00000116124.2 | 0 | NA | NA | NA | NA | NA | #VALUE! |
| ENSMUSG00000116125.2 | 0 | NA | NA | NA | NA | NA | #VALUE! |
| ENSMUSG00000116127.2 | 0 | NA | NA | NA | NA | NA | #VALUE! |
| ENSMUSG00000116129.2 | 0 | NA | NA | NA | NA | NA | #VALUE! |
| ENSMUSG00000116131.2 | 0 | NA | NA | NA | NA | NA | #VALUE! |
| ENSMUSG00000116132.2 | 0 | NA | NA | NA | NA | NA | #VALUE! |
| ENSMUSG00000116136.2 | 0 | NA | NA | NA | NA | NA | #VALUE! |
| ENSMUSG00000116138.2 | 34.70808 | 0.248196 | 1.694941 | 0.146434 | NA | NA | #VALUE! |
| ENSMUSG00000116140.2 | 0 | NA | NA | NA | NA | NA | #VALUE! |
| ENSMUSG00000116141.2 | 0.630079 | -2.90291 | 3.975047 | -0.73028 | 0.465217 | NA | #VALUE! |
| ENSMUSG00000116145.2 | 0 | NA | NA | NA | NA | NA | #VALUE! |
| ENSMUSG00000116149.2 | 0 | NA | NA | NA | NA | NA | #VALUE! |
| ENSMUSG00000116151.2 | 0 | NA | NA | NA | NA | NA | #VALUE! |
| ENSMUSG00000116155.2 | 0 | NA | NA | NA | NA | NA | #VALUE! |
| ENSMUSG00000116157.2 | 0 | NA | NA | NA | NA | NA | #VALUE! |

| | | | | | | | |
|---|---|---|---|---|---|---|---|
| ENSMUSG00000116158.2 | 0 | NA | NA | NA | NA | NA | #VALUE! |
| ENSMUSG00000116160.2 | 0 | NA | NA | NA | NA | NA | #VALUE! |
| ENSMUSG00000116162.2 | 0 | NA | NA | NA | NA | NA | #VALUE! |
| ENSMUSG00000116164.2 | 7.793525 | 6.446892 | 3.536872 | 1.822766 | 0.068339 | NA | #VALUE! |
| ENSMUSG00000116165.2 | 0 | NA | NA | NA | NA | NA | #VALUE! |
| ENSMUSG00000116166.2 | 0 | NA | NA | NA | NA | NA | #VALUE! |
| ENSMUSG00000116167.2 | 0 | NA | NA | NA | NA | NA | #VALUE! |
| ENSMUSG00000116168.2 | 0 | NA | NA | NA | NA | NA | #VALUE! |
| ENSMUSG00000116170.2 | 0 | NA | NA | NA | NA | NA | #VALUE! |
| ENSMUSG00000116171.2 | 0 | NA | NA | NA | NA | NA | #VALUE! |
| ENSMUSG00000116173.2 | 0 | NA | NA | NA | NA | NA | #VALUE! |
| ENSMUSG00000116174.2 | 0 | NA | NA | NA | NA | NA | #VALUE! |
| ENSMUSG00000116178.2 | 0 | NA | NA | NA | NA | NA | #VALUE! |
| ENSMUSG00000116179.2 | 0 | NA | NA | NA | NA | NA | #VALUE! |
| ENSMUSG00000116180.2 | 0 | NA | NA | NA | NA | NA | #VALUE! |
| ENSMUSG00000116182.2 | 0 | NA | NA | NA | NA | NA | #VALUE! |
| ENSMUSG00000116183.2 | 0 | NA | NA | NA | NA | NA | #VALUE! |
| ENSMUSG00000116184.2 | 0 | NA | NA | NA | NA | NA | #VALUE! |
| ENSMUSG00000116185.2 | 0 | NA | NA | NA | NA | NA | #VALUE! |
| ENSMUSG00000116187.2 | 0 | NA | NA | NA | NA | NA | #VALUE! |
| ENSMUSG00000116188.3 | 0 | NA | NA | NA | NA | NA | #VALUE! |
| ENSMUSG00000116190.2 | 0 | NA | NA | NA | NA | NA | #VALUE! |
| ENSMUSG00000116191.2 | 0 | NA | NA | NA | NA | NA | #VALUE! |
| ENSMUSG00000116192.2 | 0 | NA | NA | NA | NA | NA | #VALUE! |
| ENSMUSG00000116194.2 | 0 | NA | NA | NA | NA | NA | #VALUE! |
| ENSMUSG00000116197.2 | 0 | NA | NA | NA | NA | NA | #VALUE! |
| ENSMUSG00000116199.2 | 0 | NA | NA | NA | NA | NA | #VALUE! |
| ENSMUSG00000116202.2 | 0 | NA | NA | NA | NA | NA | #VALUE! |
| ENSMUSG00000116204.2 | 0 | NA | NA | NA | NA | NA | #VALUE! |
| ENSMUSG00000116207.2 | 0 | NA | NA | NA | NA | NA | #VALUE! |
| ENSMUSG00000116210.2 | 0 | NA | NA | NA | NA | NA | #VALUE! |
| ENSMUSG00000116212.2 | 0 | NA | NA | NA | NA | NA | #VALUE! |
| ENSMUSG00000116213.2 | 0 | NA | NA | NA | NA | NA | #VALUE! |
| ENSMUSG00000116215.2 | 0 | NA | NA | NA | NA | NA | #VALUE! |
| ENSMUSG00000116217.2 | 0 | NA | NA | NA | NA | NA | #VALUE! |
| ENSMUSG00000116219.2 | 8.486223 | 6.579094 | 3.768352 | 1.745881 | NA | NA | #VALUE! |
| ENSMUSG00000116220.2 | 0 | NA | NA | NA | NA | NA | #VALUE! |
| ENSMUSG00000116222.2 | 0 | NA | NA | NA | NA | NA | #VALUE! |
| ENSMUSG00000116223.2 | 0 | NA | NA | NA | NA | NA | #VALUE! |
| ENSMUSG00000116226.2 | 0 | NA | NA | NA | NA | NA | #VALUE! |
| ENSMUSG00000116227.2 | 0 | NA | NA | NA | NA | NA | #VALUE! |
| ENSMUSG00000116229.2 | 0 | NA | NA | NA | NA | NA | #VALUE! |
| ENSMUSG00000116231.2 | 0 | NA | NA | NA | NA | NA | #VALUE! |
| ENSMUSG00000116234.2 | 0 | NA | NA | NA | NA | NA | #VALUE! |

| | | | | | | | |
|---|---|---|---|---|---|---|---|
| ENSMUSG00000116237.2 | 0 | NA | NA | NA | NA | NA | #VALUE! |
| ENSMUSG00000116238.2 | 0 | NA | NA | NA | NA | NA | #VALUE! |
| ENSMUSG00000116239.2 | 0 | NA | NA | NA | NA | NA | #VALUE! |
| ENSMUSG00000116241.2 | 0 | NA | NA | NA | NA | NA | #VALUE! |
| ENSMUSG00000116242.2 | 0 | NA | NA | NA | NA | NA | #VALUE! |
| ENSMUSG00000116243.2 | 0 | NA | NA | NA | NA | NA | #VALUE! |
| ENSMUSG00000116244.2 | 0 | NA | NA | NA | NA | NA | #VALUE! |
| ENSMUSG00000116245.2 | 0 | NA | NA | NA | NA | NA | #VALUE! |
| ENSMUSG00000116249.2 | 0 | NA | NA | NA | NA | NA | #VALUE! |
| ENSMUSG00000116250.2 | 0 | NA | NA | NA | NA | NA | #VALUE! |
| ENSMUSG00000116253.2 | 0 | NA | NA | NA | NA | NA | #VALUE! |
| ENSMUSG00000116255.2 | 0 | NA | NA | NA | NA | NA | #VALUE! |
| ENSMUSG00000116256.2 | 0 | NA | NA | NA | NA | NA | #VALUE! |
| ENSMUSG00000116260.2 | 0 | NA | NA | NA | NA | NA | #VALUE! |
| ENSMUSG00000116262.2 | 0 | NA | NA | NA | NA | NA | #VALUE! |
| ENSMUSG00000116263.2 | 0 | NA | NA | NA | NA | NA | #VALUE! |
| ENSMUSG00000116265.2 | 0 | NA | NA | NA | NA | NA | #VALUE! |
| ENSMUSG00000116270.2 | 0 | NA | NA | NA | NA | NA | #VALUE! |
| ENSMUSG00000116272.2 | 0 | NA | NA | NA | NA | NA | #VALUE! |
| ENSMUSG00000116274.2 | 0 | NA | NA | NA | NA | NA | #VALUE! |
| ENSMUSG00000116275.2 | 0 | NA | NA | NA | NA | NA | #VALUE! |
| ENSMUSG00000116280.2 | 0 | NA | NA | NA | NA | NA | #VALUE! |
| ENSMUSG00000116283.2 | 0 | NA | NA | NA | NA | NA | #VALUE! |
| ENSMUSG00000116287.2 | 0 | NA | NA | NA | NA | NA | #VALUE! |
| ENSMUSG00000116288.2 | 0 | NA | NA | NA | NA | NA | #VALUE! |
| ENSMUSG00000116289.2 | 0 | NA | NA | NA | NA | NA | #VALUE! |
| ENSMUSG00000116290.2 | 0 | NA | NA | NA | NA | NA | #VALUE! |
| ENSMUSG00000116291.2 | 0 | NA | NA | NA | NA | NA | #VALUE! |
| ENSMUSG00000116292.2 | 0 | NA | NA | NA | NA | NA | #VALUE! |
| ENSMUSG00000116294.2 | 0 | NA | NA | NA | NA | NA | #VALUE! |
| ENSMUSG00000116295.2 | 0 | NA | NA | NA | NA | NA | #VALUE! |
| ENSMUSG00000116296.2 | 0 | NA | NA | NA | NA | NA | #VALUE! |
| ENSMUSG00000116297.2 | 0 | NA | NA | NA | NA | NA | #VALUE! |
| ENSMUSG00000116298.2 | 0 | NA | NA | NA | NA | NA | #VALUE! |
| ENSMUSG00000116299.2 | 0 | NA | NA | NA | NA | NA | #VALUE! |
| ENSMUSG00000116300.2 | 0 | NA | NA | NA | NA | NA | #VALUE! |
| ENSMUSG00000116301.2 | 0 | NA | NA | NA | NA | NA | #VALUE! |
| ENSMUSG00000116303.2 | 0 | NA | NA | NA | NA | NA | #VALUE! |
| ENSMUSG00000116304.2 | 0 | NA | NA | NA | NA | NA | #VALUE! |
| ENSMUSG00000116305.2 | 0 | NA | NA | NA | NA | NA | #VALUE! |
| ENSMUSG00000116307.2 | 0 | NA | NA | NA | NA | NA | #VALUE! |
| ENSMUSG00000116308.2 | 0 | NA | NA | NA | NA | NA | #VALUE! |
| ENSMUSG00000116309.2 | 0 | NA | NA | NA | NA | NA | #VALUE! |
| ENSMUSG00000116311.2 | 0 | NA | NA | NA | NA | NA | #VALUE! |

| | | | | | | | |
|---|---|---|---|---|---|---|---|
| ENSMUSG00000116314.2 | 0 | NA | NA | NA | NA | NA | #VALUE! |
| ENSMUSG00000116317.2 | 0 | NA | NA | NA | NA | NA | #VALUE! |
| ENSMUSG00000116319.2 | 0 | NA | NA | NA | NA | NA | #VALUE! |
| ENSMUSG00000116321.2 | 0 | NA | NA | NA | NA | NA | #VALUE! |
| ENSMUSG00000116323.2 | 0 | NA | NA | NA | NA | NA | #VALUE! |
| ENSMUSG00000116324.2 | 0 | NA | NA | NA | NA | NA | #VALUE! |
| ENSMUSG00000116325.2 | 0 | NA | NA | NA | NA | NA | #VALUE! |
| ENSMUSG00000116331.2 | 0 | NA | NA | NA | NA | NA | #VALUE! |
| ENSMUSG00000116335.2 | 0 | NA | NA | NA | NA | NA | #VALUE! |
| ENSMUSG00000116336.3 | 0 | NA | NA | NA | NA | NA | #VALUE! |
| ENSMUSG00000116338.2 | 0 | NA | NA | NA | NA | NA | #VALUE! |
| ENSMUSG00000116339.2 | 0 | NA | NA | NA | NA | NA | #VALUE! |
| ENSMUSG00000116341.2 | 0.673134 | 2.918386 | 3.972531 | 0.734641 | 0.462558 | NA | #VALUE! |
| ENSMUSG00000116343.2 | 0 | NA | NA | NA | NA | NA | #VALUE! |
| ENSMUSG00000116345.2 | 0 | NA | NA | NA | NA | NA | #VALUE! |
| ENSMUSG00000116347.2 | 0 | NA | NA | NA | NA | NA | #VALUE! |
| ENSMUSG00000116348.2 | 0 | NA | NA | NA | NA | NA | #VALUE! |
| ENSMUSG00000116350.2 | 0 | NA | NA | NA | NA | NA | #VALUE! |
| ENSMUSG00000116352.2 | 0 | NA | NA | NA | NA | NA | #VALUE! |
| ENSMUSG00000116353.2 | 0 | NA | NA | NA | NA | NA | #VALUE! |
| ENSMUSG00000116354.2 | 3.915782 | -5.49628 | 3.91821 | -1.40275 | 0.16069 | NA | #VALUE! |
| ENSMUSG00000116355.2 | 0 | NA | NA | NA | NA | NA | #VALUE! |
| ENSMUSG00000116356.2 | 0 | NA | NA | NA | NA | NA | #VALUE! |
| ENSMUSG00000116358.2 | 0 | NA | NA | NA | NA | NA | #VALUE! |
| ENSMUSG00000116360.2 | 0 | NA | NA | NA | NA | NA | #VALUE! |
| ENSMUSG00000116365.2 | 0 | NA | NA | NA | NA | NA | #VALUE! |
| ENSMUSG00000116366.2 | 0 | NA | NA | NA | NA | NA | #VALUE! |
| ENSMUSG00000116367.2 | 0 | NA | NA | NA | NA | NA | #VALUE! |
| ENSMUSG00000116368.2 | 0 | NA | NA | NA | NA | NA | #VALUE! |
| ENSMUSG00000116371.2 | 0 | NA | NA | NA | NA | NA | #VALUE! |
| ENSMUSG00000116376.2 | 0 | NA | NA | NA | NA | NA | #VALUE! |
| ENSMUSG00000116378.2 | 0 | NA | NA | NA | NA | NA | #VALUE! |
| ENSMUSG00000116380.2 | 0 | NA | NA | NA | NA | NA | #VALUE! |
| ENSMUSG00000116383.2 | 0 | NA | NA | NA | NA | NA | #VALUE! |
| ENSMUSG00000116385.2 | 0 | NA | NA | NA | NA | NA | #VALUE! |
| ENSMUSG00000116386.2 | 0 | NA | NA | NA | NA | NA | #VALUE! |
| ENSMUSG00000116388.2 | 0 | NA | NA | NA | NA | NA | #VALUE! |
| ENSMUSG00000116389.2 | 0 | NA | NA | NA | NA | NA | #VALUE! |
| ENSMUSG00000116395.2 | 0 | NA | NA | NA | NA | NA | #VALUE! |
| ENSMUSG00000116396.2 | 0 | NA | NA | NA | NA | NA | #VALUE! |
| ENSMUSG00000116397.2 | 0.932552 | 3.421684 | 3.953399 | 0.865504 | 0.386762 | NA | #VALUE! |
| ENSMUSG00000116398.2 | 0 | NA | NA | NA | NA | NA | #VALUE! |
| ENSMUSG00000116400.2 | 0 | NA | NA | NA | NA | NA | #VALUE! |
| ENSMUSG00000116402.2 | 0.735092 | -3.12016 | 3.965612 | -0.7868 | 0.431397 | NA | #VALUE! |

| | | | | | | | |
|---|---|---|---|---|---|---|---|
| ENSMUSG00000116403.2 | 0 | NA | NA | NA | NA | NA | #VALUE! |
| ENSMUSG00000116404.2 | 0 | NA | NA | NA | NA | NA | #VALUE! |
| ENSMUSG00000116406.2 | 0 | NA | NA | NA | NA | NA | #VALUE! |
| ENSMUSG00000116409.2 | 0 | NA | NA | NA | NA | NA | #VALUE! |
| ENSMUSG00000116412.2 | 0 | NA | NA | NA | NA | NA | #VALUE! |
| ENSMUSG00000116413.2 | 0 | NA | NA | NA | NA | NA | #VALUE! |
| ENSMUSG00000116414.3 | 0 | NA | NA | NA | NA | NA | #VALUE! |
| ENSMUSG00000116416.2 | 0 | NA | NA | NA | NA | NA | #VALUE! |
| ENSMUSG00000116419.2 | 0 | NA | NA | NA | NA | NA | #VALUE! |
| ENSMUSG00000116423.2 | 0 | NA | NA | NA | NA | NA | #VALUE! |
| ENSMUSG00000116424.2 | 0 | NA | NA | NA | NA | NA | #VALUE! |
| ENSMUSG00000116426.2 | 0 | NA | NA | NA | NA | NA | #VALUE! |
| ENSMUSG00000116428.2 | 0 | NA | NA | NA | NA | NA | #VALUE! |
| ENSMUSG00000116429.2 | 0 | NA | NA | NA | NA | NA | #VALUE! |
| ENSMUSG00000116433.2 | 0 | NA | NA | NA | NA | NA | #VALUE! |
| ENSMUSG00000116435.2 | 0 | NA | NA | NA | NA | NA | #VALUE! |
| ENSMUSG00000116437.2 | 0 | NA | NA | NA | NA | NA | #VALUE! |
| ENSMUSG00000116438.2 | 0 | NA | NA | NA | NA | NA | #VALUE! |
| ENSMUSG00000116439.2 | 0 | NA | NA | NA | NA | NA | #VALUE! |
| ENSMUSG00000116442.2 | 0 | NA | NA | NA | NA | NA | #VALUE! |
| ENSMUSG00000116445.3 | 0 | NA | NA | NA | NA | NA | #VALUE! |
| ENSMUSG00000116449.2 | 0 | NA | NA | NA | NA | NA | #VALUE! |
| ENSMUSG00000116450.2 | 0 | NA | NA | NA | NA | NA | #VALUE! |
| ENSMUSG00000116455.2 | 0 | NA | NA | NA | NA | NA | #VALUE! |
| ENSMUSG00000116461.2 | 0 | NA | NA | NA | NA | NA | #VALUE! |
| ENSMUSG00000116464.2 | 0 | NA | NA | NA | NA | NA | #VALUE! |
| ENSMUSG00000116467.2 | 0 | NA | NA | NA | NA | NA | #VALUE! |
| ENSMUSG00000116469.2 | 0 | NA | NA | NA | NA | NA | #VALUE! |
| ENSMUSG00000116472.2 | 0 | NA | NA | NA | NA | NA | #VALUE! |
| ENSMUSG00000116475.2 | 0 | NA | NA | NA | NA | NA | #VALUE! |
| ENSMUSG00000116478.2 | 0 | NA | NA | NA | NA | NA | #VALUE! |
| ENSMUSG00000116480.2 | 0 | NA | NA | NA | NA | NA | #VALUE! |
| ENSMUSG00000116481.2 | 0 | NA | NA | NA | NA | NA | #VALUE! |
| ENSMUSG00000116482.2 | 0 | NA | NA | NA | NA | NA | #VALUE! |
| ENSMUSG00000116483.2 | 0 | NA | NA | NA | NA | NA | #VALUE! |
| ENSMUSG00000116490.2 | 0 | NA | NA | NA | NA | NA | #VALUE! |
| ENSMUSG00000116491.2 | 0 | NA | NA | NA | NA | NA | #VALUE! |
| ENSMUSG00000116493.2 | 0 | NA | NA | NA | NA | NA | #VALUE! |
| ENSMUSG00000116494.2 | 0 | NA | NA | NA | NA | NA | #VALUE! |
| ENSMUSG00000116498.2 | 0 | NA | NA | NA | NA | NA | #VALUE! |
| ENSMUSG00000116499.2 | 0 | NA | NA | NA | NA | NA | #VALUE! |
| ENSMUSG00000116500.2 | 0 | NA | NA | NA | NA | NA | #VALUE! |
| ENSMUSG00000116504.2 | 0 | NA | NA | NA | NA | NA | #VALUE! |
| ENSMUSG00000116514.2 | 0 | NA | NA | NA | NA | NA | #VALUE! |

| | | | | | | | |
|---|---|---|---|---|---|---|---|
| ENSMUSG00000116516.2 | 0 | NA | NA | NA | NA | NA | #VALUE! |
| ENSMUSG00000116519.2 | 0 | NA | NA | NA | NA | NA | #VALUE! |
| ENSMUSG00000116520.2 | 0 | NA | NA | NA | NA | NA | #VALUE! |
| ENSMUSG00000116521.2 | 0 | NA | NA | NA | NA | NA | #VALUE! |
| ENSMUSG00000116525.2 | 0 | NA | NA | NA | NA | NA | #VALUE! |
| ENSMUSG00000116528.2 | 0 | NA | NA | NA | NA | NA | #VALUE! |
| ENSMUSG00000116529.2 | 0 | NA | NA | NA | NA | NA | #VALUE! |
| ENSMUSG00000116530.2 | 0 | NA | NA | NA | NA | NA | #VALUE! |
| ENSMUSG00000116531.2 | 0 | NA | NA | NA | NA | NA | #VALUE! |
| ENSMUSG00000116532.2 | 0 | NA | NA | NA | NA | NA | #VALUE! |
| ENSMUSG00000116533.2 | 0 | NA | NA | NA | NA | NA | #VALUE! |
| ENSMUSG00000116534.2 | 0 | NA | NA | NA | NA | NA | #VALUE! |
| ENSMUSG00000116535.2 | 0 | NA | NA | NA | NA | NA | #VALUE! |
| ENSMUSG00000116536.2 | 0 | NA | NA | NA | NA | NA | #VALUE! |
| ENSMUSG00000116537.2 | 0 | NA | NA | NA | NA | NA | #VALUE! |
| ENSMUSG00000116538.2 | 0 | NA | NA | NA | NA | NA | #VALUE! |
| ENSMUSG00000116539.2 | 0 | NA | NA | NA | NA | NA | #VALUE! |
| ENSMUSG00000116540.2 | 0.336567 | 1.944925 | 4.029936 | 0.482619 | 0.629366 | NA | #VALUE! |
| ENSMUSG00000116542.2 | 0 | NA | NA | NA | NA | NA | #VALUE! |
| ENSMUSG00000116543.2 | 0 | NA | NA | NA | NA | NA | #VALUE! |
| ENSMUSG00000116544.2 | 0 | NA | NA | NA | NA | NA | #VALUE! |
| ENSMUSG00000116545.2 | 0 | NA | NA | NA | NA | NA | #VALUE! |
| ENSMUSG00000116546.2 | 0 | NA | NA | NA | NA | NA | #VALUE! |
| ENSMUSG00000116547.2 | 0 | NA | NA | NA | NA | NA | #VALUE! |
| ENSMUSG00000116548.2 | 0 | NA | NA | NA | NA | NA | #VALUE! |
| ENSMUSG00000116549.2 | 0 | NA | NA | NA | NA | NA | #VALUE! |
| ENSMUSG00000116550.2 | 0.195789 | -1.05581 | 4.073302 | -0.2592 | 0.795479 | NA | #VALUE! |
| ENSMUSG00000116551.2 | 0 | NA | NA | NA | NA | NA | #VALUE! |
| ENSMUSG00000116552.2 | 0 | NA | NA | NA | NA | NA | #VALUE! |
| ENSMUSG00000116553.2 | 0 | NA | NA | NA | NA | NA | #VALUE! |
| ENSMUSG00000116554.2 | 0 | NA | NA | NA | NA | NA | #VALUE! |
| ENSMUSG00000116555.2 | 0 | NA | NA | NA | NA | NA | #VALUE! |
| ENSMUSG00000116556.2 | 0 | NA | NA | NA | NA | NA | #VALUE! |
| ENSMUSG00000116557.2 | 0 | NA | NA | NA | NA | NA | #VALUE! |
| ENSMUSG00000116558.2 | 0 | NA | NA | NA | NA | NA | #VALUE! |
| ENSMUSG00000116559.2 | 0 | NA | NA | NA | NA | NA | #VALUE! |
| ENSMUSG00000116560.2 | 0 | NA | NA | NA | NA | NA | #VALUE! |
| ENSMUSG00000116561.2 | 0 | NA | NA | NA | NA | NA | #VALUE! |
| ENSMUSG00000116562.2 | 1.162915 | -1.12607 | 3.905789 | -0.28831 | 0.773111 | NA | #VALUE! |
| ENSMUSG00000116563.2 | 0 | NA | NA | NA | NA | NA | #VALUE! |
| ENSMUSG00000116565.2 | 0 | NA | NA | NA | NA | NA | #VALUE! |
| ENSMUSG00000116566.2 | 0 | NA | NA | NA | NA | NA | #VALUE! |
| ENSMUSG00000116567.2 | 0 | NA | NA | NA | NA | NA | #VALUE! |
| ENSMUSG00000116568.2 | 0 | NA | NA | NA | NA | NA | #VALUE! |

| | | | | | | | |
|---|---|---|---|---|---|---|---|
| ENSMUSG00000116569.2 | 0 | NA | NA | NA | NA | NA | #VALUE! |
| ENSMUSG00000116570.2 | 0 | NA | NA | NA | NA | NA | #VALUE! |
| ENSMUSG00000116572.2 | 0 | NA | NA | NA | NA | NA | #VALUE! |
| ENSMUSG00000116573.2 | 0 | NA | NA | NA | NA | NA | #VALUE! |
| ENSMUSG00000116574.2 | 0 | NA | NA | NA | NA | NA | #VALUE! |
| ENSMUSG00000116575.2 | 0 | NA | NA | NA | NA | NA | #VALUE! |
| ENSMUSG00000116576.2 | 0 | NA | NA | NA | NA | NA | #VALUE! |
| ENSMUSG00000116578.2 | 0 | NA | NA | NA | NA | NA | #VALUE! |
| ENSMUSG00000116579.2 | 0 | NA | NA | NA | NA | NA | #VALUE! |
| ENSMUSG00000116580.2 | 0 | NA | NA | NA | NA | NA | #VALUE! |
| ENSMUSG00000116581.2 | 0 | NA | NA | NA | NA | NA | #VALUE! |
| ENSMUSG00000116582.2 | 0 | NA | NA | NA | NA | NA | #VALUE! |
| ENSMUSG00000116583.2 | 0 | NA | NA | NA | NA | NA | #VALUE! |
| ENSMUSG00000116585.2 | 0 | NA | NA | NA | NA | NA | #VALUE! |
| ENSMUSG00000116586.2 | 0 | NA | NA | NA | NA | NA | #VALUE! |
| ENSMUSG00000116587.2 | 0 | NA | NA | NA | NA | NA | #VALUE! |
| ENSMUSG00000116588.2 | 0 | NA | NA | NA | NA | NA | #VALUE! |
| ENSMUSG00000116589.2 | 0 | NA | NA | NA | NA | NA | #VALUE! |
| ENSMUSG00000116590.2 | 8.020808 | 6.493814 | 3.427043 | 1.894874 | 0.058109 | NA | #VALUE! |
| ENSMUSG00000116591.2 | 0 | NA | NA | NA | NA | NA | #VALUE! |
| ENSMUSG00000116592.2 | 0.105013 | -0.97324 | 4.076293 | -0.23876 | 0.811295 | NA | #VALUE! |
| ENSMUSG00000116593.2 | 0 | NA | NA | NA | NA | NA | #VALUE! |
| ENSMUSG00000116594.2 | 0 | NA | NA | NA | NA | NA | #VALUE! |
| ENSMUSG00000116595.2 | 0 | NA | NA | NA | NA | NA | #VALUE! |
| ENSMUSG00000116596.2 | 0 | NA | NA | NA | NA | NA | #VALUE! |
| ENSMUSG00000116597.2 | 0.735092 | -3.12016 | 3.965612 | -0.7868 | 0.431397 | NA | #VALUE! |
| ENSMUSG00000116598.2 | 0 | NA | NA | NA | NA | NA | #VALUE! |
| ENSMUSG00000116599.2 | 0 | NA | NA | NA | NA | NA | #VALUE! |
| ENSMUSG00000116600.2 | 0 | NA | NA | NA | NA | NA | #VALUE! |
| ENSMUSG00000116601.2 | 0 | NA | NA | NA | NA | NA | #VALUE! |
| ENSMUSG00000116603.2 | 0 | NA | NA | NA | NA | NA | #VALUE! |
| ENSMUSG00000116604.2 | 0 | NA | NA | NA | NA | NA | #VALUE! |
| ENSMUSG00000116605.2 | 0 | NA | NA | NA | NA | NA | #VALUE! |
| ENSMUSG00000116606.2 | 0 | NA | NA | NA | NA | NA | #VALUE! |
| ENSMUSG00000116607.2 | 0 | NA | NA | NA | NA | NA | #VALUE! |
| ENSMUSG00000116608.2 | 0 | NA | NA | NA | NA | NA | #VALUE! |
| ENSMUSG00000116609.2 | 0 | NA | NA | NA | NA | NA | #VALUE! |
| ENSMUSG00000116610.2 | 0 | NA | NA | NA | NA | NA | #VALUE! |
| ENSMUSG00000116611.2 | 0 | NA | NA | NA | NA | NA | #VALUE! |
| ENSMUSG00000116612.2 | 0 | NA | NA | NA | NA | NA | #VALUE! |
| ENSMUSG00000116614.2 | 0 | NA | NA | NA | NA | NA | #VALUE! |
| ENSMUSG00000116615.2 | 0 | NA | NA | NA | NA | NA | #VALUE! |
| ENSMUSG00000116616.2 | 0 | NA | NA | NA | NA | NA | #VALUE! |
| ENSMUSG00000116617.2 | 0 | NA | NA | NA | NA | NA | #VALUE! |

| | | | | | | | |
|---|---|---|---|---|---|---|---|
| ENSMUSG00000116618.2 | 0 | NA | NA | NA | NA | NA | #VALUE! |
| ENSMUSG00000116619.2 | 0 | NA | NA | NA | NA | NA | #VALUE! |
| ENSMUSG00000116620.2 | 0.204111 | -1.05581 | 4.073302 | -0.2592 | 0.795479 | NA | #VALUE! |
| ENSMUSG00000116621.2 | 0.204111 | -1.05581 | 4.073302 | -0.2592 | 0.795479 | NA | #VALUE! |
| ENSMUSG00000116622.2 | 0 | NA | NA | NA | NA | NA | #VALUE! |
| ENSMUSG00000116623.2 | 0 | NA | NA | NA | NA | NA | #VALUE! |
| ENSMUSG00000116624.2 | 0 | NA | NA | NA | NA | NA | #VALUE! |
| ENSMUSG00000116625.2 | 0 | NA | NA | NA | NA | NA | #VALUE! |
| ENSMUSG00000116626.2 | 0 | NA | NA | NA | NA | NA | #VALUE! |
| ENSMUSG00000116627.2 | 0 | NA | NA | NA | NA | NA | #VALUE! |
| ENSMUSG00000116628.2 | 0 | NA | NA | NA | NA | NA | #VALUE! |
| ENSMUSG00000116629.2 | 0 | NA | NA | NA | NA | NA | #VALUE! |
| ENSMUSG00000116630.2 | 0 | NA | NA | NA | NA | NA | #VALUE! |
| ENSMUSG00000116631.2 | 0 | NA | NA | NA | NA | NA | #VALUE! |
| ENSMUSG00000116632.2 | 0 | NA | NA | NA | NA | NA | #VALUE! |
| ENSMUSG00000116633.2 | 0 | NA | NA | NA | NA | NA | #VALUE! |
| ENSMUSG00000116634.2 | 0 | NA | NA | NA | NA | NA | #VALUE! |
| ENSMUSG00000116635.2 | 0 | NA | NA | NA | NA | NA | #VALUE! |
| ENSMUSG00000116636.2 | 0 | NA | NA | NA | NA | NA | #VALUE! |
| ENSMUSG00000116637.2 | 0.429822 | 2.285488 | 4.007814 | 0.570258 | 0.568503 | NA | #VALUE! |
| ENSMUSG00000116638.2 | 0 | NA | NA | NA | NA | NA | #VALUE! |
| ENSMUSG00000116639.2 | 0 | NA | NA | NA | NA | NA | #VALUE! |
| ENSMUSG00000116640.2 | 0.245976 | 1.027628 | 4.067126 | 0.252667 | 0.800526 | NA | #VALUE! |
| ENSMUSG00000116641.2 | 0 | NA | NA | NA | NA | NA | #VALUE! |
| ENSMUSG00000116642.2 | 0 | NA | NA | NA | NA | NA | #VALUE! |
| ENSMUSG00000116643.3 | 0 | NA | NA | NA | NA | NA | #VALUE! |
| ENSMUSG00000116644.2 | 0 | NA | NA | NA | NA | NA | #VALUE! |
| ENSMUSG00000116645.2 | 0 | NA | NA | NA | NA | NA | #VALUE! |
| ENSMUSG00000116646.2 | 0 | NA | NA | NA | NA | NA | #VALUE! |
| ENSMUSG00000116647.2 | 0 | NA | NA | NA | NA | NA | #VALUE! |
| ENSMUSG00000116648.2 | 0 | NA | NA | NA | NA | NA | #VALUE! |
| ENSMUSG00000116649.2 | 0 | NA | NA | NA | NA | NA | #VALUE! |
| ENSMUSG00000116650.2 | 0 | NA | NA | NA | NA | NA | #VALUE! |
| ENSMUSG00000116651.2 | 0 | NA | NA | NA | NA | NA | #VALUE! |
| ENSMUSG00000116652.3 | 0 | NA | NA | NA | NA | NA | #VALUE! |
| ENSMUSG00000116653.2 | 0 | NA | NA | NA | NA | NA | #VALUE! |
| ENSMUSG00000116654.2 | 0 | NA | NA | NA | NA | NA | #VALUE! |
| ENSMUSG00000116655.2 | 0 | NA | NA | NA | NA | NA | #VALUE! |
| ENSMUSG00000116656.2 | 0 | NA | NA | NA | NA | NA | #VALUE! |
| ENSMUSG00000116657.2 | 0 | NA | NA | NA | NA | NA | #VALUE! |
| ENSMUSG00000116658.2 | 0 | NA | NA | NA | NA | NA | #VALUE! |
| ENSMUSG00000116659.2 | 0 | NA | NA | NA | NA | NA | #VALUE! |
| ENSMUSG00000116660.2 | 0 | NA | NA | NA | NA | NA | #VALUE! |
| ENSMUSG00000116661.2 | 0 | NA | NA | NA | NA | NA | #VALUE! |

| | | | | | | | |
|---|---|---|---|---|---|---|---|
| ENSMUSG00000116662.2 | 0 | NA | NA | NA | NA | NA | #VALUE! |
| ENSMUSG00000116663.2 | 0 | NA | NA | NA | NA | NA | #VALUE! |
| ENSMUSG00000116664.2 | 0 | NA | NA | NA | NA | NA | #VALUE! |
| ENSMUSG00000116665.2 | 0 | NA | NA | NA | NA | NA | #VALUE! |
| ENSMUSG00000116666.2 | 0 | NA | NA | NA | NA | NA | #VALUE! |
| ENSMUSG00000116667.2 | 0 | NA | NA | NA | NA | NA | #VALUE! |
| ENSMUSG00000116668.2 | 0 | NA | NA | NA | NA | NA | #VALUE! |
| ENSMUSG00000116669.2 | 0 | NA | NA | NA | NA | NA | #VALUE! |
| ENSMUSG00000116670.2 | 0 | NA | NA | NA | NA | NA | #VALUE! |
| ENSMUSG00000116671.2 | 0 | NA | NA | NA | NA | NA | #VALUE! |
| ENSMUSG00000116672.2 | 0 | NA | NA | NA | NA | NA | #VALUE! |
| ENSMUSG00000116673.2 | 2.355969 | 4.72622 | 3.925784 | 1.203892 | 0.228631 | NA | #VALUE! |
| ENSMUSG00000116674.2 | 0 | NA | NA | NA | NA | NA | #VALUE! |
| ENSMUSG00000116675.2 | 0 | NA | NA | NA | NA | NA | #VALUE! |
| ENSMUSG00000116676.2 | 0 | NA | NA | NA | NA | NA | #VALUE! |
| ENSMUSG00000116677.2 | 0 | NA | NA | NA | NA | NA | #VALUE! |
| ENSMUSG00000116678.2 | 0 | NA | NA | NA | NA | NA | #VALUE! |
| ENSMUSG00000116679.2 | 0 | NA | NA | NA | NA | NA | #VALUE! |
| ENSMUSG00000116680.2 | 0 | NA | NA | NA | NA | NA | #VALUE! |
| ENSMUSG00000116681.2 | 0 | NA | NA | NA | NA | NA | #VALUE! |
| ENSMUSG00000116682.2 | 0 | NA | NA | NA | NA | NA | #VALUE! |
| ENSMUSG00000116684.2 | 0 | NA | NA | NA | NA | NA | #VALUE! |
| ENSMUSG00000116685.2 | 0 | NA | NA | NA | NA | NA | #VALUE! |
| ENSMUSG00000116686.2 | 0 | NA | NA | NA | NA | NA | #VALUE! |
| ENSMUSG00000116687.2 | 0 | NA | NA | NA | NA | NA | #VALUE! |
| ENSMUSG00000116688.3 | 0 | NA | NA | NA | NA | NA | #VALUE! |
| ENSMUSG00000116689.2 | 0 | NA | NA | NA | NA | NA | #VALUE! |
| ENSMUSG00000116690.2 | 0 | NA | NA | NA | NA | NA | #VALUE! |
| ENSMUSG00000116691.2 | 0 | NA | NA | NA | NA | NA | #VALUE! |
| ENSMUSG00000116692.2 | 0 | NA | NA | NA | NA | NA | #VALUE! |
| ENSMUSG00000116693.2 | 0 | NA | NA | NA | NA | NA | #VALUE! |
| ENSMUSG00000116694.2 | 3.338049 | 1.597381 | 3.741398 | 0.426948 | 0.669417 | NA | #VALUE! |
| ENSMUSG00000116695.2 | 0 | NA | NA | NA | NA | NA | #VALUE! |
| ENSMUSG00000116696.2 | 0 | NA | NA | NA | NA | NA | #VALUE! |
| ENSMUSG00000116697.2 | 0 | NA | NA | NA | NA | NA | #VALUE! |
| ENSMUSG00000116698.2 | 0 | NA | NA | NA | NA | NA | #VALUE! |
| ENSMUSG00000116699.2 | 0 | NA | NA | NA | NA | NA | #VALUE! |
| ENSMUSG00000116700.2 | 0 | NA | NA | NA | NA | NA | #VALUE! |
| ENSMUSG00000116701.2 | 0 | NA | NA | NA | NA | NA | #VALUE! |
| ENSMUSG00000116702.2 | 0 | NA | NA | NA | NA | NA | #VALUE! |
| ENSMUSG00000116703.2 | 0 | NA | NA | NA | NA | NA | #VALUE! |
| ENSMUSG00000116704.2 | 0 | NA | NA | NA | NA | NA | #VALUE! |
| ENSMUSG00000116705.2 | 0 | NA | NA | NA | NA | NA | #VALUE! |
| ENSMUSG00000116706.2 | 0 | NA | NA | NA | NA | NA | #VALUE! |

| | | | | | | | |
|---|---|---|---|---|---|---|---|
| ENSMUSG00000116707.2 | 0 | NA | NA | NA | NA | NA | #VALUE! |
| ENSMUSG00000116708.2 | 0 | NA | NA | NA | NA | NA | #VALUE! |
| ENSMUSG00000116709.2 | 0 | NA | NA | NA | NA | NA | #VALUE! |
| ENSMUSG00000116710.2 | 0 | NA | NA | NA | NA | NA | #VALUE! |
| ENSMUSG00000116711.2 | 0 | NA | NA | NA | NA | NA | #VALUE! |
| ENSMUSG00000116712.2 | 0 | NA | NA | NA | NA | NA | #VALUE! |
| ENSMUSG00000116713.2 | 0 | NA | NA | NA | NA | NA | #VALUE! |
| ENSMUSG00000116714.2 | 0 | NA | NA | NA | NA | NA | #VALUE! |
| ENSMUSG00000116715.2 | 0 | NA | NA | NA | NA | NA | #VALUE! |
| ENSMUSG00000116716.2 | 0 | NA | NA | NA | NA | NA | #VALUE! |
| ENSMUSG00000116717.2 | 0 | NA | NA | NA | NA | NA | #VALUE! |
| ENSMUSG00000116718.2 | 45.00439 | 0.021254 | 1.833235 | 0.011594 | NA | NA | #VALUE! |
| ENSMUSG00000116719.2 | 0 | NA | NA | NA | NA | NA | #VALUE! |
| ENSMUSG00000116720.2 | 0 | NA | NA | NA | NA | NA | #VALUE! |
| ENSMUSG00000116721.2 | 0 | NA | NA | NA | NA | NA | #VALUE! |
| ENSMUSG00000116722.2 | 0 | NA | NA | NA | NA | NA | #VALUE! |
| ENSMUSG00000116723.2 | 0 | NA | NA | NA | NA | NA | #VALUE! |
| ENSMUSG00000116725.2 | 0 | NA | NA | NA | NA | NA | #VALUE! |
| ENSMUSG00000116727.2 | 0 | NA | NA | NA | NA | NA | #VALUE! |
| ENSMUSG00000116728.2 | 0 | NA | NA | NA | NA | NA | #VALUE! |
| ENSMUSG00000116729.2 | 1.020554 | -3.54462 | 3.950745 | -0.8972 | 0.36961 | NA | #VALUE! |
| ENSMUSG00000116730.2 | 0 | NA | NA | NA | NA | NA | #VALUE! |
| ENSMUSG00000116731.2 | 0 | NA | NA | NA | NA | NA | #VALUE! |
| ENSMUSG00000116732.2 | 0 | NA | NA | NA | NA | NA | #VALUE! |
| ENSMUSG00000116733.2 | 0 | NA | NA | NA | NA | NA | #VALUE! |
| ENSMUSG00000116734.2 | 0 | NA | NA | NA | NA | NA | #VALUE! |
| ENSMUSG00000116735.2 | 0 | NA | NA | NA | NA | NA | #VALUE! |
| ENSMUSG00000116736.2 | 0 | NA | NA | NA | NA | NA | #VALUE! |
| ENSMUSG00000116737.2 | 0 | NA | NA | NA | NA | NA | #VALUE! |
| ENSMUSG00000116738.2 | 0 | NA | NA | NA | NA | NA | #VALUE! |
| ENSMUSG00000116739.2 | 1.025807 | 3.556426 | 3.949287 | 0.900523 | 0.367842 | NA | #VALUE! |
| ENSMUSG00000116740.2 | 0 | NA | NA | NA | NA | NA | #VALUE! |
| ENSMUSG00000116741.2 | 0 | NA | NA | NA | NA | NA | #VALUE! |
| ENSMUSG00000116742.2 | 0 | NA | NA | NA | NA | NA | #VALUE! |
| ENSMUSG00000116743.2 | 0 | NA | NA | NA | NA | NA | #VALUE! |
| ENSMUSG00000116744.2 | 0 | NA | NA | NA | NA | NA | #VALUE! |
| ENSMUSG00000116745.2 | 0 | NA | NA | NA | NA | NA | #VALUE! |
| ENSMUSG00000116746.2 | 0 | NA | NA | NA | NA | NA | #VALUE! |
| ENSMUSG00000116747.2 | 0 | NA | NA | NA | NA | NA | #VALUE! |
| ENSMUSG00000116748.2 | 0 | NA | NA | NA | NA | NA | #VALUE! |
| ENSMUSG00000116749.2 | 0 | NA | NA | NA | NA | NA | #VALUE! |
| ENSMUSG00000116750.2 | 0 | NA | NA | NA | NA | NA | #VALUE! |
| ENSMUSG00000116751.2 | 0 | NA | NA | NA | NA | NA | #VALUE! |
| ENSMUSG00000116752.2 | 0 | NA | NA | NA | NA | NA | #VALUE! |

| | | | | | | | |
|---|---|---|---|---|---|---|---|
| ENSMUSG00000116753.2 | 0 | NA | NA | NA | NA | NA | #VALUE! |
| ENSMUSG00000116754.2 | 0 | NA | NA | NA | NA | NA | #VALUE! |
| ENSMUSG00000116755.2 | 0 | NA | NA | NA | NA | NA | #VALUE! |
| ENSMUSG00000116756.2 | 0 | NA | NA | NA | NA | NA | #VALUE! |
| ENSMUSG00000116757.2 | 1.054292 | 1.293203 | 3.935181 | 0.328626 | 0.742438 | NA | #VALUE! |
| ENSMUSG00000116758.2 | 0 | NA | NA | NA | NA | NA | #VALUE! |
| ENSMUSG00000116759.2 | 0 | NA | NA | NA | NA | NA | #VALUE! |
| ENSMUSG00000116760.2 | 0 | NA | NA | NA | NA | NA | #VALUE! |
| ENSMUSG00000116761.2 | 0 | NA | NA | NA | NA | NA | #VALUE! |
| ENSMUSG00000116762.2 | 0 | NA | NA | NA | NA | NA | #VALUE! |
| ENSMUSG00000116763.2 | 0 | NA | NA | NA | NA | NA | #VALUE! |
| ENSMUSG00000116764.2 | 0 | NA | NA | NA | NA | NA | #VALUE! |
| ENSMUSG00000116765.2 | 0 | NA | NA | NA | NA | NA | #VALUE! |
| ENSMUSG00000116766.2 | 0 | NA | NA | NA | NA | NA | #VALUE! |
| ENSMUSG00000116767.2 | 0 | NA | NA | NA | NA | NA | #VALUE! |
| ENSMUSG00000116768.2 | 0 | NA | NA | NA | NA | NA | #VALUE! |
| ENSMUSG00000116769.2 | 0 | NA | NA | NA | NA | NA | #VALUE! |
| ENSMUSG00000116770.2 | 0 | NA | NA | NA | NA | NA | #VALUE! |
| ENSMUSG00000116771.2 | 0 | NA | NA | NA | NA | NA | #VALUE! |
| ENSMUSG00000116772.2 | 0 | NA | NA | NA | NA | NA | #VALUE! |
| ENSMUSG00000116773.2 | 0 | NA | NA | NA | NA | NA | #VALUE! |
| ENSMUSG00000116774.2 | 0 | NA | NA | NA | NA | NA | #VALUE! |
| ENSMUSG00000116775.2 | 0 | NA | NA | NA | NA | NA | #VALUE! |
| ENSMUSG00000116776.2 | 0 | NA | NA | NA | NA | NA | #VALUE! |
| ENSMUSG00000116777.2 | 0 | NA | NA | NA | NA | NA | #VALUE! |
| ENSMUSG00000116778.2 | 0 | NA | NA | NA | NA | NA | #VALUE! |
| ENSMUSG00000116779.2 | 0 | NA | NA | NA | NA | NA | #VALUE! |
| ENSMUSG00000116780.2 | 0 | NA | NA | NA | NA | NA | #VALUE! |
| ENSMUSG00000116781.2 | 0 | NA | NA | NA | NA | NA | #VALUE! |
| ENSMUSG00000116782.2 | 0 | NA | NA | NA | NA | NA | #VALUE! |
| ENSMUSG00000116783.2 | 0 | NA | NA | NA | NA | NA | #VALUE! |
| ENSMUSG00000116784.2 | 0 | NA | NA | NA | NA | NA | #VALUE! |
| ENSMUSG00000116785.2 | 0 | NA | NA | NA | NA | NA | #VALUE! |
| ENSMUSG00000116786.2 | 0 | NA | NA | NA | NA | NA | #VALUE! |
| ENSMUSG00000116788.2 | 0 | NA | NA | NA | NA | NA | #VALUE! |
| ENSMUSG00000116789.2 | 0 | NA | NA | NA | NA | NA | #VALUE! |
| ENSMUSG00000116790.2 | 0 | NA | NA | NA | NA | NA | #VALUE! |
| ENSMUSG00000116791.2 | 0 | NA | NA | NA | NA | NA | #VALUE! |
| ENSMUSG00000116792.2 | 0 | NA | NA | NA | NA | NA | #VALUE! |
| ENSMUSG00000116793.2 | 0 | NA | NA | NA | NA | NA | #VALUE! |
| ENSMUSG00000116794.2 | 0 | NA | NA | NA | NA | NA | #VALUE! |
| ENSMUSG00000116795.2 | 0 | NA | NA | NA | NA | NA | #VALUE! |
| ENSMUSG00000116796.2 | 0 | NA | NA | NA | NA | NA | #VALUE! |
| ENSMUSG00000116797.2 | 0 | NA | NA | NA | NA | NA | #VALUE! |

| | | | | | | | |
|---|---|---|---|---|---|---|---|
| ENSMUSG00000116798.2 | 0 | NA | NA | NA | NA | NA | #VALUE! |
| ENSMUSG00000116799.2 | 0 | NA | NA | NA | NA | NA | #VALUE! |
| ENSMUSG00000116800.2 | 0 | NA | NA | NA | NA | NA | #VALUE! |
| ENSMUSG00000116801.2 | 0 | NA | NA | NA | NA | NA | #VALUE! |
| ENSMUSG00000116802.2 | 6.104689 | 2.430613 | 3.488068 | 0.696836 | 0.485905 | NA | #VALUE! |
| ENSMUSG00000116803.2 | 0 | NA | NA | NA | NA | NA | #VALUE! |
| ENSMUSG00000116804.2 | 0 | NA | NA | NA | NA | NA | #VALUE! |
| ENSMUSG00000116805.2 | 0 | NA | NA | NA | NA | NA | #VALUE! |
| ENSMUSG00000116806.2 | 0 | NA | NA | NA | NA | NA | #VALUE! |
| ENSMUSG00000116807.2 | 0 | NA | NA | NA | NA | NA | #VALUE! |
| ENSMUSG00000116808.2 | 0 | NA | NA | NA | NA | NA | #VALUE! |
| ENSMUSG00000116809.2 | 0 | NA | NA | NA | NA | NA | #VALUE! |
| ENSMUSG00000116810.2 | 12.88385 | -2.0416 | 2.462328 | -0.82913 | NA | NA | #VALUE! |
| ENSMUSG00000116811.2 | 0 | NA | NA | NA | NA | NA | #VALUE! |
| ENSMUSG00000116812.2 | 0 | NA | NA | NA | NA | NA | #VALUE! |
| ENSMUSG00000116813.2 | 0 | NA | NA | NA | NA | NA | #VALUE! |
| ENSMUSG00000116814.2 | 0 | NA | NA | NA | NA | NA | #VALUE! |
| ENSMUSG00000116815.2 | 0 | NA | NA | NA | NA | NA | #VALUE! |
| ENSMUSG00000116816.2 | 0 | NA | NA | NA | NA | NA | #VALUE! |
| ENSMUSG00000116817.2 | 0 | NA | NA | NA | NA | NA | #VALUE! |
| ENSMUSG00000116818.2 | 0 | NA | NA | NA | NA | NA | #VALUE! |
| ENSMUSG00000116819.2 | 0 | NA | NA | NA | NA | NA | #VALUE! |
| ENSMUSG00000116820.2 | 0 | NA | NA | NA | NA | NA | #VALUE! |
| ENSMUSG00000116821.2 | 0 | NA | NA | NA | NA | NA | #VALUE! |
| ENSMUSG00000116822.2 | 0 | NA | NA | NA | NA | NA | #VALUE! |
| ENSMUSG00000116823.2 | 0 | NA | NA | NA | NA | NA | #VALUE! |
| ENSMUSG00000116824.2 | 0 | NA | NA | NA | NA | NA | #VALUE! |
| ENSMUSG00000116825.2 | 0.168284 | 1.027628 | 4.067126 | 0.252667 | 0.800526 | NA | #VALUE! |
| ENSMUSG00000116826.2 | 0 | NA | NA | NA | NA | NA | #VALUE! |
| ENSMUSG00000116827.2 | 0 | NA | NA | NA | NA | NA | #VALUE! |
| ENSMUSG00000116828.2 | 0 | NA | NA | NA | NA | NA | #VALUE! |
| ENSMUSG00000116829.2 | 0 | NA | NA | NA | NA | NA | #VALUE! |
| ENSMUSG00000116830.2 | 0 | NA | NA | NA | NA | NA | #VALUE! |
| ENSMUSG00000116831.2 | 1.866574 | -4.43493 | 3.93059 | -1.12831 | 0.259189 | NA | #VALUE! |
| ENSMUSG00000116832.2 | 0 | NA | NA | NA | NA | NA | #VALUE! |
| ENSMUSG00000116833.2 | 0 | NA | NA | NA | NA | NA | #VALUE! |
| ENSMUSG00000116834.2 | 0 | NA | NA | NA | NA | NA | #VALUE! |
| ENSMUSG00000116837.3 | 0 | NA | NA | NA | NA | NA | #VALUE! |
| ENSMUSG00000116838.2 | 0.783156 | -3.1627 | 3.963922 | -0.79787 | 0.424945 | NA | #VALUE! |
| ENSMUSG00000116839.2 | 0 | NA | NA | NA | NA | NA | #VALUE! |
| ENSMUSG00000116840.2 | 0 | NA | NA | NA | NA | NA | #VALUE! |
| ENSMUSG00000116841.2 | 0 | NA | NA | NA | NA | NA | #VALUE! |
| ENSMUSG00000116842.2 | 0 | NA | NA | NA | NA | NA | #VALUE! |
| ENSMUSG00000116843.2 | 0 | NA | NA | NA | NA | NA | #VALUE! |

| | | | | | | | |
|---|---|---|---|---|---|---|---|
| ENSMUSG00000116845.2 | 1.120042 | 2.56892 | 3.946587 | 0.650922 | 0.515097 | NA | #VALUE! |
| ENSMUSG00000116846.2 | 0.279766 | 1.808031 | 4.036658 | 0.447903 | 0.654223 | NA | #VALUE! |
| ENSMUSG00000116847.2 | 0 | NA | NA | NA | NA | NA | #VALUE! |
| ENSMUSG00000116848.2 | 0 | NA | NA | NA | NA | NA | #VALUE! |
| ENSMUSG00000116849.2 | 0 | NA | NA | NA | NA | NA | #VALUE! |
| ENSMUSG00000116850.2 | 0 | NA | NA | NA | NA | NA | #VALUE! |
| ENSMUSG00000116851.2 | 0 | NA | NA | NA | NA | NA | #VALUE! |
| ENSMUSG00000116852.2 | 3.387647 | 5.254058 | 3.919976 | 1.340329 | 0.180138 | NA | #VALUE! |
| ENSMUSG00000116853.2 | 0 | NA | NA | NA | NA | NA | #VALUE! |
| ENSMUSG00000116854.2 | 0 | NA | NA | NA | NA | NA | #VALUE! |
| ENSMUSG00000116855.2 | 0 | NA | NA | NA | NA | NA | #VALUE! |
| ENSMUSG00000116857.3 | 0 | NA | NA | NA | NA | NA | #VALUE! |
| ENSMUSG00000116859.2 | 0 | NA | NA | NA | NA | NA | #VALUE! |
| ENSMUSG00000116860.2 | 0 | NA | NA | NA | NA | NA | #VALUE! |
| ENSMUSG00000116861.2 | 0.300802 | -1.80634 | 4.050033 | -0.44601 | 0.655592 | NA | #VALUE! |
| ENSMUSG00000116862.2 | 0 | NA | NA | NA | NA | NA | #VALUE! |
| ENSMUSG00000116863.2 | 0 | NA | NA | NA | NA | NA | #VALUE! |
| ENSMUSG00000116864.2 | 0 | NA | NA | NA | NA | NA | #VALUE! |
| ENSMUSG00000116865.2 | 0 | NA | NA | NA | NA | NA | #VALUE! |
| ENSMUSG00000116866.2 | 0 | NA | NA | NA | NA | NA | #VALUE! |
| ENSMUSG00000116867.2 | 0 | NA | NA | NA | NA | NA | #VALUE! |
| ENSMUSG00000116868.2 | 0 | NA | NA | NA | NA | NA | #VALUE! |
| ENSMUSG00000116870.2 | 0 | NA | NA | NA | NA | NA | #VALUE! |
| ENSMUSG00000116871.2 | 0 | NA | NA | NA | NA | NA | #VALUE! |
| ENSMUSG00000116872.2 | 0 | NA | NA | NA | NA | NA | #VALUE! |
| ENSMUSG00000116873.2 | 0 | NA | NA | NA | NA | NA | #VALUE! |
| ENSMUSG00000116874.2 | 0 | NA | NA | NA | NA | NA | #VALUE! |
| ENSMUSG00000116875.2 | 0 | NA | NA | NA | NA | NA | #VALUE! |
| ENSMUSG00000116876.2 | 0 | NA | NA | NA | NA | NA | #VALUE! |
| ENSMUSG00000116877.2 | 0 | NA | NA | NA | NA | NA | #VALUE! |
| ENSMUSG00000116878.2 | 0 | NA | NA | NA | NA | NA | #VALUE! |
| ENSMUSG00000116879.2 | 0 | NA | NA | NA | NA | NA | #VALUE! |
| ENSMUSG00000116880.2 | 0 | NA | NA | NA | NA | NA | #VALUE! |
| ENSMUSG00000116881.2 | 0 | NA | NA | NA | NA | NA | #VALUE! |
| ENSMUSG00000116882.2 | 0 | NA | NA | NA | NA | NA | #VALUE! |
| ENSMUSG00000116883.2 | 0 | NA | NA | NA | NA | NA | #VALUE! |
| ENSMUSG00000116884.2 | 0 | NA | NA | NA | NA | NA | #VALUE! |
| ENSMUSG00000116885.2 | 0 | NA | NA | NA | NA | NA | #VALUE! |
| ENSMUSG00000116886.2 | 0 | NA | NA | NA | NA | NA | #VALUE! |
| ENSMUSG00000116887.2 | 0 | NA | NA | NA | NA | NA | #VALUE! |
| ENSMUSG00000116888.2 | 0 | NA | NA | NA | NA | NA | #VALUE! |
| ENSMUSG00000116889.2 | 0 | NA | NA | NA | NA | NA | #VALUE! |
| ENSMUSG00000116890.2 | 0 | NA | NA | NA | NA | NA | #VALUE! |
| ENSMUSG00000116891.2 | 0 | NA | NA | NA | NA | NA | #VALUE! |

| | | | | | | | |
|---|---|---|---|---|---|---|---|
| ENSMUSG00000116892.2 | 0 | NA | NA | NA | NA | NA | #VALUE! |
| ENSMUSG00000116893.2 | 0 | NA | NA | NA | NA | NA | #VALUE! |
| ENSMUSG00000116894.2 | 0 | NA | NA | NA | NA | NA | #VALUE! |
| ENSMUSG00000116895.2 | 0 | NA | NA | NA | NA | NA | #VALUE! |
| ENSMUSG00000116896.2 | 0 | NA | NA | NA | NA | NA | #VALUE! |
| ENSMUSG00000116897.2 | 0 | NA | NA | NA | NA | NA | #VALUE! |
| ENSMUSG00000116898.2 | 2.704401 | 4.936946 | 3.92321 | 1.258394 | 0.208249 | NA | #VALUE! |
| ENSMUSG00000116899.2 | 0 | NA | NA | NA | NA | NA | #VALUE! |
| ENSMUSG00000116900.2 | 0 | NA | NA | NA | NA | NA | #VALUE! |
| ENSMUSG00000116901.2 | 0 | NA | NA | NA | NA | NA | #VALUE! |
| ENSMUSG00000116902.2 | 0 | NA | NA | NA | NA | NA | #VALUE! |
| ENSMUSG00000116904.2 | 0 | NA | NA | NA | NA | NA | #VALUE! |
| ENSMUSG00000116905.2 | 0 | NA | NA | NA | NA | NA | #VALUE! |
| ENSMUSG00000116906.2 | 0 | NA | NA | NA | NA | NA | #VALUE! |
| ENSMUSG00000116907.2 | 0 | NA | NA | NA | NA | NA | #VALUE! |
| ENSMUSG00000116908.2 | 7.572759 | 6.410662 | 3.9127 | 1.638424 | 0.101333 | NA | #VALUE! |
| ENSMUSG00000116909.2 | 0 | NA | NA | NA | NA | NA | #VALUE! |
| ENSMUSG00000116910.2 | 0 | NA | NA | NA | NA | NA | #VALUE! |
| ENSMUSG00000116911.2 | 0 | NA | NA | NA | NA | NA | #VALUE! |
| ENSMUSG00000116912.2 | 0 | NA | NA | NA | NA | NA | #VALUE! |
| ENSMUSG00000116913.2 | 0 | NA | NA | NA | NA | NA | #VALUE! |
| ENSMUSG00000116914.2 | 0 | NA | NA | NA | NA | NA | #VALUE! |
| ENSMUSG00000116915.2 | 0 | NA | NA | NA | NA | NA | #VALUE! |
| ENSMUSG00000116916.2 | 0 | NA | NA | NA | NA | NA | #VALUE! |
| ENSMUSG00000116917.2 | 0 | NA | NA | NA | NA | NA | #VALUE! |
| ENSMUSG00000116918.2 | 0 | NA | NA | NA | NA | NA | #VALUE! |
| ENSMUSG00000116919.2 | 0 | NA | NA | NA | NA | NA | #VALUE! |
| ENSMUSG00000116920.2 | 0 | NA | NA | NA | NA | NA | #VALUE! |
| ENSMUSG00000116921.2 | 0 | NA | NA | NA | NA | NA | #VALUE! |
| ENSMUSG00000116922.2 | 0 | NA | NA | NA | NA | NA | #VALUE! |
| ENSMUSG00000116923.2 | 0 | NA | NA | NA | NA | NA | #VALUE! |
| ENSMUSG00000116924.2 | 0 | NA | NA | NA | NA | NA | #VALUE! |
| ENSMUSG00000116925.2 | 0 | NA | NA | NA | NA | NA | #VALUE! |
| ENSMUSG00000116926.2 | 0 | NA | NA | NA | NA | NA | #VALUE! |
| ENSMUSG00000116927.2 | 0 | NA | NA | NA | NA | NA | #VALUE! |
| ENSMUSG00000116928.2 | 0 | NA | NA | NA | NA | NA | #VALUE! |
| ENSMUSG00000116929.2 | 0 | NA | NA | NA | NA | NA | #VALUE! |
| ENSMUSG00000116930.2 | 0 | NA | NA | NA | NA | NA | #VALUE! |
| ENSMUSG00000116931.2 | 0 | NA | NA | NA | NA | NA | #VALUE! |
| ENSMUSG00000116932.2 | 0 | NA | NA | NA | NA | NA | #VALUE! |
| ENSMUSG00000116933.2 | 0 | NA | NA | NA | NA | NA | #VALUE! |
| ENSMUSG00000116934.2 | 0 | NA | NA | NA | NA | NA | #VALUE! |
| ENSMUSG00000116935.2 | 0 | NA | NA | NA | NA | NA | #VALUE! |
| ENSMUSG00000116936.2 | 0 | NA | NA | NA | NA | NA | #VALUE! |

| | | | | | | | |
|---|---|---|---|---|---|---|---|
| ENSMUSG00000116937.2 | 0 | NA | NA | NA | NA | NA | #VALUE! |
| ENSMUSG00000116938.2 | 0 | NA | NA | NA | NA | NA | #VALUE! |
| ENSMUSG00000116939.2 | 0 | NA | NA | NA | NA | NA | #VALUE! |
| ENSMUSG00000116940.2 | 0 | NA | NA | NA | NA | NA | #VALUE! |
| ENSMUSG00000116941.2 | 0 | NA | NA | NA | NA | NA | #VALUE! |
| ENSMUSG00000116942.2 | 0 | NA | NA | NA | NA | NA | #VALUE! |
| ENSMUSG00000116943.2 | 0 | NA | NA | NA | NA | NA | #VALUE! |
| ENSMUSG00000116944.2 | 0 | NA | NA | NA | NA | NA | #VALUE! |
| ENSMUSG00000116945.2 | 0 | NA | NA | NA | NA | NA | #VALUE! |
| ENSMUSG00000116946.2 | 0 | NA | NA | NA | NA | NA | #VALUE! |
| ENSMUSG00000116947.2 | 0 | NA | NA | NA | NA | NA | #VALUE! |
| ENSMUSG00000116948.2 | 0 | NA | NA | NA | NA | NA | #VALUE! |
| ENSMUSG00000116949.2 | 0 | NA | NA | NA | NA | NA | #VALUE! |
| ENSMUSG00000116950.2 | 0 | NA | NA | NA | NA | NA | #VALUE! |
| ENSMUSG00000116951.2 | 0 | NA | NA | NA | NA | NA | #VALUE! |
| ENSMUSG00000116952.2 | 0 | NA | NA | NA | NA | NA | #VALUE! |
| ENSMUSG00000116953.2 | 0.105013 | -0.97324 | 4.076293 | -0.23876 | 0.811295 | NA | #VALUE! |
| ENSMUSG00000116954.2 | 0 | NA | NA | NA | NA | NA | #VALUE! |
| ENSMUSG00000116955.2 | 0 | NA | NA | NA | NA | NA | #VALUE! |
| ENSMUSG00000116956.2 | 0 | NA | NA | NA | NA | NA | #VALUE! |
| ENSMUSG00000116957.2 | 0 | NA | NA | NA | NA | NA | #VALUE! |
| ENSMUSG00000116958.2 | 0 | NA | NA | NA | NA | NA | #VALUE! |
| ENSMUSG00000116959.2 | 0 | NA | NA | NA | NA | NA | #VALUE! |
| ENSMUSG00000116960.2 | 0 | NA | NA | NA | NA | NA | #VALUE! |
| ENSMUSG00000116961.2 | 0 | NA | NA | NA | NA | NA | #VALUE! |
| ENSMUSG00000116962.2 | 0 | NA | NA | NA | NA | NA | #VALUE! |
| ENSMUSG00000116963.2 | 0 | NA | NA | NA | NA | NA | #VALUE! |
| ENSMUSG00000116964.2 | 0 | NA | NA | NA | NA | NA | #VALUE! |
| ENSMUSG00000116965.2 | 0 | NA | NA | NA | NA | NA | #VALUE! |
| ENSMUSG00000116966.2 | 0 | NA | NA | NA | NA | NA | #VALUE! |
| ENSMUSG00000116967.2 | 0 | NA | NA | NA | NA | NA | #VALUE! |
| ENSMUSG00000116968.2 | 0 | NA | NA | NA | NA | NA | #VALUE! |
| ENSMUSG00000116969.2 | 0 | NA | NA | NA | NA | NA | #VALUE! |
| ENSMUSG00000116970.2 | 0 | NA | NA | NA | NA | NA | #VALUE! |
| ENSMUSG00000116971.2 | 0 | NA | NA | NA | NA | NA | #VALUE! |
| ENSMUSG00000116972.2 | 0.168284 | 1.027628 | 4.067126 | 0.252667 | 0.800526 | NA | #VALUE! |
| ENSMUSG00000116973.2 | 0 | NA | NA | NA | NA | NA | #VALUE! |
| ENSMUSG00000116974.2 | 0 | NA | NA | NA | NA | NA | #VALUE! |
| ENSMUSG00000116975.2 | 0 | NA | NA | NA | NA | NA | #VALUE! |
| ENSMUSG00000116976.2 | 0 | NA | NA | NA | NA | NA | #VALUE! |
| ENSMUSG00000116977.2 | 0 | NA | NA | NA | NA | NA | #VALUE! |
| ENSMUSG00000116978.2 | 0 | NA | NA | NA | NA | NA | #VALUE! |
| ENSMUSG00000116979.2 | 0 | NA | NA | NA | NA | NA | #VALUE! |
| ENSMUSG00000116980.2 | 0 | NA | NA | NA | NA | NA | #VALUE! |

| | | | | | | | |
|---|---|---|---|---|---|---|---|
| ENSMUSG00000116981.2 | 0 | NA | NA | NA | NA | NA | #VALUE! |
| ENSMUSG00000116982.2 | 0 | NA | NA | NA | NA | NA | #VALUE! |
| ENSMUSG00000116983.2 | 0 | NA | NA | NA | NA | NA | #VALUE! |
| ENSMUSG00000116984.2 | 0 | NA | NA | NA | NA | NA | #VALUE! |
| ENSMUSG00000116985.2 | 0 | NA | NA | NA | NA | NA | #VALUE! |
| ENSMUSG00000116986.2 | 0 | NA | NA | NA | NA | NA | #VALUE! |
| ENSMUSG00000116987.2 | 0 | NA | NA | NA | NA | NA | #VALUE! |
| ENSMUSG00000116988.2 | 0 | NA | NA | NA | NA | NA | #VALUE! |
| ENSMUSG00000116989.2 | 0 | NA | NA | NA | NA | NA | #VALUE! |
| ENSMUSG00000116990.2 | 0 | NA | NA | NA | NA | NA | #VALUE! |
| ENSMUSG00000116991.2 | 0 | NA | NA | NA | NA | NA | #VALUE! |
| ENSMUSG00000116992.2 | 0 | NA | NA | NA | NA | NA | #VALUE! |
| ENSMUSG00000116994.2 | 0 | NA | NA | NA | NA | NA | #VALUE! |
| ENSMUSG00000116995.2 | 6.835422 | 6.258026 | 3.595857 | 1.740343 | 0.081799 | NA | #VALUE! |
| ENSMUSG00000116996.2 | 0 | NA | NA | NA | NA | NA | #VALUE! |
| ENSMUSG00000116997.2 | 0 | NA | NA | NA | NA | NA | #VALUE! |
| ENSMUSG00000116998.2 | 0 | NA | NA | NA | NA | NA | #VALUE! |
| ENSMUSG00000116999.2 | 0 | NA | NA | NA | NA | NA | #VALUE! |
| ENSMUSG00000117000.2 | 0 | NA | NA | NA | NA | NA | #VALUE! |
| ENSMUSG00000117001.2 | 0 | NA | NA | NA | NA | NA | #VALUE! |
| ENSMUSG00000117002.2 | 0 | NA | NA | NA | NA | NA | #VALUE! |
| ENSMUSG00000117003.2 | 0 | NA | NA | NA | NA | NA | #VALUE! |
| ENSMUSG00000117004.2 | 0 | NA | NA | NA | NA | NA | #VALUE! |
| ENSMUSG00000117005.2 | 0 | NA | NA | NA | NA | NA | #VALUE! |
| ENSMUSG00000117006.2 | 0 | NA | NA | NA | NA | NA | #VALUE! |
| ENSMUSG00000117007.2 | 0 | NA | NA | NA | NA | NA | #VALUE! |
| ENSMUSG00000117008.2 | 0 | NA | NA | NA | NA | NA | #VALUE! |
| ENSMUSG00000117009.2 | 0 | NA | NA | NA | NA | NA | #VALUE! |
| ENSMUSG00000117010.2 | 0 | NA | NA | NA | NA | NA | #VALUE! |
| ENSMUSG00000117011.2 | 0 | NA | NA | NA | NA | NA | #VALUE! |
| ENSMUSG00000117012.2 | 0 | NA | NA | NA | NA | NA | #VALUE! |
| ENSMUSG00000117013.2 | 0 | NA | NA | NA | NA | NA | #VALUE! |
| ENSMUSG00000117014.2 | 0 | NA | NA | NA | NA | NA | #VALUE! |
| ENSMUSG00000117015.2 | 0 | NA | NA | NA | NA | NA | #VALUE! |
| ENSMUSG00000117016.2 | 0 | NA | NA | NA | NA | NA | #VALUE! |
| ENSMUSG00000117017.2 | 0 | NA | NA | NA | NA | NA | #VALUE! |
| ENSMUSG00000117018.2 | 0 | NA | NA | NA | NA | NA | #VALUE! |
| ENSMUSG00000117019.2 | 0 | NA | NA | NA | NA | NA | #VALUE! |
| ENSMUSG00000117020.2 | 0 | NA | NA | NA | NA | NA | #VALUE! |
| ENSMUSG00000117021.2 | 0 | NA | NA | NA | NA | NA | #VALUE! |
| ENSMUSG00000117022.2 | 0 | NA | NA | NA | NA | NA | #VALUE! |
| ENSMUSG00000117023.2 | 0 | NA | NA | NA | NA | NA | #VALUE! |
| ENSMUSG00000117024.2 | 0 | NA | NA | NA | NA | NA | #VALUE! |
| ENSMUSG00000117025.2 | 0 | NA | NA | NA | NA | NA | #VALUE! |

| | | | | | | | |
|---|---|---|---|---|---|---|---|
| ENSMUSG00000117026.2 | 0 | NA | NA | NA | NA | NA | #VALUE! |
| ENSMUSG00000117027.2 | 0 | NA | NA | NA | NA | NA | #VALUE! |
| ENSMUSG00000117028.2 | 0 | NA | NA | NA | NA | NA | #VALUE! |
| ENSMUSG00000117029.2 | 0 | NA | NA | NA | NA | NA | #VALUE! |
| ENSMUSG00000117030.2 | 0 | NA | NA | NA | NA | NA | #VALUE! |
| ENSMUSG00000117031.2 | 0 | NA | NA | NA | NA | NA | #VALUE! |
| ENSMUSG00000117033.2 | 0 | NA | NA | NA | NA | NA | #VALUE! |
| ENSMUSG00000117035.2 | 0 | NA | NA | NA | NA | NA | #VALUE! |
| ENSMUSG00000117036.2 | 0 | NA | NA | NA | NA | NA | #VALUE! |
| ENSMUSG00000117037.2 | 0 | NA | NA | NA | NA | NA | #VALUE! |
| ENSMUSG00000117038.2 | 0 | NA | NA | NA | NA | NA | #VALUE! |
| ENSMUSG00000117039.2 | 0 | NA | NA | NA | NA | NA | #VALUE! |
| ENSMUSG00000117040.2 | 0 | NA | NA | NA | NA | NA | #VALUE! |
| ENSMUSG00000117041.2 | 0 | NA | NA | NA | NA | NA | #VALUE! |
| ENSMUSG00000117042.2 | 0 | NA | NA | NA | NA | NA | #VALUE! |
| ENSMUSG00000117043.2 | 0 | NA | NA | NA | NA | NA | #VALUE! |
| ENSMUSG00000117044.2 | 0 | NA | NA | NA | NA | NA | #VALUE! |
| ENSMUSG00000117045.2 | 0 | NA | NA | NA | NA | NA | #VALUE! |
| ENSMUSG00000117046.2 | 0 | NA | NA | NA | NA | NA | #VALUE! |
| ENSMUSG00000117047.2 | 0 | NA | NA | NA | NA | NA | #VALUE! |
| ENSMUSG00000117048.2 | 0 | NA | NA | NA | NA | NA | #VALUE! |
| ENSMUSG00000117049.2 | 0.093255 | 0.868093 | 4.072464 | 0.213162 | 0.831201 | NA | #VALUE! |
| ENSMUSG00000117050.2 | 0 | NA | NA | NA | NA | NA | #VALUE! |
| ENSMUSG00000117051.2 | 0 | NA | NA | NA | NA | NA | #VALUE! |
| ENSMUSG00000117053.2 | 0 | NA | NA | NA | NA | NA | #VALUE! |
| ENSMUSG00000117054.2 | 0 | NA | NA | NA | NA | NA | #VALUE! |
| ENSMUSG00000117055.3 | 0 | NA | NA | NA | NA | NA | #VALUE! |
| ENSMUSG00000117056.2 | 0 | NA | NA | NA | NA | NA | #VALUE! |
| ENSMUSG00000117057.2 | 0 | NA | NA | NA | NA | NA | #VALUE! |
| ENSMUSG00000117058.2 | 0 | NA | NA | NA | NA | NA | #VALUE! |
| ENSMUSG00000117059.2 | 0 | NA | NA | NA | NA | NA | #VALUE! |
| ENSMUSG00000117060.2 | 0.105013 | -0.97324 | 4.076293 | -0.23876 | 0.811295 | NA | #VALUE! |
| ENSMUSG00000117061.2 | 0 | NA | NA | NA | NA | NA | #VALUE! |
| ENSMUSG00000117062.2 | 0 | NA | NA | NA | NA | NA | #VALUE! |
| ENSMUSG00000117063.2 | 0 | NA | NA | NA | NA | NA | #VALUE! |
| ENSMUSG00000117064.2 | 0 | NA | NA | NA | NA | NA | #VALUE! |
| ENSMUSG00000117065.2 | 0 | NA | NA | NA | NA | NA | #VALUE! |
| ENSMUSG00000117066.2 | 0 | NA | NA | NA | NA | NA | #VALUE! |
| ENSMUSG00000117067.2 | 0 | NA | NA | NA | NA | NA | #VALUE! |
| ENSMUSG00000117068.3 | 0 | NA | NA | NA | NA | NA | #VALUE! |
| ENSMUSG00000117069.2 | 0 | NA | NA | NA | NA | NA | #VALUE! |
| ENSMUSG00000117070.2 | 0 | NA | NA | NA | NA | NA | #VALUE! |
| ENSMUSG00000117071.2 | 0 | NA | NA | NA | NA | NA | #VALUE! |
| ENSMUSG00000117072.2 | 0 | NA | NA | NA | NA | NA | #VALUE! |

| | | | | | | | |
|---|---|---|---|---|---|---|---|
| ENSMUSG00000117073.2 | 0 | NA | NA | NA | NA | NA | #VALUE! |
| ENSMUSG00000117074.2 | 0 | NA | NA | NA | NA | NA | #VALUE! |
| ENSMUSG00000117075.2 | 0 | NA | NA | NA | NA | NA | #VALUE! |
| ENSMUSG00000117076.2 | 0 | NA | NA | NA | NA | NA | #VALUE! |
| ENSMUSG00000117077.2 | 0 | NA | NA | NA | NA | NA | #VALUE! |
| ENSMUSG00000117078.2 | 0 | NA | NA | NA | NA | NA | #VALUE! |
| ENSMUSG00000117079.2 | 0 | NA | NA | NA | NA | NA | #VALUE! |
| ENSMUSG00000117080.2 | 0 | NA | NA | NA | NA | NA | #VALUE! |
| ENSMUSG00000117081.2 | 0 | NA | NA | NA | NA | NA | #VALUE! |
| ENSMUSG00000117082.2 | 0 | NA | NA | NA | NA | NA | #VALUE! |
| ENSMUSG00000117083.2 | 0 | NA | NA | NA | NA | NA | #VALUE! |
| ENSMUSG00000117084.2 | 27.0773 | -0.15757 | 2.630967 | -0.05989 | NA | NA | #VALUE! |
| ENSMUSG00000117085.2 | 0 | NA | NA | NA | NA | NA | #VALUE! |
| ENSMUSG00000117086.2 | 0 | NA | NA | NA | NA | NA | #VALUE! |
| ENSMUSG00000117088.2 | 0 | NA | NA | NA | NA | NA | #VALUE! |
| ENSMUSG00000117089.2 | 0 | NA | NA | NA | NA | NA | #VALUE! |
| ENSMUSG00000117090.2 | 0 | NA | NA | NA | NA | NA | #VALUE! |
| ENSMUSG00000117091.2 | 0 | NA | NA | NA | NA | NA | #VALUE! |
| ENSMUSG00000117092.2 | 0 | NA | NA | NA | NA | NA | #VALUE! |
| ENSMUSG00000117093.2 | 0 | NA | NA | NA | NA | NA | #VALUE! |
| ENSMUSG00000117094.2 | 0 | NA | NA | NA | NA | NA | #VALUE! |
| ENSMUSG00000117095.2 | 0 | NA | NA | NA | NA | NA | #VALUE! |
| ENSMUSG00000117096.2 | 0 | NA | NA | NA | NA | NA | #VALUE! |
| ENSMUSG00000117098.2 | 0 | NA | NA | NA | NA | NA | #VALUE! |
| ENSMUSG00000117099.2 | 0 | NA | NA | NA | NA | NA | #VALUE! |
| ENSMUSG00000117100.2 | 0 | NA | NA | NA | NA | NA | #VALUE! |
| ENSMUSG00000117101.2 | 0 | NA | NA | NA | NA | NA | #VALUE! |
| ENSMUSG00000117102.2 | 0 | NA | NA | NA | NA | NA | #VALUE! |
| ENSMUSG00000117103.2 | 0 | NA | NA | NA | NA | NA | #VALUE! |
| ENSMUSG00000117104.2 | 0 | NA | NA | NA | NA | NA | #VALUE! |
| ENSMUSG00000117106.2 | 0 | NA | NA | NA | NA | NA | #VALUE! |
| ENSMUSG00000117107.2 | 0 | NA | NA | NA | NA | NA | #VALUE! |
| ENSMUSG00000117108.2 | 0 | NA | NA | NA | NA | NA | #VALUE! |
| ENSMUSG00000117109.2 | 0 | NA | NA | NA | NA | NA | #VALUE! |
| ENSMUSG00000117110.2 | 0 | NA | NA | NA | NA | NA | #VALUE! |
| ENSMUSG00000117111.2 | 0 | NA | NA | NA | NA | NA | #VALUE! |
| ENSMUSG00000117112.2 | 0 | NA | NA | NA | NA | NA | #VALUE! |
| ENSMUSG00000117113.2 | 3.05105 | -0.13279 | 3.786264 | -0.03507 | 0.972023 | NA | #VALUE! |
| ENSMUSG00000117114.2 | 0 | NA | NA | NA | NA | NA | #VALUE! |
| ENSMUSG00000117115.2 | 0 | NA | NA | NA | NA | NA | #VALUE! |
| ENSMUSG00000117116.2 | 0 | NA | NA | NA | NA | NA | #VALUE! |
| ENSMUSG00000117117.2 | 0 | NA | NA | NA | NA | NA | #VALUE! |
| ENSMUSG00000117118.2 | 0 | NA | NA | NA | NA | NA | #VALUE! |
| ENSMUSG00000117119.2 | 0 | NA | NA | NA | NA | NA | #VALUE! |

| | | | | | | | |
|---|---|---|---|---|---|---|---|
| ENSMUSG00000117120.2 | 0 | NA | NA | NA | NA | NA | #VALUE! |
| ENSMUSG00000117121.2 | 0 | NA | NA | NA | NA | NA | #VALUE! |
| ENSMUSG00000117122.2 | 0 | NA | NA | NA | NA | NA | #VALUE! |
| ENSMUSG00000117123.2 | 3.357187 | 5.246639 | 3.920044 | 1.338413 | 0.180762 | NA | #VALUE! |
| ENSMUSG00000117124.2 | 0 | NA | NA | NA | NA | NA | #VALUE! |
| ENSMUSG00000117125.2 | 0 | NA | NA | NA | NA | NA | #VALUE! |
| ENSMUSG00000117126.2 | 0 | NA | NA | NA | NA | NA | #VALUE! |
| ENSMUSG00000117127.2 | 0 | NA | NA | NA | NA | NA | #VALUE! |
| ENSMUSG00000117128.2 | 0 | NA | NA | NA | NA | NA | #VALUE! |
| ENSMUSG00000117129.2 | 0 | NA | NA | NA | NA | NA | #VALUE! |
| ENSMUSG00000117130.2 | 0 | NA | NA | NA | NA | NA | #VALUE! |
| ENSMUSG00000117131.2 | 0 | NA | NA | NA | NA | NA | #VALUE! |
| ENSMUSG00000117132.2 | 0 | NA | NA | NA | NA | NA | #VALUE! |
| ENSMUSG00000117133.2 | 0 | NA | NA | NA | NA | NA | #VALUE! |
| ENSMUSG00000117134.2 | 0 | NA | NA | NA | NA | NA | #VALUE! |
| ENSMUSG00000117135.2 | 0 | NA | NA | NA | NA | NA | #VALUE! |
| ENSMUSG00000117136.2 | 0 | NA | NA | NA | NA | NA | #VALUE! |
| ENSMUSG00000117137.2 | 0 | NA | NA | NA | NA | NA | #VALUE! |
| ENSMUSG00000117138.2 | 0 | NA | NA | NA | NA | NA | #VALUE! |
| ENSMUSG00000117139.2 | 0 | NA | NA | NA | NA | NA | #VALUE! |
| ENSMUSG00000117140.2 | 0 | NA | NA | NA | NA | NA | #VALUE! |
| ENSMUSG00000117141.2 | 0 | NA | NA | NA | NA | NA | #VALUE! |
| ENSMUSG00000117142.2 | 0 | NA | NA | NA | NA | NA | #VALUE! |
| ENSMUSG00000117143.2 | 0 | NA | NA | NA | NA | NA | #VALUE! |
| ENSMUSG00000117145.2 | 4.698938 | -5.75981 | 3.916303 | -1.47073 | 0.141365 | NA | #VALUE! |
| ENSMUSG00000117146.3 | 0 | NA | NA | NA | NA | NA | #VALUE! |
| ENSMUSG00000117147.2 | 0 | NA | NA | NA | NA | NA | #VALUE! |
| ENSMUSG00000117148.2 | 0 | NA | NA | NA | NA | NA | #VALUE! |
| ENSMUSG00000117149.2 | 0 | NA | NA | NA | NA | NA | #VALUE! |
| ENSMUSG00000117150.2 | 0 | NA | NA | NA | NA | NA | #VALUE! |
| ENSMUSG00000117151.2 | 0 | NA | NA | NA | NA | NA | #VALUE! |
| ENSMUSG00000117154.2 | 0 | NA | NA | NA | NA | NA | #VALUE! |
| ENSMUSG00000117156.2 | 0 | NA | NA | NA | NA | NA | #VALUE! |
| ENSMUSG00000117157.2 | 0 | NA | NA | NA | NA | NA | #VALUE! |
| ENSMUSG00000117158.2 | 0 | NA | NA | NA | NA | NA | #VALUE! |
| ENSMUSG00000117159.2 | 0 | NA | NA | NA | NA | NA | #VALUE! |
| ENSMUSG00000117160.2 | 0 | NA | NA | NA | NA | NA | #VALUE! |
| ENSMUSG00000117161.2 | 0 | NA | NA | NA | NA | NA | #VALUE! |
| ENSMUSG00000117162.2 | 0 | NA | NA | NA | NA | NA | #VALUE! |
| ENSMUSG00000117163.2 | 0 | NA | NA | NA | NA | NA | #VALUE! |
| ENSMUSG00000117164.2 | 0 | NA | NA | NA | NA | NA | #VALUE! |
| ENSMUSG00000117165.2 | 0 | NA | NA | NA | NA | NA | #VALUE! |
| ENSMUSG00000117166.2 | 0 | NA | NA | NA | NA | NA | #VALUE! |
| ENSMUSG00000117167.2 | 0.093255 | 0.868093 | 4.072464 | 0.213162 | 0.831201 | NA | #VALUE! |

| | | | | | | | |
|---|---|---|---|---|---|---|---|
| ENSMUSG00000117168.2 | 0 | NA | NA | NA | NA | NA | #VALUE! |
| ENSMUSG00000117169.3 | 0 | NA | NA | NA | NA | NA | #VALUE! |
| ENSMUSG00000117170.2 | 0 | NA | NA | NA | NA | NA | #VALUE! |
| ENSMUSG00000117171.2 | 0 | NA | NA | NA | NA | NA | #VALUE! |
| ENSMUSG00000117172.2 | 0 | NA | NA | NA | NA | NA | #VALUE! |
| ENSMUSG00000117174.2 | 0 | NA | NA | NA | NA | NA | #VALUE! |
| ENSMUSG00000117175.2 | 0 | NA | NA | NA | NA | NA | #VALUE! |
| ENSMUSG00000117176.2 | 0 | NA | NA | NA | NA | NA | #VALUE! |
| ENSMUSG00000117177.2 | 0 | NA | NA | NA | NA | NA | #VALUE! |
| ENSMUSG00000117178.2 | 0 | NA | NA | NA | NA | NA | #VALUE! |
| ENSMUSG00000117179.2 | 0 | NA | NA | NA | NA | NA | #VALUE! |
| ENSMUSG00000117180.2 | 0 | NA | NA | NA | NA | NA | #VALUE! |
| ENSMUSG00000117181.2 | 0 | NA | NA | NA | NA | NA | #VALUE! |
| ENSMUSG00000117182.2 | 0 | NA | NA | NA | NA | NA | #VALUE! |
| ENSMUSG00000117183.2 | 1.398828 | 3.99633 | 3.938205 | 1.014759 | 0.310221 | NA | #VALUE! |
| ENSMUSG00000117184.2 | 0 | NA | NA | NA | NA | NA | #VALUE! |
| ENSMUSG00000117185.2 | 0 | NA | NA | NA | NA | NA | #VALUE! |
| ENSMUSG00000117186.2 | 0 | NA | NA | NA | NA | NA | #VALUE! |
| ENSMUSG00000117187.2 | 4.428538 | 0.717733 | 3.374692 | 0.212681 | 0.831576 | NA | #VALUE! |
| ENSMUSG00000117188.2 | 0 | NA | NA | NA | NA | NA | #VALUE! |
| ENSMUSG00000117189.2 | 0 | NA | NA | NA | NA | NA | #VALUE! |
| ENSMUSG00000117190.2 | 0 | NA | NA | NA | NA | NA | #VALUE! |
| ENSMUSG00000117191.2 | 0 | NA | NA | NA | NA | NA | #VALUE! |
| ENSMUSG00000117192.2 | 0 | NA | NA | NA | NA | NA | #VALUE! |
| ENSMUSG00000117193.2 | 0 | NA | NA | NA | NA | NA | #VALUE! |
| ENSMUSG00000117194.2 | 0 | NA | NA | NA | NA | NA | #VALUE! |
| ENSMUSG00000117195.2 | 0 | NA | NA | NA | NA | NA | #VALUE! |
| ENSMUSG00000117196.2 | 0 | NA | NA | NA | NA | NA | #VALUE! |
| ENSMUSG00000117197.2 | 0 | NA | NA | NA | NA | NA | #VALUE! |
| ENSMUSG00000117198.2 | 0 | NA | NA | NA | NA | NA | #VALUE! |
| ENSMUSG00000117199.2 | 0 | NA | NA | NA | NA | NA | #VALUE! |
| ENSMUSG00000117200.2 | 0 | NA | NA | NA | NA | NA | #VALUE! |
| ENSMUSG00000117201.2 | 0 | NA | NA | NA | NA | NA | #VALUE! |
| ENSMUSG00000117202.2 | 0 | NA | NA | NA | NA | NA | #VALUE! |
| ENSMUSG00000117203.2 | 0 | NA | NA | NA | NA | NA | #VALUE! |
| ENSMUSG00000117204.2 | 0 | NA | NA | NA | NA | NA | #VALUE! |
| ENSMUSG00000117205.2 | 0 | NA | NA | NA | NA | NA | #VALUE! |
| ENSMUSG00000117206.2 | 0 | NA | NA | NA | NA | NA | #VALUE! |
| ENSMUSG00000117207.2 | 0 | NA | NA | NA | NA | NA | #VALUE! |
| ENSMUSG00000117208.2 | 0 | NA | NA | NA | NA | NA | #VALUE! |
| ENSMUSG00000117209.2 | 0 | NA | NA | NA | NA | NA | #VALUE! |
| ENSMUSG00000117210.2 | 0 | NA | NA | NA | NA | NA | #VALUE! |
| ENSMUSG00000117211.2 | 0 | NA | NA | NA | NA | NA | #VALUE! |
| ENSMUSG00000117213.2 | 0 | NA | NA | NA | NA | NA | #VALUE! |

| | | | | | | | |
|---|---|---|---|---|---|---|---|
| ENSMUSG00000117214.2 | 0 | NA | NA | NA | NA | NA | #VALUE! |
| ENSMUSG00000117215.2 | 0 | NA | NA | NA | NA | NA | #VALUE! |
| ENSMUSG00000117216.2 | 0 | NA | NA | NA | NA | NA | #VALUE! |
| ENSMUSG00000117217.2 | 0 | NA | NA | NA | NA | NA | #VALUE! |
| ENSMUSG00000117218.2 | 0 | NA | NA | NA | NA | NA | #VALUE! |
| ENSMUSG00000117219.2 | 0 | NA | NA | NA | NA | NA | #VALUE! |
| ENSMUSG00000117220.2 | 0 | NA | NA | NA | NA | NA | #VALUE! |
| ENSMUSG00000117221.2 | 0 | NA | NA | NA | NA | NA | #VALUE! |
| ENSMUSG00000117222.2 | 0 | NA | NA | NA | NA | NA | #VALUE! |
| ENSMUSG00000117223.2 | 0 | NA | NA | NA | NA | NA | #VALUE! |
| ENSMUSG00000117224.2 | 0 | NA | NA | NA | NA | NA | #VALUE! |
| ENSMUSG00000117225.2 | 0 | NA | NA | NA | NA | NA | #VALUE! |
| ENSMUSG00000117227.2 | 0 | NA | NA | NA | NA | NA | #VALUE! |
| ENSMUSG00000117228.2 | 0 | NA | NA | NA | NA | NA | #VALUE! |
| ENSMUSG00000117229.2 | 0 | NA | NA | NA | NA | NA | #VALUE! |
| ENSMUSG00000117230.2 | 0.168284 | 1.027628 | 4.067126 | 0.252667 | 0.800526 | NA | #VALUE! |
| ENSMUSG00000117231.2 | 0.105013 | -0.97324 | 4.076293 | -0.23876 | 0.811295 | NA | #VALUE! |
| ENSMUSG00000117232.2 | 0 | NA | NA | NA | NA | NA | #VALUE! |
| ENSMUSG00000117233.2 | 0 | NA | NA | NA | NA | NA | #VALUE! |
| ENSMUSG00000117234.2 | 4.91951 | 5.781601 | 3.915938 | 1.476428 | 0.139829 | NA | #VALUE! |
| ENSMUSG00000117235.2 | 0 | NA | NA | NA | NA | NA | #VALUE! |
| ENSMUSG00000117236.2 | 0 | NA | NA | NA | NA | NA | #VALUE! |
| ENSMUSG00000117237.2 | 0 | NA | NA | NA | NA | NA | #VALUE! |
| ENSMUSG00000117238.2 | 0 | NA | NA | NA | NA | NA | #VALUE! |
| ENSMUSG00000117239.2 | 0 | NA | NA | NA | NA | NA | #VALUE! |
| ENSMUSG00000117240.2 | 0 | NA | NA | NA | NA | NA | #VALUE! |
| ENSMUSG00000117241.2 | 0 | NA | NA | NA | NA | NA | #VALUE! |
| ENSMUSG00000117243.2 | 0 | NA | NA | NA | NA | NA | #VALUE! |
| ENSMUSG00000117244.2 | 0 | NA | NA | NA | NA | NA | #VALUE! |
| ENSMUSG00000117245.2 | 0 | NA | NA | NA | NA | NA | #VALUE! |
| ENSMUSG00000117246.2 | 0 | NA | NA | NA | NA | NA | #VALUE! |
| ENSMUSG00000117247.2 | 0 | NA | NA | NA | NA | NA | #VALUE! |
| ENSMUSG00000117248.2 | 0 | NA | NA | NA | NA | NA | #VALUE! |
| ENSMUSG00000117249.2 | 0 | NA | NA | NA | NA | NA | #VALUE! |
| ENSMUSG00000117250.2 | 0 | NA | NA | NA | NA | NA | #VALUE! |
| ENSMUSG00000117251.2 | 0 | NA | NA | NA | NA | NA | #VALUE! |
| ENSMUSG00000117252.2 | 0.204111 | -1.05581 | 4.073302 | -0.2592 | 0.795479 | NA | #VALUE! |
| ENSMUSG00000117253.2 | 0 | NA | NA | NA | NA | NA | #VALUE! |
| ENSMUSG00000117254.2 | 0 | NA | NA | NA | NA | NA | #VALUE! |
| ENSMUSG00000117255.2 | 0 | NA | NA | NA | NA | NA | #VALUE! |
| ENSMUSG00000117256.2 | 0 | NA | NA | NA | NA | NA | #VALUE! |
| ENSMUSG00000117257.2 | 0 | NA | NA | NA | NA | NA | #VALUE! |
| ENSMUSG00000117258.2 | 0 | NA | NA | NA | NA | NA | #VALUE! |
| ENSMUSG00000117259.2 | 0 | NA | NA | NA | NA | NA | #VALUE! |

| | | | | | | | |
|---|---|---|---|---|---|---|---|
| ENSMUSG00000117260.2 | 0 | NA | NA | NA | NA | NA | #VALUE! |
| ENSMUSG00000117262.2 | 0 | NA | NA | NA | NA | NA | #VALUE! |
| ENSMUSG00000117263.2 | 0 | NA | NA | NA | NA | NA | #VALUE! |
| ENSMUSG00000117264.2 | 0 | NA | NA | NA | NA | NA | #VALUE! |
| ENSMUSG00000117265.2 | 0 | NA | NA | NA | NA | NA | #VALUE! |
| ENSMUSG00000117266.2 | 0 | NA | NA | NA | NA | NA | #VALUE! |
| ENSMUSG00000117267.2 | 0 | NA | NA | NA | NA | NA | #VALUE! |
| ENSMUSG00000117268.2 | 0 | NA | NA | NA | NA | NA | #VALUE! |
| ENSMUSG00000117269.2 | 0 | NA | NA | NA | NA | NA | #VALUE! |
| ENSMUSG00000117270.2 | 0 | NA | NA | NA | NA | NA | #VALUE! |
| ENSMUSG00000117271.2 | 0 | NA | NA | NA | NA | NA | #VALUE! |
| ENSMUSG00000117272.2 | 0 | NA | NA | NA | NA | NA | #VALUE! |
| ENSMUSG00000117273.2 | 0 | NA | NA | NA | NA | NA | #VALUE! |
| ENSMUSG00000117274.2 | 0 | NA | NA | NA | NA | NA | #VALUE! |
| ENSMUSG00000117275.2 | 0 | NA | NA | NA | NA | NA | #VALUE! |
| ENSMUSG00000117276.2 | 0 | NA | NA | NA | NA | NA | #VALUE! |
| ENSMUSG00000117277.2 | 0 | NA | NA | NA | NA | NA | #VALUE! |
| ENSMUSG00000117278.2 | 0 | NA | NA | NA | NA | NA | #VALUE! |
| ENSMUSG00000117279.2 | 0 | NA | NA | NA | NA | NA | #VALUE! |
| ENSMUSG00000117280.2 | 0 | NA | NA | NA | NA | NA | #VALUE! |
| ENSMUSG00000117281.2 | 0 | NA | NA | NA | NA | NA | #VALUE! |
| ENSMUSG00000117282.2 | 0 | NA | NA | NA | NA | NA | #VALUE! |
| ENSMUSG00000117283.2 | 0 | NA | NA | NA | NA | NA | #VALUE! |
| ENSMUSG00000117285.2 | 0 | NA | NA | NA | NA | NA | #VALUE! |
| ENSMUSG00000117286.2 | 0 | NA | NA | NA | NA | NA | #VALUE! |
| ENSMUSG00000117287.2 | 0 | NA | NA | NA | NA | NA | #VALUE! |
| ENSMUSG00000117288.2 | 0 | NA | NA | NA | NA | NA | #VALUE! |
| ENSMUSG00000117289.2 | 0 | NA | NA | NA | NA | NA | #VALUE! |
| ENSMUSG00000117290.2 | 0 | NA | NA | NA | NA | NA | #VALUE! |
| ENSMUSG00000117291.2 | 0 | NA | NA | NA | NA | NA | #VALUE! |
| ENSMUSG00000117292.2 | 0 | NA | NA | NA | NA | NA | #VALUE! |
| ENSMUSG00000117293.2 | 0 | NA | NA | NA | NA | NA | #VALUE! |
| ENSMUSG00000117294.2 | 2.520315 | -4.8742 | 3.924358 | -1.24204 | 0.214223 | NA | #VALUE! |
| ENSMUSG00000117295.2 | 0 | NA | NA | NA | NA | NA | #VALUE! |
| ENSMUSG00000117296.2 | 0 | NA | NA | NA | NA | NA | #VALUE! |
| ENSMUSG00000117297.2 | 0 | NA | NA | NA | NA | NA | #VALUE! |
| ENSMUSG00000117298.2 | 0 | NA | NA | NA | NA | NA | #VALUE! |
| ENSMUSG00000117299.2 | 0 | NA | NA | NA | NA | NA | #VALUE! |
| ENSMUSG00000117300.2 | 0 | NA | NA | NA | NA | NA | #VALUE! |
| ENSMUSG00000117301.2 | 0 | NA | NA | NA | NA | NA | #VALUE! |
| ENSMUSG00000117302.2 | 0 | NA | NA | NA | NA | NA | #VALUE! |
| ENSMUSG00000117303.2 | 0 | NA | NA | NA | NA | NA | #VALUE! |
| ENSMUSG00000117304.2 | 0 | NA | NA | NA | NA | NA | #VALUE! |
| ENSMUSG00000117305.2 | 0 | NA | NA | NA | NA | NA | #VALUE! |

| | | | | | | | |
|---|---|---|---|---|---|---|---|
| ENSMUSG00000117306.2 | 0 | NA | NA | NA | NA | NA | #VALUE! |
| ENSMUSG00000117307.2 | 0 | NA | NA | NA | NA | NA | #VALUE! |
| ENSMUSG00000117308.2 | 0 | NA | NA | NA | NA | NA | #VALUE! |
| ENSMUSG00000117309.2 | 0 | NA | NA | NA | NA | NA | #VALUE! |
| ENSMUSG00000117310.3 | 0 | NA | NA | NA | NA | NA | #VALUE! |
| ENSMUSG00000117311.2 | 0 | NA | NA | NA | NA | NA | #VALUE! |
| ENSMUSG00000117312.2 | 0 | NA | NA | NA | NA | NA | #VALUE! |
| ENSMUSG00000117313.2 | 0 | NA | NA | NA | NA | NA | #VALUE! |
| ENSMUSG00000117314.2 | 0 | NA | NA | NA | NA | NA | #VALUE! |
| ENSMUSG00000117315.2 | 0 | NA | NA | NA | NA | NA | #VALUE! |
| ENSMUSG00000117316.2 | 0 | NA | NA | NA | NA | NA | #VALUE! |
| ENSMUSG00000117317.2 | 0 | NA | NA | NA | NA | NA | #VALUE! |
| ENSMUSG00000117318.2 | 0 | NA | NA | NA | NA | NA | #VALUE! |
| ENSMUSG00000117319.2 | 0 | NA | NA | NA | NA | NA | #VALUE! |
| ENSMUSG00000117320.2 | 0 | NA | NA | NA | NA | NA | #VALUE! |
| ENSMUSG00000117321.2 | 0 | NA | NA | NA | NA | NA | #VALUE! |
| ENSMUSG00000117322.2 | 1.682835 | 4.240781 | 3.933339 | 1.078163 | 0.280961 | NA | #VALUE! |
| ENSMUSG00000117323.2 | 0 | NA | NA | NA | NA | NA | #VALUE! |
| ENSMUSG00000117324.2 | 0 | NA | NA | NA | NA | NA | #VALUE! |
| ENSMUSG00000117325.2 | 0 | NA | NA | NA | NA | NA | #VALUE! |
| ENSMUSG00000117326.2 | 0 | NA | NA | NA | NA | NA | #VALUE! |
| ENSMUSG00000117328.2 | 0 | NA | NA | NA | NA | NA | #VALUE! |
| ENSMUSG00000117329.2 | 0 | NA | NA | NA | NA | NA | #VALUE! |
| ENSMUSG00000117330.2 | 0 | NA | NA | NA | NA | NA | #VALUE! |
| ENSMUSG00000117331.2 | 0 | NA | NA | NA | NA | NA | #VALUE! |
| ENSMUSG00000117332.2 | 0 | NA | NA | NA | NA | NA | #VALUE! |
| ENSMUSG00000117333.2 | 0 | NA | NA | NA | NA | NA | #VALUE! |
| ENSMUSG00000117334.2 | 0 | NA | NA | NA | NA | NA | #VALUE! |
| ENSMUSG00000117335.2 | 0 | NA | NA | NA | NA | NA | #VALUE! |
| ENSMUSG00000117336.2 | 0 | NA | NA | NA | NA | NA | #VALUE! |
| ENSMUSG00000117337.2 | 0 | NA | NA | NA | NA | NA | #VALUE! |
| ENSMUSG00000117338.2 | 0 | NA | NA | NA | NA | NA | #VALUE! |
| ENSMUSG00000117340.2 | 0 | NA | NA | NA | NA | NA | #VALUE! |
| ENSMUSG00000117341.2 | 0 | NA | NA | NA | NA | NA | #VALUE! |
| ENSMUSG00000117342.2 | 0 | NA | NA | NA | NA | NA | #VALUE! |
| ENSMUSG00000117343.2 | 0 | NA | NA | NA | NA | NA | #VALUE! |
| ENSMUSG00000117344.2 | 0 | NA | NA | NA | NA | NA | #VALUE! |
| ENSMUSG00000117345.2 | 0.585206 | 2.680529 | 3.984084 | 0.672809 | 0.501069 | NA | #VALUE! |
| ENSMUSG00000117346.2 | 0 | NA | NA | NA | NA | NA | #VALUE! |
| ENSMUSG00000117347.2 | 0 | NA | NA | NA | NA | NA | #VALUE! |
| ENSMUSG00000117348.2 | 0 | NA | NA | NA | NA | NA | #VALUE! |
| ENSMUSG00000117349.2 | 0 | NA | NA | NA | NA | NA | #VALUE! |
| ENSMUSG00000117350.2 | 0 | NA | NA | NA | NA | NA | #VALUE! |
| ENSMUSG00000117351.2 | 0 | NA | NA | NA | NA | NA | #VALUE! |

| | | | | | | | |
|---|---|---|---|---|---|---|---|
| ENSMUSG00000117352.2 | 0 | NA | NA | NA | NA | NA | #VALUE! |
| ENSMUSG00000117353.2 | 0 | NA | NA | NA | NA | NA | #VALUE! |
| ENSMUSG00000117354.2 | 0 | NA | NA | NA | NA | NA | #VALUE! |
| ENSMUSG00000117355.2 | 0 | NA | NA | NA | NA | NA | #VALUE! |
| ENSMUSG00000117356.2 | 0 | NA | NA | NA | NA | NA | #VALUE! |
| ENSMUSG00000117357.2 | 0 | NA | NA | NA | NA | NA | #VALUE! |
| ENSMUSG00000117358.2 | 0 | NA | NA | NA | NA | NA | #VALUE! |
| ENSMUSG00000117359.2 | 0 | NA | NA | NA | NA | NA | #VALUE! |
| ENSMUSG00000117360.2 | 0 | NA | NA | NA | NA | NA | #VALUE! |
| ENSMUSG00000117361.2 | 0 | NA | NA | NA | NA | NA | #VALUE! |
| ENSMUSG00000117362.2 | 0 | NA | NA | NA | NA | NA | #VALUE! |
| ENSMUSG00000117363.2 | 0 | NA | NA | NA | NA | NA | #VALUE! |
| ENSMUSG00000117364.2 | 0 | NA | NA | NA | NA | NA | #VALUE! |
| ENSMUSG00000117365.2 | 0 | NA | NA | NA | NA | NA | #VALUE! |
| ENSMUSG00000117366.2 | 0 | NA | NA | NA | NA | NA | #VALUE! |
| ENSMUSG00000117367.2 | 0 | NA | NA | NA | NA | NA | #VALUE! |
| ENSMUSG00000117368.2 | 0 | NA | NA | NA | NA | NA | #VALUE! |
| ENSMUSG00000117369.2 | 0 | NA | NA | NA | NA | NA | #VALUE! |
| ENSMUSG00000117370.2 | 0 | NA | NA | NA | NA | NA | #VALUE! |
| ENSMUSG00000117371.2 | 0 | NA | NA | NA | NA | NA | #VALUE! |
| ENSMUSG00000117372.2 | 0 | NA | NA | NA | NA | NA | #VALUE! |
| ENSMUSG00000117373.2 | 0 | NA | NA | NA | NA | NA | #VALUE! |
| ENSMUSG00000117374.2 | 0 | NA | NA | NA | NA | NA | #VALUE! |
| ENSMUSG00000117375.2 | 0 | NA | NA | NA | NA | NA | #VALUE! |
| ENSMUSG00000117376.2 | 0 | NA | NA | NA | NA | NA | #VALUE! |
| ENSMUSG00000117377.2 | 0 | NA | NA | NA | NA | NA | #VALUE! |
| ENSMUSG00000117378.2 | 0 | NA | NA | NA | NA | NA | #VALUE! |
| ENSMUSG00000117379.2 | 0 | NA | NA | NA | NA | NA | #VALUE! |
| ENSMUSG00000117380.2 | 0 | NA | NA | NA | NA | NA | #VALUE! |
| ENSMUSG00000117381.2 | 0 | NA | NA | NA | NA | NA | #VALUE! |
| ENSMUSG00000117382.2 | 0 | NA | NA | NA | NA | NA | #VALUE! |
| ENSMUSG00000117383.2 | 0 | NA | NA | NA | NA | NA | #VALUE! |
| ENSMUSG00000117384.2 | 0 | NA | NA | NA | NA | NA | #VALUE! |
| ENSMUSG00000117385.2 | 0 | NA | NA | NA | NA | NA | #VALUE! |
| ENSMUSG00000117386.2 | 0 | NA | NA | NA | NA | NA | #VALUE! |
| ENSMUSG00000117387.2 | 0 | NA | NA | NA | NA | NA | #VALUE! |
| ENSMUSG00000117388.2 | 0 | NA | NA | NA | NA | NA | #VALUE! |
| ENSMUSG00000117389.2 | 0 | NA | NA | NA | NA | NA | #VALUE! |
| ENSMUSG00000117390.2 | 0 | NA | NA | NA | NA | NA | #VALUE! |
| ENSMUSG00000117391.2 | 0 | NA | NA | NA | NA | NA | #VALUE! |
| ENSMUSG00000117392.2 | 0 | NA | NA | NA | NA | NA | #VALUE! |
| ENSMUSG00000117393.2 | 0 | NA | NA | NA | NA | NA | #VALUE! |
| ENSMUSG00000117394.2 | 0 | NA | NA | NA | NA | NA | #VALUE! |
| ENSMUSG00000117395.2 | 0 | NA | NA | NA | NA | NA | #VALUE! |

| | | | | | | | |
|---|---|---|---|---|---|---|---|
| ENSMUSG00000117396.2 | 0 | NA | NA | NA | NA | NA | #VALUE! |
| ENSMUSG00000117397.2 | 0 | NA | NA | NA | NA | NA | #VALUE! |
| ENSMUSG00000117398.2 | 0 | NA | NA | NA | NA | NA | #VALUE! |
| ENSMUSG00000117399.2 | 0 | NA | NA | NA | NA | NA | #VALUE! |
| ENSMUSG00000117400.2 | 0 | NA | NA | NA | NA | NA | #VALUE! |
| ENSMUSG00000117401.2 | 1.174735 | -3.75246 | 3.944889 | -0.95122 | 0.341493 | NA | #VALUE! |
| ENSMUSG00000117402.2 | 0 | NA | NA | NA | NA | NA | #VALUE! |
| ENSMUSG00000117404.2 | 0 | NA | NA | NA | NA | NA | #VALUE! |
| ENSMUSG00000117405.2 | 0.105013 | -0.97324 | 4.076293 | -0.23876 | 0.811295 | NA | #VALUE! |
| ENSMUSG00000117406.2 | 8.984013 | -1.55669 | 2.69662 | -0.57727 | 0.563755 | NA | #VALUE! |
| ENSMUSG00000117407.2 | 0 | NA | NA | NA | NA | NA | #VALUE! |
| ENSMUSG00000117408.2 | 0 | NA | NA | NA | NA | NA | #VALUE! |
| ENSMUSG00000117409.2 | 0 | NA | NA | NA | NA | NA | #VALUE! |
| ENSMUSG00000117410.2 | 0 | NA | NA | NA | NA | NA | #VALUE! |
| ENSMUSG00000117411.2 | 0 | NA | NA | NA | NA | NA | #VALUE! |
| ENSMUSG00000117412.2 | 0 | NA | NA | NA | NA | NA | #VALUE! |
| ENSMUSG00000117413.2 | 0 | NA | NA | NA | NA | NA | #VALUE! |
| ENSMUSG00000117414.2 | 0 | NA | NA | NA | NA | NA | #VALUE! |
| ENSMUSG00000117415.2 | 0 | NA | NA | NA | NA | NA | #VALUE! |
| ENSMUSG00000117416.2 | 0 | NA | NA | NA | NA | NA | #VALUE! |
| ENSMUSG00000117417.2 | 0 | NA | NA | NA | NA | NA | #VALUE! |
| ENSMUSG00000117418.2 | 7.439986 | -6.42369 | 3.91279 | -1.64172 | 0.100649 | NA | #VALUE! |
| ENSMUSG00000117419.2 | 0 | NA | NA | NA | NA | NA | #VALUE! |
| ENSMUSG00000117420.2 | 0 | NA | NA | NA | NA | NA | #VALUE! |
| ENSMUSG00000117421.2 | 0 | NA | NA | NA | NA | NA | #VALUE! |
| ENSMUSG00000117422.2 | 0 | NA | NA | NA | NA | NA | #VALUE! |
| ENSMUSG00000117423.2 | 0 | NA | NA | NA | NA | NA | #VALUE! |
| ENSMUSG00000117424.2 | 0 | NA | NA | NA | NA | NA | #VALUE! |
| ENSMUSG00000117425.2 | 0 | NA | NA | NA | NA | NA | #VALUE! |
| ENSMUSG00000117426.2 | 0 | NA | NA | NA | NA | NA | #VALUE! |
| ENSMUSG00000117427.2 | 0 | NA | NA | NA | NA | NA | #VALUE! |
| ENSMUSG00000117428.2 | 0 | NA | NA | NA | NA | NA | #VALUE! |
| ENSMUSG00000117429.2 | 0 | NA | NA | NA | NA | NA | #VALUE! |
| ENSMUSG00000117430.2 | 0 | NA | NA | NA | NA | NA | #VALUE! |
| ENSMUSG00000117431.2 | 0 | NA | NA | NA | NA | NA | #VALUE! |
| ENSMUSG00000117432.2 | 0 | NA | NA | NA | NA | NA | #VALUE! |
| ENSMUSG00000117433.2 | 0 | NA | NA | NA | NA | NA | #VALUE! |
| ENSMUSG00000117434.2 | 0 | NA | NA | NA | NA | NA | #VALUE! |
| ENSMUSG00000117435.2 | 0 | NA | NA | NA | NA | NA | #VALUE! |
| ENSMUSG00000117436.2 | 0 | NA | NA | NA | NA | NA | #VALUE! |
| ENSMUSG00000117437.2 | 0 | NA | NA | NA | NA | NA | #VALUE! |
| ENSMUSG00000117438.2 | 0.279766 | 1.808031 | 4.036658 | 0.447903 | 0.654223 | NA | #VALUE! |
| ENSMUSG00000117439.2 | 0 | NA | NA | NA | NA | NA | #VALUE! |
| ENSMUSG00000117440.2 | 0 | NA | NA | NA | NA | NA | #VALUE! |

| | | | | | | | |
|---|---|---|---|---|---|---|---|
| ENSMUSG00000117441.2 | 0 | NA | NA | NA | NA | NA | #VALUE! |
| ENSMUSG00000117442.2 | 0 | NA | NA | NA | NA | NA | #VALUE! |
| ENSMUSG00000117443.2 | 0 | NA | NA | NA | NA | NA | #VALUE! |
| ENSMUSG00000117444.2 | 0 | NA | NA | NA | NA | NA | #VALUE! |
| ENSMUSG00000117445.2 | 0 | NA | NA | NA | NA | NA | #VALUE! |
| ENSMUSG00000117446.2 | 0 | NA | NA | NA | NA | NA | #VALUE! |
| ENSMUSG00000117447.2 | 0 | NA | NA | NA | NA | NA | #VALUE! |
| ENSMUSG00000117448.2 | 0 | NA | NA | NA | NA | NA | #VALUE! |
| ENSMUSG00000117449.2 | 0 | NA | NA | NA | NA | NA | #VALUE! |
| ENSMUSG00000117450.2 | 0 | NA | NA | NA | NA | NA | #VALUE! |
| ENSMUSG00000117451.2 | 0 | NA | NA | NA | NA | NA | #VALUE! |
| ENSMUSG00000117452.2 | 0 | NA | NA | NA | NA | NA | #VALUE! |
| ENSMUSG00000117453.2 | 0 | NA | NA | NA | NA | NA | #VALUE! |
| ENSMUSG00000117454.2 | 0 | NA | NA | NA | NA | NA | #VALUE! |
| ENSMUSG00000117455.2 | 0 | NA | NA | NA | NA | NA | #VALUE! |
| ENSMUSG00000117456.2 | 0 | NA | NA | NA | NA | NA | #VALUE! |
| ENSMUSG00000117457.2 | 0 | NA | NA | NA | NA | NA | #VALUE! |
| ENSMUSG00000117458.2 | 0.89897 | -2.30704 | 3.953647 | -0.58352 | 0.559542 | NA | #VALUE! |
| ENSMUSG00000117459.2 | 0 | NA | NA | NA | NA | NA | #VALUE! |
| ENSMUSG00000117460.2 | 0 | NA | NA | NA | NA | NA | #VALUE! |
| ENSMUSG00000117461.2 | 0 | NA | NA | NA | NA | NA | #VALUE! |
| ENSMUSG00000117462.2 | 0 | NA | NA | NA | NA | NA | #VALUE! |
| ENSMUSG00000117463.2 | 0 | NA | NA | NA | NA | NA | #VALUE! |
| ENSMUSG00000117464.2 | 0 | NA | NA | NA | NA | NA | #VALUE! |
| ENSMUSG00000117465.2 | 0 | NA | NA | NA | NA | NA | #VALUE! |
| ENSMUSG00000117466.2 | 0 | NA | NA | NA | NA | NA | #VALUE! |
| ENSMUSG00000117467.2 | 0 | NA | NA | NA | NA | NA | #VALUE! |
| ENSMUSG00000117468.2 | 0 | NA | NA | NA | NA | NA | #VALUE! |
| ENSMUSG00000117469.2 | 0 | NA | NA | NA | NA | NA | #VALUE! |
| ENSMUSG00000117470.2 | 0 | NA | NA | NA | NA | NA | #VALUE! |
| ENSMUSG00000117471.2 | 0 | NA | NA | NA | NA | NA | #VALUE! |
| ENSMUSG00000117472.2 | 0 | NA | NA | NA | NA | NA | #VALUE! |
| ENSMUSG00000117473.2 | 0 | NA | NA | NA | NA | NA | #VALUE! |
| ENSMUSG00000117474.2 | 0 | NA | NA | NA | NA | NA | #VALUE! |
| ENSMUSG00000117475.2 | 0 | NA | NA | NA | NA | NA | #VALUE! |
| ENSMUSG00000117477.2 | 0 | NA | NA | NA | NA | NA | #VALUE! |
| ENSMUSG00000117478.2 | 0 | NA | NA | NA | NA | NA | #VALUE! |
| ENSMUSG00000117479.2 | 0 | NA | NA | NA | NA | NA | #VALUE! |
| ENSMUSG00000117480.2 | 0 | NA | NA | NA | NA | NA | #VALUE! |
| ENSMUSG00000117481.2 | 0 | NA | NA | NA | NA | NA | #VALUE! |
| ENSMUSG00000117482.2 | 0 | NA | NA | NA | NA | NA | #VALUE! |
| ENSMUSG00000117483.2 | 0 | NA | NA | NA | NA | NA | #VALUE! |
| ENSMUSG00000117484.2 | 0 | NA | NA | NA | NA | NA | #VALUE! |
| ENSMUSG00000117485.2 | 5.3356 | 4.831926 | 3.600648 | 1.34196 | 0.179609 | NA | #VALUE! |

| | | | | | | | |
|---|---|---|---|---|---|---|---|
| ENSMUSG00000117486.2 | 0 | NA | NA | NA | NA | NA | #VALUE! |
| ENSMUSG00000117487.2 | 0 | NA | NA | NA | NA | NA | #VALUE! |
| ENSMUSG00000117488.2 | 0 | NA | NA | NA | NA | NA | #VALUE! |
| ENSMUSG00000117489.2 | 0 | NA | NA | NA | NA | NA | #VALUE! |
| ENSMUSG00000117490.2 | 0 | NA | NA | NA | NA | NA | #VALUE! |
| ENSMUSG00000117491.2 | 0 | NA | NA | NA | NA | NA | #VALUE! |
| ENSMUSG00000117492.2 | 0 | NA | NA | NA | NA | NA | #VALUE! |
| ENSMUSG00000117493.2 | 0 | NA | NA | NA | NA | NA | #VALUE! |
| ENSMUSG00000117494.2 | 0 | NA | NA | NA | NA | NA | #VALUE! |
| ENSMUSG00000117495.3 | 0 | NA | NA | NA | NA | NA | #VALUE! |
| ENSMUSG00000117496.2 | 0 | NA | NA | NA | NA | NA | #VALUE! |
| ENSMUSG00000117497.2 | 0 | NA | NA | NA | NA | NA | #VALUE! |
| ENSMUSG00000117498.2 | 0 | NA | NA | NA | NA | NA | #VALUE! |
| ENSMUSG00000117499.2 | 8.332037 | 2.127107 | 3.082418 | 0.690078 | 0.490145 | NA | #VALUE! |
| ENSMUSG00000117500.2 | 0 | NA | NA | NA | NA | NA | #VALUE! |
| ENSMUSG00000117501.2 | 0 | NA | NA | NA | NA | NA | #VALUE! |
| ENSMUSG00000117502.2 | 0 | NA | NA | NA | NA | NA | #VALUE! |
| ENSMUSG00000117503.2 | 0 | NA | NA | NA | NA | NA | #VALUE! |
| ENSMUSG00000117504.2 | 0 | NA | NA | NA | NA | NA | #VALUE! |
| ENSMUSG00000117505.2 | 0 | NA | NA | NA | NA | NA | #VALUE! |
| ENSMUSG00000117506.2 | 0 | NA | NA | NA | NA | NA | #VALUE! |
| ENSMUSG00000117507.2 | 0 | NA | NA | NA | NA | NA | #VALUE! |
| ENSMUSG00000117508.2 | 0 | NA | NA | NA | NA | NA | #VALUE! |
| ENSMUSG00000117509.2 | 0 | NA | NA | NA | NA | NA | #VALUE! |
| ENSMUSG00000117510.2 | 0 | NA | NA | NA | NA | NA | #VALUE! |
| ENSMUSG00000117511.2 | 0 | NA | NA | NA | NA | NA | #VALUE! |
| ENSMUSG00000117512.2 | 0 | NA | NA | NA | NA | NA | #VALUE! |
| ENSMUSG00000117513.2 | 0 | NA | NA | NA | NA | NA | #VALUE! |
| ENSMUSG00000117514.2 | 0 | NA | NA | NA | NA | NA | #VALUE! |
| ENSMUSG00000117515.2 | 0.3999 | -2.17566 | 4.018831 | -0.54137 | 0.588256 | NA | #VALUE! |
| ENSMUSG00000117516.2 | 0 | NA | NA | NA | NA | NA | #VALUE! |
| ENSMUSG00000117517.2 | 0 | NA | NA | NA | NA | NA | #VALUE! |
| ENSMUSG00000117518.2 | 0 | NA | NA | NA | NA | NA | #VALUE! |
| ENSMUSG00000117519.2 | 0 | NA | NA | NA | NA | NA | #VALUE! |
| ENSMUSG00000117520.2 | 0 | NA | NA | NA | NA | NA | #VALUE! |
| ENSMUSG00000117521.2 | 0 | NA | NA | NA | NA | NA | #VALUE! |
| ENSMUSG00000117522.2 | 0 | NA | NA | NA | NA | NA | #VALUE! |
| ENSMUSG00000117523.2 | 0 | NA | NA | NA | NA | NA | #VALUE! |
| ENSMUSG00000117524.2 | 0 | NA | NA | NA | NA | NA | #VALUE! |
| ENSMUSG00000117525.2 | 0 | NA | NA | NA | NA | NA | #VALUE! |
| ENSMUSG00000117526.2 | 0 | NA | NA | NA | NA | NA | #VALUE! |
| ENSMUSG00000117527.2 | 0.261539 | 1.68833 | 4.043024 | 0.417591 | 0.676246 | NA | #VALUE! |
| ENSMUSG00000117528.2 | 0 | NA | NA | NA | NA | NA | #VALUE! |
| ENSMUSG00000117529.2 | 0 | NA | NA | NA | NA | NA | #VALUE! |

| | | | | | | | |
|---|---|---|---|---|---|---|---|
| ENSMUSG00000117530.2 | 0 | NA | NA | NA | NA | NA | #VALUE! |
| ENSMUSG00000117531.2 | 0 | NA | NA | NA | NA | NA | #VALUE! |
| ENSMUSG00000117532.2 | 0 | NA | NA | NA | NA | NA | #VALUE! |
| ENSMUSG00000117533.2 | 0 | NA | NA | NA | NA | NA | #VALUE! |
| ENSMUSG00000117534.2 | 0 | NA | NA | NA | NA | NA | #VALUE! |
| ENSMUSG00000117535.2 | 0 | NA | NA | NA | NA | NA | #VALUE! |
| ENSMUSG00000117536.2 | 0 | NA | NA | NA | NA | NA | #VALUE! |
| ENSMUSG00000117537.2 | 0 | NA | NA | NA | NA | NA | #VALUE! |
| ENSMUSG00000117538.2 | 0 | NA | NA | NA | NA | NA | #VALUE! |
| ENSMUSG00000117539.2 | 0 | NA | NA | NA | NA | NA | #VALUE! |
| ENSMUSG00000117540.2 | 0 | NA | NA | NA | NA | NA | #VALUE! |
| ENSMUSG00000117541.2 | 0 | NA | NA | NA | NA | NA | #VALUE! |
| ENSMUSG00000117542.2 | 0 | NA | NA | NA | NA | NA | #VALUE! |
| ENSMUSG00000117543.2 | 0 | NA | NA | NA | NA | NA | #VALUE! |
| ENSMUSG00000117544.2 | 0 | NA | NA | NA | NA | NA | #VALUE! |
| ENSMUSG00000117545.2 | 8.368541 | -6.59301 | 3.815764 | -1.72783 | 0.084018 | NA | #VALUE! |
| ENSMUSG00000117546.2 | 0 | NA | NA | NA | NA | NA | #VALUE! |
| ENSMUSG00000117547.2 | 0 | NA | NA | NA | NA | NA | #VALUE! |
| ENSMUSG00000117548.2 | 0 | NA | NA | NA | NA | NA | #VALUE! |
| ENSMUSG00000117549.2 | 0 | NA | NA | NA | NA | NA | #VALUE! |
| ENSMUSG00000117550.2 | 0 | NA | NA | NA | NA | NA | #VALUE! |
| ENSMUSG00000117551.2 | 0 | NA | NA | NA | NA | NA | #VALUE! |
| ENSMUSG00000117552.2 | 0 | NA | NA | NA | NA | NA | #VALUE! |
| ENSMUSG00000117553.2 | 0 | NA | NA | NA | NA | NA | #VALUE! |
| ENSMUSG00000117554.2 | 0 | NA | NA | NA | NA | NA | #VALUE! |
| ENSMUSG00000117555.2 | 0 | NA | NA | NA | NA | NA | #VALUE! |
| ENSMUSG00000117556.2 | 0 | NA | NA | NA | NA | NA | #VALUE! |
| ENSMUSG00000117557.2 | 0 | NA | NA | NA | NA | NA | #VALUE! |
| ENSMUSG00000117558.2 | 0 | NA | NA | NA | NA | NA | #VALUE! |
| ENSMUSG00000117559.2 | 0 | NA | NA | NA | NA | NA | #VALUE! |
| ENSMUSG00000117560.2 | 0 | NA | NA | NA | NA | NA | #VALUE! |
| ENSMUSG00000117561.2 | 0 | NA | NA | NA | NA | NA | #VALUE! |
| ENSMUSG00000117562.3 | 0 | NA | NA | NA | NA | NA | #VALUE! |
| ENSMUSG00000117563.2 | 0 | NA | NA | NA | NA | NA | #VALUE! |
| ENSMUSG00000117564.2 | 0 | NA | NA | NA | NA | NA | #VALUE! |
| ENSMUSG00000117565.2 | 0 | NA | NA | NA | NA | NA | #VALUE! |
| ENSMUSG00000117566.2 | 0 | NA | NA | NA | NA | NA | #VALUE! |
| ENSMUSG00000117567.2 | 0 | NA | NA | NA | NA | NA | #VALUE! |
| ENSMUSG00000117568.2 | 0 | NA | NA | NA | NA | NA | #VALUE! |
| ENSMUSG00000117569.2 | 0 | NA | NA | NA | NA | NA | #VALUE! |
| ENSMUSG00000117570.2 | 0 | NA | NA | NA | NA | NA | #VALUE! |
| ENSMUSG00000117571.2 | 0 | NA | NA | NA | NA | NA | #VALUE! |
| ENSMUSG00000117572.2 | 0 | NA | NA | NA | NA | NA | #VALUE! |
| ENSMUSG00000117573.2 | 0 | NA | NA | NA | NA | NA | #VALUE! |

| | | | | | | | |
|---|---|---|---|---|---|---|---|
| ENSMUSG00000117574.2 | 0 | NA | NA | NA | NA | NA | #VALUE! |
| ENSMUSG00000117575.2 | 0 | NA | NA | NA | NA | NA | #VALUE! |
| ENSMUSG00000117576.2 | 0 | NA | NA | NA | NA | NA | #VALUE! |
| ENSMUSG00000117577.2 | 0 | NA | NA | NA | NA | NA | #VALUE! |
| ENSMUSG00000117578.2 | 0.093255 | 0.868093 | 4.072464 | 0.213162 | 0.831201 | NA | #VALUE! |
| ENSMUSG00000117579.2 | 0 | NA | NA | NA | NA | NA | #VALUE! |
| ENSMUSG00000117580.2 | 0 | NA | NA | NA | NA | NA | #VALUE! |
| ENSMUSG00000117581.2 | 0 | NA | NA | NA | NA | NA | #VALUE! |
| ENSMUSG00000117582.2 | 0 | NA | NA | NA | NA | NA | #VALUE! |
| ENSMUSG00000117583.2 | 0 | NA | NA | NA | NA | NA | #VALUE! |
| ENSMUSG00000117584.2 | 0 | NA | NA | NA | NA | NA | #VALUE! |
| ENSMUSG00000117585.2 | 0 | NA | NA | NA | NA | NA | #VALUE! |
| ENSMUSG00000117586.2 | 0 | NA | NA | NA | NA | NA | #VALUE! |
| ENSMUSG00000117587.2 | 0 | NA | NA | NA | NA | NA | #VALUE! |
| ENSMUSG00000117588.2 | 0 | NA | NA | NA | NA | NA | #VALUE! |
| ENSMUSG00000117589.2 | 0 | NA | NA | NA | NA | NA | #VALUE! |
| ENSMUSG00000117590.2 | 0 | NA | NA | NA | NA | NA | #VALUE! |
| ENSMUSG00000117591.2 | 0 | NA | NA | NA | NA | NA | #VALUE! |
| ENSMUSG00000117592.2 | 0 | NA | NA | NA | NA | NA | #VALUE! |
| ENSMUSG00000117593.2 | 0 | NA | NA | NA | NA | NA | #VALUE! |
| ENSMUSG00000117594.2 | 0 | NA | NA | NA | NA | NA | #VALUE! |
| ENSMUSG00000117595.2 | 0 | NA | NA | NA | NA | NA | #VALUE! |
| ENSMUSG00000117596.2 | 0 | NA | NA | NA | NA | NA | #VALUE! |
| ENSMUSG00000117597.2 | 2.940368 | -5.09511 | 3.921868 | -1.29915 | 0.193891 | NA | #VALUE! |
| ENSMUSG00000117598.2 | 0 | NA | NA | NA | NA | NA | #VALUE! |
| ENSMUSG00000117599.2 | 0.093255 | 0.868093 | 4.072464 | 0.213162 | 0.831201 | NA | #VALUE! |
| ENSMUSG00000117600.2 | 0 | NA | NA | NA | NA | NA | #VALUE! |
| ENSMUSG00000117601.2 | 0 | NA | NA | NA | NA | NA | #VALUE! |
| ENSMUSG00000117602.2 | 0 | NA | NA | NA | NA | NA | #VALUE! |
| ENSMUSG00000117603.2 | 0 | NA | NA | NA | NA | NA | #VALUE! |
| ENSMUSG00000117604.2 | 0 | NA | NA | NA | NA | NA | #VALUE! |
| ENSMUSG00000117605.2 | 0 | NA | NA | NA | NA | NA | #VALUE! |
| ENSMUSG00000117606.2 | 0 | NA | NA | NA | NA | NA | #VALUE! |
| ENSMUSG00000117607.2 | 0 | NA | NA | NA | NA | NA | #VALUE! |
| ENSMUSG00000117608.2 | 0 | NA | NA | NA | NA | NA | #VALUE! |
| ENSMUSG00000117609.2 | 0 | NA | NA | NA | NA | NA | #VALUE! |
| ENSMUSG00000117610.2 | 0 | NA | NA | NA | NA | NA | #VALUE! |
| ENSMUSG00000117611.2 | 0 | NA | NA | NA | NA | NA | #VALUE! |
| ENSMUSG00000117612.2 | 0 | NA | NA | NA | NA | NA | #VALUE! |
| ENSMUSG00000117613.2 | 0 | NA | NA | NA | NA | NA | #VALUE! |
| ENSMUSG00000117614.2 | 0 | NA | NA | NA | NA | NA | #VALUE! |
| ENSMUSG00000117615.2 | 0 | NA | NA | NA | NA | NA | #VALUE! |
| ENSMUSG00000117616.2 | 0 | NA | NA | NA | NA | NA | #VALUE! |
| ENSMUSG00000117617.2 | 0 | NA | NA | NA | NA | NA | #VALUE! |

| | | | | | | | |
|---|---|---|---|---|---|---|---|
| ENSMUSG00000117618.2 | 0 | NA | NA | NA | NA | NA | #VALUE! |
| ENSMUSG00000117619.2 | 0 | NA | NA | NA | NA | NA | #VALUE! |
| ENSMUSG00000117620.2 | 0 | NA | NA | NA | NA | NA | #VALUE! |
| ENSMUSG00000117622.2 | 0 | NA | NA | NA | NA | NA | #VALUE! |
| ENSMUSG00000117623.2 | 0 | NA | NA | NA | NA | NA | #VALUE! |
| ENSMUSG00000117624.2 | 0 | NA | NA | NA | NA | NA | #VALUE! |
| ENSMUSG00000117625.2 | 0 | NA | NA | NA | NA | NA | #VALUE! |
| ENSMUSG00000117626.2 | 0 | NA | NA | NA | NA | NA | #VALUE! |
| ENSMUSG00000117627.2 | 0 | NA | NA | NA | NA | NA | #VALUE! |
| ENSMUSG00000117628.2 | 0 | NA | NA | NA | NA | NA | #VALUE! |
| ENSMUSG00000117629.2 | 0 | NA | NA | NA | NA | NA | #VALUE! |
| ENSMUSG00000117630.2 | 0 | NA | NA | NA | NA | NA | #VALUE! |
| ENSMUSG00000117631.2 | 0 | NA | NA | NA | NA | NA | #VALUE! |
| ENSMUSG00000117632.2 | 0 | NA | NA | NA | NA | NA | #VALUE! |
| ENSMUSG00000117633.2 | 0 | NA | NA | NA | NA | NA | #VALUE! |
| ENSMUSG00000117634.2 | 0 | NA | NA | NA | NA | NA | #VALUE! |
| ENSMUSG00000117635.2 | 0 | NA | NA | NA | NA | NA | #VALUE! |
| ENSMUSG00000117636.2 | 0 | NA | NA | NA | NA | NA | #VALUE! |
| ENSMUSG00000117637.2 | 0 | NA | NA | NA | NA | NA | #VALUE! |
| ENSMUSG00000117638.2 | 0 | NA | NA | NA | NA | NA | #VALUE! |
| ENSMUSG00000117639.2 | 0 | NA | NA | NA | NA | NA | #VALUE! |
| ENSMUSG00000117640.2 | 0 | NA | NA | NA | NA | NA | #VALUE! |
| ENSMUSG00000117641.2 | 0 | NA | NA | NA | NA | NA | #VALUE! |
| ENSMUSG00000117642.2 | 3.443657 | 5.264174 | 3.919884 | 1.342941 | 0.179291 | NA | #VALUE! |
| ENSMUSG00000117643.2 | 0 | NA | NA | NA | NA | NA | #VALUE! |
| ENSMUSG00000117644.2 | 0 | NA | NA | NA | NA | NA | #VALUE! |
| ENSMUSG00000117645.2 | 0 | NA | NA | NA | NA | NA | #VALUE! |
| ENSMUSG00000117646.2 | 0 | NA | NA | NA | NA | NA | #VALUE! |
| ENSMUSG00000117647.2 | 0 | NA | NA | NA | NA | NA | #VALUE! |
| ENSMUSG00000117648.2 | 3.730208 | 5.397701 | 3.918728 | 1.377412 | 0.168385 | NA | #VALUE! |
| ENSMUSG00000117649.2 | 0 | NA | NA | NA | NA | NA | #VALUE! |
| ENSMUSG00000117650.2 | 0 | NA | NA | NA | NA | NA | #VALUE! |
| ENSMUSG00000117651.2 | 0 | NA | NA | NA | NA | NA | #VALUE! |
| ENSMUSG00000117652.2 | 0 | NA | NA | NA | NA | NA | #VALUE! |
| ENSMUSG00000117653.2 | 0 | NA | NA | NA | NA | NA | #VALUE! |
| ENSMUSG00000117654.2 | 0 | NA | NA | NA | NA | NA | #VALUE! |
| ENSMUSG00000117655.2 | 0 | NA | NA | NA | NA | NA | #VALUE! |
| ENSMUSG00000117656.2 | 0 | NA | NA | NA | NA | NA | #VALUE! |
| ENSMUSG00000117657.2 | 0 | NA | NA | NA | NA | NA | #VALUE! |
| ENSMUSG00000117658.2 | 0 | NA | NA | NA | NA | NA | #VALUE! |
| ENSMUSG00000117659.2 | 0 | NA | NA | NA | NA | NA | #VALUE! |
| ENSMUSG00000117660.2 | 0 | NA | NA | NA | NA | NA | #VALUE! |
| ENSMUSG00000117661.2 | 0 | NA | NA | NA | NA | NA | #VALUE! |
| ENSMUSG00000117663.2 | 0 | NA | NA | NA | NA | NA | #VALUE! |

| | | | | | | | |
|---|---|---|---|---|---|---|---|
| ENSMUSG00000117664.2 | 0 | NA | NA | NA | NA | NA | #VALUE! |
| ENSMUSG00000117665.2 | 0 | NA | NA | NA | NA | NA | #VALUE! |
| ENSMUSG00000117666.2 | 0 | NA | NA | NA | NA | NA | #VALUE! |
| ENSMUSG00000117667.2 | 0 | NA | NA | NA | NA | NA | #VALUE! |
| ENSMUSG00000117669.2 | 0 | NA | NA | NA | NA | NA | #VALUE! |
| ENSMUSG00000117670.2 | 0 | NA | NA | NA | NA | NA | #VALUE! |
| ENSMUSG00000117671.2 | 0 | NA | NA | NA | NA | NA | #VALUE! |
| ENSMUSG00000117673.2 | 0 | NA | NA | NA | NA | NA | #VALUE! |
| ENSMUSG00000117674.2 | 0 | NA | NA | NA | NA | NA | #VALUE! |
| ENSMUSG00000117675.2 | 0 | NA | NA | NA | NA | NA | #VALUE! |
| ENSMUSG00000117676.2 | 0 | NA | NA | NA | NA | NA | #VALUE! |
| ENSMUSG00000117677.2 | 0 | NA | NA | NA | NA | NA | #VALUE! |
| ENSMUSG00000117678.2 | 0 | NA | NA | NA | NA | NA | #VALUE! |
| ENSMUSG00000117679.2 | 8.58246 | 6.591238 | 3.779215 | 1.744076 | 0.081146 | NA | #VALUE! |
| ENSMUSG00000117680.2 | 0 | NA | NA | NA | NA | NA | #VALUE! |
| ENSMUSG00000117681.2 | 0 | NA | NA | NA | NA | NA | #VALUE! |
| ENSMUSG00000117682.2 | 0 | NA | NA | NA | NA | NA | #VALUE! |
| ENSMUSG00000117683.2 | 0 | NA | NA | NA | NA | NA | #VALUE! |
| ENSMUSG00000117684.2 | 0 | NA | NA | NA | NA | NA | #VALUE! |
| ENSMUSG00000117685.2 | 0 | NA | NA | NA | NA | NA | #VALUE! |
| ENSMUSG00000117686.2 | 0 | NA | NA | NA | NA | NA | #VALUE! |
| ENSMUSG00000117687.2 | 0 | NA | NA | NA | NA | NA | #VALUE! |
| ENSMUSG00000117688.2 | 0 | NA | NA | NA | NA | NA | #VALUE! |
| ENSMUSG00000117689.2 | 0 | NA | NA | NA | NA | NA | #VALUE! |
| ENSMUSG00000117690.2 | 0 | NA | NA | NA | NA | NA | #VALUE! |
| ENSMUSG00000117691.2 | 0 | NA | NA | NA | NA | NA | #VALUE! |
| ENSMUSG00000117692.2 | 0 | NA | NA | NA | NA | NA | #VALUE! |
| ENSMUSG00000117693.2 | 0 | NA | NA | NA | NA | NA | #VALUE! |
| ENSMUSG00000117694.2 | 0 | NA | NA | NA | NA | NA | #VALUE! |
| ENSMUSG00000117695.2 | 0 | NA | NA | NA | NA | NA | #VALUE! |
| ENSMUSG00000117696.2 | 0 | NA | NA | NA | NA | NA | #VALUE! |
| ENSMUSG00000117697.2 | 0 | NA | NA | NA | NA | NA | #VALUE! |
| ENSMUSG00000117698.2 | 0.978946 | -3.48749 | 3.952505 | -0.88235 | 0.377588 | NA | #VALUE! |
| ENSMUSG00000117699.2 | 0 | NA | NA | NA | NA | NA | #VALUE! |
| ENSMUSG00000117700.2 | 0 | NA | NA | NA | NA | NA | #VALUE! |
| ENSMUSG00000117701.2 | 0 | NA | NA | NA | NA | NA | #VALUE! |
| ENSMUSG00000117702.2 | 0 | NA | NA | NA | NA | NA | #VALUE! |
| ENSMUSG00000117703.2 | 0 | NA | NA | NA | NA | NA | #VALUE! |
| ENSMUSG00000117704.2 | 0 | NA | NA | NA | NA | NA | #VALUE! |
| ENSMUSG00000117705.2 | 0 | NA | NA | NA | NA | NA | #VALUE! |
| ENSMUSG00000117706.2 | 0 | NA | NA | NA | NA | NA | #VALUE! |
| ENSMUSG00000117707.2 | 0 | NA | NA | NA | NA | NA | #VALUE! |
| ENSMUSG00000117708.2 | 0 | NA | NA | NA | NA | NA | #VALUE! |
| ENSMUSG00000117709.2 | 0 | NA | NA | NA | NA | NA | #VALUE! |

| | | | | | | | |
|---|---|---|---|---|---|---|---|
| ENSMUSG00000117710.2 | 0 | NA | NA | NA | NA | NA | #VALUE! |
| ENSMUSG00000117711.2 | 0.390621 | 0.386159 | 4.04914 | 0.095368 | 0.924022 | NA | #VALUE! |
| ENSMUSG00000117712.2 | 0 | NA | NA | NA | NA | NA | #VALUE! |
| ENSMUSG00000117713.2 | 0 | NA | NA | NA | NA | NA | #VALUE! |
| ENSMUSG00000117714.2 | 0 | NA | NA | NA | NA | NA | #VALUE! |
| ENSMUSG00000117715.2 | 0 | NA | NA | NA | NA | NA | #VALUE! |
| ENSMUSG00000117717.2 | 0 | NA | NA | NA | NA | NA | #VALUE! |
| ENSMUSG00000117718.2 | 0 | NA | NA | NA | NA | NA | #VALUE! |
| ENSMUSG00000117719.2 | 0 | NA | NA | NA | NA | NA | #VALUE! |
| ENSMUSG00000117720.2 | 0 | NA | NA | NA | NA | NA | #VALUE! |
| ENSMUSG00000117721.2 | 0 | NA | NA | NA | NA | NA | #VALUE! |
| ENSMUSG00000117722.2 | 0 | NA | NA | NA | NA | NA | #VALUE! |
| ENSMUSG00000117723.2 | 0 | NA | NA | NA | NA | NA | #VALUE! |
| ENSMUSG00000117724.2 | 0 | NA | NA | NA | NA | NA | #VALUE! |
| ENSMUSG00000117725.2 | 4.489844 | 2.342124 | 2.981861 | 0.785457 | 0.432186 | NA | #VALUE! |
| ENSMUSG00000117726.2 | 0 | NA | NA | NA | NA | NA | #VALUE! |
| ENSMUSG00000117727.2 | 0 | NA | NA | NA | NA | NA | #VALUE! |
| ENSMUSG00000117728.2 | 0 | NA | NA | NA | NA | NA | #VALUE! |
| ENSMUSG00000117729.2 | 0 | NA | NA | NA | NA | NA | #VALUE! |
| ENSMUSG00000117730.2 | 0 | NA | NA | NA | NA | NA | #VALUE! |
| ENSMUSG00000117731.2 | 0 | NA | NA | NA | NA | NA | #VALUE! |
| ENSMUSG00000117732.2 | 0 | NA | NA | NA | NA | NA | #VALUE! |
| ENSMUSG00000117733.2 | 0 | NA | NA | NA | NA | NA | #VALUE! |
| ENSMUSG00000117734.2 | 0 | NA | NA | NA | NA | NA | #VALUE! |
| ENSMUSG00000117735.2 | 0 | NA | NA | NA | NA | NA | #VALUE! |
| ENSMUSG00000117736.2 | 0 | NA | NA | NA | NA | NA | #VALUE! |
| ENSMUSG00000117737.2 | 0 | NA | NA | NA | NA | NA | #VALUE! |
| ENSMUSG00000117738.2 | 1.346268 | 3.918821 | 3.939925 | 0.994644 | 0.31991 | NA | #VALUE! |
| ENSMUSG00000117739.2 | 0 | NA | NA | NA | NA | NA | #VALUE! |
| ENSMUSG00000117740.2 | 0 | NA | NA | NA | NA | NA | #VALUE! |
| ENSMUSG00000117741.2 | 0 | NA | NA | NA | NA | NA | #VALUE! |
| ENSMUSG00000117742.2 | 0 | NA | NA | NA | NA | NA | #VALUE! |
| ENSMUSG00000117743.2 | 0 | NA | NA | NA | NA | NA | #VALUE! |
| ENSMUSG00000117744.2 | 0 | NA | NA | NA | NA | NA | #VALUE! |
| ENSMUSG00000117745.2 | 0 | NA | NA | NA | NA | NA | #VALUE! |
| ENSMUSG00000117746.2 | 0 | NA | NA | NA | NA | NA | #VALUE! |
| ENSMUSG00000117747.2 | 0 | NA | NA | NA | NA | NA | #VALUE! |
| ENSMUSG00000117748.2 | 0 | NA | NA | NA | NA | NA | #VALUE! |
| ENSMUSG00000117749.2 | 0 | NA | NA | NA | NA | NA | #VALUE! |
| ENSMUSG00000117750.2 | 0 | NA | NA | NA | NA | NA | #VALUE! |
| ENSMUSG00000117751.2 | 0 | NA | NA | NA | NA | NA | #VALUE! |
| ENSMUSG00000117752.2 | 0 | NA | NA | NA | NA | NA | #VALUE! |
| ENSMUSG00000117753.2 | 0 | NA | NA | NA | NA | NA | #VALUE! |
| ENSMUSG00000117754.2 | 0 | NA | NA | NA | NA | NA | #VALUE! |

| | | | | | | | |
|---|---|---|---|---|---|---|---|
| ENSMUSG00000117755.2 | 0 | NA | NA | NA | NA | NA | #VALUE! |
| ENSMUSG00000117756.2 | 0 | NA | NA | NA | NA | NA | #VALUE! |
| ENSMUSG00000117757.2 | 0 | NA | NA | NA | NA | NA | #VALUE! |
| ENSMUSG00000117758.2 | 0 | NA | NA | NA | NA | NA | #VALUE! |
| ENSMUSG00000117759.2 | 0 | NA | NA | NA | NA | NA | #VALUE! |
| ENSMUSG00000117760.2 | 0 | NA | NA | NA | NA | NA | #VALUE! |
| ENSMUSG00000117761.2 | 0 | NA | NA | NA | NA | NA | #VALUE! |
| ENSMUSG00000117762.2 | 0 | NA | NA | NA | NA | NA | #VALUE! |
| ENSMUSG00000117763.2 | 0 | NA | NA | NA | NA | NA | #VALUE! |
| ENSMUSG00000117764.2 | 0 | NA | NA | NA | NA | NA | #VALUE! |
| ENSMUSG00000117765.2 | 0 | NA | NA | NA | NA | NA | #VALUE! |
| ENSMUSG00000117766.2 | 0 | NA | NA | NA | NA | NA | #VALUE! |
| ENSMUSG00000117767.2 | 0 | NA | NA | NA | NA | NA | #VALUE! |
| ENSMUSG00000117768.2 | 0 | NA | NA | NA | NA | NA | #VALUE! |
| ENSMUSG00000117769.2 | 0 | NA | NA | NA | NA | NA | #VALUE! |
| ENSMUSG00000117770.2 | 0 | NA | NA | NA | NA | NA | #VALUE! |
| ENSMUSG00000117771.2 | 0 | NA | NA | NA | NA | NA | #VALUE! |
| ENSMUSG00000117772.2 | 0 | NA | NA | NA | NA | NA | #VALUE! |
| ENSMUSG00000117773.2 | 0 | NA | NA | NA | NA | NA | #VALUE! |
| ENSMUSG00000117774.2 | 0 | NA | NA | NA | NA | NA | #VALUE! |
| ENSMUSG00000117775.2 | 0 | NA | NA | NA | NA | NA | #VALUE! |
| ENSMUSG00000117777.2 | 0 | NA | NA | NA | NA | NA | #VALUE! |
| ENSMUSG00000117778.2 | 0 | NA | NA | NA | NA | NA | #VALUE! |
| ENSMUSG00000117779.2 | 0.168284 | 1.027628 | 4.067126 | 0.252667 | 0.800526 | NA | #VALUE! |
| ENSMUSG00000117781.2 | 0 | NA | NA | NA | NA | NA | #VALUE! |
| ENSMUSG00000117782.2 | 0 | NA | NA | NA | NA | NA | #VALUE! |
| ENSMUSG00000117783.2 | 0 | NA | NA | NA | NA | NA | #VALUE! |
| ENSMUSG00000117784.2 | 0 | NA | NA | NA | NA | NA | #VALUE! |
| ENSMUSG00000117785.2 | 0 | NA | NA | NA | NA | NA | #VALUE! |
| ENSMUSG00000117786.2 | 0 | NA | NA | NA | NA | NA | #VALUE! |
| ENSMUSG00000117787.2 | 0 | NA | NA | NA | NA | NA | #VALUE! |
| ENSMUSG00000117788.2 | 0 | NA | NA | NA | NA | NA | #VALUE! |
| ENSMUSG00000117789.2 | 0 | NA | NA | NA | NA | NA | #VALUE! |
| ENSMUSG00000117790.2 | 0 | NA | NA | NA | NA | NA | #VALUE! |
| ENSMUSG00000117791.2 | 0 | NA | NA | NA | NA | NA | #VALUE! |
| ENSMUSG00000117792.2 | 0 | NA | NA | NA | NA | NA | #VALUE! |
| ENSMUSG00000117793.2 | 0 | NA | NA | NA | NA | NA | #VALUE! |
| ENSMUSG00000117794.2 | 0 | NA | NA | NA | NA | NA | #VALUE! |
| ENSMUSG00000117795.2 | 0 | NA | NA | NA | NA | NA | #VALUE! |
| ENSMUSG00000117796.2 | 0 | NA | NA | NA | NA | NA | #VALUE! |
| ENSMUSG00000117797.2 | 0 | NA | NA | NA | NA | NA | #VALUE! |
| ENSMUSG00000117798.2 | 0 | NA | NA | NA | NA | NA | #VALUE! |
| ENSMUSG00000117799.2 | 0 | NA | NA | NA | NA | NA | #VALUE! |
| ENSMUSG00000117800.2 | 0 | NA | NA | NA | NA | NA | #VALUE! |

| | | | | | | | |
|---|---|---|---|---|---|---|---|
| ENSMUSG00000117801.2 | 0 | NA | NA | NA | NA | NA | #VALUE! |
| ENSMUSG00000117802.2 | 0 | NA | NA | NA | NA | NA | #VALUE! |
| ENSMUSG00000117803.2 | 0 | NA | NA | NA | NA | NA | #VALUE! |
| ENSMUSG00000117804.2 | 0.810674 | -1.42782 | 3.951856 | -0.3613 | 0.717872 | NA | #VALUE! |
| ENSMUSG00000117805.2 | 0 | NA | NA | NA | NA | NA | #VALUE! |
| ENSMUSG00000117806.2 | 0 | NA | NA | NA | NA | NA | #VALUE! |
| ENSMUSG00000117807.2 | 0 | NA | NA | NA | NA | NA | #VALUE! |
| ENSMUSG00000117808.2 | 0 | NA | NA | NA | NA | NA | #VALUE! |
| ENSMUSG00000117809.2 | 0 | NA | NA | NA | NA | NA | #VALUE! |
| ENSMUSG00000117810.2 | 0.168284 | 1.027628 | 4.067126 | 0.252667 | 0.800526 | NA | #VALUE! |
| ENSMUSG00000117811.2 | 0 | NA | NA | NA | NA | NA | #VALUE! |
| ENSMUSG00000117812.2 | 0 | NA | NA | NA | NA | NA | #VALUE! |
| ENSMUSG00000117813.2 | 0 | NA | NA | NA | NA | NA | #VALUE! |
| ENSMUSG00000117814.2 | 8.065771 | -3.66602 | 3.023417 | -1.21254 | 0.225304 | NA | #VALUE! |
| ENSMUSG00000117815.2 | 0 | NA | NA | NA | NA | NA | #VALUE! |
| ENSMUSG00000117816.2 | 0 | NA | NA | NA | NA | NA | #VALUE! |
| ENSMUSG00000117817.2 | 0 | NA | NA | NA | NA | NA | #VALUE! |
| ENSMUSG00000117818.2 | 0.195789 | -1.05581 | 4.073302 | -0.2592 | 0.795479 | NA | #VALUE! |
| ENSMUSG00000117819.2 | 0 | NA | NA | NA | NA | NA | #VALUE! |
| ENSMUSG00000117820.2 | 0 | NA | NA | NA | NA | NA | #VALUE! |
| ENSMUSG00000117821.2 | 0 | NA | NA | NA | NA | NA | #VALUE! |
| ENSMUSG00000117822.2 | 0 | NA | NA | NA | NA | NA | #VALUE! |
| ENSMUSG00000117823.2 | 0.245976 | 1.027628 | 4.067126 | 0.252667 | 0.800526 | NA | #VALUE! |
| ENSMUSG00000117824.2 | 0 | NA | NA | NA | NA | NA | #VALUE! |
| ENSMUSG00000117825.2 | 0 | NA | NA | NA | NA | NA | #VALUE! |
| ENSMUSG00000117826.2 | 0 | NA | NA | NA | NA | NA | #VALUE! |
| ENSMUSG00000117827.2 | 0 | NA | NA | NA | NA | NA | #VALUE! |
| ENSMUSG00000117828.2 | 0 | NA | NA | NA | NA | NA | #VALUE! |
| ENSMUSG00000117829.2 | 0 | NA | NA | NA | NA | NA | #VALUE! |
| ENSMUSG00000117830.2 | 0 | NA | NA | NA | NA | NA | #VALUE! |
| ENSMUSG00000117831.2 | 0 | NA | NA | NA | NA | NA | #VALUE! |
| ENSMUSG00000117832.2 | 0 | NA | NA | NA | NA | NA | #VALUE! |
| ENSMUSG00000117833.2 | 0 | NA | NA | NA | NA | NA | #VALUE! |
| ENSMUSG00000117834.2 | 0 | NA | NA | NA | NA | NA | #VALUE! |
| ENSMUSG00000117835.2 | 0 | NA | NA | NA | NA | NA | #VALUE! |
| ENSMUSG00000117836.2 | 0 | NA | NA | NA | NA | NA | #VALUE! |
| ENSMUSG00000117837.2 | 0 | NA | NA | NA | NA | NA | #VALUE! |
| ENSMUSG00000117838.2 | 0 | NA | NA | NA | NA | NA | #VALUE! |
| ENSMUSG00000117839.2 | 0 | NA | NA | NA | NA | NA | #VALUE! |
| ENSMUSG00000117840.2 | 0 | NA | NA | NA | NA | NA | #VALUE! |
| ENSMUSG00000117841.2 | 0 | NA | NA | NA | NA | NA | #VALUE! |
| ENSMUSG00000117842.2 | 0 | NA | NA | NA | NA | NA | #VALUE! |
| ENSMUSG00000117843.2 | 0 | NA | NA | NA | NA | NA | #VALUE! |
| ENSMUSG00000117844.2 | 0.279766 | 1.808031 | 4.036658 | 0.447903 | 0.654223 | NA | #VALUE! |

| | | | | | | | |
|---|---|---|---|---|---|---|---|
| ENSMUSG00000117845.2 | 0 | NA | NA | NA | NA | NA | #VALUE! |
| ENSMUSG00000117846.2 | 0 | NA | NA | NA | NA | NA | #VALUE! |
| ENSMUSG00000117847.2 | 0 | NA | NA | NA | NA | NA | #VALUE! |
| ENSMUSG00000117848.2 | 0 | NA | NA | NA | NA | NA | #VALUE! |
| ENSMUSG00000117849.2 | 0 | NA | NA | NA | NA | NA | #VALUE! |
| ENSMUSG00000117850.2 | 0 | NA | NA | NA | NA | NA | #VALUE! |
| ENSMUSG00000117851.2 | 0 | NA | NA | NA | NA | NA | #VALUE! |
| ENSMUSG00000117852.2 | 0 | NA | NA | NA | NA | NA | #VALUE! |
| ENSMUSG00000117853.2 | 0 | NA | NA | NA | NA | NA | #VALUE! |
| ENSMUSG00000117854.2 | 0 | NA | NA | NA | NA | NA | #VALUE! |
| ENSMUSG00000117855.2 | 0 | NA | NA | NA | NA | NA | #VALUE! |
| ENSMUSG00000117856.2 | 0 | NA | NA | NA | NA | NA | #VALUE! |
| ENSMUSG00000117857.2 | 0 | NA | NA | NA | NA | NA | #VALUE! |
| ENSMUSG00000117858.2 | 0 | NA | NA | NA | NA | NA | #VALUE! |
| ENSMUSG00000117859.2 | 0 | NA | NA | NA | NA | NA | #VALUE! |
| ENSMUSG00000117860.2 | 0 | NA | NA | NA | NA | NA | #VALUE! |
| ENSMUSG00000117861.2 | 0 | NA | NA | NA | NA | NA | #VALUE! |
| ENSMUSG00000117862.2 | 0 | NA | NA | NA | NA | NA | #VALUE! |
| ENSMUSG00000117863.2 | 0 | NA | NA | NA | NA | NA | #VALUE! |
| ENSMUSG00000117864.2 | 0 | NA | NA | NA | NA | NA | #VALUE! |
| ENSMUSG00000117865.2 | 0 | NA | NA | NA | NA | NA | #VALUE! |
| ENSMUSG00000117866.2 | 0 | NA | NA | NA | NA | NA | #VALUE! |
| ENSMUSG00000117867.2 | 0 | NA | NA | NA | NA | NA | #VALUE! |
| ENSMUSG00000117868.2 | 0 | NA | NA | NA | NA | NA | #VALUE! |
| ENSMUSG00000117870.2 | 0 | NA | NA | NA | NA | NA | #VALUE! |
| ENSMUSG00000117871.2 | 0 | NA | NA | NA | NA | NA | #VALUE! |
| ENSMUSG00000117872.2 | 0.571289 | 1.516851 | 3.992912 | 0.379886 | 0.70403 | NA | #VALUE! |
| ENSMUSG00000117873.2 | 0 | NA | NA | NA | NA | NA | #VALUE! |
| ENSMUSG00000117874.2 | 0 | NA | NA | NA | NA | NA | #VALUE! |
| ENSMUSG00000117875.2 | 0 | NA | NA | NA | NA | NA | #VALUE! |
| ENSMUSG00000117876.2 | 0 | NA | NA | NA | NA | NA | #VALUE! |
| ENSMUSG00000117877.2 | 0.093255 | 0.868093 | 4.072464 | 0.213162 | 0.831201 | NA | #VALUE! |
| ENSMUSG00000117878.2 | 0 | NA | NA | NA | NA | NA | #VALUE! |
| ENSMUSG00000117880.2 | 0 | NA | NA | NA | NA | NA | #VALUE! |
| ENSMUSG00000117881.2 | 0 | NA | NA | NA | NA | NA | #VALUE! |
| ENSMUSG00000117882.2 | 0 | NA | NA | NA | NA | NA | #VALUE! |
| ENSMUSG00000117883.2 | 0 | NA | NA | NA | NA | NA | #VALUE! |
| ENSMUSG00000117884.2 | 0.168284 | 1.027628 | 4.067126 | 0.252667 | 0.800526 | NA | #VALUE! |
| ENSMUSG00000117885.2 | 0 | NA | NA | NA | NA | NA | #VALUE! |
| ENSMUSG00000117886.2 | 0 | NA | NA | NA | NA | NA | #VALUE! |
| ENSMUSG00000117887.2 | 0 | NA | NA | NA | NA | NA | #VALUE! |
| ENSMUSG00000117888.2 | 0 | NA | NA | NA | NA | NA | #VALUE! |
| ENSMUSG00000117889.2 | 0 | NA | NA | NA | NA | NA | #VALUE! |
| ENSMUSG00000117890.2 | 0 | NA | NA | NA | NA | NA | #VALUE! |

| | | | | | | | |
|---|---|---|---|---|---|---|---|
| ENSMUSG00000117891.2 | 0 | NA | NA | NA | NA | NA | #VALUE! |
| ENSMUSG00000117892.2 | 0 | NA | NA | NA | NA | NA | #VALUE! |
| ENSMUSG00000117893.2 | 0 | NA | NA | NA | NA | NA | #VALUE! |
| ENSMUSG00000117894.2 | 0 | NA | NA | NA | NA | NA | #VALUE! |
| ENSMUSG00000117895.2 | 0 | NA | NA | NA | NA | NA | #VALUE! |
| ENSMUSG00000117896.2 | 0 | NA | NA | NA | NA | NA | #VALUE! |
| ENSMUSG00000117897.2 | 0 | NA | NA | NA | NA | NA | #VALUE! |
| ENSMUSG00000117898.2 | 0 | NA | NA | NA | NA | NA | #VALUE! |
| ENSMUSG00000117899.2 | 0 | NA | NA | NA | NA | NA | #VALUE! |
| ENSMUSG00000117900.2 | 0 | NA | NA | NA | NA | NA | #VALUE! |
| ENSMUSG00000117901.2 | 0 | NA | NA | NA | NA | NA | #VALUE! |
| ENSMUSG00000117902.2 | 0 | NA | NA | NA | NA | NA | #VALUE! |
| ENSMUSG00000117903.2 | 0 | NA | NA | NA | NA | NA | #VALUE! |
| ENSMUSG00000117904.2 | 0 | NA | NA | NA | NA | NA | #VALUE! |
| ENSMUSG00000117905.2 | 0 | NA | NA | NA | NA | NA | #VALUE! |
| ENSMUSG00000117906.2 | 0 | NA | NA | NA | NA | NA | #VALUE! |
| ENSMUSG00000117907.2 | 0 | NA | NA | NA | NA | NA | #VALUE! |
| ENSMUSG00000117908.2 | 0 | NA | NA | NA | NA | NA | #VALUE! |
| ENSMUSG00000117909.2 | 0 | NA | NA | NA | NA | NA | #VALUE! |
| ENSMUSG00000117910.2 | 0 | NA | NA | NA | NA | NA | #VALUE! |
| ENSMUSG00000117911.2 | 0 | NA | NA | NA | NA | NA | #VALUE! |
| ENSMUSG00000117912.2 | 0 | NA | NA | NA | NA | NA | #VALUE! |
| ENSMUSG00000117913.2 | 0 | NA | NA | NA | NA | NA | #VALUE! |
| ENSMUSG00000117914.2 | 0 | NA | NA | NA | NA | NA | #VALUE! |
| ENSMUSG00000117915.2 | 0 | NA | NA | NA | NA | NA | #VALUE! |
| ENSMUSG00000117916.2 | 0 | NA | NA | NA | NA | NA | #VALUE! |
| ENSMUSG00000117917.2 | 0 | NA | NA | NA | NA | NA | #VALUE! |
| ENSMUSG00000117918.2 | 0 | NA | NA | NA | NA | NA | #VALUE! |
| ENSMUSG00000117919.2 | 0 | NA | NA | NA | NA | NA | #VALUE! |
| ENSMUSG00000117920.2 | 0 | NA | NA | NA | NA | NA | #VALUE! |
| ENSMUSG00000117921.2 | 0 | NA | NA | NA | NA | NA | #VALUE! |
| ENSMUSG00000117922.2 | 0 | NA | NA | NA | NA | NA | #VALUE! |
| ENSMUSG00000117923.2 | 0 | NA | NA | NA | NA | NA | #VALUE! |
| ENSMUSG00000117925.2 | 0 | NA | NA | NA | NA | NA | #VALUE! |
| ENSMUSG00000117926.2 | 0 | NA | NA | NA | NA | NA | #VALUE! |
| ENSMUSG00000117927.2 | 0 | NA | NA | NA | NA | NA | #VALUE! |
| ENSMUSG00000117928.2 | 0 | NA | NA | NA | NA | NA | #VALUE! |
| ENSMUSG00000117929.2 | 0 | NA | NA | NA | NA | NA | #VALUE! |
| ENSMUSG00000117930.2 | 0 | NA | NA | NA | NA | NA | #VALUE! |
| ENSMUSG00000117931.2 | 0 | NA | NA | NA | NA | NA | #VALUE! |
| ENSMUSG00000117932.2 | 0 | NA | NA | NA | NA | NA | #VALUE! |
| ENSMUSG00000117933.2 | 0 | NA | NA | NA | NA | NA | #VALUE! |
| ENSMUSG00000117934.2 | 0 | NA | NA | NA | NA | NA | #VALUE! |
| ENSMUSG00000117935.2 | 0 | NA | NA | NA | NA | NA | #VALUE! |

| | | | | | | | |
|---|---|---|---|---|---|---|---|
| ENSMUSG00000117936.2 | 0 | NA | NA | NA | NA | NA | #VALUE! |
| ENSMUSG00000117937.2 | 0 | NA | NA | NA | NA | NA | #VALUE! |
| ENSMUSG00000117938.2 | 0 | NA | NA | NA | NA | NA | #VALUE! |
| ENSMUSG00000117939.2 | 0 | NA | NA | NA | NA | NA | #VALUE! |
| ENSMUSG00000117940.2 | 0 | NA | NA | NA | NA | NA | #VALUE! |
| ENSMUSG00000117941.2 | 0 | NA | NA | NA | NA | NA | #VALUE! |
| ENSMUSG00000117942.2 | 0 | NA | NA | NA | NA | NA | #VALUE! |
| ENSMUSG00000117943.2 | 0 | NA | NA | NA | NA | NA | #VALUE! |
| ENSMUSG00000117944.2 | 0 | NA | NA | NA | NA | NA | #VALUE! |
| ENSMUSG00000117945.2 | 0 | NA | NA | NA | NA | NA | #VALUE! |
| ENSMUSG00000117946.2 | 0 | NA | NA | NA | NA | NA | #VALUE! |
| ENSMUSG00000117947.2 | 0 | NA | NA | NA | NA | NA | #VALUE! |
| ENSMUSG00000117948.2 | 0 | NA | NA | NA | NA | NA | #VALUE! |
| ENSMUSG00000117949.2 | 0 | NA | NA | NA | NA | NA | #VALUE! |
| ENSMUSG00000117950.2 | 0 | NA | NA | NA | NA | NA | #VALUE! |
| ENSMUSG00000117951.2 | 0 | NA | NA | NA | NA | NA | #VALUE! |
| ENSMUSG00000117952.2 | 0 | NA | NA | NA | NA | NA | #VALUE! |
| ENSMUSG00000117953.2 | 0 | NA | NA | NA | NA | NA | #VALUE! |
| ENSMUSG00000117955.2 | 0 | NA | NA | NA | NA | NA | #VALUE! |
| ENSMUSG00000117956.2 | 0 | NA | NA | NA | NA | NA | #VALUE! |
| ENSMUSG00000117957.2 | 0 | NA | NA | NA | NA | NA | #VALUE! |
| ENSMUSG00000117958.2 | 0 | NA | NA | NA | NA | NA | #VALUE! |
| ENSMUSG00000117959.2 | 0 | NA | NA | NA | NA | NA | #VALUE! |
| ENSMUSG00000117960.2 | 0 | NA | NA | NA | NA | NA | #VALUE! |
| ENSMUSG00000117961.2 | 0 | NA | NA | NA | NA | NA | #VALUE! |
| ENSMUSG00000117963.2 | 0 | NA | NA | NA | NA | NA | #VALUE! |
| ENSMUSG00000117964.2 | 0 | NA | NA | NA | NA | NA | #VALUE! |
| ENSMUSG00000117965.2 | 0 | NA | NA | NA | NA | NA | #VALUE! |
| ENSMUSG00000117966.2 | 0 | NA | NA | NA | NA | NA | #VALUE! |
| ENSMUSG00000117967.2 | 0 | NA | NA | NA | NA | NA | #VALUE! |
| ENSMUSG00000117968.2 | 0 | NA | NA | NA | NA | NA | #VALUE! |
| ENSMUSG00000117969.2 | 0 | NA | NA | NA | NA | NA | #VALUE! |
| ENSMUSG00000117970.2 | 0 | NA | NA | NA | NA | NA | #VALUE! |
| ENSMUSG00000117971.2 | 0 | NA | NA | NA | NA | NA | #VALUE! |
| ENSMUSG00000117972.2 | 0 | NA | NA | NA | NA | NA | #VALUE! |
| ENSMUSG00000117973.2 | 0 | NA | NA | NA | NA | NA | #VALUE! |
| ENSMUSG00000117974.2 | 0 | NA | NA | NA | NA | NA | #VALUE! |
| ENSMUSG00000117975.2 | 0 | NA | NA | NA | NA | NA | #VALUE! |
| ENSMUSG00000117976.2 | 0 | NA | NA | NA | NA | NA | #VALUE! |
| ENSMUSG00000117977.2 | 0 | NA | NA | NA | NA | NA | #VALUE! |
| ENSMUSG00000117978.2 | 0 | NA | NA | NA | NA | NA | #VALUE! |
| ENSMUSG00000117979.2 | 0 | NA | NA | NA | NA | NA | #VALUE! |
| ENSMUSG00000117980.2 | 0 | NA | NA | NA | NA | NA | #VALUE! |
| ENSMUSG00000117981.2 | 0 | NA | NA | NA | NA | NA | #VALUE! |

| | | | | | | | |
|---|---|---|---|---|---|---|---|
| ENSMUSG00000117982.2 | 0 | NA | NA | NA | NA | NA | #VALUE! |
| ENSMUSG00000117983.2 | 0 | NA | NA | NA | NA | NA | #VALUE! |
| ENSMUSG00000117984.2 | 0 | NA | NA | NA | NA | NA | #VALUE! |
| ENSMUSG00000117985.2 | 0 | NA | NA | NA | NA | NA | #VALUE! |
| ENSMUSG00000117986.2 | 0 | NA | NA | NA | NA | NA | #VALUE! |
| ENSMUSG00000117987.2 | 0 | NA | NA | NA | NA | NA | #VALUE! |
| ENSMUSG00000117988.2 | 0 | NA | NA | NA | NA | NA | #VALUE! |
| ENSMUSG00000117989.2 | 0 | NA | NA | NA | NA | NA | #VALUE! |
| ENSMUSG00000117990.2 | 0 | NA | NA | NA | NA | NA | #VALUE! |
| ENSMUSG00000117991.2 | 0 | NA | NA | NA | NA | NA | #VALUE! |
| ENSMUSG00000117992.2 | 0 | NA | NA | NA | NA | NA | #VALUE! |
| ENSMUSG00000117993.2 | 0 | NA | NA | NA | NA | NA | #VALUE! |
| ENSMUSG00000117994.2 | 0 | NA | NA | NA | NA | NA | #VALUE! |
| ENSMUSG00000117995.2 | 0 | NA | NA | NA | NA | NA | #VALUE! |
| ENSMUSG00000117996.2 | 0 | NA | NA | NA | NA | NA | #VALUE! |
| ENSMUSG00000117998.2 | 0 | NA | NA | NA | NA | NA | #VALUE! |
| ENSMUSG00000117999.2 | 0 | NA | NA | NA | NA | NA | #VALUE! |
| ENSMUSG00000118000.2 | 0 | NA | NA | NA | NA | NA | #VALUE! |
| ENSMUSG00000118001.2 | 0 | NA | NA | NA | NA | NA | #VALUE! |
| ENSMUSG00000118002.2 | 0 | NA | NA | NA | NA | NA | #VALUE! |
| ENSMUSG00000118003.2 | 0 | NA | NA | NA | NA | NA | #VALUE! |
| ENSMUSG00000118004.2 | 0 | NA | NA | NA | NA | NA | #VALUE! |
| ENSMUSG00000118005.2 | 0 | NA | NA | NA | NA | NA | #VALUE! |
| ENSMUSG00000118006.2 | 0 | NA | NA | NA | NA | NA | #VALUE! |
| ENSMUSG00000118007.2 | 0 | NA | NA | NA | NA | NA | #VALUE! |
| ENSMUSG00000118008.2 | 0 | NA | NA | NA | NA | NA | #VALUE! |
| ENSMUSG00000118009.2 | 0 | NA | NA | NA | NA | NA | #VALUE! |
| ENSMUSG00000118010.2 | 0 | NA | NA | NA | NA | NA | #VALUE! |
| ENSMUSG00000118011.2 | 0 | NA | NA | NA | NA | NA | #VALUE! |
| ENSMUSG00000118012.2 | 0 | NA | NA | NA | NA | NA | #VALUE! |
| ENSMUSG00000118013.2 | 0 | NA | NA | NA | NA | NA | #VALUE! |
| ENSMUSG00000118014.2 | 0 | NA | NA | NA | NA | NA | #VALUE! |
| ENSMUSG00000118015.2 | 0 | NA | NA | NA | NA | NA | #VALUE! |
| ENSMUSG00000118016.2 | 0 | NA | NA | NA | NA | NA | #VALUE! |
| ENSMUSG00000118018.2 | 0 | NA | NA | NA | NA | NA | #VALUE! |
| ENSMUSG00000118020.2 | 0 | NA | NA | NA | NA | NA | #VALUE! |
| ENSMUSG00000118021.2 | 0 | NA | NA | NA | NA | NA | #VALUE! |
| ENSMUSG00000118022.2 | 0 | NA | NA | NA | NA | NA | #VALUE! |
| ENSMUSG00000118023.2 | 0 | NA | NA | NA | NA | NA | #VALUE! |
| ENSMUSG00000118024.2 | 0 | NA | NA | NA | NA | NA | #VALUE! |
| ENSMUSG00000118025.2 | 0 | NA | NA | NA | NA | NA | #VALUE! |
| ENSMUSG00000118026.2 | 0 | NA | NA | NA | NA | NA | #VALUE! |
| ENSMUSG00000118027.2 | 0 | NA | NA | NA | NA | NA | #VALUE! |
| ENSMUSG00000118028.2 | 0 | NA | NA | NA | NA | NA | #VALUE! |

| | | | | | | | |
|---|---|---|---|---|---|---|---|
| ENSMUSG00000118029.2 | 0 | NA | NA | NA | NA | NA | #VALUE! |
| ENSMUSG00000118030.2 | 0.466276 | 2.449889 | 3.997179 | 0.612904 | 0.53994 | NA | #VALUE! |
| ENSMUSG00000118031.2 | 0 | NA | NA | NA | NA | NA | #VALUE! |
| ENSMUSG00000118032.2 | 0 | NA | NA | NA | NA | NA | #VALUE! |
| ENSMUSG00000118033.2 | 0 | NA | NA | NA | NA | NA | #VALUE! |
| ENSMUSG00000118034.2 | 0 | NA | NA | NA | NA | NA | #VALUE! |
| ENSMUSG00000118035.2 | 0 | NA | NA | NA | NA | NA | #VALUE! |
| ENSMUSG00000118036.2 | 0 | NA | NA | NA | NA | NA | #VALUE! |
| ENSMUSG00000118037.2 | 0 | NA | NA | NA | NA | NA | #VALUE! |
| ENSMUSG00000118038.2 | 0 | NA | NA | NA | NA | NA | #VALUE! |
| ENSMUSG00000118039.2 | 0 | NA | NA | NA | NA | NA | #VALUE! |
| ENSMUSG00000118040.2 | 0 | NA | NA | NA | NA | NA | #VALUE! |
| ENSMUSG00000118041.2 | 0 | NA | NA | NA | NA | NA | #VALUE! |
| ENSMUSG00000118042.2 | 0 | NA | NA | NA | NA | NA | #VALUE! |
| ENSMUSG00000118043.2 | 0 | NA | NA | NA | NA | NA | #VALUE! |
| ENSMUSG00000118044.2 | 0 | NA | NA | NA | NA | NA | #VALUE! |
| ENSMUSG00000118045.2 | 0 | NA | NA | NA | NA | NA | #VALUE! |
| ENSMUSG00000118046.2 | 0 | NA | NA | NA | NA | NA | #VALUE! |
| ENSMUSG00000118047.2 | 0 | NA | NA | NA | NA | NA | #VALUE! |
| ENSMUSG00000118048.2 | 0 | NA | NA | NA | NA | NA | #VALUE! |
| ENSMUSG00000118049.2 | 0 | NA | NA | NA | NA | NA | #VALUE! |
| ENSMUSG00000118050.2 | 0 | NA | NA | NA | NA | NA | #VALUE! |
| ENSMUSG00000118051.2 | 0 | NA | NA | NA | NA | NA | #VALUE! |
| ENSMUSG00000118053.2 | 0 | NA | NA | NA | NA | NA | #VALUE! |
| ENSMUSG00000118054.2 | 0 | NA | NA | NA | NA | NA | #VALUE! |
| ENSMUSG00000118055.2 | 0 | NA | NA | NA | NA | NA | #VALUE! |
| ENSMUSG00000118056.2 | 0 | NA | NA | NA | NA | NA | #VALUE! |
| ENSMUSG00000118057.2 | 0 | NA | NA | NA | NA | NA | #VALUE! |
| ENSMUSG00000118058.2 | 0 | NA | NA | NA | NA | NA | #VALUE! |
| ENSMUSG00000118059.2 | 0 | NA | NA | NA | NA | NA | #VALUE! |
| ENSMUSG00000118060.2 | 0 | NA | NA | NA | NA | NA | #VALUE! |
| ENSMUSG00000118061.2 | 0 | NA | NA | NA | NA | NA | #VALUE! |
| ENSMUSG00000118062.2 | 0 | NA | NA | NA | NA | NA | #VALUE! |
| ENSMUSG00000118063.2 | 0 | NA | NA | NA | NA | NA | #VALUE! |
| ENSMUSG00000118064.2 | 0 | NA | NA | NA | NA | NA | #VALUE! |
| ENSMUSG00000118065.2 | 0 | NA | NA | NA | NA | NA | #VALUE! |
| ENSMUSG00000118066.2 | 0 | NA | NA | NA | NA | NA | #VALUE! |
| ENSMUSG00000118067.2 | 0 | NA | NA | NA | NA | NA | #VALUE! |
| ENSMUSG00000118068.2 | 0 | NA | NA | NA | NA | NA | #VALUE! |
| ENSMUSG00000118070.2 | 0 | NA | NA | NA | NA | NA | #VALUE! |
| ENSMUSG00000118071.2 | 1.475853 | 4.029233 | 3.937502 | 1.023297 | 0.306167 | NA | #VALUE! |
| ENSMUSG00000118072.2 | 0 | NA | NA | NA | NA | NA | #VALUE! |
| ENSMUSG00000118073.2 | 0 | NA | NA | NA | NA | NA | #VALUE! |
| ENSMUSG00000118074.2 | 0.105013 | -0.97324 | 4.076293 | -0.23876 | 0.811295 | NA | #VALUE! |

| | | | | | | | |
|---|---|---|---|---|---|---|---|
| ENSMUSG00000118075.2 | 0 | NA | NA | NA | NA | NA | #VALUE! |
| ENSMUSG00000118076.2 | 0 | NA | NA | NA | NA | NA | #VALUE! |
| ENSMUSG00000118077.2 | 0 | NA | NA | NA | NA | NA | #VALUE! |
| ENSMUSG00000118078.2 | 0 | NA | NA | NA | NA | NA | #VALUE! |
| ENSMUSG00000118079.2 | 0 | NA | NA | NA | NA | NA | #VALUE! |
| ENSMUSG00000118080.2 | 0 | NA | NA | NA | NA | NA | #VALUE! |
| ENSMUSG00000118081.2 | 0 | NA | NA | NA | NA | NA | #VALUE! |
| ENSMUSG00000118082.2 | 0 | NA | NA | NA | NA | NA | #VALUE! |
| ENSMUSG00000118083.2 | 0 | NA | NA | NA | NA | NA | #VALUE! |
| ENSMUSG00000118084.2 | 2.705731 | 4.913666 | 3.923477 | 1.252375 | 0.210433 | NA | #VALUE! |
| ENSMUSG00000118085.2 | 0 | NA | NA | NA | NA | NA | #VALUE! |
| ENSMUSG00000118086.2 | 0 | NA | NA | NA | NA | NA | #VALUE! |
| ENSMUSG00000118087.2 | 3.132626 | -5.17361 | 3.921071 | -1.31944 | 0.187023 | NA | #VALUE! |
| ENSMUSG00000118088.2 | 0 | NA | NA | NA | NA | NA | #VALUE! |
| ENSMUSG00000118089.2 | 0 | NA | NA | NA | NA | NA | #VALUE! |
| ENSMUSG00000118090.2 | 0 | NA | NA | NA | NA | NA | #VALUE! |
| ENSMUSG00000118091.2 | 0 | NA | NA | NA | NA | NA | #VALUE! |
| ENSMUSG00000118092.2 | 0 | NA | NA | NA | NA | NA | #VALUE! |
| ENSMUSG00000118093.2 | 0 | NA | NA | NA | NA | NA | #VALUE! |
| ENSMUSG00000118094.2 | 0 | NA | NA | NA | NA | NA | #VALUE! |
| ENSMUSG00000118096.2 | 0 | NA | NA | NA | NA | NA | #VALUE! |
| ENSMUSG00000118097.2 | 0 | NA | NA | NA | NA | NA | #VALUE! |
| ENSMUSG00000118098.2 | 0 | NA | NA | NA | NA | NA | #VALUE! |
| ENSMUSG00000118099.2 | 0 | NA | NA | NA | NA | NA | #VALUE! |
| ENSMUSG00000118100.2 | 0.466276 | 2.449889 | 3.997179 | 0.612904 | 0.53994 | NA | #VALUE! |
| ENSMUSG00000118101.2 | 0 | NA | NA | NA | NA | NA | #VALUE! |
| ENSMUSG00000118102.2 | 0 | NA | NA | NA | NA | NA | #VALUE! |
| ENSMUSG00000118103.2 | 0 | NA | NA | NA | NA | NA | #VALUE! |
| ENSMUSG00000118104.2 | 0 | NA | NA | NA | NA | NA | #VALUE! |
| ENSMUSG00000118105.2 | 0 | NA | NA | NA | NA | NA | #VALUE! |
| ENSMUSG00000118106.2 | 0 | NA | NA | NA | NA | NA | #VALUE! |
| ENSMUSG00000118107.2 | 0 | NA | NA | NA | NA | NA | #VALUE! |
| ENSMUSG00000118108.2 | 0 | NA | NA | NA | NA | NA | #VALUE! |
| ENSMUSG00000118109.2 | 0 | NA | NA | NA | NA | NA | #VALUE! |
| ENSMUSG00000118110.2 | 0 | NA | NA | NA | NA | NA | #VALUE! |
| ENSMUSG00000118111.2 | 0 | NA | NA | NA | NA | NA | #VALUE! |
| ENSMUSG00000118112.2 | 0 | NA | NA | NA | NA | NA | #VALUE! |
| ENSMUSG00000118113.2 | 0 | NA | NA | NA | NA | NA | #VALUE! |
| ENSMUSG00000118114.2 | 0 | NA | NA | NA | NA | NA | #VALUE! |
| ENSMUSG00000118115.2 | 0 | NA | NA | NA | NA | NA | #VALUE! |
| ENSMUSG00000118116.2 | 0 | NA | NA | NA | NA | NA | #VALUE! |
| ENSMUSG00000118117.2 | 0 | NA | NA | NA | NA | NA | #VALUE! |
| ENSMUSG00000118118.2 | 0 | NA | NA | NA | NA | NA | #VALUE! |
| ENSMUSG00000118119.2 | 0 | NA | NA | NA | NA | NA | #VALUE! |

| | | | | | | | |
|---|---|---|---|---|---|---|---|
| ENSMUSG00000118120.2 | 0 | NA | NA | NA | NA | NA | #VALUE! |
| ENSMUSG00000118121.2 | 0 | NA | NA | NA | NA | NA | #VALUE! |
| ENSMUSG00000118122.2 | 0 | NA | NA | NA | NA | NA | #VALUE! |
| ENSMUSG00000118123.2 | 0 | NA | NA | NA | NA | NA | #VALUE! |
| ENSMUSG00000118124.2 | 0 | NA | NA | NA | NA | NA | #VALUE! |
| ENSMUSG00000118125.2 | 0 | NA | NA | NA | NA | NA | #VALUE! |
| ENSMUSG00000118126.2 | 0 | NA | NA | NA | NA | NA | #VALUE! |
| ENSMUSG00000118127.2 | 0 | NA | NA | NA | NA | NA | #VALUE! |
| ENSMUSG00000118128.2 | 0 | NA | NA | NA | NA | NA | #VALUE! |
| ENSMUSG00000118129.2 | 0 | NA | NA | NA | NA | NA | #VALUE! |
| ENSMUSG00000118130.2 | 0 | NA | NA | NA | NA | NA | #VALUE! |
| ENSMUSG00000118131.2 | 0 | NA | NA | NA | NA | NA | #VALUE! |
| ENSMUSG00000118132.2 | 0 | NA | NA | NA | NA | NA | #VALUE! |
| ENSMUSG00000118133.2 | 0.441765 | 0.012025 | 4.059931 | 0.002962 | 0.997637 | NA | #VALUE! |
| ENSMUSG00000118134.2 | 0.841418 | 3.240539 | 3.959551 | 0.818411 | 0.413123 | NA | #VALUE! |
| ENSMUSG00000118135.2 | 0 | NA | NA | NA | NA | NA | #VALUE! |
| ENSMUSG00000118136.2 | 0 | NA | NA | NA | NA | NA | #VALUE! |
| ENSMUSG00000118137.2 | 0 | NA | NA | NA | NA | NA | #VALUE! |
| ENSMUSG00000118138.2 | 0 | NA | NA | NA | NA | NA | #VALUE! |
| ENSMUSG00000118139.2 | 0 | NA | NA | NA | NA | NA | #VALUE! |
| ENSMUSG00000118140.2 | 0 | NA | NA | NA | NA | NA | #VALUE! |
| ENSMUSG00000118141.2 | 0 | NA | NA | NA | NA | NA | #VALUE! |
| ENSMUSG00000118142.2 | 0 | NA | NA | NA | NA | NA | #VALUE! |
| ENSMUSG00000118143.2 | 0.195789 | -1.05581 | 4.073302 | -0.2592 | 0.795479 | NA | #VALUE! |
| ENSMUSG00000118144.2 | 0 | NA | NA | NA | NA | NA | #VALUE! |
| ENSMUSG00000118145.2 | 0 | NA | NA | NA | NA | NA | #VALUE! |
| ENSMUSG00000118146.2 | 0 | NA | NA | NA | NA | NA | #VALUE! |
| ENSMUSG00000118147.2 | 0 | NA | NA | NA | NA | NA | #VALUE! |
| ENSMUSG00000118148.2 | 0 | NA | NA | NA | NA | NA | #VALUE! |
| ENSMUSG00000118149.2 | 0 | NA | NA | NA | NA | NA | #VALUE! |
| ENSMUSG00000118150.2 | 0 | NA | NA | NA | NA | NA | #VALUE! |
| ENSMUSG00000118151.2 | 0 | NA | NA | NA | NA | NA | #VALUE! |
| ENSMUSG00000118152.2 | 0 | NA | NA | NA | NA | NA | #VALUE! |
| ENSMUSG00000118153.2 | 0 | NA | NA | NA | NA | NA | #VALUE! |
| ENSMUSG00000118154.2 | 0 | NA | NA | NA | NA | NA | #VALUE! |
| ENSMUSG00000118156.2 | 0 | NA | NA | NA | NA | NA | #VALUE! |
| ENSMUSG00000118157.2 | 0 | NA | NA | NA | NA | NA | #VALUE! |
| ENSMUSG00000118158.2 | 0 | NA | NA | NA | NA | NA | #VALUE! |
| ENSMUSG00000118159.2 | 0 | NA | NA | NA | NA | NA | #VALUE! |
| ENSMUSG00000118160.2 | 0 | NA | NA | NA | NA | NA | #VALUE! |
| ENSMUSG00000118161.2 | 0 | NA | NA | NA | NA | NA | #VALUE! |
| ENSMUSG00000118162.2 | 0 | NA | NA | NA | NA | NA | #VALUE! |
| ENSMUSG00000118163.2 | 0 | NA | NA | NA | NA | NA | #VALUE! |
| ENSMUSG00000118164.2 | 0 | NA | NA | NA | NA | NA | #VALUE! |

| | | | | | | | |
|---|---|---|---|---|---|---|---|
| ENSMUSG00000118165.2 | 0 | NA | NA | NA | NA | NA | #VALUE! |
| ENSMUSG00000118166.2 | 0 | NA | NA | NA | NA | NA | #VALUE! |
| ENSMUSG00000118167.2 | 0 | NA | NA | NA | NA | NA | #VALUE! |
| ENSMUSG00000118168.2 | 0 | NA | NA | NA | NA | NA | #VALUE! |
| ENSMUSG00000118169.2 | 0 | NA | NA | NA | NA | NA | #VALUE! |
| ENSMUSG00000118170.2 | 0 | NA | NA | NA | NA | NA | #VALUE! |
| ENSMUSG00000118171.2 | 0 | NA | NA | NA | NA | NA | #VALUE! |
| ENSMUSG00000118172.2 | 0 | NA | NA | NA | NA | NA | #VALUE! |
| ENSMUSG00000118173.2 | 0 | NA | NA | NA | NA | NA | #VALUE! |
| ENSMUSG00000118174.2 | 0 | NA | NA | NA | NA | NA | #VALUE! |
| ENSMUSG00000118175.2 | 0 | NA | NA | NA | NA | NA | #VALUE! |
| ENSMUSG00000118176.2 | 0 | NA | NA | NA | NA | NA | #VALUE! |
| ENSMUSG00000118177.2 | 0 | NA | NA | NA | NA | NA | #VALUE! |
| ENSMUSG00000118178.2 | 0 | NA | NA | NA | NA | NA | #VALUE! |
| ENSMUSG00000118179.2 | 0 | NA | NA | NA | NA | NA | #VALUE! |
| ENSMUSG00000118180.2 | 0 | NA | NA | NA | NA | NA | #VALUE! |
| ENSMUSG00000118181.2 | 0 | NA | NA | NA | NA | NA | #VALUE! |
| ENSMUSG00000118182.2 | 0 | NA | NA | NA | NA | NA | #VALUE! |
| ENSMUSG00000118183.2 | 0 | NA | NA | NA | NA | NA | #VALUE! |
| ENSMUSG00000118184.2 | 0 | NA | NA | NA | NA | NA | #VALUE! |
| ENSMUSG00000118185.2 | 0 | NA | NA | NA | NA | NA | #VALUE! |
| ENSMUSG00000118186.2 | 0 | NA | NA | NA | NA | NA | #VALUE! |
| ENSMUSG00000118187.2 | 0 | NA | NA | NA | NA | NA | #VALUE! |
| ENSMUSG00000118188.2 | 0 | NA | NA | NA | NA | NA | #VALUE! |
| ENSMUSG00000118189.2 | 0 | NA | NA | NA | NA | NA | #VALUE! |
| ENSMUSG00000118190.2 | 0 | NA | NA | NA | NA | NA | #VALUE! |
| ENSMUSG00000118191.2 | 0 | NA | NA | NA | NA | NA | #VALUE! |
| ENSMUSG00000118192.2 | 0 | NA | NA | NA | NA | NA | #VALUE! |
| ENSMUSG00000118193.2 | 0 | NA | NA | NA | NA | NA | #VALUE! |
| ENSMUSG00000118194.2 | 0 | NA | NA | NA | NA | NA | #VALUE! |
| ENSMUSG00000118195.2 | 0 | NA | NA | NA | NA | NA | #VALUE! |
| ENSMUSG00000118196.2 | 0 | NA | NA | NA | NA | NA | #VALUE! |
| ENSMUSG00000118197.2 | 1.92544 | -3.49268 | 3.922839 | -0.89035 | 0.373281 | NA | #VALUE! |
| ENSMUSG00000118198.2 | 0 | NA | NA | NA | NA | NA | #VALUE! |
| ENSMUSG00000118199.2 | 0 | NA | NA | NA | NA | NA | #VALUE! |
| ENSMUSG00000118200.2 | 0 | NA | NA | NA | NA | NA | #VALUE! |
| ENSMUSG00000118201.2 | 0 | NA | NA | NA | NA | NA | #VALUE! |
| ENSMUSG00000118202.2 | 0 | NA | NA | NA | NA | NA | #VALUE! |
| ENSMUSG00000118203.2 | 0 | NA | NA | NA | NA | NA | #VALUE! |
| ENSMUSG00000118204.2 | 0 | NA | NA | NA | NA | NA | #VALUE! |
| ENSMUSG00000118205.2 | 0 | NA | NA | NA | NA | NA | #VALUE! |
| ENSMUSG00000118206.2 | 0 | NA | NA | NA | NA | NA | #VALUE! |
| ENSMUSG00000118207.2 | 0 | NA | NA | NA | NA | NA | #VALUE! |
| ENSMUSG00000118208.2 | 0 | NA | NA | NA | NA | NA | #VALUE! |

| | | | | | | | |
|---|---|---|---|---|---|---|---|
| ENSMUSG00000118209.2 | 0 | NA | NA | NA | NA | NA | #VALUE! |
| ENSMUSG00000118210.2 | 0 | NA | NA | NA | NA | NA | #VALUE! |
| ENSMUSG00000118211.2 | 0 | NA | NA | NA | NA | NA | #VALUE! |
| ENSMUSG00000118212.2 | 0 | NA | NA | NA | NA | NA | #VALUE! |
| ENSMUSG00000118213.2 | 0 | NA | NA | NA | NA | NA | #VALUE! |
| ENSMUSG00000118214.2 | 0 | NA | NA | NA | NA | NA | #VALUE! |
| ENSMUSG00000118215.2 | 0 | NA | NA | NA | NA | NA | #VALUE! |
| ENSMUSG00000118216.2 | 0 | NA | NA | NA | NA | NA | #VALUE! |
| ENSMUSG00000118217.2 | 0 | NA | NA | NA | NA | NA | #VALUE! |
| ENSMUSG00000118218.2 | 0 | NA | NA | NA | NA | NA | #VALUE! |
| ENSMUSG00000118219.2 | 0 | NA | NA | NA | NA | NA | #VALUE! |
| ENSMUSG00000118220.2 | 0.105013 | -0.97324 | 4.076293 | -0.23876 | 0.811295 | NA | #VALUE! |
| ENSMUSG00000118221.2 | 0 | NA | NA | NA | NA | NA | #VALUE! |
| ENSMUSG00000118222.2 | 0 | NA | NA | NA | NA | NA | #VALUE! |
| ENSMUSG00000118223.2 | 0 | NA | NA | NA | NA | NA | #VALUE! |
| ENSMUSG00000118224.2 | 0 | NA | NA | NA | NA | NA | #VALUE! |
| ENSMUSG00000118225.2 | 0 | NA | NA | NA | NA | NA | #VALUE! |
| ENSMUSG00000118226.2 | 0 | NA | NA | NA | NA | NA | #VALUE! |
| ENSMUSG00000118227.2 | 0 | NA | NA | NA | NA | NA | #VALUE! |
| ENSMUSG00000118228.2 | 0 | NA | NA | NA | NA | NA | #VALUE! |
| ENSMUSG00000118229.2 | 0 | NA | NA | NA | NA | NA | #VALUE! |
| ENSMUSG00000118230.2 | 0 | NA | NA | NA | NA | NA | #VALUE! |
| ENSMUSG00000118231.2 | 0 | NA | NA | NA | NA | NA | #VALUE! |
| ENSMUSG00000118232.2 | 0 | NA | NA | NA | NA | NA | #VALUE! |
| ENSMUSG00000118233.2 | 0 | NA | NA | NA | NA | NA | #VALUE! |
| ENSMUSG00000118234.2 | 0 | NA | NA | NA | NA | NA | #VALUE! |
| ENSMUSG00000118235.2 | 0 | NA | NA | NA | NA | NA | #VALUE! |
| ENSMUSG00000118236.2 | 0 | NA | NA | NA | NA | NA | #VALUE! |
| ENSMUSG00000118237.2 | 0 | NA | NA | NA | NA | NA | #VALUE! |
| ENSMUSG00000118238.2 | 0 | NA | NA | NA | NA | NA | #VALUE! |
| ENSMUSG00000118239.2 | 3.328415 | -5.26129 | 3.920229 | -1.34209 | 0.179567 | NA | #VALUE! |
| ENSMUSG00000118240.2 | 0 | NA | NA | NA | NA | NA | #VALUE! |
| ENSMUSG00000118241.2 | 0 | NA | NA | NA | NA | NA | #VALUE! |
| ENSMUSG00000118242.2 | 0 | NA | NA | NA | NA | NA | #VALUE! |
| ENSMUSG00000118243.2 | 1.260158 | -3.88425 | 3.941584 | -0.98545 | 0.324401 | NA | #VALUE! |
| ENSMUSG00000118244.2 | 0 | NA | NA | NA | NA | NA | #VALUE! |
| ENSMUSG00000118245.2 | 0 | NA | NA | NA | NA | NA | #VALUE! |
| ENSMUSG00000118246.2 | 0 | NA | NA | NA | NA | NA | #VALUE! |
| ENSMUSG00000118247.2 | 0 | NA | NA | NA | NA | NA | #VALUE! |
| ENSMUSG00000118248.2 | 0 | NA | NA | NA | NA | NA | #VALUE! |
| ENSMUSG00000118249.2 | 0 | NA | NA | NA | NA | NA | #VALUE! |
| ENSMUSG00000118250.2 | 0 | NA | NA | NA | NA | NA | #VALUE! |
| ENSMUSG00000118251.2 | 0 | NA | NA | NA | NA | NA | #VALUE! |
| ENSMUSG00000118252.2 | 0 | NA | NA | NA | NA | NA | #VALUE! |

| | | | | | | | |
|---|---|---|---|---|---|---|---|
| ENSMUSG00000118253.2 | 0 | NA | NA | NA | NA | NA | #VALUE! |
| ENSMUSG00000118254.2 | 0 | NA | NA | NA | NA | NA | #VALUE! |
| ENSMUSG00000118255.2 | 0 | NA | NA | NA | NA | NA | #VALUE! |
| ENSMUSG00000118256.2 | 0 | NA | NA | NA | NA | NA | #VALUE! |
| ENSMUSG00000118257.2 | 0 | NA | NA | NA | NA | NA | #VALUE! |
| ENSMUSG00000118258.2 | 0 | NA | NA | NA | NA | NA | #VALUE! |
| ENSMUSG00000118259.2 | 0 | NA | NA | NA | NA | NA | #VALUE! |
| ENSMUSG00000118260.2 | 0 | NA | NA | NA | NA | NA | #VALUE! |
| ENSMUSG00000118261.2 | 0 | NA | NA | NA | NA | NA | #VALUE! |
| ENSMUSG00000118262.2 | 0 | NA | NA | NA | NA | NA | #VALUE! |
| ENSMUSG00000118263.2 | 0 | NA | NA | NA | NA | NA | #VALUE! |
| ENSMUSG00000118264.2 | 0.457328 | 0.67272 | 4.035785 | 0.166689 | 0.867615 | NA | #VALUE! |
| ENSMUSG00000118265.2 | 0 | NA | NA | NA | NA | NA | #VALUE! |
| ENSMUSG00000118266.2 | 0 | NA | NA | NA | NA | NA | #VALUE! |
| ENSMUSG00000118267.2 | 0 | NA | NA | NA | NA | NA | #VALUE! |
| ENSMUSG00000118268.2 | 0 | NA | NA | NA | NA | NA | #VALUE! |
| ENSMUSG00000118269.2 | 0 | NA | NA | NA | NA | NA | #VALUE! |
| ENSMUSG00000118270.2 | 0 | NA | NA | NA | NA | NA | #VALUE! |
| ENSMUSG00000118271.2 | 0 | NA | NA | NA | NA | NA | #VALUE! |
| ENSMUSG00000118272.2 | 0 | NA | NA | NA | NA | NA | #VALUE! |
| ENSMUSG00000118273.2 | 0 | NA | NA | NA | NA | NA | #VALUE! |
| ENSMUSG00000118274.2 | 0 | NA | NA | NA | NA | NA | #VALUE! |
| ENSMUSG00000118275.2 | 0 | NA | NA | NA | NA | NA | #VALUE! |
| ENSMUSG00000118276.2 | 0 | NA | NA | NA | NA | NA | #VALUE! |
| ENSMUSG00000118277.2 | 0 | NA | NA | NA | NA | NA | #VALUE! |
| ENSMUSG00000118278.2 | 0 | NA | NA | NA | NA | NA | #VALUE! |
| ENSMUSG00000118279.2 | 0 | NA | NA | NA | NA | NA | #VALUE! |
| ENSMUSG00000118280.2 | 0 | NA | NA | NA | NA | NA | #VALUE! |
| ENSMUSG00000118281.2 | 0 | NA | NA | NA | NA | NA | #VALUE! |
| ENSMUSG00000118282.2 | 0 | NA | NA | NA | NA | NA | #VALUE! |
| ENSMUSG00000118283.2 | 0 | NA | NA | NA | NA | NA | #VALUE! |
| ENSMUSG00000118284.2 | 0 | NA | NA | NA | NA | NA | #VALUE! |
| ENSMUSG00000118285.2 | 0.168284 | 1.027628 | 4.067126 | 0.252667 | 0.800526 | NA | #VALUE! |
| ENSMUSG00000118286.2 | 0 | NA | NA | NA | NA | NA | #VALUE! |
| ENSMUSG00000118287.2 | 0 | NA | NA | NA | NA | NA | #VALUE! |
| ENSMUSG00000118288.2 | 0 | NA | NA | NA | NA | NA | #VALUE! |
| ENSMUSG00000118289.2 | 0 | NA | NA | NA | NA | NA | #VALUE! |
| ENSMUSG00000118290.2 | 0 | NA | NA | NA | NA | NA | #VALUE! |
| ENSMUSG00000118291.2 | 0 | NA | NA | NA | NA | NA | #VALUE! |
| ENSMUSG00000118292.2 | 0 | NA | NA | NA | NA | NA | #VALUE! |
| ENSMUSG00000118293.2 | 0 | NA | NA | NA | NA | NA | #VALUE! |
| ENSMUSG00000118294.2 | 0 | NA | NA | NA | NA | NA | #VALUE! |
| ENSMUSG00000118295.2 | 0 | NA | NA | NA | NA | NA | #VALUE! |
| ENSMUSG00000118296.2 | 0 | NA | NA | NA | NA | NA | #VALUE! |

| | | | | | | | |
|---|---|---|---|---|---|---|---|
| ENSMUSG00000118297.2 | 0 | NA | NA | NA | NA | NA | #VALUE! |
| ENSMUSG00000118298.2 | 0 | NA | NA | NA | NA | NA | #VALUE! |
| ENSMUSG00000118299.2 | 0.309124 | -1.83395 | 4.047982 | -0.45305 | 0.650511 | NA | #VALUE! |
| ENSMUSG00000118300.2 | 0 | NA | NA | NA | NA | NA | #VALUE! |
| ENSMUSG00000118301.2 | 0 | NA | NA | NA | NA | NA | #VALUE! |
| ENSMUSG00000118302.2 | 0 | NA | NA | NA | NA | NA | #VALUE! |
| ENSMUSG00000118303.2 | 0 | NA | NA | NA | NA | NA | #VALUE! |
| ENSMUSG00000118304.2 | 0 | NA | NA | NA | NA | NA | #VALUE! |
| ENSMUSG00000118305.2 | 0 | NA | NA | NA | NA | NA | #VALUE! |
| ENSMUSG00000118306.2 | 0 | NA | NA | NA | NA | NA | #VALUE! |
| ENSMUSG00000118307.2 | 0 | NA | NA | NA | NA | NA | #VALUE! |
| ENSMUSG00000118308.2 | 0 | NA | NA | NA | NA | NA | #VALUE! |
| ENSMUSG00000118309.2 | 0 | NA | NA | NA | NA | NA | #VALUE! |
| ENSMUSG00000118310.2 | 0 | NA | NA | NA | NA | NA | #VALUE! |
| ENSMUSG00000118311.2 | 0 | NA | NA | NA | NA | NA | #VALUE! |
| ENSMUSG00000118312.2 | 0 | NA | NA | NA | NA | NA | #VALUE! |
| ENSMUSG00000118313.2 | 0 | NA | NA | NA | NA | NA | #VALUE! |
| ENSMUSG00000118314.2 | 0 | NA | NA | NA | NA | NA | #VALUE! |
| ENSMUSG00000118315.2 | 0 | NA | NA | NA | NA | NA | #VALUE! |
| ENSMUSG00000118316.2 | 0 | NA | NA | NA | NA | NA | #VALUE! |
| ENSMUSG00000118317.2 | 0 | NA | NA | NA | NA | NA | #VALUE! |
| ENSMUSG00000118318.2 | 0 | NA | NA | NA | NA | NA | #VALUE! |
| ENSMUSG00000118319.2 | 0 | NA | NA | NA | NA | NA | #VALUE! |
| ENSMUSG00000118320.2 | 0 | NA | NA | NA | NA | NA | #VALUE! |
| ENSMUSG00000118321.2 | 0 | NA | NA | NA | NA | NA | #VALUE! |
| ENSMUSG00000118322.2 | 0 | NA | NA | NA | NA | NA | #VALUE! |
| ENSMUSG00000118324.2 | 0 | NA | NA | NA | NA | NA | #VALUE! |
| ENSMUSG00000118325.2 | 0 | NA | NA | NA | NA | NA | #VALUE! |
| ENSMUSG00000118326.2 | 0 | NA | NA | NA | NA | NA | #VALUE! |
| ENSMUSG00000118327.2 | 0 | NA | NA | NA | NA | NA | #VALUE! |
| ENSMUSG00000118328.2 | 0 | NA | NA | NA | NA | NA | #VALUE! |
| ENSMUSG00000118329.2 | 0 | NA | NA | NA | NA | NA | #VALUE! |
| ENSMUSG00000118330.2 | 0 | NA | NA | NA | NA | NA | #VALUE! |
| ENSMUSG00000118331.2 | 0 | NA | NA | NA | NA | NA | #VALUE! |
| ENSMUSG00000118332.2 | 0 | NA | NA | NA | NA | NA | #VALUE! |
| ENSMUSG00000118333.2 | 0 | NA | NA | NA | NA | NA | #VALUE! |
| ENSMUSG00000118334.2 | 0 | NA | NA | NA | NA | NA | #VALUE! |
| ENSMUSG00000118335.2 | 0 | NA | NA | NA | NA | NA | #VALUE! |
| ENSMUSG00000118336.2 | 0 | NA | NA | NA | NA | NA | #VALUE! |
| ENSMUSG00000118337.2 | 0 | NA | NA | NA | NA | NA | #VALUE! |
| ENSMUSG00000118338.2 | 0 | NA | NA | NA | NA | NA | #VALUE! |
| ENSMUSG00000118339.2 | 0 | NA | NA | NA | NA | NA | #VALUE! |
| ENSMUSG00000118340.2 | 0 | NA | NA | NA | NA | NA | #VALUE! |
| ENSMUSG00000118341.2 | 0 | NA | NA | NA | NA | NA | #VALUE! |

| | | | | | | | |
|---|---|---|---|---|---|---|---|
| ENSMUSG00000118342.2 | 0 | NA | NA | NA | NA | NA | #VALUE! |
| ENSMUSG00000118343.2 | 0 | NA | NA | NA | NA | NA | #VALUE! |
| ENSMUSG00000118344.2 | 0 | NA | NA | NA | NA | NA | #VALUE! |
| ENSMUSG00000118345.2 | 0 | NA | NA | NA | NA | NA | #VALUE! |
| ENSMUSG00000118347.2 | 0 | NA | NA | NA | NA | NA | #VALUE! |
| ENSMUSG00000118348.2 | 0 | NA | NA | NA | NA | NA | #VALUE! |
| ENSMUSG00000118349.2 | 0 | NA | NA | NA | NA | NA | #VALUE! |
| ENSMUSG00000118350.2 | 0 | NA | NA | NA | NA | NA | #VALUE! |
| ENSMUSG00000118351.2 | 0 | NA | NA | NA | NA | NA | #VALUE! |
| ENSMUSG00000118352.2 | 0 | NA | NA | NA | NA | NA | #VALUE! |
| ENSMUSG00000118353.2 | 0.939203 | -3.45767 | 3.953451 | -0.87459 | 0.381795 | NA | #VALUE! |
| ENSMUSG00000118354.2 | 0 | NA | NA | NA | NA | NA | #VALUE! |
| ENSMUSG00000118355.2 | 0 | NA | NA | NA | NA | NA | #VALUE! |
| ENSMUSG00000118356.2 | 0 | NA | NA | NA | NA | NA | #VALUE! |
| ENSMUSG00000118357.2 | 0 | NA | NA | NA | NA | NA | #VALUE! |
| ENSMUSG00000118358.2 | 0 | NA | NA | NA | NA | NA | #VALUE! |
| ENSMUSG00000118360.2 | 0 | NA | NA | NA | NA | NA | #VALUE! |
| ENSMUSG00000118361.2 | 0 | NA | NA | NA | NA | NA | #VALUE! |
| ENSMUSG00000118362.2 | 0 | NA | NA | NA | NA | NA | #VALUE! |
| ENSMUSG00000118363.2 | 0 | NA | NA | NA | NA | NA | #VALUE! |
| ENSMUSG00000118364.2 | 0 | NA | NA | NA | NA | NA | #VALUE! |
| ENSMUSG00000118365.2 | 0 | NA | NA | NA | NA | NA | #VALUE! |
| ENSMUSG00000118367.2 | 0 | NA | NA | NA | NA | NA | #VALUE! |
| ENSMUSG00000118368.2 | 0 | NA | NA | NA | NA | NA | #VALUE! |
| ENSMUSG00000118369.2 | 0 | NA | NA | NA | NA | NA | #VALUE! |
| ENSMUSG00000118370.2 | 0 | NA | NA | NA | NA | NA | #VALUE! |
| ENSMUSG00000118371.2 | 0 | NA | NA | NA | NA | NA | #VALUE! |
| ENSMUSG00000118372.2 | 0 | NA | NA | NA | NA | NA | #VALUE! |
| ENSMUSG00000118373.2 | 0 | NA | NA | NA | NA | NA | #VALUE! |
| ENSMUSG00000118374.2 | 0 | NA | NA | NA | NA | NA | #VALUE! |
| ENSMUSG00000118375.2 | 0 | NA | NA | NA | NA | NA | #VALUE! |
| ENSMUSG00000118376.2 | 0 | NA | NA | NA | NA | NA | #VALUE! |
| ENSMUSG00000118377.2 | 0 | NA | NA | NA | NA | NA | #VALUE! |
| ENSMUSG00000118378.2 | 0 | NA | NA | NA | NA | NA | #VALUE! |
| ENSMUSG00000118379.2 | 0 | NA | NA | NA | NA | NA | #VALUE! |
| ENSMUSG00000118380.2 | 0 | NA | NA | NA | NA | NA | #VALUE! |
| ENSMUSG00000118381.2 | 0 | NA | NA | NA | NA | NA | #VALUE! |
| ENSMUSG00000118382.2 | 0.841418 | 3.240539 | 3.959551 | 0.818411 | 0.413123 | NA | #VALUE! |
| ENSMUSG00000118383.2 | 0 | NA | NA | NA | NA | NA | #VALUE! |
| ENSMUSG00000118384.2 | 0 | NA | NA | NA | NA | NA | #VALUE! |
| ENSMUSG00000118385.2 | 0 | NA | NA | NA | NA | NA | #VALUE! |
| ENSMUSG00000118386.2 | 0 | NA | NA | NA | NA | NA | #VALUE! |
| ENSMUSG00000118387.2 | 0 | NA | NA | NA | NA | NA | #VALUE! |
| ENSMUSG00000118388.2 | 0 | NA | NA | NA | NA | NA | #VALUE! |

| | | | | | | | |
|---|---|---|---|---|---|---|---|
| ENSMUSG00000118389.2 | 0 | NA | NA | NA | NA | NA | #VALUE! |
| ENSMUSG00000118390.2 | 0 | NA | NA | NA | NA | NA | #VALUE! |
| ENSMUSG00000118391.2 | 0 | NA | NA | NA | NA | NA | #VALUE! |
| ENSMUSG00000118392.2 | 0 | NA | NA | NA | NA | NA | #VALUE! |
| ENSMUSG00000118393.2 | 0 | NA | NA | NA | NA | NA | #VALUE! |
| ENSMUSG00000118394.2 | 0 | NA | NA | NA | NA | NA | #VALUE! |
| ENSMUSG00000118395.2 | 0 | NA | NA | NA | NA | NA | #VALUE! |
| ENSMUSG00000118396.3 | 0 | NA | NA | NA | NA | NA | #VALUE! |
| ENSMUSG00000118397.2 | 0 | NA | NA | NA | NA | NA | #VALUE! |
| ENSMUSG00000118398.2 | 0 | NA | NA | NA | NA | NA | #VALUE! |
| ENSMUSG00000118399.2 | 0 | NA | NA | NA | NA | NA | #VALUE! |
| ENSMUSG00000118400.2 | 0.300802 | -1.80634 | 4.050033 | -0.44601 | 0.655592 | NA | #VALUE! |
| ENSMUSG00000118401.2 | 0 | NA | NA | NA | NA | NA | #VALUE! |
| ENSMUSG00000118402.2 | 0 | NA | NA | NA | NA | NA | #VALUE! |
| ENSMUSG00000118403.2 | 0 | NA | NA | NA | NA | NA | #VALUE! |
| ENSMUSG00000118404.2 | 0 | NA | NA | NA | NA | NA | #VALUE! |
| ENSMUSG00000118406.2 | 0 | NA | NA | NA | NA | NA | #VALUE! |
| ENSMUSG00000118407.2 | 0 | NA | NA | NA | NA | NA | #VALUE! |
| ENSMUSG00000118408.2 | 0 | NA | NA | NA | NA | NA | #VALUE! |
| ENSMUSG00000118409.2 | 0 | NA | NA | NA | NA | NA | #VALUE! |
| ENSMUSG00000118410.2 | 0 | NA | NA | NA | NA | NA | #VALUE! |
| ENSMUSG00000118411.2 | 0 | NA | NA | NA | NA | NA | #VALUE! |
| ENSMUSG00000118412.2 | 0 | NA | NA | NA | NA | NA | #VALUE! |
| ENSMUSG00000118413.2 | 0 | NA | NA | NA | NA | NA | #VALUE! |
| ENSMUSG00000118414.2 | 0 | NA | NA | NA | NA | NA | #VALUE! |
| ENSMUSG00000118415.2 | 0 | NA | NA | NA | NA | NA | #VALUE! |
| ENSMUSG00000118416.2 | 0 | NA | NA | NA | NA | NA | #VALUE! |
| ENSMUSG00000118417.2 | 0 | NA | NA | NA | NA | NA | #VALUE! |
| ENSMUSG00000118418.2 | 0 | NA | NA | NA | NA | NA | #VALUE! |
| ENSMUSG00000118419.2 | 0 | NA | NA | NA | NA | NA | #VALUE! |
| ENSMUSG00000118420.2 | 0 | NA | NA | NA | NA | NA | #VALUE! |
| ENSMUSG00000118421.2 | 0 | NA | NA | NA | NA | NA | #VALUE! |
| ENSMUSG00000118422.2 | 0 | NA | NA | NA | NA | NA | #VALUE! |
| ENSMUSG00000118423.2 | 0 | NA | NA | NA | NA | NA | #VALUE! |
| ENSMUSG00000118424.2 | 0 | NA | NA | NA | NA | NA | #VALUE! |
| ENSMUSG00000118425.2 | 0 | NA | NA | NA | NA | NA | #VALUE! |
| ENSMUSG00000118426.2 | 0 | NA | NA | NA | NA | NA | #VALUE! |
| ENSMUSG00000118427.2 | 0 | NA | NA | NA | NA | NA | #VALUE! |
| ENSMUSG00000118428.2 | 0 | NA | NA | NA | NA | NA | #VALUE! |
| ENSMUSG00000118429.2 | 0 | NA | NA | NA | NA | NA | #VALUE! |
| ENSMUSG00000118430.2 | 0 | NA | NA | NA | NA | NA | #VALUE! |
| ENSMUSG00000118431.2 | 0 | NA | NA | NA | NA | NA | #VALUE! |
| ENSMUSG00000118432.2 | 0 | NA | NA | NA | NA | NA | #VALUE! |
| ENSMUSG00000118433.2 | 0 | NA | NA | NA | NA | NA | #VALUE! |

| | | | | | | | |
|---|---|---|---|---|---|---|---|
| ENSMUSG00000118434.2 | 0 | NA | NA | NA | NA | NA | #VALUE! |
| ENSMUSG00000118435.3 | 0 | NA | NA | NA | NA | NA | #VALUE! |
| ENSMUSG00000118436.2 | 0 | NA | NA | NA | NA | NA | #VALUE! |
| ENSMUSG00000118437.2 | 0 | NA | NA | NA | NA | NA | #VALUE! |
| ENSMUSG00000118438.2 | 0 | NA | NA | NA | NA | NA | #VALUE! |
| ENSMUSG00000118439.2 | 0 | NA | NA | NA | NA | NA | #VALUE! |
| ENSMUSG00000118440.2 | 0 | NA | NA | NA | NA | NA | #VALUE! |
| ENSMUSG00000118441.2 | 0 | NA | NA | NA | NA | NA | #VALUE! |
| ENSMUSG00000118442.2 | 0 | NA | NA | NA | NA | NA | #VALUE! |
| ENSMUSG00000118443.2 | 0 | NA | NA | NA | NA | NA | #VALUE! |
| ENSMUSG00000118444.2 | 0 | NA | NA | NA | NA | NA | #VALUE! |
| ENSMUSG00000118445.2 | 0 | NA | NA | NA | NA | NA | #VALUE! |
| ENSMUSG00000118447.2 | 0 | NA | NA | NA | NA | NA | #VALUE! |
| ENSMUSG00000118448.2 | 0 | NA | NA | NA | NA | NA | #VALUE! |
| ENSMUSG00000118449.2 | 2.424263 | 0.131437 | 3.800731 | 0.034582 | 0.972413 | NA | #VALUE! |
| ENSMUSG00000118450.2 | 0 | NA | NA | NA | NA | NA | #VALUE! |
| ENSMUSG00000118451.2 | 0 | NA | NA | NA | NA | NA | #VALUE! |
| ENSMUSG00000118452.2 | 0 | NA | NA | NA | NA | NA | #VALUE! |
| ENSMUSG00000118453.2 | 0 | NA | NA | NA | NA | NA | #VALUE! |
| ENSMUSG00000118454.2 | 0 | NA | NA | NA | NA | NA | #VALUE! |
| ENSMUSG00000118456.2 | 0 | NA | NA | NA | NA | NA | #VALUE! |
| ENSMUSG00000118457.2 | 0 | NA | NA | NA | NA | NA | #VALUE! |
| ENSMUSG00000118458.2 | 0 | NA | NA | NA | NA | NA | #VALUE! |
| ENSMUSG00000118459.2 | 0 | NA | NA | NA | NA | NA | #VALUE! |
| ENSMUSG00000118460.2 | 0 | NA | NA | NA | NA | NA | #VALUE! |
| ENSMUSG00000118461.2 | 0 | NA | NA | NA | NA | NA | #VALUE! |
| ENSMUSG00000118462.2 | 0 | NA | NA | NA | NA | NA | #VALUE! |
| ENSMUSG00000118463.2 | 0 | NA | NA | NA | NA | NA | #VALUE! |
| ENSMUSG00000118464.2 | 0 | NA | NA | NA | NA | NA | #VALUE! |
| ENSMUSG00000118465.2 | 0 | NA | NA | NA | NA | NA | #VALUE! |
| ENSMUSG00000118466.2 | 0 | NA | NA | NA | NA | NA | #VALUE! |
| ENSMUSG00000118467.2 | 0 | NA | NA | NA | NA | NA | #VALUE! |
| ENSMUSG00000118468.2 | 0 | NA | NA | NA | NA | NA | #VALUE! |
| ENSMUSG00000118469.2 | 0 | NA | NA | NA | NA | NA | #VALUE! |
| ENSMUSG00000118470.2 | 0 | NA | NA | NA | NA | NA | #VALUE! |
| ENSMUSG00000118471.2 | 0 | NA | NA | NA | NA | NA | #VALUE! |
| ENSMUSG00000118472.2 | 0 | NA | NA | NA | NA | NA | #VALUE! |
| ENSMUSG00000118473.2 | 0 | NA | NA | NA | NA | NA | #VALUE! |
| ENSMUSG00000118474.2 | 0 | NA | NA | NA | NA | NA | #VALUE! |
| ENSMUSG00000118475.2 | 0 | NA | NA | NA | NA | NA | #VALUE! |
| ENSMUSG00000118476.2 | 0 | NA | NA | NA | NA | NA | #VALUE! |
| ENSMUSG00000118477.2 | 0 | NA | NA | NA | NA | NA | #VALUE! |
| ENSMUSG00000118478.2 | 0 | NA | NA | NA | NA | NA | #VALUE! |
| ENSMUSG00000118479.2 | 0 | NA | NA | NA | NA | NA | #VALUE! |

| | | | | | | | |
|---|---|---|---|---|---|---|---|
| ENSMUSG00000118480.2 | 0 | NA | NA | NA | NA | NA | #VALUE! |
| ENSMUSG00000118481.2 | 0 | NA | NA | NA | NA | NA | #VALUE! |
| ENSMUSG00000118482.2 | 0 | NA | NA | NA | NA | NA | #VALUE! |
| ENSMUSG00000118483.2 | 0 | NA | NA | NA | NA | NA | #VALUE! |
| ENSMUSG00000118484.2 | 0 | NA | NA | NA | NA | NA | #VALUE! |
| ENSMUSG00000118485.2 | 0 | NA | NA | NA | NA | NA | #VALUE! |
| ENSMUSG00000118486.2 | 0 | NA | NA | NA | NA | NA | #VALUE! |
| ENSMUSG00000118487.2 | 0 | NA | NA | NA | NA | NA | #VALUE! |
| ENSMUSG00000118488.2 | 0 | NA | NA | NA | NA | NA | #VALUE! |
| ENSMUSG00000118489.2 | 0 | NA | NA | NA | NA | NA | #VALUE! |
| ENSMUSG00000118490.2 | 0 | NA | NA | NA | NA | NA | #VALUE! |
| ENSMUSG00000118491.2 | 0 | NA | NA | NA | NA | NA | #VALUE! |
| ENSMUSG00000118492.2 | 0 | NA | NA | NA | NA | NA | #VALUE! |
| ENSMUSG00000118493.2 | 0 | NA | NA | NA | NA | NA | #VALUE! |
| ENSMUSG00000118494.2 | 0 | NA | NA | NA | NA | NA | #VALUE! |
| ENSMUSG00000118495.2 | 0 | NA | NA | NA | NA | NA | #VALUE! |
| ENSMUSG00000118496.2 | 0 | NA | NA | NA | NA | NA | #VALUE! |
| ENSMUSG00000118497.2 | 0 | NA | NA | NA | NA | NA | #VALUE! |
| ENSMUSG00000118498.3 | 0 | NA | NA | NA | NA | NA | #VALUE! |
| ENSMUSG00000118499.2 | 0 | NA | NA | NA | NA | NA | #VALUE! |
| ENSMUSG00000118500.2 | 0 | NA | NA | NA | NA | NA | #VALUE! |
| ENSMUSG00000118501.2 | 0 | NA | NA | NA | NA | NA | #VALUE! |
| ENSMUSG00000118502.2 | 0 | NA | NA | NA | NA | NA | #VALUE! |
| ENSMUSG00000118503.2 | 0 | NA | NA | NA | NA | NA | #VALUE! |
| ENSMUSG00000118504.2 | 0 | NA | NA | NA | NA | NA | #VALUE! |
| ENSMUSG00000118505.2 | 0 | NA | NA | NA | NA | NA | #VALUE! |
| ENSMUSG00000118507.2 | 0 | NA | NA | NA | NA | NA | #VALUE! |
| ENSMUSG00000118508.2 | 0 | NA | NA | NA | NA | NA | #VALUE! |
| ENSMUSG00000118509.2 | 0 | NA | NA | NA | NA | NA | #VALUE! |
| ENSMUSG00000118510.2 | 0 | NA | NA | NA | NA | NA | #VALUE! |
| ENSMUSG00000118511.2 | 0 | NA | NA | NA | NA | NA | #VALUE! |
| ENSMUSG00000118512.2 | 0 | NA | NA | NA | NA | NA | #VALUE! |
| ENSMUSG00000118513.2 | 0 | NA | NA | NA | NA | NA | #VALUE! |
| ENSMUSG00000118514.2 | 0 | NA | NA | NA | NA | NA | #VALUE! |
| ENSMUSG00000118515.2 | 0 | NA | NA | NA | NA | NA | #VALUE! |
| ENSMUSG00000118517.2 | 0 | NA | NA | NA | NA | NA | #VALUE! |
| ENSMUSG00000118518.2 | 0 | NA | NA | NA | NA | NA | #VALUE! |
| ENSMUSG00000118519.2 | 0 | NA | NA | NA | NA | NA | #VALUE! |
| ENSMUSG00000118520.2 | 0 | NA | NA | NA | NA | NA | #VALUE! |
| ENSMUSG00000118521.2 | 0 | NA | NA | NA | NA | NA | #VALUE! |
| ENSMUSG00000118522.2 | 0 | NA | NA | NA | NA | NA | #VALUE! |
| ENSMUSG00000118523.2 | 2.692536 | 4.918862 | 3.923417 | 1.253719 | 0.209944 | NA | #VALUE! |
| ENSMUSG00000118524.2 | 0 | NA | NA | NA | NA | NA | #VALUE! |
| ENSMUSG00000118525.2 | 0 | NA | NA | NA | NA | NA | #VALUE! |

| | | | | | | | |
|---|---|---|---|---|---|---|---|
| ENSMUSG00000118526.3 | 0 | NA | NA | NA | NA | NA | #VALUE! |
| ENSMUSG00000118527.2 | 0 | NA | NA | NA | NA | NA | #VALUE! |
| ENSMUSG00000118528.2 | 0 | NA | NA | NA | NA | NA | #VALUE! |
| ENSMUSG00000118529.2 | 0 | NA | NA | NA | NA | NA | #VALUE! |
| ENSMUSG00000118530.2 | 0 | NA | NA | NA | NA | NA | #VALUE! |
| ENSMUSG00000118531.2 | 0 | NA | NA | NA | NA | NA | #VALUE! |
| ENSMUSG00000118532.2 | 0 | NA | NA | NA | NA | NA | #VALUE! |
| ENSMUSG00000118533.2 | 0 | NA | NA | NA | NA | NA | #VALUE! |
| ENSMUSG00000118534.2 | 0 | NA | NA | NA | NA | NA | #VALUE! |
| ENSMUSG00000118535.2 | 0 | NA | NA | NA | NA | NA | #VALUE! |
| ENSMUSG00000118536.2 | 0 | NA | NA | NA | NA | NA | #VALUE! |
| ENSMUSG00000118538.2 | 0 | NA | NA | NA | NA | NA | #VALUE! |
| ENSMUSG00000118539.2 | 0 | NA | NA | NA | NA | NA | #VALUE! |
| ENSMUSG00000118540.2 | 0 | NA | NA | NA | NA | NA | #VALUE! |
| ENSMUSG00000118541.2 | 0 | NA | NA | NA | NA | NA | #VALUE! |
| ENSMUSG00000118542.2 | 0 | NA | NA | NA | NA | NA | #VALUE! |
| ENSMUSG00000118543.2 | 0 | NA | NA | NA | NA | NA | #VALUE! |
| ENSMUSG00000118544.2 | 0 | NA | NA | NA | NA | NA | #VALUE! |
| ENSMUSG00000118545.2 | 0 | NA | NA | NA | NA | NA | #VALUE! |
| ENSMUSG00000118546.2 | 0 | NA | NA | NA | NA | NA | #VALUE! |
| ENSMUSG00000118547.2 | 0 | NA | NA | NA | NA | NA | #VALUE! |
| ENSMUSG00000118548.2 | 0 | NA | NA | NA | NA | NA | #VALUE! |
| ENSMUSG00000118549.2 | 0 | NA | NA | NA | NA | NA | #VALUE! |
| ENSMUSG00000118550.2 | 0 | NA | NA | NA | NA | NA | #VALUE! |
| ENSMUSG00000118551.2 | 0 | NA | NA | NA | NA | NA | #VALUE! |
| ENSMUSG00000118552.2 | 0 | NA | NA | NA | NA | NA | #VALUE! |
| ENSMUSG00000118553.2 | 0 | NA | NA | NA | NA | NA | #VALUE! |
| ENSMUSG00000118554.2 | 0 | NA | NA | NA | NA | NA | #VALUE! |
| ENSMUSG00000118555.2 | 0 | NA | NA | NA | NA | NA | #VALUE! |
| ENSMUSG00000118556.2 | 0 | NA | NA | NA | NA | NA | #VALUE! |
| ENSMUSG00000118557.2 | 0 | NA | NA | NA | NA | NA | #VALUE! |
| ENSMUSG00000118558.2 | 0 | NA | NA | NA | NA | NA | #VALUE! |
| ENSMUSG00000118559.3 | 0 | NA | NA | NA | NA | NA | #VALUE! |
| ENSMUSG00000118560.2 | 0 | NA | NA | NA | NA | NA | #VALUE! |
| ENSMUSG00000118561.2 | 0 | NA | NA | NA | NA | NA | #VALUE! |
| ENSMUSG00000118562.2 | 0 | NA | NA | NA | NA | NA | #VALUE! |
| ENSMUSG00000118563.2 | 0 | NA | NA | NA | NA | NA | #VALUE! |
| ENSMUSG00000118565.2 | 0 | NA | NA | NA | NA | NA | #VALUE! |
| ENSMUSG00000118566.2 | 0 | NA | NA | NA | NA | NA | #VALUE! |
| ENSMUSG00000118568.2 | 0 | NA | NA | NA | NA | NA | #VALUE! |
| ENSMUSG00000118569.2 | 0 | NA | NA | NA | NA | NA | #VALUE! |
| ENSMUSG00000118570.2 | 0 | NA | NA | NA | NA | NA | #VALUE! |
| ENSMUSG00000118571.2 | 0 | NA | NA | NA | NA | NA | #VALUE! |
| ENSMUSG00000118572.2 | 0 | NA | NA | NA | NA | NA | #VALUE! |

| | | | | | | | |
|---|---|---|---|---|---|---|---|
| ENSMUSG00000118573.2 | 0 | NA | NA | NA | NA | NA | #VALUE! |
| ENSMUSG00000118574.2 | 0 | NA | NA | NA | NA | NA | #VALUE! |
| ENSMUSG00000118576.2 | 0 | NA | NA | NA | NA | NA | #VALUE! |
| ENSMUSG00000118578.2 | 0 | NA | NA | NA | NA | NA | #VALUE! |
| ENSMUSG00000118579.2 | 0 | NA | NA | NA | NA | NA | #VALUE! |
| ENSMUSG00000118580.2 | 0 | NA | NA | NA | NA | NA | #VALUE! |
| ENSMUSG00000118581.2 | 0 | NA | NA | NA | NA | NA | #VALUE! |
| ENSMUSG00000118582.2 | 0 | NA | NA | NA | NA | NA | #VALUE! |
| ENSMUSG00000118583.2 | 0 | NA | NA | NA | NA | NA | #VALUE! |
| ENSMUSG00000118584.2 | 0 | NA | NA | NA | NA | NA | #VALUE! |
| ENSMUSG00000118585.2 | 0 | NA | NA | NA | NA | NA | #VALUE! |
| ENSMUSG00000118586.2 | 0 | NA | NA | NA | NA | NA | #VALUE! |
| ENSMUSG00000118587.2 | 0 | NA | NA | NA | NA | NA | #VALUE! |
| ENSMUSG00000118588.2 | 0 | NA | NA | NA | NA | NA | #VALUE! |
| ENSMUSG00000118589.2 | 0 | NA | NA | NA | NA | NA | #VALUE! |
| ENSMUSG00000118590.2 | 0 | NA | NA | NA | NA | NA | #VALUE! |
| ENSMUSG00000118591.2 | 0 | NA | NA | NA | NA | NA | #VALUE! |
| ENSMUSG00000118592.2 | 0 | NA | NA | NA | NA | NA | #VALUE! |
| ENSMUSG00000118593.2 | 0 | NA | NA | NA | NA | NA | #VALUE! |
| ENSMUSG00000118594.2 | 0 | NA | NA | NA | NA | NA | #VALUE! |
| ENSMUSG00000118595.2 | 0 | NA | NA | NA | NA | NA | #VALUE! |
| ENSMUSG00000118596.2 | 0 | NA | NA | NA | NA | NA | #VALUE! |
| ENSMUSG00000118597.2 | 0 | NA | NA | NA | NA | NA | #VALUE! |
| ENSMUSG00000118598.2 | 0 | NA | NA | NA | NA | NA | #VALUE! |
| ENSMUSG00000118599.2 | 0 | NA | NA | NA | NA | NA | #VALUE! |
| ENSMUSG00000118600.2 | 0 | NA | NA | NA | NA | NA | #VALUE! |
| ENSMUSG00000118601.2 | 0 | NA | NA | NA | NA | NA | #VALUE! |
| ENSMUSG00000118602.2 | 0 | NA | NA | NA | NA | NA | #VALUE! |
| ENSMUSG00000118603.2 | 0 | NA | NA | NA | NA | NA | #VALUE! |
| ENSMUSG00000118604.2 | 0 | NA | NA | NA | NA | NA | #VALUE! |
| ENSMUSG00000118605.2 | 0 | NA | NA | NA | NA | NA | #VALUE! |
| ENSMUSG00000118606.2 | 0 | NA | NA | NA | NA | NA | #VALUE! |
| ENSMUSG00000118607.2 | 0 | NA | NA | NA | NA | NA | #VALUE! |
| ENSMUSG00000118608.2 | 0 | NA | NA | NA | NA | NA | #VALUE! |
| ENSMUSG00000118609.2 | 0 | NA | NA | NA | NA | NA | #VALUE! |
| ENSMUSG00000118610.2 | 0 | NA | NA | NA | NA | NA | #VALUE! |
| ENSMUSG00000118611.2 | 0 | NA | NA | NA | NA | NA | #VALUE! |
| ENSMUSG00000118612.2 | 0 | NA | NA | NA | NA | NA | #VALUE! |
| ENSMUSG00000118613.2 | 0 | NA | NA | NA | NA | NA | #VALUE! |
| ENSMUSG00000118614.2 | 0 | NA | NA | NA | NA | NA | #VALUE! |
| ENSMUSG00000118615.2 | 0 | NA | NA | NA | NA | NA | #VALUE! |
| ENSMUSG00000118616.2 | 0 | NA | NA | NA | NA | NA | #VALUE! |
| ENSMUSG00000118617.2 | 0 | NA | NA | NA | NA | NA | #VALUE! |
| ENSMUSG00000118618.2 | 0 | NA | NA | NA | NA | NA | #VALUE! |

| | | | | | | | |
|---|---|---|---|---|---|---|---|
| ENSMUSG00000118619.2 | 0 | NA | NA | NA | NA | NA | #VALUE! |
| ENSMUSG00000118620.2 | 0 | NA | NA | NA | NA | NA | #VALUE! |
| ENSMUSG00000118621.2 | 0 | NA | NA | NA | NA | NA | #VALUE! |
| ENSMUSG00000118622.2 | 0 | NA | NA | NA | NA | NA | #VALUE! |
| ENSMUSG00000118623.2 | 0 | NA | NA | NA | NA | NA | #VALUE! |
| ENSMUSG00000118624.2 | 0 | NA | NA | NA | NA | NA | #VALUE! |
| ENSMUSG00000118625.2 | 0 | NA | NA | NA | NA | NA | #VALUE! |
| ENSMUSG00000118626.2 | 0 | NA | NA | NA | NA | NA | #VALUE! |
| ENSMUSG00000118627.2 | 0 | NA | NA | NA | NA | NA | #VALUE! |
| ENSMUSG00000118628.2 | 0 | NA | NA | NA | NA | NA | #VALUE! |
| ENSMUSG00000118629.2 | 0 | NA | NA | NA | NA | NA | #VALUE! |
| ENSMUSG00000118630.2 | 0 | NA | NA | NA | NA | NA | #VALUE! |
| ENSMUSG00000118631.2 | 7.960319 | -6.52083 | 3.880481 | -1.68042 | 0.092876 | NA | #VALUE! |
| ENSMUSG00000118632.2 | 0 | NA | NA | NA | NA | NA | #VALUE! |
| ENSMUSG00000118633.2 | 0 | NA | NA | NA | NA | NA | #VALUE! |
| ENSMUSG00000118634.2 | 0 | NA | NA | NA | NA | NA | #VALUE! |
| ENSMUSG00000118635.2 | 0 | NA | NA | NA | NA | NA | #VALUE! |
| ENSMUSG00000118636.2 | 0 | NA | NA | NA | NA | NA | #VALUE! |
| ENSMUSG00000118637.2 | 0 | NA | NA | NA | NA | NA | #VALUE! |
| ENSMUSG00000118638.2 | 0 | NA | NA | NA | NA | NA | #VALUE! |
| ENSMUSG00000118639.2 | 0 | NA | NA | NA | NA | NA | #VALUE! |
| ENSMUSG00000118640.2 | 0 | NA | NA | NA | NA | NA | #VALUE! |
| ENSMUSG00000118641.2 | 0 | NA | NA | NA | NA | NA | #VALUE! |
| ENSMUSG00000118642.2 | 0 | NA | NA | NA | NA | NA | #VALUE! |
| ENSMUSG00000118644.2 | 0 | NA | NA | NA | NA | NA | #VALUE! |
| ENSMUSG00000118645.2 | 0 | NA | NA | NA | NA | NA | #VALUE! |
| ENSMUSG00000118646.2 | 0 | NA | NA | NA | NA | NA | #VALUE! |
| ENSMUSG00000118647.2 | 0 | NA | NA | NA | NA | NA | #VALUE! |
| ENSMUSG00000118648.2 | 0 | NA | NA | NA | NA | NA | #VALUE! |
| ENSMUSG00000118649.2 | 0 | NA | NA | NA | NA | NA | #VALUE! |
| ENSMUSG00000118650.2 | 0 | NA | NA | NA | NA | NA | #VALUE! |
| ENSMUSG00000118651.2 | 0 | NA | NA | NA | NA | NA | #VALUE! |
| ENSMUSG00000118652.2 | 0 | NA | NA | NA | NA | NA | #VALUE! |
| ENSMUSG00000118653.2 | 2.091057 | 2.07894 | 3.867285 | 0.537571 | 0.590873 | NA | #VALUE! |
| ENSMUSG00000118654.2 | 0 | NA | NA | NA | NA | NA | #VALUE! |
| ENSMUSG00000118656.2 | 0 | NA | NA | NA | NA | NA | #VALUE! |
| ENSMUSG00000118657.2 | 0 | NA | NA | NA | NA | NA | #VALUE! |
| ENSMUSG00000118658.2 | 0 | NA | NA | NA | NA | NA | #VALUE! |
| ENSMUSG00000118659.2 | 0 | NA | NA | NA | NA | NA | #VALUE! |
| ENSMUSG00000118660.1 | 0 | NA | NA | NA | NA | NA | #VALUE! |
| ENSMUSG00000118661.1 | 0 | NA | NA | NA | NA | NA | #VALUE! |
| ENSMUSG00000118662.2 | 0.816443 | -3.21914 | 3.961753 | -0.81255 | 0.416474 | NA | #VALUE! |
| ENSMUSG00000118663.1 | 9.888331 | 0.168401 | 2.615084 | 0.064396 | 0.948655 | NA | #VALUE! |
| ENSMUSG00000118666.1 | 0 | NA | NA | NA | NA | NA | #VALUE! |

| | | | | | | | |
|---|---|---|---|---|---|---|---|
| ENSMUSG00000118667.1 | 0.168284 | 1.027628 | 4.067126 | 0.252667 | 0.800526 | NA | #VALUE! |
| ENSMUSG00000118668.1 | 0 | NA | NA | NA | NA | NA | #VALUE! |
| ENSMUSG00000118670.1 | 0 | NA | NA | NA | NA | NA | #VALUE! |
| ENSMUSG00000118672.1 | 0 | NA | NA | NA | NA | NA | #VALUE! |
| ENSMUSG00000118674.1 | 0 | NA | NA | NA | NA | NA | #VALUE! |
| ENSMUSG00000118675.1 | 0 | NA | NA | NA | NA | NA | #VALUE! |
| ENSMUSG00000118676.1 | 0 | NA | NA | NA | NA | NA | #VALUE! |
| ENSMUSG00000118677.1 | 0 | NA | NA | NA | NA | NA | #VALUE! |
| ENSMUSG00000118678.1 | 0 | NA | NA | NA | NA | NA | #VALUE! |
| ENSMUSG00000118680.1 | 0 | NA | NA | NA | NA | NA | #VALUE! |
| ENSMUSG00000118681.1 | 0 | NA | NA | NA | NA | NA | #VALUE! |
| ENSMUSG00000118687.1 | 0 | NA | NA | NA | NA | NA | #VALUE! |
| ENSMUSG00000118688.1 | 0 | NA | NA | NA | NA | NA | #VALUE! |
| ENSMUSG00000118690.1 | 0 | NA | NA | NA | NA | NA | #VALUE! |
| ENSMUSG00000118692.1 | 0 | NA | NA | NA | NA | NA | #VALUE! |
| ENSMUSG00000118694.2 | 0 | NA | NA | NA | NA | NA | #VALUE! |
| ENSMUSG00000118695.1 | 0 | NA | NA | NA | NA | NA | #VALUE! |
| ENSMUSG00000118696.1 | 0 | NA | NA | NA | NA | NA | #VALUE! |
| ENSMUSG00000118697.1 | 0 | NA | NA | NA | NA | NA | #VALUE! |
| ENSMUSG00000118698.1 | 0 | NA | NA | NA | NA | NA | #VALUE! |
| ENSMUSG00000118699.1 | 0 | NA | NA | NA | NA | NA | #VALUE! |
| ENSMUSG00000118700.1 | 0 | NA | NA | NA | NA | NA | #VALUE! |
| ENSMUSG00000118701.2 | 0 | NA | NA | NA | NA | NA | #VALUE! |
| ENSMUSG00000118703.1 | 0 | NA | NA | NA | NA | NA | #VALUE! |
| ENSMUSG00000118706.1 | 0 | NA | NA | NA | NA | NA | #VALUE! |
| ENSMUSG00000118708.1 | 0 | NA | NA | NA | NA | NA | #VALUE! |
| ENSMUSG00000118710.1 | 0 | NA | NA | NA | NA | NA | #VALUE! |
| ENSMUSG00000118711.1 | 0 | NA | NA | NA | NA | NA | #VALUE! |
| ENSMUSG00000118712.1 | 0 | NA | NA | NA | NA | NA | #VALUE! |
| ENSMUSG00000118714.1 | 0 | NA | NA | NA | NA | NA | #VALUE! |
| ENSMUSG00000118715.1 | 0 | NA | NA | NA | NA | NA | #VALUE! |
| ENSMUSG00000118716.1 | 0 | NA | NA | NA | NA | NA | #VALUE! |
| ENSMUSG00000118717.1 | 0 | NA | NA | NA | NA | NA | #VALUE! |
| ENSMUSG00000118718.1 | 0 | NA | NA | NA | NA | NA | #VALUE! |
| ENSMUSG00000118719.1 | 0 | NA | NA | NA | NA | NA | #VALUE! |
| ENSMUSG00000118720.1 | 0 | NA | NA | NA | NA | NA | #VALUE! |
| ENSMUSG00000118721.1 | 0 | NA | NA | NA | NA | NA | #VALUE! |
| ENSMUSG00000118723.1 | 0 | NA | NA | NA | NA | NA | #VALUE! |
| ENSMUSG00000118725.1 | 0 | NA | NA | NA | NA | NA | #VALUE! |
| ENSMUSG00000118726.1 | 0 | NA | NA | NA | NA | NA | #VALUE! |
| ENSMUSG00000118727.1 | 0 | NA | NA | NA | NA | NA | #VALUE! |
| ENSMUSG00000118728.1 | 0 | NA | NA | NA | NA | NA | #VALUE! |
| ENSMUSG00000118729.1 | 0 | NA | NA | NA | NA | NA | #VALUE! |
| ENSMUSG00000118730.1 | 0 | NA | NA | NA | NA | NA | #VALUE! |

| | | | | | | | |
|---|---|---|---|---|---|---|---|
| ENSMUSG00000118731.1 | 0 | NA | NA | NA | NA | NA | #VALUE! |
| ENSMUSG00000118732.1 | 0 | NA | NA | NA | NA | NA | #VALUE! |
| ENSMUSG00000118733.2 | 0 | NA | NA | NA | NA | NA | #VALUE! |
| ENSMUSG00000118734.1 | 0 | NA | NA | NA | NA | NA | #VALUE! |
| ENSMUSG00000118735.1 | 0 | NA | NA | NA | NA | NA | #VALUE! |
| ENSMUSG00000118736.2 | 0 | NA | NA | NA | NA | NA | #VALUE! |
| ENSMUSG00000118737.1 | 0 | NA | NA | NA | NA | NA | #VALUE! |
| ENSMUSG00000118738.1 | 0 | NA | NA | NA | NA | NA | #VALUE! |
| ENSMUSG00000118741.1 | 0 | NA | NA | NA | NA | NA | #VALUE! |
| ENSMUSG00000118743.1 | 0 | NA | NA | NA | NA | NA | #VALUE! |
| ENSMUSG00000118744.1 | 0 | NA | NA | NA | NA | NA | #VALUE! |
| ENSMUSG00000118746.1 | 0 | NA | NA | NA | NA | NA | #VALUE! |
| ENSMUSG00000118747.1 | 0 | NA | NA | NA | NA | NA | #VALUE! |
| ENSMUSG00000118748.1 | 0 | NA | NA | NA | NA | NA | #VALUE! |
| ENSMUSG00000118751.1 | 0 | NA | NA | NA | NA | NA | #VALUE! |
| ENSMUSG00000118753.1 | 0 | NA | NA | NA | NA | NA | #VALUE! |
| ENSMUSG00000118755.1 | 0 | NA | NA | NA | NA | NA | #VALUE! |
| ENSMUSG00000118756.1 | 0 | NA | NA | NA | NA | NA | #VALUE! |
| ENSMUSG00000118757.1 | 0 | NA | NA | NA | NA | NA | #VALUE! |
| ENSMUSG00000118759.1 | 0 | NA | NA | NA | NA | NA | #VALUE! |
| ENSMUSG00000118760.1 | 0 | NA | NA | NA | NA | NA | #VALUE! |
| ENSMUSG00000118761.1 | 0 | NA | NA | NA | NA | NA | #VALUE! |
| ENSMUSG00000118762.1 | 0 | NA | NA | NA | NA | NA | #VALUE! |
| ENSMUSG00000118763.1 | 0 | NA | NA | NA | NA | NA | #VALUE! |
| ENSMUSG00000118764.1 | 0 | NA | NA | NA | NA | NA | #VALUE! |
| ENSMUSG00000118768.1 | 0 | NA | NA | NA | NA | NA | #VALUE! |
| ENSMUSG00000118769.1 | 0 | NA | NA | NA | NA | NA | #VALUE! |
| ENSMUSG00000118770.1 | 0 | NA | NA | NA | NA | NA | #VALUE! |
| ENSMUSG00000118772.1 | 0 | NA | NA | NA | NA | NA | #VALUE! |
| ENSMUSG00000118773.1 | 0 | NA | NA | NA | NA | NA | #VALUE! |
| ENSMUSG00000118776.2 | 0 | NA | NA | NA | NA | NA | #VALUE! |
| ENSMUSG00000118777.1 | 0 | NA | NA | NA | NA | NA | #VALUE! |
| ENSMUSG00000118778.1 | 0 | NA | NA | NA | NA | NA | #VALUE! |
| ENSMUSG00000118779.2 | 0 | NA | NA | NA | NA | NA | #VALUE! |
| ENSMUSG00000118780.1 | 0 | NA | NA | NA | NA | NA | #VALUE! |
| ENSMUSG00000118781.1 | 0 | NA | NA | NA | NA | NA | #VALUE! |
| ENSMUSG00000118782.1 | 0 | NA | NA | NA | NA | NA | #VALUE! |
| ENSMUSG00000118786.1 | 0 | NA | NA | NA | NA | NA | #VALUE! |
| ENSMUSG00000118787.1 | 0 | NA | NA | NA | NA | NA | #VALUE! |
| ENSMUSG00000118788.1 | 0 | NA | NA | NA | NA | NA | #VALUE! |
| ENSMUSG00000118789.2 | 0 | NA | NA | NA | NA | NA | #VALUE! |
| ENSMUSG00000118790.1 | 3.469883 | -5.32046 | 3.91969 | -1.35737 | 0.174664 | NA | #VALUE! |
| ENSMUSG00000118792.1 | 0 | NA | NA | NA | NA | NA | #VALUE! |
| ENSMUSG00000118793.1 | 0 | NA | NA | NA | NA | NA | #VALUE! |

| | | | | | | | |
|---|---|---|---|---|---|---|---|
| ENSMUSG00000118795.2 | 0 | NA | NA | NA | NA | NA | #VALUE! |
| ENSMUSG00000118796.1 | 0 | NA | NA | NA | NA | NA | #VALUE! |
| ENSMUSG00000118798.1 | 0 | NA | NA | NA | NA | NA | #VALUE! |
| ENSMUSG00000118799.1 | 0 | NA | NA | NA | NA | NA | #VALUE! |
| ENSMUSG00000118803.1 | 0 | NA | NA | NA | NA | NA | #VALUE! |
| ENSMUSG00000118804.1 | 0 | NA | NA | NA | NA | NA | #VALUE! |
| ENSMUSG00000118807.1 | 0 | NA | NA | NA | NA | NA | #VALUE! |
| ENSMUSG00000118809.1 | 0 | NA | NA | NA | NA | NA | #VALUE! |
| ENSMUSG00000118810.1 | 0 | NA | NA | NA | NA | NA | #VALUE! |
| ENSMUSG00000118811.1 | 0 | NA | NA | NA | NA | NA | #VALUE! |
| ENSMUSG00000118813.1 | 0 | NA | NA | NA | NA | NA | #VALUE! |
| ENSMUSG00000118814.1 | 0 | NA | NA | NA | NA | NA | #VALUE! |
| ENSMUSG00000118815.1 | 0 | NA | NA | NA | NA | NA | #VALUE! |
| ENSMUSG00000118816.1 | 0 | NA | NA | NA | NA | NA | #VALUE! |
| ENSMUSG00000118820.1 | 0 | NA | NA | NA | NA | NA | #VALUE! |
| ENSMUSG00000118824.1 | 0 | NA | NA | NA | NA | NA | #VALUE! |
| ENSMUSG00000118825.1 | 0 | NA | NA | NA | NA | NA | #VALUE! |
| ENSMUSG00000118827.1 | 0 | NA | NA | NA | NA | NA | #VALUE! |
| ENSMUSG00000118828.1 | 0 | NA | NA | NA | NA | NA | #VALUE! |
| ENSMUSG00000118829.1 | 0 | NA | NA | NA | NA | NA | #VALUE! |
| ENSMUSG00000118832.1 | 0 | NA | NA | NA | NA | NA | #VALUE! |
| ENSMUSG00000118833.1 | 0 | NA | NA | NA | NA | NA | #VALUE! |
| ENSMUSG00000118834.1 | 0 | NA | NA | NA | NA | NA | #VALUE! |
| ENSMUSG00000118836.1 | 0 | NA | NA | NA | NA | NA | #VALUE! |
| ENSMUSG00000118837.1 | 0 | NA | NA | NA | NA | NA | #VALUE! |
| ENSMUSG00000118838.1 | 0 | NA | NA | NA | NA | NA | #VALUE! |
| ENSMUSG00000118839.1 | 1.622033 | 0.745622 | 3.847106 | 0.193814 | 0.846322 | NA | #VALUE! |
| ENSMUSG00000118841.1 | 0 | NA | NA | NA | NA | NA | #VALUE! |
| ENSMUSG00000118842.1 | 0 | NA | NA | NA | NA | NA | #VALUE! |
| ENSMUSG00000118845.1 | 0 | NA | NA | NA | NA | NA | #VALUE! |
| ENSMUSG00000118846.1 | 0 | NA | NA | NA | NA | NA | #VALUE! |
| ENSMUSG00000118848.1 | 0 | NA | NA | NA | NA | NA | #VALUE! |
| ENSMUSG00000118852.1 | 0 | NA | NA | NA | NA | NA | #VALUE! |
| ENSMUSG00000118853.2 | 0 | NA | NA | NA | NA | NA | #VALUE! |
| ENSMUSG00000118855.1 | 0 | NA | NA | NA | NA | NA | #VALUE! |
| ENSMUSG00000118856.1 | 0 | NA | NA | NA | NA | NA | #VALUE! |
| ENSMUSG00000118857.1 | 0 | NA | NA | NA | NA | NA | #VALUE! |
| ENSMUSG00000118858.1 | 0 | NA | NA | NA | NA | NA | #VALUE! |
| ENSMUSG00000118859.1 | 0 | NA | NA | NA | NA | NA | #VALUE! |
| ENSMUSG00000118860.1 | 0 | NA | NA | NA | NA | NA | #VALUE! |
| ENSMUSG00000118861.1 | 0 | NA | NA | NA | NA | NA | #VALUE! |
| ENSMUSG00000118864.1 | 0 | NA | NA | NA | NA | NA | #VALUE! |
| ENSMUSG00000118865.1 | 0 | NA | NA | NA | NA | NA | #VALUE! |
| ENSMUSG00000118866.1 | 0 | NA | NA | NA | NA | NA | #VALUE! |

| | | | | | | | |
|---|---|---|---|---|---|---|---|
| ENSMUSG00000118867.1 | 0 | NA | NA | NA | NA | NA | #VALUE! |
| ENSMUSG00000118868.1 | 0 | NA | NA | NA | NA | NA | #VALUE! |
| ENSMUSG00000118870.1 | 0 | NA | NA | NA | NA | NA | #VALUE! |
| ENSMUSG00000118871.1 | 0 | NA | NA | NA | NA | NA | #VALUE! |
| ENSMUSG00000118872.2 | 0 | NA | NA | NA | NA | NA | #VALUE! |
| ENSMUSG00000118873.1 | 0 | NA | NA | NA | NA | NA | #VALUE! |
| ENSMUSG00000118876.1 | 0 | NA | NA | NA | NA | NA | #VALUE! |
| ENSMUSG00000118877.1 | 0 | NA | NA | NA | NA | NA | #VALUE! |
| ENSMUSG00000118878.1 | 0 | NA | NA | NA | NA | NA | #VALUE! |
| ENSMUSG00000118879.1 | 0 | NA | NA | NA | NA | NA | #VALUE! |
| ENSMUSG00000118880.1 | 0 | NA | NA | NA | NA | NA | #VALUE! |
| ENSMUSG00000118881.1 | 0 | NA | NA | NA | NA | NA | #VALUE! |
| ENSMUSG00000118883.1 | 0 | NA | NA | NA | NA | NA | #VALUE! |
| ENSMUSG00000118885.1 | 0 | NA | NA | NA | NA | NA | #VALUE! |
| ENSMUSG00000118886.1 | 0 | NA | NA | NA | NA | NA | #VALUE! |
| ENSMUSG00000118888.1 | 0 | NA | NA | NA | NA | NA | #VALUE! |
| ENSMUSG00000118889.1 | 0 | NA | NA | NA | NA | NA | #VALUE! |
| ENSMUSG00000118890.1 | 0 | NA | NA | NA | NA | NA | #VALUE! |
| ENSMUSG00000118894.1 | 0 | NA | NA | NA | NA | NA | #VALUE! |
| ENSMUSG00000118897.1 | 0 | NA | NA | NA | NA | NA | #VALUE! |
| ENSMUSG00000118898.1 | 0 | NA | NA | NA | NA | NA | #VALUE! |
| ENSMUSG00000118899.1 | 0 | NA | NA | NA | NA | NA | #VALUE! |
| ENSMUSG00000118900.1 | 0 | NA | NA | NA | NA | NA | #VALUE! |
| ENSMUSG00000118901.1 | 0 | NA | NA | NA | NA | NA | #VALUE! |
| ENSMUSG00000118902.1 | 0 | NA | NA | NA | NA | NA | #VALUE! |
| ENSMUSG00000118903.1 | 0 | NA | NA | NA | NA | NA | #VALUE! |
| ENSMUSG00000118904.1 | 0 | NA | NA | NA | NA | NA | #VALUE! |
| ENSMUSG00000118905.1 | 0 | NA | NA | NA | NA | NA | #VALUE! |
| ENSMUSG00000118906.1 | 0 | NA | NA | NA | NA | NA | #VALUE! |
| ENSMUSG00000118907.1 | 0 | NA | NA | NA | NA | NA | #VALUE! |
| ENSMUSG00000118908.1 | 0 | NA | NA | NA | NA | NA | #VALUE! |
| ENSMUSG00000118909.1 | 0 | NA | NA | NA | NA | NA | #VALUE! |
| ENSMUSG00000118910.1 | 0 | NA | NA | NA | NA | NA | #VALUE! |
| ENSMUSG00000118911.1 | 0 | NA | NA | NA | NA | NA | #VALUE! |
| ENSMUSG00000118912.1 | 0 | NA | NA | NA | NA | NA | #VALUE! |
| ENSMUSG00000118913.1 | 0 | NA | NA | NA | NA | NA | #VALUE! |
| ENSMUSG00000118914.1 | 0 | NA | NA | NA | NA | NA | #VALUE! |
| ENSMUSG00000118915.1 | 0 | NA | NA | NA | NA | NA | #VALUE! |
| ENSMUSG00000118916.1 | 0 | NA | NA | NA | NA | NA | #VALUE! |
| ENSMUSG00000118917.1 | 0 | NA | NA | NA | NA | NA | #VALUE! |
| ENSMUSG00000118918.1 | 0 | NA | NA | NA | NA | NA | #VALUE! |
| ENSMUSG00000118919.1 | 0 | NA | NA | NA | NA | NA | #VALUE! |
| ENSMUSG00000118920.1 | 0 | NA | NA | NA | NA | NA | #VALUE! |
| ENSMUSG00000118922.1 | 0 | NA | NA | NA | NA | NA | #VALUE! |

| | | | | | | | |
|---|---|---|---|---|---|---|---|
| ENSMUSG00000118923.1 | 0 | NA | NA | NA | NA | NA | #VALUE! |
| ENSMUSG00000118924.1 | 0 | NA | NA | NA | NA | NA | #VALUE! |
| ENSMUSG00000118925.1 | 0 | NA | NA | NA | NA | NA | #VALUE! |
| ENSMUSG00000118926.1 | 0 | NA | NA | NA | NA | NA | #VALUE! |
| ENSMUSG00000118928.1 | 0 | NA | NA | NA | NA | NA | #VALUE! |
| ENSMUSG00000118930.1 | 0 | NA | NA | NA | NA | NA | #VALUE! |
| ENSMUSG00000118933.1 | 0 | NA | NA | NA | NA | NA | #VALUE! |
| ENSMUSG00000118934.1 | 0 | NA | NA | NA | NA | NA | #VALUE! |
| ENSMUSG00000118936.1 | 0 | NA | NA | NA | NA | NA | #VALUE! |
| ENSMUSG00000118937.1 | 0 | NA | NA | NA | NA | NA | #VALUE! |
| ENSMUSG00000118939.1 | 0 | NA | NA | NA | NA | NA | #VALUE! |
| ENSMUSG00000118941.1 | 0 | NA | NA | NA | NA | NA | #VALUE! |
| ENSMUSG00000118943.1 | 0 | NA | NA | NA | NA | NA | #VALUE! |
| ENSMUSG00000118945.1 | 0 | NA | NA | NA | NA | NA | #VALUE! |
| ENSMUSG00000118946.1 | 0 | NA | NA | NA | NA | NA | #VALUE! |
| ENSMUSG00000118948.1 | 0 | NA | NA | NA | NA | NA | #VALUE! |
| ENSMUSG00000118952.1 | 0 | NA | NA | NA | NA | NA | #VALUE! |
| ENSMUSG00000118955.1 | 0 | NA | NA | NA | NA | NA | #VALUE! |
| ENSMUSG00000118956.1 | 0 | NA | NA | NA | NA | NA | #VALUE! |
| ENSMUSG00000118957.1 | 0 | NA | NA | NA | NA | NA | #VALUE! |
| ENSMUSG00000118960.1 | 0 | NA | NA | NA | NA | NA | #VALUE! |
| ENSMUSG00000118963.1 | 0 | NA | NA | NA | NA | NA | #VALUE! |
| ENSMUSG00000118965.1 | 0 | NA | NA | NA | NA | NA | #VALUE! |
| ENSMUSG00000118966.1 | 0 | NA | NA | NA | NA | NA | #VALUE! |
| ENSMUSG00000118967.1 | 0 | NA | NA | NA | NA | NA | #VALUE! |
| ENSMUSG00000118968.1 | 0 | NA | NA | NA | NA | NA | #VALUE! |
| ENSMUSG00000118969.1 | 0 | NA | NA | NA | NA | NA | #VALUE! |
| ENSMUSG00000118970.1 | 0 | NA | NA | NA | NA | NA | #VALUE! |
| ENSMUSG00000118971.1 | 0 | NA | NA | NA | NA | NA | #VALUE! |
| ENSMUSG00000118972.1 | 0 | NA | NA | NA | NA | NA | #VALUE! |
| ENSMUSG00000118975.1 | 0 | NA | NA | NA | NA | NA | #VALUE! |
| ENSMUSG00000118976.1 | 0 | NA | NA | NA | NA | NA | #VALUE! |
| ENSMUSG00000118977.1 | 0 | NA | NA | NA | NA | NA | #VALUE! |
| ENSMUSG00000118978.1 | 0 | NA | NA | NA | NA | NA | #VALUE! |
| ENSMUSG00000118979.1 | 0 | NA | NA | NA | NA | NA | #VALUE! |
| ENSMUSG00000118981.1 | 0 | NA | NA | NA | NA | NA | #VALUE! |
| ENSMUSG00000118983.1 | 0 | NA | NA | NA | NA | NA | #VALUE! |
| ENSMUSG00000118984.1 | 0 | NA | NA | NA | NA | NA | #VALUE! |
| ENSMUSG00000118991.1 | 0 | NA | NA | NA | NA | NA | #VALUE! |
| ENSMUSG00000118995.1 | 0 | NA | NA | NA | NA | NA | #VALUE! |
| ENSMUSG00000118996.1 | 0 | NA | NA | NA | NA | NA | #VALUE! |
| ENSMUSG00000118997.1 | 0 | NA | NA | NA | NA | NA | #VALUE! |
| ENSMUSG00000118998.1 | 0 | NA | NA | NA | NA | NA | #VALUE! |
| ENSMUSG00000118999.1 | 0 | NA | NA | NA | NA | NA | #VALUE! |

| | | | | | | | |
|---|---|---|---|---|---|---|---|
| ENSMUSG00000119001.1 | 0 | NA | NA | NA | NA | NA | #VALUE! |
| ENSMUSG00000119002.1 | 0 | NA | NA | NA | NA | NA | #VALUE! |
| ENSMUSG00000119004.1 | 0 | NA | NA | NA | NA | NA | #VALUE! |
| ENSMUSG00000119005.1 | 0 | NA | NA | NA | NA | NA | #VALUE! |
| ENSMUSG00000119006.1 | 0 | NA | NA | NA | NA | NA | #VALUE! |
| ENSMUSG00000119009.1 | 0 | NA | NA | NA | NA | NA | #VALUE! |
| ENSMUSG00000119010.1 | 0 | NA | NA | NA | NA | NA | #VALUE! |
| ENSMUSG00000119011.1 | 0 | NA | NA | NA | NA | NA | #VALUE! |
| ENSMUSG00000119012.1 | 0 | NA | NA | NA | NA | NA | #VALUE! |
| ENSMUSG00000119013.1 | 0 | NA | NA | NA | NA | NA | #VALUE! |
| ENSMUSG00000119014.1 | 0 | NA | NA | NA | NA | NA | #VALUE! |
| ENSMUSG00000119015.1 | 0 | NA | NA | NA | NA | NA | #VALUE! |
| ENSMUSG00000119019.1 | 0 | NA | NA | NA | NA | NA | #VALUE! |
| ENSMUSG00000119020.1 | 0 | NA | NA | NA | NA | NA | #VALUE! |
| ENSMUSG00000119021.1 | 0 | NA | NA | NA | NA | NA | #VALUE! |
| ENSMUSG00000119023.1 | 0 | NA | NA | NA | NA | NA | #VALUE! |
| ENSMUSG00000119024.1 | 0 | NA | NA | NA | NA | NA | #VALUE! |
| ENSMUSG00000119025.1 | 0 | NA | NA | NA | NA | NA | #VALUE! |
| ENSMUSG00000119029.1 | 0 | NA | NA | NA | NA | NA | #VALUE! |
| ENSMUSG00000119030.1 | 0 | NA | NA | NA | NA | NA | #VALUE! |
| ENSMUSG00000119031.1 | 0 | NA | NA | NA | NA | NA | #VALUE! |
| ENSMUSG00000119032.1 | 0 | NA | NA | NA | NA | NA | #VALUE! |
| ENSMUSG00000119033.2 | 0 | NA | NA | NA | NA | NA | #VALUE! |
| ENSMUSG00000119035.1 | 0 | NA | NA | NA | NA | NA | #VALUE! |
| ENSMUSG00000119036.1 | 0 | NA | NA | NA | NA | NA | #VALUE! |
| ENSMUSG00000119038.1 | 0 | NA | NA | NA | NA | NA | #VALUE! |
| ENSMUSG00000119039.1 | 0 | NA | NA | NA | NA | NA | #VALUE! |
| ENSMUSG00000119040.1 | 0 | NA | NA | NA | NA | NA | #VALUE! |
| ENSMUSG00000119041.1 | 0 | NA | NA | NA | NA | NA | #VALUE! |
| ENSMUSG00000119042.1 | 0 | NA | NA | NA | NA | NA | #VALUE! |
| ENSMUSG00000119044.1 | 0 | NA | NA | NA | NA | NA | #VALUE! |
| ENSMUSG00000119045.1 | 0 | NA | NA | NA | NA | NA | #VALUE! |
| ENSMUSG00000119047.1 | 0 | NA | NA | NA | NA | NA | #VALUE! |
| ENSMUSG00000119048.1 | 0 | NA | NA | NA | NA | NA | #VALUE! |
| ENSMUSG00000119049.1 | 0 | NA | NA | NA | NA | NA | #VALUE! |
| ENSMUSG00000119051.1 | 0 | NA | NA | NA | NA | NA | #VALUE! |
| ENSMUSG00000119052.1 | 0 | NA | NA | NA | NA | NA | #VALUE! |
| ENSMUSG00000119053.1 | 0 | NA | NA | NA | NA | NA | #VALUE! |
| ENSMUSG00000119055.1 | 0 | NA | NA | NA | NA | NA | #VALUE! |
| ENSMUSG00000119056.2 | 0 | NA | NA | NA | NA | NA | #VALUE! |
| ENSMUSG00000119057.1 | 0 | NA | NA | NA | NA | NA | #VALUE! |
| ENSMUSG00000119058.1 | 0 | NA | NA | NA | NA | NA | #VALUE! |
| ENSMUSG00000119061.1 | 0 | NA | NA | NA | NA | NA | #VALUE! |
| ENSMUSG00000119062.1 | 0 | NA | NA | NA | NA | NA | #VALUE! |

| | | | | | | | |
|---|---|---|---|---|---|---|---|
| ENSMUSG00000119063.1 | 0 | NA | NA | NA | NA | NA | #VALUE! |
| ENSMUSG00000119065.1 | 0 | NA | NA | NA | NA | NA | #VALUE! |
| ENSMUSG00000119068.1 | 0 | NA | NA | NA | NA | NA | #VALUE! |
| ENSMUSG00000119069.1 | 0 | NA | NA | NA | NA | NA | #VALUE! |
| ENSMUSG00000119071.1 | 0 | NA | NA | NA | NA | NA | #VALUE! |
| ENSMUSG00000119072.1 | 0 | NA | NA | NA | NA | NA | #VALUE! |
| ENSMUSG00000119073.1 | 0 | NA | NA | NA | NA | NA | #VALUE! |
| ENSMUSG00000119074.1 | 0 | NA | NA | NA | NA | NA | #VALUE! |
| ENSMUSG00000119078.1 | 0 | NA | NA | NA | NA | NA | #VALUE! |
| ENSMUSG00000119079.1 | 0 | NA | NA | NA | NA | NA | #VALUE! |
| ENSMUSG00000119080.1 | 0 | NA | NA | NA | NA | NA | #VALUE! |
| ENSMUSG00000119082.1 | 0 | NA | NA | NA | NA | NA | #VALUE! |
| ENSMUSG00000119084.1 | 0 | NA | NA | NA | NA | NA | #VALUE! |
| ENSMUSG00000119085.1 | 0 | NA | NA | NA | NA | NA | #VALUE! |
| ENSMUSG00000119086.1 | 0 | NA | NA | NA | NA | NA | #VALUE! |
| ENSMUSG00000119087.1 | 0 | NA | NA | NA | NA | NA | #VALUE! |
| ENSMUSG00000119088.1 | 0 | NA | NA | NA | NA | NA | #VALUE! |
| ENSMUSG00000119089.1 | 0 | NA | NA | NA | NA | NA | #VALUE! |
| ENSMUSG00000119090.1 | 0 | NA | NA | NA | NA | NA | #VALUE! |
| ENSMUSG00000119091.1 | 0 | NA | NA | NA | NA | NA | #VALUE! |
| ENSMUSG00000119092.1 | 0 | NA | NA | NA | NA | NA | #VALUE! |
| ENSMUSG00000119093.1 | 0 | NA | NA | NA | NA | NA | #VALUE! |
| ENSMUSG00000119095.1 | 0 | NA | NA | NA | NA | NA | #VALUE! |
| ENSMUSG00000119097.1 | 0 | NA | NA | NA | NA | NA | #VALUE! |
| ENSMUSG00000119098.1 | 0 | NA | NA | NA | NA | NA | #VALUE! |
| ENSMUSG00000119100.1 | 0 | NA | NA | NA | NA | NA | #VALUE! |
| ENSMUSG00000119101.1 | 0 | NA | NA | NA | NA | NA | #VALUE! |
| ENSMUSG00000119104.1 | 0 | NA | NA | NA | NA | NA | #VALUE! |
| ENSMUSG00000119105.1 | 0 | NA | NA | NA | NA | NA | #VALUE! |
| ENSMUSG00000119106.1 | 0 | NA | NA | NA | NA | NA | #VALUE! |
| ENSMUSG00000119108.2 | 0 | NA | NA | NA | NA | NA | #VALUE! |
| ENSMUSG00000119111.1 | 0 | NA | NA | NA | NA | NA | #VALUE! |
| ENSMUSG00000119112.1 | 0 | NA | NA | NA | NA | NA | #VALUE! |
| ENSMUSG00000119114.2 | 0 | NA | NA | NA | NA | NA | #VALUE! |
| ENSMUSG00000119115.1 | 0 | NA | NA | NA | NA | NA | #VALUE! |
| ENSMUSG00000119117.1 | 0 | NA | NA | NA | NA | NA | #VALUE! |
| ENSMUSG00000119118.1 | 0 | NA | NA | NA | NA | NA | #VALUE! |
| ENSMUSG00000119119.1 | 0 | NA | NA | NA | NA | NA | #VALUE! |
| ENSMUSG00000119121.1 | 0 | NA | NA | NA | NA | NA | #VALUE! |
| ENSMUSG00000119122.1 | 0 | NA | NA | NA | NA | NA | #VALUE! |
| ENSMUSG00000119124.1 | 0 | NA | NA | NA | NA | NA | #VALUE! |
| ENSMUSG00000119125.1 | 0 | NA | NA | NA | NA | NA | #VALUE! |
| ENSMUSG00000119126.1 | 0 | NA | NA | NA | NA | NA | #VALUE! |
| ENSMUSG00000119127.1 | 0 | NA | NA | NA | NA | NA | #VALUE! |

| | | | | | | | |
|---|---|---|---|---|---|---|---|
| ENSMUSG00000119128.2 | 0 | NA | NA | NA | NA | NA | #VALUE! |
| ENSMUSG00000119129.1 | 0 | NA | NA | NA | NA | NA | #VALUE! |
| ENSMUSG00000119130.1 | 0 | NA | NA | NA | NA | NA | #VALUE! |
| ENSMUSG00000119131.1 | 0 | NA | NA | NA | NA | NA | #VALUE! |
| ENSMUSG00000119132.1 | 1.025807 | 3.556426 | 3.949287 | 0.900523 | 0.367842 | NA | #VALUE! |
| ENSMUSG00000119133.1 | 0 | NA | NA | NA | NA | NA | #VALUE! |
| ENSMUSG00000119136.1 | 0 | NA | NA | NA | NA | NA | #VALUE! |
| ENSMUSG00000119137.1 | 0 | NA | NA | NA | NA | NA | #VALUE! |
| ENSMUSG00000119139.1 | 0 | NA | NA | NA | NA | NA | #VALUE! |
| ENSMUSG00000119140.2 | 0 | NA | NA | NA | NA | NA | #VALUE! |
| ENSMUSG00000119141.1 | 0 | NA | NA | NA | NA | NA | #VALUE! |
| ENSMUSG00000119142.1 | 0 | NA | NA | NA | NA | NA | #VALUE! |
| ENSMUSG00000119143.1 | 0 | NA | NA | NA | NA | NA | #VALUE! |
| ENSMUSG00000119144.1 | 0 | NA | NA | NA | NA | NA | #VALUE! |
| ENSMUSG00000119145.1 | 0 | NA | NA | NA | NA | NA | #VALUE! |
| ENSMUSG00000119148.1 | 0 | NA | NA | NA | NA | NA | #VALUE! |
| ENSMUSG00000119149.1 | 0 | NA | NA | NA | NA | NA | #VALUE! |
| ENSMUSG00000119150.1 | 0 | NA | NA | NA | NA | NA | #VALUE! |
| ENSMUSG00000119153.1 | 0 | NA | NA | NA | NA | NA | #VALUE! |
| ENSMUSG00000119155.1 | 0 | NA | NA | NA | NA | NA | #VALUE! |
| ENSMUSG00000119157.1 | 0 | NA | NA | NA | NA | NA | #VALUE! |
| ENSMUSG00000119159.1 | 0 | NA | NA | NA | NA | NA | #VALUE! |
| ENSMUSG00000119160.1 | 0 | NA | NA | NA | NA | NA | #VALUE! |
| ENSMUSG00000119161.1 | 0 | NA | NA | NA | NA | NA | #VALUE! |
| ENSMUSG00000119162.1 | 0 | NA | NA | NA | NA | NA | #VALUE! |
| ENSMUSG00000119163.1 | 0 | NA | NA | NA | NA | NA | #VALUE! |
| ENSMUSG00000119166.1 | 0 | NA | NA | NA | NA | NA | #VALUE! |
| ENSMUSG00000119168.1 | 0 | NA | NA | NA | NA | NA | #VALUE! |
| ENSMUSG00000119170.1 | 0 | NA | NA | NA | NA | NA | #VALUE! |
| ENSMUSG00000119171.1 | 0 | NA | NA | NA | NA | NA | #VALUE! |
| ENSMUSG00000119172.1 | 0 | NA | NA | NA | NA | NA | #VALUE! |
| ENSMUSG00000119173.1 | 0 | NA | NA | NA | NA | NA | #VALUE! |
| ENSMUSG00000119174.1 | 0 | NA | NA | NA | NA | NA | #VALUE! |
| ENSMUSG00000119175.1 | 0 | NA | NA | NA | NA | NA | #VALUE! |
| ENSMUSG00000119176.1 | 0 | NA | NA | NA | NA | NA | #VALUE! |
| ENSMUSG00000119179.1 | 0 | NA | NA | NA | NA | NA | #VALUE! |
| ENSMUSG00000119180.1 | 0 | NA | NA | NA | NA | NA | #VALUE! |
| ENSMUSG00000119181.1 | 0 | NA | NA | NA | NA | NA | #VALUE! |
| ENSMUSG00000119183.1 | 0 | NA | NA | NA | NA | NA | #VALUE! |
| ENSMUSG00000119184.1 | 0 | NA | NA | NA | NA | NA | #VALUE! |
| ENSMUSG00000119186.1 | 0 | NA | NA | NA | NA | NA | #VALUE! |
| ENSMUSG00000119190.1 | 0 | NA | NA | NA | NA | NA | #VALUE! |
| ENSMUSG00000119191.1 | 0 | NA | NA | NA | NA | NA | #VALUE! |
| ENSMUSG00000119192.1 | 0 | NA | NA | NA | NA | NA | #VALUE! |

| | | | | | | | |
|---|---|---|---|---|---|---|---|
| ENSMUSG00000119194.1 | 0 | NA | NA | NA | NA | NA | #VALUE! |
| ENSMUSG00000119195.1 | 0 | NA | NA | NA | NA | NA | #VALUE! |
| ENSMUSG00000119196.1 | 0 | NA | NA | NA | NA | NA | #VALUE! |
| ENSMUSG00000119197.1 | 0 | NA | NA | NA | NA | NA | #VALUE! |
| ENSMUSG00000119199.1 | 0 | NA | NA | NA | NA | NA | #VALUE! |
| ENSMUSG00000119200.1 | 0 | NA | NA | NA | NA | NA | #VALUE! |
| ENSMUSG00000119201.1 | 0 | NA | NA | NA | NA | NA | #VALUE! |
| ENSMUSG00000119204.1 | 0 | NA | NA | NA | NA | NA | #VALUE! |
| ENSMUSG00000119205.1 | 0 | NA | NA | NA | NA | NA | #VALUE! |
| ENSMUSG00000119206.1 | 0 | NA | NA | NA | NA | NA | #VALUE! |
| ENSMUSG00000119207.1 | 0 | NA | NA | NA | NA | NA | #VALUE! |
| ENSMUSG00000119208.1 | 0 | NA | NA | NA | NA | NA | #VALUE! |
| ENSMUSG00000119210.1 | 0 | NA | NA | NA | NA | NA | #VALUE! |
| ENSMUSG00000119211.1 | 0 | NA | NA | NA | NA | NA | #VALUE! |
| ENSMUSG00000119213.1 | 0 | NA | NA | NA | NA | NA | #VALUE! |
| ENSMUSG00000119214.1 | 0 | NA | NA | NA | NA | NA | #VALUE! |
| ENSMUSG00000119215.1 | 0 | NA | NA | NA | NA | NA | #VALUE! |
| ENSMUSG00000119216.1 | 0 | NA | NA | NA | NA | NA | #VALUE! |
| ENSMUSG00000119217.1 | 0 | NA | NA | NA | NA | NA | #VALUE! |
| ENSMUSG00000119219.1 | 0 | NA | NA | NA | NA | NA | #VALUE! |
| ENSMUSG00000119221.1 | 0 | NA | NA | NA | NA | NA | #VALUE! |
| ENSMUSG00000119223.1 | 0 | NA | NA | NA | NA | NA | #VALUE! |
| ENSMUSG00000119224.1 | 0 | NA | NA | NA | NA | NA | #VALUE! |
| ENSMUSG00000119225.1 | 0 | NA | NA | NA | NA | NA | #VALUE! |
| ENSMUSG00000119226.1 | 0 | NA | NA | NA | NA | NA | #VALUE! |
| ENSMUSG00000119227.1 | 0 | NA | NA | NA | NA | NA | #VALUE! |
| ENSMUSG00000119228.1 | 0 | NA | NA | NA | NA | NA | #VALUE! |
| ENSMUSG00000119229.1 | 0 | NA | NA | NA | NA | NA | #VALUE! |
| ENSMUSG00000119232.1 | 0 | NA | NA | NA | NA | NA | #VALUE! |
| ENSMUSG00000119233.1 | 0 | NA | NA | NA | NA | NA | #VALUE! |
| ENSMUSG00000119234.1 | 0 | NA | NA | NA | NA | NA | #VALUE! |
| ENSMUSG00000119236.1 | 0 | NA | NA | NA | NA | NA | #VALUE! |
| ENSMUSG00000119238.1 | 0 | NA | NA | NA | NA | NA | #VALUE! |
| ENSMUSG00000119239.1 | 0 | NA | NA | NA | NA | NA | #VALUE! |
| ENSMUSG00000119240.1 | 0 | NA | NA | NA | NA | NA | #VALUE! |
| ENSMUSG00000119241.1 | 0 | NA | NA | NA | NA | NA | #VALUE! |
| ENSMUSG00000119242.1 | 0 | NA | NA | NA | NA | NA | #VALUE! |
| ENSMUSG00000119244.1 | 0 | NA | NA | NA | NA | NA | #VALUE! |
| ENSMUSG00000119247.1 | 0 | NA | NA | NA | NA | NA | #VALUE! |
| ENSMUSG00000119250.1 | 0 | NA | NA | NA | NA | NA | #VALUE! |
| ENSMUSG00000119251.1 | 0 | NA | NA | NA | NA | NA | #VALUE! |
| ENSMUSG00000119254.1 | 0 | NA | NA | NA | NA | NA | #VALUE! |
| ENSMUSG00000119255.2 | 0 | NA | NA | NA | NA | NA | #VALUE! |
| ENSMUSG00000119256.1 | 0 | NA | NA | NA | NA | NA | #VALUE! |

| | | | | | | | |
|---|---|---|---|---|---|---|---|
| ENSMUSG00000119257.1 | 0 | NA | NA | NA | NA | NA | #VALUE! |
| ENSMUSG00000119258.1 | 0 | NA | NA | NA | NA | NA | #VALUE! |
| ENSMUSG00000119259.1 | 0 | NA | NA | NA | NA | NA | #VALUE! |
| ENSMUSG00000119260.1 | 0 | NA | NA | NA | NA | NA | #VALUE! |
| ENSMUSG00000119262.1 | 0 | NA | NA | NA | NA | NA | #VALUE! |
| ENSMUSG00000119263.1 | 0 | NA | NA | NA | NA | NA | #VALUE! |
| ENSMUSG00000119265.2 | 0 | NA | NA | NA | NA | NA | #VALUE! |
| ENSMUSG00000119267.1 | 0 | NA | NA | NA | NA | NA | #VALUE! |
| ENSMUSG00000119268.1 | 0 | NA | NA | NA | NA | NA | #VALUE! |
| ENSMUSG00000119269.1 | 0 | NA | NA | NA | NA | NA | #VALUE! |
| ENSMUSG00000119270.1 | 0 | NA | NA | NA | NA | NA | #VALUE! |
| ENSMUSG00000119271.1 | 0 | NA | NA | NA | NA | NA | #VALUE! |
| ENSMUSG00000119272.1 | 0 | NA | NA | NA | NA | NA | #VALUE! |
| ENSMUSG00000119273.1 | 0 | NA | NA | NA | NA | NA | #VALUE! |
| ENSMUSG00000119274.1 | 0 | NA | NA | NA | NA | NA | #VALUE! |
| ENSMUSG00000119275.1 | 0 | NA | NA | NA | NA | NA | #VALUE! |
| ENSMUSG00000119277.1 | 0 | NA | NA | NA | NA | NA | #VALUE! |
| ENSMUSG00000119278.1 | 0 | NA | NA | NA | NA | NA | #VALUE! |
| ENSMUSG00000119279.1 | 0 | NA | NA | NA | NA | NA | #VALUE! |
| ENSMUSG00000119280.1 | 0 | NA | NA | NA | NA | NA | #VALUE! |
| ENSMUSG00000119282.1 | 0 | NA | NA | NA | NA | NA | #VALUE! |
| ENSMUSG00000119285.1 | 0 | NA | NA | NA | NA | NA | #VALUE! |
| ENSMUSG00000119286.1 | 0 | NA | NA | NA | NA | NA | #VALUE! |
| ENSMUSG00000119287.1 | 0 | NA | NA | NA | NA | NA | #VALUE! |
| ENSMUSG00000119288.1 | 0 | NA | NA | NA | NA | NA | #VALUE! |
| ENSMUSG00000119290.1 | 0 | NA | NA | NA | NA | NA | #VALUE! |
| ENSMUSG00000119291.1 | 0 | NA | NA | NA | NA | NA | #VALUE! |
| ENSMUSG00000119292.1 | 0 | NA | NA | NA | NA | NA | #VALUE! |
| ENSMUSG00000119293.1 | 0 | NA | NA | NA | NA | NA | #VALUE! |
| ENSMUSG00000119294.1 | 0 | NA | NA | NA | NA | NA | #VALUE! |
| ENSMUSG00000119295.1 | 0 | NA | NA | NA | NA | NA | #VALUE! |
| ENSMUSG00000119296.1 | 0 | NA | NA | NA | NA | NA | #VALUE! |
| ENSMUSG00000119298.1 | 0 | NA | NA | NA | NA | NA | #VALUE! |
| ENSMUSG00000119299.1 | 0 | NA | NA | NA | NA | NA | #VALUE! |
| ENSMUSG00000119304.1 | 0 | NA | NA | NA | NA | NA | #VALUE! |
| ENSMUSG00000119306.1 | 0 | NA | NA | NA | NA | NA | #VALUE! |
| ENSMUSG00000119307.1 | 0 | NA | NA | NA | NA | NA | #VALUE! |
| ENSMUSG00000119308.1 | 0 | NA | NA | NA | NA | NA | #VALUE! |
| ENSMUSG00000119310.1 | 0 | NA | NA | NA | NA | NA | #VALUE! |
| ENSMUSG00000119311.1 | 0 | NA | NA | NA | NA | NA | #VALUE! |
| ENSMUSG00000119312.1 | 0 | NA | NA | NA | NA | NA | #VALUE! |
| ENSMUSG00000119314.1 | 0 | NA | NA | NA | NA | NA | #VALUE! |
| ENSMUSG00000119316.1 | 0 | NA | NA | NA | NA | NA | #VALUE! |
| ENSMUSG00000119317.1 | 0 | NA | NA | NA | NA | NA | #VALUE! |

| | | | | | | | |
|---|---|---|---|---|---|---|---|
| ENSMUSG00000119318.1 | 0 | NA | NA | NA | NA | NA | #VALUE! |
| ENSMUSG00000119320.1 | 0 | NA | NA | NA | NA | NA | #VALUE! |
| ENSMUSG00000119321.1 | 0 | NA | NA | NA | NA | NA | #VALUE! |
| ENSMUSG00000119322.1 | 0 | NA | NA | NA | NA | NA | #VALUE! |
| ENSMUSG00000119323.1 | 0 | NA | NA | NA | NA | NA | #VALUE! |
| ENSMUSG00000119324.1 | 0 | NA | NA | NA | NA | NA | #VALUE! |
| ENSMUSG00000119325.1 | 0 | NA | NA | NA | NA | NA | #VALUE! |
| ENSMUSG00000119326.1 | 0 | NA | NA | NA | NA | NA | #VALUE! |
| ENSMUSG00000119327.1 | 0 | NA | NA | NA | NA | NA | #VALUE! |
| ENSMUSG00000119329.1 | 0 | NA | NA | NA | NA | NA | #VALUE! |
| ENSMUSG00000119331.1 | 0 | NA | NA | NA | NA | NA | #VALUE! |
| ENSMUSG00000119332.1 | 0 | NA | NA | NA | NA | NA | #VALUE! |
| ENSMUSG00000119334.1 | 0 | NA | NA | NA | NA | NA | #VALUE! |
| ENSMUSG00000119336.1 | 0 | NA | NA | NA | NA | NA | #VALUE! |
| ENSMUSG00000119338.1 | 0 | NA | NA | NA | NA | NA | #VALUE! |
| ENSMUSG00000119339.1 | 0 | NA | NA | NA | NA | NA | #VALUE! |
| ENSMUSG00000119342.1 | 0 | NA | NA | NA | NA | NA | #VALUE! |
| ENSMUSG00000119343.1 | 0 | NA | NA | NA | NA | NA | #VALUE! |
| ENSMUSG00000119344.1 | 0 | NA | NA | NA | NA | NA | #VALUE! |
| ENSMUSG00000119345.1 | 0 | NA | NA | NA | NA | NA | #VALUE! |
| ENSMUSG00000119346.1 | 0 | NA | NA | NA | NA | NA | #VALUE! |
| ENSMUSG00000119347.1 | 0 | NA | NA | NA | NA | NA | #VALUE! |
| ENSMUSG00000119350.1 | 0 | NA | NA | NA | NA | NA | #VALUE! |
| ENSMUSG00000119351.1 | 0 | NA | NA | NA | NA | NA | #VALUE! |
| ENSMUSG00000119352.1 | 0 | NA | NA | NA | NA | NA | #VALUE! |
| ENSMUSG00000119353.1 | 0 | NA | NA | NA | NA | NA | #VALUE! |
| ENSMUSG00000119355.1 | 0 | NA | NA | NA | NA | NA | #VALUE! |
| ENSMUSG00000119356.1 | 0 | NA | NA | NA | NA | NA | #VALUE! |
| ENSMUSG00000119357.1 | 0 | NA | NA | NA | NA | NA | #VALUE! |
| ENSMUSG00000119358.1 | 0 | NA | NA | NA | NA | NA | #VALUE! |
| ENSMUSG00000119359.1 | 0 | NA | NA | NA | NA | NA | #VALUE! |
| ENSMUSG00000119360.1 | 0 | NA | NA | NA | NA | NA | #VALUE! |
| ENSMUSG00000119361.1 | 0 | NA | NA | NA | NA | NA | #VALUE! |
| ENSMUSG00000119363.1 | 0 | NA | NA | NA | NA | NA | #VALUE! |
| ENSMUSG00000119364.1 | 0 | NA | NA | NA | NA | NA | #VALUE! |
| ENSMUSG00000119367.1 | 0 | NA | NA | NA | NA | NA | #VALUE! |
| ENSMUSG00000119368.1 | 0 | NA | NA | NA | NA | NA | #VALUE! |
| ENSMUSG00000119371.1 | 0 | NA | NA | NA | NA | NA | #VALUE! |
| ENSMUSG00000119373.1 | 0 | NA | NA | NA | NA | NA | #VALUE! |
| ENSMUSG00000119374.1 | 0 | NA | NA | NA | NA | NA | #VALUE! |
| ENSMUSG00000119375.1 | 0 | NA | NA | NA | NA | NA | #VALUE! |
| ENSMUSG00000119376.1 | 0 | NA | NA | NA | NA | NA | #VALUE! |
| ENSMUSG00000119377.1 | 0 | NA | NA | NA | NA | NA | #VALUE! |
| ENSMUSG00000119378.1 | 0 | NA | NA | NA | NA | NA | #VALUE! |

| | | | | | | | |
|---|---|---|---|---|---|---|---|
| ENSMUSG00000119380.1 | 0 | NA | NA | NA | NA | NA | #VALUE! |
| ENSMUSG00000119381.1 | 0 | NA | NA | NA | NA | NA | #VALUE! |
| ENSMUSG00000119383.1 | 0 | NA | NA | NA | NA | NA | #VALUE! |
| ENSMUSG00000119385.1 | 0 | NA | NA | NA | NA | NA | #VALUE! |
| ENSMUSG00000119386.1 | 0 | NA | NA | NA | NA | NA | #VALUE! |
| ENSMUSG00000119387.1 | 0 | NA | NA | NA | NA | NA | #VALUE! |
| ENSMUSG00000119388.1 | 0 | NA | NA | NA | NA | NA | #VALUE! |
| ENSMUSG00000119391.1 | 0 | NA | NA | NA | NA | NA | #VALUE! |
| ENSMUSG00000119394.1 | 0 | NA | NA | NA | NA | NA | #VALUE! |
| ENSMUSG00000119395.1 | 0 | NA | NA | NA | NA | NA | #VALUE! |
| ENSMUSG00000119396.1 | 0 | NA | NA | NA | NA | NA | #VALUE! |
| ENSMUSG00000119397.1 | 0 | NA | NA | NA | NA | NA | #VALUE! |
| ENSMUSG00000119398.1 | 0 | NA | NA | NA | NA | NA | #VALUE! |
| ENSMUSG00000119399.1 | 0 | NA | NA | NA | NA | NA | #VALUE! |
| ENSMUSG00000119402.1 | 0 | NA | NA | NA | NA | NA | #VALUE! |
| ENSMUSG00000119403.1 | 0 | NA | NA | NA | NA | NA | #VALUE! |
| ENSMUSG00000119404.1 | 0 | NA | NA | NA | NA | NA | #VALUE! |
| ENSMUSG00000119405.1 | 0 | NA | NA | NA | NA | NA | #VALUE! |
| ENSMUSG00000119409.1 | 0 | NA | NA | NA | NA | NA | #VALUE! |
| ENSMUSG00000119410.1 | 0 | NA | NA | NA | NA | NA | #VALUE! |
| ENSMUSG00000119411.1 | 0 | NA | NA | NA | NA | NA | #VALUE! |
| ENSMUSG00000119412.1 | 0 | NA | NA | NA | NA | NA | #VALUE! |
| ENSMUSG00000119413.1 | 0 | NA | NA | NA | NA | NA | #VALUE! |
| ENSMUSG00000119414.1 | 0 | NA | NA | NA | NA | NA | #VALUE! |
| ENSMUSG00000119415.1 | 0 | NA | NA | NA | NA | NA | #VALUE! |
| ENSMUSG00000119417.2 | 0 | NA | NA | NA | NA | NA | #VALUE! |
| ENSMUSG00000119426.1 | 0 | NA | NA | NA | NA | NA | #VALUE! |
| ENSMUSG00000119427.1 | 0 | NA | NA | NA | NA | NA | #VALUE! |
| ENSMUSG00000119428.1 | 0 | NA | NA | NA | NA | NA | #VALUE! |
| ENSMUSG00000119430.1 | 0 | NA | NA | NA | NA | NA | #VALUE! |
| ENSMUSG00000119431.2 | 0 | NA | NA | NA | NA | NA | #VALUE! |
| ENSMUSG00000119432.1 | 0 | NA | NA | NA | NA | NA | #VALUE! |
| ENSMUSG00000119433.1 | 0 | NA | NA | NA | NA | NA | #VALUE! |
| ENSMUSG00000119434.1 | 0 | NA | NA | NA | NA | NA | #VALUE! |
| ENSMUSG00000119436.1 | 0 | NA | NA | NA | NA | NA | #VALUE! |
| ENSMUSG00000119440.1 | 0 | NA | NA | NA | NA | NA | #VALUE! |
| ENSMUSG00000119441.1 | 0 | NA | NA | NA | NA | NA | #VALUE! |
| ENSMUSG00000119442.1 | 0 | NA | NA | NA | NA | NA | #VALUE! |
| ENSMUSG00000119444.1 | 0 | NA | NA | NA | NA | NA | #VALUE! |
| ENSMUSG00000119445.1 | 0 | NA | NA | NA | NA | NA | #VALUE! |
| ENSMUSG00000119446.1 | 0 | NA | NA | NA | NA | NA | #VALUE! |
| ENSMUSG00000119447.1 | 0 | NA | NA | NA | NA | NA | #VALUE! |
| ENSMUSG00000119449.1 | 0 | NA | NA | NA | NA | NA | #VALUE! |
| ENSMUSG00000119450.1 | 0 | NA | NA | NA | NA | NA | #VALUE! |

| | | | | | | | |
|---|---|---|---|---|---|---|---|
| ENSMUSG00000119451.1 | 0 | NA | NA | NA | NA | NA | #VALUE! |
| ENSMUSG00000119452.1 | 0 | NA | NA | NA | NA | NA | #VALUE! |
| ENSMUSG00000119456.1 | 0 | NA | NA | NA | NA | NA | #VALUE! |
| ENSMUSG00000119457.1 | 0 | NA | NA | NA | NA | NA | #VALUE! |
| ENSMUSG00000119458.1 | 0 | NA | NA | NA | NA | NA | #VALUE! |
| ENSMUSG00000119459.1 | 0 | NA | NA | NA | NA | NA | #VALUE! |
| ENSMUSG00000119461.1 | 0 | NA | NA | NA | NA | NA | #VALUE! |
| ENSMUSG00000119462.1 | 0 | NA | NA | NA | NA | NA | #VALUE! |
| ENSMUSG00000119463.1 | 0 | NA | NA | NA | NA | NA | #VALUE! |
| ENSMUSG00000119464.1 | 0 | NA | NA | NA | NA | NA | #VALUE! |
| ENSMUSG00000119465.1 | 0 | NA | NA | NA | NA | NA | #VALUE! |
| ENSMUSG00000119467.1 | 0 | NA | NA | NA | NA | NA | #VALUE! |
| ENSMUSG00000119468.1 | 0 | NA | NA | NA | NA | NA | #VALUE! |
| ENSMUSG00000119469.1 | 0 | NA | NA | NA | NA | NA | #VALUE! |
| ENSMUSG00000119470.1 | 0 | NA | NA | NA | NA | NA | #VALUE! |
| ENSMUSG00000119471.1 | 0 | NA | NA | NA | NA | NA | #VALUE! |
| ENSMUSG00000119473.1 | 0 | NA | NA | NA | NA | NA | #VALUE! |
| ENSMUSG00000119475.1 | 0 | NA | NA | NA | NA | NA | #VALUE! |
| ENSMUSG00000119476.1 | 0 | NA | NA | NA | NA | NA | #VALUE! |
| ENSMUSG00000119477.1 | 0 | NA | NA | NA | NA | NA | #VALUE! |
| ENSMUSG00000119481.1 | 0 | NA | NA | NA | NA | NA | #VALUE! |
| ENSMUSG00000119482.1 | 0 | NA | NA | NA | NA | NA | #VALUE! |
| ENSMUSG00000119485.1 | 0 | NA | NA | NA | NA | NA | #VALUE! |
| ENSMUSG00000119486.1 | 0 | NA | NA | NA | NA | NA | #VALUE! |
| ENSMUSG00000119487.1 | 0 | NA | NA | NA | NA | NA | #VALUE! |
| ENSMUSG00000119488.2 | 0 | NA | NA | NA | NA | NA | #VALUE! |
| ENSMUSG00000119490.1 | 0 | NA | NA | NA | NA | NA | #VALUE! |
| ENSMUSG00000119491.1 | 0 | NA | NA | NA | NA | NA | #VALUE! |
| ENSMUSG00000119493.1 | 0 | NA | NA | NA | NA | NA | #VALUE! |
| ENSMUSG00000119496.1 | 0 | NA | NA | NA | NA | NA | #VALUE! |
| ENSMUSG00000119497.1 | 0 | NA | NA | NA | NA | NA | #VALUE! |
| ENSMUSG00000119498.1 | 0 | NA | NA | NA | NA | NA | #VALUE! |
| ENSMUSG00000119499.1 | 0 | NA | NA | NA | NA | NA | #VALUE! |
| ENSMUSG00000119500.1 | 0 | NA | NA | NA | NA | NA | #VALUE! |
| ENSMUSG00000119501.1 | 0 | NA | NA | NA | NA | NA | #VALUE! |
| ENSMUSG00000119502.2 | 0 | NA | NA | NA | NA | NA | #VALUE! |
| ENSMUSG00000119503.1 | 0 | NA | NA | NA | NA | NA | #VALUE! |
| ENSMUSG00000119504.1 | 0 | NA | NA | NA | NA | NA | #VALUE! |
| ENSMUSG00000119505.1 | 0 | NA | NA | NA | NA | NA | #VALUE! |
| ENSMUSG00000119506.1 | 0 | NA | NA | NA | NA | NA | #VALUE! |
| ENSMUSG00000119508.1 | 0 | NA | NA | NA | NA | NA | #VALUE! |
| ENSMUSG00000119509.1 | 0 | NA | NA | NA | NA | NA | #VALUE! |
| ENSMUSG00000119510.1 | 0 | NA | NA | NA | NA | NA | #VALUE! |
| ENSMUSG00000119511.1 | 0 | NA | NA | NA | NA | NA | #VALUE! |

| | | | | | | | |
|---|---|---|---|---|---|---|---|
| ENSMUSG00000119513.1 | 0 | NA | NA | NA | NA | NA | #VALUE! |
| ENSMUSG00000119515.1 | 0 | NA | NA | NA | NA | NA | #VALUE! |
| ENSMUSG00000119516.1 | 0 | NA | NA | NA | NA | NA | #VALUE! |
| ENSMUSG00000119520.1 | 1.890236 | -4.46257 | 3.93014 | -1.13547 | 0.256177 | NA | #VALUE! |
| ENSMUSG00000119521.1 | 0 | NA | NA | NA | NA | NA | #VALUE! |
| ENSMUSG00000119523.1 | 0 | NA | NA | NA | NA | NA | #VALUE! |
| ENSMUSG00000119524.1 | 0 | NA | NA | NA | NA | NA | #VALUE! |
| ENSMUSG00000119525.1 | 0 | NA | NA | NA | NA | NA | #VALUE! |
| ENSMUSG00000119528.1 | 0 | NA | NA | NA | NA | NA | #VALUE! |
| ENSMUSG00000119529.1 | 0 | NA | NA | NA | NA | NA | #VALUE! |
| ENSMUSG00000119530.1 | 0 | NA | NA | NA | NA | NA | #VALUE! |
| ENSMUSG00000119532.1 | 0 | NA | NA | NA | NA | NA | #VALUE! |
| ENSMUSG00000119538.1 | 0 | NA | NA | NA | NA | NA | #VALUE! |
| ENSMUSG00000119539.1 | 0 | NA | NA | NA | NA | NA | #VALUE! |
| ENSMUSG00000119542.1 | 0 | NA | NA | NA | NA | NA | #VALUE! |
| ENSMUSG00000119543.1 | 0 | NA | NA | NA | NA | NA | #VALUE! |
| ENSMUSG00000119544.2 | 0 | NA | NA | NA | NA | NA | #VALUE! |
| ENSMUSG00000119546.1 | 0 | NA | NA | NA | NA | NA | #VALUE! |
| ENSMUSG00000119547.1 | 0 | NA | NA | NA | NA | NA | #VALUE! |
| ENSMUSG00000119548.1 | 0 | NA | NA | NA | NA | NA | #VALUE! |
| ENSMUSG00000119550.1 | 0 | NA | NA | NA | NA | NA | #VALUE! |
| ENSMUSG00000119552.1 | 0 | NA | NA | NA | NA | NA | #VALUE! |
| ENSMUSG00000119553.1 | 0 | NA | NA | NA | NA | NA | #VALUE! |
| ENSMUSG00000119554.1 | 0 | NA | NA | NA | NA | NA | #VALUE! |
| ENSMUSG00000119555.1 | 0 | NA | NA | NA | NA | NA | #VALUE! |
| ENSMUSG00000119556.1 | 0 | NA | NA | NA | NA | NA | #VALUE! |
| ENSMUSG00000119558.1 | 0 | NA | NA | NA | NA | NA | #VALUE! |
| ENSMUSG00000119559.1 | 0 | NA | NA | NA | NA | NA | #VALUE! |
| ENSMUSG00000119560.1 | 0 | NA | NA | NA | NA | NA | #VALUE! |
| ENSMUSG00000119561.1 | 0 | NA | NA | NA | NA | NA | #VALUE! |
| ENSMUSG00000119562.1 | 0 | NA | NA | NA | NA | NA | #VALUE! |
| ENSMUSG00000119563.1 | 0 | NA | NA | NA | NA | NA | #VALUE! |
| ENSMUSG00000119566.1 | 0 | NA | NA | NA | NA | NA | #VALUE! |
| ENSMUSG00000119567.1 | 0 | NA | NA | NA | NA | NA | #VALUE! |
| ENSMUSG00000119568.1 | 0 | NA | NA | NA | NA | NA | #VALUE! |
| ENSMUSG00000119569.1 | 0 | NA | NA | NA | NA | NA | #VALUE! |
| ENSMUSG00000119570.1 | 0 | NA | NA | NA | NA | NA | #VALUE! |
| ENSMUSG00000119572.1 | 0 | NA | NA | NA | NA | NA | #VALUE! |
| ENSMUSG00000119573.1 | 0 | NA | NA | NA | NA | NA | #VALUE! |
| ENSMUSG00000119576.1 | 0 | NA | NA | NA | NA | NA | #VALUE! |
| ENSMUSG00000119577.1 | 0 | NA | NA | NA | NA | NA | #VALUE! |
| ENSMUSG00000119578.1 | 0 | NA | NA | NA | NA | NA | #VALUE! |
| ENSMUSG00000119579.1 | 0 | NA | NA | NA | NA | NA | #VALUE! |
| ENSMUSG00000119580.1 | 0 | NA | NA | NA | NA | NA | #VALUE! |

| | | | | | | | |
|---|---|---|---|---|---|---|---|
| ENSMUSG00000119581.1 | 0 | NA | NA | NA | NA | NA | #VALUE! |
| ENSMUSG00000119582.1 | 0 | NA | NA | NA | NA | NA | #VALUE! |
| ENSMUSG00000119586.1 | 0 | NA | NA | NA | NA | NA | #VALUE! |
| ENSMUSG00000119587.1 | 0 | NA | NA | NA | NA | NA | #VALUE! |
| ENSMUSG00000119588.1 | 0 | NA | NA | NA | NA | NA | #VALUE! |
| ENSMUSG00000119589.1 | 0 | NA | NA | NA | NA | NA | #VALUE! |
| ENSMUSG00000119593.1 | 0 | NA | NA | NA | NA | NA | #VALUE! |
| ENSMUSG00000119594.1 | 0 | NA | NA | NA | NA | NA | #VALUE! |
| ENSMUSG00000119598.1 | 0 | NA | NA | NA | NA | NA | #VALUE! |
| ENSMUSG00000119601.1 | 0 | NA | NA | NA | NA | NA | #VALUE! |
| ENSMUSG00000119602.1 | 0 | NA | NA | NA | NA | NA | #VALUE! |
| ENSMUSG00000119603.1 | 0 | NA | NA | NA | NA | NA | #VALUE! |
| ENSMUSG00000119606.1 | 0 | NA | NA | NA | NA | NA | #VALUE! |
| ENSMUSG00000119608.1 | 0 | NA | NA | NA | NA | NA | #VALUE! |
| ENSMUSG00000119609.2 | 0 | NA | NA | NA | NA | NA | #VALUE! |
| ENSMUSG00000119610.1 | 0 | NA | NA | NA | NA | NA | #VALUE! |
| ENSMUSG00000119611.1 | 0 | NA | NA | NA | NA | NA | #VALUE! |
| ENSMUSG00000119612.1 | 0 | NA | NA | NA | NA | NA | #VALUE! |
| ENSMUSG00000119614.1 | 0 | NA | NA | NA | NA | NA | #VALUE! |
| ENSMUSG00000119615.1 | 0 | NA | NA | NA | NA | NA | #VALUE! |
| ENSMUSG00000119616.1 | 0 | NA | NA | NA | NA | NA | #VALUE! |
| ENSMUSG00000119617.1 | 0 | NA | NA | NA | NA | NA | #VALUE! |
| ENSMUSG00000119619.1 | 0 | NA | NA | NA | NA | NA | #VALUE! |
| ENSMUSG00000119624.1 | 0 | NA | NA | NA | NA | NA | #VALUE! |
| ENSMUSG00000119625.1 | 0 | NA | NA | NA | NA | NA | #VALUE! |
| ENSMUSG00000119626.1 | 0 | NA | NA | NA | NA | NA | #VALUE! |
| ENSMUSG00000119629.1 | 0 | NA | NA | NA | NA | NA | #VALUE! |
| ENSMUSG00000119631.2 | 0 | NA | NA | NA | NA | NA | #VALUE! |
| ENSMUSG00000119633.1 | 0 | NA | NA | NA | NA | NA | #VALUE! |
| ENSMUSG00000119635.1 | 0 | NA | NA | NA | NA | NA | #VALUE! |
| ENSMUSG00000119636.1 | 0 | NA | NA | NA | NA | NA | #VALUE! |
| ENSMUSG00000119637.1 | 0 | NA | NA | NA | NA | NA | #VALUE! |
| ENSMUSG00000119638.1 | 0 | NA | NA | NA | NA | NA | #VALUE! |
| ENSMUSG00000119639.1 | 0 | NA | NA | NA | NA | NA | #VALUE! |
| ENSMUSG00000119640.1 | 0 | NA | NA | NA | NA | NA | #VALUE! |
| ENSMUSG00000119641.1 | 0 | NA | NA | NA | NA | NA | #VALUE! |
| ENSMUSG00000119642.1 | 0 | NA | NA | NA | NA | NA | #VALUE! |
| ENSMUSG00000119643.1 | 0 | NA | NA | NA | NA | NA | #VALUE! |
| ENSMUSG00000119647.1 | 0 | NA | NA | NA | NA | NA | #VALUE! |
| ENSMUSG00000119648.1 | 0 | NA | NA | NA | NA | NA | #VALUE! |
| ENSMUSG00000119649.1 | 0 | NA | NA | NA | NA | NA | #VALUE! |
| ENSMUSG00000119653.1 | 0 | NA | NA | NA | NA | NA | #VALUE! |
| ENSMUSG00000119654.1 | 0 | NA | NA | NA | NA | NA | #VALUE! |
| ENSMUSG00000119655.1 | 0 | NA | NA | NA | NA | NA | #VALUE! |

| | | | | | | | |
|---|---|---|---|---|---|---|---|
| ENSMUSG00000119656.1 | 0 | NA | NA | NA | NA | NA | #VALUE! |
| ENSMUSG00000119657.1 | 0 | NA | NA | NA | NA | NA | #VALUE! |
| ENSMUSG00000119658.1 | 0 | NA | NA | NA | NA | NA | #VALUE! |
| ENSMUSG00000119659.1 | 0 | NA | NA | NA | NA | NA | #VALUE! |
| ENSMUSG00000119660.1 | 0 | NA | NA | NA | NA | NA | #VALUE! |
| ENSMUSG00000119661.1 | 0 | NA | NA | NA | NA | NA | #VALUE! |
| ENSMUSG00000119662.1 | 0 | NA | NA | NA | NA | NA | #VALUE! |
| ENSMUSG00000119665.1 | 0 | NA | NA | NA | NA | NA | #VALUE! |
| ENSMUSG00000119666.1 | 0 | NA | NA | NA | NA | NA | #VALUE! |
| ENSMUSG00000119667.1 | 0 | NA | NA | NA | NA | NA | #VALUE! |
| ENSMUSG00000119668.1 | 0 | NA | NA | NA | NA | NA | #VALUE! |
| ENSMUSG00000119670.1 | 0 | NA | NA | NA | NA | NA | #VALUE! |
| ENSMUSG00000119672.1 | 0 | NA | NA | NA | NA | NA | #VALUE! |
| ENSMUSG00000119673.1 | 0 | NA | NA | NA | NA | NA | #VALUE! |
| ENSMUSG00000119677.1 | 0 | NA | NA | NA | NA | NA | #VALUE! |
| ENSMUSG00000119678.1 | 0 | NA | NA | NA | NA | NA | #VALUE! |
| ENSMUSG00000119679.1 | 0 | NA | NA | NA | NA | NA | #VALUE! |
| ENSMUSG00000119680.1 | 0 | NA | NA | NA | NA | NA | #VALUE! |
| ENSMUSG00000119681.1 | 0 | NA | NA | NA | NA | NA | #VALUE! |
| ENSMUSG00000119682.1 | 0 | NA | NA | NA | NA | NA | #VALUE! |
| ENSMUSG00000119684.1 | 0 | NA | NA | NA | NA | NA | #VALUE! |
| ENSMUSG00000119685.1 | 0 | NA | NA | NA | NA | NA | #VALUE! |
| ENSMUSG00000119686.1 | 0 | NA | NA | NA | NA | NA | #VALUE! |
| ENSMUSG00000119687.1 | 0 | NA | NA | NA | NA | NA | #VALUE! |
| ENSMUSG00000119688.1 | 0 | NA | NA | NA | NA | NA | #VALUE! |
| ENSMUSG00000119689.1 | 0 | NA | NA | NA | NA | NA | #VALUE! |
| ENSMUSG00000119692.2 | 0 | NA | NA | NA | NA | NA | #VALUE! |
| ENSMUSG00000119694.1 | 0 | NA | NA | NA | NA | NA | #VALUE! |
| ENSMUSG00000119695.1 | 0 | NA | NA | NA | NA | NA | #VALUE! |
| ENSMUSG00000119696.1 | 0 | NA | NA | NA | NA | NA | #VALUE! |
| ENSMUSG00000119700.1 | 0 | NA | NA | NA | NA | NA | #VALUE! |
| ENSMUSG00000119701.1 | 0 | NA | NA | NA | NA | NA | #VALUE! |
| ENSMUSG00000119705.1 | 0 | NA | NA | NA | NA | NA | #VALUE! |
| ENSMUSG00000119706.1 | 0 | NA | NA | NA | NA | NA | #VALUE! |
| ENSMUSG00000119708.1 | 0 | NA | NA | NA | NA | NA | #VALUE! |
| ENSMUSG00000119710.1 | 0 | NA | NA | NA | NA | NA | #VALUE! |
| ENSMUSG00000119711.1 | 0 | NA | NA | NA | NA | NA | #VALUE! |
| ENSMUSG00000119713.1 | 0 | NA | NA | NA | NA | NA | #VALUE! |
| ENSMUSG00000119714.1 | 0 | NA | NA | NA | NA | NA | #VALUE! |
| ENSMUSG00000119719.1 | 0 | NA | NA | NA | NA | NA | #VALUE! |
| ENSMUSG00000119721.2 | 0 | NA | NA | NA | NA | NA | #VALUE! |
| ENSMUSG00000119722.1 | 0 | NA | NA | NA | NA | NA | #VALUE! |
| ENSMUSG00000119723.1 | 0 | NA | NA | NA | NA | NA | #VALUE! |
| ENSMUSG00000119724.1 | 0 | NA | NA | NA | NA | NA | #VALUE! |

| | | | | | | | |
|---|---|---|---|---|---|---|---|
| ENSMUSG00000119726.1 | 0 | NA | NA | NA | NA | NA | #VALUE! |
| ENSMUSG00000119728.1 | 0 | NA | NA | NA | NA | NA | #VALUE! |
| ENSMUSG00000119730.1 | 0 | NA | NA | NA | NA | NA | #VALUE! |
| ENSMUSG00000119731.1 | 0 | NA | NA | NA | NA | NA | #VALUE! |
| ENSMUSG00000119732.1 | 0 | NA | NA | NA | NA | NA | #VALUE! |
| ENSMUSG00000119733.1 | 0 | NA | NA | NA | NA | NA | #VALUE! |
| ENSMUSG00000119734.1 | 0 | NA | NA | NA | NA | NA | #VALUE! |
| ENSMUSG00000119735.1 | 0 | NA | NA | NA | NA | NA | #VALUE! |
| ENSMUSG00000119736.1 | 0 | NA | NA | NA | NA | NA | #VALUE! |
| ENSMUSG00000119737.1 | 0 | NA | NA | NA | NA | NA | #VALUE! |
| ENSMUSG00000119738.1 | 0 | NA | NA | NA | NA | NA | #VALUE! |
| ENSMUSG00000119739.1 | 0 | NA | NA | NA | NA | NA | #VALUE! |
| ENSMUSG00000119740.1 | 0 | NA | NA | NA | NA | NA | #VALUE! |
| ENSMUSG00000119741.1 | 0 | NA | NA | NA | NA | NA | #VALUE! |
| ENSMUSG00000119742.1 | 0 | NA | NA | NA | NA | NA | #VALUE! |
| ENSMUSG00000119743.1 | 0 | NA | NA | NA | NA | NA | #VALUE! |
| ENSMUSG00000119744.1 | 0 | NA | NA | NA | NA | NA | #VALUE! |
| ENSMUSG00000119745.1 | 0 | NA | NA | NA | NA | NA | #VALUE! |
| ENSMUSG00000119746.1 | 0 | NA | NA | NA | NA | NA | #VALUE! |
| ENSMUSG00000119747.1 | 0 | NA | NA | NA | NA | NA | #VALUE! |
| ENSMUSG00000119748.1 | 0 | NA | NA | NA | NA | NA | #VALUE! |
| ENSMUSG00000119750.1 | 0 | NA | NA | NA | NA | NA | #VALUE! |
| ENSMUSG00000119751.1 | 0 | NA | NA | NA | NA | NA | #VALUE! |
| ENSMUSG00000119752.1 | 0 | NA | NA | NA | NA | NA | #VALUE! |
| ENSMUSG00000119753.1 | 0 | NA | NA | NA | NA | NA | #VALUE! |
| ENSMUSG00000119754.1 | 0 | NA | NA | NA | NA | NA | #VALUE! |
| ENSMUSG00000119755.1 | 0 | NA | NA | NA | NA | NA | #VALUE! |
| ENSMUSG00000119756.1 | 0 | NA | NA | NA | NA | NA | #VALUE! |
| ENSMUSG00000119757.2 | 0 | NA | NA | NA | NA | NA | #VALUE! |
| ENSMUSG00000119759.1 | 0 | NA | NA | NA | NA | NA | #VALUE! |
| ENSMUSG00000119760.1 | 0 | NA | NA | NA | NA | NA | #VALUE! |
| ENSMUSG00000119761.1 | 0 | NA | NA | NA | NA | NA | #VALUE! |
| ENSMUSG00000119762.1 | 0 | NA | NA | NA | NA | NA | #VALUE! |
| ENSMUSG00000119763.1 | 0 | NA | NA | NA | NA | NA | #VALUE! |
| ENSMUSG00000119764.1 | 0 | NA | NA | NA | NA | NA | #VALUE! |
| ENSMUSG00000119765.1 | 0 | NA | NA | NA | NA | NA | #VALUE! |
| ENSMUSG00000119766.1 | 0 | NA | NA | NA | NA | NA | #VALUE! |
| ENSMUSG00000119767.1 | 0 | NA | NA | NA | NA | NA | #VALUE! |
| ENSMUSG00000119769.1 | 0 | NA | NA | NA | NA | NA | #VALUE! |
| ENSMUSG00000119770.2 | 0 | NA | NA | NA | NA | NA | #VALUE! |
| ENSMUSG00000119771.1 | 0 | NA | NA | NA | NA | NA | #VALUE! |
| ENSMUSG00000119773.1 | 0 | NA | NA | NA | NA | NA | #VALUE! |
| ENSMUSG00000119774.1 | 0 | NA | NA | NA | NA | NA | #VALUE! |
| ENSMUSG00000119776.1 | 14.28775 | -7.36505 | 3.229818 | -2.28033 | NA | NA | #VALUE! |

| | | | | | | | |
|---|---|---|---|---|---|---|---|
| ENSMUSG00000119777.1 | 0 | NA | NA | NA | NA | NA | #VALUE! |
| ENSMUSG00000119778.1 | 0 | NA | NA | NA | NA | NA | #VALUE! |
| ENSMUSG00000119779.1 | 0 | NA | NA | NA | NA | NA | #VALUE! |
| ENSMUSG00000119780.1 | 0 | NA | NA | NA | NA | NA | #VALUE! |
| ENSMUSG00000119782.1 | 0 | NA | NA | NA | NA | NA | #VALUE! |
| ENSMUSG00000119783.1 | 0 | NA | NA | NA | NA | NA | #VALUE! |
| ENSMUSG00000119785.1 | 0 | NA | NA | NA | NA | NA | #VALUE! |
| ENSMUSG00000119786.1 | 0 | NA | NA | NA | NA | NA | #VALUE! |
| ENSMUSG00000119789.1 | 0 | NA | NA | NA | NA | NA | #VALUE! |
| ENSMUSG00000119790.1 | 0 | NA | NA | NA | NA | NA | #VALUE! |
| ENSMUSG00000119791.1 | 0 | NA | NA | NA | NA | NA | #VALUE! |
| ENSMUSG00000119793.1 | 0 | NA | NA | NA | NA | NA | #VALUE! |
| ENSMUSG00000119795.1 | 0 | NA | NA | NA | NA | NA | #VALUE! |
| ENSMUSG00000119797.1 | 0 | NA | NA | NA | NA | NA | #VALUE! |
| ENSMUSG00000119799.1 | 0 | NA | NA | NA | NA | NA | #VALUE! |
| ENSMUSG00000119800.1 | 0 | NA | NA | NA | NA | NA | #VALUE! |
| ENSMUSG00000119801.1 | 0 | NA | NA | NA | NA | NA | #VALUE! |
| ENSMUSG00000119802.1 | 0 | NA | NA | NA | NA | NA | #VALUE! |
| ENSMUSG00000119803.1 | 0 | NA | NA | NA | NA | NA | #VALUE! |
| ENSMUSG00000119804.2 | 0 | NA | NA | NA | NA | NA | #VALUE! |
| ENSMUSG00000119805.1 | 0 | NA | NA | NA | NA | NA | #VALUE! |
| ENSMUSG00000119806.1 | 0 | NA | NA | NA | NA | NA | #VALUE! |
| ENSMUSG00000119807.1 | 0 | NA | NA | NA | NA | NA | #VALUE! |
| ENSMUSG00000119808.1 | 0 | NA | NA | NA | NA | NA | #VALUE! |
| ENSMUSG00000119809.1 | 0 | NA | NA | NA | NA | NA | #VALUE! |
| ENSMUSG00000119810.1 | 0 | NA | NA | NA | NA | NA | #VALUE! |
| ENSMUSG00000119815.1 | 0 | NA | NA | NA | NA | NA | #VALUE! |
| ENSMUSG00000119816.1 | 0 | NA | NA | NA | NA | NA | #VALUE! |
| ENSMUSG00000119817.1 | 0 | NA | NA | NA | NA | NA | #VALUE! |
| ENSMUSG00000119819.1 | 0 | NA | NA | NA | NA | NA | #VALUE! |
| ENSMUSG00000119820.1 | 0 | NA | NA | NA | NA | NA | #VALUE! |
| ENSMUSG00000119822.1 | 0 | NA | NA | NA | NA | NA | #VALUE! |
| ENSMUSG00000119823.1 | 0 | NA | NA | NA | NA | NA | #VALUE! |
| ENSMUSG00000119825.1 | 0 | NA | NA | NA | NA | NA | #VALUE! |
| ENSMUSG00000119826.1 | 0 | NA | NA | NA | NA | NA | #VALUE! |
| ENSMUSG00000119827.1 | 0 | NA | NA | NA | NA | NA | #VALUE! |
| ENSMUSG00000119828.1 | 0 | NA | NA | NA | NA | NA | #VALUE! |
| ENSMUSG00000119829.2 | 0 | NA | NA | NA | NA | NA | #VALUE! |
| ENSMUSG00000119831.1 | 0 | NA | NA | NA | NA | NA | #VALUE! |
| ENSMUSG00000119833.1 | 0 | NA | NA | NA | NA | NA | #VALUE! |
| ENSMUSG00000119835.1 | 0 | NA | NA | NA | NA | NA | #VALUE! |
| ENSMUSG00000119837.1 | 0 | NA | NA | NA | NA | NA | #VALUE! |
| ENSMUSG00000119838.1 | 0 | NA | NA | NA | NA | NA | #VALUE! |
| ENSMUSG00000119841.1 | 0 | NA | NA | NA | NA | NA | #VALUE! |

| | | | | | | | |
|---|---|---|---|---|---|---|---|
| ENSMUSG00000119843.1 | 0 | NA | NA | NA | NA | NA | #VALUE! |
| ENSMUSG00000119844.1 | 0 | NA | NA | NA | NA | NA | #VALUE! |
| ENSMUSG00000119845.1 | 0 | NA | NA | NA | NA | NA | #VALUE! |
| ENSMUSG00000119846.1 | 0 | NA | NA | NA | NA | NA | #VALUE! |
| ENSMUSG00000119847.1 | 0 | NA | NA | NA | NA | NA | #VALUE! |
| ENSMUSG00000119848.1 | 0 | NA | NA | NA | NA | NA | #VALUE! |
| ENSMUSG00000119849.1 | 0 | NA | NA | NA | NA | NA | #VALUE! |
| ENSMUSG00000119851.1 | 0 | NA | NA | NA | NA | NA | #VALUE! |
| ENSMUSG00000119855.1 | 0 | NA | NA | NA | NA | NA | #VALUE! |
| ENSMUSG00000119857.1 | 0 | NA | NA | NA | NA | NA | #VALUE! |
| ENSMUSG00000119858.1 | 0 | NA | NA | NA | NA | NA | #VALUE! |
| ENSMUSG00000119859.1 | 0 | NA | NA | NA | NA | NA | #VALUE! |
| ENSMUSG00000119860.1 | 0 | NA | NA | NA | NA | NA | #VALUE! |
| ENSMUSG00000119861.1 | 0 | NA | NA | NA | NA | NA | #VALUE! |
| ENSMUSG00000119862.1 | 0.245976 | 1.027628 | 4.067126 | 0.252667 | 0.800526 | NA | #VALUE! |
| ENSMUSG00000119863.1 | 0 | NA | NA | NA | NA | NA | #VALUE! |
| ENSMUSG00000119864.1 | 0 | NA | NA | NA | NA | NA | #VALUE! |
| ENSMUSG00000119866.1 | 0 | NA | NA | NA | NA | NA | #VALUE! |
| ENSMUSG00000119867.1 | 0 | NA | NA | NA | NA | NA | #VALUE! |
| ENSMUSG00000119869.1 | 0 | NA | NA | NA | NA | NA | #VALUE! |
| ENSMUSG00000119872.1 | 0 | NA | NA | NA | NA | NA | #VALUE! |
| ENSMUSG00000119873.1 | 0 | NA | NA | NA | NA | NA | #VALUE! |
| ENSMUSG00000119874.1 | 0 | NA | NA | NA | NA | NA | #VALUE! |
| ENSMUSG00000119876.1 | 0 | NA | NA | NA | NA | NA | #VALUE! |
| ENSMUSG00000119877.1 | 0 | NA | NA | NA | NA | NA | #VALUE! |
| ENSMUSG00000119878.1 | 0 | NA | NA | NA | NA | NA | #VALUE! |
| ENSMUSG00000119880.1 | 0 | NA | NA | NA | NA | NA | #VALUE! |
| ENSMUSG00000119882.1 | 0 | NA | NA | NA | NA | NA | #VALUE! |
| ENSMUSG00000119883.1 | 0 | NA | NA | NA | NA | NA | #VALUE! |
| ENSMUSG00000119884.1 | 0 | NA | NA | NA | NA | NA | #VALUE! |
| ENSMUSG00000119885.1 | 0 | NA | NA | NA | NA | NA | #VALUE! |
| ENSMUSG00000119886.1 | 0 | NA | NA | NA | NA | NA | #VALUE! |
| ENSMUSG00000119887.1 | 0 | NA | NA | NA | NA | NA | #VALUE! |
| ENSMUSG00000119891.1 | 0 | NA | NA | NA | NA | NA | #VALUE! |
| ENSMUSG00000119892.1 | 0 | NA | NA | NA | NA | NA | #VALUE! |
| ENSMUSG00000119893.1 | 0 | NA | NA | NA | NA | NA | #VALUE! |
| ENSMUSG00000119894.1 | 0 | NA | NA | NA | NA | NA | #VALUE! |
| ENSMUSG00000119895.1 | 0 | NA | NA | NA | NA | NA | #VALUE! |
| ENSMUSG00000119897.1 | 0 | NA | NA | NA | NA | NA | #VALUE! |
| ENSMUSG00000119899.1 | 0 | NA | NA | NA | NA | NA | #VALUE! |
| ENSMUSG00000119901.1 | 0 | NA | NA | NA | NA | NA | #VALUE! |
| ENSMUSG00000119903.1 | 0 | NA | NA | NA | NA | NA | #VALUE! |
| ENSMUSG00000119904.1 | 0 | NA | NA | NA | NA | NA | #VALUE! |
| ENSMUSG00000119905.1 | 0 | NA | NA | NA | NA | NA | #VALUE! |

| | | | | | | | |
|---|---|---|---|---|---|---|---|
| ENSMUSG00000119906.1 | 0 | NA | NA | NA | NA | NA | #VALUE! |
| ENSMUSG00000119907.1 | 0 | NA | NA | NA | NA | NA | #VALUE! |
| ENSMUSG00000119908.1 | 0 | NA | NA | NA | NA | NA | #VALUE! |
| ENSMUSG00000119912.1 | 0 | NA | NA | NA | NA | NA | #VALUE! |
| ENSMUSG00000119913.1 | 0 | NA | NA | NA | NA | NA | #VALUE! |
| ENSMUSG00000119917.1 | 0 | NA | NA | NA | NA | NA | #VALUE! |
| ENSMUSG00000119919.1 | 0 | NA | NA | NA | NA | NA | #VALUE! |
| ENSMUSG00000119920.1 | 0 | NA | NA | NA | NA | NA | #VALUE! |
| ENSMUSG00000119921.1 | 0 | NA | NA | NA | NA | NA | #VALUE! |
| ENSMUSG00000119922.1 | 0 | NA | NA | NA | NA | NA | #VALUE! |
| ENSMUSG00000119925.1 | 0 | NA | NA | NA | NA | NA | #VALUE! |
| ENSMUSG00000119926.1 | 0 | NA | NA | NA | NA | NA | #VALUE! |
| ENSMUSG00000119927.1 | 0 | NA | NA | NA | NA | NA | #VALUE! |
| ENSMUSG00000119928.1 | 0 | NA | NA | NA | NA | NA | #VALUE! |
| ENSMUSG00000119929.1 | 0 | NA | NA | NA | NA | NA | #VALUE! |
| ENSMUSG00000119930.1 | 0 | NA | NA | NA | NA | NA | #VALUE! |
| ENSMUSG00000119931.1 | 0 | NA | NA | NA | NA | NA | #VALUE! |
| ENSMUSG00000119932.1 | 0 | NA | NA | NA | NA | NA | #VALUE! |
| ENSMUSG00000119936.1 | 0 | NA | NA | NA | NA | NA | #VALUE! |
| ENSMUSG00000119937.1 | 0 | NA | NA | NA | NA | NA | #VALUE! |
| ENSMUSG00000119938.1 | 0 | NA | NA | NA | NA | NA | #VALUE! |
| ENSMUSG00001074846.1 | 0 | NA | NA | NA | NA | NA | #VALUE! |
| ENSMUSG00001118660.1 | 0 | NA | NA | NA | NA | NA | #VALUE! |
| ENSMUSG00001118661.1 | 0 | NA | NA | NA | NA | NA | #VALUE! |
| ENSMUSG00001118662.1 | 0 | NA | NA | NA | NA | NA | #VALUE! |
| ENSMUSG00001118663.1 | 0 | NA | NA | NA | NA | NA | #VALUE! |
| ENSMUSG00001118664.1 | 0 | NA | NA | NA | NA | NA | #VALUE! |
| ENSMUSG00001118666.1 | 0 | NA | NA | NA | NA | NA | #VALUE! |
| ENSMUSG00001118668.1 | 0 | NA | NA | NA | NA | NA | #VALUE! |
| ENSMUSG00001118669.1 | 0 | NA | NA | NA | NA | NA | #VALUE! |
| ENSMUSG00001118671.1 | 0 | NA | NA | NA | NA | NA | #VALUE! |
| ENSMUSG00002074846.1 | 0 | NA | NA | NA | NA | NA | #VALUE! |
| ENSMUSG00002074847.1 | 0 | NA | NA | NA | NA | NA | #VALUE! |
| ENSMUSG00002074848.1 | 0 | NA | NA | NA | NA | NA | #VALUE! |
| ENSMUSG00002074849.1 | 0 | NA | NA | NA | NA | NA | #VALUE! |
| ENSMUSG00002074850.1 | 0 | NA | NA | NA | NA | NA | #VALUE! |
| ENSMUSG00002074851.1 | 0 | NA | NA | NA | NA | NA | #VALUE! |
| ENSMUSG00002074852.1 | 0 | NA | NA | NA | NA | NA | #VALUE! |
| ENSMUSG00002074853.1 | 0 | NA | NA | NA | NA | NA | #VALUE! |
| ENSMUSG00002074854.1 | 0 | NA | NA | NA | NA | NA | #VALUE! |
| ENSMUSG00002074855.1 | 0 | NA | NA | NA | NA | NA | #VALUE! |
| ENSMUSG00002074856.1 | 0 | NA | NA | NA | NA | NA | #VALUE! |
| ENSMUSG00002074857.1 | 1.428775 | -4.03418 | 3.938169 | -1.02438 | 0.305656 | NA | #VALUE! |
| ENSMUSG00002074858.1 | 0 | NA | NA | NA | NA | NA | #VALUE! |

| | | | | | | | |
|---|---|---|---|---|---|---|---|
| ENSMUSG00002074859.1 | 0 | NA | NA | NA | NA | NA | #VALUE! |
| ENSMUSG00002074860.1 | 0 | NA | NA | NA | NA | NA | #VALUE! |
| ENSMUSG00002074861.1 | 0 | NA | NA | NA | NA | NA | #VALUE! |
| ENSMUSG00002074862.1 | 0 | NA | NA | NA | NA | NA | #VALUE! |
| ENSMUSG00002074863.1 | 0 | NA | NA | NA | NA | NA | #VALUE! |
| ENSMUSG00002074864.1 | 0 | NA | NA | NA | NA | NA | #VALUE! |
| ENSMUSG00002074865.1 | 0 | NA | NA | NA | NA | NA | #VALUE! |
| ENSMUSG00002074866.1 | 0 | NA | NA | NA | NA | NA | #VALUE! |
| ENSMUSG00002074867.1 | 0 | NA | NA | NA | NA | NA | #VALUE! |
| ENSMUSG00002074868.1 | 0 | NA | NA | NA | NA | NA | #VALUE! |
| ENSMUSG00002074869.1 | 0 | NA | NA | NA | NA | NA | #VALUE! |
| ENSMUSG00002074870.1 | 0 | NA | NA | NA | NA | NA | #VALUE! |
| ENSMUSG00002074871.1 | 0 | NA | NA | NA | NA | NA | #VALUE! |
| ENSMUSG00002074872.1 | 0 | NA | NA | NA | NA | NA | #VALUE! |
| ENSMUSG00002074873.1 | 0 | NA | NA | NA | NA | NA | #VALUE! |
| ENSMUSG00002074874.1 | 0 | NA | NA | NA | NA | NA | #VALUE! |
| ENSMUSG00002074875.1 | 0 | NA | NA | NA | NA | NA | #VALUE! |
| ENSMUSG00002074876.1 | 0 | NA | NA | NA | NA | NA | #VALUE! |
| ENSMUSG00002074877.1 | 0 | NA | NA | NA | NA | NA | #VALUE! |
| ENSMUSG00002074878.1 | 0 | NA | NA | NA | NA | NA | #VALUE! |
| ENSMUSG00002074879.1 | 0 | NA | NA | NA | NA | NA | #VALUE! |
| ENSMUSG00002074880.1 | 0 | NA | NA | NA | NA | NA | #VALUE! |
| ENSMUSG00002074881.1 | 0 | NA | NA | NA | NA | NA | #VALUE! |
| ENSMUSG00002074882.1 | 0 | NA | NA | NA | NA | NA | #VALUE! |
| ENSMUSG00002074883.1 | 0 | NA | NA | NA | NA | NA | #VALUE! |
| ENSMUSG00002074884.1 | 0 | NA | NA | NA | NA | NA | #VALUE! |
| ENSMUSG00002074885.1 | 0 | NA | NA | NA | NA | NA | #VALUE! |
| ENSMUSG00002074886.1 | 0 | NA | NA | NA | NA | NA | #VALUE! |
| ENSMUSG00002074887.1 | 0 | NA | NA | NA | NA | NA | #VALUE! |
| ENSMUSG00002074888.1 | 0 | NA | NA | NA | NA | NA | #VALUE! |
| ENSMUSG00002074889.1 | 0 | NA | NA | NA | NA | NA | #VALUE! |
| ENSMUSG00002074890.1 | 0 | NA | NA | NA | NA | NA | #VALUE! |
| ENSMUSG00002074891.1 | 0 | NA | NA | NA | NA | NA | #VALUE! |
| ENSMUSG00002074892.1 | 0 | NA | NA | NA | NA | NA | #VALUE! |
| ENSMUSG00002074893.1 | 0 | NA | NA | NA | NA | NA | #VALUE! |
| ENSMUSG00002074894.1 | 0 | NA | NA | NA | NA | NA | #VALUE! |
| ENSMUSG00002074895.1 | 0 | NA | NA | NA | NA | NA | #VALUE! |
| ENSMUSG00002074896.1 | 0 | NA | NA | NA | NA | NA | #VALUE! |
| ENSMUSG00002074897.1 | 0 | NA | NA | NA | NA | NA | #VALUE! |
| ENSMUSG00002074898.1 | 0 | NA | NA | NA | NA | NA | #VALUE! |
| ENSMUSG00002074899.1 | 0 | NA | NA | NA | NA | NA | #VALUE! |
| ENSMUSG00002074900.1 | 0 | NA | NA | NA | NA | NA | #VALUE! |
| ENSMUSG00002074901.1 | 0 | NA | NA | NA | NA | NA | #VALUE! |
| ENSMUSG00002074902.1 | 0 | NA | NA | NA | NA | NA | #VALUE! |

| | | | | | | | |
|---|---|---|---|---|---|---|---|
| ENSMUSG00002074903.1 | 0 | NA | NA | NA | NA | NA | #VALUE! |
| ENSMUSG00002074904.1 | 0 | NA | NA | NA | NA | NA | #VALUE! |
| ENSMUSG00002074905.1 | 0 | NA | NA | NA | NA | NA | #VALUE! |
| ENSMUSG00002074906.1 | 0 | NA | NA | NA | NA | NA | #VALUE! |
| ENSMUSG00002074907.1 | 0 | NA | NA | NA | NA | NA | #VALUE! |
| ENSMUSG00002074908.1 | 0 | NA | NA | NA | NA | NA | #VALUE! |
| ENSMUSG00002074909.1 | 0 | NA | NA | NA | NA | NA | #VALUE! |
| ENSMUSG00002074910.1 | 0 | NA | NA | NA | NA | NA | #VALUE! |
| ENSMUSG00002074911.1 | 0 | NA | NA | NA | NA | NA | #VALUE! |
| ENSMUSG00002074912.1 | 0 | NA | NA | NA | NA | NA | #VALUE! |
| ENSMUSG00002074913.1 | 0 | NA | NA | NA | NA | NA | #VALUE! |
| ENSMUSG00002074914.1 | 0 | NA | NA | NA | NA | NA | #VALUE! |
| ENSMUSG00002074915.1 | 0 | NA | NA | NA | NA | NA | #VALUE! |
| ENSMUSG00002074916.1 | 0 | NA | NA | NA | NA | NA | #VALUE! |
| ENSMUSG00002074917.1 | 0 | NA | NA | NA | NA | NA | #VALUE! |
| ENSMUSG00002074918.1 | 0 | NA | NA | NA | NA | NA | #VALUE! |
| ENSMUSG00002074919.1 | 0 | NA | NA | NA | NA | NA | #VALUE! |
| ENSMUSG00002074920.1 | 0 | NA | NA | NA | NA | NA | #VALUE! |
| ENSMUSG00002074921.1 | 0 | NA | NA | NA | NA | NA | #VALUE! |
| ENSMUSG00002074922.1 | 0 | NA | NA | NA | NA | NA | #VALUE! |
| ENSMUSG00002074923.1 | 0 | NA | NA | NA | NA | NA | #VALUE! |
| ENSMUSG00002074924.1 | 0 | NA | NA | NA | NA | NA | #VALUE! |
| ENSMUSG00002074925.1 | 0 | NA | NA | NA | NA | NA | #VALUE! |
| ENSMUSG00002074926.1 | 0 | NA | NA | NA | NA | NA | #VALUE! |
| ENSMUSG00002074927.1 | 0 | NA | NA | NA | NA | NA | #VALUE! |
| ENSMUSG00002074928.1 | 0 | NA | NA | NA | NA | NA | #VALUE! |
| ENSMUSG00002074929.1 | 0 | NA | NA | NA | NA | NA | #VALUE! |
| ENSMUSG00002074930.1 | 0 | NA | NA | NA | NA | NA | #VALUE! |
| ENSMUSG00002074931.1 | 0 | NA | NA | NA | NA | NA | #VALUE! |
| ENSMUSG00002074932.1 | 0 | NA | NA | NA | NA | NA | #VALUE! |
| ENSMUSG00002074933.1 | 0 | NA | NA | NA | NA | NA | #VALUE! |
| ENSMUSG00002074934.1 | 0 | NA | NA | NA | NA | NA | #VALUE! |
| ENSMUSG00002074935.1 | 0 | NA | NA | NA | NA | NA | #VALUE! |
| ENSMUSG00002074936.1 | 0 | NA | NA | NA | NA | NA | #VALUE! |
| ENSMUSG00002074937.1 | 0 | NA | NA | NA | NA | NA | #VALUE! |
| ENSMUSG00002074938.1 | 0 | NA | NA | NA | NA | NA | #VALUE! |
| ENSMUSG00002074939.1 | 0 | NA | NA | NA | NA | NA | #VALUE! |
| ENSMUSG00002074940.1 | 0 | NA | NA | NA | NA | NA | #VALUE! |
| ENSMUSG00002074941.1 | 0 | NA | NA | NA | NA | NA | #VALUE! |
| ENSMUSG00002074942.1 | 0 | NA | NA | NA | NA | NA | #VALUE! |
| ENSMUSG00002074943.1 | 0 | NA | NA | NA | NA | NA | #VALUE! |
| ENSMUSG00002074944.1 | 0 | NA | NA | NA | NA | NA | #VALUE! |
| ENSMUSG00002074945.1 | 0 | NA | NA | NA | NA | NA | #VALUE! |
| ENSMUSG00002074946.1 | 0 | NA | NA | NA | NA | NA | #VALUE! |

| | | | | | | | |
|---|---|---|---|---|---|---|---|
| ENSMUSG00002074947.1 | 0 | NA | NA | NA | NA | NA | #VALUE! |
| ENSMUSG00002074948.1 | 0 | NA | NA | NA | NA | NA | #VALUE! |
| ENSMUSG00002074949.1 | 0 | NA | NA | NA | NA | NA | #VALUE! |
| ENSMUSG00002074950.1 | 0 | NA | NA | NA | NA | NA | #VALUE! |
| ENSMUSG00002074951.1 | 0 | NA | NA | NA | NA | NA | #VALUE! |
| ENSMUSG00002074952.1 | 0 | NA | NA | NA | NA | NA | #VALUE! |
| ENSMUSG00002074953.1 | 0 | NA | NA | NA | NA | NA | #VALUE! |
| ENSMUSG00002074954.1 | 0 | NA | NA | NA | NA | NA | #VALUE! |
| ENSMUSG00002074955.1 | 0 | NA | NA | NA | NA | NA | #VALUE! |
| ENSMUSG00002074956.1 | 0 | NA | NA | NA | NA | NA | #VALUE! |
| ENSMUSG00002074957.1 | 0 | NA | NA | NA | NA | NA | #VALUE! |
| ENSMUSG00002074958.1 | 0 | NA | NA | NA | NA | NA | #VALUE! |
| ENSMUSG00002074959.1 | 0 | NA | NA | NA | NA | NA | #VALUE! |
| ENSMUSG00002074960.1 | 0 | NA | NA | NA | NA | NA | #VALUE! |
| ENSMUSG00002074961.1 | 0 | NA | NA | NA | NA | NA | #VALUE! |
| ENSMUSG00002074962.1 | 0 | NA | NA | NA | NA | NA | #VALUE! |
| ENSMUSG00002074963.1 | 0 | NA | NA | NA | NA | NA | #VALUE! |
| ENSMUSG00002074964.1 | 0 | NA | NA | NA | NA | NA | #VALUE! |
| ENSMUSG00002074965.1 | 0 | NA | NA | NA | NA | NA | #VALUE! |
| ENSMUSG00002074966.1 | 0 | NA | NA | NA | NA | NA | #VALUE! |
| ENSMUSG00002074967.1 | 0 | NA | NA | NA | NA | NA | #VALUE! |
| ENSMUSG00002074968.1 | 0 | NA | NA | NA | NA | NA | #VALUE! |
| ENSMUSG00002074969.1 | 0 | NA | NA | NA | NA | NA | #VALUE! |
| ENSMUSG00002074970.1 | 0 | NA | NA | NA | NA | NA | #VALUE! |
| ENSMUSG00002074971.1 | 0 | NA | NA | NA | NA | NA | #VALUE! |
| ENSMUSG00002074972.1 | 0 | NA | NA | NA | NA | NA | #VALUE! |
| ENSMUSG00002074973.1 | 0 | NA | NA | NA | NA | NA | #VALUE! |
| ENSMUSG00002074974.1 | 0 | NA | NA | NA | NA | NA | #VALUE! |
| ENSMUSG00002074975.1 | 0 | NA | NA | NA | NA | NA | #VALUE! |
| ENSMUSG00002074976.1 | 0 | NA | NA | NA | NA | NA | #VALUE! |
| ENSMUSG00002074977.1 | 0 | NA | NA | NA | NA | NA | #VALUE! |
| ENSMUSG00002074978.1 | 0 | NA | NA | NA | NA | NA | #VALUE! |
| ENSMUSG00002074979.1 | 0 | NA | NA | NA | NA | NA | #VALUE! |
| ENSMUSG00002074980.1 | 0 | NA | NA | NA | NA | NA | #VALUE! |
| ENSMUSG00002074981.1 | 0 | NA | NA | NA | NA | NA | #VALUE! |
| ENSMUSG00002074982.1 | 0 | NA | NA | NA | NA | NA | #VALUE! |
| ENSMUSG00002074983.1 | 0 | NA | NA | NA | NA | NA | #VALUE! |
| ENSMUSG00002074984.1 | 0 | NA | NA | NA | NA | NA | #VALUE! |
| ENSMUSG00002074985.1 | 0 | NA | NA | NA | NA | NA | #VALUE! |
| ENSMUSG00002074986.1 | 0 | NA | NA | NA | NA | NA | #VALUE! |
| ENSMUSG00002074987.1 | 0 | NA | NA | NA | NA | NA | #VALUE! |
| ENSMUSG00002074988.1 | 4.830604 | -5.80779 | 3.915992 | -1.4831 | 0.138049 | NA | #VALUE! |
| ENSMUSG00002074989.1 | 0 | NA | NA | NA | NA | NA | #VALUE! |
| ENSMUSG00002074990.1 | 0 | NA | NA | NA | NA | NA | #VALUE! |

| | | | | | | | |
|---|---|---|---|---|---|---|---|
| ENSMUSG00002074991.1 | 0 | NA | NA | NA | NA | NA | #VALUE! |
| ENSMUSG00002074992.1 | 0 | NA | NA | NA | NA | NA | #VALUE! |
| ENSMUSG00002074993.1 | 0 | NA | NA | NA | NA | NA | #VALUE! |
| ENSMUSG00002074994.1 | 0 | NA | NA | NA | NA | NA | #VALUE! |
| ENSMUSG00002074995.1 | 0 | NA | NA | NA | NA | NA | #VALUE! |
| ENSMUSG00002074996.1 | 0 | NA | NA | NA | NA | NA | #VALUE! |
| ENSMUSG00002074997.1 | 0 | NA | NA | NA | NA | NA | #VALUE! |
| ENSMUSG00002074998.1 | 0 | NA | NA | NA | NA | NA | #VALUE! |
| ENSMUSG00002074999.1 | 0 | NA | NA | NA | NA | NA | #VALUE! |
| ENSMUSG00002075000.1 | 0 | NA | NA | NA | NA | NA | #VALUE! |
| ENSMUSG00002075001.1 | 0 | NA | NA | NA | NA | NA | #VALUE! |
| ENSMUSG00002075002.1 | 0 | NA | NA | NA | NA | NA | #VALUE! |
| ENSMUSG00002075003.1 | 0 | NA | NA | NA | NA | NA | #VALUE! |
| ENSMUSG00002075004.1 | 0 | NA | NA | NA | NA | NA | #VALUE! |
| ENSMUSG00002075005.1 | 0 | NA | NA | NA | NA | NA | #VALUE! |
| ENSMUSG00002075006.1 | 0 | NA | NA | NA | NA | NA | #VALUE! |
| ENSMUSG00002075007.1 | 0 | NA | NA | NA | NA | NA | #VALUE! |
| ENSMUSG00002075008.1 | 0 | NA | NA | NA | NA | NA | #VALUE! |
| ENSMUSG00002075009.1 | 0 | NA | NA | NA | NA | NA | #VALUE! |
| ENSMUSG00002075010.1 | 0 | NA | NA | NA | NA | NA | #VALUE! |
| ENSMUSG00002075011.1 | 0 | NA | NA | NA | NA | NA | #VALUE! |
| ENSMUSG00002075012.1 | 0 | NA | NA | NA | NA | NA | #VALUE! |
| ENSMUSG00002075013.1 | 0 | NA | NA | NA | NA | NA | #VALUE! |
| ENSMUSG00002075014.1 | 0 | NA | NA | NA | NA | NA | #VALUE! |
| ENSMUSG00002075015.1 | 0 | NA | NA | NA | NA | NA | #VALUE! |
| ENSMUSG00002075016.1 | 0 | NA | NA | NA | NA | NA | #VALUE! |
| ENSMUSG00002075017.1 | 0 | NA | NA | NA | NA | NA | #VALUE! |
| ENSMUSG00002075018.1 | 0 | NA | NA | NA | NA | NA | #VALUE! |
| ENSMUSG00002075019.1 | 0 | NA | NA | NA | NA | NA | #VALUE! |
| ENSMUSG00002075020.1 | 0 | NA | NA | NA | NA | NA | #VALUE! |
| ENSMUSG00002075021.1 | 0 | NA | NA | NA | NA | NA | #VALUE! |
| ENSMUSG00002075022.1 | 0 | NA | NA | NA | NA | NA | #VALUE! |
| ENSMUSG00002075023.1 | 0 | NA | NA | NA | NA | NA | #VALUE! |
| ENSMUSG00002075024.1 | 0 | NA | NA | NA | NA | NA | #VALUE! |
| ENSMUSG00002075025.1 | 0 | NA | NA | NA | NA | NA | #VALUE! |
| ENSMUSG00002075026.1 | 0 | NA | NA | NA | NA | NA | #VALUE! |
| ENSMUSG00002075027.1 | 0 | NA | NA | NA | NA | NA | #VALUE! |
| ENSMUSG00002075028.1 | 0 | NA | NA | NA | NA | NA | #VALUE! |
| ENSMUSG00002075029.1 | 0 | NA | NA | NA | NA | NA | #VALUE! |
| ENSMUSG00002075030.1 | 0 | NA | NA | NA | NA | NA | #VALUE! |
| ENSMUSG00002075031.1 | 0 | NA | NA | NA | NA | NA | #VALUE! |
| ENSMUSG00002075032.1 | 0 | NA | NA | NA | NA | NA | #VALUE! |
| ENSMUSG00002075033.1 | 0 | NA | NA | NA | NA | NA | #VALUE! |
| ENSMUSG00002075034.1 | 0 | NA | NA | NA | NA | NA | #VALUE! |

| | | | | | | | |
|---|---|---|---|---|---|---|---|
| ENSMUSG00002075035.1 | 0 | NA | NA | NA | NA | NA | #VALUE! |
| ENSMUSG00002075036.1 | 0 | NA | NA | NA | NA | NA | #VALUE! |
| ENSMUSG00002075037.1 | 0 | NA | NA | NA | NA | NA | #VALUE! |
| ENSMUSG00002075038.1 | 0 | NA | NA | NA | NA | NA | #VALUE! |
| ENSMUSG00002075039.1 | 0 | NA | NA | NA | NA | NA | #VALUE! |
| ENSMUSG00002075040.1 | 0 | NA | NA | NA | NA | NA | #VALUE! |
| ENSMUSG00002075041.1 | 0 | NA | NA | NA | NA | NA | #VALUE! |
| ENSMUSG00002075042.1 | 0 | NA | NA | NA | NA | NA | #VALUE! |
| ENSMUSG00002075043.1 | 0 | NA | NA | NA | NA | NA | #VALUE! |
| ENSMUSG00002075044.1 | 0 | NA | NA | NA | NA | NA | #VALUE! |
| ENSMUSG00002075045.1 | 0 | NA | NA | NA | NA | NA | #VALUE! |
| ENSMUSG00002075046.1 | 0 | NA | NA | NA | NA | NA | #VALUE! |
| ENSMUSG00002075047.1 | 0 | NA | NA | NA | NA | NA | #VALUE! |
| ENSMUSG00002075048.1 | 0 | NA | NA | NA | NA | NA | #VALUE! |
| ENSMUSG00002075049.1 | 0 | NA | NA | NA | NA | NA | #VALUE! |
| ENSMUSG00002075050.1 | 0 | NA | NA | NA | NA | NA | #VALUE! |
| ENSMUSG00002075051.1 | 0 | NA | NA | NA | NA | NA | #VALUE! |
| ENSMUSG00002075052.1 | 0 | NA | NA | NA | NA | NA | #VALUE! |
| ENSMUSG00002075053.1 | 0 | NA | NA | NA | NA | NA | #VALUE! |
| ENSMUSG00002075054.1 | 0 | NA | NA | NA | NA | NA | #VALUE! |
| ENSMUSG00002075055.1 | 0 | NA | NA | NA | NA | NA | #VALUE! |
| ENSMUSG00002075056.1 | 0 | NA | NA | NA | NA | NA | #VALUE! |
| ENSMUSG00002075057.1 | 0 | NA | NA | NA | NA | NA | #VALUE! |
| ENSMUSG00002075058.1 | 0 | NA | NA | NA | NA | NA | #VALUE! |
| ENSMUSG00002075059.1 | 0 | NA | NA | NA | NA | NA | #VALUE! |
| ENSMUSG00002075060.1 | 0 | NA | NA | NA | NA | NA | #VALUE! |
| ENSMUSG00002075061.1 | 0 | NA | NA | NA | NA | NA | #VALUE! |
| ENSMUSG00002075062.1 | 0 | NA | NA | NA | NA | NA | #VALUE! |
| ENSMUSG00002075063.1 | 0 | NA | NA | NA | NA | NA | #VALUE! |
| ENSMUSG00002075064.1 | 0 | NA | NA | NA | NA | NA | #VALUE! |
| ENSMUSG00002075065.1 | 0 | NA | NA | NA | NA | NA | #VALUE! |
| ENSMUSG00002075066.1 | 0 | NA | NA | NA | NA | NA | #VALUE! |
| ENSMUSG00002075067.1 | 0 | NA | NA | NA | NA | NA | #VALUE! |
| ENSMUSG00002075068.1 | 0 | NA | NA | NA | NA | NA | #VALUE! |
| ENSMUSG00002075069.1 | 0 | NA | NA | NA | NA | NA | #VALUE! |
| ENSMUSG00002075070.1 | 0 | NA | NA | NA | NA | NA | #VALUE! |
| ENSMUSG00002075071.1 | 0 | NA | NA | NA | NA | NA | #VALUE! |
| ENSMUSG00002075072.1 | 0 | NA | NA | NA | NA | NA | #VALUE! |
| ENSMUSG00002075073.1 | 0 | NA | NA | NA | NA | NA | #VALUE! |
| ENSMUSG00002075074.1 | 0 | NA | NA | NA | NA | NA | #VALUE! |
| ENSMUSG00002075075.1 | 0 | NA | NA | NA | NA | NA | #VALUE! |
| ENSMUSG00002075076.1 | 0 | NA | NA | NA | NA | NA | #VALUE! |
| ENSMUSG00002075077.1 | 0 | NA | NA | NA | NA | NA | #VALUE! |
| ENSMUSG00002075078.1 | 0 | NA | NA | NA | NA | NA | #VALUE! |

| | | | | | | | |
|---|---|---|---|---|---|---|---|
| ENSMUSG00002075079.1 | 0 | NA | NA | NA | NA | NA | #VALUE! |
| ENSMUSG00002075080.1 | 0 | NA | NA | NA | NA | NA | #VALUE! |
| ENSMUSG00002075081.1 | 0 | NA | NA | NA | NA | NA | #VALUE! |
| ENSMUSG00002075082.1 | 0 | NA | NA | NA | NA | NA | #VALUE! |
| ENSMUSG00002075083.1 | 0 | NA | NA | NA | NA | NA | #VALUE! |
| ENSMUSG00002075084.1 | 0 | NA | NA | NA | NA | NA | #VALUE! |
| ENSMUSG00002075085.1 | 0 | NA | NA | NA | NA | NA | #VALUE! |
| ENSMUSG00002075086.1 | 0 | NA | NA | NA | NA | NA | #VALUE! |
| ENSMUSG00002075087.1 | 0 | NA | NA | NA | NA | NA | #VALUE! |
| ENSMUSG00002075088.1 | 0 | NA | NA | NA | NA | NA | #VALUE! |
| ENSMUSG00002075089.1 | 0 | NA | NA | NA | NA | NA | #VALUE! |
| ENSMUSG00002075090.1 | 0 | NA | NA | NA | NA | NA | #VALUE! |
| ENSMUSG00002075091.1 | 0 | NA | NA | NA | NA | NA | #VALUE! |
| ENSMUSG00002075092.1 | 0 | NA | NA | NA | NA | NA | #VALUE! |
| ENSMUSG00002075093.1 | 0 | NA | NA | NA | NA | NA | #VALUE! |
| ENSMUSG00002075094.1 | 0 | NA | NA | NA | NA | NA | #VALUE! |
| ENSMUSG00002075095.1 | 0 | NA | NA | NA | NA | NA | #VALUE! |
| ENSMUSG00002075096.1 | 0 | NA | NA | NA | NA | NA | #VALUE! |
| ENSMUSG00002075097.1 | 0 | NA | NA | NA | NA | NA | #VALUE! |
| ENSMUSG00002075098.1 | 0 | NA | NA | NA | NA | NA | #VALUE! |
| ENSMUSG00002075099.1 | 0 | NA | NA | NA | NA | NA | #VALUE! |
| ENSMUSG00002075100.1 | 0 | NA | NA | NA | NA | NA | #VALUE! |
| ENSMUSG00002075101.1 | 0 | NA | NA | NA | NA | NA | #VALUE! |
| ENSMUSG00002075102.1 | 0 | NA | NA | NA | NA | NA | #VALUE! |
| ENSMUSG00002075103.1 | 0 | NA | NA | NA | NA | NA | #VALUE! |
| ENSMUSG00002075104.1 | 0 | NA | NA | NA | NA | NA | #VALUE! |
| ENSMUSG00002075105.1 | 0 | NA | NA | NA | NA | NA | #VALUE! |
| ENSMUSG00002075106.1 | 0 | NA | NA | NA | NA | NA | #VALUE! |
| ENSMUSG00002075107.1 | 0 | NA | NA | NA | NA | NA | #VALUE! |
| ENSMUSG00002075108.1 | 0 | NA | NA | NA | NA | NA | #VALUE! |
| ENSMUSG00002075109.1 | 0 | NA | NA | NA | NA | NA | #VALUE! |
| ENSMUSG00002075110.1 | 0 | NA | NA | NA | NA | NA | #VALUE! |
| ENSMUSG00002075111.1 | 0 | NA | NA | NA | NA | NA | #VALUE! |
| ENSMUSG00002075112.1 | 0 | NA | NA | NA | NA | NA | #VALUE! |
| ENSMUSG00002075113.1 | 0 | NA | NA | NA | NA | NA | #VALUE! |
| ENSMUSG00002075114.1 | 0 | NA | NA | NA | NA | NA | #VALUE! |
| ENSMUSG00002075115.1 | 0 | NA | NA | NA | NA | NA | #VALUE! |
| ENSMUSG00002075116.1 | 0 | NA | NA | NA | NA | NA | #VALUE! |
| ENSMUSG00002075117.1 | 0 | NA | NA | NA | NA | NA | #VALUE! |
| ENSMUSG00002075118.1 | 0 | NA | NA | NA | NA | NA | #VALUE! |
| ENSMUSG00002075119.1 | 0 | NA | NA | NA | NA | NA | #VALUE! |
| ENSMUSG00002075120.1 | 0 | NA | NA | NA | NA | NA | #VALUE! |
| ENSMUSG00002075121.1 | 0 | NA | NA | NA | NA | NA | #VALUE! |
| ENSMUSG00002075122.1 | 0 | NA | NA | NA | NA | NA | #VALUE! |

| | | | | | | | |
|---|---|---|---|---|---|---|---|
| ENSMUSG00002075123.1 | 0 | NA | NA | NA | NA | NA | #VALUE! |
| ENSMUSG00002075124.1 | 0 | NA | NA | NA | NA | NA | #VALUE! |
| ENSMUSG00002075125.1 | 0 | NA | NA | NA | NA | NA | #VALUE! |
| ENSMUSG00002075126.1 | 0 | NA | NA | NA | NA | NA | #VALUE! |
| ENSMUSG00002075127.1 | 0 | NA | NA | NA | NA | NA | #VALUE! |
| ENSMUSG00002075128.1 | 0 | NA | NA | NA | NA | NA | #VALUE! |
| ENSMUSG00002075129.1 | 0 | NA | NA | NA | NA | NA | #VALUE! |
| ENSMUSG00002075130.1 | 0 | NA | NA | NA | NA | NA | #VALUE! |
| ENSMUSG00002075131.1 | 0 | NA | NA | NA | NA | NA | #VALUE! |
| ENSMUSG00002075132.1 | 0 | NA | NA | NA | NA | NA | #VALUE! |
| ENSMUSG00002075133.1 | 0 | NA | NA | NA | NA | NA | #VALUE! |
| ENSMUSG00002075134.1 | 0 | NA | NA | NA | NA | NA | #VALUE! |
| ENSMUSG00002075135.1 | 0 | NA | NA | NA | NA | NA | #VALUE! |
| ENSMUSG00002075136.1 | 0 | NA | NA | NA | NA | NA | #VALUE! |
| ENSMUSG00002075137.1 | 0 | NA | NA | NA | NA | NA | #VALUE! |
| ENSMUSG00002075138.1 | 0 | NA | NA | NA | NA | NA | #VALUE! |
| ENSMUSG00002075139.1 | 0 | NA | NA | NA | NA | NA | #VALUE! |
| ENSMUSG00002075140.1 | 0 | NA | NA | NA | NA | NA | #VALUE! |
| ENSMUSG00002075141.1 | 0 | NA | NA | NA | NA | NA | #VALUE! |
| ENSMUSG00002075142.1 | 0 | NA | NA | NA | NA | NA | #VALUE! |
| ENSMUSG00002075143.1 | 0 | NA | NA | NA | NA | NA | #VALUE! |
| ENSMUSG00002075144.1 | 0 | NA | NA | NA | NA | NA | #VALUE! |
| ENSMUSG00002075145.1 | 0 | NA | NA | NA | NA | NA | #VALUE! |
| ENSMUSG00002075146.1 | 0 | NA | NA | NA | NA | NA | #VALUE! |
| ENSMUSG00002075147.1 | 0 | NA | NA | NA | NA | NA | #VALUE! |
| ENSMUSG00002075148.1 | 0 | NA | NA | NA | NA | NA | #VALUE! |
| ENSMUSG00002075149.1 | 0 | NA | NA | NA | NA | NA | #VALUE! |
| ENSMUSG00002075150.1 | 0 | NA | NA | NA | NA | NA | #VALUE! |
| ENSMUSG00002075151.1 | 0 | NA | NA | NA | NA | NA | #VALUE! |
| ENSMUSG00002075152.1 | 0 | NA | NA | NA | NA | NA | #VALUE! |
| ENSMUSG00002075153.1 | 0 | NA | NA | NA | NA | NA | #VALUE! |
| ENSMUSG00002075154.1 | 0 | NA | NA | NA | NA | NA | #VALUE! |
| ENSMUSG00002075155.1 | 0 | NA | NA | NA | NA | NA | #VALUE! |
| ENSMUSG00002075156.1 | 0 | NA | NA | NA | NA | NA | #VALUE! |
| ENSMUSG00002075157.1 | 0 | NA | NA | NA | NA | NA | #VALUE! |
| ENSMUSG00002075158.1 | 0 | NA | NA | NA | NA | NA | #VALUE! |
| ENSMUSG00002075159.1 | 0 | NA | NA | NA | NA | NA | #VALUE! |
| ENSMUSG00002075160.1 | 0 | NA | NA | NA | NA | NA | #VALUE! |
| ENSMUSG00002075161.1 | 0 | NA | NA | NA | NA | NA | #VALUE! |
| ENSMUSG00002075162.1 | 0 | NA | NA | NA | NA | NA | #VALUE! |
| ENSMUSG00002075163.1 | 0 | NA | NA | NA | NA | NA | #VALUE! |
| ENSMUSG00002075164.1 | 0 | NA | NA | NA | NA | NA | #VALUE! |
| ENSMUSG00002075165.1 | 0 | NA | NA | NA | NA | NA | #VALUE! |
| ENSMUSG00002075166.1 | 0 | NA | NA | NA | NA | NA | #VALUE! |

| | | | | | | | |
|---|---|---|---|---|---|---|---|
| ENSMUSG00002075167.1 | 0 | NA | NA | NA | NA | NA | #VALUE! |
| ENSMUSG00002075168.1 | 0 | NA | NA | NA | NA | NA | #VALUE! |
| ENSMUSG00002075169.1 | 0 | NA | NA | NA | NA | NA | #VALUE! |
| ENSMUSG00002075170.1 | 0 | NA | NA | NA | NA | NA | #VALUE! |
| ENSMUSG00002075171.1 | 0 | NA | NA | NA | NA | NA | #VALUE! |
| ENSMUSG00002075172.1 | 0 | NA | NA | NA | NA | NA | #VALUE! |
| ENSMUSG00002075173.1 | 0 | NA | NA | NA | NA | NA | #VALUE! |
| ENSMUSG00002075174.1 | 0 | NA | NA | NA | NA | NA | #VALUE! |
| ENSMUSG00002075175.1 | 0 | NA | NA | NA | NA | NA | #VALUE! |
| ENSMUSG00002075176.1 | 0 | NA | NA | NA | NA | NA | #VALUE! |
| ENSMUSG00002075177.1 | 0 | NA | NA | NA | NA | NA | #VALUE! |
| ENSMUSG00002075178.1 | 0 | NA | NA | NA | NA | NA | #VALUE! |
| ENSMUSG00002075179.1 | 0 | NA | NA | NA | NA | NA | #VALUE! |
| ENSMUSG00002075180.1 | 0 | NA | NA | NA | NA | NA | #VALUE! |
| ENSMUSG00002075181.1 | 0 | NA | NA | NA | NA | NA | #VALUE! |
| ENSMUSG00002075182.1 | 0 | NA | NA | NA | NA | NA | #VALUE! |
| ENSMUSG00002075183.1 | 0 | NA | NA | NA | NA | NA | #VALUE! |
| ENSMUSG00002075184.1 | 0 | NA | NA | NA | NA | NA | #VALUE! |
| ENSMUSG00002075185.1 | 0 | NA | NA | NA | NA | NA | #VALUE! |
| ENSMUSG00002075186.1 | 0 | NA | NA | NA | NA | NA | #VALUE! |
| ENSMUSG00002075187.1 | 0 | NA | NA | NA | NA | NA | #VALUE! |
| ENSMUSG00002075188.1 | 0 | NA | NA | NA | NA | NA | #VALUE! |
| ENSMUSG00002075189.1 | 0 | NA | NA | NA | NA | NA | #VALUE! |
| ENSMUSG00002075190.1 | 0 | NA | NA | NA | NA | NA | #VALUE! |
| ENSMUSG00002075191.1 | 0 | NA | NA | NA | NA | NA | #VALUE! |
| ENSMUSG00002075192.1 | 0 | NA | NA | NA | NA | NA | #VALUE! |
| ENSMUSG00002075193.1 | 0 | NA | NA | NA | NA | NA | #VALUE! |
| ENSMUSG00002075194.1 | 0 | NA | NA | NA | NA | NA | #VALUE! |
| ENSMUSG00002075195.1 | 0 | NA | NA | NA | NA | NA | #VALUE! |
| ENSMUSG00002075196.1 | 0 | NA | NA | NA | NA | NA | #VALUE! |
| ENSMUSG00002075197.1 | 0 | NA | NA | NA | NA | NA | #VALUE! |
| ENSMUSG00002075198.1 | 0 | NA | NA | NA | NA | NA | #VALUE! |
| ENSMUSG00002075199.1 | 0 | NA | NA | NA | NA | NA | #VALUE! |
| ENSMUSG00002075200.1 | 0 | NA | NA | NA | NA | NA | #VALUE! |
| ENSMUSG00002075201.1 | 0 | NA | NA | NA | NA | NA | #VALUE! |
| ENSMUSG00002075202.1 | 0 | NA | NA | NA | NA | NA | #VALUE! |
| ENSMUSG00002075203.1 | 0 | NA | NA | NA | NA | NA | #VALUE! |
| ENSMUSG00002075204.1 | 0 | NA | NA | NA | NA | NA | #VALUE! |
| ENSMUSG00002075205.1 | 0 | NA | NA | NA | NA | NA | #VALUE! |
| ENSMUSG00002075206.1 | 0 | NA | NA | NA | NA | NA | #VALUE! |
| ENSMUSG00002075207.1 | 0 | NA | NA | NA | NA | NA | #VALUE! |
| ENSMUSG00002075208.1 | 0 | NA | NA | NA | NA | NA | #VALUE! |
| ENSMUSG00002075209.1 | 0 | NA | NA | NA | NA | NA | #VALUE! |
| ENSMUSG00002075210.1 | 0 | NA | NA | NA | NA | NA | #VALUE! |

| | | | | | | | |
|---|---|---|---|---|---|---|---|
| ENSMUSG00002075211.1 | 0 | NA | NA | NA | NA | NA | #VALUE! |
| ENSMUSG00002075212.1 | 0 | NA | NA | NA | NA | NA | #VALUE! |
| ENSMUSG00002075213.1 | 0 | NA | NA | NA | NA | NA | #VALUE! |
| ENSMUSG00002075214.1 | 0 | NA | NA | NA | NA | NA | #VALUE! |
| ENSMUSG00002075215.1 | 0 | NA | NA | NA | NA | NA | #VALUE! |
| ENSMUSG00002075216.1 | 0 | NA | NA | NA | NA | NA | #VALUE! |
| ENSMUSG00002075217.1 | 0 | NA | NA | NA | NA | NA | #VALUE! |
| ENSMUSG00002075218.1 | 0 | NA | NA | NA | NA | NA | #VALUE! |
| ENSMUSG00002075219.1 | 0 | NA | NA | NA | NA | NA | #VALUE! |
| ENSMUSG00002075220.1 | 0 | NA | NA | NA | NA | NA | #VALUE! |
| ENSMUSG00002075221.1 | 0 | NA | NA | NA | NA | NA | #VALUE! |
| ENSMUSG00002075222.1 | 0 | NA | NA | NA | NA | NA | #VALUE! |
| ENSMUSG00002075223.1 | 0 | NA | NA | NA | NA | NA | #VALUE! |
| ENSMUSG00002075224.1 | 0 | NA | NA | NA | NA | NA | #VALUE! |
| ENSMUSG00002075225.1 | 0 | NA | NA | NA | NA | NA | #VALUE! |
| ENSMUSG00002075226.1 | 0 | NA | NA | NA | NA | NA | #VALUE! |
| ENSMUSG00002075227.1 | 0 | NA | NA | NA | NA | NA | #VALUE! |
| ENSMUSG00002075228.1 | 0 | NA | NA | NA | NA | NA | #VALUE! |
| ENSMUSG00002075229.1 | 0 | NA | NA | NA | NA | NA | #VALUE! |
| ENSMUSG00002075230.1 | 0 | NA | NA | NA | NA | NA | #VALUE! |
| ENSMUSG00002075231.1 | 0 | NA | NA | NA | NA | NA | #VALUE! |
| ENSMUSG00002075232.1 | 0 | NA | NA | NA | NA | NA | #VALUE! |
| ENSMUSG00002075233.1 | 0 | NA | NA | NA | NA | NA | #VALUE! |
| ENSMUSG00002075234.1 | 0 | NA | NA | NA | NA | NA | #VALUE! |
| ENSMUSG00002075235.1 | 0 | NA | NA | NA | NA | NA | #VALUE! |
| ENSMUSG00002075236.1 | 0 | NA | NA | NA | NA | NA | #VALUE! |
| ENSMUSG00002075237.1 | 0 | NA | NA | NA | NA | NA | #VALUE! |
| ENSMUSG00002075238.1 | 0 | NA | NA | NA | NA | NA | #VALUE! |
| ENSMUSG00002075239.1 | 0 | NA | NA | NA | NA | NA | #VALUE! |
| ENSMUSG00002075240.1 | 0 | NA | NA | NA | NA | NA | #VALUE! |
| ENSMUSG00002075241.1 | 0 | NA | NA | NA | NA | NA | #VALUE! |
| ENSMUSG00002075242.1 | 0 | NA | NA | NA | NA | NA | #VALUE! |
| ENSMUSG00002075243.1 | 0 | NA | NA | NA | NA | NA | #VALUE! |
| ENSMUSG00002075244.1 | 0 | NA | NA | NA | NA | NA | #VALUE! |
| ENSMUSG00002075245.1 | 0 | NA | NA | NA | NA | NA | #VALUE! |
| ENSMUSG00002075246.1 | 0 | NA | NA | NA | NA | NA | #VALUE! |
| ENSMUSG00002075247.1 | 0 | NA | NA | NA | NA | NA | #VALUE! |
| ENSMUSG00002075248.1 | 0 | NA | NA | NA | NA | NA | #VALUE! |
| ENSMUSG00002075249.1 | 0 | NA | NA | NA | NA | NA | #VALUE! |
| ENSMUSG00002075250.1 | 0 | NA | NA | NA | NA | NA | #VALUE! |
| ENSMUSG00002075251.1 | 0 | NA | NA | NA | NA | NA | #VALUE! |
| ENSMUSG00002075252.1 | 0 | NA | NA | NA | NA | NA | #VALUE! |
| ENSMUSG00002075253.1 | 0 | NA | NA | NA | NA | NA | #VALUE! |
| ENSMUSG00002075254.1 | 0 | NA | NA | NA | NA | NA | #VALUE! |

| | | | | | | | |
|---|---|---|---|---|---|---|---|
| ENSMUSG00002075255.1 | 0 | NA | NA | NA | NA | NA | #VALUE! |
| ENSMUSG00002075256.1 | 0 | NA | NA | NA | NA | NA | #VALUE! |
| ENSMUSG00002075257.1 | 0 | NA | NA | NA | NA | NA | #VALUE! |
| ENSMUSG00002075258.1 | 0 | NA | NA | NA | NA | NA | #VALUE! |
| ENSMUSG00002075259.1 | 0 | NA | NA | NA | NA | NA | #VALUE! |
| ENSMUSG00002075260.1 | 0 | NA | NA | NA | NA | NA | #VALUE! |
| ENSMUSG00002075261.1 | 0 | NA | NA | NA | NA | NA | #VALUE! |
| ENSMUSG00002075262.1 | 0 | NA | NA | NA | NA | NA | #VALUE! |
| ENSMUSG00002075263.1 | 0 | NA | NA | NA | NA | NA | #VALUE! |
| ENSMUSG00002075264.1 | 0 | NA | NA | NA | NA | NA | #VALUE! |
| ENSMUSG00002075265.1 | 0 | NA | NA | NA | NA | NA | #VALUE! |
| ENSMUSG00002075266.1 | 0 | NA | NA | NA | NA | NA | #VALUE! |
| ENSMUSG00002075267.1 | 0 | NA | NA | NA | NA | NA | #VALUE! |
| ENSMUSG00002075268.1 | 0 | NA | NA | NA | NA | NA | #VALUE! |
| ENSMUSG00002075269.1 | 0 | NA | NA | NA | NA | NA | #VALUE! |
| ENSMUSG00002075270.1 | 0 | NA | NA | NA | NA | NA | #VALUE! |
| ENSMUSG00002075271.1 | 0 | NA | NA | NA | NA | NA | #VALUE! |
| ENSMUSG00002075272.1 | 0 | NA | NA | NA | NA | NA | #VALUE! |
| ENSMUSG00002075273.1 | 0 | NA | NA | NA | NA | NA | #VALUE! |
| ENSMUSG00002075274.1 | 0 | NA | NA | NA | NA | NA | #VALUE! |
| ENSMUSG00002075275.1 | 0 | NA | NA | NA | NA | NA | #VALUE! |
| ENSMUSG00002075276.1 | 0 | NA | NA | NA | NA | NA | #VALUE! |
| ENSMUSG00002075277.1 | 0 | NA | NA | NA | NA | NA | #VALUE! |
| ENSMUSG00002075278.1 | 0 | NA | NA | NA | NA | NA | #VALUE! |
| ENSMUSG00002075279.1 | 0 | NA | NA | NA | NA | NA | #VALUE! |
| ENSMUSG00002075280.1 | 0 | NA | NA | NA | NA | NA | #VALUE! |
| ENSMUSG00002075281.1 | 0 | NA | NA | NA | NA | NA | #VALUE! |
| ENSMUSG00002075282.1 | 0 | NA | NA | NA | NA | NA | #VALUE! |
| ENSMUSG00002075283.1 | 0 | NA | NA | NA | NA | NA | #VALUE! |
| ENSMUSG00002075284.1 | 0 | NA | NA | NA | NA | NA | #VALUE! |
| ENSMUSG00002075285.1 | 0 | NA | NA | NA | NA | NA | #VALUE! |
| ENSMUSG00002075286.1 | 0 | NA | NA | NA | NA | NA | #VALUE! |
| ENSMUSG00002075287.1 | 0 | NA | NA | NA | NA | NA | #VALUE! |
| ENSMUSG00002075288.1 | 0 | NA | NA | NA | NA | NA | #VALUE! |
| ENSMUSG00002075289.1 | 0 | NA | NA | NA | NA | NA | #VALUE! |
| ENSMUSG00002075290.1 | 0 | NA | NA | NA | NA | NA | #VALUE! |
| ENSMUSG00002075291.1 | 0 | NA | NA | NA | NA | NA | #VALUE! |
| ENSMUSG00002075292.1 | 0 | NA | NA | NA | NA | NA | #VALUE! |
| ENSMUSG00002075293.1 | 0 | NA | NA | NA | NA | NA | #VALUE! |
| ENSMUSG00002075294.1 | 0 | NA | NA | NA | NA | NA | #VALUE! |
| ENSMUSG00002075295.1 | 0 | NA | NA | NA | NA | NA | #VALUE! |
| ENSMUSG00002075296.1 | 0 | NA | NA | NA | NA | NA | #VALUE! |
| ENSMUSG00002075297.1 | 0 | NA | NA | NA | NA | NA | #VALUE! |
| ENSMUSG00002075298.1 | 0 | NA | NA | NA | NA | NA | #VALUE! |

| | | | | | | | |
|---|---|---|---|---|---|---|---|
| ENSMUSG00002075299.1 | 0 | NA | NA | NA | NA | NA | #VALUE! |
| ENSMUSG00002075300.1 | 0 | NA | NA | NA | NA | NA | #VALUE! |
| ENSMUSG00002075301.1 | 0 | NA | NA | NA | NA | NA | #VALUE! |
| ENSMUSG00002075302.1 | 0 | NA | NA | NA | NA | NA | #VALUE! |
| ENSMUSG00002075303.1 | 0 | NA | NA | NA | NA | NA | #VALUE! |
| ENSMUSG00002075304.1 | 0 | NA | NA | NA | NA | NA | #VALUE! |
| ENSMUSG00002075305.1 | 0 | NA | NA | NA | NA | NA | #VALUE! |
| ENSMUSG00002075306.1 | 0 | NA | NA | NA | NA | NA | #VALUE! |
| ENSMUSG00002075307.1 | 0 | NA | NA | NA | NA | NA | #VALUE! |
| ENSMUSG00002075308.1 | 0 | NA | NA | NA | NA | NA | #VALUE! |
| ENSMUSG00002075309.1 | 0 | NA | NA | NA | NA | NA | #VALUE! |
| ENSMUSG00002075310.1 | 0 | NA | NA | NA | NA | NA | #VALUE! |
| ENSMUSG00002075311.1 | 0 | NA | NA | NA | NA | NA | #VALUE! |
| ENSMUSG00002075312.1 | 0 | NA | NA | NA | NA | NA | #VALUE! |
| ENSMUSG00002075313.1 | 0 | NA | NA | NA | NA | NA | #VALUE! |
| ENSMUSG00002075314.1 | 0 | NA | NA | NA | NA | NA | #VALUE! |
| ENSMUSG00002075315.1 | 0 | NA | NA | NA | NA | NA | #VALUE! |
| ENSMUSG00002075316.1 | 0 | NA | NA | NA | NA | NA | #VALUE! |
| ENSMUSG00002075317.1 | 0 | NA | NA | NA | NA | NA | #VALUE! |
| ENSMUSG00002075318.1 | 0 | NA | NA | NA | NA | NA | #VALUE! |
| ENSMUSG00002075319.1 | 0 | NA | NA | NA | NA | NA | #VALUE! |
| ENSMUSG00002075320.1 | 0 | NA | NA | NA | NA | NA | #VALUE! |
| ENSMUSG00002075321.1 | 0 | NA | NA | NA | NA | NA | #VALUE! |
| ENSMUSG00002075322.1 | 0 | NA | NA | NA | NA | NA | #VALUE! |
| ENSMUSG00002075323.1 | 0 | NA | NA | NA | NA | NA | #VALUE! |
| ENSMUSG00002075324.1 | 0 | NA | NA | NA | NA | NA | #VALUE! |
| ENSMUSG00002075325.1 | 0 | NA | NA | NA | NA | NA | #VALUE! |
| ENSMUSG00002075326.1 | 0 | NA | NA | NA | NA | NA | #VALUE! |
| ENSMUSG00002075327.1 | 0 | NA | NA | NA | NA | NA | #VALUE! |
| ENSMUSG00002075328.1 | 0 | NA | NA | NA | NA | NA | #VALUE! |
| ENSMUSG00002075329.1 | 0 | NA | NA | NA | NA | NA | #VALUE! |
| ENSMUSG00002075330.1 | 0 | NA | NA | NA | NA | NA | #VALUE! |
| ENSMUSG00002075331.1 | 0 | NA | NA | NA | NA | NA | #VALUE! |
| ENSMUSG00002075332.1 | 0 | NA | NA | NA | NA | NA | #VALUE! |
| ENSMUSG00002075333.1 | 0 | NA | NA | NA | NA | NA | #VALUE! |
| ENSMUSG00002075334.1 | 0 | NA | NA | NA | NA | NA | #VALUE! |
| ENSMUSG00002075335.1 | 0 | NA | NA | NA | NA | NA | #VALUE! |
| ENSMUSG00002075336.1 | 0 | NA | NA | NA | NA | NA | #VALUE! |
| ENSMUSG00002075337.1 | 0 | NA | NA | NA | NA | NA | #VALUE! |
| ENSMUSG00002075338.1 | 0 | NA | NA | NA | NA | NA | #VALUE! |
| ENSMUSG00002075339.1 | 0 | NA | NA | NA | NA | NA | #VALUE! |
| ENSMUSG00002075340.1 | 0 | NA | NA | NA | NA | NA | #VALUE! |
| ENSMUSG00002075341.1 | 0 | NA | NA | NA | NA | NA | #VALUE! |
| ENSMUSG00002075342.1 | 0 | NA | NA | NA | NA | NA | #VALUE! |

| | | | | | | | |
|---|---|---|---|---|---|---|---|
| ENSMUSG00002075343.1 | 0 | NA | NA | NA | NA | NA | #VALUE! |
| ENSMUSG00002075344.1 | 0 | NA | NA | NA | NA | NA | #VALUE! |
| ENSMUSG00002075345.1 | 0 | NA | NA | NA | NA | NA | #VALUE! |
| ENSMUSG00002075346.1 | 0 | NA | NA | NA | NA | NA | #VALUE! |
| ENSMUSG00002075347.1 | 0 | NA | NA | NA | NA | NA | #VALUE! |
| ENSMUSG00002075348.1 | 0 | NA | NA | NA | NA | NA | #VALUE! |
| ENSMUSG00002075349.1 | 0 | NA | NA | NA | NA | NA | #VALUE! |
| ENSMUSG00002075350.1 | 0 | NA | NA | NA | NA | NA | #VALUE! |
| ENSMUSG00002075351.1 | 0 | NA | NA | NA | NA | NA | #VALUE! |
| ENSMUSG00002075352.1 | 0 | NA | NA | NA | NA | NA | #VALUE! |
| ENSMUSG00002075353.1 | 0 | NA | NA | NA | NA | NA | #VALUE! |
| ENSMUSG00002075354.1 | 0 | NA | NA | NA | NA | NA | #VALUE! |
| ENSMUSG00002075355.1 | 0 | NA | NA | NA | NA | NA | #VALUE! |
| ENSMUSG00002075356.1 | 0 | NA | NA | NA | NA | NA | #VALUE! |
| ENSMUSG00002075357.1 | 0 | NA | NA | NA | NA | NA | #VALUE! |
| ENSMUSG00002075358.1 | 0 | NA | NA | NA | NA | NA | #VALUE! |
| ENSMUSG00002075359.1 | 0 | NA | NA | NA | NA | NA | #VALUE! |
| ENSMUSG00002075360.1 | 0 | NA | NA | NA | NA | NA | #VALUE! |
| ENSMUSG00002075361.1 | 0 | NA | NA | NA | NA | NA | #VALUE! |
| ENSMUSG00002075362.1 | 0 | NA | NA | NA | NA | NA | #VALUE! |
| ENSMUSG00002075363.1 | 0 | NA | NA | NA | NA | NA | #VALUE! |
| ENSMUSG00002075364.1 | 0 | NA | NA | NA | NA | NA | #VALUE! |
| ENSMUSG00002075365.1 | 0 | NA | NA | NA | NA | NA | #VALUE! |
| ENSMUSG00002075366.1 | 0 | NA | NA | NA | NA | NA | #VALUE! |
| ENSMUSG00002075367.1 | 0 | NA | NA | NA | NA | NA | #VALUE! |
| ENSMUSG00002075368.1 | 0 | NA | NA | NA | NA | NA | #VALUE! |
| ENSMUSG00002075369.1 | 0 | NA | NA | NA | NA | NA | #VALUE! |
| ENSMUSG00002075370.1 | 0 | NA | NA | NA | NA | NA | #VALUE! |
| ENSMUSG00002075371.1 | 0 | NA | NA | NA | NA | NA | #VALUE! |
| ENSMUSG00002075372.1 | 0 | NA | NA | NA | NA | NA | #VALUE! |
| ENSMUSG00002075373.1 | 0 | NA | NA | NA | NA | NA | #VALUE! |
| ENSMUSG00002075374.1 | 0 | NA | NA | NA | NA | NA | #VALUE! |
| ENSMUSG00002075375.1 | 0 | NA | NA | NA | NA | NA | #VALUE! |
| ENSMUSG00002075376.1 | 0 | NA | NA | NA | NA | NA | #VALUE! |
| ENSMUSG00002075377.1 | 0 | NA | NA | NA | NA | NA | #VALUE! |
| ENSMUSG00002075378.1 | 0 | NA | NA | NA | NA | NA | #VALUE! |
| ENSMUSG00002075379.1 | 0 | NA | NA | NA | NA | NA | #VALUE! |
| ENSMUSG00002075380.1 | 0 | NA | NA | NA | NA | NA | #VALUE! |
| ENSMUSG00002075381.1 | 0 | NA | NA | NA | NA | NA | #VALUE! |
| ENSMUSG00002075382.1 | 0 | NA | NA | NA | NA | NA | #VALUE! |
| ENSMUSG00002075383.1 | 0 | NA | NA | NA | NA | NA | #VALUE! |
| ENSMUSG00002075384.1 | 0 | NA | NA | NA | NA | NA | #VALUE! |
| ENSMUSG00002075385.1 | 0 | NA | NA | NA | NA | NA | #VALUE! |
| ENSMUSG00002075386.1 | 0 | NA | NA | NA | NA | NA | #VALUE! |

| | | | | | | | |
|---|---|---|---|---|---|---|---|
| ENSMUSG00002075387.1 | 0 | NA | NA | NA | NA | NA | #VALUE! |
| ENSMUSG00002075388.1 | 0 | NA | NA | NA | NA | NA | #VALUE! |
| ENSMUSG00002075389.1 | 0 | NA | NA | NA | NA | NA | #VALUE! |
| ENSMUSG00002075390.1 | 0 | NA | NA | NA | NA | NA | #VALUE! |
| ENSMUSG00002075391.1 | 0 | NA | NA | NA | NA | NA | #VALUE! |
| ENSMUSG00002075392.1 | 0 | NA | NA | NA | NA | NA | #VALUE! |
| ENSMUSG00002075393.1 | 0 | NA | NA | NA | NA | NA | #VALUE! |
| ENSMUSG00002075394.1 | 0 | NA | NA | NA | NA | NA | #VALUE! |
| ENSMUSG00002075395.1 | 0 | NA | NA | NA | NA | NA | #VALUE! |
| ENSMUSG00002075396.1 | 0 | NA | NA | NA | NA | NA | #VALUE! |
| ENSMUSG00002075397.1 | 0 | NA | NA | NA | NA | NA | #VALUE! |
| ENSMUSG00002075398.1 | 0 | NA | NA | NA | NA | NA | #VALUE! |
| ENSMUSG00002075399.1 | 0 | NA | NA | NA | NA | NA | #VALUE! |
| ENSMUSG00002075400.1 | 0 | NA | NA | NA | NA | NA | #VALUE! |
| ENSMUSG00002075401.1 | 0 | NA | NA | NA | NA | NA | #VALUE! |
| ENSMUSG00002075402.1 | 0 | NA | NA | NA | NA | NA | #VALUE! |
| ENSMUSG00002075403.1 | 0 | NA | NA | NA | NA | NA | #VALUE! |
| ENSMUSG00002075404.1 | 0 | NA | NA | NA | NA | NA | #VALUE! |
| ENSMUSG00002075405.1 | 0 | NA | NA | NA | NA | NA | #VALUE! |
| ENSMUSG00002075406.1 | 0 | NA | NA | NA | NA | NA | #VALUE! |
| ENSMUSG00002075407.1 | 0 | NA | NA | NA | NA | NA | #VALUE! |
| ENSMUSG00002075408.1 | 0 | NA | NA | NA | NA | NA | #VALUE! |
| ENSMUSG00002075409.1 | 0 | NA | NA | NA | NA | NA | #VALUE! |
| ENSMUSG00002075410.1 | 0 | NA | NA | NA | NA | NA | #VALUE! |
| ENSMUSG00002075411.1 | 0 | NA | NA | NA | NA | NA | #VALUE! |
| ENSMUSG00002075412.1 | 0 | NA | NA | NA | NA | NA | #VALUE! |
| ENSMUSG00002075413.1 | 0 | NA | NA | NA | NA | NA | #VALUE! |
| ENSMUSG00002075414.1 | 0 | NA | NA | NA | NA | NA | #VALUE! |
| ENSMUSG00002075415.1 | 0 | NA | NA | NA | NA | NA | #VALUE! |
| ENSMUSG00002075416.1 | 0 | NA | NA | NA | NA | NA | #VALUE! |
| ENSMUSG00002075417.1 | 0 | NA | NA | NA | NA | NA | #VALUE! |
| ENSMUSG00002075418.1 | 0 | NA | NA | NA | NA | NA | #VALUE! |
| ENSMUSG00002075419.1 | 0 | NA | NA | NA | NA | NA | #VALUE! |
| ENSMUSG00002075420.1 | 0 | NA | NA | NA | NA | NA | #VALUE! |
| ENSMUSG00002075421.1 | 0 | NA | NA | NA | NA | NA | #VALUE! |
| ENSMUSG00002075422.1 | 0 | NA | NA | NA | NA | NA | #VALUE! |
| ENSMUSG00002075423.1 | 0 | NA | NA | NA | NA | NA | #VALUE! |
| ENSMUSG00002075424.1 | 0 | NA | NA | NA | NA | NA | #VALUE! |
| ENSMUSG00002075425.1 | 0 | NA | NA | NA | NA | NA | #VALUE! |
| ENSMUSG00002075426.1 | 0 | NA | NA | NA | NA | NA | #VALUE! |
| ENSMUSG00002075427.1 | 0 | NA | NA | NA | NA | NA | #VALUE! |
| ENSMUSG00002075428.1 | 0 | NA | NA | NA | NA | NA | #VALUE! |
| ENSMUSG00002075429.1 | 0 | NA | NA | NA | NA | NA | #VALUE! |
| ENSMUSG00002075430.1 | 0 | NA | NA | NA | NA | NA | #VALUE! |

| | | | | | | | |
|---|---|---|---|---|---|---|---|
| ENSMUSG00002075431.1 | 0 | NA | NA | NA | NA | NA | #VALUE! |
| ENSMUSG00002075432.1 | 0 | NA | NA | NA | NA | NA | #VALUE! |
| ENSMUSG00002075433.1 | 0 | NA | NA | NA | NA | NA | #VALUE! |
| ENSMUSG00002075434.1 | 0 | NA | NA | NA | NA | NA | #VALUE! |
| ENSMUSG00002075435.1 | 0 | NA | NA | NA | NA | NA | #VALUE! |
| ENSMUSG00002075436.1 | 0 | NA | NA | NA | NA | NA | #VALUE! |
| ENSMUSG00002075437.1 | 0 | NA | NA | NA | NA | NA | #VALUE! |
| ENSMUSG00002075438.1 | 0 | NA | NA | NA | NA | NA | #VALUE! |
| ENSMUSG00002075439.1 | 0 | NA | NA | NA | NA | NA | #VALUE! |
| ENSMUSG00002075440.1 | 0 | NA | NA | NA | NA | NA | #VALUE! |
| ENSMUSG00002075441.1 | 0 | NA | NA | NA | NA | NA | #VALUE! |
| ENSMUSG00002075442.1 | 0 | NA | NA | NA | NA | NA | #VALUE! |
| ENSMUSG00002075443.1 | 0 | NA | NA | NA | NA | NA | #VALUE! |
| ENSMUSG00002075444.1 | 0 | NA | NA | NA | NA | NA | #VALUE! |
| ENSMUSG00002075445.1 | 0 | NA | NA | NA | NA | NA | #VALUE! |
| ENSMUSG00002075446.1 | 0 | NA | NA | NA | NA | NA | #VALUE! |
| ENSMUSG00002075447.1 | 0 | NA | NA | NA | NA | NA | #VALUE! |
| ENSMUSG00002075448.1 | 0 | NA | NA | NA | NA | NA | #VALUE! |
| ENSMUSG00002075449.1 | 0 | NA | NA | NA | NA | NA | #VALUE! |
| ENSMUSG00002075450.1 | 0 | NA | NA | NA | NA | NA | #VALUE! |
| ENSMUSG00002075451.1 | 0 | NA | NA | NA | NA | NA | #VALUE! |
| ENSMUSG00002075452.1 | 0 | NA | NA | NA | NA | NA | #VALUE! |
| ENSMUSG00002075453.1 | 0 | NA | NA | NA | NA | NA | #VALUE! |
| ENSMUSG00002075454.1 | 0 | NA | NA | NA | NA | NA | #VALUE! |
| ENSMUSG00002075455.1 | 0 | NA | NA | NA | NA | NA | #VALUE! |
| ENSMUSG00002075456.1 | 0 | NA | NA | NA | NA | NA | #VALUE! |
| ENSMUSG00002075457.1 | 0 | NA | NA | NA | NA | NA | #VALUE! |
| ENSMUSG00002075458.1 | 0 | NA | NA | NA | NA | NA | #VALUE! |
| ENSMUSG00002075459.1 | 0 | NA | NA | NA | NA | NA | #VALUE! |
| ENSMUSG00002075460.1 | 0 | NA | NA | NA | NA | NA | #VALUE! |
| ENSMUSG00002075461.1 | 0 | NA | NA | NA | NA | NA | #VALUE! |
| ENSMUSG00002075462.1 | 0 | NA | NA | NA | NA | NA | #VALUE! |
| ENSMUSG00002075463.1 | 0 | NA | NA | NA | NA | NA | #VALUE! |
| ENSMUSG00002075464.1 | 0 | NA | NA | NA | NA | NA | #VALUE! |
| ENSMUSG00002075465.1 | 0 | NA | NA | NA | NA | NA | #VALUE! |
| ENSMUSG00002075466.1 | 0 | NA | NA | NA | NA | NA | #VALUE! |
| ENSMUSG00002075467.1 | 0 | NA | NA | NA | NA | NA | #VALUE! |
| ENSMUSG00002075468.1 | 0 | NA | NA | NA | NA | NA | #VALUE! |
| ENSMUSG00002075469.1 | 0 | NA | NA | NA | NA | NA | #VALUE! |
| ENSMUSG00002075470.1 | 0 | NA | NA | NA | NA | NA | #VALUE! |
| ENSMUSG00002075471.1 | 0 | NA | NA | NA | NA | NA | #VALUE! |
| ENSMUSG00002075472.1 | 0 | NA | NA | NA | NA | NA | #VALUE! |
| ENSMUSG00002075473.1 | 0 | NA | NA | NA | NA | NA | #VALUE! |
| ENSMUSG00002075474.1 | 0 | NA | NA | NA | NA | NA | #VALUE! |

| | | | | | | | |
|---|---|---|---|---|---|---|---|
| ENSMUSG00002075475.1 | 0 | NA | NA | NA | NA | NA | #VALUE! |
| ENSMUSG00002075476.1 | 0 | NA | NA | NA | NA | NA | #VALUE! |
| ENSMUSG00002075477.1 | 0 | NA | NA | NA | NA | NA | #VALUE! |
| ENSMUSG00002075478.1 | 0 | NA | NA | NA | NA | NA | #VALUE! |
| ENSMUSG00002075479.1 | 0 | NA | NA | NA | NA | NA | #VALUE! |
| ENSMUSG00002075480.1 | 0 | NA | NA | NA | NA | NA | #VALUE! |
| ENSMUSG00002075481.1 | 0 | NA | NA | NA | NA | NA | #VALUE! |
| ENSMUSG00002075482.1 | 0 | NA | NA | NA | NA | NA | #VALUE! |
| ENSMUSG00002075483.1 | 0 | NA | NA | NA | NA | NA | #VALUE! |
| ENSMUSG00002075484.1 | 0 | NA | NA | NA | NA | NA | #VALUE! |
| ENSMUSG00002075485.1 | 0 | NA | NA | NA | NA | NA | #VALUE! |
| ENSMUSG00002075486.1 | 0.297366 | -0.14751 | 4.065279 | -0.03629 | 0.971054 | NA | #VALUE! |
| ENSMUSG00002075487.1 | 0 | NA | NA | NA | NA | NA | #VALUE! |
| ENSMUSG00002075488.1 | 0 | NA | NA | NA | NA | NA | #VALUE! |
| ENSMUSG00002075489.1 | 0 | NA | NA | NA | NA | NA | #VALUE! |
| ENSMUSG00002075490.1 | 0 | NA | NA | NA | NA | NA | #VALUE! |
| ENSMUSG00002075491.1 | 0 | NA | NA | NA | NA | NA | #VALUE! |
| ENSMUSG00002075492.1 | 0 | NA | NA | NA | NA | NA | #VALUE! |
| ENSMUSG00002075493.1 | 0 | NA | NA | NA | NA | NA | #VALUE! |
| ENSMUSG00002075494.1 | 0 | NA | NA | NA | NA | NA | #VALUE! |
| ENSMUSG00002075495.1 | 0 | NA | NA | NA | NA | NA | #VALUE! |
| ENSMUSG00002075496.1 | 0 | NA | NA | NA | NA | NA | #VALUE! |
| ENSMUSG00002075497.1 | 0 | NA | NA | NA | NA | NA | #VALUE! |
| ENSMUSG00002075498.1 | 0 | NA | NA | NA | NA | NA | #VALUE! |
| ENSMUSG00002075499.1 | 0 | NA | NA | NA | NA | NA | #VALUE! |
| ENSMUSG00002075500.1 | 0 | NA | NA | NA | NA | NA | #VALUE! |
| ENSMUSG00002075501.1 | 0 | NA | NA | NA | NA | NA | #VALUE! |
| ENSMUSG00002075502.1 | 0 | NA | NA | NA | NA | NA | #VALUE! |
| ENSMUSG00002075503.1 | 0 | NA | NA | NA | NA | NA | #VALUE! |
| ENSMUSG00002075504.1 | 0 | NA | NA | NA | NA | NA | #VALUE! |
| ENSMUSG00002075505.1 | 0 | NA | NA | NA | NA | NA | #VALUE! |
| ENSMUSG00002075506.1 | 0 | NA | NA | NA | NA | NA | #VALUE! |
| ENSMUSG00002075507.1 | 0 | NA | NA | NA | NA | NA | #VALUE! |
| ENSMUSG00002075508.1 | 0 | NA | NA | NA | NA | NA | #VALUE! |
| ENSMUSG00002075509.1 | 0 | NA | NA | NA | NA | NA | #VALUE! |
| ENSMUSG00002075510.1 | 0 | NA | NA | NA | NA | NA | #VALUE! |
| ENSMUSG00002075511.1 | 0 | NA | NA | NA | NA | NA | #VALUE! |
| ENSMUSG00002075512.1 | 0 | NA | NA | NA | NA | NA | #VALUE! |
| ENSMUSG00002075513.1 | 0 | NA | NA | NA | NA | NA | #VALUE! |
| ENSMUSG00002075514.1 | 0 | NA | NA | NA | NA | NA | #VALUE! |
| ENSMUSG00002075515.1 | 0 | NA | NA | NA | NA | NA | #VALUE! |
| ENSMUSG00002075516.1 | 0 | NA | NA | NA | NA | NA | #VALUE! |
| ENSMUSG00002075517.1 | 0 | NA | NA | NA | NA | NA | #VALUE! |
| ENSMUSG00002075518.1 | 0 | NA | NA | NA | NA | NA | #VALUE! |

| | | | | | | | |
|---|---|---|---|---|---|---|---|
| ENSMUSG00002075519.1 | 0 | NA | NA | NA | NA | NA | #VALUE! |
| ENSMUSG00002075520.1 | 0 | NA | NA | NA | NA | NA | #VALUE! |
| ENSMUSG00002075521.1 | 0 | NA | NA | NA | NA | NA | #VALUE! |
| ENSMUSG00002075522.1 | 0 | NA | NA | NA | NA | NA | #VALUE! |
| ENSMUSG00002075523.1 | 0 | NA | NA | NA | NA | NA | #VALUE! |
| ENSMUSG00002075524.1 | 0 | NA | NA | NA | NA | NA | #VALUE! |
| ENSMUSG00002075525.1 | 0 | NA | NA | NA | NA | NA | #VALUE! |
| ENSMUSG00002075526.1 | 0 | NA | NA | NA | NA | NA | #VALUE! |
| ENSMUSG00002075527.1 | 0 | NA | NA | NA | NA | NA | #VALUE! |
| ENSMUSG00002075528.1 | 0 | NA | NA | NA | NA | NA | #VALUE! |
| ENSMUSG00002075529.1 | 0 | NA | NA | NA | NA | NA | #VALUE! |
| ENSMUSG00002075530.1 | 0 | NA | NA | NA | NA | NA | #VALUE! |
| ENSMUSG00002075531.1 | 0 | NA | NA | NA | NA | NA | #VALUE! |
| ENSMUSG00002075532.1 | 0 | NA | NA | NA | NA | NA | #VALUE! |
| ENSMUSG00002075533.1 | 0 | NA | NA | NA | NA | NA | #VALUE! |
| ENSMUSG00002075534.1 | 0 | NA | NA | NA | NA | NA | #VALUE! |
| ENSMUSG00002075535.1 | 0 | NA | NA | NA | NA | NA | #VALUE! |
| ENSMUSG00002075536.1 | 0 | NA | NA | NA | NA | NA | #VALUE! |
| ENSMUSG00002075537.1 | 0 | NA | NA | NA | NA | NA | #VALUE! |
| ENSMUSG00002075538.1 | 0 | NA | NA | NA | NA | NA | #VALUE! |
| ENSMUSG00002075539.1 | 0 | NA | NA | NA | NA | NA | #VALUE! |
| ENSMUSG00002075540.1 | 0 | NA | NA | NA | NA | NA | #VALUE! |
| ENSMUSG00002075541.1 | 0 | NA | NA | NA | NA | NA | #VALUE! |
| ENSMUSG00002075542.1 | 0 | NA | NA | NA | NA | NA | #VALUE! |
| ENSMUSG00002075543.1 | 0 | NA | NA | NA | NA | NA | #VALUE! |
| ENSMUSG00002075544.1 | 0 | NA | NA | NA | NA | NA | #VALUE! |
| ENSMUSG00002075545.1 | 0 | NA | NA | NA | NA | NA | #VALUE! |
| ENSMUSG00002075546.1 | 0 | NA | NA | NA | NA | NA | #VALUE! |
| ENSMUSG00002075547.1 | 0 | NA | NA | NA | NA | NA | #VALUE! |
| ENSMUSG00002075548.1 | 0 | NA | NA | NA | NA | NA | #VALUE! |
| ENSMUSG00002075549.1 | 0 | NA | NA | NA | NA | NA | #VALUE! |
| ENSMUSG00002075550.1 | 0 | NA | NA | NA | NA | NA | #VALUE! |
| ENSMUSG00002075551.1 | 0 | NA | NA | NA | NA | NA | #VALUE! |
| ENSMUSG00002075552.1 | 0 | NA | NA | NA | NA | NA | #VALUE! |
| ENSMUSG00002075553.1 | 0 | NA | NA | NA | NA | NA | #VALUE! |
| ENSMUSG00002075554.1 | 0 | NA | NA | NA | NA | NA | #VALUE! |
| ENSMUSG00002075555.1 | 0 | NA | NA | NA | NA | NA | #VALUE! |
| ENSMUSG00002075556.1 | 0 | NA | NA | NA | NA | NA | #VALUE! |
| ENSMUSG00002075557.1 | 0 | NA | NA | NA | NA | NA | #VALUE! |
| ENSMUSG00002075558.1 | 0 | NA | NA | NA | NA | NA | #VALUE! |
| ENSMUSG00002075559.1 | 0 | NA | NA | NA | NA | NA | #VALUE! |
| ENSMUSG00002075560.1 | 0 | NA | NA | NA | NA | NA | #VALUE! |
| ENSMUSG00002075561.1 | 0 | NA | NA | NA | NA | NA | #VALUE! |
| ENSMUSG00002075562.1 | 0 | NA | NA | NA | NA | NA | #VALUE! |

| | | | | | | | |
|---|---|---|---|---|---|---|---|
| ENSMUSG00002075563.1 | 0 | NA | NA | NA | NA | NA | #VALUE! |
| ENSMUSG00002075564.1 | 0 | NA | NA | NA | NA | NA | #VALUE! |
| ENSMUSG00002075565.1 | 0 | NA | NA | NA | NA | NA | #VALUE! |
| ENSMUSG00002075566.1 | 0 | NA | NA | NA | NA | NA | #VALUE! |
| ENSMUSG00002075567.1 | 0 | NA | NA | NA | NA | NA | #VALUE! |
| ENSMUSG00002075568.1 | 0 | NA | NA | NA | NA | NA | #VALUE! |
| ENSMUSG00002075569.1 | 0 | NA | NA | NA | NA | NA | #VALUE! |
| ENSMUSG00002075570.1 | 0 | NA | NA | NA | NA | NA | #VALUE! |
| ENSMUSG00002075571.1 | 0 | NA | NA | NA | NA | NA | #VALUE! |
| ENSMUSG00002075572.1 | 0 | NA | NA | NA | NA | NA | #VALUE! |
| ENSMUSG00002075573.1 | 0 | NA | NA | NA | NA | NA | #VALUE! |
| ENSMUSG00002075574.1 | 0 | NA | NA | NA | NA | NA | #VALUE! |
| ENSMUSG00002075575.1 | 0 | NA | NA | NA | NA | NA | #VALUE! |
| ENSMUSG00002075576.1 | 0 | NA | NA | NA | NA | NA | #VALUE! |
| ENSMUSG00002075577.1 | 0 | NA | NA | NA | NA | NA | #VALUE! |
| ENSMUSG00002075578.1 | 0 | NA | NA | NA | NA | NA | #VALUE! |
| ENSMUSG00002075579.1 | 0 | NA | NA | NA | NA | NA | #VALUE! |
| ENSMUSG00002075580.1 | 0 | NA | NA | NA | NA | NA | #VALUE! |
| ENSMUSG00002075581.1 | 0 | NA | NA | NA | NA | NA | #VALUE! |
| ENSMUSG00002075582.1 | 0 | NA | NA | NA | NA | NA | #VALUE! |
| ENSMUSG00002075583.1 | 0 | NA | NA | NA | NA | NA | #VALUE! |
| ENSMUSG00002075584.1 | 0 | NA | NA | NA | NA | NA | #VALUE! |
| ENSMUSG00002075585.1 | 0 | NA | NA | NA | NA | NA | #VALUE! |
| ENSMUSG00002075586.1 | 0 | NA | NA | NA | NA | NA | #VALUE! |
| ENSMUSG00002075587.1 | 0 | NA | NA | NA | NA | NA | #VALUE! |
| ENSMUSG00002075588.1 | 0 | NA | NA | NA | NA | NA | #VALUE! |
| ENSMUSG00002075589.1 | 0 | NA | NA | NA | NA | NA | #VALUE! |
| ENSMUSG00002075590.1 | 0 | NA | NA | NA | NA | NA | #VALUE! |
| ENSMUSG00002075591.1 | 0 | NA | NA | NA | NA | NA | #VALUE! |
| ENSMUSG00002075592.1 | 0 | NA | NA | NA | NA | NA | #VALUE! |
| ENSMUSG00002075593.1 | 0 | NA | NA | NA | NA | NA | #VALUE! |
| ENSMUSG00002075594.1 | 0 | NA | NA | NA | NA | NA | #VALUE! |
| ENSMUSG00002075595.1 | 0 | NA | NA | NA | NA | NA | #VALUE! |
| ENSMUSG00002075596.1 | 0 | NA | NA | NA | NA | NA | #VALUE! |
| ENSMUSG00002075597.1 | 0 | NA | NA | NA | NA | NA | #VALUE! |
| ENSMUSG00002075598.1 | 0 | NA | NA | NA | NA | NA | #VALUE! |
| ENSMUSG00002075599.1 | 0 | NA | NA | NA | NA | NA | #VALUE! |
| ENSMUSG00002075600.1 | 0 | NA | NA | NA | NA | NA | #VALUE! |
| ENSMUSG00002075601.1 | 0 | NA | NA | NA | NA | NA | #VALUE! |
| ENSMUSG00002075602.1 | 0 | NA | NA | NA | NA | NA | #VALUE! |
| ENSMUSG00002075603.1 | 0 | NA | NA | NA | NA | NA | #VALUE! |
| ENSMUSG00002075604.1 | 0 | NA | NA | NA | NA | NA | #VALUE! |
| ENSMUSG00002075605.1 | 0 | NA | NA | NA | NA | NA | #VALUE! |
| ENSMUSG00002075606.1 | 0 | NA | NA | NA | NA | NA | #VALUE! |

| | | | | | | | |
|---|---|---|---|---|---|---|---|
| ENSMUSG00002075607.1 | 0 | NA | NA | NA | NA | NA | #VALUE! |
| ENSMUSG00002075608.1 | 0 | NA | NA | NA | NA | NA | #VALUE! |
| ENSMUSG00002075609.1 | 0 | NA | NA | NA | NA | NA | #VALUE! |
| ENSMUSG00002075610.1 | 0 | NA | NA | NA | NA | NA | #VALUE! |
| ENSMUSG00002075611.1 | 0 | NA | NA | NA | NA | NA | #VALUE! |
| ENSMUSG00002075612.1 | 0 | NA | NA | NA | NA | NA | #VALUE! |
| ENSMUSG00002075613.1 | 0 | NA | NA | NA | NA | NA | #VALUE! |
| ENSMUSG00002075614.1 | 0 | NA | NA | NA | NA | NA | #VALUE! |
| ENSMUSG00002075615.1 | 0 | NA | NA | NA | NA | NA | #VALUE! |
| ENSMUSG00002075616.1 | 0 | NA | NA | NA | NA | NA | #VALUE! |
| ENSMUSG00002075617.1 | 0 | NA | NA | NA | NA | NA | #VALUE! |
| ENSMUSG00002075618.1 | 0 | NA | NA | NA | NA | NA | #VALUE! |
| ENSMUSG00002075619.1 | 0 | NA | NA | NA | NA | NA | #VALUE! |
| ENSMUSG00002075620.1 | 0 | NA | NA | NA | NA | NA | #VALUE! |
| ENSMUSG00002075621.1 | 0 | NA | NA | NA | NA | NA | #VALUE! |
| ENSMUSG00002075622.1 | 0 | NA | NA | NA | NA | NA | #VALUE! |
| ENSMUSG00002075623.1 | 0 | NA | NA | NA | NA | NA | #VALUE! |
| ENSMUSG00002075624.1 | 0 | NA | NA | NA | NA | NA | #VALUE! |
| ENSMUSG00002075625.1 | 0 | NA | NA | NA | NA | NA | #VALUE! |
| ENSMUSG00002075626.1 | 0 | NA | NA | NA | NA | NA | #VALUE! |
| ENSMUSG00002075627.1 | 0 | NA | NA | NA | NA | NA | #VALUE! |
| ENSMUSG00002075628.1 | 0 | NA | NA | NA | NA | NA | #VALUE! |
| ENSMUSG00002075629.1 | 0 | NA | NA | NA | NA | NA | #VALUE! |
| ENSMUSG00002075630.1 | 0 | NA | NA | NA | NA | NA | #VALUE! |
| ENSMUSG00002075631.1 | 0 | NA | NA | NA | NA | NA | #VALUE! |
| ENSMUSG00002075632.1 | 0 | NA | NA | NA | NA | NA | #VALUE! |
| ENSMUSG00002075633.1 | 0 | NA | NA | NA | NA | NA | #VALUE! |
| ENSMUSG00002075634.1 | 0 | NA | NA | NA | NA | NA | #VALUE! |
| ENSMUSG00002075635.1 | 0 | NA | NA | NA | NA | NA | #VALUE! |
| ENSMUSG00002075636.1 | 0 | NA | NA | NA | NA | NA | #VALUE! |
| ENSMUSG00002075637.1 | 0 | NA | NA | NA | NA | NA | #VALUE! |
| ENSMUSG00002075638.1 | 0 | NA | NA | NA | NA | NA | #VALUE! |
| ENSMUSG00002075639.1 | 0 | NA | NA | NA | NA | NA | #VALUE! |
| ENSMUSG00002075640.1 | 0 | NA | NA | NA | NA | NA | #VALUE! |
| ENSMUSG00002075641.1 | 0 | NA | NA | NA | NA | NA | #VALUE! |
| ENSMUSG00002075642.1 | 0 | NA | NA | NA | NA | NA | #VALUE! |
| ENSMUSG00002075643.1 | 0 | NA | NA | NA | NA | NA | #VALUE! |
| ENSMUSG00002075644.1 | 0 | NA | NA | NA | NA | NA | #VALUE! |
| ENSMUSG00002075645.1 | 0 | NA | NA | NA | NA | NA | #VALUE! |
| ENSMUSG00002075646.1 | 0 | NA | NA | NA | NA | NA | #VALUE! |
| ENSMUSG00002075647.1 | 0 | NA | NA | NA | NA | NA | #VALUE! |
| ENSMUSG00002075648.1 | 0 | NA | NA | NA | NA | NA | #VALUE! |
| ENSMUSG00002075649.1 | 0 | NA | NA | NA | NA | NA | #VALUE! |
| ENSMUSG00002075650.1 | 0 | NA | NA | NA | NA | NA | #VALUE! |

| | | | | | | | |
|---|---|---|---|---|---|---|---|
| ENSMUSG00002075651.1 | 0 | NA | NA | NA | NA | NA | #VALUE! |
| ENSMUSG00002075652.1 | 0 | NA | NA | NA | NA | NA | #VALUE! |
| ENSMUSG00002075653.1 | 0 | NA | NA | NA | NA | NA | #VALUE! |
| ENSMUSG00002075654.1 | 0 | NA | NA | NA | NA | NA | #VALUE! |
| ENSMUSG00002075655.1 | 0 | NA | NA | NA | NA | NA | #VALUE! |
| ENSMUSG00002075656.1 | 0 | NA | NA | NA | NA | NA | #VALUE! |
| ENSMUSG00002075657.1 | 0 | NA | NA | NA | NA | NA | #VALUE! |
| ENSMUSG00002075658.1 | 0 | NA | NA | NA | NA | NA | #VALUE! |
| ENSMUSG00002075659.1 | 0 | NA | NA | NA | NA | NA | #VALUE! |
| ENSMUSG00002075660.1 | 0 | NA | NA | NA | NA | NA | #VALUE! |
| ENSMUSG00002075661.1 | 0 | NA | NA | NA | NA | NA | #VALUE! |
| ENSMUSG00002075662.1 | 0 | NA | NA | NA | NA | NA | #VALUE! |
| ENSMUSG00002075663.1 | 0 | NA | NA | NA | NA | NA | #VALUE! |
| ENSMUSG00002075664.1 | 0 | NA | NA | NA | NA | NA | #VALUE! |
| ENSMUSG00002075665.1 | 0 | NA | NA | NA | NA | NA | #VALUE! |
| ENSMUSG00002075666.1 | 0 | NA | NA | NA | NA | NA | #VALUE! |
| ENSMUSG00002075667.1 | 0 | NA | NA | NA | NA | NA | #VALUE! |
| ENSMUSG00002075668.1 | 0 | NA | NA | NA | NA | NA | #VALUE! |
| ENSMUSG00002075669.1 | 0 | NA | NA | NA | NA | NA | #VALUE! |
| ENSMUSG00002075670.1 | 0 | NA | NA | NA | NA | NA | #VALUE! |
| ENSMUSG00002075671.1 | 0 | NA | NA | NA | NA | NA | #VALUE! |
| ENSMUSG00002075672.1 | 0 | NA | NA | NA | NA | NA | #VALUE! |
| ENSMUSG00002075673.1 | 0 | NA | NA | NA | NA | NA | #VALUE! |
| ENSMUSG00002075674.1 | 0 | NA | NA | NA | NA | NA | #VALUE! |
| ENSMUSG00002075675.1 | 0 | NA | NA | NA | NA | NA | #VALUE! |
| ENSMUSG00002075676.1 | 0 | NA | NA | NA | NA | NA | #VALUE! |
| ENSMUSG00002075677.1 | 0 | NA | NA | NA | NA | NA | #VALUE! |
| ENSMUSG00002075678.1 | 0 | NA | NA | NA | NA | NA | #VALUE! |
| ENSMUSG00002075679.1 | 0 | NA | NA | NA | NA | NA | #VALUE! |
| ENSMUSG00002075680.1 | 0 | NA | NA | NA | NA | NA | #VALUE! |
| ENSMUSG00002075681.1 | 0 | NA | NA | NA | NA | NA | #VALUE! |
| ENSMUSG00002075682.1 | 0 | NA | NA | NA | NA | NA | #VALUE! |
| ENSMUSG00002075683.1 | 0 | NA | NA | NA | NA | NA | #VALUE! |
| ENSMUSG00002075684.1 | 0 | NA | NA | NA | NA | NA | #VALUE! |
| ENSMUSG00002075685.1 | 0 | NA | NA | NA | NA | NA | #VALUE! |
| ENSMUSG00002075686.1 | 0 | NA | NA | NA | NA | NA | #VALUE! |
| ENSMUSG00002075687.1 | 0 | NA | NA | NA | NA | NA | #VALUE! |
| ENSMUSG00002075688.1 | 0 | NA | NA | NA | NA | NA | #VALUE! |
| ENSMUSG00002075689.1 | 0 | NA | NA | NA | NA | NA | #VALUE! |
| ENSMUSG00002075690.1 | 0 | NA | NA | NA | NA | NA | #VALUE! |
| ENSMUSG00002075691.1 | 0 | NA | NA | NA | NA | NA | #VALUE! |
| ENSMUSG00002075692.1 | 0 | NA | NA | NA | NA | NA | #VALUE! |
| ENSMUSG00002075693.1 | 0 | NA | NA | NA | NA | NA | #VALUE! |
| ENSMUSG00002075694.1 | 0 | NA | NA | NA | NA | NA | #VALUE! |

| | | | | | | | |
|---|---|---|---|---|---|---|---|
| ENSMUSG00002075695.1 | 0 | NA | NA | NA | NA | NA | #VALUE! |
| ENSMUSG00002075696.1 | 0 | NA | NA | NA | NA | NA | #VALUE! |
| ENSMUSG00002075697.1 | 0 | NA | NA | NA | NA | NA | #VALUE! |
| ENSMUSG00002075698.1 | 0 | NA | NA | NA | NA | NA | #VALUE! |
| ENSMUSG00002075699.1 | 0 | NA | NA | NA | NA | NA | #VALUE! |
| ENSMUSG00002075700.1 | 0 | NA | NA | NA | NA | NA | #VALUE! |
| ENSMUSG00002075701.1 | 0 | NA | NA | NA | NA | NA | #VALUE! |
| ENSMUSG00002075702.1 | 0 | NA | NA | NA | NA | NA | #VALUE! |
| ENSMUSG00002075703.1 | 0 | NA | NA | NA | NA | NA | #VALUE! |
| ENSMUSG00002075704.1 | 0 | NA | NA | NA | NA | NA | #VALUE! |
| ENSMUSG00002075705.1 | 0 | NA | NA | NA | NA | NA | #VALUE! |
| ENSMUSG00002075706.1 | 0 | NA | NA | NA | NA | NA | #VALUE! |
| ENSMUSG00002075707.1 | 0 | NA | NA | NA | NA | NA | #VALUE! |
| ENSMUSG00002075708.1 | 0 | NA | NA | NA | NA | NA | #VALUE! |
| ENSMUSG00002075709.1 | 0 | NA | NA | NA | NA | NA | #VALUE! |
| ENSMUSG00002075710.1 | 0 | NA | NA | NA | NA | NA | #VALUE! |
| ENSMUSG00002075711.1 | 0 | NA | NA | NA | NA | NA | #VALUE! |
| ENSMUSG00002075712.1 | 0 | NA | NA | NA | NA | NA | #VALUE! |
| ENSMUSG00002075713.1 | 0 | NA | NA | NA | NA | NA | #VALUE! |
| ENSMUSG00002075714.1 | 0 | NA | NA | NA | NA | NA | #VALUE! |
| ENSMUSG00002075715.1 | 0 | NA | NA | NA | NA | NA | #VALUE! |
| ENSMUSG00002075716.1 | 0 | NA | NA | NA | NA | NA | #VALUE! |
| ENSMUSG00002075717.1 | 0 | NA | NA | NA | NA | NA | #VALUE! |
| ENSMUSG00002075718.1 | 0 | NA | NA | NA | NA | NA | #VALUE! |
| ENSMUSG00002075719.1 | 0 | NA | NA | NA | NA | NA | #VALUE! |
| ENSMUSG00002075720.1 | 0 | NA | NA | NA | NA | NA | #VALUE! |
| ENSMUSG00002075721.1 | 0 | NA | NA | NA | NA | NA | #VALUE! |
| ENSMUSG00002075722.1 | 0 | NA | NA | NA | NA | NA | #VALUE! |
| ENSMUSG00002075723.1 | 0 | NA | NA | NA | NA | NA | #VALUE! |
| ENSMUSG00002075724.1 | 0 | NA | NA | NA | NA | NA | #VALUE! |
| ENSMUSG00002075725.1 | 0 | NA | NA | NA | NA | NA | #VALUE! |
| ENSMUSG00002075726.1 | 0 | NA | NA | NA | NA | NA | #VALUE! |
| ENSMUSG00002075727.1 | 0 | NA | NA | NA | NA | NA | #VALUE! |
| ENSMUSG00002075728.1 | 0 | NA | NA | NA | NA | NA | #VALUE! |
| ENSMUSG00002075729.1 | 0 | NA | NA | NA | NA | NA | #VALUE! |
| ENSMUSG00002075730.1 | 0 | NA | NA | NA | NA | NA | #VALUE! |
| ENSMUSG00002075731.1 | 0 | NA | NA | NA | NA | NA | #VALUE! |
| ENSMUSG00002075732.1 | 0 | NA | NA | NA | NA | NA | #VALUE! |
| ENSMUSG00002075733.1 | 0 | NA | NA | NA | NA | NA | #VALUE! |
| ENSMUSG00002075734.1 | 0 | NA | NA | NA | NA | NA | #VALUE! |
| ENSMUSG00002075735.1 | 0 | NA | NA | NA | NA | NA | #VALUE! |
| ENSMUSG00002075736.1 | 0 | NA | NA | NA | NA | NA | #VALUE! |
| ENSMUSG00002075737.1 | 0 | NA | NA | NA | NA | NA | #VALUE! |
| ENSMUSG00002075738.1 | 0 | NA | NA | NA | NA | NA | #VALUE! |

| | | | | | | | |
|---|---|---|---|---|---|---|---|
| ENSMUSG00002075739.1 | 0 | NA | NA | NA | NA | NA | #VALUE! |
| ENSMUSG00002075740.1 | 0 | NA | NA | NA | NA | NA | #VALUE! |
| ENSMUSG00002075741.1 | 0 | NA | NA | NA | NA | NA | #VALUE! |
| ENSMUSG00002075742.1 | 0 | NA | NA | NA | NA | NA | #VALUE! |
| ENSMUSG00002075743.1 | 0 | NA | NA | NA | NA | NA | #VALUE! |
| ENSMUSG00002075744.1 | 0 | NA | NA | NA | NA | NA | #VALUE! |
| ENSMUSG00002075745.1 | 0 | NA | NA | NA | NA | NA | #VALUE! |
| ENSMUSG00002075746.1 | 0 | NA | NA | NA | NA | NA | #VALUE! |
| ENSMUSG00002075747.1 | 0 | NA | NA | NA | NA | NA | #VALUE! |
| ENSMUSG00002075748.1 | 0 | NA | NA | NA | NA | NA | #VALUE! |
| ENSMUSG00002075749.1 | 0 | NA | NA | NA | NA | NA | #VALUE! |
| ENSMUSG00002075750.1 | 0 | NA | NA | NA | NA | NA | #VALUE! |
| ENSMUSG00002075751.1 | 0 | NA | NA | NA | NA | NA | #VALUE! |
| ENSMUSG00002075752.1 | 0 | NA | NA | NA | NA | NA | #VALUE! |
| ENSMUSG00002075753.1 | 0 | NA | NA | NA | NA | NA | #VALUE! |
| ENSMUSG00002075754.1 | 0 | NA | NA | NA | NA | NA | #VALUE! |
| ENSMUSG00002075755.1 | 0 | NA | NA | NA | NA | NA | #VALUE! |
| ENSMUSG00002075756.1 | 0 | NA | NA | NA | NA | NA | #VALUE! |
| ENSMUSG00002075757.1 | 0 | NA | NA | NA | NA | NA | #VALUE! |
| ENSMUSG00002075758.1 | 0 | NA | NA | NA | NA | NA | #VALUE! |
| ENSMUSG00002075759.1 | 0 | NA | NA | NA | NA | NA | #VALUE! |
| ENSMUSG00002075760.1 | 0 | NA | NA | NA | NA | NA | #VALUE! |
| ENSMUSG00002075761.1 | 0 | NA | NA | NA | NA | NA | #VALUE! |
| ENSMUSG00002075762.1 | 0 | NA | NA | NA | NA | NA | #VALUE! |
| ENSMUSG00002075763.1 | 0 | NA | NA | NA | NA | NA | #VALUE! |
| ENSMUSG00002075764.1 | 0 | NA | NA | NA | NA | NA | #VALUE! |
| ENSMUSG00002075765.1 | 0 | NA | NA | NA | NA | NA | #VALUE! |
| ENSMUSG00002075766.1 | 0 | NA | NA | NA | NA | NA | #VALUE! |
| ENSMUSG00002075767.1 | 0 | NA | NA | NA | NA | NA | #VALUE! |
| ENSMUSG00002075768.1 | 0 | NA | NA | NA | NA | NA | #VALUE! |
| ENSMUSG00002075769.1 | 0 | NA | NA | NA | NA | NA | #VALUE! |
| ENSMUSG00002075770.1 | 0 | NA | NA | NA | NA | NA | #VALUE! |
| ENSMUSG00002075771.1 | 0 | NA | NA | NA | NA | NA | #VALUE! |
| ENSMUSG00002075772.1 | 0 | NA | NA | NA | NA | NA | #VALUE! |
| ENSMUSG00002075773.1 | 0 | NA | NA | NA | NA | NA | #VALUE! |
| ENSMUSG00002075774.1 | 0 | NA | NA | NA | NA | NA | #VALUE! |
| ENSMUSG00002075775.1 | 0 | NA | NA | NA | NA | NA | #VALUE! |
| ENSMUSG00002075776.1 | 0 | NA | NA | NA | NA | NA | #VALUE! |
| ENSMUSG00002075777.1 | 0 | NA | NA | NA | NA | NA | #VALUE! |
| ENSMUSG00002075778.1 | 0 | NA | NA | NA | NA | NA | #VALUE! |
| ENSMUSG00002075779.1 | 0 | NA | NA | NA | NA | NA | #VALUE! |
| ENSMUSG00002075780.1 | 0 | NA | NA | NA | NA | NA | #VALUE! |
| ENSMUSG00002075781.1 | 0 | NA | NA | NA | NA | NA | #VALUE! |
| ENSMUSG00002075782.1 | 0 | NA | NA | NA | NA | NA | #VALUE! |

| | | | | | | | |
|---|---|---|---|---|---|---|---|
| ENSMUSG00002075783.1 | 0 | NA | NA | NA | NA | NA | #VALUE! |
| ENSMUSG00002075784.1 | 0 | NA | NA | NA | NA | NA | #VALUE! |
| ENSMUSG00002075785.1 | 0 | NA | NA | NA | NA | NA | #VALUE! |
| ENSMUSG00002075786.1 | 0 | NA | NA | NA | NA | NA | #VALUE! |
| ENSMUSG00002075787.1 | 0 | NA | NA | NA | NA | NA | #VALUE! |
| ENSMUSG00002075788.1 | 0 | NA | NA | NA | NA | NA | #VALUE! |
| ENSMUSG00002075789.1 | 0 | NA | NA | NA | NA | NA | #VALUE! |
| ENSMUSG00002075790.1 | 0 | NA | NA | NA | NA | NA | #VALUE! |
| ENSMUSG00002075791.1 | 0 | NA | NA | NA | NA | NA | #VALUE! |
| ENSMUSG00002075792.1 | 0 | NA | NA | NA | NA | NA | #VALUE! |
| ENSMUSG00002075793.1 | 0 | NA | NA | NA | NA | NA | #VALUE! |
| ENSMUSG00002075794.1 | 0 | NA | NA | NA | NA | NA | #VALUE! |
| ENSMUSG00002075796.1 | 0 | NA | NA | NA | NA | NA | #VALUE! |
| ENSMUSG00002075797.1 | 0 | NA | NA | NA | NA | NA | #VALUE! |
| ENSMUSG00002075798.1 | 0 | NA | NA | NA | NA | NA | #VALUE! |
| ENSMUSG00002075799.1 | 0 | NA | NA | NA | NA | NA | #VALUE! |
| ENSMUSG00002075800.1 | 0 | NA | NA | NA | NA | NA | #VALUE! |
| ENSMUSG00002075801.1 | 0 | NA | NA | NA | NA | NA | #VALUE! |
| ENSMUSG00002075802.1 | 0 | NA | NA | NA | NA | NA | #VALUE! |
| ENSMUSG00002075803.1 | 0 | NA | NA | NA | NA | NA | #VALUE! |
| ENSMUSG00002075804.1 | 0 | NA | NA | NA | NA | NA | #VALUE! |
| ENSMUSG00002075805.1 | 0 | NA | NA | NA | NA | NA | #VALUE! |
| ENSMUSG00002075806.1 | 0 | NA | NA | NA | NA | NA | #VALUE! |
| ENSMUSG00002075807.1 | 0 | NA | NA | NA | NA | NA | #VALUE! |
| ENSMUSG00002075808.1 | 0 | NA | NA | NA | NA | NA | #VALUE! |
| ENSMUSG00002075809.1 | 0 | NA | NA | NA | NA | NA | #VALUE! |
| ENSMUSG00002075810.1 | 0 | NA | NA | NA | NA | NA | #VALUE! |
| ENSMUSG00002075811.1 | 0 | NA | NA | NA | NA | NA | #VALUE! |
| ENSMUSG00002075812.1 | 0 | NA | NA | NA | NA | NA | #VALUE! |
| ENSMUSG00002075813.1 | 0 | NA | NA | NA | NA | NA | #VALUE! |
| ENSMUSG00002075814.1 | 0 | NA | NA | NA | NA | NA | #VALUE! |
| ENSMUSG00002075815.1 | 0 | NA | NA | NA | NA | NA | #VALUE! |
| ENSMUSG00002075816.1 | 0 | NA | NA | NA | NA | NA | #VALUE! |
| ENSMUSG00002075817.1 | 0 | NA | NA | NA | NA | NA | #VALUE! |
| ENSMUSG00002075818.1 | 0 | NA | NA | NA | NA | NA | #VALUE! |
| ENSMUSG00002075819.1 | 0 | NA | NA | NA | NA | NA | #VALUE! |
| ENSMUSG00002075820.1 | 0 | NA | NA | NA | NA | NA | #VALUE! |
| ENSMUSG00002075821.1 | 0 | NA | NA | NA | NA | NA | #VALUE! |
| ENSMUSG00002075822.1 | 0 | NA | NA | NA | NA | NA | #VALUE! |
| ENSMUSG00002075823.1 | 0 | NA | NA | NA | NA | NA | #VALUE! |
| ENSMUSG00002075824.1 | 0 | NA | NA | NA | NA | NA | #VALUE! |
| ENSMUSG00002075825.1 | 0 | NA | NA | NA | NA | NA | #VALUE! |
| ENSMUSG00002075826.1 | 0 | NA | NA | NA | NA | NA | #VALUE! |
| ENSMUSG00002075827.1 | 0 | NA | NA | NA | NA | NA | #VALUE! |

| | | | | | | | |
|---|---|---|---|---|---|---|---|
| ENSMUSG00002075828.1 | 0 | NA | NA | NA | NA | NA | #VALUE! |
| ENSMUSG00002075829.1 | 0 | NA | NA | NA | NA | NA | #VALUE! |
| ENSMUSG00002075830.1 | 0 | NA | NA | NA | NA | NA | #VALUE! |
| ENSMUSG00002075831.1 | 0 | NA | NA | NA | NA | NA | #VALUE! |
| ENSMUSG00002075832.1 | 0 | NA | NA | NA | NA | NA | #VALUE! |
| ENSMUSG00002075833.1 | 0 | NA | NA | NA | NA | NA | #VALUE! |
| ENSMUSG00002075834.1 | 0 | NA | NA | NA | NA | NA | #VALUE! |
| ENSMUSG00002075835.1 | 0 | NA | NA | NA | NA | NA | #VALUE! |
| ENSMUSG00002075836.1 | 0 | NA | NA | NA | NA | NA | #VALUE! |
| ENSMUSG00002075837.1 | 0 | NA | NA | NA | NA | NA | #VALUE! |
| ENSMUSG00002075838.1 | 0 | NA | NA | NA | NA | NA | #VALUE! |
| ENSMUSG00002075839.1 | 0 | NA | NA | NA | NA | NA | #VALUE! |
| ENSMUSG00002075840.1 | 0 | NA | NA | NA | NA | NA | #VALUE! |
| ENSMUSG00002075841.1 | 0 | NA | NA | NA | NA | NA | #VALUE! |
| ENSMUSG00002075842.1 | 0 | NA | NA | NA | NA | NA | #VALUE! |
| ENSMUSG00002075843.1 | 0 | NA | NA | NA | NA | NA | #VALUE! |
| ENSMUSG00002075844.1 | 0 | NA | NA | NA | NA | NA | #VALUE! |
| ENSMUSG00002075845.1 | 0 | NA | NA | NA | NA | NA | #VALUE! |
| ENSMUSG00002075846.1 | 0 | NA | NA | NA | NA | NA | #VALUE! |
| ENSMUSG00002075847.1 | 0 | NA | NA | NA | NA | NA | #VALUE! |
| ENSMUSG00002075848.1 | 0 | NA | NA | NA | NA | NA | #VALUE! |
| ENSMUSG00002075849.1 | 0 | NA | NA | NA | NA | NA | #VALUE! |
| ENSMUSG00002075850.1 | 0 | NA | NA | NA | NA | NA | #VALUE! |
| ENSMUSG00002075851.1 | 0 | NA | NA | NA | NA | NA | #VALUE! |
| ENSMUSG00002075852.1 | 0 | NA | NA | NA | NA | NA | #VALUE! |
| ENSMUSG00002075853.1 | 0 | NA | NA | NA | NA | NA | #VALUE! |
| ENSMUSG00002075854.1 | 0 | NA | NA | NA | NA | NA | #VALUE! |
| ENSMUSG00002075855.1 | 0 | NA | NA | NA | NA | NA | #VALUE! |
| ENSMUSG00002075856.1 | 0 | NA | NA | NA | NA | NA | #VALUE! |
| ENSMUSG00002075857.1 | 0 | NA | NA | NA | NA | NA | #VALUE! |
| ENSMUSG00002075858.1 | 0 | NA | NA | NA | NA | NA | #VALUE! |
| ENSMUSG00002075859.1 | 0 | NA | NA | NA | NA | NA | #VALUE! |
| ENSMUSG00002075860.1 | 0 | NA | NA | NA | NA | NA | #VALUE! |
| ENSMUSG00002075861.1 | 0 | NA | NA | NA | NA | NA | #VALUE! |
| ENSMUSG00002075862.1 | 0 | NA | NA | NA | NA | NA | #VALUE! |
| ENSMUSG00002075863.1 | 0 | NA | NA | NA | NA | NA | #VALUE! |
| ENSMUSG00002075864.1 | 0 | NA | NA | NA | NA | NA | #VALUE! |
| ENSMUSG00002075865.1 | 0 | NA | NA | NA | NA | NA | #VALUE! |
| ENSMUSG00002075866.1 | 0 | NA | NA | NA | NA | NA | #VALUE! |
| ENSMUSG00002075867.1 | 0 | NA | NA | NA | NA | NA | #VALUE! |
| ENSMUSG00002075868.1 | 0 | NA | NA | NA | NA | NA | #VALUE! |
| ENSMUSG00002075869.1 | 0 | NA | NA | NA | NA | NA | #VALUE! |
| ENSMUSG00002075870.1 | 0 | NA | NA | NA | NA | NA | #VALUE! |
| ENSMUSG00002075871.1 | 0 | NA | NA | NA | NA | NA | #VALUE! |

| | | | | | | | |
|---|---|---|---|---|---|---|---|
| ENSMUSG00002075872.1 | 0 | NA | NA | NA | NA | NA | #VALUE! |
| ENSMUSG00002075873.1 | 0 | NA | NA | NA | NA | NA | #VALUE! |
| ENSMUSG00002075874.1 | 0 | NA | NA | NA | NA | NA | #VALUE! |
| ENSMUSG00002075875.1 | 0 | NA | NA | NA | NA | NA | #VALUE! |
| ENSMUSG00002075876.1 | 0 | NA | NA | NA | NA | NA | #VALUE! |
| ENSMUSG00002075877.1 | 0 | NA | NA | NA | NA | NA | #VALUE! |
| ENSMUSG00002075878.1 | 0 | NA | NA | NA | NA | NA | #VALUE! |
| ENSMUSG00002075879.1 | 0 | NA | NA | NA | NA | NA | #VALUE! |
| ENSMUSG00002075880.1 | 0 | NA | NA | NA | NA | NA | #VALUE! |
| ENSMUSG00002075881.1 | 0 | NA | NA | NA | NA | NA | #VALUE! |
| ENSMUSG00002075882.1 | 0 | NA | NA | NA | NA | NA | #VALUE! |
| ENSMUSG00002075883.1 | 0 | NA | NA | NA | NA | NA | #VALUE! |
| ENSMUSG00002075884.1 | 0 | NA | NA | NA | NA | NA | #VALUE! |
| ENSMUSG00002075885.1 | 0 | NA | NA | NA | NA | NA | #VALUE! |
| ENSMUSG00002075886.1 | 0 | NA | NA | NA | NA | NA | #VALUE! |
| ENSMUSG00002075887.1 | 0 | NA | NA | NA | NA | NA | #VALUE! |
| ENSMUSG00002075888.1 | 0 | NA | NA | NA | NA | NA | #VALUE! |
| ENSMUSG00002075889.1 | 0 | NA | NA | NA | NA | NA | #VALUE! |
| ENSMUSG00002075890.1 | 0 | NA | NA | NA | NA | NA | #VALUE! |
| ENSMUSG00002075891.1 | 0 | NA | NA | NA | NA | NA | #VALUE! |
| ENSMUSG00002075892.1 | 0 | NA | NA | NA | NA | NA | #VALUE! |
| ENSMUSG00002075893.1 | 0 | NA | NA | NA | NA | NA | #VALUE! |
| ENSMUSG00002075894.1 | 0 | NA | NA | NA | NA | NA | #VALUE! |
| ENSMUSG00002075895.1 | 0 | NA | NA | NA | NA | NA | #VALUE! |
| ENSMUSG00002075896.1 | 0 | NA | NA | NA | NA | NA | #VALUE! |
| ENSMUSG00002075897.1 | 0 | NA | NA | NA | NA | NA | #VALUE! |
| ENSMUSG00002075898.1 | 0 | NA | NA | NA | NA | NA | #VALUE! |
| ENSMUSG00002075899.1 | 0 | NA | NA | NA | NA | NA | #VALUE! |
| ENSMUSG00002075900.1 | 0 | NA | NA | NA | NA | NA | #VALUE! |
| ENSMUSG00002075901.1 | 0 | NA | NA | NA | NA | NA | #VALUE! |
| ENSMUSG00002075902.1 | 0 | NA | NA | NA | NA | NA | #VALUE! |
| ENSMUSG00002075903.1 | 0 | NA | NA | NA | NA | NA | #VALUE! |
| ENSMUSG00002075904.1 | 0 | NA | NA | NA | NA | NA | #VALUE! |
| ENSMUSG00002075905.1 | 0 | NA | NA | NA | NA | NA | #VALUE! |
| ENSMUSG00002075906.1 | 0 | NA | NA | NA | NA | NA | #VALUE! |
| ENSMUSG00002075907.1 | 0 | NA | NA | NA | NA | NA | #VALUE! |
| ENSMUSG00002075908.1 | 0 | NA | NA | NA | NA | NA | #VALUE! |
| ENSMUSG00002075909.1 | 0 | NA | NA | NA | NA | NA | #VALUE! |
| ENSMUSG00002075910.1 | 0 | NA | NA | NA | NA | NA | #VALUE! |
| ENSMUSG00002075911.1 | 0 | NA | NA | NA | NA | NA | #VALUE! |
| ENSMUSG00002075912.1 | 0 | NA | NA | NA | NA | NA | #VALUE! |
| ENSMUSG00002075913.1 | 0 | NA | NA | NA | NA | NA | #VALUE! |
| ENSMUSG00002075914.1 | 0 | NA | NA | NA | NA | NA | #VALUE! |
| ENSMUSG00002075915.1 | 0 | NA | NA | NA | NA | NA | #VALUE! |

| | | | | | | | |
|---|---|---|---|---|---|---|---|
| ENSMUSG00002075916.1 | 0 | NA | NA | NA | NA | NA | #VALUE! |
| ENSMUSG00002075917.1 | 0 | NA | NA | NA | NA | NA | #VALUE! |
| ENSMUSG00002075918.1 | 0 | NA | NA | NA | NA | NA | #VALUE! |
| ENSMUSG00002075919.1 | 0 | NA | NA | NA | NA | NA | #VALUE! |
| ENSMUSG00002075920.1 | 0 | NA | NA | NA | NA | NA | #VALUE! |
| ENSMUSG00002075921.1 | 0 | NA | NA | NA | NA | NA | #VALUE! |
| ENSMUSG00002075922.1 | 0 | NA | NA | NA | NA | NA | #VALUE! |
| ENSMUSG00002075923.1 | 0 | NA | NA | NA | NA | NA | #VALUE! |
| ENSMUSG00002075924.1 | 0 | NA | NA | NA | NA | NA | #VALUE! |
| ENSMUSG00002075925.1 | 0 | NA | NA | NA | NA | NA | #VALUE! |
| ENSMUSG00002075926.1 | 0 | NA | NA | NA | NA | NA | #VALUE! |
| ENSMUSG00002075927.1 | 0 | NA | NA | NA | NA | NA | #VALUE! |
| ENSMUSG00002075928.1 | 0 | NA | NA | NA | NA | NA | #VALUE! |
| ENSMUSG00002075929.1 | 0 | NA | NA | NA | NA | NA | #VALUE! |
| ENSMUSG00002075930.1 | 0 | NA | NA | NA | NA | NA | #VALUE! |
| ENSMUSG00002075931.1 | 0 | NA | NA | NA | NA | NA | #VALUE! |
| ENSMUSG00002075932.1 | 0 | NA | NA | NA | NA | NA | #VALUE! |
| ENSMUSG00002075933.1 | 0 | NA | NA | NA | NA | NA | #VALUE! |
| ENSMUSG00002075934.1 | 0 | NA | NA | NA | NA | NA | #VALUE! |
| ENSMUSG00002075935.1 | 0 | NA | NA | NA | NA | NA | #VALUE! |
| ENSMUSG00002075936.1 | 0 | NA | NA | NA | NA | NA | #VALUE! |
| ENSMUSG00002075937.1 | 0 | NA | NA | NA | NA | NA | #VALUE! |
| ENSMUSG00002075938.1 | 0 | NA | NA | NA | NA | NA | #VALUE! |
| ENSMUSG00002075939.1 | 0 | NA | NA | NA | NA | NA | #VALUE! |
| ENSMUSG00002075940.1 | 0 | NA | NA | NA | NA | NA | #VALUE! |
| ENSMUSG00002075941.1 | 0 | NA | NA | NA | NA | NA | #VALUE! |
| ENSMUSG00002075942.1 | 0 | NA | NA | NA | NA | NA | #VALUE! |
| ENSMUSG00002075943.1 | 0 | NA | NA | NA | NA | NA | #VALUE! |
| ENSMUSG00002075944.1 | 0 | NA | NA | NA | NA | NA | #VALUE! |
| ENSMUSG00002075945.1 | 0 | NA | NA | NA | NA | NA | #VALUE! |
| ENSMUSG00002075946.1 | 0 | NA | NA | NA | NA | NA | #VALUE! |
| ENSMUSG00002075947.1 | 0 | NA | NA | NA | NA | NA | #VALUE! |
| ENSMUSG00002075948.1 | 0 | NA | NA | NA | NA | NA | #VALUE! |
| ENSMUSG00002075949.1 | 0 | NA | NA | NA | NA | NA | #VALUE! |
| ENSMUSG00002075950.1 | 0 | NA | NA | NA | NA | NA | #VALUE! |
| ENSMUSG00002075951.1 | 0 | NA | NA | NA | NA | NA | #VALUE! |
| ENSMUSG00002075952.1 | 0 | NA | NA | NA | NA | NA | #VALUE! |
| ENSMUSG00002075953.1 | 0 | NA | NA | NA | NA | NA | #VALUE! |
| ENSMUSG00002075954.1 | 0 | NA | NA | NA | NA | NA | #VALUE! |
| ENSMUSG00002075955.1 | 0 | NA | NA | NA | NA | NA | #VALUE! |
| ENSMUSG00002075956.1 | 0 | NA | NA | NA | NA | NA | #VALUE! |
| ENSMUSG00002075957.1 | 0 | NA | NA | NA | NA | NA | #VALUE! |
| ENSMUSG00002075958.1 | 0 | NA | NA | NA | NA | NA | #VALUE! |
| ENSMUSG00002075959.1 | 0 | NA | NA | NA | NA | NA | #VALUE! |

| | | | | | | | |
|---|---|---|---|---|---|---|---|
| ENSMUSG00002075960.1 | 0 | NA | NA | NA | NA | NA | #VALUE! |
| ENSMUSG00002075961.1 | 0 | NA | NA | NA | NA | NA | #VALUE! |
| ENSMUSG00002075962.1 | 0 | NA | NA | NA | NA | NA | #VALUE! |
| ENSMUSG00002075963.1 | 0 | NA | NA | NA | NA | NA | #VALUE! |
| ENSMUSG00002075964.1 | 0 | NA | NA | NA | NA | NA | #VALUE! |
| ENSMUSG00002075965.1 | 0 | NA | NA | NA | NA | NA | #VALUE! |
| ENSMUSG00002075966.1 | 0 | NA | NA | NA | NA | NA | #VALUE! |
| ENSMUSG00002075967.1 | 0.093255 | 0.868093 | 4.072464 | 0.213162 | 0.831201 | NA | #VALUE! |
| ENSMUSG00002075968.1 | 0 | NA | NA | NA | NA | NA | #VALUE! |
| ENSMUSG00002075969.1 | 0 | NA | NA | NA | NA | NA | #VALUE! |
| ENSMUSG00002075970.1 | 0 | NA | NA | NA | NA | NA | #VALUE! |
| ENSMUSG00002075971.1 | 0 | NA | NA | NA | NA | NA | #VALUE! |
| ENSMUSG00002075972.1 | 0 | NA | NA | NA | NA | NA | #VALUE! |
| ENSMUSG00002075973.1 | 0 | NA | NA | NA | NA | NA | #VALUE! |
| ENSMUSG00002075974.1 | 0 | NA | NA | NA | NA | NA | #VALUE! |
| ENSMUSG00002075975.1 | 0 | NA | NA | NA | NA | NA | #VALUE! |
| ENSMUSG00002075976.1 | 0 | NA | NA | NA | NA | NA | #VALUE! |
| ENSMUSG00002075977.1 | 0 | NA | NA | NA | NA | NA | #VALUE! |
| ENSMUSG00002075978.1 | 0 | NA | NA | NA | NA | NA | #VALUE! |
| ENSMUSG00002075979.1 | 0 | NA | NA | NA | NA | NA | #VALUE! |
| ENSMUSG00002075980.1 | 0 | NA | NA | NA | NA | NA | #VALUE! |
| ENSMUSG00002075981.1 | 0 | NA | NA | NA | NA | NA | #VALUE! |
| ENSMUSG00002075982.1 | 0 | NA | NA | NA | NA | NA | #VALUE! |
| ENSMUSG00002075983.1 | 0 | NA | NA | NA | NA | NA | #VALUE! |
| ENSMUSG00002075984.1 | 0 | NA | NA | NA | NA | NA | #VALUE! |
| ENSMUSG00002075985.1 | 0 | NA | NA | NA | NA | NA | #VALUE! |
| ENSMUSG00002075986.1 | 0 | NA | NA | NA | NA | NA | #VALUE! |
| ENSMUSG00002075987.1 | 0 | NA | NA | NA | NA | NA | #VALUE! |
| ENSMUSG00002075988.1 | 0 | NA | NA | NA | NA | NA | #VALUE! |
| ENSMUSG00002075989.1 | 0 | NA | NA | NA | NA | NA | #VALUE! |
| ENSMUSG00002075990.1 | 0 | NA | NA | NA | NA | NA | #VALUE! |
| ENSMUSG00002075991.1 | 0 | NA | NA | NA | NA | NA | #VALUE! |
| ENSMUSG00002075992.1 | 0 | NA | NA | NA | NA | NA | #VALUE! |
| ENSMUSG00002075993.1 | 0 | NA | NA | NA | NA | NA | #VALUE! |
| ENSMUSG00002075994.1 | 0 | NA | NA | NA | NA | NA | #VALUE! |
| ENSMUSG00002075995.1 | 0 | NA | NA | NA | NA | NA | #VALUE! |
| ENSMUSG00002075996.1 | 0 | NA | NA | NA | NA | NA | #VALUE! |
| ENSMUSG00002075997.1 | 0 | NA | NA | NA | NA | NA | #VALUE! |
| ENSMUSG00002075998.1 | 0 | NA | NA | NA | NA | NA | #VALUE! |
| ENSMUSG00002075999.1 | 0 | NA | NA | NA | NA | NA | #VALUE! |
| ENSMUSG00002076000.1 | 0 | NA | NA | NA | NA | NA | #VALUE! |
| ENSMUSG00002076001.1 | 0 | NA | NA | NA | NA | NA | #VALUE! |
| ENSMUSG00002076002.1 | 0 | NA | NA | NA | NA | NA | #VALUE! |
| ENSMUSG00002076003.1 | 0 | NA | NA | NA | NA | NA | #VALUE! |

| | | | | | | | |
|---|---|---|---|---|---|---|---|
| ENSMUSG00002076004.1 | 0 | NA | NA | NA | NA | NA | #VALUE! |
| ENSMUSG00002076005.1 | 0 | NA | NA | NA | NA | NA | #VALUE! |
| ENSMUSG00002076006.1 | 0 | NA | NA | NA | NA | NA | #VALUE! |
| ENSMUSG00002076007.1 | 0 | NA | NA | NA | NA | NA | #VALUE! |
| ENSMUSG00002076008.1 | 0 | NA | NA | NA | NA | NA | #VALUE! |
| ENSMUSG00002076009.1 | 0 | NA | NA | NA | NA | NA | #VALUE! |
| ENSMUSG00002076010.1 | 0 | NA | NA | NA | NA | NA | #VALUE! |
| ENSMUSG00002076011.1 | 0 | NA | NA | NA | NA | NA | #VALUE! |
| ENSMUSG00002076012.1 | 0 | NA | NA | NA | NA | NA | #VALUE! |
| ENSMUSG00002076013.1 | 0 | NA | NA | NA | NA | NA | #VALUE! |
| ENSMUSG00002076014.1 | 0 | NA | NA | NA | NA | NA | #VALUE! |
| ENSMUSG00002076015.1 | 0 | NA | NA | NA | NA | NA | #VALUE! |
| ENSMUSG00002076016.1 | 0 | NA | NA | NA | NA | NA | #VALUE! |
| ENSMUSG00002076017.1 | 0 | NA | NA | NA | NA | NA | #VALUE! |
| ENSMUSG00002076018.1 | 0 | NA | NA | NA | NA | NA | #VALUE! |
| ENSMUSG00002076019.1 | 0 | NA | NA | NA | NA | NA | #VALUE! |
| ENSMUSG00002076020.1 | 0 | NA | NA | NA | NA | NA | #VALUE! |
| ENSMUSG00002076021.1 | 0 | NA | NA | NA | NA | NA | #VALUE! |
| ENSMUSG00002076022.1 | 0 | NA | NA | NA | NA | NA | #VALUE! |
| ENSMUSG00002076023.1 | 0 | NA | NA | NA | NA | NA | #VALUE! |
| ENSMUSG00002076024.1 | 0 | NA | NA | NA | NA | NA | #VALUE! |
| ENSMUSG00002076025.1 | 0 | NA | NA | NA | NA | NA | #VALUE! |
| ENSMUSG00002076026.1 | 3.524204 | -5.34395 | 3.919482 | -1.36343 | 0.172746 | NA | #VALUE! |
| ENSMUSG00002076027.1 | 0 | NA | NA | NA | NA | NA | #VALUE! |
| ENSMUSG00002076028.1 | 0 | NA | NA | NA | NA | NA | #VALUE! |
| ENSMUSG00002076029.1 | 0 | NA | NA | NA | NA | NA | #VALUE! |
| ENSMUSG00002076030.1 | 0 | NA | NA | NA | NA | NA | #VALUE! |
| ENSMUSG00002076031.1 | 0 | NA | NA | NA | NA | NA | #VALUE! |
| ENSMUSG00002076032.1 | 0 | NA | NA | NA | NA | NA | #VALUE! |
| ENSMUSG00002076033.1 | 0 | NA | NA | NA | NA | NA | #VALUE! |
| ENSMUSG00002076034.1 | 0 | NA | NA | NA | NA | NA | #VALUE! |
| ENSMUSG00002076035.1 | 0 | NA | NA | NA | NA | NA | #VALUE! |
| ENSMUSG00002076036.1 | 0 | NA | NA | NA | NA | NA | #VALUE! |
| ENSMUSG00002076037.1 | 0 | NA | NA | NA | NA | NA | #VALUE! |
| ENSMUSG00002076038.1 | 0 | NA | NA | NA | NA | NA | #VALUE! |
| ENSMUSG00002076039.1 | 0 | NA | NA | NA | NA | NA | #VALUE! |
| ENSMUSG00002076040.1 | 0 | NA | NA | NA | NA | NA | #VALUE! |
| ENSMUSG00002076041.1 | 0 | NA | NA | NA | NA | NA | #VALUE! |
| ENSMUSG00002076042.1 | 0 | NA | NA | NA | NA | NA | #VALUE! |
| ENSMUSG00002076043.1 | 0 | NA | NA | NA | NA | NA | #VALUE! |
| ENSMUSG00002076044.1 | 0 | NA | NA | NA | NA | NA | #VALUE! |
| ENSMUSG00002076045.1 | 0 | NA | NA | NA | NA | NA | #VALUE! |
| ENSMUSG00002076046.1 | 0 | NA | NA | NA | NA | NA | #VALUE! |
| ENSMUSG00002076047.1 | 0 | NA | NA | NA | NA | NA | #VALUE! |

| | | | | | | | |
|---|---|---|---|---|---|---|---|
| ENSMUSG00002076048.1 | 0 | NA | NA | NA | NA | NA | #VALUE! |
| ENSMUSG00002076049.1 | 0 | NA | NA | NA | NA | NA | #VALUE! |
| ENSMUSG00002076050.1 | 0 | NA | NA | NA | NA | NA | #VALUE! |
| ENSMUSG00002076051.1 | 0 | NA | NA | NA | NA | NA | #VALUE! |
| ENSMUSG00002076052.1 | 0 | NA | NA | NA | NA | NA | #VALUE! |
| ENSMUSG00002076053.1 | 0 | NA | NA | NA | NA | NA | #VALUE! |
| ENSMUSG00002076054.1 | 0 | NA | NA | NA | NA | NA | #VALUE! |
| ENSMUSG00002076055.1 | 0 | NA | NA | NA | NA | NA | #VALUE! |
| ENSMUSG00002076056.1 | 0 | NA | NA | NA | NA | NA | #VALUE! |
| ENSMUSG00002076057.1 | 0 | NA | NA | NA | NA | NA | #VALUE! |
| ENSMUSG00002076058.1 | 0 | NA | NA | NA | NA | NA | #VALUE! |
| ENSMUSG00002076059.1 | 0 | NA | NA | NA | NA | NA | #VALUE! |
| ENSMUSG00002076060.1 | 0 | NA | NA | NA | NA | NA | #VALUE! |
| ENSMUSG00002076061.1 | 0 | NA | NA | NA | NA | NA | #VALUE! |
| ENSMUSG00002076062.1 | 0 | NA | NA | NA | NA | NA | #VALUE! |
| ENSMUSG00002076063.1 | 0 | NA | NA | NA | NA | NA | #VALUE! |
| ENSMUSG00002076064.1 | 0 | NA | NA | NA | NA | NA | #VALUE! |
| ENSMUSG00002076065.1 | 0 | NA | NA | NA | NA | NA | #VALUE! |
| ENSMUSG00002076066.1 | 0 | NA | NA | NA | NA | NA | #VALUE! |
| ENSMUSG00002076067.1 | 0 | NA | NA | NA | NA | NA | #VALUE! |
| ENSMUSG00002076068.1 | 0 | NA | NA | NA | NA | NA | #VALUE! |
| ENSMUSG00002076069.1 | 0 | NA | NA | NA | NA | NA | #VALUE! |
| ENSMUSG00002076070.1 | 0 | NA | NA | NA | NA | NA | #VALUE! |
| ENSMUSG00002076071.1 | 0 | NA | NA | NA | NA | NA | #VALUE! |
| ENSMUSG00002076072.1 | 0 | NA | NA | NA | NA | NA | #VALUE! |
| ENSMUSG00002076073.1 | 0 | NA | NA | NA | NA | NA | #VALUE! |
| ENSMUSG00002076074.1 | 0 | NA | NA | NA | NA | NA | #VALUE! |
| ENSMUSG00002076075.1 | 0 | NA | NA | NA | NA | NA | #VALUE! |
| ENSMUSG00002076076.1 | 0 | NA | NA | NA | NA | NA | #VALUE! |
| ENSMUSG00002076077.1 | 0 | NA | NA | NA | NA | NA | #VALUE! |
| ENSMUSG00002076078.1 | 0 | NA | NA | NA | NA | NA | #VALUE! |
| ENSMUSG00002076079.1 | 0 | NA | NA | NA | NA | NA | #VALUE! |
| ENSMUSG00002076080.1 | 0 | NA | NA | NA | NA | NA | #VALUE! |
| ENSMUSG00002076081.1 | 0 | NA | NA | NA | NA | NA | #VALUE! |
| ENSMUSG00002076082.1 | 0 | NA | NA | NA | NA | NA | #VALUE! |
| ENSMUSG00002076083.1 | 0 | NA | NA | NA | NA | NA | #VALUE! |
| ENSMUSG00002076084.1 | 0 | NA | NA | NA | NA | NA | #VALUE! |
| ENSMUSG00002076085.1 | 0 | NA | NA | NA | NA | NA | #VALUE! |
| ENSMUSG00002076086.1 | 0 | NA | NA | NA | NA | NA | #VALUE! |
| ENSMUSG00002076087.1 | 0 | NA | NA | NA | NA | NA | #VALUE! |
| ENSMUSG00002076088.1 | 0 | NA | NA | NA | NA | NA | #VALUE! |
| ENSMUSG00002076089.1 | 0 | NA | NA | NA | NA | NA | #VALUE! |
| ENSMUSG00002076090.1 | 0 | NA | NA | NA | NA | NA | #VALUE! |
| ENSMUSG00002076091.1 | 0 | NA | NA | NA | NA | NA | #VALUE! |

| | | | | | | | |
|---|---|---|---|---|---|---|---|
| ENSMUSG00002076092.1 | 0 | NA | NA | NA | NA | NA | #VALUE! |
| ENSMUSG00002076093.1 | 0 | NA | NA | NA | NA | NA | #VALUE! |
| ENSMUSG00002076094.1 | 0 | NA | NA | NA | NA | NA | #VALUE! |
| ENSMUSG00002076095.1 | 0 | NA | NA | NA | NA | NA | #VALUE! |
| ENSMUSG00002076096.1 | 0 | NA | NA | NA | NA | NA | #VALUE! |
| ENSMUSG00002076097.1 | 0 | NA | NA | NA | NA | NA | #VALUE! |
| ENSMUSG00002076098.1 | 0 | NA | NA | NA | NA | NA | #VALUE! |
| ENSMUSG00002076099.1 | 0 | NA | NA | NA | NA | NA | #VALUE! |
| ENSMUSG00002076100.1 | 0 | NA | NA | NA | NA | NA | #VALUE! |
| ENSMUSG00002076101.1 | 0 | NA | NA | NA | NA | NA | #VALUE! |
| ENSMUSG00002076102.1 | 0 | NA | NA | NA | NA | NA | #VALUE! |
| ENSMUSG00002076103.1 | 0 | NA | NA | NA | NA | NA | #VALUE! |
| ENSMUSG00002076104.1 | 0 | NA | NA | NA | NA | NA | #VALUE! |
| ENSMUSG00002076105.1 | 0 | NA | NA | NA | NA | NA | #VALUE! |
| ENSMUSG00002076106.1 | 0 | NA | NA | NA | NA | NA | #VALUE! |
| ENSMUSG00002076107.1 | 0 | NA | NA | NA | NA | NA | #VALUE! |
| ENSMUSG00002076108.1 | 0 | NA | NA | NA | NA | NA | #VALUE! |
| ENSMUSG00002076109.1 | 0 | NA | NA | NA | NA | NA | #VALUE! |
| ENSMUSG00002076110.1 | 0 | NA | NA | NA | NA | NA | #VALUE! |
| ENSMUSG00002076111.1 | 0 | NA | NA | NA | NA | NA | #VALUE! |
| ENSMUSG00002076112.1 | 0 | NA | NA | NA | NA | NA | #VALUE! |
| ENSMUSG00002076113.1 | 0 | NA | NA | NA | NA | NA | #VALUE! |
| ENSMUSG00002076114.1 | 0 | NA | NA | NA | NA | NA | #VALUE! |
| ENSMUSG00002076115.1 | 0 | NA | NA | NA | NA | NA | #VALUE! |
| ENSMUSG00002076116.1 | 0 | NA | NA | NA | NA | NA | #VALUE! |
| ENSMUSG00002076117.1 | 0 | NA | NA | NA | NA | NA | #VALUE! |
| ENSMUSG00002076118.1 | 0 | NA | NA | NA | NA | NA | #VALUE! |
| ENSMUSG00002076119.1 | 0 | NA | NA | NA | NA | NA | #VALUE! |
| ENSMUSG00002076120.1 | 0 | NA | NA | NA | NA | NA | #VALUE! |
| ENSMUSG00002076121.1 | 0 | NA | NA | NA | NA | NA | #VALUE! |
| ENSMUSG00002076122.1 | 0 | NA | NA | NA | NA | NA | #VALUE! |
| ENSMUSG00002076123.1 | 0 | NA | NA | NA | NA | NA | #VALUE! |
| ENSMUSG00002076124.1 | 0 | NA | NA | NA | NA | NA | #VALUE! |
| ENSMUSG00002076125.1 | 0 | NA | NA | NA | NA | NA | #VALUE! |
| ENSMUSG00002076126.1 | 0 | NA | NA | NA | NA | NA | #VALUE! |
| ENSMUSG00002076127.1 | 0 | NA | NA | NA | NA | NA | #VALUE! |
| ENSMUSG00002076128.1 | 0 | NA | NA | NA | NA | NA | #VALUE! |
| ENSMUSG00002076129.1 | 0 | NA | NA | NA | NA | NA | #VALUE! |
| ENSMUSG00002076130.1 | 0 | NA | NA | NA | NA | NA | #VALUE! |
| ENSMUSG00002076131.1 | 0 | NA | NA | NA | NA | NA | #VALUE! |
| ENSMUSG00002076132.1 | 0 | NA | NA | NA | NA | NA | #VALUE! |
| ENSMUSG00002076133.1 | 0 | NA | NA | NA | NA | NA | #VALUE! |
| ENSMUSG00002076134.1 | 0 | NA | NA | NA | NA | NA | #VALUE! |
| ENSMUSG00002076135.1 | 0 | NA | NA | NA | NA | NA | #VALUE! |

| | | | | | | | |
|---|---|---|---|---|---|---|---|
| ENSMUSG00002076136.1 | 0 | NA | NA | NA | NA | NA | #VALUE! |
| ENSMUSG00002076137.1 | 0 | NA | NA | NA | NA | NA | #VALUE! |
| ENSMUSG00002076138.1 | 0 | NA | NA | NA | NA | NA | #VALUE! |
| ENSMUSG00002076139.1 | 0 | NA | NA | NA | NA | NA | #VALUE! |
| ENSMUSG00002076140.1 | 0 | NA | NA | NA | NA | NA | #VALUE! |
| ENSMUSG00002076141.1 | 0 | NA | NA | NA | NA | NA | #VALUE! |
| ENSMUSG00002076142.1 | 0 | NA | NA | NA | NA | NA | #VALUE! |
| ENSMUSG00002076143.1 | 0 | NA | NA | NA | NA | NA | #VALUE! |
| ENSMUSG00002076144.1 | 0 | NA | NA | NA | NA | NA | #VALUE! |
| ENSMUSG00002076145.1 | 0 | NA | NA | NA | NA | NA | #VALUE! |
| ENSMUSG00002076146.1 | 0 | NA | NA | NA | NA | NA | #VALUE! |
| ENSMUSG00002076147.1 | 0 | NA | NA | NA | NA | NA | #VALUE! |
| ENSMUSG00002076148.1 | 0 | NA | NA | NA | NA | NA | #VALUE! |
| ENSMUSG00002076149.1 | 0 | NA | NA | NA | NA | NA | #VALUE! |
| ENSMUSG00002076150.1 | 0 | NA | NA | NA | NA | NA | #VALUE! |
| ENSMUSG00002076151.1 | 0 | NA | NA | NA | NA | NA | #VALUE! |
| ENSMUSG00002076152.1 | 0 | NA | NA | NA | NA | NA | #VALUE! |
| ENSMUSG00002076153.1 | 0 | NA | NA | NA | NA | NA | #VALUE! |
| ENSMUSG00002076154.1 | 0 | NA | NA | NA | NA | NA | #VALUE! |
| ENSMUSG00002076155.1 | 0 | NA | NA | NA | NA | NA | #VALUE! |
| ENSMUSG00002076156.1 | 0 | NA | NA | NA | NA | NA | #VALUE! |
| ENSMUSG00002076157.1 | 0 | NA | NA | NA | NA | NA | #VALUE! |
| ENSMUSG00002076158.1 | 0 | NA | NA | NA | NA | NA | #VALUE! |
| ENSMUSG00002076159.1 | 0 | NA | NA | NA | NA | NA | #VALUE! |
| ENSMUSG00002076160.1 | 0 | NA | NA | NA | NA | NA | #VALUE! |
| ENSMUSG00002076161.1 | 3.646231 | 4.270293 | 3.91222 | 1.091527 | 0.275041 | NA | #VALUE! |
| ENSMUSG00002076162.1 | 0 | NA | NA | NA | NA | NA | #VALUE! |
| ENSMUSG00002076163.1 | 0 | NA | NA | NA | NA | NA | #VALUE! |
| ENSMUSG00002076164.1 | 0 | NA | NA | NA | NA | NA | #VALUE! |
| ENSMUSG00002076165.1 | 0 | NA | NA | NA | NA | NA | #VALUE! |
| ENSMUSG00002076166.1 | 0 | NA | NA | NA | NA | NA | #VALUE! |
| ENSMUSG00002076167.1 | 0 | NA | NA | NA | NA | NA | #VALUE! |
| ENSMUSG00002076168.1 | 0 | NA | NA | NA | NA | NA | #VALUE! |
| ENSMUSG00002076169.1 | 0 | NA | NA | NA | NA | NA | #VALUE! |
| ENSMUSG00002076170.1 | 0 | NA | NA | NA | NA | NA | #VALUE! |
| ENSMUSG00002076171.1 | 0 | NA | NA | NA | NA | NA | #VALUE! |
| ENSMUSG00002076172.1 | 0 | NA | NA | NA | NA | NA | #VALUE! |
| ENSMUSG00002076173.1 | 0 | NA | NA | NA | NA | NA | #VALUE! |
| ENSMUSG00002076174.1 | 0 | NA | NA | NA | NA | NA | #VALUE! |
| ENSMUSG00002076175.1 | 0 | NA | NA | NA | NA | NA | #VALUE! |
| ENSMUSG00002076176.1 | 0 | NA | NA | NA | NA | NA | #VALUE! |
| ENSMUSG00002076177.1 | 0 | NA | NA | NA | NA | NA | #VALUE! |
| ENSMUSG00002076178.1 | 0 | NA | NA | NA | NA | NA | #VALUE! |
| ENSMUSG00002076179.1 | 0 | NA | NA | NA | NA | NA | #VALUE! |

| | | | | | | | |
|---|---|---|---|---|---|---|---|
| ENSMUSG00002076180.1 | 0 | NA | NA | NA | NA | NA | #VALUE! |
| ENSMUSG00002076181.1 | 0 | NA | NA | NA | NA | NA | #VALUE! |
| ENSMUSG00002076182.1 | 0 | NA | NA | NA | NA | NA | #VALUE! |
| ENSMUSG00002076183.1 | 0 | NA | NA | NA | NA | NA | #VALUE! |
| ENSMUSG00002076184.1 | 0 | NA | NA | NA | NA | NA | #VALUE! |
| ENSMUSG00002076185.1 | 0 | NA | NA | NA | NA | NA | #VALUE! |
| ENSMUSG00002076186.1 | 0 | NA | NA | NA | NA | NA | #VALUE! |
| ENSMUSG00002076187.1 | 0 | NA | NA | NA | NA | NA | #VALUE! |
| ENSMUSG00002076188.1 | 0 | NA | NA | NA | NA | NA | #VALUE! |
| ENSMUSG00002076189.1 | 0 | NA | NA | NA | NA | NA | #VALUE! |
| ENSMUSG00002076190.1 | 0 | NA | NA | NA | NA | NA | #VALUE! |
| ENSMUSG00002076191.1 | 0 | NA | NA | NA | NA | NA | #VALUE! |
| ENSMUSG00002076192.1 | 0 | NA | NA | NA | NA | NA | #VALUE! |
| ENSMUSG00002076193.1 | 0 | NA | NA | NA | NA | NA | #VALUE! |
| ENSMUSG00002076194.1 | 0 | NA | NA | NA | NA | NA | #VALUE! |
| ENSMUSG00002076195.1 | 0 | NA | NA | NA | NA | NA | #VALUE! |
| ENSMUSG00002076196.1 | 0 | NA | NA | NA | NA | NA | #VALUE! |
| ENSMUSG00002076197.1 | 0 | NA | NA | NA | NA | NA | #VALUE! |
| ENSMUSG00002076198.1 | 0 | NA | NA | NA | NA | NA | #VALUE! |
| ENSMUSG00002076199.1 | 0 | NA | NA | NA | NA | NA | #VALUE! |
| ENSMUSG00002076200.1 | 0 | NA | NA | NA | NA | NA | #VALUE! |
| ENSMUSG00002076201.1 | 0 | NA | NA | NA | NA | NA | #VALUE! |
| ENSMUSG00002076202.1 | 0 | NA | NA | NA | NA | NA | #VALUE! |
| ENSMUSG00002076203.1 | 0 | NA | NA | NA | NA | NA | #VALUE! |
| ENSMUSG00002076204.1 | 0 | NA | NA | NA | NA | NA | #VALUE! |
| ENSMUSG00002076205.1 | 0 | NA | NA | NA | NA | NA | #VALUE! |
| ENSMUSG00002076206.1 | 0 | NA | NA | NA | NA | NA | #VALUE! |
| ENSMUSG00002076207.1 | 0 | NA | NA | NA | NA | NA | #VALUE! |
| ENSMUSG00002076208.1 | 0 | NA | NA | NA | NA | NA | #VALUE! |
| ENSMUSG00002076209.1 | 0 | NA | NA | NA | NA | NA | #VALUE! |
| ENSMUSG00002076210.1 | 0 | NA | NA | NA | NA | NA | #VALUE! |
| ENSMUSG00002076211.1 | 0 | NA | NA | NA | NA | NA | #VALUE! |
| ENSMUSG00002076212.1 | 0 | NA | NA | NA | NA | NA | #VALUE! |
| ENSMUSG00002076213.1 | 0 | NA | NA | NA | NA | NA | #VALUE! |
| ENSMUSG00002076214.1 | 0 | NA | NA | NA | NA | NA | #VALUE! |
| ENSMUSG00002076215.1 | 0 | NA | NA | NA | NA | NA | #VALUE! |
| ENSMUSG00002076216.1 | 0 | NA | NA | NA | NA | NA | #VALUE! |
| ENSMUSG00002076217.1 | 0 | NA | NA | NA | NA | NA | #VALUE! |
| ENSMUSG00002076218.1 | 0 | NA | NA | NA | NA | NA | #VALUE! |
| ENSMUSG00002076219.1 | 0 | NA | NA | NA | NA | NA | #VALUE! |
| ENSMUSG00002076220.1 | 0.309124 | -1.83395 | 4.047982 | -0.45305 | 0.650511 | NA | #VALUE! |
| ENSMUSG00002076221.1 | 0 | NA | NA | NA | NA | NA | #VALUE! |
| ENSMUSG00002076222.1 | 0 | NA | NA | NA | NA | NA | #VALUE! |
| ENSMUSG00002076223.1 | 0 | NA | NA | NA | NA | NA | #VALUE! |

| | | | | | | | |
|---|---|---|---|---|---|---|---|
| ENSMUSG00002076224.1 | 0 | NA | NA | NA | NA | NA | #VALUE! |
| ENSMUSG00002076225.1 | 0 | NA | NA | NA | NA | NA | #VALUE! |
| ENSMUSG00002076226.1 | 0 | NA | NA | NA | NA | NA | #VALUE! |
| ENSMUSG00002076227.1 | 0 | NA | NA | NA | NA | NA | #VALUE! |
| ENSMUSG00002076228.1 | 0 | NA | NA | NA | NA | NA | #VALUE! |
| ENSMUSG00002076229.1 | 0 | NA | NA | NA | NA | NA | #VALUE! |
| ENSMUSG00002076230.1 | 0 | NA | NA | NA | NA | NA | #VALUE! |
| ENSMUSG00002076231.1 | 0 | NA | NA | NA | NA | NA | #VALUE! |
| ENSMUSG00002076232.1 | 0 | NA | NA | NA | NA | NA | #VALUE! |
| ENSMUSG00002076233.1 | 0 | NA | NA | NA | NA | NA | #VALUE! |
| ENSMUSG00002076234.1 | 0 | NA | NA | NA | NA | NA | #VALUE! |
| ENSMUSG00002076235.1 | 0 | NA | NA | NA | NA | NA | #VALUE! |
| ENSMUSG00002076236.1 | 0 | NA | NA | NA | NA | NA | #VALUE! |
| ENSMUSG00002076237.1 | 0 | NA | NA | NA | NA | NA | #VALUE! |
| ENSMUSG00002076238.1 | 0 | NA | NA | NA | NA | NA | #VALUE! |
| ENSMUSG00002076239.1 | 0 | NA | NA | NA | NA | NA | #VALUE! |
| ENSMUSG00002076240.1 | 0.204111 | -1.05581 | 4.073302 | -0.2592 | 0.795479 | NA | #VALUE! |
| ENSMUSG00002076241.1 | 0 | NA | NA | NA | NA | NA | #VALUE! |
| ENSMUSG00002076242.1 | 0 | NA | NA | NA | NA | NA | #VALUE! |
| ENSMUSG00002076243.1 | 0 | NA | NA | NA | NA | NA | #VALUE! |
| ENSMUSG00002076244.1 | 0 | NA | NA | NA | NA | NA | #VALUE! |
| ENSMUSG00002076245.1 | 0 | NA | NA | NA | NA | NA | #VALUE! |
| ENSMUSG00002076246.1 | 0 | NA | NA | NA | NA | NA | #VALUE! |
| ENSMUSG00002076247.1 | 0 | NA | NA | NA | NA | NA | #VALUE! |
| ENSMUSG00002076248.1 | 0 | NA | NA | NA | NA | NA | #VALUE! |
| ENSMUSG00002076249.1 | 0 | NA | NA | NA | NA | NA | #VALUE! |
| ENSMUSG00002076250.1 | 0 | NA | NA | NA | NA | NA | #VALUE! |
| ENSMUSG00002076251.1 | 0 | NA | NA | NA | NA | NA | #VALUE! |
| ENSMUSG00002076252.1 | 0 | NA | NA | NA | NA | NA | #VALUE! |
| ENSMUSG00002076253.1 | 0 | NA | NA | NA | NA | NA | #VALUE! |
| ENSMUSG00002076254.1 | 0 | NA | NA | NA | NA | NA | #VALUE! |
| ENSMUSG00002076255.1 | 0 | NA | NA | NA | NA | NA | #VALUE! |
| ENSMUSG00002076256.1 | 0 | NA | NA | NA | NA | NA | #VALUE! |
| ENSMUSG00002076257.1 | 0 | NA | NA | NA | NA | NA | #VALUE! |
| ENSMUSG00002076258.1 | 0 | NA | NA | NA | NA | NA | #VALUE! |
| ENSMUSG00002076259.1 | 0 | NA | NA | NA | NA | NA | #VALUE! |
| ENSMUSG00002076260.1 | 0 | NA | NA | NA | NA | NA | #VALUE! |
| ENSMUSG00002076261.1 | 0 | NA | NA | NA | NA | NA | #VALUE! |
| ENSMUSG00002076262.1 | 0 | NA | NA | NA | NA | NA | #VALUE! |
| ENSMUSG00002076263.1 | 0 | NA | NA | NA | NA | NA | #VALUE! |
| ENSMUSG00002076264.1 | 0 | NA | NA | NA | NA | NA | #VALUE! |
| ENSMUSG00002076265.1 | 0 | NA | NA | NA | NA | NA | #VALUE! |
| ENSMUSG00002076266.1 | 0 | NA | NA | NA | NA | NA | #VALUE! |
| ENSMUSG00002076267.1 | 0 | NA | NA | NA | NA | NA | #VALUE! |

| | | | | | | | |
|---|---|---|---|---|---|---|---|
| ENSMUSG00002076268.1 | 0 | NA | NA | NA | NA | NA | #VALUE! |
| ENSMUSG00002076269.1 | 0 | NA | NA | NA | NA | NA | #VALUE! |
| ENSMUSG00002076270.1 | 0 | NA | NA | NA | NA | NA | #VALUE! |
| ENSMUSG00002076271.1 | 0 | NA | NA | NA | NA | NA | #VALUE! |
| ENSMUSG00002076272.1 | 0 | NA | NA | NA | NA | NA | #VALUE! |
| ENSMUSG00002076273.1 | 0 | NA | NA | NA | NA | NA | #VALUE! |
| ENSMUSG00002076274.1 | 0 | NA | NA | NA | NA | NA | #VALUE! |
| ENSMUSG00002076275.1 | 0 | NA | NA | NA | NA | NA | #VALUE! |
| ENSMUSG00002076276.1 | 0 | NA | NA | NA | NA | NA | #VALUE! |
| ENSMUSG00002076277.1 | 0 | NA | NA | NA | NA | NA | #VALUE! |
| ENSMUSG00002076278.1 | 0 | NA | NA | NA | NA | NA | #VALUE! |
| ENSMUSG00002076279.1 | 0 | NA | NA | NA | NA | NA | #VALUE! |
| ENSMUSG00002076280.1 | 0 | NA | NA | NA | NA | NA | #VALUE! |
| ENSMUSG00002076281.1 | 0 | NA | NA | NA | NA | NA | #VALUE! |
| ENSMUSG00002076282.1 | 0 | NA | NA | NA | NA | NA | #VALUE! |
| ENSMUSG00002076283.1 | 0 | NA | NA | NA | NA | NA | #VALUE! |
| ENSMUSG00002076284.1 | 0 | NA | NA | NA | NA | NA | #VALUE! |
| ENSMUSG00002076285.1 | 0 | NA | NA | NA | NA | NA | #VALUE! |
| ENSMUSG00002076286.1 | 0 | NA | NA | NA | NA | NA | #VALUE! |
| ENSMUSG00002076287.1 | 0 | NA | NA | NA | NA | NA | #VALUE! |
| ENSMUSG00002076288.1 | 0 | NA | NA | NA | NA | NA | #VALUE! |
| ENSMUSG00002076289.1 | 0 | NA | NA | NA | NA | NA | #VALUE! |
| ENSMUSG00002076290.1 | 0 | NA | NA | NA | NA | NA | #VALUE! |
| ENSMUSG00002076291.1 | 0 | NA | NA | NA | NA | NA | #VALUE! |
| ENSMUSG00002076292.1 | 0 | NA | NA | NA | NA | NA | #VALUE! |
| ENSMUSG00002076293.1 | 0 | NA | NA | NA | NA | NA | #VALUE! |
| ENSMUSG00002076294.1 | 0 | NA | NA | NA | NA | NA | #VALUE! |
| ENSMUSG00002076295.1 | 0 | NA | NA | NA | NA | NA | #VALUE! |
| ENSMUSG00002076296.1 | 0 | NA | NA | NA | NA | NA | #VALUE! |
| ENSMUSG00002076297.1 | 0 | NA | NA | NA | NA | NA | #VALUE! |
| ENSMUSG00002076298.1 | 0 | NA | NA | NA | NA | NA | #VALUE! |
| ENSMUSG00002076299.1 | 0 | NA | NA | NA | NA | NA | #VALUE! |
| ENSMUSG00002076300.1 | 0 | NA | NA | NA | NA | NA | #VALUE! |
| ENSMUSG00002076301.1 | 0 | NA | NA | NA | NA | NA | #VALUE! |
| ENSMUSG00002076302.1 | 0 | NA | NA | NA | NA | NA | #VALUE! |
| ENSMUSG00002076303.1 | 0 | NA | NA | NA | NA | NA | #VALUE! |
| ENSMUSG00002076304.1 | 0 | NA | NA | NA | NA | NA | #VALUE! |
| ENSMUSG00002076305.1 | 0 | NA | NA | NA | NA | NA | #VALUE! |
| ENSMUSG00002076306.1 | 0 | NA | NA | NA | NA | NA | #VALUE! |
| ENSMUSG00002076307.1 | 0 | NA | NA | NA | NA | NA | #VALUE! |
| ENSMUSG00002076308.1 | 0 | NA | NA | NA | NA | NA | #VALUE! |
| ENSMUSG00002076309.1 | 0 | NA | NA | NA | NA | NA | #VALUE! |
| ENSMUSG00002076310.1 | 0 | NA | NA | NA | NA | NA | #VALUE! |
| ENSMUSG00002076311.1 | 0 | NA | NA | NA | NA | NA | #VALUE! |

| | | | | | | | |
|---|---|---|---|---|---|---|---|
| ENSMUSG00002076312.1 | 0 | NA | NA | NA | NA | NA | #VALUE! |
| ENSMUSG00002076313.1 | 0 | NA | NA | NA | NA | NA | #VALUE! |
| ENSMUSG00002076314.1 | 0 | NA | NA | NA | NA | NA | #VALUE! |
| ENSMUSG00002076315.1 | 0 | NA | NA | NA | NA | NA | #VALUE! |
| ENSMUSG00002076316.1 | 0 | NA | NA | NA | NA | NA | #VALUE! |
| ENSMUSG00002076317.1 | 0 | NA | NA | NA | NA | NA | #VALUE! |
| ENSMUSG00002076318.1 | 0 | NA | NA | NA | NA | NA | #VALUE! |
| ENSMUSG00002076319.1 | 0 | NA | NA | NA | NA | NA | #VALUE! |
| ENSMUSG00002076320.1 | 0 | NA | NA | NA | NA | NA | #VALUE! |
| ENSMUSG00002076321.1 | 0 | NA | NA | NA | NA | NA | #VALUE! |
| ENSMUSG00002076322.1 | 0 | NA | NA | NA | NA | NA | #VALUE! |
| ENSMUSG00002076323.1 | 0 | NA | NA | NA | NA | NA | #VALUE! |
| ENSMUSG00002076324.1 | 0 | NA | NA | NA | NA | NA | #VALUE! |
| ENSMUSG00002076325.1 | 0 | NA | NA | NA | NA | NA | #VALUE! |
| ENSMUSG00002076326.1 | 0 | NA | NA | NA | NA | NA | #VALUE! |
| ENSMUSG00002076327.1 | 0 | NA | NA | NA | NA | NA | #VALUE! |
| ENSMUSG00002076328.1 | 0 | NA | NA | NA | NA | NA | #VALUE! |
| ENSMUSG00002076329.1 | 0 | NA | NA | NA | NA | NA | #VALUE! |
| ENSMUSG00002076330.1 | 0 | NA | NA | NA | NA | NA | #VALUE! |
| ENSMUSG00002076331.1 | 0 | NA | NA | NA | NA | NA | #VALUE! |
| ENSMUSG00002076332.1 | 0 | NA | NA | NA | NA | NA | #VALUE! |
| ENSMUSG00002076333.1 | 0 | NA | NA | NA | NA | NA | #VALUE! |
| ENSMUSG00002076334.1 | 0 | NA | NA | NA | NA | NA | #VALUE! |
| ENSMUSG00002076335.1 | 0 | NA | NA | NA | NA | NA | #VALUE! |
| ENSMUSG00002076336.1 | 0 | NA | NA | NA | NA | NA | #VALUE! |
| ENSMUSG00002076337.1 | 0 | NA | NA | NA | NA | NA | #VALUE! |
| ENSMUSG00002076338.1 | 0 | NA | NA | NA | NA | NA | #VALUE! |
| ENSMUSG00002076339.1 | 0 | NA | NA | NA | NA | NA | #VALUE! |
| ENSMUSG00002076340.1 | 0 | NA | NA | NA | NA | NA | #VALUE! |
| ENSMUSG00002076341.1 | 0 | NA | NA | NA | NA | NA | #VALUE! |
| ENSMUSG00002076342.1 | 0 | NA | NA | NA | NA | NA | #VALUE! |
| ENSMUSG00002076343.1 | 0 | NA | NA | NA | NA | NA | #VALUE! |
| ENSMUSG00002076344.1 | 0 | NA | NA | NA | NA | NA | #VALUE! |
| ENSMUSG00002076345.1 | 0 | NA | NA | NA | NA | NA | #VALUE! |
| ENSMUSG00002076346.1 | 0 | NA | NA | NA | NA | NA | #VALUE! |
| ENSMUSG00002076347.1 | 0 | NA | NA | NA | NA | NA | #VALUE! |
| ENSMUSG00002076348.1 | 0 | NA | NA | NA | NA | NA | #VALUE! |
| ENSMUSG00002076349.1 | 0 | NA | NA | NA | NA | NA | #VALUE! |
| ENSMUSG00002076350.1 | 0 | NA | NA | NA | NA | NA | #VALUE! |
| ENSMUSG00002076351.1 | 0 | NA | NA | NA | NA | NA | #VALUE! |
| ENSMUSG00002076352.1 | 0 | NA | NA | NA | NA | NA | #VALUE! |
| ENSMUSG00002076353.1 | 0 | NA | NA | NA | NA | NA | #VALUE! |
| ENSMUSG00002076354.1 | 0 | NA | NA | NA | NA | NA | #VALUE! |
| ENSMUSG00002076355.1 | 0 | NA | NA | NA | NA | NA | #VALUE! |

| | | | | | | | |
|---|---|---|---|---|---|---|---|
| ENSMUSG00002076356.1 | 0 | NA | NA | NA | NA | NA | #VALUE! |
| ENSMUSG00002076357.1 | 0 | NA | NA | NA | NA | NA | #VALUE! |
| ENSMUSG00002076358.1 | 0 | NA | NA | NA | NA | NA | #VALUE! |
| ENSMUSG00002076359.1 | 0 | NA | NA | NA | NA | NA | #VALUE! |
| ENSMUSG00002076360.1 | 0 | NA | NA | NA | NA | NA | #VALUE! |
| ENSMUSG00002076361.1 | 0 | NA | NA | NA | NA | NA | #VALUE! |
| ENSMUSG00002076362.1 | 0 | NA | NA | NA | NA | NA | #VALUE! |
| ENSMUSG00002076363.1 | 0 | NA | NA | NA | NA | NA | #VALUE! |
| ENSMUSG00002076364.1 | 0 | NA | NA | NA | NA | NA | #VALUE! |
| ENSMUSG00002076365.1 | 0 | NA | NA | NA | NA | NA | #VALUE! |
| ENSMUSG00002076366.1 | 0 | NA | NA | NA | NA | NA | #VALUE! |
| ENSMUSG00002076367.1 | 0 | NA | NA | NA | NA | NA | #VALUE! |
| ENSMUSG00002076368.1 | 0 | NA | NA | NA | NA | NA | #VALUE! |
| ENSMUSG00002076369.1 | 0 | NA | NA | NA | NA | NA | #VALUE! |
| ENSMUSG00002076370.1 | 0 | NA | NA | NA | NA | NA | #VALUE! |
| ENSMUSG00002076371.1 | 0 | NA | NA | NA | NA | NA | #VALUE! |
| ENSMUSG00002076372.1 | 0 | NA | NA | NA | NA | NA | #VALUE! |
| ENSMUSG00002076373.1 | 0 | NA | NA | NA | NA | NA | #VALUE! |
| ENSMUSG00002076374.1 | 0 | NA | NA | NA | NA | NA | #VALUE! |
| ENSMUSG00002076375.1 | 0 | NA | NA | NA | NA | NA | #VALUE! |
| ENSMUSG00002076376.1 | 0 | NA | NA | NA | NA | NA | #VALUE! |
| ENSMUSG00002076377.1 | 0 | NA | NA | NA | NA | NA | #VALUE! |
| ENSMUSG00002076378.1 | 0 | NA | NA | NA | NA | NA | #VALUE! |
| ENSMUSG00002076379.1 | 0 | NA | NA | NA | NA | NA | #VALUE! |
| ENSMUSG00002076380.1 | 0 | NA | NA | NA | NA | NA | #VALUE! |
| ENSMUSG00002076381.1 | 0 | NA | NA | NA | NA | NA | #VALUE! |
| ENSMUSG00002076382.1 | 0 | NA | NA | NA | NA | NA | #VALUE! |
| ENSMUSG00002076383.1 | 0 | NA | NA | NA | NA | NA | #VALUE! |
| ENSMUSG00002076384.1 | 0 | NA | NA | NA | NA | NA | #VALUE! |
| ENSMUSG00002076385.1 | 0 | NA | NA | NA | NA | NA | #VALUE! |
| ENSMUSG00002076386.1 | 0 | NA | NA | NA | NA | NA | #VALUE! |
| ENSMUSG00002076387.1 | 0 | NA | NA | NA | NA | NA | #VALUE! |
| ENSMUSG00002076388.1 | 0 | NA | NA | NA | NA | NA | #VALUE! |
| ENSMUSG00002076389.1 | 0 | NA | NA | NA | NA | NA | #VALUE! |
| ENSMUSG00002076390.1 | 0 | NA | NA | NA | NA | NA | #VALUE! |
| ENSMUSG00002076391.1 | 0 | NA | NA | NA | NA | NA | #VALUE! |
| ENSMUSG00002076392.1 | 0 | NA | NA | NA | NA | NA | #VALUE! |
| ENSMUSG00002076393.1 | 0 | NA | NA | NA | NA | NA | #VALUE! |
| ENSMUSG00002076394.1 | 0 | NA | NA | NA | NA | NA | #VALUE! |
| ENSMUSG00002076395.1 | 0 | NA | NA | NA | NA | NA | #VALUE! |
| ENSMUSG00002076396.1 | 0 | NA | NA | NA | NA | NA | #VALUE! |
| ENSMUSG00002076397.1 | 0 | NA | NA | NA | NA | NA | #VALUE! |
| ENSMUSG00002076398.1 | 0 | NA | NA | NA | NA | NA | #VALUE! |
| ENSMUSG00002076399.1 | 0 | NA | NA | NA | NA | NA | #VALUE! |

| | | | | | | | |
|---|---|---|---|---|---|---|---|
| ENSMUSG00002076400.1 | 0 | NA | NA | NA | NA | NA | #VALUE! |
| ENSMUSG00002076401.1 | 0 | NA | NA | NA | NA | NA | #VALUE! |
| ENSMUSG00002076402.1 | 0 | NA | NA | NA | NA | NA | #VALUE! |
| ENSMUSG00002076403.1 | 0 | NA | NA | NA | NA | NA | #VALUE! |
| ENSMUSG00002076404.1 | 0 | NA | NA | NA | NA | NA | #VALUE! |
| ENSMUSG00002076405.1 | 0 | NA | NA | NA | NA | NA | #VALUE! |
| ENSMUSG00002076406.1 | 0 | NA | NA | NA | NA | NA | #VALUE! |
| ENSMUSG00002076407.1 | 0 | NA | NA | NA | NA | NA | #VALUE! |
| ENSMUSG00002076408.1 | 0 | NA | NA | NA | NA | NA | #VALUE! |
| ENSMUSG00002076409.1 | 0 | NA | NA | NA | NA | NA | #VALUE! |
| ENSMUSG00002076410.1 | 0 | NA | NA | NA | NA | NA | #VALUE! |
| ENSMUSG00002076411.1 | 0 | NA | NA | NA | NA | NA | #VALUE! |
| ENSMUSG00002076412.1 | 0 | NA | NA | NA | NA | NA | #VALUE! |
| ENSMUSG00002076413.1 | 0 | NA | NA | NA | NA | NA | #VALUE! |
| ENSMUSG00002076414.1 | 0 | NA | NA | NA | NA | NA | #VALUE! |
| ENSMUSG00002076415.1 | 0 | NA | NA | NA | NA | NA | #VALUE! |
| ENSMUSG00002076416.1 | 0 | NA | NA | NA | NA | NA | #VALUE! |
| ENSMUSG00002076417.1 | 0 | NA | NA | NA | NA | NA | #VALUE! |
| ENSMUSG00002076418.1 | 0 | NA | NA | NA | NA | NA | #VALUE! |
| ENSMUSG00002076419.1 | 0 | NA | NA | NA | NA | NA | #VALUE! |
| ENSMUSG00002076420.1 | 0 | NA | NA | NA | NA | NA | #VALUE! |
| ENSMUSG00002076421.1 | 0 | NA | NA | NA | NA | NA | #VALUE! |
| ENSMUSG00002076422.1 | 0 | NA | NA | NA | NA | NA | #VALUE! |
| ENSMUSG00002076423.1 | 0 | NA | NA | NA | NA | NA | #VALUE! |
| ENSMUSG00002076424.1 | 0 | NA | NA | NA | NA | NA | #VALUE! |
| ENSMUSG00002076425.1 | 0 | NA | NA | NA | NA | NA | #VALUE! |
| ENSMUSG00002076426.1 | 0 | NA | NA | NA | NA | NA | #VALUE! |
| ENSMUSG00002076427.1 | 0 | NA | NA | NA | NA | NA | #VALUE! |
| ENSMUSG00002076428.1 | 0 | NA | NA | NA | NA | NA | #VALUE! |
| ENSMUSG00002076429.1 | 0 | NA | NA | NA | NA | NA | #VALUE! |
| ENSMUSG00002076430.1 | 0 | NA | NA | NA | NA | NA | #VALUE! |
| ENSMUSG00002076431.1 | 0 | NA | NA | NA | NA | NA | #VALUE! |
| ENSMUSG00002076432.1 | 0 | NA | NA | NA | NA | NA | #VALUE! |
| ENSMUSG00002076433.1 | 0 | NA | NA | NA | NA | NA | #VALUE! |
| ENSMUSG00002076434.1 | 0 | NA | NA | NA | NA | NA | #VALUE! |
| ENSMUSG00002076435.1 | 0 | NA | NA | NA | NA | NA | #VALUE! |
| ENSMUSG00002076436.1 | 0 | NA | NA | NA | NA | NA | #VALUE! |
| ENSMUSG00002076437.1 | 0 | NA | NA | NA | NA | NA | #VALUE! |
| ENSMUSG00002076438.1 | 0 | NA | NA | NA | NA | NA | #VALUE! |
| ENSMUSG00002076439.1 | 0 | NA | NA | NA | NA | NA | #VALUE! |
| ENSMUSG00002076440.1 | 0 | NA | NA | NA | NA | NA | #VALUE! |
| ENSMUSG00002076441.1 | 0 | NA | NA | NA | NA | NA | #VALUE! |
| ENSMUSG00002076442.1 | 0 | NA | NA | NA | NA | NA | #VALUE! |
| ENSMUSG00002076443.1 | 0 | NA | NA | NA | NA | NA | #VALUE! |

| | | | | | | | |
|---|---|---|---|---|---|---|---|
| ENSMUSG00002076444.1 | 0 | NA | NA | NA | NA | NA | #VALUE! |
| ENSMUSG00002076445.1 | 0 | NA | NA | NA | NA | NA | #VALUE! |
| ENSMUSG00002076446.1 | 0 | NA | NA | NA | NA | NA | #VALUE! |
| ENSMUSG00002076447.1 | 0 | NA | NA | NA | NA | NA | #VALUE! |
| ENSMUSG00002076448.1 | 0 | NA | NA | NA | NA | NA | #VALUE! |
| ENSMUSG00002076449.1 | 0 | NA | NA | NA | NA | NA | #VALUE! |
| ENSMUSG00002076450.1 | 0 | NA | NA | NA | NA | NA | #VALUE! |
| ENSMUSG00002076451.1 | 0 | NA | NA | NA | NA | NA | #VALUE! |
| ENSMUSG00002076452.1 | 0 | NA | NA | NA | NA | NA | #VALUE! |
| ENSMUSG00002076453.1 | 0 | NA | NA | NA | NA | NA | #VALUE! |
| ENSMUSG00002076454.1 | 0 | NA | NA | NA | NA | NA | #VALUE! |
| ENSMUSG00002076455.1 | 0 | NA | NA | NA | NA | NA | #VALUE! |
| ENSMUSG00002076456.1 | 0 | NA | NA | NA | NA | NA | #VALUE! |
| ENSMUSG00002076457.1 | 0 | NA | NA | NA | NA | NA | #VALUE! |
| ENSMUSG00002076458.1 | 0 | NA | NA | NA | NA | NA | #VALUE! |
| ENSMUSG00002076459.1 | 0 | NA | NA | NA | NA | NA | #VALUE! |
| ENSMUSG00002076460.1 | 0 | NA | NA | NA | NA | NA | #VALUE! |
| ENSMUSG00002076461.1 | 0 | NA | NA | NA | NA | NA | #VALUE! |
| ENSMUSG00002076462.1 | 0 | NA | NA | NA | NA | NA | #VALUE! |
| ENSMUSG00002076463.1 | 0 | NA | NA | NA | NA | NA | #VALUE! |
| ENSMUSG00002076464.1 | 0 | NA | NA | NA | NA | NA | #VALUE! |
| ENSMUSG00002076465.1 | 0 | NA | NA | NA | NA | NA | #VALUE! |
| ENSMUSG00002076466.1 | 0 | NA | NA | NA | NA | NA | #VALUE! |
| ENSMUSG00002076467.1 | 0 | NA | NA | NA | NA | NA | #VALUE! |
| ENSMUSG00002076468.1 | 0 | NA | NA | NA | NA | NA | #VALUE! |
| ENSMUSG00002076469.1 | 0 | NA | NA | NA | NA | NA | #VALUE! |
| ENSMUSG00002076470.1 | 0 | NA | NA | NA | NA | NA | #VALUE! |
| ENSMUSG00002076471.1 | 0 | NA | NA | NA | NA | NA | #VALUE! |
| ENSMUSG00002076472.1 | 0 | NA | NA | NA | NA | NA | #VALUE! |
| ENSMUSG00002076473.1 | 0 | NA | NA | NA | NA | NA | #VALUE! |
| ENSMUSG00002076474.1 | 0 | NA | NA | NA | NA | NA | #VALUE! |
| ENSMUSG00002076475.1 | 0 | NA | NA | NA | NA | NA | #VALUE! |
| ENSMUSG00002076476.1 | 0 | NA | NA | NA | NA | NA | #VALUE! |
| ENSMUSG00002076477.1 | 0 | NA | NA | NA | NA | NA | #VALUE! |
| ENSMUSG00002076478.1 | 0 | NA | NA | NA | NA | NA | #VALUE! |
| ENSMUSG00002076479.1 | 0 | NA | NA | NA | NA | NA | #VALUE! |
| ENSMUSG00002076480.1 | 0 | NA | NA | NA | NA | NA | #VALUE! |
| ENSMUSG00002076481.1 | 0 | NA | NA | NA | NA | NA | #VALUE! |
| ENSMUSG00002076482.1 | 0 | NA | NA | NA | NA | NA | #VALUE! |
| ENSMUSG00002076483.1 | 0 | NA | NA | NA | NA | NA | #VALUE! |
| ENSMUSG00002076484.1 | 0 | NA | NA | NA | NA | NA | #VALUE! |
| ENSMUSG00002076485.1 | 0 | NA | NA | NA | NA | NA | #VALUE! |
| ENSMUSG00002076486.1 | 0 | NA | NA | NA | NA | NA | #VALUE! |
| ENSMUSG00002076487.1 | 0 | NA | NA | NA | NA | NA | #VALUE! |

| | | | | | | | |
|---|---|---|---|---|---|---|---|
| ENSMUSG00002076488.1 | 0 | NA | NA | NA | NA | NA | #VALUE! |
| ENSMUSG00002076489.1 | 0 | NA | NA | NA | NA | NA | #VALUE! |
| ENSMUSG00002076490.1 | 0 | NA | NA | NA | NA | NA | #VALUE! |
| ENSMUSG00002076491.1 | 0 | NA | NA | NA | NA | NA | #VALUE! |
| ENSMUSG00002076492.1 | 0 | NA | NA | NA | NA | NA | #VALUE! |
| ENSMUSG00002076493.1 | 0 | NA | NA | NA | NA | NA | #VALUE! |
| ENSMUSG00002076494.1 | 0 | NA | NA | NA | NA | NA | #VALUE! |
| ENSMUSG00002076495.1 | 0 | NA | NA | NA | NA | NA | #VALUE! |
| ENSMUSG00002076496.1 | 0 | NA | NA | NA | NA | NA | #VALUE! |
| ENSMUSG00002076497.1 | 0 | NA | NA | NA | NA | NA | #VALUE! |
| ENSMUSG00002076498.1 | 0 | NA | NA | NA | NA | NA | #VALUE! |
| ENSMUSG00002076499.1 | 0 | NA | NA | NA | NA | NA | #VALUE! |
| ENSMUSG00002076500.1 | 0 | NA | NA | NA | NA | NA | #VALUE! |
| ENSMUSG00002076501.1 | 0 | NA | NA | NA | NA | NA | #VALUE! |
| ENSMUSG00002076502.1 | 0 | NA | NA | NA | NA | NA | #VALUE! |
| ENSMUSG00002076503.1 | 0 | NA | NA | NA | NA | NA | #VALUE! |
| ENSMUSG00002076504.1 | 0 | NA | NA | NA | NA | NA | #VALUE! |
| ENSMUSG00002076505.1 | 0 | NA | NA | NA | NA | NA | #VALUE! |
| ENSMUSG00002076506.1 | 0 | NA | NA | NA | NA | NA | #VALUE! |
| ENSMUSG00002076507.1 | 0 | NA | NA | NA | NA | NA | #VALUE! |
| ENSMUSG00002076508.1 | 0 | NA | NA | NA | NA | NA | #VALUE! |
| ENSMUSG00002076509.1 | 0 | NA | NA | NA | NA | NA | #VALUE! |
| ENSMUSG00002076510.1 | 0 | NA | NA | NA | NA | NA | #VALUE! |
| ENSMUSG00002076511.1 | 0 | NA | NA | NA | NA | NA | #VALUE! |
| ENSMUSG00002076512.1 | 0 | NA | NA | NA | NA | NA | #VALUE! |
| ENSMUSG00002076513.1 | 0 | NA | NA | NA | NA | NA | #VALUE! |
| ENSMUSG00002076514.1 | 0 | NA | NA | NA | NA | NA | #VALUE! |
| ENSMUSG00002076515.1 | 0 | NA | NA | NA | NA | NA | #VALUE! |
| ENSMUSG00002076516.1 | 0 | NA | NA | NA | NA | NA | #VALUE! |
| ENSMUSG00002076517.1 | 0 | NA | NA | NA | NA | NA | #VALUE! |
| ENSMUSG00002076518.1 | 0 | NA | NA | NA | NA | NA | #VALUE! |
| ENSMUSG00002076519.1 | 0 | NA | NA | NA | NA | NA | #VALUE! |
| ENSMUSG00002076520.1 | 0 | NA | NA | NA | NA | NA | #VALUE! |
| ENSMUSG00002076521.1 | 0 | NA | NA | NA | NA | NA | #VALUE! |
| ENSMUSG00002076522.1 | 0 | NA | NA | NA | NA | NA | #VALUE! |
| ENSMUSG00002076523.1 | 0 | NA | NA | NA | NA | NA | #VALUE! |
| ENSMUSG00002076524.1 | 0 | NA | NA | NA | NA | NA | #VALUE! |
| ENSMUSG00002076525.1 | 0 | NA | NA | NA | NA | NA | #VALUE! |
| ENSMUSG00002076526.1 | 0 | NA | NA | NA | NA | NA | #VALUE! |
| ENSMUSG00002076527.1 | 0 | NA | NA | NA | NA | NA | #VALUE! |
| ENSMUSG00002076528.1 | 0 | NA | NA | NA | NA | NA | #VALUE! |
| ENSMUSG00002076529.1 | 0 | NA | NA | NA | NA | NA | #VALUE! |
| ENSMUSG00002076530.1 | 0 | NA | NA | NA | NA | NA | #VALUE! |
| ENSMUSG00002076531.1 | 0 | NA | NA | NA | NA | NA | #VALUE! |

| | | | | | | | |
|---|---|---|---|---|---|---|---|
| ENSMUSG00002076532.1 | 0 | NA | NA | NA | NA | NA | #VALUE! |
| ENSMUSG00002076533.1 | 0 | NA | NA | NA | NA | NA | #VALUE! |
| ENSMUSG00002076534.1 | 0 | NA | NA | NA | NA | NA | #VALUE! |
| ENSMUSG00002076535.1 | 0 | NA | NA | NA | NA | NA | #VALUE! |
| ENSMUSG00002076536.1 | 0 | NA | NA | NA | NA | NA | #VALUE! |
| ENSMUSG00002076537.1 | 0 | NA | NA | NA | NA | NA | #VALUE! |
| ENSMUSG00002076538.1 | 0 | NA | NA | NA | NA | NA | #VALUE! |
| ENSMUSG00002076539.1 | 0 | NA | NA | NA | NA | NA | #VALUE! |
| ENSMUSG00002076540.1 | 0 | NA | NA | NA | NA | NA | #VALUE! |
| ENSMUSG00002076541.1 | 0 | NA | NA | NA | NA | NA | #VALUE! |
| ENSMUSG00002076542.1 | 0 | NA | NA | NA | NA | NA | #VALUE! |
| ENSMUSG00002076543.1 | 0 | NA | NA | NA | NA | NA | #VALUE! |
| ENSMUSG00002076544.1 | 0 | NA | NA | NA | NA | NA | #VALUE! |
| ENSMUSG00002076545.1 | 0 | NA | NA | NA | NA | NA | #VALUE! |
| ENSMUSG00002076546.1 | 0 | NA | NA | NA | NA | NA | #VALUE! |
| ENSMUSG00002076547.1 | 0 | NA | NA | NA | NA | NA | #VALUE! |
| ENSMUSG00002076548.1 | 0 | NA | NA | NA | NA | NA | #VALUE! |
| ENSMUSG00002076549.1 | 0 | NA | NA | NA | NA | NA | #VALUE! |
| ENSMUSG00002076550.1 | 0 | NA | NA | NA | NA | NA | #VALUE! |
| ENSMUSG00002076551.1 | 0 | NA | NA | NA | NA | NA | #VALUE! |
| ENSMUSG00002076552.1 | 0 | NA | NA | NA | NA | NA | #VALUE! |
| ENSMUSG00002076553.1 | 0 | NA | NA | NA | NA | NA | #VALUE! |
| ENSMUSG00002076554.1 | 0 | NA | NA | NA | NA | NA | #VALUE! |
| ENSMUSG00002076555.1 | 0 | NA | NA | NA | NA | NA | #VALUE! |
| ENSMUSG00002076556.1 | 0 | NA | NA | NA | NA | NA | #VALUE! |
| ENSMUSG00002076557.1 | 0 | NA | NA | NA | NA | NA | #VALUE! |
| ENSMUSG00002076558.1 | 0 | NA | NA | NA | NA | NA | #VALUE! |
| ENSMUSG00002076559.1 | 0 | NA | NA | NA | NA | NA | #VALUE! |
| ENSMUSG00002076560.1 | 0 | NA | NA | NA | NA | NA | #VALUE! |
| ENSMUSG00002076561.1 | 0 | NA | NA | NA | NA | NA | #VALUE! |
| ENSMUSG00002076562.1 | 0 | NA | NA | NA | NA | NA | #VALUE! |
| ENSMUSG00002076563.1 | 0 | NA | NA | NA | NA | NA | #VALUE! |
| ENSMUSG00002076564.1 | 0 | NA | NA | NA | NA | NA | #VALUE! |
| ENSMUSG00002076565.1 | 0 | NA | NA | NA | NA | NA | #VALUE! |
| ENSMUSG00002076566.1 | 0 | NA | NA | NA | NA | NA | #VALUE! |
| ENSMUSG00002076567.1 | 0 | NA | NA | NA | NA | NA | #VALUE! |
| ENSMUSG00002076568.1 | 0 | NA | NA | NA | NA | NA | #VALUE! |
| ENSMUSG00002076569.1 | 0 | NA | NA | NA | NA | NA | #VALUE! |
| ENSMUSG00002076570.1 | 0 | NA | NA | NA | NA | NA | #VALUE! |
| ENSMUSG00002076571.1 | 0 | NA | NA | NA | NA | NA | #VALUE! |
| ENSMUSG00002076572.1 | 0 | NA | NA | NA | NA | NA | #VALUE! |
| ENSMUSG00002076573.1 | 0 | NA | NA | NA | NA | NA | #VALUE! |
| ENSMUSG00002076574.1 | 0 | NA | NA | NA | NA | NA | #VALUE! |
| ENSMUSG00002076575.1 | 0 | NA | NA | NA | NA | NA | #VALUE! |

| | | | | | | | |
|---|---|---|---|---|---|---|---|
| ENSMUSG00002076576.1 | 0 | NA | NA | NA | NA | NA | #VALUE! |
| ENSMUSG00002076577.1 | 0 | NA | NA | NA | NA | NA | #VALUE! |
| ENSMUSG00002076578.1 | 0 | NA | NA | NA | NA | NA | #VALUE! |
| ENSMUSG00002076579.1 | 0 | NA | NA | NA | NA | NA | #VALUE! |
| ENSMUSG00002076580.1 | 0 | NA | NA | NA | NA | NA | #VALUE! |
| ENSMUSG00002076581.1 | 0 | NA | NA | NA | NA | NA | #VALUE! |
| ENSMUSG00002076582.1 | 0 | NA | NA | NA | NA | NA | #VALUE! |
| ENSMUSG00002076583.1 | 0 | NA | NA | NA | NA | NA | #VALUE! |
| ENSMUSG00002076584.1 | 0 | NA | NA | NA | NA | NA | #VALUE! |
| ENSMUSG00002076585.1 | 0 | NA | NA | NA | NA | NA | #VALUE! |
| ENSMUSG00002076586.1 | 0 | NA | NA | NA | NA | NA | #VALUE! |
| ENSMUSG00002076587.1 | 0 | NA | NA | NA | NA | NA | #VALUE! |
| ENSMUSG00002076588.1 | 0 | NA | NA | NA | NA | NA | #VALUE! |
| ENSMUSG00002076589.1 | 0 | NA | NA | NA | NA | NA | #VALUE! |
| ENSMUSG00002076590.1 | 0 | NA | NA | NA | NA | NA | #VALUE! |
| ENSMUSG00002076591.1 | 0 | NA | NA | NA | NA | NA | #VALUE! |
| ENSMUSG00002076592.1 | 0 | NA | NA | NA | NA | NA | #VALUE! |
| ENSMUSG00002076593.1 | 0 | NA | NA | NA | NA | NA | #VALUE! |
| ENSMUSG00002076594.1 | 0 | NA | NA | NA | NA | NA | #VALUE! |
| ENSMUSG00002076595.1 | 0 | NA | NA | NA | NA | NA | #VALUE! |
| ENSMUSG00002076596.1 | 0 | NA | NA | NA | NA | NA | #VALUE! |
| ENSMUSG00002076597.1 | 0 | NA | NA | NA | NA | NA | #VALUE! |
| ENSMUSG00002076598.1 | 0 | NA | NA | NA | NA | NA | #VALUE! |
| ENSMUSG00002076599.1 | 0 | NA | NA | NA | NA | NA | #VALUE! |
| ENSMUSG00002076600.1 | 0 | NA | NA | NA | NA | NA | #VALUE! |
| ENSMUSG00002076601.1 | 0 | NA | NA | NA | NA | NA | #VALUE! |
| ENSMUSG00002076602.1 | 0 | NA | NA | NA | NA | NA | #VALUE! |
| ENSMUSG00002076603.1 | 0 | NA | NA | NA | NA | NA | #VALUE! |
| ENSMUSG00002076604.1 | 0 | NA | NA | NA | NA | NA | #VALUE! |
| ENSMUSG00002076605.1 | 0 | NA | NA | NA | NA | NA | #VALUE! |
| ENSMUSG00002076606.1 | 0 | NA | NA | NA | NA | NA | #VALUE! |
| ENSMUSG00002076607.1 | 0 | NA | NA | NA | NA | NA | #VALUE! |
| ENSMUSG00002076608.1 | 0 | NA | NA | NA | NA | NA | #VALUE! |
| ENSMUSG00002076609.1 | 0 | NA | NA | NA | NA | NA | #VALUE! |
| ENSMUSG00002076610.1 | 0 | NA | NA | NA | NA | NA | #VALUE! |
| ENSMUSG00002076611.1 | 0 | NA | NA | NA | NA | NA | #VALUE! |
| ENSMUSG00002076612.1 | 0 | NA | NA | NA | NA | NA | #VALUE! |
| ENSMUSG00002076613.1 | 0 | NA | NA | NA | NA | NA | #VALUE! |
| ENSMUSG00002076614.1 | 0 | NA | NA | NA | NA | NA | #VALUE! |
| ENSMUSG00002076615.1 | 0 | NA | NA | NA | NA | NA | #VALUE! |
| ENSMUSG00002076616.1 | 0 | NA | NA | NA | NA | NA | #VALUE! |
| ENSMUSG00002076617.1 | 0 | NA | NA | NA | NA | NA | #VALUE! |
| ENSMUSG00002076618.1 | 0 | NA | NA | NA | NA | NA | #VALUE! |
| ENSMUSG00002076619.1 | 0 | NA | NA | NA | NA | NA | #VALUE! |

| | | | | | | | |
|---|---|---|---|---|---|---|---|
| ENSMUSG00002076620.1 | 0 | NA | NA | NA | NA | NA | #VALUE! |
| ENSMUSG00002076621.1 | 0 | NA | NA | NA | NA | NA | #VALUE! |
| ENSMUSG00002076622.1 | 0 | NA | NA | NA | NA | NA | #VALUE! |
| ENSMUSG00002076623.1 | 0 | NA | NA | NA | NA | NA | #VALUE! |
| ENSMUSG00002076624.1 | 0 | NA | NA | NA | NA | NA | #VALUE! |
| ENSMUSG00002076625.1 | 0 | NA | NA | NA | NA | NA | #VALUE! |
| ENSMUSG00002076626.1 | 0 | NA | NA | NA | NA | NA | #VALUE! |
| ENSMUSG00002076627.1 | 0 | NA | NA | NA | NA | NA | #VALUE! |
| ENSMUSG00002076628.1 | 0 | NA | NA | NA | NA | NA | #VALUE! |
| ENSMUSG00002076629.1 | 0 | NA | NA | NA | NA | NA | #VALUE! |
| ENSMUSG00002076630.1 | 0 | NA | NA | NA | NA | NA | #VALUE! |
| ENSMUSG00002076631.1 | 0 | NA | NA | NA | NA | NA | #VALUE! |
| ENSMUSG00002076632.1 | 0 | NA | NA | NA | NA | NA | #VALUE! |
| ENSMUSG00002076633.1 | 0 | NA | NA | NA | NA | NA | #VALUE! |
| ENSMUSG00002076634.1 | 0 | NA | NA | NA | NA | NA | #VALUE! |
| ENSMUSG00002076635.1 | 0 | NA | NA | NA | NA | NA | #VALUE! |
| ENSMUSG00002076636.1 | 0 | NA | NA | NA | NA | NA | #VALUE! |
| ENSMUSG00002076637.1 | 0 | NA | NA | NA | NA | NA | #VALUE! |
| ENSMUSG00002076638.1 | 0 | NA | NA | NA | NA | NA | #VALUE! |
| ENSMUSG00002076639.1 | 0 | NA | NA | NA | NA | NA | #VALUE! |
| ENSMUSG00002076640.1 | 0 | NA | NA | NA | NA | NA | #VALUE! |
| ENSMUSG00002076641.1 | 0 | NA | NA | NA | NA | NA | #VALUE! |
| ENSMUSG00002076642.1 | 0 | NA | NA | NA | NA | NA | #VALUE! |
| ENSMUSG00002076643.1 | 0 | NA | NA | NA | NA | NA | #VALUE! |
| ENSMUSG00002076644.1 | 0 | NA | NA | NA | NA | NA | #VALUE! |
| ENSMUSG00002076645.1 | 0 | NA | NA | NA | NA | NA | #VALUE! |
| ENSMUSG00002076646.1 | 0 | NA | NA | NA | NA | NA | #VALUE! |
| ENSMUSG00002076647.1 | 0 | NA | NA | NA | NA | NA | #VALUE! |
| ENSMUSG00002076648.1 | 0 | NA | NA | NA | NA | NA | #VALUE! |
| ENSMUSG00002076649.1 | 0 | NA | NA | NA | NA | NA | #VALUE! |
| ENSMUSG00002076650.1 | 0 | NA | NA | NA | NA | NA | #VALUE! |
| ENSMUSG00002076651.1 | 0 | NA | NA | NA | NA | NA | #VALUE! |
| ENSMUSG00002076652.1 | 0 | NA | NA | NA | NA | NA | #VALUE! |
| ENSMUSG00002076653.1 | 0 | NA | NA | NA | NA | NA | #VALUE! |
| ENSMUSG00002076654.1 | 0 | NA | NA | NA | NA | NA | #VALUE! |
| ENSMUSG00002076655.1 | 0 | NA | NA | NA | NA | NA | #VALUE! |
| ENSMUSG00002076656.1 | 0 | NA | NA | NA | NA | NA | #VALUE! |
| ENSMUSG00002076657.1 | 0 | NA | NA | NA | NA | NA | #VALUE! |
| ENSMUSG00002076658.1 | 0 | NA | NA | NA | NA | NA | #VALUE! |
| ENSMUSG00002076659.1 | 0 | NA | NA | NA | NA | NA | #VALUE! |
| ENSMUSG00002076660.1 | 0 | NA | NA | NA | NA | NA | #VALUE! |
| ENSMUSG00002076661.1 | 0 | NA | NA | NA | NA | NA | #VALUE! |
| ENSMUSG00002076662.1 | 0 | NA | NA | NA | NA | NA | #VALUE! |
| ENSMUSG00002076663.1 | 0 | NA | NA | NA | NA | NA | #VALUE! |

| | | | | | | | |
|---|---|---|---|---|---|---|---|
| ENSMUSG00002076664.1 | 0 | NA | NA | NA | NA | NA | #VALUE! |
| ENSMUSG00002076665.1 | 0 | NA | NA | NA | NA | NA | #VALUE! |
| ENSMUSG00002076666.1 | 0 | NA | NA | NA | NA | NA | #VALUE! |
| ENSMUSG00002076667.1 | 0 | NA | NA | NA | NA | NA | #VALUE! |
| ENSMUSG00002076668.1 | 0 | NA | NA | NA | NA | NA | #VALUE! |
| ENSMUSG00002076669.1 | 0 | NA | NA | NA | NA | NA | #VALUE! |
| ENSMUSG00002076670.1 | 0 | NA | NA | NA | NA | NA | #VALUE! |
| ENSMUSG00002076671.1 | 0 | NA | NA | NA | NA | NA | #VALUE! |
| ENSMUSG00002076672.1 | 0 | NA | NA | NA | NA | NA | #VALUE! |
| ENSMUSG00002076673.1 | 0 | NA | NA | NA | NA | NA | #VALUE! |
| ENSMUSG00002076674.1 | 0 | NA | NA | NA | NA | NA | #VALUE! |
| ENSMUSG00002076675.1 | 0 | NA | NA | NA | NA | NA | #VALUE! |
| ENSMUSG00002076676.1 | 0 | NA | NA | NA | NA | NA | #VALUE! |
| ENSMUSG00002076677.1 | 0 | NA | NA | NA | NA | NA | #VALUE! |
| ENSMUSG00002076678.1 | 0 | NA | NA | NA | NA | NA | #VALUE! |
| ENSMUSG00002076679.1 | 0 | NA | NA | NA | NA | NA | #VALUE! |
| ENSMUSG00002076680.1 | 0 | NA | NA | NA | NA | NA | #VALUE! |
| ENSMUSG00002076681.1 | 0 | NA | NA | NA | NA | NA | #VALUE! |
| ENSMUSG00002076682.1 | 0 | NA | NA | NA | NA | NA | #VALUE! |
| ENSMUSG00002076683.1 | 0 | NA | NA | NA | NA | NA | #VALUE! |
| ENSMUSG00002076684.1 | 0 | NA | NA | NA | NA | NA | #VALUE! |
| ENSMUSG00002076685.1 | 0 | NA | NA | NA | NA | NA | #VALUE! |
| ENSMUSG00002076686.1 | 0 | NA | NA | NA | NA | NA | #VALUE! |
| ENSMUSG00002076687.1 | 0 | NA | NA | NA | NA | NA | #VALUE! |
| ENSMUSG00002076688.1 | 0 | NA | NA | NA | NA | NA | #VALUE! |
| ENSMUSG00002076689.1 | 0 | NA | NA | NA | NA | NA | #VALUE! |
| ENSMUSG00002076690.1 | 0 | NA | NA | NA | NA | NA | #VALUE! |
| ENSMUSG00002076691.1 | 0 | NA | NA | NA | NA | NA | #VALUE! |
| ENSMUSG00002076692.1 | 0 | NA | NA | NA | NA | NA | #VALUE! |
| ENSMUSG00002076693.1 | 0 | NA | NA | NA | NA | NA | #VALUE! |
| ENSMUSG00002076694.1 | 0 | NA | NA | NA | NA | NA | #VALUE! |
| ENSMUSG00002076695.1 | 0 | NA | NA | NA | NA | NA | #VALUE! |
| ENSMUSG00002076696.1 | 0 | NA | NA | NA | NA | NA | #VALUE! |
| ENSMUSG00002076697.1 | 0 | NA | NA | NA | NA | NA | #VALUE! |
| ENSMUSG00002076698.1 | 0 | NA | NA | NA | NA | NA | #VALUE! |
| ENSMUSG00002076699.1 | 0 | NA | NA | NA | NA | NA | #VALUE! |
| ENSMUSG00002076700.1 | 0 | NA | NA | NA | NA | NA | #VALUE! |
| ENSMUSG00002076701.1 | 0 | NA | NA | NA | NA | NA | #VALUE! |
| ENSMUSG00002076702.1 | 0 | NA | NA | NA | NA | NA | #VALUE! |
| ENSMUSG00002076703.1 | 0 | NA | NA | NA | NA | NA | #VALUE! |
| ENSMUSG00002076704.1 | 0 | NA | NA | NA | NA | NA | #VALUE! |
| ENSMUSG00002076705.1 | 0 | NA | NA | NA | NA | NA | #VALUE! |
| ENSMUSG00002076706.1 | 0 | NA | NA | NA | NA | NA | #VALUE! |
| ENSMUSG00002076707.1 | 0 | NA | NA | NA | NA | NA | #VALUE! |

| | | | | | | | |
|---|---|---|---|---|---|---|---|
| ENSMUSG00002076708.1 | 0 | NA | NA | NA | NA | NA | #VALUE! |
| ENSMUSG00002076709.1 | 0 | NA | NA | NA | NA | NA | #VALUE! |
| ENSMUSG00002076710.1 | 0 | NA | NA | NA | NA | NA | #VALUE! |
| ENSMUSG00002076711.1 | 0 | NA | NA | NA | NA | NA | #VALUE! |
| ENSMUSG00002076712.1 | 0 | NA | NA | NA | NA | NA | #VALUE! |
| ENSMUSG00002076713.1 | 0 | NA | NA | NA | NA | NA | #VALUE! |
| ENSMUSG00002076714.1 | 0 | NA | NA | NA | NA | NA | #VALUE! |
| ENSMUSG00002076715.1 | 0 | NA | NA | NA | NA | NA | #VALUE! |
| ENSMUSG00002076716.1 | 0 | NA | NA | NA | NA | NA | #VALUE! |
| ENSMUSG00002076717.1 | 0 | NA | NA | NA | NA | NA | #VALUE! |
| ENSMUSG00002076718.1 | 0 | NA | NA | NA | NA | NA | #VALUE! |
| ENSMUSG00002076719.1 | 0 | NA | NA | NA | NA | NA | #VALUE! |
| ENSMUSG00002076720.1 | 0 | NA | NA | NA | NA | NA | #VALUE! |
| ENSMUSG00002076721.1 | 0 | NA | NA | NA | NA | NA | #VALUE! |
| ENSMUSG00002076722.1 | 0 | NA | NA | NA | NA | NA | #VALUE! |
| ENSMUSG00002076723.1 | 0 | NA | NA | NA | NA | NA | #VALUE! |
| ENSMUSG00002076724.1 | 0 | NA | NA | NA | NA | NA | #VALUE! |
| ENSMUSG00002076725.1 | 0 | NA | NA | NA | NA | NA | #VALUE! |
| ENSMUSG00002076726.1 | 0 | NA | NA | NA | NA | NA | #VALUE! |
| ENSMUSG00002076727.1 | 0 | NA | NA | NA | NA | NA | #VALUE! |
| ENSMUSG00002076728.1 | 0 | NA | NA | NA | NA | NA | #VALUE! |
| ENSMUSG00002076729.1 | 0 | NA | NA | NA | NA | NA | #VALUE! |
| ENSMUSG00002076730.1 | 0 | NA | NA | NA | NA | NA | #VALUE! |
| ENSMUSG00002076731.1 | 0 | NA | NA | NA | NA | NA | #VALUE! |
| ENSMUSG00002076732.1 | 0 | NA | NA | NA | NA | NA | #VALUE! |
| ENSMUSG00002076733.1 | 0 | NA | NA | NA | NA | NA | #VALUE! |
| ENSMUSG00002076734.1 | 0 | NA | NA | NA | NA | NA | #VALUE! |
| ENSMUSG00002076735.1 | 0 | NA | NA | NA | NA | NA | #VALUE! |
| ENSMUSG00002076736.1 | 0 | NA | NA | NA | NA | NA | #VALUE! |
| ENSMUSG00002076737.1 | 0 | NA | NA | NA | NA | NA | #VALUE! |
| ENSMUSG00002076738.1 | 0 | NA | NA | NA | NA | NA | #VALUE! |
| ENSMUSG00002076739.1 | 0 | NA | NA | NA | NA | NA | #VALUE! |
| ENSMUSG00002076740.1 | 0 | NA | NA | NA | NA | NA | #VALUE! |
| ENSMUSG00002076741.1 | 0 | NA | NA | NA | NA | NA | #VALUE! |
| ENSMUSG00002076742.1 | 0 | NA | NA | NA | NA | NA | #VALUE! |
| ENSMUSG00002076743.1 | 0 | NA | NA | NA | NA | NA | #VALUE! |
| ENSMUSG00002076744.1 | 0 | NA | NA | NA | NA | NA | #VALUE! |
| ENSMUSG00002076745.1 | 0 | NA | NA | NA | NA | NA | #VALUE! |
| ENSMUSG00002076746.1 | 0 | NA | NA | NA | NA | NA | #VALUE! |
| ENSMUSG00002076747.1 | 0 | NA | NA | NA | NA | NA | #VALUE! |
| ENSMUSG00002076748.1 | 0 | NA | NA | NA | NA | NA | #VALUE! |
| ENSMUSG00002076749.1 | 0 | NA | NA | NA | NA | NA | #VALUE! |
| ENSMUSG00002076750.1 | 0 | NA | NA | NA | NA | NA | #VALUE! |
| ENSMUSG00002076751.1 | 0 | NA | NA | NA | NA | NA | #VALUE! |

| | | | | | | | |
|---|---|---|---|---|---|---|---|
| ENSMUSG00002076752.1 | 0 | NA | NA | NA | NA | NA | #VALUE! |
| ENSMUSG00002076753.1 | 0 | NA | NA | NA | NA | NA | #VALUE! |
| ENSMUSG00002076754.1 | 0 | NA | NA | NA | NA | NA | #VALUE! |
| ENSMUSG00002076755.1 | 0 | NA | NA | NA | NA | NA | #VALUE! |
| ENSMUSG00002076756.1 | 0 | NA | NA | NA | NA | NA | #VALUE! |
| ENSMUSG00002076757.1 | 0 | NA | NA | NA | NA | NA | #VALUE! |
| ENSMUSG00002076758.1 | 0 | NA | NA | NA | NA | NA | #VALUE! |
| ENSMUSG00002076759.1 | 0 | NA | NA | NA | NA | NA | #VALUE! |
| ENSMUSG00002076760.1 | 0 | NA | NA | NA | NA | NA | #VALUE! |
| ENSMUSG00002076761.1 | 0 | NA | NA | NA | NA | NA | #VALUE! |
| ENSMUSG00002076762.1 | 0 | NA | NA | NA | NA | NA | #VALUE! |
| ENSMUSG00002076763.1 | 0 | NA | NA | NA | NA | NA | #VALUE! |
| ENSMUSG00002076764.1 | 0 | NA | NA | NA | NA | NA | #VALUE! |
| ENSMUSG00002076765.1 | 0 | NA | NA | NA | NA | NA | #VALUE! |
| ENSMUSG00002076766.1 | 0 | NA | NA | NA | NA | NA | #VALUE! |
| ENSMUSG00002076767.1 | 0 | NA | NA | NA | NA | NA | #VALUE! |
| ENSMUSG00002076768.1 | 0 | NA | NA | NA | NA | NA | #VALUE! |
| ENSMUSG00002076769.1 | 0 | NA | NA | NA | NA | NA | #VALUE! |
| ENSMUSG00002076770.1 | 0 | NA | NA | NA | NA | NA | #VALUE! |
| ENSMUSG00002076771.1 | 0 | NA | NA | NA | NA | NA | #VALUE! |
| ENSMUSG00002076772.1 | 0 | NA | NA | NA | NA | NA | #VALUE! |
| ENSMUSG00002076773.1 | 0 | NA | NA | NA | NA | NA | #VALUE! |
| ENSMUSG00002076774.1 | 0 | NA | NA | NA | NA | NA | #VALUE! |
| ENSMUSG00002076775.1 | 0 | NA | NA | NA | NA | NA | #VALUE! |
| ENSMUSG00002076776.1 | 0 | NA | NA | NA | NA | NA | #VALUE! |
| ENSMUSG00002076777.1 | 0 | NA | NA | NA | NA | NA | #VALUE! |
| ENSMUSG00002076778.1 | 0 | NA | NA | NA | NA | NA | #VALUE! |
| ENSMUSG00002076779.1 | 0 | NA | NA | NA | NA | NA | #VALUE! |
| ENSMUSG00002076780.1 | 0 | NA | NA | NA | NA | NA | #VALUE! |
| ENSMUSG00002076781.1 | 0 | NA | NA | NA | NA | NA | #VALUE! |
| ENSMUSG00002076782.1 | 0 | NA | NA | NA | NA | NA | #VALUE! |
| ENSMUSG00002076783.1 | 0 | NA | NA | NA | NA | NA | #VALUE! |
| ENSMUSG00002076784.1 | 0 | NA | NA | NA | NA | NA | #VALUE! |
| ENSMUSG00002076785.1 | 0 | NA | NA | NA | NA | NA | #VALUE! |
| ENSMUSG00002076786.1 | 0 | NA | NA | NA | NA | NA | #VALUE! |
| ENSMUSG00002076787.1 | 0 | NA | NA | NA | NA | NA | #VALUE! |
| ENSMUSG00002076788.1 | 0 | NA | NA | NA | NA | NA | #VALUE! |
| ENSMUSG00002076789.1 | 0 | NA | NA | NA | NA | NA | #VALUE! |
| ENSMUSG00002076790.1 | 0 | NA | NA | NA | NA | NA | #VALUE! |
| ENSMUSG00002076791.1 | 0 | NA | NA | NA | NA | NA | #VALUE! |
| ENSMUSG00002076792.1 | 0 | NA | NA | NA | NA | NA | #VALUE! |
| ENSMUSG00002076793.1 | 0 | NA | NA | NA | NA | NA | #VALUE! |
| ENSMUSG00002076794.1 | 0 | NA | NA | NA | NA | NA | #VALUE! |
| ENSMUSG00002076795.1 | 0 | NA | NA | NA | NA | NA | #VALUE! |

| | | | | | | | |
|---|---|---|---|---|---|---|---|
| ENSMUSG00002076796.1 | 0 | NA | NA | NA | NA | NA | #VALUE! |
| ENSMUSG00002076797.1 | 0 | NA | NA | NA | NA | NA | #VALUE! |
| ENSMUSG00002076798.1 | 0.204111 | -1.05581 | 4.073302 | -0.2592 | 0.795479 | NA | #VALUE! |
| ENSMUSG00002076799.1 | 0 | NA | NA | NA | NA | NA | #VALUE! |
| ENSMUSG00002076800.1 | 0 | NA | NA | NA | NA | NA | #VALUE! |
| ENSMUSG00002076801.1 | 0 | NA | NA | NA | NA | NA | #VALUE! |
| ENSMUSG00002076802.1 | 0 | NA | NA | NA | NA | NA | #VALUE! |
| ENSMUSG00002076803.1 | 0 | NA | NA | NA | NA | NA | #VALUE! |
| ENSMUSG00002076804.1 | 0 | NA | NA | NA | NA | NA | #VALUE! |
| ENSMUSG00002076805.1 | 0 | NA | NA | NA | NA | NA | #VALUE! |
| ENSMUSG00002076806.1 | 0 | NA | NA | NA | NA | NA | #VALUE! |
| ENSMUSG00002076807.1 | 0 | NA | NA | NA | NA | NA | #VALUE! |
| ENSMUSG00002076808.1 | 0 | NA | NA | NA | NA | NA | #VALUE! |
| ENSMUSG00002076809.1 | 0 | NA | NA | NA | NA | NA | #VALUE! |
| ENSMUSG00002076810.1 | 0 | NA | NA | NA | NA | NA | #VALUE! |
| ENSMUSG00002076811.1 | 0 | NA | NA | NA | NA | NA | #VALUE! |
| ENSMUSG00002076812.1 | 0 | NA | NA | NA | NA | NA | #VALUE! |
| ENSMUSG00002076813.1 | 0 | NA | NA | NA | NA | NA | #VALUE! |
| ENSMUSG00002076814.1 | 0 | NA | NA | NA | NA | NA | #VALUE! |
| ENSMUSG00002076815.1 | 0 | NA | NA | NA | NA | NA | #VALUE! |
| ENSMUSG00002076816.1 | 0 | NA | NA | NA | NA | NA | #VALUE! |
| ENSMUSG00002076817.1 | 0 | NA | NA | NA | NA | NA | #VALUE! |
| ENSMUSG00002076818.1 | 0 | NA | NA | NA | NA | NA | #VALUE! |
| ENSMUSG00002076819.1 | 0 | NA | NA | NA | NA | NA | #VALUE! |
| ENSMUSG00002076820.1 | 0 | NA | NA | NA | NA | NA | #VALUE! |
| ENSMUSG00002076821.1 | 0 | NA | NA | NA | NA | NA | #VALUE! |
| ENSMUSG00002076822.1 | 0 | NA | NA | NA | NA | NA | #VALUE! |
| ENSMUSG00002076823.1 | 0 | NA | NA | NA | NA | NA | #VALUE! |
| ENSMUSG00002076824.1 | 0 | NA | NA | NA | NA | NA | #VALUE! |
| ENSMUSG00002076825.1 | 0 | NA | NA | NA | NA | NA | #VALUE! |
| ENSMUSG00002076826.1 | 0 | NA | NA | NA | NA | NA | #VALUE! |
| ENSMUSG00002076827.1 | 0 | NA | NA | NA | NA | NA | #VALUE! |
| ENSMUSG00002076828.1 | 0 | NA | NA | NA | NA | NA | #VALUE! |
| ENSMUSG00002076829.1 | 0 | NA | NA | NA | NA | NA | #VALUE! |
| ENSMUSG00002076830.1 | 0 | NA | NA | NA | NA | NA | #VALUE! |
| ENSMUSG00002076831.1 | 0 | NA | NA | NA | NA | NA | #VALUE! |
| ENSMUSG00002076832.1 | 0 | NA | NA | NA | NA | NA | #VALUE! |
| ENSMUSG00002076833.1 | 0 | NA | NA | NA | NA | NA | #VALUE! |
| ENSMUSG00002076834.1 | 0 | NA | NA | NA | NA | NA | #VALUE! |
| ENSMUSG00002076835.1 | 0 | NA | NA | NA | NA | NA | #VALUE! |
| ENSMUSG00002076836.1 | 0 | NA | NA | NA | NA | NA | #VALUE! |
| ENSMUSG00002076837.1 | 0 | NA | NA | NA | NA | NA | #VALUE! |
| ENSMUSG00002076838.1 | 0 | NA | NA | NA | NA | NA | #VALUE! |
| ENSMUSG00002076839.1 | 0 | NA | NA | NA | NA | NA | #VALUE! |

| | | | | | | | |
|---|---|---|---|---|---|---|---|
| ENSMUSG00002076840.1 | 0 | NA | NA | NA | NA | NA | #VALUE! |
| ENSMUSG00002076841.1 | 0 | NA | NA | NA | NA | NA | #VALUE! |
| ENSMUSG00002076842.1 | 0 | NA | NA | NA | NA | NA | #VALUE! |
| ENSMUSG00002076843.1 | 0 | NA | NA | NA | NA | NA | #VALUE! |
| ENSMUSG00002076844.1 | 0 | NA | NA | NA | NA | NA | #VALUE! |
| ENSMUSG00002076845.1 | 0 | NA | NA | NA | NA | NA | #VALUE! |
| ENSMUSG00002076846.1 | 0 | NA | NA | NA | NA | NA | #VALUE! |
| ENSMUSG00002076847.1 | 0 | NA | NA | NA | NA | NA | #VALUE! |
| ENSMUSG00002076848.1 | 0 | NA | NA | NA | NA | NA | #VALUE! |
| ENSMUSG00002076849.1 | 0 | NA | NA | NA | NA | NA | #VALUE! |
| ENSMUSG00002076850.1 | 0 | NA | NA | NA | NA | NA | #VALUE! |
| ENSMUSG00002076851.1 | 0 | NA | NA | NA | NA | NA | #VALUE! |
| ENSMUSG00002076852.1 | 0 | NA | NA | NA | NA | NA | #VALUE! |
| ENSMUSG00002076853.1 | 0 | NA | NA | NA | NA | NA | #VALUE! |
| ENSMUSG00002076854.1 | 0 | NA | NA | NA | NA | NA | #VALUE! |
| ENSMUSG00002076855.1 | 0 | NA | NA | NA | NA | NA | #VALUE! |
| ENSMUSG00002076856.1 | 0 | NA | NA | NA | NA | NA | #VALUE! |
| ENSMUSG00002076857.1 | 0 | NA | NA | NA | NA | NA | #VALUE! |
| ENSMUSG00002076858.1 | 0 | NA | NA | NA | NA | NA | #VALUE! |
| ENSMUSG00002076859.1 | 0 | NA | NA | NA | NA | NA | #VALUE! |
| ENSMUSG00002076860.1 | 0 | NA | NA | NA | NA | NA | #VALUE! |
| ENSMUSG00002076861.1 | 0 | NA | NA | NA | NA | NA | #VALUE! |
| ENSMUSG00002076862.1 | 0 | NA | NA | NA | NA | NA | #VALUE! |
| ENSMUSG00002076863.1 | 0 | NA | NA | NA | NA | NA | #VALUE! |
| ENSMUSG00002076864.1 | 0 | NA | NA | NA | NA | NA | #VALUE! |
| ENSMUSG00002076865.1 | 0 | NA | NA | NA | NA | NA | #VALUE! |
| ENSMUSG00002076866.1 | 0 | NA | NA | NA | NA | NA | #VALUE! |
| ENSMUSG00002076867.1 | 0 | NA | NA | NA | NA | NA | #VALUE! |
| ENSMUSG00002076868.1 | 0 | NA | NA | NA | NA | NA | #VALUE! |
| ENSMUSG00002076869.1 | 0 | NA | NA | NA | NA | NA | #VALUE! |
| ENSMUSG00002076870.1 | 0 | NA | NA | NA | NA | NA | #VALUE! |
| ENSMUSG00002076871.1 | 0 | NA | NA | NA | NA | NA | #VALUE! |
| ENSMUSG00002076872.1 | 0 | NA | NA | NA | NA | NA | #VALUE! |
| ENSMUSG00002076873.1 | 0 | NA | NA | NA | NA | NA | #VALUE! |
| ENSMUSG00002076874.1 | 0 | NA | NA | NA | NA | NA | #VALUE! |
| ENSMUSG00002076875.1 | 0 | NA | NA | NA | NA | NA | #VALUE! |
| ENSMUSG00002076876.1 | 0 | NA | NA | NA | NA | NA | #VALUE! |
| ENSMUSG00002076877.1 | 0 | NA | NA | NA | NA | NA | #VALUE! |
| ENSMUSG00002076878.1 | 0 | NA | NA | NA | NA | NA | #VALUE! |
| ENSMUSG00002076879.1 | 0 | NA | NA | NA | NA | NA | #VALUE! |
| ENSMUSG00002076880.1 | 0 | NA | NA | NA | NA | NA | #VALUE! |
| ENSMUSG00002076881.1 | 0 | NA | NA | NA | NA | NA | #VALUE! |
| ENSMUSG00002076882.1 | 0 | NA | NA | NA | NA | NA | #VALUE! |
| ENSMUSG00002076883.1 | 0 | NA | NA | NA | NA | NA | #VALUE! |

| | | | | | | | |
|---|---|---|---|---|---|---|---|
| ENSMUSG00002076884.1 | 0 | NA | NA | NA | NA | NA | #VALUE! |
| ENSMUSG00002076885.1 | 0 | NA | NA | NA | NA | NA | #VALUE! |
| ENSMUSG00002076886.1 | 0 | NA | NA | NA | NA | NA | #VALUE! |
| ENSMUSG00002076887.1 | 0 | NA | NA | NA | NA | NA | #VALUE! |
| ENSMUSG00002076888.1 | 0 | NA | NA | NA | NA | NA | #VALUE! |
| ENSMUSG00002076889.1 | 0 | NA | NA | NA | NA | NA | #VALUE! |
| ENSMUSG00002076890.1 | 0 | NA | NA | NA | NA | NA | #VALUE! |
| ENSMUSG00002076891.1 | 0 | NA | NA | NA | NA | NA | #VALUE! |
| ENSMUSG00002076892.1 | 0 | NA | NA | NA | NA | NA | #VALUE! |
| ENSMUSG00002076893.1 | 0 | NA | NA | NA | NA | NA | #VALUE! |
| ENSMUSG00002076894.1 | 0 | NA | NA | NA | NA | NA | #VALUE! |
| ENSMUSG00002076895.1 | 0 | NA | NA | NA | NA | NA | #VALUE! |
| ENSMUSG00002076896.1 | 0 | NA | NA | NA | NA | NA | #VALUE! |
| ENSMUSG00002076897.1 | 0 | NA | NA | NA | NA | NA | #VALUE! |
| ENSMUSG00002076898.1 | 0 | NA | NA | NA | NA | NA | #VALUE! |
| ENSMUSG00002076899.1 | 0 | NA | NA | NA | NA | NA | #VALUE! |
| ENSMUSG00002076900.1 | 0 | NA | NA | NA | NA | NA | #VALUE! |
| ENSMUSG00002076901.1 | 0 | NA | NA | NA | NA | NA | #VALUE! |
| ENSMUSG00002076902.1 | 0 | NA | NA | NA | NA | NA | #VALUE! |
| ENSMUSG00002076903.1 | 0 | NA | NA | NA | NA | NA | #VALUE! |
| ENSMUSG00002076904.1 | 0 | NA | NA | NA | NA | NA | #VALUE! |
| ENSMUSG00002076905.1 | 0 | NA | NA | NA | NA | NA | #VALUE! |
| ENSMUSG00002076906.1 | 0 | NA | NA | NA | NA | NA | #VALUE! |
| ENSMUSG00002076907.1 | 0 | NA | NA | NA | NA | NA | #VALUE! |
| ENSMUSG00002076908.1 | 0 | NA | NA | NA | NA | NA | #VALUE! |
| ENSMUSG00002076909.1 | 0 | NA | NA | NA | NA | NA | #VALUE! |
| ENSMUSG00002076910.1 | 0 | NA | NA | NA | NA | NA | #VALUE! |
| ENSMUSG00002076911.1 | 0 | NA | NA | NA | NA | NA | #VALUE! |
| ENSMUSG00002076912.1 | 0 | NA | NA | NA | NA | NA | #VALUE! |
| ENSMUSG00002076913.1 | 0 | NA | NA | NA | NA | NA | #VALUE! |
| ENSMUSG00002076914.1 | 0 | NA | NA | NA | NA | NA | #VALUE! |
| ENSMUSG00002076915.1 | 0 | NA | NA | NA | NA | NA | #VALUE! |
| ENSMUSG00002076916.1 | 0 | NA | NA | NA | NA | NA | #VALUE! |
| ENSMUSG00002076917.1 | 0 | NA | NA | NA | NA | NA | #VALUE! |
| ENSMUSG00002076918.1 | 0 | NA | NA | NA | NA | NA | #VALUE! |
| ENSMUSG00002076919.1 | 0 | NA | NA | NA | NA | NA | #VALUE! |
| ENSMUSG00002076920.1 | 0 | NA | NA | NA | NA | NA | #VALUE! |
| ENSMUSG00002076921.1 | 0 | NA | NA | NA | NA | NA | #VALUE! |
| ENSMUSG00002076922.1 | 0 | NA | NA | NA | NA | NA | #VALUE! |
| ENSMUSG00002076923.1 | 0 | NA | NA | NA | NA | NA | #VALUE! |
| ENSMUSG00002076924.1 | 0 | NA | NA | NA | NA | NA | #VALUE! |
| ENSMUSG00002076925.1 | 0 | NA | NA | NA | NA | NA | #VALUE! |
| ENSMUSG00002076926.1 | 0 | NA | NA | NA | NA | NA | #VALUE! |
| ENSMUSG00002076927.1 | 0 | NA | NA | NA | NA | NA | #VALUE! |

| | | | | | | | |
|---|---|---|---|---|---|---|---|
| ENSMUSG00002076928.1 | 0 | NA | NA | NA | NA | NA | #VALUE! |
| ENSMUSG00002076929.1 | 0 | NA | NA | NA | NA | NA | #VALUE! |
| ENSMUSG00002076930.1 | 0 | NA | NA | NA | NA | NA | #VALUE! |
| ENSMUSG00002076931.1 | 0 | NA | NA | NA | NA | NA | #VALUE! |
| ENSMUSG00002076932.1 | 0 | NA | NA | NA | NA | NA | #VALUE! |
| ENSMUSG00002076933.1 | 0 | NA | NA | NA | NA | NA | #VALUE! |
| ENSMUSG00002076934.1 | 0 | NA | NA | NA | NA | NA | #VALUE! |
| ENSMUSG00002076935.1 | 0 | NA | NA | NA | NA | NA | #VALUE! |
| ENSMUSG00002076936.1 | 0 | NA | NA | NA | NA | NA | #VALUE! |
| ENSMUSG00002076937.1 | 0 | NA | NA | NA | NA | NA | #VALUE! |
| ENSMUSG00002076938.1 | 0 | NA | NA | NA | NA | NA | #VALUE! |
| ENSMUSG00002076939.1 | 0 | NA | NA | NA | NA | NA | #VALUE! |
| ENSMUSG00002076940.1 | 0 | NA | NA | NA | NA | NA | #VALUE! |
| ENSMUSG00002076941.1 | 0 | NA | NA | NA | NA | NA | #VALUE! |
| ENSMUSG00002076942.1 | 0 | NA | NA | NA | NA | NA | #VALUE! |
| ENSMUSG00002076943.1 | 0 | NA | NA | NA | NA | NA | #VALUE! |
| ENSMUSG00002076944.1 | 0 | NA | NA | NA | NA | NA | #VALUE! |
| ENSMUSG00002076945.1 | 0 | NA | NA | NA | NA | NA | #VALUE! |
| ENSMUSG00002076946.1 | 0 | NA | NA | NA | NA | NA | #VALUE! |
| ENSMUSG00002076947.1 | 0 | NA | NA | NA | NA | NA | #VALUE! |
| ENSMUSG00002076948.1 | 0 | NA | NA | NA | NA | NA | #VALUE! |
| ENSMUSG00002076949.1 | 0 | NA | NA | NA | NA | NA | #VALUE! |
| ENSMUSG00002076950.1 | 0 | NA | NA | NA | NA | NA | #VALUE! |
| ENSMUSG00002076951.1 | 0 | NA | NA | NA | NA | NA | #VALUE! |
| ENSMUSG00002076952.1 | 0 | NA | NA | NA | NA | NA | #VALUE! |
| ENSMUSG00002076953.1 | 0 | NA | NA | NA | NA | NA | #VALUE! |
| ENSMUSG00002076954.1 | 0 | NA | NA | NA | NA | NA | #VALUE! |
| ENSMUSG00002076955.1 | 0 | NA | NA | NA | NA | NA | #VALUE! |
| ENSMUSG00002076956.1 | 0 | NA | NA | NA | NA | NA | #VALUE! |
| ENSMUSG00002076957.1 | 0 | NA | NA | NA | NA | NA | #VALUE! |
| ENSMUSG00002076958.1 | 0 | NA | NA | NA | NA | NA | #VALUE! |
| ENSMUSG00002076959.1 | 0 | NA | NA | NA | NA | NA | #VALUE! |
| ENSMUSG00002076960.1 | 0 | NA | NA | NA | NA | NA | #VALUE! |
| ENSMUSG00002076961.1 | 0 | NA | NA | NA | NA | NA | #VALUE! |
| ENSMUSG00002076962.1 | 0 | NA | NA | NA | NA | NA | #VALUE! |
| ENSMUSG00002076963.1 | 0 | NA | NA | NA | NA | NA | #VALUE! |
| ENSMUSG00002076964.1 | 0 | NA | NA | NA | NA | NA | #VALUE! |
| ENSMUSG00002076965.1 | 0 | NA | NA | NA | NA | NA | #VALUE! |
| ENSMUSG00002076966.1 | 0 | NA | NA | NA | NA | NA | #VALUE! |
| ENSMUSG00002076967.1 | 0 | NA | NA | NA | NA | NA | #VALUE! |
| ENSMUSG00002076968.1 | 0 | NA | NA | NA | NA | NA | #VALUE! |
| ENSMUSG00002076969.1 | 0 | NA | NA | NA | NA | NA | #VALUE! |
| ENSMUSG00002076970.1 | 0 | NA | NA | NA | NA | NA | #VALUE! |
| ENSMUSG00002076971.1 | 0 | NA | NA | NA | NA | NA | #VALUE! |

| | | | | | | | |
|---|---|---|---|---|---|---|---|
| ENSMUSG00002076972.1 | 0 | NA | NA | NA | NA | NA | #VALUE! |
| ENSMUSG00002076973.1 | 0 | NA | NA | NA | NA | NA | #VALUE! |
| ENSMUSG00002076974.1 | 0 | NA | NA | NA | NA | NA | #VALUE! |
| ENSMUSG00002076975.1 | 0 | NA | NA | NA | NA | NA | #VALUE! |
| ENSMUSG00002076976.1 | 0 | NA | NA | NA | NA | NA | #VALUE! |
| ENSMUSG00002076977.1 | 0 | NA | NA | NA | NA | NA | #VALUE! |
| ENSMUSG00002076978.1 | 0 | NA | NA | NA | NA | NA | #VALUE! |
| ENSMUSG00002076979.1 | 0 | NA | NA | NA | NA | NA | #VALUE! |
| ENSMUSG00002076980.1 | 0 | NA | NA | NA | NA | NA | #VALUE! |
| ENSMUSG00002076981.1 | 0 | NA | NA | NA | NA | NA | #VALUE! |
| ENSMUSG00002076982.1 | 0 | NA | NA | NA | NA | NA | #VALUE! |
| ENSMUSG00002076983.1 | 0 | NA | NA | NA | NA | NA | #VALUE! |
| ENSMUSG00002076984.1 | 0 | NA | NA | NA | NA | NA | #VALUE! |
| ENSMUSG00002076985.1 | 0 | NA | NA | NA | NA | NA | #VALUE! |
| ENSMUSG00002076986.1 | 0 | NA | NA | NA | NA | NA | #VALUE! |
| ENSMUSG00002076987.1 | 0 | NA | NA | NA | NA | NA | #VALUE! |
| ENSMUSG00002076988.1 | 0 | NA | NA | NA | NA | NA | #VALUE! |
| ENSMUSG00002076989.1 | 0 | NA | NA | NA | NA | NA | #VALUE! |
| ENSMUSG00002076990.1 | 0 | NA | NA | NA | NA | NA | #VALUE! |
| ENSMUSG00002076991.1 | 0 | NA | NA | NA | NA | NA | #VALUE! |
| ENSMUSG00002076992.1 | 0 | NA | NA | NA | NA | NA | #VALUE! |
| __too_low_aQual | 0 | NA | NA | NA | NA | NA | |
| __not_aligned | 0 | NA | NA | NA | NA | NA | |
| __alignment_not_unique | 0 | NA | NA | NA | NA | NA | |

j
