## Supplemental Data 2 for "Liver-specific Mettl14 deletion induces nuclear heterotypia and dysregulates RNA export machinery"

Table: Gene sets enriched in phenotype *Mettl14 (3 samples)* ***[plain text format]***

| | Shorthand Name for Figure | GS<br>follow link to MSigDB | GS DETAILS | SIZE | ES | NES | NOM p-val | FDR q-val | FWER p-val | RANK AT MAX | LEADING EDGE |
|---|---|---|---|---|---|---|---|---|---|---|---|
| 3 | Signature upregulated genes: invasive hepatoblastoma | CAIRO_LIVER_DEVELOPMENT_UP | Details ... | 90 | 0.34 | 1.61 | 0 | 0.902 | 0.968 | 1236 | tags=23%, list=12%, signal=26% |
| 2 | c-Myc targets: hepatocyte proliferation and regeneration | YU_MYC_TARGETS_UP | Details ... | 27 | 0.44 | 1.66 | 0.016 | 0.983 | 0.931 | 481 | tags=22%, list=5%, signal=23% |
| 10 | | GAUSSMANN_MLL_AF4_FUSION_TARGETS_D_UP | Details ... | 18 | 0.44 | 1.46 | 0.075 | 0.693 | 1 | 1050 | tags=28%, list=10%, signal=31% |
| 4 | signature upregulated HNF1A targets: hepatocellular carcinoma state | SERVITJA_LIVER_HNF1A_TARGETS_UP | Details ... | 89 | 0.33 | 1.6 | 0 | 0.748 | 0.978 | 1062 | tags=31%, list=11%, signal=35% |
| 7 | Sphingolipid metabolism | REACTOME_SPHINGOLIPID_METABOLISM | Details ... | 54 | 0.34 | 1.5 | 0 | 0.791 | 0.998 | 562 | tags=17%, list=6%, signal=18% |
| 1 | Fatty acid metabolism | WP_FATTY_ACID_BIOSYNTHESIS | Details ... | 21 | 0.49 | 1.7 | 0.013 | 1 | 0.871 | 1692 | tags=38%, list=17%, signal=46% |
| 6 | Mitochondrial fatty acid beta oxidation | REACTOME_MITOCHONDRIAL_FATTY_ACID_BETA_OXIDATION | Details ... | 34 | 0.39 | 1.51 | 0.021 | 0.835 | 0.997 | 1282 | tags=26%, list=13%, signal=30% |
| 19 | PPAR signaling pathway | WP_PPAR_SIGNALING_PATHWAY | Details ... | 58 | 0.3 | 1.31 | 0.061 | 0.821 | 1 | 1507 | tags=29%, list=15%, signal=34% |
| 9 | signature upregulated HFN4A targets: hepatocellular carcinoma | OHGUCHI_LIVER_HNF4A_TARGETS_UP | Details ... | 27 | 0.4 | 1.48 | 0.066 | 0.676 | 0.999 | 1275 | tags=37%, list=13%, signal=42% |
| 40 | | REACTOME_FATTY_ACID_METABOLISM | | 119 | 0.24 | 1.16 | 0.085 | 0.837 | 1 | 1302 | tags=24%, list=13%, signal=27% |
| 11 | | WP_SPHINGOLIPID_METABOLISM_OVERVIEW | Details ... | 16 | 0.44 | 1.38 | 0.107 | 0.963 | 1 | 486 | tags=25%, list=5%, signal=26% |
| 12 | | WP_SPHINGOLIPID_METABOLISM_INTEGRATED_PATHWAY | Details ... | 16 | 0.44 | 1.38 | 0.11 | 0.887 | 1 | 486 | tags=25%, list=5%, signal=26% |
| 14 | | REACTOME_SPHINGOLIPID_DE_NOVO_BIOSYNTHESIS | Details ... | 15 | 0.42 | 1.35 | 0.113 | 0.885 | 1 | 486 | tags=20%, list=5%, signal=21% |
| 28 | | WENG_POR_TARGETS_LIVER_UP | | 46 | 0.28 | 1.22 | 0.133 | 0.897 | 1 | 662 | tags=17%, list=7%, signal=19% |
| 22 | Classic adipogenic targets of PPARG | WANG_CLASSIC_ADIPOGENIC_TARGETS_OF_PPARG | | 21 | 0.37 | 1.29 | 0.143 | 0.786 | 1 | 1412 | tags=38%, list=14%, signal=44% |
| 56 | Nucleotide di and triphosphate conversion | REACTOME_INTERCONVERSION_OF_NUCLEOTIDE_DI_AND_TRIPHOSPHATES | | 19 | 0.32 | 1.08 | 0.352 | 0.88 | 1 | 399 | tags=16%, list=4%, signal=16% |
| 79 | RNA polymerase III transcription | REACTOME_RNA_POLYMERASE_III_TRANSCRIPTION | | 31 | 0.26 | 0.98 | 0.474 | 0.924 | 1 | 1273 | tags=29%, list=13%, signal=33% |
| 90 | Metabolism of Nucleotides | REACTOME_METABOLISM_OF_NUCLEOTIDES | | 72 | 0.2 | 0.94 | 0.611 | 0.949 | 1 | 1618 | tags=24%, list=16%, signal=28% |
| 112 | Nucleotide salvage | REACTOME_NUCLEOTIDE_SALVAGE | | 17 | 0.26 | 0.85 | 0.711 | 0.971 | 1 | 1200 | tags=24%, list=12%, signal=27% |
| 135 | Trans-lesion synthesis by PolH: DNA damage repair | REACTOME_TRANSLESION_SYNTHESIS_BY_POLH | | 17 | 0.24 | 0.74 | 0.8 | 0.992 | 1 | 991 | tags=18%, list=10%, signal=20% |
| 129 | rRNA processing in nucleolus and cytosol | REACTOME_MAJOR_PATHWAY_OF_RRNA_PROCESSING_IN_THE_NUCLEOLUS_AND_CYTOSOL | | 167 | 0.15 | 0.78 | 1 | 0.975 | 1 | 1524 | tags=17%, list=15%, signal=20% |
| 20 | | REACTOME_RECRUITMENT_OF_NUMA_TO_MITOTIC_CENTROSOMES | Details ... | 64 | 0.3 | 1.31 | 0.051 | 0.785 | 1 | 628 | tags=16%, list=6%, signal=17% |
| 44 | | REACTOME_REGULATION_OF_PLK1_ACTIVITY_AT_G2_M_TRANSITION | | 66 | 0.26 | 1.15 | 0.177 | 0.799 | 1 | 991 | tags=17%, list=10%, signal=18% |
| 46 | | REACTOME_MISCELLANEOUS_TRANSPORT_AND_BINDING_EVENTS | | 16 | 0.36 | 1.15 | 0.249 | 0.769 | 1 | 707 | tags=19%, list=7%, signal=20% |
| 29 | | LAZARO_GENETIC_MOUSE_MODEL_HIGH_GRADE_LARGE_CELL_NEUROENDOCRINE_LUNG_CARCINOMA_DN | | 222 | 0.21 | 1.21 | 0 | 0.895 | 1 | 1541 | tags=24%, list=15%, signal=28% |
| 5 | | MCCLUNG_COCAINE_REWARD_5D | Details ... | 38 | 0.4 | 1.59 | 0.013 | 0.62 | 0.981 | 349 | tags=16%, list=3%, signal=16% |
| 15 | | BOYLAN_MULTIPLE_MYELOMA_PCA3_UP | Details ... | 58 | 0.31 | 1.35 | 0.029 | 0.861 | 1 | 1398 | tags=31%, list=14%, signal=36% |
| 13 | | REACTOME_CARDIAC_CONDUCTION | Details ... | 29 | 0.37 | 1.38 | 0.051 | 0.843 | 1 | 1604 | tags=31%, list=16%, signal=37% |
| 8 | | CUI_TCF21_TARGETS_DN | Details ... | 19 | 0.45 | 1.49 | 0.058 | 0.742 | 0.999 | 1150 | tags=37%, list=11%, signal=41% |
| 16 | | REACTOME_DOWNREGULATION_OF_TGF_BETA_RECEPTOR_SIGNALING | Details ... | 15 | 0.41 | 1.34 | 0.097 | 0.841 | 1 | 991 | tags=27%, list=10%, signal=30% |
| 24 | | WENG_POR_TARGETS_GLOBAL_DN | | 23 | 0.35 | 1.28 | 0.104 | 0.777 | 1 | 1302 | tags=39%, list=13%, signal=45% |
| 21 | | REACTOME_SYNTHESIS_OF_IP3_AND_IP4_IN_THE_CYTOSOL | | 15 | 0.4 | 1.3 | 0.115 | 0.8 | 1 | 1025 | tags=33%, list=10%, signal=37% |
| 17 | | WENG_POR_DOSAGE | Details ... | 22 | 0.39 | 1.33 | 0.122 | 0.838 | 1 | 599 | tags=27%, list=6%, signal=29% |
| 27 | | REACTOME_AURKA_ACTIVATION_BY_TPX2 | | 53 | 0.29 | 1.23 | 0.122 | 0.877 | 1 | 504 | tags=13%, list=5%, signal=14% |
| 18 | | GAUSSMANN_MLL_AF4_FUSION_TARGETS_F_DN | Details ... | 21 | 0.39 | 1.32 | 0.128 | 0.836 | 1 | 779 | tags=29%, list=8%, signal=31% |
| 47 | | SANSOM_APC_TARGETS_REQUIRE_MYC | | 155 | 0.22 | 1.15 | 0.143 | 0.754 | 1 | 1314 | tags=21%, list=13%, signal=23% |
| 26 | | REACTOME_SYNTHESIS_OF_PC | | 18 | 0.38 | 1.27 | 0.145 | 0.75 | 1 | 179 | tags=11%, list=2%, signal=11% |
| 34 | | MCCLUNG_CREB1_TARGETS_DN | | 40 | 0.3 | 1.19 | 0.155 | 0.865 | 1 | 95 | tags=10%, list=1%, signal=10% |
| 23 | | ZHENG_IL22_SIGNALING_DN | | 19 | 0.37 | 1.28 | 0.162 | 0.8 | 1 | 1368 | tags=26%, list=14%, signal=30% |
| 33 | | BURTON_ADIPOGENESIS_PEAK_AT_8HR | | 31 | 0.31 | 1.19 | 0.172 | 0.856 | 1 | 704 | tags=19%, list=7%, signal=21% |
| 39 | | CHEBOTAEV_GR_TARGETS_DN | | 58 | 0.26 | 1.17 | 0.177 | 0.797 | 1 | 664 | tags=19%, list=7%, signal=20% |
| 25 | | WP_EICOSANOID_METABOLISM_VIA_LIPOXYGENASES_LOX | | 17 | 0.39 | 1.27 | 0.18 | 0.777 | 1 | 615 | tags=24%, list=6%, signal=25% |
| 32 | | HEVNER_CORTEX_CAUDAL_SUBVENTRICULAR_ZONE | | 19 | 0.35 | 1.2 | 0.186 | 0.866 | 1 | 1045 | tags=21%, list=10%, signal=23% |
| 36 | | REACTOME_G_PROTEIN_MEDIATED_EVENTS | | 22 | 0.33 | 1.18 | 0.186 | 0.826 | 1 | 1025 | tags=27%, list=10%, signal=30% |
| 30 | | WANG_ADIPOGENIC_GENES_REPRESSED_BY_SIRT1 | | 22 | 0.34 | 1.2 | 0.189 | 0.899 | 1 | 610 | tags=18%, list=6%, signal=19% |
| 38 | | REACTOME_GLYCOSPHINGOLIPID_CATABOLISM | | 25 | 0.33 | 1.18 | 0.197 | 0.817 | 1 | 562 | tags=16%, list=6%, signal=17% |
| 41 | | ZHENG_IL22_SIGNALING_UP | | 33 | 0.3 | 1.16 | 0.2 | 0.819 | 1 | 292 | tags=12%, list=3%, signal=12% |
| 35 | | WP_CHOLESTEROL_METABOLISM_WITH_BLOCH_AND_KANDUTSCH_RUSSELL_PATHWAYS | | 47 | 0.28 | 1.18 | 0.203 | 0.849 | 1 | 567 | tags=17%, list=6%, signal=18% |
| 31 | | REACTOME_METABOLISM_OF_COFACTORS | | 15 | 0.38 | 1.2 | 0.216 | 0.877 | 1 | 382 | tags=20%, list=4%, signal=21% |
| 37 | | TSENG_ADIPOGENIC_POTENTIAL_UP | | 18 | 0.35 | 1.18 | 0.239 | 0.817 | 1 | 1395 | tags=44%, list=14%, signal=51% |
| 43 | | REACTOME_FORMATION_OF_THE_BETA_CATENIN_TCF_TRANSACTIVATING_COMPLEX | | 17 | 0.36 | 1.16 | 0.244 | 0.791 | 1 | 909 | tags=35%, list=9%, signal=39% |
| 48 | | ZHANG_GATA6_TARGETS_DN | | 24 | 0.32 | 1.15 | 0.246 | 0.743 | 1 | 1147 | tags=29%, list=11%, signal=33% |
| 52 | | WP_PRIMARY_FOCAL_SEGMENTAL_GLOMERULOSCLEROSIS_FSGS | | 45 | 0.26 | 1.1 | 0.254 | 0.872 | 1 | 364 | tags=9%, list=4%, signal=9% |
| 57 | | YAO_TEMPORAL_RESPONSE_TO_PROGESTERONE_CLUSTER_11 | | 92 | 0.22 | 1.07 | 0.26 | 0.912 | 1 | 1412 | tags=18%, list=14%, signal=21% |
| 49 | | HU_GENOTOXIN_ACTION_DIRECT_VS_INDIRECT_4HR | | 23 | 0.33 | 1.14 | 0.273 | 0.754 | 1 | 1 | tags=4%, list=0%, signal=4% |
| 45 | | BIOCARTA_CCR5_PATHWAY | | 15 | 0.37 | 1.15 | 0.294 | 0.785 | 1 | 1287 | tags=40%, list=13%, signal=46% |
| 51 | | MCCLUNG_DELTA_FOSB_TARGETS_2WK | | 23 | 0.32 | 1.11 | 0.297 | 0.824 | 1 | 1190 | tags=30%, list=12%, signal=34% |
| 53 | | REACTOME_FATTY_ACYL_COA_BIOSYNTHESIS | | 25 | 0.3 | 1.09 | 0.304 | 0.892 | 1 | 1154 | tags=28%, list=11%, signal=32% |
| 42 | | REACTOME_AZATHIOPRINE_ADME | | 17 | 0.37 | 1.16 | 0.308 | 0.805 | 1 | 1598 | tags=41%, list=16%, signal=49% |
| 54 | | TERAMOTO_OPN_TARGETS_CLUSTER_6 | | 16 | 0.34 | 1.08 | 0.313 | 0.899 | 1 | 471 | tags=25%, list=5%, signal=26% |
| 59 | | WORSCHECH_TUMOR_REJECTION_UP | | 24 | 0.29 | 1.05 | 0.313 | 0.943 | 1 | 757 | tags=17%, list=7%, signal=18% |
| 50 | | NADLER_OBESITY_DN | | 45 | 0.27 | 1.12 | 0.323 | 0.824 | 1 | 848 | tags=24%, list=8%, signal=27% |
| 55 | | REACTOME_DNA_DAMAGE_TELOMERE_STRESS_INDUCED_SENESCENCE | | 16 | 0.36 | 1.08 | 0.326 | 0.896 | 1 | 257 | tags=13%, list=3%, signal=13% |
| 58 | | REACTOME_SIGNAL_AMPLIFICATION | | 19 | 0.31 | 1.05 | 0.368 | 0.943 | 1 | 1425 | tags=47%, list=14%, signal=55% |
| 60 | | REACTOME_OPIOID_SIGNALLING | | 37 | 0.26 | 1.04 | 0.368 | 0.982 | 1 | 1025 | tags=27%, list=10%, signal=30% |
| 61 | | HEVNER_INTERMEDIATE_ZONE_AND_UP_POSTMITOTIC_NEURONS | | 21 | 0.31 | 1.03 | 0.389 | 1 | 1 | 689 | tags=14%, list=7%, signal=15% |
| 62 | | WP_GPCRS_ODORANT | | 17 | 0.32 | 1.02 | 0.398 | 1 | 1 | 2394 | tags=53%, list=24%, signal=69% |
| 65 | | WP_PURINE_METABOLISM | | 108 | 0.2 | 1.01 | 0.404 | 1 | 1 | 1618 | tags=26%, list=16%, signal=31% |
| 68 | | GROSS_HYPOXIA_VIA_ELK3_ONLY_DN | | 31 | 0.26 | 1 | 0.435 | 0.986 | 1 | 1287 | tags=26%, list=13%, signal=29% |
| 67 | | WENG_POR_TARGETS_GLOBAL_UP | | 22 | 0.29 | 1.01 | 0.436 | 0.998 | 1 | 848 | tags=23%, list=8%, signal=25% |
| 70 | | RAMPON_ENRICHED_LEARNING_ENVIRONMENT_LATE_UP | | 15 | 0.32 | 1 | 0.44 | 0.976 | 1 | 440 | tags=13%, list=4%, signal=14% |
| 85 | | REACTOME_IKK_COMPLEX_RECRUITMENT_MEDIATED_BY_RIP1 | | 18 | 0.3 | 0.96 | 0.441 | 0.917 | 1 | 991 | tags=17%, list=10%, signal=18% |
| 69 | | NADLER_HYPERGLYCEMIA_AT_OBESITY | | 38 | 0.25 | 1 | 0.451 | 0.978 | 1 | 1575 | tags=32%, list=16%, signal=37% |
| 66 | | BERENJENO_TRANSFORMED_BY_RHOA_FOREVER_UP | | 19 | 0.3 | 1.01 | 0.452 | 1 | 1 | 593 | tags=21%, list=6%, signal=22% |
| 77 | | WP_MITOCHONDRIAL_LONG_CHAIN_FATTY_ACID_BETA_OXIDATION | | 16 | 0.3 | 0.99 | 0.453 | 0.933 | 1 | 610 | tags=19%, list=6%, signal=20% |
| 71 | | SERVITJA_ISLET_HNF1A_TARGETS_UP | | 88 | 0.21 | 1 | 0.459 | 0.965 | 1 | 1233 | tags=22%, list=12%, signal=24% |
| 72 | | REACTOME_GLYCOSPHINGOLIPID_METABOLISM | | 31 | 0.27 | 1 | 0.46 | 0.956 | 1 | 562 | tags=13%, list=6%, signal=14% |
| 73 | | REACTOME_TP53_REGULATES_TRANSCRIPTION_OF_CELL_CYCLE_GENES | | 23 | 0.28 | 0.99 | 0.462 | 0.959 | 1 | 193 | tags=13%, list=2%, signal=13% |
| 74 | | HOLLERN_SQUAMOUS_BREAST_TUMOR | | 38 | 0.25 | 0.99 | 0.463 | 0.956 | 1 | 1716 | tags=34%, list=17%, signal=41% |
| 75 | | MARSHALL_VIRAL_INFECTION_RESPONSE_DN | | 17 | 0.31 | 0.99 | 0.463 | 0.946 | 1 | 1280 | tags=29%, list=13%, signal=34% |
| 83 | | REACTOME_SIGNALING_BY_HIPPO | | 19 | 0.29 | 0.97 | 0.465 | 0.915 | 1 | 172 | tags=11%, list=2%, signal=11% |
| 63 | | BOYLAN_MULTIPLE_MYELOMA_C_DN | | 43 | 0.24 | 1.02 | 0.466 | 0.987 | 1 | 1270 | tags=28%, list=13%, signal=32% |
| 64 | | REACTOME_MITOTIC_PROMETAPHASE | | 129 | 0.2 | 1.02 | 0.468 | 0.977 | 1 | 825 | tags=13%, list=8%, signal=14% |
| 80 | | HEVNER_CORTICAL_PLATE_POSTMITOTIC_PROJECTION_NEURONS | | 52 | 0.23 | 0.98 | 0.468 | 0.924 | 1 | 101 | tags=6%, list=1%, signal=6% |
| 81 | | BIOCARTA_EDG1_PATHWAY | | 17 | 0.3 | 0.97 | 0.468 | 0.93 | 1 | 555 | tags=18%, list=5%, signal=19% |
| 82 | | RAO_BOUND_BY_SALL4_ISOFORM_A | | 101 | 0.2 | 0.97 | 0.475 | 0.924 | 1 | 938 | tags=14%, list=9%, signal=15% |
| 84 | | DARWICHE_PAPILLOMA_PROGRESSION_RISK | | 37 | 0.24 | 0.97 | 0.494 | 0.914 | 1 | 1092 | tags=24%, list=11%, signal=27% |
| 89 | | REACTOME_SYNTHESIS_OF_VERY_LONG_CHAIN_FATTY_ACYL_COAS | | 15 | 0.31 | 0.94 | 0.494 | 0.957 | 1 | 993 | tags=27%, list=10%, signal=30% |
| 78 | | REACTOME_RAC3_GTPASE_CYCLE | | 61 | 0.22 | 0.98 | 0.508 | 0.93 | 1 | 1104 | tags=16%, list=11%, signal=18% |
| 76 | | REACTOME_ACTIVATION_OF_ATR_IN_RESPONSE_TO_REPLICATION_STRESS | | 23 | 0.29 | 0.99 | 0.522 | 0.939 | 1 | 1115 | tags=26%, list=11%, signal=29% |
| 87 | | REACTOME_CA2_PATHWAY | | 25 | 0.26 | 0.95 | 0.542 | 0.936 | 1 | 1133 | tags=28%, list=11%, signal=31% |
| 86 | | NABA_ECM_AFFILIATED | | 52 | 0.22 | 0.96 | 0.571 | 0.937 | 1 | 498 | tags=12%, list=5%, signal=12% |
| 91 | | WP_EICOSANOID_METABOLISM_VIA_CYCLOOXYGENASES_COX | | 16 | 0.29 | 0.91 | 0.581 | 1 | 1 | 1721 | tags=38%, list=17%, signal=45% |
| 88 | | REACTOME_MUSCLE_CONTRACTION | | 51 | 0.22 | 0.94 | 0.59 | 0.957 | 1 | 1604 | tags=24%, list=16%, signal=28% |
| 101 | | HOFFMANN_PRE_BI_TO_LARGE_PRE_BII_LYMPHOCYTE_UP | | 15 | 0.28 | 0.88 | 0.6 | 0.996 | 1 | 2199 | tags=53%, list=22%, signal=68% |
| 97 | | DAWSON_METHYLATED_IN_LYMPHOMA_TCL1 | | 21 | 0.26 | 0.9 | 0.607 | 1 | 1 | 7464 | tags=100%, list=74%, signal=382% |
| 94 | | GERHOLD_ADIPOGENESIS_UP | | 42 | 0.22 | 0.91 | 0.608 | 1 | 1 | 1154 | tags=24%, list=11%, signal=27% |
| 93 | | REACTOME_TELOMERE_C_STRAND_LAGGING_STRAND_SYNTHESIS | | 23 | 0.26 | 0.91 | 0.619 | 1 | 1 | 1114 | tags=22%, list=11%, signal=24% |
| 105 | | BIOCARTA_G1_PATHWAY | | 17 | 0.27 | 0.88 | 0.62 | 0.972 | 1 | 652 | tags=18%, list=6%, signal=19% |
| 104 | | BIOCARTA_CASPASE_PATHWAY | | 17 | 0.27 | 0.88 | 0.623 | 0.976 | 1 | 2126 | tags=53%, list=21%, signal=67% |
| 96 | | SCHLINGEMANN_SKIN_CARCINOGENESIS_TPA_DN | | 17 | 0.28 | 0.9 | 0.624 | 1 | 1 | 607 | tags=18%, list=6%, signal=19% |
| 99 | | WP_GLUTATHIONE_METABOLISM | | 15 | 0.28 | 0.89 | 0.628 | 0.997 | 1 | 520 | tags=13%, list=5%, signal=14% |
| 100 | | ISHIDA_E2F_TARGETS | | 31 | 0.24 | 0.89 | 0.629 | 1 | 1 | 346 | tags=13%, list=3%, signal=13% |
| 111 | | MATZUK_SPERMATID_DIFFERENTIATION | | 19 | 0.25 | 0.86 | 0.636 | 0.967 | 1 | 533 | tags=11%, list=5%, signal=11% |
| 95 | | REACTOME_BASE_EXCISION_REPAIR | | 25 | 0.25 | 0.9 | 0.644 | 1 | 1 | 516 | tags=12%, list=5%, signal=13% |
| 98 | | REACTOME_PEROXISOMAL_LIPID_METABOLISM | | 24 | 0.26 | 0.89 | 0.651 | 0.998 | 1 | 1765 | tags=42%, list=17%, signal=50% |
| 107 | | WP_TRIACYLGLYCERIDE_SYNTHESIS | | 18 | 0.26 | 0.88 | 0.651 | 0.964 | 1 | 1213 | tags=22%, list=12%, signal=25% |
| 103 | | LIANG_HEMATOPOIESIS_STEM_CELL_NUMBER_SMALL_VS_HUGE_UP | | 32 | 0.24 | 0.88 | 0.658 | 0.978 | 1 | 1520 | tags=25%, list=15%, signal=29% |
| 114 | | REACTOME_RESOLUTION_OF_D_LOOP_STRUCTURES | | 16 | 0.28 | 0.85 | 0.659 | 0.957 | 1 | 2053 | tags=44%, list=20%, signal=55% |
| 119 | | REACTOME_FANCONI_ANEMIA_PATHWAY | | 20 | 0.24 | 0.83 | 0.672 | 0.972 | 1 | 1065 | tags=20%, list=11%, signal=22% |
| 110 | | KAMMINGA_EZH2_TARGETS | | 29 | 0.23 | 0.86 | 0.674 | 0.968 | 1 | 316 | tags=10%, list=3%, signal=11% |
| 106 | | REACTOME_SYNTHESIS_OF_SUBSTRATES_IN_N_GLYCAN_BIOSYTHESIS | | 38 | 0.22 | 0.88 | 0.675 | 0.971 | 1 | 874 | tags=18%, list=9%, signal=20% |
| 92 | | WP_CYTOPLASMIC_RIBOSOMAL_PROTEINS | | 77 | 0.2 | 0.91 | 0.688 | 1 | 1 | 6994 | tags=96%, list=69%, [illegible] |
| 124 | | REACTOME_POLYMERASE_SWITCHING_ON_THE_C_STRAND_OF_THE_TELOMERE | | 17 | 0.25 | 0.81 | 0.702 | 0.962 | 1 | 1114 | tags=24%, list=11%, signal=26% |
| 115 | | REACTOME_CHONDROITIN_SULFATE_DERMATAN_SULFATE_METABOLISM | | 26 | 0.23 | 0.85 | 0.706 | 0.954 | 1 | 646 | tags=12%, list=6%, signal=12% |
| 116 | | REACTOME_VITAMIN_B5_PANTOTHENATE_METABOLISM | | 15 | 0.26 | 0.84 | 0.715 | 0.977 | 1 | 1154 | tags=27%, list=11%, signal=30% |
| 113 | | REACTOME_HEPARAN_SULFATE_HEPARIN_HS_GAG_METABOLISM | | 29 | 0.23 | 0.85 | 0.718 | 0.964 | 1 | 695 | tags=17%, list=7%, signal=18% |
| 121 | | REACTOME_ION_HOMEOSTASIS | | 21 | 0.24 | 0.83 | 0.726 | 0.959 | 1 | 1505 | tags=29%, list=15%, signal=34% |
| 122 | | REACTOME_PURINE_CATABOLISM | | 15 | 0.26 | 0.82 | 0.73 | 0.964 | 1 | 7463 | tags=100%, list=74%, signal=382% |
| 118 | | REACTOME_GLUTATHIONE_CONJUGATION | | 29 | 0.23 | 0.83 | 0.735 | 0.969 | 1 | 848 | tags=17%, list=8%, signal=19% |
| 117 | | SHETH_LIVER_CANCER_VS_TXNIP_LOSS_PAM5 | | 35 | 0.21 | 0.83 | 0.736 | 0.973 | 1 | 1535 | tags=29%, list=15%, signal=34% |
| 109 | | REACTOME_RND1_GTPASE_CYCLE | | 29 | 0.22 | 0.86 | 0.741 | 0.976 | 1 | 1620 | tags=31%, list=16%, signal=37% |
| 108 | | COATES_MACROPHAGE_M1_VS_M2_UP | | 64 | 0.19 | 0.87 | 0.755 | 0.96 | 1 | 890 | tags=14%, list=9%, signal=15% |
| 123 | | REACTOME_ION_TRANSPORT_BY_P_TYPE_ATPASES | | 25 | 0.22 | 0.82 | 0.759 | 0.962 | 1 | 1505 | tags=28%, list=15%, signal=33% |
| 102 | | BURTON_ADIPOGENESIS_3 | | 59 | 0.19 | 0.88 | 0.794 | 0.987 | 1 | 1443 | tags=20%, list=14%, signal=24% |
| 134 | | BIOCARTA_HDAC_PATHWAY | | 17 | 0.23 | 0.74 | 0.795 | 0.997 | 1 | 1687 | tags=29%, list=17%, signal=35% |
| 126 | | REACTOME_HDR_THROUGH_HOMOLOGOUS_RECOMBINATION_HRR | | 33 | 0.21 | 0.81 | 0.799 | 0.957 | 1 | 2053 | tags=33%, list=20%, signal=42% |
| 125 | | WP_FATTY_ACID_BETA_OXIDATION | | 29 | 0.21 | 0.81 | 0.801 | 0.956 | 1 | 1028 | tags=17%, list=10%, signal=19% |
| 128 | | MIKKELSEN_ES_ICP_WITH_H3K4ME3_AND_H3K27ME3 | | 24 | 0.22 | 0.79 | 0.802 | 0.979 | 1 | 469 | tags=13%, list=5%, signal=13% |
| 131 | | SCHRAETS_MLL_TARGETS_DN | | 21 | 0.22 | 0.76 | 0.833 | 0.993 | 1 | 1198 | tags=19%, list=12%, signal=22% |
| 133 | | MEISSNER_BRAIN_HCP_WITH_H3K27ME3 | | 35 | 0.2 | 0.74 | 0.834 | 1 | 1 | 1126 | tags=17%, list=11%, signal=19% |
| 136 | | REACTOME_HSF1_DEPENDENT_TRANSACTIVATION | | 17 | 0.23 | 0.74 | 0.836 | 0.985 | 1 | 1294 | tags=24%, list=13%, signal=27% |
| 120 | | REACTOME_NONSENSE_MEDIATED_DECAY_NMD_INDEPENDENT_OF_THE_EXON_JUNCTION_COMPLEX_EJC | | 86 | 0.17 | 0.83 | 0.864 | 0.967 | 1 | 8350 | tags=100%, list=83%, signal=572% |
| 138 | | REACTOME_METABOLISM_OF_PORPHYRINS | | 25 | 0.2 | 0.71 | 0.872 | 1 | 1 | 1674 | tags=32%, list=17%, signal=38% |
| 130 | | CHEBOTAEV_GR_TARGETS_UP | | 45 | 0.18 | 0.77 | 0.883 | 0.986 | 1 | 984 | tags=13%, list=10%, signal=15% |
| 127 | | LEIN_PONS_MARKERS | | 41 | 0.19 | 0.79 | 0.887 | 0.979 | 1 | 1056 | tags=15%, list=10%, signal=16% |
| 132 | | REACTOME_PKMTS_METHYLATE_HISTONE_LYSINES | | 37 | 0.19 | 0.76 | 0.89 | 0.99 | 1 | 1392 | tags=24%, list=14%, signal=28% |
| 139 | | REACTOME_TRANSLESION_SYNTHESIS_BY_POLK | | 16 | 0.22 | 0.7 | 0.897 | 1 | 1 | 991 | tags=19%, list=10%, signal=21% |
| 137 | | WP_ALZHEIMER_39_S_DISEASE | | 50 | 0.17 | 0.74 | 0.916 | 0.979 | 1 | 762 | tags=10%, list=8%, signal=11% |
| 142 | | REACTOME_CYTOSOLIC_SULFONATION_OF_SMALL_MOLECULES | | 15 | 0.2 | 0.61 | 0.935 | 1 | 1 | 589 | tags=13%, list=6%, signal=14% |
| 140 | | BRIDEAU_IMPRINTED_GENES | | 27 | 0.18 | 0.67 | 0.949 | 1 | 1 | 945 | tags=15%, list=9%, signal=16% |
| 141 | | WP_STATIN_PATHWAY | | 18 | 0.19 | 0.62 | 0.962 | 1 | 1 | 1464 | tags=22%, list=14%, signal=26% |
| 143 | | REACTOME_GAP_FILLING_DNA_REPAIR_SYNTHESIS_AND_LIGATION_IN_GG_NER | | 17 | 0.18 | 0.6 | 0.962 | 1 | 1 | 1114 | tags=18%, list=11%, signal=20% |
| 145 | | REACTOME_CONDENSATION_OF_PROPHASE_CHROMOSOMES | | 15 | 0.19 | 0.57 | 0.974 | 1 | 1 | 2418 | tags=47%, list=24%, signal=61% |
| 144 | | REACTOME_RUNX1_REGULATES_GENES_INVOLVED_IN_MEGAKARYOCYTE_DIFFERENTIATION_AND_PLATELET_FUNCTION | | 25 | 0.16 | 0.59 | 0.985 | 1 | 1 | 1747 | tags=28%, list=17%, signal=34% |
| 149 | | REACTOME_ONCOGENE_INDUCED_SENESCENCE | | 15 | 0.17 | 0.53 | 0.988 | 1 | 1 | 1930 | tags=27%, list=19%, signal=33% |
| 147 | | JIANG_AGING_CEREBRAL_CORTEX_UP | | 28 | 0.15 | 0.54 | 0.989 | 1 | 1 | 1280 | tags=14%, list=13%, signal=16% |
| 150 | | REACTOME_INTRINSIC_PATHWAY_FOR_APOPTOSIS | | 32 | 0.14 | 0.52 | 0.994 | 1 | 1 | 1657 | tags=22%, list=16%, signal=26% |
| 146 | | REACTOME_SULFUR_AMINO_ACID_METABOLISM | | 24 | 0.16 | 0.55 | 0.995 | 1 | 1 | 7432 | tags=96%, list=74%, [illegible] |
| 148 | | WP_TOLL_LIKE_RECEPTOR_SIGNALING | | 26 | 0.15 | 0.54 | 1 | 1 | 1 | 8586 | tags=100%, list=85%, signal=665% |
| 151 | | REACTOME_RECOGNITION_OF_DNA_DAMAGE_BY_PCNA_CONTAINING_REPLICATION_COMPLEX | | 23 | 0.14 | 0.49 | 1 | 1 | 1 | 8707 | tags=100%, list=86%, signal=723% |
| 152 | | REACTOME_INTERFERON_ALPHA_BETA_SIGNALING | | 17 | 0.14 | 0.46 | 1 | 0.999 | 1 | 5719 | tags=88%, list=57%, [illegible] |

Table: Gene sets enriched in phenotype *WT (3 samples)* ***[plain text format]***

| | Shorthand name for figure | GS follow link to MSigDB | GS DETAILS | SIZE | ES | NES | NOM p-val | FDR q-val | FWER p-val | RANK AT MAX | LEADING EDGE | |
|---|---|---|---|---|---|---|---|---|---|---|---|---|
| 1 | signature downregulated HNF1A targets: hepatocellular carcinoma state | SERVITIA_LIVER_HNF1A_TARGETS_DN | Details ... | 169 | -0.53 | -1.98 | 0 | 0.03 | 0.029 | 2604 | tags=51%, list=26%, | |
| 2 | Complement activation: liver regeneration and autoimmunity | WP_COMPLEMENT_ACTIVATION_CLASSICAL_PATHWAY | Details ... | 15 | -0.72 | -1.85 | 0 | 0.125 | 0.221 | 976 | tags=60%, list=10%, | |
| 5 | Integrin cell-surface interactions: disruption of hepatocyte polarity leading to widespread fibrosis | REACTOME_INTEGRIN_CELL_SURFACE_INTERACTIONS | Details ... | 36 | -0.58 | -1.78 | 0 | 0.166 | 0.552 | 1953 | tags=50%, list=19%, | |
| 6 | Assembly of collagen fibrils | REACTOME_ASSEMBLY_OF_COLLAGEN_FIBRILS_AND_OTHER_MULTIMERIC_STRUCTURES | Details ... | 19 | -0.66 | -1.75 | 0.003 | 0.221 | 0.715 | 1810 | tags=58%, list=18%, | |
| 7 | ERBB4 signaling: Liver injury response defects | REACTOME_SIGNALING_BY_ERBB4 | Details ... | 25 | -0.61 | -1.75 | 0.004 | 0.194 | 0.721 | 2416 | tags=48%, list=24%, | |
| 8 | IL6 pathway: impaired liver regeneration | BIOCARTA_IL6_PATHWAY | Details ... | 18 | -0.64 | -1.74 | 0 | 0.185 | 0.759 | 2648 | tags=72%, list=26%, | |
| 9 | MET signaling: Hepatocyte growth factor receptor | REACTOME_SIGNALING_BY_MET | Details ... | 49 | -0.54 | -1.74 | 0.003 | 0.172 | 0.776 | 2597 | tags=47%, list=26%, | |
| 19 | Metabolism of steroid hormones | REACTOME_METABOLISM_OF_STEROID_HORMONES | Details ... | 19 | -0.62 | -1.68 | 0.001 | 0.171 | 0.955 | 1281 | tags=37%, list=13%, | |
| 110 | Lysosome vesicle biogenesis | REACTOME_LYSOSOME_VESICLE_BIOGENESIS | | 23 | -0.52 | -1.47 | 0.047 | 0.32 | 1 | 1336 | tags=35%, list=13%, | |
| 119 | Regulation of Insulin-like Growth Factor (IGF) transport and uptake by Insulin-like Growth Factor Binding Proteins (IGFBPs | REACTOME_REGULATION_OF_INSULIN_LIKE_GROWTH_FACTOR_IGF_TRANSPORT_AND_UPTAKE_BY_INSULIN_LIKE_GROWTH_FACTOR_BINDING_PROTEINS_IGFBPS | | 76 | -0.42 | -1.46 | 0.026 | 0.327 | 1 | 2988 | tags=49%, list=30%, signal=69% | |
| 239 | Focal adhesion PI3K/Akt/mTOR Signaling Pathway: dysregulation of autophagy in liver injury progression | WP_FOCAL_ADHESION_PI3K_AKT_MTOR_SIGNALING_PATHWAY | | 167 | -0.36 | -1.34 | 0.026 | 0.41 | 1 | 2745 | tags=37%, list=27%, | |
| 11 | signature downregulated genes: invasive and metastatic hepatoblastoma | CAIRO_LIVER_DEVELOPMENT_DN | Details ... | 190 | -0.46 | -1.73 | 0 | 0.167 | 0.831 | 3437 | tags=54%, list=34%, | |
| 16 | signature downregulated HFN4A targets: hepatocellular carcinoma | OHGUCHI_LIVER_HNF4A_TARGETS_DN | Details ... | 190 | -0.46 | -1.71 | 0 | 0.147 | 0.906 | 2604 | tags=45%, list=26%, | |
| 43 | PTEN regulation | REACTOME_PTEN_REGULATION | | 97 | -0.45 | -1.6 | 0 | 0.208 | 0.999 | 2710 | tags=41%, list=27%, | |
| 287 | Recycling of bile acids and salts | REACTOME_RECYCLING_OF_BILE_ACIDS_AND_SALTS | | 15 | -0.52 | -1.31 | 0.143 | 0.428 | 1 | 2653 | tags=47%, list=26%, | |
| 366 | Transport of bile salts | REACTOME_TRANSPORT_OF_BILE_SALTS_AND_ORGANIC_ACIDS_METAL_IONS_AND_AMINE_COMPOUNDS | | 34 | -0.41 | -1.25 | 0.155 | 0.493 | 1 | 2378 | tags=38%, list=24%, signal=50% | |
| 440 | Synthesis of bile acid salts via 7 alpha hydroxycholesterol | REACTOME_SYNTHESIS_OF_BILE_ACIDS_AND_BILE_SALTS_VIA_7ALPHA_HYDROXYCHOLESTEROL | | 20 | -0.43 | -1.2 | 0.22 | 0.548 | 1 | 3437 | tags=45%, list=34%, signal=68% | |
| 574 | Bile acid and bile salt metabolism | REACTOME_BILE_ACID_AND_BILE_SALT_METABOLISM | | 34 | -0.37 | -1.13 | 0.3 | 0.635 | 1 | 3437 | tags=44%, list=34%, | |
| 39 | mRNA decay by 3' to 5' exoribonuclease | REACTOME_MRNA_DECAY_BY_3_TO_5_EXORIBONUCLEASE | | 15 | -0.63 | -1.61 | 0.016 | 0.201 | 0.998 | 356 | tags=27%, list=4%, signal=28% | |
| 51 | Deadenylation dependent mRNA decay | REACTOME_DEADENYLATION_DEPENDENT_MRNA_DECAY | | 49 | -0.48 | -1.56 | 0.008 | 0.271 | 1 | 3006 | tags=47%, list=30%, | |
| 64 | Deadenylation of mRNA | REACTOME_DEADENYLATION_OF_MRNA | | 21 | -0.55 | -1.53 | 0.031 | 0.297 | 1 | 2993 | tags=57%, list=30%, | |
| 278 | Auf1 Hnrnp D0 binding: degradation of mRNAs | REACTOME_AUF1_HNRNP_D0_BINDS_AND_DESTABILIZES_MRNA | | 52 | -0.4 | -1.32 | 0.093 | 0.425 | 1 | 3600 | tags=52%, list=36%, | |
| 303 | Processing of intronless pre-mRNA | REACTOME_PROCESSING_OF_INTRONLESS_PRE_MRNAS | | 18 | -0.49 | -1.3 | 0.165 | 0.445 | 1 | 2029 | tags=39%, list=20%, | |
| 362 | Processing of capped intron containing pre-mRNA | REACTOME_PROCESSING_OF_CAPPED_INTRON_CONTAINING_PRE_MRNA | | 245 | -0.33 | -1.25 | 0.041 | 0.493 | 1 | 3999 | tags=46%, list=40%, | |
| 399 | AU-rich element binding proteins: mRNA destabilization | REACTOME_REGULATION_OF_MRNA_STABILITY_BY_PROTEINS_THAT_BIND_AU_RICH_ELEMENTS | | 79 | -0.35 | -1.24 | 0.157 | 0.5 | 1 | 3600 | tags=46%, list=36%, signal=70% | |
| 414 | RNA Polymerase I transcription | REACTOME_RNA_POLYMERASE_I_TRANSCRIPTION | | 41 | -0.39 | -1.22 | 0.182 | 0.53 | 1 | 1585 | tags=29%, list=16%, | Loss of ERBB4 related to increased liver injury and HBV chronicity https://www.ncbi.nlm.nih.gov/pmc/articles/PMC4826259/ |
| 447 | mRNA splicing | REACTOME_MRNA_SPLICING | | 186 | -0.32 | -1.2 | 0.106 | 0.552 | 1 | 3970 | tags=45%, list=39%, | |
| 500 | RNA polymerase II transcription termination | REACTOME_RNA_POLYMERASE_II_TRANSCRIPTION_TERMINATION | | 56 | -0.35 | -1.17 | 0.234 | 0.587 | 1 | 2222 | tags=29%, list=22%, | |
| 521 | Sumoylation of RNA binding proteins | REACTOME_SUMOYLATION_OF_RNA_BINDING_PROTEINS | | 36 | -0.38 | -1.15 | 0.289 | 0.607 | 1 | 1967 | tags=28%, list=19%, | |
| 551 | mRNA processing | WP_MRNA_PROCESSING | | 358 | -0.29 | -1.14 | 0.144 | 0.616 | 1 | 3540 | tags=39%, list=35%, | |
| 23 | Proteasome degradation | WP_PROTEASOME_DEGRADATION | | 47 | -0.52 | -1.67 | 0 | 0.165 | 0.971 | 2710 | tags=51%, list=27%, | |
| 80 | Circadian clock peripheral genes | UEDA_PERIFERAL_CLOCK | | 167 | -0.41 | -1.51 | 0 | 0.305 | 1 | 2416 | tags=38%, list=24%, | |
| 101 | Antigen processing: Ubiquitination & Proteasome degradation | REACTOME_ANTIGEN_PROCESSING_UBIQUITINATION_PROTEASOME_DEGRADATI | | 233 | -0.39 | -1.49 | 0 | 0.282 | 1 | 2751 | tags=36%, list=27%, | |
| 24 | Degradation of extracellular matrix | REACTOME_DEGRADATION_OF_THE_EXTRACELLULAR_MATRIX | | 40 | -0.53 | -1.66 | 0.002 | 0.179 | 0.977 | 2121 | tags=48%, list=21%, | |
| 40 | Inflammatory response | NEMETH_INFLAMMATORY_RESPONSE_LPS_UP | | 70 | -0.47 | -1.61 | 0.003 | 0.206 | 0.999 | 1685 | tags=26%, list=17%, | |
| 63 | Gap junction trafficking and regulation: hepatocyte cell cycle regulation | REACTOME_GAP_JUNCTION_TRAFFICKING_AND_REGULATION | | 21 | -0.55 | -1.54 | 0.031 | 0.286 | 1 | 999 | tags=33%, list=10%, | |
| 100 | Adaptive immune system | REACTOME_ADAPTIVE_IMMUNE_SYSTEM | | 450 | -0.38 | -1.49 | 0 | 0.279 | 1 | 2751 | tags=37%, list=27%, | |
| 102 | Collagen degradation | REACTOME_COLLAGEN_DEGRADATION | | 20 | -0.55 | -1.49 | 0.036 | 0.28 | 1 | 1621 | tags=45%, list=16%, | |
| 133 | Degradation of mitotic proteins in late mitosis / early G1 phase | REACTOME_APC_C_CDH1_MEDIATED_DEGRADATION_OF_CDC20_AND_OTHER_APC_C_CDH1_TARGETED_PROTEINS_IN_LATE_MITOSIS_EARLY_G1 | | 65 | -0.43 | -1.44 | 0.033 | 0.35 | 1 | 2710 | tags=42%, list=27%, signal=56% | |
| 382 | G1-S phase transition | REACTOME_MITOTIC_G1_PHASE_AND_G1_S_TRANSITION | | 109 | -0.35 | -1.25 | 0.108 | 0.491 | 1 | 2710 | tags=38%, list=27%, | |
| 469 | Regulation of lipid metabolism by PPAR alpha | REACTOME_REGULATION_OF_LIPID_METABOLISM_BY_PPARALPHA | | 15 | -0.47 | -1.19 | 0.269 | 0.568 | 1 | 755 | tags=20%, list=7%, signal=22% | |
| 10 | Lif signaling | ABBUD_LIF_SIGNALING_1_UP | Details ... | 33 | -0.57 | -1.74 | 0 | 0.164 | 0.797 | 2597 | tags=55%, list=26%, | |
| 17 | Notch signaling: inhibition of hepatic stellate cell activation to myofibroblasts | REACTOME_SIGNALING_BY_NOTCH | Details ... | 28 | -0.58 | -1.69 | 0.006 | 0.174 | 0.945 | 784 | tags=25%, list=8%, signal=27% | |
| 26 | EGF pathway: amelioration of liver fibrosis | BIOCARTA_EGF_PATHWAY | | 25 | -0.57 | -1.65 | 0.006 | 0.198 | 0.986 | 2648 | tags=60%, list=26%, | |
| 65 | RTK signaling | REACTOME_SIGNALING_BY_RECEPTOR_TYROSINE_KINASES | | 267 | -0.4 | -1.53 | 0 | 0.293 | 1 | 2655 | tags=37%, list=26%, | |
| 74 | Pluripotency associated mechanisms | WP_MECHANISMS_ASSOCIATED_WITH_PLURIPOTENCY | | 201 | -0.4 | -1.52 | 0 | 0.284 | 1 | 2111 | tags=32%, list=21%, | |
| 79 | ABC protein family mediated transport: bile salt transport | REACTOME_ABC_FAMILY_PROTEINS_MEDIATED_TRANSPORT | | 82 | -0.44 | -1.51 | 0.008 | 0.298 | 1 | 3048 | tags=48%, list=30%, | |
| 103 | Cargo targeting to cilia: Hedgehog signaling and liver development | REACTOME_VXPX_CARGO_TARGETING_TO_CILIUM | | 15 | -0.58 | -1.49 | 0.039 | 0.279 | 1 | 2419 | tags=60%, list=24%, | |
| 97 | MAPK family signaling cascades | REACTOME_MAPK_FAMILY_SIGNALING_CASCADES | | 183 | -0.4 | -1.49 | 0 | 0.286 | 1 | 2735 | tags=39%, list=27%, | |
| 159 | Deubiquitination: inhibition of hepatic stellate cell activation to myofibroblasts | REACTOME_DEUBIQUITINATION | | 189 | -0.38 | -1.41 | 0.004 | 0.372 | 1 | 2783 | tags=35%, list=28%, | |
| 153 | Intracellular signaling by second messangers | REACTOME_INTRACELLULAR_SIGNALING_BY_SECOND_MESSENGERS | | 178 | -0.38 | -1.41 | 0.006 | 0.364 | 1 | 2726 | tags=38%, list=27%, | |
| 228 | Hedgehog ligand biogenesis | REACTOME_HEDGEHOG_LIGAND_BIOGENESIS | | 54 | -0.41 | -1.35 | 0.067 | 0.405 | 1 | 3701 | tags=52%, list=37%, | |
| 343 | Hedgehog on state | REACTOME_HEDGEHOG_ON_STATE | | 63 | -0.37 | -1.27 | 0.112 | 0.475 | 1 | 3600 | tags=48%, list=36%, | |
| 435 | Vesicle mediated transport: hepatocellular bile secretion | REACTOME_VESICLE_MEDIATED_TRANSPORT | | 430 | -0.31 | -1.21 | 0.032 | 0.538 | 1 | 3596 | tags=42%, list=36%, | |
| 591 | signaling by hedgehog | REACTOME_SIGNALING_BY_HEDGEHOG | | 94 | -0.32 | -1.11 | 0.304 | 0.66 | 1 | 3600 | tags=44%, list=36%, | |
| 47 | | REACTOME_REGULATION_OF_PTEN_STABILITY_AND_ACTIVITY | | 63 | -0.47 | -1.58 | 0 | 0.235 | 1 | 2710 | tags=43%, list=27%, | |
| 3 | | REACTOME_COMPLEMENT_CASCADE | Details ... | 37 | -0.59 | -1.82 | 0 | 0.137 | 0.341 | 2395 | tags=59%, list=24%, | |
| 216 | | PILON_KLF1_TARGETS_UP | | 390 | -0.35 | -1.36 | 0 | 0.407 | 1 | 2442 | tags=32%, list=24%, | |
| 196 | | MARTORIATI_MDM4_TARGETS_FETAL_LIVER_DN | | 430 | -0.35 | -1.37 | 0.001 | 0.403 | 1 | 2820 | tags=35%, list=28%, | |
| 309 | | MARTINEZ_TP53_TARGETS_UP | | 442 | -0.33 | -1.29 | 0.001 | 0.448 | 1 | 3044 | tags=35%, list=30%, | |
| 4 | | WANG_RESPONSE_TO_BEXAROTENE_DN | Details ... | 17 | -0.69 | -1.8 | 0.001 | 0.169 | 0.487 | 1264 | tags=47%, list=13%, | |
| 44 | | GROSS_HYPOXIA_VIA_ELK3_AND_HIF1A_DN | | 88 | -0.45 | -1.6 | 0.001 | 0.21 | 0.999 | 3687 | tags=55%, list=37%, | |
| 34 | | BREDEMEYER_RAG_SIGNALING_NOT_VIA_ATM_DN | | 39 | -0.52 | -1.62 | 0.006 | 0.201 | 0.995 | 1591 | tags=31%, list=16%, | |
| 304 | | REACTOME_TRANSPORT_OF_SMALL_MOLECULES | | 377 | -0.33 | -1.3 | 0.008 | 0.445 | 1 | 3065 | tags=37%, list=30%, | |
| 49 | | WUNDER_INFLAMMATORY_RESPONSE_AND_CHOLESTEROL_UP | | 51 | -0.48 | -1.57 | 0.01 | 0.25 | 1 | 2126 | tags=41%, list=21%, | |
| 141 | | REACTOME_REGULATION_OF_RUNX2_EXPRESSION_AND_ACTIVITY | | 50 | -0.44 | -1.42 | 0.039 | 0.361 | 1 | 3600 | tags=52%, list=36%, | |
| 136 | | REACTOME_UBIQUITIN_MEDIATED_DEGRADATION_OF_PHOSPHORYLATED_CDC25 | | 46 | -0.45 | -1.43 | 0.039 | 0.349 | 1 | 2710 | tags=43%, list=27%, | |
| 183 | | REACTOME_ACTIVATION_OF_APC_C_AND_APC_C_CDC20_MEDIATED_DEGRADATION_OF_MITOTIC_PROTEINS | | 66 | -0.41 | -1.38 | 0.041 | 0.398 | 1 | 2710 | tags=41%, list=27%, signal=56% | |
| 147 | | SHETH_LIVER_CANCER_VS_TXNIP_LOSS_PAM3 | | 43 | -0.45 | -1.42 | 0.044 | 0.363 | 1 | 1817 | tags=40%, list=18%, | |
| 205 | | REACTOME_TRANSCRIPTIONAL_REGULATION_BY_RUNX2 | | 61 | -0.41 | -1.36 | 0.047 | 0.408 | 1 | 2710 | tags=41%, list=27%, | |
| 210 | | REACTOME_PINK1_PRKN_MEDIATED_MITOPHAGY | | 20 | -0.5 | -1.36 | 0.112 | 0.412 | 1 | 882 | tags=20%, list=9%, signal=22% | |
| 456 | | REACTOME_HEDGEHOG_OFF_STATE | | 76 | -0.35 | -1.19 | 0.203 | 0.569 | 1 | 3600 | tags=45%, list=36%, | |
| 596 | | REACTOME_GENE_SILENCING_BY_RNA | | 36 | -0.36 | -1.11 | 0.329 | 0.657 | 1 | 1967 | tags=25%, list=19%, | |
| 624 | | REACTOME_ACTIVATION_OF_THE_MRNA_UPON_BINDING_OF_THE_CAP_BINDING_COMPLEX_AND_EIFS_AND_SUBSEQUENT_BINDING_TO_43S | | 57 | -0.33 | -1.09 | 0.334 | 0.684 | 1 | 4335 | tags=47%, list=43%, signal=83% | |
| 660 | | REACTOME_TRANSPORT_OF_MATURE_MRNAS_DERIVED_FROM_INTRONLESS_TRANSCRIPTS | | 37 | -0.35 | -1.07 | 0.38 | 0.712 | 1 | 2029 | tags=27%, list=20%, signal=34% | |
| 709 | | YAUCH_HEDGEHOG_SIGNALING_PARACRINE_DN | | 64 | -0.31 | -1.04 | 0.418 | 0.737 | 1 | 1844 | tags=22%, list=18%, | |
| 719 | | REACTOME_SYNTHESIS_OF_BILE_ACIDS_AND_BILE_SALTS | | 26 | -0.36 | -1.04 | 0.424 | 0.738 | 1 | 1192 | tags=19%, list=12%, | |
| 808 | | REACTOME_RNA_POLYMERASE_II_PRE_TRANSCRIPTION_EVENTS | | 68 | -0.29 | -0.99 | 0.481 | 0.779 | 1 | 3907 | tags=44%, list=39%, | |
| 853 | | WP_DYSREGULATED_MIRNA_TARGETING_IN_INSULIN_PI3K_AKT_SIGNALING | | 23 | -0.34 | -0.97 | 0.514 | 0.802 | 1 | 1621 | tags=30%, list=16%, | |
| 858 | | REACTOME_PROCESSING_OF_CAPPED_INTRONLESS_PRE_MRNA | | 25 | -0.33 | -0.97 | 0.533 | 0.801 | 1 | 2029 | tags=28%, list=20%, | |
| 859 | | REACTOME_TRISTETRAPROLIN_TTP_ZFP36_BINDS_AND_DESTABILIZES_MRNA | | 16 | -0.37 | -0.97 | 0.541 | 0.801 | 1 | 2655 | tags=38%, list=26%, | |
| 863 | | REACTOME_FORMATION_OF_RNA_POL_II_ELONGATION_COMPLEX | | 48 | -0.3 | -0.96 | 0.546 | 0.803 | 1 | 4385 | tags=52%, list=43%, | |
| 947 | | REACTOME_MRNA_CAPPING | | 26 | -0.32 | -0.92 | 0.606 | 0.833 | 1 | 633 | tags=12%, list=6%, signal=12% | |
| 964 | | REACTOME_TRANSCRIPTIONAL_REGULATION_BY_SMALL_RNAS | | 30 | -0.31 | -0.91 | 0.606 | 0.837 | 1 | 5345 | tags=70%, list=53%, | |
| 961 | | REACTOME_BUTYRATE_RESPONSE_FACTOR_1_BRF1_BINDS_AND_DESTABILIZES | | 16 | -0.35 | -0.91 | 0.614 | 0.836 | 1 | 1193 | tags=19%, list=12%, | |
| 988 | | REACTOME_POSITIVE_EPIGENETIC_REGULATION_OF_RRNA_EXPRESSION | | 26 | -0.31 | -0.89 | 0.624 | 0.859 | 1 | 3987 | tags=58%, list=39%, | |
| 1036 | | REACTOME_MRNA_SPLICING_MINOR_PATHWAY | | 45 | -0.27 | -0.86 | 0.696 | 0.875 | 1 | 3704 | tags=38%, list=37%, | |
| 1058 | | ULE_SPLICING_VIA_NOVA2 | | 27 | -0.29 | -0.84 | 0.708 | 0.896 | 1 | 2818 | tags=33%, list=28%, | |
| 1078 | | REACTOME_RNA_POLYMERASE_III_TRANSCRIPTION_INITIATION_FROM_TYPE_3_PROMOTER | | 24 | -0.29 | -0.82 | 0.728 | 0.901 | 1 | 775 | tags=17%, list=8%, signal=18% | |
| 1096 | | REACTOME_RNA_POLYMERASE_III_TRANSCRIPTION_INITIATION_FROM_TYPE_1_PROMOTER | | 25 | -0.28 | -0.81 | 0.745 | 0.907 | 1 | 775 | tags=16%, list=8%, signal=17% | |
| 1090 | | YAUCH_HEDGEHOG_SIGNALING_PARACRINE_UP | | 60 | -0.24 | -0.81 | 0.774 | 0.909 | 1 | 3224 | tags=33%, list=32%, | |
| 1081 | | REACTOME_RNA_POLYMERASE_II_TRANSCRIBES_SNRNA_GENES | | 67 | -0.24 | -0.82 | 0.774 | 0.902 | 1 | 3315 | tags=34%, list=33%, | |
| 1120 | | REACTOME_RNA_POLYMERASE_II_PROMOTER_ESCAPE | | 41 | -0.25 | -0.79 | 0.804 | 0.915 | 1 | 712 | tags=12%, list=7%, signal=13% | |
| 1205 | | REACTOME_TRANSPORT_OF_THE_SLBP_DEPENDANT_MATURE_MRNA | | 31 | -0.23 | -0.69 | 0.907 | 0.965 | 1 | 5134 | tags=58%, list=51%, | |
| 1242 | | WP_SPLICING_FACTOR_NOVA_REGULATED_SYNAPTIC_PROTEINS | | 26 | -0.22 | -0.63 | 0.933 | 0.979 | 1 | 2902 | tags=42%, list=29%, | |
| 1267 | | REACTOME_FGFR2_ALTERNATIVE_SPLICING | | 16 | -0.19 | -0.5 | 0.977 | 0.997 | 1 | 3315 | tags=31%, list=33%, | |
| 20 | | ICHIBA_GRAFT_VERSUS_HOST_DISEASE_D7_UP | Details ... | 107 | -0.47 | -1.68 | 0 | 0.175 | 0.963 | 2651 | tags=45%, list=26%, | |
| 38 | | GROSS_HYPOXIA_VIA_ELK3_DN | | 112 | -0.45 | -1.61 | 0 | 0.203 | 0.998 | 2354 | tags=38%, list=23%, | |
| 68 | | REACTOME_CLASS_I_MHC_MEDIATED_ANTIGEN_PROCESSING_PRESENTATION | | 270 | -0.4 | -1.53 | 0 | 0.289 | 1 | 2537 | tags=35%, list=25%, | |
| 15 | | HOWLIN_CITED1_TARGETS_1_UP | Details ... | 17 | -0.66 | -1.71 | 0.001 | 0.15 | 0.893 | 2155 | tags=59%, list=21%, | |
| 225 | | MARTINEZ_RB1_TARGETS_UP | | 468 | -0.35 | -1.35 | 0.001 | 0.406 | 1 | 2568 | tags=31%, list=25%, | |
| 21 | | REACTOME_ANTIGEN_PROCESSING_CROSS_PRESENTATION | | 68 | -0.5 | -1.68 | 0.002 | 0.168 | 0.965 | 2710 | tags=50%, list=27%, | |
| 27 | | YAO_TEMPORAL_RESPONSE_TO_PROGESTERONE_CLUSTER_0 | | 60 | -0.49 | -1.65 | 0.002 | 0.193 | 0.987 | 965 | tags=33%, list=10%, | |
| 86 | | MARKEY_RB1_ACUTE_LOF_DN | | 183 | -0.4 | -1.5 | 0.002 | 0.293 | 1 | 2718 | tags=39%, list=27%, | |
| 42 | | REACTOME_METABOLISM_OF_POLYAMINES | | 54 | -0.49 | -1.6 | 0.003 | 0.21 | 0.999 | 2710 | tags=48%, list=27%, | |
| 77 | | GAVIN_FOXP3_TARGETS_CLUSTER_T4 | | 77 | -0.44 | -1.52 | 0.003 | 0.292 | 1 | 3616 | tags=57%, list=36%, | |
| 132 | | REACTOME_NEDDYLATION | | 178 | -0.38 | -1.44 | 0.003 | 0.351 | 1 | 2969 | tags=38%, list=29%, | |
| 13 | | RUAN_RESPONSE_TO_TNF_TROGLITAZONE_UP | Details ... | 16 | -0.65 | -1.72 | 0.004 | 0.158 | 0.865 | 952 | tags=38%, list=9%, signal=41% | |
| 14 | | WP_BLOOD_CLOTTING_CASCADE | Details ... | 15 | -0.65 | -1.71 | 0.004 | 0.157 | 0.886 | 2395 | tags=73%, list=24%, | |
| 46 | | BOYLAN_MULTIPLE_MYELOMA_PCA1_UP | | 63 | -0.47 | -1.58 | 0.004 | 0.239 | 1 | 2082 | tags=40%, list=21%, | |
| 71 | | REACTOME_EXTRACELLULAR_MATRIX_ORGANIZATION | | 100 | -0.43 | -1.53 | 0.004 | 0.283 | 1 | 3853 | tags=61%, list=38%, | |
| 137 | | ZHENG_FOXP3_TARGETS_IN_THYMUS_UP | | 166 | -0.38 | -1.43 | 0.004 | 0.352 | 1 | 2112 | tags=33%, list=21%, | |
| 140 | | PASQUALUCCI_LYMPHOMA_BY_GC_STAGE_UP | | 202 | -0.38 | -1.43 | 0.004 | 0.361 | 1 | 3173 | tags=43%, list=31%, | |
| 25 | | GUILLAUMOND_KLF10_TARGETS_UP | | 30 | -0.55 | -1.66 | 0.005 | 0.182 | 0.979 | 2553 | tags=57%, list=25%, | |
| 33 | | BIOCARTA_INTRINSIC_PATHWAY | | 19 | -0.61 | -1.63 | 0.005 | 0.2 | 0.995 | 1953 | tags=58%, list=19%, | |
| 59 | | REACTOME_DOWNSTREAM_TCR_SIGNALING | | 73 | -0.45 | -1.55 | 0.005 | 0.265 | 1 | 2710 | tags=42%, list=27%, | |
| 81 | | GUO_TARGETS_OF_IRS1_AND_IRS2 | | 87 | -0.43 | -1.51 | 0.005 | 0.303 | 1 | 3433 | tags=52%, list=34%, | |
| 117 | | GRAESSMANN_RESPONSE_TO_MC_AND_SERUM_DEPRIVATION_UP | | 147 | -0.39 | -1.46 | 0.005 | 0.33 | 1 | 2137 | tags=33%, list=21%, | |
| 12 | | REACTOME_RHOBTB1_GTPASE_CYCLE | Details ... | 21 | -0.62 | -1.72 | 0.006 | 0.167 | 0.862 | 2370 | tags=48%, list=23%, | |
| 18 | | MA_MYELOID_DIFFERENTIATION_UP | Details ... | 29 | -0.58 | -1.69 | 0.006 | 0.173 | 0.951 | 1825 | tags=34%, list=18%, | |
| 126 | | BROWN_MYELOID_CELL_DEVELOPMENT_UP | | 110 | -0.4 | -1.45 | 0.006 | 0.34 | 1 | 3253 | tags=44%, list=32%, | |
| 48 | | KHETCHOUMIAN_TRIM24_TARGETS_UP | | 37 | -0.51 | -1.58 | 0.007 | 0.249 | 1 | 2314 | tags=43%, list=23%, | |
| 30 | | YAO_TEMPORAL_RESPONSE_TO_PROGESTERONE_CLUSTER_5 | | 21 | -0.61 | -1.64 | 0.008 | 0.19 | 0.992 | 2030 | tags=48%, list=20%, | |
| 31 | | REACTOME_ELASTIC_FIBRE_FORMATION | | 18 | -0.62 | -1.63 | 0.008 | 0.198 | 0.992 | 2120 | tags=61%, list=21%, | |
| 53 | | WP_DELTA_NOTCH_SIGNALING_PATHWAY | | 56 | -0.47 | -1.56 | 0.008 | 0.269 | 1 | 1685 | tags=32%, list=17%, | |
| 125 | | REACTOME_UB_SPECIFIC_PROCESSING_PROTEASES | | 136 | -0.4 | -1.45 | 0.009 | 0.342 | 1 | 2768 | tags=38%, list=27%, | |
| 131 | | GAUSSMANN_MLL_AF4_FUSION_TARGETS_C_UP | | 127 | -0.39 | -1.44 | 0.009 | 0.352 | 1 | 2198 | tags=31%, list=22%, | |
| 22 | | REACTOME_ER_PHAGOSOME_PATHWAY | | 17 | -0.63 | -1.67 | 0.01 | 0.17 | 0.971 | 2520 | tags=65%, list=25%, | |
| 29 | | BIOCARTA_AGR_PATHWAY | | 19 | -0.61 | -1.64 | 0.01 | 0.183 | 0.988 | 1345 | tags=42%, list=13%, | |
| 41 | | NADLER_OBESITY_UP | | 51 | -0.48 | -1.61 | 0.01 | 0.203 | 0.999 | 2718 | tags=49%, list=27%, | |
| 92 | | REACTOME_TCR_SIGNALING | | 84 | -0.44 | -1.5 | 0.01 | 0.295 | 1 | 2710 | tags=43%, list=27%, | |
| 98 | | BRUINS_UVC_RESPONSE_MIDDLE | | 73 | -0.44 | -1.49 | 0.01 | 0.284 | 1 | 2171 | tags=38%, list=21%, | |
| 107 | | REACTOME_PCP_CE_PATHWAY | | 72 | -0.44 | -1.48 | 0.01 | 0.303 | 1 | 2834 | tags=43%, list=28%, | |
| 37 | | CUI_TCF21_TARGETS_UP | | 20 | -0.59 | -1.62 | 0.011 | 0.202 | 0.998 | 688 | tags=40%, list=7%, signal=43% | |
| 45 | | REACTOME_PLATELET_AGGREGATION_PLUG_FORMATION | | 25 | -0.55 | -1.6 | 0.011 | 0.215 | 1 | 2652 | tags=52%, list=26%, | |
| 104 | | ICHIBA_GRAFT_VERSUS_HOST_DISEASE_35D_DN | | 56 | -0.45 | -1.49 | 0.011 | 0.277 | 1 | 1799 | tags=30%, list=18%, | |
| 122 | | WP_EGFR1_SIGNALING_PATHWAY | | 132 | -0.4 | -1.45 | 0.011 | 0.332 | 1 | 2875 | tags=39%, list=28%, | |
| 201 | | REACTOME_METABOLISM_OF_AMINO_ACIDS_AND_DERIVATIVES | | 201 | -0.36 | -1.37 | 0.011 | 0.406 | 1 | 3677 | tags=45%, list=36%, | |
| 377 | | GRAESSMANN_APOPTOSIS_BY_SERUM_DEPRIVATION_UP | | 388 | -0.32 | -1.25 | 0.011 | 0.49 | 1 | 2547 | tags=29%, list=25%, | |
| 35 | | REACTOME_ECM_PROTEOGLYCANS | | 19 | -0.6 | -1.62 | 0.012 | 0.199 | 0.995 | 1811 | tags=53%, list=18%, | |
| 54 | | HAN_JNK_SINGALING_DN | | 30 | -0.52 | -1.56 | 0.012 | 0.264 | 1 | 949 | tags=30%, list=9%, signal=33% | |
| 58 | | REACTOME_RHOBTB_GTPASE_CYCLE | | 32 | -0.53 | -1.55 | 0.012 | 0.259 | 1 | 1321 | tags=31%, list=13%, | |
| 32 | | KOHOUTEK_CCNT1_TARGETS | | 25 | -0.57 | -1.63 | 0.013 | 0.206 | 0.994 | 1426 | tags=40%, list=14%, | |
| 28 | | REACTOME_NON_INTEGRIN_MEMBRANE_ECM_INTERACTIONS | | 16 | -0.63 | -1.65 | 0.014 | 0.187 | 0.987 | 1849 | tags=56%, list=18%, | |
| 56 | | WP_TYPE_II_INTERFERON_SIGNALING_IFNG | | 21 | -0.57 | -1.56 | 0.014 | 0.262 | 1 | 1367 | tags=38%, list=14%, | |
| 145 | | HUANG_GATA2_TARGETS_UP | | 104 | -0.4 | -1.42 | 0.014 | 0.366 | 1 | 2635 | tags=37%, list=26%, | |
| 170 | | REACTOME_CELLULAR_RESPONSE_TO_CHEMICAL_STRESS | | 145 | -0.38 | -1.4 | 0.014 | 0.369 | 1 | 2710 | tags=35%, list=27%, | |
| 36 | | OUYANG_PROSTATE_CANCER_PROGRESSION_UP | | 19 | -0.6 | -1.62 | 0.015 | 0.197 | 0.996 | 625 | tags=32%, list=6%, signal=34% | |
| 94 | | REACTOME_G_ALPHA_Q_SIGNALLING_EVENTS | | 66 | -0.44 | -1.49 | 0.015 | 0.292 | 1 | 1031 | tags=23%, list=10%, | |
| 96 | | REACTOME_DEGRADATION_OF_AXIN | | 52 | -0.45 | -1.49 | 0.015 | 0.287 | 1 | 2710 | tags=40%, list=27%, | |
| 148 | | REACTOME_FC_EPSILON_RECEPTOR_FCERI_SIGNALING | | 98 | -0.4 | -1.42 | 0.016 | 0.365 | 1 | 2902 | tags=45%, list=29%, | |
| 217 | | RAO_BOUND_BY_SALL4 | | 147 | -0.37 | -1.36 | 0.016 | 0.408 | 1 | 2341 | tags=32%, list=23%, | |
| 60 | | REACTOME_SIGNALING_BY_ERBB2 | | 32 | -0.52 | -1.55 | 0.017 | 0.261 | 1 | 2539 | tags=41%, list=25%, | |
| 69 | | BIOCARTA_TPO_PATHWAY | | 23 | -0.54 | -1.53 | 0.017 | 0.286 | 1 | 2648 | tags=57%, list=26%, | |
| 75 | | YAO_TEMPORAL_RESPONSE_TO_PROGESTERONE_CLUSTER_1 | | 51 | -0.46 | -1.52 | 0.017 | 0.28 | 1 | 1911 | tags=33%, list=19%, | |
| 127 | | REACTOME_UCH_PROTEINASES | | 72 | -0.43 | -1.44 | 0.017 | 0.339 | 1 | 3600 | tags=50%, list=36%, | |
| 361 | | MARTINEZ_RB1_AND_TP53_TARGETS_UP | | 443 | -0.32 | -1.25 | 0.017 | 0.492 | 1 | 3044 | tags=34%, list=30%, | |
| 52 | | REACTOME_SIGNALING_BY_PDGF | | 35 | -0.51 | -1.56 | 0.018 | 0.271 | 1 | 2597 | tags=51%, list=26%, | |
| 95 | | ZHONG_SECRETOME_OF_LUNG_CANCER_AND_ENDOTHELIUM | | 60 | -0.44 | -1.49 | 0.018 | 0.29 | 1 | 4012 | tags=57%, list=40%, | |
| 62 | | REACTOME_ANTIGEN_PRESENTATION_FOLDING_ASSEMBLY_AND_PEPTIDE_LOADING_OF_CLASS_I_MHC | | 24 | -0.55 | -1.55 | 0.019 | 0.26 | 1 | 1914 | tags=42%, list=19%, signal=51% | |
| 66 | | BIOCARTA_TFF_PATHWAY | | 18 | -0.58 | -1.53 | 0.019 | 0.289 | 1 | 1213 | tags=33%, list=12%, | |
| 105 | | ZENG_GU_ICB_CONTROL_METAGENE_31_PRECICTIVE_ICB_RESISTANCE | | 46 | -0.46 | -1.48 | 0.019 | 0.299 | 1 | 2557 | tags=39%, list=25%, | |
| 249 | | DURCHDEWALD_SKIN_CARCINOGENESIS_DN | | 191 | -0.35 | -1.33 | 0.019 | 0.414 | 1 | 3272 | tags=39%, list=32%, | |
| 380 | | MARTINEZ_RB1_AND_TP53_TARGETS_DN | | 375 | -0.32 | -1.25 | 0.019 | 0.492 | 1 | 3044 | tags=36%, list=30%, | |
| 89 | | HESS_TARGETS_OF_HOXA9_AND_MEIS1_UP | | 49 | -0.47 | -1.5 | 0.02 | 0.296 | 1 | 1216 | tags=20%, list=12%, | |
| 154 | | IVANOVA_HEMATOPOIESIS_STEM_CELL | | 105 | -0.4 | -1.41 | 0.02 | 0.363 | 1 | 1977 | tags=39%, list=20%, | |
| 231 | | KARLSSON_TGFB1_TARGETS_DN | | 167 | -0.36 | -1.35 | 0.02 | 0.408 | 1 | 2901 | tags=35%, list=29%, | |
| 57 | | REACTOME_RHOU_GTPASE_CYCLE | | 31 | -0.52 | -1.56 | 0.021 | 0.26 | 1 | 783 | tags=29%, list=8%, signal=31% | |
| 67 | | MCDOWELL_ACUTE_LUNG_INJURY_UP | | 34 | -0.51 | -1.53 | 0.021 | 0.291 | 1 | 2314 | tags=44%, list=23%, | |
| 50 | | BIOCARTA_IL2_PATHWAY | | 15 | -0.6 | -1.57 | 0.022 | 0.269 | 1 | 2648 | tags=60%, list=26%, | |
| 90 | | WP_IL_7_SIGNALING_PATHWAY | | 37 | -0.49 | -1.5 | 0.022 | 0.296 | 1 | 799 | tags=27%, list=8%, signal=29% | |
| 108 | | LEE_AGING_NEOCORTEX_UP | | 58 | -0.44 | -1.47 | 0.022 | 0.314 | 1 | 2762 | tags=38%, list=27%, | |
| 61 | | SHIN_B_CELL_LYMPHOMA_CLUSTER_2 | | 16 | -0.6 | -1.55 | 0.023 | 0.26 | 1 | 1267 | tags=38%, list=13%, | |
| 111 | | REACTOME_REGULATION_OF_RAS_BY_GAPS | | 57 | -0.44 | -1.47 | 0.023 | 0.317 | 1 | 2735 | tags=42%, list=27%, | |
| 190 | | WP_FOCAL_ADHESION | | 108 | -0.38 | -1.38 | 0.023 | 0.4 | 1 | 2745 | tags=37%, list=27%, | |
| 185 | | LANDIS_ERBB2_BREAST_TUMORS_324_UP | | 121 | -0.38 | -1.38 | 0.024 | 0.396 | 1 | 3241 | tags=40%, list=32%, | |
| 374 | | ZHENG_BOUND_BY_FOXP3 | | 346 | -0.32 | -1.25 | 0.024 | 0.491 | 1 | 2725 | tags=31%, list=27%, | |
| 55 | | REACTOME_MET_PROMOTES_CELL_MOTILITY | | 20 | -0.58 | -1.56 | 0.025 | 0.262 | 1 | 2579 | tags=60%, list=26%, | |
| 83 | | REACTOME_ENDOGENOUS_STEROLS | | 15 | -0.6 | -1.5 | 0.025 | 0.299 | 1 | 1375 | tags=47%, list=14%, | |
| 93 | | REACTOME_FORMATION_OF_FIBRIN_CLOT_CLOTTING_CASCADE | | 25 | -0.53 | -1.5 | 0.025 | 0.292 | 1 | 2540 | tags=56%, list=25%, | |
| 99 | | REACTOME_PEPTIDE_LIGAND_BINDING_RECEPTORS | | 33 | -0.49 | -1.49 | 0.025 | 0.282 | 1 | 2713 | tags=45%, list=27%, | |
| 112 | | LEE_AGING_CEREBELLUM_UP | | 61 | -0.44 | -1.46 | 0.025 | 0.325 | 1 | 2919 | tags=41%, list=29%, | |
| 176 | | ZENG_GU_ICB_CONTROL_METAGENE_42_PRECICTIVE_ICB_RESISTANCE | | 120 | -0.38 | -1.38 | 0.025 | 0.405 | 1 | 2817 | tags=35%, list=28%, | |
| 70 | | HAN_JNK_SINGALING_UP | | 29 | -0.51 | -1.53 | 0.026 | 0.284 | 1 | 884 | tags=28%, list=9%, signal=30% | |
| 78 | | REACTOME_EPH_EPHRIN_SIGNALING | | 38 | -0.49 | -1.51 | 0.026 | 0.301 | 1 | 2654 | tags=42%, list=26%, | |
| 142 | | REACTOME_SIGNALING_BY_THE_B_CELL_RECEPTOR_BCR | | 85 | -0.41 | -1.42 | 0.026 | 0.365 | 1 | 2894 | tags=41%, list=29%, | |
| 84 | | BILANGES_SERUM_SENSITIVE_VIA_TSC2 | | 26 | -0.52 | -1.5 | 0.027 | 0.298 | 1 | 1885 | tags=42%, list=19%, | |
| 88 | | WANG_TARGETS_OF_MLL_CBP_FUSION_UP | | 31 | -0.5 | -1.5 | 0.027 | 0.295 | 1 | 2150 | tags=48%, list=21%, | |
| 158 | | REACTOME_FCERI_MEDIATED_NF_KB_ACTIVATION | | 69 | -0.41 | -1.41 | 0.027 | 0.364 | 1 | 2710 | tags=42%, list=27%, | |
| 87 | | REACTOME_DAP12_INTERACTIONS | | 25 | -0.54 | -1.5 | 0.028 | 0.292 | 1 | 2520 | tags=56%, list=25%, | |
| 91 | | LEIN_NEURON_MARKERS | | 24 | -0.53 | -1.5 | 0.028 | 0.294 | 1 | 2408 | tags=46%, list=24%, | |
| 181 | | REACTOME_DEATH_RECEPTOR_SIGNALING | | 102 | -0.39 | -1.38 | 0.028 | 0.398 | 1 | 2654 | tags=35%, list=26%, | |
| 130 | | WP_IL_5_SIGNALING_PATHWAY | | 50 | -0.44 | -1.44 | 0.029 | 0.353 | 1 | 2597 | tags=44%, list=26%, | |
| 157 | | KIM_GLIS2_TARGETS_UP | | 57 | -0.43 | -1.41 | 0.029 | 0.363 | 1 | 2534 | tags=47%, list=25%, | |
| 73 | | BIOCARTA_GLEEVEC_PATHWAY | | 21 | -0.56 | -1.52 | 0.03 | 0.285 | 1 | 2579 | tags=48%, list=26%, | |
| 82 | | KUNINGER_IGF1_VS_PDGFB_TARGETS_DN | | 25 | -0.53 | -1.51 | 0.031 | 0.302 | 1 | 907 | tags=28%, list=9%, signal=31% | |
| 109 | | REACTOME_N_GLYCAN_TRIMMING_IN_THE_ER_AND_CALNEXIN_CALRETICULIN_C | | 15 | -0.58 | -1.47 | 0.031 | 0.319 | 1 | 1684 | tags=40%, list=17%, | |
| 411 | | SWEET_LUNG_CANCER_KRAS_UP | | 362 | -0.32 | -1.22 | 0.031 | 0.525 | 1 | 2550 | tags=30%, list=25%, | |
| 149 | | REACTOME_CELL_SURFACE_INTERACTIONS_AT_THE_VASCULAR_WALL | | 55 | -0.43 | -1.42 | 0.032 | 0.37 | 1 | 2646 | tags=40%, list=26%, | |
| 262 | | MCBRYAN_PUBERTAL_BREAST_4_5WK_UP | | 144 | -0.36 | -1.32 | 0.032 | 0.422 | 1 | 1679 | tags=26%, list=17%, | |
| 323 | | RAMALHO_STEMNESS_UP | | 183 | -0.34 | -1.28 | 0.032 | 0.466 | 1 | 3372 | tags=39%, list=33%, | |
| 115 | | HOWLIN_PUBERTAL_MAMMARY_GLAND | | 38 | -0.48 | -1.46 | 0.033 | 0.332 | 1 | 1026 | tags=21%, list=10%, | |
| 155 | | REACTOME_KEAP1_NFE2L2_PATHWAY | | 73 | -0.41 | -1.41 | 0.033 | 0.36 | 1 | 2710 | tags=40%, list=27%, | |
| 409 | | REACTOME_HEMOSTASIS | | 302 | -0.32 | -1.23 | 0.033 | 0.519 | 1 | 2713 | tags=33%, list=27%, | |
| 72 | | BERENJENO_ROCK_SIGNALING_NOT_VIA_RHOA_UP | | 16 | -0.59 | -1.52 | 0.034 | 0.286 | 1 | 2774 | tags=63%, list=27%, | |
| 120 | | BYSTROEM_CORRELATED_WITH_IL5_DN | | 46 | -0.45 | -1.45 | 0.034 | 0.334 | 1 | 2985 | tags=41%, list=30%, | |
| 106 | | BIOCARTA_HER2_PATHWAY | | 18 | -0.56 | -1.48 | 0.035 | 0.303 | 1 | 2597 | tags=50%, list=26%, | |
| 161 | | REACTOME_CDK_MEDIATED_PHOSPHORYLATION_AND_REMOVAL_OF_CDC6 | | 64 | -0.42 | -1.41 | 0.035 | 0.372 | 1 | 2710 | tags=41%, list=27%, | |
| 139 | | REACTOME_REGULATION_OF_RUNX3_EXPRESSION_AND_ACTIVITY | | 50 | -0.44 | -1.43 | 0.036 | 0.36 | 1 | 2710 | tags=42%, list=27%, | |
| 113 | | APRELIKOVA_BRCA1_TARGETS | | 37 | -0.48 | -1.46 | 0.037 | 0.334 | 1 | 1770 | tags=35%, list=18%, | |
| 123 | | WP_TRANSLATION_FACTORS | | 45 | -0.46 | -1.45 | 0.037 | 0.339 | 1 | 3250 | tags=53%, list=32%, | |
| 114 | | REACTOME_HDACS_DEACETYLATE_HISTONES | | 21 | -0.53 | -1.46 | 0.038 | 0.334 | 1 | 3647 | tags=67%, list=36%, | |
| 118 | | LIN_NPAS4_TARGETS_DN | | 34 | -0.47 | -1.46 | 0.038 | 0.327 | 1 | 3358 | tags=53%, list=33%, | |
| 193 | | REACTOME_ACTIVATION_OF_NF_KAPPAB_IN_B_CELLS | | 61 | -0.41 | -1.37 | 0.038 | 0.399 | 1 | 3600 | tags=49%, list=36%, | |
| 251 | | REACTOME_NERVOUS_SYSTEM_DEVELOPMENT | | 147 | -0.36 | -1.33 | 0.038 | 0.412 | 1 | 2369 | tags=31%, list=23%, | |
| 85 | | REACTOME_MOLECULES_ASSOCIATED_WITH_ELASTIC_FIBRES | | 16 | -0.58 | -1.5 | 0.039 | 0.295 | 1 | 2120 | tags=56%, list=21%, | |

| | | | | | | | | | | | | |
|---|---|---|---|---|---|---|---|---|---|---|---|---|
| 135 | | | ZHENG_FOXP3_TARGETS_IN_T_LYMPHOCYTE_DN | | 31 | -0.48 | -1.43 | 0.039 | 0.35 | 1 | 2895 | tags=45%, list=29%, |
| 156 | | | REACTOME_CROSS_PRESENTATION_OF_SOLUBLE_EXOGENOUS_ANTIGENS_ENDOSOMES | | 44 | -0.44 | -1.41 | 0.039 | 0.358 | 1 | 2710 | tags=45%, list=27%, signal=62% |
| 116 | | | REACTOME_EGFR_DOWNREGULATION | | 19 | -0.54 | -1.46 | 0.04 | 0.33 | 1 | 458 | tags=16%, list=5%, signal=17% |
| 169 | | | HUMMERICH_SKIN_CANCER_PROGRESSION_UP | | 64 | -0.42 | -1.4 | 0.04 | 0.371 | 1 | 3648 | tags=53%, list=36%, |
| 299 | | | REACTOME_SIGNALING_BY_INTERLEUKINS | | 175 | -0.35 | -1.3 | 0.04 | 0.434 | 1 | 2935 | tags=38%, list=29%, |
| 121 | | | MCBRYAN_PUBERTAL_TGFB1_TARGETS_DN | | 42 | -0.46 | -1.45 | 0.043 | 0.333 | 1 | 1044 | tags=21%, list=10%, |
| 209 | | | ZENG_GU_ICB_CONTROL_METAGENE_25_PRECICTIVE_ICB_RESPONSE | | 83 | -0.39 | -1.36 | 0.044 | 0.41 | 1 | 2578 | tags=40%, list=26%, |
| 76 | | | REACTOME_EPH_EPHRIN_MEDIATED_REPULSION_OF_CELLS | | 15 | -0.6 | -1.52 | 0.045 | 0.284 | 1 | 908 | tags=33%, list=9%, signal=37% |
| 168 | | | REACTOME_CELL_CELL_COMMUNICATION | | 54 | -0.43 | -1.4 | 0.045 | 0.368 | 1 | 2707 | tags=41%, list=27%, |
| 194 | | | REACTOME_DEGRADATION_OF_BETA_CATENIN_BY_THE_DESTRUCTION_COMPLE | | 71 | -0.4 | -1.37 | 0.046 | 0.398 | 1 | 2710 | tags=38%, list=27%, |
| 247 | | | ZHENG_GLIOBLASTOMA_PLASTICITY_UP | | 125 | -0.37 | -1.34 | 0.046 | 0.414 | 1 | 2660 | tags=38%, list=26%, |
| 358 | | | GOLDRATH_ANTIGEN_RESPONSE | | 239 | -0.33 | -1.26 | 0.046 | 0.485 | 1 | 2546 | tags=31%, list=25%, |
| 296 | | | BRUINS_UVC_RESPONSE_VIA_TP53_GROUP_D | | 152 | -0.35 | -1.3 | 0.048 | 0.438 | 1 | 3456 | tags=42%, list=34%, |
| 195 | | | BIOCARTA_MAPK_PATHWAY | | 55 | -0.42 | -1.37 | 0.049 | 0.4 | 1 | 4488 | tags=67%, list=44%, |
| 283 | | | ZHONG_SECRETOME_OF_LUNG_CANCER_AND_FIBROBLAST | | 112 | -0.36 | -1.31 | 0.049 | 0.429 | 1 | 4096 | tags=47%, list=41%, |
| 165 | | | LANDIS_BREAST_CANCER_PROGRESSION_UP | | 37 | -0.46 | -1.4 | 0.05 | 0.368 | 1 | 2363 | tags=41%, list=23%, |
| 187 | | | JACKSON_DNMT1_TARGETS_UP | | 61 | -0.41 | -1.38 | 0.05 | 0.396 | 1 | 3160 | tags=48%, list=31%, |
| 215 | | | REACTOME_FACTORS_INVOLVED_IN_MEGAKARYOCYTE_DEVELOPMENT_AND_PLATELET_PRODUCTION | | 65 | -0.41 | -1.36 | 0.05 | 0.408 | 1 | 3495 | tags=48%, list=35%, signal=72% |
| 221 | | | YAO_HOXA10_TARGETS_VIA_PROGESTERONE_UP | | 58 | -0.41 | -1.35 | 0.05 | 0.405 | 1 | 1190 | tags=24%, list=12%, |
| 164 | | | YAO_TEMPORAL_RESPONSE_TO_PROGESTERONE_CLUSTER_16 | | 42 | -0.44 | -1.4 | 0.051 | 0.369 | 1 | 2507 | tags=43%, list=25%, |
| 290 | | | SANSOM_APC_TARGETS | | 113 | -0.36 | -1.31 | 0.051 | 0.435 | 1 | 3597 | tags=47%, list=36%, |
| 230 | | | REACTOME_MHC_CLASS_II_ANTIGEN_PRESENTATION | | 71 | -0.4 | -1.35 | 0.052 | 0.409 | 1 | 3581 | tags=49%, list=35%, |
| 413 | | | REACTOME_CYTOKINE_SIGNALING_IN_IMMUNE_SYSTEM | | 285 | -0.32 | -1.22 | 0.052 | 0.531 | 1 | 2935 | tags=35%, list=29%, |
| 124 | | | STARK_HYPPOCAMPUS_22Q11_DELETION_DN | | 18 | -0.54 | -1.45 | 0.053 | 0.338 | 1 | 2266 | tags=50%, list=22%, |
| 452 | | | MARTINEZ_TP53_TARGETS_DN | | 380 | -0.31 | -1.19 | 0.053 | 0.568 | 1 | 3044 | tags=35%, list=30%, |
| 138 | | | REACTOME_SIGNALING_BY_EGFR | | 33 | -0.47 | -1.43 | 0.055 | 0.353 | 1 | 1524 | tags=24%, list=15%, |
| 204 | | | REACTOME_STABILIZATION_OF_P53 | | 52 | -0.41 | -1.36 | 0.055 | 0.409 | 1 | 2710 | tags=40%, list=27%, |
| 235 | | | YAO_TEMPORAL_RESPONSE_TO_PROGESTERONE_CLUSTER_9 | | 62 | -0.4 | -1.34 | 0.055 | 0.407 | 1 | 2430 | tags=34%, list=24%, |
| 175 | | | LEIN_MEDULLA_MARKERS | | 44 | -0.44 | -1.39 | 0.056 | 0.403 | 1 | 2061 | tags=34%, list=20%, |
| 188 | | | REACTOME_SELECTIVE_AUTOPHAGY | | 48 | -0.43 | -1.38 | 0.056 | 0.394 | 1 | 2728 | tags=42%, list=27%, |
| 192 | | | REACTOME_DEGRADATION_OF_DVL | | 53 | -0.42 | -1.38 | 0.056 | 0.397 | 1 | 3676 | tags=53%, list=36%, |
| 128 | | | REACTOME_BINDING_AND_UPTAKE_OF_LIGANDS_BY_SCAVENGER_RECEPTORS | | 26 | -0.49 | -1.44 | 0.057 | 0.355 | 1 | 1756 | tags=38%, list=17%, |
| 172 | | | REACTOME_CELL_JUNCTION_ORGANIZATION | | 43 | -0.44 | -1.4 | 0.057 | 0.377 | 1 | 2306 | tags=37%, list=23%, |
| 178 | | | REACTOME_P75_NTR_RECEPTOR_MEDIATED_SIGNALLING | | 56 | -0.42 | -1.38 | 0.057 | 0.401 | 1 | 2654 | tags=38%, list=26%, |
| 134 | | | REACTOME_INTEGRIN_SIGNALING | | 22 | -0.51 | -1.43 | 0.058 | 0.35 | 1 | 2652 | tags=45%, list=26%, |
| 318 | | | PLASARI_TGFB1_TARGETS_10HR_DN | | 164 | -0.35 | -1.28 | 0.058 | 0.462 | 1 | 1004 | tags=20%, list=10%, |
| 184 | | | MCBRYAN_PUBERTAL_BREAST_6_7WK_DN | | 49 | -0.43 | -1.38 | 0.059 | 0.397 | 1 | 1335 | tags=29%, list=13%, |
| 202 | | | WP_INTEGRIN_MEDIATED_CELL_ADHESION | | 61 | -0.41 | -1.37 | 0.059 | 0.405 | 1 | 2817 | tags=39%, list=28%, |
| 167 | | | WP_OXIDATIVE_DAMAGE_RESPONSE | | 27 | -0.48 | -1.4 | 0.06 | 0.366 | 1 | 976 | tags=26%, list=10%, |
| 311 | | | REACTOME_TRANSPORT_TO_THE_GOLGI_AND_SUBSEQUENT_MODIFICATION | | 138 | -0.35 | -1.29 | 0.06 | 0.452 | 1 | 3596 | tags=43%, list=36%, |
| 150 | | | REACTOME_NCAM_SIGNALING_FOR_NEURITE_OUT_GROWTH | | 20 | -0.52 | -1.42 | 0.061 | 0.368 | 1 | 1621 | tags=35%, list=16%, |
| 163 | | | STEARMAN_LUNG_CANCER_EARLY_VS_LATE_DN | | 45 | -0.44 | -1.4 | 0.061 | 0.371 | 1 | 949 | tags=27%, list=9%, signal=29% |
| 229 | | | REACTOME_SIGNALING_BY_NTRK1_TRKA | | 46 | -0.42 | -1.35 | 0.061 | 0.409 | 1 | 2834 | tags=43%, list=28%, |
| 324 | | | GROSS_HYPOXIA_VIA_ELK3_UP | | 158 | -0.34 | -1.28 | 0.061 | 0.468 | 1 | 2313 | tags=29%, list=23%, |
| 375 | | | BOYLAN_MULTIPLE_MYELOMA_C_D_DN | | 154 | -0.34 | -1.25 | 0.061 | 0.49 | 1 | 3354 | tags=44%, list=33%, |
| 144 | | | LA_MEN1_TARGETS | | 15 | -0.56 | -1.42 | 0.062 | 0.368 | 1 | 2033 | tags=40%, list=20%, |
| 180 | | | BERENJENO_ROCK_SIGNALING_NOT_VIA_RHOA_DN | | 43 | -0.44 | -1.38 | 0.062 | 0.398 | 1 | 2621 | tags=42%, list=26%, |
| 281 | | | ZENG_GU_ICB_CONTROL_METAGENE_44_PRECICTIVE_ICB_RESPONSE | | 117 | -0.36 | -1.31 | 0.062 | 0.43 | 1 | 1406 | tags=22%, list=14%, |
| 351 | | | REACTOME_SIGNALING_BY_GPCR | | 170 | -0.34 | -1.26 | 0.062 | 0.485 | 1 | 2654 | tags=32%, list=26%, |
| 410 | | | DELACROIX_RARG_BOUND_MEF | | 244 | -0.32 | -1.23 | 0.062 | 0.518 | 1 | 2143 | tags=27%, list=21%, |
| 146 | | | REACTOME_RUNX1_INTERACTS_WITH_CO_FACTORS_WHOSE_PRECISE_EFFECT_ON_RUNX1_TARGETS_IS_NOT_KNOWN | | 25 | -0.49 | -1.42 | 0.063 | 0.365 | 1 | 1530 | tags=36%, list=15%, signal=42% |
| 151 | | | KASLER_HDAC7_TARGETS_2_DN | | 21 | -0.51 | -1.41 | 0.063 | 0.366 | 1 | 3192 | tags=48%, list=32%, |
| 276 | | | REACTOME_DNA_REPLICATION_PRE_INITIATION | | 88 | -0.37 | -1.32 | 0.063 | 0.427 | 1 | 2710 | tags=38%, list=27%, |
| 166 | | | BIOCARTA_PDGF_PATHWAY | | 26 | -0.5 | -1.4 | 0.064 | 0.368 | 1 | 2648 | tags=58%, list=26%, |
| 208 | | | SEKI_INFLAMMATORY_RESPONSE_LPS_UP | | 45 | -0.42 | -1.36 | 0.064 | 0.412 | 1 | 3432 | tags=51%, list=34%, |
| 129 | | | REACTOME_DOWNREGULATION_OF_ERBB2_SIGNALING | | 15 | -0.56 | -1.44 | 0.065 | 0.354 | 1 | 148 | tags=20%, list=1%, signal=20% |
| 162 | | | GREENBAUM_E2A_TARGETS_UP | | 16 | -0.55 | -1.4 | 0.065 | 0.372 | 1 | 2004 | tags=31%, list=20%, |
| 353 | | | REACTOME_DEVELOPMENTAL_BIOLOGY | | 184 | -0.34 | -1.26 | 0.066 | 0.486 | 1 | 3113 | tags=38%, list=31%, |
| 143 | | | GROSS_HYPOXIA_VIA_ELK3_ONLY_UP | | 23 | -0.5 | -1.42 | 0.067 | 0.364 | 1 | 2197 | tags=48%, list=22%, |
| 233 | | | REACTOME_GLI3_IS_PROCESSED_TO_GLI3R_BY_THE_PROTEASOME | | 54 | -0.41 | -1.35 | 0.068 | 0.407 | 1 | 3600 | tags=50%, list=36%, |
| 284 | | | FOSTER_KDM1A_TARGETS_UP | | 94 | -0.37 | -1.31 | 0.068 | 0.427 | 1 | 2745 | tags=38%, list=27%, |
| 320 | | | ZHANG_BREAST_CANCER_PROGENITORS_DN | | 102 | -0.36 | -1.28 | 0.069 | 0.462 | 1 | 2769 | tags=34%, list=27%, |
| 152 | | | BIOCARTA_EPO_PATHWAY | | 17 | -0.54 | -1.41 | 0.07 | 0.366 | 1 | 2648 | tags=53%, list=26%, |
| 198 | | | REACTOME_COSTIMULATION_BY_THE_CD28_FAMILY | | 34 | -0.45 | -1.37 | 0.07 | 0.4 | 1 | 2652 | tags=44%, list=26%, |
| 261 | | | REACTOME_THE_ROLE_OF_GTSE1_IN_G2_M_PROGRESSION_AFTER_G2_CHECKP | | 59 | -0.4 | -1.33 | 0.07 | 0.423 | 1 | 2710 | tags=42%, list=27%, |
| 253 | | | ZEMEK_IMMUNE_CHECKPOINT_BLOCKADE_OVARIAN_CANCER_RENCA_UP | | 51 | -0.42 | -1.33 | 0.072 | 0.411 | 1 | 2869 | tags=45%, list=28%, |
| 334 | | | MCBRYAN_PUBERTAL_BREAST_5_6WK_DN | | 114 | -0.36 | -1.27 | 0.072 | 0.472 | 1 | 3721 | tags=48%, list=37%, |
| 213 | | | GAVIN_FOXP3_TARGETS_CLUSTER_P2 | | 41 | -0.44 | -1.36 | 0.073 | 0.41 | 1 | 2297 | tags=39%, list=23%, |
| 246 | | | REACTOME_NUCLEAR_EVENTS_MEDIATED_BY_NFE2L2 | | 54 | -0.41 | -1.34 | 0.074 | 0.415 | 1 | 3600 | tags=48%, list=36%, |
| 255 | | | REACTOME_DOWNSTREAM_SIGNALING_EVENTS_OF_B_CELL_RECEPTOR_BCR | | 74 | -0.39 | -1.33 | 0.074 | 0.421 | 1 | 2894 | tags=41%, list=29%, |
| 298 | | | TSENG_IRS1_TARGETS_DN | | 93 | -0.37 | -1.3 | 0.074 | 0.435 | 1 | 2477 | tags=38%, list=25%, |
| 289 | | | REACTOME_C_TYPE_LECTIN_RECEPTORS_CLRS | | 91 | -0.37 | -1.31 | 0.075 | 0.431 | 1 | 3287 | tags=46%, list=33%, |
| 171 | | | STEARMAN_TUMOR_FIELD_EFFECT_UP | | 30 | -0.46 | -1.4 | 0.076 | 0.368 | 1 | 2082 | tags=40%, list=21%, |
| 174 | | | BRACHAT_RESPONSE_TO_CAMPTOTHECIN_UP | | 20 | -0.51 | -1.39 | 0.076 | 0.402 | 1 | 1132 | tags=30%, list=11%, |
| 244 | | | REACTOME_ASYMMETRIC_LOCALIZATION_OF_PCP_PROTEINS | | 55 | -0.41 | -1.34 | 0.076 | 0.416 | 1 | 3676 | tags=51%, list=36%, |
| 245 | | | CASTELLANO_NRAS_TARGETS_UP | | 58 | -0.4 | -1.34 | 0.076 | 0.414 | 1 | 2955 | tags=43%, list=29%, |
| 266 | | | WP_ESC_PLURIPOTENCY_PATHWAYS | | 63 | -0.39 | -1.32 | 0.076 | 0.424 | 1 | 998 | tags=21%, list=10%, |
| 160 | | | LEE_AGING_MUSCLE_UP | | 26 | -0.48 | -1.41 | 0.077 | 0.373 | 1 | 1619 | tags=27%, list=16%, |
| 302 | | | RASHI_RESPONSE_TO_IONIZING_RADIATION_5 | | 94 | -0.37 | -1.3 | 0.078 | 0.441 | 1 | 3096 | tags=40%, list=31%, |
| 237 | | | ZEMEK_IMMUNE_CHECKPOINT_BLOCKADE_OVARIAN_CANCER_OVERLAP_UP | | 50 | -0.42 | -1.34 | 0.079 | 0.407 | 1 | 2869 | tags=46%, list=28%, |
| 238 | | | IKEDA_MIR1_TARGETS_UP | | 49 | -0.41 | -1.34 | 0.081 | 0.406 | 1 | 2037 | tags=35%, list=20%, |
| 482 | | | NABA_MATRISOME | | 325 | -0.31 | -1.18 | 0.081 | 0.582 | 1 | 3068 | tags=41%, list=30%, |
| 173 | | | REACTOME_CARGO_CONCENTRATION_IN_THE_ER | | 25 | -0.48 | -1.39 | 0.082 | 0.378 | 1 | 3581 | tags=52%, list=35%, |
| 186 | | | JOHANSSON_BRAIN_CANCER_EARLY_VS_LATE_DN | | 36 | -0.44 | -1.38 | 0.082 | 0.397 | 1 | 2722 | tags=42%, list=27%, |
| 291 | | | REACTOME_CARGO_RECOGNITION_FOR_CLATHRIN_MEDIATED_ENDOCYTOSIS | | 70 | -0.38 | -1.31 | 0.082 | 0.435 | 1 | 3113 | tags=41%, list=31%, |
| 352 | | | CHYLA_CBFA2T3_TARGETS_DN | | 127 | -0.35 | -1.26 | 0.082 | 0.484 | 1 | 3353 | tags=41%, list=33%, |
| 232 | | | LEE_TARGETS_OF_PTCH1_AND_SUFU_UP | | 36 | -0.44 | -1.35 | 0.083 | 0.407 | 1 | 2672 | tags=44%, list=26%, |
| 301 | | | REACTOME_APC_C_MEDIATED_DEGRADATION_OF_CELL_CYCLE_PROTEINS | | 74 | -0.38 | -1.3 | 0.083 | 0.433 | 1 | 2710 | tags=38%, list=27%, |
| 331 | | | REACTOME_MACROAUTOPHAGY | | 99 | -0.36 | -1.28 | 0.083 | 0.469 | 1 | 2743 | tags=35%, list=27%, |
| 243 | | | CHIARADONNA_NEOPLASTIC_TRANSFORMATION_KRAS_CDC25_DN | | 37 | -0.43 | -1.34 | 0.084 | 0.416 | 1 | 2669 | tags=46%, list=26%, |
| 257 | | | BURTON_ADIPOGENESIS_11 | | 48 | -0.41 | -1.33 | 0.084 | 0.42 | 1 | 2306 | tags=31%, list=23%, |
| 212 | | | LEE_AGING_MUSCLE_DN | | 31 | -0.46 | -1.36 | 0.085 | 0.411 | 1 | 2203 | tags=45%, list=22%, |
| 236 | | | BIOCARTA_KERATINOCYTE_PATHWAY | | 37 | -0.44 | -1.34 | 0.085 | 0.408 | 1 | 807 | tags=22%, list=8%, signal=23% |
| 275 | | | REACTOME_MAPK6_MAPK4_SIGNALING | | 63 | -0.39 | -1.32 | 0.085 | 0.425 | 1 | 2710 | tags=41%, list=27%, |
| 407 | | | WANG_TUMOR_INVASIVENESS_DN | | 184 | -0.33 | -1.23 | 0.085 | 0.52 | 1 | 4724 | tags=53%, list=47%, |
| 223 | | | ZHENG_GLIOBLASTOMA_PLASTICITY_DN | | 29 | -0.46 | -1.35 | 0.086 | 0.404 | 1 | 2089 | tags=52%, list=21%, |
| 248 | | | MCCLUNG_COCAIN_REWARD_4WK | | 46 | -0.41 | -1.34 | 0.086 | 0.413 | 1 | 2323 | tags=43%, list=23%, |
| 345 | | | REACTOME_CLATHRIN_MEDIATED_ENDOCYTOSIS | | 100 | -0.36 | -1.27 | 0.086 | 0.476 | 1 | 2890 | tags=39%, list=29%, |
| 514 | | | REACTOME_CELLULAR_RESPONSES_TO_STIMULI | | 371 | -0.3 | -1.16 | 0.087 | 0.594 | 1 | 3728 | tags=41%, list=37%, |
| 177 | | | GROSS_ELK3_TARGETS_UP | | 21 | -0.5 | -1.38 | 0.088 | 0.403 | 1 | 1927 | tags=33%, list=19%, |
| 206 | | | BIOCARTA_NUCLEARRS_PATHWAY | | 25 | -0.47 | -1.36 | 0.088 | 0.409 | 1 | 3052 | tags=48%, list=30%, |
| 260 | | | MARSON_FOXP3_TARGETS_UP | | 53 | -0.4 | -1.33 | 0.088 | 0.424 | 1 | 3046 | tags=45%, list=30%, |
| 288 | | | REACTOME_TRANSPORT_OF_MATURE_TRANSCRIPT_TO_CYTOPLASM | | 69 | -0.39 | -1.31 | 0.088 | 0.427 | 1 | 2222 | tags=30%, list=22%, |
| 207 | | | WP_IL_9_SIGNALING_PATHWAY | | 18 | -0.51 | -1.36 | 0.089 | 0.411 | 1 | 2597 | tags=50%, list=26%, |
| 234 | | | JIANG_AGING_CEREBRAL_CORTEX_DN | | 37 | -0.43 | -1.35 | 0.09 | 0.406 | 1 | 4208 | tags=51%, list=42%, |
| 182 | | | BAUS_TFF2_TARGETS_UP | | 23 | -0.49 | -1.38 | 0.091 | 0.398 | 1 | 2649 | tags=61%, list=26%, |
| 219 | | | REACTOME_EPHB_MEDIATED_FORWARD_SIGNALING | | 25 | -0.48 | -1.35 | 0.091 | 0.407 | 1 | 2451 | tags=36%, list=24%, |
| 280 | | | MORI_PRE_BI_LYMPHOCYTE_DN | | 64 | -0.39 | -1.31 | 0.091 | 0.429 | 1 | 2335 | tags=39%, list=23%, |
| 328 | | | NABA_CORE_MATRISOME | | 95 | -0.36 | -1.28 | 0.091 | 0.469 | 1 | 2745 | tags=40%, list=27%, |
| 191 | | | BIOCARTA_NGF_PATHWAY | | 16 | -0.52 | -1.38 | 0.092 | 0.398 | 1 | 2648 | tags=50%, list=26%, |
| 199 | | | REACTOME_FCERI_MEDIATED_MAPK_ACTIVATION | | 20 | -0.5 | -1.37 | 0.092 | 0.4 | 1 | 3699 | tags=70%, list=37%, |
| 200 | | | LEE_AGING_NEOCORTEX_DN | | 35 | -0.45 | -1.37 | 0.092 | 0.398 | 1 | 2593 | tags=34%, list=26%, |
| 218 | | | WP_ONE_CARBON_METABOLISM | | 24 | -0.47 | -1.35 | 0.092 | 0.409 | 1 | 3582 | tags=50%, list=35%, |
| 240 | | | HOUSTIS_ROS | | 33 | -0.45 | -1.34 | 0.092 | 0.414 | 1 | 4692 | tags=70%, list=46%, |
| 386 | | | ICHIBA_GRAFT_VERSUS_HOST_DISEASE_35D_UP | | 116 | -0.35 | -1.24 | 0.092 | 0.491 | 1 | 2530 | tags=36%, list=25%, |
| 400 | | | GRAESSMANN_APOPTOSIS_BY_SERUM_DEPRIVATION_DN | | 139 | -0.34 | -1.23 | 0.092 | 0.509 | 1 | 2620 | tags=32%, list=26%, |
| 241 | | | LEE_AGING_CEREBELLUM_DN | | 37 | -0.44 | -1.34 | 0.093 | 0.412 | 1 | 1761 | tags=32%, list=17%, |
| 267 | | | REACTOME_CELL_DEATH_SIGNALLING_VIA_NRAGE_NRIF_AND_NADE | | 44 | -0.41 | -1.32 | 0.093 | 0.423 | 1 | 2654 | tags=39%, list=26%, |
| 509 | | | WANG_TUMOR_INVASIVENESS_UP | | 340 | -0.3 | -1.16 | 0.093 | 0.596 | 1 | 4011 | tags=40%, list=40%, |
| 279 | | | WP_ALPHA_6_BETA_4_INTEGRIN_SIGNALING_PATHWAY | | 42 | -0.42 | -1.31 | 0.094 | 0.426 | 1 | 1213 | tags=24%, list=12%, |
| 344 | | | REACTOME_ASSEMBLY_OF_THE_PRE_REPLICATIVE_COMPLEX | | 77 | -0.37 | -1.27 | 0.094 | 0.476 | 1 | 2710 | tags=38%, list=27%, |
| 189 | | | STEGER_ADIPOGENESIS_UP | | 16 | -0.52 | -1.38 | 0.095 | 0.393 | 1 | 952 | tags=31%, list=9%, signal=34% |
| 203 | | | BIOCARTA_MET_PATHWAY | | 28 | -0.47 | -1.36 | 0.095 | 0.407 | 1 | 2597 | tags=39%, list=26%, |
| 314 | | | REACTOME_SCF_SKP2_MEDIATED_DEGRADATION_OF_P27_P21 | | 55 | -0.39 | -1.29 | 0.095 | 0.453 | 1 | 2710 | tags=40%, list=27%, |
| 342 | | | INGRAM_SHH_TARGETS_UP | | 93 | -0.36 | -1.27 | 0.095 | 0.472 | 1 | 2726 | tags=32%, list=27%, |
| 179 | | | REACTOME_SUMOYLATION_OF_INTRACELLULAR_RECEPTORS | | 19 | -0.51 | -1.38 | 0.096 | 0.399 | 1 | 3052 | tags=53%, list=30%, |
| 211 | | | REACTOME_SIGNAL_TRANSDUCTION_BY_L1 | | 17 | -0.52 | -1.36 | 0.096 | 0.41 | 1 | 4509 | tags=82%, list=45%, |
| 294 | | | REACTOME_CYCLIN_A_CDK2_ASSOCIATED_EVENTS_AT_S_PHASE_ENTRY | | 63 | -0.39 | -1.3 | 0.096 | 0.439 | 1 | 2710 | tags=38%, list=27%, |
| 242 | | | BIOCARTA_ECM_PATHWAY | | 19 | -0.5 | -1.34 | 0.098 | 0.411 | 1 | 1213 | tags=26%, list=12%, |
| 286 | | | BILANGES_RAPAMYCIN_SENSITIVE_VIA_TSC1_AND_TSC2 | | 63 | -0.39 | -1.31 | 0.098 | 0.427 | 1 | 2225 | tags=29%, list=22%, |
| 305 | | | ZENG_GU_ICB_CONTROL_METAGENE_8_PRECICTIVE_ICB_RESPONSE | | 57 | -0.39 | -1.3 | 0.098 | 0.446 | 1 | 1949 | tags=28%, list=19%, |
| 306 | | | GAVIN_FOXP3_TARGETS_CLUSTER_P7 | | 51 | -0.4 | -1.29 | 0.099 | 0.446 | 1 | 3342 | tags=47%, list=33%, |
| 197 | | | REACTOME_THROMBIN_SIGNALLING_THROUGH_PROTEINASE_ACTIVATED_RECEPTORS_PARS | | 17 | -0.52 | -1.37 | 0.1 | 0.402 | 1 | 804 | tags=24%, list=8%, signal=26% |
| 227 | | | REACTOME_COMMON_PATHWAY_OF_FIBRIN_CLOT_FORMATION | | 15 | -0.53 | -1.35 | 0.1 | 0.403 | 1 | 3530 | tags=73%, list=35%, |
| 254 | | | POS_HISTAMINE_RESPONSE_NETWORK | | 20 | -0.49 | -1.33 | 0.101 | 0.413 | 1 | 2519 | tags=50%, list=25%, |
| 297 | | | CHESLER_BRAIN_QTL_CIS | | 52 | -0.4 | -1.3 | 0.101 | 0.436 | 1 | 2128 | tags=29%, list=21%, |
| 513 | | | BOCHKIS_FOXA2_TARGETS | | 383 | -0.3 | -1.16 | 0.101 | 0.594 | 1 | 3472 | tags=38%, list=34%, |
| 224 | | | RASHI_NFKB1_TARGETS | | 15 | -0.53 | -1.35 | 0.102 | 0.404 | 1 | 2621 | tags=47%, list=26%, |
| 264 | | | BORLAK_LIVER_CANCER_EGF_UP | | 38 | -0.43 | -1.32 | 0.102 | 0.423 | 1 | 2651 | tags=47%, list=26%, |
| 347 | | | GAVIN_FOXP3_TARGETS_CLUSTER_T7 | | 87 | -0.36 | -1.26 | 0.102 | 0.481 | 1 | 2635 | tags=29%, list=26%, |
| 359 | | | REACTOME_METABOLISM_OF_STEROIDS | | 85 | -0.36 | -1.26 | 0.102 | 0.488 | 1 | 3437 | tags=45%, list=34%, |
| 222 | | | REACTOME_INTERLEUKIN_2_FAMILY_SIGNALING | | 17 | -0.52 | -1.35 | 0.103 | 0.406 | 1 | 2597 | tags=53%, list=26%, |
| 250 | | | BIOCARTA_P38MAPK_PATHWAY | | 25 | -0.47 | -1.33 | 0.104 | 0.413 | 1 | 4410 | tags=72%, list=44%, |
| 271 | | | MARSON_FOXP3_TARGETS_DN | | 38 | -0.43 | -1.32 | 0.105 | 0.426 | 1 | 2649 | tags=37%, list=26%, |
| 424 | | | TORCHIA_TARGETS_OF_EWSR1_FLI1_FUSION_UP | | 173 | -0.33 | -1.22 | 0.105 | 0.527 | 1 | 1572 | tags=23%, list=16%, |
| 325 | | | ZAMORA_NOS2_TARGETS_DN | | 71 | -0.38 | -1.28 | 0.107 | 0.468 | 1 | 3406 | tags=45%, list=34%, |
| 330 | | | REACTOME_TRANSCRIPTIONAL_REGULATION_BY_RUNX3 | | 67 | -0.38 | -1.28 | 0.107 | 0.468 | 1 | 2710 | tags=37%, list=27%, |
| 357 | | | REACTOME_BETA_CATENIN_INDEPENDENT_WNT_SIGNALING | | 94 | -0.36 | -1.26 | 0.107 | 0.486 | 1 | 2834 | tags=37%, list=28%, |
| 392 | | | NABA_ECM_REGULATORS | | 106 | -0.35 | -1.24 | 0.107 | 0.497 | 1 | 3068 | tags=46%, list=30%, |
| 270 | | | BURTON_ADIPOGENESIS_4 | | 35 | -0.43 | -1.32 | 0.108 | 0.424 | 1 | 1946 | tags=34%, list=19%, |
| 537 | | | REACTOME_MEMBRANE_TRAFFICKING | | 405 | -0.3 | -1.15 | 0.108 | 0.607 | 1 | 3596 | tags=41%, list=36%, |
| 263 | | | MCDOWELL_ACUTE_LUNG_INJURY_DN | | 32 | -0.45 | -1.32 | 0.109 | 0.424 | 1 | 1340 | tags=31%, list=13%, |
| 327 | | | QI_PLASMACYTOMA_DN | | 68 | -0.38 | -1.28 | 0.109 | 0.468 | 1 | 3330 | tags=46%, list=33%, |
| 312 | | | REACTOME_GAMMA_CARBOXYLATION_HYPUSINYLATION_HYDROXYLATION_AND_ARYLSULFATASE_ACTIVATION | | 44 | -0.41 | -1.29 | 0.11 | 0.451 | 1 | 3630 | tags=52%, list=36%, signal=81% |
| 341 | | | REACTOME_TNFR2_NON_CANONICAL_NF_KB_PATHWAY | | 61 | -0.38 | -1.27 | 0.11 | 0.473 | 1 | 3600 | tags=49%, list=36%, |
| 226 | | | GAZIN_EPIGENETIC_SILENCING_BY_KRAS | | 16 | -0.53 | -1.35 | 0.111 | 0.405 | 1 | 1547 | tags=31%, list=15%, |
| 497 | | | MONNIER_POSTRADIATION_TUMOR_ESCAPE_DN | | 304 | -0.3 | -1.17 | 0.111 | 0.588 | 1 | 2670 | tags=31%, list=26%, |
| 220 | | | REACTOME_GPVI_MEDIATED_ACTIVATION_CASCADE | | 20 | -0.5 | -1.35 | 0.112 | 0.406 | 1 | 2241 | tags=45%, list=22%, |
| 259 | | | REACTOME_ISG15_ANTIVIRAL_MECHANISM | | 27 | -0.45 | -1.33 | 0.113 | 0.423 | 1 | 3585 | tags=52%, list=35%, |
| 256 | | | MIKKELSEN_ES_HCP_WITH_H3_UNMETHYLATED | | 15 | -0.51 | -1.33 | 0.114 | 0.421 | 1 | 3065 | tags=60%, list=30%, |
| 214 | | | REACTOME_PARACETAMOL_ADME | | 21 | -0.49 | -1.36 | 0.115 | 0.409 | 1 | 2442 | tags=33%, list=24%, |
| 417 | | | REACTOME_SUMOYLATION | | 121 | -0.34 | -1.22 | 0.115 | 0.531 | 1 | 2170 | tags=27%, list=21%, |
| 370 | | | GROSS_HYPOXIA_VIA_ELK3_AND_HIF1A_UP | | 109 | -0.35 | -1.25 | 0.116 | 0.492 | 1 | 2938 | tags=37%, list=29%, |
| 252 | | | REACTOME_CLASS_B_2_SECRETIN_FAMILY_RECEPTORS | | 17 | -0.51 | -1.33 | 0.117 | 0.412 | 1 | 521 | tags=18%, list=5%, signal=19% |
| 258 | | | WP_WHITE_FAT_CELL_DIFFERENTIATION | | 20 | -0.48 | -1.33 | 0.118 | 0.424 | 1 | 4116 | tags=65%, list=41%, |
| 282 | | | REACTOME_FLT3_SIGNALING | | 25 | -0.46 | -1.31 | 0.119 | 0.428 | 1 | 604 | tags=20%, list=6%, signal=21% |
| 384 | | | GAUSSMANN_MLL_AF4_FUSION_TARGETS_F_UP | | 102 | -0.35 | -1.25 | 0.119 | 0.491 | 1 | 3061 | tags=39%, list=30%, |
| 437 | | | YAO_TEMPORAL_RESPONSE_TO_PROGESTERONE_CLUSTER_17 | | 174 | -0.32 | -1.21 | 0.119 | 0.543 | 1 | 3368 | tags=36%, list=33%, |
| 395 | | | GERY_CEBP_TARGETS | | 94 | -0.35 | -1.24 | 0.12 | 0.495 | 1 | 2690 | tags=36%, list=27%, |
| 273 | | | HEVNER_CORTEX_CAUDAL_VENTRICULAR_ZONE | | 26 | -0.46 | -1.32 | 0.121 | 0.427 | 1 | 1723 | tags=31%, list=17%, |
| 316 | | | REACTOME_DECTIN_1_MEDIATED_NONCANONICAL_NF_KB_SIGNALING | | 52 | -0.39 | -1.29 | 0.121 | 0.46 | 1 | 2710 | tags=40%, list=27%, |
| 315 | | | MODY_HIPPOCAMPUS_POSTNATAL | | 34 | -0.43 | -1.29 | 0.122 | 0.461 | 1 | 3585 | tags=44%, list=35%, |
| 268 | | | REACTOME_SIGNALING_BY_SCF_KIT | | 24 | -0.47 | -1.32 | 0.123 | 0.423 | 1 | 2597 | tags=42%, list=26%, |
| 313 | | | REACTOME_TRANS_GOLGI_NETWORK_VESICLE_BUDDING | | 56 | -0.39 | -1.29 | 0.123 | 0.452 | 1 | 2061 | tags=34%, list=20%, |
| 272 | | | BURTON_ADIPOGENESIS_1 | | 25 | -0.46 | -1.32 | 0.124 | 0.426 | 1 | 2089 | tags=32%, list=21%, |
| 365 | | | MIKKELSEN_MEF_LCP_WITH_H3K4ME3 | | 71 | -0.37 | -1.25 | 0.125 | 0.495 | 1 | 3172 | tags=49%, list=31%, |
| 265 | | | REACTOME_CD28_CO_STIMULATION | | 20 | -0.49 | -1.32 | 0.126 | 0.425 | 1 | 2552 | tags=45%, list=25%, |
| 355 | | | HUMMERICH_SKIN_CANCER_PROGRESSION_DN | | 66 | -0.37 | -1.26 | 0.126 | 0.486 | 1 | 2044 | tags=29%, list=20%, |
| 308 | | | RASHI_RESPONSE_TO_IONIZING_RADIATION_3 | | 33 | -0.43 | -1.29 | 0.127 | 0.449 | 1 | 1563 | tags=30%, list=15%, |
| 348 | | | SAFFORD_T_LYMPHOCYTE_ANERGY | | 46 | -0.39 | -1.26 | 0.127 | 0.48 | 1 | 627 | tags=15%, list=6%, signal=16% |
| 402 | | | REACTOME_CLEC7A_DECTIN_1_SIGNALING | | 80 | -0.35 | -1.23 | 0.128 | 0.518 | 1 | 2894 | tags=41%, list=29%, |
| 408 | | | MA_RAT_AGING_DN | | 117 | -0.34 | -1.23 | 0.129 | 0.519 | 1 | 2137 | tags=30%, list=21%, |
| 422 | | | BYSTRYKH_HEMATOPOIESIS_STEM_CELL_AND_BRAIN_QTL_TRANS | | 114 | -0.34 | -1.22 | 0.129 | 0.529 | 1 | 3111 | tags=39%, list=31%, |
| 292 | | | BIOCARTA_INSULIN_PATHWAY | | 18 | -0.49 | -1.31 | 0.13 | 0.434 | 1 | 2648 | tags=50%, list=26%, |
| 360 | | | YAO_TEMPORAL_RESPONSE_TO_PROGESTERONE_CLUSTER_7 | | 63 | -0.37 | -1.26 | 0.13 | 0.488 | 1 | 2796 | tags=38%, list=28%, |
| 397 | | | REACTOME_CELLULAR_RESPONSE_TO_HYPOXIA | | 65 | -0.37 | -1.24 | 0.13 | 0.501 | 1 | 2749 | tags=35%, list=27%, |
| 269 | | | ONO_FOXP3_TARGETS_DN | | 16 | -0.5 | -1.32 | 0.131 | 0.422 | 1 | 2577 | tags=56%, list=26%, |
| 285 | | | HOFFMANN_IMMATURE_TO_MATURE_B_LYMPHOCYTE_UP | | 19 | -0.48 | -1.31 | 0.131 | 0.426 | 1 | 2919 | tags=53%, list=29%, |
| 319 | | | LEIN_MIDBRAIN_MARKERS | | 32 | -0.44 | -1.28 | 0.133 | 0.463 | 1 | 1309 | tags=22%, list=13%, |
| 431 | | | MCBRYAN_PUBERTAL_BREAST_3_4WK_UP | | 124 | -0.33 | -1.21 | 0.133 | 0.531 | 1 | 2651 | tags=35%, list=26%, |
| 434 | | | REACTOME_SEPARATION_OF_SISTER_CHROMATIDS | | 136 | -0.33 | -1.21 | 0.133 | 0.535 | 1 | 2710 | tags=36%, list=27%, |
| 453 | | | REACTOME_ASPARAGINE_N_LINKED_GLYCOSYLATION | | 201 | -0.32 | -1.19 | 0.133 | 0.567 | 1 | 3596 | tags=41%, list=36%, |
| 317 | | | REACTOME_SUMOYLATION_OF_TRANSCRIPTION_COFACTORS | | 28 | -0.44 | -1.29 | 0.136 | 0.46 | 1 | 2091 | tags=36%, list=21%, |
| 293 | | | BRACHAT_RESPONSE_TO_METHOTREXATE_DN | | 22 | -0.48 | -1.3 | 0.138 | 0.439 | 1 | 2068 | tags=36%, list=20%, |
| 335 | | | REACTOME_CYTOPROTECTION_BY_HMOX1 | | 46 | -0.4 | -1.27 | 0.138 | 0.472 | 1 | 3728 | tags=46%, list=37%, |
| 338 | | | RASHI_RESPONSE_TO_IONIZING_RADIATION_1 | | 33 | -0.42 | -1.27 | 0.138 | 0.474 | 1 | 2503 | tags=36%, list=25%, |
| 428 | | | REACTOME_G1_S_TRANSITION | | 90 | -0.35 | -1.22 | 0.14 | 0.527 | 1 | 2710 | tags=36%, list=27%, |
| 555 | | | SCHAEFFER_PROSTATE_DEVELOPMENT_6HR_DN | | 372 | -0.29 | -1.14 | 0.14 | 0.619 | 1 | 3313 | tags=37%, list=33%, |
| 321 | | | SCHAEFFER_PROSTATE_DEVELOPMENT_AND_CANCER_BOX4_DN | | 31 | -0.43 | -1.28 | 0.141 | 0.464 | 1 | 2503 | tags=42%, list=25%, |
| 340 | | | MORI_PLASMA_CELL_UP | | 45 | -0.4 | -1.27 | 0.141 | 0.473 | 1 | 3176 | tags=40%, list=31%, |
| 356 | | | REACTOME_SIGNALING_BY_NTRKS | | 54 | -0.39 | -1.26 | 0.141 | 0.485 | 1 | 2655 | tags=39%, list=26%, |
| 426 | | | REACTOME_SLC_MEDIATED_TRANSMEMBRANE_TRANSPORT | | 107 | -0.34 | -1.22 | 0.141 | 0.527 | 1 | 2417 | tags=33%, list=24%, |
| 307 | | | BURTON_ADIPOGENESIS_PEAK_AT_2HR | | 28 | -0.44 | -1.29 | 0.142 | 0.45 | 1 | 3021 | tags=39%, list=30%, |
| 329 | | | LIAN_LIPA_TARGETS_3M | | 40 | -0.4 | -1.28 | 0.142 | 0.469 | 1 | 3482 | tags=48%, list=34%, |
| 396 | | | REACTOME_SUMOYLATION_OF_DNA_DAMAGE_RESPONSE_AND_REPAIR_PROTEI | | 58 | -0.37 | -1.24 | 0.142 | 0.496 | 1 | 2091 | tags=29%, list=21%, |
| 332 | | | REACTOME_NUCLEAR_RECEPTOR_TRANSCRIPTION_PATHWAY | | 28 | -0.43 | -1.28 | 0.143 | 0.469 | 1 | 3052 | tags=50%, list=30%, |
| 322 | | | REACTOME_MAPK_TARGETS_NUCLEAR_EVENTS_MEDIATED_BY_MAP_KINASES | | 23 | -0.45 | -1.28 | 0.144 | 0.466 | 1 | 4697 | tags=83%, list=47%, |
| 406 | | | ZENG_GU_ICB_CONTROL_METAGENE_14_PRECICTIVE_ICB_RESPONSE | | 67 | -0.37 | -1.23 | 0.144 | 0.519 | 1 | 2073 | tags=30%, list=21%, |
| 436 | | | REACTOME_TRANSCRIPTIONAL_REGULATION_BY_RUNX1 | | 113 | -0.34 | -1.21 | 0.144 | 0.541 | 1 | 2710 | tags=35%, list=27%, |
| 277 | | | WP_MICROGLIA_PATHOGEN_PHAGOCYTOSIS_PATHWAY | | 23 | -0.46 | -1.32 | 0.145 | 0.425 | 1 | 2451 | tags=48%, list=24%, |
| 300 | | | REACTOME_CARGO_TRAFFICKING_TO_THE_PERICILIARY_MEMBRANE | | 27 | -0.45 | -1.3 | 0.145 | 0.433 | 1 | 2836 | tags=48%, list=28%, |
| 274 | | | REACTOME_RHO_GTPASES_ACTIVATE_NADPH_OXIDASES | | 17 | -0.5 | -1.32 | 0.146 | 0.426 | 1 | 2149 | tags=47%, list=21%, |
| 398 | | | REACTOME_AMINO_ACIDS_REGULATE_MTORC1 | | 40 | -0.4 | -1.24 | 0.148 | 0.5 | 1 | 2436 | tags=35%, list=24%, |
| 295 | | | REACTOME_DAP12_SIGNALING | | 16 | -0.5 | -1.3 | 0.15 | 0.438 | 1 | 2369 | tags=44%, list=23%, |
| 310 | | | REACTOME_INTRINSIC_PATHWAY_OF_FIBRIN_CLOT_FORMATION | | 16 | -0.5 | -1.29 | 0.151 | 0.45 | 1 | 4545 | tags=94%, list=45%, |
| 523 | | | KUMAR_TARGETS_OF_MLL_AF9_FUSION | | 261 | -0.3 | -1.15 | 0.153 | 0.607 | 1 | 2598 | tags=28%, list=26%, |
| 333 | | | WP_RETINOL_METABOLISM | | 25 | -0.44 | -1.27 | 0.154 | 0.472 | 1 | 1219 | tags=24%, list=12%, |
| 372 | | | VANDESLUIS_COMMD1_TARGETS_GROUP_3_UP | | 47 | -0.39 | -1.25 | 0.154 | 0.491 | 1 | 638 | tags=15%, list=6%, signal=16% |
| 403 | | | KATSANOU_ELAVL1_TARGETS_DN | | 69 | -0.36 | -1.23 | 0.155 | 0.517 | 1 | 1988 | tags=30%, list=20%, |
| 326 | | | TERAO_AOX4_TARGETS_SKIN_UP | | 20 | -0.46 | -1.28 | 0.156 | 0.467 | 1 | 2364 | tags=35%, list=23%, |
| 337 | | | REACTOME_REGULATION_OF_PTEN_GENE_TRANSCRIPTION | | 32 | -0.42 | -1.27 | 0.157 | 0.475 | 1 | 3069 | tags=44%, list=30%, |
| 378 | | | REACTOME_SIGNALING_BY_INSULIN_RECEPTOR | | 47 | -0.39 | -1.25 | 0.157 | 0.49 | 1 | 3023 | tags=38%, list=30%, |
| 557 | | | MILI_PSEUDOPODIA_CHEMOTAXIS_DN | | 326 | -0.3 | -1.14 | 0.158 | 0.621 | 1 | 3496 | tags=37%, list=35%, |
| 390 | | | LIM_MAMMARY_LUMINAL_MATURE_UP | | 59 | -0.37 | -1.24 | 0.16 | 0.494 | 1 | 1431 | tags=25%, list=14%, |

| | | | | | | | | | | | | |
|---|---|---|---|---|---|---|---|---|---|---|---|---|
| 415 | | | BILANGES_SERUM_SENSITIVE_GENES | | 52 | -0.38 | -1.22 | 0.16 | 0.529 | 1 | 2716 | tags=35%, list=27%, |
| 336 | | | MULLIGAN_NTF3_SIGNALING_VIA_INSR_AND_IGF1R_UP | | 19 | -0.47 | -1.27 | 0.161 | 0.473 | 1 | 3610 | tags=58%, list=36%, |
| 383 | | | WP_EXERCISE_INDUCED_CIRCADIAN_REGULATION | | 42 | -0.39 | -1.25 | 0.161 | 0.491 | 1 | 2391 | tags=40%, list=24%, |
| 394 | | | BURTON_ADIPOGENESIS_7 | | 38 | -0.4 | -1.24 | 0.161 | 0.496 | 1 | 3785 | tags=58%, list=37%, |
| 421 | | | REACTOME_INTERLEUKIN_1_FAMILY_SIGNALING | | 93 | -0.35 | -1.22 | 0.161 | 0.53 | 1 | 3654 | tags=44%, list=36%, |
| 350 | | | BIOCARTA_ERK_PATHWAY | | 22 | -0.45 | -1.26 | 0.163 | 0.486 | 1 | 2597 | tags=41%, list=26%, |
| 387 | | | TSENG_ADIPOGENIC_POTENTIAL_DN | | 34 | -0.4 | -1.24 | 0.163 | 0.491 | 1 | 1381 | tags=29%, list=14%, |
| 582 | | | MARTINEZ_RB1_TARGETS_DN | | 371 | -0.29 | -1.12 | 0.163 | 0.649 | 1 | 3037 | tags=35%, list=30%, |
| 354 | | | WP_NOTCH_SIGNALING_PATHWAY | | 28 | -0.44 | -1.26 | 0.164 | 0.486 | 1 | 2342 | tags=29%, list=23%, |
| 419 | | | SCHAEFFER_PROSTATE_DEVELOPMENT_12HR_UP | | 70 | -0.36 | -1.22 | 0.164 | 0.53 | 1 | 2518 | tags=37%, list=25%, |
| 373 | | | WANG_RESPONSE_TO_BEXAROTENE_UP | | 18 | -0.47 | -1.25 | 0.166 | 0.49 | 1 | 1921 | tags=44%, list=19%, |
| 518 | | | VANOEVELEN_MYOGENESIS_SIN3A_TARGETS | | 171 | -0.31 | -1.16 | 0.167 | 0.603 | 1 | 3056 | tags=33%, list=30%, |
| 346 | | | RIZ_ERYTHROID_DIFFERENTIATION_12HR | | 23 | -0.45 | -1.26 | 0.168 | 0.482 | 1 | 2314 | tags=39%, list=23%, |
| 349 | | | VANASSE_BCL2_TARGETS_UP | | 21 | -0.46 | -1.26 | 0.168 | 0.479 | 1 | 3196 | tags=52%, list=32%, |
| 376 | | | REACTOME_L1CAM_INTERACTIONS | | 44 | -0.39 | -1.25 | 0.168 | 0.489 | 1 | 1997 | tags=32%, list=20%, |
| 368 | | | REACTOME_DNA_DOUBLE_STRAND_BREAK_RESPONSE | | 38 | -0.41 | -1.25 | 0.169 | 0.492 | 1 | 292 | tags=13%, list=3%, signal=13% |
| 339 | | | LANDIS_ERBB2_BREAST_TUMORS_65_UP | | 20 | -0.47 | -1.27 | 0.172 | 0.473 | 1 | 2368 | tags=40%, list=23%, |
| 369 | | | SHIN_B_CELL_LYMPHOMA_CLUSTER_8 | | 20 | -0.46 | -1.25 | 0.172 | 0.491 | 1 | 1685 | tags=35%, list=17%, |
| 367 | | | CHESLER_BRAIN_HIGHEST_GENETIC_VARIANCE | | 29 | -0.42 | -1.25 | 0.175 | 0.492 | 1 | 2978 | tags=45%, list=29%, |
| 450 | | | REACTOME_SWITCHING_OF_ORIGINS_TO_A_POST_REPLICATIVE_STATE | | 81 | -0.35 | -1.2 | 0.176 | 0.562 | 1 | 2710 | tags=36%, list=27%, |
| 459 | | | NABA_ECM_GLYCOPROTEINS | | 71 | -0.35 | -1.19 | 0.177 | 0.569 | 1 | 2745 | tags=41%, list=27%, |
| 391 | | | YU_MYC_TARGETS_DN | | 40 | -0.4 | -1.24 | 0.179 | 0.495 | 1 | 2846 | tags=43%, list=28%, |
| 418 | | | QUINTENS_EMBRYONIC_BRAIN_RESPONSE_TO_IR | | 50 | -0.37 | -1.22 | 0.18 | 0.531 | 1 | 2106 | tags=32%, list=21%, |
| 488 | | | REACTOME_ER_TO_GOLGI_ANTEROGRADE_TRANSPORT | | 120 | -0.33 | -1.18 | 0.181 | 0.577 | 1 | 3596 | tags=42%, list=36%, |
| 393 | | | REACTOME_INITIAL_TRIGGERING_OF_COMPLEMENT | | 16 | -0.48 | -1.24 | 0.183 | 0.497 | 1 | 976 | tags=31%, list=10%, |
| 576 | | | FOSTER_TOLERANT_MACROPHAGE_DN | | 302 | -0.29 | -1.12 | 0.183 | 0.64 | 1 | 2876 | tags=33%, list=28%, |
| 427 | | | REACTOME_GOLGI_ASSOCIATED_VESICLE_BIOGENESIS | | 43 | -0.39 | -1.22 | 0.184 | 0.528 | 1 | 2061 | tags=35%, list=20%, |
| 532 | | | LI_INDUCED_T_TO_NATURAL_KILLER_UP | | 191 | -0.31 | -1.15 | 0.184 | 0.608 | 1 | 2774 | tags=36%, list=27%, |
| 505 | | | MCBRYAN_PUBERTAL_TGFB1_TARGETS_UP | | 133 | -0.32 | -1.16 | 0.185 | 0.593 | 1 | 1626 | tags=24%, list=16%, |
| 423 | | | REACTOME_SIGNALING_BY_PTK6 | | 32 | -0.41 | -1.22 | 0.186 | 0.528 | 1 | 1165 | tags=22%, list=12%, |
| 404 | | | REACTOME_INSULIN_RECEPTOR_SIGNALLING_CASCADE | | 26 | -0.42 | -1.23 | 0.187 | 0.516 | 1 | 2726 | tags=42%, list=27%, |
| 472 | | | MARKEY_RB1_CHRONIC_LOF_DN | | 82 | -0.34 | -1.18 | 0.187 | 0.579 | 1 | 3136 | tags=37%, list=31%, |
| 528 | | | REACTOME_PLATELET_ACTIVATION_SIGNALING_AND_AGGREGATION | | 169 | -0.31 | -1.15 | 0.187 | 0.607 | 1 | 2713 | tags=34%, list=27%, |
| 363 | | | WANG_IMMORTALIZED_BY_HOXA9_AND_MEIS1_UP | | 19 | -0.46 | -1.25 | 0.188 | 0.493 | 1 | 1267 | tags=32%, list=13%, |
| 487 | | | WP_IL_6_SIGNALING_PATHWAY | | 80 | -0.34 | -1.18 | 0.188 | 0.578 | 1 | 2416 | tags=31%, list=24%, |
| 485 | | | CHIARADONNA_NEOPLASTIC_TRANSFORMATION_KRAS_DN | | 112 | -0.32 | -1.18 | 0.19 | 0.579 | 1 | 905 | tags=17%, list=9%, signal=18% |
| 438 | | | REACTOME_CYTOCHROME_P450_ARRANGED_BY_SUBSTRATE_TYPE | | 38 | -0.39 | -1.21 | 0.191 | 0.547 | 1 | 1658 | tags=34%, list=16%, |
| 495 | | | REACTOME_S_PHASE | | 123 | -0.33 | -1.17 | 0.191 | 0.589 | 1 | 1713 | tags=24%, list=17%, |
| 364 | | | WP_GPCRS_CLASS_A_RHODOPSIN_LIKE | | 17 | -0.47 | -1.25 | 0.192 | 0.496 | 1 | 668 | tags=18%, list=7%, signal=19% |
| 371 | | | REACTOME_MTORC1_MEDIATED_SIGNALLING | | 21 | -0.45 | -1.25 | 0.192 | 0.491 | 1 | 2655 | tags=48%, list=26%, |
| 381 | | | OUYANG_PROSTATE_CANCER_MARKERS | | 16 | -0.48 | -1.25 | 0.193 | 0.491 | 1 | 2857 | tags=44%, list=28%, |
| 433 | | | SASAI_RESISTANCE_TO_NEOPLASTIC_TRANSFROMATION | | 30 | -0.41 | -1.21 | 0.193 | 0.535 | 1 | 2214 | tags=33%, list=22%, |
| 483 | | | PLASARI_TGFB1_SIGNALING_VIA_NFIC_1HR_DN | | 71 | -0.35 | -1.18 | 0.193 | 0.581 | 1 | 1890 | tags=28%, list=19%, |
| 444 | | | REACTOME_GPCR_LIGAND_BINDING | | 65 | -0.36 | -1.2 | 0.195 | 0.549 | 1 | 1337 | tags=20%, list=13%, |
| 446 | | | LEE_CALORIE_RESTRICTION_NEOCORTEX_UP | | 42 | -0.38 | -1.2 | 0.195 | 0.553 | 1 | 2695 | tags=33%, list=27%, |
| 468 | | | PANGAS_TUMOR_SUPPRESSION_BY_SMAD1_AND_SMAD5_UP | | 79 | -0.34 | -1.19 | 0.195 | 0.569 | 1 | 2203 | tags=25%, list=22%, |
| 593 | | | IVANOVA_HEMATOPOIESIS_STEM_CELL_AND_PROGENITOR | | 349 | -0.29 | -1.11 | 0.196 | 0.658 | 1 | 2661 | tags=30%, list=26%, |
| 464 | | | REACTOME_NEGATIVE_REGULATION_OF_THE_PI3K_AKT_NETWORK | | 58 | -0.36 | -1.19 | 0.198 | 0.567 | 1 | 2534 | tags=34%, list=25%, |
| 379 | | | REACTOME_TRANSCRIPTIONAL_REGULATION_BY_E2F6 | | 18 | -0.47 | -1.25 | 0.199 | 0.49 | 1 | 1267 | tags=28%, list=13%, |
| 389 | | | REACTOME_NEGATIVE_REGULATION_OF_MET_ACTIVITY | | 17 | -0.47 | -1.24 | 0.199 | 0.492 | 1 | 2539 | tags=35%, list=25%, |
| 385 | | | NABA_MATRISOME_PRIMARY_METASTATIC_LUNG_TUMOR | | 21 | -0.45 | -1.24 | 0.201 | 0.491 | 1 | 2395 | tags=48%, list=24%, |
| 405 | | | SHIN_B_CELL_LYMPHOMA_CLUSTER_7 | | 19 | -0.46 | -1.23 | 0.201 | 0.517 | 1 | 2024 | tags=32%, list=20%, |
| 602 | | | IVANOVA_HEMATOPOIESIS_LATE_PROGENITOR | | 364 | -0.29 | -1.11 | 0.201 | 0.657 | 1 | 2559 | tags=27%, list=25%, |
| 481 | | | WP_REGULATION_OF_ACTIN_CYTOSKELETON | | 87 | -0.33 | -1.18 | 0.203 | 0.583 | 1 | 2652 | tags=33%, list=26%, |
| 486 | | | REACTOME_PROTEIN_UBIQUITINATION | | 51 | -0.36 | -1.18 | 0.204 | 0.579 | 1 | 3726 | tags=41%, list=37%, |
| 507 | | | IVANOVA_HEMATOPOIESIS_STEM_CELL_LONG_TERM | | 107 | -0.33 | -1.16 | 0.204 | 0.597 | 1 | 2019 | tags=29%, list=20%, |
| 432 | | | WP_G13_SIGNALING_PATHWAY | | 27 | -0.41 | -1.21 | 0.205 | 0.534 | 1 | 2322 | tags=30%, list=23%, |
| 484 | | | MA_RAT_AGING_UP | | 74 | -0.34 | -1.18 | 0.205 | 0.58 | 1 | 3796 | tags=46%, list=38%, |
| 388 | | | BIOCARTA_VDR_PATHWAY | | 19 | -0.46 | -1.24 | 0.206 | 0.493 | 1 | 1078 | tags=32%, list=11%, |
| 443 | | | REACTOME_CLASS_A_1_RHODOPSIN_LIKE_RECEPTORS | | 47 | -0.37 | -1.2 | 0.206 | 0.55 | 1 | 2713 | tags=36%, list=27%, |
| 420 | | | REACTOME_CELL_CELL_JUNCTION_ORGANIZATION | | 26 | -0.42 | -1.22 | 0.207 | 0.531 | 1 | 3645 | tags=54%, list=36%, |
| 448 | | | BOYLAN_MULTIPLE_MYELOMA_C_UP | | 36 | -0.39 | -1.2 | 0.207 | 0.558 | 1 | 1136 | tags=19%, list=11%, |
| 498 | | | QI_HYPOXIA | | 86 | -0.34 | -1.17 | 0.207 | 0.588 | 1 | 3165 | tags=38%, list=31%, |
| 425 | | | REACTOME_PROTEIN_HYDROXYLATION | | 15 | -0.48 | -1.22 | 0.212 | 0.526 | 1 | 3416 | tags=40%, list=34%, |
| 412 | | | CHEN_PDGF_TARGETS | | 17 | -0.47 | -1.22 | 0.213 | 0.525 | 1 | 3298 | tags=59%, list=33%, |
| 462 | | | WP_TGF_BETA_SIGNALING_PATHWAY | | 38 | -0.38 | -1.19 | 0.213 | 0.568 | 1 | 868 | tags=21%, list=9%, signal=23% |
| 465 | | | ZHONG_SECRETOME_OF_LUNG_CANCER_AND_MACROPHAGE | | 64 | -0.35 | -1.19 | 0.213 | 0.566 | 1 | 3685 | tags=42%, list=36%, |
| 430 | | | REACTOME_MITOPHAGY | | 26 | -0.42 | -1.21 | 0.214 | 0.531 | 1 | 1530 | tags=27%, list=15%, |
| 525 | | | MIKKELSEN_MCV6_LCP_WITH_H3K4ME3 | | 73 | -0.34 | -1.15 | 0.214 | 0.61 | 1 | 3172 | tags=40%, list=31%, |
| 460 | | | REACTOME_G_ALPHA_S_SIGNALLING_EVENTS | | 29 | -0.41 | -1.19 | 0.215 | 0.568 | 1 | 1615 | tags=24%, list=16%, |
| 441 | | | REACTOME_RHOBTB2_GTPASE_CYCLE | | 21 | -0.44 | -1.2 | 0.216 | 0.548 | 1 | 2682 | tags=38%, list=27%, |
| 478 | | | REACTOME_PLASMA_LIPOPROTEIN_ASSEMBLY_REMODELING_AND_CLEARANCE | | 56 | -0.36 | -1.18 | 0.216 | 0.582 | 1 | 3126 | tags=41%, list=31%, |
| 489 | | | RASHI_RESPONSE_TO_IONIZING_RADIATION_6 | | 56 | -0.36 | -1.18 | 0.217 | 0.579 | 1 | 3548 | tags=52%, list=35%, |
| 502 | | | LI_INDUCED_T_TO_NATURAL_KILLER_DN | | 85 | -0.34 | -1.17 | 0.217 | 0.588 | 1 | 2466 | tags=32%, list=24%, |
| 449 | | | JIANG_AGING_HYPOTHALAMUS_UP | | 47 | -0.37 | -1.2 | 0.221 | 0.559 | 1 | 2742 | tags=30%, list=27%, |
| 475 | | | GUO_HEX_TARGETS_UP | | 58 | -0.36 | -1.18 | 0.221 | 0.578 | 1 | 2408 | tags=33%, list=24%, |
| 401 | | | REACTOME_FOXO_MEDIATED_TRANSCRIPTION | | 16 | -0.48 | -1.23 | 0.222 | 0.514 | 1 | 2655 | tags=38%, list=26%, |
| 517 | | | REACTOME_ORC1_REMOVAL_FROM_CHROMATIN | | 63 | -0.35 | -1.16 | 0.225 | 0.598 | 1 | 2710 | tags=37%, list=27%, |
| 439 | | | REACTOME_COLLAGEN_FORMATION | | 31 | -0.4 | -1.21 | 0.226 | 0.545 | 1 | 3820 | tags=68%, list=38%, |
| 416 | | | HU_GENOTOXIC_DAMAGE_4HR | | 21 | -0.44 | -1.22 | 0.228 | 0.53 | 1 | 2209 | tags=38%, list=22%, |
| 429 | | | REACTOME_SMAD2_SMAD3_SMAD4_HETEROTRIMER_REGULATES_TRANSCRIPTI | | 23 | -0.43 | -1.21 | 0.228 | 0.529 | 1 | 750 | tags=22%, list=7%, signal=23% |
| 458 | | | QI_HYPOXIA_TARGETS_OF_HIF1A_AND_FOXA2 | | 20 | -0.43 | -1.19 | 0.228 | 0.571 | 1 | 1615 | tags=25%, list=16%, |
| 559 | | | REACTOME_CHROMATIN_MODIFYING_ENZYMES | | 127 | -0.31 | -1.13 | 0.229 | 0.625 | 1 | 3105 | tags=35%, list=31%, |
| 445 | | | REACTOME_REGULATION_OF_TP53_ACTIVITY_THROUGH_ACETYLATION | | 27 | -0.41 | -1.2 | 0.23 | 0.553 | 1 | 4034 | tags=56%, list=40%, |
| 451 | | | REACTOME_IRS_MEDIATED_SIGNALLING | | 21 | -0.44 | -1.19 | 0.23 | 0.566 | 1 | 2726 | tags=43%, list=27%, |
| 463 | | | WP_MECP2_AND_ASSOCIATED_RETT_SYNDROME | | 27 | -0.41 | -1.19 | 0.231 | 0.567 | 1 | 1616 | tags=30%, list=16%, |
| 473 | | | REACTOME_TRANSCRIPTIONAL_ACTIVITY_OF_SMAD2_SMAD3_SMAD4_HETEROT | | 34 | -0.39 | -1.18 | 0.232 | 0.578 | 1 | 1626 | tags=24%, list=16%, |
| 476 | | | MEISSNER_NPC_HCP_WITH_H3K4ME3_AND_H3K27ME3 | | 35 | -0.39 | -1.18 | 0.234 | 0.58 | 1 | 2871 | tags=43%, list=28%, |
| 457 | | | REACTOME_LDL_CLEARANCE | | 17 | -0.45 | -1.19 | 0.235 | 0.57 | 1 | 1465 | tags=35%, list=15%, |
| 554 | | | REACTOME_DNA_REPLICATION | | 107 | -0.32 | -1.14 | 0.235 | 0.62 | 1 | 1799 | tags=25%, list=18%, |
| 455 | | | REACTOME_ADHERENS_JUNCTIONS_INTERACTIONS | | 22 | -0.43 | -1.19 | 0.237 | 0.566 | 1 | 3645 | tags=59%, list=36%, |
| 546 | | | REACTOME_TOLL_LIKE_RECEPTOR_CASCADES | | 111 | -0.32 | -1.14 | 0.237 | 0.615 | 1 | 2902 | tags=36%, list=29%, |
| 516 | | | WP_IL_3_SIGNALING_PATHWAY | | 72 | -0.34 | -1.16 | 0.239 | 0.596 | 1 | 1367 | tags=19%, list=14%, |
| 577 | | | MARKEY_RB1_ACUTE_LOF_UP | | 167 | -0.3 | -1.12 | 0.241 | 0.641 | 1 | 3006 | tags=31%, list=30%, |
| 470 | | | REACTOME_HATS_ACETYLATE_HISTONES | | 32 | -0.39 | -1.19 | 0.243 | 0.57 | 1 | 1787 | tags=31%, list=18%, |
| 503 | | | BREDEMEYER_RAG_SIGNALING_NOT_VIA_ATM_UP | | 44 | -0.37 | -1.17 | 0.243 | 0.59 | 1 | 2392 | tags=30%, list=24%, |
| 512 | | | REACTOME_G1_S_DNA_DAMAGE_CHECKPOINTS | | 55 | -0.36 | -1.16 | 0.243 | 0.595 | 1 | 2710 | tags=38%, list=27%, |
| 494 | | | REACTOME_E3_UBIQUITIN_LIGASES_UBIQUITINATE_TARGET_PROTEINS | | 35 | -0.39 | -1.17 | 0.244 | 0.588 | 1 | 3706 | tags=43%, list=37%, |
| 508 | | | CHANGOLKAR_H2AFY_TARGETS_DN | | 40 | -0.37 | -1.16 | 0.244 | 0.596 | 1 | 3209 | tags=45%, list=32%, |
| 570 | | | REACTOME_SIGNALING_BY_WNT | | 170 | -0.3 | -1.13 | 0.244 | 0.634 | 1 | 2710 | tags=32%, list=27%, |
| 520 | | | MORI_MATURE_B_LYMPHOCYTE_UP | | 64 | -0.34 | -1.16 | 0.249 | 0.606 | 1 | 3806 | tags=50%, list=38%, |
| 527 | | | REACTOME_COPI_MEDIATED_ANTEROGRADE_TRANSPORT | | 75 | -0.34 | -1.15 | 0.249 | 0.608 | 1 | 3715 | tags=43%, list=37%, |
| 467 | | | WELCH_GATA1_TARGETS | | 16 | -0.45 | -1.19 | 0.25 | 0.57 | 1 | 3385 | tags=56%, list=34%, |
| 471 | | | REACTOME_COLLAGEN_BIOSYNTHESIS_AND_MODIFYING_ENZYMES | | 27 | -0.4 | -1.18 | 0.251 | 0.579 | 1 | 1653 | tags=37%, list=16%, |
| 474 | | | GROSS_ELK3_TARGETS_DN | | 21 | -0.44 | -1.18 | 0.251 | 0.578 | 1 | 3116 | tags=48%, list=31%, |
| 511 | | | CHIARADONNA_NEOPLASTIC_TRANSFORMATION_KRAS_CDC25_UP | | 38 | -0.37 | -1.16 | 0.252 | 0.596 | 1 | 2654 | tags=39%, list=26%, |
| 579 | | | GOTZMANN_EPITHELIAL_TO_MESENCHYMAL_TRANSITION_DN | | 163 | -0.3 | -1.12 | 0.252 | 0.643 | 1 | 3131 | tags=36%, list=31%, |
| 442 | | | REACTOME_CONVERSION_FROM_APC_C_CDC20_TO_APC_C_CDH1_IN_LATE_ANA | | 17 | -0.46 | -1.2 | 0.253 | 0.55 | 1 | 1626 | tags=35%, list=16%, |
| 530 | | | ICHIBA_GRAFT_VERSUS_HOST_DISEASE_D7_DN | | 41 | -0.37 | -1.15 | 0.253 | 0.608 | 1 | 928 | tags=22%, list=9%, signal=24% |
| 493 | | | REACTOME_SIGNALING_BY_FGFR1 | | 29 | -0.4 | -1.17 | 0.254 | 0.585 | 1 | 3997 | tags=55%, list=40%, |
| 504 | | | WANG_TARGETS_OF_MLL_CBP_FUSION_DN | | 42 | -0.37 | -1.17 | 0.254 | 0.591 | 1 | 3563 | tags=50%, list=35%, |
| 548 | | | WP_ADIPOGENESIS_GENES | | 84 | -0.33 | -1.14 | 0.254 | 0.614 | 1 | 1030 | tags=21%, list=10%, |
| 575 | | | ZENG_GU_ICB_CONTROL_METAGENE_3_PRECICTIVE_ICB_RESPONSE | | 107 | -0.31 | -1.13 | 0.254 | 0.634 | 1 | 2662 | tags=35%, list=26%, |
| 492 | | | REACTOME_PREDNISONE_ADME | | 15 | -0.45 | -1.17 | 0.255 | 0.585 | 1 | 3599 | tags=47%, list=36%, |
| 543 | | | VANLOO_SP3_TARGETS_DN | | 66 | -0.34 | -1.15 | 0.256 | 0.606 | 1 | 1847 | tags=26%, list=18%, |
| 547 | | | GAUSSMANN_MLL_AF4_FUSION_TARGETS_E_UP | | 51 | -0.35 | -1.14 | 0.257 | 0.615 | 1 | 3548 | tags=51%, list=35%, |
| 461 | | | XU_CREBBP_TARGETS_UP | | 16 | -0.46 | -1.19 | 0.258 | 0.569 | 1 | 2778 | tags=44%, list=28%, |
| 466 | | | BOYLAN_MULTIPLE_MYELOMA_C_CLUSTER_UP | | 27 | -0.41 | -1.19 | 0.258 | 0.571 | 1 | 996 | tags=22%, list=10%, |
| 585 | | | REACTOME_TCF_DEPENDENT_SIGNALING_IN_RESPONSE_TO_WNT | | 122 | -0.31 | -1.12 | 0.258 | 0.649 | 1 | 2710 | tags=34%, list=27%, |
| 454 | | | BIOCARTA_AT1R_PATHWAY | | 23 | -0.42 | -1.19 | 0.259 | 0.568 | 1 | 1213 | tags=30%, list=12%, |
| 538 | | | GOLDRATH_IMMUNE_MEMORY | | 52 | -0.36 | -1.15 | 0.259 | 0.607 | 1 | 1136 | tags=19%, list=11%, |
| 480 | | | BIOCARTA_FCER1_PATHWAY | | 29 | -0.4 | -1.18 | 0.261 | 0.582 | 1 | 2800 | tags=45%, list=28%, |
| 549 | | | BURTON_ADIPOGENESIS_9 | | 68 | -0.34 | -1.14 | 0.262 | 0.614 | 1 | 3004 | tags=41%, list=30%, |
| 566 | | | RIZ_ERYTHROID_DIFFERENTIATION | | 57 | -0.34 | -1.13 | 0.262 | 0.63 | 1 | 1591 | tags=25%, list=16%, |
| 501 | | | REACTOME_SIGNALING_BY_TYPE_1_INSULIN_LIKE_GROWTH_FACTOR_1_RECEPTOR_IGF1R | | 24 | -0.41 | -1.17 | 0.263 | 0.588 | 1 | 2726 | tags=42%, list=27%, signal=57% |
| 529 | | | TERAO_AOX4_TARGETS_HG_UP | | 24 | -0.41 | -1.15 | 0.263 | 0.608 | 1 | 2722 | tags=42%, list=27%, |
| 563 | | | GROSS_HYPOXIA_VIA_HIF1A_DN | | 77 | -0.33 | -1.13 | 0.263 | 0.631 | 1 | 3345 | tags=39%, list=33%, |
| 477 | | | BIOCARTA_CXCR4_PATHWAY | | 18 | -0.44 | -1.18 | 0.265 | 0.582 | 1 | 783 | tags=22%, list=8%, signal=24% |
| 544 | | | REACTOME_IRON_UPTAKE_AND_TRANSPORT | | 44 | -0.36 | -1.15 | 0.265 | 0.605 | 1 | 3445 | tags=43%, list=34%, |
| 510 | | | REACTOME_INSULIN_RECEPTOR_RECYCLING | | 22 | -0.42 | -1.16 | 0.266 | 0.595 | 1 | 4704 | tags=64%, list=47%, |
| 479 | | | BIOCARTA_MTOR_PATHWAY | | 19 | -0.44 | -1.18 | 0.267 | 0.583 | 1 | 3997 | tags=58%, list=40%, |
| 539 | | | REACTOME_NUCLEAR_ENVELOPE_BREAKDOWN | | 41 | -0.37 | -1.15 | 0.267 | 0.606 | 1 | 1602 | tags=24%, list=16%, |
| 499 | | | TERAO_AOX4_TARGETS_SKIN_DN | | 16 | -0.45 | -1.17 | 0.268 | 0.588 | 1 | 1311 | tags=25%, list=13%, |
| 533 | | | REACTOME_NEGATIVE_REGULATION_OF_MAPK_PATHWAY | | 32 | -0.39 | -1.15 | 0.268 | 0.608 | 1 | 4651 | tags=66%, list=46%, |
| 534 | | | IKEDA_MIR133_TARGETS_UP | | 41 | -0.36 | -1.15 | 0.268 | 0.609 | 1 | 5098 | tags=66%, list=50%, |
| 490 | | | VILIMAS_NOTCH1_TARGETS_UP | | 15 | -0.46 | -1.18 | 0.269 | 0.579 | 1 | 2335 | tags=47%, list=23%, |
| 515 | | | WP_P38_MAPK_SIGNALING_PATHWAY | | 28 | -0.4 | -1.16 | 0.269 | 0.596 | 1 | 2746 | tags=43%, list=27%, |
| 641 | | | MATSUDA_NATURAL_KILLER_DIFFERENTIATION | | 321 | -0.28 | -1.08 | 0.269 | 0.692 | 1 | 2576 | tags=28%, list=26%, |
| 619 | | | PASINI_SUZ12_TARGETS_DN | | 229 | -0.29 | -1.1 | 0.27 | 0.674 | 1 | 2707 | tags=30%, list=27%, |
| 491 | | | REACTOME_GPER1_SIGNALING | | 15 | -0.46 | -1.17 | 0.271 | 0.582 | 1 | 1615 | tags=33%, list=16%, |
| 496 | | | REACTOME_SIGNALING_BY_CSF3_G_CSF | | 18 | -0.44 | -1.17 | 0.271 | 0.589 | 1 | 2749 | tags=39%, list=27%, |
| 569 | | | REACTOME_SIGNALING_BY_TGFB_FAMILY_MEMBERS | | 77 | -0.33 | -1.13 | 0.272 | 0.632 | 1 | 1635 | tags=22%, list=16%, |
| 553 | | | REACTOME_IMMUNOREGULATORY_INTERACTIONS_BETWEEN_A_LYMPHOID_AND_A_NON_LYMPHOID_CELL | | 36 | -0.37 | -1.14 | 0.273 | 0.618 | 1 | 2520 | tags=47%, list=25%, signal=63% |
| 597 | | | REACTOME_MITOTIC_METAPHASE_AND_ANAPHASE | | 170 | -0.3 | -1.11 | 0.273 | 0.657 | 1 | 2710 | tags=32%, list=27%, |
| 531 | | | REACTOME_REGULATION_OF_TNFR1_SIGNALING | | 41 | -0.37 | -1.15 | 0.275 | 0.607 | 1 | 2563 | tags=32%, list=25%, |
| 622 | | | NABA_MATRISOME_ASSOCIATED | | 230 | -0.29 | -1.09 | 0.275 | 0.681 | 1 | 3195 | tags=42%, list=32%, |
| 564 | | | REACTOME_SIGNALING_BY_VEGF | | 77 | -0.33 | -1.13 | 0.277 | 0.63 | 1 | 2387 | tags=31%, list=24%, |
| 545 | | | REACTOME_TRANSPORT_OF_INORGANIC_CATIONS_ANIONS_AND_AMINO_ACIDS_OLIGOPEPTIDES | | 45 | -0.36 | -1.14 | 0.28 | 0.615 | 1 | 2012 | tags=31%, list=20%, signal=39% |
| 542 | | | REACTOME_SIGNALING_BY_BMP | | 22 | -0.42 | -1.15 | 0.282 | 0.604 | 1 | 1635 | tags=27%, list=16%, |
| 595 | | | REACTOME_METABOLISM_OF_WATER_SOLUBLE_VITAMINS_AND_COFACTORS | | 91 | -0.32 | -1.11 | 0.282 | 0.658 | 1 | 3163 | tags=36%, list=31%, |
| 603 | | | STARK_PREFRONTAL_CORTEX_22Q11_DELETION_UP | | 139 | -0.3 | -1.11 | 0.282 | 0.66 | 1 | 2735 | tags=34%, list=27%, |
| 608 | | | WANG_MLL_TARGETS | | 139 | -0.3 | -1.1 | 0.282 | 0.665 | 1 | 2288 | tags=29%, list=23%, |
| 506 | | | RUAN_RESPONSE_TO_TROGLITAZONE_UP | | 18 | -0.44 | -1.16 | 0.283 | 0.596 | 1 | 135 | tags=11%, list=1%, signal=11% |
| 561 | | | WP_NEURAL_CREST_DIFFERENTIATION | | 42 | -0.36 | -1.13 | 0.283 | 0.628 | 1 | 2707 | tags=33%, list=27%, |
| 610 | | | FOSTER_KDM1A_TARGETS_DN | | 131 | -0.3 | -1.1 | 0.285 | 0.666 | 1 | 1934 | tags=24%, list=19%, |
| 524 | | | WP_NUCLEAR_RECEPTORS_IN_LIPID_METABOLISM_AND_TOXICITY | | 24 | -0.4 | -1.15 | 0.286 | 0.609 | 1 | 3052 | tags=42%, list=30%, |
| 584 | | | RAY_TUMORIGENESIS_BY_ERBB2_CDC25A_UP | | 93 | -0.32 | -1.12 | 0.286 | 0.649 | 1 | 1921 | tags=26%, list=19%, |
| 635 | | | IVANOVA_HEMATOPOIESIS_MATURE_CELL | | 219 | -0.29 | -1.09 | 0.287 | 0.692 | 1 | 2363 | tags=29%, list=23%, |
| 522 | | | REACTOME_NRAGE_SIGNALS_DEATH_THROUGH_JNK | | 29 | -0.39 | -1.15 | 0.288 | 0.607 | 1 | 3854 | tags=59%, list=38%, |
| 587 | | | REACTOME_RHOA_GTPASE_CYCLE | | 90 | -0.32 | -1.12 | 0.288 | 0.654 | 1 | 3061 | tags=39%, list=30%, |
| 562 | | | MILI_PSEUDOPODIA_CHEMOTAXIS_UP | | 71 | -0.33 | -1.13 | 0.29 | 0.63 | 1 | 3461 | tags=45%, list=34%, |
| 617 | | | ZENG_GU_POST_ICB_METAGENE_41 | | 156 | -0.3 | -1.1 | 0.29 | 0.672 | 1 | 3379 | tags=39%, list=33%, |
| 535 | | | REACTOME_METHYLATION | | 15 | -0.45 | -1.15 | 0.291 | 0.609 | 1 | 2759 | tags=33%, list=27%, |
| 552 | | | REACTOME_ASPIRIN_ADME | | 20 | -0.42 | -1.14 | 0.291 | 0.618 | 1 | 2442 | tags=35%, list=24%, |
| 526 | | | REACTOME_INHIBITION_OF_THE_PROTEOLYTIC_ACTIVITY_OF_APC_C_REQUIRED_FOR_THE_ONSET_OF_ANAPHASE_BY_MITOTIC_SPINDLE_CHECKPOINT_COMPO | | 17 | -0.44 | -1.15 | 0.292 | 0.609 | 1 | 1626 | tags=29%, list=16%, signal=35% |
| 572 | | | GALINDO_IMMUNE_RESPONSE_TO_ENTEROTOXIN | | 53 | -0.34 | -1.13 | 0.292 | 0.634 | 1 | 912 | tags=21%, list=9%, signal=23% |
| 669 | | | REACTOME_SIGNALING_BY_RHO_GTPASES_MIRO_GTPASES_AND_RHOBTB3 | | 428 | -0.28 | -1.07 | 0.292 | 0.712 | 1 | 2700 | tags=32%, list=27%, |
| 550 | | | BIOCARTA_INTEGRIN_PATHWAY | | 27 | -0.39 | -1.14 | 0.294 | 0.613 | 1 | 3585 | tags=48%, list=35%, |
| 540 | | | BERENJENO_TRANSFORMED_BY_RHOA_REVERSIBLY_DN | | 23 | -0.41 | -1.15 | 0.295 | 0.605 | 1 | 1937 | tags=30%, list=19%, |
| 519 | | | NABA_MATRISOME_METASTATIC_LUNG_LYMPH_NODE_METASTASIS | | 19 | -0.43 | -1.16 | 0.296 | 0.603 | 1 | 1879 | tags=53%, list=19%, |
| 536 | | | REACTOME_METABOLISM_OF_NITRIC_OXIDE_NOS3_ACTIVATION_AND_REGULATI | | 15 | -0.45 | -1.15 | 0.296 | 0.608 | 1 | 1208 | tags=33%, list=12%, |
| 565 | | | REACTOME_REGULATION_OF_TLR_BY_ENDOGENOUS_LIGAND | | 15 | -0.44 | -1.13 | 0.296 | 0.63 | 1 | 1953 | tags=47%, list=19%, |
| 567 | | | HU_GENOTOXIC_DAMAGE_24HR | | 29 | -0.39 | -1.13 | 0.296 | 0.629 | 1 | 2374 | tags=24%, list=24%, |
| 625 | | | WP_MAPK_SIGNALING_PATHWAY | | 107 | -0.3 | -1.09 | 0.299 | 0.684 | 1 | 908 | tags=14%, list=9%, signal=15% |
| 581 | | | BIOCARTA_IL2RB_PATHWAY | | 25 | -0.39 | -1.12 | 0.3 | 0.643 | 1 | 2579 | tags=36%, list=26%, |
| 541 | | | REACTOME_CELL_EXTRACELLULAR_MATRIX_INTERACTIONS | | 17 | -0.44 | -1.15 | 0.304 | 0.604 | 1 | 2177 | tags=35%, list=22%, |
| 600 | | | WP_COMPREHENSIVE_IL_17A_SIGNALING | | 74 | -0.33 | -1.11 | 0.304 | 0.657 | 1 | 1607 | tags=26%, list=16%, |
| 560 | | | LEE_CALORIE_RESTRICTION_MUSCLE_UP | | 26 | -0.4 | -1.13 | 0.305 | 0.626 | 1 | 3356 | tags=42%, list=33%, |
| 580 | | | WP_FIBRIN_COMPLEMENT_RECEPTOR_3_SIGNALING_PATHWAY | | 31 | -0.38 | -1.12 | 0.305 | 0.644 | 1 | 2102 | tags=29%, list=21%, |
| 556 | | | SCHLINGEMANN_SKIN_CARCINOGENESIS_TPA_UP | | 26 | -0.39 | -1.14 | 0.306 | 0.621 | 1 | 3260 | tags=38%, list=32%, |
| 558 | | | MIKKELSEN_MCV6_HCP_WITH_H3K27ME3 | | 43 | -0.36 | -1.13 | 0.306 | 0.625 | 1 | 1605 | tags=28%, list=16%, |
| 573 | | | BIOCARTA_GSK3_PATHWAY | | 19 | -0.42 | -1.13 | 0.31 | 0.633 | 1 | 579 | tags=16%, list=6%, signal=17% |
| 589 | | | WP_APOPTOSIS | | 57 | -0.34 | -1.11 | 0.31 | 0.656 | 1 | 3270 | tags=39%, list=32%, |
| 616 | | | WP_ENDOCHONDRAL_OSSIFICATION | | 36 | -0.35 | -1.1 | 0.31 | 0.67 | 1 | 1593 | tags=28%, list=16%, |
| 640 | | | YAMAZAKI_TCEB3_TARGETS_DN | | 146 | -0.29 | -1.08 | 0.312 | 0.691 | 1 | 3341 | tags=38%, list=33%, |
| 592 | | | WP_ESTROGEN_SIGNALING | | 64 | -0.33 | -1.11 | 0.314 | 0.659 | 1 | 1327 | tags=17%, list=13%, |
| 633 | | | SHETH_LIVER_CANCER_VS_TXNIP_LOSS_PAM1 | | 158 | -0.29 | -1.09 | 0.314 | 0.69 | 1 | 2080 | tags=25%, list=21%, |
| 598 | | | WP_KIT_RECEPTOR_SIGNALING_PATHWAY | | 49 | -0.34 | -1.11 | 0.316 | 0.656 | 1 | 2735 | tags=33%, list=27%, |
| 571 | | | REACTOME_RAF_INDEPENDENT_MAPK1_3_ACTIVATION | | 16 | -0.43 | -1.13 | 0.317 | 0.635 | 1 | 1583 | tags=31%, list=16%, |
| 594 | | | LIAN_LIPA_TARGETS_6M | | 47 | -0.35 | -1.11 | 0.318 | 0.658 | 1 | 2737 | tags=36%, list=27%, |
| 586 | | | REACTOME_PLASMA_LIPOPROTEIN_CLEARANCE | | 30 | -0.37 | -1.12 | 0.32 | 0.649 | 1 | 3126 | tags=47%, list=31%, |
| 590 | | | BIOCARTA_IGF1_PATHWAY | | 19 | -0.41 | -1.11 | 0.321 | 0.661 | 1 | 2648 | tags=42%, list=26%, |
| 605 | | | CHANGOLKAR_H2AFY_TARGETS_UP | | 43 | -0.35 | -1.11 | 0.322 | 0.66 | 1 | 3692 | tags=44%, list=37%, |
| 628 | | | CHIARADONNA_NEOPLASTIC_TRANSFORMATION_KRAS_UP | | 94 | -0.31 | -1.09 | 0.322 | 0.691 | 1 | 1989 | tags=22%, list=20%, |
| 614 | | | WP_SELENIUM_METABOLISM_SELENOPROTEINS | | 40 | -0.35 | -1.1 | 0.323 | 0.666 | 1 | 644 | tags=15%, list=6%, signal=16% |
| 631 | | | WP_MYOMETRIAL_RELAXATION_AND_CONTRACTION_PATHWAYS | | 74 | -0.32 | -1.09 | 0.325 | 0.691 | 1 | 1817 | tags=24%, list=18%, |
| 613 | | | REACTOME_SUMOYLATION_OF_CHROMATIN_ORGANIZATION_PROTEINS | | 42 | -0.35 | -1.1 | 0.326 | 0.667 | 1 | 2091 | tags=26%, list=21%, |
| 604 | | | REACTOME_SIGNALING_BY_FGFR | | 48 | -0.34 | -1.11 | 0.328 | 0.661 | 1 | 3997 | tags=50%, list=40%, |
| 646 | | | REACTOME_REGULATION_OF_TP53_ACTIVITY | | 117 | -0.3 | -1.08 | 0.328 | 0.694 | 1 | 3190 | tags=38%, list=32%, |
| 599 | | | WP_DRAVET_SYNDROME_SCN1A_A1783V_POINT_MUTATION_MODEL | | 29 | -0.38 | -1.11 | 0.329 | 0.655 | 1 | 1596 | tags=24%, list=16%, |
| 578 | | | KAMMINGA_SENESCENCE | | 26 | -0.39 | -1.12 | 0.331 | 0.64 | 1 | 2900 | tags=35%, list=29%, |
| 606 | | | KAMIKUBO_MYELOID_CEBPA_NETWORK | | 22 | -0.39 | -1.11 | 0.332 | 0.663 | 1 | 1156 | tags=23%, list=11%, |
| 568 | | | PAL_PRMT5_TARGETS_DN | | 16 | -0.43 | -1.13 | 0.333 | 0.629 | 1 | 1833 | tags=38%, list=18%, |
| 583 | | | XU_CREBBP_TARGETS_DN | | 23 | -0.4 | -1.12 | 0.335 | 0.648 | 1 | 1612 | tags=30%, list=16%, |
| 620 | | | BYSTRYKH_HEMATOPOIESIS_STEM_CELL_AND_BRAIN_QTL_CIS | | 63 | -0.33 | -1.1 | 0.336 | 0.676 | 1 | 2358 | tags=33%, list=23%, |
| 607 | | | BILANGES_RAPAMYCIN_SENSITIVE_GENES | | 27 | -0.38 | -1.1 | 0.337 | 0.664 | 1 | 1882 | tags=26%, list=19%, |
| 639 | | | REACTOME_RUNX1_REGULATES_TRANSCRIPTION_OF_GENES_INVOLVED_IN_DIFFERENTIATION_OF_HSCS | | 58 | -0.33 | -1.08 | 0.338 | 0.691 | 1 | 2710 | tags=36%, list=27%, signal=49% |
| 621 | | | REACTOME_CHAPERONIN_MEDIATED_PROTEIN_FOLDING | | 22 | -0.39 | -1.1 | 0.34 | 0.675 | 1 | 2928 | tags=41%, list=29%, |
| 655 | | | REACTOME_INTRA_GOLGI_AND_RETROGRADE_GOLGI_TO_ER_TRAFFIC | | 133 | -0.29 | -1.07 | 0.34 | 0.708 | 1 | 3537 | tags=41%, list=35%, |
| 642 | | | BRUNO_HEMATOPOIESIS | | 49 | -0.33 | -1.08 | 0.341 | 0.691 | 1 | 1345 | tags=20%, list=13%, |
| 601 | | | BIOCARTA_TOLL_PATHWAY | | 21 | -0.41 | -1.11 | 0.342 | 0.657 | 1 | 3699 | tags=52%, list=37%, |
| 611 | | | CAIRO_PML_TARGETS_BOUND_BY_MYC_UP | | 19 | -0.41 | -1.1 | 0.343 | 0.667 | 1 | 796 | tags=16%, list=8%, signal=17% |
| 637 | | | HESS_TARGETS_OF_HOXA9_AND_MEIS1_DN | | 56 | -0.33 | -1.09 | 0.343 | 0.69 | 1 | 3433 | tags=41%, list=34%, |
| 647 | | | WP_INSULIN_SIGNALING | | 114 | -0.3 | -1.08 | 0.343 | 0.694 | 1 | 2743 | tags=33%, list=27%, |
| 588 | | | REACTOME_PHOSPHORYLATION_OF_THE_APC_C | | 17 | -0.42 | -1.11 | 0.346 | 0.657 | 1 | 1626 | tags=35%, list=16%, |
| 623 | | | BIOCARTA_BCR_PATHWAY | | 23 | -0.39 | -1.09 | 0.346 | 0.681 | 1 | 625 | tags=22%, list=6%, signal=23% |
| 654 | | | YAMAZAKI_TCEB3_TARGETS_UP | | 125 | -0.29 | -1.07 | 0.347 | 0.709 | 1 | 3409 | tags=34%, list=34%, |
| 615 | | | LIM_MAMMARY_LUMINAL_PROGENITOR_UP | | 22 | -0.39 | -1.1 | 0.348 | 0.667 | 1 | 2182 | tags=36%, list=22%, |
| 630 | | | RAMALHO_STEMNESS_DN | | 44 | -0.34 | -1.09 | 0.348 | 0.692 | 1 | 3505 | tags=50%, list=35%, |
| 629 | | | GOBERT_CORE_OLIGODENDROCYTE_DIFFERENTIATION | | 24 | -0.38 | -1.09 | 0.349 | 0.69 | 1 | 3780 | tags=50%, list=37%, |
| 665 | | | WP_CHEMOKINE_SIGNALING_PATHWAY | | 91 | -0.3 | -1.07 | 0.349 | 0.713 | 1 | 2652 | tags=31%, list=26%, |
| 686 | | | ZHANG_BREAST_CANCER_PROGENITORS_UP | | 340 | -0.27 | -1.06 | 0.349 | 0.723 | 1 | 2502 | tags=28%, list=25%, |
| 612 | | | REACTOME_MITOCHONDRIAL_CALCIUM_ION_TRANSPORT | | 16 | -0.42 | -1.1 | 0.35 | 0.668 | 1 | 4343 | tags=56%, list=43%, |
| 638 | | | HOFFMANN_LARGE_TO_SMALL_PRE_BII_LYMPHOCYTE_DN | | 36 | -0.35 | -1.09 | 0.35 | 0.691 | 1 | 3368 | tags=42%, list=33%, |

| | | | | | | | | | | | |
|---|---|---|---|---|---|---|---|---|---|---|---|
| 618 | | BRACHAT_RESPONSE_TO_METHOTREXATE_UP | | 17 | -0.42 | -1.1 | 0.352 | 0.671 | 1 | 515 | tags=18%, list=5%, signal=19% |
| 636 | | RASHI_RESPONSE_TO_IONIZING_RADIATION_4 | | 39 | -0.36 | -1.09 | 0.352 | 0.691 | 1 | 1937 | tags=26%, list=19%, |
| 627 | | GAVIN_FOXP3_TARGETS_CLUSTER_P4 | | 57 | -0.33 | -1.09 | 0.353 | 0.69 | 1 | 3682 | tags=49%, list=36%, |
| 652 | | GOLDRATH_HOMEOSTATIC_PROLIFERATION | | 131 | -0.3 | -1.07 | 0.355 | 0.711 | 1 | 2590 | tags=28%, list=26%, |
| 703 | | GRAESSMANN_RESPONSE_TO_MC_AND_DOXORUBICIN_UP | | 417 | -0.27 | -1.05 | 0.356 | 0.733 | 1 | 2966 | tags=32%, list=29%, |
| 683 | | REACTOME_BIOLOGICAL_OXIDATIONS | | 144 | -0.29 | -1.06 | 0.36 | 0.721 | 1 | 2929 | tags=32%, list=29%, |
| 626 | | REACTOME_MTOR_SIGNALLING | | 36 | -0.35 | -1.09 | 0.362 | 0.686 | 1 | 2743 | tags=36%, list=27%, |
| 609 | | REACTOME_ERK_MAPK_TARGETS | | 15 | -0.43 | -1.1 | 0.364 | 0.665 | 1 | 4651 | tags=80%, list=46%, |
| 632 | | REACTOME_DDX58_IFIH1_MEDIATED_INDUCTION_OF_INTERFERON_ALPHA_BETA | | 28 | -0.37 | -1.09 | 0.364 | 0.69 | 1 | 3101 | tags=43%, list=31%, |
| 648 | | MORI_LARGE_PRE_BII_LYMPHOCYTE_UP | | 58 | -0.33 | -1.08 | 0.364 | 0.701 | 1 | 1591 | tags=26%, list=16%, |
| 644 | | REACTOME_MITOCHONDRIAL_BIOGENESIS | | 25 | -0.38 | -1.08 | 0.365 | 0.691 | 1 | 4131 | tags=40%, list=41%, |
| 657 | | MORI_SMALL_PRE_BII_LYMPHOCYTE_DN | | 63 | -0.32 | -1.07 | 0.365 | 0.714 | 1 | 3898 | tags=56%, list=39%, |
| 662 | | STEARMAN_LUNG_CANCER_EARLY_VS_LATE_UP | | 115 | -0.3 | -1.07 | 0.366 | 0.713 | 1 | 1807 | tags=25%, list=18%, |
| 653 | | REACTOME_MAP2K_AND_MAPK_ACTIVATION | | 30 | -0.36 | -1.07 | 0.367 | 0.71 | 1 | 3636 | tags=43%, list=36%, |
| 656 | | GUILLAUMOND_KLF10_TARGETS_DN | | 30 | -0.36 | -1.07 | 0.367 | 0.71 | 1 | 2195 | tags=33%, list=22%, |
| 682 | | BRUINS_UVC_RESPONSE_EARLY_LATE | | 228 | -0.28 | -1.06 | 0.367 | 0.722 | 1 | 3273 | tags=36%, list=32%, |
| 710 | | MONNIER_POSTRADIATION_TUMOR_ESCAPE_UP | | 329 | -0.27 | -1.04 | 0.367 | 0.737 | 1 | 2701 | tags=28%, list=27%, |
| 650 | | NABA_SECRETED_FACTORS | | 72 | -0.32 | -1.08 | 0.368 | 0.7 | 1 | 2621 | tags=36%, list=26%, |
| 664 | | LEIN_CHOROID_PLEXUS_MARKERS | | 60 | -0.32 | -1.07 | 0.368 | 0.712 | 1 | 600 | tags=12%, list=6%, signal=12% |
| 680 | | REACTOME_G2_M_CHECKPOINTS | | 108 | -0.29 | -1.06 | 0.368 | 0.718 | 1 | 2710 | tags=31%, list=27%, |
| 684 | | WP_AMINO_ACID_METABOLISM | | 82 | -0.3 | -1.06 | 0.369 | 0.721 | 1 | 3589 | tags=39%, list=36%, |
| 634 | | HEVNER_CORTEX_BASAL_INTERMEDIATE_PROGENITOR_CELLS | | 32 | -0.37 | -1.09 | 0.37 | 0.69 | 1 | 2006 | tags=28%, list=20%, |
| 645 | | WORSCHECH_TUMOR_EVASION_AND_TOLEROGENICITY_UP | | 15 | -0.42 | -1.08 | 0.37 | 0.692 | 1 | 1458 | tags=40%, list=14%, |
| 666 | | GOTZMANN_EPITHELIAL_TO_MESENCHYMAL_TRANSITION_UP | | 47 | -0.34 | -1.07 | 0.37 | 0.713 | 1 | 2707 | tags=45%, list=27%, |
| 674 | | MCBRYAN_PUBERTAL_BREAST_4_5WK_DN | | 151 | -0.29 | -1.06 | 0.37 | 0.715 | 1 | 1800 | tags=21%, list=18%, |
| 679 | | INGRAM_SHH_TARGETS_DN | | 37 | -0.34 | -1.06 | 0.371 | 0.717 | 1 | 1721 | tags=30%, list=17%, |
| 661 | | REACTOME_RHOB_GTPASE_CYCLE | | 45 | -0.33 | -1.07 | 0.372 | 0.713 | 1 | 2553 | tags=38%, list=25%, |
| 663 | | REACTOME_GLYOXYLATE_METABOLISM_AND_GLYCINE_DEGRADATION | | 28 | -0.36 | -1.07 | 0.373 | 0.713 | 1 | 5881 | tags=79%, list=58%, |
| 643 | | MORI_IMMATURE_B_LYMPHOCYTE_UP | | 35 | -0.35 | -1.08 | 0.374 | 0.69 | 1 | 2335 | tags=37%, list=23%, |
| 649 | | MARTIN_NFKB_TARGETS_UP | | 28 | -0.37 | -1.08 | 0.374 | 0.7 | 1 | 2262 | tags=36%, list=22%, |
| 651 | | REACTOME_SEALING_OF_THE_NUCLEAR_ENVELOPE_NE_BY_ESCRT_III | | 18 | -0.41 | -1.07 | 0.381 | 0.712 | 1 | 3336 | tags=50%, list=33%, |
| 673 | | REACTOME_G_ALPHA_12_13_SIGNALLING_EVENTS | | 40 | -0.34 | -1.06 | 0.381 | 0.716 | 1 | 2654 | tags=33%, list=26%, |
| 689 | | REACTOME_PHASE_II_CONJUGATION_OF_COMPOUNDS | | 75 | -0.31 | -1.06 | 0.382 | 0.728 | 1 | 2759 | tags=28%, list=27%, |
| 670 | | REACTOME_TNFR1_INDUCED_PROAPOPTOTIC_SIGNALING | | 19 | -0.4 | -1.07 | 0.386 | 0.711 | 1 | 2563 | tags=42%, list=25%, |
| 685 | | WP_MAPK_CASCADE | | 22 | -0.38 | -1.06 | 0.386 | 0.721 | 1 | 625 | tags=18%, list=6%, signal=19% |
| 672 | | BIOCARTA_PYK2_PATHWAY | | 24 | -0.38 | -1.07 | 0.387 | 0.715 | 1 | 1213 | tags=25%, list=12%, |
| 658 | | REACTOME_RND3_GTPASE_CYCLE | | 29 | -0.36 | -1.07 | 0.388 | 0.714 | 1 | 3410 | tags=48%, list=34%, |
| 678 | | LIU_OVARIAN_CANCER_TUMORS_AND_XENOGRAFTS_XDGS_UP | | 21 | -0.39 | -1.06 | 0.388 | 0.718 | 1 | 3103 | tags=38%, list=31%, |
| 659 | | VANASSE_BCL2_TARGETS_DN | | 34 | -0.35 | -1.07 | 0.389 | 0.713 | 1 | 2782 | tags=32%, list=28%, |
| 681 | | REACTOME_MYD88_MAL_TIRAP_CASCADE_INITIATED_ON_PLASMA_MEMBRANE | | 66 | -0.31 | -1.06 | 0.391 | 0.721 | 1 | 3724 | tags=45%, list=37%, |
| 696 | | IWANAGA_CARCINOGENESIS_BY_KRAS_PTEN_UP | | 121 | -0.29 | -1.05 | 0.391 | 0.728 | 1 | 2994 | tags=33%, list=30%, |
| 698 | | WP_SPINAL_CORD_INJURY | | 48 | -0.32 | -1.05 | 0.391 | 0.729 | 1 | 4051 | tags=50%, list=40%, |
| 692 | | REACTOME_RESPONSE_TO_ELEVATED_PLATELET_CYTOSOLIC_CA2 | | 93 | -0.3 | -1.05 | 0.392 | 0.727 | 1 | 3028 | tags=39%, list=30%, |
| 702 | | WP_MICRORNAS_IN_CARDIOMYOCYTE_HYPERTROPHY | | 58 | -0.31 | -1.05 | 0.393 | 0.731 | 1 | 2800 | tags=33%, list=28%, |
| 675 | | SHETH_LIVER_CANCER_VS_TXNIP_LOSS_PAM6 | | 33 | -0.35 | -1.06 | 0.396 | 0.715 | 1 | 2654 | tags=39%, list=26%, |
| 690 | | MODY_HIPPOCAMPUS_PRENATAL | | 39 | -0.34 | -1.06 | 0.396 | 0.727 | 1 | 4290 | tags=49%, list=42%, |
| 701 | | REACTOME_SIGNALING_BY_TGF_BETA_RECEPTOR_COMPLEX | | 61 | -0.31 | -1.05 | 0.396 | 0.73 | 1 | 1626 | tags=21%, list=16%, |
| 671 | | BIOCARTA_HCMV_PATHWAY | | 16 | -0.42 | -1.07 | 0.398 | 0.716 | 1 | 644 | tags=19%, list=6%, signal=20% |
| 695 | | KASLER_HDAC7_TARGETS_1_UP | | 125 | -0.29 | -1.05 | 0.399 | 0.728 | 1 | 2729 | tags=34%, list=27%, |
| 668 | | LIAN_NEUTROPHIL_GRANULE_CONSTITUENTS | | 16 | -0.41 | -1.07 | 0.401 | 0.713 | 1 | 3581 | tags=50%, list=35%, |
| 706 | | GAUSSMANN_MLL_AF4_FUSION_TARGETS_G_UP | | 116 | -0.29 | -1.05 | 0.401 | 0.739 | 1 | 2262 | tags=27%, list=22%, |
| 708 | | MATZUK_SPERMATOZOA | | 53 | -0.31 | -1.04 | 0.403 | 0.738 | 1 | 3832 | tags=53%, list=38%, |
| 700 | | BOYLAN_MULTIPLE_MYELOMA_PCA3_DN | | 46 | -0.33 | -1.05 | 0.404 | 0.731 | 1 | 4274 | tags=52%, list=42%, |
| 713 | | REACTOME_COPII_MEDIATED_VESICLE_TRANSPORT | | 58 | -0.31 | -1.04 | 0.405 | 0.736 | 1 | 4091 | tags=50%, list=41%, |
| 667 | | REACTOME_XENOBIOTICS | | 16 | -0.41 | -1.07 | 0.406 | 0.712 | 1 | 1658 | tags=31%, list=16%, |
| 676 | | REACTOME_SIALIC_ACID_METABOLISM | | 16 | -0.41 | -1.06 | 0.407 | 0.718 | 1 | 3595 | tags=56%, list=36%, |
| 677 | | BIOCARTA_NFKB_PATHWAY | | 18 | -0.39 | -1.06 | 0.408 | 0.718 | 1 | 3180 | tags=44%, list=31%, |
| 694 | | REACTOME_DISASSEMBLY_OF_THE_DESTRUCTION_COMPLEX_AND_RECRUITMENT_OF_AXIN_TO_THE_MEMBRANE | | 22 | -0.38 | -1.05 | 0.408 | 0.729 | 1 | 4557 | tags=59%, list=45%, signal=107% |
| 717 | | BRACHAT_RESPONSE_TO_CAMPTOTHECIN_DN | | 37 | -0.34 | -1.04 | 0.408 | 0.735 | 1 | 2177 | tags=27%, list=22%, |
| 688 | | REACTOME_SIGNALING_BY_FGFR3 | | 25 | -0.37 | -1.06 | 0.41 | 0.728 | 1 | 3997 | tags=52%, list=40%, |
| 697 | | RIZ_ERYTHROID_DIFFERENTIATION_CCNE1 | | 32 | -0.36 | -1.05 | 0.411 | 0.728 | 1 | 2245 | tags=38%, list=22%, |
| 712 | | REACTOME_AGGREPHAGY | | 22 | -0.37 | -1.04 | 0.411 | 0.735 | 1 | 2728 | tags=36%, list=27%, |
| 724 | | REACTOME_RHO_GTPASE_EFFECTORS | | 160 | -0.28 | -1.04 | 0.411 | 0.739 | 1 | 2655 | tags=33%, list=26%, |
| 727 | | HOLLERN_EMT_BREAST_TUMOR_DN | | 65 | -0.31 | -1.04 | 0.412 | 0.741 | 1 | 1235 | tags=20%, list=12%, |
| 687 | | REACTOME_RECYCLING_PATHWAY_OF_L1 | | 22 | -0.38 | -1.06 | 0.414 | 0.729 | 1 | 1389 | tags=27%, list=14%, |
| 726 | | MCBRYAN_PUBERTAL_BREAST_6_7WK_UP | | 158 | -0.28 | -1.04 | 0.414 | 0.742 | 1 | 3241 | tags=37%, list=32%, |
| 699 | | BYSTRYKH_HEMATOPOIESIS_STEM_CELL_QTL_CIS | | 108 | -0.3 | -1.05 | 0.415 | 0.732 | 1 | 2358 | tags=25%, list=23%, |
| 721 | | BIOCARTA_IGF1MTOR_PATHWAY | | 18 | -0.39 | -1.04 | 0.417 | 0.741 | 1 | 2004 | tags=28%, list=20%, |
| 714 | | REACTOME_O_LINKED_GLYCOSYLATION | | 32 | -0.34 | -1.04 | 0.418 | 0.735 | 1 | 2393 | tags=34%, list=24%, |
| 716 | | FOSTER_TOLERANT_MACROPHAGE_UP | | 110 | -0.29 | -1.04 | 0.418 | 0.736 | 1 | 1379 | tags=19%, list=14%, |
| 691 | | JEPSEN_SMRT_TARGETS | | 21 | -0.38 | -1.05 | 0.419 | 0.726 | 1 | 3522 | tags=57%, list=35%, |
| 705 | | WP_SIDS_SUSCEPTIBILITY_PATHWAYS | | 22 | -0.37 | -1.05 | 0.419 | 0.74 | 1 | 4403 | tags=64%, list=44%, |
| 739 | | REACTOME_RMTS_METHYLATE_HISTONE_ARGININES | | 29 | -0.35 | -1.03 | 0.42 | 0.754 | 1 | 1284 | tags=21%, list=13%, |
| 693 | | LIANG_HEMATOPOIESIS_STEM_CELL_NUMBER_LARGE_VS_TINY_UP | | 26 | -0.36 | -1.05 | 0.421 | 0.728 | 1 | 1891 | tags=31%, list=19%, |
| 707 | | WP_FOLIC_ACID_NETWORK | | 17 | -0.4 | -1.04 | 0.422 | 0.738 | 1 | 4256 | tags=59%, list=42%, |
| 729 | | MCCLUNG_CREB1_TARGETS_UP | | 57 | -0.31 | -1.04 | 0.423 | 0.741 | 1 | 2445 | tags=28%, list=24%, |
| 720 | | LEIN_OLIGODENDROCYTE_MARKERS | | 43 | -0.33 | -1.04 | 0.425 | 0.737 | 1 | 2883 | tags=47%, list=29%, |
| 740 | | GAUSSMANN_MLL_AF4_FUSION_TARGETS_A_UP | | 132 | -0.28 | -1.03 | 0.425 | 0.754 | 1 | 3037 | tags=33%, list=30%, |
| 734 | | BIOCARTA_WNT_PATHWAY | | 19 | -0.37 | -1.03 | 0.426 | 0.746 | 1 | 1530 | tags=21%, list=15%, |
| 711 | | WP_OVARIAN_INFERTILITY | | 15 | -0.41 | -1.04 | 0.427 | 0.736 | 1 | 2444 | tags=47%, list=24%, |
| 723 | | REACTOME_APC_CDC20_MEDIATED_DEGRADATION_OF_NEK2A | | 21 | -0.38 | -1.04 | 0.427 | 0.74 | 1 | 1626 | tags=24%, list=16%, |
| 756 | | LEE_CALORIE_RESTRICTION_NEOCORTEX_DN | | 46 | -0.32 | -1.02 | 0.427 | 0.764 | 1 | 1390 | tags=20%, list=14%, |
| 743 | | LANG_MYB_FAMILY_TARGETS | | 26 | -0.36 | -1.03 | 0.428 | 0.756 | 1 | 3014 | tags=38%, list=30%, |
| 704 | | REACTOME_SIGNALING_BY_FGFR4 | | 26 | -0.36 | -1.05 | 0.429 | 0.736 | 1 | 3997 | tags=50%, list=40%, |
| 741 | | REACTOME_REGULATION_OF_APC_C_ACTIVATORS_BETWEEN_G1_S_AND_EARLY_ANAPHASE | | 28 | -0.35 | -1.03 | 0.429 | 0.753 | 1 | 1626 | tags=25%, list=16%, signal=30% |
| 731 | | REACTOME_RHOV_GTPASE_CYCLE | | 27 | -0.35 | -1.03 | 0.43 | 0.743 | 1 | 779 | tags=19%, list=8%, signal=20% |
| 736 | | REACTOME_EPIGENETIC_REGULATION_OF_GENE_EXPRESSION | | 78 | -0.29 | -1.03 | 0.43 | 0.752 | 1 | 4010 | tags=49%, list=40%, |
| 732 | | REACTOME_EXTRA_NUCLEAR_ESTROGEN_SIGNALING | | 41 | -0.33 | -1.03 | 0.431 | 0.746 | 1 | 1547 | tags=27%, list=15%, |
| 728 | | BIOCARTA_CTCF_PATHWAY | | 17 | -0.4 | -1.04 | 0.433 | 0.741 | 1 | 817 | tags=24%, list=8%, signal=26% |
| 730 | | BIOCARTA_MTA3_PATHWAY | | 15 | -0.4 | -1.04 | 0.433 | 0.742 | 1 | 3336 | tags=47%, list=33%, |
| 733 | | BANG_VERTEPORFIN_ENDOMETRIAL_CANCER_CELLS_UP | | 16 | -0.4 | -1.03 | 0.433 | 0.747 | 1 | 4734 | tags=81%, list=47%, |
| 722 | | MIKKELSEN_ES_LCP_WITH_H3K4ME3 | | 48 | -0.32 | -1.04 | 0.435 | 0.74 | 1 | 1465 | tags=25%, list=15%, |
| 715 | | BIOCARTA_NTHI_PATHWAY | | 19 | -0.39 | -1.04 | 0.436 | 0.736 | 1 | 4488 | tags=63%, list=44%, |
| 746 | | JECHLINGER_EPITHELIAL_TO_MESENCHYMAL_TRANSITION_DN | | 45 | -0.32 | -1.02 | 0.438 | 0.757 | 1 | 2459 | tags=33%, list=24%, |
| 757 | | REACTOME_CHOLESTEROL_BIOSYNTHESIS | | 25 | -0.35 | -1.02 | 0.438 | 0.763 | 1 | 3663 | tags=52%, list=36%, |
| 751 | | MARTORIATI_MDM4_TARGETS_NEUROEPITHELIUM_UP | | 112 | -0.28 | -1.02 | 0.441 | 0.76 | 1 | 4096 | tags=47%, list=41%, |
| 758 | | MORI_SMALL_PRE_BII_LYMPHOCYTE_UP | | 49 | -0.31 | -1.02 | 0.441 | 0.763 | 1 | 2823 | tags=31%, list=28%, |
| 744 | | REACTOME_VASOPRESSIN_REGULATES_RENAL_WATER_HOMEOSTASIS_VIA_AQUAPORINS | | 16 | -0.39 | -1.03 | 0.444 | 0.755 | 1 | 466 | tags=13%, list=5%, signal=13% |
| 748 | | REACTOME_PHASE_I_FUNCTIONALIZATION_OF_COMPOUNDS | | 67 | -0.3 | -1.02 | 0.444 | 0.756 | 1 | 1835 | tags=27%, list=18%, |
| 725 | | MILI_PSEUDOPODIA | | 40 | -0.34 | -1.04 | 0.446 | 0.742 | 1 | 3458 | tags=48%, list=34%, |
| 749 | | REACTOME_RNA_POLYMERASE_I_TRANSCRIPTION_TERMINATION | | 26 | -0.35 | -1.02 | 0.446 | 0.756 | 1 | 1002 | tags=23%, list=10%, |
| 754 | | BIOCARTA_ARENRF2_PATHWAY | | 19 | -0.38 | -1.02 | 0.446 | 0.764 | 1 | 3809 | tags=63%, list=38%, |
| 737 | | ZENG_GU_ICB_CONTROL_METAGENE_21_PRECICTIVE_ICB_RESISTANCE | | 128 | -0.28 | -1.03 | 0.447 | 0.752 | 1 | 1682 | tags=18%, list=17%, |
| 738 | | WP_FAS_PATHWAY_AND_STRESS_INDUCTION_OF_HSP_REGULATION | | 28 | -0.35 | -1.03 | 0.447 | 0.754 | 1 | 625 | tags=14%, list=6%, signal=15% |
| 750 | | REACTOME_NEUROTRANSMITTER_RECEPTORS_AND_POSTSYNAPTIC_SIGNAL_TRANSMISSION | | 43 | -0.32 | -1.02 | 0.447 | 0.757 | 1 | 1561 | tags=21%, list=15%, signal=25% |
| 764 | | WP_GLYCOLYSIS_AND_GLUCONEOGENESIS | | 31 | -0.35 | -1.02 | 0.447 | 0.765 | 1 | 4130 | tags=48%, list=41%, |
| 768 | | MCBRYAN_PUBERTAL_BREAST_5_6WK_UP | | 93 | -0.29 | -1.01 | 0.447 | 0.767 | 1 | 1769 | tags=27%, list=18%, |
| 747 | | REACTOME_DETOXIFICATION_OF_REACTIVE_OXYGEN_SPECIES | | 28 | -0.35 | -1.02 | 0.449 | 0.757 | 1 | 4222 | tags=50%, list=42%, |
| 718 | | ABBUD_LIF_SIGNALING_1_DN | | 20 | -0.38 | -1.04 | 0.45 | 0.737 | 1 | 2347 | tags=35%, list=23%, |
| 765 | | SWEET_LUNG_CANCER_KRAS_DN | | 271 | -0.27 | -1.02 | 0.45 | 0.765 | 1 | 1991 | tags=22%, list=20%, |
| 742 | | REACTOME_KINESINS | | 20 | -0.37 | -1.03 | 0.451 | 0.753 | 1 | 3457 | tags=50%, list=34%, |
| 755 | | REACTOME_VEGFR2_MEDIATED_VASCULAR_PERMEABILITY | | 23 | -0.37 | -1.02 | 0.451 | 0.764 | 1 | 1596 | tags=35%, list=16%, |
| 753 | | REACTOME_COPI_INDEPENDENT_GOLGI_TO_ER_RETROGRADE_TRAFFIC | | 34 | -0.33 | -1.02 | 0.452 | 0.764 | 1 | 3801 | tags=50%, list=38%, |
| 745 | | DELASERNA_MYOD_TARGETS_DN | | 40 | -0.33 | -1.03 | 0.455 | 0.756 | 1 | 1980 | tags=25%, list=20%, |
| 769 | | GERHOLD_ADIPOGENESIS_DN | | 53 | -0.31 | -1.01 | 0.459 | 0.769 | 1 | 1473 | tags=19%, list=15%, |
| 793 | | WP_OXIDATIVE_STRESS_RESPONSE | | 24 | -0.35 | -1 | 0.459 | 0.775 | 1 | 4403 | tags=54%, list=44%, |
| 770 | | BERENJENO_TRANSFORMED_BY_RHOA_DN | | 330 | -0.26 | -1.01 | 0.461 | 0.77 | 1 | 3027 | tags=34%, list=30%, |
| 761 | | MCCLUNG_DELTA_FOSB_TARGETS_8WK | | 30 | -0.34 | -1.02 | 0.462 | 0.767 | 1 | 3089 | tags=50%, list=31%, |
| 752 | | REACTOME_APC_C_CDC20_MEDIATED_DEGRADATION_OF_CYCLIN_B | | 21 | -0.38 | -1.02 | 0.463 | 0.763 | 1 | 1626 | tags=29%, list=16%, |
| 762 | | REACTOME_AQUAPORIN_MEDIATED_TRANSPORT | | 19 | -0.38 | -1.02 | 0.463 | 0.766 | 1 | 810 | tags=16%, list=8%, signal=17% |
| 771 | | REACTOME_ION_CHANNEL_TRANSPORT | | 72 | -0.3 | -1.01 | 0.464 | 0.769 | 1 | 2191 | tags=22%, list=22%, |
| 786 | | REACTOME_INOSITOL_PHOSPHATE_METABOLISM | | 30 | -0.33 | -1 | 0.464 | 0.778 | 1 | 1499 | tags=27%, list=15%, |
| 735 | | REACTOME_ACTIVATION_OF_KAINATE_RECEPTORS_UPON_GLUTAMATE_BINDIN | | 16 | -0.4 | -1.03 | 0.465 | 0.748 | 1 | 51 | tags=13%, list=1%, signal=13% |
| 775 | | REACTOME_TOLL_LIKE_RECEPTOR_4_TLR4_CASCADE | | 88 | -0.29 | -1.01 | 0.466 | 0.77 | 1 | 3724 | tags=43%, list=37%, |
| 799 | | HOFFMANN_IMMATURE_TO_MATURE_B_LYMPHOCYTE_DN | | 34 | -0.33 | -1 | 0.467 | 0.777 | 1 | 3429 | tags=41%, list=34%, |
| 759 | | PASQUALUCCI_LYMPHOMA_BY_GC_STAGE_DN | | 108 | -0.28 | -1.02 | 0.468 | 0.765 | 1 | 2827 | tags=29%, list=28%, |
| 783 | | REACTOME_RHOG_GTPASE_CYCLE | | 56 | -0.31 | -1.01 | 0.468 | 0.774 | 1 | 3693 | tags=48%, list=37%, |
| 763 | | MIKKELSEN_IPS_LCP_WITH_H3K4ME3 | | 64 | -0.3 | -1.02 | 0.47 | 0.766 | 1 | 1465 | tags=22%, list=15%, |
| 772 | | IRITANI_MAD1_TARGETS_DN | | 42 | -0.32 | -1.01 | 0.47 | 0.769 | 1 | 1864 | tags=19%, list=18%, |
| 760 | | WP_P53_SIGNALING | | 45 | -0.32 | -1.02 | 0.471 | 0.765 | 1 | 1944 | tags=24%, list=19%, |
| 781 | | WP_WNT_SIGNALING_PATHWAY_WP403 | | 32 | -0.33 | -1.01 | 0.471 | 0.774 | 1 | 908 | tags=19%, list=9%, signal=21% |
| 774 | | BROWN_MYELOID_CELL_DEVELOPMENT_DN | | 96 | -0.28 | -1.01 | 0.472 | 0.771 | 1 | 1147 | tags=15%, list=11%, |
| 778 | | REACTOME_ROS_AND_RNS_PRODUCTION_IN_PHAGOCYTES | | 22 | -0.37 | -1.01 | 0.472 | 0.775 | 1 | 3023 | tags=36%, list=30%, |
| 767 | | BIOCARTA_STRESS_PATHWAY | | 21 | -0.37 | -1.01 | 0.473 | 0.768 | 1 | 4697 | tags=67%, list=47%, |
| 790 | | WP_G1_TO_S_CELL_CYCLE_CONTROL | | 43 | -0.32 | -1 | 0.473 | 0.776 | 1 | 1944 | tags=26%, list=19%, |
| 789 | | REACTOME_SYNTHESIS_OF_PIPS_AT_THE_PLASMA_MEMBRANE | | 31 | -0.34 | -1 | 0.475 | 0.777 | 1 | 2241 | tags=29%, list=22%, |
| 766 | | REACTOME_NUCLEAR_EVENTS_KINASE_AND_TRANSCRIPTION_FACTOR_ACTIVAT | | 20 | -0.38 | -1.01 | 0.476 | 0.765 | 1 | 4697 | tags=75%, list=47%, |
| 779 | | KONG_E2F3_TARGETS | | 44 | -0.32 | -1.01 | 0.476 | 0.775 | 1 | 796 | tags=11%, list=8%, signal=12% |
| 782 | | JECHLINGER_EPITHELIAL_TO_MESENCHYMAL_TRANSITION_UP | | 47 | -0.31 | -1.01 | 0.479 | 0.773 | 1 | 3120 | tags=45%, list=31%, |
| 773 | | BIOCARTA_BIOPEPTIDES_PATHWAY | | 27 | -0.34 | -1.01 | 0.48 | 0.769 | 1 | 3585 | tags=52%, list=35%, |
| 777 | | HEVNER_TELENCEPHALON_MICROGLIA | | 18 | -0.38 | -1.01 | 0.48 | 0.775 | 1 | 328 | tags=17%, list=3%, signal=17% |
| 795 | | REACTOME_RHOF_GTPASE_CYCLE | | 29 | -0.33 | -1 | 0.48 | 0.776 | 1 | 1461 | tags=31%, list=14%, |
| 780 | | WP_WNT_SIGNALING_PATHWAY_WP539 | | 68 | -0.3 | -1.01 | 0.481 | 0.774 | 1 | 1556 | tags=22%, list=15%, |
| 785 | | LIANG_HEMATOPOIESIS_STEM_CELL_NUMBER_LARGE_VS_TINY_DN | | 28 | -0.34 | -1 | 0.481 | 0.776 | 1 | 971 | tags=21%, list=10%, |
| 810 | | HU_GENOTOXIN_ACTION_DIRECT_VS_INDIRECT_24HR | | 37 | -0.32 | -0.99 | 0.481 | 0.779 | 1 | 3526 | tags=38%, list=35%, |
| 791 | | SARTIPY_NORMAL_AT_INSULIN_RESISTANCE_DN | | 15 | -0.39 | -1 | 0.483 | 0.775 | 1 | 3225 | tags=53%, list=32%, |
| 798 | | BIOCARTA_RAS_PATHWAY | | 17 | -0.38 | -1 | 0.483 | 0.776 | 1 | 471 | tags=12%, list=5%, signal=12% |
| 815 | | WP_EPO_RECEPTOR_SIGNALING | | 20 | -0.36 | -0.99 | 0.483 | 0.778 | 1 | 2597 | tags=35%, list=26%, |
| 784 | | LIM_MAMMARY_LUMINAL_MATURE_DN | | 63 | -0.29 | -1 | 0.484 | 0.775 | 1 | 1000 | tags=17%, list=10%, |
| 801 | | REACTOME_TNF_SIGNALING | | 50 | -0.3 | -1 | 0.485 | 0.776 | 1 | 2563 | tags=28%, list=25%, |
| 800 | | REACTOME_CELL_CYCLE_CHECKPOINTS | | 187 | -0.26 | -1 | 0.486 | 0.776 | 1 | 2710 | tags=31%, list=27%, |
| 811 | | BOYLAN_MULTIPLE_MYELOMA_D_UP | | 62 | -0.3 | -0.99 | 0.486 | 0.778 | 1 | 1717 | tags=24%, list=17%, |
| 776 | | CHYLA_CBFA2T3_TARGETS_UP | | 214 | -0.27 | -1.01 | 0.487 | 0.775 | 1 | 3402 | tags=36%, list=34%, |
| 796 | | REACTOME_NEUTROPHIL_DEGRANULATION | | 336 | -0.26 | -1 | 0.487 | 0.775 | 1 | 2869 | tags=31%, list=28%, |
| 788 | | COATES_MACROPHAGE_M1_VS_M2_DN | | 61 | -0.3 | -1 | 0.488 | 0.777 | 1 | 1652 | tags=21%, list=16%, |
| 817 | | UEDA_CENTRAL_CLOCK | | 74 | -0.29 | -0.99 | 0.488 | 0.779 | 1 | 1301 | tags=16%, list=13%, |
| 787 | | AFFAR_YY1_TARGETS_UP | | 88 | -0.29 | -1 | 0.489 | 0.777 | 1 | 3119 | tags=38%, list=31%, |
| 816 | | REACTOME_GLUCONEOGENESIS | | 24 | -0.35 | -0.99 | 0.489 | 0.778 | 1 | 4096 | tags=50%, list=41%, |
| 819 | | WP_TRYPTOPHAN_METABOLISM | | 33 | -0.33 | -0.99 | 0.49 | 0.783 | 1 | 1001 | tags=15%, list=10%, |
| 794 | | PIONTEK_PKD1_TARGETS_UP | | 34 | -0.33 | -1 | 0.491 | 0.776 | 1 | 2915 | tags=41%, list=29%, |
| 806 | | VIETOR_IFRD1_TARGETS | | 19 | -0.37 | -1 | 0.494 | 0.775 | 1 | 2814 | tags=37%, list=28%, |
| 831 | | REACTOME_RIPK1_MEDIATED_REGULATED_NECROSIS | | 26 | -0.34 | -0.98 | 0.496 | 0.79 | 1 | 3270 | tags=38%, list=32%, |
| 807 | | DARWICHE_PAPILLOMA_RISK_LOW_DN | | 105 | -0.28 | -1 | 0.497 | 0.774 | 1 | 2064 | tags=25%, list=20%, |
| 830 | | LIANG_HEMATOPOIESIS_STEM_CELL_NUMBER_SMALL_VS_HUGE_DN | | 23 | -0.35 | -0.98 | 0.497 | 0.791 | 1 | 1891 | tags=26%, list=19%, |
| 826 | | WENG_POR_TARGETS_LIVER_DN | | 22 | -0.35 | -0.98 | 0.499 | 0.786 | 1 | 1264 | tags=18%, list=13%, |
| 829 | | REACTOME_RAS_PROCESSING | | 18 | -0.37 | -0.98 | 0.499 | 0.791 | 1 | 1300 | tags=22%, list=13%, |
| 818 | | REACTOME_RAB_GEFS_EXCHANGE_GTP_FOR_GDP_ON_RABS | | 69 | -0.29 | -0.99 | 0.5 | 0.779 | 1 | 1324 | tags=19%, list=13%, |
| 812 | | WP_G_PROTEIN_SIGNALING_PATHWAYS | | 49 | -0.31 | -0.99 | 0.501 | 0.781 | 1 | 807 | tags=14%, list=8%, signal=15% |
| 802 | | VARELA_ZMPSTE24_TARGETS_UP | | 34 | -0.33 | -1 | 0.502 | 0.777 | 1 | 1769 | tags=29%, list=18%, |
| 804 | | HOFFMANN_LARGE_TO_SMALL_PRE_BII_LYMPHOCYTE_UP | | 88 | -0.28 | -1 | 0.502 | 0.777 | 1 | 1799 | tags=20%, list=18%, |
| 803 | | DELACROIX_RAR_BOUND_ES | | 281 | -0.26 | -1 | 0.503 | 0.777 | 1 | 1890 | tags=22%, list=19%, |
| 825 | | REACTOME_NEUROTRANSMITTER_RELEASE_CYCLE | | 21 | -0.36 | -0.99 | 0.505 | 0.787 | 1 | 2389 | tags=33%, list=24%, |
| 792 | | HEVNER_CORTEX_RADIAL_GLIA_PROGENITORS | | 255 | -0.26 | -1 | 0.506 | 0.775 | 1 | 2163 | tags=22%, list=21%, |
| 836 | | MIKKELSEN_MEF_HCP_WITH_H3K27ME3 | | 64 | -0.29 | -0.98 | 0.508 | 0.79 | 1 | 3096 | tags=38%, list=31%, |
| 851 | | REACTOME_ACTIVATION_OF_THE_PRE_REPLICATIVE_COMPLEX | | 24 | -0.35 | -0.97 | 0.509 | 0.796 | 1 | 1799 | tags=29%, list=18%, |
| 820 | | WP_EUKARYOTIC_TRANSCRIPTION_INITIATION | | 38 | -0.32 | -0.99 | 0.51 | 0.786 | 1 | 1327 | tags=18%, list=13%, |
| 809 | | REACTOME_GOLGI_TO_ER_RETROGRADE_TRANSPORT | | 85 | -0.28 | -0.99 | 0.511 | 0.779 | 1 | 4065 | tags=47%, list=40%, |
| 839 | | REACTOME_PLASMA_LIPOPROTEIN_REMODELING | | 25 | -0.34 | -0.98 | 0.511 | 0.795 | 1 | 1574 | tags=24%, list=16%, |
| 822 | | BURTON_ADIPOGENESIS_PEAK_AT_16HR | | 29 | -0.34 | -0.99 | 0.512 | 0.785 | 1 | 2794 | tags=38%, list=28%, |
| 824 | | REACTOME_TOLL_LIKE_RECEPTOR_9_TLR9_CASCADE | | 72 | -0.29 | -0.99 | 0.512 | 0.785 | 1 | 3724 | tags=44%, list=37%, |
| 797 | | REACTOME_RHO_GTPASE_CYCLE | | 297 | -0.26 | -1 | 0.513 | 0.776 | 1 | 3377 | tags=37%, list=33%, |
| 805 | | REACTOME_TRANSCRIPTIONAL_REGULATION_BY_TP53 | | 239 | -0.26 | -1 | 0.513 | 0.776 | 1 | 3914 | tags=43%, list=39%, |
| 827 | | SANSOM_APC_TARGETS_UP | | 86 | -0.28 | -0.98 | 0.513 | 0.786 | 1 | 2761 | tags=34%, list=27%, |
| 813 | | REACTOME_GLUCAGON_LIKE_PEPTIDE_1_GLP1_REGULATES_INSULIN_SECRETIO | | 15 | -0.39 | -0.99 | 0.514 | 0.78 | 1 | 1247 | tags=20%, list=12%, |
| 828 | | REACTOME_RETROGRADE_TRANSPORT_AT_THE_TRANS_GOLGI_NETWORK | | 41 | -0.32 | -0.98 | 0.514 | 0.79 | 1 | 4070 | tags=51%, list=40%, |
| 832 | | REACTOME_TBC_RABGAPS | | 26 | -0.34 | -0.98 | 0.515 | 0.79 | 1 | 3914 | tags=58%, list=39%, |
| 823 | | IVANOVA_HEMATOPOIESIS_INTERMEDIATE_PROGENITOR | | 110 | -0.28 | -0.99 | 0.516 | 0.785 | 1 | 3305 | tags=35%, list=33%, |
| 834 | | BIOCARTA_CHEMICAL_PATHWAY | | 17 | -0.38 | -0.98 | 0.516 | 0.79 | 1 | 573 | tags=18%, list=6%, signal=19% |
| 837 | | REACTOME_MYD88_INDEPENDENT_TLR4_CASCADE | | 79 | -0.28 | -0.98 | 0.516 | 0.79 | 1 | 3724 | tags=44%, list=37%, |
| 838 | | REACTOME_RAF_ACTIVATION | | 29 | -0.33 | -0.98 | 0.516 | 0.79 | 1 | 4557 | tags=62%, list=45%, |
| 840 | | WOTTON_RUNX_TARGETS_DN | | 17 | -0.38 | -0.98 | 0.518 | 0.794 | 1 | 978 | tags=18%, list=10%, |
| 848 | | REACTOME_NUCLEAR_ENVELOPE_NE_REASSEMBLY | | 52 | -0.3 | -0.97 | 0.521 | 0.796 | 1 | 999 | tags=17%, list=10%, |
| 821 | | REACTOME_ESR_MEDIATED_SIGNALING | | 85 | -0.28 | -0.99 | 0.522 | 0.785 | 1 | 1573 | tags=21%, list=16%, |
| 875 | | REACTOME_MITOTIC_PROPHASE | | 64 | -0.28 | -0.96 | 0.522 | 0.801 | 1 | 5211 | tags=63%, list=52%, |
| 814 | | REACTOME_RAB_REGULATION_OF_TRAFFICKING | | 88 | -0.28 | -0.99 | 0.523 | 0.779 | 1 | 4546 | tags=58%, list=45%, |
| 845 | | REACTOME_FORMATION_OF_ATP_BY_CHEMIOSMOTIC_COUPLING | | 15 | -0.38 | -0.97 | 0.524 | 0.794 | 1 | 5948 | tags=73%, list=59%, |
| 833 | | PASINI_SUZ12_TARGETS_UP | | 50 | -0.3 | -0.98 | 0.525 | 0.789 | 1 | 1864 | tags=28%, list=18%, |
| 841 | | MORI_PRE_BI_LYMPHOCYTE_UP | | 50 | -0.3 | -0.98 | 0.527 | 0.794 | 1 | 1828 | tags=22%, list=18%, |
| 867 | | BOYLAN_MULTIPLE_MYELOMA_D_CLUSTER_UP | | 18 | -0.36 | -0.96 | 0.528 | 0.801 | 1 | 1717 | tags=28%, list=17%, |
| 869 | | REACTOME_SENESCENCE_ASSOCIATED_SECRETORY_PHENOTYPE_SASP | | 32 | -0.32 | -0.96 | 0.528 | 0.8 | 1 | 1944 | tags=28%, list=19%, |
| 862 | | SEKI_INFLAMMATORY_RESPONSE_LPS_DN | | 16 | -0.37 | -0.96 | 0.529 | 0.804 | 1 | 2135 | tags=25%, list=21%, |
| 852 | | REACTOME_COPI_DEPENDENT_GOLGI_TO_ER_RETROGRADE_TRAFFIC | | 58 | -0.29 | -0.97 | 0.53 | 0.796 | 1 | 3537 | tags=41%, list=35%, |
| 872 | | REACTOME_REGULATED_NECROSIS | | 41 | -0.31 | -0.96 | 0.53 | 0.802 | 1 | 3270 | tags=39%, list=32%, |
| 849 | | LEE_CALORIE_RESTRICTION_MUSCLE_DN | | 37 | -0.32 | -0.97 | 0.532 | 0.795 | 1 | 4473 | tags=59%, list=44%, |
| 844 | | YAO_TEMPORAL_RESPONSE_TO_PROGESTERONE_CLUSTER_14 | | 125 | -0.27 | -0.98 | 0.533 | 0.794 | 1 | 2139 | tags=27%, list=21%, |
| 847 | | REACTOME_OVARIAN_TUMOR_DOMAIN_PROTEASES | | 25 | -0.35 | -0.97 | 0.533 | 0.794 | 1 | 2525 | tags=28%, list=25%, |
| 870 | | REACTOME_FCERI_MEDIATED_CA_2_MOBILIZATION | | 16 | -0.37 | -0.96 | 0.533 | 0.8 | 1 | 2800 | tags=50%, list=28%, |
| 843 | | REACTOME_G_ALPHA_I_SIGNALLING_EVENTS | | 74 | -0.28 | -0.98 | 0.534 | 0.795 | 1 | 2540 | tags=30%, list=25%, |
| 885 | | REACTOME_OXIDATIVE_STRESS_INDUCED_SENESCENCE | | 29 | -0.32 | -0.95 | 0.536 | 0.813 | 1 | 4329 | tags=55%, list=43%, |
| 866 | | REACTOME_FCGAMMA_RECEPTOR_FCGR_DEPENDENT_PHAGOCYTOSIS | | 56 | -0.29 | -0.96 | 0.54 | 0.802 | 1 | 2478 | tags=32%, list=25%, |
| 892 | | WP_NOVEL_JUN_DMP1_PATHWAY | | 21 | -0.35 | -0.95 | 0.541 | 0.814 | 1 | 625 | tags=14%, list=6%, signal=15% |
| 854 | | WP_ERBB_SIGNALING_PATHWAY | | 29 | -0.33 | -0.97 | 0.542 | 0.802 | 1 | 2416 | tags=38%, list=24%, |
| 884 | | MIKKELSEN_MEF_ICP_WITH_H3K27ME3 | | 30 | -0.32 | -0.95 | 0.542 | 0.814 | 1 | 1975 | tags=30%, list=20%, |
| 880 | | REACTOME_SIGNALING_BY_ROBO_RECEPTORS | | 15 | -0.38 | -0.96 | 0.544 | 0.804 | 1 | 2172 | tags=33%, list=22%, |
| 871 | | SERVITJA_ISLET_HNF1A_TARGETS_DN | | 67 | -0.29 | -0.96 | 0.545 | 0.802 | 1 | 2669 | tags=30%, list=26%, |
| 879 | | REACTOME_HDMS_DEMETHYLATE_HISTONES | | 23 | -0.34 | -0.96 | 0.545 | 0.803 | 1 | 1746 | tags=26%, list=17%, |
| 883 | | REACTOME_REGULATION_OF_TP53_DEGRADATION | | 30 | -0.32 | -0.95 | 0.545 | 0.809 | 1 | 1696 | tags=27%, list=17%, |
| 890 | | DELACROIX_RAR_TARGETS_UP | | 33 | -0.31 | -0.95 | 0.545 | 0.814 | 1 | 1440 | tags=21%, list=14%, |
| 846 | | BAKKER_FOXO3_TARGETS_DN | | 116 | -0.27 | -0.97 | 0.547 | 0.795 | 1 | 3330 | tags=36%, list=33%, |
| 864 | | REACTOME_ARACHIDONIC_ACID_METABOLISM | | 24 | -0.34 | -0.96 | 0.547 | 0.802 | 1 | 2799 | tags=38%, list=28%, |
| 882 | | BURTON_ADIPOGENESIS_10 | | 29 | -0.32 | -0.95 | 0.547 | 0.809 | 1 | 1879 | tags=34%, list=19%, |
| 856 | | REACTOME_PEROXISOMAL_PROTEIN_IMPORT | | 59 | -0.29 | -0.97 | 0.548 | 0.803 | 1 | 2983 | tags=34%, list=30%, |
| 888 | | ZENG_GU_POST_ICB_METAGENE_39 | | 33 | -0.31 | -0.95 | 0.548 | 0.814 | 1 | 2684 | tags=33%, list=27%, |
| 895 | | WP_OXIDATION_BY_CYTOCHROME_P450 | | 23 | -0.34 | -0.95 | 0.548 | 0.815 | 1 | 1374 | tags=26%, list=14%, |

| | | | | | | | | | | | |
|---|---|---|---|---|---|---|---|---|---|---|---|
| 855 | | REACTOME_SYNTHESIS_OF_PA | | 18 | -0.36 | -0.97 | 0.55 | 0.803 | 1 | 4421 | tags=61%, list=44%, |
| 897 | | REACTOME_NUCLEOTIDE_BINDING_DOMAIN_LEUCINE_RICH_REPEAT_CONTAINING_RECEPTOR_NLR_SIGNALING_PATHWAYS | | 30 | -0.32 | -0.95 | 0.55 | 0.817 | 1 | 2525 | tags=30%, list=25%, signal=40% |
| 868 | | SAUL_SEN_MAYO | | 58 | -0.29 | -0.96 | 0.552 | 0.8 | 1 | 2621 | tags=29%, list=26%, |
| 873 | | SCHAEFFER_PROSTATE_DEVELOPMENT_12HR_DN | | 21 | -0.35 | -0.96 | 0.553 | 0.802 | 1 | 804 | tags=19%, list=8%, signal=21% |
| 876 | | ZENG_GU_ICB_CONTROL_METAGENE_19_PRECICTIVE_ICB_RESPONSE | | 84 | -0.27 | -0.96 | 0.555 | 0.801 | 1 | 2380 | tags=25%, list=24%, |
| 886 | | LEIN_CEREBELLUM_MARKERS | | 41 | -0.3 | -0.95 | 0.555 | 0.813 | 1 | 3402 | tags=46%, list=34%, |
| 899 | | BIOCARTA_CREB_PATHWAY | | 17 | -0.36 | -0.95 | 0.556 | 0.815 | 1 | 573 | tags=18%, list=6%, signal=19% |
| 835 | | REACTOME_M_PHASE | | 263 | -0.26 | -0.98 | 0.559 | 0.791 | 1 | 2710 | tags=29%, list=27%, |
| 860 | | DARWICHE_SKIN_TUMOR_PROMOTER_UP | | 92 | -0.28 | -0.97 | 0.559 | 0.802 | 1 | 2180 | tags=27%, list=22%, |
| 902 | | WP_CHOLESTEROL_BIOSYNTHESIS | | 15 | -0.37 | -0.94 | 0.559 | 0.817 | 1 | 1672 | tags=33%, list=17%, |
| 914 | | BOYLAN_MULTIPLE_MYELOMA_D_CLUSTER_DN | | 23 | -0.33 | -0.94 | 0.559 | 0.825 | 1 | 1489 | tags=26%, list=15%, |
| 865 | | BIOCARTA_GH_PATHWAY | | 22 | -0.34 | -0.96 | 0.56 | 0.802 | 1 | 2369 | tags=36%, list=23%, |
| 915 | | BIOCARTA_TCR_PATHWAY | | 30 | -0.31 | -0.94 | 0.56 | 0.824 | 1 | 625 | tags=17%, list=6%, signal=18% |
| 891 | | ZHENG_FOXP3_TARGETS_UP | | 20 | -0.34 | -0.95 | 0.561 | 0.813 | 1 | 4301 | tags=45%, list=43%, |
| 904 | | BIOCARTA_EIF4_PATHWAY | | 21 | -0.35 | -0.94 | 0.562 | 0.817 | 1 | 3845 | tags=57%, list=38%, |
| 861 | | HOLLERN_EMT_BREAST_TUMOR_UP | | 78 | -0.28 | -0.97 | 0.563 | 0.803 | 1 | 2696 | tags=32%, list=27%, |
| 901 | | BREDEMEYER_RAG_SIGNALING_VIA_ATM_NOT_VIA_NFKB_UP | | 34 | -0.31 | -0.95 | 0.563 | 0.815 | 1 | 1504 | tags=26%, list=15%, |
| 903 | | REACTOME_TRANSFERRIN_ENDOCYTOSIS_AND_RECYCLING | | 24 | -0.33 | -0.94 | 0.563 | 0.817 | 1 | 4704 | tags=58%, list=47%, |
| 935 | | SAENZ_DETOX_PATHWAY_AND_CARCINOGENESIS_DN | | 17 | -0.35 | -0.93 | 0.566 | 0.825 | 1 | 4790 | tags=59%, list=47%, |
| 907 | | CLASPER_LYMPHATIC_VESSELS_DURING_METASTASIS_DN | | 18 | -0.35 | -0.94 | 0.567 | 0.818 | 1 | 1849 | tags=33%, list=18%, |
| 898 | | LEE_TARGETS_OF_PTCH1_AND_SUFU_DN | | 32 | -0.31 | -0.95 | 0.568 | 0.816 | 1 | 2268 | tags=34%, list=22%, |
| 929 | | REACTOME_REGULATION_OF_INSULIN_SECRETION | | 30 | -0.32 | -0.93 | 0.57 | 0.825 | 1 | 1247 | tags=20%, list=12%, |
| 932 | | REACTOME_REGULATION_OF_TP53_ACTIVITY_THROUGH_METHYLATION | | 16 | -0.36 | -0.93 | 0.57 | 0.826 | 1 | 4034 | tags=56%, list=40%, |
| 912 | | BILANGES_SERUM_RESPONSE_TRANSLATION | | 26 | -0.33 | -0.94 | 0.572 | 0.821 | 1 | 4996 | tags=62%, list=49%, |
| 850 | | TORCHIA_TARGETS_OF_EWSR1_FLI1_FUSION_DN | | 228 | -0.26 | -0.97 | 0.573 | 0.796 | 1 | 2478 | tags=27%, list=25%, |
| 877 | | DURAND_STROMA_S_UP | | 147 | -0.26 | -0.96 | 0.573 | 0.802 | 1 | 2578 | tags=30%, list=26%, |
| 911 | | REACTOME_FORMATION_OF_WDR5_CONTAINING_HISTONE_MODIFYING_COMPLE | | 34 | -0.31 | -0.94 | 0.573 | 0.821 | 1 | 1475 | tags=24%, list=15%, |
| 927 | | PLASARI_TGFB1_SIGNALING_VIA_NFIC_1HR_UP | | 21 | -0.34 | -0.93 | 0.574 | 0.826 | 1 | 1375 | tags=24%, list=14%, |
| 900 | | REACTOME_SEMAPHORIN_INTERACTIONS | | 30 | -0.32 | -0.95 | 0.576 | 0.816 | 1 | 1370 | tags=20%, list=14%, |
| 905 | | REACTOME_STIMULI_SENSING_CHANNELS | | 34 | -0.31 | -0.94 | 0.577 | 0.816 | 1 | 228 | tags=9%, list=2%, signal=9% |
| 874 | | DARWICHE_SQUAMOUS_CELL_CARCINOMA_UP | | 102 | -0.27 | -0.96 | 0.578 | 0.801 | 1 | 3648 | tags=41%, list=36%, |
| 889 | | DARWICHE_PAPILLOMA_RISK_HIGH_UP | | 93 | -0.27 | -0.95 | 0.579 | 0.814 | 1 | 3648 | tags=40%, list=36%, |
| 941 | | WP_NUCLEAR_RECEPTORS | | 24 | -0.32 | -0.92 | 0.579 | 0.833 | 1 | 3052 | tags=42%, list=30%, |
| 906 | | REACTOME_TGF_BETA_RECEPTOR_SIGNALING_ACTIVATES_SMADS | | 31 | -0.31 | -0.94 | 0.581 | 0.817 | 1 | 1564 | tags=23%, list=15%, |
| 918 | | YAO_TEMPORAL_RESPONSE_TO_PROGESTERONE_CLUSTER_6 | | 59 | -0.28 | -0.93 | 0.581 | 0.824 | 1 | 2044 | tags=25%, list=20%, |
| 937 | | BIOCARTA_TEL_PATHWAY | | 17 | -0.35 | -0.92 | 0.581 | 0.83 | 1 | 178 | tags=12%, list=2%, signal=12% |
| 910 | | REACTOME_DOWNREGULATION_OF_SMAD2_3_SMAD4_TRANSCRIPTIONAL_ACTIVI | | 20 | -0.34 | -0.94 | 0.582 | 0.821 | 1 | 1626 | tags=20%, list=16%, |
| 925 | | REACTOME_RHOC_GTPASE_CYCLE | | 49 | -0.28 | -0.93 | 0.582 | 0.826 | 1 | 2553 | tags=31%, list=25%, |
| 928 | | REACTOME_RHOD_GTPASE_CYCLE | | 39 | -0.29 | -0.93 | 0.582 | 0.825 | 1 | 3663 | tags=49%, list=36%, |
| 936 | | REACTOME_RHO_GTPASES_ACTIVATE_IQGAPS | | 20 | -0.34 | -0.93 | 0.583 | 0.827 | 1 | 1208 | tags=25%, list=12%, |
| 955 | | MCBRYAN_PUBERTAL_BREAST_3_4WK_DN | | 25 | -0.32 | -0.91 | 0.584 | 0.835 | 1 | 1932 | tags=24%, list=19%, |
| 893 | | LIN_NPAS4_TARGETS_UP | | 109 | -0.26 | -0.95 | 0.585 | 0.814 | 1 | 2559 | tags=28%, list=25%, |
| 916 | | MIKKELSEN_NPC_HCP_WITH_H3K27ME3 | | 51 | -0.29 | -0.94 | 0.588 | 0.823 | 1 | 1949 | tags=29%, list=19%, |
| 931 | | GRAESSMANN_RESPONSE_TO_MC_AND_SERUM_DEPRIVATION_DN | | 51 | -0.28 | -0.93 | 0.588 | 0.825 | 1 | 1269 | tags=18%, list=13%, |
| 887 | | IKEDA_MIR30_TARGETS_UP | | 100 | -0.27 | -0.95 | 0.589 | 0.813 | 1 | 2054 | tags=24%, list=20%, |
| 926 | | REACTOME_INTERLEUKIN_3_INTERLEUKIN_5_AND_GM_CSF_SIGNALING | | 28 | -0.31 | -0.93 | 0.589 | 0.827 | 1 | 2579 | tags=29%, list=26%, |
| 919 | | REACTOME_SIGNALING_BY_FGFR2 | | 40 | -0.29 | -0.93 | 0.591 | 0.825 | 1 | 3997 | tags=45%, list=40%, |
| 954 | | BERENJENO_TRANSFORMED_BY_RHOA_FOREVER_DN | | 22 | -0.33 | -0.91 | 0.592 | 0.836 | 1 | 2306 | tags=32%, list=23%, |
| 924 | | REACTOME_SIGNALLING_TO_ERKS | | 19 | -0.35 | -0.93 | 0.593 | 0.825 | 1 | 4406 | tags=58%, list=44%, |
| 959 | | REACTOME_PROTEIN_PROTEIN_INTERACTIONS_AT_SYNAPSES | | 25 | -0.31 | -0.91 | 0.595 | 0.836 | 1 | 1348 | tags=16%, list=13%, |
| 963 | | BIOCARTA_IL1R_PATHWAY | | 21 | -0.33 | -0.91 | 0.595 | 0.838 | 1 | 3253 | tags=38%, list=32%, |
| 842 | | MILI_PSEUDOPODIA_HAPTOTAXIS_UP | | 482 | -0.25 | -0.98 | 0.596 | 0.795 | 1 | 3053 | tags=28%, list=30%, |
| 943 | | SHIN_B_CELL_LYMPHOMA_CLUSTER_3 | | 16 | -0.35 | -0.92 | 0.596 | 0.835 | 1 | 4918 | tags=75%, list=49%, |
| 971 | | BIOCARTA_PML_PATHWAY | | 15 | -0.35 | -0.9 | 0.597 | 0.846 | 1 | 5496 | tags=67%, list=54%, |
| 909 | | CHIARADONNA_NEOPLASTIC_TRANSFORMATION_CDC25_DN | | 115 | -0.26 | -0.94 | 0.598 | 0.822 | 1 | 2053 | tags=23%, list=20%, |
| 878 | | RAY_TUMORIGENESIS_BY_ERBB2_CDC25A_DN | | 197 | -0.26 | -0.96 | 0.601 | 0.801 | 1 | 3228 | tags=36%, list=32%, |
| 894 | | IGLESIAS_E2F_TARGETS_UP | | 121 | -0.26 | -0.95 | 0.601 | 0.815 | 1 | 2955 | tags=33%, list=29%, |
| 951 | | REACTOME_VISUAL_PHOTOTRANSDUCTION | | 47 | -0.28 | -0.92 | 0.601 | 0.834 | 1 | 3771 | tags=40%, list=37%, |
| 934 | | MORI_EMU_MYC_LYMPHOMA_BY_ONSET_TIME_UP | | 72 | -0.27 | -0.93 | 0.602 | 0.826 | 1 | 3794 | tags=42%, list=38%, |
| 857 | | IVANOVA_HEMATOPOIESIS_EARLY_PROGENITOR | | 349 | -0.25 | -0.97 | 0.605 | 0.802 | 1 | 2492 | tags=25%, list=25%, |
| 939 | | HORTON_SREBF_TARGETS | | 24 | -0.33 | -0.92 | 0.607 | 0.833 | 1 | 1672 | tags=25%, list=17%, |
| 980 | | REACTOME_CLASS_I_PEROXISOMAL_MEMBRANE_PROTEIN_IMPORT | | 17 | -0.34 | -0.9 | 0.607 | 0.845 | 1 | 2488 | tags=35%, list=25%, |
| 944 | | REACTOME_PRC2_METHYLATES_HISTONES_AND_DNA | | 20 | -0.33 | -0.92 | 0.609 | 0.834 | 1 | 4232 | tags=45%, list=42%, |
| 985 | | WAKABAYASHI_ADIPOGENESIS_PPARG_BOUND_36HR | | 22 | -0.32 | -0.9 | 0.609 | 0.849 | 1 | 1824 | tags=23%, list=18%, |
| 956 | | BOYLAN_MULTIPLE_MYELOMA_D_DN | | 55 | -0.28 | -0.91 | 0.61 | 0.835 | 1 | 2026 | tags=24%, list=20%, |
| 953 | | REACTOME_TP53_REGULATES_METABOLIC_GENES | | 61 | -0.28 | -0.92 | 0.611 | 0.835 | 1 | 2743 | tags=30%, list=27%, |
| 965 | | BOYLAN_MULTIPLE_MYELOMA_C_CLUSTER_DN | | 24 | -0.32 | -0.91 | 0.612 | 0.837 | 1 | 1784 | tags=33%, list=18%, |
| 923 | | BURTON_ADIPOGENESIS_8 | | 54 | -0.29 | -0.93 | 0.613 | 0.823 | 1 | 4792 | tags=57%, list=47%, |
| 922 | | SHETH_LIVER_CANCER_VS_TXNIP_LOSS_PAM2 | | 93 | -0.27 | -0.93 | 0.615 | 0.824 | 1 | 796 | tags=13%, list=8%, signal=14% |
| 940 | | REACTOME_RESOLUTION_OF_SISTER_CHROMATID_COHESION | | 76 | -0.26 | -0.92 | 0.615 | 0.833 | 1 | 2406 | tags=30%, list=24%, |
| 952 | | REACTOME_CDC42_GTPASE_CYCLE | | 57 | -0.28 | -0.92 | 0.616 | 0.836 | 1 | 3693 | tags=46%, list=37%, |
| 973 | | HOFFMANN_PRE_BI_TO_LARGE_PRE_BII_LYMPHOCYTE_DN | | 47 | -0.28 | -0.9 | 0.618 | 0.846 | 1 | 3511 | tags=43%, list=35%, |
| 976 | | REACTOME_RHO_GTPASES_ACTIVATE_PKNS | | 16 | -0.35 | -0.9 | 0.618 | 0.845 | 1 | 4984 | tags=50%, list=49%, |
| 981 | | WP_GLUTATHIONE_AND_ONE_CARBON_METABOLISM | | 25 | -0.31 | -0.9 | 0.618 | 0.844 | 1 | 4222 | tags=48%, list=42%, |
| 974 | | REACTOME_TELOMERE_EXTENSION_BY_TELOMERASE | | 15 | -0.35 | -0.9 | 0.619 | 0.845 | 1 | 4513 | tags=67%, list=45%, |
| 969 | | DELACROIX_RAR_TARGETS_DN | | 17 | -0.34 | -0.91 | 0.62 | 0.84 | 1 | 608 | tags=12%, list=6%, signal=12% |
| 948 | | WP_METAPATHWAY_BIOTRANSFORMATION | | 76 | -0.27 | -0.92 | 0.625 | 0.832 | 1 | 2489 | tags=25%, list=25%, |
| 917 | | MEISSNER_NPC_HCP_WITH_H3_UNMETHYLATED | | 153 | -0.25 | -0.93 | 0.626 | 0.824 | 1 | 3150 | tags=37%, list=31%, |
| 967 | | WP_IL_2_SIGNALING_PATHWAY | | 62 | -0.27 | -0.91 | 0.626 | 0.837 | 1 | 1367 | tags=18%, list=14%, |
| 930 | | PLASARI_TGFB1_TARGETS_10HR_UP | | 90 | -0.26 | -0.93 | 0.628 | 0.825 | 1 | 1864 | tags=27%, list=18%, |
| 1000 | | REACTOME_RHO_GTPASES_ACTIVATE_WASPS_AND_WAVES | | 28 | -0.3 | -0.88 | 0.629 | 0.865 | 1 | 2478 | tags=36%, list=25%, |
| 993 | | JIANG_AGING_HYPOTHALAMUS_DN | | 28 | -0.3 | -0.89 | 0.63 | 0.857 | 1 | 4215 | tags=50%, list=42%, |
| 1007 | | BYSTRYKH_HEMATOPOIESIS_STEM_CELL_SCP2_QTL_TRANS | | 17 | -0.33 | -0.88 | 0.631 | 0.866 | 1 | 640 | tags=18%, list=6%, signal=19% |
| 982 | | REACTOME_INTRA_GOLGI_TRAFFIC | | 30 | -0.31 | -0.9 | 0.633 | 0.844 | 1 | 3537 | tags=43%, list=35%, |
| 987 | | BIOCARTA_CERAMIDE_PATHWAY | | 21 | -0.33 | -0.89 | 0.634 | 0.859 | 1 | 5085 | tags=71%, list=50%, |
| 977 | | DELASERNA_MYOD_TARGETS_UP | | 49 | -0.28 | -0.9 | 0.635 | 0.844 | 1 | 3290 | tags=37%, list=33%, |
| 881 | | REACTOME_CELL_CYCLE | | 412 | -0.25 | -0.96 | 0.636 | 0.805 | 1 | 2710 | tags=28%, list=27%, |
| 950 | | REACTOME_TRANSMISSION_ACROSS_CHEMICAL_SYNAPSES | | 68 | -0.27 | -0.92 | 0.636 | 0.835 | 1 | 2389 | tags=25%, list=24%, |
| 938 | | REACTOME_MITOTIC_SPINDLE_CHECKPOINT | | 73 | -0.27 | -0.92 | 0.637 | 0.834 | 1 | 2406 | tags=30%, list=24%, |
| 1003 | | HOFFMANN_SMALL_PRE_BII_TO_IMMATURE_B_LYMPHOCYTE_DN | | 23 | -0.31 | -0.88 | 0.637 | 0.866 | 1 | 1834 | tags=26%, list=18%, |
| 972 | | REACTOME_ESTROGEN_DEPENDENT_GENE_EXPRESSION | | 42 | -0.29 | -0.9 | 0.638 | 0.846 | 1 | 1934 | tags=21%, list=19%, |
| 997 | | SARTIPY_NORMAL_AT_INSULIN_RESISTANCE_UP | | 23 | -0.32 | -0.88 | 0.638 | 0.863 | 1 | 935 | tags=17%, list=9%, signal=19% |
| 979 | | REACTOME_ANCHORING_OF_THE_BASAL_BODY_TO_THE_PLASMA_MEMBRANE | | 66 | -0.27 | -0.9 | 0.639 | 0.844 | 1 | 1191 | tags=17%, list=12%, |
| 1004 | | SCHMIDT_POR_TARGETS_IN_LIMB_BUD_UP | | 25 | -0.31 | -0.88 | 0.643 | 0.866 | 1 | 1672 | tags=28%, list=17%, |
| 946 | | LE_EGR2_TARGETS_DN | | 72 | -0.27 | -0.92 | 0.644 | 0.833 | 1 | 2831 | tags=32%, list=28%, |
| 1006 | | WP_GLYCEROLIPIDS_AND_GLYCEROPHOSPHOLIPIDS_WP4345 | | 17 | -0.33 | -0.88 | 0.645 | 0.865 | 1 | 876 | tags=18%, list=9%, signal=19% |
| 1010 | | REACTOME_PROTEIN_METHYLATION | | 17 | -0.33 | -0.87 | 0.645 | 0.869 | 1 | 2318 | tags=35%, list=23%, |
| 968 | | REACTOME_PI_METABOLISM | | 50 | -0.28 | -0.91 | 0.647 | 0.838 | 1 | 1636 | tags=20%, list=16%, |
| 996 | | REACTOME_DOWNSTREAM_SIGNAL_TRANSDUCTION | | 22 | -0.31 | -0.88 | 0.648 | 0.861 | 1 | 2597 | tags=36%, list=26%, |
| 958 | | REACTOME_ANTIVIRAL_MECHANISM_BY_IFN_STIMULATED_GENES | | 71 | -0.27 | -0.91 | 0.65 | 0.835 | 1 | 3654 | tags=41%, list=36%, |
| 970 | | REACTOME_PROGRAMMED_CELL_DEATH | | 102 | -0.25 | -0.9 | 0.65 | 0.843 | 1 | 3270 | tags=33%, list=32%, |
| 994 | | REACTOME_METABOLISM_OF_FAT_SOLUBLE_VITAMINS | | 33 | -0.29 | -0.89 | 0.651 | 0.86 | 1 | 3771 | tags=39%, list=37%, |
| 1019 | | BIOCARTA_PTEN_PATHWAY | | 16 | -0.33 | -0.87 | 0.651 | 0.873 | 1 | 1547 | tags=25%, list=15%, |
| 1005 | | REACTOME_SUMOYLATION_OF_SUMOYLATION_PROTEINS | | 28 | -0.3 | -0.88 | 0.652 | 0.865 | 1 | 5134 | tags=68%, list=51%, |
| 1013 | | WP_GLYCEROLIPIDS_AND_GLYCEROPHOSPHOLIPIDS_WP5296 | | 17 | -0.33 | -0.87 | 0.652 | 0.87 | 1 | 876 | tags=18%, list=9%, signal=19% |
| 975 | | WP_OXIDATIVE_STRESS_AND_REDOX_PATHWAY | | 69 | -0.26 | -0.9 | 0.655 | 0.844 | 1 | 1234 | tags=14%, list=12%, |
| 990 | | REACTOME_NEGATIVE_REGULATION_OF_FGFR3_SIGNALING | | 17 | -0.34 | -0.89 | 0.655 | 0.858 | 1 | 3997 | tags=53%, list=40%, |
| 1035 | | REACTOME_GLYCOGEN_METABOLISM | | 15 | -0.34 | -0.86 | 0.656 | 0.875 | 1 | 1208 | tags=27%, list=12%, |
| 1039 | | REACTOME_NEGATIVE_REGULATION_OF_FGFR2_SIGNALING | | 16 | -0.32 | -0.85 | 0.656 | 0.881 | 1 | 3997 | tags=50%, list=40%, |
| 1028 | | MORI_EMU_MYC_LYMPHOMA_BY_ONSET_TIME_DN | | 17 | -0.33 | -0.86 | 0.657 | 0.875 | 1 | 2649 | tags=29%, list=26%, |
| 1023 | | PLASARI_TGFB1_SIGNALING_VIA_NFIC_10HR_DN | | 16 | -0.33 | -0.87 | 0.658 | 0.873 | 1 | 1688 | tags=25%, list=17%, |
| 896 | | LIM_MAMMARY_STEM_CELL_DN | | 300 | -0.25 | -0.95 | 0.659 | 0.815 | 1 | 3371 | tags=36%, list=33%, |
| 1001 | | GAVIN_FOXP3_TARGETS_CLUSTER_P6 | | 44 | -0.28 | -0.88 | 0.661 | 0.866 | 1 | 1799 | tags=20%, list=18%, |
| 1002 | | REACTOME_NEGATIVE_REGULATION_OF_FGFR4_SIGNALING | | 18 | -0.33 | -0.88 | 0.661 | 0.867 | 1 | 3997 | tags=50%, list=40%, |
| 962 | | REACTOME_RHO_GTPASES_ACTIVATE_FORMINS | | 84 | -0.26 | -0.91 | 0.662 | 0.837 | 1 | 2406 | tags=29%, list=24%, |
| 957 | | SCHAEFFER_PROSTATE_DEVELOPMENT_6HR_UP | | 114 | -0.25 | -0.91 | 0.664 | 0.835 | 1 | 1182 | tags=15%, list=12%, |
| 1012 | | WP_GPCRS_NON_ODORANT | | 39 | -0.28 | -0.87 | 0.664 | 0.871 | 1 | 2806 | tags=28%, list=28%, |
| 992 | | REACTOME_GLUCOSE_METABOLISM | | 59 | -0.27 | -0.89 | 0.665 | 0.858 | 1 | 4397 | tags=47%, list=44%, |
| 983 | | REACTOME_NUCLEOTIDE_EXCISION_REPAIR | | 93 | -0.26 | -0.9 | 0.669 | 0.844 | 1 | 3918 | tags=41%, list=39%, |
| 998 | | REACTOME_INTERLEUKIN_17_SIGNALING | | 49 | -0.27 | -0.88 | 0.669 | 0.863 | 1 | 4697 | tags=59%, list=47%, |
| 1016 | | REACTOME_PKR_MEDIATED_SIGNALING | | 45 | -0.27 | -0.87 | 0.67 | 0.869 | 1 | 3101 | tags=38%, list=31%, |
| 1027 | | WP_HEART_DEVELOPMENT | | 24 | -0.31 | -0.86 | 0.67 | 0.875 | 1 | 750 | tags=13%, list=7%, signal=13% |
| 908 | | RAO_BOUND_BY_SALL4_ISOFORM_B | | 330 | -0.24 | -0.94 | 0.671 | 0.818 | 1 | 2882 | tags=27%, list=29%, |
| 945 | | QI_PLASMACYTOMA_UP | | 140 | -0.25 | -0.92 | 0.671 | 0.834 | 1 | 3558 | tags=35%, list=35%, |
| 1025 | | REACTOME_PEPTIDE_HORMONE_METABOLISM | | 32 | -0.29 | -0.87 | 0.672 | 0.873 | 1 | 3830 | tags=56%, list=38%, |
| 960 | | REACTOME_METABOLISM_OF_VITAMINS_AND_COFACTORS | | 138 | -0.25 | -0.91 | 0.673 | 0.837 | 1 | 3163 | tags=32%, list=31%, |
| 1033 | | REACTOME_SUMOYLATION_OF_DNA_REPLICATION_PROTEINS | | 33 | -0.29 | -0.86 | 0.676 | 0.874 | 1 | 2091 | tags=24%, list=21%, |
| 1011 | | MIKKELSEN_NPC_HCP_WITH_H3K4ME3_AND_H3K27ME3 | | 43 | -0.28 | -0.87 | 0.677 | 0.869 | 1 | 2745 | tags=40%, list=27%, |
| 1015 | | MEISSNER_NPC_HCP_WITH_H3K4ME2_AND_H3K27ME3 | | 52 | -0.27 | -0.87 | 0.677 | 0.87 | 1 | 2391 | tags=35%, list=24%, |
| 933 | | SCHAEFFER_PROSTATE_DEVELOPMENT_48HR_UP | | 255 | -0.24 | -0.93 | 0.679 | 0.826 | 1 | 2593 | tags=27%, list=26%, |
| 1034 | | REACTOME_SMOOTH_MUSCLE_CONTRACTION | | 20 | -0.31 | -0.86 | 0.68 | 0.875 | 1 | 1403 | tags=20%, list=14%, |
| 1022 | | BAKKER_FOXO3_TARGETS_UP | | 47 | -0.27 | -0.87 | 0.681 | 0.874 | 1 | 2650 | tags=34%, list=26%, |
| 1060 | | REACTOME_BRANCHED_CHAIN_AMINO_ACID_CATABOLISM | | 19 | -0.31 | -0.84 | 0.685 | 0.895 | 1 | 4944 | tags=58%, list=49%, |
| 921 | | SHETH_LIVER_CANCER_VS_TXNIP_LOSS_PAM4 | | 255 | -0.24 | -0.93 | 0.686 | 0.824 | 1 | 2389 | tags=23%, list=24%, |
| 984 | | DARWICHE_PAPILLOMA_RISK_HIGH_DN | | 110 | -0.25 | -0.9 | 0.686 | 0.848 | 1 | 3534 | tags=39%, list=35%, |
| 1029 | | REACTOME_CYCLIN_D_ASSOCIATED_EVENTS_IN_G1 | | 35 | -0.28 | -0.86 | 0.686 | 0.876 | 1 | 2577 | tags=29%, list=26%, |
| 1040 | | HEVNER_TELENCEPHALON_MENINGEAL_CELLS | | 25 | -0.3 | -0.85 | 0.686 | 0.88 | 1 | 2545 | tags=40%, list=25%, |
| 966 | | ZENG_GU_POST_ICB_METAGENE_31 | | 118 | -0.25 | -0.91 | 0.687 | 0.837 | 1 | 3023 | tags=35%, list=30%, |
| 1043 | | REACTOME_TRANSPORT_OF_VITAMINS_NUCLEOSIDES_AND_RELATED_MOLECUL | | 25 | -0.3 | -0.85 | 0.687 | 0.879 | 1 | 3423 | tags=40%, list=34%, |
| 1026 | | MA_MYELOID_DIFFERENTIATION_DN | | 29 | -0.29 | -0.86 | 0.688 | 0.874 | 1 | 2649 | tags=38%, list=26%, |
| 913 | | MIKKELSEN_ES_ICP_WITH_H3K4ME3 | | 354 | -0.24 | -0.94 | 0.691 | 0.82 | 1 | 2388 | tags=25%, list=24%, |
| 1050 | | REACTOME_RHOH_GTPASE_CYCLE | | 29 | -0.29 | -0.84 | 0.693 | 0.891 | 1 | 2652 | tags=38%, list=26%, |
| 995 | | CHIARADONNA_NEOPLASTIC_TRANSFORMATION_CDC25_UP | | 87 | -0.25 | -0.89 | 0.695 | 0.859 | 1 | 3372 | tags=34%, list=33%, |
| 986 | | GAVIN_FOXP3_TARGETS_CLUSTER_P3 | | 99 | -0.25 | -0.89 | 0.697 | 0.854 | 1 | 1291 | tags=16%, list=13%, |
| 1045 | | REACTOME_NONHOMOLOGOUS_END_JOINING_NHEJ | | 32 | -0.28 | -0.85 | 0.698 | 0.879 | 1 | 4834 | tags=53%, list=48%, |
| 1041 | | REACTOME_TP53_REGULATES_TRANSCRIPTION_OF_DNA_REPAIR_GENES | | 39 | -0.27 | -0.85 | 0.703 | 0.88 | 1 | 4385 | tags=49%, list=43%, |
| 989 | | LANDIS_ERBB2_BREAST_TUMORS_324_DN | | 125 | -0.24 | -0.89 | 0.705 | 0.858 | 1 | 2578 | tags=26%, list=26%, |
| 991 | | DARWICHE_SKIN_TUMOR_PROMOTER_DN | | 111 | -0.25 | -0.89 | 0.705 | 0.858 | 1 | 4061 | tags=44%, list=40%, |
| 920 | | BRUINS_UVC_RESPONSE_VIA_TP53_GROUP_A | | 436 | -0.24 | -0.93 | 0.706 | 0.824 | 1 | 3519 | tags=38%, list=35%, |
| 1032 | | REACTOME_CHROMOSOME_MAINTENANCE | | 60 | -0.26 | -0.86 | 0.71 | 0.875 | 1 | 4091 | tags=47%, list=41%, |
| 1063 | | REACTOME_DEACTIVATION_OF_THE_BETA_CATENIN_TRANSACTIVATING_COMPL | | 25 | -0.29 | -0.83 | 0.71 | 0.896 | 1 | 2563 | tags=24%, list=25%, |
| 1046 | | HUANG_GATA2_TARGETS_DN | | 42 | -0.27 | -0.85 | 0.712 | 0.883 | 1 | 1403 | tags=21%, list=14%, |
| 1055 | | YAO_TEMPORAL_RESPONSE_TO_PROGESTERONE_CLUSTER_8 | | 36 | -0.27 | -0.84 | 0.712 | 0.893 | 1 | 3454 | tags=39%, list=34%, |
| 1038 | | REACTOME_CELLULAR_SENESCENCE | | 68 | -0.26 | -0.86 | 0.713 | 0.88 | 1 | 4069 | tags=46%, list=40%, |
| 1067 | | STEGER_ADIPOGENESIS_DN | | 15 | -0.32 | -0.83 | 0.717 | 0.896 | 1 | 509 | tags=13%, list=5%, signal=14% |
| 1037 | | REACTOME_DRUG_ADME | | 66 | -0.25 | -0.86 | 0.719 | 0.876 | 1 | 3599 | tags=32%, list=36%, |
| 1024 | | LANDIS_BREAST_CANCER_PROGRESSION_DN | | 57 | -0.26 | -0.87 | 0.72 | 0.873 | 1 | 2253 | tags=25%, list=22%, |
| 1048 | | REACTOME_PROCESSING_OF_DNA_DOUBLE_STRAND_BREAK_ENDS | | 43 | -0.27 | -0.85 | 0.72 | 0.884 | 1 | 3898 | tags=40%, list=39%, |
| 1076 | | ONGUSAHA_TP53_TARGETS | | 27 | -0.29 | -0.82 | 0.72 | 0.902 | 1 | 1187 | tags=22%, list=12%, |
| 999 | | ZENG_GU_POST_ICB_METAGENE_22 | | 116 | -0.24 | -0.88 | 0.722 | 0.864 | 1 | 4051 | tags=39%, list=40%, |
| 1066 | | REACTOME_SYNTHESIS_OF_ACTIVE_UBIQUITIN_ROLES_OF_E1_AND_E2_ENZYME | | 26 | -0.28 | -0.83 | 0.722 | 0.896 | 1 | 2228 | tags=23%, list=22%, |
| 1071 | | REACTOME_TAK1_DEPENDENT_IKK_AND_NF_KAPPA_B_ACTIVATION | | 29 | -0.29 | -0.83 | 0.722 | 0.9 | 1 | 3453 | tags=38%, list=34%, |
| 1047 | | BIOCARTA_HIVNEF_PATHWAY | | 47 | -0.26 | -0.85 | 0.723 | 0.884 | 1 | 3270 | tags=32%, list=32%, |
| 1069 | | REACTOME_DNA_DAMAGE_RECOGNITION_IN_GG_NER | | 34 | -0.28 | -0.83 | 0.723 | 0.897 | 1 | 4508 | tags=50%, list=45%, |
| 1021 | | DURAND_STROMA_NS_UP | | 89 | -0.25 | -0.87 | 0.725 | 0.874 | 1 | 2530 | tags=26%, list=25%, |
| 1044 | | YAO_TEMPORAL_RESPONSE_TO_PROGESTERONE_CLUSTER_10 | | 65 | -0.25 | -0.85 | 0.725 | 0.88 | 1 | 1836 | tags=22%, list=18%, |
| 1079 | | REACTOME_CITRIC_ACID_CYCLE_TCA_CYCLE | | 19 | -0.31 | -0.82 | 0.725 | 0.902 | 1 | 5476 | tags=63%, list=54%, |
| 1088 | | KAMIKUBO_MYELOID_MN1_NETWORK | | 17 | -0.31 | -0.81 | 0.725 | 0.909 | 1 | 5132 | tags=76%, list=51%, |
| 942 | | LAZARO_GENETIC_MOUSE_MODEL_HIGH_GRADE_SMALL_CELL_NEUROENDOCRINE_LUNG_CARCINOMA_UP | | 290 | -0.24 | -0.92 | 0.726 | 0.836 | 1 | 2437 | tags=24%, list=24%, signal=31% |
| 1018 | | DARWICHE_SQUAMOUS_CELL_CARCINOMA_DN | | 101 | -0.25 | -0.87 | 0.729 | 0.871 | 1 | 3822 | tags=42%, list=38%, |
| 1014 | | REACTOME_PROTEIN_LOCALIZATION | | 90 | -0.25 | -0.87 | 0.73 | 0.87 | 1 | 3256 | tags=32%, list=32%, |
| 1051 | | GUO_HEX_TARGETS_DN | | 51 | -0.26 | -0.84 | 0.73 | 0.891 | 1 | 425 | tags=8%, list=4%, signal=8% |
| 1102 | | REACTOME_METALLOPROTEASE_DUBS | | 17 | -0.31 | -0.81 | 0.73 | 0.905 | 1 | 4511 | tags=59%, list=45%, |
| 1056 | | MORI_IMMATURE_B_LYMPHOCYTE_DN | | 58 | -0.25 | -0.84 | 0.731 | 0.895 | 1 | 1799 | tags=21%, list=18%, |
| 978 | | REACTOME_METABOLISM_OF_CARBOHYDRATES | | 167 | -0.24 | -0.9 | 0.732 | 0.843 | 1 | 4023 | tags=41%, list=40%, |
| 1098 | | REACTOME_G_PROTEIN_BETA_GAMMA_SIGNALLING | | 16 | -0.31 | -0.81 | 0.732 | 0.907 | 1 | 1547 | tags=19%, list=15%, |
| 1054 | | BRUINS_UVC_RESPONSE_VIA_TP53_GROUP_C | | 52 | -0.26 | -0.84 | 0.735 | 0.891 | 1 | 1624 | tags=21%, list=16%, |
| 1065 | | REACTOME_RAB_GERANYLGERANYLATION | | 39 | -0.27 | -0.83 | 0.737 | 0.896 | 1 | 1460 | tags=18%, list=14%, |
| 1074 | | REACTOME_ACTIVATION_OF_BH3_ONLY_PROTEINS | | 16 | -0.32 | -0.82 | 0.737 | 0.903 | 1 | 5108 | tags=63%, list=51%, |
| 1082 | | MIKKELSEN_MEF_LCP_WITH_H3K27ME3 | | 16 | -0.31 | -0.82 | 0.737 | 0.902 | 1 | 473 | tags=13%, list=5%, signal=13% |
| 1009 | | REACTOME_ORGANELLE_BIOGENESIS_AND_MAINTENANCE | | 149 | -0.24 | -0.88 | 0.738 | 0.866 | 1 | 2419 | tags=25%, list=24%, |
| 1042 | | REACTOME_DNA_DOUBLE_STRAND_BREAK_REPAIR | | 88 | -0.24 | -0.85 | 0.739 | 0.88 | 1 | 3629 | tags=34%, list=36%, |
| 1059 | | WP_CALCIUM_REGULATION_IN_CARDIAC_CELLS | | 56 | -0.26 | -0.84 | 0.739 | 0.896 | 1 | 1879 | tags=20%, list=19%, |
| 1061 | | BURTON_ADIPOGENESIS_PEAK_AT_24HR | | 29 | -0.28 | -0.83 | 0.741 | 0.897 | 1 | 3068 | tags=38%, list=30%, |
| 1073 | | WP_OXIDATIVE_PHOSPHORYLATION | | 47 | -0.26 | -0.83 | 0.741 | 0.9 | 1 | 5948 | tags=62%, list=59%, |
| 1077 | | REACTOME_TRANSLESION_SYNTHESIS_BY_Y_FAMILY_DNA_POLYMERASES_BYPASSES_LESIONS_ON_DNA_TEMPLATE | | 31 | -0.28 | -0.82 | 0.744 | 0.901 | 1 | 1417 | tags=19%, list=14%, signal=22% |
| 1091 | | WP_LUNG_FIBROSIS | | 26 | -0.28 | -0.81 | 0.744 | 0.908 | 1 | 3330 | tags=46%, list=33%, |
| 1080 | | REACTOME_POST_TRANSLATIONAL_MODIFICATION_SYNTHESIS_OF_GPI_ANCHORED_PROTEINS | | 36 | -0.27 | -0.82 | 0.747 | 0.901 | 1 | 3995 | tags=50%, list=40%, signal=82% |
| 1093 | | REACTOME_DEPOSITION_OF_NEW_CENPA_CONTAINING_NUCLEOSOMES_AT_THE_CENTROMERE | | 24 | -0.29 | -0.81 | 0.747 | 0.907 | 1 | 3982 | tags=46%, list=39%, signal=75% |
| 1106 | | REACTOME_POSTMITOTIC_NUCLEAR_PORE_COMPLEX_NPC_REFORMATION | | 18 | -0.3 | -0.81 | 0.747 | 0.904 | 1 | 986 | tags=17%, list=10%, |
| 1083 | | REACTOME_SENSORY_PERCEPTION | | 48 | -0.26 | -0.82 | 0.748 | 0.904 | 1 | 2224 | tags=23%, list=22%, |
| 1084 | | BURTON_ADIPOGENESIS_PEAK_AT_0HR | | 44 | -0.26 | -0.82 | 0.748 | 0.904 | 1 | 2716 | tags=27%, list=27%, |
| 1109 | | REACTOME_NOD1_2_SIGNALING_PATHWAY | | 25 | -0.28 | -0.8 | 0.749 | 0.905 | 1 | 2525 | tags=28%, list=25%, |
| 1020 | | REACTOME_SIGNALING_BY_NUCLEAR_RECEPTORS | | 119 | -0.24 | -0.87 | 0.75 | 0.874 | 1 | 1573 | tags=18%, list=16%, |
| 1114 | | REACTOME_CYCLIN_A_B1_B2_ASSOCIATED_EVENTS_DURING_G2_M_TRANSITIO | | 16 | -0.31 | -0.8 | 0.75 | 0.913 | 1 | 2677 | tags=31%, list=27%, |
| 949 | | WAKABAYASHI_ADIPOGENESIS_PPARG_BOUND_8D | | 497 | -0.24 | -0.92 | 0.752 | 0.832 | 1 | 2140 | tags=21%, list=21%, |
| 1130 | | MARCINIAK_ER_STRESS_RESPONSE_VIA_CHOP | | 16 | -0.31 | -0.79 | 0.752 | 0.914 | 1 | 1465 | tags=19%, list=15%, |
| 1075 | | DURCHDEWALD_SKIN_CARCINOGENESIS_UP | | 54 | -0.25 | -0.82 | 0.756 | 0.903 | 1 | 1993 | tags=20%, list=20%, |
| 1087 | | REACTOME_APOPTOTIC_EXECUTION_PHASE | | 36 | -0.26 | -0.81 | 0.756 | 0.909 | 1 | 3225 | tags=33%, list=32%, |
| 1094 | | BURTON_ADIPOGENESIS_12 | | 27 | -0.28 | -0.81 | 0.757 | 0.908 | 1 | 4738 | tags=63%, list=47%, |
| 1008 | | REACTOME_DNA_REPAIR | | 197 | -0.23 | -0.88 | 0.758 | 0.866 | 1 | 4049 | tags=41%, list=40%, |
| 1062 | | REACTOME_FORMATION_OF_TC_NER_PRE_INCISION_COMPLEX | | 50 | -0.26 | -0.83 | 0.759 | 0.897 | 1 | 4508 | tags=50%, list=45%, |
| 1100 | | REACTOME_CYTOSOLIC_SENSORS_OF_PATHOGEN_ASSOCIATED_DNA | | 27 | -0.28 | -0.81 | 0.759 | 0.906 | 1 | 1841 | tags=22%, list=18%, |
| 1089 | | MODY_HIPPOCAMPUS_NEONATAL | | 24 | -0.29 | -0.81 | 0.76 | 0.909 | 1 | 3584 | tags=46%, list=35%, |
| 1111 | | STARK_HYPPOCAMPUS_22Q11_DELETION_UP | | 32 | -0.27 | -0.8 | 0.76 | 0.908 | 1 | 3476 | tags=44%, list=34%, |
| 1121 | | REACTOME_KERATAN_SULFATE_KERATIN_METABOLISM | | 16 | -0.31 | -0.79 | 0.761 | 0.915 | 1 | 3326 | tags=38%, list=33%, |
| 1049 | | DARWICHE_PAPILLOMA_RISK_LOW_UP | | 98 | -0.24 | -0.85 | 0.762 | 0.883 | 1 | 3648 | tags=40%, list=36%, |
| 1068 | | REACTOME_GLOBAL_GENOME_NUCLEOTIDE_EXCISION_REPAIR_GG_NER | | 68 | -0.24 | -0.83 | 0.762 | 0.897 | 1 | 3567 | tags=32%, list=35%, |
| 1117 | | WP_FACTORS_AND_PATHWAYS_AFFECTING_INSULIN_LIKE_GROWTH_FACTOR_IGF1_AKT_SIGNALING | | 25 | -0.27 | -0.79 | 0.764 | 0.915 | 1 | 3749 | tags=44%, list=37%, signal=70% |
| 1017 | | SCHAEFFER_PROSTATE_DEVELOPMENT_48HR_DN | | 185 | -0.24 | -0.87 | 0.765 | 0.871 | 1 | 2281 | tags=23%, list=23%, |
| 1057 | | CHEN_LIVER_METABOLISM_QTL_CIS | | 77 | -0.24 | -0.84 | 0.766 | 0.894 | 1 | 3352 | tags=34%, list=33%, |
| 1070 | | YAO_TEMPORAL_RESPONSE_TO_PROGESTERONE_CLUSTER_2 | | 39 | -0.27 | -0.83 | 0.768 | 0.9 | 1 | 677 | tags=13%, list=7%, signal=14% |
| 1053 | | RASHI_RESPONSE_TO_IONIZING_RADIATION_2 | | 89 | -0.24 | -0.84 | 0.769 | 0.89 | 1 | 2621 | tags=29%, list=26%, |
| 1134 | | WP_TCA_CYCLE | | 29 | -0.26 | -0.78 | 0.769 | 0.915 | 1 | 4993 | tags=48%, list=49%, |
| 1110 | | REACTOME_TELOMERE_MAINTENANCE | | 36 | -0.26 | -0.8 | 0.77 | 0.906 | 1 | 4542 | tags=56%, list=45%, |
| 1142 | | REACTOME_RET_SIGNALING | | 17 | -0.29 | -0.77 | 0.772 | 0.92 | 1 | 4107 | tags=53%, list=41%, |
| 1064 | | PANGAS_TUMOR_SUPPRESSION_BY_SMAD1_AND_SMAD5_DN | | 94 | -0.24 | -0.83 | 0.773 | 0.896 | 1 | 2476 | tags=27%, list=25%, |
| 1085 | | CADWELL_ATG16L1_TARGETS_UP | | 59 | -0.25 | -0.82 | 0.781 | 0.904 | 1 | 1966 | tags=22%, list=19%, |
| 1132 | | MCMURRAY_TP53_HRAS_COOPERATION_RESPONSE_DN | | 28 | -0.27 | -0.79 | 0.781 | 0.914 | 1 | 3732 | tags=46%, list=37%, |
| 1138 | | CHESLER_BRAIN_HIGHEST_EXPRESSION | | 33 | -0.26 | -0.78 | 0.782 | 0.919 | 1 | 4411 | tags=42%, list=44%, |
| 1118 | | WP_PARKINSON_39_S_DISEASE | | 23 | -0.28 | -0.79 | 0.783 | 0.916 | 1 | 428 | tags=9%, list=4%, signal=9% |
| 1145 | | BIOCARTA_IGF1R_PATHWAY | | 19 | -0.29 | -0.77 | 0.783 | 0.921 | 1 | 1213 | tags=16%, list=12%, |
| 1031 | | YAO_TEMPORAL_RESPONSE_TO_PROGESTERONE_CLUSTER_13 | | 163 | -0.23 | -0.86 | 0.786 | 0.875 | 1 | 3070 | tags=26%, list=30%, |

| | | | | | | | | | | | |
|---|---|---|---|---|---|---|---|---|---|---|---|
| 1115 | | JOHANSSON_GLIOMAGENESIS_BY_PDGFB_UP | | 46 | -0.25 | -0.8 | 0.786 | 0.912 | 1 | 3225 | tags=37%, list=32%, |
| 1144 | | SCHAEFFER_SOX9_TARGETS_IN_PROSTATE_DEVELOPMENT_DN | | 31 | -0.26 | -0.77 | 0.786 | 0.921 | 1 | 3897 | tags=45%, list=39%, |
| 1097 | | REACTOME_SNRNP_ASSEMBLY | | 44 | -0.26 | -0.81 | 0.787 | 0.907 | 1 | 3383 | tags=32%, list=33%, |
| 1137 | | BIOCARTA_FAS_PATHWAY | | 24 | -0.28 | -0.78 | 0.787 | 0.918 | 1 | 625 | tags=13%, list=6%, signal=13% |
| 1099 | | CADWELL_ATG16L1_TARGETS_DN | | 52 | -0.25 | -0.81 | 0.788 | 0.907 | 1 | 4196 | tags=46%, list=42%, |
| 1104 | | REACTOME_AMPLIFICATION_OF_SIGNAL_FROM_THE_KINETOCHORES | | 58 | -0.24 | -0.81 | 0.788 | 0.904 | 1 | 2571 | tags=31%, list=25%, |
| 1155 | | REACTOME_ACTIVATION_OF_NMDA_RECEPTORS_AND_POSTSYNAPTIC_EVENTS | | 15 | -0.3 | -0.76 | 0.789 | 0.935 | 1 | 51 | tags=7%, list=1%, signal=7% |
| 1133 | | REACTOME_SUMOYLATION_OF_UBIQUITINYLATION_PROTEINS | | 33 | -0.26 | -0.78 | 0.79 | 0.914 | 1 | 4962 | tags=58%, list=49%, |
| 1149 | | BIOCARTA_BAD_PATHWAY | | 18 | -0.29 | -0.76 | 0.79 | 0.927 | 1 | 4840 | tags=61%, list=48%, |
| 1123 | | MATZUK_SPERMATOCYTE | | 36 | -0.25 | -0.79 | 0.792 | 0.914 | 1 | 2484 | tags=28%, list=25%, |
| 1151 | | GAUSSMANN_MLL_AF4_FUSION_TARGETS_B_UP | | 17 | -0.29 | -0.76 | 0.792 | 0.932 | 1 | 3752 | tags=53%, list=37%, |
| 1086 | | KATSANOU_ELAVL1_TARGETS_UP | | 83 | -0.23 | -0.81 | 0.793 | 0.909 | 1 | 1937 | tags=22%, list=19%, |
| 1107 | | REACTOME_TRANSCRIPTION_COUPLED_NUCLEOTIDE_EXCISION_REPAIR_TC_NER | | 67 | -0.24 | -0.81 | 0.795 | 0.904 | 1 | 3918 | tags=39%, list=39%, |
| 1072 | | IWANAGA_CARCINOGENESIS_BY_KRAS_UP | | 105 | -0.23 | -0.83 | 0.799 | 0.899 | 1 | 2195 | tags=22%, list=22%, |
| 1136 | | REACTOME_G2_M_DNA_DAMAGE_CHECKPOINT | | 48 | -0.24 | -0.78 | 0.804 | 0.916 | 1 | 4034 | tags=40%, list=40%, |
| 1152 | | REACTOME_PYRUVATE_METABOLISM | | 18 | -0.28 | -0.76 | 0.805 | 0.935 | 1 | 3884 | tags=44%, list=38%, |
| 1141 | | REACTOME_PLATELET_HOMEOSTASIS | | 36 | -0.25 | -0.78 | 0.807 | 0.92 | 1 | 2430 | tags=28%, list=24%, |
| 1175 | | WP_PROSTAGLANDIN_SYNTHESIS_AND_REGULATION | | 15 | -0.29 | -0.74 | 0.81 | 0.942 | 1 | 2430 | tags=27%, list=24%, |
| 1140 | | REACTOME_GLYCOLYSIS | | 44 | -0.24 | -0.78 | 0.811 | 0.919 | 1 | 5158 | tags=61%, list=51%, |
| 1153 | | REACTOME_ENERGY_DEPENDENT_REGULATION_OF_MTOR_BY_LKB1_AMPK | | 24 | -0.27 | -0.76 | 0.812 | 0.934 | 1 | 2743 | tags=29%, list=27%, |
| 1170 | | SCHRAETS_MLL_TARGETS_UP | | 19 | -0.27 | -0.74 | 0.814 | 0.941 | 1 | 3604 | tags=58%, list=36%, |
| 1150 | | MORI_LARGE_PRE_BII_LYMPHOCYTE_DN | | 35 | -0.25 | -0.76 | 0.815 | 0.932 | 1 | 2478 | tags=34%, list=25%, |
| 1171 | | REACTOME_RESOLUTION_OF_AP_SITES_VIA_THE_MULTIPLE_NUCLEOTIDE_PATCH_REPLACEMENT_PATHWAY | | 19 | -0.27 | -0.74 | 0.817 | 0.943 | 1 | 3262 | tags=32%, list=32%, signal=47% |
| 1135 | | HOWLIN_CITED1_TARGETS_1_DN | | 28 | -0.27 | -0.78 | 0.818 | 0.915 | 1 | 2138 | tags=29%, list=21%, |
| 1129 | | REACTOME_APOPTOSIS | | 76 | -0.23 | -0.79 | 0.821 | 0.911 | 1 | 3270 | tags=33%, list=32%, |
| 1157 | | BIOCARTA_FMLP_PATHWAY | | 24 | -0.27 | -0.75 | 0.822 | 0.937 | 1 | 217 | tags=8%, list=2%, signal=8% |
| 1159 | | MIKKELSEN_NPC_LCP_WITH_H3K4ME3 | | 31 | -0.25 | -0.75 | 0.823 | 0.937 | 1 | 1370 | tags=23%, list=14%, |
| 1172 | | REACTOME_RHO_GTPASES_ACTIVATE_PAKS | | 15 | -0.29 | -0.74 | 0.823 | 0.943 | 1 | 2627 | tags=33%, list=26%, |
| 1030 | | CUI_TCF21_TARGETS_2_UP | | 232 | -0.23 | -0.86 | 0.824 | 0.876 | 1 | 1621 | tags=17%, list=16%, |
| 1112 | | TSENG_IRS1_TARGETS_UP | | 78 | -0.24 | -0.8 | 0.825 | 0.91 | 1 | 1899 | tags=23%, list=19%, |
| 1163 | | WP_IL_1_SIGNALING_PATHWAY | | 32 | -0.25 | -0.74 | 0.828 | 0.942 | 1 | 3585 | tags=38%, list=35%, |
| 1095 | | REACTOME_PHOSPHOLIPID_METABOLISM | | 117 | -0.22 | -0.81 | 0.829 | 0.908 | 1 | 1865 | tags=20%, list=18%, |
| 1108 | | REACTOME_RAC1_GTPASE_CYCLE | | 110 | -0.22 | -0.81 | 0.829 | 0.903 | 1 | 2658 | tags=30%, list=26%, |
| 1156 | | MIKKELSEN_IPS_ICP_WITH_H3K4ME3_AND_H327ME3 | | 28 | -0.26 | -0.75 | 0.829 | 0.934 | 1 | 2442 | tags=32%, list=24%, |
| 1154 | | REACTOME_TNFR1_INDUCED_NF_KAPPA_B_SIGNALING_PATHWAY | | 29 | -0.26 | -0.76 | 0.83 | 0.934 | 1 | 2563 | tags=28%, list=25%, |
| 1092 | | WAKABAYASHI_ADIPOGENESIS_PPARG_RXRA_BOUND_WITH_H4K20ME1_MARK | | 132 | -0.22 | -0.81 | 0.831 | 0.908 | 1 | 2731 | tags=24%, list=27%, |
| 1105 | | REACTOME_INTERFERON_SIGNALING | | 96 | -0.23 | -0.81 | 0.833 | 0.903 | 1 | 2597 | tags=25%, list=26%, |
| 1122 | | REACTOME_NEURONAL_SYSTEM | | 93 | -0.23 | -0.79 | 0.834 | 0.914 | 1 | 2389 | tags=22%, list=24%, |
| 1160 | | BIOCARTA_GPCR_PATHWAY | | 22 | -0.27 | -0.75 | 0.834 | 0.938 | 1 | 625 | tags=14%, list=6%, signal=15% |
| 1116 | | REACTOME_RESPIRATORY_ELECTRON_TRANSPORT_ATP_SYNTHESIS_BY_CHEMIOSMOTIC_COUPLING_AND_HEAT_PRODUCTION_BY_UNCOUPLING_PROTEINS | | 110 | -0.22 | -0.79 | 0.835 | 0.915 | 1 | 4638 | tags=39%, list=46%, signal=71% |
| 1143 | | REACTOME_HOMOLOGY_DIRECTED_REPAIR | | 67 | -0.23 | -0.77 | 0.835 | 0.92 | 1 | 4026 | tags=39%, list=40%, |
| 1165 | | REACTOME_APOPTOTIC_CLEAVAGE_OF_CELLULAR_PROTEINS | | 29 | -0.25 | -0.74 | 0.836 | 0.942 | 1 | 2660 | tags=28%, list=26%, |
| 1187 | | HARRIS_BRAIN_CANCER_PROGENITORS | | 18 | -0.27 | -0.73 | 0.836 | 0.944 | 1 | 3964 | tags=61%, list=39%, |
| 1173 | | GAUSSMANN_MLL_AF4_FUSION_TARGETS_G_DN | | 18 | -0.27 | -0.74 | 0.837 | 0.943 | 1 | 569 | tags=11%, list=6%, signal=12% |
| 1181 | | MATZUK_MALE_REPRODUCTION_SERTOLI | | 16 | -0.28 | -0.73 | 0.839 | 0.941 | 1 | 3246 | tags=31%, list=32%, |
| 1169 | | REACTOME_NUCLEAR_PORE_COMPLEX_NPC_DISASSEMBLY | | 28 | -0.25 | -0.74 | 0.84 | 0.941 | 1 | 1967 | tags=21%, list=19%, |
| 1188 | | BYSTROEM_CORRELATED_WITH_IL5_UP | | 24 | -0.25 | -0.72 | 0.84 | 0.945 | 1 | 4403 | tags=54%, list=44%, |
| 1101 | | REACTOME_EUKARYOTIC_TRANSLATION_INITIATION | | 112 | -0.23 | -0.81 | 0.841 | 0.906 | 1 | 5310 | tags=53%, list=53%, |
| 1119 | | REACTOME_RESPIRATORY_ELECTRON_TRANSPORT | | 92 | -0.22 | -0.79 | 0.841 | 0.915 | 1 | 4917 | tags=43%, list=49%, |
| 1139 | | BURTON_ADIPOGENESIS_2 | | 49 | -0.24 | -0.78 | 0.841 | 0.92 | 1 | 1615 | tags=18%, list=16%, |
| 1148 | | HOFFMANN_SMALL_PRE_BII_TO_IMMATURE_B_LYMPHOCYTE_UP | | 43 | -0.24 | -0.77 | 0.841 | 0.927 | 1 | 2578 | tags=35%, list=26%, |
| 1125 | | IWANAGA_CARCINOGENESIS_BY_KRAS_DN | | 75 | -0.23 | -0.79 | 0.842 | 0.912 | 1 | 2831 | tags=24%, list=28%, |
| 1164 | | ELLWOOD_MYC_TARGETS_DN | | 34 | -0.25 | -0.74 | 0.844 | 0.943 | 1 | 2841 | tags=26%, list=28%, |
| 1128 | | MARKEY_RB1_CHRONIC_LOF_UP | | 70 | -0.23 | -0.79 | 0.845 | 0.912 | 1 | 1672 | tags=16%, list=17%, |
| 1180 | | REACTOME_REGULATION_OF_SIGNALING_BY_CBL | | 17 | -0.28 | -0.73 | 0.845 | 0.942 | 1 | 2579 | tags=29%, list=26%, |
| 1184 | | REACTOME_DNA_STRAND_ELONGATION | | 19 | -0.27 | -0.73 | 0.845 | 0.943 | 1 | 3262 | tags=37%, list=32%, |
| 1185 | | HOLLERN_MICROACINAR_BREAST_TUMOR_UP | | 26 | -0.25 | -0.73 | 0.848 | 0.942 | 1 | 1106 | tags=12%, list=11%, |
| 1146 | | YAO_TEMPORAL_RESPONSE_TO_PROGESTERONE_CLUSTER_12 | | 70 | -0.23 | -0.77 | 0.85 | 0.925 | 1 | 2586 | tags=20%, list=26%, |
| 1178 | | PLASARI_TGFB1_SIGNALING_VIA_NFIC_10HR_UP | | 32 | -0.25 | -0.73 | 0.854 | 0.943 | 1 | 3504 | tags=44%, list=35%, |
| 1177 | | WP_TYROBP_CAUSAL_NETWORK_IN_MICROGLIA | | 38 | -0.24 | -0.73 | 0.855 | 0.941 | 1 | 2074 | tags=21%, list=21%, |
| 1052 | | MIKKELSEN_NPC_ICP_WITH_H3K4ME3 | | 264 | -0.22 | -0.84 | 0.861 | 0.891 | 1 | 2051 | tags=20%, list=20%, |
| 1189 | | BIOCARTA_VIP_PATHWAY | | 17 | -0.28 | -0.71 | 0.861 | 0.952 | 1 | 5108 | tags=71%, list=51%, |
| 1200 | | WP_MITOCHONDRIAL_GENE_EXPRESSION | | 17 | -0.26 | -0.69 | 0.866 | 0.963 | 1 | 997 | tags=24%, list=10%, |
| 1179 | | ZAMORA_NOS2_TARGETS_UP | | 48 | -0.23 | -0.73 | 0.867 | 0.942 | 1 | 3262 | tags=33%, list=32%, |
| 1168 | | REACTOME_INTEGRATION_OF_ENERGY_METABOLISM | | 42 | -0.24 | -0.74 | 0.868 | 0.942 | 1 | 1247 | tags=14%, list=12%, |
| 1194 | | REACTOME_RND2_GTPASE_CYCLE | | 28 | -0.24 | -0.7 | 0.872 | 0.962 | 1 | 3614 | tags=46%, list=36%, |
| 1191 | | REACTOME_NICOTINATE_METABOLISM | | 23 | -0.25 | -0.71 | 0.876 | 0.958 | 1 | 1117 | tags=13%, list=11%, |
| 1214 | | IKEDA_MIR30_TARGETS_DN | | 15 | -0.27 | -0.67 | 0.879 | 0.972 | 1 | 937 | tags=13%, list=9%, signal=15% |
| 1195 | | RUAN_RESPONSE_TO_TNF_TROGLITAZONE_DN | | 29 | -0.24 | -0.7 | 0.88 | 0.963 | 1 | 1860 | tags=17%, list=18%, |
| 1199 | | BIOCARTA_AKT_PATHWAY | | 18 | -0.26 | -0.7 | 0.883 | 0.963 | 1 | 1711 | tags=22%, list=17%, |
| 1196 | | REACTOME_DUAL_INCISION_IN_GG_NER | | 34 | -0.23 | -0.7 | 0.884 | 0.963 | 1 | 1576 | tags=15%, list=16%, |
| 1113 | | REACTOME_THE_CITRIC_ACID_TCA_CYCLE_AND_RESPIRATORY_ELECTRON_TRANSPORT | | 149 | -0.22 | -0.8 | 0.885 | 0.913 | 1 | 4638 | tags=40%, list=46%, signal=73% |
| 1127 | | LAZARO_GENETIC_MOUSE_MODEL_HIGH_GRADE_LARGE_CELL_NEUROENDOCRINE_LUNG_CARCINOMA_UP | | 176 | -0.21 | -0.79 | 0.886 | 0.912 | 1 | 2163 | tags=21%, list=21%, signal=26% |
| 1182 | | REACTOME_BIOSYNTHESIS_OF_THE_N_GLYCAN_PRECURSOR_DOLICHOL_LIPID_LINKED_OLIGOSACCHARIDE_LLO_AND_TRANSFER_TO_A_NASCENT_PROTEIN | | 49 | -0.23 | -0.73 | 0.886 | 0.942 | 1 | 1661 | tags=18%, list=16%, signal=22% |
| 1166 | | REACTOME_GLYCEROPHOSPHOLIPID_BIOSYNTHESIS | | 68 | -0.22 | -0.74 | 0.888 | 0.941 | 1 | 3770 | tags=43%, list=37%, |
| 1203 | | REACTOME_DNA_DAMAGE_BYPASS | | 38 | -0.22 | -0.69 | 0.889 | 0.963 | 1 | 1417 | tags=16%, list=14%, |
| 1124 | | AFFAR_YY1_TARGETS_DN | | 136 | -0.22 | -0.79 | 0.89 | 0.913 | 1 | 1825 | tags=18%, list=18%, |
| 1161 | | WP_ELECTRON_TRANSPORT_CHAIN | | 81 | -0.21 | -0.75 | 0.892 | 0.94 | 1 | 4593 | tags=36%, list=45%, |
| 1216 | | REACTOME_REGULATION_OF_FZD_BY_UBIQUITINATION | | 15 | -0.26 | -0.67 | 0.893 | 0.973 | 1 | 3977 | tags=47%, list=39%, |
| 1183 | | REACTOME_REGULATION_OF_TP53_ACTIVITY_THROUGH_PHOSPHORYLATION | | 63 | -0.22 | -0.73 | 0.894 | 0.943 | 1 | 3190 | tags=32%, list=32%, |
| 1186 | | RUAN_RESPONSE_TO_TNF_DN | | 63 | -0.22 | -0.73 | 0.894 | 0.944 | 1 | 2018 | tags=16%, list=20%, |
| 1224 | | REACTOME_O_LINKED_GLYCOSYLATION_OF_MUCINS | | 17 | -0.25 | -0.66 | 0.895 | 0.972 | 1 | 2393 | tags=29%, list=24%, |
| 1190 | | LE_EGR2_TARGETS_UP | | 65 | -0.21 | -0.71 | 0.896 | 0.957 | 1 | 1907 | tags=18%, list=19%, |
| 1158 | | KARLSSON_TGFB1_TARGETS_UP | | 110 | -0.21 | -0.75 | 0.897 | 0.937 | 1 | 3871 | tags=38%, list=38%, |
| 1211 | | REACTOME_INTRAFLAGELLAR_TRANSPORT | | 36 | -0.22 | -0.67 | 0.897 | 0.972 | 1 | 2309 | tags=28%, list=23%, |
| 1131 | | SANSOM_APC_MYC_TARGETS | | 183 | -0.21 | -0.79 | 0.898 | 0.914 | 1 | 3694 | tags=39%, list=37%, |
| 1192 | | REACTOME_RAC2_GTPASE_CYCLE | | 60 | -0.21 | -0.71 | 0.898 | 0.957 | 1 | 3663 | tags=43%, list=36%, |
| 1207 | | KUNINGER_IGF1_VS_PDGFB_TARGETS_UP | | 29 | -0.23 | -0.68 | 0.898 | 0.969 | 1 | 2416 | tags=24%, list=24%, |
| 1206 | | REACTOME_INITIATION_OF_NUCLEAR_ENVELOPE_NE_REFORMATION | | 16 | -0.26 | -0.68 | 0.899 | 0.969 | 1 | 5006 | tags=69%, list=50%, |
| 1193 | | HEVNER_CORTICAL_PLATE_POSTMITOTIC_NEURONS | | 49 | -0.22 | -0.71 | 0.9 | 0.957 | 1 | 3782 | tags=37%, list=37%, |
| 1197 | | REACTOME_HSP90_CHAPERONE_CYCLE_FOR_STEROID_HORMONE_RECEPTORS_SHR_IN_THE_PRESENCE_OF_LIGAND | | 38 | -0.22 | -0.7 | 0.9 | 0.963 | 1 | 3625 | tags=42%, list=36%, signal=65% |
| 1147 | | REACTOME_MITOTIC_G2_G2_M_PHASES | | 139 | -0.21 | -0.77 | 0.901 | 0.926 | 1 | 2710 | tags=27%, list=27%, |
| 1208 | | BIOCARTA_DEATH_PATHWAY | | 23 | -0.24 | -0.68 | 0.901 | 0.972 | 1 | 3180 | tags=35%, list=31%, |
| 1176 | | BOYLAN_MULTIPLE_MYELOMA_C_D_UP | | 93 | -0.21 | -0.74 | 0.902 | 0.941 | 1 | 545 | tags=6%, list=5%, signal=7% |
| 1204 | | BREDEMEYER_RAG_SIGNALING_VIA_ATM_NOT_VIA_NFKB_DN | | 34 | -0.22 | -0.69 | 0.902 | 0.965 | 1 | 3433 | tags=35%, list=34%, |
| 1209 | | BIOCARTA_NFAT_PATHWAY | | 31 | -0.22 | -0.68 | 0.902 | 0.972 | 1 | 478 | tags=10%, list=5%, signal=10% |
| 1198 | | WP_ONE_CARBON_METABOLISM_AND_RELATED_PATHWAYS | | 42 | -0.22 | -0.7 | 0.904 | 0.963 | 1 | 4236 | tags=40%, list=42%, |
| 1215 | | MORI_PLASMA_CELL_DN | | 21 | -0.24 | -0.67 | 0.904 | 0.973 | 1 | 5320 | tags=67%, list=53%, |
| 1221 | | BIOCARTA_MITOCHONDRIA_PATHWAY | | 16 | -0.26 | -0.66 | 0.904 | 0.974 | 1 | 4977 | tags=69%, list=49%, |
| 1230 | | BRACHAT_RESPONSE_TO_CISPLATIN | | 20 | -0.24 | -0.65 | 0.909 | 0.975 | 1 | 3222 | tags=35%, list=32%, |
| 1213 | | WP_GLYCOGEN_METABOLISM | | 26 | -0.23 | -0.67 | 0.91 | 0.973 | 1 | 3038 | tags=35%, list=30%, |
| 1228 | | PLASARI_TGFB1_TARGETS_1HR_UP | | 15 | -0.25 | -0.65 | 0.91 | 0.973 | 1 | 119 | tags=7%, list=1%, signal=7% |
| 1222 | | REACTOME_ACTIVATED_TAK1_MEDIATES_P38_MAPK_ACTIVATION | | 19 | -0.25 | -0.66 | 0.911 | 0.973 | 1 | 2689 | tags=26%, list=27%, |
| 1212 | | REACTOME_PYRUVATE_METABOLISM_AND_CITRIC_ACID_TCA_CYCLE | | 39 | -0.21 | -0.67 | 0.912 | 0.973 | 1 | 4605 | tags=44%, list=46%, |
| 1226 | | REACTOME_TERMINATION_OF_TRANSLESION_DNA_SYNTHESIS | | 24 | -0.23 | -0.66 | 0.913 | 0.972 | 1 | 4049 | tags=38%, list=40%, |
| 1235 | | BIOCARTA_TNFR1_PATHWAY | | 24 | -0.22 | -0.64 | 0.913 | 0.979 | 1 | 2660 | tags=25%, list=26%, |
| 1229 | | SIMBULAN_PARP1_TARGETS_UP | | 22 | -0.23 | -0.65 | 0.915 | 0.976 | 1 | 1291 | tags=23%, list=13%, |
| 1218 | | REACTOME_ENDOSOMAL_SORTING_COMPLEX_REQUIRED_FOR_TRANSPORT_ES | | 28 | -0.23 | -0.67 | 0.917 | 0.972 | 1 | 4376 | tags=43%, list=43%, |
| 1240 | | WP_SELENIUM_MICRONUTRIENT_NETWORK | | 15 | -0.25 | -0.63 | 0.919 | 0.98 | 1 | 4256 | tags=47%, list=42%, |
| 1201 | | BILANGES_SERUM_AND_RAPAMYCIN_SENSITIVE_GENES | | 61 | -0.21 | -0.69 | 0.922 | 0.963 | 1 | 5601 | tags=52%, list=55%, |
| 1223 | | WP_ID_SIGNALING_PATHWAY | | 32 | -0.22 | -0.66 | 0.922 | 0.972 | 1 | 2894 | tags=31%, list=29%, |
| 1234 | | REACTOME_PYROPTOSIS | | 15 | -0.25 | -0.64 | 0.922 | 0.978 | 1 | 3253 | tags=40%, list=32%, |
| 1227 | | LANDIS_ERBB2_BREAST_TUMORS_65_DN | | 29 | -0.22 | -0.65 | 0.923 | 0.973 | 1 | 1859 | tags=17%, list=18%, |
| 1231 | | BIOCARTA_RACCYCD_PATHWAY | | 21 | -0.23 | -0.64 | 0.923 | 0.977 | 1 | 471 | tags=10%, list=5%, signal=10% |
| 1238 | | REACTOME_JNK_C_JUN_KINASES_PHOSPHORYLATION_AND_ACTIVATION_MEDIATED_BY_ACTIVATED_HUMAN_TAK1 | | 19 | -0.23 | -0.63 | 0.924 | 0.979 | 1 | 2102 | tags=21%, list=21%, signal=27% |
| 1241 | | REACTOME_PCNA_DEPENDENT_LONG_PATCH_BASE_EXCISION_REPAIR | | 16 | -0.24 | -0.63 | 0.924 | 0.979 | 1 | 3898 | tags=38%, list=39%, |
| 1243 | | WP_SIGNAL_TRANSDUCTION_OF_S1P_RECEPTOR | | 16 | -0.24 | -0.62 | 0.929 | 0.98 | 1 | 1771 | tags=19%, list=18%, |
| 1237 | | REACTOME_NUCLEOTIDE_CATABOLISM | | 26 | -0.22 | -0.63 | 0.93 | 0.979 | 1 | 826 | tags=8%, list=8%, signal=8% |
| 1232 | | NEMETH_INFLAMMATORY_RESPONSE_LPS_DN | | 20 | -0.23 | -0.64 | 0.934 | 0.977 | 1 | 1384 | tags=15%, list=14%, |
| 1162 | | MEISSNER_NPC_HCP_WITH_H3K4ME2 | | 156 | -0.2 | -0.75 | 0.935 | 0.94 | 1 | 2084 | tags=21%, list=21%, |
| 1103 | | BERENJENO_TRANSFORMED_BY_RHOA_UP | | 400 | -0.21 | -0.81 | 0.94 | 0.905 | 1 | 2106 | tags=20%, list=21%, |
| 1126 | | GOBERT_OLIGODENDROCYTE_DIFFERENTIATION_UP | | 323 | -0.21 | -0.79 | 0.94 | 0.912 | 1 | 2421 | tags=23%, list=24%, |
| 1219 | | REACTOME_DUAL_INCISION_IN_TC_NER | | 56 | -0.2 | -0.66 | 0.94 | 0.973 | 1 | 3918 | tags=38%, list=39%, |
| 1245 | | HEVNER_INTERMEDIATE_ZONE_AND_UP_POSTMITOTIC_PROJECTION_NEURONS | | 28 | -0.21 | -0.61 | 0.942 | 0.985 | 1 | 4007 | tags=32%, list=40%, |
| 1248 | | HEVNER_CORTEX_APICAL_AND_BASAL_INTERMEDIATE_PROGENITOR_CELLS | | 30 | -0.2 | -0.6 | 0.944 | 0.985 | 1 | 3899 | tags=37%, list=39%, |
| 1220 | | WP_WNT_SIGNALING_PATHWAY_AND_PLURIPOTENCY | | 56 | -0.2 | -0.66 | 0.945 | 0.974 | 1 | 4112 | tags=41%, list=41%, |
| 1249 | | REACTOME_SIGNALING_BY_RETINOIC_ACID | | 28 | -0.21 | -0.6 | 0.945 | 0.985 | 1 | 3514 | tags=32%, list=35%, |
| 1233 | | REACTOME_RHOQ_GTPASE_CYCLE | | 42 | -0.2 | -0.64 | 0.946 | 0.976 | 1 | 2728 | tags=31%, list=27%, |
| 1252 | | REACTOME_LAGGING_STRAND_SYNTHESIS | | 17 | -0.22 | -0.59 | 0.949 | 0.987 | 1 | 3262 | tags=35%, list=32%, |
| 1251 | | WP_WNT_SIGNALING_IN_KIDNEY_DISEASE | | 22 | -0.22 | -0.6 | 0.953 | 0.984 | 1 | 4029 | tags=41%, list=40%, |
| 1167 | | REACTOME_TRANSLATION | | 216 | -0.2 | -0.74 | 0.956 | 0.941 | 1 | 4392 | tags=38%, list=43%, |
| 1225 | | REACTOME_CELLULAR_RESPONSE_TO_HEAT_STRESS | | 71 | -0.19 | -0.66 | 0.959 | 0.972 | 1 | 5236 | tags=56%, list=52%, |
| 1174 | | LIM_MAMMARY_STEM_CELL_UP | | 221 | -0.2 | -0.74 | 0.961 | 0.942 | 1 | 1980 | tags=19%, list=20%, |
| 1255 | | YAO_TEMPORAL_RESPONSE_TO_PROGESTERONE_CLUSTER_15 | | 19 | -0.22 | -0.58 | 0.964 | 0.989 | 1 | 1980 | tags=21%, list=20%, |
| 1202 | | PAL_PRMT5_TARGETS_UP | | 163 | -0.19 | -0.69 | 0.968 | 0.963 | 1 | 3703 | tags=34%, list=37%, |
| 1250 | | REACTOME_FORMATION_OF_THE_EARLY_ELONGATION_COMPLEX | | 29 | -0.2 | -0.6 | 0.968 | 0.985 | 1 | 4385 | tags=41%, list=43%, |
| 1257 | | KOHOUTEK_CCNT2_TARGETS | | 27 | -0.2 | -0.57 | 0.97 | 0.989 | 1 | 3326 | tags=33%, list=33%, |
| 1253 | | VARELA_ZMPSTE24_TARGETS_DN | | 45 | -0.19 | -0.59 | 0.971 | 0.988 | 1 | 2452 | tags=22%, list=24%, |
| 1259 | | SANSOM_WNT_PATHWAY_REQUIRE_MYC | | 42 | -0.18 | -0.56 | 0.979 | 0.992 | 1 | 3732 | tags=38%, list=37%, |
| 1256 | | REACTOME_FORMATION_OF_INCISION_COMPLEX_IN_GG_NER | | 38 | -0.18 | -0.58 | 0.98 | 0.989 | 1 | 2091 | tags=16%, list=21%, |
| 1269 | | MADAN_DPPA4_TARGETS | | 17 | -0.19 | -0.49 | 0.982 | 0.997 | 1 | 260 | tags=6%, list=3%, signal=6% |
| 1254 | | REACTOME_REGULATION_OF_HSF1_MEDIATED_HEAT_SHOCK_RESPONSE | | 56 | -0.18 | -0.58 | 0.983 | 0.99 | 1 | 5236 | tags=55%, list=52%, |
| 1247 | | REACTOME_GLYCOSAMINOGLYCAN_METABOLISM | | 63 | -0.18 | -0.61 | 0.984 | 0.984 | 1 | 4023 | tags=38%, list=40%, |
| 1246 | | BURTON_ADIPOGENESIS_5 | | 107 | -0.17 | -0.61 | 0.986 | 0.984 | 1 | 3271 | tags=31%, list=32%, |
| 1236 | | WAKABAYASHI_ADIPOGENESIS_PPARG_RXRA_BOUND_36HR | | 116 | -0.18 | -0.63 | 0.989 | 0.978 | 1 | 2222 | tags=19%, list=22%, |
| 1261 | | LANDIS_ERBB2_BREAST_PRENEOPLASTIC_DN | | 51 | -0.17 | -0.55 | 0.991 | 0.993 | 1 | 2018 | tags=18%, list=20%, |
| 1244 | | MARTORIATI_MDM4_TARGETS_FETAL_LIVER_UP | | 150 | -0.17 | -0.61 | 0.993 | 0.983 | 1 | 2276 | tags=17%, list=23%, |
| 1260 | | MARTORIATI_MDM4_TARGETS_NEUROEPITHELIUM_DN | | 71 | -0.16 | -0.55 | 0.993 | 0.994 | 1 | 2794 | tags=24%, list=28%, |
| 1262 | | GAUSSMANN_MLL_AF4_FUSION_TARGETS_A_DN | | 48 | -0.17 | -0.54 | 0.993 | 0.994 | 1 | 3353 | tags=33%, list=33%, |
| 1264 | | MORI_MATURE_B_LYMPHOCYTE_DN | | 53 | -0.16 | -0.52 | 0.993 | 0.997 | 1 | 4380 | tags=42%, list=43%, |
| 1268 | | LEIN_ASTROCYTE_MARKERS | | 28 | -0.17 | -0.49 | 0.994 | 0.997 | 1 | 1723 | tags=14%, list=17%, |
| 1270 | | REACTOME_HDR_THROUGH_SINGLE_STRAND_ANNEALING_SSA | | 23 | -0.17 | -0.47 | 0.995 | 0.997 | 1 | 4026 | tags=39%, list=40%, |
| 1239 | | IWANAGA_CARCINOGENESIS_BY_KRAS_PTEN_DN | | 217 | -0.17 | -0.63 | 0.996 | 0.98 | 1 | 3570 | tags=34%, list=35%, |
| 1265 | | MIKKELSEN_MEF_HCP_WITH_H3_UNMETHYLATED | | 33 | -0.17 | -0.51 | 0.996 | 0.998 | 1 | 4925 | tags=58%, list=49%, |
| 1258 | | REACTOME_NONSENSE_MEDIATED_DECAY_NMD | | 102 | -0.16 | -0.57 | 0.998 | 0.991 | 1 | 5602 | tags=54%, list=55%, |
| 1266 | | REACTOME_MITOCHONDRIAL_TRANSLATION | | 87 | -0.14 | -0.5 | 0.998 | 0.998 | 1 | 3700 | tags=32%, list=37%, |
| 1210 | | STARK_PREFRONTAL_CORTEX_22Q11_DELETION_DN | | 455 | -0.17 | -0.67 | 0.999 | 0.972 | 1 | 4020 | tags=35%, list=40%, |
| 1217 | | REACTOME_METABOLISM_OF_LIPIDS | | 399 | -0.17 | -0.67 | 0.999 | 0.973 | 1 | 3571 | tags=33%, list=35%, |
| 1263 | | BURTON_ADIPOGENESIS_6 | | 164 | -0.14 | -0.54 | 1 | 0.994 | 1 | 3465 | tags=24%, list=34%, |
| 1271 | | REACTOME_COMPLEX_I_BIOGENESIS | | 55 | -0.13 | -0.44 | 1 | 0.998 | 1 | 5526 | tags=53%, list=55%, |
